# Probabilistic graph-based model uncovers previously unseen druggable vulnerabilities in major solid cancers

**DOI:** 10.1101/2024.06.04.597409

**Authors:** Ying Zhu, Stephanie T. Schmidt, Li Zhao, Chunjie Jiang, Patrizio Di Micco, Costas Mitsopoulos, Andrew Futreal, Bissan Al-Lazikani

## Abstract

Over half cancer patients lack safe, effective, targeted therapies despite abundant molecular profiling data. Statistically recurrent cancer drivers have provided fertile ground for drug discovery where they exist. But in rare, complex, and heterogeneous cancers, strong driver signals are elusive. Moreover, therapeutically exploitable molecular vulnerabilities extend beyond classical drivers. Here we describe a novel, integrative, generalizable graph-based, cooperativity-led Markov chain model, A_3_D_3_a’s MVP (Adaptive AI-Augmented Drug Discovery and Development Molecular Vulnerability Picker), to identify and prioritize key druggable molecular vulnerabilities in cancer. The algorithm exploits cooperativity of weak signals within a cancer molecular network to enhance the signal of true molecular vulnerabilities. We apply A_3_D_3_a’s MVP to 19 solid cancer types and demonstrate that it outperforms standard approaches for target hypothesis generation by >3-fold as benchmarked against cell line genetic perturbation and drug screening data. Importantly, we demonstrate its ability to identify non-driver druggable vulnerabilities and highlight 43 novel or emergent druggable targets for these tumors.

## Introduction

Advances in molecular profiling technologies in the post genomic era have served cancer through the discovery of driver genes that were successfully exploited for the discovery of novel therapeutics^1, 2^. Indeed, oncology has seen the greatest growth in novel, first-in-class drugs since the maturation of the Human Genome Project^3^. Despite these successes, 32% of cancer patients do not achieve 5-year relative survival^4^ and 53% do not achieve 10 years. The successes in molecular-profiling-driven drug discovery have been cases with clear, statistically differentiated, druggable driver genes. Meanwhile, cancers that continue to lack lasting, effective treatments tend to be rare cancers and heterogeneous complex cancers such as those that recur after developing drug resistance^5, 6^. For these 53% of patients, the signal of key molecular vulnerabilities is often drowned by the noise of heterogeneity or small numbers of samples.

Harnessing network biology as a tool to understand disease-development is a well-established field both in oncology and other diseases. We previously investigated graph-based patterns in biological networks and identified distinct graph behaviors predictive of cancer targets^7^; and demonstrated that graph-based environmental perturbations can strongly predict individual patient drug responses in lung^8^. Both studies indicated the potential to use inherent graph-patterns and signal cooperativity to uncover hidden patterns. Others have shown success when utilizing transcriptional networks and identifying ‘Master Regulators’ from transcriptomic and genomic data combined within a network-based model^9^. Recently, Geneformer^10^, a transfer learning model developed from single-cell transcriptome data, was applied to perform disease modeling of cardiomyopathy. While hugely informative to understand the biology, these methods rely on strong recurrent signals of disease genes in specific cancers and thus will miss weaker signals in heterogeneous, complex, and rare cancers.

A key misconception is that that cancer drug targets must be disease drivers. Almost half cancer drugs (e.g., PARP and aromatase inhibitors) target non-driver proteins^3^ that influence or cooperate with driver pathways. Therefore, to maximally impact cancers of unmet need we need to consider all druggable, biologically impactful molecular vulnerabilities – drivers, or non-drivers. Further, to accelerate drug discovery in these areas of need, we must prioritize those impactful vulnerabilities that are also technically tractable^11^. We previously combined pharmacological annotation and machine-learning-based druggability assessment to objectively prioritize cancer genes for drug discovery^11, 12^.

To date, there is no generalizable, extensible mathematical framework that is able to integrate noisy, multimodal *<u>patient</u>* data to identify the most likely druggable molecular vulnerabilities regardless of their mutation or expression signal. Yet, such a method is urgently needed to accelerate the next generation of targeted therapeutics to address the unmet needs of the >50% cancer patients with rare, complex, and heterogeneous disease. This method must apply to noisy patient data rather than derive from idealized and largely artificial cell line data.

Here we describe **A_3_D_3_a’s MVP** (**M**olecular **V**ulnerability **P**icker), a novel, integrative, graph-based cooperativity-led Markov chain model to exploit cooperativity of weak signals across a molecular interaction network to highlight and prioritize molecular vulnerabilities regardless of their mutation frequencies or driver status. We further integrate it with AI-enabled pharmacological and druggability annotation from canSAR.ai^13^. We apply A_3_D_3_a’s MVP to major solid cancers and demonstrate its capability to identify genuine molecular vulnerabilities beyond obvious known targets. We make A_3_D_3_a’s MVP publicly available to the research community.

## Results

### A_3_D_3_a’s MVP model architecture and optimization

A_3_D_3_a’s MVP applies a cooperativity-led Markov Chain model (MCm) to highlight regions of empirically derived protein-interaction networks most important for signal transmission. We hypothesize that our approach increases the signal-to-noise ratio of actual druggable molecular vulnerabilities relevant to specified cancer contexts and can identify vulnerabilities regardless of their alteration status. We assume: 1) each node fully transmits its signal to itself or its neighbors; 2) cooperativity remains within each node’s first-degree neighbors. Though gross assumptions, they are a useful start for the model. The goal is to rank nodes by signal accumulation and enable prioritization of vulnerabilities for experimental validation towards drug discovery.

First, we analyzed DNA and RNA data from 15 TCGA solid tumor studies (Supplementary Table 1) to identify “seed” genes whose corresponding proteins were used to initiate networks for each cancer. In this initial step, we applied a more permissive recurrence threshold for inclusion of genes than is typical bioinformatics approaches. This is to capture any genes that may carry a weak, but biologically important signal (Online Methods and Supplementary Table 2). The DNA seed genes were selected from mutation, copy number variation, and structural variation events as described (Online Methods). Tumor and normal RNA expression data were additionally available for 11 of 15 cancers. For these cancers, we added differentially expressed genes that are strongly predictive of tumor versus normal phenotypes using a support vector machine classifier (Online Methods).

Using these seeds, we query a background human interactome of 471,200 interactions between 18,614 proteins created using curated, experimentally determined protein interactions from canSAR^13^, KEGG^14^, and HI-union^15^. This returns fragmented networks connecting the seeds. Next, we connected the fragments into larger networks by incorporating non-seed proteins from the background interactome that were most selectively connected with the seeds (Fig. 1A, Online Methods). The resulting networks consist of 636.0±4.7 proteins (DNA data alone) and 653.8±0.3 proteins (combined DNA and RNA) across the 15 cancers. These networks form the starting graph for the mathematical model.

**Figure 1.**
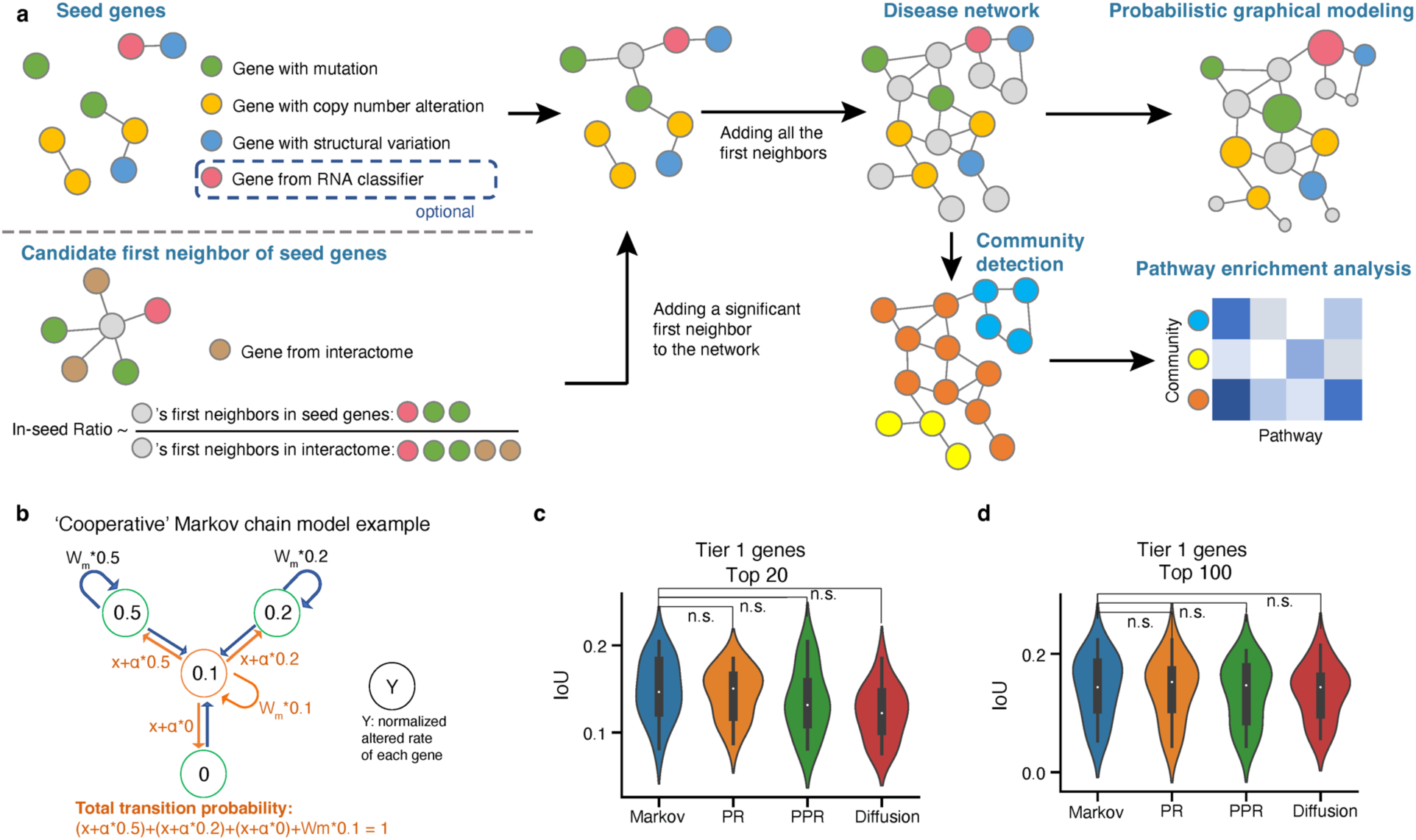
Schematics of model architecture and model selection. **a.** Schematics of probabilistic graphical modeling of the disease network. **b.** Illustration of the transition biased Markov chain model that is employed by A3D3a’s MVP. **c-d.** Intersection over union (IoU) of the top 20 genes (c) or top 100 genes (d) ranked by model (A3D3a’s MVP, PR, PPR or raw diffusion) and Tier 1 genes of every cancer type. 15 cancer types were included in the analysis. Comparison between optimized models of A3D3a’s MVP, PR, PPR and raw diffusion by one-sided Rank-sum test, p-values were adjusted by Benjamini/Hochberg method (n.s.: p_adj>=0.05).

We use frequency of genomic and/or expression alteration, f_alteration_, as each node’s initial state. We encode f_alteration_ into the transition probability of each node’s self-loop and edges to adjacent nodes (Fig. 1B). Nodes are ranked by their final score at steady state, representing their likelihood of being a vulnerability (Online Methods). We optimized three parameters – W_m_ (weighting of nodes’ self-loop), α (cooperativity factor), and N (number of top-ranked non-seed genes) – by maximizing the average rank of the intersection over union of the top 20 genes predicted in each indication and the Cancer Gene Census’s^16^ Tier 1 genes filtered by hotspot mutations^17, 18^ (Fig. S1, Online Methods).

We evaluated MCm, PageRank^19^, Personalized PageRank^20^, and Raw Diffusion^21, 22^ algorithms for use in A_3_D_3_a’s MVP; we observed similar performance across models from DNA data (Fig.1C-D, Fig. S1) and DNA and RNA data (Fig. S2). Since study bias is inherent to empirical protein-interaction data^7^; well-studied proteins are more likely to have known interaction partners. To mitigate this, we normalize against background interaction rates and implement the optimized model least susceptible to study bias, which was a MCm (Fig. S2, S3). Across indications, A_3_D_3_a’s MVP provides a different prioritization of genes than f_alteration_. We find modest overlap (24%±13% genes shared) between the top 20 genes ranked by f_alteration_ and A_3_D_3_a’s MVP (Fig. 2A). Moreover, the top 20 genes returned by A_3_D_3_a’s MVP contain more Tier 1 genes (p=1.5×10^-5^, rank-sum test) than those by f_alteration_ (Fig. 2B).

**Figure 2.**
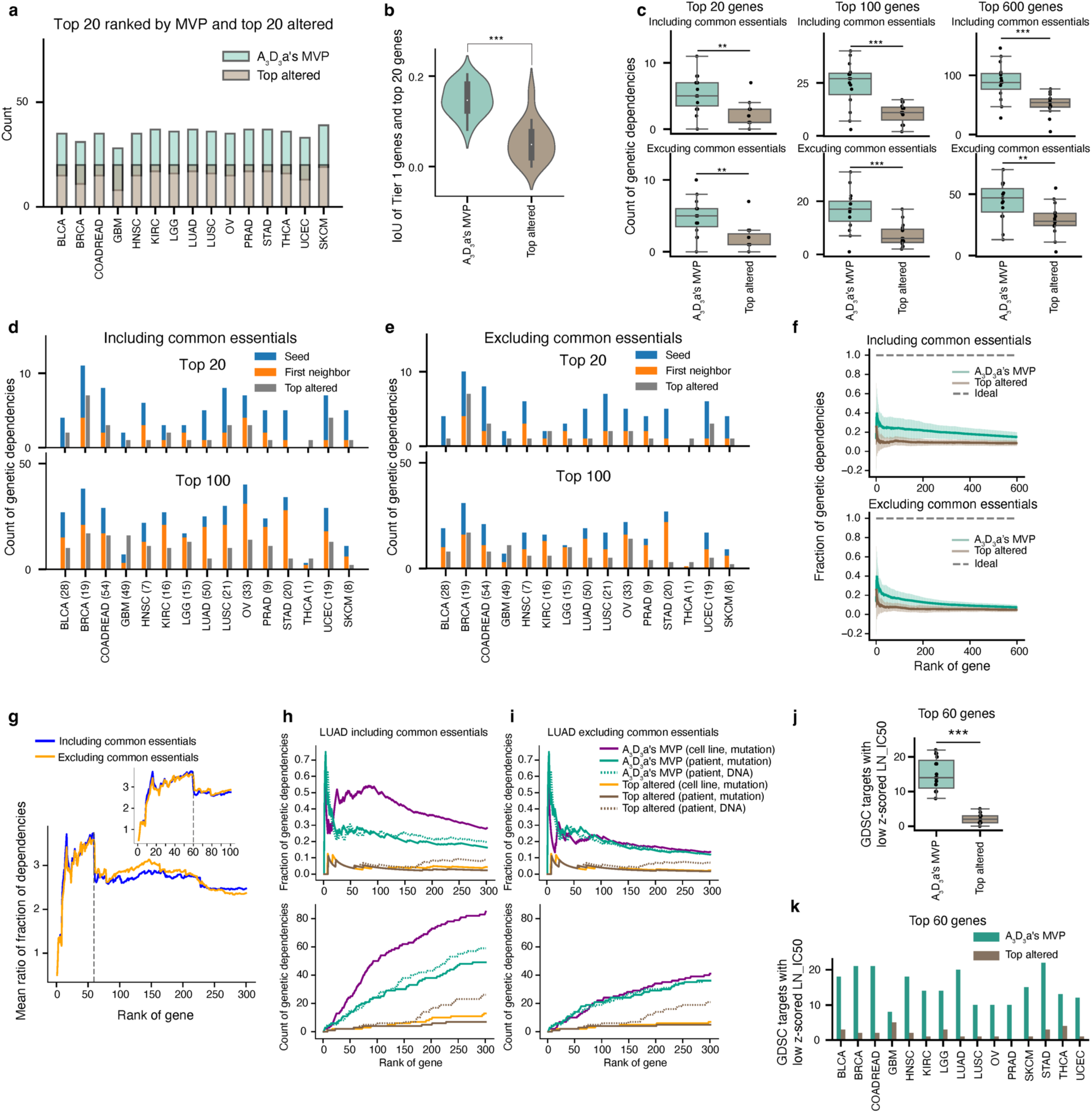
Model validation by Depmap genetic dependencies and GDSC targets. **a.** The intersection of top 20 ranked genes by A3D3a’s MVP and top 20 altered genes for each cancer type. **b.** Intersection over union (IoU) of Tier 1 genes and top 20 ranked genes by A3D3a’s MVP or top 20 altered genes. (n = 15 cancer types). **c.** Comparison of count of genetic dependencies (including common essentials) pooled from all Depmap cell lines of each cancer type among top 20/100/600 ranked genes by A3D3a’s MVP and top altered genes (top panel). (Bottom panel) the genetic dependencies did not include the common essentials). (n = 15 cancer types). **d.** Count of genetic dependencies (including common essentials) pooled from all Depmap cell lines of each cancer type among top 20 ranked genes by A3D3a’s MVP and top 20 altered genes (top panel). The same analysis was performed for top 100 ranked or altered genes (bottom panel). For the top ranked genes, the proportion of seed genes and first neighbors were shown. **e.** The same analysis as in d. except that the genetic dependencies did not include common essentials. **f**. Mean and standard deviation of fraction of genetic dependencies among top ranked genes by A3D3a’s MVP and top altered genes across 15 cancer types. (Top) genetic dependencies include common essentials. (Bottom) genetic dependencies do not include common essentials. **g.** Mean ratio of fraction of genetic dependencies between top ranked genes by A3D3a’s MVP and top altered genes. Gray dashed line indicates the sharp decline. Inset is a zoom-in view of the same figure. **h.** Fraction (top) and count (bottom) of genetic dependencies among top ranked genes by A3D3a’s MVP and top altered genes in LUAD. Purple solid line: model constructed using seed from mutation data of cell lines. Green solid line: model constructed using seed from mutation data of patients. Green dotted line: model constructed using seed genes from DNA data (mutation, CNV, SV) of patients. Top altered genes from mutation data of cell lines and patients are plotted in orange and brown solid lines, respectively. Top altered genes from DNA data of patients are plotted in brown dotted line. We find that networks from cell line profiling significantly outperform networks from patient data as shown by the mean and standard deviation of fraction recovered: 0.40 ± 0.09 for A3D3a’s MVP applied to cell line data, 0.23 ± 0.07 for A3D3a’s MVP applied to patient mutation data, and 0.25 ± 0.07 for A3D3a’s MVP applied to patient DNA data across the top 300 ranked genes. **i.** The same plots as in j. except that the genetic dependencies do not include common essentials. When excluding common essentials, A3D3a’s MVP maintains its enhanced performance against frequency-based prioritization over the top 300 ranked genes (p = 8.9 × 10^-51^ Wilcoxon sign-rank test), but no additional value is gained from generating the disease networks from cell line profiling data. **j.** Group comparison of genes with low z-score of LN_IC50 (<-1.5) among top 60 ranked genes by A3D3a’s MVP and top 60 altered genes. **k.** Comparisons of genes with low z-score of LN_IC50 (<-1.5) among top 60 ranked genes by A3D3a’s MVP and top 60 altered genes for each cancer type. **a-k.** seed genes from DNA data.

### Validation of A_3_D_3_a’s MVP in cancer cell lines

To systematically evaluate A_3_D_3_a’s MVP performance in prioritizing molecular vulnerabilities, we use large-scale, experimental data from DepMap^23^ and the Genomics of Drug Sensitivity in Cancer (GDSC)^24^. These resources are the most objective, high-throughput datasets of cell-line-validated dependencies and are invaluable for benchmarking against standard approaches. For both DepMap and GDSC, we pooled cell lines by broad histological indication to ensure sufficient numbers for analysis. It is important to note that cell line models and large-scale growth inhibition assays have limitations and cannot fully capture the complexity of patient tumors (Supplementary Information). Accordingly, A_3_D_3_a’s MVP is neither trained nor optimized using cell line dependencies. We merely use them as a standardized experimental validation baseline for assessing performance.

We examined the top 20, 100, and 600 genes returned by A_3_D_3_a’s MVP to assess how many affected 2D cell growth. Using DepMap, we defined genetic dependencies as those with Chronos scores^25^ < -1 in cell lines from the appropriate histological type. A_3_D_3_a’s MVP identified both cancer-specific and “common essential” genes (CEs: widely observed dependencies^26^). Many CEs are targets of FDA-approved drugs and/or are cancer drivers under pharmaceutical development, e.g., MTOR^27^, CHEK1^28^, and ATR^29^. Hence, we included these CEs and report results with and without them (Fig. S4). Using DNA data, we found that A_3_D_3_a’s MVP identified more dependencies in the top 20, 100, and 600 genes whether including (p = 1.3×10^-3^, 9.1×10^-4^, and 6.7×10^-4^, rank-sum test) or excluding CEs (p = 1.4×10^-3^, 9.1×10^-4^, and 7.9×10^-3^, rank-sum test) (Figure 2C, Table 1). This trend held when RNA data was added (Fig. S5).

**Table 1.** Depmap validation of top 20 ranked genes by A3D3a’s MVP (DNA as seed) and top 20 altered genes from DNA data.

| Tumor type | Method | Count of genetic dependencies | Percent of genetic dependencies (%) | Genetic dependencies | Common essentials |
| --- | --- | --- | --- | --- | --- |
| BLCA | A<sub>3</sub>D<sub>3</sub>a's MVP | 4 | 20 | CREBBP, E2F3, EP300, PIK3CA |  |
|  | Top altered | 2 | 10 | PFDN2, PIK3CA | PFDN2 |
| BRCA | A<sub>3</sub>D<sub>3</sub>a's MVP | 11 | 55 | AKT1, BRCA1, CCND1, EGFR, EP300, ERBB2, ESR1, MYC, PIK3CA, RAD21, UBC | BRCA1 |
|  | Top altered | 7 | 35 | CCND1, EIF3H, FADD, MYC, PIK3CA, RAD21, UTP23 |  |
| COADREAD | A<sub>3</sub>D<sub>3</sub>a's MVP | 8 | 40 | APC, CTNNB1, ELAVL1, KRAS, MAPK1, PIK3CA, SMAD4, TCF7L2 |  |
|  | Top altered | 3 | 15 | APC, KRAS, PIK3CA |  |
| GBM | A<sub>3</sub>D<sub>3</sub>a's MVP | 2 | 10 | PDGFRA, PIK3CA |  |
|  | Top altered | 1 | 5 | SEC61G |  |
| HNSC | A<sub>3</sub>D<sub>3</sub>a's MVP | 6 | 30 | CCND1, CTNNB1, EGFR, HUWE1, MYC, TP63 | BRCA1 |
|  | Top altered | 1 | 5 | CCND1 |  |
| KIRC | A<sub>3</sub>D<sub>3</sub>a's MVP | 3 | 15 | ACTB, MTOR, VHL | MTOR |
|  | Top altered | 2 | 10 | DDX41, VHL |  |
| LGG | A<sub>3</sub>D<sub>3</sub>a's MVP | 3 | 15 | MYC, PTPN11, RELA |  |
|  | Top altered | 1 | 5 | SEC61G | ATRX |
| LUAD | A<sub>3</sub>D<sub>3</sub>a's MVP | 5 | 25 | CTNNB1, EGFR, KRAS, MYC, PIK3CA |  |
|  | Top altered | 1 | 5 | KRAS |  |
| LUSC | A<sub>3</sub>D<sub>3</sub>a's MVP | 8 | 40 | ACTL6A, AP2M1, CREBBP, CUL3, EGFR, PIK3CA, SOX2, UBC | ACTL6A |
|  | Top altered | 3 | 15 | AP2M1, EIF2B5, SOX2 | EIF2B5 |
| OV | A<sub>3</sub>D<sub>3</sub>a's MVP | 7 | 35 | CDK1, CDK2, EGFR, MYC, PCNA, PTK2, PUF60 | BRCA1, CDK1, PCNA |
|  | Top altered | 3 | 15 | EFR3A, MYC, PTK2 |  |
| PRAD | A<sub>3</sub>D<sub>3</sub>a's MVP | 5 | 25 | EP300, ERG, FOXA1, MYC, SNW1 | HSPA8, SNW1 |
|  | Top altered | 1 | 5 | ERG |  |
| STAD | A<sub>3</sub>D<sub>3</sub>a's MVP | 5 | 25 | CTNNB1, EGFR, KRAS, MYC, PIK3CA |  |
|  | Top altered | 0 | 0 |  |  |
| THCA | A<sub>3</sub>D<sub>3</sub>a's MVP | 0 | 0 |  |  |
|  | Top altered | 1 | 5 | EIF1AX | EHMT1 |
| UCEC | A<sub>3</sub>D<sub>3</sub>a's MVP | 7 | 35 | CHD4, FGFR2, KRAS, MYC, PIK3CA, PPP2R1A, UBC | PPP2R1A |
|  | Top altered | 4 | 20 | CHD4, CTCF, KMT2D, PIK3CA | CTCF |
| SKCM | A<sub>3</sub>D<sub>3</sub>a's MVP | 5 | 25 | BRAF, CCND1, MAPK1, NRAS, RAC1 | GRB2, RAC1 |
|  | Top altered | 1 | 5 | BRAF |  |

Since A_3_D_3_a’s MVP identifies vulnerabilities regardless of alteration status, we determined the number of top-ranked genes that were non-seeds, i.e., those highlighted by A_3_D_3_a’s MVP due to their role in network communication rather than f_alteration_. With DNA data, we find in the top 20 genes that at least one non-seed was a dependency in relevant cell lines for 12 of 15 indications (range: 1-4, average: 2±1.1 including CEs; range: 1-4, average: 1.8±0.9 excluding CEs) (Fig. 2D-E). In the top 100 genes, we find non-seeds were dependencies across all indications (range: 2-31, mean: 16.7±8). When including RNA data, we observed marginal impact by these additional seeds (4 out of 11 with improvement, Fig. S5C).

Next, we assessed the fraction of genes that were dependencies at each rank. A_3_D_3_a’s MVP consistently returns a greater fraction of dependencies (R_mvp_) than f_alteration_ (R_f_) (p=0, rank-sum test, Fig. 2F), which was consistent across seed types (Fig. S5-8). We then plotted the ratio R_mvp_/R_f_ against rank (Fig. 2G). For the first 8 genes by rank, we found R_mvp_/R_f_ was less than 1.5, indicating that A_3_D_3_a’s MVP offers little additional value where genes are frequently mutated. The benefits of our cooperative model become evident after this point: R_mvp_/R_f_ remains around or above 3 until rank point ∼60 when it drops again to below 3-fold. Thus, the greatest value of A_3_D_3_a’s MVP is in its top 60 genes (Fig. S9).

Cancer cell lines are inherently limited in representing real tumors. Thus, it is difficult to assess how many vulnerabilities predicted by A_3_D_3_a’s MVP are algorithm false positives versus genuine vulnerabilities not represented by the 2D cancer cell line assays. To approximate this, we tested the performance baseline if molecular profiling data input into A_3_D_3_a’s MVP came directly from the experimental model, thus reducing (but not fully eliminating) the impact of the 2D experimental system’s limitations. We selected lung adenocarcinoma (LUAD) due to the large number of cell lines (50) available and their relative consistency with patient data^30^. We constructed graphs using LUAD cell line molecular profiling data (MVP_cells_) and compared the dependency retrieval metrics to the graphs constructed using patient data above (MVP_patients_). The area under the curve (AUC) between the MVP_cells_ and the MVP_patients_, is 99% of the AUC between the MVP_patients_ and the patient baseline (alteration curve; Fig. 2H lower panel). Moreover, while MVP_cells_ returned nearly double the dependencies as MVP_patients_ with CEs but roughly the same without them (Fig. 2H-2I). This is consistent with CEs being important for 2D cell growth^31, 32^. These results demonstrate both that nearly half the signal is lost due to differences between the cancer cell lines and patient tumors (Supplementary Information).

Pharmacological perturbation provides an orthogonal view that overcomes some limitations of genetic knock-out (Online Methods), so we further assessed the performance of A_3_D_3_a’s MVP in identifying true drug targets using the GDSC^24^. For each gene in the top 20 and top 100 genes by A_3_D_3_a’s MVP or f_alteration_, we used GDSC-provided mappings to identify drugs targeting its corresponding protein. We excluded promiscuous drugs with > 4 targets and consider a gene a ‘hit’ if at least one of its drugs had z-scored ln(IC50 values) < -1.5 in relevant cancer cell lines (Supplementary Table 3).

As with genetic dependencies, we find that A_3_D_3_a’s MVP prioritized more targets whose drugs were active in relevant cell lines than f_alteration_ (p=3.4×10^-6^ for top 20 genes; 3.1×10^-6^ for top 100 genes, rank-sum test), both in individual indications and in aggregate (Fig. 2J-K, S10, Table 2). The results remained consistent when networks were constructed from combined DNA and RNA data (Fig. S11, Supplementary Table 4).

**Table 2.** GDSC validation of top 20 ranked genes by A3D3a’s MVP (DNA as seed) and top 20 altered genes from DNA data.

| Tumor type | Method | GDSC target count | GDSC target count with low z-scored LN IC50 | Percent of top 20 genes with low z-scored LN IC50 (%) | GDSC targets among top 20 with low z-scored LN IC50 |
| --- | --- | --- | --- | --- | --- |
| BLCA | A3D3a's MVP | 9 | 9 | 45 | ATM,CREBBP,EP300,ERBB2,ERBB3,ESR1,FGFR3,PIK3CA,TP53 |
|  | Top altered | 2 | 2 | 10 | PIK3CA,TP53 |
| BRCA | A3D3a's MVP | 7 | 7 | 35 | AKT1,EGFR,EP300,ERBB2,ESR1,PIK3CA,TP53 |
|  | Top altered | 2 | 2 | 10 | PIK3CA,TP53 |
| COADREAD | A3D3a's MVP | 8 | 6 | 30 | ATM,ESR1,MAPK1,PIK3CA,PIK3CB,SRC |
|  | Top altered | 3 | 1 | 5 | PIK3CA |
| GBM | A3D3a's MVP | 4 | 3 | 15 | EGFR,PDGFRA,PIK3CA |
|  | Top altered | 2 | 1 | 5 | EGFR |
| HNSC | A3D3a's MVP | 7 | 7 | 35 | CREBBP,EGFR,EP300,ESR1,NTRK1,PIK3CA,TP53 |
|  | Top altered | 2 | 2 | 10 | PIK3CA,TP53 |
| KIRC | A3D3a's MVP | 11 | 10 | 50 | ATM,EP300,ESR1,FGFR4,HDAC1,HSP90AA1,IGF1R,MTOR,PIK3CA,SMARCA4 |
|  | Top altered | 0 | 0 | 0 |  |
| LGG | A3D3a's MVP | 9 | 6 | 30 | EGFR,HSF1,HSP90AA1,JAK2,NTRK1,PIK3CA |
|  | Top altered | 4 | 2 | 10 | EGFR,PIK3CA |
| LUAD | A3D3a's MVP | 12 | 10 | 50 | ATM,EGFR,ERBB2,HSP90AA1,KRAS,MAPK1,MET,PIK3CA,SMARCA4,SRC |
|  | Top altered | 2 | 1 | 5 | KRAS |
| LUSC | A3D3a's MVP | 5 | 4 | 20 | CREBBP,EGFR,ESR1,PIK3CA |
|  | Top altered | 1 | 0 | 0 |  |
| OV | A3D3a's MVP | 5 | 4 | 20 | CDK1,CDK2,EGFR,PTK2 |
|  | Top altered | 1 | 1 | 5 | PTK2 |
| PRAD | A3D3a's MVP | 9 | 6 | 30 | AR,ATM,EP300,ESR1,GSK3B,PIK3CA |
|  | Top altered | 1 | 0 | 0 |  |
| SKCM | A3D3a's MVP | 7 | 7 | 35 | BRAF,EGFR,EP300,MAPK1,MAPK3,RAC1,TP53 |
|  | Top altered | 1 | 1 | 5 | BRAF |
| STAD | A3D3a's MVP | 10 | 8 | 40 | ATM,EGFR,ERBB2,ERBB3,ERBB4,ESR1,PIK3CA, SRC |
|  | Top altered | 1 | 0 | 0 |  |
| THCA | A3D3a's MVP | 10 | 8 | 40 | AKT1,AKT2,ATM,BRAF,GSK3B,MAPK1,MAPK3,RET |
|  | Top altered | 5 | 3 | 15 | BRAF,NTRK1,RET |
| UCEC | A3D3a's MVP | 8 | 5 | 25 | EGFR,EP300,ESR1,FGFR2,PIK3CA |
|  | Top altered | 2 | 1 | 5 | PIK3CA |

### A_3_D_3_a’s MVP can rescue clinically relevant genes even if not mutated

The hypothesis for A_3_D_3_a’s MVP cooperativity model is that it should identify disease-important genes even if they had no or little mutational or differential expression patterns. To test this, we identified 36 distinct (54 gene-cancer pairs) Clinically Relevant Genes (CRGs) from oncoKB^33^ (levels 1-3) and combined them with the expert-curated FDA-approved 64 distinct genes (133 gene-cancer pairs) drug targets from cansar.ai^3, 13^. Together, this resulted in 85 distinct (168 gene-cancer pairs) genes that are either efficacy targets or clinical biomarkers for the cancers in this analysis (Supplementary Table 5). First, we examined A_3_D_3_a’s MVP performance from the patient data. On average in the top 60 ranked genes across indications, A_3_D_3_a’s MVP identified 69% of genes of oncoKB level 1 compared to 34% by f_alteration_ and found 70% of genes with FDA approved drugs, clinical evidence, or part of standard of care (oncoKB levels 1-3) compared to 25% by f_alteration_ (Supplementary Table 5). A_3_D_3_a’s MVP recapitulated 19% of pharmacological targets (canSAR.ai) in the top_60_ compared to 3% by f_alteration_. Those not in the top 60 genes by A_3_D_3_a’s MVP included biologically non-specific targets (e.g., HDACs, topoisomerases) and immunotherapeutic targets distal to disease pathways.

To test whether A_3_D_3_a’s MVP’s cooperativity can identify latent clinically relevant molecular vulnerabilities (i.e., those unaltered in patients), we simulated indication-specific networks as if the major CRGs were not mutated (Fig. 3A). For example, in breast cancer, we set f_alteration_ of PIK3CA or ERBB2 to zero. We then constructed graphs, ranking nodes using A_3_D_3_a’s MVP as before to test whether signal cooperativity modeled by A_3_D_3_a’s MVP was sufficient to rescue these CRGs as molecular vulnerabilities (Fig. S12).

**Figure 3.**
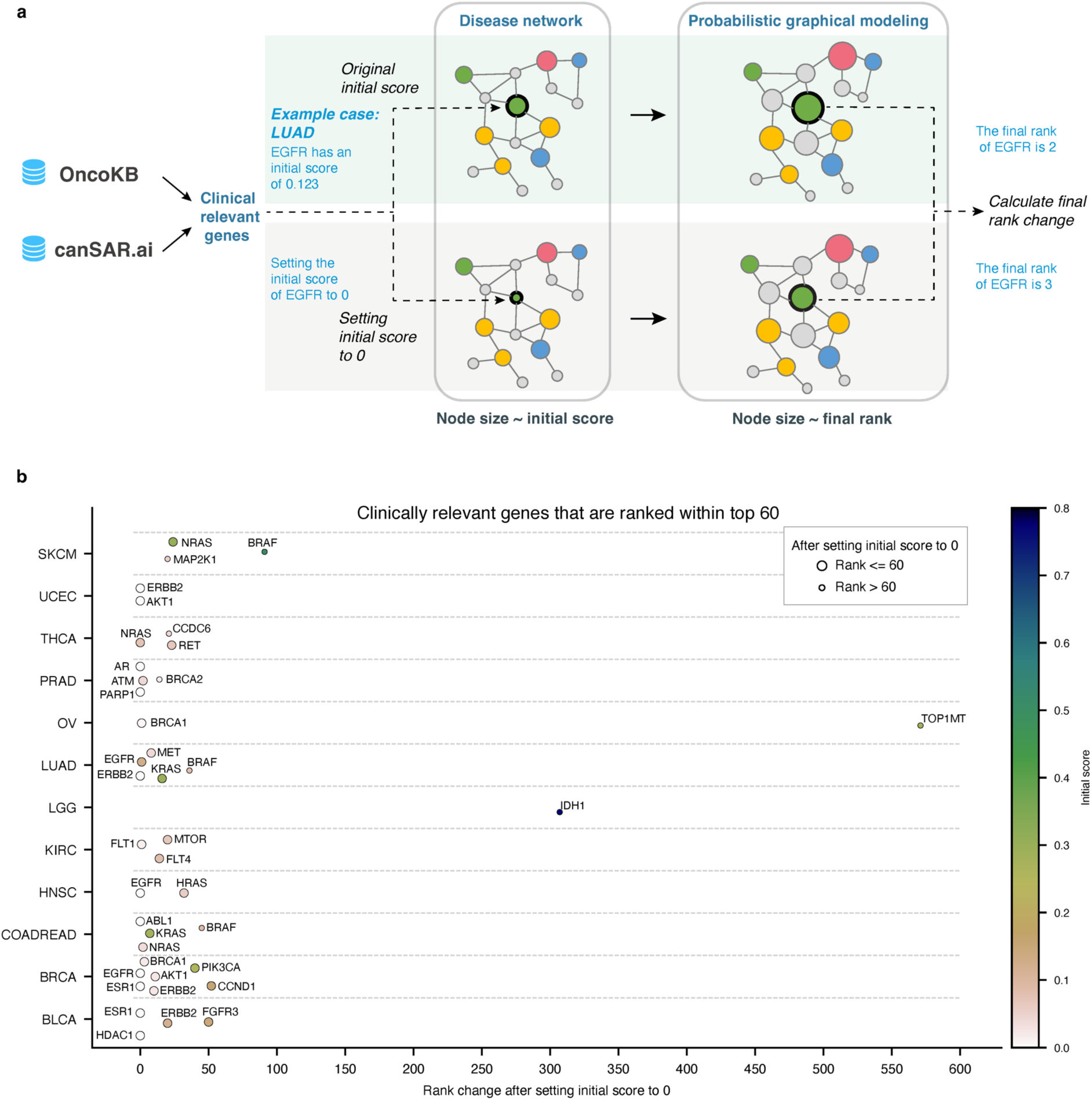
Initial score perturbation and model properties investigation. **a.** Illustration of the workflow of the initial score perturbation by setting the initial score of the clinically relevant genes obtained from OncoKB or canSAR.ai to zero and calculate the rank change predicted by A3D3a’s MVP model. **b.** Rank change after setting the initial score to zero for the clinically relevant genes that were ranked among top 60 by the original A3D3a’s MVP model for each cancer types. Top1mt as a putative target of cytotoxic drugs is included for comparison with other clinically relevant targets.

Of the 39 gene-cancer combinations (26 distinct genes), 32 (21 distinct genes) remain in the top 60 genes (Fig. 3B). A_3_D_3_a’s MVP rescues 82% of gene-cancer pairs (81% of distinct genes). In fact, signal cooperativity modeled by A_3_D_3_a’s MVP was sufficiently strong to maintain the ranking within ≤10 places 21 gene-cancer pairs (14 distinct genes). One notable exception, where the rank changes dramatically from 34 to 341, is IDH1 in Low Grade Glioma (LGG). Upon inspection, IDH1’s high rank was primarily due to its high f_alteration_. When f_alteration_ was set to zero, IDH1 was not rescued by our model. One might speculate that this is due to IDH1 acting indirectly in LGG via its oncometabolite, D-2-Hydroxyglutarate, which then spreads and affects many pathways and biological processes^34^, rather than LGG-specific pathways. Another exception, TOP1MT, although not a CRG, serves as a target for cytotoxic drugs, included here for comparison with CRGs. Full results are provided in Supplementary Table 6 and underscore the strength of A_3_D_3_a’s MVP in highlighting CRGs.

### Case study: A_3_D_3_a’s MVP to common cancer types

We examined the top 60 genes returned by A_3_D_3_a’s MVP for Lung Adenocarcinomas, the most common subtype of non-small cell lung cancer (NSCLC) comprising 55% of all new lung cancer diagnoses^35^. Onto the A_3_D_3_a’s MVP network (Fig. 4A) we map the genes that are genetic dependencies and/or GDSC targets, common essential genes, or have drugs; tool compounds or are ‘druggable’. Networks for all 15 cancers including are in the Supplementary Information (Fig. S13-14) along with summaries of genetic dependencies observed in each (Supplementary Table 7).

**Figure 4.**
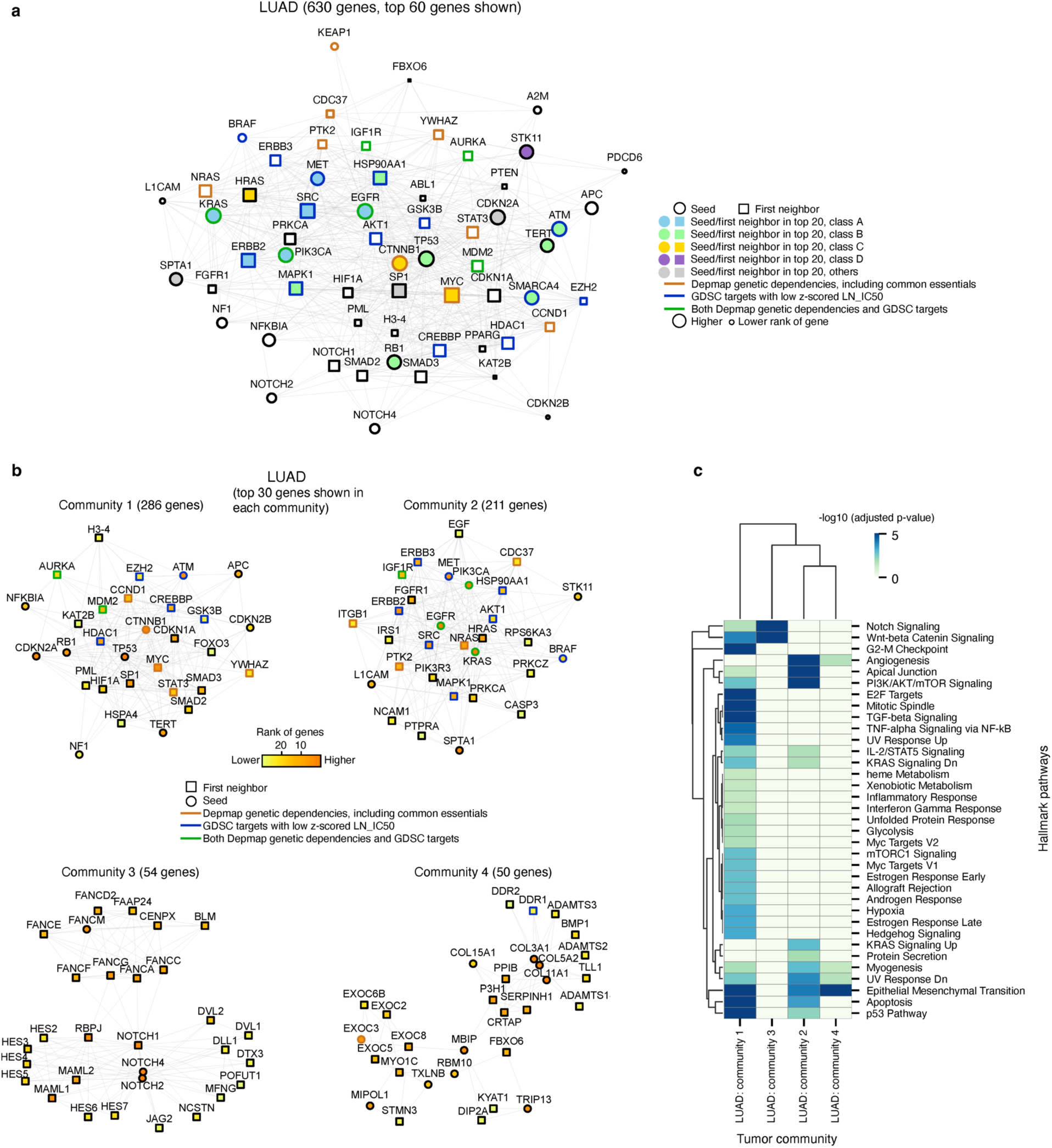
Comparison of model predictions for LUAD. **a.** The largest connected graph of the top 60 ranked genes by A3D3a’s MVP using DNA as seed for LUAD were displayed. Top 20 ranked genes were highlighted in blue (class A), green (class B), yellow (class C), purple (class D), and gray (others) face colors. Depmap genetic dependencies (including common essentials) were labeled in orange edge color. GDSC targets with low z-scored LN_IC50 were labeled in blue edge color. The genes that were both genetic dependency and GDSC targets with low z-scored LN_IC50 were labeled in green edge color. **b.** Top four communities of the LUAD disease network. The largest connected graph of the top 30 ranked genes by A3D3a’s MVP of each community were shown. **c.** Over-representation analysis showed the significantly enriched pathways of each community of LUAD.

Among the top 20 ranked genes, EGFR, KRAS, ERBB2, and MET are targets of FDA-approved drugs for NSCLC. Interestingly, ERBB2 and MYC are not sufficiently altered in the patient data to count as ‘drivers’ or seeds, yet were retrieved and ranked highly by A_3_D_3_a’s MVP. We observed that MYC is a genetic dependency in LUAD cell lines^23^ with MYC copy number gain reported as an independent poor-prognostic factor of patient survival in LUAD^36^. We note that because MYC’s rate of copy number gain was low (8.4% of patients) in the TCGA cohort used, MYC was not a seed gene; despite this, A_3_D_3_a’s MVP brought it into the network as a non-seed gene. Taken together, the presence of approved drug targets, genetic dependencies, and prognostic biomarkers among the genes prioritized by A_3_D_3_a’s MVP underscore the utility of the model developed.

To map the biological processes within the LUAD network, we performed community detection to identify subnetworks (Fig. 4B) and annotated them by the enrichment of cancer hallmark pathways (Fig. 4C). For ease of visualization, we show the largest connected graph of the top 30 genes from A3D3a’s MVP for the top four communities found in LUAD. The largest community (community 1) has 286 genes with associations including G2-M checkpoint, p53 pathway, mitotic spindle, and TGF-beta signaling. The second largest community (community 2) has 211 genes with associations including KRAS signaling UP, angiogenesis, and PI3K/AKT/mTOR signaling (Fig. 4C). Community 3 (54 genes) and community 4 (50 genes) are associated with notch signaling and the epithelial mesenchymal transition, respectively. By applying community detection to our network representations of cancer, we can enable new biological insights anchored in the unique vulnerabilities identified by our cooperativity-led approach.

### Application of A_3_D_3_a’s MVP to difficult-to-treat cancers

The cancers analyzed above all mostly common and relatively well-understood cancers. Moreover, although A_3_D_3_a’s MVP was not ‘trained’ on the above indications, known genes from them were used to help optimize early parameters of the algorithm (Online Method). We examined whether A_3_D_3_a’s MVP is generalizable outside these common cancers and especially whether it can identify molecular vulnerabilities in four difficult-to-treat cancers (Fig. S15, Supplementary Table 8), including pancreatic adenocarcinoma (PAAD) and esophageal adenocarcinoma (ESCA), which have the lowest 5-year relative survival rates of all cancers at 12% and 21%, respectively^4^. We found that across indications, A_3_D_3_a’s MVP returned more genetic dependencies in its top 60 genes compared to f_alteration:_ 10.75±1.92 (average ± standard deviation) versus 6.25±1.92 (p<0.05, rank-sum test) (Fig. 5A-C). A_3_D_3_a’s MVP identified more targets with active drugs in relevant cell lines (10.75±1.09) than f_alteration_ (0.5±0.5) (p<0.05, rank-sum test) (Fig. 5D). Network views of the top 60 ranked genes in ESCA and PAAD are shown (Fig. 5E, 5F). Taken together, even when applied to difficult indications, A_3_D_3_a’s MVP highlights important vulnerabilities, which can pave the way for new treatment options in hard-to-treat indications.

**Figure 5.**
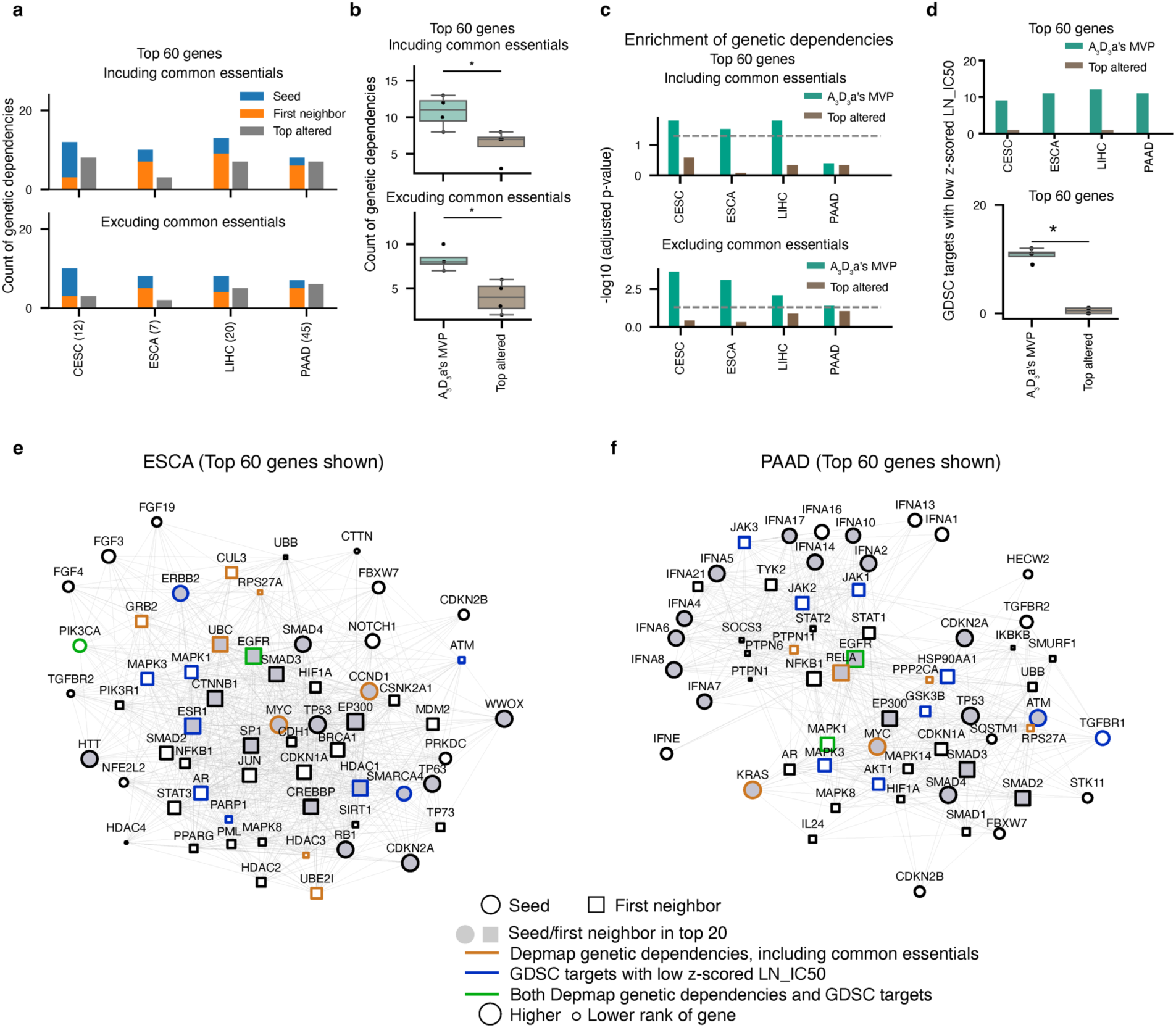
Model predictions for cancer types in the test set. **a.** Count of Depmap genetic dependencies among top 60 genes ranked by A3D3a’s MVP model using DNA as seed or by altered rate for each cancer type. Pooled genetic dependencies of all cell lines of each cancer type were considered. **b.** Group comparison of count of genetic dependencies among top 60 genes ranked by A3D3a’s MVP model and by altered rate across cancer types in a. **c.** Enrichment of genetic dependencies among top 60 ranked genes or top 60 altered genes in each cancer type computed by hypergeometric test. An adjusted p-value less than 0.05 was considered significantly enriched. Dashed gray horizontal line represented an adjusted p-value at 0.05. y axis was plotted as the negative log10 of the adjusted p-value. **d.** GDSC targets with low z-scored LN of IC50 among top 60 ranked genes and top 60 altered genes. **e.** Group comparison of the GDSC targets with low z-scored LN of IC50 among top 60 ranked genes across cancer types in d. Group comparisons in b and d by two-sided rank-sum test, *: p<0.05. **e.** and **f.** Network view of the largest connected graph of top 60 ranked genes by A3D3a’s MVP for ESCA and PAAD, respectively. Top 20 ranked genes were highlighted in gray face colors. Depmap genetic dependencies (including common essentials) were labeled in orange edge color. GDSC targets with low z-scored LN_IC50 were labeled in blue edge color. The genes that were both genetic dependency and GDSC targets with low z-scored LN_IC50 were labeled in green edge color.

### Novel therapeutic opportunities identified by A_3_D_3_a’s MVP

The integrative pharmacological annotation from canSAR.ai^7, 11–13^ with A_3_D_3_a’s MVP provides for the first time a seamless tool to identify targets with both biological and technical promise for drug discovery. As an illustration, we examined the combined, pharmacologically annotated 162 genes from the top 20 ranked genes in all the 15 cancers. We label the genes based on their proximity to the clinic as above, finding 94 across the four classes. We additionally annotate whether they are hits in relevant cell lines in the DepMap or GDSC databases (Fig. 6A, Supplementary Table 9).

**Figure 6.**
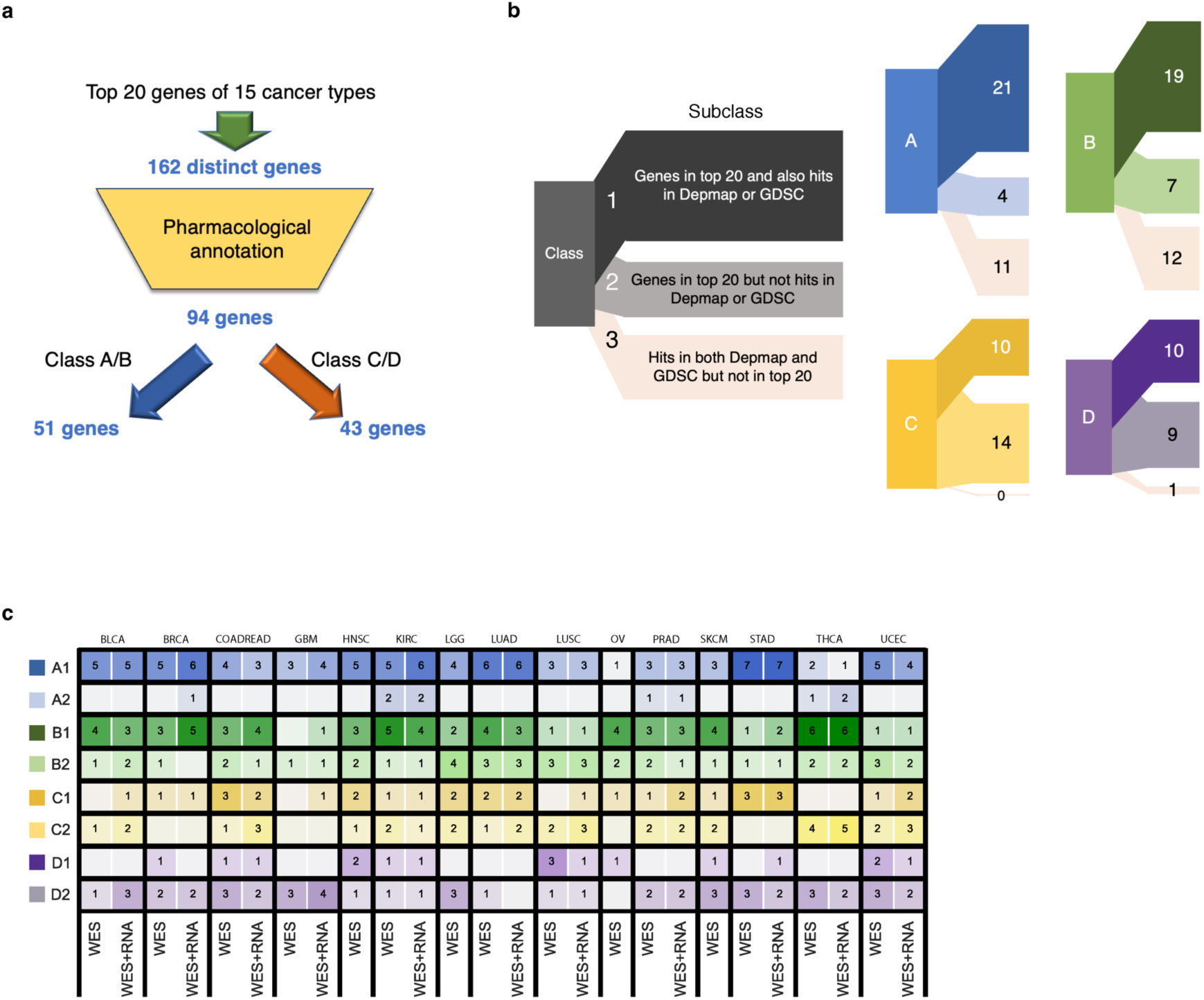
Summary of druggable opportunities across indications. **a.** The number of genes from the top 20 ranked genes by A3D3a’s MVP whose corresponding protein falls into one of the four druggability classes. **b.** For genes whose corresponding protein in each druggability class, they were further separated in three subclasses: 1. Genes in top 20 and also hits in Depmap or GDSC targets for at least one indication; 2. Genes in top 20 but not hits in either Depmap or GDSC targets for any of the indications; 3. Genes are hits in both Depmap and GDSC targets but not in top 20 for any of the indications. **c.** Number of genes ranked in top 20 by A3D3a’s MVP of classes A, B, C, and D and subclasses 1 or 2 for individual indications for DNA-only and DNA+RNA models. The subclass-1 indicates the genes that are among top 20 and also hits in Depmap or GDSC targets for that indication, and the subclass-2 indicates the genes that are among top 20 but not hits in Depmap or GDSC targets for that indication.

Of the 162 genes, 94 are druggable, amenable to modulation by a small molecule or biotherapeutic (Fig. 6B). Of these 94 genes, 60 were also hits in DepMap, GDSC, or both for the appropriate cell lines in at least one cancer. 34 genes were not hits in either cell line database, despite ranking in the top 20. 14 of these 34 were brought into the network as loss events: nonsense mutations, copy number deletion in more than 2% of patients, or low RNA expression (Supplementary Table 10). The other 20 may be irrelevant genes or genes that may be important but not observable in limited growth inhibition assays in 2D cell lines. For example, among these 20 genes, STAT3 is known to be associated with colorectal cancer proliferation^37^ and treatment resistance^38^. Finally, 24 genes were hits in both databases but not ranked in the top 20 by A_3_D_3_a’s MVP. (Fig. 6A). 17 of these 24 ranked in the top 60.

Of the 94 druggable genes, 51 are already targeted by FDA-approved or clinical stage drugs. 43 are either in earlier stages of discovery or have no public modulators available. This latter set present potential novel targets for therapeutics in these cancers. Figure 6C summarizes these per cancer type and reports how many targets were identified by DNA or combined DNA and RNA data. Finally, with the increasing interest in covalent modulators as drugs^39^, we used canSAR.ai to identify druggable pockets with exposed cysteine (C), lysine (K), and tyrosine (Y) amino acids which may potentially be targetable with covalent inhibitors. We found that 55 proteins were brought into the networks as gain events or latent vulnerabilities and also had at least one of these residues accessible within their primary druggable pocket (Supplementary Table 10, Supplementary Table 11).

## Discussion

To discover the next generation of drugs for cancer, we need to better identify novel, pharmaceutically targetable molecular vulnerabilities, regardless of their driver status, from noisy, heterogenous patient data. A_3_D_3_a’s MVP, an integrative, generalizable graph-based, cooperativity led Markov-chain model to identify molecular vulnerabilities from multimodal patient data. We demonstrated that it outperforms standard frequency-based approaches by 3-fold at identifying true molecular vulnerabilities as confirmed in large-scale genetic knockout and drug sensitivity screens. We demonstrate the model’s ability to identify druggable ‘latent’ vulnerabilities – molecules that are important for signaling and survival of a specific cancer that are not themselves mutated or altered. We use A_3_D_3_a’s MVP as a framework to map pharmaceutically relevant information and target druggability from canSAR.ai. We demonstrate the power of this approach at retrieving known drug targets and propose novel targets that could, with additional validation, be exploited for the development of new cancer therapies. A_3_D_3_a’s MVP sets the foundation for accelerating target discovery by prioritizing the most likely druggable Achilles’ heels for cancer.

## Methods

### Processing of DNA data from the TCGA

Genomic data from solid cancers in the TCGA PanCancer studies were downloaded from cBioPortal (https://github.com/cBioPortal/datahub/tree/master/public). Cancer indications were ranked in descending order by cohort size. The top fifteen were used for model optimization and benchmarking; these included Breast invasive carcinoma (BRCA), Colorectal cancer (COADREAD), Glioblastoma multiforme (GBM), Ovarian serous cystadenocarcinoma (OV), Lung adenocarcinoma (LUAD), Uterine corpus endometrial cancer (UCEC), Head and neck squamous cell carcinoma (HNSC), Brain lower grade glioma (LGG), Kidney renal clear cell carcinoma (KIRC), Thyroid carcinoma (THCA), Prostate adenocarcinoma (PRAD), Lung squamous cell carcinoma (LUSC), Skin cutaneous melanoma (SKCM), Stomach adenocarcinoma (STAD), and Bladder urothelial carcinoma (BLCA). Four indications were used as an independent test set and included Cervical squamous cell carcinoma and endocervical adenocarcinoma (CESC), Esophageal carcinoma (ESCA), Liver hepatocellular carcinoma (LIHC), and Pancreatic adenocarcinoma (PAAD). The indications in the test set were selected since they had the next highest cohort sizes and had DepMap data from at least 7 cell lines from the corresponding indication. Only primary tumor samples were analyzed except in the case of SKCM for which most of the samples were metastatic samples.

These data were analyzed as follows: Mutsig2CV^1^ with default settings was used to identify potential drivers from all somatic variants identified from whole exome sequencing, including single nucleotide variants (SNVs) and indels. To select mutations with impact on protein function, all intronic, intergenic and silent variants were excluded; exonic variants annotated as ‘LOW’ for IMPACT were removed; and variants annotated as ‘PASS’ for FILTER were retained as described^2^. Somatic copy number variation (CNV) data were obtained from Affymetrix SNP arrays by TCGA. Somatic copy number profiles were created based on putative copy number information generated using GISTIC 2.0^3^ CNVs meeting the threshold either for deep deletions (less than -2) or high-level amplifications (greater than 2) were retained. Structural variation (SV) frequency data (percentage of samples with at least one structural variation event) for all indications in the TCGA pancancer studies^2^ were downloaded from cBioPortal^4^ directly. Samples with somatic mutation, copy number, and structural variation data were included in downstream analysis, and the alteration frequency for each gene was calculated for mutation, copy number and structural variation data modality individually. An overall DNA alteration frequency was calculated for each gene as the maximum altered frequency among mutation, copy number, and structure variation modalities of that gene and employed in validation process by checking the number of hits in the DepMap and GDSC datasets among top altered genes.

### Processing of DNA data from DepMap cell line profiling

Somatic mutation data from cell lines profiled in DepMap were accessed from the DepMap portal through the ‘CCLE_mutation_2020Q3.txt’ file; these data were aligned to the hg19 reference genome matching the alignment used in the TCGA data analyzed^2^. LUAD cell lines were selected by filtering using the Model ID parameter. MutSig2CV was used for driver gene identification and applied to all somatic mutations identified in LUAD cell lines. Duplicated and silent variants were removed. Alteration frequency of each gene was then calculated by dividing the number of LUAD cell lines mutated in a given gene by the total number of LUAD cell lines.

### Processing of RNA-seq data from TCGA and GTEx

Raw read counts were obtained from the Cancer Genome Atlas (TCGA) and Genotype-Tissue Expression (GTEx) datasets, as indicated in Table S1. Eleven indications with no less than five samples in each of the following classes were included in the analysis: ‘TCGA tumor’, ‘TCGA normal-adjacent to tumor (NAT)’, and ‘GTEx normal’. These indications included BLCA, BRCA, COADREAD, GBM, KIRC, LUAD, LUSC, PRAD, STAD, THCA, and UCEC. Genes detected in less than 10% of samples in either the TCGA or GTEx were removed, and only genes detected in both TCGA and GTEx were considered for downstream analysis.

ComBat-seq was used to perform batch effect correction; each indication was processed separately^5^. For each indication, samples from the TCGA and GTEx were regarded as being from two different batches; ‘TCGA tumor’ samples were set as one group, while ‘GTEx normal’ and ‘TCGA NAT’ samples were assigned to another group. The resulting expression matrix underwent transformation and normalization using DESeq2 by established methods^6^.

### Interactome construction

The interactome was constructed from the canSAR^7^, KEGG^8^ and HI-union^9^ databases.

canSAR protein interaction data were downloaded from canSAR.ai (https://cansar.ai/cpat). Interactions with a provided confidence score greater than or equal to 0.2 were retained. Interactions whose type were designated as ‘low-confidence complexation’ or ‘drug synergy’ were excluded, while interactions whose type were ‘complex’, ‘direct’, ‘reaction’, ‘signaling’, or ‘transcriptional’ were included. Multiple interactions connecting any two proteins were replaced with a single interaction, condensing 550,270 interactions to 419,705 interactions. These interactions connecting 18,096 proteins were brought into the interactome from canSAR.

KEGG data were accessed through MD Anderson’s institutional repository. Interactions were parsed and extracted from xml files using the R KEGGgraph package^10^. 65,924 unique interactions between 6,357 proteins were incorporated into the interactome and were of the following types: 18,896 ECrel interactions (enzyme-enzyme relation, indicating two enzymes catalyzing successive reaction steps), 5,264 GErel interactions (gene expression interaction, indicating interaction of transcription factor and target gene), 738 PCrel interactions (protein-compound interaction), and 41,026 PPrel interactions (protein-protein interaction, such as binding and modification).

HI-union data were downloaded from the human reference interactome (HuRI) database^9^. 63,800 unique interactions between 9,050 proteins from HI-union were brought into the interactome.

Data from these three sources were combined and multiple interactions between any two proteins were replaced with a single interaction. With each protein serving as a node and each interaction as an edge, the resulting interactome contains 18,614 protein nodes connected by 471,200 edges.

### Selection of seed genes for networks

Networks were generated for each indication using proteins whose corresponding genes were altered and/or differentially expressed in that indication. These genes are referred to as ‘seed genes’; the process used to select them is described below:

Seed genes derived from DNA data were selected from mutation, CNV, and SV events. For mutation data, genes were ranked by the geometric average of the ranking by alteration frequency of genes in descending order, and the ranking by p-values determined by MutSig2CV for each gene were then arranged in ascending order. The top 50 genes provided by the final ranking were selected as seed genes. For CNV data, the top 50 genes ranked by alteration frequency in descending order were selected as the seed genes. For SV data, the top 50 genes ranked by alteration frequency in descending order that had SV events in more than 2 percent of patients were chosen as seed genes. TTN and MUC16 genes were finally excluded from the DNA seed for the downstream analysis because their extensive protein size tends to result in a high susceptibility to mutations.

Seed genes derived from RNA-seq were selected from genes differentially expressed between tumor and normal samples. A regularized linear SVM classifier based on elastic net regularization was created to further filtering gene features by classifying samples as tumor or normal. The model was implemented using Python’s sklearn package^11^. The data were split in a stratified fashion into a training set consisting of 70% of the data and a test set consisting of 30% of the data.

For the training data, differential expression analysis between tumor and normal samples was performed by the rank sum test. False discovery rate (FDR) was calculated by Benjamini and Hochberg’s method. Genes with a FDR < 0.05 and an absolute log2 fold change (abs_log2_FC) > 1 were used as input features for the classification model. The maximum number of features was limited to 2500 which were selected by sorting by FDR (ascending) and abs_log2_FC (descending). For indications with imbalanced class labels, Synthetic Minority Oversampling Technique (SMOTE) was used to synthesize new samples for the minority class. After this data augmentation step, features were scaled to values between 0 and 1 in the training set by min-max scaling.

The scaler generated from the training set was applied to the test set. Five-fold cross-validation was employed for hyperparameter tuning with grid search. Stochastic gradient descent was used during training. Validation and test accuracy were monitored to avoid overfitting or underfitting. After model tuning, features with non-zero coefficients were selected as seed genes. The average number of seed genes selected from RNA-seq data across indications was 22 with a standard deviation of 8.

Seed genes derived from mutation data only were selected from the mutation events of DNA data. Genes were sorted by the geometric average of ranking by alteration frequency in descending order, and the ranking by p-values determined by MutSig2CV for each gene were then arranged in ascending order. The top 100 genes provided by the final ranking were selected as the seed genes.

### Selection of non-seed genes for networks

Networks for each indication were augmented by the inclusion of ‘non-seed’ genes from the interactome to bring together disconnected portions of the network.

A first neighbor of a given gene is defined as another gene directly connected to it. To begin with, all first neighbors of seed genes were retrieved from the interactome as candidate non-seed genes. If a candidate non-seed gene has more than two first neighbors in the seed list, or if it has only one first neighbor and that first neighbor is in the seed list, the candidate non-seed gene was included in the following analysis. For each candidate non-seed gene *A*, the in-seed ratio (ISR) representing the normalized fraction of its first neighbors in the original seed list was computed as follows:

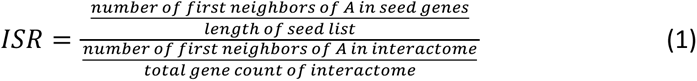

Next, whether the ISR occurred by chance was evaluated using a permutation test. The interactome was randomly permuted 10,000 times to generate new seed lists of the same size as the original seed list; then how often the observed ISR could be achieved by chance was computed (empirical p-value). p-values were corrected by multiple hypothesis testing by Benjamini and Hochberg’s method and genes with FDR < 0.05 were retained. First neighbors were ranked by the FDR (ascending) and the number of neighbors in seed (descending). The top *N* ranked first neighbors were included in the graphical modeling of the disease network. *N* was optimized together with other parameters as explained below in the “Tier 1 genes and model parameter optimization” section.

### Graphical modeling

The largest connected graph of the disease network was used to perform graphical modeling by applying a Markov chain model.

Markov chains provide stochastic models in which the probability of each event is dependent only on its immediately prior state. Computationally efficient because of their memorylessness, Markov chain models^12^, have been applied to a variety of biological questions, including molecular phylogenetics in bioinformatics^13^, stochastic chemical kinetics in systems biology^14^, and modeling of cortical micro-circuits in neuroscience^15^. In particular, Markov chain models have important features that lend themselves well to our question of interest. First, altered signal travels in a probabilistic manner within the network and finally reaches a stable state. We hypothesize that, when modelling the cooperativity of signals in a set of adjacent nodes, those nodes with highly accumulated signal at the stable state are likely to be therapeutic vulnerabilities. Second, Markov chain models are explainable and controllable, thus lending themselves well to interpreting results and providing clear insights into the underlying data.

The Markov chain model used in A_3_D_3_a’s MVP was written in Python and defined as follows:

For a graph with *n* nodes and an adjacency matrix *A* with zeros on its diagonal, let ***x***(*t*) be the variable describing the state of all the nodes at step *t*:

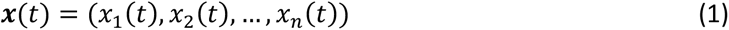

The state transition follows:

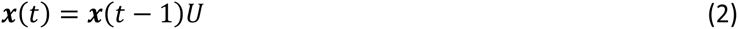

*U* is the transition matrix of size *n*-by-*n*.

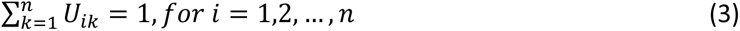

*U*_*ik*_ is the transition probability from node *i* to node *k*.

If the altered rate (initial score) of nodes is ***f*** = (*f*_1_, *f*_2_, …, *f*_*n*_), the normalized altered rate ***p*** = (*p*_1_, *p*_2_, …, *p*_*n*_) is:

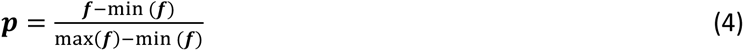

The transition matrix can be set as:

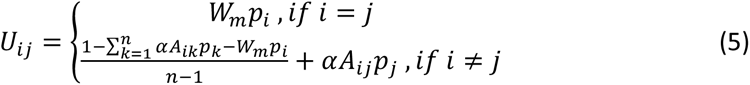

Which satisfies equation (3). *W*_*m*_ is the weight parameter on the self-loop of nodes of the Markov chain model, and *α* is the cooperativity factor describing transition tendency toward the altered nodes. The choice of *W*_*m*_ and *α* should guarantee that *U*_*i*j_ > 0, for *i*, *j* = 1, 2,…,*n*.

If ***x***(*t*) = ***x***(*t* − 1) + ***ε***, where ***ε*** = (*ε*_1_, *ε*_2_, …, *ε*_*n*_) satisfies *ε*_*i*_ < 10^−8^ for every *i* in 1,2 *,…, n,* the model was regarded as having converged and the state transition was stopped. Otherwise, the model was stopped after reaching 10,000 iterations. The final state ***x***(*t*) of the nodes was taken as the final score output of the model.

The initial score of a gene from the DNA seed was determined by the altered rate of that gene. The initial score of a gene from the RNA seed varied depending on whether the gene is up-regulated or down-regulated in tumors. For an up-regulated gene, the initial score was determined by the fraction of tumor samples exhibiting expression levels higher than the top 1% (99th percentile) of the normal group. Conversely, for a down-regulated gene, the score was established by the fraction of tumor samples showing expression levels lower than the bottom 1% (1st percentile) of the normal group. If a gene is both DNA seed and RNA seed, its initial score is taken as the maximum of its DNA seed initial score and RNA seed initial score. The initial score of non-seed gene was set to 0.

For comparison, Page Rank^16^ and Personalized Page Rank^17,18^) models were implemented using the Python networkx package^19^. The ‘personalization’ parameter of nodes in the Personalized Page Rank model was set as the altered rate **f** of nodes. Classical Diffusion models on graphs^20^ were implemented using the diffuStats^21^ package in R . It computes the classical Gaussian diffusion kernel that involves matrix exponentiation, and it has a ‘bandwidth’ parameter σ^2^ that controls the extent of spreading. The initial score of nodes in the Classical Diffusion model was the altered rate **f** of nodes.

### Tier 1 genes and model parameter optimization

Tier 1 genes from the Cancer Gene Census, 578 genes in total, were downloaded from the COSMIC database^22^. Tier 1 genes for each of the 15 TCGA indications were identified by filtering the Tier 1 genes to only those with at least one hotspot mutation in a specified indication.

Cancer hotspot mutations were downloaded from https://www.cancerhotspots.org ^23^.

The intersection over union (IoU) of the top 20 ranked genes returned by graphical models and the number of Tier 1 genes for each indication was used to optimize model parameters. The model parameters corresponding to the highest average rank of IoU across indications were selected. IoU was used to perform parameter optimization by grid search for all models considered: Markov chain, Page Rank, Personalized Page Rank, and Raw Diffusion models.

For Markov chain models (MCm), the parameters tuned were *W*_*m*_, *α*, and *N* (number of top ranked non-seed genes). The optimal MCm derived from DNA data only has α =0.1, W_m_ of 0.5, and N =550, and that derived from DNA and RNA-seq data has an α = 0.1, W_m_ =0.4, and N =550.

For Page Rank and Personalized Page Rank models, *α* (damping parameter), and *N* (number of top ranked non-seed genes) were tuned. For Raw Diffusion models, *σ*^2^ (bandwidth parameter controlling extent of the spreading) and *N* (number of top ranked non-seed genes) were tuned.

For each model type, models using different parameter settings were ranked by the IoU of Tier 1 genes among the top 20 genes predicted in each indication. The model parameters providing the highest average rank across indications were selected, and the same model parameters were applied across indications, including the 15 used in model optimization and the 4 used as an independent test set.

### Study bias estimation

Data on publications and their association with specified genes were downloaded from NCBI (https://ftp.ncbi.nlm.nih.gov/gene/DATA/gene2pubmed.gz) on Aug-9-2022. The number of publications corresponding to each gene was calculated and then used to compute the Spearman correlation with the rank of genes predicted by Markov chain, Page Rank, Personalized Page Rank, and Raw Diffusion models.

### Model validation by DepMap and GDSC data

We used genetic and pharmacological perturbation data in cancer cell lines from DepMap^24^ (22Q4) and the Genomics of Drug Sensitivity in Cancer (GDSC)^25^, respectively, to evaluate our results. In DepMap, genetic knock-out via CRISPR removes the entire gene and thus impacts all its interactions, rather than just modulating its activity. In contrast, in the GDSC, drugs can focus on the specific site of activity (e.g., the catalytic site of an enzyme) but are more prone to non-selective effects as many drugs show polypharmacology. The use of both genetic and pharmacological perturbation as orthogonal tools enhances confidence in observed experimental results.

The DepMap public 22Q4 datasets including common essentials, CRISPR gene effect, and indication-specific cell line information were downloaded from the DepMap portal. 1,247 Common essential genes were obtained from ‘Achilles’ common essential controls^24^ which drew upon two studies^26,27^. DepMap genetic dependencies in each indication were defined as those with a Chronos score less than -1 in the cell lines corresponding to that indication^28^. The union of genetic dependencies across all cell lines corresponding to a given indication was used to evaluate the results provided by A_3_D_3_a’s MVP for that indication.

The July 2022 release of the Genomics of Drug Sensitivity in Cancer (GDSC) database was downloaded for both GDSC1 and GDSC2^25^. Data from drugs applied to all cell lines in GDSC1 and GDSC2 were retained, and the z-scored natural logarithm of fitted IC50 values (LN_IC50) was used to estimate the drug sensitivity in each cell line. Genes that listed by the GDSC as putative targets of a drug with a z-scored LN_IC50 value < -1.5 in cell lines corresponding to particular indication were regarded as targets. Drugs with more than four distinct putative targets were excluded.

The fraction of DepMap genetic dependencies and GDSC targets among top-ranked genes provided by A_3_D_3_a’s MVP were used to validate the model performance and were compared to rankings returned by alteration frequency alone.

### Recapitulation of clinically relevant genes by A_3_D_3_a’s MVP

Clinically relevant genes were collected from oncoKB^29^ and mechanistic targets of FDA-approved drugs as curated in cansar.ai^7,30^ ). From oncoKB, genes from level 1 (FDA-approved drugs), level 2 (Standard care) and level 3 (Clinical evidence) were included for each cancer type.

### canSAR druggability annotation and pocket accessibility analysis

Proteins corresponding to top-ranked genes were annotated using the CPAT tool from cansar.ai^7^ to determine associated compounds and druggability based on a manual curation of small molecule drugs. The following classification was used:

- Class A: target of FDA-approved drug
- Class B: target of investigational drug
- Class C: target of discovery-stage drug with reported bioactivity <100nM for IC50/Ki/Kd,
- Class D: 3D-ligandable structures available.

To identify potentially accessible protein pockets within these proteins, relative solvent accessibility (RSA) was calculated using DSSP^31^ by dividing solvent accessibility by the maximum solvent accessibility of each chain. A threshold of 0.25 was used; greater than was regarded as exposed and less than as buried. Accordingly, pockets containing at least one accessible cysteine, lysine, or tyrosine residue were counted, with protein chains with a Jaccard distance > 0.5 assigned as being representations of the same pocket.

### Statistical analysis

The Wilcoxon Rank Sum Test was performed to test the equality of means of two sample groups; p < 0.05 was used as the significance threshold. Spearman correlations were calculated to analyze study bias estimation. Hypergeometric testing was performed as part of pathway enrichment analysis and enrichment analysis of genetic dependencies. The Benjamini-Hochberg method was used for multiple hypothesis correction.

### Community detection and pathway enrichment analysis

Community detection was performed using the Louvain method with resolution set to 0.7; this approach was implemented using the Python networkx package^19^.

Pathway enrichment was performed through overrepresentation analysis using the hypergeometric test as implemented by the Python gseapy^32^ package.

### Code availability

The codebase for A3D3a’s MVP is publicly available at https://github.com/YingZ-A3D3a/A3D3a_MVP with the MIT License.

