## Supplementary figures for "Probabilistic graph-based model uncovers previously unseen druggable vulnerabilities in major solid cancers"

Supp. Fig. 1. Model selection, DNA as seed

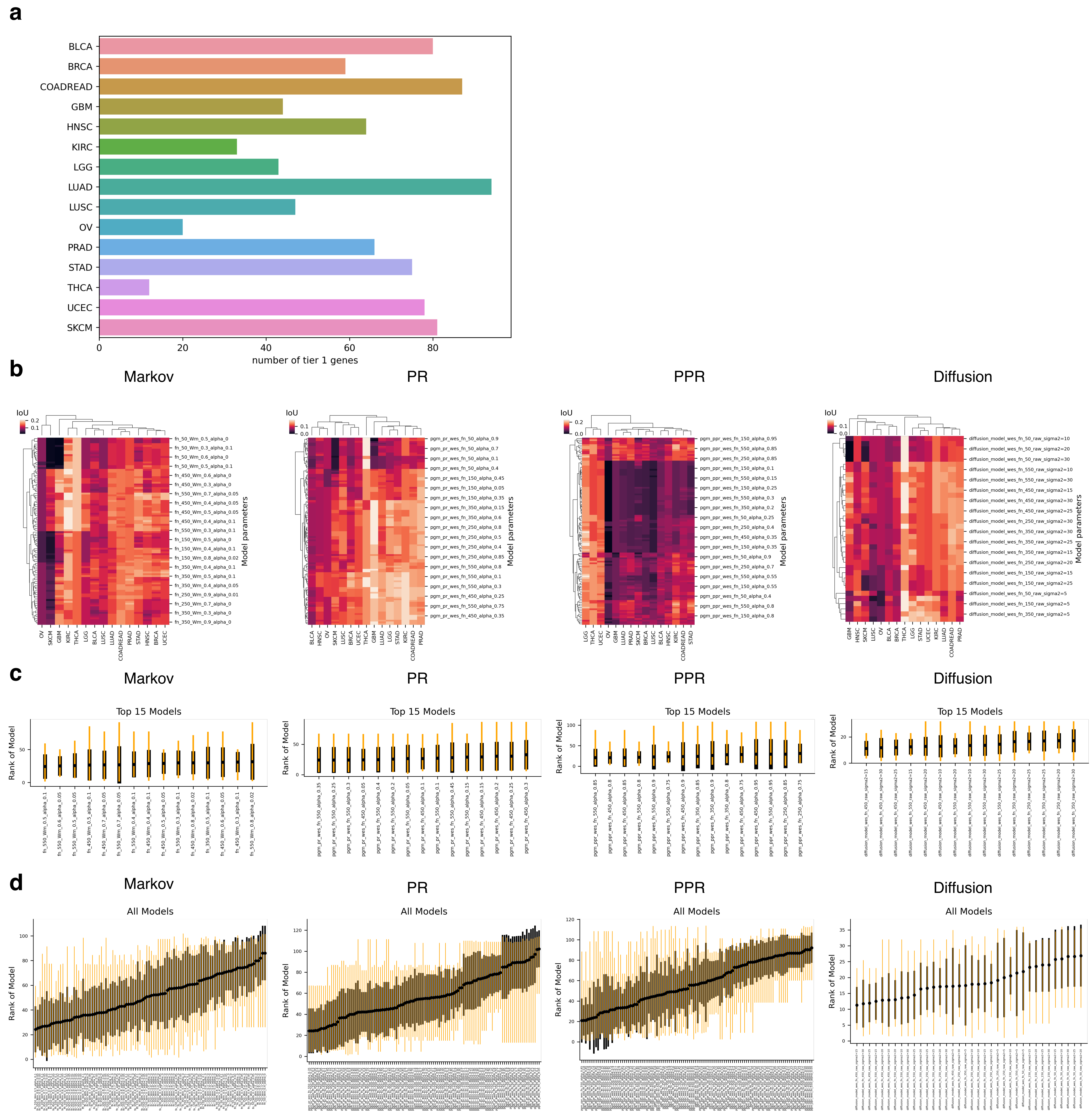


**Supp.Fig.1.** Tier 1 genes and model selection (DNA as seed). **a.** Number of Tier 1 genes in each cancer type filtered by hotspot mutations. **b-d.** Model parameter optimization for Markov chain, PR, PPR and classical diffusion models. **b.** Heatmap of IoU of Tier 1 genes among the top 20 ranked genes for each model. **c.** For each cancer type, the models were ranked by the IoU of Tier 1 genes among the top 20 genes predicted by the models. Top 15 models with highest average rank across cancer types were plotted for Markov chain, PR, PPR and classical diffusion models. Mean and standard deviation of model rank across cancer types were indicated by dot and black bars, respectively. **d.** The average rank across cancer types for all models was plotted for Markov chain, PR, PPR and classical diffusion models.

### Supp. Fig. 2. Model selection, DNA+RNA as seed

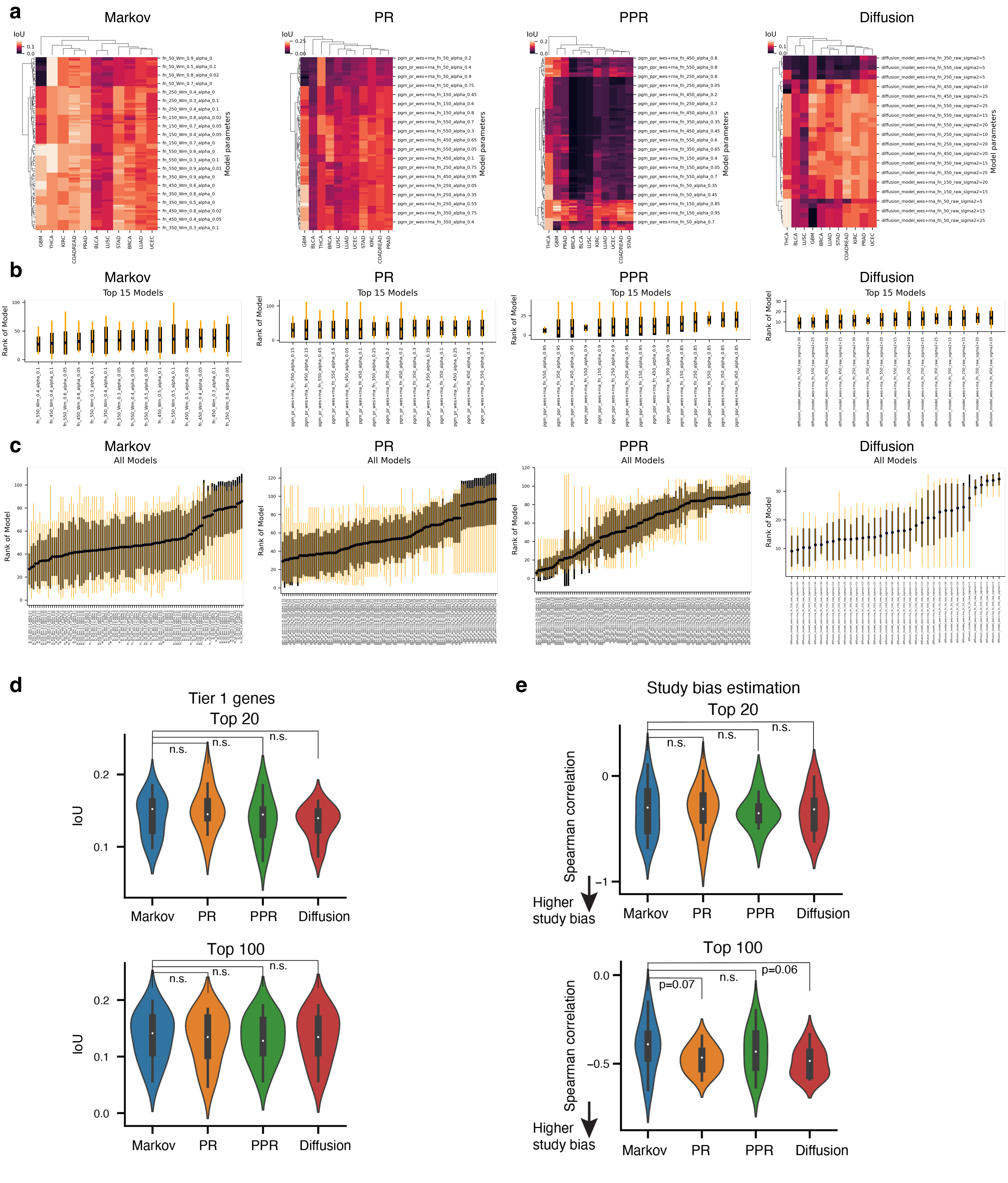


**Supp.Fig.2.** Model selection and comparison (DNA+RNA as seed). **a-c.** Model parameter optimization for Markov chain, PR, PPR and classical diffusion models. **a.** Heatmap of IoU of Tier 1 genes among the top 20 ranked genes for each model. **b.** For each cancer type, the models were ranked by the IoU of Tier 1 genes among the top 20 genes predicted by the models. Top 15 models with highest average rank across cancer types were plotted for Markov chain, PR, PPR and classical diffusion models. Mean and standard deviation of model rank across cancer types were indicated by dot and black bars, respectively. **c.** The average rank across cancer types for all models was plotted for Markov chain, PR, PPR and classical diffusion models. **d.** Model comparisons of IoU of Tier 1 genes among top 20 ranked genes (top) and top 100 ranked genes (bottom). **e.** Study bias estimation between models (right) by computing the Spearman correlation of the rank and the number of publications for the top 20 ranked genes (top), and for the top 100 ranked genes (bottom). One-sided rank-sum test was performed for group comparison.

Supp. Fig. 3. Model properties and study bias, DNA as seed

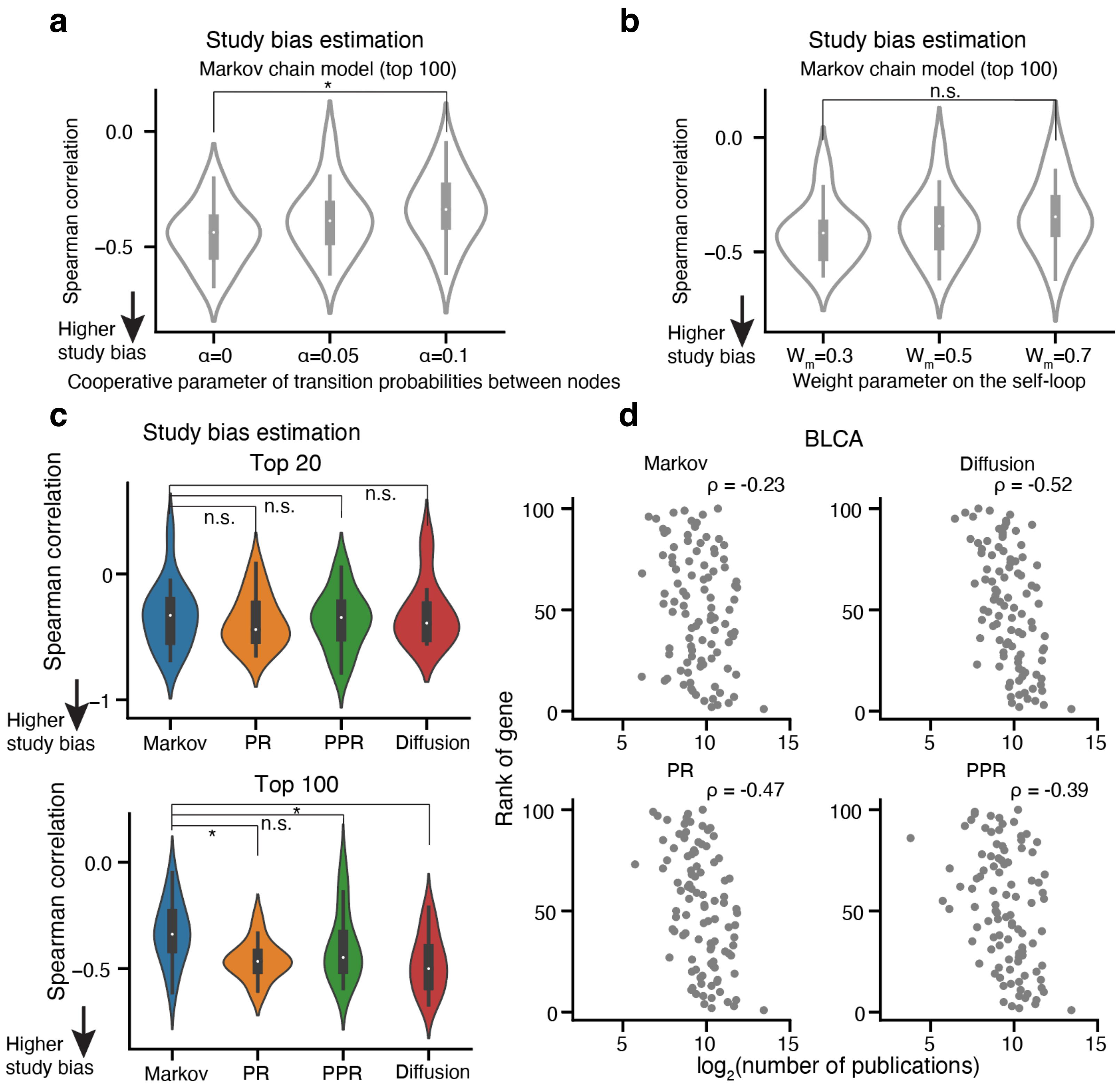


**Supp.Fig.3** Model properties and study bias (DNA as seed). **a.** The influence of the bias parameter α on the study bias. Study bias estimation by computing the Spearman correlation of the model output score and the $\log_2$(number of publications) of each gene among the top 100 ranked for 15 cancer types. $W_m$ is set at 0.5 and number of first neighbors is set at 550. **b.** The influence of the weight parameter $W_m$ on the study bias. Study bias calculation is the same as c. α is set at 0.05 and number of first neighbors is set at 550. **c.** Study bias estimation by computing the Spearman correlation of the model output score and the $\log_2$(number of publications) of each gene among the top 20 ranked (top panel), and top 100 ranked (bottom panel) for 15 cancer types. Comparison between optimized models of $A_3D_3$a's MVP, PR, PPR and raw diffusion by one-sided Rank-sum test, p-values were adjusted by Benjamini/Hochberg method (n.s.: p_adj>=0.05, *: p_adj<0.05). **d.** Correlation of rank of top 100 ranked genes with the $\log_2$(number of publications) of different models for BLCA as an example case.

Supp. Fig. 4. Depmap validation of top ranked genes using DNA as seed

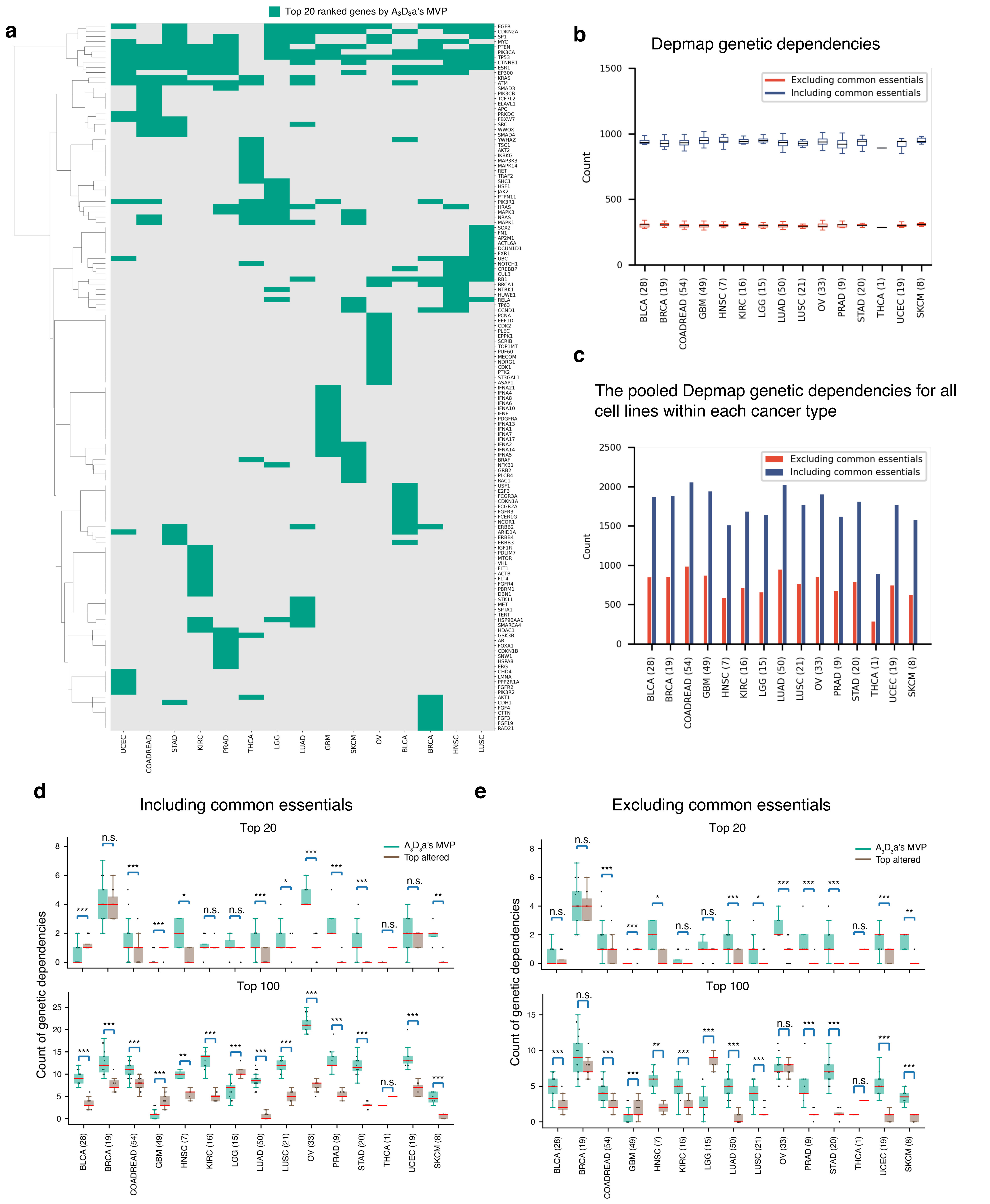


**Supp.Fig.4.** Depmap validation of top ranked genes using DNA as seed. **a.** Top 20 ranked genes of each cancer type by $A_3D_3$a's MVP model. **b.** Depmap genetic dependencies and common essentials for individual cell lines of each cancer type. **c.** The pooled Depmap genetic dependencies for all cell lines within each cancer type. **d.** Count of Depmap genetic dependencies of individual cancer cell lines (including common essentials) among top 20 or top 100 ranked genes by $A_3D_3$a's MVP model vs. by altered rate. **e.** Count of Depmap genetic dependencies of individual cancer cell lines (excluding common essentials) among top 20 ranked genes by $A_3D_3$a's MVP model vs. by altered rate. Group comparisons by two-sided rank-sum test. *: p<0.05, **: p<0.01, ***: p<0.001, n.s.: not significant.

### Supp. Fig. 5. Depmap validation of top ranked genes using DNA+RNA as seed

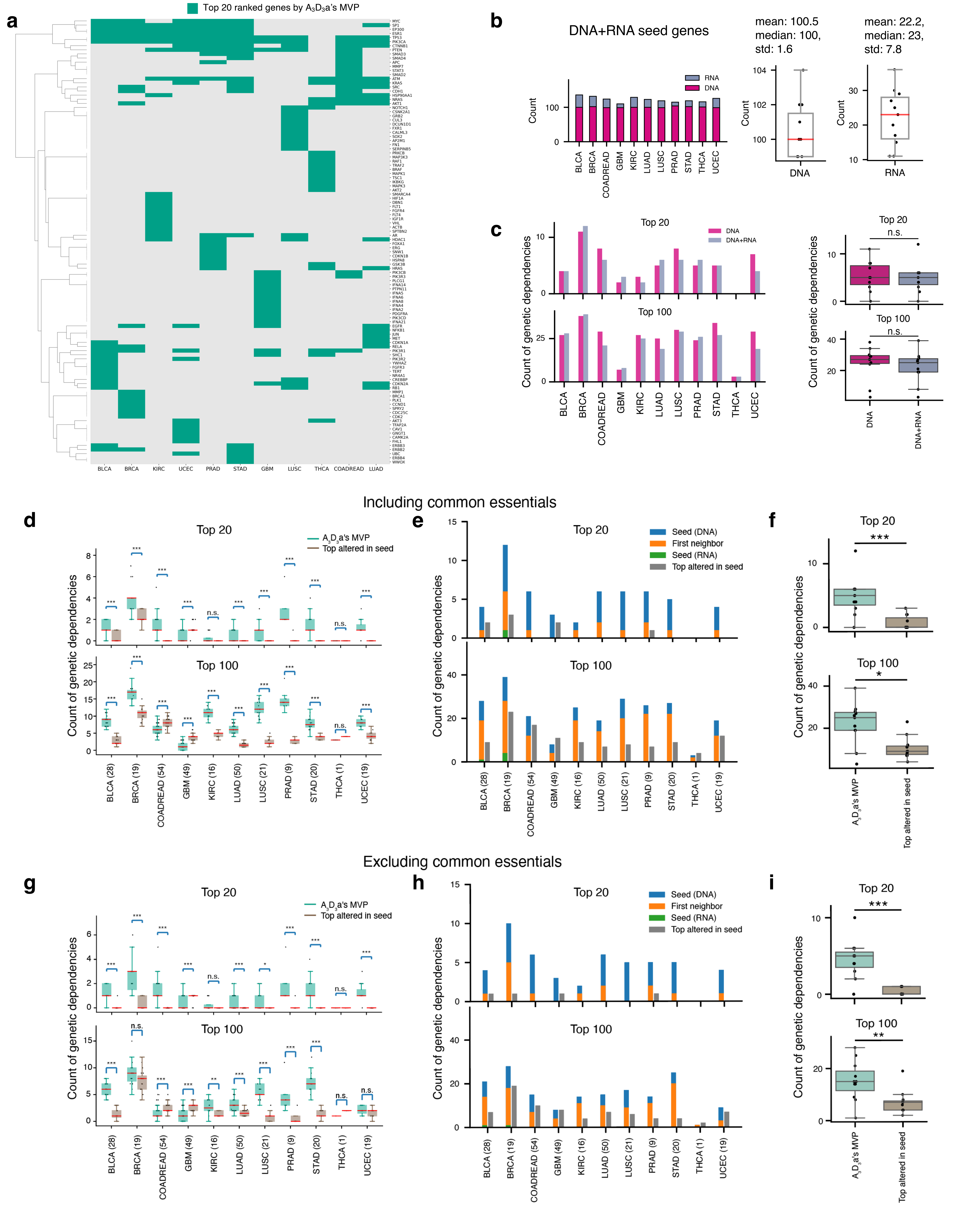


**Supp.Fig. 5.** Depmap validation of top ranked genes using DNA+RNA as seed. **a.** Top 20 ranked genes of $A_3D_3a$'s MVP for each cancer type using DNA+RNA as seed. **b.** (Left panel) Seed gene composition. (Right panel) distribution of seed from DNA and seed from RNA. **c.** (Left panel) count of genetic dependencies of top ranked genes using DNA as seed versus using DNA+RNA as seed. (Right panel) group comparison of the data in c. **d.** Count of genetic dependencies of individual cancer cell lines among top 20 or 100 ranked genes by $A_3D_3a$'s MVP vs. among top altered genes in seed. **e.** Count of pooled genetic dependencies of all cell lines of each cancer type among top ranked genes by $A_3D_3a$'s MVP vs. top altered seed genes. **f.** Group comparison of the data in e. **g-i.** The same analyses as in c-e except that common essentials were excluded. Group comparisons by two-sided rank-sum test. *: p<0.05, **: p<0.01, ***: p<0.001, n.s.: not significant.

Supp. Fig. 6. Depmap genetic dependencies among top ranked genes by $A_3D_3a$'s MVP (DNA as seed) and top altered genes of DNA

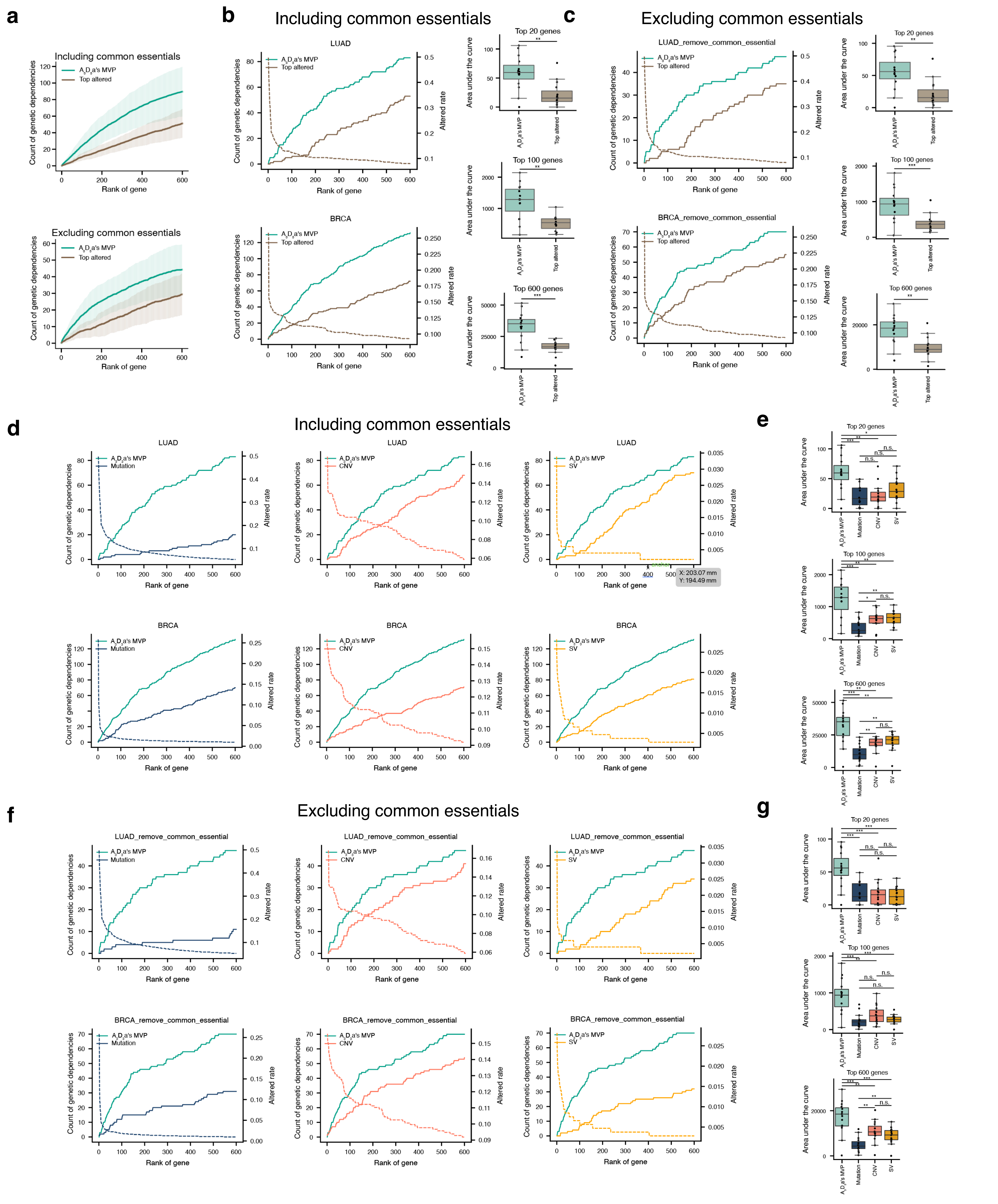


**Supp.Fig.6.** Comparison of count of Depmap genetic dependencies among top ranked genes by $A_3D_3a$'s MVP model using DNA as seed and by altered rate. **a.** Mean and standard deviation of count of genetic dependencies (including common essentials) among the top ranked genes by $A_3D_3a$'s MVP and top altered genes across 15 cancer types. Genetic dependencies did (top) and did not include common essentials (bottom). **b.** (Left panel) Count of genetic dependencies (common essentials included) among top ranked genes by $A_3D_3a$'s MVP (green) and top altered genes in DNA (brown) for LUAD and BRCA. Dashed brown lines showed the altered rate of the genes. (Right panel) The area under the curve of the count of genetic dependencies as a function of top ranked or top altered genes as shown in the left panel across 15 cancer types. **c.** the same plots as in a. except that the genetic dependencies do not include the common essentials. **d.** Count of genetic dependencies (common essentials included) among top ranked genes by $A_3D_3a$'s MVP (green) and top altered genes in mutation (blue), CNV (red) and SV (orange), respectively. Dashed lines showed the altered rate of the genes. **e.** The area under the curve of the count of genetic dependencies as a function of top ranked or top altered genes. **f.** The same plots as in d. except that the genetic dependencies do not include the common essentials. **g.** The area under the curve of the count of genetic dependencies as a function of top ranked or top altered genes as shown in e across 15 cancer types. Group comparisons in a, b, d and f by two-sided rank-sum test. *: p<0.05, **: p<0.01, ***: p<0.001, n.s.: not significant.

### Supp. Fig. 7. Comparison of Depmap genetic dependencies among top ranked genes by $A_3D_3$a's MVP (DNA+RNA as seed) with top altered genes

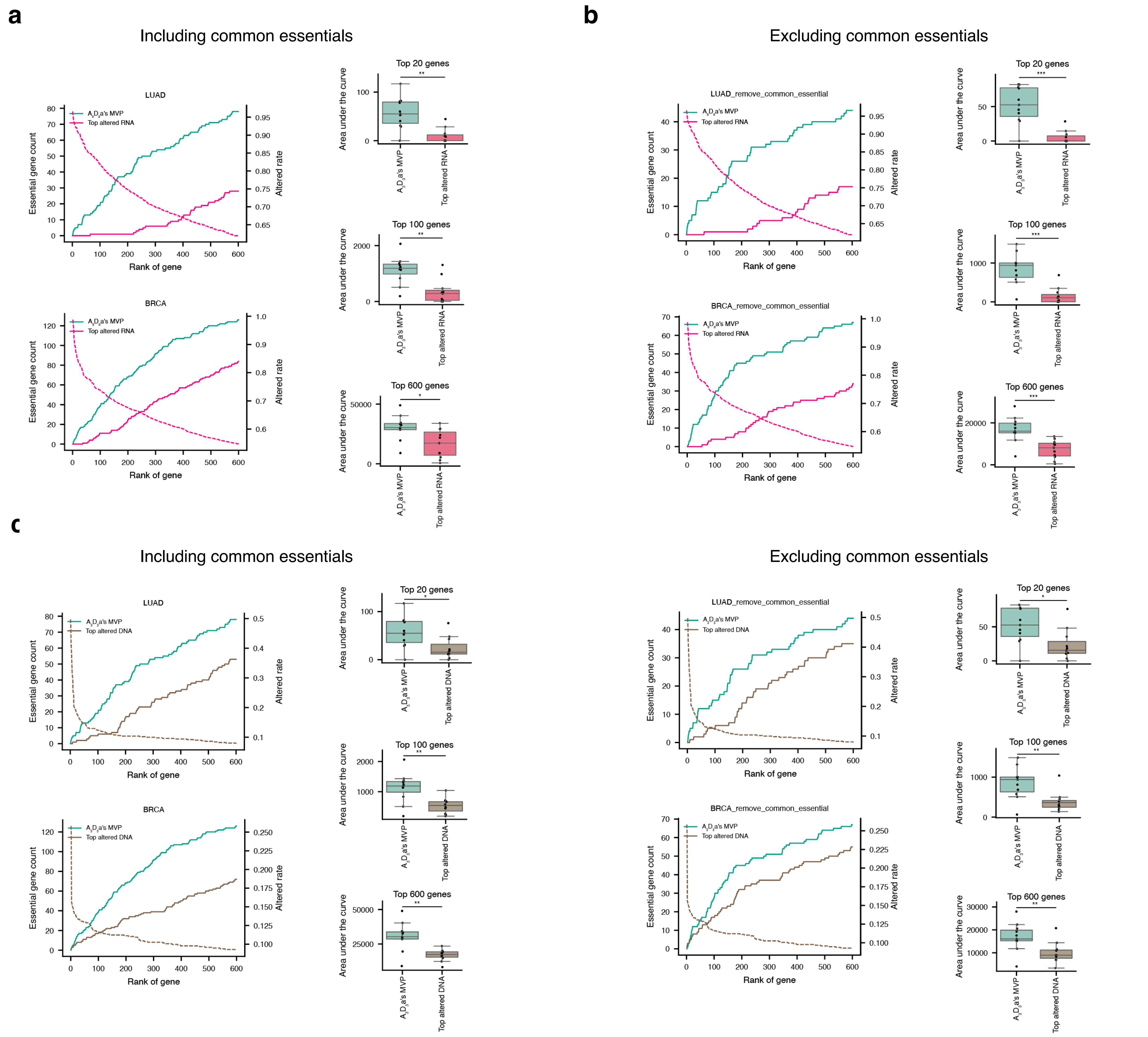


**Supp.Fig.7.** Comparison of Depmap genetic dependencies among top ranked genes by $A_3D_3$a's MVP (DNA+RNA as seed) with top altered genes. **a.** (Left panel) count of genetic dependencies (common essentials included) among top ranked genes by $A_3D_3$a's MVP (green) and top altered genes in RNA (pink). Dashed pink lines showed the altered rate of the genes. (Right panel) the area under the curve of the count of genetic dependencies as a function of top ranked or top altered genes as shown in the left panels across 11 cancer types. **b.** The same analyses as a. except that the genetic dependencies did not include the common essentials. **c.** (Left panel) count of genetic dependencies (common essentials included) among top ranked genes by $A_3D_3$a's MVP (green) and top altered genes in DNA (brown). Dashed brown lines showed the altered rate of the genes. (Right panel) the area under the curve of the count of genetic dependencies as a function of top ranked or top altered genes as shown in the left panels across 11 cancer types. **d.** The same analyses as c. except that the genetic dependencies did not include the common essentials.

### Supp. Fig. 8. Depmap essentials among top ranked genes by MVP (DNA as seed) and MVP (mutation as seed)

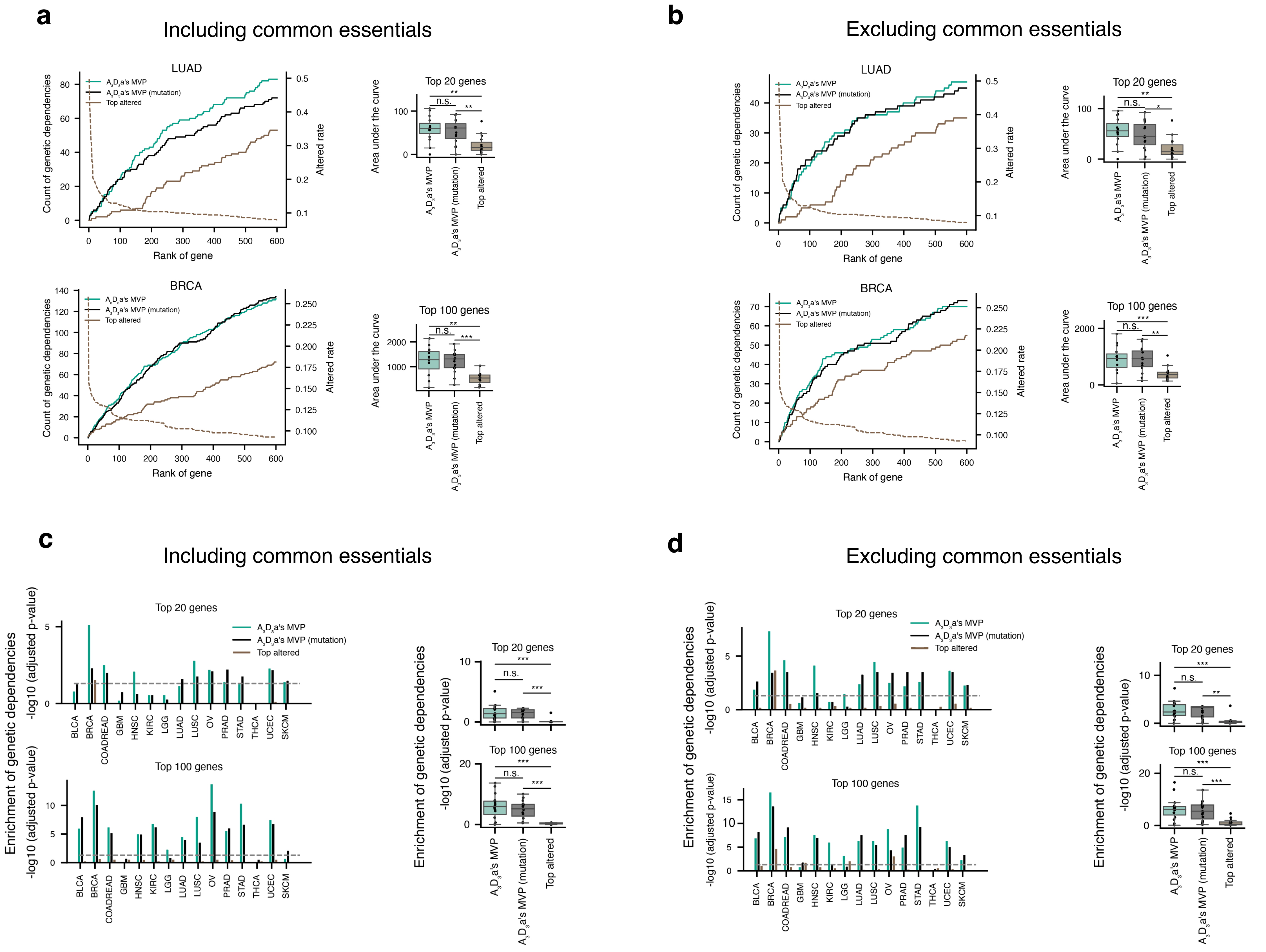


**Supp.Fig.8.** Comparison of count of Depmap genetic dependencies among top ranked genes by $A_3D_3$a's MVP (DNA as seed) and by $A_3D_3$a's MVP (mutation as seed). **a.** Count of genetic dependencies (common essentials included) among top ranked genes by $A_3D_3$a's MVP (DNA as seed, including mutation, CNV and SV, green), by $A_3D_3$a's MVP (only using mutation data as seed genes, black), and top altered genes (brown). Dashed brown lines showed the altered rate. Right panels are the area under the curve of the count of the genetic dependencies as a function of top ranked genes as shown in the left panels across 15 cancer types. **b.** the same plots as in a. except that the genetic dependencies do not include the common essentials. **c.** Enrichment analysis of genetic dependencies (common essential genes included) among top ranked genes by hypergeometric test for each cancer type. Gray horizontal dashed lines indicate the adjusted p-value of 0.05 of the hypergeometric test. Right panels are the group comparisons of the data in the left panels. **d.** the same plots as in c. except that the genetic dependencies do not include the common essentials. Group comparisons in a-d by two-sided rank-sum test. *: $p<0.05$, **: $p<0.01$, ***: $p<0.001$, n.s.: not significant.

Supp. Fig. 9. Depmap genetic dependencies among top 60 ranked genes, DNA as seed

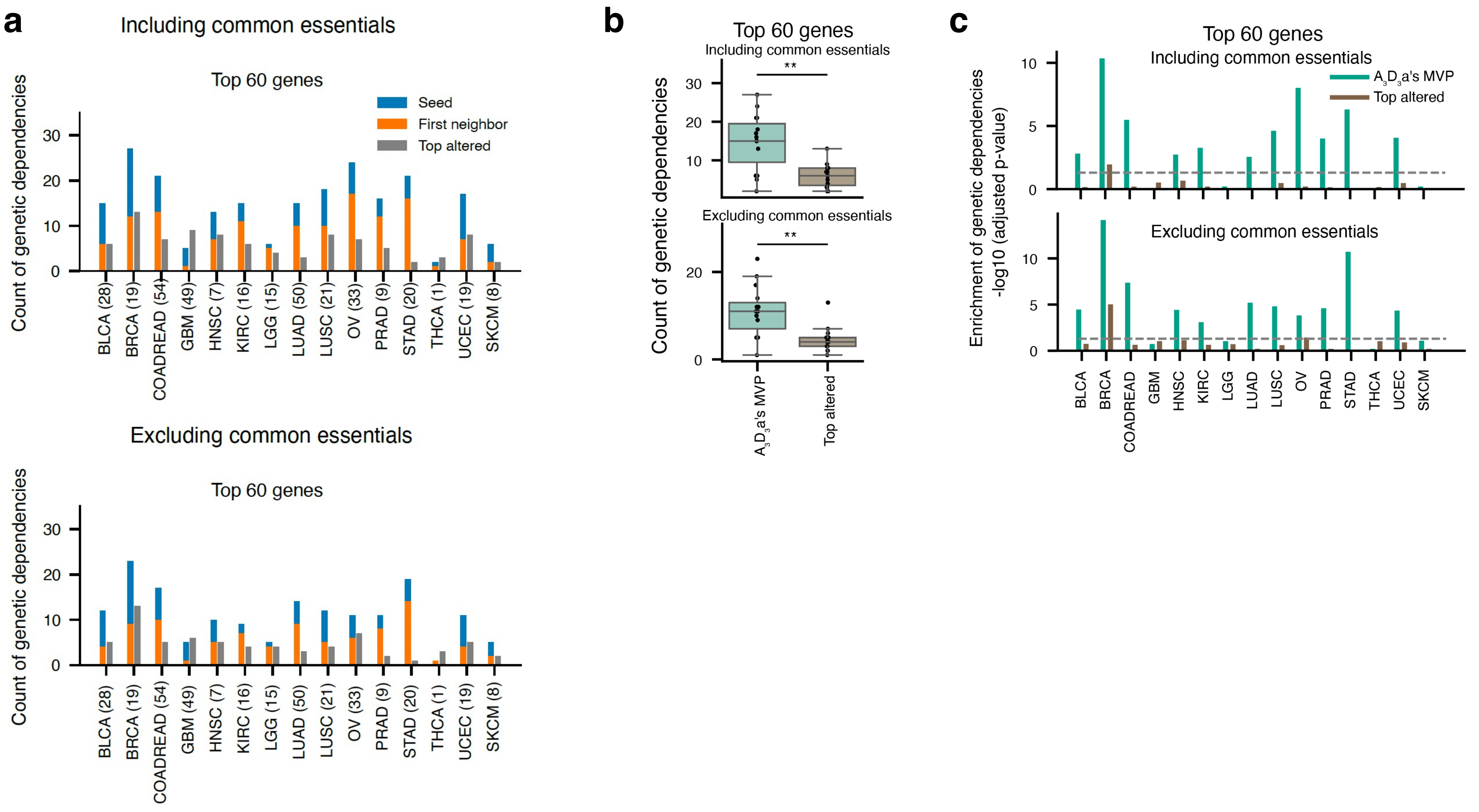


**Supp.Fig.9.** Depmap genetic dependencies among top 60 ranked genes. **a.** Count of Depmap genetic dependencies among top 60 genes ranked by $A_3D_3$a's MVP model using DNA as seed or by altered rate for each cancer type. Pooled genetic dependencies of all cell lines of each cancer type were considered. **b.** Comparison of count of genetic dependencies (including common essentials) pooled from all Depmap cell lines of each cancer type among top 60 ranked genes by $A_3D_3$a's MVP and top altered genes (top panel). (Bottom panel) the genetic dependencies did not include the common essentials. (n = 15 cancer types). **c.** Enrichment of genetic dependencies (including common essentials) among top 60 ranked genes in each cancer type computed by hypergeometric test. An adjusted p-value less than 0.05 was considered significantly enriched. Dashed gray horizontal line represented an adjusted p-value at 0.05. y axis was plotted as the negative $\log_{10}$ of the adjusted p-value.

Supp. Fig. 10. GDSC validation of top ranked genes using DNA as seed

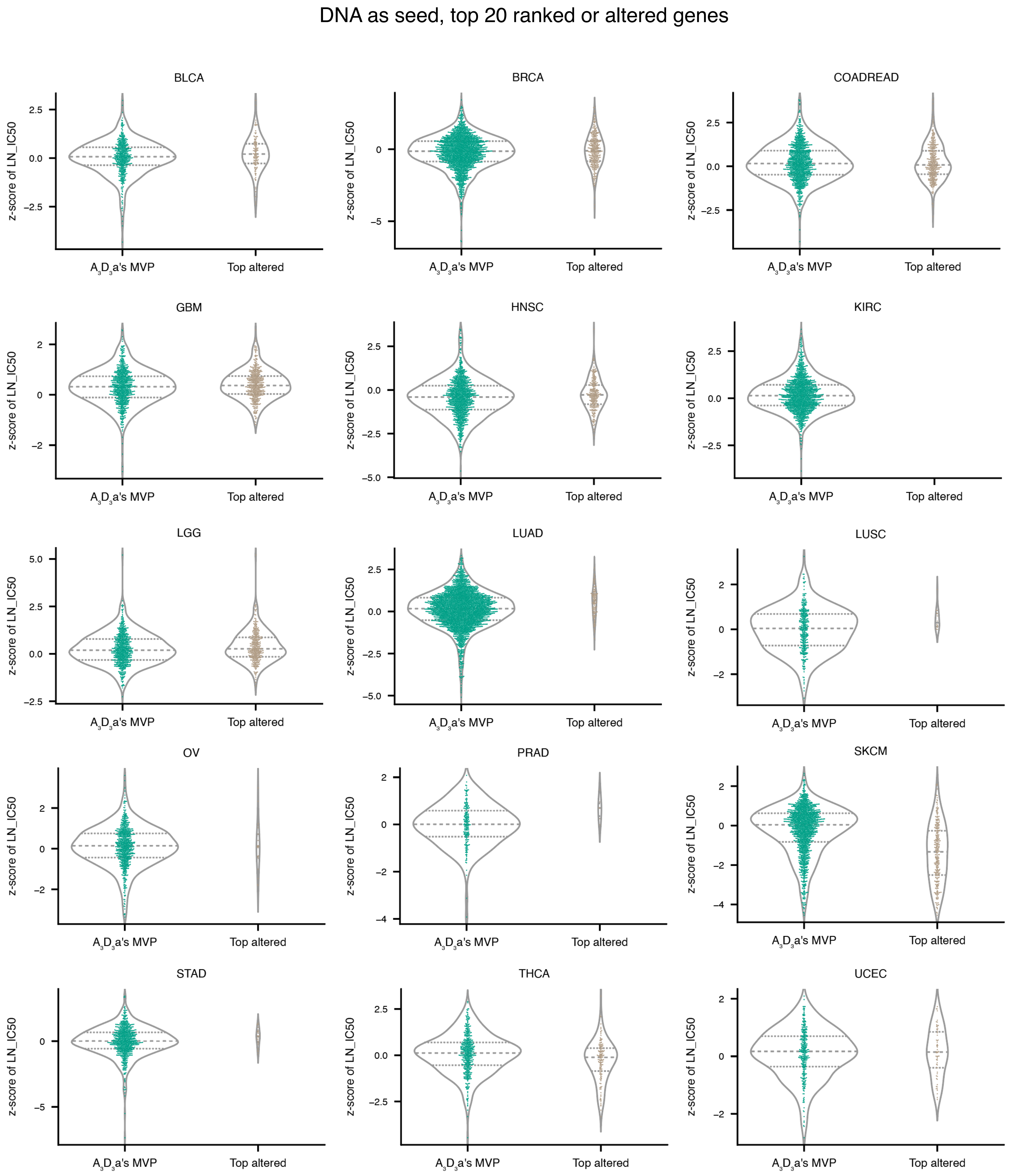


**Supp.Fig.10.** GDSC validation of top ranked genes using DNA as seed. Comparison of z-score of LN_IC50 of dose response of GDSC targets among top 20 ranked genes by $A_3D_3$a's MVP using DNA as seed and top 20 altered genes in seed for each cancer type. Dotted horizontal lines represent the 25$^{th}$ and 75$^{th}$ percentiles, and the dashed line represent the median.

Supp. Fig. 11. GDSC validation of top ranked genes using DNA+RNA as seed

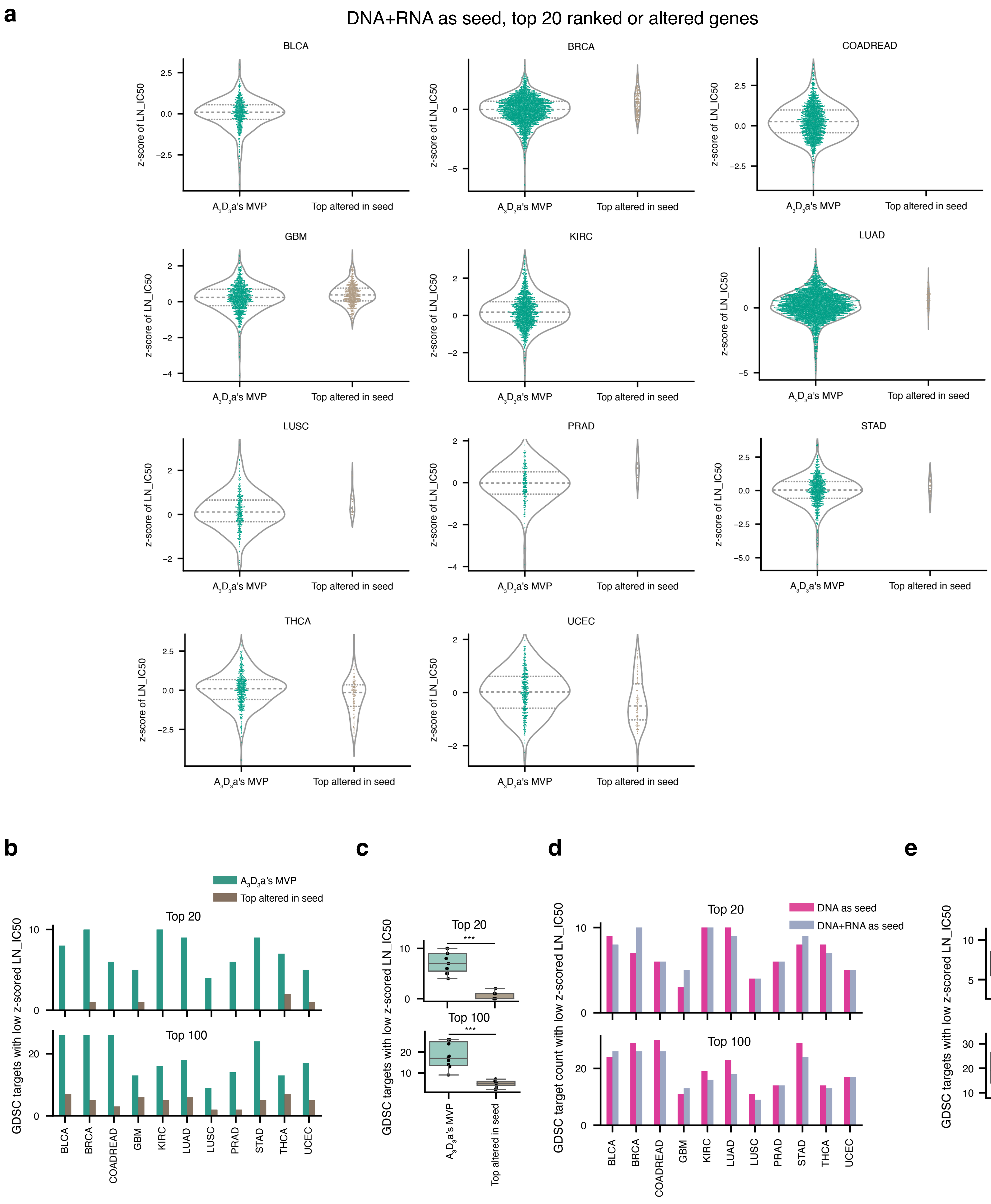


**Supp.Fig.11.** GDSC validation of top ranked genes using DNA+RNA as seed. **a.** Comparison of z-score of LN_IC50 of dose response of GDSC targets among top 20 ranked genes by $A_3D_3$a's MVP using DNA+RNA as seed and top 20 altered genes in seed for each cancer type. Dotted horizontal lines represent the 25$^{th}$ and 75$^{th}$ percentiles, and the dashed line represent the median. **b.** Comparisons of genes with low z-score of LN_IC50 (<-1.5) among top 20 ranked genes by $A_3D_3$a's MVP using DNA+RNA as seed and top 20 altered genes in seed . **c.** Group comparison of data in b. **d.** Comparisons of genes with low z-score of LN_IC50 (<-1.5) among top 20 ranked genes by $A_3D_3$a's MVP using DNA as seed or using DNA+RNA as seed. **e.** Group comparison of data in d.

### Supp. Fig. 12. Setting initial score of known driver genes to zero

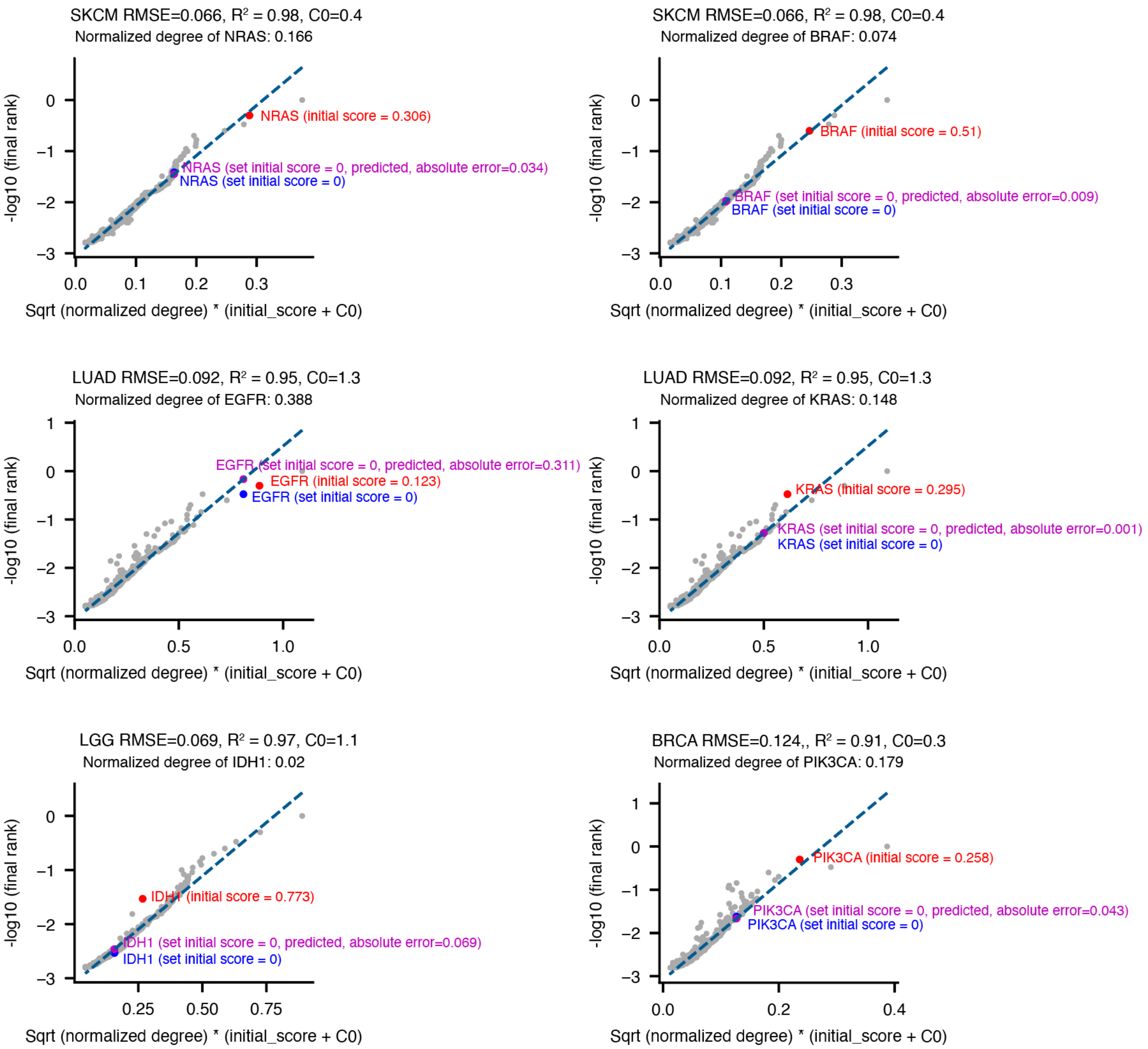


**Supp.Fig.12**. Final rank change after setting initial score of known driver genes to zero (DNA as seed). –log10 (final rank) can be fitted by a linear regression to the multiplication of square root of normalized degree of a given gene in the disease network and the initial score of that gene added by a constant. The constant was tuned for each cancer type specifically by minimizing the root mean squared error (RMSE) of the prediction of linear regression model and the –log10 (final rank). Four example cancer types, SKCM , LUAD, LGG and BRCA were shown. The actual initial score of the driver gene was shown in red. The output of the $A_3D_3$a's MVP model after setting the initial score of the driver gene was shown in blue. The prediction by the linear regression model for setting the initial score of the driver gene was shown in purple.

### Supp. Fig. 13. Network view of the top ranked genes by $A_3D_3a$'s MVP using DNA as seed for the major cancer types

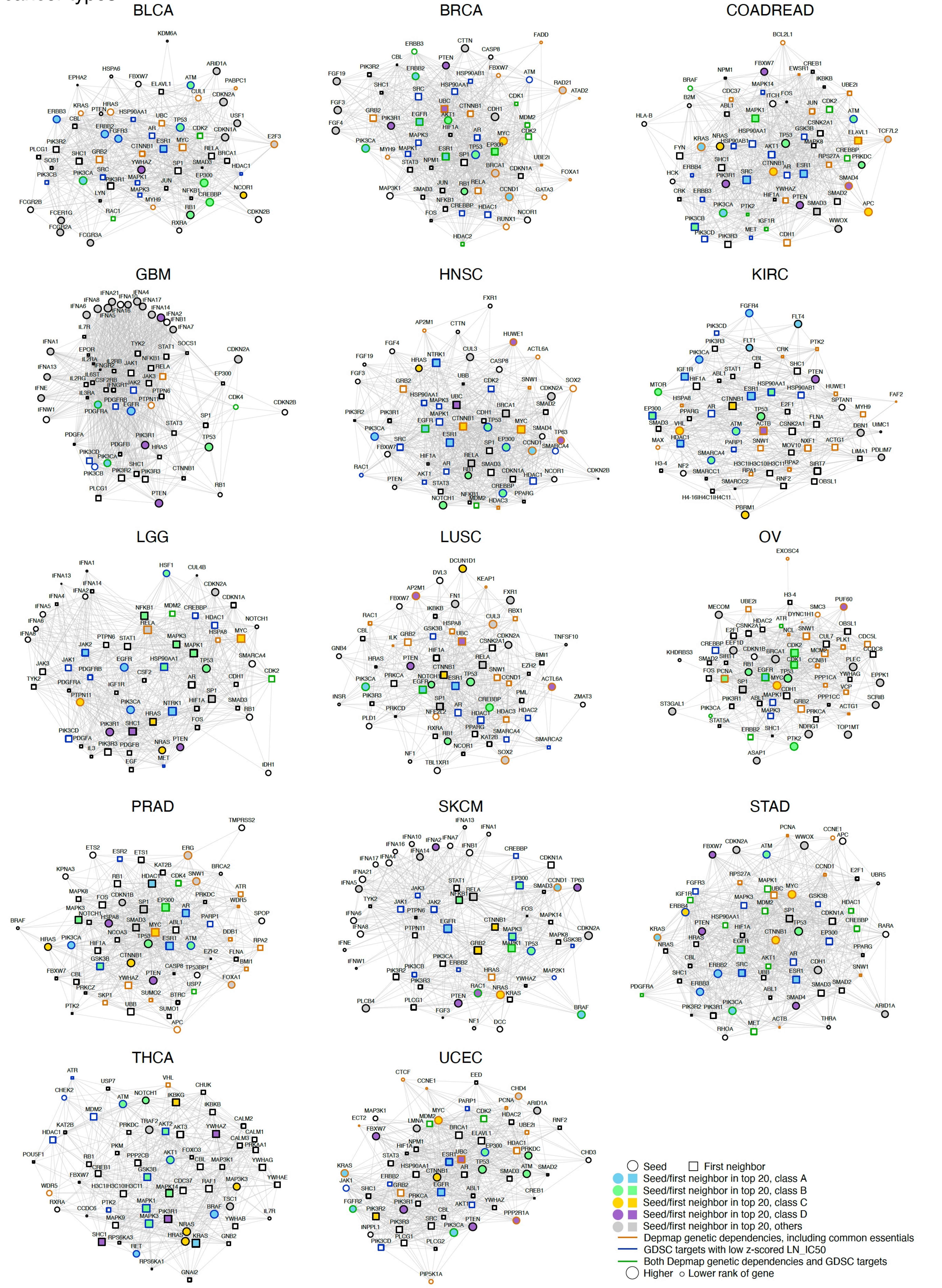


**Supp.Fig.13.** Network view of the top ranked genes by $A_3D_3a$'s MVP using DNA as seed for the major cancer types. The largest connected graph of the top 60 ranked genes by $A_3D_3a$'s MVP for each cancer type was displayed. Top 20 ranked genes were highlighted in blue (class A), green (class B), yellow (class C), purple (class D), and gray (others) face colors. Depmap genetic dependencies (including common essentials) were labeled in orange edge color. GDSC targets with low z-scored LN_IC50 were labeled in blue edge color. The genes that were both genetic dependency and GDSC targets with low z-scored LN_IC50 were labeled in green edge color.

Supp. Fig. 14. Network view of the top ranked genes by $A_3D_3a$'s MVP using DNA+RNA as seed for the major cancer types

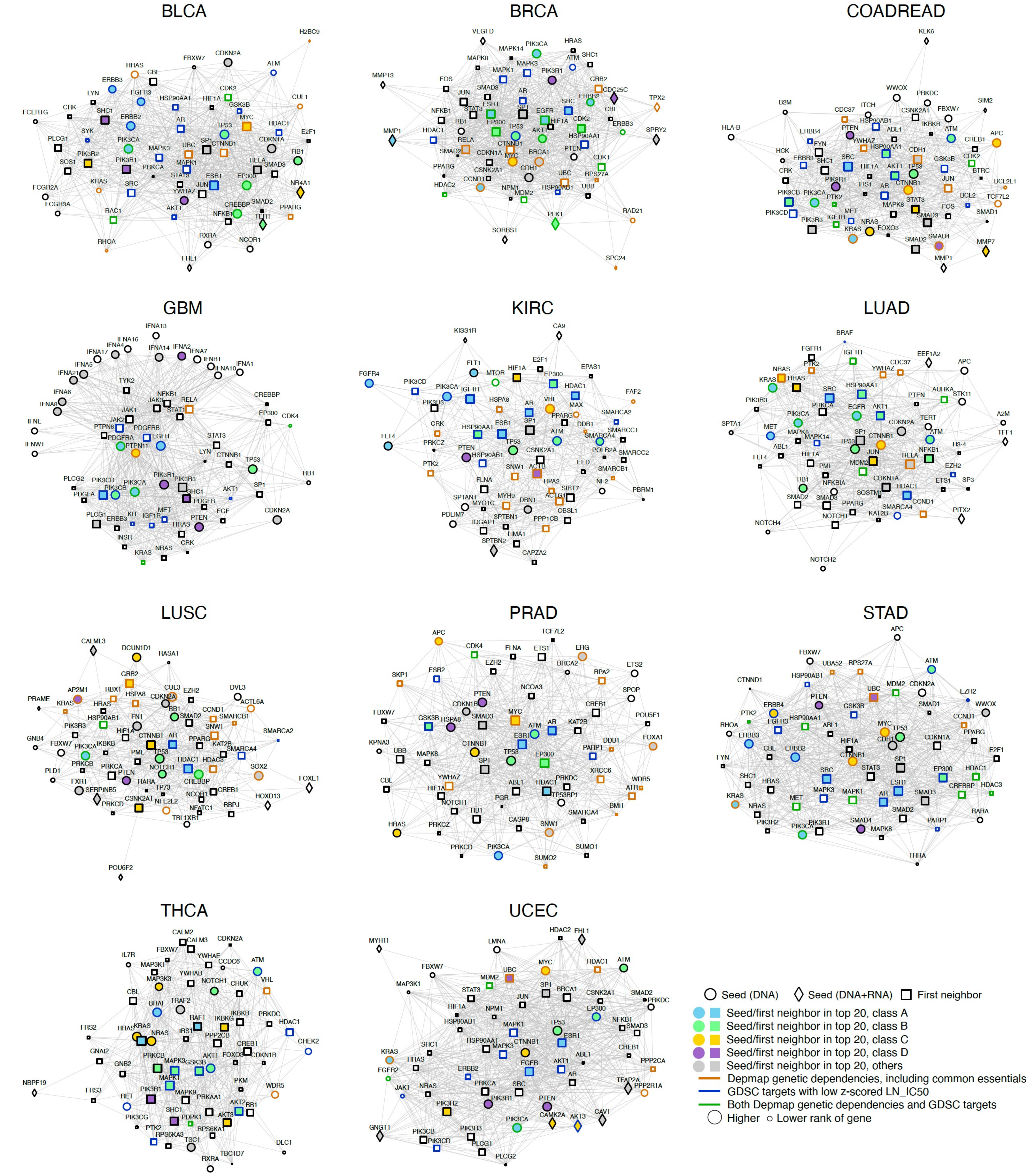


**Supp.Fig.14.** Network view of the top ranked genes by $A_3D_3a$'s MVP using DNA+RNA as seed for the major cancer types. The largest connected graph of the top 60 ranked genes by $A_3D_3a$'s MVP for each cancer type was displayed. Top 20 ranked genes were highlighted in blue (class A), green (class B), yellow (class C), purple (class D), and gray (others) face colors. Depmap genetic dependencies (including common essentials) were labeled in orange edge color. GDSC targets with low z-scored LN_IC50 were labeled in blue edge color. The genes that were both genetic dependency and GDSC targets with low z-scored LN_IC50 were labeled in green edge color.

Supp. Fig. 15. Depmap validation of cancer type in the test set

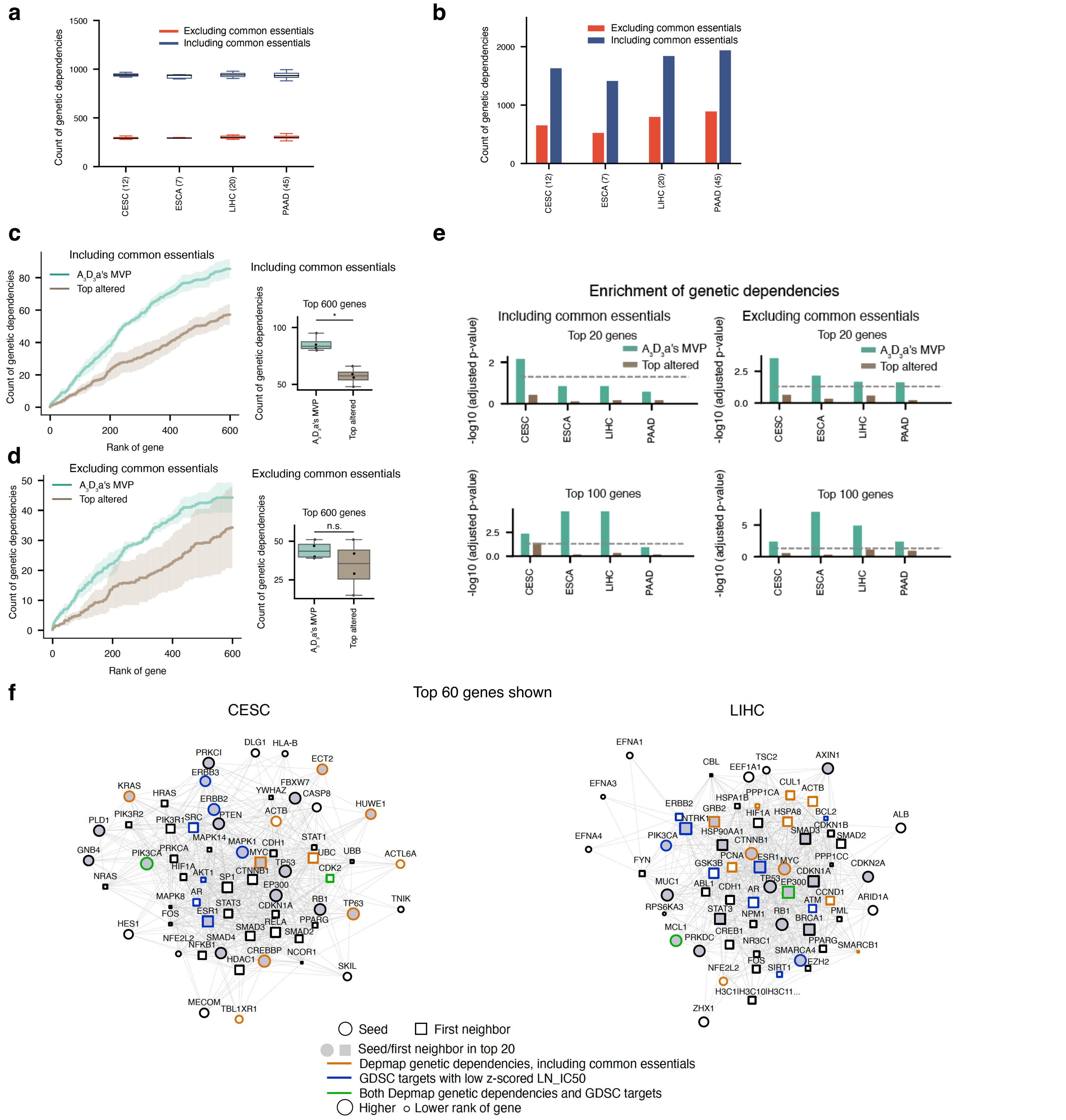


**Supp.Fig.15.** Depmap validation of cancer types in the test set. **a.** Depmap genetic dependencies and common essentials for individual cell lines of each cancer type. **b.** The pooled Depmap genetic dependencies for all cell lines within each cancer type. **c.** (Left panel) mean and standard deviation of count of genetic dependencies (including common essentials) among the top ranked genes by A$_3$D$_3$a's MVP using DNA as seed and top altered genes across cancer types. (Right panel) Group comparison of the count of genetic dependencies (including common essentials) among the top 600 ranked genes by A$_3$D$_3$a's MVP using DNA as seed and top altered genes across cancer types. **d.** The same analyses as e. except that the genetic dependencies did not include the common essentials. **e.** Enrichment of genetic dependencies among top 20 or 100 ranked genes in each cancer type computed by hypergeometric test. An adjusted p-value less than 0.05 was considered significantly enriched. Dashed gray horizontal line represented an adjusted p-value at 0.05. y axis was plotted as the negative log$_{10}$ of the adjusted p-value. **f.** Network view of the top 60 ranked genes by A$_3$D$_3$a's MVP using DNA as seed for the CESC and LIHC. The largest connected graph of the top 60 ranked genes by A$_3$D$_3$a's MVP for each cancer type was displayed. Top 20 ranked genes were highlighted in gray face colors. Depmap genetic dependencies (including common essentials) were labeled in orange edge color. GDSC targets with low z-scored LN_IC50 were labeled in blue edge color. The genes that were both genetic dependency and GDSC targets with low z-scored LN_IC50 were labeled in green edge color. Group comparisons in c and d by two-sided rank-sum test. *: p<0.05, n.s.: not significant.

### Supp. Fig. 16. Compare the prioritized genes by MVP and by Pacini et al. Cancer Cell paper

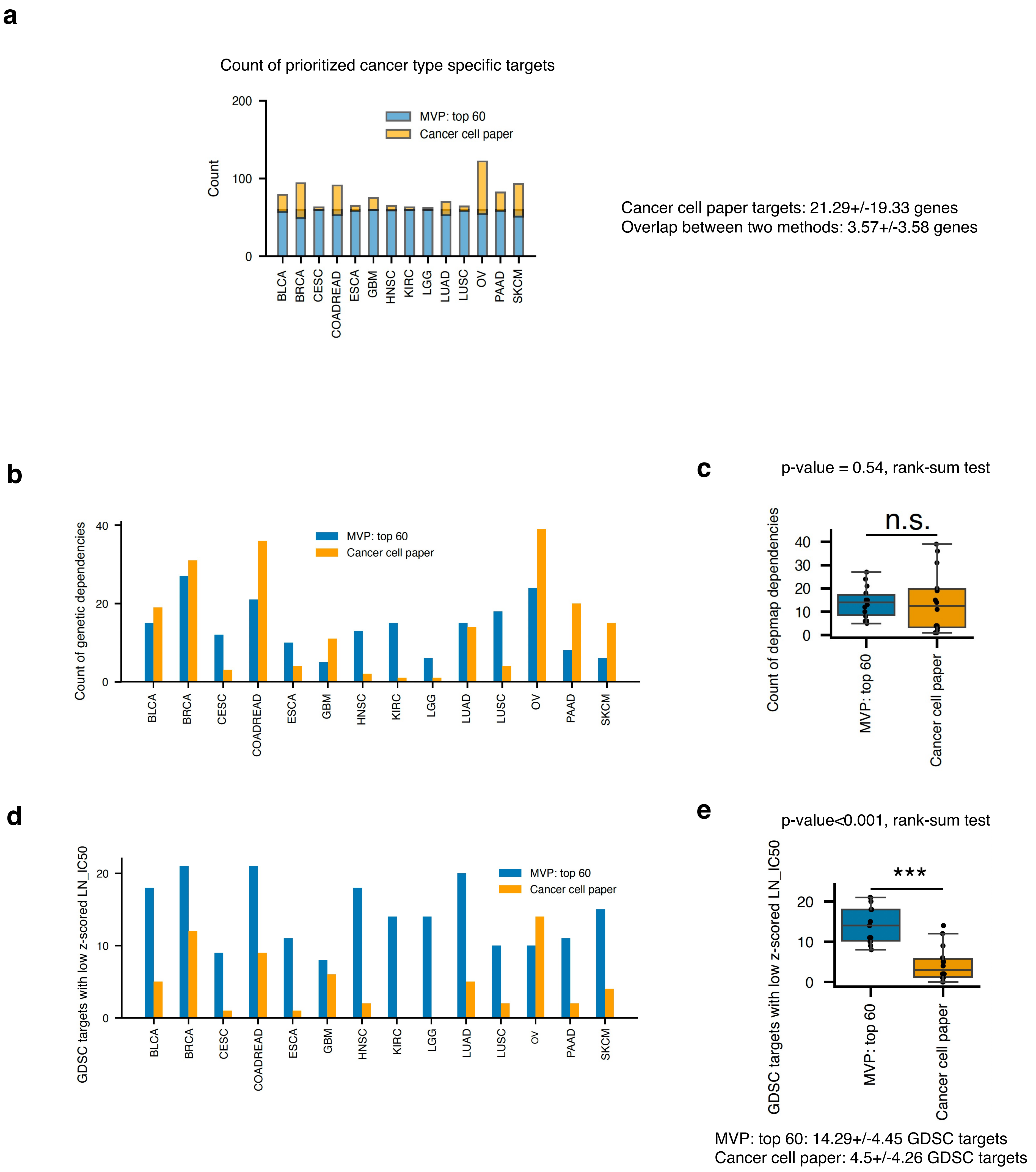


**Supp.Fig.16.** Comparison of the top ranked genes by MVP and by Pacini et al. Cancer Cell paper. (The Cancer Cell paper can be downloaded from https://doi.org/10.1016/j.ccell.2023.12.016). **a.** Count of prioritized genes and the overlap between them of MVP and Cancer Cell paper for the 14 cancer types common in both studies. **b.** Count of depmap dependencies for each cancer type among the prioritized genes of the two studies. **c.** Boxplot of the count data in b. Group comparison by the rank-sum test. **c.** Count of GDSC targets with low z-scored LN_IC50 (<-1.5) for each cancer type among the prioritized genes of the two studies. **d.** Boxplot of the count data in c. Group comparison by the rank-sum test.
