## Supplementary material for "Probabilistic graph-based model uncovers previously unseen druggable vulnerabilities in major solid cancers": supp table 1

| Name | Abbreviation | All Samples From Pancancer Atlas | Selected Tumor Sample Type | Samples with Mutation and CNA | Samples with SV | Tumor RNA | NAT RNA | GTEX RNA | GTEX tissue type |
|---|---|---|---|---|---|---|---|---|---|
| Breast Invasive Carcinoma (TCGA, PanCancer Atlas) | BRCA | 1084 | primary | 1034 | 1066 | 1108 | 113 | 459 | breast_mammary_tissue |
| Colorectal Adenocarcinoma (TCGA, PanCancer Atlas) | COAD,READ | 594 | primary | 581 | 534 | 638 | 51 | 779 | colon sigmoid and tranverse |
| Glioblastoma Multiforme (TCGA, PanCancer Atlas) | GBM | 592 | primary | 378 | 390 | 168 | 5 | 2483 | all 12 brain section |
| Ovarian Serous Cystadenocarcinoma (TCGA, PanCancer Atlas) | OV | 585 | primary | 564 | 521 | 381 | 0 | | |
| Lung Adenocarcinoma (TCGA, PanCancer Atlas) | LUAD | 566 | primary | 502 | 502 | 530 | 59 | 578 | lung |
| Uterine Corpus Endometrial Carcinoma (TCGA, PanCancer Atlas) | UCEC | 529 | primary | 408 | 517 | 550 | 35 | 142 | uterus |
| Head and Neck Squamous Cell Carcinoma (TCGA, PanCancer Atlas) | HNSC | 523 | primary | 517 | 515 | 504 | 44 | | |
| Brain Lower Grade Glioma (TCGA, PanCancer Atlas) | LGG | 514 | primary | 510 | 514 | 532 | 0 | | |
| Kidney Renal Clear Cell Carcinoma (TCGA, PanCancer Atlas) | KIRC | 512 | primary | 468 | 402 | 537 | 72 | 85 | kidney cortex |
| Thyroid Carcinoma (TCGA, PanCancer Atlas) | THCA | 500 | primary | 494 | 489 | 512 | 59 | 653 | thyroid |
| Prostate Adenocarcinoma (TCGA, PanCancer Atlas) | PRAD | 494 | primary | 489 | 494 | 501 | 52 | 245 | prostate |
| Lung Squamous Cell Carcinoma (TCGA, PanCancer Atlas) | LUSC | 487 | primary | 469 | 484 | 501 | 49 | 578 | lung |
| Skin Cutaneous Melanoma (TCGA, PanCancer Atlas) | SKCM | 448 | Met | 363 | 363 | 472 | 1 | | |
| Stomach Adenocarcinoma (TCGA, PanCancer Atlas) | STAD | 440 | primary | 436 | 436 | 375 | 32 | 359 | stomach |
| Bladder Urothelial Carcinoma (TCGA, PanCancer Atlas) | BLCA | 411 | primary | 408 | 410 | 409 | 19 | 21 | bladder |
| Liver Hepatocellular Carcinoma (TCGA, PanCancer Atlas) | LIHC | 372 | primary | 366 | 366 | | | | |
| Cervical Squamous Cell Carcinoma (TCGA, PanCancer Atlas) | CESC | 297 | primary | 291 | 291 | | | | |
| Kidney Renal Papillary Cell Carcinoma (TCGA, PanCancer Atlas) | KIRP | 283 | | 283 | | | | | |
| Sarcoma (TCGA, PanCancer Atlas) | SARC | 255 | | 253 | | | | | |
| Acute Myeloid Leukemia (TCGA, PanCancer Atlas) | LAML | 200 | | 190 | | | | | |
| Pancreatic Adenocarcinoma (TCGA, PanCancer Atlas) | PAAD | 184 | primary | 176 | 184 | | | | |
| Esophageal Adenocarcinoma (TCGA, PanCancer Atlas) | ESCA | 182 | primary | 174 | 182 | | | | |
| Pheochromocytoma and Paraganglioma (TCGA, PanCancer Atlas) | PCPG | 178 | | 161 | | | | | |
| Testicular Germ Cell Tumors (TCGA, PanCancer Atlas) | TGCT | 149 | | 149 | | | | | |
| Thymoma (TCGA, PanCancer Atlas) | THYM | 123 | | 123 | | | | | |
| Adrenocortical Carcinoma (TCGA, PanCancer Atlas) | ACC | 92 | | 89 | | | | | |
| Mesothelioma (TCGA, PanCancer Atlas) | MESO | 87 | | 83 | | | | | |
| Uveal Melanoma (TCGA, PanCancer Atlas) | UVM | 80 | | 80 | | | | | |
| Kidney Chromophobe (TCGA, PanCancer Atlas) | KICH | 65 | | 65 | | | | | |
| Uterine Carcinosarcoma (TCGA, PanCancer Atlas) | UCS | 57 | | 56 | | | | | |
| Diffuse Large B-Cell Lymphoma (TCGA, PanCancer Atlas) | DLBC | 48 | | 48 | | | | | |
| Cholangiocarcinoma (TCGA, PanCancer Atlas) | CHOL | 36 | | 36 | | | | | |
