## Supplementary material for "Probabilistic graph-based model uncovers previously unseen druggable vulnerabilities in major solid cancers": supp table 2

| Cancer | Gene | Source |
|---|---|---|
| BLCA | CDKN2A | CNV |
| BLCA | CDKN2B | CNV |
| BLCA | MTAP | CNV |
| BLCA | DMRTA1 | CNV |
| BLCA | CDKAL1 | CNV |
| BLCA | ARHGAP30 | CNV |
| BLCA | PFDN2 | CNV |
| BLCA | KLHDC9 | CNV |
| BLCA | PVRL4 | CNV |
| BLCA | TSTD1 | CNV |
| BLCA | DEDD | CNV |
| BLCA | NIT1 | CNV |
| BLCA | USF1 | CNV |
| BLCA | F11R | CNV |
| BLCA | ITLN2 | CNV |
| BLCA | UFC1 | CNV |
| BLCA | ADAMTS4 | CNV |
| BLCA | NR1I3 | CNV |
| BLCA | PPOX | CNV |
| BLCA | B4GALT3 | CNV |
| BLCA | APOA2 | CNV |
| BLCA | PCP4L1 | CNV |
| BLCA | NDUFS2 | CNV |
| BLCA | TOMM40L | CNV |
| BLCA | FCER1G | CNV |
| BLCA | USP21 | CNV |
| BLCA | SOX4 | CNV |
| BLCA | ZNF706 | CNV |
| BLCA | MPZ | CNV |
| BLCA | SDHC | CNV |
| BLCA | YWHAZ | CNV |
| BLCA | E2F3 | CNV |
| BLCA | ITLN1 | CNV |
| BLCA | CD244 | CNV |
| BLCA | C1orf192 | CNV |
| BLCA | PABPC1 | CNV |
| BLCA | GRHL2 | CNV |
| BLCA | RNF19A | CNV |
| BLCA | ANKRD46 | CNV |
| BLCA | SNX31 | CNV |
| BLCA | SPAG1 | CNV |
| BLCA | LY9 | CNV |

| | | |
|---|---|---|
| BLCA | FCGR2C | CNV |
| BLCA | NCALD | CNV |
| BLCA | HSPA6 | CNV |
| BLCA | FCGR3A | CNV |
| BLCA | FCGR2B | CNV |
| BLCA | FCGR3B | CNV |
| BLCA | FCGR2A | CNV |
| BLCA | NOS1AP | CNV |
| BLCA | TP53 | Mutation |
| BLCA | KDM6A | Mutation |
| BLCA | PIK3CA | Mutation |
| BLCA | ARID1A | Mutation |
| BLCA | RB1 | Mutation |
| BLCA | ELF3 | Mutation |
| BLCA | STAG2 | Mutation |
| BLCA | EP300 | Mutation |
| BLCA | FGFR3 | Mutation |
| BLCA | CREBBP | Mutation |
| BLCA | CDKN2A | Mutation |
| BLCA | CDKN1A | Mutation |
| BLCA | ATM | Mutation |
| BLCA | RHOB | Mutation |
| BLCA | ERBB2 | Mutation |
| BLCA | TSC1 | Mutation |
| BLCA | SPTAN1 | Mutation |
| BLCA | ERCC2 | Mutation |
| BLCA | FAT1 | Mutation |
| BLCA | ASXL2 | Mutation |
| BLCA | FBXW7 | Mutation |
| BLCA | SYNE1 | Mutation |
| BLCA | ERBB3 | Mutation |
| BLCA | ARID2 | Mutation |
| BLCA | RXRA | Mutation |
| BLCA | NFE2L2 | Mutation |
| BLCA | COL7A1 | Mutation |
| BLCA | RHOA | Mutation |
| BLCA | MACF1 | Mutation |
| BLCA | PARD3 | Mutation |
| BLCA | C3orf70 | Mutation |
| BLCA | EPHA2 | Mutation |
| BLCA | AHR | Mutation |
| BLCA | MYH9 | Mutation |
| BLCA | ASXL1 | Mutation |

| | | |
|---|---|---|
| BLCA | HRAS | Mutation |
| BLCA | PSIP1 | Mutation |
| BLCA | SF3B1 | Mutation |
| BLCA | NCOR1 | Mutation |
| BLCA | ARID1B | Mutation |
| BLCA | KRAS | Mutation |
| BLCA | NF1 | Mutation |
| BLCA | PTEN | Mutation |
| BLCA | ARHGAP35 | Mutation |
| BLCA | TRRAP | Mutation |
| BLCA | CUL1 | Mutation |
| BLCA | CHD2 | Mutation |
| BLCA | HMCN1 | Mutation |
| BLCA | PHF3 | Mutation |
| BLCA | KLF5 | Mutation |
| BLCA | CDKAL1 | SV |
| BLCA | TACC3 | SV |
| BRCA | MYC | CNV |
| BRCA | CCND1 | CNV |
| BRCA | ORAOV1 | CNV |
| BRCA | POU5F1B | CNV |
| BRCA | FGF19 | CNV |
| BRCA | FGF4 | CNV |
| BRCA | FGF3 | CNV |
| BRCA | ANO1 | CNV |
| BRCA | FADD | CNV |
| BRCA | PPFIA1 | CNV |
| BRCA | EIF3H | CNV |
| BRCA | TRPS1 | CNV |
| BRCA | CTTN | CNV |
| BRCA | SHANK2 | CNV |
| BRCA | UTP23 | CNV |
| BRCA | SLC30A8 | CNV |
| BRCA | SAMD12 | CNV |
| BRCA | RAD21 | CNV |
| BRCA | FAM84B | CNV |
| BRCA | FER1L6 | CNV |
| BRCA | ANXA13 | CNV |
| BRCA | NSMCE2 | CNV |
| BRCA | MED30 | CNV |
| BRCA | FAM91A1 | CNV |
| BRCA | TRIB1 | CNV |
| BRCA | KIAA0196 | CNV |

| | | |
|---|---|---|
| BRCA | EXT1 | CNV |
| BRCA | ERLIN2 | CNV |
| BRCA | KLHL38 | CNV |
| BRCA | MAL2 | CNV |
| BRCA | COLEC10 | CNV |
| BRCA | TNFRSF11B | CNV |
| BRCA | GPR124 | CNV |
| BRCA | TMEM65 | CNV |
| BRCA | ZNF703 | CNV |
| BRCA | MYEOV | CNV |
| BRCA | MTSS1 | CNV |
| BRCA | ATAD2 | CNV |
| BRCA | RNF139 | CNV |
| BRCA | ASAP1 | CNV |
| BRCA | TATDN1 | CNV |
| BRCA | COL14A1 | CNV |
| BRCA | PROSC | CNV |
| BRCA | NDUFB9 | CNV |
| BRCA | NOV | CNV |
| BRCA | TAF2 | CNV |
| BRCA | TRMT12 | CNV |
| BRCA | FAM83A | CNV |
| BRCA | SNTB1 | CNV |
| BRCA | CSMD3 | CNV |
| BRCA | PIK3CA | Mutation |
| BRCA | TP53 | Mutation |
| BRCA | CDH1 | Mutation |
| BRCA | GATA3 | Mutation |
| BRCA | MAP3K1 | Mutation |
| BRCA | PTEN | Mutation |
| BRCA | NCOR1 | Mutation |
| BRCA | MAP2K4 | Mutation |
| BRCA | ARID1A | Mutation |
| BRCA | NF1 | Mutation |
| BRCA | RUNX1 | Mutation |
| BRCA | FOXA1 | Mutation |
| BRCA | ERBB2 | Mutation |
| BRCA | RB1 | Mutation |
| BRCA | TBX3 | Mutation |
| BRCA | PIK3R1 | Mutation |
| BRCA | CBFB | Mutation |
| BRCA | AKT1 | Mutation |
| BRCA | CTCF | Mutation |

| | | |
|---|---|---|
| BRCA | BRCA1 | Mutation |
| BRCA | CHD4 | Mutation |
| BRCA | KDM6A | Mutation |
| BRCA | SF3B1 | Mutation |
| BRCA | BRCA2 | Mutation |
| BRCA | MYH9 | Mutation |
| BRCA | SYNE1 | Mutation |
| BRCA | MUC17 | Mutation |
| BRCA | SPEN | Mutation |
| BRCA | FBXW7 | Mutation |
| BRCA | NIPBL | Mutation |
| BRCA | MEN1 | Mutation |
| BRCA | DIAPH1 | Mutation |
| BRCA | GPS2 | Mutation |
| BRCA | PHKA2 | Mutation |
| BRCA | CASP8 | Mutation |
| BRCA | EYS | Mutation |
| BRCA | ABCA7 | Mutation |
| BRCA | ERBB3 | Mutation |
| BRCA | TTN | Mutation |
| BRCA | ATP10B | Mutation |
| BRCA | MYB | Mutation |
| BRCA | SPTA1 | Mutation |
| BRCA | BAP1 | Mutation |
| BRCA | ANKHD1 | Mutation |
| BRCA | COL6A6 | Mutation |
| BRCA | HCFC2 | Mutation |
| BRCA | DMD | Mutation |
| BRCA | HIVEP1 | Mutation |
| BRCA | ATM | Mutation |
| BRCA | ZMYM3 | Mutation |
| BRCA | SHANK2 | SV |
| BRCA | BCAS3 | SV |
| BRCA | FBXL20 | SV |
| COADREAD | MACROD2 | CNV |
| COADREAD | WWOX | CNV |
| COADREAD | PLAGL2 | CNV |
| COADREAD | TM9SF4 | CNV |
| COADREAD | POFUT1 | CNV |
| COADREAD | ASXL1 | CNV |
| COADREAD | KIF3B | CNV |
| COADREAD | HCK | CNV |
| COADREAD | DEFB119 | CNV |

| | | |
|---|---|---|
| COADREAD | FOXS1 | CNV |
| COADREAD | HM13 | CNV |
| COADREAD | DEFB121 | CNV |
| COADREAD | DEFB118 | CNV |
| COADREAD | C20orf112 | CNV |
| COADREAD | PDRG1 | CNV |
| COADREAD | DEFB116 | CNV |
| COADREAD | DEFB115 | CNV |
| COADREAD | MYLK2 | CNV |
| COADREAD | TPX2 | CNV |
| COADREAD | TTLL9 | CNV |
| COADREAD | XKR7 | CNV |
| COADREAD | DEFB123 | CNV |
| COADREAD | ITCH | CNV |
| COADREAD | DEFB124 | CNV |
| COADREAD | COX4I2 | CNV |
| COADREAD | ID1 | CNV |
| COADREAD | REM1 | CNV |
| COADREAD | DUSP15 | CNV |
| COADREAD | AHCY | CNV |
| COADREAD | NFS1 | CNV |
| COADREAD | MAP1LC3A | CNV |
| COADREAD | DYNLRB1 | CNV |
| COADREAD | PIGU | CNV |
| COADREAD | RBM39 | CNV |
| COADREAD | EIF2S2 | CNV |
| COADREAD | DLGAP4 | CNV |
| COADREAD | PHF20 | CNV |
| COADREAD | ROMO1 | CNV |
| COADREAD | BCL2L1 | CNV |
| COADREAD | PKIG | CNV |
| COADREAD | EPB41L1 | CNV |
| COADREAD | NCOA6 | CNV |
| COADREAD | C20orf203 | CNV |
| COADREAD | FITM2 | CNV |
| COADREAD | TP53INP2 | CNV |
| COADREAD | TTPAL | CNV |
| COADREAD | SCAND1 | CNV |
| COADREAD | RALY | CNV |
| COADREAD | ASIP | CNV |
| COADREAD | COMMD7 | CNV |
| COADREAD | APC | Mutation |
| COADREAD | TP53 | Mutation |

| | | |
|---|---|---|
| COADREAD | KRAS | Mutation |
| COADREAD | FBXW7 | Mutation |
| COADREAD | ARID1A | Mutation |
| COADREAD | SOX9 | Mutation |
| COADREAD | PIK3CA | Mutation |
| COADREAD | SMAD4 | Mutation |
| COADREAD | RNF43 | Mutation |
| COADREAD | ATM | Mutation |
| COADREAD | TCF7L2 | Mutation |
| COADREAD | MBD6 | Mutation |
| COADREAD | BRAF | Mutation |
| COADREAD | PTEN | Mutation |
| COADREAD | ZFP36L2 | Mutation |
| COADREAD | BMPR2 | Mutation |
| COADREAD | DOCK3 | Mutation |
| COADREAD | BCL9L | Mutation |
| COADREAD | DYSF | Mutation |
| COADREAD | ACVR2A | Mutation |
| COADREAD | BCL9 | Mutation |
| COADREAD | ZNF469 | Mutation |
| COADREAD | RYR2 | Mutation |
| COADREAD | CTNNB1 | Mutation |
| COADREAD | CIC | Mutation |
| COADREAD | PIK3R1 | Mutation |
| COADREAD | FLNB | Mutation |
| COADREAD | ASXL1 | Mutation |
| COADREAD | HLA-B | Mutation |
| COADREAD | NRAS | Mutation |
| COADREAD | PCBP1 | Mutation |
| COADREAD | LARP4B | Mutation |
| COADREAD | ANK3 | Mutation |
| COADREAD | PRKDC | Mutation |
| COADREAD | CSMD3 | Mutation |
| COADREAD | B2M | Mutation |
| COADREAD | DNAH12 | Mutation |
| COADREAD | PLEKHA6 | Mutation |
| COADREAD | BCORL1 | Mutation |
| COADREAD | SYNE1 | Mutation |
| COADREAD | ZBTB20 | Mutation |
| COADREAD | TTN | Mutation |
| COADREAD | NFASC | Mutation |
| COADREAD | DNAH11 | Mutation |
| COADREAD | CHD3 | Mutation |

| | | |
|---|---|---|
| COADREAD | EYS | Mutation |
| COADREAD | MECOM | Mutation |
| COADREAD | ARHGAP5 | Mutation |
| COADREAD | MVK | Mutation |
| COADREAD | FHOD3 | Mutation |
| GBM | CDKN2A | CNV |
| GBM | CDKN2B | CNV |
| GBM | EGFR | CNV |
| GBM | MTAP | CNV |
| GBM | SEC61G | CNV |
| GBM | LANCL2 | CNV |
| GBM | DMRTA1 | CNV |
| GBM | IFNE | CNV |
| GBM | IFNA1 | CNV |
| GBM | IFNA8 | CNV |
| GBM | IFNA13 | CNV |
| GBM | IFNA2 | CNV |
| GBM | KLHL9 | CNV |
| GBM | IFNA6 | CNV |
| GBM | IFNA5 | CNV |
| GBM | VOPP1 | CNV |
| GBM | VSTM2A | CNV |
| GBM | IFNA14 | CNV |
| GBM | IFNA17 | CNV |
| GBM | IFNA4 | CNV |
| GBM | IFNA10 | CNV |
| GBM | IFNA16 | CNV |
| GBM | IFNA7 | CNV |
| GBM | IFNA21 | CNV |
| GBM | IFNW1 | CNV |
| GBM | ELAVL2 | CNV |
| GBM | IFNB1 | CNV |
| GBM | PTPLAD2 | CNV |
| GBM | 9-Mar | CNV |
| GBM | CDK4 | CNV |
| GBM | TSPAN31 | CNV |
| GBM | AGAP2 | CNV |
| GBM | CYP27B1 | CNV |
| GBM | 14-Sep | CNV |
| GBM | METTL1 | CNV |
| GBM | TSFM | CNV |
| GBM | PDGFRA | CNV |
| GBM | OS9 | CNV |

| | | |
|---|---|---|
| GBM | GSX2 | CNV |
| GBM | GBAS | CNV |
| GBM | ZNF713 | CNV |
| GBM | AVIL | CNV |
| GBM | CHIC2 | CNV |
| GBM | PSPH | CNV |
| GBM | MRPS17 | CNV |
| GBM | CTDSP2 | CNV |
| GBM | MLLT3 | CNV |
| GBM | TUSC1 | CNV |
| GBM | CCT6A | CNV |
| GBM | SUMF2 | CNV |
| GBM | PTEN | Mutation |
| GBM | TP53 | Mutation |
| GBM | PIK3R1 | Mutation |
| GBM | NF1 | Mutation |
| GBM | EGFR | Mutation |
| GBM | RB1 | Mutation |
| GBM | PIK3CA | Mutation |
| GBM | IDH1 | Mutation |
| GBM | ATRX | Mutation |
| GBM | STAG2 | Mutation |
| GBM | KEL | Mutation |
| GBM | LZTR1 | Mutation |
| GBM | CHD8 | Mutation |
| GBM | MUC17 | Mutation |
| GBM | F5 | Mutation |
| GBM | QKI | Mutation |
| GBM | BRAF | Mutation |
| GBM | FLG | Mutation |
| GBM | PTPN11 | Mutation |
| GBM | SYNE1 | Mutation |
| GBM | RPL5 | Mutation |
| GBM | LRP2 | Mutation |
| GBM | ASTL | Mutation |
| GBM | DEPDC5 | Mutation |
| GBM | COL1A2 | Mutation |
| GBM | FRMD7 | Mutation |
| GBM | UGT2A3 | Mutation |
| GBM | NOTCH2 | Mutation |
| GBM | CDKN2C | Mutation |
| GBM | ACRC | Mutation |
| GBM | SLC26A3 | Mutation |

| | | |
|---|---|---|
| GBM | DAO | Mutation |
| GBM | FAM120B | Mutation |
| GBM | TTN | Mutation |
| GBM | TCF12 | Mutation |
| GBM | PIK3CB | Mutation |
| GBM | PRPF40B | Mutation |
| GBM | RIMS2 | Mutation |
| GBM | TEX15 | Mutation |
| GBM | FBN2 | Mutation |
| GBM | IL18RAP | Mutation |
| GBM | EXOC2 | Mutation |
| GBM | MYOF | Mutation |
| GBM | AHNAK2 | Mutation |
| GBM | MYH11 | Mutation |
| GBM | PIK3C2G | Mutation |
| GBM | KRT34 | Mutation |
| GBM | MAP3K1 | Mutation |
| GBM | GABRA6 | Mutation |
| GBM | PDGFRA | Mutation |
| GBM | EGFR | SV |
| GBM | SEPTIN14 | SV |
| GBM | TSFM | SV |
| GBM | OS9 | SV |
| GBM | SEC61G | SV |
| HNSC | CDKN2A | CNV |
| HNSC | CDKN2B | CNV |
| HNSC | PPFIA1 | CNV |
| HNSC | FADD | CNV |
| HNSC | CTTN | CNV |
| HNSC | ANO1 | CNV |
| HNSC | SHANK2 | CNV |
| HNSC | FGF3 | CNV |
| HNSC | FGF4 | CNV |
| HNSC | FGF19 | CNV |
| HNSC | CCND1 | CNV |
| HNSC | ORAOV1 | CNV |
| HNSC | CSMD1 | CNV |
| HNSC | MYEOV | CNV |
| HNSC | TP63 | CNV |
| HNSC | TPCN2 | CNV |
| HNSC | LEPREL1 | CNV |
| HNSC | SOX2 | CNV |
| HNSC | ZMAT3 | CNV |

| | | |
|---|---|---|
| HNSC | PIK3CA | CNV |
| HNSC | LAMP3 | CNV |
| HNSC | DCUN1D1 | CNV |
| HNSC | KCNMB3 | CNV |
| HNSC | TPRG1 | CNV |
| HNSC | ATP11B | CNV |
| HNSC | MCCC1 | CNV |
| HNSC | NAALADL2 | CNV |
| HNSC | MCF2L2 | CNV |
| HNSC | ACTL6A | CNV |
| HNSC | USP13 | CNV |
| HNSC | GNB4 | CNV |
| HNSC | MFN1 | CNV |
| HNSC | NDUFB5 | CNV |
| HNSC | FXR1 | CNV |
| HNSC | DNAJC19 | CNV |
| HNSC | ZNF639 | CNV |
| HNSC | MRPL47 | CNV |
| HNSC | PEX5L | CNV |
| HNSC | AP2M1 | CNV |
| HNSC | DVL3 | CNV |
| HNSC | PTPRD | CNV |
| HNSC | YEATS2 | CNV |
| HNSC | HTR3E | CNV |
| HNSC | KLHL24 | CNV |
| HNSC | TBL1XR1 | CNV |
| HNSC | HTR3C | CNV |
| HNSC | KCNMB2 | CNV |
| HNSC | MAP6D1 | CNV |
| HNSC | HTR3D | CNV |
| HNSC | ABCC5 | CNV |
| HNSC | TP53 | Mutation |
| HNSC | CDKN2A | Mutation |
| HNSC | FAT1 | Mutation |
| HNSC | NSD1 | Mutation |
| HNSC | NOTCH1 | Mutation |
| HNSC | CASP8 | Mutation |
| HNSC | PIK3CA | Mutation |
| HNSC | EP300 | Mutation |
| HNSC | TGFBR2 | Mutation |
| HNSC | FBXW7 | Mutation |
| HNSC | HUWE1 | Mutation |
| HNSC | EPHA2 | Mutation |

| | | |
|---|---|---|
| HNSC | HLA-B | Mutation |
| HNSC | HRAS | Mutation |
| HNSC | NFE2L2 | Mutation |
| HNSC | CREBBP | Mutation |
| HNSC | ZNF750 | Mutation |
| HNSC | KDM6A | Mutation |
| HNSC | HLA-A | Mutation |
| HNSC | DYSF | Mutation |
| HNSC | RASA1 | Mutation |
| HNSC | RAC1 | Mutation |
| HNSC | MYH9 | Mutation |
| HNSC | CTCF | Mutation |
| HNSC | TTN | Mutation |
| HNSC | RB1 | Mutation |
| HNSC | SMARCA4 | Mutation |
| HNSC | KEAP1 | Mutation |
| HNSC | NOTCH2 | Mutation |
| HNSC | SYNE1 | Mutation |
| HNSC | DNAH5 | Mutation |
| HNSC | PTEN | Mutation |
| HNSC | ASXL3 | Mutation |
| HNSC | SEMA5A | Mutation |
| HNSC | SMAD4 | Mutation |
| HNSC | TANC1 | Mutation |
| HNSC | NCOR1 | Mutation |
| HNSC | AKAP9 | Mutation |
| HNSC | FAT2 | Mutation |
| HNSC | RHOA | Mutation |
| HNSC | AK5 | Mutation |
| HNSC | CUL3 | Mutation |
| HNSC | NAA25 | Mutation |
| HNSC | FNBP4 | Mutation |
| HNSC | ASXL1 | Mutation |
| HNSC | PCDH9 | Mutation |
| HNSC | ARID2 | Mutation |
| HNSC | GIGYF2 | Mutation |
| HNSC | MUC16 | Mutation |
| HNSC | NUMA1 | Mutation |
| KIRC | FAM193B | CNV |
| KIRC | RAB24 | CNV |
| KIRC | OR4F16 | CNV |
| KIRC | SLC34A1 | CNV |
| KIRC | UIMC1 | CNV |

| | | |
|---|---|---|
| KIRC | RGS14 | CNV |
| KIRC | ZNF346 | CNV |
| KIRC | MXD3 | CNV |
| KIRC | PRELID1 | CNV |
| KIRC | ERGIC1 | CNV |
| KIRC | GRK6 | CNV |
| KIRC | PDLIM7 | CNV |
| KIRC | DBN1 | CNV |
| KIRC | NKX2-5 | CNV |
| KIRC | HK3 | CNV |
| KIRC | LMAN2 | CNV |
| KIRC | NEURL1B | CNV |
| KIRC | PFN3 | CNV |
| KIRC | STC2 | CNV |
| KIRC | FGFR4 | CNV |
| KIRC | DOK3 | CNV |
| KIRC | OR4F29 | CNV |
| KIRC | DDX41 | CNV |
| KIRC | B4GALT7 | CNV |
| KIRC | TMED9 | CNV |
| KIRC | BNIP1 | CNV |
| KIRC | PRR7 | CNV |
| KIRC | DUSP1 | CNV |
| KIRC | BTNL8 | CNV |
| KIRC | F12 | CNV |
| KIRC | NSD1 | CNV |
| KIRC | FAM153C | CNV |
| KIRC | C5orf45 | CNV |
| KIRC | PROP1 | CNV |
| KIRC | GFPT2 | CNV |
| KIRC | TRIM7 | CNV |
| KIRC | KIAA1191 | CNV |
| KIRC | TRIM41 | CNV |
| KIRC | DRD1 | CNV |
| KIRC | BOD1 | CNV |
| KIRC | ZNF879 | CNV |
| KIRC | RNF44 | CNV |
| KIRC | MAML1 | CNV |
| KIRC | COL23A1 | CNV |
| KIRC | FLT4 | CNV |
| KIRC | CNOT6 | CNV |
| KIRC | CLK4 | CNV |
| KIRC | FAF2 | CNV |

| | | |
|---|---|---|
| KIRC | HNRNPAB | CNV |
| KIRC | ZNF354B | CNV |
| KIRC | VHL | Mutation |
| KIRC | SETD2 | Mutation |
| KIRC | PBRM1 | Mutation |
| KIRC | BAP1 | Mutation |
| KIRC | MTOR | Mutation |
| KIRC | KDM5C | Mutation |
| KIRC | PTEN | Mutation |
| KIRC | ARID1A | Mutation |
| KIRC | ATM | Mutation |
| KIRC | TP53 | Mutation |
| KIRC | SMARCA4 | Mutation |
| KIRC | RTTN | Mutation |
| KIRC | SPEN | Mutation |
| KIRC | ZNF800 | Mutation |
| KIRC | PIK3CA | Mutation |
| KIRC | NF2 | Mutation |
| KIRC | DENND4A | Mutation |
| KIRC | UBR1 | Mutation |
| KIRC | STAG2 | Mutation |
| KIRC | SDAD1 | Mutation |
| KIRC | ACLY | Mutation |
| KIRC | HMCN1 | Mutation |
| KIRC | KIF1A | Mutation |
| KIRC | SDR16C5 | Mutation |
| KIRC | MGA | Mutation |
| KIRC | CDK12 | Mutation |
| KIRC | PCK1 | Mutation |
| KIRC | ARHGAP35 | Mutation |
| KIRC | NRIP1 | Mutation |
| KIRC | PRPF8 | Mutation |
| KIRC | KIF13B | Mutation |
| KIRC | MAX | Mutation |
| KIRC | OTOF | Mutation |
| KIRC | DST | Mutation |
| KIRC | TGM5 | Mutation |
| KIRC | FLT1 | Mutation |
| KIRC | ROCK1 | Mutation |
| KIRC | PCDHGB3 | Mutation |
| KIRC | AHNAK2 | Mutation |
| KIRC | SPHKAP | Mutation |
| KIRC | SLK | Mutation |

| | | |
|---|---|---|
| KIRC | COL6A6 | Mutation |
| KIRC | ATG7 | Mutation |
| KIRC | SPTAN1 | Mutation |
| KIRC | RNF43 | Mutation |
| KIRC | COL5A3 | Mutation |
| KIRC | MUC17 | Mutation |
| KIRC | OPTC | Mutation |
| KIRC | NFASC | Mutation |
| KIRC | MYH2 | Mutation |
| LGG | CDKN2B | CNV |
| LGG | CDKN2A | CNV |
| LGG | MTAP | CNV |
| LGG | EGFR | CNV |
| LGG | DMRTA1 | CNV |
| LGG | SEC61G | CNV |
| LGG | IFNE | CNV |
| LGG | IFNA1 | CNV |
| LGG | IFNA8 | CNV |
| LGG | ZNF251 | CNV |
| LGG | ZNF250 | CNV |
| LGG | PARP11 | CNV |
| LGG | FBXL6 | CNV |
| LGG | IFNA6 | CNV |
| LGG | IFNA5 | CNV |
| LGG | SCRIB | CNV |
| LGG | MAPK15 | CNV |
| LGG | FOXH1 | CNV |
| LGG | RPL8 | CNV |
| LGG | ADCK5 | CNV |
| LGG | SLC39A4 | CNV |
| LGG | GPT | CNV |
| LGG | EPPK1 | CNV |
| LGG | KIFC2 | CNV |
| LGG | DGAT1 | CNV |
| LGG | ZNF517 | CNV |
| LGG | EXOSC4 | CNV |
| LGG | ZNF7 | CNV |
| LGG | LRRC24 | CNV |
| LGG | MFSD3 | CNV |
| LGG | SHARPIN | CNV |
| LGG | COMMD5 | CNV |
| LGG | OPLAH | CNV |
| LGG | KLHL9 | CNV |

| | | |
|---|---|---|
| LGG | IFNA2 | CNV |
| LGG | CYHR1 | CNV |
| LGG | C8orf33 | CNV |
| LGG | GRINA | CNV |
| LGG | HSF1 | CNV |
| LGG | PPP1R16A | CNV |
| LGG | GPAA1 | CNV |
| LGG | IFNA13 | CNV |
| LGG | ZNF16 | CNV |
| LGG | RECQL4 | CNV |
| LGG | CPSF1 | CNV |
| LGG | PARP10 | CNV |
| LGG | CYC1 | CNV |
| LGG | ZNF707 | CNV |
| LGG | FAM83H | CNV |
| LGG | GSDMD | CNV |
| LGG | IDH1 | Mutation |
| LGG | TP53 | Mutation |
| LGG | ATRX | Mutation |
| LGG | CIC | Mutation |
| LGG | NOTCH1 | Mutation |
| LGG | FUBP1 | Mutation |
| LGG | EGFR | Mutation |
| LGG | NF1 | Mutation |
| LGG | PIK3R1 | Mutation |
| LGG | PIK3CA | Mutation |
| LGG | SMARCA4 | Mutation |
| LGG | IDH2 | Mutation |
| LGG | ARID1A | Mutation |
| LGG | NIPBL | Mutation |
| LGG | PTEN | Mutation |
| LGG | ZBTB20 | Mutation |
| LGG | TCF12 | Mutation |
| LGG | ZNF292 | Mutation |
| LGG | ARID2 | Mutation |
| LGG | SETD2 | Mutation |
| LGG | MYH8 | Mutation |
| LGG | DNMT3A | Mutation |
| LGG | CUL4B | Mutation |
| LGG | PTPN11 | Mutation |
| LGG | KEL | Mutation |
| LGG | CNOT1 | Mutation |
| LGG | RB1 | Mutation |

| | | |
|---|---|---|
| LGG | BCOR | Mutation |
| LGG | KRT15 | Mutation |
| LGG | PDE8A | Mutation |
| LGG | R3HDM1 | Mutation |
| LGG | DOCK5 | Mutation |
| LGG | TLR7 | Mutation |
| LGG | ARID1B | Mutation |
| LGG | MYH11 | Mutation |
| LGG | SLC6A4 | Mutation |
| LGG | SELE | Mutation |
| LGG | FHOD1 | Mutation |
| LGG | NRAS | Mutation |
| LGG | SLC6A3 | Mutation |
| LGG | OGT | Mutation |
| LGG | C4BPA | Mutation |
| LGG | RIPK4 | Mutation |
| LGG | FER1L6 | Mutation |
| LGG | PFKL | Mutation |
| LGG | HCFC1 | Mutation |
| LGG | ANKRD17 | Mutation |
| LGG | WWC3 | Mutation |
| LGG | COL6A3 | Mutation |
| LGG | MYOCD | Mutation |
| LUAD | CDKN2A | CNV |
| LUAD | CDKN2B | CNV |
| LUAD | CLPTM1L | CNV |
| LUAD | SFTA3 | CNV |
| LUAD | NKX2-1 | CNV |
| LUAD | MBIP | CNV |
| LUAD | TRIP13 | CNV |
| LUAD | SLC9A3 | CNV |
| LUAD | C5orf55 | CNV |
| LUAD | PDCD6 | CNV |
| LUAD | SLC6A18 | CNV |
| LUAD | TERT | CNV |
| LUAD | EXOC3 | CNV |
| LUAD | AHRR | CNV |
| LUAD | SDHA | CNV |
| LUAD | CEP72 | CNV |
| LUAD | SLC6A19 | CNV |
| LUAD | TPPP | CNV |
| LUAD | ZDHHC11 | CNV |
| LUAD | BRD9 | CNV |

| | | |
|---|---|---|
| LUAD | LRRC14B | CNV |
| LUAD | PLEKHG4B | CNV |
| LUAD | NKD2 | CNV |
| LUAD | SLC12A7 | CNV |
| LUAD | CCDC127 | CNV |
| LUAD | SLC6A3 | CNV |
| LUAD | SLC25A21 | CNV |
| LUAD | NDUFS6 | CNV |
| LUAD | MRPL36 | CNV |
| LUAD | LPCAT1 | CNV |
| LUAD | IRX4 | CNV |
| LUAD | PAX9 | CNV |
| LUAD | NKX2-8 | CNV |
| LUAD | MTAP | CNV |
| LUAD | RALGAPA1 | CNV |
| LUAD | FAM105B | CNV |
| LUAD | BRMS1L | CNV |
| LUAD | FAM105A | CNV |
| LUAD | TRIO | CNV |
| LUAD | ANKH | CNV |
| LUAD | C5orf38 | CNV |
| LUAD | INSM2 | CNV |
| LUAD | IRX2 | CNV |
| LUAD | NFKBIA | CNV |
| LUAD | DNAH5 | CNV |
| LUAD | NOTCH2 | CNV |
| LUAD | KIAA0391 | CNV |
| LUAD | MIPOL1 | CNV |
| LUAD | DMRTA1 | CNV |
| LUAD | PSMA6 | CNV |
| LUAD | TP53 | Mutation |
| LUAD | KRAS | Mutation |
| LUAD | KEAP1 | Mutation |
| LUAD | EGFR | Mutation |
| LUAD | STK11 | Mutation |
| LUAD | NF1 | Mutation |
| LUAD | SMARCA4 | Mutation |
| LUAD | RBM10 | Mutation |
| LUAD | MGA | Mutation |
| LUAD | BRAF | Mutation |
| LUAD | ATM | Mutation |
| LUAD | SETD2 | Mutation |
| LUAD | ARID1A | Mutation |

| | | |
|---|---|---|
| LUAD | RB1 | Mutation |
| LUAD | COL5A2 | Mutation |
| LUAD | MYH7 | Mutation |
| LUAD | DST | Mutation |
| LUAD | FBN2 | Mutation |
| LUAD | PIK3CA | Mutation |
| LUAD | COL3A1 | Mutation |
| LUAD | SPTA1 | Mutation |
| LUAD | SYNE2 | Mutation |
| LUAD | COL11A1 | Mutation |
| LUAD | ARID2 | Mutation |
| LUAD | MET | Mutation |
| LUAD | ITGAX | Mutation |
| LUAD | FHOD3 | Mutation |
| LUAD | CDKN2A | Mutation |
| LUAD | VCAN | Mutation |
| LUAD | FANCM | Mutation |
| LUAD | L1CAM | Mutation |
| LUAD | COL15A1 | Mutation |
| LUAD | LAMA4 | Mutation |
| LUAD | NOTCH4 | Mutation |
| LUAD | TXLNB | Mutation |
| LUAD | FAM71A | Mutation |
| LUAD | APC | Mutation |
| LUAD | TNN | Mutation |
| LUAD | LRP1B | Mutation |
| LUAD | CSMD3 | Mutation |
| LUAD | DNMT3A | Mutation |
| LUAD | TGFBR3 | Mutation |
| LUAD | HEATR5B | Mutation |
| LUAD | TTN | Mutation |
| LUAD | CTNNB1 | Mutation |
| LUAD | FRMPD1 | Mutation |
| LUAD | A2M | Mutation |
| LUAD | HYDIN | Mutation |
| LUAD | MUC16 | Mutation |
| LUAD | LUM | Mutation |
| LUAD | SFTPB | SV |
| LUSC | MCF2L2 | CNV |
| LUSC | ATP11B | CNV |
| LUSC | MCCC1 | CNV |
| LUSC | DCUN1D1 | CNV |
| LUSC | SOX2 | CNV |

| | | |
|---|---|---|
| LUSC | B3GNT5 | CNV |
| LUSC | LAMP3 | CNV |
| LUSC | PARL | CNV |
| LUSC | HTR3D | CNV |
| LUSC | YEATS2 | CNV |
| LUSC | ABCC5 | CNV |
| LUSC | HTR3E | CNV |
| LUSC | KLHL6 | CNV |
| LUSC | KLHL24 | CNV |
| LUSC | DVL3 | CNV |
| LUSC | HTR3C | CNV |
| LUSC | MAP6D1 | CNV |
| LUSC | AP2M1 | CNV |
| LUSC | EIF2B5 | CNV |
| LUSC | FXR1 | CNV |
| LUSC | DNAJC19 | CNV |
| LUSC | ZNF639 | CNV |
| LUSC | PEX5L | CNV |
| LUSC | KCNMB3 | CNV |
| LUSC | PIK3CA | CNV |
| LUSC | TTC14 | CNV |
| LUSC | KCNMB2 | CNV |
| LUSC | ZMAT3 | CNV |
| LUSC | USP13 | CNV |
| LUSC | GNB4 | CNV |
| LUSC | CCDC39 | CNV |
| LUSC | MFN1 | CNV |
| LUSC | ACTL6A | CNV |
| LUSC | NDUFB5 | CNV |
| LUSC | MRPL47 | CNV |
| LUSC | NAALADL2 | CNV |
| LUSC | ABCF3 | CNV |
| LUSC | TNFSF10 | CNV |
| LUSC | TMEM212 | CNV |
| LUSC | PLD1 | CNV |
| LUSC | FNDC3B | CNV |
| LUSC | MAGEF1 | CNV |
| LUSC | TBL1XR1 | CNV |
| LUSC | GHSR | CNV |
| LUSC | EPHB3 | CNV |
| LUSC | NCEH1 | CNV |
| LUSC | VPS8 | CNV |
| LUSC | VWA5B2 | CNV |

| | | |
|---|---|---|
| LUSC | CHRD | CNV |
| LUSC | ALG3 | CNV |
| LUSC | TP53 | Mutation |
| LUSC | CDKN2A | Mutation |
| LUSC | FAT1 | Mutation |
| LUSC | NFE2L2 | Mutation |
| LUSC | PTEN | Mutation |
| LUSC | NF1 | Mutation |
| LUSC | PIK3CA | Mutation |
| LUSC | RB1 | Mutation |
| LUSC | RYR1 | Mutation |
| LUSC | KEAP1 | Mutation |
| LUSC | NOTCH1 | Mutation |
| LUSC | RASA1 | Mutation |
| LUSC | ARID1A | Mutation |
| LUSC | ARHGAP35 | Mutation |
| LUSC | KDM6A | Mutation |
| LUSC | CUL3 | Mutation |
| LUSC | PTPRC | Mutation |
| LUSC | DOCK10 | Mutation |
| LUSC | NSD1 | Mutation |
| LUSC | FBXW7 | Mutation |
| LUSC | ACACB | Mutation |
| LUSC | CREBBP | Mutation |
| LUSC | ALS2 | Mutation |
| LUSC | RYR3 | Mutation |
| LUSC | APOB | Mutation |
| LUSC | LCT | Mutation |
| LUSC | IBTK | Mutation |
| LUSC | CLIP1 | Mutation |
| LUSC | ADAMTS12 | Mutation |
| LUSC | FN1 | Mutation |
| LUSC | TTN | Mutation |
| LUSC | NR1H4 | Mutation |
| LUSC | DRD3 | Mutation |
| LUSC | KCNG4 | Mutation |
| LUSC | ZNF236 | Mutation |
| LUSC | FLG | Mutation |
| LUSC | SELP | Mutation |
| LUSC | CYP11B1 | Mutation |
| LUSC | SLC28A1 | Mutation |
| LUSC | TTPA | Mutation |
| LUSC | COL22A1 | Mutation |

| | | |
|---|---|---|
| LUSC | CYBB | Mutation |
| LUSC | MYH7 | Mutation |
| LUSC | COL12A1 | Mutation |
| LUSC | COL15A1 | Mutation |
| LUSC | KLF5 | Mutation |
| LUSC | TYK2 | Mutation |
| LUSC | INSR | Mutation |
| LUSC | CSMD3 | Mutation |
| LUSC | GRID2 | Mutation |
| OV | MYC | CNV |
| OV | GSDMC | CNV |
| OV | POU5F1B | CNV |
| OV | ASAP1 | CNV |
| OV | FAM49B | CNV |
| OV | ADCY8 | CNV |
| OV | FAM84B | CNV |
| OV | EFR3A | CNV |
| OV | KCNQ3 | CNV |
| OV | LRRC6 | CNV |
| OV | OC90 | CNV |
| OV | HHLA1 | CNV |
| OV | PHF20L1 | CNV |
| OV | TG | CNV |
| OV | TMEM71 | CNV |
| OV | ST3GAL1 | CNV |
| OV | NDRG1 | CNV |
| OV | ZFAT | CNV |
| OV | WISP1 | CNV |
| OV | SLA | CNV |
| OV | TRAPPC9 | CNV |
| OV | KHDRBS3 | CNV |
| OV | DENND3 | CNV |
| OV | PTK2 | CNV |
| OV | SQLE | CNV |
| OV | TRIB1 | CNV |
| OV | MECOM | CNV |
| OV | NSMCE2 | CNV |
| OV | ZNF572 | CNV |
| OV | SLC45A4 | CNV |
| OV | ZC3H3 | CNV |
| OV | KIAA0196 | CNV |
| OV | EEF1D | CNV |
| OV | GSDMD | CNV |

| | | |
|---|---|---|
| OV | MAFA | CNV |
| OV | NAPRT1 | CNV |
| OV | CHRAC1 | CNV |
| OV | MTSS1 | CNV |
| OV | OPLAH | CNV |
| OV | SPATC1 | CNV |
| OV | GPAA1 | CNV |
| OV | SCRIB | CNV |
| OV | PUF60 | CNV |
| OV | PLEC | CNV |
| OV | EPPK1 | CNV |
| OV | TOP1MT | CNV |
| OV | COL22A1 | CNV |
| OV | NRBP2 | CNV |
| OV | EXOSC4 | CNV |
| OV | RHPN1 | CNV |
| OV | TP53 | Mutation |
| OV | BRCA1 | Mutation |
| OV | NF1 | Mutation |
| OV | RB1 | Mutation |
| OV | PPM1F | Mutation |
| OV | MYF5 | Mutation |
| OV | PTPN4 | Mutation |
| OV | TMC1 | Mutation |
| OV | CDK12 | Mutation |
| OV | PCDHA2 | Mutation |
| OV | DYNC1H1 | Mutation |
| OV | KRT4 | Mutation |
| OV | PHACTR3 | Mutation |
| OV | LAMC1 | Mutation |
| OV | SVIL | Mutation |
| OV | AHNAK | Mutation |
| OV | NPAS3 | Mutation |
| OV | TRMT1 | Mutation |
| OV | ABHD11 | Mutation |
| OV | MYOC | Mutation |
| OV | ARHGEF7 | Mutation |
| OV | MYH4 | Mutation |
| OV | PLAU | Mutation |
| OV | PIGO | Mutation |
| OV | SMC3 | Mutation |
| OV | PCDHGC5 | Mutation |
| OV | CLCN6 | Mutation |

| | | |
|---|---|---|
| OV | IL21R | Mutation |
| OV | SBNO1 | Mutation |
| OV | ADAMTS13 | Mutation |
| OV | BCR | Mutation |
| OV | IGSF10 | Mutation |
| OV | DNAH8 | Mutation |
| OV | AP1B1 | Mutation |
| OV | PLD2 | Mutation |
| OV | PKD2L1 | Mutation |
| OV | CASR | Mutation |
| OV | TBX21 | Mutation |
| OV | ABCC3 | Mutation |
| OV | MAGEA11 | Mutation |
| OV | TOP2A | Mutation |
| OV | ATP1A2 | Mutation |
| OV | CDKN1B | Mutation |
| OV | DSN1 | Mutation |
| OV | NOTCH4 | Mutation |
| OV | DNAH3 | Mutation |
| OV | PIK3CA | Mutation |
| OV | HIPK1 | Mutation |
| OV | SETD1A | Mutation |
| OV | TTN | Mutation |
| PRAD | PTEN | CNV |
| PRAD | RNLS | CNV |
| PRAD | ATAD1 | CNV |
| PRAD | TMPRSS2 | CNV |
| PRAD | LIPJ | CNV |
| PRAD | LRCH1 | CNV |
| PRAD | ZNF292 | CNV |
| PRAD | ERG | CNV |
| PRAD | HTR2A | CNV |
| PRAD | WDFY2 | CNV |
| PRAD | RNASEH2B | CNV |
| PRAD | KPNA3 | CNV |
| PRAD | PHF11 | CNV |
| PRAD | EBPL | CNV |
| PRAD | EPHA7 | CNV |
| PRAD | ARL11 | CNV |
| PRAD | KCNRG | CNV |
| PRAD | CSMD3 | CNV |
| PRAD | TRIM13 | CNV |
| PRAD | DLEU7 | CNV |

| | | |
|---|---|---|
| PRAD | RCBTB1 | CNV |
| PRAD | CAB39L | CNV |
| PRAD | SIAH3 | CNV |
| PRAD | INTS6 | CNV |
| PRAD | CCDC70 | CNV |
| PRAD | DHRS12 | CNV |
| PRAD | ZC3H13 | CNV |
| PRAD | SERPINE3 | CNV |
| PRAD | ATP7B | CNV |
| PRAD | LCP1 | CNV |
| PRAD | SETDB2 | CNV |
| PRAD | ESD | CNV |
| PRAD | NEK3 | CNV |
| PRAD | CPB2 | CNV |
| PRAD | ENOX1 | CNV |
| PRAD | DSCAM | CNV |
| PRAD | ALG11 | CNV |
| PRAD | MX1 | CNV |
| PRAD | ANKRD6 | CNV |
| PRAD | FAM124A | CNV |
| PRAD | CASP8AP2 | CNV |
| PRAD | MX2 | CNV |
| PRAD | MDN1 | CNV |
| PRAD | UTP14C | CNV |
| PRAD | NUDT15 | CNV |
| PRAD | SPERT | CNV |
| PRAD | NEK5 | CNV |
| PRAD | ETS2 | CNV |
| PRAD | FNDC3A | CNV |
| PRAD | SUCLA2 | CNV |
| PRAD | TP53 | Mutation |
| PRAD | SPOP | Mutation |
| PRAD | FOXA1 | Mutation |
| PRAD | PTEN | Mutation |
| PRAD | ATM | Mutation |
| PRAD | CTNNB1 | Mutation |
| PRAD | PIK3CA | Mutation |
| PRAD | APC | Mutation |
| PRAD | CDK12 | Mutation |
| PRAD | ZFHX3 | Mutation |
| PRAD | KDM6A | Mutation |
| PRAD | LAMA3 | Mutation |
| PRAD | NTM | Mutation |

| | | |
|---|---|---|
| PRAD | CDKN1B | Mutation |
| PRAD | MED12 | Mutation |
| PRAD | HSPA8 | Mutation |
| PRAD | ZMYM3 | Mutation |
| PRAD | TP53BP1 | Mutation |
| PRAD | SMARCA1 | Mutation |
| PRAD | BRAF | Mutation |
| PRAD | IDH1 | Mutation |
| PRAD | CDKL2 | Mutation |
| PRAD | DNAH7 | Mutation |
| PRAD | BRCA2 | Mutation |
| PRAD | RNF43 | Mutation |
| PRAD | ERF | Mutation |
| PRAD | AADACL4 | Mutation |
| PRAD | FBN1 | Mutation |
| PRAD | CASZ1 | Mutation |
| PRAD | NRXN3 | Mutation |
| PRAD | COL11A1 | Mutation |
| PRAD | RYBP | Mutation |
| PRAD | KRT25 | Mutation |
| PRAD | RNF31 | Mutation |
| PRAD | COL6A3 | Mutation |
| PRAD | NCKAP1L | Mutation |
| PRAD | BTBD11 | Mutation |
| PRAD | SALL1 | Mutation |
| PRAD | GRIA1 | Mutation |
| PRAD | LILRB5 | Mutation |
| PRAD | HOXB13 | Mutation |
| PRAD | KLHL18 | Mutation |
| PRAD | USP28 | Mutation |
| PRAD | TTN | Mutation |
| PRAD | IL6ST | Mutation |
| PRAD | AFG3L2 | Mutation |
| PRAD | PTPRC | Mutation |
| PRAD | HRAS | Mutation |
| PRAD | ETV3 | Mutation |
| PRAD | SNW1 | Mutation |
| PRAD | TMPRSS2 | SV |
| PRAD | ERG | SV |
| PRAD | SLC45A3 | SV |
| PRAD | ACP3 | SV |
| PRAD | ETV4 | SV |
| PRAD | ETV1 | SV |

| | | |
|---|---|---|
| PRAD | PLPP1 | SV |
| SKCM | CDKN2A | CNV |
| SKCM | CDKN2B | CNV |
| SKCM | MTAP | CNV |
| SKCM | DMRTA1 | CNV |
| SKCM | IFNE | CNV |
| SKCM | IFNA8 | CNV |
| SKCM | IFNA2 | CNV |
| SKCM | IFNA1 | CNV |
| SKCM | IFNA6 | CNV |
| SKCM | IFNA13 | CNV |
| SKCM | IFNA17 | CNV |
| SKCM | KLHL9 | CNV |
| SKCM | IFNA14 | CNV |
| SKCM | IFNA7 | CNV |
| SKCM | IFNA4 | CNV |
| SKCM | IFNA5 | CNV |
| SKCM | IFNA10 | CNV |
| SKCM | IFNA16 | CNV |
| SKCM | ELAVL2 | CNV |
| SKCM | IFNA21 | CNV |
| SKCM | IFNW1 | CNV |
| SKCM | PTEN | CNV |
| SKCM | IFNB1 | CNV |
| SKCM | PTPLAD2 | CNV |
| SKCM | MITF | CNV |
| SKCM | CCND1 | CNV |
| SKCM | NOTCH2 | CNV |
| SKCM | TERT | CNV |
| SKCM | TUSC1 | CNV |
| SKCM | SLC12A7 | CNV |
| SKCM | SLC6A18 | CNV |
| SKCM | FGF3 | CNV |
| SKCM | SLC6A19 | CNV |
| SKCM | NKD2 | CNV |
| SKCM | SHANK2 | CNV |
| SKCM | MLLT3 | CNV |
| SKCM | FGF19 | CNV |
| SKCM | FGF4 | CNV |
| SKCM | ORAOV1 | CNV |
| SKCM | ZNF697 | CNV |
| SKCM | FRMD4B | CNV |
| SKCM | ANO1 | CNV |

| | | |
|---|---|---|
| SKCM | MYEOV | CNV |
| SKCM | PPFIA1 | CNV |
| SKCM | REG4 | CNV |
| SKCM | ADAM30 | CNV |
| SKCM | NBPF7 | CNV |
| SKCM | PHGDH | CNV |
| SKCM | HMGCS2 | CNV |
| SKCM | C5orf55 | CNV |
| SKCM | BRAF | Mutation |
| SKCM | TP53 | Mutation |
| SKCM | NRAS | Mutation |
| SKCM | CDKN2A | Mutation |
| SKCM | XIRP2 | Mutation |
| SKCM | THSD7B | Mutation |
| SKCM | CSMD3 | Mutation |
| SKCM | NF1 | Mutation |
| SKCM | DNAH3 | Mutation |
| SKCM | PCLO | Mutation |
| SKCM | DSP | Mutation |
| SKCM | STAB2 | Mutation |
| SKCM | PAPPA2 | Mutation |
| SKCM | MAGEC1 | Mutation |
| SKCM | PTEN | Mutation |
| SKCM | ARID2 | Mutation |
| SKCM | C6 | Mutation |
| SKCM | CNTNAP2 | Mutation |
| SKCM | RAC1 | Mutation |
| SKCM | KCNB2 | Mutation |
| SKCM | DCC | Mutation |
| SKCM | PCDH15 | Mutation |
| SKCM | PPP6C | Mutation |
| SKCM | FAM135B | Mutation |
| SKCM | TMC5 | Mutation |
| SKCM | CNTN5 | Mutation |
| SKCM | PTPRT | Mutation |
| SKCM | MECOM | Mutation |
| SKCM | MYH1 | Mutation |
| SKCM | TPTE | Mutation |
| SKCM | SPAG17 | Mutation |
| SKCM | MYH7 | Mutation |
| SKCM | KIAA1109 | Mutation |
| SKCM | ASTN1 | Mutation |
| SKCM | SPHKAP | Mutation |

| | | |
|---|---|---|
| SKCM | PLCB4 | Mutation |
| SKCM | MYO5B | Mutation |
| SKCM | PDE1A | Mutation |
| SKCM | DDX3X | Mutation |
| SKCM | SELP | Mutation |
| SKCM | SCN10A | Mutation |
| SKCM | TTN | Mutation |
| SKCM | APOB | Mutation |
| SKCM | MAP2K1 | Mutation |
| SKCM | PTPRB | Mutation |
| SKCM | FAM83B | Mutation |
| SKCM | DSCAM | Mutation |
| SKCM | TRRAP | Mutation |
| SKCM | PARM1 | Mutation |
| SKCM | TP63 | Mutation |
| STAD | WWOX | CNV |
| STAD | PDE4D | CNV |
| STAD | PTPRD | CNV |
| STAD | GRB7 | CNV |
| STAD | ERBB2 | CNV |
| STAD | NAALADL2 | CNV |
| STAD | PGAP3 | CNV |
| STAD | DMD | CNV |
| STAD | IKZF3 | CNV |
| STAD | PNMT | CNV |
| STAD | TCAP | CNV |
| STAD | IMMP2L | CNV |
| STAD | MACROD2 | CNV |
| STAD | GMDS | CNV |
| STAD | STARD3 | CNV |
| STAD | MYC | CNV |
| STAD | FHIT | CNV |
| STAD | CDKN2A | CNV |
| STAD | PARK2 | CNV |
| STAD | CCNE1 | CNV |
| STAD | POU5F1B | CNV |
| STAD | PPP1R1B | CNV |
| STAD | CDKN2B | CNV |
| STAD | C19orf12 | CNV |
| STAD | NEUROD2 | CNV |
| STAD | PLEKHF1 | CNV |
| STAD | MTAP | CNV |
| STAD | MED24 | CNV |

| | | |
|---|---|---|
| STAD | CSF3 | CNV |
| STAD | PSMD3 | CNV |
| STAD | FAM84B | CNV |
| STAD | GSDMA | CNV |
| STAD | WIPF2 | CNV |
| STAD | ZPBP2 | CNV |
| STAD | GSDMB | CNV |
| STAD | GATA6 | CNV |
| STAD | POP4 | CNV |
| STAD | CDK12 | CNV |
| STAD | VSTM2B | CNV |
| STAD | THRA | CNV |
| STAD | ORMDL3 | CNV |
| STAD | IGFBP4 | CNV |
| STAD | TNS4 | CNV |
| STAD | RARA | CNV |
| STAD | CDC6 | CNV |
| STAD | NSMCE2 | CNV |
| STAD | TRIB1 | CNV |
| STAD | NR1D1 | CNV |
| STAD | UQCRFS1 | CNV |
| STAD | CASC3 | CNV |
| STAD | TP53 | Mutation |
| STAD | ARID1A | Mutation |
| STAD | ACVR2A | Mutation |
| STAD | RNF43 | Mutation |
| STAD | KRAS | Mutation |
| STAD | PIK3CA | Mutation |
| STAD | DOCK3 | Mutation |
| STAD | PTEN | Mutation |
| STAD | CDH1 | Mutation |
| STAD | ZBTB20 | Mutation |
| STAD | FBXW7 | Mutation |
| STAD | UBR5 | Mutation |
| STAD | LARP4B | Mutation |
| STAD | RPL22 | Mutation |
| STAD | ZFHX4 | Mutation |
| STAD | COL12A1 | Mutation |
| STAD | LRP1B | Mutation |
| STAD | APC | Mutation |
| STAD | SMAD4 | Mutation |
| STAD | KLF3 | Mutation |
| STAD | PLEKHA6 | Mutation |

| | | |
|---|---|---|
| STAD | RHOA | Mutation |
| STAD | MUC16 | Mutation |
| STAD | MUC6 | Mutation |
| STAD | PGM5 | Mutation |
| STAD | CTCF | Mutation |
| STAD | BCOR | Mutation |
| STAD | HLA-B | Mutation |
| STAD | ATM | Mutation |
| STAD | TTK | Mutation |
| STAD | CIC | Mutation |
| STAD | CTNND1 | Mutation |
| STAD | TTN | Mutation |
| STAD | ESRP1 | Mutation |
| STAD | FHOD3 | Mutation |
| STAD | CTNNB1 | Mutation |
| STAD | CUBN | Mutation |
| STAD | RIMS2 | Mutation |
| STAD | ERBB4 | Mutation |
| STAD | SACS | Mutation |
| STAD | ERBB3 | Mutation |
| STAD | BTBD11 | Mutation |
| STAD | MAP2K7 | Mutation |
| STAD | SPTA1 | Mutation |
| STAD | XIRP2 | Mutation |
| STAD | BMPR2 | Mutation |
| STAD | ARID2 | Mutation |
| STAD | KIAA0195 | Mutation |
| STAD | HMCN1 | Mutation |
| STAD | GTF3C1 | Mutation |
| STAD | CLDN18 | SV |
| STAD | ARHGAP26 | SV |
| THCA | CACNA1B | CNV |
| THCA | FAM157B | CNV |
| THCA | EHMT1 | CNV |
| THCA | ASTN2 | CNV |
| THCA | FCN1 | CNV |
| THCA | LAMC3 | CNV |
| THCA | CCDC6 | CNV |
| THCA | NOTCH1 | CNV |
| THCA | FIBCD1 | CNV |
| THCA | SEC16A | CNV |
| THCA | COL5A1 | CNV |
| THCA | FCN2 | CNV |

| | | |
|---|---|---|
| THCA | C9orf163 | CNV |
| THCA | ZER1 | CNV |
| THCA | XKR3 | CNV |
| THCA | WDR5 | CNV |
| THCA | VBP1 | CNV |
| THCA | VAMP7 | CNV |
| THCA | SLC2A6 | CNV |
| THCA | USP20 | CNV |
| THCA | ZDHHC12 | CNV |
| THCA | WDR34 | CNV |
| THCA | TNFSF8 | CNV |
| THCA | ZMYND19 | CNV |
| THCA | UCK1 | CNV |
| THCA | OBP2A | CNV |
| THCA | OBP2B | CNV |
| THCA | ODF2 | CNV |
| THCA | TRUB2 | CNV |
| THCA | TTF1 | CNV |
| THCA | TSC1 | CNV |
| THCA | URM1 | CNV |
| THCA | UAP1L1 | CNV |
| THCA | SLC16A9 | CNV |
| THCA | UBAC1 | CNV |
| THCA | TMEM8C | CNV |
| THCA | TMEM141 | CNV |
| THCA | TPRN | CNV |
| THCA | TOR1B | CNV |
| THCA | TRAF2 | CNV |
| THCA | TOR1A | CNV |
| THCA | SLC34A3 | CNV |
| THCA | RXRA | CNV |
| THCA | SLC25A25 | CNV |
| THCA | SNAPC4 | CNV |
| THCA | DDX31 | CNV |
| THCA | DPP7 | CNV |
| THCA | DOLPP1 | CNV |
| THCA | DOLK | CNV |
| THCA | DNM1 | CNV |
| THCA | BRAF | Mutation |
| THCA | NRAS | Mutation |
| THCA | HRAS | Mutation |
| THCA | EIF1AX | Mutation |
| THCA | AKT1 | Mutation |

| | | |
|---|---|---|
| THCA | NUP93 | Mutation |
| THCA | PPM1D | Mutation |
| THCA | SPTA1 | Mutation |
| THCA | ATM | Mutation |
| THCA | CHEK2 | Mutation |
| THCA | DNMT3A | Mutation |
| THCA | JMJD1C | Mutation |
| THCA | IL7R | Mutation |
| THCA | OR56A1 | Mutation |
| THCA | GBF1 | Mutation |
| THCA | RPRD1A | Mutation |
| THCA | MAP3K3 | Mutation |
| THCA | USP9X | Mutation |
| THCA | TG | Mutation |
| THCA | GRM6 | Mutation |
| THCA | TBC1D7 | Mutation |
| THCA | PLK2 | Mutation |
| THCA | CSMD3 | Mutation |
| THCA | FAM3C | Mutation |
| THCA | TRIP11 | Mutation |
| THCA | SIGLEC6 | Mutation |
| THCA | ZFHX3 | Mutation |
| THCA | MTMR12 | Mutation |
| THCA | KDM2B | Mutation |
| THCA | SUOX | Mutation |
| THCA | TCF7L1 | Mutation |
| THCA | ABHD4 | Mutation |
| THCA | BNC2 | Mutation |
| THCA | SCAP | Mutation |
| THCA | LRRC19 | Mutation |
| THCA | KDM2A | Mutation |
| THCA | EFCAB1 | Mutation |
| THCA | BACH1 | Mutation |
| THCA | PIK3R5 | Mutation |
| THCA | SLCO2B1 | Mutation |
| THCA | PHKA2 | Mutation |
| THCA | RMND1 | Mutation |
| THCA | SHROOM2 | Mutation |
| THCA | VPS13D | Mutation |
| THCA | POLR1B | Mutation |
| THCA | ARID2 | Mutation |
| THCA | SLC34A2 | Mutation |
| THCA | GIMAP6 | Mutation |

| | | |
|---|---|---|
| THCA | TTN | Mutation |
| THCA | DLC1 | Mutation |
| THCA | RET | SV |
| THCA | CCDC6 | SV |
| THCA | BRAF | SV |
| UCEC | MECOM | CNV |
| UCEC | MYC | CNV |
| UCEC | FNDC3B | CNV |
| UCEC | NCEH1 | CNV |
| UCEC | MYNN | CNV |
| UCEC | UBQLN4 | CNV |
| UCEC | ARHGEF2 | CNV |
| UCEC | RIT1 | CNV |
| UCEC | KIAA0907 | CNV |
| UCEC | LRRC34 | CNV |
| UCEC | CCNE1 | CNV |
| UCEC | GHSR | CNV |
| UCEC | SSR2 | CNV |
| UCEC | RAB25 | CNV |
| UCEC | MEX3A | CNV |
| UCEC | LMNA | CNV |
| UCEC | RXFP4 | CNV |
| UCEC | TNFSF10 | CNV |
| UCEC | LRRC31 | CNV |
| UCEC | ZNF687 | CNV |
| UCEC | SYT11 | CNV |
| UCEC | RFX5 | CNV |
| UCEC | PI4KB | CNV |
| UCEC | LRRIQ4 | CNV |
| UCEC | ECT2 | CNV |
| UCEC | GON4L | CNV |
| UCEC | LPP | CNV |
| UCEC | SAMD7 | CNV |
| UCEC | POGZ | CNV |
| UCEC | SEMA4A | CNV |
| UCEC | PSMD4 | CNV |
| UCEC | PRUNE | CNV |
| UCEC | NAALADL2 | CNV |
| UCEC | GPR160 | CNV |
| UCEC | CDC42SE1 | CNV |
| UCEC | APOD | CNV |
| UCEC | SLC25A44 | CNV |
| UCEC | SPATA16 | CNV |

| | | |
|---|---|---|
| UCEC | MUC20 | CNV |
| UCEC | TNK2 | CNV |
| UCEC | TFRC | CNV |
| UCEC | PIP5K1A | CNV |
| UCEC | PIK3CA | CNV |
| UCEC | BNIPL | CNV |
| UCEC | SELENBP1 | CNV |
| UCEC | ACAP2 | CNV |
| UCEC | C1orf56 | CNV |
| UCEC | SEC62 | CNV |
| UCEC | GABPB2 | CNV |
| UCEC | PSMB4 | CNV |
| UCEC | PTEN | Mutation |
| UCEC | ARID1A | Mutation |
| UCEC | TP53 | Mutation |
| UCEC | PIK3R1 | Mutation |
| UCEC | CTCF | Mutation |
| UCEC | ARHGAP35 | Mutation |
| UCEC | ZFHX3 | Mutation |
| UCEC | PIK3CA | Mutation |
| UCEC | KRAS | Mutation |
| UCEC | CTNNB1 | Mutation |
| UCEC | RNF43 | Mutation |
| UCEC | CHD4 | Mutation |
| UCEC | ATM | Mutation |
| UCEC | NSD1 | Mutation |
| UCEC | JAK1 | Mutation |
| UCEC | INPPL1 | Mutation |
| UCEC | NIPBL | Mutation |
| UCEC | FBXW7 | Mutation |
| UCEC | ARID5B | Mutation |
| UCEC | CHD3 | Mutation |
| UCEC | MGA | Mutation |
| UCEC | FGFR2 | Mutation |
| UCEC | DOCK3 | Mutation |
| UCEC | RPL22 | Mutation |
| UCEC | MAP3K1 | Mutation |
| UCEC | FAT1 | Mutation |
| UCEC | DNAH7 | Mutation |
| UCEC | ESRP1 | Mutation |
| UCEC | PPP2R1A | Mutation |
| UCEC | ANK3 | Mutation |
| UCEC | ZMYM2 | Mutation |

| | | |
|---|---|---|
| UCEC | BCORL1 | Mutation |
| UCEC | NF1 | Mutation |
| UCEC | SVIL | Mutation |
| UCEC | ACVR2A | Mutation |
| UCEC | PTCH1 | Mutation |
| UCEC | BCOR | Mutation |
| UCEC | EP300 | Mutation |
| UCEC | EYS | Mutation |
| UCEC | POLE | Mutation |
| UCEC | TCERG1 | Mutation |
| UCEC | RBM27 | Mutation |
| UCEC | ARID4B | Mutation |
| UCEC | PRKDC | Mutation |
| UCEC | ZNF292 | Mutation |
| UCEC | PCLO | Mutation |
| UCEC | CSMD3 | Mutation |
| UCEC | TTC3 | Mutation |
| UCEC | DOCK11 | Mutation |
| UCEC | MSH3 | Mutation |
