## Supplementary material for "Probabilistic graph-based model uncovers previously unseen druggable vulnerabilities in major solid cancers": supp table 3

| method | gene | z_score_LN_IC5( | tumor_type | cell_line | drug | method | gene | z_score_LN_IC50 | tumor_type | cell_line | drug |
|---|---|---|---|---|---|---|---|---|---|---|---|
| MVP_top60_DNA_seed | ERBB2 | -4.438821 | BLCA | DSH1 | Lapatinib | MVP_top60_DNA+RNA_seed | ERBB2 | -4.438821 | BLCA | DSH1 | Lapatinib |
| MVP_top60_DNA_seed | ERBB2 | -4.325046 | BLCA | DSH1 | PF-00299804 | MVP_top60_DNA+RNA_seed | ERBB2 | -4.325046 | BLCA | DSH1 | PF-00299804 |
| MVP_top60_DNA_seed | FGFR3 | -3.328411 | BLCA | RT-112 | PD173074 | MVP_top60_DNA+RNA_seed | FGFR3 | -3.328411 | BLCA | RT-112 | PD173074 |
| MVP_top60_DNA_seed | ERBB2 | -3.254309 | BLCA | DSH1 | CP724714 | MVP_top60_DNA+RNA_seed | ERBB2 | -3.254309 | BLCA | DSH1 | CP724714 |
| MVP_top60_DNA_seed | ERBB2 | -2.742925 | BLCA | DSH1 | Afatinib | MVP_top60_DNA+RNA_seed | ERBB2 | -2.742925 | BLCA | DSH1 | Afatinib |
| MVP_top60_DNA_seed | ERBB2 | -2.707938 | BLCA | DSH1 | CI-1033 | MVP_top60_DNA+RNA_seed | ERBB2 | -2.707938 | BLCA | DSH1 | CI-1033 |
| MVP_top60_DNA_seed | CDK2 | -2.612901 | BLCA | T24 | AZD5438 | MVP_top60_DNA+RNA_seed | CDK2 | -2.612901 | BLCA | T24 | AZD5438 |
| MVP_top60_DNA_seed | EP300 | -2.606148 | BLCA | BFTC-905 | I-CBP112 | MVP_top60_DNA+RNA_seed | EP300 | -2.606148 | BLCA | BFTC-905 | I-CBP112 |
| MVP_top60_DNA_seed | CREBBP | -2.606148 | BLCA | BFTC-905 | I-CBP112 | MVP_top60_DNA+RNA_seed | CREBBP | -2.606148 | BLCA | BFTC-905 | I-CBP112 |
| MVP_top60_DNA_seed | RAC1 | -2.524394 | BLCA | DSH1 | EHT-1864 | MVP_top60_DNA+RNA_seed | RAC1 | -2.524394 | BLCA | DSH1 | EHT-1864 |
| MVP_top60_DNA_seed | FGFR3 | -2.484826 | BLCA | RT-112 | AZD4547 | MVP_top60_DNA+RNA_seed | FGFR3 | -2.484826 | BLCA | RT-112 | AZD4547 |
| MVP_top60_DNA_seed | FGFR3 | -2.467491 | BLCA | RT4 | AZD4547 | MVP_top60_DNA+RNA_seed | FGFR3 | -2.467491 | BLCA | RT4 | AZD4547 |
| MVP_top60_DNA_seed | PIK3CA | -2.363975 | BLCA | CAL-29 | AZD8835 | MVP_top60_DNA+RNA_seed | PIK3CA | -2.363975 | BLCA | CAL-29 | AZD8835 |
| MVP_top60_DNA_seed | FGFR3 | -2.295108 | BLCA | RT-112 | FGFR_3831 | MVP_top60_DNA+RNA_seed | FGFR3 | -2.295108 | BLCA | RT-112 | FGFR_3831 |
| MVP_top60_DNA_seed | AR | -2.288176 | BLCA | KU-19-19 | Bicalutamide | MVP_top60_DNA+RNA_seed | AKT1 | -2.291563 | BLCA | DSH1 | AKT inhibitor VIII |
| MVP_top60_DNA_seed | PIK3CA | -2.256468 | BLCA | CAL-29 | PI3Ka_4409 | MVP_top60_DNA+RNA_seed | AR | -2.288176 | BLCA | KU-19-19 | Bicalutamide |
| MVP_top60_DNA_seed | MAPK1 | -2.237887 | BLCA | KU-19-19 | VX-11e | MVP_top60_DNA+RNA_seed | PIK3CA | -2.256468 | BLCA | CAL-29 | PI3Ka_4409 |
| MVP_top60_DNA_seed | AR | -2.183409 | BLCA | SW780 | Bicalutamide | MVP_top60_DNA+RNA_seed | MAPK1 | -2.237887 | BLCA | KU-19-19 | VX-11e |
| MVP_top60_DNA_seed | PIK3CB | -2.181479 | BLCA | SW1710 | AZD6482 | MVP_top60_DNA+RNA_seed | AR | -2.183409 | BLCA | SW780 | Bicalutamide |
| MVP_top60_DNA_seed | ERBB2 | -2.088312 | BLCA | DSH1 | Afatinib | MVP_top60_DNA+RNA_seed | ERBB2 | -2.088312 | BLCA | DSH1 | Afatinib |
| MVP_top60_DNA_seed | AR | -2.080597 | BLCA | BFTC-905 | Bicalutamide | MVP_top60_DNA+RNA_seed | AR | -2.080597 | BLCA | BFTC-905 | Bicalutamide |
| MVP_top60_DNA_seed | SRC | -2.053829 | BLCA | LB831-BLC | WH-4-023 | MVP_top60_DNA+RNA_seed | SRC | -2.053829 | BLCA | LB831-BLC | WH-4-023 |
| MVP_top60_DNA_seed | HSP90AA1 | -1.980061 | BLCA | T24 | Elesclomol | MVP_top60_DNA+RNA_seed | HSP90AA1 | -1.980061 | BLCA | T24 | Elesclomol |
| MVP_top60_DNA_seed | PIK3CB | -1.963598 | BLCA | J82 | AZD6482 | MVP_top60_DNA+RNA_seed | GSK3B | -1.972706 | BLCA | BFTC-905 | SB216763 |
| MVP_top60_DNA_seed | CDK2 | -1.939887 | BLCA | KU-19-19 | AZD5438 | MVP_top60_DNA+RNA_seed | CDK2 | -1.939887 | BLCA | KU-19-19 | AZD5438 |
| MVP_top60_DNA_seed | FGFR3 | -1.8724 | BLCA | RT-112 | AZD4547 | MVP_top60_DNA+RNA_seed | FGFR3 | -1.8724 | BLCA | RT-112 | AZD4547 |
| MVP_top60_DNA_seed | CDK2 | -1.854325 | BLCA | BFTC-905 | AZD5438 | MVP_top60_DNA+RNA_seed | GSK3B | -1.855615 | BLCA | SW1710 | CHIR-99021 |
| MVP_top60_DNA_seed | PIK3CB | -1.842964 | BLCA | SW1710 | AZD8186 | MVP_top60_DNA+RNA_seed | CDK2 | -1.854325 | BLCA | BFTC-905 | AZD5438 |
| MVP_top60_DNA_seed | ESR1 | -1.840861 | BLCA | UM-UC-3 | Fulvestrant | MVP_top60_DNA+RNA_seed | ESR1 | -1.840861 | BLCA | UM-UC-3 | Fulvestrant |
| MVP_top60_DNA_seed | AR | -1.811428 | BLCA | 647-V | AZD3514 | MVP_top60_DNA+RNA_seed | GSK3B | -1.825552 | BLCA | LB831-BLC | SB216763 |
| MVP_top60_DNA_seed | MAPK1 | -1.796703 | BLCA | RT4 | VX-11e | MVP_top60_DNA+RNA_seed | AR | -1.811428 | BLCA | 647-V | AZD3514 |
| MVP_top60_DNA_seed | AR | -1.745904 | BLCA | SW780 | Bicalutamide | MVP_top60_DNA+RNA_seed | MAPK1 | -1.796703 | BLCA | RT4 | VX-11e |
| MVP_top60_DNA_seed | ATM | -1.693137 | BLCA | 639-V | KU-55933 | MVP_top60_DNA+RNA_seed | GSK3B | -1.757738 | BLCA | 5637 | SB216763 |
| MVP_top60_DNA_seed | ESR1 | -1.682553 | BLCA | BFTC-905 | Fulvestrant | MVP_top60_DNA+RNA_seed | AR | -1.745904 | BLCA | SW780 | Bicalutamide |
| MVP_top60_DNA_seed | RAC1 | -1.681755 | BLCA | 647-V | EHT-1864 | MVP_top60_DNA+RNA_seed | AKT1 | -1.741462 | BLCA | SW1710 | AKT inhibitor VIII |
| MVP_top60_DNA_seed | HSP90AA1 | -1.678847 | BLCA | CAL-29 | Tanespimycin | MVP_top60_DNA+RNA_seed | ATM | -1.693137 | BLCA | 639-V | KU-55933 |
| MVP_top60_DNA_seed | AR | -1.616688 | BLCA | KU-19-19 | AZD3514 | MVP_top60_DNA+RNA_seed | AKT1 | -1.682623 | BLCA | CAL-29 | BAY AKT1 |
| MVP_top60_DNA_seed | ERBB2 | -1.589285 | BLCA | 5637 | CUDC-101 | MVP_top60_DNA+RNA_seed | ESR1 | -1.682553 | BLCA | BFTC-905 | Fulvestrant |
| MVP_top60_DNA_seed | PIK3CA | -1.581022 | BLCA | BFTC-905 | AZD8835 | MVP_top60_DNA+RNA_seed | RAC1 | -1.681755 | BLCA | 647-V | EHT-1864 |

| | | | | | | | | | | | |
|---|---|---|---|---|---|---|---|---|---|---|---|
| MVP_top60_DNA_seed | AR | -1.577378 | BLCA | UM-UC-3 | Bicalutamide | MVP_top60_DNA+RNA_seed | HSP90AA1 | -1.678847 | BLCA | CAL-29 | Tanespimycin |
| MVP_top60_DNA_seed | HSP90AA1 | -1.564901 | BLCA | DSH1 | CCT-018159 | MVP_top60_DNA+RNA_seed | AR | -1.616688 | BLCA | KU-19-19 | AZD3514 |
| MVP_top60_DNA_seed | PIK3CB | -1.542007 | BLCA | T24 | AZD8186 | MVP_top60_DNA+RNA_seed | ERBB2 | -1.589285 | BLCA | 5637 | CUDC-101 |
| MVP_top60_DNA_seed | FGFR3 | -1.522641 | BLCA | SW780 | PD173074 | MVP_top60_DNA+RNA_seed | PIK3CA | -1.581022 | BLCA | BFTC-905 | AZD8835 |
| MVP_top60_DNA_seed | AR | -1.515838 | BLCA | CAL-29 | AZD3514 | MVP_top60_DNA+RNA_seed | AR | -1.577378 | BLCA | UM-UC-3 | Bicalutamide |
| MVP_top60_DNA_seed | HSP90AA1 | -1.513749 | BLCA | SW1710 | Luminespib | MVP_top60_DNA+RNA_seed | SYK | -1.574792 | BLCA | LB831-BLC | BAY-61-3606 |
| MVP_top60_DNA_seed | PIK3CB | -1.510381 | BLCA | CAL-29 | AZD6482 | MVP_top60_DNA+RNA_seed | HSP90AA1 | -1.564901 | BLCA | DSH1 | CCT-018159 |
| MVP_top60_DNA_seed | CDK2 | -1.505427 | BLCA | LB831-BLC | Seliciclib | MVP_top60_DNA+RNA_seed | GSK3B | -1.524438 | BLCA | LB831-BLC | CHIR-99021 |
| MVP_top60_DNA_seed | PIK3CA | -1.503629 | BLCA | BFTC-905 | PI3Ka_4409 | MVP_top60_DNA+RNA_seed | FGFR3 | -1.522641 | BLCA | SW780 | PD173074 |
| MVP_top60_DNA_seed | CDK2 | -1.498968 | BLCA | UM-UC-3 | AZD5438 | MVP_top60_DNA+RNA_seed | AR | -1.515838 | BLCA | CAL-29 | AZD3514 |
| MVP_top60_DNA_seed | HDAC1 | -1.495752 | BLCA | RT-112 | Dacinostat | MVP_top60_DNA+RNA_seed | HSP90AA1 | -1.513749 | BLCA | SW1710 | Luminespib |
| MVP_top60_DNA_seed | ESR1 | -1.493015 | BLCA | HT-1376 | Tamoxifen | MVP_top60_DNA+RNA_seed | CDK2 | -1.505427 | BLCA | LB831-BLC | Seliciclib |
| MVP_top60_DNA_seed | ATM | -1.478445 | BLCA | 639-V | KU-60019 | MVP_top60_DNA+RNA_seed | PIK3CA | -1.503629 | BLCA | BFTC-905 | PI3Ka_4409 |
| MVP_top60_DNA_seed | HSP90AA1 | -1.448314 | BLCA | CAL-29 | Elesclomol | MVP_top60_DNA+RNA_seed | CDK2 | -1.498968 | BLCA | UM-UC-3 | AZD5438 |
| MVP_top60_DNA_seed | HSP90AA1 | -1.445142 | BLCA | KU-19-19 | CCT-018159 | MVP_top60_DNA+RNA_seed | HDAC1 | -1.495752 | BLCA | RT-112 | Dacinostat |
| MVP_top60_DNA_seed | ESR1 | -1.430641 | BLCA | J82 | Tamoxifen | MVP_top60_DNA+RNA_seed | ESR1 | -1.493015 | BLCA | HT-1376 | Tamoxifen |
| MVP_top60_DNA_seed | HDAC1 | -1.42959 | BLCA | SW1710 | Dacinostat | MVP_top60_DNA+RNA_seed | ATM | -1.478445 | BLCA | 639-V | KU-60019 |
| MVP_top60_DNA_seed | PIK3CB | -1.414083 | BLCA | 639-V | AZD6482 | MVP_top60_DNA+RNA_seed | GSK3B | -1.460044 | BLCA | RT-112 | CHIR-99021 |
| MVP_top60_DNA_seed | HSP90AA1 | -1.41335 | BLCA | SW780 | Tanespimycin | MVP_top60_DNA+RNA_seed | HSP90AA1 | -1.448314 | BLCA | CAL-29 | Elesclomol |
| MVP_top60_DNA_seed | HSP90AA1 | -1.409313 | BLCA | RT-112 | Luminespib | MVP_top60_DNA+RNA_seed | HSP90AA1 | -1.445142 | BLCA | KU-19-19 | CCT-018159 |
| MVP_top60_DNA_seed | HSP90AA1 | -1.394994 | BLCA | 639-V | Luminespib | MVP_top60_DNA+RNA_seed | GSK3B | -1.444594 | BLCA | SW1710 | SB216763 |
| MVP_top60_DNA_seed | ATM | -1.376261 | BLCA | DSH1 | KU-60019 | MVP_top60_DNA+RNA_seed | GSK3B | -1.439964 | BLCA | KU-19-19 | CHIR-99021 |
| MVP_top60_DNA_seed | HSP90AA1 | -1.375886 | BLCA | DSH1 | Tanespimycin | MVP_top60_DNA+RNA_seed | ESR1 | -1.430641 | BLCA | J82 | Tamoxifen |
| MVP_top60_DNA_seed | ESR1 | -1.369715 | BLCA | SW780 | Tamoxifen | MVP_top60_DNA+RNA_seed | HDAC1 | -1.42959 | BLCA | SW1710 | Dacinostat |
| MVP_top60_DNA_seed | SRC | -1.36019 | BLCA | RT4 | NVP-BHG712 | MVP_top60_DNA+RNA_seed | HSP90AA1 | -1.41335 | BLCA | SW780 | Tanespimycin |
| MVP_top60_DNA_seed | SRC | -1.319527 | BLCA | 5637 | Bosutinib | MVP_top60_DNA+RNA_seed | HSP90AA1 | -1.409313 | BLCA | RT-112 | Luminespib |
| MVP_top60_DNA_seed | HSP90AA1 | -1.315604 | BLCA | DSH1 | SNX-2112 | MVP_top60_DNA+RNA_seed | HSP90AA1 | -1.394994 | BLCA | 639-V | Luminespib |
| MVP_top60_DNA_seed | HDAC1 | -1.305161 | BLCA | J82 | JQ12 | MVP_top60_DNA+RNA_seed | GSK3B | -1.376858 | BLCA | SW1710 | CHIR-99021 |
| MVP_top60_DNA_seed | EP300 | -1.304192 | BLCA | SW780 | I-CBP112 | MVP_top60_DNA+RNA_seed | ATM | -1.376261 | BLCA | DSH1 | KU-60019 |
| MVP_top60_DNA_seed | CREBBP | -1.304192 | BLCA | SW780 | I-CBP112 | MVP_top60_DNA+RNA_seed | HSP90AA1 | -1.375886 | BLCA | DSH1 | Tanespimycin |
| MVP_top60_DNA_seed | ATM | -1.290857 | BLCA | KU-19-19 | KU-60019 | MVP_top60_DNA+RNA_seed | ESR1 | -1.369715 | BLCA | SW780 | Tamoxifen |
| MVP_top60_DNA_seed | MAPK1 | -1.266642 | BLCA | BFTC-905 | VX-11e | MVP_top60_DNA+RNA_seed | SRC | -1.36019 | BLCA | RT4 | NVP-BHG712 |
| MVP_top60_DNA_seed | RAC1 | -1.26211 | BLCA | T24 | EHT-1864 | MVP_top60_DNA+RNA_seed | SRC | -1.319527 | BLCA | 5637 | Bosutinib |
| MVP_top60_DNA_seed | HDAC1 | -1.261335 | BLCA | DSH1 | JQ12 | MVP_top60_DNA+RNA_seed | HSP90AA1 | -1.315604 | BLCA | DSH1 | SNX-2112 |
| MVP_top60_DNA_seed | HDAC1 | -1.260079 | BLCA | RT-112 | JQ12 | MVP_top60_DNA+RNA_seed | GSK3B | -1.308247 | BLCA | KU-19-19 | AZD7969 |
| MVP_top60_DNA_seed | ERBB2 | -1.242668 | BLCA | DSH1 | AZD8931 | MVP_top60_DNA+RNA_seed | HDAC1 | -1.305161 | BLCA | J82 | JQ12 |
| MVP_top60_DNA_seed | ERBB3 | -1.242668 | BLCA | DSH1 | AZD8931 | MVP_top60_DNA+RNA_seed | EP300 | -1.304192 | BLCA | SW780 | I-CBP112 |
| MVP_top60_DNA_seed | HSP90AA1 | -1.23511 | BLCA | 5637 | Tanespimycin | MVP_top60_DNA+RNA_seed | CREBBP | -1.304192 | BLCA | SW780 | I-CBP112 |
| MVP_top60_DNA_seed | ATM | -1.225021 | BLCA | BFTC-905 | KU-55933 | MVP_top60_DNA+RNA_seed | ATM | -1.290857 | BLCA | KU-19-19 | KU-60019 |
| MVP_top60_DNA_seed | RAC1 | -1.224489 | BLCA | SW1710 | EHT-1864 | MVP_top60_DNA+RNA_seed | SYK | -1.269576 | BLCA | J82 | BAY-61-3606 |
| MVP_top60_DNA_seed | HSP90AA1 | -1.205194 | BLCA | UM-UC-3 | CCT-018159 | MVP_top60_DNA+RNA_seed | MAPK1 | -1.266642 | BLCA | BFTC-905 | VX-11e |

| | | | | | | | | | | | |
|---|---|---|---|---|---|---|---|---|---|---|---|
| MVP_top60_DNA_seed | ATM | -1.199163 | BLCA | SW1710 | KU-55933 | MVP_top60_DNA+RNA_seed | RAC1 | -1.26211 | BLCA | T24 | EHT-1864 |
| MVP_top60_DNA_seed | FGFR3 | -1.196634 | BLCA | RT4 | FGFR_3831 | MVP_top60_DNA+RNA_seed | HDAC1 | -1.261335 | BLCA | DSH1 | JQ12 |
| MVP_top60_DNA_seed | HSP90AA1 | -1.180781 | BLCA | TCCSUP | Tanespimycin | MVP_top60_DNA+RNA_seed | HDAC1 | -1.260079 | BLCA | RT-112 | JQ12 |
| MVP_top60_DNA_seed | HDAC1 | -1.179582 | BLCA | DSH1 | Entinostat | MVP_top60_DNA+RNA_seed | AKT1 | -1.251451 | BLCA | SW780 | AKT inhibitor VIII |
| MVP_top60_DNA_seed | HDAC1 | -1.173169 | BLCA | SW1710 | JQ12 | MVP_top60_DNA+RNA_seed | ERBB2 | -1.242668 | BLCA | DSH1 | AZD8931 |
| MVP_top60_DNA_seed | HSP90AA1 | -1.160563 | BLCA | SW1710 | Elesclomol | MVP_top60_DNA+RNA_seed | ERBB3 | -1.242668 | BLCA | DSH1 | AZD8931 |
| MVP_top60_DNA_seed | HSP90AA1 | -1.160202 | BLCA | KU-19-19 | Elesclomol | MVP_top60_DNA+RNA_seed | GSK3B | -1.239198 | BLCA | 639-V | SB216763 |
| MVP_top60_DNA_seed | FGFR3 | -1.157505 | BLCA | CAL-29 | FGFR_3831 | MVP_top60_DNA+RNA_seed | HSP90AA1 | -1.23511 | BLCA | 5637 | Tanespimycin |
| MVP_top60_DNA_seed | CDK2 | -1.152268 | BLCA | CAL-29 | AZD5438 | MVP_top60_DNA+RNA_seed | ATM | -1.225021 | BLCA | BFTC-905 | KU-55933 |
| MVP_top60_DNA_seed | FGFR3 | -1.133878 | BLCA | BFTC-905 | FGFR_3831 | MVP_top60_DNA+RNA_seed | RAC1 | -1.224489 | BLCA | SW1710 | EHT-1864 |
| MVP_top60_DNA_seed | ATM | -1.126226 | BLCA | 5637 | KU-60019 | MVP_top60_DNA+RNA_seed | HSP90AA1 | -1.205194 | BLCA | UM-UC-3 | CCT-018159 |
| MVP_top60_DNA_seed | AR | -1.110713 | BLCA | TCCSUP | Bicalutamide | MVP_top60_DNA+RNA_seed | ATM | -1.199163 | BLCA | SW1710 | KU-55933 |
| MVP_top60_DNA_seed | ESR1 | -1.109265 | BLCA | HT-1197 | Tamoxifen | MVP_top60_DNA+RNA_seed | FGFR3 | -1.196634 | BLCA | RT4 | FGFR_3831 |
| MVP_top60_DNA_seed | ERBB2 | -1.101271 | BLCA | 5637 | PF-00299804 | MVP_top60_DNA+RNA_seed | GSK3B | -1.19099 | BLCA | DSH1 | CHIR-99021 |
| MVP_top60_DNA_seed | HSP90AA1 | -1.094539 | BLCA | T24 | CCT-018159 | MVP_top60_DNA+RNA_seed | GSK3B | -1.185545 | BLCA | CAL-29 | SB216763 |
| MVP_top60_DNA_seed | HDAC1 | -1.085001 | BLCA | RT4 | AR-42 | MVP_top60_DNA+RNA_seed | HSP90AA1 | -1.180781 | BLCA | TCCSUP | Tanespimycin |
| MVP_top60_DNA_seed | HSP90AA1 | -1.084309 | BLCA | UM-UC-3 | SNX-2112 | MVP_top60_DNA+RNA_seed | HDAC1 | -1.179582 | BLCA | DSH1 | Entinostat |
| MVP_top60_DNA_seed | SRC | -1.082232 | BLCA | T24 | Bosutinib | MVP_top60_DNA+RNA_seed | GSK3B | -1.175102 | BLCA | BFTC-905 | AZD7969 |
| MVP_top60_DNA_seed | FGFR3 | -1.072767 | BLCA | SW780 | AZD4547 | MVP_top60_DNA+RNA_seed | HDAC1 | -1.173169 | BLCA | SW1710 | JQ12 |
| MVP_top60_DNA_seed | PIK3CA | -1.067843 | BLCA | HT-1197 | AZD8835 | MVP_top60_DNA+RNA_seed | HSP90AA1 | -1.160563 | BLCA | SW1710 | Elesclomol |
| MVP_top60_DNA_seed | RAC1 | -1.064962 | BLCA | 5637 | EHT-1864 | MVP_top60_DNA+RNA_seed | HSP90AA1 | -1.160202 | BLCA | KU-19-19 | Elesclomol |
| MVP_top60_DNA_seed | MAPK1 | -1.06321 | BLCA | UM-UC-3 | VX-11e | MVP_top60_DNA+RNA_seed | FGFR3 | -1.157505 | BLCA | CAL-29 | FGFR_3831 |
| MVP_top60_DNA_seed | ESR1 | -1.061734 | BLCA | 5637 | Fulvestrant | MVP_top60_DNA+RNA_seed | CDK2 | -1.152268 | BLCA | CAL-29 | AZD5438 |
| MVP_top60_DNA_seed | HSP90AA1 | -1.056867 | BLCA | 5637 | CCT-018159 | MVP_top60_DNA+RNA_seed | AKT1 | -1.138382 | BLCA | T24 | MK-2206 |
| MVP_top60_DNA_seed | ATM | -1.048221 | BLCA | CAL-29 | KU-60019 | MVP_top60_DNA+RNA_seed | FGFR3 | -1.133878 | BLCA | BFTC-905 | FGFR_3831 |
| MVP_top60_DNA_seed | ATM | -1.04677 | BLCA | UM-UC-3 | KU-55933 | MVP_top60_DNA+RNA_seed | AKT1 | -1.126506 | BLCA | CAL-29 | AKT inhibitor VIII |
| MVP_top60_DNA_seed | RAC1 | -1.033788 | BLCA | UM-UC-3 | EHT-1864 | MVP_top60_DNA+RNA_seed | ATM | -1.126226 | BLCA | 5637 | KU-60019 |
| MVP_top60_DNA_seed | ATM | -1.029138 | BLCA | KU-19-19 | KU-55933 | MVP_top60_DNA+RNA_seed | GSK3B | -1.111157 | BLCA | TCCSUP | SB216763 |
| MVP_top60_DNA_seed | HSP90AA1 | -1.026715 | BLCA | RT-112 | SNX-2112 | MVP_top60_DNA+RNA_seed | AR | -1.110713 | BLCA | TCCSUP | Bicalutamide |
| MVP_top60_DNA_seed | FGFR3 | -1.0258 | BLCA | CAL-29 | AZD4547 | MVP_top60_DNA+RNA_seed | ESR1 | -1.109265 | BLCA | HT-1197 | Tamoxifen |
| MVP_top60_DNA_seed | CDK2 | -1.018856 | BLCA | SW1710 | AZD5438 | MVP_top60_DNA+RNA_seed | ERBB2 | -1.101271 | BLCA | 5637 | PF-00299804 |
| MVP_top60_DNA_seed | ATM | -1.016126 | BLCA | RT4 | CP466722 | MVP_top60_DNA+RNA_seed | HSP90AA1 | -1.094539 | BLCA | T24 | CCT-018159 |
| MVP_top60_DNA_seed | HSP90AA1 | -1.011196 | BLCA | 647-V | Elesclomol | MVP_top60_DNA+RNA_seed | HDAC1 | -1.085001 | BLCA | RT4 | AR-42 |
| MVP_top60_DNA_seed | ERBB2 | -1.010918 | BLCA | UM-UC-3 | CP724714 | MVP_top60_DNA+RNA_seed | HSP90AA1 | -1.084309 | BLCA | UM-UC-3 | SNX-2112 |
| MVP_top60_DNA_seed | HDAC1 | -1.00867 | BLCA | 5637 | Belinostat | MVP_top60_DNA+RNA_seed | SRC | -1.082232 | BLCA | T24 | Bosutinib |
| MVP_top60_DNA_seed | PIK3CA | -0.997563 | BLCA | RT-112 | PI3Ka_4409 | MVP_top60_DNA+RNA_seed | AKT1 | -1.07599 | BLCA | CAL-29 | MK-2206 |
| MVP_top60_DNA_seed | AR | -0.988827 | BLCA | 639-V | Bicalutamide | MVP_top60_DNA+RNA_seed | FGFR3 | -1.072767 | BLCA | SW780 | AZD4547 |
| MVP_top60_DNA_seed | HDAC1 | -0.972542 | BLCA | 5637 | AR-42 | MVP_top60_DNA+RNA_seed | PIK3CA | -1.067843 | BLCA | HT-1197 | AZD8835 |
| MVP_top60_DNA_seed | ATM | -0.967163 | BLCA | BFTC-905 | KU-60019 | MVP_top60_DNA+RNA_seed | RAC1 | -1.064962 | BLCA | 5637 | EHT-1864 |
| MVP_top60_DNA_seed | HSP90AA1 | -0.966633 | BLCA | RT-112 | CCT-018159 | MVP_top60_DNA+RNA_seed | MAPK1 | -1.06321 | BLCA | UM-UC-3 | VX-11e |
| MVP_top60_DNA_seed | SRC | -0.95912 | BLCA | HT-1376 | Bosutinib | MVP_top60_DNA+RNA_seed | ESR1 | -1.061734 | BLCA | 5637 | Fulvestrant |

| | | | | | | | | | | | |
|---|---|---|---|---|---|---|---|---|---|---|---|
| MVP_top60_DNA_seed | CDK2 | -0.956892 | BLCA | TCCSUP | AZD5438 | MVP_top60_DNA+RNA_seed | SYK | -1.0612 | BLCA | 639-V | BAY-61-3606 |
| MVP_top60_DNA_seed | ERBB2 | -0.954745 | BLCA | KU-19-19 | Afatinib | MVP_top60_DNA+RNA_seed | HSP90AA1 | -1.056867 | BLCA | 5637 | CCT-018159 |
| MVP_top60_DNA_seed | FGFR3 | -0.950371 | BLCA | TCCSUP | AZD4547 | MVP_top60_DNA+RNA_seed | ATM | -1.048221 | BLCA | CAL-29 | KU-60019 |
| MVP_top60_DNA_seed | ERBB2 | -0.94915 | BLCA | 5637 | Afatinib | MVP_top60_DNA+RNA_seed | ATM | -1.04677 | BLCA | UM-UC-3 | KU-55933 |
| MVP_top60_DNA_seed | FGFR3 | -0.9455 | BLCA | KU-19-19 | AZD4547 | MVP_top60_DNA+RNA_seed | SYK | -1.034625 | BLCA | SW1710 | BAY-61-3606 |
| MVP_top60_DNA_seed | ESR1 | -0.9383 | BLCA | CAL-29 | Tamoxifen | MVP_top60_DNA+RNA_seed | RAC1 | -1.033788 | BLCA | UM-UC-3 | EHT-1864 |
| MVP_top60_DNA_seed | ESR1 | -0.934027 | BLCA | KU-19-19 | Fulvestrant | MVP_top60_DNA+RNA_seed | ATM | -1.029138 | BLCA | KU-19-19 | KU-55933 |
| MVP_top60_DNA_seed | ERBB2 | -0.932357 | BLCA | CAL-29 | Afatinib | MVP_top60_DNA+RNA_seed | HSP90AA1 | -1.026715 | BLCA | RT-112 | SNX-2112 |
| MVP_top60_DNA_seed | FGFR3 | -0.924806 | BLCA | 5637 | FGFR_3831 | MVP_top60_DNA+RNA_seed | FGFR3 | -1.0258 | BLCA | CAL-29 | AZD4547 |
| MVP_top60_DNA_seed | FGFR3 | -0.924536 | BLCA | KU-19-19 | FGFR_3831 | MVP_top60_DNA+RNA_seed | CDK2 | -1.018856 | BLCA | SW1710 | AZD5438 |
| MVP_top60_DNA_seed | HSP90AA1 | -0.901971 | BLCA | 639-V | CCT-018159 | MVP_top60_DNA+RNA_seed | ATM | -1.016126 | BLCA | RT4 | CP466722 |
| MVP_top60_DNA_seed | ERBB2 | -0.899297 | BLCA | CAL-29 | Afatinib | MVP_top60_DNA+RNA_seed | GSK3B | -1.013155 | BLCA | DSH1 | SB216763 |
| MVP_top60_DNA_seed | HSP90AA1 | -0.893812 | BLCA | 639-V | Elesclomol | MVP_top60_DNA+RNA_seed | HSP90AA1 | -1.011196 | BLCA | 647-V | Elesclomol |
| MVP_top60_DNA_seed | ATM | -0.890909 | BLCA | KU-19-19 | torin2 | MVP_top60_DNA+RNA_seed | ERBB2 | -1.010918 | BLCA | UM-UC-3 | CP724714 |
| MVP_top60_DNA_seed | FGFR3 | -0.877443 | BLCA | 639-V | FGFR_3831 | MVP_top60_DNA+RNA_seed | HDAC1 | -1.00867 | BLCA | 5637 | Belinostat |
| MVP_top60_DNA_seed | FGFR3 | -0.874883 | BLCA | LB831-BLC | AZD4547 | MVP_top60_DNA+RNA_seed | PIK3CA | -0.997563 | BLCA | RT-112 | PI3Ka_4409 |
| MVP_top60_DNA_seed | CDK2 | -0.871595 | BLCA | BFTC-905 | PHA-793887 | MVP_top60_DNA+RNA_seed | GSK3B | -0.99266 | BLCA | RT-112 | KIN001-042 |
| MVP_top60_DNA_seed | HDAC1 | -0.866688 | BLCA | 5637 | CAY10603 | MVP_top60_DNA+RNA_seed | AR | -0.988827 | BLCA | 639-V | Bicalutamide |
| MVP_top60_DNA_seed | MAPK1 | -0.864832 | BLCA | DSH1 | FR-180204 | MVP_top60_DNA+RNA_seed | GSK3B | -0.980675 | BLCA | SW1710 | AZD7969 |
| MVP_top60_DNA_seed | MAPK3 | -0.864832 | BLCA | DSH1 | FR-180204 | MVP_top60_DNA+RNA_seed | HDAC1 | -0.972542 | BLCA | 5637 | AR-42 |
| MVP_top60_DNA_seed | ERBB2 | -0.864677 | BLCA | 5637 | AZD8931 | MVP_top60_DNA+RNA_seed | ATM | -0.967163 | BLCA | BFTC-905 | KU-60019 |
| MVP_top60_DNA_seed | ERBB3 | -0.864677 | BLCA | 5637 | AZD8931 | MVP_top60_DNA+RNA_seed | HSP90AA1 | -0.966633 | BLCA | RT-112 | CCT-018159 |
| MVP_top60_DNA_seed | HDAC1 | -0.849471 | BLCA | DSH1 | Tubastatin A | MVP_top60_DNA+RNA_seed | SRC | -0.95912 | BLCA | HT-1376 | Bosutinib |
| MVP_top60_DNA_seed | PIK3CB | -0.848089 | BLCA | SW780 | AZD8186 | MVP_top60_DNA+RNA_seed | CDK2 | -0.956892 | BLCA | TCCSUP | AZD5438 |
| MVP_top60_DNA_seed | HSP90AA1 | -0.84521 | BLCA | BFTC-905 | Luminespib | MVP_top60_DNA+RNA_seed | ERBB2 | -0.954745 | BLCA | KU-19-19 | Afatinib |
| MVP_top60_DNA_seed | HDAC1 | -0.842454 | BLCA | DSH1 | AR-42 | MVP_top60_DNA+RNA_seed | FGFR3 | -0.950371 | BLCA | TCCSUP | AZD4547 |
| MVP_top60_DNA_seed | FGFR3 | -0.832065 | BLCA | SW780 | FGFR_3831 | MVP_top60_DNA+RNA_seed | ERBB2 | -0.94915 | BLCA | 5637 | Afatinib |
| MVP_top60_DNA_seed | HSP90AA1 | -0.831803 | BLCA | HT-1376 | Tanespimycin | MVP_top60_DNA+RNA_seed | FGFR3 | -0.9455 | BLCA | KU-19-19 | AZD4547 |
| MVP_top60_DNA_seed | ATM | -0.831072 | BLCA | SW1710 | KU-60019 | MVP_top60_DNA+RNA_seed | ESR1 | -0.9383 | BLCA | CAL-29 | Tamoxifen |
| MVP_top60_DNA_seed | HDAC1 | -0.808963 | BLCA | RT4 | Tubastatin A | MVP_top60_DNA+RNA_seed | AKT1 | -0.937897 | BLCA | DSH1 | AKT inhibitor VIII |
| MVP_top60_DNA_seed | ERBB2 | -0.802694 | BLCA | DSH1 | CUDC-101 | MVP_top60_DNA+RNA_seed | ESR1 | -0.934027 | BLCA | KU-19-19 | Fulvestrant |
| MVP_top60_DNA_seed | CDK2 | -0.795987 | BLCA | 5637 | AZD5438 | MVP_top60_DNA+RNA_seed | ERBB2 | -0.932357 | BLCA | CAL-29 | Afatinib |
| MVP_top60_DNA_seed | SRC | -0.794606 | BLCA | UM-UC-3 | NVP-BHG712 | MVP_top60_DNA+RNA_seed | GSK3B | -0.930535 | BLCA | RT4 | CHIR-99021 |
| MVP_top60_DNA_seed | FGFR3 | -0.778905 | BLCA | J82 | PD173074 | MVP_top60_DNA+RNA_seed | FGFR3 | -0.924806 | BLCA | 5637 | FGFR_3831 |
| MVP_top60_DNA_seed | SRC | -0.77268 | BLCA | KU-19-19 | HG-6-71-01 | MVP_top60_DNA+RNA_seed | FGFR3 | -0.924536 | BLCA | KU-19-19 | FGFR_3831 |
| MVP_top60_DNA_seed | HDAC1 | -0.772117 | BLCA | RT4 | CAY10603 | MVP_top60_DNA+RNA_seed | HSP90AA1 | -0.901971 | BLCA | 639-V | CCT-018159 |
| MVP_top60_DNA_seed | RAC1 | -0.766762 | BLCA | KU-19-19 | EHT-1864 | MVP_top60_DNA+RNA_seed | ERBB2 | -0.899297 | BLCA | CAL-29 | Afatinib |
| MVP_top60_DNA_seed | FGFR3 | -0.765011 | BLCA | DSH1 | AZD4547 | MVP_top60_DNA+RNA_seed | GSK3B | -0.898562 | BLCA | T24 | CHIR-99021 |
| MVP_top60_DNA_seed | HSP90AA1 | -0.759682 | BLCA | J82 | Elesclomol | MVP_top60_DNA+RNA_seed | HSP90AA1 | -0.893812 | BLCA | 639-V | Elesclomol |
| MVP_top60_DNA_seed | ERBB2 | -0.758431 | BLCA | 5637 | CI-1033 | MVP_top60_DNA+RNA_seed | AKT1 | -0.891537 | BLCA | TCCSUP | AKT inhibitor VIII |
| MVP_top60_DNA_seed | PIK3CB | -0.75273 | BLCA | LB831-BLC | TGX221 | MVP_top60_DNA+RNA_seed | AKT1 | -0.890988 | BLCA | RT-112 | MK-2206 |

| | | | | | | | | | | | |
|---|---|---|---|---|---|---|---|---|---|---|---|
| MVP_top60_DNA_seed | ERBB2 | -0.748832 | BLCA | 5637 | Afatinib | MVP_top60_DNA+RNA_seed | ATM | -0.890909 | BLCA | KU-19-19 | torin2 |
| MVP_top60_DNA_seed | HSP90AA1 | -0.74246 | BLCA | RT-112 | Tanespimycin | MVP_top60_DNA+RNA_seed | FGFR3 | -0.877443 | BLCA | 639-V | FGFR_3831 |
| MVP_top60_DNA_seed | HSP90AA1 | -0.735504 | BLCA | UM-UC-3 | Elesclomol | MVP_top60_DNA+RNA_seed | FGFR3 | -0.874883 | BLCA | LB831-BLC | AZD4547 |
| MVP_top60_DNA_seed | HSP90AA1 | -0.733889 | BLCA | RT4 | SNX-2112 | MVP_top60_DNA+RNA_seed | CDK2 | -0.871595 | BLCA | BFTC-905 | PHA-793887 |
| MVP_top60_DNA_seed | ATM | -0.727612 | BLCA | 5637 | KU-55933 | MVP_top60_DNA+RNA_seed | GSK3B | -0.870229 | BLCA | 639-V | AZD7969 |
| MVP_top60_DNA_seed | ESR1 | -0.724177 | BLCA | CAL-29 | Fulvestrant | MVP_top60_DNA+RNA_seed | HDAC1 | -0.866688 | BLCA | 5637 | CAY10603 |
| MVP_top60_DNA_seed | MAPK1 | -0.724107 | BLCA | UM-UC-3 | FR-180204 | MVP_top60_DNA+RNA_seed | MAPK1 | -0.864832 | BLCA | DSH1 | FR-180204 |
| MVP_top60_DNA_seed | MAPK3 | -0.724107 | BLCA | UM-UC-3 | FR-180204 | MVP_top60_DNA+RNA_seed | MAPK3 | -0.864832 | BLCA | DSH1 | FR-180204 |
| MVP_top60_DNA_seed | ATM | -0.720967 | BLCA | BFTC-905 | CP466722 | MVP_top60_DNA+RNA_seed | ERBB2 | -0.864677 | BLCA | 5637 | AZD8931 |
| MVP_top60_DNA_seed | FGFR3 | -0.717796 | BLCA | 639-V | PD173074 | MVP_top60_DNA+RNA_seed | ERBB3 | -0.864677 | BLCA | 5637 | AZD8931 |
| MVP_top60_DNA_seed | AR | -0.714716 | BLCA | BFTC-905 | Bicalutamide | MVP_top60_DNA+RNA_seed | HDAC1 | -0.849471 | BLCA | DSH1 | Tubastatin A |
| MVP_top60_DNA_seed | MAPK1 | -0.712393 | BLCA | SW1710 | FR-180204 | MVP_top60_DNA+RNA_seed | SYK | -0.848167 | BLCA | BFTC-905 | BAY-61-3606 |
| MVP_top60_DNA_seed | MAPK3 | -0.712393 | BLCA | SW1710 | FR-180204 | MVP_top60_DNA+RNA_seed | HSP90AA1 | -0.84521 | BLCA | BFTC-905 | Luminespib |
| MVP_top60_DNA_seed | HSP90AA1 | -0.700782 | BLCA | LB831-BLC | Tanespimycin | MVP_top60_DNA+RNA_seed | HDAC1 | -0.842454 | BLCA | DSH1 | AR-42 |
| MVP_top60_DNA_seed | HSP90AA1 | -0.700131 | BLCA | SW780 | CCT-018159 | MVP_top60_DNA+RNA_seed | GSK3B | -0.833715 | BLCA | T24 | AZD7969 |
| MVP_top60_DNA_seed | AR | -0.696803 | BLCA | HT-1376 | Bicalutamide | MVP_top60_DNA+RNA_seed | FGFR3 | -0.832065 | BLCA | SW780 | FGFR_3831 |
| MVP_top60_DNA_seed | AR | -0.69572 | BLCA | 5637 | Bicalutamide | MVP_top60_DNA+RNA_seed | HSP90AA1 | -0.831803 | BLCA | HT-1376 | Tanespimycin |
| MVP_top60_DNA_seed | HSP90AA1 | -0.69275 | BLCA | RT-112 | Elesclomol | MVP_top60_DNA+RNA_seed | AKT1 | -0.831636 | BLCA | BFTC-905 | AKT inhibitor VIII |
| MVP_top60_DNA_seed | HDAC1 | -0.692167 | BLCA | DSH1 | Belinostat | MVP_top60_DNA+RNA_seed | ATM | -0.831072 | BLCA | SW1710 | KU-60019 |
| MVP_top60_DNA_seed | HSP90AA1 | -0.688518 | BLCA | HT-1197 | Luminespib | MVP_top60_DNA+RNA_seed | HDAC1 | -0.808963 | BLCA | RT4 | Tubastatin A |
| MVP_top60_DNA_seed | PIK3CB | -0.686329 | BLCA | UM-UC-3 | AZD8186 | MVP_top60_DNA+RNA_seed | ERBB2 | -0.802694 | BLCA | DSH1 | CUDC-101 |
| MVP_top60_DNA_seed | ERBB2 | -0.681252 | BLCA | RT4 | CUDC-101 | MVP_top60_DNA+RNA_seed | CDK2 | -0.795987 | BLCA | 5637 | AZD5438 |
| MVP_top60_DNA_seed | HSP90AA1 | -0.678584 | BLCA | J82 | Luminespib | MVP_top60_DNA+RNA_seed | SRC | -0.794606 | BLCA | UM-UC-3 | NVP-BHG712 |
| MVP_top60_DNA_seed | HDAC1 | -0.671066 | BLCA | 5637 | Dacinostat | MVP_top60_DNA+RNA_seed | FGFR3 | -0.778905 | BLCA | J82 | PD173074 |
| MVP_top60_DNA_seed | ESR1 | -0.670112 | BLCA | SW1710 | Fulvestrant | MVP_top60_DNA+RNA_seed | AKT1 | -0.774524 | BLCA | RT-112 | AKT inhibitor VIII |
| MVP_top60_DNA_seed | RAC1 | -0.669924 | BLCA | HT-1376 | EHT-1864 | MVP_top60_DNA+RNA_seed | SRC | -0.77268 | BLCA | KU-19-19 | HG-6-71-01 |
| MVP_top60_DNA_seed | CDK2 | -0.668988 | BLCA | 639-V | AZD5438 | MVP_top60_DNA+RNA_seed | HDAC1 | -0.772117 | BLCA | RT4 | CAY10603 |
| MVP_top60_DNA_seed | HDAC1 | -0.667546 | BLCA | 5637 | JQ12 | MVP_top60_DNA+RNA_seed | RAC1 | -0.766762 | BLCA | KU-19-19 | EHT-1864 |
| MVP_top60_DNA_seed | HDAC1 | -0.665691 | BLCA | BFTC-905 | JQ12 | MVP_top60_DNA+RNA_seed | FGFR3 | -0.765011 | BLCA | DSH1 | AZD4547 |
| MVP_top60_DNA_seed | HSP90AA1 | -0.663418 | BLCA | 5637 | Luminespib | MVP_top60_DNA+RNA_seed | AKT1 | -0.760402 | BLCA | J82 | AKT inhibitor VIII |
| MVP_top60_DNA_seed | HDAC1 | -0.658994 | BLCA | 5637 | Tubastatin A | MVP_top60_DNA+RNA_seed | HSP90AA1 | -0.759682 | BLCA | J82 | Elesclomol |
| MVP_top60_DNA_seed | ATM | -0.658293 | BLCA | HT-1376 | KU-55933 | MVP_top60_DNA+RNA_seed | ERBB2 | -0.758431 | BLCA | 5637 | CI-1033 |
| MVP_top60_DNA_seed | HSP90AA1 | -0.657866 | BLCA | TCCSUP | Elesclomol | MVP_top60_DNA+RNA_seed | ERBB2 | -0.748832 | BLCA | 5637 | Afatinib |
| MVP_top60_DNA_seed | ERBB2 | -0.657314 | BLCA | CAL-29 | AZD8931 | MVP_top60_DNA+RNA_seed | HSP90AA1 | -0.74246 | BLCA | RT-112 | Tanespimycin |
| MVP_top60_DNA_seed | ERBB3 | -0.657314 | BLCA | CAL-29 | AZD8931 | MVP_top60_DNA+RNA_seed | AKT1 | -0.742079 | BLCA | SW780 | BAY AKT1 |
| MVP_top60_DNA_seed | ATM | -0.641694 | BLCA | 647-V | KU-55933 | MVP_top60_DNA+RNA_seed | HSP90AA1 | -0.735504 | BLCA | UM-UC-3 | Elesclomol |
| MVP_top60_DNA_seed | HDAC1 | -0.635339 | BLCA | RT4 | Belinostat | MVP_top60_DNA+RNA_seed | HSP90AA1 | -0.733889 | BLCA | RT4 | SNX-2112 |
| MVP_top60_DNA_seed | AR | -0.632205 | BLCA | SW1710 | AZD3514 | MVP_top60_DNA+RNA_seed | ATM | -0.727612 | BLCA | 5637 | KU-55933 |
| MVP_top60_DNA_seed | EP300 | -0.631944 | BLCA | LB831-BLC | I-CBP112 | MVP_top60_DNA+RNA_seed | ESR1 | -0.724177 | BLCA | CAL-29 | Fulvestrant |
| MVP_top60_DNA_seed | CREBBP | -0.631944 | BLCA | LB831-BLC | I-CBP112 | MVP_top60_DNA+RNA_seed | MAPK1 | -0.724107 | BLCA | UM-UC-3 | FR-180204 |
| MVP_top60_DNA_seed | FGFR3 | -0.631871 | BLCA | SW1710 | FGFR_3831 | MVP_top60_DNA+RNA_seed | MAPK3 | -0.724107 | BLCA | UM-UC-3 | FR-180204 |

| | | | | | | | | | | | |
|---|---|---|---|---|---|---|---|---|---|---|---|
| MVP_top60_DNA_seed | SRC | -0.629658 | BLCA | KU-19-19 | NVP-BHG712 | MVP_top60_DNA+RNA_seed | ATM | -0.720967 | BLCA | BFTC-905 | CP466722 |
| MVP_top60_DNA_seed | ATM | -0.626703 | BLCA | UM-UC-3 | CP466722 | MVP_top60_DNA+RNA_seed | FGFR3 | -0.717796 | BLCA | 639-V | PD173074 |
| MVP_top60_DNA_seed | PIK3CA | -0.625709 | BLCA | RT-112 | AZD8835 | MVP_top60_DNA+RNA_seed | AR | -0.714716 | BLCA | BFTC-905 | Bicalutamide |
| MVP_top60_DNA_seed | PIK3CB | -0.62266 | BLCA | 647-V | AZD8186 | MVP_top60_DNA+RNA_seed | MAPK1 | -0.712393 | BLCA | SW1710 | FR-180204 |
| MVP_top60_DNA_seed | SRC | -0.621414 | BLCA | UM-UC-3 | Bosutinib | MVP_top60_DNA+RNA_seed | MAPK3 | -0.712393 | BLCA | SW1710 | FR-180204 |
| MVP_top60_DNA_seed | SRC | -0.615669 | BLCA | SW1710 | NVP-BHG712 | MVP_top60_DNA+RNA_seed | AKT1 | -0.704688 | BLCA | SW1710 | MK-2206 |
| MVP_top60_DNA_seed | ESR1 | -0.614965 | BLCA | 5637 | Tamoxifen | MVP_top60_DNA+RNA_seed | HSP90AA1 | -0.700782 | BLCA | LB831-BLC | Tanespimycin |
| MVP_top60_DNA_seed | ATM | -0.613331 | BLCA | UM-UC-3 | KU-60019 | MVP_top60_DNA+RNA_seed | HSP90AA1 | -0.700131 | BLCA | SW780 | CCT-018159 |
| MVP_top60_DNA_seed | SRC | -0.612926 | BLCA | CAL-29 | Bosutinib | MVP_top60_DNA+RNA_seed | GSK3B | -0.699159 | BLCA | UM-UC-3 | CHIR-99021 |
| MVP_top60_DNA_seed | FGFR3 | -0.605211 | BLCA | 5637 | AZD4547 | MVP_top60_DNA+RNA_seed | AR | -0.696803 | BLCA | HT-1376 | Bicalutamide |
| MVP_top60_DNA_seed | PIK3CA | -0.57933 | BLCA | TCCSUP | AZD8835 | MVP_top60_DNA+RNA_seed | AR | -0.69572 | BLCA | 5637 | Bicalutamide |
| MVP_top60_DNA_seed | HSP90AA1 | -0.576004 | BLCA | 639-V | SNX-2112 | MVP_top60_DNA+RNA_seed | HSP90AA1 | -0.69275 | BLCA | RT-112 | Elesclomol |
| MVP_top60_DNA_seed | ERBB2 | -0.573449 | BLCA | RT4 | CI-1033 | MVP_top60_DNA+RNA_seed | HDAC1 | -0.692167 | BLCA | DSH1 | Belinostat |
| MVP_top60_DNA_seed | PIK3CB | -0.567239 | BLCA | UM-UC-3 | AZD6482 | MVP_top60_DNA+RNA_seed | HSP90AA1 | -0.688518 | BLCA | HT-1197 | Luminespib |
| MVP_top60_DNA_seed | FGFR3 | -0.564687 | BLCA | RT4 | PD173074 | MVP_top60_DNA+RNA_seed | ERBB2 | -0.681252 | BLCA | RT4 | CUDC-101 |
| MVP_top60_DNA_seed | AR | -0.564587 | BLCA | UM-UC-3 | Bicalutamide | MVP_top60_DNA+RNA_seed | AKT1 | -0.678804 | BLCA | 647-V | AKT inhibitor VIII |
| MVP_top60_DNA_seed | ERBB2 | -0.561997 | BLCA | RT4 | CP724714 | MVP_top60_DNA+RNA_seed | HSP90AA1 | -0.678584 | BLCA | J82 | Luminespib |
| MVP_top60_DNA_seed | FGFR3 | -0.5617 | BLCA | 5637 | AZD4547 | MVP_top60_DNA+RNA_seed | HDAC1 | -0.671066 | BLCA | 5637 | Dacinostat |
| MVP_top60_DNA_seed | HDAC1 | -0.560452 | BLCA | TCCSUP | JQ12 | MVP_top60_DNA+RNA_seed | ESR1 | -0.670112 | BLCA | SW1710 | Fulvestrant |
| MVP_top60_DNA_seed | ERBB2 | -0.557493 | BLCA | CAL-29 | CI-1033 | MVP_top60_DNA+RNA_seed | RAC1 | -0.669924 | BLCA | HT-1376 | EHT-1864 |
| MVP_top60_DNA_seed | PIK3CB | -0.544892 | BLCA | 647-V | AZD6482 | MVP_top60_DNA+RNA_seed | CDK2 | -0.668988 | BLCA | 639-V | AZD5438 |
| MVP_top60_DNA_seed | HSP90AA1 | -0.544765 | BLCA | CAL-29 | SNX-2112 | MVP_top60_DNA+RNA_seed | HDAC1 | -0.667546 | BLCA | 5637 | JQ12 |
| MVP_top60_DNA_seed | HSP90AA1 | -0.542981 | BLCA | 639-V | Tanespimycin | MVP_top60_DNA+RNA_seed | HDAC1 | -0.665691 | BLCA | BFTC-905 | JQ12 |
| MVP_top60_DNA_seed | SRC | -0.542338 | BLCA | DSH1 | Bosutinib | MVP_top60_DNA+RNA_seed | HSP90AA1 | -0.663418 | BLCA | 5637 | Luminespib |
| MVP_top60_DNA_seed | ERBB2 | -0.538975 | BLCA | HT-1376 | Afatinib | MVP_top60_DNA+RNA_seed | HDAC1 | -0.658994 | BLCA | 5637 | Tubastatin A |
| MVP_top60_DNA_seed | HSP90AA1 | -0.534086 | BLCA | UM-UC-3 | Luminespib | MVP_top60_DNA+RNA_seed | ATM | -0.658293 | BLCA | HT-1376 | KU-55933 |
| MVP_top60_DNA_seed | HDAC1 | -0.531103 | BLCA | UM-UC-3 | Tubastatin A | MVP_top60_DNA+RNA_seed | HSP90AA1 | -0.657866 | BLCA | TCCSUP | Elesclomol |
| MVP_top60_DNA_seed | HDAC1 | -0.52919 | BLCA | HT-1197 | Dacinostat | MVP_top60_DNA+RNA_seed | ERBB2 | -0.657314 | BLCA | CAL-29 | AZD8931 |
| MVP_top60_DNA_seed | MAPK1 | -0.525094 | BLCA | BFTC-905 | FR-180204 | MVP_top60_DNA+RNA_seed | ERBB3 | -0.657314 | BLCA | CAL-29 | AZD8931 |
| MVP_top60_DNA_seed | MAPK3 | -0.525094 | BLCA | BFTC-905 | FR-180204 | MVP_top60_DNA+RNA_seed | ATM | -0.641694 | BLCA | 647-V | KU-55933 |
| MVP_top60_DNA_seed | PIK3CA | -0.521185 | BLCA | SW1710 | AZD8835 | MVP_top60_DNA+RNA_seed | HDAC1 | -0.635339 | BLCA | RT4 | Belinostat |
| MVP_top60_DNA_seed | PIK3CB | -0.516605 | BLCA | CAL-29 | AZD8186 | MVP_top60_DNA+RNA_seed | AR | -0.632205 | BLCA | SW1710 | AZD3514 |
| MVP_top60_DNA_seed | HDAC1 | -0.494043 | BLCA | RT-112 | Tubastatin A | MVP_top60_DNA+RNA_seed | EP300 | -0.631944 | BLCA | LB831-BLC | I-CBP112 |
| MVP_top60_DNA_seed | ATM | -0.48821 | BLCA | CAL-29 | KU-55933 | MVP_top60_DNA+RNA_seed | CREBBP | -0.631944 | BLCA | LB831-BLC | I-CBP112 |
| MVP_top60_DNA_seed | CDK2 | -0.487247 | BLCA | 647-V | AZD5438 | MVP_top60_DNA+RNA_seed | FGFR3 | -0.631871 | BLCA | SW1710 | FGFR_3831 |
| MVP_top60_DNA_seed | MAPK1 | -0.48396 | BLCA | RT4 | FR-180204 | MVP_top60_DNA+RNA_seed | SRC | -0.629658 | BLCA | KU-19-19 | NVP-BHG712 |
| MVP_top60_DNA_seed | MAPK3 | -0.48396 | BLCA | RT4 | FR-180204 | MVP_top60_DNA+RNA_seed | ATM | -0.626703 | BLCA | UM-UC-3 | CP466722 |
| MVP_top60_DNA_seed | PIK3CA | -0.475212 | BLCA | 5637 | PI3Ka_4409 | MVP_top60_DNA+RNA_seed | PIK3CA | -0.625709 | BLCA | RT-112 | AZD8835 |
| MVP_top60_DNA_seed | ESR1 | -0.474641 | BLCA | SW1710 | Tamoxifen | MVP_top60_DNA+RNA_seed | SRC | -0.621414 | BLCA | UM-UC-3 | Bosutinib |
| MVP_top60_DNA_seed | ESR1 | -0.469291 | BLCA | 647-V | Tamoxifen | MVP_top60_DNA+RNA_seed | SRC | -0.615669 | BLCA | SW1710 | NVP-BHG712 |
| MVP_top60_DNA_seed | SRC | -0.469089 | BLCA | 5637 | NVP-BHG712 | MVP_top60_DNA+RNA_seed | ESR1 | -0.614965 | BLCA | 5637 | Tamoxifen |

| | | | | | | | | | | | |
|---|---|---|---|---|---|---|---|---|---|---|---|
| MVP_top60_DNA_seed | MAPK1 | -0.468498 | BLCA | RT-112 | FR-180204 | MVP_top60_DNA+RNA_seed | ATM | -0.613331 | BLCA | UM-UC-3 | KU-60019 |
| MVP_top60_DNA_seed | MAPK3 | -0.468498 | BLCA | RT-112 | FR-180204 | MVP_top60_DNA+RNA_seed | SRC | -0.612926 | BLCA | CAL-29 | Bosutinib |
| MVP_top60_DNA_seed | HDAC1 | -0.465922 | BLCA | BFTC-905 | Dacinostat | MVP_top60_DNA+RNA_seed | AKT1 | -0.612252 | BLCA | LB831-BLC | A-443654 |
| MVP_top60_DNA_seed | ATM | -0.465896 | BLCA | 5637 | CP466722 | MVP_top60_DNA+RNA_seed | AKT1 | -0.607351 | BLCA | BFTC-905 | BAY AKT1 |
| MVP_top60_DNA_seed | PIK3CB | -0.463004 | BLCA | BFTC-905 | AZD8186 | MVP_top60_DNA+RNA_seed | FGFR3 | -0.605211 | BLCA | 5637 | AZD4547 |
| MVP_top60_DNA_seed | ERBB2 | -0.461972 | BLCA | SW780 | Afatinib | MVP_top60_DNA+RNA_seed | AKT1 | -0.601039 | BLCA | 639-V | AKT inhibitor VIII |
| MVP_top60_DNA_seed | HDAC1 | -0.459474 | BLCA | DSH1 | CAY10603 | MVP_top60_DNA+RNA_seed | GSK3B | -0.591195 | BLCA | 647-V | AZD7969 |
| MVP_top60_DNA_seed | AR | -0.45165 | BLCA | HT-1197 | Bicalutamide | MVP_top60_DNA+RNA_seed | GSK3B | -0.58388 | BLCA | KU-19-19 | CHIR-99021 |
| MVP_top60_DNA_seed | PIK3CB | -0.447244 | BLCA | BFTC-905 | AZD6482 | MVP_top60_DNA+RNA_seed | PIK3CA | -0.57933 | BLCA | TCCSUP | AZD8835 |
| MVP_top60_DNA_seed | FGFR3 | -0.445267 | BLCA | UM-UC-3 | PD173074 | MVP_top60_DNA+RNA_seed | HSP90AA1 | -0.576004 | BLCA | 639-V | SNX-2112 |
| MVP_top60_DNA_seed | AR | -0.440359 | BLCA | SW780 | AZD3514 | MVP_top60_DNA+RNA_seed | SYK | -0.575483 | BLCA | RT4 | BAY-61-3606 |
| MVP_top60_DNA_seed | SRC | -0.43792 | BLCA | BFTC-905 | NVP-BHG712 | MVP_top60_DNA+RNA_seed | ERBB2 | -0.573449 | BLCA | RT4 | CI-1033 |
| MVP_top60_DNA_seed | HSP90AA1 | -0.430859 | BLCA | J82 | Tanespimycin | MVP_top60_DNA+RNA_seed | GSK3B | -0.572213 | BLCA | KU-19-19 | KIN001-042 |
| MVP_top60_DNA_seed | HSP90AA1 | -0.430173 | BLCA | BFTC-905 | Elesclomol | MVP_top60_DNA+RNA_seed | FGFR3 | -0.564687 | BLCA | RT4 | PD173074 |
| MVP_top60_DNA_seed | HSP90AA1 | -0.427552 | BLCA | CAL-29 | CCT-018159 | MVP_top60_DNA+RNA_seed | AR | -0.564587 | BLCA | UM-UC-3 | Bicalutamide |
| MVP_top60_DNA_seed | ERBB2 | -0.421078 | BLCA | UM-UC-3 | CI-1033 | MVP_top60_DNA+RNA_seed | ERBB2 | -0.561997 | BLCA | RT4 | CP724714 |
| MVP_top60_DNA_seed | FGFR3 | -0.419476 | BLCA | SW1710 | AZD4547 | MVP_top60_DNA+RNA_seed | FGFR3 | -0.5617 | BLCA | 5637 | AZD4547 |
| MVP_top60_DNA_seed | ERBB2 | -0.415892 | BLCA | HT-1376 | Afatinib | MVP_top60_DNA+RNA_seed | HDAC1 | -0.560452 | BLCA | TCCSUP | JQ12 |
| MVP_top60_DNA_seed | SRC | -0.408333 | BLCA | SW780 | Bosutinib | MVP_top60_DNA+RNA_seed | GSK3B | -0.559871 | BLCA | CAL-29 | CHIR-99021 |
| MVP_top60_DNA_seed | HSP90AA1 | -0.404858 | BLCA | TCCSUP | SNX-2112 | MVP_top60_DNA+RNA_seed | ERBB2 | -0.557493 | BLCA | CAL-29 | CI-1033 |
| MVP_top60_DNA_seed | HDAC1 | -0.402258 | BLCA | LB831-BLC | Belinostat | MVP_top60_DNA+RNA_seed | HSP90AA1 | -0.544765 | BLCA | CAL-29 | SNX-2112 |
| MVP_top60_DNA_seed | ERBB2 | -0.40167 | BLCA | CAL-29 | CUDC-101 | MVP_top60_DNA+RNA_seed | HSP90AA1 | -0.542981 | BLCA | 639-V | Tanespimycin |
| MVP_top60_DNA_seed | RAC1 | -0.396124 | BLCA | BFTC-905 | EHT-1864 | MVP_top60_DNA+RNA_seed | GSK3B | -0.542709 | BLCA | UM-UC-3 | AZD7969 |
| MVP_top60_DNA_seed | HSP90AA1 | -0.385703 | BLCA | BFTC-905 | SNX-2112 | MVP_top60_DNA+RNA_seed | SRC | -0.542338 | BLCA | DSH1 | Bosutinib |
| MVP_top60_DNA_seed | HSP90AA1 | -0.384152 | BLCA | SW1710 | Tanespimycin | MVP_top60_DNA+RNA_seed | GSK3B | -0.541612 | BLCA | 5637 | AZD7969 |
| MVP_top60_DNA_seed | HDAC1 | -0.380506 | BLCA | DSH1 | Dacinostat | MVP_top60_DNA+RNA_seed | ERBB2 | -0.538975 | BLCA | HT-1376 | Afatinib |
| MVP_top60_DNA_seed | SRC | -0.379895 | BLCA | SW1710 | Bosutinib | MVP_top60_DNA+RNA_seed | GSK3B | -0.534461 | BLCA | BFTC-905 | CHIR-99021 |
| MVP_top60_DNA_seed | ERBB2 | -0.37621 | BLCA | 647-V | Afatinib | MVP_top60_DNA+RNA_seed | HSP90AA1 | -0.534086 | BLCA | UM-UC-3 | Luminespib |
| MVP_top60_DNA_seed | ATM | -0.367496 | BLCA | SW1710 | CP466722 | MVP_top60_DNA+RNA_seed | GSK3B | -0.532854 | BLCA | T24 | CHIR-99021 |
| MVP_top60_DNA_seed | PIK3CA | -0.367211 | BLCA | 639-V | AZD8835 | MVP_top60_DNA+RNA_seed | HDAC1 | -0.531103 | BLCA | UM-UC-3 | Tubastatin A |
| MVP_top60_DNA_seed | HDAC1 | -0.361213 | BLCA | RT-112 | Belinostat | MVP_top60_DNA+RNA_seed | HDAC1 | -0.52919 | BLCA | HT-1197 | Dacinostat |
| MVP_top60_DNA_seed | ESR1 | -0.359325 | BLCA | HT-1376 | Fulvestrant | MVP_top60_DNA+RNA_seed | MAPK1 | -0.525094 | BLCA | BFTC-905 | FR-180204 |
| MVP_top60_DNA_seed | HDAC1 | -0.352598 | BLCA | 639-V | Dacinostat | MVP_top60_DNA+RNA_seed | MAPK3 | -0.525094 | BLCA | BFTC-905 | FR-180204 |
| MVP_top60_DNA_seed | ATM | -0.349145 | BLCA | LB831-BLC | KU-60019 | MVP_top60_DNA+RNA_seed | PIK3CA | -0.521185 | BLCA | SW1710 | AZD8835 |
| MVP_top60_DNA_seed | HDAC1 | -0.348338 | BLCA | RT-112 | AR-42 | MVP_top60_DNA+RNA_seed | AKT1 | -0.514432 | BLCA | 5637 | GSK690693 |
| MVP_top60_DNA_seed | HSP90AA1 | -0.345031 | BLCA | DSH1 | Elesclomol | MVP_top60_DNA+RNA_seed | GSK3B | -0.499208 | BLCA | RT4 | KIN001-042 |
| MVP_top60_DNA_seed | ERBB2 | -0.343765 | BLCA | BFTC-905 | AZD8931 | MVP_top60_DNA+RNA_seed | HDAC1 | -0.494043 | BLCA | RT-112 | Tubastatin A |
| MVP_top60_DNA_seed | ERBB3 | -0.343765 | BLCA | BFTC-905 | AZD8931 | MVP_top60_DNA+RNA_seed | ATM | -0.48821 | BLCA | CAL-29 | KU-55933 |
| MVP_top60_DNA_seed | ERBB2 | -0.34311 | BLCA | RT-112 | CUDC-101 | MVP_top60_DNA+RNA_seed | CDK2 | -0.487247 | BLCA | 647-V | AZD5438 |
| MVP_top60_DNA_seed | ERBB2 | -0.341824 | BLCA | SW1710 | AZD8931 | MVP_top60_DNA+RNA_seed | MAPK1 | -0.48396 | BLCA | RT4 | FR-180204 |
| MVP_top60_DNA_seed | ERBB3 | -0.341824 | BLCA | SW1710 | AZD8931 | MVP_top60_DNA+RNA_seed | MAPK3 | -0.48396 | BLCA | RT4 | FR-180204 |

| | | | | | | | | | | | |
|---|---|---|---|---|---|---|---|---|---|---|---|
| MVP_top60_DNA_seed | PIK3CB | -0.340728 | BLCA | LB831-BLC | AZD6482 | MVP_top60_DNA+RNA_seed | PIK3CA | -0.475212 | BLCA | 5637 | PI3Ka_4409 |
| MVP_top60_DNA_seed | FGFR3 | -0.340278 | BLCA | SW1710 | AZD4547 | MVP_top60_DNA+RNA_seed | ESR1 | -0.474641 | BLCA | SW1710 | Tamoxifen |
| MVP_top60_DNA_seed | AR | -0.338001 | BLCA | LB831-BLC | Bicalutamide | MVP_top60_DNA+RNA_seed | ESR1 | -0.469291 | BLCA | 647-V | Tamoxifen |
| MVP_top60_DNA_seed | MAPK1 | -0.337749 | BLCA | T24 | FR-180204 | MVP_top60_DNA+RNA_seed | SRC | -0.469089 | BLCA | 5637 | NVP-BHG712 |
| MVP_top60_DNA_seed | MAPK3 | -0.337749 | BLCA | T24 | FR-180204 | MVP_top60_DNA+RNA_seed | MAPK1 | -0.468498 | BLCA | RT-112 | FR-180204 |
| MVP_top60_DNA_seed | CDK2 | -0.337426 | BLCA | UM-UC-3 | PHA-793887 | MVP_top60_DNA+RNA_seed | MAPK3 | -0.468498 | BLCA | RT-112 | FR-180204 |
| MVP_top60_DNA_seed | MAPK1 | -0.327587 | BLCA | 5637 | FR-180204 | MVP_top60_DNA+RNA_seed | HDAC1 | -0.465922 | BLCA | BFTC-905 | Dacinostat |
| MVP_top60_DNA_seed | MAPK3 | -0.327587 | BLCA | 5637 | FR-180204 | MVP_top60_DNA+RNA_seed | ATM | -0.465896 | BLCA | 5637 | CP466722 |
| MVP_top60_DNA_seed | AR | -0.321946 | BLCA | 639-V | AZD3514 | MVP_top60_DNA+RNA_seed | ERBB2 | -0.461972 | BLCA | SW780 | Afatinib |
| MVP_top60_DNA_seed | HDAC1 | -0.321691 | BLCA | KU-19-19 | AR-42 | MVP_top60_DNA+RNA_seed | HDAC1 | -0.459474 | BLCA | DSH1 | CAY10603 |
| MVP_top60_DNA_seed | CDK2 | -0.321306 | BLCA | RT-112 | PHA-793887 | MVP_top60_DNA+RNA_seed | AKT1 | -0.453968 | BLCA | CAL-29 | GSK690693 |
| MVP_top60_DNA_seed | ERBB2 | -0.310446 | BLCA | BFTC-905 | Afatinib | MVP_top60_DNA+RNA_seed | AR | -0.45165 | BLCA | HT-1197 | Bicalutamide |
| MVP_top60_DNA_seed | HSP90AA1 | -0.308167 | BLCA | LB831-BLC | Luminespib | MVP_top60_DNA+RNA_seed | AKT1 | -0.450326 | BLCA | SW1710 | BAY AKT1 |
| MVP_top60_DNA_seed | ERBB2 | -0.305638 | BLCA | 647-V | AZD8931 | MVP_top60_DNA+RNA_seed | AKT1 | -0.44527 | BLCA | CAL-29 | AKT inhibitor VIII |
| MVP_top60_DNA_seed | ERBB3 | -0.305638 | BLCA | 647-V | AZD8931 | MVP_top60_DNA+RNA_seed | FGFR3 | -0.445267 | BLCA | UM-UC-3 | PD173074 |
| MVP_top60_DNA_seed | HSP90AA1 | -0.304812 | BLCA | T24 | Tanespimycin | MVP_top60_DNA+RNA_seed | AR | -0.440359 | BLCA | SW780 | AZD3514 |
| MVP_top60_DNA_seed | ESR1 | -0.302726 | BLCA | KU-19-19 | Tamoxifen | MVP_top60_DNA+RNA_seed | AKT1 | -0.440034 | BLCA | RT-112 | BAY AKT1 |
| MVP_top60_DNA_seed | ERBB2 | -0.302246 | BLCA | HT-1376 | AZD8931 | MVP_top60_DNA+RNA_seed | SRC | -0.43792 | BLCA | BFTC-905 | NVP-BHG712 |
| MVP_top60_DNA_seed | ERBB3 | -0.302246 | BLCA | HT-1376 | AZD8931 | MVP_top60_DNA+RNA_seed | AKT1 | -0.432145 | BLCA | BFTC-905 | AKT inhibitor VIII |
| MVP_top60_DNA_seed | PIK3CA | -0.302125 | BLCA | SW780 | PI3Ka_4409 | MVP_top60_DNA+RNA_seed | HSP90AA1 | -0.430859 | BLCA | J82 | Tanespimycin |
| MVP_top60_DNA_seed | ERBB2 | -0.301657 | BLCA | HT-1376 | PF-00299804 | MVP_top60_DNA+RNA_seed | HSP90AA1 | -0.430173 | BLCA | BFTC-905 | Elesclomol |
| MVP_top60_DNA_seed | ATM | -0.298965 | BLCA | 647-V | KU-60019 | MVP_top60_DNA+RNA_seed | HSP90AA1 | -0.427552 | BLCA | CAL-29 | CCT-018159 |
| MVP_top60_DNA_seed | PIK3CA | -0.295805 | BLCA | LB831-BLC | AZD8835 | MVP_top60_DNA+RNA_seed | SYK | -0.426475 | BLCA | 647-V | BAY-61-3606 |
| MVP_top60_DNA_seed | HSP90AA1 | -0.288093 | BLCA | 647-V | Luminespib | MVP_top60_DNA+RNA_seed | ERBB2 | -0.421078 | BLCA | UM-UC-3 | CI-1033 |
| MVP_top60_DNA_seed | FGFR3 | -0.281437 | BLCA | SW1710 | PD173074 | MVP_top60_DNA+RNA_seed | FGFR3 | -0.419476 | BLCA | SW1710 | AZD4547 |
| MVP_top60_DNA_seed | HSP90AA1 | -0.279167 | BLCA | TCCSUP | CCT-018159 | MVP_top60_DNA+RNA_seed | ERBB2 | -0.415892 | BLCA | HT-1376 | Afatinib |
| MVP_top60_DNA_seed | HDAC1 | -0.275553 | BLCA | HT-1197 | Belinostat | MVP_top60_DNA+RNA_seed | SRC | -0.408333 | BLCA | SW780 | Bosutinib |
| MVP_top60_DNA_seed | HSP90AA1 | -0.275021 | BLCA | SW780 | Elesclomol | MVP_top60_DNA+RNA_seed | HSP90AA1 | -0.404858 | BLCA | TCCSUP | SNX-2112 |
| MVP_top60_DNA_seed | ERBB2 | -0.274014 | BLCA | RT4 | AZD8931 | MVP_top60_DNA+RNA_seed | AKT1 | -0.402547 | BLCA | DSH1 | A-443654 |
| MVP_top60_DNA_seed | ERBB3 | -0.274014 | BLCA | RT4 | AZD8931 | MVP_top60_DNA+RNA_seed | HDAC1 | -0.402258 | BLCA | LB831-BLC | Belinostat |
| MVP_top60_DNA_seed | FGFR3 | -0.269176 | BLCA | TCCSUP | FGFR_3831 | MVP_top60_DNA+RNA_seed | ERBB2 | -0.40167 | BLCA | CAL-29 | CUDC-101 |
| MVP_top60_DNA_seed | ATM | -0.265803 | BLCA | SW780 | KU-60019 | MVP_top60_DNA+RNA_seed | RAC1 | -0.396124 | BLCA | BFTC-905 | EHT-1864 |
| MVP_top60_DNA_seed | AR | -0.257535 | BLCA | 5637 | Bicalutamide | MVP_top60_DNA+RNA_seed | AKT1 | -0.389026 | BLCA | RT-112 | GSK690693 |
| MVP_top60_DNA_seed | HSP90AA1 | -0.253879 | BLCA | HT-1197 | CCT-018159 | MVP_top60_DNA+RNA_seed | HSP90AA1 | -0.385703 | BLCA | BFTC-905 | SNX-2112 |
| MVP_top60_DNA_seed | SRC | -0.253838 | BLCA | KU-19-19 | Bosutinib | MVP_top60_DNA+RNA_seed | HSP90AA1 | -0.384152 | BLCA | SW1710 | Tanespimycin |
| MVP_top60_DNA_seed | ERBB2 | -0.251038 | BLCA | SW780 | AZD8931 | MVP_top60_DNA+RNA_seed | HDAC1 | -0.380506 | BLCA | DSH1 | Dacinostat |
| MVP_top60_DNA_seed | ERBB3 | -0.251038 | BLCA | SW780 | AZD8931 | MVP_top60_DNA+RNA_seed | SRC | -0.379895 | BLCA | SW1710 | Bosutinib |
| MVP_top60_DNA_seed | HSP90AA1 | -0.249839 | BLCA | 5637 | Elesclomol | MVP_top60_DNA+RNA_seed | ERBB2 | -0.37621 | BLCA | 647-V | Afatinib |
| MVP_top60_DNA_seed | HSP90AA1 | -0.249534 | BLCA | LB831-BLC | Elesclomol | MVP_top60_DNA+RNA_seed | GSK3B | -0.373137 | BLCA | T24 | SB216763 |
| MVP_top60_DNA_seed | PIK3CB | -0.236967 | BLCA | 5637 | AZD8186 | MVP_top60_DNA+RNA_seed | GSK3B | -0.372554 | BLCA | CAL-29 | CHIR-99021 |
| MVP_top60_DNA_seed | HDAC1 | -0.234497 | BLCA | SW1710 | Belinostat | MVP_top60_DNA+RNA_seed | ATM | -0.367496 | BLCA | SW1710 | CP466722 |

| | | | | | | | | | | | |
|---|---|---|---|---|---|---|---|---|---|---|---|
| MVP_top60_DNA_seed | ERBB2 | -0.234315 | BLCA | 5637 | CP724714 | MVP_top60_DNA+RNA_seed | PIK3CA | -0.367211 | BLCA | 639-V | AZD8835 |
| MVP_top60_DNA_seed | AR | -0.233886 | BLCA | DSH1 | Bicalutamide | MVP_top60_DNA+RNA_seed | HDAC1 | -0.361213 | BLCA | RT-112 | Belinostat |
| MVP_top60_DNA_seed | SRC | -0.232832 | BLCA | 647-V | Bosutinib | MVP_top60_DNA+RNA_seed | ESR1 | -0.359325 | BLCA | HT-1376 | Fulvestrant |
| MVP_top60_DNA_seed | ATM | -0.232808 | BLCA | 647-V | CP466722 | MVP_top60_DNA+RNA_seed | HDAC1 | -0.352598 | BLCA | 639-V | Dacinostat |
| MVP_top60_DNA_seed | PIK3CA | -0.227711 | BLCA | HT-1197 | PI3Ka_4409 | MVP_top60_DNA+RNA_seed | ATM | -0.349145 | BLCA | LB831-BLC | KU-60019 |
| MVP_top60_DNA_seed | PIK3CA | -0.226616 | BLCA | 647-V | PI3Ka_4409 | MVP_top60_DNA+RNA_seed | HDAC1 | -0.348338 | BLCA | RT-112 | AR-42 |
| MVP_top60_DNA_seed | PIK3CB | -0.219449 | BLCA | 639-V | AZD8186 | MVP_top60_DNA+RNA_seed | HSP90AA1 | -0.345031 | BLCA | DSH1 | Elesclomol |
| MVP_top60_DNA_seed | HDAC1 | -0.218998 | BLCA | KU-19-19 | Belinostat | MVP_top60_DNA+RNA_seed | ERBB2 | -0.343765 | BLCA | BFTC-905 | AZD8931 |
| MVP_top60_DNA_seed | ERBB2 | -0.209528 | BLCA | 647-V | CI-1033 | MVP_top60_DNA+RNA_seed | ERBB3 | -0.343765 | BLCA | BFTC-905 | AZD8931 |
| MVP_top60_DNA_seed | CDK2 | -0.208471 | BLCA | J82 | AZD5438 | MVP_top60_DNA+RNA_seed | ERBB2 | -0.34311 | BLCA | RT-112 | CUDC-101 |
| MVP_top60_DNA_seed | HSP90AA1 | -0.207432 | BLCA | 5637 | SNX-2112 | MVP_top60_DNA+RNA_seed | ERBB2 | -0.341824 | BLCA | SW1710 | AZD8931 |
| MVP_top60_DNA_seed | SRC | -0.205708 | BLCA | RT-112 | Bosutinib | MVP_top60_DNA+RNA_seed | ERBB3 | -0.341824 | BLCA | SW1710 | AZD8931 |
| MVP_top60_DNA_seed | AR | -0.204971 | BLCA | DSH1 | AZD3514 | MVP_top60_DNA+RNA_seed | FGFR3 | -0.340278 | BLCA | SW1710 | AZD4547 |
| MVP_top60_DNA_seed | ERBB2 | -0.2026 | BLCA | 639-V | CI-1033 | MVP_top60_DNA+RNA_seed | AR | -0.338001 | BLCA | LB831-BLC | Bicalutamide |
| MVP_top60_DNA_seed | MAPK1 | -0.196632 | BLCA | 5637 | VX-11e | MVP_top60_DNA+RNA_seed | MAPK1 | -0.337749 | BLCA | T24 | FR-180204 |
| MVP_top60_DNA_seed | ERBB2 | -0.194179 | BLCA | 639-V | CP724714 | MVP_top60_DNA+RNA_seed | MAPK3 | -0.337749 | BLCA | T24 | FR-180204 |
| MVP_top60_DNA_seed | HSP90AA1 | -0.194016 | BLCA | DSH1 | Luminespib | MVP_top60_DNA+RNA_seed | CDK2 | -0.337426 | BLCA | UM-UC-3 | PHA-793887 |
| MVP_top60_DNA_seed | HSP90AA1 | -0.190025 | BLCA | BFTC-905 | Tanespimycin | MVP_top60_DNA+RNA_seed | MAPK1 | -0.327587 | BLCA | 5637 | FR-180204 |
| MVP_top60_DNA_seed | AR | -0.188814 | BLCA | SW1710 | Bicalutamide | MVP_top60_DNA+RNA_seed | MAPK3 | -0.327587 | BLCA | 5637 | FR-180204 |
| MVP_top60_DNA_seed | ATM | -0.188418 | BLCA | HT-1376 | KU-60019 | MVP_top60_DNA+RNA_seed | AR | -0.321946 | BLCA | 639-V | AZD3514 |
| MVP_top60_DNA_seed | AR | -0.186299 | BLCA | 639-V | Bicalutamide | MVP_top60_DNA+RNA_seed | HDAC1 | -0.321691 | BLCA | KU-19-19 | AR-42 |
| MVP_top60_DNA_seed | HDAC1 | -0.183451 | BLCA | UM-UC-3 | Belinostat | MVP_top60_DNA+RNA_seed | CDK2 | -0.321306 | BLCA | RT-112 | PHA-793887 |
| MVP_top60_DNA_seed | ERBB2 | -0.179867 | BLCA | RT4 | PF-00299804 | MVP_top60_DNA+RNA_seed | GSK3B | -0.317313 | BLCA | HT-1197 | SB216763 |
| MVP_top60_DNA_seed | HDAC1 | -0.177665 | BLCA | HT-1197 | JQ12 | MVP_top60_DNA+RNA_seed | ERBB2 | -0.310446 | BLCA | BFTC-905 | Afatinib |
| MVP_top60_DNA_seed | HDAC1 | -0.173597 | BLCA | SW1710 | AR-42 | MVP_top60_DNA+RNA_seed | HSP90AA1 | -0.308167 | BLCA | LB831-BLC | Luminespib |
| MVP_top60_DNA_seed | HSP90AA1 | -0.172612 | BLCA | SW1710 | SNX-2112 | MVP_top60_DNA+RNA_seed | ERBB2 | -0.305638 | BLCA | 647-V | AZD8931 |
| MVP_top60_DNA_seed | ERBB2 | -0.17012 | BLCA | BFTC-905 | CI-1033 | MVP_top60_DNA+RNA_seed | ERBB3 | -0.305638 | BLCA | 647-V | AZD8931 |
| MVP_top60_DNA_seed | FGFR3 | -0.166442 | BLCA | SW780 | AZD4547 | MVP_top60_DNA+RNA_seed | HSP90AA1 | -0.304812 | BLCA | T24 | Tanespimycin |
| MVP_top60_DNA_seed | ERBB2 | -0.164055 | BLCA | RT-112 | AZD8931 | MVP_top60_DNA+RNA_seed | ESR1 | -0.302726 | BLCA | KU-19-19 | Tamoxifen |
| MVP_top60_DNA_seed | ERBB3 | -0.164055 | BLCA | RT-112 | AZD8931 | MVP_top60_DNA+RNA_seed | ERBB2 | -0.302246 | BLCA | HT-1376 | AZD8931 |
| MVP_top60_DNA_seed | ERBB2 | -0.159624 | BLCA | 647-V | Afatinib | MVP_top60_DNA+RNA_seed | ERBB3 | -0.302246 | BLCA | HT-1376 | AZD8931 |
| MVP_top60_DNA_seed | ERBB2 | -0.158559 | BLCA | CAL-29 | CP724714 | MVP_top60_DNA+RNA_seed | PIK3CA | -0.302125 | BLCA | SW780 | PI3Ka_4409 |
| MVP_top60_DNA_seed | ATM | -0.158345 | BLCA | J82 | KU-55933 | MVP_top60_DNA+RNA_seed | ERBB2 | -0.301657 | BLCA | HT-1376 | PF-00299804 |
| MVP_top60_DNA_seed | FGFR3 | -0.158158 | BLCA | J82 | AZD4547 | MVP_top60_DNA+RNA_seed | ATM | -0.298965 | BLCA | 647-V | KU-60019 |
| MVP_top60_DNA_seed | EP300 | -0.156056 | BLCA | J82 | I-CBP112 | MVP_top60_DNA+RNA_seed | AKT1 | -0.29773 | BLCA | RT-112 | AKT inhibitor VIII |
| MVP_top60_DNA_seed | CREBBP | -0.156056 | BLCA | J82 | I-CBP112 | MVP_top60_DNA+RNA_seed | PIK3CA | -0.295805 | BLCA | LB831-BLC | AZD8835 |
| MVP_top60_DNA_seed | PIK3CB | -0.151697 | BLCA | TCCSUP | AZD6482 | MVP_top60_DNA+RNA_seed | HSP90AA1 | -0.288093 | BLCA | 647-V | Luminespib |
| MVP_top60_DNA_seed | ERBB2 | -0.151256 | BLCA | 639-V | AZD8931 | MVP_top60_DNA+RNA_seed | GSK3B | -0.283072 | BLCA | LB831-BLC | CHIR-99021 |
| MVP_top60_DNA_seed | ERBB3 | -0.151256 | BLCA | 639-V | AZD8931 | MVP_top60_DNA+RNA_seed | FGFR3 | -0.281437 | BLCA | SW1710 | PD173074 |
| MVP_top60_DNA_seed | ESR1 | -0.150683 | BLCA | RT-112 | Fulvestrant | MVP_top60_DNA+RNA_seed | HSP90AA1 | -0.279167 | BLCA | TCCSUP | CCT-018159 |
| MVP_top60_DNA_seed | EP300 | -0.150236 | BLCA | 639-V | I-CBP112 | MVP_top60_DNA+RNA_seed | HDAC1 | -0.275553 | BLCA | HT-1197 | Belinostat |

| | | | | | | | | | | | |
|---|---|---|---|---|---|---|---|---|---|---|---|
| MVP_top60_DNA_seed | CREBBP | -0.150236 | BLCA | 639-V | I-CBP112 | MVP_top60_DNA+RNA_seed | HSP90AA1 | -0.275021 | BLCA | SW780 | Elesclomol |
| MVP_top60_DNA_seed | HDAC1 | -0.140899 | BLCA | RT-112 | CAY10603 | MVP_top60_DNA+RNA_seed | ERBB2 | -0.274014 | BLCA | RT4 | AZD8931 |
| MVP_top60_DNA_seed | MAPK1 | -0.137653 | BLCA | SW1710 | VX-11e | MVP_top60_DNA+RNA_seed | ERBB3 | -0.274014 | BLCA | RT4 | AZD8931 |
| MVP_top60_DNA_seed | MAPK1 | -0.137387 | BLCA | CAL-29 | FR-180204 | MVP_top60_DNA+RNA_seed | FGFR3 | -0.269176 | BLCA | TCCSUP | FGFR_3831 |
| MVP_top60_DNA_seed | MAPK3 | -0.137387 | BLCA | CAL-29 | FR-180204 | MVP_top60_DNA+RNA_seed | ATM | -0.265803 | BLCA | SW780 | KU-60019 |
| MVP_top60_DNA_seed | AR | -0.130993 | BLCA | J82 | AZD3514 | MVP_top60_DNA+RNA_seed | GSK3B | -0.257777 | BLCA | HT-1376 | KIN001-042 |
| MVP_top60_DNA_seed | PIK3CB | -0.125699 | BLCA | HT-1376 | AZD8186 | MVP_top60_DNA+RNA_seed | AR | -0.257535 | BLCA | 5637 | Bicalutamide |
| MVP_top60_DNA_seed | FGFR3 | -0.121367 | BLCA | TCCSUP | PD173074 | MVP_top60_DNA+RNA_seed | GSK3B | -0.254205 | BLCA | UM-UC-3 | SB216763 |
| MVP_top60_DNA_seed | ERBB2 | -0.116867 | BLCA | BFTC-905 | CUDC-101 | MVP_top60_DNA+RNA_seed | HSP90AA1 | -0.253879 | BLCA | HT-1197 | CCT-018159 |
| MVP_top60_DNA_seed | EP300 | -0.115471 | BLCA | 5637 | I-CBP112 | MVP_top60_DNA+RNA_seed | SRC | -0.253838 | BLCA | KU-19-19 | Bosutinib |
| MVP_top60_DNA_seed | CREBBP | -0.115471 | BLCA | 5637 | I-CBP112 | MVP_top60_DNA+RNA_seed | GSK3B | -0.253007 | BLCA | UM-UC-3 | KIN001-042 |
| MVP_top60_DNA_seed | HDAC1 | -0.113782 | BLCA | HT-1197 | AR-42 | MVP_top60_DNA+RNA_seed | AKT1 | -0.252239 | BLCA | J82 | BAY AKT1 |
| MVP_top60_DNA_seed | HSP90AA1 | -0.109124 | BLCA | UM-UC-3 | Tanespimycin | MVP_top60_DNA+RNA_seed | ERBB2 | -0.251038 | BLCA | SW780 | AZD8931 |
| MVP_top60_DNA_seed | SRC | -0.107021 | BLCA | BFTC-905 | Bosutinib | MVP_top60_DNA+RNA_seed | ERBB3 | -0.251038 | BLCA | SW780 | AZD8931 |
| MVP_top60_DNA_seed | ATM | -0.106696 | BLCA | T24 | KU-60019 | MVP_top60_DNA+RNA_seed | HSP90AA1 | -0.249839 | BLCA | 5637 | Elesclomol |
| MVP_top60_DNA_seed | AR | -0.102666 | BLCA | HT-1376 | AZD3514 | MVP_top60_DNA+RNA_seed | HSP90AA1 | -0.249534 | BLCA | LB831-BLC | Elesclomol |
| MVP_top60_DNA_seed | ERBB2 | -0.097358 | BLCA | LB831-BLC | CUDC-101 | MVP_top60_DNA+RNA_seed | AKT1 | -0.245482 | BLCA | HT-1197 | BAY AKT1 |
| MVP_top60_DNA_seed | ERBB2 | -0.0953 | BLCA | RT-112 | CI-1033 | MVP_top60_DNA+RNA_seed | GSK3B | -0.241863 | BLCA | 5637 | KIN001-042 |
| MVP_top60_DNA_seed | PIK3CB | -0.094715 | BLCA | SW780 | AZD6482 | MVP_top60_DNA+RNA_seed | GSK3B | -0.241592 | BLCA | J82 | CHIR-99021 |
| MVP_top60_DNA_seed | HDAC1 | -0.091872 | BLCA | CAL-29 | AR-42 | MVP_top60_DNA+RNA_seed | HDAC1 | -0.234497 | BLCA | SW1710 | Belinostat |
| MVP_top60_DNA_seed | AR | -0.090396 | BLCA | TCCSUP | AZD3514 | MVP_top60_DNA+RNA_seed | ERBB2 | -0.234315 | BLCA | 5637 | CP724714 |
| MVP_top60_DNA_seed | ERBB2 | -0.088 | BLCA | 639-V | PF-00299804 | MVP_top60_DNA+RNA_seed | AR | -0.233886 | BLCA | DSH1 | Bicalutamide |
| MVP_top60_DNA_seed | HDAC1 | -0.087833 | BLCA | CAL-29 | CAY10603 | MVP_top60_DNA+RNA_seed | SRC | -0.232832 | BLCA | 647-V | Bosutinib |
| MVP_top60_DNA_seed | HSP90AA1 | -0.087304 | BLCA | J82 | CCT-018159 | MVP_top60_DNA+RNA_seed | ATM | -0.232808 | BLCA | 647-V | CP466722 |
| MVP_top60_DNA_seed | FGFR3 | -0.086873 | BLCA | LB831-BLC | FGFR_3831 | MVP_top60_DNA+RNA_seed | GSK3B | -0.227977 | BLCA | J82 | AZD7969 |
| MVP_top60_DNA_seed | ERBB2 | -0.086413 | BLCA | UM-UC-3 | PF-00299804 | MVP_top60_DNA+RNA_seed | PIK3CA | -0.227711 | BLCA | HT-1197 | PI3Ka_4409 |
| MVP_top60_DNA_seed | HDAC1 | -0.084414 | BLCA | BFTC-905 | Belinostat | MVP_top60_DNA+RNA_seed | GSK3B | -0.227657 | BLCA | 5637 | CHIR-99021 |
| MVP_top60_DNA_seed | PIK3CB | -0.079745 | BLCA | J82 | AZD8186 | MVP_top60_DNA+RNA_seed | PIK3CA | -0.226616 | BLCA | 647-V | PI3Ka_4409 |
| MVP_top60_DNA_seed | HDAC1 | -0.075407 | BLCA | UM-UC-3 | CAY10603 | MVP_top60_DNA+RNA_seed | GSK3B | -0.222144 | BLCA | TCCSUP | CHIR-99021 |
| MVP_top60_DNA_seed | SRC | -0.070686 | BLCA | RT-112 | NVP-BHG712 | MVP_top60_DNA+RNA_seed | GSK3B | -0.220462 | BLCA | RT4 | CHIR-99021 |
| MVP_top60_DNA_seed | FGFR3 | -0.069224 | BLCA | BFTC-905 | AZD4547 | MVP_top60_DNA+RNA_seed | HDAC1 | -0.218998 | BLCA | KU-19-19 | Belinostat |
| MVP_top60_DNA_seed | ATM | -0.065788 | BLCA | RT-112 | KU-60019 | MVP_top60_DNA+RNA_seed | GSK3B | -0.209674 | BLCA | 647-V | SB216763 |
| MVP_top60_DNA_seed | ATM | -0.0563 | BLCA | J82 | KU-60019 | MVP_top60_DNA+RNA_seed | ERBB2 | -0.209528 | BLCA | 647-V | CI-1033 |
| MVP_top60_DNA_seed | EP300 | -0.056293 | BLCA | UM-UC-3 | I-CBP112 | MVP_top60_DNA+RNA_seed | AKT1 | -0.208629 | BLCA | DSH1 | GSK690693 |
| MVP_top60_DNA_seed | CREBBP | -0.056293 | BLCA | UM-UC-3 | I-CBP112 | MVP_top60_DNA+RNA_seed | CDK2 | -0.208471 | BLCA | J82 | AZD5438 |
| MVP_top60_DNA_seed | ERBB2 | -0.053534 | BLCA | UM-UC-3 | CUDC-101 | MVP_top60_DNA+RNA_seed | HSP90AA1 | -0.207432 | BLCA | 5637 | SNX-2112 |
| MVP_top60_DNA_seed | ERBB2 | -0.053279 | BLCA | SW1710 | CUDC-101 | MVP_top60_DNA+RNA_seed | SRC | -0.205708 | BLCA | RT-112 | Bosutinib |
| MVP_top60_DNA_seed | HDAC1 | -0.045825 | BLCA | LB831-BLC | CAY10603 | MVP_top60_DNA+RNA_seed | AR | -0.204971 | BLCA | DSH1 | AZD3514 |
| MVP_top60_DNA_seed | ERBB2 | -0.045528 | BLCA | 639-V | Afatinib | MVP_top60_DNA+RNA_seed | GSK3B | -0.204719 | BLCA | TCCSUP | KIN001-042 |
| MVP_top60_DNA_seed | HDAC1 | -0.044736 | BLCA | SW1710 | CAY10603 | MVP_top60_DNA+RNA_seed | ERBB2 | -0.2026 | BLCA | 639-V | CI-1033 |
| MVP_top60_DNA_seed | HDAC1 | -0.042739 | BLCA | TCCSUP | Dacinostat | MVP_top60_DNA+RNA_seed | GSK3B | -0.198619 | BLCA | BFTC-905 | CHIR-99021 |

| | | | | | | | | | | | |
|---|---|---|---|---|---|---|---|---|---|---|---|
| MVP_top60_DNA_seed | CDK2 | -0.041383 | BLCA | 5637 | PHA-793887 | MVP_top60_DNA+RNA_seed | MAPK1 | -0.196632 | BLCA | 5637 | VX-11e |
| MVP_top60_DNA_seed | ERBB2 | -0.039289 | BLCA | SW1710 | Afatinib | MVP_top60_DNA+RNA_seed | ERBB2 | -0.194179 | BLCA | 639-V | CP724714 |
| MVP_top60_DNA_seed | ESR1 | -0.036355 | BLCA | 639-V | Tamoxifen | MVP_top60_DNA+RNA_seed | HSP90AA1 | -0.194016 | BLCA | DSH1 | Luminespib |
| MVP_top60_DNA_seed | CDK2 | -0.033633 | BLCA | 647-V | PHA-793887 | MVP_top60_DNA+RNA_seed | HSP90AA1 | -0.190025 | BLCA | BFTC-905 | Tanespimycin |
| MVP_top60_DNA_seed | EP300 | -0.031533 | BLCA | SW1710 | I-CBP112 | MVP_top60_DNA+RNA_seed | AR | -0.188814 | BLCA | SW1710 | Bicalutamide |
| MVP_top60_DNA_seed | CREBBP | -0.031533 | BLCA | SW1710 | I-CBP112 | MVP_top60_DNA+RNA_seed | ATM | -0.188418 | BLCA | HT-1376 | KU-60019 |
| MVP_top60_DNA_seed | HSP90AA1 | -0.029601 | BLCA | T24 | Luminespib | MVP_top60_DNA+RNA_seed | AKT1 | -0.187703 | BLCA | DSH1 | MK-2206 |
| MVP_top60_DNA_seed | ERBB2 | -0.02186 | BLCA | HT-1376 | CI-1033 | MVP_top60_DNA+RNA_seed | AR | -0.186299 | BLCA | 639-V | Bicalutamide |
| MVP_top60_DNA_seed | RAC1 | -0.021736 | BLCA | SW780 | EHT-1864 | MVP_top60_DNA+RNA_seed | HDAC1 | -0.183451 | BLCA | UM-UC-3 | Belinostat |
| MVP_top60_DNA_seed | HDAC1 | -0.020237 | BLCA | TCCSUP | Belinostat | MVP_top60_DNA+RNA_seed | ERBB2 | -0.179867 | BLCA | RT4 | PF-00299804 |
| MVP_top60_DNA_seed | HDAC1 | -0.018529 | BLCA | LB831-BLC | AR-42 | MVP_top60_DNA+RNA_seed | GSK3B | -0.177962 | BLCA | UM-UC-3 | CHIR-99021 |
| MVP_top60_DNA_seed | HDAC1 | -0.015948 | BLCA | LB831-BLC | Entinostat | MVP_top60_DNA+RNA_seed | HDAC1 | -0.177665 | BLCA | HT-1197 | JQ12 |
| MVP_top60_DNA_seed | AR | -0.015503 | BLCA | CAL-29 | Bicalutamide | MVP_top60_DNA+RNA_seed | AKT1 | -0.17555 | BLCA | SW1710 | GSK690693 |
| MVP_top60_DNA_seed | ERBB2 | -0.015376 | BLCA | RT-112 | Afatinib | MVP_top60_DNA+RNA_seed | AKT1 | -0.175255 | BLCA | SW1710 | AKT inhibitor VIII |
| MVP_top60_DNA_seed | HDAC1 | -0.012599 | BLCA | BFTC-905 | CAY10603 | MVP_top60_DNA+RNA_seed | HDAC1 | -0.173597 | BLCA | SW1710 | AR-42 |
| MVP_top60_DNA_seed | HSP90AA1 | -0.011266 | BLCA | SW1710 | CCT-018159 | MVP_top60_DNA+RNA_seed | HSP90AA1 | -0.172612 | BLCA | SW1710 | SNX-2112 |
| MVP_top60_DNA_seed | ERBB2 | -0.011232 | BLCA | BFTC-905 | PF-00299804 | MVP_top60_DNA+RNA_seed | ERBB2 | -0.17012 | BLCA | BFTC-905 | CI-1033 |
| MVP_top60_DNA_seed | ERBB2 | -0.008406 | BLCA | HT-1197 | CI-1033 | MVP_top60_DNA+RNA_seed | GSK3B | -0.168516 | BLCA | KU-19-19 | SB216763 |
| MVP_top60_DNA_seed | HDAC1 | -0.007912 | BLCA | KU-19-19 | CAY10603 | MVP_top60_DNA+RNA_seed | FGFR3 | -0.166442 | BLCA | SW780 | AZD4547 |
| MVP_top60_DNA_seed | AR | -0.005619 | BLCA | 647-V | Bicalutamide | MVP_top60_DNA+RNA_seed | AKT1 | -0.166319 | BLCA | LB831-BLC | AKT inhibitor VIII |
| MVP_top60_DNA_seed | FGFR3 | -0.005326 | BLCA | 647-V | AZD4547 | MVP_top60_DNA+RNA_seed | ERBB2 | -0.164055 | BLCA | RT-112 | AZD8931 |
| MVP_top60_DNA_seed | FGFR3 | -0.004722 | BLCA | LB831-BLC | AZD4547 | MVP_top60_DNA+RNA_seed | ERBB3 | -0.164055 | BLCA | RT-112 | AZD8931 |
| MVP_top60_DNA_seed | HSP90AA1 | -0.000209 | BLCA | RT4 | Tanespimycin | MVP_top60_DNA+RNA_seed | ERBB2 | -0.159624 | BLCA | 647-V | Afatinib |
| MVP_top60_DNA_seed | ESR1 | 0.002839 | BLCA | UM-UC-3 | Tamoxifen | MVP_top60_DNA+RNA_seed | ERBB2 | -0.158559 | BLCA | CAL-29 | CP724714 |
| MVP_top60_DNA_seed | MAPK1 | 0.003524 | BLCA | T24 | VX-11e | MVP_top60_DNA+RNA_seed | ATM | -0.158345 | BLCA | J82 | KU-55933 |
| MVP_top60_DNA_seed | ERBB2 | 0.004606 | BLCA | BFTC-905 | CP724714 | MVP_top60_DNA+RNA_seed | FGFR3 | -0.158158 | BLCA | J82 | AZD4547 |
| MVP_top60_DNA_seed | AR | 0.005566 | BLCA | RT4 | AZD3514 | MVP_top60_DNA+RNA_seed | EP300 | -0.156056 | BLCA | J82 | I-CBP112 |
| MVP_top60_DNA_seed | ESR1 | 0.005904 | BLCA | LB831-BLC | Fulvestrant | MVP_top60_DNA+RNA_seed | CREBBP | -0.156056 | BLCA | J82 | I-CBP112 |
| MVP_top60_DNA_seed | ERBB2 | 0.00736 | BLCA | KU-19-19 | CI-1033 | MVP_top60_DNA+RNA_seed | AKT1 | -0.152953 | BLCA | 639-V | MK-2206 |
| MVP_top60_DNA_seed | ERBB2 | 0.01718 | BLCA | SW1710 | CP724714 | MVP_top60_DNA+RNA_seed | ERBB2 | -0.151256 | BLCA | 639-V | AZD8931 |
| MVP_top60_DNA_seed | PIK3CB | 0.018024 | BLCA | DSH1 | AZD6482 | MVP_top60_DNA+RNA_seed | ERBB3 | -0.151256 | BLCA | 639-V | AZD8931 |
| MVP_top60_DNA_seed | ERBB2 | 0.026348 | BLCA | 639-V | Afatinib | MVP_top60_DNA+RNA_seed | ESR1 | -0.150683 | BLCA | RT-112 | Fulvestrant |
| MVP_top60_DNA_seed | SRC | 0.031539 | BLCA | DSH1 | Saracatinib | MVP_top60_DNA+RNA_seed | EP300 | -0.150236 | BLCA | 639-V | I-CBP112 |
| MVP_top60_DNA_seed | HDAC1 | 0.032543 | BLCA | 647-V | Dacinostat | MVP_top60_DNA+RNA_seed | CREBBP | -0.150236 | BLCA | 639-V | I-CBP112 |
| MVP_top60_DNA_seed | CDK2 | 0.033175 | BLCA | DSH1 | Seliciclib | MVP_top60_DNA+RNA_seed | HDAC1 | -0.140899 | BLCA | RT-112 | CAY10603 |
| MVP_top60_DNA_seed | HDAC1 | 0.034249 | BLCA | CAL-29 | Dacinostat | MVP_top60_DNA+RNA_seed | MAPK1 | -0.137653 | BLCA | SW1710 | VX-11e |
| MVP_top60_DNA_seed | HDAC1 | 0.041985 | BLCA | SW780 | JQ12 | MVP_top60_DNA+RNA_seed | MAPK1 | -0.137387 | BLCA | CAL-29 | FR-180204 |
| MVP_top60_DNA_seed | ATM | 0.043332 | BLCA | T24 | CP466722 | MVP_top60_DNA+RNA_seed | MAPK3 | -0.137387 | BLCA | CAL-29 | FR-180204 |
| MVP_top60_DNA_seed | ATM | 0.046629 | BLCA | HT-1376 | CP466722 | MVP_top60_DNA+RNA_seed | SYK | -0.133351 | BLCA | UM-UC-3 | BAY-61-3606 |
| MVP_top60_DNA_seed | RAC1 | 0.048684 | BLCA | 639-V | EHT-1864 | MVP_top60_DNA+RNA_seed | AR | -0.130993 | BLCA | J82 | AZD3514 |
| MVP_top60_DNA_seed | ERBB2 | 0.049261 | BLCA | 647-V | PF-00299804 | MVP_top60_DNA+RNA_seed | FGFR3 | -0.121367 | BLCA | TCCSUP | PD173074 |

| | | | | | | | | | | | |
|---|---|---|---|---|---|---|---|---|---|---|---|
| MVP_top60_DNA_seed | PIK3CA | 0.049539 | BLCA | SW1710 | PI3Ka_4409 | MVP_top60_DNA+RNA_seed | ERBB2 | -0.116867 | BLCA | BFTC-905 | CUDC-101 |
| MVP_top60_DNA_seed | ERBB2 | 0.050835 | BLCA | UM-UC-3 | AZD8931 | MVP_top60_DNA+RNA_seed | EP300 | -0.115471 | BLCA | 5637 | I-CBP112 |
| MVP_top60_DNA_seed | ERBB3 | 0.050835 | BLCA | UM-UC-3 | AZD8931 | MVP_top60_DNA+RNA_seed | CREBBP | -0.115471 | BLCA | 5637 | I-CBP112 |
| MVP_top60_DNA_seed | PIK3CA | 0.050942 | BLCA | J82 | PI3Ka_4409 | MVP_top60_DNA+RNA_seed | AKT1 | -0.114217 | BLCA | 647-V | BAY AKT1 |
| MVP_top60_DNA_seed | ATM | 0.053801 | BLCA | LB831-BLC | torin2 | MVP_top60_DNA+RNA_seed | HDAC1 | -0.113782 | BLCA | HT-1197 | AR-42 |
| MVP_top60_DNA_seed | ESR1 | 0.055445 | BLCA | BFTC-905 | Tamoxifen | MVP_top60_DNA+RNA_seed | GSK3B | -0.112884 | BLCA | DSH1 | AZD7969 |
| MVP_top60_DNA_seed | ERBB2 | 0.059361 | BLCA | SW1710 | CI-1033 | MVP_top60_DNA+RNA_seed | HSP90AA1 | -0.109124 | BLCA | UM-UC-3 | Tanespimycin |
| MVP_top60_DNA_seed | HDAC1 | 0.070256 | BLCA | CAL-29 | Belinostat | MVP_top60_DNA+RNA_seed | SRC | -0.107021 | BLCA | BFTC-905 | Bosutinib |
| MVP_top60_DNA_seed | HDAC1 | 0.075413 | BLCA | HT-1197 | CAY10603 | MVP_top60_DNA+RNA_seed | ATM | -0.106696 | BLCA | T24 | KU-60019 |
| MVP_top60_DNA_seed | ERBB2 | 0.075794 | BLCA | KU-19-19 | CUDC-101 | MVP_top60_DNA+RNA_seed | AR | -0.102666 | BLCA | HT-1376 | AZD3514 |
| MVP_top60_DNA_seed | AR | 0.078502 | BLCA | J82 | Bicalutamide | MVP_top60_DNA+RNA_seed | GSK3B | -0.098456 | BLCA | CAL-29 | AZD7969 |
| MVP_top60_DNA_seed | ERBB2 | 0.080014 | BLCA | SW1710 | Afatinib | MVP_top60_DNA+RNA_seed | ERBB2 | -0.097358 | BLCA | LB831-BLC | CUDC-101 |
| MVP_top60_DNA_seed | ERBB2 | 0.080143 | BLCA | TCCSUP | AZD8931 | MVP_top60_DNA+RNA_seed | ERBB2 | -0.0953 | BLCA | RT-112 | CI-1033 |
| MVP_top60_DNA_seed | ERBB3 | 0.080143 | BLCA | TCCSUP | AZD8931 | MVP_top60_DNA+RNA_seed | AKT1 | -0.094724 | BLCA | 5637 | AKT inhibitor VIII |
| MVP_top60_DNA_seed | HDAC1 | 0.088324 | BLCA | LB831-BLC | JQ12 | MVP_top60_DNA+RNA_seed | HDAC1 | -0.091872 | BLCA | CAL-29 | AR-42 |
| MVP_top60_DNA_seed | FGFR3 | 0.089341 | BLCA | HT-1197 | AZD4547 | MVP_top60_DNA+RNA_seed | AR | -0.090396 | BLCA | TCCSUP | AZD3514 |
| MVP_top60_DNA_seed | AR | 0.090211 | BLCA | T24 | Bicalutamide | MVP_top60_DNA+RNA_seed | GSK3B | -0.089379 | BLCA | J82 | CHIR-99021 |
| MVP_top60_DNA_seed | HDAC1 | 0.092252 | BLCA | KU-19-19 | Dacinostat | MVP_top60_DNA+RNA_seed | ERBB2 | -0.088 | BLCA | 639-V | PF-00299804 |
| MVP_top60_DNA_seed | PIK3CB | 0.092891 | BLCA | T24 | AZD6482 | MVP_top60_DNA+RNA_seed | HDAC1 | -0.087833 | BLCA | CAL-29 | CAY10603 |
| MVP_top60_DNA_seed | ATM | 0.093754 | BLCA | DSH1 | CP466722 | MVP_top60_DNA+RNA_seed | HSP90AA1 | -0.087304 | BLCA | J82 | CCT-018159 |
| MVP_top60_DNA_seed | SRC | 0.101669 | BLCA | 639-V | Bosutinib | MVP_top60_DNA+RNA_seed | FGFR3 | -0.086873 | BLCA | LB831-BLC | FGFR_3831 |
| MVP_top60_DNA_seed | PIK3CB | 0.102349 | BLCA | LB831-BLC | AZD8186 | MVP_top60_DNA+RNA_seed | ERBB2 | -0.086413 | BLCA | UM-UC-3 | PF-00299804 |
| MVP_top60_DNA_seed | ERBB2 | 0.103219 | BLCA | KU-19-19 | CP724714 | MVP_top60_DNA+RNA_seed | HDAC1 | -0.084414 | BLCA | BFTC-905 | Belinostat |
| MVP_top60_DNA_seed | FGFR3 | 0.103477 | BLCA | CAL-29 | AZD4547 | MVP_top60_DNA+RNA_seed | HDAC1 | -0.075407 | BLCA | UM-UC-3 | CAY10603 |
| MVP_top60_DNA_seed | EP300 | 0.10791 | BLCA | HT-1197 | I-CBP112 | MVP_top60_DNA+RNA_seed | SYK | -0.071346 | BLCA | HT-1376 | BAY-61-3606 |
| MVP_top60_DNA_seed | CREBBP | 0.10791 | BLCA | HT-1197 | I-CBP112 | MVP_top60_DNA+RNA_seed | SRC | -0.070686 | BLCA | RT-112 | NVP-BHG712 |
| MVP_top60_DNA_seed | HDAC1 | 0.10976 | BLCA | BFTC-905 | AR-42 | MVP_top60_DNA+RNA_seed | FGFR3 | -0.069224 | BLCA | BFTC-905 | AZD4547 |
| MVP_top60_DNA_seed | PIK3CA | 0.120483 | BLCA | DSH1 | AZD8835 | MVP_top60_DNA+RNA_seed | ATM | -0.065788 | BLCA | RT-112 | KU-60019 |
| MVP_top60_DNA_seed | ATM | 0.121958 | BLCA | LB831-BLC | KU-55933 | MVP_top60_DNA+RNA_seed | ATM | -0.0563 | BLCA | J82 | KU-60019 |
| MVP_top60_DNA_seed | PIK3CA | 0.123523 | BLCA | 5637 | AZD8835 | MVP_top60_DNA+RNA_seed | EP300 | -0.056293 | BLCA | UM-UC-3 | I-CBP112 |
| MVP_top60_DNA_seed | HDAC1 | 0.125089 | BLCA | KU-19-19 | JQ12 | MVP_top60_DNA+RNA_seed | CREBBP | -0.056293 | BLCA | UM-UC-3 | I-CBP112 |
| MVP_top60_DNA_seed | FGFR3 | 0.130293 | BLCA | HT-1376 | AZD4547 | MVP_top60_DNA+RNA_seed | ERBB2 | -0.053534 | BLCA | UM-UC-3 | CUDC-101 |
| MVP_top60_DNA_seed | PIK3CB | 0.131687 | BLCA | TCCSUP | AZD8186 | MVP_top60_DNA+RNA_seed | ERBB2 | -0.053279 | BLCA | SW1710 | CUDC-101 |
| MVP_top60_DNA_seed | ERBB2 | 0.132366 | BLCA | CAL-29 | PF-00299804 | MVP_top60_DNA+RNA_seed | HDAC1 | -0.045825 | BLCA | LB831-BLC | CAY10603 |
| MVP_top60_DNA_seed | FGFR3 | 0.13299 | BLCA | J82 | FGFR_3831 | MVP_top60_DNA+RNA_seed | ERBB2 | -0.045528 | BLCA | 639-V | Afatinib |
| MVP_top60_DNA_seed | HDAC1 | 0.134787 | BLCA | UM-UC-3 | AR-42 | MVP_top60_DNA+RNA_seed | HDAC1 | -0.044736 | BLCA | SW1710 | CAY10603 |
| MVP_top60_DNA_seed | CDK2 | 0.13613 | BLCA | LB831-BLC | PHA-793887 | MVP_top60_DNA+RNA_seed | HDAC1 | -0.042739 | BLCA | TCCSUP | Dacinostat |
| MVP_top60_DNA_seed | ESR1 | 0.14097 | BLCA | HT-1197 | Fulvestrant | MVP_top60_DNA+RNA_seed | CDK2 | -0.041383 | BLCA | 5637 | PHA-793887 |
| MVP_top60_DNA_seed | CDK2 | 0.148468 | BLCA | KU-19-19 | PHA-793887 | MVP_top60_DNA+RNA_seed | ERBB2 | -0.039289 | BLCA | SW1710 | Afatinib |
| MVP_top60_DNA_seed | HDAC1 | 0.15055 | BLCA | TCCSUP | CAY10603 | MVP_top60_DNA+RNA_seed | ESR1 | -0.036355 | BLCA | 639-V | Tamoxifen |
| MVP_top60_DNA_seed | FGFR3 | 0.151513 | BLCA | 639-V | AZD4547 | MVP_top60_DNA+RNA_seed | CDK2 | -0.033633 | BLCA | 647-V | PHA-793887 |

| | | | | | | | | | | | |
|---|---|---|---|---|---|---|---|---|---|---|---|
| MVP_top60_DNA_seed | ERBB2 | 0.153305 | BLCA | T24 | PF-00299804 | MVP_top60_DNA+RNA_seed | SYK | -0.032905 | BLCA | 5637 | BAY-61-3606 |
| MVP_top60_DNA_seed | AR | 0.154262 | BLCA | RT-112 | Bicalutamide | MVP_top60_DNA+RNA_seed | EP300 | -0.031533 | BLCA | SW1710 | I-CBP112 |
| MVP_top60_DNA_seed | CDK2 | 0.154787 | BLCA | 639-V | PHA-793887 | MVP_top60_DNA+RNA_seed | CREBBP | -0.031533 | BLCA | SW1710 | I-CBP112 |
| MVP_top60_DNA_seed | AR | 0.156358 | BLCA | J82 | Bicalutamide | MVP_top60_DNA+RNA_seed | HSP90AA1 | -0.029601 | BLCA | T24 | Luminespib |
| MVP_top60_DNA_seed | ERBB2 | 0.171571 | BLCA | KU-19-19 | AZD8931 | MVP_top60_DNA+RNA_seed | ERBB2 | -0.02186 | BLCA | HT-1376 | CI-1033 |
| MVP_top60_DNA_seed | ERBB3 | 0.171571 | BLCA | KU-19-19 | AZD8931 | MVP_top60_DNA+RNA_seed | RAC1 | -0.021736 | BLCA | SW780 | EHT-1864 |
| MVP_top60_DNA_seed | FGFR3 | 0.172147 | BLCA | UM-UC-3 | FGFR_3831 | MVP_top60_DNA+RNA_seed | HDAC1 | -0.020237 | BLCA | TCCSUP | Belinostat |
| MVP_top60_DNA_seed | MAPK1 | 0.176751 | BLCA | SW780 | VX-11e | MVP_top60_DNA+RNA_seed | HDAC1 | -0.018529 | BLCA | LB831-BLC | AR-42 |
| MVP_top60_DNA_seed | RAC1 | 0.178272 | BLCA | LB831-BLC | EHT-1864 | MVP_top60_DNA+RNA_seed | HDAC1 | -0.015948 | BLCA | LB831-BLC | Entinostat |
| MVP_top60_DNA_seed | HDAC1 | 0.18213 | BLCA | TCCSUP | AR-42 | MVP_top60_DNA+RNA_seed | AR | -0.015503 | BLCA | CAL-29 | Bicalutamide |
| MVP_top60_DNA_seed | FGFR3 | 0.184439 | BLCA | LB831-BLC | PD173074 | MVP_top60_DNA+RNA_seed | ERBB2 | -0.015376 | BLCA | RT-112 | Afatinib |
| MVP_top60_DNA_seed | SRC | 0.188404 | BLCA | J82 | Bosutinib | MVP_top60_DNA+RNA_seed | HDAC1 | -0.012599 | BLCA | BFTC-905 | CAY10603 |
| MVP_top60_DNA_seed | ERBB2 | 0.18863 | BLCA | LB831-BLC | CI-1033 | MVP_top60_DNA+RNA_seed | HSP90AA1 | -0.011266 | BLCA | SW1710 | CCT-018159 |
| MVP_top60_DNA_seed | FGFR3 | 0.189791 | BLCA | DSH1 | PD173074 | MVP_top60_DNA+RNA_seed | ERBB2 | -0.011232 | BLCA | BFTC-905 | PF-00299804 |
| MVP_top60_DNA_seed | ERBB2 | 0.190569 | BLCA | TCCSUP | CI-1033 | MVP_top60_DNA+RNA_seed | ERBB2 | -0.008406 | BLCA | HT-1197 | CI-1033 |
| MVP_top60_DNA_seed | RAC1 | 0.195741 | BLCA | HT-1197 | EHT-1864 | MVP_top60_DNA+RNA_seed | HDAC1 | -0.007912 | BLCA | KU-19-19 | CAY10603 |
| MVP_top60_DNA_seed | ATM | 0.20477 | BLCA | TCCSUP | KU-60019 | MVP_top60_DNA+RNA_seed | GSK3B | -0.006204 | BLCA | 639-V | CHIR-99021 |
| MVP_top60_DNA_seed | FGFR3 | 0.20775 | BLCA | DSH1 | AZD4547 | MVP_top60_DNA+RNA_seed | AR | -0.005619 | BLCA | 647-V | Bicalutamide |
| MVP_top60_DNA_seed | ERBB2 | 0.20964 | BLCA | LB831-BLC | Lapatinib | MVP_top60_DNA+RNA_seed | FGFR3 | -0.005326 | BLCA | 647-V | AZD4547 |
| MVP_top60_DNA_seed | EP300 | 0.212048 | BLCA | DSH1 | I-CBP112 | MVP_top60_DNA+RNA_seed | AKT1 | -0.005232 | BLCA | UM-UC-3 | AKT inhibitor VIII |
| MVP_top60_DNA_seed | CREBBP | 0.212048 | BLCA | DSH1 | I-CBP112 | MVP_top60_DNA+RNA_seed | FGFR3 | -0.004722 | BLCA | LB831-BLC | AZD4547 |
| MVP_top60_DNA_seed | SRC | 0.212184 | BLCA | LB831-BLC | HG-6-71-01 | MVP_top60_DNA+RNA_seed | HSP90AA1 | -0.000209 | BLCA | RT4 | Tanespimycin |
| MVP_top60_DNA_seed | PIK3CA | 0.212412 | BLCA | DSH1 | PI3Ka_4409 | MVP_top60_DNA+RNA_seed | ESR1 | 0.002839 | BLCA | UM-UC-3 | Tamoxifen |
| MVP_top60_DNA_seed | HSP90AA1 | 0.214774 | BLCA | HT-1197 | Elesclomol | MVP_top60_DNA+RNA_seed | AKT1 | 0.002971 | BLCA | RT4 | AKT inhibitor VIII |
| MVP_top60_DNA_seed | PIK3CA | 0.217414 | BLCA | T24 | AZD8835 | MVP_top60_DNA+RNA_seed | MAPK1 | 0.003524 | BLCA | T24 | VX-11e |
| MVP_top60_DNA_seed | AR | 0.222905 | BLCA | CAL-29 | Bicalutamide | MVP_top60_DNA+RNA_seed | ERBB2 | 0.004606 | BLCA | BFTC-905 | CP724714 |
| MVP_top60_DNA_seed | MAPK1 | 0.225432 | BLCA | RT-112 | VX-11e | MVP_top60_DNA+RNA_seed | AR | 0.005566 | BLCA | RT4 | AZD3514 |
| MVP_top60_DNA_seed | HDAC1 | 0.227031 | BLCA | 647-V | JQ12 | MVP_top60_DNA+RNA_seed | ESR1 | 0.005904 | BLCA | LB831-BLC | Fulvestrant |
| MVP_top60_DNA_seed | MAPK1 | 0.230195 | BLCA | 639-V | FR-180204 | MVP_top60_DNA+RNA_seed | ERBB2 | 0.00736 | BLCA | KU-19-19 | CI-1033 |
| MVP_top60_DNA_seed | MAPK3 | 0.230195 | BLCA | 639-V | FR-180204 | MVP_top60_DNA+RNA_seed | GSK3B | 0.014993 | BLCA | 639-V | KIN001-042 |
| MVP_top60_DNA_seed | PIK3CA | 0.230402 | BLCA | 639-V | PI3Ka_4409 | MVP_top60_DNA+RNA_seed | GSK3B | 0.016266 | BLCA | 639-V | CHIR-99021 |
| MVP_top60_DNA_seed | ERBB2 | 0.239817 | BLCA | J82 | AZD8931 | MVP_top60_DNA+RNA_seed | ERBB2 | 0.01718 | BLCA | SW1710 | CP724714 |
| MVP_top60_DNA_seed | ERBB3 | 0.239817 | BLCA | J82 | AZD8931 | MVP_top60_DNA+RNA_seed | ERBB2 | 0.026348 | BLCA | 639-V | Afatinib |
| MVP_top60_DNA_seed | SRC | 0.240198 | BLCA | 647-V | NVP-BHG712 | MVP_top60_DNA+RNA_seed | SRC | 0.031539 | BLCA | DSH1 | Saracatinib |
| MVP_top60_DNA_seed | FGFR3 | 0.242725 | BLCA | UM-UC-3 | AZD4547 | MVP_top60_DNA+RNA_seed | HDAC1 | 0.032543 | BLCA | 647-V | Dacinostat |
| MVP_top60_DNA_seed | FGFR3 | 0.243276 | BLCA | UM-UC-3 | AZD4547 | MVP_top60_DNA+RNA_seed | CDK2 | 0.033175 | BLCA | DSH1 | Seliciclib |
| MVP_top60_DNA_seed | ERBB2 | 0.244047 | BLCA | T24 | AZD8931 | MVP_top60_DNA+RNA_seed | HDAC1 | 0.034249 | BLCA | CAL-29 | Dacinostat |
| MVP_top60_DNA_seed | ERBB3 | 0.244047 | BLCA | T24 | AZD8931 | MVP_top60_DNA+RNA_seed | HDAC1 | 0.041985 | BLCA | SW780 | JQ12 |
| MVP_top60_DNA_seed | CDK2 | 0.246186 | BLCA | RT-112 | AZD5438 | MVP_top60_DNA+RNA_seed | ATM | 0.043332 | BLCA | T24 | CP466722 |
| MVP_top60_DNA_seed | ERBB2 | 0.249308 | BLCA | UM-UC-3 | Afatinib | MVP_top60_DNA+RNA_seed | AKT1 | 0.043799 | BLCA | KU-19-19 | MK-2206 |
| MVP_top60_DNA_seed | HSP90AA1 | 0.255261 | BLCA | BFTC-905 | CCT-018159 | MVP_top60_DNA+RNA_seed | ATM | 0.046629 | BLCA | HT-1376 | CP466722 |

| | | | | | | | | | | | |
|---|---|---|---|---|---|---|---|---|---|---|---|
| MVP_top60_DNA_seed | ESR1 | 0.262134 | BLCA | TCCSUP | Fulvestrant | MVP_top60_DNA+RNA_seed | RAC1 | 0.048684 | BLCA | 639-V | EHT-1864 |
| MVP_top60_DNA_seed | AR | 0.269045 | BLCA | 5637 | AZD3514 | MVP_top60_DNA+RNA_seed | ERBB2 | 0.049261 | BLCA | 647-V | PF-00299804 |
| MVP_top60_DNA_seed | ATM | 0.269272 | BLCA | RT-112 | KU-55933 | MVP_top60_DNA+RNA_seed | PIK3CA | 0.049539 | BLCA | SW1710 | PI3Ka_4409 |
| MVP_top60_DNA_seed | FGFR3 | 0.27548 | BLCA | HT-1376 | AZD4547 | MVP_top60_DNA+RNA_seed | ERBB2 | 0.050835 | BLCA | UM-UC-3 | AZD8931 |
| MVP_top60_DNA_seed | HSP90AA1 | 0.275767 | BLCA | 647-V | SNX-2112 | MVP_top60_DNA+RNA_seed | ERBB3 | 0.050835 | BLCA | UM-UC-3 | AZD8931 |
| MVP_top60_DNA_seed | ATM | 0.276932 | BLCA | CAL-29 | CP466722 | MVP_top60_DNA+RNA_seed | PIK3CA | 0.050942 | BLCA | J82 | PI3Ka_4409 |
| MVP_top60_DNA_seed | AR | 0.277381 | BLCA | HT-1197 | AZD3514 | MVP_top60_DNA+RNA_seed | GSK3B | 0.052863 | BLCA | CAL-29 | KIN001-042 |
| MVP_top60_DNA_seed | CDK2 | 0.278442 | BLCA | DSH1 | PHA-793887 | MVP_top60_DNA+RNA_seed | ATM | 0.053801 | BLCA | LB831-BLC | torin2 |
| MVP_top60_DNA_seed | ATM | 0.280053 | BLCA | SW780 | KU-55933 | MVP_top60_DNA+RNA_seed | GSK3B | 0.055128 | BLCA | BFTC-905 | KIN001-042 |
| MVP_top60_DNA_seed | HDAC1 | 0.282295 | BLCA | BFTC-905 | Tubastatin A | MVP_top60_DNA+RNA_seed | ESR1 | 0.055445 | BLCA | BFTC-905 | Tamoxifen |
| MVP_top60_DNA_seed | SRC | 0.290119 | BLCA | LB831-BLC | Saracatinib | MVP_top60_DNA+RNA_seed | ERBB2 | 0.059361 | BLCA | SW1710 | CI-1033 |
| MVP_top60_DNA_seed | FGFR3 | 0.290289 | BLCA | KU-19-19 | PD173074 | MVP_top60_DNA+RNA_seed | HDAC1 | 0.070256 | BLCA | CAL-29 | Belinostat |
| MVP_top60_DNA_seed | MAPK1 | 0.290837 | BLCA | LB831-BLC | FR-180204 | MVP_top60_DNA+RNA_seed | HDAC1 | 0.075413 | BLCA | HT-1197 | CAY10603 |
| MVP_top60_DNA_seed | MAPK3 | 0.290837 | BLCA | LB831-BLC | FR-180204 | MVP_top60_DNA+RNA_seed | ERBB2 | 0.075794 | BLCA | KU-19-19 | CUDC-101 |
| MVP_top60_DNA_seed | FGFR3 | 0.29394 | BLCA | 647-V | FGFR_3831 | MVP_top60_DNA+RNA_seed | AR | 0.078502 | BLCA | J82 | Bicalutamide |
| MVP_top60_DNA_seed | CDK2 | 0.294887 | BLCA | T24 | PHA-793887 | MVP_top60_DNA+RNA_seed | ERBB2 | 0.080014 | BLCA | SW1710 | Afatinib |
| MVP_top60_DNA_seed | PIK3CB | 0.297701 | BLCA | HT-1197 | AZD8186 | MVP_top60_DNA+RNA_seed | ERBB2 | 0.080143 | BLCA | TCCSUP | AZD8931 |
| MVP_top60_DNA_seed | RAC1 | 0.302502 | BLCA | J82 | EHT-1864 | MVP_top60_DNA+RNA_seed | ERBB3 | 0.080143 | BLCA | TCCSUP | AZD8931 |
| MVP_top60_DNA_seed | ERBB2 | 0.303676 | BLCA | J82 | Afatinib | MVP_top60_DNA+RNA_seed | AKT1 | 0.085794 | BLCA | TCCSUP | BAY AKT1 |
| MVP_top60_DNA_seed | ERBB2 | 0.304186 | BLCA | T24 | CUDC-101 | MVP_top60_DNA+RNA_seed | HDAC1 | 0.088324 | BLCA | LB831-BLC | JQ12 |
| MVP_top60_DNA_seed | PIK3CA | 0.306539 | BLCA | UM-UC-3 | AZD8835 | MVP_top60_DNA+RNA_seed | AKT1 | 0.089081 | BLCA | UM-UC-3 | GSK690693 |
| MVP_top60_DNA_seed | FGFR3 | 0.310781 | BLCA | KU-19-19 | AZD4547 | MVP_top60_DNA+RNA_seed | FGFR3 | 0.089341 | BLCA | HT-1197 | AZD4547 |
| MVP_top60_DNA_seed | CDK2 | 0.314928 | BLCA | RT4 | PHA-793887 | MVP_top60_DNA+RNA_seed | AR | 0.090211 | BLCA | T24 | Bicalutamide |
| MVP_top60_DNA_seed | HDAC1 | 0.317185 | BLCA | T24 | Tubastatin A | MVP_top60_DNA+RNA_seed | HDAC1 | 0.092252 | BLCA | KU-19-19 | Dacinostat |
| MVP_top60_DNA_seed | ERBB2 | 0.318815 | BLCA | LB831-BLC | Afatinib | MVP_top60_DNA+RNA_seed | ATM | 0.093754 | BLCA | DSH1 | CP466722 |
| MVP_top60_DNA_seed | HDAC1 | 0.327679 | BLCA | J82 | Dacinostat | MVP_top60_DNA+RNA_seed | SRC | 0.101669 | BLCA | 639-V | Bosutinib |
| MVP_top60_DNA_seed | FGFR3 | 0.328257 | BLCA | 5637 | PD173074 | MVP_top60_DNA+RNA_seed | ERBB2 | 0.103219 | BLCA | KU-19-19 | CP724714 |
| MVP_top60_DNA_seed | ERBB2 | 0.328999 | BLCA | RT-112 | CP724714 | MVP_top60_DNA+RNA_seed | FGFR3 | 0.103477 | BLCA | CAL-29 | AZD4547 |
| MVP_top60_DNA_seed | AR | 0.332016 | BLCA | DSH1 | Bicalutamide | MVP_top60_DNA+RNA_seed | EP300 | 0.10791 | BLCA | HT-1197 | I-CBP112 |
| MVP_top60_DNA_seed | FGFR3 | 0.334738 | BLCA | BFTC-905 | PD173074 | MVP_top60_DNA+RNA_seed | CREBBP | 0.10791 | BLCA | HT-1197 | I-CBP112 |
| MVP_top60_DNA_seed | ERBB2 | 0.338084 | BLCA | RT-112 | PF-00299804 | MVP_top60_DNA+RNA_seed | HDAC1 | 0.10976 | BLCA | BFTC-905 | AR-42 |
| MVP_top60_DNA_seed | RAC1 | 0.340131 | BLCA | CAL-29 | EHT-1864 | MVP_top60_DNA+RNA_seed | GSK3B | 0.113053 | BLCA | LB831-BLC | KIN001-042 |
| MVP_top60_DNA_seed | ERBB2 | 0.341313 | BLCA | J82 | Afatinib | MVP_top60_DNA+RNA_seed | PIK3CA | 0.120483 | BLCA | DSH1 | AZD8835 |
| MVP_top60_DNA_seed | ERBB2 | 0.346372 | BLCA | UM-UC-3 | Afatinib | MVP_top60_DNA+RNA_seed | ATM | 0.121958 | BLCA | LB831-BLC | KU-55933 |
| MVP_top60_DNA_seed | HDAC1 | 0.349277 | BLCA | SW1710 | Tubastatin A | MVP_top60_DNA+RNA_seed | GSK3B | 0.123165 | BLCA | HT-1197 | AZD7969 |
| MVP_top60_DNA_seed | EP300 | 0.351883 | BLCA | CAL-29 | I-CBP112 | MVP_top60_DNA+RNA_seed | PIK3CA | 0.123523 | BLCA | 5637 | AZD8835 |
| MVP_top60_DNA_seed | CREBBP | 0.351883 | BLCA | CAL-29 | I-CBP112 | MVP_top60_DNA+RNA_seed | HDAC1 | 0.125089 | BLCA | KU-19-19 | JQ12 |
| MVP_top60_DNA_seed | FGFR3 | 0.35798 | BLCA | TCCSUP | AZD4547 | MVP_top60_DNA+RNA_seed | GSK3B | 0.125728 | BLCA | LB831-BLC | AZD7969 |
| MVP_top60_DNA_seed | HDAC1 | 0.363134 | BLCA | T24 | AR-42 | MVP_top60_DNA+RNA_seed | FGFR3 | 0.130293 | BLCA | HT-1376 | AZD4547 |
| MVP_top60_DNA_seed | HDAC1 | 0.369948 | BLCA | T24 | Belinostat | MVP_top60_DNA+RNA_seed | ERBB2 | 0.132366 | BLCA | CAL-29 | PF-00299804 |
| MVP_top60_DNA_seed | ESR1 | 0.370063 | BLCA | DSH1 | Tamoxifen | MVP_top60_DNA+RNA_seed | FGFR3 | 0.13299 | BLCA | J82 | FGFR_3831 |

| | | | | | | | | | | | |
|---|---|---|---|---|---|---|---|---|---|---|---|
| MVP_top60_DNA_seed | ERBB2 | 0.373692 | BLCA | SW1710 | PF-00299804 | MVP_top60_DNA+RNA_seed | HDAC1 | 0.134787 | BLCA | UM-UC-3 | AR-42 |
| MVP_top60_DNA_seed | MAPK1 | 0.374806 | BLCA | CAL-29 | VX-11e | MVP_top60_DNA+RNA_seed | CDK2 | 0.13613 | BLCA | LB831-BLC | PHA-793887 |
| MVP_top60_DNA_seed | ESR1 | 0.378035 | BLCA | J82 | Fulvestrant | MVP_top60_DNA+RNA_seed | ESR1 | 0.14097 | BLCA | HT-1197 | Fulvestrant |
| MVP_top60_DNA_seed | PIK3CA | 0.378449 | BLCA | 647-V | AZD8835 | MVP_top60_DNA+RNA_seed | AKT1 | 0.147947 | BLCA | BFTC-905 | MK-2206 |
| MVP_top60_DNA_seed | PIK3CA | 0.379359 | BLCA | LB831-BLC | PI3Ka_4409 | MVP_top60_DNA+RNA_seed | CDK2 | 0.148468 | BLCA | KU-19-19 | PHA-793887 |
| MVP_top60_DNA_seed | ERBB2 | 0.380391 | BLCA | 639-V | CUDC-101 | MVP_top60_DNA+RNA_seed | GSK3B | 0.14915 | BLCA | SW1710 | KIN001-042 |
| MVP_top60_DNA_seed | AR | 0.381983 | BLCA | SW1710 | Bicalutamide | MVP_top60_DNA+RNA_seed | HDAC1 | 0.15055 | BLCA | TCCSUP | CAY10603 |
| MVP_top60_DNA_seed | ERBB2 | 0.385041 | BLCA | TCCSUP | Afatinib | MVP_top60_DNA+RNA_seed | FGFR3 | 0.151513 | BLCA | 639-V | AZD4547 |
| MVP_top60_DNA_seed | ERBB2 | 0.388544 | BLCA | SW780 | Afatinib | MVP_top60_DNA+RNA_seed | ERBB2 | 0.153305 | BLCA | T24 | PF-00299804 |
| MVP_top60_DNA_seed | CDK2 | 0.390177 | BLCA | HT-1197 | AZD5438 | MVP_top60_DNA+RNA_seed | AR | 0.154262 | BLCA | RT-112 | Bicalutamide |
| MVP_top60_DNA_seed | HDAC1 | 0.393925 | BLCA | T24 | CAY10603 | MVP_top60_DNA+RNA_seed | CDK2 | 0.154787 | BLCA | 639-V | PHA-793887 |
| MVP_top60_DNA_seed | CDK2 | 0.394048 | BLCA | LB831-BLC | AZD5438 | MVP_top60_DNA+RNA_seed | AR | 0.156358 | BLCA | J82 | Bicalutamide |
| MVP_top60_DNA_seed | ERBB2 | 0.394126 | BLCA | T24 | Afatinib | MVP_top60_DNA+RNA_seed | ERBB2 | 0.171571 | BLCA | KU-19-19 | AZD8931 |
| MVP_top60_DNA_seed | ERBB2 | 0.394816 | BLCA | 647-V | CP724714 | MVP_top60_DNA+RNA_seed | ERBB3 | 0.171571 | BLCA | KU-19-19 | AZD8931 |
| MVP_top60_DNA_seed | FGFR3 | 0.40049 | BLCA | T24 | AZD4547 | MVP_top60_DNA+RNA_seed | AKT1 | 0.171944 | BLCA | 5637 | MK-2206 |
| MVP_top60_DNA_seed | EP300 | 0.410601 | BLCA | T24 | I-CBP112 | MVP_top60_DNA+RNA_seed | FGFR3 | 0.172147 | BLCA | UM-UC-3 | FGFR_3831 |
| MVP_top60_DNA_seed | CREBBP | 0.410601 | BLCA | T24 | I-CBP112 | MVP_top60_DNA+RNA_seed | AKT1 | 0.172759 | BLCA | HT-1376 | AKT inhibitor VIII |
| MVP_top60_DNA_seed | MAPK1 | 0.411417 | BLCA | DSH1 | VX-11e | MVP_top60_DNA+RNA_seed | AKT1 | 0.17531 | BLCA | SW780 | MK-2206 |
| MVP_top60_DNA_seed | PIK3CB | 0.419708 | BLCA | RT-112 | AZD6482 | MVP_top60_DNA+RNA_seed | MAPK1 | 0.176751 | BLCA | SW780 | VX-11e |
| MVP_top60_DNA_seed | AR | 0.424763 | BLCA | RT-112 | AZD3514 | MVP_top60_DNA+RNA_seed | RAC1 | 0.178272 | BLCA | LB831-BLC | EHT-1864 |
| MVP_top60_DNA_seed | HDAC1 | 0.426475 | BLCA | 639-V | Tubastatin A | MVP_top60_DNA+RNA_seed | GSK3B | 0.179014 | BLCA | 5637 | CHIR-99021 |
| MVP_top60_DNA_seed | MAPK1 | 0.428655 | BLCA | 639-V | VX-11e | MVP_top60_DNA+RNA_seed | HDAC1 | 0.18213 | BLCA | TCCSUP | AR-42 |
| MVP_top60_DNA_seed | HSP90AA1 | 0.432685 | BLCA | 647-V | CCT-018159 | MVP_top60_DNA+RNA_seed | FGFR3 | 0.184439 | BLCA | LB831-BLC | PD173074 |
| MVP_top60_DNA_seed | ATM | 0.434257 | BLCA | TCCSUP | KU-55933 | MVP_top60_DNA+RNA_seed | SRC | 0.188404 | BLCA | J82 | Bosutinib |
| MVP_top60_DNA_seed | ERBB2 | 0.436909 | BLCA | TCCSUP | CUDC-101 | MVP_top60_DNA+RNA_seed | ERBB2 | 0.18863 | BLCA | LB831-BLC | CI-1033 |
| MVP_top60_DNA_seed | PIK3CB | 0.439983 | BLCA | HT-1197 | AZD6482 | MVP_top60_DNA+RNA_seed | FGFR3 | 0.189791 | BLCA | DSH1 | PD173074 |
| MVP_top60_DNA_seed | SRC | 0.442002 | BLCA | CAL-29 | NVP-BHG712 | MVP_top60_DNA+RNA_seed | ERBB2 | 0.190569 | BLCA | TCCSUP | CI-1033 |
| MVP_top60_DNA_seed | ESR1 | 0.44238 | BLCA | LB831-BLC | Tamoxifen | MVP_top60_DNA+RNA_seed | RAC1 | 0.195741 | BLCA | HT-1197 | EHT-1864 |
| MVP_top60_DNA_seed | HDAC1 | 0.445041 | BLCA | 639-V | CAY10603 | MVP_top60_DNA+RNA_seed | GSK3B | 0.196065 | BLCA | RT4 | AZD7969 |
| MVP_top60_DNA_seed | ERBB2 | 0.449358 | BLCA | BFTC-905 | Afatinib | MVP_top60_DNA+RNA_seed | AKT1 | 0.198713 | BLCA | UM-UC-3 | BAY AKT1 |
| MVP_top60_DNA_seed | HDAC1 | 0.449822 | BLCA | 639-V | Belinostat | MVP_top60_DNA+RNA_seed | ATM | 0.20477 | BLCA | TCCSUP | KU-60019 |
| MVP_top60_DNA_seed | EP300 | 0.458008 | BLCA | 647-V | I-CBP112 | MVP_top60_DNA+RNA_seed | FGFR3 | 0.20775 | BLCA | DSH1 | AZD4547 |
| MVP_top60_DNA_seed | CREBBP | 0.458008 | BLCA | 647-V | I-CBP112 | MVP_top60_DNA+RNA_seed | ERBB2 | 0.20964 | BLCA | LB831-BLC | Lapatinib |
| MVP_top60_DNA_seed | ERBB2 | 0.459341 | BLCA | LB831-BLC | AZD8931 | MVP_top60_DNA+RNA_seed | EP300 | 0.212048 | BLCA | DSH1 | I-CBP112 |
| MVP_top60_DNA_seed | ERBB3 | 0.459341 | BLCA | LB831-BLC | AZD8931 | MVP_top60_DNA+RNA_seed | CREBBP | 0.212048 | BLCA | DSH1 | I-CBP112 |
| MVP_top60_DNA_seed | AR | 0.463076 | BLCA | LB831-BLC | Bicalutamide | MVP_top60_DNA+RNA_seed | SRC | 0.212184 | BLCA | LB831-BLC | HG-6-71-01 |
| MVP_top60_DNA_seed | CDK2 | 0.473515 | BLCA | CAL-29 | PHA-793887 | MVP_top60_DNA+RNA_seed | PIK3CA | 0.212412 | BLCA | DSH1 | PI3Ka_4409 |
| MVP_top60_DNA_seed | HDAC1 | 0.473914 | BLCA | LB831-BLC | Dacinostat | MVP_top60_DNA+RNA_seed | HSP90AA1 | 0.214774 | BLCA | HT-1197 | Elesclomol |
| MVP_top60_DNA_seed | FGFR3 | 0.475546 | BLCA | J82 | AZD4547 | MVP_top60_DNA+RNA_seed | PIK3CA | 0.217414 | BLCA | T24 | AZD8835 |
| MVP_top60_DNA_seed | HSP90AA1 | 0.478719 | BLCA | KU-19-19 | Luminespib | MVP_top60_DNA+RNA_seed | AR | 0.222905 | BLCA | CAL-29 | Bicalutamide |
| MVP_top60_DNA_seed | EP300 | 0.480541 | BLCA | RT-112 | I-CBP112 | MVP_top60_DNA+RNA_seed | MAPK1 | 0.225432 | BLCA | RT-112 | VX-11e |

| | | | | | | | | | | | |
|---|---|---|---|---|---|---|---|---|---|---|---|
| MVP_top60_DNA_seed | CREBBP | 0.480541 | BLCA | RT-112 | I-CBP112 | MVP_top60_DNA+RNA_seed | HDAC1 | 0.227031 | BLCA | 647-V | JQ12 |
| MVP_top60_DNA_seed | PIK3CA | 0.485958 | BLCA | J82 | AZD8835 | MVP_top60_DNA+RNA_seed | MAPK1 | 0.230195 | BLCA | 639-V | FR-180204 |
| MVP_top60_DNA_seed | ERBB2 | 0.489032 | BLCA | SW780 | CP724714 | MVP_top60_DNA+RNA_seed | MAPK3 | 0.230195 | BLCA | 639-V | FR-180204 |
| MVP_top60_DNA_seed | HDAC1 | 0.489722 | BLCA | LB831-BLC | Tubastatin A | MVP_top60_DNA+RNA_seed | PIK3CA | 0.230402 | BLCA | 639-V | PI3Ka_4409 |
| MVP_top60_DNA_seed | FGFR3 | 0.507279 | BLCA | CAL-29 | PD173074 | MVP_top60_DNA+RNA_seed | ERBB2 | 0.239817 | BLCA | J82 | AZD8931 |
| MVP_top60_DNA_seed | ATM | 0.5075 | BLCA | 639-V | CP466722 | MVP_top60_DNA+RNA_seed | ERBB3 | 0.239817 | BLCA | J82 | AZD8931 |
| MVP_top60_DNA_seed | ERBB2 | 0.510722 | BLCA | HT-1197 | PF-00299804 | MVP_top60_DNA+RNA_seed | SRC | 0.240198 | BLCA | 647-V | NVP-BHG712 |
| MVP_top60_DNA_seed | MAPK1 | 0.511499 | BLCA | KU-19-19 | FR-180204 | MVP_top60_DNA+RNA_seed | AKT1 | 0.242338 | BLCA | UM-UC-3 | AKT inhibitor VIII |
| MVP_top60_DNA_seed | MAPK3 | 0.511499 | BLCA | KU-19-19 | FR-180204 | MVP_top60_DNA+RNA_seed | FGFR3 | 0.242725 | BLCA | UM-UC-3 | AZD4547 |
| MVP_top60_DNA_seed | RAC1 | 0.52361 | BLCA | RT4 | EHT-1864 | MVP_top60_DNA+RNA_seed | FGFR3 | 0.243276 | BLCA | UM-UC-3 | AZD4547 |
| MVP_top60_DNA_seed | SRC | 0.529177 | BLCA | T24 | NVP-BHG712 | MVP_top60_DNA+RNA_seed | ERBB2 | 0.244047 | BLCA | T24 | AZD8931 |
| MVP_top60_DNA_seed | ERBB2 | 0.532934 | BLCA | T24 | CI-1033 | MVP_top60_DNA+RNA_seed | ERBB3 | 0.244047 | BLCA | T24 | AZD8931 |
| MVP_top60_DNA_seed | RAC1 | 0.541463 | BLCA | TCCSUP | EHT-1864 | MVP_top60_DNA+RNA_seed | CDK2 | 0.246186 | BLCA | RT-112 | AZD5438 |
| MVP_top60_DNA_seed | HDAC1 | 0.543064 | BLCA | 639-V | JQ12 | MVP_top60_DNA+RNA_seed | ERBB2 | 0.249308 | BLCA | UM-UC-3 | Afatinib |
| MVP_top60_DNA_seed | FGFR3 | 0.545051 | BLCA | T24 | AZD4547 | MVP_top60_DNA+RNA_seed | HSP90AA1 | 0.255261 | BLCA | BFTC-905 | CCT-018159 |
| MVP_top60_DNA_seed | FGFR3 | 0.548163 | BLCA | T24 | PD173074 | MVP_top60_DNA+RNA_seed | AKT1 | 0.255487 | BLCA | 5637 | BAY AKT1 |
| MVP_top60_DNA_seed | HSP90AA1 | 0.548901 | BLCA | HT-1376 | CCT-018159 | MVP_top60_DNA+RNA_seed | ESR1 | 0.262134 | BLCA | TCCSUP | Fulvestrant |
| MVP_top60_DNA_seed | FGFR3 | 0.554717 | BLCA | 639-V | AZD4547 | MVP_top60_DNA+RNA_seed | AKT1 | 0.267143 | BLCA | LB831-BLC | BAY AKT1 |
| MVP_top60_DNA_seed | AR | 0.55611 | BLCA | 647-V | Bicalutamide | MVP_top60_DNA+RNA_seed | AR | 0.269045 | BLCA | 5637 | AZD3514 |
| MVP_top60_DNA_seed | ERBB2 | 0.562585 | BLCA | T24 | CP724714 | MVP_top60_DNA+RNA_seed | ATM | 0.269272 | BLCA | RT-112 | KU-55933 |
| MVP_top60_DNA_seed | HDAC1 | 0.568724 | BLCA | HT-1376 | Dacinostat | MVP_top60_DNA+RNA_seed | GSK3B | 0.274457 | BLCA | SW780 | KIN001-042 |
| MVP_top60_DNA_seed | PIK3CB | 0.570484 | BLCA | 5637 | AZD6482 | MVP_top60_DNA+RNA_seed | FGFR3 | 0.27548 | BLCA | HT-1376 | AZD4547 |
| MVP_top60_DNA_seed | ERBB2 | 0.574242 | BLCA | LB831-BLC | CP724714 | MVP_top60_DNA+RNA_seed | HSP90AA1 | 0.275767 | BLCA | 647-V | SNX-2112 |
| MVP_top60_DNA_seed | HDAC1 | 0.591335 | BLCA | LB831-BLC | Parthenolide | MVP_top60_DNA+RNA_seed | ATM | 0.276932 | BLCA | CAL-29 | CP466722 |
| MVP_top60_DNA_seed | HSP90AA1 | 0.600068 | BLCA | LB831-BLC | CCT-018159 | MVP_top60_DNA+RNA_seed | AR | 0.277381 | BLCA | HT-1197 | AZD3514 |
| MVP_top60_DNA_seed | ESR1 | 0.605609 | BLCA | T24 | Tamoxifen | MVP_top60_DNA+RNA_seed | CDK2 | 0.278442 | BLCA | DSH1 | PHA-793887 |
| MVP_top60_DNA_seed | HSP90AA1 | 0.607712 | BLCA | HT-1197 | Tanespimycin | MVP_top60_DNA+RNA_seed | ATM | 0.280053 | BLCA | SW780 | KU-55933 |
| MVP_top60_DNA_seed | ATM | 0.612735 | BLCA | RT4 | KU-60019 | MVP_top60_DNA+RNA_seed | HDAC1 | 0.282295 | BLCA | BFTC-905 | Tubastatin A |
| MVP_top60_DNA_seed | HSP90AA1 | 0.614559 | BLCA | KU-19-19 | SNX-2112 | MVP_top60_DNA+RNA_seed | SRC | 0.290119 | BLCA | LB831-BLC | Saracatinib |
| MVP_top60_DNA_seed | ESR1 | 0.615755 | BLCA | 647-V | Fulvestrant | MVP_top60_DNA+RNA_seed | FGFR3 | 0.290289 | BLCA | KU-19-19 | PD173074 |
| MVP_top60_DNA_seed | HSP90AA1 | 0.61851 | BLCA | HT-1376 | Elesclomol | MVP_top60_DNA+RNA_seed | MAPK1 | 0.290837 | BLCA | LB831-BLC | FR-180204 |
| MVP_top60_DNA_seed | ESR1 | 0.620536 | BLCA | RT-112 | Tamoxifen | MVP_top60_DNA+RNA_seed | MAPK3 | 0.290837 | BLCA | LB831-BLC | FR-180204 |
| MVP_top60_DNA_seed | HSP90AA1 | 0.623416 | BLCA | HT-1376 | Luminespib | MVP_top60_DNA+RNA_seed | FGFR3 | 0.29394 | BLCA | 647-V | FGFR_3831 |
| MVP_top60_DNA_seed | ERBB2 | 0.628933 | BLCA | 647-V | CUDC-101 | MVP_top60_DNA+RNA_seed | CDK2 | 0.294887 | BLCA | T24 | PHA-793887 |
| MVP_top60_DNA_seed | HDAC1 | 0.629172 | BLCA | 647-V | CAY10603 | MVP_top60_DNA+RNA_seed | GSK3B | 0.299235 | BLCA | SW780 | AZD7969 |
| MVP_top60_DNA_seed | AR | 0.635087 | BLCA | LB831-BLC | AZD3514 | MVP_top60_DNA+RNA_seed | GSK3B | 0.30034 | BLCA | TCCSUP | AZD7969 |
| MVP_top60_DNA_seed | CDK2 | 0.637335 | BLCA | SW780 | AZD5438 | MVP_top60_DNA+RNA_seed | RAC1 | 0.302502 | BLCA | J82 | EHT-1864 |
| MVP_top60_DNA_seed | HSP90AA1 | 0.643852 | BLCA | T24 | SNX-2112 | MVP_top60_DNA+RNA_seed | ERBB2 | 0.303676 | BLCA | J82 | Afatinib |
| MVP_top60_DNA_seed | ESR1 | 0.647181 | BLCA | DSH1 | Fulvestrant | MVP_top60_DNA+RNA_seed | ERBB2 | 0.304186 | BLCA | T24 | CUDC-101 |
| MVP_top60_DNA_seed | EP300 | 0.653108 | BLCA | RT4 | I-CBP112 | MVP_top60_DNA+RNA_seed | PIK3CA | 0.306539 | BLCA | UM-UC-3 | AZD8835 |
| MVP_top60_DNA_seed | CREBBP | 0.653108 | BLCA | RT4 | I-CBP112 | MVP_top60_DNA+RNA_seed | AKT1 | 0.310573 | BLCA | J82 | AKT inhibitor VIII |

| | | | | | | | | | | | |
|---|---|---|---|---|---|---|---|---|---|---|---|
| MVP_top60_DNA_seed | ERBB2 | 0.655604 | BLCA | TCCSUP | PF-00299804 | MVP_top60_DNA+RNA_seed | FGFR3 | 0.310781 | BLCA | KU-19-19 | AZD4547 |
| MVP_top60_DNA_seed | FGFR3 | 0.658264 | BLCA | HT-1197 | AZD4547 | MVP_top60_DNA+RNA_seed | CDK2 | 0.314928 | BLCA | RT4 | PHA-793887 |
| MVP_top60_DNA_seed | FGFR3 | 0.659795 | BLCA | HT-1376 | FGFR_3831 | MVP_top60_DNA+RNA_seed | HDAC1 | 0.317185 | BLCA | T24 | Tubastatin A |
| MVP_top60_DNA_seed | MAPK1 | 0.660416 | BLCA | HT-1197 | VX-11e | MVP_top60_DNA+RNA_seed | ERBB2 | 0.318815 | BLCA | LB831-BLC | Afatinib |
| MVP_top60_DNA_seed | HDAC1 | 0.661461 | BLCA | J82 | Tubastatin A | MVP_top60_DNA+RNA_seed | HDAC1 | 0.327679 | BLCA | J82 | Dacinostat |
| MVP_top60_DNA_seed | AR | 0.663194 | BLCA | T24 | Bicalutamide | MVP_top60_DNA+RNA_seed | FGFR3 | 0.328257 | BLCA | 5637 | PD173074 |
| MVP_top60_DNA_seed | PIK3CA | 0.670531 | BLCA | HT-1376 | AZD8835 | MVP_top60_DNA+RNA_seed | ERBB2 | 0.328999 | BLCA | RT-112 | CP724714 |
| MVP_top60_DNA_seed | HDAC1 | 0.670655 | BLCA | UM-UC-3 | Dacinostat | MVP_top60_DNA+RNA_seed | AKT1 | 0.330672 | BLCA | 647-V | AKT inhibitor VIII |
| MVP_top60_DNA_seed | CDK2 | 0.675658 | BLCA | RT4 | AZD5438 | MVP_top60_DNA+RNA_seed | AR | 0.332016 | BLCA | DSH1 | Bicalutamide |
| MVP_top60_DNA_seed | AR | 0.68006 | BLCA | RT-112 | Bicalutamide | MVP_top60_DNA+RNA_seed | FGFR3 | 0.334738 | BLCA | BFTC-905 | PD173074 |
| MVP_top60_DNA_seed | ERBB2 | 0.701632 | BLCA | SW780 | PF-00299804 | MVP_top60_DNA+RNA_seed | ERBB2 | 0.338084 | BLCA | RT-112 | PF-00299804 |
| MVP_top60_DNA_seed | ERBB2 | 0.702484 | BLCA | J82 | CP724714 | MVP_top60_DNA+RNA_seed | RAC1 | 0.340131 | BLCA | CAL-29 | EHT-1864 |
| MVP_top60_DNA_seed | HDAC1 | 0.706485 | BLCA | J82 | Belinostat | MVP_top60_DNA+RNA_seed | ERBB2 | 0.341313 | BLCA | J82 | Afatinib |
| MVP_top60_DNA_seed | CDK2 | 0.707769 | BLCA | HT-1197 | PHA-793887 | MVP_top60_DNA+RNA_seed | GSK3B | 0.345328 | BLCA | T24 | KIN001-042 |
| MVP_top60_DNA_seed | FGFR3 | 0.709857 | BLCA | 647-V | AZD4547 | MVP_top60_DNA+RNA_seed | ERBB2 | 0.346372 | BLCA | UM-UC-3 | Afatinib |
| MVP_top60_DNA_seed | HDAC1 | 0.711897 | BLCA | KU-19-19 | Tubastatin A | MVP_top60_DNA+RNA_seed | HDAC1 | 0.349277 | BLCA | SW1710 | Tubastatin A |
| MVP_top60_DNA_seed | SRC | 0.713668 | BLCA | LB831-BLC | Bosutinib | MVP_top60_DNA+RNA_seed | EP300 | 0.351883 | BLCA | CAL-29 | I-CBP112 |
| MVP_top60_DNA_seed | MAPK1 | 0.713982 | BLCA | TCCSUP | FR-180204 | MVP_top60_DNA+RNA_seed | CREBBP | 0.351883 | BLCA | CAL-29 | I-CBP112 |
| MVP_top60_DNA_seed | MAPK3 | 0.713982 | BLCA | TCCSUP | FR-180204 | MVP_top60_DNA+RNA_seed | FGFR3 | 0.35798 | BLCA | TCCSUP | AZD4547 |
| MVP_top60_DNA_seed | MAPK1 | 0.716533 | BLCA | LB831-BLC | VX-11e | MVP_top60_DNA+RNA_seed | HDAC1 | 0.363134 | BLCA | T24 | AR-42 |
| MVP_top60_DNA_seed | PIK3CB | 0.725715 | BLCA | DSH1 | TGX221 | MVP_top60_DNA+RNA_seed | HDAC1 | 0.369948 | BLCA | T24 | Belinostat |
| MVP_top60_DNA_seed | HDAC1 | 0.729354 | BLCA | HT-1376 | JQ12 | MVP_top60_DNA+RNA_seed | ESR1 | 0.370063 | BLCA | DSH1 | Tamoxifen |
| MVP_top60_DNA_seed | ERBB2 | 0.731252 | BLCA | LB831-BLC | Afatinib | MVP_top60_DNA+RNA_seed | ERBB2 | 0.373692 | BLCA | SW1710 | PF-00299804 |
| MVP_top60_DNA_seed | ERBB2 | 0.733738 | BLCA | KU-19-19 | PF-00299804 | MVP_top60_DNA+RNA_seed | MAPK1 | 0.374806 | BLCA | CAL-29 | VX-11e |
| MVP_top60_DNA_seed | ERBB2 | 0.741394 | BLCA | KU-19-19 | Afatinib | MVP_top60_DNA+RNA_seed | ESR1 | 0.378035 | BLCA | J82 | Fulvestrant |
| MVP_top60_DNA_seed | FGFR3 | 0.746584 | BLCA | DSH1 | FGFR_3831 | MVP_top60_DNA+RNA_seed | PIK3CA | 0.378449 | BLCA | 647-V | AZD8835 |
| MVP_top60_DNA_seed | HDAC1 | 0.754019 | BLCA | CAL-29 | Tubastatin A | MVP_top60_DNA+RNA_seed | PIK3CA | 0.379359 | BLCA | LB831-BLC | PI3Ka_4409 |
| MVP_top60_DNA_seed | PIK3CA | 0.758114 | BLCA | KU-19-19 | AZD8835 | MVP_top60_DNA+RNA_seed | ERBB2 | 0.380391 | BLCA | 639-V | CUDC-101 |
| MVP_top60_DNA_seed | ERBB2 | 0.759855 | BLCA | J82 | PF-00299804 | MVP_top60_DNA+RNA_seed | AR | 0.381983 | BLCA | SW1710 | Bicalutamide |
| MVP_top60_DNA_seed | PIK3CB | 0.760959 | BLCA | HT-1376 | AZD6482 | MVP_top60_DNA+RNA_seed | ERBB2 | 0.385041 | BLCA | TCCSUP | Afatinib |
| MVP_top60_DNA_seed | FGFR3 | 0.765492 | BLCA | 647-V | PD173074 | MVP_top60_DNA+RNA_seed | GSK3B | 0.38636 | BLCA | J82 | KIN001-042 |
| MVP_top60_DNA_seed | HDAC1 | 0.767131 | BLCA | RT4 | Dacinostat | MVP_top60_DNA+RNA_seed | AKT1 | 0.388401 | BLCA | HT-1376 | BAY AKT1 |
| MVP_top60_DNA_seed | HDAC1 | 0.775161 | BLCA | 639-V | AR-42 | MVP_top60_DNA+RNA_seed | ERBB2 | 0.388544 | BLCA | SW780 | Afatinib |
| MVP_top60_DNA_seed | SRC | 0.7796 | BLCA | DSH1 | WH-4-023 | MVP_top60_DNA+RNA_seed | CDK2 | 0.390177 | BLCA | HT-1197 | AZD5438 |
| MVP_top60_DNA_seed | ERBB2 | 0.783366 | BLCA | T24 | Afatinib | MVP_top60_DNA+RNA_seed | SYK | 0.393408 | BLCA | T24 | BAY-61-3606 |
| MVP_top60_DNA_seed | ATM | 0.784378 | BLCA | LB831-BLC | CP466722 | MVP_top60_DNA+RNA_seed | HDAC1 | 0.393925 | BLCA | T24 | CAY10603 |
| MVP_top60_DNA_seed | ERBB2 | 0.799432 | BLCA | HT-1197 | CUDC-101 | MVP_top60_DNA+RNA_seed | CDK2 | 0.394048 | BLCA | LB831-BLC | AZD5438 |
| MVP_top60_DNA_seed | PIK3CA | 0.811732 | BLCA | UM-UC-3 | PI3Ka_4409 | MVP_top60_DNA+RNA_seed | ERBB2 | 0.394126 | BLCA | T24 | Afatinib |
| MVP_top60_DNA_seed | EP300 | 0.818554 | BLCA | TCCSUP | I-CBP112 | MVP_top60_DNA+RNA_seed | ERBB2 | 0.394816 | BLCA | 647-V | CP724714 |
| MVP_top60_DNA_seed | CREBBP | 0.818554 | BLCA | TCCSUP | I-CBP112 | MVP_top60_DNA+RNA_seed | SYK | 0.39889 | BLCA | CAL-29 | BAY-61-3606 |
| MVP_top60_DNA_seed | ESR1 | 0.829707 | BLCA | RT4 | Tamoxifen | MVP_top60_DNA+RNA_seed | FGFR3 | 0.40049 | BLCA | T24 | AZD4547 |

| | | | | | | | | | | | |
|---|---|---|---|---|---|---|---|---|---|---|---|
| MVP_top60_DNA_seed | SRC | 0.830927 | BLCA | LB831-BLC | NVP-BHG712 | MVP_top60_DNA+RNA_seed | EP300 | 0.410601 | BLCA | T24 | I-CBP112 |
| MVP_top60_DNA_seed | PIK3CA | 0.831832 | BLCA | HT-1376 | PI3Ka_4409 | MVP_top60_DNA+RNA_seed | CREBBP | 0.410601 | BLCA | T24 | I-CBP112 |
| MVP_top60_DNA_seed | ERBB2 | 0.832581 | BLCA | SW780 | CI-1033 | MVP_top60_DNA+RNA_seed | MAPK1 | 0.411417 | BLCA | DSH1 | VX-11e |
| MVP_top60_DNA_seed | PIK3CA | 0.840047 | BLCA | RT4 | AZD8835 | MVP_top60_DNA+RNA_seed | AR | 0.424763 | BLCA | RT-112 | AZD3514 |
| MVP_top60_DNA_seed | AR | 0.840871 | BLCA | TCCSUP | Bicalutamide | MVP_top60_DNA+RNA_seed | HDAC1 | 0.426475 | BLCA | 639-V | Tubastatin A |
| MVP_top60_DNA_seed | ATM | 0.843254 | BLCA | DSH1 | KU-55933 | MVP_top60_DNA+RNA_seed | MAPK1 | 0.428655 | BLCA | 639-V | VX-11e |
| MVP_top60_DNA_seed | ESR1 | 0.844687 | BLCA | TCCSUP | Tamoxifen | MVP_top60_DNA+RNA_seed | HSP90AA1 | 0.432685 | BLCA | 647-V | CCT-018159 |
| MVP_top60_DNA_seed | ERBB2 | 0.845006 | BLCA | J82 | CI-1033 | MVP_top60_DNA+RNA_seed | ATM | 0.434257 | BLCA | TCCSUP | KU-55933 |
| MVP_top60_DNA_seed | HDAC1 | 0.860853 | BLCA | CAL-29 | JQ12 | MVP_top60_DNA+RNA_seed | ERBB2 | 0.436909 | BLCA | TCCSUP | CUDC-101 |
| MVP_top60_DNA_seed | FGFR3 | 0.868491 | BLCA | T24 | FGFR_3831 | MVP_top60_DNA+RNA_seed | SRC | 0.442002 | BLCA | CAL-29 | NVP-BHG712 |
| MVP_top60_DNA_seed | ATM | 0.869617 | BLCA | RT-112 | CP466722 | MVP_top60_DNA+RNA_seed | ESR1 | 0.44238 | BLCA | LB831-BLC | Tamoxifen |
| MVP_top60_DNA_seed | CDK2 | 0.877469 | BLCA | SW780 | PHA-793887 | MVP_top60_DNA+RNA_seed | HDAC1 | 0.445041 | BLCA | 639-V | CAY10603 |
| MVP_top60_DNA_seed | MAPK1 | 0.878392 | BLCA | J82 | FR-180204 | MVP_top60_DNA+RNA_seed | ERBB2 | 0.449358 | BLCA | BFTC-905 | Afatinib |
| MVP_top60_DNA_seed | MAPK3 | 0.878392 | BLCA | J82 | FR-180204 | MVP_top60_DNA+RNA_seed | HDAC1 | 0.449822 | BLCA | 639-V | Belinostat |
| MVP_top60_DNA_seed | ERBB2 | 0.878812 | BLCA | LB831-BLC | PF-00299804 | MVP_top60_DNA+RNA_seed | EP300 | 0.458008 | BLCA | 647-V | I-CBP112 |
| MVP_top60_DNA_seed | CDK2 | 0.87978 | BLCA | HT-1376 | AZD5438 | MVP_top60_DNA+RNA_seed | CREBBP | 0.458008 | BLCA | 647-V | I-CBP112 |
| MVP_top60_DNA_seed | HDAC1 | 0.882037 | BLCA | 647-V | Tubastatin A | MVP_top60_DNA+RNA_seed | ERBB2 | 0.459341 | BLCA | LB831-BLC | AZD8931 |
| MVP_top60_DNA_seed | ERBB2 | 0.891491 | BLCA | HT-1197 | Afatinib | MVP_top60_DNA+RNA_seed | ERBB3 | 0.459341 | BLCA | LB831-BLC | AZD8931 |
| MVP_top60_DNA_seed | SRC | 0.897409 | BLCA | SW780 | NVP-BHG712 | MVP_top60_DNA+RNA_seed | AR | 0.463076 | BLCA | LB831-BLC | Bicalutamide |
| MVP_top60_DNA_seed | ERBB2 | 0.899072 | BLCA | RT4 | Afatinib | MVP_top60_DNA+RNA_seed | CDK2 | 0.473515 | BLCA | CAL-29 | PHA-793887 |
| MVP_top60_DNA_seed | ERBB2 | 0.901325 | BLCA | RT-112 | Afatinib | MVP_top60_DNA+RNA_seed | HDAC1 | 0.473914 | BLCA | LB831-BLC | Dacinostat |
| MVP_top60_DNA_seed | SRC | 0.902829 | BLCA | 639-V | NVP-BHG712 | MVP_top60_DNA+RNA_seed | FGFR3 | 0.475546 | BLCA | J82 | AZD4547 |
| MVP_top60_DNA_seed | HDAC1 | 0.908418 | BLCA | DSH1 | Parthenolide | MVP_top60_DNA+RNA_seed | HSP90AA1 | 0.478719 | BLCA | KU-19-19 | Luminespib |
| MVP_top60_DNA_seed | ATM | 0.913186 | BLCA | KU-19-19 | CP466722 | MVP_top60_DNA+RNA_seed | EP300 | 0.480541 | BLCA | RT-112 | I-CBP112 |
| MVP_top60_DNA_seed | HDAC1 | 0.920123 | BLCA | HT-1197 | Tubastatin A | MVP_top60_DNA+RNA_seed | CREBBP | 0.480541 | BLCA | RT-112 | I-CBP112 |
| MVP_top60_DNA_seed | ERBB2 | 0.921234 | BLCA | HT-1197 | AZD8931 | MVP_top60_DNA+RNA_seed | PIK3CA | 0.485958 | BLCA | J82 | AZD8835 |
| MVP_top60_DNA_seed | ERBB3 | 0.921234 | BLCA | HT-1197 | AZD8931 | MVP_top60_DNA+RNA_seed | AKT1 | 0.487781 | BLCA | 639-V | AKT inhibitor VIII |
| MVP_top60_DNA_seed | HSP90AA1 | 0.93104 | BLCA | J82 | SNX-2112 | MVP_top60_DNA+RNA_seed | ERBB2 | 0.489032 | BLCA | SW780 | CP724714 |
| MVP_top60_DNA_seed | ATM | 0.932838 | BLCA | HT-1197 | CP466722 | MVP_top60_DNA+RNA_seed | HDAC1 | 0.489722 | BLCA | LB831-BLC | Tubastatin A |
| MVP_top60_DNA_seed | PIK3CA | 0.936587 | BLCA | SW780 | AZD8835 | MVP_top60_DNA+RNA_seed | AKT1 | 0.492095 | BLCA | BFTC-905 | GSK690693 |
| MVP_top60_DNA_seed | ERBB2 | 0.942534 | BLCA | HT-1197 | CP724714 | MVP_top60_DNA+RNA_seed | AKT1 | 0.501236 | BLCA | T24 | GSK690693 |
| MVP_top60_DNA_seed | MAPK1 | 0.944128 | BLCA | 647-V | VX-11e | MVP_top60_DNA+RNA_seed | FGFR3 | 0.507279 | BLCA | CAL-29 | PD173074 |
| MVP_top60_DNA_seed | CDK2 | 0.964557 | BLCA | DSH1 | AZD5438 | MVP_top60_DNA+RNA_seed | ATM | 0.5075 | BLCA | 639-V | CP466722 |
| MVP_top60_DNA_seed | SRC | 0.971585 | BLCA | TCCSUP | Bosutinib | MVP_top60_DNA+RNA_seed | ERBB2 | 0.510722 | BLCA | HT-1197 | PF-00299804 |
| MVP_top60_DNA_seed | PIK3CA | 0.983085 | BLCA | T24 | PI3Ka_4409 | MVP_top60_DNA+RNA_seed | MAPK1 | 0.511499 | BLCA | KU-19-19 | FR-180204 |
| MVP_top60_DNA_seed | HDAC1 | 0.98705 | BLCA | T24 | JQ12 | MVP_top60_DNA+RNA_seed | MAPK3 | 0.511499 | BLCA | KU-19-19 | FR-180204 |
| MVP_top60_DNA_seed | HDAC1 | 0.989847 | BLCA | 647-V | AR-42 | MVP_top60_DNA+RNA_seed | AKT1 | 0.517201 | BLCA | J82 | GSK690693 |
| MVP_top60_DNA_seed | AR | 0.990879 | BLCA | RT4 | Bicalutamide | MVP_top60_DNA+RNA_seed | AKT1 | 0.517447 | BLCA | T24 | BAY AKT1 |
| MVP_top60_DNA_seed | MAPK1 | 1.002308 | BLCA | SW780 | FR-180204 | MVP_top60_DNA+RNA_seed | GSK3B | 0.518503 | BLCA | J82 | SB216763 |
| MVP_top60_DNA_seed | MAPK3 | 1.002308 | BLCA | SW780 | FR-180204 | MVP_top60_DNA+RNA_seed | RAC1 | 0.52361 | BLCA | RT4 | EHT-1864 |
| MVP_top60_DNA_seed | ERBB2 | 1.003533 | BLCA | TCCSUP | CP724714 | MVP_top60_DNA+RNA_seed | AKT1 | 0.526465 | BLCA | 639-V | GSK690693 |

| | | | | | | | | | | | |
|---|---|---|---|---|---|---|---|---|---|---|---|
| MVP_top60_DNA_seed | HDAC1 | 1.004622 | BLCA | 647-V | Belinostat | MVP_top60_DNA+RNA_seed | SRC | 0.529177 | BLCA | T24 | NVP-BHG712 |
| MVP_top60_DNA_seed | HDAC1 | 1.018985 | BLCA | TCCSUP | Tubastatin A | MVP_top60_DNA+RNA_seed | GSK3B | 0.532341 | BLCA | SW780 | CHIR-99021 |
| MVP_top60_DNA_seed | ERBB2 | 1.023381 | BLCA | TCCSUP | Afatinib | MVP_top60_DNA+RNA_seed | ERBB2 | 0.532934 | BLCA | T24 | CI-1033 |
| MVP_top60_DNA_seed | HDAC1 | 1.038622 | BLCA | SW780 | Tubastatin A | MVP_top60_DNA+RNA_seed | AKT1 | 0.535225 | BLCA | KU-19-19 | GSK690693 |
| MVP_top60_DNA_seed | HDAC1 | 1.043258 | BLCA | T24 | Dacinostat | MVP_top60_DNA+RNA_seed | GSK3B | 0.535633 | BLCA | HT-1376 | CHIR-99021 |
| MVP_top60_DNA_seed | MAPK1 | 1.04827 | BLCA | J82 | VX-11e | MVP_top60_DNA+RNA_seed | RAC1 | 0.541463 | BLCA | TCCSUP | EHT-1864 |
| MVP_top60_DNA_seed | ATM | 1.05073 | BLCA | HT-1197 | KU-55933 | MVP_top60_DNA+RNA_seed | HDAC1 | 0.543064 | BLCA | 639-V | JQ12 |
| MVP_top60_DNA_seed | PIK3CB | 1.058871 | BLCA | KU-19-19 | AZD8186 | MVP_top60_DNA+RNA_seed | FGFR3 | 0.545051 | BLCA | T24 | AZD4547 |
| MVP_top60_DNA_seed | ATM | 1.059921 | BLCA | TCCSUP | CP466722 | MVP_top60_DNA+RNA_seed | FGFR3 | 0.548163 | BLCA | T24 | PD173074 |
| MVP_top60_DNA_seed | AR | 1.062939 | BLCA | UM-UC-3 | AZD3514 | MVP_top60_DNA+RNA_seed | HSP90AA1 | 0.548901 | BLCA | HT-1376 | CCT-018159 |
| MVP_top60_DNA_seed | HDAC1 | 1.064198 | BLCA | J82 | CAY10603 | MVP_top60_DNA+RNA_seed | SYK | 0.550639 | BLCA | TCCSUP | BAY-61-3606 |
| MVP_top60_DNA_seed | SRC | 1.066257 | BLCA | HT-1197 | Bosutinib | MVP_top60_DNA+RNA_seed | FGFR3 | 0.554717 | BLCA | 639-V | AZD4547 |
| MVP_top60_DNA_seed | PIK3CB | 1.069444 | BLCA | RT4 | AZD8186 | MVP_top60_DNA+RNA_seed | AR | 0.55611 | BLCA | 647-V | Bicalutamide |
| MVP_top60_DNA_seed | CDK2 | 1.073477 | BLCA | SW1710 | PHA-793887 | MVP_top60_DNA+RNA_seed | SYK | 0.559285 | BLCA | HT-1197 | BAY-61-3606 |
| MVP_top60_DNA_seed | RAC1 | 1.078141 | BLCA | RT-112 | EHT-1864 | MVP_top60_DNA+RNA_seed | ERBB2 | 0.562585 | BLCA | T24 | CP724714 |
| MVP_top60_DNA_seed | HSP90AA1 | 1.098969 | BLCA | KU-19-19 | Tanespimycin | MVP_top60_DNA+RNA_seed | HDAC1 | 0.568724 | BLCA | HT-1376 | Dacinostat |
| MVP_top60_DNA_seed | FGFR3 | 1.099201 | BLCA | HT-1376 | PD173074 | MVP_top60_DNA+RNA_seed | ERBB2 | 0.574242 | BLCA | LB831-BLC | CP724714 |
| MVP_top60_DNA_seed | PIK3CA | 1.110391 | BLCA | KU-19-19 | PI3Ka_4409 | MVP_top60_DNA+RNA_seed | GSK3B | 0.580351 | BLCA | 647-V | KIN001-042 |
| MVP_top60_DNA_seed | ERBB2 | 1.114642 | BLCA | HT-1376 | CP724714 | MVP_top60_DNA+RNA_seed | HDAC1 | 0.591335 | BLCA | LB831-BLC | Parthenolide |
| MVP_top60_DNA_seed | PIK3CB | 1.133734 | BLCA | KU-19-19 | AZD6482 | MVP_top60_DNA+RNA_seed | GSK3B | 0.592192 | BLCA | 647-V | CHIR-99021 |
| MVP_top60_DNA_seed | MAPK1 | 1.134826 | BLCA | 647-V | FR-180204 | MVP_top60_DNA+RNA_seed | HSP90AA1 | 0.600068 | BLCA | LB831-BLC | CCT-018159 |
| MVP_top60_DNA_seed | MAPK3 | 1.134826 | BLCA | 647-V | FR-180204 | MVP_top60_DNA+RNA_seed | ESR1 | 0.605609 | BLCA | T24 | Tamoxifen |
| MVP_top60_DNA_seed | AR | 1.151308 | BLCA | HT-1197 | Bicalutamide | MVP_top60_DNA+RNA_seed | HSP90AA1 | 0.607712 | BLCA | HT-1197 | Tanespimycin |
| MVP_top60_DNA_seed | ERBB2 | 1.152584 | BLCA | RT4 | Afatinib | MVP_top60_DNA+RNA_seed | ATM | 0.612735 | BLCA | RT4 | KU-60019 |
| MVP_top60_DNA_seed | ESR1 | 1.161236 | BLCA | SW780 | Fulvestrant | MVP_top60_DNA+RNA_seed | HSP90AA1 | 0.614559 | BLCA | KU-19-19 | SNX-2112 |
| MVP_top60_DNA_seed | ESR1 | 1.175275 | BLCA | RT4 | Fulvestrant | MVP_top60_DNA+RNA_seed | ESR1 | 0.615755 | BLCA | 647-V | Fulvestrant |
| MVP_top60_DNA_seed | SRC | 1.186874 | BLCA | DSH1 | NVP-BHG712 | MVP_top60_DNA+RNA_seed | HSP90AA1 | 0.61851 | BLCA | HT-1376 | Elesclomol |
| MVP_top60_DNA_seed | ATM | 1.197124 | BLCA | T24 | KU-55933 | MVP_top60_DNA+RNA_seed | ESR1 | 0.620536 | BLCA | RT-112 | Tamoxifen |
| MVP_top60_DNA_seed | ERBB2 | 1.205992 | BLCA | HT-1197 | Afatinib | MVP_top60_DNA+RNA_seed | HSP90AA1 | 0.623416 | BLCA | HT-1376 | Luminespib |
| MVP_top60_DNA_seed | PIK3CB | 1.212955 | BLCA | DSH1 | AZD8186 | MVP_top60_DNA+RNA_seed | ERBB2 | 0.628933 | BLCA | 647-V | CUDC-101 |
| MVP_top60_DNA_seed | HDAC1 | 1.216008 | BLCA | UM-UC-3 | JQ12 | MVP_top60_DNA+RNA_seed | HDAC1 | 0.629172 | BLCA | 647-V | CAY10603 |
| MVP_top60_DNA_seed | MAPK1 | 1.218684 | BLCA | TCCSUP | VX-11e | MVP_top60_DNA+RNA_seed | AR | 0.635087 | BLCA | LB831-BLC | AZD3514 |
| MVP_top60_DNA_seed | HSP90AA1 | 1.219865 | BLCA | 647-V | Tanespimycin | MVP_top60_DNA+RNA_seed | CDK2 | 0.637335 | BLCA | SW780 | AZD5438 |
| MVP_top60_DNA_seed | SRC | 1.220215 | BLCA | J82 | NVP-BHG712 | MVP_top60_DNA+RNA_seed | GSK3B | 0.641527 | BLCA | DSH1 | KIN001-042 |
| MVP_top60_DNA_seed | SRC | 1.223682 | BLCA | RT4 | Bosutinib | MVP_top60_DNA+RNA_seed | HSP90AA1 | 0.643852 | BLCA | T24 | SNX-2112 |
| MVP_top60_DNA_seed | ERBB2 | 1.227829 | BLCA | J82 | CUDC-101 | MVP_top60_DNA+RNA_seed | ESR1 | 0.647181 | BLCA | DSH1 | Fulvestrant |
| MVP_top60_DNA_seed | CDK2 | 1.272512 | BLCA | J82 | PHA-793887 | MVP_top60_DNA+RNA_seed | EP300 | 0.653108 | BLCA | RT4 | I-CBP112 |
| MVP_top60_DNA_seed | HDAC1 | 1.273782 | BLCA | HT-1376 | Tubastatin A | MVP_top60_DNA+RNA_seed | CREBBP | 0.653108 | BLCA | RT4 | I-CBP112 |
| MVP_top60_DNA_seed | HDAC1 | 1.293242 | BLCA | J82 | AR-42 | MVP_top60_DNA+RNA_seed | ERBB2 | 0.655604 | BLCA | TCCSUP | PF-00299804 |
| MVP_top60_DNA_seed | FGFR3 | 1.331466 | BLCA | HT-1197 | PD173074 | MVP_top60_DNA+RNA_seed | FGFR3 | 0.658264 | BLCA | HT-1197 | AZD4547 |
| MVP_top60_DNA_seed | HDAC1 | 1.333382 | BLCA | HT-1376 | Belinostat | MVP_top60_DNA+RNA_seed | FGFR3 | 0.659795 | BLCA | HT-1376 | FGFR_3831 |

| | | | | | | | | | | | |
|---|---|---|---|---|---|---|---|---|---|---|---|
| MVP_top60_DNA_seed | MAPK1 | 1.334568 | BLCA | HT-1376 | FR-180204 | MVP_top60_DNA+RNA_seed | MAPK1 | 0.660416 | BLCA | HT-1197 | VX-11e |
| MVP_top60_DNA_seed | MAPK3 | 1.334568 | BLCA | HT-1376 | FR-180204 | MVP_top60_DNA+RNA_seed | HDAC1 | 0.661461 | BLCA | J82 | Tubastatin A |
| MVP_top60_DNA_seed | MAPK1 | 1.334672 | BLCA | HT-1197 | FR-180204 | MVP_top60_DNA+RNA_seed | AKT1 | 0.662963 | BLCA | KU-19-19 | AKT inhibitor VIII |
| MVP_top60_DNA_seed | MAPK3 | 1.334672 | BLCA | HT-1197 | FR-180204 | MVP_top60_DNA+RNA_seed | AR | 0.663194 | BLCA | T24 | Bicalutamide |
| MVP_top60_DNA_seed | SRC | 1.344916 | BLCA | TCCSUP | NVP-BHG712 | MVP_top60_DNA+RNA_seed | PIK3CA | 0.670531 | BLCA | HT-1376 | AZD8835 |
| MVP_top60_DNA_seed | MAPK1 | 1.346925 | BLCA | HT-1376 | VX-11e | MVP_top60_DNA+RNA_seed | HDAC1 | 0.670655 | BLCA | UM-UC-3 | Dacinostat |
| MVP_top60_DNA_seed | AR | 1.355719 | BLCA | T24 | AZD3514 | MVP_top60_DNA+RNA_seed | CDK2 | 0.675658 | BLCA | RT4 | AZD5438 |
| MVP_top60_DNA_seed | AR | 1.358262 | BLCA | HT-1376 | Bicalutamide | MVP_top60_DNA+RNA_seed | AR | 0.68006 | BLCA | RT-112 | Bicalutamide |
| MVP_top60_DNA_seed | HSP90AA1 | 1.362939 | BLCA | HT-1197 | SNX-2112 | MVP_top60_DNA+RNA_seed | AKT1 | 0.680569 | BLCA | LB831-BLC | GSK690693 |
| MVP_top60_DNA_seed | AR | 1.363457 | BLCA | RT4 | Bicalutamide | MVP_top60_DNA+RNA_seed | GSK3B | 0.68592 | BLCA | HT-1376 | CHIR-99021 |
| MVP_top60_DNA_seed | ATM | 1.383265 | BLCA | RT4 | KU-55933 | MVP_top60_DNA+RNA_seed | AKT1 | 0.688368 | BLCA | 647-V | GSK690693 |
| MVP_top60_DNA_seed | CDK2 | 1.386626 | BLCA | TCCSUP | PHA-793887 | MVP_top60_DNA+RNA_seed | AKT1 | 0.693244 | BLCA | 647-V | MK-2206 |
| MVP_top60_DNA_seed | SRC | 1.427045 | BLCA | HT-1197 | NVP-BHG712 | MVP_top60_DNA+RNA_seed | ERBB2 | 0.701632 | BLCA | SW780 | PF-00299804 |
| MVP_top60_DNA_seed | SRC | 1.468831 | BLCA | HT-1376 | NVP-BHG712 | MVP_top60_DNA+RNA_seed | ERBB2 | 0.702484 | BLCA | J82 | CP724714 |
| MVP_top60_DNA_seed | CDK2 | 1.47139 | BLCA | HT-1376 | PHA-793887 | MVP_top60_DNA+RNA_seed | HDAC1 | 0.706485 | BLCA | J82 | Belinostat |
| MVP_top60_DNA_seed | HDAC1 | 1.527156 | BLCA | RT4 | JQ12 | MVP_top60_DNA+RNA_seed | CDK2 | 0.707769 | BLCA | HT-1197 | PHA-793887 |
| MVP_top60_DNA_seed | PIK3CB | 1.532403 | BLCA | RT4 | AZD6482 | MVP_top60_DNA+RNA_seed | FGFR3 | 0.709857 | BLCA | 647-V | AZD4547 |
| MVP_top60_DNA_seed | EP300 | 1.592007 | BLCA | HT-1376 | I-CBP112 | MVP_top60_DNA+RNA_seed | HDAC1 | 0.711897 | BLCA | KU-19-19 | Tubastatin A |
| MVP_top60_DNA_seed | CREBBP | 1.592007 | BLCA | HT-1376 | I-CBP112 | MVP_top60_DNA+RNA_seed | SRC | 0.713668 | BLCA | LB831-BLC | Bosutinib |
| MVP_top60_DNA_seed | HSP90AA1 | 1.59846 | BLCA | SW780 | SNX-2112 | MVP_top60_DNA+RNA_seed | MAPK1 | 0.713982 | BLCA | TCCSUP | FR-180204 |
| MVP_top60_DNA_seed | ERBB2 | 1.651326 | BLCA | SW780 | CUDC-101 | MVP_top60_DNA+RNA_seed | MAPK3 | 0.713982 | BLCA | TCCSUP | FR-180204 |
| MVP_top60_DNA_seed | ERBB2 | 1.656002 | BLCA | HT-1376 | CUDC-101 | MVP_top60_DNA+RNA_seed | MAPK1 | 0.716533 | BLCA | LB831-BLC | VX-11e |
| MVP_top60_DNA_seed | HDAC1 | 1.65752 | BLCA | SW780 | CAY10603 | MVP_top60_DNA+RNA_seed | AKT1 | 0.721057 | BLCA | DSH1 | BAY AKT1 |
| MVP_top60_DNA_seed | HSP90AA1 | 1.711511 | BLCA | RT4 | CCT-018159 | MVP_top60_DNA+RNA_seed | HDAC1 | 0.729354 | BLCA | HT-1376 | JQ12 |
| MVP_top60_DNA_seed | HDAC1 | 1.759039 | BLCA | HT-1376 | CAY10603 | MVP_top60_DNA+RNA_seed | ERBB2 | 0.731252 | BLCA | LB831-BLC | Afatinib |
| MVP_top60_DNA_seed | ATM | 1.785292 | BLCA | J82 | CP466722 | MVP_top60_DNA+RNA_seed | ERBB2 | 0.733738 | BLCA | KU-19-19 | PF-00299804 |
| MVP_top60_DNA_seed | FGFR3 | 1.805235 | BLCA | HT-1197 | FGFR_3831 | MVP_top60_DNA+RNA_seed | ERBB2 | 0.741394 | BLCA | KU-19-19 | Afatinib |
| MVP_top60_DNA_seed | HDAC1 | 1.818339 | BLCA | HT-1376 | AR-42 | MVP_top60_DNA+RNA_seed | FGFR3 | 0.746584 | BLCA | DSH1 | FGFR_3831 |
| MVP_top60_DNA_seed | PIK3CA | 1.846686 | BLCA | RT4 | PI3Ka_4409 | MVP_top60_DNA+RNA_seed | SYK | 0.751473 | BLCA | RT-112 | BAY-61-3606 |
| MVP_top60_DNA_seed | HDAC1 | 1.855702 | BLCA | SW780 | AR-42 | MVP_top60_DNA+RNA_seed | HDAC1 | 0.754019 | BLCA | CAL-29 | Tubastatin A |
| MVP_top60_DNA_seed | HSP90AA1 | 1.896371 | BLCA | HT-1376 | SNX-2112 | MVP_top60_DNA+RNA_seed | PIK3CA | 0.758114 | BLCA | KU-19-19 | AZD8835 |
| MVP_top60_DNA_seed | ATM | 1.932395 | BLCA | HT-1197 | KU-60019 | MVP_top60_DNA+RNA_seed | ERBB2 | 0.759855 | BLCA | J82 | PF-00299804 |
| MVP_top60_DNA_seed | HSP90AA1 | 1.936929 | BLCA | RT4 | Elesclomol | MVP_top60_DNA+RNA_seed | GSK3B | 0.761388 | BLCA | RT-112 | SB216763 |
| MVP_top60_DNA_seed | HDAC1 | 1.976456 | BLCA | SW780 | Dacinostat | MVP_top60_DNA+RNA_seed | FGFR3 | 0.765492 | BLCA | 647-V | PD173074 |
| MVP_top60_DNA_seed | HDAC1 | 2.114746 | BLCA | SW780 | Belinostat | MVP_top60_DNA+RNA_seed | HDAC1 | 0.767131 | BLCA | RT4 | Dacinostat |
| MVP_top60_DNA_seed | HSP90AA1 | 2.3012 | BLCA | TCCSUP | Luminespib | MVP_top60_DNA+RNA_seed | AKT1 | 0.768035 | BLCA | LB831-BLC | MK-2206 |
| MVP_top60_DNA_seed | ATM | 2.355221 | BLCA | SW780 | CP466722 | MVP_top60_DNA+RNA_seed | HDAC1 | 0.775161 | BLCA | 639-V | AR-42 |
| MVP_top60_DNA_seed | HSP90AA1 | 2.39341 | BLCA | CAL-29 | Luminespib | MVP_top60_DNA+RNA_seed | GSK3B | 0.776275 | BLCA | HT-1376 | AZD7969 |
| MVP_top60_DNA_seed | HSP90AA1 | 2.489461 | BLCA | SW780 | Luminespib | MVP_top60_DNA+RNA_seed | SRC | 0.7796 | BLCA | DSH1 | WH-4-023 |
| MVP_top60_DNA_seed | HSP90AA1 | 2.923546 | BLCA | RT4 | Luminespib | MVP_top60_DNA+RNA_seed | ERBB2 | 0.783366 | BLCA | T24 | Afatinib |
| MVP_top60_DNA_seed | MAPK1 | -3.08539 | BLCA | KU-19-19 | VX-11e | MVP_top60_DNA+RNA_seed | ATM | 0.784378 | BLCA | LB831-BLC | CP466722 |

| | | | | | | | | | | | |
|---|---|---|---|---|---|---|---|---|---|---|---|
| MVP_top60_DNA_seed | ERBB2 | -2.945931 | BLCA | DSH1 | Afatinib | MVP_top60_DNA+RNA_seed | ERBB2 | 0.799432 | BLCA | HT-1197 | CUDC-101 |
| MVP_top60_DNA_seed | FGFR3 | -2.672072 | BLCA | RT-112 | PD173074 | MVP_top60_DNA+RNA_seed | AKT1 | 0.808711 | BLCA | HT-1197 | MK-2206 |
| MVP_top60_DNA_seed | FGFR3 | -2.555222 | BLCA | RT-112 | AZD4547 | MVP_top60_DNA+RNA_seed | PIK3CA | 0.811732 | BLCA | UM-UC-3 | PI3Ka_4409 |
| MVP_top60_DNA_seed | ERBB2 | -2.538815 | BLCA | DSH1 | Lapatinib | MVP_top60_DNA+RNA_seed | GSK3B | 0.817255 | BLCA | DSH1 | CHIR-99021 |
| MVP_top60_DNA_seed | MAPK1 | -2.499572 | BLCA | KU-19-19 | Ulixertinib | MVP_top60_DNA+RNA_seed | EP300 | 0.818554 | BLCA | TCCSUP | I-CBP112 |
| MVP_top60_DNA_seed | MAPK3 | -2.499572 | BLCA | KU-19-19 | Ulixertinib | MVP_top60_DNA+RNA_seed | CREBBP | 0.818554 | BLCA | TCCSUP | I-CBP112 |
| MVP_top60_DNA_seed | MAPK1 | -2.327525 | BLCA | KU-19-19 | SCH772984 | MVP_top60_DNA+RNA_seed | AKT1 | 0.81943 | BLCA | LB831-BLC | AKT inhibitor VIII |
| MVP_top60_DNA_seed | MAPK3 | -2.327525 | BLCA | KU-19-19 | SCH772984 | MVP_top60_DNA+RNA_seed | ESR1 | 0.829707 | BLCA | RT4 | Tamoxifen |
| MVP_top60_DNA_seed | ERBB2 | -2.184704 | BLCA | DSH1 | Sapitinib | MVP_top60_DNA+RNA_seed | SRC | 0.830927 | BLCA | LB831-BLC | NVP-BHG712 |
| MVP_top60_DNA_seed | ERBB3 | -2.184704 | BLCA | DSH1 | Sapitinib | MVP_top60_DNA+RNA_seed | PIK3CA | 0.831832 | BLCA | HT-1376 | PI3Ka_4409 |
| MVP_top60_DNA_seed | MAPK1 | -2.053093 | BLCA | KU-19-19 | Ulixertinib | MVP_top60_DNA+RNA_seed | AKT1 | 0.831914 | BLCA | 5637 | AKT inhibitor VIII |
| MVP_top60_DNA_seed | MAPK3 | -2.053093 | BLCA | KU-19-19 | Ulixertinib | MVP_top60_DNA+RNA_seed | ERBB2 | 0.832581 | BLCA | SW780 | CI-1033 |
| MVP_top60_DNA_seed | PIK3CA | -1.984079 | BLCA | CAL-29 | Alpelisib | MVP_top60_DNA+RNA_seed | PIK3CA | 0.840047 | BLCA | RT4 | AZD8835 |
| MVP_top60_DNA_seed | FGFR3 | -1.797738 | BLCA | RT4 | PD173074 | MVP_top60_DNA+RNA_seed | AR | 0.840871 | BLCA | TCCSUP | Bicalutamide |
| MVP_top60_DNA_seed | TP53 | -1.730724 | BLCA | BFTC-905 | MIRA-1 | MVP_top60_DNA+RNA_seed | GSK3B | 0.842794 | BLCA | 647-V | CHIR-99021 |
| MVP_top60_DNA_seed | MAPK1 | -1.720698 | BLCA | BFTC-905 | Ulixertinib | MVP_top60_DNA+RNA_seed | ATM | 0.843254 | BLCA | DSH1 | KU-55933 |
| MVP_top60_DNA_seed | MAPK3 | -1.720698 | BLCA | BFTC-905 | Ulixertinib | MVP_top60_DNA+RNA_seed | ESR1 | 0.844687 | BLCA | TCCSUP | Tamoxifen |
| MVP_top60_DNA_seed | MAPK1 | -1.711996 | BLCA | DSH1 | Ulixertinib | MVP_top60_DNA+RNA_seed | ERBB2 | 0.845006 | BLCA | J82 | CI-1033 |
| MVP_top60_DNA_seed | MAPK3 | -1.711996 | BLCA | DSH1 | Ulixertinib | MVP_top60_DNA+RNA_seed | HDAC1 | 0.860853 | BLCA | CAL-29 | JQ12 |
| MVP_top60_DNA_seed | HDAC1 | -1.658672 | BLCA | KU-19-19 | Romidepsin | MVP_top60_DNA+RNA_seed | FGFR3 | 0.868491 | BLCA | T24 | FGFR_3831 |
| MVP_top60_DNA_seed | SRC | -1.602043 | BLCA | BFTC-905 | Bosutinib | MVP_top60_DNA+RNA_seed | ATM | 0.869617 | BLCA | RT-112 | CP466722 |
| MVP_top60_DNA_seed | HSP90AA1 | -1.578527 | BLCA | DSH1 | Tanespimycin | MVP_top60_DNA+RNA_seed | AKT1 | 0.873048 | BLCA | SW780 | AKT inhibitor VIII |
| MVP_top60_DNA_seed | FGFR3 | -1.568543 | BLCA | RT4 | AZD4547 | MVP_top60_DNA+RNA_seed | CDK2 | 0.877469 | BLCA | SW780 | PHA-793887 |
| MVP_top60_DNA_seed | MAPK1 | -1.521053 | BLCA | BFTC-905 | SCH772984 | MVP_top60_DNA+RNA_seed | MAPK1 | 0.878392 | BLCA | J82 | FR-180204 |
| MVP_top60_DNA_seed | MAPK3 | -1.521053 | BLCA | BFTC-905 | SCH772984 | MVP_top60_DNA+RNA_seed | MAPK3 | 0.878392 | BLCA | J82 | FR-180204 |
| MVP_top60_DNA_seed | ERBB2 | -1.500321 | BLCA | CAL-29 | Sapitinib | MVP_top60_DNA+RNA_seed | ERBB2 | 0.878812 | BLCA | LB831-BLC | PF-00299804 |
| MVP_top60_DNA_seed | ERBB3 | -1.500321 | BLCA | CAL-29 | Sapitinib | MVP_top60_DNA+RNA_seed | CDK2 | 0.87978 | BLCA | HT-1376 | AZD5438 |
| MVP_top60_DNA_seed | KRAS | -1.451969 | BLCA | KU-19-19 | KRAS (G12C) Inhibitor-1 | MVP_top60_DNA+RNA_seed | HDAC1 | 0.882037 | BLCA | 647-V | Tubastatin A |
| MVP_top60_DNA_seed | HSP90AA1 | -1.449154 | BLCA | CAL-29 | Tanespimycin | MVP_top60_DNA+RNA_seed | AKT1 | 0.882481 | BLCA | SW780 | GSK690693 |
| MVP_top60_DNA_seed | HSP90AA1 | -1.347454 | BLCA | RT4 | Tanespimycin | MVP_top60_DNA+RNA_seed | ERBB2 | 0.891491 | BLCA | HT-1197 | Afatinib |
| MVP_top60_DNA_seed | ESR1 | -1.319957 | BLCA | BFTC-905 | GDC0810 | MVP_top60_DNA+RNA_seed | SRC | 0.897409 | BLCA | SW780 | NVP-BHG712 |
| MVP_top60_DNA_seed | ESR1 | -1.311481 | BLCA | BFTC-905 | Tamoxifen | MVP_top60_DNA+RNA_seed | GSK3B | 0.898258 | BLCA | SW780 | SB216763 |
| MVP_top60_DNA_seed | CDK2 | -1.311034 | BLCA | KU-19-19 | AZD5438 | MVP_top60_DNA+RNA_seed | ERBB2 | 0.899072 | BLCA | RT4 | Afatinib |
| MVP_top60_DNA_seed | ESR1 | -1.258992 | BLCA | HT-1197 | Tamoxifen | MVP_top60_DNA+RNA_seed | AKT1 | 0.899598 | BLCA | RT4 | GSK690693 |
| MVP_top60_DNA_seed | ERBB2 | -1.252804 | BLCA | 5637 | Sapitinib | MVP_top60_DNA+RNA_seed | ERBB2 | 0.901325 | BLCA | RT-112 | Afatinib |
| MVP_top60_DNA_seed | ERBB3 | -1.252804 | BLCA | 5637 | Sapitinib | MVP_top60_DNA+RNA_seed | SRC | 0.902829 | BLCA | 639-V | NVP-BHG712 |
| MVP_top60_DNA_seed | FGFR3 | -1.251877 | BLCA | SW780 | AZD4547 | MVP_top60_DNA+RNA_seed | HDAC1 | 0.908418 | BLCA | DSH1 | Parthenolide |
| MVP_top60_DNA_seed | FGFR3 | -1.242566 | BLCA | J82 | AZD4547 | MVP_top60_DNA+RNA_seed | ATM | 0.913186 | BLCA | KU-19-19 | CP466722 |
| MVP_top60_DNA_seed | HSP90AA1 | -1.214951 | BLCA | TCCSUP | Tanespimycin | MVP_top60_DNA+RNA_seed | HDAC1 | 0.920123 | BLCA | HT-1197 | Tubastatin A |
| MVP_top60_DNA_seed | EP300 | -1.208234 | BLCA | BFTC-905 | SGC-CBP30 | MVP_top60_DNA+RNA_seed | ERBB2 | 0.921234 | BLCA | HT-1197 | AZD8931 |
| MVP_top60_DNA_seed | CREBBP | -1.208234 | BLCA | BFTC-905 | SGC-CBP30 | MVP_top60_DNA+RNA_seed | ERBB3 | 0.921234 | BLCA | HT-1197 | AZD8931 |

| | | | | | | | | | | | |
|---|---|---|---|---|---|---|---|---|---|---|---|
| MVP_top60_DNA_seed | HSP90AA1 | -1.178451 | BLCA | BFTC-905 | Tanespimycin | MVP_top60_DNA+RNA_seed | AKT1 | 0.927205 | BLCA | 639-V | BAY AKT1 |
| MVP_top60_DNA_seed | PIK3CA | -1.147597 | BLCA | CAL-29 | GNE-317 | MVP_top60_DNA+RNA_seed | HSP90AA1 | 0.93104 | BLCA | J82 | SNX-2112 |
| MVP_top60_DNA_seed | RAC1 | -1.146802 | BLCA | T24 | EHT-1864 | MVP_top60_DNA+RNA_seed | ATM | 0.932838 | BLCA | HT-1197 | CP466722 |
| MVP_top60_DNA_seed | ESR1 | -1.127363 | BLCA | DSH1 | Tamoxifen | MVP_top60_DNA+RNA_seed | PIK3CA | 0.936587 | BLCA | SW780 | AZD8835 |
| MVP_top60_DNA_seed | ERBB2 | -1.126995 | BLCA | 5637 | Afatinib | MVP_top60_DNA+RNA_seed | AKT1 | 0.936991 | BLCA | TCCSUP | GSK690693 |
| MVP_top60_DNA_seed | PIK3CA | -1.094133 | BLCA | CAL-29 | Buparlisib | MVP_top60_DNA+RNA_seed | ERBB2 | 0.942534 | BLCA | HT-1197 | CP724714 |
| MVP_top60_DNA_seed | PIK3CB | -1.094133 | BLCA | CAL-29 | Buparlisib | MVP_top60_DNA+RNA_seed | MAPK1 | 0.944128 | BLCA | 647-V | VX-11e |
| MVP_top60_DNA_seed | CDK2 | -1.069996 | BLCA | KU-19-19 | Dinaciclib | MVP_top60_DNA+RNA_seed | AKT1 | 0.954369 | BLCA | T24 | AKT inhibitor VIII |
| MVP_top60_DNA_seed | FGFR3 | -1.064624 | BLCA | SW780 | PD173074 | MVP_top60_DNA+RNA_seed | AKT1 | 0.955762 | BLCA | TCCSUP | MK-2206 |
| MVP_top60_DNA_seed | EP300 | -1.03655 | BLCA | BFTC-905 | CPI-637 | MVP_top60_DNA+RNA_seed | AKT1 | 0.963504 | BLCA | HT-1376 | MK-2206 |
| MVP_top60_DNA_seed | MAPK1 | -1.033519 | BLCA | BFTC-905 | Ulixertinib | MVP_top60_DNA+RNA_seed | CDK2 | 0.964557 | BLCA | DSH1 | AZD5438 |
| MVP_top60_DNA_seed | MAPK3 | -1.033519 | BLCA | BFTC-905 | Ulixertinib | MVP_top60_DNA+RNA_seed | SRC | 0.971585 | BLCA | TCCSUP | Bosutinib |
| MVP_top60_DNA_seed | MAPK1 | -1.021434 | BLCA | BFTC-905 | VX-11e | MVP_top60_DNA+RNA_seed | GSK3B | 0.971983 | BLCA | HT-1197 | CHIR-99021 |
| MVP_top60_DNA_seed | KRAS | -0.992505 | BLCA | DSH1 | KRAS (G12C) Inhibitor-12 | MVP_top60_DNA+RNA_seed | GSK3B | 0.972491 | BLCA | RT-112 | CHIR-99021 |
| MVP_top60_DNA_seed | MAPK1 | -0.982858 | BLCA | DSH1 | SCH772984 | MVP_top60_DNA+RNA_seed | PIK3CA | 0.983085 | BLCA | T24 | PI3Ka_4409 |
| MVP_top60_DNA_seed | MAPK3 | -0.982858 | BLCA | DSH1 | SCH772984 | MVP_top60_DNA+RNA_seed | AKT1 | 0.983347 | BLCA | T24 | AKT inhibitor VIII |
| MVP_top60_DNA_seed | EP300 | -0.963364 | BLCA | RT4 | SGC-CBP30 | MVP_top60_DNA+RNA_seed | HDAC1 | 0.98705 | BLCA | T24 | JQ12 |
| MVP_top60_DNA_seed | CREBBP | -0.963364 | BLCA | RT4 | SGC-CBP30 | MVP_top60_DNA+RNA_seed | HDAC1 | 0.989847 | BLCA | 647-V | AR-42 |
| MVP_top60_DNA_seed | RAC1 | -0.963187 | BLCA | RT4 | EHT-1864 | MVP_top60_DNA+RNA_seed | AR | 0.990879 | BLCA | RT4 | Bicalutamide |
| MVP_top60_DNA_seed | ATM | -0.962046 | BLCA | BFTC-905 | KU-55933 | MVP_top60_DNA+RNA_seed | MAPK1 | 1.002308 | BLCA | SW780 | FR-180204 |
| MVP_top60_DNA_seed | ESR1 | -0.960303 | BLCA | KU-19-19 | GDC0810 | MVP_top60_DNA+RNA_seed | MAPK3 | 1.002308 | BLCA | SW780 | FR-180204 |
| MVP_top60_DNA_seed | HSP90AA1 | -0.957798 | BLCA | 639-V | Luminespib | MVP_top60_DNA+RNA_seed | ERBB2 | 1.003533 | BLCA | TCCSUP | CP724714 |
| MVP_top60_DNA_seed | HSP90AA1 | -0.9546 | BLCA | KU-19-19 | Luminespib | MVP_top60_DNA+RNA_seed | HDAC1 | 1.004622 | BLCA | 647-V | Belinostat |
| MVP_top60_DNA_seed | SRC | -0.938558 | BLCA | DSH1 | Bosutinib | MVP_top60_DNA+RNA_seed | HDAC1 | 1.018985 | BLCA | TCCSUP | Tubastatin A |
| MVP_top60_DNA_seed | MAPK1 | -0.929273 | BLCA | DSH1 | VX-11e | MVP_top60_DNA+RNA_seed | ERBB2 | 1.023381 | BLCA | TCCSUP | Afatinib |
| MVP_top60_DNA_seed | FGFR3 | -0.923771 | BLCA | 639-V | AZD4547 | MVP_top60_DNA+RNA_seed | HDAC1 | 1.038622 | BLCA | SW780 | Tubastatin A |
| MVP_top60_DNA_seed | HDAC1 | -0.90454 | BLCA | LB831-BLC | Romidepsin | MVP_top60_DNA+RNA_seed | HDAC1 | 1.043258 | BLCA | T24 | Dacinostat |
| MVP_top60_DNA_seed | PIK3CA | -0.884619 | BLCA | CAL-29 | AZD8186 | MVP_top60_DNA+RNA_seed | MAPK1 | 1.04827 | BLCA | J82 | VX-11e |
| MVP_top60_DNA_seed | PIK3CB | -0.884619 | BLCA | CAL-29 | AZD8186 | MVP_top60_DNA+RNA_seed | ATM | 1.05073 | BLCA | HT-1197 | KU-55933 |
| MVP_top60_DNA_seed | HSP90AA1 | -0.88273 | BLCA | UM-UC-3 | Luminespib | MVP_top60_DNA+RNA_seed | AKT1 | 1.059757 | BLCA | HT-1376 | GSK690693 |
| MVP_top60_DNA_seed | EP300 | -0.879263 | BLCA | SW1710 | SGC-CBP30 | MVP_top60_DNA+RNA_seed | ATM | 1.059921 | BLCA | TCCSUP | CP466722 |
| MVP_top60_DNA_seed | CREBBP | -0.879263 | BLCA | SW1710 | SGC-CBP30 | MVP_top60_DNA+RNA_seed | AR | 1.062939 | BLCA | UM-UC-3 | AZD3514 |
| MVP_top60_DNA_seed | CDK2 | -0.851637 | BLCA | DSH1 | Dinaciclib | MVP_top60_DNA+RNA_seed | HDAC1 | 1.064198 | BLCA | J82 | CAY10603 |
| MVP_top60_DNA_seed | KRAS | -0.841657 | BLCA | 639-V | KRAS (G12C) Inhibitor-12 | MVP_top60_DNA+RNA_seed | SRC | 1.066257 | BLCA | HT-1197 | Bosutinib |
| MVP_top60_DNA_seed | ESR1 | -0.840238 | BLCA | 5637 | GDC0810 | MVP_top60_DNA+RNA_seed | GSK3B | 1.067595 | BLCA | HT-1197 | CHIR-99021 |
| MVP_top60_DNA_seed | PIK3CA | -0.837054 | BLCA | BFTC-905 | Buparlisib | MVP_top60_DNA+RNA_seed | CDK2 | 1.073477 | BLCA | SW1710 | PHA-793887 |
| MVP_top60_DNA_seed | PIK3CB | -0.837054 | BLCA | BFTC-905 | Buparlisib | MVP_top60_DNA+RNA_seed | RAC1 | 1.078141 | BLCA | RT-112 | EHT-1864 |
| MVP_top60_DNA_seed | ATM | -0.822909 | BLCA | 647-V | KU-55933 | MVP_top60_DNA+RNA_seed | AKT1 | 1.091105 | BLCA | HT-1197 | GSK690693 |
| MVP_top60_DNA_seed | HSP90AA1 | -0.821776 | BLCA | 639-V | Tanespimycin | MVP_top60_DNA+RNA_seed | AKT1 | 1.098725 | BLCA | HT-1376 | AKT inhibitor VIII |
| MVP_top60_DNA_seed | ESR1 | -0.818628 | BLCA | HT-1376 | GDC0810 | MVP_top60_DNA+RNA_seed | HSP90AA1 | 1.098969 | BLCA | KU-19-19 | Tanespimycin |
| MVP_top60_DNA_seed | MAPK1 | -0.803411 | BLCA | RT-112 | Ulixertinib | MVP_top60_DNA+RNA_seed | AKT1 | 1.098994 | BLCA | HT-1197 | AKT inhibitor VIII |

| | | | | | | | | | | | |
|---|---|---|---|---|---|---|---|---|---|---|---|
| MVP_top60_DNA_seed | MAPK3 | -0.803411 | BLCA | RT-112 | Ulixertinib | MVP_top60_DNA+RNA_seed | FGFR3 | 1.099201 | BLCA | HT-1376 | PD173074 |
| MVP_top60_DNA_seed | PIK3CA | -0.799839 | BLCA | BFTC-905 | Alpelisib | MVP_top60_DNA+RNA_seed | PIK3CA | 1.110391 | BLCA | KU-19-19 | PI3Ka_4409 |
| MVP_top60_DNA_seed | ERBB2 | -0.795081 | BLCA | 5637 | Lapatinib | MVP_top60_DNA+RNA_seed | ERBB2 | 1.114642 | BLCA | HT-1376 | CP724714 |
| MVP_top60_DNA_seed | ESR1 | -0.788135 | BLCA | HT-1376 | Tamoxifen | MVP_top60_DNA+RNA_seed | GSK3B | 1.114646 | BLCA | SW780 | CHIR-99021 |
| MVP_top60_DNA_seed | MAPK1 | -0.786778 | BLCA | RT-112 | SCH772984 | MVP_top60_DNA+RNA_seed | GSK3B | 1.115586 | BLCA | HT-1376 | SB216763 |
| MVP_top60_DNA_seed | MAPK3 | -0.786778 | BLCA | RT-112 | SCH772984 | MVP_top60_DNA+RNA_seed | MAPK1 | 1.134826 | BLCA | 647-V | FR-180204 |
| MVP_top60_DNA_seed | PIK3CA | -0.777686 | BLCA | 5637 | Buparlisib | MVP_top60_DNA+RNA_seed | MAPK3 | 1.134826 | BLCA | 647-V | FR-180204 |
| MVP_top60_DNA_seed | PIK3CB | -0.777686 | BLCA | 5637 | Buparlisib | MVP_top60_DNA+RNA_seed | AR | 1.151308 | BLCA | HT-1197 | Bicalutamide |
| MVP_top60_DNA_seed | CDK2 | -0.760694 | BLCA | RT-112 | AZD5438 | MVP_top60_DNA+RNA_seed | ERBB2 | 1.152584 | BLCA | RT4 | Afatinib |
| MVP_top60_DNA_seed | RAC1 | -0.75836 | BLCA | DSH1 | EHT-1864 | MVP_top60_DNA+RNA_seed | ESR1 | 1.161236 | BLCA | SW780 | Fulvestrant |
| MVP_top60_DNA_seed | ESR1 | -0.752726 | BLCA | CAL-29 | Tamoxifen | MVP_top60_DNA+RNA_seed | ESR1 | 1.175275 | BLCA | RT4 | Fulvestrant |
| MVP_top60_DNA_seed | RAC1 | -0.747393 | BLCA | 639-V | EHT-1864 | MVP_top60_DNA+RNA_seed | SRC | 1.186874 | BLCA | DSH1 | NVP-BHG712 |
| MVP_top60_DNA_seed | MAPK1 | -0.747032 | BLCA | RT4 | VX-11e | MVP_top60_DNA+RNA_seed | GSK3B | 1.18976 | BLCA | TCCSUP | CHIR-99021 |
| MVP_top60_DNA_seed | CDK2 | -0.738271 | BLCA | RT-112 | Dinaciclib | MVP_top60_DNA+RNA_seed | ATM | 1.197124 | BLCA | T24 | KU-55933 |
| MVP_top60_DNA_seed | ESR1 | -0.698797 | BLCA | 639-V | Tamoxifen | MVP_top60_DNA+RNA_seed | ERBB2 | 1.205992 | BLCA | HT-1197 | Afatinib |
| MVP_top60_DNA_seed | ERBB2 | -0.696024 | BLCA | CAL-29 | Lapatinib | MVP_top60_DNA+RNA_seed | HDAC1 | 1.216008 | BLCA | UM-UC-3 | JQ12 |
| MVP_top60_DNA_seed | PIK3CA | -0.69325 | BLCA | HT-1197 | Buparlisib | MVP_top60_DNA+RNA_seed | MAPK1 | 1.218684 | BLCA | TCCSUP | VX-11e |
| MVP_top60_DNA_seed | PIK3CB | -0.69325 | BLCA | HT-1197 | Buparlisib | MVP_top60_DNA+RNA_seed | HSP90AA1 | 1.219865 | BLCA | 647-V | Tanespimycin |
| MVP_top60_DNA_seed | HSP90AA1 | -0.692026 | BLCA | KU-19-19 | Tanespimycin | MVP_top60_DNA+RNA_seed | SRC | 1.220215 | BLCA | J82 | NVP-BHG712 |
| MVP_top60_DNA_seed | ESR1 | -0.689053 | BLCA | UM-UC-3 | GDC0810 | MVP_top60_DNA+RNA_seed | SRC | 1.223682 | BLCA | RT4 | Bosutinib |
| MVP_top60_DNA_seed | HDAC1 | -0.684011 | BLCA | KU-19-19 | Entinostat | MVP_top60_DNA+RNA_seed | ERBB2 | 1.227829 | BLCA | J82 | CUDC-101 |
| MVP_top60_DNA_seed | KRAS | -0.678566 | BLCA | CAL-29 | KRAS (G12C) Inhibitor-1 | MVP_top60_DNA+RNA_seed | AKT1 | 1.235147 | BLCA | HT-1197 | AKT inhibitor VIII |
| MVP_top60_DNA_seed | PIK3CA | -0.670616 | BLCA | RT-112 | Alpelisib | MVP_top60_DNA+RNA_seed | AKT1 | 1.266484 | BLCA | RT4 | BAY AKT1 |
| MVP_top60_DNA_seed | MAPK1 | -0.658453 | BLCA | RT4 | SCH772984 | MVP_top60_DNA+RNA_seed | CDK2 | 1.272512 | BLCA | J82 | PHA-793887 |
| MVP_top60_DNA_seed | MAPK3 | -0.658453 | BLCA | RT4 | SCH772984 | MVP_top60_DNA+RNA_seed | HDAC1 | 1.273782 | BLCA | HT-1376 | Tubastatin A |
| MVP_top60_DNA_seed | SRC | -0.656227 | BLCA | 5637 | Bosutinib | MVP_top60_DNA+RNA_seed | HDAC1 | 1.293242 | BLCA | J82 | AR-42 |
| MVP_top60_DNA_seed | HDAC1 | -0.639754 | BLCA | 5637 | Entinostat | MVP_top60_DNA+RNA_seed | AKT1 | 1.324218 | BLCA | KU-19-19 | AKT inhibitor VIII |
| MVP_top60_DNA_seed | KRAS | -0.626387 | BLCA | BFTC-905 | KRAS (G12C) Inhibitor-1 | MVP_top60_DNA+RNA_seed | FGFR3 | 1.331466 | BLCA | HT-1197 | PD173074 |
| MVP_top60_DNA_seed | CDK2 | -0.620205 | BLCA | BFTC-905 | AZD5438 | MVP_top60_DNA+RNA_seed | HDAC1 | 1.333382 | BLCA | HT-1376 | Belinostat |
| MVP_top60_DNA_seed | ERBB2 | -0.602856 | BLCA | CAL-29 | Afatinib | MVP_top60_DNA+RNA_seed | MAPK1 | 1.334568 | BLCA | HT-1376 | FR-180204 |
| MVP_top60_DNA_seed | ATM | -0.602001 | BLCA | 639-V | KU-55933 | MVP_top60_DNA+RNA_seed | MAPK3 | 1.334568 | BLCA | HT-1376 | FR-180204 |
| MVP_top60_DNA_seed | HSP90AA1 | -0.600517 | BLCA | BFTC-905 | Luminespib | MVP_top60_DNA+RNA_seed | MAPK1 | 1.334672 | BLCA | HT-1197 | FR-180204 |
| MVP_top60_DNA_seed | HSP90AA1 | -0.587736 | BLCA | DSH1 | Luminespib | MVP_top60_DNA+RNA_seed | MAPK3 | 1.334672 | BLCA | HT-1197 | FR-180204 |
| MVP_top60_DNA_seed | CDK2 | -0.587022 | BLCA | CAL-29 | Dinaciclib | MVP_top60_DNA+RNA_seed | SRC | 1.344916 | BLCA | TCCSUP | NVP-BHG712 |
| MVP_top60_DNA_seed | MAPK1 | -0.582567 | BLCA | HT-1197 | Ulixertinib | MVP_top60_DNA+RNA_seed | MAPK1 | 1.346925 | BLCA | HT-1376 | VX-11e |
| MVP_top60_DNA_seed | MAPK3 | -0.582567 | BLCA | HT-1197 | Ulixertinib | MVP_top60_DNA+RNA_seed | AR | 1.355719 | BLCA | T24 | AZD3514 |
| MVP_top60_DNA_seed | HSP90AA1 | -0.579553 | BLCA | RT-112 | Luminespib | MVP_top60_DNA+RNA_seed | AR | 1.358262 | BLCA | HT-1376 | Bicalutamide |
| MVP_top60_DNA_seed | EP300 | -0.567635 | BLCA | KU-19-19 | SGC-CBP30 | MVP_top60_DNA+RNA_seed | HSP90AA1 | 1.362939 | BLCA | HT-1197 | SNX-2112 |
| MVP_top60_DNA_seed | CREBBP | -0.567635 | BLCA | KU-19-19 | SGC-CBP30 | MVP_top60_DNA+RNA_seed | AR | 1.363457 | BLCA | RT4 | Bicalutamide |
| MVP_top60_DNA_seed | PIK3CA | -0.562267 | BLCA | DSH1 | GNE-317 | MVP_top60_DNA+RNA_seed | ATM | 1.383265 | BLCA | RT4 | KU-55933 |
| MVP_top60_DNA_seed | RAC1 | -0.554754 | BLCA | CAL-29 | EHT-1864 | MVP_top60_DNA+RNA_seed | CDK2 | 1.386626 | BLCA | TCCSUP | PHA-793887 |

| | | | | | | | | | | | |
|---|---|---|---|---|---|---|---|---|---|---|---|
| MVP_top60_DNA_seed | KRAS | -0.550227 | BLCA | 5637 | KRAS (G12C) Inhibitor-12 | MVP_top60_DNA+RNA_seed | AKT1 | 1.398668 | BLCA | RT4 | AKT inhibitor VIII |
| MVP_top60_DNA_seed | CDK2 | -0.538674 | BLCA | LB831-BLC | Dinaciclib | MVP_top60_DNA+RNA_seed | SRC | 1.427045 | BLCA | HT-1197 | NVP-BHG712 |
| MVP_top60_DNA_seed | KRAS | -0.533955 | BLCA | RT4 | KRAS (G12C) Inhibitor-12 | MVP_top60_DNA+RNA_seed | SYK | 1.457517 | BLCA | KU-19-19 | BAY-61-3606 |
| MVP_top60_DNA_seed | PIK3CA | -0.533608 | BLCA | HT-1197 | GNE-317 | MVP_top60_DNA+RNA_seed | SRC | 1.468831 | BLCA | HT-1376 | NVP-BHG712 |
| MVP_top60_DNA_seed | HDAC1 | -0.532909 | BLCA | BFTC-905 | Romidepsin | MVP_top60_DNA+RNA_seed | CDK2 | 1.47139 | BLCA | HT-1376 | PHA-793887 |
| MVP_top60_DNA_seed | ESR1 | -0.522591 | BLCA | SW1710 | Tamoxifen | MVP_top60_DNA+RNA_seed | SYK | 1.518029 | BLCA | DSH1 | BAY-61-3606 |
| MVP_top60_DNA_seed | HDAC1 | -0.522475 | BLCA | DSH1 | PCI-34051 | MVP_top60_DNA+RNA_seed | HDAC1 | 1.527156 | BLCA | RT4 | JQ12 |
| MVP_top60_DNA_seed | SRC | -0.517813 | BLCA | KU-19-19 | Bosutinib | MVP_top60_DNA+RNA_seed | EP300 | 1.592007 | BLCA | HT-1376 | I-CBP112 |
| MVP_top60_DNA_seed | CDK2 | -0.51229 | BLCA | BFTC-905 | Dinaciclib | MVP_top60_DNA+RNA_seed | CREBBP | 1.592007 | BLCA | HT-1376 | I-CBP112 |
| MVP_top60_DNA_seed | PIK3CA | -0.481732 | BLCA | KU-19-19 | Buparlisib | MVP_top60_DNA+RNA_seed | HSP90AA1 | 1.59846 | BLCA | SW780 | SNX-2112 |
| MVP_top60_DNA_seed | PIK3CB | -0.481732 | BLCA | KU-19-19 | Buparlisib | MVP_top60_DNA+RNA_seed | ERBB2 | 1.651326 | BLCA | SW780 | CUDC-101 |
| MVP_top60_DNA_seed | PIK3CA | -0.476776 | BLCA | KU-19-19 | GNE-317 | MVP_top60_DNA+RNA_seed | GSK3B | 1.652269 | BLCA | HT-1197 | KIN001-042 |
| MVP_top60_DNA_seed | RAC1 | -0.475895 | BLCA | SW780 | EHT-1864 | MVP_top60_DNA+RNA_seed | ERBB2 | 1.656002 | BLCA | HT-1376 | CUDC-101 |
| MVP_top60_DNA_seed | EP300 | -0.459899 | BLCA | DSH1 | SGC-CBP30 | MVP_top60_DNA+RNA_seed | HDAC1 | 1.65752 | BLCA | SW780 | CAY10603 |
| MVP_top60_DNA_seed | CREBBP | -0.459899 | BLCA | DSH1 | SGC-CBP30 | MVP_top60_DNA+RNA_seed | SYK | 1.672616 | BLCA | SW780 | BAY-61-3606 |
| MVP_top60_DNA_seed | MAPK1 | -0.458567 | BLCA | RT-112 | Ulixertinib | MVP_top60_DNA+RNA_seed | HSP90AA1 | 1.711511 | BLCA | RT4 | CCT-018159 |
| MVP_top60_DNA_seed | MAPK3 | -0.458567 | BLCA | RT-112 | Ulixertinib | MVP_top60_DNA+RNA_seed | HDAC1 | 1.759039 | BLCA | HT-1376 | CAY10603 |
| MVP_top60_DNA_seed | EP300 | -0.45414 | BLCA | SW780 | SGC-CBP30 | MVP_top60_DNA+RNA_seed | ATM | 1.785292 | BLCA | J82 | CP466722 |
| MVP_top60_DNA_seed | CREBBP | -0.45414 | BLCA | SW780 | SGC-CBP30 | MVP_top60_DNA+RNA_seed | AKT1 | 1.787245 | BLCA | TCCSUP | AKT inhibitor VIII |
| MVP_top60_DNA_seed | HSP90AA1 | -0.447393 | BLCA | LB831-BLC | Luminespib | MVP_top60_DNA+RNA_seed | FGFR3 | 1.805235 | BLCA | HT-1197 | FGFR_3831 |
| MVP_top60_DNA_seed | HDAC1 | -0.422774 | BLCA | RT-112 | Romidepsin | MVP_top60_DNA+RNA_seed | HDAC1 | 1.818339 | BLCA | HT-1376 | AR-42 |
| MVP_top60_DNA_seed | FGFR3 | -0.40624 | BLCA | 647-V | AZD4547 | MVP_top60_DNA+RNA_seed | AKT1 | 1.83911 | BLCA | RT4 | MK-2206 |
| MVP_top60_DNA_seed | RAC1 | -0.404138 | BLCA | BFTC-905 | EHT-1864 | MVP_top60_DNA+RNA_seed | PIK3CA | 1.846686 | BLCA | RT4 | PI3Ka_4409 |
| MVP_top60_DNA_seed | HSP90AA1 | -0.403736 | BLCA | 5637 | Tanespimycin | MVP_top60_DNA+RNA_seed | HDAC1 | 1.855702 | BLCA | SW780 | AR-42 |
| MVP_top60_DNA_seed | FGFR3 | -0.401204 | BLCA | CAL-29 | AZD4547 | MVP_top60_DNA+RNA_seed | HSP90AA1 | 1.896371 | BLCA | HT-1376 | SNX-2112 |
| MVP_top60_DNA_seed | HSP90AA1 | -0.383208 | BLCA | RT-112 | Tanespimycin | MVP_top60_DNA+RNA_seed | ATM | 1.932395 | BLCA | HT-1197 | KU-60019 |
| MVP_top60_DNA_seed | MAPK1 | -0.381047 | BLCA | LB831-BLC | Ulixertinib | MVP_top60_DNA+RNA_seed | HSP90AA1 | 1.936929 | BLCA | RT4 | Elesclomol |
| MVP_top60_DNA_seed | MAPK3 | -0.381047 | BLCA | LB831-BLC | Ulixertinib | MVP_top60_DNA+RNA_seed | HDAC1 | 1.976456 | BLCA | SW780 | Dacinostat |
| MVP_top60_DNA_seed | HDAC1 | -0.378929 | BLCA | KU-19-19 | PCI-34051 | MVP_top60_DNA+RNA_seed | GSK3B | 2.015226 | BLCA | RT-112 | AZD7969 |
| MVP_top60_DNA_seed | MAPK1 | -0.337577 | BLCA | DSH1 | Ulixertinib | MVP_top60_DNA+RNA_seed | HDAC1 | 2.114746 | BLCA | SW780 | Belinostat |
| MVP_top60_DNA_seed | MAPK3 | -0.337577 | BLCA | DSH1 | Ulixertinib | MVP_top60_DNA+RNA_seed | HSP90AA1 | 2.3012 | BLCA | TCCSUP | Luminespib |
| MVP_top60_DNA_seed | PIK3CA | -0.335329 | BLCA | 647-V | AZD8186 | MVP_top60_DNA+RNA_seed | ATM | 2.355221 | BLCA | SW780 | CP466722 |
| MVP_top60_DNA_seed | PIK3CB | -0.335329 | BLCA | 647-V | AZD8186 | MVP_top60_DNA+RNA_seed | HSP90AA1 | 2.39341 | BLCA | CAL-29 | Luminespib |
| MVP_top60_DNA_seed | ESR1 | -0.33484 | BLCA | DSH1 | GDC0810 | MVP_top60_DNA+RNA_seed | HSP90AA1 | 2.489461 | BLCA | SW780 | Luminespib |
| MVP_top60_DNA_seed | EP300 | -0.327979 | BLCA | KU-19-19 | CPI-637 | MVP_top60_DNA+RNA_seed | HSP90AA1 | 2.923546 | BLCA | RT4 | Luminespib |
| MVP_top60_DNA_seed | CDK2 | -0.319533 | BLCA | RT4 | AZD5438 | MVP_top60_DNA+RNA_seed | MAPK1 | -3.08539 | BLCA | KU-19-19 | VX-11e |
| MVP_top60_DNA_seed | HDAC1 | -0.31773 | BLCA | 5637 | Romidepsin | MVP_top60_DNA+RNA_seed | ERBB2 | -2.945931 | BLCA | DSH1 | Afatinib |
| MVP_top60_DNA_seed | CDK2 | -0.315001 | BLCA | RT4 | Dinaciclib | MVP_top60_DNA+RNA_seed | FGFR3 | -2.672072 | BLCA | RT-112 | PD173074 |
| MVP_top60_DNA_seed | FGFR3 | -0.314594 | BLCA | 5637 | AZD4547 | MVP_top60_DNA+RNA_seed | FGFR3 | -2.555222 | BLCA | RT-112 | AZD4547 |
| MVP_top60_DNA_seed | PIK3CA | -0.311178 | BLCA | 639-V | AZD8186 | MVP_top60_DNA+RNA_seed | ERBB2 | -2.538815 | BLCA | DSH1 | Lapatinib |
| MVP_top60_DNA_seed | PIK3CB | -0.311178 | BLCA | 639-V | AZD8186 | MVP_top60_DNA+RNA_seed | MAPK1 | -2.499572 | BLCA | KU-19-19 | Ulixertinib |

| | | | | | | | | | | | |
|---|---|---|---|---|---|---|---|---|---|---|---|
| MVP_top60_DNA_seed | MAPK1 | -0.306001 | BLCA | RT-112 | VX-11e | MVP_top60_DNA+RNA_seed | MAPK3 | -2.499572 | BLCA | KU-19-19 | Ulixertinib |
| MVP_top60_DNA_seed | ATM | -0.302883 | BLCA | 5637 | KU-55933 | MVP_top60_DNA+RNA_seed | MAPK1 | -2.327525 | BLCA | KU-19-19 | SCH772984 |
| MVP_top60_DNA_seed | PIK3CA | -0.297558 | BLCA | 639-V | Buparlisib | MVP_top60_DNA+RNA_seed | MAPK3 | -2.327525 | BLCA | KU-19-19 | SCH772984 |
| MVP_top60_DNA_seed | PIK3CB | -0.297558 | BLCA | 639-V | Buparlisib | MVP_top60_DNA+RNA_seed | ERBB2 | -2.184704 | BLCA | DSH1 | Sapitinib |
| MVP_top60_DNA_seed | CDK2 | -0.292461 | BLCA | LB831-BLC | AZD5438 | MVP_top60_DNA+RNA_seed | ERBB3 | -2.184704 | BLCA | DSH1 | Sapitinib |
| MVP_top60_DNA_seed | MAPK1 | -0.288714 | BLCA | CAL-29 | SCH772984 | MVP_top60_DNA+RNA_seed | SYK | -2.111682 | BLCA | DSH1 | PRT062607 |
| MVP_top60_DNA_seed | MAPK3 | -0.288714 | BLCA | CAL-29 | SCH772984 | MVP_top60_DNA+RNA_seed | MAPK1 | -2.053093 | BLCA | KU-19-19 | Ulixertinib |
| MVP_top60_DNA_seed | PIK3CA | -0.28342 | BLCA | RT-112 | GNE-317 | MVP_top60_DNA+RNA_seed | MAPK3 | -2.053093 | BLCA | KU-19-19 | Ulixertinib |
| MVP_top60_DNA_seed | EP300 | -0.281952 | BLCA | 639-V | CPI-637 | MVP_top60_DNA+RNA_seed | PIK3CA | -1.984079 | BLCA | CAL-29 | Alpelisib |
| MVP_top60_DNA_seed | MAPK1 | -0.274744 | BLCA | CAL-29 | VX-11e | MVP_top60_DNA+RNA_seed | FGFR3 | -1.797738 | BLCA | RT4 | PD173074 |
| MVP_top60_DNA_seed | EP300 | -0.273802 | BLCA | J82 | SGC-CBP30 | MVP_top60_DNA+RNA_seed | TP53 | -1.730724 | BLCA | BFTC-905 | MIRA-1 |
| MVP_top60_DNA_seed | CREBBP | -0.273802 | BLCA | J82 | SGC-CBP30 | MVP_top60_DNA+RNA_seed | MAPK1 | -1.720698 | BLCA | BFTC-905 | Ulixertinib |
| MVP_top60_DNA_seed | PIK3CA | -0.273018 | BLCA | BFTC-905 | GNE-317 | MVP_top60_DNA+RNA_seed | MAPK3 | -1.720698 | BLCA | BFTC-905 | Ulixertinib |
| MVP_top60_DNA_seed | ESR1 | -0.271128 | BLCA | RT4 | Tamoxifen | MVP_top60_DNA+RNA_seed | MAPK1 | -1.711996 | BLCA | DSH1 | Ulixertinib |
| MVP_top60_DNA_seed | PIK3CA | -0.270785 | BLCA | TCCSUP | GNE-317 | MVP_top60_DNA+RNA_seed | MAPK3 | -1.711996 | BLCA | DSH1 | Ulixertinib |
| MVP_top60_DNA_seed | CDK2 | -0.263229 | BLCA | 639-V | Dinaciclib | MVP_top60_DNA+RNA_seed | HDAC1 | -1.658672 | BLCA | KU-19-19 | Romidepsin |
| MVP_top60_DNA_seed | MAPK1 | -0.261527 | BLCA | 5637 | Ulixertinib | MVP_top60_DNA+RNA_seed | SRC | -1.602043 | BLCA | BFTC-905 | Bosutinib |
| MVP_top60_DNA_seed | MAPK3 | -0.261527 | BLCA | 5637 | Ulixertinib | MVP_top60_DNA+RNA_seed | HSP90AA1 | -1.578527 | BLCA | DSH1 | Tanespimycin |
| MVP_top60_DNA_seed | ESR1 | -0.259256 | BLCA | CAL-29 | GDC0810 | MVP_top60_DNA+RNA_seed | FGFR3 | -1.568543 | BLCA | RT4 | AZD4547 |
| MVP_top60_DNA_seed | ESR1 | -0.253103 | BLCA | KU-19-19 | Tamoxifen | MVP_top60_DNA+RNA_seed | MAPK1 | -1.521053 | BLCA | BFTC-905 | SCH772984 |
| MVP_top60_DNA_seed | EP300 | -0.250773 | BLCA | TCCSUP | SGC-CBP30 | MVP_top60_DNA+RNA_seed | MAPK3 | -1.521053 | BLCA | BFTC-905 | SCH772984 |
| MVP_top60_DNA_seed | CREBBP | -0.250773 | BLCA | TCCSUP | SGC-CBP30 | MVP_top60_DNA+RNA_seed | GSK3B | -1.503833 | BLCA | KU-19-19 | SB216763 |
| MVP_top60_DNA_seed | ERBB2 | -0.249731 | BLCA | BFTC-905 | Afatinib | MVP_top60_DNA+RNA_seed | ERBB2 | -1.500321 | BLCA | CAL-29 | Sapitinib |
| MVP_top60_DNA_seed | HDAC1 | -0.249172 | BLCA | RT4 | PCI-34051 | MVP_top60_DNA+RNA_seed | ERBB3 | -1.500321 | BLCA | CAL-29 | Sapitinib |
| MVP_top60_DNA_seed | MAPK1 | -0.233154 | BLCA | RT4 | Ulixertinib | MVP_top60_DNA+RNA_seed | KRAS | -1.451969 | BLCA | KU-19-19 | (RAS (G12C) Inhibitor-1 |
| MVP_top60_DNA_seed | MAPK3 | -0.233154 | BLCA | RT4 | Ulixertinib | MVP_top60_DNA+RNA_seed | HSP90AA1 | -1.449154 | BLCA | CAL-29 | Tanespimycin |
| MVP_top60_DNA_seed | HSP90AA1 | -0.230539 | BLCA | LB831-BLC | Tanespimycin | MVP_top60_DNA+RNA_seed | HSP90AA1 | -1.347454 | BLCA | RT4 | Tanespimycin |
| MVP_top60_DNA_seed | RAC1 | -0.226074 | BLCA | RT-112 | EHT-1864 | MVP_top60_DNA+RNA_seed | ESR1 | -1.319957 | BLCA | BFTC-905 | GDC0810 |
| MVP_top60_DNA_seed | PIK3CA | -0.22432 | BLCA | UM-UC-3 | GNE-317 | MVP_top60_DNA+RNA_seed | ESR1 | -1.311481 | BLCA | BFTC-905 | Tamoxifen |
| MVP_top60_DNA_seed | ATM | -0.222383 | BLCA | LB831-BLC | KU-55933 | MVP_top60_DNA+RNA_seed | CDK2 | -1.311034 | BLCA | KU-19-19 | AZD5438 |
| MVP_top60_DNA_seed | HDAC1 | -0.219178 | BLCA | BFTC-905 | PCI-34051 | MVP_top60_DNA+RNA_seed | ESR1 | -1.258992 | BLCA | HT-1197 | Tamoxifen |
| MVP_top60_DNA_seed | TP53 | -0.214167 | BLCA | 639-V | MIRA-1 | MVP_top60_DNA+RNA_seed | ERBB2 | -1.252804 | BLCA | 5637 | Sapitinib |
| MVP_top60_DNA_seed | CDK2 | -0.205094 | BLCA | 639-V | AZD5438 | MVP_top60_DNA+RNA_seed | ERBB3 | -1.252804 | BLCA | 5637 | Sapitinib |
| MVP_top60_DNA_seed | PIK3CA | -0.202548 | BLCA | HT-1197 | AZD8186 | MVP_top60_DNA+RNA_seed | FGFR3 | -1.251877 | BLCA | SW780 | AZD4547 |
| MVP_top60_DNA_seed | PIK3CB | -0.202548 | BLCA | HT-1197 | AZD8186 | MVP_top60_DNA+RNA_seed | FGFR3 | -1.242566 | BLCA | J82 | AZD4547 |
| MVP_top60_DNA_seed | ATM | -0.195009 | BLCA | KU-19-19 | KU-55933 | MVP_top60_DNA+RNA_seed | HSP90AA1 | -1.214951 | BLCA | TCCSUP | Tanespimycin |
| MVP_top60_DNA_seed | EP300 | -0.180177 | BLCA | DSH1 | CPI-637 | MVP_top60_DNA+RNA_seed | EP300 | -1.208234 | BLCA | BFTC-905 | SGC-CBP30 |
| MVP_top60_DNA_seed | ERBB2 | -0.179974 | BLCA | BFTC-905 | Lapatinib | MVP_top60_DNA+RNA_seed | CREBBP | -1.208234 | BLCA | BFTC-905 | SGC-CBP30 |
| MVP_top60_DNA_seed | KRAS | -0.17804 | BLCA | RT-112 | (RAS (G12C) Inhibitor-12 | MVP_top60_DNA+RNA_seed | HSP90AA1 | -1.178451 | BLCA | BFTC-905 | Tanespimycin |
| MVP_top60_DNA_seed | MAPK1 | -0.17267 | BLCA | 5637 | SCH772984 | MVP_top60_DNA+RNA_seed | PIK3CA | -1.147597 | BLCA | CAL-29 | GNE-317 |
| MVP_top60_DNA_seed | MAPK3 | -0.17267 | BLCA | 5637 | SCH772984 | MVP_top60_DNA+RNA_seed | RAC1 | -1.146802 | BLCA | T24 | EHT-1864 |

| | | | | | | | | | | | |
|---|---|---|---|---|---|---|---|---|---|---|---|
| MVP_top60_DNA_seed | PIK3CA | -0.167433 | BLCA | RT-112 | Buparlisib | MVP_top60_DNA+RNA_seed | AKT1 | -1.129855 | BLCA | CAL-29 | GSK2110183B |
| MVP_top60_DNA_seed | PIK3CB | -0.167433 | BLCA | RT-112 | Buparlisib | MVP_top60_DNA+RNA_seed | ESR1 | -1.127363 | BLCA | DSH1 | Tamoxifen |
| MVP_top60_DNA_seed | ESR1 | -0.164304 | BLCA | RT-112 | Tamoxifen | MVP_top60_DNA+RNA_seed | ERBB2 | -1.126995 | BLCA | 5637 | Afatinib |
| MVP_top60_DNA_seed | HDAC1 | -0.157222 | BLCA | CAL-29 | Entinostat | MVP_top60_DNA+RNA_seed | SYK | -1.123493 | BLCA | KU-19-19 | PRT062607 |
| MVP_top60_DNA_seed | RAC1 | -0.146832 | BLCA | SW1710 | EHT-1864 | MVP_top60_DNA+RNA_seed | PIK3CA | -1.094133 | BLCA | CAL-29 | Buparlisib |
| MVP_top60_DNA_seed | PIK3CA | -0.141735 | BLCA | UM-UC-3 | AZD8186 | MVP_top60_DNA+RNA_seed | CDK2 | -1.069996 | BLCA | KU-19-19 | Dinaciclib |
| MVP_top60_DNA_seed | PIK3CB | -0.141735 | BLCA | UM-UC-3 | AZD8186 | MVP_top60_DNA+RNA_seed | FGFR3 | -1.064624 | BLCA | SW780 | PD173074 |
| MVP_top60_DNA_seed | TP53 | -0.130501 | BLCA | RT-112 | MIRA-1 | MVP_top60_DNA+RNA_seed | EP300 | -1.03655 | BLCA | BFTC-905 | CPI-637 |
| MVP_top60_DNA_seed | FGFR3 | -0.12048 | BLCA | LB831-BLC | PD173074 | MVP_top60_DNA+RNA_seed | MAPK1 | -1.033519 | BLCA | BFTC-905 | Ulixertinib |
| MVP_top60_DNA_seed | FGFR3 | -0.120234 | BLCA | LB831-BLC | AZD4547 | MVP_top60_DNA+RNA_seed | MAPK3 | -1.033519 | BLCA | BFTC-905 | Ulixertinib |
| MVP_top60_DNA_seed | PIK3CA | -0.11969 | BLCA | 647-V | GNE-317 | MVP_top60_DNA+RNA_seed | AKT1 | -1.026716 | BLCA | LB831-BLC | AT13148 |
| MVP_top60_DNA_seed | ESR1 | -0.118755 | BLCA | 647-V | Tamoxifen | MVP_top60_DNA+RNA_seed | TERT | -1.023961 | BLCA | 639-V | BIBR-1532 |
| MVP_top60_DNA_seed | PIK3CA | -0.110066 | BLCA | TCCSUP | Buparlisib | MVP_top60_DNA+RNA_seed | MAPK1 | -1.021434 | BLCA | BFTC-905 | VX-11e |
| MVP_top60_DNA_seed | PIK3CB | -0.110066 | BLCA | TCCSUP | Buparlisib | MVP_top60_DNA+RNA_seed | AKT1 | -1.018623 | BLCA | HT-1197 | Uprosertib |
| MVP_top60_DNA_seed | CDK2 | -0.103417 | BLCA | DSH1 | AZD5438 | MVP_top60_DNA+RNA_seed | TERT | -1.011537 | BLCA | DSH1 | BIBR-1532 |
| MVP_top60_DNA_seed | ESR1 | -0.102027 | BLCA | J82 | GDC0810 | MVP_top60_DNA+RNA_seed | KRAS | -0.992505 | BLCA | DSH1 | KRAS (G12C) Inhibitor-1 |
| MVP_top60_DNA_seed | PIK3CA | -0.101739 | BLCA | BFTC-905 | AZD8186 | MVP_top60_DNA+RNA_seed | MAPK1 | -0.982858 | BLCA | DSH1 | SCH772984 |
| MVP_top60_DNA_seed | PIK3CB | -0.101739 | BLCA | BFTC-905 | AZD8186 | MVP_top60_DNA+RNA_seed | MAPK3 | -0.982858 | BLCA | DSH1 | SCH772984 |
| MVP_top60_DNA_seed | MAPK1 | -0.092468 | BLCA | 639-V | Ulixertinib | MVP_top60_DNA+RNA_seed | AKT1 | -0.98006 | BLCA | DSH1 | Uprosertib |
| MVP_top60_DNA_seed | MAPK3 | -0.092468 | BLCA | 639-V | Ulixertinib | MVP_top60_DNA+RNA_seed | AKT1 | -0.963439 | BLCA | HT-1197 | Ipatasertib |
| MVP_top60_DNA_seed | FGFR3 | -0.092348 | BLCA | KU-19-19 | PD173074 | MVP_top60_DNA+RNA_seed | EP300 | -0.963364 | BLCA | RT4 | SGC-CBP30 |
| MVP_top60_DNA_seed | SRC | -0.089659 | BLCA | HT-1197 | Bosutinib | MVP_top60_DNA+RNA_seed | CREBBP | -0.963364 | BLCA | RT4 | SGC-CBP30 |
| MVP_top60_DNA_seed | PIK3CA | -0.087925 | BLCA | LB831-BLC | Alpelisib | MVP_top60_DNA+RNA_seed | RAC1 | -0.963187 | BLCA | RT4 | EHT-1864 |
| MVP_top60_DNA_seed | HSP90AA1 | -0.083219 | BLCA | UM-UC-3 | Tanespimycin | MVP_top60_DNA+RNA_seed | ATM | -0.962046 | BLCA | BFTC-905 | KU-55933 |
| MVP_top60_DNA_seed | ESR1 | -0.080712 | BLCA | 639-V | GDC0810 | MVP_top60_DNA+RNA_seed | ESR1 | -0.960303 | BLCA | KU-19-19 | GDC0810 |
| MVP_top60_DNA_seed | ERBB2 | -0.074152 | BLCA | BFTC-905 | Sapitinib | MVP_top60_DNA+RNA_seed | HSP90AA1 | -0.957798 | BLCA | 639-V | Luminespib |
| MVP_top60_DNA_seed | ERBB3 | -0.074152 | BLCA | BFTC-905 | Sapitinib | MVP_top60_DNA+RNA_seed | HSP90AA1 | -0.9546 | BLCA | KU-19-19 | Luminespib |
| MVP_top60_DNA_seed | PIK3CA | -0.070269 | BLCA | DSH1 | Buparlisib | MVP_top60_DNA+RNA_seed | AKT1 | -0.942744 | BLCA | DSH1 | MK-2206 |
| MVP_top60_DNA_seed | PIK3CB | -0.070269 | BLCA | DSH1 | Buparlisib | MVP_top60_DNA+RNA_seed | SRC | -0.938558 | BLCA | DSH1 | Bosutinib |
| MVP_top60_DNA_seed | CDK2 | -0.069245 | BLCA | RT-112 | MK-8776 | MVP_top60_DNA+RNA_seed | MAPK1 | -0.929273 | BLCA | DSH1 | VX-11e |
| MVP_top60_DNA_seed | CDK2 | -0.069157 | BLCA | DSH1 | MK-8776 | MVP_top60_DNA+RNA_seed | FGFR3 | -0.923771 | BLCA | 639-V | AZD4547 |
| MVP_top60_DNA_seed | ERBB2 | -0.063729 | BLCA | 647-V | Afatinib | MVP_top60_DNA+RNA_seed | AKT1 | -0.917154 | BLCA | CAL-29 | Uprosertib |
| MVP_top60_DNA_seed | FGFR3 | -0.061481 | BLCA | 639-V | PD173074 | MVP_top60_DNA+RNA_seed | HDAC1 | -0.90454 | BLCA | LB831-BLC | Romidepsin |
| MVP_top60_DNA_seed | RAC1 | -0.052606 | BLCA | KU-19-19 | EHT-1864 | MVP_top60_DNA+RNA_seed | TERT | -0.902284 | BLCA | HT-1376 | BIBR-1532 |
| MVP_top60_DNA_seed | FGFR3 | -0.043731 | BLCA | DSH1 | PD173074 | MVP_top60_DNA+RNA_seed | PIK3CA | -0.884619 | BLCA | CAL-29 | AZD8186 |
| MVP_top60_DNA_seed | HSP90AA1 | -0.041544 | BLCA | 5637 | Luminespib | MVP_top60_DNA+RNA_seed | HSP90AA1 | -0.88273 | BLCA | UM-UC-3 | Luminespib |
| MVP_top60_DNA_seed | AR | -0.040754 | BLCA | T24 | Bicalutamide | MVP_top60_DNA+RNA_seed | EP300 | -0.879263 | BLCA | SW1710 | SGC-CBP30 |
| MVP_top60_DNA_seed | ERBB2 | -0.030175 | BLCA | SW1710 | Lapatinib | MVP_top60_DNA+RNA_seed | CREBBP | -0.879263 | BLCA | SW1710 | SGC-CBP30 |
| MVP_top60_DNA_seed | ERBB2 | -0.026496 | BLCA | 639-V | Afatinib | MVP_top60_DNA+RNA_seed | AKT1 | -0.868757 | BLCA | CAL-29 | MK-2206 |
| MVP_top60_DNA_seed | PIK3CA | -0.013456 | BLCA | 647-V | Buparlisib | MVP_top60_DNA+RNA_seed | CDK2 | -0.851637 | BLCA | DSH1 | Dinaciclib |
| MVP_top60_DNA_seed | PIK3CB | -0.013456 | BLCA | 647-V | Buparlisib | MVP_top60_DNA+RNA_seed | KRAS | -0.841657 | BLCA | 639-V | KRAS (G12C) Inhibitor-1 |

| | | | | | | | | | | | |
|---|---|---|---|---|---|---|---|---|---|---|---|
| MVP_top60_DNA_seed | MAPK1 | -0.011556 | BLCA | SW780 | Ulixertinib | MVP_top60_DNA+RNA_seed | ESR1 | -0.840238 | BLCA | 5637 | GDC0810 |
| MVP_top60_DNA_seed | MAPK3 | -0.011556 | BLCA | SW780 | Ulixertinib | MVP_top60_DNA+RNA_seed | PIK3CA | -0.837054 | BLCA | BFTC-905 | Buparlisib |
| MVP_top60_DNA_seed | PIK3CA | -0.010391 | BLCA | 5637 | GNE-317 | MVP_top60_DNA+RNA_seed | AKT1 | -0.834798 | BLCA | HT-1197 | Afuresertib |
| MVP_top60_DNA_seed | RAC1 | -0.009372 | BLCA | LB831-BLC | EHT-1864 | MVP_top60_DNA+RNA_seed | ATM | -0.822909 | BLCA | 647-V | KU-55933 |
| MVP_top60_DNA_seed | PIK3CA | -0.005178 | BLCA | HT-1197 | Alpelisib | MVP_top60_DNA+RNA_seed | HSP90AA1 | -0.821776 | BLCA | 639-V | Tanespimycin |
| MVP_top60_DNA_seed | HSP90AA1 | -0.003759 | BLCA | T24 | Luminespib | MVP_top60_DNA+RNA_seed | ESR1 | -0.818628 | BLCA | HT-1376 | GDC0810 |
| MVP_top60_DNA_seed | RAC1 | -0.002287 | BLCA | 647-V | EHT-1864 | MVP_top60_DNA+RNA_seed | AKT1 | -0.804174 | BLCA | CAL-29 | AT13148 |
| MVP_top60_DNA_seed | EP300 | 0.003175 | BLCA | 639-V | SGC-CBP30 | MVP_top60_DNA+RNA_seed | MAPK1 | -0.803411 | BLCA | RT-112 | Ulixertinib |
| MVP_top60_DNA_seed | CREBBP | 0.003175 | BLCA | 639-V | SGC-CBP30 | MVP_top60_DNA+RNA_seed | MAPK3 | -0.803411 | BLCA | RT-112 | Ulixertinib |
| MVP_top60_DNA_seed | CDK2 | 0.005737 | BLCA | CAL-29 | AZD5438 | MVP_top60_DNA+RNA_seed | PIK3CA | -0.799839 | BLCA | BFTC-905 | Alpelisib |
| MVP_top60_DNA_seed | HDAC1 | 0.006348 | BLCA | TCCSUP | Entinostat | MVP_top60_DNA+RNA_seed | ERBB2 | -0.795081 | BLCA | 5637 | Lapatinib |
| MVP_top60_DNA_seed | EP300 | 0.007679 | BLCA | RT4 | CPI-637 | MVP_top60_DNA+RNA_seed | AKT1 | -0.789461 | BLCA | SW1710 | AT13148 |
| MVP_top60_DNA_seed | PIK3CA | 0.010687 | BLCA | 5637 | Alpelisib | MVP_top60_DNA+RNA_seed | ESR1 | -0.788135 | BLCA | HT-1376 | Tamoxifen |
| MVP_top60_DNA_seed | ERBB2 | 0.01589 | BLCA | RT4 | Lapatinib | MVP_top60_DNA+RNA_seed | MAPK1 | -0.786778 | BLCA | RT-112 | SCH772984 |
| MVP_top60_DNA_seed | ESR1 | 0.017008 | BLCA | 5637 | Tamoxifen | MVP_top60_DNA+RNA_seed | MAPK3 | -0.786778 | BLCA | RT-112 | SCH772984 |
| MVP_top60_DNA_seed | PIK3CA | 0.019742 | BLCA | UM-UC-3 | Buparlisib | MVP_top60_DNA+RNA_seed | PIK3CA | -0.777686 | BLCA | 5637 | Buparlisib |
| MVP_top60_DNA_seed | PIK3CB | 0.019742 | BLCA | UM-UC-3 | Buparlisib | MVP_top60_DNA+RNA_seed | TERT | -0.77393 | BLCA | HT-1197 | BIBR-1532 |
| MVP_top60_DNA_seed | MAPK1 | 0.019759 | BLCA | 5637 | VX-11e | MVP_top60_DNA+RNA_seed | CDK2 | -0.760694 | BLCA | RT-112 | AZD5438 |
| MVP_top60_DNA_seed | HDAC1 | 0.032682 | BLCA | CAL-29 | Romidepsin | MVP_top60_DNA+RNA_seed | RAC1 | -0.75836 | BLCA | DSH1 | EHT-1864 |
| MVP_top60_DNA_seed | ESR1 | 0.039152 | BLCA | UM-UC-3 | Tamoxifen | MVP_top60_DNA+RNA_seed | ESR1 | -0.752726 | BLCA | CAL-29 | Tamoxifen |
| MVP_top60_DNA_seed | SRC | 0.039951 | BLCA | HT-1376 | Bosutinib | MVP_top60_DNA+RNA_seed | AKT1 | -0.751268 | BLCA | CAL-29 | Uprosertib |
| MVP_top60_DNA_seed | TP53 | 0.039981 | BLCA | KU-19-19 | MIRA-1 | MVP_top60_DNA+RNA_seed | RAC1 | -0.747393 | BLCA | 639-V | EHT-1864 |
| MVP_top60_DNA_seed | CDK2 | 0.04224 | BLCA | SW780 | AZD5438 | MVP_top60_DNA+RNA_seed | MAPK1 | -0.747032 | BLCA | RT4 | VX-11e |
| MVP_top60_DNA_seed | HDAC1 | 0.042626 | BLCA | LB831-BLC | PCI-34051 | MVP_top60_DNA+RNA_seed | CDK2 | -0.738271 | BLCA | RT-112 | Dinaciclib |
| MVP_top60_DNA_seed | HDAC1 | 0.043913 | BLCA | DSH1 | Romidepsin | MVP_top60_DNA+RNA_seed | ESR1 | -0.698797 | BLCA | 639-V | Tamoxifen |
| MVP_top60_DNA_seed | SRC | 0.057773 | BLCA | CAL-29 | Bosutinib | MVP_top60_DNA+RNA_seed | ERBB2 | -0.696024 | BLCA | CAL-29 | Lapatinib |
| MVP_top60_DNA_seed | ERBB2 | 0.065332 | BLCA | 647-V | Sapitinib | MVP_top60_DNA+RNA_seed | PIK3CA | -0.69325 | BLCA | HT-1197 | Buparlisib |
| MVP_top60_DNA_seed | ERBB3 | 0.065332 | BLCA | 647-V | Sapitinib | MVP_top60_DNA+RNA_seed | HSP90AA1 | -0.692026 | BLCA | KU-19-19 | Tanespimycin |
| MVP_top60_DNA_seed | ERBB2 | 0.066548 | BLCA | RT4 | Sapitinib | MVP_top60_DNA+RNA_seed | ESR1 | -0.689053 | BLCA | UM-UC-3 | GDC0810 |
| MVP_top60_DNA_seed | ERBB3 | 0.066548 | BLCA | RT4 | Sapitinib | MVP_top60_DNA+RNA_seed | HDAC1 | -0.684011 | BLCA | KU-19-19 | Entinostat |
| MVP_top60_DNA_seed | CDK2 | 0.069504 | BLCA | SW780 | Dinaciclib | MVP_top60_DNA+RNA_seed | KRAS | -0.678566 | BLCA | CAL-29 | KRAS (G12C) Inhibitor-1 |
| MVP_top60_DNA_seed | PIK3CA | 0.070222 | BLCA | DSH1 | Alpelisib | MVP_top60_DNA+RNA_seed | PIK3CA | -0.670616 | BLCA | RT-112 | Alpelisib |
| MVP_top60_DNA_seed | FGFR3 | 0.07172 | BLCA | BFTC-905 | PD173074 | MVP_top60_DNA+RNA_seed | MAPK1 | -0.658453 | BLCA | RT4 | SCH772984 |
| MVP_top60_DNA_seed | FGFR3 | 0.072193 | BLCA | UM-UC-3 | AZD4547 | MVP_top60_DNA+RNA_seed | MAPK3 | -0.658453 | BLCA | RT4 | SCH772984 |
| MVP_top60_DNA_seed | PIK3CA | 0.077521 | BLCA | 639-V | GNE-317 | MVP_top60_DNA+RNA_seed | SRC | -0.656227 | BLCA | 5637 | Bosutinib |
| MVP_top60_DNA_seed | TP53 | 0.082077 | BLCA | CAL-29 | MIRA-1 | MVP_top60_DNA+RNA_seed | HDAC1 | -0.639754 | BLCA | 5637 | Entinostat |
| MVP_top60_DNA_seed | MAPK1 | 0.082733 | BLCA | SW780 | SCH772984 | MVP_top60_DNA+RNA_seed | AKT1 | -0.63809 | BLCA | DSH1 | Ipatasertib |
| MVP_top60_DNA_seed | MAPK3 | 0.082733 | BLCA | SW780 | SCH772984 | MVP_top60_DNA+RNA_seed | KRAS | -0.626387 | BLCA | BFTC-905 | KRAS (G12C) Inhibitor-1 |
| MVP_top60_DNA_seed | ESR1 | 0.092829 | BLCA | SW1710 | GDC0810 | MVP_top60_DNA+RNA_seed | CDK2 | -0.620205 | BLCA | BFTC-905 | AZD5438 |
| MVP_top60_DNA_seed | PIK3CA | 0.095867 | BLCA | SW1710 | Buparlisib | MVP_top60_DNA+RNA_seed | ERBB2 | -0.602856 | BLCA | CAL-29 | Afatinib |
| MVP_top60_DNA_seed | PIK3CB | 0.095867 | BLCA | SW1710 | Buparlisib | MVP_top60_DNA+RNA_seed | ATM | -0.602001 | BLCA | 639-V | KU-55933 |

| | | | | | | | | | | | |
|---|---|---|---|---|---|---|---|---|---|---|---|
| MVP_top60_DNA_seed | SRC | 0.104374 | BLCA | SW1710 | Bosutinib | MVP_top60_DNA+RNA_seed | HSP90AA1 | -0.600517 | BLCA | BFTC-905 | Luminespib |
| MVP_top60_DNA_seed | CDK2 | 0.107665 | BLCA | LB831-BLC | MK-8776 | MVP_top60_DNA+RNA_seed | HSP90AA1 | -0.587736 | BLCA | DSH1 | Luminespib |
| MVP_top60_DNA_seed | ATM | 0.111065 | BLCA | DSH1 | KU-55933 | MVP_top60_DNA+RNA_seed | CDK2 | -0.587022 | BLCA | CAL-29 | Dinaciclib |
| MVP_top60_DNA_seed | FGFR3 | 0.114756 | BLCA | SW1710 | AZD4547 | MVP_top60_DNA+RNA_seed | MAPK1 | -0.582567 | BLCA | HT-1197 | Ulixertinib |
| MVP_top60_DNA_seed | HDAC1 | 0.11586 | BLCA | 647-V | Romidepsin | MVP_top60_DNA+RNA_seed | MAPK3 | -0.582567 | BLCA | HT-1197 | Ulixertinib |
| MVP_top60_DNA_seed | HSP90AA1 | 0.118117 | BLCA | 647-V | Tanespimycin | MVP_top60_DNA+RNA_seed | HSP90AA1 | -0.579553 | BLCA | RT-112 | Luminespib |
| MVP_top60_DNA_seed | HDAC1 | 0.120188 | BLCA | 647-V | PCI-34051 | MVP_top60_DNA+RNA_seed | SYK | -0.574401 | BLCA | BFTC-905 | PRT062607 |
| MVP_top60_DNA_seed | MAPK1 | 0.122208 | BLCA | CAL-29 | Ulixertinib | MVP_top60_DNA+RNA_seed | EP300 | -0.567635 | BLCA | KU-19-19 | SGC-CBP30 |
| MVP_top60_DNA_seed | MAPK3 | 0.122208 | BLCA | CAL-29 | Ulixertinib | MVP_top60_DNA+RNA_seed | CREBBP | -0.567635 | BLCA | KU-19-19 | SGC-CBP30 |
| MVP_top60_DNA_seed | EP300 | 0.126142 | BLCA | 5637 | SGC-CBP30 | MVP_top60_DNA+RNA_seed | AKT1 | -0.562278 | BLCA | DSH1 | Uprosertib |
| MVP_top60_DNA_seed | CREBBP | 0.126142 | BLCA | 5637 | SGC-CBP30 | MVP_top60_DNA+RNA_seed | PIK3CA | -0.562267 | BLCA | DSH1 | GNE-317 |
| MVP_top60_DNA_seed | EP300 | 0.129522 | BLCA | SW1710 | CPI-637 | MVP_top60_DNA+RNA_seed | RAC1 | -0.554754 | BLCA | CAL-29 | EHT-1864 |
| MVP_top60_DNA_seed | ATM | 0.140815 | BLCA | UM-UC-3 | KU-55933 | MVP_top60_DNA+RNA_seed | KRAS | -0.550227 | BLCA | 5637 | KRAS (G12C) Inhibitor-1 |
| MVP_top60_DNA_seed | FGFR3 | 0.141684 | BLCA | J82 | PD173074 | MVP_top60_DNA+RNA_seed | CDK2 | -0.538674 | BLCA | LB831-BLC | Dinaciclib |
| MVP_top60_DNA_seed | MAPK1 | 0.142674 | BLCA | SW780 | VX-11e | MVP_top60_DNA+RNA_seed | KRAS | -0.533955 | BLCA | RT4 | KRAS (G12C) Inhibitor-1 |
| MVP_top60_DNA_seed | ERBB2 | 0.142791 | BLCA | KU-19-19 | Lapatinib | MVP_top60_DNA+RNA_seed | PIK3CA | -0.533608 | BLCA | HT-1197 | GNE-317 |
| MVP_top60_DNA_seed | MAPK1 | 0.147848 | BLCA | SW1710 | Ulixertinib | MVP_top60_DNA+RNA_seed | HDAC1 | -0.532909 | BLCA | BFTC-905 | Romidepsin |
| MVP_top60_DNA_seed | MAPK3 | 0.147848 | BLCA | SW1710 | Ulixertinib | MVP_top60_DNA+RNA_seed | ESR1 | -0.522591 | BLCA | SW1710 | Tamoxifen |
| MVP_top60_DNA_seed | FGFR3 | 0.149376 | BLCA | CAL-29 | PD173074 | MVP_top60_DNA+RNA_seed | HDAC1 | -0.522475 | BLCA | DSH1 | PCI-34051 |
| MVP_top60_DNA_seed | HDAC1 | 0.152653 | BLCA | HT-1197 | Romidepsin | MVP_top60_DNA+RNA_seed | TERT | -0.520061 | BLCA | 647-V | BIBR-1532 |
| MVP_top60_DNA_seed | RAC1 | 0.159193 | BLCA | UM-UC-3 | EHT-1864 | MVP_top60_DNA+RNA_seed | SRC | -0.517813 | BLCA | KU-19-19 | Bosutinib |
| MVP_top60_DNA_seed | EP300 | 0.159683 | BLCA | CAL-29 | SGC-CBP30 | MVP_top60_DNA+RNA_seed | SYK | -0.517734 | BLCA | BFTC-905 | Entospletinib |
| MVP_top60_DNA_seed | CREBBP | 0.159683 | BLCA | CAL-29 | SGC-CBP30 | MVP_top60_DNA+RNA_seed | AKT1 | -0.513118 | BLCA | RT4 | AT13148 |
| MVP_top60_DNA_seed | EP300 | 0.162684 | BLCA | HT-1376 | CPI-637 | MVP_top60_DNA+RNA_seed | CDK2 | -0.51229 | BLCA | BFTC-905 | Dinaciclib |
| MVP_top60_DNA_seed | EP300 | 0.163524 | BLCA | 647-V | CPI-637 | MVP_top60_DNA+RNA_seed | AKT1 | -0.496974 | BLCA | CAL-29 | Ipatasertib |
| MVP_top60_DNA_seed | HSP90AA1 | 0.163691 | BLCA | T24 | Tanespimycin | MVP_top60_DNA+RNA_seed | AKT1 | -0.483977 | BLCA | J82 | AT13148 |
| MVP_top60_DNA_seed | PIK3CA | 0.16847 | BLCA | SW780 | GNE-317 | MVP_top60_DNA+RNA_seed | PIK3CA | -0.481732 | BLCA | KU-19-19 | Buparlisib |
| MVP_top60_DNA_seed | MAPK1 | 0.16983 | BLCA | CAL-29 | Ulixertinib | MVP_top60_DNA+RNA_seed | PIK3CA | -0.476776 | BLCA | KU-19-19 | GNE-317 |
| MVP_top60_DNA_seed | MAPK3 | 0.16983 | BLCA | CAL-29 | Ulixertinib | MVP_top60_DNA+RNA_seed | RAC1 | -0.475895 | BLCA | SW780 | EHT-1864 |
| MVP_top60_DNA_seed | TP53 | 0.170108 | BLCA | UM-UC-3 | MIRA-1 | MVP_top60_DNA+RNA_seed | SYK | -0.462184 | BLCA | KU-19-19 | Entospletinib |
| MVP_top60_DNA_seed | HDAC1 | 0.172076 | BLCA | RT-112 | Entinostat | MVP_top60_DNA+RNA_seed | EP300 | -0.459899 | BLCA | DSH1 | SGC-CBP30 |
| MVP_top60_DNA_seed | MAPK1 | 0.186558 | BLCA | SW1710 | SCH772984 | MVP_top60_DNA+RNA_seed | CREBBP | -0.459899 | BLCA | DSH1 | SGC-CBP30 |
| MVP_top60_DNA_seed | MAPK3 | 0.186558 | BLCA | SW1710 | SCH772984 | MVP_top60_DNA+RNA_seed | MAPK1 | -0.458567 | BLCA | RT-112 | Ulixertinib |
| MVP_top60_DNA_seed | FGFR3 | 0.194693 | BLCA | BFTC-905 | AZD4547 | MVP_top60_DNA+RNA_seed | MAPK3 | -0.458567 | BLCA | RT-112 | Ulixertinib |
| MVP_top60_DNA_seed | ERBB2 | 0.19596 | BLCA | KU-19-19 | Afatinib | MVP_top60_DNA+RNA_seed | AKT1 | -0.456477 | BLCA | KU-19-19 | Uprosertib |
| MVP_top60_DNA_seed | HSP90AA1 | 0.200704 | BLCA | SW780 | Tanespimycin | MVP_top60_DNA+RNA_seed | EP300 | -0.45414 | BLCA | SW780 | SGC-CBP30 |
| MVP_top60_DNA_seed | FGFR3 | 0.201579 | BLCA | 647-V | PD173074 | MVP_top60_DNA+RNA_seed | CREBBP | -0.45414 | BLCA | SW780 | SGC-CBP30 |
| MVP_top60_DNA_seed | CDK2 | 0.201717 | BLCA | 647-V | Dinaciclib | MVP_top60_DNA+RNA_seed | HSP90AA1 | -0.447393 | BLCA | LB831-BLC | Luminespib |
| MVP_top60_DNA_seed | HSP90AA1 | 0.202892 | BLCA | SW1710 | Tanespimycin | MVP_top60_DNA+RNA_seed | SYK | -0.446466 | BLCA | RT-112 | PRT062607 |
| MVP_top60_DNA_seed | CDK2 | 0.205152 | BLCA | KU-19-19 | MK-8776 | MVP_top60_DNA+RNA_seed | AKT1 | -0.435484 | BLCA | KU-19-19 | AT13148 |
| MVP_top60_DNA_seed | TP53 | 0.209842 | BLCA | DSH1 | MIRA-1 | MVP_top60_DNA+RNA_seed | AKT1 | -0.423251 | BLCA | DSH1 | Afuresertib |

| | | | | | | | | | | | |
|---|---|---|---|---|---|---|---|---|---|---|---|
| MVP_top60_DNA_seed | HDAC1 | 0.211596 | BLCA | SW1710 | PCI-34051 | MVP_top60_DNA+RNA_seed | HDAC1 | -0.422774 | BLCA | RT-112 | Romidepsin |
| MVP_top60_DNA_seed | MAPK1 | 0.216558 | BLCA | 639-V | Ulixertinib | MVP_top60_DNA+RNA_seed | TERT | -0.410889 | BLCA | SW780 | BIBR-1532 |
| MVP_top60_DNA_seed | MAPK3 | 0.216558 | BLCA | 639-V | Ulixertinib | MVP_top60_DNA+RNA_seed | FGFR3 | -0.40624 | BLCA | 647-V | AZD4547 |
| MVP_top60_DNA_seed | EP300 | 0.219325 | BLCA | LB831-BLC | CPI-637 | MVP_top60_DNA+RNA_seed | RAC1 | -0.404138 | BLCA | BFTC-905 | EHT-1864 |
| MVP_top60_DNA_seed | PIK3CA | 0.220832 | BLCA | SW1710 | GNE-317 | MVP_top60_DNA+RNA_seed | HSP90AA1 | -0.403736 | BLCA | 5637 | Tanespimycin |
| MVP_top60_DNA_seed | FGFR3 | 0.225976 | BLCA | KU-19-19 | AZD4547 | MVP_top60_DNA+RNA_seed | SYK | -0.403604 | BLCA | RT-112 | Entospletinib |
| MVP_top60_DNA_seed | HDAC1 | 0.235783 | BLCA | 639-V | PCI-34051 | MVP_top60_DNA+RNA_seed | FGFR3 | -0.401204 | BLCA | CAL-29 | AZD4547 |
| MVP_top60_DNA_seed | RAC1 | 0.241462 | BLCA | 5637 | EHT-1864 | MVP_top60_DNA+RNA_seed | AKT1 | -0.38569 | BLCA | SW1710 | AZD5363 |
| MVP_top60_DNA_seed | HSP90AA1 | 0.247158 | BLCA | SW1710 | Luminespib | MVP_top60_DNA+RNA_seed | AKT1 | -0.385281 | BLCA | CAL-29 | Afuresertib |
| MVP_top60_DNA_seed | SRC | 0.252369 | BLCA | 639-V | Bosutinib | MVP_top60_DNA+RNA_seed | HSP90AA1 | -0.383208 | BLCA | RT-112 | Tanespimycin |
| MVP_top60_DNA_seed | ESR1 | 0.254018 | BLCA | 647-V | GDC0810 | MVP_top60_DNA+RNA_seed | MAPK1 | -0.381047 | BLCA | LB831-BLC | Ulixertinib |
| MVP_top60_DNA_seed | PIK3CA | 0.255993 | BLCA | LB831-BLC | GNE-317 | MVP_top60_DNA+RNA_seed | MAPK3 | -0.381047 | BLCA | LB831-BLC | Ulixertinib |
| MVP_top60_DNA_seed | SRC | 0.270978 | BLCA | T24 | Bosutinib | MVP_top60_DNA+RNA_seed | HDAC1 | -0.378929 | BLCA | KU-19-19 | PCI-34051 |
| MVP_top60_DNA_seed | ERBB2 | 0.28053 | BLCA | 639-V | Sapitinib | MVP_top60_DNA+RNA_seed | SYK | -0.37714 | BLCA | SW1710 | Entospletinib |
| MVP_top60_DNA_seed | ERBB3 | 0.28053 | BLCA | 639-V | Sapitinib | MVP_top60_DNA+RNA_seed | AKT1 | -0.373692 | BLCA | HT-1197 | GSK2110183B |
| MVP_top60_DNA_seed | HDAC1 | 0.283604 | BLCA | 639-V | Romidepsin | MVP_top60_DNA+RNA_seed | AKT1 | -0.351272 | BLCA | 647-V | AT13148 |
| MVP_top60_DNA_seed | FGFR3 | 0.29218 | BLCA | SW1710 | PD173074 | MVP_top60_DNA+RNA_seed | MAPK1 | -0.337577 | BLCA | DSH1 | Ulixertinib |
| MVP_top60_DNA_seed | CDK2 | 0.297523 | BLCA | 647-V | AZD5438 | MVP_top60_DNA+RNA_seed | MAPK3 | -0.337577 | BLCA | DSH1 | Ulixertinib |
| MVP_top60_DNA_seed | MAPK1 | 0.31246 | BLCA | 647-V | Ulixertinib | MVP_top60_DNA+RNA_seed | PIK3CA | -0.335329 | BLCA | 647-V | AZD8186 |
| MVP_top60_DNA_seed | MAPK3 | 0.31246 | BLCA | 647-V | Ulixertinib | MVP_top60_DNA+RNA_seed | ESR1 | -0.33484 | BLCA | DSH1 | GDC0810 |
| MVP_top60_DNA_seed | HSP90AA1 | 0.320718 | BLCA | 647-V | Luminespib | MVP_top60_DNA+RNA_seed | EP300 | -0.327979 | BLCA | KU-19-19 | CPI-637 |
| MVP_top60_DNA_seed | FGFR3 | 0.326347 | BLCA | DSH1 | AZD4547 | MVP_top60_DNA+RNA_seed | AKT1 | -0.327574 | BLCA | HT-1197 | AZD5363 |
| MVP_top60_DNA_seed | MAPK1 | 0.334225 | BLCA | RT4 | Ulixertinib | MVP_top60_DNA+RNA_seed | TERT | -0.327465 | BLCA | RT-112 | BIBR-1532 |
| MVP_top60_DNA_seed | MAPK3 | 0.334225 | BLCA | RT4 | Ulixertinib | MVP_top60_DNA+RNA_seed | AKT1 | -0.322344 | BLCA | BFTC-905 | MK-2206 |
| MVP_top60_DNA_seed | KRAS | 0.335212 | BLCA | T24 | KRAS (G12C) Inhibitor-1[illegible] | MVP_top60_DNA+RNA_seed | CDK2 | -0.319533 | BLCA | RT4 | AZD5438 |
| MVP_top60_DNA_seed | PIK3CA | 0.336452 | BLCA | LB831-BLC | AZD8186 | MVP_top60_DNA+RNA_seed | HDAC1 | -0.31773 | BLCA | 5637 | Romidepsin |
| MVP_top60_DNA_seed | PIK3CB | 0.336452 | BLCA | LB831-BLC | AZD8186 | MVP_top60_DNA+RNA_seed | SYK | -0.316387 | BLCA | DSH1 | Entospletinib |
| MVP_top60_DNA_seed | ESR1 | 0.338352 | BLCA | HT-1197 | GDC0810 | MVP_top60_DNA+RNA_seed | CDK2 | -0.315001 | BLCA | RT4 | Dinaciclib |
| MVP_top60_DNA_seed | SRC | 0.342638 | BLCA | UM-UC-3 | Bosutinib | MVP_top60_DNA+RNA_seed | FGFR3 | -0.314594 | BLCA | 5637 | AZD4547 |
| MVP_top60_DNA_seed | MAPK1 | 0.343628 | BLCA | 639-V | SCH772984 | MVP_top60_DNA+RNA_seed | PIK3CA | -0.311178 | BLCA | 639-V | AZD8186 |
| MVP_top60_DNA_seed | MAPK3 | 0.343628 | BLCA | 639-V | SCH772984 | MVP_top60_DNA+RNA_seed | AKT1 | -0.308006 | BLCA | RT-112 | Uprosertib |
| MVP_top60_DNA_seed | EP300 | 0.343881 | BLCA | SW780 | CPI-637 | MVP_top60_DNA+RNA_seed | MAPK1 | -0.306001 | BLCA | RT-112 | VX-11e |
| MVP_top60_DNA_seed | PIK3CA | 0.349232 | BLCA | SW1710 | AZD8186 | MVP_top60_DNA+RNA_seed | ATM | -0.302883 | BLCA | 5637 | KU-55933 |
| MVP_top60_DNA_seed | PIK3CB | 0.349232 | BLCA | SW1710 | AZD8186 | MVP_top60_DNA+RNA_seed | GSK3B | -0.299059 | BLCA | 5637 | SB216763 |
| MVP_top60_DNA_seed | CDK2 | 0.35568 | BLCA | SW1710 | AZD5438 | MVP_top60_DNA+RNA_seed | PIK3CA | -0.297558 | BLCA | 639-V | Buparlisib |
| MVP_top60_DNA_seed | FGFR3 | 0.363163 | BLCA | TCCSUP | AZD4547 | MVP_top60_DNA+RNA_seed | CDK2 | -0.292461 | BLCA | LB831-BLC | AZD5438 |
| MVP_top60_DNA_seed | HDAC1 | 0.363548 | BLCA | SW780 | Entinostat | MVP_top60_DNA+RNA_seed | MAPK1 | -0.288714 | BLCA | CAL-29 | SCH772984 |
| MVP_top60_DNA_seed | CDK2 | 0.376723 | BLCA | 5637 | MK-8776 | MVP_top60_DNA+RNA_seed | MAPK3 | -0.288714 | BLCA | CAL-29 | SCH772984 |
| MVP_top60_DNA_seed | PIK3CA | 0.380056 | BLCA | DSH1 | AZD8186 | MVP_top60_DNA+RNA_seed | AKT1 | -0.286014 | BLCA | DSH1 | AT13148 |
| MVP_top60_DNA_seed | PIK3CB | 0.380056 | BLCA | DSH1 | AZD8186 | MVP_top60_DNA+RNA_seed | PIK3CA | -0.28342 | BLCA | RT-112 | GNE-317 |
| MVP_top60_DNA_seed | TP53 | 0.382708 | BLCA | 647-V | MIRA-1 | MVP_top60_DNA+RNA_seed | EP300 | -0.281952 | BLCA | 639-V | CPI-637 |

| | | | | | | | | | | | |
|---|---|---|---|---|---|---|---|---|---|---|---|
| MVP_top60_DNA_seed | MAPK1 | 0.385978 | BLCA | 647-V | SCH772984 | MVP_top60_DNA+RNA_seed | MAPK1 | -0.274744 | BLCA | CAL-29 | VX-11e |
| MVP_top60_DNA_seed | MAPK3 | 0.385978 | BLCA | 647-V | SCH772984 | MVP_top60_DNA+RNA_seed | EP300 | -0.273802 | BLCA | J82 | SGC-CBP30 |
| MVP_top60_DNA_seed | SRC | 0.397445 | BLCA | RT-112 | Bosutinib | MVP_top60_DNA+RNA_seed | CREBBP | -0.273802 | BLCA | J82 | SGC-CBP30 |
| MVP_top60_DNA_seed | CDK2 | 0.397772 | BLCA | HT-1376 | MK-8776 | MVP_top60_DNA+RNA_seed | PIK3CA | -0.273018 | BLCA | BFTC-905 | GNE-317 |
| MVP_top60_DNA_seed | KRAS | 0.399391 | BLCA | SW780 | KRAS (G12C) Inhibitor-1 | MVP_top60_DNA+RNA_seed | ESR1 | -0.271128 | BLCA | RT4 | Tamoxifen |
| MVP_top60_DNA_seed | ATM | 0.400572 | BLCA | CAL-29 | KU-55933 | MVP_top60_DNA+RNA_seed | PIK3CA | -0.270785 | BLCA | TCCSUP | GNE-317 |
| MVP_top60_DNA_seed | PIK3CA | 0.404888 | BLCA | 5637 | AZD8186 | MVP_top60_DNA+RNA_seed | CDK2 | -0.263229 | BLCA | 639-V | Dinaciclib |
| MVP_top60_DNA_seed | PIK3CB | 0.404888 | BLCA | 5637 | AZD8186 | MVP_top60_DNA+RNA_seed | MAPK1 | -0.261527 | BLCA | 5637 | Ulixertinib |
| MVP_top60_DNA_seed | HDAC1 | 0.413044 | BLCA | HT-1197 | Entinostat | MVP_top60_DNA+RNA_seed | MAPK3 | -0.261527 | BLCA | 5637 | Ulixertinib |
| MVP_top60_DNA_seed | HDAC1 | 0.413164 | BLCA | DSH1 | Entinostat | MVP_top60_DNA+RNA_seed | ESR1 | -0.259256 | BLCA | CAL-29 | GDC0810 |
| MVP_top60_DNA_seed | CDK2 | 0.414545 | BLCA | BFTC-905 | MK-8776 | MVP_top60_DNA+RNA_seed | SYK | -0.25889 | BLCA | CAL-29 | Entospletinib |
| MVP_top60_DNA_seed | FGFR3 | 0.416917 | BLCA | 5637 | PD173074 | MVP_top60_DNA+RNA_seed | ESR1 | -0.253103 | BLCA | KU-19-19 | Tamoxifen |
| MVP_top60_DNA_seed | TP53 | 0.417231 | BLCA | SW1710 | MIRA-1 | MVP_top60_DNA+RNA_seed | EP300 | -0.250773 | BLCA | TCCSUP | SGC-CBP30 |
| MVP_top60_DNA_seed | CDK2 | 0.420146 | BLCA | SW1710 | Dinaciclib | MVP_top60_DNA+RNA_seed | CREBBP | -0.250773 | BLCA | TCCSUP | SGC-CBP30 |
| MVP_top60_DNA_seed | HSP90AA1 | 0.423604 | BLCA | T24 | CCT-018159 | MVP_top60_DNA+RNA_seed | ERBB2 | -0.249731 | BLCA | BFTC-905 | Afatinib |
| MVP_top60_DNA_seed | RAC1 | 0.423903 | BLCA | HT-1197 | EHT-1864 | MVP_top60_DNA+RNA_seed | HDAC1 | -0.249172 | BLCA | RT4 | PCI-34051 |
| MVP_top60_DNA_seed | ERBB2 | 0.424951 | BLCA | UM-UC-3 | Sapitinib | MVP_top60_DNA+RNA_seed | MAPK1 | -0.233154 | BLCA | RT4 | Ulixertinib |
| MVP_top60_DNA_seed | ERBB3 | 0.424951 | BLCA | UM-UC-3 | Sapitinib | MVP_top60_DNA+RNA_seed | MAPK3 | -0.233154 | BLCA | RT4 | Ulixertinib |
| MVP_top60_DNA_seed | CDK2 | 0.428728 | BLCA | J82 | MK-8776 | MVP_top60_DNA+RNA_seed | HSP90AA1 | -0.230539 | BLCA | LB831-BLC | Tanespimycin |
| MVP_top60_DNA_seed | PIK3CA | 0.435335 | BLCA | HT-1376 | AZD8186 | MVP_top60_DNA+RNA_seed | RAC1 | -0.226074 | BLCA | RT-112 | EHT-1864 |
| MVP_top60_DNA_seed | PIK3CB | 0.435335 | BLCA | HT-1376 | AZD8186 | MVP_top60_DNA+RNA_seed | PIK3CA | -0.22432 | BLCA | UM-UC-3 | GNE-317 |
| MVP_top60_DNA_seed | ERBB2 | 0.441624 | BLCA | RT-112 | Afatinib | MVP_top60_DNA+RNA_seed | ATM | -0.222383 | BLCA | LB831-BLC | KU-55933 |
| MVP_top60_DNA_seed | ERBB2 | 0.442474 | BLCA | RT-112 | Sapitinib | MVP_top60_DNA+RNA_seed | GSK3B | -0.221771 | BLCA | HT-1197 | SB216763 |
| MVP_top60_DNA_seed | ERBB3 | 0.442474 | BLCA | RT-112 | Sapitinib | MVP_top60_DNA+RNA_seed | HDAC1 | -0.219178 | BLCA | BFTC-905 | PCI-34051 |
| MVP_top60_DNA_seed | PIK3CA | 0.44544 | BLCA | SW780 | Buparlisib | MVP_top60_DNA+RNA_seed | AKT1 | -0.217535 | BLCA | DSH1 | GSK2110183B |
| MVP_top60_DNA_seed | PIK3CB | 0.44544 | BLCA | SW780 | Buparlisib | MVP_top60_DNA+RNA_seed | TP53 | -0.214167 | BLCA | 639-V | MIRA-1 |
| MVP_top60_DNA_seed | ESR1 | 0.446831 | BLCA | SW780 | Tamoxifen | MVP_top60_DNA+RNA_seed | SYK | -0.212842 | BLCA | LB831-BLC | PRT062607 |
| MVP_top60_DNA_seed | ERBB2 | 0.446839 | BLCA | RT4 | Afatinib | MVP_top60_DNA+RNA_seed | CDK2 | -0.205094 | BLCA | 639-V | AZD5438 |
| MVP_top60_DNA_seed | EP300 | 0.457723 | BLCA | CAL-29 | CPI-637 | MVP_top60_DNA+RNA_seed | PIK3CA | -0.202548 | BLCA | HT-1197 | AZD8186 |
| MVP_top60_DNA_seed | HDAC1 | 0.458762 | BLCA | BFTC-905 | Entinostat | MVP_top60_DNA+RNA_seed | ATM | -0.195009 | BLCA | KU-19-19 | KU-55933 |
| MVP_top60_DNA_seed | ATM | 0.459207 | BLCA | RT4 | KU-55933 | MVP_top60_DNA+RNA_seed | SYK | -0.188305 | BLCA | LB831-BLC | Entospletinib |
| MVP_top60_DNA_seed | CDK2 | 0.460578 | BLCA | T24 | MK-8776 | MVP_top60_DNA+RNA_seed | GSK3B | -0.187494 | BLCA | 647-V | SB216763 |
| MVP_top60_DNA_seed | HDAC1 | 0.462096 | BLCA | SW780 | PCI-34051 | MVP_top60_DNA+RNA_seed | EP300 | -0.180177 | BLCA | DSH1 | CPI-637 |
| MVP_top60_DNA_seed | ESR1 | 0.464937 | BLCA | TCCSUP | GDC0810 | MVP_top60_DNA+RNA_seed | ERBB2 | -0.179974 | BLCA | BFTC-905 | Lapatinib |
| MVP_top60_DNA_seed | ATM | 0.468521 | BLCA | RT-112 | KU-55933 | MVP_top60_DNA+RNA_seed | AKT1 | -0.178759 | BLCA | UM-UC-3 | AZD5363 |
| MVP_top60_DNA_seed | ERBB2 | 0.470151 | BLCA | LB831-BLC | Sapitinib | MVP_top60_DNA+RNA_seed | KRAS | -0.17804 | BLCA | RT-112 | (RAS (G12C) Inhibitor-1 |
| MVP_top60_DNA_seed | ERBB3 | 0.470151 | BLCA | LB831-BLC | Sapitinib | MVP_top60_DNA+RNA_seed | AKT1 | -0.173619 | BLCA | HT-1197 | Uprosertib |
| MVP_top60_DNA_seed | EP300 | 0.474652 | BLCA | 5637 | CPI-637 | MVP_top60_DNA+RNA_seed | MAPK1 | -0.17267 | BLCA | 5637 | SCH772984 |
| MVP_top60_DNA_seed | HDAC1 | 0.478699 | BLCA | TCCSUP | Romidepsin | MVP_top60_DNA+RNA_seed | MAPK3 | -0.17267 | BLCA | 5637 | SCH772984 |
| MVP_top60_DNA_seed | TP53 | 0.480846 | BLCA | T24 | MIRA-1 | MVP_top60_DNA+RNA_seed | AKT1 | -0.171751 | BLCA | CAL-29 | AZD5363 |
| MVP_top60_DNA_seed | ERBB2 | 0.482534 | BLCA | HT-1376 | Sapitinib | MVP_top60_DNA+RNA_seed | PIK3CA | -0.167433 | BLCA | RT-112 | Buparlisib |

| | | | | | | | | | | | |
|---|---|---|---|---|---|---|---|---|---|---|---|
| MVP_top60_DNA_seed | ERBB3 | 0.482534 | BLCA | HT-1376 | Sapitinib | MVP_top60_DNA+RNA_seed | ESR1 | -0.164304 | BLCA | RT-112 | Tamoxifen |
| MVP_top60_DNA_seed | ESR1 | 0.488574 | BLCA | RT4 | GDC0810 | MVP_top60_DNA+RNA_seed | AKT1 | -0.160602 | BLCA | KU-19-19 | Afuresertib |
| MVP_top60_DNA_seed | FGFR3 | 0.488856 | BLCA | HT-1376 | AZD4547 | MVP_top60_DNA+RNA_seed | HDAC1 | -0.157222 | BLCA | CAL-29 | Entinostat |
| MVP_top60_DNA_seed | CDK2 | 0.489465 | BLCA | HT-1197 | Dinaciclib | MVP_top60_DNA+RNA_seed | GSK3B | -0.15395 | BLCA | 639-V | SB216763 |
| MVP_top60_DNA_seed | PIK3CA | 0.491344 | BLCA | RT4 | Buparlisib | MVP_top60_DNA+RNA_seed | AKT1 | -0.15172 | BLCA | DSH1 | AZD5363 |
| MVP_top60_DNA_seed | PIK3CB | 0.491344 | BLCA | RT4 | Buparlisib | MVP_top60_DNA+RNA_seed | RAC1 | -0.146832 | BLCA | SW1710 | EHT-1864 |
| MVP_top60_DNA_seed | ESR1 | 0.496316 | BLCA | RT-112 | GDC0810 | MVP_top60_DNA+RNA_seed | PIK3CA | -0.141735 | BLCA | UM-UC-3 | AZD8186 |
| MVP_top60_DNA_seed | HDAC1 | 0.50013 | BLCA | RT-112 | PCI-34051 | MVP_top60_DNA+RNA_seed | TP53 | -0.130501 | BLCA | RT-112 | MIRA-1 |
| MVP_top60_DNA_seed | PIK3CA | 0.505059 | BLCA | TCCSUP | Alpelisib | MVP_top60_DNA+RNA_seed | AKT1 | -0.128144 | BLCA | KU-19-19 | Ipatasertib |
| MVP_top60_DNA_seed | ERBB2 | 0.508525 | BLCA | RT-112 | Lapatinib | MVP_top60_DNA+RNA_seed | FGFR3 | -0.12048 | BLCA | LB831-BLC | PD173074 |
| MVP_top60_DNA_seed | CDK2 | 0.514899 | BLCA | 5637 | Dinaciclib | MVP_top60_DNA+RNA_seed | FGFR3 | -0.120234 | BLCA | LB831-BLC | AZD4547 |
| MVP_top60_DNA_seed | MAPK1 | 0.515561 | BLCA | 647-V | Ulixertinib | MVP_top60_DNA+RNA_seed | PIK3CA | -0.11969 | BLCA | 647-V | GNE-317 |
| MVP_top60_DNA_seed | MAPK3 | 0.515561 | BLCA | 647-V | Ulixertinib | MVP_top60_DNA+RNA_seed | ESR1 | -0.118755 | BLCA | 647-V | Tamoxifen |
| MVP_top60_DNA_seed | TP53 | 0.520183 | BLCA | SW780 | MIRA-1 | MVP_top60_DNA+RNA_seed | AKT1 | -0.110779 | BLCA | KU-19-19 | GSK2110183B |
| MVP_top60_DNA_seed | TP53 | 0.520968 | BLCA | RT4 | MIRA-1 | MVP_top60_DNA+RNA_seed | PIK3CA | -0.110066 | BLCA | TCCSUP | Buparlisib |
| MVP_top60_DNA_seed | HSP90AA1 | 0.521502 | BLCA | TCCSUP | Luminespib | MVP_top60_DNA+RNA_seed | AKT1 | -0.108392 | BLCA | LB831-BLC | Uprosertib |
| MVP_top60_DNA_seed | ESR1 | 0.522589 | BLCA | TCCSUP | Tamoxifen | MVP_top60_DNA+RNA_seed | CDK2 | -0.103417 | BLCA | DSH1 | AZD5438 |
| MVP_top60_DNA_seed | CDK2 | 0.522928 | BLCA | TCCSUP | MK-8776 | MVP_top60_DNA+RNA_seed | ESR1 | -0.102027 | BLCA | J82 | GDC0810 |
| MVP_top60_DNA_seed | HDAC1 | 0.526095 | BLCA | T24 | Entinostat | MVP_top60_DNA+RNA_seed | PIK3CA | -0.101739 | BLCA | BFTC-905 | AZD8186 |
| MVP_top60_DNA_seed | CDK2 | 0.527036 | BLCA | T24 | Dinaciclib | MVP_top60_DNA+RNA_seed | MAPK1 | -0.092468 | BLCA | 639-V | Ulixertinib |
| MVP_top60_DNA_seed | HSP90AA1 | 0.5282 | BLCA | HT-1376 | Tanespimycin | MVP_top60_DNA+RNA_seed | MAPK3 | -0.092468 | BLCA | 639-V | Ulixertinib |
| MVP_top60_DNA_seed | HDAC1 | 0.528543 | BLCA | T24 | PCI-34051 | MVP_top60_DNA+RNA_seed | FGFR3 | -0.092348 | BLCA | KU-19-19 | PD173074 |
| MVP_top60_DNA_seed | CDK2 | 0.531484 | BLCA | 5637 | AZD5438 | MVP_top60_DNA+RNA_seed | SRC | -0.089659 | BLCA | HT-1197 | Bosutinib |
| MVP_top60_DNA_seed | ERBB2 | 0.532001 | BLCA | T24 | Sapitinib | MVP_top60_DNA+RNA_seed | PIK3CA | -0.087925 | BLCA | LB831-BLC | Alpelisib |
| MVP_top60_DNA_seed | ERBB3 | 0.532001 | BLCA | T24 | Sapitinib | MVP_top60_DNA+RNA_seed | HSP90AA1 | -0.083219 | BLCA | UM-UC-3 | Tanespimycin |
| MVP_top60_DNA_seed | HDAC1 | 0.536889 | BLCA | 647-V | Entinostat | MVP_top60_DNA+RNA_seed | ESR1 | -0.080712 | BLCA | 639-V | GDC0810 |
| MVP_top60_DNA_seed | MAPK1 | 0.537592 | BLCA | LB831-BLC | SCH772984 | MVP_top60_DNA+RNA_seed | ERBB2 | -0.074152 | BLCA | BFTC-905 | Sapitinib |
| MVP_top60_DNA_seed | MAPK3 | 0.537592 | BLCA | LB831-BLC | SCH772984 | MVP_top60_DNA+RNA_seed | ERBB3 | -0.074152 | BLCA | BFTC-905 | Sapitinib |
| MVP_top60_DNA_seed | EP300 | 0.543802 | BLCA | RT-112 | SGC-CBP30 | MVP_top60_DNA+RNA_seed | AKT1 | -0.07177 | BLCA | BFTC-905 | Ipatasertib |
| MVP_top60_DNA_seed | CREBBP | 0.543802 | BLCA | RT-112 | SGC-CBP30 | MVP_top60_DNA+RNA_seed | PIK3CA | -0.070269 | BLCA | DSH1 | Buparlisib |
| MVP_top60_DNA_seed | HSP90AA1 | 0.551154 | BLCA | HT-1376 | Luminespib | MVP_top60_DNA+RNA_seed | CDK2 | -0.069245 | BLCA | RT-112 | MK-8776 |
| MVP_top60_DNA_seed | HDAC1 | 0.554861 | BLCA | TCCSUP | PCI-34051 | MVP_top60_DNA+RNA_seed | CDK2 | -0.069157 | BLCA | DSH1 | MK-8776 |
| MVP_top60_DNA_seed | ESR1 | 0.555564 | BLCA | LB831-BLC | Tamoxifen | MVP_top60_DNA+RNA_seed | ERBB2 | -0.063729 | BLCA | 647-V | Afatinib |
| MVP_top60_DNA_seed | PIK3CA | 0.557464 | BLCA | RT-112 | AZD8186 | MVP_top60_DNA+RNA_seed | FGFR3 | -0.061481 | BLCA | 639-V | PD173074 |
| MVP_top60_DNA_seed | PIK3CB | 0.557464 | BLCA | RT-112 | AZD8186 | MVP_top60_DNA+RNA_seed | RAC1 | -0.052606 | BLCA | KU-19-19 | EHT-1864 |
| MVP_top60_DNA_seed | EP300 | 0.572369 | BLCA | J82 | CPI-637 | MVP_top60_DNA+RNA_seed | FGFR3 | -0.043731 | BLCA | DSH1 | PD173074 |
| MVP_top60_DNA_seed | CDK2 | 0.583289 | BLCA | CAL-29 | MK-8776 | MVP_top60_DNA+RNA_seed | HSP90AA1 | -0.041544 | BLCA | 5637 | Luminespib |
| MVP_top60_DNA_seed | SRC | 0.591106 | BLCA | 647-V | Bosutinib | MVP_top60_DNA+RNA_seed | AR | -0.040754 | BLCA | T24 | Bicalutamide |
| MVP_top60_DNA_seed | MAPK1 | 0.593041 | BLCA | HT-1376 | Ulixertinib | MVP_top60_DNA+RNA_seed | TERT | -0.031656 | BLCA | 5637 | BIBR-1532 |
| MVP_top60_DNA_seed | MAPK3 | 0.593041 | BLCA | HT-1376 | Ulixertinib | MVP_top60_DNA+RNA_seed | GSK3B | -0.03108 | BLCA | CAL-29 | SB216763 |
| MVP_top60_DNA_seed | MAPK1 | 0.59314 | BLCA | 5637 | Ulixertinib | MVP_top60_DNA+RNA_seed | ERBB2 | -0.030175 | BLCA | SW1710 | Lapatinib |

| | | | | | | | | | | | |
|---|---|---|---|---|---|---|---|---|---|---|---|
| MVP_top60_DNA_seed | MAPK3 | 0.59314 | BLCA | 5637 | Ulixertinib | MVP_top60_DNA+RNA_seed | ERBB2 | -0.026496 | BLCA | 639-V | Afatinib |
| MVP_top60_DNA_seed | PIK3CA | 0.599658 | BLCA | 639-V | Alpelisib | MVP_top60_DNA+RNA_seed | GSK3B | -0.021785 | BLCA | RT4 | SB216763 |
| MVP_top60_DNA_seed | MAPK1 | 0.603553 | BLCA | SW780 | Ulixertinib | MVP_top60_DNA+RNA_seed | GSK3B | -0.01428 | BLCA | BFTC-905 | SB216763 |
| MVP_top60_DNA_seed | MAPK3 | 0.603553 | BLCA | SW780 | Ulixertinib | MVP_top60_DNA+RNA_seed | PIK3CA | -0.013456 | BLCA | 647-V | Buparlisib |
| MVP_top60_DNA_seed | HDAC1 | 0.609223 | BLCA | HT-1197 | PCI-34051 | MVP_top60_DNA+RNA_seed | MAPK1 | -0.011556 | BLCA | SW780 | Ulixertinib |
| MVP_top60_DNA_seed | PIK3CB | 0.615372 | BLCA | T24 | AZD6482 | MVP_top60_DNA+RNA_seed | MAPK3 | -0.011556 | BLCA | SW780 | Ulixertinib |
| MVP_top60_DNA_seed | PIK3CA | 0.61902 | BLCA | LB831-BLC | Buparlisib | MVP_top60_DNA+RNA_seed | PIK3CA | -0.010391 | BLCA | 5637 | GNE-317 |
| MVP_top60_DNA_seed | PIK3CB | 0.61902 | BLCA | LB831-BLC | Buparlisib | MVP_top60_DNA+RNA_seed | RAC1 | -0.009372 | BLCA | LB831-BLC | EHT-1864 |
| MVP_top60_DNA_seed | ERBB2 | 0.623149 | BLCA | LB831-BLC | Afatinib | MVP_top60_DNA+RNA_seed | AKT1 | -0.00531 | BLCA | SW1710 | Uprosertib |
| MVP_top60_DNA_seed | ERBB2 | 0.626589 | BLCA | KU-19-19 | Sapitinib | MVP_top60_DNA+RNA_seed | PIK3CA | -0.005178 | BLCA | HT-1197 | Alpelisib |
| MVP_top60_DNA_seed | ERBB3 | 0.626589 | BLCA | KU-19-19 | Sapitinib | MVP_top60_DNA+RNA_seed | HSP90AA1 | -0.003759 | BLCA | T24 | Luminespib |
| MVP_top60_DNA_seed | HDAC1 | 0.64124 | BLCA | SW1710 | Entinostat | MVP_top60_DNA+RNA_seed | RAC1 | -0.002287 | BLCA | 647-V | EHT-1864 |
| MVP_top60_DNA_seed | KRAS | 0.649053 | BLCA | J82 | KRAS (G12C) Inhibitor-12 | MVP_top60_DNA+RNA_seed | AKT1 | -0.000916 | BLCA | LB831-BLC | AZD5363 |
| MVP_top60_DNA_seed | FGFR3 | 0.650917 | BLCA | HT-1197 | PD173074 | MVP_top60_DNA+RNA_seed | EP300 | 0.003175 | BLCA | 639-V | SGC-CBP30 |
| MVP_top60_DNA_seed | ERBB2 | 0.661388 | BLCA | SW1710 | Sapitinib | MVP_top60_DNA+RNA_seed | CREBBP | 0.003175 | BLCA | 639-V | SGC-CBP30 |
| MVP_top60_DNA_seed | ERBB3 | 0.661388 | BLCA | SW1710 | Sapitinib | MVP_top60_DNA+RNA_seed | CDK2 | 0.005737 | BLCA | CAL-29 | AZD5438 |
| MVP_top60_DNA_seed | HSP90AA1 | 0.669108 | BLCA | CAL-29 | Luminespib | MVP_top60_DNA+RNA_seed | HDAC1 | 0.006348 | BLCA | TCCSUP | Entinostat |
| MVP_top60_DNA_seed | TP53 | 0.670544 | BLCA | 5637 | MIRA-1 | MVP_top60_DNA+RNA_seed | EP300 | 0.007679 | BLCA | RT4 | CPI-637 |
| MVP_top60_DNA_seed | PIK3CA | 0.672557 | BLCA | TCCSUP | AZD8186 | MVP_top60_DNA+RNA_seed | PIK3CA | 0.010687 | BLCA | 5637 | Alpelisib |
| MVP_top60_DNA_seed | PIK3CB | 0.672557 | BLCA | TCCSUP | AZD8186 | MVP_top60_DNA+RNA_seed | ERBB2 | 0.01589 | BLCA | RT4 | Lapatinib |
| MVP_top60_DNA_seed | MAPK1 | 0.674563 | BLCA | T24 | VX-11e | MVP_top60_DNA+RNA_seed | ESR1 | 0.017008 | BLCA | 5637 | Tamoxifen |
| MVP_top60_DNA_seed | HDAC1 | 0.678048 | BLCA | CAL-29 | PCI-34051 | MVP_top60_DNA+RNA_seed | AKT1 | 0.017193 | BLCA | KU-19-19 | AZD5363 |
| MVP_top60_DNA_seed | ERBB2 | 0.689029 | BLCA | HT-1376 | Lapatinib | MVP_top60_DNA+RNA_seed | PIK3CA | 0.019742 | BLCA | UM-UC-3 | Buparlisib |
| MVP_top60_DNA_seed | ESR1 | 0.69589 | BLCA | LB831-BLC | GDC0810 | MVP_top60_DNA+RNA_seed | MAPK1 | 0.019759 | BLCA | 5637 | VX-11e |
| MVP_top60_DNA_seed | KRAS | 0.69703 | BLCA | 647-V | KRAS (G12C) Inhibitor-12 | MVP_top60_DNA+RNA_seed | AKT1 | 0.030888 | BLCA | LB831-BLC | Ipatasertib |
| MVP_top60_DNA_seed | TP53 | 0.706531 | BLCA | HT-1197 | MIRA-1 | MVP_top60_DNA+RNA_seed | HDAC1 | 0.032682 | BLCA | CAL-29 | Romidepsin |
| MVP_top60_DNA_seed | ERBB2 | 0.707956 | BLCA | TCCSUP | Sapitinib | MVP_top60_DNA+RNA_seed | ESR1 | 0.039152 | BLCA | UM-UC-3 | Tamoxifen |
| MVP_top60_DNA_seed | ERBB3 | 0.707956 | BLCA | TCCSUP | Sapitinib | MVP_top60_DNA+RNA_seed | SRC | 0.039951 | BLCA | HT-1376 | Bosutinib |
| MVP_top60_DNA_seed | MAPK1 | 0.715589 | BLCA | J82 | Ulixertinib | MVP_top60_DNA+RNA_seed | TP53 | 0.039981 | BLCA | KU-19-19 | MIRA-1 |
| MVP_top60_DNA_seed | MAPK3 | 0.715589 | BLCA | J82 | Ulixertinib | MVP_top60_DNA+RNA_seed | CDK2 | 0.04224 | BLCA | SW780 | AZD5438 |
| MVP_top60_DNA_seed | MAPK1 | 0.71866 | BLCA | 639-V | VX-11e | MVP_top60_DNA+RNA_seed | HDAC1 | 0.042626 | BLCA | LB831-BLC | PCI-34051 |
| MVP_top60_DNA_seed | TP53 | 0.720144 | BLCA | TCCSUP | MIRA-1 | MVP_top60_DNA+RNA_seed | HDAC1 | 0.043913 | BLCA | DSH1 | Romidepsin |
| MVP_top60_DNA_seed | MAPK1 | 0.725801 | BLCA | UM-UC-3 | SCH772984 | MVP_top60_DNA+RNA_seed | AKT1 | 0.044309 | BLCA | BFTC-905 | Afuresertib |
| MVP_top60_DNA_seed | MAPK3 | 0.725801 | BLCA | UM-UC-3 | SCH772984 | MVP_top60_DNA+RNA_seed | AKT1 | 0.0549 | BLCA | 639-V | Ipatasertib |
| MVP_top60_DNA_seed | ERBB2 | 0.726094 | BLCA | 639-V | Lapatinib | MVP_top60_DNA+RNA_seed | SRC | 0.057773 | BLCA | CAL-29 | Bosutinib |
| MVP_top60_DNA_seed | FGFR3 | 0.728333 | BLCA | T24 | PD173074 | MVP_top60_DNA+RNA_seed | GSK3B | 0.05915 | BLCA | TCCSUP | SB216763 |
| MVP_top60_DNA_seed | TP53 | 0.736355 | BLCA | LB831-BLC | MIRA-1 | MVP_top60_DNA+RNA_seed | ERBB2 | 0.065332 | BLCA | 647-V | Sapitinib |
| MVP_top60_DNA_seed | EP300 | 0.736911 | BLCA | 647-V | SGC-CBP30 | MVP_top60_DNA+RNA_seed | ERBB3 | 0.065332 | BLCA | 647-V | Sapitinib |
| MVP_top60_DNA_seed | CREBBP | 0.736911 | BLCA | 647-V | SGC-CBP30 | MVP_top60_DNA+RNA_seed | ERBB2 | 0.066548 | BLCA | RT4 | Sapitinib |
| MVP_top60_DNA_seed | ERBB2 | 0.757283 | BLCA | SW1710 | Afatinib | MVP_top60_DNA+RNA_seed | ERBB3 | 0.066548 | BLCA | RT4 | Sapitinib |
| MVP_top60_DNA_seed | CDK2 | 0.76688 | BLCA | 647-V | MK-8776 | MVP_top60_DNA+RNA_seed | CDK2 | 0.069504 | BLCA | SW780 | Dinaciclib |

| | | | | | | | | | | | |
|---|---|---|---|---|---|---|---|---|---|---|---|
| MVP_top60_DNA_seed | PIK3CA | 0.770447 | BLCA | RT4 | Alpelisib | MVP_top60_DNA+RNA_seed | PIK3CA | 0.070222 | BLCA | DSH1 | Alpelisib |
| MVP_top60_DNA_seed | EP300 | 0.772096 | BLCA | LB831-BLC | SGC-CBP30 | MVP_top60_DNA+RNA_seed | FGFR3 | 0.07172 | BLCA | BFTC-905 | PD173074 |
| MVP_top60_DNA_seed | CREBBP | 0.772096 | BLCA | LB831-BLC | SGC-CBP30 | MVP_top60_DNA+RNA_seed | FGFR3 | 0.072193 | BLCA | UM-UC-3 | AZD4547 |
| MVP_top60_DNA_seed | CDK2 | 0.773827 | BLCA | 639-V | MK-8776 | MVP_top60_DNA+RNA_seed | PIK3CA | 0.077521 | BLCA | 639-V | GNE-317 |
| MVP_top60_DNA_seed | ATM | 0.777271 | BLCA | SW1710 | KU-55933 | MVP_top60_DNA+RNA_seed | AKT1 | 0.078109 | BLCA | RT-112 | Afuresertib |
| MVP_top60_DNA_seed | TP53 | 0.778608 | BLCA | J82 | MIRA-1 | MVP_top60_DNA+RNA_seed | TP53 | 0.082077 | BLCA | CAL-29 | MIRA-1 |
| MVP_top60_DNA_seed | FGFR3 | 0.784985 | BLCA | HT-1197 | AZD4547 | MVP_top60_DNA+RNA_seed | MAPK1 | 0.082733 | BLCA | SW780 | SCH772984 |
| MVP_top60_DNA_seed | ERBB2 | 0.786125 | BLCA | UM-UC-3 | Afatinib | MVP_top60_DNA+RNA_seed | MAPK3 | 0.082733 | BLCA | SW780 | SCH772984 |
| MVP_top60_DNA_seed | PIK3CA | 0.807117 | BLCA | T24 | AZD8186 | MVP_top60_DNA+RNA_seed | AKT1 | 0.091251 | BLCA | SW1710 | Ipatasertib |
| MVP_top60_DNA_seed | PIK3CB | 0.807117 | BLCA | T24 | AZD8186 | MVP_top60_DNA+RNA_seed | ESR1 | 0.092829 | BLCA | SW1710 | GDC0810 |
| MVP_top60_DNA_seed | MAPK1 | 0.810133 | BLCA | 647-V | VX-11e | MVP_top60_DNA+RNA_seed | PIK3CA | 0.095867 | BLCA | SW1710 | Buparlisib |
| MVP_top60_DNA_seed | ERBB2 | 0.812624 | BLCA | HT-1197 | Afatinib | MVP_top60_DNA+RNA_seed | GSK3B | 0.100865 | BLCA | UM-UC-3 | SB216763 |
| MVP_top60_DNA_seed | ERBB2 | 0.814626 | BLCA | 647-V | Lapatinib | MVP_top60_DNA+RNA_seed | SRC | 0.104374 | BLCA | SW1710 | Bosutinib |
| MVP_top60_DNA_seed | ESR1 | 0.818888 | BLCA | J82 | Tamoxifen | MVP_top60_DNA+RNA_seed | CDK2 | 0.107665 | BLCA | LB831-BLC | MK-8776 |
| MVP_top60_DNA_seed | MAPK1 | 0.821146 | BLCA | HT-1376 | Ulixertinib | MVP_top60_DNA+RNA_seed | ATM | 0.111065 | BLCA | DSH1 | KU-55933 |
| MVP_top60_DNA_seed | MAPK3 | 0.821146 | BLCA | HT-1376 | Ulixertinib | MVP_top60_DNA+RNA_seed | TERT | 0.113943 | BLCA | KU-19-19 | BIBR-1532 |
| MVP_top60_DNA_seed | ATM | 0.843566 | BLCA | T24 | KU-55933 | MVP_top60_DNA+RNA_seed | FGFR3 | 0.114756 | BLCA | SW1710 | AZD4547 |
| MVP_top60_DNA_seed | RAC1 | 0.85142 | BLCA | HT-1376 | EHT-1864 | MVP_top60_DNA+RNA_seed | HDAC1 | 0.11586 | BLCA | 647-V | Romidepsin |
| MVP_top60_DNA_seed | HSP90AA1 | 0.852704 | BLCA | T24 | Elesclomol | MVP_top60_DNA+RNA_seed | TERT | 0.117448 | BLCA | SW1710 | BIBR-1532 |
| MVP_top60_DNA_seed | PIK3CA | 0.864754 | BLCA | KU-19-19 | AZD8186 | MVP_top60_DNA+RNA_seed | HSP90AA1 | 0.118117 | BLCA | 647-V | Tanespimycin |
| MVP_top60_DNA_seed | PIK3CB | 0.864754 | BLCA | KU-19-19 | AZD8186 | MVP_top60_DNA+RNA_seed | HDAC1 | 0.120188 | BLCA | 647-V | PCI-34051 |
| MVP_top60_DNA_seed | SRC | 0.868372 | BLCA | RT4 | Bosutinib | MVP_top60_DNA+RNA_seed | AKT1 | 0.121163 | BLCA | SW1710 | Afuresertib |
| MVP_top60_DNA_seed | EP300 | 0.883701 | BLCA | RT-112 | CPI-637 | MVP_top60_DNA+RNA_seed | MAPK1 | 0.122208 | BLCA | CAL-29 | Ulixertinib |
| MVP_top60_DNA_seed | HDAC1 | 0.886583 | BLCA | J82 | Entinostat | MVP_top60_DNA+RNA_seed | MAPK3 | 0.122208 | BLCA | CAL-29 | Ulixertinib |
| MVP_top60_DNA_seed | MAPK1 | 0.886949 | BLCA | UM-UC-3 | VX-11e | MVP_top60_DNA+RNA_seed | EP300 | 0.126142 | BLCA | 5637 | SGC-CBP30 |
| MVP_top60_DNA_seed | CDK2 | 0.904883 | BLCA | UM-UC-3 | MK-8776 | MVP_top60_DNA+RNA_seed | CREBBP | 0.126142 | BLCA | 5637 | SGC-CBP30 |
| MVP_top60_DNA_seed | SRC | 0.908303 | BLCA | LB831-BLC | Bosutinib | MVP_top60_DNA+RNA_seed | TERT | 0.129428 | BLCA | BFTC-905 | BIBR-1532 |
| MVP_top60_DNA_seed | ERBB2 | 0.923167 | BLCA | SW780 | Lapatinib | MVP_top60_DNA+RNA_seed | EP300 | 0.129522 | BLCA | SW1710 | CPI-637 |
| MVP_top60_DNA_seed | HDAC1 | 0.923567 | BLCA | J82 | PCI-34051 | MVP_top60_DNA+RNA_seed | ATM | 0.140815 | BLCA | UM-UC-3 | KU-55933 |
| MVP_top60_DNA_seed | RAC1 | 0.925929 | BLCA | TCCSUP | EHT-1864 | MVP_top60_DNA+RNA_seed | FGFR3 | 0.141684 | BLCA | J82 | PD173074 |
| MVP_top60_DNA_seed | PIK3CA | 0.933356 | BLCA | T24 | GNE-317 | MVP_top60_DNA+RNA_seed | MAPK1 | 0.142674 | BLCA | SW780 | VX-11e |
| MVP_top60_DNA_seed | MAPK1 | 0.943222 | BLCA | TCCSUP | Ulixertinib | MVP_top60_DNA+RNA_seed | ERBB2 | 0.142791 | BLCA | KU-19-19 | Lapatinib |
| MVP_top60_DNA_seed | MAPK3 | 0.943222 | BLCA | TCCSUP | Ulixertinib | MVP_top60_DNA+RNA_seed | MAPK1 | 0.147848 | BLCA | SW1710 | Ulixertinib |
| MVP_top60_DNA_seed | ESR1 | 0.954019 | BLCA | T24 | GDC0810 | MVP_top60_DNA+RNA_seed | MAPK3 | 0.147848 | BLCA | SW1710 | Ulixertinib |
| MVP_top60_DNA_seed | KRAS | 0.957842 | BLCA | HT-1197 | KRAS (G12C) Inhibitor-1[illegible] | MVP_top60_DNA+RNA_seed | FGFR3 | 0.149376 | BLCA | CAL-29 | PD173074 |
| MVP_top60_DNA_seed | ERBB2 | 0.957884 | BLCA | HT-1197 | Sapitinib | MVP_top60_DNA+RNA_seed | SYK | 0.150809 | BLCA | 5637 | PRT062607 |
| MVP_top60_DNA_seed | ERBB3 | 0.957884 | BLCA | HT-1197 | Sapitinib | MVP_top60_DNA+RNA_seed | HDAC1 | 0.152653 | BLCA | HT-1197 | Romidepsin |
| MVP_top60_DNA_seed | EP300 | 0.95824 | BLCA | TCCSUP | CPI-637 | MVP_top60_DNA+RNA_seed | AKT1 | 0.157412 | BLCA | BFTC-905 | Uprosertib |
| MVP_top60_DNA_seed | HSP90AA1 | 0.971267 | BLCA | HT-1197 | Tanespimycin | MVP_top60_DNA+RNA_seed | RAC1 | 0.159193 | BLCA | UM-UC-3 | EHT-1864 |
| MVP_top60_DNA_seed | ATM | 0.972707 | BLCA | SW780 | KU-55933 | MVP_top60_DNA+RNA_seed | EP300 | 0.159683 | BLCA | CAL-29 | SGC-CBP30 |
| MVP_top60_DNA_seed | ATM | 0.97483 | BLCA | TCCSUP | KU-55933 | MVP_top60_DNA+RNA_seed | CREBBP | 0.159683 | BLCA | CAL-29 | SGC-CBP30 |

| | | | | | | | | | | | |
|---|---|---|---|---|---|---|---|---|---|---|---|
| MVP_top60_DNA_seed | CDK2 | 0.982875 | BLCA | RT4 | MK-8776 | MVP_top60_DNA+RNA_seed | EP300 | 0.162684 | BLCA | HT-1376 | CPI-637 |
| MVP_top60_DNA_seed | TP53 | 0.988377 | BLCA | HT-1376 | MIRA-1 | MVP_top60_DNA+RNA_seed | EP300 | 0.163524 | BLCA | 647-V | CPI-637 |
| MVP_top60_DNA_seed | MAPK1 | 0.990776 | BLCA | T24 | Ulixertinib | MVP_top60_DNA+RNA_seed | HSP90AA1 | 0.163691 | BLCA | T24 | Tanespimycin |
| MVP_top60_DNA_seed | MAPK3 | 0.990776 | BLCA | T24 | Ulixertinib | MVP_top60_DNA+RNA_seed | TERT | 0.165043 | BLCA | CAL-29 | BIBR-1532 |
| MVP_top60_DNA_seed | MAPK1 | 0.995933 | BLCA | HT-1376 | SCH772984 | MVP_top60_DNA+RNA_seed | SYK | 0.166649 | BLCA | RT4 | Entospletinib |
| MVP_top60_DNA_seed | MAPK3 | 0.995933 | BLCA | HT-1376 | SCH772984 | MVP_top60_DNA+RNA_seed | SYK | 0.166801 | BLCA | SW780 | Entospletinib |
| MVP_top60_DNA_seed | MAPK1 | 0.998349 | BLCA | T24 | SCH772984 | MVP_top60_DNA+RNA_seed | PIK3CA | 0.16847 | BLCA | SW780 | GNE-317 |
| MVP_top60_DNA_seed | MAPK3 | 0.998349 | BLCA | T24 | SCH772984 | MVP_top60_DNA+RNA_seed | MAPK1 | 0.16983 | BLCA | CAL-29 | Ulixertinib |
| MVP_top60_DNA_seed | MAPK1 | 1.003831 | BLCA | SW1710 | Ulixertinib | MVP_top60_DNA+RNA_seed | MAPK3 | 0.16983 | BLCA | CAL-29 | Ulixertinib |
| MVP_top60_DNA_seed | MAPK3 | 1.003831 | BLCA | SW1710 | Ulixertinib | MVP_top60_DNA+RNA_seed | SYK | 0.16993 | BLCA | 639-V | Entospletinib |
| MVP_top60_DNA_seed | HDAC1 | 1.005992 | BLCA | LB831-BLC | Entinostat | MVP_top60_DNA+RNA_seed | TP53 | 0.170108 | BLCA | UM-UC-3 | MIRA-1 |
| MVP_top60_DNA_seed | ESR1 | 1.007142 | BLCA | SW780 | GDC0810 | MVP_top60_DNA+RNA_seed | HDAC1 | 0.172076 | BLCA | RT-112 | Entinostat |
| MVP_top60_DNA_seed | EP300 | 1.009242 | BLCA | UM-UC-3 | CPI-637 | MVP_top60_DNA+RNA_seed | AKT1 | 0.182792 | BLCA | RT-112 | GSK2110183B |
| MVP_top60_DNA_seed | MAPK1 | 1.022648 | BLCA | J82 | VX-11e | MVP_top60_DNA+RNA_seed | MAPK1 | 0.186558 | BLCA | SW1710 | SCH772984 |
| MVP_top60_DNA_seed | PIK3CA | 1.024332 | BLCA | J82 | AZD8186 | MVP_top60_DNA+RNA_seed | MAPK3 | 0.186558 | BLCA | SW1710 | SCH772984 |
| MVP_top60_DNA_seed | PIK3CB | 1.024332 | BLCA | J82 | AZD8186 | MVP_top60_DNA+RNA_seed | AKT1 | 0.188141 | BLCA | RT-112 | MK-2206 |
| MVP_top60_DNA_seed | KRAS | 1.033816 | BLCA | SW1710 | KRAS (G12C) Inhibitor-12 | MVP_top60_DNA+RNA_seed | FGFR3 | 0.194693 | BLCA | BFTC-905 | AZD4547 |
| MVP_top60_DNA_seed | CDK2 | 1.04764 | BLCA | SW1710 | MK-8776 | MVP_top60_DNA+RNA_seed | ERBB2 | 0.19596 | BLCA | KU-19-19 | Afatinib |
| MVP_top60_DNA_seed | PIK3CA | 1.077254 | BLCA | KU-19-19 | Alpelisib | MVP_top60_DNA+RNA_seed | SYK | 0.198086 | BLCA | 5637 | Entospletinib |
| MVP_top60_DNA_seed | CDK2 | 1.078331 | BLCA | TCCSUP | Dinaciclib | MVP_top60_DNA+RNA_seed | HSP90AA1 | 0.200704 | BLCA | SW780 | Tanespimycin |
| MVP_top60_DNA_seed | CDK2 | 1.078401 | BLCA | SW780 | MK-8776 | MVP_top60_DNA+RNA_seed | AKT1 | 0.200817 | BLCA | 639-V | AZD5363 |
| MVP_top60_DNA_seed | MAPK1 | 1.091312 | BLCA | LB831-BLC | Ulixertinib | MVP_top60_DNA+RNA_seed | FGFR3 | 0.201579 | BLCA | 647-V | PD173074 |
| MVP_top60_DNA_seed | MAPK3 | 1.091312 | BLCA | LB831-BLC | Ulixertinib | MVP_top60_DNA+RNA_seed | CDK2 | 0.201717 | BLCA | 647-V | Dinaciclib |
| MVP_top60_DNA_seed | ERBB2 | 1.0985 | BLCA | J82 | Sapitinib | MVP_top60_DNA+RNA_seed | AKT1 | 0.202145 | BLCA | 5637 | Uprosertib |
| MVP_top60_DNA_seed | ERBB3 | 1.0985 | BLCA | J82 | Sapitinib | MVP_top60_DNA+RNA_seed | HSP90AA1 | 0.202892 | BLCA | SW1710 | Tanespimycin |
| MVP_top60_DNA_seed | HDAC1 | 1.100254 | BLCA | 5637 | PCI-34051 | MVP_top60_DNA+RNA_seed | CDK2 | 0.205152 | BLCA | KU-19-19 | MK-8776 |
| MVP_top60_DNA_seed | PIK3CA | 1.103866 | BLCA | T24 | Alpelisib | MVP_top60_DNA+RNA_seed | TP53 | 0.209842 | BLCA | DSH1 | MIRA-1 |
| MVP_top60_DNA_seed | MAPK1 | 1.104656 | BLCA | SW1710 | VX-11e | MVP_top60_DNA+RNA_seed | HDAC1 | 0.211596 | BLCA | SW1710 | PCI-34051 |
| MVP_top60_DNA_seed | PIK3CA | 1.110764 | BLCA | RT4 | GNE-317 | MVP_top60_DNA+RNA_seed | MAPK1 | 0.216558 | BLCA | 639-V | Ulixertinib |
| MVP_top60_DNA_seed | MAPK1 | 1.111692 | BLCA | J82 | SCH772984 | MVP_top60_DNA+RNA_seed | MAPK3 | 0.216558 | BLCA | 639-V | Ulixertinib |
| MVP_top60_DNA_seed | MAPK3 | 1.111692 | BLCA | J82 | SCH772984 | MVP_top60_DNA+RNA_seed | EP300 | 0.219325 | BLCA | LB831-BLC | CPI-637 |
| MVP_top60_DNA_seed | PIK3CA | 1.13296 | BLCA | SW1710 | Alpelisib | MVP_top60_DNA+RNA_seed | PIK3CA | 0.220832 | BLCA | SW1710 | GNE-317 |
| MVP_top60_DNA_seed | KRAS | 1.133926 | BLCA | LB831-BLC | KRAS (G12C) Inhibitor-12 | MVP_top60_DNA+RNA_seed | FGFR3 | 0.225976 | BLCA | KU-19-19 | AZD4547 |
| MVP_top60_DNA_seed | ERBB2 | 1.137594 | BLCA | LB831-BLC | Lapatinib | MVP_top60_DNA+RNA_seed | HDAC1 | 0.235783 | BLCA | 639-V | PCI-34051 |
| MVP_top60_DNA_seed | FGFR3 | 1.141148 | BLCA | UM-UC-3 | PD173074 | MVP_top60_DNA+RNA_seed | RAC1 | 0.241462 | BLCA | 5637 | EHT-1864 |
| MVP_top60_DNA_seed | ERBB2 | 1.143819 | BLCA | J82 | Lapatinib | MVP_top60_DNA+RNA_seed | HSP90AA1 | 0.247158 | BLCA | SW1710 | Luminespib |
| MVP_top60_DNA_seed | MAPK1 | 1.150021 | BLCA | LB831-BLC | VX-11e | MVP_top60_DNA+RNA_seed | AKT1 | 0.248396 | BLCA | KU-19-19 | Uprosertib |
| MVP_top60_DNA_seed | MAPK1 | 1.159215 | BLCA | HT-1197 | Ulixertinib | MVP_top60_DNA+RNA_seed | SRC | 0.252369 | BLCA | 639-V | Bosutinib |
| MVP_top60_DNA_seed | MAPK3 | 1.159215 | BLCA | HT-1197 | Ulixertinib | MVP_top60_DNA+RNA_seed | ESR1 | 0.254018 | BLCA | 647-V | GDC0810 |
| MVP_top60_DNA_seed | SRC | 1.161011 | BLCA | SW780 | Bosutinib | MVP_top60_DNA+RNA_seed | PIK3CA | 0.255993 | BLCA | LB831-BLC | GNE-317 |
| MVP_top60_DNA_seed | ERBB2 | 1.171975 | BLCA | J82 | Afatinib | MVP_top60_DNA+RNA_seed | SYK | 0.259331 | BLCA | 647-V | Entospletinib |

| | | | | | | | | | | | |
|---|---|---|---|---|---|---|---|---|---|---|---|
| MVP_top60_DNA_seed | ATM | 1.190943 | BLCA | J82 | KU-55933 | MVP_top60_DNA+RNA_seed | GSK3B | 0.264377 | BLCA | RT-112 | SB216763 |
| MVP_top60_DNA_seed | ERBB2 | 1.197309 | BLCA | SW780 | Sapitinib | MVP_top60_DNA+RNA_seed | SRC | 0.270978 | BLCA | T24 | Bosutinib |
| MVP_top60_DNA_seed | ERBB3 | 1.197309 | BLCA | SW780 | Sapitinib | MVP_top60_DNA+RNA_seed | AKT1 | 0.274371 | BLCA | SW1710 | MK-2206 |
| MVP_top60_DNA_seed | ERBB2 | 1.211017 | BLCA | UM-UC-3 | Lapatinib | MVP_top60_DNA+RNA_seed | AKT1 | 0.277424 | BLCA | RT4 | Uprosertib |
| MVP_top60_DNA_seed | CDK2 | 1.237529 | BLCA | TCCSUP | AZD5438 | MVP_top60_DNA+RNA_seed | ERBB2 | 0.28053 | BLCA | 639-V | Sapitinib |
| MVP_top60_DNA_seed | SRC | 1.238383 | BLCA | J82 | Bosutinib | MVP_top60_DNA+RNA_seed | ERBB3 | 0.28053 | BLCA | 639-V | Sapitinib |
| MVP_top60_DNA_seed | CDK2 | 1.241545 | BLCA | HT-1197 | MK-8776 | MVP_top60_DNA+RNA_seed | HDAC1 | 0.283604 | BLCA | 639-V | Romidepsin |
| MVP_top60_DNA_seed | ATM | 1.25491 | BLCA | HT-1197 | KU-55933 | MVP_top60_DNA+RNA_seed | FGFR3 | 0.29218 | BLCA | SW1710 | PD173074 |
| MVP_top60_DNA_seed | PIK3CA | 1.261823 | BLCA | 647-V | Alpelisib | MVP_top60_DNA+RNA_seed | AKT1 | 0.295047 | BLCA | BFTC-905 | AT13148 |
| MVP_top60_DNA_seed | ERBB2 | 1.262369 | BLCA | SW780 | Afatinib | MVP_top60_DNA+RNA_seed | CDK2 | 0.297523 | BLCA | 647-V | AZD5438 |
| MVP_top60_DNA_seed | MAPK1 | 1.263677 | BLCA | UM-UC-3 | Ulixertinib | MVP_top60_DNA+RNA_seed | MAPK1 | 0.31246 | BLCA | 647-V | Ulixertinib |
| MVP_top60_DNA_seed | MAPK3 | 1.263677 | BLCA | UM-UC-3 | Ulixertinib | MVP_top60_DNA+RNA_seed | MAPK3 | 0.31246 | BLCA | 647-V | Ulixertinib |
| MVP_top60_DNA_seed | ATM | 1.269779 | BLCA | HT-1376 | KU-55933 | MVP_top60_DNA+RNA_seed | TERT | 0.318581 | BLCA | T24 | BIBR-1532 |
| MVP_top60_DNA_seed | PIK3CA | 1.29491 | BLCA | UM-UC-3 | Alpelisib | MVP_top60_DNA+RNA_seed | HSP90AA1 | 0.320718 | BLCA | 647-V | Luminespib |
| MVP_top60_DNA_seed | SRC | 1.296924 | BLCA | TCCSUP | Bosutinib | MVP_top60_DNA+RNA_seed | FGFR3 | 0.326347 | BLCA | DSH1 | AZD4547 |
| MVP_top60_DNA_seed | HDAC1 | 1.297524 | BLCA | SW1710 | Romidepsin | MVP_top60_DNA+RNA_seed | MAPK1 | 0.334225 | BLCA | RT4 | Ulixertinib |
| MVP_top60_DNA_seed | HSP90AA1 | 1.303137 | BLCA | HT-1197 | Luminespib | MVP_top60_DNA+RNA_seed | MAPK3 | 0.334225 | BLCA | RT4 | Ulixertinib |
| MVP_top60_DNA_seed | MAPK1 | 1.303322 | BLCA | HT-1197 | SCH772984 | MVP_top60_DNA+RNA_seed | KRAS | 0.335212 | BLCA | T24 | :RAS (G12C) Inhibitor-1 |
| MVP_top60_DNA_seed | MAPK3 | 1.303322 | BLCA | HT-1197 | SCH772984 | MVP_top60_DNA+RNA_seed | PIK3CA | 0.336452 | BLCA | LB831-BLC | AZD8186 |
| MVP_top60_DNA_seed | PIK3CA | 1.309147 | BLCA | J82 | Alpelisib | MVP_top60_DNA+RNA_seed | ESR1 | 0.338352 | BLCA | HT-1197 | GDC0810 |
| MVP_top60_DNA_seed | ERBB2 | 1.311765 | BLCA | TCCSUP | Afatinib | MVP_top60_DNA+RNA_seed | SRC | 0.342638 | BLCA | UM-UC-3 | Bosutinib |
| MVP_top60_DNA_seed | CDK2 | 1.322245 | BLCA | HT-1197 | AZD5438 | MVP_top60_DNA+RNA_seed | MAPK1 | 0.343628 | BLCA | 639-V | SCH772984 |
| MVP_top60_DNA_seed | ERBB2 | 1.332755 | BLCA | T24 | Lapatinib | MVP_top60_DNA+RNA_seed | MAPK3 | 0.343628 | BLCA | 639-V | SCH772984 |
| MVP_top60_DNA_seed | KRAS | 1.344899 | BLCA | TCCSUP | KRAS (G12C) Inhibitor-1: | MVP_top60_DNA+RNA_seed | EP300 | 0.343881 | BLCA | SW780 | CPI-637 |
| MVP_top60_DNA_seed | HDAC1 | 1.357084 | BLCA | SW780 | Romidepsin | MVP_top60_DNA+RNA_seed | AKT1 | 0.344699 | BLCA | BFTC-905 | GSK2110183B |
| MVP_top60_DNA_seed | HDAC1 | 1.39272 | BLCA | 639-V | Entinostat | MVP_top60_DNA+RNA_seed | GSK3B | 0.348467 | BLCA | HT-1376 | SB216763 |
| MVP_top60_DNA_seed | PIK3CA | 1.405059 | BLCA | SW780 | Alpelisib | MVP_top60_DNA+RNA_seed | GSK3B | 0.348599 | BLCA | J82 | SB216763 |
| MVP_top60_DNA_seed | ERBB2 | 1.405351 | BLCA | HT-1376 | Afatinib | MVP_top60_DNA+RNA_seed | PIK3CA | 0.349232 | BLCA | SW1710 | AZD8186 |
| MVP_top60_DNA_seed | FGFR3 | 1.427875 | BLCA | HT-1376 | PD173074 | MVP_top60_DNA+RNA_seed | CDK2 | 0.35568 | BLCA | SW1710 | AZD5438 |
| MVP_top60_DNA_seed | ERBB2 | 1.483823 | BLCA | HT-1197 | Lapatinib | MVP_top60_DNA+RNA_seed | AKT1 | 0.363026 | BLCA | UM-UC-3 | Ipatasertib |
| MVP_top60_DNA_seed | ESR1 | 1.491866 | BLCA | T24 | Tamoxifen | MVP_top60_DNA+RNA_seed | FGFR3 | 0.363163 | BLCA | TCCSUP | AZD4547 |
| MVP_top60_DNA_seed | EP300 | 1.51147 | BLCA | HT-1197 | SGC-CBP30 | MVP_top60_DNA+RNA_seed | HDAC1 | 0.363548 | BLCA | SW780 | Entinostat |
| MVP_top60_DNA_seed | CREBBP | 1.51147 | BLCA | HT-1197 | SGC-CBP30 | MVP_top60_DNA+RNA_seed | AKT1 | 0.372025 | BLCA | UM-UC-3 | AT13148 |
| MVP_top60_DNA_seed | MAPK1 | 1.518474 | BLCA | UM-UC-3 | Ulixertinib | MVP_top60_DNA+RNA_seed | CDK2 | 0.376723 | BLCA | 5637 | MK-8776 |
| MVP_top60_DNA_seed | MAPK3 | 1.518474 | BLCA | UM-UC-3 | Ulixertinib | MVP_top60_DNA+RNA_seed | PIK3CA | 0.380056 | BLCA | DSH1 | AZD8186 |
| MVP_top60_DNA_seed | MAPK1 | 1.563748 | BLCA | J82 | Ulixertinib | MVP_top60_DNA+RNA_seed | TP53 | 0.382708 | BLCA | 647-V | MIRA-1 |
| MVP_top60_DNA_seed | MAPK3 | 1.563748 | BLCA | J82 | Ulixertinib | MVP_top60_DNA+RNA_seed | MAPK1 | 0.385978 | BLCA | 647-V | SCH772984 |
| MVP_top60_DNA_seed | MAPK1 | 1.565737 | BLCA | TCCSUP | Ulixertinib | MVP_top60_DNA+RNA_seed | MAPK3 | 0.385978 | BLCA | 647-V | SCH772984 |
| MVP_top60_DNA_seed | MAPK3 | 1.565737 | BLCA | TCCSUP | Ulixertinib | MVP_top60_DNA+RNA_seed | SYK | 0.396566 | BLCA | TCCSUP | Entospletinib |
| MVP_top60_DNA_seed | HDAC1 | 1.565886 | BLCA | RT4 | Entinostat | MVP_top60_DNA+RNA_seed | SRC | 0.397445 | BLCA | RT-112 | Bosutinib |
| MVP_top60_DNA_seed | MAPK1 | 1.618588 | BLCA | HT-1197 | VX-11e | MVP_top60_DNA+RNA_seed | CDK2 | 0.397772 | BLCA | HT-1376 | MK-8776 |

| | | | | | | | | | | | |
|---|---|---|---|---|---|---|---|---|---|---|---|
| MVP_top60_DNA_seed | HSP90AA1 | 1.649132 | BLCA | J82 | Tanespimycin | MVP_top60_DNA+RNA_seed | KRAS | 0.399391 | BLCA | SW780 | KRAS (G12C) Inhibitor-1 |
| MVP_top60_DNA_seed | PIK3CA | 1.680588 | BLCA | T24 | Buparlisib | MVP_top60_DNA+RNA_seed | ATM | 0.400572 | BLCA | CAL-29 | KU-55933 |
| MVP_top60_DNA_seed | PIK3CB | 1.680588 | BLCA | T24 | Buparlisib | MVP_top60_DNA+RNA_seed | AKT1 | 0.403904 | BLCA | 647-V | AZD5363 |
| MVP_top60_DNA_seed | MAPK1 | 1.699284 | BLCA | TCCSUP | SCH772984 | MVP_top60_DNA+RNA_seed | PIK3CA | 0.404888 | BLCA | 5637 | AZD8186 |
| MVP_top60_DNA_seed | MAPK3 | 1.699284 | BLCA | TCCSUP | SCH772984 | MVP_top60_DNA+RNA_seed | HDAC1 | 0.413044 | BLCA | HT-1197 | Entinostat |
| MVP_top60_DNA_seed | PIK3CA | 1.706332 | BLCA | RT4 | AZD8186 | MVP_top60_DNA+RNA_seed | HDAC1 | 0.413164 | BLCA | DSH1 | Entinostat |
| MVP_top60_DNA_seed | PIK3CB | 1.706332 | BLCA | RT4 | AZD8186 | MVP_top60_DNA+RNA_seed | CDK2 | 0.414545 | BLCA | BFTC-905 | MK-8776 |
| MVP_top60_DNA_seed | PIK3CA | 1.715415 | BLCA | HT-1376 | GNE-317 | MVP_top60_DNA+RNA_seed | SYK | 0.414892 | BLCA | SW780 | PRT062607 |
| MVP_top60_DNA_seed | PIK3CA | 1.716928 | BLCA | SW780 | AZD8186 | MVP_top60_DNA+RNA_seed | FGFR3 | 0.416917 | BLCA | 5637 | PD173074 |
| MVP_top60_DNA_seed | PIK3CB | 1.716928 | BLCA | SW780 | AZD8186 | MVP_top60_DNA+RNA_seed | TP53 | 0.417231 | BLCA | SW1710 | MIRA-1 |
| MVP_top60_DNA_seed | PIK3CA | 1.756588 | BLCA | HT-1376 | Alpelisib | MVP_top60_DNA+RNA_seed | CDK2 | 0.420146 | BLCA | SW1710 | Dinaciclib |
| MVP_top60_DNA_seed | CDK2 | 1.765449 | BLCA | J82 | AZD5438 | MVP_top60_DNA+RNA_seed | AKT1 | 0.420243 | BLCA | BFTC-905 | Uprosertib |
| MVP_top60_DNA_seed | ERBB2 | 1.799722 | BLCA | T24 | Afatinib | MVP_top60_DNA+RNA_seed | HSP90AA1 | 0.423604 | BLCA | T24 | CCT-018159 |
| MVP_top60_DNA_seed | FGFR3 | 1.805725 | BLCA | T24 | AZD4547 | MVP_top60_DNA+RNA_seed | RAC1 | 0.423903 | BLCA | HT-1197 | EHT-1864 |
| MVP_top60_DNA_seed | FGFR3 | 1.826144 | BLCA | TCCSUP | PD173074 | MVP_top60_DNA+RNA_seed | AKT1 | 0.424804 | BLCA | RT-112 | Ipatasertib |
| MVP_top60_DNA_seed | PIK3CA | 1.951983 | BLCA | J82 | Buparlisib | MVP_top60_DNA+RNA_seed | ERBB2 | 0.424951 | BLCA | UM-UC-3 | Sapitinib |
| MVP_top60_DNA_seed | PIK3CB | 1.951983 | BLCA | J82 | Buparlisib | MVP_top60_DNA+RNA_seed | ERBB3 | 0.424951 | BLCA | UM-UC-3 | Sapitinib |
| MVP_top60_DNA_seed | HSP90AA1 | 1.968166 | BLCA | SW780 | Luminespib | MVP_top60_DNA+RNA_seed | CDK2 | 0.428728 | BLCA | J82 | MK-8776 |
| MVP_top60_DNA_seed | HSP90AA1 | 2.011633 | BLCA | J82 | Luminespib | MVP_top60_DNA+RNA_seed | AKT1 | 0.435136 | BLCA | BFTC-905 | AZD5363 |
| MVP_top60_DNA_seed | HDAC1 | 2.059134 | BLCA | RT4 | Romidepsin | MVP_top60_DNA+RNA_seed | PIK3CA | 0.435335 | BLCA | HT-1376 | AZD8186 |
| MVP_top60_DNA_seed | ERBB2 | 2.060922 | BLCA | TCCSUP | Lapatinib | MVP_top60_DNA+RNA_seed | ERBB2 | 0.441624 | BLCA | RT-112 | Afatinib |
| MVP_top60_DNA_seed | HSP90AA1 | 2.071933 | BLCA | RT4 | Luminespib | MVP_top60_DNA+RNA_seed | ERBB2 | 0.442474 | BLCA | RT-112 | Sapitinib |
| MVP_top60_DNA_seed | RAC1 | 2.108204 | BLCA | J82 | EHT-1864 | MVP_top60_DNA+RNA_seed | ERBB3 | 0.442474 | BLCA | RT-112 | Sapitinib |
| MVP_top60_DNA_seed | MAPK1 | 2.116787 | BLCA | HT-1376 | VX-11e | MVP_top60_DNA+RNA_seed | PIK3CA | 0.44544 | BLCA | SW780 | Buparlisib |
| MVP_top60_DNA_seed | MAPK1 | 2.238414 | BLCA | TCCSUP | VX-11e | MVP_top60_DNA+RNA_seed | AKT1 | 0.445614 | BLCA | 5637 | AZD5363 |
| MVP_top60_DNA_seed | PIK3CA | 2.716316 | BLCA | HT-1376 | Buparlisib | MVP_top60_DNA+RNA_seed | GSK3B | 0.445743 | BLCA | SW1710 | SB216763 |
| MVP_top60_DNA_seed | PIK3CB | 2.716316 | BLCA | HT-1376 | Buparlisib | MVP_top60_DNA+RNA_seed | AKT1 | 0.44611 | BLCA | TCCSUP | GSK2110183B |
| MVP_top60_DNA_seed | CDK2 | 2.72311 | BLCA | HT-1376 | AZD5438 | MVP_top60_DNA+RNA_seed | ESR1 | 0.446831 | BLCA | SW780 | Tamoxifen |
| MVP_top60_DNA_seed | CDK2 | 2.922617 | BLCA | J82 | Dinaciclib | MVP_top60_DNA+RNA_seed | ERBB2 | 0.446839 | BLCA | RT4 | Afatinib |
| MVP_top60_DNA_seed | PIK3CA | 2.971117 | BLCA | J82 | GNE-317 | MVP_top60_DNA+RNA_seed | AKT1 | 0.448445 | BLCA | T24 | AT13148 |
| MVP_top60_DNA_seed | HDAC1 | 3.488344 | BLCA | J82 | Romidepsin | MVP_top60_DNA+RNA_seed | AKT1 | 0.449686 | BLCA | LB831-BLC | Afuresertib |
| MVP_top60_DNA_seed | ESR1 | -6.409629 | BRCA | Hs-578-T | Fulvestrant | MVP_top60_DNA+RNA_seed | AKT1 | 0.457008 | BLCA | TCCSUP | Uprosertib |
| MVP_top60_DNA_seed | ERBB2 | -6.352842 | BRCA | HCC2218 | CP724714 | MVP_top60_DNA+RNA_seed | EP300 | 0.457723 | BLCA | CAL-29 | CPI-637 |
| MVP_top60_DNA_seed | ERBB2 | -5.634798 | BRCA | BT-474 | CP724714 | MVP_top60_DNA+RNA_seed | HDAC1 | 0.458762 | BLCA | BFTC-905 | Entinostat |
| MVP_top60_DNA_seed | ERBB2 | -4.8899 | BRCA | OCUB-M | Lapatinib | MVP_top60_DNA+RNA_seed | ATM | 0.459207 | BLCA | RT4 | KU-55933 |
| MVP_top60_DNA_seed | EGFR | -4.8899 | BRCA | OCUB-M | Lapatinib | MVP_top60_DNA+RNA_seed | CDK2 | 0.460578 | BLCA | T24 | MK-8776 |
| MVP_top60_DNA_seed | ERBB2 | -4.736905 | BRCA | HCC2218 | Lapatinib | MVP_top60_DNA+RNA_seed | AKT1 | 0.461441 | BLCA | 639-V | Uprosertib |
| MVP_top60_DNA_seed | EGFR | -4.736905 | BRCA | HCC2218 | Lapatinib | MVP_top60_DNA+RNA_seed | HDAC1 | 0.462096 | BLCA | SW780 | PCI-34051 |
| MVP_top60_DNA_seed | ERBB2 | -4.54251 | BRCA | BT-474 | PF-00299804 | MVP_top60_DNA+RNA_seed | ESR1 | 0.464937 | BLCA | TCCSUP | GDC0810 |
| MVP_top60_DNA_seed | EGFR | -4.54251 | BRCA | BT-474 | PF-00299804 | MVP_top60_DNA+RNA_seed | ATM | 0.468521 | BLCA | RT-112 | KU-55933 |
| MVP_top60_DNA_seed | ESR1 | -4.456821 | BRCA | CAMA-1 | Fulvestrant | MVP_top60_DNA+RNA_seed | ERBB2 | 0.470151 | BLCA | LB831-BLC | Sapitinib |

| | | | | | | | | | | | |
|---|---|---|---|---|---|---|---|---|---|---|---|
| MVP_top60_DNA_seed | ERBB2 | -4.30329 | BRCA | MDA-MB-175-VII | CP724714 | MVP_top60_DNA+RNA_seed | ERBB3 | 0.470151 | BLCA | LB831-BLC | Sapitinib |
| MVP_top60_DNA_seed | EGFR | -4.118491 | BRCA | BT-474 | AST-1306 | MVP_top60_DNA+RNA_seed | EP300 | 0.474652 | BLCA | 5637 | CPI-637 |
| MVP_top60_DNA_seed | ERBB2 | -4.04302 | BRCA | AU565 | PF-00299804 | MVP_top60_DNA+RNA_seed | HDAC1 | 0.478699 | BLCA | TCCSUP | Romidepsin |
| MVP_top60_DNA_seed | EGFR | -4.04302 | BRCA | AU565 | PF-00299804 | MVP_top60_DNA+RNA_seed | TP53 | 0.480846 | BLCA | T24 | MIRA-1 |
| MVP_top60_DNA_seed | ERBB2 | -3.927648 | BRCA | BT-474 | CI-1033 | MVP_top60_DNA+RNA_seed | ERBB2 | 0.482534 | BLCA | HT-1376 | Sapitinib |
| MVP_top60_DNA_seed | EGFR | -3.927648 | BRCA | BT-474 | CI-1033 | MVP_top60_DNA+RNA_seed | ERBB3 | 0.482534 | BLCA | HT-1376 | Sapitinib |
| MVP_top60_DNA_seed | ERBB2 | -3.805563 | BRCA | OCUB-M | CP724714 | MVP_top60_DNA+RNA_seed | AKT1 | 0.482751 | BLCA | 639-V | Afuresertib |
| MVP_top60_DNA_seed | EGFR | -3.767342 | BRCA | AU565 | AST-1306 | MVP_top60_DNA+RNA_seed | TERT | 0.486949 | BLCA | RT4 | BIBR-1532 |
| MVP_top60_DNA_seed | ESR1 | -3.509161 | BRCA | T47D | Tamoxifen | MVP_top60_DNA+RNA_seed | AKT1 | 0.487589 | BLCA | LB831-BLC | GSK2110183B |
| MVP_top60_DNA_seed | ERBB2 | -3.449584 | BRCA | AU565 | CI-1033 | MVP_top60_DNA+RNA_seed | ESR1 | 0.488574 | BLCA | RT4 | GDC0810 |
| MVP_top60_DNA_seed | EGFR | -3.449584 | BRCA | AU565 | CI-1033 | MVP_top60_DNA+RNA_seed | FGFR3 | 0.488856 | BLCA | HT-1376 | AZD4547 |
| MVP_top60_DNA_seed | AKT1 | -3.350027 | BRCA | OCUB-M | GSK690693 | MVP_top60_DNA+RNA_seed | CDK2 | 0.489465 | BLCA | HT-1197 | Dinaciclib |
| MVP_top60_DNA_seed | MAPK1 | -3.312718 | BRCA | DU-4475 | VX-11e | MVP_top60_DNA+RNA_seed | PIK3CA | 0.491344 | BLCA | RT4 | Buparlisib |
| MVP_top60_DNA_seed | ERBB2 | -3.307809 | BRCA | MDA-MB-175-VII | PF-00299804 | MVP_top60_DNA+RNA_seed | ESR1 | 0.496316 | BLCA | RT-112 | GDC0810 |
| MVP_top60_DNA_seed | EGFR | -3.307809 | BRCA | MDA-MB-175-VII | PF-00299804 | MVP_top60_DNA+RNA_seed | HDAC1 | 0.50013 | BLCA | RT-112 | PCI-34051 |
| MVP_top60_DNA_seed | ESR1 | -3.268441 | BRCA | MDA-MB-361 | Fulvestrant | MVP_top60_DNA+RNA_seed | GSK3B | 0.502878 | BLCA | SW780 | SB216763 |
| MVP_top60_DNA_seed | ERBB2 | -3.245884 | BRCA | HCC2218 | PF-00299804 | MVP_top60_DNA+RNA_seed | PIK3CA | 0.505059 | BLCA | TCCSUP | Alpelisib |
| MVP_top60_DNA_seed | EGFR | -3.245884 | BRCA | HCC2218 | PF-00299804 | MVP_top60_DNA+RNA_seed | ERBB2 | 0.508525 | BLCA | RT-112 | Lapatinib |
| MVP_top60_DNA_seed | ERBB2 | -3.213861 | BRCA | BT-474 | Lapatinib | MVP_top60_DNA+RNA_seed | AKT1 | 0.509973 | BLCA | RT-112 | Uprosertib |
| MVP_top60_DNA_seed | EGFR | -3.213861 | BRCA | BT-474 | Lapatinib | MVP_top60_DNA+RNA_seed | CDK2 | 0.514899 | BLCA | 5637 | Dinaciclib |
| MVP_top60_DNA_seed | EGFR | -3.106854 | BRCA | HCC2218 | AST-1306 | MVP_top60_DNA+RNA_seed | MAPK1 | 0.515561 | BLCA | 647-V | Ulixertinib |
| MVP_top60_DNA_seed | AKT1 | -3.102256 | BRCA | EVSA-T | BAY AKT1 | MVP_top60_DNA+RNA_seed | MAPK3 | 0.515561 | BLCA | 647-V | Ulixertinib |
| MVP_top60_DNA_seed | AKT1 | -3.048792 | BRCA | EFM-19 | AKT inhibitor VIII | MVP_top60_DNA+RNA_seed | AKT1 | 0.516403 | BLCA | UM-UC-3 | MK-2206 |
| MVP_top60_DNA_seed | ERBB2 | -3.044946 | BRCA | HCC1419 | CP724714 | MVP_top60_DNA+RNA_seed | SYK | 0.516863 | BLCA | J82 | Entospletinib |
| MVP_top60_DNA_seed | ERBB2 | -3.042142 | BRCA | HCC2218 | Afatinib | MVP_top60_DNA+RNA_seed | TP53 | 0.520183 | BLCA | SW780 | MIRA-1 |
| MVP_top60_DNA_seed | EGFR | -3.042142 | BRCA | HCC2218 | Afatinib | MVP_top60_DNA+RNA_seed | TP53 | 0.520968 | BLCA | RT4 | MIRA-1 |
| MVP_top60_DNA_seed | ERBB2 | -3.006821 | BRCA | AU565 | CP724714 | MVP_top60_DNA+RNA_seed | HSP90AA1 | 0.521502 | BLCA | TCCSUP | Luminespib |
| MVP_top60_DNA_seed | EGFR | -2.982644 | BRCA | MDA-MB-175-VII | Pelitinib | MVP_top60_DNA+RNA_seed | ESR1 | 0.522589 | BLCA | TCCSUP | Tamoxifen |
| MVP_top60_DNA_seed | ERBB2 | -2.939075 | BRCA | HCC1419 | PF-00299804 | MVP_top60_DNA+RNA_seed | CDK2 | 0.522928 | BLCA | TCCSUP | MK-8776 |
| MVP_top60_DNA_seed | EGFR | -2.939075 | BRCA | HCC1419 | PF-00299804 | MVP_top60_DNA+RNA_seed | HDAC1 | 0.526095 | BLCA | T24 | Entinostat |
| MVP_top60_DNA_seed | EGFR | -2.903137 | BRCA | HCC1419 | AST-1306 | MVP_top60_DNA+RNA_seed | CDK2 | 0.527036 | BLCA | T24 | Dinaciclib |
| MVP_top60_DNA_seed | AKT1 | -2.871968 | BRCA | MDA-MB-361 | AKT inhibitor VIII | MVP_top60_DNA+RNA_seed | HSP90AA1 | 0.5282 | BLCA | HT-1376 | Tanespimycin |
| MVP_top60_DNA_seed | EGFR | -2.869472 | BRCA | MDA-MB-175-VII | AST-1306 | MVP_top60_DNA+RNA_seed | HDAC1 | 0.528543 | BLCA | T24 | PCI-34051 |
| MVP_top60_DNA_seed | ERBB2 | -2.831997 | BRCA | HCC2218 | Afatinib | MVP_top60_DNA+RNA_seed | CDK2 | 0.531484 | BLCA | 5637 | AZD5438 |
| MVP_top60_DNA_seed | EGFR | -2.831997 | BRCA | HCC2218 | Afatinib | MVP_top60_DNA+RNA_seed | ERBB2 | 0.532001 | BLCA | T24 | Sapitinib |
| MVP_top60_DNA_seed | ERBB2 | -2.816371 | BRCA | AU565 | Afatinib | MVP_top60_DNA+RNA_seed | ERBB3 | 0.532001 | BLCA | T24 | Sapitinib |
| MVP_top60_DNA_seed | EGFR | -2.816371 | BRCA | AU565 | Afatinib | MVP_top60_DNA+RNA_seed | AKT1 | 0.535776 | BLCA | RT-112 | AT13148 |
| MVP_top60_DNA_seed | AKT1 | -2.815989 | BRCA | EVSA-T | MK-2206 | MVP_top60_DNA+RNA_seed | HDAC1 | 0.536889 | BLCA | 647-V | Entinostat |
| MVP_top60_DNA_seed | ERBB2 | -2.797545 | BRCA | EFM-192A | Afatinib | MVP_top60_DNA+RNA_seed | MAPK1 | 0.537592 | BLCA | LB831-BLC | SCH772984 |
| MVP_top60_DNA_seed | EGFR | -2.797545 | BRCA | EFM-192A | Afatinib | MVP_top60_DNA+RNA_seed | MAPK3 | 0.537592 | BLCA | LB831-BLC | SCH772984 |
| MVP_top60_DNA_seed | AKT1 | -2.795246 | BRCA | OCUB-M | BAY AKT1 | MVP_top60_DNA+RNA_seed | AKT1 | 0.538528 | BLCA | RT4 | GSK2110183B |

| | | | | | | | | | | | |
|---|---|---|---|---|---|---|---|---|---|---|---|
| MVP_top60_DNA_seed | ERBB2 | -2.779453 | BRCA | MDA-MB-175-VII | CI-1033 | MVP_top60_DNA+RNA_seed | EP300 | 0.543802 | BLCA | RT-112 | SGC-CBP30 |
| MVP_top60_DNA_seed | EGFR | -2.779453 | BRCA | MDA-MB-175-VII | CI-1033 | MVP_top60_DNA+RNA_seed | CREBBP | 0.543802 | BLCA | RT-112 | SGC-CBP30 |
| MVP_top60_DNA_seed | EGFR | -2.760498 | BRCA | OCUB-M | AST-1306 | MVP_top60_DNA+RNA_seed | AKT1 | 0.548216 | BLCA | KU-19-19 | MK-2206 |
| MVP_top60_DNA_seed | ERBB2 | -2.741944 | BRCA | MDA-MB-175-VII | Afatinib | MVP_top60_DNA+RNA_seed | SYK | 0.550016 | BLCA | 639-V | PRT062607 |
| MVP_top60_DNA_seed | EGFR | -2.741944 | BRCA | MDA-MB-175-VII | Afatinib | MVP_top60_DNA+RNA_seed | HSP90AA1 | 0.551154 | BLCA | HT-1376 | Luminespib |
| MVP_top60_DNA_seed | AR | -2.677455 | BRCA | HCC1599 | AZD3514 | MVP_top60_DNA+RNA_seed | HDAC1 | 0.554861 | BLCA | TCCSUP | PCI-34051 |
| MVP_top60_DNA_seed | ERBB2 | -2.665276 | BRCA | MDA-MB-175-VII | Afatinib | MVP_top60_DNA+RNA_seed | ESR1 | 0.555564 | BLCA | LB831-BLC | Tamoxifen |
| MVP_top60_DNA_seed | EGFR | -2.665276 | BRCA | MDA-MB-175-VII | Afatinib | MVP_top60_DNA+RNA_seed | PIK3CA | 0.557464 | BLCA | RT-112 | AZD8186 |
| MVP_top60_DNA_seed | EGFR | -2.650893 | BRCA | MDA-MB-468 | Cetuximab | MVP_top60_DNA+RNA_seed | AKT1 | 0.558315 | BLCA | 647-V | Uprosertib |
| MVP_top60_DNA_seed | AR | -2.642482 | BRCA | CAMA-1 | Bicalutamide | MVP_top60_DNA+RNA_seed | EP300 | 0.572369 | BLCA | J82 | CPI-637 |
| MVP_top60_DNA_seed | ERBB2 | -2.628485 | BRCA | OCUB-M | PF-00299804 | MVP_top60_DNA+RNA_seed | AKT1 | 0.576319 | BLCA | HT-1197 | MK-2206 |
| MVP_top60_DNA_seed | EGFR | -2.628485 | BRCA | OCUB-M | PF-00299804 | MVP_top60_DNA+RNA_seed | AKT1 | 0.581651 | BLCA | 639-V | GSK2110183B |
| MVP_top60_DNA_seed | ERBB2 | -2.555571 | BRCA | BT-474 | Afatinib | MVP_top60_DNA+RNA_seed | GSK3B | 0.58282 | BLCA | DSH1 | SB216763 |
| MVP_top60_DNA_seed | EGFR | -2.555571 | BRCA | BT-474 | Afatinib | MVP_top60_DNA+RNA_seed | CDK2 | 0.583289 | BLCA | CAL-29 | MK-8776 |
| MVP_top60_DNA_seed | PIK3CA | -2.548715 | BRCA | OCUB-M | PI3Ka_4409 | MVP_top60_DNA+RNA_seed | GSK3B | 0.591009 | BLCA | LB831-BLC | SB216763 |
| MVP_top60_DNA_seed | AKT1 | -2.54498 | BRCA | T47D | AKT inhibitor VIII | MVP_top60_DNA+RNA_seed | SRC | 0.591106 | BLCA | 647-V | Bosutinib |
| MVP_top60_DNA_seed | CDK1 | -2.537566 | BRCA | MDA-MB-468 | RO-3306 | MVP_top60_DNA+RNA_seed | AKT1 | 0.592416 | BLCA | HT-1197 | AT13148 |
| MVP_top60_DNA_seed | AKT1 | -2.526133 | BRCA | HCC70 | AKT inhibitor VIII | MVP_top60_DNA+RNA_seed | AKT1 | 0.592433 | BLCA | 647-V | MK-2206 |
| MVP_top60_DNA_seed | ERBB2 | -2.499472 | BRCA | OCUB-M | CI-1033 | MVP_top60_DNA+RNA_seed | MAPK1 | 0.593041 | BLCA | HT-1376 | Ulixertinib |
| MVP_top60_DNA_seed | EGFR | -2.499472 | BRCA | OCUB-M | CI-1033 | MVP_top60_DNA+RNA_seed | MAPK3 | 0.593041 | BLCA | HT-1376 | Ulixertinib |
| MVP_top60_DNA_seed | ERBB2 | -2.454956 | BRCA | HCC1419 | CI-1033 | MVP_top60_DNA+RNA_seed | MAPK1 | 0.59314 | BLCA | 5637 | Ulixertinib |
| MVP_top60_DNA_seed | EGFR | -2.454956 | BRCA | HCC1419 | CI-1033 | MVP_top60_DNA+RNA_seed | MAPK3 | 0.59314 | BLCA | 5637 | Ulixertinib |
| MVP_top60_DNA_seed | ESR1 | -2.449334 | BRCA | EFM-19 | Fulvestrant | MVP_top60_DNA+RNA_seed | PIK3CA | 0.599658 | BLCA | 639-V | Alpelisib |
| MVP_top60_DNA_seed | AKT1 | -2.446538 | BRCA | MDA-MB-175-VII | MK-2206 | MVP_top60_DNA+RNA_seed | MAPK1 | 0.603553 | BLCA | SW780 | Ulixertinib |
| MVP_top60_DNA_seed | ERBB2 | -2.42065 | BRCA | MDA-MB-453 | CP724714 | MVP_top60_DNA+RNA_seed | MAPK3 | 0.603553 | BLCA | SW780 | Ulixertinib |
| MVP_top60_DNA_seed | EGFR | -2.41971 | BRCA | BT-474 | Pelitinib | MVP_top60_DNA+RNA_seed | HDAC1 | 0.609223 | BLCA | HT-1197 | PCI-34051 |
| MVP_top60_DNA_seed | AKT1 | -2.397081 | BRCA | CAL-148 | GSK690693 | MVP_top60_DNA+RNA_seed | PIK3CA | 0.61902 | BLCA | LB831-BLC | Buparlisib |
| MVP_top60_DNA_seed | ERBB2 | -2.39683 | BRCA | HCC2218 | CI-1033 | MVP_top60_DNA+RNA_seed | ERBB2 | 0.623149 | BLCA | LB831-BLC | Afatinib |
| MVP_top60_DNA_seed | EGFR | -2.39683 | BRCA | HCC2218 | CI-1033 | MVP_top60_DNA+RNA_seed | ERBB2 | 0.626589 | BLCA | KU-19-19 | Sapitinib |
| MVP_top60_DNA_seed | CDK1 | -2.394293 | BRCA | Hs-578-T | RO-3306 | MVP_top60_DNA+RNA_seed | ERBB3 | 0.626589 | BLCA | KU-19-19 | Sapitinib |
| MVP_top60_DNA_seed | AKT1 | -2.378894 | BRCA | MDA-MB-330 | AKT inhibitor VIII | MVP_top60_DNA+RNA_seed | AKT1 | 0.63092 | BLCA | RT-112 | AZD5363 |
| MVP_top60_DNA_seed | PIK3CA | -2.357662 | BRCA | MDA-MB-453 | PI3Ka_4409 | MVP_top60_DNA+RNA_seed | AKT1 | 0.632431 | BLCA | SW780 | MK-2206 |
| MVP_top60_DNA_seed | ERBB2 | -2.35215 | BRCA | UACC-893 | Afatinib | MVP_top60_DNA+RNA_seed | HDAC1 | 0.64124 | BLCA | SW1710 | Entinostat |
| MVP_top60_DNA_seed | EGFR | -2.35215 | BRCA | UACC-893 | Afatinib | MVP_top60_DNA+RNA_seed | AKT1 | 0.645267 | BLCA | J82 | Ipatasertib |
| MVP_top60_DNA_seed | EGFR | -2.333524 | BRCA | OCUB-M | Pelitinib | MVP_top60_DNA+RNA_seed | KRAS | 0.649053 | BLCA | J82 | KRAS (G12C) Inhibitor-1 |
| MVP_top60_DNA_seed | AKT1 | -2.312486 | BRCA | OCUB-M | AKT inhibitor VIII | MVP_top60_DNA+RNA_seed | AKT1 | 0.649413 | BLCA | SW1710 | GSK2110183B |
| MVP_top60_DNA_seed | ESR1 | -2.285577 | BRCA | HCC1187 | Tamoxifen | MVP_top60_DNA+RNA_seed | FGFR3 | 0.650917 | BLCA | HT-1197 | PD173074 |
| MVP_top60_DNA_seed | ESR1 | -2.276771 | BRCA | MCF7 | Tamoxifen | MVP_top60_DNA+RNA_seed | ERBB2 | 0.661388 | BLCA | SW1710 | Sapitinib |
| MVP_top60_DNA_seed | ERBB2 | -2.275965 | BRCA | AU565 | Afatinib | MVP_top60_DNA+RNA_seed | ERBB3 | 0.661388 | BLCA | SW1710 | Sapitinib |
| MVP_top60_DNA_seed | EGFR | -2.275965 | BRCA | AU565 | Afatinib | MVP_top60_DNA+RNA_seed | AKT1 | 0.662226 | BLCA | J82 | Uprosertib |
| MVP_top60_DNA_seed | AKT1 | -2.255889 | BRCA | HCC70 | MK-2206 | MVP_top60_DNA+RNA_seed | AKT1 | 0.666149 | BLCA | J82 | Afuresertib |

| | | | | | | | | | | | |
|---|---|---|---|---|---|---|---|---|---|---|---|
| MVP_top60_DNA_seed | ERBB2 | -2.239042 | BRCA | ZR-75-30 | Afatinib | MVP_top60_DNA+RNA_seed | HSP90AA1 | 0.669108 | BLCA | CAL-29 | Luminespib |
| MVP_top60_DNA_seed | EGFR | -2.239042 | BRCA | ZR-75-30 | Afatinib | MVP_top60_DNA+RNA_seed | TP53 | 0.670544 | BLCA | 5637 | MIRA-1 |
| MVP_top60_DNA_seed | AKT1 | -2.234574 | BRCA | EFM-19 | MK-2206 | MVP_top60_DNA+RNA_seed | AKT1 | 0.670994 | BLCA | SW780 | AT13148 |
| MVP_top60_DNA_seed | AKT1 | -2.219478 | BRCA | ZR-75-30 | AKT inhibitor VIII | MVP_top60_DNA+RNA_seed | PIK3CA | 0.672557 | BLCA | TCCSUP | AZD8186 |
| MVP_top60_DNA_seed | AKT1 | -2.217692 | BRCA | AU565 | AKT inhibitor VIII | MVP_top60_DNA+RNA_seed | MAPK1 | 0.674563 | BLCA | T24 | VX-11e |
| MVP_top60_DNA_seed | AKT1 | -2.217372 | BRCA | ZR-75-30 | AKT inhibitor VIII | MVP_top60_DNA+RNA_seed | HDAC1 | 0.678048 | BLCA | CAL-29 | PCI-34051 |
| MVP_top60_DNA_seed | ERBB2 | -2.167766 | BRCA | UACC-893 | CP724714 | MVP_top60_DNA+RNA_seed | AKT1 | 0.686049 | BLCA | 639-V | MK-2206 |
| MVP_top60_DNA_seed | ERBB2 | -2.160443 | BRCA | BT-474 | Afatinib | MVP_top60_DNA+RNA_seed | ERBB2 | 0.689029 | BLCA | HT-1376 | Lapatinib |
| MVP_top60_DNA_seed | EGFR | -2.160443 | BRCA | BT-474 | Afatinib | MVP_top60_DNA+RNA_seed | ESR1 | 0.69589 | BLCA | LB831-BLC | GDC0810 |
| MVP_top60_DNA_seed | CDK1 | -2.155946 | BRCA | EVSA-T | RO-3306 | MVP_top60_DNA+RNA_seed | KRAS | 0.69703 | BLCA | 647-V | KRAS (G12C) Inhibitor-1 |
| MVP_top60_DNA_seed | MAPK3 | -2.155364 | BRCA | HCC70 | FR-180204 | MVP_top60_DNA+RNA_seed | TP53 | 0.706531 | BLCA | HT-1197 | MIRA-1 |
| MVP_top60_DNA_seed | MAPK1 | -2.155364 | BRCA | HCC70 | FR-180204 | MVP_top60_DNA+RNA_seed | ERBB2 | 0.707956 | BLCA | TCCSUP | Sapitinib |
| MVP_top60_DNA_seed | ERBB2 | -2.14666 | BRCA | MDA-MB-175-VII | AZD8931 | MVP_top60_DNA+RNA_seed | ERBB3 | 0.707956 | BLCA | TCCSUP | Sapitinib |
| MVP_top60_DNA_seed | EGFR | -2.14666 | BRCA | MDA-MB-175-VII | AZD8931 | MVP_top60_DNA+RNA_seed | MAPK1 | 0.715589 | BLCA | J82 | Ulixertinib |
| MVP_top60_DNA_seed | ERBB3 | -2.14666 | BRCA | MDA-MB-175-VII | AZD8931 | MVP_top60_DNA+RNA_seed | MAPK3 | 0.715589 | BLCA | J82 | Ulixertinib |
| MVP_top60_DNA_seed | ERBB2 | -2.129764 | BRCA | HCC1599 | AZD8931 | MVP_top60_DNA+RNA_seed | TERT | 0.718228 | BLCA | UM-UC-3 | BIBR-1532 |
| MVP_top60_DNA_seed | EGFR | -2.129764 | BRCA | HCC1599 | AZD8931 | MVP_top60_DNA+RNA_seed | MAPK1 | 0.71866 | BLCA | 639-V | VX-11e |
| MVP_top60_DNA_seed | ERBB3 | -2.129764 | BRCA | HCC1599 | AZD8931 | MVP_top60_DNA+RNA_seed | TP53 | 0.720144 | BLCA | TCCSUP | MIRA-1 |
| MVP_top60_DNA_seed | ERBB2 | -2.128122 | BRCA | OCUB-M | Afatinib | MVP_top60_DNA+RNA_seed | MAPK1 | 0.725801 | BLCA | UM-UC-3 | SCH772984 |
| MVP_top60_DNA_seed | EGFR | -2.128122 | BRCA | OCUB-M | Afatinib | MVP_top60_DNA+RNA_seed | MAPK3 | 0.725801 | BLCA | UM-UC-3 | SCH772984 |
| MVP_top60_DNA_seed | AKT1 | -2.112252 | BRCA | T47D | MK-2206 | MVP_top60_DNA+RNA_seed | ERBB2 | 0.726094 | BLCA | 639-V | Lapatinib |
| MVP_top60_DNA_seed | EGFR | -2.09853 | BRCA | HCC202 | AST-1306 | MVP_top60_DNA+RNA_seed | FGFR3 | 0.728333 | BLCA | T24 | PD173074 |
| MVP_top60_DNA_seed | AKT1 | -2.097631 | BRCA | OCUB-M | A-443654 | MVP_top60_DNA+RNA_seed | AKT1 | 0.732163 | BLCA | UM-UC-3 | Uprosertib |
| MVP_top60_DNA_seed | PIK3CA | -2.077815 | BRCA | MDA-MB-453 | AZD8835 | MVP_top60_DNA+RNA_seed | TP53 | 0.736355 | BLCA | LB831-BLC | MIRA-1 |
| MVP_top60_DNA_seed | ERBB2 | -2.072793 | BRCA | UACC-893 | Afatinib | MVP_top60_DNA+RNA_seed | EP300 | 0.736911 | BLCA | 647-V | SGC-CBP30 |
| MVP_top60_DNA_seed | EGFR | -2.072793 | BRCA | UACC-893 | Afatinib | MVP_top60_DNA+RNA_seed | CREBBP | 0.736911 | BLCA | 647-V | SGC-CBP30 |
| MVP_top60_DNA_seed | AKT1 | -2.045806 | BRCA | MDA-MB-415 | BAY AKT1 | MVP_top60_DNA+RNA_seed | AKT1 | 0.743589 | BLCA | J82 | AZD5363 |
| MVP_top60_DNA_seed | HSP90AA1 | -2.039732 | BRCA | MDA-MB-468 | Elesclomol | MVP_top60_DNA+RNA_seed | SYK | 0.749309 | BLCA | SW1710 | PRT062607 |
| MVP_top60_DNA_seed | HSP90AB1 | -2.039732 | BRCA | MDA-MB-468 | Elesclomol | MVP_top60_DNA+RNA_seed | ERBB2 | 0.757283 | BLCA | SW1710 | Afatinib |
| MVP_top60_DNA_seed | ERBB2 | -2.036727 | BRCA | HCC1954 | CP724714 | MVP_top60_DNA+RNA_seed | CDK2 | 0.76688 | BLCA | 647-V | MK-8776 |
| MVP_top60_DNA_seed | AKT1 | -2.033369 | BRCA | MDA-MB-361 | BAY AKT1 | MVP_top60_DNA+RNA_seed | PIK3CA | 0.770447 | BLCA | RT4 | Alpelisib |
| MVP_top60_DNA_seed | PIK3CA | -2.033158 | BRCA | HCC202 | PI3Ka_4409 | MVP_top60_DNA+RNA_seed | SYK | 0.770644 | BLCA | RT4 | PRT062607 |
| MVP_top60_DNA_seed | ERBB2 | -2.033014 | BRCA | ZR-75-30 | Afatinib | MVP_top60_DNA+RNA_seed | AKT1 | 0.772087 | BLCA | 647-V | GSK2110183B |
| MVP_top60_DNA_seed | EGFR | -2.033014 | BRCA | ZR-75-30 | Afatinib | MVP_top60_DNA+RNA_seed | EP300 | 0.772096 | BLCA | LB831-BLC | SGC-CBP30 |
| MVP_top60_DNA_seed | PIK3CA | -2.01571 | BRCA | MDA-MB-361 | PI3Ka_4409 | MVP_top60_DNA+RNA_seed | CREBBP | 0.772096 | BLCA | LB831-BLC | SGC-CBP30 |
| MVP_top60_DNA_seed | AKT1 | -1.997287 | BRCA | T47D | GSK690693 | MVP_top60_DNA+RNA_seed | CDK2 | 0.773827 | BLCA | 639-V | MK-8776 |
| MVP_top60_DNA_seed | PIK3CA | -1.993881 | BRCA | MDA-MB-361 | AZD8835 | MVP_top60_DNA+RNA_seed | ATM | 0.777271 | BLCA | SW1710 | KU-55933 |
| MVP_top60_DNA_seed | EGFR | -1.978114 | BRCA | HCC2218 | Pelitinib | MVP_top60_DNA+RNA_seed | TP53 | 0.778608 | BLCA | J82 | MIRA-1 |
| MVP_top60_DNA_seed | AR | -1.968538 | BRCA | DU-4475 | Bicalutamide | MVP_top60_DNA+RNA_seed | FGFR3 | 0.784985 | BLCA | HT-1197 | AZD4547 |
| MVP_top60_DNA_seed | AKT1 | -1.967086 | BRCA | UACC-893 | GSK690693 | MVP_top60_DNA+RNA_seed | GSK3B | 0.785001 | BLCA | T24 | SB216763 |
| MVP_top60_DNA_seed | AKT1 | -1.962525 | BRCA | BT-474 | GSK690693 | MVP_top60_DNA+RNA_seed | ERBB2 | 0.786125 | BLCA | UM-UC-3 | Afatinib |

| | | | | | | | | | | | |
|---|---|---|---|---|---|---|---|---|---|---|---|
| MVP_top60_DNA_seed | EGFR | -1.961958 | BRCA | UACC-893 | AST-1306 | MVP_top60_DNA+RNA_seed | SYK | 0.795258 | BLCA | CAL-29 | PRT062607 |
| MVP_top60_DNA_seed | ESR1 | -1.960994 | BRCA | HCC1500 | Fulvestrant | MVP_top60_DNA+RNA_seed | SYK | 0.797459 | BLCA | HT-1197 | Entospletinib |
| MVP_top60_DNA_seed | ERBB2 | -1.94645 | BRCA | UACC-893 | PF-00299804 | MVP_top60_DNA+RNA_seed | PIK3CA | 0.807117 | BLCA | T24 | AZD8186 |
| MVP_top60_DNA_seed | EGFR | -1.94645 | BRCA | UACC-893 | PF-00299804 | MVP_top60_DNA+RNA_seed | MAPK1 | 0.810133 | BLCA | 647-V | VX-11e |
| MVP_top60_DNA_seed | PIK3CA | -1.923421 | BRCA | HCC1954 | PI3Ka_4409 | MVP_top60_DNA+RNA_seed | ERBB2 | 0.812624 | BLCA | HT-1197 | Afatinib |
| MVP_top60_DNA_seed | ERBB2 | -1.895571 | BRCA | HCC2157 | Afatinib | MVP_top60_DNA+RNA_seed | ERBB2 | 0.814626 | BLCA | 647-V | Lapatinib |
| MVP_top60_DNA_seed | EGFR | -1.895571 | BRCA | HCC2157 | Afatinib | MVP_top60_DNA+RNA_seed | ESR1 | 0.818888 | BLCA | J82 | Tamoxifen |
| MVP_top60_DNA_seed | HSP90AA1 | -1.892612 | BRCA | CAL-51 | Elesclomol | MVP_top60_DNA+RNA_seed | MAPK1 | 0.821146 | BLCA | HT-1376 | Ulixertinib |
| MVP_top60_DNA_seed | HSP90AB1 | -1.892612 | BRCA | CAL-51 | Elesclomol | MVP_top60_DNA+RNA_seed | MAPK3 | 0.821146 | BLCA | HT-1376 | Ulixertinib |
| MVP_top60_DNA_seed | PIK3CA | -1.888956 | BRCA | MDA-MB-330 | AZD8835 | MVP_top60_DNA+RNA_seed | AKT1 | 0.821533 | BLCA | 5637 | GSK2110183B |
| MVP_top60_DNA_seed | AKT1 | -1.881377 | BRCA | OCUB-M | AKT inhibitor VIII | MVP_top60_DNA+RNA_seed | ATM | 0.843566 | BLCA | T24 | KU-55933 |
| MVP_top60_DNA_seed | ERBB2 | -1.872608 | BRCA | HCC202 | CI-1033 | MVP_top60_DNA+RNA_seed | AKT1 | 0.846677 | BLCA | 647-V | Uprosertib |
| MVP_top60_DNA_seed | EGFR | -1.872608 | BRCA | HCC202 | CI-1033 | MVP_top60_DNA+RNA_seed | RAC1 | 0.85142 | BLCA | HT-1376 | EHT-1864 |
| MVP_top60_DNA_seed | AKT1 | -1.85086 | BRCA | HCC70 | AKT inhibitor VIII | MVP_top60_DNA+RNA_seed | HSP90AA1 | 0.852704 | BLCA | T24 | Elesclomol |
| MVP_top60_DNA_seed | ERBB2 | -1.848209 | BRCA | UACC-893 | CI-1033 | MVP_top60_DNA+RNA_seed | TERT | 0.853057 | BLCA | LB831-BLC | BIBR-1532 |
| MVP_top60_DNA_seed | EGFR | -1.848209 | BRCA | UACC-893 | CI-1033 | MVP_top60_DNA+RNA_seed | TERT | 0.853201 | BLCA | TCCSUP | BIBR-1532 |
| MVP_top60_DNA_seed | AKT1 | -1.83964 | BRCA | YMB-1-E | GSK690693 | MVP_top60_DNA+RNA_seed | PIK3CA | 0.864754 | BLCA | KU-19-19 | AZD8186 |
| MVP_top60_DNA_seed | EGFR | -1.831664 | BRCA | UACC-893 | Pelitinib | MVP_top60_DNA+RNA_seed | AKT1 | 0.867568 | BLCA | 647-V | Ipatasertib |
| MVP_top60_DNA_seed | PIK3CA | -1.819452 | BRCA | OCUB-M | AZD8835 | MVP_top60_DNA+RNA_seed | SRC | 0.868372 | BLCA | RT4 | Bosutinib |
| MVP_top60_DNA_seed | ERBB2 | -1.811388 | BRCA | HCC202 | Afatinib | MVP_top60_DNA+RNA_seed | SYK | 0.868935 | BLCA | TCCSUP | PRT062607 |
| MVP_top60_DNA_seed | EGFR | -1.811388 | BRCA | HCC202 | Afatinib | MVP_top60_DNA+RNA_seed | AKT1 | 0.881831 | BLCA | TCCSUP | Afuresertib |
| MVP_top60_DNA_seed | AKT1 | -1.804262 | BRCA | MFM-223 | BAY AKT1 | MVP_top60_DNA+RNA_seed | EP300 | 0.883701 | BLCA | RT-112 | CPI-637 |
| MVP_top60_DNA_seed | AKT1 | -1.799965 | BRCA | HCC202 | BAY AKT1 | MVP_top60_DNA+RNA_seed | HDAC1 | 0.886583 | BLCA | J82 | Entinostat |
| MVP_top60_DNA_seed | ESR1 | -1.798991 | BRCA | HCC1428 | Fulvestrant | MVP_top60_DNA+RNA_seed | MAPK1 | 0.886949 | BLCA | UM-UC-3 | VX-11e |
| MVP_top60_DNA_seed | HSP90AA1 | -1.790851 | BRCA | MDA-MB-361 | Tanespimycin | MVP_top60_DNA+RNA_seed | AKT1 | 0.90383 | BLCA | J82 | GSK2110183B |
| MVP_top60_DNA_seed | HSP90AB1 | -1.790851 | BRCA | MDA-MB-361 | Tanespimycin | MVP_top60_DNA+RNA_seed | CDK2 | 0.904883 | BLCA | UM-UC-3 | MK-8776 |
| MVP_top60_DNA_seed | AKT1 | -1.788657 | BRCA | MDA-MB-415 | AKT inhibitor VIII | MVP_top60_DNA+RNA_seed | SRC | 0.908303 | BLCA | LB831-BLC | Bosutinib |
| MVP_top60_DNA_seed | ESR1 | -1.77589 | BRCA | BT-474 | Fulvestrant | MVP_top60_DNA+RNA_seed | GSK3B | 0.915374 | BLCA | T24 | CHIR-99021 |
| MVP_top60_DNA_seed | EGFR | -1.755145 | BRCA | CAL-85-1 | Pelitinib | MVP_top60_DNA+RNA_seed | ERBB2 | 0.923167 | BLCA | SW780 | Lapatinib |
| MVP_top60_DNA_seed | AKT1 | -1.753459 | BRCA | BT-474 | AKT inhibitor VIII | MVP_top60_DNA+RNA_seed | HDAC1 | 0.923567 | BLCA | J82 | PCI-34051 |
| MVP_top60_DNA_seed | HSP90AA1 | -1.749076 | BRCA | MDA-MB-231 | CCT-018159 | MVP_top60_DNA+RNA_seed | RAC1 | 0.925929 | BLCA | TCCSUP | EHT-1864 |
| MVP_top60_DNA_seed | HSP90AB1 | -1.749076 | BRCA | MDA-MB-231 | CCT-018159 | MVP_top60_DNA+RNA_seed | AKT1 | 0.927534 | BLCA | TCCSUP | AZD5363 |
| MVP_top60_DNA_seed | HSP90AA1 | -1.746968 | BRCA | CAL-51 | Tanespimycin | MVP_top60_DNA+RNA_seed | PIK3CA | 0.933356 | BLCA | T24 | GNE-317 |
| MVP_top60_DNA_seed | HSP90AB1 | -1.746968 | BRCA | CAL-51 | Tanespimycin | MVP_top60_DNA+RNA_seed | MAPK1 | 0.943222 | BLCA | TCCSUP | Ulixertinib |
| MVP_top60_DNA_seed | PIK3CA | -1.72137 | BRCA | HCC1599 | PI3Ka_4409 | MVP_top60_DNA+RNA_seed | MAPK3 | 0.943222 | BLCA | TCCSUP | Ulixertinib |
| MVP_top60_DNA_seed | AKT1 | -1.699765 | BRCA | ZR-75-30 | GSK690693 | MVP_top60_DNA+RNA_seed | AKT1 | 0.949045 | BLCA | SW1710 | Uprosertib |
| MVP_top60_DNA_seed | AKT1 | -1.696774 | BRCA | HCC70 | BAY AKT1 | MVP_top60_DNA+RNA_seed | ESR1 | 0.954019 | BLCA | T24 | GDC0810 |
| MVP_top60_DNA_seed | EGFR | -1.684412 | BRCA | AU565 | Pelitinib | MVP_top60_DNA+RNA_seed | KRAS | 0.957842 | BLCA | HT-1197 | RAS (G12C) Inhibitor-1 |
| MVP_top60_DNA_seed | ERBB2 | -1.683543 | BRCA | EFM-192A | Afatinib | MVP_top60_DNA+RNA_seed | ERBB2 | 0.957884 | BLCA | HT-1197 | Sapitinib |
| MVP_top60_DNA_seed | EGFR | -1.683543 | BRCA | EFM-192A | Afatinib | MVP_top60_DNA+RNA_seed | ERBB3 | 0.957884 | BLCA | HT-1197 | Sapitinib |
| MVP_top60_DNA_seed | AKT1 | -1.682431 | BRCA | HCC1419 | MK-2206 | MVP_top60_DNA+RNA_seed | EP300 | 0.95824 | BLCA | TCCSUP | CPI-637 |

| | | | | | | | | | | | |
|---|---|---|---|---|---|---|---|---|---|---|---|
| MVP_top60_DNA_seed | AKT1 | -1.663806 | BRCA | T47D | AKT inhibitor VIII | MVP_top60_DNA+RNA_seed | SYK | 0.959906 | BLCA | HT-1197 | PRT062607 |
| MVP_top60_DNA_seed | MDM2 | -1.662462 | BRCA | DU-4475 | Nutlin-3a (-) | MVP_top60_DNA+RNA_seed | AKT1 | 0.963379 | BLCA | RT4 | MK-2206 |
| MVP_top60_DNA_seed | ATM | -1.656866 | BRCA | HCC70 | KU-55933 | MVP_top60_DNA+RNA_seed | HSP90AA1 | 0.971267 | BLCA | HT-1197 | Tanespimycin |
| MVP_top60_DNA_seed | AKT1 | -1.65644 | BRCA | MDA-MB-361 | MK-2206 | MVP_top60_DNA+RNA_seed | ATM | 0.972707 | BLCA | SW780 | KU-55933 |
| MVP_top60_DNA_seed | ATM | -1.646731 | BRCA | HCC2157 | KU-55933 | MVP_top60_DNA+RNA_seed | ATM | 0.97483 | BLCA | TCCSUP | KU-55933 |
| MVP_top60_DNA_seed | AKT1 | -1.64532 | BRCA | HCC1954 | GSK690693 | MVP_top60_DNA+RNA_seed | AKT1 | 0.976965 | BLCA | SW780 | GSK2110183B |
| MVP_top60_DNA_seed | AKT1 | -1.644698 | BRCA | EFM-19 | BAY AKT1 | MVP_top60_DNA+RNA_seed | AKT1 | 0.981496 | BLCA | 5637 | MK-2206 |
| MVP_top60_DNA_seed | ESR1 | -1.636771 | BRCA | OCUB-M | Fulvestrant | MVP_top60_DNA+RNA_seed | CDK2 | 0.982875 | BLCA | RT4 | MK-8776 |
| MVP_top60_DNA_seed | AKT1 | -1.632491 | BRCA | MDA-MB-330 | MK-2206 | MVP_top60_DNA+RNA_seed | TP53 | 0.988377 | BLCA | HT-1376 | MIRA-1 |
| MVP_top60_DNA_seed | SRC | -1.627486 | BRCA | DU-4475 | NVP-BHG712 | MVP_top60_DNA+RNA_seed | MAPK1 | 0.990776 | BLCA | T24 | Ulixertinib |
| MVP_top60_DNA_seed | EGFR | -1.624663 | BRCA | BT-20 | Cetuximab | MVP_top60_DNA+RNA_seed | MAPK3 | 0.990776 | BLCA | T24 | Ulixertinib |
| MVP_top60_DNA_seed | ATM | -1.614636 | BRCA | EVSA-T | KU-55933 | MVP_top60_DNA+RNA_seed | AKT1 | 0.993103 | BLCA | LB831-BLC | Uprosertib |
| MVP_top60_DNA_seed | CDK2 | -1.608597 | BRCA | HCC70 | PHA-793887 | MVP_top60_DNA+RNA_seed | MAPK1 | 0.995933 | BLCA | HT-1376 | SCH772984 |
| MVP_top60_DNA_seed | HDAC1 | -1.602832 | BRCA | EFM-19 | JQ12 | MVP_top60_DNA+RNA_seed | MAPK3 | 0.995933 | BLCA | HT-1376 | SCH772984 |
| MVP_top60_DNA_seed | HDAC2 | -1.602832 | BRCA | EFM-19 | JQ12 | MVP_top60_DNA+RNA_seed | MAPK1 | 0.998349 | BLCA | T24 | SCH772984 |
| MVP_top60_DNA_seed | AKT1 | -1.591133 | BRCA | MDA-MB-330 | AKT inhibitor VIII | MVP_top60_DNA+RNA_seed | MAPK3 | 0.998349 | BLCA | T24 | SCH772984 |
| MVP_top60_DNA_seed | ESR1 | -1.58436 | BRCA | EVSA-T | Fulvestrant | MVP_top60_DNA+RNA_seed | MAPK1 | 1.003831 | BLCA | SW1710 | Ulixertinib |
| MVP_top60_DNA_seed | AKT1 | -1.575452 | BRCA | MDA-MB-453 | BAY AKT1 | MVP_top60_DNA+RNA_seed | MAPK3 | 1.003831 | BLCA | SW1710 | Ulixertinib |
| MVP_top60_DNA_seed | HSP90AA1 | -1.563298 | BRCA | CAL-51 | CCT-018159 | MVP_top60_DNA+RNA_seed | HDAC1 | 1.005992 | BLCA | LB831-BLC | Entinostat |
| MVP_top60_DNA_seed | HSP90AB1 | -1.563298 | BRCA | CAL-51 | CCT-018159 | MVP_top60_DNA+RNA_seed | ESR1 | 1.007142 | BLCA | SW780 | GDC0810 |
| MVP_top60_DNA_seed | HDAC1 | -1.562873 | BRCA | CAL-85-1 | JQ12 | MVP_top60_DNA+RNA_seed | EP300 | 1.009242 | BLCA | UM-UC-3 | CPI-637 |
| MVP_top60_DNA_seed | HDAC2 | -1.562873 | BRCA | CAL-85-1 | JQ12 | MVP_top60_DNA+RNA_seed | MAPK1 | 1.022648 | BLCA | J82 | VX-11e |
| MVP_top60_DNA_seed | ERBB2 | -1.562326 | BRCA | HCC1954 | Afatinib | MVP_top60_DNA+RNA_seed | PIK3CA | 1.024332 | BLCA | J82 | AZD8186 |
| MVP_top60_DNA_seed | EGFR | -1.562326 | BRCA | HCC1954 | Afatinib | MVP_top60_DNA+RNA_seed | AKT1 | 1.02633 | BLCA | HT-1376 | Afuresertib |
| MVP_top60_DNA_seed | AKT1 | -1.555749 | BRCA | T47D | BAY AKT1 | MVP_top60_DNA+RNA_seed | AKT1 | 1.030586 | BLCA | 5637 | Ipatasertib |
| MVP_top60_DNA_seed | ERBB2 | -1.552322 | BRCA | MDA-MB-330 | Afatinib | MVP_top60_DNA+RNA_seed | AKT1 | 1.032042 | BLCA | T24 | Afuresertib |
| MVP_top60_DNA_seed | EGFR | -1.552322 | BRCA | MDA-MB-330 | Afatinib | MVP_top60_DNA+RNA_seed | AKT1 | 1.032414 | BLCA | UM-UC-3 | Afuresertib |
| MVP_top60_DNA_seed | AR | -1.551632 | BRCA | OCUB-M | AZD3514 | MVP_top60_DNA+RNA_seed | KRAS | 1.033816 | BLCA | SW1710 | KRAS (G12C) Inhibitor-1 |
| MVP_top60_DNA_seed | AKT1 | -1.543239 | BRCA | HCC38 | AKT inhibitor VIII | MVP_top60_DNA+RNA_seed | AKT1 | 1.035223 | BLCA | 647-V | Afuresertib |
| MVP_top60_DNA_seed | HSP90AA1 | -1.533125 | BRCA | MCF7 | Luminespib | MVP_top60_DNA+RNA_seed | AKT1 | 1.036531 | BLCA | LB831-BLC | MK-2206 |
| MVP_top60_DNA_seed | HSP90AB1 | -1.533125 | BRCA | MCF7 | Luminespib | MVP_top60_DNA+RNA_seed | TERT | 1.043692 | BLCA | J82 | BIBR-1532 |
| MVP_top60_DNA_seed | AKT1 | -1.528884 | BRCA | CAL-148 | A-443654 | MVP_top60_DNA+RNA_seed | CDK2 | 1.04764 | BLCA | SW1710 | MK-8776 |
| MVP_top60_DNA_seed | HSP90AA1 | -1.527797 | BRCA | MDA-MB-175-VII | Tanespimycin | MVP_top60_DNA+RNA_seed | PIK3CA | 1.077254 | BLCA | KU-19-19 | Alpelisib |
| MVP_top60_DNA_seed | HSP90AB1 | -1.527797 | BRCA | MDA-MB-175-VII | Tanespimycin | MVP_top60_DNA+RNA_seed | CDK2 | 1.078331 | BLCA | TCCSUP | Dinaciclib |
| MVP_top60_DNA_seed | AKT1 | -1.520264 | BRCA | MDA-MB-453 | GSK690693 | MVP_top60_DNA+RNA_seed | CDK2 | 1.078401 | BLCA | SW780 | MK-8776 |
| MVP_top60_DNA_seed | HDAC1 | -1.499309 | BRCA | MDA-MB-453 | Belinostat | MVP_top60_DNA+RNA_seed | MAPK1 | 1.091312 | BLCA | LB831-BLC | Ulixertinib |
| MVP_top60_DNA_seed | HDAC1 | -1.497798 | BRCA | MDA-MB-453 | AR-42 | MVP_top60_DNA+RNA_seed | MAPK3 | 1.091312 | BLCA | LB831-BLC | Ulixertinib |
| MVP_top60_DNA_seed | ESR1 | -1.497497 | BRCA | T47D | Fulvestrant | MVP_top60_DNA+RNA_seed | AKT1 | 1.093093 | BLCA | T24 | Ipatasertib |
| MVP_top60_DNA_seed | AKT1 | -1.492177 | BRCA | MDA-MB-453 | AKT inhibitor VIII | MVP_top60_DNA+RNA_seed | ERBB2 | 1.0985 | BLCA | J82 | Sapitinib |
| MVP_top60_DNA_seed | HSP90AA1 | -1.490882 | BRCA | BT-474 | Tanespimycin | MVP_top60_DNA+RNA_seed | ERBB3 | 1.0985 | BLCA | J82 | Sapitinib |
| MVP_top60_DNA_seed | HSP90AB1 | -1.490882 | BRCA | BT-474 | Tanespimycin | MVP_top60_DNA+RNA_seed | HDAC1 | 1.100254 | BLCA | 5637 | PCI-34051 |

| | | | | | | | | | | | |
|---|---|---|---|---|---|---|---|---|---|---|---|
| MVP_top60_DNA_seed | AKT1 | -1.48983 | BRCA | EFM-19 | AKT inhibitor VIII | MVP_top60_DNA+RNA_seed | PIK3CA | 1.103866 | BLCA | T24 | Alpelisib |
| MVP_top60_DNA_seed | AKT1 | -1.487882 | BRCA | EFM-192A | MK-2206 | MVP_top60_DNA+RNA_seed | MAPK1 | 1.104656 | BLCA | SW1710 | VX-11e |
| MVP_top60_DNA_seed | AKT1 | -1.487501 | BRCA | HCC2218 | AKT inhibitor VIII | MVP_top60_DNA+RNA_seed | AKT1 | 1.105274 | BLCA | 5637 | AT13148 |
| MVP_top60_DNA_seed | EGFR | -1.487398 | BRCA | HCC2218 | Gefitinib | MVP_top60_DNA+RNA_seed | PIK3CA | 1.110764 | BLCA | RT4 | GNE-317 |
| MVP_top60_DNA_seed | ESR1 | -1.480426 | BRCA | MDA-MB-361 | Tamoxifen | MVP_top60_DNA+RNA_seed | MAPK1 | 1.111692 | BLCA | J82 | SCH772984 |
| MVP_top60_DNA_seed | ERBB2 | -1.463658 | BRCA | AU565 | AZD8931 | MVP_top60_DNA+RNA_seed | MAPK3 | 1.111692 | BLCA | J82 | SCH772984 |
| MVP_top60_DNA_seed | EGFR | -1.463658 | BRCA | AU565 | AZD8931 | MVP_top60_DNA+RNA_seed | PIK3CA | 1.13296 | BLCA | SW1710 | Alpelisib |
| MVP_top60_DNA_seed | ERBB3 | -1.463658 | BRCA | AU565 | AZD8931 | MVP_top60_DNA+RNA_seed | KRAS | 1.133926 | BLCA | LB831-BLC | KRAS (G12C) Inhibitor-1 |
| MVP_top60_DNA_seed | AKT1 | -1.46047 | BRCA | MDA-MB-415 | MK-2206 | MVP_top60_DNA+RNA_seed | ERBB2 | 1.137594 | BLCA | LB831-BLC | Lapatinib |
| MVP_top60_DNA_seed | EP300 | -1.459423 | BRCA | JIMT-1 | I-CBP112 | MVP_top60_DNA+RNA_seed | FGFR3 | 1.141148 | BLCA | UM-UC-3 | PD173074 |
| MVP_top60_DNA_seed | CREBBP | -1.459423 | BRCA | JIMT-1 | I-CBP112 | MVP_top60_DNA+RNA_seed | ERBB2 | 1.143819 | BLCA | J82 | Lapatinib |
| MVP_top60_DNA_seed | AKT1 | -1.459332 | BRCA | HCC1569 | AKT inhibitor VIII | MVP_top60_DNA+RNA_seed | MAPK1 | 1.150021 | BLCA | LB831-BLC | VX-11e |
| MVP_top60_DNA_seed | EGFR | -1.453118 | BRCA | ZR-75-30 | Pelitinib | MVP_top60_DNA+RNA_seed | MAPK1 | 1.159215 | BLCA | HT-1197 | Ulixertinib |
| MVP_top60_DNA_seed | HSP90AA1 | -1.4471 | BRCA | MCF7 | Tanespimycin | MVP_top60_DNA+RNA_seed | MAPK3 | 1.159215 | BLCA | HT-1197 | Ulixertinib |
| MVP_top60_DNA_seed | HSP90AB1 | -1.4471 | BRCA | MCF7 | Tanespimycin | MVP_top60_DNA+RNA_seed | SRC | 1.161011 | BLCA | SW780 | Bosutinib |
| MVP_top60_DNA_seed | ERBB2 | -1.442501 | BRCA | MDA-MB-453 | CUDC-101 | MVP_top60_DNA+RNA_seed | SYK | 1.166608 | BLCA | 647-V | PRT062607 |
| MVP_top60_DNA_seed | EGFR | -1.442501 | BRCA | MDA-MB-453 | CUDC-101 | MVP_top60_DNA+RNA_seed | ERBB2 | 1.171975 | BLCA | J82 | Afatinib |
| MVP_top60_DNA_seed | ATM | -1.440139 | BRCA | CAL-51 | KU-60019 | MVP_top60_DNA+RNA_seed | AKT1 | 1.178829 | BLCA | 639-V | AT13148 |
| MVP_top60_DNA_seed | PIK3CA | -1.437506 | BRCA | JIMT-1 | AZD8835 | MVP_top60_DNA+RNA_seed | AKT1 | 1.179464 | BLCA | SW780 | Ipatasertib |
| MVP_top60_DNA_seed | AKT1 | -1.431509 | BRCA | HCC1954 | BAY AKT1 | MVP_top60_DNA+RNA_seed | AKT1 | 1.181698 | BLCA | J82 | MK-2206 |
| MVP_top60_DNA_seed | ATM | -1.430587 | BRCA | OCUB-M | KU-60019 | MVP_top60_DNA+RNA_seed | AKT1 | 1.186752 | BLCA | 5637 | Afuresertib |
| MVP_top60_DNA_seed | AKT1 | -1.423091 | BRCA | HCC1599 | AKT inhibitor VIII | MVP_top60_DNA+RNA_seed | ATM | 1.190943 | BLCA | J82 | KU-55933 |
| MVP_top60_DNA_seed | AKT1 | -1.421005 | BRCA | EVSA-T | AKT inhibitor VIII | MVP_top60_DNA+RNA_seed | ERBB2 | 1.197309 | BLCA | SW780 | Sapitinib |
| MVP_top60_DNA_seed | HDAC1 | -1.414216 | BRCA | CAL-85-1 | Dacinostat | MVP_top60_DNA+RNA_seed | ERBB3 | 1.197309 | BLCA | SW780 | Sapitinib |
| MVP_top60_DNA_seed | ERBB2 | -1.413423 | BRCA | ZR-75-30 | CP724714 | MVP_top60_DNA+RNA_seed | ERBB2 | 1.211017 | BLCA | UM-UC-3 | Lapatinib |
| MVP_top60_DNA_seed | AKT1 | -1.410322 | BRCA | OCUB-M | MK-2206 | MVP_top60_DNA+RNA_seed | SYK | 1.220589 | BLCA | T24 | PRT062607 |
| MVP_top60_DNA_seed | HDAC1 | -1.407702 | BRCA | MCF7 | JQ12 | MVP_top60_DNA+RNA_seed | CDK2 | 1.237529 | BLCA | TCCSUP | AZD5438 |
| MVP_top60_DNA_seed | HDAC2 | -1.407702 | BRCA | MCF7 | JQ12 | MVP_top60_DNA+RNA_seed | SRC | 1.238383 | BLCA | J82 | Bosutinib |
| MVP_top60_DNA_seed | HDAC1 | -1.406492 | BRCA | MDA-MB-361 | JQ12 | MVP_top60_DNA+RNA_seed | CDK2 | 1.241545 | BLCA | HT-1197 | MK-8776 |
| MVP_top60_DNA_seed | HDAC2 | -1.406492 | BRCA | MDA-MB-361 | JQ12 | MVP_top60_DNA+RNA_seed | ATM | 1.25491 | BLCA | HT-1197 | KU-55933 |
| MVP_top60_DNA_seed | AR | -1.3957 | BRCA | OCUB-M | Bicalutamide | MVP_top60_DNA+RNA_seed | PIK3CA | 1.261823 | BLCA | 647-V | Alpelisib |
| MVP_top60_DNA_seed | AR | -1.395492 | BRCA | MDA-MB-453 | Bicalutamide | MVP_top60_DNA+RNA_seed | ERBB2 | 1.262369 | BLCA | SW780 | Afatinib |
| MVP_top60_DNA_seed | HSP90AA1 | -1.391886 | BRCA | EVSA-T | Tanespimycin | MVP_top60_DNA+RNA_seed | AKT1 | 1.262599 | BLCA | SW780 | Afuresertib |
| MVP_top60_DNA_seed | HSP90AB1 | -1.391886 | BRCA | EVSA-T | Tanespimycin | MVP_top60_DNA+RNA_seed | MAPK1 | 1.263677 | BLCA | UM-UC-3 | Ulixertinib |
| MVP_top60_DNA_seed | EGFR | -1.383862 | BRCA | BT-549 | Pelitinib | MVP_top60_DNA+RNA_seed | MAPK3 | 1.263677 | BLCA | UM-UC-3 | Ulixertinib |
| MVP_top60_DNA_seed | HDAC1 | -1.372223 | BRCA | MDA-MB-453 | Tubastatin A | MVP_top60_DNA+RNA_seed | AKT1 | 1.266405 | BLCA | J82 | Uprosertib |
| MVP_top60_DNA_seed | ERBB2 | -1.356386 | BRCA | HDQ-P1 | Afatinib | MVP_top60_DNA+RNA_seed | ATM | 1.269779 | BLCA | HT-1376 | KU-55933 |
| MVP_top60_DNA_seed | EGFR | -1.356386 | BRCA | HDQ-P1 | Afatinib | MVP_top60_DNA+RNA_seed | AKT1 | 1.271487 | BLCA | HT-1376 | Uprosertib |
| MVP_top60_DNA_seed | CDK1 | -1.352117 | BRCA | MCF7 | RO-3306 | MVP_top60_DNA+RNA_seed | AKT1 | 1.27278 | BLCA | 639-V | Uprosertib |
| MVP_top60_DNA_seed | ERBB2 | -1.346121 | BRCA | HCC1419 | Afatinib | MVP_top60_DNA+RNA_seed | AKT1 | 1.276749 | BLCA | HT-1376 | MK-2206 |
| MVP_top60_DNA_seed | EGFR | -1.346121 | BRCA | HCC1419 | Afatinib | MVP_top60_DNA+RNA_seed | PIK3CA | 1.29491 | BLCA | UM-UC-3 | Alpelisib |

| | | | | | | | | | | | |
|---|---|---|---|---|---|---|---|---|---|---|---|
| MVP_top60_DNA_seed | AKT1 | -1.345647 | BRCA | YMB-1-E | AKT inhibitor VIII | MVP_top60_DNA+RNA_seed | SRC | 1.296924 | BLCA | TCCSUP | Bosutinib |
| MVP_top60_DNA_seed | EGFR | -1.342716 | BRCA | MDA-MB-468 | Gefitinib | MVP_top60_DNA+RNA_seed | HDAC1 | 1.297524 | BLCA | SW1710 | Romidepsin |
| MVP_top60_DNA_seed | HSP90AA1 | -1.339572 | BRCA | MDA-MB-330 | Tanespimycin | MVP_top60_DNA+RNA_seed | HSP90AA1 | 1.303137 | BLCA | HT-1197 | Luminespib |
| MVP_top60_DNA_seed | HSP90AB1 | -1.339572 | BRCA | MDA-MB-330 | Tanespimycin | MVP_top60_DNA+RNA_seed | MAPK1 | 1.303322 | BLCA | HT-1197 | SCH772984 |
| MVP_top60_DNA_seed | MDM2 | -1.339111 | BRCA | MDA-MB-231 | Serdemetan | MVP_top60_DNA+RNA_seed | MAPK3 | 1.303322 | BLCA | HT-1197 | SCH772984 |
| MVP_top60_DNA_seed | MDM2 | -1.332638 | BRCA | MCF7 | Nutlin-3a (-) | MVP_top60_DNA+RNA_seed | PIK3CA | 1.309147 | BLCA | J82 | Alpelisib |
| MVP_top60_DNA_seed | AKT1 | -1.332366 | BRCA | HCC1569 | AKT inhibitor VIII | MVP_top60_DNA+RNA_seed | ERBB2 | 1.311765 | BLCA | TCCSUP | Afatinib |
| MVP_top60_DNA_seed | HSP90AA1 | -1.331114 | BRCA | AU565 | CCT-018159 | MVP_top60_DNA+RNA_seed | AKT1 | 1.319851 | BLCA | SW780 | Uprosertib |
| MVP_top60_DNA_seed | HSP90AB1 | -1.331114 | BRCA | AU565 | CCT-018159 | MVP_top60_DNA+RNA_seed | CDK2 | 1.322245 | BLCA | HT-1197 | AZD5438 |
| MVP_top60_DNA_seed | HSP90AA1 | -1.331059 | BRCA | OCUB-M | CCT-018159 | MVP_top60_DNA+RNA_seed | AKT1 | 1.323849 | BLCA | HT-1376 | AZD5363 |
| MVP_top60_DNA_seed | HSP90AB1 | -1.331059 | BRCA | OCUB-M | CCT-018159 | MVP_top60_DNA+RNA_seed | ERBB2 | 1.332755 | BLCA | T24 | Lapatinib |
| MVP_top60_DNA_seed | EGFR | -1.328804 | BRCA | MDA-MB-175-VII | Gefitinib | MVP_top60_DNA+RNA_seed | KRAS | 1.344899 | BLCA | TCCSUP | (RAS (G12C) Inhibitor-1 |
| MVP_top60_DNA_seed | EGFR | -1.326821 | BRCA | EFM-192A | AST-1306 | MVP_top60_DNA+RNA_seed | HDAC1 | 1.357084 | BLCA | SW780 | Romidepsin |
| MVP_top60_DNA_seed | ATM | -1.322394 | BRCA | Hs-578-T | KU-55933 | MVP_top60_DNA+RNA_seed | AKT1 | 1.358885 | BLCA | HT-1376 | Uprosertib |
| MVP_top60_DNA_seed | AKT1 | -1.318777 | BRCA | HCC1569 | GSK690693 | MVP_top60_DNA+RNA_seed | AKT1 | 1.372699 | BLCA | SW780 | AZD5363 |
| MVP_top60_DNA_seed | AKT1 | -1.312031 | BRCA | MDA-MB-468 | AKT inhibitor VIII | MVP_top60_DNA+RNA_seed | HDAC1 | 1.39272 | BLCA | 639-V | Entinostat |
| MVP_top60_DNA_seed | PIK3CA | -1.311004 | BRCA | EFM-19 | PI3Ka_4409 | MVP_top60_DNA+RNA_seed | PIK3CA | 1.405059 | BLCA | SW780 | Alpelisib |
| MVP_top60_DNA_seed | PIK3CA | -1.288787 | BRCA | EFM-192A | PI3Ka_4409 | MVP_top60_DNA+RNA_seed | ERBB2 | 1.405351 | BLCA | HT-1376 | Afatinib |
| MVP_top60_DNA_seed | ESR1 | -1.285457 | BRCA | AU565 | Fulvestrant | MVP_top60_DNA+RNA_seed | FGFR3 | 1.427875 | BLCA | HT-1376 | PD173074 |
| MVP_top60_DNA_seed | ERBB2 | -1.285071 | BRCA | MDA-MB-468 | Afatinib | MVP_top60_DNA+RNA_seed | AKT1 | 1.454679 | BLCA | TCCSUP | AT13148 |
| MVP_top60_DNA_seed | EGFR | -1.285071 | BRCA | MDA-MB-468 | Afatinib | MVP_top60_DNA+RNA_seed | AKT1 | 1.461819 | BLCA | 5637 | Uprosertib |
| MVP_top60_DNA_seed | EGFR | -1.280387 | BRCA | MDA-MB-330 | AST-1306 | MVP_top60_DNA+RNA_seed | AKT1 | 1.480396 | BLCA | RT4 | Ipatasertib |
| MVP_top60_DNA_seed | HSP90AA1 | -1.278815 | BRCA | DU-4475 | SNX-2112 | MVP_top60_DNA+RNA_seed | ERBB2 | 1.483823 | BLCA | HT-1197 | Lapatinib |
| MVP_top60_DNA_seed | HSP90AB1 | -1.278815 | BRCA | DU-4475 | SNX-2112 | MVP_top60_DNA+RNA_seed | AKT1 | 1.486132 | BLCA | SW780 | Uprosertib |
| MVP_top60_DNA_seed | AKT1 | -1.278181 | BRCA | HCC1599 | GSK690693 | MVP_top60_DNA+RNA_seed | SYK | 1.488834 | BLCA | T24 | Entospletinib |
| MVP_top60_DNA_seed | HSP90AA1 | -1.275733 | BRCA | CAL-51 | Luminespib | MVP_top60_DNA+RNA_seed | ESR1 | 1.491866 | BLCA | T24 | Tamoxifen |
| MVP_top60_DNA_seed | HSP90AB1 | -1.275733 | BRCA | CAL-51 | Luminespib | MVP_top60_DNA+RNA_seed | EP300 | 1.51147 | BLCA | HT-1197 | SGC-CBP30 |
| MVP_top60_DNA_seed | EGFR | -1.274644 | BRCA | MDA-MB-468 | Pelitinib | MVP_top60_DNA+RNA_seed | CREBBP | 1.51147 | BLCA | HT-1197 | SGC-CBP30 |
| MVP_top60_DNA_seed | HDAC1 | -1.274228 | BRCA | HCC1428 | Dacinostat | MVP_top60_DNA+RNA_seed | MAPK1 | 1.518474 | BLCA | UM-UC-3 | Ulixertinib |
| MVP_top60_DNA_seed | ERBB2 | -1.26613 | BRCA | HCC202 | PF-00299804 | MVP_top60_DNA+RNA_seed | MAPK3 | 1.518474 | BLCA | UM-UC-3 | Ulixertinib |
| MVP_top60_DNA_seed | EGFR | -1.26613 | BRCA | HCC202 | PF-00299804 | MVP_top60_DNA+RNA_seed | AKT1 | 1.54701 | BLCA | HT-1376 | AT13148 |
| MVP_top60_DNA_seed | HDAC1 | -1.265904 | BRCA | EVSA-T | Parthenolide | MVP_top60_DNA+RNA_seed | AKT1 | 1.559286 | BLCA | T24 | MK-2206 |
| MVP_top60_DNA_seed | AKT1 | -1.255299 | BRCA | CAL-51 | AKT inhibitor VIII | MVP_top60_DNA+RNA_seed | MAPK1 | 1.563748 | BLCA | J82 | Ulixertinib |
| MVP_top60_DNA_seed | HSP90AA1 | -1.253933 | BRCA | HCC2157 | CCT-018159 | MVP_top60_DNA+RNA_seed | MAPK3 | 1.563748 | BLCA | J82 | Ulixertinib |
| MVP_top60_DNA_seed | HSP90AB1 | -1.253933 | BRCA | HCC2157 | CCT-018159 | MVP_top60_DNA+RNA_seed | MAPK1 | 1.565737 | BLCA | TCCSUP | Ulixertinib |
| MVP_top60_DNA_seed | ATM | -1.251797 | BRCA | HCC1500 | CP466722 | MVP_top60_DNA+RNA_seed | MAPK3 | 1.565737 | BLCA | TCCSUP | Ulixertinib |
| MVP_top60_DNA_seed | ERBB2 | -1.245807 | BRCA | HCC202 | Afatinib | MVP_top60_DNA+RNA_seed | HDAC1 | 1.565886 | BLCA | RT4 | Entinostat |
| MVP_top60_DNA_seed | EGFR | -1.245807 | BRCA | HCC202 | Afatinib | MVP_top60_DNA+RNA_seed | AKT1 | 1.579811 | BLCA | RT4 | Uprosertib |
| MVP_top60_DNA_seed | ERBB2 | -1.243057 | BRCA | HCC1599 | PF-00299804 | MVP_top60_DNA+RNA_seed | AKT1 | 1.583184 | BLCA | T24 | AZD5363 |
| MVP_top60_DNA_seed | EGFR | -1.243057 | BRCA | HCC1599 | PF-00299804 | MVP_top60_DNA+RNA_seed | AKT1 | 1.585922 | BLCA | TCCSUP | Ipatasertib |
| MVP_top60_DNA_seed | ERBB2 | -1.24242 | BRCA | HCC1419 | Afatinib | MVP_top60_DNA+RNA_seed | MAPK1 | 1.618588 | BLCA | HT-1197 | VX-11e |

| | | | | | | | | | | | |
|---|---|---|---|---|---|---|---|---|---|---|---|
| MVP_top60_DNA_seed | EGFR | -1.24242 | BRCA | HCC1419 | Afatinib | MVP_top60_DNA+RNA_seed | AKT1 | 1.621279 | BLCA | TCCSUP | MK-2206 |
| MVP_top60_DNA_seed | MDM2 | -1.24201 | BRCA | BT-549 | Serdemetan | MVP_top60_DNA+RNA_seed | AKT1 | 1.622517 | BLCA | TCCSUP | Uprosertib |
| MVP_top60_DNA_seed | EGFR | -1.241509 | BRCA | MDA-MB-453 | AST-1306 | MVP_top60_DNA+RNA_seed | AKT1 | 1.644244 | BLCA | RT4 | AZD5363 |
| MVP_top60_DNA_seed | AKT1 | -1.240709 | BRCA | MFM-223 | MK-2206 | MVP_top60_DNA+RNA_seed | AKT1 | 1.645548 | BLCA | HT-1376 | Ipatasertib |
| MVP_top60_DNA_seed | ESR1 | -1.23885 | BRCA | BT-549 | Fulvestrant | MVP_top60_DNA+RNA_seed | HSP90AA1 | 1.649132 | BLCA | J82 | Tanespimycin |
| MVP_top60_DNA_seed | AKT1 | -1.228396 | BRCA | UACC-893 | AKT inhibitor VIII | MVP_top60_DNA+RNA_seed | PIK3CA | 1.680588 | BLCA | T24 | Buparlisib |
| MVP_top60_DNA_seed | ERBB2 | -1.226681 | BRCA | Hs-578-T | PF-00299804 | MVP_top60_DNA+RNA_seed | AKT1 | 1.682355 | BLCA | RT4 | Afuresertib |
| MVP_top60_DNA_seed | EGFR | -1.226681 | BRCA | Hs-578-T | PF-00299804 | MVP_top60_DNA+RNA_seed | MAPK1 | 1.699284 | BLCA | TCCSUP | SCH772984 |
| MVP_top60_DNA_seed | EGFR | -1.220525 | BRCA | HCC1419 | Pelitinib | MVP_top60_DNA+RNA_seed | MAPK3 | 1.699284 | BLCA | TCCSUP | SCH772984 |
| MVP_top60_DNA_seed | AKT1 | -1.21847 | BRCA | HCC1954 | AKT inhibitor VIII | MVP_top60_DNA+RNA_seed | PIK3CA | 1.706332 | BLCA | RT4 | AZD8186 |
| MVP_top60_DNA_seed | ATM | -1.215249 | BRCA | MDA-MB-468 | KU-55933 | MVP_top60_DNA+RNA_seed | PIK3CA | 1.715415 | BLCA | HT-1376 | GNE-317 |
| MVP_top60_DNA_seed | AKT1 | -1.21461 | BRCA | BT-474 | AKT inhibitor VIII | MVP_top60_DNA+RNA_seed | PIK3CA | 1.716928 | BLCA | SW780 | AZD8186 |
| MVP_top60_DNA_seed | AR | -1.212017 | BRCA | MCF7 | Bicalutamide | MVP_top60_DNA+RNA_seed | PIK3CA | 1.756588 | BLCA | HT-1376 | Alpelisib |
| MVP_top60_DNA_seed | AKT1 | -1.203389 | BRCA | AU565 | AKT inhibitor VIII | MVP_top60_DNA+RNA_seed | CDK2 | 1.765449 | BLCA | J82 | AZD5438 |
| MVP_top60_DNA_seed | EGFR | -1.201192 | BRCA | CAL-85-1 | Cetuximab | MVP_top60_DNA+RNA_seed | SYK | 1.780851 | BLCA | J82 | PRT062607 |
| MVP_top60_DNA_seed | PIK3CA | -1.199455 | BRCA | BT-20 | AZD8835 | MVP_top60_DNA+RNA_seed | ERBB2 | 1.799722 | BLCA | T24 | Afatinib |
| MVP_top60_DNA_seed | HDAC1 | -1.19842 | BRCA | DU-4475 | Entinostat | MVP_top60_DNA+RNA_seed | FGFR3 | 1.805725 | BLCA | T24 | AZD4547 |
| MVP_top60_DNA_seed | PIK3CA | -1.194497 | BRCA | EFM-19 | AZD8835 | MVP_top60_DNA+RNA_seed | FGFR3 | 1.826144 | BLCA | TCCSUP | PD173074 |
| MVP_top60_DNA_seed | AKT1 | -1.188487 | BRCA | HCC1419 | AKT inhibitor VIII | MVP_top60_DNA+RNA_seed | AKT1 | 1.90821 | BLCA | T24 | Uprosertib |
| MVP_top60_DNA_seed | AKT1 | -1.186063 | BRCA | HCC1419 | AKT inhibitor VIII | MVP_top60_DNA+RNA_seed | PIK3CA | 1.951983 | BLCA | J82 | Buparlisib |
| MVP_top60_DNA_seed | ATM | -1.18475 | BRCA | ZR-75-30 | CP466722 | MVP_top60_DNA+RNA_seed | HSP90AA1 | 1.968166 | BLCA | SW780 | Luminespib |
| MVP_top60_DNA_seed | ATM | -1.178105 | BRCA | MDA-MB-231 | KU-60019 | MVP_top60_DNA+RNA_seed | HSP90AA1 | 2.011633 | BLCA | J82 | Luminespib |
| MVP_top60_DNA_seed | AKT1 | -1.166147 | BRCA | MCF7 | BAY AKT1 | MVP_top60_DNA+RNA_seed | HDAC1 | 2.059134 | BLCA | RT4 | Romidepsin |
| MVP_top60_DNA_seed | ATM | -1.163912 | BRCA | CAL-51 | KU-55933 | MVP_top60_DNA+RNA_seed | ERBB2 | 2.060922 | BLCA | TCCSUP | Lapatinib |
| MVP_top60_DNA_seed | PIK3CA | -1.156943 | BRCA | HCC1599 | AZD8835 | MVP_top60_DNA+RNA_seed | HSP90AA1 | 2.071933 | BLCA | RT4 | Luminespib |
| MVP_top60_DNA_seed | HSP90AA1 | -1.155473 | BRCA | HCC2218 | Tanespimycin | MVP_top60_DNA+RNA_seed | RAC1 | 2.108204 | BLCA | J82 | EHT-1864 |
| MVP_top60_DNA_seed | HSP90AB1 | -1.155473 | BRCA | HCC2218 | Tanespimycin | MVP_top60_DNA+RNA_seed | MAPK1 | 2.116787 | BLCA | HT-1376 | VX-11e |
| MVP_top60_DNA_seed | HSP90AA1 | -1.151911 | BRCA | CAL-85-1 | Luminespib | MVP_top60_DNA+RNA_seed | MAPK1 | 2.238414 | BLCA | TCCSUP | VX-11e |
| MVP_top60_DNA_seed | HSP90AB1 | -1.151911 | BRCA | CAL-85-1 | Luminespib | MVP_top60_DNA+RNA_seed | PIK3CA | 2.716316 | BLCA | HT-1376 | Buparlisib |
| MVP_top60_DNA_seed | CDK2 | -1.151067 | BRCA | HCC1599 | PHA-793887 | MVP_top60_DNA+RNA_seed | CDK2 | 2.72311 | BLCA | HT-1376 | AZD5438 |
| MVP_top60_DNA_seed | AKT1 | -1.149798 | BRCA | MDA-MB-361 | AKT inhibitor VIII | MVP_top60_DNA+RNA_seed | CDK2 | 2.922617 | BLCA | J82 | Dinaciclib |
| MVP_top60_DNA_seed | ERBB2 | -1.146226 | BRCA | CAL-85-1 | Afatinib | MVP_top60_DNA+RNA_seed | PIK3CA | 2.971117 | BLCA | J82 | GNE-317 |
| MVP_top60_DNA_seed | EGFR | -1.146226 | BRCA | CAL-85-1 | Afatinib | MVP_top60_DNA+RNA_seed | HDAC1 | 3.488344 | BLCA | J82 | Romidepsin |
| MVP_top60_DNA_seed | CDK2 | -1.144685 | BRCA | MDA-MB-468 | PHA-793887 | MVP_top60_DNA+RNA_seed | ESR1 | -6.409629 | BRCA | Hs-578-T | Fulvestrant |
| MVP_top60_DNA_seed | HSP90AA1 | -1.141385 | BRCA | OCUB-M | SNX-2112 | MVP_top60_DNA+RNA_seed | ERBB2 | -6.352842 | BRCA | HCC2218 | CP724714 |
| MVP_top60_DNA_seed | HSP90AB1 | -1.141385 | BRCA | OCUB-M | SNX-2112 | MVP_top60_DNA+RNA_seed | ERBB2 | -5.634798 | BRCA | BT-474 | CP724714 |
| MVP_top60_DNA_seed | AR | -1.138431 | BRCA | OCUB-M | Bicalutamide | MVP_top60_DNA+RNA_seed | EGFR | -4.8899 | BRCA | OCUB-M | Lapatinib |
| MVP_top60_DNA_seed | PIK3CA | -1.1376 | BRCA | UACC-893 | PI3Ka_4409 | MVP_top60_DNA+RNA_seed | ERBB2 | -4.8899 | BRCA | OCUB-M | Lapatinib |
| MVP_top60_DNA_seed | CDK1 | -1.136599 | BRCA | AU565 | RO-3306 | MVP_top60_DNA+RNA_seed | EGFR | -4.736905 | BRCA | HCC2218 | Lapatinib |
| MVP_top60_DNA_seed | HSP90AA1 | -1.134675 | BRCA | MDA-MB-453 | SNX-2112 | MVP_top60_DNA+RNA_seed | ERBB2 | -4.736905 | BRCA | HCC2218 | Lapatinib |
| MVP_top60_DNA_seed | HSP90AB1 | -1.134675 | BRCA | MDA-MB-453 | SNX-2112 | MVP_top60_DNA+RNA_seed | EGFR | -4.54251 | BRCA | BT-474 | PF-00299804 |

| | | | | | | | | | | | |
|---|---|---|---|---|---|---|---|---|---|---|---|
| MVP_top60_DNA_seed | ERBB2 | -1.132719 | BRCA | MDA-MB-453 | CI-1033 | MVP_top60_DNA+RNA_seed | ERBB2 | -4.54251 | BRCA | BT-474 | PF-00299804 |
| MVP_top60_DNA_seed | EGFR | -1.132719 | BRCA | MDA-MB-453 | CI-1033 | MVP_top60_DNA+RNA_seed | ESR1 | -4.456821 | BRCA | CAMA-1 | Fulvestrant |
| MVP_top60_DNA_seed | CDK2 | -1.131387 | BRCA | HCC1187 | PHA-793887 | MVP_top60_DNA+RNA_seed | ERBB2 | -4.30329 | BRCA | MDA-MB-175-VII | CP724714 |
| MVP_top60_DNA_seed | PIK3CA | -1.130069 | BRCA | T47D | PI3Ka_4409 | MVP_top60_DNA+RNA_seed | EGFR | -4.118491 | BRCA | BT-474 | AST-1306 |
| MVP_top60_DNA_seed | ATM | -1.126387 | BRCA | CAL-85-1 | CP466722 | MVP_top60_DNA+RNA_seed | EGFR | -4.04302 | BRCA | AU565 | PF-00299804 |
| MVP_top60_DNA_seed | HDAC1 | -1.124987 | BRCA | MDA-MB-330 | JQ12 | MVP_top60_DNA+RNA_seed | ERBB2 | -4.04302 | BRCA | AU565 | PF-00299804 |
| MVP_top60_DNA_seed | HDAC2 | -1.124987 | BRCA | MDA-MB-330 | JQ12 | MVP_top60_DNA+RNA_seed | EGFR | -3.927648 | BRCA | BT-474 | CI-1033 |
| MVP_top60_DNA_seed | PIK3CA | -1.12156 | BRCA | MCF7 | AZD8835 | MVP_top60_DNA+RNA_seed | ERBB2 | -3.927648 | BRCA | BT-474 | CI-1033 |
| MVP_top60_DNA_seed | HDAC1 | -1.119708 | BRCA | DU-4475 | Dacinostat | MVP_top60_DNA+RNA_seed | ERBB2 | -3.805563 | BRCA | OCUB-M | CP724714 |
| MVP_top60_DNA_seed | HDAC1 | -1.108087 | BRCA | OCUB-M | Dacinostat | MVP_top60_DNA+RNA_seed | EGFR | -3.767342 | BRCA | AU565 | AST-1306 |
| MVP_top60_DNA_seed | ERBB2 | -1.104227 | BRCA | DU-4475 | CP724714 | MVP_top60_DNA+RNA_seed | ESR1 | -3.509161 | BRCA | T47D | Tamoxifen |
| MVP_top60_DNA_seed | ESR1 | -1.102562 | BRCA | MDA-MB-157 | Fulvestrant | MVP_top60_DNA+RNA_seed | EGFR | -3.449584 | BRCA | AU565 | CI-1033 |
| MVP_top60_DNA_seed | ERBB2 | -1.101357 | BRCA | HCC1187 | PF-00299804 | MVP_top60_DNA+RNA_seed | ERBB2 | -3.449584 | BRCA | AU565 | CI-1033 |
| MVP_top60_DNA_seed | EGFR | -1.101357 | BRCA | HCC1187 | PF-00299804 | MVP_top60_DNA+RNA_seed | AKT1 | -3.350027 | BRCA | OCUB-M | GSK690693 |
| MVP_top60_DNA_seed | ATM | -1.10092 | BRCA | T47D | KU-60019 | MVP_top60_DNA+RNA_seed | MAPK1 | -3.312718 | BRCA | DU-4475 | VX-11e |
| MVP_top60_DNA_seed | EGFR | -1.099539 | BRCA | EFM-19 | AST-1306 | MVP_top60_DNA+RNA_seed | EGFR | -3.307809 | BRCA | MDA-MB-175-VII | PF-00299804 |
| MVP_top60_DNA_seed | ERBB2 | -1.099158 | BRCA | MDA-MB-468 | PF-00299804 | MVP_top60_DNA+RNA_seed | ERBB2 | -3.307809 | BRCA | MDA-MB-175-VII | PF-00299804 |
| MVP_top60_DNA_seed | EGFR | -1.099158 | BRCA | MDA-MB-468 | PF-00299804 | MVP_top60_DNA+RNA_seed | ESR1 | -3.268441 | BRCA | MDA-MB-361 | Fulvestrant |
| MVP_top60_DNA_seed | AKT1 | -1.093649 | BRCA | HCC2157 | MK-2206 | MVP_top60_DNA+RNA_seed | EGFR | -3.245884 | BRCA | HCC2218 | PF-00299804 |
| MVP_top60_DNA_seed | HSP90AA1 | -1.090695 | BRCA | EFM-192A | Tanespimycin | MVP_top60_DNA+RNA_seed | ERBB2 | -3.245884 | BRCA | HCC2218 | PF-00299804 |
| MVP_top60_DNA_seed | HSP90AB1 | -1.090695 | BRCA | EFM-192A | Tanespimycin | MVP_top60_DNA+RNA_seed | EGFR | -3.213861 | BRCA | BT-474 | Lapatinib |
| MVP_top60_DNA_seed | HDAC1 | -1.08089 | BRCA | HCC2218 | Dacinostat | MVP_top60_DNA+RNA_seed | ERBB2 | -3.213861 | BRCA | BT-474 | Lapatinib |
| MVP_top60_DNA_seed | CDK1 | -1.078041 | BRCA | JIMT-1 | RO-3306 | MVP_top60_DNA+RNA_seed | EGFR | -3.106854 | BRCA | HCC2218 | AST-1306 |
| MVP_top60_DNA_seed | HDAC1 | -1.076885 | BRCA | MDA-MB-157 | Dacinostat | MVP_top60_DNA+RNA_seed | AKT1 | -3.102256 | BRCA | EVSA-T | BAY AKT1 |
| MVP_top60_DNA_seed | ERBB2 | -1.074989 | BRCA | BT-549 | CP724714 | MVP_top60_DNA+RNA_seed | AKT1 | -3.048792 | BRCA | EFM-19 | AKT inhibitor VIII |
| MVP_top60_DNA_seed | AR | -1.074078 | BRCA | AU565 | Bicalutamide | MVP_top60_DNA+RNA_seed | ERBB2 | -3.044946 | BRCA | HCC1419 | CP724714 |
| MVP_top60_DNA_seed | PIK3CA | -1.072273 | BRCA | JIMT-1 | PI3Ka_4409 | MVP_top60_DNA+RNA_seed | EGFR | -3.042142 | BRCA | HCC2218 | Afatinib |
| MVP_top60_DNA_seed | EGFR | -1.070307 | BRCA | HCC1954 | AST-1306 | MVP_top60_DNA+RNA_seed | ERBB2 | -3.042142 | BRCA | HCC2218 | Afatinib |
| MVP_top60_DNA_seed | AKT1 | -1.067314 | BRCA | HCC2157 | GSK690693 | MVP_top60_DNA+RNA_seed | ERBB2 | -3.006821 | BRCA | AU565 | CP724714 |
| MVP_top60_DNA_seed | MDM2 | -1.066872 | BRCA | CAL-51 | Nutlin-3a (-) | MVP_top60_DNA+RNA_seed | EGFR | -2.982644 | BRCA | MDA-MB-175-VII | Pelitinib |
| MVP_top60_DNA_seed | AKT1 | -1.063205 | BRCA | UACC-893 | AKT inhibitor VIII | MVP_top60_DNA+RNA_seed | EGFR | -2.939075 | BRCA | HCC1419 | PF-00299804 |
| MVP_top60_DNA_seed | HSP90AA1 | -1.063194 | BRCA | JIMT-1 | Tanespimycin | MVP_top60_DNA+RNA_seed | ERBB2 | -2.939075 | BRCA | HCC1419 | PF-00299804 |
| MVP_top60_DNA_seed | HSP90AB1 | -1.063194 | BRCA | JIMT-1 | Tanespimycin | MVP_top60_DNA+RNA_seed | EGFR | -2.903137 | BRCA | HCC1419 | AST-1306 |
| MVP_top60_DNA_seed | HSP90AA1 | -1.059699 | BRCA | BT-549 | Elesclomol | MVP_top60_DNA+RNA_seed | AKT1 | -2.871968 | BRCA | MDA-MB-361 | AKT inhibitor VIII |
| MVP_top60_DNA_seed | HSP90AB1 | -1.059699 | BRCA | BT-549 | Elesclomol | MVP_top60_DNA+RNA_seed | EGFR | -2.869472 | BRCA | MDA-MB-175-VII | AST-1306 |
| MVP_top60_DNA_seed | HSP90AA1 | -1.055238 | BRCA | Hs-578-T | Tanespimycin | MVP_top60_DNA+RNA_seed | EGFR | -2.831997 | BRCA | HCC2218 | Afatinib |
| MVP_top60_DNA_seed | HSP90AB1 | -1.055238 | BRCA | Hs-578-T | Tanespimycin | MVP_top60_DNA+RNA_seed | ERBB2 | -2.831997 | BRCA | HCC2218 | Afatinib |
| MVP_top60_DNA_seed | AKT1 | -1.044714 | BRCA | EVSA-T | GSK690693 | MVP_top60_DNA+RNA_seed | EGFR | -2.816371 | BRCA | AU565 | Afatinib |
| MVP_top60_DNA_seed | HDAC1 | -1.043514 | BRCA | OCUB-M | Parthenolide | MVP_top60_DNA+RNA_seed | ERBB2 | -2.816371 | BRCA | AU565 | Afatinib |
| MVP_top60_DNA_seed | HDAC1 | -1.043264 | BRCA | UACC-812 | Tubastatin A | MVP_top60_DNA+RNA_seed | AKT1 | -2.815989 | BRCA | EVSA-T | MK-2206 |
| MVP_top60_DNA_seed | HSP90AA1 | -1.04278 | BRCA | MRK-nu-1 | Luminespib | MVP_top60_DNA+RNA_seed | EGFR | -2.797545 | BRCA | EFM-192A | Afatinib |

| | | | | | | | | | | | |
|---|---|---|---|---|---|---|---|---|---|---|---|
| MVP_top60_DNA_seed | HSP90AB1 | -1.04278 | BRCA | MRK-nu-1 | Luminespib | MVP_top60_DNA+RNA_seed | ERBB2 | -2.797545 | BRCA | EFM-192A | Afatinib |
| MVP_top60_DNA_seed | CDK2 | -1.042589 | BRCA | DU-4475 | PHA-793887 | MVP_top60_DNA+RNA_seed | AKT1 | -2.795246 | BRCA | OCUB-M | BAY AKT1 |
| MVP_top60_DNA_seed | ERBB2 | -1.041564 | BRCA | MDA-MB-468 | AZD8931 | MVP_top60_DNA+RNA_seed | EGFR | -2.779453 | BRCA | MDA-MB-175-VII | CI-1033 |
| MVP_top60_DNA_seed | EGFR | -1.041564 | BRCA | MDA-MB-468 | AZD8931 | MVP_top60_DNA+RNA_seed | ERBB2 | -2.779453 | BRCA | MDA-MB-175-VII | CI-1033 |
| MVP_top60_DNA_seed | ERBB3 | -1.041564 | BRCA | MDA-MB-468 | AZD8931 | MVP_top60_DNA+RNA_seed | EGFR | -2.760498 | BRCA | OCUB-M | AST-1306 |
| MVP_top60_DNA_seed | AKT1 | -1.041496 | BRCA | BT-474 | BAY AKT1 | MVP_top60_DNA+RNA_seed | EGFR | -2.741944 | BRCA | MDA-MB-175-VII | Afatinib |
| MVP_top60_DNA_seed | HDAC1 | -1.032349 | BRCA | HCC1599 | AR-42 | MVP_top60_DNA+RNA_seed | ERBB2 | -2.741944 | BRCA | MDA-MB-175-VII | Afatinib |
| MVP_top60_DNA_seed | ESR1 | -1.031006 | BRCA | OCUB-M | Tamoxifen | MVP_top60_DNA+RNA_seed | AR | -2.677455 | BRCA | HCC1599 | AZD3514 |
| MVP_top60_DNA_seed | AKT1 | -1.026894 | BRCA | BT-20 | AKT inhibitor VIII | MVP_top60_DNA+RNA_seed | EGFR | -2.665276 | BRCA | MDA-MB-175-VII | Afatinib |
| MVP_top60_DNA_seed | HSP90AA1 | -1.02336 | BRCA | DU-4475 | Luminespib | MVP_top60_DNA+RNA_seed | ERBB2 | -2.665276 | BRCA | MDA-MB-175-VII | Afatinib |
| MVP_top60_DNA_seed | HSP90AB1 | -1.02336 | BRCA | DU-4475 | Luminespib | MVP_top60_DNA+RNA_seed | EGFR | -2.650893 | BRCA | MDA-MB-468 | Cetuximab |
| MVP_top60_DNA_seed | EGFR | -1.019592 | BRCA | HCC1599 | Pelitinib | MVP_top60_DNA+RNA_seed | AR | -2.642482 | BRCA | CAMA-1 | Bicalutamide |
| MVP_top60_DNA_seed | HDAC1 | -1.016586 | BRCA | MDA-MB-453 | CAY10603 | MVP_top60_DNA+RNA_seed | EGFR | -2.628485 | BRCA | OCUB-M | PF-00299804 |
| MVP_top60_DNA_seed | EGFR | -1.015866 | BRCA | HDQ-P1 | Gefitinib | MVP_top60_DNA+RNA_seed | ERBB2 | -2.628485 | BRCA | OCUB-M | PF-00299804 |
| MVP_top60_DNA_seed | ESR1 | -1.003676 | BRCA | EVSA-T | Tamoxifen | MVP_top60_DNA+RNA_seed | EGFR | -2.555571 | BRCA | BT-474 | Afatinib |
| MVP_top60_DNA_seed | ERBB2 | -0.996338 | BRCA | MDA-MB-330 | PF-00299804 | MVP_top60_DNA+RNA_seed | ERBB2 | -2.555571 | BRCA | BT-474 | Afatinib |
| MVP_top60_DNA_seed | EGFR | -0.996338 | BRCA | MDA-MB-330 | PF-00299804 | MVP_top60_DNA+RNA_seed | PIK3CA | -2.548715 | BRCA | OCUB-M | PI3Ka_4409 |
| MVP_top60_DNA_seed | ATM | -0.995295 | BRCA | MCF7 | KU-60019 | MVP_top60_DNA+RNA_seed | AKT1 | -2.54498 | BRCA | T47D | AKT inhibitor VIII |
| MVP_top60_DNA_seed | ESR1 | -0.990266 | BRCA | BT-549 | Tamoxifen | MVP_top60_DNA+RNA_seed | CDK1 | -2.537566 | BRCA | MDA-MB-468 | RO-3306 |
| MVP_top60_DNA_seed | HDAC1 | -0.987847 | BRCA | HCC1599 | Tubastatin A | MVP_top60_DNA+RNA_seed | AKT1 | -2.526133 | BRCA | HCC70 | AKT inhibitor VIII |
| MVP_top60_DNA_seed | EGFR | -0.986708 | BRCA | HCC2157 | Erlotinib | MVP_top60_DNA+RNA_seed | EGFR | -2.499472 | BRCA | OCUB-M | CI-1033 |
| MVP_top60_DNA_seed | AKT1 | -0.985059 | BRCA | BT-549 | GSK690693 | MVP_top60_DNA+RNA_seed | ERBB2 | -2.499472 | BRCA | OCUB-M | CI-1033 |
| MVP_top60_DNA_seed | SRC | -0.984937 | BRCA | MFM-223 | HG-6-71-01 | MVP_top60_DNA+RNA_seed | EGFR | -2.454956 | BRCA | HCC1419 | CI-1033 |
| MVP_top60_DNA_seed | ERBB2 | -0.983487 | BRCA | OCUB-M | CUDC-101 | MVP_top60_DNA+RNA_seed | ERBB2 | -2.454956 | BRCA | HCC1419 | CI-1033 |
| MVP_top60_DNA_seed | EGFR | -0.983487 | BRCA | OCUB-M | CUDC-101 | MVP_top60_DNA+RNA_seed | ESR1 | -2.449334 | BRCA | EFM-19 | Fulvestrant |
| MVP_top60_DNA_seed | AR | -0.974125 | BRCA | HCC1500 | AZD3514 | MVP_top60_DNA+RNA_seed | AKT1 | -2.446538 | BRCA | MDA-MB-175-VII | MK-2206 |
| MVP_top60_DNA_seed | AKT1 | -0.971857 | BRCA | HCC1187 | BAY AKT1 | MVP_top60_DNA+RNA_seed | ERBB2 | -2.42065 | BRCA | MDA-MB-453 | CP724714 |
| MVP_top60_DNA_seed | SRC | -0.966973 | BRCA | JIMT-1 | Bosutinib | MVP_top60_DNA+RNA_seed | EGFR | -2.41971 | BRCA | BT-474 | Pelitinib |
| MVP_top60_DNA_seed | AR | -0.966711 | BRCA | HCC2157 | AZD3514 | MVP_top60_DNA+RNA_seed | AKT1 | -2.397081 | BRCA | CAL-148 | GSK690693 |
| MVP_top60_DNA_seed | ERBB2 | -0.965679 | BRCA | DU-4475 | CI-1033 | MVP_top60_DNA+RNA_seed | EGFR | -2.39683 | BRCA | HCC2218 | CI-1033 |
| MVP_top60_DNA_seed | EGFR | -0.965679 | BRCA | DU-4475 | CI-1033 | MVP_top60_DNA+RNA_seed | ERBB2 | -2.39683 | BRCA | HCC2218 | CI-1033 |
| MVP_top60_DNA_seed | AKT1 | -0.964751 | BRCA | HCC2157 | AKT inhibitor VIII | MVP_top60_DNA+RNA_seed | CDK1 | -2.394293 | BRCA | Hs-578-T | RO-3306 |
| MVP_top60_DNA_seed | EGFR | -0.963921 | BRCA | Hs-578-T | AST-1306 | MVP_top60_DNA+RNA_seed | AKT1 | -2.378894 | BRCA | MDA-MB-330 | AKT inhibitor VIII |
| MVP_top60_DNA_seed | EGFR | -0.963661 | BRCA | T47D | Pelitinib | MVP_top60_DNA+RNA_seed | PIK3CA | -2.357662 | BRCA | MDA-MB-453 | PI3Ka_4409 |
| MVP_top60_DNA_seed | PIK3CA | -0.962684 | BRCA | EVSA-T | PI3Ka_4409 | MVP_top60_DNA+RNA_seed | EGFR | -2.35215 | BRCA | UACC-893 | Afatinib |
| MVP_top60_DNA_seed | ERBB2 | -0.957423 | BRCA | HCC1187 | CI-1033 | MVP_top60_DNA+RNA_seed | ERBB2 | -2.35215 | BRCA | UACC-893 | Afatinib |
| MVP_top60_DNA_seed | EGFR | -0.957423 | BRCA | HCC1187 | CI-1033 | MVP_top60_DNA+RNA_seed | EGFR | -2.333524 | BRCA | OCUB-M | Pelitinib |
| MVP_top60_DNA_seed | ERBB2 | -0.957203 | BRCA | HCC1599 | CUDC-101 | MVP_top60_DNA+RNA_seed | AKT1 | -2.312486 | BRCA | OCUB-M | AKT inhibitor VIII |
| MVP_top60_DNA_seed | EGFR | -0.957203 | BRCA | HCC1599 | CUDC-101 | MVP_top60_DNA+RNA_seed | ESR1 | -2.285577 | BRCA | HCC1187 | Tamoxifen |
| MVP_top60_DNA_seed | HSP90AA1 | -0.956587 | BRCA | HCC1599 | SNX-2112 | MVP_top60_DNA+RNA_seed | ESR1 | -2.276771 | BRCA | MCF7 | Tamoxifen |
| MVP_top60_DNA_seed | HSP90AB1 | -0.956587 | BRCA | HCC1599 | SNX-2112 | MVP_top60_DNA+RNA_seed | EGFR | -2.275965 | BRCA | AU565 | Afatinib |

| | | | | | | | | | | | |
|---|---|---|---|---|---|---|---|---|---|---|---|
| MVP_top60_DNA_seed | CDK2 | -0.953106 | BRCA | HCC70 | AZD5438 | MVP_top60_DNA+RNA_seed | ERBB2 | -2.275965 | BRCA | AU565 | Afatinib |
| MVP_top60_DNA_seed | HDAC1 | -0.951777 | BRCA | AU565 | Dacinostat | MVP_top60_DNA+RNA_seed | AKT1 | -2.255889 | BRCA | HCC70 | MK-2206 |
| MVP_top60_DNA_seed | ATM | -0.950399 | BRCA | MCF7 | KU-55933 | MVP_top60_DNA+RNA_seed | EGFR | -2.239042 | BRCA | ZR-75-30 | Afatinib |
| MVP_top60_DNA_seed | HDAC1 | -0.946798 | BRCA | HCC1599 | Entinostat | MVP_top60_DNA+RNA_seed | ERBB2 | -2.239042 | BRCA | ZR-75-30 | Afatinib |
| MVP_top60_DNA_seed | ERBB2 | -0.945299 | BRCA | MDA-MB-468 | CP724714 | MVP_top60_DNA+RNA_seed | AKT1 | -2.234574 | BRCA | EFM-19 | MK-2206 |
| MVP_top60_DNA_seed | EGFR | -0.943089 | BRCA | HCC70 | Gefitinib | MVP_top60_DNA+RNA_seed | AKT1 | -2.219478 | BRCA | ZR-75-30 | AKT inhibitor VIII |
| MVP_top60_DNA_seed | PIK3CA | -0.937617 | BRCA | HCC1954 | AZD8835 | MVP_top60_DNA+RNA_seed | AKT1 | -2.217692 | BRCA | AU565 | AKT inhibitor VIII |
| MVP_top60_DNA_seed | HSP90AA1 | -0.931812 | BRCA | MCF7 | CCT-018159 | MVP_top60_DNA+RNA_seed | AKT1 | -2.217372 | BRCA | ZR-75-30 | AKT inhibitor VIII |
| MVP_top60_DNA_seed | HSP90AB1 | -0.931812 | BRCA | MCF7 | CCT-018159 | MVP_top60_DNA+RNA_seed | ERBB2 | -2.167766 | BRCA | UACC-893 | CP724714 |
| MVP_top60_DNA_seed | ERBB2 | -0.931348 | BRCA | MDA-MB-361 | CP724714 | MVP_top60_DNA+RNA_seed | EGFR | -2.160443 | BRCA | BT-474 | Afatinib |
| MVP_top60_DNA_seed | AKT1 | -0.931273 | BRCA | MFM-223 | GSK690693 | MVP_top60_DNA+RNA_seed | ERBB2 | -2.160443 | BRCA | BT-474 | Afatinib |
| MVP_top60_DNA_seed | HDAC1 | -0.929575 | BRCA | HCC1599 | CAY10603 | MVP_top60_DNA+RNA_seed | CDK1 | -2.155946 | BRCA | EVSA-T | RO-3306 |
| MVP_top60_DNA_seed | ERBB2 | -0.926688 | BRCA | OCUB-M | AZD8931 | MVP_top60_DNA+RNA_seed | MAPK3 | -2.155364 | BRCA | HCC70 | FR-180204 |
| MVP_top60_DNA_seed | EGFR | -0.926688 | BRCA | OCUB-M | AZD8931 | MVP_top60_DNA+RNA_seed | MAPK1 | -2.155364 | BRCA | HCC70 | FR-180204 |
| MVP_top60_DNA_seed | ERBB3 | -0.926688 | BRCA | OCUB-M | AZD8931 | MVP_top60_DNA+RNA_seed | EGFR | -2.14666 | BRCA | MDA-MB-175-VII | AZD8931 |
| MVP_top60_DNA_seed | ERBB2 | -0.925262 | BRCA | MDA-MB-468 | CI-1033 | MVP_top60_DNA+RNA_seed | ERBB2 | -2.14666 | BRCA | MDA-MB-175-VII | AZD8931 |
| MVP_top60_DNA_seed | EGFR | -0.925262 | BRCA | MDA-MB-468 | CI-1033 | MVP_top60_DNA+RNA_seed | ERBB3 | -2.14666 | BRCA | MDA-MB-175-VII | AZD8931 |
| MVP_top60_DNA_seed | ERBB2 | -0.923674 | BRCA | HCC38 | CI-1033 | MVP_top60_DNA+RNA_seed | EGFR | -2.129764 | BRCA | HCC1599 | AZD8931 |
| MVP_top60_DNA_seed | EGFR | -0.923674 | BRCA | HCC38 | CI-1033 | MVP_top60_DNA+RNA_seed | ERBB2 | -2.129764 | BRCA | HCC1599 | AZD8931 |
| MVP_top60_DNA_seed | EGFR | -0.921856 | BRCA | AU565 | Gefitinib | MVP_top60_DNA+RNA_seed | ERBB3 | -2.129764 | BRCA | HCC1599 | AZD8931 |
| MVP_top60_DNA_seed | ATM | -0.920671 | BRCA | BT-549 | CP466722 | MVP_top60_DNA+RNA_seed | EGFR | -2.128122 | BRCA | OCUB-M | Afatinib |
| MVP_top60_DNA_seed | MDM2 | -0.920505 | BRCA | AU565 | Serdemetan | MVP_top60_DNA+RNA_seed | ERBB2 | -2.128122 | BRCA | OCUB-M | Afatinib |
| MVP_top60_DNA_seed | HDAC1 | -0.918874 | BRCA | HCC1599 | Dacinostat | MVP_top60_DNA+RNA_seed | AKT1 | -2.112252 | BRCA | T47D | MK-2206 |
| MVP_top60_DNA_seed | ERBB2 | -0.916199 | BRCA | EFM-192A | PF-00299804 | MVP_top60_DNA+RNA_seed | EGFR | -2.09853 | BRCA | HCC202 | AST-1306 |
| MVP_top60_DNA_seed | EGFR | -0.916199 | BRCA | EFM-192A | PF-00299804 | MVP_top60_DNA+RNA_seed | AKT1 | -2.097631 | BRCA | OCUB-M | A-443654 |
| MVP_top60_DNA_seed | ESR1 | -0.915395 | BRCA | HCC1569 | Tamoxifen | MVP_top60_DNA+RNA_seed | PIK3CA | -2.077815 | BRCA | MDA-MB-453 | AZD8835 |
| MVP_top60_DNA_seed | ATM | -0.91518 | BRCA | OCUB-M | CP466722 | MVP_top60_DNA+RNA_seed | EGFR | -2.072793 | BRCA | UACC-893 | Afatinib |
| MVP_top60_DNA_seed | EGFR | -0.912772 | BRCA | BT-474 | Gefitinib | MVP_top60_DNA+RNA_seed | ERBB2 | -2.072793 | BRCA | UACC-893 | Afatinib |
| MVP_top60_DNA_seed | HSP90AA1 | -0.910938 | BRCA | HCC1937 | Elesclomol | MVP_top60_DNA+RNA_seed | AKT1 | -2.045806 | BRCA | MDA-MB-415 | BAY AKT1 |
| MVP_top60_DNA_seed | HSP90AB1 | -0.910938 | BRCA | HCC1937 | Elesclomol | MVP_top60_DNA+RNA_seed | HSP90AA1 | -2.039732 | BRCA | MDA-MB-468 | Elesclomol |
| MVP_top60_DNA_seed | ERBB2 | -0.910203 | BRCA | HCC2157 | Lapatinib | MVP_top60_DNA+RNA_seed | HSP90AB1 | -2.039732 | BRCA | MDA-MB-468 | Elesclomol |
| MVP_top60_DNA_seed | EGFR | -0.910203 | BRCA | HCC2157 | Lapatinib | MVP_top60_DNA+RNA_seed | ERBB2 | -2.036727 | BRCA | HCC1954 | CP724714 |
| MVP_top60_DNA_seed | CDK1 | -0.908468 | BRCA | HCC1395 | RO-3306 | MVP_top60_DNA+RNA_seed | AKT1 | -2.033369 | BRCA | MDA-MB-361 | BAY AKT1 |
| MVP_top60_DNA_seed | HDAC1 | -0.907799 | BRCA | MRK-nu-1 | Parthenolide | MVP_top60_DNA+RNA_seed | PIK3CA | -2.033158 | BRCA | HCC202 | PI3Ka_4409 |
| MVP_top60_DNA_seed | CDK2 | -0.905273 | BRCA | HCC1599 | Seliciclib | MVP_top60_DNA+RNA_seed | EGFR | -2.033014 | BRCA | ZR-75-30 | Afatinib |
| MVP_top60_DNA_seed | AKT1 | -0.905204 | BRCA | Hs-578-T | MK-2206 | MVP_top60_DNA+RNA_seed | ERBB2 | -2.033014 | BRCA | ZR-75-30 | Afatinib |
| MVP_top60_DNA_seed | HDAC1 | -0.903705 | BRCA | HCC2157 | Dacinostat | MVP_top60_DNA+RNA_seed | PIK3CA | -2.01571 | BRCA | MDA-MB-361 | PI3Ka_4409 |
| MVP_top60_DNA_seed | ERBB2 | -0.903416 | BRCA | EFM-192A | CI-1033 | MVP_top60_DNA+RNA_seed | AKT1 | -1.997287 | BRCA | T47D | GSK690693 |
| MVP_top60_DNA_seed | EGFR | -0.903416 | BRCA | EFM-192A | CI-1033 | MVP_top60_DNA+RNA_seed | PIK3CA | -1.993881 | BRCA | MDA-MB-361 | AZD8835 |
| MVP_top60_DNA_seed | ERBB2 | -0.899524 | BRCA | COLO-824 | PF-00299804 | MVP_top60_DNA+RNA_seed | EGFR | -1.978114 | BRCA | HCC2218 | Pelitinib |
| MVP_top60_DNA_seed | EGFR | -0.899524 | BRCA | COLO-824 | PF-00299804 | MVP_top60_DNA+RNA_seed | AR | -1.968538 | BRCA | DU-4475 | Bicalutamide |

| | | | | | | | | | | | |
|---|---|---|---|---|---|---|---|---|---|---|---|
| MVP_top60_DNA_seed | MAPK3 | -0.898571 | BRCA | DU-4475 | FR-180204 | MVP_top60_DNA+RNA_seed | AKT1 | -1.967086 | BRCA | UACC-893 | GSK690693 |
| MVP_top60_DNA_seed | MAPK1 | -0.898571 | BRCA | DU-4475 | FR-180204 | MVP_top60_DNA+RNA_seed | AKT1 | -1.962525 | BRCA | BT-474 | GSK690693 |
| MVP_top60_DNA_seed | PIK3CA | -0.895769 | BRCA | BT-20 | PI3Ka_4409 | MVP_top60_DNA+RNA_seed | EGFR | -1.961958 | BRCA | UACC-893 | AST-1306 |
| MVP_top60_DNA_seed | ERBB2 | -0.894579 | BRCA | T47D | CUDC-101 | MVP_top60_DNA+RNA_seed | ESR1 | -1.960994 | BRCA | HCC1500 | Fulvestrant |
| MVP_top60_DNA_seed | EGFR | -0.894579 | BRCA | T47D | CUDC-101 | MVP_top60_DNA+RNA_seed | EGFR | -1.94645 | BRCA | UACC-893 | PF-00299804 |
| MVP_top60_DNA_seed | EGFR | -0.893788 | BRCA | HCC38 | Cetuximab | MVP_top60_DNA+RNA_seed | ERBB2 | -1.94645 | BRCA | UACC-893 | PF-00299804 |
| MVP_top60_DNA_seed | PIK3CA | -0.8937 | BRCA | BT-474 | PI3Ka_4409 | MVP_top60_DNA+RNA_seed | PIK3CA | -1.923421 | BRCA | HCC1954 | PI3Ka_4409 |
| MVP_top60_DNA_seed | HSP90AA1 | -0.892775 | BRCA | CAL-148 | SNX-2112 | MVP_top60_DNA+RNA_seed | EGFR | -1.895571 | BRCA | HCC2157 | Afatinib |
| MVP_top60_DNA_seed | HSP90AB1 | -0.892775 | BRCA | CAL-148 | SNX-2112 | MVP_top60_DNA+RNA_seed | ERBB2 | -1.895571 | BRCA | HCC2157 | Afatinib |
| MVP_top60_DNA_seed | ERBB2 | -0.89228 | BRCA | BT-474 | AZD8931 | MVP_top60_DNA+RNA_seed | HSP90AA1 | -1.892612 | BRCA | CAL-51 | Elesclomol |
| MVP_top60_DNA_seed | EGFR | -0.89228 | BRCA | BT-474 | AZD8931 | MVP_top60_DNA+RNA_seed | HSP90AB1 | -1.892612 | BRCA | CAL-51 | Elesclomol |
| MVP_top60_DNA_seed | ERBB3 | -0.89228 | BRCA | BT-474 | AZD8931 | MVP_top60_DNA+RNA_seed | PIK3CA | -1.888956 | BRCA | MDA-MB-330 | AZD8835 |
| MVP_top60_DNA_seed | CDK2 | -0.891753 | BRCA | CAL-51 | AZD5438 | MVP_top60_DNA+RNA_seed | AKT1 | -1.881377 | BRCA | OCUB-M | AKT inhibitor VIII |
| MVP_top60_DNA_seed | CDK2 | -0.88982 | BRCA | OCUB-M | PHA-793887 | MVP_top60_DNA+RNA_seed | EGFR | -1.872608 | BRCA | HCC202 | CI-1033 |
| MVP_top60_DNA_seed | CDK2 | -0.887057 | BRCA | HCC1187 | AZD5438 | MVP_top60_DNA+RNA_seed | ERBB2 | -1.872608 | BRCA | HCC202 | CI-1033 |
| MVP_top60_DNA_seed | HDAC1 | -0.886035 | BRCA | HCC1419 | JQ12 | MVP_top60_DNA+RNA_seed | AKT1 | -1.85086 | BRCA | HCC70 | AKT inhibitor VIII |
| MVP_top60_DNA_seed | HDAC2 | -0.886035 | BRCA | HCC1419 | JQ12 | MVP_top60_DNA+RNA_seed | EGFR | -1.848209 | BRCA | UACC-893 | CI-1033 |
| MVP_top60_DNA_seed | AKT1 | -0.885571 | BRCA | MDA-MB-468 | GSK690693 | MVP_top60_DNA+RNA_seed | ERBB2 | -1.848209 | BRCA | UACC-893 | CI-1033 |
| MVP_top60_DNA_seed | ERBB2 | -0.882029 | BRCA | MDA-MB-361 | Afatinib | MVP_top60_DNA+RNA_seed | AKT1 | -1.83964 | BRCA | YMB-1-E | GSK690693 |
| MVP_top60_DNA_seed | EGFR | -0.882029 | BRCA | MDA-MB-361 | Afatinib | MVP_top60_DNA+RNA_seed | EGFR | -1.831664 | BRCA | UACC-893 | Pelitinib |
| MVP_top60_DNA_seed | AR | -0.881055 | BRCA | CAL-51 | Bicalutamide | MVP_top60_DNA+RNA_seed | PIK3CA | -1.819452 | BRCA | OCUB-M | AZD8835 |
| MVP_top60_DNA_seed | EGFR | -0.880323 | BRCA | MDA-MB-468 | AST-1306 | MVP_top60_DNA+RNA_seed | EGFR | -1.811388 | BRCA | HCC202 | Afatinib |
| MVP_top60_DNA_seed | AKT1 | -0.877569 | BRCA | BT-20 | GSK690693 | MVP_top60_DNA+RNA_seed | ERBB2 | -1.811388 | BRCA | HCC202 | Afatinib |
| MVP_top60_DNA_seed | ERBB2 | -0.873298 | BRCA | MDA-MB-330 | Afatinib | MVP_top60_DNA+RNA_seed | AKT1 | -1.804262 | BRCA | MFM-223 | BAY AKT1 |
| MVP_top60_DNA_seed | EGFR | -0.873298 | BRCA | MDA-MB-330 | Afatinib | MVP_top60_DNA+RNA_seed | AKT1 | -1.799965 | BRCA | HCC202 | BAY AKT1 |
| MVP_top60_DNA_seed | ERBB2 | -0.870149 | BRCA | UACC-812 | Lapatinib | MVP_top60_DNA+RNA_seed | ESR1 | -1.798991 | BRCA | HCC1428 | Fulvestrant |
| MVP_top60_DNA_seed | EGFR | -0.870149 | BRCA | UACC-812 | Lapatinib | MVP_top60_DNA+RNA_seed | HSP90AA1 | -1.790851 | BRCA | MDA-MB-361 | Tanespimycin |
| MVP_top60_DNA_seed | CDK2 | -0.869125 | BRCA | BT-549 | AZD5438 | MVP_top60_DNA+RNA_seed | HSP90AB1 | -1.790851 | BRCA | MDA-MB-361 | Tanespimycin |
| MVP_top60_DNA_seed | HSP90AA1 | -0.869099 | BRCA | MDA-MB-436 | Elesclomol | MVP_top60_DNA+RNA_seed | AKT1 | -1.788657 | BRCA | MDA-MB-415 | AKT inhibitor VIII |
| MVP_top60_DNA_seed | HSP90AB1 | -0.869099 | BRCA | MDA-MB-436 | Elesclomol | MVP_top60_DNA+RNA_seed | ESR1 | -1.77589 | BRCA | BT-474 | Fulvestrant |
| MVP_top60_DNA_seed | AR | -0.865078 | BRCA | EFM-19 | AZD3514 | MVP_top60_DNA+RNA_seed | EGFR | -1.755145 | BRCA | CAL-85-1 | Pelitinib |
| MVP_top60_DNA_seed | ERBB2 | -0.862815 | BRCA | EVSA-T | CUDC-101 | MVP_top60_DNA+RNA_seed | AKT1 | -1.753459 | BRCA | BT-474 | AKT inhibitor VIII |
| MVP_top60_DNA_seed | EGFR | -0.862815 | BRCA | EVSA-T | CUDC-101 | MVP_top60_DNA+RNA_seed | HSP90AA1 | -1.749076 | BRCA | MDA-MB-231 | CCT-018159 |
| MVP_top60_DNA_seed | EGFR | -0.862629 | BRCA | HCC1954 | Pelitinib | MVP_top60_DNA+RNA_seed | HSP90AB1 | -1.749076 | BRCA | MDA-MB-231 | CCT-018159 |
| MVP_top60_DNA_seed | ERBB2 | -0.862256 | BRCA | MDA-MB-330 | CI-1033 | MVP_top60_DNA+RNA_seed | HSP90AA1 | -1.746968 | BRCA | CAL-51 | Tanespimycin |
| MVP_top60_DNA_seed | EGFR | -0.862256 | BRCA | MDA-MB-330 | CI-1033 | MVP_top60_DNA+RNA_seed | HSP90AB1 | -1.746968 | BRCA | CAL-51 | Tanespimycin |
| MVP_top60_DNA_seed | HDAC1 | -0.858594 | BRCA | MDA-MB-157 | JQ12 | MVP_top60_DNA+RNA_seed | PIK3CA | -1.72137 | BRCA | HCC1599 | PI3Ka_4409 |
| MVP_top60_DNA_seed | HDAC2 | -0.858594 | BRCA | MDA-MB-157 | JQ12 | MVP_top60_DNA+RNA_seed | AKT1 | -1.699765 | BRCA | ZR-75-30 | GSK690693 |
| MVP_top60_DNA_seed | AKT1 | -0.857239 | BRCA | MDA-MB-175-VII | AKT inhibitor VIII | MVP_top60_DNA+RNA_seed | AKT1 | -1.696774 | BRCA | HCC70 | BAY AKT1 |
| MVP_top60_DNA_seed | HDAC1 | -0.857238 | BRCA | HCC202 | Dacinostat | MVP_top60_DNA+RNA_seed | EGFR | -1.684412 | BRCA | AU565 | Pelitinib |
| MVP_top60_DNA_seed | AKT1 | -0.856328 | BRCA | JIMT-1 | MK-2206 | MVP_top60_DNA+RNA_seed | EGFR | -1.683543 | BRCA | EFM-192A | Afatinib |

| | | | | | | | | | | | |
|---|---|---|---|---|---|---|---|---|---|---|---|
| MVP_top60_DNA_seed | HSP90AA1 | -0.855657 | BRCA | DU-4475 | Tanespimycin | MVP_top60_DNA+RNA_seed | ERBB2 | -1.683543 | BRCA | EFM-192A | Afatinib |
| MVP_top60_DNA_seed | HSP90AB1 | -0.855657 | BRCA | DU-4475 | Tanespimycin | MVP_top60_DNA+RNA_seed | AKT1 | -1.682431 | BRCA | HCC1419 | MK-2206 |
| MVP_top60_DNA_seed | HSP90AA1 | -0.855371 | BRCA | JIMT-1 | Luminespib | MVP_top60_DNA+RNA_seed | AKT1 | -1.663806 | BRCA | T47D | AKT inhibitor VIII |
| MVP_top60_DNA_seed | HSP90AB1 | -0.855371 | BRCA | JIMT-1 | Luminespib | MVP_top60_DNA+RNA_seed | MDM2 | -1.662462 | BRCA | DU-4475 | Nutlin-3a (-) |
| MVP_top60_DNA_seed | ESR1 | -0.855114 | BRCA | HCC1428 | Tamoxifen | MVP_top60_DNA+RNA_seed | ATM | -1.656866 | BRCA | HCC70 | KU-55933 |
| MVP_top60_DNA_seed | PIK3CA | -0.852086 | BRCA | UACC-893 | AZD8835 | MVP_top60_DNA+RNA_seed | AKT1 | -1.65644 | BRCA | MDA-MB-361 | MK-2206 |
| MVP_top60_DNA_seed | ERBB2 | -0.852045 | BRCA | DU-4475 | PF-00299804 | MVP_top60_DNA+RNA_seed | ATM | -1.646731 | BRCA | HCC2157 | KU-55933 |
| MVP_top60_DNA_seed | EGFR | -0.852045 | BRCA | DU-4475 | PF-00299804 | MVP_top60_DNA+RNA_seed | AKT1 | -1.64532 | BRCA | HCC1954 | GSK690693 |
| MVP_top60_DNA_seed | HSP90AA1 | -0.85197 | BRCA | HCC1187 | CCT-018159 | MVP_top60_DNA+RNA_seed | AKT1 | -1.644698 | BRCA | EFM-19 | BAY AKT1 |
| MVP_top60_DNA_seed | HSP90AB1 | -0.85197 | BRCA | HCC1187 | CCT-018159 | MVP_top60_DNA+RNA_seed | ESR1 | -1.636771 | BRCA | OCUB-M | Fulvestrant |
| MVP_top60_DNA_seed | HDAC1 | -0.850372 | BRCA | EVSA-T | Tubastatin A | MVP_top60_DNA+RNA_seed | AKT1 | -1.632491 | BRCA | MDA-MB-330 | MK-2206 |
| MVP_top60_DNA_seed | CDK1 | -0.844429 | BRCA | MDA-MB-330 | RO-3306 | MVP_top60_DNA+RNA_seed | SRC | -1.627486 | BRCA | DU-4475 | NVP-BHG712 |
| MVP_top60_DNA_seed | AR | -0.836762 | BRCA | HCC1954 | Bicalutamide | MVP_top60_DNA+RNA_seed | EGFR | -1.624663 | BRCA | BT-20 | Cetuximab |
| MVP_top60_DNA_seed | MAPK1 | -0.834528 | BRCA | CAL-85-1 | VX-11e | MVP_top60_DNA+RNA_seed | ATM | -1.614636 | BRCA | EVSA-T | KU-55933 |
| MVP_top60_DNA_seed | ERBB2 | -0.834036 | BRCA | Hs-578-T | CI-1033 | MVP_top60_DNA+RNA_seed | CDK2 | -1.608597 | BRCA | HCC70 | PHA-793887 |
| MVP_top60_DNA_seed | EGFR | -0.834036 | BRCA | Hs-578-T | CI-1033 | MVP_top60_DNA+RNA_seed | HDAC1 | -1.602832 | BRCA | EFM-19 | JQ12 |
| MVP_top60_DNA_seed | PIK3CA | -0.832895 | BRCA | MFM-223 | AZD8835 | MVP_top60_DNA+RNA_seed | HDAC2 | -1.602832 | BRCA | EFM-19 | JQ12 |
| MVP_top60_DNA_seed | ERBB2 | -0.829601 | BRCA | HCC1569 | Afatinib | MVP_top60_DNA+RNA_seed | AKT1 | -1.591133 | BRCA | MDA-MB-330 | AKT inhibitor VIII |
| MVP_top60_DNA_seed | EGFR | -0.829601 | BRCA | HCC1569 | Afatinib | MVP_top60_DNA+RNA_seed | ESR1 | -1.58436 | BRCA | EVSA-T | Fulvestrant |
| MVP_top60_DNA_seed | HSP90AA1 | -0.828051 | BRCA | Hs-578-T | Elesclomol | MVP_top60_DNA+RNA_seed | AKT1 | -1.575452 | BRCA | MDA-MB-453 | BAY AKT1 |
| MVP_top60_DNA_seed | HSP90AB1 | -0.828051 | BRCA | Hs-578-T | Elesclomol | MVP_top60_DNA+RNA_seed | HSP90AA1 | -1.563298 | BRCA | CAL-51 | CCT-018159 |
| MVP_top60_DNA_seed | ERBB2 | -0.827642 | BRCA | CAL-148 | CUDC-101 | MVP_top60_DNA+RNA_seed | HSP90AB1 | -1.563298 | BRCA | CAL-51 | CCT-018159 |
| MVP_top60_DNA_seed | EGFR | -0.827642 | BRCA | CAL-148 | CUDC-101 | MVP_top60_DNA+RNA_seed | HDAC1 | -1.562873 | BRCA | CAL-85-1 | JQ12 |
| MVP_top60_DNA_seed | ERBB2 | -0.827082 | BRCA | EFM-192A | AZD8931 | MVP_top60_DNA+RNA_seed | HDAC2 | -1.562873 | BRCA | CAL-85-1 | JQ12 |
| MVP_top60_DNA_seed | EGFR | -0.827082 | BRCA | EFM-192A | AZD8931 | MVP_top60_DNA+RNA_seed | EGFR | -1.562326 | BRCA | HCC1954 | Afatinib |
| MVP_top60_DNA_seed | ERBB3 | -0.827082 | BRCA | EFM-192A | AZD8931 | MVP_top60_DNA+RNA_seed | ERBB2 | -1.562326 | BRCA | HCC1954 | Afatinib |
| MVP_top60_DNA_seed | ATM | -0.82688 | BRCA | HCC1419 | CP466722 | MVP_top60_DNA+RNA_seed | AKT1 | -1.555749 | BRCA | T47D | BAY AKT1 |
| MVP_top60_DNA_seed | ERBB2 | -0.824701 | BRCA | BT-20 | Afatinib | MVP_top60_DNA+RNA_seed | EGFR | -1.552322 | BRCA | MDA-MB-330 | Afatinib |
| MVP_top60_DNA_seed | EGFR | -0.824701 | BRCA | BT-20 | Afatinib | MVP_top60_DNA+RNA_seed | ERBB2 | -1.552322 | BRCA | MDA-MB-330 | Afatinib |
| MVP_top60_DNA_seed | ATM | -0.823608 | BRCA | HCC2157 | KU-60019 | MVP_top60_DNA+RNA_seed | AR | -1.551632 | BRCA | OCUB-M | AZD3514 |
| MVP_top60_DNA_seed | ESR1 | -0.823213 | BRCA | HCC1954 | Tamoxifen | MVP_top60_DNA+RNA_seed | AKT1 | -1.543239 | BRCA | HCC38 | AKT inhibitor VIII |
| MVP_top60_DNA_seed | CDK2 | -0.822768 | BRCA | HCC2218 | PHA-793887 | MVP_top60_DNA+RNA_seed | HSP90AA1 | -1.533125 | BRCA | MCF7 | Luminespib |
| MVP_top60_DNA_seed | HDAC1 | -0.821157 | BRCA | EVSA-T | AR-42 | MVP_top60_DNA+RNA_seed | HSP90AB1 | -1.533125 | BRCA | MCF7 | Luminespib |
| MVP_top60_DNA_seed | AKT1 | -0.820497 | BRCA | MDA-MB-175-VII | GSK690693 | MVP_top60_DNA+RNA_seed | AKT1 | -1.528884 | BRCA | CAL-148 | A-443654 |
| MVP_top60_DNA_seed | ERBB2 | -0.817278 | BRCA | UACC-812 | CP724714 | MVP_top60_DNA+RNA_seed | HSP90AA1 | -1.527797 | BRCA | MDA-MB-175-VII | Tanespimycin |
| MVP_top60_DNA_seed | EGFR | -0.816214 | BRCA | DU-4475 | Gefitinib | MVP_top60_DNA+RNA_seed | HSP90AB1 | -1.527797 | BRCA | MDA-MB-175-VII | Tanespimycin |
| MVP_top60_DNA_seed | AKT1 | -0.813966 | BRCA | AU565 | MK-2206 | MVP_top60_DNA+RNA_seed | AKT1 | -1.520264 | BRCA | MDA-MB-453 | GSK690693 |
| MVP_top60_DNA_seed | MDM2 | -0.8133 | BRCA | MDA-MB-361 | Serdemetan | MVP_top60_DNA+RNA_seed | HDAC1 | -1.499309 | BRCA | MDA-MB-453 | Belinostat |
| MVP_top60_DNA_seed | AKT1 | -0.812343 | BRCA | HCC2218 | GSK690693 | MVP_top60_DNA+RNA_seed | HDAC1 | -1.497798 | BRCA | MDA-MB-453 | AR-42 |
| MVP_top60_DNA_seed | HSP90AA1 | -0.804905 | BRCA | HCC1954 | CCT-018159 | MVP_top60_DNA+RNA_seed | ESR1 | -1.497497 | BRCA | T47D | Fulvestrant |
| MVP_top60_DNA_seed | HSP90AB1 | -0.804905 | BRCA | HCC1954 | CCT-018159 | MVP_top60_DNA+RNA_seed | AKT1 | -1.492177 | BRCA | MDA-MB-453 | AKT inhibitor VIII |

| | | | | | | | | | | | |
|---|---|---|---|---|---|---|---|---|---|---|---|
| MVP_top60_DNA_seed | HDAC1 | -0.802507 | BRCA | OCUB-M | CAY10603 | MVP_top60_DNA+RNA_seed | HSP90AA1 | -1.490882 | BRCA | BT-474 | Tanespimycin |
| MVP_top60_DNA_seed | AKT1 | -0.801952 | BRCA | HCC1599 | A-443654 | MVP_top60_DNA+RNA_seed | HSP90AB1 | -1.490882 | BRCA | BT-474 | Tanespimycin |
| MVP_top60_DNA_seed | HSP90AA1 | -0.79718 | BRCA | EFM-192A | Luminespib | MVP_top60_DNA+RNA_seed | AKT1 | -1.48983 | BRCA | EFM-19 | AKT inhibitor VIII |
| MVP_top60_DNA_seed | HSP90AB1 | -0.79718 | BRCA | EFM-192A | Luminespib | MVP_top60_DNA+RNA_seed | AKT1 | -1.487882 | BRCA | EFM-192A | MK-2206 |
| MVP_top60_DNA_seed | ESR1 | -0.796847 | BRCA | JIMT-1 | Tamoxifen | MVP_top60_DNA+RNA_seed | AKT1 | -1.487501 | BRCA | HCC2218 | AKT inhibitor VIII |
| MVP_top60_DNA_seed | AKT1 | -0.796096 | BRCA | MDA-MB-453 | AKT inhibitor VIII | MVP_top60_DNA+RNA_seed | EGFR | -1.487398 | BRCA | HCC2218 | Gefitinib |
| MVP_top60_DNA_seed | HDAC1 | -0.795522 | BRCA | OCUB-M | Tubastatin A | MVP_top60_DNA+RNA_seed | ESR1 | -1.480426 | BRCA | MDA-MB-361 | Tamoxifen |
| MVP_top60_DNA_seed | EGFR | -0.792913 | BRCA | MDA-MB-361 | Pelitinib | MVP_top60_DNA+RNA_seed | EGFR | -1.463658 | BRCA | AU565 | AZD8931 |
| MVP_top60_DNA_seed | MDM2 | -0.791316 | BRCA | DU-4475 | Serdemetan | MVP_top60_DNA+RNA_seed | ERBB2 | -1.463658 | BRCA | AU565 | AZD8931 |
| MVP_top60_DNA_seed | ATM | -0.787646 | BRCA | HCC1569 | CP466722 | MVP_top60_DNA+RNA_seed | ERBB3 | -1.463658 | BRCA | AU565 | AZD8931 |
| MVP_top60_DNA_seed | HDAC1 | -0.787546 | BRCA | MDA-MB-361 | Dacinostat | MVP_top60_DNA+RNA_seed | AKT1 | -1.46047 | BRCA | MDA-MB-415 | MK-2206 |
| MVP_top60_DNA_seed | ERBB2 | -0.787435 | BRCA | ZR-75-30 | CUDC-101 | MVP_top60_DNA+RNA_seed | EP300 | -1.459423 | BRCA | JIMT-1 | I-CBP112 |
| MVP_top60_DNA_seed | EGFR | -0.787435 | BRCA | ZR-75-30 | CUDC-101 | MVP_top60_DNA+RNA_seed | AKT1 | -1.459332 | BRCA | HCC1569 | AKT inhibitor VIII |
| MVP_top60_DNA_seed | PIK3CA | -0.787269 | BRCA | MCF7 | PI3Ka_4409 | MVP_top60_DNA+RNA_seed | EGFR | -1.453118 | BRCA | ZR-75-30 | Pelitinib |
| MVP_top60_DNA_seed | HSP90AA1 | -0.785228 | BRCA | MDA-MB-157 | Tanespimycin | MVP_top60_DNA+RNA_seed | HSP90AA1 | -1.4471 | BRCA | MCF7 | Tanespimycin |
| MVP_top60_DNA_seed | HSP90AB1 | -0.785228 | BRCA | MDA-MB-157 | Tanespimycin | MVP_top60_DNA+RNA_seed | HSP90AB1 | -1.4471 | BRCA | MCF7 | Tanespimycin |
| MVP_top60_DNA_seed | EGFR | -0.783679 | BRCA | HCC38 | AST-1306 | MVP_top60_DNA+RNA_seed | EGFR | -1.442501 | BRCA | MDA-MB-453 | CUDC-101 |
| MVP_top60_DNA_seed | PIK3CA | -0.782566 | BRCA | EFM-192A | AZD8835 | MVP_top60_DNA+RNA_seed | ERBB2 | -1.442501 | BRCA | MDA-MB-453 | CUDC-101 |
| MVP_top60_DNA_seed | ERBB2 | -0.781674 | BRCA | UACC-893 | AZD8931 | MVP_top60_DNA+RNA_seed | ATM | -1.440139 | BRCA | CAL-51 | KU-60019 |
| MVP_top60_DNA_seed | EGFR | -0.781674 | BRCA | UACC-893 | AZD8931 | MVP_top60_DNA+RNA_seed | PIK3CA | -1.437506 | BRCA | JIMT-1 | AZD8835 |
| MVP_top60_DNA_seed | ERBB3 | -0.781674 | BRCA | UACC-893 | AZD8931 | MVP_top60_DNA+RNA_seed | AKT1 | -1.431509 | BRCA | HCC1954 | BAY AKT1 |
| MVP_top60_DNA_seed | HDAC1 | -0.781366 | BRCA | EFM-192A | JQ12 | MVP_top60_DNA+RNA_seed | ATM | -1.430587 | BRCA | OCUB-M | KU-60019 |
| MVP_top60_DNA_seed | HDAC2 | -0.781366 | BRCA | EFM-192A | JQ12 | MVP_top60_DNA+RNA_seed | AKT1 | -1.423091 | BRCA | HCC1599 | AKT inhibitor VIII |
| MVP_top60_DNA_seed | HDAC1 | -0.781025 | BRCA | EFM-19 | Tubastatin A | MVP_top60_DNA+RNA_seed | AKT1 | -1.421005 | BRCA | EVSA-T | AKT inhibitor VIII |
| MVP_top60_DNA_seed | HDAC1 | -0.778595 | BRCA | HCC1599 | Belinostat | MVP_top60_DNA+RNA_seed | HDAC1 | -1.414216 | BRCA | CAL-85-1 | Dacinostat |
| MVP_top60_DNA_seed | ERBB2 | -0.778568 | BRCA | HCC1954 | AZD8931 | MVP_top60_DNA+RNA_seed | ERBB2 | -1.413423 | BRCA | ZR-75-30 | CP724714 |
| MVP_top60_DNA_seed | EGFR | -0.778568 | BRCA | HCC1954 | AZD8931 | MVP_top60_DNA+RNA_seed | AKT1 | -1.410322 | BRCA | OCUB-M | MK-2206 |
| MVP_top60_DNA_seed | ERBB3 | -0.778568 | BRCA | HCC1954 | AZD8931 | MVP_top60_DNA+RNA_seed | HDAC1 | -1.407702 | BRCA | MCF7 | JQ12 |
| MVP_top60_DNA_seed | PIK3CA | -0.778535 | BRCA | BT-474 | AZD8835 | MVP_top60_DNA+RNA_seed | HDAC2 | -1.407702 | BRCA | MCF7 | JQ12 |
| MVP_top60_DNA_seed | AR | -0.778391 | BRCA | BT-549 | Bicalutamide | MVP_top60_DNA+RNA_seed | HDAC1 | -1.406492 | BRCA | MDA-MB-361 | JQ12 |
| MVP_top60_DNA_seed | HSP90AA1 | -0.777219 | BRCA | CAMA-1 | Tanespimycin | MVP_top60_DNA+RNA_seed | HDAC2 | -1.406492 | BRCA | MDA-MB-361 | JQ12 |
| MVP_top60_DNA_seed | HSP90AB1 | -0.777219 | BRCA | CAMA-1 | Tanespimycin | MVP_top60_DNA+RNA_seed | AR | -1.3957 | BRCA | OCUB-M | Bicalutamide |
| MVP_top60_DNA_seed | HDAC1 | -0.776846 | BRCA | MCF7 | Dacinostat | MVP_top60_DNA+RNA_seed | AR | -1.395492 | BRCA | MDA-MB-453 | Bicalutamide |
| MVP_top60_DNA_seed | AKT1 | -0.775694 | BRCA | HCC70 | GSK690693 | MVP_top60_DNA+RNA_seed | HSP90AA1 | -1.391886 | BRCA | EVSA-T | Tanespimycin |
| MVP_top60_DNA_seed | ERBB2 | -0.773581 | BRCA | MDA-MB-468 | Afatinib | MVP_top60_DNA+RNA_seed | HSP90AB1 | -1.391886 | BRCA | EVSA-T | Tanespimycin |
| MVP_top60_DNA_seed | EGFR | -0.773581 | BRCA | MDA-MB-468 | Afatinib | MVP_top60_DNA+RNA_seed | EGFR | -1.383862 | BRCA | BT-549 | Pelitinib |
| MVP_top60_DNA_seed | EP300 | -0.773089 | BRCA | BT-474 | I-CBP112 | MVP_top60_DNA+RNA_seed | HDAC1 | -1.372223 | BRCA | MDA-MB-453 | Tubastatin A |
| MVP_top60_DNA_seed | CREBBP | -0.773089 | BRCA | BT-474 | I-CBP112 | MVP_top60_DNA+RNA_seed | EGFR | -1.356386 | BRCA | HDQ-P1 | Afatinib |
| MVP_top60_DNA_seed | ERBB2 | -0.772035 | BRCA | UACC-812 | CI-1033 | MVP_top60_DNA+RNA_seed | ERBB2 | -1.356386 | BRCA | HDQ-P1 | Afatinib |
| MVP_top60_DNA_seed | EGFR | -0.772035 | BRCA | UACC-812 | CI-1033 | MVP_top60_DNA+RNA_seed | CDK1 | -1.352117 | BRCA | MCF7 | RO-3306 |
| MVP_top60_DNA_seed | ATM | -0.770453 | BRCA | AU565 | KU-60019 | MVP_top60_DNA+RNA_seed | EGFR | -1.346121 | BRCA | HCC1419 | Afatinib |

| | | | | | | | | | | | |
|---|---|---|---|---|---|---|---|---|---|---|---|
| MVP_top60_DNA_seed | CDK2 | -0.769496 | BRCA | MRK-nu-1 | Seliciclib | MVP_top60_DNA+RNA_seed | ERBB2 | -1.346121 | BRCA | HCC1419 | Afatinib |
| MVP_top60_DNA_seed | PIK3CA | -0.767928 | BRCA | HCC202 | AZD8835 | MVP_top60_DNA+RNA_seed | AKT1 | -1.345647 | BRCA | YMB-1-E | AKT inhibitor VIII |
| MVP_top60_DNA_seed | EGFR | -0.765775 | BRCA | BT-20 | Gefitinib | MVP_top60_DNA+RNA_seed | EGFR | -1.342716 | BRCA | MDA-MB-468 | Gefitinib |
| MVP_top60_DNA_seed | CDK2 | -0.765333 | BRCA | Hs-578-T | AZD5438 | MVP_top60_DNA+RNA_seed | HSP90AA1 | -1.339572 | BRCA | MDA-MB-330 | Tanespimycin |
| MVP_top60_DNA_seed | ESR1 | -0.764831 | BRCA | MDA-MB-468 | Fulvestrant | MVP_top60_DNA+RNA_seed | HSP90AB1 | -1.339572 | BRCA | MDA-MB-330 | Tanespimycin |
| MVP_top60_DNA_seed | CDK1 | -0.763281 | BRCA | HCC1428 | RO-3306 | MVP_top60_DNA+RNA_seed | MDM2 | -1.339111 | BRCA | MDA-MB-231 | Serdemetan |
| MVP_top60_DNA_seed | AR | -0.760143 | BRCA | MRK-nu-1 | Bicalutamide | MVP_top60_DNA+RNA_seed | MDM2 | -1.332638 | BRCA | MCF7 | Nutlin-3a (-) |
| MVP_top60_DNA_seed | HDAC1 | -0.756349 | BRCA | EVSA-T | Belinostat | MVP_top60_DNA+RNA_seed | AKT1 | -1.332366 | BRCA | HCC1569 | AKT inhibitor VIII |
| MVP_top60_DNA_seed | AKT1 | -0.752134 | BRCA | HCC2218 | MK-2206 | MVP_top60_DNA+RNA_seed | HSP90AA1 | -1.331114 | BRCA | AU565 | CCT-018159 |
| MVP_top60_DNA_seed | HSP90AA1 | -0.7521 | BRCA | EVSA-T | CCT-018159 | MVP_top60_DNA+RNA_seed | HSP90AB1 | -1.331114 | BRCA | AU565 | CCT-018159 |
| MVP_top60_DNA_seed | HSP90AB1 | -0.7521 | BRCA | EVSA-T | CCT-018159 | MVP_top60_DNA+RNA_seed | HSP90AA1 | -1.331059 | BRCA | OCUB-M | CCT-018159 |
| MVP_top60_DNA_seed | HDAC1 | -0.74829 | BRCA | HCC1428 | JQ12 | MVP_top60_DNA+RNA_seed | HSP90AB1 | -1.331059 | BRCA | OCUB-M | CCT-018159 |
| MVP_top60_DNA_seed | HDAC2 | -0.74829 | BRCA | HCC1428 | JQ12 | MVP_top60_DNA+RNA_seed | EGFR | -1.328804 | BRCA | MDA-MB-175-VII | Gefitinib |
| MVP_top60_DNA_seed | CDK2 | -0.747896 | BRCA | HCC2157 | PHA-793887 | MVP_top60_DNA+RNA_seed | EGFR | -1.326821 | BRCA | EFM-192A | AST-1306 |
| MVP_top60_DNA_seed | HSP90AA1 | -0.74642 | BRCA | COLO-824 | Tanespimycin | MVP_top60_DNA+RNA_seed | MAPK14 | -1.322826 | BRCA | OCUB-M | TAK-715 |
| MVP_top60_DNA_seed | HSP90AB1 | -0.74642 | BRCA | COLO-824 | Tanespimycin | MVP_top60_DNA+RNA_seed | ATM | -1.322394 | BRCA | Hs-578-T | KU-55933 |
| MVP_top60_DNA_seed | ERBB2 | -0.74533 | BRCA | EFM-19 | CUDC-101 | MVP_top60_DNA+RNA_seed | AKT1 | -1.318777 | BRCA | HCC1569 | GSK690693 |
| MVP_top60_DNA_seed | EGFR | -0.74533 | BRCA | EFM-19 | CUDC-101 | MVP_top60_DNA+RNA_seed | AKT1 | -1.312031 | BRCA | MDA-MB-468 | AKT inhibitor VIII |
| MVP_top60_DNA_seed | ERBB2 | -0.744696 | BRCA | HCC1599 | CI-1033 | MVP_top60_DNA+RNA_seed | PIK3CA | -1.311004 | BRCA | EFM-19 | PI3Ka_4409 |
| MVP_top60_DNA_seed | EGFR | -0.744696 | BRCA | HCC1599 | CI-1033 | MVP_top60_DNA+RNA_seed | PIK3CA | -1.288787 | BRCA | EFM-192A | PI3Ka_4409 |
| MVP_top60_DNA_seed | AKT1 | -0.743904 | BRCA | CAL-148 | AKT inhibitor VIII | MVP_top60_DNA+RNA_seed | ESR1 | -1.285457 | BRCA | AU565 | Fulvestrant |
| MVP_top60_DNA_seed | ESR1 | -0.743047 | BRCA | COLO-824 | Fulvestrant | MVP_top60_DNA+RNA_seed | EGFR | -1.285071 | BRCA | MDA-MB-468 | Afatinib |
| MVP_top60_DNA_seed | SRC | -0.741454 | BRCA | UACC-812 | Saracatinib | MVP_top60_DNA+RNA_seed | ERBB2 | -1.285071 | BRCA | MDA-MB-468 | Afatinib |
| MVP_top60_DNA_seed | ERBB2 | -0.738983 | BRCA | BT-20 | AZD8931 | MVP_top60_DNA+RNA_seed | EGFR | -1.280387 | BRCA | MDA-MB-330 | AST-1306 |
| MVP_top60_DNA_seed | EGFR | -0.738983 | BRCA | BT-20 | AZD8931 | MVP_top60_DNA+RNA_seed | HSP90AA1 | -1.278815 | BRCA | DU-4475 | SNX-2112 |
| MVP_top60_DNA_seed | ERBB3 | -0.738983 | BRCA | BT-20 | AZD8931 | MVP_top60_DNA+RNA_seed | HSP90AB1 | -1.278815 | BRCA | DU-4475 | SNX-2112 |
| MVP_top60_DNA_seed | AKT1 | -0.738526 | BRCA | CAMA-1 | GSK690693 | MVP_top60_DNA+RNA_seed | AKT1 | -1.278181 | BRCA | HCC1599 | GSK690693 |
| MVP_top60_DNA_seed | AKT1 | -0.737176 | BRCA | MDA-MB-415 | AKT inhibitor VIII | MVP_top60_DNA+RNA_seed | HSP90AA1 | -1.275733 | BRCA | CAL-51 | Luminespib |
| MVP_top60_DNA_seed | ERBB2 | -0.734953 | BRCA | CAL-85-1 | AZD8931 | MVP_top60_DNA+RNA_seed | HSP90AB1 | -1.275733 | BRCA | CAL-51 | Luminespib |
| MVP_top60_DNA_seed | EGFR | -0.734953 | BRCA | CAL-85-1 | AZD8931 | MVP_top60_DNA+RNA_seed | EGFR | -1.274644 | BRCA | MDA-MB-468 | Pelitinib |
| MVP_top60_DNA_seed | ERBB3 | -0.734953 | BRCA | CAL-85-1 | AZD8931 | MVP_top60_DNA+RNA_seed | HDAC1 | -1.274228 | BRCA | HCC1428 | Dacinostat |
| MVP_top60_DNA_seed | ERBB2 | -0.729207 | BRCA | MDA-MB-361 | CUDC-101 | MVP_top60_DNA+RNA_seed | EGFR | -1.26613 | BRCA | HCC202 | PF-00299804 |
| MVP_top60_DNA_seed | EGFR | -0.729207 | BRCA | MDA-MB-361 | CUDC-101 | MVP_top60_DNA+RNA_seed | ERBB2 | -1.26613 | BRCA | HCC202 | PF-00299804 |
| MVP_top60_DNA_seed | HDAC1 | -0.727561 | BRCA | CAL-148 | Belinostat | MVP_top60_DNA+RNA_seed | HDAC1 | -1.265904 | BRCA | EVSA-T | Parthenolide |
| MVP_top60_DNA_seed | HDAC1 | -0.727364 | BRCA | T47D | Belinostat | MVP_top60_DNA+RNA_seed | PLK1 | -1.265636 | BRCA | COLO-824 | PLK_6522 |
| MVP_top60_DNA_seed | AR | -0.726063 | BRCA | HCC1187 | Bicalutamide | MVP_top60_DNA+RNA_seed | AKT1 | -1.255299 | BRCA | CAL-51 | AKT inhibitor VIII |
| MVP_top60_DNA_seed | AR | -0.724022 | BRCA | HCC1569 | Bicalutamide | MVP_top60_DNA+RNA_seed | HSP90AA1 | -1.253933 | BRCA | HCC2157 | CCT-018159 |
| MVP_top60_DNA_seed | EGFR | -0.723274 | BRCA | MRK-nu-1 | AST-1306 | MVP_top60_DNA+RNA_seed | HSP90AB1 | -1.253933 | BRCA | HCC2157 | CCT-018159 |
| MVP_top60_DNA_seed | EGFR | -0.723214 | BRCA | COLO-824 | AST-1306 | MVP_top60_DNA+RNA_seed | ATM | -1.251797 | BRCA | HCC1500 | CP466722 |
| MVP_top60_DNA_seed | HSP90AA1 | -0.72126 | BRCA | BT-549 | Tanespimycin | MVP_top60_DNA+RNA_seed | EGFR | -1.245807 | BRCA | HCC202 | Afatinib |
| MVP_top60_DNA_seed | HSP90AB1 | -0.72126 | BRCA | BT-549 | Tanespimycin | MVP_top60_DNA+RNA_seed | ERBB2 | -1.245807 | BRCA | HCC202 | Afatinib |

| | | | | | | | | | | | |
|---|---|---|---|---|---|---|---|---|---|---|---|
| MVP_top60_DNA_seed | HDAC1 | -0.717399 | BRCA | HCC38 | Dacinostat | MVP_top60_DNA+RNA_seed | EGFR | -1.243057 | BRCA | HCC1599 | PF-00299804 |
| MVP_top60_DNA_seed | HDAC1 | -0.713041 | BRCA | OCUB-M | Belinostat | MVP_top60_DNA+RNA_seed | ERBB2 | -1.243057 | BRCA | HCC1599 | PF-00299804 |
| MVP_top60_DNA_seed | AR | -0.712505 | BRCA | HCC1187 | Bicalutamide | MVP_top60_DNA+RNA_seed | EGFR | -1.24242 | BRCA | HCC1419 | Afatinib |
| MVP_top60_DNA_seed | HDAC1 | -0.703683 | BRCA | HCC202 | Belinostat | MVP_top60_DNA+RNA_seed | ERBB2 | -1.24242 | BRCA | HCC1419 | Afatinib |
| MVP_top60_DNA_seed | EGFR | -0.703436 | BRCA | ZR-75-30 | AST-1306 | MVP_top60_DNA+RNA_seed | MDM2 | -1.24201 | BRCA | BT-549 | Serdemetan |
| MVP_top60_DNA_seed | ERBB2 | -0.701954 | BRCA | HCC70 | Afatinib | MVP_top60_DNA+RNA_seed | EGFR | -1.241509 | BRCA | MDA-MB-453 | AST-1306 |
| MVP_top60_DNA_seed | EGFR | -0.701954 | BRCA | HCC70 | Afatinib | MVP_top60_DNA+RNA_seed | AKT1 | -1.240709 | BRCA | MFM-223 | MK-2206 |
| MVP_top60_DNA_seed | ERBB2 | -0.700818 | BRCA | HCC202 | CUDC-101 | MVP_top60_DNA+RNA_seed | ESR1 | -1.23885 | BRCA | BT-549 | Fulvestrant |
| MVP_top60_DNA_seed | EGFR | -0.700818 | BRCA | HCC202 | CUDC-101 | MVP_top60_DNA+RNA_seed | AKT1 | -1.228396 | BRCA | UACC-893 | AKT inhibitor VIII |
| MVP_top60_DNA_seed | AKT1 | -0.700671 | BRCA | CAL-85-1 | AKT inhibitor VIII | MVP_top60_DNA+RNA_seed | EGFR | -1.226681 | BRCA | Hs-578-T | PF-00299804 |
| MVP_top60_DNA_seed | HDAC1 | -0.698216 | BRCA | HCC202 | AR-42 | MVP_top60_DNA+RNA_seed | ERBB2 | -1.226681 | BRCA | Hs-578-T | PF-00299804 |
| MVP_top60_DNA_seed | AR | -0.697133 | BRCA | HCC1599 | Bicalutamide | MVP_top60_DNA+RNA_seed | EGFR | -1.220525 | BRCA | HCC1419 | Pelitinib |
| MVP_top60_DNA_seed | EP300 | -0.697074 | BRCA | OCUB-M | I-CBP112 | MVP_top60_DNA+RNA_seed | AKT1 | -1.21847 | BRCA | HCC1954 | AKT inhibitor VIII |
| MVP_top60_DNA_seed | CREBBP | -0.697074 | BRCA | OCUB-M | I-CBP112 | MVP_top60_DNA+RNA_seed | ATM | -1.215249 | BRCA | MDA-MB-468 | KU-55933 |
| MVP_top60_DNA_seed | AKT1 | -0.696958 | BRCA | HCC202 | AKT inhibitor VIII | MVP_top60_DNA+RNA_seed | AKT1 | -1.21461 | BRCA | BT-474 | AKT inhibitor VIII |
| MVP_top60_DNA_seed | ERBB2 | -0.694728 | BRCA | MRK-nu-1 | CI-1033 | MVP_top60_DNA+RNA_seed | AR | -1.212017 | BRCA | MCF7 | Bicalutamide |
| MVP_top60_DNA_seed | EGFR | -0.694728 | BRCA | MRK-nu-1 | CI-1033 | MVP_top60_DNA+RNA_seed | AKT1 | -1.203389 | BRCA | AU565 | AKT inhibitor VIII |
| MVP_top60_DNA_seed | ERBB2 | -0.693747 | BRCA | ZR-75-30 | PF-00299804 | MVP_top60_DNA+RNA_seed | EGFR | -1.201192 | BRCA | CAL-85-1 | Cetuximab |
| MVP_top60_DNA_seed | EGFR | -0.693747 | BRCA | ZR-75-30 | PF-00299804 | MVP_top60_DNA+RNA_seed | PIK3CA | -1.199455 | BRCA | BT-20 | AZD8835 |
| MVP_top60_DNA_seed | AKT1 | -0.692199 | BRCA | HCC1187 | GSK690693 | MVP_top60_DNA+RNA_seed | HDAC1 | -1.19842 | BRCA | DU-4475 | Entinostat |
| MVP_top60_DNA_seed | HSP90AA1 | -0.692142 | BRCA | MDA-MB-436 | Tanespimycin | MVP_top60_DNA+RNA_seed | PIK3CA | -1.194497 | BRCA | EFM-19 | AZD8835 |
| MVP_top60_DNA_seed | HSP90AB1 | -0.692142 | BRCA | MDA-MB-436 | Tanespimycin | MVP_top60_DNA+RNA_seed | AKT1 | -1.188487 | BRCA | HCC1419 | AKT inhibitor VIII |
| MVP_top60_DNA_seed | ERBB2 | -0.688979 | BRCA | CAL-148 | CI-1033 | MVP_top60_DNA+RNA_seed | AKT1 | -1.186063 | BRCA | HCC1419 | AKT inhibitor VIII |
| MVP_top60_DNA_seed | EGFR | -0.688979 | BRCA | CAL-148 | CI-1033 | MVP_top60_DNA+RNA_seed | ATM | -1.18475 | BRCA | ZR-75-30 | CP466722 |
| MVP_top60_DNA_seed | ESR1 | -0.688846 | BRCA | HCC1954 | Fulvestrant | MVP_top60_DNA+RNA_seed | PLK1 | -1.179885 | BRCA | OCUB-M | BI-2536 |
| MVP_top60_DNA_seed | MAPK3 | -0.685212 | BRCA | CAL-85-1 | FR-180204 | MVP_top60_DNA+RNA_seed | ATM | -1.178105 | BRCA | MDA-MB-231 | KU-60019 |
| MVP_top60_DNA_seed | MAPK1 | -0.685212 | BRCA | CAL-85-1 | FR-180204 | MVP_top60_DNA+RNA_seed | PLK1 | -1.173377 | BRCA | AU565 | PLK_6522 |
| MVP_top60_DNA_seed | ATM | -0.684548 | BRCA | Hs-578-T | KU-60019 | MVP_top60_DNA+RNA_seed | AKT1 | -1.166147 | BRCA | MCF7 | BAY AKT1 |
| MVP_top60_DNA_seed | ATM | -0.683833 | BRCA | HCC1569 | KU-55933 | MVP_top60_DNA+RNA_seed | ATM | -1.163912 | BRCA | CAL-51 | KU-55933 |
| MVP_top60_DNA_seed | AKT1 | -0.681203 | BRCA | HCC2218 | BAY AKT1 | MVP_top60_DNA+RNA_seed | PIK3CA | -1.156943 | BRCA | HCC1599 | AZD8835 |
| MVP_top60_DNA_seed | HDAC1 | -0.679711 | BRCA | HCC2218 | Entinostat | MVP_top60_DNA+RNA_seed | HSP90AA1 | -1.155473 | BRCA | HCC2218 | Tanespimycin |
| MVP_top60_DNA_seed | HSP90AA1 | -0.67693 | BRCA | MDA-MB-361 | Elesclomol | MVP_top60_DNA+RNA_seed | HSP90AB1 | -1.155473 | BRCA | HCC2218 | Tanespimycin |
| MVP_top60_DNA_seed | HSP90AB1 | -0.67693 | BRCA | MDA-MB-361 | Elesclomol | MVP_top60_DNA+RNA_seed | HSP90AA1 | -1.151911 | BRCA | CAL-85-1 | Luminespib |
| MVP_top60_DNA_seed | ERBB2 | -0.673998 | BRCA | EFM-19 | PF-00299804 | MVP_top60_DNA+RNA_seed | HSP90AB1 | -1.151911 | BRCA | CAL-85-1 | Luminespib |
| MVP_top60_DNA_seed | EGFR | -0.673998 | BRCA | EFM-19 | PF-00299804 | MVP_top60_DNA+RNA_seed | CDK2 | -1.151067 | BRCA | HCC1599 | PHA-793887 |
| MVP_top60_DNA_seed | HDAC1 | -0.673952 | BRCA | MDA-MB-361 | AR-42 | MVP_top60_DNA+RNA_seed | AKT1 | -1.149798 | BRCA | MDA-MB-361 | AKT inhibitor VIII |
| MVP_top60_DNA_seed | EGFR | -0.672527 | BRCA | MDA-MB-361 | Gefitinib | MVP_top60_DNA+RNA_seed | EGFR | -1.146226 | BRCA | CAL-85-1 | Afatinib |
| MVP_top60_DNA_seed | AR | -0.67137 | BRCA | MCF7 | AZD3514 | MVP_top60_DNA+RNA_seed | ERBB2 | -1.146226 | BRCA | CAL-85-1 | Afatinib |
| MVP_top60_DNA_seed | SRC | -0.670324 | BRCA | OCUB-M | NVP-BHG712 | MVP_top60_DNA+RNA_seed | CDK2 | -1.144685 | BRCA | MDA-MB-468 | PHA-793887 |
| MVP_top60_DNA_seed | ESR1 | -0.669935 | BRCA | HCC1187 | Fulvestrant | MVP_top60_DNA+RNA_seed | HSP90AA1 | -1.141385 | BRCA | OCUB-M | SNX-2112 |
| MVP_top60_DNA_seed | ATM | -0.669773 | BRCA | MDA-MB-468 | CP466722 | MVP_top60_DNA+RNA_seed | HSP90AB1 | -1.141385 | BRCA | OCUB-M | SNX-2112 |

| | | | | | | | | | | | |
|---|---|---|---|---|---|---|---|---|---|---|---|
| MVP_top60_DNA_seed | EGFR | -0.669623 | BRCA | MDA-MB-157 | Gefitinib | MVP_top60_DNA+RNA_seed | AR | -1.138431 | BRCA | OCUB-M | Bicalutamide |
| MVP_top60_DNA_seed | EGFR | -0.666571 | BRCA | JIMT-1 | Cetuximab | MVP_top60_DNA+RNA_seed | PIK3CA | -1.1376 | BRCA | UACC-893 | PI3Ka_4409 |
| MVP_top60_DNA_seed | HSP90AA1 | -0.665852 | BRCA | JIMT-1 | Elesclomol | MVP_top60_DNA+RNA_seed | CDK1 | -1.136599 | BRCA | AU565 | RO-3306 |
| MVP_top60_DNA_seed | HSP90AB1 | -0.665852 | BRCA | JIMT-1 | Elesclomol | MVP_top60_DNA+RNA_seed | HSP90AA1 | -1.134675 | BRCA | MDA-MB-453 | SNX-2112 |
| MVP_top60_DNA_seed | HSP90AA1 | -0.665215 | BRCA | CAL-120 | Elesclomol | MVP_top60_DNA+RNA_seed | HSP90AB1 | -1.134675 | BRCA | MDA-MB-453 | SNX-2112 |
| MVP_top60_DNA_seed | HSP90AB1 | -0.665215 | BRCA | CAL-120 | Elesclomol | MVP_top60_DNA+RNA_seed | EGFR | -1.132719 | BRCA | MDA-MB-453 | CI-1033 |
| MVP_top60_DNA_seed | ESR1 | -0.661274 | BRCA | BT-483 | Fulvestrant | MVP_top60_DNA+RNA_seed | ERBB2 | -1.132719 | BRCA | MDA-MB-453 | CI-1033 |
| MVP_top60_DNA_seed | ESR1 | -0.660956 | BRCA | HCC1937 | Fulvestrant | MVP_top60_DNA+RNA_seed | CDK2 | -1.131387 | BRCA | HCC1187 | PHA-793887 |
| MVP_top60_DNA_seed | ERBB2 | -0.659806 | BRCA | CAL-85-1 | CP724714 | MVP_top60_DNA+RNA_seed | PIK3CA | -1.130069 | BRCA | T47D | PI3Ka_4409 |
| MVP_top60_DNA_seed | MDM2 | -0.657863 | BRCA | HCC1954 | Nutlin-3a (-) | MVP_top60_DNA+RNA_seed | ATM | -1.126387 | BRCA | CAL-85-1 | CP466722 |
| MVP_top60_DNA_seed | ERBB2 | -0.657423 | BRCA | BT-483 | CI-1033 | MVP_top60_DNA+RNA_seed | HDAC1 | -1.124987 | BRCA | MDA-MB-330 | JQ12 |
| MVP_top60_DNA_seed | EGFR | -0.657423 | BRCA | BT-483 | CI-1033 | MVP_top60_DNA+RNA_seed | HDAC2 | -1.124987 | BRCA | MDA-MB-330 | JQ12 |
| MVP_top60_DNA_seed | EP300 | -0.654789 | BRCA | CAMA-1 | I-CBP112 | MVP_top60_DNA+RNA_seed | PIK3CA | -1.12156 | BRCA | MCF7 | AZD8835 |
| MVP_top60_DNA_seed | CREBBP | -0.654789 | BRCA | CAMA-1 | I-CBP112 | MVP_top60_DNA+RNA_seed | HDAC1 | -1.119708 | BRCA | DU-4475 | Dacinostat |
| MVP_top60_DNA_seed | CDK2 | -0.654369 | BRCA | HCC2218 | Seliciclib | MVP_top60_DNA+RNA_seed | PLK1 | -1.116155 | BRCA | OCUB-M | GW843682X |
| MVP_top60_DNA_seed | CDK2 | -0.654038 | BRCA | HCC1954 | PHA-793887 | MVP_top60_DNA+RNA_seed | HDAC1 | -1.108087 | BRCA | OCUB-M | Dacinostat |
| MVP_top60_DNA_seed | AKT1 | -0.646971 | BRCA | HCC1937 | AKT inhibitor VIII | MVP_top60_DNA+RNA_seed | ERBB2 | -1.104227 | BRCA | DU-4475 | CP724714 |
| MVP_top60_DNA_seed | HDAC1 | -0.64648 | BRCA | ZR-75-30 | Dacinostat | MVP_top60_DNA+RNA_seed | ESR1 | -1.102562 | BRCA | MDA-MB-157 | Fulvestrant |
| MVP_top60_DNA_seed | HDAC1 | -0.646383 | BRCA | OCUB-M | AR-42 | MVP_top60_DNA+RNA_seed | EGFR | -1.101357 | BRCA | HCC1187 | PF-00299804 |
| MVP_top60_DNA_seed | HDAC1 | -0.64417 | BRCA | T47D | Dacinostat | MVP_top60_DNA+RNA_seed | ERBB2 | -1.101357 | BRCA | HCC1187 | PF-00299804 |
| MVP_top60_DNA_seed | ESR1 | -0.644129 | BRCA | EFM-192A | Fulvestrant | MVP_top60_DNA+RNA_seed | ATM | -1.10092 | BRCA | T47D | KU-60019 |
| MVP_top60_DNA_seed | AR | -0.642307 | BRCA | MDA-MB-415 | Bicalutamide | MVP_top60_DNA+RNA_seed | EGFR | -1.099539 | BRCA | EFM-19 | AST-1306 |
| MVP_top60_DNA_seed | EGFR | -0.641862 | BRCA | CAL-120 | Pelitinib | MVP_top60_DNA+RNA_seed | EGFR | -1.099158 | BRCA | MDA-MB-468 | PF-00299804 |
| MVP_top60_DNA_seed | ERBB2 | -0.641509 | BRCA | AU565 | CUDC-101 | MVP_top60_DNA+RNA_seed | ERBB2 | -1.099158 | BRCA | MDA-MB-468 | PF-00299804 |
| MVP_top60_DNA_seed | EGFR | -0.641509 | BRCA | AU565 | CUDC-101 | MVP_top60_DNA+RNA_seed | AKT1 | -1.093649 | BRCA | HCC2157 | MK-2206 |
| MVP_top60_DNA_seed | AR | -0.639064 | BRCA | UACC-893 | Bicalutamide | MVP_top60_DNA+RNA_seed | HSP90AA1 | -1.090695 | BRCA | EFM-192A | Tanespimycin |
| MVP_top60_DNA_seed | HDAC1 | -0.638339 | BRCA | ZR-75-30 | Belinostat | MVP_top60_DNA+RNA_seed | HSP90AB1 | -1.090695 | BRCA | EFM-192A | Tanespimycin |
| MVP_top60_DNA_seed | AKT1 | -0.637138 | BRCA | CAMA-1 | BAY AKT1 | MVP_top60_DNA+RNA_seed | MAPK14 | -1.084629 | BRCA | DU-4475 | TAK-715 |
| MVP_top60_DNA_seed | ERBB2 | -0.636479 | BRCA | HCC38 | PF-00299804 | MVP_top60_DNA+RNA_seed | HDAC1 | -1.08089 | BRCA | HCC2218 | Dacinostat |
| MVP_top60_DNA_seed | EGFR | -0.636479 | BRCA | HCC38 | PF-00299804 | MVP_top60_DNA+RNA_seed | CDK1 | -1.078041 | BRCA | JIMT-1 | RO-3306 |
| MVP_top60_DNA_seed | HSP90AA1 | -0.636017 | BRCA | ZR-75-30 | Elesclomol | MVP_top60_DNA+RNA_seed | HDAC1 | -1.076885 | BRCA | MDA-MB-157 | Dacinostat |
| MVP_top60_DNA_seed | HSP90AB1 | -0.636017 | BRCA | ZR-75-30 | Elesclomol | MVP_top60_DNA+RNA_seed | ERBB2 | -1.074989 | BRCA | BT-549 | CP724714 |
| MVP_top60_DNA_seed | AKT1 | -0.635331 | BRCA | HCC1569 | BAY AKT1 | MVP_top60_DNA+RNA_seed | AR | -1.074078 | BRCA | AU565 | Bicalutamide |
| MVP_top60_DNA_seed | HDAC1 | -0.635159 | BRCA | MRK-nu-1 | Tubastatin A | MVP_top60_DNA+RNA_seed | PIK3CA | -1.072273 | BRCA | JIMT-1 | PI3Ka_4409 |
| MVP_top60_DNA_seed | ATM | -0.633036 | BRCA | HCC202 | CP466722 | MVP_top60_DNA+RNA_seed | EGFR | -1.070307 | BRCA | HCC1954 | AST-1306 |
| MVP_top60_DNA_seed | HDAC1 | -0.632153 | BRCA | T47D | AR-42 | MVP_top60_DNA+RNA_seed | AKT1 | -1.067314 | BRCA | HCC2157 | GSK690693 |
| MVP_top60_DNA_seed | HSP90AA1 | -0.630822 | BRCA | HCC38 | SNX-2112 | MVP_top60_DNA+RNA_seed | MDM2 | -1.066872 | BRCA | CAL-51 | Nutlin-3a (-) |
| MVP_top60_DNA_seed | HSP90AB1 | -0.630822 | BRCA | HCC38 | SNX-2112 | MVP_top60_DNA+RNA_seed | AKT1 | -1.063205 | BRCA | UACC-893 | AKT inhibitor VIII |
| MVP_top60_DNA_seed | ERBB2 | -0.629646 | BRCA | HCC1187 | CP724714 | MVP_top60_DNA+RNA_seed | HSP90AA1 | -1.063194 | BRCA | JIMT-1 | Tanespimycin |
| MVP_top60_DNA_seed | ERBB2 | -0.629127 | BRCA | HCC1569 | Afatinib | MVP_top60_DNA+RNA_seed | HSP90AB1 | -1.063194 | BRCA | JIMT-1 | Tanespimycin |
| MVP_top60_DNA_seed | EGFR | -0.629127 | BRCA | HCC1569 | Afatinib | MVP_top60_DNA+RNA_seed | HSP90AA1 | -1.059699 | BRCA | BT-549 | Elesclomol |

| | | | | | | | | | | | |
|---|---|---|---|---|---|---|---|---|---|---|---|
| MVP_top60_DNA_seed | ATM | -0.628853 | BRCA | COLO-824 | KU-60019 | MVP_top60_DNA+RNA_seed | HSP90AB1 | -1.059699 | BRCA | BT-549 | Elesclomol |
| MVP_top60_DNA_seed | AKT1 | -0.62829 | BRCA | HCC1187 | AKT inhibitor VIII | MVP_top60_DNA+RNA_seed | MAPK14 | -1.05643 | BRCA | HCC1500 | TAK-715 |
| MVP_top60_DNA_seed | HSP90AA1 | -0.628036 | BRCA | T47D | Elesclomol | MVP_top60_DNA+RNA_seed | MAPK14 | -1.055777 | BRCA | HCC2218 | TAK-715 |
| MVP_top60_DNA_seed | HSP90AB1 | -0.628036 | BRCA | T47D | Elesclomol | MVP_top60_DNA+RNA_seed | HSP90AA1 | -1.055238 | BRCA | Hs-578-T | Tanespimycin |
| MVP_top60_DNA_seed | AKT1 | -0.627952 | BRCA | JIMT-1 | BAY AKT1 | MVP_top60_DNA+RNA_seed | HSP90AB1 | -1.055238 | BRCA | Hs-578-T | Tanespimycin |
| MVP_top60_DNA_seed | ATM | -0.626746 | BRCA | MFM-223 | torin2 | MVP_top60_DNA+RNA_seed | AKT1 | -1.044714 | BRCA | EVSA-T | GSK690693 |
| MVP_top60_DNA_seed | ERBB2 | -0.624557 | BRCA | EVSA-T | AZD8931 | MVP_top60_DNA+RNA_seed | HDAC1 | -1.043514 | BRCA | OCUB-M | Parthenolide |
| MVP_top60_DNA_seed | EGFR | -0.624557 | BRCA | EVSA-T | AZD8931 | MVP_top60_DNA+RNA_seed | HDAC1 | -1.043264 | BRCA | UACC-812 | Tubastatin A |
| MVP_top60_DNA_seed | ERBB3 | -0.624557 | BRCA | EVSA-T | AZD8931 | MVP_top60_DNA+RNA_seed | HSP90AA1 | -1.04278 | BRCA | MRK-nu-1 | Luminespib |
| MVP_top60_DNA_seed | ESR1 | -0.624056 | BRCA | JIMT-1 | Fulvestrant | MVP_top60_DNA+RNA_seed | HSP90AB1 | -1.04278 | BRCA | MRK-nu-1 | Luminespib |
| MVP_top60_DNA_seed | AKT1 | -0.622998 | BRCA | MDA-MB-330 | BAY AKT1 | MVP_top60_DNA+RNA_seed | CDK2 | -1.042589 | BRCA | DU-4475 | PHA-793887 |
| MVP_top60_DNA_seed | ERBB2 | -0.622722 | BRCA | HCC2157 | AZD8931 | MVP_top60_DNA+RNA_seed | EGFR | -1.041564 | BRCA | MDA-MB-468 | AZD8931 |
| MVP_top60_DNA_seed | EGFR | -0.622722 | BRCA | HCC2157 | AZD8931 | MVP_top60_DNA+RNA_seed | ERBB2 | -1.041564 | BRCA | MDA-MB-468 | AZD8931 |
| MVP_top60_DNA_seed | ERBB3 | -0.622722 | BRCA | HCC2157 | AZD8931 | MVP_top60_DNA+RNA_seed | ERBB3 | -1.041564 | BRCA | MDA-MB-468 | AZD8931 |
| MVP_top60_DNA_seed | AR | -0.622551 | BRCA | EVSA-T | Bicalutamide | MVP_top60_DNA+RNA_seed | AKT1 | -1.041496 | BRCA | BT-474 | BAY AKT1 |
| MVP_top60_DNA_seed | HDAC1 | -0.622258 | BRCA | EVSA-T | CAY10603 | MVP_top60_DNA+RNA_seed | HDAC1 | -1.032349 | BRCA | HCC1599 | AR-42 |
| MVP_top60_DNA_seed | AR | -0.621906 | BRCA | MDA-MB-468 | Bicalutamide | MVP_top60_DNA+RNA_seed | ESR1 | -1.031006 | BRCA | OCUB-M | Tamoxifen |
| MVP_top60_DNA_seed | AR | -0.618247 | BRCA | MDA-MB-468 | AZD3514 | MVP_top60_DNA+RNA_seed | AKT1 | -1.026894 | BRCA | BT-20 | AKT inhibitor VIII |
| MVP_top60_DNA_seed | AKT1 | -0.614502 | BRCA | BT-549 | AKT inhibitor VIII | MVP_top60_DNA+RNA_seed | HSP90AA1 | -1.02336 | BRCA | DU-4475 | Luminespib |
| MVP_top60_DNA_seed | ERBB2 | -0.613571 | BRCA | MDA-MB-330 | AZD8931 | MVP_top60_DNA+RNA_seed | HSP90AB1 | -1.02336 | BRCA | DU-4475 | Luminespib |
| MVP_top60_DNA_seed | EGFR | -0.613571 | BRCA | MDA-MB-330 | AZD8931 | MVP_top60_DNA+RNA_seed | EGFR | -1.019592 | BRCA | HCC1599 | Pelitinib |
| MVP_top60_DNA_seed | ERBB3 | -0.613571 | BRCA | MDA-MB-330 | AZD8931 | MVP_top60_DNA+RNA_seed | HDAC1 | -1.016586 | BRCA | MDA-MB-453 | CAY10603 |
| MVP_top60_DNA_seed | AKT1 | -0.611586 | BRCA | DU-4475 | AKT inhibitor VIII | MVP_top60_DNA+RNA_seed | EGFR | -1.015866 | BRCA | HDQ-P1 | Gefitinib |
| MVP_top60_DNA_seed | ERBB2 | -0.607462 | BRCA | HCC1419 | AZD8931 | MVP_top60_DNA+RNA_seed | ESR1 | -1.003676 | BRCA | EVSA-T | Tamoxifen |
| MVP_top60_DNA_seed | EGFR | -0.607462 | BRCA | HCC1419 | AZD8931 | MVP_top60_DNA+RNA_seed | EGFR | -0.996338 | BRCA | MDA-MB-330 | PF-00299804 |
| MVP_top60_DNA_seed | ERBB3 | -0.607462 | BRCA | HCC1419 | AZD8931 | MVP_top60_DNA+RNA_seed | ERBB2 | -0.996338 | BRCA | MDA-MB-330 | PF-00299804 |
| MVP_top60_DNA_seed | SRC | -0.60714 | BRCA | HCC1599 | NVP-BHG712 | MVP_top60_DNA+RNA_seed | ATM | -0.995295 | BRCA | MCF7 | KU-60019 |
| MVP_top60_DNA_seed | ERBB2 | -0.605689 | BRCA | JIMT-1 | Afatinib | MVP_top60_DNA+RNA_seed | ESR1 | -0.990266 | BRCA | BT-549 | Tamoxifen |
| MVP_top60_DNA_seed | EGFR | -0.605689 | BRCA | JIMT-1 | Afatinib | MVP_top60_DNA+RNA_seed | HDAC1 | -0.987847 | BRCA | HCC1599 | Tubastatin A |
| MVP_top60_DNA_seed | HDAC1 | -0.60563 | BRCA | HCC38 | Tubastatin A | MVP_top60_DNA+RNA_seed | EGFR | -0.986708 | BRCA | HCC2157 | Erlotinib |
| MVP_top60_DNA_seed | ESR1 | -0.605107 | BRCA | HCC1569 | Fulvestrant | MVP_top60_DNA+RNA_seed | AKT1 | -0.985059 | BRCA | BT-549 | GSK690693 |
| MVP_top60_DNA_seed | HDAC1 | -0.597841 | BRCA | DU-4475 | AR-42 | MVP_top60_DNA+RNA_seed | SRC | -0.984937 | BRCA | MFM-223 | HG-6-71-01 |
| MVP_top60_DNA_seed | MAPK3 | -0.59783 | BRCA | MDA-MB-175-VII | FR-180204 | MVP_top60_DNA+RNA_seed | EGFR | -0.983487 | BRCA | OCUB-M | CUDC-101 |
| MVP_top60_DNA_seed | MAPK1 | -0.59783 | BRCA | MDA-MB-175-VII | FR-180204 | MVP_top60_DNA+RNA_seed | ERBB2 | -0.983487 | BRCA | OCUB-M | CUDC-101 |
| MVP_top60_DNA_seed | AR | -0.597128 | BRCA | T47D | Bicalutamide | MVP_top60_DNA+RNA_seed | AR | -0.974125 | BRCA | HCC1500 | AZD3514 |
| MVP_top60_DNA_seed | EGFR | -0.595857 | BRCA | MDA-MB-361 | AST-1306 | MVP_top60_DNA+RNA_seed | AKT1 | -0.971857 | BRCA | HCC1187 | BAY AKT1 |
| MVP_top60_DNA_seed | AKT1 | -0.593677 | BRCA | MDA-MB-175-VII | AKT inhibitor VIII | MVP_top60_DNA+RNA_seed | PLK1 | -0.969456 | BRCA | BT-549 | PLK_6522 |
| MVP_top60_DNA_seed | ATM | -0.593621 | BRCA | EFM-19 | CP466722 | MVP_top60_DNA+RNA_seed | SRC | -0.966973 | BRCA | JIMT-1 | Bosutinib |
| MVP_top60_DNA_seed | HSP90AA1 | -0.593513 | BRCA | MDA-MB-468 | SNX-2112 | MVP_top60_DNA+RNA_seed | AR | -0.966711 | BRCA | HCC2157 | AZD3514 |
| MVP_top60_DNA_seed | HSP90AB1 | -0.593513 | BRCA | MDA-MB-468 | SNX-2112 | MVP_top60_DNA+RNA_seed | EGFR | -0.965679 | BRCA | DU-4475 | CI-1033 |
| MVP_top60_DNA_seed | ERBB2 | -0.592496 | BRCA | EVSA-T | Lapatinib | MVP_top60_DNA+RNA_seed | ERBB2 | -0.965679 | BRCA | DU-4475 | CI-1033 |

| | | | | | | | | | | | |
|---|---|---|---|---|---|---|---|---|---|---|---|
| MVP_top60_DNA_seed | EGFR | -0.592496 | BRCA | EVSA-T | Lapatinib | MVP_top60_DNA+RNA_seed | AKT1 | -0.964751 | BRCA | HCC2157 | AKT inhibitor VIII |
| MVP_top60_DNA_seed | ERBB2 | -0.591098 | BRCA | COLO-824 | CP724714 | MVP_top60_DNA+RNA_seed | EGFR | -0.963921 | BRCA | Hs-578-T | AST-1306 |
| MVP_top60_DNA_seed | HDAC1 | -0.589299 | BRCA | CAL-148 | AR-42 | MVP_top60_DNA+RNA_seed | EGFR | -0.963661 | BRCA | T47D | Pelitinib |
| MVP_top60_DNA_seed | ERBB2 | -0.588791 | BRCA | JIMT-1 | AZD8931 | MVP_top60_DNA+RNA_seed | PIK3CA | -0.962684 | BRCA | EVSA-T | PI3Ka_4409 |
| MVP_top60_DNA_seed | EGFR | -0.588791 | BRCA | JIMT-1 | AZD8931 | MVP_top60_DNA+RNA_seed | EGFR | -0.957423 | BRCA | HCC1187 | CI-1033 |
| MVP_top60_DNA_seed | ERBB3 | -0.588791 | BRCA | JIMT-1 | AZD8931 | MVP_top60_DNA+RNA_seed | ERBB2 | -0.957423 | BRCA | HCC1187 | CI-1033 |
| MVP_top60_DNA_seed | HSP90AA1 | -0.588493 | BRCA | MFM-223 | Elesclomol | MVP_top60_DNA+RNA_seed | EGFR | -0.957203 | BRCA | HCC1599 | CUDC-101 |
| MVP_top60_DNA_seed | HSP90AB1 | -0.588493 | BRCA | MFM-223 | Elesclomol | MVP_top60_DNA+RNA_seed | ERBB2 | -0.957203 | BRCA | HCC1599 | CUDC-101 |
| MVP_top60_DNA_seed | AKT1 | -0.587493 | BRCA | HCC1937 | GSK690693 | MVP_top60_DNA+RNA_seed | HSP90AA1 | -0.956587 | BRCA | HCC1599 | SNX-2112 |
| MVP_top60_DNA_seed | AKT1 | -0.586046 | BRCA | MCF7 | MK-2206 | MVP_top60_DNA+RNA_seed | HSP90AB1 | -0.956587 | BRCA | HCC1599 | SNX-2112 |
| MVP_top60_DNA_seed | EGFR | -0.584069 | BRCA | JIMT-1 | Gefitinib | MVP_top60_DNA+RNA_seed | CDK2 | -0.953106 | BRCA | HCC70 | AZD5438 |
| MVP_top60_DNA_seed | HSP90AA1 | -0.583879 | BRCA | MDA-MB-361 | SNX-2112 | MVP_top60_DNA+RNA_seed | HDAC1 | -0.951777 | BRCA | AU565 | Dacinostat |
| MVP_top60_DNA_seed | HSP90AB1 | -0.583879 | BRCA | MDA-MB-361 | SNX-2112 | MVP_top60_DNA+RNA_seed | ATM | -0.950399 | BRCA | MCF7 | KU-55933 |
| MVP_top60_DNA_seed | ERBB2 | -0.582703 | BRCA | MDA-MB-453 | Afatinib | MVP_top60_DNA+RNA_seed | PLK1 | -0.948112 | BRCA | MFM-223 | PLK_6522 |
| MVP_top60_DNA_seed | EGFR | -0.582703 | BRCA | MDA-MB-453 | Afatinib | MVP_top60_DNA+RNA_seed | HDAC1 | -0.946798 | BRCA | HCC1599 | Entinostat |
| MVP_top60_DNA_seed | AR | -0.57956 | BRCA | HCC38 | Bicalutamide | MVP_top60_DNA+RNA_seed | ERBB2 | -0.945299 | BRCA | MDA-MB-468 | CP724714 |
| MVP_top60_DNA_seed | ATM | -0.575105 | BRCA | HCC1187 | CP466722 | MVP_top60_DNA+RNA_seed | EGFR | -0.943089 | BRCA | HCC70 | Gefitinib |
| MVP_top60_DNA_seed | PIK3CA | -0.574712 | BRCA | HCC1187 | AZD8835 | MVP_top60_DNA+RNA_seed | PIK3CA | -0.937617 | BRCA | HCC1954 | AZD8835 |
| MVP_top60_DNA_seed | ERBB2 | -0.573468 | BRCA | HCC1500 | PF-00299804 | MVP_top60_DNA+RNA_seed | HSP90AA1 | -0.931812 | BRCA | MCF7 | CCT-018159 |
| MVP_top60_DNA_seed | EGFR | -0.573468 | BRCA | HCC1500 | PF-00299804 | MVP_top60_DNA+RNA_seed | HSP90AB1 | -0.931812 | BRCA | MCF7 | CCT-018159 |
| MVP_top60_DNA_seed | PIK3CA | -0.573288 | BRCA | T47D | AZD8835 | MVP_top60_DNA+RNA_seed | ERBB2 | -0.931348 | BRCA | MDA-MB-361 | CP724714 |
| MVP_top60_DNA_seed | EGFR | -0.571153 | BRCA | Hs-578-T | Cetuximab | MVP_top60_DNA+RNA_seed | AKT1 | -0.931273 | BRCA | MFM-223 | GSK690693 |
| MVP_top60_DNA_seed | HSP90AA1 | -0.569932 | BRCA | HCC1419 | Tanespimycin | MVP_top60_DNA+RNA_seed | HDAC1 | -0.929575 | BRCA | HCC1599 | CAY10603 |
| MVP_top60_DNA_seed | HSP90AB1 | -0.569932 | BRCA | HCC1419 | Tanespimycin | MVP_top60_DNA+RNA_seed | EGFR | -0.926688 | BRCA | OCUB-M | AZD8931 |
| MVP_top60_DNA_seed | HSP90AA1 | -0.569817 | BRCA | MCF7 | Elesclomol | MVP_top60_DNA+RNA_seed | ERBB2 | -0.926688 | BRCA | OCUB-M | AZD8931 |
| MVP_top60_DNA_seed | HSP90AB1 | -0.569817 | BRCA | MCF7 | Elesclomol | MVP_top60_DNA+RNA_seed | ERBB3 | -0.926688 | BRCA | OCUB-M | AZD8931 |
| MVP_top60_DNA_seed | PIK3CA | -0.568687 | BRCA | MFM-223 | PI3Ka_4409 | MVP_top60_DNA+RNA_seed | EGFR | -0.925262 | BRCA | MDA-MB-468 | CI-1033 |
| MVP_top60_DNA_seed | AKT1 | -0.56628 | BRCA | EFM-192A | BAY AKT1 | MVP_top60_DNA+RNA_seed | ERBB2 | -0.925262 | BRCA | MDA-MB-468 | CI-1033 |
| MVP_top60_DNA_seed | PIK3CA | -0.565849 | BRCA | HCC1187 | PI3Ka_4409 | MVP_top60_DNA+RNA_seed | EGFR | -0.923674 | BRCA | HCC38 | CI-1033 |
| MVP_top60_DNA_seed | EGFR | -0.565733 | BRCA | OCUB-M | Cetuximab | MVP_top60_DNA+RNA_seed | ERBB2 | -0.923674 | BRCA | HCC38 | CI-1033 |
| MVP_top60_DNA_seed | ATM | -0.565203 | BRCA | AU565 | KU-55933 | MVP_top60_DNA+RNA_seed | EGFR | -0.921856 | BRCA | AU565 | Gefitinib |
| MVP_top60_DNA_seed | ERBB2 | -0.564873 | BRCA | ZR-75-30 | AZD8931 | MVP_top60_DNA+RNA_seed | ATM | -0.920671 | BRCA | BT-549 | CP466722 |
| MVP_top60_DNA_seed | EGFR | -0.564873 | BRCA | ZR-75-30 | AZD8931 | MVP_top60_DNA+RNA_seed | MDM2 | -0.920505 | BRCA | AU565 | Serdemetan |
| MVP_top60_DNA_seed | ERBB3 | -0.564873 | BRCA | ZR-75-30 | AZD8931 | MVP_top60_DNA+RNA_seed | HDAC1 | -0.918874 | BRCA | HCC1599 | Dacinostat |
| MVP_top60_DNA_seed | HDAC1 | -0.564583 | BRCA | EFM-192A | Dacinostat | MVP_top60_DNA+RNA_seed | EGFR | -0.916199 | BRCA | EFM-192A | PF-00299804 |
| MVP_top60_DNA_seed | HDAC1 | -0.564211 | BRCA | EFM-19 | Dacinostat | MVP_top60_DNA+RNA_seed | ERBB2 | -0.916199 | BRCA | EFM-192A | PF-00299804 |
| MVP_top60_DNA_seed | EGFR | -0.562484 | BRCA | MDA-MB-415 | Cetuximab | MVP_top60_DNA+RNA_seed | ESR1 | -0.915395 | BRCA | HCC1569 | Tamoxifen |
| MVP_top60_DNA_seed | ATM | -0.562424 | BRCA | HCC70 | CP466722 | MVP_top60_DNA+RNA_seed | ATM | -0.91518 | BRCA | OCUB-M | CP466722 |
| MVP_top60_DNA_seed | ERBB2 | -0.56204 | BRCA | MRK-nu-1 | PF-00299804 | MVP_top60_DNA+RNA_seed | EGFR | -0.912772 | BRCA | BT-474 | Gefitinib |
| MVP_top60_DNA_seed | EGFR | -0.56204 | BRCA | MRK-nu-1 | PF-00299804 | MVP_top60_DNA+RNA_seed | HSP90AA1 | -0.910938 | BRCA | HCC1937 | Elesclomol |
| MVP_top60_DNA_seed | EP300 | -0.560005 | BRCA | MDA-MB-231 | I-CBP112 | MVP_top60_DNA+RNA_seed | HSP90AB1 | -0.910938 | BRCA | HCC1937 | Elesclomol |

| | | | | | | | | | | | |
|---|---|---|---|---|---|---|---|---|---|---|---|
| MVP_top60_DNA_seed | CREBBP | -0.560005 | BRCA | MDA-MB-231 | I-CBP112 | MVP_top60_DNA+RNA_seed | EGFR | -0.910203 | BRCA | HCC2157 | Lapatinib |
| MVP_top60_DNA_seed | HSP90AA1 | -0.558557 | BRCA | HCC1569 | CCT-018159 | MVP_top60_DNA+RNA_seed | ERBB2 | -0.910203 | BRCA | HCC2157 | Lapatinib |
| MVP_top60_DNA_seed | HSP90AB1 | -0.558557 | BRCA | HCC1569 | CCT-018159 | MVP_top60_DNA+RNA_seed | CDK1 | -0.908468 | BRCA | HCC1395 | RO-3306 |
| MVP_top60_DNA_seed | ERBB2 | -0.556597 | BRCA | HCC1569 | CI-1033 | MVP_top60_DNA+RNA_seed | HDAC1 | -0.907799 | BRCA | MRK-nu-1 | Parthenolide |
| MVP_top60_DNA_seed | EGFR | -0.556597 | BRCA | HCC1569 | CI-1033 | MVP_top60_DNA+RNA_seed | CDK2 | -0.905273 | BRCA | HCC1599 | Seliciclib |
| MVP_top60_DNA_seed | AKT1 | -0.55622 | BRCA | EVSA-T | A-443654 | MVP_top60_DNA+RNA_seed | AKT1 | -0.905204 | BRCA | Hs-578-T | MK-2206 |
| MVP_top60_DNA_seed | HSP90AA1 | -0.554147 | BRCA | BT-474 | Elesclomol | MVP_top60_DNA+RNA_seed | HDAC1 | -0.903705 | BRCA | HCC2157 | Dacinostat |
| MVP_top60_DNA_seed | HSP90AB1 | -0.554147 | BRCA | BT-474 | Elesclomol | MVP_top60_DNA+RNA_seed | EGFR | -0.903416 | BRCA | EFM-192A | CI-1033 |
| MVP_top60_DNA_seed | HSP90AA1 | -0.550969 | BRCA | OCUB-M | Luminespib | MVP_top60_DNA+RNA_seed | ERBB2 | -0.903416 | BRCA | EFM-192A | CI-1033 |
| MVP_top60_DNA_seed | HSP90AB1 | -0.550969 | BRCA | OCUB-M | Luminespib | MVP_top60_DNA+RNA_seed | EGFR | -0.899524 | BRCA | COLO-824 | PF-00299804 |
| MVP_top60_DNA_seed | AKT1 | -0.550963 | BRCA | MDA-MB-175-VII | BAY AKT1 | MVP_top60_DNA+RNA_seed | ERBB2 | -0.899524 | BRCA | COLO-824 | PF-00299804 |
| MVP_top60_DNA_seed | HDAC1 | -0.549887 | BRCA | HCC70 | CAY10603 | MVP_top60_DNA+RNA_seed | MAPK3 | -0.898571 | BRCA | DU-4475 | FR-180204 |
| MVP_top60_DNA_seed | AR | -0.54888 | BRCA | CAL-148 | Bicalutamide | MVP_top60_DNA+RNA_seed | MAPK1 | -0.898571 | BRCA | DU-4475 | FR-180204 |
| MVP_top60_DNA_seed | EGFR | -0.546664 | BRCA | HCC2157 | AST-1306 | MVP_top60_DNA+RNA_seed | PIK3CA | -0.895769 | BRCA | BT-20 | PI3Ka_4409 |
| MVP_top60_DNA_seed | MDM2 | -0.546559 | BRCA | HCC2218 | Nutlin-3a (-) | MVP_top60_DNA+RNA_seed | EGFR | -0.894579 | BRCA | T47D | CUDC-101 |
| MVP_top60_DNA_seed | ERBB2 | -0.542227 | BRCA | HCC2157 | CI-1033 | MVP_top60_DNA+RNA_seed | ERBB2 | -0.894579 | BRCA | T47D | CUDC-101 |
| MVP_top60_DNA_seed | EGFR | -0.542227 | BRCA | HCC2157 | CI-1033 | MVP_top60_DNA+RNA_seed | EGFR | -0.893788 | BRCA | HCC38 | Cetuximab |
| MVP_top60_DNA_seed | ERBB2 | -0.540024 | BRCA | HCC1954 | PF-00299804 | MVP_top60_DNA+RNA_seed | PIK3CA | -0.8937 | BRCA | BT-474 | PI3Ka_4409 |
| MVP_top60_DNA_seed | EGFR | -0.540024 | BRCA | HCC1954 | PF-00299804 | MVP_top60_DNA+RNA_seed | HSP90AA1 | -0.892775 | BRCA | CAL-148 | SNX-2112 |
| MVP_top60_DNA_seed | EGFR | -0.539556 | BRCA | HDQ-P1 | Cetuximab | MVP_top60_DNA+RNA_seed | HSP90AB1 | -0.892775 | BRCA | CAL-148 | SNX-2112 |
| MVP_top60_DNA_seed | ATM | -0.537684 | BRCA | HCC1187 | KU-55933 | MVP_top60_DNA+RNA_seed | EGFR | -0.89228 | BRCA | BT-474 | AZD8931 |
| MVP_top60_DNA_seed | AKT1 | -0.536812 | BRCA | HCC1395 | MK-2206 | MVP_top60_DNA+RNA_seed | ERBB2 | -0.89228 | BRCA | BT-474 | AZD8931 |
| MVP_top60_DNA_seed | HDAC1 | -0.536721 | BRCA | ZR-75-30 | JQ12 | MVP_top60_DNA+RNA_seed | ERBB3 | -0.89228 | BRCA | BT-474 | AZD8931 |
| MVP_top60_DNA_seed | HDAC2 | -0.536721 | BRCA | ZR-75-30 | JQ12 | MVP_top60_DNA+RNA_seed | CDK2 | -0.891753 | BRCA | CAL-51 | AZD5438 |
| MVP_top60_DNA_seed | AR | -0.535914 | BRCA | EVSA-T | AZD3514 | MVP_top60_DNA+RNA_seed | CDK2 | -0.88982 | BRCA | OCUB-M | PHA-793887 |
| MVP_top60_DNA_seed | ERBB2 | -0.532808 | BRCA | HCC2157 | PF-00299804 | MVP_top60_DNA+RNA_seed | CDK2 | -0.887057 | BRCA | HCC1187 | AZD5438 |
| MVP_top60_DNA_seed | EGFR | -0.532808 | BRCA | HCC2157 | PF-00299804 | MVP_top60_DNA+RNA_seed | HDAC1 | -0.886035 | BRCA | HCC1419 | JQ12 |
| MVP_top60_DNA_seed | EGFR | -0.531315 | BRCA | MDA-MB-175-VII | Cetuximab | MVP_top60_DNA+RNA_seed | HDAC2 | -0.886035 | BRCA | HCC1419 | JQ12 |
| MVP_top60_DNA_seed | HDAC1 | -0.531225 | BRCA | CAMA-1 | JQ12 | MVP_top60_DNA+RNA_seed | AKT1 | -0.885571 | BRCA | MDA-MB-468 | GSK690693 |
| MVP_top60_DNA_seed | HDAC2 | -0.531225 | BRCA | CAMA-1 | JQ12 | MVP_top60_DNA+RNA_seed | EGFR | -0.882029 | BRCA | MDA-MB-361 | Afatinib |
| MVP_top60_DNA_seed | AKT1 | -0.531059 | BRCA | MDA-MB-361 | GSK690693 | MVP_top60_DNA+RNA_seed | ERBB2 | -0.882029 | BRCA | MDA-MB-361 | Afatinib |
| MVP_top60_DNA_seed | AR | -0.530111 | BRCA | CAL-51 | Bicalutamide | MVP_top60_DNA+RNA_seed | AR | -0.881055 | BRCA | CAL-51 | Bicalutamide |
| MVP_top60_DNA_seed | HDAC1 | -0.526599 | BRCA | MDA-MB-231 | JQ12 | MVP_top60_DNA+RNA_seed | EGFR | -0.880323 | BRCA | MDA-MB-468 | AST-1306 |
| MVP_top60_DNA_seed | HDAC2 | -0.526599 | BRCA | MDA-MB-231 | JQ12 | MVP_top60_DNA+RNA_seed | AKT1 | -0.877569 | BRCA | BT-20 | GSK690693 |
| MVP_top60_DNA_seed | EGFR | -0.526292 | BRCA | HCC1187 | AST-1306 | MVP_top60_DNA+RNA_seed | EGFR | -0.873298 | BRCA | MDA-MB-330 | Afatinib |
| MVP_top60_DNA_seed | ESR1 | -0.52464 | BRCA | HCC1500 | Tamoxifen | MVP_top60_DNA+RNA_seed | ERBB2 | -0.873298 | BRCA | MDA-MB-330 | Afatinib |
| MVP_top60_DNA_seed | HDAC1 | -0.52226 | BRCA | CAL-148 | Tubastatin A | MVP_top60_DNA+RNA_seed | EGFR | -0.870149 | BRCA | UACC-812 | Lapatinib |
| MVP_top60_DNA_seed | ERBB2 | -0.520793 | BRCA | HCC1187 | AZD8931 | MVP_top60_DNA+RNA_seed | ERBB2 | -0.870149 | BRCA | UACC-812 | Lapatinib |
| MVP_top60_DNA_seed | EGFR | -0.520793 | BRCA | HCC1187 | AZD8931 | MVP_top60_DNA+RNA_seed | PLK1 | -0.869562 | BRCA | MDA-MB-361 | PLK_6522 |
| MVP_top60_DNA_seed | ERBB3 | -0.520793 | BRCA | HCC1187 | AZD8931 | MVP_top60_DNA+RNA_seed | CDK2 | -0.869125 | BRCA | BT-549 | AZD5438 |
| MVP_top60_DNA_seed | CDK1 | -0.519871 | BRCA | HCC38 | RO-3306 | MVP_top60_DNA+RNA_seed | HSP90AA1 | -0.869099 | BRCA | MDA-MB-436 | Elesclomol |

| | | | | | | | | | | | |
|---|---|---|---|---|---|---|---|---|---|---|---|
| MVP_top60_DNA_seed | ATM | -0.519858 | BRCA | MDA-MB-330 | CP466722 | MVP_top60_DNA+RNA_seed | HSP90AB1 | -0.869099 | BRCA | MDA-MB-436 | Elesclomol |
| MVP_top60_DNA_seed | ERBB2 | -0.519458 | BRCA | MDA-MB-361 | PF-00299804 | MVP_top60_DNA+RNA_seed | AR | -0.865078 | BRCA | EFM-19 | AZD3514 |
| MVP_top60_DNA_seed | EGFR | -0.519458 | BRCA | MDA-MB-361 | PF-00299804 | MVP_top60_DNA+RNA_seed | EGFR | -0.862815 | BRCA | EVSA-T | CUDC-101 |
| MVP_top60_DNA_seed | ERBB2 | -0.514997 | BRCA | COLO-824 | CI-1033 | MVP_top60_DNA+RNA_seed | ERBB2 | -0.862815 | BRCA | EVSA-T | CUDC-101 |
| MVP_top60_DNA_seed | EGFR | -0.514997 | BRCA | COLO-824 | CI-1033 | MVP_top60_DNA+RNA_seed | EGFR | -0.862629 | BRCA | HCC1954 | Pelitinib |
| MVP_top60_DNA_seed | EGFR | -0.513168 | BRCA | MDA-MB-157 | AST-1306 | MVP_top60_DNA+RNA_seed | EGFR | -0.862256 | BRCA | MDA-MB-330 | CI-1033 |
| MVP_top60_DNA_seed | ATM | -0.51309 | BRCA | MDA-MB-157 | KU-60019 | MVP_top60_DNA+RNA_seed | ERBB2 | -0.862256 | BRCA | MDA-MB-330 | CI-1033 |
| MVP_top60_DNA_seed | HSP90AA1 | -0.509269 | BRCA | MDA-MB-231 | Elesclomol | MVP_top60_DNA+RNA_seed | HDAC1 | -0.858594 | BRCA | MDA-MB-157 | JQ12 |
| MVP_top60_DNA_seed | HSP90AB1 | -0.509269 | BRCA | MDA-MB-231 | Elesclomol | MVP_top60_DNA+RNA_seed | HDAC2 | -0.858594 | BRCA | MDA-MB-157 | JQ12 |
| MVP_top60_DNA_seed | MDM2 | -0.509041 | BRCA | UACC-893 | Serdemetan | MVP_top60_DNA+RNA_seed | PLK1 | -0.857366 | BRCA | OCUB-M | PLK_6522 |
| MVP_top60_DNA_seed | MAPK3 | -0.508185 | BRCA | BT-474 | FR-180204 | MVP_top60_DNA+RNA_seed | AKT1 | -0.857239 | BRCA | MDA-MB-175-VII | AKT inhibitor VIII |
| MVP_top60_DNA_seed | MAPK1 | -0.508185 | BRCA | BT-474 | FR-180204 | MVP_top60_DNA+RNA_seed | HDAC1 | -0.857238 | BRCA | HCC202 | Dacinostat |
| MVP_top60_DNA_seed | ATM | -0.50763 | BRCA | HCC1954 | CP466722 | MVP_top60_DNA+RNA_seed | AKT1 | -0.856328 | BRCA | JIMT-1 | MK-2206 |
| MVP_top60_DNA_seed | ATM | -0.505788 | BRCA | T47D | CP466722 | MVP_top60_DNA+RNA_seed | HSP90AA1 | -0.855657 | BRCA | DU-4475 | Tanespimycin |
| MVP_top60_DNA_seed | ESR1 | -0.505314 | BRCA | DU-4475 | Tamoxifen | MVP_top60_DNA+RNA_seed | HSP90AB1 | -0.855657 | BRCA | DU-4475 | Tanespimycin |
| MVP_top60_DNA_seed | CDK2 | -0.501561 | BRCA | DU-4475 | Seliciclib | MVP_top60_DNA+RNA_seed | HSP90AA1 | -0.855371 | BRCA | JIMT-1 | Luminespib |
| MVP_top60_DNA_seed | HDAC1 | -0.500232 | BRCA | DU-4475 | Tubastatin A | MVP_top60_DNA+RNA_seed | HSP90AB1 | -0.855371 | BRCA | JIMT-1 | Luminespib |
| MVP_top60_DNA_seed | MAPK3 | -0.499002 | BRCA | HCC1599 | FR-180204 | MVP_top60_DNA+RNA_seed | ESR1 | -0.855114 | BRCA | HCC1428 | Tamoxifen |
| MVP_top60_DNA_seed | MAPK1 | -0.499002 | BRCA | HCC1599 | FR-180204 | MVP_top60_DNA+RNA_seed | PLK1 | -0.853843 | BRCA | MDA-MB-468 | PLK_6522 |
| MVP_top60_DNA_seed | EGFR | -0.498948 | BRCA | MFM-223 | AST-1306 | MVP_top60_DNA+RNA_seed | PIK3CA | -0.852086 | BRCA | UACC-893 | AZD8835 |
| MVP_top60_DNA_seed | HSP90AA1 | -0.498653 | BRCA | HCC1419 | Elesclomol | MVP_top60_DNA+RNA_seed | EGFR | -0.852045 | BRCA | DU-4475 | PF-00299804 |
| MVP_top60_DNA_seed | HSP90AB1 | -0.498653 | BRCA | HCC1419 | Elesclomol | MVP_top60_DNA+RNA_seed | ERBB2 | -0.852045 | BRCA | DU-4475 | PF-00299804 |
| MVP_top60_DNA_seed | CDK1 | -0.497766 | BRCA | OCUB-M | RO-3306 | MVP_top60_DNA+RNA_seed | HSP90AA1 | -0.85197 | BRCA | HCC1187 | CCT-018159 |
| MVP_top60_DNA_seed | EGFR | -0.496611 | BRCA | HCC1569 | AST-1306 | MVP_top60_DNA+RNA_seed | HSP90AB1 | -0.85197 | BRCA | HCC1187 | CCT-018159 |
| MVP_top60_DNA_seed | HSP90AA1 | -0.495431 | BRCA | MDA-MB-361 | Luminespib | MVP_top60_DNA+RNA_seed | MAPK14 | -0.851816 | BRCA | MDA-MB-468 | TAK-715 |
| MVP_top60_DNA_seed | HSP90AB1 | -0.495431 | BRCA | MDA-MB-361 | Luminespib | MVP_top60_DNA+RNA_seed | HDAC1 | -0.850372 | BRCA | EVSA-T | Tubastatin A |
| MVP_top60_DNA_seed | HDAC1 | -0.495061 | BRCA | HCC70 | Belinostat | MVP_top60_DNA+RNA_seed | CDK1 | -0.844429 | BRCA | MDA-MB-330 | RO-3306 |
| MVP_top60_DNA_seed | ESR1 | -0.49031 | BRCA | MDA-MB-415 | Fulvestrant | MVP_top60_DNA+RNA_seed | AR | -0.836762 | BRCA | HCC1954 | Bicalutamide |
| MVP_top60_DNA_seed | HDAC1 | -0.489166 | BRCA | DU-4475 | Belinostat | MVP_top60_DNA+RNA_seed | MAPK1 | -0.834528 | BRCA | CAL-85-1 | VX-11e |
| MVP_top60_DNA_seed | AKT1 | -0.488589 | BRCA | BT-20 | BAY AKT1 | MVP_top60_DNA+RNA_seed | EGFR | -0.834036 | BRCA | Hs-578-T | CI-1033 |
| MVP_top60_DNA_seed | MDM2 | -0.487753 | BRCA | HCC1954 | Serdemetan | MVP_top60_DNA+RNA_seed | ERBB2 | -0.834036 | BRCA | Hs-578-T | CI-1033 |
| MVP_top60_DNA_seed | CDK2 | -0.487203 | BRCA | ZR-75-30 | PHA-793887 | MVP_top60_DNA+RNA_seed | PLK1 | -0.833838 | BRCA | HCC1599 | GW843682X |
| MVP_top60_DNA_seed | ERBB2 | -0.485702 | BRCA | OCUB-M | Afatinib | MVP_top60_DNA+RNA_seed | PIK3CA | -0.832895 | BRCA | MFM-223 | AZD8835 |
| MVP_top60_DNA_seed | EGFR | -0.485702 | BRCA | OCUB-M | Afatinib | MVP_top60_DNA+RNA_seed | EGFR | -0.829601 | BRCA | HCC1569 | Afatinib |
| MVP_top60_DNA_seed | HSP90AA1 | -0.484939 | BRCA | HCC38 | Luminespib | MVP_top60_DNA+RNA_seed | ERBB2 | -0.829601 | BRCA | HCC1569 | Afatinib |
| MVP_top60_DNA_seed | HSP90AB1 | -0.484939 | BRCA | HCC38 | Luminespib | MVP_top60_DNA+RNA_seed | HSP90AA1 | -0.828051 | BRCA | Hs-578-T | Elesclomol |
| MVP_top60_DNA_seed | HSP90AA1 | -0.484702 | BRCA | COLO-824 | SNX-2112 | MVP_top60_DNA+RNA_seed | HSP90AB1 | -0.828051 | BRCA | Hs-578-T | Elesclomol |
| MVP_top60_DNA_seed | HSP90AB1 | -0.484702 | BRCA | COLO-824 | SNX-2112 | MVP_top60_DNA+RNA_seed | EGFR | -0.827642 | BRCA | CAL-148 | CUDC-101 |
| MVP_top60_DNA_seed | ERBB2 | -0.484012 | BRCA | HDQ-P1 | CI-1033 | MVP_top60_DNA+RNA_seed | ERBB2 | -0.827642 | BRCA | CAL-148 | CUDC-101 |
| MVP_top60_DNA_seed | EGFR | -0.484012 | BRCA | HDQ-P1 | CI-1033 | MVP_top60_DNA+RNA_seed | EGFR | -0.827082 | BRCA | EFM-192A | AZD8931 |
| MVP_top60_DNA_seed | HSP90AA1 | -0.483835 | BRCA | HCC1395 | Elesclomol | MVP_top60_DNA+RNA_seed | ERBB2 | -0.827082 | BRCA | EFM-192A | AZD8931 |

| | | | | | | | | | | | |
|---|---|---|---|---|---|---|---|---|---|---|---|
| MVP_top60_DNA_seed | HSP90AB1 | -0.483835 | BRCA | HCC1395 | Elesclomol | MVP_top60_DNA+RNA_seed | ERBB3 | -0.827082 | BRCA | EFM-192A | AZD8931 |
| MVP_top60_DNA_seed | ERBB2 | -0.483653 | BRCA | HCC1569 | PF-00299804 | MVP_top60_DNA+RNA_seed | ATM | -0.82688 | BRCA | HCC1419 | CP466722 |
| MVP_top60_DNA_seed | EGFR | -0.483653 | BRCA | HCC1569 | PF-00299804 | MVP_top60_DNA+RNA_seed | EGFR | -0.824701 | BRCA | BT-20 | Afatinib |
| MVP_top60_DNA_seed | ATM | -0.483183 | BRCA | UACC-812 | CP466722 | MVP_top60_DNA+RNA_seed | ERBB2 | -0.824701 | BRCA | BT-20 | Afatinib |
| MVP_top60_DNA_seed | HSP90AA1 | -0.483061 | BRCA | HCC2157 | Tanespimycin | MVP_top60_DNA+RNA_seed | ATM | -0.823608 | BRCA | HCC2157 | KU-60019 |
| MVP_top60_DNA_seed | HSP90AB1 | -0.483061 | BRCA | HCC2157 | Tanespimycin | MVP_top60_DNA+RNA_seed | ESR1 | -0.823213 | BRCA | HCC1954 | Tamoxifen |
| MVP_top60_DNA_seed | AR | -0.482859 | BRCA | CAL-120 | Bicalutamide | MVP_top60_DNA+RNA_seed | CDK2 | -0.822768 | BRCA | HCC2218 | PHA-793887 |
| MVP_top60_DNA_seed | HSP90AA1 | -0.481917 | BRCA | BT-483 | Elesclomol | MVP_top60_DNA+RNA_seed | PLK1 | -0.821775 | BRCA | HCC1187 | PLK_6522 |
| MVP_top60_DNA_seed | HSP90AB1 | -0.481917 | BRCA | BT-483 | Elesclomol | MVP_top60_DNA+RNA_seed | HDAC1 | -0.821157 | BRCA | EVSA-T | AR-42 |
| MVP_top60_DNA_seed | ERBB2 | -0.48186 | BRCA | DU-4475 | Afatinib | MVP_top60_DNA+RNA_seed | AKT1 | -0.820497 | BRCA | MDA-MB-175-VII | GSK690693 |
| MVP_top60_DNA_seed | EGFR | -0.48186 | BRCA | DU-4475 | Afatinib | MVP_top60_DNA+RNA_seed | ERBB2 | -0.817278 | BRCA | UACC-812 | CP724714 |
| MVP_top60_DNA_seed | ATM | -0.481795 | BRCA | MDA-MB-175-VII | CP466722 | MVP_top60_DNA+RNA_seed | MAPK14 | -0.817041 | BRCA | CAL-85-1 | TAK-715 |
| MVP_top60_DNA_seed | ATM | -0.479321 | BRCA | HCC1599 | CP466722 | MVP_top60_DNA+RNA_seed | EGFR | -0.816214 | BRCA | DU-4475 | Gefitinib |
| MVP_top60_DNA_seed | ESR1 | -0.479239 | BRCA | EFM-19 | Tamoxifen | MVP_top60_DNA+RNA_seed | AKT1 | -0.813966 | BRCA | AU565 | MK-2206 |
| MVP_top60_DNA_seed | AR | -0.474902 | BRCA | JIMT-1 | Bicalutamide | MVP_top60_DNA+RNA_seed | MDM2 | -0.8133 | BRCA | MDA-MB-361 | Serdemetan |
| MVP_top60_DNA_seed | CDK2 | -0.473969 | BRCA | MRK-nu-1 | AZD5438 | MVP_top60_DNA+RNA_seed | AKT1 | -0.812343 | BRCA | HCC2218 | GSK690693 |
| MVP_top60_DNA_seed | AR | -0.473498 | BRCA | CAL-120 | AZD3514 | MVP_top60_DNA+RNA_seed | HSP90AA1 | -0.804905 | BRCA | HCC1954 | CCT-018159 |
| MVP_top60_DNA_seed | ERBB2 | -0.470583 | BRCA | CAL-85-1 | CI-1033 | MVP_top60_DNA+RNA_seed | HSP90AB1 | -0.804905 | BRCA | HCC1954 | CCT-018159 |
| MVP_top60_DNA_seed | EGFR | -0.470583 | BRCA | CAL-85-1 | CI-1033 | MVP_top60_DNA+RNA_seed | HDAC1 | -0.802507 | BRCA | OCUB-M | CAY10603 |
| MVP_top60_DNA_seed | AR | -0.467232 | BRCA | HCC70 | Bicalutamide | MVP_top60_DNA+RNA_seed | AKT1 | -0.801952 | BRCA | HCC1599 | A-443654 |
| MVP_top60_DNA_seed | HSP90AA1 | -0.465864 | BRCA | HCC2157 | Elesclomol | MVP_top60_DNA+RNA_seed | HSP90AA1 | -0.79718 | BRCA | EFM-192A | Luminespib |
| MVP_top60_DNA_seed | HSP90AB1 | -0.465864 | BRCA | HCC2157 | Elesclomol | MVP_top60_DNA+RNA_seed | HSP90AB1 | -0.79718 | BRCA | EFM-192A | Luminespib |
| MVP_top60_DNA_seed | HSP90AA1 | -0.464356 | BRCA | AU565 | Tanespimycin | MVP_top60_DNA+RNA_seed | ESR1 | -0.796847 | BRCA | JIMT-1 | Tamoxifen |
| MVP_top60_DNA_seed | HSP90AB1 | -0.464356 | BRCA | AU565 | Tanespimycin | MVP_top60_DNA+RNA_seed | PLK1 | -0.796427 | BRCA | CAL-120 | PLK_6522 |
| MVP_top60_DNA_seed | ERBB2 | -0.46147 | BRCA | HCC1954 | CI-1033 | MVP_top60_DNA+RNA_seed | AKT1 | -0.796096 | BRCA | MDA-MB-453 | AKT inhibitor VIII |
| MVP_top60_DNA_seed | EGFR | -0.46147 | BRCA | HCC1954 | CI-1033 | MVP_top60_DNA+RNA_seed | HDAC1 | -0.795522 | BRCA | OCUB-M | Tubastatin A |
| MVP_top60_DNA_seed | HDAC1 | -0.461138 | BRCA | HCC1428 | AR-42 | MVP_top60_DNA+RNA_seed | EGFR | -0.792913 | BRCA | MDA-MB-361 | Pelitinib |
| MVP_top60_DNA_seed | HDAC1 | -0.461089 | BRCA | DU-4475 | Parthenolide | MVP_top60_DNA+RNA_seed | MDM2 | -0.791316 | BRCA | DU-4475 | Serdemetan |
| MVP_top60_DNA_seed | HDAC1 | -0.460676 | BRCA | ZR-75-30 | AR-42 | MVP_top60_DNA+RNA_seed | ATM | -0.787646 | BRCA | HCC1569 | CP466722 |
| MVP_top60_DNA_seed | HDAC1 | -0.459233 | BRCA | T47D | Tubastatin A | MVP_top60_DNA+RNA_seed | HDAC1 | -0.787546 | BRCA | MDA-MB-361 | Dacinostat |
| MVP_top60_DNA_seed | ERBB2 | -0.457829 | BRCA | CAL-85-1 | PF-00299804 | MVP_top60_DNA+RNA_seed | EGFR | -0.787435 | BRCA | ZR-75-30 | CUDC-101 |
| MVP_top60_DNA_seed | EGFR | -0.457829 | BRCA | CAL-85-1 | PF-00299804 | MVP_top60_DNA+RNA_seed | ERBB2 | -0.787435 | BRCA | ZR-75-30 | CUDC-101 |
| MVP_top60_DNA_seed | HDAC1 | -0.457802 | BRCA | MDA-MB-330 | Dacinostat | MVP_top60_DNA+RNA_seed | PIK3CA | -0.787269 | BRCA | MCF7 | PI3Ka_4409 |
| MVP_top60_DNA_seed | ERBB2 | -0.457104 | BRCA | MDA-MB-453 | PF-00299804 | MVP_top60_DNA+RNA_seed | HSP90AA1 | -0.785228 | BRCA | MDA-MB-157 | Tanespimycin |
| MVP_top60_DNA_seed | EGFR | -0.457104 | BRCA | MDA-MB-453 | PF-00299804 | MVP_top60_DNA+RNA_seed | HSP90AB1 | -0.785228 | BRCA | MDA-MB-157 | Tanespimycin |
| MVP_top60_DNA_seed | AKT1 | -0.455335 | BRCA | EFM-19 | GSK690693 | MVP_top60_DNA+RNA_seed | EGFR | -0.783679 | BRCA | HCC38 | AST-1306 |
| MVP_top60_DNA_seed | SRC | -0.455244 | BRCA | BT-549 | Bosutinib | MVP_top60_DNA+RNA_seed | PIK3CA | -0.782566 | BRCA | EFM-192A | AZD8835 |
| MVP_top60_DNA_seed | HSP90AA1 | -0.454606 | BRCA | HCC1569 | Luminespib | MVP_top60_DNA+RNA_seed | EGFR | -0.781674 | BRCA | UACC-893 | AZD8931 |
| MVP_top60_DNA_seed | HSP90AB1 | -0.454606 | BRCA | HCC1569 | Luminespib | MVP_top60_DNA+RNA_seed | ERBB2 | -0.781674 | BRCA | UACC-893 | AZD8931 |
| MVP_top60_DNA_seed | ERBB2 | -0.454469 | BRCA | MDA-MB-361 | CI-1033 | MVP_top60_DNA+RNA_seed | ERBB3 | -0.781674 | BRCA | UACC-893 | AZD8931 |
| MVP_top60_DNA_seed | EGFR | -0.454469 | BRCA | MDA-MB-361 | CI-1033 | MVP_top60_DNA+RNA_seed | HDAC1 | -0.781366 | BRCA | EFM-192A | JQ12 |

| | | | | | | | | | | | |
|---|---|---|---|---|---|---|---|---|---|---|---|
| MVP_top60_DNA_seed | ATM | -0.453857 | BRCA | DU-4475 | KU-55933 | MVP_top60_DNA+RNA_seed | HDAC2 | -0.781366 | BRCA | EFM-192A | JQ12 |
| MVP_top60_DNA_seed | PIK3CA | -0.453369 | BRCA | MDA-MB-330 | PI3Ka_4409 | MVP_top60_DNA+RNA_seed | HDAC1 | -0.781025 | BRCA | EFM-19 | Tubastatin A |
| MVP_top60_DNA_seed | AR | -0.453109 | BRCA | HCC1954 | Bicalutamide | MVP_top60_DNA+RNA_seed | HDAC1 | -0.778595 | BRCA | HCC1599 | Belinostat |
| MVP_top60_DNA_seed | HSP90AA1 | -0.452918 | BRCA | BT-474 | SNX-2112 | MVP_top60_DNA+RNA_seed | EGFR | -0.778568 | BRCA | HCC1954 | AZD8931 |
| MVP_top60_DNA_seed | HSP90AB1 | -0.452918 | BRCA | BT-474 | SNX-2112 | MVP_top60_DNA+RNA_seed | ERBB2 | -0.778568 | BRCA | HCC1954 | AZD8931 |
| MVP_top60_DNA_seed | CDK2 | -0.452877 | BRCA | MDA-MB-468 | AZD5438 | MVP_top60_DNA+RNA_seed | ERBB3 | -0.778568 | BRCA | HCC1954 | AZD8931 |
| MVP_top60_DNA_seed | ESR1 | -0.450343 | BRCA | MFM-223 | Fulvestrant | MVP_top60_DNA+RNA_seed | PIK3CA | -0.778535 | BRCA | BT-474 | AZD8835 |
| MVP_top60_DNA_seed | AKT1 | -0.447618 | BRCA | HCC2218 | A-443654 | MVP_top60_DNA+RNA_seed | AR | -0.778391 | BRCA | BT-549 | Bicalutamide |
| MVP_top60_DNA_seed | HDAC1 | -0.447284 | BRCA | HCC1419 | Dacinostat | MVP_top60_DNA+RNA_seed | HSP90AA1 | -0.777219 | BRCA | CAMA-1 | Tanespimycin |
| MVP_top60_DNA_seed | HDAC1 | -0.44226 | BRCA | AU565 | Belinostat | MVP_top60_DNA+RNA_seed | HSP90AB1 | -0.777219 | BRCA | CAMA-1 | Tanespimycin |
| MVP_top60_DNA_seed | ERBB2 | -0.441241 | BRCA | HCC2157 | Afatinib | MVP_top60_DNA+RNA_seed | HDAC1 | -0.776846 | BRCA | MCF7 | Dacinostat |
| MVP_top60_DNA_seed | EGFR | -0.441241 | BRCA | HCC2157 | Afatinib | MVP_top60_DNA+RNA_seed | AKT1 | -0.775694 | BRCA | HCC70 | GSK690693 |
| MVP_top60_DNA_seed | PIK3CA | -0.43744 | BRCA | EVSA-T | AZD8835 | MVP_top60_DNA+RNA_seed | EGFR | -0.773581 | BRCA | MDA-MB-468 | Afatinib |
| MVP_top60_DNA_seed | HDAC1 | -0.436725 | BRCA | MDA-MB-330 | AR-42 | MVP_top60_DNA+RNA_seed | ERBB2 | -0.773581 | BRCA | MDA-MB-468 | Afatinib |
| MVP_top60_DNA_seed | HDAC1 | -0.435708 | BRCA | CAL-148 | CAY10603 | MVP_top60_DNA+RNA_seed | EP300 | -0.773089 | BRCA | BT-474 | I-CBP112 |
| MVP_top60_DNA_seed | HSP90AA1 | -0.4352 | BRCA | HCC2218 | CCT-018159 | MVP_top60_DNA+RNA_seed | EGFR | -0.772035 | BRCA | UACC-812 | CI-1033 |
| MVP_top60_DNA_seed | HSP90AB1 | -0.4352 | BRCA | HCC2218 | CCT-018159 | MVP_top60_DNA+RNA_seed | ERBB2 | -0.772035 | BRCA | UACC-812 | CI-1033 |
| MVP_top60_DNA_seed | MAPK3 | -0.435194 | BRCA | OCUB-M | FR-180204 | MVP_top60_DNA+RNA_seed | ATM | -0.770453 | BRCA | AU565 | KU-60019 |
| MVP_top60_DNA_seed | MAPK1 | -0.435194 | BRCA | OCUB-M | FR-180204 | MVP_top60_DNA+RNA_seed | CDK2 | -0.769496 | BRCA | MRK-nu-1 | Seliciclib |
| MVP_top60_DNA_seed | EP300 | -0.432685 | BRCA | HCC2157 | I-CBP112 | MVP_top60_DNA+RNA_seed | PIK3CA | -0.767928 | BRCA | HCC202 | AZD8835 |
| MVP_top60_DNA_seed | CREBBP | -0.432685 | BRCA | HCC2157 | I-CBP112 | MVP_top60_DNA+RNA_seed | EGFR | -0.765775 | BRCA | BT-20 | Gefitinib |
| MVP_top60_DNA_seed | HDAC1 | -0.432422 | BRCA | AU565 | AR-42 | MVP_top60_DNA+RNA_seed | CDK2 | -0.765333 | BRCA | Hs-578-T | AZD5438 |
| MVP_top60_DNA_seed | EP300 | -0.432125 | BRCA | HCC1428 | I-CBP112 | MVP_top60_DNA+RNA_seed | ESR1 | -0.764831 | BRCA | MDA-MB-468 | Fulvestrant |
| MVP_top60_DNA_seed | CREBBP | -0.432125 | BRCA | HCC1428 | I-CBP112 | MVP_top60_DNA+RNA_seed | CDK1 | -0.763281 | BRCA | HCC1428 | RO-3306 |
| MVP_top60_DNA_seed | AKT1 | -0.427629 | BRCA | BT-20 | AKT inhibitor VIII | MVP_top60_DNA+RNA_seed | AR | -0.760143 | BRCA | MRK-nu-1 | Bicalutamide |
| MVP_top60_DNA_seed | ERBB2 | -0.426082 | BRCA | HCC70 | AZD8931 | MVP_top60_DNA+RNA_seed | HDAC1 | -0.756349 | BRCA | EVSA-T | Belinostat |
| MVP_top60_DNA_seed | EGFR | -0.426082 | BRCA | HCC70 | AZD8931 | MVP_top60_DNA+RNA_seed | AKT1 | -0.752134 | BRCA | HCC2218 | MK-2206 |
| MVP_top60_DNA_seed | ERBB3 | -0.426082 | BRCA | HCC70 | AZD8931 | MVP_top60_DNA+RNA_seed | HSP90AA1 | -0.7521 | BRCA | EVSA-T | CCT-018159 |
| MVP_top60_DNA_seed | AKT1 | -0.424946 | BRCA | HCC1954 | AKT inhibitor VIII | MVP_top60_DNA+RNA_seed | HSP90AB1 | -0.7521 | BRCA | EVSA-T | CCT-018159 |
| MVP_top60_DNA_seed | ERBB2 | -0.424329 | BRCA | HDQ-P1 | PF-00299804 | MVP_top60_DNA+RNA_seed | HDAC1 | -0.74829 | BRCA | HCC1428 | JQ12 |
| MVP_top60_DNA_seed | EGFR | -0.424329 | BRCA | HDQ-P1 | PF-00299804 | MVP_top60_DNA+RNA_seed | HDAC2 | -0.74829 | BRCA | HCC1428 | JQ12 |
| MVP_top60_DNA_seed | ATM | -0.42349 | BRCA | EFM-19 | KU-55933 | MVP_top60_DNA+RNA_seed | CDK2 | -0.747896 | BRCA | HCC2157 | PHA-793887 |
| MVP_top60_DNA_seed | ATM | -0.423416 | BRCA | DU-4475 | CP466722 | MVP_top60_DNA+RNA_seed | HSP90AA1 | -0.74642 | BRCA | COLO-824 | Tanespimycin |
| MVP_top60_DNA_seed | SRC | -0.422841 | BRCA | HCC1937 | Bosutinib | MVP_top60_DNA+RNA_seed | HSP90AB1 | -0.74642 | BRCA | COLO-824 | Tanespimycin |
| MVP_top60_DNA_seed | CDK1 | -0.417586 | BRCA | DU-4475 | RO-3306 | MVP_top60_DNA+RNA_seed | EGFR | -0.74533 | BRCA | EFM-19 | CUDC-101 |
| MVP_top60_DNA_seed | ERBB2 | -0.415889 | BRCA | HCC1143 | AZD8931 | MVP_top60_DNA+RNA_seed | ERBB2 | -0.74533 | BRCA | EFM-19 | CUDC-101 |
| MVP_top60_DNA_seed | EGFR | -0.415889 | BRCA | HCC1143 | AZD8931 | MVP_top60_DNA+RNA_seed | EGFR | -0.744696 | BRCA | HCC1599 | CI-1033 |
| MVP_top60_DNA_seed | ERBB3 | -0.415889 | BRCA | HCC1143 | AZD8931 | MVP_top60_DNA+RNA_seed | ERBB2 | -0.744696 | BRCA | HCC1599 | CI-1033 |
| MVP_top60_DNA_seed | SRC | -0.415427 | BRCA | HCC1599 | Saracatinib | MVP_top60_DNA+RNA_seed | AKT1 | -0.743904 | BRCA | CAL-148 | AKT inhibitor VIII |
| MVP_top60_DNA_seed | EGFR | -0.413221 | BRCA | HCC1187 | Pelitinib | MVP_top60_DNA+RNA_seed | ESR1 | -0.743047 | BRCA | COLO-824 | Fulvestrant |
| MVP_top60_DNA_seed | ERBB2 | -0.412765 | BRCA | HCC1395 | PF-00299804 | MVP_top60_DNA+RNA_seed | SRC | -0.741454 | BRCA | UACC-812 | Saracatinib |

| | | | | | | | | | | | |
|---|---|---|---|---|---|---|---|---|---|---|---|
| MVP_top60_DNA_seed | EGFR | -0.412765 | BRCA | HCC1395 | PF-00299804 | MVP_top60_DNA+RNA_seed | EGFR | -0.738983 | BRCA | BT-20 | AZD8931 |
| MVP_top60_DNA_seed | ATM | -0.405786 | BRCA | CAL-51 | CP466722 | MVP_top60_DNA+RNA_seed | ERBB2 | -0.738983 | BRCA | BT-20 | AZD8931 |
| MVP_top60_DNA_seed | EGFR | -0.405305 | BRCA | HCC1395 | AST-1306 | MVP_top60_DNA+RNA_seed | ERBB3 | -0.738983 | BRCA | BT-20 | AZD8931 |
| MVP_top60_DNA_seed | ERBB2 | -0.404112 | BRCA | CAMA-1 | CI-1033 | MVP_top60_DNA+RNA_seed | AKT1 | -0.738526 | BRCA | CAMA-1 | GSK690693 |
| MVP_top60_DNA_seed | EGFR | -0.404112 | BRCA | CAMA-1 | CI-1033 | MVP_top60_DNA+RNA_seed | AKT1 | -0.737176 | BRCA | MDA-MB-415 | AKT inhibitor VIII |
| MVP_top60_DNA_seed | HDAC1 | -0.401732 | BRCA | HCC1500 | AR-42 | MVP_top60_DNA+RNA_seed | EGFR | -0.734953 | BRCA | CAL-85-1 | AZD8931 |
| MVP_top60_DNA_seed | EGFR | -0.400758 | BRCA | BT-483 | AST-1306 | MVP_top60_DNA+RNA_seed | ERBB2 | -0.734953 | BRCA | CAL-85-1 | AZD8931 |
| MVP_top60_DNA_seed | MDM2 | -0.399239 | BRCA | JIMT-1 | Serdemetan | MVP_top60_DNA+RNA_seed | ERBB3 | -0.734953 | BRCA | CAL-85-1 | AZD8931 |
| MVP_top60_DNA_seed | HSP90AA1 | -0.398809 | BRCA | MFM-223 | Tanespimycin | MVP_top60_DNA+RNA_seed | PLK1 | -0.733683 | BRCA | HCC1599 | BI-2536 |
| MVP_top60_DNA_seed | HSP90AB1 | -0.398809 | BRCA | MFM-223 | Tanespimycin | MVP_top60_DNA+RNA_seed | EGFR | -0.729207 | BRCA | MDA-MB-361 | CUDC-101 |
| MVP_top60_DNA_seed | AKT1 | -0.398249 | BRCA | BT-474 | MK-2206 | MVP_top60_DNA+RNA_seed | ERBB2 | -0.729207 | BRCA | MDA-MB-361 | CUDC-101 |
| MVP_top60_DNA_seed | HSP90AA1 | -0.397831 | BRCA | MDA-MB-453 | CCT-018159 | MVP_top60_DNA+RNA_seed | HDAC1 | -0.727561 | BRCA | CAL-148 | Belinostat |
| MVP_top60_DNA_seed | HSP90AB1 | -0.397831 | BRCA | MDA-MB-453 | CCT-018159 | MVP_top60_DNA+RNA_seed | HDAC1 | -0.727364 | BRCA | T47D | Belinostat |
| MVP_top60_DNA_seed | HDAC1 | -0.396341 | BRCA | CAL-51 | JQ12 | MVP_top60_DNA+RNA_seed | AR | -0.726063 | BRCA | HCC1187 | Bicalutamide |
| MVP_top60_DNA_seed | HDAC2 | -0.396341 | BRCA | CAL-51 | JQ12 | MVP_top60_DNA+RNA_seed | AR | -0.724022 | BRCA | HCC1569 | Bicalutamide |
| MVP_top60_DNA_seed | HSP90AA1 | -0.391822 | BRCA | DU-4475 | CCT-018159 | MVP_top60_DNA+RNA_seed | EGFR | -0.723274 | BRCA | MRK-nu-1 | AST-1306 |
| MVP_top60_DNA_seed | HSP90AB1 | -0.391822 | BRCA | DU-4475 | CCT-018159 | MVP_top60_DNA+RNA_seed | EGFR | -0.723214 | BRCA | COLO-824 | AST-1306 |
| MVP_top60_DNA_seed | MDM2 | -0.389164 | BRCA | CAL-51 | Serdemetan | MVP_top60_DNA+RNA_seed | HSP90AA1 | -0.72126 | BRCA | BT-549 | Tanespimycin |
| MVP_top60_DNA_seed | EGFR | -0.38752 | BRCA | HCC1599 | AST-1306 | MVP_top60_DNA+RNA_seed | HSP90AB1 | -0.72126 | BRCA | BT-549 | Tanespimycin |
| MVP_top60_DNA_seed | HDAC1 | -0.386741 | BRCA | OCUB-M | Entinostat | MVP_top60_DNA+RNA_seed | MAPK14 | -0.718252 | BRCA | HCC1187 | TAK-715 |
| MVP_top60_DNA_seed | ERBB2 | -0.386319 | BRCA | MDA-MB-361 | AZD8931 | MVP_top60_DNA+RNA_seed | HDAC1 | -0.717399 | BRCA | HCC38 | Dacinostat |
| MVP_top60_DNA_seed | EGFR | -0.386319 | BRCA | MDA-MB-361 | AZD8931 | MVP_top60_DNA+RNA_seed | HDAC1 | -0.713041 | BRCA | OCUB-M | Belinostat |
| MVP_top60_DNA_seed | ERBB3 | -0.386319 | BRCA | MDA-MB-361 | AZD8931 | MVP_top60_DNA+RNA_seed | AR | -0.712505 | BRCA | HCC1187 | Bicalutamide |
| MVP_top60_DNA_seed | EP300 | -0.385811 | BRCA | COLO-824 | I-CBP112 | MVP_top60_DNA+RNA_seed | HDAC1 | -0.703683 | BRCA | HCC202 | Belinostat |
| MVP_top60_DNA_seed | CREBBP | -0.385811 | BRCA | COLO-824 | I-CBP112 | MVP_top60_DNA+RNA_seed | EGFR | -0.703436 | BRCA | ZR-75-30 | AST-1306 |
| MVP_top60_DNA_seed | ESR1 | -0.385659 | BRCA | DU-4475 | Fulvestrant | MVP_top60_DNA+RNA_seed | EGFR | -0.701954 | BRCA | HCC70 | Afatinib |
| MVP_top60_DNA_seed | CDK2 | -0.382457 | BRCA | BT-549 | PHA-793887 | MVP_top60_DNA+RNA_seed | ERBB2 | -0.701954 | BRCA | HCC70 | Afatinib |
| MVP_top60_DNA_seed | EGFR | -0.382117 | BRCA | CAL-148 | AST-1306 | MVP_top60_DNA+RNA_seed | EGFR | -0.700818 | BRCA | HCC202 | CUDC-101 |
| MVP_top60_DNA_seed | ERBB2 | -0.380465 | BRCA | HCC1428 | CUDC-101 | MVP_top60_DNA+RNA_seed | ERBB2 | -0.700818 | BRCA | HCC202 | CUDC-101 |
| MVP_top60_DNA_seed | EGFR | -0.380465 | BRCA | HCC1428 | CUDC-101 | MVP_top60_DNA+RNA_seed | AKT1 | -0.700671 | BRCA | CAL-85-1 | AKT inhibitor VIII |
| MVP_top60_DNA_seed | HDAC1 | -0.376742 | BRCA | HCC2157 | JQ12 | MVP_top60_DNA+RNA_seed | HDAC1 | -0.698216 | BRCA | HCC202 | AR-42 |
| MVP_top60_DNA_seed | HDAC2 | -0.376742 | BRCA | HCC2157 | JQ12 | MVP_top60_DNA+RNA_seed | PLK1 | -0.698038 | BRCA | EFM-19 | PLK_6522 |
| MVP_top60_DNA_seed | ESR1 | -0.374809 | BRCA | BT-20 | Fulvestrant | MVP_top60_DNA+RNA_seed | AR | -0.697133 | BRCA | HCC1599 | Bicalutamide |
| MVP_top60_DNA_seed | ATM | -0.372726 | BRCA | CAL-120 | KU-60019 | MVP_top60_DNA+RNA_seed | EP300 | -0.697074 | BRCA | OCUB-M | I-CBP112 |
| MVP_top60_DNA_seed | HDAC1 | -0.371402 | BRCA | HCC1954 | JQ12 | MVP_top60_DNA+RNA_seed | AKT1 | -0.696958 | BRCA | HCC202 | AKT inhibitor VIII |
| MVP_top60_DNA_seed | HDAC2 | -0.371402 | BRCA | HCC1954 | JQ12 | MVP_top60_DNA+RNA_seed | EGFR | -0.694728 | BRCA | MRK-nu-1 | CI-1033 |
| MVP_top60_DNA_seed | HDAC1 | -0.371322 | BRCA | MDA-MB-361 | Belinostat | MVP_top60_DNA+RNA_seed | ERBB2 | -0.694728 | BRCA | MRK-nu-1 | CI-1033 |
| MVP_top60_DNA_seed | HDAC1 | -0.370033 | BRCA | HCC1599 | JQ12 | MVP_top60_DNA+RNA_seed | EGFR | -0.693747 | BRCA | ZR-75-30 | PF-00299804 |
| MVP_top60_DNA_seed | HDAC2 | -0.370033 | BRCA | HCC1599 | JQ12 | MVP_top60_DNA+RNA_seed | ERBB2 | -0.693747 | BRCA | ZR-75-30 | PF-00299804 |
| MVP_top60_DNA_seed | HDAC1 | -0.369541 | BRCA | AU565 | JQ12 | MVP_top60_DNA+RNA_seed | AKT1 | -0.692199 | BRCA | HCC1187 | GSK690693 |
| MVP_top60_DNA_seed | HDAC2 | -0.369541 | BRCA | AU565 | JQ12 | MVP_top60_DNA+RNA_seed | HSP90AA1 | -0.692142 | BRCA | MDA-MB-436 | Tanespimycin |

| | | | | | | | | | | | |
|---|---|---|---|---|---|---|---|---|---|---|---|
| MVP_top60_DNA_seed | CDK1 | -0.36943 | BRCA | HCC1419 | RO-3306 | MVP_top60_DNA+RNA_seed | HSP90AB1 | -0.692142 | BRCA | MDA-MB-436 | Tanespimycin |
| MVP_top60_DNA_seed | ERBB2 | -0.368993 | BRCA | HCC202 | AZD8931 | MVP_top60_DNA+RNA_seed | PLK1 | -0.689839 | BRCA | HCC38 | PLK_6522 |
| MVP_top60_DNA_seed | EGFR | -0.368993 | BRCA | HCC202 | AZD8931 | MVP_top60_DNA+RNA_seed | EGFR | -0.688979 | BRCA | CAL-148 | CI-1033 |
| MVP_top60_DNA_seed | ERBB3 | -0.368993 | BRCA | HCC202 | AZD8931 | MVP_top60_DNA+RNA_seed | ERBB2 | -0.688979 | BRCA | CAL-148 | CI-1033 |
| MVP_top60_DNA_seed | ATM | -0.363855 | BRCA | MDA-MB-415 | KU-55933 | MVP_top60_DNA+RNA_seed | ESR1 | -0.688846 | BRCA | HCC1954 | Fulvestrant |
| MVP_top60_DNA_seed | MAPK1 | -0.362996 | BRCA | HCC70 | VX-11e | MVP_top60_DNA+RNA_seed | MAPK3 | -0.685212 | BRCA | CAL-85-1 | FR-180204 |
| MVP_top60_DNA_seed | AKT1 | -0.362291 | BRCA | MDA-MB-468 | MK-2206 | MVP_top60_DNA+RNA_seed | MAPK1 | -0.685212 | BRCA | CAL-85-1 | FR-180204 |
| MVP_top60_DNA_seed | AKT1 | -0.361687 | BRCA | DU-4475 | AKT inhibitor VIII | MVP_top60_DNA+RNA_seed | ATM | -0.684548 | BRCA | Hs-578-T | KU-60019 |
| MVP_top60_DNA_seed | SRC | -0.359665 | BRCA | MDA-MB-231 | NVP-BHG712 | MVP_top60_DNA+RNA_seed | ATM | -0.683833 | BRCA | HCC1569 | KU-55933 |
| MVP_top60_DNA_seed | HSP90AA1 | -0.352942 | BRCA | CAL-120 | SNX-2112 | MVP_top60_DNA+RNA_seed | AKT1 | -0.681203 | BRCA | HCC2218 | BAY AKT1 |
| MVP_top60_DNA_seed | HSP90AB1 | -0.352942 | BRCA | CAL-120 | SNX-2112 | MVP_top60_DNA+RNA_seed | HDAC1 | -0.679711 | BRCA | HCC2218 | Entinostat |
| MVP_top60_DNA_seed | HSP90AA1 | -0.35281 | BRCA | HCC1937 | Tanespimycin | MVP_top60_DNA+RNA_seed | HSP90AA1 | -0.67693 | BRCA | MDA-MB-361 | Elesclomol |
| MVP_top60_DNA_seed | HSP90AB1 | -0.35281 | BRCA | HCC1937 | Tanespimycin | MVP_top60_DNA+RNA_seed | HSP90AB1 | -0.67693 | BRCA | MDA-MB-361 | Elesclomol |
| MVP_top60_DNA_seed | AKT1 | -0.351138 | BRCA | COLO-824 | BAY AKT1 | MVP_top60_DNA+RNA_seed | EGFR | -0.673998 | BRCA | EFM-19 | PF-00299804 |
| MVP_top60_DNA_seed | ATM | -0.349512 | BRCA | AU565 | CP466722 | MVP_top60_DNA+RNA_seed | ERBB2 | -0.673998 | BRCA | EFM-19 | PF-00299804 |
| MVP_top60_DNA_seed | MDM2 | -0.349217 | BRCA | HCC70 | Nutlin-3a (-) | MVP_top60_DNA+RNA_seed | HDAC1 | -0.673952 | BRCA | MDA-MB-361 | AR-42 |
| MVP_top60_DNA_seed | CDK2 | -0.346458 | BRCA | CAL-148 | Seliciclib | MVP_top60_DNA+RNA_seed | EGFR | -0.672527 | BRCA | MDA-MB-361 | Gefitinib |
| MVP_top60_DNA_seed | AR | -0.344532 | BRCA | Hs-578-T | Bicalutamide | MVP_top60_DNA+RNA_seed | AR | -0.67137 | BRCA | MCF7 | AZD3514 |
| MVP_top60_DNA_seed | EGFR | -0.343659 | BRCA | HCC1569 | Pelitinib | MVP_top60_DNA+RNA_seed | SRC | -0.670324 | BRCA | OCUB-M | NVP-BHG712 |
| MVP_top60_DNA_seed | ERBB2 | -0.341543 | BRCA | HCC1937 | PF-00299804 | MVP_top60_DNA+RNA_seed | ESR1 | -0.669935 | BRCA | HCC1187 | Fulvestrant |
| MVP_top60_DNA_seed | EGFR | -0.341543 | BRCA | HCC1937 | PF-00299804 | MVP_top60_DNA+RNA_seed | ATM | -0.669773 | BRCA | MDA-MB-468 | CP466722 |
| MVP_top60_DNA_seed | HDAC1 | -0.339472 | BRCA | HCC1428 | Belinostat | MVP_top60_DNA+RNA_seed | EGFR | -0.669623 | BRCA | MDA-MB-157 | Gefitinib |
| MVP_top60_DNA_seed | HDAC1 | -0.336182 | BRCA | CAL-148 | Entinostat | MVP_top60_DNA+RNA_seed | EGFR | -0.666571 | BRCA | JIMT-1 | Cetuximab |
| MVP_top60_DNA_seed | EGFR | -0.331544 | BRCA | DU-4475 | AST-1306 | MVP_top60_DNA+RNA_seed | HSP90AA1 | -0.665852 | BRCA | JIMT-1 | Elesclomol |
| MVP_top60_DNA_seed | SRC | -0.330301 | BRCA | HCC1187 | NVP-BHG712 | MVP_top60_DNA+RNA_seed | HSP90AB1 | -0.665852 | BRCA | JIMT-1 | Elesclomol |
| MVP_top60_DNA_seed | HSP90AA1 | -0.327699 | BRCA | CAL-120 | Luminespib | MVP_top60_DNA+RNA_seed | HSP90AA1 | -0.665215 | BRCA | CAL-120 | Elesclomol |
| MVP_top60_DNA_seed | HSP90AB1 | -0.327699 | BRCA | CAL-120 | Luminespib | MVP_top60_DNA+RNA_seed | HSP90AB1 | -0.665215 | BRCA | CAL-120 | Elesclomol |
| MVP_top60_DNA_seed | HDAC1 | -0.327204 | BRCA | EFM-19 | CAY10603 | MVP_top60_DNA+RNA_seed | ESR1 | -0.661274 | BRCA | BT-483 | Fulvestrant |
| MVP_top60_DNA_seed | HSP90AA1 | -0.326974 | BRCA | MDA-MB-453 | Elesclomol | MVP_top60_DNA+RNA_seed | ESR1 | -0.660956 | BRCA | HCC1937 | Fulvestrant |
| MVP_top60_DNA_seed | HSP90AB1 | -0.326974 | BRCA | MDA-MB-453 | Elesclomol | MVP_top60_DNA+RNA_seed | ERBB2 | -0.659806 | BRCA | CAL-85-1 | CP724714 |
| MVP_top60_DNA_seed | ERBB2 | -0.326213 | BRCA | COLO-824 | AZD8931 | MVP_top60_DNA+RNA_seed | MDM2 | -0.657863 | BRCA | HCC1954 | Nutlin-3a (-) |
| MVP_top60_DNA_seed | EGFR | -0.326213 | BRCA | COLO-824 | AZD8931 | MVP_top60_DNA+RNA_seed | EGFR | -0.657423 | BRCA | BT-483 | CI-1033 |
| MVP_top60_DNA_seed | ERBB3 | -0.326213 | BRCA | COLO-824 | AZD8931 | MVP_top60_DNA+RNA_seed | ERBB2 | -0.657423 | BRCA | BT-483 | CI-1033 |
| MVP_top60_DNA_seed | HSP90AA1 | -0.32541 | BRCA | MRK-nu-1 | SNX-2112 | MVP_top60_DNA+RNA_seed | EP300 | -0.654789 | BRCA | CAMA-1 | I-CBP112 |
| MVP_top60_DNA_seed | HSP90AB1 | -0.32541 | BRCA | MRK-nu-1 | SNX-2112 | MVP_top60_DNA+RNA_seed | CDK2 | -0.654369 | BRCA | HCC2218 | Seliciclib |
| MVP_top60_DNA_seed | CDK2 | -0.323705 | BRCA | MDA-MB-453 | PHA-793887 | MVP_top60_DNA+RNA_seed | CDK2 | -0.654038 | BRCA | HCC1954 | PHA-793887 |
| MVP_top60_DNA_seed | MDM2 | -0.322278 | BRCA | MDA-MB-175-VII | Nutlin-3a (-) | MVP_top60_DNA+RNA_seed | PLK1 | -0.653011 | BRCA | HCC1143 | PLK_6522 |
| MVP_top60_DNA_seed | HDAC1 | -0.317886 | BRCA | CAL-120 | CAY10603 | MVP_top60_DNA+RNA_seed | AKT1 | -0.646971 | BRCA | HCC1937 | AKT inhibitor VIII |
| MVP_top60_DNA_seed | AKT1 | -0.317256 | BRCA | HCC1419 | GSK690693 | MVP_top60_DNA+RNA_seed | HDAC1 | -0.64648 | BRCA | ZR-75-30 | Dacinostat |
| MVP_top60_DNA_seed | ERBB2 | -0.316624 | BRCA | BT-549 | CI-1033 | MVP_top60_DNA+RNA_seed | HDAC1 | -0.646383 | BRCA | OCUB-M | AR-42 |
| MVP_top60_DNA_seed | EGFR | -0.316624 | BRCA | BT-549 | CI-1033 | MVP_top60_DNA+RNA_seed | HDAC1 | -0.64417 | BRCA | T47D | Dacinostat |

| | | | | | | | | | | | |
|---|---|---|---|---|---|---|---|---|---|---|---|
| MVP_top60_DNA_seed | HDAC1 | -0.316158 | BRCA | CAL-148 | Dacinostat | MVP_top60_DNA+RNA_seed | ESR1 | -0.644129 | BRCA | EFM-192A | Fulvestrant |
| MVP_top60_DNA_seed | ESR1 | -0.315186 | BRCA | EFM-192A | Tamoxifen | MVP_top60_DNA+RNA_seed | AR | -0.642307 | BRCA | MDA-MB-415 | Bicalutamide |
| MVP_top60_DNA_seed | ERBB2 | -0.315164 | BRCA | HCC38 | CUDC-101 | MVP_top60_DNA+RNA_seed | EGFR | -0.641862 | BRCA | CAL-120 | Pelitinib |
| MVP_top60_DNA_seed | EGFR | -0.315164 | BRCA | HCC38 | CUDC-101 | MVP_top60_DNA+RNA_seed | EGFR | -0.641509 | BRCA | AU565 | CUDC-101 |
| MVP_top60_DNA_seed | ERBB2 | -0.312949 | BRCA | MDA-MB-453 | AZD8931 | MVP_top60_DNA+RNA_seed | ERBB2 | -0.641509 | BRCA | AU565 | CUDC-101 |
| MVP_top60_DNA_seed | EGFR | -0.312949 | BRCA | MDA-MB-453 | AZD8931 | MVP_top60_DNA+RNA_seed | AR | -0.639064 | BRCA | UACC-893 | Bicalutamide |
| MVP_top60_DNA_seed | ERBB3 | -0.312949 | BRCA | MDA-MB-453 | AZD8931 | MVP_top60_DNA+RNA_seed | HDAC1 | -0.638339 | BRCA | ZR-75-30 | Belinostat |
| MVP_top60_DNA_seed | HSP90AA1 | -0.309917 | BRCA | HCC1143 | Tanespimycin | MVP_top60_DNA+RNA_seed | PLK1 | -0.637338 | BRCA | MRK-nu-1 | PLK_6522 |
| MVP_top60_DNA_seed | HSP90AB1 | -0.309917 | BRCA | HCC1143 | Tanespimycin | MVP_top60_DNA+RNA_seed | AKT1 | -0.637138 | BRCA | CAMA-1 | BAY AKT1 |
| MVP_top60_DNA_seed | HSP90AA1 | -0.309789 | BRCA | BT-549 | CCT-018159 | MVP_top60_DNA+RNA_seed | EGFR | -0.636479 | BRCA | HCC38 | PF-00299804 |
| MVP_top60_DNA_seed | HSP90AB1 | -0.309789 | BRCA | BT-549 | CCT-018159 | MVP_top60_DNA+RNA_seed | ERBB2 | -0.636479 | BRCA | HCC38 | PF-00299804 |
| MVP_top60_DNA_seed | MAPK1 | -0.309321 | BRCA | OCUB-M | VX-11e | MVP_top60_DNA+RNA_seed | HSP90AA1 | -0.636017 | BRCA | ZR-75-30 | Elesclomol |
| MVP_top60_DNA_seed | AKT1 | -0.309279 | BRCA | CAMA-1 | AKT inhibitor VIII | MVP_top60_DNA+RNA_seed | HSP90AB1 | -0.636017 | BRCA | ZR-75-30 | Elesclomol |
| MVP_top60_DNA_seed | HSP90AA1 | -0.309029 | BRCA | MDA-MB-330 | Luminespib | MVP_top60_DNA+RNA_seed | AKT1 | -0.635331 | BRCA | HCC1569 | BAY AKT1 |
| MVP_top60_DNA_seed | HSP90AB1 | -0.309029 | BRCA | MDA-MB-330 | Luminespib | MVP_top60_DNA+RNA_seed | HDAC1 | -0.635159 | BRCA | MRK-nu-1 | Tubastatin A |
| MVP_top60_DNA_seed | ERBB2 | -0.307356 | BRCA | T47D | Afatinib | MVP_top60_DNA+RNA_seed | ATM | -0.633036 | BRCA | HCC202 | CP466722 |
| MVP_top60_DNA_seed | EGFR | -0.307356 | BRCA | T47D | Afatinib | MVP_top60_DNA+RNA_seed | HDAC1 | -0.632153 | BRCA | T47D | AR-42 |
| MVP_top60_DNA_seed | MDM2 | -0.30579 | BRCA | HCC38 | Serdemetan | MVP_top60_DNA+RNA_seed | HSP90AA1 | -0.630822 | BRCA | HCC38 | SNX-2112 |
| MVP_top60_DNA_seed | EGFR | -0.304055 | BRCA | HDQ-P1 | AST-1306 | MVP_top60_DNA+RNA_seed | HSP90AB1 | -0.630822 | BRCA | HCC38 | SNX-2112 |
| MVP_top60_DNA_seed | HDAC1 | -0.303572 | BRCA | HCC70 | Tubastatin A | MVP_top60_DNA+RNA_seed | ERBB2 | -0.629646 | BRCA | HCC1187 | CP724714 |
| MVP_top60_DNA_seed | HDAC1 | -0.302652 | BRCA | HCC1500 | Dacinostat | MVP_top60_DNA+RNA_seed | EGFR | -0.629127 | BRCA | HCC1569 | Afatinib |
| MVP_top60_DNA_seed | HDAC1 | -0.30263 | BRCA | EVSA-T | Entinostat | MVP_top60_DNA+RNA_seed | ERBB2 | -0.629127 | BRCA | HCC1569 | Afatinib |
| MVP_top60_DNA_seed | HSP90AA1 | -0.301063 | BRCA | HCC38 | Tanespimycin | MVP_top60_DNA+RNA_seed | ATM | -0.628853 | BRCA | COLO-824 | KU-60019 |
| MVP_top60_DNA_seed | HSP90AB1 | -0.301063 | BRCA | HCC38 | Tanespimycin | MVP_top60_DNA+RNA_seed | AKT1 | -0.62829 | BRCA | HCC1187 | AKT inhibitor VIII |
| MVP_top60_DNA_seed | AKT1 | -0.30011 | BRCA | MDA-MB-468 | AKT inhibitor VIII | MVP_top60_DNA+RNA_seed | HSP90AA1 | -0.628036 | BRCA | T47D | Elesclomol |
| MVP_top60_DNA_seed | EGFR | -0.299922 | BRCA | HCC70 | AST-1306 | MVP_top60_DNA+RNA_seed | HSP90AB1 | -0.628036 | BRCA | T47D | Elesclomol |
| MVP_top60_DNA_seed | PIK3CA | -0.297693 | BRCA | MDA-MB-175-VII | AZD8835 | MVP_top60_DNA+RNA_seed | AKT1 | -0.627952 | BRCA | JIMT-1 | BAY AKT1 |
| MVP_top60_DNA_seed | MDM2 | -0.296172 | BRCA | HCC1569 | Serdemetan | MVP_top60_DNA+RNA_seed | ATM | -0.626746 | BRCA | MFM-223 | torin2 |
| MVP_top60_DNA_seed | ERBB2 | -0.295236 | BRCA | T47D | CP724714 | MVP_top60_DNA+RNA_seed | EGFR | -0.624557 | BRCA | EVSA-T | AZD8931 |
| MVP_top60_DNA_seed | HDAC1 | -0.292521 | BRCA | CAMA-1 | Dacinostat | MVP_top60_DNA+RNA_seed | ERBB2 | -0.624557 | BRCA | EVSA-T | AZD8931 |
| MVP_top60_DNA_seed | HDAC1 | -0.291465 | BRCA | ZR-75-30 | CAY10603 | MVP_top60_DNA+RNA_seed | ERBB3 | -0.624557 | BRCA | EVSA-T | AZD8931 |
| MVP_top60_DNA_seed | ATM | -0.289568 | BRCA | HCC1428 | KU-60019 | MVP_top60_DNA+RNA_seed | ESR1 | -0.624056 | BRCA | JIMT-1 | Fulvestrant |
| MVP_top60_DNA_seed | HSP90AA1 | -0.288799 | BRCA | BT-474 | CCT-018159 | MVP_top60_DNA+RNA_seed | AKT1 | -0.622998 | BRCA | MDA-MB-330 | BAY AKT1 |
| MVP_top60_DNA_seed | HSP90AB1 | -0.288799 | BRCA | BT-474 | CCT-018159 | MVP_top60_DNA+RNA_seed | EGFR | -0.622722 | BRCA | HCC2157 | AZD8931 |
| MVP_top60_DNA_seed | ERBB2 | -0.288053 | BRCA | DU-4475 | CUDC-101 | MVP_top60_DNA+RNA_seed | ERBB2 | -0.622722 | BRCA | HCC2157 | AZD8931 |
| MVP_top60_DNA_seed | EGFR | -0.288053 | BRCA | DU-4475 | CUDC-101 | MVP_top60_DNA+RNA_seed | ERBB3 | -0.622722 | BRCA | HCC2157 | AZD8931 |
| MVP_top60_DNA_seed | HSP90AA1 | -0.287209 | BRCA | MDA-MB-453 | Tanespimycin | MVP_top60_DNA+RNA_seed | AR | -0.622551 | BRCA | EVSA-T | Bicalutamide |
| MVP_top60_DNA_seed | HSP90AB1 | -0.287209 | BRCA | MDA-MB-453 | Tanespimycin | MVP_top60_DNA+RNA_seed | HDAC1 | -0.622258 | BRCA | EVSA-T | CAY10603 |
| MVP_top60_DNA_seed | EGFR | -0.287018 | BRCA | HCC2157 | Gefitinib | MVP_top60_DNA+RNA_seed | AR | -0.621906 | BRCA | MDA-MB-468 | Bicalutamide |
| MVP_top60_DNA_seed | HDAC1 | -0.286241 | BRCA | T47D | CAY10603 | MVP_top60_DNA+RNA_seed | AR | -0.618247 | BRCA | MDA-MB-468 | AZD3514 |
| MVP_top60_DNA_seed | AR | -0.285949 | BRCA | COLO-824 | Bicalutamide | MVP_top60_DNA+RNA_seed | AKT1 | -0.614502 | BRCA | BT-549 | AKT inhibitor VIII |

| | | | | | | | | | | | |
|---|---|---|---|---|---|---|---|---|---|---|---|
| MVP_top60_DNA_seed | AR | -0.285626 | BRCA | HCC2218 | AZD3514 | MVP_top60_DNA+RNA_seed | EGFR | -0.613571 | BRCA | MDA-MB-330 | AZD8931 |
| MVP_top60_DNA_seed | AR | -0.284682 | BRCA | DU-4475 | Bicalutamide | MVP_top60_DNA+RNA_seed | ERBB2 | -0.613571 | BRCA | MDA-MB-330 | AZD8931 |
| MVP_top60_DNA_seed | CDK2 | -0.283028 | BRCA | T47D | PHA-793887 | MVP_top60_DNA+RNA_seed | ERBB3 | -0.613571 | BRCA | MDA-MB-330 | AZD8931 |
| MVP_top60_DNA_seed | EP300 | -0.282905 | BRCA | DU-4475 | I-CBP112 | MVP_top60_DNA+RNA_seed | AKT1 | -0.611586 | BRCA | DU-4475 | AKT inhibitor VIII |
| MVP_top60_DNA_seed | CREBBP | -0.282905 | BRCA | DU-4475 | I-CBP112 | MVP_top60_DNA+RNA_seed | EGFR | -0.607462 | BRCA | HCC1419 | AZD8931 |
| MVP_top60_DNA_seed | HDAC1 | -0.281786 | BRCA | HCC202 | CAY10603 | MVP_top60_DNA+RNA_seed | ERBB2 | -0.607462 | BRCA | HCC1419 | AZD8931 |
| MVP_top60_DNA_seed | MDM2 | -0.279105 | BRCA | MRK-nu-1 | Nutlin-3a (-) | MVP_top60_DNA+RNA_seed | ERBB3 | -0.607462 | BRCA | HCC1419 | AZD8931 |
| MVP_top60_DNA_seed | HDAC1 | -0.274384 | BRCA | HCC1569 | Dacinostat | MVP_top60_DNA+RNA_seed | SRC | -0.60714 | BRCA | HCC1599 | NVP-BHG712 |
| MVP_top60_DNA_seed | HSP90AA1 | -0.272654 | BRCA | ZR-75-30 | Luminespib | MVP_top60_DNA+RNA_seed | EGFR | -0.605689 | BRCA | JIMT-1 | Afatinib |
| MVP_top60_DNA_seed | HSP90AB1 | -0.272654 | BRCA | ZR-75-30 | Luminespib | MVP_top60_DNA+RNA_seed | ERBB2 | -0.605689 | BRCA | JIMT-1 | Afatinib |
| MVP_top60_DNA_seed | ERBB2 | -0.27155 | BRCA | MDA-MB-157 | Afatinib | MVP_top60_DNA+RNA_seed | HDAC1 | -0.60563 | BRCA | HCC38 | Tubastatin A |
| MVP_top60_DNA_seed | EGFR | -0.27155 | BRCA | MDA-MB-157 | Afatinib | MVP_top60_DNA+RNA_seed | ESR1 | -0.605107 | BRCA | HCC1569 | Fulvestrant |
| MVP_top60_DNA_seed | AR | -0.270686 | BRCA | MDA-MB-453 | AZD3514 | MVP_top60_DNA+RNA_seed | HDAC1 | -0.597841 | BRCA | DU-4475 | AR-42 |
| MVP_top60_DNA_seed | SRC | -0.270308 | BRCA | HDQ-P1 | Bosutinib | MVP_top60_DNA+RNA_seed | MAPK3 | -0.59783 | BRCA | MDA-MB-175-VII | FR-180204 |
| MVP_top60_DNA_seed | MDM2 | -0.269769 | BRCA | MDA-MB-157 | Nutlin-3a (-) | MVP_top60_DNA+RNA_seed | MAPK1 | -0.59783 | BRCA | MDA-MB-175-VII | FR-180204 |
| MVP_top60_DNA_seed | AR | -0.268955 | BRCA | CAL-85-1 | AZD3514 | MVP_top60_DNA+RNA_seed | AR | -0.597128 | BRCA | T47D | Bicalutamide |
| MVP_top60_DNA_seed | AKT1 | -0.267296 | BRCA | CAL-51 | BAY AKT1 | MVP_top60_DNA+RNA_seed | EGFR | -0.595857 | BRCA | MDA-MB-361 | AST-1306 |
| MVP_top60_DNA_seed | AKT1 | -0.265996 | BRCA | CAL-51 | GSK690693 | MVP_top60_DNA+RNA_seed | AKT1 | -0.593677 | BRCA | MDA-MB-175-VII | AKT inhibitor VIII |
| MVP_top60_DNA_seed | ERBB2 | -0.264912 | BRCA | COLO-824 | Afatinib | MVP_top60_DNA+RNA_seed | ATM | -0.593621 | BRCA | EFM-19 | CP466722 |
| MVP_top60_DNA_seed | EGFR | -0.264912 | BRCA | COLO-824 | Afatinib | MVP_top60_DNA+RNA_seed | HSP90AA1 | -0.593513 | BRCA | MDA-MB-468 | SNX-2112 |
| MVP_top60_DNA_seed | AKT1 | -0.263242 | BRCA | CAL-85-1 | GSK690693 | MVP_top60_DNA+RNA_seed | HSP90AB1 | -0.593513 | BRCA | MDA-MB-468 | SNX-2112 |
| MVP_top60_DNA_seed | ERBB2 | -0.262769 | BRCA | HCC1569 | CP724714 | MVP_top60_DNA+RNA_seed | EGFR | -0.592496 | BRCA | EVSA-T | Lapatinib |
| MVP_top60_DNA_seed | SRC | -0.262509 | BRCA | CAL-120 | NVP-BHG712 | MVP_top60_DNA+RNA_seed | ERBB2 | -0.592496 | BRCA | EVSA-T | Lapatinib |
| MVP_top60_DNA_seed | ESR1 | -0.262397 | BRCA | CAL-51 | Fulvestrant | MVP_top60_DNA+RNA_seed | ERBB2 | -0.591098 | BRCA | COLO-824 | CP724714 |
| MVP_top60_DNA_seed | AKT1 | -0.253143 | BRCA | CAL-85-1 | AKT inhibitor VIII | MVP_top60_DNA+RNA_seed | HDAC1 | -0.589299 | BRCA | CAL-148 | AR-42 |
| MVP_top60_DNA_seed | ERBB2 | -0.252589 | BRCA | HCC1500 | CP724714 | MVP_top60_DNA+RNA_seed | EGFR | -0.588791 | BRCA | JIMT-1 | AZD8931 |
| MVP_top60_DNA_seed | ERBB2 | -0.252435 | BRCA | HCC1187 | Afatinib | MVP_top60_DNA+RNA_seed | ERBB2 | -0.588791 | BRCA | JIMT-1 | AZD8931 |
| MVP_top60_DNA_seed | EGFR | -0.252435 | BRCA | HCC1187 | Afatinib | MVP_top60_DNA+RNA_seed | ERBB3 | -0.588791 | BRCA | JIMT-1 | AZD8931 |
| MVP_top60_DNA_seed | ATM | -0.251995 | BRCA | BT-20 | CP466722 | MVP_top60_DNA+RNA_seed | HSP90AA1 | -0.588493 | BRCA | MFM-223 | Elesclomol |
| MVP_top60_DNA_seed | SRC | -0.251688 | BRCA | MDA-MB-361 | Bosutinib | MVP_top60_DNA+RNA_seed | HSP90AB1 | -0.588493 | BRCA | MFM-223 | Elesclomol |
| MVP_top60_DNA_seed | ATM | -0.251051 | BRCA | EFM-192A | CP466722 | MVP_top60_DNA+RNA_seed | AKT1 | -0.587493 | BRCA | HCC1937 | GSK690693 |
| MVP_top60_DNA_seed | ERBB2 | -0.250836 | BRCA | MDA-MB-361 | Afatinib | MVP_top60_DNA+RNA_seed | AKT1 | -0.586046 | BRCA | MCF7 | MK-2206 |
| MVP_top60_DNA_seed | EGFR | -0.250836 | BRCA | MDA-MB-361 | Afatinib | MVP_top60_DNA+RNA_seed | EGFR | -0.584069 | BRCA | JIMT-1 | Gefitinib |
| MVP_top60_DNA_seed | AR | -0.250694 | BRCA | MDA-MB-436 | AZD3514 | MVP_top60_DNA+RNA_seed | HSP90AA1 | -0.583879 | BRCA | MDA-MB-361 | SNX-2112 |
| MVP_top60_DNA_seed | MAPK1 | -0.250553 | BRCA | MDA-MB-175-VII | VX-11e | MVP_top60_DNA+RNA_seed | HSP90AB1 | -0.583879 | BRCA | MDA-MB-361 | SNX-2112 |
| MVP_top60_DNA_seed | SRC | -0.24862 | BRCA | MDA-MB-468 | Bosutinib | MVP_top60_DNA+RNA_seed | EGFR | -0.582703 | BRCA | MDA-MB-453 | Afatinib |
| MVP_top60_DNA_seed | MAPK3 | -0.24839 | BRCA | MDA-MB-361 | FR-180204 | MVP_top60_DNA+RNA_seed | ERBB2 | -0.582703 | BRCA | MDA-MB-453 | Afatinib |
| MVP_top60_DNA_seed | MAPK1 | -0.24839 | BRCA | MDA-MB-361 | FR-180204 | MVP_top60_DNA+RNA_seed | AR | -0.57956 | BRCA | HCC38 | Bicalutamide |
| MVP_top60_DNA_seed | HSP90AA1 | -0.247247 | BRCA | HCC1937 | CCT-018159 | MVP_top60_DNA+RNA_seed | PLK1 | -0.578569 | BRCA | JIMT-1 | PLK_6522 |
| MVP_top60_DNA_seed | HSP90AB1 | -0.247247 | BRCA | HCC1937 | CCT-018159 | MVP_top60_DNA+RNA_seed | PLK1 | -0.576057 | BRCA | HCC1395 | PLK_6522 |
| MVP_top60_DNA_seed | AKT1 | -0.247109 | BRCA | HCC1187 | MK-2206 | MVP_top60_DNA+RNA_seed | ATM | -0.575105 | BRCA | HCC1187 | CP466722 |

| | | | | | | | | | | | |
|---|---|---|---|---|---|---|---|---|---|---|---|
| MVP_top60_DNA_seed | HDAC1 | -0.246923 | BRCA | EFM-19 | AR-42 | MVP_top60_DNA+RNA_seed | PIK3CA | -0.574712 | BRCA | HCC1187 | AZD8835 |
| MVP_top60_DNA_seed | ERBB2 | -0.244686 | BRCA | EFM-19 | CI-1033 | MVP_top60_DNA+RNA_seed | EGFR | -0.573468 | BRCA | HCC1500 | PF-00299804 |
| MVP_top60_DNA_seed | EGFR | -0.244686 | BRCA | EFM-19 | CI-1033 | MVP_top60_DNA+RNA_seed | ERBB2 | -0.573468 | BRCA | HCC1500 | PF-00299804 |
| MVP_top60_DNA_seed | HDAC1 | -0.243026 | BRCA | HCC2157 | Entinostat | MVP_top60_DNA+RNA_seed | PIK3CA | -0.573288 | BRCA | T47D | AZD8835 |
| MVP_top60_DNA_seed | EGFR | -0.242339 | BRCA | HCC70 | Cetuximab | MVP_top60_DNA+RNA_seed | EGFR | -0.571153 | BRCA | Hs-578-T | Cetuximab |
| MVP_top60_DNA_seed | PIK3CA | -0.24184 | BRCA | AU565 | AZD8835 | MVP_top60_DNA+RNA_seed | HSP90AA1 | -0.569932 | BRCA | HCC1419 | Tanespimycin |
| MVP_top60_DNA_seed | AKT1 | -0.240728 | BRCA | HCC38 | AKT inhibitor VIII | MVP_top60_DNA+RNA_seed | HSP90AB1 | -0.569932 | BRCA | HCC1419 | Tanespimycin |
| MVP_top60_DNA_seed | ERBB2 | -0.240265 | BRCA | BT-20 | Afatinib | MVP_top60_DNA+RNA_seed | HSP90AA1 | -0.569817 | BRCA | MCF7 | Elesclomol |
| MVP_top60_DNA_seed | EGFR | -0.240265 | BRCA | BT-20 | Afatinib | MVP_top60_DNA+RNA_seed | HSP90AB1 | -0.569817 | BRCA | MCF7 | Elesclomol |
| MVP_top60_DNA_seed | HDAC1 | -0.239537 | BRCA | MDA-MB-361 | CAY10603 | MVP_top60_DNA+RNA_seed | PIK3CA | -0.568687 | BRCA | MFM-223 | PI3Ka_4409 |
| MVP_top60_DNA_seed | ERBB2 | -0.238543 | BRCA | HCC1428 | PF-00299804 | MVP_top60_DNA+RNA_seed | AKT1 | -0.56628 | BRCA | EFM-192A | BAY AKT1 |
| MVP_top60_DNA_seed | EGFR | -0.238543 | BRCA | HCC1428 | PF-00299804 | MVP_top60_DNA+RNA_seed | PIK3CA | -0.565849 | BRCA | HCC1187 | PI3Ka_4409 |
| MVP_top60_DNA_seed | HSP90AA1 | -0.237874 | BRCA | BT-20 | CCT-018159 | MVP_top60_DNA+RNA_seed | EGFR | -0.565733 | BRCA | OCUB-M | Cetuximab |
| MVP_top60_DNA_seed | HSP90AB1 | -0.237874 | BRCA | BT-20 | CCT-018159 | MVP_top60_DNA+RNA_seed | ATM | -0.565203 | BRCA | AU565 | KU-55933 |
| MVP_top60_DNA_seed | ATM | -0.2371 | BRCA | HCC38 | CP466722 | MVP_top60_DNA+RNA_seed | EGFR | -0.564873 | BRCA | ZR-75-30 | AZD8931 |
| MVP_top60_DNA_seed | AKT1 | -0.234712 | BRCA | HCC2218 | AKT inhibitor VIII | MVP_top60_DNA+RNA_seed | ERBB2 | -0.564873 | BRCA | ZR-75-30 | AZD8931 |
| MVP_top60_DNA_seed | PIK3CA | -0.234688 | BRCA | MDA-MB-175-VII | PI3Ka_4409 | MVP_top60_DNA+RNA_seed | ERBB3 | -0.564873 | BRCA | ZR-75-30 | AZD8931 |
| MVP_top60_DNA_seed | EP300 | -0.233121 | BRCA | BT-483 | I-CBP112 | MVP_top60_DNA+RNA_seed | HDAC1 | -0.564583 | BRCA | EFM-192A | Dacinostat |
| MVP_top60_DNA_seed | CREBBP | -0.233121 | BRCA | BT-483 | I-CBP112 | MVP_top60_DNA+RNA_seed | HDAC1 | -0.564211 | BRCA | EFM-19 | Dacinostat |
| MVP_top60_DNA_seed | HDAC1 | -0.232503 | BRCA | HCC1500 | Belinostat | MVP_top60_DNA+RNA_seed | EGFR | -0.562484 | BRCA | MDA-MB-415 | Cetuximab |
| MVP_top60_DNA_seed | HDAC1 | -0.231314 | BRCA | BT-474 | CAY10603 | MVP_top60_DNA+RNA_seed | ATM | -0.562424 | BRCA | HCC70 | CP466722 |
| MVP_top60_DNA_seed | EGFR | -0.22943 | BRCA | BT-549 | AST-1306 | MVP_top60_DNA+RNA_seed | EGFR | -0.56204 | BRCA | MRK-nu-1 | PF-00299804 |
| MVP_top60_DNA_seed | MDM2 | -0.228918 | BRCA | T47D | Serdemetan | MVP_top60_DNA+RNA_seed | ERBB2 | -0.56204 | BRCA | MRK-nu-1 | PF-00299804 |
| MVP_top60_DNA_seed | HDAC1 | -0.227297 | BRCA | HCC202 | Tubastatin A | MVP_top60_DNA+RNA_seed | EP300 | -0.560005 | BRCA | MDA-MB-231 | I-CBP112 |
| MVP_top60_DNA_seed | ERBB2 | -0.224574 | BRCA | HCC1500 | CI-1033 | MVP_top60_DNA+RNA_seed | HSP90AA1 | -0.558557 | BRCA | HCC1569 | CCT-018159 |
| MVP_top60_DNA_seed | EGFR | -0.224574 | BRCA | HCC1500 | CI-1033 | MVP_top60_DNA+RNA_seed | HSP90AB1 | -0.558557 | BRCA | HCC1569 | CCT-018159 |
| MVP_top60_DNA_seed | ATM | -0.220896 | BRCA | JIMT-1 | CP466722 | MVP_top60_DNA+RNA_seed | EGFR | -0.556597 | BRCA | HCC1569 | CI-1033 |
| MVP_top60_DNA_seed | HDAC1 | -0.216818 | BRCA | MDA-MB-330 | Belinostat | MVP_top60_DNA+RNA_seed | ERBB2 | -0.556597 | BRCA | HCC1569 | CI-1033 |
| MVP_top60_DNA_seed | AKT1 | -0.216371 | BRCA | COLO-824 | AKT inhibitor VIII | MVP_top60_DNA+RNA_seed | AKT1 | -0.55622 | BRCA | EVSA-T | A-443654 |
| MVP_top60_DNA_seed | AKT1 | -0.214062 | BRCA | HCC1395 | AKT inhibitor VIII | MVP_top60_DNA+RNA_seed | HSP90AA1 | -0.554147 | BRCA | BT-474 | Elesclomol |
| MVP_top60_DNA_seed | MDM2 | -0.212422 | BRCA | HCC1937 | Serdemetan | MVP_top60_DNA+RNA_seed | HSP90AB1 | -0.554147 | BRCA | BT-474 | Elesclomol |
| MVP_top60_DNA_seed | AKT1 | -0.211224 | BRCA | MDA-MB-453 | MK-2206 | MVP_top60_DNA+RNA_seed | HSP90AA1 | -0.550969 | BRCA | OCUB-M | Luminespib |
| MVP_top60_DNA_seed | AKT1 | -0.209807 | BRCA | EFM-192A | AKT inhibitor VIII | MVP_top60_DNA+RNA_seed | HSP90AB1 | -0.550969 | BRCA | OCUB-M | Luminespib |
| MVP_top60_DNA_seed | ERBB2 | -0.208009 | BRCA | BT-549 | PF-00299804 | MVP_top60_DNA+RNA_seed | AKT1 | -0.550963 | BRCA | MDA-MB-175-VII | BAY AKT1 |
| MVP_top60_DNA_seed | EGFR | -0.208009 | BRCA | BT-549 | PF-00299804 | MVP_top60_DNA+RNA_seed | HDAC1 | -0.549887 | BRCA | HCC70 | CAY10603 |
| MVP_top60_DNA_seed | CDK1 | -0.207887 | BRCA | HCC70 | RO-3306 | MVP_top60_DNA+RNA_seed | AR | -0.54888 | BRCA | CAL-148 | Bicalutamide |
| MVP_top60_DNA_seed | ESR1 | -0.206923 | BRCA | HCC1599 | Fulvestrant | MVP_top60_DNA+RNA_seed | EGFR | -0.546664 | BRCA | HCC2157 | AST-1306 |
| MVP_top60_DNA_seed | MDM2 | -0.20646 | BRCA | CAL-120 | Serdemetan | MVP_top60_DNA+RNA_seed | MDM2 | -0.546559 | BRCA | HCC2218 | Nutlin-3a (-) |
| MVP_top60_DNA_seed | HSP90AA1 | -0.205534 | BRCA | BT-474 | Luminespib | MVP_top60_DNA+RNA_seed | EGFR | -0.542227 | BRCA | HCC2157 | CI-1033 |
| MVP_top60_DNA_seed | HSP90AB1 | -0.205534 | BRCA | BT-474 | Luminespib | MVP_top60_DNA+RNA_seed | ERBB2 | -0.542227 | BRCA | HCC2157 | CI-1033 |
| MVP_top60_DNA_seed | ESR1 | -0.205499 | BRCA | HCC70 | Fulvestrant | MVP_top60_DNA+RNA_seed | EGFR | -0.540024 | BRCA | HCC1954 | PF-00299804 |

| | | | | | | | | | | | |
|---|---|---|---|---|---|---|---|---|---|---|---|
| MVP_top60_DNA_seed | CDK2 | -0.205035 | BRCA | OCUB-M | Seliciclib | MVP_top60_DNA+RNA_seed | ERBB2 | -0.540024 | BRCA | HCC1954 | PF-00299804 |
| MVP_top60_DNA_seed | HSP90AA1 | -0.204579 | BRCA | CAL-148 | Luminespib | MVP_top60_DNA+RNA_seed | EGFR | -0.539556 | BRCA | HDQ-P1 | Cetuximab |
| MVP_top60_DNA_seed | HSP90AB1 | -0.204579 | BRCA | CAL-148 | Luminespib | MVP_top60_DNA+RNA_seed | ATM | -0.537684 | BRCA | HCC1187 | KU-55933 |
| MVP_top60_DNA_seed | ERBB2 | -0.204165 | BRCA | HCC70 | CUDC-101 | MVP_top60_DNA+RNA_seed | AKT1 | -0.536812 | BRCA | HCC1395 | MK-2206 |
| MVP_top60_DNA_seed | EGFR | -0.204165 | BRCA | HCC70 | CUDC-101 | MVP_top60_DNA+RNA_seed | HDAC1 | -0.536721 | BRCA | ZR-75-30 | JQ12 |
| MVP_top60_DNA_seed | CDK2 | -0.203436 | BRCA | MDA-MB-231 | AZD5438 | MVP_top60_DNA+RNA_seed | HDAC2 | -0.536721 | BRCA | ZR-75-30 | JQ12 |
| MVP_top60_DNA_seed | HDAC1 | -0.202709 | BRCA | MDA-MB-330 | Tubastatin A | MVP_top60_DNA+RNA_seed | AR | -0.535914 | BRCA | EVSA-T | AZD3514 |
| MVP_top60_DNA_seed | SRC | -0.202072 | BRCA | MRK-nu-1 | NVP-BHG712 | MVP_top60_DNA+RNA_seed | EGFR | -0.532808 | BRCA | HCC2157 | PF-00299804 |
| MVP_top60_DNA_seed | HDAC1 | -0.201216 | BRCA | JIMT-1 | JQ12 | MVP_top60_DNA+RNA_seed | ERBB2 | -0.532808 | BRCA | HCC2157 | PF-00299804 |
| MVP_top60_DNA_seed | HDAC2 | -0.201216 | BRCA | JIMT-1 | JQ12 | MVP_top60_DNA+RNA_seed | EGFR | -0.531315 | BRCA | MDA-MB-175-VII | Cetuximab |
| MVP_top60_DNA_seed | EGFR | -0.201204 | BRCA | CAMA-1 | AST-1306 | MVP_top60_DNA+RNA_seed | HDAC1 | -0.531225 | BRCA | CAMA-1 | JQ12 |
| MVP_top60_DNA_seed | ESR1 | -0.200337 | BRCA | MDA-MB-436 | Fulvestrant | MVP_top60_DNA+RNA_seed | HDAC2 | -0.531225 | BRCA | CAMA-1 | JQ12 |
| MVP_top60_DNA_seed | HDAC1 | -0.200168 | BRCA | MDA-MB-453 | Dacinostat | MVP_top60_DNA+RNA_seed | AKT1 | -0.531059 | BRCA | MDA-MB-361 | GSK690693 |
| MVP_top60_DNA_seed | HDAC1 | -0.198542 | BRCA | CAMA-1 | CAY10603 | MVP_top60_DNA+RNA_seed | AR | -0.530111 | BRCA | CAL-51 | Bicalutamide |
| MVP_top60_DNA_seed | AR | -0.198342 | BRCA | MDA-MB-453 | Bicalutamide | MVP_top60_DNA+RNA_seed | HDAC1 | -0.526599 | BRCA | MDA-MB-231 | JQ12 |
| MVP_top60_DNA_seed | AR | -0.196671 | BRCA | MDA-MB-415 | AZD3514 | MVP_top60_DNA+RNA_seed | HDAC2 | -0.526599 | BRCA | MDA-MB-231 | JQ12 |
| MVP_top60_DNA_seed | EGFR | -0.196267 | BRCA | HCC1937 | AST-1306 | MVP_top60_DNA+RNA_seed | EGFR | -0.526292 | BRCA | HCC1187 | AST-1306 |
| MVP_top60_DNA_seed | CDK2 | -0.193133 | BRCA | CAL-85-1 | PHA-793887 | MVP_top60_DNA+RNA_seed | MAPK14 | -0.525091 | BRCA | BT-474 | TAK-715 |
| MVP_top60_DNA_seed | HDAC1 | -0.191114 | BRCA | AU565 | CAY10603 | MVP_top60_DNA+RNA_seed | ESR1 | -0.52464 | BRCA | HCC1500 | Tamoxifen |
| MVP_top60_DNA_seed | EGFR | -0.189242 | BRCA | HCC1937 | Pelitinib | MVP_top60_DNA+RNA_seed | HDAC1 | -0.52226 | BRCA | CAL-148 | Tubastatin A |
| MVP_top60_DNA_seed | HDAC1 | -0.188971 | BRCA | HCC38 | Belinostat | MVP_top60_DNA+RNA_seed | EGFR | -0.520793 | BRCA | HCC1187 | AZD8931 |
| MVP_top60_DNA_seed | EGFR | -0.188355 | BRCA | JIMT-1 | Pelitinib | MVP_top60_DNA+RNA_seed | ERBB2 | -0.520793 | BRCA | HCC1187 | AZD8931 |
| MVP_top60_DNA_seed | PIK3CA | -0.185494 | BRCA | HCC1569 | PI3Ka_4409 | MVP_top60_DNA+RNA_seed | ERBB3 | -0.520793 | BRCA | HCC1187 | AZD8931 |
| MVP_top60_DNA_seed | ESR1 | -0.184403 | BRCA | HDQ-P1 | Fulvestrant | MVP_top60_DNA+RNA_seed | CDK1 | -0.519871 | BRCA | HCC38 | RO-3306 |
| MVP_top60_DNA_seed | PIK3CA | -0.182922 | BRCA | CAL-51 | PI3Ka_4409 | MVP_top60_DNA+RNA_seed | ATM | -0.519858 | BRCA | MDA-MB-330 | CP466722 |
| MVP_top60_DNA_seed | EGFR | -0.182163 | BRCA | HCC1937 | Gefitinib | MVP_top60_DNA+RNA_seed | EGFR | -0.519458 | BRCA | MDA-MB-361 | PF-00299804 |
| MVP_top60_DNA_seed | ERBB2 | -0.181836 | BRCA | HCC1395 | AZD8931 | MVP_top60_DNA+RNA_seed | ERBB2 | -0.519458 | BRCA | MDA-MB-361 | PF-00299804 |
| MVP_top60_DNA_seed | EGFR | -0.181836 | BRCA | HCC1395 | AZD8931 | MVP_top60_DNA+RNA_seed | PLK1 | -0.516691 | BRCA | CAL-51 | PLK_6522 |
| MVP_top60_DNA_seed | ERBB3 | -0.181836 | BRCA | HCC1395 | AZD8931 | MVP_top60_DNA+RNA_seed | EGFR | -0.514997 | BRCA | COLO-824 | CI-1033 |
| MVP_top60_DNA_seed | MAPK3 | -0.181278 | BRCA | HDQ-P1 | FR-180204 | MVP_top60_DNA+RNA_seed | ERBB2 | -0.514997 | BRCA | COLO-824 | CI-1033 |
| MVP_top60_DNA_seed | MAPK1 | -0.181278 | BRCA | HDQ-P1 | FR-180204 | MVP_top60_DNA+RNA_seed | EGFR | -0.513168 | BRCA | MDA-MB-157 | AST-1306 |
| MVP_top60_DNA_seed | ATM | -0.179992 | BRCA | HCC1428 | CP466722 | MVP_top60_DNA+RNA_seed | ATM | -0.51309 | BRCA | MDA-MB-157 | KU-60019 |
| MVP_top60_DNA_seed | ESR1 | -0.178356 | BRCA | HCC1395 | Fulvestrant | MVP_top60_DNA+RNA_seed | HSP90AA1 | -0.509269 | BRCA | MDA-MB-231 | Elesclomol |
| MVP_top60_DNA_seed | CDK2 | -0.177967 | BRCA | CAL-120 | AZD5438 | MVP_top60_DNA+RNA_seed | HSP90AB1 | -0.509269 | BRCA | MDA-MB-231 | Elesclomol |
| MVP_top60_DNA_seed | AR | -0.177442 | BRCA | HCC70 | Bicalutamide | MVP_top60_DNA+RNA_seed | MDM2 | -0.509041 | BRCA | UACC-893 | Serdemetan |
| MVP_top60_DNA_seed | ERBB2 | -0.176416 | BRCA | DU-4475 | Lapatinib | MVP_top60_DNA+RNA_seed | MAPK3 | -0.508185 | BRCA | BT-474 | FR-180204 |
| MVP_top60_DNA_seed | EGFR | -0.176416 | BRCA | DU-4475 | Lapatinib | MVP_top60_DNA+RNA_seed | MAPK1 | -0.508185 | BRCA | BT-474 | FR-180204 |
| MVP_top60_DNA_seed | MAPK1 | -0.174389 | BRCA | EVSA-T | VX-11e | MVP_top60_DNA+RNA_seed | ATM | -0.50763 | BRCA | HCC1954 | CP466722 |
| MVP_top60_DNA_seed | ERBB2 | -0.173906 | BRCA | HCC1500 | CUDC-101 | MVP_top60_DNA+RNA_seed | ATM | -0.505788 | BRCA | T47D | CP466722 |
| MVP_top60_DNA_seed | EGFR | -0.173906 | BRCA | HCC1500 | CUDC-101 | MVP_top60_DNA+RNA_seed | ESR1 | -0.505314 | BRCA | DU-4475 | Tamoxifen |
| MVP_top60_DNA_seed | HDAC1 | -0.173043 | BRCA | BT-474 | Tubastatin A | MVP_top60_DNA+RNA_seed | CDK2 | -0.501561 | BRCA | DU-4475 | Seliciclib |

| | | | | | | | | | | | |
|---|---|---|---|---|---|---|---|---|---|---|---|
| MVP_top60_DNA_seed | EGFR | -0.172309 | BRCA | CAL-85-1 | AST-1306 | MVP_top60_DNA+RNA_seed | HDAC1 | -0.500232 | BRCA | DU-4475 | Tubastatin A |
| MVP_top60_DNA_seed | AKT1 | -0.171418 | BRCA | BT-20 | MK-2206 | MVP_top60_DNA+RNA_seed | MAPK3 | -0.499002 | BRCA | HCC1599 | FR-180204 |
| MVP_top60_DNA_seed | HDAC1 | -0.171175 | BRCA | HCC1187 | Parthenolide | MVP_top60_DNA+RNA_seed | MAPK1 | -0.499002 | BRCA | HCC1599 | FR-180204 |
| MVP_top60_DNA_seed | PIK3CA | -0.171098 | BRCA | CAL-120 | AZD8835 | MVP_top60_DNA+RNA_seed | EGFR | -0.498948 | BRCA | MFM-223 | AST-1306 |
| MVP_top60_DNA_seed | EGFR | -0.170635 | BRCA | HCC1500 | AST-1306 | MVP_top60_DNA+RNA_seed | HSP90AA1 | -0.498653 | BRCA | HCC1419 | Elesclomol |
| MVP_top60_DNA_seed | AKT1 | -0.166268 | BRCA | JIMT-1 | AKT inhibitor VIII | MVP_top60_DNA+RNA_seed | HSP90AB1 | -0.498653 | BRCA | HCC1419 | Elesclomol |
| MVP_top60_DNA_seed | HDAC1 | -0.164273 | BRCA | HCC70 | JQ12 | MVP_top60_DNA+RNA_seed | CDK1 | -0.497766 | BRCA | OCUB-M | RO-3306 |
| MVP_top60_DNA_seed | HDAC2 | -0.164273 | BRCA | HCC70 | JQ12 | MVP_top60_DNA+RNA_seed | EGFR | -0.496611 | BRCA | HCC1569 | AST-1306 |
| MVP_top60_DNA_seed | HDAC1 | -0.162591 | BRCA | HCC1569 | CAY10603 | MVP_top60_DNA+RNA_seed | HSP90AA1 | -0.495431 | BRCA | MDA-MB-361 | Luminespib |
| MVP_top60_DNA_seed | AKT1 | -0.159768 | BRCA | MCF7 | AKT inhibitor VIII | MVP_top60_DNA+RNA_seed | HSP90AB1 | -0.495431 | BRCA | MDA-MB-361 | Luminespib |
| MVP_top60_DNA_seed | ATM | -0.159495 | BRCA | CAL-120 | CP466722 | MVP_top60_DNA+RNA_seed | HDAC1 | -0.495061 | BRCA | HCC70 | Belinostat |
| MVP_top60_DNA_seed | ERBB2 | -0.159472 | BRCA | HCC70 | CI-1033 | MVP_top60_DNA+RNA_seed | ESR1 | -0.49031 | BRCA | MDA-MB-415 | Fulvestrant |
| MVP_top60_DNA_seed | EGFR | -0.159472 | BRCA | HCC70 | CI-1033 | MVP_top60_DNA+RNA_seed | HDAC1 | -0.489166 | BRCA | DU-4475 | Belinostat |
| MVP_top60_DNA_seed | EP300 | -0.158065 | BRCA | MDA-MB-361 | I-CBP112 | MVP_top60_DNA+RNA_seed | AKT1 | -0.488589 | BRCA | BT-20 | BAY AKT1 |
| MVP_top60_DNA_seed | CREBBP | -0.158065 | BRCA | MDA-MB-361 | I-CBP112 | MVP_top60_DNA+RNA_seed | MDM2 | -0.487753 | BRCA | HCC1954 | Serdemetan |
| MVP_top60_DNA_seed | ATM | -0.158058 | BRCA | MDA-MB-231 | CP466722 | MVP_top60_DNA+RNA_seed | CDK2 | -0.487203 | BRCA | ZR-75-30 | PHA-793887 |
| MVP_top60_DNA_seed | MAPK3 | -0.156945 | BRCA | HCC202 | FR-180204 | MVP_top60_DNA+RNA_seed | EGFR | -0.485702 | BRCA | OCUB-M | Afatinib |
| MVP_top60_DNA_seed | MAPK1 | -0.156945 | BRCA | HCC202 | FR-180204 | MVP_top60_DNA+RNA_seed | ERBB2 | -0.485702 | BRCA | OCUB-M | Afatinib |
| MVP_top60_DNA_seed | HDAC1 | -0.155937 | BRCA | ZR-75-30 | Tubastatin A | MVP_top60_DNA+RNA_seed | HSP90AA1 | -0.484939 | BRCA | HCC38 | Luminespib |
| MVP_top60_DNA_seed | EGFR | -0.152968 | BRCA | OCUB-M | Gefitinib | MVP_top60_DNA+RNA_seed | HSP90AB1 | -0.484939 | BRCA | HCC38 | Luminespib |
| MVP_top60_DNA_seed | SRC | -0.15267 | BRCA | MDA-MB-157 | Bosutinib | MVP_top60_DNA+RNA_seed | HSP90AA1 | -0.484702 | BRCA | COLO-824 | SNX-2112 |
| MVP_top60_DNA_seed | AR | -0.151122 | BRCA | DU-4475 | AZD3514 | MVP_top60_DNA+RNA_seed | HSP90AB1 | -0.484702 | BRCA | COLO-824 | SNX-2112 |
| MVP_top60_DNA_seed | PIK3CA | -0.14942 | BRCA | CAL-85-1 | PI3Ka_4409 | MVP_top60_DNA+RNA_seed | EGFR | -0.484012 | BRCA | HDQ-P1 | CI-1033 |
| MVP_top60_DNA_seed | PIK3CA | -0.149209 | BRCA | AU565 | PI3Ka_4409 | MVP_top60_DNA+RNA_seed | ERBB2 | -0.484012 | BRCA | HDQ-P1 | CI-1033 |
| MVP_top60_DNA_seed | HDAC1 | -0.148313 | BRCA | EFM-192A | Tubastatin A | MVP_top60_DNA+RNA_seed | HSP90AA1 | -0.483835 | BRCA | HCC1395 | Elesclomol |
| MVP_top60_DNA_seed | ERBB2 | -0.146968 | BRCA | HCC1143 | Afatinib | MVP_top60_DNA+RNA_seed | HSP90AB1 | -0.483835 | BRCA | HCC1395 | Elesclomol |
| MVP_top60_DNA_seed | EGFR | -0.146968 | BRCA | HCC1143 | Afatinib | MVP_top60_DNA+RNA_seed | EGFR | -0.483653 | BRCA | HCC1569 | PF-00299804 |
| MVP_top60_DNA_seed | EP300 | -0.146959 | BRCA | EVSA-T | I-CBP112 | MVP_top60_DNA+RNA_seed | ERBB2 | -0.483653 | BRCA | HCC1569 | PF-00299804 |
| MVP_top60_DNA_seed | CREBBP | -0.146959 | BRCA | EVSA-T | I-CBP112 | MVP_top60_DNA+RNA_seed | ATM | -0.483183 | BRCA | UACC-812 | CP466722 |
| MVP_top60_DNA_seed | MAPK1 | -0.146919 | BRCA | BT-474 | VX-11e | MVP_top60_DNA+RNA_seed | HSP90AA1 | -0.483061 | BRCA | HCC2157 | Tanespimycin |
| MVP_top60_DNA_seed | EGFR | -0.143765 | BRCA | HCC1500 | Pelitinib | MVP_top60_DNA+RNA_seed | HSP90AB1 | -0.483061 | BRCA | HCC2157 | Tanespimycin |
| MVP_top60_DNA_seed | HSP90AA1 | -0.142816 | BRCA | EVSA-T | Elesclomol | MVP_top60_DNA+RNA_seed | AR | -0.482859 | BRCA | CAL-120 | Bicalutamide |
| MVP_top60_DNA_seed | HSP90AB1 | -0.142816 | BRCA | EVSA-T | Elesclomol | MVP_top60_DNA+RNA_seed | HSP90AA1 | -0.481917 | BRCA | BT-483 | Elesclomol |
| MVP_top60_DNA_seed | EP300 | -0.141759 | BRCA | HCC2218 | I-CBP112 | MVP_top60_DNA+RNA_seed | HSP90AB1 | -0.481917 | BRCA | BT-483 | Elesclomol |
| MVP_top60_DNA_seed | CREBBP | -0.141759 | BRCA | HCC2218 | I-CBP112 | MVP_top60_DNA+RNA_seed | EGFR | -0.48186 | BRCA | DU-4475 | Afatinib |
| MVP_top60_DNA_seed | HSP90AA1 | -0.141117 | BRCA | COLO-824 | CCT-018159 | MVP_top60_DNA+RNA_seed | ERBB2 | -0.48186 | BRCA | DU-4475 | Afatinib |
| MVP_top60_DNA_seed | HSP90AB1 | -0.141117 | BRCA | COLO-824 | CCT-018159 | MVP_top60_DNA+RNA_seed | ATM | -0.481795 | BRCA | MDA-MB-175-VII | CP466722 |
| MVP_top60_DNA_seed | ERBB2 | -0.140173 | BRCA | HCC38 | AZD8931 | MVP_top60_DNA+RNA_seed | MAPK14 | -0.479322 | BRCA | MDA-MB-453 | TAK-715 |
| MVP_top60_DNA_seed | EGFR | -0.140173 | BRCA | HCC38 | AZD8931 | MVP_top60_DNA+RNA_seed | ATM | -0.479321 | BRCA | HCC1599 | CP466722 |
| MVP_top60_DNA_seed | ERBB3 | -0.140173 | BRCA | HCC38 | AZD8931 | MVP_top60_DNA+RNA_seed | ESR1 | -0.479239 | BRCA | EFM-19 | Tamoxifen |
| MVP_top60_DNA_seed | HDAC1 | -0.140055 | BRCA | MDA-MB-231 | AR-42 | MVP_top60_DNA+RNA_seed | AR | -0.474902 | BRCA | JIMT-1 | Bicalutamide |

| | | | | | | | | | | | |
|---|---|---|---|---|---|---|---|---|---|---|---|
| MVP_top60_DNA_seed | HSP90AA1 | -0.139833 | BRCA | BT-20 | Elesclomol | MVP_top60_DNA+RNA_seed | CDK2 | -0.473969 | BRCA | MRK-nu-1 | AZD5438 |
| MVP_top60_DNA_seed | HSP90AB1 | -0.139833 | BRCA | BT-20 | Elesclomol | MVP_top60_DNA+RNA_seed | AR | -0.473498 | BRCA | CAL-120 | AZD3514 |
| MVP_top60_DNA_seed | EP300 | -0.139638 | BRCA | HCC1569 | I-CBP112 | MVP_top60_DNA+RNA_seed | EGFR | -0.470583 | BRCA | CAL-85-1 | CI-1033 |
| MVP_top60_DNA_seed | CREBBP | -0.139638 | BRCA | HCC1569 | I-CBP112 | MVP_top60_DNA+RNA_seed | ERBB2 | -0.470583 | BRCA | CAL-85-1 | CI-1033 |
| MVP_top60_DNA_seed | ERBB2 | -0.139142 | BRCA | CAMA-1 | CUDC-101 | MVP_top60_DNA+RNA_seed | AR | -0.467232 | BRCA | HCC70 | Bicalutamide |
| MVP_top60_DNA_seed | EGFR | -0.139142 | BRCA | CAMA-1 | CUDC-101 | MVP_top60_DNA+RNA_seed | HSP90AA1 | -0.465864 | BRCA | HCC2157 | Elesclomol |
| MVP_top60_DNA_seed | HDAC1 | -0.139049 | BRCA | JIMT-1 | Dacinostat | MVP_top60_DNA+RNA_seed | HSP90AB1 | -0.465864 | BRCA | HCC2157 | Elesclomol |
| MVP_top60_DNA_seed | HSP90AA1 | -0.138766 | BRCA | HCC70 | Tanespimycin | MVP_top60_DNA+RNA_seed | HSP90AA1 | -0.464356 | BRCA | AU565 | Tanespimycin |
| MVP_top60_DNA_seed | HSP90AB1 | -0.138766 | BRCA | HCC70 | Tanespimycin | MVP_top60_DNA+RNA_seed | HSP90AB1 | -0.464356 | BRCA | AU565 | Tanespimycin |
| MVP_top60_DNA_seed | CDK1 | -0.137951 | BRCA | HCC1954 | RO-3306 | MVP_top60_DNA+RNA_seed | MAPK14 | -0.463549 | BRCA | CAL-51 | TAK-715 |
| MVP_top60_DNA_seed | ATM | -0.137945 | BRCA | MDA-MB-468 | KU-60019 | MVP_top60_DNA+RNA_seed | EGFR | -0.46147 | BRCA | HCC1954 | CI-1033 |
| MVP_top60_DNA_seed | AR | -0.137819 | BRCA | CAMA-1 | Bicalutamide | MVP_top60_DNA+RNA_seed | ERBB2 | -0.46147 | BRCA | HCC1954 | CI-1033 |
| MVP_top60_DNA_seed | ERBB2 | -0.137279 | BRCA | HCC2218 | AZD8931 | MVP_top60_DNA+RNA_seed | HDAC1 | -0.461138 | BRCA | HCC1428 | AR-42 |
| MVP_top60_DNA_seed | EGFR | -0.137279 | BRCA | HCC2218 | AZD8931 | MVP_top60_DNA+RNA_seed | HDAC1 | -0.461089 | BRCA | DU-4475 | Parthenolide |
| MVP_top60_DNA_seed | ERBB3 | -0.137279 | BRCA | HCC2218 | AZD8931 | MVP_top60_DNA+RNA_seed | HDAC1 | -0.460676 | BRCA | ZR-75-30 | AR-42 |
| MVP_top60_DNA_seed | ATM | -0.136084 | BRCA | HDQ-P1 | CP466722 | MVP_top60_DNA+RNA_seed | HDAC1 | -0.459233 | BRCA | T47D | Tubastatin A |
| MVP_top60_DNA_seed | ATM | -0.134713 | BRCA | BT-474 | CP466722 | MVP_top60_DNA+RNA_seed | EGFR | -0.457829 | BRCA | CAL-85-1 | PF-00299804 |
| MVP_top60_DNA_seed | CDK2 | -0.133968 | BRCA | EVSA-T | PHA-793887 | MVP_top60_DNA+RNA_seed | ERBB2 | -0.457829 | BRCA | CAL-85-1 | PF-00299804 |
| MVP_top60_DNA_seed | ERBB2 | -0.133964 | BRCA | CAMA-1 | PF-00299804 | MVP_top60_DNA+RNA_seed | HDAC1 | -0.457802 | BRCA | MDA-MB-330 | Dacinostat |
| MVP_top60_DNA_seed | EGFR | -0.133964 | BRCA | CAMA-1 | PF-00299804 | MVP_top60_DNA+RNA_seed | EGFR | -0.457104 | BRCA | MDA-MB-453 | PF-00299804 |
| MVP_top60_DNA_seed | EGFR | -0.133527 | BRCA | DU-4475 | Pelitinib | MVP_top60_DNA+RNA_seed | ERBB2 | -0.457104 | BRCA | MDA-MB-453 | PF-00299804 |
| MVP_top60_DNA_seed | MAPK3 | -0.132932 | BRCA | HCC1187 | FR-180204 | MVP_top60_DNA+RNA_seed | AKT1 | -0.455335 | BRCA | EFM-19 | GSK690693 |
| MVP_top60_DNA_seed | MAPK1 | -0.132932 | BRCA | HCC1187 | FR-180204 | MVP_top60_DNA+RNA_seed | SRC | -0.455244 | BRCA | BT-549 | Bosutinib |
| MVP_top60_DNA_seed | HDAC1 | -0.132579 | BRCA | MFM-223 | AR-42 | MVP_top60_DNA+RNA_seed | HSP90AA1 | -0.454606 | BRCA | HCC1569 | Luminespib |
| MVP_top60_DNA_seed | EP300 | -0.132404 | BRCA | MDA-MB-468 | I-CBP112 | MVP_top60_DNA+RNA_seed | HSP90AB1 | -0.454606 | BRCA | HCC1569 | Luminespib |
| MVP_top60_DNA_seed | CREBBP | -0.132404 | BRCA | MDA-MB-468 | I-CBP112 | MVP_top60_DNA+RNA_seed | EGFR | -0.454469 | BRCA | MDA-MB-361 | CI-1033 |
| MVP_top60_DNA_seed | ATM | -0.131959 | BRCA | BT-483 | CP466722 | MVP_top60_DNA+RNA_seed | ERBB2 | -0.454469 | BRCA | MDA-MB-361 | CI-1033 |
| MVP_top60_DNA_seed | ESR1 | -0.131703 | BRCA | CAL-120 | Tamoxifen | MVP_top60_DNA+RNA_seed | ATM | -0.453857 | BRCA | DU-4475 | KU-55933 |
| MVP_top60_DNA_seed | HDAC1 | -0.131661 | BRCA | MFM-223 | Belinostat | MVP_top60_DNA+RNA_seed | PIK3CA | -0.453369 | BRCA | MDA-MB-330 | PI3Ka_4409 |
| MVP_top60_DNA_seed | MDM2 | -0.128592 | BRCA | OCUB-M | Nutlin-3a (-) | MVP_top60_DNA+RNA_seed | AR | -0.453109 | BRCA | HCC1954 | Bicalutamide |
| MVP_top60_DNA_seed | EGFR | -0.12757 | BRCA | HCC1428 | AST-1306 | MVP_top60_DNA+RNA_seed | HSP90AA1 | -0.452918 | BRCA | BT-474 | SNX-2112 |
| MVP_top60_DNA_seed | ERBB2 | -0.127057 | BRCA | EVSA-T | PF-00299804 | MVP_top60_DNA+RNA_seed | HSP90AB1 | -0.452918 | BRCA | BT-474 | SNX-2112 |
| MVP_top60_DNA_seed | EGFR | -0.127057 | BRCA | EVSA-T | PF-00299804 | MVP_top60_DNA+RNA_seed | CDK2 | -0.452877 | BRCA | MDA-MB-468 | AZD5438 |
| MVP_top60_DNA_seed | EGFR | -0.126935 | BRCA | HCC38 | Pelitinib | MVP_top60_DNA+RNA_seed | ESR1 | -0.450343 | BRCA | MFM-223 | Fulvestrant |
| MVP_top60_DNA_seed | ERBB2 | -0.124578 | BRCA | HCC70 | PF-00299804 | MVP_top60_DNA+RNA_seed | AKT1 | -0.447618 | BRCA | HCC2218 | A-443654 |
| MVP_top60_DNA_seed | EGFR | -0.124578 | BRCA | HCC70 | PF-00299804 | MVP_top60_DNA+RNA_seed | HDAC1 | -0.447284 | BRCA | HCC1419 | Dacinostat |
| MVP_top60_DNA_seed | HDAC1 | -0.123964 | BRCA | MDA-MB-361 | Tubastatin A | MVP_top60_DNA+RNA_seed | HDAC1 | -0.44226 | BRCA | AU565 | Belinostat |
| MVP_top60_DNA_seed | HSP90AA1 | -0.123933 | BRCA | BT-20 | Tanespimycin | MVP_top60_DNA+RNA_seed | EGFR | -0.441241 | BRCA | HCC2157 | Afatinib |
| MVP_top60_DNA_seed | HSP90AB1 | -0.123933 | BRCA | BT-20 | Tanespimycin | MVP_top60_DNA+RNA_seed | ERBB2 | -0.441241 | BRCA | HCC2157 | Afatinib |
| MVP_top60_DNA_seed | HDAC1 | -0.123044 | BRCA | HCC2218 | CAY10603 | MVP_top60_DNA+RNA_seed | PIK3CA | -0.43744 | BRCA | EVSA-T | AZD8835 |
| MVP_top60_DNA_seed | AKT1 | -0.122759 | BRCA | HCC1395 | BAY AKT1 | MVP_top60_DNA+RNA_seed | HDAC1 | -0.436725 | BRCA | MDA-MB-330 | AR-42 |

| | | | | | | | | | | | |
|---|---|---|---|---|---|---|---|---|---|---|---|
| MVP_top60_DNA_seed | ERBB2 | -0.121454 | BRCA | HCC1569 | CUDC-101 | MVP_top60_DNA+RNA_seed | MAPK14 | -0.436125 | BRCA | HCC1599 | TAK-715 |
| MVP_top60_DNA_seed | EGFR | -0.121454 | BRCA | HCC1569 | CUDC-101 | MVP_top60_DNA+RNA_seed | HDAC1 | -0.435708 | BRCA | CAL-148 | CAY10603 |
| MVP_top60_DNA_seed | AR | -0.120436 | BRCA | MDA-MB-330 | Bicalutamide | MVP_top60_DNA+RNA_seed | HSP90AA1 | -0.4352 | BRCA | HCC2218 | CCT-018159 |
| MVP_top60_DNA_seed | HSP90AA1 | -0.119617 | BRCA | HCC202 | SNX-2112 | MVP_top60_DNA+RNA_seed | HSP90AB1 | -0.4352 | BRCA | HCC2218 | CCT-018159 |
| MVP_top60_DNA_seed | HSP90AB1 | -0.119617 | BRCA | HCC202 | SNX-2112 | MVP_top60_DNA+RNA_seed | MAPK3 | -0.435194 | BRCA | OCUB-M | FR-180204 |
| MVP_top60_DNA_seed | AR | -0.118924 | BRCA | MDA-MB-231 | AZD3514 | MVP_top60_DNA+RNA_seed | MAPK1 | -0.435194 | BRCA | OCUB-M | FR-180204 |
| MVP_top60_DNA_seed | EGFR | -0.118894 | BRCA | CAL-51 | Gefitinib | MVP_top60_DNA+RNA_seed | EP300 | -0.432685 | BRCA | HCC2157 | I-CBP112 |
| MVP_top60_DNA_seed | EGFR | -0.118836 | BRCA | MDA-MB-231 | AST-1306 | MVP_top60_DNA+RNA_seed | HDAC1 | -0.432422 | BRCA | AU565 | AR-42 |
| MVP_top60_DNA_seed | ATM | -0.116796 | BRCA | MFM-223 | KU-55933 | MVP_top60_DNA+RNA_seed | EP300 | -0.432125 | BRCA | HCC1428 | I-CBP112 |
| MVP_top60_DNA_seed | ERBB2 | -0.116294 | BRCA | T47D | Afatinib | MVP_top60_DNA+RNA_seed | AKT1 | -0.427629 | BRCA | BT-20 | AKT inhibitor VIII |
| MVP_top60_DNA_seed | EGFR | -0.116294 | BRCA | T47D | Afatinib | MVP_top60_DNA+RNA_seed | EGFR | -0.426082 | BRCA | HCC70 | AZD8931 |
| MVP_top60_DNA_seed | AR | -0.116142 | BRCA | MDA-MB-361 | Bicalutamide | MVP_top60_DNA+RNA_seed | ERBB2 | -0.426082 | BRCA | HCC70 | AZD8931 |
| MVP_top60_DNA_seed | HDAC1 | -0.11562 | BRCA | BT-474 | Entinostat | MVP_top60_DNA+RNA_seed | ERBB3 | -0.426082 | BRCA | HCC70 | AZD8931 |
| MVP_top60_DNA_seed | SRC | -0.115369 | BRCA | EFM-19 | NVP-BHG712 | MVP_top60_DNA+RNA_seed | PLK1 | -0.425524 | BRCA | EVSA-T | PLK_6522 |
| MVP_top60_DNA_seed | AR | -0.114835 | BRCA | JIMT-1 | AZD3514 | MVP_top60_DNA+RNA_seed | MAPK14 | -0.425075 | BRCA | HCC1954 | TAK-715 |
| MVP_top60_DNA_seed | MDM2 | -0.113512 | BRCA | MDA-MB-468 | Serdemetan | MVP_top60_DNA+RNA_seed | AKT1 | -0.424946 | BRCA | HCC1954 | AKT inhibitor VIII |
| MVP_top60_DNA_seed | HSP90AA1 | -0.113122 | BRCA | EFM-19 | Tanespimycin | MVP_top60_DNA+RNA_seed | EGFR | -0.424329 | BRCA | HDQ-P1 | PF-00299804 |
| MVP_top60_DNA_seed | HSP90AB1 | -0.113122 | BRCA | EFM-19 | Tanespimycin | MVP_top60_DNA+RNA_seed | ERBB2 | -0.424329 | BRCA | HDQ-P1 | PF-00299804 |
| MVP_top60_DNA_seed | HSP90AA1 | -0.112393 | BRCA | MDA-MB-468 | Luminespib | MVP_top60_DNA+RNA_seed | ATM | -0.42349 | BRCA | EFM-19 | KU-55933 |
| MVP_top60_DNA_seed | HSP90AB1 | -0.112393 | BRCA | MDA-MB-468 | Luminespib | MVP_top60_DNA+RNA_seed | ATM | -0.423416 | BRCA | DU-4475 | CP466722 |
| MVP_top60_DNA_seed | ATM | -0.112313 | BRCA | MDA-MB-453 | CP466722 | MVP_top60_DNA+RNA_seed | SRC | -0.422841 | BRCA | HCC1937 | Bosutinib |
| MVP_top60_DNA_seed | PIK3CA | -0.111097 | BRCA | HCC2218 | PI3Ka_4409 | MVP_top60_DNA+RNA_seed | PLK1 | -0.420428 | BRCA | HCC1954 | PLK_6522 |
| MVP_top60_DNA_seed | MDM2 | -0.110434 | BRCA | MDA-MB-157 | Serdemetan | MVP_top60_DNA+RNA_seed | CDK1 | -0.417586 | BRCA | DU-4475 | RO-3306 |
| MVP_top60_DNA_seed | HSP90AA1 | -0.110167 | BRCA | MDA-MB-175-VII | CCT-018159 | MVP_top60_DNA+RNA_seed | EGFR | -0.415889 | BRCA | HCC1143 | AZD8931 |
| MVP_top60_DNA_seed | HSP90AB1 | -0.110167 | BRCA | MDA-MB-175-VII | CCT-018159 | MVP_top60_DNA+RNA_seed | ERBB2 | -0.415889 | BRCA | HCC1143 | AZD8931 |
| MVP_top60_DNA_seed | HSP90AA1 | -0.110099 | BRCA | CAL-120 | CCT-018159 | MVP_top60_DNA+RNA_seed | ERBB3 | -0.415889 | BRCA | HCC1143 | AZD8931 |
| MVP_top60_DNA_seed | HSP90AB1 | -0.110099 | BRCA | CAL-120 | CCT-018159 | MVP_top60_DNA+RNA_seed | SRC | -0.415427 | BRCA | HCC1599 | Saracatinib |
| MVP_top60_DNA_seed | ERBB2 | -0.110048 | BRCA | HDQ-P1 | Afatinib | MVP_top60_DNA+RNA_seed | EGFR | -0.413221 | BRCA | HCC1187 | Pelitinib |
| MVP_top60_DNA_seed | EGFR | -0.110048 | BRCA | HDQ-P1 | Afatinib | MVP_top60_DNA+RNA_seed | EGFR | -0.412765 | BRCA | HCC1395 | PF-00299804 |
| MVP_top60_DNA_seed | HSP90AA1 | -0.109325 | BRCA | UACC-893 | Tanespimycin | MVP_top60_DNA+RNA_seed | ERBB2 | -0.412765 | BRCA | HCC1395 | PF-00299804 |
| MVP_top60_DNA_seed | HSP90AB1 | -0.109325 | BRCA | UACC-893 | Tanespimycin | MVP_top60_DNA+RNA_seed | PLK1 | -0.408664 | BRCA | DU-4475 | GW843682X |
| MVP_top60_DNA_seed | SRC | -0.108857 | BRCA | EVSA-T | NVP-BHG712 | MVP_top60_DNA+RNA_seed | ATM | -0.405786 | BRCA | CAL-51 | CP466722 |
| MVP_top60_DNA_seed | ERBB2 | -0.104941 | BRCA | MCF7 | CI-1033 | MVP_top60_DNA+RNA_seed | EGFR | -0.405305 | BRCA | HCC1395 | AST-1306 |
| MVP_top60_DNA_seed | EGFR | -0.104941 | BRCA | MCF7 | CI-1033 | MVP_top60_DNA+RNA_seed | EGFR | -0.404112 | BRCA | CAMA-1 | CI-1033 |
| MVP_top60_DNA_seed | HSP90AA1 | -0.102275 | BRCA | HCC1937 | Luminespib | MVP_top60_DNA+RNA_seed | ERBB2 | -0.404112 | BRCA | CAMA-1 | CI-1033 |
| MVP_top60_DNA_seed | HSP90AB1 | -0.102275 | BRCA | HCC1937 | Luminespib | MVP_top60_DNA+RNA_seed | HDAC1 | -0.401732 | BRCA | HCC1500 | AR-42 |
| MVP_top60_DNA_seed | MAPK3 | -0.099685 | BRCA | BT-549 | FR-180204 | MVP_top60_DNA+RNA_seed | EGFR | -0.400758 | BRCA | BT-483 | AST-1306 |
| MVP_top60_DNA_seed | MAPK1 | -0.099685 | BRCA | BT-549 | FR-180204 | MVP_top60_DNA+RNA_seed | MDM2 | -0.399239 | BRCA | JIMT-1 | Serdemetan |
| MVP_top60_DNA_seed | CDK2 | -0.095464 | BRCA | DU-4475 | AZD5438 | MVP_top60_DNA+RNA_seed | HSP90AA1 | -0.398809 | BRCA | MFM-223 | Tanespimycin |
| MVP_top60_DNA_seed | HDAC1 | -0.09419 | BRCA | HCC1569 | Belinostat | MVP_top60_DNA+RNA_seed | HSP90AB1 | -0.398809 | BRCA | MFM-223 | Tanespimycin |
| MVP_top60_DNA_seed | HDAC1 | -0.093054 | BRCA | OCUB-M | JQ12 | MVP_top60_DNA+RNA_seed | AKT1 | -0.398249 | BRCA | BT-474 | MK-2206 |

| | | | | | | | | | | | |
|---|---|---|---|---|---|---|---|---|---|---|---|
| MVP_top60_DNA_seed | HDAC2 | -0.093054 | BRCA | OCUB-M | JQ12 | MVP_top60_DNA+RNA_seed | HSP90AA1 | -0.397831 | BRCA | MDA-MB-453 | CCT-018159 |
| MVP_top60_DNA_seed | CDK2 | -0.090015 | BRCA | HCC1937 | PHA-793887 | MVP_top60_DNA+RNA_seed | HSP90AB1 | -0.397831 | BRCA | MDA-MB-453 | CCT-018159 |
| MVP_top60_DNA_seed | HDAC1 | -0.088683 | BRCA | CAMA-1 | Belinostat | MVP_top60_DNA+RNA_seed | HDAC1 | -0.396341 | BRCA | CAL-51 | JQ12 |
| MVP_top60_DNA_seed | HDAC1 | -0.087323 | BRCA | HCC38 | AR-42 | MVP_top60_DNA+RNA_seed | HDAC2 | -0.396341 | BRCA | CAL-51 | JQ12 |
| MVP_top60_DNA_seed | HDAC1 | -0.086792 | BRCA | HCC1569 | AR-42 | MVP_top60_DNA+RNA_seed | HSP90AA1 | -0.391822 | BRCA | DU-4475 | CCT-018159 |
| MVP_top60_DNA_seed | AKT1 | -0.085986 | BRCA | CAL-120 | BAY AKT1 | MVP_top60_DNA+RNA_seed | HSP90AB1 | -0.391822 | BRCA | DU-4475 | CCT-018159 |
| MVP_top60_DNA_seed | CDK2 | -0.085464 | BRCA | HCC1187 | Seliciclib | MVP_top60_DNA+RNA_seed | MDM2 | -0.389164 | BRCA | CAL-51 | Serdemetan |
| MVP_top60_DNA_seed | CDK2 | -0.085366 | BRCA | HCC38 | PHA-793887 | MVP_top60_DNA+RNA_seed | EGFR | -0.38752 | BRCA | HCC1599 | AST-1306 |
| MVP_top60_DNA_seed | ATM | -0.084442 | BRCA | CAMA-1 | CP466722 | MVP_top60_DNA+RNA_seed | HDAC1 | -0.386741 | BRCA | OCUB-M | Entinostat |
| MVP_top60_DNA_seed | MDM2 | -0.0826 | BRCA | HCC1395 | Serdemetan | MVP_top60_DNA+RNA_seed | EGFR | -0.386319 | BRCA | MDA-MB-361 | AZD8931 |
| MVP_top60_DNA_seed | HSP90AA1 | -0.080607 | BRCA | MDA-MB-361 | CCT-018159 | MVP_top60_DNA+RNA_seed | ERBB2 | -0.386319 | BRCA | MDA-MB-361 | AZD8931 |
| MVP_top60_DNA_seed | HSP90AB1 | -0.080607 | BRCA | MDA-MB-361 | CCT-018159 | MVP_top60_DNA+RNA_seed | ERBB3 | -0.386319 | BRCA | MDA-MB-361 | AZD8931 |
| MVP_top60_DNA_seed | ERBB2 | -0.079762 | BRCA | MDA-MB-175-VII | CUDC-101 | MVP_top60_DNA+RNA_seed | EP300 | -0.385811 | BRCA | COLO-824 | I-CBP112 |
| MVP_top60_DNA_seed | EGFR | -0.079762 | BRCA | MDA-MB-175-VII | CUDC-101 | MVP_top60_DNA+RNA_seed | ESR1 | -0.385659 | BRCA | DU-4475 | Fulvestrant |
| MVP_top60_DNA_seed | AKT1 | -0.079275 | BRCA | CAL-120 | AKT inhibitor VIII | MVP_top60_DNA+RNA_seed | CDK2 | -0.382457 | BRCA | BT-549 | PHA-793887 |
| MVP_top60_DNA_seed | EGFR | -0.077216 | BRCA | COLO-824 | Pelitinib | MVP_top60_DNA+RNA_seed | EGFR | -0.382117 | BRCA | CAL-148 | AST-1306 |
| MVP_top60_DNA_seed | HSP90AA1 | -0.073555 | BRCA | HDQ-P1 | CCT-018159 | MVP_top60_DNA+RNA_seed | EGFR | -0.380465 | BRCA | HCC1428 | CUDC-101 |
| MVP_top60_DNA_seed | HSP90AB1 | -0.073555 | BRCA | HDQ-P1 | CCT-018159 | MVP_top60_DNA+RNA_seed | ERBB2 | -0.380465 | BRCA | HCC1428 | CUDC-101 |
| MVP_top60_DNA_seed | HSP90AA1 | -0.073326 | BRCA | MDA-MB-157 | CCT-018159 | MVP_top60_DNA+RNA_seed | HDAC1 | -0.376742 | BRCA | HCC2157 | JQ12 |
| MVP_top60_DNA_seed | HSP90AB1 | -0.073326 | BRCA | MDA-MB-157 | CCT-018159 | MVP_top60_DNA+RNA_seed | HDAC2 | -0.376742 | BRCA | HCC2157 | JQ12 |
| MVP_top60_DNA_seed | CDK2 | -0.07315 | BRCA | HCC1143 | AZD5438 | MVP_top60_DNA+RNA_seed | ESR1 | -0.374809 | BRCA | BT-20 | Fulvestrant |
| MVP_top60_DNA_seed | EP300 | -0.073097 | BRCA | CAL-51 | I-CBP112 | MVP_top60_DNA+RNA_seed | ATM | -0.372726 | BRCA | CAL-120 | KU-60019 |
| MVP_top60_DNA_seed | CREBBP | -0.073097 | BRCA | CAL-51 | I-CBP112 | MVP_top60_DNA+RNA_seed | HDAC1 | -0.371402 | BRCA | HCC1954 | JQ12 |
| MVP_top60_DNA_seed | EGFR | -0.071951 | BRCA | BT-20 | Pelitinib | MVP_top60_DNA+RNA_seed | HDAC2 | -0.371402 | BRCA | HCC1954 | JQ12 |
| MVP_top60_DNA_seed | ATM | -0.071417 | BRCA | JIMT-1 | KU-60019 | MVP_top60_DNA+RNA_seed | HDAC1 | -0.371322 | BRCA | MDA-MB-361 | Belinostat |
| MVP_top60_DNA_seed | AR | -0.071413 | BRCA | EFM-192A | AZD3514 | MVP_top60_DNA+RNA_seed | HDAC1 | -0.370033 | BRCA | HCC1599 | JQ12 |
| MVP_top60_DNA_seed | SRC | -0.069612 | BRCA | EVSA-T | Saracatinib | MVP_top60_DNA+RNA_seed | HDAC2 | -0.370033 | BRCA | HCC1599 | JQ12 |
| MVP_top60_DNA_seed | ATM | -0.066741 | BRCA | MDA-MB-157 | KU-55933 | MVP_top60_DNA+RNA_seed | HDAC1 | -0.369541 | BRCA | AU565 | JQ12 |
| MVP_top60_DNA_seed | HDAC1 | -0.064702 | BRCA | HCC1500 | Tubastatin A | MVP_top60_DNA+RNA_seed | HDAC2 | -0.369541 | BRCA | AU565 | JQ12 |
| MVP_top60_DNA_seed | ERBB2 | -0.063861 | BRCA | JIMT-1 | Afatinib | MVP_top60_DNA+RNA_seed | CDK1 | -0.36943 | BRCA | HCC1419 | RO-3306 |
| MVP_top60_DNA_seed | EGFR | -0.063861 | BRCA | JIMT-1 | Afatinib | MVP_top60_DNA+RNA_seed | EGFR | -0.368993 | BRCA | HCC202 | AZD8931 |
| MVP_top60_DNA_seed | HSP90AA1 | -0.062509 | BRCA | HCC2157 | Luminespib | MVP_top60_DNA+RNA_seed | ERBB2 | -0.368993 | BRCA | HCC202 | AZD8931 |
| MVP_top60_DNA_seed | HSP90AB1 | -0.062509 | BRCA | HCC2157 | Luminespib | MVP_top60_DNA+RNA_seed | ERBB3 | -0.368993 | BRCA | HCC202 | AZD8931 |
| MVP_top60_DNA_seed | ATM | -0.059343 | BRCA | JIMT-1 | KU-55933 | MVP_top60_DNA+RNA_seed | ATM | -0.363855 | BRCA | MDA-MB-415 | KU-55933 |
| MVP_top60_DNA_seed | ERBB2 | -0.058118 | BRCA | CAL-51 | Afatinib | MVP_top60_DNA+RNA_seed | MAPK1 | -0.362996 | BRCA | HCC70 | VX-11e |
| MVP_top60_DNA_seed | EGFR | -0.058118 | BRCA | CAL-51 | Afatinib | MVP_top60_DNA+RNA_seed | AKT1 | -0.362291 | BRCA | MDA-MB-468 | MK-2206 |
| MVP_top60_DNA_seed | ERBB2 | -0.057841 | BRCA | MDA-MB-330 | CUDC-101 | MVP_top60_DNA+RNA_seed | MAPK14 | -0.361725 | BRCA | CAL-148 | TAK-715 |
| MVP_top60_DNA_seed | EGFR | -0.057841 | BRCA | MDA-MB-330 | CUDC-101 | MVP_top60_DNA+RNA_seed | AKT1 | -0.361687 | BRCA | DU-4475 | AKT inhibitor VIII |
| MVP_top60_DNA_seed | PIK3CA | -0.057415 | BRCA | HCC70 | PI3Ka_4409 | MVP_top60_DNA+RNA_seed | SRC | -0.359665 | BRCA | MDA-MB-231 | NVP-BHG712 |
| MVP_top60_DNA_seed | AR | -0.056003 | BRCA | HCC1187 | AZD3514 | MVP_top60_DNA+RNA_seed | HSP90AA1 | -0.352942 | BRCA | CAL-120 | SNX-2112 |
| MVP_top60_DNA_seed | HDAC1 | -0.055193 | BRCA | EFM-19 | Belinostat | MVP_top60_DNA+RNA_seed | HSP90AB1 | -0.352942 | BRCA | CAL-120 | SNX-2112 |

| | | | | | | | | | | | |
|---|---|---|---|---|---|---|---|---|---|---|---|
| MVP_top60_DNA_seed | AKT1 | -0.054143 | BRCA | MDA-MB-468 | BAY AKT1 | MVP_top60_DNA+RNA_seed | HSP90AA1 | -0.35281 | BRCA | HCC1937 | Tanespimycin |
| MVP_top60_DNA_seed | HDAC1 | -0.052786 | BRCA | HCC38 | CAY10603 | MVP_top60_DNA+RNA_seed | HSP90AB1 | -0.35281 | BRCA | HCC1937 | Tanespimycin |
| MVP_top60_DNA_seed | ATM | -0.051489 | BRCA | HCC1569 | KU-60019 | MVP_top60_DNA+RNA_seed | AKT1 | -0.351138 | BRCA | COLO-824 | BAY AKT1 |
| MVP_top60_DNA_seed | MDM2 | -0.050959 | BRCA | Hs-578-T | Serdemetan | MVP_top60_DNA+RNA_seed | ATM | -0.349512 | BRCA | AU565 | CP466722 |
| MVP_top60_DNA_seed | MDM2 | -0.046605 | BRCA | MCF7 | Serdemetan | MVP_top60_DNA+RNA_seed | MDM2 | -0.349217 | BRCA | HCC70 | Nutlin-3a (-) |
| MVP_top60_DNA_seed | MDM2 | -0.042679 | BRCA | MDA-MB-436 | Serdemetan | MVP_top60_DNA+RNA_seed | CDK2 | -0.346458 | BRCA | CAL-148 | Seliciclib |
| MVP_top60_DNA_seed | HSP90AA1 | -0.040835 | BRCA | MDA-MB-330 | SNX-2112 | MVP_top60_DNA+RNA_seed | AR | -0.344532 | BRCA | Hs-578-T | Bicalutamide |
| MVP_top60_DNA_seed | HSP90AB1 | -0.040835 | BRCA | MDA-MB-330 | SNX-2112 | MVP_top60_DNA+RNA_seed | EGFR | -0.343659 | BRCA | HCC1569 | Pelitinib |
| MVP_top60_DNA_seed | HDAC1 | -0.039788 | BRCA | MRK-nu-1 | Entinostat | MVP_top60_DNA+RNA_seed | EGFR | -0.341543 | BRCA | HCC1937 | PF-00299804 |
| MVP_top60_DNA_seed | ERBB2 | -0.03723 | BRCA | MDA-MB-157 | CUDC-101 | MVP_top60_DNA+RNA_seed | ERBB2 | -0.341543 | BRCA | HCC1937 | PF-00299804 |
| MVP_top60_DNA_seed | EGFR | -0.03723 | BRCA | MDA-MB-157 | CUDC-101 | MVP_top60_DNA+RNA_seed | HDAC1 | -0.339472 | BRCA | HCC1428 | Belinostat |
| MVP_top60_DNA_seed | ERBB2 | -0.03415 | BRCA | CAL-51 | CI-1033 | MVP_top60_DNA+RNA_seed | HDAC1 | -0.336182 | BRCA | CAL-148 | Entinostat |
| MVP_top60_DNA_seed | EGFR | -0.03415 | BRCA | CAL-51 | CI-1033 | MVP_top60_DNA+RNA_seed | MAPK14 | -0.334738 | BRCA | HCC202 | TAK-715 |
| MVP_top60_DNA_seed | HDAC1 | -0.033859 | BRCA | DU-4475 | JQ12 | MVP_top60_DNA+RNA_seed | PLK1 | -0.334662 | BRCA | HCC1187 | GW843682X |
| MVP_top60_DNA_seed | HDAC2 | -0.033859 | BRCA | DU-4475 | JQ12 | MVP_top60_DNA+RNA_seed | EGFR | -0.331544 | BRCA | DU-4475 | AST-1306 |
| MVP_top60_DNA_seed | AKT1 | -0.032697 | BRCA | CAMA-1 | MK-2206 | MVP_top60_DNA+RNA_seed | SRC | -0.330301 | BRCA | HCC1187 | NVP-BHG712 |
| MVP_top60_DNA_seed | ATM | -0.030413 | BRCA | HCC1937 | KU-60019 | MVP_top60_DNA+RNA_seed | HSP90AA1 | -0.327699 | BRCA | CAL-120 | Luminespib |
| MVP_top60_DNA_seed | HSP90AA1 | -0.030141 | BRCA | MDA-MB-415 | Tanespimycin | MVP_top60_DNA+RNA_seed | HSP90AB1 | -0.327699 | BRCA | CAL-120 | Luminespib |
| MVP_top60_DNA_seed | HSP90AB1 | -0.030141 | BRCA | MDA-MB-415 | Tanespimycin | MVP_top60_DNA+RNA_seed | HDAC1 | -0.327204 | BRCA | EFM-19 | CAY10603 |
| MVP_top60_DNA_seed | AKT1 | -0.029633 | BRCA | CAL-148 | AKT inhibitor VIII | MVP_top60_DNA+RNA_seed | HSP90AA1 | -0.326974 | BRCA | MDA-MB-453 | Elesclomol |
| MVP_top60_DNA_seed | ERBB2 | -0.027313 | BRCA | MFM-223 | CUDC-101 | MVP_top60_DNA+RNA_seed | HSP90AB1 | -0.326974 | BRCA | MDA-MB-453 | Elesclomol |
| MVP_top60_DNA_seed | EGFR | -0.027313 | BRCA | MFM-223 | CUDC-101 | MVP_top60_DNA+RNA_seed | EGFR | -0.326213 | BRCA | COLO-824 | AZD8931 |
| MVP_top60_DNA_seed | ESR1 | -0.027226 | BRCA | MDA-MB-453 | Fulvestrant | MVP_top60_DNA+RNA_seed | ERBB2 | -0.326213 | BRCA | COLO-824 | AZD8931 |
| MVP_top60_DNA_seed | AKT1 | -0.026829 | BRCA | BT-549 | BAY AKT1 | MVP_top60_DNA+RNA_seed | ERBB3 | -0.326213 | BRCA | COLO-824 | AZD8931 |
| MVP_top60_DNA_seed | ATM | -0.02663 | BRCA | UACC-893 | torin2 | MVP_top60_DNA+RNA_seed | HSP90AA1 | -0.32541 | BRCA | MRK-nu-1 | SNX-2112 |
| MVP_top60_DNA_seed | AR | -0.026229 | BRCA | MFM-223 | Bicalutamide | MVP_top60_DNA+RNA_seed | HSP90AB1 | -0.32541 | BRCA | MRK-nu-1 | SNX-2112 |
| MVP_top60_DNA_seed | MDM2 | -0.024945 | BRCA | HCC1187 | Serdemetan | MVP_top60_DNA+RNA_seed | CDK2 | -0.323705 | BRCA | MDA-MB-453 | PHA-793887 |
| MVP_top60_DNA_seed | ATM | -0.024624 | BRCA | HCC38 | KU-55933 | MVP_top60_DNA+RNA_seed | MDM2 | -0.322278 | BRCA | MDA-MB-175-VII | Nutlin-3a (-) |
| MVP_top60_DNA_seed | ATM | -0.024072 | BRCA | BT-549 | KU-60019 | MVP_top60_DNA+RNA_seed | HDAC1 | -0.317886 | BRCA | CAL-120 | CAY10603 |
| MVP_top60_DNA_seed | HSP90AA1 | -0.023728 | BRCA | MDA-MB-175-VII | SNX-2112 | MVP_top60_DNA+RNA_seed | AKT1 | -0.317256 | BRCA | HCC1419 | GSK690693 |
| MVP_top60_DNA_seed | HSP90AB1 | -0.023728 | BRCA | MDA-MB-175-VII | SNX-2112 | MVP_top60_DNA+RNA_seed | EGFR | -0.316624 | BRCA | BT-549 | CI-1033 |
| MVP_top60_DNA_seed | EGFR | -0.022716 | BRCA | MDA-MB-453 | Pelitinib | MVP_top60_DNA+RNA_seed | ERBB2 | -0.316624 | BRCA | BT-549 | CI-1033 |
| MVP_top60_DNA_seed | ERBB2 | -0.022567 | BRCA | BT-483 | CP724714 | MVP_top60_DNA+RNA_seed | HDAC1 | -0.316158 | BRCA | CAL-148 | Dacinostat |
| MVP_top60_DNA_seed | PIK3CA | -0.021242 | BRCA | HCC1143 | AZD8835 | MVP_top60_DNA+RNA_seed | ESR1 | -0.315186 | BRCA | EFM-192A | Tamoxifen |
| MVP_top60_DNA_seed | ERBB2 | -0.021151 | BRCA | CAL-51 | CP724714 | MVP_top60_DNA+RNA_seed | EGFR | -0.315164 | BRCA | HCC38 | CUDC-101 |
| MVP_top60_DNA_seed | SRC | -0.021041 | BRCA | HCC1954 | Bosutinib | MVP_top60_DNA+RNA_seed | ERBB2 | -0.315164 | BRCA | HCC38 | CUDC-101 |
| MVP_top60_DNA_seed | HDAC1 | -0.020495 | BRCA | HCC2218 | Tubastatin A | MVP_top60_DNA+RNA_seed | EGFR | -0.312949 | BRCA | MDA-MB-453 | AZD8931 |
| MVP_top60_DNA_seed | ERBB2 | -0.019741 | BRCA | COLO-824 | CUDC-101 | MVP_top60_DNA+RNA_seed | ERBB2 | -0.312949 | BRCA | MDA-MB-453 | AZD8931 |
| MVP_top60_DNA_seed | EGFR | -0.019741 | BRCA | COLO-824 | CUDC-101 | MVP_top60_DNA+RNA_seed | ERBB3 | -0.312949 | BRCA | MDA-MB-453 | AZD8931 |
| MVP_top60_DNA_seed | ERBB2 | -0.017937 | BRCA | EVSA-T | Afatinib | MVP_top60_DNA+RNA_seed | HSP90AA1 | -0.309917 | BRCA | HCC1143 | Tanespimycin |
| MVP_top60_DNA_seed | EGFR | -0.017937 | BRCA | EVSA-T | Afatinib | MVP_top60_DNA+RNA_seed | HSP90AB1 | -0.309917 | BRCA | HCC1143 | Tanespimycin |

| | | | | | | | | | | | |
|---|---|---|---|---|---|---|---|---|---|---|---|
| MVP_top60_DNA_seed | AKT1 | -0.017303 | BRCA | Hs-578-T | BAY AKT1 | MVP_top60_DNA+RNA_seed | HSP90AA1 | -0.309789 | BRCA | BT-549 | CCT-018159 |
| MVP_top60_DNA_seed | AR | -0.016785 | BRCA | HCC202 | AZD3514 | MVP_top60_DNA+RNA_seed | HSP90AB1 | -0.309789 | BRCA | BT-549 | CCT-018159 |
| MVP_top60_DNA_seed | AKT1 | -0.016601 | BRCA | MFM-223 | A-443654 | MVP_top60_DNA+RNA_seed | MAPK1 | -0.309321 | BRCA | OCUB-M | VX-11e |
| MVP_top60_DNA_seed | AKT1 | -0.014649 | BRCA | DU-4475 | A-443654 | MVP_top60_DNA+RNA_seed | AKT1 | -0.309279 | BRCA | CAMA-1 | AKT inhibitor VIII |
| MVP_top60_DNA_seed | HDAC1 | -0.012392 | BRCA | BT-474 | Dacinostat | MVP_top60_DNA+RNA_seed | HSP90AA1 | -0.309029 | BRCA | MDA-MB-330 | Luminespib |
| MVP_top60_DNA_seed | HDAC1 | -0.01202 | BRCA | HCC2157 | Tubastatin A | MVP_top60_DNA+RNA_seed | HSP90AB1 | -0.309029 | BRCA | MDA-MB-330 | Luminespib |
| MVP_top60_DNA_seed | EP300 | -0.01052 | BRCA | HCC1500 | I-CBP112 | MVP_top60_DNA+RNA_seed | EGFR | -0.307356 | BRCA | T47D | Afatinib |
| MVP_top60_DNA_seed | CREBBP | -0.01052 | BRCA | HCC1500 | I-CBP112 | MVP_top60_DNA+RNA_seed | ERBB2 | -0.307356 | BRCA | T47D | Afatinib |
| MVP_top60_DNA_seed | AKT1 | -0.009745 | BRCA | CAMA-1 | AKT inhibitor VIII | MVP_top60_DNA+RNA_seed | MDM2 | -0.30579 | BRCA | HCC38 | Serdemetan |
| MVP_top60_DNA_seed | HDAC1 | -0.00784 | BRCA | EVSA-T | Dacinostat | MVP_top60_DNA+RNA_seed | EGFR | -0.304055 | BRCA | HDQ-P1 | AST-1306 |
| MVP_top60_DNA_seed | EP300 | -0.006419 | BRCA | HCC1187 | I-CBP112 | MVP_top60_DNA+RNA_seed | HDAC1 | -0.303572 | BRCA | HCC70 | Tubastatin A |
| MVP_top60_DNA_seed | CREBBP | -0.006419 | BRCA | HCC1187 | I-CBP112 | MVP_top60_DNA+RNA_seed | HDAC1 | -0.302652 | BRCA | HCC1500 | Dacinostat |
| MVP_top60_DNA_seed | HDAC1 | -0.004466 | BRCA | DU-4475 | CAY10603 | MVP_top60_DNA+RNA_seed | HDAC1 | -0.30263 | BRCA | EVSA-T | Entinostat |
| MVP_top60_DNA_seed | HDAC1 | -0.003002 | BRCA | COLO-824 | Belinostat | MVP_top60_DNA+RNA_seed | HSP90AA1 | -0.301063 | BRCA | HCC38 | Tanespimycin |
| MVP_top60_DNA_seed | HDAC1 | -0.002936 | BRCA | T47D | JQ12 | MVP_top60_DNA+RNA_seed | HSP90AB1 | -0.301063 | BRCA | HCC38 | Tanespimycin |
| MVP_top60_DNA_seed | HDAC2 | -0.002936 | BRCA | T47D | JQ12 | MVP_top60_DNA+RNA_seed | AKT1 | -0.30011 | BRCA | MDA-MB-468 | AKT inhibitor VIII |
| MVP_top60_DNA_seed | SRC | -0.00118 | BRCA | OCUB-M | WH-4-023 | MVP_top60_DNA+RNA_seed | EGFR | -0.299922 | BRCA | HCC70 | AST-1306 |
| MVP_top60_DNA_seed | EGFR | 0.000901 | BRCA | MDA-MB-231 | Cetuximab | MVP_top60_DNA+RNA_seed | PIK3CA | -0.297693 | BRCA | MDA-MB-175-VII | AZD8835 |
| MVP_top60_DNA_seed | HDAC1 | 0.000929 | BRCA | MFM-223 | CAY10603 | MVP_top60_DNA+RNA_seed | MDM2 | -0.296172 | BRCA | HCC1569 | Serdemetan |
| MVP_top60_DNA_seed | HSP90AA1 | 0.001793 | BRCA | ZR-75-30 | SNX-2112 | MVP_top60_DNA+RNA_seed | ERBB2 | -0.295236 | BRCA | T47D | CP724714 |
| MVP_top60_DNA_seed | HSP90AB1 | 0.001793 | BRCA | ZR-75-30 | SNX-2112 | MVP_top60_DNA+RNA_seed | PLK1 | -0.292947 | BRCA | HCC1187 | BI-2536 |
| MVP_top60_DNA_seed | ERBB2 | 0.002719 | BRCA | CAL-120 | AZD8931 | MVP_top60_DNA+RNA_seed | HDAC1 | -0.292521 | BRCA | CAMA-1 | Dacinostat |
| MVP_top60_DNA_seed | EGFR | 0.002719 | BRCA | CAL-120 | AZD8931 | MVP_top60_DNA+RNA_seed | HDAC1 | -0.291465 | BRCA | ZR-75-30 | CAY10603 |
| MVP_top60_DNA_seed | ERBB3 | 0.002719 | BRCA | CAL-120 | AZD8931 | MVP_top60_DNA+RNA_seed | ATM | -0.289568 | BRCA | HCC1428 | KU-60019 |
| MVP_top60_DNA_seed | AKT1 | 0.002886 | BRCA | MFM-223 | AKT inhibitor VIII | MVP_top60_DNA+RNA_seed | HSP90AA1 | -0.288799 | BRCA | BT-474 | CCT-018159 |
| MVP_top60_DNA_seed | ATM | 0.003691 | BRCA | UACC-893 | KU-60019 | MVP_top60_DNA+RNA_seed | HSP90AB1 | -0.288799 | BRCA | BT-474 | CCT-018159 |
| MVP_top60_DNA_seed | AKT1 | 0.006228 | BRCA | HCC202 | MK-2206 | MVP_top60_DNA+RNA_seed | EGFR | -0.288053 | BRCA | DU-4475 | CUDC-101 |
| MVP_top60_DNA_seed | MAPK3 | 0.006984 | BRCA | EVSA-T | FR-180204 | MVP_top60_DNA+RNA_seed | ERBB2 | -0.288053 | BRCA | DU-4475 | CUDC-101 |
| MVP_top60_DNA_seed | MAPK1 | 0.006984 | BRCA | EVSA-T | FR-180204 | MVP_top60_DNA+RNA_seed | HSP90AA1 | -0.287209 | BRCA | MDA-MB-453 | Tanespimycin |
| MVP_top60_DNA_seed | HDAC1 | 0.00842 | BRCA | HCC1419 | AR-42 | MVP_top60_DNA+RNA_seed | HSP90AB1 | -0.287209 | BRCA | MDA-MB-453 | Tanespimycin |
| MVP_top60_DNA_seed | ERBB2 | 0.008665 | BRCA | BT-20 | CI-1033 | MVP_top60_DNA+RNA_seed | EGFR | -0.287018 | BRCA | HCC2157 | Gefitinib |
| MVP_top60_DNA_seed | EGFR | 0.008665 | BRCA | BT-20 | CI-1033 | MVP_top60_DNA+RNA_seed | HDAC1 | -0.286241 | BRCA | T47D | CAY10603 |
| MVP_top60_DNA_seed | HSP90AA1 | 0.008874 | BRCA | MDA-MB-468 | CCT-018159 | MVP_top60_DNA+RNA_seed | AR | -0.285949 | BRCA | COLO-824 | Bicalutamide |
| MVP_top60_DNA_seed | HSP90AB1 | 0.008874 | BRCA | MDA-MB-468 | CCT-018159 | MVP_top60_DNA+RNA_seed | AR | -0.285626 | BRCA | HCC2218 | AZD3514 |
| MVP_top60_DNA_seed | AKT1 | 0.009825 | BRCA | AU565 | GSK690693 | MVP_top60_DNA+RNA_seed | AR | -0.284682 | BRCA | DU-4475 | Bicalutamide |
| MVP_top60_DNA_seed | MAPK3 | 0.010996 | BRCA | AU565 | FR-180204 | MVP_top60_DNA+RNA_seed | CDK2 | -0.283028 | BRCA | T47D | PHA-793887 |
| MVP_top60_DNA_seed | MAPK1 | 0.010996 | BRCA | AU565 | FR-180204 | MVP_top60_DNA+RNA_seed | EP300 | -0.282905 | BRCA | DU-4475 | I-CBP112 |
| MVP_top60_DNA_seed | ERBB2 | 0.014621 | BRCA | MDA-MB-330 | CP724714 | MVP_top60_DNA+RNA_seed | HDAC1 | -0.281786 | BRCA | HCC202 | CAY10603 |
| MVP_top60_DNA_seed | ATM | 0.014622 | BRCA | HCC1954 | KU-60019 | MVP_top60_DNA+RNA_seed | MDM2 | -0.279105 | BRCA | MRK-nu-1 | Nutlin-3a (-) |
| MVP_top60_DNA_seed | SRC | 0.016492 | BRCA | MDA-MB-175-VII | NVP-BHG712 | MVP_top60_DNA+RNA_seed | HDAC1 | -0.274384 | BRCA | HCC1569 | Dacinostat |
| MVP_top60_DNA_seed | AKT1 | 0.016839 | BRCA | MDA-MB-436 | BAY AKT1 | MVP_top60_DNA+RNA_seed | HSP90AA1 | -0.272654 | BRCA | ZR-75-30 | Luminespib |

| | | | | | | | | | | | |
|---|---|---|---|---|---|---|---|---|---|---|---|
| MVP_top60_DNA_seed | PIK3CA | 0.017213 | BRCA | HCC2157 | AZD8835 | MVP_top60_DNA+RNA_seed | HSP90AB1 | -0.272654 | BRCA | ZR-75-30 | Luminespib |
| MVP_top60_DNA_seed | SRC | 0.018149 | BRCA | HCC1187 | Bosutinib | MVP_top60_DNA+RNA_seed | PLK1 | -0.271905 | BRCA | MCF7 | PLK_6522 |
| MVP_top60_DNA_seed | ERBB2 | 0.018477 | BRCA | MDA-MB-231 | CP724714 | MVP_top60_DNA+RNA_seed | EGFR | -0.27155 | BRCA | MDA-MB-157 | Afatinib |
| MVP_top60_DNA_seed | AKT1 | 0.020005 | BRCA | HCC1937 | AKT inhibitor VIII | MVP_top60_DNA+RNA_seed | ERBB2 | -0.27155 | BRCA | MDA-MB-157 | Afatinib |
| MVP_top60_DNA_seed | MDM2 | 0.021967 | BRCA | HCC70 | Serdemetan | MVP_top60_DNA+RNA_seed | AR | -0.270686 | BRCA | MDA-MB-453 | AZD3514 |
| MVP_top60_DNA_seed | SRC | 0.022424 | BRCA | DU-4475 | Bosutinib | MVP_top60_DNA+RNA_seed | SRC | -0.270308 | BRCA | HDQ-P1 | Bosutinib |
| MVP_top60_DNA_seed | ERBB2 | 0.022436 | BRCA | CAL-51 | AZD8931 | MVP_top60_DNA+RNA_seed | MDM2 | -0.269769 | BRCA | MDA-MB-157 | Nutlin-3a (-) |
| MVP_top60_DNA_seed | EGFR | 0.022436 | BRCA | CAL-51 | AZD8931 | MVP_top60_DNA+RNA_seed | AR | -0.268955 | BRCA | CAL-85-1 | AZD3514 |
| MVP_top60_DNA_seed | ERBB3 | 0.022436 | BRCA | CAL-51 | AZD8931 | MVP_top60_DNA+RNA_seed | AKT1 | -0.267296 | BRCA | CAL-51 | BAY AKT1 |
| MVP_top60_DNA_seed | HSP90AA1 | 0.022726 | BRCA | JIMT-1 | CCT-018159 | MVP_top60_DNA+RNA_seed | AKT1 | -0.265996 | BRCA | CAL-51 | GSK690693 |
| MVP_top60_DNA_seed | HSP90AB1 | 0.022726 | BRCA | JIMT-1 | CCT-018159 | MVP_top60_DNA+RNA_seed | EGFR | -0.264912 | BRCA | COLO-824 | Afatinib |
| MVP_top60_DNA_seed | HDAC1 | 0.023683 | BRCA | HCC70 | Dacinostat | MVP_top60_DNA+RNA_seed | ERBB2 | -0.264912 | BRCA | COLO-824 | Afatinib |
| MVP_top60_DNA_seed | SRC | 0.024022 | BRCA | EVSA-T | WH-4-023 | MVP_top60_DNA+RNA_seed | AKT1 | -0.263242 | BRCA | CAL-85-1 | GSK690693 |
| MVP_top60_DNA_seed | SRC | 0.025152 | BRCA | UACC-893 | NVP-BHG712 | MVP_top60_DNA+RNA_seed | ERBB2 | -0.262769 | BRCA | HCC1569 | CP724714 |
| MVP_top60_DNA_seed | HDAC1 | 0.02672 | BRCA | CAL-51 | Dacinostat | MVP_top60_DNA+RNA_seed | SRC | -0.262509 | BRCA | CAL-120 | NVP-BHG712 |
| MVP_top60_DNA_seed | AKT1 | 0.027052 | BRCA | BT-483 | AKT inhibitor VIII | MVP_top60_DNA+RNA_seed | ESR1 | -0.262397 | BRCA | CAL-51 | Fulvestrant |
| MVP_top60_DNA_seed | ERBB2 | 0.027379 | BRCA | MFM-223 | CI-1033 | MVP_top60_DNA+RNA_seed | PLK1 | -0.253283 | BRCA | MDA-MB-231 | PLK_6522 |
| MVP_top60_DNA_seed | EGFR | 0.027379 | BRCA | MFM-223 | CI-1033 | MVP_top60_DNA+RNA_seed | AKT1 | -0.253143 | BRCA | CAL-85-1 | AKT inhibitor VIII |
| MVP_top60_DNA_seed | AR | 0.02763 | BRCA | Hs-578-T | AZD3514 | MVP_top60_DNA+RNA_seed | ERBB2 | -0.252589 | BRCA | HCC1500 | CP724714 |
| MVP_top60_DNA_seed | SRC | 0.028744 | BRCA | MRK-nu-1 | WH-4-023 | MVP_top60_DNA+RNA_seed | EGFR | -0.252435 | BRCA | HCC1187 | Afatinib |
| MVP_top60_DNA_seed | HDAC1 | 0.030211 | BRCA | MDA-MB-157 | CAY10603 | MVP_top60_DNA+RNA_seed | ERBB2 | -0.252435 | BRCA | HCC1187 | Afatinib |
| MVP_top60_DNA_seed | AR | 0.030327 | BRCA | BT-549 | AZD3514 | MVP_top60_DNA+RNA_seed | ATM | -0.251995 | BRCA | BT-20 | CP466722 |
| MVP_top60_DNA_seed | ERBB2 | 0.030356 | BRCA | MDA-MB-231 | CUDC-101 | MVP_top60_DNA+RNA_seed | SRC | -0.251688 | BRCA | MDA-MB-361 | Bosutinib |
| MVP_top60_DNA_seed | EGFR | 0.030356 | BRCA | MDA-MB-231 | CUDC-101 | MVP_top60_DNA+RNA_seed | ATM | -0.251051 | BRCA | EFM-192A | CP466722 |
| MVP_top60_DNA_seed | EGFR | 0.031371 | BRCA | CAL-148 | Erlotinib | MVP_top60_DNA+RNA_seed | EGFR | -0.250836 | BRCA | MDA-MB-361 | Afatinib |
| MVP_top60_DNA_seed | HSP90AA1 | 0.032538 | BRCA | HCC70 | Luminespib | MVP_top60_DNA+RNA_seed | ERBB2 | -0.250836 | BRCA | MDA-MB-361 | Afatinib |
| MVP_top60_DNA_seed | HSP90AB1 | 0.032538 | BRCA | HCC70 | Luminespib | MVP_top60_DNA+RNA_seed | AR | -0.250694 | BRCA | MDA-MB-436 | AZD3514 |
| MVP_top60_DNA_seed | HSP90AA1 | 0.036266 | BRCA | HCC1187 | Elesclomol | MVP_top60_DNA+RNA_seed | MAPK1 | -0.250553 | BRCA | MDA-MB-175-VII | VX-11e |
| MVP_top60_DNA_seed | HSP90AB1 | 0.036266 | BRCA | HCC1187 | Elesclomol | MVP_top60_DNA+RNA_seed | SRC | -0.24862 | BRCA | MDA-MB-468 | Bosutinib |
| MVP_top60_DNA_seed | EGFR | 0.03818 | BRCA | EFM-192A | Cetuximab | MVP_top60_DNA+RNA_seed | MAPK3 | -0.24839 | BRCA | MDA-MB-361 | FR-180204 |
| MVP_top60_DNA_seed | HDAC1 | 0.041152 | BRCA | COLO-824 | CAY10603 | MVP_top60_DNA+RNA_seed | MAPK1 | -0.24839 | BRCA | MDA-MB-361 | FR-180204 |
| MVP_top60_DNA_seed | HDAC1 | 0.042391 | BRCA | HCC1428 | CAY10603 | MVP_top60_DNA+RNA_seed | HSP90AA1 | -0.247247 | BRCA | HCC1937 | CCT-018159 |
| MVP_top60_DNA_seed | HDAC1 | 0.043389 | BRCA | HCC70 | AR-42 | MVP_top60_DNA+RNA_seed | HSP90AB1 | -0.247247 | BRCA | HCC1937 | CCT-018159 |
| MVP_top60_DNA_seed | PIK3CA | 0.04396 | BRCA | HCC1143 | PI3Ka_4409 | MVP_top60_DNA+RNA_seed | AKT1 | -0.247109 | BRCA | HCC1187 | MK-2206 |
| MVP_top60_DNA_seed | AKT1 | 0.04507 | BRCA | JIMT-1 | AKT inhibitor VIII | MVP_top60_DNA+RNA_seed | HDAC1 | -0.246923 | BRCA | EFM-19 | AR-42 |
| MVP_top60_DNA_seed | AR | 0.045406 | BRCA | MDA-MB-468 | Bicalutamide | MVP_top60_DNA+RNA_seed | EGFR | -0.244686 | BRCA | EFM-19 | CI-1033 |
| MVP_top60_DNA_seed | MDM2 | 0.045815 | BRCA | HCC2157 | Serdemetan | MVP_top60_DNA+RNA_seed | ERBB2 | -0.244686 | BRCA | EFM-19 | CI-1033 |
| MVP_top60_DNA_seed | MAPK1 | 0.046108 | BRCA | MDA-MB-468 | VX-11e | MVP_top60_DNA+RNA_seed | HDAC1 | -0.243026 | BRCA | HCC2157 | Entinostat |
| MVP_top60_DNA_seed | ATM | 0.046754 | BRCA | UACC-893 | CP466722 | MVP_top60_DNA+RNA_seed | EGFR | -0.242339 | BRCA | HCC70 | Cetuximab |
| MVP_top60_DNA_seed | AKT1 | 0.046837 | BRCA | MDA-MB-157 | MK-2206 | MVP_top60_DNA+RNA_seed | PIK3CA | -0.24184 | BRCA | AU565 | AZD8835 |
| MVP_top60_DNA_seed | HSP90AA1 | 0.047191 | BRCA | HCC1599 | Luminespib | MVP_top60_DNA+RNA_seed | AKT1 | -0.240728 | BRCA | HCC38 | AKT inhibitor VIII |

| | | | | | | | | | | | |
|---|---|---|---|---|---|---|---|---|---|---|---|
| MVP_top60_DNA_seed | HSP90AB1 | 0.047191 | BRCA | HCC1599 | Luminespib | MVP_top60_DNA+RNA_seed | EGFR | -0.240265 | BRCA | BT-20 | Afatinib |
| MVP_top60_DNA_seed | MDM2 | 0.047969 | BRCA | MRK-nu-1 | Serdemetan | MVP_top60_DNA+RNA_seed | ERBB2 | -0.240265 | BRCA | BT-20 | Afatinib |
| MVP_top60_DNA_seed | MDM2 | 0.048309 | BRCA | HCC1569 | Nutlin-3a (-) | MVP_top60_DNA+RNA_seed | HDAC1 | -0.239537 | BRCA | MDA-MB-361 | CAY10603 |
| MVP_top60_DNA_seed | AKT1 | 0.048326 | BRCA | HCC1937 | MK-2206 | MVP_top60_DNA+RNA_seed | EGFR | -0.238543 | BRCA | HCC1428 | PF-00299804 |
| MVP_top60_DNA_seed | ERBB2 | 0.051118 | BRCA | CAL-120 | Afatinib | MVP_top60_DNA+RNA_seed | ERBB2 | -0.238543 | BRCA | HCC1428 | PF-00299804 |
| MVP_top60_DNA_seed | EGFR | 0.051118 | BRCA | CAL-120 | Afatinib | MVP_top60_DNA+RNA_seed | HSP90AA1 | -0.237874 | BRCA | BT-20 | CCT-018159 |
| MVP_top60_DNA_seed | HSP90AA1 | 0.053462 | BRCA | COLO-824 | Luminespib | MVP_top60_DNA+RNA_seed | HSP90AB1 | -0.237874 | BRCA | BT-20 | CCT-018159 |
| MVP_top60_DNA_seed | HSP90AB1 | 0.053462 | BRCA | COLO-824 | Luminespib | MVP_top60_DNA+RNA_seed | ATM | -0.2371 | BRCA | HCC38 | CP466722 |
| MVP_top60_DNA_seed | MAPK3 | 0.055312 | BRCA | EFM-192A | FR-180204 | MVP_top60_DNA+RNA_seed | AKT1 | -0.234712 | BRCA | HCC2218 | AKT inhibitor VIII |
| MVP_top60_DNA_seed | MAPK1 | 0.055312 | BRCA | EFM-192A | FR-180204 | MVP_top60_DNA+RNA_seed | PIK3CA | -0.234688 | BRCA | MDA-MB-175-VII | PI3Ka_4409 |
| MVP_top60_DNA_seed | HDAC1 | 0.055563 | BRCA | MDA-MB-330 | CAY10603 | MVP_top60_DNA+RNA_seed | EP300 | -0.233121 | BRCA | BT-483 | I-CBP112 |
| MVP_top60_DNA_seed | ATM | 0.05617 | BRCA | MRK-nu-1 | KU-60019 | MVP_top60_DNA+RNA_seed | HDAC1 | -0.232503 | BRCA | HCC1500 | Belinostat |
| MVP_top60_DNA_seed | ATM | 0.056716 | BRCA | HCC1395 | KU-60019 | MVP_top60_DNA+RNA_seed | HDAC1 | -0.231314 | BRCA | BT-474 | CAY10603 |
| MVP_top60_DNA_seed | CDK2 | 0.056791 | BRCA | HCC1395 | AZD5438 | MVP_top60_DNA+RNA_seed | EGFR | -0.22943 | BRCA | BT-549 | AST-1306 |
| MVP_top60_DNA_seed | MDM2 | 0.057521 | BRCA | EFM-192A | Nutlin-3a (-) | MVP_top60_DNA+RNA_seed | MDM2 | -0.228918 | BRCA | T47D | Serdemetan |
| MVP_top60_DNA_seed | ERBB2 | 0.057974 | BRCA | BT-483 | PF-00299804 | MVP_top60_DNA+RNA_seed | HDAC1 | -0.227297 | BRCA | HCC202 | Tubastatin A |
| MVP_top60_DNA_seed | EGFR | 0.057974 | BRCA | BT-483 | PF-00299804 | MVP_top60_DNA+RNA_seed | EGFR | -0.224574 | BRCA | HCC1500 | CI-1033 |
| MVP_top60_DNA_seed | HSP90AA1 | 0.058874 | BRCA | HCC1143 | Elesclomol | MVP_top60_DNA+RNA_seed | ERBB2 | -0.224574 | BRCA | HCC1500 | CI-1033 |
| MVP_top60_DNA_seed | HSP90AB1 | 0.058874 | BRCA | HCC1143 | Elesclomol | MVP_top60_DNA+RNA_seed | ATM | -0.220896 | BRCA | JIMT-1 | CP466722 |
| MVP_top60_DNA_seed | HSP90AA1 | 0.060412 | BRCA | UACC-893 | Luminespib | MVP_top60_DNA+RNA_seed | HDAC1 | -0.216818 | BRCA | MDA-MB-330 | Belinostat |
| MVP_top60_DNA_seed | HSP90AB1 | 0.060412 | BRCA | UACC-893 | Luminespib | MVP_top60_DNA+RNA_seed | AKT1 | -0.216371 | BRCA | COLO-824 | AKT inhibitor VIII |
| MVP_top60_DNA_seed | AKT1 | 0.062104 | BRCA | HCC1599 | AKT inhibitor VIII | MVP_top60_DNA+RNA_seed | AKT1 | -0.214062 | BRCA | HCC1395 | AKT inhibitor VIII |
| MVP_top60_DNA_seed | PIK3CA | 0.062903 | BRCA | MDA-MB-415 | PI3Ka_4409 | MVP_top60_DNA+RNA_seed | MDM2 | -0.212422 | BRCA | HCC1937 | Serdemetan |
| MVP_top60_DNA_seed | MDM2 | 0.066885 | BRCA | CAMA-1 | Serdemetan | MVP_top60_DNA+RNA_seed | AKT1 | -0.211224 | BRCA | MDA-MB-453 | MK-2206 |
| MVP_top60_DNA_seed | SRC | 0.067162 | BRCA | OCUB-M | Saracatinib | MVP_top60_DNA+RNA_seed | AKT1 | -0.209807 | BRCA | EFM-192A | AKT inhibitor VIII |
| MVP_top60_DNA_seed | HDAC1 | 0.067624 | BRCA | MRK-nu-1 | JQ12 | MVP_top60_DNA+RNA_seed | EGFR | -0.208009 | BRCA | BT-549 | PF-00299804 |
| MVP_top60_DNA_seed | HDAC2 | 0.067624 | BRCA | MRK-nu-1 | JQ12 | MVP_top60_DNA+RNA_seed | ERBB2 | -0.208009 | BRCA | BT-549 | PF-00299804 |
| MVP_top60_DNA_seed | CDK2 | 0.068255 | BRCA | YMB-1-E | PHA-793887 | MVP_top60_DNA+RNA_seed | CDK1 | -0.207887 | BRCA | HCC70 | RO-3306 |
| MVP_top60_DNA_seed | ERBB2 | 0.069502 | BRCA | MCF7 | AZD8931 | MVP_top60_DNA+RNA_seed | ESR1 | -0.206923 | BRCA | HCC1599 | Fulvestrant |
| MVP_top60_DNA_seed | EGFR | 0.069502 | BRCA | MCF7 | AZD8931 | MVP_top60_DNA+RNA_seed | MDM2 | -0.20646 | BRCA | CAL-120 | Serdemetan |
| MVP_top60_DNA_seed | ERBB3 | 0.069502 | BRCA | MCF7 | AZD8931 | MVP_top60_DNA+RNA_seed | HSP90AA1 | -0.205534 | BRCA | BT-474 | Luminespib |
| MVP_top60_DNA_seed | ESR1 | 0.069655 | BRCA | MDA-MB-415 | Tamoxifen | MVP_top60_DNA+RNA_seed | HSP90AB1 | -0.205534 | BRCA | BT-474 | Luminespib |
| MVP_top60_DNA_seed | PIK3CA | 0.070522 | BRCA | HCC2157 | PI3Ka_4409 | MVP_top60_DNA+RNA_seed | ESR1 | -0.205499 | BRCA | HCC70 | Fulvestrant |
| MVP_top60_DNA_seed | AKT1 | 0.071425 | BRCA | CAL-51 | AKT inhibitor VIII | MVP_top60_DNA+RNA_seed | CDK2 | -0.205035 | BRCA | OCUB-M | Seliciclib |
| MVP_top60_DNA_seed | ESR1 | 0.072287 | BRCA | MDA-MB-330 | Fulvestrant | MVP_top60_DNA+RNA_seed | HSP90AA1 | -0.204579 | BRCA | CAL-148 | Luminespib |
| MVP_top60_DNA_seed | AKT1 | 0.072316 | BRCA | ZR-75-30 | MK-2206 | MVP_top60_DNA+RNA_seed | HSP90AB1 | -0.204579 | BRCA | CAL-148 | Luminespib |
| MVP_top60_DNA_seed | SRC | 0.07244 | BRCA | JIMT-1 | NVP-BHG712 | MVP_top60_DNA+RNA_seed | EGFR | -0.204165 | BRCA | HCC70 | CUDC-101 |
| MVP_top60_DNA_seed | CDK1 | 0.074795 | BRCA | EFM-192A | RO-3306 | MVP_top60_DNA+RNA_seed | ERBB2 | -0.204165 | BRCA | HCC70 | CUDC-101 |
| MVP_top60_DNA_seed | CDK2 | 0.07494 | BRCA | MCF7 | AZD5438 | MVP_top60_DNA+RNA_seed | CDK2 | -0.203436 | BRCA | MDA-MB-231 | AZD5438 |
| MVP_top60_DNA_seed | AKT1 | 0.075811 | BRCA | HCC2157 | AKT inhibitor VIII | MVP_top60_DNA+RNA_seed | HDAC1 | -0.202709 | BRCA | MDA-MB-330 | Tubastatin A |
| MVP_top60_DNA_seed | PIK3CA | 0.077335 | BRCA | COLO-824 | AZD8835 | MVP_top60_DNA+RNA_seed | SRC | -0.202072 | BRCA | MRK-nu-1 | NVP-BHG712 |

| | | | | | | | | | | | |
|---|---|---|---|---|---|---|---|---|---|---|---|
| MVP_top60_DNA_seed | HSP90AA1 | 0.077646 | BRCA | HCC1428 | CCT-018159 | MVP_top60_DNA+RNA_seed | HDAC1 | -0.201216 | BRCA | JIMT-1 | JQ12 |
| MVP_top60_DNA_seed | HSP90AB1 | 0.077646 | BRCA | HCC1428 | CCT-018159 | MVP_top60_DNA+RNA_seed | HDAC2 | -0.201216 | BRCA | JIMT-1 | JQ12 |
| MVP_top60_DNA_seed | HSP90AA1 | 0.077779 | BRCA | CAL-120 | Tanespimycin | MVP_top60_DNA+RNA_seed | EGFR | -0.201204 | BRCA | CAMA-1 | AST-1306 |
| MVP_top60_DNA_seed | HSP90AB1 | 0.077779 | BRCA | CAL-120 | Tanespimycin | MVP_top60_DNA+RNA_seed | ESR1 | -0.200337 | BRCA | MDA-MB-436 | Fulvestrant |
| MVP_top60_DNA_seed | ATM | 0.079098 | BRCA | MDA-MB-175-VII | KU-60019 | MVP_top60_DNA+RNA_seed | HDAC1 | -0.200168 | BRCA | MDA-MB-453 | Dacinostat |
| MVP_top60_DNA_seed | AKT1 | 0.07947 | BRCA | MDA-MB-436 | MK-2206 | MVP_top60_DNA+RNA_seed | HDAC1 | -0.198542 | BRCA | CAMA-1 | CAY10603 |
| MVP_top60_DNA_seed | EGFR | 0.081099 | BRCA | HCC1143 | Gefitinib | MVP_top60_DNA+RNA_seed | AR | -0.198342 | BRCA | MDA-MB-453 | Bicalutamide |
| MVP_top60_DNA_seed | AKT1 | 0.081173 | BRCA | DU-4475 | GSK690693 | MVP_top60_DNA+RNA_seed | PLK1 | -0.197318 | BRCA | CAL-148 | BI-2536 |
| MVP_top60_DNA_seed | MDM2 | 0.081959 | BRCA | MDA-MB-330 | Nutlin-3a (-) | MVP_top60_DNA+RNA_seed | AR | -0.196671 | BRCA | MDA-MB-415 | AZD3514 |
| MVP_top60_DNA_seed | AKT1 | 0.083383 | BRCA | UACC-812 | GSK690693 | MVP_top60_DNA+RNA_seed | EGFR | -0.196267 | BRCA | HCC1937 | AST-1306 |
| MVP_top60_DNA_seed | ATM | 0.086015 | BRCA | MDA-MB-415 | torin2 | MVP_top60_DNA+RNA_seed | CDK2 | -0.193133 | BRCA | CAL-85-1 | PHA-793887 |
| MVP_top60_DNA_seed | MDM2 | 0.086929 | BRCA | CAL-120 | Nutlin-3a (-) | MVP_top60_DNA+RNA_seed | HDAC1 | -0.191114 | BRCA | AU565 | CAY10603 |
| MVP_top60_DNA_seed | ERBB2 | 0.087979 | BRCA | CAMA-1 | Afatinib | MVP_top60_DNA+RNA_seed | PLK1 | -0.18996 | BRCA | CAL-148 | GW843682X |
| MVP_top60_DNA_seed | EGFR | 0.087979 | BRCA | CAMA-1 | Afatinib | MVP_top60_DNA+RNA_seed | EGFR | -0.189242 | BRCA | HCC1937 | Pelitinib |
| MVP_top60_DNA_seed | EGFR | 0.088177 | BRCA | MCF7 | Gefitinib | MVP_top60_DNA+RNA_seed | HDAC1 | -0.188971 | BRCA | HCC38 | Belinostat |
| MVP_top60_DNA_seed | MDM2 | 0.089254 | BRCA | HCC1143 | Serdemetan | MVP_top60_DNA+RNA_seed | EGFR | -0.188355 | BRCA | JIMT-1 | Pelitinib |
| MVP_top60_DNA_seed | ERBB2 | 0.089457 | BRCA | MCF7 | PF-00299804 | MVP_top60_DNA+RNA_seed | PIK3CA | -0.185494 | BRCA | HCC1569 | PI3Ka_4409 |
| MVP_top60_DNA_seed | EGFR | 0.089457 | BRCA | MCF7 | PF-00299804 | MVP_top60_DNA+RNA_seed | ESR1 | -0.184403 | BRCA | HDQ-P1 | Fulvestrant |
| MVP_top60_DNA_seed | PIK3CA | 0.089588 | BRCA | BT-549 | PI3Ka_4409 | MVP_top60_DNA+RNA_seed | PIK3CA | -0.182922 | BRCA | CAL-51 | PI3Ka_4409 |
| MVP_top60_DNA_seed | AR | 0.090976 | BRCA | MDA-MB-436 | Bicalutamide | MVP_top60_DNA+RNA_seed | MAPK14 | -0.182244 | BRCA | CAMA-1 | TAK-715 |
| MVP_top60_DNA_seed | ERBB2 | 0.091137 | BRCA | MDA-MB-231 | PF-00299804 | MVP_top60_DNA+RNA_seed | EGFR | -0.182163 | BRCA | HCC1937 | Gefitinib |
| MVP_top60_DNA_seed | EGFR | 0.091137 | BRCA | MDA-MB-231 | PF-00299804 | MVP_top60_DNA+RNA_seed | EGFR | -0.181836 | BRCA | HCC1395 | AZD8931 |
| MVP_top60_DNA_seed | ATM | 0.091322 | BRCA | HCC38 | KU-60019 | MVP_top60_DNA+RNA_seed | ERBB2 | -0.181836 | BRCA | HCC1395 | AZD8931 |
| MVP_top60_DNA_seed | HDAC1 | 0.091611 | BRCA | HCC1500 | CAY10603 | MVP_top60_DNA+RNA_seed | ERBB3 | -0.181836 | BRCA | HCC1395 | AZD8931 |
| MVP_top60_DNA_seed | AKT1 | 0.09276 | BRCA | MRK-nu-1 | A-443654 | MVP_top60_DNA+RNA_seed | MAPK14 | -0.181781 | BRCA | MFM-223 | TAK-715 |
| MVP_top60_DNA_seed | AR | 0.094804 | BRCA | HCC1500 | Bicalutamide | MVP_top60_DNA+RNA_seed | MAPK3 | -0.181278 | BRCA | HDQ-P1 | FR-180204 |
| MVP_top60_DNA_seed | MAPK1 | 0.094908 | BRCA | HCC1599 | VX-11e | MVP_top60_DNA+RNA_seed | MAPK1 | -0.181278 | BRCA | HDQ-P1 | FR-180204 |
| MVP_top60_DNA_seed | AR | 0.095665 | BRCA | T47D | AZD3514 | MVP_top60_DNA+RNA_seed | ATM | -0.179992 | BRCA | HCC1428 | CP466722 |
| MVP_top60_DNA_seed | PIK3CA | 0.096349 | BRCA | HCC1569 | AZD8835 | MVP_top60_DNA+RNA_seed | ESR1 | -0.178356 | BRCA | HCC1395 | Fulvestrant |
| MVP_top60_DNA_seed | MAPK3 | 0.096423 | BRCA | UACC-812 | FR-180204 | MVP_top60_DNA+RNA_seed | CDK2 | -0.177967 | BRCA | CAL-120 | AZD5438 |
| MVP_top60_DNA_seed | MAPK1 | 0.096423 | BRCA | UACC-812 | FR-180204 | MVP_top60_DNA+RNA_seed | AR | -0.177442 | BRCA | HCC70 | Bicalutamide |
| MVP_top60_DNA_seed | HSP90AA1 | 0.097434 | BRCA | AU565 | Elesclomol | MVP_top60_DNA+RNA_seed | EGFR | -0.176416 | BRCA | DU-4475 | Lapatinib |
| MVP_top60_DNA_seed | HSP90AB1 | 0.097434 | BRCA | AU565 | Elesclomol | MVP_top60_DNA+RNA_seed | ERBB2 | -0.176416 | BRCA | DU-4475 | Lapatinib |
| MVP_top60_DNA_seed | MAPK1 | 0.09773 | BRCA | UACC-893 | VX-11e | MVP_top60_DNA+RNA_seed | MAPK1 | -0.174389 | BRCA | EVSA-T | VX-11e |
| MVP_top60_DNA_seed | ERBB2 | 0.098501 | BRCA | CAMA-1 | CP724714 | MVP_top60_DNA+RNA_seed | EGFR | -0.173906 | BRCA | HCC1500 | CUDC-101 |
| MVP_top60_DNA_seed | ATM | 0.099442 | BRCA | OCUB-M | KU-55933 | MVP_top60_DNA+RNA_seed | ERBB2 | -0.173906 | BRCA | HCC1500 | CUDC-101 |
| MVP_top60_DNA_seed | EGFR | 0.100288 | BRCA | HCC1954 | Cetuximab | MVP_top60_DNA+RNA_seed | HDAC1 | -0.173043 | BRCA | BT-474 | Tubastatin A |
| MVP_top60_DNA_seed | HSP90AA1 | 0.100619 | BRCA | HCC1419 | Luminespib | MVP_top60_DNA+RNA_seed | EGFR | -0.172309 | BRCA | CAL-85-1 | AST-1306 |
| MVP_top60_DNA_seed | HSP90AB1 | 0.100619 | BRCA | HCC1419 | Luminespib | MVP_top60_DNA+RNA_seed | AKT1 | -0.171418 | BRCA | BT-20 | MK-2206 |
| MVP_top60_DNA_seed | HDAC1 | 0.103185 | BRCA | EFM-192A | CAY10603 | MVP_top60_DNA+RNA_seed | HDAC1 | -0.171175 | BRCA | HCC1187 | Parthenolide |
| MVP_top60_DNA_seed | ERBB2 | 0.10463 | BRCA | HCC1937 | CI-1033 | MVP_top60_DNA+RNA_seed | PIK3CA | -0.171098 | BRCA | CAL-120 | AZD8835 |

| | | | | | | | | | | | |
|---|---|---|---|---|---|---|---|---|---|---|---|
| MVP_top60_DNA_seed | EGFR | 0.10463 | BRCA | HCC1937 | CI-1033 | MVP_top60_DNA+RNA_seed | EGFR | -0.170635 | BRCA | HCC1500 | AST-1306 |
| MVP_top60_DNA_seed | ERBB2 | 0.106027 | BRCA | MFM-223 | PF-00299804 | MVP_top60_DNA+RNA_seed | AKT1 | -0.166268 | BRCA | JIMT-1 | AKT inhibitor VIII |
| MVP_top60_DNA_seed | EGFR | 0.106027 | BRCA | MFM-223 | PF-00299804 | MVP_top60_DNA+RNA_seed | HDAC1 | -0.164273 | BRCA | HCC70 | JQ12 |
| MVP_top60_DNA_seed | AKT1 | 0.106134 | BRCA | HCC1937 | BAY AKT1 | MVP_top60_DNA+RNA_seed | HDAC2 | -0.164273 | BRCA | HCC70 | JQ12 |
| MVP_top60_DNA_seed | EP300 | 0.108224 | BRCA | MDA-MB-453 | I-CBP112 | MVP_top60_DNA+RNA_seed | HDAC1 | -0.162591 | BRCA | HCC1569 | CAY10603 |
| MVP_top60_DNA_seed | CREBBP | 0.108224 | BRCA | MDA-MB-453 | I-CBP112 | MVP_top60_DNA+RNA_seed | AKT1 | -0.159768 | BRCA | MCF7 | AKT inhibitor VIII |
| MVP_top60_DNA_seed | HSP90AA1 | 0.109041 | BRCA | COLO-824 | Elesclomol | MVP_top60_DNA+RNA_seed | ATM | -0.159495 | BRCA | CAL-120 | CP466722 |
| MVP_top60_DNA_seed | HSP90AB1 | 0.109041 | BRCA | COLO-824 | Elesclomol | MVP_top60_DNA+RNA_seed | EGFR | -0.159472 | BRCA | HCC70 | CI-1033 |
| MVP_top60_DNA_seed | HDAC1 | 0.10997 | BRCA | COLO-824 | AR-42 | MVP_top60_DNA+RNA_seed | ERBB2 | -0.159472 | BRCA | HCC70 | CI-1033 |
| MVP_top60_DNA_seed | HDAC1 | 0.111111 | BRCA | CAL-120 | Belinostat | MVP_top60_DNA+RNA_seed | EP300 | -0.158065 | BRCA | MDA-MB-361 | I-CBP112 |
| MVP_top60_DNA_seed | HDAC1 | 0.112541 | BRCA | EFM-192A | AR-42 | MVP_top60_DNA+RNA_seed | ATM | -0.158058 | BRCA | MDA-MB-231 | CP466722 |
| MVP_top60_DNA_seed | CDK2 | 0.114245 | BRCA | JIMT-1 | AZD5438 | MVP_top60_DNA+RNA_seed | MAPK14 | -0.157427 | BRCA | EVSA-T | TAK-715 |
| MVP_top60_DNA_seed | ERBB2 | 0.114469 | BRCA | HCC1395 | CI-1033 | MVP_top60_DNA+RNA_seed | MAPK3 | -0.156945 | BRCA | HCC202 | FR-180204 |
| MVP_top60_DNA_seed | EGFR | 0.114469 | BRCA | HCC1395 | CI-1033 | MVP_top60_DNA+RNA_seed | MAPK1 | -0.156945 | BRCA | HCC202 | FR-180204 |
| MVP_top60_DNA_seed | HDAC1 | 0.114589 | BRCA | CAL-120 | Tubastatin A | MVP_top60_DNA+RNA_seed | HDAC1 | -0.155937 | BRCA | ZR-75-30 | Tubastatin A |
| MVP_top60_DNA_seed | ESR1 | 0.115618 | BRCA | MDA-MB-468 | Tamoxifen | MVP_top60_DNA+RNA_seed | EGFR | -0.152968 | BRCA | OCUB-M | Gefitinib |
| MVP_top60_DNA_seed | MDM2 | 0.116123 | BRCA | HCC1419 | Serdemetan | MVP_top60_DNA+RNA_seed | SRC | -0.15267 | BRCA | MDA-MB-157 | Bosutinib |
| MVP_top60_DNA_seed | ERBB2 | 0.116567 | BRCA | HCC1937 | CUDC-101 | MVP_top60_DNA+RNA_seed | AR | -0.151122 | BRCA | DU-4475 | AZD3514 |
| MVP_top60_DNA_seed | EGFR | 0.116567 | BRCA | HCC1937 | CUDC-101 | MVP_top60_DNA+RNA_seed | PIK3CA | -0.14942 | BRCA | CAL-85-1 | PI3Ka_4409 |
| MVP_top60_DNA_seed | EGFR | 0.1175 | BRCA | HCC1937 | Cetuximab | MVP_top60_DNA+RNA_seed | PIK3CA | -0.149209 | BRCA | AU565 | PI3Ka_4409 |
| MVP_top60_DNA_seed | HDAC1 | 0.121783 | BRCA | HCC1937 | Dacinostat | MVP_top60_DNA+RNA_seed | HDAC1 | -0.148313 | BRCA | EFM-192A | Tubastatin A |
| MVP_top60_DNA_seed | EGFR | 0.122589 | BRCA | CAL-51 | Cetuximab | MVP_top60_DNA+RNA_seed | EGFR | -0.146968 | BRCA | HCC1143 | Afatinib |
| MVP_top60_DNA_seed | AKT1 | 0.123049 | BRCA | HCC1500 | AKT inhibitor VIII | MVP_top60_DNA+RNA_seed | ERBB2 | -0.146968 | BRCA | HCC1143 | Afatinib |
| MVP_top60_DNA_seed | ERBB2 | 0.126355 | BRCA | CAL-148 | CP724714 | MVP_top60_DNA+RNA_seed | EP300 | -0.146959 | BRCA | EVSA-T | I-CBP112 |
| MVP_top60_DNA_seed | HDAC1 | 0.126956 | BRCA | UACC-812 | AR-42 | MVP_top60_DNA+RNA_seed | MAPK1 | -0.146919 | BRCA | BT-474 | VX-11e |
| MVP_top60_DNA_seed | ERBB2 | 0.130992 | BRCA | MDA-MB-436 | PF-00299804 | MVP_top60_DNA+RNA_seed | EGFR | -0.143765 | BRCA | HCC1500 | Pelitinib |
| MVP_top60_DNA_seed | EGFR | 0.130992 | BRCA | MDA-MB-436 | PF-00299804 | MVP_top60_DNA+RNA_seed | HSP90AA1 | -0.142816 | BRCA | EVSA-T | Elesclomol |
| MVP_top60_DNA_seed | AR | 0.131472 | BRCA | MCF7 | Bicalutamide | MVP_top60_DNA+RNA_seed | HSP90AB1 | -0.142816 | BRCA | EVSA-T | Elesclomol |
| MVP_top60_DNA_seed | SRC | 0.132356 | BRCA | CAL-85-1 | NVP-BHG712 | MVP_top60_DNA+RNA_seed | EP300 | -0.141759 | BRCA | HCC2218 | I-CBP112 |
| MVP_top60_DNA_seed | HDAC1 | 0.133469 | BRCA | MDA-MB-231 | Dacinostat | MVP_top60_DNA+RNA_seed | HSP90AA1 | -0.141117 | BRCA | COLO-824 | CCT-018159 |
| MVP_top60_DNA_seed | MDM2 | 0.134263 | BRCA | HCC1500 | Nutlin-3a (-) | MVP_top60_DNA+RNA_seed | HSP90AB1 | -0.141117 | BRCA | COLO-824 | CCT-018159 |
| MVP_top60_DNA_seed | ERBB2 | 0.134345 | BRCA | MDA-MB-415 | PF-00299804 | MVP_top60_DNA+RNA_seed | EGFR | -0.140173 | BRCA | HCC38 | AZD8931 |
| MVP_top60_DNA_seed | EGFR | 0.134345 | BRCA | MDA-MB-415 | PF-00299804 | MVP_top60_DNA+RNA_seed | ERBB2 | -0.140173 | BRCA | HCC38 | AZD8931 |
| MVP_top60_DNA_seed | ERBB2 | 0.134791 | BRCA | MDA-MB-436 | Afatinib | MVP_top60_DNA+RNA_seed | ERBB3 | -0.140173 | BRCA | HCC38 | AZD8931 |
| MVP_top60_DNA_seed | EGFR | 0.134791 | BRCA | MDA-MB-436 | Afatinib | MVP_top60_DNA+RNA_seed | HDAC1 | -0.140055 | BRCA | MDA-MB-231 | AR-42 |
| MVP_top60_DNA_seed | ERBB2 | 0.135771 | BRCA | HCC2218 | CUDC-101 | MVP_top60_DNA+RNA_seed | HSP90AA1 | -0.139833 | BRCA | BT-20 | Elesclomol |
| MVP_top60_DNA_seed | EGFR | 0.135771 | BRCA | HCC2218 | CUDC-101 | MVP_top60_DNA+RNA_seed | HSP90AB1 | -0.139833 | BRCA | BT-20 | Elesclomol |
| MVP_top60_DNA_seed | ERBB2 | 0.136246 | BRCA | MDA-MB-436 | CI-1033 | MVP_top60_DNA+RNA_seed | EP300 | -0.139638 | BRCA | HCC1569 | I-CBP112 |
| MVP_top60_DNA_seed | EGFR | 0.136246 | BRCA | MDA-MB-436 | CI-1033 | MVP_top60_DNA+RNA_seed | EGFR | -0.139142 | BRCA | CAMA-1 | CUDC-101 |
| MVP_top60_DNA_seed | CDK2 | 0.136873 | BRCA | AU565 | PHA-793887 | MVP_top60_DNA+RNA_seed | ERBB2 | -0.139142 | BRCA | CAMA-1 | CUDC-101 |
| MVP_top60_DNA_seed | PIK3CA | 0.140748 | BRCA | COLO-824 | PI3Ka_4409 | MVP_top60_DNA+RNA_seed | HDAC1 | -0.139049 | BRCA | JIMT-1 | Dacinostat |

| | | | | | | | | | | | |
|---|---|---|---|---|---|---|---|---|---|---|---|
| MVP_top60_DNA_seed | MDM2 | 0.141089 | BRCA | COLO-824 | Serdemetan | MVP_top60_DNA+RNA_seed | HSP90AA1 | -0.138766 | BRCA | HCC70 | Tanespimycin |
| MVP_top60_DNA_seed | AKT1 | 0.141542 | BRCA | BT-474 | A-443654 | MVP_top60_DNA+RNA_seed | HSP90AB1 | -0.138766 | BRCA | HCC70 | Tanespimycin |
| MVP_top60_DNA_seed | HSP90AA1 | 0.141877 | BRCA | CAMA-1 | Luminespib | MVP_top60_DNA+RNA_seed | CDK1 | -0.137951 | BRCA | HCC1954 | RO-3306 |
| MVP_top60_DNA_seed | HSP90AB1 | 0.141877 | BRCA | CAMA-1 | Luminespib | MVP_top60_DNA+RNA_seed | ATM | -0.137945 | BRCA | MDA-MB-468 | KU-60019 |
| MVP_top60_DNA_seed | CDK2 | 0.143741 | BRCA | BT-474 | PHA-793887 | MVP_top60_DNA+RNA_seed | AR | -0.137819 | BRCA | CAMA-1 | Bicalutamide |
| MVP_top60_DNA_seed | AKT1 | 0.14389 | BRCA | JIMT-1 | GSK690693 | MVP_top60_DNA+RNA_seed | PLK1 | -0.137538 | BRCA | Hs-578-T | PLK_6522 |
| MVP_top60_DNA_seed | HSP90AA1 | 0.144053 | BRCA | EFM-19 | Elesclomol | MVP_top60_DNA+RNA_seed | EGFR | -0.137279 | BRCA | HCC2218 | AZD8931 |
| MVP_top60_DNA_seed | HSP90AB1 | 0.144053 | BRCA | EFM-19 | Elesclomol | MVP_top60_DNA+RNA_seed | ERBB2 | -0.137279 | BRCA | HCC2218 | AZD8931 |
| MVP_top60_DNA_seed | ERBB2 | 0.144631 | BRCA | HDQ-P1 | CUDC-101 | MVP_top60_DNA+RNA_seed | ERBB3 | -0.137279 | BRCA | HCC2218 | AZD8931 |
| MVP_top60_DNA_seed | EGFR | 0.144631 | BRCA | HDQ-P1 | CUDC-101 | MVP_top60_DNA+RNA_seed | ATM | -0.136084 | BRCA | HDQ-P1 | CP466722 |
| MVP_top60_DNA_seed | SRC | 0.144917 | BRCA | BT-549 | NVP-BHG712 | MVP_top60_DNA+RNA_seed | ATM | -0.134713 | BRCA | BT-474 | CP466722 |
| MVP_top60_DNA_seed | HDAC1 | 0.145114 | BRCA | MDA-MB-157 | Belinostat | MVP_top60_DNA+RNA_seed | CDK2 | -0.133968 | BRCA | EVSA-T | PHA-793887 |
| MVP_top60_DNA_seed | ERBB2 | 0.146357 | BRCA | HCC1599 | CP724714 | MVP_top60_DNA+RNA_seed | EGFR | -0.133964 | BRCA | CAMA-1 | PF-00299804 |
| MVP_top60_DNA_seed | AKT1 | 0.146771 | BRCA | HCC2157 | BAY AKT1 | MVP_top60_DNA+RNA_seed | ERBB2 | -0.133964 | BRCA | CAMA-1 | PF-00299804 |
| MVP_top60_DNA_seed | HDAC1 | 0.1476 | BRCA | MDA-MB-415 | JQ12 | MVP_top60_DNA+RNA_seed | EGFR | -0.133527 | BRCA | DU-4475 | Pelitinib |
| MVP_top60_DNA_seed | HDAC2 | 0.1476 | BRCA | MDA-MB-415 | JQ12 | MVP_top60_DNA+RNA_seed | MAPK3 | -0.132932 | BRCA | HCC1187 | FR-180204 |
| MVP_top60_DNA_seed | SRC | 0.147974 | BRCA | OCUB-M | Bosutinib | MVP_top60_DNA+RNA_seed | MAPK1 | -0.132932 | BRCA | HCC1187 | FR-180204 |
| MVP_top60_DNA_seed | MDM2 | 0.147997 | BRCA | OCUB-M | Serdemetan | MVP_top60_DNA+RNA_seed | HDAC1 | -0.132579 | BRCA | MFM-223 | AR-42 |
| MVP_top60_DNA_seed | ERBB2 | 0.150503 | BRCA | MDA-MB-231 | CI-1033 | MVP_top60_DNA+RNA_seed | EP300 | -0.132404 | BRCA | MDA-MB-468 | I-CBP112 |
| MVP_top60_DNA_seed | EGFR | 0.150503 | BRCA | MDA-MB-231 | CI-1033 | MVP_top60_DNA+RNA_seed | ATM | -0.131959 | BRCA | BT-483 | CP466722 |
| MVP_top60_DNA_seed | AR | 0.151776 | BRCA | HCC1428 | AZD3514 | MVP_top60_DNA+RNA_seed | ESR1 | -0.131703 | BRCA | CAL-120 | Tamoxifen |
| MVP_top60_DNA_seed | EGFR | 0.155196 | BRCA | MDA-MB-436 | AST-1306 | MVP_top60_DNA+RNA_seed | HDAC1 | -0.131661 | BRCA | MFM-223 | Belinostat |
| MVP_top60_DNA_seed | SRC | 0.159669 | BRCA | MDA-MB-361 | NVP-BHG712 | MVP_top60_DNA+RNA_seed | MDM2 | -0.128592 | BRCA | OCUB-M | Nutlin-3a (-) |
| MVP_top60_DNA_seed | MAPK3 | 0.164286 | BRCA | MDA-MB-468 | FR-180204 | MVP_top60_DNA+RNA_seed | EGFR | -0.12757 | BRCA | HCC1428 | AST-1306 |
| MVP_top60_DNA_seed | MAPK1 | 0.164286 | BRCA | MDA-MB-468 | FR-180204 | MVP_top60_DNA+RNA_seed | EGFR | -0.127057 | BRCA | EVSA-T | PF-00299804 |
| MVP_top60_DNA_seed | AR | 0.164707 | BRCA | EVSA-T | Bicalutamide | MVP_top60_DNA+RNA_seed | ERBB2 | -0.127057 | BRCA | EVSA-T | PF-00299804 |
| MVP_top60_DNA_seed | SRC | 0.165582 | BRCA | HCC70 | NVP-BHG712 | MVP_top60_DNA+RNA_seed | EGFR | -0.126935 | BRCA | HCC38 | Pelitinib |
| MVP_top60_DNA_seed | ATM | 0.166204 | BRCA | EFM-192A | KU-60019 | MVP_top60_DNA+RNA_seed | EGFR | -0.124578 | BRCA | HCC70 | PF-00299804 |
| MVP_top60_DNA_seed | ERBB2 | 0.167667 | BRCA | EVSA-T | CP724714 | MVP_top60_DNA+RNA_seed | ERBB2 | -0.124578 | BRCA | HCC70 | PF-00299804 |
| MVP_top60_DNA_seed | EGFR | 0.169938 | BRCA | OCUB-M | Erlotinib | MVP_top60_DNA+RNA_seed | HDAC1 | -0.123964 | BRCA | MDA-MB-361 | Tubastatin A |
| MVP_top60_DNA_seed | ERBB2 | 0.171885 | BRCA | HCC1937 | Afatinib | MVP_top60_DNA+RNA_seed | HSP90AA1 | -0.123933 | BRCA | BT-20 | Tanespimycin |
| MVP_top60_DNA_seed | EGFR | 0.171885 | BRCA | HCC1937 | Afatinib | MVP_top60_DNA+RNA_seed | HSP90AB1 | -0.123933 | BRCA | BT-20 | Tanespimycin |
| MVP_top60_DNA_seed | HDAC1 | 0.172948 | BRCA | HCC1937 | CAY10603 | MVP_top60_DNA+RNA_seed | HDAC1 | -0.123044 | BRCA | HCC2218 | CAY10603 |
| MVP_top60_DNA_seed | ATM | 0.174024 | BRCA | HCC1428 | KU-55933 | MVP_top60_DNA+RNA_seed | AKT1 | -0.122759 | BRCA | HCC1395 | BAY AKT1 |
| MVP_top60_DNA_seed | MDM2 | 0.175351 | BRCA | HCC1187 | Nutlin-3a (-) | MVP_top60_DNA+RNA_seed | EGFR | -0.121454 | BRCA | HCC1569 | CUDC-101 |
| MVP_top60_DNA_seed | MDM2 | 0.175657 | BRCA | CAMA-1 | Nutlin-3a (-) | MVP_top60_DNA+RNA_seed | ERBB2 | -0.121454 | BRCA | HCC1569 | CUDC-101 |
| MVP_top60_DNA_seed | ERBB2 | 0.178231 | BRCA | MDA-MB-453 | Afatinib | MVP_top60_DNA+RNA_seed | AR | -0.120436 | BRCA | MDA-MB-330 | Bicalutamide |
| MVP_top60_DNA_seed | EGFR | 0.178231 | BRCA | MDA-MB-453 | Afatinib | MVP_top60_DNA+RNA_seed | HSP90AA1 | -0.119617 | BRCA | HCC202 | SNX-2112 |
| MVP_top60_DNA_seed | SRC | 0.18019 | BRCA | HCC1500 | Bosutinib | MVP_top60_DNA+RNA_seed | HSP90AB1 | -0.119617 | BRCA | HCC202 | SNX-2112 |
| MVP_top60_DNA_seed | HSP90AA1 | 0.180214 | BRCA | ZR-75-30 | Tanespimycin | MVP_top60_DNA+RNA_seed | AR | -0.118924 | BRCA | MDA-MB-231 | AZD3514 |
| MVP_top60_DNA_seed | HSP90AB1 | 0.180214 | BRCA | ZR-75-30 | Tanespimycin | MVP_top60_DNA+RNA_seed | EGFR | -0.118894 | BRCA | CAL-51 | Gefitinib |

| | | | | | | | | | | | |
|---|---|---|---|---|---|---|---|---|---|---|---|
| MVP_top60_DNA_seed | ATM | 0.182381 | BRCA | HCC1937 | CP466722 | MVP_top60_DNA+RNA_seed | EGFR | -0.118836 | BRCA | MDA-MB-231 | AST-1306 |
| MVP_top60_DNA_seed | EGFR | 0.182819 | BRCA | HCC1500 | Gefitinib | MVP_top60_DNA+RNA_seed | ATM | -0.116796 | BRCA | MFM-223 | KU-55933 |
| MVP_top60_DNA_seed | HDAC1 | 0.182972 | BRCA | HCC1187 | Tubastatin A | MVP_top60_DNA+RNA_seed | EGFR | -0.116294 | BRCA | T47D | Afatinib |
| MVP_top60_DNA_seed | ERBB2 | 0.18347 | BRCA | CAL-51 | PF-00299804 | MVP_top60_DNA+RNA_seed | ERBB2 | -0.116294 | BRCA | T47D | Afatinib |
| MVP_top60_DNA_seed | EGFR | 0.18347 | BRCA | CAL-51 | PF-00299804 | MVP_top60_DNA+RNA_seed | AR | -0.116142 | BRCA | MDA-MB-361 | Bicalutamide |
| MVP_top60_DNA_seed | AKT1 | 0.18445 | BRCA | DU-4475 | BAY AKT1 | MVP_top60_DNA+RNA_seed | HDAC1 | -0.11562 | BRCA | BT-474 | Entinostat |
| MVP_top60_DNA_seed | ERBB2 | 0.185998 | BRCA | UACC-812 | PF-00299804 | MVP_top60_DNA+RNA_seed | SRC | -0.115369 | BRCA | EFM-19 | NVP-BHG712 |
| MVP_top60_DNA_seed | EGFR | 0.185998 | BRCA | UACC-812 | PF-00299804 | MVP_top60_DNA+RNA_seed | AR | -0.114835 | BRCA | JIMT-1 | AZD3514 |
| MVP_top60_DNA_seed | HSP90AA1 | 0.186117 | BRCA | HDQ-P1 | Tanespimycin | MVP_top60_DNA+RNA_seed | MDM2 | -0.113512 | BRCA | MDA-MB-468 | Serdemetan |
| MVP_top60_DNA_seed | HSP90AB1 | 0.186117 | BRCA | HDQ-P1 | Tanespimycin | MVP_top60_DNA+RNA_seed | HSP90AA1 | -0.113122 | BRCA | EFM-19 | Tanespimycin |
| MVP_top60_DNA_seed | HSP90AA1 | 0.186202 | BRCA | EVSA-T | SNX-2112 | MVP_top60_DNA+RNA_seed | HSP90AB1 | -0.113122 | BRCA | EFM-19 | Tanespimycin |
| MVP_top60_DNA_seed | HSP90AB1 | 0.186202 | BRCA | EVSA-T | SNX-2112 | MVP_top60_DNA+RNA_seed | HSP90AA1 | -0.112393 | BRCA | MDA-MB-468 | Luminespib |
| MVP_top60_DNA_seed | ATM | 0.187041 | BRCA | HCC1500 | KU-55933 | MVP_top60_DNA+RNA_seed | HSP90AB1 | -0.112393 | BRCA | MDA-MB-468 | Luminespib |
| MVP_top60_DNA_seed | ESR1 | 0.187775 | BRCA | MDA-MB-231 | Tamoxifen | MVP_top60_DNA+RNA_seed | ATM | -0.112313 | BRCA | MDA-MB-453 | CP466722 |
| MVP_top60_DNA_seed | CDK2 | 0.188939 | BRCA | HCC1500 | PHA-793887 | MVP_top60_DNA+RNA_seed | PIK3CA | -0.111097 | BRCA | HCC2218 | PI3Ka_4409 |
| MVP_top60_DNA_seed | HDAC1 | 0.189323 | BRCA | HCC1599 | Parthenolide | MVP_top60_DNA+RNA_seed | MDM2 | -0.110434 | BRCA | MDA-MB-157 | Serdemetan |
| MVP_top60_DNA_seed | MAPK3 | 0.189665 | BRCA | CAL-148 | FR-180204 | MVP_top60_DNA+RNA_seed | HSP90AA1 | -0.110167 | BRCA | MDA-MB-175-VII | CCT-018159 |
| MVP_top60_DNA_seed | MAPK1 | 0.189665 | BRCA | CAL-148 | FR-180204 | MVP_top60_DNA+RNA_seed | HSP90AB1 | -0.110167 | BRCA | MDA-MB-175-VII | CCT-018159 |
| MVP_top60_DNA_seed | ERBB2 | 0.191482 | BRCA | MCF7 | Afatinib | MVP_top60_DNA+RNA_seed | HSP90AA1 | -0.110099 | BRCA | CAL-120 | CCT-018159 |
| MVP_top60_DNA_seed | EGFR | 0.191482 | BRCA | MCF7 | Afatinib | MVP_top60_DNA+RNA_seed | HSP90AB1 | -0.110099 | BRCA | CAL-120 | CCT-018159 |
| MVP_top60_DNA_seed | ERBB2 | 0.191675 | BRCA | Hs-578-T | AZD8931 | MVP_top60_DNA+RNA_seed | EGFR | -0.110048 | BRCA | HDQ-P1 | Afatinib |
| MVP_top60_DNA_seed | EGFR | 0.191675 | BRCA | Hs-578-T | AZD8931 | MVP_top60_DNA+RNA_seed | ERBB2 | -0.110048 | BRCA | HDQ-P1 | Afatinib |
| MVP_top60_DNA_seed | ERBB3 | 0.191675 | BRCA | Hs-578-T | AZD8931 | MVP_top60_DNA+RNA_seed | HSP90AA1 | -0.109325 | BRCA | UACC-893 | Tanespimycin |
| MVP_top60_DNA_seed | EGFR | 0.194689 | BRCA | MDA-MB-361 | Cetuximab | MVP_top60_DNA+RNA_seed | HSP90AB1 | -0.109325 | BRCA | UACC-893 | Tanespimycin |
| MVP_top60_DNA_seed | HDAC1 | 0.194952 | BRCA | UACC-812 | Belinostat | MVP_top60_DNA+RNA_seed | SRC | -0.108857 | BRCA | EVSA-T | NVP-BHG712 |
| MVP_top60_DNA_seed | PIK3CA | 0.195146 | BRCA | CAL-120 | PI3Ka_4409 | MVP_top60_DNA+RNA_seed | EGFR | -0.104941 | BRCA | MCF7 | CI-1033 |
| MVP_top60_DNA_seed | SRC | 0.196157 | BRCA | HCC2218 | Bosutinib | MVP_top60_DNA+RNA_seed | ERBB2 | -0.104941 | BRCA | MCF7 | CI-1033 |
| MVP_top60_DNA_seed | HDAC1 | 0.196217 | BRCA | HCC1937 | Belinostat | MVP_top60_DNA+RNA_seed | HSP90AA1 | -0.102275 | BRCA | HCC1937 | Luminespib |
| MVP_top60_DNA_seed | ERBB2 | 0.196219 | BRCA | CAL-120 | CI-1033 | MVP_top60_DNA+RNA_seed | HSP90AB1 | -0.102275 | BRCA | HCC1937 | Luminespib |
| MVP_top60_DNA_seed | EGFR | 0.196219 | BRCA | CAL-120 | CI-1033 | MVP_top60_DNA+RNA_seed | MAPK3 | -0.099685 | BRCA | BT-549 | FR-180204 |
| MVP_top60_DNA_seed | HDAC1 | 0.196222 | BRCA | MDA-MB-175-VII | AR-42 | MVP_top60_DNA+RNA_seed | MAPK1 | -0.099685 | BRCA | BT-549 | FR-180204 |
| MVP_top60_DNA_seed | AR | 0.197116 | BRCA | HCC2157 | Bicalutamide | MVP_top60_DNA+RNA_seed | PLK1 | -0.096615 | BRCA | MDA-MB-436 | PLK_6522 |
| MVP_top60_DNA_seed | AKT1 | 0.199236 | BRCA | MDA-MB-157 | AKT inhibitor VIII | MVP_top60_DNA+RNA_seed | CDK2 | -0.095464 | BRCA | DU-4475 | AZD5438 |
| MVP_top60_DNA_seed | SRC | 0.200347 | BRCA | BT-474 | NVP-BHG712 | MVP_top60_DNA+RNA_seed | PLK1 | -0.095186 | BRCA | DU-4475 | BI-2536 |
| MVP_top60_DNA_seed | ATM | 0.202256 | BRCA | HCC1500 | KU-60019 | MVP_top60_DNA+RNA_seed | HDAC1 | -0.09419 | BRCA | HCC1569 | Belinostat |
| MVP_top60_DNA_seed | PIK3CA | 0.20947 | BRCA | DU-4475 | AZD8835 | MVP_top60_DNA+RNA_seed | HDAC1 | -0.093054 | BRCA | OCUB-M | JQ12 |
| MVP_top60_DNA_seed | AKT1 | 0.209578 | BRCA | HCC2157 | A-443654 | MVP_top60_DNA+RNA_seed | HDAC2 | -0.093054 | BRCA | OCUB-M | JQ12 |
| MVP_top60_DNA_seed | ERBB2 | 0.210319 | BRCA | MDA-MB-157 | AZD8931 | MVP_top60_DNA+RNA_seed | CDK2 | -0.090015 | BRCA | HCC1937 | PHA-793887 |
| MVP_top60_DNA_seed | EGFR | 0.210319 | BRCA | MDA-MB-157 | AZD8931 | MVP_top60_DNA+RNA_seed | HDAC1 | -0.088683 | BRCA | CAMA-1 | Belinostat |
| MVP_top60_DNA_seed | ERBB3 | 0.210319 | BRCA | MDA-MB-157 | AZD8931 | MVP_top60_DNA+RNA_seed | HDAC1 | -0.087323 | BRCA | HCC38 | AR-42 |
| MVP_top60_DNA_seed | AKT1 | 0.212269 | BRCA | HCC1500 | GSK690693 | MVP_top60_DNA+RNA_seed | HDAC1 | -0.086792 | BRCA | HCC1569 | AR-42 |

| | | | | | | | | | | | |
|---|---|---|---|---|---|---|---|---|---|---|---|
| MVP_top60_DNA_seed | CDK2 | 0.213091 | BRCA | MDA-MB-157 | AZD5438 | MVP_top60_DNA+RNA_seed | AKT1 | -0.085986 | BRCA | CAL-120 | BAY AKT1 |
| MVP_top60_DNA_seed | ESR1 | 0.213694 | BRCA | CAL-51 | Tamoxifen | MVP_top60_DNA+RNA_seed | CDK2 | -0.085464 | BRCA | HCC1187 | Seliciclib |
| MVP_top60_DNA_seed | CDK2 | 0.214181 | BRCA | HCC1569 | PHA-793887 | MVP_top60_DNA+RNA_seed | CDK2 | -0.085366 | BRCA | HCC38 | PHA-793887 |
| MVP_top60_DNA_seed | HDAC1 | 0.215845 | BRCA | MRK-nu-1 | CAY10603 | MVP_top60_DNA+RNA_seed | ATM | -0.084442 | BRCA | CAMA-1 | CP466722 |
| MVP_top60_DNA_seed | HSP90AA1 | 0.215933 | BRCA | HCC70 | SNX-2112 | MVP_top60_DNA+RNA_seed | MDM2 | -0.0826 | BRCA | HCC1395 | Serdemetan |
| MVP_top60_DNA_seed | HSP90AB1 | 0.215933 | BRCA | HCC70 | SNX-2112 | MVP_top60_DNA+RNA_seed | HSP90AA1 | -0.080607 | BRCA | MDA-MB-361 | CCT-018159 |
| MVP_top60_DNA_seed | ERBB2 | 0.215937 | BRCA | HCC1569 | AZD8931 | MVP_top60_DNA+RNA_seed | HSP90AB1 | -0.080607 | BRCA | MDA-MB-361 | CCT-018159 |
| MVP_top60_DNA_seed | EGFR | 0.215937 | BRCA | HCC1569 | AZD8931 | MVP_top60_DNA+RNA_seed | EGFR | -0.079762 | BRCA | MDA-MB-175-VII | CUDC-101 |
| MVP_top60_DNA_seed | ERBB3 | 0.215937 | BRCA | HCC1569 | AZD8931 | MVP_top60_DNA+RNA_seed | ERBB2 | -0.079762 | BRCA | MDA-MB-175-VII | CUDC-101 |
| MVP_top60_DNA_seed | ERBB2 | 0.21707 | BRCA | EFM-192A | CUDC-101 | MVP_top60_DNA+RNA_seed | AKT1 | -0.079275 | BRCA | CAL-120 | AKT inhibitor VIII |
| MVP_top60_DNA_seed | EGFR | 0.21707 | BRCA | EFM-192A | CUDC-101 | MVP_top60_DNA+RNA_seed | EGFR | -0.077216 | BRCA | COLO-824 | Pelitinib |
| MVP_top60_DNA_seed | HDAC1 | 0.217211 | BRCA | MDA-MB-157 | AR-42 | MVP_top60_DNA+RNA_seed | HSP90AA1 | -0.073555 | BRCA | HDQ-P1 | CCT-018159 |
| MVP_top60_DNA_seed | ESR1 | 0.218847 | BRCA | CAL-120 | Fulvestrant | MVP_top60_DNA+RNA_seed | HSP90AB1 | -0.073555 | BRCA | HDQ-P1 | CCT-018159 |
| MVP_top60_DNA_seed | ERBB2 | 0.219009 | BRCA | BT-549 | Afatinib | MVP_top60_DNA+RNA_seed | HSP90AA1 | -0.073326 | BRCA | MDA-MB-157 | CCT-018159 |
| MVP_top60_DNA_seed | EGFR | 0.219009 | BRCA | BT-549 | Afatinib | MVP_top60_DNA+RNA_seed | HSP90AB1 | -0.073326 | BRCA | MDA-MB-157 | CCT-018159 |
| MVP_top60_DNA_seed | CDK2 | 0.219732 | BRCA | HCC1599 | AZD5438 | MVP_top60_DNA+RNA_seed | CDK2 | -0.07315 | BRCA | HCC1143 | AZD5438 |
| MVP_top60_DNA_seed | CDK2 | 0.220263 | BRCA | BT-20 | PHA-793887 | MVP_top60_DNA+RNA_seed | EP300 | -0.073097 | BRCA | CAL-51 | I-CBP112 |
| MVP_top60_DNA_seed | ERBB2 | 0.223009 | BRCA | BT-549 | AZD8931 | MVP_top60_DNA+RNA_seed | EGFR | -0.071951 | BRCA | BT-20 | Pelitinib |
| MVP_top60_DNA_seed | EGFR | 0.223009 | BRCA | BT-549 | AZD8931 | MVP_top60_DNA+RNA_seed | ATM | -0.071417 | BRCA | JIMT-1 | KU-60019 |
| MVP_top60_DNA_seed | ERBB3 | 0.223009 | BRCA | BT-549 | AZD8931 | MVP_top60_DNA+RNA_seed | AR | -0.071413 | BRCA | EFM-192A | AZD3514 |
| MVP_top60_DNA_seed | HDAC1 | 0.223073 | BRCA | HDQ-P1 | CAY10603 | MVP_top60_DNA+RNA_seed | SRC | -0.069612 | BRCA | EVSA-T | Saracatinib |
| MVP_top60_DNA_seed | ERBB2 | 0.223207 | BRCA | HCC70 | Afatinib | MVP_top60_DNA+RNA_seed | ATM | -0.066741 | BRCA | MDA-MB-157 | KU-55933 |
| MVP_top60_DNA_seed | EGFR | 0.223207 | BRCA | HCC70 | Afatinib | MVP_top60_DNA+RNA_seed | HDAC1 | -0.064702 | BRCA | HCC1500 | Tubastatin A |
| MVP_top60_DNA_seed | EGFR | 0.223422 | BRCA | MDA-MB-453 | Cetuximab | MVP_top60_DNA+RNA_seed | EGFR | -0.063861 | BRCA | JIMT-1 | Afatinib |
| MVP_top60_DNA_seed | ERBB2 | 0.224232 | BRCA | DU-4475 | AZD8931 | MVP_top60_DNA+RNA_seed | ERBB2 | -0.063861 | BRCA | JIMT-1 | Afatinib |
| MVP_top60_DNA_seed | EGFR | 0.224232 | BRCA | DU-4475 | AZD8931 | MVP_top60_DNA+RNA_seed | HSP90AA1 | -0.062509 | BRCA | HCC2157 | Luminespib |
| MVP_top60_DNA_seed | ERBB3 | 0.224232 | BRCA | DU-4475 | AZD8931 | MVP_top60_DNA+RNA_seed | HSP90AB1 | -0.062509 | BRCA | HCC2157 | Luminespib |
| MVP_top60_DNA_seed | HDAC1 | 0.22814 | BRCA | HCC2218 | AR-42 | MVP_top60_DNA+RNA_seed | ATM | -0.059343 | BRCA | JIMT-1 | KU-55933 |
| MVP_top60_DNA_seed | HDAC1 | 0.228217 | BRCA | HDQ-P1 | Belinostat | MVP_top60_DNA+RNA_seed | EGFR | -0.058118 | BRCA | CAL-51 | Afatinib |
| MVP_top60_DNA_seed | ERBB2 | 0.228485 | BRCA | CAL-148 | PF-00299804 | MVP_top60_DNA+RNA_seed | ERBB2 | -0.058118 | BRCA | CAL-51 | Afatinib |
| MVP_top60_DNA_seed | EGFR | 0.228485 | BRCA | CAL-148 | PF-00299804 | MVP_top60_DNA+RNA_seed | EGFR | -0.057841 | BRCA | MDA-MB-330 | CUDC-101 |
| MVP_top60_DNA_seed | ERBB2 | 0.230246 | BRCA | HCC1143 | Afatinib | MVP_top60_DNA+RNA_seed | ERBB2 | -0.057841 | BRCA | MDA-MB-330 | CUDC-101 |
| MVP_top60_DNA_seed | EGFR | 0.230246 | BRCA | HCC1143 | Afatinib | MVP_top60_DNA+RNA_seed | PIK3CA | -0.057415 | BRCA | HCC70 | PI3Ka_4409 |
| MVP_top60_DNA_seed | HDAC1 | 0.230496 | BRCA | AU565 | Tubastatin A | MVP_top60_DNA+RNA_seed | AR | -0.056003 | BRCA | HCC1187 | AZD3514 |
| MVP_top60_DNA_seed | ERBB2 | 0.231426 | BRCA | HCC1599 | Lapatinib | MVP_top60_DNA+RNA_seed | HDAC1 | -0.055193 | BRCA | EFM-19 | Belinostat |
| MVP_top60_DNA_seed | EGFR | 0.231426 | BRCA | HCC1599 | Lapatinib | MVP_top60_DNA+RNA_seed | AKT1 | -0.054143 | BRCA | MDA-MB-468 | BAY AKT1 |
| MVP_top60_DNA_seed | HSP90AA1 | 0.231471 | BRCA | HCC2218 | SNX-2112 | MVP_top60_DNA+RNA_seed | MAPK14 | -0.053791 | BRCA | HCC70 | TAK-715 |
| MVP_top60_DNA_seed | HSP90AB1 | 0.231471 | BRCA | HCC2218 | SNX-2112 | MVP_top60_DNA+RNA_seed | HDAC1 | -0.052786 | BRCA | HCC38 | CAY10603 |
| MVP_top60_DNA_seed | ATM | 0.232119 | BRCA | HCC1937 | KU-55933 | MVP_top60_DNA+RNA_seed | ATM | -0.051489 | BRCA | HCC1569 | KU-60019 |
| MVP_top60_DNA_seed | EGFR | 0.23466 | BRCA | HCC1954 | Gefitinib | MVP_top60_DNA+RNA_seed | MDM2 | -0.050959 | BRCA | Hs-578-T | Serdemetan |
| MVP_top60_DNA_seed | AR | 0.237601 | BRCA | HCC202 | Bicalutamide | MVP_top60_DNA+RNA_seed | MDM2 | -0.046605 | BRCA | MCF7 | Serdemetan |

| | | | | | | | | | | | |
|---|---|---|---|---|---|---|---|---|---|---|---|
| MVP_top60_DNA_seed | AKT1 | 0.240572 | BRCA | CAL-51 | MK-2206 | MVP_top60_DNA+RNA_seed | MDM2 | -0.042679 | BRCA | MDA-MB-436 | Serdemetan |
| MVP_top60_DNA_seed | EGFR | 0.243617 | BRCA | HCC1599 | Erlotinib | MVP_top60_DNA+RNA_seed | HSP90AA1 | -0.040835 | BRCA | MDA-MB-330 | SNX-2112 |
| MVP_top60_DNA_seed | AKT1 | 0.243661 | BRCA | HCC1395 | GSK690693 | MVP_top60_DNA+RNA_seed | HSP90AB1 | -0.040835 | BRCA | MDA-MB-330 | SNX-2112 |
| MVP_top60_DNA_seed | AKT1 | 0.246166 | BRCA | HCC1954 | MK-2206 | MVP_top60_DNA+RNA_seed | HDAC1 | -0.039788 | BRCA | MRK-nu-1 | Entinostat |
| MVP_top60_DNA_seed | EGFR | 0.246834 | BRCA | MCF7 | AST-1306 | MVP_top60_DNA+RNA_seed | EGFR | -0.03723 | BRCA | MDA-MB-157 | CUDC-101 |
| MVP_top60_DNA_seed | ATM | 0.250078 | BRCA | HCC1187 | KU-60019 | MVP_top60_DNA+RNA_seed | ERBB2 | -0.03723 | BRCA | MDA-MB-157 | CUDC-101 |
| MVP_top60_DNA_seed | AR | 0.250352 | BRCA | HDQ-P1 | AZD3514 | MVP_top60_DNA+RNA_seed | EGFR | -0.03415 | BRCA | CAL-51 | CI-1033 |
| MVP_top60_DNA_seed | AR | 0.250388 | BRCA | JIMT-1 | Bicalutamide | MVP_top60_DNA+RNA_seed | ERBB2 | -0.03415 | BRCA | CAL-51 | CI-1033 |
| MVP_top60_DNA_seed | ERBB2 | 0.250815 | BRCA | HCC1954 | Afatinib | MVP_top60_DNA+RNA_seed | HDAC1 | -0.033859 | BRCA | DU-4475 | JQ12 |
| MVP_top60_DNA_seed | EGFR | 0.250815 | BRCA | HCC1954 | Afatinib | MVP_top60_DNA+RNA_seed | HDAC2 | -0.033859 | BRCA | DU-4475 | JQ12 |
| MVP_top60_DNA_seed | MAPK1 | 0.254507 | BRCA | MDA-MB-231 | VX-11e | MVP_top60_DNA+RNA_seed | AKT1 | -0.032697 | BRCA | CAMA-1 | MK-2206 |
| MVP_top60_DNA_seed | AKT1 | 0.254693 | BRCA | MDA-MB-231 | AKT inhibitor VIII | MVP_top60_DNA+RNA_seed | ATM | -0.030413 | BRCA | HCC1937 | KU-60019 |
| MVP_top60_DNA_seed | AR | 0.260478 | BRCA | MDA-MB-231 | Bicalutamide | MVP_top60_DNA+RNA_seed | HSP90AA1 | -0.030141 | BRCA | MDA-MB-415 | Tanespimycin |
| MVP_top60_DNA_seed | HSP90AA1 | 0.260855 | BRCA | HCC2218 | Luminespib | MVP_top60_DNA+RNA_seed | HSP90AB1 | -0.030141 | BRCA | MDA-MB-415 | Tanespimycin |
| MVP_top60_DNA_seed | HSP90AB1 | 0.260855 | BRCA | HCC2218 | Luminespib | MVP_top60_DNA+RNA_seed | AKT1 | -0.029633 | BRCA | CAL-148 | AKT inhibitor VIII |
| MVP_top60_DNA_seed | CDK2 | 0.262919 | BRCA | EVSA-T | AZD5438 | MVP_top60_DNA+RNA_seed | EGFR | -0.027313 | BRCA | MFM-223 | CUDC-101 |
| MVP_top60_DNA_seed | PIK3CA | 0.263913 | BRCA | Hs-578-T | PI3Ka_4409 | MVP_top60_DNA+RNA_seed | ERBB2 | -0.027313 | BRCA | MFM-223 | CUDC-101 |
| MVP_top60_DNA_seed | AKT1 | 0.26403 | BRCA | HCC1419 | BAY AKT1 | MVP_top60_DNA+RNA_seed | ESR1 | -0.027226 | BRCA | MDA-MB-453 | Fulvestrant |
| MVP_top60_DNA_seed | ERBB2 | 0.265165 | BRCA | CAMA-1 | AZD8931 | MVP_top60_DNA+RNA_seed | AKT1 | -0.026829 | BRCA | BT-549 | BAY AKT1 |
| MVP_top60_DNA_seed | EGFR | 0.265165 | BRCA | CAMA-1 | AZD8931 | MVP_top60_DNA+RNA_seed | ATM | -0.02663 | BRCA | UACC-893 | torin2 |
| MVP_top60_DNA_seed | ERBB3 | 0.265165 | BRCA | CAMA-1 | AZD8931 | MVP_top60_DNA+RNA_seed | AR | -0.026229 | BRCA | MFM-223 | Bicalutamide |
| MVP_top60_DNA_seed | ERBB2 | 0.265168 | BRCA | CAL-51 | Afatinib | MVP_top60_DNA+RNA_seed | MDM2 | -0.024945 | BRCA | HCC1187 | Serdemetan |
| MVP_top60_DNA_seed | EGFR | 0.265168 | BRCA | CAL-51 | Afatinib | MVP_top60_DNA+RNA_seed | ATM | -0.024624 | BRCA | HCC38 | KU-55933 |
| MVP_top60_DNA_seed | ERBB2 | 0.265947 | BRCA | EVSA-T | CI-1033 | MVP_top60_DNA+RNA_seed | ATM | -0.024072 | BRCA | BT-549 | KU-60019 |
| MVP_top60_DNA_seed | EGFR | 0.265947 | BRCA | EVSA-T | CI-1033 | MVP_top60_DNA+RNA_seed | HSP90AA1 | -0.023728 | BRCA | MDA-MB-175-VII | SNX-2112 |
| MVP_top60_DNA_seed | EGFR | 0.266082 | BRCA | EVSA-T | Erlotinib | MVP_top60_DNA+RNA_seed | HSP90AB1 | -0.023728 | BRCA | MDA-MB-175-VII | SNX-2112 |
| MVP_top60_DNA_seed | HDAC1 | 0.266171 | BRCA | HCC2157 | Belinostat | MVP_top60_DNA+RNA_seed | EGFR | -0.022716 | BRCA | MDA-MB-453 | Pelitinib |
| MVP_top60_DNA_seed | MDM2 | 0.266786 | BRCA | MDA-MB-453 | Serdemetan | MVP_top60_DNA+RNA_seed | ERBB2 | -0.022567 | BRCA | BT-483 | CP724714 |
| MVP_top60_DNA_seed | ERBB2 | 0.267458 | BRCA | HCC1500 | Afatinib | MVP_top60_DNA+RNA_seed | PIK3CA | -0.021242 | BRCA | HCC1143 | AZD8835 |
| MVP_top60_DNA_seed | EGFR | 0.267458 | BRCA | HCC1500 | Afatinib | MVP_top60_DNA+RNA_seed | ERBB2 | -0.021151 | BRCA | CAL-51 | CP724714 |
| MVP_top60_DNA_seed | HSP90AA1 | 0.268208 | BRCA | HCC1569 | Tanespimycin | MVP_top60_DNA+RNA_seed | SRC | -0.021041 | BRCA | HCC1954 | Bosutinib |
| MVP_top60_DNA_seed | HSP90AB1 | 0.268208 | BRCA | HCC1569 | Tanespimycin | MVP_top60_DNA+RNA_seed | HDAC1 | -0.020495 | BRCA | HCC2218 | Tubastatin A |
| MVP_top60_DNA_seed | PIK3CA | 0.273674 | BRCA | MDA-MB-231 | AZD8835 | MVP_top60_DNA+RNA_seed | EGFR | -0.019741 | BRCA | COLO-824 | CUDC-101 |
| MVP_top60_DNA_seed | EGFR | 0.275647 | BRCA | CAL-51 | AST-1306 | MVP_top60_DNA+RNA_seed | ERBB2 | -0.019741 | BRCA | COLO-824 | CUDC-101 |
| MVP_top60_DNA_seed | ATM | 0.27759 | BRCA | MFM-223 | CP466722 | MVP_top60_DNA+RNA_seed | EGFR | -0.017937 | BRCA | EVSA-T | Afatinib |
| MVP_top60_DNA_seed | AR | 0.278215 | BRCA | MDA-MB-157 | Bicalutamide | MVP_top60_DNA+RNA_seed | ERBB2 | -0.017937 | BRCA | EVSA-T | Afatinib |
| MVP_top60_DNA_seed | ERBB2 | 0.278677 | BRCA | MDA-MB-468 | CUDC-101 | MVP_top60_DNA+RNA_seed | AKT1 | -0.017303 | BRCA | Hs-578-T | BAY AKT1 |
| MVP_top60_DNA_seed | EGFR | 0.278677 | BRCA | MDA-MB-468 | CUDC-101 | MVP_top60_DNA+RNA_seed | AR | -0.016785 | BRCA | HCC202 | AZD3514 |
| MVP_top60_DNA_seed | ATM | 0.27932 | BRCA | BT-20 | KU-60019 | MVP_top60_DNA+RNA_seed | AKT1 | -0.016601 | BRCA | MFM-223 | A-443654 |
| MVP_top60_DNA_seed | MDM2 | 0.283597 | BRCA | UACC-893 | Nutlin-3a (-) | MVP_top60_DNA+RNA_seed | AKT1 | -0.014649 | BRCA | DU-4475 | A-443654 |
| MVP_top60_DNA_seed | AKT1 | 0.284119 | BRCA | HCC1143 | MK-2206 | MVP_top60_DNA+RNA_seed | HDAC1 | -0.012392 | BRCA | BT-474 | Dacinostat |

| | | | | | | | | | | | |
|---|---|---|---|---|---|---|---|---|---|---|---|
| MVP_top60_DNA_seed | ERBB2 | 0.284801 | BRCA | MCF7 | CP724714 | MVP_top60_DNA+RNA_seed | MAPK14 | -0.012185 | BRCA | ZR-75-30 | TAK-715 |
| MVP_top60_DNA_seed | HDAC1 | 0.284886 | BRCA | MDA-MB-468 | CAY10603 | MVP_top60_DNA+RNA_seed | MAPK14 | -0.012089 | BRCA | HCC1569 | TAK-715 |
| MVP_top60_DNA_seed | ESR1 | 0.2857 | BRCA | HCC38 | Fulvestrant | MVP_top60_DNA+RNA_seed | HDAC1 | -0.01202 | BRCA | HCC2157 | Tubastatin A |
| MVP_top60_DNA_seed | HDAC1 | 0.289726 | BRCA | MDA-MB-468 | Tubastatin A | MVP_top60_DNA+RNA_seed | EP300 | -0.01052 | BRCA | HCC1500 | I-CBP112 |
| MVP_top60_DNA_seed | AR | 0.289937 | BRCA | COLO-824 | AZD3514 | MVP_top60_DNA+RNA_seed | AKT1 | -0.009745 | BRCA | CAMA-1 | AKT inhibitor VIII |
| MVP_top60_DNA_seed | EGFR | 0.290741 | BRCA | CAL-120 | Gefitinib | MVP_top60_DNA+RNA_seed | HDAC1 | -0.00784 | BRCA | EVSA-T | Dacinostat |
| MVP_top60_DNA_seed | ESR1 | 0.291541 | BRCA | HCC1143 | Fulvestrant | MVP_top60_DNA+RNA_seed | EP300 | -0.006419 | BRCA | HCC1187 | I-CBP112 |
| MVP_top60_DNA_seed | MAPK3 | 0.291552 | BRCA | HCC38 | FR-180204 | MVP_top60_DNA+RNA_seed | HDAC1 | -0.004466 | BRCA | DU-4475 | CAY10603 |
| MVP_top60_DNA_seed | MAPK1 | 0.291552 | BRCA | HCC38 | FR-180204 | MVP_top60_DNA+RNA_seed | HDAC1 | -0.003002 | BRCA | COLO-824 | Belinostat |
| MVP_top60_DNA_seed | HSP90AA1 | 0.291787 | BRCA | HCC1428 | Tanespimycin | MVP_top60_DNA+RNA_seed | HDAC1 | -0.002936 | BRCA | T47D | JQ12 |
| MVP_top60_DNA_seed | HSP90AB1 | 0.291787 | BRCA | HCC1428 | Tanespimycin | MVP_top60_DNA+RNA_seed | HDAC2 | -0.002936 | BRCA | T47D | JQ12 |
| MVP_top60_DNA_seed | CDK1 | 0.29219 | BRCA | CAL-120 | RO-3306 | MVP_top60_DNA+RNA_seed | SRC | -0.00118 | BRCA | OCUB-M | WH-4-023 |
| MVP_top60_DNA_seed | MAPK1 | 0.292472 | BRCA | ZR-75-30 | VX-11e | MVP_top60_DNA+RNA_seed | MAPK14 | 0.000561 | BRCA | COLO-824 | TAK-715 |
| MVP_top60_DNA_seed | HSP90AA1 | 0.293983 | BRCA | MDA-MB-231 | Luminespib | MVP_top60_DNA+RNA_seed | EGFR | 0.000901 | BRCA | MDA-MB-231 | Cetuximab |
| MVP_top60_DNA_seed | HSP90AB1 | 0.293983 | BRCA | MDA-MB-231 | Luminespib | MVP_top60_DNA+RNA_seed | HDAC1 | 0.000929 | BRCA | MFM-223 | CAY10603 |
| MVP_top60_DNA_seed | SRC | 0.295214 | BRCA | HCC1599 | WH-4-023 | MVP_top60_DNA+RNA_seed | HSP90AA1 | 0.001793 | BRCA | ZR-75-30 | SNX-2112 |
| MVP_top60_DNA_seed | AR | 0.3003 | BRCA | HDQ-P1 | Bicalutamide | MVP_top60_DNA+RNA_seed | HSP90AB1 | 0.001793 | BRCA | ZR-75-30 | SNX-2112 |
| MVP_top60_DNA_seed | ERBB2 | 0.301572 | BRCA | MRK-nu-1 | AZD8931 | MVP_top60_DNA+RNA_seed | EGFR | 0.002719 | BRCA | CAL-120 | AZD8931 |
| MVP_top60_DNA_seed | EGFR | 0.301572 | BRCA | MRK-nu-1 | AZD8931 | MVP_top60_DNA+RNA_seed | ERBB2 | 0.002719 | BRCA | CAL-120 | AZD8931 |
| MVP_top60_DNA_seed | ERBB3 | 0.301572 | BRCA | MRK-nu-1 | AZD8931 | MVP_top60_DNA+RNA_seed | ERBB3 | 0.002719 | BRCA | CAL-120 | AZD8931 |
| MVP_top60_DNA_seed | SRC | 0.302479 | BRCA | T47D | NVP-BHG712 | MVP_top60_DNA+RNA_seed | AKT1 | 0.002886 | BRCA | MFM-223 | AKT inhibitor VIII |
| MVP_top60_DNA_seed | AR | 0.30375 | BRCA | MDA-MB-361 | AZD3514 | MVP_top60_DNA+RNA_seed | ATM | 0.003691 | BRCA | UACC-893 | KU-60019 |
| MVP_top60_DNA_seed | HDAC1 | 0.3063 | BRCA | CAL-120 | AR-42 | MVP_top60_DNA+RNA_seed | AKT1 | 0.006228 | BRCA | HCC202 | MK-2206 |
| MVP_top60_DNA_seed | ERBB2 | 0.306594 | BRCA | BT-20 | CUDC-101 | MVP_top60_DNA+RNA_seed | MAPK3 | 0.006984 | BRCA | EVSA-T | FR-180204 |
| MVP_top60_DNA_seed | EGFR | 0.306594 | BRCA | BT-20 | CUDC-101 | MVP_top60_DNA+RNA_seed | MAPK1 | 0.006984 | BRCA | EVSA-T | FR-180204 |
| MVP_top60_DNA_seed | MAPK3 | 0.311931 | BRCA | HCC1569 | FR-180204 | MVP_top60_DNA+RNA_seed | HDAC1 | 0.00842 | BRCA | HCC1419 | AR-42 |
| MVP_top60_DNA_seed | MAPK1 | 0.311931 | BRCA | HCC1569 | FR-180204 | MVP_top60_DNA+RNA_seed | EGFR | 0.008665 | BRCA | BT-20 | CI-1033 |
| MVP_top60_DNA_seed | CDK1 | 0.314311 | BRCA | HCC1187 | RO-3306 | MVP_top60_DNA+RNA_seed | ERBB2 | 0.008665 | BRCA | BT-20 | CI-1033 |
| MVP_top60_DNA_seed | SRC | 0.314802 | BRCA | HCC1187 | WH-4-023 | MVP_top60_DNA+RNA_seed | HSP90AA1 | 0.008874 | BRCA | MDA-MB-468 | CCT-018159 |
| MVP_top60_DNA_seed | ERBB2 | 0.317858 | BRCA | HCC38 | CP724714 | MVP_top60_DNA+RNA_seed | HSP90AB1 | 0.008874 | BRCA | MDA-MB-468 | CCT-018159 |
| MVP_top60_DNA_seed | ERBB2 | 0.319282 | BRCA | HCC1428 | CP724714 | MVP_top60_DNA+RNA_seed | AKT1 | 0.009825 | BRCA | AU565 | GSK690693 |
| MVP_top60_DNA_seed | ERBB2 | 0.31949 | BRCA | MFM-223 | AZD8931 | MVP_top60_DNA+RNA_seed | MAPK3 | 0.010996 | BRCA | AU565 | FR-180204 |
| MVP_top60_DNA_seed | EGFR | 0.31949 | BRCA | MFM-223 | AZD8931 | MVP_top60_DNA+RNA_seed | MAPK1 | 0.010996 | BRCA | AU565 | FR-180204 |
| MVP_top60_DNA_seed | ERBB3 | 0.31949 | BRCA | MFM-223 | AZD8931 | MVP_top60_DNA+RNA_seed | ERBB2 | 0.014621 | BRCA | MDA-MB-330 | CP724714 |
| MVP_top60_DNA_seed | EGFR | 0.320176 | BRCA | MDA-MB-415 | AST-1306 | MVP_top60_DNA+RNA_seed | ATM | 0.014622 | BRCA | HCC1954 | KU-60019 |
| MVP_top60_DNA_seed | AKT1 | 0.320529 | BRCA | HCC1428 | AKT inhibitor VIII | MVP_top60_DNA+RNA_seed | SRC | 0.016492 | BRCA | MDA-MB-175-VII | NVP-BHG712 |
| MVP_top60_DNA_seed | SRC | 0.322569 | BRCA | MDA-MB-175-VII | Bosutinib | MVP_top60_DNA+RNA_seed | AKT1 | 0.016839 | BRCA | MDA-MB-436 | BAY AKT1 |
| MVP_top60_DNA_seed | HSP90AA1 | 0.322793 | BRCA | HCC70 | Elesclomol | MVP_top60_DNA+RNA_seed | PIK3CA | 0.017213 | BRCA | HCC2157 | AZD8835 |
| MVP_top60_DNA_seed | HSP90AB1 | 0.322793 | BRCA | HCC70 | Elesclomol | MVP_top60_DNA+RNA_seed | SRC | 0.018149 | BRCA | HCC1187 | Bosutinib |
| MVP_top60_DNA_seed | HSP90AA1 | 0.323507 | BRCA | DU-4475 | Elesclomol | MVP_top60_DNA+RNA_seed | PLK1 | 0.018303 | BRCA | MDA-MB-453 | PLK_6522 |
| MVP_top60_DNA_seed | HSP90AB1 | 0.323507 | BRCA | DU-4475 | Elesclomol | MVP_top60_DNA+RNA_seed | ERBB2 | 0.018477 | BRCA | MDA-MB-231 | CP724714 |

| | | | | | | | | | | | |
|---|---|---|---|---|---|---|---|---|---|---|---|
| MVP_top60_DNA_seed | HDAC1 | 0.326547 | BRCA | CAL-120 | Dacinostat | MVP_top60_DNA+RNA_seed | AKT1 | 0.020005 | BRCA | HCC1937 | AKT inhibitor VIII |
| MVP_top60_DNA_seed | ERBB2 | 0.329717 | BRCA | HCC1187 | CUDC-101 | MVP_top60_DNA+RNA_seed | MDM2 | 0.021967 | BRCA | HCC70 | Serdemetan |
| MVP_top60_DNA_seed | EGFR | 0.329717 | BRCA | HCC1187 | CUDC-101 | MVP_top60_DNA+RNA_seed | SRC | 0.022424 | BRCA | DU-4475 | Bosutinib |
| MVP_top60_DNA_seed | HDAC1 | 0.330276 | BRCA | HCC1500 | JQ12 | MVP_top60_DNA+RNA_seed | EGFR | 0.022436 | BRCA | CAL-51 | AZD8931 |
| MVP_top60_DNA_seed | HDAC2 | 0.330276 | BRCA | HCC1500 | JQ12 | MVP_top60_DNA+RNA_seed | ERBB2 | 0.022436 | BRCA | CAL-51 | AZD8931 |
| MVP_top60_DNA_seed | SRC | 0.331932 | BRCA | UACC-812 | NVP-BHG712 | MVP_top60_DNA+RNA_seed | ERBB3 | 0.022436 | BRCA | CAL-51 | AZD8931 |
| MVP_top60_DNA_seed | SRC | 0.333426 | BRCA | DU-4475 | WH-4-023 | MVP_top60_DNA+RNA_seed | HSP90AA1 | 0.022726 | BRCA | JIMT-1 | CCT-018159 |
| MVP_top60_DNA_seed | ERBB2 | 0.334683 | BRCA | HDQ-P1 | AZD8931 | MVP_top60_DNA+RNA_seed | HSP90AB1 | 0.022726 | BRCA | JIMT-1 | CCT-018159 |
| MVP_top60_DNA_seed | EGFR | 0.334683 | BRCA | HDQ-P1 | AZD8931 | MVP_top60_DNA+RNA_seed | HDAC1 | 0.023683 | BRCA | HCC70 | Dacinostat |
| MVP_top60_DNA_seed | ERBB3 | 0.334683 | BRCA | HDQ-P1 | AZD8931 | MVP_top60_DNA+RNA_seed | SRC | 0.024022 | BRCA | EVSA-T | WH-4-023 |
| MVP_top60_DNA_seed | EGFR | 0.335152 | BRCA | HCC1187 | Cetuximab | MVP_top60_DNA+RNA_seed | SRC | 0.025152 | BRCA | UACC-893 | NVP-BHG712 |
| MVP_top60_DNA_seed | AKT1 | 0.336009 | BRCA | MDA-MB-330 | GSK690693 | MVP_top60_DNA+RNA_seed | HDAC1 | 0.02672 | BRCA | CAL-51 | Dacinostat |
| MVP_top60_DNA_seed | AR | 0.338021 | BRCA | CAL-51 | AZD3514 | MVP_top60_DNA+RNA_seed | AKT1 | 0.027052 | BRCA | BT-483 | AKT inhibitor VIII |
| MVP_top60_DNA_seed | HSP90AA1 | 0.33898 | BRCA | T47D | SNX-2112 | MVP_top60_DNA+RNA_seed | EGFR | 0.027379 | BRCA | MFM-223 | CI-1033 |
| MVP_top60_DNA_seed | HSP90AB1 | 0.33898 | BRCA | T47D | SNX-2112 | MVP_top60_DNA+RNA_seed | ERBB2 | 0.027379 | BRCA | MFM-223 | CI-1033 |
| MVP_top60_DNA_seed | EGFR | 0.339472 | BRCA | HCC1569 | Cetuximab | MVP_top60_DNA+RNA_seed | AR | 0.02763 | BRCA | Hs-578-T | AZD3514 |
| MVP_top60_DNA_seed | CDK1 | 0.340687 | BRCA | MRK-nu-1 | RO-3306 | MVP_top60_DNA+RNA_seed | SRC | 0.028744 | BRCA | MRK-nu-1 | WH-4-023 |
| MVP_top60_DNA_seed | HDAC1 | 0.341897 | BRCA | HCC2157 | CAY10603 | MVP_top60_DNA+RNA_seed | HDAC1 | 0.030211 | BRCA | MDA-MB-157 | CAY10603 |
| MVP_top60_DNA_seed | HDAC1 | 0.342318 | BRCA | EFM-192A | Belinostat | MVP_top60_DNA+RNA_seed | AR | 0.030327 | BRCA | BT-549 | AZD3514 |
| MVP_top60_DNA_seed | ERBB2 | 0.34598 | BRCA | CAL-120 | CUDC-101 | MVP_top60_DNA+RNA_seed | EGFR | 0.030356 | BRCA | MDA-MB-231 | CUDC-101 |
| MVP_top60_DNA_seed | EGFR | 0.34598 | BRCA | CAL-120 | CUDC-101 | MVP_top60_DNA+RNA_seed | ERBB2 | 0.030356 | BRCA | MDA-MB-231 | CUDC-101 |
| MVP_top60_DNA_seed | AKT1 | 0.347618 | BRCA | EVSA-T | AKT inhibitor VIII | MVP_top60_DNA+RNA_seed | EGFR | 0.031371 | BRCA | CAL-148 | Erlotinib |
| MVP_top60_DNA_seed | ATM | 0.348123 | BRCA | MDA-MB-436 | CP466722 | MVP_top60_DNA+RNA_seed | HSP90AA1 | 0.032538 | BRCA | HCC70 | Luminespib |
| MVP_top60_DNA_seed | AKT1 | 0.348169 | BRCA | UACC-812 | AKT inhibitor VIII | MVP_top60_DNA+RNA_seed | HSP90AB1 | 0.032538 | BRCA | HCC70 | Luminespib |
| MVP_top60_DNA_seed | HDAC1 | 0.349991 | BRCA | CAMA-1 | Tubastatin A | MVP_top60_DNA+RNA_seed | HSP90AA1 | 0.036266 | BRCA | HCC1187 | Elesclomol |
| MVP_top60_DNA_seed | CDK2 | 0.351101 | BRCA | HCC2157 | AZD5438 | MVP_top60_DNA+RNA_seed | HSP90AB1 | 0.036266 | BRCA | HCC1187 | Elesclomol |
| MVP_top60_DNA_seed | HSP90AA1 | 0.351219 | BRCA | HCC1428 | Elesclomol | MVP_top60_DNA+RNA_seed | EGFR | 0.03818 | BRCA | EFM-192A | Cetuximab |
| MVP_top60_DNA_seed | HSP90AB1 | 0.351219 | BRCA | HCC1428 | Elesclomol | MVP_top60_DNA+RNA_seed | HDAC1 | 0.041152 | BRCA | COLO-824 | CAY10603 |
| MVP_top60_DNA_seed | ERBB2 | 0.352074 | BRCA | COLO-824 | Afatinib | MVP_top60_DNA+RNA_seed | HDAC1 | 0.042391 | BRCA | HCC1428 | CAY10603 |
| MVP_top60_DNA_seed | EGFR | 0.352074 | BRCA | COLO-824 | Afatinib | MVP_top60_DNA+RNA_seed | HDAC1 | 0.043389 | BRCA | HCC70 | AR-42 |
| MVP_top60_DNA_seed | MAPK1 | 0.35362 | BRCA | JIMT-1 | VX-11e | MVP_top60_DNA+RNA_seed | PIK3CA | 0.04396 | BRCA | HCC1143 | PI3Ka_4409 |
| MVP_top60_DNA_seed | SRC | 0.35421 | BRCA | HCC1187 | Saracatinib | MVP_top60_DNA+RNA_seed | AKT1 | 0.04507 | BRCA | JIMT-1 | AKT inhibitor VIII |
| MVP_top60_DNA_seed | EGFR | 0.354893 | BRCA | HCC1395 | Cetuximab | MVP_top60_DNA+RNA_seed | AR | 0.045406 | BRCA | MDA-MB-468 | Bicalutamide |
| MVP_top60_DNA_seed | HSP90AA1 | 0.356661 | BRCA | AU565 | Luminespib | MVP_top60_DNA+RNA_seed | MDM2 | 0.045815 | BRCA | HCC2157 | Serdemetan |
| MVP_top60_DNA_seed | HSP90AB1 | 0.356661 | BRCA | AU565 | Luminespib | MVP_top60_DNA+RNA_seed | MAPK1 | 0.046108 | BRCA | MDA-MB-468 | VX-11e |
| MVP_top60_DNA_seed | PIK3CA | 0.357184 | BRCA | BT-483 | PI3Ka_4409 | MVP_top60_DNA+RNA_seed | ATM | 0.046754 | BRCA | UACC-893 | CP466722 |
| MVP_top60_DNA_seed | AKT1 | 0.360402 | BRCA | HCC1395 | AKT inhibitor VIII | MVP_top60_DNA+RNA_seed | AKT1 | 0.046837 | BRCA | MDA-MB-157 | MK-2206 |
| MVP_top60_DNA_seed | MAPK3 | 0.360534 | BRCA | ZR-75-30 | FR-180204 | MVP_top60_DNA+RNA_seed | HSP90AA1 | 0.047191 | BRCA | HCC1599 | Luminespib |
| MVP_top60_DNA_seed | MAPK1 | 0.360534 | BRCA | ZR-75-30 | FR-180204 | MVP_top60_DNA+RNA_seed | HSP90AB1 | 0.047191 | BRCA | HCC1599 | Luminespib |
| MVP_top60_DNA_seed | EGFR | 0.362866 | BRCA | DU-4475 | Erlotinib | MVP_top60_DNA+RNA_seed | MDM2 | 0.047969 | BRCA | MRK-nu-1 | Serdemetan |
| MVP_top60_DNA_seed | HSP90AA1 | 0.363172 | BRCA | CAL-85-1 | Elesclomol | MVP_top60_DNA+RNA_seed | MDM2 | 0.048309 | BRCA | HCC1569 | Nutlin-3a (-) |

| | | | | | | | | | | | |
|---|---|---|---|---|---|---|---|---|---|---|---|
| MVP_top60_DNA_seed | HSP90AB1 | 0.363172 | BRCA | CAL-85-1 | Elesclomol | MVP_top60_DNA+RNA_seed | AKT1 | 0.048326 | BRCA | HCC1937 | MK-2206 |
| MVP_top60_DNA_seed | CDK2 | 0.363205 | BRCA | MRK-nu-1 | PHA-793887 | MVP_top60_DNA+RNA_seed | PLK1 | 0.04863 | BRCA | EVSA-T | BI-2536 |
| MVP_top60_DNA_seed | EGFR | 0.366594 | BRCA | EVSA-T | Cetuximab | MVP_top60_DNA+RNA_seed | EGFR | 0.051118 | BRCA | CAL-120 | Afatinib |
| MVP_top60_DNA_seed | HSP90AA1 | 0.36707 | BRCA | HCC70 | CCT-018159 | MVP_top60_DNA+RNA_seed | ERBB2 | 0.051118 | BRCA | CAL-120 | Afatinib |
| MVP_top60_DNA_seed | HSP90AB1 | 0.36707 | BRCA | HCC70 | CCT-018159 | MVP_top60_DNA+RNA_seed | HSP90AA1 | 0.053462 | BRCA | COLO-824 | Luminespib |
| MVP_top60_DNA_seed | HSP90AA1 | 0.37156 | BRCA | HCC38 | CCT-018159 | MVP_top60_DNA+RNA_seed | HSP90AB1 | 0.053462 | BRCA | COLO-824 | Luminespib |
| MVP_top60_DNA_seed | HSP90AB1 | 0.37156 | BRCA | HCC38 | CCT-018159 | MVP_top60_DNA+RNA_seed | MAPK3 | 0.055312 | BRCA | EFM-192A | FR-180204 |
| MVP_top60_DNA_seed | EGFR | 0.371745 | BRCA | CAL-120 | AST-1306 | MVP_top60_DNA+RNA_seed | MAPK1 | 0.055312 | BRCA | EFM-192A | FR-180204 |
| MVP_top60_DNA_seed | HDAC1 | 0.372329 | BRCA | MDA-MB-231 | CAY10603 | MVP_top60_DNA+RNA_seed | HDAC1 | 0.055563 | BRCA | MDA-MB-330 | CAY10603 |
| MVP_top60_DNA_seed | CDK2 | 0.375242 | BRCA | CAMA-1 | PHA-793887 | MVP_top60_DNA+RNA_seed | ATM | 0.05617 | BRCA | MRK-nu-1 | KU-60019 |
| MVP_top60_DNA_seed | AR | 0.375664 | BRCA | Hs-578-T | Bicalutamide | MVP_top60_DNA+RNA_seed | ATM | 0.056716 | BRCA | HCC1395 | KU-60019 |
| MVP_top60_DNA_seed | SRC | 0.376331 | BRCA | MFM-223 | NVP-BHG712 | MVP_top60_DNA+RNA_seed | CDK2 | 0.056791 | BRCA | HCC1395 | AZD5438 |
| MVP_top60_DNA_seed | AR | 0.377115 | BRCA | COLO-824 | Bicalutamide | MVP_top60_DNA+RNA_seed | MDM2 | 0.057521 | BRCA | EFM-192A | Nutlin-3a (-) |
| MVP_top60_DNA_seed | ERBB2 | 0.379639 | BRCA | MDA-MB-157 | Afatinib | MVP_top60_DNA+RNA_seed | EGFR | 0.057974 | BRCA | BT-483 | PF-00299804 |
| MVP_top60_DNA_seed | EGFR | 0.379639 | BRCA | MDA-MB-157 | Afatinib | MVP_top60_DNA+RNA_seed | ERBB2 | 0.057974 | BRCA | BT-483 | PF-00299804 |
| MVP_top60_DNA_seed | AKT1 | 0.380008 | BRCA | HCC1569 | MK-2206 | MVP_top60_DNA+RNA_seed | HSP90AA1 | 0.058874 | BRCA | HCC1143 | Elesclomol |
| MVP_top60_DNA_seed | HSP90AA1 | 0.380733 | BRCA | HCC202 | Luminespib | MVP_top60_DNA+RNA_seed | HSP90AB1 | 0.058874 | BRCA | HCC1143 | Elesclomol |
| MVP_top60_DNA_seed | HSP90AB1 | 0.380733 | BRCA | HCC202 | Luminespib | MVP_top60_DNA+RNA_seed | HSP90AA1 | 0.060412 | BRCA | UACC-893 | Luminespib |
| MVP_top60_DNA_seed | HSP90AA1 | 0.382742 | BRCA | MFM-223 | SNX-2112 | MVP_top60_DNA+RNA_seed | HSP90AB1 | 0.060412 | BRCA | UACC-893 | Luminespib |
| MVP_top60_DNA_seed | HSP90AB1 | 0.382742 | BRCA | MFM-223 | SNX-2112 | MVP_top60_DNA+RNA_seed | AKT1 | 0.062104 | BRCA | HCC1599 | AKT inhibitor VIII |
| MVP_top60_DNA_seed | ERBB2 | 0.382975 | BRCA | HCC1187 | Lapatinib | MVP_top60_DNA+RNA_seed | PIK3CA | 0.062903 | BRCA | MDA-MB-415 | PI3Ka_4409 |
| MVP_top60_DNA_seed | EGFR | 0.382975 | BRCA | HCC1187 | Lapatinib | MVP_top60_DNA+RNA_seed | MDM2 | 0.066885 | BRCA | CAMA-1 | Serdemetan |
| MVP_top60_DNA_seed | ATM | 0.383024 | BRCA | HCC2218 | CP466722 | MVP_top60_DNA+RNA_seed | SRC | 0.067162 | BRCA | OCUB-M | Saracatinib |
| MVP_top60_DNA_seed | AR | 0.383444 | BRCA | CAL-85-1 | Bicalutamide | MVP_top60_DNA+RNA_seed | HDAC1 | 0.067624 | BRCA | MRK-nu-1 | JQ12 |
| MVP_top60_DNA_seed | CDK2 | 0.383499 | BRCA | AU565 | AZD5438 | MVP_top60_DNA+RNA_seed | HDAC2 | 0.067624 | BRCA | MRK-nu-1 | JQ12 |
| MVP_top60_DNA_seed | AR | 0.384808 | BRCA | MDA-MB-361 | Bicalutamide | MVP_top60_DNA+RNA_seed | CDK2 | 0.068255 | BRCA | YMB-1-E | PHA-793887 |
| MVP_top60_DNA_seed | AKT1 | 0.385717 | BRCA | HCC1187 | A-443654 | MVP_top60_DNA+RNA_seed | EGFR | 0.069502 | BRCA | MCF7 | AZD8931 |
| MVP_top60_DNA_seed | CDK2 | 0.385996 | BRCA | MDA-MB-330 | PHA-793887 | MVP_top60_DNA+RNA_seed | ERBB2 | 0.069502 | BRCA | MCF7 | AZD8931 |
| MVP_top60_DNA_seed | CDK1 | 0.387682 | BRCA | MDA-MB-157 | RO-3306 | MVP_top60_DNA+RNA_seed | ERBB3 | 0.069502 | BRCA | MCF7 | AZD8931 |
| MVP_top60_DNA_seed | EP300 | 0.388149 | BRCA | MCF7 | I-CBP112 | MVP_top60_DNA+RNA_seed | ESR1 | 0.069655 | BRCA | MDA-MB-415 | Tamoxifen |
| MVP_top60_DNA_seed | CREBBP | 0.388149 | BRCA | MCF7 | I-CBP112 | MVP_top60_DNA+RNA_seed | PIK3CA | 0.070522 | BRCA | HCC2157 | PI3Ka_4409 |
| MVP_top60_DNA_seed | ERBB2 | 0.39012 | BRCA | CAL-148 | Lapatinib | MVP_top60_DNA+RNA_seed | AKT1 | 0.071425 | BRCA | CAL-51 | AKT inhibitor VIII |
| MVP_top60_DNA_seed | EGFR | 0.39012 | BRCA | CAL-148 | Lapatinib | MVP_top60_DNA+RNA_seed | ESR1 | 0.072287 | BRCA | MDA-MB-330 | Fulvestrant |
| MVP_top60_DNA_seed | AR | 0.391131 | BRCA | MDA-MB-157 | Bicalutamide | MVP_top60_DNA+RNA_seed | AKT1 | 0.072316 | BRCA | ZR-75-30 | MK-2206 |
| MVP_top60_DNA_seed | ERBB2 | 0.392644 | BRCA | YMB-1-E | CP724714 | MVP_top60_DNA+RNA_seed | SRC | 0.07244 | BRCA | JIMT-1 | NVP-BHG712 |
| MVP_top60_DNA_seed | HDAC1 | 0.394204 | BRCA | JIMT-1 | CAY10603 | MVP_top60_DNA+RNA_seed | CDK1 | 0.074795 | BRCA | EFM-192A | RO-3306 |
| MVP_top60_DNA_seed | EGFR | 0.39458 | BRCA | HCC1187 | Erlotinib | MVP_top60_DNA+RNA_seed | CDK2 | 0.07494 | BRCA | MCF7 | AZD5438 |
| MVP_top60_DNA_seed | MAPK1 | 0.394598 | BRCA | HCC1187 | VX-11e | MVP_top60_DNA+RNA_seed | AKT1 | 0.075811 | BRCA | HCC2157 | AKT inhibitor VIII |
| MVP_top60_DNA_seed | AR | 0.396533 | BRCA | HCC1428 | Bicalutamide | MVP_top60_DNA+RNA_seed | PIK3CA | 0.077335 | BRCA | COLO-824 | AZD8835 |
| MVP_top60_DNA_seed | EGFR | 0.397457 | BRCA | DU-4475 | Cetuximab | MVP_top60_DNA+RNA_seed | HSP90AA1 | 0.077646 | BRCA | HCC1428 | CCT-018159 |
| MVP_top60_DNA_seed | ERBB2 | 0.398169 | BRCA | MDA-MB-231 | Afatinib | MVP_top60_DNA+RNA_seed | HSP90AB1 | 0.077646 | BRCA | HCC1428 | CCT-018159 |

| | | | | | | | | | | | |
|---|---|---|---|---|---|---|---|---|---|---|---|
| MVP_top60_DNA_seed | EGFR | 0.398169 | BRCA | MDA-MB-231 | Afatinib | MVP_top60_DNA+RNA_seed | HSP90AA1 | 0.077779 | BRCA | CAL-120 | Tanespimycin |
| MVP_top60_DNA_seed | ATM | 0.400196 | BRCA | EFM-192A | KU-55933 | MVP_top60_DNA+RNA_seed | HSP90AB1 | 0.077779 | BRCA | CAL-120 | Tanespimycin |
| MVP_top60_DNA_seed | CDK2 | 0.406485 | BRCA | HCC1569 | AZD5438 | MVP_top60_DNA+RNA_seed | ATM | 0.079098 | BRCA | MDA-MB-175-VII | KU-60019 |
| MVP_top60_DNA_seed | EGFR | 0.40687 | BRCA | BT-483 | Cetuximab | MVP_top60_DNA+RNA_seed | AKT1 | 0.07947 | BRCA | MDA-MB-436 | MK-2206 |
| MVP_top60_DNA_seed | ERBB2 | 0.407315 | BRCA | MDA-MB-231 | AZD8931 | MVP_top60_DNA+RNA_seed | EGFR | 0.081099 | BRCA | HCC1143 | Gefitinib |
| MVP_top60_DNA_seed | EGFR | 0.407315 | BRCA | MDA-MB-231 | AZD8931 | MVP_top60_DNA+RNA_seed | AKT1 | 0.081173 | BRCA | DU-4475 | GSK690693 |
| MVP_top60_DNA_seed | ERBB3 | 0.407315 | BRCA | MDA-MB-231 | AZD8931 | MVP_top60_DNA+RNA_seed | MDM2 | 0.081959 | BRCA | MDA-MB-330 | Nutlin-3a (-) |
| MVP_top60_DNA_seed | ERBB2 | 0.407404 | BRCA | CAL-85-1 | Afatinib | MVP_top60_DNA+RNA_seed | AKT1 | 0.083383 | BRCA | UACC-812 | GSK690693 |
| MVP_top60_DNA_seed | EGFR | 0.407404 | BRCA | CAL-85-1 | Afatinib | MVP_top60_DNA+RNA_seed | ATM | 0.086015 | BRCA | MDA-MB-415 | torin2 |
| MVP_top60_DNA_seed | HSP90AA1 | 0.40808 | BRCA | HDQ-P1 | Luminespib | MVP_top60_DNA+RNA_seed | MDM2 | 0.086929 | BRCA | CAL-120 | Nutlin-3a (-) |
| MVP_top60_DNA_seed | HSP90AB1 | 0.40808 | BRCA | HDQ-P1 | Luminespib | MVP_top60_DNA+RNA_seed | EGFR | 0.087979 | BRCA | CAMA-1 | Afatinib |
| MVP_top60_DNA_seed | ERBB2 | 0.410087 | BRCA | Hs-578-T | CUDC-101 | MVP_top60_DNA+RNA_seed | ERBB2 | 0.087979 | BRCA | CAMA-1 | Afatinib |
| MVP_top60_DNA_seed | EGFR | 0.410087 | BRCA | Hs-578-T | CUDC-101 | MVP_top60_DNA+RNA_seed | EGFR | 0.088177 | BRCA | MCF7 | Gefitinib |
| MVP_top60_DNA_seed | HDAC1 | 0.411973 | BRCA | HCC38 | JQ12 | MVP_top60_DNA+RNA_seed | MDM2 | 0.089254 | BRCA | HCC1143 | Serdemetan |
| MVP_top60_DNA_seed | HDAC2 | 0.411973 | BRCA | HCC38 | JQ12 | MVP_top60_DNA+RNA_seed | EGFR | 0.089457 | BRCA | MCF7 | PF-00299804 |
| MVP_top60_DNA_seed | CDK2 | 0.412647 | BRCA | UACC-893 | PHA-793887 | MVP_top60_DNA+RNA_seed | ERBB2 | 0.089457 | BRCA | MCF7 | PF-00299804 |
| MVP_top60_DNA_seed | AR | 0.412676 | BRCA | MDA-MB-157 | AZD3514 | MVP_top60_DNA+RNA_seed | PIK3CA | 0.089588 | BRCA | BT-549 | PI3Ka_4409 |
| MVP_top60_DNA_seed | EGFR | 0.412689 | BRCA | YMB-1-E | AST-1306 | MVP_top60_DNA+RNA_seed | MAPK14 | 0.090355 | BRCA | AU565 | TAK-715 |
| MVP_top60_DNA_seed | AKT1 | 0.413122 | BRCA | CAL-120 | MK-2206 | MVP_top60_DNA+RNA_seed | AR | 0.090976 | BRCA | MDA-MB-436 | Bicalutamide |
| MVP_top60_DNA_seed | ERBB2 | 0.416688 | BRCA | MDA-MB-415 | AZD8931 | MVP_top60_DNA+RNA_seed | EGFR | 0.091137 | BRCA | MDA-MB-231 | PF-00299804 |
| MVP_top60_DNA_seed | EGFR | 0.416688 | BRCA | MDA-MB-415 | AZD8931 | MVP_top60_DNA+RNA_seed | ERBB2 | 0.091137 | BRCA | MDA-MB-231 | PF-00299804 |
| MVP_top60_DNA_seed | ERBB3 | 0.416688 | BRCA | MDA-MB-415 | AZD8931 | MVP_top60_DNA+RNA_seed | ATM | 0.091322 | BRCA | HCC38 | KU-60019 |
| MVP_top60_DNA_seed | CDK2 | 0.417262 | BRCA | MDA-MB-436 | AZD5438 | MVP_top60_DNA+RNA_seed | HDAC1 | 0.091611 | BRCA | HCC1500 | CAY10603 |
| MVP_top60_DNA_seed | EGFR | 0.419688 | BRCA | MCF7 | Cetuximab | MVP_top60_DNA+RNA_seed | AKT1 | 0.09276 | BRCA | MRK-nu-1 | A-443654 |
| MVP_top60_DNA_seed | ERBB2 | 0.419985 | BRCA | HCC1937 | AZD8931 | MVP_top60_DNA+RNA_seed | AR | 0.094804 | BRCA | HCC1500 | Bicalutamide |
| MVP_top60_DNA_seed | EGFR | 0.419985 | BRCA | HCC1937 | AZD8931 | MVP_top60_DNA+RNA_seed | MAPK1 | 0.094908 | BRCA | HCC1599 | VX-11e |
| MVP_top60_DNA_seed | ERBB3 | 0.419985 | BRCA | HCC1937 | AZD8931 | MVP_top60_DNA+RNA_seed | AR | 0.095665 | BRCA | T47D | AZD3514 |
| MVP_top60_DNA_seed | HDAC1 | 0.420588 | BRCA | HCC1954 | Dacinostat | MVP_top60_DNA+RNA_seed | PIK3CA | 0.096349 | BRCA | HCC1569 | AZD8835 |
| MVP_top60_DNA_seed | AKT1 | 0.421063 | BRCA | MDA-MB-436 | AKT inhibitor VIII | MVP_top60_DNA+RNA_seed | MAPK3 | 0.096423 | BRCA | UACC-812 | FR-180204 |
| MVP_top60_DNA_seed | ERBB2 | 0.424701 | BRCA | CAL-120 | Afatinib | MVP_top60_DNA+RNA_seed | MAPK1 | 0.096423 | BRCA | UACC-812 | FR-180204 |
| MVP_top60_DNA_seed | EGFR | 0.424701 | BRCA | CAL-120 | Afatinib | MVP_top60_DNA+RNA_seed | HSP90AA1 | 0.097434 | BRCA | AU565 | Elesclomol |
| MVP_top60_DNA_seed | SRC | 0.427708 | BRCA | MFM-223 | Bosutinib | MVP_top60_DNA+RNA_seed | HSP90AB1 | 0.097434 | BRCA | AU565 | Elesclomol |
| MVP_top60_DNA_seed | CDK2 | 0.429085 | BRCA | COLO-824 | PHA-793887 | MVP_top60_DNA+RNA_seed | MAPK1 | 0.09773 | BRCA | UACC-893 | VX-11e |
| MVP_top60_DNA_seed | CDK2 | 0.429125 | BRCA | EVSA-T | Seliciclib | MVP_top60_DNA+RNA_seed | ERBB2 | 0.098501 | BRCA | CAMA-1 | CP724714 |
| MVP_top60_DNA_seed | ATM | 0.430449 | BRCA | MDA-MB-175-VII | KU-55933 | MVP_top60_DNA+RNA_seed | ATM | 0.099442 | BRCA | OCUB-M | KU-55933 |
| MVP_top60_DNA_seed | ATM | 0.431319 | BRCA | UACC-893 | KU-55933 | MVP_top60_DNA+RNA_seed | EGFR | 0.100288 | BRCA | HCC1954 | Cetuximab |
| MVP_top60_DNA_seed | CDK2 | 0.431438 | BRCA | CAL-148 | PHA-793887 | MVP_top60_DNA+RNA_seed | HSP90AA1 | 0.100619 | BRCA | HCC1419 | Luminespib |
| MVP_top60_DNA_seed | ERBB2 | 0.431992 | BRCA | CAL-85-1 | CUDC-101 | MVP_top60_DNA+RNA_seed | HSP90AB1 | 0.100619 | BRCA | HCC1419 | Luminespib |
| MVP_top60_DNA_seed | EGFR | 0.431992 | BRCA | CAL-85-1 | CUDC-101 | MVP_top60_DNA+RNA_seed | HDAC1 | 0.103185 | BRCA | EFM-192A | CAY10603 |
| MVP_top60_DNA_seed | HSP90AA1 | 0.432539 | BRCA | CAL-85-1 | SNX-2112 | MVP_top60_DNA+RNA_seed | EGFR | 0.10463 | BRCA | HCC1937 | CI-1033 |
| MVP_top60_DNA_seed | HSP90AB1 | 0.432539 | BRCA | CAL-85-1 | SNX-2112 | MVP_top60_DNA+RNA_seed | ERBB2 | 0.10463 | BRCA | HCC1937 | CI-1033 |

| | | | | | | | | | | | |
|---|---|---|---|---|---|---|---|---|---|---|---|
| MVP_top60_DNA_seed | MDM2 | 0.433099 | BRCA | T47D | Nutlin-3a (-) | MVP_top60_DNA+RNA_seed | EGFR | 0.106027 | BRCA | MFM-223 | PF-00299804 |
| MVP_top60_DNA_seed | HSP90AA1 | 0.436866 | BRCA | HCC1954 | Tanespimycin | MVP_top60_DNA+RNA_seed | ERBB2 | 0.106027 | BRCA | MFM-223 | PF-00299804 |
| MVP_top60_DNA_seed | HSP90AB1 | 0.436866 | BRCA | HCC1954 | Tanespimycin | MVP_top60_DNA+RNA_seed | AKT1 | 0.106134 | BRCA | HCC1937 | BAY AKT1 |
| MVP_top60_DNA_seed | HDAC1 | 0.436938 | BRCA | HDQ-P1 | Tubastatin A | MVP_top60_DNA+RNA_seed | EP300 | 0.108224 | BRCA | MDA-MB-453 | I-CBP112 |
| MVP_top60_DNA_seed | ATM | 0.437909 | BRCA | MFM-223 | KU-60019 | MVP_top60_DNA+RNA_seed | HSP90AA1 | 0.109041 | BRCA | COLO-824 | Elesclomol |
| MVP_top60_DNA_seed | AR | 0.439375 | BRCA | EFM-19 | Bicalutamide | MVP_top60_DNA+RNA_seed | HSP90AB1 | 0.109041 | BRCA | COLO-824 | Elesclomol |
| MVP_top60_DNA_seed | HSP90AA1 | 0.442632 | BRCA | Hs-578-T | CCT-018159 | MVP_top60_DNA+RNA_seed | HDAC1 | 0.10997 | BRCA | COLO-824 | AR-42 |
| MVP_top60_DNA_seed | HSP90AB1 | 0.442632 | BRCA | Hs-578-T | CCT-018159 | MVP_top60_DNA+RNA_seed | HDAC1 | 0.111111 | BRCA | CAL-120 | Belinostat |
| MVP_top60_DNA_seed | HDAC1 | 0.445017 | BRCA | HCC1428 | Tubastatin A | MVP_top60_DNA+RNA_seed | HDAC1 | 0.112541 | BRCA | EFM-192A | AR-42 |
| MVP_top60_DNA_seed | HDAC1 | 0.445725 | BRCA | HDQ-P1 | AR-42 | MVP_top60_DNA+RNA_seed | CDK2 | 0.114245 | BRCA | JIMT-1 | AZD5438 |
| MVP_top60_DNA_seed | ERBB2 | 0.448714 | BRCA | JIMT-1 | CP724714 | MVP_top60_DNA+RNA_seed | EGFR | 0.114469 | BRCA | HCC1395 | CI-1033 |
| MVP_top60_DNA_seed | CDK1 | 0.449258 | BRCA | MDA-MB-361 | RO-3306 | MVP_top60_DNA+RNA_seed | ERBB2 | 0.114469 | BRCA | HCC1395 | CI-1033 |
| MVP_top60_DNA_seed | CDK1 | 0.450365 | BRCA | MDA-MB-453 | RO-3306 | MVP_top60_DNA+RNA_seed | HDAC1 | 0.114589 | BRCA | CAL-120 | Tubastatin A |
| MVP_top60_DNA_seed | SRC | 0.451443 | BRCA | UACC-893 | Bosutinib | MVP_top60_DNA+RNA_seed | ESR1 | 0.115618 | BRCA | MDA-MB-468 | Tamoxifen |
| MVP_top60_DNA_seed | AR | 0.451459 | BRCA | MFM-223 | Bicalutamide | MVP_top60_DNA+RNA_seed | MDM2 | 0.116123 | BRCA | HCC1419 | Serdemetan |
| MVP_top60_DNA_seed | HDAC1 | 0.452287 | BRCA | MRK-nu-1 | Dacinostat | MVP_top60_DNA+RNA_seed | EGFR | 0.116567 | BRCA | HCC1937 | CUDC-101 |
| MVP_top60_DNA_seed | HDAC1 | 0.455539 | BRCA | CAL-148 | JQ12 | MVP_top60_DNA+RNA_seed | ERBB2 | 0.116567 | BRCA | HCC1937 | CUDC-101 |
| MVP_top60_DNA_seed | HDAC2 | 0.455539 | BRCA | CAL-148 | JQ12 | MVP_top60_DNA+RNA_seed | EGFR | 0.1175 | BRCA | HCC1937 | Cetuximab |
| MVP_top60_DNA_seed | MDM2 | 0.457127 | BRCA | EFM-192A | Serdemetan | MVP_top60_DNA+RNA_seed | HDAC1 | 0.121783 | BRCA | HCC1937 | Dacinostat |
| MVP_top60_DNA_seed | AKT1 | 0.457567 | BRCA | DU-4475 | MK-2206 | MVP_top60_DNA+RNA_seed | EGFR | 0.122589 | BRCA | CAL-51 | Cetuximab |
| MVP_top60_DNA_seed | HDAC1 | 0.458625 | BRCA | HCC1937 | AR-42 | MVP_top60_DNA+RNA_seed | AKT1 | 0.123049 | BRCA | HCC1500 | AKT inhibitor VIII |
| MVP_top60_DNA_seed | EGFR | 0.459879 | BRCA | EVSA-T | AST-1306 | MVP_top60_DNA+RNA_seed | ERBB2 | 0.126355 | BRCA | CAL-148 | CP724714 |
| MVP_top60_DNA_seed | ERBB2 | 0.460134 | BRCA | HCC2157 | CP724714 | MVP_top60_DNA+RNA_seed | HDAC1 | 0.126956 | BRCA | UACC-812 | AR-42 |
| MVP_top60_DNA_seed | EGFR | 0.461271 | BRCA | CAL-51 | Pelitinib | MVP_top60_DNA+RNA_seed | EGFR | 0.130992 | BRCA | MDA-MB-436 | PF-00299804 |
| MVP_top60_DNA_seed | CDK1 | 0.464012 | BRCA | CAL-51 | RO-3306 | MVP_top60_DNA+RNA_seed | ERBB2 | 0.130992 | BRCA | MDA-MB-436 | PF-00299804 |
| MVP_top60_DNA_seed | ATM | 0.46592 | BRCA | MDA-MB-453 | KU-60019 | MVP_top60_DNA+RNA_seed | AR | 0.131472 | BRCA | MCF7 | Bicalutamide |
| MVP_top60_DNA_seed | HDAC1 | 0.466444 | BRCA | JIMT-1 | Belinostat | MVP_top60_DNA+RNA_seed | SRC | 0.132356 | BRCA | CAL-85-1 | NVP-BHG712 |
| MVP_top60_DNA_seed | AR | 0.469028 | BRCA | HCC38 | AZD3514 | MVP_top60_DNA+RNA_seed | HDAC1 | 0.133469 | BRCA | MDA-MB-231 | Dacinostat |
| MVP_top60_DNA_seed | CDK2 | 0.470625 | BRCA | JIMT-1 | PHA-793887 | MVP_top60_DNA+RNA_seed | MDM2 | 0.134263 | BRCA | HCC1500 | Nutlin-3a (-) |
| MVP_top60_DNA_seed | HDAC1 | 0.471323 | BRCA | HCC1419 | Tubastatin A | MVP_top60_DNA+RNA_seed | EGFR | 0.134345 | BRCA | MDA-MB-415 | PF-00299804 |
| MVP_top60_DNA_seed | HSP90AA1 | 0.471802 | BRCA | CAL-85-1 | CCT-018159 | MVP_top60_DNA+RNA_seed | ERBB2 | 0.134345 | BRCA | MDA-MB-415 | PF-00299804 |
| MVP_top60_DNA_seed | HSP90AB1 | 0.471802 | BRCA | CAL-85-1 | CCT-018159 | MVP_top60_DNA+RNA_seed | EGFR | 0.134791 | BRCA | MDA-MB-436 | Afatinib |
| MVP_top60_DNA_seed | ERBB2 | 0.473766 | BRCA | HCC1937 | Afatinib | MVP_top60_DNA+RNA_seed | ERBB2 | 0.134791 | BRCA | MDA-MB-436 | Afatinib |
| MVP_top60_DNA_seed | EGFR | 0.473766 | BRCA | HCC1937 | Afatinib | MVP_top60_DNA+RNA_seed | EGFR | 0.135771 | BRCA | HCC2218 | CUDC-101 |
| MVP_top60_DNA_seed | HSP90AA1 | 0.474087 | BRCA | T47D | CCT-018159 | MVP_top60_DNA+RNA_seed | ERBB2 | 0.135771 | BRCA | HCC2218 | CUDC-101 |
| MVP_top60_DNA_seed | HSP90AB1 | 0.474087 | BRCA | T47D | CCT-018159 | MVP_top60_DNA+RNA_seed | EGFR | 0.136246 | BRCA | MDA-MB-436 | CI-1033 |
| MVP_top60_DNA_seed | HSP90AA1 | 0.475484 | BRCA | BT-483 | Tanespimycin | MVP_top60_DNA+RNA_seed | ERBB2 | 0.136246 | BRCA | MDA-MB-436 | CI-1033 |
| MVP_top60_DNA_seed | HSP90AB1 | 0.475484 | BRCA | BT-483 | Tanespimycin | MVP_top60_DNA+RNA_seed | CDK2 | 0.136873 | BRCA | AU565 | PHA-793887 |
| MVP_top60_DNA_seed | ESR1 | 0.477032 | BRCA | HCC2157 | Tamoxifen | MVP_top60_DNA+RNA_seed | PIK3CA | 0.140748 | BRCA | COLO-824 | PI3Ka_4409 |
| MVP_top60_DNA_seed | ERBB2 | 0.477275 | BRCA | MDA-MB-415 | CI-1033 | MVP_top60_DNA+RNA_seed | MDM2 | 0.141089 | BRCA | COLO-824 | Serdemetan |
| MVP_top60_DNA_seed | EGFR | 0.477275 | BRCA | MDA-MB-415 | CI-1033 | MVP_top60_DNA+RNA_seed | AKT1 | 0.141542 | BRCA | BT-474 | A-443654 |

| | | | | | | | | | | | |
|---|---|---|---|---|---|---|---|---|---|---|---|
| MVP_top60_DNA_seed | HDAC1 | 0.480472 | BRCA | HCC1187 | CAY10603 | MVP_top60_DNA+RNA_seed | HSP90AA1 | 0.141877 | BRCA | CAMA-1 | Luminespib |
| MVP_top60_DNA_seed | CDK2 | 0.480921 | BRCA | BT-483 | PHA-793887 | MVP_top60_DNA+RNA_seed | HSP90AB1 | 0.141877 | BRCA | CAMA-1 | Luminespib |
| MVP_top60_DNA_seed | AR | 0.482647 | BRCA | EFM-192A | Bicalutamide | MVP_top60_DNA+RNA_seed | CDK2 | 0.143741 | BRCA | BT-474 | PHA-793887 |
| MVP_top60_DNA_seed | ERBB2 | 0.483363 | BRCA | HCC1419 | CUDC-101 | MVP_top60_DNA+RNA_seed | AKT1 | 0.14389 | BRCA | JIMT-1 | GSK690693 |
| MVP_top60_DNA_seed | EGFR | 0.483363 | BRCA | HCC1419 | CUDC-101 | MVP_top60_DNA+RNA_seed | HSP90AA1 | 0.144053 | BRCA | EFM-19 | Elesclomol |
| MVP_top60_DNA_seed | CDK1 | 0.485959 | BRCA | COLO-824 | RO-3306 | MVP_top60_DNA+RNA_seed | HSP90AB1 | 0.144053 | BRCA | EFM-19 | Elesclomol |
| MVP_top60_DNA_seed | CDK1 | 0.490931 | BRCA | HCC1937 | RO-3306 | MVP_top60_DNA+RNA_seed | EGFR | 0.144631 | BRCA | HDQ-P1 | CUDC-101 |
| MVP_top60_DNA_seed | ATM | 0.491327 | BRCA | EVSA-T | KU-60019 | MVP_top60_DNA+RNA_seed | ERBB2 | 0.144631 | BRCA | HDQ-P1 | CUDC-101 |
| MVP_top60_DNA_seed | CDK2 | 0.491604 | BRCA | COLO-824 | AZD5438 | MVP_top60_DNA+RNA_seed | SRC | 0.144917 | BRCA | BT-549 | NVP-BHG712 |
| MVP_top60_DNA_seed | ERBB2 | 0.493138 | BRCA | YMB-1-E | CI-1033 | MVP_top60_DNA+RNA_seed | HDAC1 | 0.145114 | BRCA | MDA-MB-157 | Belinostat |
| MVP_top60_DNA_seed | EGFR | 0.493138 | BRCA | YMB-1-E | CI-1033 | MVP_top60_DNA+RNA_seed | ERBB2 | 0.146357 | BRCA | HCC1599 | CP724714 |
| MVP_top60_DNA_seed | ERBB2 | 0.49404 | BRCA | YMB-1-E | PF-00299804 | MVP_top60_DNA+RNA_seed | AKT1 | 0.146771 | BRCA | HCC2157 | BAY AKT1 |
| MVP_top60_DNA_seed | EGFR | 0.49404 | BRCA | YMB-1-E | PF-00299804 | MVP_top60_DNA+RNA_seed | HDAC1 | 0.1476 | BRCA | MDA-MB-415 | JQ12 |
| MVP_top60_DNA_seed | PIK3CA | 0.494257 | BRCA | HDQ-P1 | PI3Ka_4409 | MVP_top60_DNA+RNA_seed | HDAC2 | 0.1476 | BRCA | MDA-MB-415 | JQ12 |
| MVP_top60_DNA_seed | SRC | 0.494855 | BRCA | HCC38 | NVP-BHG712 | MVP_top60_DNA+RNA_seed | SRC | 0.147974 | BRCA | OCUB-M | Bosutinib |
| MVP_top60_DNA_seed | MAPK1 | 0.497419 | BRCA | EFM-192A | VX-11e | MVP_top60_DNA+RNA_seed | MDM2 | 0.147997 | BRCA | OCUB-M | Serdemetan |
| MVP_top60_DNA_seed | SRC | 0.498405 | BRCA | COLO-824 | NVP-BHG712 | MVP_top60_DNA+RNA_seed | EGFR | 0.150503 | BRCA | MDA-MB-231 | CI-1033 |
| MVP_top60_DNA_seed | ATM | 0.502609 | BRCA | CAL-85-1 | KU-60019 | MVP_top60_DNA+RNA_seed | ERBB2 | 0.150503 | BRCA | MDA-MB-231 | CI-1033 |
| MVP_top60_DNA_seed | SRC | 0.502828 | BRCA | CAL-148 | Saracatinib | MVP_top60_DNA+RNA_seed | AR | 0.151776 | BRCA | HCC1428 | AZD3514 |
| MVP_top60_DNA_seed | ATM | 0.503081 | BRCA | MDA-MB-361 | KU-60019 | MVP_top60_DNA+RNA_seed | EGFR | 0.155196 | BRCA | MDA-MB-436 | AST-1306 |
| MVP_top60_DNA_seed | HDAC1 | 0.503395 | BRCA | UACC-893 | Dacinostat | MVP_top60_DNA+RNA_seed | SRC | 0.159669 | BRCA | MDA-MB-361 | NVP-BHG712 |
| MVP_top60_DNA_seed | ERBB2 | 0.50455 | BRCA | T47D | PF-00299804 | MVP_top60_DNA+RNA_seed | MAPK3 | 0.164286 | BRCA | MDA-MB-468 | FR-180204 |
| MVP_top60_DNA_seed | EGFR | 0.50455 | BRCA | T47D | PF-00299804 | MVP_top60_DNA+RNA_seed | MAPK1 | 0.164286 | BRCA | MDA-MB-468 | FR-180204 |
| MVP_top60_DNA_seed | AR | 0.50526 | BRCA | HCC1395 | Bicalutamide | MVP_top60_DNA+RNA_seed | AR | 0.164707 | BRCA | EVSA-T | Bicalutamide |
| MVP_top60_DNA_seed | ATM | 0.506636 | BRCA | HCC1143 | KU-60019 | MVP_top60_DNA+RNA_seed | SRC | 0.165582 | BRCA | HCC70 | NVP-BHG712 |
| MVP_top60_DNA_seed | HDAC1 | 0.50914 | BRCA | COLO-824 | Dacinostat | MVP_top60_DNA+RNA_seed | ATM | 0.166204 | BRCA | EFM-192A | KU-60019 |
| MVP_top60_DNA_seed | MDM2 | 0.509296 | BRCA | MDA-MB-361 | Nutlin-3a (-) | MVP_top60_DNA+RNA_seed | ERBB2 | 0.167667 | BRCA | EVSA-T | CP724714 |
| MVP_top60_DNA_seed | HDAC1 | 0.509605 | BRCA | HCC1569 | Tubastatin A | MVP_top60_DNA+RNA_seed | EGFR | 0.169938 | BRCA | OCUB-M | Erlotinib |
| MVP_top60_DNA_seed | AKT1 | 0.510231 | BRCA | CAL-120 | GSK690693 | MVP_top60_DNA+RNA_seed | EGFR | 0.171885 | BRCA | HCC1937 | Afatinib |
| MVP_top60_DNA_seed | EGFR | 0.51134 | BRCA | HCC1428 | Cetuximab | MVP_top60_DNA+RNA_seed | ERBB2 | 0.171885 | BRCA | HCC1937 | Afatinib |
| MVP_top60_DNA_seed | ERBB2 | 0.512721 | BRCA | BT-474 | CUDC-101 | MVP_top60_DNA+RNA_seed | HDAC1 | 0.172948 | BRCA | HCC1937 | CAY10603 |
| MVP_top60_DNA_seed | EGFR | 0.512721 | BRCA | BT-474 | CUDC-101 | MVP_top60_DNA+RNA_seed | ATM | 0.174024 | BRCA | HCC1428 | KU-55933 |
| MVP_top60_DNA_seed | PIK3CA | 0.513118 | BRCA | HCC2218 | AZD8835 | MVP_top60_DNA+RNA_seed | MDM2 | 0.175351 | BRCA | HCC1187 | Nutlin-3a (-) |
| MVP_top60_DNA_seed | HDAC1 | 0.513258 | BRCA | CAMA-1 | AR-42 | MVP_top60_DNA+RNA_seed | MDM2 | 0.175657 | BRCA | CAMA-1 | Nutlin-3a (-) |
| MVP_top60_DNA_seed | PIK3CA | 0.51515 | BRCA | DU-4475 | PI3Ka_4409 | MVP_top60_DNA+RNA_seed | EGFR | 0.178231 | BRCA | MDA-MB-453 | Afatinib |
| MVP_top60_DNA_seed | HDAC1 | 0.51526 | BRCA | HCC1419 | Belinostat | MVP_top60_DNA+RNA_seed | ERBB2 | 0.178231 | BRCA | MDA-MB-453 | Afatinib |
| MVP_top60_DNA_seed | HSP90AA1 | 0.516196 | BRCA | OCUB-M | Elesclomol | MVP_top60_DNA+RNA_seed | SRC | 0.18019 | BRCA | HCC1500 | Bosutinib |
| MVP_top60_DNA_seed | HSP90AB1 | 0.516196 | BRCA | OCUB-M | Elesclomol | MVP_top60_DNA+RNA_seed | HSP90AA1 | 0.180214 | BRCA | ZR-75-30 | Tanespimycin |
| MVP_top60_DNA_seed | HSP90AA1 | 0.516744 | BRCA | BT-549 | SNX-2112 | MVP_top60_DNA+RNA_seed | HSP90AB1 | 0.180214 | BRCA | ZR-75-30 | Tanespimycin |
| MVP_top60_DNA_seed | HSP90AB1 | 0.516744 | BRCA | BT-549 | SNX-2112 | MVP_top60_DNA+RNA_seed | ATM | 0.182381 | BRCA | HCC1937 | CP466722 |
| MVP_top60_DNA_seed | HSP90AA1 | 0.517006 | BRCA | EVSA-T | Luminespib | MVP_top60_DNA+RNA_seed | EGFR | 0.182819 | BRCA | HCC1500 | Gefitinib |

| | | | | | | | | | | | |
|---|---|---|---|---|---|---|---|---|---|---|---|
| MVP_top60_DNA_seed | HSP90AB1 | 0.517006 | BRCA | EVSA-T | Luminespib | MVP_top60_DNA+RNA_seed | HDAC1 | 0.182972 | BRCA | HCC1187 | Tubastatin A |
| MVP_top60_DNA_seed | HDAC1 | 0.518816 | BRCA | CAL-85-1 | Belinostat | MVP_top60_DNA+RNA_seed | EGFR | 0.18347 | BRCA | CAL-51 | PF-00299804 |
| MVP_top60_DNA_seed | ERBB2 | 0.520022 | BRCA | BT-483 | AZD8931 | MVP_top60_DNA+RNA_seed | ERBB2 | 0.18347 | BRCA | CAL-51 | PF-00299804 |
| MVP_top60_DNA_seed | EGFR | 0.520022 | BRCA | BT-483 | AZD8931 | MVP_top60_DNA+RNA_seed | AKT1 | 0.18445 | BRCA | DU-4475 | BAY AKT1 |
| MVP_top60_DNA_seed | ERBB3 | 0.520022 | BRCA | BT-483 | AZD8931 | MVP_top60_DNA+RNA_seed | EGFR | 0.185998 | BRCA | UACC-812 | PF-00299804 |
| MVP_top60_DNA_seed | EGFR | 0.520231 | BRCA | CAL-120 | Cetuximab | MVP_top60_DNA+RNA_seed | ERBB2 | 0.185998 | BRCA | UACC-812 | PF-00299804 |
| MVP_top60_DNA_seed | HSP90AA1 | 0.521592 | BRCA | HCC1937 | SNX-2112 | MVP_top60_DNA+RNA_seed | HSP90AA1 | 0.186117 | BRCA | HDQ-P1 | Tanespimycin |
| MVP_top60_DNA_seed | HSP90AB1 | 0.521592 | BRCA | HCC1937 | SNX-2112 | MVP_top60_DNA+RNA_seed | HSP90AB1 | 0.186117 | BRCA | HDQ-P1 | Tanespimycin |
| MVP_top60_DNA_seed | ERBB2 | 0.522645 | BRCA | BT-20 | PF-00299804 | MVP_top60_DNA+RNA_seed | HSP90AA1 | 0.186202 | BRCA | EVSA-T | SNX-2112 |
| MVP_top60_DNA_seed | EGFR | 0.522645 | BRCA | BT-20 | PF-00299804 | MVP_top60_DNA+RNA_seed | HSP90AB1 | 0.186202 | BRCA | EVSA-T | SNX-2112 |
| MVP_top60_DNA_seed | EGFR | 0.523688 | BRCA | T47D | AST-1306 | MVP_top60_DNA+RNA_seed | ATM | 0.187041 | BRCA | HCC1500 | KU-55933 |
| MVP_top60_DNA_seed | ERBB2 | 0.524095 | BRCA | DU-4475 | Afatinib | MVP_top60_DNA+RNA_seed | ESR1 | 0.187775 | BRCA | MDA-MB-231 | Tamoxifen |
| MVP_top60_DNA_seed | EGFR | 0.524095 | BRCA | DU-4475 | Afatinib | MVP_top60_DNA+RNA_seed | CDK2 | 0.188939 | BRCA | HCC1500 | PHA-793887 |
| MVP_top60_DNA_seed | HDAC1 | 0.524606 | BRCA | CAL-85-1 | CAY10603 | MVP_top60_DNA+RNA_seed | HDAC1 | 0.189323 | BRCA | HCC1599 | Parthenolide |
| MVP_top60_DNA_seed | AR | 0.52493 | BRCA | MDA-MB-231 | Bicalutamide | MVP_top60_DNA+RNA_seed | MAPK3 | 0.189665 | BRCA | CAL-148 | FR-180204 |
| MVP_top60_DNA_seed | HSP90AA1 | 0.526635 | BRCA | AU565 | SNX-2112 | MVP_top60_DNA+RNA_seed | MAPK1 | 0.189665 | BRCA | CAL-148 | FR-180204 |
| MVP_top60_DNA_seed | HSP90AB1 | 0.526635 | BRCA | AU565 | SNX-2112 | MVP_top60_DNA+RNA_seed | EGFR | 0.191482 | BRCA | MCF7 | Afatinib |
| MVP_top60_DNA_seed | HSP90AA1 | 0.52781 | BRCA | HCC202 | Elesclomol | MVP_top60_DNA+RNA_seed | ERBB2 | 0.191482 | BRCA | MCF7 | Afatinib |
| MVP_top60_DNA_seed | HSP90AB1 | 0.52781 | BRCA | HCC202 | Elesclomol | MVP_top60_DNA+RNA_seed | EGFR | 0.191675 | BRCA | Hs-578-T | AZD8931 |
| MVP_top60_DNA_seed | HDAC1 | 0.529142 | BRCA | BT-20 | Belinostat | MVP_top60_DNA+RNA_seed | ERBB2 | 0.191675 | BRCA | Hs-578-T | AZD8931 |
| MVP_top60_DNA_seed | ERBB2 | 0.529395 | BRCA | EFM-19 | AZD8931 | MVP_top60_DNA+RNA_seed | ERBB3 | 0.191675 | BRCA | Hs-578-T | AZD8931 |
| MVP_top60_DNA_seed | EGFR | 0.529395 | BRCA | EFM-19 | AZD8931 | MVP_top60_DNA+RNA_seed | EGFR | 0.194689 | BRCA | MDA-MB-361 | Cetuximab |
| MVP_top60_DNA_seed | ERBB3 | 0.529395 | BRCA | EFM-19 | AZD8931 | MVP_top60_DNA+RNA_seed | HDAC1 | 0.194952 | BRCA | UACC-812 | Belinostat |
| MVP_top60_DNA_seed | ERBB2 | 0.530013 | BRCA | MDA-MB-231 | Afatinib | MVP_top60_DNA+RNA_seed | PIK3CA | 0.195146 | BRCA | CAL-120 | PI3Ka_4409 |
| MVP_top60_DNA_seed | EGFR | 0.530013 | BRCA | MDA-MB-231 | Afatinib | MVP_top60_DNA+RNA_seed | SRC | 0.196157 | BRCA | HCC2218 | Bosutinib |
| MVP_top60_DNA_seed | SRC | 0.530179 | BRCA | HCC1395 | Bosutinib | MVP_top60_DNA+RNA_seed | HDAC1 | 0.196217 | BRCA | HCC1937 | Belinostat |
| MVP_top60_DNA_seed | AR | 0.53045 | BRCA | BT-474 | Bicalutamide | MVP_top60_DNA+RNA_seed | EGFR | 0.196219 | BRCA | CAL-120 | CI-1033 |
| MVP_top60_DNA_seed | ERBB2 | 0.53203 | BRCA | HCC1428 | CI-1033 | MVP_top60_DNA+RNA_seed | ERBB2 | 0.196219 | BRCA | CAL-120 | CI-1033 |
| MVP_top60_DNA_seed | EGFR | 0.53203 | BRCA | HCC1428 | CI-1033 | MVP_top60_DNA+RNA_seed | HDAC1 | 0.196222 | BRCA | MDA-MB-175-VII | AR-42 |
| MVP_top60_DNA_seed | EGFR | 0.532044 | BRCA | MDA-MB-330 | Gefitinib | MVP_top60_DNA+RNA_seed | AR | 0.197116 | BRCA | HCC2157 | Bicalutamide |
| MVP_top60_DNA_seed | MAPK3 | 0.533184 | BRCA | MDA-MB-453 | FR-180204 | MVP_top60_DNA+RNA_seed | AKT1 | 0.199236 | BRCA | MDA-MB-157 | AKT inhibitor VIII |
| MVP_top60_DNA_seed | MAPK1 | 0.533184 | BRCA | MDA-MB-453 | FR-180204 | MVP_top60_DNA+RNA_seed | SRC | 0.200347 | BRCA | BT-474 | NVP-BHG712 |
| MVP_top60_DNA_seed | AR | 0.535518 | BRCA | MDA-MB-175-VII | Bicalutamide | MVP_top60_DNA+RNA_seed | ATM | 0.202256 | BRCA | HCC1500 | KU-60019 |
| MVP_top60_DNA_seed | AKT1 | 0.536015 | BRCA | BT-549 | AKT inhibitor VIII | MVP_top60_DNA+RNA_seed | PIK3CA | 0.20947 | BRCA | DU-4475 | AZD8835 |
| MVP_top60_DNA_seed | ERBB2 | 0.537272 | BRCA | EFM-19 | Afatinib | MVP_top60_DNA+RNA_seed | AKT1 | 0.209578 | BRCA | HCC2157 | A-443654 |
| MVP_top60_DNA_seed | EGFR | 0.537272 | BRCA | EFM-19 | Afatinib | MVP_top60_DNA+RNA_seed | EGFR | 0.210319 | BRCA | MDA-MB-157 | AZD8931 |
| MVP_top60_DNA_seed | AR | 0.537986 | BRCA | HCC1143 | Bicalutamide | MVP_top60_DNA+RNA_seed | ERBB2 | 0.210319 | BRCA | MDA-MB-157 | AZD8931 |
| MVP_top60_DNA_seed | HDAC1 | 0.541957 | BRCA | BT-474 | Belinostat | MVP_top60_DNA+RNA_seed | ERBB3 | 0.210319 | BRCA | MDA-MB-157 | AZD8931 |
| MVP_top60_DNA_seed | HDAC1 | 0.543264 | BRCA | MDA-MB-175-VII | Belinostat | MVP_top60_DNA+RNA_seed | AKT1 | 0.212269 | BRCA | HCC1500 | GSK690693 |
| MVP_top60_DNA_seed | EGFR | 0.544561 | BRCA | MDA-MB-231 | Pelitinib | MVP_top60_DNA+RNA_seed | CDK2 | 0.213091 | BRCA | MDA-MB-157 | AZD5438 |
| MVP_top60_DNA_seed | CDK1 | 0.545483 | BRCA | BT-20 | RO-3306 | MVP_top60_DNA+RNA_seed | ESR1 | 0.213694 | BRCA | CAL-51 | Tamoxifen |

| | | | | | | | | | | | |
|---|---|---|---|---|---|---|---|---|---|---|---|
| MVP_top60_DNA_seed | ESR1 | 0.546651 | BRCA | HCC2157 | Fulvestrant | MVP_top60_DNA+RNA_seed | CDK2 | 0.214181 | BRCA | HCC1569 | PHA-793887 |
| MVP_top60_DNA_seed | ERBB2 | 0.548275 | BRCA | BT-549 | CUDC-101 | MVP_top60_DNA+RNA_seed | PLK1 | 0.215164 | BRCA | MRK-nu-1 | BI-2536 |
| MVP_top60_DNA_seed | EGFR | 0.548275 | BRCA | BT-549 | CUDC-101 | MVP_top60_DNA+RNA_seed | HDAC1 | 0.215845 | BRCA | MRK-nu-1 | CAY10603 |
| MVP_top60_DNA_seed | EP300 | 0.549712 | BRCA | HCC70 | I-CBP112 | MVP_top60_DNA+RNA_seed | HSP90AA1 | 0.215933 | BRCA | HCC70 | SNX-2112 |
| MVP_top60_DNA_seed | CREBBP | 0.549712 | BRCA | HCC70 | I-CBP112 | MVP_top60_DNA+RNA_seed | HSP90AB1 | 0.215933 | BRCA | HCC70 | SNX-2112 |
| MVP_top60_DNA_seed | EGFR | 0.55082 | BRCA | HCC1187 | Gefitinib | MVP_top60_DNA+RNA_seed | EGFR | 0.215937 | BRCA | HCC1569 | AZD8931 |
| MVP_top60_DNA_seed | PIK3CA | 0.551516 | BRCA | HCC1419 | AZD8835 | MVP_top60_DNA+RNA_seed | ERBB2 | 0.215937 | BRCA | HCC1569 | AZD8931 |
| MVP_top60_DNA_seed | ATM | 0.5518 | BRCA | MDA-MB-330 | KU-55933 | MVP_top60_DNA+RNA_seed | ERBB3 | 0.215937 | BRCA | HCC1569 | AZD8931 |
| MVP_top60_DNA_seed | HDAC1 | 0.551837 | BRCA | MDA-MB-175-VII | JQ12 | MVP_top60_DNA+RNA_seed | EGFR | 0.21707 | BRCA | EFM-192A | CUDC-101 |
| MVP_top60_DNA_seed | HDAC2 | 0.551837 | BRCA | MDA-MB-175-VII | JQ12 | MVP_top60_DNA+RNA_seed | ERBB2 | 0.21707 | BRCA | EFM-192A | CUDC-101 |
| MVP_top60_DNA_seed | MAPK1 | 0.552745 | BRCA | HCC202 | VX-11e | MVP_top60_DNA+RNA_seed | HDAC1 | 0.217211 | BRCA | MDA-MB-157 | AR-42 |
| MVP_top60_DNA_seed | EGFR | 0.553932 | BRCA | HCC38 | Gefitinib | MVP_top60_DNA+RNA_seed | ESR1 | 0.218847 | BRCA | CAL-120 | Fulvestrant |
| MVP_top60_DNA_seed | ESR1 | 0.554975 | BRCA | MDA-MB-330 | Tamoxifen | MVP_top60_DNA+RNA_seed | EGFR | 0.219009 | BRCA | BT-549 | Afatinib |
| MVP_top60_DNA_seed | SRC | 0.555526 | BRCA | EFM-192A | Bosutinib | MVP_top60_DNA+RNA_seed | ERBB2 | 0.219009 | BRCA | BT-549 | Afatinib |
| MVP_top60_DNA_seed | HDAC1 | 0.555635 | BRCA | UACC-812 | CAY10603 | MVP_top60_DNA+RNA_seed | CDK2 | 0.219732 | BRCA | HCC1599 | AZD5438 |
| MVP_top60_DNA_seed | CDK2 | 0.556501 | BRCA | HCC1954 | AZD5438 | MVP_top60_DNA+RNA_seed | CDK2 | 0.220263 | BRCA | BT-20 | PHA-793887 |
| MVP_top60_DNA_seed | HDAC1 | 0.55672 | BRCA | CAL-85-1 | AR-42 | MVP_top60_DNA+RNA_seed | EGFR | 0.223009 | BRCA | BT-549 | AZD8931 |
| MVP_top60_DNA_seed | AKT1 | 0.559417 | BRCA | HDQ-P1 | GSK690693 | MVP_top60_DNA+RNA_seed | ERBB2 | 0.223009 | BRCA | BT-549 | AZD8931 |
| MVP_top60_DNA_seed | SRC | 0.560439 | BRCA | MDA-MB-436 | Bosutinib | MVP_top60_DNA+RNA_seed | ERBB3 | 0.223009 | BRCA | BT-549 | AZD8931 |
| MVP_top60_DNA_seed | AR | 0.562717 | BRCA | HCC1937 | AZD3514 | MVP_top60_DNA+RNA_seed | HDAC1 | 0.223073 | BRCA | HDQ-P1 | CAY10603 |
| MVP_top60_DNA_seed | ERBB2 | 0.563051 | BRCA | MDA-MB-157 | PF-00299804 | MVP_top60_DNA+RNA_seed | EGFR | 0.223207 | BRCA | HCC70 | Afatinib |
| MVP_top60_DNA_seed | EGFR | 0.563051 | BRCA | MDA-MB-157 | PF-00299804 | MVP_top60_DNA+RNA_seed | ERBB2 | 0.223207 | BRCA | HCC70 | Afatinib |
| MVP_top60_DNA_seed | HSP90AA1 | 0.564069 | BRCA | HCC1569 | SNX-2112 | MVP_top60_DNA+RNA_seed | EGFR | 0.223422 | BRCA | MDA-MB-453 | Cetuximab |
| MVP_top60_DNA_seed | HSP90AB1 | 0.564069 | BRCA | HCC1569 | SNX-2112 | MVP_top60_DNA+RNA_seed | EGFR | 0.224232 | BRCA | DU-4475 | AZD8931 |
| MVP_top60_DNA_seed | EP300 | 0.564155 | BRCA | MDA-MB-157 | I-CBP112 | MVP_top60_DNA+RNA_seed | ERBB2 | 0.224232 | BRCA | DU-4475 | AZD8931 |
| MVP_top60_DNA_seed | CREBBP | 0.564155 | BRCA | MDA-MB-157 | I-CBP112 | MVP_top60_DNA+RNA_seed | ERBB3 | 0.224232 | BRCA | DU-4475 | AZD8931 |
| MVP_top60_DNA_seed | HSP90AA1 | 0.564583 | BRCA | EFM-192A | CCT-018159 | MVP_top60_DNA+RNA_seed | HDAC1 | 0.22814 | BRCA | HCC2218 | AR-42 |
| MVP_top60_DNA_seed | HSP90AB1 | 0.564583 | BRCA | EFM-192A | CCT-018159 | MVP_top60_DNA+RNA_seed | HDAC1 | 0.228217 | BRCA | HDQ-P1 | Belinostat |
| MVP_top60_DNA_seed | MAPK1 | 0.569103 | BRCA | COLO-824 | VX-11e | MVP_top60_DNA+RNA_seed | EGFR | 0.228485 | BRCA | CAL-148 | PF-00299804 |
| MVP_top60_DNA_seed | EP300 | 0.569813 | BRCA | Hs-578-T | I-CBP112 | MVP_top60_DNA+RNA_seed | ERBB2 | 0.228485 | BRCA | CAL-148 | PF-00299804 |
| MVP_top60_DNA_seed | CREBBP | 0.569813 | BRCA | Hs-578-T | I-CBP112 | MVP_top60_DNA+RNA_seed | MAPK14 | 0.228814 | BRCA | BT-549 | TAK-715 |
| MVP_top60_DNA_seed | AKT1 | 0.570828 | BRCA | HCC38 | GSK690693 | MVP_top60_DNA+RNA_seed | EGFR | 0.230246 | BRCA | HCC1143 | Afatinib |
| MVP_top60_DNA_seed | HDAC1 | 0.57103 | BRCA | MRK-nu-1 | Belinostat | MVP_top60_DNA+RNA_seed | ERBB2 | 0.230246 | BRCA | HCC1143 | Afatinib |
| MVP_top60_DNA_seed | MAPK1 | 0.573084 | BRCA | MRK-nu-1 | VX-11e | MVP_top60_DNA+RNA_seed | HDAC1 | 0.230496 | BRCA | AU565 | Tubastatin A |
| MVP_top60_DNA_seed | EGFR | 0.573091 | BRCA | BT-474 | Erlotinib | MVP_top60_DNA+RNA_seed | EGFR | 0.231426 | BRCA | HCC1599 | Lapatinib |
| MVP_top60_DNA_seed | ATM | 0.575337 | BRCA | HCC1419 | KU-60019 | MVP_top60_DNA+RNA_seed | ERBB2 | 0.231426 | BRCA | HCC1599 | Lapatinib |
| MVP_top60_DNA_seed | HDAC1 | 0.57712 | BRCA | UACC-812 | Parthenolide | MVP_top60_DNA+RNA_seed | HSP90AA1 | 0.231471 | BRCA | HCC2218 | SNX-2112 |
| MVP_top60_DNA_seed | EGFR | 0.577188 | BRCA | UACC-812 | Pelitinib | MVP_top60_DNA+RNA_seed | HSP90AB1 | 0.231471 | BRCA | HCC2218 | SNX-2112 |
| MVP_top60_DNA_seed | HDAC1 | 0.577867 | BRCA | MDA-MB-436 | CAY10603 | MVP_top60_DNA+RNA_seed | ATM | 0.232119 | BRCA | HCC1937 | KU-55933 |
| MVP_top60_DNA_seed | HDAC1 | 0.579872 | BRCA | CAL-85-1 | Tubastatin A | MVP_top60_DNA+RNA_seed | EGFR | 0.23466 | BRCA | HCC1954 | Gefitinib |
| MVP_top60_DNA_seed | SRC | 0.580742 | BRCA | HCC2218 | NVP-BHG712 | MVP_top60_DNA+RNA_seed | AR | 0.237601 | BRCA | HCC202 | Bicalutamide |

| | | | | | | | | | | | |
|---|---|---|---|---|---|---|---|---|---|---|---|
| MVP_top60_DNA_seed | AKT1 | 0.58148 | BRCA | Hs-578-T | AKT inhibitor VIII | MVP_top60_DNA+RNA_seed | AKT1 | 0.240572 | BRCA | CAL-51 | MK-2206 |
| MVP_top60_DNA_seed | ATM | 0.581752 | BRCA | MDA-MB-361 | KU-55933 | MVP_top60_DNA+RNA_seed | EGFR | 0.243617 | BRCA | HCC1599 | Erlotinib |
| MVP_top60_DNA_seed | CDK1 | 0.582603 | BRCA | MDA-MB-231 | RO-3306 | MVP_top60_DNA+RNA_seed | AKT1 | 0.243661 | BRCA | HCC1395 | GSK690693 |
| MVP_top60_DNA_seed | HDAC1 | 0.583193 | BRCA | YMB-1-E | CAY10603 | MVP_top60_DNA+RNA_seed | AKT1 | 0.246166 | BRCA | HCC1954 | MK-2206 |
| MVP_top60_DNA_seed | SRC | 0.585664 | BRCA | AU565 | NVP-BHG712 | MVP_top60_DNA+RNA_seed | MAPK14 | 0.246168 | BRCA | HCC38 | TAK-715 |
| MVP_top60_DNA_seed | PIK3CA | 0.586171 | BRCA | CAL-85-1 | AZD8835 | MVP_top60_DNA+RNA_seed | EGFR | 0.246834 | BRCA | MCF7 | AST-1306 |
| MVP_top60_DNA_seed | HDAC1 | 0.586438 | BRCA | COLO-824 | Tubastatin A | MVP_top60_DNA+RNA_seed | ATM | 0.250078 | BRCA | HCC1187 | KU-60019 |
| MVP_top60_DNA_seed | CDK2 | 0.587705 | BRCA | HDQ-P1 | AZD5438 | MVP_top60_DNA+RNA_seed | AR | 0.250352 | BRCA | HDQ-P1 | AZD3514 |
| MVP_top60_DNA_seed | CDK1 | 0.589198 | BRCA | MDA-MB-175-VII | RO-3306 | MVP_top60_DNA+RNA_seed | AR | 0.250388 | BRCA | JIMT-1 | Bicalutamide |
| MVP_top60_DNA_seed | CDK2 | 0.589889 | BRCA | MFM-223 | AZD5438 | MVP_top60_DNA+RNA_seed | EGFR | 0.250815 | BRCA | HCC1954 | Afatinib |
| MVP_top60_DNA_seed | ATM | 0.58995 | BRCA | COLO-824 | CP466722 | MVP_top60_DNA+RNA_seed | ERBB2 | 0.250815 | BRCA | HCC1954 | Afatinib |
| MVP_top60_DNA_seed | CDK2 | 0.591078 | BRCA | HCC202 | AZD5438 | MVP_top60_DNA+RNA_seed | MAPK1 | 0.254507 | BRCA | MDA-MB-231 | VX-11e |
| MVP_top60_DNA_seed | HDAC1 | 0.592524 | BRCA | HCC2218 | JQ12 | MVP_top60_DNA+RNA_seed | AKT1 | 0.254693 | BRCA | MDA-MB-231 | AKT inhibitor VIII |
| MVP_top60_DNA_seed | HDAC2 | 0.592524 | BRCA | HCC2218 | JQ12 | MVP_top60_DNA+RNA_seed | AR | 0.260478 | BRCA | MDA-MB-231 | Bicalutamide |
| MVP_top60_DNA_seed | ERBB2 | 0.596476 | BRCA | CAMA-1 | Afatinib | MVP_top60_DNA+RNA_seed | HSP90AA1 | 0.260855 | BRCA | HCC2218 | Luminespib |
| MVP_top60_DNA_seed | EGFR | 0.596476 | BRCA | CAMA-1 | Afatinib | MVP_top60_DNA+RNA_seed | HSP90AB1 | 0.260855 | BRCA | HCC2218 | Luminespib |
| MVP_top60_DNA_seed | EGFR | 0.598134 | BRCA | HDQ-P1 | Pelitinib | MVP_top60_DNA+RNA_seed | CDK2 | 0.262919 | BRCA | EVSA-T | AZD5438 |
| MVP_top60_DNA_seed | ERBB2 | 0.59822 | BRCA | MRK-nu-1 | CUDC-101 | MVP_top60_DNA+RNA_seed | PIK3CA | 0.263913 | BRCA | Hs-578-T | PI3Ka_4409 |
| MVP_top60_DNA_seed | EGFR | 0.59822 | BRCA | MRK-nu-1 | CUDC-101 | MVP_top60_DNA+RNA_seed | AKT1 | 0.26403 | BRCA | HCC1419 | BAY AKT1 |
| MVP_top60_DNA_seed | EGFR | 0.600228 | BRCA | MDA-MB-157 | Cetuximab | MVP_top60_DNA+RNA_seed | EGFR | 0.265165 | BRCA | CAMA-1 | AZD8931 |
| MVP_top60_DNA_seed | ERBB2 | 0.602708 | BRCA | EFM-192A | CP724714 | MVP_top60_DNA+RNA_seed | ERBB2 | 0.265165 | BRCA | CAMA-1 | AZD8931 |
| MVP_top60_DNA_seed | HDAC1 | 0.604339 | BRCA | HCC2157 | Parthenolide | MVP_top60_DNA+RNA_seed | ERBB3 | 0.265165 | BRCA | CAMA-1 | AZD8931 |
| MVP_top60_DNA_seed | AKT1 | 0.604704 | BRCA | MDA-MB-415 | GSK690693 | MVP_top60_DNA+RNA_seed | EGFR | 0.265168 | BRCA | CAL-51 | Afatinib |
| MVP_top60_DNA_seed | SRC | 0.605145 | BRCA | HCC2157 | Bosutinib | MVP_top60_DNA+RNA_seed | ERBB2 | 0.265168 | BRCA | CAL-51 | Afatinib |
| MVP_top60_DNA_seed | PIK3CA | 0.605227 | BRCA | HDQ-P1 | AZD8835 | MVP_top60_DNA+RNA_seed | EGFR | 0.265947 | BRCA | EVSA-T | CI-1033 |
| MVP_top60_DNA_seed | SRC | 0.608014 | BRCA | MDA-MB-468 | NVP-BHG712 | MVP_top60_DNA+RNA_seed | ERBB2 | 0.265947 | BRCA | EVSA-T | CI-1033 |
| MVP_top60_DNA_seed | ESR1 | 0.609086 | BRCA | AU565 | Tamoxifen | MVP_top60_DNA+RNA_seed | EGFR | 0.266082 | BRCA | EVSA-T | Erlotinib |
| MVP_top60_DNA_seed | SRC | 0.609227 | BRCA | HCC1428 | Bosutinib | MVP_top60_DNA+RNA_seed | HDAC1 | 0.266171 | BRCA | HCC2157 | Belinostat |
| MVP_top60_DNA_seed | SRC | 0.61239 | BRCA | HCC1569 | NVP-BHG712 | MVP_top60_DNA+RNA_seed | MDM2 | 0.266786 | BRCA | MDA-MB-453 | Serdemetan |
| MVP_top60_DNA_seed | HDAC1 | 0.614849 | BRCA | MDA-MB-175-VII | CAY10603 | MVP_top60_DNA+RNA_seed | EGFR | 0.267458 | BRCA | HCC1500 | Afatinib |
| MVP_top60_DNA_seed | AKT1 | 0.614997 | BRCA | COLO-824 | GSK690693 | MVP_top60_DNA+RNA_seed | ERBB2 | 0.267458 | BRCA | HCC1500 | Afatinib |
| MVP_top60_DNA_seed | MAPK3 | 0.615108 | BRCA | JIMT-1 | FR-180204 | MVP_top60_DNA+RNA_seed | HSP90AA1 | 0.268208 | BRCA | HCC1569 | Tanespimycin |
| MVP_top60_DNA_seed | MAPK1 | 0.615108 | BRCA | JIMT-1 | FR-180204 | MVP_top60_DNA+RNA_seed | HSP90AB1 | 0.268208 | BRCA | HCC1569 | Tanespimycin |
| MVP_top60_DNA_seed | HSP90AA1 | 0.615237 | BRCA | JIMT-1 | SNX-2112 | MVP_top60_DNA+RNA_seed | PIK3CA | 0.273674 | BRCA | MDA-MB-231 | AZD8835 |
| MVP_top60_DNA_seed | HSP90AB1 | 0.615237 | BRCA | JIMT-1 | SNX-2112 | MVP_top60_DNA+RNA_seed | EGFR | 0.275647 | BRCA | CAL-51 | AST-1306 |
| MVP_top60_DNA_seed | AR | 0.618493 | BRCA | HCC1419 | Bicalutamide | MVP_top60_DNA+RNA_seed | ATM | 0.27759 | BRCA | MFM-223 | CP466722 |
| MVP_top60_DNA_seed | EGFR | 0.618902 | BRCA | HCC202 | Cetuximab | MVP_top60_DNA+RNA_seed | AR | 0.278215 | BRCA | MDA-MB-157 | Bicalutamide |
| MVP_top60_DNA_seed | ATM | 0.62006 | BRCA | EVSA-T | CP466722 | MVP_top60_DNA+RNA_seed | EGFR | 0.278677 | BRCA | MDA-MB-468 | CUDC-101 |
| MVP_top60_DNA_seed | ERBB2 | 0.622148 | BRCA | MFM-223 | CP724714 | MVP_top60_DNA+RNA_seed | ERBB2 | 0.278677 | BRCA | MDA-MB-468 | CUDC-101 |
| MVP_top60_DNA_seed | ESR1 | 0.622334 | BRCA | MDA-MB-453 | Tamoxifen | MVP_top60_DNA+RNA_seed | ATM | 0.27932 | BRCA | BT-20 | KU-60019 |
| MVP_top60_DNA_seed | MAPK1 | 0.623426 | BRCA | MDA-MB-453 | VX-11e | MVP_top60_DNA+RNA_seed | MAPK14 | 0.280379 | BRCA | MCF7 | TAK-715 |

| | | | | | | | | | | | |
|---|---|---|---|---|---|---|---|---|---|---|---|
| MVP_top60_DNA_seed | ATM | 0.624342 | BRCA | HCC1143 | KU-55933 | MVP_top60_DNA+RNA_seed | MDM2 | 0.283597 | BRCA | UACC-893 | Nutlin-3a (-) |
| MVP_top60_DNA_seed | CDK2 | 0.625674 | BRCA | CAL-51 | PHA-793887 | MVP_top60_DNA+RNA_seed | AKT1 | 0.284119 | BRCA | HCC1143 | MK-2206 |
| MVP_top60_DNA_seed | SRC | 0.626035 | BRCA | BT-474 | Saracatinib | MVP_top60_DNA+RNA_seed | ERBB2 | 0.284801 | BRCA | MCF7 | CP724714 |
| MVP_top60_DNA_seed | AKT1 | 0.626182 | BRCA | EFM-192A | AKT inhibitor VIII | MVP_top60_DNA+RNA_seed | HDAC1 | 0.284886 | BRCA | MDA-MB-468 | CAY10603 |
| MVP_top60_DNA_seed | MAPK3 | 0.627238 | BRCA | HCC1419 | FR-180204 | MVP_top60_DNA+RNA_seed | ESR1 | 0.2857 | BRCA | HCC38 | Fulvestrant |
| MVP_top60_DNA_seed | MAPK1 | 0.627238 | BRCA | HCC1419 | FR-180204 | MVP_top60_DNA+RNA_seed | PLK1 | 0.287728 | BRCA | BT-20 | PLK_6522 |
| MVP_top60_DNA_seed | HDAC1 | 0.628216 | BRCA | UACC-893 | Belinostat | MVP_top60_DNA+RNA_seed | HDAC1 | 0.289726 | BRCA | MDA-MB-468 | Tubastatin A |
| MVP_top60_DNA_seed | EP300 | 0.628665 | BRCA | HCC38 | I-CBP112 | MVP_top60_DNA+RNA_seed | AR | 0.289937 | BRCA | COLO-824 | AZD3514 |
| MVP_top60_DNA_seed | CREBBP | 0.628665 | BRCA | HCC38 | I-CBP112 | MVP_top60_DNA+RNA_seed | EGFR | 0.290741 | BRCA | CAL-120 | Gefitinib |
| MVP_top60_DNA_seed | ATM | 0.628962 | BRCA | T47D | KU-55933 | MVP_top60_DNA+RNA_seed | ESR1 | 0.291541 | BRCA | HCC1143 | Fulvestrant |
| MVP_top60_DNA_seed | HDAC1 | 0.630533 | BRCA | MDA-MB-231 | Tubastatin A | MVP_top60_DNA+RNA_seed | MAPK3 | 0.291552 | BRCA | HCC38 | FR-180204 |
| MVP_top60_DNA_seed | EGFR | 0.630657 | BRCA | HCC1419 | Gefitinib | MVP_top60_DNA+RNA_seed | MAPK1 | 0.291552 | BRCA | HCC38 | FR-180204 |
| MVP_top60_DNA_seed | EGFR | 0.631566 | BRCA | Hs-578-T | Gefitinib | MVP_top60_DNA+RNA_seed | HSP90AA1 | 0.291787 | BRCA | HCC1428 | Tanespimycin |
| MVP_top60_DNA_seed | ATM | 0.632667 | BRCA | EFM-19 | KU-60019 | MVP_top60_DNA+RNA_seed | HSP90AB1 | 0.291787 | BRCA | HCC1428 | Tanespimycin |
| MVP_top60_DNA_seed | HDAC1 | 0.6329 | BRCA | UACC-893 | CAY10603 | MVP_top60_DNA+RNA_seed | CDK1 | 0.29219 | BRCA | CAL-120 | RO-3306 |
| MVP_top60_DNA_seed | HDAC1 | 0.633283 | BRCA | EVSA-T | JQ12 | MVP_top60_DNA+RNA_seed | MAPK1 | 0.292472 | BRCA | ZR-75-30 | VX-11e |
| MVP_top60_DNA_seed | HDAC2 | 0.633283 | BRCA | EVSA-T | JQ12 | MVP_top60_DNA+RNA_seed | HSP90AA1 | 0.293983 | BRCA | MDA-MB-231 | Luminespib |
| MVP_top60_DNA_seed | MDM2 | 0.633327 | BRCA | MDA-MB-436 | Nutlin-3a (-) | MVP_top60_DNA+RNA_seed | HSP90AB1 | 0.293983 | BRCA | MDA-MB-231 | Luminespib |
| MVP_top60_DNA_seed | ERBB2 | 0.634073 | BRCA | BT-20 | CP724714 | MVP_top60_DNA+RNA_seed | SRC | 0.295214 | BRCA | HCC1599 | WH-4-023 |
| MVP_top60_DNA_seed | HSP90AA1 | 0.634147 | BRCA | MDA-MB-436 | Luminespib | MVP_top60_DNA+RNA_seed | AR | 0.3003 | BRCA | HDQ-P1 | Bicalutamide |
| MVP_top60_DNA_seed | HSP90AB1 | 0.634147 | BRCA | MDA-MB-436 | Luminespib | MVP_top60_DNA+RNA_seed | EGFR | 0.301572 | BRCA | MRK-nu-1 | AZD8931 |
| MVP_top60_DNA_seed | MDM2 | 0.634267 | BRCA | MDA-MB-468 | Nutlin-3a (-) | MVP_top60_DNA+RNA_seed | ERBB2 | 0.301572 | BRCA | MRK-nu-1 | AZD8931 |
| MVP_top60_DNA_seed | EGFR | 0.635143 | BRCA | CAL-148 | Pelitinib | MVP_top60_DNA+RNA_seed | ERBB3 | 0.301572 | BRCA | MRK-nu-1 | AZD8931 |
| MVP_top60_DNA_seed | EGFR | 0.635787 | BRCA | EVSA-T | Gefitinib | MVP_top60_DNA+RNA_seed | SRC | 0.302479 | BRCA | T47D | NVP-BHG712 |
| MVP_top60_DNA_seed | HDAC1 | 0.638395 | BRCA | HCC1395 | Tubastatin A | MVP_top60_DNA+RNA_seed | AR | 0.30375 | BRCA | MDA-MB-361 | AZD3514 |
| MVP_top60_DNA_seed | SRC | 0.639092 | BRCA | AU565 | Bosutinib | MVP_top60_DNA+RNA_seed | HDAC1 | 0.3063 | BRCA | CAL-120 | AR-42 |
| MVP_top60_DNA_seed | HDAC1 | 0.63959 | BRCA | Hs-578-T | CAY10603 | MVP_top60_DNA+RNA_seed | EGFR | 0.306594 | BRCA | BT-20 | CUDC-101 |
| MVP_top60_DNA_seed | HDAC1 | 0.640473 | BRCA | BT-20 | CAY10603 | MVP_top60_DNA+RNA_seed | ERBB2 | 0.306594 | BRCA | BT-20 | CUDC-101 |
| MVP_top60_DNA_seed | HDAC1 | 0.642319 | BRCA | BT-483 | Dacinostat | MVP_top60_DNA+RNA_seed | MAPK3 | 0.311931 | BRCA | HCC1569 | FR-180204 |
| MVP_top60_DNA_seed | HDAC1 | 0.645582 | BRCA | BT-474 | AR-42 | MVP_top60_DNA+RNA_seed | MAPK1 | 0.311931 | BRCA | HCC1569 | FR-180204 |
| MVP_top60_DNA_seed | AKT1 | 0.645613 | BRCA | HCC1428 | AKT inhibitor VIII | MVP_top60_DNA+RNA_seed | CDK1 | 0.314311 | BRCA | HCC1187 | RO-3306 |
| MVP_top60_DNA_seed | AKT1 | 0.645868 | BRCA | BT-549 | MK-2206 | MVP_top60_DNA+RNA_seed | SRC | 0.314802 | BRCA | HCC1187 | WH-4-023 |
| MVP_top60_DNA_seed | HDAC1 | 0.645925 | BRCA | HCC2157 | AR-42 | MVP_top60_DNA+RNA_seed | ERBB2 | 0.317858 | BRCA | HCC38 | CP724714 |
| MVP_top60_DNA_seed | SRC | 0.646892 | BRCA | MDA-MB-453 | Bosutinib | MVP_top60_DNA+RNA_seed | ERBB2 | 0.319282 | BRCA | HCC1428 | CP724714 |
| MVP_top60_DNA_seed | ESR1 | 0.649359 | BRCA | MRK-nu-1 | Tamoxifen | MVP_top60_DNA+RNA_seed | EGFR | 0.31949 | BRCA | MFM-223 | AZD8931 |
| MVP_top60_DNA_seed | HDAC1 | 0.650665 | BRCA | UACC-893 | JQ12 | MVP_top60_DNA+RNA_seed | ERBB2 | 0.31949 | BRCA | MFM-223 | AZD8931 |
| MVP_top60_DNA_seed | HDAC2 | 0.650665 | BRCA | UACC-893 | JQ12 | MVP_top60_DNA+RNA_seed | ERBB3 | 0.31949 | BRCA | MFM-223 | AZD8931 |
| MVP_top60_DNA_seed | PIK3CA | 0.650719 | BRCA | HCC70 | AZD8835 | MVP_top60_DNA+RNA_seed | EGFR | 0.320176 | BRCA | MDA-MB-415 | AST-1306 |
| MVP_top60_DNA_seed | HSP90AA1 | 0.652526 | BRCA | HCC2218 | Elesclomol | MVP_top60_DNA+RNA_seed | AKT1 | 0.320529 | BRCA | HCC1428 | AKT inhibitor VIII |
| MVP_top60_DNA_seed | HSP90AB1 | 0.652526 | BRCA | HCC2218 | Elesclomol | MVP_top60_DNA+RNA_seed | MAPK14 | 0.322519 | BRCA | BT-20 | TAK-715 |
| MVP_top60_DNA_seed | SRC | 0.653248 | BRCA | MDA-MB-231 | Bosutinib | MVP_top60_DNA+RNA_seed | SRC | 0.322569 | BRCA | MDA-MB-175-VII | Bosutinib |

| | | | | | | | | | | | |
|---|---|---|---|---|---|---|---|---|---|---|---|
| MVP_top60_DNA_seed | EGFR | 0.653912 | BRCA | HCC1428 | Gefitinib | MVP_top60_DNA+RNA_seed | HSP90AA1 | 0.322793 | BRCA | HCC70 | Elesclomol |
| MVP_top60_DNA_seed | MAPK3 | 0.656137 | BRCA | MDA-MB-231 | FR-180204 | MVP_top60_DNA+RNA_seed | HSP90AB1 | 0.322793 | BRCA | HCC70 | Elesclomol |
| MVP_top60_DNA_seed | MAPK1 | 0.656137 | BRCA | MDA-MB-231 | FR-180204 | MVP_top60_DNA+RNA_seed | HSP90AA1 | 0.323507 | BRCA | DU-4475 | Elesclomol |
| MVP_top60_DNA_seed | SRC | 0.657088 | BRCA | EVSA-T | Bosutinib | MVP_top60_DNA+RNA_seed | HSP90AB1 | 0.323507 | BRCA | DU-4475 | Elesclomol |
| MVP_top60_DNA_seed | HSP90AA1 | 0.658284 | BRCA | UACC-893 | SNX-2112 | MVP_top60_DNA+RNA_seed | HDAC1 | 0.326547 | BRCA | CAL-120 | Dacinostat |
| MVP_top60_DNA_seed | HSP90AB1 | 0.658284 | BRCA | UACC-893 | SNX-2112 | MVP_top60_DNA+RNA_seed | EGFR | 0.329717 | BRCA | HCC1187 | CUDC-101 |
| MVP_top60_DNA_seed | ERBB2 | 0.658619 | BRCA | HCC1937 | CP724714 | MVP_top60_DNA+RNA_seed | ERBB2 | 0.329717 | BRCA | HCC1187 | CUDC-101 |
| MVP_top60_DNA_seed | SRC | 0.661681 | BRCA | BT-20 | Bosutinib | MVP_top60_DNA+RNA_seed | HDAC1 | 0.330276 | BRCA | HCC1500 | JQ12 |
| MVP_top60_DNA_seed | HDAC1 | 0.661751 | BRCA | MDA-MB-415 | Dacinostat | MVP_top60_DNA+RNA_seed | HDAC2 | 0.330276 | BRCA | HCC1500 | JQ12 |
| MVP_top60_DNA_seed | ESR1 | 0.66269 | BRCA | MDA-MB-157 | Tamoxifen | MVP_top60_DNA+RNA_seed | SRC | 0.331932 | BRCA | UACC-812 | NVP-BHG712 |
| MVP_top60_DNA_seed | HSP90AA1 | 0.663046 | BRCA | MDA-MB-415 | Elesclomol | MVP_top60_DNA+RNA_seed | SRC | 0.333426 | BRCA | DU-4475 | WH-4-023 |
| MVP_top60_DNA_seed | HSP90AB1 | 0.663046 | BRCA | MDA-MB-415 | Elesclomol | MVP_top60_DNA+RNA_seed | EGFR | 0.334683 | BRCA | HDQ-P1 | AZD8931 |
| MVP_top60_DNA_seed | ATM | 0.663625 | BRCA | CAL-85-1 | KU-55933 | MVP_top60_DNA+RNA_seed | ERBB2 | 0.334683 | BRCA | HDQ-P1 | AZD8931 |
| MVP_top60_DNA_seed | HDAC1 | 0.665461 | BRCA | BT-549 | Belinostat | MVP_top60_DNA+RNA_seed | ERBB3 | 0.334683 | BRCA | HDQ-P1 | AZD8931 |
| MVP_top60_DNA_seed | SRC | 0.668118 | BRCA | MRK-nu-1 | Saracatinib | MVP_top60_DNA+RNA_seed | EGFR | 0.335152 | BRCA | HCC1187 | Cetuximab |
| MVP_top60_DNA_seed | CDK2 | 0.669238 | BRCA | HDQ-P1 | PHA-793887 | MVP_top60_DNA+RNA_seed | AKT1 | 0.336009 | BRCA | MDA-MB-330 | GSK690693 |
| MVP_top60_DNA_seed | MAPK1 | 0.669275 | BRCA | MDA-MB-361 | VX-11e | MVP_top60_DNA+RNA_seed | AR | 0.338021 | BRCA | CAL-51 | AZD3514 |
| MVP_top60_DNA_seed | ERBB2 | 0.669388 | BRCA | CAL-120 | PF-00299804 | MVP_top60_DNA+RNA_seed | HSP90AA1 | 0.33898 | BRCA | T47D | SNX-2112 |
| MVP_top60_DNA_seed | EGFR | 0.669388 | BRCA | CAL-120 | PF-00299804 | MVP_top60_DNA+RNA_seed | HSP90AB1 | 0.33898 | BRCA | T47D | SNX-2112 |
| MVP_top60_DNA_seed | ESR1 | 0.671297 | BRCA | HCC70 | Tamoxifen | MVP_top60_DNA+RNA_seed | EGFR | 0.339472 | BRCA | HCC1569 | Cetuximab |
| MVP_top60_DNA_seed | ESR1 | 0.671432 | BRCA | HCC38 | Tamoxifen | MVP_top60_DNA+RNA_seed | CDK1 | 0.340687 | BRCA | MRK-nu-1 | RO-3306 |
| MVP_top60_DNA_seed | MDM2 | 0.671566 | BRCA | JIMT-1 | Nutlin-3a (-) | MVP_top60_DNA+RNA_seed | HDAC1 | 0.341897 | BRCA | HCC2157 | CAY10603 |
| MVP_top60_DNA_seed | SRC | 0.671946 | BRCA | HCC1937 | NVP-BHG712 | MVP_top60_DNA+RNA_seed | HDAC1 | 0.342318 | BRCA | EFM-192A | Belinostat |
| MVP_top60_DNA_seed | ERBB2 | 0.672107 | BRCA | JIMT-1 | CUDC-101 | MVP_top60_DNA+RNA_seed | EGFR | 0.34598 | BRCA | CAL-120 | CUDC-101 |
| MVP_top60_DNA_seed | EGFR | 0.672107 | BRCA | JIMT-1 | CUDC-101 | MVP_top60_DNA+RNA_seed | ERBB2 | 0.34598 | BRCA | CAL-120 | CUDC-101 |
| MVP_top60_DNA_seed | ERBB2 | 0.672211 | BRCA | MDA-MB-436 | AZD8931 | MVP_top60_DNA+RNA_seed | AKT1 | 0.347618 | BRCA | EVSA-T | AKT inhibitor VIII |
| MVP_top60_DNA_seed | EGFR | 0.672211 | BRCA | MDA-MB-436 | AZD8931 | MVP_top60_DNA+RNA_seed | ATM | 0.348123 | BRCA | MDA-MB-436 | CP466722 |
| MVP_top60_DNA_seed | ERBB3 | 0.672211 | BRCA | MDA-MB-436 | AZD8931 | MVP_top60_DNA+RNA_seed | AKT1 | 0.348169 | BRCA | UACC-812 | AKT inhibitor VIII |
| MVP_top60_DNA_seed | SRC | 0.673371 | BRCA | HCC202 | NVP-BHG712 | MVP_top60_DNA+RNA_seed | HDAC1 | 0.349991 | BRCA | CAMA-1 | Tubastatin A |
| MVP_top60_DNA_seed | PIK3CA | 0.674489 | BRCA | BT-549 | AZD8835 | MVP_top60_DNA+RNA_seed | CDK2 | 0.351101 | BRCA | HCC2157 | AZD5438 |
| MVP_top60_DNA_seed | MDM2 | 0.674932 | BRCA | EVSA-T | Nutlin-3a (-) | MVP_top60_DNA+RNA_seed | HSP90AA1 | 0.351219 | BRCA | HCC1428 | Elesclomol |
| MVP_top60_DNA_seed | ERBB2 | 0.676459 | BRCA | HCC1187 | Afatinib | MVP_top60_DNA+RNA_seed | HSP90AB1 | 0.351219 | BRCA | HCC1428 | Elesclomol |
| MVP_top60_DNA_seed | EGFR | 0.676459 | BRCA | HCC1187 | Afatinib | MVP_top60_DNA+RNA_seed | EGFR | 0.352074 | BRCA | COLO-824 | Afatinib |
| MVP_top60_DNA_seed | PIK3CA | 0.676875 | BRCA | HCC1419 | PI3Ka_4409 | MVP_top60_DNA+RNA_seed | ERBB2 | 0.352074 | BRCA | COLO-824 | Afatinib |
| MVP_top60_DNA_seed | HDAC1 | 0.676988 | BRCA | HCC1395 | Dacinostat | MVP_top60_DNA+RNA_seed | MAPK1 | 0.35362 | BRCA | JIMT-1 | VX-11e |
| MVP_top60_DNA_seed | EP300 | 0.679417 | BRCA | CAL-85-1 | I-CBP112 | MVP_top60_DNA+RNA_seed | SRC | 0.35421 | BRCA | HCC1187 | Saracatinib |
| MVP_top60_DNA_seed | CREBBP | 0.679417 | BRCA | CAL-85-1 | I-CBP112 | MVP_top60_DNA+RNA_seed | EGFR | 0.354893 | BRCA | HCC1395 | Cetuximab |
| MVP_top60_DNA_seed | HDAC1 | 0.680194 | BRCA | MRK-nu-1 | AR-42 | MVP_top60_DNA+RNA_seed | HSP90AA1 | 0.356661 | BRCA | AU565 | Luminespib |
| MVP_top60_DNA_seed | HSP90AA1 | 0.681098 | BRCA | HCC1428 | Luminespib | MVP_top60_DNA+RNA_seed | HSP90AB1 | 0.356661 | BRCA | AU565 | Luminespib |
| MVP_top60_DNA_seed | HSP90AB1 | 0.681098 | BRCA | HCC1428 | Luminespib | MVP_top60_DNA+RNA_seed | PIK3CA | 0.357184 | BRCA | BT-483 | PI3Ka_4409 |
| MVP_top60_DNA_seed | SRC | 0.681715 | BRCA | CAL-120 | Bosutinib | MVP_top60_DNA+RNA_seed | AKT1 | 0.360402 | BRCA | HCC1395 | AKT inhibitor VIII |

| | | | | | | | | | | | |
|---|---|---|---|---|---|---|---|---|---|---|---|
| MVP_top60_DNA_seed | ERBB2 | 0.682007 | BRCA | T47D | CI-1033 | MVP_top60_DNA+RNA_seed | MAPK3 | 0.360534 | BRCA | ZR-75-30 | FR-180204 |
| MVP_top60_DNA_seed | EGFR | 0.682007 | BRCA | T47D | CI-1033 | MVP_top60_DNA+RNA_seed | MAPK1 | 0.360534 | BRCA | ZR-75-30 | FR-180204 |
| MVP_top60_DNA_seed | AR | 0.682349 | BRCA | BT-474 | AZD3514 | MVP_top60_DNA+RNA_seed | EGFR | 0.362866 | BRCA | DU-4475 | Erlotinib |
| MVP_top60_DNA_seed | ATM | 0.683365 | BRCA | HCC1395 | CP466722 | MVP_top60_DNA+RNA_seed | HSP90AA1 | 0.363172 | BRCA | CAL-85-1 | Elesclomol |
| MVP_top60_DNA_seed | HDAC1 | 0.684444 | BRCA | BT-549 | Dacinostat | MVP_top60_DNA+RNA_seed | HSP90AB1 | 0.363172 | BRCA | CAL-85-1 | Elesclomol |
| MVP_top60_DNA_seed | HSP90AA1 | 0.684676 | BRCA | MDA-MB-436 | SNX-2112 | MVP_top60_DNA+RNA_seed | CDK2 | 0.363205 | BRCA | MRK-nu-1 | PHA-793887 |
| MVP_top60_DNA_seed | HSP90AB1 | 0.684676 | BRCA | MDA-MB-436 | SNX-2112 | MVP_top60_DNA+RNA_seed | EGFR | 0.366594 | BRCA | EVSA-T | Cetuximab |
| MVP_top60_DNA_seed | EP300 | 0.686821 | BRCA | MFM-223 | I-CBP112 | MVP_top60_DNA+RNA_seed | HSP90AA1 | 0.36707 | BRCA | HCC70 | CCT-018159 |
| MVP_top60_DNA_seed | CREBBP | 0.686821 | BRCA | MFM-223 | I-CBP112 | MVP_top60_DNA+RNA_seed | HSP90AB1 | 0.36707 | BRCA | HCC70 | CCT-018159 |
| MVP_top60_DNA_seed | HSP90AA1 | 0.687342 | BRCA | CAMA-1 | SNX-2112 | MVP_top60_DNA+RNA_seed | HSP90AA1 | 0.37156 | BRCA | HCC38 | CCT-018159 |
| MVP_top60_DNA_seed | HSP90AB1 | 0.687342 | BRCA | CAMA-1 | SNX-2112 | MVP_top60_DNA+RNA_seed | HSP90AB1 | 0.37156 | BRCA | HCC38 | CCT-018159 |
| MVP_top60_DNA_seed | ERBB2 | 0.690179 | BRCA | HCC1428 | Afatinib | MVP_top60_DNA+RNA_seed | EGFR | 0.371745 | BRCA | CAL-120 | AST-1306 |
| MVP_top60_DNA_seed | EGFR | 0.690179 | BRCA | HCC1428 | Afatinib | MVP_top60_DNA+RNA_seed | HDAC1 | 0.372329 | BRCA | MDA-MB-231 | CAY10603 |
| MVP_top60_DNA_seed | EGFR | 0.690557 | BRCA | HCC1428 | Pelitinib | MVP_top60_DNA+RNA_seed | PLK1 | 0.374765 | BRCA | EVSA-T | GW843682X |
| MVP_top60_DNA_seed | SRC | 0.69163 | BRCA | HCC1500 | NVP-BHG712 | MVP_top60_DNA+RNA_seed | CDK2 | 0.375242 | BRCA | CAMA-1 | PHA-793887 |
| MVP_top60_DNA_seed | CDK2 | 0.692347 | BRCA | MDA-MB-453 | AZD5438 | MVP_top60_DNA+RNA_seed | AR | 0.375664 | BRCA | Hs-578-T | Bicalutamide |
| MVP_top60_DNA_seed | HDAC1 | 0.695362 | BRCA | HCC1187 | Entinostat | MVP_top60_DNA+RNA_seed | SRC | 0.376331 | BRCA | MFM-223 | NVP-BHG712 |
| MVP_top60_DNA_seed | ERBB2 | 0.6967 | BRCA | EVSA-T | Afatinib | MVP_top60_DNA+RNA_seed | AR | 0.377115 | BRCA | COLO-824 | Bicalutamide |
| MVP_top60_DNA_seed | EGFR | 0.6967 | BRCA | EVSA-T | Afatinib | MVP_top60_DNA+RNA_seed | EGFR | 0.379639 | BRCA | MDA-MB-157 | Afatinib |
| MVP_top60_DNA_seed | ERBB2 | 0.700015 | BRCA | HCC1500 | Afatinib | MVP_top60_DNA+RNA_seed | ERBB2 | 0.379639 | BRCA | MDA-MB-157 | Afatinib |
| MVP_top60_DNA_seed | EGFR | 0.700015 | BRCA | HCC1500 | Afatinib | MVP_top60_DNA+RNA_seed | AKT1 | 0.380008 | BRCA | HCC1569 | MK-2206 |
| MVP_top60_DNA_seed | HDAC1 | 0.700463 | BRCA | CAL-51 | Tubastatin A | MVP_top60_DNA+RNA_seed | HSP90AA1 | 0.380733 | BRCA | HCC202 | Luminespib |
| MVP_top60_DNA_seed | HDAC1 | 0.700607 | BRCA | YMB-1-E | Belinostat | MVP_top60_DNA+RNA_seed | HSP90AB1 | 0.380733 | BRCA | HCC202 | Luminespib |
| MVP_top60_DNA_seed | AR | 0.702264 | BRCA | UACC-812 | Bicalutamide | MVP_top60_DNA+RNA_seed | HSP90AA1 | 0.382742 | BRCA | MFM-223 | SNX-2112 |
| MVP_top60_DNA_seed | MAPK3 | 0.705806 | BRCA | HCC1937 | FR-180204 | MVP_top60_DNA+RNA_seed | HSP90AB1 | 0.382742 | BRCA | MFM-223 | SNX-2112 |
| MVP_top60_DNA_seed | MAPK1 | 0.705806 | BRCA | HCC1937 | FR-180204 | MVP_top60_DNA+RNA_seed | EGFR | 0.382975 | BRCA | HCC1187 | Lapatinib |
| MVP_top60_DNA_seed | AR | 0.70582 | BRCA | BT-549 | Bicalutamide | MVP_top60_DNA+RNA_seed | ERBB2 | 0.382975 | BRCA | HCC1187 | Lapatinib |
| MVP_top60_DNA_seed | SRC | 0.70602 | BRCA | CAL-51 | NVP-BHG712 | MVP_top60_DNA+RNA_seed | ATM | 0.383024 | BRCA | HCC2218 | CP466722 |
| MVP_top60_DNA_seed | SRC | 0.706497 | BRCA | HCC1419 | NVP-BHG712 | MVP_top60_DNA+RNA_seed | AR | 0.383444 | BRCA | CAL-85-1 | Bicalutamide |
| MVP_top60_DNA_seed | MAPK3 | 0.706987 | BRCA | MRK-nu-1 | FR-180204 | MVP_top60_DNA+RNA_seed | CDK2 | 0.383499 | BRCA | AU565 | AZD5438 |
| MVP_top60_DNA_seed | MAPK1 | 0.706987 | BRCA | MRK-nu-1 | FR-180204 | MVP_top60_DNA+RNA_seed | AR | 0.384808 | BRCA | MDA-MB-361 | Bicalutamide |
| MVP_top60_DNA_seed | AR | 0.70817 | BRCA | HCC1569 | AZD3514 | MVP_top60_DNA+RNA_seed | AKT1 | 0.385717 | BRCA | HCC1187 | A-443654 |
| MVP_top60_DNA_seed | ERBB2 | 0.70822 | BRCA | HCC38 | Afatinib | MVP_top60_DNA+RNA_seed | CDK2 | 0.385996 | BRCA | MDA-MB-330 | PHA-793887 |
| MVP_top60_DNA_seed | EGFR | 0.70822 | BRCA | HCC38 | Afatinib | MVP_top60_DNA+RNA_seed | CDK1 | 0.387682 | BRCA | MDA-MB-157 | RO-3306 |
| MVP_top60_DNA_seed | HDAC1 | 0.708266 | BRCA | JIMT-1 | AR-42 | MVP_top60_DNA+RNA_seed | EP300 | 0.388149 | BRCA | MCF7 | I-CBP112 |
| MVP_top60_DNA_seed | MDM2 | 0.708748 | BRCA | MDA-MB-231 | Nutlin-3a (-) | MVP_top60_DNA+RNA_seed | EGFR | 0.39012 | BRCA | CAL-148 | Lapatinib |
| MVP_top60_DNA_seed | AKT1 | 0.709522 | BRCA | COLO-824 | AKT inhibitor VIII | MVP_top60_DNA+RNA_seed | ERBB2 | 0.39012 | BRCA | CAL-148 | Lapatinib |
| MVP_top60_DNA_seed | CDK2 | 0.709921 | BRCA | OCUB-M | AZD5438 | MVP_top60_DNA+RNA_seed | AR | 0.391131 | BRCA | MDA-MB-157 | Bicalutamide |
| MVP_top60_DNA_seed | HSP90AA1 | 0.711067 | BRCA | UACC-893 | CCT-018159 | MVP_top60_DNA+RNA_seed | ERBB2 | 0.392644 | BRCA | YMB-1-E | CP724714 |
| MVP_top60_DNA_seed | HSP90AB1 | 0.711067 | BRCA | UACC-893 | CCT-018159 | MVP_top60_DNA+RNA_seed | HDAC1 | 0.394204 | BRCA | JIMT-1 | CAY10603 |
| MVP_top60_DNA_seed | HSP90AA1 | 0.711759 | BRCA | ZR-75-30 | CCT-018159 | MVP_top60_DNA+RNA_seed | EGFR | 0.39458 | BRCA | HCC1187 | Erlotinib |

| | | | | | | | | | | | |
|---|---|---|---|---|---|---|---|---|---|---|---|
| MVP_top60_DNA_seed | HSP90AB1 | 0.711759 | BRCA | ZR-75-30 | CCT-018159 | MVP_top60_DNA+RNA_seed | MAPK1 | 0.394598 | BRCA | HCC1187 | VX-11e |
| MVP_top60_DNA_seed | HDAC1 | 0.712204 | BRCA | HCC1419 | CAY10603 | MVP_top60_DNA+RNA_seed | AR | 0.396533 | BRCA | HCC1428 | Bicalutamide |
| MVP_top60_DNA_seed | HSP90AA1 | 0.713635 | BRCA | YMB-1-E | SNX-2112 | MVP_top60_DNA+RNA_seed | EGFR | 0.397457 | BRCA | DU-4475 | Cetuximab |
| MVP_top60_DNA_seed | HSP90AB1 | 0.713635 | BRCA | YMB-1-E | SNX-2112 | MVP_top60_DNA+RNA_seed | EGFR | 0.398169 | BRCA | MDA-MB-231 | Afatinib |
| MVP_top60_DNA_seed | HSP90AA1 | 0.714584 | BRCA | EFM-192A | SNX-2112 | MVP_top60_DNA+RNA_seed | ERBB2 | 0.398169 | BRCA | MDA-MB-231 | Afatinib |
| MVP_top60_DNA_seed | HSP90AB1 | 0.714584 | BRCA | EFM-192A | SNX-2112 | MVP_top60_DNA+RNA_seed | ATM | 0.400196 | BRCA | EFM-192A | KU-55933 |
| MVP_top60_DNA_seed | SRC | 0.715428 | BRCA | ZR-75-30 | NVP-BHG712 | MVP_top60_DNA+RNA_seed | CDK2 | 0.406485 | BRCA | HCC1569 | AZD5438 |
| MVP_top60_DNA_seed | CDK1 | 0.715652 | BRCA | T47D | RO-3306 | MVP_top60_DNA+RNA_seed | EGFR | 0.40687 | BRCA | BT-483 | Cetuximab |
| MVP_top60_DNA_seed | EGFR | 0.718217 | BRCA | UACC-812 | AST-1306 | MVP_top60_DNA+RNA_seed | EGFR | 0.407315 | BRCA | MDA-MB-231 | AZD8931 |
| MVP_top60_DNA_seed | CDK2 | 0.718726 | BRCA | MDA-MB-175-VII | PHA-793887 | MVP_top60_DNA+RNA_seed | ERBB2 | 0.407315 | BRCA | MDA-MB-231 | AZD8931 |
| MVP_top60_DNA_seed | ERBB2 | 0.719779 | BRCA | HCC38 | Afatinib | MVP_top60_DNA+RNA_seed | ERBB3 | 0.407315 | BRCA | MDA-MB-231 | AZD8931 |
| MVP_top60_DNA_seed | EGFR | 0.719779 | BRCA | HCC38 | Afatinib | MVP_top60_DNA+RNA_seed | EGFR | 0.407404 | BRCA | CAL-85-1 | Afatinib |
| MVP_top60_DNA_seed | ERBB2 | 0.720678 | BRCA | MRK-nu-1 | Lapatinib | MVP_top60_DNA+RNA_seed | ERBB2 | 0.407404 | BRCA | CAL-85-1 | Afatinib |
| MVP_top60_DNA_seed | EGFR | 0.720678 | BRCA | MRK-nu-1 | Lapatinib | MVP_top60_DNA+RNA_seed | HSP90AA1 | 0.40808 | BRCA | HDQ-P1 | Luminespib |
| MVP_top60_DNA_seed | ATM | 0.721892 | BRCA | YMB-1-E | CP466722 | MVP_top60_DNA+RNA_seed | HSP90AB1 | 0.40808 | BRCA | HDQ-P1 | Luminespib |
| MVP_top60_DNA_seed | EP300 | 0.721992 | BRCA | MRK-nu-1 | I-CBP112 | MVP_top60_DNA+RNA_seed | EGFR | 0.410087 | BRCA | Hs-578-T | CUDC-101 |
| MVP_top60_DNA_seed | CREBBP | 0.721992 | BRCA | MRK-nu-1 | I-CBP112 | MVP_top60_DNA+RNA_seed | ERBB2 | 0.410087 | BRCA | Hs-578-T | CUDC-101 |
| MVP_top60_DNA_seed | MAPK1 | 0.722424 | BRCA | CAL-148 | VX-11e | MVP_top60_DNA+RNA_seed | HDAC1 | 0.411973 | BRCA | HCC38 | JQ12 |
| MVP_top60_DNA_seed | SRC | 0.722521 | BRCA | DU-4475 | Saracatinib | MVP_top60_DNA+RNA_seed | HDAC2 | 0.411973 | BRCA | HCC38 | JQ12 |
| MVP_top60_DNA_seed | HDAC1 | 0.723137 | BRCA | BT-549 | CAY10603 | MVP_top60_DNA+RNA_seed | CDK2 | 0.412647 | BRCA | UACC-893 | PHA-793887 |
| MVP_top60_DNA_seed | EGFR | 0.723789 | BRCA | HCC1419 | Cetuximab | MVP_top60_DNA+RNA_seed | AR | 0.412676 | BRCA | MDA-MB-157 | AZD3514 |
| MVP_top60_DNA_seed | SRC | 0.724489 | BRCA | YMB-1-E | NVP-BHG712 | MVP_top60_DNA+RNA_seed | EGFR | 0.412689 | BRCA | YMB-1-E | AST-1306 |
| MVP_top60_DNA_seed | ERBB2 | 0.726607 | BRCA | MDA-MB-436 | Afatinib | MVP_top60_DNA+RNA_seed | AKT1 | 0.413122 | BRCA | CAL-120 | MK-2206 |
| MVP_top60_DNA_seed | EGFR | 0.726607 | BRCA | MDA-MB-436 | Afatinib | MVP_top60_DNA+RNA_seed | EGFR | 0.416688 | BRCA | MDA-MB-415 | AZD8931 |
| MVP_top60_DNA_seed | ATM | 0.727851 | BRCA | MDA-MB-415 | CP466722 | MVP_top60_DNA+RNA_seed | ERBB2 | 0.416688 | BRCA | MDA-MB-415 | AZD8931 |
| MVP_top60_DNA_seed | CDK2 | 0.728712 | BRCA | HCC1937 | AZD5438 | MVP_top60_DNA+RNA_seed | ERBB3 | 0.416688 | BRCA | MDA-MB-415 | AZD8931 |
| MVP_top60_DNA_seed | ESR1 | 0.731603 | BRCA | Hs-578-T | Tamoxifen | MVP_top60_DNA+RNA_seed | CDK2 | 0.417262 | BRCA | MDA-MB-436 | AZD5438 |
| MVP_top60_DNA_seed | ATM | 0.732635 | BRCA | CAL-120 | KU-55933 | MVP_top60_DNA+RNA_seed | EGFR | 0.419688 | BRCA | MCF7 | Cetuximab |
| MVP_top60_DNA_seed | HDAC1 | 0.733702 | BRCA | COLO-824 | JQ12 | MVP_top60_DNA+RNA_seed | EGFR | 0.419985 | BRCA | HCC1937 | AZD8931 |
| MVP_top60_DNA_seed | HDAC2 | 0.733702 | BRCA | COLO-824 | JQ12 | MVP_top60_DNA+RNA_seed | ERBB2 | 0.419985 | BRCA | HCC1937 | AZD8931 |
| MVP_top60_DNA_seed | SRC | 0.733831 | BRCA | EFM-192A | NVP-BHG712 | MVP_top60_DNA+RNA_seed | ERBB3 | 0.419985 | BRCA | HCC1937 | AZD8931 |
| MVP_top60_DNA_seed | ERBB2 | 0.736135 | BRCA | MDA-MB-436 | CUDC-101 | MVP_top60_DNA+RNA_seed | HDAC1 | 0.420588 | BRCA | HCC1954 | Dacinostat |
| MVP_top60_DNA_seed | EGFR | 0.736135 | BRCA | MDA-MB-436 | CUDC-101 | MVP_top60_DNA+RNA_seed | AKT1 | 0.421063 | BRCA | MDA-MB-436 | AKT inhibitor VIII |
| MVP_top60_DNA_seed | AR | 0.738074 | BRCA | MRK-nu-1 | Bicalutamide | MVP_top60_DNA+RNA_seed | EGFR | 0.424701 | BRCA | CAL-120 | Afatinib |
| MVP_top60_DNA_seed | AR | 0.73952 | BRCA | BT-20 | Bicalutamide | MVP_top60_DNA+RNA_seed | ERBB2 | 0.424701 | BRCA | CAL-120 | Afatinib |
| MVP_top60_DNA_seed | MAPK3 | 0.740058 | BRCA | HCC1428 | FR-180204 | MVP_top60_DNA+RNA_seed | SRC | 0.427708 | BRCA | MFM-223 | Bosutinib |
| MVP_top60_DNA_seed | MAPK1 | 0.740058 | BRCA | HCC1428 | FR-180204 | MVP_top60_DNA+RNA_seed | CDK2 | 0.429085 | BRCA | COLO-824 | PHA-793887 |
| MVP_top60_DNA_seed | ERBB2 | 0.740161 | BRCA | HCC70 | CP724714 | MVP_top60_DNA+RNA_seed | CDK2 | 0.429125 | BRCA | EVSA-T | Seliciclib |
| MVP_top60_DNA_seed | HSP90AA1 | 0.741213 | BRCA | HCC1419 | SNX-2112 | MVP_top60_DNA+RNA_seed | ATM | 0.430449 | BRCA | MDA-MB-175-VII | KU-55933 |
| MVP_top60_DNA_seed | HSP90AB1 | 0.741213 | BRCA | HCC1419 | SNX-2112 | MVP_top60_DNA+RNA_seed | MAPK14 | 0.431141 | BRCA | UACC-812 | TAK-715 |
| MVP_top60_DNA_seed | HDAC1 | 0.74274 | BRCA | CAL-148 | Parthenolide | MVP_top60_DNA+RNA_seed | ATM | 0.431319 | BRCA | UACC-893 | KU-55933 |

| | | | | | | | | | | | |
|---|---|---|---|---|---|---|---|---|---|---|---|
| MVP_top60_DNA_seed | EGFR | 0.74285 | BRCA | UACC-893 | Cetuximab | MVP_top60_DNA+RNA_seed | CDK2 | 0.431438 | BRCA | CAL-148 | PHA-793887 |
| MVP_top60_DNA_seed | SRC | 0.743804 | BRCA | HCC2157 | Saracatinib | MVP_top60_DNA+RNA_seed | EGFR | 0.431992 | BRCA | CAL-85-1 | CUDC-101 |
| MVP_top60_DNA_seed | AR | 0.744404 | BRCA | HCC202 | Bicalutamide | MVP_top60_DNA+RNA_seed | ERBB2 | 0.431992 | BRCA | CAL-85-1 | CUDC-101 |
| MVP_top60_DNA_seed | ATM | 0.7454 | BRCA | MDA-MB-361 | CP466722 | MVP_top60_DNA+RNA_seed | HSP90AA1 | 0.432539 | BRCA | CAL-85-1 | SNX-2112 |
| MVP_top60_DNA_seed | EGFR | 0.747787 | BRCA | CAMA-1 | Pelitinib | MVP_top60_DNA+RNA_seed | HSP90AB1 | 0.432539 | BRCA | CAL-85-1 | SNX-2112 |
| MVP_top60_DNA_seed | EGFR | 0.749414 | BRCA | BT-549 | Cetuximab | MVP_top60_DNA+RNA_seed | MDM2 | 0.433099 | BRCA | T47D | Nutlin-3a (-) |
| MVP_top60_DNA_seed | MAPK3 | 0.749428 | BRCA | MDA-MB-330 | FR-180204 | MVP_top60_DNA+RNA_seed | MAPK14 | 0.43529 | BRCA | JIMT-1 | TAK-715 |
| MVP_top60_DNA_seed | MAPK1 | 0.749428 | BRCA | MDA-MB-330 | FR-180204 | MVP_top60_DNA+RNA_seed | HSP90AA1 | 0.436866 | BRCA | HCC1954 | Tanespimycin |
| MVP_top60_DNA_seed | SRC | 0.750777 | BRCA | CAL-148 | NVP-BHG712 | MVP_top60_DNA+RNA_seed | HSP90AB1 | 0.436866 | BRCA | HCC1954 | Tanespimycin |
| MVP_top60_DNA_seed | SRC | 0.753952 | BRCA | ZR-75-30 | Bosutinib | MVP_top60_DNA+RNA_seed | HDAC1 | 0.436938 | BRCA | HDQ-P1 | Tubastatin A |
| MVP_top60_DNA_seed | PIK3CA | 0.754134 | BRCA | MDA-MB-468 | PI3Ka_4409 | MVP_top60_DNA+RNA_seed | ATM | 0.437909 | BRCA | MFM-223 | KU-60019 |
| MVP_top60_DNA_seed | CDK2 | 0.755465 | BRCA | HCC2157 | Seliciclib | MVP_top60_DNA+RNA_seed | AR | 0.439375 | BRCA | EFM-19 | Bicalutamide |
| MVP_top60_DNA_seed | MAPK1 | 0.756585 | BRCA | T47D | VX-11e | MVP_top60_DNA+RNA_seed | HSP90AA1 | 0.442632 | BRCA | Hs-578-T | CCT-018159 |
| MVP_top60_DNA_seed | EP300 | 0.756949 | BRCA | HCC1954 | I-CBP112 | MVP_top60_DNA+RNA_seed | HSP90AB1 | 0.442632 | BRCA | Hs-578-T | CCT-018159 |
| MVP_top60_DNA_seed | CREBBP | 0.756949 | BRCA | HCC1954 | I-CBP112 | MVP_top60_DNA+RNA_seed | HDAC1 | 0.445017 | BRCA | HCC1428 | Tubastatin A |
| MVP_top60_DNA_seed | EP300 | 0.757292 | BRCA | MDA-MB-415 | I-CBP112 | MVP_top60_DNA+RNA_seed | HDAC1 | 0.445725 | BRCA | HDQ-P1 | AR-42 |
| MVP_top60_DNA_seed | CREBBP | 0.757292 | BRCA | MDA-MB-415 | I-CBP112 | MVP_top60_DNA+RNA_seed | ERBB2 | 0.448714 | BRCA | JIMT-1 | CP724714 |
| MVP_top60_DNA_seed | AR | 0.757631 | BRCA | MRK-nu-1 | AZD3514 | MVP_top60_DNA+RNA_seed | CDK1 | 0.449258 | BRCA | MDA-MB-361 | RO-3306 |
| MVP_top60_DNA_seed | AR | 0.759007 | BRCA | AU565 | AZD3514 | MVP_top60_DNA+RNA_seed | CDK1 | 0.450365 | BRCA | MDA-MB-453 | RO-3306 |
| MVP_top60_DNA_seed | SRC | 0.761482 | BRCA | EFM-19 | Bosutinib | MVP_top60_DNA+RNA_seed | SRC | 0.451443 | BRCA | UACC-893 | Bosutinib |
| MVP_top60_DNA_seed | HDAC1 | 0.761535 | BRCA | MDA-MB-468 | AR-42 | MVP_top60_DNA+RNA_seed | AR | 0.451459 | BRCA | MFM-223 | Bicalutamide |
| MVP_top60_DNA_seed | ERBB2 | 0.761735 | BRCA | HCC1954 | CUDC-101 | MVP_top60_DNA+RNA_seed | HDAC1 | 0.452287 | BRCA | MRK-nu-1 | Dacinostat |
| MVP_top60_DNA_seed | EGFR | 0.761735 | BRCA | HCC1954 | CUDC-101 | MVP_top60_DNA+RNA_seed | HDAC1 | 0.455539 | BRCA | CAL-148 | JQ12 |
| MVP_top60_DNA_seed | HDAC1 | 0.762701 | BRCA | HCC1187 | AR-42 | MVP_top60_DNA+RNA_seed | HDAC2 | 0.455539 | BRCA | CAL-148 | JQ12 |
| MVP_top60_DNA_seed | HSP90AA1 | 0.762805 | BRCA | MFM-223 | CCT-018159 | MVP_top60_DNA+RNA_seed | MDM2 | 0.457127 | BRCA | EFM-192A | Serdemetan |
| MVP_top60_DNA_seed | HSP90AB1 | 0.762805 | BRCA | MFM-223 | CCT-018159 | MVP_top60_DNA+RNA_seed | AKT1 | 0.457567 | BRCA | DU-4475 | MK-2206 |
| MVP_top60_DNA_seed | SRC | 0.7645 | BRCA | Hs-578-T | Bosutinib | MVP_top60_DNA+RNA_seed | HDAC1 | 0.458625 | BRCA | HCC1937 | AR-42 |
| MVP_top60_DNA_seed | HDAC1 | 0.764697 | BRCA | MDA-MB-436 | JQ12 | MVP_top60_DNA+RNA_seed | MAPK14 | 0.458704 | BRCA | HCC2157 | TAK-715 |
| MVP_top60_DNA_seed | HDAC2 | 0.764697 | BRCA | MDA-MB-436 | JQ12 | MVP_top60_DNA+RNA_seed | MAPK14 | 0.459822 | BRCA | HCC1428 | TAK-715 |
| MVP_top60_DNA_seed | AKT1 | 0.765361 | BRCA | MCF7 | GSK690693 | MVP_top60_DNA+RNA_seed | EGFR | 0.459879 | BRCA | EVSA-T | AST-1306 |
| MVP_top60_DNA_seed | EGFR | 0.768046 | BRCA | BT-20 | AST-1306 | MVP_top60_DNA+RNA_seed | ERBB2 | 0.460134 | BRCA | HCC2157 | CP724714 |
| MVP_top60_DNA_seed | SRC | 0.768971 | BRCA | BT-20 | NVP-BHG712 | MVP_top60_DNA+RNA_seed | EGFR | 0.461271 | BRCA | CAL-51 | Pelitinib |
| MVP_top60_DNA_seed | HDAC1 | 0.771997 | BRCA | HCC1500 | Parthenolide | MVP_top60_DNA+RNA_seed | CDK1 | 0.464012 | BRCA | CAL-51 | RO-3306 |
| MVP_top60_DNA_seed | SRC | 0.772994 | BRCA | BT-474 | HG-6-71-01 | MVP_top60_DNA+RNA_seed | ATM | 0.46592 | BRCA | MDA-MB-453 | KU-60019 |
| MVP_top60_DNA_seed | MAPK3 | 0.773322 | BRCA | MCF7 | FR-180204 | MVP_top60_DNA+RNA_seed | HDAC1 | 0.466444 | BRCA | JIMT-1 | Belinostat |
| MVP_top60_DNA_seed | MAPK1 | 0.773322 | BRCA | MCF7 | FR-180204 | MVP_top60_DNA+RNA_seed | AR | 0.469028 | BRCA | HCC38 | AZD3514 |
| MVP_top60_DNA_seed | HDAC1 | 0.774583 | BRCA | MDA-MB-157 | Tubastatin A | MVP_top60_DNA+RNA_seed | CDK2 | 0.470625 | BRCA | JIMT-1 | PHA-793887 |
| MVP_top60_DNA_seed | ERBB2 | 0.776247 | BRCA | Hs-578-T | Afatinib | MVP_top60_DNA+RNA_seed | HDAC1 | 0.471323 | BRCA | HCC1419 | Tubastatin A |
| MVP_top60_DNA_seed | EGFR | 0.776247 | BRCA | Hs-578-T | Afatinib | MVP_top60_DNA+RNA_seed | HSP90AA1 | 0.471802 | BRCA | CAL-85-1 | CCT-018159 |
| MVP_top60_DNA_seed | MDM2 | 0.779843 | BRCA | HCC1428 | Serdemetan | MVP_top60_DNA+RNA_seed | HSP90AB1 | 0.471802 | BRCA | CAL-85-1 | CCT-018159 |
| MVP_top60_DNA_seed | EGFR | 0.780746 | BRCA | HCC2157 | Cetuximab | MVP_top60_DNA+RNA_seed | EGFR | 0.473766 | BRCA | HCC1937 | Afatinib |

| | | | | | | | | | | | |
|---|---|---|---|---|---|---|---|---|---|---|---|
| MVP_top60_DNA_seed | ERBB2 | 0.782216 | BRCA | MFM-223 | Afatinib | MVP_top60_DNA+RNA_seed | ERBB2 | 0.473766 | BRCA | HCC1937 | Afatinib |
| MVP_top60_DNA_seed | EGFR | 0.782216 | BRCA | MFM-223 | Afatinib | MVP_top60_DNA+RNA_seed | HSP90AA1 | 0.474087 | BRCA | T47D | CCT-018159 |
| MVP_top60_DNA_seed | CDK2 | 0.787585 | BRCA | HCC1419 | PHA-793887 | MVP_top60_DNA+RNA_seed | HSP90AB1 | 0.474087 | BRCA | T47D | CCT-018159 |
| MVP_top60_DNA_seed | HDAC1 | 0.788546 | BRCA | BT-20 | AR-42 | MVP_top60_DNA+RNA_seed | HSP90AA1 | 0.475484 | BRCA | BT-483 | Tanespimycin |
| MVP_top60_DNA_seed | AR | 0.788998 | BRCA | HCC2157 | Bicalutamide | MVP_top60_DNA+RNA_seed | HSP90AB1 | 0.475484 | BRCA | BT-483 | Tanespimycin |
| MVP_top60_DNA_seed | AR | 0.789313 | BRCA | HCC1500 | Bicalutamide | MVP_top60_DNA+RNA_seed | ESR1 | 0.477032 | BRCA | HCC2157 | Tamoxifen |
| MVP_top60_DNA_seed | EGFR | 0.792312 | BRCA | EFM-192A | Gefitinib | MVP_top60_DNA+RNA_seed | EGFR | 0.477275 | BRCA | MDA-MB-415 | CI-1033 |
| MVP_top60_DNA_seed | MAPK1 | 0.794154 | BRCA | CAMA-1 | VX-11e | MVP_top60_DNA+RNA_seed | ERBB2 | 0.477275 | BRCA | MDA-MB-415 | CI-1033 |
| MVP_top60_DNA_seed | ATM | 0.794938 | BRCA | MDA-MB-453 | KU-55933 | MVP_top60_DNA+RNA_seed | HDAC1 | 0.480472 | BRCA | HCC1187 | CAY10603 |
| MVP_top60_DNA_seed | SRC | 0.795733 | BRCA | MDA-MB-415 | HG-6-71-01 | MVP_top60_DNA+RNA_seed | CDK2 | 0.480921 | BRCA | BT-483 | PHA-793887 |
| MVP_top60_DNA_seed | SRC | 0.798275 | BRCA | CAL-148 | WH-4-023 | MVP_top60_DNA+RNA_seed | AR | 0.482647 | BRCA | EFM-192A | Bicalutamide |
| MVP_top60_DNA_seed | MAPK3 | 0.801731 | BRCA | MDA-MB-157 | FR-180204 | MVP_top60_DNA+RNA_seed | EGFR | 0.483363 | BRCA | HCC1419 | CUDC-101 |
| MVP_top60_DNA_seed | MAPK1 | 0.801731 | BRCA | MDA-MB-157 | FR-180204 | MVP_top60_DNA+RNA_seed | ERBB2 | 0.483363 | BRCA | HCC1419 | CUDC-101 |
| MVP_top60_DNA_seed | MAPK3 | 0.804182 | BRCA | CAL-51 | FR-180204 | MVP_top60_DNA+RNA_seed | CDK1 | 0.485959 | BRCA | COLO-824 | RO-3306 |
| MVP_top60_DNA_seed | MAPK1 | 0.804182 | BRCA | CAL-51 | FR-180204 | MVP_top60_DNA+RNA_seed | CDK1 | 0.490931 | BRCA | HCC1937 | RO-3306 |
| MVP_top60_DNA_seed | ERBB2 | 0.805516 | BRCA | MDA-MB-415 | Afatinib | MVP_top60_DNA+RNA_seed | ATM | 0.491327 | BRCA | EVSA-T | KU-60019 |
| MVP_top60_DNA_seed | EGFR | 0.805516 | BRCA | MDA-MB-415 | Afatinib | MVP_top60_DNA+RNA_seed | CDK2 | 0.491604 | BRCA | COLO-824 | AZD5438 |
| MVP_top60_DNA_seed | EGFR | 0.805539 | BRCA | HCC202 | Gefitinib | MVP_top60_DNA+RNA_seed | EGFR | 0.493138 | BRCA | YMB-1-E | CI-1033 |
| MVP_top60_DNA_seed | AR | 0.809381 | BRCA | MDA-MB-330 | Bicalutamide | MVP_top60_DNA+RNA_seed | ERBB2 | 0.493138 | BRCA | YMB-1-E | CI-1033 |
| MVP_top60_DNA_seed | AR | 0.810886 | BRCA | EFM-192A | Bicalutamide | MVP_top60_DNA+RNA_seed | EGFR | 0.49404 | BRCA | YMB-1-E | PF-00299804 |
| MVP_top60_DNA_seed | HDAC1 | 0.811737 | BRCA | BT-549 | Tubastatin A | MVP_top60_DNA+RNA_seed | ERBB2 | 0.49404 | BRCA | YMB-1-E | PF-00299804 |
| MVP_top60_DNA_seed | MDM2 | 0.812224 | BRCA | HCC1937 | Nutlin-3a (-) | MVP_top60_DNA+RNA_seed | PIK3CA | 0.494257 | BRCA | HDQ-P1 | PI3Ka_4409 |
| MVP_top60_DNA_seed | AR | 0.814415 | BRCA | HCC1937 | Bicalutamide | MVP_top60_DNA+RNA_seed | SRC | 0.494855 | BRCA | HCC38 | NVP-BHG712 |
| MVP_top60_DNA_seed | HSP90AA1 | 0.815104 | BRCA | HDQ-P1 | SNX-2112 | MVP_top60_DNA+RNA_seed | MAPK1 | 0.497419 | BRCA | EFM-192A | VX-11e |
| MVP_top60_DNA_seed | HSP90AB1 | 0.815104 | BRCA | HDQ-P1 | SNX-2112 | MVP_top60_DNA+RNA_seed | SRC | 0.498405 | BRCA | COLO-824 | NVP-BHG712 |
| MVP_top60_DNA_seed | EGFR | 0.815608 | BRCA | CAMA-1 | Gefitinib | MVP_top60_DNA+RNA_seed | ATM | 0.502609 | BRCA | CAL-85-1 | KU-60019 |
| MVP_top60_DNA_seed | SRC | 0.815843 | BRCA | HCC1500 | Saracatinib | MVP_top60_DNA+RNA_seed | SRC | 0.502828 | BRCA | CAL-148 | Saracatinib |
| MVP_top60_DNA_seed | ERBB2 | 0.817017 | BRCA | UACC-893 | CUDC-101 | MVP_top60_DNA+RNA_seed | ATM | 0.503081 | BRCA | MDA-MB-361 | KU-60019 |
| MVP_top60_DNA_seed | EGFR | 0.817017 | BRCA | UACC-893 | CUDC-101 | MVP_top60_DNA+RNA_seed | HDAC1 | 0.503395 | BRCA | UACC-893 | Dacinostat |
| MVP_top60_DNA_seed | CDK2 | 0.818316 | BRCA | MFM-223 | PHA-793887 | MVP_top60_DNA+RNA_seed | EGFR | 0.50455 | BRCA | T47D | PF-00299804 |
| MVP_top60_DNA_seed | CDK1 | 0.818427 | BRCA | MDA-MB-436 | RO-3306 | MVP_top60_DNA+RNA_seed | ERBB2 | 0.50455 | BRCA | T47D | PF-00299804 |
| MVP_top60_DNA_seed | HSP90AA1 | 0.818776 | BRCA | HCC202 | Tanespimycin | MVP_top60_DNA+RNA_seed | AR | 0.50526 | BRCA | HCC1395 | Bicalutamide |
| MVP_top60_DNA_seed | HSP90AB1 | 0.818776 | BRCA | HCC202 | Tanespimycin | MVP_top60_DNA+RNA_seed | ATM | 0.506636 | BRCA | HCC1143 | KU-60019 |
| MVP_top60_DNA_seed | CDK2 | 0.819318 | BRCA | MDA-MB-157 | PHA-793887 | MVP_top60_DNA+RNA_seed | HDAC1 | 0.50914 | BRCA | COLO-824 | Dacinostat |
| MVP_top60_DNA_seed | EGFR | 0.820375 | BRCA | HCC1500 | Cetuximab | MVP_top60_DNA+RNA_seed | MDM2 | 0.509296 | BRCA | MDA-MB-361 | Nutlin-3a (-) |
| MVP_top60_DNA_seed | MAPK1 | 0.821476 | BRCA | CAL-51 | VX-11e | MVP_top60_DNA+RNA_seed | HDAC1 | 0.509605 | BRCA | HCC1569 | Tubastatin A |
| MVP_top60_DNA_seed | ATM | 0.823726 | BRCA | COLO-824 | KU-55933 | MVP_top60_DNA+RNA_seed | AKT1 | 0.510231 | BRCA | CAL-120 | GSK690693 |
| MVP_top60_DNA_seed | HSP90AA1 | 0.825108 | BRCA | MDA-MB-453 | Luminespib | MVP_top60_DNA+RNA_seed | EGFR | 0.51134 | BRCA | HCC1428 | Cetuximab |
| MVP_top60_DNA_seed | HSP90AB1 | 0.825108 | BRCA | MDA-MB-453 | Luminespib | MVP_top60_DNA+RNA_seed | EGFR | 0.512721 | BRCA | BT-474 | CUDC-101 |
| MVP_top60_DNA_seed | EGFR | 0.825797 | BRCA | HCC70 | Pelitinib | MVP_top60_DNA+RNA_seed | ERBB2 | 0.512721 | BRCA | BT-474 | CUDC-101 |
| MVP_top60_DNA_seed | HDAC1 | 0.827368 | BRCA | HCC1569 | JQ12 | MVP_top60_DNA+RNA_seed | PIK3CA | 0.513118 | BRCA | HCC2218 | AZD8835 |

| | | | | | | | | | | | |
|---|---|---|---|---|---|---|---|---|---|---|---|
| MVP_top60_DNA_seed | HDAC2 | 0.827368 | BRCA | HCC1569 | JQ12 | MVP_top60_DNA+RNA_seed | HDAC1 | 0.513258 | BRCA | CAMA-1 | AR-42 |
| MVP_top60_DNA_seed | HDAC1 | 0.827447 | BRCA | HCC1937 | Tubastatin A | MVP_top60_DNA+RNA_seed | PIK3CA | 0.51515 | BRCA | DU-4475 | PI3Ka_4409 |
| MVP_top60_DNA_seed | HDAC1 | 0.82824 | BRCA | Hs-578-T | AR-42 | MVP_top60_DNA+RNA_seed | HDAC1 | 0.51526 | BRCA | HCC1419 | Belinostat |
| MVP_top60_DNA_seed | EP300 | 0.828722 | BRCA | BT-549 | I-CBP112 | MVP_top60_DNA+RNA_seed | HSP90AA1 | 0.516196 | BRCA | OCUB-M | Elesclomol |
| MVP_top60_DNA_seed | CREBBP | 0.828722 | BRCA | BT-549 | I-CBP112 | MVP_top60_DNA+RNA_seed | HSP90AB1 | 0.516196 | BRCA | OCUB-M | Elesclomol |
| MVP_top60_DNA_seed | AKT1 | 0.828966 | BRCA | MDA-MB-157 | GSK690693 | MVP_top60_DNA+RNA_seed | HSP90AA1 | 0.516744 | BRCA | BT-549 | SNX-2112 |
| MVP_top60_DNA_seed | EGFR | 0.829146 | BRCA | MRK-nu-1 | Cetuximab | MVP_top60_DNA+RNA_seed | HSP90AB1 | 0.516744 | BRCA | BT-549 | SNX-2112 |
| MVP_top60_DNA_seed | HDAC1 | 0.829223 | BRCA | Hs-578-T | Belinostat | MVP_top60_DNA+RNA_seed | HSP90AA1 | 0.517006 | BRCA | EVSA-T | Luminespib |
| MVP_top60_DNA_seed | EGFR | 0.829341 | BRCA | EFM-192A | Pelitinib | MVP_top60_DNA+RNA_seed | HSP90AB1 | 0.517006 | BRCA | EVSA-T | Luminespib |
| MVP_top60_DNA_seed | ERBB2 | 0.833459 | BRCA | MRK-nu-1 | Afatinib | MVP_top60_DNA+RNA_seed | HDAC1 | 0.518816 | BRCA | CAL-85-1 | Belinostat |
| MVP_top60_DNA_seed | EGFR | 0.833459 | BRCA | MRK-nu-1 | Afatinib | MVP_top60_DNA+RNA_seed | EGFR | 0.520022 | BRCA | BT-483 | AZD8931 |
| MVP_top60_DNA_seed | ERBB2 | 0.83371 | BRCA | HCC2157 | CUDC-101 | MVP_top60_DNA+RNA_seed | ERBB2 | 0.520022 | BRCA | BT-483 | AZD8931 |
| MVP_top60_DNA_seed | EGFR | 0.83371 | BRCA | HCC2157 | CUDC-101 | MVP_top60_DNA+RNA_seed | ERBB3 | 0.520022 | BRCA | BT-483 | AZD8931 |
| MVP_top60_DNA_seed | MAPK1 | 0.835356 | BRCA | HCC38 | VX-11e | MVP_top60_DNA+RNA_seed | EGFR | 0.520231 | BRCA | CAL-120 | Cetuximab |
| MVP_top60_DNA_seed | HDAC1 | 0.839258 | BRCA | BT-474 | JQ12 | MVP_top60_DNA+RNA_seed | HSP90AA1 | 0.521592 | BRCA | HCC1937 | SNX-2112 |
| MVP_top60_DNA_seed | HDAC2 | 0.839258 | BRCA | BT-474 | JQ12 | MVP_top60_DNA+RNA_seed | HSP90AB1 | 0.521592 | BRCA | HCC1937 | SNX-2112 |
| MVP_top60_DNA_seed | SRC | 0.84048 | BRCA | CAMA-1 | NVP-BHG712 | MVP_top60_DNA+RNA_seed | EGFR | 0.522645 | BRCA | BT-20 | PF-00299804 |
| MVP_top60_DNA_seed | HDAC1 | 0.840586 | BRCA | MDA-MB-175-VII | Dacinostat | MVP_top60_DNA+RNA_seed | ERBB2 | 0.522645 | BRCA | BT-20 | PF-00299804 |
| MVP_top60_DNA_seed | ERBB2 | 0.841687 | BRCA | MDA-MB-157 | CP724714 | MVP_top60_DNA+RNA_seed | EGFR | 0.523688 | BRCA | T47D | AST-1306 |
| MVP_top60_DNA_seed | AR | 0.842606 | BRCA | HCC38 | Bicalutamide | MVP_top60_DNA+RNA_seed | EGFR | 0.524095 | BRCA | DU-4475 | Afatinib |
| MVP_top60_DNA_seed | ESR1 | 0.843092 | BRCA | MDA-MB-231 | Fulvestrant | MVP_top60_DNA+RNA_seed | ERBB2 | 0.524095 | BRCA | DU-4475 | Afatinib |
| MVP_top60_DNA_seed | ATM | 0.843967 | BRCA | MDA-MB-436 | KU-60019 | MVP_top60_DNA+RNA_seed | HDAC1 | 0.524606 | BRCA | CAL-85-1 | CAY10603 |
| MVP_top60_DNA_seed | MAPK3 | 0.844382 | BRCA | YMB-1-E | FR-180204 | MVP_top60_DNA+RNA_seed | AR | 0.52493 | BRCA | MDA-MB-231 | Bicalutamide |
| MVP_top60_DNA_seed | MAPK1 | 0.844382 | BRCA | YMB-1-E | FR-180204 | MVP_top60_DNA+RNA_seed | HSP90AA1 | 0.526635 | BRCA | AU565 | SNX-2112 |
| MVP_top60_DNA_seed | HDAC1 | 0.844563 | BRCA | MDA-MB-436 | Belinostat | MVP_top60_DNA+RNA_seed | HSP90AB1 | 0.526635 | BRCA | AU565 | SNX-2112 |
| MVP_top60_DNA_seed | HDAC1 | 0.845404 | BRCA | HCC2218 | Belinostat | MVP_top60_DNA+RNA_seed | HSP90AA1 | 0.52781 | BRCA | HCC202 | Elesclomol |
| MVP_top60_DNA_seed | EGFR | 0.845407 | BRCA | MDA-MB-231 | Gefitinib | MVP_top60_DNA+RNA_seed | HSP90AB1 | 0.52781 | BRCA | HCC202 | Elesclomol |
| MVP_top60_DNA_seed | AKT1 | 0.84634 | BRCA | MDA-MB-157 | AKT inhibitor VIII | MVP_top60_DNA+RNA_seed | HDAC1 | 0.529142 | BRCA | BT-20 | Belinostat |
| MVP_top60_DNA_seed | AR | 0.851371 | BRCA | HCC1419 | Bicalutamide | MVP_top60_DNA+RNA_seed | EGFR | 0.529395 | BRCA | EFM-19 | AZD8931 |
| MVP_top60_DNA_seed | ERBB2 | 0.851706 | BRCA | MFM-223 | Afatinib | MVP_top60_DNA+RNA_seed | ERBB2 | 0.529395 | BRCA | EFM-19 | AZD8931 |
| MVP_top60_DNA_seed | EGFR | 0.851706 | BRCA | MFM-223 | Afatinib | MVP_top60_DNA+RNA_seed | ERBB3 | 0.529395 | BRCA | EFM-19 | AZD8931 |
| MVP_top60_DNA_seed | SRC | 0.851957 | BRCA | MDA-MB-436 | NVP-BHG712 | MVP_top60_DNA+RNA_seed | EGFR | 0.530013 | BRCA | MDA-MB-231 | Afatinib |
| MVP_top60_DNA_seed | MAPK1 | 0.852481 | BRCA | HCC1500 | VX-11e | MVP_top60_DNA+RNA_seed | ERBB2 | 0.530013 | BRCA | MDA-MB-231 | Afatinib |
| MVP_top60_DNA_seed | CDK2 | 0.85286 | BRCA | CAL-120 | PHA-793887 | MVP_top60_DNA+RNA_seed | SRC | 0.530179 | BRCA | HCC1395 | Bosutinib |
| MVP_top60_DNA_seed | MAPK3 | 0.853616 | BRCA | COLO-824 | FR-180204 | MVP_top60_DNA+RNA_seed | AR | 0.53045 | BRCA | BT-474 | Bicalutamide |
| MVP_top60_DNA_seed | MAPK1 | 0.853616 | BRCA | COLO-824 | FR-180204 | MVP_top60_DNA+RNA_seed | EGFR | 0.53203 | BRCA | HCC1428 | CI-1033 |
| MVP_top60_DNA_seed | HSP90AA1 | 0.854733 | BRCA | HCC1500 | Tanespimycin | MVP_top60_DNA+RNA_seed | ERBB2 | 0.53203 | BRCA | HCC1428 | CI-1033 |
| MVP_top60_DNA_seed | HSP90AB1 | 0.854733 | BRCA | HCC1500 | Tanespimycin | MVP_top60_DNA+RNA_seed | EGFR | 0.532044 | BRCA | MDA-MB-330 | Gefitinib |
| MVP_top60_DNA_seed | HDAC1 | 0.85709 | BRCA | MCF7 | CAY10603 | MVP_top60_DNA+RNA_seed | MAPK3 | 0.533184 | BRCA | MDA-MB-453 | FR-180204 |
| MVP_top60_DNA_seed | HSP90AA1 | 0.859483 | BRCA | OCUB-M | Tanespimycin | MVP_top60_DNA+RNA_seed | MAPK1 | 0.533184 | BRCA | MDA-MB-453 | FR-180204 |
| MVP_top60_DNA_seed | HSP90AB1 | 0.859483 | BRCA | OCUB-M | Tanespimycin | MVP_top60_DNA+RNA_seed | AR | 0.535518 | BRCA | MDA-MB-175-VII | Bicalutamide |

| | | | | | | | | | | | |
|---|---|---|---|---|---|---|---|---|---|---|---|
| MVP_top60_DNA_seed | ERBB2 | 0.860178 | BRCA | MCF7 | Afatinib | MVP_top60_DNA+RNA_seed | AKT1 | 0.536015 | BRCA | BT-549 | AKT inhibitor VIII |
| MVP_top60_DNA_seed | EGFR | 0.860178 | BRCA | MCF7 | Afatinib | MVP_top60_DNA+RNA_seed | EGFR | 0.537272 | BRCA | EFM-19 | Afatinib |
| MVP_top60_DNA_seed | EP300 | 0.860822 | BRCA | CAL-120 | I-CBP112 | MVP_top60_DNA+RNA_seed | ERBB2 | 0.537272 | BRCA | EFM-19 | Afatinib |
| MVP_top60_DNA_seed | CREBBP | 0.860822 | BRCA | CAL-120 | I-CBP112 | MVP_top60_DNA+RNA_seed | AR | 0.537986 | BRCA | HCC1143 | Bicalutamide |
| MVP_top60_DNA_seed | HSP90AA1 | 0.861234 | BRCA | T47D | Luminespib | MVP_top60_DNA+RNA_seed | HDAC1 | 0.541957 | BRCA | BT-474 | Belinostat |
| MVP_top60_DNA_seed | HSP90AB1 | 0.861234 | BRCA | T47D | Luminespib | MVP_top60_DNA+RNA_seed | HDAC1 | 0.543264 | BRCA | MDA-MB-175-VII | Belinostat |
| MVP_top60_DNA_seed | ESR1 | 0.861865 | BRCA | HDQ-P1 | Tamoxifen | MVP_top60_DNA+RNA_seed | EGFR | 0.544561 | BRCA | MDA-MB-231 | Pelitinib |
| MVP_top60_DNA_seed | AKT1 | 0.865286 | BRCA | MDA-MB-231 | GSK690693 | MVP_top60_DNA+RNA_seed | CDK1 | 0.545483 | BRCA | BT-20 | RO-3306 |
| MVP_top60_DNA_seed | CDK1 | 0.86555 | BRCA | HCC1569 | RO-3306 | MVP_top60_DNA+RNA_seed | ESR1 | 0.546651 | BRCA | HCC2157 | Fulvestrant |
| MVP_top60_DNA_seed | HSP90AA1 | 0.866896 | BRCA | MDA-MB-157 | Elesclomol | MVP_top60_DNA+RNA_seed | EGFR | 0.548275 | BRCA | BT-549 | CUDC-101 |
| MVP_top60_DNA_seed | HSP90AB1 | 0.866896 | BRCA | MDA-MB-157 | Elesclomol | MVP_top60_DNA+RNA_seed | ERBB2 | 0.548275 | BRCA | BT-549 | CUDC-101 |
| MVP_top60_DNA_seed | HSP90AA1 | 0.868122 | BRCA | MDA-MB-415 | Luminespib | MVP_top60_DNA+RNA_seed | MAPK14 | 0.549016 | BRCA | T47D | TAK-715 |
| MVP_top60_DNA_seed | HSP90AB1 | 0.868122 | BRCA | MDA-MB-415 | Luminespib | MVP_top60_DNA+RNA_seed | EP300 | 0.549712 | BRCA | HCC70 | I-CBP112 |
| MVP_top60_DNA_seed | MAPK3 | 0.870383 | BRCA | T47D | FR-180204 | MVP_top60_DNA+RNA_seed | EGFR | 0.55082 | BRCA | HCC1187 | Gefitinib |
| MVP_top60_DNA_seed | MAPK1 | 0.870383 | BRCA | T47D | FR-180204 | MVP_top60_DNA+RNA_seed | PIK3CA | 0.551516 | BRCA | HCC1419 | AZD8835 |
| MVP_top60_DNA_seed | EGFR | 0.8716 | BRCA | COLO-824 | Gefitinib | MVP_top60_DNA+RNA_seed | ATM | 0.5518 | BRCA | MDA-MB-330 | KU-55933 |
| MVP_top60_DNA_seed | MDM2 | 0.871691 | BRCA | EVSA-T | Serdemetan | MVP_top60_DNA+RNA_seed | HDAC1 | 0.551837 | BRCA | MDA-MB-175-VII | JQ12 |
| MVP_top60_DNA_seed | AKT1 | 0.871972 | BRCA | Hs-578-T | GSK690693 | MVP_top60_DNA+RNA_seed | HDAC2 | 0.551837 | BRCA | MDA-MB-175-VII | JQ12 |
| MVP_top60_DNA_seed | CDK2 | 0.872436 | BRCA | HCC1428 | PHA-793887 | MVP_top60_DNA+RNA_seed | MAPK1 | 0.552745 | BRCA | HCC202 | VX-11e |
| MVP_top60_DNA_seed | ATM | 0.873695 | BRCA | MDA-MB-157 | CP466722 | MVP_top60_DNA+RNA_seed | MAPK14 | 0.553797 | BRCA | MDA-MB-175-VII | TAK-715 |
| MVP_top60_DNA_seed | ERBB2 | 0.875026 | BRCA | BT-549 | Afatinib | MVP_top60_DNA+RNA_seed | EGFR | 0.553932 | BRCA | HCC38 | Gefitinib |
| MVP_top60_DNA_seed | EGFR | 0.875026 | BRCA | BT-549 | Afatinib | MVP_top60_DNA+RNA_seed | ESR1 | 0.554975 | BRCA | MDA-MB-330 | Tamoxifen |
| MVP_top60_DNA_seed | EGFR | 0.876027 | BRCA | MFM-223 | Pelitinib | MVP_top60_DNA+RNA_seed | SRC | 0.555526 | BRCA | EFM-192A | Bosutinib |
| MVP_top60_DNA_seed | HSP90AA1 | 0.882227 | BRCA | MDA-MB-175-VII | Luminespib | MVP_top60_DNA+RNA_seed | HDAC1 | 0.555635 | BRCA | UACC-812 | CAY10603 |
| MVP_top60_DNA_seed | HSP90AB1 | 0.882227 | BRCA | MDA-MB-175-VII | Luminespib | MVP_top60_DNA+RNA_seed | CDK2 | 0.556501 | BRCA | HCC1954 | AZD5438 |
| MVP_top60_DNA_seed | EGFR | 0.885029 | BRCA | EVSA-T | Pelitinib | MVP_top60_DNA+RNA_seed | HDAC1 | 0.55672 | BRCA | CAL-85-1 | AR-42 |
| MVP_top60_DNA_seed | HDAC1 | 0.8854 | BRCA | MCF7 | Tubastatin A | MVP_top60_DNA+RNA_seed | AKT1 | 0.559417 | BRCA | HDQ-P1 | GSK690693 |
| MVP_top60_DNA_seed | ERBB2 | 0.888675 | BRCA | Hs-578-T | CP724714 | MVP_top60_DNA+RNA_seed | PLK1 | 0.560414 | BRCA | MDA-MB-157 | PLK_6522 |
| MVP_top60_DNA_seed | HSP90AA1 | 0.889434 | BRCA | BT-549 | Luminespib | MVP_top60_DNA+RNA_seed | SRC | 0.560439 | BRCA | MDA-MB-436 | Bosutinib |
| MVP_top60_DNA_seed | HSP90AB1 | 0.889434 | BRCA | BT-549 | Luminespib | MVP_top60_DNA+RNA_seed | AR | 0.562717 | BRCA | HCC1937 | AZD3514 |
| MVP_top60_DNA_seed | MAPK1 | 0.88952 | BRCA | CAL-120 | VX-11e | MVP_top60_DNA+RNA_seed | EGFR | 0.563051 | BRCA | MDA-MB-157 | PF-00299804 |
| MVP_top60_DNA_seed | MDM2 | 0.890229 | BRCA | HCC2157 | Nutlin-3a (-) | MVP_top60_DNA+RNA_seed | ERBB2 | 0.563051 | BRCA | MDA-MB-157 | PF-00299804 |
| MVP_top60_DNA_seed | MAPK3 | 0.891714 | BRCA | CAL-120 | FR-180204 | MVP_top60_DNA+RNA_seed | HSP90AA1 | 0.564069 | BRCA | HCC1569 | SNX-2112 |
| MVP_top60_DNA_seed | MAPK1 | 0.891714 | BRCA | CAL-120 | FR-180204 | MVP_top60_DNA+RNA_seed | HSP90AB1 | 0.564069 | BRCA | HCC1569 | SNX-2112 |
| MVP_top60_DNA_seed | HDAC1 | 0.892382 | BRCA | MDA-MB-436 | Tubastatin A | MVP_top60_DNA+RNA_seed | EP300 | 0.564155 | BRCA | MDA-MB-157 | I-CBP112 |
| MVP_top60_DNA_seed | SRC | 0.892723 | BRCA | HCC2218 | Saracatinib | MVP_top60_DNA+RNA_seed | HSP90AA1 | 0.564583 | BRCA | EFM-192A | CCT-018159 |
| MVP_top60_DNA_seed | ESR1 | 0.893809 | BRCA | COLO-824 | Tamoxifen | MVP_top60_DNA+RNA_seed | HSP90AB1 | 0.564583 | BRCA | EFM-192A | CCT-018159 |
| MVP_top60_DNA_seed | HDAC1 | 0.893912 | BRCA | JIMT-1 | Tubastatin A | MVP_top60_DNA+RNA_seed | MAPK1 | 0.569103 | BRCA | COLO-824 | VX-11e |
| MVP_top60_DNA_seed | HSP90AA1 | 0.897371 | BRCA | HCC1395 | Tanespimycin | MVP_top60_DNA+RNA_seed | EP300 | 0.569813 | BRCA | Hs-578-T | I-CBP112 |
| MVP_top60_DNA_seed | HSP90AB1 | 0.897371 | BRCA | HCC1395 | Tanespimycin | MVP_top60_DNA+RNA_seed | AKT1 | 0.570828 | BRCA | HCC38 | GSK690693 |
| MVP_top60_DNA_seed | AKT1 | 0.89809 | BRCA | HCC1428 | GSK690693 | MVP_top60_DNA+RNA_seed | HDAC1 | 0.57103 | BRCA | MRK-nu-1 | Belinostat |

| | | | | | | | | | | | |
|---|---|---|---|---|---|---|---|---|---|---|---|
| MVP_top60_DNA_seed | HDAC1 | 0.898834 | BRCA | YMB-1-E | Tubastatin A | MVP_top60_DNA+RNA_seed | MAPK1 | 0.573084 | BRCA | MRK-nu-1 | VX-11e |
| MVP_top60_DNA_seed | CDK2 | 0.902109 | BRCA | MDA-MB-361 | PHA-793887 | MVP_top60_DNA+RNA_seed | EGFR | 0.573091 | BRCA | BT-474 | Erlotinib |
| MVP_top60_DNA_seed | AR | 0.902566 | BRCA | HCC1569 | Bicalutamide | MVP_top60_DNA+RNA_seed | ATM | 0.575337 | BRCA | HCC1419 | KU-60019 |
| MVP_top60_DNA_seed | MAPK1 | 0.902579 | BRCA | MDA-MB-157 | VX-11e | MVP_top60_DNA+RNA_seed | HDAC1 | 0.57712 | BRCA | UACC-812 | Parthenolide |
| MVP_top60_DNA_seed | EGFR | 0.903784 | BRCA | COLO-824 | Cetuximab | MVP_top60_DNA+RNA_seed | EGFR | 0.577188 | BRCA | UACC-812 | Pelitinib |
| MVP_top60_DNA_seed | EGFR | 0.903969 | BRCA | MDA-MB-330 | Pelitinib | MVP_top60_DNA+RNA_seed | HDAC1 | 0.577867 | BRCA | MDA-MB-436 | CAY10603 |
| MVP_top60_DNA_seed | AKT1 | 0.904081 | BRCA | MRK-nu-1 | GSK690693 | MVP_top60_DNA+RNA_seed | HDAC1 | 0.579872 | BRCA | CAL-85-1 | Tubastatin A |
| MVP_top60_DNA_seed | HDAC1 | 0.904316 | BRCA | MDA-MB-468 | Dacinostat | MVP_top60_DNA+RNA_seed | SRC | 0.580742 | BRCA | HCC2218 | NVP-BHG712 |
| MVP_top60_DNA_seed | EP300 | 0.906945 | BRCA | HCC202 | I-CBP112 | MVP_top60_DNA+RNA_seed | AKT1 | 0.58148 | BRCA | Hs-578-T | AKT inhibitor VIII |
| MVP_top60_DNA_seed | CREBBP | 0.906945 | BRCA | HCC202 | I-CBP112 | MVP_top60_DNA+RNA_seed | ATM | 0.581752 | BRCA | MDA-MB-361 | KU-55933 |
| MVP_top60_DNA_seed | AR | 0.907571 | BRCA | CAL-120 | Bicalutamide | MVP_top60_DNA+RNA_seed | CDK1 | 0.582603 | BRCA | MDA-MB-231 | RO-3306 |
| MVP_top60_DNA_seed | ATM | 0.908144 | BRCA | CAMA-1 | KU-60019 | MVP_top60_DNA+RNA_seed | PLK1 | 0.582692 | BRCA | DU-4475 | PLK_6522 |
| MVP_top60_DNA_seed | SRC | 0.909176 | BRCA | MCF7 | NVP-BHG712 | MVP_top60_DNA+RNA_seed | HDAC1 | 0.583193 | BRCA | YMB-1-E | CAY10603 |
| MVP_top60_DNA_seed | MDM2 | 0.911452 | BRCA | COLO-824 | Nutlin-3a (-) | MVP_top60_DNA+RNA_seed | SRC | 0.585664 | BRCA | AU565 | NVP-BHG712 |
| MVP_top60_DNA_seed | AKT1 | 0.912439 | BRCA | HCC1187 | AKT inhibitor VIII | MVP_top60_DNA+RNA_seed | PIK3CA | 0.586171 | BRCA | CAL-85-1 | AZD8835 |
| MVP_top60_DNA_seed | HSP90AA1 | 0.913598 | BRCA | BT-20 | SNX-2112 | MVP_top60_DNA+RNA_seed | HDAC1 | 0.586438 | BRCA | COLO-824 | Tubastatin A |
| MVP_top60_DNA_seed | HSP90AB1 | 0.913598 | BRCA | BT-20 | SNX-2112 | MVP_top60_DNA+RNA_seed | CDK2 | 0.587705 | BRCA | HDQ-P1 | AZD5438 |
| MVP_top60_DNA_seed | SRC | 0.914616 | BRCA | COLO-824 | WH-4-023 | MVP_top60_DNA+RNA_seed | CDK1 | 0.589198 | BRCA | MDA-MB-175-VII | RO-3306 |
| MVP_top60_DNA_seed | SRC | 0.914661 | BRCA | BT-474 | WH-4-023 | MVP_top60_DNA+RNA_seed | CDK2 | 0.589889 | BRCA | MFM-223 | AZD5438 |
| MVP_top60_DNA_seed | CDK2 | 0.915436 | BRCA | MDA-MB-231 | PHA-793887 | MVP_top60_DNA+RNA_seed | ATM | 0.58995 | BRCA | COLO-824 | CP466722 |
| MVP_top60_DNA_seed | EGFR | 0.915618 | BRCA | MDA-MB-453 | Gefitinib | MVP_top60_DNA+RNA_seed | CDK2 | 0.591078 | BRCA | HCC202 | AZD5438 |
| MVP_top60_DNA_seed | CDK2 | 0.915737 | BRCA | CAL-85-1 | AZD5438 | MVP_top60_DNA+RNA_seed | HDAC1 | 0.592524 | BRCA | HCC2218 | JQ12 |
| MVP_top60_DNA_seed | EGFR | 0.916803 | BRCA | MDA-MB-436 | Cetuximab | MVP_top60_DNA+RNA_seed | HDAC2 | 0.592524 | BRCA | HCC2218 | JQ12 |
| MVP_top60_DNA_seed | EGFR | 0.917078 | BRCA | CAL-85-1 | Gefitinib | MVP_top60_DNA+RNA_seed | EGFR | 0.596476 | BRCA | CAMA-1 | Afatinib |
| MVP_top60_DNA_seed | AKT1 | 0.91751 | BRCA | MDA-MB-231 | AKT inhibitor VIII | MVP_top60_DNA+RNA_seed | ERBB2 | 0.596476 | BRCA | CAMA-1 | Afatinib |
| MVP_top60_DNA_seed | CDK2 | 0.917545 | BRCA | MDA-MB-361 | AZD5438 | MVP_top60_DNA+RNA_seed | EGFR | 0.598134 | BRCA | HDQ-P1 | Pelitinib |
| MVP_top60_DNA_seed | ESR1 | 0.918374 | BRCA | BT-474 | Tamoxifen | MVP_top60_DNA+RNA_seed | EGFR | 0.59822 | BRCA | MRK-nu-1 | CUDC-101 |
| MVP_top60_DNA_seed | HDAC1 | 0.919348 | BRCA | MDA-MB-468 | JQ12 | MVP_top60_DNA+RNA_seed | ERBB2 | 0.59822 | BRCA | MRK-nu-1 | CUDC-101 |
| MVP_top60_DNA_seed | HDAC2 | 0.919348 | BRCA | MDA-MB-468 | JQ12 | MVP_top60_DNA+RNA_seed | EGFR | 0.600228 | BRCA | MDA-MB-157 | Cetuximab |
| MVP_top60_DNA_seed | MAPK3 | 0.922607 | BRCA | HCC1500 | FR-180204 | MVP_top60_DNA+RNA_seed | ERBB2 | 0.602708 | BRCA | EFM-192A | CP724714 |
| MVP_top60_DNA_seed | MAPK1 | 0.922607 | BRCA | HCC1500 | FR-180204 | MVP_top60_DNA+RNA_seed | HDAC1 | 0.604339 | BRCA | HCC2157 | Parthenolide |
| MVP_top60_DNA_seed | MDM2 | 0.923737 | BRCA | AU565 | Nutlin-3a (-) | MVP_top60_DNA+RNA_seed | AKT1 | 0.604704 | BRCA | MDA-MB-415 | GSK690693 |
| MVP_top60_DNA_seed | HDAC1 | 0.927028 | BRCA | MDA-MB-468 | Belinostat | MVP_top60_DNA+RNA_seed | SRC | 0.605145 | BRCA | HCC2157 | Bosutinib |
| MVP_top60_DNA_seed | ERBB2 | 0.927758 | BRCA | EFM-19 | CP724714 | MVP_top60_DNA+RNA_seed | PIK3CA | 0.605227 | BRCA | HDQ-P1 | AZD8835 |
| MVP_top60_DNA_seed | ATM | 0.929542 | BRCA | MDA-MB-231 | KU-55933 | MVP_top60_DNA+RNA_seed | SRC | 0.608014 | BRCA | MDA-MB-468 | NVP-BHG712 |
| MVP_top60_DNA_seed | MDM2 | 0.929939 | BRCA | BT-483 | Serdemetan | MVP_top60_DNA+RNA_seed | ESR1 | 0.609086 | BRCA | AU565 | Tamoxifen |
| MVP_top60_DNA_seed | MAPK3 | 0.930072 | BRCA | UACC-893 | FR-180204 | MVP_top60_DNA+RNA_seed | SRC | 0.609227 | BRCA | HCC1428 | Bosutinib |
| MVP_top60_DNA_seed | MAPK1 | 0.930072 | BRCA | UACC-893 | FR-180204 | MVP_top60_DNA+RNA_seed | SRC | 0.61239 | BRCA | HCC1569 | NVP-BHG712 |
| MVP_top60_DNA_seed | MAPK3 | 0.930932 | BRCA | HCC2157 | FR-180204 | MVP_top60_DNA+RNA_seed | HDAC1 | 0.614849 | BRCA | MDA-MB-175-VII | CAY10603 |
| MVP_top60_DNA_seed | MAPK1 | 0.930932 | BRCA | HCC2157 | FR-180204 | MVP_top60_DNA+RNA_seed | AKT1 | 0.614997 | BRCA | COLO-824 | GSK690693 |
| MVP_top60_DNA_seed | HDAC1 | 0.931174 | BRCA | MDA-MB-175-VII | Tubastatin A | MVP_top60_DNA+RNA_seed | MAPK3 | 0.615108 | BRCA | JIMT-1 | FR-180204 |

| | | | | | | | | | | | |
|---|---|---|---|---|---|---|---|---|---|---|---|
| MVP_top60_DNA_seed | HDAC1 | 0.931531 | BRCA | MFM-223 | Tubastatin A | MVP_top60_DNA+RNA_seed | MAPK1 | 0.615108 | BRCA | JIMT-1 | FR-180204 |
| MVP_top60_DNA_seed | AKT1 | 0.93287 | BRCA | HCC38 | BAY AKT1 | MVP_top60_DNA+RNA_seed | HSP90AA1 | 0.615237 | BRCA | JIMT-1 | SNX-2112 |
| MVP_top60_DNA_seed | AR | 0.933287 | BRCA | HCC1395 | Bicalutamide | MVP_top60_DNA+RNA_seed | HSP90AB1 | 0.615237 | BRCA | JIMT-1 | SNX-2112 |
| MVP_top60_DNA_seed | ERBB2 | 0.934991 | BRCA | HCC1428 | AZD8931 | MVP_top60_DNA+RNA_seed | AR | 0.618493 | BRCA | HCC1419 | Bicalutamide |
| MVP_top60_DNA_seed | EGFR | 0.934991 | BRCA | HCC1428 | AZD8931 | MVP_top60_DNA+RNA_seed | EGFR | 0.618902 | BRCA | HCC202 | Cetuximab |
| MVP_top60_DNA_seed | ERBB3 | 0.934991 | BRCA | HCC1428 | AZD8931 | MVP_top60_DNA+RNA_seed | MAPK14 | 0.619317 | BRCA | HCC1419 | TAK-715 |
| MVP_top60_DNA_seed | PIK3CA | 0.936188 | BRCA | MDA-MB-415 | AZD8835 | MVP_top60_DNA+RNA_seed | ATM | 0.62006 | BRCA | EVSA-T | CP466722 |
| MVP_top60_DNA_seed | MDM2 | 0.936859 | BRCA | MDA-MB-453 | Nutlin-3a (-) | MVP_top60_DNA+RNA_seed | ERBB2 | 0.622148 | BRCA | MFM-223 | CP724714 |
| MVP_top60_DNA_seed | EGFR | 0.937526 | BRCA | HCC2218 | Erlotinib | MVP_top60_DNA+RNA_seed | ESR1 | 0.622334 | BRCA | MDA-MB-453 | Tamoxifen |
| MVP_top60_DNA_seed | EP300 | 0.937796 | BRCA | MDA-MB-175-VII | I-CBP112 | MVP_top60_DNA+RNA_seed | MAPK1 | 0.623426 | BRCA | MDA-MB-453 | VX-11e |
| MVP_top60_DNA_seed | CREBBP | 0.937796 | BRCA | MDA-MB-175-VII | I-CBP112 | MVP_top60_DNA+RNA_seed | ATM | 0.624342 | BRCA | HCC1143 | KU-55933 |
| MVP_top60_DNA_seed | ERBB2 | 0.937899 | BRCA | MFM-223 | Lapatinib | MVP_top60_DNA+RNA_seed | CDK2 | 0.625674 | BRCA | CAL-51 | PHA-793887 |
| MVP_top60_DNA_seed | EGFR | 0.937899 | BRCA | MFM-223 | Lapatinib | MVP_top60_DNA+RNA_seed | SRC | 0.626035 | BRCA | BT-474 | Saracatinib |
| MVP_top60_DNA_seed | SRC | 0.938497 | BRCA | HCC2157 | WH-4-023 | MVP_top60_DNA+RNA_seed | AKT1 | 0.626182 | BRCA | EFM-192A | AKT inhibitor VIII |
| MVP_top60_DNA_seed | HDAC1 | 0.9386 | BRCA | UACC-893 | Tubastatin A | MVP_top60_DNA+RNA_seed | MAPK3 | 0.627238 | BRCA | HCC1419 | FR-180204 |
| MVP_top60_DNA_seed | ESR1 | 0.942013 | BRCA | HCC2218 | Tamoxifen | MVP_top60_DNA+RNA_seed | MAPK1 | 0.627238 | BRCA | HCC1419 | FR-180204 |
| MVP_top60_DNA_seed | EP300 | 0.942172 | BRCA | EFM-192A | I-CBP112 | MVP_top60_DNA+RNA_seed | HDAC1 | 0.628216 | BRCA | UACC-893 | Belinostat |
| MVP_top60_DNA_seed | CREBBP | 0.942172 | BRCA | EFM-192A | I-CBP112 | MVP_top60_DNA+RNA_seed | EP300 | 0.628665 | BRCA | HCC38 | I-CBP112 |
| MVP_top60_DNA_seed | MAPK3 | 0.944657 | BRCA | HCC2218 | FR-180204 | MVP_top60_DNA+RNA_seed | ATM | 0.628962 | BRCA | T47D | KU-55933 |
| MVP_top60_DNA_seed | MAPK1 | 0.944657 | BRCA | HCC2218 | FR-180204 | MVP_top60_DNA+RNA_seed | HDAC1 | 0.630533 | BRCA | MDA-MB-231 | Tubastatin A |
| MVP_top60_DNA_seed | ATM | 0.945688 | BRCA | MDA-MB-436 | KU-55933 | MVP_top60_DNA+RNA_seed | EGFR | 0.630657 | BRCA | HCC1419 | Gefitinib |
| MVP_top60_DNA_seed | HSP90AA1 | 0.945909 | BRCA | HCC1500 | SNX-2112 | MVP_top60_DNA+RNA_seed | EGFR | 0.631566 | BRCA | Hs-578-T | Gefitinib |
| MVP_top60_DNA_seed | HSP90AB1 | 0.945909 | BRCA | HCC1500 | SNX-2112 | MVP_top60_DNA+RNA_seed | ATM | 0.632667 | BRCA | EFM-19 | KU-60019 |
| MVP_top60_DNA_seed | HSP90AA1 | 0.946881 | BRCA | MRK-nu-1 | CCT-018159 | MVP_top60_DNA+RNA_seed | HDAC1 | 0.6329 | BRCA | UACC-893 | CAY10603 |
| MVP_top60_DNA_seed | HSP90AB1 | 0.946881 | BRCA | MRK-nu-1 | CCT-018159 | MVP_top60_DNA+RNA_seed | HDAC1 | 0.633283 | BRCA | EVSA-T | JQ12 |
| MVP_top60_DNA_seed | MDM2 | 0.947786 | BRCA | HCC1428 | Nutlin-3a (-) | MVP_top60_DNA+RNA_seed | HDAC2 | 0.633283 | BRCA | EVSA-T | JQ12 |
| MVP_top60_DNA_seed | ERBB2 | 0.947969 | BRCA | COLO-824 | Lapatinib | MVP_top60_DNA+RNA_seed | MDM2 | 0.633327 | BRCA | MDA-MB-436 | Nutlin-3a (-) |
| MVP_top60_DNA_seed | EGFR | 0.947969 | BRCA | COLO-824 | Lapatinib | MVP_top60_DNA+RNA_seed | ERBB2 | 0.634073 | BRCA | BT-20 | CP724714 |
| MVP_top60_DNA_seed | HDAC1 | 0.948571 | BRCA | HCC1187 | Belinostat | MVP_top60_DNA+RNA_seed | HSP90AA1 | 0.634147 | BRCA | MDA-MB-436 | Luminespib |
| MVP_top60_DNA_seed | HSP90AA1 | 0.949756 | BRCA | HCC38 | Elesclomol | MVP_top60_DNA+RNA_seed | HSP90AB1 | 0.634147 | BRCA | MDA-MB-436 | Luminespib |
| MVP_top60_DNA_seed | HSP90AB1 | 0.949756 | BRCA | HCC38 | Elesclomol | MVP_top60_DNA+RNA_seed | MDM2 | 0.634267 | BRCA | MDA-MB-468 | Nutlin-3a (-) |
| MVP_top60_DNA_seed | AR | 0.949852 | BRCA | CAL-85-1 | Bicalutamide | MVP_top60_DNA+RNA_seed | EGFR | 0.635143 | BRCA | CAL-148 | Pelitinib |
| MVP_top60_DNA_seed | EP300 | 0.949904 | BRCA | HCC1937 | I-CBP112 | MVP_top60_DNA+RNA_seed | EGFR | 0.635787 | BRCA | EVSA-T | Gefitinib |
| MVP_top60_DNA_seed | CREBBP | 0.949904 | BRCA | HCC1937 | I-CBP112 | MVP_top60_DNA+RNA_seed | HDAC1 | 0.638395 | BRCA | HCC1395 | Tubastatin A |
| MVP_top60_DNA_seed | ERBB2 | 0.950042 | BRCA | MRK-nu-1 | CP724714 | MVP_top60_DNA+RNA_seed | SRC | 0.639092 | BRCA | AU565 | Bosutinib |
| MVP_top60_DNA_seed | SRC | 0.950429 | BRCA | CAL-51 | Bosutinib | MVP_top60_DNA+RNA_seed | HDAC1 | 0.63959 | BRCA | Hs-578-T | CAY10603 |
| MVP_top60_DNA_seed | AR | 0.954621 | BRCA | HCC70 | AZD3514 | MVP_top60_DNA+RNA_seed | HDAC1 | 0.640473 | BRCA | BT-20 | CAY10603 |
| MVP_top60_DNA_seed | AKT1 | 0.954682 | BRCA | EFM-192A | GSK690693 | MVP_top60_DNA+RNA_seed | HDAC1 | 0.642319 | BRCA | BT-483 | Dacinostat |
| MVP_top60_DNA_seed | HDAC1 | 0.955723 | BRCA | MDA-MB-436 | AR-42 | MVP_top60_DNA+RNA_seed | HDAC1 | 0.645582 | BRCA | BT-474 | AR-42 |
| MVP_top60_DNA_seed | ERBB2 | 0.956597 | BRCA | YMB-1-E | CUDC-101 | MVP_top60_DNA+RNA_seed | AKT1 | 0.645613 | BRCA | HCC1428 | AKT inhibitor VIII |
| MVP_top60_DNA_seed | EGFR | 0.956597 | BRCA | YMB-1-E | CUDC-101 | MVP_top60_DNA+RNA_seed | AKT1 | 0.645868 | BRCA | BT-549 | MK-2206 |

| | | | | | | | | | | | |
|---|---|---|---|---|---|---|---|---|---|---|---|
| MVP_top60_DNA_seed | CDK1 | 0.956686 | BRCA | HCC2157 | RO-3306 | MVP_top60_DNA+RNA_seed | HDAC1 | 0.645925 | BRCA | HCC2157 | AR-42 |
| MVP_top60_DNA_seed | AR | 0.957918 | BRCA | HCC1428 | Bicalutamide | MVP_top60_DNA+RNA_seed | SRC | 0.646892 | BRCA | MDA-MB-453 | Bosutinib |
| MVP_top60_DNA_seed | HDAC1 | 0.959736 | BRCA | BT-549 | AR-42 | MVP_top60_DNA+RNA_seed | ESR1 | 0.649359 | BRCA | MRK-nu-1 | Tamoxifen |
| MVP_top60_DNA_seed | HSP90AA1 | 0.960136 | BRCA | HCC1500 | Luminespib | MVP_top60_DNA+RNA_seed | HDAC1 | 0.650665 | BRCA | UACC-893 | JQ12 |
| MVP_top60_DNA_seed | HSP90AB1 | 0.960136 | BRCA | HCC1500 | Luminespib | MVP_top60_DNA+RNA_seed | HDAC2 | 0.650665 | BRCA | UACC-893 | JQ12 |
| MVP_top60_DNA_seed | HSP90AA1 | 0.960632 | BRCA | MDA-MB-231 | SNX-2112 | MVP_top60_DNA+RNA_seed | PIK3CA | 0.650719 | BRCA | HCC70 | AZD8835 |
| MVP_top60_DNA_seed | HSP90AB1 | 0.960632 | BRCA | MDA-MB-231 | SNX-2112 | MVP_top60_DNA+RNA_seed | HSP90AA1 | 0.652526 | BRCA | HCC2218 | Elesclomol |
| MVP_top60_DNA_seed | EGFR | 0.963089 | BRCA | BT-474 | Cetuximab | MVP_top60_DNA+RNA_seed | HSP90AB1 | 0.652526 | BRCA | HCC2218 | Elesclomol |
| MVP_top60_DNA_seed | ATM | 0.963402 | BRCA | BT-474 | KU-60019 | MVP_top60_DNA+RNA_seed | SRC | 0.653248 | BRCA | MDA-MB-231 | Bosutinib |
| MVP_top60_DNA_seed | ERBB2 | 0.964885 | BRCA | HDQ-P1 | CP724714 | MVP_top60_DNA+RNA_seed | EGFR | 0.653912 | BRCA | HCC1428 | Gefitinib |
| MVP_top60_DNA_seed | ATM | 0.966125 | BRCA | MCF7 | CP466722 | MVP_top60_DNA+RNA_seed | MAPK3 | 0.656137 | BRCA | MDA-MB-231 | FR-180204 |
| MVP_top60_DNA_seed | ATM | 0.968245 | BRCA | Hs-578-T | CP466722 | MVP_top60_DNA+RNA_seed | MAPK1 | 0.656137 | BRCA | MDA-MB-231 | FR-180204 |
| MVP_top60_DNA_seed | HDAC1 | 0.970554 | BRCA | BT-20 | Tubastatin A | MVP_top60_DNA+RNA_seed | SRC | 0.657088 | BRCA | EVSA-T | Bosutinib |
| MVP_top60_DNA_seed | MDM2 | 0.970658 | BRCA | BT-483 | Nutlin-3a (-) | MVP_top60_DNA+RNA_seed | HSP90AA1 | 0.658284 | BRCA | UACC-893 | SNX-2112 |
| MVP_top60_DNA_seed | PIK3CA | 0.972751 | BRCA | Hs-578-T | AZD8835 | MVP_top60_DNA+RNA_seed | HSP90AB1 | 0.658284 | BRCA | UACC-893 | SNX-2112 |
| MVP_top60_DNA_seed | ESR1 | 0.978747 | BRCA | HCC202 | Fulvestrant | MVP_top60_DNA+RNA_seed | ERBB2 | 0.658619 | BRCA | HCC1937 | CP724714 |
| MVP_top60_DNA_seed | ERBB2 | 0.979905 | BRCA | Hs-578-T | Afatinib | MVP_top60_DNA+RNA_seed | SRC | 0.661681 | BRCA | BT-20 | Bosutinib |
| MVP_top60_DNA_seed | EGFR | 0.979905 | BRCA | Hs-578-T | Afatinib | MVP_top60_DNA+RNA_seed | HDAC1 | 0.661751 | BRCA | MDA-MB-415 | Dacinostat |
| MVP_top60_DNA_seed | MAPK3 | 0.981035 | BRCA | Hs-578-T | FR-180204 | MVP_top60_DNA+RNA_seed | ESR1 | 0.66269 | BRCA | MDA-MB-157 | Tamoxifen |
| MVP_top60_DNA_seed | MAPK1 | 0.981035 | BRCA | Hs-578-T | FR-180204 | MVP_top60_DNA+RNA_seed | HSP90AA1 | 0.663046 | BRCA | MDA-MB-415 | Elesclomol |
| MVP_top60_DNA_seed | MDM2 | 0.981959 | BRCA | HCC1419 | Nutlin-3a (-) | MVP_top60_DNA+RNA_seed | HSP90AB1 | 0.663046 | BRCA | MDA-MB-415 | Elesclomol |
| MVP_top60_DNA_seed | SRC | 0.982827 | BRCA | MCF7 | Bosutinib | MVP_top60_DNA+RNA_seed | ATM | 0.663625 | BRCA | CAL-85-1 | KU-55933 |
| MVP_top60_DNA_seed | EP300 | 0.982974 | BRCA | BT-20 | I-CBP112 | MVP_top60_DNA+RNA_seed | HDAC1 | 0.665461 | BRCA | BT-549 | Belinostat |
| MVP_top60_DNA_seed | CREBBP | 0.982974 | BRCA | BT-20 | I-CBP112 | MVP_top60_DNA+RNA_seed | SRC | 0.668118 | BRCA | MRK-nu-1 | Saracatinib |
| MVP_top60_DNA_seed | CDK2 | 0.984427 | BRCA | HCC38 | AZD5438 | MVP_top60_DNA+RNA_seed | CDK2 | 0.669238 | BRCA | HDQ-P1 | PHA-793887 |
| MVP_top60_DNA_seed | PIK3CA | 0.987493 | BRCA | HCC1395 | AZD8835 | MVP_top60_DNA+RNA_seed | MAPK1 | 0.669275 | BRCA | MDA-MB-361 | VX-11e |
| MVP_top60_DNA_seed | ATM | 0.991155 | BRCA | CAL-148 | CP466722 | MVP_top60_DNA+RNA_seed | EGFR | 0.669388 | BRCA | CAL-120 | PF-00299804 |
| MVP_top60_DNA_seed | MAPK1 | 0.991266 | BRCA | MCF7 | VX-11e | MVP_top60_DNA+RNA_seed | ERBB2 | 0.669388 | BRCA | CAL-120 | PF-00299804 |
| MVP_top60_DNA_seed | HDAC1 | 0.993604 | BRCA | CAL-120 | JQ12 | MVP_top60_DNA+RNA_seed | ESR1 | 0.671297 | BRCA | HCC70 | Tamoxifen |
| MVP_top60_DNA_seed | HDAC2 | 0.993604 | BRCA | CAL-120 | JQ12 | MVP_top60_DNA+RNA_seed | ESR1 | 0.671432 | BRCA | HCC38 | Tamoxifen |
| MVP_top60_DNA_seed | AKT1 | 0.994011 | BRCA | HCC1428 | MK-2206 | MVP_top60_DNA+RNA_seed | MDM2 | 0.671566 | BRCA | JIMT-1 | Nutlin-3a (-) |
| MVP_top60_DNA_seed | EGFR | 0.994099 | BRCA | UACC-812 | Erlotinib | MVP_top60_DNA+RNA_seed | SRC | 0.671946 | BRCA | HCC1937 | NVP-BHG712 |
| MVP_top60_DNA_seed | SRC | 0.995394 | BRCA | HCC1395 | NVP-BHG712 | MVP_top60_DNA+RNA_seed | EGFR | 0.672107 | BRCA | JIMT-1 | CUDC-101 |
| MVP_top60_DNA_seed | PIK3CA | 0.997241 | BRCA | ZR-75-30 | AZD8835 | MVP_top60_DNA+RNA_seed | ERBB2 | 0.672107 | BRCA | JIMT-1 | CUDC-101 |
| MVP_top60_DNA_seed | ERBB2 | 0.998925 | BRCA | MRK-nu-1 | Afatinib | MVP_top60_DNA+RNA_seed | EGFR | 0.672211 | BRCA | MDA-MB-436 | AZD8931 |
| MVP_top60_DNA_seed | EGFR | 0.998925 | BRCA | MRK-nu-1 | Afatinib | MVP_top60_DNA+RNA_seed | ERBB2 | 0.672211 | BRCA | MDA-MB-436 | AZD8931 |
| MVP_top60_DNA_seed | AKT1 | 1.00147 | BRCA | CAL-120 | AKT inhibitor VIII | MVP_top60_DNA+RNA_seed | ERBB3 | 0.672211 | BRCA | MDA-MB-436 | AZD8931 |
| MVP_top60_DNA_seed | MAPK1 | 1.003265 | BRCA | AU565 | VX-11e | MVP_top60_DNA+RNA_seed | MAPK14 | 0.672998 | BRCA | UACC-893 | TAK-715 |
| MVP_top60_DNA_seed | ERBB2 | 1.004373 | BRCA | HCC1395 | Afatinib | MVP_top60_DNA+RNA_seed | SRC | 0.673371 | BRCA | HCC202 | NVP-BHG712 |
| MVP_top60_DNA_seed | EGFR | 1.004373 | BRCA | HCC1395 | Afatinib | MVP_top60_DNA+RNA_seed | PIK3CA | 0.674489 | BRCA | BT-549 | AZD8835 |
| MVP_top60_DNA_seed | HDAC1 | 1.00563 | BRCA | UACC-893 | AR-42 | MVP_top60_DNA+RNA_seed | MDM2 | 0.674932 | BRCA | EVSA-T | Nutlin-3a (-) |

| | | | | | | | | | | | |
|---|---|---|---|---|---|---|---|---|---|---|---|
| MVP_top60_DNA_seed | ERBB2 | 1.006389 | BRCA | HCC1428 | Afatinib | MVP_top60_DNA+RNA_seed | EGFR | 0.676459 | BRCA | HCC1187 | Afatinib |
| MVP_top60_DNA_seed | EGFR | 1.006389 | BRCA | HCC1428 | Afatinib | MVP_top60_DNA+RNA_seed | ERBB2 | 0.676459 | BRCA | HCC1187 | Afatinib |
| MVP_top60_DNA_seed | HSP90AA1 | 1.010285 | BRCA | CAMA-1 | CCT-018159 | MVP_top60_DNA+RNA_seed | PIK3CA | 0.676875 | BRCA | HCC1419 | PI3Ka_4409 |
| MVP_top60_DNA_seed | HSP90AB1 | 1.010285 | BRCA | CAMA-1 | CCT-018159 | MVP_top60_DNA+RNA_seed | HDAC1 | 0.676988 | BRCA | HCC1395 | Dacinostat |
| MVP_top60_DNA_seed | HDAC1 | 1.010608 | BRCA | HCC1954 | AR-42 | MVP_top60_DNA+RNA_seed | EP300 | 0.679417 | BRCA | CAL-85-1 | I-CBP112 |
| MVP_top60_DNA_seed | HDAC1 | 1.011245 | BRCA | Hs-578-T | Tubastatin A | MVP_top60_DNA+RNA_seed | HDAC1 | 0.680194 | BRCA | MRK-nu-1 | AR-42 |
| MVP_top60_DNA_seed | PIK3CA | 1.012949 | BRCA | HCC38 | PI3Ka_4409 | MVP_top60_DNA+RNA_seed | HSP90AA1 | 0.681098 | BRCA | HCC1428 | Luminespib |
| MVP_top60_DNA_seed | MDM2 | 1.013197 | BRCA | CAL-85-1 | Nutlin-3a (-) | MVP_top60_DNA+RNA_seed | HSP90AB1 | 0.681098 | BRCA | HCC1428 | Luminespib |
| MVP_top60_DNA_seed | SRC | 1.014293 | BRCA | COLO-824 | Saracatinib | MVP_top60_DNA+RNA_seed | SRC | 0.681715 | BRCA | CAL-120 | Bosutinib |
| MVP_top60_DNA_seed | MDM2 | 1.014353 | BRCA | Hs-578-T | Nutlin-3a (-) | MVP_top60_DNA+RNA_seed | EGFR | 0.682007 | BRCA | T47D | CI-1033 |
| MVP_top60_DNA_seed | EGFR | 1.014678 | BRCA | HCC1395 | Pelitinib | MVP_top60_DNA+RNA_seed | ERBB2 | 0.682007 | BRCA | T47D | CI-1033 |
| MVP_top60_DNA_seed | EGFR | 1.015577 | BRCA | MRK-nu-1 | Erlotinib | MVP_top60_DNA+RNA_seed | AR | 0.682349 | BRCA | BT-474 | AZD3514 |
| MVP_top60_DNA_seed | EGFR | 1.018191 | BRCA | MFM-223 | Erlotinib | MVP_top60_DNA+RNA_seed | ATM | 0.683365 | BRCA | HCC1395 | CP466722 |
| MVP_top60_DNA_seed | EGFR | 1.021568 | BRCA | MDA-MB-330 | Cetuximab | MVP_top60_DNA+RNA_seed | HDAC1 | 0.684444 | BRCA | BT-549 | Dacinostat |
| MVP_top60_DNA_seed | HDAC1 | 1.023373 | BRCA | YMB-1-E | AR-42 | MVP_top60_DNA+RNA_seed | HSP90AA1 | 0.684676 | BRCA | MDA-MB-436 | SNX-2112 |
| MVP_top60_DNA_seed | AR | 1.023938 | BRCA | UACC-893 | Bicalutamide | MVP_top60_DNA+RNA_seed | HSP90AB1 | 0.684676 | BRCA | MDA-MB-436 | SNX-2112 |
| MVP_top60_DNA_seed | MDM2 | 1.025946 | BRCA | HCC38 | Nutlin-3a (-) | MVP_top60_DNA+RNA_seed | EP300 | 0.686821 | BRCA | MFM-223 | I-CBP112 |
| MVP_top60_DNA_seed | AKT1 | 1.030633 | BRCA | COLO-824 | MK-2206 | MVP_top60_DNA+RNA_seed | HSP90AA1 | 0.687342 | BRCA | CAMA-1 | SNX-2112 |
| MVP_top60_DNA_seed | ESR1 | 1.031725 | BRCA | MDA-MB-175-VII | Tamoxifen | MVP_top60_DNA+RNA_seed | HSP90AB1 | 0.687342 | BRCA | CAMA-1 | SNX-2112 |
| MVP_top60_DNA_seed | ERBB2 | 1.032075 | BRCA | ZR-75-30 | CI-1033 | MVP_top60_DNA+RNA_seed | EGFR | 0.690179 | BRCA | HCC1428 | Afatinib |
| MVP_top60_DNA_seed | EGFR | 1.032075 | BRCA | ZR-75-30 | CI-1033 | MVP_top60_DNA+RNA_seed | ERBB2 | 0.690179 | BRCA | HCC1428 | Afatinib |
| MVP_top60_DNA_seed | CDK1 | 1.032567 | BRCA | HCC1143 | RO-3306 | MVP_top60_DNA+RNA_seed | EGFR | 0.690557 | BRCA | HCC1428 | Pelitinib |
| MVP_top60_DNA_seed | ATM | 1.032779 | BRCA | HCC1395 | KU-55933 | MVP_top60_DNA+RNA_seed | SRC | 0.69163 | BRCA | HCC1500 | NVP-BHG712 |
| MVP_top60_DNA_seed | AKT1 | 1.036376 | BRCA | HCC1143 | BAY AKT1 | MVP_top60_DNA+RNA_seed | CDK2 | 0.692347 | BRCA | MDA-MB-453 | AZD5438 |
| MVP_top60_DNA_seed | CDK1 | 1.03844 | BRCA | HCC1500 | RO-3306 | MVP_top60_DNA+RNA_seed | HDAC1 | 0.695362 | BRCA | HCC1187 | Entinostat |
| MVP_top60_DNA_seed | AKT1 | 1.038997 | BRCA | Hs-578-T | AKT inhibitor VIII | MVP_top60_DNA+RNA_seed | EGFR | 0.6967 | BRCA | EVSA-T | Afatinib |
| MVP_top60_DNA_seed | ATM | 1.040636 | BRCA | HDQ-P1 | KU-55933 | MVP_top60_DNA+RNA_seed | ERBB2 | 0.6967 | BRCA | EVSA-T | Afatinib |
| MVP_top60_DNA_seed | ESR1 | 1.042918 | BRCA | HCC1937 | Tamoxifen | MVP_top60_DNA+RNA_seed | EGFR | 0.700015 | BRCA | HCC1500 | Afatinib |
| MVP_top60_DNA_seed | HSP90AA1 | 1.044194 | BRCA | HCC1954 | SNX-2112 | MVP_top60_DNA+RNA_seed | ERBB2 | 0.700015 | BRCA | HCC1500 | Afatinib |
| MVP_top60_DNA_seed | HSP90AB1 | 1.044194 | BRCA | HCC1954 | SNX-2112 | MVP_top60_DNA+RNA_seed | HDAC1 | 0.700463 | BRCA | CAL-51 | Tubastatin A |
| MVP_top60_DNA_seed | MDM2 | 1.048777 | BRCA | HCC1500 | Serdemetan | MVP_top60_DNA+RNA_seed | HDAC1 | 0.700607 | BRCA | YMB-1-E | Belinostat |
| MVP_top60_DNA_seed | HDAC1 | 1.052385 | BRCA | HCC1954 | Tubastatin A | MVP_top60_DNA+RNA_seed | AR | 0.702264 | BRCA | UACC-812 | Bicalutamide |
| MVP_top60_DNA_seed | EGFR | 1.052745 | BRCA | COLO-824 | Erlotinib | MVP_top60_DNA+RNA_seed | MAPK3 | 0.705806 | BRCA | HCC1937 | FR-180204 |
| MVP_top60_DNA_seed | ESR1 | 1.053103 | BRCA | CAMA-1 | Tamoxifen | MVP_top60_DNA+RNA_seed | MAPK1 | 0.705806 | BRCA | HCC1937 | FR-180204 |
| MVP_top60_DNA_seed | AR | 1.053818 | BRCA | CAMA-1 | AZD3514 | MVP_top60_DNA+RNA_seed | AR | 0.70582 | BRCA | BT-549 | Bicalutamide |
| MVP_top60_DNA_seed | ESR1 | 1.055534 | BRCA | BT-20 | Tamoxifen | MVP_top60_DNA+RNA_seed | SRC | 0.70602 | BRCA | CAL-51 | NVP-BHG712 |
| MVP_top60_DNA_seed | EGFR | 1.056015 | BRCA | MDA-MB-436 | Gefitinib | MVP_top60_DNA+RNA_seed | SRC | 0.706497 | BRCA | HCC1419 | NVP-BHG712 |
| MVP_top60_DNA_seed | MDM2 | 1.056745 | BRCA | BT-474 | Nutlin-3a (-) | MVP_top60_DNA+RNA_seed | MAPK3 | 0.706987 | BRCA | MRK-nu-1 | FR-180204 |
| MVP_top60_DNA_seed | AKT1 | 1.061477 | BRCA | HDQ-P1 | MK-2206 | MVP_top60_DNA+RNA_seed | MAPK1 | 0.706987 | BRCA | MRK-nu-1 | FR-180204 |
| MVP_top60_DNA_seed | ERBB2 | 1.061921 | BRCA | MDA-MB-157 | CI-1033 | MVP_top60_DNA+RNA_seed | AR | 0.70817 | BRCA | HCC1569 | AZD3514 |
| MVP_top60_DNA_seed | EGFR | 1.061921 | BRCA | MDA-MB-157 | CI-1033 | MVP_top60_DNA+RNA_seed | EGFR | 0.70822 | BRCA | HCC38 | Afatinib |

| | | | | | | | | | | | |
|---|---|---|---|---|---|---|---|---|---|---|---|
| MVP_top60_DNA_seed | HSP90AA1 | 1.064246 | BRCA | BT-20 | Luminespib | MVP_top60_DNA+RNA_seed | ERBB2 | 0.70822 | BRCA | HCC38 | Afatinib |
| MVP_top60_DNA_seed | HSP90AB1 | 1.064246 | BRCA | BT-20 | Luminespib | MVP_top60_DNA+RNA_seed | HDAC1 | 0.708266 | BRCA | JIMT-1 | AR-42 |
| MVP_top60_DNA_seed | CDK2 | 1.06477 | BRCA | HCC1395 | PHA-793887 | MVP_top60_DNA+RNA_seed | MDM2 | 0.708748 | BRCA | MDA-MB-231 | Nutlin-3a (-) |
| MVP_top60_DNA_seed | EP300 | 1.065074 | BRCA | HDQ-P1 | I-CBP112 | MVP_top60_DNA+RNA_seed | AKT1 | 0.709522 | BRCA | COLO-824 | AKT inhibitor VIII |
| MVP_top60_DNA_seed | CREBBP | 1.065074 | BRCA | HDQ-P1 | I-CBP112 | MVP_top60_DNA+RNA_seed | CDK2 | 0.709921 | BRCA | OCUB-M | AZD5438 |
| MVP_top60_DNA_seed | AKT1 | 1.065861 | BRCA | CAL-85-1 | BAY AKT1 | MVP_top60_DNA+RNA_seed | HSP90AA1 | 0.711067 | BRCA | UACC-893 | CCT-018159 |
| MVP_top60_DNA_seed | MAPK3 | 1.067461 | BRCA | HCC1954 | FR-180204 | MVP_top60_DNA+RNA_seed | HSP90AB1 | 0.711067 | BRCA | UACC-893 | CCT-018159 |
| MVP_top60_DNA_seed | MAPK1 | 1.067461 | BRCA | HCC1954 | FR-180204 | MVP_top60_DNA+RNA_seed | HSP90AA1 | 0.711759 | BRCA | ZR-75-30 | CCT-018159 |
| MVP_top60_DNA_seed | AR | 1.073449 | BRCA | HDQ-P1 | Bicalutamide | MVP_top60_DNA+RNA_seed | HSP90AB1 | 0.711759 | BRCA | ZR-75-30 | CCT-018159 |
| MVP_top60_DNA_seed | SRC | 1.075631 | BRCA | HCC1143 | Bosutinib | MVP_top60_DNA+RNA_seed | HDAC1 | 0.712204 | BRCA | HCC1419 | CAY10603 |
| MVP_top60_DNA_seed | HSP90AA1 | 1.078525 | BRCA | UACC-893 | Elesclomol | MVP_top60_DNA+RNA_seed | HSP90AA1 | 0.713635 | BRCA | YMB-1-E | SNX-2112 |
| MVP_top60_DNA_seed | HSP90AB1 | 1.078525 | BRCA | UACC-893 | Elesclomol | MVP_top60_DNA+RNA_seed | HSP90AB1 | 0.713635 | BRCA | YMB-1-E | SNX-2112 |
| MVP_top60_DNA_seed | PIK3CA | 1.082358 | BRCA | HCC38 | AZD8835 | MVP_top60_DNA+RNA_seed | HSP90AA1 | 0.714584 | BRCA | EFM-192A | SNX-2112 |
| MVP_top60_DNA_seed | MDM2 | 1.083206 | BRCA | MFM-223 | Nutlin-3a (-) | MVP_top60_DNA+RNA_seed | HSP90AB1 | 0.714584 | BRCA | EFM-192A | SNX-2112 |
| MVP_top60_DNA_seed | MAPK1 | 1.084861 | BRCA | HCC2157 | VX-11e | MVP_top60_DNA+RNA_seed | SRC | 0.715428 | BRCA | ZR-75-30 | NVP-BHG712 |
| MVP_top60_DNA_seed | MDM2 | 1.085644 | BRCA | EFM-19 | Nutlin-3a (-) | MVP_top60_DNA+RNA_seed | CDK1 | 0.715652 | BRCA | T47D | RO-3306 |
| MVP_top60_DNA_seed | MAPK1 | 1.089046 | BRCA | YMB-1-E | VX-11e | MVP_top60_DNA+RNA_seed | EGFR | 0.718217 | BRCA | UACC-812 | AST-1306 |
| MVP_top60_DNA_seed | HSP90AA1 | 1.091441 | BRCA | HCC1143 | CCT-018159 | MVP_top60_DNA+RNA_seed | CDK2 | 0.718726 | BRCA | MDA-MB-175-VII | PHA-793887 |
| MVP_top60_DNA_seed | HSP90AB1 | 1.091441 | BRCA | HCC1143 | CCT-018159 | MVP_top60_DNA+RNA_seed | EGFR | 0.719779 | BRCA | HCC38 | Afatinib |
| MVP_top60_DNA_seed | ERBB2 | 1.091944 | BRCA | MDA-MB-436 | CP724714 | MVP_top60_DNA+RNA_seed | ERBB2 | 0.719779 | BRCA | HCC38 | Afatinib |
| MVP_top60_DNA_seed | HDAC1 | 1.092439 | BRCA | MDA-MB-453 | JQ12 | MVP_top60_DNA+RNA_seed | EGFR | 0.720678 | BRCA | MRK-nu-1 | Lapatinib |
| MVP_top60_DNA_seed | HDAC2 | 1.092439 | BRCA | MDA-MB-453 | JQ12 | MVP_top60_DNA+RNA_seed | ERBB2 | 0.720678 | BRCA | MRK-nu-1 | Lapatinib |
| MVP_top60_DNA_seed | EGFR | 1.09424 | BRCA | MRK-nu-1 | Gefitinib | MVP_top60_DNA+RNA_seed | ATM | 0.721892 | BRCA | YMB-1-E | CP466722 |
| MVP_top60_DNA_seed | CDK2 | 1.094768 | BRCA | EFM-192A | AZD5438 | MVP_top60_DNA+RNA_seed | EP300 | 0.721992 | BRCA | MRK-nu-1 | I-CBP112 |
| MVP_top60_DNA_seed | ERBB2 | 1.096387 | BRCA | HCC1395 | Afatinib | MVP_top60_DNA+RNA_seed | MAPK1 | 0.722424 | BRCA | CAL-148 | VX-11e |
| MVP_top60_DNA_seed | EGFR | 1.096387 | BRCA | HCC1395 | Afatinib | MVP_top60_DNA+RNA_seed | SRC | 0.722521 | BRCA | DU-4475 | Saracatinib |
| MVP_top60_DNA_seed | MDM2 | 1.098218 | BRCA | HCC1143 | Nutlin-3a (-) | MVP_top60_DNA+RNA_seed | HDAC1 | 0.723137 | BRCA | BT-549 | CAY10603 |
| MVP_top60_DNA_seed | ERBB2 | 1.098459 | BRCA | MDA-MB-415 | Afatinib | MVP_top60_DNA+RNA_seed | EGFR | 0.723789 | BRCA | HCC1419 | Cetuximab |
| MVP_top60_DNA_seed | EGFR | 1.098459 | BRCA | MDA-MB-415 | Afatinib | MVP_top60_DNA+RNA_seed | SRC | 0.724489 | BRCA | YMB-1-E | NVP-BHG712 |
| MVP_top60_DNA_seed | MAPK1 | 1.099008 | BRCA | HCC1428 | VX-11e | MVP_top60_DNA+RNA_seed | EGFR | 0.726607 | BRCA | MDA-MB-436 | Afatinib |
| MVP_top60_DNA_seed | ATM | 1.10329 | BRCA | MRK-nu-1 | CP466722 | MVP_top60_DNA+RNA_seed | ERBB2 | 0.726607 | BRCA | MDA-MB-436 | Afatinib |
| MVP_top60_DNA_seed | SRC | 1.108298 | BRCA | MDA-MB-330 | Bosutinib | MVP_top60_DNA+RNA_seed | ATM | 0.727851 | BRCA | MDA-MB-415 | CP466722 |
| MVP_top60_DNA_seed | EGFR | 1.110495 | BRCA | CAMA-1 | Cetuximab | MVP_top60_DNA+RNA_seed | CDK2 | 0.728712 | BRCA | HCC1937 | AZD5438 |
| MVP_top60_DNA_seed | MAPK1 | 1.112239 | BRCA | Hs-578-T | VX-11e | MVP_top60_DNA+RNA_seed | ESR1 | 0.731603 | BRCA | Hs-578-T | Tamoxifen |
| MVP_top60_DNA_seed | ESR1 | 1.113248 | BRCA | UACC-893 | Tamoxifen | MVP_top60_DNA+RNA_seed | ATM | 0.732635 | BRCA | CAL-120 | KU-55933 |
| MVP_top60_DNA_seed | ERBB2 | 1.114931 | BRCA | HCC1395 | CP724714 | MVP_top60_DNA+RNA_seed | HDAC1 | 0.733702 | BRCA | COLO-824 | JQ12 |
| MVP_top60_DNA_seed | SRC | 1.11809 | BRCA | CAMA-1 | Bosutinib | MVP_top60_DNA+RNA_seed | HDAC2 | 0.733702 | BRCA | COLO-824 | JQ12 |
| MVP_top60_DNA_seed | HDAC1 | 1.119004 | BRCA | BT-549 | JQ12 | MVP_top60_DNA+RNA_seed | SRC | 0.733831 | BRCA | EFM-192A | NVP-BHG712 |
| MVP_top60_DNA_seed | HDAC2 | 1.119004 | BRCA | BT-549 | JQ12 | MVP_top60_DNA+RNA_seed | EGFR | 0.736135 | BRCA | MDA-MB-436 | CUDC-101 |
| MVP_top60_DNA_seed | SRC | 1.120187 | BRCA | HCC2218 | WH-4-023 | MVP_top60_DNA+RNA_seed | ERBB2 | 0.736135 | BRCA | MDA-MB-436 | CUDC-101 |
| MVP_top60_DNA_seed | AKT1 | 1.123569 | BRCA | COLO-824 | A-443654 | MVP_top60_DNA+RNA_seed | AR | 0.738074 | BRCA | MRK-nu-1 | Bicalutamide |

| | | | | | | | | | | | |
|---|---|---|---|---|---|---|---|---|---|---|---|
| MVP_top60_DNA_seed | EP300 | 1.123824 | BRCA | HCC1419 | I-CBP112 | MVP_top60_DNA+RNA_seed | AR | 0.73952 | BRCA | BT-20 | Bicalutamide |
| MVP_top60_DNA_seed | CREBBP | 1.123824 | BRCA | HCC1419 | I-CBP112 | MVP_top60_DNA+RNA_seed | MAPK3 | 0.740058 | BRCA | HCC1428 | FR-180204 |
| MVP_top60_DNA_seed | MDM2 | 1.124313 | BRCA | CAL-85-1 | Serdemetan | MVP_top60_DNA+RNA_seed | MAPK1 | 0.740058 | BRCA | HCC1428 | FR-180204 |
| MVP_top60_DNA_seed | HSP90AA1 | 1.124441 | BRCA | CAL-51 | SNX-2112 | MVP_top60_DNA+RNA_seed | ERBB2 | 0.740161 | BRCA | HCC70 | CP724714 |
| MVP_top60_DNA_seed | HSP90AB1 | 1.124441 | BRCA | CAL-51 | SNX-2112 | MVP_top60_DNA+RNA_seed | HSP90AA1 | 0.741213 | BRCA | HCC1419 | SNX-2112 |
| MVP_top60_DNA_seed | ERBB2 | 1.126794 | BRCA | EFM-19 | Afatinib | MVP_top60_DNA+RNA_seed | HSP90AB1 | 0.741213 | BRCA | HCC1419 | SNX-2112 |
| MVP_top60_DNA_seed | EGFR | 1.126794 | BRCA | EFM-19 | Afatinib | MVP_top60_DNA+RNA_seed | HDAC1 | 0.74274 | BRCA | CAL-148 | Parthenolide |
| MVP_top60_DNA_seed | AKT1 | 1.128053 | BRCA | MDA-MB-231 | BAY AKT1 | MVP_top60_DNA+RNA_seed | EGFR | 0.74285 | BRCA | UACC-893 | Cetuximab |
| MVP_top60_DNA_seed | SRC | 1.129413 | BRCA | MFM-223 | WH-4-023 | MVP_top60_DNA+RNA_seed | SRC | 0.743804 | BRCA | HCC2157 | Saracatinib |
| MVP_top60_DNA_seed | ESR1 | 1.130416 | BRCA | ZR-75-30 | Fulvestrant | MVP_top60_DNA+RNA_seed | AR | 0.744404 | BRCA | HCC202 | Bicalutamide |
| MVP_top60_DNA_seed | AR | 1.135081 | BRCA | HCC1143 | AZD3514 | MVP_top60_DNA+RNA_seed | ATM | 0.7454 | BRCA | MDA-MB-361 | CP466722 |
| MVP_top60_DNA_seed | HSP90AA1 | 1.142179 | BRCA | CAL-85-1 | Tanespimycin | MVP_top60_DNA+RNA_seed | EGFR | 0.747787 | BRCA | CAMA-1 | Pelitinib |
| MVP_top60_DNA_seed | HSP90AB1 | 1.142179 | BRCA | CAL-85-1 | Tanespimycin | MVP_top60_DNA+RNA_seed | EGFR | 0.749414 | BRCA | BT-549 | Cetuximab |
| MVP_top60_DNA_seed | MDM2 | 1.144594 | BRCA | MFM-223 | Serdemetan | MVP_top60_DNA+RNA_seed | MAPK3 | 0.749428 | BRCA | MDA-MB-330 | FR-180204 |
| MVP_top60_DNA_seed | MAPK1 | 1.14544 | BRCA | HCC1569 | VX-11e | MVP_top60_DNA+RNA_seed | MAPK1 | 0.749428 | BRCA | MDA-MB-330 | FR-180204 |
| MVP_top60_DNA_seed | EGFR | 1.14782 | BRCA | MFM-223 | Cetuximab | MVP_top60_DNA+RNA_seed | SRC | 0.750777 | BRCA | CAL-148 | NVP-BHG712 |
| MVP_top60_DNA_seed | SRC | 1.148501 | BRCA | BT-474 | Bosutinib | MVP_top60_DNA+RNA_seed | SRC | 0.753952 | BRCA | ZR-75-30 | Bosutinib |
| MVP_top60_DNA_seed | EGFR | 1.150186 | BRCA | UACC-893 | Gefitinib | MVP_top60_DNA+RNA_seed | PIK3CA | 0.754134 | BRCA | MDA-MB-468 | PI3Ka_4409 |
| MVP_top60_DNA_seed | PIK3CA | 1.15106 | BRCA | BT-483 | AZD8835 | MVP_top60_DNA+RNA_seed | CDK2 | 0.755465 | BRCA | HCC2157 | Seliciclib |
| MVP_top60_DNA_seed | CDK1 | 1.15205 | BRCA | BT-549 | RO-3306 | MVP_top60_DNA+RNA_seed | MAPK1 | 0.756585 | BRCA | T47D | VX-11e |
| MVP_top60_DNA_seed | ESR1 | 1.156381 | BRCA | MRK-nu-1 | Fulvestrant | MVP_top60_DNA+RNA_seed | EP300 | 0.756949 | BRCA | HCC1954 | I-CBP112 |
| MVP_top60_DNA_seed | PIK3CA | 1.15663 | BRCA | MDA-MB-468 | AZD8835 | MVP_top60_DNA+RNA_seed | EP300 | 0.757292 | BRCA | MDA-MB-415 | I-CBP112 |
| MVP_top60_DNA_seed | ATM | 1.158436 | BRCA | HCC202 | KU-60019 | MVP_top60_DNA+RNA_seed | AR | 0.757631 | BRCA | MRK-nu-1 | AZD3514 |
| MVP_top60_DNA_seed | MAPK1 | 1.158625 | BRCA | HCC2218 | VX-11e | MVP_top60_DNA+RNA_seed | AR | 0.759007 | BRCA | AU565 | AZD3514 |
| MVP_top60_DNA_seed | CDK1 | 1.165613 | BRCA | EFM-19 | RO-3306 | MVP_top60_DNA+RNA_seed | SRC | 0.761482 | BRCA | EFM-19 | Bosutinib |
| MVP_top60_DNA_seed | MAPK3 | 1.169895 | BRCA | BT-20 | FR-180204 | MVP_top60_DNA+RNA_seed | HDAC1 | 0.761535 | BRCA | MDA-MB-468 | AR-42 |
| MVP_top60_DNA_seed | MAPK1 | 1.169895 | BRCA | BT-20 | FR-180204 | MVP_top60_DNA+RNA_seed | EGFR | 0.761735 | BRCA | HCC1954 | CUDC-101 |
| MVP_top60_DNA_seed | CDK1 | 1.173316 | BRCA | UACC-893 | RO-3306 | MVP_top60_DNA+RNA_seed | ERBB2 | 0.761735 | BRCA | HCC1954 | CUDC-101 |
| MVP_top60_DNA_seed | AR | 1.173383 | BRCA | MDA-MB-175-VII | Bicalutamide | MVP_top60_DNA+RNA_seed | HDAC1 | 0.762701 | BRCA | HCC1187 | AR-42 |
| MVP_top60_DNA_seed | HSP90AA1 | 1.173827 | BRCA | CAMA-1 | Elesclomol | MVP_top60_DNA+RNA_seed | HSP90AA1 | 0.762805 | BRCA | MFM-223 | CCT-018159 |
| MVP_top60_DNA_seed | HSP90AB1 | 1.173827 | BRCA | CAMA-1 | Elesclomol | MVP_top60_DNA+RNA_seed | HSP90AB1 | 0.762805 | BRCA | MFM-223 | CCT-018159 |
| MVP_top60_DNA_seed | PIK3CA | 1.175306 | BRCA | CAL-51 | AZD8835 | MVP_top60_DNA+RNA_seed | SRC | 0.7645 | BRCA | Hs-578-T | Bosutinib |
| MVP_top60_DNA_seed | ERBB2 | 1.176434 | BRCA | CAL-120 | CP724714 | MVP_top60_DNA+RNA_seed | HDAC1 | 0.764697 | BRCA | MDA-MB-436 | JQ12 |
| MVP_top60_DNA_seed | EP300 | 1.176935 | BRCA | HCC1395 | I-CBP112 | MVP_top60_DNA+RNA_seed | HDAC2 | 0.764697 | BRCA | MDA-MB-436 | JQ12 |
| MVP_top60_DNA_seed | CREBBP | 1.176935 | BRCA | HCC1395 | I-CBP112 | MVP_top60_DNA+RNA_seed | AKT1 | 0.765361 | BRCA | MCF7 | GSK690693 |
| MVP_top60_DNA_seed | MDM2 | 1.177493 | BRCA | BT-20 | Serdemetan | MVP_top60_DNA+RNA_seed | EGFR | 0.768046 | BRCA | BT-20 | AST-1306 |
| MVP_top60_DNA_seed | AKT1 | 1.178983 | BRCA | HDQ-P1 | AKT inhibitor VIII | MVP_top60_DNA+RNA_seed | SRC | 0.768971 | BRCA | BT-20 | NVP-BHG712 |
| MVP_top60_DNA_seed | MAPK1 | 1.179384 | BRCA | HCC1954 | VX-11e | MVP_top60_DNA+RNA_seed | PLK1 | 0.771832 | BRCA | BT-474 | BI-2536 |
| MVP_top60_DNA_seed | MDM2 | 1.182363 | BRCA | BT-549 | Nutlin-3a (-) | MVP_top60_DNA+RNA_seed | HDAC1 | 0.771997 | BRCA | HCC1500 | Parthenolide |
| MVP_top60_DNA_seed | EP300 | 1.186063 | BRCA | EFM-19 | I-CBP112 | MVP_top60_DNA+RNA_seed | SRC | 0.772994 | BRCA | BT-474 | HG-6-71-01 |
| MVP_top60_DNA_seed | CREBBP | 1.186063 | BRCA | EFM-19 | I-CBP112 | MVP_top60_DNA+RNA_seed | MAPK3 | 0.773322 | BRCA | MCF7 | FR-180204 |

| | | | | | | | | | | | |
|---|---|---|---|---|---|---|---|---|---|---|---|
| MVP_top60_DNA_seed | AR | 1.187736 | BRCA | HCC1937 | Bicalutamide | MVP_top60_DNA+RNA_seed | MAPK1 | 0.773322 | BRCA | MCF7 | FR-180204 |
| MVP_top60_DNA_seed | HDAC1 | 1.18868 | BRCA | BT-483 | JQ12 | MVP_top60_DNA+RNA_seed | HDAC1 | 0.774583 | BRCA | MDA-MB-157 | Tubastatin A |
| MVP_top60_DNA_seed | HDAC2 | 1.18868 | BRCA | BT-483 | JQ12 | MVP_top60_DNA+RNA_seed | EGFR | 0.776247 | BRCA | Hs-578-T | Afatinib |
| MVP_top60_DNA_seed | SRC | 1.188852 | BRCA | MDA-MB-157 | NVP-BHG712 | MVP_top60_DNA+RNA_seed | ERBB2 | 0.776247 | BRCA | Hs-578-T | Afatinib |
| MVP_top60_DNA_seed | EGFR | 1.190486 | BRCA | HCC1143 | Cetuximab | MVP_top60_DNA+RNA_seed | MDM2 | 0.779843 | BRCA | HCC1428 | Serdemetan |
| MVP_top60_DNA_seed | MAPK1 | 1.192202 | BRCA | UACC-812 | VX-11e | MVP_top60_DNA+RNA_seed | EGFR | 0.780746 | BRCA | HCC2157 | Cetuximab |
| MVP_top60_DNA_seed | HDAC1 | 1.195305 | BRCA | HCC1395 | Belinostat | MVP_top60_DNA+RNA_seed | EGFR | 0.782216 | BRCA | MFM-223 | Afatinib |
| MVP_top60_DNA_seed | SRC | 1.198467 | BRCA | MDA-MB-453 | NVP-BHG712 | MVP_top60_DNA+RNA_seed | ERBB2 | 0.782216 | BRCA | MFM-223 | Afatinib |
| MVP_top60_DNA_seed | ATM | 1.198858 | BRCA | DU-4475 | KU-60019 | MVP_top60_DNA+RNA_seed | PLK1 | 0.782784 | BRCA | HCC2218 | GW843682X |
| MVP_top60_DNA_seed | MAPK1 | 1.200177 | BRCA | HCC1395 | VX-11e | MVP_top60_DNA+RNA_seed | CDK2 | 0.787585 | BRCA | HCC1419 | PHA-793887 |
| MVP_top60_DNA_seed | AKT1 | 1.201319 | BRCA | MDA-MB-436 | GSK690693 | MVP_top60_DNA+RNA_seed | HDAC1 | 0.788546 | BRCA | BT-20 | AR-42 |
| MVP_top60_DNA_seed | CDK2 | 1.20265 | BRCA | BT-20 | AZD5438 | MVP_top60_DNA+RNA_seed | AR | 0.788998 | BRCA | HCC2157 | Bicalutamide |
| MVP_top60_DNA_seed | HDAC1 | 1.202879 | BRCA | COLO-824 | Parthenolide | MVP_top60_DNA+RNA_seed | AR | 0.789313 | BRCA | HCC1500 | Bicalutamide |
| MVP_top60_DNA_seed | MDM2 | 1.205442 | BRCA | BT-474 | Serdemetan | MVP_top60_DNA+RNA_seed | EGFR | 0.792312 | BRCA | EFM-192A | Gefitinib |
| MVP_top60_DNA_seed | MDM2 | 1.206361 | BRCA | HCC1395 | Nutlin-3a (-) | MVP_top60_DNA+RNA_seed | MAPK1 | 0.794154 | BRCA | CAMA-1 | VX-11e |
| MVP_top60_DNA_seed | HDAC1 | 1.209938 | BRCA | HCC1500 | Entinostat | MVP_top60_DNA+RNA_seed | ATM | 0.794938 | BRCA | MDA-MB-453 | KU-55933 |
| MVP_top60_DNA_seed | HSP90AA1 | 1.21152 | BRCA | EFM-192A | Elesclomol | MVP_top60_DNA+RNA_seed | SRC | 0.795733 | BRCA | MDA-MB-415 | HG-6-71-01 |
| MVP_top60_DNA_seed | HSP90AB1 | 1.21152 | BRCA | EFM-192A | Elesclomol | MVP_top60_DNA+RNA_seed | SRC | 0.798275 | BRCA | CAL-148 | WH-4-023 |
| MVP_top60_DNA_seed | SRC | 1.212273 | BRCA | HCC70 | Bosutinib | MVP_top60_DNA+RNA_seed | MAPK3 | 0.801731 | BRCA | MDA-MB-157 | FR-180204 |
| MVP_top60_DNA_seed | PIK3CA | 1.217205 | BRCA | HCC1937 | PI3Ka_4409 | MVP_top60_DNA+RNA_seed | MAPK1 | 0.801731 | BRCA | MDA-MB-157 | FR-180204 |
| MVP_top60_DNA_seed | MAPK1 | 1.217349 | BRCA | BT-549 | VX-11e | MVP_top60_DNA+RNA_seed | MAPK3 | 0.804182 | BRCA | CAL-51 | FR-180204 |
| MVP_top60_DNA_seed | HDAC1 | 1.218545 | BRCA | HCC1187 | JQ12 | MVP_top60_DNA+RNA_seed | MAPK1 | 0.804182 | BRCA | CAL-51 | FR-180204 |
| MVP_top60_DNA_seed | HDAC2 | 1.218545 | BRCA | HCC1187 | JQ12 | MVP_top60_DNA+RNA_seed | EGFR | 0.805516 | BRCA | MDA-MB-415 | Afatinib |
| MVP_top60_DNA_seed | MAPK3 | 1.218793 | BRCA | MFM-223 | FR-180204 | MVP_top60_DNA+RNA_seed | ERBB2 | 0.805516 | BRCA | MDA-MB-415 | Afatinib |
| MVP_top60_DNA_seed | MAPK1 | 1.218793 | BRCA | MFM-223 | FR-180204 | MVP_top60_DNA+RNA_seed | EGFR | 0.805539 | BRCA | HCC202 | Gefitinib |
| MVP_top60_DNA_seed | CDK2 | 1.221172 | BRCA | MCF7 | PHA-793887 | MVP_top60_DNA+RNA_seed | AR | 0.809381 | BRCA | MDA-MB-330 | Bicalutamide |
| MVP_top60_DNA_seed | ATM | 1.221655 | BRCA | BT-483 | KU-55933 | MVP_top60_DNA+RNA_seed | AR | 0.810886 | BRCA | EFM-192A | Bicalutamide |
| MVP_top60_DNA_seed | AR | 1.222609 | BRCA | BT-20 | Bicalutamide | MVP_top60_DNA+RNA_seed | HDAC1 | 0.811737 | BRCA | BT-549 | Tubastatin A |
| MVP_top60_DNA_seed | HSP90AA1 | 1.224278 | BRCA | BT-483 | SNX-2112 | MVP_top60_DNA+RNA_seed | MDM2 | 0.812224 | BRCA | HCC1937 | Nutlin-3a (-) |
| MVP_top60_DNA_seed | HSP90AB1 | 1.224278 | BRCA | BT-483 | SNX-2112 | MVP_top60_DNA+RNA_seed | AR | 0.814415 | BRCA | HCC1937 | Bicalutamide |
| MVP_top60_DNA_seed | HDAC1 | 1.22688 | BRCA | HCC1954 | CAY10603 | MVP_top60_DNA+RNA_seed | HSP90AA1 | 0.815104 | BRCA | HDQ-P1 | SNX-2112 |
| MVP_top60_DNA_seed | CDK2 | 1.226965 | BRCA | MDA-MB-330 | AZD5438 | MVP_top60_DNA+RNA_seed | HSP90AB1 | 0.815104 | BRCA | HDQ-P1 | SNX-2112 |
| MVP_top60_DNA_seed | HSP90AA1 | 1.227944 | BRCA | UACC-812 | Luminespib | MVP_top60_DNA+RNA_seed | EGFR | 0.815608 | BRCA | CAMA-1 | Gefitinib |
| MVP_top60_DNA_seed | HSP90AB1 | 1.227944 | BRCA | UACC-812 | Luminespib | MVP_top60_DNA+RNA_seed | SRC | 0.815843 | BRCA | HCC1500 | Saracatinib |
| MVP_top60_DNA_seed | EGFR | 1.232463 | BRCA | BT-549 | Gefitinib | MVP_top60_DNA+RNA_seed | EGFR | 0.817017 | BRCA | UACC-893 | CUDC-101 |
| MVP_top60_DNA_seed | SRC | 1.23678 | BRCA | HCC2157 | NVP-BHG712 | MVP_top60_DNA+RNA_seed | ERBB2 | 0.817017 | BRCA | UACC-893 | CUDC-101 |
| MVP_top60_DNA_seed | ATM | 1.236821 | BRCA | MRK-nu-1 | KU-55933 | MVP_top60_DNA+RNA_seed | PLK1 | 0.818127 | BRCA | HCC1569 | PLK_6522 |
| MVP_top60_DNA_seed | MAPK1 | 1.238212 | BRCA | MFM-223 | VX-11e | MVP_top60_DNA+RNA_seed | CDK2 | 0.818316 | BRCA | MFM-223 | PHA-793887 |
| MVP_top60_DNA_seed | CDK2 | 1.240101 | BRCA | MDA-MB-436 | PHA-793887 | MVP_top60_DNA+RNA_seed | CDK1 | 0.818427 | BRCA | MDA-MB-436 | RO-3306 |
| MVP_top60_DNA_seed | AKT1 | 1.242022 | BRCA | MRK-nu-1 | AKT inhibitor VIII | MVP_top60_DNA+RNA_seed | HSP90AA1 | 0.818776 | BRCA | HCC202 | Tanespimycin |
| MVP_top60_DNA_seed | HDAC1 | 1.242255 | BRCA | HCC1954 | Belinostat | MVP_top60_DNA+RNA_seed | HSP90AB1 | 0.818776 | BRCA | HCC202 | Tanespimycin |

| | | | | | | | | | | | |
|---|---|---|---|---|---|---|---|---|---|---|---|
| MVP_top60_DNA_seed | MDM2 | 1.244282 | BRCA | BT-20 | Nutlin-3a (-) | MVP_top60_DNA+RNA_seed | CDK2 | 0.819318 | BRCA | MDA-MB-157 | PHA-793887 |
| MVP_top60_DNA_seed | MDM2 | 1.245407 | BRCA | HDQ-P1 | Nutlin-3a (-) | MVP_top60_DNA+RNA_seed | EGFR | 0.820375 | BRCA | HCC1500 | Cetuximab |
| MVP_top60_DNA_seed | SRC | 1.250813 | BRCA | MRK-nu-1 | Bosutinib | MVP_top60_DNA+RNA_seed | MAPK1 | 0.821476 | BRCA | CAL-51 | VX-11e |
| MVP_top60_DNA_seed | ESR1 | 1.251149 | BRCA | HCC1395 | Tamoxifen | MVP_top60_DNA+RNA_seed | ATM | 0.823726 | BRCA | COLO-824 | KU-55933 |
| MVP_top60_DNA_seed | AR | 1.251462 | BRCA | MDA-MB-330 | AZD3514 | MVP_top60_DNA+RNA_seed | HSP90AA1 | 0.825108 | BRCA | MDA-MB-453 | Luminespib |
| MVP_top60_DNA_seed | AR | 1.252344 | BRCA | UACC-893 | AZD3514 | MVP_top60_DNA+RNA_seed | HSP90AB1 | 0.825108 | BRCA | MDA-MB-453 | Luminespib |
| MVP_top60_DNA_seed | EGFR | 1.256459 | BRCA | MCF7 | Pelitinib | MVP_top60_DNA+RNA_seed | EGFR | 0.825797 | BRCA | HCC70 | Pelitinib |
| MVP_top60_DNA_seed | MAPK1 | 1.25669 | BRCA | HCC1937 | VX-11e | MVP_top60_DNA+RNA_seed | HDAC1 | 0.827368 | BRCA | HCC1569 | JQ12 |
| MVP_top60_DNA_seed | CDK1 | 1.258323 | BRCA | CAMA-1 | RO-3306 | MVP_top60_DNA+RNA_seed | HDAC2 | 0.827368 | BRCA | HCC1569 | JQ12 |
| MVP_top60_DNA_seed | PIK3CA | 1.260121 | BRCA | MDA-MB-231 | PI3Ka_4409 | MVP_top60_DNA+RNA_seed | HDAC1 | 0.827447 | BRCA | HCC1937 | Tubastatin A |
| MVP_top60_DNA_seed | CDK2 | 1.26254 | BRCA | EFM-19 | PHA-793887 | MVP_top60_DNA+RNA_seed | HDAC1 | 0.82824 | BRCA | Hs-578-T | AR-42 |
| MVP_top60_DNA_seed | SRC | 1.269252 | BRCA | Hs-578-T | NVP-BHG712 | MVP_top60_DNA+RNA_seed | EP300 | 0.828722 | BRCA | BT-549 | I-CBP112 |
| MVP_top60_DNA_seed | HSP90AA1 | 1.270252 | BRCA | HCC1187 | Luminespib | MVP_top60_DNA+RNA_seed | AKT1 | 0.828966 | BRCA | MDA-MB-157 | GSK690693 |
| MVP_top60_DNA_seed | HSP90AB1 | 1.270252 | BRCA | HCC1187 | Luminespib | MVP_top60_DNA+RNA_seed | EGFR | 0.829146 | BRCA | MRK-nu-1 | Cetuximab |
| MVP_top60_DNA_seed | HSP90AA1 | 1.271272 | BRCA | MDA-MB-175-VII | Elesclomol | MVP_top60_DNA+RNA_seed | HDAC1 | 0.829223 | BRCA | Hs-578-T | Belinostat |
| MVP_top60_DNA_seed | HSP90AB1 | 1.271272 | BRCA | MDA-MB-175-VII | Elesclomol | MVP_top60_DNA+RNA_seed | EGFR | 0.829341 | BRCA | EFM-192A | Pelitinib |
| MVP_top60_DNA_seed | AKT1 | 1.271434 | BRCA | HDQ-P1 | AKT inhibitor VIII | MVP_top60_DNA+RNA_seed | EGFR | 0.833459 | BRCA | MRK-nu-1 | Afatinib |
| MVP_top60_DNA_seed | CDK1 | 1.274383 | BRCA | BT-483 | RO-3306 | MVP_top60_DNA+RNA_seed | ERBB2 | 0.833459 | BRCA | MRK-nu-1 | Afatinib |
| MVP_top60_DNA_seed | SRC | 1.274686 | BRCA | MDA-MB-330 | NVP-BHG712 | MVP_top60_DNA+RNA_seed | EGFR | 0.83371 | BRCA | HCC2157 | CUDC-101 |
| MVP_top60_DNA_seed | SRC | 1.276089 | BRCA | HCC38 | Bosutinib | MVP_top60_DNA+RNA_seed | ERBB2 | 0.83371 | BRCA | HCC2157 | CUDC-101 |
| MVP_top60_DNA_seed | AR | 1.278358 | BRCA | HCC2218 | Bicalutamide | MVP_top60_DNA+RNA_seed | MAPK1 | 0.835356 | BRCA | HCC38 | VX-11e |
| MVP_top60_DNA_seed | SRC | 1.278701 | BRCA | ZR-75-30 | HG-6-71-01 | MVP_top60_DNA+RNA_seed | HDAC1 | 0.839258 | BRCA | BT-474 | JQ12 |
| MVP_top60_DNA_seed | HDAC1 | 1.279063 | BRCA | MDA-MB-415 | CAY10603 | MVP_top60_DNA+RNA_seed | HDAC2 | 0.839258 | BRCA | BT-474 | JQ12 |
| MVP_top60_DNA_seed | SRC | 1.281685 | BRCA | COLO-824 | Bosutinib | MVP_top60_DNA+RNA_seed | SRC | 0.84048 | BRCA | CAMA-1 | NVP-BHG712 |
| MVP_top60_DNA_seed | AKT1 | 1.285223 | BRCA | BT-483 | AKT inhibitor VIII | MVP_top60_DNA+RNA_seed | HDAC1 | 0.840586 | BRCA | MDA-MB-175-VII | Dacinostat |
| MVP_top60_DNA_seed | EGFR | 1.285272 | BRCA | HCC1569 | Gefitinib | MVP_top60_DNA+RNA_seed | ERBB2 | 0.841687 | BRCA | MDA-MB-157 | CP724714 |
| MVP_top60_DNA_seed | MAPK1 | 1.28531 | BRCA | BT-20 | VX-11e | MVP_top60_DNA+RNA_seed | AR | 0.842606 | BRCA | HCC38 | Bicalutamide |
| MVP_top60_DNA_seed | ATM | 1.286046 | BRCA | ZR-75-30 | KU-60019 | MVP_top60_DNA+RNA_seed | ESR1 | 0.843092 | BRCA | MDA-MB-231 | Fulvestrant |
| MVP_top60_DNA_seed | AKT1 | 1.29213 | BRCA | MCF7 | AKT inhibitor VIII | MVP_top60_DNA+RNA_seed | ATM | 0.843967 | BRCA | MDA-MB-436 | KU-60019 |
| MVP_top60_DNA_seed | ATM | 1.292581 | BRCA | BT-474 | torin2 | MVP_top60_DNA+RNA_seed | MAPK3 | 0.844382 | BRCA | YMB-1-E | FR-180204 |
| MVP_top60_DNA_seed | HSP90AA1 | 1.294714 | BRCA | HCC1500 | Elesclomol | MVP_top60_DNA+RNA_seed | MAPK1 | 0.844382 | BRCA | YMB-1-E | FR-180204 |
| MVP_top60_DNA_seed | HSP90AB1 | 1.294714 | BRCA | HCC1500 | Elesclomol | MVP_top60_DNA+RNA_seed | HDAC1 | 0.844563 | BRCA | MDA-MB-436 | Belinostat |
| MVP_top60_DNA_seed | SRC | 1.29646 | BRCA | HCC1428 | NVP-BHG712 | MVP_top60_DNA+RNA_seed | HDAC1 | 0.845404 | BRCA | HCC2218 | Belinostat |
| MVP_top60_DNA_seed | MAPK1 | 1.296682 | BRCA | MDA-MB-330 | VX-11e | MVP_top60_DNA+RNA_seed | EGFR | 0.845407 | BRCA | MDA-MB-231 | Gefitinib |
| MVP_top60_DNA_seed | ESR1 | 1.298138 | BRCA | CAL-85-1 | Tamoxifen | MVP_top60_DNA+RNA_seed | AKT1 | 0.84634 | BRCA | MDA-MB-157 | AKT inhibitor VIII |
| MVP_top60_DNA_seed | MAPK3 | 1.29992 | BRCA | CAMA-1 | FR-180204 | MVP_top60_DNA+RNA_seed | AR | 0.851371 | BRCA | HCC1419 | Bicalutamide |
| MVP_top60_DNA_seed | MAPK1 | 1.29992 | BRCA | CAMA-1 | FR-180204 | MVP_top60_DNA+RNA_seed | EGFR | 0.851706 | BRCA | MFM-223 | Afatinib |
| MVP_top60_DNA_seed | ESR1 | 1.305586 | BRCA | HCC1143 | Tamoxifen | MVP_top60_DNA+RNA_seed | ERBB2 | 0.851706 | BRCA | MFM-223 | Afatinib |
| MVP_top60_DNA_seed | AR | 1.307055 | BRCA | MDA-MB-175-VII | AZD3514 | MVP_top60_DNA+RNA_seed | SRC | 0.851957 | BRCA | MDA-MB-436 | NVP-BHG712 |
| MVP_top60_DNA_seed | CDK2 | 1.309309 | BRCA | EFM-19 | AZD5438 | MVP_top60_DNA+RNA_seed | MAPK1 | 0.852481 | BRCA | HCC1500 | VX-11e |
| MVP_top60_DNA_seed | CDK2 | 1.312185 | BRCA | Hs-578-T | PHA-793887 | MVP_top60_DNA+RNA_seed | CDK2 | 0.85286 | BRCA | CAL-120 | PHA-793887 |

| | | | | | | | | | | | |
|---|---|---|---|---|---|---|---|---|---|---|---|
| MVP_top60_DNA_seed | MAPK1 | 1.312262 | BRCA | HDQ-P1 | VX-11e | MVP_top60_DNA+RNA_seed | MAPK3 | 0.853616 | BRCA | COLO-824 | FR-180204 |
| MVP_top60_DNA_seed | CDK1 | 1.314105 | BRCA | MDA-MB-415 | RO-3306 | MVP_top60_DNA+RNA_seed | MAPK1 | 0.853616 | BRCA | COLO-824 | FR-180204 |
| MVP_top60_DNA_seed | EGFR | 1.315048 | BRCA | MFM-223 | Gefitinib | MVP_top60_DNA+RNA_seed | HSP90AA1 | 0.854733 | BRCA | HCC1500 | Tanespimycin |
| MVP_top60_DNA_seed | SRC | 1.316657 | BRCA | UACC-893 | HG-6-71-01 | MVP_top60_DNA+RNA_seed | HSP90AB1 | 0.854733 | BRCA | HCC1500 | Tanespimycin |
| MVP_top60_DNA_seed | AKT1 | 1.316765 | BRCA | HCC1428 | BAY AKT1 | MVP_top60_DNA+RNA_seed | MAPK14 | 0.855899 | BRCA | EFM-192A | TAK-715 |
| MVP_top60_DNA_seed | PIK3CA | 1.317085 | BRCA | MDA-MB-157 | AZD8835 | MVP_top60_DNA+RNA_seed | HDAC1 | 0.85709 | BRCA | MCF7 | CAY10603 |
| MVP_top60_DNA_seed | HSP90AA1 | 1.317554 | BRCA | MDA-MB-468 | Tanespimycin | MVP_top60_DNA+RNA_seed | HSP90AA1 | 0.859483 | BRCA | OCUB-M | Tanespimycin |
| MVP_top60_DNA_seed | HSP90AB1 | 1.317554 | BRCA | MDA-MB-468 | Tanespimycin | MVP_top60_DNA+RNA_seed | HSP90AB1 | 0.859483 | BRCA | OCUB-M | Tanespimycin |
| MVP_top60_DNA_seed | HDAC1 | 1.318724 | BRCA | BT-20 | Dacinostat | MVP_top60_DNA+RNA_seed | EGFR | 0.860178 | BRCA | MCF7 | Afatinib |
| MVP_top60_DNA_seed | SRC | 1.319888 | BRCA | HCC1954 | NVP-BHG712 | MVP_top60_DNA+RNA_seed | ERBB2 | 0.860178 | BRCA | MCF7 | Afatinib |
| MVP_top60_DNA_seed | HDAC1 | 1.319932 | BRCA | HCC1395 | JQ12 | MVP_top60_DNA+RNA_seed | EP300 | 0.860822 | BRCA | CAL-120 | I-CBP112 |
| MVP_top60_DNA_seed | HDAC2 | 1.319932 | BRCA | HCC1395 | JQ12 | MVP_top60_DNA+RNA_seed | HSP90AA1 | 0.861234 | BRCA | T47D | Luminespib |
| MVP_top60_DNA_seed | PIK3CA | 1.320549 | BRCA | HCC1428 | AZD8835 | MVP_top60_DNA+RNA_seed | HSP90AB1 | 0.861234 | BRCA | T47D | Luminespib |
| MVP_top60_DNA_seed | ERBB2 | 1.321515 | BRCA | MDA-MB-415 | CP724714 | MVP_top60_DNA+RNA_seed | ESR1 | 0.861865 | BRCA | HDQ-P1 | Tamoxifen |
| MVP_top60_DNA_seed | EGFR | 1.322106 | BRCA | EFM-19 | Cetuximab | MVP_top60_DNA+RNA_seed | AKT1 | 0.865286 | BRCA | MDA-MB-231 | GSK690693 |
| MVP_top60_DNA_seed | ESR1 | 1.324028 | BRCA | MDA-MB-175-VII | Fulvestrant | MVP_top60_DNA+RNA_seed | CDK1 | 0.86555 | BRCA | HCC1569 | RO-3306 |
| MVP_top60_DNA_seed | AKT1 | 1.325025 | BRCA | CAL-85-1 | MK-2206 | MVP_top60_DNA+RNA_seed | HSP90AA1 | 0.866896 | BRCA | MDA-MB-157 | Elesclomol |
| MVP_top60_DNA_seed | SRC | 1.329337 | BRCA | MFM-223 | Saracatinib | MVP_top60_DNA+RNA_seed | HSP90AB1 | 0.866896 | BRCA | MDA-MB-157 | Elesclomol |
| MVP_top60_DNA_seed | CDK2 | 1.330978 | BRCA | UACC-812 | Seliciclib | MVP_top60_DNA+RNA_seed | HSP90AA1 | 0.868122 | BRCA | MDA-MB-415 | Luminespib |
| MVP_top60_DNA_seed | HDAC1 | 1.335948 | BRCA | HCC1395 | CAY10603 | MVP_top60_DNA+RNA_seed | HSP90AB1 | 0.868122 | BRCA | MDA-MB-415 | Luminespib |
| MVP_top60_DNA_seed | MDM2 | 1.337037 | BRCA | HDQ-P1 | Serdemetan | MVP_top60_DNA+RNA_seed | MAPK3 | 0.870383 | BRCA | T47D | FR-180204 |
| MVP_top60_DNA_seed | PIK3CA | 1.344327 | BRCA | MDA-MB-436 | AZD8835 | MVP_top60_DNA+RNA_seed | MAPK1 | 0.870383 | BRCA | T47D | FR-180204 |
| MVP_top60_DNA_seed | MDM2 | 1.346095 | BRCA | ZR-75-30 | Nutlin-3a (-) | MVP_top60_DNA+RNA_seed | EGFR | 0.8716 | BRCA | COLO-824 | Gefitinib |
| MVP_top60_DNA_seed | AR | 1.347654 | BRCA | MDA-MB-415 | Bicalutamide | MVP_top60_DNA+RNA_seed | MDM2 | 0.871691 | BRCA | EVSA-T | Serdemetan |
| MVP_top60_DNA_seed | SRC | 1.349832 | BRCA | HCC1569 | Bosutinib | MVP_top60_DNA+RNA_seed | AKT1 | 0.871972 | BRCA | Hs-578-T | GSK690693 |
| MVP_top60_DNA_seed | ATM | 1.350035 | BRCA | HDQ-P1 | KU-60019 | MVP_top60_DNA+RNA_seed | CDK2 | 0.872436 | BRCA | HCC1428 | PHA-793887 |
| MVP_top60_DNA_seed | CDK1 | 1.351208 | BRCA | BT-474 | RO-3306 | MVP_top60_DNA+RNA_seed | ATM | 0.873695 | BRCA | MDA-MB-157 | CP466722 |
| MVP_top60_DNA_seed | CDK2 | 1.353857 | BRCA | EFM-192A | PHA-793887 | MVP_top60_DNA+RNA_seed | EGFR | 0.875026 | BRCA | BT-549 | Afatinib |
| MVP_top60_DNA_seed | HDAC1 | 1.358523 | BRCA | MFM-223 | Dacinostat | MVP_top60_DNA+RNA_seed | ERBB2 | 0.875026 | BRCA | BT-549 | Afatinib |
| MVP_top60_DNA_seed | MAPK3 | 1.359593 | BRCA | HCC1395 | FR-180204 | MVP_top60_DNA+RNA_seed | EGFR | 0.876027 | BRCA | MFM-223 | Pelitinib |
| MVP_top60_DNA_seed | MAPK1 | 1.359593 | BRCA | HCC1395 | FR-180204 | MVP_top60_DNA+RNA_seed | PLK1 | 0.87786 | BRCA | HCC1937 | PLK_6522 |
| MVP_top60_DNA_seed | EGFR | 1.361535 | BRCA | T47D | Cetuximab | MVP_top60_DNA+RNA_seed | HSP90AA1 | 0.882227 | BRCA | MDA-MB-175-VII | Luminespib |
| MVP_top60_DNA_seed | CDK2 | 1.362343 | BRCA | UACC-812 | PHA-793887 | MVP_top60_DNA+RNA_seed | HSP90AB1 | 0.882227 | BRCA | MDA-MB-175-VII | Luminespib |
| MVP_top60_DNA_seed | AKT1 | 1.363286 | BRCA | MRK-nu-1 | AKT inhibitor VIII | MVP_top60_DNA+RNA_seed | EGFR | 0.885029 | BRCA | EVSA-T | Pelitinib |
| MVP_top60_DNA_seed | MAPK3 | 1.363836 | BRCA | MDA-MB-436 | FR-180204 | MVP_top60_DNA+RNA_seed | HDAC1 | 0.8854 | BRCA | MCF7 | Tubastatin A |
| MVP_top60_DNA_seed | MAPK1 | 1.363836 | BRCA | MDA-MB-436 | FR-180204 | MVP_top60_DNA+RNA_seed | ERBB2 | 0.888675 | BRCA | Hs-578-T | CP724714 |
| MVP_top60_DNA_seed | EGFR | 1.367886 | BRCA | HCC2157 | Pelitinib | MVP_top60_DNA+RNA_seed | HSP90AA1 | 0.889434 | BRCA | BT-549 | Luminespib |
| MVP_top60_DNA_seed | HDAC1 | 1.373532 | BRCA | MFM-223 | Parthenolide | MVP_top60_DNA+RNA_seed | HSP90AB1 | 0.889434 | BRCA | BT-549 | Luminespib |
| MVP_top60_DNA_seed | HDAC1 | 1.374431 | BRCA | BT-483 | CAY10603 | MVP_top60_DNA+RNA_seed | MAPK1 | 0.88952 | BRCA | CAL-120 | VX-11e |
| MVP_top60_DNA_seed | CDK1 | 1.381748 | BRCA | HDQ-P1 | RO-3306 | MVP_top60_DNA+RNA_seed | MDM2 | 0.890229 | BRCA | HCC2157 | Nutlin-3a (-) |
| MVP_top60_DNA_seed | AR | 1.384759 | BRCA | EFM-19 | Bicalutamide | MVP_top60_DNA+RNA_seed | MAPK3 | 0.891714 | BRCA | CAL-120 | FR-180204 |

| | | | | | | | | | | | |
|---|---|---|---|---|---|---|---|---|---|---|---|
| MVP_top60_DNA_seed | SRC | 1.385285 | BRCA | CAL-85-1 | Bosutinib | MVP_top60_DNA+RNA_seed | MAPK1 | 0.891714 | BRCA | CAL-120 | FR-180204 |
| MVP_top60_DNA_seed | PIK3CA | 1.386546 | BRCA | HCC1428 | PI3Ka_4409 | MVP_top60_DNA+RNA_seed | HDAC1 | 0.892382 | BRCA | MDA-MB-436 | Tubastatin A |
| MVP_top60_DNA_seed | EGFR | 1.392698 | BRCA | YMB-1-E | Pelitinib | MVP_top60_DNA+RNA_seed | SRC | 0.892723 | BRCA | HCC2218 | Saracatinib |
| MVP_top60_DNA_seed | HDAC1 | 1.392781 | BRCA | MDA-MB-415 | Tubastatin A | MVP_top60_DNA+RNA_seed | ESR1 | 0.893809 | BRCA | COLO-824 | Tamoxifen |
| MVP_top60_DNA_seed | AR | 1.394125 | BRCA | HCC1419 | AZD3514 | MVP_top60_DNA+RNA_seed | HDAC1 | 0.893912 | BRCA | JIMT-1 | Tubastatin A |
| MVP_top60_DNA_seed | EP300 | 1.395697 | BRCA | MDA-MB-436 | I-CBP112 | MVP_top60_DNA+RNA_seed | HSP90AA1 | 0.897371 | BRCA | HCC1395 | Tanespimycin |
| MVP_top60_DNA_seed | CREBBP | 1.395697 | BRCA | MDA-MB-436 | I-CBP112 | MVP_top60_DNA+RNA_seed | HSP90AB1 | 0.897371 | BRCA | HCC1395 | Tanespimycin |
| MVP_top60_DNA_seed | AKT1 | 1.396284 | BRCA | BT-483 | GSK690693 | MVP_top60_DNA+RNA_seed | AKT1 | 0.89809 | BRCA | HCC1428 | GSK690693 |
| MVP_top60_DNA_seed | MAPK1 | 1.396348 | BRCA | HCC1419 | VX-11e | MVP_top60_DNA+RNA_seed | HDAC1 | 0.898834 | BRCA | YMB-1-E | Tubastatin A |
| MVP_top60_DNA_seed | CDK2 | 1.39664 | BRCA | COLO-824 | Seliciclib | MVP_top60_DNA+RNA_seed | CDK2 | 0.902109 | BRCA | MDA-MB-361 | PHA-793887 |
| MVP_top60_DNA_seed | ERBB2 | 1.39735 | BRCA | MCF7 | CUDC-101 | MVP_top60_DNA+RNA_seed | AR | 0.902566 | BRCA | HCC1569 | Bicalutamide |
| MVP_top60_DNA_seed | EGFR | 1.39735 | BRCA | MCF7 | CUDC-101 | MVP_top60_DNA+RNA_seed | MAPK1 | 0.902579 | BRCA | MDA-MB-157 | VX-11e |
| MVP_top60_DNA_seed | ATM | 1.397527 | BRCA | HCC1954 | KU-55933 | MVP_top60_DNA+RNA_seed | EGFR | 0.903784 | BRCA | COLO-824 | Cetuximab |
| MVP_top60_DNA_seed | EGFR | 1.398585 | BRCA | MDA-MB-157 | Pelitinib | MVP_top60_DNA+RNA_seed | EGFR | 0.903969 | BRCA | MDA-MB-330 | Pelitinib |
| MVP_top60_DNA_seed | MAPK3 | 1.405187 | BRCA | EFM-19 | FR-180204 | MVP_top60_DNA+RNA_seed | AKT1 | 0.904081 | BRCA | MRK-nu-1 | GSK690693 |
| MVP_top60_DNA_seed | MAPK1 | 1.405187 | BRCA | EFM-19 | FR-180204 | MVP_top60_DNA+RNA_seed | HDAC1 | 0.904316 | BRCA | MDA-MB-468 | Dacinostat |
| MVP_top60_DNA_seed | EGFR | 1.405752 | BRCA | Hs-578-T | Pelitinib | MVP_top60_DNA+RNA_seed | EP300 | 0.906945 | BRCA | HCC202 | I-CBP112 |
| MVP_top60_DNA_seed | SRC | 1.407518 | BRCA | T47D | Bosutinib | MVP_top60_DNA+RNA_seed | AR | 0.907571 | BRCA | CAL-120 | Bicalutamide |
| MVP_top60_DNA_seed | HDAC1 | 1.418512 | BRCA | CAL-51 | CAY10603 | MVP_top60_DNA+RNA_seed | ATM | 0.908144 | BRCA | CAMA-1 | KU-60019 |
| MVP_top60_DNA_seed | ESR1 | 1.423801 | BRCA | MDA-MB-436 | Tamoxifen | MVP_top60_DNA+RNA_seed | SRC | 0.909176 | BRCA | MCF7 | NVP-BHG712 |
| MVP_top60_DNA_seed | HDAC1 | 1.426478 | BRCA | BT-474 | Parthenolide | MVP_top60_DNA+RNA_seed | MDM2 | 0.911452 | BRCA | COLO-824 | Nutlin-3a (-) |
| MVP_top60_DNA_seed | HSP90AA1 | 1.42892 | BRCA | HCC1428 | SNX-2112 | MVP_top60_DNA+RNA_seed | AKT1 | 0.912439 | BRCA | HCC1187 | AKT inhibitor VIII |
| MVP_top60_DNA_seed | HSP90AB1 | 1.42892 | BRCA | HCC1428 | SNX-2112 | MVP_top60_DNA+RNA_seed | HSP90AA1 | 0.913598 | BRCA | BT-20 | SNX-2112 |
| MVP_top60_DNA_seed | CDK1 | 1.430098 | BRCA | CAL-85-1 | RO-3306 | MVP_top60_DNA+RNA_seed | HSP90AB1 | 0.913598 | BRCA | BT-20 | SNX-2112 |
| MVP_top60_DNA_seed | EGFR | 1.433212 | BRCA | MRK-nu-1 | Pelitinib | MVP_top60_DNA+RNA_seed | SRC | 0.914616 | BRCA | COLO-824 | WH-4-023 |
| MVP_top60_DNA_seed | CDK2 | 1.436536 | BRCA | HCC2218 | AZD5438 | MVP_top60_DNA+RNA_seed | SRC | 0.914661 | BRCA | BT-474 | WH-4-023 |
| MVP_top60_DNA_seed | SRC | 1.443748 | BRCA | MDA-MB-415 | NVP-BHG712 | MVP_top60_DNA+RNA_seed | CDK2 | 0.915436 | BRCA | MDA-MB-231 | PHA-793887 |
| MVP_top60_DNA_seed | AR | 1.449382 | BRCA | MDA-MB-436 | Bicalutamide | MVP_top60_DNA+RNA_seed | MAPK14 | 0.915581 | BRCA | YMB-1-E | TAK-715 |
| MVP_top60_DNA_seed | PIK3CA | 1.450804 | BRCA | ZR-75-30 | PI3Ka_4409 | MVP_top60_DNA+RNA_seed | EGFR | 0.915618 | BRCA | MDA-MB-453 | Gefitinib |
| MVP_top60_DNA_seed | AR | 1.451524 | BRCA | MFM-223 | AZD3514 | MVP_top60_DNA+RNA_seed | CDK2 | 0.915737 | BRCA | CAL-85-1 | AZD5438 |
| MVP_top60_DNA_seed | MAPK1 | 1.45488 | BRCA | EFM-19 | VX-11e | MVP_top60_DNA+RNA_seed | EGFR | 0.916803 | BRCA | MDA-MB-436 | Cetuximab |
| MVP_top60_DNA_seed | AR | 1.460438 | BRCA | ZR-75-30 | Bicalutamide | MVP_top60_DNA+RNA_seed | EGFR | 0.917078 | BRCA | CAL-85-1 | Gefitinib |
| MVP_top60_DNA_seed | CDK2 | 1.464954 | BRCA | BT-474 | Seliciclib | MVP_top60_DNA+RNA_seed | PLK1 | 0.917323 | BRCA | HCC2218 | BI-2536 |
| MVP_top60_DNA_seed | SRC | 1.46543 | BRCA | HDQ-P1 | NVP-BHG712 | MVP_top60_DNA+RNA_seed | AKT1 | 0.91751 | BRCA | MDA-MB-231 | AKT inhibitor VIII |
| MVP_top60_DNA_seed | ESR1 | 1.470021 | BRCA | HCC202 | Tamoxifen | MVP_top60_DNA+RNA_seed | CDK2 | 0.917545 | BRCA | MDA-MB-361 | AZD5438 |
| MVP_top60_DNA_seed | ESR1 | 1.471784 | BRCA | HCC1419 | Tamoxifen | MVP_top60_DNA+RNA_seed | ESR1 | 0.918374 | BRCA | BT-474 | Tamoxifen |
| MVP_top60_DNA_seed | HSP90AA1 | 1.472149 | BRCA | Hs-578-T | SNX-2112 | MVP_top60_DNA+RNA_seed | HDAC1 | 0.919348 | BRCA | MDA-MB-468 | JQ12 |
| MVP_top60_DNA_seed | HSP90AB1 | 1.472149 | BRCA | Hs-578-T | SNX-2112 | MVP_top60_DNA+RNA_seed | HDAC2 | 0.919348 | BRCA | MDA-MB-468 | JQ12 |
| MVP_top60_DNA_seed | MDM2 | 1.477896 | BRCA | MDA-MB-415 | Nutlin-3a (-) | MVP_top60_DNA+RNA_seed | MAPK3 | 0.922607 | BRCA | HCC1500 | FR-180204 |
| MVP_top60_DNA_seed | AKT1 | 1.479658 | BRCA | MDA-MB-157 | BAY AKT1 | MVP_top60_DNA+RNA_seed | MAPK1 | 0.922607 | BRCA | HCC1500 | FR-180204 |
| MVP_top60_DNA_seed | PIK3CA | 1.479792 | BRCA | HCC1937 | AZD8835 | MVP_top60_DNA+RNA_seed | MDM2 | 0.923737 | BRCA | AU565 | Nutlin-3a (-) |

| | | | | | | | | | | | |
|---|---|---|---|---|---|---|---|---|---|---|---|
| MVP_top60_DNA_seed | PIK3CA | 1.482499 | BRCA | MDA-MB-157 | PI3Ka_4409 | MVP_top60_DNA+RNA_seed | MAPK14 | 0.925797 | BRCA | CAL-120 | TAK-715 |
| MVP_top60_DNA_seed | HSP90AA1 | 1.483976 | BRCA | HCC1395 | SNX-2112 | MVP_top60_DNA+RNA_seed | HDAC1 | 0.927028 | BRCA | MDA-MB-468 | Belinostat |
| MVP_top60_DNA_seed | HSP90AB1 | 1.483976 | BRCA | HCC1395 | SNX-2112 | MVP_top60_DNA+RNA_seed | ERBB2 | 0.927758 | BRCA | EFM-19 | CP724714 |
| MVP_top60_DNA_seed | ATM | 1.491857 | BRCA | HCC2218 | KU-55933 | MVP_top60_DNA+RNA_seed | ATM | 0.929542 | BRCA | MDA-MB-231 | KU-55933 |
| MVP_top60_DNA_seed | MAPK1 | 1.495213 | BRCA | MDA-MB-436 | VX-11e | MVP_top60_DNA+RNA_seed | MDM2 | 0.929939 | BRCA | BT-483 | Serdemetan |
| MVP_top60_DNA_seed | AKT1 | 1.500272 | BRCA | BT-483 | MK-2206 | MVP_top60_DNA+RNA_seed | MAPK3 | 0.930072 | BRCA | UACC-893 | FR-180204 |
| MVP_top60_DNA_seed | AKT1 | 1.500373 | BRCA | HDQ-P1 | BAY AKT1 | MVP_top60_DNA+RNA_seed | MAPK1 | 0.930072 | BRCA | UACC-893 | FR-180204 |
| MVP_top60_DNA_seed | HDAC1 | 1.502827 | BRCA | HCC1937 | JQ12 | MVP_top60_DNA+RNA_seed | MAPK3 | 0.930932 | BRCA | HCC2157 | FR-180204 |
| MVP_top60_DNA_seed | HDAC2 | 1.502827 | BRCA | HCC1937 | JQ12 | MVP_top60_DNA+RNA_seed | MAPK1 | 0.930932 | BRCA | HCC2157 | FR-180204 |
| MVP_top60_DNA_seed | MDM2 | 1.508596 | BRCA | HCC202 | Nutlin-3a (-) | MVP_top60_DNA+RNA_seed | HDAC1 | 0.931174 | BRCA | MDA-MB-175-VII | Tubastatin A |
| MVP_top60_DNA_seed | HSP90AA1 | 1.515681 | BRCA | EFM-19 | SNX-2112 | MVP_top60_DNA+RNA_seed | HDAC1 | 0.931531 | BRCA | MFM-223 | Tubastatin A |
| MVP_top60_DNA_seed | HSP90AB1 | 1.515681 | BRCA | EFM-19 | SNX-2112 | MVP_top60_DNA+RNA_seed | AKT1 | 0.93287 | BRCA | HCC38 | BAY AKT1 |
| MVP_top60_DNA_seed | EGFR | 1.517091 | BRCA | BT-483 | Gefitinib | MVP_top60_DNA+RNA_seed | AR | 0.933287 | BRCA | HCC1395 | Bicalutamide |
| MVP_top60_DNA_seed | MAPK1 | 1.522008 | BRCA | MDA-MB-415 | VX-11e | MVP_top60_DNA+RNA_seed | EGFR | 0.934991 | BRCA | HCC1428 | AZD8931 |
| MVP_top60_DNA_seed | EGFR | 1.523588 | BRCA | HCC2218 | Cetuximab | MVP_top60_DNA+RNA_seed | ERBB2 | 0.934991 | BRCA | HCC1428 | AZD8931 |
| MVP_top60_DNA_seed | ATM | 1.527524 | BRCA | BT-20 | KU-55933 | MVP_top60_DNA+RNA_seed | ERBB3 | 0.934991 | BRCA | HCC1428 | AZD8931 |
| MVP_top60_DNA_seed | ERBB2 | 1.531003 | BRCA | MDA-MB-415 | CUDC-101 | MVP_top60_DNA+RNA_seed | PIK3CA | 0.936188 | BRCA | MDA-MB-415 | AZD8835 |
| MVP_top60_DNA_seed | EGFR | 1.531003 | BRCA | MDA-MB-415 | CUDC-101 | MVP_top60_DNA+RNA_seed | MDM2 | 0.936859 | BRCA | MDA-MB-453 | Nutlin-3a (-) |
| MVP_top60_DNA_seed | ERBB2 | 1.531241 | BRCA | HCC1395 | CUDC-101 | MVP_top60_DNA+RNA_seed | EGFR | 0.937526 | BRCA | HCC2218 | Erlotinib |
| MVP_top60_DNA_seed | EGFR | 1.531241 | BRCA | HCC1395 | CUDC-101 | MVP_top60_DNA+RNA_seed | EP300 | 0.937796 | BRCA | MDA-MB-175-VII | I-CBP112 |
| MVP_top60_DNA_seed | HDAC1 | 1.542562 | BRCA | CAL-51 | AR-42 | MVP_top60_DNA+RNA_seed | EGFR | 0.937899 | BRCA | MFM-223 | Lapatinib |
| MVP_top60_DNA_seed | EP300 | 1.543025 | BRCA | HCC1143 | I-CBP112 | MVP_top60_DNA+RNA_seed | ERBB2 | 0.937899 | BRCA | MFM-223 | Lapatinib |
| MVP_top60_DNA_seed | CREBBP | 1.543025 | BRCA | HCC1143 | I-CBP112 | MVP_top60_DNA+RNA_seed | SRC | 0.938497 | BRCA | HCC2157 | WH-4-023 |
| MVP_top60_DNA_seed | HSP90AA1 | 1.543495 | BRCA | HCC1187 | Tanespimycin | MVP_top60_DNA+RNA_seed | HDAC1 | 0.9386 | BRCA | UACC-893 | Tubastatin A |
| MVP_top60_DNA_seed | HSP90AB1 | 1.543495 | BRCA | HCC1187 | Tanespimycin | MVP_top60_DNA+RNA_seed | ESR1 | 0.942013 | BRCA | HCC2218 | Tamoxifen |
| MVP_top60_DNA_seed | HDAC1 | 1.544541 | BRCA | MCF7 | AR-42 | MVP_top60_DNA+RNA_seed | EP300 | 0.942172 | BRCA | EFM-192A | I-CBP112 |
| MVP_top60_DNA_seed | EGFR | 1.544867 | BRCA | EFM-19 | Pelitinib | MVP_top60_DNA+RNA_seed | MAPK3 | 0.944657 | BRCA | HCC2218 | FR-180204 |
| MVP_top60_DNA_seed | HSP90AA1 | 1.545609 | BRCA | EFM-19 | CCT-018159 | MVP_top60_DNA+RNA_seed | MAPK1 | 0.944657 | BRCA | HCC2218 | FR-180204 |
| MVP_top60_DNA_seed | HSP90AB1 | 1.545609 | BRCA | EFM-19 | CCT-018159 | MVP_top60_DNA+RNA_seed | ATM | 0.945688 | BRCA | MDA-MB-436 | KU-55933 |
| MVP_top60_DNA_seed | AR | 1.549658 | BRCA | BT-474 | Bicalutamide | MVP_top60_DNA+RNA_seed | HSP90AA1 | 0.945909 | BRCA | HCC1500 | SNX-2112 |
| MVP_top60_DNA_seed | CDK1 | 1.550917 | BRCA | MFM-223 | RO-3306 | MVP_top60_DNA+RNA_seed | HSP90AB1 | 0.945909 | BRCA | HCC1500 | SNX-2112 |
| MVP_top60_DNA_seed | AKT1 | 1.556124 | BRCA | UACC-812 | AKT inhibitor VIII | MVP_top60_DNA+RNA_seed | HSP90AA1 | 0.946881 | BRCA | MRK-nu-1 | CCT-018159 |
| MVP_top60_DNA_seed | ERBB2 | 1.557459 | BRCA | CAL-51 | CUDC-101 | MVP_top60_DNA+RNA_seed | HSP90AB1 | 0.946881 | BRCA | MRK-nu-1 | CCT-018159 |
| MVP_top60_DNA_seed | EGFR | 1.557459 | BRCA | CAL-51 | CUDC-101 | MVP_top60_DNA+RNA_seed | MDM2 | 0.947786 | BRCA | HCC1428 | Nutlin-3a (-) |
| MVP_top60_DNA_seed | AR | 1.569871 | BRCA | HCC1954 | AZD3514 | MVP_top60_DNA+RNA_seed | EGFR | 0.947969 | BRCA | COLO-824 | Lapatinib |
| MVP_top60_DNA_seed | HDAC1 | 1.572447 | BRCA | CAL-51 | Belinostat | MVP_top60_DNA+RNA_seed | ERBB2 | 0.947969 | BRCA | COLO-824 | Lapatinib |
| MVP_top60_DNA_seed | CDK1 | 1.574741 | BRCA | HCC2218 | RO-3306 | MVP_top60_DNA+RNA_seed | HDAC1 | 0.948571 | BRCA | HCC1187 | Belinostat |
| MVP_top60_DNA_seed | CDK2 | 1.575152 | BRCA | HCC1428 | AZD5438 | MVP_top60_DNA+RNA_seed | HSP90AA1 | 0.949756 | BRCA | HCC38 | Elesclomol |
| MVP_top60_DNA_seed | AKT1 | 1.57811 | BRCA | MRK-nu-1 | BAY AKT1 | MVP_top60_DNA+RNA_seed | HSP90AB1 | 0.949756 | BRCA | HCC38 | Elesclomol |
| MVP_top60_DNA_seed | HSP90AA1 | 1.579625 | BRCA | MDA-MB-157 | Luminespib | MVP_top60_DNA+RNA_seed | AR | 0.949852 | BRCA | CAL-85-1 | Bicalutamide |
| MVP_top60_DNA_seed | HSP90AB1 | 1.579625 | BRCA | MDA-MB-157 | Luminespib | MVP_top60_DNA+RNA_seed | EP300 | 0.949904 | BRCA | HCC1937 | I-CBP112 |

| | | | | | | | | | | | |
|---|---|---|---|---|---|---|---|---|---|---|---|
| MVP_top60_DNA_seed | ERBB2 | 1.580409 | BRCA | BT-483 | Afatinib | MVP_top60_DNA+RNA_seed | ERBB2 | 0.950042 | BRCA | MRK-nu-1 | CP724714 |
| MVP_top60_DNA_seed | EGFR | 1.580409 | BRCA | BT-483 | Afatinib | MVP_top60_DNA+RNA_seed | SRC | 0.950429 | BRCA | CAL-51 | Bosutinib |
| MVP_top60_DNA_seed | AKT1 | 1.58138 | BRCA | HCC1500 | BAY AKT1 | MVP_top60_DNA+RNA_seed | AR | 0.954621 | BRCA | HCC70 | AZD3514 |
| MVP_top60_DNA_seed | ATM | 1.58686 | BRCA | HCC2157 | CP466722 | MVP_top60_DNA+RNA_seed | AKT1 | 0.954682 | BRCA | EFM-192A | GSK690693 |
| MVP_top60_DNA_seed | CDK1 | 1.587102 | BRCA | ZR-75-30 | RO-3306 | MVP_top60_DNA+RNA_seed | HDAC1 | 0.955723 | BRCA | MDA-MB-436 | AR-42 |
| MVP_top60_DNA_seed | HSP90AA1 | 1.589529 | BRCA | HCC1395 | CCT-018159 | MVP_top60_DNA+RNA_seed | MAPK14 | 0.956238 | BRCA | MDA-MB-231 | TAK-715 |
| MVP_top60_DNA_seed | HSP90AB1 | 1.589529 | BRCA | HCC1395 | CCT-018159 | MVP_top60_DNA+RNA_seed | EGFR | 0.956597 | BRCA | YMB-1-E | CUDC-101 |
| MVP_top60_DNA_seed | CDK2 | 1.589824 | BRCA | MDA-MB-415 | PHA-793887 | MVP_top60_DNA+RNA_seed | ERBB2 | 0.956597 | BRCA | YMB-1-E | CUDC-101 |
| MVP_top60_DNA_seed | HDAC1 | 1.591187 | BRCA | UACC-812 | Entinostat | MVP_top60_DNA+RNA_seed | CDK1 | 0.956686 | BRCA | HCC2157 | RO-3306 |
| MVP_top60_DNA_seed | SRC | 1.593186 | BRCA | HCC1419 | Bosutinib | MVP_top60_DNA+RNA_seed | MAPK14 | 0.957127 | BRCA | HCC1937 | TAK-715 |
| MVP_top60_DNA_seed | AKT1 | 1.593829 | BRCA | MFM-223 | AKT inhibitor VIII | MVP_top60_DNA+RNA_seed | AR | 0.957918 | BRCA | HCC1428 | Bicalutamide |
| MVP_top60_DNA_seed | ESR1 | 1.600764 | BRCA | UACC-893 | Fulvestrant | MVP_top60_DNA+RNA_seed | HDAC1 | 0.959736 | BRCA | BT-549 | AR-42 |
| MVP_top60_DNA_seed | EGFR | 1.611809 | BRCA | EFM-19 | Gefitinib | MVP_top60_DNA+RNA_seed | HSP90AA1 | 0.960136 | BRCA | HCC1500 | Luminespib |
| MVP_top60_DNA_seed | CDK2 | 1.612211 | BRCA | BT-474 | AZD5438 | MVP_top60_DNA+RNA_seed | HSP90AB1 | 0.960136 | BRCA | HCC1500 | Luminespib |
| MVP_top60_DNA_seed | AKT1 | 1.616176 | BRCA | HCC1500 | AKT inhibitor VIII | MVP_top60_DNA+RNA_seed | HSP90AA1 | 0.960632 | BRCA | MDA-MB-231 | SNX-2112 |
| MVP_top60_DNA_seed | AKT1 | 1.618654 | BRCA | MDA-MB-231 | MK-2206 | MVP_top60_DNA+RNA_seed | HSP90AB1 | 0.960632 | BRCA | MDA-MB-231 | SNX-2112 |
| MVP_top60_DNA_seed | EP300 | 1.625018 | BRCA | T47D | I-CBP112 | MVP_top60_DNA+RNA_seed | EGFR | 0.963089 | BRCA | BT-474 | Cetuximab |
| MVP_top60_DNA_seed | CREBBP | 1.625018 | BRCA | T47D | I-CBP112 | MVP_top60_DNA+RNA_seed | ATM | 0.963402 | BRCA | BT-474 | KU-60019 |
| MVP_top60_DNA_seed | MAPK3 | 1.628262 | BRCA | MDA-MB-415 | FR-180204 | MVP_top60_DNA+RNA_seed | ERBB2 | 0.964885 | BRCA | HDQ-P1 | CP724714 |
| MVP_top60_DNA_seed | MAPK1 | 1.628262 | BRCA | MDA-MB-415 | FR-180204 | MVP_top60_DNA+RNA_seed | ATM | 0.966125 | BRCA | MCF7 | CP466722 |
| MVP_top60_DNA_seed | EGFR | 1.629093 | BRCA | MDA-MB-415 | Gefitinib | MVP_top60_DNA+RNA_seed | ATM | 0.968245 | BRCA | Hs-578-T | CP466722 |
| MVP_top60_DNA_seed | ATM | 1.635209 | BRCA | BT-474 | KU-55933 | MVP_top60_DNA+RNA_seed | PLK1 | 0.970128 | BRCA | HDQ-P1 | PLK_6522 |
| MVP_top60_DNA_seed | AR | 1.64456 | BRCA | HCC1395 | AZD3514 | MVP_top60_DNA+RNA_seed | HDAC1 | 0.970554 | BRCA | BT-20 | Tubastatin A |
| MVP_top60_DNA_seed | HDAC1 | 1.651381 | BRCA | MDA-MB-436 | Dacinostat | MVP_top60_DNA+RNA_seed | MDM2 | 0.970658 | BRCA | BT-483 | Nutlin-3a (-) |
| MVP_top60_DNA_seed | HDAC1 | 1.651663 | BRCA | MCF7 | Belinostat | MVP_top60_DNA+RNA_seed | PIK3CA | 0.972751 | BRCA | Hs-578-T | AZD8835 |
| MVP_top60_DNA_seed | EGFR | 1.659673 | BRCA | MDA-MB-415 | Pelitinib | MVP_top60_DNA+RNA_seed | ESR1 | 0.978747 | BRCA | HCC202 | Fulvestrant |
| MVP_top60_DNA_seed | HDAC1 | 1.660249 | BRCA | MDA-MB-415 | Belinostat | MVP_top60_DNA+RNA_seed | EGFR | 0.979905 | BRCA | Hs-578-T | Afatinib |
| MVP_top60_DNA_seed | ESR1 | 1.663496 | BRCA | CAL-85-1 | Fulvestrant | MVP_top60_DNA+RNA_seed | ERBB2 | 0.979905 | BRCA | Hs-578-T | Afatinib |
| MVP_top60_DNA_seed | ERBB2 | 1.667051 | BRCA | UACC-812 | CUDC-101 | MVP_top60_DNA+RNA_seed | MAPK3 | 0.981035 | BRCA | Hs-578-T | FR-180204 |
| MVP_top60_DNA_seed | EGFR | 1.667051 | BRCA | UACC-812 | CUDC-101 | MVP_top60_DNA+RNA_seed | MAPK1 | 0.981035 | BRCA | Hs-578-T | FR-180204 |
| MVP_top60_DNA_seed | ERBB2 | 1.669678 | BRCA | T47D | AZD8931 | MVP_top60_DNA+RNA_seed | MDM2 | 0.981959 | BRCA | HCC1419 | Nutlin-3a (-) |
| MVP_top60_DNA_seed | EGFR | 1.669678 | BRCA | T47D | AZD8931 | MVP_top60_DNA+RNA_seed | SRC | 0.982827 | BRCA | MCF7 | Bosutinib |
| MVP_top60_DNA_seed | ERBB3 | 1.669678 | BRCA | T47D | AZD8931 | MVP_top60_DNA+RNA_seed | EP300 | 0.982974 | BRCA | BT-20 | I-CBP112 |
| MVP_top60_DNA_seed | HDAC1 | 1.672678 | BRCA | HCC1395 | AR-42 | MVP_top60_DNA+RNA_seed | CDK2 | 0.984427 | BRCA | HCC38 | AZD5438 |
| MVP_top60_DNA_seed | HSP90AA1 | 1.681601 | BRCA | MDA-MB-231 | Tanespimycin | MVP_top60_DNA+RNA_seed | PIK3CA | 0.987493 | BRCA | HCC1395 | AZD8835 |
| MVP_top60_DNA_seed | HSP90AB1 | 1.681601 | BRCA | MDA-MB-231 | Tanespimycin | MVP_top60_DNA+RNA_seed | ATM | 0.991155 | BRCA | CAL-148 | CP466722 |
| MVP_top60_DNA_seed | HSP90AA1 | 1.681777 | BRCA | HDQ-P1 | Elesclomol | MVP_top60_DNA+RNA_seed | MAPK1 | 0.991266 | BRCA | MCF7 | VX-11e |
| MVP_top60_DNA_seed | HSP90AB1 | 1.681777 | BRCA | HDQ-P1 | Elesclomol | MVP_top60_DNA+RNA_seed | HDAC1 | 0.993604 | BRCA | CAL-120 | JQ12 |
| MVP_top60_DNA_seed | PIK3CA | 1.682452 | BRCA | HCC1500 | AZD8835 | MVP_top60_DNA+RNA_seed | HDAC2 | 0.993604 | BRCA | CAL-120 | JQ12 |
| MVP_top60_DNA_seed | CDK1 | 1.686275 | BRCA | HCC202 | RO-3306 | MVP_top60_DNA+RNA_seed | AKT1 | 0.994011 | BRCA | HCC1428 | MK-2206 |
| MVP_top60_DNA_seed | CDK2 | 1.688732 | BRCA | MFM-223 | Seliciclib | MVP_top60_DNA+RNA_seed | EGFR | 0.994099 | BRCA | UACC-812 | Erlotinib |

| | | | | | | | | | | | |
|---|---|---|---|---|---|---|---|---|---|---|---|
| MVP_top60_DNA_seed | MDM2 | 1.689582 | BRCA | EFM-19 | Serdemetan | MVP_top60_DNA+RNA_seed | SRC | 0.995394 | BRCA | HCC1395 | NVP-BHG712 |
| MVP_top60_DNA_seed | ATM | 1.693808 | BRCA | HCC70 | KU-60019 | MVP_top60_DNA+RNA_seed | PIK3CA | 0.997241 | BRCA | ZR-75-30 | AZD8835 |
| MVP_top60_DNA_seed | ATM | 1.698489 | BRCA | CAMA-1 | KU-55933 | MVP_top60_DNA+RNA_seed | EGFR | 0.998925 | BRCA | MRK-nu-1 | Afatinib |
| MVP_top60_DNA_seed | CDK2 | 1.701804 | BRCA | T47D | AZD5438 | MVP_top60_DNA+RNA_seed | ERBB2 | 0.998925 | BRCA | MRK-nu-1 | Afatinib |
| MVP_top60_DNA_seed | HDAC1 | 1.705346 | BRCA | BT-483 | Tubastatin A | MVP_top60_DNA+RNA_seed | AKT1 | 1.00147 | BRCA | CAL-120 | AKT inhibitor VIII |
| MVP_top60_DNA_seed | ERBB2 | 1.721437 | BRCA | HCC1500 | AZD8931 | MVP_top60_DNA+RNA_seed | MAPK1 | 1.003265 | BRCA | AU565 | VX-11e |
| MVP_top60_DNA_seed | EGFR | 1.721437 | BRCA | HCC1500 | AZD8931 | MVP_top60_DNA+RNA_seed | EGFR | 1.004373 | BRCA | HCC1395 | Afatinib |
| MVP_top60_DNA_seed | ERBB3 | 1.721437 | BRCA | HCC1500 | AZD8931 | MVP_top60_DNA+RNA_seed | ERBB2 | 1.004373 | BRCA | HCC1395 | Afatinib |
| MVP_top60_DNA_seed | EGFR | 1.725387 | BRCA | MDA-MB-436 | Pelitinib | MVP_top60_DNA+RNA_seed | HDAC1 | 1.00563 | BRCA | UACC-893 | AR-42 |
| MVP_top60_DNA_seed | SRC | 1.727797 | BRCA | MDA-MB-415 | Bosutinib | MVP_top60_DNA+RNA_seed | EGFR | 1.006389 | BRCA | HCC1428 | Afatinib |
| MVP_top60_DNA_seed | ERBB2 | 1.737899 | BRCA | BT-483 | Afatinib | MVP_top60_DNA+RNA_seed | ERBB2 | 1.006389 | BRCA | HCC1428 | Afatinib |
| MVP_top60_DNA_seed | EGFR | 1.737899 | BRCA | BT-483 | Afatinib | MVP_top60_DNA+RNA_seed | MAPK14 | 1.007654 | BRCA | EFM-19 | TAK-715 |
| MVP_top60_DNA_seed | EGFR | 1.738674 | BRCA | HCC1395 | Gefitinib | MVP_top60_DNA+RNA_seed | HSP90AA1 | 1.010285 | BRCA | CAMA-1 | CCT-018159 |
| MVP_top60_DNA_seed | ESR1 | 1.73933 | BRCA | HCC2218 | Fulvestrant | MVP_top60_DNA+RNA_seed | HSP90AB1 | 1.010285 | BRCA | CAMA-1 | CCT-018159 |
| MVP_top60_DNA_seed | HSP90AA1 | 1.744701 | BRCA | HCC1954 | Luminespib | MVP_top60_DNA+RNA_seed | HDAC1 | 1.010608 | BRCA | HCC1954 | AR-42 |
| MVP_top60_DNA_seed | HSP90AB1 | 1.744701 | BRCA | HCC1954 | Luminespib | MVP_top60_DNA+RNA_seed | HDAC1 | 1.011245 | BRCA | Hs-578-T | Tubastatin A |
| MVP_top60_DNA_seed | ATM | 1.748778 | BRCA | HCC1419 | KU-55933 | MVP_top60_DNA+RNA_seed | PIK3CA | 1.012949 | BRCA | HCC38 | PI3Ka_4409 |
| MVP_top60_DNA_seed | HSP90AA1 | 1.760544 | BRCA | T47D | Tanespimycin | MVP_top60_DNA+RNA_seed | MDM2 | 1.013197 | BRCA | CAL-85-1 | Nutlin-3a (-) |
| MVP_top60_DNA_seed | HSP90AB1 | 1.760544 | BRCA | T47D | Tanespimycin | MVP_top60_DNA+RNA_seed | SRC | 1.014293 | BRCA | COLO-824 | Saracatinib |
| MVP_top60_DNA_seed | HSP90AA1 | 1.763084 | BRCA | HCC1419 | CCT-018159 | MVP_top60_DNA+RNA_seed | MDM2 | 1.014353 | BRCA | Hs-578-T | Nutlin-3a (-) |
| MVP_top60_DNA_seed | HSP90AB1 | 1.763084 | BRCA | HCC1419 | CCT-018159 | MVP_top60_DNA+RNA_seed | EGFR | 1.014678 | BRCA | HCC1395 | Pelitinib |
| MVP_top60_DNA_seed | HSP90AA1 | 1.76859 | BRCA | HCC1500 | CCT-018159 | MVP_top60_DNA+RNA_seed | EGFR | 1.015577 | BRCA | MRK-nu-1 | Erlotinib |
| MVP_top60_DNA_seed | HSP90AB1 | 1.76859 | BRCA | HCC1500 | CCT-018159 | MVP_top60_DNA+RNA_seed | EGFR | 1.018191 | BRCA | MFM-223 | Erlotinib |
| MVP_top60_DNA_seed | AR | 1.773014 | BRCA | HCC2218 | Bicalutamide | MVP_top60_DNA+RNA_seed | EGFR | 1.021568 | BRCA | MDA-MB-330 | Cetuximab |
| MVP_top60_DNA_seed | SRC | 1.782302 | BRCA | HCC202 | Bosutinib | MVP_top60_DNA+RNA_seed | HDAC1 | 1.023373 | BRCA | YMB-1-E | AR-42 |
| MVP_top60_DNA_seed | EGFR | 1.790184 | BRCA | BT-483 | Pelitinib | MVP_top60_DNA+RNA_seed | AR | 1.023938 | BRCA | UACC-893 | Bicalutamide |
| MVP_top60_DNA_seed | MDM2 | 1.79149 | BRCA | HCC202 | Serdemetan | MVP_top60_DNA+RNA_seed | MDM2 | 1.025946 | BRCA | HCC38 | Nutlin-3a (-) |
| MVP_top60_DNA_seed | ESR1 | 1.792802 | BRCA | HCC1419 | Fulvestrant | MVP_top60_DNA+RNA_seed | PLK1 | 1.026712 | BRCA | HCC2157 | BI-2536 |
| MVP_top60_DNA_seed | AKT1 | 1.797169 | BRCA | HCC38 | MK-2206 | MVP_top60_DNA+RNA_seed | AKT1 | 1.030633 | BRCA | COLO-824 | MK-2206 |
| MVP_top60_DNA_seed | AKT1 | 1.801117 | BRCA | MRK-nu-1 | MK-2206 | MVP_top60_DNA+RNA_seed | ESR1 | 1.031725 | BRCA | MDA-MB-175-VII | Tamoxifen |
| MVP_top60_DNA_seed | EGFR | 1.804296 | BRCA | T47D | Gefitinib | MVP_top60_DNA+RNA_seed | EGFR | 1.032075 | BRCA | ZR-75-30 | CI-1033 |
| MVP_top60_DNA_seed | AKT1 | 1.80809 | BRCA | HCC1500 | MK-2206 | MVP_top60_DNA+RNA_seed | ERBB2 | 1.032075 | BRCA | ZR-75-30 | CI-1033 |
| MVP_top60_DNA_seed | HSP90AA1 | 1.810166 | BRCA | MDA-MB-157 | SNX-2112 | MVP_top60_DNA+RNA_seed | CDK1 | 1.032567 | BRCA | HCC1143 | RO-3306 |
| MVP_top60_DNA_seed | HSP90AB1 | 1.810166 | BRCA | MDA-MB-157 | SNX-2112 | MVP_top60_DNA+RNA_seed | ATM | 1.032779 | BRCA | HCC1395 | KU-55933 |
| MVP_top60_DNA_seed | AR | 1.816611 | BRCA | T47D | Bicalutamide | MVP_top60_DNA+RNA_seed | AKT1 | 1.036376 | BRCA | HCC1143 | BAY AKT1 |
| MVP_top60_DNA_seed | HSP90AA1 | 1.819943 | BRCA | HCC1395 | Luminespib | MVP_top60_DNA+RNA_seed | CDK1 | 1.03844 | BRCA | HCC1500 | RO-3306 |
| MVP_top60_DNA_seed | HSP90AB1 | 1.819943 | BRCA | HCC1395 | Luminespib | MVP_top60_DNA+RNA_seed | AKT1 | 1.038997 | BRCA | Hs-578-T | AKT inhibitor VIII |
| MVP_top60_DNA_seed | MAPK1 | 1.820287 | BRCA | BT-483 | VX-11e | MVP_top60_DNA+RNA_seed | ATM | 1.040636 | BRCA | HDQ-P1 | KU-55933 |
| MVP_top60_DNA_seed | CDK2 | 1.822255 | BRCA | HCC1500 | AZD5438 | MVP_top60_DNA+RNA_seed | ESR1 | 1.042918 | BRCA | HCC1937 | Tamoxifen |
| MVP_top60_DNA_seed | MDM2 | 1.841397 | BRCA | ZR-75-30 | Serdemetan | MVP_top60_DNA+RNA_seed | HSP90AA1 | 1.044194 | BRCA | HCC1954 | SNX-2112 |
| MVP_top60_DNA_seed | ESR1 | 1.844901 | BRCA | ZR-75-30 | Tamoxifen | MVP_top60_DNA+RNA_seed | HSP90AB1 | 1.044194 | BRCA | HCC1954 | SNX-2112 |

| | | | | | | | | | | | |
|---|---|---|---|---|---|---|---|---|---|---|---|
| MVP_top60_DNA_seed | CDK2 | 1.852959 | BRCA | UACC-893 | AZD5438 | MVP_top60_DNA+RNA_seed | MDM2 | 1.048777 | BRCA | HCC1500 | Serdemetan |
| MVP_top60_DNA_seed | HSP90AA1 | 1.855709 | BRCA | MDA-MB-436 | CCT-018159 | MVP_top60_DNA+RNA_seed | HDAC1 | 1.052385 | BRCA | HCC1954 | Tubastatin A |
| MVP_top60_DNA_seed | HSP90AB1 | 1.855709 | BRCA | MDA-MB-436 | CCT-018159 | MVP_top60_DNA+RNA_seed | EGFR | 1.052745 | BRCA | COLO-824 | Erlotinib |
| MVP_top60_DNA_seed | HDAC1 | 1.855711 | BRCA | Hs-578-T | JQ12 | MVP_top60_DNA+RNA_seed | ESR1 | 1.053103 | BRCA | CAMA-1 | Tamoxifen |
| MVP_top60_DNA_seed | HDAC2 | 1.855711 | BRCA | Hs-578-T | JQ12 | MVP_top60_DNA+RNA_seed | AR | 1.053818 | BRCA | CAMA-1 | AZD3514 |
| MVP_top60_DNA_seed | HDAC1 | 1.867294 | BRCA | BT-20 | JQ12 | MVP_top60_DNA+RNA_seed | ESR1 | 1.055534 | BRCA | BT-20 | Tamoxifen |
| MVP_top60_DNA_seed | HDAC2 | 1.867294 | BRCA | BT-20 | JQ12 | MVP_top60_DNA+RNA_seed | EGFR | 1.056015 | BRCA | MDA-MB-436 | Gefitinib |
| MVP_top60_DNA_seed | HDAC1 | 1.869746 | BRCA | MFM-223 | JQ12 | MVP_top60_DNA+RNA_seed | MDM2 | 1.056745 | BRCA | BT-474 | Nutlin-3a (-) |
| MVP_top60_DNA_seed | HDAC2 | 1.869746 | BRCA | MFM-223 | JQ12 | MVP_top60_DNA+RNA_seed | AKT1 | 1.061477 | BRCA | HDQ-P1 | MK-2206 |
| MVP_top60_DNA_seed | MAPK3 | 1.874762 | BRCA | BT-483 | FR-180204 | MVP_top60_DNA+RNA_seed | EGFR | 1.061921 | BRCA | MDA-MB-157 | CI-1033 |
| MVP_top60_DNA_seed | MAPK1 | 1.874762 | BRCA | BT-483 | FR-180204 | MVP_top60_DNA+RNA_seed | ERBB2 | 1.061921 | BRCA | MDA-MB-157 | CI-1033 |
| MVP_top60_DNA_seed | HSP90AA1 | 1.875977 | BRCA | MRK-nu-1 | Tanespimycin | MVP_top60_DNA+RNA_seed | HSP90AA1 | 1.064246 | BRCA | BT-20 | Luminespib |
| MVP_top60_DNA_seed | HSP90AB1 | 1.875977 | BRCA | MRK-nu-1 | Tanespimycin | MVP_top60_DNA+RNA_seed | HSP90AB1 | 1.064246 | BRCA | BT-20 | Luminespib |
| MVP_top60_DNA_seed | ATM | 1.886238 | BRCA | HCC202 | KU-55933 | MVP_top60_DNA+RNA_seed | CDK2 | 1.06477 | BRCA | HCC1395 | PHA-793887 |
| MVP_top60_DNA_seed | PIK3CA | 1.886279 | BRCA | MRK-nu-1 | PI3Ka_4409 | MVP_top60_DNA+RNA_seed | EP300 | 1.065074 | BRCA | HDQ-P1 | I-CBP112 |
| MVP_top60_DNA_seed | HSP90AA1 | 1.901656 | BRCA | MCF7 | SNX-2112 | MVP_top60_DNA+RNA_seed | AKT1 | 1.065861 | BRCA | CAL-85-1 | BAY AKT1 |
| MVP_top60_DNA_seed | HSP90AB1 | 1.901656 | BRCA | MCF7 | SNX-2112 | MVP_top60_DNA+RNA_seed | MAPK3 | 1.067461 | BRCA | HCC1954 | FR-180204 |
| MVP_top60_DNA_seed | PIK3CA | 1.903586 | BRCA | MRK-nu-1 | AZD8835 | MVP_top60_DNA+RNA_seed | MAPK1 | 1.067461 | BRCA | HCC1954 | FR-180204 |
| MVP_top60_DNA_seed | HSP90AA1 | 1.905592 | BRCA | MDA-MB-415 | SNX-2112 | MVP_top60_DNA+RNA_seed | AR | 1.073449 | BRCA | HDQ-P1 | Bicalutamide |
| MVP_top60_DNA_seed | HSP90AB1 | 1.905592 | BRCA | MDA-MB-415 | SNX-2112 | MVP_top60_DNA+RNA_seed | SRC | 1.075631 | BRCA | HCC1143 | Bosutinib |
| MVP_top60_DNA_seed | CDK2 | 1.908081 | BRCA | MDA-MB-415 | AZD5438 | MVP_top60_DNA+RNA_seed | HSP90AA1 | 1.078525 | BRCA | UACC-893 | Elesclomol |
| MVP_top60_DNA_seed | HDAC1 | 1.919463 | BRCA | HDQ-P1 | Dacinostat | MVP_top60_DNA+RNA_seed | HSP90AB1 | 1.078525 | BRCA | UACC-893 | Elesclomol |
| MVP_top60_DNA_seed | HDAC1 | 1.923513 | BRCA | HCC2218 | Parthenolide | MVP_top60_DNA+RNA_seed | PIK3CA | 1.082358 | BRCA | HCC38 | AZD8835 |
| MVP_top60_DNA_seed | HSP90AA1 | 1.955992 | BRCA | HCC2157 | SNX-2112 | MVP_top60_DNA+RNA_seed | PLK1 | 1.083181 | BRCA | COLO-824 | GW843682X |
| MVP_top60_DNA_seed | HSP90AB1 | 1.955992 | BRCA | HCC2157 | SNX-2112 | MVP_top60_DNA+RNA_seed | MDM2 | 1.083206 | BRCA | MFM-223 | Nutlin-3a (-) |
| MVP_top60_DNA_seed | PIK3CA | 1.962095 | BRCA | HCC1500 | PI3Ka_4409 | MVP_top60_DNA+RNA_seed | MAPK1 | 1.084861 | BRCA | HCC2157 | VX-11e |
| MVP_top60_DNA_seed | HDAC1 | 1.975689 | BRCA | MDA-MB-415 | AR-42 | MVP_top60_DNA+RNA_seed | MDM2 | 1.085644 | BRCA | EFM-19 | Nutlin-3a (-) |
| MVP_top60_DNA_seed | SRC | 1.989396 | BRCA | BT-483 | Bosutinib | MVP_top60_DNA+RNA_seed | MAPK1 | 1.089046 | BRCA | YMB-1-E | VX-11e |
| MVP_top60_DNA_seed | SRC | 2.000384 | BRCA | BT-483 | NVP-BHG712 | MVP_top60_DNA+RNA_seed | HSP90AA1 | 1.091441 | BRCA | HCC1143 | CCT-018159 |
| MVP_top60_DNA_seed | HSP90AA1 | 2.02296 | BRCA | MRK-nu-1 | Elesclomol | MVP_top60_DNA+RNA_seed | HSP90AB1 | 1.091441 | BRCA | HCC1143 | CCT-018159 |
| MVP_top60_DNA_seed | HSP90AB1 | 2.02296 | BRCA | MRK-nu-1 | Elesclomol | MVP_top60_DNA+RNA_seed | ERBB2 | 1.091944 | BRCA | MDA-MB-436 | CP724714 |
| MVP_top60_DNA_seed | HDAC1 | 2.024309 | BRCA | UACC-812 | JQ12 | MVP_top60_DNA+RNA_seed | HDAC1 | 1.092439 | BRCA | MDA-MB-453 | JQ12 |
| MVP_top60_DNA_seed | HDAC2 | 2.024309 | BRCA | UACC-812 | JQ12 | MVP_top60_DNA+RNA_seed | HDAC2 | 1.092439 | BRCA | MDA-MB-453 | JQ12 |
| MVP_top60_DNA_seed | ATM | 2.042981 | BRCA | ZR-75-30 | KU-55933 | MVP_top60_DNA+RNA_seed | EGFR | 1.09424 | BRCA | MRK-nu-1 | Gefitinib |
| MVP_top60_DNA_seed | PIK3CA | 2.047064 | BRCA | HCC1395 | PI3Ka_4409 | MVP_top60_DNA+RNA_seed | CDK2 | 1.094768 | BRCA | EFM-192A | AZD5438 |
| MVP_top60_DNA_seed | HSP90AA1 | 2.052114 | BRCA | UACC-812 | SNX-2112 | MVP_top60_DNA+RNA_seed | MAPK14 | 1.09606 | BRCA | MDA-MB-361 | TAK-715 |
| MVP_top60_DNA_seed | HSP90AB1 | 2.052114 | BRCA | UACC-812 | SNX-2112 | MVP_top60_DNA+RNA_seed | EGFR | 1.096387 | BRCA | HCC1395 | Afatinib |
| MVP_top60_DNA_seed | CDK2 | 2.053182 | BRCA | CAMA-1 | AZD5438 | MVP_top60_DNA+RNA_seed | ERBB2 | 1.096387 | BRCA | HCC1395 | Afatinib |
| MVP_top60_DNA_seed | HDAC1 | 2.060857 | BRCA | HDQ-P1 | JQ12 | MVP_top60_DNA+RNA_seed | MDM2 | 1.098218 | BRCA | HCC1143 | Nutlin-3a (-) |
| MVP_top60_DNA_seed | HDAC2 | 2.060857 | BRCA | HDQ-P1 | JQ12 | MVP_top60_DNA+RNA_seed | EGFR | 1.098459 | BRCA | MDA-MB-415 | Afatinib |
| MVP_top60_DNA_seed | EGFR | 2.075293 | BRCA | ZR-75-30 | Gefitinib | MVP_top60_DNA+RNA_seed | ERBB2 | 1.098459 | BRCA | MDA-MB-415 | Afatinib |

| | | | | | | | | | | | |
|---|---|---|---|---|---|---|---|---|---|---|---|
| MVP_top60_DNA_seed | HSP90AA1 | 2.100611 | BRCA | HCC202 | CCT-018159 | MVP_top60_DNA+RNA_seed | MAPK1 | 1.099008 | BRCA | HCC1428 | VX-11e |
| MVP_top60_DNA_seed | HSP90AB1 | 2.100611 | BRCA | HCC202 | CCT-018159 | MVP_top60_DNA+RNA_seed | ATM | 1.10329 | BRCA | MRK-nu-1 | CP466722 |
| MVP_top60_DNA_seed | HSP90AA1 | 2.102781 | BRCA | MFM-223 | Luminespib | MVP_top60_DNA+RNA_seed | SRC | 1.108298 | BRCA | MDA-MB-330 | Bosutinib |
| MVP_top60_DNA_seed | HSP90AB1 | 2.102781 | BRCA | MFM-223 | Luminespib | MVP_top60_DNA+RNA_seed | EGFR | 1.110495 | BRCA | CAMA-1 | Cetuximab |
| MVP_top60_DNA_seed | MDM2 | 2.135298 | BRCA | HCC2218 | Serdemetan | MVP_top60_DNA+RNA_seed | MAPK1 | 1.112239 | BRCA | Hs-578-T | VX-11e |
| MVP_top60_DNA_seed | HDAC1 | 2.165719 | BRCA | COLO-824 | Entinostat | MVP_top60_DNA+RNA_seed | ESR1 | 1.113248 | BRCA | UACC-893 | Tamoxifen |
| MVP_top60_DNA_seed | ATM | 2.172968 | BRCA | BT-483 | KU-60019 | MVP_top60_DNA+RNA_seed | ERBB2 | 1.114931 | BRCA | HCC1395 | CP724714 |
| MVP_top60_DNA_seed | PIK3CA | 2.180415 | BRCA | CAMA-1 | AZD8835 | MVP_top60_DNA+RNA_seed | SRC | 1.11809 | BRCA | CAMA-1 | Bosutinib |
| MVP_top60_DNA_seed | HSP90AA1 | 2.268071 | BRCA | HCC1569 | Elesclomol | MVP_top60_DNA+RNA_seed | HDAC1 | 1.119004 | BRCA | BT-549 | JQ12 |
| MVP_top60_DNA_seed | HSP90AB1 | 2.268071 | BRCA | HCC1569 | Elesclomol | MVP_top60_DNA+RNA_seed | HDAC2 | 1.119004 | BRCA | BT-549 | JQ12 |
| MVP_top60_DNA_seed | MDM2 | 2.282995 | BRCA | MDA-MB-175-VII | Serdemetan | MVP_top60_DNA+RNA_seed | SRC | 1.120187 | BRCA | HCC2218 | WH-4-023 |
| MVP_top60_DNA_seed | PIK3CA | 2.290532 | BRCA | CAMA-1 | PI3Ka_4409 | MVP_top60_DNA+RNA_seed | AKT1 | 1.123569 | BRCA | COLO-824 | A-443654 |
| MVP_top60_DNA_seed | PIK3CA | 2.332789 | BRCA | MDA-MB-436 | PI3Ka_4409 | MVP_top60_DNA+RNA_seed | EP300 | 1.123824 | BRCA | HCC1419 | I-CBP112 |
| MVP_top60_DNA_seed | ESR1 | 2.342078 | BRCA | BT-483 | Tamoxifen | MVP_top60_DNA+RNA_seed | MDM2 | 1.124313 | BRCA | CAL-85-1 | Serdemetan |
| MVP_top60_DNA_seed | ESR1 | 2.378933 | BRCA | MFM-223 | Tamoxifen | MVP_top60_DNA+RNA_seed | HSP90AA1 | 1.124441 | BRCA | CAL-51 | SNX-2112 |
| MVP_top60_DNA_seed | HDAC1 | 2.379139 | BRCA | MFM-223 | Entinostat | MVP_top60_DNA+RNA_seed | HSP90AB1 | 1.124441 | BRCA | CAL-51 | SNX-2112 |
| MVP_top60_DNA_seed | CDK2 | 2.407839 | BRCA | ZR-75-30 | AZD5438 | MVP_top60_DNA+RNA_seed | EGFR | 1.126794 | BRCA | EFM-19 | Afatinib |
| MVP_top60_DNA_seed | AKT1 | 2.444972 | BRCA | BT-483 | BAY AKT1 | MVP_top60_DNA+RNA_seed | ERBB2 | 1.126794 | BRCA | EFM-19 | Afatinib |
| MVP_top60_DNA_seed | HSP90AA1 | 2.448277 | BRCA | HCC1954 | Elesclomol | MVP_top60_DNA+RNA_seed | AKT1 | 1.128053 | BRCA | MDA-MB-231 | BAY AKT1 |
| MVP_top60_DNA_seed | HSP90AB1 | 2.448277 | BRCA | HCC1954 | Elesclomol | MVP_top60_DNA+RNA_seed | SRC | 1.129413 | BRCA | MFM-223 | WH-4-023 |
| MVP_top60_DNA_seed | AR | 2.513702 | BRCA | ZR-75-30 | AZD3514 | MVP_top60_DNA+RNA_seed | ESR1 | 1.130416 | BRCA | ZR-75-30 | Fulvestrant |
| MVP_top60_DNA_seed | ATM | 2.584383 | BRCA | HCC2218 | KU-60019 | MVP_top60_DNA+RNA_seed | AR | 1.135081 | BRCA | HCC1143 | AZD3514 |
| MVP_top60_DNA_seed | CDK2 | 2.633355 | BRCA | MDA-MB-175-VII | AZD5438 | MVP_top60_DNA+RNA_seed | HSP90AA1 | 1.142179 | BRCA | CAL-85-1 | Tanespimycin |
| MVP_top60_DNA_seed | HSP90AA1 | 2.640077 | BRCA | Hs-578-T | Luminespib | MVP_top60_DNA+RNA_seed | HSP90AB1 | 1.142179 | BRCA | CAL-85-1 | Tanespimycin |
| MVP_top60_DNA_seed | HSP90AB1 | 2.640077 | BRCA | Hs-578-T | Luminespib | MVP_top60_DNA+RNA_seed | MDM2 | 1.144594 | BRCA | MFM-223 | Serdemetan |
| MVP_top60_DNA_seed | AR | 2.667127 | BRCA | BT-483 | AZD3514 | MVP_top60_DNA+RNA_seed | MAPK1 | 1.14544 | BRCA | HCC1569 | VX-11e |
| MVP_top60_DNA_seed | CDK2 | 2.692984 | BRCA | HCC1419 | AZD5438 | MVP_top60_DNA+RNA_seed | EGFR | 1.14782 | BRCA | MFM-223 | Cetuximab |
| MVP_top60_DNA_seed | HDAC1 | 2.711435 | BRCA | BT-483 | Belinostat | MVP_top60_DNA+RNA_seed | SRC | 1.148501 | BRCA | BT-474 | Bosutinib |
| MVP_top60_DNA_seed | AR | 2.773452 | BRCA | BT-483 | Bicalutamide | MVP_top60_DNA+RNA_seed | EGFR | 1.150186 | BRCA | UACC-893 | Gefitinib |
| MVP_top60_DNA_seed | HDAC1 | 2.794843 | BRCA | HCC1187 | Dacinostat | MVP_top60_DNA+RNA_seed | PIK3CA | 1.15106 | BRCA | BT-483 | AZD8835 |
| MVP_top60_DNA_seed | AR | 2.819253 | BRCA | BT-483 | Bicalutamide | MVP_top60_DNA+RNA_seed | CDK1 | 1.15205 | BRCA | BT-549 | RO-3306 |
| MVP_top60_DNA_seed | HDAC1 | 2.839094 | BRCA | UACC-812 | Dacinostat | MVP_top60_DNA+RNA_seed | MAPK14 | 1.153392 | BRCA | MDA-MB-330 | TAK-715 |
| MVP_top60_DNA_seed | HSP90AA1 | 2.975847 | BRCA | BT-483 | CCT-018159 | MVP_top60_DNA+RNA_seed | ESR1 | 1.156381 | BRCA | MRK-nu-1 | Fulvestrant |
| MVP_top60_DNA_seed | HSP90AB1 | 2.975847 | BRCA | BT-483 | CCT-018159 | MVP_top60_DNA+RNA_seed | PIK3CA | 1.15663 | BRCA | MDA-MB-468 | AZD8835 |
| MVP_top60_DNA_seed | ATM | 3.125079 | BRCA | ZR-75-30 | torin2 | MVP_top60_DNA+RNA_seed | ATM | 1.158436 | BRCA | HCC202 | KU-60019 |
| MVP_top60_DNA_seed | ERBB2 | 3.424488 | BRCA | BT-483 | CUDC-101 | MVP_top60_DNA+RNA_seed | MAPK1 | 1.158625 | BRCA | HCC2218 | VX-11e |
| MVP_top60_DNA_seed | EGFR | 3.424488 | BRCA | BT-483 | CUDC-101 | MVP_top60_DNA+RNA_seed | PLK1 | 1.158757 | BRCA | MDA-MB-415 | PLK_6522 |
| MVP_top60_DNA_seed | HDAC1 | 3.455672 | BRCA | BT-483 | AR-42 | MVP_top60_DNA+RNA_seed | CDK1 | 1.165613 | BRCA | EFM-19 | RO-3306 |
| MVP_top60_DNA_seed | HSP90AA1 | 3.512918 | BRCA | BT-483 | Luminespib | MVP_top60_DNA+RNA_seed | MAPK3 | 1.169895 | BRCA | BT-20 | FR-180204 |
| MVP_top60_DNA_seed | HSP90AB1 | 3.512918 | BRCA | BT-483 | Luminespib | MVP_top60_DNA+RNA_seed | MAPK1 | 1.169895 | BRCA | BT-20 | FR-180204 |
| MVP_top60_DNA_seed | CDK2 | 3.717065 | BRCA | BT-483 | AZD5438 | MVP_top60_DNA+RNA_seed | PLK1 | 1.172116 | BRCA | COLO-824 | BI-2536 |

| | | | | | | | | | | | |
|---|---|---|---|---|---|---|---|---|---|---|---|
| MVP_top60_DNA_seed | HDAC1 | 4.486456 | BRCA | Hs-578-T | Dacinostat | MVP_top60_DNA+RNA_seed | CDK1 | 1.173316 | BRCA | UACC-893 | RO-3306 |
| MVP_top60_DNA_seed | PIK3CA | -4.087508 | BRCA | YMB-1-E | AZD8186 | MVP_top60_DNA+RNA_seed | AR | 1.173383 | BRCA | MDA-MB-175-VII | Bicalutamide |
| MVP_top60_DNA_seed | ATM | -3.853213 | BRCA | HCC2157 | KU-55933 | MVP_top60_DNA+RNA_seed | HSP90AA1 | 1.173827 | BRCA | CAMA-1 | Elesclomol |
| MVP_top60_DNA_seed | PIK3CA | -3.846124 | BRCA | EVSA-T | AZD8186 | MVP_top60_DNA+RNA_seed | HSP90AB1 | 1.173827 | BRCA | CAMA-1 | Elesclomol |
| MVP_top60_DNA_seed | ERBB2 | -3.754 | BRCA | MDA-MB-175-VII | Afatinib | MVP_top60_DNA+RNA_seed | PIK3CA | 1.175306 | BRCA | CAL-51 | AZD8835 |
| MVP_top60_DNA_seed | EGFR | -3.754 | BRCA | MDA-MB-175-VII | Afatinib | MVP_top60_DNA+RNA_seed | ERBB2 | 1.176434 | BRCA | CAL-120 | CP724714 |
| MVP_top60_DNA_seed | MAPK3 | -3.557454 | BRCA | DU-4475 | SCH772984 | MVP_top60_DNA+RNA_seed | EP300 | 1.176935 | BRCA | HCC1395 | I-CBP112 |
| MVP_top60_DNA_seed | MAPK1 | -3.557454 | BRCA | DU-4475 | SCH772984 | MVP_top60_DNA+RNA_seed | MDM2 | 1.177493 | BRCA | BT-20 | Serdemetan |
| MVP_top60_DNA_seed | ERBB2 | -3.497705 | BRCA | HCC2218 | Afatinib | MVP_top60_DNA+RNA_seed | AKT1 | 1.178983 | BRCA | HDQ-P1 | AKT inhibitor VIII |
| MVP_top60_DNA_seed | EGFR | -3.497705 | BRCA | HCC2218 | Afatinib | MVP_top60_DNA+RNA_seed | MAPK1 | 1.179384 | BRCA | HCC1954 | VX-11e |
| MVP_top60_DNA_seed | PIK3CA | -3.378294 | BRCA | MDA-MB-415 | AZD8186 | MVP_top60_DNA+RNA_seed | MDM2 | 1.182363 | BRCA | BT-549 | Nutlin-3a (-) |
| MVP_top60_DNA_seed | AKT1 | -3.302738 | BRCA | CAL-148 | Ipatasertib | MVP_top60_DNA+RNA_seed | EP300 | 1.186063 | BRCA | EFM-19 | I-CBP112 |
| MVP_top60_DNA_seed | AKT1 | -3.282229 | BRCA | EVSA-T | GSK2110183B | MVP_top60_DNA+RNA_seed | AR | 1.187736 | BRCA | HCC1937 | Bicalutamide |
| MVP_top60_DNA_seed | AKT1 | -3.259805 | BRCA | CAL-148 | Afuresertib | MVP_top60_DNA+RNA_seed | HDAC1 | 1.18868 | BRCA | BT-483 | JQ12 |
| MVP_top60_DNA_seed | ERBB2 | -3.16649 | BRCA | UACC-893 | Afatinib | MVP_top60_DNA+RNA_seed | HDAC2 | 1.18868 | BRCA | BT-483 | JQ12 |
| MVP_top60_DNA_seed | EGFR | -3.16649 | BRCA | UACC-893 | Afatinib | MVP_top60_DNA+RNA_seed | SRC | 1.188852 | BRCA | MDA-MB-157 | NVP-BHG712 |
| MVP_top60_DNA_seed | ERBB2 | -3.158051 | BRCA | MDA-MB-175-VII | Lapatinib | MVP_top60_DNA+RNA_seed | EGFR | 1.190486 | BRCA | HCC1143 | Cetuximab |
| MVP_top60_DNA_seed | EGFR | -3.158051 | BRCA | MDA-MB-175-VII | Lapatinib | MVP_top60_DNA+RNA_seed | MAPK1 | 1.192202 | BRCA | UACC-812 | VX-11e |
| MVP_top60_DNA_seed | AKT1 | -3.141626 | BRCA | CAL-148 | AZD5363 | MVP_top60_DNA+RNA_seed | HDAC1 | 1.195305 | BRCA | HCC1395 | Belinostat |
| MVP_top60_DNA_seed | AKT1 | -3.138923 | BRCA | CAL-148 | Uprosertib | MVP_top60_DNA+RNA_seed | SRC | 1.198467 | BRCA | MDA-MB-453 | NVP-BHG712 |
| MVP_top60_DNA_seed | ERBB2 | -3.13007 | BRCA | MDA-MB-175-VII | Sapitinib | MVP_top60_DNA+RNA_seed | ATM | 1.198858 | BRCA | DU-4475 | KU-60019 |
| MVP_top60_DNA_seed | EGFR | -3.13007 | BRCA | MDA-MB-175-VII | Sapitinib | MVP_top60_DNA+RNA_seed | MAPK1 | 1.200177 | BRCA | HCC1395 | VX-11e |
| MVP_top60_DNA_seed | ERBB3 | -3.13007 | BRCA | MDA-MB-175-VII | Sapitinib | MVP_top60_DNA+RNA_seed | AKT1 | 1.201319 | BRCA | MDA-MB-436 | GSK690693 |
| MVP_top60_DNA_seed | AKT1 | -3.113381 | BRCA | OCUB-M | Uprosertib | MVP_top60_DNA+RNA_seed | CDK2 | 1.20265 | BRCA | BT-20 | AZD5438 |
| MVP_top60_DNA_seed | MAPK1 | -3.094224 | BRCA | DU-4475 | VX-11e | MVP_top60_DNA+RNA_seed | HDAC1 | 1.202879 | BRCA | COLO-824 | Parthenolide |
| MVP_top60_DNA_seed | AKT1 | -3.011198 | BRCA | OCUB-M | Afuresertib | MVP_top60_DNA+RNA_seed | MDM2 | 1.205442 | BRCA | BT-474 | Serdemetan |
| MVP_top60_DNA_seed | AKT1 | -2.964343 | BRCA | EVSA-T | Afuresertib | MVP_top60_DNA+RNA_seed | MDM2 | 1.206361 | BRCA | HCC1395 | Nutlin-3a (-) |
| MVP_top60_DNA_seed | EGFR | -2.963037 | BRCA | MDA-MB-175-VII | Gefitinib | MVP_top60_DNA+RNA_seed | HDAC1 | 1.209938 | BRCA | HCC1500 | Entinostat |
| MVP_top60_DNA_seed | AKT1 | -2.904853 | BRCA | OCUB-M | GSK2110183B | MVP_top60_DNA+RNA_seed | HSP90AA1 | 1.21152 | BRCA | EFM-192A | Elesclomol |
| MVP_top60_DNA_seed | AKT1 | -2.891728 | BRCA | CAL-148 | Uprosertib | MVP_top60_DNA+RNA_seed | HSP90AB1 | 1.21152 | BRCA | EFM-192A | Elesclomol |
| MVP_top60_DNA_seed | EGFR | -2.870454 | BRCA | HCC2218 | Osimertinib | MVP_top60_DNA+RNA_seed | SRC | 1.212273 | BRCA | HCC70 | Bosutinib |
| MVP_top60_DNA_seed | AKT1 | -2.836144 | BRCA | EVSA-T | AZD5363 | MVP_top60_DNA+RNA_seed | PIK3CA | 1.217205 | BRCA | HCC1937 | PI3Ka_4409 |
| MVP_top60_DNA_seed | MAPK3 | -2.82499 | BRCA | DU-4475 | Ulixertinib | MVP_top60_DNA+RNA_seed | MAPK1 | 1.217349 | BRCA | BT-549 | VX-11e |
| MVP_top60_DNA_seed | MAPK1 | -2.82499 | BRCA | DU-4475 | Ulixertinib | MVP_top60_DNA+RNA_seed | HDAC1 | 1.218545 | BRCA | HCC1187 | JQ12 |
| MVP_top60_DNA_seed | AKT1 | -2.81292 | BRCA | MDA-MB-415 | Ipatasertib | MVP_top60_DNA+RNA_seed | HDAC2 | 1.218545 | BRCA | HCC1187 | JQ12 |
| MVP_top60_DNA_seed | AKT1 | -2.789178 | BRCA | OCUB-M | Ipatasertib | MVP_top60_DNA+RNA_seed | MAPK3 | 1.218793 | BRCA | MFM-223 | FR-180204 |
| MVP_top60_DNA_seed | EGFR | -2.775513 | BRCA | MDA-MB-175-VII | Osimertinib | MVP_top60_DNA+RNA_seed | MAPK1 | 1.218793 | BRCA | MFM-223 | FR-180204 |
| MVP_top60_DNA_seed | AKT1 | -2.754233 | BRCA | OCUB-M | Uprosertib | MVP_top60_DNA+RNA_seed | CDK2 | 1.221172 | BRCA | MCF7 | PHA-793887 |
| MVP_top60_DNA_seed | CDK2 | -2.74532 | BRCA | HCC2218 | MK-8776 | MVP_top60_DNA+RNA_seed | ATM | 1.221655 | BRCA | BT-483 | KU-55933 |
| MVP_top60_DNA_seed | EGFR | -2.709433 | BRCA | HDQ-P1 | AZD3759 | MVP_top60_DNA+RNA_seed | AR | 1.222609 | BRCA | BT-20 | Bicalutamide |
| MVP_top60_DNA_seed | AKT1 | -2.692292 | BRCA | OCUB-M | AZD5363 | MVP_top60_DNA+RNA_seed | HSP90AA1 | 1.224278 | BRCA | BT-483 | SNX-2112 |

| | | | | | | | | | | | |
|---|---|---|---|---|---|---|---|---|---|---|---|
| MVP_top60_DNA_seed | AKT1 | -2.666864 | BRCA | EVSA-T | Ipatasertib | MVP_top60_DNA+RNA_seed | HSP90AB1 | 1.224278 | BRCA | BT-483 | SNX-2112 |
| MVP_top60_DNA_seed | ESR1 | -2.666749 | BRCA | MCF7 | GDC0810 | MVP_top60_DNA+RNA_seed | HDAC1 | 1.22688 | BRCA | HCC1954 | CAY10603 |
| MVP_top60_DNA_seed | ERBB2 | -2.653491 | BRCA | HCC2218 | Lapatinib | MVP_top60_DNA+RNA_seed | CDK2 | 1.226965 | BRCA | MDA-MB-330 | AZD5438 |
| MVP_top60_DNA_seed | EGFR | -2.653491 | BRCA | HCC2218 | Lapatinib | MVP_top60_DNA+RNA_seed | PLK1 | 1.227898 | BRCA | MFM-223 | GW843682X |
| MVP_top60_DNA_seed | ERBB2 | -2.611412 | BRCA | MDA-MB-330 | Afatinib | MVP_top60_DNA+RNA_seed | HSP90AA1 | 1.227944 | BRCA | UACC-812 | Luminespib |
| MVP_top60_DNA_seed | EGFR | -2.611412 | BRCA | MDA-MB-330 | Afatinib | MVP_top60_DNA+RNA_seed | HSP90AB1 | 1.227944 | BRCA | UACC-812 | Luminespib |
| MVP_top60_DNA_seed | AKT1 | -2.602613 | BRCA | YMB-1-E | MK-2206 | MVP_top60_DNA+RNA_seed | EGFR | 1.232463 | BRCA | BT-549 | Gefitinib |
| MVP_top60_DNA_seed | ERBB2 | -2.580583 | BRCA | AU565 | Lapatinib | MVP_top60_DNA+RNA_seed | SRC | 1.23678 | BRCA | HCC2157 | NVP-BHG712 |
| MVP_top60_DNA_seed | EGFR | -2.580583 | BRCA | AU565 | Lapatinib | MVP_top60_DNA+RNA_seed | ATM | 1.236821 | BRCA | MRK-nu-1 | KU-55933 |
| MVP_top60_DNA_seed | EGFR | -2.550187 | BRCA | UACC-893 | Osimertinib | MVP_top60_DNA+RNA_seed | MAPK1 | 1.238212 | BRCA | MFM-223 | VX-11e |
| MVP_top60_DNA_seed | AKT1 | -2.527069 | BRCA | EVSA-T | MK-2206 | MVP_top60_DNA+RNA_seed | CDK2 | 1.240101 | BRCA | MDA-MB-436 | PHA-793887 |
| MVP_top60_DNA_seed | AKT1 | -2.51042 | BRCA | HCC2157 | GSK2110183B | MVP_top60_DNA+RNA_seed | AKT1 | 1.242022 | BRCA | MRK-nu-1 | AKT inhibitor VIII |
| MVP_top60_DNA_seed | ESR1 | -2.502182 | BRCA | CAMA-1 | GDC0810 | MVP_top60_DNA+RNA_seed | HDAC1 | 1.242255 | BRCA | HCC1954 | Belinostat |
| MVP_top60_DNA_seed | PIK3CA | -2.478395 | BRCA | MDA-MB-361 | Alpelisib | MVP_top60_DNA+RNA_seed | MDM2 | 1.244282 | BRCA | BT-20 | Nutlin-3a (-) |
| MVP_top60_DNA_seed | PIK3CA | -2.467028 | BRCA | MDA-MB-330 | Alpelisib | MVP_top60_DNA+RNA_seed | MDM2 | 1.245407 | BRCA | HDQ-P1 | Nutlin-3a (-) |
| MVP_top60_DNA_seed | MAPK3 | -2.458065 | BRCA | DU-4475 | Ulixertinib | MVP_top60_DNA+RNA_seed | MAPK14 | 1.248142 | BRCA | MDA-MB-415 | TAK-715 |
| MVP_top60_DNA_seed | MAPK1 | -2.458065 | BRCA | DU-4475 | Ulixertinib | MVP_top60_DNA+RNA_seed | SRC | 1.250813 | BRCA | MRK-nu-1 | Bosutinib |
| MVP_top60_DNA_seed | AKT1 | -2.454941 | BRCA | HCC2157 | AZD5363 | MVP_top60_DNA+RNA_seed | ESR1 | 1.251149 | BRCA | HCC1395 | Tamoxifen |
| MVP_top60_DNA_seed | AKT1 | -2.448408 | BRCA | EVSA-T | Uprosertib | MVP_top60_DNA+RNA_seed | AR | 1.251462 | BRCA | MDA-MB-330 | AZD3514 |
| MVP_top60_DNA_seed | AKT1 | -2.434601 | BRCA | EVSA-T | Uprosertib | MVP_top60_DNA+RNA_seed | AR | 1.252344 | BRCA | UACC-893 | AZD3514 |
| MVP_top60_DNA_seed | ERBB2 | -2.433481 | BRCA | UACC-893 | Lapatinib | MVP_top60_DNA+RNA_seed | EGFR | 1.256459 | BRCA | MCF7 | Pelitinib |
| MVP_top60_DNA_seed | EGFR | -2.433481 | BRCA | UACC-893 | Lapatinib | MVP_top60_DNA+RNA_seed | MAPK1 | 1.25669 | BRCA | HCC1937 | VX-11e |
| MVP_top60_DNA_seed | AKT1 | -2.413609 | BRCA | HCC2157 | Afuresertib | MVP_top60_DNA+RNA_seed | CDK1 | 1.258323 | BRCA | CAMA-1 | RO-3306 |
| MVP_top60_DNA_seed | EGFR | -2.391917 | BRCA | MDA-MB-330 | Osimertinib | MVP_top60_DNA+RNA_seed | PIK3CA | 1.260121 | BRCA | MDA-MB-231 | PI3Ka_4409 |
| MVP_top60_DNA_seed | AKT1 | -2.377263 | BRCA | YMB-1-E | Ipatasertib | MVP_top60_DNA+RNA_seed | CDK2 | 1.26254 | BRCA | EFM-19 | PHA-793887 |
| MVP_top60_DNA_seed | PIK3CA | -2.349952 | BRCA | OCUB-M | Alpelisib | MVP_top60_DNA+RNA_seed | SRC | 1.269252 | BRCA | Hs-578-T | NVP-BHG712 |
| MVP_top60_DNA_seed | AKT1 | -2.345353 | BRCA | CAMA-1 | Afuresertib | MVP_top60_DNA+RNA_seed | HSP90AA1 | 1.270252 | BRCA | HCC1187 | Luminespib |
| MVP_top60_DNA_seed | TP53 | -2.331311 | BRCA | HCC1599 | MIRA-1 | MVP_top60_DNA+RNA_seed | HSP90AB1 | 1.270252 | BRCA | HCC1187 | Luminespib |
| MVP_top60_DNA_seed | MAPK3 | -2.323983 | BRCA | MDA-MB-453 | Ulixertinib | MVP_top60_DNA+RNA_seed | HSP90AA1 | 1.271272 | BRCA | MDA-MB-175-VII | Elesclomol |
| MVP_top60_DNA_seed | MAPK1 | -2.323983 | BRCA | MDA-MB-453 | Ulixertinib | MVP_top60_DNA+RNA_seed | HSP90AB1 | 1.271272 | BRCA | MDA-MB-175-VII | Elesclomol |
| MVP_top60_DNA_seed | MAPK3 | -2.288881 | BRCA | MDA-MB-468 | Ulixertinib | MVP_top60_DNA+RNA_seed | AKT1 | 1.271434 | BRCA | HDQ-P1 | AKT inhibitor VIII |
| MVP_top60_DNA_seed | MAPK1 | -2.288881 | BRCA | MDA-MB-468 | Ulixertinib | MVP_top60_DNA+RNA_seed | CDK1 | 1.274383 | BRCA | BT-483 | RO-3306 |
| MVP_top60_DNA_seed | AKT1 | -2.284351 | BRCA | MDA-MB-415 | AZD5363 | MVP_top60_DNA+RNA_seed | MAPK14 | 1.274441 | BRCA | MDA-MB-436 | TAK-715 |
| MVP_top60_DNA_seed | AKT1 | -2.279725 | BRCA | OCUB-M | MK-2206 | MVP_top60_DNA+RNA_seed | SRC | 1.274686 | BRCA | MDA-MB-330 | NVP-BHG712 |
| MVP_top60_DNA_seed | AKT1 | -2.278051 | BRCA | CAMA-1 | Ipatasertib | MVP_top60_DNA+RNA_seed | SRC | 1.276089 | BRCA | HCC38 | Bosutinib |
| MVP_top60_DNA_seed | ERBB2 | -2.264811 | BRCA | ZR-75-30 | Afatinib | MVP_top60_DNA+RNA_seed | AR | 1.278358 | BRCA | HCC2218 | Bicalutamide |
| MVP_top60_DNA_seed | EGFR | -2.264811 | BRCA | ZR-75-30 | Afatinib | MVP_top60_DNA+RNA_seed | SRC | 1.278701 | BRCA | ZR-75-30 | HG-6-71-01 |
| MVP_top60_DNA_seed | EGFR | -2.262981 | BRCA | HDQ-P1 | Osimertinib | MVP_top60_DNA+RNA_seed | HDAC1 | 1.279063 | BRCA | MDA-MB-415 | CAY10603 |
| MVP_top60_DNA_seed | ESR1 | -2.250288 | BRCA | EFM-19 | GDC0810 | MVP_top60_DNA+RNA_seed | SRC | 1.281685 | BRCA | COLO-824 | Bosutinib |
| MVP_top60_DNA_seed | AKT1 | -2.214992 | BRCA | MDA-MB-415 | Afuresertib | MVP_top60_DNA+RNA_seed | AKT1 | 1.285223 | BRCA | BT-483 | AKT inhibitor VIII |
| MVP_top60_DNA_seed | PIK3CA | -2.200339 | BRCA | UACC-893 | Alpelisib | MVP_top60_DNA+RNA_seed | EGFR | 1.285272 | BRCA | HCC1569 | Gefitinib |

| | | | | | | | | | | | |
|---|---|---|---|---|---|---|---|---|---|---|---|
| MVP_top60_DNA_seed | PIK3CA | -2.198675 | BRCA | MDA-MB-453 | Alpelisib | MVP_top60_DNA+RNA_seed | MAPK1 | 1.28531 | BRCA | BT-20 | VX-11e |
| MVP_top60_DNA_seed | AKT1 | -2.181131 | BRCA | UACC-893 | AZD5363 | MVP_top60_DNA+RNA_seed | ATM | 1.286046 | BRCA | ZR-75-30 | KU-60019 |
| MVP_top60_DNA_seed | AKT1 | -2.168426 | BRCA | UACC-893 | Uprosertib | MVP_top60_DNA+RNA_seed | AKT1 | 1.29213 | BRCA | MCF7 | AKT inhibitor VIII |
| MVP_top60_DNA_seed | AKT1 | -2.162809 | BRCA | HCC1569 | GSK2110183B | MVP_top60_DNA+RNA_seed | ATM | 1.292581 | BRCA | BT-474 | torin2 |
| MVP_top60_DNA_seed | AKT1 | -2.153509 | BRCA | MDA-MB-453 | GSK2110183B | MVP_top60_DNA+RNA_seed | HSP90AA1 | 1.294714 | BRCA | HCC1500 | Elesclomol |
| MVP_top60_DNA_seed | ERBB2 | -2.126294 | BRCA | HCC2157 | Lapatinib | MVP_top60_DNA+RNA_seed | HSP90AB1 | 1.294714 | BRCA | HCC1500 | Elesclomol |
| MVP_top60_DNA_seed | EGFR | -2.126294 | BRCA | HCC2157 | Lapatinib | MVP_top60_DNA+RNA_seed | SRC | 1.29646 | BRCA | HCC1428 | NVP-BHG712 |
| MVP_top60_DNA_seed | AKT1 | -2.118229 | BRCA | UACC-893 | MK-2206 | MVP_top60_DNA+RNA_seed | MAPK1 | 1.296682 | BRCA | MDA-MB-330 | VX-11e |
| MVP_top60_DNA_seed | AKT1 | -2.115307 | BRCA | MDA-MB-415 | Uprosertib | MVP_top60_DNA+RNA_seed | ESR1 | 1.298138 | BRCA | CAL-85-1 | Tamoxifen |
| MVP_top60_DNA_seed | ERBB2 | -2.114903 | BRCA | ZR-75-30 | Lapatinib | MVP_top60_DNA+RNA_seed | MAPK3 | 1.29992 | BRCA | CAMA-1 | FR-180204 |
| MVP_top60_DNA_seed | EGFR | -2.114903 | BRCA | ZR-75-30 | Lapatinib | MVP_top60_DNA+RNA_seed | MAPK1 | 1.29992 | BRCA | CAMA-1 | FR-180204 |
| MVP_top60_DNA_seed | ERBB2 | -2.112493 | BRCA | AU565 | Afatinib | MVP_top60_DNA+RNA_seed | ESR1 | 1.305586 | BRCA | HCC1143 | Tamoxifen |
| MVP_top60_DNA_seed | EGFR | -2.112493 | BRCA | AU565 | Afatinib | MVP_top60_DNA+RNA_seed | AR | 1.307055 | BRCA | MDA-MB-175-VII | AZD3514 |
| MVP_top60_DNA_seed | PIK3CA | -2.109527 | BRCA | HCC70 | AZD8186 | MVP_top60_DNA+RNA_seed | CDK2 | 1.309309 | BRCA | EFM-19 | AZD5438 |
| MVP_top60_DNA_seed | AKT1 | -2.10873 | BRCA | CAL-148 | MK-2206 | MVP_top60_DNA+RNA_seed | PLK1 | 1.309873 | BRCA | CAL-85-1 | PLK_6522 |
| MVP_top60_DNA_seed | AKT1 | -2.090514 | BRCA | MDA-MB-361 | Uprosertib | MVP_top60_DNA+RNA_seed | CDK2 | 1.312185 | BRCA | Hs-578-T | PHA-793887 |
| MVP_top60_DNA_seed | AKT1 | -2.087608 | BRCA | MDA-MB-415 | GSK2110183B | MVP_top60_DNA+RNA_seed | MAPK1 | 1.312262 | BRCA | HDQ-P1 | VX-11e |
| MVP_top60_DNA_seed | EGFR | -2.086603 | BRCA | AU565 | Osimertinib | MVP_top60_DNA+RNA_seed | CDK1 | 1.314105 | BRCA | MDA-MB-415 | RO-3306 |
| MVP_top60_DNA_seed | AKT1 | -2.085156 | BRCA | MDA-MB-453 | Uprosertib | MVP_top60_DNA+RNA_seed | EGFR | 1.315048 | BRCA | MFM-223 | Gefitinib |
| MVP_top60_DNA_seed | MDM2 | -2.073491 | BRCA | DU-4475 | Nutlin-3a (-) | MVP_top60_DNA+RNA_seed | SRC | 1.316657 | BRCA | UACC-893 | HG-6-71-01 |
| MVP_top60_DNA_seed | AKT1 | -2.06845 | BRCA | YMB-1-E | AZD5363 | MVP_top60_DNA+RNA_seed | AKT1 | 1.316765 | BRCA | HCC1428 | BAY AKT1 |
| MVP_top60_DNA_seed | ERBB2 | -2.058312 | BRCA | UACC-893 | Sapitinib | MVP_top60_DNA+RNA_seed | PIK3CA | 1.317085 | BRCA | MDA-MB-157 | AZD8835 |
| MVP_top60_DNA_seed | EGFR | -2.058312 | BRCA | UACC-893 | Sapitinib | MVP_top60_DNA+RNA_seed | HSP90AA1 | 1.317554 | BRCA | MDA-MB-468 | Tanespimycin |
| MVP_top60_DNA_seed | ERBB3 | -2.058312 | BRCA | UACC-893 | Sapitinib | MVP_top60_DNA+RNA_seed | HSP90AB1 | 1.317554 | BRCA | MDA-MB-468 | Tanespimycin |
| MVP_top60_DNA_seed | AKT1 | -2.053109 | BRCA | HCC2157 | Ipatasertib | MVP_top60_DNA+RNA_seed | HDAC1 | 1.318724 | BRCA | BT-20 | Dacinostat |
| MVP_top60_DNA_seed | AKT1 | -2.046083 | BRCA | UACC-893 | Afuresertib | MVP_top60_DNA+RNA_seed | SRC | 1.319888 | BRCA | HCC1954 | NVP-BHG712 |
| MVP_top60_DNA_seed | HSP90AA1 | -2.032451 | BRCA | MDA-MB-453 | Tanespimycin | MVP_top60_DNA+RNA_seed | HDAC1 | 1.319932 | BRCA | HCC1395 | JQ12 |
| MVP_top60_DNA_seed | HSP90AB1 | -2.032451 | BRCA | MDA-MB-453 | Tanespimycin | MVP_top60_DNA+RNA_seed | HDAC2 | 1.319932 | BRCA | HCC1395 | JQ12 |
| MVP_top60_DNA_seed | TP53 | -2.017852 | BRCA | MRK-nu-1 | MIRA-1 | MVP_top60_DNA+RNA_seed | PIK3CA | 1.320549 | BRCA | HCC1428 | AZD8835 |
| MVP_top60_DNA_seed | ERBB2 | -2.007439 | BRCA | HDQ-P1 | Sapitinib | MVP_top60_DNA+RNA_seed | ERBB2 | 1.321515 | BRCA | MDA-MB-415 | CP724714 |
| MVP_top60_DNA_seed | EGFR | -2.007439 | BRCA | HDQ-P1 | Sapitinib | MVP_top60_DNA+RNA_seed | EGFR | 1.322106 | BRCA | EFM-19 | Cetuximab |
| MVP_top60_DNA_seed | ERBB3 | -2.007439 | BRCA | HDQ-P1 | Sapitinib | MVP_top60_DNA+RNA_seed | ESR1 | 1.324028 | BRCA | MDA-MB-175-VII | Fulvestrant |
| MVP_top60_DNA_seed | AKT1 | -1.996394 | BRCA | MDA-MB-415 | MK-2206 | MVP_top60_DNA+RNA_seed | AKT1 | 1.325025 | BRCA | CAL-85-1 | MK-2206 |
| MVP_top60_DNA_seed | AKT1 | -1.996282 | BRCA | CAMA-1 | AZD5363 | MVP_top60_DNA+RNA_seed | SRC | 1.329337 | BRCA | MFM-223 | Saracatinib |
| MVP_top60_DNA_seed | ATM | -1.990948 | BRCA | MRK-nu-1 | KU-55933 | MVP_top60_DNA+RNA_seed | CDK2 | 1.330978 | BRCA | UACC-812 | Seliciclib |
| MVP_top60_DNA_seed | AKT1 | -1.988244 | BRCA | HCC1569 | Uprosertib | MVP_top60_DNA+RNA_seed | MAPK14 | 1.335513 | BRCA | HCC1395 | TAK-715 |
| MVP_top60_DNA_seed | PIK3CA | -1.979001 | BRCA | HCC2157 | AZD8186 | MVP_top60_DNA+RNA_seed | HDAC1 | 1.335948 | BRCA | HCC1395 | CAY10603 |
| MVP_top60_DNA_seed | AKT1 | -1.97538 | BRCA | YMB-1-E | Afuresertib | MVP_top60_DNA+RNA_seed | MDM2 | 1.337037 | BRCA | HDQ-P1 | Serdemetan |
| MVP_top60_DNA_seed | ERBB2 | -1.965332 | BRCA | BT-474 | Lapatinib | MVP_top60_DNA+RNA_seed | PLK1 | 1.340377 | BRCA | MFM-223 | BI-2536 |
| MVP_top60_DNA_seed | EGFR | -1.965332 | BRCA | BT-474 | Lapatinib | MVP_top60_DNA+RNA_seed | PIK3CA | 1.344327 | BRCA | MDA-MB-436 | AZD8835 |
| MVP_top60_DNA_seed | AKT1 | -1.963085 | BRCA | YMB-1-E | Uprosertib | MVP_top60_DNA+RNA_seed | MDM2 | 1.346095 | BRCA | ZR-75-30 | Nutlin-3a (-) |

| | | | | | | | | | | | |
|---|---|---|---|---|---|---|---|---|---|---|---|
| MVP_top60_DNA_seed | AKT1 | -1.95807 | BRCA | MDA-MB-453 | MK-2206 | MVP_top60_DNA+RNA_seed | AR | 1.347654 | BRCA | MDA-MB-415 | Bicalutamide |
| MVP_top60_DNA_seed | ESR1 | -1.956708 | BRCA | T47D | GDC0810 | MVP_top60_DNA+RNA_seed | SRC | 1.349832 | BRCA | HCC1569 | Bosutinib |
| MVP_top60_DNA_seed | AKT1 | -1.942434 | BRCA | UACC-893 | Ipatasertib | MVP_top60_DNA+RNA_seed | ATM | 1.350035 | BRCA | HDQ-P1 | KU-60019 |
| MVP_top60_DNA_seed | EGFR | -1.938209 | BRCA | MDA-MB-175-VII | Erlotinib | MVP_top60_DNA+RNA_seed | CDK1 | 1.351208 | BRCA | BT-474 | RO-3306 |
| MVP_top60_DNA_seed | EGFR | -1.935551 | BRCA | HCC2157 | Gefitinib | MVP_top60_DNA+RNA_seed | CDK2 | 1.353857 | BRCA | EFM-192A | PHA-793887 |
| MVP_top60_DNA_seed | EGFR | -1.932837 | BRCA | HDQ-P1 | Gefitinib | MVP_top60_DNA+RNA_seed | PLK1 | 1.357253 | BRCA | BT-474 | GW843682X |
| MVP_top60_DNA_seed | AKT1 | -1.930333 | BRCA | HCC1954 | GSK2110183B | MVP_top60_DNA+RNA_seed | HDAC1 | 1.358523 | BRCA | MFM-223 | Dacinostat |
| MVP_top60_DNA_seed | ATM | -1.924923 | BRCA | Hs-578-T | KU-55933 | MVP_top60_DNA+RNA_seed | MAPK3 | 1.359593 | BRCA | HCC1395 | FR-180204 |
| MVP_top60_DNA_seed | MDM2 | -1.919876 | BRCA | CAL-148 | Nutlin-3a (-) | MVP_top60_DNA+RNA_seed | MAPK1 | 1.359593 | BRCA | HCC1395 | FR-180204 |
| MVP_top60_DNA_seed | HSP90AA1 | -1.914909 | BRCA | MDA-MB-361 | Tanespimycin | MVP_top60_DNA+RNA_seed | EGFR | 1.361535 | BRCA | T47D | Cetuximab |
| MVP_top60_DNA_seed | HSP90AB1 | -1.914909 | BRCA | MDA-MB-361 | Tanespimycin | MVP_top60_DNA+RNA_seed | CDK2 | 1.362343 | BRCA | UACC-812 | PHA-793887 |
| MVP_top60_DNA_seed | HDAC1 | -1.913332 | BRCA | DU-4475 | Entinostat | MVP_top60_DNA+RNA_seed | AKT1 | 1.363286 | BRCA | MRK-nu-1 | AKT inhibitor VIII |
| MVP_top60_DNA_seed | EP300 | -1.912318 | BRCA | CAMA-1 | SGC-CBP30 | MVP_top60_DNA+RNA_seed | MAPK3 | 1.363836 | BRCA | MDA-MB-436 | FR-180204 |
| MVP_top60_DNA_seed | CREBBP | -1.912318 | BRCA | CAMA-1 | SGC-CBP30 | MVP_top60_DNA+RNA_seed | MAPK1 | 1.363836 | BRCA | MDA-MB-436 | FR-180204 |
| MVP_top60_DNA_seed | EGFR | -1.906474 | BRCA | OCUB-M | Osimertinib | MVP_top60_DNA+RNA_seed | EGFR | 1.367886 | BRCA | HCC2157 | Pelitinib |
| MVP_top60_DNA_seed | HSP90AA1 | -1.896893 | BRCA | MDA-MB-330 | Tanespimycin | MVP_top60_DNA+RNA_seed | HDAC1 | 1.373532 | BRCA | MFM-223 | Parthenolide |
| MVP_top60_DNA_seed | HSP90AB1 | -1.896893 | BRCA | MDA-MB-330 | Tanespimycin | MVP_top60_DNA+RNA_seed | HDAC1 | 1.374431 | BRCA | BT-483 | CAY10603 |
| MVP_top60_DNA_seed | MAPK3 | -1.894407 | BRCA | MRK-nu-1 | Ulixertinib | MVP_top60_DNA+RNA_seed | CDK1 | 1.381748 | BRCA | HDQ-P1 | RO-3306 |
| MVP_top60_DNA_seed | MAPK1 | -1.894407 | BRCA | MRK-nu-1 | Ulixertinib | MVP_top60_DNA+RNA_seed | AR | 1.384759 | BRCA | EFM-19 | Bicalutamide |
| MVP_top60_DNA_seed | AKT1 | -1.893713 | BRCA | HCC1569 | Afuresertib | MVP_top60_DNA+RNA_seed | SRC | 1.385285 | BRCA | CAL-85-1 | Bosutinib |
| MVP_top60_DNA_seed | AKT1 | -1.890909 | BRCA | CAL-148 | AT13148 | MVP_top60_DNA+RNA_seed | PIK3CA | 1.386546 | BRCA | HCC1428 | PI3Ka_4409 |
| MVP_top60_DNA_seed | PIK3CA | -1.88702 | BRCA | MDA-MB-330 | AZD8186 | MVP_top60_DNA+RNA_seed | MAPK14 | 1.389415 | BRCA | MDA-MB-157 | TAK-715 |
| MVP_top60_DNA_seed | PIK3CA | -1.882454 | BRCA | CAMA-1 | AZD8186 | MVP_top60_DNA+RNA_seed | EGFR | 1.392698 | BRCA | YMB-1-E | Pelitinib |
| MVP_top60_DNA_seed | AKT1 | -1.877085 | BRCA | BT-20 | GSK2110183B | MVP_top60_DNA+RNA_seed | HDAC1 | 1.392781 | BRCA | MDA-MB-415 | Tubastatin A |
| MVP_top60_DNA_seed | AKT1 | -1.869376 | BRCA | HCC70 | Ipatasertib | MVP_top60_DNA+RNA_seed | AR | 1.394125 | BRCA | HCC1419 | AZD3514 |
| MVP_top60_DNA_seed | AKT1 | -1.868896 | BRCA | HCC2157 | Uprosertib | MVP_top60_DNA+RNA_seed | EP300 | 1.395697 | BRCA | MDA-MB-436 | I-CBP112 |
| MVP_top60_DNA_seed | AKT1 | -1.850647 | BRCA | HCC1569 | Ipatasertib | MVP_top60_DNA+RNA_seed | AKT1 | 1.396284 | BRCA | BT-483 | GSK690693 |
| MVP_top60_DNA_seed | ERBB2 | -1.837248 | BRCA | AU565 | Sapitinib | MVP_top60_DNA+RNA_seed | MAPK1 | 1.396348 | BRCA | HCC1419 | VX-11e |
| MVP_top60_DNA_seed | EGFR | -1.837248 | BRCA | AU565 | Sapitinib | MVP_top60_DNA+RNA_seed | CDK2 | 1.39664 | BRCA | COLO-824 | Seliciclib |
| MVP_top60_DNA_seed | ERBB3 | -1.837248 | BRCA | AU565 | Sapitinib | MVP_top60_DNA+RNA_seed | EGFR | 1.39735 | BRCA | MCF7 | CUDC-101 |
| MVP_top60_DNA_seed | HDAC1 | -1.822743 | BRCA | CAL-148 | Entinostat | MVP_top60_DNA+RNA_seed | ERBB2 | 1.39735 | BRCA | MCF7 | CUDC-101 |
| MVP_top60_DNA_seed | ERBB2 | -1.813625 | BRCA | MDA-MB-330 | Lapatinib | MVP_top60_DNA+RNA_seed | ATM | 1.397527 | BRCA | HCC1954 | KU-55933 |
| MVP_top60_DNA_seed | EGFR | -1.813625 | BRCA | MDA-MB-330 | Lapatinib | MVP_top60_DNA+RNA_seed | EGFR | 1.398585 | BRCA | MDA-MB-157 | Pelitinib |
| MVP_top60_DNA_seed | ESR1 | -1.79809 | BRCA | HCC1419 | GDC0810 | MVP_top60_DNA+RNA_seed | MAPK3 | 1.405187 | BRCA | EFM-19 | FR-180204 |
| MVP_top60_DNA_seed | EGFR | -1.783818 | BRCA | EFM-192A | Osimertinib | MVP_top60_DNA+RNA_seed | MAPK1 | 1.405187 | BRCA | EFM-19 | FR-180204 |
| MVP_top60_DNA_seed | EGFR | -1.777703 | BRCA | HDQ-P1 | Erlotinib | MVP_top60_DNA+RNA_seed | EGFR | 1.405752 | BRCA | Hs-578-T | Pelitinib |
| MVP_top60_DNA_seed | AKT1 | -1.768701 | BRCA | HCC70 | AZD5363 | MVP_top60_DNA+RNA_seed | SRC | 1.407518 | BRCA | T47D | Bosutinib |
| MVP_top60_DNA_seed | ESR1 | -1.762729 | BRCA | HCC2157 | Tamoxifen | MVP_top60_DNA+RNA_seed | PLK1 | 1.415635 | BRCA | HCC2157 | GW843682X |
| MVP_top60_DNA_seed | AKT1 | -1.756361 | BRCA | HCC70 | Uprosertib | MVP_top60_DNA+RNA_seed | HDAC1 | 1.418512 | BRCA | CAL-51 | CAY10603 |
| MVP_top60_DNA_seed | AKT1 | -1.744543 | BRCA | MDA-MB-415 | Uprosertib | MVP_top60_DNA+RNA_seed | ESR1 | 1.423801 | BRCA | MDA-MB-436 | Tamoxifen |
| MVP_top60_DNA_seed | CDK2 | -1.742885 | BRCA | MDA-MB-361 | MK-8776 | MVP_top60_DNA+RNA_seed | HDAC1 | 1.426478 | BRCA | BT-474 | Parthenolide |

| | | | | | | | | | | | |
|---|---|---|---|---|---|---|---|---|---|---|---|
| MVP_top60_DNA_seed | ERBB2 | -1.734971 | BRCA | BT-474 | Afatinib | MVP_top60_DNA+RNA_seed | HSP90AA1 | 1.42892 | BRCA | HCC1428 | SNX-2112 |
| MVP_top60_DNA_seed | EGFR | -1.734971 | BRCA | BT-474 | Afatinib | MVP_top60_DNA+RNA_seed | HSP90AB1 | 1.42892 | BRCA | HCC1428 | SNX-2112 |
| MVP_top60_DNA_seed | AKT1 | -1.733572 | BRCA | AU565 | AT13148 | MVP_top60_DNA+RNA_seed | CDK1 | 1.430098 | BRCA | CAL-85-1 | RO-3306 |
| MVP_top60_DNA_seed | HDAC1 | -1.730747 | BRCA | HCC2157 | Entinostat | MVP_top60_DNA+RNA_seed | EGFR | 1.433212 | BRCA | MRK-nu-1 | Pelitinib |
| MVP_top60_DNA_seed | PIK3CA | -1.729419 | BRCA | CAL-148 | Alpelisib | MVP_top60_DNA+RNA_seed | MAPK14 | 1.434403 | BRCA | MRK-nu-1 | TAK-715 |
| MVP_top60_DNA_seed | AKT1 | -1.723778 | BRCA | HCC70 | MK-2206 | MVP_top60_DNA+RNA_seed | CDK2 | 1.436536 | BRCA | HCC2218 | AZD5438 |
| MVP_top60_DNA_seed | ERBB2 | -1.706325 | BRCA | OCUB-M | Afatinib | MVP_top60_DNA+RNA_seed | SRC | 1.443748 | BRCA | MDA-MB-415 | NVP-BHG712 |
| MVP_top60_DNA_seed | EGFR | -1.706325 | BRCA | OCUB-M | Afatinib | MVP_top60_DNA+RNA_seed | AR | 1.449382 | BRCA | MDA-MB-436 | Bicalutamide |
| MVP_top60_DNA_seed | AKT1 | -1.705594 | BRCA | HCC2157 | MK-2206 | MVP_top60_DNA+RNA_seed | PIK3CA | 1.450804 | BRCA | ZR-75-30 | PI3Ka_4409 |
| MVP_top60_DNA_seed | EP300 | -1.70371 | BRCA | CAL-148 | CPI-637 | MVP_top60_DNA+RNA_seed | AR | 1.451524 | BRCA | MFM-223 | AZD3514 |
| MVP_top60_DNA_seed | CDK1 | -1.692986 | BRCA | MRK-nu-1 | RO-3306 | MVP_top60_DNA+RNA_seed | MAPK1 | 1.45488 | BRCA | EFM-19 | VX-11e |
| MVP_top60_DNA_seed | AKT1 | -1.689145 | BRCA | AU565 | Uprosertib | MVP_top60_DNA+RNA_seed | AR | 1.460438 | BRCA | ZR-75-30 | Bicalutamide |
| MVP_top60_DNA_seed | AKT1 | -1.681967 | BRCA | MDA-MB-175-VII | Afuresertib | MVP_top60_DNA+RNA_seed | CDK2 | 1.464954 | BRCA | BT-474 | Seliciclib |
| MVP_top60_DNA_seed | PIK3CA | -1.675579 | BRCA | BT-20 | Alpelisib | MVP_top60_DNA+RNA_seed | SRC | 1.46543 | BRCA | HDQ-P1 | NVP-BHG712 |
| MVP_top60_DNA_seed | PIK3CA | -1.673389 | BRCA | OCUB-M | GNE-317 | MVP_top60_DNA+RNA_seed | PLK1 | 1.465972 | BRCA | HCC1599 | PLK_6522 |
| MVP_top60_DNA_seed | EGFR | -1.655282 | BRCA | AU565 | Gefitinib | MVP_top60_DNA+RNA_seed | ESR1 | 1.470021 | BRCA | HCC202 | Tamoxifen |
| MVP_top60_DNA_seed | MAPK3 | -1.650711 | BRCA | MDA-MB-157 | Ulixertinib | MVP_top60_DNA+RNA_seed | ESR1 | 1.471784 | BRCA | HCC1419 | Tamoxifen |
| MVP_top60_DNA_seed | MAPK1 | -1.650711 | BRCA | MDA-MB-157 | Ulixertinib | MVP_top60_DNA+RNA_seed | HSP90AA1 | 1.472149 | BRCA | Hs-578-T | SNX-2112 |
| MVP_top60_DNA_seed | CDK1 | -1.642671 | BRCA | DU-4475 | RO-3306 | MVP_top60_DNA+RNA_seed | HSP90AB1 | 1.472149 | BRCA | Hs-578-T | SNX-2112 |
| MVP_top60_DNA_seed | AKT1 | -1.636872 | BRCA | HCC1569 | AZD5363 | MVP_top60_DNA+RNA_seed | MDM2 | 1.477896 | BRCA | MDA-MB-415 | Nutlin-3a (-) |
| MVP_top60_DNA_seed | AKT1 | -1.629614 | BRCA | MFM-223 | Afuresertib | MVP_top60_DNA+RNA_seed | AKT1 | 1.479658 | BRCA | MDA-MB-157 | BAY AKT1 |
| MVP_top60_DNA_seed | ERBB2 | -1.622725 | BRCA | EFM-192A | Lapatinib | MVP_top60_DNA+RNA_seed | PIK3CA | 1.479792 | BRCA | HCC1937 | AZD8835 |
| MVP_top60_DNA_seed | EGFR | -1.622725 | BRCA | EFM-192A | Lapatinib | MVP_top60_DNA+RNA_seed | PLK1 | 1.482361 | BRCA | CAMA-1 | PLK_6522 |
| MVP_top60_DNA_seed | AKT1 | -1.61593 | BRCA | MDA-MB-415 | AT13148 | MVP_top60_DNA+RNA_seed | PIK3CA | 1.482499 | BRCA | MDA-MB-157 | PI3Ka_4409 |
| MVP_top60_DNA_seed | AKT1 | -1.609161 | BRCA | CAMA-1 | GSK2110183B | MVP_top60_DNA+RNA_seed | HSP90AA1 | 1.483976 | BRCA | HCC1395 | SNX-2112 |
| MVP_top60_DNA_seed | PIK3CA | -1.607008 | BRCA | EFM-19 | Alpelisib | MVP_top60_DNA+RNA_seed | HSP90AB1 | 1.483976 | BRCA | HCC1395 | SNX-2112 |
| MVP_top60_DNA_seed | ERBB2 | -1.597645 | BRCA | BT-474 | Sapitinib | MVP_top60_DNA+RNA_seed | ATM | 1.491857 | BRCA | HCC2218 | KU-55933 |
| MVP_top60_DNA_seed | EGFR | -1.597645 | BRCA | BT-474 | Sapitinib | MVP_top60_DNA+RNA_seed | MAPK1 | 1.495213 | BRCA | MDA-MB-436 | VX-11e |
| MVP_top60_DNA_seed | ERBB3 | -1.597645 | BRCA | BT-474 | Sapitinib | MVP_top60_DNA+RNA_seed | AKT1 | 1.500272 | BRCA | BT-483 | MK-2206 |
| MVP_top60_DNA_seed | AKT1 | -1.585314 | BRCA | MDA-MB-453 | Uprosertib | MVP_top60_DNA+RNA_seed | AKT1 | 1.500373 | BRCA | HDQ-P1 | BAY AKT1 |
| MVP_top60_DNA_seed | EGFR | -1.584241 | BRCA | ZR-75-30 | Osimertinib | MVP_top60_DNA+RNA_seed | HDAC1 | 1.502827 | BRCA | HCC1937 | JQ12 |
| MVP_top60_DNA_seed | CDK2 | -1.574131 | BRCA | HCC1395 | MK-8776 | MVP_top60_DNA+RNA_seed | HDAC2 | 1.502827 | BRCA | HCC1937 | JQ12 |
| MVP_top60_DNA_seed | MAPK3 | -1.573528 | BRCA | CAL-85-1 | Ulixertinib | MVP_top60_DNA+RNA_seed | MDM2 | 1.508596 | BRCA | HCC202 | Nutlin-3a (-) |
| MVP_top60_DNA_seed | MAPK1 | -1.573528 | BRCA | CAL-85-1 | Ulixertinib | MVP_top60_DNA+RNA_seed | HSP90AA1 | 1.515681 | BRCA | EFM-19 | SNX-2112 |
| MVP_top60_DNA_seed | AKT1 | -1.569484 | BRCA | MDA-MB-175-VII | Uprosertib | MVP_top60_DNA+RNA_seed | HSP90AB1 | 1.515681 | BRCA | EFM-19 | SNX-2112 |
| MVP_top60_DNA_seed | HDAC1 | -1.559249 | BRCA | MDA-MB-330 | Romidepsin | MVP_top60_DNA+RNA_seed | EGFR | 1.517091 | BRCA | BT-483 | Gefitinib |
| MVP_top60_DNA_seed | HDAC2 | -1.559249 | BRCA | MDA-MB-330 | Romidepsin | MVP_top60_DNA+RNA_seed | MAPK1 | 1.522008 | BRCA | MDA-MB-415 | VX-11e |
| MVP_top60_DNA_seed | CDK2 | -1.558595 | BRCA | DU-4475 | AZD5438 | MVP_top60_DNA+RNA_seed | EGFR | 1.523588 | BRCA | HCC2218 | Cetuximab |
| MVP_top60_DNA_seed | MDM2 | -1.551871 | BRCA | DU-4475 | Serdemetan | MVP_top60_DNA+RNA_seed | ATM | 1.527524 | BRCA | BT-20 | KU-55933 |
| MVP_top60_DNA_seed | AKT1 | -1.551098 | BRCA | MDA-MB-361 | GSK2110183B | MVP_top60_DNA+RNA_seed | EGFR | 1.531003 | BRCA | MDA-MB-415 | CUDC-101 |
| MVP_top60_DNA_seed | PIK3CA | -1.544994 | BRCA | MRK-nu-1 | GNE-317 | MVP_top60_DNA+RNA_seed | ERBB2 | 1.531003 | BRCA | MDA-MB-415 | CUDC-101 |

| | | | | | | | | | | | |
|---|---|---|---|---|---|---|---|---|---|---|---|
| MVP_top60_DNA_seed | ATM | -1.538073 | BRCA | EVSA-T | KU-55933 | MVP_top60_DNA+RNA_seed | EGFR | 1.531241 | BRCA | HCC1395 | CUDC-101 |
| MVP_top60_DNA_seed | AKT1 | -1.524496 | BRCA | MDA-MB-175-VII | Ipatasertib | MVP_top60_DNA+RNA_seed | ERBB2 | 1.531241 | BRCA | HCC1395 | CUDC-101 |
| MVP_top60_DNA_seed | AKT1 | -1.517368 | BRCA | CAMA-1 | Uprosertib | MVP_top60_DNA+RNA_seed | HDAC1 | 1.542562 | BRCA | CAL-51 | AR-42 |
| MVP_top60_DNA_seed | EP300 | -1.517151 | BRCA | MDA-MB-453 | CPI-637 | MVP_top60_DNA+RNA_seed | EP300 | 1.543025 | BRCA | HCC1143 | I-CBP112 |
| MVP_top60_DNA_seed | ATM | -1.516846 | BRCA | MDA-MB-415 | KU-55933 | MVP_top60_DNA+RNA_seed | HSP90AA1 | 1.543495 | BRCA | HCC1187 | Tanespimycin |
| MVP_top60_DNA_seed | ATM | -1.510452 | BRCA | DU-4475 | KU-55933 | MVP_top60_DNA+RNA_seed | HSP90AB1 | 1.543495 | BRCA | HCC1187 | Tanespimycin |
| MVP_top60_DNA_seed | ERBB2 | -1.506965 | BRCA | OCUB-M | Sapitinib | MVP_top60_DNA+RNA_seed | HDAC1 | 1.544541 | BRCA | MCF7 | AR-42 |
| MVP_top60_DNA_seed | EGFR | -1.506965 | BRCA | OCUB-M | Sapitinib | MVP_top60_DNA+RNA_seed | EGFR | 1.544867 | BRCA | EFM-19 | Pelitinib |
| MVP_top60_DNA_seed | ERBB3 | -1.506965 | BRCA | OCUB-M | Sapitinib | MVP_top60_DNA+RNA_seed | HSP90AA1 | 1.545609 | BRCA | EFM-19 | CCT-018159 |
| MVP_top60_DNA_seed | EGFR | -1.50638 | BRCA | MDA-MB-330 | AZD3759 | MVP_top60_DNA+RNA_seed | HSP90AB1 | 1.545609 | BRCA | EFM-19 | CCT-018159 |
| MVP_top60_DNA_seed | ERBB2 | -1.504361 | BRCA | HCC2218 | Sapitinib | MVP_top60_DNA+RNA_seed | AR | 1.549658 | BRCA | BT-474 | Bicalutamide |
| MVP_top60_DNA_seed | EGFR | -1.504361 | BRCA | HCC2218 | Sapitinib | MVP_top60_DNA+RNA_seed | CDK1 | 1.550917 | BRCA | MFM-223 | RO-3306 |
| MVP_top60_DNA_seed | ERBB3 | -1.504361 | BRCA | HCC2218 | Sapitinib | MVP_top60_DNA+RNA_seed | AKT1 | 1.556124 | BRCA | UACC-812 | AKT inhibitor VIII |
| MVP_top60_DNA_seed | HSP90AA1 | -1.504166 | BRCA | COLO-824 | Tanespimycin | MVP_top60_DNA+RNA_seed | EGFR | 1.557459 | BRCA | CAL-51 | CUDC-101 |
| MVP_top60_DNA_seed | HSP90AB1 | -1.504166 | BRCA | COLO-824 | Tanespimycin | MVP_top60_DNA+RNA_seed | ERBB2 | 1.557459 | BRCA | CAL-51 | CUDC-101 |
| MVP_top60_DNA_seed | ESR1 | -1.503186 | BRCA | COLO-824 | Tamoxifen | MVP_top60_DNA+RNA_seed | AR | 1.569871 | BRCA | HCC1954 | AZD3514 |
| MVP_top60_DNA_seed | AKT1 | -1.502818 | BRCA | MDA-MB-175-VII | MK-2206 | MVP_top60_DNA+RNA_seed | HDAC1 | 1.572447 | BRCA | CAL-51 | Belinostat |
| MVP_top60_DNA_seed | AKT1 | -1.501786 | BRCA | MFM-223 | Uprosertib | MVP_top60_DNA+RNA_seed | CDK1 | 1.574741 | BRCA | HCC2218 | RO-3306 |
| MVP_top60_DNA_seed | ERBB2 | -1.486205 | BRCA | OCUB-M | Lapatinib | MVP_top60_DNA+RNA_seed | CDK2 | 1.575152 | BRCA | HCC1428 | AZD5438 |
| MVP_top60_DNA_seed | EGFR | -1.486205 | BRCA | OCUB-M | Lapatinib | MVP_top60_DNA+RNA_seed | PLK1 | 1.575889 | BRCA | HCC1428 | PLK_6522 |
| MVP_top60_DNA_seed | AKT1 | -1.476894 | BRCA | MFM-223 | AZD5363 | MVP_top60_DNA+RNA_seed | MAPK14 | 1.577486 | BRCA | Hs-578-T | TAK-715 |
| MVP_top60_DNA_seed | MDM2 | -1.474321 | BRCA | MRK-nu-1 | Nutlin-3a (-) | MVP_top60_DNA+RNA_seed | AKT1 | 1.57811 | BRCA | MRK-nu-1 | BAY AKT1 |
| MVP_top60_DNA_seed | HSP90AA1 | -1.474037 | BRCA | CAL-51 | Luminespib | MVP_top60_DNA+RNA_seed | HSP90AA1 | 1.579625 | BRCA | MDA-MB-157 | Luminespib |
| MVP_top60_DNA_seed | HSP90AB1 | -1.474037 | BRCA | CAL-51 | Luminespib | MVP_top60_DNA+RNA_seed | HSP90AB1 | 1.579625 | BRCA | MDA-MB-157 | Luminespib |
| MVP_top60_DNA_seed | HDAC1 | -1.47315 | BRCA | EVSA-T | Entinostat | MVP_top60_DNA+RNA_seed | EGFR | 1.580409 | BRCA | BT-483 | Afatinib |
| MVP_top60_DNA_seed | HDAC1 | -1.472168 | BRCA | HCC1599 | PCI-34051 | MVP_top60_DNA+RNA_seed | ERBB2 | 1.580409 | BRCA | BT-483 | Afatinib |
| MVP_top60_DNA_seed | HSP90AA1 | -1.469543 | BRCA | DU-4475 | Tanespimycin | MVP_top60_DNA+RNA_seed | AKT1 | 1.58138 | BRCA | HCC1500 | BAY AKT1 |
| MVP_top60_DNA_seed | HSP90AB1 | -1.469543 | BRCA | DU-4475 | Tanespimycin | MVP_top60_DNA+RNA_seed | ATM | 1.58686 | BRCA | HCC2157 | CP466722 |
| MVP_top60_DNA_seed | HSP90AA1 | -1.465141 | BRCA | MDA-MB-330 | CCT-018159 | MVP_top60_DNA+RNA_seed | CDK1 | 1.587102 | BRCA | ZR-75-30 | RO-3306 |
| MVP_top60_DNA_seed | HSP90AB1 | -1.465141 | BRCA | MDA-MB-330 | CCT-018159 | MVP_top60_DNA+RNA_seed | HSP90AA1 | 1.589529 | BRCA | HCC1395 | CCT-018159 |
| MVP_top60_DNA_seed | AKT1 | -1.463847 | BRCA | MFM-223 | Ipatasertib | MVP_top60_DNA+RNA_seed | HSP90AB1 | 1.589529 | BRCA | HCC1395 | CCT-018159 |
| MVP_top60_DNA_seed | AKT1 | -1.459284 | BRCA | CAMA-1 | Uprosertib | MVP_top60_DNA+RNA_seed | CDK2 | 1.589824 | BRCA | MDA-MB-415 | PHA-793887 |
| MVP_top60_DNA_seed | CDK1 | -1.454586 | BRCA | OCUB-M | RO-3306 | MVP_top60_DNA+RNA_seed | HDAC1 | 1.591187 | BRCA | UACC-812 | Entinostat |
| MVP_top60_DNA_seed | PIK3CA | -1.452375 | BRCA | MDA-MB-415 | GNE-317 | MVP_top60_DNA+RNA_seed | SRC | 1.593186 | BRCA | HCC1419 | Bosutinib |
| MVP_top60_DNA_seed | ERBB2 | -1.451084 | BRCA | MDA-MB-468 | Sapitinib | MVP_top60_DNA+RNA_seed | AKT1 | 1.593829 | BRCA | MFM-223 | AKT inhibitor VIII |
| MVP_top60_DNA_seed | EGFR | -1.451084 | BRCA | MDA-MB-468 | Sapitinib | MVP_top60_DNA+RNA_seed | ESR1 | 1.600764 | BRCA | UACC-893 | Fulvestrant |
| MVP_top60_DNA_seed | ERBB3 | -1.451084 | BRCA | MDA-MB-468 | Sapitinib | MVP_top60_DNA+RNA_seed | MAPK14 | 1.607179 | BRCA | HDQ-P1 | TAK-715 |
| MVP_top60_DNA_seed | HDAC1 | -1.449642 | BRCA | MDA-MB-468 | PCI-34051 | MVP_top60_DNA+RNA_seed | EGFR | 1.611809 | BRCA | EFM-19 | Gefitinib |
| MVP_top60_DNA_seed | HDAC1 | -1.4418 | BRCA | HCC1599 | Entinostat | MVP_top60_DNA+RNA_seed | CDK2 | 1.612211 | BRCA | BT-474 | AZD5438 |
| MVP_top60_DNA_seed | AKT1 | -1.43768 | BRCA | T47D | Ipatasertib | MVP_top60_DNA+RNA_seed | AKT1 | 1.616176 | BRCA | HCC1500 | AKT inhibitor VIII |
| MVP_top60_DNA_seed | ERBB2 | -1.432996 | BRCA | ZR-75-30 | Sapitinib | MVP_top60_DNA+RNA_seed | AKT1 | 1.618654 | BRCA | MDA-MB-231 | MK-2206 |

| | | | | | | | | | | | |
|---|---|---|---|---|---|---|---|---|---|---|---|
| MVP_top60_DNA_seed | EGFR | -1.432996 | BRCA | ZR-75-30 | Sapitinib | MVP_top60_DNA+RNA_seed | EP300 | 1.625018 | BRCA | T47D | I-CBP112 |
| MVP_top60_DNA_seed | ERBB3 | -1.432996 | BRCA | ZR-75-30 | Sapitinib | MVP_top60_DNA+RNA_seed | MAPK3 | 1.628262 | BRCA | MDA-MB-415 | FR-180204 |
| MVP_top60_DNA_seed | AKT1 | -1.42518 | BRCA | HCC70 | Uprosertib | MVP_top60_DNA+RNA_seed | MAPK1 | 1.628262 | BRCA | MDA-MB-415 | FR-180204 |
| MVP_top60_DNA_seed | PIK3CA | -1.424528 | BRCA | HCC2157 | GNE-317 | MVP_top60_DNA+RNA_seed | EGFR | 1.629093 | BRCA | MDA-MB-415 | Gefitinib |
| MVP_top60_DNA_seed | CDK1 | -1.423225 | BRCA | CAL-148 | RO-3306 | MVP_top60_DNA+RNA_seed | ATM | 1.635209 | BRCA | BT-474 | KU-55933 |
| MVP_top60_DNA_seed | EGFR | -1.421015 | BRCA | BT-474 | Osimertinib | MVP_top60_DNA+RNA_seed | AR | 1.64456 | BRCA | HCC1395 | AZD3514 |
| MVP_top60_DNA_seed | AKT1 | -1.410435 | BRCA | BT-20 | Uprosertib | MVP_top60_DNA+RNA_seed | HDAC1 | 1.651381 | BRCA | MDA-MB-436 | Dacinostat |
| MVP_top60_DNA_seed | ERBB2 | -1.409896 | BRCA | MDA-MB-330 | Sapitinib | MVP_top60_DNA+RNA_seed | HDAC1 | 1.651663 | BRCA | MCF7 | Belinostat |
| MVP_top60_DNA_seed | EGFR | -1.409896 | BRCA | MDA-MB-330 | Sapitinib | MVP_top60_DNA+RNA_seed | EGFR | 1.659673 | BRCA | MDA-MB-415 | Pelitinib |
| MVP_top60_DNA_seed | ERBB3 | -1.409896 | BRCA | MDA-MB-330 | Sapitinib | MVP_top60_DNA+RNA_seed | HDAC1 | 1.660249 | BRCA | MDA-MB-415 | Belinostat |
| MVP_top60_DNA_seed | AKT1 | -1.40714 | BRCA | MFM-223 | GSK2110183B | MVP_top60_DNA+RNA_seed | ESR1 | 1.663496 | BRCA | CAL-85-1 | Fulvestrant |
| MVP_top60_DNA_seed | AKT1 | -1.405907 | BRCA | MRK-nu-1 | AT13148 | MVP_top60_DNA+RNA_seed | EGFR | 1.667051 | BRCA | UACC-812 | CUDC-101 |
| MVP_top60_DNA_seed | ESR1 | -1.402278 | BRCA | EVSA-T | GDC0810 | MVP_top60_DNA+RNA_seed | ERBB2 | 1.667051 | BRCA | UACC-812 | CUDC-101 |
| MVP_top60_DNA_seed | PIK3CA | -1.401092 | BRCA | HCC1954 | Alpelisib | MVP_top60_DNA+RNA_seed | EGFR | 1.669678 | BRCA | T47D | AZD8931 |
| MVP_top60_DNA_seed | AKT1 | -1.400544 | BRCA | CAL-51 | Ipatasertib | MVP_top60_DNA+RNA_seed | ERBB2 | 1.669678 | BRCA | T47D | AZD8931 |
| MVP_top60_DNA_seed | PIK3CA | -1.397942 | BRCA | MDA-MB-175-VII | GNE-317 | MVP_top60_DNA+RNA_seed | ERBB3 | 1.669678 | BRCA | T47D | AZD8931 |
| MVP_top60_DNA_seed | MDM2 | -1.396027 | BRCA | CAL-148 | Serdemetan | MVP_top60_DNA+RNA_seed | HDAC1 | 1.672678 | BRCA | HCC1395 | AR-42 |
| MVP_top60_DNA_seed | ATM | -1.393523 | BRCA | HCC70 | KU-55933 | MVP_top60_DNA+RNA_seed | HSP90AA1 | 1.681601 | BRCA | MDA-MB-231 | Tanespimycin |
| MVP_top60_DNA_seed | HDAC1 | -1.392804 | BRCA | OCUB-M | PCI-34051 | MVP_top60_DNA+RNA_seed | HSP90AB1 | 1.681601 | BRCA | MDA-MB-231 | Tanespimycin |
| MVP_top60_DNA_seed | ERBB2 | -1.390275 | BRCA | UACC-812 | Lapatinib | MVP_top60_DNA+RNA_seed | HSP90AA1 | 1.681777 | BRCA | HDQ-P1 | Elesclomol |
| MVP_top60_DNA_seed | EGFR | -1.390275 | BRCA | UACC-812 | Lapatinib | MVP_top60_DNA+RNA_seed | HSP90AB1 | 1.681777 | BRCA | HDQ-P1 | Elesclomol |
| MVP_top60_DNA_seed | AKT1 | -1.388491 | BRCA | HCC70 | GSK2110183B | MVP_top60_DNA+RNA_seed | PIK3CA | 1.682452 | BRCA | HCC1500 | AZD8835 |
| MVP_top60_DNA_seed | AKT1 | -1.380943 | BRCA | EFM-19 | Ipatasertib | MVP_top60_DNA+RNA_seed | CDK1 | 1.686275 | BRCA | HCC202 | RO-3306 |
| MVP_top60_DNA_seed | AKT1 | -1.375726 | BRCA | MFM-223 | Uprosertib | MVP_top60_DNA+RNA_seed | CDK2 | 1.688732 | BRCA | MFM-223 | Seliciclib |
| MVP_top60_DNA_seed | EGFR | -1.374025 | BRCA | HCC1419 | Osimertinib | MVP_top60_DNA+RNA_seed | MDM2 | 1.689582 | BRCA | EFM-19 | Serdemetan |
| MVP_top60_DNA_seed | AKT1 | -1.372254 | BRCA | HCC70 | Afuresertib | MVP_top60_DNA+RNA_seed | ATM | 1.693808 | BRCA | HCC70 | KU-60019 |
| MVP_top60_DNA_seed | PIK3CA | -1.37047 | BRCA | MDA-MB-361 | GNE-317 | MVP_top60_DNA+RNA_seed | ATM | 1.698489 | BRCA | CAMA-1 | KU-55933 |
| MVP_top60_DNA_seed | ERBB2 | -1.367107 | BRCA | UACC-812 | Sapitinib | MVP_top60_DNA+RNA_seed | CDK2 | 1.701804 | BRCA | T47D | AZD5438 |
| MVP_top60_DNA_seed | EGFR | -1.367107 | BRCA | UACC-812 | Sapitinib | MVP_top60_DNA+RNA_seed | HDAC1 | 1.705346 | BRCA | BT-483 | Tubastatin A |
| MVP_top60_DNA_seed | ERBB3 | -1.367107 | BRCA | UACC-812 | Sapitinib | MVP_top60_DNA+RNA_seed | EGFR | 1.721437 | BRCA | HCC1500 | AZD8931 |
| MVP_top60_DNA_seed | ERBB2 | -1.357323 | BRCA | HDQ-P1 | Lapatinib | MVP_top60_DNA+RNA_seed | ERBB2 | 1.721437 | BRCA | HCC1500 | AZD8931 |
| MVP_top60_DNA_seed | EGFR | -1.357323 | BRCA | HDQ-P1 | Lapatinib | MVP_top60_DNA+RNA_seed | ERBB3 | 1.721437 | BRCA | HCC1500 | AZD8931 |
| MVP_top60_DNA_seed | HDAC1 | -1.354088 | BRCA | MDA-MB-361 | Entinostat | MVP_top60_DNA+RNA_seed | EGFR | 1.725387 | BRCA | MDA-MB-436 | Pelitinib |
| MVP_top60_DNA_seed | ERBB2 | -1.352 | BRCA | DU-4475 | Lapatinib | MVP_top60_DNA+RNA_seed | PLK1 | 1.72722 | BRCA | MDA-MB-330 | PLK_6522 |
| MVP_top60_DNA_seed | EGFR | -1.352 | BRCA | DU-4475 | Lapatinib | MVP_top60_DNA+RNA_seed | SRC | 1.727797 | BRCA | MDA-MB-415 | Bosutinib |
| MVP_top60_DNA_seed | ESR1 | -1.348401 | BRCA | MDA-MB-415 | GDC0810 | MVP_top60_DNA+RNA_seed | EGFR | 1.737899 | BRCA | BT-483 | Afatinib |
| MVP_top60_DNA_seed | HDAC1 | -1.345298 | BRCA | Hs-578-T | Romidepsin | MVP_top60_DNA+RNA_seed | ERBB2 | 1.737899 | BRCA | BT-483 | Afatinib |
| MVP_top60_DNA_seed | HDAC2 | -1.345298 | BRCA | Hs-578-T | Romidepsin | MVP_top60_DNA+RNA_seed | EGFR | 1.738674 | BRCA | HCC1395 | Gefitinib |
| MVP_top60_DNA_seed | EGFR | -1.339172 | BRCA | UACC-812 | Osimertinib | MVP_top60_DNA+RNA_seed | ESR1 | 1.73933 | BRCA | HCC2218 | Fulvestrant |
| MVP_top60_DNA_seed | ESR1 | -1.328841 | BRCA | EVSA-T | Tamoxifen | MVP_top60_DNA+RNA_seed | HSP90AA1 | 1.744701 | BRCA | HCC1954 | Luminespib |
| MVP_top60_DNA_seed | MDM2 | -1.328609 | BRCA | HCC1569 | Serdemetan | MVP_top60_DNA+RNA_seed | HSP90AB1 | 1.744701 | BRCA | HCC1954 | Luminespib |

| | | | | | | | | | | | |
|---|---|---|---|---|---|---|---|---|---|---|---|
| MVP_top60_DNA_seed | AKT1 | -1.323851 | BRCA | MDA-MB-175-VII | AZD5363 | MVP_top60_DNA+RNA_seed | ATM | 1.748778 | BRCA | HCC1419 | KU-55933 |
| MVP_top60_DNA_seed | CDK1 | -1.320518 | BRCA | HCC2157 | RO-3306 | MVP_top60_DNA+RNA_seed | PLK1 | 1.753902 | BRCA | T47D | PLK_6522 |
| MVP_top60_DNA_seed | PIK3CA | -1.319474 | BRCA | EVSA-T | GNE-317 | MVP_top60_DNA+RNA_seed | HSP90AA1 | 1.760544 | BRCA | T47D | Tanespimycin |
| MVP_top60_DNA_seed | HDAC1 | -1.317187 | BRCA | DU-4475 | Romidepsin | MVP_top60_DNA+RNA_seed | HSP90AB1 | 1.760544 | BRCA | T47D | Tanespimycin |
| MVP_top60_DNA_seed | HDAC2 | -1.317187 | BRCA | DU-4475 | Romidepsin | MVP_top60_DNA+RNA_seed | HSP90AA1 | 1.763084 | BRCA | HCC1419 | CCT-018159 |
| MVP_top60_DNA_seed | ATM | -1.316938 | BRCA | OCUB-M | KU-55933 | MVP_top60_DNA+RNA_seed | HSP90AB1 | 1.763084 | BRCA | HCC1419 | CCT-018159 |
| MVP_top60_DNA_seed | CDK2 | -1.315471 | BRCA | HCC1599 | MK-8776 | MVP_top60_DNA+RNA_seed | HSP90AA1 | 1.76859 | BRCA | HCC1500 | CCT-018159 |
| MVP_top60_DNA_seed | AKT1 | -1.315157 | BRCA | HCC1569 | AT13148 | MVP_top60_DNA+RNA_seed | HSP90AB1 | 1.76859 | BRCA | HCC1500 | CCT-018159 |
| MVP_top60_DNA_seed | ERBB2 | -1.309951 | BRCA | HCC2157 | Sapitinib | MVP_top60_DNA+RNA_seed | AR | 1.773014 | BRCA | HCC2218 | Bicalutamide |
| MVP_top60_DNA_seed | EGFR | -1.309951 | BRCA | HCC2157 | Sapitinib | MVP_top60_DNA+RNA_seed | SRC | 1.782302 | BRCA | HCC202 | Bosutinib |
| MVP_top60_DNA_seed | ERBB3 | -1.309951 | BRCA | HCC2157 | Sapitinib | MVP_top60_DNA+RNA_seed | EGFR | 1.790184 | BRCA | BT-483 | Pelitinib |
| MVP_top60_DNA_seed | CDK2 | -1.306551 | BRCA | HCC2218 | Dinaciclib | MVP_top60_DNA+RNA_seed | MAPK14 | 1.790881 | BRCA | BT-483 | TAK-715 |
| MVP_top60_DNA_seed | CDK1 | -1.306551 | BRCA | HCC2218 | Dinaciclib | MVP_top60_DNA+RNA_seed | MDM2 | 1.79149 | BRCA | HCC202 | Serdemetan |
| MVP_top60_DNA_seed | AKT1 | -1.306424 | BRCA | MDA-MB-361 | MK-2206 | MVP_top60_DNA+RNA_seed | ESR1 | 1.792802 | BRCA | HCC1419 | Fulvestrant |
| MVP_top60_DNA_seed | PIK3CA | -1.305722 | BRCA | T47D | Alpelisib | MVP_top60_DNA+RNA_seed | AKT1 | 1.797169 | BRCA | HCC38 | MK-2206 |
| MVP_top60_DNA_seed | ERBB2 | -1.304562 | BRCA | EFM-192A | Sapitinib | MVP_top60_DNA+RNA_seed | AKT1 | 1.801117 | BRCA | MRK-nu-1 | MK-2206 |
| MVP_top60_DNA_seed | EGFR | -1.304562 | BRCA | EFM-192A | Sapitinib | MVP_top60_DNA+RNA_seed | EGFR | 1.804296 | BRCA | T47D | Gefitinib |
| MVP_top60_DNA_seed | ERBB3 | -1.304562 | BRCA | EFM-192A | Sapitinib | MVP_top60_DNA+RNA_seed | AKT1 | 1.80809 | BRCA | HCC1500 | MK-2206 |
| MVP_top60_DNA_seed | PIK3CA | -1.30157 | BRCA | HDQ-P1 | GNE-317 | MVP_top60_DNA+RNA_seed | HSP90AA1 | 1.810166 | BRCA | MDA-MB-157 | SNX-2112 |
| MVP_top60_DNA_seed | PIK3CA | -1.298818 | BRCA | HCC2157 | Buparlisib | MVP_top60_DNA+RNA_seed | HSP90AB1 | 1.810166 | BRCA | MDA-MB-157 | SNX-2112 |
| MVP_top60_DNA_seed | AKT1 | -1.296755 | BRCA | Hs-578-T | GSK2110183B | MVP_top60_DNA+RNA_seed | AR | 1.816611 | BRCA | T47D | Bicalutamide |
| MVP_top60_DNA_seed | AKT1 | -1.290018 | BRCA | MDA-MB-330 | Uprosertib | MVP_top60_DNA+RNA_seed | HSP90AA1 | 1.819943 | BRCA | HCC1395 | Luminespib |
| MVP_top60_DNA_seed | EGFR | -1.287227 | BRCA | MDA-MB-468 | AZD3759 | MVP_top60_DNA+RNA_seed | HSP90AB1 | 1.819943 | BRCA | HCC1395 | Luminespib |
| MVP_top60_DNA_seed | ESR1 | -1.286758 | BRCA | OCUB-M | GDC0810 | MVP_top60_DNA+RNA_seed | MAPK1 | 1.820287 | BRCA | BT-483 | VX-11e |
| MVP_top60_DNA_seed | ERBB2 | -1.286186 | BRCA | HCC2157 | Afatinib | MVP_top60_DNA+RNA_seed | CDK2 | 1.822255 | BRCA | HCC1500 | AZD5438 |
| MVP_top60_DNA_seed | EGFR | -1.286186 | BRCA | HCC2157 | Afatinib | MVP_top60_DNA+RNA_seed | PLK1 | 1.827528 | BRCA | HCC202 | PLK_6522 |
| MVP_top60_DNA_seed | AKT1 | -1.284561 | BRCA | MDA-MB-453 | Afuresertib | MVP_top60_DNA+RNA_seed | MDM2 | 1.841397 | BRCA | ZR-75-30 | Serdemetan |
| MVP_top60_DNA_seed | PIK3CA | -1.282233 | BRCA | MDA-MB-330 | GNE-317 | MVP_top60_DNA+RNA_seed | ESR1 | 1.844901 | BRCA | ZR-75-30 | Tamoxifen |
| MVP_top60_DNA_seed | CDK1 | -1.28002 | BRCA | Hs-578-T | RO-3306 | MVP_top60_DNA+RNA_seed | CDK2 | 1.852959 | BRCA | UACC-893 | AZD5438 |
| MVP_top60_DNA_seed | ERBB2 | -1.277934 | BRCA | MDA-MB-453 | Lapatinib | MVP_top60_DNA+RNA_seed | HSP90AA1 | 1.855709 | BRCA | MDA-MB-436 | CCT-018159 |
| MVP_top60_DNA_seed | EGFR | -1.277934 | BRCA | MDA-MB-453 | Lapatinib | MVP_top60_DNA+RNA_seed | HSP90AB1 | 1.855709 | BRCA | MDA-MB-436 | CCT-018159 |
| MVP_top60_DNA_seed | HSP90AA1 | -1.276484 | BRCA | CAL-51 | Tanespimycin | MVP_top60_DNA+RNA_seed | HDAC1 | 1.855711 | BRCA | Hs-578-T | JQ12 |
| MVP_top60_DNA_seed | HSP90AB1 | -1.276484 | BRCA | CAL-51 | Tanespimycin | MVP_top60_DNA+RNA_seed | HDAC2 | 1.855711 | BRCA | Hs-578-T | JQ12 |
| MVP_top60_DNA_seed | AKT1 | -1.275226 | BRCA | Hs-578-T | Uprosertib | MVP_top60_DNA+RNA_seed | HDAC1 | 1.867294 | BRCA | BT-20 | JQ12 |
| MVP_top60_DNA_seed | EP300 | -1.262581 | BRCA | COLO-824 | CPI-637 | MVP_top60_DNA+RNA_seed | HDAC2 | 1.867294 | BRCA | BT-20 | JQ12 |
| MVP_top60_DNA_seed | SRC | -1.259319 | BRCA | HDQ-P1 | Bosutinib | MVP_top60_DNA+RNA_seed | HDAC1 | 1.869746 | BRCA | MFM-223 | JQ12 |
| MVP_top60_DNA_seed | ERBB2 | -1.251343 | BRCA | HCC1419 | Lapatinib | MVP_top60_DNA+RNA_seed | HDAC2 | 1.869746 | BRCA | MFM-223 | JQ12 |
| MVP_top60_DNA_seed | EGFR | -1.251343 | BRCA | HCC1419 | Lapatinib | MVP_top60_DNA+RNA_seed | MAPK3 | 1.874762 | BRCA | BT-483 | FR-180204 |
| MVP_top60_DNA_seed | AKT1 | -1.251025 | BRCA | CAL-51 | Afuresertib | MVP_top60_DNA+RNA_seed | MAPK1 | 1.874762 | BRCA | BT-483 | FR-180204 |
| MVP_top60_DNA_seed | ESR1 | -1.244882 | BRCA | T47D | Tamoxifen | MVP_top60_DNA+RNA_seed | HSP90AA1 | 1.875977 | BRCA | MRK-nu-1 | Tanespimycin |
| MVP_top60_DNA_seed | PIK3CA | -1.243567 | BRCA | HDQ-P1 | AZD8186 | MVP_top60_DNA+RNA_seed | HSP90AB1 | 1.875977 | BRCA | MRK-nu-1 | Tanespimycin |

| | | | | | | | | | | | |
|---|---|---|---|---|---|---|---|---|---|---|---|
| MVP_top60_DNA_seed | CDK2 | -1.242961 | BRCA | MDA-MB-157 | MK-8776 | MVP_top60_DNA+RNA_seed | ATM | 1.886238 | BRCA | HCC202 | KU-55933 |
| MVP_top60_DNA_seed | PIK3CA | -1.240029 | BRCA | EFM-19 | GNE-317 | MVP_top60_DNA+RNA_seed | PIK3CA | 1.886279 | BRCA | MRK-nu-1 | PI3Ka_4409 |
| MVP_top60_DNA_seed | PIK3CA | -1.236731 | BRCA | CAL-51 | GNE-317 | MVP_top60_DNA+RNA_seed | HSP90AA1 | 1.901656 | BRCA | MCF7 | SNX-2112 |
| MVP_top60_DNA_seed | AKT1 | -1.234477 | BRCA | HCC1419 | Ipatasertib | MVP_top60_DNA+RNA_seed | HSP90AB1 | 1.901656 | BRCA | MCF7 | SNX-2112 |
| MVP_top60_DNA_seed | ESR1 | -1.231812 | BRCA | HCC1395 | Tamoxifen | MVP_top60_DNA+RNA_seed | PIK3CA | 1.903586 | BRCA | MRK-nu-1 | AZD8835 |
| MVP_top60_DNA_seed | HDAC1 | -1.231066 | BRCA | YMB-1-E | PCI-34051 | MVP_top60_DNA+RNA_seed | HSP90AA1 | 1.905592 | BRCA | MDA-MB-415 | SNX-2112 |
| MVP_top60_DNA_seed | PIK3CA | -1.223717 | BRCA | EVSA-T | Buparlisib | MVP_top60_DNA+RNA_seed | HSP90AB1 | 1.905592 | BRCA | MDA-MB-415 | SNX-2112 |
| MVP_top60_DNA_seed | AKT1 | -1.223013 | BRCA | HDQ-P1 | AT13148 | MVP_top60_DNA+RNA_seed | CDK2 | 1.908081 | BRCA | MDA-MB-415 | AZD5438 |
| MVP_top60_DNA_seed | EGFR | -1.218025 | BRCA | HCC1569 | Osimertinib | MVP_top60_DNA+RNA_seed | PLK1 | 1.919401 | BRCA | UACC-893 | PLK_6522 |
| MVP_top60_DNA_seed | PIK3CA | -1.217177 | BRCA | BT-474 | Alpelisib | MVP_top60_DNA+RNA_seed | HDAC1 | 1.919463 | BRCA | HDQ-P1 | Dacinostat |
| MVP_top60_DNA_seed | PIK3CA | -1.207284 | BRCA | UACC-893 | GNE-317 | MVP_top60_DNA+RNA_seed | HDAC1 | 1.923513 | BRCA | HCC2218 | Parthenolide |
| MVP_top60_DNA_seed | HDAC1 | -1.207032 | BRCA | HCC1569 | PCI-34051 | MVP_top60_DNA+RNA_seed | HSP90AA1 | 1.955992 | BRCA | HCC2157 | SNX-2112 |
| MVP_top60_DNA_seed | ERBB2 | -1.206512 | BRCA | MDA-MB-468 | Lapatinib | MVP_top60_DNA+RNA_seed | HSP90AB1 | 1.955992 | BRCA | HCC2157 | SNX-2112 |
| MVP_top60_DNA_seed | EGFR | -1.206512 | BRCA | MDA-MB-468 | Lapatinib | MVP_top60_DNA+RNA_seed | PIK3CA | 1.962095 | BRCA | HCC1500 | PI3Ka_4409 |
| MVP_top60_DNA_seed | MAPK1 | -1.203858 | BRCA | HDQ-P1 | VX-11e | MVP_top60_DNA+RNA_seed | HDAC1 | 1.975689 | BRCA | MDA-MB-415 | AR-42 |
| MVP_top60_DNA_seed | HDAC1 | -1.202038 | BRCA | MDA-MB-330 | Entinostat | MVP_top60_DNA+RNA_seed | SRC | 1.989396 | BRCA | BT-483 | Bosutinib |
| MVP_top60_DNA_seed | HSP90AA1 | -1.201796 | BRCA | DU-4475 | Luminespib | MVP_top60_DNA+RNA_seed | SRC | 2.000384 | BRCA | BT-483 | NVP-BHG712 |
| MVP_top60_DNA_seed | HSP90AB1 | -1.201796 | BRCA | DU-4475 | Luminespib | MVP_top60_DNA+RNA_seed | PLK1 | 2.016825 | BRCA | HCC1500 | PLK_6522 |
| MVP_top60_DNA_seed | AKT1 | -1.193382 | BRCA | HCC1395 | Ipatasertib | MVP_top60_DNA+RNA_seed | HSP90AA1 | 2.02296 | BRCA | MRK-nu-1 | Elesclomol |
| MVP_top60_DNA_seed | AKT1 | -1.190906 | BRCA | MDA-MB-330 | Ipatasertib | MVP_top60_DNA+RNA_seed | HSP90AB1 | 2.02296 | BRCA | MRK-nu-1 | Elesclomol |
| MVP_top60_DNA_seed | AKT1 | -1.190831 | BRCA | Hs-578-T | Ipatasertib | MVP_top60_DNA+RNA_seed | HDAC1 | 2.024309 | BRCA | UACC-812 | JQ12 |
| MVP_top60_DNA_seed | CDK2 | -1.189686 | BRCA | HCC1569 | MK-8776 | MVP_top60_DNA+RNA_seed | HDAC2 | 2.024309 | BRCA | UACC-812 | JQ12 |
| MVP_top60_DNA_seed | TP53 | -1.185478 | BRCA | HCC2157 | MIRA-1 | MVP_top60_DNA+RNA_seed | ATM | 2.042981 | BRCA | ZR-75-30 | KU-55933 |
| MVP_top60_DNA_seed | ERBB2 | -1.173941 | BRCA | MDA-MB-361 | Sapitinib | MVP_top60_DNA+RNA_seed | PIK3CA | 2.047064 | BRCA | HCC1395 | PI3Ka_4409 |
| MVP_top60_DNA_seed | EGFR | -1.173941 | BRCA | MDA-MB-361 | Sapitinib | MVP_top60_DNA+RNA_seed | HSP90AA1 | 2.052114 | BRCA | UACC-812 | SNX-2112 |
| MVP_top60_DNA_seed | ERBB3 | -1.173941 | BRCA | MDA-MB-361 | Sapitinib | MVP_top60_DNA+RNA_seed | HSP90AB1 | 2.052114 | BRCA | UACC-812 | SNX-2112 |
| MVP_top60_DNA_seed | CDK2 | -1.172171 | BRCA | HCC1954 | Dinaciclib | MVP_top60_DNA+RNA_seed | CDK2 | 2.053182 | BRCA | CAMA-1 | AZD5438 |
| MVP_top60_DNA_seed | CDK1 | -1.172171 | BRCA | HCC1954 | Dinaciclib | MVP_top60_DNA+RNA_seed | HDAC1 | 2.060857 | BRCA | HDQ-P1 | JQ12 |
| MVP_top60_DNA_seed | ESR1 | -1.17002 | BRCA | AU565 | GDC0810 | MVP_top60_DNA+RNA_seed | HDAC2 | 2.060857 | BRCA | HDQ-P1 | JQ12 |
| MVP_top60_DNA_seed | ERBB2 | -1.168457 | BRCA | HCC1419 | Sapitinib | MVP_top60_DNA+RNA_seed | EGFR | 2.075293 | BRCA | ZR-75-30 | Gefitinib |
| MVP_top60_DNA_seed | EGFR | -1.168457 | BRCA | HCC1419 | Sapitinib | MVP_top60_DNA+RNA_seed | HSP90AA1 | 2.100611 | BRCA | HCC202 | CCT-018159 |
| MVP_top60_DNA_seed | ERBB3 | -1.168457 | BRCA | HCC1419 | Sapitinib | MVP_top60_DNA+RNA_seed | HSP90AB1 | 2.100611 | BRCA | HCC202 | CCT-018159 |
| MVP_top60_DNA_seed | HSP90AA1 | -1.159436 | BRCA | HCC2218 | CCT-018159 | MVP_top60_DNA+RNA_seed | HSP90AA1 | 2.102781 | BRCA | MFM-223 | Luminespib |
| MVP_top60_DNA_seed | HSP90AB1 | -1.159436 | BRCA | HCC2218 | CCT-018159 | MVP_top60_DNA+RNA_seed | HSP90AB1 | 2.102781 | BRCA | MFM-223 | Luminespib |
| MVP_top60_DNA_seed | TP53 | -1.155661 | BRCA | HCC1419 | MIRA-1 | MVP_top60_DNA+RNA_seed | MDM2 | 2.135298 | BRCA | HCC2218 | Serdemetan |
| MVP_top60_DNA_seed | AKT1 | -1.154297 | BRCA | MDA-MB-361 | Uprosertib | MVP_top60_DNA+RNA_seed | HDAC1 | 2.165719 | BRCA | COLO-824 | Entinostat |
| MVP_top60_DNA_seed | AKT1 | -1.152108 | BRCA | MDA-MB-361 | Ipatasertib | MVP_top60_DNA+RNA_seed | ATM | 2.172968 | BRCA | BT-483 | KU-60019 |
| MVP_top60_DNA_seed | AKT1 | -1.151762 | BRCA | YMB-1-E | AT13148 | MVP_top60_DNA+RNA_seed | PIK3CA | 2.180415 | BRCA | CAMA-1 | AZD8835 |
| MVP_top60_DNA_seed | AKT1 | -1.148076 | BRCA | HCC1395 | Afuresertib | MVP_top60_DNA+RNA_seed | HSP90AA1 | 2.268071 | BRCA | HCC1569 | Elesclomol |
| MVP_top60_DNA_seed | AKT1 | -1.143101 | BRCA | MDA-MB-453 | Ipatasertib | MVP_top60_DNA+RNA_seed | HSP90AB1 | 2.268071 | BRCA | HCC1569 | Elesclomol |
| MVP_top60_DNA_seed | AKT1 | -1.137307 | BRCA | EFM-19 | GSK2110183B | MVP_top60_DNA+RNA_seed | MDM2 | 2.282995 | BRCA | MDA-MB-175-VII | Serdemetan |

| | | | | | | | | | | | |
|---|---|---|---|---|---|---|---|---|---|---|---|
| MVP_top60_DNA_seed | CDK1 | -1.136104 | BRCA | MDA-MB-468 | RO-3306 | MVP_top60_DNA+RNA_seed | PIK3CA | 2.290532 | BRCA | CAMA-1 | PI3Ka_4409 |
| MVP_top60_DNA_seed | HDAC1 | -1.135274 | BRCA | MDA-MB-453 | Romidepsin | MVP_top60_DNA+RNA_seed | PLK1 | 2.296477 | BRCA | HCC70 | PLK_6522 |
| MVP_top60_DNA_seed | HDAC2 | -1.135274 | BRCA | MDA-MB-453 | Romidepsin | MVP_top60_DNA+RNA_seed | PIK3CA | 2.332789 | BRCA | MDA-MB-436 | PI3Ka_4409 |
| MVP_top60_DNA_seed | PIK3CA | -1.128218 | BRCA | MDA-MB-175-VII | Alpelisib | MVP_top60_DNA+RNA_seed | ESR1 | 2.342078 | BRCA | BT-483 | Tamoxifen |
| MVP_top60_DNA_seed | AKT1 | -1.123818 | BRCA | MDA-MB-330 | GSK2110183B | MVP_top60_DNA+RNA_seed | ESR1 | 2.378933 | BRCA | MFM-223 | Tamoxifen |
| MVP_top60_DNA_seed | CDK2 | -1.115391 | BRCA | MDA-MB-468 | Dinaciclib | MVP_top60_DNA+RNA_seed | HDAC1 | 2.379139 | BRCA | MFM-223 | Entinostat |
| MVP_top60_DNA_seed | CDK1 | -1.115391 | BRCA | MDA-MB-468 | Dinaciclib | MVP_top60_DNA+RNA_seed | PLK1 | 2.396567 | BRCA | HCC2218 | PLK_6522 |
| MVP_top60_DNA_seed | EGFR | -1.109276 | BRCA | MDA-MB-453 | Gefitinib | MVP_top60_DNA+RNA_seed | CDK2 | 2.407839 | BRCA | ZR-75-30 | AZD5438 |
| MVP_top60_DNA_seed | AKT1 | -1.108911 | BRCA | MDA-MB-330 | AZD5363 | MVP_top60_DNA+RNA_seed | AKT1 | 2.444972 | BRCA | BT-483 | BAY AKT1 |
| MVP_top60_DNA_seed | HSP90AA1 | -1.105739 | BRCA | MDA-MB-330 | Elesclomol | MVP_top60_DNA+RNA_seed | HSP90AA1 | 2.448277 | BRCA | HCC1954 | Elesclomol |
| MVP_top60_DNA_seed | HSP90AB1 | -1.105739 | BRCA | MDA-MB-330 | Elesclomol | MVP_top60_DNA+RNA_seed | HSP90AB1 | 2.448277 | BRCA | HCC1954 | Elesclomol |
| MVP_top60_DNA_seed | CDK2 | -1.102498 | BRCA | HCC1187 | Dinaciclib | MVP_top60_DNA+RNA_seed | PLK1 | 2.488824 | BRCA | HCC1419 | PLK_6522 |
| MVP_top60_DNA_seed | CDK1 | -1.102498 | BRCA | HCC1187 | Dinaciclib | MVP_top60_DNA+RNA_seed | PLK1 | 2.498751 | BRCA | BT-474 | PLK_6522 |
| MVP_top60_DNA_seed | AKT1 | -1.100187 | BRCA | BT-20 | Ipatasertib | MVP_top60_DNA+RNA_seed | AR | 2.513702 | BRCA | ZR-75-30 | AZD3514 |
| MVP_top60_DNA_seed | AKT1 | -1.100014 | BRCA | EFM-19 | MK-2206 | MVP_top60_DNA+RNA_seed | ATM | 2.584383 | BRCA | HCC2218 | KU-60019 |
| MVP_top60_DNA_seed | PIK3CA | -1.098943 | BRCA | CAL-148 | GNE-317 | MVP_top60_DNA+RNA_seed | PLK1 | 2.618082 | BRCA | MDA-MB-175-VII | PLK_6522 |
| MVP_top60_DNA_seed | MDM2 | -1.097828 | BRCA | HCC1500 | Nutlin-3a (-) | MVP_top60_DNA+RNA_seed | CDK2 | 2.633355 | BRCA | MDA-MB-175-VII | AZD5438 |
| MVP_top60_DNA_seed | ESR1 | -1.094082 | BRCA | CAMA-1 | Tamoxifen | MVP_top60_DNA+RNA_seed | HSP90AA1 | 2.640077 | BRCA | Hs-578-T | Luminespib |
| MVP_top60_DNA_seed | ATM | -1.087495 | BRCA | AU565 | KU-55933 | MVP_top60_DNA+RNA_seed | HSP90AB1 | 2.640077 | BRCA | Hs-578-T | Luminespib |
| MVP_top60_DNA_seed | EP300 | -1.087417 | BRCA | HCC1937 | SGC-CBP30 | MVP_top60_DNA+RNA_seed | PLK1 | 2.642613 | BRCA | ZR-75-30 | PLK_6522 |
| MVP_top60_DNA_seed | CREBBP | -1.087417 | BRCA | HCC1937 | SGC-CBP30 | MVP_top60_DNA+RNA_seed | PLK1 | 2.659119 | BRCA | BT-483 | PLK_6522 |
| MVP_top60_DNA_seed | EGFR | -1.087029 | BRCA | MDA-MB-468 | Gefitinib | MVP_top60_DNA+RNA_seed | AR | 2.667127 | BRCA | BT-483 | AZD3514 |
| MVP_top60_DNA_seed | AR | -1.083394 | BRCA | HCC2218 | Bicalutamide | MVP_top60_DNA+RNA_seed | CDK2 | 2.692984 | BRCA | HCC1419 | AZD5438 |
| MVP_top60_DNA_seed | CDK1 | -1.083037 | BRCA | EVSA-T | RO-3306 | MVP_top60_DNA+RNA_seed | HDAC1 | 2.711435 | BRCA | BT-483 | Belinostat |
| MVP_top60_DNA_seed | MAPK1 | -1.078733 | BRCA | HCC1569 | VX-11e | MVP_top60_DNA+RNA_seed | AR | 2.773452 | BRCA | BT-483 | Bicalutamide |
| MVP_top60_DNA_seed | PIK3CA | -1.078618 | BRCA | HCC1395 | AZD8186 | MVP_top60_DNA+RNA_seed | HDAC1 | 2.794843 | BRCA | HCC1187 | Dacinostat |
| MVP_top60_DNA_seed | HSP90AA1 | -1.077325 | BRCA | MDA-MB-361 | Luminespib | MVP_top60_DNA+RNA_seed | AR | 2.819253 | BRCA | BT-483 | Bicalutamide |
| MVP_top60_DNA_seed | HSP90AB1 | -1.077325 | BRCA | MDA-MB-361 | Luminespib | MVP_top60_DNA+RNA_seed | HDAC1 | 2.839094 | BRCA | UACC-812 | Dacinostat |
| MVP_top60_DNA_seed | PIK3CA | -1.076969 | BRCA | CAMA-1 | GNE-317 | MVP_top60_DNA+RNA_seed | HSP90AA1 | 2.975847 | BRCA | BT-483 | CCT-018159 |
| MVP_top60_DNA_seed | ESR1 | -1.07538 | BRCA | DU-4475 | Tamoxifen | MVP_top60_DNA+RNA_seed | HSP90AB1 | 2.975847 | BRCA | BT-483 | CCT-018159 |
| MVP_top60_DNA_seed | PIK3CA | -1.074375 | BRCA | T47D | GNE-317 | MVP_top60_DNA+RNA_seed | ATM | 3.125079 | BRCA | ZR-75-30 | torin2 |
| MVP_top60_DNA_seed | PIK3CA | -1.073624 | BRCA | HCC1599 | Alpelisib | MVP_top60_DNA+RNA_seed | EGFR | 3.424488 | BRCA | BT-483 | CUDC-101 |
| MVP_top60_DNA_seed | AKT1 | -1.071297 | BRCA | Hs-578-T | MK-2206 | MVP_top60_DNA+RNA_seed | ERBB2 | 3.424488 | BRCA | BT-483 | CUDC-101 |
| MVP_top60_DNA_seed | HSP90AA1 | -1.07054 | BRCA | MDA-MB-453 | Luminespib | MVP_top60_DNA+RNA_seed | HDAC1 | 3.455672 | BRCA | BT-483 | AR-42 |
| MVP_top60_DNA_seed | HSP90AB1 | -1.07054 | BRCA | MDA-MB-453 | Luminespib | MVP_top60_DNA+RNA_seed | HSP90AA1 | 3.512918 | BRCA | BT-483 | Luminespib |
| MVP_top60_DNA_seed | AKT1 | -1.062409 | BRCA | HCC1954 | Uprosertib | MVP_top60_DNA+RNA_seed | HSP90AB1 | 3.512918 | BRCA | BT-483 | Luminespib |
| MVP_top60_DNA_seed | HSP90AA1 | -1.061235 | BRCA | CAL-148 | Tanespimycin | MVP_top60_DNA+RNA_seed | CDK2 | 3.717065 | BRCA | BT-483 | AZD5438 |
| MVP_top60_DNA_seed | HSP90AB1 | -1.061235 | BRCA | CAL-148 | Tanespimycin | MVP_top60_DNA+RNA_seed | HDAC1 | 4.486456 | BRCA | Hs-578-T | Dacinostat |
| MVP_top60_DNA_seed | AKT1 | -1.056711 | BRCA | T47D | GSK2110183B | MVP_top60_DNA+RNA_seed | PIK3CA | -4.087508 | BRCA | YMB-1-E | AZD8186 |
| MVP_top60_DNA_seed | HDAC1 | -1.054149 | BRCA | CAL-51 | PCI-34051 | MVP_top60_DNA+RNA_seed | ATM | -3.853213 | BRCA | HCC2157 | KU-55933 |
| MVP_top60_DNA_seed | ESR1 | -1.05175 | BRCA | MCF7 | Tamoxifen | MVP_top60_DNA+RNA_seed | PIK3CA | -3.846124 | BRCA | EVSA-T | AZD8186 |

| | | | | | | | | | | | |
|---|---|---|---|---|---|---|---|---|---|---|---|
| MVP_top60_DNA_seed | HSP90AA1 | -1.051622 | BRCA | UACC-893 | Tanespimycin | MVP_top60_DNA+RNA_seed | EGFR | -3.754 | BRCA | MDA-MB-175-VII | Afatinib |
| MVP_top60_DNA_seed | HSP90AB1 | -1.051622 | BRCA | UACC-893 | Tanespimycin | MVP_top60_DNA+RNA_seed | ERBB2 | -3.754 | BRCA | MDA-MB-175-VII | Afatinib |
| MVP_top60_DNA_seed | AKT1 | -1.050404 | BRCA | Hs-578-T | AZD5363 | MVP_top60_DNA+RNA_seed | MAPK3 | -3.557454 | BRCA | DU-4475 | SCH772984 |
| MVP_top60_DNA_seed | HSP90AA1 | -1.043814 | BRCA | Hs-578-T | Tanespimycin | MVP_top60_DNA+RNA_seed | MAPK1 | -3.557454 | BRCA | DU-4475 | SCH772984 |
| MVP_top60_DNA_seed | HSP90AB1 | -1.043814 | BRCA | Hs-578-T | Tanespimycin | MVP_top60_DNA+RNA_seed | EGFR | -3.497705 | BRCA | HCC2218 | Afatinib |
| MVP_top60_DNA_seed | CDK2 | -1.039464 | BRCA | HCC1187 | AZD5438 | MVP_top60_DNA+RNA_seed | ERBB2 | -3.497705 | BRCA | HCC2218 | Afatinib |
| MVP_top60_DNA_seed | AKT1 | -1.038614 | BRCA | CAMA-1 | MK-2206 | MVP_top60_DNA+RNA_seed | PIK3CA | -3.378294 | BRCA | MDA-MB-415 | AZD8186 |
| MVP_top60_DNA_seed | AKT1 | -1.037668 | BRCA | EFM-19 | Uprosertib | MVP_top60_DNA+RNA_seed | AKT1 | -3.302738 | BRCA | CAL-148 | Ipatasertib |
| MVP_top60_DNA_seed | MAPK3 | -1.036445 | BRCA | MDA-MB-436 | Ulixertinib | MVP_top60_DNA+RNA_seed | AKT1 | -3.282229 | BRCA | EVSA-T | GSK2110183B |
| MVP_top60_DNA_seed | MAPK1 | -1.036445 | BRCA | MDA-MB-436 | Ulixertinib | MVP_top60_DNA+RNA_seed | AKT1 | -3.259805 | BRCA | CAL-148 | Afuresertib |
| MVP_top60_DNA_seed | AKT1 | -1.034662 | BRCA | MDA-MB-330 | Afuresertib | MVP_top60_DNA+RNA_seed | EGFR | -3.16649 | BRCA | UACC-893 | Afatinib |
| MVP_top60_DNA_seed | AKT1 | -1.028377 | BRCA | BT-20 | AZD5363 | MVP_top60_DNA+RNA_seed | ERBB2 | -3.16649 | BRCA | UACC-893 | Afatinib |
| MVP_top60_DNA_seed | PIK3CA | -1.026325 | BRCA | CAL-51 | Buparlisib | MVP_top60_DNA+RNA_seed | EGFR | -3.158051 | BRCA | MDA-MB-175-VII | Lapatinib |
| MVP_top60_DNA_seed | TP53 | -1.02495 | BRCA | OCUB-M | MIRA-1 | MVP_top60_DNA+RNA_seed | ERBB2 | -3.158051 | BRCA | MDA-MB-175-VII | Lapatinib |
| MVP_top60_DNA_seed | PIK3CA | -1.022631 | BRCA | EFM-19 | AZD8186 | MVP_top60_DNA+RNA_seed | AKT1 | -3.141626 | BRCA | CAL-148 | AZD5363 |
| MVP_top60_DNA_seed | PIK3CA | -1.019348 | BRCA | MDA-MB-361 | Buparlisib | MVP_top60_DNA+RNA_seed | AKT1 | -3.138923 | BRCA | CAL-148 | Uprosertib |
| MVP_top60_DNA_seed | PIK3CA | -1.018997 | BRCA | MDA-MB-175-VII | Buparlisib | MVP_top60_DNA+RNA_seed | EGFR | -3.13007 | BRCA | MDA-MB-175-VII | Sapitinib |
| MVP_top60_DNA_seed | ESR1 | -1.018263 | BRCA | DU-4475 | GDC0810 | MVP_top60_DNA+RNA_seed | ERBB2 | -3.13007 | BRCA | MDA-MB-175-VII | Sapitinib |
| MVP_top60_DNA_seed | PIK3CA | -1.016139 | BRCA | HCC1569 | Alpelisib | MVP_top60_DNA+RNA_seed | ERBB3 | -3.13007 | BRCA | MDA-MB-175-VII | Sapitinib |
| MVP_top60_DNA_seed | AKT1 | -1.013224 | BRCA | CAL-85-1 | GSK2110183B | MVP_top60_DNA+RNA_seed | AKT1 | -3.113381 | BRCA | OCUB-M | Uprosertib |
| MVP_top60_DNA_seed | CDK2 | -1.007732 | BRCA | MDA-MB-468 | AZD5438 | MVP_top60_DNA+RNA_seed | MAPK1 | -3.094224 | BRCA | DU-4475 | VX-11e |
| MVP_top60_DNA_seed | CDK2 | -1.007576 | BRCA | HCC2157 | Dinaciclib | MVP_top60_DNA+RNA_seed | AKT1 | -3.011198 | BRCA | OCUB-M | Afuresertib |
| MVP_top60_DNA_seed | CDK1 | -1.007576 | BRCA | HCC2157 | Dinaciclib | MVP_top60_DNA+RNA_seed | AKT1 | -2.964343 | BRCA | EVSA-T | Afuresertib |
| MVP_top60_DNA_seed | AKT1 | -1.007023 | BRCA | COLO-824 | AT13148 | MVP_top60_DNA+RNA_seed | EGFR | -2.963037 | BRCA | MDA-MB-175-VII | Gefitinib |
| MVP_top60_DNA_seed | ESR1 | -1.005566 | BRCA | MDA-MB-361 | GDC0810 | MVP_top60_DNA+RNA_seed | AKT1 | -2.904853 | BRCA | OCUB-M | GSK2110183B |
| MVP_top60_DNA_seed | AKT1 | -0.997542 | BRCA | HCC1569 | MK-2206 | MVP_top60_DNA+RNA_seed | AKT1 | -2.891728 | BRCA | CAL-148 | Uprosertib |
| MVP_top60_DNA_seed | EGFR | -0.997178 | BRCA | MDA-MB-330 | Gefitinib | MVP_top60_DNA+RNA_seed | EGFR | -2.870454 | BRCA | HCC2218 | Osimertinib |
| MVP_top60_DNA_seed | HDAC1 | -0.996057 | BRCA | AU565 | Entinostat | MVP_top60_DNA+RNA_seed | AKT1 | -2.836144 | BRCA | EVSA-T | AZD5363 |
| MVP_top60_DNA_seed | EP300 | -0.99594 | BRCA | MDA-MB-157 | CPI-637 | MVP_top60_DNA+RNA_seed | MAPK3 | -2.82499 | BRCA | DU-4475 | Ulixertinib |
| MVP_top60_DNA_seed | EP300 | -0.993573 | BRCA | HCC1419 | SGC-CBP30 | MVP_top60_DNA+RNA_seed | MAPK1 | -2.82499 | BRCA | DU-4475 | Ulixertinib |
| MVP_top60_DNA_seed | CREBBP | -0.993573 | BRCA | HCC1419 | SGC-CBP30 | MVP_top60_DNA+RNA_seed | AKT1 | -2.81292 | BRCA | MDA-MB-415 | Ipatasertib |
| MVP_top60_DNA_seed | AKT1 | -0.990002 | BRCA | BT-549 | AT13148 | MVP_top60_DNA+RNA_seed | AKT1 | -2.789178 | BRCA | OCUB-M | Ipatasertib |
| MVP_top60_DNA_seed | HDAC1 | -0.989535 | BRCA | HCC70 | Entinostat | MVP_top60_DNA+RNA_seed | EGFR | -2.775513 | BRCA | MDA-MB-175-VII | Osimertinib |
| MVP_top60_DNA_seed | AKT1 | -0.982903 | BRCA | COLO-824 | Uprosertib | MVP_top60_DNA+RNA_seed | AKT1 | -2.754233 | BRCA | OCUB-M | Uprosertib |
| MVP_top60_DNA_seed | EP300 | -0.982896 | BRCA | OCUB-M | SGC-CBP30 | MVP_top60_DNA+RNA_seed | CDK2 | -2.74532 | BRCA | HCC2218 | MK-8776 |
| MVP_top60_DNA_seed | CREBBP | -0.982896 | BRCA | OCUB-M | SGC-CBP30 | MVP_top60_DNA+RNA_seed | EGFR | -2.709433 | BRCA | HDQ-P1 | AZD3759 |
| MVP_top60_DNA_seed | CDK2 | -0.981077 | BRCA | Hs-578-T | MK-8776 | MVP_top60_DNA+RNA_seed | AKT1 | -2.692292 | BRCA | OCUB-M | AZD5363 |
| MVP_top60_DNA_seed | AKT1 | -0.980322 | BRCA | CAL-51 | AZD5363 | MVP_top60_DNA+RNA_seed | AKT1 | -2.666864 | BRCA | EVSA-T | Ipatasertib |
| MVP_top60_DNA_seed | AKT1 | -0.979968 | BRCA | CAMA-1 | AT13148 | MVP_top60_DNA+RNA_seed | ESR1 | -2.666749 | BRCA | MCF7 | GDC0810 |
| MVP_top60_DNA_seed | HSP90AA1 | -0.977994 | BRCA | EVSA-T | Tanespimycin | MVP_top60_DNA+RNA_seed | EGFR | -2.653491 | BRCA | HCC2218 | Lapatinib |
| MVP_top60_DNA_seed | HSP90AB1 | -0.977994 | BRCA | EVSA-T | Tanespimycin | MVP_top60_DNA+RNA_seed | ERBB2 | -2.653491 | BRCA | HCC2218 | Lapatinib |

| | | | | | | | | | | | |
|---|---|---|---|---|---|---|---|---|---|---|---|
| MVP_top60_DNA_seed | MDM2 | -0.976296 | BRCA | CAL-51 | Nutlin-3a (-) | MVP_top60_DNA+RNA_seed | EGFR | -2.611412 | BRCA | MDA-MB-330 | Afatinib |
| MVP_top60_DNA_seed | ATM | -0.975642 | BRCA | HCC1395 | KU-55933 | MVP_top60_DNA+RNA_seed | ERBB2 | -2.611412 | BRCA | MDA-MB-330 | Afatinib |
| MVP_top60_DNA_seed | EGFR | -0.973906 | BRCA | COLO-824 | Gefitinib | MVP_top60_DNA+RNA_seed | AKT1 | -2.602613 | BRCA | YMB-1-E | MK-2206 |
| MVP_top60_DNA_seed | CDK2 | -0.973722 | BRCA | MRK-nu-1 | AZD5438 | MVP_top60_DNA+RNA_seed | EGFR | -2.580583 | BRCA | AU565 | Lapatinib |
| MVP_top60_DNA_seed | AKT1 | -0.969774 | BRCA | MDA-MB-175-VII | Uprosertib | MVP_top60_DNA+RNA_seed | ERBB2 | -2.580583 | BRCA | AU565 | Lapatinib |
| MVP_top60_DNA_seed | ATM | -0.962891 | BRCA | BT-549 | KU-55933 | MVP_top60_DNA+RNA_seed | EGFR | -2.550187 | BRCA | UACC-893 | Osimertinib |
| MVP_top60_DNA_seed | CDK2 | -0.962587 | BRCA | HCC1954 | AZD5438 | MVP_top60_DNA+RNA_seed | AKT1 | -2.527069 | BRCA | EVSA-T | MK-2206 |
| MVP_top60_DNA_seed | EGFR | -0.962319 | BRCA | DU-4475 | Gefitinib | MVP_top60_DNA+RNA_seed | AKT1 | -2.51042 | BRCA | HCC2157 | GSK2110183B |
| MVP_top60_DNA_seed | ESR1 | -0.960347 | BRCA | HCC1500 | GDC0810 | MVP_top60_DNA+RNA_seed | ESR1 | -2.502182 | BRCA | CAMA-1 | GDC0810 |
| MVP_top60_DNA_seed | ERBB2 | -0.95876 | BRCA | MDA-MB-468 | Afatinib | MVP_top60_DNA+RNA_seed | PIK3CA | -2.478395 | BRCA | MDA-MB-361 | Alpelisib |
| MVP_top60_DNA_seed | EGFR | -0.95876 | BRCA | MDA-MB-468 | Afatinib | MVP_top60_DNA+RNA_seed | PIK3CA | -2.467028 | BRCA | MDA-MB-330 | Alpelisib |
| MVP_top60_DNA_seed | CDK2 | -0.957054 | BRCA | EVSA-T | Dinaciclib | MVP_top60_DNA+RNA_seed | MAPK3 | -2.458065 | BRCA | DU-4475 | Ulixertinib |
| MVP_top60_DNA_seed | CDK1 | -0.957054 | BRCA | EVSA-T | Dinaciclib | MVP_top60_DNA+RNA_seed | MAPK1 | -2.458065 | BRCA | DU-4475 | Ulixertinib |
| MVP_top60_DNA_seed | AKT1 | -0.956436 | BRCA | HCC1395 | MK-2206 | MVP_top60_DNA+RNA_seed | AKT1 | -2.454941 | BRCA | HCC2157 | AZD5363 |
| MVP_top60_DNA_seed | AKT1 | -0.950649 | BRCA | HCC1187 | AT13148 | MVP_top60_DNA+RNA_seed | AKT1 | -2.448408 | BRCA | EVSA-T | Uprosertib |
| MVP_top60_DNA_seed | AKT1 | -0.945921 | BRCA | COLO-824 | Ipatasertib | MVP_top60_DNA+RNA_seed | AKT1 | -2.434601 | BRCA | EVSA-T | Uprosertib |
| MVP_top60_DNA_seed | AKT1 | -0.942142 | BRCA | HCC1395 | Uprosertib | MVP_top60_DNA+RNA_seed | EGFR | -2.433481 | BRCA | UACC-893 | Lapatinib |
| MVP_top60_DNA_seed | HSP90AA1 | -0.94195 | BRCA | CAL-148 | Luminespib | MVP_top60_DNA+RNA_seed | ERBB2 | -2.433481 | BRCA | UACC-893 | Lapatinib |
| MVP_top60_DNA_seed | HSP90AB1 | -0.94195 | BRCA | CAL-148 | Luminespib | MVP_top60_DNA+RNA_seed | AKT1 | -2.413609 | BRCA | HCC2157 | Afuresertib |
| MVP_top60_DNA_seed | MDM2 | -0.940786 | BRCA | OCUB-M | Serdemetan | MVP_top60_DNA+RNA_seed | EGFR | -2.391917 | BRCA | MDA-MB-330 | Osimertinib |
| MVP_top60_DNA_seed | MDM2 | -0.939178 | BRCA | MCF7 | Nutlin-3a (-) | MVP_top60_DNA+RNA_seed | AKT1 | -2.377263 | BRCA | YMB-1-E | Ipatasertib |
| MVP_top60_DNA_seed | ERBB2 | -0.93287 | BRCA | EVSA-T | Lapatinib | MVP_top60_DNA+RNA_seed | PIK3CA | -2.349952 | BRCA | OCUB-M | Alpelisib |
| MVP_top60_DNA_seed | EGFR | -0.93287 | BRCA | EVSA-T | Lapatinib | MVP_top60_DNA+RNA_seed | AKT1 | -2.345353 | BRCA | CAMA-1 | Afuresertib |
| MVP_top60_DNA_seed | PIK3CA | -0.932406 | BRCA | HDQ-P1 | Buparlisib | MVP_top60_DNA+RNA_seed | TP53 | -2.331311 | BRCA | HCC1599 | MIRA-1 |
| MVP_top60_DNA_seed | AKT1 | -0.932132 | BRCA | MDA-MB-453 | AT13148 | MVP_top60_DNA+RNA_seed | MAPK3 | -2.323983 | BRCA | MDA-MB-453 | Ulixertinib |
| MVP_top60_DNA_seed | PIK3CA | -0.931641 | BRCA | AU565 | Alpelisib | MVP_top60_DNA+RNA_seed | MAPK1 | -2.323983 | BRCA | MDA-MB-453 | Ulixertinib |
| MVP_top60_DNA_seed | ERBB2 | -0.9296 | BRCA | HCC1954 | Lapatinib | MVP_top60_DNA+RNA_seed | MAPK3 | -2.288881 | BRCA | MDA-MB-468 | Ulixertinib |
| MVP_top60_DNA_seed | EGFR | -0.9296 | BRCA | HCC1954 | Lapatinib | MVP_top60_DNA+RNA_seed | MAPK1 | -2.288881 | BRCA | MDA-MB-468 | Ulixertinib |
| MVP_top60_DNA_seed | PIK3CA | -0.927964 | BRCA | T47D | AZD8186 | MVP_top60_DNA+RNA_seed | AKT1 | -2.284351 | BRCA | MDA-MB-415 | AZD5363 |
| MVP_top60_DNA_seed | HDAC1 | -0.922488 | BRCA | HCC1500 | Entinostat | MVP_top60_DNA+RNA_seed | AKT1 | -2.279725 | BRCA | OCUB-M | MK-2206 |
| MVP_top60_DNA_seed | EGFR | -0.92219 | BRCA | HCC2157 | Osimertinib | MVP_top60_DNA+RNA_seed | AKT1 | -2.278051 | BRCA | CAMA-1 | Ipatasertib |
| MVP_top60_DNA_seed | SRC | -0.921268 | BRCA | MDA-MB-361 | Bosutinib | MVP_top60_DNA+RNA_seed | EGFR | -2.264811 | BRCA | ZR-75-30 | Afatinib |
| MVP_top60_DNA_seed | AKT1 | -0.92097 | BRCA | T47D | Afuresertib | MVP_top60_DNA+RNA_seed | ERBB2 | -2.264811 | BRCA | ZR-75-30 | Afatinib |
| MVP_top60_DNA_seed | ESR1 | -0.920678 | BRCA | MDA-MB-415 | Tamoxifen | MVP_top60_DNA+RNA_seed | EGFR | -2.262981 | BRCA | HDQ-P1 | Osimertinib |
| MVP_top60_DNA_seed | TP53 | -0.920382 | BRCA | MDA-MB-415 | MIRA-1 | MVP_top60_DNA+RNA_seed | ESR1 | -2.250288 | BRCA | EFM-19 | GDC0810 |
| MVP_top60_DNA_seed | HSP90AA1 | -0.919906 | BRCA | OCUB-M | Luminespib | MVP_top60_DNA+RNA_seed | AKT1 | -2.214992 | BRCA | MDA-MB-415 | Afuresertib |
| MVP_top60_DNA_seed | HSP90AB1 | -0.919906 | BRCA | OCUB-M | Luminespib | MVP_top60_DNA+RNA_seed | PIK3CA | -2.200339 | BRCA | UACC-893 | Alpelisib |
| MVP_top60_DNA_seed | PIK3CA | -0.917627 | BRCA | CAL-120 | GNE-317 | MVP_top60_DNA+RNA_seed | PIK3CA | -2.198675 | BRCA | MDA-MB-453 | Alpelisib |
| MVP_top60_DNA_seed | AKT1 | -0.916813 | BRCA | OCUB-M | AT13148 | MVP_top60_DNA+RNA_seed | AKT1 | -2.181131 | BRCA | UACC-893 | AZD5363 |
| MVP_top60_DNA_seed | MAPK3 | -0.915602 | BRCA | HCC1143 | Ulixertinib | MVP_top60_DNA+RNA_seed | AKT1 | -2.168426 | BRCA | UACC-893 | Uprosertib |
| MVP_top60_DNA_seed | MAPK1 | -0.915602 | BRCA | HCC1143 | Ulixertinib | MVP_top60_DNA+RNA_seed | AKT1 | -2.162809 | BRCA | HCC1569 | GSK2110183B |

| | | | | | | | | | | | |
|---|---|---|---|---|---|---|---|---|---|---|---|
| MVP_top60_DNA_seed | ERBB2 | -0.915335 | BRCA | COLO-824 | Lapatinib | MVP_top60_DNA+RNA_seed | AKT1 | -2.153509 | BRCA | MDA-MB-453 | GSK2110183B |
| MVP_top60_DNA_seed | EGFR | -0.915335 | BRCA | COLO-824 | Lapatinib | MVP_top60_DNA+RNA_seed | EGFR | -2.126294 | BRCA | HCC2157 | Lapatinib |
| MVP_top60_DNA_seed | SRC | -0.915074 | BRCA | HCC2157 | Bosutinib | MVP_top60_DNA+RNA_seed | ERBB2 | -2.126294 | BRCA | HCC2157 | Lapatinib |
| MVP_top60_DNA_seed | EP300 | -0.909177 | BRCA | EVSA-T | SGC-CBP30 | MVP_top60_DNA+RNA_seed | AKT1 | -2.118229 | BRCA | UACC-893 | MK-2206 |
| MVP_top60_DNA_seed | CREBBP | -0.909177 | BRCA | EVSA-T | SGC-CBP30 | MVP_top60_DNA+RNA_seed | AKT1 | -2.115307 | BRCA | MDA-MB-415 | Uprosertib |
| MVP_top60_DNA_seed | AKT1 | -0.90838 | BRCA | HCC1954 | AZD5363 | MVP_top60_DNA+RNA_seed | EGFR | -2.114903 | BRCA | ZR-75-30 | Lapatinib |
| MVP_top60_DNA_seed | ERBB2 | -0.906452 | BRCA | HCC1143 | Sapitinib | MVP_top60_DNA+RNA_seed | ERBB2 | -2.114903 | BRCA | ZR-75-30 | Lapatinib |
| MVP_top60_DNA_seed | EGFR | -0.906452 | BRCA | HCC1143 | Sapitinib | MVP_top60_DNA+RNA_seed | EGFR | -2.112493 | BRCA | AU565 | Afatinib |
| MVP_top60_DNA_seed | ERBB3 | -0.906452 | BRCA | HCC1143 | Sapitinib | MVP_top60_DNA+RNA_seed | ERBB2 | -2.112493 | BRCA | AU565 | Afatinib |
| MVP_top60_DNA_seed | EGFR | -0.903875 | BRCA | OCUB-M | AZD3759 | MVP_top60_DNA+RNA_seed | PIK3CA | -2.109527 | BRCA | HCC70 | AZD8186 |
| MVP_top60_DNA_seed | ESR1 | -0.901197 | BRCA | COLO-824 | GDC0810 | MVP_top60_DNA+RNA_seed | AKT1 | -2.10873 | BRCA | CAL-148 | MK-2206 |
| MVP_top60_DNA_seed | CDK1 | -0.896132 | BRCA | BT-549 | RO-3306 | MVP_top60_DNA+RNA_seed | AKT1 | -2.090514 | BRCA | MDA-MB-361 | Uprosertib |
| MVP_top60_DNA_seed | ESR1 | -0.895712 | BRCA | MRK-nu-1 | GDC0810 | MVP_top60_DNA+RNA_seed | AKT1 | -2.087608 | BRCA | MDA-MB-415 | GSK2110183B |
| MVP_top60_DNA_seed | AKT1 | -0.894297 | BRCA | CAL-51 | Uprosertib | MVP_top60_DNA+RNA_seed | EGFR | -2.086603 | BRCA | AU565 | Osimertinib |
| MVP_top60_DNA_seed | PIK3CA | -0.892798 | BRCA | T47D | Buparlisib | MVP_top60_DNA+RNA_seed | AKT1 | -2.085156 | BRCA | MDA-MB-453 | Uprosertib |
| MVP_top60_DNA_seed | ERBB2 | -0.892471 | BRCA | MDA-MB-361 | Afatinib | MVP_top60_DNA+RNA_seed | MDM2 | -2.073491 | BRCA | DU-4475 | Nutlin-3a (-) |
| MVP_top60_DNA_seed | EGFR | -0.892471 | BRCA | MDA-MB-361 | Afatinib | MVP_top60_DNA+RNA_seed | AKT1 | -2.06845 | BRCA | YMB-1-E | AZD5363 |
| MVP_top60_DNA_seed | CDK2 | -0.891599 | BRCA | HCC2157 | AZD5438 | MVP_top60_DNA+RNA_seed | EGFR | -2.058312 | BRCA | UACC-893 | Sapitinib |
| MVP_top60_DNA_seed | ESR1 | -0.891472 | BRCA | HCC1187 | GDC0810 | MVP_top60_DNA+RNA_seed | ERBB2 | -2.058312 | BRCA | UACC-893 | Sapitinib |
| MVP_top60_DNA_seed | ESR1 | -0.886636 | BRCA | HDQ-P1 | Tamoxifen | MVP_top60_DNA+RNA_seed | ERBB3 | -2.058312 | BRCA | UACC-893 | Sapitinib |
| MVP_top60_DNA_seed | HSP90AA1 | -0.883579 | BRCA | HCC2157 | Tanespimycin | MVP_top60_DNA+RNA_seed | AKT1 | -2.053109 | BRCA | HCC2157 | Ipatasertib |
| MVP_top60_DNA_seed | HSP90AB1 | -0.883579 | BRCA | HCC2157 | Tanespimycin | MVP_top60_DNA+RNA_seed | AKT1 | -2.046083 | BRCA | UACC-893 | Afuresertib |
| MVP_top60_DNA_seed | HDAC1 | -0.883106 | BRCA | UACC-893 | PCI-34051 | MVP_top60_DNA+RNA_seed | HSP90AA1 | -2.032451 | BRCA | MDA-MB-453 | Tanespimycin |
| MVP_top60_DNA_seed | PIK3CA | -0.882716 | BRCA | AU565 | AZD8186 | MVP_top60_DNA+RNA_seed | HSP90AB1 | -2.032451 | BRCA | MDA-MB-453 | Tanespimycin |
| MVP_top60_DNA_seed | AKT1 | -0.881891 | BRCA | COLO-824 | MK-2206 | MVP_top60_DNA+RNA_seed | TP53 | -2.017852 | BRCA | MRK-nu-1 | MIRA-1 |
| MVP_top60_DNA_seed | PIK3CA | -0.880401 | BRCA | Hs-578-T | AZD8186 | MVP_top60_DNA+RNA_seed | EGFR | -2.007439 | BRCA | HDQ-P1 | Sapitinib |
| MVP_top60_DNA_seed | SRC | -0.879599 | BRCA | HCC2218 | Bosutinib | MVP_top60_DNA+RNA_seed | ERBB2 | -2.007439 | BRCA | HDQ-P1 | Sapitinib |
| MVP_top60_DNA_seed | EP300 | -0.878249 | BRCA | MRK-nu-1 | CPI-637 | MVP_top60_DNA+RNA_seed | ERBB3 | -2.007439 | BRCA | HDQ-P1 | Sapitinib |
| MVP_top60_DNA_seed | AKT1 | -0.872927 | BRCA | EVSA-T | AT13148 | MVP_top60_DNA+RNA_seed | AKT1 | -1.996394 | BRCA | MDA-MB-415 | MK-2206 |
| MVP_top60_DNA_seed | ESR1 | -0.872757 | BRCA | AU565 | Tamoxifen | MVP_top60_DNA+RNA_seed | AKT1 | -1.996282 | BRCA | CAMA-1 | AZD5363 |
| MVP_top60_DNA_seed | AKT1 | -0.8665 | BRCA | Hs-578-T | AT13148 | MVP_top60_DNA+RNA_seed | ATM | -1.990948 | BRCA | MRK-nu-1 | KU-55933 |
| MVP_top60_DNA_seed | AKT1 | -0.861728 | BRCA | COLO-824 | AZD5363 | MVP_top60_DNA+RNA_seed | AKT1 | -1.988244 | BRCA | HCC1569 | Uprosertib |
| MVP_top60_DNA_seed | EGFR | -0.860086 | BRCA | DU-4475 | Osimertinib | MVP_top60_DNA+RNA_seed | PIK3CA | -1.979001 | BRCA | HCC2157 | AZD8186 |
| MVP_top60_DNA_seed | AKT1 | -0.856041 | BRCA | HCC1954 | AT13148 | MVP_top60_DNA+RNA_seed | AKT1 | -1.97538 | BRCA | YMB-1-E | Afuresertib |
| MVP_top60_DNA_seed | PIK3CA | -0.853556 | BRCA | MCF7 | Alpelisib | MVP_top60_DNA+RNA_seed | EGFR | -1.965332 | BRCA | BT-474 | Lapatinib |
| MVP_top60_DNA_seed | PIK3CA | -0.850261 | BRCA | EFM-192A | Alpelisib | MVP_top60_DNA+RNA_seed | ERBB2 | -1.965332 | BRCA | BT-474 | Lapatinib |
| MVP_top60_DNA_seed | ESR1 | -0.848816 | BRCA | OCUB-M | Tamoxifen | MVP_top60_DNA+RNA_seed | AKT1 | -1.963085 | BRCA | YMB-1-E | Uprosertib |
| MVP_top60_DNA_seed | EP300 | -0.841965 | BRCA | BT-20 | SGC-CBP30 | MVP_top60_DNA+RNA_seed | AKT1 | -1.95807 | BRCA | MDA-MB-453 | MK-2206 |
| MVP_top60_DNA_seed | CREBBP | -0.841965 | BRCA | BT-20 | SGC-CBP30 | MVP_top60_DNA+RNA_seed | ESR1 | -1.956708 | BRCA | T47D | GDC0810 |
| MVP_top60_DNA_seed | PIK3CA | -0.840747 | BRCA | MDA-MB-361 | AZD8186 | MVP_top60_DNA+RNA_seed | AKT1 | -1.942434 | BRCA | UACC-893 | Ipatasertib |
| MVP_top60_DNA_seed | MAPK3 | -0.837772 | BRCA | HDQ-P1 | Ulixertinib | MVP_top60_DNA+RNA_seed | EGFR | -1.938209 | BRCA | MDA-MB-175-VII | Erlotinib |

| | | | | | | | | | | | |
|---|---|---|---|---|---|---|---|---|---|---|---|
| MVP_top60_DNA_seed | MAPK1 | -0.837772 | BRCA | HDQ-P1 | Ulixertinib | MVP_top60_DNA+RNA_seed | EGFR | -1.935551 | BRCA | HCC2157 | Gefitinib |
| MVP_top60_DNA_seed | MAPK3 | -0.836915 | BRCA | UACC-893 | Ulixertinib | MVP_top60_DNA+RNA_seed | EGFR | -1.932837 | BRCA | HDQ-P1 | Gefitinib |
| MVP_top60_DNA_seed | MAPK1 | -0.836915 | BRCA | UACC-893 | Ulixertinib | MVP_top60_DNA+RNA_seed | AKT1 | -1.930333 | BRCA | HCC1954 | GSK2110183B |
| MVP_top60_DNA_seed | AKT1 | -0.835386 | BRCA | AU565 | GSK2110183B | MVP_top60_DNA+RNA_seed | ATM | -1.924923 | BRCA | Hs-578-T | KU-55933 |
| MVP_top60_DNA_seed | AKT1 | -0.832701 | BRCA | EFM-19 | Uprosertib | MVP_top60_DNA+RNA_seed | MDM2 | -1.919876 | BRCA | CAL-148 | Nutlin-3a (-) |
| MVP_top60_DNA_seed | MAPK3 | -0.83266 | BRCA | MDA-MB-231 | SCH772984 | MVP_top60_DNA+RNA_seed | HSP90AA1 | -1.914909 | BRCA | MDA-MB-361 | Tanespimycin |
| MVP_top60_DNA_seed | MAPK1 | -0.83266 | BRCA | MDA-MB-231 | SCH772984 | MVP_top60_DNA+RNA_seed | HSP90AB1 | -1.914909 | BRCA | MDA-MB-361 | Tanespimycin |
| MVP_top60_DNA_seed | AKT1 | -0.830537 | BRCA | MFM-223 | MK-2206 | MVP_top60_DNA+RNA_seed | HDAC1 | -1.913332 | BRCA | DU-4475 | Entinostat |
| MVP_top60_DNA_seed | ESR1 | -0.826148 | BRCA | CAL-148 | GDC0810 | MVP_top60_DNA+RNA_seed | EP300 | -1.912318 | BRCA | CAMA-1 | SGC-CBP30 |
| MVP_top60_DNA_seed | HSP90AA1 | -0.825778 | BRCA | HCC2157 | Luminespib | MVP_top60_DNA+RNA_seed | EGFR | -1.906474 | BRCA | OCUB-M | Osimertinib |
| MVP_top60_DNA_seed | HSP90AB1 | -0.825778 | BRCA | HCC2157 | Luminespib | MVP_top60_DNA+RNA_seed | HSP90AA1 | -1.896893 | BRCA | MDA-MB-330 | Tanespimycin |
| MVP_top60_DNA_seed | EP300 | -0.825162 | BRCA | HDQ-P1 | SGC-CBP30 | MVP_top60_DNA+RNA_seed | HSP90AB1 | -1.896893 | BRCA | MDA-MB-330 | Tanespimycin |
| MVP_top60_DNA_seed | CREBBP | -0.825162 | BRCA | HDQ-P1 | SGC-CBP30 | MVP_top60_DNA+RNA_seed | MAPK3 | -1.894407 | BRCA | MRK-nu-1 | Ulixertinib |
| MVP_top60_DNA_seed | CDK2 | -0.823937 | BRCA | HCC70 | MK-8776 | MVP_top60_DNA+RNA_seed | MAPK1 | -1.894407 | BRCA | MRK-nu-1 | Ulixertinib |
| MVP_top60_DNA_seed | AKT1 | -0.822013 | BRCA | CAL-51 | GSK2110183B | MVP_top60_DNA+RNA_seed | AKT1 | -1.893713 | BRCA | HCC1569 | Afuresertib |
| MVP_top60_DNA_seed | PIK3CA | -0.819784 | BRCA | COLO-824 | AZD8186 | MVP_top60_DNA+RNA_seed | AKT1 | -1.890909 | BRCA | CAL-148 | AT13148 |
| MVP_top60_DNA_seed | AKT1 | -0.81715 | BRCA | MDA-MB-453 | AZD5363 | MVP_top60_DNA+RNA_seed | PIK3CA | -1.88702 | BRCA | MDA-MB-330 | AZD8186 |
| MVP_top60_DNA_seed | CDK2 | -0.81652 | BRCA | HCC38 | MK-8776 | MVP_top60_DNA+RNA_seed | PIK3CA | -1.882454 | BRCA | CAMA-1 | AZD8186 |
| MVP_top60_DNA_seed | MAPK3 | -0.816422 | BRCA | MDA-MB-175-VII | Ulixertinib | MVP_top60_DNA+RNA_seed | AKT1 | -1.877085 | BRCA | BT-20 | GSK2110183B |
| MVP_top60_DNA_seed | MAPK1 | -0.816422 | BRCA | MDA-MB-175-VII | Ulixertinib | MVP_top60_DNA+RNA_seed | AKT1 | -1.869376 | BRCA | HCC70 | Ipatasertib |
| MVP_top60_DNA_seed | CDK2 | -0.815706 | BRCA | HCC70 | Dinaciclib | MVP_top60_DNA+RNA_seed | AKT1 | -1.868896 | BRCA | HCC2157 | Uprosertib |
| MVP_top60_DNA_seed | CDK1 | -0.815706 | BRCA | HCC70 | Dinaciclib | MVP_top60_DNA+RNA_seed | AKT1 | -1.850647 | BRCA | HCC1569 | Ipatasertib |
| MVP_top60_DNA_seed | PIK3CA | -0.809161 | BRCA | MRK-nu-1 | Buparlisib | MVP_top60_DNA+RNA_seed | EGFR | -1.837248 | BRCA | AU565 | Sapitinib |
| MVP_top60_DNA_seed | ERBB2 | -0.805849 | BRCA | HDQ-P1 | Afatinib | MVP_top60_DNA+RNA_seed | ERBB2 | -1.837248 | BRCA | AU565 | Sapitinib |
| MVP_top60_DNA_seed | EGFR | -0.805849 | BRCA | HDQ-P1 | Afatinib | MVP_top60_DNA+RNA_seed | ERBB3 | -1.837248 | BRCA | AU565 | Sapitinib |
| MVP_top60_DNA_seed | ATM | -0.802533 | BRCA | MDA-MB-468 | KU-55933 | MVP_top60_DNA+RNA_seed | HDAC1 | -1.822743 | BRCA | CAL-148 | Entinostat |
| MVP_top60_DNA_seed | ERBB2 | -0.800196 | BRCA | BT-20 | Sapitinib | MVP_top60_DNA+RNA_seed | EGFR | -1.813625 | BRCA | MDA-MB-330 | Lapatinib |
| MVP_top60_DNA_seed | EGFR | -0.800196 | BRCA | BT-20 | Sapitinib | MVP_top60_DNA+RNA_seed | ERBB2 | -1.813625 | BRCA | MDA-MB-330 | Lapatinib |
| MVP_top60_DNA_seed | ERBB3 | -0.800196 | BRCA | BT-20 | Sapitinib | MVP_top60_DNA+RNA_seed | ESR1 | -1.79809 | BRCA | HCC1419 | GDC0810 |
| MVP_top60_DNA_seed | HSP90AA1 | -0.79664 | BRCA | COLO-824 | Luminespib | MVP_top60_DNA+RNA_seed | EGFR | -1.783818 | BRCA | EFM-192A | Osimertinib |
| MVP_top60_DNA_seed | HSP90AB1 | -0.79664 | BRCA | COLO-824 | Luminespib | MVP_top60_DNA+RNA_seed | EGFR | -1.777703 | BRCA | HDQ-P1 | Erlotinib |
| MVP_top60_DNA_seed | CDK2 | -0.786787 | BRCA | MDA-MB-330 | Dinaciclib | MVP_top60_DNA+RNA_seed | AKT1 | -1.768701 | BRCA | HCC70 | AZD5363 |
| MVP_top60_DNA_seed | CDK1 | -0.786787 | BRCA | MDA-MB-330 | Dinaciclib | MVP_top60_DNA+RNA_seed | ESR1 | -1.762729 | BRCA | HCC2157 | Tamoxifen |
| MVP_top60_DNA_seed | AKT1 | -0.785847 | BRCA | HCC1395 | AZD5363 | MVP_top60_DNA+RNA_seed | AKT1 | -1.756361 | BRCA | HCC70 | Uprosertib |
| MVP_top60_DNA_seed | ESR1 | -0.78522 | BRCA | MDA-MB-157 | GDC0810 | MVP_top60_DNA+RNA_seed | AKT1 | -1.744543 | BRCA | MDA-MB-415 | Uprosertib |
| MVP_top60_DNA_seed | ERBB2 | -0.780079 | BRCA | JIMT-1 | Sapitinib | MVP_top60_DNA+RNA_seed | CDK2 | -1.742885 | BRCA | MDA-MB-361 | MK-8776 |
| MVP_top60_DNA_seed | EGFR | -0.780079 | BRCA | JIMT-1 | Sapitinib | MVP_top60_DNA+RNA_seed | EGFR | -1.734971 | BRCA | BT-474 | Afatinib |
| MVP_top60_DNA_seed | ERBB3 | -0.780079 | BRCA | JIMT-1 | Sapitinib | MVP_top60_DNA+RNA_seed | ERBB2 | -1.734971 | BRCA | BT-474 | Afatinib |
| MVP_top60_DNA_seed | HDAC1 | -0.774694 | BRCA | HCC1569 | Romidepsin | MVP_top60_DNA+RNA_seed | AKT1 | -1.733572 | BRCA | AU565 | AT13148 |
| MVP_top60_DNA_seed | HDAC2 | -0.774694 | BRCA | HCC1569 | Romidepsin | MVP_top60_DNA+RNA_seed | HDAC1 | -1.730747 | BRCA | HCC2157 | Entinostat |
| MVP_top60_DNA_seed | ERBB2 | -0.774536 | BRCA | HCC1569 | Lapatinib | MVP_top60_DNA+RNA_seed | PIK3CA | -1.729419 | BRCA | CAL-148 | Alpelisib |

| | | | | | | | | | | | |
|---|---|---|---|---|---|---|---|---|---|---|---|
| MVP_top60_DNA_seed | EGFR | -0.774536 | BRCA | HCC1569 | Lapatinib | MVP_top60_DNA+RNA_seed | AKT1 | -1.723778 | BRCA | HCC70 | MK-2206 |
| MVP_top60_DNA_seed | PIK3CA | -0.772239 | BRCA | OCUB-M | Buparlisib | MVP_top60_DNA+RNA_seed | EGFR | -1.706325 | BRCA | OCUB-M | Afatinib |
| MVP_top60_DNA_seed | MDM2 | -0.770785 | BRCA | YMB-1-E | Nutlin-3a (-) | MVP_top60_DNA+RNA_seed | ERBB2 | -1.706325 | BRCA | OCUB-M | Afatinib |
| MVP_top60_DNA_seed | EP300 | -0.767298 | BRCA | MDA-MB-453 | SGC-CBP30 | MVP_top60_DNA+RNA_seed | AKT1 | -1.705594 | BRCA | HCC2157 | MK-2206 |
| MVP_top60_DNA_seed | CREBBP | -0.767298 | BRCA | MDA-MB-453 | SGC-CBP30 | MVP_top60_DNA+RNA_seed | EP300 | -1.70371 | BRCA | CAL-148 | CPI-637 |
| MVP_top60_DNA_seed | HSP90AA1 | -0.767242 | BRCA | MRK-nu-1 | Tanespimycin | MVP_top60_DNA+RNA_seed | CDK1 | -1.692986 | BRCA | MRK-nu-1 | RO-3306 |
| MVP_top60_DNA_seed | HSP90AB1 | -0.767242 | BRCA | MRK-nu-1 | Tanespimycin | MVP_top60_DNA+RNA_seed | AKT1 | -1.689145 | BRCA | AU565 | Uprosertib |
| MVP_top60_DNA_seed | AKT1 | -0.766869 | BRCA | HCC202 | MK-2206 | MVP_top60_DNA+RNA_seed | PLK1 | -1.686304 | BRCA | MRK-nu-1 | BI-2536 |
| MVP_top60_DNA_seed | CDK1 | -0.764868 | BRCA | COLO-824 | RO-3306 | MVP_top60_DNA+RNA_seed | AKT1 | -1.681967 | BRCA | MDA-MB-175-VII | Afuresertib |
| MVP_top60_DNA_seed | EGFR | -0.762929 | BRCA | MDA-MB-468 | Osimertinib | MVP_top60_DNA+RNA_seed | PIK3CA | -1.675579 | BRCA | BT-20 | Alpelisib |
| MVP_top60_DNA_seed | AKT1 | -0.762558 | BRCA | DU-4475 | Uprosertib | MVP_top60_DNA+RNA_seed | PIK3CA | -1.673389 | BRCA | OCUB-M | GNE-317 |
| MVP_top60_DNA_seed | AKT1 | -0.762364 | BRCA | MCF7 | Uprosertib | MVP_top60_DNA+RNA_seed | EGFR | -1.655282 | BRCA | AU565 | Gefitinib |
| MVP_top60_DNA_seed | ERBB2 | -0.75585 | BRCA | HCC1143 | Lapatinib | MVP_top60_DNA+RNA_seed | MAPK3 | -1.650711 | BRCA | MDA-MB-157 | Ulixertinib |
| MVP_top60_DNA_seed | EGFR | -0.75585 | BRCA | HCC1143 | Lapatinib | MVP_top60_DNA+RNA_seed | MAPK1 | -1.650711 | BRCA | MDA-MB-157 | Ulixertinib |
| MVP_top60_DNA_seed | MDM2 | -0.755338 | BRCA | AU565 | Serdemetan | MVP_top60_DNA+RNA_seed | CDK1 | -1.642671 | BRCA | DU-4475 | RO-3306 |
| MVP_top60_DNA_seed | SRC | -0.753389 | BRCA | DU-4475 | Bosutinib | MVP_top60_DNA+RNA_seed | AKT1 | -1.636872 | BRCA | HCC1569 | AZD5363 |
| MVP_top60_DNA_seed | EGFR | -0.751663 | BRCA | DU-4475 | Erlotinib | MVP_top60_DNA+RNA_seed | AKT1 | -1.629614 | BRCA | MFM-223 | Afuresertib |
| MVP_top60_DNA_seed | CDK2 | -0.750382 | BRCA | MDA-MB-231 | AZD5438 | MVP_top60_DNA+RNA_seed | EGFR | -1.622725 | BRCA | EFM-192A | Lapatinib |
| MVP_top60_DNA_seed | AKT1 | -0.74949 | BRCA | MDA-MB-330 | MK-2206 | MVP_top60_DNA+RNA_seed | ERBB2 | -1.622725 | BRCA | EFM-192A | Lapatinib |
| MVP_top60_DNA_seed | CDK2 | -0.749482 | BRCA | MRK-nu-1 | Dinaciclib | MVP_top60_DNA+RNA_seed | AKT1 | -1.61593 | BRCA | MDA-MB-415 | AT13148 |
| MVP_top60_DNA_seed | CDK1 | -0.749482 | BRCA | MRK-nu-1 | Dinaciclib | MVP_top60_DNA+RNA_seed | AKT1 | -1.609161 | BRCA | CAMA-1 | GSK2110183B |
| MVP_top60_DNA_seed | AKT1 | -0.746946 | BRCA | HDQ-P1 | Uprosertib | MVP_top60_DNA+RNA_seed | PIK3CA | -1.607008 | BRCA | EFM-19 | Alpelisib |
| MVP_top60_DNA_seed | ESR1 | -0.74636 | BRCA | MDA-MB-361 | Tamoxifen | MVP_top60_DNA+RNA_seed | EGFR | -1.597645 | BRCA | BT-474 | Sapitinib |
| MVP_top60_DNA_seed | PIK3CA | -0.745503 | BRCA | HCC1143 | AZD8186 | MVP_top60_DNA+RNA_seed | ERBB2 | -1.597645 | BRCA | BT-474 | Sapitinib |
| MVP_top60_DNA_seed | PIK3CA | -0.743736 | BRCA | Hs-578-T | Buparlisib | MVP_top60_DNA+RNA_seed | ERBB3 | -1.597645 | BRCA | BT-474 | Sapitinib |
| MVP_top60_DNA_seed | PIK3CA | -0.741553 | BRCA | MDA-MB-468 | AZD8186 | MVP_top60_DNA+RNA_seed | AKT1 | -1.585314 | BRCA | MDA-MB-453 | Uprosertib |
| MVP_top60_DNA_seed | CDK2 | -0.738308 | BRCA | Hs-578-T | AZD5438 | MVP_top60_DNA+RNA_seed | EGFR | -1.584241 | BRCA | ZR-75-30 | Osimertinib |
| MVP_top60_DNA_seed | AKT1 | -0.737288 | BRCA | UACC-893 | AT13148 | MVP_top60_DNA+RNA_seed | CDK2 | -1.574131 | BRCA | HCC1395 | MK-8776 |
| MVP_top60_DNA_seed | PIK3CA | -0.73116 | BRCA | JIMT-1 | GNE-317 | MVP_top60_DNA+RNA_seed | MAPK3 | -1.573528 | BRCA | CAL-85-1 | Ulixertinib |
| MVP_top60_DNA_seed | PIK3CA | -0.730409 | BRCA | Hs-578-T | Alpelisib | MVP_top60_DNA+RNA_seed | MAPK1 | -1.573528 | BRCA | CAL-85-1 | Ulixertinib |
| MVP_top60_DNA_seed | CDK2 | -0.72988 | BRCA | OCUB-M | Dinaciclib | MVP_top60_DNA+RNA_seed | AKT1 | -1.569484 | BRCA | MDA-MB-175-VII | Uprosertib |
| MVP_top60_DNA_seed | CDK1 | -0.72988 | BRCA | OCUB-M | Dinaciclib | MVP_top60_DNA+RNA_seed | HDAC1 | -1.559249 | BRCA | MDA-MB-330 | Romidepsin |
| MVP_top60_DNA_seed | CDK2 | -0.727971 | BRCA | MDA-MB-157 | Dinaciclib | MVP_top60_DNA+RNA_seed | HDAC2 | -1.559249 | BRCA | MDA-MB-330 | Romidepsin |
| MVP_top60_DNA_seed | CDK1 | -0.727971 | BRCA | MDA-MB-157 | Dinaciclib | MVP_top60_DNA+RNA_seed | CDK2 | -1.558595 | BRCA | DU-4475 | AZD5438 |
| MVP_top60_DNA_seed | AKT1 | -0.726847 | BRCA | AU565 | AZD5363 | MVP_top60_DNA+RNA_seed | MDM2 | -1.551871 | BRCA | DU-4475 | Serdemetan |
| MVP_top60_DNA_seed | AKT1 | -0.726761 | BRCA | AU565 | Uprosertib | MVP_top60_DNA+RNA_seed | AKT1 | -1.551098 | BRCA | MDA-MB-361 | GSK2110183B |
| MVP_top60_DNA_seed | MDM2 | -0.724274 | BRCA | COLO-824 | Serdemetan | MVP_top60_DNA+RNA_seed | PIK3CA | -1.544994 | BRCA | MRK-nu-1 | GNE-317 |
| MVP_top60_DNA_seed | AKT1 | -0.722234 | BRCA | EFM-19 | Afuresertib | MVP_top60_DNA+RNA_seed | ATM | -1.538073 | BRCA | EVSA-T | KU-55933 |
| MVP_top60_DNA_seed | ERBB2 | -0.719294 | BRCA | JIMT-1 | Lapatinib | MVP_top60_DNA+RNA_seed | AKT1 | -1.524496 | BRCA | MDA-MB-175-VII | Ipatasertib |
| MVP_top60_DNA_seed | EGFR | -0.719294 | BRCA | JIMT-1 | Lapatinib | MVP_top60_DNA+RNA_seed | AKT1 | -1.517368 | BRCA | CAMA-1 | Uprosertib |
| MVP_top60_DNA_seed | EGFR | -0.717571 | BRCA | CAL-148 | Osimertinib | MVP_top60_DNA+RNA_seed | EP300 | -1.517151 | BRCA | MDA-MB-453 | CPI-637 |

| | | | | | | | | | | | |
|---|---|---|---|---|---|---|---|---|---|---|---|
| MVP_top60_DNA_seed | EGFR | -0.716149 | BRCA | MDA-MB-468 | Erlotinib | MVP_top60_DNA+RNA_seed | ATM | -1.516846 | BRCA | MDA-MB-415 | KU-55933 |
| MVP_top60_DNA_seed | HSP90AA1 | -0.716111 | BRCA | MDA-MB-231 | Luminespib | MVP_top60_DNA+RNA_seed | ATM | -1.510452 | BRCA | DU-4475 | KU-55933 |
| MVP_top60_DNA_seed | HSP90AB1 | -0.716111 | BRCA | MDA-MB-231 | Luminespib | MVP_top60_DNA+RNA_seed | EGFR | -1.506965 | BRCA | OCUB-M | Sapitinib |
| MVP_top60_DNA_seed | CDK2 | -0.714111 | BRCA | HCC70 | AZD5438 | MVP_top60_DNA+RNA_seed | ERBB2 | -1.506965 | BRCA | OCUB-M | Sapitinib |
| MVP_top60_DNA_seed | AKT1 | -0.713725 | BRCA | AU565 | Afuresertib | MVP_top60_DNA+RNA_seed | ERBB3 | -1.506965 | BRCA | OCUB-M | Sapitinib |
| MVP_top60_DNA_seed | EGFR | -0.711807 | BRCA | AU565 | Erlotinib | MVP_top60_DNA+RNA_seed | EGFR | -1.50638 | BRCA | MDA-MB-330 | AZD3759 |
| MVP_top60_DNA_seed | EGFR | -0.708757 | BRCA | UACC-893 | Erlotinib | MVP_top60_DNA+RNA_seed | EGFR | -1.504361 | BRCA | HCC2218 | Sapitinib |
| MVP_top60_DNA_seed | MDM2 | -0.70811 | BRCA | UACC-812 | Nutlin-3a (-) | MVP_top60_DNA+RNA_seed | ERBB2 | -1.504361 | BRCA | HCC2218 | Sapitinib |
| MVP_top60_DNA_seed | EP300 | -0.707949 | BRCA | AU565 | SGC-CBP30 | MVP_top60_DNA+RNA_seed | ERBB3 | -1.504361 | BRCA | HCC2218 | Sapitinib |
| MVP_top60_DNA_seed | CREBBP | -0.707949 | BRCA | AU565 | SGC-CBP30 | MVP_top60_DNA+RNA_seed | HSP90AA1 | -1.504166 | BRCA | COLO-824 | Tanespimycin |
| MVP_top60_DNA_seed | EGFR | -0.707066 | BRCA | COLO-824 | Osimertinib | MVP_top60_DNA+RNA_seed | HSP90AB1 | -1.504166 | BRCA | COLO-824 | Tanespimycin |
| MVP_top60_DNA_seed | PIK3CA | -0.704781 | BRCA | CAL-148 | AZD8186 | MVP_top60_DNA+RNA_seed | ESR1 | -1.503186 | BRCA | COLO-824 | Tamoxifen |
| MVP_top60_DNA_seed | AKT1 | -0.703104 | BRCA | MDA-MB-361 | Afuresertib | MVP_top60_DNA+RNA_seed | AKT1 | -1.502818 | BRCA | MDA-MB-175-VII | MK-2206 |
| MVP_top60_DNA_seed | PIK3CA | -0.702855 | BRCA | HCC1395 | Buparlisib | MVP_top60_DNA+RNA_seed | AKT1 | -1.501786 | BRCA | MFM-223 | Uprosertib |
| MVP_top60_DNA_seed | HDAC1 | -0.699312 | BRCA | CAL-85-1 | PCI-34051 | MVP_top60_DNA+RNA_seed | EGFR | -1.486205 | BRCA | OCUB-M | Lapatinib |
| MVP_top60_DNA_seed | HSP90AA1 | -0.695131 | BRCA | JIMT-1 | Luminespib | MVP_top60_DNA+RNA_seed | ERBB2 | -1.486205 | BRCA | OCUB-M | Lapatinib |
| MVP_top60_DNA_seed | HSP90AB1 | -0.695131 | BRCA | JIMT-1 | Luminespib | MVP_top60_DNA+RNA_seed | AKT1 | -1.476894 | BRCA | MFM-223 | AZD5363 |
| MVP_top60_DNA_seed | AKT1 | -0.692554 | BRCA | COLO-824 | Uprosertib | MVP_top60_DNA+RNA_seed | MDM2 | -1.474321 | BRCA | MRK-nu-1 | Nutlin-3a (-) |
| MVP_top60_DNA_seed | PIK3CA | -0.691065 | BRCA | EFM-192A | GNE-317 | MVP_top60_DNA+RNA_seed | HSP90AA1 | -1.474037 | BRCA | CAL-51 | Luminespib |
| MVP_top60_DNA_seed | AKT1 | -0.690475 | BRCA | AU565 | Ipatasertib | MVP_top60_DNA+RNA_seed | HSP90AB1 | -1.474037 | BRCA | CAL-51 | Luminespib |
| MVP_top60_DNA_seed | CDK2 | -0.689955 | BRCA | DU-4475 | Dinaciclib | MVP_top60_DNA+RNA_seed | HDAC1 | -1.47315 | BRCA | EVSA-T | Entinostat |
| MVP_top60_DNA_seed | CDK1 | -0.689955 | BRCA | DU-4475 | Dinaciclib | MVP_top60_DNA+RNA_seed | HDAC1 | -1.472168 | BRCA | HCC1599 | PCI-34051 |
| MVP_top60_DNA_seed | AKT1 | -0.686996 | BRCA | HCC1954 | Ipatasertib | MVP_top60_DNA+RNA_seed | HSP90AA1 | -1.469543 | BRCA | DU-4475 | Tanespimycin |
| MVP_top60_DNA_seed | AKT1 | -0.686102 | BRCA | AU565 | MK-2206 | MVP_top60_DNA+RNA_seed | HSP90AB1 | -1.469543 | BRCA | DU-4475 | Tanespimycin |
| MVP_top60_DNA_seed | ATM | -0.684788 | BRCA | HCC1419 | KU-55933 | MVP_top60_DNA+RNA_seed | HSP90AA1 | -1.465141 | BRCA | MDA-MB-330 | CCT-018159 |
| MVP_top60_DNA_seed | ERBB2 | -0.679345 | BRCA | HCC1395 | Lapatinib | MVP_top60_DNA+RNA_seed | HSP90AB1 | -1.465141 | BRCA | MDA-MB-330 | CCT-018159 |
| MVP_top60_DNA_seed | EGFR | -0.679345 | BRCA | HCC1395 | Lapatinib | MVP_top60_DNA+RNA_seed | AKT1 | -1.463847 | BRCA | MFM-223 | Ipatasertib |
| MVP_top60_DNA_seed | HDAC1 | -0.678338 | BRCA | CAL-85-1 | Romidepsin | MVP_top60_DNA+RNA_seed | AKT1 | -1.459284 | BRCA | CAMA-1 | Uprosertib |
| MVP_top60_DNA_seed | HDAC2 | -0.678338 | BRCA | CAL-85-1 | Romidepsin | MVP_top60_DNA+RNA_seed | CDK1 | -1.454586 | BRCA | OCUB-M | RO-3306 |
| MVP_top60_DNA_seed | HDAC1 | -0.677763 | BRCA | MDA-MB-453 | Entinostat | MVP_top60_DNA+RNA_seed | PIK3CA | -1.452375 | BRCA | MDA-MB-415 | GNE-317 |
| MVP_top60_DNA_seed | PIK3CA | -0.67704 | BRCA | UACC-812 | AZD8186 | MVP_top60_DNA+RNA_seed | EGFR | -1.451084 | BRCA | MDA-MB-468 | Sapitinib |
| MVP_top60_DNA_seed | ERBB2 | -0.675895 | BRCA | HCC1954 | Sapitinib | MVP_top60_DNA+RNA_seed | ERBB2 | -1.451084 | BRCA | MDA-MB-468 | Sapitinib |
| MVP_top60_DNA_seed | EGFR | -0.675895 | BRCA | HCC1954 | Sapitinib | MVP_top60_DNA+RNA_seed | ERBB3 | -1.451084 | BRCA | MDA-MB-468 | Sapitinib |
| MVP_top60_DNA_seed | ERBB3 | -0.675895 | BRCA | HCC1954 | Sapitinib | MVP_top60_DNA+RNA_seed | HDAC1 | -1.449642 | BRCA | MDA-MB-468 | PCI-34051 |
| MVP_top60_DNA_seed | CDK2 | -0.668885 | BRCA | CAL-51 | Dinaciclib | MVP_top60_DNA+RNA_seed | HDAC1 | -1.4418 | BRCA | HCC1599 | Entinostat |
| MVP_top60_DNA_seed | CDK1 | -0.668885 | BRCA | CAL-51 | Dinaciclib | MVP_top60_DNA+RNA_seed | AKT1 | -1.43768 | BRCA | T47D | Ipatasertib |
| MVP_top60_DNA_seed | AKT1 | -0.668123 | BRCA | BT-20 | Uprosertib | MVP_top60_DNA+RNA_seed | EGFR | -1.432996 | BRCA | ZR-75-30 | Sapitinib |
| MVP_top60_DNA_seed | ESR1 | -0.666552 | BRCA | MDA-MB-453 | Tamoxifen | MVP_top60_DNA+RNA_seed | ERBB2 | -1.432996 | BRCA | ZR-75-30 | Sapitinib |
| MVP_top60_DNA_seed | HSP90AA1 | -0.663581 | BRCA | HCC1419 | Tanespimycin | MVP_top60_DNA+RNA_seed | ERBB3 | -1.432996 | BRCA | ZR-75-30 | Sapitinib |
| MVP_top60_DNA_seed | HSP90AB1 | -0.663581 | BRCA | HCC1419 | Tanespimycin | MVP_top60_DNA+RNA_seed | AKT1 | -1.42518 | BRCA | HCC70 | Uprosertib |
| MVP_top60_DNA_seed | PIK3CA | -0.660445 | BRCA | EFM-192A | Buparlisib | MVP_top60_DNA+RNA_seed | PIK3CA | -1.424528 | BRCA | HCC2157 | GNE-317 |

| | | | | | | | | | | | |
|---|---|---|---|---|---|---|---|---|---|---|---|
| MVP_top60_DNA_seed | AKT1 | -0.656801 | BRCA | MDA-MB-468 | Afuresertib | MVP_top60_DNA+RNA_seed | CDK1 | -1.423225 | BRCA | CAL-148 | RO-3306 |
| MVP_top60_DNA_seed | HDAC1 | -0.654747 | BRCA | HCC1428 | Entinostat | MVP_top60_DNA+RNA_seed | EGFR | -1.421015 | BRCA | BT-474 | Osimertinib |
| MVP_top60_DNA_seed | ERBB2 | -0.654271 | BRCA | BT-20 | Lapatinib | MVP_top60_DNA+RNA_seed | AKT1 | -1.410435 | BRCA | BT-20 | Uprosertib |
| MVP_top60_DNA_seed | EGFR | -0.654271 | BRCA | BT-20 | Lapatinib | MVP_top60_DNA+RNA_seed | EGFR | -1.409896 | BRCA | MDA-MB-330 | Sapitinib |
| MVP_top60_DNA_seed | PIK3CA | -0.653864 | BRCA | AU565 | GNE-317 | MVP_top60_DNA+RNA_seed | ERBB2 | -1.409896 | BRCA | MDA-MB-330 | Sapitinib |
| MVP_top60_DNA_seed | HDAC1 | -0.653387 | BRCA | MDA-MB-231 | Romidepsin | MVP_top60_DNA+RNA_seed | ERBB3 | -1.409896 | BRCA | MDA-MB-330 | Sapitinib |
| MVP_top60_DNA_seed | HDAC2 | -0.653387 | BRCA | MDA-MB-231 | Romidepsin | MVP_top60_DNA+RNA_seed | AKT1 | -1.40714 | BRCA | MFM-223 | GSK2110183B |
| MVP_top60_DNA_seed | AKT1 | -0.653351 | BRCA | HDQ-P1 | AZD5363 | MVP_top60_DNA+RNA_seed | AKT1 | -1.405907 | BRCA | MRK-nu-1 | AT13148 |
| MVP_top60_DNA_seed | PIK3CA | -0.652526 | BRCA | MDA-MB-330 | Buparlisib | MVP_top60_DNA+RNA_seed | ESR1 | -1.402278 | BRCA | EVSA-T | GDC0810 |
| MVP_top60_DNA_seed | PIK3CA | -0.652473 | BRCA | UACC-812 | Alpelisib | MVP_top60_DNA+RNA_seed | PIK3CA | -1.401092 | BRCA | HCC1954 | Alpelisib |
| MVP_top60_DNA_seed | CDK2 | -0.651474 | BRCA | HCC1937 | MK-8776 | MVP_top60_DNA+RNA_seed | AKT1 | -1.400544 | BRCA | CAL-51 | Ipatasertib |
| MVP_top60_DNA_seed | EGFR | -0.65124 | BRCA | COLO-824 | Erlotinib | MVP_top60_DNA+RNA_seed | PIK3CA | -1.397942 | BRCA | MDA-MB-175-VII | GNE-317 |
| MVP_top60_DNA_seed | EP300 | -0.648117 | BRCA | Hs-578-T | CPI-637 | MVP_top60_DNA+RNA_seed | MDM2 | -1.396027 | BRCA | CAL-148 | Serdemetan |
| MVP_top60_DNA_seed | TP53 | -0.647937 | BRCA | EVSA-T | MIRA-1 | MVP_top60_DNA+RNA_seed | ATM | -1.393523 | BRCA | HCC70 | KU-55933 |
| MVP_top60_DNA_seed | ERBB2 | -0.643898 | BRCA | MDA-MB-453 | Sapitinib | MVP_top60_DNA+RNA_seed | HDAC1 | -1.392804 | BRCA | OCUB-M | PCI-34051 |
| MVP_top60_DNA_seed | EGFR | -0.643898 | BRCA | MDA-MB-453 | Sapitinib | MVP_top60_DNA+RNA_seed | EGFR | -1.390275 | BRCA | UACC-812 | Lapatinib |
| MVP_top60_DNA_seed | ERBB3 | -0.643898 | BRCA | MDA-MB-453 | Sapitinib | MVP_top60_DNA+RNA_seed | ERBB2 | -1.390275 | BRCA | UACC-812 | Lapatinib |
| MVP_top60_DNA_seed | AKT1 | -0.643539 | BRCA | HCC1395 | Uprosertib | MVP_top60_DNA+RNA_seed | AKT1 | -1.388491 | BRCA | HCC70 | GSK2110183B |
| MVP_top60_DNA_seed | HSP90AA1 | -0.64115 | BRCA | MDA-MB-330 | Luminespib | MVP_top60_DNA+RNA_seed | AKT1 | -1.380943 | BRCA | EFM-19 | Ipatasertib |
| MVP_top60_DNA_seed | HSP90AB1 | -0.64115 | BRCA | MDA-MB-330 | Luminespib | MVP_top60_DNA+RNA_seed | AKT1 | -1.375726 | BRCA | MFM-223 | Uprosertib |
| MVP_top60_DNA_seed | ESR1 | -0.637152 | BRCA | CAL-85-1 | GDC0810 | MVP_top60_DNA+RNA_seed | EGFR | -1.374025 | BRCA | HCC1419 | Osimertinib |
| MVP_top60_DNA_seed | HSP90AA1 | -0.635658 | BRCA | CAL-85-1 | Luminespib | MVP_top60_DNA+RNA_seed | AKT1 | -1.372254 | BRCA | HCC70 | Afuresertib |
| MVP_top60_DNA_seed | HSP90AB1 | -0.635658 | BRCA | CAL-85-1 | Luminespib | MVP_top60_DNA+RNA_seed | PIK3CA | -1.37047 | BRCA | MDA-MB-361 | GNE-317 |
| MVP_top60_DNA_seed | EGFR | -0.63156 | BRCA | HCC1143 | Erlotinib | MVP_top60_DNA+RNA_seed | EGFR | -1.367107 | BRCA | UACC-812 | Sapitinib |
| MVP_top60_DNA_seed | EP300 | -0.629102 | BRCA | OCUB-M | CPI-637 | MVP_top60_DNA+RNA_seed | ERBB2 | -1.367107 | BRCA | UACC-812 | Sapitinib |
| MVP_top60_DNA_seed | EGFR | -0.628902 | BRCA | EVSA-T | Erlotinib | MVP_top60_DNA+RNA_seed | ERBB3 | -1.367107 | BRCA | UACC-812 | Sapitinib |
| MVP_top60_DNA_seed | ESR1 | -0.626137 | BRCA | Hs-578-T | Tamoxifen | MVP_top60_DNA+RNA_seed | EGFR | -1.357323 | BRCA | HDQ-P1 | Lapatinib |
| MVP_top60_DNA_seed | AKT1 | -0.625503 | BRCA | HCC1419 | GSK2110183B | MVP_top60_DNA+RNA_seed | ERBB2 | -1.357323 | BRCA | HDQ-P1 | Lapatinib |
| MVP_top60_DNA_seed | EP300 | -0.6246 | BRCA | HCC1419 | CPI-637 | MVP_top60_DNA+RNA_seed | HDAC1 | -1.354088 | BRCA | MDA-MB-361 | Entinostat |
| MVP_top60_DNA_seed | EGFR | -0.62256 | BRCA | HCC1419 | Gefitinib | MVP_top60_DNA+RNA_seed | EGFR | -1.352 | BRCA | DU-4475 | Lapatinib |
| MVP_top60_DNA_seed | AKT1 | -0.621252 | BRCA | T47D | AZD5363 | MVP_top60_DNA+RNA_seed | ERBB2 | -1.352 | BRCA | DU-4475 | Lapatinib |
| MVP_top60_DNA_seed | AKT1 | -0.61829 | BRCA | Hs-578-T | Uprosertib | MVP_top60_DNA+RNA_seed | ESR1 | -1.348401 | BRCA | MDA-MB-415 | GDC0810 |
| MVP_top60_DNA_seed | CDK1 | -0.618099 | BRCA | CAL-51 | RO-3306 | MVP_top60_DNA+RNA_seed | HDAC1 | -1.345298 | BRCA | Hs-578-T | Romidepsin |
| MVP_top60_DNA_seed | AKT1 | -0.617231 | BRCA | UACC-812 | MK-2206 | MVP_top60_DNA+RNA_seed | HDAC2 | -1.345298 | BRCA | Hs-578-T | Romidepsin |
| MVP_top60_DNA_seed | HDAC1 | -0.615121 | BRCA | BT-474 | Entinostat | MVP_top60_DNA+RNA_seed | EGFR | -1.339172 | BRCA | UACC-812 | Osimertinib |
| MVP_top60_DNA_seed | AKT1 | -0.614668 | BRCA | MCF7 | Uprosertib | MVP_top60_DNA+RNA_seed | ESR1 | -1.328841 | BRCA | EVSA-T | Tamoxifen |
| MVP_top60_DNA_seed | AKT1 | -0.612365 | BRCA | CAL-51 | AT13148 | MVP_top60_DNA+RNA_seed | MDM2 | -1.328609 | BRCA | HCC1569 | Serdemetan |
| MVP_top60_DNA_seed | EGFR | -0.610433 | BRCA | HCC2157 | Erlotinib | MVP_top60_DNA+RNA_seed | AKT1 | -1.323851 | BRCA | MDA-MB-175-VII | AZD5363 |
| MVP_top60_DNA_seed | AKT1 | -0.608807 | BRCA | HDQ-P1 | Afuresertib | MVP_top60_DNA+RNA_seed | CDK1 | -1.320518 | BRCA | HCC2157 | RO-3306 |
| MVP_top60_DNA_seed | PIK3CA | -0.607739 | BRCA | Hs-578-T | GNE-317 | MVP_top60_DNA+RNA_seed | PIK3CA | -1.319474 | BRCA | EVSA-T | GNE-317 |
| MVP_top60_DNA_seed | CDK2 | -0.606584 | BRCA | Hs-578-T | Dinaciclib | MVP_top60_DNA+RNA_seed | HDAC1 | -1.317187 | BRCA | DU-4475 | Romidepsin |

| | | | | | | | | | | | |
|---|---|---|---|---|---|---|---|---|---|---|---|
| MVP_top60_DNA_seed | CDK1 | -0.606584 | BRCA | Hs-578-T | Dinaciclib | MVP_top60_DNA+RNA_seed | HDAC2 | -1.317187 | BRCA | DU-4475 | Romidepsin |
| MVP_top60_DNA_seed | HSP90AA1 | -0.604727 | BRCA | HDQ-P1 | Tanespimycin | MVP_top60_DNA+RNA_seed | ATM | -1.316938 | BRCA | OCUB-M | KU-55933 |
| MVP_top60_DNA_seed | HSP90AB1 | -0.604727 | BRCA | HDQ-P1 | Tanespimycin | MVP_top60_DNA+RNA_seed | CDK2 | -1.315471 | BRCA | HCC1599 | MK-8776 |
| MVP_top60_DNA_seed | PIK3CA | -0.601721 | BRCA | COLO-824 | Alpelisib | MVP_top60_DNA+RNA_seed | AKT1 | -1.315157 | BRCA | HCC1569 | AT13148 |
| MVP_top60_DNA_seed | EGFR | -0.600252 | BRCA | HCC1143 | AZD3759 | MVP_top60_DNA+RNA_seed | EGFR | -1.309951 | BRCA | HCC2157 | Sapitinib |
| MVP_top60_DNA_seed | HDAC1 | -0.599259 | BRCA | HCC1419 | Romidepsin | MVP_top60_DNA+RNA_seed | ERBB2 | -1.309951 | BRCA | HCC2157 | Sapitinib |
| MVP_top60_DNA_seed | HDAC2 | -0.599259 | BRCA | HCC1419 | Romidepsin | MVP_top60_DNA+RNA_seed | ERBB3 | -1.309951 | BRCA | HCC2157 | Sapitinib |
| MVP_top60_DNA_seed | HDAC1 | -0.598366 | BRCA | EFM-192A | Entinostat | MVP_top60_DNA+RNA_seed | CDK2 | -1.306551 | BRCA | HCC2218 | Dinaciclib |
| MVP_top60_DNA_seed | CDK2 | -0.597811 | BRCA | AU565 | MK-8776 | MVP_top60_DNA+RNA_seed | CDK1 | -1.306551 | BRCA | HCC2218 | Dinaciclib |
| MVP_top60_DNA_seed | HSP90AA1 | -0.597687 | BRCA | MFM-223 | Tanespimycin | MVP_top60_DNA+RNA_seed | AKT1 | -1.306424 | BRCA | MDA-MB-361 | MK-2206 |
| MVP_top60_DNA_seed | HSP90AB1 | -0.597687 | BRCA | MFM-223 | Tanespimycin | MVP_top60_DNA+RNA_seed | PLK1 | -1.306051 | BRCA | Hs-578-T | BI-2536 |
| MVP_top60_DNA_seed | PIK3CA | -0.596498 | BRCA | DU-4475 | Buparlisib | MVP_top60_DNA+RNA_seed | PIK3CA | -1.305722 | BRCA | T47D | Alpelisib |
| MVP_top60_DNA_seed | PIK3CA | -0.595752 | BRCA | HCC1143 | Buparlisib | MVP_top60_DNA+RNA_seed | EGFR | -1.304562 | BRCA | EFM-192A | Sapitinib |
| MVP_top60_DNA_seed | AKT1 | -0.593354 | BRCA | HCC1599 | Uprosertib | MVP_top60_DNA+RNA_seed | ERBB2 | -1.304562 | BRCA | EFM-192A | Sapitinib |
| MVP_top60_DNA_seed | AKT1 | -0.593165 | BRCA | BT-474 | Uprosertib | MVP_top60_DNA+RNA_seed | ERBB3 | -1.304562 | BRCA | EFM-192A | Sapitinib |
| MVP_top60_DNA_seed | PIK3CA | -0.592111 | BRCA | MDA-MB-175-VII | AZD8186 | MVP_top60_DNA+RNA_seed | PIK3CA | -1.30157 | BRCA | HDQ-P1 | GNE-317 |
| MVP_top60_DNA_seed | ESR1 | -0.590756 | BRCA | MDA-MB-231 | GDC0810 | MVP_top60_DNA+RNA_seed | PLK1 | -1.299101 | BRCA | OCUB-M | BI-2536 |
| MVP_top60_DNA_seed | EP300 | -0.5902 | BRCA | DU-4475 | SGC-CBP30 | MVP_top60_DNA+RNA_seed | PIK3CA | -1.298818 | BRCA | HCC2157 | Buparlisib |
| MVP_top60_DNA_seed | CREBBP | -0.5902 | BRCA | DU-4475 | SGC-CBP30 | MVP_top60_DNA+RNA_seed | AKT1 | -1.296755 | BRCA | Hs-578-T | GSK2110183B |
| MVP_top60_DNA_seed | HDAC1 | -0.589309 | BRCA | MDA-MB-468 | Romidepsin | MVP_top60_DNA+RNA_seed | AKT1 | -1.290018 | BRCA | MDA-MB-330 | Uprosertib |
| MVP_top60_DNA_seed | HDAC2 | -0.589309 | BRCA | MDA-MB-468 | Romidepsin | MVP_top60_DNA+RNA_seed | EGFR | -1.287227 | BRCA | MDA-MB-468 | AZD3759 |
| MVP_top60_DNA_seed | ATM | -0.58849 | BRCA | CAMA-1 | KU-55933 | MVP_top60_DNA+RNA_seed | ESR1 | -1.286758 | BRCA | OCUB-M | GDC0810 |
| MVP_top60_DNA_seed | CDK1 | -0.583503 | BRCA | AU565 | RO-3306 | MVP_top60_DNA+RNA_seed | EGFR | -1.286186 | BRCA | HCC2157 | Afatinib |
| MVP_top60_DNA_seed | AKT1 | -0.576975 | BRCA | HCC1937 | GSK2110183B | MVP_top60_DNA+RNA_seed | ERBB2 | -1.286186 | BRCA | HCC2157 | Afatinib |
| MVP_top60_DNA_seed | MAPK1 | -0.576253 | BRCA | MDA-MB-231 | VX-11e | MVP_top60_DNA+RNA_seed | AKT1 | -1.284561 | BRCA | MDA-MB-453 | Afuresertib |
| MVP_top60_DNA_seed | EGFR | -0.574657 | BRCA | HCC2157 | AZD3759 | MVP_top60_DNA+RNA_seed | PIK3CA | -1.282233 | BRCA | MDA-MB-330 | GNE-317 |
| MVP_top60_DNA_seed | HDAC1 | -0.571774 | BRCA | HCC1954 | Romidepsin | MVP_top60_DNA+RNA_seed | CDK1 | -1.28002 | BRCA | Hs-578-T | RO-3306 |
| MVP_top60_DNA_seed | HDAC2 | -0.571774 | BRCA | HCC1954 | Romidepsin | MVP_top60_DNA+RNA_seed | EGFR | -1.277934 | BRCA | MDA-MB-453 | Lapatinib |
| MVP_top60_DNA_seed | PIK3CA | -0.57039 | BRCA | HCC1569 | AZD8186 | MVP_top60_DNA+RNA_seed | ERBB2 | -1.277934 | BRCA | MDA-MB-453 | Lapatinib |
| MVP_top60_DNA_seed | PIK3CA | -0.568248 | BRCA | CAL-148 | Buparlisib | MVP_top60_DNA+RNA_seed | HSP90AA1 | -1.276484 | BRCA | CAL-51 | Tanespimycin |
| MVP_top60_DNA_seed | HSP90AA1 | -0.567335 | BRCA | EVSA-T | Luminespib | MVP_top60_DNA+RNA_seed | HSP90AB1 | -1.276484 | BRCA | CAL-51 | Tanespimycin |
| MVP_top60_DNA_seed | HSP90AB1 | -0.567335 | BRCA | EVSA-T | Luminespib | MVP_top60_DNA+RNA_seed | AKT1 | -1.275226 | BRCA | Hs-578-T | Uprosertib |
| MVP_top60_DNA_seed | AKT1 | -0.567116 | BRCA | MDA-MB-231 | AT13148 | MVP_top60_DNA+RNA_seed | EP300 | -1.262581 | BRCA | COLO-824 | CPI-637 |
| MVP_top60_DNA_seed | HSP90AA1 | -0.564846 | BRCA | HDQ-P1 | Luminespib | MVP_top60_DNA+RNA_seed | SRC | -1.259319 | BRCA | HDQ-P1 | Bosutinib |
| MVP_top60_DNA_seed | HSP90AB1 | -0.564846 | BRCA | HDQ-P1 | Luminespib | MVP_top60_DNA+RNA_seed | EGFR | -1.251343 | BRCA | HCC1419 | Lapatinib |
| MVP_top60_DNA_seed | HSP90AA1 | -0.563338 | BRCA | JIMT-1 | Tanespimycin | MVP_top60_DNA+RNA_seed | ERBB2 | -1.251343 | BRCA | HCC1419 | Lapatinib |
| MVP_top60_DNA_seed | HSP90AB1 | -0.563338 | BRCA | JIMT-1 | Tanespimycin | MVP_top60_DNA+RNA_seed | AKT1 | -1.251025 | BRCA | CAL-51 | Afuresertib |
| MVP_top60_DNA_seed | AKT1 | -0.562517 | BRCA | EFM-19 | AZD5363 | MVP_top60_DNA+RNA_seed | ESR1 | -1.244882 | BRCA | T47D | Tamoxifen |
| MVP_top60_DNA_seed | SRC | -0.5615 | BRCA | MRK-nu-1 | Bosutinib | MVP_top60_DNA+RNA_seed | PIK3CA | -1.243567 | BRCA | HDQ-P1 | AZD8186 |
| MVP_top60_DNA_seed | ATM | -0.554212 | BRCA | HCC1954 | KU-55933 | MVP_top60_DNA+RNA_seed | CDK2 | -1.242961 | BRCA | MDA-MB-157 | MK-8776 |
| MVP_top60_DNA_seed | ERBB2 | -0.553837 | BRCA | MCF7 | Lapatinib | MVP_top60_DNA+RNA_seed | PIK3CA | -1.240029 | BRCA | EFM-19 | GNE-317 |

| | | | | | | | | | | | |
|---|---|---|---|---|---|---|---|---|---|---|---|
| MVP_top60_DNA_seed | EGFR | -0.553837 | BRCA | MCF7 | Lapatinib | MVP_top60_DNA+RNA_seed | PLK1 | -1.238342 | BRCA | HCC1599 | BI-2536 |
| MVP_top60_DNA_seed | CDK2 | -0.550668 | BRCA | CAL-51 | AZD5438 | MVP_top60_DNA+RNA_seed | PIK3CA | -1.236731 | BRCA | CAL-51 | GNE-317 |
| MVP_top60_DNA_seed | HDAC1 | -0.550593 | BRCA | OCUB-M | Entinostat | MVP_top60_DNA+RNA_seed | AKT1 | -1.234477 | BRCA | HCC1419 | Ipatasertib |
| MVP_top60_DNA_seed | HDAC1 | -0.548279 | BRCA | EVSA-T | PCI-34051 | MVP_top60_DNA+RNA_seed | ESR1 | -1.231812 | BRCA | HCC1395 | Tamoxifen |
| MVP_top60_DNA_seed | ERBB2 | -0.544447 | BRCA | EFM-192A | Afatinib | MVP_top60_DNA+RNA_seed | HDAC1 | -1.231066 | BRCA | YMB-1-E | PCI-34051 |
| MVP_top60_DNA_seed | EGFR | -0.544447 | BRCA | EFM-192A | Afatinib | MVP_top60_DNA+RNA_seed | PIK3CA | -1.223717 | BRCA | EVSA-T | Buparlisib |
| MVP_top60_DNA_seed | HSP90AA1 | -0.543351 | BRCA | HCC1569 | Tanespimycin | MVP_top60_DNA+RNA_seed | AKT1 | -1.223013 | BRCA | HDQ-P1 | AT13148 |
| MVP_top60_DNA_seed | HSP90AB1 | -0.543351 | BRCA | HCC1569 | Tanespimycin | MVP_top60_DNA+RNA_seed | EGFR | -1.218025 | BRCA | HCC1569 | Osimertinib |
| MVP_top60_DNA_seed | EGFR | -0.541206 | BRCA | MDA-MB-361 | AZD3759 | MVP_top60_DNA+RNA_seed | PIK3CA | -1.217177 | BRCA | BT-474 | Alpelisib |
| MVP_top60_DNA_seed | AKT1 | -0.539013 | BRCA | Hs-578-T | Afuresertib | MVP_top60_DNA+RNA_seed | PIK3CA | -1.207284 | BRCA | UACC-893 | GNE-317 |
| MVP_top60_DNA_seed | ERBB2 | -0.538031 | BRCA | MDA-MB-415 | Lapatinib | MVP_top60_DNA+RNA_seed | HDAC1 | -1.207032 | BRCA | HCC1569 | PCI-34051 |
| MVP_top60_DNA_seed | EGFR | -0.538031 | BRCA | MDA-MB-415 | Lapatinib | MVP_top60_DNA+RNA_seed | EGFR | -1.206512 | BRCA | MDA-MB-468 | Lapatinib |
| MVP_top60_DNA_seed | MDM2 | -0.537312 | BRCA | HCC2218 | Serdemetan | MVP_top60_DNA+RNA_seed | ERBB2 | -1.206512 | BRCA | MDA-MB-468 | Lapatinib |
| MVP_top60_DNA_seed | EGFR | -0.533864 | BRCA | MDA-MB-361 | Osimertinib | MVP_top60_DNA+RNA_seed | MAPK1 | -1.203858 | BRCA | HDQ-P1 | VX-11e |
| MVP_top60_DNA_seed | ERBB2 | -0.533007 | BRCA | CAL-85-1 | Sapitinib | MVP_top60_DNA+RNA_seed | HDAC1 | -1.202038 | BRCA | MDA-MB-330 | Entinostat |
| MVP_top60_DNA_seed | EGFR | -0.533007 | BRCA | CAL-85-1 | Sapitinib | MVP_top60_DNA+RNA_seed | HSP90AA1 | -1.201796 | BRCA | DU-4475 | Luminespib |
| MVP_top60_DNA_seed | ERBB3 | -0.533007 | BRCA | CAL-85-1 | Sapitinib | MVP_top60_DNA+RNA_seed | HSP90AB1 | -1.201796 | BRCA | DU-4475 | Luminespib |
| MVP_top60_DNA_seed | AKT1 | -0.528852 | BRCA | MRK-nu-1 | Uprosertib | MVP_top60_DNA+RNA_seed | AKT1 | -1.193382 | BRCA | HCC1395 | Ipatasertib |
| MVP_top60_DNA_seed | TP53 | -0.527752 | BRCA | HCC70 | MIRA-1 | MVP_top60_DNA+RNA_seed | AKT1 | -1.190906 | BRCA | MDA-MB-330 | Ipatasertib |
| MVP_top60_DNA_seed | EGFR | -0.526819 | BRCA | AU565 | AZD3759 | MVP_top60_DNA+RNA_seed | AKT1 | -1.190831 | BRCA | Hs-578-T | Ipatasertib |
| MVP_top60_DNA_seed | AKT1 | -0.525991 | BRCA | HCC1954 | Afuresertib | MVP_top60_DNA+RNA_seed | CDK2 | -1.189686 | BRCA | HCC1569 | MK-8776 |
| MVP_top60_DNA_seed | CDK1 | -0.525477 | BRCA | HCC1500 | RO-3306 | MVP_top60_DNA+RNA_seed | TP53 | -1.185478 | BRCA | HCC2157 | MIRA-1 |
| MVP_top60_DNA_seed | ERBB2 | -0.524376 | BRCA | HCC1569 | Sapitinib | MVP_top60_DNA+RNA_seed | EGFR | -1.173941 | BRCA | MDA-MB-361 | Sapitinib |
| MVP_top60_DNA_seed | EGFR | -0.524376 | BRCA | HCC1569 | Sapitinib | MVP_top60_DNA+RNA_seed | ERBB2 | -1.173941 | BRCA | MDA-MB-361 | Sapitinib |
| MVP_top60_DNA_seed | ERBB3 | -0.524376 | BRCA | HCC1569 | Sapitinib | MVP_top60_DNA+RNA_seed | ERBB3 | -1.173941 | BRCA | MDA-MB-361 | Sapitinib |
| MVP_top60_DNA_seed | MAPK1 | -0.523588 | BRCA | MDA-MB-453 | VX-11e | MVP_top60_DNA+RNA_seed | CDK2 | -1.172171 | BRCA | HCC1954 | Dinaciclib |
| MVP_top60_DNA_seed | ESR1 | -0.521907 | BRCA | HCC1419 | Tamoxifen | MVP_top60_DNA+RNA_seed | CDK1 | -1.172171 | BRCA | HCC1954 | Dinaciclib |
| MVP_top60_DNA_seed | CDK2 | -0.51965 | BRCA | MDA-MB-361 | Dinaciclib | MVP_top60_DNA+RNA_seed | ESR1 | -1.17002 | BRCA | AU565 | GDC0810 |
| MVP_top60_DNA_seed | CDK1 | -0.51965 | BRCA | MDA-MB-361 | Dinaciclib | MVP_top60_DNA+RNA_seed | EGFR | -1.168457 | BRCA | HCC1419 | Sapitinib |
| MVP_top60_DNA_seed | EP300 | -0.516204 | BRCA | MDA-MB-231 | CPI-637 | MVP_top60_DNA+RNA_seed | ERBB2 | -1.168457 | BRCA | HCC1419 | Sapitinib |
| MVP_top60_DNA_seed | AKT1 | -0.516089 | BRCA | T47D | MK-2206 | MVP_top60_DNA+RNA_seed | ERBB3 | -1.168457 | BRCA | HCC1419 | Sapitinib |
| MVP_top60_DNA_seed | PIK3CA | -0.515105 | BRCA | AU565 | Buparlisib | MVP_top60_DNA+RNA_seed | HSP90AA1 | -1.159436 | BRCA | HCC2218 | CCT-018159 |
| MVP_top60_DNA_seed | AKT1 | -0.514955 | BRCA | EFM-192A | Uprosertib | MVP_top60_DNA+RNA_seed | HSP90AB1 | -1.159436 | BRCA | HCC2218 | CCT-018159 |
| MVP_top60_DNA_seed | PIK3CA | -0.514386 | BRCA | HCC1143 | GNE-317 | MVP_top60_DNA+RNA_seed | TP53 | -1.155661 | BRCA | HCC1419 | MIRA-1 |
| MVP_top60_DNA_seed | AKT1 | -0.512371 | BRCA | DU-4475 | GSK2110183B | MVP_top60_DNA+RNA_seed | AKT1 | -1.154297 | BRCA | MDA-MB-361 | Uprosertib |
| MVP_top60_DNA_seed | HDAC1 | -0.511999 | BRCA | AU565 | PCI-34051 | MVP_top60_DNA+RNA_seed | AKT1 | -1.152108 | BRCA | MDA-MB-361 | Ipatasertib |
| MVP_top60_DNA_seed | EGFR | -0.511941 | BRCA | MDA-MB-175-VII | AZD3759 | MVP_top60_DNA+RNA_seed | AKT1 | -1.151762 | BRCA | YMB-1-E | AT13148 |
| MVP_top60_DNA_seed | AKT1 | -0.50826 | BRCA | HDQ-P1 | Ipatasertib | MVP_top60_DNA+RNA_seed | AKT1 | -1.148076 | BRCA | HCC1395 | Afuresertib |
| MVP_top60_DNA_seed | AKT1 | -0.505243 | BRCA | MDA-MB-361 | AZD5363 | MVP_top60_DNA+RNA_seed | AKT1 | -1.143101 | BRCA | MDA-MB-453 | Ipatasertib |
| MVP_top60_DNA_seed | PIK3CA | -0.503639 | BRCA | HCC70 | GNE-317 | MVP_top60_DNA+RNA_seed | AKT1 | -1.137307 | BRCA | EFM-19 | GSK2110183B |
| MVP_top60_DNA_seed | CDK1 | -0.502564 | BRCA | HCC1428 | RO-3306 | MVP_top60_DNA+RNA_seed | CDK1 | -1.136104 | BRCA | MDA-MB-468 | RO-3306 |

| | | | | | | | | | | | |
|---|---|---|---|---|---|---|---|---|---|---|---|
| MVP_top60_DNA_seed | HSP90AA1 | -0.50009 | BRCA | UACC-893 | Luminespib | MVP_top60_DNA+RNA_seed | HDAC1 | -1.135274 | BRCA | MDA-MB-453 | Romidepsin |
| MVP_top60_DNA_seed | HSP90AB1 | -0.50009 | BRCA | UACC-893 | Luminespib | MVP_top60_DNA+RNA_seed | HDAC2 | -1.135274 | BRCA | MDA-MB-453 | Romidepsin |
| MVP_top60_DNA_seed | TP53 | -0.499832 | BRCA | CAL-148 | MIRA-1 | MVP_top60_DNA+RNA_seed | PIK3CA | -1.128218 | BRCA | MDA-MB-175-VII | Alpelisib |
| MVP_top60_DNA_seed | CDK2 | -0.496807 | BRCA | BT-20 | AZD5438 | MVP_top60_DNA+RNA_seed | AKT1 | -1.123818 | BRCA | MDA-MB-330 | GSK2110183B |
| MVP_top60_DNA_seed | ATM | -0.49617 | BRCA | YMB-1-E | KU-55933 | MVP_top60_DNA+RNA_seed | CDK2 | -1.115391 | BRCA | MDA-MB-468 | Dinaciclib |
| MVP_top60_DNA_seed | CDK1 | -0.494962 | BRCA | BT-20 | RO-3306 | MVP_top60_DNA+RNA_seed | CDK1 | -1.115391 | BRCA | MDA-MB-468 | Dinaciclib |
| MVP_top60_DNA_seed | HDAC1 | -0.494453 | BRCA | HCC2157 | Romidepsin | MVP_top60_DNA+RNA_seed | EGFR | -1.109276 | BRCA | MDA-MB-453 | Gefitinib |
| MVP_top60_DNA_seed | HDAC2 | -0.494453 | BRCA | HCC2157 | Romidepsin | MVP_top60_DNA+RNA_seed | AKT1 | -1.108911 | BRCA | MDA-MB-330 | AZD5363 |
| MVP_top60_DNA_seed | HSP90AA1 | -0.490478 | BRCA | Hs-578-T | Luminespib | MVP_top60_DNA+RNA_seed | HSP90AA1 | -1.105739 | BRCA | MDA-MB-330 | Elesclomol |
| MVP_top60_DNA_seed | HSP90AB1 | -0.490478 | BRCA | Hs-578-T | Luminespib | MVP_top60_DNA+RNA_seed | HSP90AB1 | -1.105739 | BRCA | MDA-MB-330 | Elesclomol |
| MVP_top60_DNA_seed | ESR1 | -0.486566 | BRCA | HCC1500 | Tamoxifen | MVP_top60_DNA+RNA_seed | CDK2 | -1.102498 | BRCA | HCC1187 | Dinaciclib |
| MVP_top60_DNA_seed | PIK3CA | -0.486361 | BRCA | HCC1954 | Buparlisib | MVP_top60_DNA+RNA_seed | CDK1 | -1.102498 | BRCA | HCC1187 | Dinaciclib |
| MVP_top60_DNA_seed | EP300 | -0.484749 | BRCA | CAMA-1 | CPI-637 | MVP_top60_DNA+RNA_seed | AKT1 | -1.100187 | BRCA | BT-20 | Ipatasertib |
| MVP_top60_DNA_seed | ERBB2 | -0.484072 | BRCA | CAL-51 | Lapatinib | MVP_top60_DNA+RNA_seed | AKT1 | -1.100014 | BRCA | EFM-19 | MK-2206 |
| MVP_top60_DNA_seed | EGFR | -0.484072 | BRCA | CAL-51 | Lapatinib | MVP_top60_DNA+RNA_seed | PIK3CA | -1.098943 | BRCA | CAL-148 | GNE-317 |
| MVP_top60_DNA_seed | AKT1 | -0.482557 | BRCA | HCC1395 | AT13148 | MVP_top60_DNA+RNA_seed | MDM2 | -1.097828 | BRCA | HCC1500 | Nutlin-3a (-) |
| MVP_top60_DNA_seed | ATM | -0.481969 | BRCA | COLO-824 | KU-55933 | MVP_top60_DNA+RNA_seed | ESR1 | -1.094082 | BRCA | CAMA-1 | Tamoxifen |
| MVP_top60_DNA_seed | AKT1 | -0.478356 | BRCA | HCC1419 | Uprosertib | MVP_top60_DNA+RNA_seed | ATM | -1.087495 | BRCA | AU565 | KU-55933 |
| MVP_top60_DNA_seed | MDM2 | -0.477942 | BRCA | EVSA-T | Serdemetan | MVP_top60_DNA+RNA_seed | EP300 | -1.087417 | BRCA | HCC1937 | SGC-CBP30 |
| MVP_top60_DNA_seed | HSP90AA1 | -0.476288 | BRCA | HCC2218 | Elesclomol | MVP_top60_DNA+RNA_seed | EGFR | -1.087029 | BRCA | MDA-MB-468 | Gefitinib |
| MVP_top60_DNA_seed | HSP90AB1 | -0.476288 | BRCA | HCC2218 | Elesclomol | MVP_top60_DNA+RNA_seed | AR | -1.083394 | BRCA | HCC2218 | Bicalutamide |
| MVP_top60_DNA_seed | CDK2 | -0.473378 | BRCA | MDA-MB-231 | Dinaciclib | MVP_top60_DNA+RNA_seed | CDK1 | -1.083037 | BRCA | EVSA-T | RO-3306 |
| MVP_top60_DNA_seed | CDK1 | -0.473378 | BRCA | MDA-MB-231 | Dinaciclib | MVP_top60_DNA+RNA_seed | MAPK1 | -1.078733 | BRCA | HCC1569 | VX-11e |
| MVP_top60_DNA_seed | EGFR | -0.473315 | BRCA | HCC1143 | Osimertinib | MVP_top60_DNA+RNA_seed | PIK3CA | -1.078618 | BRCA | HCC1395 | AZD8186 |
| MVP_top60_DNA_seed | AKT1 | -0.473254 | BRCA | MCF7 | AT13148 | MVP_top60_DNA+RNA_seed | HSP90AA1 | -1.077325 | BRCA | MDA-MB-361 | Luminespib |
| MVP_top60_DNA_seed | EP300 | -0.470434 | BRCA | EFM-192A | SGC-CBP30 | MVP_top60_DNA+RNA_seed | HSP90AB1 | -1.077325 | BRCA | MDA-MB-361 | Luminespib |
| MVP_top60_DNA_seed | CREBBP | -0.470434 | BRCA | EFM-192A | SGC-CBP30 | MVP_top60_DNA+RNA_seed | PIK3CA | -1.076969 | BRCA | CAMA-1 | GNE-317 |
| MVP_top60_DNA_seed | PIK3CA | -0.464579 | BRCA | MDA-MB-231 | GNE-317 | MVP_top60_DNA+RNA_seed | ESR1 | -1.07538 | BRCA | DU-4475 | Tamoxifen |
| MVP_top60_DNA_seed | EGFR | -0.461709 | BRCA | HCC1954 | AZD3759 | MVP_top60_DNA+RNA_seed | PIK3CA | -1.074375 | BRCA | T47D | GNE-317 |
| MVP_top60_DNA_seed | CDK1 | -0.458783 | BRCA | HCC1395 | RO-3306 | MVP_top60_DNA+RNA_seed | PIK3CA | -1.073624 | BRCA | HCC1599 | Alpelisib |
| MVP_top60_DNA_seed | TP53 | -0.454454 | BRCA | DU-4475 | MIRA-1 | MVP_top60_DNA+RNA_seed | AKT1 | -1.071297 | BRCA | Hs-578-T | MK-2206 |
| MVP_top60_DNA_seed | EP300 | -0.451755 | BRCA | BT-549 | CPI-637 | MVP_top60_DNA+RNA_seed | HSP90AA1 | -1.07054 | BRCA | MDA-MB-453 | Luminespib |
| MVP_top60_DNA_seed | CDK2 | -0.448916 | BRCA | MDA-MB-157 | AZD5438 | MVP_top60_DNA+RNA_seed | HSP90AB1 | -1.07054 | BRCA | MDA-MB-453 | Luminespib |
| MVP_top60_DNA_seed | MAPK3 | -0.448886 | BRCA | UACC-893 | SCH772984 | MVP_top60_DNA+RNA_seed | PLK1 | -1.068707 | BRCA | HCC2157 | BI-2536 |
| MVP_top60_DNA_seed | MAPK1 | -0.448886 | BRCA | UACC-893 | SCH772984 | MVP_top60_DNA+RNA_seed | AKT1 | -1.062409 | BRCA | HCC1954 | Uprosertib |
| MVP_top60_DNA_seed | EGFR | -0.444722 | BRCA | EFM-192A | Erlotinib | MVP_top60_DNA+RNA_seed | HSP90AA1 | -1.061235 | BRCA | CAL-148 | Tanespimycin |
| MVP_top60_DNA_seed | HSP90AA1 | -0.441942 | BRCA | CAL-85-1 | Tanespimycin | MVP_top60_DNA+RNA_seed | HSP90AB1 | -1.061235 | BRCA | CAL-148 | Tanespimycin |
| MVP_top60_DNA_seed | HSP90AB1 | -0.441942 | BRCA | CAL-85-1 | Tanespimycin | MVP_top60_DNA+RNA_seed | AKT1 | -1.056711 | BRCA | T47D | GSK2110183B |
| MVP_top60_DNA_seed | PIK3CA | -0.441574 | BRCA | COLO-824 | Buparlisib | MVP_top60_DNA+RNA_seed | HDAC1 | -1.054149 | BRCA | CAL-51 | PCI-34051 |
| MVP_top60_DNA_seed | PIK3CA | -0.441497 | BRCA | HCC1143 | Alpelisib | MVP_top60_DNA+RNA_seed | ESR1 | -1.05175 | BRCA | MCF7 | Tamoxifen |
| MVP_top60_DNA_seed | ESR1 | -0.440353 | BRCA | HCC1954 | Tamoxifen | MVP_top60_DNA+RNA_seed | HSP90AA1 | -1.051622 | BRCA | UACC-893 | Tanespimycin |

| | | | | | | | | | | | |
|---|---|---|---|---|---|---|---|---|---|---|---|
| MVP_top60_DNA_seed | AKT1 | -0.438643 | BRCA | BT-474 | Ipatasertib | MVP_top60_DNA+RNA_seed | HSP90AB1 | -1.051622 | BRCA | UACC-893 | Tanespimycin |
| MVP_top60_DNA_seed | PIK3CA | -0.438056 | BRCA | HCC1954 | GNE-317 | MVP_top60_DNA+RNA_seed | AKT1 | -1.050404 | BRCA | Hs-578-T | AZD5363 |
| MVP_top60_DNA_seed | PIK3CA | -0.43752 | BRCA | MCF7 | Buparlisib | MVP_top60_DNA+RNA_seed | HSP90AA1 | -1.043814 | BRCA | Hs-578-T | Tanespimycin |
| MVP_top60_DNA_seed | HSP90AA1 | -0.437013 | BRCA | MDA-MB-175-VII | Luminespib | MVP_top60_DNA+RNA_seed | HSP90AB1 | -1.043814 | BRCA | Hs-578-T | Tanespimycin |
| MVP_top60_DNA_seed | HSP90AB1 | -0.437013 | BRCA | MDA-MB-175-VII | Luminespib | MVP_top60_DNA+RNA_seed | CDK2 | -1.039464 | BRCA | HCC1187 | AZD5438 |
| MVP_top60_DNA_seed | PIK3CA | -0.436057 | BRCA | HDQ-P1 | Alpelisib | MVP_top60_DNA+RNA_seed | AKT1 | -1.038614 | BRCA | CAMA-1 | MK-2206 |
| MVP_top60_DNA_seed | ATM | -0.435736 | BRCA | CAL-85-1 | KU-55933 | MVP_top60_DNA+RNA_seed | AKT1 | -1.037668 | BRCA | EFM-19 | Uprosertib |
| MVP_top60_DNA_seed | HSP90AA1 | -0.434775 | BRCA | HCC38 | Tanespimycin | MVP_top60_DNA+RNA_seed | MAPK3 | -1.036445 | BRCA | MDA-MB-436 | Ulixertinib |
| MVP_top60_DNA_seed | HSP90AB1 | -0.434775 | BRCA | HCC38 | Tanespimycin | MVP_top60_DNA+RNA_seed | MAPK1 | -1.036445 | BRCA | MDA-MB-436 | Ulixertinib |
| MVP_top60_DNA_seed | AKT1 | -0.434648 | BRCA | HCC1599 | Afuresertib | MVP_top60_DNA+RNA_seed | AKT1 | -1.034662 | BRCA | MDA-MB-330 | Afuresertib |
| MVP_top60_DNA_seed | MAPK3 | -0.433042 | BRCA | UACC-893 | Ulixertinib | MVP_top60_DNA+RNA_seed | AKT1 | -1.028377 | BRCA | BT-20 | AZD5363 |
| MVP_top60_DNA_seed | MAPK1 | -0.433042 | BRCA | UACC-893 | Ulixertinib | MVP_top60_DNA+RNA_seed | PIK3CA | -1.026325 | BRCA | CAL-51 | Buparlisib |
| MVP_top60_DNA_seed | ATM | -0.432764 | BRCA | BT-20 | KU-55933 | MVP_top60_DNA+RNA_seed | TP53 | -1.02495 | BRCA | OCUB-M | MIRA-1 |
| MVP_top60_DNA_seed | AKT1 | -0.429136 | BRCA | HCC2157 | AT13148 | MVP_top60_DNA+RNA_seed | PIK3CA | -1.022631 | BRCA | EFM-19 | AZD8186 |
| MVP_top60_DNA_seed | ESR1 | -0.428431 | BRCA | MDA-MB-468 | Tamoxifen | MVP_top60_DNA+RNA_seed | PIK3CA | -1.019348 | BRCA | MDA-MB-361 | Buparlisib |
| MVP_top60_DNA_seed | AKT1 | -0.425352 | BRCA | EFM-192A | AZD5363 | MVP_top60_DNA+RNA_seed | PIK3CA | -1.018997 | BRCA | MDA-MB-175-VII | Buparlisib |
| MVP_top60_DNA_seed | MDM2 | -0.424858 | BRCA | YMB-1-E | Serdemetan | MVP_top60_DNA+RNA_seed | ESR1 | -1.018263 | BRCA | DU-4475 | GDC0810 |
| MVP_top60_DNA_seed | ATM | -0.423123 | BRCA | T47D | KU-55933 | MVP_top60_DNA+RNA_seed | PIK3CA | -1.016139 | BRCA | HCC1569 | Alpelisib |
| MVP_top60_DNA_seed | AKT1 | -0.422735 | BRCA | BT-474 | MK-2206 | MVP_top60_DNA+RNA_seed | AKT1 | -1.013224 | BRCA | CAL-85-1 | GSK2110183B |
| MVP_top60_DNA_seed | AKT1 | -0.422422 | BRCA | COLO-824 | Afuresertib | MVP_top60_DNA+RNA_seed | CDK2 | -1.007732 | BRCA | MDA-MB-468 | AZD5438 |
| MVP_top60_DNA_seed | SRC | -0.42217 | BRCA | CAL-85-1 | Bosutinib | MVP_top60_DNA+RNA_seed | CDK2 | -1.007576 | BRCA | HCC2157 | Dinaciclib |
| MVP_top60_DNA_seed | EGFR | -0.418449 | BRCA | HCC1937 | Erlotinib | MVP_top60_DNA+RNA_seed | CDK1 | -1.007576 | BRCA | HCC2157 | Dinaciclib |
| MVP_top60_DNA_seed | PIK3CA | -0.413514 | BRCA | EFM-192A | AZD8186 | MVP_top60_DNA+RNA_seed | AKT1 | -1.007023 | BRCA | COLO-824 | AT13148 |
| MVP_top60_DNA_seed | PIK3CA | -0.410153 | BRCA | YMB-1-E | GNE-317 | MVP_top60_DNA+RNA_seed | ESR1 | -1.005566 | BRCA | MDA-MB-361 | GDC0810 |
| MVP_top60_DNA_seed | ERBB2 | -0.406105 | BRCA | COLO-824 | Afatinib | MVP_top60_DNA+RNA_seed | AKT1 | -0.997542 | BRCA | HCC1569 | MK-2206 |
| MVP_top60_DNA_seed | EGFR | -0.406105 | BRCA | COLO-824 | Afatinib | MVP_top60_DNA+RNA_seed | EGFR | -0.997178 | BRCA | MDA-MB-330 | Gefitinib |
| MVP_top60_DNA_seed | ESR1 | -0.404312 | BRCA | HCC70 | GDC0810 | MVP_top60_DNA+RNA_seed | HDAC1 | -0.996057 | BRCA | AU565 | Entinostat |
| MVP_top60_DNA_seed | PIK3CA | -0.402635 | BRCA | EFM-19 | Buparlisib | MVP_top60_DNA+RNA_seed | EP300 | -0.99594 | BRCA | MDA-MB-157 | CPI-637 |
| MVP_top60_DNA_seed | EGFR | -0.401897 | BRCA | HCC1954 | Osimertinib | MVP_top60_DNA+RNA_seed | EP300 | -0.993573 | BRCA | HCC1419 | SGC-CBP30 |
| MVP_top60_DNA_seed | TP53 | -0.401514 | BRCA | AU565 | MIRA-1 | MVP_top60_DNA+RNA_seed | AKT1 | -0.990002 | BRCA | BT-549 | AT13148 |
| MVP_top60_DNA_seed | CDK2 | -0.401164 | BRCA | CAL-85-1 | AZD5438 | MVP_top60_DNA+RNA_seed | HDAC1 | -0.989535 | BRCA | HCC70 | Entinostat |
| MVP_top60_DNA_seed | AKT1 | -0.400745 | BRCA | HCC1500 | AT13148 | MVP_top60_DNA+RNA_seed | AKT1 | -0.982903 | BRCA | COLO-824 | Uprosertib |
| MVP_top60_DNA_seed | CDK2 | -0.397833 | BRCA | MFM-223 | Dinaciclib | MVP_top60_DNA+RNA_seed | EP300 | -0.982896 | BRCA | OCUB-M | SGC-CBP30 |
| MVP_top60_DNA_seed | CDK1 | -0.397833 | BRCA | MFM-223 | Dinaciclib | MVP_top60_DNA+RNA_seed | CDK2 | -0.981077 | BRCA | Hs-578-T | MK-8776 |
| MVP_top60_DNA_seed | EGFR | -0.39758 | BRCA | HCC1500 | AZD3759 | MVP_top60_DNA+RNA_seed | AKT1 | -0.980322 | BRCA | CAL-51 | AZD5363 |
| MVP_top60_DNA_seed | EGFR | -0.39627 | BRCA | MRK-nu-1 | AZD3759 | MVP_top60_DNA+RNA_seed | AKT1 | -0.979968 | BRCA | CAMA-1 | AT13148 |
| MVP_top60_DNA_seed | AKT1 | -0.391369 | BRCA | BT-20 | Afuresertib | MVP_top60_DNA+RNA_seed | HSP90AA1 | -0.977994 | BRCA | EVSA-T | Tanespimycin |
| MVP_top60_DNA_seed | CDK2 | -0.388864 | BRCA | BT-20 | Dinaciclib | MVP_top60_DNA+RNA_seed | HSP90AB1 | -0.977994 | BRCA | EVSA-T | Tanespimycin |
| MVP_top60_DNA_seed | CDK1 | -0.388864 | BRCA | BT-20 | Dinaciclib | MVP_top60_DNA+RNA_seed | MDM2 | -0.976296 | BRCA | CAL-51 | Nutlin-3a (-) |
| MVP_top60_DNA_seed | HDAC1 | -0.386675 | BRCA | MRK-nu-1 | Entinostat | MVP_top60_DNA+RNA_seed | ATM | -0.975642 | BRCA | HCC1395 | KU-55933 |
| MVP_top60_DNA_seed | ERBB2 | -0.386482 | BRCA | UACC-812 | Afatinib | MVP_top60_DNA+RNA_seed | EGFR | -0.973906 | BRCA | COLO-824 | Gefitinib |

| | | | | | | | | | | | |
|---|---|---|---|---|---|---|---|---|---|---|---|
| MVP_top60_DNA_seed | EGFR | -0.386482 | BRCA | UACC-812 | Afatinib | MVP_top60_DNA+RNA_seed | CDK2 | -0.973722 | BRCA | MRK-nu-1 | AZD5438 |
| MVP_top60_DNA_seed | ERBB2 | -0.384797 | BRCA | MDA-MB-361 | Lapatinib | MVP_top60_DNA+RNA_seed | AKT1 | -0.969774 | BRCA | MDA-MB-175-VII | Uprosertib |
| MVP_top60_DNA_seed | EGFR | -0.384797 | BRCA | MDA-MB-361 | Lapatinib | MVP_top60_DNA+RNA_seed | ATM | -0.962891 | BRCA | BT-549 | KU-55933 |
| MVP_top60_DNA_seed | AKT1 | -0.384099 | BRCA | HCC2218 | AT13148 | MVP_top60_DNA+RNA_seed | CDK2 | -0.962587 | BRCA | HCC1954 | AZD5438 |
| MVP_top60_DNA_seed | ERBB2 | -0.383961 | BRCA | HCC1599 | Lapatinib | MVP_top60_DNA+RNA_seed | EGFR | -0.962319 | BRCA | DU-4475 | Gefitinib |
| MVP_top60_DNA_seed | EGFR | -0.383961 | BRCA | HCC1599 | Lapatinib | MVP_top60_DNA+RNA_seed | ESR1 | -0.960347 | BRCA | HCC1500 | GDC0810 |
| MVP_top60_DNA_seed | AKT1 | -0.383738 | BRCA | BT-20 | MK-2206 | MVP_top60_DNA+RNA_seed | EGFR | -0.95876 | BRCA | MDA-MB-468 | Afatinib |
| MVP_top60_DNA_seed | HDAC1 | -0.383662 | BRCA | T47D | PCI-34051 | MVP_top60_DNA+RNA_seed | ERBB2 | -0.95876 | BRCA | MDA-MB-468 | Afatinib |
| MVP_top60_DNA_seed | EGFR | -0.382351 | BRCA | T47D | Osimertinib | MVP_top60_DNA+RNA_seed | CDK2 | -0.957054 | BRCA | EVSA-T | Dinaciclib |
| MVP_top60_DNA_seed | MDM2 | -0.382284 | BRCA | HCC1395 | Serdemetan | MVP_top60_DNA+RNA_seed | CDK1 | -0.957054 | BRCA | EVSA-T | Dinaciclib |
| MVP_top60_DNA_seed | EP300 | -0.3819 | BRCA | HCC1569 | SGC-CBP30 | MVP_top60_DNA+RNA_seed | AKT1 | -0.956436 | BRCA | HCC1395 | MK-2206 |
| MVP_top60_DNA_seed | CREBBP | -0.3819 | BRCA | HCC1569 | SGC-CBP30 | MVP_top60_DNA+RNA_seed | PLK1 | -0.952557 | BRCA | CAL-51 | BI-2536 |
| MVP_top60_DNA_seed | ERBB2 | -0.380733 | BRCA | HCC1937 | Lapatinib | MVP_top60_DNA+RNA_seed | AKT1 | -0.950649 | BRCA | HCC1187 | AT13148 |
| MVP_top60_DNA_seed | EGFR | -0.380733 | BRCA | HCC1937 | Lapatinib | MVP_top60_DNA+RNA_seed | AKT1 | -0.945921 | BRCA | COLO-824 | Ipatasertib |
| MVP_top60_DNA_seed | HDAC1 | -0.379507 | BRCA | COLO-824 | Entinostat | MVP_top60_DNA+RNA_seed | AKT1 | -0.942142 | BRCA | HCC1395 | Uprosertib |
| MVP_top60_DNA_seed | HSP90AA1 | -0.375534 | BRCA | MDA-MB-468 | Luminespib | MVP_top60_DNA+RNA_seed | HSP90AA1 | -0.94195 | BRCA | CAL-148 | Luminespib |
| MVP_top60_DNA_seed | HSP90AB1 | -0.375534 | BRCA | MDA-MB-468 | Luminespib | MVP_top60_DNA+RNA_seed | HSP90AB1 | -0.94195 | BRCA | CAL-148 | Luminespib |
| MVP_top60_DNA_seed | ESR1 | -0.373597 | BRCA | HCC1143 | Tamoxifen | MVP_top60_DNA+RNA_seed | MDM2 | -0.940786 | BRCA | OCUB-M | Serdemetan |
| MVP_top60_DNA_seed | CDK2 | -0.372683 | BRCA | AU565 | Dinaciclib | MVP_top60_DNA+RNA_seed | MDM2 | -0.939178 | BRCA | MCF7 | Nutlin-3a (-) |
| MVP_top60_DNA_seed | CDK1 | -0.372683 | BRCA | AU565 | Dinaciclib | MVP_top60_DNA+RNA_seed | PLK1 | -0.935033 | BRCA | COLO-824 | BI-2536 |
| MVP_top60_DNA_seed | PIK3CA | -0.372092 | BRCA | ZR-75-30 | Alpelisib | MVP_top60_DNA+RNA_seed | EGFR | -0.93287 | BRCA | EVSA-T | Lapatinib |
| MVP_top60_DNA_seed | AKT1 | -0.371385 | BRCA | HCC1954 | Uprosertib | MVP_top60_DNA+RNA_seed | ERBB2 | -0.93287 | BRCA | EVSA-T | Lapatinib |
| MVP_top60_DNA_seed | CDK2 | -0.370665 | BRCA | HCC1569 | AZD5438 | MVP_top60_DNA+RNA_seed | PIK3CA | -0.932406 | BRCA | HDQ-P1 | Buparlisib |
| MVP_top60_DNA_seed | PIK3CA | -0.37003 | BRCA | HCC1599 | Buparlisib | MVP_top60_DNA+RNA_seed | AKT1 | -0.932132 | BRCA | MDA-MB-453 | AT13148 |
| MVP_top60_DNA_seed | ESR1 | -0.369648 | BRCA | EFM-19 | Tamoxifen | MVP_top60_DNA+RNA_seed | PIK3CA | -0.931641 | BRCA | AU565 | Alpelisib |
| MVP_top60_DNA_seed | MAPK3 | -0.369141 | BRCA | MDA-MB-231 | Ulixertinib | MVP_top60_DNA+RNA_seed | EGFR | -0.9296 | BRCA | HCC1954 | Lapatinib |
| MVP_top60_DNA_seed | MAPK1 | -0.369141 | BRCA | MDA-MB-231 | Ulixertinib | MVP_top60_DNA+RNA_seed | ERBB2 | -0.9296 | BRCA | HCC1954 | Lapatinib |
| MVP_top60_DNA_seed | EP300 | -0.368969 | BRCA | Hs-578-T | SGC-CBP30 | MVP_top60_DNA+RNA_seed | PIK3CA | -0.927964 | BRCA | T47D | AZD8186 |
| MVP_top60_DNA_seed | CREBBP | -0.368969 | BRCA | Hs-578-T | SGC-CBP30 | MVP_top60_DNA+RNA_seed | HDAC1 | -0.922488 | BRCA | HCC1500 | Entinostat |
| MVP_top60_DNA_seed | HDAC1 | -0.367144 | BRCA | HCC2218 | Entinostat | MVP_top60_DNA+RNA_seed | EGFR | -0.92219 | BRCA | HCC2157 | Osimertinib |
| MVP_top60_DNA_seed | MAPK3 | -0.366678 | BRCA | CAL-51 | Ulixertinib | MVP_top60_DNA+RNA_seed | SRC | -0.921268 | BRCA | MDA-MB-361 | Bosutinib |
| MVP_top60_DNA_seed | MAPK1 | -0.366678 | BRCA | CAL-51 | Ulixertinib | MVP_top60_DNA+RNA_seed | AKT1 | -0.92097 | BRCA | T47D | Afuresertib |
| MVP_top60_DNA_seed | MAPK3 | -0.365342 | BRCA | MDA-MB-175-VII | SCH772984 | MVP_top60_DNA+RNA_seed | ESR1 | -0.920678 | BRCA | MDA-MB-415 | Tamoxifen |
| MVP_top60_DNA_seed | MAPK1 | -0.365342 | BRCA | MDA-MB-175-VII | SCH772984 | MVP_top60_DNA+RNA_seed | TP53 | -0.920382 | BRCA | MDA-MB-415 | MIRA-1 |
| MVP_top60_DNA_seed | MAPK3 | -0.364407 | BRCA | CAL-85-1 | SCH772984 | MVP_top60_DNA+RNA_seed | HSP90AA1 | -0.919906 | BRCA | OCUB-M | Luminespib |
| MVP_top60_DNA_seed | MAPK1 | -0.364407 | BRCA | CAL-85-1 | SCH772984 | MVP_top60_DNA+RNA_seed | HSP90AB1 | -0.919906 | BRCA | OCUB-M | Luminespib |
| MVP_top60_DNA_seed | AKT1 | -0.363355 | BRCA | HCC1500 | Ipatasertib | MVP_top60_DNA+RNA_seed | PIK3CA | -0.917627 | BRCA | CAL-120 | GNE-317 |
| MVP_top60_DNA_seed | MDM2 | -0.361414 | BRCA | MRK-nu-1 | Serdemetan | MVP_top60_DNA+RNA_seed | AKT1 | -0.916813 | BRCA | OCUB-M | AT13148 |
| MVP_top60_DNA_seed | ERBB2 | -0.358688 | BRCA | MDA-MB-231 | Lapatinib | MVP_top60_DNA+RNA_seed | MAPK3 | -0.915602 | BRCA | HCC1143 | Ulixertinib |
| MVP_top60_DNA_seed | EGFR | -0.358688 | BRCA | MDA-MB-231 | Lapatinib | MVP_top60_DNA+RNA_seed | MAPK1 | -0.915602 | BRCA | HCC1143 | Ulixertinib |
| MVP_top60_DNA_seed | HDAC1 | -0.356851 | BRCA | EVSA-T | Romidepsin | MVP_top60_DNA+RNA_seed | EGFR | -0.915335 | BRCA | COLO-824 | Lapatinib |

| | | | | | | | | | | | |
|---|---|---|---|---|---|---|---|---|---|---|---|
| MVP_top60_DNA_seed | HDAC2 | -0.356851 | BRCA | EVSA-T | Romidepsin | MVP_top60_DNA+RNA_seed | ERBB2 | -0.915335 | BRCA | COLO-824 | Lapatinib |
| MVP_top60_DNA_seed | EGFR | -0.355947 | BRCA | MDA-MB-330 | Erlotinib | MVP_top60_DNA+RNA_seed | SRC | -0.915074 | BRCA | HCC2157 | Bosutinib |
| MVP_top60_DNA_seed | ERBB2 | -0.355762 | BRCA | Hs-578-T | Lapatinib | MVP_top60_DNA+RNA_seed | EP300 | -0.909177 | BRCA | EVSA-T | SGC-CBP30 |
| MVP_top60_DNA_seed | EGFR | -0.355762 | BRCA | Hs-578-T | Lapatinib | MVP_top60_DNA+RNA_seed | AKT1 | -0.90838 | BRCA | HCC1954 | AZD5363 |
| MVP_top60_DNA_seed | AKT1 | -0.35495 | BRCA | EFM-192A | Ipatasertib | MVP_top60_DNA+RNA_seed | EGFR | -0.906452 | BRCA | HCC1143 | Sapitinib |
| MVP_top60_DNA_seed | AKT1 | -0.354806 | BRCA | HCC1954 | MK-2206 | MVP_top60_DNA+RNA_seed | ERBB2 | -0.906452 | BRCA | HCC1143 | Sapitinib |
| MVP_top60_DNA_seed | EP300 | -0.353751 | BRCA | CAL-51 | SGC-CBP30 | MVP_top60_DNA+RNA_seed | ERBB3 | -0.906452 | BRCA | HCC1143 | Sapitinib |
| MVP_top60_DNA_seed | CREBBP | -0.353751 | BRCA | CAL-51 | SGC-CBP30 | MVP_top60_DNA+RNA_seed | EGFR | -0.903875 | BRCA | OCUB-M | AZD3759 |
| MVP_top60_DNA_seed | EGFR | -0.351396 | BRCA | MDA-MB-361 | Erlotinib | MVP_top60_DNA+RNA_seed | ESR1 | -0.901197 | BRCA | COLO-824 | GDC0810 |
| MVP_top60_DNA_seed | HDAC1 | -0.347376 | BRCA | EFM-192A | Romidepsin | MVP_top60_DNA+RNA_seed | CDK1 | -0.896132 | BRCA | BT-549 | RO-3306 |
| MVP_top60_DNA_seed | HDAC2 | -0.347376 | BRCA | EFM-192A | Romidepsin | MVP_top60_DNA+RNA_seed | ESR1 | -0.895712 | BRCA | MRK-nu-1 | GDC0810 |
| MVP_top60_DNA_seed | AKT1 | -0.345408 | BRCA | EFM-192A | GSK2110183B | MVP_top60_DNA+RNA_seed | AKT1 | -0.894297 | BRCA | CAL-51 | Uprosertib |
| MVP_top60_DNA_seed | CDK2 | -0.344741 | BRCA | CAL-85-1 | Dinaciclib | MVP_top60_DNA+RNA_seed | PIK3CA | -0.892798 | BRCA | T47D | Buparlisib |
| MVP_top60_DNA_seed | CDK1 | -0.344741 | BRCA | CAL-85-1 | Dinaciclib | MVP_top60_DNA+RNA_seed | EGFR | -0.892471 | BRCA | MDA-MB-361 | Afatinib |
| MVP_top60_DNA_seed | MAPK3 | -0.339309 | BRCA | HCC70 | SCH772984 | MVP_top60_DNA+RNA_seed | ERBB2 | -0.892471 | BRCA | MDA-MB-361 | Afatinib |
| MVP_top60_DNA_seed | MAPK1 | -0.339309 | BRCA | HCC70 | SCH772984 | MVP_top60_DNA+RNA_seed | CDK2 | -0.891599 | BRCA | HCC2157 | AZD5438 |
| MVP_top60_DNA_seed | HSP90AA1 | -0.338967 | BRCA | UACC-812 | Luminespib | MVP_top60_DNA+RNA_seed | ESR1 | -0.891472 | BRCA | HCC1187 | GDC0810 |
| MVP_top60_DNA_seed | HSP90AB1 | -0.338967 | BRCA | UACC-812 | Luminespib | MVP_top60_DNA+RNA_seed | ESR1 | -0.886636 | BRCA | HDQ-P1 | Tamoxifen |
| MVP_top60_DNA_seed | EGFR | -0.337079 | BRCA | HCC1954 | Erlotinib | MVP_top60_DNA+RNA_seed | HSP90AA1 | -0.883579 | BRCA | HCC2157 | Tanespimycin |
| MVP_top60_DNA_seed | CDK2 | -0.333141 | BRCA | MRK-nu-1 | MK-8776 | MVP_top60_DNA+RNA_seed | HSP90AB1 | -0.883579 | BRCA | HCC2157 | Tanespimycin |
| MVP_top60_DNA_seed | PIK3CA | -0.332338 | BRCA | HCC70 | Buparlisib | MVP_top60_DNA+RNA_seed | HDAC1 | -0.883106 | BRCA | UACC-893 | PCI-34051 |
| MVP_top60_DNA_seed | EGFR | -0.33203 | BRCA | UACC-893 | AZD3759 | MVP_top60_DNA+RNA_seed | PIK3CA | -0.882716 | BRCA | AU565 | AZD8186 |
| MVP_top60_DNA_seed | AKT1 | -0.330614 | BRCA | MDA-MB-468 | Ipatasertib | MVP_top60_DNA+RNA_seed | AKT1 | -0.881891 | BRCA | COLO-824 | MK-2206 |
| MVP_top60_DNA_seed | EGFR | -0.330414 | BRCA | MDA-MB-453 | Osimertinib | MVP_top60_DNA+RNA_seed | PIK3CA | -0.880401 | BRCA | Hs-578-T | AZD8186 |
| MVP_top60_DNA_seed | CDK1 | -0.330299 | BRCA | MDA-MB-453 | RO-3306 | MVP_top60_DNA+RNA_seed | SRC | -0.879599 | BRCA | HCC2218 | Bosutinib |
| MVP_top60_DNA_seed | PIK3CA | -0.32986 | BRCA | MFM-223 | Alpelisib | MVP_top60_DNA+RNA_seed | EP300 | -0.878249 | BRCA | MRK-nu-1 | CPI-637 |
| MVP_top60_DNA_seed | ESR1 | -0.329342 | BRCA | HCC1187 | Tamoxifen | MVP_top60_DNA+RNA_seed | AKT1 | -0.872927 | BRCA | EVSA-T | AT13148 |
| MVP_top60_DNA_seed | AKT1 | -0.329262 | BRCA | MDA-MB-468 | Uprosertib | MVP_top60_DNA+RNA_seed | ESR1 | -0.872757 | BRCA | AU565 | Tamoxifen |
| MVP_top60_DNA_seed | PIK3CA | -0.328263 | BRCA | CAL-85-1 | GNE-317 | MVP_top60_DNA+RNA_seed | AKT1 | -0.8665 | BRCA | Hs-578-T | AT13148 |
| MVP_top60_DNA_seed | HSP90AA1 | -0.326785 | BRCA | HCC2218 | Tanespimycin | MVP_top60_DNA+RNA_seed | AKT1 | -0.861728 | BRCA | COLO-824 | AZD5363 |
| MVP_top60_DNA_seed | HSP90AB1 | -0.326785 | BRCA | HCC2218 | Tanespimycin | MVP_top60_DNA+RNA_seed | EGFR | -0.860086 | BRCA | DU-4475 | Osimertinib |
| MVP_top60_DNA_seed | MAPK3 | -0.324759 | BRCA | HCC2157 | Ulixertinib | MVP_top60_DNA+RNA_seed | AKT1 | -0.856041 | BRCA | HCC1954 | AT13148 |
| MVP_top60_DNA_seed | MAPK1 | -0.324759 | BRCA | HCC2157 | Ulixertinib | MVP_top60_DNA+RNA_seed | PIK3CA | -0.853556 | BRCA | MCF7 | Alpelisib |
| MVP_top60_DNA_seed | AKT1 | -0.322016 | BRCA | HCC1599 | MK-2206 | MVP_top60_DNA+RNA_seed | PIK3CA | -0.850261 | BRCA | EFM-192A | Alpelisib |
| MVP_top60_DNA_seed | PIK3CA | -0.321277 | BRCA | MDA-MB-453 | Buparlisib | MVP_top60_DNA+RNA_seed | ESR1 | -0.848816 | BRCA | OCUB-M | Tamoxifen |
| MVP_top60_DNA_seed | PIK3CA | -0.320262 | BRCA | BT-20 | Buparlisib | MVP_top60_DNA+RNA_seed | EP300 | -0.841965 | BRCA | BT-20 | SGC-CBP30 |
| MVP_top60_DNA_seed | EGFR | -0.317185 | BRCA | DU-4475 | AZD3759 | MVP_top60_DNA+RNA_seed | PIK3CA | -0.840747 | BRCA | MDA-MB-361 | AZD8186 |
| MVP_top60_DNA_seed | EGFR | -0.316982 | BRCA | UACC-893 | Gefitinib | MVP_top60_DNA+RNA_seed | MAPK3 | -0.837772 | BRCA | HDQ-P1 | Ulixertinib |
| MVP_top60_DNA_seed | SRC | -0.313141 | BRCA | HCC1395 | Bosutinib | MVP_top60_DNA+RNA_seed | MAPK1 | -0.837772 | BRCA | HDQ-P1 | Ulixertinib |
| MVP_top60_DNA_seed | AKT1 | -0.312627 | BRCA | BT-549 | MK-2206 | MVP_top60_DNA+RNA_seed | MAPK3 | -0.836915 | BRCA | UACC-893 | Ulixertinib |
| MVP_top60_DNA_seed | PIK3CA | -0.312548 | BRCA | BT-20 | GNE-317 | MVP_top60_DNA+RNA_seed | MAPK1 | -0.836915 | BRCA | UACC-893 | Ulixertinib |

| | | | | | | | | | | | |
|---|---|---|---|---|---|---|---|---|---|---|---|
| MVP_top60_DNA_seed | EGFR | -0.311841 | BRCA | MDA-MB-415 | Gefitinib | MVP_top60_DNA+RNA_seed | AKT1 | -0.835386 | BRCA | AU565 | GSK2110183B |
| MVP_top60_DNA_seed | EGFR | -0.311725 | BRCA | T47D | AZD3759 | MVP_top60_DNA+RNA_seed | AKT1 | -0.832701 | BRCA | EFM-19 | Uprosertib |
| MVP_top60_DNA_seed | AKT1 | -0.310426 | BRCA | MDA-MB-175-VII | AT13148 | MVP_top60_DNA+RNA_seed | MAPK3 | -0.83266 | BRCA | MDA-MB-231 | SCH772984 |
| MVP_top60_DNA_seed | EGFR | -0.309153 | BRCA | EVSA-T | AZD3759 | MVP_top60_DNA+RNA_seed | MAPK1 | -0.83266 | BRCA | MDA-MB-231 | SCH772984 |
| MVP_top60_DNA_seed | ATM | -0.307615 | BRCA | MDA-MB-231 | KU-55933 | MVP_top60_DNA+RNA_seed | AKT1 | -0.830537 | BRCA | MFM-223 | MK-2206 |
| MVP_top60_DNA_seed | AKT1 | -0.305241 | BRCA | BT-549 | Ipatasertib | MVP_top60_DNA+RNA_seed | ESR1 | -0.826148 | BRCA | CAL-148 | GDC0810 |
| MVP_top60_DNA_seed | EP300 | -0.301915 | BRCA | MDA-MB-415 | CPI-637 | MVP_top60_DNA+RNA_seed | HSP90AA1 | -0.825778 | BRCA | HCC2157 | Luminespib |
| MVP_top60_DNA_seed | AKT1 | -0.295996 | BRCA | DU-4475 | Uprosertib | MVP_top60_DNA+RNA_seed | HSP90AB1 | -0.825778 | BRCA | HCC2157 | Luminespib |
| MVP_top60_DNA_seed | EGFR | -0.295966 | BRCA | MDA-MB-415 | Osimertinib | MVP_top60_DNA+RNA_seed | EP300 | -0.825162 | BRCA | HDQ-P1 | SGC-CBP30 |
| MVP_top60_DNA_seed | MAPK3 | -0.289547 | BRCA | HCC1395 | Ulixertinib | MVP_top60_DNA+RNA_seed | CDK2 | -0.823937 | BRCA | HCC70 | MK-8776 |
| MVP_top60_DNA_seed | MAPK1 | -0.289547 | BRCA | HCC1395 | Ulixertinib | MVP_top60_DNA+RNA_seed | AKT1 | -0.822013 | BRCA | CAL-51 | GSK2110183B |
| MVP_top60_DNA_seed | AKT1 | -0.286204 | BRCA | HCC2157 | Uprosertib | MVP_top60_DNA+RNA_seed | PIK3CA | -0.819784 | BRCA | COLO-824 | AZD8186 |
| MVP_top60_DNA_seed | HDAC1 | -0.286013 | BRCA | EFM-19 | PCI-34051 | MVP_top60_DNA+RNA_seed | AKT1 | -0.81715 | BRCA | MDA-MB-453 | AZD5363 |
| MVP_top60_DNA_seed | ESR1 | -0.283836 | BRCA | BT-20 | GDC0810 | MVP_top60_DNA+RNA_seed | CDK2 | -0.81652 | BRCA | HCC38 | MK-8776 |
| MVP_top60_DNA_seed | AKT1 | -0.280625 | BRCA | HCC1143 | Ipatasertib | MVP_top60_DNA+RNA_seed | MAPK3 | -0.816422 | BRCA | MDA-MB-175-VII | Ulixertinib |
| MVP_top60_DNA_seed | AKT1 | -0.280149 | BRCA | HCC1187 | GSK2110183B | MVP_top60_DNA+RNA_seed | MAPK1 | -0.816422 | BRCA | MDA-MB-175-VII | Ulixertinib |
| MVP_top60_DNA_seed | AKT1 | -0.278866 | BRCA | HCC1419 | AZD5363 | MVP_top60_DNA+RNA_seed | CDK2 | -0.815706 | BRCA | HCC70 | Dinaciclib |
| MVP_top60_DNA_seed | AKT1 | -0.277893 | BRCA | MRK-nu-1 | AZD5363 | MVP_top60_DNA+RNA_seed | CDK1 | -0.815706 | BRCA | HCC70 | Dinaciclib |
| MVP_top60_DNA_seed | EGFR | -0.276886 | BRCA | HCC1419 | Erlotinib | MVP_top60_DNA+RNA_seed | PIK3CA | -0.809161 | BRCA | MRK-nu-1 | Buparlisib |
| MVP_top60_DNA_seed | CDK2 | -0.276467 | BRCA | AU565 | AZD5438 | MVP_top60_DNA+RNA_seed | EGFR | -0.805849 | BRCA | HDQ-P1 | Afatinib |
| MVP_top60_DNA_seed | CDK2 | -0.273945 | BRCA | BT-549 | MK-8776 | MVP_top60_DNA+RNA_seed | ERBB2 | -0.805849 | BRCA | HDQ-P1 | Afatinib |
| MVP_top60_DNA_seed | ERBB2 | -0.271284 | BRCA | HCC1937 | Sapitinib | MVP_top60_DNA+RNA_seed | ATM | -0.802533 | BRCA | MDA-MB-468 | KU-55933 |
| MVP_top60_DNA_seed | EGFR | -0.271284 | BRCA | HCC1937 | Sapitinib | MVP_top60_DNA+RNA_seed | EGFR | -0.800196 | BRCA | BT-20 | Sapitinib |
| MVP_top60_DNA_seed | ERBB3 | -0.271284 | BRCA | HCC1937 | Sapitinib | MVP_top60_DNA+RNA_seed | ERBB2 | -0.800196 | BRCA | BT-20 | Sapitinib |
| MVP_top60_DNA_seed | HDAC1 | -0.268851 | BRCA | HCC1143 | Entinostat | MVP_top60_DNA+RNA_seed | ERBB3 | -0.800196 | BRCA | BT-20 | Sapitinib |
| MVP_top60_DNA_seed | CDK1 | -0.268381 | BRCA | MDA-MB-415 | RO-3306 | MVP_top60_DNA+RNA_seed | HSP90AA1 | -0.79664 | BRCA | COLO-824 | Luminespib |
| MVP_top60_DNA_seed | AKT1 | -0.265106 | BRCA | BT-474 | GSK2110183B | MVP_top60_DNA+RNA_seed | HSP90AB1 | -0.79664 | BRCA | COLO-824 | Luminespib |
| MVP_top60_DNA_seed | ESR1 | -0.26497 | BRCA | EFM-192A | GDC0810 | MVP_top60_DNA+RNA_seed | CDK2 | -0.786787 | BRCA | MDA-MB-330 | Dinaciclib |
| MVP_top60_DNA_seed | CDK2 | -0.26356 | BRCA | EFM-192A | Dinaciclib | MVP_top60_DNA+RNA_seed | CDK1 | -0.786787 | BRCA | MDA-MB-330 | Dinaciclib |
| MVP_top60_DNA_seed | CDK1 | -0.26356 | BRCA | EFM-192A | Dinaciclib | MVP_top60_DNA+RNA_seed | AKT1 | -0.785847 | BRCA | HCC1395 | AZD5363 |
| MVP_top60_DNA_seed | PIK3CA | -0.261162 | BRCA | JIMT-1 | Alpelisib | MVP_top60_DNA+RNA_seed | ESR1 | -0.78522 | BRCA | MDA-MB-157 | GDC0810 |
| MVP_top60_DNA_seed | MAPK3 | -0.259304 | BRCA | HCC1569 | SCH772984 | MVP_top60_DNA+RNA_seed | PLK1 | -0.780478 | BRCA | EVSA-T | BI-2536 |
| MVP_top60_DNA_seed | MAPK1 | -0.259304 | BRCA | HCC1569 | SCH772984 | MVP_top60_DNA+RNA_seed | EGFR | -0.780079 | BRCA | JIMT-1 | Sapitinib |
| MVP_top60_DNA_seed | CDK2 | -0.259041 | BRCA | DU-4475 | MK-8776 | MVP_top60_DNA+RNA_seed | ERBB2 | -0.780079 | BRCA | JIMT-1 | Sapitinib |
| MVP_top60_DNA_seed | SRC | -0.257781 | BRCA | HCC1569 | Bosutinib | MVP_top60_DNA+RNA_seed | ERBB3 | -0.780079 | BRCA | JIMT-1 | Sapitinib |
| MVP_top60_DNA_seed | EGFR | -0.257672 | BRCA | HCC1143 | Gefitinib | MVP_top60_DNA+RNA_seed | HDAC1 | -0.774694 | BRCA | HCC1569 | Romidepsin |
| MVP_top60_DNA_seed | AKT1 | -0.254692 | BRCA | UACC-812 | AT13148 | MVP_top60_DNA+RNA_seed | HDAC2 | -0.774694 | BRCA | HCC1569 | Romidepsin |
| MVP_top60_DNA_seed | HDAC1 | -0.25232 | BRCA | DU-4475 | PCI-34051 | MVP_top60_DNA+RNA_seed | EGFR | -0.774536 | BRCA | HCC1569 | Lapatinib |
| MVP_top60_DNA_seed | AKT1 | -0.250284 | BRCA | HCC1937 | Ipatasertib | MVP_top60_DNA+RNA_seed | ERBB2 | -0.774536 | BRCA | HCC1569 | Lapatinib |
| MVP_top60_DNA_seed | HSP90AA1 | -0.249708 | BRCA | BT-549 | Luminespib | MVP_top60_DNA+RNA_seed | PIK3CA | -0.772239 | BRCA | OCUB-M | Buparlisib |
| MVP_top60_DNA_seed | HSP90AB1 | -0.249708 | BRCA | BT-549 | Luminespib | MVP_top60_DNA+RNA_seed | MDM2 | -0.770785 | BRCA | YMB-1-E | Nutlin-3a (-) |

| | | | | | | | | | | | |
|---|---|---|---|---|---|---|---|---|---|---|---|
| MVP_top60_DNA_seed | CDK2 | -0.249661 | BRCA | OCUB-M | AZD5438 | MVP_top60_DNA+RNA_seed | EP300 | -0.767298 | BRCA | MDA-MB-453 | SGC-CBP30 |
| MVP_top60_DNA_seed | AKT1 | -0.249464 | BRCA | BT-549 | Afuresertib | MVP_top60_DNA+RNA_seed | HSP90AA1 | -0.767242 | BRCA | MRK-nu-1 | Tanespimycin |
| MVP_top60_DNA_seed | HDAC1 | -0.246658 | BRCA | Hs-578-T | Entinostat | MVP_top60_DNA+RNA_seed | HSP90AB1 | -0.767242 | BRCA | MRK-nu-1 | Tanespimycin |
| MVP_top60_DNA_seed | ESR1 | -0.246657 | BRCA | HCC38 | Tamoxifen | MVP_top60_DNA+RNA_seed | AKT1 | -0.766869 | BRCA | HCC202 | MK-2206 |
| MVP_top60_DNA_seed | CDK2 | -0.245615 | BRCA | YMB-1-E | MK-8776 | MVP_top60_DNA+RNA_seed | CDK1 | -0.764868 | BRCA | COLO-824 | RO-3306 |
| MVP_top60_DNA_seed | ERBB2 | -0.244856 | BRCA | HCC1500 | Afatinib | MVP_top60_DNA+RNA_seed | EGFR | -0.762929 | BRCA | MDA-MB-468 | Osimertinib |
| MVP_top60_DNA_seed | EGFR | -0.244856 | BRCA | HCC1500 | Afatinib | MVP_top60_DNA+RNA_seed | AKT1 | -0.762558 | BRCA | DU-4475 | Uprosertib |
| MVP_top60_DNA_seed | AKT1 | -0.242575 | BRCA | HCC70 | AT13148 | MVP_top60_DNA+RNA_seed | AKT1 | -0.762364 | BRCA | MCF7 | Uprosertib |
| MVP_top60_DNA_seed | HDAC1 | -0.241981 | BRCA | HCC2218 | PCI-34051 | MVP_top60_DNA+RNA_seed | EGFR | -0.75585 | BRCA | HCC1143 | Lapatinib |
| MVP_top60_DNA_seed | ERBB2 | -0.241773 | BRCA | HCC1599 | Sapitinib | MVP_top60_DNA+RNA_seed | ERBB2 | -0.75585 | BRCA | HCC1143 | Lapatinib |
| MVP_top60_DNA_seed | EGFR | -0.241773 | BRCA | HCC1599 | Sapitinib | MVP_top60_DNA+RNA_seed | MDM2 | -0.755338 | BRCA | AU565 | Serdemetan |
| MVP_top60_DNA_seed | ERBB3 | -0.241773 | BRCA | HCC1599 | Sapitinib | MVP_top60_DNA+RNA_seed | SRC | -0.753389 | BRCA | DU-4475 | Bosutinib |
| MVP_top60_DNA_seed | AKT1 | -0.241393 | BRCA | MDA-MB-468 | AT13148 | MVP_top60_DNA+RNA_seed | EGFR | -0.751663 | BRCA | DU-4475 | Erlotinib |
| MVP_top60_DNA_seed | MAPK3 | -0.240852 | BRCA | EVSA-T | Ulixertinib | MVP_top60_DNA+RNA_seed | CDK2 | -0.750382 | BRCA | MDA-MB-231 | AZD5438 |
| MVP_top60_DNA_seed | MAPK1 | -0.240852 | BRCA | EVSA-T | Ulixertinib | MVP_top60_DNA+RNA_seed | AKT1 | -0.74949 | BRCA | MDA-MB-330 | MK-2206 |
| MVP_top60_DNA_seed | AKT1 | -0.238961 | BRCA | BT-474 | Afuresertib | MVP_top60_DNA+RNA_seed | CDK2 | -0.749482 | BRCA | MRK-nu-1 | Dinaciclib |
| MVP_top60_DNA_seed | PIK3CA | -0.238073 | BRCA | CAL-85-1 | Alpelisib | MVP_top60_DNA+RNA_seed | CDK1 | -0.749482 | BRCA | MRK-nu-1 | Dinaciclib |
| MVP_top60_DNA_seed | EP300 | -0.237628 | BRCA | MDA-MB-468 | SGC-CBP30 | MVP_top60_DNA+RNA_seed | AKT1 | -0.746946 | BRCA | HDQ-P1 | Uprosertib |
| MVP_top60_DNA_seed | CREBBP | -0.237628 | BRCA | MDA-MB-468 | SGC-CBP30 | MVP_top60_DNA+RNA_seed | ESR1 | -0.74636 | BRCA | MDA-MB-361 | Tamoxifen |
| MVP_top60_DNA_seed | CDK2 | -0.236229 | BRCA | MDA-MB-361 | AZD5438 | MVP_top60_DNA+RNA_seed | PIK3CA | -0.745503 | BRCA | HCC1143 | AZD8186 |
| MVP_top60_DNA_seed | SRC | -0.235273 | BRCA | MDA-MB-468 | Bosutinib | MVP_top60_DNA+RNA_seed | PIK3CA | -0.743736 | BRCA | Hs-578-T | Buparlisib |
| MVP_top60_DNA_seed | ATM | -0.234404 | BRCA | HCC1937 | KU-55933 | MVP_top60_DNA+RNA_seed | PIK3CA | -0.741553 | BRCA | MDA-MB-468 | AZD8186 |
| MVP_top60_DNA_seed | HSP90AA1 | -0.231679 | BRCA | HCC1599 | Elesclomol | MVP_top60_DNA+RNA_seed | CDK2 | -0.738308 | BRCA | Hs-578-T | AZD5438 |
| MVP_top60_DNA_seed | HSP90AB1 | -0.231679 | BRCA | HCC1599 | Elesclomol | MVP_top60_DNA+RNA_seed | AKT1 | -0.737288 | BRCA | UACC-893 | AT13148 |
| MVP_top60_DNA_seed | HDAC1 | -0.227067 | BRCA | EFM-19 | Romidepsin | MVP_top60_DNA+RNA_seed | PIK3CA | -0.73116 | BRCA | JIMT-1 | GNE-317 |
| MVP_top60_DNA_seed | HDAC2 | -0.227067 | BRCA | EFM-19 | Romidepsin | MVP_top60_DNA+RNA_seed | PIK3CA | -0.730409 | BRCA | Hs-578-T | Alpelisib |
| MVP_top60_DNA_seed | MAPK1 | -0.223007 | BRCA | MDA-MB-415 | VX-11e | MVP_top60_DNA+RNA_seed | CDK2 | -0.72988 | BRCA | OCUB-M | Dinaciclib |
| MVP_top60_DNA_seed | EGFR | -0.22275 | BRCA | HCC1187 | Erlotinib | MVP_top60_DNA+RNA_seed | CDK1 | -0.72988 | BRCA | OCUB-M | Dinaciclib |
| MVP_top60_DNA_seed | EGFR | -0.22114 | BRCA | HCC1937 | Gefitinib | MVP_top60_DNA+RNA_seed | CDK2 | -0.727971 | BRCA | MDA-MB-157 | Dinaciclib |
| MVP_top60_DNA_seed | AKT1 | -0.217162 | BRCA | DU-4475 | MK-2206 | MVP_top60_DNA+RNA_seed | CDK1 | -0.727971 | BRCA | MDA-MB-157 | Dinaciclib |
| MVP_top60_DNA_seed | HSP90AA1 | -0.214866 | BRCA | YMB-1-E | Elesclomol | MVP_top60_DNA+RNA_seed | AKT1 | -0.726847 | BRCA | AU565 | AZD5363 |
| MVP_top60_DNA_seed | HSP90AB1 | -0.214866 | BRCA | YMB-1-E | Elesclomol | MVP_top60_DNA+RNA_seed | AKT1 | -0.726761 | BRCA | AU565 | Uprosertib |
| MVP_top60_DNA_seed | PIK3CA | -0.212837 | BRCA | HCC1937 | AZD8186 | MVP_top60_DNA+RNA_seed | MDM2 | -0.724274 | BRCA | COLO-824 | Serdemetan |
| MVP_top60_DNA_seed | CDK1 | -0.212401 | BRCA | HCC70 | RO-3306 | MVP_top60_DNA+RNA_seed | AKT1 | -0.722234 | BRCA | EFM-19 | Afuresertib |
| MVP_top60_DNA_seed | AKT1 | -0.211479 | BRCA | UACC-812 | Afuresertib | MVP_top60_DNA+RNA_seed | EGFR | -0.719294 | BRCA | JIMT-1 | Lapatinib |
| MVP_top60_DNA_seed | AKT1 | -0.210983 | BRCA | MDA-MB-468 | AZD5363 | MVP_top60_DNA+RNA_seed | ERBB2 | -0.719294 | BRCA | JIMT-1 | Lapatinib |
| MVP_top60_DNA_seed | PIK3CA | -0.208771 | BRCA | MFM-223 | GNE-317 | MVP_top60_DNA+RNA_seed | EGFR | -0.717571 | BRCA | CAL-148 | Osimertinib |
| MVP_top60_DNA_seed | PIK3CA | -0.208619 | BRCA | HCC1569 | GNE-317 | MVP_top60_DNA+RNA_seed | EGFR | -0.716149 | BRCA | MDA-MB-468 | Erlotinib |
| MVP_top60_DNA_seed | SRC | -0.206993 | BRCA | OCUB-M | Bosutinib | MVP_top60_DNA+RNA_seed | HSP90AA1 | -0.716111 | BRCA | MDA-MB-231 | Luminespib |
| MVP_top60_DNA_seed | AKT1 | -0.206989 | BRCA | MRK-nu-1 | Uprosertib | MVP_top60_DNA+RNA_seed | HSP90AB1 | -0.716111 | BRCA | MDA-MB-231 | Luminespib |
| MVP_top60_DNA_seed | CDK2 | -0.206906 | BRCA | MDA-MB-453 | Dinaciclib | MVP_top60_DNA+RNA_seed | CDK2 | -0.714111 | BRCA | HCC70 | AZD5438 |

| | | | | | | | | | | | |
|---|---|---|---|---|---|---|---|---|---|---|---|
| MVP_top60_DNA_seed | CDK1 | -0.206906 | BRCA | MDA-MB-453 | Dinaciclib | MVP_top60_DNA+RNA_seed | AKT1 | -0.713725 | BRCA | AU565 | Afuresertib |
| MVP_top60_DNA_seed | AKT1 | -0.205739 | BRCA | COLO-824 | GSK2110183B | MVP_top60_DNA+RNA_seed | EGFR | -0.711807 | BRCA | AU565 | Erlotinib |
| MVP_top60_DNA_seed | TP53 | -0.205086 | BRCA | HCC1187 | MIRA-1 | MVP_top60_DNA+RNA_seed | PLK1 | -0.711158 | BRCA | HCC1599 | MK-1775 |
| MVP_top60_DNA_seed | MDM2 | -0.201917 | BRCA | HCC1419 | Serdemetan | MVP_top60_DNA+RNA_seed | EGFR | -0.708757 | BRCA | UACC-893 | Erlotinib |
| MVP_top60_DNA_seed | ESR1 | -0.201884 | BRCA | HCC1954 | GDC0810 | MVP_top60_DNA+RNA_seed | MDM2 | -0.70811 | BRCA | UACC-812 | Nutlin-3a (-) |
| MVP_top60_DNA_seed | HSP90AA1 | -0.201748 | BRCA | HCC1569 | Luminespib | MVP_top60_DNA+RNA_seed | EP300 | -0.707949 | BRCA | AU565 | SGC-CBP30 |
| MVP_top60_DNA_seed | HSP90AB1 | -0.201748 | BRCA | HCC1569 | Luminespib | MVP_top60_DNA+RNA_seed | EGFR | -0.707066 | BRCA | COLO-824 | Osimertinib |
| MVP_top60_DNA_seed | HSP90AA1 | -0.201721 | BRCA | HCC1937 | Tanespimycin | MVP_top60_DNA+RNA_seed | PIK3CA | -0.704781 | BRCA | CAL-148 | AZD8186 |
| MVP_top60_DNA_seed | HSP90AB1 | -0.201721 | BRCA | HCC1937 | Tanespimycin | MVP_top60_DNA+RNA_seed | AKT1 | -0.703104 | BRCA | MDA-MB-361 | Afuresertib |
| MVP_top60_DNA_seed | TP53 | -0.201608 | BRCA | HCC1500 | MIRA-1 | MVP_top60_DNA+RNA_seed | PIK3CA | -0.702855 | BRCA | HCC1395 | Buparlisib |
| MVP_top60_DNA_seed | AKT1 | -0.197573 | BRCA | MCF7 | Ipatasertib | MVP_top60_DNA+RNA_seed | PLK1 | -0.700967 | BRCA | JIMT-1 | BI-2536 |
| MVP_top60_DNA_seed | HDAC1 | -0.196695 | BRCA | UACC-893 | Entinostat | MVP_top60_DNA+RNA_seed | HDAC1 | -0.699312 | BRCA | CAL-85-1 | PCI-34051 |
| MVP_top60_DNA_seed | ERBB2 | -0.196205 | BRCA | MDA-MB-157 | Sapitinib | MVP_top60_DNA+RNA_seed | PLK1 | -0.699233 | BRCA | DU-4475 | BI-2536 |
| MVP_top60_DNA_seed | EGFR | -0.196205 | BRCA | MDA-MB-157 | Sapitinib | MVP_top60_DNA+RNA_seed | HSP90AA1 | -0.695131 | BRCA | JIMT-1 | Luminespib |
| MVP_top60_DNA_seed | ERBB3 | -0.196205 | BRCA | MDA-MB-157 | Sapitinib | MVP_top60_DNA+RNA_seed | HSP90AB1 | -0.695131 | BRCA | JIMT-1 | Luminespib |
| MVP_top60_DNA_seed | HSP90AA1 | -0.194768 | BRCA | MDA-MB-175-VII | Tanespimycin | MVP_top60_DNA+RNA_seed | AKT1 | -0.692554 | BRCA | COLO-824 | Uprosertib |
| MVP_top60_DNA_seed | HSP90AB1 | -0.194768 | BRCA | MDA-MB-175-VII | Tanespimycin | MVP_top60_DNA+RNA_seed | PIK3CA | -0.691065 | BRCA | EFM-192A | GNE-317 |
| MVP_top60_DNA_seed | EGFR | -0.192831 | BRCA | HCC1937 | Osimertinib | MVP_top60_DNA+RNA_seed | AKT1 | -0.690475 | BRCA | AU565 | Ipatasertib |
| MVP_top60_DNA_seed | ERBB2 | -0.191157 | BRCA | MDA-MB-453 | Afatinib | MVP_top60_DNA+RNA_seed | CDK2 | -0.689955 | BRCA | DU-4475 | Dinaciclib |
| MVP_top60_DNA_seed | EGFR | -0.191157 | BRCA | MDA-MB-453 | Afatinib | MVP_top60_DNA+RNA_seed | CDK1 | -0.689955 | BRCA | DU-4475 | Dinaciclib |
| MVP_top60_DNA_seed | CDK1 | -0.190001 | BRCA | MDA-MB-231 | RO-3306 | MVP_top60_DNA+RNA_seed | AKT1 | -0.686996 | BRCA | HCC1954 | Ipatasertib |
| MVP_top60_DNA_seed | ESR1 | -0.187918 | BRCA | HCC2157 | GDC0810 | MVP_top60_DNA+RNA_seed | AKT1 | -0.686102 | BRCA | AU565 | MK-2206 |
| MVP_top60_DNA_seed | EGFR | -0.187262 | BRCA | T47D | Erlotinib | MVP_top60_DNA+RNA_seed | ATM | -0.684788 | BRCA | HCC1419 | KU-55933 |
| MVP_top60_DNA_seed | EGFR | -0.185884 | BRCA | HCC2218 | Erlotinib | MVP_top60_DNA+RNA_seed | EGFR | -0.679345 | BRCA | HCC1395 | Lapatinib |
| MVP_top60_DNA_seed | HDAC1 | -0.185616 | BRCA | MDA-MB-231 | Entinostat | MVP_top60_DNA+RNA_seed | ERBB2 | -0.679345 | BRCA | HCC1395 | Lapatinib |
| MVP_top60_DNA_seed | ERBB2 | -0.185169 | BRCA | T47D | Sapitinib | MVP_top60_DNA+RNA_seed | HDAC1 | -0.678338 | BRCA | CAL-85-1 | Romidepsin |
| MVP_top60_DNA_seed | EGFR | -0.185169 | BRCA | T47D | Sapitinib | MVP_top60_DNA+RNA_seed | HDAC2 | -0.678338 | BRCA | CAL-85-1 | Romidepsin |
| MVP_top60_DNA_seed | ERBB3 | -0.185169 | BRCA | T47D | Sapitinib | MVP_top60_DNA+RNA_seed | HDAC1 | -0.677763 | BRCA | MDA-MB-453 | Entinostat |
| MVP_top60_DNA_seed | ESR1 | -0.184272 | BRCA | EFM-192A | Tamoxifen | MVP_top60_DNA+RNA_seed | PIK3CA | -0.67704 | BRCA | UACC-812 | AZD8186 |
| MVP_top60_DNA_seed | AKT1 | -0.183703 | BRCA | HCC1937 | Uprosertib | MVP_top60_DNA+RNA_seed | EGFR | -0.675895 | BRCA | HCC1954 | Sapitinib |
| MVP_top60_DNA_seed | HDAC1 | -0.178024 | BRCA | BT-549 | Entinostat | MVP_top60_DNA+RNA_seed | ERBB2 | -0.675895 | BRCA | HCC1954 | Sapitinib |
| MVP_top60_DNA_seed | PIK3CA | -0.175846 | BRCA | UACC-812 | GNE-317 | MVP_top60_DNA+RNA_seed | ERBB3 | -0.675895 | BRCA | HCC1954 | Sapitinib |
| MVP_top60_DNA_seed | MAPK3 | -0.175576 | BRCA | Hs-578-T | SCH772984 | MVP_top60_DNA+RNA_seed | CDK2 | -0.668885 | BRCA | CAL-51 | Dinaciclib |
| MVP_top60_DNA_seed | MAPK1 | -0.175576 | BRCA | Hs-578-T | SCH772984 | MVP_top60_DNA+RNA_seed | CDK1 | -0.668885 | BRCA | CAL-51 | Dinaciclib |
| MVP_top60_DNA_seed | PIK3CA | -0.175299 | BRCA | UACC-893 | AZD8186 | MVP_top60_DNA+RNA_seed | AKT1 | -0.668123 | BRCA | BT-20 | Uprosertib |
| MVP_top60_DNA_seed | HDAC1 | -0.173626 | BRCA | HDQ-P1 | Romidepsin | MVP_top60_DNA+RNA_seed | ESR1 | -0.666552 | BRCA | MDA-MB-453 | Tamoxifen |
| MVP_top60_DNA_seed | HDAC2 | -0.173626 | BRCA | HDQ-P1 | Romidepsin | MVP_top60_DNA+RNA_seed | HSP90AA1 | -0.663581 | BRCA | HCC1419 | Tanespimycin |
| MVP_top60_DNA_seed | EGFR | -0.173574 | BRCA | HCC1569 | Gefitinib | MVP_top60_DNA+RNA_seed | HSP90AB1 | -0.663581 | BRCA | HCC1419 | Tanespimycin |
| MVP_top60_DNA_seed | CDK1 | -0.171546 | BRCA | CAMA-1 | RO-3306 | MVP_top60_DNA+RNA_seed | PIK3CA | -0.660445 | BRCA | EFM-192A | Buparlisib |
| MVP_top60_DNA_seed | PIK3CA | -0.168908 | BRCA | MDA-MB-453 | AZD8186 | MVP_top60_DNA+RNA_seed | AKT1 | -0.656801 | BRCA | MDA-MB-468 | Afuresertib |
| MVP_top60_DNA_seed | AKT1 | -0.16772 | BRCA | DU-4475 | AT13148 | MVP_top60_DNA+RNA_seed | HDAC1 | -0.654747 | BRCA | HCC1428 | Entinostat |

| | | | | | | | | | | | |
|---|---|---|---|---|---|---|---|---|---|---|---|
| MVP_top60_DNA_seed | TP53 | -0.167296 | BRCA | BT-20 | MIRA-1 | MVP_top60_DNA+RNA_seed | EGFR | -0.654271 | BRCA | BT-20 | Lapatinib |
| MVP_top60_DNA_seed | TP53 | -0.164861 | BRCA | MDA-MB-453 | MIRA-1 | MVP_top60_DNA+RNA_seed | ERBB2 | -0.654271 | BRCA | BT-20 | Lapatinib |
| MVP_top60_DNA_seed | EGFR | -0.163452 | BRCA | CAL-148 | Erlotinib | MVP_top60_DNA+RNA_seed | PIK3CA | -0.653864 | BRCA | AU565 | GNE-317 |
| MVP_top60_DNA_seed | ESR1 | -0.161042 | BRCA | CAL-51 | Tamoxifen | MVP_top60_DNA+RNA_seed | HDAC1 | -0.653387 | BRCA | MDA-MB-231 | Romidepsin |
| MVP_top60_DNA_seed | HDAC1 | -0.159195 | BRCA | OCUB-M | Romidepsin | MVP_top60_DNA+RNA_seed | HDAC2 | -0.653387 | BRCA | MDA-MB-231 | Romidepsin |
| MVP_top60_DNA_seed | HDAC2 | -0.159195 | BRCA | OCUB-M | Romidepsin | MVP_top60_DNA+RNA_seed | AKT1 | -0.653351 | BRCA | HDQ-P1 | AZD5363 |
| MVP_top60_DNA_seed | EP300 | -0.158123 | BRCA | BT-474 | CPI-637 | MVP_top60_DNA+RNA_seed | PIK3CA | -0.652526 | BRCA | MDA-MB-330 | Buparlisib |
| MVP_top60_DNA_seed | MDM2 | -0.15666 | BRCA | MDA-MB-175-VII | Nutlin-3a (-) | MVP_top60_DNA+RNA_seed | PIK3CA | -0.652473 | BRCA | UACC-812 | Alpelisib |
| MVP_top60_DNA_seed | EGFR | -0.152889 | BRCA | HCC1569 | AZD3759 | MVP_top60_DNA+RNA_seed | CDK2 | -0.651474 | BRCA | HCC1937 | MK-8776 |
| MVP_top60_DNA_seed | MAPK1 | -0.150985 | BRCA | CAL-51 | VX-11e | MVP_top60_DNA+RNA_seed | EGFR | -0.65124 | BRCA | COLO-824 | Erlotinib |
| MVP_top60_DNA_seed | ERBB2 | -0.148272 | BRCA | EVSA-T | Sapitinib | MVP_top60_DNA+RNA_seed | EP300 | -0.648117 | BRCA | Hs-578-T | CPI-637 |
| MVP_top60_DNA_seed | EGFR | -0.148272 | BRCA | EVSA-T | Sapitinib | MVP_top60_DNA+RNA_seed | TP53 | -0.647937 | BRCA | EVSA-T | MIRA-1 |
| MVP_top60_DNA_seed | ERBB3 | -0.148272 | BRCA | EVSA-T | Sapitinib | MVP_top60_DNA+RNA_seed | EGFR | -0.643898 | BRCA | MDA-MB-453 | Sapitinib |
| MVP_top60_DNA_seed | PIK3CA | -0.147878 | BRCA | MDA-MB-468 | GNE-317 | MVP_top60_DNA+RNA_seed | ERBB2 | -0.643898 | BRCA | MDA-MB-453 | Sapitinib |
| MVP_top60_DNA_seed | ERBB2 | -0.147212 | BRCA | T47D | Lapatinib | MVP_top60_DNA+RNA_seed | ERBB3 | -0.643898 | BRCA | MDA-MB-453 | Sapitinib |
| MVP_top60_DNA_seed | EGFR | -0.147212 | BRCA | T47D | Lapatinib | MVP_top60_DNA+RNA_seed | AKT1 | -0.643539 | BRCA | HCC1395 | Uprosertib |
| MVP_top60_DNA_seed | PIK3CA | -0.142764 | BRCA | CAL-51 | Alpelisib | MVP_top60_DNA+RNA_seed | HSP90AA1 | -0.64115 | BRCA | MDA-MB-330 | Luminespib |
| MVP_top60_DNA_seed | ERBB2 | -0.142341 | BRCA | HCC70 | Lapatinib | MVP_top60_DNA+RNA_seed | HSP90AB1 | -0.64115 | BRCA | MDA-MB-330 | Luminespib |
| MVP_top60_DNA_seed | EGFR | -0.142341 | BRCA | HCC70 | Lapatinib | MVP_top60_DNA+RNA_seed | ESR1 | -0.637152 | BRCA | CAL-85-1 | GDC0810 |
| MVP_top60_DNA_seed | AKT1 | -0.14134 | BRCA | CAL-51 | Uprosertib | MVP_top60_DNA+RNA_seed | HSP90AA1 | -0.635658 | BRCA | CAL-85-1 | Luminespib |
| MVP_top60_DNA_seed | MAPK1 | -0.140194 | BRCA | COLO-824 | VX-11e | MVP_top60_DNA+RNA_seed | HSP90AB1 | -0.635658 | BRCA | CAL-85-1 | Luminespib |
| MVP_top60_DNA_seed | ERBB2 | -0.140165 | BRCA | DU-4475 | Sapitinib | MVP_top60_DNA+RNA_seed | EGFR | -0.63156 | BRCA | HCC1143 | Erlotinib |
| MVP_top60_DNA_seed | EGFR | -0.140165 | BRCA | DU-4475 | Sapitinib | MVP_top60_DNA+RNA_seed | EP300 | -0.629102 | BRCA | OCUB-M | CPI-637 |
| MVP_top60_DNA_seed | ERBB3 | -0.140165 | BRCA | DU-4475 | Sapitinib | MVP_top60_DNA+RNA_seed | EGFR | -0.628902 | BRCA | EVSA-T | Erlotinib |
| MVP_top60_DNA_seed | MDM2 | -0.139946 | BRCA | HCC70 | Serdemetan | MVP_top60_DNA+RNA_seed | ESR1 | -0.626137 | BRCA | Hs-578-T | Tamoxifen |
| MVP_top60_DNA_seed | AKT1 | -0.139743 | BRCA | HCC1419 | MK-2206 | MVP_top60_DNA+RNA_seed | AKT1 | -0.625503 | BRCA | HCC1419 | GSK2110183B |
| MVP_top60_DNA_seed | AKT1 | -0.138311 | BRCA | DU-4475 | Ipatasertib | MVP_top60_DNA+RNA_seed | EP300 | -0.6246 | BRCA | HCC1419 | CPI-637 |
| MVP_top60_DNA_seed | MAPK1 | -0.135935 | BRCA | HCC1954 | VX-11e | MVP_top60_DNA+RNA_seed | EGFR | -0.62256 | BRCA | HCC1419 | Gefitinib |
| MVP_top60_DNA_seed | ATM | -0.13526 | BRCA | EFM-19 | KU-55933 | MVP_top60_DNA+RNA_seed | AKT1 | -0.621252 | BRCA | T47D | AZD5363 |
| MVP_top60_DNA_seed | EP300 | -0.134227 | BRCA | MRK-nu-1 | SGC-CBP30 | MVP_top60_DNA+RNA_seed | AKT1 | -0.61829 | BRCA | Hs-578-T | Uprosertib |
| MVP_top60_DNA_seed | CREBBP | -0.134227 | BRCA | MRK-nu-1 | SGC-CBP30 | MVP_top60_DNA+RNA_seed | CDK1 | -0.618099 | BRCA | CAL-51 | RO-3306 |
| MVP_top60_DNA_seed | HSP90AA1 | -0.134212 | BRCA | MDA-MB-415 | Tanespimycin | MVP_top60_DNA+RNA_seed | AKT1 | -0.617231 | BRCA | UACC-812 | MK-2206 |
| MVP_top60_DNA_seed | HSP90AB1 | -0.134212 | BRCA | MDA-MB-415 | Tanespimycin | MVP_top60_DNA+RNA_seed | HDAC1 | -0.615121 | BRCA | BT-474 | Entinostat |
| MVP_top60_DNA_seed | AKT1 | -0.133466 | BRCA | HCC1599 | AZD5363 | MVP_top60_DNA+RNA_seed | AKT1 | -0.614668 | BRCA | MCF7 | Uprosertib |
| MVP_top60_DNA_seed | PIK3CA | -0.132617 | BRCA | JIMT-1 | Buparlisib | MVP_top60_DNA+RNA_seed | AKT1 | -0.612365 | BRCA | CAL-51 | AT13148 |
| MVP_top60_DNA_seed | ATM | -0.132384 | BRCA | HCC1143 | KU-55933 | MVP_top60_DNA+RNA_seed | EGFR | -0.610433 | BRCA | HCC2157 | Erlotinib |
| MVP_top60_DNA_seed | EP300 | -0.132295 | BRCA | T47D | CPI-637 | MVP_top60_DNA+RNA_seed | AKT1 | -0.608807 | BRCA | HDQ-P1 | Afuresertib |
| MVP_top60_DNA_seed | PIK3CA | -0.13028 | BRCA | YMB-1-E | Buparlisib | MVP_top60_DNA+RNA_seed | PIK3CA | -0.607739 | BRCA | Hs-578-T | GNE-317 |
| MVP_top60_DNA_seed | TP53 | -0.128637 | BRCA | HCC2218 | MIRA-1 | MVP_top60_DNA+RNA_seed | CDK2 | -0.606584 | BRCA | Hs-578-T | Dinaciclib |
| MVP_top60_DNA_seed | ERBB2 | -0.127189 | BRCA | CAL-85-1 | Lapatinib | MVP_top60_DNA+RNA_seed | CDK1 | -0.606584 | BRCA | Hs-578-T | Dinaciclib |
| MVP_top60_DNA_seed | EGFR | -0.127189 | BRCA | CAL-85-1 | Lapatinib | MVP_top60_DNA+RNA_seed | HSP90AA1 | -0.604727 | BRCA | HDQ-P1 | Tanespimycin |

| | | | | | | | | | | | |
|---|---|---|---|---|---|---|---|---|---|---|---|
| MVP_top60_DNA_seed | ESR1 | -0.127146 | BRCA | JIMT-1 | Tamoxifen | MVP_top60_DNA+RNA_seed | HSP90AB1 | -0.604727 | BRCA | HDQ-P1 | Tanespimycin |
| MVP_top60_DNA_seed | HDAC1 | -0.126853 | BRCA | MDA-MB-361 | Romidepsin | MVP_top60_DNA+RNA_seed | PIK3CA | -0.601721 | BRCA | COLO-824 | Alpelisib |
| MVP_top60_DNA_seed | HDAC2 | -0.126853 | BRCA | MDA-MB-361 | Romidepsin | MVP_top60_DNA+RNA_seed | EGFR | -0.600252 | BRCA | HCC1143 | AZD3759 |
| MVP_top60_DNA_seed | ATM | -0.126464 | BRCA | HCC38 | KU-55933 | MVP_top60_DNA+RNA_seed | HDAC1 | -0.599259 | BRCA | HCC1419 | Romidepsin |
| MVP_top60_DNA_seed | AKT1 | -0.126359 | BRCA | MDA-MB-468 | GSK2110183B | MVP_top60_DNA+RNA_seed | HDAC2 | -0.599259 | BRCA | HCC1419 | Romidepsin |
| MVP_top60_DNA_seed | AKT1 | -0.126093 | BRCA | EFM-192A | Uprosertib | MVP_top60_DNA+RNA_seed | HDAC1 | -0.598366 | BRCA | EFM-192A | Entinostat |
| MVP_top60_DNA_seed | AKT1 | -0.124824 | BRCA | EFM-192A | Afuresertib | MVP_top60_DNA+RNA_seed | CDK2 | -0.597811 | BRCA | AU565 | MK-8776 |
| MVP_top60_DNA_seed | HSP90AA1 | -0.124607 | BRCA | MDA-MB-415 | Luminespib | MVP_top60_DNA+RNA_seed | HSP90AA1 | -0.597687 | BRCA | MFM-223 | Tanespimycin |
| MVP_top60_DNA_seed | HSP90AB1 | -0.124607 | BRCA | MDA-MB-415 | Luminespib | MVP_top60_DNA+RNA_seed | HSP90AB1 | -0.597687 | BRCA | MFM-223 | Tanespimycin |
| MVP_top60_DNA_seed | ATM | -0.123973 | BRCA | HCC2218 | KU-55933 | MVP_top60_DNA+RNA_seed | PIK3CA | -0.596498 | BRCA | DU-4475 | Buparlisib |
| MVP_top60_DNA_seed | EP300 | -0.123753 | BRCA | HCC2157 | SGC-CBP30 | MVP_top60_DNA+RNA_seed | PIK3CA | -0.595752 | BRCA | HCC1143 | Buparlisib |
| MVP_top60_DNA_seed | CREBBP | -0.123753 | BRCA | HCC2157 | SGC-CBP30 | MVP_top60_DNA+RNA_seed | AKT1 | -0.593354 | BRCA | HCC1599 | Uprosertib |
| MVP_top60_DNA_seed | PIK3CA | -0.122867 | BRCA | HCC1187 | AZD8186 | MVP_top60_DNA+RNA_seed | AKT1 | -0.593165 | BRCA | BT-474 | Uprosertib |
| MVP_top60_DNA_seed | EGFR | -0.117539 | BRCA | CAL-51 | Erlotinib | MVP_top60_DNA+RNA_seed | PIK3CA | -0.592111 | BRCA | MDA-MB-175-VII | AZD8186 |
| MVP_top60_DNA_seed | ERBB2 | -0.113196 | BRCA | HCC38 | Lapatinib | MVP_top60_DNA+RNA_seed | ESR1 | -0.590756 | BRCA | MDA-MB-231 | GDC0810 |
| MVP_top60_DNA_seed | EGFR | -0.113196 | BRCA | HCC38 | Lapatinib | MVP_top60_DNA+RNA_seed | EP300 | -0.5902 | BRCA | DU-4475 | SGC-CBP30 |
| MVP_top60_DNA_seed | CDK2 | -0.111853 | BRCA | EVSA-T | AZD5438 | MVP_top60_DNA+RNA_seed | HDAC1 | -0.589309 | BRCA | MDA-MB-468 | Romidepsin |
| MVP_top60_DNA_seed | CDK1 | -0.111581 | BRCA | JIMT-1 | RO-3306 | MVP_top60_DNA+RNA_seed | HDAC2 | -0.589309 | BRCA | MDA-MB-468 | Romidepsin |
| MVP_top60_DNA_seed | HSP90AA1 | -0.110684 | BRCA | HCC1954 | Tanespimycin | MVP_top60_DNA+RNA_seed | ATM | -0.58849 | BRCA | CAMA-1 | KU-55933 |
| MVP_top60_DNA_seed | HSP90AB1 | -0.110684 | BRCA | HCC1954 | Tanespimycin | MVP_top60_DNA+RNA_seed | CDK1 | -0.583503 | BRCA | AU565 | RO-3306 |
| MVP_top60_DNA_seed | HSP90AA1 | -0.109998 | BRCA | HCC1599 | Luminespib | MVP_top60_DNA+RNA_seed | AKT1 | -0.576975 | BRCA | HCC1937 | GSK2110183B |
| MVP_top60_DNA_seed | HSP90AB1 | -0.109998 | BRCA | HCC1599 | Luminespib | MVP_top60_DNA+RNA_seed | MAPK1 | -0.576253 | BRCA | MDA-MB-231 | VX-11e |
| MVP_top60_DNA_seed | PIK3CA | -0.105804 | BRCA | MDA-MB-453 | GNE-317 | MVP_top60_DNA+RNA_seed | EGFR | -0.574657 | BRCA | HCC2157 | AZD3759 |
| MVP_top60_DNA_seed | CDK1 | -0.104339 | BRCA | UACC-893 | RO-3306 | MVP_top60_DNA+RNA_seed | HDAC1 | -0.571774 | BRCA | HCC1954 | Romidepsin |
| MVP_top60_DNA_seed | HSP90AA1 | -0.104295 | BRCA | MCF7 | Tanespimycin | MVP_top60_DNA+RNA_seed | HDAC2 | -0.571774 | BRCA | HCC1954 | Romidepsin |
| MVP_top60_DNA_seed | HSP90AB1 | -0.104295 | BRCA | MCF7 | Tanespimycin | MVP_top60_DNA+RNA_seed | PIK3CA | -0.57039 | BRCA | HCC1569 | AZD8186 |
| MVP_top60_DNA_seed | AKT1 | -0.10414 | BRCA | EFM-192A | MK-2206 | MVP_top60_DNA+RNA_seed | PIK3CA | -0.568248 | BRCA | CAL-148 | Buparlisib |
| MVP_top60_DNA_seed | SRC | -0.100852 | BRCA | MDA-MB-231 | Bosutinib | MVP_top60_DNA+RNA_seed | HSP90AA1 | -0.567335 | BRCA | EVSA-T | Luminespib |
| MVP_top60_DNA_seed | CDK1 | -0.10033 | BRCA | HDQ-P1 | RO-3306 | MVP_top60_DNA+RNA_seed | HSP90AB1 | -0.567335 | BRCA | EVSA-T | Luminespib |
| MVP_top60_DNA_seed | AKT1 | -0.097581 | BRCA | BT-549 | AZD5363 | MVP_top60_DNA+RNA_seed | AKT1 | -0.567116 | BRCA | MDA-MB-231 | AT13148 |
| MVP_top60_DNA_seed | PIK3CA | -0.096342 | BRCA | HCC2218 | Alpelisib | MVP_top60_DNA+RNA_seed | HSP90AA1 | -0.564846 | BRCA | HDQ-P1 | Luminespib |
| MVP_top60_DNA_seed | ERBB2 | -0.094247 | BRCA | Hs-578-T | Afatinib | MVP_top60_DNA+RNA_seed | HSP90AB1 | -0.564846 | BRCA | HDQ-P1 | Luminespib |
| MVP_top60_DNA_seed | EGFR | -0.094247 | BRCA | Hs-578-T | Afatinib | MVP_top60_DNA+RNA_seed | HSP90AA1 | -0.563338 | BRCA | JIMT-1 | Tanespimycin |
| MVP_top60_DNA_seed | SRC | -0.093435 | BRCA | EVSA-T | Bosutinib | MVP_top60_DNA+RNA_seed | HSP90AB1 | -0.563338 | BRCA | JIMT-1 | Tanespimycin |
| MVP_top60_DNA_seed | PIK3CA | -0.09236 | BRCA | HCC70 | Alpelisib | MVP_top60_DNA+RNA_seed | AKT1 | -0.562517 | BRCA | EFM-19 | AZD5363 |
| MVP_top60_DNA_seed | CDK1 | -0.090985 | BRCA | MCF7 | RO-3306 | MVP_top60_DNA+RNA_seed | SRC | -0.5615 | BRCA | MRK-nu-1 | Bosutinib |
| MVP_top60_DNA_seed | MDM2 | -0.089748 | BRCA | BT-20 | Serdemetan | MVP_top60_DNA+RNA_seed | ATM | -0.554212 | BRCA | HCC1954 | KU-55933 |
| MVP_top60_DNA_seed | EGFR | -0.087398 | BRCA | HCC70 | Erlotinib | MVP_top60_DNA+RNA_seed | EGFR | -0.553837 | BRCA | MCF7 | Lapatinib |
| MVP_top60_DNA_seed | PIK3CA | -0.086838 | BRCA | DU-4475 | GNE-317 | MVP_top60_DNA+RNA_seed | ERBB2 | -0.553837 | BRCA | MCF7 | Lapatinib |
| MVP_top60_DNA_seed | EGFR | -0.085134 | BRCA | BT-20 | AZD3759 | MVP_top60_DNA+RNA_seed | CDK2 | -0.550668 | BRCA | CAL-51 | AZD5438 |
| MVP_top60_DNA_seed | MAPK3 | -0.082166 | BRCA | BT-20 | SCH772984 | MVP_top60_DNA+RNA_seed | HDAC1 | -0.550593 | BRCA | OCUB-M | Entinostat |

| | | | | | | | | | | | |
|---|---|---|---|---|---|---|---|---|---|---|---|
| MVP_top60_DNA_seed | MAPK1 | -0.082166 | BRCA | BT-20 | SCH772984 | MVP_top60_DNA+RNA_seed | HDAC1 | -0.548279 | BRCA | EVSA-T | PCI-34051 |
| MVP_top60_DNA_seed | EGFR | -0.081593 | BRCA | MDA-MB-361 | Gefitinib | MVP_top60_DNA+RNA_seed | EGFR | -0.544447 | BRCA | EFM-192A | Afatinib |
| MVP_top60_DNA_seed | MAPK3 | -0.080819 | BRCA | HDQ-P1 | SCH772984 | MVP_top60_DNA+RNA_seed | ERBB2 | -0.544447 | BRCA | EFM-192A | Afatinib |
| MVP_top60_DNA_seed | MAPK1 | -0.080819 | BRCA | HDQ-P1 | SCH772984 | MVP_top60_DNA+RNA_seed | HSP90AA1 | -0.543351 | BRCA | HCC1569 | Tanespimycin |
| MVP_top60_DNA_seed | HSP90AA1 | -0.079782 | BRCA | MCF7 | Luminespib | MVP_top60_DNA+RNA_seed | HSP90AB1 | -0.543351 | BRCA | HCC1569 | Tanespimycin |
| MVP_top60_DNA_seed | HSP90AB1 | -0.079782 | BRCA | MCF7 | Luminespib | MVP_top60_DNA+RNA_seed | EGFR | -0.541206 | BRCA | MDA-MB-361 | AZD3759 |
| MVP_top60_DNA_seed | HDAC1 | -0.078202 | BRCA | CAL-51 | Entinostat | MVP_top60_DNA+RNA_seed | AKT1 | -0.539013 | BRCA | Hs-578-T | Afuresertib |
| MVP_top60_DNA_seed | PIK3CA | -0.077314 | BRCA | HCC2157 | Alpelisib | MVP_top60_DNA+RNA_seed | EGFR | -0.538031 | BRCA | MDA-MB-415 | Lapatinib |
| MVP_top60_DNA_seed | HSP90AA1 | -0.074845 | BRCA | HCC1419 | Luminespib | MVP_top60_DNA+RNA_seed | ERBB2 | -0.538031 | BRCA | MDA-MB-415 | Lapatinib |
| MVP_top60_DNA_seed | HSP90AB1 | -0.074845 | BRCA | HCC1419 | Luminespib | MVP_top60_DNA+RNA_seed | MDM2 | -0.537312 | BRCA | HCC2218 | Serdemetan |
| MVP_top60_DNA_seed | ESR1 | -0.071202 | BRCA | HCC1937 | GDC0810 | MVP_top60_DNA+RNA_seed | EGFR | -0.533864 | BRCA | MDA-MB-361 | Osimertinib |
| MVP_top60_DNA_seed | AKT1 | -0.071095 | BRCA | MCF7 | MK-2206 | MVP_top60_DNA+RNA_seed | EGFR | -0.533007 | BRCA | CAL-85-1 | Sapitinib |
| MVP_top60_DNA_seed | HSP90AA1 | -0.066654 | BRCA | AU565 | Tanespimycin | MVP_top60_DNA+RNA_seed | ERBB2 | -0.533007 | BRCA | CAL-85-1 | Sapitinib |
| MVP_top60_DNA_seed | HSP90AB1 | -0.066654 | BRCA | AU565 | Tanespimycin | MVP_top60_DNA+RNA_seed | ERBB3 | -0.533007 | BRCA | CAL-85-1 | Sapitinib |
| MVP_top60_DNA_seed | PIK3CA | -0.064745 | BRCA | MDA-MB-415 | Buparlisib | MVP_top60_DNA+RNA_seed | AKT1 | -0.528852 | BRCA | MRK-nu-1 | Uprosertib |
| MVP_top60_DNA_seed | AKT1 | -0.063491 | BRCA | EFM-19 | AT13148 | MVP_top60_DNA+RNA_seed | TP53 | -0.527752 | BRCA | HCC70 | MIRA-1 |
| MVP_top60_DNA_seed | ERBB2 | -0.062007 | BRCA | YMB-1-E | Lapatinib | MVP_top60_DNA+RNA_seed | EGFR | -0.526819 | BRCA | AU565 | AZD3759 |
| MVP_top60_DNA_seed | EGFR | -0.062007 | BRCA | YMB-1-E | Lapatinib | MVP_top60_DNA+RNA_seed | AKT1 | -0.525991 | BRCA | HCC1954 | Afuresertib |
| MVP_top60_DNA_seed | ATM | -0.059547 | BRCA | MDA-MB-361 | KU-55933 | MVP_top60_DNA+RNA_seed | PLK1 | -0.525683 | BRCA | CAL-148 | MK-1775 |
| MVP_top60_DNA_seed | HSP90AA1 | -0.057207 | BRCA | HCC38 | Luminespib | MVP_top60_DNA+RNA_seed | CDK1 | -0.525477 | BRCA | HCC1500 | RO-3306 |
| MVP_top60_DNA_seed | HSP90AB1 | -0.057207 | BRCA | HCC38 | Luminespib | MVP_top60_DNA+RNA_seed | PLK1 | -0.524674 | BRCA | Hs-578-T | MK-1775 |
| MVP_top60_DNA_seed | ERBB2 | -0.053825 | BRCA | HCC1569 | Afatinib | MVP_top60_DNA+RNA_seed | EGFR | -0.524376 | BRCA | HCC1569 | Sapitinib |
| MVP_top60_DNA_seed | EGFR | -0.053825 | BRCA | HCC1569 | Afatinib | MVP_top60_DNA+RNA_seed | ERBB2 | -0.524376 | BRCA | HCC1569 | Sapitinib |
| MVP_top60_DNA_seed | MAPK3 | -0.053441 | BRCA | HCC1569 | Ulixertinib | MVP_top60_DNA+RNA_seed | ERBB3 | -0.524376 | BRCA | HCC1569 | Sapitinib |
| MVP_top60_DNA_seed | MAPK1 | -0.053441 | BRCA | HCC1569 | Ulixertinib | MVP_top60_DNA+RNA_seed | MAPK1 | -0.523588 | BRCA | MDA-MB-453 | VX-11e |
| MVP_top60_DNA_seed | MAPK3 | -0.053414 | BRCA | HCC1419 | Ulixertinib | MVP_top60_DNA+RNA_seed | ESR1 | -0.521907 | BRCA | HCC1419 | Tamoxifen |
| MVP_top60_DNA_seed | MAPK1 | -0.053414 | BRCA | HCC1419 | Ulixertinib | MVP_top60_DNA+RNA_seed | CDK2 | -0.51965 | BRCA | MDA-MB-361 | Dinaciclib |
| MVP_top60_DNA_seed | AKT1 | -0.052553 | BRCA | UACC-812 | Ipatasertib | MVP_top60_DNA+RNA_seed | CDK1 | -0.51965 | BRCA | MDA-MB-361 | Dinaciclib |
| MVP_top60_DNA_seed | EGFR | -0.052387 | BRCA | MRK-nu-1 | Erlotinib | MVP_top60_DNA+RNA_seed | EP300 | -0.516204 | BRCA | MDA-MB-231 | CPI-637 |
| MVP_top60_DNA_seed | AKT1 | -0.049565 | BRCA | HCC1937 | Uprosertib | MVP_top60_DNA+RNA_seed | AKT1 | -0.516089 | BRCA | T47D | MK-2206 |
| MVP_top60_DNA_seed | AKT1 | -0.048284 | BRCA | EFM-192A | AT13148 | MVP_top60_DNA+RNA_seed | PIK3CA | -0.515105 | BRCA | AU565 | Buparlisib |
| MVP_top60_DNA_seed | MDM2 | -0.047509 | BRCA | T47D | Nutlin-3a (-) | MVP_top60_DNA+RNA_seed | AKT1 | -0.514955 | BRCA | EFM-192A | Uprosertib |
| MVP_top60_DNA_seed | AKT1 | -0.046702 | BRCA | HDQ-P1 | Uprosertib | MVP_top60_DNA+RNA_seed | PIK3CA | -0.514386 | BRCA | HCC1143 | GNE-317 |
| MVP_top60_DNA_seed | MAPK3 | -0.044559 | BRCA | HCC1143 | SCH772984 | MVP_top60_DNA+RNA_seed | AKT1 | -0.512371 | BRCA | DU-4475 | GSK2110183B |
| MVP_top60_DNA_seed | MAPK1 | -0.044559 | BRCA | HCC1143 | SCH772984 | MVP_top60_DNA+RNA_seed | HDAC1 | -0.511999 | BRCA | AU565 | PCI-34051 |
| MVP_top60_DNA_seed | AKT1 | -0.042614 | BRCA | HCC38 | AZD5363 | MVP_top60_DNA+RNA_seed | EGFR | -0.511941 | BRCA | MDA-MB-175-VII | AZD3759 |
| MVP_top60_DNA_seed | CDK2 | -0.040349 | BRCA | CAL-51 | MK-8776 | MVP_top60_DNA+RNA_seed | AKT1 | -0.50826 | BRCA | HDQ-P1 | Ipatasertib |
| MVP_top60_DNA_seed | ATM | -0.039755 | BRCA | HCC1187 | KU-55933 | MVP_top60_DNA+RNA_seed | AKT1 | -0.505243 | BRCA | MDA-MB-361 | AZD5363 |
| MVP_top60_DNA_seed | AKT1 | -0.038662 | BRCA | MDA-MB-361 | AT13148 | MVP_top60_DNA+RNA_seed | PIK3CA | -0.503639 | BRCA | HCC70 | GNE-317 |
| MVP_top60_DNA_seed | CDK1 | -0.038444 | BRCA | HCC1569 | RO-3306 | MVP_top60_DNA+RNA_seed | PLK1 | -0.502995 | BRCA | DU-4475 | MK-1775 |
| MVP_top60_DNA_seed | EGFR | -0.03659 | BRCA | HCC70 | AZD3759 | MVP_top60_DNA+RNA_seed | CDK1 | -0.502564 | BRCA | HCC1428 | RO-3306 |

| | | | | | | | | | | | |
|---|---|---|---|---|---|---|---|---|---|---|---|
| MVP_top60_DNA_seed | HSP90AA1 | -0.034519 | BRCA | HCC2218 | Luminespib | MVP_top60_DNA+RNA_seed | HSP90AA1 | -0.50009 | BRCA | UACC-893 | Luminespib |
| MVP_top60_DNA_seed | HSP90AB1 | -0.034519 | BRCA | HCC2218 | Luminespib | MVP_top60_DNA+RNA_seed | HSP90AB1 | -0.50009 | BRCA | UACC-893 | Luminespib |
| MVP_top60_DNA_seed | AKT1 | -0.032897 | BRCA | MDA-MB-330 | AT13148 | MVP_top60_DNA+RNA_seed | TP53 | -0.499832 | BRCA | CAL-148 | MIRA-1 |
| MVP_top60_DNA_seed | HDAC1 | -0.031421 | BRCA | CAMA-1 | Entinostat | MVP_top60_DNA+RNA_seed | CDK2 | -0.496807 | BRCA | BT-20 | AZD5438 |
| MVP_top60_DNA_seed | ESR1 | -0.031292 | BRCA | UACC-812 | Tamoxifen | MVP_top60_DNA+RNA_seed | ATM | -0.49617 | BRCA | YMB-1-E | KU-55933 |
| MVP_top60_DNA_seed | MAPK3 | -0.028843 | BRCA | HCC2218 | SCH772984 | MVP_top60_DNA+RNA_seed | CDK1 | -0.494962 | BRCA | BT-20 | RO-3306 |
| MVP_top60_DNA_seed | MAPK1 | -0.028843 | BRCA | HCC2218 | SCH772984 | MVP_top60_DNA+RNA_seed | HDAC1 | -0.494453 | BRCA | HCC2157 | Romidepsin |
| MVP_top60_DNA_seed | ESR1 | -0.026363 | BRCA | BT-549 | Tamoxifen | MVP_top60_DNA+RNA_seed | HDAC2 | -0.494453 | BRCA | HCC2157 | Romidepsin |
| MVP_top60_DNA_seed | AKT1 | -0.026331 | BRCA | CAL-51 | MK-2206 | MVP_top60_DNA+RNA_seed | HSP90AA1 | -0.490478 | BRCA | Hs-578-T | Luminespib |
| MVP_top60_DNA_seed | CDK2 | -0.025684 | BRCA | EFM-19 | Dinaciclib | MVP_top60_DNA+RNA_seed | HSP90AB1 | -0.490478 | BRCA | Hs-578-T | Luminespib |
| MVP_top60_DNA_seed | CDK1 | -0.025684 | BRCA | EFM-19 | Dinaciclib | MVP_top60_DNA+RNA_seed | ESR1 | -0.486566 | BRCA | HCC1500 | Tamoxifen |
| MVP_top60_DNA_seed | CDK2 | -0.02457 | BRCA | MDA-MB-415 | Dinaciclib | MVP_top60_DNA+RNA_seed | PIK3CA | -0.486361 | BRCA | HCC1954 | Buparlisib |
| MVP_top60_DNA_seed | CDK1 | -0.02457 | BRCA | MDA-MB-415 | Dinaciclib | MVP_top60_DNA+RNA_seed | EP300 | -0.484749 | BRCA | CAMA-1 | CPI-637 |
| MVP_top60_DNA_seed | EGFR | -0.023951 | BRCA | MRK-nu-1 | Gefitinib | MVP_top60_DNA+RNA_seed | EGFR | -0.484072 | BRCA | CAL-51 | Lapatinib |
| MVP_top60_DNA_seed | HSP90AA1 | -0.022439 | BRCA | HCC1937 | Luminespib | MVP_top60_DNA+RNA_seed | ERBB2 | -0.484072 | BRCA | CAL-51 | Lapatinib |
| MVP_top60_DNA_seed | HSP90AB1 | -0.022439 | BRCA | HCC1937 | Luminespib | MVP_top60_DNA+RNA_seed | AKT1 | -0.482557 | BRCA | HCC1395 | AT13148 |
| MVP_top60_DNA_seed | HDAC1 | -0.022102 | BRCA | HCC1954 | Entinostat | MVP_top60_DNA+RNA_seed | ATM | -0.481969 | BRCA | COLO-824 | KU-55933 |
| MVP_top60_DNA_seed | EGFR | -0.021254 | BRCA | CAL-51 | Gefitinib | MVP_top60_DNA+RNA_seed | AKT1 | -0.478356 | BRCA | HCC1419 | Uprosertib |
| MVP_top60_DNA_seed | SRC | -0.020417 | BRCA | BT-20 | Bosutinib | MVP_top60_DNA+RNA_seed | MDM2 | -0.477942 | BRCA | EVSA-T | Serdemetan |
| MVP_top60_DNA_seed | HDAC1 | -0.018234 | BRCA | CAMA-1 | PCI-34051 | MVP_top60_DNA+RNA_seed | HSP90AA1 | -0.476288 | BRCA | HCC2218 | Elesclomol |
| MVP_top60_DNA_seed | ATM | -0.017561 | BRCA | HDQ-P1 | KU-55933 | MVP_top60_DNA+RNA_seed | HSP90AB1 | -0.476288 | BRCA | HCC2218 | Elesclomol |
| MVP_top60_DNA_seed | ERBB2 | -0.016339 | BRCA | BT-549 | Lapatinib | MVP_top60_DNA+RNA_seed | CDK2 | -0.473378 | BRCA | MDA-MB-231 | Dinaciclib |
| MVP_top60_DNA_seed | EGFR | -0.016339 | BRCA | BT-549 | Lapatinib | MVP_top60_DNA+RNA_seed | CDK1 | -0.473378 | BRCA | MDA-MB-231 | Dinaciclib |
| MVP_top60_DNA_seed | EGFR | -0.016185 | BRCA | EVSA-T | Gefitinib | MVP_top60_DNA+RNA_seed | EGFR | -0.473315 | BRCA | HCC1143 | Osimertinib |
| MVP_top60_DNA_seed | AKT1 | -0.015569 | BRCA | T47D | Uprosertib | MVP_top60_DNA+RNA_seed | AKT1 | -0.473254 | BRCA | MCF7 | AT13148 |
| MVP_top60_DNA_seed | ERBB2 | -0.015539 | BRCA | MRK-nu-1 | Afatinib | MVP_top60_DNA+RNA_seed | EP300 | -0.470434 | BRCA | EFM-192A | SGC-CBP30 |
| MVP_top60_DNA_seed | EGFR | -0.015539 | BRCA | MRK-nu-1 | Afatinib | MVP_top60_DNA+RNA_seed | PIK3CA | -0.464579 | BRCA | MDA-MB-231 | GNE-317 |
| MVP_top60_DNA_seed | PIK3CA | -0.014331 | BRCA | BT-549 | GNE-317 | MVP_top60_DNA+RNA_seed | EGFR | -0.461709 | BRCA | HCC1954 | AZD3759 |
| MVP_top60_DNA_seed | AKT1 | -0.01307 | BRCA | HCC1937 | MK-2206 | MVP_top60_DNA+RNA_seed | CDK1 | -0.458783 | BRCA | HCC1395 | RO-3306 |
| MVP_top60_DNA_seed | AKT1 | -0.010066 | BRCA | HCC2218 | AZD5363 | MVP_top60_DNA+RNA_seed | TP53 | -0.454454 | BRCA | DU-4475 | MIRA-1 |
| MVP_top60_DNA_seed | CDK2 | -0.007275 | BRCA | HDQ-P1 | AZD5438 | MVP_top60_DNA+RNA_seed | EP300 | -0.451755 | BRCA | BT-549 | CPI-637 |
| MVP_top60_DNA_seed | PIK3CA | -0.006998 | BRCA | MCF7 | AZD8186 | MVP_top60_DNA+RNA_seed | CDK2 | -0.448916 | BRCA | MDA-MB-157 | AZD5438 |
| MVP_top60_DNA_seed | HSP90AA1 | -0.006935 | BRCA | HCC1500 | Tanespimycin | MVP_top60_DNA+RNA_seed | MAPK3 | -0.448886 | BRCA | UACC-893 | SCH772984 |
| MVP_top60_DNA_seed | HSP90AB1 | -0.006935 | BRCA | HCC1500 | Tanespimycin | MVP_top60_DNA+RNA_seed | MAPK1 | -0.448886 | BRCA | UACC-893 | SCH772984 |
| MVP_top60_DNA_seed | EGFR | -0.006254 | BRCA | EFM-19 | Osimertinib | MVP_top60_DNA+RNA_seed | EGFR | -0.444722 | BRCA | EFM-192A | Erlotinib |
| MVP_top60_DNA_seed | ERBB2 | -0.006149 | BRCA | EFM-19 | Sapitinib | MVP_top60_DNA+RNA_seed | HSP90AA1 | -0.441942 | BRCA | CAL-85-1 | Tanespimycin |
| MVP_top60_DNA_seed | EGFR | -0.006149 | BRCA | EFM-19 | Sapitinib | MVP_top60_DNA+RNA_seed | HSP90AB1 | -0.441942 | BRCA | CAL-85-1 | Tanespimycin |
| MVP_top60_DNA_seed | ERBB3 | -0.006149 | BRCA | EFM-19 | Sapitinib | MVP_top60_DNA+RNA_seed | PIK3CA | -0.441574 | BRCA | COLO-824 | Buparlisib |
| MVP_top60_DNA_seed | ESR1 | -0.00481 | BRCA | MDA-MB-436 | Tamoxifen | MVP_top60_DNA+RNA_seed | PIK3CA | -0.441497 | BRCA | HCC1143 | Alpelisib |
| MVP_top60_DNA_seed | MAPK3 | -0.004705 | BRCA | MDA-MB-468 | Ulixertinib | MVP_top60_DNA+RNA_seed | ESR1 | -0.440353 | BRCA | HCC1954 | Tamoxifen |
| MVP_top60_DNA_seed | MAPK1 | -0.004705 | BRCA | MDA-MB-468 | Ulixertinib | MVP_top60_DNA+RNA_seed | AKT1 | -0.438643 | BRCA | BT-474 | Ipatasertib |

| | | | | | | | | | | | |
|---|---|---|---|---|---|---|---|---|---|---|---|
| MVP_top60_DNA_seed | PIK3CA | -0.003606 | BRCA | COLO-824 | GNE-317 | MVP_top60_DNA+RNA_seed | PIK3CA | -0.438056 | BRCA | HCC1954 | GNE-317 |
| MVP_top60_DNA_seed | PIK3CA | -0.003436 | BRCA | EVSA-T | Alpelisib | MVP_top60_DNA+RNA_seed | PIK3CA | -0.43752 | BRCA | MCF7 | Buparlisib |
| MVP_top60_DNA_seed | EP300 | -0.002939 | BRCA | JIMT-1 | SGC-CBP30 | MVP_top60_DNA+RNA_seed | HSP90AA1 | -0.437013 | BRCA | MDA-MB-175-VII | Luminespib |
| MVP_top60_DNA_seed | CREBBP | -0.002939 | BRCA | JIMT-1 | SGC-CBP30 | MVP_top60_DNA+RNA_seed | HSP90AB1 | -0.437013 | BRCA | MDA-MB-175-VII | Luminespib |
| MVP_top60_DNA_seed | HDAC1 | -0.002552 | BRCA | CAL-85-1 | Entinostat | MVP_top60_DNA+RNA_seed | PIK3CA | -0.436057 | BRCA | HDQ-P1 | Alpelisib |
| MVP_top60_DNA_seed | MDM2 | -0.002526 | BRCA | AU565 | Nutlin-3a (-) | MVP_top60_DNA+RNA_seed | ATM | -0.435736 | BRCA | CAL-85-1 | KU-55933 |
| MVP_top60_DNA_seed | ERBB2 | 0.001844 | BRCA | CAL-148 | Afatinib | MVP_top60_DNA+RNA_seed | HSP90AA1 | -0.434775 | BRCA | HCC38 | Tanespimycin |
| MVP_top60_DNA_seed | EGFR | 0.001844 | BRCA | CAL-148 | Afatinib | MVP_top60_DNA+RNA_seed | HSP90AB1 | -0.434775 | BRCA | HCC38 | Tanespimycin |
| MVP_top60_DNA_seed | EGFR | 0.002988 | BRCA | OCUB-M | Gefitinib | MVP_top60_DNA+RNA_seed | AKT1 | -0.434648 | BRCA | HCC1599 | Afuresertib |
| MVP_top60_DNA_seed | AKT1 | 0.003273 | BRCA | HCC2218 | Uprosertib | MVP_top60_DNA+RNA_seed | MAPK3 | -0.433042 | BRCA | UACC-893 | Ulixertinib |
| MVP_top60_DNA_seed | SRC | 0.005662 | BRCA | HCC1419 | Bosutinib | MVP_top60_DNA+RNA_seed | MAPK1 | -0.433042 | BRCA | UACC-893 | Ulixertinib |
| MVP_top60_DNA_seed | HSP90AA1 | 0.005988 | BRCA | MRK-nu-1 | Luminespib | MVP_top60_DNA+RNA_seed | ATM | -0.432764 | BRCA | BT-20 | KU-55933 |
| MVP_top60_DNA_seed | HSP90AB1 | 0.005988 | BRCA | MRK-nu-1 | Luminespib | MVP_top60_DNA+RNA_seed | AKT1 | -0.429136 | BRCA | HCC2157 | AT13148 |
| MVP_top60_DNA_seed | EGFR | 0.006975 | BRCA | HCC1419 | AZD3759 | MVP_top60_DNA+RNA_seed | ESR1 | -0.428431 | BRCA | MDA-MB-468 | Tamoxifen |
| MVP_top60_DNA_seed | HDAC1 | 0.010375 | BRCA | JIMT-1 | Romidepsin | MVP_top60_DNA+RNA_seed | AKT1 | -0.425352 | BRCA | EFM-192A | AZD5363 |
| MVP_top60_DNA_seed | HDAC2 | 0.010375 | BRCA | JIMT-1 | Romidepsin | MVP_top60_DNA+RNA_seed | MDM2 | -0.424858 | BRCA | YMB-1-E | Serdemetan |
| MVP_top60_DNA_seed | TP53 | 0.010507 | BRCA | MDA-MB-330 | MIRA-1 | MVP_top60_DNA+RNA_seed | ATM | -0.423123 | BRCA | T47D | KU-55933 |
| MVP_top60_DNA_seed | EGFR | 0.015833 | BRCA | HCC70 | Gefitinib | MVP_top60_DNA+RNA_seed | AKT1 | -0.422735 | BRCA | BT-474 | MK-2206 |
| MVP_top60_DNA_seed | MAPK1 | 0.017338 | BRCA | EVSA-T | VX-11e | MVP_top60_DNA+RNA_seed | AKT1 | -0.422422 | BRCA | COLO-824 | Afuresertib |
| MVP_top60_DNA_seed | HDAC1 | 0.017551 | BRCA | MDA-MB-157 | Romidepsin | MVP_top60_DNA+RNA_seed | SRC | -0.42217 | BRCA | CAL-85-1 | Bosutinib |
| MVP_top60_DNA_seed | HDAC2 | 0.017551 | BRCA | MDA-MB-157 | Romidepsin | MVP_top60_DNA+RNA_seed | EGFR | -0.418449 | BRCA | HCC1937 | Erlotinib |
| MVP_top60_DNA_seed | SRC | 0.017835 | BRCA | CAL-148 | Bosutinib | MVP_top60_DNA+RNA_seed | PIK3CA | -0.413514 | BRCA | EFM-192A | AZD8186 |
| MVP_top60_DNA_seed | ESR1 | 0.019116 | BRCA | HCC70 | Tamoxifen | MVP_top60_DNA+RNA_seed | PIK3CA | -0.410153 | BRCA | YMB-1-E | GNE-317 |
| MVP_top60_DNA_seed | AKT1 | 0.020388 | BRCA | CAL-85-1 | Ipatasertib | MVP_top60_DNA+RNA_seed | PLK1 | -0.406585 | BRCA | CAL-51 | MK-1775 |
| MVP_top60_DNA_seed | HSP90AA1 | 0.021405 | BRCA | CAL-120 | Luminespib | MVP_top60_DNA+RNA_seed | EGFR | -0.406105 | BRCA | COLO-824 | Afatinib |
| MVP_top60_DNA_seed | HSP90AB1 | 0.021405 | BRCA | CAL-120 | Luminespib | MVP_top60_DNA+RNA_seed | ERBB2 | -0.406105 | BRCA | COLO-824 | Afatinib |
| MVP_top60_DNA_seed | AKT1 | 0.021467 | BRCA | MDA-MB-468 | MK-2206 | MVP_top60_DNA+RNA_seed | ESR1 | -0.404312 | BRCA | HCC70 | GDC0810 |
| MVP_top60_DNA_seed | EGFR | 0.02231 | BRCA | CAMA-1 | Osimertinib | MVP_top60_DNA+RNA_seed | PIK3CA | -0.402635 | BRCA | EFM-19 | Buparlisib |
| MVP_top60_DNA_seed | HSP90AA1 | 0.024163 | BRCA | HCC1569 | CCT-018159 | MVP_top60_DNA+RNA_seed | EGFR | -0.401897 | BRCA | HCC1954 | Osimertinib |
| MVP_top60_DNA_seed | HSP90AB1 | 0.024163 | BRCA | HCC1569 | CCT-018159 | MVP_top60_DNA+RNA_seed | TP53 | -0.401514 | BRCA | AU565 | MIRA-1 |
| MVP_top60_DNA_seed | MDM2 | 0.024757 | BRCA | ZR-75-30 | Serdemetan | MVP_top60_DNA+RNA_seed | CDK2 | -0.401164 | BRCA | CAL-85-1 | AZD5438 |
| MVP_top60_DNA_seed | EGFR | 0.025478 | BRCA | CAL-51 | Osimertinib | MVP_top60_DNA+RNA_seed | AKT1 | -0.400745 | BRCA | HCC1500 | AT13148 |
| MVP_top60_DNA_seed | HSP90AA1 | 0.026115 | BRCA | YMB-1-E | Tanespimycin | MVP_top60_DNA+RNA_seed | CDK2 | -0.397833 | BRCA | MFM-223 | Dinaciclib |
| MVP_top60_DNA_seed | HSP90AB1 | 0.026115 | BRCA | YMB-1-E | Tanespimycin | MVP_top60_DNA+RNA_seed | CDK1 | -0.397833 | BRCA | MFM-223 | Dinaciclib |
| MVP_top60_DNA_seed | MDM2 | 0.02657 | BRCA | MDA-MB-415 | Nutlin-3a (-) | MVP_top60_DNA+RNA_seed | EGFR | -0.39758 | BRCA | HCC1500 | AZD3759 |
| MVP_top60_DNA_seed | MAPK3 | 0.026659 | BRCA | EVSA-T | Ulixertinib | MVP_top60_DNA+RNA_seed | EGFR | -0.39627 | BRCA | MRK-nu-1 | AZD3759 |
| MVP_top60_DNA_seed | MAPK1 | 0.026659 | BRCA | EVSA-T | Ulixertinib | MVP_top60_DNA+RNA_seed | AKT1 | -0.391369 | BRCA | BT-20 | Afuresertib |
| MVP_top60_DNA_seed | AKT1 | 0.027095 | BRCA | HCC1143 | GSK2110183B | MVP_top60_DNA+RNA_seed | CDK2 | -0.388864 | BRCA | BT-20 | Dinaciclib |
| MVP_top60_DNA_seed | PIK3CA | 0.031282 | BRCA | HCC1954 | AZD8186 | MVP_top60_DNA+RNA_seed | CDK1 | -0.388864 | BRCA | BT-20 | Dinaciclib |
| MVP_top60_DNA_seed | CDK2 | 0.031771 | BRCA | HCC1419 | MK-8776 | MVP_top60_DNA+RNA_seed | HDAC1 | -0.386675 | BRCA | MRK-nu-1 | Entinostat |
| MVP_top60_DNA_seed | HSP90AA1 | 0.035358 | BRCA | YMB-1-E | Luminespib | MVP_top60_DNA+RNA_seed | EGFR | -0.386482 | BRCA | UACC-812 | Afatinib |

| | | | | | | | | | | | |
|---|---|---|---|---|---|---|---|---|---|---|---|
| MVP_top60_DNA_seed | HSP90AB1 | 0.035358 | BRCA | YMB-1-E | Luminespib | MVP_top60_DNA+RNA_seed | ERBB2 | -0.386482 | BRCA | UACC-812 | Afatinib |
| MVP_top60_DNA_seed | CDK1 | 0.036086 | BRCA | HCC2218 | RO-3306 | MVP_top60_DNA+RNA_seed | EGFR | -0.384797 | BRCA | MDA-MB-361 | Lapatinib |
| MVP_top60_DNA_seed | MAPK3 | 0.037352 | BRCA | OCUB-M | Ulixertinib | MVP_top60_DNA+RNA_seed | ERBB2 | -0.384797 | BRCA | MDA-MB-361 | Lapatinib |
| MVP_top60_DNA_seed | MAPK1 | 0.037352 | BRCA | OCUB-M | Ulixertinib | MVP_top60_DNA+RNA_seed | AKT1 | -0.384099 | BRCA | HCC2218 | AT13148 |
| MVP_top60_DNA_seed | HSP90AA1 | 0.039856 | BRCA | HCC1143 | Luminespib | MVP_top60_DNA+RNA_seed | EGFR | -0.383961 | BRCA | HCC1599 | Lapatinib |
| MVP_top60_DNA_seed | HSP90AB1 | 0.039856 | BRCA | HCC1143 | Luminespib | MVP_top60_DNA+RNA_seed | ERBB2 | -0.383961 | BRCA | HCC1599 | Lapatinib |
| MVP_top60_DNA_seed | EGFR | 0.040789 | BRCA | HCC2218 | AZD3759 | MVP_top60_DNA+RNA_seed | AKT1 | -0.383738 | BRCA | BT-20 | MK-2206 |
| MVP_top60_DNA_seed | HDAC1 | 0.041152 | BRCA | MDA-MB-361 | PCI-34051 | MVP_top60_DNA+RNA_seed | HDAC1 | -0.383662 | BRCA | T47D | PCI-34051 |
| MVP_top60_DNA_seed | ERBB2 | 0.041701 | BRCA | JIMT-1 | Afatinib | MVP_top60_DNA+RNA_seed | EGFR | -0.382351 | BRCA | T47D | Osimertinib |
| MVP_top60_DNA_seed | EGFR | 0.041701 | BRCA | JIMT-1 | Afatinib | MVP_top60_DNA+RNA_seed | MDM2 | -0.382284 | BRCA | HCC1395 | Serdemetan |
| MVP_top60_DNA_seed | CDK2 | 0.043953 | BRCA | UACC-893 | MK-8776 | MVP_top60_DNA+RNA_seed | EP300 | -0.3819 | BRCA | HCC1569 | SGC-CBP30 |
| MVP_top60_DNA_seed | ESR1 | 0.044537 | BRCA | CAL-85-1 | Tamoxifen | MVP_top60_DNA+RNA_seed | EGFR | -0.380733 | BRCA | HCC1937 | Lapatinib |
| MVP_top60_DNA_seed | ERBB2 | 0.044667 | BRCA | DU-4475 | Afatinib | MVP_top60_DNA+RNA_seed | ERBB2 | -0.380733 | BRCA | HCC1937 | Lapatinib |
| MVP_top60_DNA_seed | EGFR | 0.044667 | BRCA | DU-4475 | Afatinib | MVP_top60_DNA+RNA_seed | HDAC1 | -0.379507 | BRCA | COLO-824 | Entinostat |
| MVP_top60_DNA_seed | AKT1 | 0.044804 | BRCA | HCC1187 | Ipatasertib | MVP_top60_DNA+RNA_seed | HSP90AA1 | -0.375534 | BRCA | MDA-MB-468 | Luminespib |
| MVP_top60_DNA_seed | PIK3CA | 0.044886 | BRCA | BT-549 | AZD8186 | MVP_top60_DNA+RNA_seed | HSP90AB1 | -0.375534 | BRCA | MDA-MB-468 | Luminespib |
| MVP_top60_DNA_seed | MDM2 | 0.045823 | BRCA | MDA-MB-453 | Serdemetan | MVP_top60_DNA+RNA_seed | ESR1 | -0.373597 | BRCA | HCC1143 | Tamoxifen |
| MVP_top60_DNA_seed | MDM2 | 0.048354 | BRCA | HCC2157 | Serdemetan | MVP_top60_DNA+RNA_seed | CDK2 | -0.372683 | BRCA | AU565 | Dinaciclib |
| MVP_top60_DNA_seed | AKT1 | 0.051383 | BRCA | HCC1937 | Afuresertib | MVP_top60_DNA+RNA_seed | CDK1 | -0.372683 | BRCA | AU565 | Dinaciclib |
| MVP_top60_DNA_seed | PIK3CA | 0.053041 | BRCA | CAMA-1 | Buparlisib | MVP_top60_DNA+RNA_seed | PIK3CA | -0.372092 | BRCA | ZR-75-30 | Alpelisib |
| MVP_top60_DNA_seed | HDAC1 | 0.053309 | BRCA | CAMA-1 | Romidepsin | MVP_top60_DNA+RNA_seed | AKT1 | -0.371385 | BRCA | HCC1954 | Uprosertib |
| MVP_top60_DNA_seed | HDAC2 | 0.053309 | BRCA | CAMA-1 | Romidepsin | MVP_top60_DNA+RNA_seed | CDK2 | -0.370665 | BRCA | HCC1569 | AZD5438 |
| MVP_top60_DNA_seed | PIK3CA | 0.05419 | BRCA | MDA-MB-468 | Buparlisib | MVP_top60_DNA+RNA_seed | PIK3CA | -0.37003 | BRCA | HCC1599 | Buparlisib |
| MVP_top60_DNA_seed | MDM2 | 0.058001 | BRCA | T47D | Serdemetan | MVP_top60_DNA+RNA_seed | ESR1 | -0.369648 | BRCA | EFM-19 | Tamoxifen |
| MVP_top60_DNA_seed | MAPK3 | 0.058565 | BRCA | CAL-51 | Ulixertinib | MVP_top60_DNA+RNA_seed | MAPK3 | -0.369141 | BRCA | MDA-MB-231 | Ulixertinib |
| MVP_top60_DNA_seed | MAPK1 | 0.058565 | BRCA | CAL-51 | Ulixertinib | MVP_top60_DNA+RNA_seed | MAPK1 | -0.369141 | BRCA | MDA-MB-231 | Ulixertinib |
| MVP_top60_DNA_seed | SRC | 0.059465 | BRCA | HCC1143 | Bosutinib | MVP_top60_DNA+RNA_seed | EP300 | -0.368969 | BRCA | Hs-578-T | SGC-CBP30 |
| MVP_top60_DNA_seed | MDM2 | 0.059967 | BRCA | HCC2157 | Nutlin-3a (-) | MVP_top60_DNA+RNA_seed | HDAC1 | -0.367144 | BRCA | HCC2218 | Entinostat |
| MVP_top60_DNA_seed | MDM2 | 0.060373 | BRCA | HCC1569 | Nutlin-3a (-) | MVP_top60_DNA+RNA_seed | MAPK3 | -0.366678 | BRCA | CAL-51 | Ulixertinib |
| MVP_top60_DNA_seed | EGFR | 0.06129 | BRCA | BT-474 | Erlotinib | MVP_top60_DNA+RNA_seed | MAPK1 | -0.366678 | BRCA | CAL-51 | Ulixertinib |
| MVP_top60_DNA_seed | MAPK3 | 0.062979 | BRCA | YMB-1-E | SCH772984 | MVP_top60_DNA+RNA_seed | MAPK3 | -0.365342 | BRCA | MDA-MB-175-VII | SCH772984 |
| MVP_top60_DNA_seed | MAPK1 | 0.062979 | BRCA | YMB-1-E | SCH772984 | MVP_top60_DNA+RNA_seed | MAPK1 | -0.365342 | BRCA | MDA-MB-175-VII | SCH772984 |
| MVP_top60_DNA_seed | EP300 | 0.062984 | BRCA | HCC70 | SGC-CBP30 | MVP_top60_DNA+RNA_seed | MAPK3 | -0.364407 | BRCA | CAL-85-1 | SCH772984 |
| MVP_top60_DNA_seed | CREBBP | 0.062984 | BRCA | HCC70 | SGC-CBP30 | MVP_top60_DNA+RNA_seed | MAPK1 | -0.364407 | BRCA | CAL-85-1 | SCH772984 |
| MVP_top60_DNA_seed | TP53 | 0.067588 | BRCA | CAL-85-1 | MIRA-1 | MVP_top60_DNA+RNA_seed | AKT1 | -0.363355 | BRCA | HCC1500 | Ipatasertib |
| MVP_top60_DNA_seed | HSP90AA1 | 0.067877 | BRCA | MDA-MB-157 | Luminespib | MVP_top60_DNA+RNA_seed | MDM2 | -0.361414 | BRCA | MRK-nu-1 | Serdemetan |
| MVP_top60_DNA_seed | HSP90AB1 | 0.067877 | BRCA | MDA-MB-157 | Luminespib | MVP_top60_DNA+RNA_seed | EGFR | -0.358688 | BRCA | MDA-MB-231 | Lapatinib |
| MVP_top60_DNA_seed | HDAC1 | 0.068065 | BRCA | MDA-MB-453 | PCI-34051 | MVP_top60_DNA+RNA_seed | ERBB2 | -0.358688 | BRCA | MDA-MB-231 | Lapatinib |
| MVP_top60_DNA_seed | MDM2 | 0.069172 | BRCA | JIMT-1 | Serdemetan | MVP_top60_DNA+RNA_seed | HDAC1 | -0.356851 | BRCA | EVSA-T | Romidepsin |
| MVP_top60_DNA_seed | HDAC1 | 0.069842 | BRCA | MDA-MB-468 | Entinostat | MVP_top60_DNA+RNA_seed | HDAC2 | -0.356851 | BRCA | EVSA-T | Romidepsin |
| MVP_top60_DNA_seed | MDM2 | 0.071249 | BRCA | HCC1187 | Nutlin-3a (-) | MVP_top60_DNA+RNA_seed | EGFR | -0.355947 | BRCA | MDA-MB-330 | Erlotinib |

| | | | | | | | | | | | |
|---|---|---|---|---|---|---|---|---|---|---|---|
| MVP_top60_DNA_seed | PIK3CA | 0.072001 | BRCA | HCC1395 | Alpelisib | MVP_top60_DNA+RNA_seed | EGFR | -0.355762 | BRCA | Hs-578-T | Lapatinib |
| MVP_top60_DNA_seed | AKT1 | 0.072133 | BRCA | HCC1599 | AT13148 | MVP_top60_DNA+RNA_seed | ERBB2 | -0.355762 | BRCA | Hs-578-T | Lapatinib |
| MVP_top60_DNA_seed | ATM | 0.073825 | BRCA | HCC1569 | KU-55933 | MVP_top60_DNA+RNA_seed | AKT1 | -0.35495 | BRCA | EFM-192A | Ipatasertib |
| MVP_top60_DNA_seed | CDK1 | 0.073852 | BRCA | T47D | RO-3306 | MVP_top60_DNA+RNA_seed | AKT1 | -0.354806 | BRCA | HCC1954 | MK-2206 |
| MVP_top60_DNA_seed | MAPK3 | 0.073975 | BRCA | MDA-MB-453 | Ulixertinib | MVP_top60_DNA+RNA_seed | EP300 | -0.353751 | BRCA | CAL-51 | SGC-CBP30 |
| MVP_top60_DNA_seed | MAPK1 | 0.073975 | BRCA | MDA-MB-453 | Ulixertinib | MVP_top60_DNA+RNA_seed | EGFR | -0.351396 | BRCA | MDA-MB-361 | Erlotinib |
| MVP_top60_DNA_seed | ERBB2 | 0.074967 | BRCA | MDA-MB-231 | Afatinib | MVP_top60_DNA+RNA_seed | PLK1 | -0.348205 | BRCA | HCC1395 | MK-1775 |
| MVP_top60_DNA_seed | EGFR | 0.074967 | BRCA | MDA-MB-231 | Afatinib | MVP_top60_DNA+RNA_seed | HDAC1 | -0.347376 | BRCA | EFM-192A | Romidepsin |
| MVP_top60_DNA_seed | HSP90AA1 | 0.075628 | BRCA | BT-20 | Tanespimycin | MVP_top60_DNA+RNA_seed | HDAC2 | -0.347376 | BRCA | EFM-192A | Romidepsin |
| MVP_top60_DNA_seed | HSP90AB1 | 0.075628 | BRCA | BT-20 | Tanespimycin | MVP_top60_DNA+RNA_seed | AKT1 | -0.345408 | BRCA | EFM-192A | GSK2110183B |
| MVP_top60_DNA_seed | MAPK3 | 0.076447 | BRCA | MDA-MB-453 | SCH772984 | MVP_top60_DNA+RNA_seed | CDK2 | -0.344741 | BRCA | CAL-85-1 | Dinaciclib |
| MVP_top60_DNA_seed | MAPK1 | 0.076447 | BRCA | MDA-MB-453 | SCH772984 | MVP_top60_DNA+RNA_seed | CDK1 | -0.344741 | BRCA | CAL-85-1 | Dinaciclib |
| MVP_top60_DNA_seed | SRC | 0.077267 | BRCA | UACC-893 | Bosutinib | MVP_top60_DNA+RNA_seed | MAPK3 | -0.339309 | BRCA | HCC70 | SCH772984 |
| MVP_top60_DNA_seed | ESR1 | 0.07979 | BRCA | HCC1569 | Tamoxifen | MVP_top60_DNA+RNA_seed | MAPK1 | -0.339309 | BRCA | HCC70 | SCH772984 |
| MVP_top60_DNA_seed | TP53 | 0.079871 | BRCA | CAMA-1 | MIRA-1 | MVP_top60_DNA+RNA_seed | HSP90AA1 | -0.338967 | BRCA | UACC-812 | Luminespib |
| MVP_top60_DNA_seed | EGFR | 0.0799 | BRCA | COLO-824 | AZD3759 | MVP_top60_DNA+RNA_seed | HSP90AB1 | -0.338967 | BRCA | UACC-812 | Luminespib |
| MVP_top60_DNA_seed | MAPK1 | 0.081773 | BRCA | UACC-893 | VX-11e | MVP_top60_DNA+RNA_seed | EGFR | -0.337079 | BRCA | HCC1954 | Erlotinib |
| MVP_top60_DNA_seed | AKT1 | 0.085434 | BRCA | ZR-75-30 | Uprosertib | MVP_top60_DNA+RNA_seed | CDK2 | -0.333141 | BRCA | MRK-nu-1 | MK-8776 |
| MVP_top60_DNA_seed | HSP90AA1 | 0.087714 | BRCA | HCC70 | Luminespib | MVP_top60_DNA+RNA_seed | PIK3CA | -0.332338 | BRCA | HCC70 | Buparlisib |
| MVP_top60_DNA_seed | HSP90AB1 | 0.087714 | BRCA | HCC70 | Luminespib | MVP_top60_DNA+RNA_seed | EGFR | -0.33203 | BRCA | UACC-893 | AZD3759 |
| MVP_top60_DNA_seed | MAPK1 | 0.088764 | BRCA | OCUB-M | VX-11e | MVP_top60_DNA+RNA_seed | AKT1 | -0.330614 | BRCA | MDA-MB-468 | Ipatasertib |
| MVP_top60_DNA_seed | ESR1 | 0.089536 | BRCA | MRK-nu-1 | Tamoxifen | MVP_top60_DNA+RNA_seed | EGFR | -0.330414 | BRCA | MDA-MB-453 | Osimertinib |
| MVP_top60_DNA_seed | MDM2 | 0.091379 | BRCA | MDA-MB-468 | Serdemetan | MVP_top60_DNA+RNA_seed | CDK1 | -0.330299 | BRCA | MDA-MB-453 | RO-3306 |
| MVP_top60_DNA_seed | AKT1 | 0.093271 | BRCA | HCC1187 | MK-2206 | MVP_top60_DNA+RNA_seed | PIK3CA | -0.32986 | BRCA | MFM-223 | Alpelisib |
| MVP_top60_DNA_seed | TP53 | 0.09638 | BRCA | HDQ-P1 | MIRA-1 | MVP_top60_DNA+RNA_seed | ESR1 | -0.329342 | BRCA | HCC1187 | Tamoxifen |
| MVP_top60_DNA_seed | AR | 0.098345 | BRCA | YMB-1-E | Bicalutamide | MVP_top60_DNA+RNA_seed | AKT1 | -0.329262 | BRCA | MDA-MB-468 | Uprosertib |
| MVP_top60_DNA_seed | AKT1 | 0.099004 | BRCA | BT-549 | Uprosertib | MVP_top60_DNA+RNA_seed | PIK3CA | -0.328263 | BRCA | CAL-85-1 | GNE-317 |
| MVP_top60_DNA_seed | CDK1 | 0.099478 | BRCA | CAL-120 | RO-3306 | MVP_top60_DNA+RNA_seed | HSP90AA1 | -0.326785 | BRCA | HCC2218 | Tanespimycin |
| MVP_top60_DNA_seed | PIK3CA | 0.103107 | BRCA | UACC-893 | Buparlisib | MVP_top60_DNA+RNA_seed | HSP90AB1 | -0.326785 | BRCA | HCC2218 | Tanespimycin |
| MVP_top60_DNA_seed | AKT1 | 0.104465 | BRCA | UACC-812 | AZD5363 | MVP_top60_DNA+RNA_seed | MAPK3 | -0.324759 | BRCA | HCC2157 | Ulixertinib |
| MVP_top60_DNA_seed | EGFR | 0.107777 | BRCA | MDA-MB-415 | AZD3759 | MVP_top60_DNA+RNA_seed | MAPK1 | -0.324759 | BRCA | HCC2157 | Ulixertinib |
| MVP_top60_DNA_seed | AKT1 | 0.108611 | BRCA | BT-474 | Uprosertib | MVP_top60_DNA+RNA_seed | AKT1 | -0.322016 | BRCA | HCC1599 | MK-2206 |
| MVP_top60_DNA_seed | HSP90AA1 | 0.108982 | BRCA | EFM-192A | Tanespimycin | MVP_top60_DNA+RNA_seed | PIK3CA | -0.321277 | BRCA | MDA-MB-453 | Buparlisib |
| MVP_top60_DNA_seed | HSP90AB1 | 0.108982 | BRCA | EFM-192A | Tanespimycin | MVP_top60_DNA+RNA_seed | PIK3CA | -0.320262 | BRCA | BT-20 | Buparlisib |
| MVP_top60_DNA_seed | AKT1 | 0.110268 | BRCA | ZR-75-30 | MK-2206 | MVP_top60_DNA+RNA_seed | EGFR | -0.317185 | BRCA | DU-4475 | AZD3759 |
| MVP_top60_DNA_seed | EGFR | 0.110414 | BRCA | BT-20 | Erlotinib | MVP_top60_DNA+RNA_seed | EGFR | -0.316982 | BRCA | UACC-893 | Gefitinib |
| MVP_top60_DNA_seed | EP300 | 0.110681 | BRCA | DU-4475 | CPI-637 | MVP_top60_DNA+RNA_seed | SRC | -0.313141 | BRCA | HCC1395 | Bosutinib |
| MVP_top60_DNA_seed | TP53 | 0.112399 | BRCA | HCC1569 | MIRA-1 | MVP_top60_DNA+RNA_seed | AKT1 | -0.312627 | BRCA | BT-549 | MK-2206 |
| MVP_top60_DNA_seed | HDAC1 | 0.113509 | BRCA | HCC1500 | PCI-34051 | MVP_top60_DNA+RNA_seed | PIK3CA | -0.312548 | BRCA | BT-20 | GNE-317 |
| MVP_top60_DNA_seed | ESR1 | 0.113837 | BRCA | HCC2218 | GDC0810 | MVP_top60_DNA+RNA_seed | EGFR | -0.311841 | BRCA | MDA-MB-415 | Gefitinib |
| MVP_top60_DNA_seed | MAPK3 | 0.115631 | BRCA | COLO-824 | SCH772984 | MVP_top60_DNA+RNA_seed | EGFR | -0.311725 | BRCA | T47D | AZD3759 |

| | | | | | | | | | | | |
|---|---|---|---|---|---|---|---|---|---|---|---|
| MVP_top60_DNA_seed | MAPK1 | 0.115631 | BRCA | COLO-824 | SCH772984 | MVP_top60_DNA+RNA_seed | AKT1 | -0.310426 | BRCA | MDA-MB-175-VII | AT13148 |
| MVP_top60_DNA_seed | EGFR | 0.116354 | BRCA | HCC2218 | Gefitinib | MVP_top60_DNA+RNA_seed | EGFR | -0.309153 | BRCA | EVSA-T | AZD3759 |
| MVP_top60_DNA_seed | ATM | 0.117551 | BRCA | MFM-223 | KU-55933 | MVP_top60_DNA+RNA_seed | ATM | -0.307615 | BRCA | MDA-MB-231 | KU-55933 |
| MVP_top60_DNA_seed | CDK2 | 0.118606 | BRCA | HCC1143 | Dinaciclib | MVP_top60_DNA+RNA_seed | AKT1 | -0.305241 | BRCA | BT-549 | Ipatasertib |
| MVP_top60_DNA_seed | CDK1 | 0.118606 | BRCA | HCC1143 | Dinaciclib | MVP_top60_DNA+RNA_seed | EP300 | -0.301915 | BRCA | MDA-MB-415 | CPI-637 |
| MVP_top60_DNA_seed | SRC | 0.119608 | BRCA | CAL-51 | Bosutinib | MVP_top60_DNA+RNA_seed | AKT1 | -0.295996 | BRCA | DU-4475 | Uprosertib |
| MVP_top60_DNA_seed | MAPK1 | 0.120396 | BRCA | MDA-MB-468 | VX-11e | MVP_top60_DNA+RNA_seed | EGFR | -0.295966 | BRCA | MDA-MB-415 | Osimertinib |
| MVP_top60_DNA_seed | ESR1 | 0.1208 | BRCA | HCC1937 | Tamoxifen | MVP_top60_DNA+RNA_seed | MAPK3 | -0.289547 | BRCA | HCC1395 | Ulixertinib |
| MVP_top60_DNA_seed | HDAC1 | 0.122261 | BRCA | HCC1569 | Entinostat | MVP_top60_DNA+RNA_seed | MAPK1 | -0.289547 | BRCA | HCC1395 | Ulixertinib |
| MVP_top60_DNA_seed | ERBB2 | 0.122275 | BRCA | HCC1143 | Afatinib | MVP_top60_DNA+RNA_seed | AKT1 | -0.286204 | BRCA | HCC2157 | Uprosertib |
| MVP_top60_DNA_seed | EGFR | 0.122275 | BRCA | HCC1143 | Afatinib | MVP_top60_DNA+RNA_seed | HDAC1 | -0.286013 | BRCA | EFM-19 | PCI-34051 |
| MVP_top60_DNA_seed | AKT1 | 0.122831 | BRCA | HCC1419 | Afuresertib | MVP_top60_DNA+RNA_seed | ESR1 | -0.283836 | BRCA | BT-20 | GDC0810 |
| MVP_top60_DNA_seed | ERBB2 | 0.123387 | BRCA | MDA-MB-415 | Sapitinib | MVP_top60_DNA+RNA_seed | AKT1 | -0.280625 | BRCA | HCC1143 | Ipatasertib |
| MVP_top60_DNA_seed | EGFR | 0.123387 | BRCA | MDA-MB-415 | Sapitinib | MVP_top60_DNA+RNA_seed | AKT1 | -0.280149 | BRCA | HCC1187 | GSK2110183B |
| MVP_top60_DNA_seed | ERBB3 | 0.123387 | BRCA | MDA-MB-415 | Sapitinib | MVP_top60_DNA+RNA_seed | AKT1 | -0.278866 | BRCA | HCC1419 | AZD5363 |
| MVP_top60_DNA_seed | CDK1 | 0.123649 | BRCA | MDA-MB-436 | RO-3306 | MVP_top60_DNA+RNA_seed | AKT1 | -0.277893 | BRCA | MRK-nu-1 | AZD5363 |
| MVP_top60_DNA_seed | TP53 | 0.124896 | BRCA | Hs-578-T | MIRA-1 | MVP_top60_DNA+RNA_seed | EGFR | -0.276886 | BRCA | HCC1419 | Erlotinib |
| MVP_top60_DNA_seed | ATM | 0.125494 | BRCA | MDA-MB-453 | KU-55933 | MVP_top60_DNA+RNA_seed | CDK2 | -0.276467 | BRCA | AU565 | AZD5438 |
| MVP_top60_DNA_seed | EP300 | 0.128188 | BRCA | MCF7 | CPI-637 | MVP_top60_DNA+RNA_seed | CDK2 | -0.273945 | BRCA | BT-549 | MK-8776 |
| MVP_top60_DNA_seed | ERBB2 | 0.128273 | BRCA | HCC38 | Sapitinib | MVP_top60_DNA+RNA_seed | EGFR | -0.271284 | BRCA | HCC1937 | Sapitinib |
| MVP_top60_DNA_seed | EGFR | 0.128273 | BRCA | HCC38 | Sapitinib | MVP_top60_DNA+RNA_seed | ERBB2 | -0.271284 | BRCA | HCC1937 | Sapitinib |
| MVP_top60_DNA_seed | ERBB3 | 0.128273 | BRCA | HCC38 | Sapitinib | MVP_top60_DNA+RNA_seed | ERBB3 | -0.271284 | BRCA | HCC1937 | Sapitinib |
| MVP_top60_DNA_seed | TP53 | 0.129468 | BRCA | JIMT-1 | MIRA-1 | MVP_top60_DNA+RNA_seed | HDAC1 | -0.268851 | BRCA | HCC1143 | Entinostat |
| MVP_top60_DNA_seed | HDAC1 | 0.132359 | BRCA | COLO-824 | PCI-34051 | MVP_top60_DNA+RNA_seed | CDK1 | -0.268381 | BRCA | MDA-MB-415 | RO-3306 |
| MVP_top60_DNA_seed | EP300 | 0.132601 | BRCA | MDA-MB-157 | SGC-CBP30 | MVP_top60_DNA+RNA_seed | AKT1 | -0.265106 | BRCA | BT-474 | GSK2110183B |
| MVP_top60_DNA_seed | CREBBP | 0.132601 | BRCA | MDA-MB-157 | SGC-CBP30 | MVP_top60_DNA+RNA_seed | ESR1 | -0.26497 | BRCA | EFM-192A | GDC0810 |
| MVP_top60_DNA_seed | ERBB2 | 0.133351 | BRCA | T47D | Afatinib | MVP_top60_DNA+RNA_seed | CDK2 | -0.26356 | BRCA | EFM-192A | Dinaciclib |
| MVP_top60_DNA_seed | EGFR | 0.133351 | BRCA | T47D | Afatinib | MVP_top60_DNA+RNA_seed | CDK1 | -0.26356 | BRCA | EFM-192A | Dinaciclib |
| MVP_top60_DNA_seed | EGFR | 0.137305 | BRCA | OCUB-M | Erlotinib | MVP_top60_DNA+RNA_seed | PIK3CA | -0.261162 | BRCA | JIMT-1 | Alpelisib |
| MVP_top60_DNA_seed | AR | 0.139365 | BRCA | HCC1569 | Bicalutamide | MVP_top60_DNA+RNA_seed | MAPK3 | -0.259304 | BRCA | HCC1569 | SCH772984 |
| MVP_top60_DNA_seed | ESR1 | 0.139713 | BRCA | MDA-MB-468 | GDC0810 | MVP_top60_DNA+RNA_seed | MAPK1 | -0.259304 | BRCA | HCC1569 | SCH772984 |
| MVP_top60_DNA_seed | EGFR | 0.140037 | BRCA | MDA-MB-436 | AZD3759 | MVP_top60_DNA+RNA_seed | CDK2 | -0.259041 | BRCA | DU-4475 | MK-8776 |
| MVP_top60_DNA_seed | CDK1 | 0.141477 | BRCA | MDA-MB-361 | RO-3306 | MVP_top60_DNA+RNA_seed | SRC | -0.257781 | BRCA | HCC1569 | Bosutinib |
| MVP_top60_DNA_seed | MDM2 | 0.141721 | BRCA | CAL-85-1 | Serdemetan | MVP_top60_DNA+RNA_seed | EGFR | -0.257672 | BRCA | HCC1143 | Gefitinib |
| MVP_top60_DNA_seed | AKT1 | 0.142988 | BRCA | BT-474 | AZD5363 | MVP_top60_DNA+RNA_seed | AKT1 | -0.254692 | BRCA | UACC-812 | AT13148 |
| MVP_top60_DNA_seed | EGFR | 0.145195 | BRCA | HCC1187 | Osimertinib | MVP_top60_DNA+RNA_seed | HDAC1 | -0.25232 | BRCA | DU-4475 | PCI-34051 |
| MVP_top60_DNA_seed | AKT1 | 0.146456 | BRCA | HCC1599 | Ipatasertib | MVP_top60_DNA+RNA_seed | AKT1 | -0.250284 | BRCA | HCC1937 | Ipatasertib |
| MVP_top60_DNA_seed | CDK2 | 0.147343 | BRCA | MDA-MB-415 | AZD5438 | MVP_top60_DNA+RNA_seed | HSP90AA1 | -0.249708 | BRCA | BT-549 | Luminespib |
| MVP_top60_DNA_seed | CDK2 | 0.149829 | BRCA | COLO-824 | AZD5438 | MVP_top60_DNA+RNA_seed | HSP90AB1 | -0.249708 | BRCA | BT-549 | Luminespib |
| MVP_top60_DNA_seed | MAPK1 | 0.150034 | BRCA | CAL-148 | VX-11e | MVP_top60_DNA+RNA_seed | CDK2 | -0.249661 | BRCA | OCUB-M | AZD5438 |
| MVP_top60_DNA_seed | AKT1 | 0.150584 | BRCA | DU-4475 | Afuresertib | MVP_top60_DNA+RNA_seed | AKT1 | -0.249464 | BRCA | BT-549 | Afuresertib |

| | | | | | | | | | | | |
|---|---|---|---|---|---|---|---|---|---|---|---|
| MVP_top60_DNA_seed | CDK2 | 0.151225 | BRCA | EVSA-T | MK-8776 | MVP_top60_DNA+RNA_seed | HDAC1 | -0.246658 | BRCA | Hs-578-T | Entinostat |
| MVP_top60_DNA_seed | HSP90AA1 | 0.152291 | BRCA | AU565 | Luminespib | MVP_top60_DNA+RNA_seed | ESR1 | -0.246657 | BRCA | HCC38 | Tamoxifen |
| MVP_top60_DNA_seed | HSP90AB1 | 0.152291 | BRCA | AU565 | Luminespib | MVP_top60_DNA+RNA_seed | CDK2 | -0.245615 | BRCA | YMB-1-E | MK-8776 |
| MVP_top60_DNA_seed | TP53 | 0.15232 | BRCA | MDA-MB-231 | MIRA-1 | MVP_top60_DNA+RNA_seed | EGFR | -0.244856 | BRCA | HCC1500 | Afatinib |
| MVP_top60_DNA_seed | AKT1 | 0.152356 | BRCA | JIMT-1 | Ipatasertib | MVP_top60_DNA+RNA_seed | ERBB2 | -0.244856 | BRCA | HCC1500 | Afatinib |
| MVP_top60_DNA_seed | MDM2 | 0.15388 | BRCA | HCC1187 | Serdemetan | MVP_top60_DNA+RNA_seed | AKT1 | -0.242575 | BRCA | HCC70 | AT13148 |
| MVP_top60_DNA_seed | MAPK3 | 0.154001 | BRCA | MDA-MB-415 | SCH772984 | MVP_top60_DNA+RNA_seed | HDAC1 | -0.241981 | BRCA | HCC2218 | PCI-34051 |
| MVP_top60_DNA_seed | MAPK1 | 0.154001 | BRCA | MDA-MB-415 | SCH772984 | MVP_top60_DNA+RNA_seed | EGFR | -0.241773 | BRCA | HCC1599 | Sapitinib |
| MVP_top60_DNA_seed | MAPK3 | 0.155134 | BRCA | CAL-51 | SCH772984 | MVP_top60_DNA+RNA_seed | ERBB2 | -0.241773 | BRCA | HCC1599 | Sapitinib |
| MVP_top60_DNA_seed | MAPK1 | 0.155134 | BRCA | CAL-51 | SCH772984 | MVP_top60_DNA+RNA_seed | ERBB3 | -0.241773 | BRCA | HCC1599 | Sapitinib |
| MVP_top60_DNA_seed | ESR1 | 0.156679 | BRCA | CAL-51 | GDC0810 | MVP_top60_DNA+RNA_seed | AKT1 | -0.241393 | BRCA | MDA-MB-468 | AT13148 |
| MVP_top60_DNA_seed | EGFR | 0.156795 | BRCA | HCC1500 | Erlotinib | MVP_top60_DNA+RNA_seed | MAPK3 | -0.240852 | BRCA | EVSA-T | Ulixertinib |
| MVP_top60_DNA_seed | EGFR | 0.157517 | BRCA | BT-474 | Gefitinib | MVP_top60_DNA+RNA_seed | MAPK1 | -0.240852 | BRCA | EVSA-T | Ulixertinib |
| MVP_top60_DNA_seed | AKT1 | 0.157924 | BRCA | HCC2218 | Ipatasertib | MVP_top60_DNA+RNA_seed | AKT1 | -0.238961 | BRCA | BT-474 | Afuresertib |
| MVP_top60_DNA_seed | HSP90AA1 | 0.16055 | BRCA | OCUB-M | Tanespimycin | MVP_top60_DNA+RNA_seed | PIK3CA | -0.238073 | BRCA | CAL-85-1 | Alpelisib |
| MVP_top60_DNA_seed | HSP90AB1 | 0.16055 | BRCA | OCUB-M | Tanespimycin | MVP_top60_DNA+RNA_seed | EP300 | -0.237628 | BRCA | MDA-MB-468 | SGC-CBP30 |
| MVP_top60_DNA_seed | EGFR | 0.16357 | BRCA | CAL-148 | Gefitinib | MVP_top60_DNA+RNA_seed | CDK2 | -0.236229 | BRCA | MDA-MB-361 | AZD5438 |
| MVP_top60_DNA_seed | MAPK3 | 0.163814 | BRCA | OCUB-M | SCH772984 | MVP_top60_DNA+RNA_seed | SRC | -0.235273 | BRCA | MDA-MB-468 | Bosutinib |
| MVP_top60_DNA_seed | MAPK1 | 0.163814 | BRCA | OCUB-M | SCH772984 | MVP_top60_DNA+RNA_seed | ATM | -0.234404 | BRCA | HCC1937 | KU-55933 |
| MVP_top60_DNA_seed | PIK3CA | 0.163942 | BRCA | CAL-51 | AZD8186 | MVP_top60_DNA+RNA_seed | HSP90AA1 | -0.231679 | BRCA | HCC1599 | Elesclomol |
| MVP_top60_DNA_seed | HDAC1 | 0.165217 | BRCA | HCC2157 | PCI-34051 | MVP_top60_DNA+RNA_seed | HSP90AB1 | -0.231679 | BRCA | HCC1599 | Elesclomol |
| MVP_top60_DNA_seed | CDK2 | 0.165858 | BRCA | MDA-MB-453 | MK-8776 | MVP_top60_DNA+RNA_seed | HDAC1 | -0.227067 | BRCA | EFM-19 | Romidepsin |
| MVP_top60_DNA_seed | EGFR | 0.169747 | BRCA | MDA-MB-453 | Erlotinib | MVP_top60_DNA+RNA_seed | HDAC2 | -0.227067 | BRCA | EFM-19 | Romidepsin |
| MVP_top60_DNA_seed | MAPK3 | 0.171978 | BRCA | CAMA-1 | Ulixertinib | MVP_top60_DNA+RNA_seed | MAPK1 | -0.223007 | BRCA | MDA-MB-415 | VX-11e |
| MVP_top60_DNA_seed | MAPK1 | 0.171978 | BRCA | CAMA-1 | Ulixertinib | MVP_top60_DNA+RNA_seed | EGFR | -0.22275 | BRCA | HCC1187 | Erlotinib |
| MVP_top60_DNA_seed | AKT1 | 0.178007 | BRCA | CAL-120 | Afuresertib | MVP_top60_DNA+RNA_seed | EGFR | -0.22114 | BRCA | HCC1937 | Gefitinib |
| MVP_top60_DNA_seed | MDM2 | 0.178802 | BRCA | Hs-578-T | Nutlin-3a (-) | MVP_top60_DNA+RNA_seed | AKT1 | -0.217162 | BRCA | DU-4475 | MK-2206 |
| MVP_top60_DNA_seed | MDM2 | 0.181521 | BRCA | MDA-MB-231 | Serdemetan | MVP_top60_DNA+RNA_seed | HSP90AA1 | -0.214866 | BRCA | YMB-1-E | Elesclomol |
| MVP_top60_DNA_seed | ERBB2 | 0.182481 | BRCA | CAL-51 | Sapitinib | MVP_top60_DNA+RNA_seed | HSP90AB1 | -0.214866 | BRCA | YMB-1-E | Elesclomol |
| MVP_top60_DNA_seed | EGFR | 0.182481 | BRCA | CAL-51 | Sapitinib | MVP_top60_DNA+RNA_seed | PIK3CA | -0.212837 | BRCA | HCC1937 | AZD8186 |
| MVP_top60_DNA_seed | ERBB3 | 0.182481 | BRCA | CAL-51 | Sapitinib | MVP_top60_DNA+RNA_seed | CDK1 | -0.212401 | BRCA | HCC70 | RO-3306 |
| MVP_top60_DNA_seed | MAPK3 | 0.185853 | BRCA | AU565 | SCH772984 | MVP_top60_DNA+RNA_seed | AKT1 | -0.211479 | BRCA | UACC-812 | Afuresertib |
| MVP_top60_DNA_seed | MAPK1 | 0.185853 | BRCA | AU565 | SCH772984 | MVP_top60_DNA+RNA_seed | AKT1 | -0.210983 | BRCA | MDA-MB-468 | AZD5363 |
| MVP_top60_DNA_seed | EGFR | 0.187275 | BRCA | HCC1187 | AZD3759 | MVP_top60_DNA+RNA_seed | PIK3CA | -0.208771 | BRCA | MFM-223 | GNE-317 |
| MVP_top60_DNA_seed | HSP90AA1 | 0.187992 | BRCA | BT-474 | Luminespib | MVP_top60_DNA+RNA_seed | PIK3CA | -0.208619 | BRCA | HCC1569 | GNE-317 |
| MVP_top60_DNA_seed | HSP90AB1 | 0.187992 | BRCA | BT-474 | Luminespib | MVP_top60_DNA+RNA_seed | SRC | -0.206993 | BRCA | OCUB-M | Bosutinib |
| MVP_top60_DNA_seed | PIK3CA | 0.19012 | BRCA | MDA-MB-436 | AZD8186 | MVP_top60_DNA+RNA_seed | AKT1 | -0.206989 | BRCA | MRK-nu-1 | Uprosertib |
| MVP_top60_DNA_seed | MDM2 | 0.192487 | BRCA | HCC1954 | Serdemetan | MVP_top60_DNA+RNA_seed | CDK2 | -0.206906 | BRCA | MDA-MB-453 | Dinaciclib |
| MVP_top60_DNA_seed | CDK1 | 0.192632 | BRCA | YMB-1-E | RO-3306 | MVP_top60_DNA+RNA_seed | CDK1 | -0.206906 | BRCA | MDA-MB-453 | Dinaciclib |
| MVP_top60_DNA_seed | AKT1 | 0.196025 | BRCA | MRK-nu-1 | GSK2110183B | MVP_top60_DNA+RNA_seed | AKT1 | -0.205739 | BRCA | COLO-824 | GSK2110183B |
| MVP_top60_DNA_seed | HDAC1 | 0.197657 | BRCA | HCC1395 | Entinostat | MVP_top60_DNA+RNA_seed | TP53 | -0.205086 | BRCA | HCC1187 | MIRA-1 |

| | | | | | | | | | | | |
|---|---|---|---|---|---|---|---|---|---|---|---|
| MVP_top60_DNA_seed | MAPK3 | 0.197716 | BRCA | HCC1500 | SCH772984 | MVP_top60_DNA+RNA_seed | PLK1 | -0.203687 | BRCA | HCC1569 | MK-1775 |
| MVP_top60_DNA_seed | MAPK1 | 0.197716 | BRCA | HCC1500 | SCH772984 | MVP_top60_DNA+RNA_seed | MDM2 | -0.201917 | BRCA | HCC1419 | Serdemetan |
| MVP_top60_DNA_seed | MAPK1 | 0.198648 | BRCA | CAL-85-1 | VX-11e | MVP_top60_DNA+RNA_seed | ESR1 | -0.201884 | BRCA | HCC1954 | GDC0810 |
| MVP_top60_DNA_seed | MAPK3 | 0.20069 | BRCA | HCC70 | Ulixertinib | MVP_top60_DNA+RNA_seed | HSP90AA1 | -0.201748 | BRCA | HCC1569 | Luminespib |
| MVP_top60_DNA_seed | MAPK1 | 0.20069 | BRCA | HCC70 | Ulixertinib | MVP_top60_DNA+RNA_seed | HSP90AB1 | -0.201748 | BRCA | HCC1569 | Luminespib |
| MVP_top60_DNA_seed | AKT1 | 0.203556 | BRCA | HCC1419 | Uprosertib | MVP_top60_DNA+RNA_seed | HSP90AA1 | -0.201721 | BRCA | HCC1937 | Tanespimycin |
| MVP_top60_DNA_seed | ERBB2 | 0.20404 | BRCA | EFM-19 | Afatinib | MVP_top60_DNA+RNA_seed | HSP90AB1 | -0.201721 | BRCA | HCC1937 | Tanespimycin |
| MVP_top60_DNA_seed | EGFR | 0.20404 | BRCA | EFM-19 | Afatinib | MVP_top60_DNA+RNA_seed | TP53 | -0.201608 | BRCA | HCC1500 | MIRA-1 |
| MVP_top60_DNA_seed | CDK2 | 0.204974 | BRCA | HCC1143 | MK-8776 | MVP_top60_DNA+RNA_seed | AKT1 | -0.197573 | BRCA | MCF7 | Ipatasertib |
| MVP_top60_DNA_seed | ERBB2 | 0.205415 | BRCA | HCC1954 | Afatinib | MVP_top60_DNA+RNA_seed | HDAC1 | -0.196695 | BRCA | UACC-893 | Entinostat |
| MVP_top60_DNA_seed | EGFR | 0.205415 | BRCA | HCC1954 | Afatinib | MVP_top60_DNA+RNA_seed | EGFR | -0.196205 | BRCA | MDA-MB-157 | Sapitinib |
| MVP_top60_DNA_seed | AKT1 | 0.206739 | BRCA | DU-4475 | AZD5363 | MVP_top60_DNA+RNA_seed | ERBB2 | -0.196205 | BRCA | MDA-MB-157 | Sapitinib |
| MVP_top60_DNA_seed | MAPK3 | 0.208441 | BRCA | HCC1419 | Ulixertinib | MVP_top60_DNA+RNA_seed | ERBB3 | -0.196205 | BRCA | MDA-MB-157 | Sapitinib |
| MVP_top60_DNA_seed | MAPK1 | 0.208441 | BRCA | HCC1419 | Ulixertinib | MVP_top60_DNA+RNA_seed | HSP90AA1 | -0.194768 | BRCA | MDA-MB-175-VII | Tanespimycin |
| MVP_top60_DNA_seed | EP300 | 0.210694 | BRCA | CAL-85-1 | SGC-CBP30 | MVP_top60_DNA+RNA_seed | HSP90AB1 | -0.194768 | BRCA | MDA-MB-175-VII | Tanespimycin |
| MVP_top60_DNA_seed | CREBBP | 0.210694 | BRCA | CAL-85-1 | SGC-CBP30 | MVP_top60_DNA+RNA_seed | EGFR | -0.192831 | BRCA | HCC1937 | Osimertinib |
| MVP_top60_DNA_seed | EGFR | 0.211014 | BRCA | HCC70 | Osimertinib | MVP_top60_DNA+RNA_seed | EGFR | -0.191157 | BRCA | MDA-MB-453 | Afatinib |
| MVP_top60_DNA_seed | CDK1 | 0.212461 | BRCA | HCC1143 | RO-3306 | MVP_top60_DNA+RNA_seed | ERBB2 | -0.191157 | BRCA | MDA-MB-453 | Afatinib |
| MVP_top60_DNA_seed | PIK3CA | 0.213926 | BRCA | MDA-MB-157 | Buparlisib | MVP_top60_DNA+RNA_seed | CDK1 | -0.190001 | BRCA | MDA-MB-231 | RO-3306 |
| MVP_top60_DNA_seed | EGFR | 0.215344 | BRCA | MDA-MB-415 | Erlotinib | MVP_top60_DNA+RNA_seed | ESR1 | -0.187918 | BRCA | HCC2157 | GDC0810 |
| MVP_top60_DNA_seed | CDK1 | 0.21569 | BRCA | HCC1599 | RO-3306 | MVP_top60_DNA+RNA_seed | EGFR | -0.187262 | BRCA | T47D | Erlotinib |
| MVP_top60_DNA_seed | ATM | 0.217352 | BRCA | MDA-MB-436 | KU-55933 | MVP_top60_DNA+RNA_seed | EGFR | -0.185884 | BRCA | HCC2218 | Erlotinib |
| MVP_top60_DNA_seed | HDAC1 | 0.218582 | BRCA | T47D | Entinostat | MVP_top60_DNA+RNA_seed | HDAC1 | -0.185616 | BRCA | MDA-MB-231 | Entinostat |
| MVP_top60_DNA_seed | TP53 | 0.221206 | BRCA | UACC-812 | MIRA-1 | MVP_top60_DNA+RNA_seed | EGFR | -0.185169 | BRCA | T47D | Sapitinib |
| MVP_top60_DNA_seed | MDM2 | 0.22449 | BRCA | MDA-MB-468 | Nutlin-3a (-) | MVP_top60_DNA+RNA_seed | ERBB2 | -0.185169 | BRCA | T47D | Sapitinib |
| MVP_top60_DNA_seed | EGFR | 0.226585 | BRCA | CAL-148 | AZD3759 | MVP_top60_DNA+RNA_seed | ERBB3 | -0.185169 | BRCA | T47D | Sapitinib |
| MVP_top60_DNA_seed | PIK3CA | 0.227552 | BRCA | CAL-85-1 | Buparlisib | MVP_top60_DNA+RNA_seed | ESR1 | -0.184272 | BRCA | EFM-192A | Tamoxifen |
| MVP_top60_DNA_seed | MDM2 | 0.230602 | BRCA | EVSA-T | Nutlin-3a (-) | MVP_top60_DNA+RNA_seed | AKT1 | -0.183703 | BRCA | HCC1937 | Uprosertib |
| MVP_top60_DNA_seed | MDM2 | 0.231145 | BRCA | CAMA-1 | Nutlin-3a (-) | MVP_top60_DNA+RNA_seed | HDAC1 | -0.178024 | BRCA | BT-549 | Entinostat |
| MVP_top60_DNA_seed | ERBB2 | 0.23241 | BRCA | HCC1395 | Afatinib | MVP_top60_DNA+RNA_seed | PIK3CA | -0.175846 | BRCA | UACC-812 | GNE-317 |
| MVP_top60_DNA_seed | EGFR | 0.23241 | BRCA | HCC1395 | Afatinib | MVP_top60_DNA+RNA_seed | MAPK3 | -0.175576 | BRCA | Hs-578-T | SCH772984 |
| MVP_top60_DNA_seed | AKT1 | 0.232639 | BRCA | BT-549 | Uprosertib | MVP_top60_DNA+RNA_seed | MAPK1 | -0.175576 | BRCA | Hs-578-T | SCH772984 |
| MVP_top60_DNA_seed | MDM2 | 0.234278 | BRCA | EFM-192A | Serdemetan | MVP_top60_DNA+RNA_seed | PIK3CA | -0.175299 | BRCA | UACC-893 | AZD8186 |
| MVP_top60_DNA_seed | TP53 | 0.238802 | BRCA | HCC1954 | MIRA-1 | MVP_top60_DNA+RNA_seed | HDAC1 | -0.173626 | BRCA | HDQ-P1 | Romidepsin |
| MVP_top60_DNA_seed | CDK2 | 0.239184 | BRCA | CAL-148 | MK-8776 | MVP_top60_DNA+RNA_seed | HDAC2 | -0.173626 | BRCA | HDQ-P1 | Romidepsin |
| MVP_top60_DNA_seed | HDAC1 | 0.240572 | BRCA | YMB-1-E | Entinostat | MVP_top60_DNA+RNA_seed | EGFR | -0.173574 | BRCA | HCC1569 | Gefitinib |
| MVP_top60_DNA_seed | EGFR | 0.241128 | BRCA | HCC1599 | Osimertinib | MVP_top60_DNA+RNA_seed | CDK1 | -0.171546 | BRCA | CAMA-1 | RO-3306 |
| MVP_top60_DNA_seed | EP300 | 0.2459 | BRCA | AU565 | CPI-637 | MVP_top60_DNA+RNA_seed | PIK3CA | -0.168908 | BRCA | MDA-MB-453 | AZD8186 |
| MVP_top60_DNA_seed | TP53 | 0.246462 | BRCA | MDA-MB-361 | MIRA-1 | MVP_top60_DNA+RNA_seed | AKT1 | -0.16772 | BRCA | DU-4475 | AT13148 |
| MVP_top60_DNA_seed | MDM2 | 0.249105 | BRCA | HCC1419 | Nutlin-3a (-) | MVP_top60_DNA+RNA_seed | TP53 | -0.167296 | BRCA | BT-20 | MIRA-1 |
| MVP_top60_DNA_seed | SRC | 0.251062 | BRCA | MDA-MB-415 | Bosutinib | MVP_top60_DNA+RNA_seed | TP53 | -0.164861 | BRCA | MDA-MB-453 | MIRA-1 |

| | | | | | | | | | | | |
|---|---|---|---|---|---|---|---|---|---|---|---|
| MVP_top60_DNA_seed | MDM2 | 0.254507 | BRCA | HCC1395 | Nutlin-3a (-) | MVP_top60_DNA+RNA_seed | EGFR | -0.163452 | BRCA | CAL-148 | Erlotinib |
| MVP_top60_DNA_seed | EGFR | 0.254665 | BRCA | CAMA-1 | AZD3759 | MVP_top60_DNA+RNA_seed | ESR1 | -0.161042 | BRCA | CAL-51 | Tamoxifen |
| MVP_top60_DNA_seed | AKT1 | 0.25478 | BRCA | MCF7 | Afuresertib | MVP_top60_DNA+RNA_seed | HDAC1 | -0.159195 | BRCA | OCUB-M | Romidepsin |
| MVP_top60_DNA_seed | MAPK3 | 0.255539 | BRCA | MDA-MB-468 | SCH772984 | MVP_top60_DNA+RNA_seed | HDAC2 | -0.159195 | BRCA | OCUB-M | Romidepsin |
| MVP_top60_DNA_seed | MAPK1 | 0.255539 | BRCA | MDA-MB-468 | SCH772984 | MVP_top60_DNA+RNA_seed | EP300 | -0.158123 | BRCA | BT-474 | CPI-637 |
| MVP_top60_DNA_seed | ERBB2 | 0.25621 | BRCA | HCC1395 | Sapitinib | MVP_top60_DNA+RNA_seed | MDM2 | -0.15666 | BRCA | MDA-MB-175-VII | Nutlin-3a (-) |
| MVP_top60_DNA_seed | EGFR | 0.25621 | BRCA | HCC1395 | Sapitinib | MVP_top60_DNA+RNA_seed | EGFR | -0.152889 | BRCA | HCC1569 | AZD3759 |
| MVP_top60_DNA_seed | ERBB3 | 0.25621 | BRCA | HCC1395 | Sapitinib | MVP_top60_DNA+RNA_seed | MAPK1 | -0.150985 | BRCA | CAL-51 | VX-11e |
| MVP_top60_DNA_seed | EP300 | 0.256755 | BRCA | EVSA-T | CPI-637 | MVP_top60_DNA+RNA_seed | EGFR | -0.148272 | BRCA | EVSA-T | Sapitinib |
| MVP_top60_DNA_seed | MAPK3 | 0.258694 | BRCA | CAL-120 | Ulixertinib | MVP_top60_DNA+RNA_seed | ERBB2 | -0.148272 | BRCA | EVSA-T | Sapitinib |
| MVP_top60_DNA_seed | MAPK1 | 0.258694 | BRCA | CAL-120 | Ulixertinib | MVP_top60_DNA+RNA_seed | ERBB3 | -0.148272 | BRCA | EVSA-T | Sapitinib |
| MVP_top60_DNA_seed | EGFR | 0.259729 | BRCA | YMB-1-E | AZD3759 | MVP_top60_DNA+RNA_seed | PIK3CA | -0.147878 | BRCA | MDA-MB-468 | GNE-317 |
| MVP_top60_DNA_seed | CDK1 | 0.260846 | BRCA | MDA-MB-175-VII | RO-3306 | MVP_top60_DNA+RNA_seed | EGFR | -0.147212 | BRCA | T47D | Lapatinib |
| MVP_top60_DNA_seed | EGFR | 0.262971 | BRCA | BT-20 | Gefitinib | MVP_top60_DNA+RNA_seed | ERBB2 | -0.147212 | BRCA | T47D | Lapatinib |
| MVP_top60_DNA_seed | EP300 | 0.263757 | BRCA | MDA-MB-361 | SGC-CBP30 | MVP_top60_DNA+RNA_seed | PIK3CA | -0.142764 | BRCA | CAL-51 | Alpelisib |
| MVP_top60_DNA_seed | CREBBP | 0.263757 | BRCA | MDA-MB-361 | SGC-CBP30 | MVP_top60_DNA+RNA_seed | EGFR | -0.142341 | BRCA | HCC70 | Lapatinib |
| MVP_top60_DNA_seed | HSP90AA1 | 0.264808 | BRCA | HCC1569 | Elesclomol | MVP_top60_DNA+RNA_seed | ERBB2 | -0.142341 | BRCA | HCC70 | Lapatinib |
| MVP_top60_DNA_seed | HSP90AB1 | 0.264808 | BRCA | HCC1569 | Elesclomol | MVP_top60_DNA+RNA_seed | AKT1 | -0.14134 | BRCA | CAL-51 | Uprosertib |
| MVP_top60_DNA_seed | CDK2 | 0.265977 | BRCA | JIMT-1 | AZD5438 | MVP_top60_DNA+RNA_seed | MAPK1 | -0.140194 | BRCA | COLO-824 | VX-11e |
| MVP_top60_DNA_seed | HDAC1 | 0.266299 | BRCA | T47D | Romidepsin | MVP_top60_DNA+RNA_seed | EGFR | -0.140165 | BRCA | DU-4475 | Sapitinib |
| MVP_top60_DNA_seed | HDAC2 | 0.266299 | BRCA | T47D | Romidepsin | MVP_top60_DNA+RNA_seed | ERBB2 | -0.140165 | BRCA | DU-4475 | Sapitinib |
| MVP_top60_DNA_seed | CDK2 | 0.266426 | BRCA | OCUB-M | MK-8776 | MVP_top60_DNA+RNA_seed | ERBB3 | -0.140165 | BRCA | DU-4475 | Sapitinib |
| MVP_top60_DNA_seed | EP300 | 0.266513 | BRCA | MDA-MB-330 | SGC-CBP30 | MVP_top60_DNA+RNA_seed | MDM2 | -0.139946 | BRCA | HCC70 | Serdemetan |
| MVP_top60_DNA_seed | CREBBP | 0.266513 | BRCA | MDA-MB-330 | SGC-CBP30 | MVP_top60_DNA+RNA_seed | AKT1 | -0.139743 | BRCA | HCC1419 | MK-2206 |
| MVP_top60_DNA_seed | EGFR | 0.26705 | BRCA | CAL-51 | AZD3759 | MVP_top60_DNA+RNA_seed | AKT1 | -0.138311 | BRCA | DU-4475 | Ipatasertib |
| MVP_top60_DNA_seed | HDAC1 | 0.268117 | BRCA | HCC70 | Romidepsin | MVP_top60_DNA+RNA_seed | MAPK1 | -0.135935 | BRCA | HCC1954 | VX-11e |
| MVP_top60_DNA_seed | HDAC2 | 0.268117 | BRCA | HCC70 | Romidepsin | MVP_top60_DNA+RNA_seed | ATM | -0.13526 | BRCA | EFM-19 | KU-55933 |
| MVP_top60_DNA_seed | PIK3CA | 0.268441 | BRCA | YMB-1-E | Alpelisib | MVP_top60_DNA+RNA_seed | EP300 | -0.134227 | BRCA | MRK-nu-1 | SGC-CBP30 |
| MVP_top60_DNA_seed | MAPK3 | 0.270589 | BRCA | AU565 | Ulixertinib | MVP_top60_DNA+RNA_seed | HSP90AA1 | -0.134212 | BRCA | MDA-MB-415 | Tanespimycin |
| MVP_top60_DNA_seed | MAPK1 | 0.270589 | BRCA | AU565 | Ulixertinib | MVP_top60_DNA+RNA_seed | HSP90AB1 | -0.134212 | BRCA | MDA-MB-415 | Tanespimycin |
| MVP_top60_DNA_seed | PIK3CA | 0.273017 | BRCA | BT-483 | Alpelisib | MVP_top60_DNA+RNA_seed | AKT1 | -0.133466 | BRCA | HCC1599 | AZD5363 |
| MVP_top60_DNA_seed | HDAC1 | 0.273117 | BRCA | CAL-51 | Romidepsin | MVP_top60_DNA+RNA_seed | PIK3CA | -0.132617 | BRCA | JIMT-1 | Buparlisib |
| MVP_top60_DNA_seed | HDAC2 | 0.273117 | BRCA | CAL-51 | Romidepsin | MVP_top60_DNA+RNA_seed | ATM | -0.132384 | BRCA | HCC1143 | KU-55933 |
| MVP_top60_DNA_seed | EGFR | 0.274221 | BRCA | HCC1954 | Gefitinib | MVP_top60_DNA+RNA_seed | EP300 | -0.132295 | BRCA | T47D | CPI-637 |
| MVP_top60_DNA_seed | PIK3CA | 0.274275 | BRCA | CAL-120 | Buparlisib | MVP_top60_DNA+RNA_seed | PIK3CA | -0.13028 | BRCA | YMB-1-E | Buparlisib |
| MVP_top60_DNA_seed | MAPK3 | 0.277716 | BRCA | MDA-MB-415 | Ulixertinib | MVP_top60_DNA+RNA_seed | PLK1 | -0.129855 | BRCA | BT-549 | MK-1775 |
| MVP_top60_DNA_seed | MAPK1 | 0.277716 | BRCA | MDA-MB-415 | Ulixertinib | MVP_top60_DNA+RNA_seed | TP53 | -0.128637 | BRCA | HCC2218 | MIRA-1 |
| MVP_top60_DNA_seed | MAPK1 | 0.279317 | BRCA | HCC1395 | VX-11e | MVP_top60_DNA+RNA_seed | EGFR | -0.127189 | BRCA | CAL-85-1 | Lapatinib |
| MVP_top60_DNA_seed | ATM | 0.279826 | BRCA | MCF7 | KU-55933 | MVP_top60_DNA+RNA_seed | ERBB2 | -0.127189 | BRCA | CAL-85-1 | Lapatinib |
| MVP_top60_DNA_seed | PIK3CA | 0.280119 | BRCA | DU-4475 | Alpelisib | MVP_top60_DNA+RNA_seed | ESR1 | -0.127146 | BRCA | JIMT-1 | Tamoxifen |
| MVP_top60_DNA_seed | ESR1 | 0.281353 | BRCA | JIMT-1 | GDC0810 | MVP_top60_DNA+RNA_seed | HDAC1 | -0.126853 | BRCA | MDA-MB-361 | Romidepsin |

| | | | | | | | | | | | |
|---|---|---|---|---|---|---|---|---|---|---|---|
| MVP_top60_DNA_seed | MAPK3 | 0.282012 | BRCA | MDA-MB-361 | Ulixertinib | MVP_top60_DNA+RNA_seed | HDAC2 | -0.126853 | BRCA | MDA-MB-361 | Romidepsin |
| MVP_top60_DNA_seed | MAPK1 | 0.282012 | BRCA | MDA-MB-361 | Ulixertinib | MVP_top60_DNA+RNA_seed | ATM | -0.126464 | BRCA | HCC38 | KU-55933 |
| MVP_top60_DNA_seed | MAPK3 | 0.282838 | BRCA | MFM-223 | Ulixertinib | MVP_top60_DNA+RNA_seed | AKT1 | -0.126359 | BRCA | MDA-MB-468 | GSK2110183B |
| MVP_top60_DNA_seed | MAPK1 | 0.282838 | BRCA | MFM-223 | Ulixertinib | MVP_top60_DNA+RNA_seed | AKT1 | -0.126093 | BRCA | EFM-192A | Uprosertib |
| MVP_top60_DNA_seed | EGFR | 0.283823 | BRCA | JIMT-1 | Erlotinib | MVP_top60_DNA+RNA_seed | AKT1 | -0.124824 | BRCA | EFM-192A | Afuresertib |
| MVP_top60_DNA_seed | PIK3CA | 0.286683 | BRCA | BT-474 | AZD8186 | MVP_top60_DNA+RNA_seed | HSP90AA1 | -0.124607 | BRCA | MDA-MB-415 | Luminespib |
| MVP_top60_DNA_seed | EGFR | 0.288612 | BRCA | YMB-1-E | Osimertinib | MVP_top60_DNA+RNA_seed | HSP90AB1 | -0.124607 | BRCA | MDA-MB-415 | Luminespib |
| MVP_top60_DNA_seed | MAPK3 | 0.289274 | BRCA | HCC2218 | Ulixertinib | MVP_top60_DNA+RNA_seed | ATM | -0.123973 | BRCA | HCC2218 | KU-55933 |
| MVP_top60_DNA_seed | MAPK1 | 0.289274 | BRCA | HCC2218 | Ulixertinib | MVP_top60_DNA+RNA_seed | EP300 | -0.123753 | BRCA | HCC2157 | SGC-CBP30 |
| MVP_top60_DNA_seed | ATM | 0.291028 | BRCA | HCC1500 | KU-55933 | MVP_top60_DNA+RNA_seed | PIK3CA | -0.122867 | BRCA | HCC1187 | AZD8186 |
| MVP_top60_DNA_seed | EP300 | 0.292118 | BRCA | MDA-MB-231 | SGC-CBP30 | MVP_top60_DNA+RNA_seed | EGFR | -0.117539 | BRCA | CAL-51 | Erlotinib |
| MVP_top60_DNA_seed | CREBBP | 0.292118 | BRCA | MDA-MB-231 | SGC-CBP30 | MVP_top60_DNA+RNA_seed | EGFR | -0.113196 | BRCA | HCC38 | Lapatinib |
| MVP_top60_DNA_seed | TP53 | 0.294139 | BRCA | MDA-MB-157 | MIRA-1 | MVP_top60_DNA+RNA_seed | ERBB2 | -0.113196 | BRCA | HCC38 | Lapatinib |
| MVP_top60_DNA_seed | ESR1 | 0.295298 | BRCA | HDQ-P1 | GDC0810 | MVP_top60_DNA+RNA_seed | CDK2 | -0.111853 | BRCA | EVSA-T | AZD5438 |
| MVP_top60_DNA_seed | MDM2 | 0.295542 | BRCA | MDA-MB-157 | Nutlin-3a (-) | MVP_top60_DNA+RNA_seed | CDK1 | -0.111581 | BRCA | JIMT-1 | RO-3306 |
| MVP_top60_DNA_seed | HSP90AA1 | 0.296061 | BRCA | BT-549 | Tanespimycin | MVP_top60_DNA+RNA_seed | HSP90AA1 | -0.110684 | BRCA | HCC1954 | Tanespimycin |
| MVP_top60_DNA_seed | HSP90AB1 | 0.296061 | BRCA | BT-549 | Tanespimycin | MVP_top60_DNA+RNA_seed | HSP90AB1 | -0.110684 | BRCA | HCC1954 | Tanespimycin |
| MVP_top60_DNA_seed | AKT1 | 0.297367 | BRCA | CAL-120 | AZD5363 | MVP_top60_DNA+RNA_seed | HSP90AA1 | -0.109998 | BRCA | HCC1599 | Luminespib |
| MVP_top60_DNA_seed | ESR1 | 0.297503 | BRCA | BT-20 | Tamoxifen | MVP_top60_DNA+RNA_seed | HSP90AB1 | -0.109998 | BRCA | HCC1599 | Luminespib |
| MVP_top60_DNA_seed | ERBB2 | 0.298848 | BRCA | HCC1428 | Lapatinib | MVP_top60_DNA+RNA_seed | PIK3CA | -0.105804 | BRCA | MDA-MB-453 | GNE-317 |
| MVP_top60_DNA_seed | EGFR | 0.298848 | BRCA | HCC1428 | Lapatinib | MVP_top60_DNA+RNA_seed | CDK1 | -0.104339 | BRCA | UACC-893 | RO-3306 |
| MVP_top60_DNA_seed | HDAC1 | 0.299565 | BRCA | MDA-MB-415 | Romidepsin | MVP_top60_DNA+RNA_seed | HSP90AA1 | -0.104295 | BRCA | MCF7 | Tanespimycin |
| MVP_top60_DNA_seed | HDAC2 | 0.299565 | BRCA | MDA-MB-415 | Romidepsin | MVP_top60_DNA+RNA_seed | HSP90AB1 | -0.104295 | BRCA | MCF7 | Tanespimycin |
| MVP_top60_DNA_seed | ERBB2 | 0.300245 | BRCA | BT-549 | Afatinib | MVP_top60_DNA+RNA_seed | AKT1 | -0.10414 | BRCA | EFM-192A | MK-2206 |
| MVP_top60_DNA_seed | EGFR | 0.300245 | BRCA | BT-549 | Afatinib | MVP_top60_DNA+RNA_seed | SRC | -0.100852 | BRCA | MDA-MB-231 | Bosutinib |
| MVP_top60_DNA_seed | ESR1 | 0.303056 | BRCA | MFM-223 | Tamoxifen | MVP_top60_DNA+RNA_seed | CDK1 | -0.10033 | BRCA | HDQ-P1 | RO-3306 |
| MVP_top60_DNA_seed | EP300 | 0.30463 | BRCA | HCC70 | CPI-637 | MVP_top60_DNA+RNA_seed | AKT1 | -0.097581 | BRCA | BT-549 | AZD5363 |
| MVP_top60_DNA_seed | ERBB2 | 0.307912 | BRCA | COLO-824 | Sapitinib | MVP_top60_DNA+RNA_seed | PIK3CA | -0.096342 | BRCA | HCC2218 | Alpelisib |
| MVP_top60_DNA_seed | EGFR | 0.307912 | BRCA | COLO-824 | Sapitinib | MVP_top60_DNA+RNA_seed | EGFR | -0.094247 | BRCA | Hs-578-T | Afatinib |
| MVP_top60_DNA_seed | ERBB3 | 0.307912 | BRCA | COLO-824 | Sapitinib | MVP_top60_DNA+RNA_seed | ERBB2 | -0.094247 | BRCA | Hs-578-T | Afatinib |
| MVP_top60_DNA_seed | MAPK3 | 0.308046 | BRCA | CAL-85-1 | Ulixertinib | MVP_top60_DNA+RNA_seed | SRC | -0.093435 | BRCA | EVSA-T | Bosutinib |
| MVP_top60_DNA_seed | MAPK1 | 0.308046 | BRCA | CAL-85-1 | Ulixertinib | MVP_top60_DNA+RNA_seed | PIK3CA | -0.09236 | BRCA | HCC70 | Alpelisib |
| MVP_top60_DNA_seed | HDAC1 | 0.308293 | BRCA | Hs-578-T | PCI-34051 | MVP_top60_DNA+RNA_seed | CDK1 | -0.090985 | BRCA | MCF7 | RO-3306 |
| MVP_top60_DNA_seed | TP53 | 0.311105 | BRCA | UACC-893 | MIRA-1 | MVP_top60_DNA+RNA_seed | MDM2 | -0.089748 | BRCA | BT-20 | Serdemetan |
| MVP_top60_DNA_seed | CDK2 | 0.311469 | BRCA | COLO-824 | Dinaciclib | MVP_top60_DNA+RNA_seed | EGFR | -0.087398 | BRCA | HCC70 | Erlotinib |
| MVP_top60_DNA_seed | CDK1 | 0.311469 | BRCA | COLO-824 | Dinaciclib | MVP_top60_DNA+RNA_seed | PIK3CA | -0.086838 | BRCA | DU-4475 | GNE-317 |
| MVP_top60_DNA_seed | ERBB2 | 0.312327 | BRCA | MDA-MB-231 | Sapitinib | MVP_top60_DNA+RNA_seed | PLK1 | -0.086005 | BRCA | BT-549 | BI-2536 |
| MVP_top60_DNA_seed | EGFR | 0.312327 | BRCA | MDA-MB-231 | Sapitinib | MVP_top60_DNA+RNA_seed | EGFR | -0.085134 | BRCA | BT-20 | AZD3759 |
| MVP_top60_DNA_seed | ERBB3 | 0.312327 | BRCA | MDA-MB-231 | Sapitinib | MVP_top60_DNA+RNA_seed | MAPK3 | -0.082166 | BRCA | BT-20 | SCH772984 |
| MVP_top60_DNA_seed | CDK1 | 0.313471 | BRCA | HCC1954 | RO-3306 | MVP_top60_DNA+RNA_seed | MAPK1 | -0.082166 | BRCA | BT-20 | SCH772984 |
| MVP_top60_DNA_seed | MAPK3 | 0.315106 | BRCA | CAL-148 | Ulixertinib | MVP_top60_DNA+RNA_seed | EGFR | -0.081593 | BRCA | MDA-MB-361 | Gefitinib |

| | | | | | | | | | | | |
|---|---|---|---|---|---|---|---|---|---|---|---|
| MVP_top60_DNA_seed | MAPK1 | 0.315106 | BRCA | CAL-148 | Ulixertinib | MVP_top60_DNA+RNA_seed | MAPK3 | -0.080819 | BRCA | HDQ-P1 | SCH772984 |
| MVP_top60_DNA_seed | PIK3CA | 0.315274 | BRCA | DU-4475 | AZD8186 | MVP_top60_DNA+RNA_seed | MAPK1 | -0.080819 | BRCA | HDQ-P1 | SCH772984 |
| MVP_top60_DNA_seed | SRC | 0.315316 | BRCA | HCC1937 | Bosutinib | MVP_top60_DNA+RNA_seed | HSP90AA1 | -0.079782 | BRCA | MCF7 | Luminespib |
| MVP_top60_DNA_seed | SRC | 0.315473 | BRCA | T47D | Bosutinib | MVP_top60_DNA+RNA_seed | HSP90AB1 | -0.079782 | BRCA | MCF7 | Luminespib |
| MVP_top60_DNA_seed | EGFR | 0.316304 | BRCA | HCC1187 | Gefitinib | MVP_top60_DNA+RNA_seed | PLK1 | -0.078954 | BRCA | HCC38 | MK-1775 |
| MVP_top60_DNA_seed | ESR1 | 0.318846 | BRCA | CAL-148 | Tamoxifen | MVP_top60_DNA+RNA_seed | HDAC1 | -0.078202 | BRCA | CAL-51 | Entinostat |
| MVP_top60_DNA_seed | PIK3CA | 0.319258 | BRCA | HCC1395 | GNE-317 | MVP_top60_DNA+RNA_seed | PIK3CA | -0.077314 | BRCA | HCC2157 | Alpelisib |
| MVP_top60_DNA_seed | EGFR | 0.319356 | BRCA | HCC1395 | Erlotinib | MVP_top60_DNA+RNA_seed | HSP90AA1 | -0.074845 | BRCA | HCC1419 | Luminespib |
| MVP_top60_DNA_seed | ERBB2 | 0.320019 | BRCA | MRK-nu-1 | Sapitinib | MVP_top60_DNA+RNA_seed | HSP90AB1 | -0.074845 | BRCA | HCC1419 | Luminespib |
| MVP_top60_DNA_seed | EGFR | 0.320019 | BRCA | MRK-nu-1 | Sapitinib | MVP_top60_DNA+RNA_seed | ESR1 | -0.071202 | BRCA | HCC1937 | GDC0810 |
| MVP_top60_DNA_seed | ERBB3 | 0.320019 | BRCA | MRK-nu-1 | Sapitinib | MVP_top60_DNA+RNA_seed | AKT1 | -0.071095 | BRCA | MCF7 | MK-2206 |
| MVP_top60_DNA_seed | AKT1 | 0.320294 | BRCA | MDA-MB-231 | Uprosertib | MVP_top60_DNA+RNA_seed | HSP90AA1 | -0.066654 | BRCA | AU565 | Tanespimycin |
| MVP_top60_DNA_seed | ERBB2 | 0.320661 | BRCA | EVSA-T | Afatinib | MVP_top60_DNA+RNA_seed | HSP90AB1 | -0.066654 | BRCA | AU565 | Tanespimycin |
| MVP_top60_DNA_seed | EGFR | 0.320661 | BRCA | EVSA-T | Afatinib | MVP_top60_DNA+RNA_seed | PIK3CA | -0.064745 | BRCA | MDA-MB-415 | Buparlisib |
| MVP_top60_DNA_seed | EP300 | 0.321388 | BRCA | MDA-MB-415 | SGC-CBP30 | MVP_top60_DNA+RNA_seed | AKT1 | -0.063491 | BRCA | EFM-19 | AT13148 |
| MVP_top60_DNA_seed | CREBBP | 0.321388 | BRCA | MDA-MB-415 | SGC-CBP30 | MVP_top60_DNA+RNA_seed | EGFR | -0.062007 | BRCA | YMB-1-E | Lapatinib |
| MVP_top60_DNA_seed | CDK2 | 0.32236 | BRCA | HCC1569 | Dinaciclib | MVP_top60_DNA+RNA_seed | ERBB2 | -0.062007 | BRCA | YMB-1-E | Lapatinib |
| MVP_top60_DNA_seed | CDK1 | 0.32236 | BRCA | HCC1569 | Dinaciclib | MVP_top60_DNA+RNA_seed | PLK1 | -0.06018 | BRCA | JIMT-1 | MK-1775 |
| MVP_top60_DNA_seed | EGFR | 0.323621 | BRCA | CAMA-1 | Gefitinib | MVP_top60_DNA+RNA_seed | ATM | -0.059547 | BRCA | MDA-MB-361 | KU-55933 |
| MVP_top60_DNA_seed | MDM2 | 0.325432 | BRCA | HCC2218 | Nutlin-3a (-) | MVP_top60_DNA+RNA_seed | HSP90AA1 | -0.057207 | BRCA | HCC38 | Luminespib |
| MVP_top60_DNA_seed | EP300 | 0.327685 | BRCA | HCC1428 | CPI-637 | MVP_top60_DNA+RNA_seed | HSP90AB1 | -0.057207 | BRCA | HCC38 | Luminespib |
| MVP_top60_DNA_seed | ESR1 | 0.328086 | BRCA | Hs-578-T | GDC0810 | MVP_top60_DNA+RNA_seed | EGFR | -0.053825 | BRCA | HCC1569 | Afatinib |
| MVP_top60_DNA_seed | MAPK3 | 0.329549 | BRCA | EFM-192A | Ulixertinib | MVP_top60_DNA+RNA_seed | ERBB2 | -0.053825 | BRCA | HCC1569 | Afatinib |
| MVP_top60_DNA_seed | MAPK1 | 0.329549 | BRCA | EFM-192A | Ulixertinib | MVP_top60_DNA+RNA_seed | MAPK3 | -0.053441 | BRCA | HCC1569 | Ulixertinib |
| MVP_top60_DNA_seed | AKT1 | 0.330373 | BRCA | CAL-85-1 | Afuresertib | MVP_top60_DNA+RNA_seed | MAPK1 | -0.053441 | BRCA | HCC1569 | Ulixertinib |
| MVP_top60_DNA_seed | HDAC1 | 0.332186 | BRCA | MRK-nu-1 | PCI-34051 | MVP_top60_DNA+RNA_seed | MAPK3 | -0.053414 | BRCA | HCC1419 | Ulixertinib |
| MVP_top60_DNA_seed | MAPK1 | 0.332632 | BRCA | MFM-223 | VX-11e | MVP_top60_DNA+RNA_seed | MAPK1 | -0.053414 | BRCA | HCC1419 | Ulixertinib |
| MVP_top60_DNA_seed | ESR1 | 0.333839 | BRCA | MDA-MB-231 | Tamoxifen | MVP_top60_DNA+RNA_seed | AKT1 | -0.052553 | BRCA | UACC-812 | Ipatasertib |
| MVP_top60_DNA_seed | MAPK1 | 0.337024 | BRCA | CAMA-1 | VX-11e | MVP_top60_DNA+RNA_seed | EGFR | -0.052387 | BRCA | MRK-nu-1 | Erlotinib |
| MVP_top60_DNA_seed | EP300 | 0.337381 | BRCA | ZR-75-30 | CPI-637 | MVP_top60_DNA+RNA_seed | AKT1 | -0.049565 | BRCA | HCC1937 | Uprosertib |
| MVP_top60_DNA_seed | MAPK3 | 0.338626 | BRCA | HCC70 | Ulixertinib | MVP_top60_DNA+RNA_seed | AKT1 | -0.048284 | BRCA | EFM-192A | AT13148 |
| MVP_top60_DNA_seed | MAPK1 | 0.338626 | BRCA | HCC70 | Ulixertinib | MVP_top60_DNA+RNA_seed | MDM2 | -0.047509 | BRCA | T47D | Nutlin-3a (-) |
| MVP_top60_DNA_seed | ERBB2 | 0.3401 | BRCA | MFM-223 | Lapatinib | MVP_top60_DNA+RNA_seed | AKT1 | -0.046702 | BRCA | HDQ-P1 | Uprosertib |
| MVP_top60_DNA_seed | EGFR | 0.3401 | BRCA | MFM-223 | Lapatinib | MVP_top60_DNA+RNA_seed | MAPK3 | -0.044559 | BRCA | HCC1143 | SCH772984 |
| MVP_top60_DNA_seed | PIK3CA | 0.342549 | BRCA | HCC1599 | GNE-317 | MVP_top60_DNA+RNA_seed | MAPK1 | -0.044559 | BRCA | HCC1143 | SCH772984 |
| MVP_top60_DNA_seed | CDK2 | 0.343294 | BRCA | HCC1419 | AZD5438 | MVP_top60_DNA+RNA_seed | AKT1 | -0.042614 | BRCA | HCC38 | AZD5363 |
| MVP_top60_DNA_seed | HSP90AA1 | 0.346003 | BRCA | EFM-192A | Luminespib | MVP_top60_DNA+RNA_seed | CDK2 | -0.040349 | BRCA | CAL-51 | MK-8776 |
| MVP_top60_DNA_seed | HSP90AB1 | 0.346003 | BRCA | EFM-192A | Luminespib | MVP_top60_DNA+RNA_seed | ATM | -0.039755 | BRCA | HCC1187 | KU-55933 |
| MVP_top60_DNA_seed | ATM | 0.346457 | BRCA | CAL-51 | KU-55933 | MVP_top60_DNA+RNA_seed | AKT1 | -0.038662 | BRCA | MDA-MB-361 | AT13148 |
| MVP_top60_DNA_seed | MAPK3 | 0.346819 | BRCA | COLO-824 | Ulixertinib | MVP_top60_DNA+RNA_seed | CDK1 | -0.038444 | BRCA | HCC1569 | RO-3306 |
| MVP_top60_DNA_seed | MAPK1 | 0.346819 | BRCA | COLO-824 | Ulixertinib | MVP_top60_DNA+RNA_seed | PLK1 | -0.037378 | BRCA | MDA-MB-453 | BI-2536 |

| | | | | | | | | | | | |
|---|---|---|---|---|---|---|---|---|---|---|---|
| MVP_top60_DNA_seed | HSP90AA1 | 0.348532 | BRCA | BT-474 | Tanespimycin | MVP_top60_DNA+RNA_seed | EGFR | -0.03659 | BRCA | HCC70 | AZD3759 |
| MVP_top60_DNA_seed | HSP90AB1 | 0.348532 | BRCA | BT-474 | Tanespimycin | MVP_top60_DNA+RNA_seed | HSP90AA1 | -0.034519 | BRCA | HCC2218 | Luminespib |
| MVP_top60_DNA_seed | PIK3CA | 0.349377 | BRCA | HCC38 | GNE-317 | MVP_top60_DNA+RNA_seed | HSP90AB1 | -0.034519 | BRCA | HCC2218 | Luminespib |
| MVP_top60_DNA_seed | MAPK3 | 0.35042 | BRCA | MDA-MB-231 | Ulixertinib | MVP_top60_DNA+RNA_seed | AKT1 | -0.032897 | BRCA | MDA-MB-330 | AT13148 |
| MVP_top60_DNA_seed | MAPK1 | 0.35042 | BRCA | MDA-MB-231 | Ulixertinib | MVP_top60_DNA+RNA_seed | HDAC1 | -0.031421 | BRCA | CAMA-1 | Entinostat |
| MVP_top60_DNA_seed | EGFR | 0.350665 | BRCA | EVSA-T | Osimertinib | MVP_top60_DNA+RNA_seed | ESR1 | -0.031292 | BRCA | UACC-812 | Tamoxifen |
| MVP_top60_DNA_seed | ERBB2 | 0.350742 | BRCA | Hs-578-T | Sapitinib | MVP_top60_DNA+RNA_seed | MAPK3 | -0.028843 | BRCA | HCC2218 | SCH772984 |
| MVP_top60_DNA_seed | EGFR | 0.350742 | BRCA | Hs-578-T | Sapitinib | MVP_top60_DNA+RNA_seed | MAPK1 | -0.028843 | BRCA | HCC2218 | SCH772984 |
| MVP_top60_DNA_seed | ERBB3 | 0.350742 | BRCA | Hs-578-T | Sapitinib | MVP_top60_DNA+RNA_seed | ESR1 | -0.026363 | BRCA | BT-549 | Tamoxifen |
| MVP_top60_DNA_seed | ERBB2 | 0.351015 | BRCA | HCC1187 | Afatinib | MVP_top60_DNA+RNA_seed | AKT1 | -0.026331 | BRCA | CAL-51 | MK-2206 |
| MVP_top60_DNA_seed | EGFR | 0.351015 | BRCA | HCC1187 | Afatinib | MVP_top60_DNA+RNA_seed | CDK2 | -0.025684 | BRCA | EFM-19 | Dinaciclib |
| MVP_top60_DNA_seed | HDAC1 | 0.355513 | BRCA | HDQ-P1 | Entinostat | MVP_top60_DNA+RNA_seed | CDK1 | -0.025684 | BRCA | EFM-19 | Dinaciclib |
| MVP_top60_DNA_seed | EGFR | 0.35555 | BRCA | HCC1937 | AZD3759 | MVP_top60_DNA+RNA_seed | CDK2 | -0.02457 | BRCA | MDA-MB-415 | Dinaciclib |
| MVP_top60_DNA_seed | EGFR | 0.355994 | BRCA | MDA-MB-231 | AZD3759 | MVP_top60_DNA+RNA_seed | CDK1 | -0.02457 | BRCA | MDA-MB-415 | Dinaciclib |
| MVP_top60_DNA_seed | MDM2 | 0.356087 | BRCA | OCUB-M | Nutlin-3a (-) | MVP_top60_DNA+RNA_seed | EGFR | -0.023951 | BRCA | MRK-nu-1 | Gefitinib |
| MVP_top60_DNA_seed | PIK3CA | 0.358545 | BRCA | HCC2218 | AZD8186 | MVP_top60_DNA+RNA_seed | HSP90AA1 | -0.022439 | BRCA | HCC1937 | Luminespib |
| MVP_top60_DNA_seed | MAPK3 | 0.361365 | BRCA | HCC1954 | Ulixertinib | MVP_top60_DNA+RNA_seed | HSP90AB1 | -0.022439 | BRCA | HCC1937 | Luminespib |
| MVP_top60_DNA_seed | MAPK1 | 0.361365 | BRCA | HCC1954 | Ulixertinib | MVP_top60_DNA+RNA_seed | HDAC1 | -0.022102 | BRCA | HCC1954 | Entinostat |
| MVP_top60_DNA_seed | MAPK3 | 0.364817 | BRCA | HCC1395 | SCH772984 | MVP_top60_DNA+RNA_seed | EGFR | -0.021254 | BRCA | CAL-51 | Gefitinib |
| MVP_top60_DNA_seed | MAPK1 | 0.364817 | BRCA | HCC1395 | SCH772984 | MVP_top60_DNA+RNA_seed | SRC | -0.020417 | BRCA | BT-20 | Bosutinib |
| MVP_top60_DNA_seed | HDAC1 | 0.365853 | BRCA | AU565 | Romidepsin | MVP_top60_DNA+RNA_seed | HDAC1 | -0.018234 | BRCA | CAMA-1 | PCI-34051 |
| MVP_top60_DNA_seed | HDAC2 | 0.365853 | BRCA | AU565 | Romidepsin | MVP_top60_DNA+RNA_seed | ATM | -0.017561 | BRCA | HDQ-P1 | KU-55933 |
| MVP_top60_DNA_seed | HSP90AA1 | 0.366832 | BRCA | BT-20 | Luminespib | MVP_top60_DNA+RNA_seed | EGFR | -0.016339 | BRCA | BT-549 | Lapatinib |
| MVP_top60_DNA_seed | HSP90AB1 | 0.366832 | BRCA | BT-20 | Luminespib | MVP_top60_DNA+RNA_seed | ERBB2 | -0.016339 | BRCA | BT-549 | Lapatinib |
| MVP_top60_DNA_seed | ERBB2 | 0.367445 | BRCA | CAMA-1 | Sapitinib | MVP_top60_DNA+RNA_seed | EGFR | -0.016185 | BRCA | EVSA-T | Gefitinib |
| MVP_top60_DNA_seed | EGFR | 0.367445 | BRCA | CAMA-1 | Sapitinib | MVP_top60_DNA+RNA_seed | AKT1 | -0.015569 | BRCA | T47D | Uprosertib |
| MVP_top60_DNA_seed | ERBB3 | 0.367445 | BRCA | CAMA-1 | Sapitinib | MVP_top60_DNA+RNA_seed | EGFR | -0.015539 | BRCA | MRK-nu-1 | Afatinib |
| MVP_top60_DNA_seed | MAPK3 | 0.36984 | BRCA | HCC1395 | Ulixertinib | MVP_top60_DNA+RNA_seed | ERBB2 | -0.015539 | BRCA | MRK-nu-1 | Afatinib |
| MVP_top60_DNA_seed | MAPK1 | 0.36984 | BRCA | HCC1395 | Ulixertinib | MVP_top60_DNA+RNA_seed | PIK3CA | -0.014331 | BRCA | BT-549 | GNE-317 |
| MVP_top60_DNA_seed | AKT1 | 0.370756 | BRCA | HCC1937 | AZD5363 | MVP_top60_DNA+RNA_seed | AKT1 | -0.01307 | BRCA | HCC1937 | MK-2206 |
| MVP_top60_DNA_seed | CDK2 | 0.373398 | BRCA | T47D | AZD5438 | MVP_top60_DNA+RNA_seed | PLK1 | -0.010128 | BRCA | MDA-MB-231 | BI-2536 |
| MVP_top60_DNA_seed | PIK3CA | 0.373971 | BRCA | BT-20 | AZD8186 | MVP_top60_DNA+RNA_seed | AKT1 | -0.010066 | BRCA | HCC2218 | AZD5363 |
| MVP_top60_DNA_seed | MAPK1 | 0.374475 | BRCA | HCC70 | VX-11e | MVP_top60_DNA+RNA_seed | CDK2 | -0.007275 | BRCA | HDQ-P1 | AZD5438 |
| MVP_top60_DNA_seed | MAPK3 | 0.377161 | BRCA | HCC1419 | SCH772984 | MVP_top60_DNA+RNA_seed | PIK3CA | -0.006998 | BRCA | MCF7 | AZD8186 |
| MVP_top60_DNA_seed | MAPK1 | 0.377161 | BRCA | HCC1419 | SCH772984 | MVP_top60_DNA+RNA_seed | HSP90AA1 | -0.006935 | BRCA | HCC1500 | Tanespimycin |
| MVP_top60_DNA_seed | SRC | 0.379868 | BRCA | AU565 | Bosutinib | MVP_top60_DNA+RNA_seed | HSP90AB1 | -0.006935 | BRCA | HCC1500 | Tanespimycin |
| MVP_top60_DNA_seed | MDM2 | 0.381394 | BRCA | HCC202 | Nutlin-3a (-) | MVP_top60_DNA+RNA_seed | EGFR | -0.006254 | BRCA | EFM-19 | Osimertinib |
| MVP_top60_DNA_seed | CDK2 | 0.38377 | BRCA | CAL-120 | Dinaciclib | MVP_top60_DNA+RNA_seed | EGFR | -0.006149 | BRCA | EFM-19 | Sapitinib |
| MVP_top60_DNA_seed | CDK1 | 0.38377 | BRCA | CAL-120 | Dinaciclib | MVP_top60_DNA+RNA_seed | ERBB2 | -0.006149 | BRCA | EFM-19 | Sapitinib |
| MVP_top60_DNA_seed | CDK1 | 0.383838 | BRCA | CAL-85-1 | RO-3306 | MVP_top60_DNA+RNA_seed | ERBB3 | -0.006149 | BRCA | EFM-19 | Sapitinib |
| MVP_top60_DNA_seed | ERBB2 | 0.383858 | BRCA | HCC70 | Afatinib | MVP_top60_DNA+RNA_seed | ESR1 | -0.00481 | BRCA | MDA-MB-436 | Tamoxifen |

| | | | | | | | | | | | |
|---|---|---|---|---|---|---|---|---|---|---|---|
| MVP_top60_DNA_seed | EGFR | 0.383858 | BRCA | HCC70 | Afatinib | MVP_top60_DNA+RNA_seed | MAPK3 | -0.004705 | BRCA | MDA-MB-468 | Ulixertinib |
| MVP_top60_DNA_seed | MDM2 | 0.38466 | BRCA | HDQ-P1 | Serdemetan | MVP_top60_DNA+RNA_seed | MAPK1 | -0.004705 | BRCA | MDA-MB-468 | Ulixertinib |
| MVP_top60_DNA_seed | ESR1 | 0.386212 | BRCA | HCC2218 | Tamoxifen | MVP_top60_DNA+RNA_seed | PIK3CA | -0.003606 | BRCA | COLO-824 | GNE-317 |
| MVP_top60_DNA_seed | CDK2 | 0.388468 | BRCA | JIMT-1 | MK-8776 | MVP_top60_DNA+RNA_seed | PIK3CA | -0.003436 | BRCA | EVSA-T | Alpelisib |
| MVP_top60_DNA_seed | ATM | 0.388842 | BRCA | HCC1599 | KU-55933 | MVP_top60_DNA+RNA_seed | EP300 | -0.002939 | BRCA | JIMT-1 | SGC-CBP30 |
| MVP_top60_DNA_seed | TP53 | 0.389817 | BRCA | CAL-51 | MIRA-1 | MVP_top60_DNA+RNA_seed | HDAC1 | -0.002552 | BRCA | CAL-85-1 | Entinostat |
| MVP_top60_DNA_seed | CDK2 | 0.390245 | BRCA | MDA-MB-330 | AZD5438 | MVP_top60_DNA+RNA_seed | MDM2 | -0.002526 | BRCA | AU565 | Nutlin-3a (-) |
| MVP_top60_DNA_seed | HSP90AA1 | 0.390655 | BRCA | T47D | Luminespib | MVP_top60_DNA+RNA_seed | EGFR | 0.001844 | BRCA | CAL-148 | Afatinib |
| MVP_top60_DNA_seed | HSP90AB1 | 0.390655 | BRCA | T47D | Luminespib | MVP_top60_DNA+RNA_seed | ERBB2 | 0.001844 | BRCA | CAL-148 | Afatinib |
| MVP_top60_DNA_seed | EP300 | 0.393664 | BRCA | JIMT-1 | CPI-637 | MVP_top60_DNA+RNA_seed | EGFR | 0.002988 | BRCA | OCUB-M | Gefitinib |
| MVP_top60_DNA_seed | TP53 | 0.393963 | BRCA | YMB-1-E | MIRA-1 | MVP_top60_DNA+RNA_seed | AKT1 | 0.003273 | BRCA | HCC2218 | Uprosertib |
| MVP_top60_DNA_seed | MAPK1 | 0.394005 | BRCA | MRK-nu-1 | VX-11e | MVP_top60_DNA+RNA_seed | SRC | 0.005662 | BRCA | HCC1419 | Bosutinib |
| MVP_top60_DNA_seed | MAPK3 | 0.395886 | BRCA | COLO-824 | Ulixertinib | MVP_top60_DNA+RNA_seed | HSP90AA1 | 0.005988 | BRCA | MRK-nu-1 | Luminespib |
| MVP_top60_DNA_seed | MAPK1 | 0.395886 | BRCA | COLO-824 | Ulixertinib | MVP_top60_DNA+RNA_seed | HSP90AB1 | 0.005988 | BRCA | MRK-nu-1 | Luminespib |
| MVP_top60_DNA_seed | PIK3CA | 0.396797 | BRCA | OCUB-M | AZD8186 | MVP_top60_DNA+RNA_seed | PLK1 | 0.006015 | BRCA | MCF7 | BI-2536 |
| MVP_top60_DNA_seed | HSP90AA1 | 0.399075 | BRCA | CAMA-1 | Luminespib | MVP_top60_DNA+RNA_seed | EGFR | 0.006975 | BRCA | HCC1419 | AZD3759 |
| MVP_top60_DNA_seed | HSP90AB1 | 0.399075 | BRCA | CAMA-1 | Luminespib | MVP_top60_DNA+RNA_seed | HDAC1 | 0.010375 | BRCA | JIMT-1 | Romidepsin |
| MVP_top60_DNA_seed | MAPK3 | 0.400136 | BRCA | T47D | SCH772984 | MVP_top60_DNA+RNA_seed | HDAC2 | 0.010375 | BRCA | JIMT-1 | Romidepsin |
| MVP_top60_DNA_seed | MAPK1 | 0.400136 | BRCA | T47D | SCH772984 | MVP_top60_DNA+RNA_seed | TP53 | 0.010507 | BRCA | MDA-MB-330 | MIRA-1 |
| MVP_top60_DNA_seed | AR | 0.401465 | BRCA | HCC1599 | Bicalutamide | MVP_top60_DNA+RNA_seed | EGFR | 0.015833 | BRCA | HCC70 | Gefitinib |
| MVP_top60_DNA_seed | MAPK3 | 0.401974 | BRCA | Hs-578-T | Ulixertinib | MVP_top60_DNA+RNA_seed | MAPK1 | 0.017338 | BRCA | EVSA-T | VX-11e |
| MVP_top60_DNA_seed | MAPK1 | 0.401974 | BRCA | Hs-578-T | Ulixertinib | MVP_top60_DNA+RNA_seed | HDAC1 | 0.017551 | BRCA | MDA-MB-157 | Romidepsin |
| MVP_top60_DNA_seed | HDAC1 | 0.402384 | BRCA | HDQ-P1 | PCI-34051 | MVP_top60_DNA+RNA_seed | HDAC2 | 0.017551 | BRCA | MDA-MB-157 | Romidepsin |
| MVP_top60_DNA_seed | MDM2 | 0.402435 | BRCA | UACC-812 | Serdemetan | MVP_top60_DNA+RNA_seed | SRC | 0.017835 | BRCA | CAL-148 | Bosutinib |
| MVP_top60_DNA_seed | PIK3CA | 0.405435 | BRCA | UACC-812 | Buparlisib | MVP_top60_DNA+RNA_seed | ESR1 | 0.019116 | BRCA | HCC70 | Tamoxifen |
| MVP_top60_DNA_seed | CDK1 | 0.405889 | BRCA | ZR-75-30 | RO-3306 | MVP_top60_DNA+RNA_seed | AKT1 | 0.020388 | BRCA | CAL-85-1 | Ipatasertib |
| MVP_top60_DNA_seed | MAPK3 | 0.40609 | BRCA | T47D | Ulixertinib | MVP_top60_DNA+RNA_seed | HSP90AA1 | 0.021405 | BRCA | CAL-120 | Luminespib |
| MVP_top60_DNA_seed | MAPK1 | 0.40609 | BRCA | T47D | Ulixertinib | MVP_top60_DNA+RNA_seed | HSP90AB1 | 0.021405 | BRCA | CAL-120 | Luminespib |
| MVP_top60_DNA_seed | MAPK3 | 0.406631 | BRCA | AU565 | Ulixertinib | MVP_top60_DNA+RNA_seed | AKT1 | 0.021467 | BRCA | MDA-MB-468 | MK-2206 |
| MVP_top60_DNA_seed | MAPK1 | 0.406631 | BRCA | AU565 | Ulixertinib | MVP_top60_DNA+RNA_seed | EGFR | 0.02231 | BRCA | CAMA-1 | Osimertinib |
| MVP_top60_DNA_seed | HSP90AA1 | 0.406809 | BRCA | HCC70 | Tanespimycin | MVP_top60_DNA+RNA_seed | HSP90AA1 | 0.024163 | BRCA | HCC1569 | CCT-018159 |
| MVP_top60_DNA_seed | HSP90AB1 | 0.406809 | BRCA | HCC70 | Tanespimycin | MVP_top60_DNA+RNA_seed | HSP90AB1 | 0.024163 | BRCA | HCC1569 | CCT-018159 |
| MVP_top60_DNA_seed | CDK2 | 0.409051 | BRCA | BT-549 | AZD5438 | MVP_top60_DNA+RNA_seed | MDM2 | 0.024757 | BRCA | ZR-75-30 | Serdemetan |
| MVP_top60_DNA_seed | HSP90AA1 | 0.413036 | BRCA | UACC-893 | Elesclomol | MVP_top60_DNA+RNA_seed | EGFR | 0.025478 | BRCA | CAL-51 | Osimertinib |
| MVP_top60_DNA_seed | HSP90AB1 | 0.413036 | BRCA | UACC-893 | Elesclomol | MVP_top60_DNA+RNA_seed | HSP90AA1 | 0.026115 | BRCA | YMB-1-E | Tanespimycin |
| MVP_top60_DNA_seed | HDAC1 | 0.41397 | BRCA | HCC1187 | Entinostat | MVP_top60_DNA+RNA_seed | HSP90AB1 | 0.026115 | BRCA | YMB-1-E | Tanespimycin |
| MVP_top60_DNA_seed | HDAC1 | 0.414933 | BRCA | MDA-MB-231 | PCI-34051 | MVP_top60_DNA+RNA_seed | MDM2 | 0.02657 | BRCA | MDA-MB-415 | Nutlin-3a (-) |
| MVP_top60_DNA_seed | MDM2 | 0.415639 | BRCA | HCC1599 | Nutlin-3a (-) | MVP_top60_DNA+RNA_seed | MAPK3 | 0.026659 | BRCA | EVSA-T | Ulixertinib |
| MVP_top60_DNA_seed | MDM2 | 0.416264 | BRCA | HCC1954 | Nutlin-3a (-) | MVP_top60_DNA+RNA_seed | MAPK1 | 0.026659 | BRCA | EVSA-T | Ulixertinib |
| MVP_top60_DNA_seed | AKT1 | 0.417909 | BRCA | HCC2218 | MK-2206 | MVP_top60_DNA+RNA_seed | AKT1 | 0.027095 | BRCA | HCC1143 | GSK2110183B |
| MVP_top60_DNA_seed | MDM2 | 0.419545 | BRCA | MCF7 | Serdemetan | MVP_top60_DNA+RNA_seed | PIK3CA | 0.031282 | BRCA | HCC1954 | AZD8186 |

| | | | | | | | | | | | |
|---|---|---|---|---|---|---|---|---|---|---|---|
| MVP_top60_DNA_seed | EGFR | 0.419947 | BRCA | MDA-MB-231 | Erlotinib | MVP_top60_DNA+RNA_seed | CDK2 | 0.031771 | BRCA | HCC1419 | MK-8776 |
| MVP_top60_DNA_seed | MAPK3 | 0.420019 | BRCA | HCC1954 | SCH772984 | MVP_top60_DNA+RNA_seed | HSP90AA1 | 0.035358 | BRCA | YMB-1-E | Luminespib |
| MVP_top60_DNA_seed | MAPK1 | 0.420019 | BRCA | HCC1954 | SCH772984 | MVP_top60_DNA+RNA_seed | HSP90AB1 | 0.035358 | BRCA | YMB-1-E | Luminespib |
| MVP_top60_DNA_seed | ERBB2 | 0.420646 | BRCA | CAL-148 | Sapitinib | MVP_top60_DNA+RNA_seed | CDK1 | 0.036086 | BRCA | HCC2218 | RO-3306 |
| MVP_top60_DNA_seed | EGFR | 0.420646 | BRCA | CAL-148 | Sapitinib | MVP_top60_DNA+RNA_seed | MAPK3 | 0.037352 | BRCA | OCUB-M | Ulixertinib |
| MVP_top60_DNA_seed | ERBB3 | 0.420646 | BRCA | CAL-148 | Sapitinib | MVP_top60_DNA+RNA_seed | MAPK1 | 0.037352 | BRCA | OCUB-M | Ulixertinib |
| MVP_top60_DNA_seed | AKT1 | 0.420836 | BRCA | HCC1187 | Afuresertib | MVP_top60_DNA+RNA_seed | HSP90AA1 | 0.039856 | BRCA | HCC1143 | Luminespib |
| MVP_top60_DNA_seed | PIK3CA | 0.422393 | BRCA | HCC1419 | AZD8186 | MVP_top60_DNA+RNA_seed | HSP90AB1 | 0.039856 | BRCA | HCC1143 | Luminespib |
| MVP_top60_DNA_seed | MDM2 | 0.423598 | BRCA | CAL-51 | Serdemetan | MVP_top60_DNA+RNA_seed | EGFR | 0.040789 | BRCA | HCC2218 | AZD3759 |
| MVP_top60_DNA_seed | HDAC1 | 0.425495 | BRCA | HCC1419 | PCI-34051 | MVP_top60_DNA+RNA_seed | HDAC1 | 0.041152 | BRCA | MDA-MB-361 | PCI-34051 |
| MVP_top60_DNA_seed | MAPK1 | 0.427494 | BRCA | HCC2157 | VX-11e | MVP_top60_DNA+RNA_seed | EGFR | 0.041701 | BRCA | JIMT-1 | Afatinib |
| MVP_top60_DNA_seed | EP300 | 0.429734 | BRCA | EFM-19 | CPI-637 | MVP_top60_DNA+RNA_seed | ERBB2 | 0.041701 | BRCA | JIMT-1 | Afatinib |
| MVP_top60_DNA_seed | HDAC1 | 0.432214 | BRCA | HCC70 | PCI-34051 | MVP_top60_DNA+RNA_seed | CDK2 | 0.043953 | BRCA | UACC-893 | MK-8776 |
| MVP_top60_DNA_seed | ESR1 | 0.432784 | BRCA | MDA-MB-157 | Tamoxifen | MVP_top60_DNA+RNA_seed | ESR1 | 0.044537 | BRCA | CAL-85-1 | Tamoxifen |
| MVP_top60_DNA_seed | MDM2 | 0.432936 | BRCA | MDA-MB-415 | Serdemetan | MVP_top60_DNA+RNA_seed | EGFR | 0.044667 | BRCA | DU-4475 | Afatinib |
| MVP_top60_DNA_seed | ERBB2 | 0.433847 | BRCA | HCC1187 | Sapitinib | MVP_top60_DNA+RNA_seed | ERBB2 | 0.044667 | BRCA | DU-4475 | Afatinib |
| MVP_top60_DNA_seed | EGFR | 0.433847 | BRCA | HCC1187 | Sapitinib | MVP_top60_DNA+RNA_seed | AKT1 | 0.044804 | BRCA | HCC1187 | Ipatasertib |
| MVP_top60_DNA_seed | ERBB3 | 0.433847 | BRCA | HCC1187 | Sapitinib | MVP_top60_DNA+RNA_seed | PIK3CA | 0.044886 | BRCA | BT-549 | AZD8186 |
| MVP_top60_DNA_seed | CDK2 | 0.434729 | BRCA | MDA-MB-453 | AZD5438 | MVP_top60_DNA+RNA_seed | MDM2 | 0.045823 | BRCA | MDA-MB-453 | Serdemetan |
| MVP_top60_DNA_seed | ESR1 | 0.436325 | BRCA | HCC1143 | GDC0810 | MVP_top60_DNA+RNA_seed | MDM2 | 0.048354 | BRCA | HCC2157 | Serdemetan |
| MVP_top60_DNA_seed | ERBB2 | 0.437476 | BRCA | MDA-MB-415 | Afatinib | MVP_top60_DNA+RNA_seed | AKT1 | 0.051383 | BRCA | HCC1937 | Afuresertib |
| MVP_top60_DNA_seed | EGFR | 0.437476 | BRCA | MDA-MB-415 | Afatinib | MVP_top60_DNA+RNA_seed | PIK3CA | 0.053041 | BRCA | CAMA-1 | Buparlisib |
| MVP_top60_DNA_seed | CDK1 | 0.438682 | BRCA | HCC38 | RO-3306 | MVP_top60_DNA+RNA_seed | HDAC1 | 0.053309 | BRCA | CAMA-1 | Romidepsin |
| MVP_top60_DNA_seed | HDAC1 | 0.438968 | BRCA | BT-474 | Romidepsin | MVP_top60_DNA+RNA_seed | HDAC2 | 0.053309 | BRCA | CAMA-1 | Romidepsin |
| MVP_top60_DNA_seed | HDAC2 | 0.438968 | BRCA | BT-474 | Romidepsin | MVP_top60_DNA+RNA_seed | PIK3CA | 0.05419 | BRCA | MDA-MB-468 | Buparlisib |
| MVP_top60_DNA_seed | EP300 | 0.439838 | BRCA | BT-474 | SGC-CBP30 | MVP_top60_DNA+RNA_seed | MDM2 | 0.058001 | BRCA | T47D | Serdemetan |
| MVP_top60_DNA_seed | CREBBP | 0.439838 | BRCA | BT-474 | SGC-CBP30 | MVP_top60_DNA+RNA_seed | MAPK3 | 0.058565 | BRCA | CAL-51 | Ulixertinib |
| MVP_top60_DNA_seed | MDM2 | 0.43996 | BRCA | COLO-824 | Nutlin-3a (-) | MVP_top60_DNA+RNA_seed | MAPK1 | 0.058565 | BRCA | CAL-51 | Ulixertinib |
| MVP_top60_DNA_seed | ESR1 | 0.442264 | BRCA | MDA-MB-436 | GDC0810 | MVP_top60_DNA+RNA_seed | SRC | 0.059465 | BRCA | HCC1143 | Bosutinib |
| MVP_top60_DNA_seed | AKT1 | 0.443013 | BRCA | HDQ-P1 | MK-2206 | MVP_top60_DNA+RNA_seed | MDM2 | 0.059967 | BRCA | HCC2157 | Nutlin-3a (-) |
| MVP_top60_DNA_seed | CDK2 | 0.445264 | BRCA | HCC38 | Dinaciclib | MVP_top60_DNA+RNA_seed | MDM2 | 0.060373 | BRCA | HCC1569 | Nutlin-3a (-) |
| MVP_top60_DNA_seed | CDK1 | 0.445264 | BRCA | HCC38 | Dinaciclib | MVP_top60_DNA+RNA_seed | EGFR | 0.06129 | BRCA | BT-474 | Erlotinib |
| MVP_top60_DNA_seed | AKT1 | 0.447848 | BRCA | MFM-223 | AT13148 | MVP_top60_DNA+RNA_seed | MAPK3 | 0.062979 | BRCA | YMB-1-E | SCH772984 |
| MVP_top60_DNA_seed | EGFR | 0.44962 | BRCA | HCC1500 | Gefitinib | MVP_top60_DNA+RNA_seed | MAPK1 | 0.062979 | BRCA | YMB-1-E | SCH772984 |
| MVP_top60_DNA_seed | MDM2 | 0.450224 | BRCA | CAMA-1 | Serdemetan | MVP_top60_DNA+RNA_seed | EP300 | 0.062984 | BRCA | HCC70 | SGC-CBP30 |
| MVP_top60_DNA_seed | CDK2 | 0.450349 | BRCA | UACC-893 | Dinaciclib | MVP_top60_DNA+RNA_seed | PLK1 | 0.067211 | BRCA | HCC1143 | MK-1775 |
| MVP_top60_DNA_seed | CDK1 | 0.450349 | BRCA | UACC-893 | Dinaciclib | MVP_top60_DNA+RNA_seed | TP53 | 0.067588 | BRCA | CAL-85-1 | MIRA-1 |
| MVP_top60_DNA_seed | EP300 | 0.451761 | BRCA | HCC1395 | CPI-637 | MVP_top60_DNA+RNA_seed | HSP90AA1 | 0.067877 | BRCA | MDA-MB-157 | Luminespib |
| MVP_top60_DNA_seed | AKT1 | 0.452331 | BRCA | HCC1143 | Afuresertib | MVP_top60_DNA+RNA_seed | HSP90AB1 | 0.067877 | BRCA | MDA-MB-157 | Luminespib |
| MVP_top60_DNA_seed | EP300 | 0.453045 | BRCA | CAL-85-1 | CPI-637 | MVP_top60_DNA+RNA_seed | HDAC1 | 0.068065 | BRCA | MDA-MB-453 | PCI-34051 |
| MVP_top60_DNA_seed | MAPK1 | 0.453514 | BRCA | HCC2218 | VX-11e | MVP_top60_DNA+RNA_seed | MDM2 | 0.069172 | BRCA | JIMT-1 | Serdemetan |

| | | | | | | | | | | | |
|---|---|---|---|---|---|---|---|---|---|---|---|
| MVP_top60_DNA_seed | SRC | 0.453599 | BRCA | Hs-578-T | Bosutinib | MVP_top60_DNA+RNA_seed | HDAC1 | 0.069842 | BRCA | MDA-MB-468 | Entinostat |
| MVP_top60_DNA_seed | TP53 | 0.453655 | BRCA | MDA-MB-468 | MIRA-1 | MVP_top60_DNA+RNA_seed | MDM2 | 0.071249 | BRCA | HCC1187 | Nutlin-3a (-) |
| MVP_top60_DNA_seed | CDK2 | 0.454282 | BRCA | BT-20 | MK-8776 | MVP_top60_DNA+RNA_seed | PIK3CA | 0.072001 | BRCA | HCC1395 | Alpelisib |
| MVP_top60_DNA_seed | CDK2 | 0.45477 | BRCA | MDA-MB-436 | MK-8776 | MVP_top60_DNA+RNA_seed | AKT1 | 0.072133 | BRCA | HCC1599 | AT13148 |
| MVP_top60_DNA_seed | HDAC1 | 0.457268 | BRCA | HCC1419 | Entinostat | MVP_top60_DNA+RNA_seed | ATM | 0.073825 | BRCA | HCC1569 | KU-55933 |
| MVP_top60_DNA_seed | AKT1 | 0.457967 | BRCA | MDA-MB-157 | AT13148 | MVP_top60_DNA+RNA_seed | CDK1 | 0.073852 | BRCA | T47D | RO-3306 |
| MVP_top60_DNA_seed | MAPK1 | 0.459014 | BRCA | HCC1187 | VX-11e | MVP_top60_DNA+RNA_seed | MAPK3 | 0.073975 | BRCA | MDA-MB-453 | Ulixertinib |
| MVP_top60_DNA_seed | PIK3CA | 0.459333 | BRCA | BT-549 | Buparlisib | MVP_top60_DNA+RNA_seed | MAPK1 | 0.073975 | BRCA | MDA-MB-453 | Ulixertinib |
| MVP_top60_DNA_seed | ERBB2 | 0.459412 | BRCA | HCC70 | Sapitinib | MVP_top60_DNA+RNA_seed | EGFR | 0.074967 | BRCA | MDA-MB-231 | Afatinib |
| MVP_top60_DNA_seed | EGFR | 0.459412 | BRCA | HCC70 | Sapitinib | MVP_top60_DNA+RNA_seed | ERBB2 | 0.074967 | BRCA | MDA-MB-231 | Afatinib |
| MVP_top60_DNA_seed | ERBB3 | 0.459412 | BRCA | HCC70 | Sapitinib | MVP_top60_DNA+RNA_seed | HSP90AA1 | 0.075628 | BRCA | BT-20 | Tanespimycin |
| MVP_top60_DNA_seed | AKT1 | 0.462422 | BRCA | JIMT-1 | MK-2206 | MVP_top60_DNA+RNA_seed | HSP90AB1 | 0.075628 | BRCA | BT-20 | Tanespimycin |
| MVP_top60_DNA_seed | MAPK1 | 0.464492 | BRCA | AU565 | VX-11e | MVP_top60_DNA+RNA_seed | MAPK3 | 0.076447 | BRCA | MDA-MB-453 | SCH772984 |
| MVP_top60_DNA_seed | MAPK1 | 0.465861 | BRCA | HCC1500 | VX-11e | MVP_top60_DNA+RNA_seed | MAPK1 | 0.076447 | BRCA | MDA-MB-453 | SCH772984 |
| MVP_top60_DNA_seed | MDM2 | 0.466177 | BRCA | UACC-893 | Nutlin-3a (-) | MVP_top60_DNA+RNA_seed | SRC | 0.077267 | BRCA | UACC-893 | Bosutinib |
| MVP_top60_DNA_seed | EGFR | 0.466315 | BRCA | EFM-192A | AZD3759 | MVP_top60_DNA+RNA_seed | ESR1 | 0.07979 | BRCA | HCC1569 | Tamoxifen |
| MVP_top60_DNA_seed | MAPK1 | 0.466883 | BRCA | BT-20 | VX-11e | MVP_top60_DNA+RNA_seed | TP53 | 0.079871 | BRCA | CAMA-1 | MIRA-1 |
| MVP_top60_DNA_seed | CDK2 | 0.46745 | BRCA | MDA-MB-468 | MK-8776 | MVP_top60_DNA+RNA_seed | EGFR | 0.0799 | BRCA | COLO-824 | AZD3759 |
| MVP_top60_DNA_seed | PIK3CA | 0.4689 | BRCA | MDA-MB-415 | Alpelisib | MVP_top60_DNA+RNA_seed | MAPK1 | 0.081773 | BRCA | UACC-893 | VX-11e |
| MVP_top60_DNA_seed | AKT1 | 0.470302 | BRCA | MDA-MB-468 | Uprosertib | MVP_top60_DNA+RNA_seed | AKT1 | 0.085434 | BRCA | ZR-75-30 | Uprosertib |
| MVP_top60_DNA_seed | HSP90AA1 | 0.471624 | BRCA | MDA-MB-157 | Tanespimycin | MVP_top60_DNA+RNA_seed | PLK1 | 0.087046 | BRCA | MDA-MB-157 | MK-1775 |
| MVP_top60_DNA_seed | HSP90AB1 | 0.471624 | BRCA | MDA-MB-157 | Tanespimycin | MVP_top60_DNA+RNA_seed | HSP90AA1 | 0.087714 | BRCA | HCC70 | Luminespib |
| MVP_top60_DNA_seed | AKT1 | 0.472093 | BRCA | HCC1143 | AZD5363 | MVP_top60_DNA+RNA_seed | HSP90AB1 | 0.087714 | BRCA | HCC70 | Luminespib |
| MVP_top60_DNA_seed | MDM2 | 0.47277 | BRCA | BT-20 | Nutlin-3a (-) | MVP_top60_DNA+RNA_seed | MAPK1 | 0.088764 | BRCA | OCUB-M | VX-11e |
| MVP_top60_DNA_seed | ESR1 | 0.476116 | BRCA | BT-474 | GDC0810 | MVP_top60_DNA+RNA_seed | ESR1 | 0.089536 | BRCA | MRK-nu-1 | Tamoxifen |
| MVP_top60_DNA_seed | MAPK3 | 0.476896 | BRCA | MFM-223 | SCH772984 | MVP_top60_DNA+RNA_seed | MDM2 | 0.091379 | BRCA | MDA-MB-468 | Serdemetan |
| MVP_top60_DNA_seed | MAPK1 | 0.476896 | BRCA | MFM-223 | SCH772984 | MVP_top60_DNA+RNA_seed | AKT1 | 0.093271 | BRCA | HCC1187 | MK-2206 |
| MVP_top60_DNA_seed | AKT1 | 0.478651 | BRCA | MDA-MB-157 | Afuresertib | MVP_top60_DNA+RNA_seed | TP53 | 0.09638 | BRCA | HDQ-P1 | MIRA-1 |
| MVP_top60_DNA_seed | CDK2 | 0.47911 | BRCA | YMB-1-E | Dinaciclib | MVP_top60_DNA+RNA_seed | AR | 0.098345 | BRCA | YMB-1-E | Bicalutamide |
| MVP_top60_DNA_seed | CDK1 | 0.47911 | BRCA | YMB-1-E | Dinaciclib | MVP_top60_DNA+RNA_seed | AKT1 | 0.099004 | BRCA | BT-549 | Uprosertib |
| MVP_top60_DNA_seed | MAPK3 | 0.479354 | BRCA | HCC1187 | Ulixertinib | MVP_top60_DNA+RNA_seed | CDK1 | 0.099478 | BRCA | CAL-120 | RO-3306 |
| MVP_top60_DNA_seed | MAPK1 | 0.479354 | BRCA | HCC1187 | Ulixertinib | MVP_top60_DNA+RNA_seed | PIK3CA | 0.103107 | BRCA | UACC-893 | Buparlisib |
| MVP_top60_DNA_seed | EP300 | 0.480464 | BRCA | HCC1187 | SGC-CBP30 | MVP_top60_DNA+RNA_seed | AKT1 | 0.104465 | BRCA | UACC-812 | AZD5363 |
| MVP_top60_DNA_seed | CREBBP | 0.480464 | BRCA | HCC1187 | SGC-CBP30 | MVP_top60_DNA+RNA_seed | EGFR | 0.107777 | BRCA | MDA-MB-415 | AZD3759 |
| MVP_top60_DNA_seed | EGFR | 0.481107 | BRCA | EFM-19 | AZD3759 | MVP_top60_DNA+RNA_seed | AKT1 | 0.108611 | BRCA | BT-474 | Uprosertib |
| MVP_top60_DNA_seed | AKT1 | 0.481963 | BRCA | HCC1143 | Uprosertib | MVP_top60_DNA+RNA_seed | HSP90AA1 | 0.108982 | BRCA | EFM-192A | Tanespimycin |
| MVP_top60_DNA_seed | ESR1 | 0.484817 | BRCA | MDA-MB-453 | GDC0810 | MVP_top60_DNA+RNA_seed | HSP90AB1 | 0.108982 | BRCA | EFM-192A | Tanespimycin |
| MVP_top60_DNA_seed | ESR1 | 0.48503 | BRCA | MDA-MB-330 | GDC0810 | MVP_top60_DNA+RNA_seed | AKT1 | 0.110268 | BRCA | ZR-75-30 | MK-2206 |
| MVP_top60_DNA_seed | MAPK3 | 0.485736 | BRCA | MRK-nu-1 | SCH772984 | MVP_top60_DNA+RNA_seed | EGFR | 0.110414 | BRCA | BT-20 | Erlotinib |
| MVP_top60_DNA_seed | MAPK1 | 0.485736 | BRCA | MRK-nu-1 | SCH772984 | MVP_top60_DNA+RNA_seed | EP300 | 0.110681 | BRCA | DU-4475 | CPI-637 |
| MVP_top60_DNA_seed | PIK3CA | 0.487822 | BRCA | CAL-85-1 | AZD8186 | MVP_top60_DNA+RNA_seed | TP53 | 0.112399 | BRCA | HCC1569 | MIRA-1 |

| | | | | | | | | | | | |
|---|---|---|---|---|---|---|---|---|---|---|---|
| MVP_top60_DNA_seed | HSP90AA1 | 0.488262 | BRCA | HCC1500 | Luminespib | MVP_top60_DNA+RNA_seed | HDAC1 | 0.113509 | BRCA | HCC1500 | PCI-34051 |
| MVP_top60_DNA_seed | HSP90AB1 | 0.488262 | BRCA | HCC1500 | Luminespib | MVP_top60_DNA+RNA_seed | ESR1 | 0.113837 | BRCA | HCC2218 | GDC0810 |
| MVP_top60_DNA_seed | EP300 | 0.488286 | BRCA | HCC1500 | SGC-CBP30 | MVP_top60_DNA+RNA_seed | MAPK3 | 0.115631 | BRCA | COLO-824 | SCH772984 |
| MVP_top60_DNA_seed | CREBBP | 0.488286 | BRCA | HCC1500 | SGC-CBP30 | MVP_top60_DNA+RNA_seed | MAPK1 | 0.115631 | BRCA | COLO-824 | SCH772984 |
| MVP_top60_DNA_seed | SRC | 0.488445 | BRCA | JIMT-1 | Bosutinib | MVP_top60_DNA+RNA_seed | EGFR | 0.116354 | BRCA | HCC2218 | Gefitinib |
| MVP_top60_DNA_seed | EGFR | 0.491244 | BRCA | UACC-812 | Gefitinib | MVP_top60_DNA+RNA_seed | ATM | 0.117551 | BRCA | MFM-223 | KU-55933 |
| MVP_top60_DNA_seed | EP300 | 0.492236 | BRCA | HCC38 | SGC-CBP30 | MVP_top60_DNA+RNA_seed | CDK2 | 0.118606 | BRCA | HCC1143 | Dinaciclib |
| MVP_top60_DNA_seed | CREBBP | 0.492236 | BRCA | HCC38 | SGC-CBP30 | MVP_top60_DNA+RNA_seed | CDK1 | 0.118606 | BRCA | HCC1143 | Dinaciclib |
| MVP_top60_DNA_seed | EGFR | 0.493629 | BRCA | HCC1599 | AZD3759 | MVP_top60_DNA+RNA_seed | SRC | 0.119608 | BRCA | CAL-51 | Bosutinib |
| MVP_top60_DNA_seed | EP300 | 0.495564 | BRCA | HDQ-P1 | CPI-637 | MVP_top60_DNA+RNA_seed | MAPK1 | 0.120396 | BRCA | MDA-MB-468 | VX-11e |
| MVP_top60_DNA_seed | MAPK3 | 0.497144 | BRCA | HCC2157 | Ulixertinib | MVP_top60_DNA+RNA_seed | ESR1 | 0.1208 | BRCA | HCC1937 | Tamoxifen |
| MVP_top60_DNA_seed | MAPK1 | 0.497144 | BRCA | HCC2157 | Ulixertinib | MVP_top60_DNA+RNA_seed | HDAC1 | 0.122261 | BRCA | HCC1569 | Entinostat |
| MVP_top60_DNA_seed | AKT1 | 0.497206 | BRCA | T47D | Uprosertib | MVP_top60_DNA+RNA_seed | EGFR | 0.122275 | BRCA | HCC1143 | Afatinib |
| MVP_top60_DNA_seed | EP300 | 0.498955 | BRCA | HCC1954 | SGC-CBP30 | MVP_top60_DNA+RNA_seed | ERBB2 | 0.122275 | BRCA | HCC1143 | Afatinib |
| MVP_top60_DNA_seed | CREBBP | 0.498955 | BRCA | HCC1954 | SGC-CBP30 | MVP_top60_DNA+RNA_seed | AKT1 | 0.122831 | BRCA | HCC1419 | Afuresertib |
| MVP_top60_DNA_seed | TP53 | 0.499196 | BRCA | T47D | MIRA-1 | MVP_top60_DNA+RNA_seed | EGFR | 0.123387 | BRCA | MDA-MB-415 | Sapitinib |
| MVP_top60_DNA_seed | ATM | 0.499748 | BRCA | CAL-148 | KU-55933 | MVP_top60_DNA+RNA_seed | ERBB2 | 0.123387 | BRCA | MDA-MB-415 | Sapitinib |
| MVP_top60_DNA_seed | MAPK3 | 0.500147 | BRCA | JIMT-1 | Ulixertinib | MVP_top60_DNA+RNA_seed | ERBB3 | 0.123387 | BRCA | MDA-MB-415 | Sapitinib |
| MVP_top60_DNA_seed | MAPK1 | 0.500147 | BRCA | JIMT-1 | Ulixertinib | MVP_top60_DNA+RNA_seed | CDK1 | 0.123649 | BRCA | MDA-MB-436 | RO-3306 |
| MVP_top60_DNA_seed | AKT1 | 0.50023 | BRCA | MDA-MB-231 | Uprosertib | MVP_top60_DNA+RNA_seed | TP53 | 0.124896 | BRCA | Hs-578-T | MIRA-1 |
| MVP_top60_DNA_seed | PIK3CA | 0.501662 | BRCA | HCC1937 | Alpelisib | MVP_top60_DNA+RNA_seed | ATM | 0.125494 | BRCA | MDA-MB-453 | KU-55933 |
| MVP_top60_DNA_seed | HDAC1 | 0.504462 | BRCA | MDA-MB-157 | Entinostat | MVP_top60_DNA+RNA_seed | EP300 | 0.128188 | BRCA | MCF7 | CPI-637 |
| MVP_top60_DNA_seed | CDK2 | 0.507491 | BRCA | CAMA-1 | AZD5438 | MVP_top60_DNA+RNA_seed | EGFR | 0.128273 | BRCA | HCC38 | Sapitinib |
| MVP_top60_DNA_seed | ERBB2 | 0.510696 | BRCA | MDA-MB-157 | Lapatinib | MVP_top60_DNA+RNA_seed | ERBB2 | 0.128273 | BRCA | HCC38 | Sapitinib |
| MVP_top60_DNA_seed | EGFR | 0.510696 | BRCA | MDA-MB-157 | Lapatinib | MVP_top60_DNA+RNA_seed | ERBB3 | 0.128273 | BRCA | HCC38 | Sapitinib |
| MVP_top60_DNA_seed | ERBB2 | 0.515466 | BRCA | CAMA-1 | Lapatinib | MVP_top60_DNA+RNA_seed | TP53 | 0.129468 | BRCA | JIMT-1 | MIRA-1 |
| MVP_top60_DNA_seed | EGFR | 0.515466 | BRCA | CAMA-1 | Lapatinib | MVP_top60_DNA+RNA_seed | HDAC1 | 0.132359 | BRCA | COLO-824 | PCI-34051 |
| MVP_top60_DNA_seed | ESR1 | 0.516456 | BRCA | BT-474 | Tamoxifen | MVP_top60_DNA+RNA_seed | EP300 | 0.132601 | BRCA | MDA-MB-157 | SGC-CBP30 |
| MVP_top60_DNA_seed | CDK1 | 0.517717 | BRCA | HCC1419 | RO-3306 | MVP_top60_DNA+RNA_seed | EGFR | 0.133351 | BRCA | T47D | Afatinib |
| MVP_top60_DNA_seed | ERBB2 | 0.518824 | BRCA | HCC1937 | Afatinib | MVP_top60_DNA+RNA_seed | ERBB2 | 0.133351 | BRCA | T47D | Afatinib |
| MVP_top60_DNA_seed | EGFR | 0.518824 | BRCA | HCC1937 | Afatinib | MVP_top60_DNA+RNA_seed | EGFR | 0.137305 | BRCA | OCUB-M | Erlotinib |
| MVP_top60_DNA_seed | SRC | 0.519776 | BRCA | HCC1500 | Bosutinib | MVP_top60_DNA+RNA_seed | AR | 0.139365 | BRCA | HCC1569 | Bicalutamide |
| MVP_top60_DNA_seed | CDK2 | 0.520274 | BRCA | MCF7 | AZD5438 | MVP_top60_DNA+RNA_seed | ESR1 | 0.139713 | BRCA | MDA-MB-468 | GDC0810 |
| MVP_top60_DNA_seed | EGFR | 0.52298 | BRCA | HCC1569 | Erlotinib | MVP_top60_DNA+RNA_seed | EGFR | 0.140037 | BRCA | MDA-MB-436 | AZD3759 |
| MVP_top60_DNA_seed | PIK3CA | 0.522996 | BRCA | MDA-MB-157 | GNE-317 | MVP_top60_DNA+RNA_seed | CDK1 | 0.141477 | BRCA | MDA-MB-361 | RO-3306 |
| MVP_top60_DNA_seed | HDAC1 | 0.52851 | BRCA | HCC1143 | Romidepsin | MVP_top60_DNA+RNA_seed | MDM2 | 0.141721 | BRCA | CAL-85-1 | Serdemetan |
| MVP_top60_DNA_seed | HDAC2 | 0.52851 | BRCA | HCC1143 | Romidepsin | MVP_top60_DNA+RNA_seed | AKT1 | 0.142988 | BRCA | BT-474 | AZD5363 |
| MVP_top60_DNA_seed | AKT1 | 0.529439 | BRCA | MDA-MB-436 | Afuresertib | MVP_top60_DNA+RNA_seed | EGFR | 0.145195 | BRCA | HCC1187 | Osimertinib |
| MVP_top60_DNA_seed | PIK3CA | 0.530461 | BRCA | HCC38 | Alpelisib | MVP_top60_DNA+RNA_seed | AKT1 | 0.146456 | BRCA | HCC1599 | Ipatasertib |
| MVP_top60_DNA_seed | AKT1 | 0.532261 | BRCA | HCC1187 | AZD5363 | MVP_top60_DNA+RNA_seed | CDK2 | 0.147343 | BRCA | MDA-MB-415 | AZD5438 |
| MVP_top60_DNA_seed | CDK2 | 0.532401 | BRCA | HCC2157 | MK-8776 | MVP_top60_DNA+RNA_seed | CDK2 | 0.149829 | BRCA | COLO-824 | AZD5438 |

| | | | | | | | | | | | |
|---|---|---|---|---|---|---|---|---|---|---|---|
| MVP_top60_DNA_seed | PIK3CA | 0.533957 | BRCA | HCC1419 | Alpelisib | MVP_top60_DNA+RNA_seed | MAPK1 | 0.150034 | BRCA | CAL-148 | VX-11e |
| MVP_top60_DNA_seed | EP300 | 0.534131 | BRCA | EFM-19 | SGC-CBP30 | MVP_top60_DNA+RNA_seed | AKT1 | 0.150584 | BRCA | DU-4475 | Afuresertib |
| MVP_top60_DNA_seed | CREBBP | 0.534131 | BRCA | EFM-19 | SGC-CBP30 | MVP_top60_DNA+RNA_seed | CDK2 | 0.151225 | BRCA | EVSA-T | MK-8776 |
| MVP_top60_DNA_seed | PIK3CA | 0.535915 | BRCA | HCC38 | AZD8186 | MVP_top60_DNA+RNA_seed | HSP90AA1 | 0.152291 | BRCA | AU565 | Luminespib |
| MVP_top60_DNA_seed | MAPK3 | 0.535941 | BRCA | EFM-19 | SCH772984 | MVP_top60_DNA+RNA_seed | HSP90AB1 | 0.152291 | BRCA | AU565 | Luminespib |
| MVP_top60_DNA_seed | MAPK1 | 0.535941 | BRCA | EFM-19 | SCH772984 | MVP_top60_DNA+RNA_seed | TP53 | 0.15232 | BRCA | MDA-MB-231 | MIRA-1 |
| MVP_top60_DNA_seed | ATM | 0.537588 | BRCA | MDA-MB-330 | KU-55933 | MVP_top60_DNA+RNA_seed | AKT1 | 0.152356 | BRCA | JIMT-1 | Ipatasertib |
| MVP_top60_DNA_seed | HDAC1 | 0.539044 | BRCA | BT-20 | PCI-34051 | MVP_top60_DNA+RNA_seed | MDM2 | 0.15388 | BRCA | HCC1187 | Serdemetan |
| MVP_top60_DNA_seed | PIK3CA | 0.540628 | BRCA | JIMT-1 | AZD8186 | MVP_top60_DNA+RNA_seed | MAPK3 | 0.154001 | BRCA | MDA-MB-415 | SCH772984 |
| MVP_top60_DNA_seed | PIK3CA | 0.541288 | BRCA | BT-549 | Alpelisib | MVP_top60_DNA+RNA_seed | MAPK1 | 0.154001 | BRCA | MDA-MB-415 | SCH772984 |
| MVP_top60_DNA_seed | ERBB2 | 0.543644 | BRCA | MRK-nu-1 | Lapatinib | MVP_top60_DNA+RNA_seed | MAPK3 | 0.155134 | BRCA | CAL-51 | SCH772984 |
| MVP_top60_DNA_seed | EGFR | 0.543644 | BRCA | MRK-nu-1 | Lapatinib | MVP_top60_DNA+RNA_seed | MAPK1 | 0.155134 | BRCA | CAL-51 | SCH772984 |
| MVP_top60_DNA_seed | HSP90AA1 | 0.545039 | BRCA | MFM-223 | Luminespib | MVP_top60_DNA+RNA_seed | ESR1 | 0.156679 | BRCA | CAL-51 | GDC0810 |
| MVP_top60_DNA_seed | HSP90AB1 | 0.545039 | BRCA | MFM-223 | Luminespib | MVP_top60_DNA+RNA_seed | EGFR | 0.156795 | BRCA | HCC1500 | Erlotinib |
| MVP_top60_DNA_seed | CDK2 | 0.545852 | BRCA | MDA-MB-231 | MK-8776 | MVP_top60_DNA+RNA_seed | EGFR | 0.157517 | BRCA | BT-474 | Gefitinib |
| MVP_top60_DNA_seed | ESR1 | 0.547848 | BRCA | UACC-812 | GDC0810 | MVP_top60_DNA+RNA_seed | AKT1 | 0.157924 | BRCA | HCC2218 | Ipatasertib |
| MVP_top60_DNA_seed | EGFR | 0.548339 | BRCA | BT-549 | Osimertinib | MVP_top60_DNA+RNA_seed | HSP90AA1 | 0.16055 | BRCA | OCUB-M | Tanespimycin |
| MVP_top60_DNA_seed | AKT1 | 0.549188 | BRCA | JIMT-1 | AZD5363 | MVP_top60_DNA+RNA_seed | HSP90AB1 | 0.16055 | BRCA | OCUB-M | Tanespimycin |
| MVP_top60_DNA_seed | CDK2 | 0.550147 | BRCA | BT-549 | Dinaciclib | MVP_top60_DNA+RNA_seed | EGFR | 0.16357 | BRCA | CAL-148 | Gefitinib |
| MVP_top60_DNA_seed | CDK1 | 0.550147 | BRCA | BT-549 | Dinaciclib | MVP_top60_DNA+RNA_seed | MAPK3 | 0.163814 | BRCA | OCUB-M | SCH772984 |
| MVP_top60_DNA_seed | AKT1 | 0.552365 | BRCA | BT-20 | AT13148 | MVP_top60_DNA+RNA_seed | MAPK1 | 0.163814 | BRCA | OCUB-M | SCH772984 |
| MVP_top60_DNA_seed | HDAC1 | 0.553209 | BRCA | HCC1500 | Romidepsin | MVP_top60_DNA+RNA_seed | PIK3CA | 0.163942 | BRCA | CAL-51 | AZD8186 |
| MVP_top60_DNA_seed | HDAC2 | 0.553209 | BRCA | HCC1500 | Romidepsin | MVP_top60_DNA+RNA_seed | HDAC1 | 0.165217 | BRCA | HCC2157 | PCI-34051 |
| MVP_top60_DNA_seed | SRC | 0.553834 | BRCA | HCC70 | Bosutinib | MVP_top60_DNA+RNA_seed | CDK2 | 0.165858 | BRCA | MDA-MB-453 | MK-8776 |
| MVP_top60_DNA_seed | AKT1 | 0.556612 | BRCA | MDA-MB-157 | AZD5363 | MVP_top60_DNA+RNA_seed | EGFR | 0.169747 | BRCA | MDA-MB-453 | Erlotinib |
| MVP_top60_DNA_seed | ERBB2 | 0.556828 | BRCA | EFM-19 | Lapatinib | MVP_top60_DNA+RNA_seed | MAPK3 | 0.171978 | BRCA | CAMA-1 | Ulixertinib |
| MVP_top60_DNA_seed | EGFR | 0.556828 | BRCA | EFM-19 | Lapatinib | MVP_top60_DNA+RNA_seed | MAPK1 | 0.171978 | BRCA | CAMA-1 | Ulixertinib |
| MVP_top60_DNA_seed | TP53 | 0.557608 | BRCA | ZR-75-30 | MIRA-1 | MVP_top60_DNA+RNA_seed | PLK1 | 0.176091 | BRCA | HCC1143 | BI-2536 |
| MVP_top60_DNA_seed | ERBB2 | 0.559887 | BRCA | HCC1599 | Afatinib | MVP_top60_DNA+RNA_seed | AKT1 | 0.178007 | BRCA | CAL-120 | Afuresertib |
| MVP_top60_DNA_seed | EGFR | 0.559887 | BRCA | HCC1599 | Afatinib | MVP_top60_DNA+RNA_seed | MDM2 | 0.178802 | BRCA | Hs-578-T | Nutlin-3a (-) |
| MVP_top60_DNA_seed | PIK3CA | 0.561574 | BRCA | MFM-223 | Buparlisib | MVP_top60_DNA+RNA_seed | MDM2 | 0.181521 | BRCA | MDA-MB-231 | Serdemetan |
| MVP_top60_DNA_seed | AKT1 | 0.561979 | BRCA | BT-549 | GSK2110183B | MVP_top60_DNA+RNA_seed | EGFR | 0.182481 | BRCA | CAL-51 | Sapitinib |
| MVP_top60_DNA_seed | MAPK3 | 0.562695 | BRCA | YMB-1-E | Ulixertinib | MVP_top60_DNA+RNA_seed | ERBB2 | 0.182481 | BRCA | CAL-51 | Sapitinib |
| MVP_top60_DNA_seed | MAPK1 | 0.562695 | BRCA | YMB-1-E | Ulixertinib | MVP_top60_DNA+RNA_seed | ERBB3 | 0.182481 | BRCA | CAL-51 | Sapitinib |
| MVP_top60_DNA_seed | PIK3CA | 0.56285 | BRCA | MDA-MB-231 | Buparlisib | MVP_top60_DNA+RNA_seed | MAPK3 | 0.185853 | BRCA | AU565 | SCH772984 |
| MVP_top60_DNA_seed | EGFR | 0.565418 | BRCA | CAL-85-1 | Gefitinib | MVP_top60_DNA+RNA_seed | MAPK1 | 0.185853 | BRCA | AU565 | SCH772984 |
| MVP_top60_DNA_seed | AKT1 | 0.565608 | BRCA | MRK-nu-1 | Ipatasertib | MVP_top60_DNA+RNA_seed | EGFR | 0.187275 | BRCA | HCC1187 | AZD3759 |
| MVP_top60_DNA_seed | HDAC1 | 0.566626 | BRCA | HCC1937 | Entinostat | MVP_top60_DNA+RNA_seed | HSP90AA1 | 0.187992 | BRCA | BT-474 | Luminespib |
| MVP_top60_DNA_seed | AKT1 | 0.568882 | BRCA | ZR-75-30 | AT13148 | MVP_top60_DNA+RNA_seed | HSP90AB1 | 0.187992 | BRCA | BT-474 | Luminespib |
| MVP_top60_DNA_seed | AKT1 | 0.570058 | BRCA | HCC1143 | MK-2206 | MVP_top60_DNA+RNA_seed | PIK3CA | 0.19012 | BRCA | MDA-MB-436 | AZD8186 |
| MVP_top60_DNA_seed | HDAC1 | 0.570452 | BRCA | MFM-223 | Romidepsin | MVP_top60_DNA+RNA_seed | MDM2 | 0.192487 | BRCA | HCC1954 | Serdemetan |

| | | | | | | | | | | | |
|---|---|---|---|---|---|---|---|---|---|---|---|
| MVP_top60_DNA_seed | HDAC2 | 0.570452 | BRCA | MFM-223 | Romidepsin | MVP_top60_DNA+RNA_seed | CDK1 | 0.192632 | BRCA | YMB-1-E | RO-3306 |
| MVP_top60_DNA_seed | EGFR | 0.570613 | BRCA | HCC1395 | AZD3759 | MVP_top60_DNA+RNA_seed | AKT1 | 0.196025 | BRCA | MRK-nu-1 | GSK2110183B |
| MVP_top60_DNA_seed | CDK2 | 0.572848 | BRCA | CAMA-1 | Dinaciclib | MVP_top60_DNA+RNA_seed | HDAC1 | 0.197657 | BRCA | HCC1395 | Entinostat |
| MVP_top60_DNA_seed | CDK1 | 0.572848 | BRCA | CAMA-1 | Dinaciclib | MVP_top60_DNA+RNA_seed | MAPK3 | 0.197716 | BRCA | HCC1500 | SCH772984 |
| MVP_top60_DNA_seed | HSP90AA1 | 0.573333 | BRCA | UACC-893 | CCT-018159 | MVP_top60_DNA+RNA_seed | MAPK1 | 0.197716 | BRCA | HCC1500 | SCH772984 |
| MVP_top60_DNA_seed | HSP90AB1 | 0.573333 | BRCA | UACC-893 | CCT-018159 | MVP_top60_DNA+RNA_seed | MAPK1 | 0.198648 | BRCA | CAL-85-1 | VX-11e |
| MVP_top60_DNA_seed | HDAC1 | 0.573789 | BRCA | HCC1954 | PCI-34051 | MVP_top60_DNA+RNA_seed | MAPK3 | 0.20069 | BRCA | HCC70 | Ulixertinib |
| MVP_top60_DNA_seed | EGFR | 0.574579 | BRCA | MDA-MB-231 | Gefitinib | MVP_top60_DNA+RNA_seed | MAPK1 | 0.20069 | BRCA | HCC70 | Ulixertinib |
| MVP_top60_DNA_seed | AKT1 | 0.575969 | BRCA | HCC1143 | AT13148 | MVP_top60_DNA+RNA_seed | AKT1 | 0.203556 | BRCA | HCC1419 | Uprosertib |
| MVP_top60_DNA_seed | CDK1 | 0.579946 | BRCA | HCC1937 | RO-3306 | MVP_top60_DNA+RNA_seed | EGFR | 0.20404 | BRCA | EFM-19 | Afatinib |
| MVP_top60_DNA_seed | AKT1 | 0.582031 | BRCA | BT-483 | Uprosertib | MVP_top60_DNA+RNA_seed | ERBB2 | 0.20404 | BRCA | EFM-19 | Afatinib |
| MVP_top60_DNA_seed | AKT1 | 0.58658 | BRCA | HDQ-P1 | GSK2110183B | MVP_top60_DNA+RNA_seed | CDK2 | 0.204974 | BRCA | HCC1143 | MK-8776 |
| MVP_top60_DNA_seed | HSP90AA1 | 0.587856 | BRCA | MDA-MB-436 | Luminespib | MVP_top60_DNA+RNA_seed | EGFR | 0.205415 | BRCA | HCC1954 | Afatinib |
| MVP_top60_DNA_seed | HSP90AB1 | 0.587856 | BRCA | MDA-MB-436 | Luminespib | MVP_top60_DNA+RNA_seed | ERBB2 | 0.205415 | BRCA | HCC1954 | Afatinib |
| MVP_top60_DNA_seed | ERBB2 | 0.591309 | BRCA | CAL-51 | Afatinib | MVP_top60_DNA+RNA_seed | AKT1 | 0.206739 | BRCA | DU-4475 | AZD5363 |
| MVP_top60_DNA_seed | EGFR | 0.591309 | BRCA | CAL-51 | Afatinib | MVP_top60_DNA+RNA_seed | PLK1 | 0.207684 | BRCA | HDQ-P1 | MK-1775 |
| MVP_top60_DNA_seed | MDM2 | 0.593548 | BRCA | BT-549 | Serdemetan | MVP_top60_DNA+RNA_seed | MAPK3 | 0.208441 | BRCA | HCC1419 | Ulixertinib |
| MVP_top60_DNA_seed | EGFR | 0.596675 | BRCA | HCC1395 | Gefitinib | MVP_top60_DNA+RNA_seed | MAPK1 | 0.208441 | BRCA | HCC1419 | Ulixertinib |
| MVP_top60_DNA_seed | TP53 | 0.597125 | BRCA | COLO-824 | MIRA-1 | MVP_top60_DNA+RNA_seed | EP300 | 0.210694 | BRCA | CAL-85-1 | SGC-CBP30 |
| MVP_top60_DNA_seed | SRC | 0.597296 | BRCA | HCC1599 | Bosutinib | MVP_top60_DNA+RNA_seed | EGFR | 0.211014 | BRCA | HCC70 | Osimertinib |
| MVP_top60_DNA_seed | EGFR | 0.599271 | BRCA | HCC38 | AZD3759 | MVP_top60_DNA+RNA_seed | CDK1 | 0.212461 | BRCA | HCC1143 | RO-3306 |
| MVP_top60_DNA_seed | MAPK3 | 0.599324 | BRCA | Hs-578-T | Ulixertinib | MVP_top60_DNA+RNA_seed | PLK1 | 0.212815 | BRCA | EVSA-T | MK-1775 |
| MVP_top60_DNA_seed | MAPK1 | 0.599324 | BRCA | Hs-578-T | Ulixertinib | MVP_top60_DNA+RNA_seed | PIK3CA | 0.213926 | BRCA | MDA-MB-157 | Buparlisib |
| MVP_top60_DNA_seed | HDAC1 | 0.600131 | BRCA | MDA-MB-436 | Romidepsin | MVP_top60_DNA+RNA_seed | EGFR | 0.215344 | BRCA | MDA-MB-415 | Erlotinib |
| MVP_top60_DNA_seed | HDAC2 | 0.600131 | BRCA | MDA-MB-436 | Romidepsin | MVP_top60_DNA+RNA_seed | CDK1 | 0.21569 | BRCA | HCC1599 | RO-3306 |
| MVP_top60_DNA_seed | EGFR | 0.600241 | BRCA | MDA-MB-231 | Osimertinib | MVP_top60_DNA+RNA_seed | ATM | 0.217352 | BRCA | MDA-MB-436 | KU-55933 |
| MVP_top60_DNA_seed | PIK3CA | 0.60026 | BRCA | ZR-75-30 | AZD8186 | MVP_top60_DNA+RNA_seed | HDAC1 | 0.218582 | BRCA | T47D | Entinostat |
| MVP_top60_DNA_seed | PIK3CA | 0.600468 | BRCA | MCF7 | GNE-317 | MVP_top60_DNA+RNA_seed | TP53 | 0.221206 | BRCA | UACC-812 | MIRA-1 |
| MVP_top60_DNA_seed | HDAC1 | 0.600637 | BRCA | MDA-MB-415 | PCI-34051 | MVP_top60_DNA+RNA_seed | MDM2 | 0.22449 | BRCA | MDA-MB-468 | Nutlin-3a (-) |
| MVP_top60_DNA_seed | HDAC1 | 0.600773 | BRCA | JIMT-1 | Entinostat | MVP_top60_DNA+RNA_seed | PLK1 | 0.224669 | BRCA | COLO-824 | MK-1775 |
| MVP_top60_DNA_seed | HDAC1 | 0.60109 | BRCA | MDA-MB-415 | Entinostat | MVP_top60_DNA+RNA_seed | EGFR | 0.226585 | BRCA | CAL-148 | AZD3759 |
| MVP_top60_DNA_seed | ERBB2 | 0.60238 | BRCA | HCC1428 | Afatinib | MVP_top60_DNA+RNA_seed | PIK3CA | 0.227552 | BRCA | CAL-85-1 | Buparlisib |
| MVP_top60_DNA_seed | EGFR | 0.60238 | BRCA | HCC1428 | Afatinib | MVP_top60_DNA+RNA_seed | MDM2 | 0.230602 | BRCA | EVSA-T | Nutlin-3a (-) |
| MVP_top60_DNA_seed | MAPK3 | 0.603121 | BRCA | MDA-MB-175-VII | Ulixertinib | MVP_top60_DNA+RNA_seed | PLK1 | 0.230955 | BRCA | MDA-MB-231 | MK-1775 |
| MVP_top60_DNA_seed | MAPK1 | 0.603121 | BRCA | MDA-MB-175-VII | Ulixertinib | MVP_top60_DNA+RNA_seed | MDM2 | 0.231145 | BRCA | CAMA-1 | Nutlin-3a (-) |
| MVP_top60_DNA_seed | MAPK1 | 0.60327 | BRCA | HCC1937 | VX-11e | MVP_top60_DNA+RNA_seed | PLK1 | 0.231566 | BRCA | MFM-223 | MK-1775 |
| MVP_top60_DNA_seed | AKT1 | 0.603419 | BRCA | HCC1500 | GSK2110183B | MVP_top60_DNA+RNA_seed | EGFR | 0.23241 | BRCA | HCC1395 | Afatinib |
| MVP_top60_DNA_seed | PIK3CA | 0.604254 | BRCA | MDA-MB-157 | Alpelisib | MVP_top60_DNA+RNA_seed | ERBB2 | 0.23241 | BRCA | HCC1395 | Afatinib |
| MVP_top60_DNA_seed | MDM2 | 0.60526 | BRCA | HCC1143 | Serdemetan | MVP_top60_DNA+RNA_seed | AKT1 | 0.232639 | BRCA | BT-549 | Uprosertib |
| MVP_top60_DNA_seed | AKT1 | 0.607114 | BRCA | MCF7 | AZD5363 | MVP_top60_DNA+RNA_seed | MDM2 | 0.234278 | BRCA | EFM-192A | Serdemetan |
| MVP_top60_DNA_seed | EGFR | 0.60732 | BRCA | MDA-MB-157 | Erlotinib | MVP_top60_DNA+RNA_seed | PLK1 | 0.237452 | BRCA | MDA-MB-468 | MK-1775 |

| | | | | | | | | | | | |
|---|---|---|---|---|---|---|---|---|---|---|---|
| MVP_top60_DNA_seed | MAPK3 | 0.610385 | BRCA | HCC1143 | Ulixertinib | MVP_top60_DNA+RNA_seed | TP53 | 0.238802 | BRCA | HCC1954 | MIRA-1 |
| MVP_top60_DNA_seed | MAPK1 | 0.610385 | BRCA | HCC1143 | Ulixertinib | MVP_top60_DNA+RNA_seed | CDK2 | 0.239184 | BRCA | CAL-148 | MK-8776 |
| MVP_top60_DNA_seed | PIK3CA | 0.611107 | BRCA | CAMA-1 | Alpelisib | MVP_top60_DNA+RNA_seed | HDAC1 | 0.240572 | BRCA | YMB-1-E | Entinostat |
| MVP_top60_DNA_seed | EP300 | 0.612053 | BRCA | BT-483 | CPI-637 | MVP_top60_DNA+RNA_seed | EGFR | 0.241128 | BRCA | HCC1599 | Osimertinib |
| MVP_top60_DNA_seed | TP53 | 0.612158 | BRCA | EFM-19 | MIRA-1 | MVP_top60_DNA+RNA_seed | EP300 | 0.2459 | BRCA | AU565 | CPI-637 |
| MVP_top60_DNA_seed | HSP90AA1 | 0.613982 | BRCA | ZR-75-30 | Tanespimycin | MVP_top60_DNA+RNA_seed | TP53 | 0.246462 | BRCA | MDA-MB-361 | MIRA-1 |
| MVP_top60_DNA_seed | HSP90AB1 | 0.613982 | BRCA | ZR-75-30 | Tanespimycin | MVP_top60_DNA+RNA_seed | MDM2 | 0.249105 | BRCA | HCC1419 | Nutlin-3a (-) |
| MVP_top60_DNA_seed | ERBB2 | 0.615211 | BRCA | CAL-85-1 | Afatinib | MVP_top60_DNA+RNA_seed | SRC | 0.251062 | BRCA | MDA-MB-415 | Bosutinib |
| MVP_top60_DNA_seed | EGFR | 0.615211 | BRCA | CAL-85-1 | Afatinib | MVP_top60_DNA+RNA_seed | MDM2 | 0.254507 | BRCA | HCC1395 | Nutlin-3a (-) |
| MVP_top60_DNA_seed | CDK2 | 0.615699 | BRCA | CAMA-1 | MK-8776 | MVP_top60_DNA+RNA_seed | EGFR | 0.254665 | BRCA | CAMA-1 | AZD3759 |
| MVP_top60_DNA_seed | MAPK3 | 0.618559 | BRCA | MDA-MB-415 | Ulixertinib | MVP_top60_DNA+RNA_seed | AKT1 | 0.25478 | BRCA | MCF7 | Afuresertib |
| MVP_top60_DNA_seed | MAPK1 | 0.618559 | BRCA | MDA-MB-415 | Ulixertinib | MVP_top60_DNA+RNA_seed | MAPK3 | 0.255539 | BRCA | MDA-MB-468 | SCH772984 |
| MVP_top60_DNA_seed | PIK3CA | 0.618825 | BRCA | HCC1187 | Buparlisib | MVP_top60_DNA+RNA_seed | MAPK1 | 0.255539 | BRCA | MDA-MB-468 | SCH772984 |
| MVP_top60_DNA_seed | AKT1 | 0.619 | BRCA | ZR-75-30 | Afuresertib | MVP_top60_DNA+RNA_seed | EGFR | 0.25621 | BRCA | HCC1395 | Sapitinib |
| MVP_top60_DNA_seed | AKT1 | 0.619932 | BRCA | MDA-MB-157 | GSK2110183B | MVP_top60_DNA+RNA_seed | ERBB2 | 0.25621 | BRCA | HCC1395 | Sapitinib |
| MVP_top60_DNA_seed | MAPK3 | 0.621142 | BRCA | CAMA-1 | Ulixertinib | MVP_top60_DNA+RNA_seed | ERBB3 | 0.25621 | BRCA | HCC1395 | Sapitinib |
| MVP_top60_DNA_seed | MAPK1 | 0.621142 | BRCA | CAMA-1 | Ulixertinib | MVP_top60_DNA+RNA_seed | EP300 | 0.256755 | BRCA | EVSA-T | CPI-637 |
| MVP_top60_DNA_seed | HDAC1 | 0.62297 | BRCA | HCC38 | Romidepsin | MVP_top60_DNA+RNA_seed | MAPK3 | 0.258694 | BRCA | CAL-120 | Ulixertinib |
| MVP_top60_DNA_seed | HDAC2 | 0.62297 | BRCA | HCC38 | Romidepsin | MVP_top60_DNA+RNA_seed | MAPK1 | 0.258694 | BRCA | CAL-120 | Ulixertinib |
| MVP_top60_DNA_seed | MDM2 | 0.623686 | BRCA | EFM-19 | Nutlin-3a (-) | MVP_top60_DNA+RNA_seed | EGFR | 0.259729 | BRCA | YMB-1-E | AZD3759 |
| MVP_top60_DNA_seed | MAPK1 | 0.628375 | BRCA | MDA-MB-330 | VX-11e | MVP_top60_DNA+RNA_seed | CDK1 | 0.260846 | BRCA | MDA-MB-175-VII | RO-3306 |
| MVP_top60_DNA_seed | SRC | 0.628974 | BRCA | BT-549 | Bosutinib | MVP_top60_DNA+RNA_seed | EGFR | 0.262971 | BRCA | BT-20 | Gefitinib |
| MVP_top60_DNA_seed | AKT1 | 0.631516 | BRCA | ZR-75-30 | Uprosertib | MVP_top60_DNA+RNA_seed | EP300 | 0.263757 | BRCA | MDA-MB-361 | SGC-CBP30 |
| MVP_top60_DNA_seed | AKT1 | 0.631719 | BRCA | MDA-MB-231 | GSK2110183B | MVP_top60_DNA+RNA_seed | HSP90AA1 | 0.264808 | BRCA | HCC1569 | Elesclomol |
| MVP_top60_DNA_seed | EP300 | 0.634874 | BRCA | UACC-812 | CPI-637 | MVP_top60_DNA+RNA_seed | HSP90AB1 | 0.264808 | BRCA | HCC1569 | Elesclomol |
| MVP_top60_DNA_seed | HDAC1 | 0.644273 | BRCA | MCF7 | Romidepsin | MVP_top60_DNA+RNA_seed | CDK2 | 0.265977 | BRCA | JIMT-1 | AZD5438 |
| MVP_top60_DNA_seed | HDAC2 | 0.644273 | BRCA | MCF7 | Romidepsin | MVP_top60_DNA+RNA_seed | HDAC1 | 0.266299 | BRCA | T47D | Romidepsin |
| MVP_top60_DNA_seed | ATM | 0.645282 | BRCA | EFM-192A | KU-55933 | MVP_top60_DNA+RNA_seed | HDAC2 | 0.266299 | BRCA | T47D | Romidepsin |
| MVP_top60_DNA_seed | MDM2 | 0.646593 | BRCA | MDA-MB-361 | Nutlin-3a (-) | MVP_top60_DNA+RNA_seed | CDK2 | 0.266426 | BRCA | OCUB-M | MK-8776 |
| MVP_top60_DNA_seed | CDK2 | 0.647108 | BRCA | EFM-192A | AZD5438 | MVP_top60_DNA+RNA_seed | EP300 | 0.266513 | BRCA | MDA-MB-330 | SGC-CBP30 |
| MVP_top60_DNA_seed | MAPK3 | 0.647704 | BRCA | CAMA-1 | SCH772984 | MVP_top60_DNA+RNA_seed | EGFR | 0.26705 | BRCA | CAL-51 | AZD3759 |
| MVP_top60_DNA_seed | MAPK1 | 0.647704 | BRCA | CAMA-1 | SCH772984 | MVP_top60_DNA+RNA_seed | HDAC1 | 0.268117 | BRCA | HCC70 | Romidepsin |
| MVP_top60_DNA_seed | AKT1 | 0.651384 | BRCA | ZR-75-30 | AZD5363 | MVP_top60_DNA+RNA_seed | HDAC2 | 0.268117 | BRCA | HCC70 | Romidepsin |
| MVP_top60_DNA_seed | EGFR | 0.651455 | BRCA | YMB-1-E | Gefitinib | MVP_top60_DNA+RNA_seed | PIK3CA | 0.268441 | BRCA | YMB-1-E | Alpelisib |
| MVP_top60_DNA_seed | MAPK3 | 0.652549 | BRCA | HCC1937 | SCH772984 | MVP_top60_DNA+RNA_seed | MAPK3 | 0.270589 | BRCA | AU565 | Ulixertinib |
| MVP_top60_DNA_seed | MAPK1 | 0.652549 | BRCA | HCC1937 | SCH772984 | MVP_top60_DNA+RNA_seed | MAPK1 | 0.270589 | BRCA | AU565 | Ulixertinib |
| MVP_top60_DNA_seed | EP300 | 0.654191 | BRCA | HCC1143 | SGC-CBP30 | MVP_top60_DNA+RNA_seed | PIK3CA | 0.273017 | BRCA | BT-483 | Alpelisib |
| MVP_top60_DNA_seed | CREBBP | 0.654191 | BRCA | HCC1143 | SGC-CBP30 | MVP_top60_DNA+RNA_seed | HDAC1 | 0.273117 | BRCA | CAL-51 | Romidepsin |
| MVP_top60_DNA_seed | CDK1 | 0.655792 | BRCA | MFM-223 | RO-3306 | MVP_top60_DNA+RNA_seed | HDAC2 | 0.273117 | BRCA | CAL-51 | Romidepsin |
| MVP_top60_DNA_seed | MAPK3 | 0.656012 | BRCA | BT-549 | SCH772984 | MVP_top60_DNA+RNA_seed | EGFR | 0.274221 | BRCA | HCC1954 | Gefitinib |
| MVP_top60_DNA_seed | MAPK1 | 0.656012 | BRCA | BT-549 | SCH772984 | MVP_top60_DNA+RNA_seed | PIK3CA | 0.274275 | BRCA | CAL-120 | Buparlisib |

| | | | | | | | | | | | |
|---|---|---|---|---|---|---|---|---|---|---|---|
| MVP_top60_DNA_seed | EGFR | 0.656821 | BRCA | EFM-192A | Gefitinib | MVP_top60_DNA+RNA_seed | MAPK3 | 0.277716 | BRCA | MDA-MB-415 | Ulixertinib |
| MVP_top60_DNA_seed | CDK2 | 0.660289 | BRCA | EFM-19 | AZD5438 | MVP_top60_DNA+RNA_seed | MAPK1 | 0.277716 | BRCA | MDA-MB-415 | Ulixertinib |
| MVP_top60_DNA_seed | ERBB2 | 0.662206 | BRCA | YMB-1-E | Sapitinib | MVP_top60_DNA+RNA_seed | MAPK1 | 0.279317 | BRCA | HCC1395 | VX-11e |
| MVP_top60_DNA_seed | EGFR | 0.662206 | BRCA | YMB-1-E | Sapitinib | MVP_top60_DNA+RNA_seed | ATM | 0.279826 | BRCA | MCF7 | KU-55933 |
| MVP_top60_DNA_seed | ERBB3 | 0.662206 | BRCA | YMB-1-E | Sapitinib | MVP_top60_DNA+RNA_seed | PIK3CA | 0.280119 | BRCA | DU-4475 | Alpelisib |
| MVP_top60_DNA_seed | MDM2 | 0.664847 | BRCA | EFM-192A | Nutlin-3a (-) | MVP_top60_DNA+RNA_seed | ESR1 | 0.281353 | BRCA | JIMT-1 | GDC0810 |
| MVP_top60_DNA_seed | HSP90AA1 | 0.666444 | BRCA | EFM-19 | Tanespimycin | MVP_top60_DNA+RNA_seed | MAPK3 | 0.282012 | BRCA | MDA-MB-361 | Ulixertinib |
| MVP_top60_DNA_seed | HSP90AB1 | 0.666444 | BRCA | EFM-19 | Tanespimycin | MVP_top60_DNA+RNA_seed | MAPK1 | 0.282012 | BRCA | MDA-MB-361 | Ulixertinib |
| MVP_top60_DNA_seed | AKT1 | 0.667264 | BRCA | JIMT-1 | Uprosertib | MVP_top60_DNA+RNA_seed | MAPK3 | 0.282838 | BRCA | MFM-223 | Ulixertinib |
| MVP_top60_DNA_seed | TP53 | 0.671396 | BRCA | EFM-192A | MIRA-1 | MVP_top60_DNA+RNA_seed | MAPK1 | 0.282838 | BRCA | MFM-223 | Ulixertinib |
| MVP_top60_DNA_seed | HSP90AA1 | 0.673884 | BRCA | EFM-19 | Luminespib | MVP_top60_DNA+RNA_seed | EGFR | 0.283823 | BRCA | JIMT-1 | Erlotinib |
| MVP_top60_DNA_seed | HSP90AB1 | 0.673884 | BRCA | EFM-19 | Luminespib | MVP_top60_DNA+RNA_seed | PIK3CA | 0.286683 | BRCA | BT-474 | AZD8186 |
| MVP_top60_DNA_seed | EGFR | 0.67533 | BRCA | JIMT-1 | Gefitinib | MVP_top60_DNA+RNA_seed | EGFR | 0.288612 | BRCA | YMB-1-E | Osimertinib |
| MVP_top60_DNA_seed | HDAC1 | 0.675533 | BRCA | HCC1428 | PCI-34051 | MVP_top60_DNA+RNA_seed | MAPK3 | 0.289274 | BRCA | HCC2218 | Ulixertinib |
| MVP_top60_DNA_seed | MAPK1 | 0.67569 | BRCA | JIMT-1 | VX-11e | MVP_top60_DNA+RNA_seed | MAPK1 | 0.289274 | BRCA | HCC2218 | Ulixertinib |
| MVP_top60_DNA_seed | AKT1 | 0.675987 | BRCA | HCC38 | Afuresertib | MVP_top60_DNA+RNA_seed | ATM | 0.291028 | BRCA | HCC1500 | KU-55933 |
| MVP_top60_DNA_seed | PIK3CA | 0.678447 | BRCA | MDA-MB-231 | Alpelisib | MVP_top60_DNA+RNA_seed | EP300 | 0.292118 | BRCA | MDA-MB-231 | SGC-CBP30 |
| MVP_top60_DNA_seed | ERBB2 | 0.679063 | BRCA | BT-20 | Afatinib | MVP_top60_DNA+RNA_seed | TP53 | 0.294139 | BRCA | MDA-MB-157 | MIRA-1 |
| MVP_top60_DNA_seed | EGFR | 0.679063 | BRCA | BT-20 | Afatinib | MVP_top60_DNA+RNA_seed | ESR1 | 0.295298 | BRCA | HDQ-P1 | GDC0810 |
| MVP_top60_DNA_seed | EGFR | 0.67979 | BRCA | MRK-nu-1 | Osimertinib | MVP_top60_DNA+RNA_seed | MDM2 | 0.295542 | BRCA | MDA-MB-157 | Nutlin-3a (-) |
| MVP_top60_DNA_seed | MAPK1 | 0.680542 | BRCA | T47D | VX-11e | MVP_top60_DNA+RNA_seed | HSP90AA1 | 0.296061 | BRCA | BT-549 | Tanespimycin |
| MVP_top60_DNA_seed | PIK3CA | 0.685429 | BRCA | MRK-nu-1 | Alpelisib | MVP_top60_DNA+RNA_seed | HSP90AB1 | 0.296061 | BRCA | BT-549 | Tanespimycin |
| MVP_top60_DNA_seed | MAPK3 | 0.688152 | BRCA | T47D | Ulixertinib | MVP_top60_DNA+RNA_seed | AKT1 | 0.297367 | BRCA | CAL-120 | AZD5363 |
| MVP_top60_DNA_seed | MAPK1 | 0.688152 | BRCA | T47D | Ulixertinib | MVP_top60_DNA+RNA_seed | ESR1 | 0.297503 | BRCA | BT-20 | Tamoxifen |
| MVP_top60_DNA_seed | ESR1 | 0.69315 | BRCA | UACC-893 | GDC0810 | MVP_top60_DNA+RNA_seed | EGFR | 0.298848 | BRCA | HCC1428 | Lapatinib |
| MVP_top60_DNA_seed | AKT1 | 0.696434 | BRCA | HCC2218 | Afuresertib | MVP_top60_DNA+RNA_seed | ERBB2 | 0.298848 | BRCA | HCC1428 | Lapatinib |
| MVP_top60_DNA_seed | AKT1 | 0.697626 | BRCA | MDA-MB-436 | Uprosertib | MVP_top60_DNA+RNA_seed | HDAC1 | 0.299565 | BRCA | MDA-MB-415 | Romidepsin |
| MVP_top60_DNA_seed | EGFR | 0.698003 | BRCA | MDA-MB-436 | Osimertinib | MVP_top60_DNA+RNA_seed | HDAC2 | 0.299565 | BRCA | MDA-MB-415 | Romidepsin |
| MVP_top60_DNA_seed | MDM2 | 0.699228 | BRCA | MDA-MB-231 | Nutlin-3a (-) | MVP_top60_DNA+RNA_seed | EGFR | 0.300245 | BRCA | BT-549 | Afatinib |
| MVP_top60_DNA_seed | MAPK3 | 0.699837 | BRCA | EVSA-T | SCH772984 | MVP_top60_DNA+RNA_seed | ERBB2 | 0.300245 | BRCA | BT-549 | Afatinib |
| MVP_top60_DNA_seed | MAPK1 | 0.699837 | BRCA | EVSA-T | SCH772984 | MVP_top60_DNA+RNA_seed | ESR1 | 0.303056 | BRCA | MFM-223 | Tamoxifen |
| MVP_top60_DNA_seed | SRC | 0.700187 | BRCA | COLO-824 | Bosutinib | MVP_top60_DNA+RNA_seed | EP300 | 0.30463 | BRCA | HCC70 | CPI-637 |
| MVP_top60_DNA_seed | HSP90AA1 | 0.700301 | BRCA | HCC1143 | Tanespimycin | MVP_top60_DNA+RNA_seed | EGFR | 0.307912 | BRCA | COLO-824 | Sapitinib |
| MVP_top60_DNA_seed | HSP90AB1 | 0.700301 | BRCA | HCC1143 | Tanespimycin | MVP_top60_DNA+RNA_seed | ERBB2 | 0.307912 | BRCA | COLO-824 | Sapitinib |
| MVP_top60_DNA_seed | EGFR | 0.701297 | BRCA | Hs-578-T | Erlotinib | MVP_top60_DNA+RNA_seed | ERBB3 | 0.307912 | BRCA | COLO-824 | Sapitinib |
| MVP_top60_DNA_seed | MAPK3 | 0.702898 | BRCA | HCC1954 | Ulixertinib | MVP_top60_DNA+RNA_seed | MAPK3 | 0.308046 | BRCA | CAL-85-1 | Ulixertinib |
| MVP_top60_DNA_seed | MAPK1 | 0.702898 | BRCA | HCC1954 | Ulixertinib | MVP_top60_DNA+RNA_seed | MAPK1 | 0.308046 | BRCA | CAL-85-1 | Ulixertinib |
| MVP_top60_DNA_seed | HSP90AA1 | 0.703344 | BRCA | MDA-MB-468 | Tanespimycin | MVP_top60_DNA+RNA_seed | HDAC1 | 0.308293 | BRCA | Hs-578-T | PCI-34051 |
| MVP_top60_DNA_seed | HSP90AB1 | 0.703344 | BRCA | MDA-MB-468 | Tanespimycin | MVP_top60_DNA+RNA_seed | TP53 | 0.311105 | BRCA | UACC-893 | MIRA-1 |
| MVP_top60_DNA_seed | MDM2 | 0.704055 | BRCA | MDA-MB-361 | Serdemetan | MVP_top60_DNA+RNA_seed | CDK2 | 0.311469 | BRCA | COLO-824 | Dinaciclib |
| MVP_top60_DNA_seed | MDM2 | 0.706376 | BRCA | MDA-MB-453 | Nutlin-3a (-) | MVP_top60_DNA+RNA_seed | CDK1 | 0.311469 | BRCA | COLO-824 | Dinaciclib |

| | | | | | | | | | | | |
|---|---|---|---|---|---|---|---|---|---|---|---|
| MVP_top60_DNA_seed | EP300 | 0.706754 | BRCA | MDA-MB-361 | CPI-637 | MVP_top60_DNA+RNA_seed | EGFR | 0.312327 | BRCA | MDA-MB-231 | Sapitinib |
| MVP_top60_DNA_seed | PIK3CA | 0.706951 | BRCA | MDA-MB-468 | Alpelisib | MVP_top60_DNA+RNA_seed | ERBB2 | 0.312327 | BRCA | MDA-MB-231 | Sapitinib |
| MVP_top60_DNA_seed | EP300 | 0.707106 | BRCA | COLO-824 | SGC-CBP30 | MVP_top60_DNA+RNA_seed | ERBB3 | 0.312327 | BRCA | MDA-MB-231 | Sapitinib |
| MVP_top60_DNA_seed | CREBBP | 0.707106 | BRCA | COLO-824 | SGC-CBP30 | MVP_top60_DNA+RNA_seed | CDK1 | 0.313471 | BRCA | HCC1954 | RO-3306 |
| MVP_top60_DNA_seed | MDM2 | 0.709502 | BRCA | UACC-893 | Serdemetan | MVP_top60_DNA+RNA_seed | MAPK3 | 0.315106 | BRCA | CAL-148 | Ulixertinib |
| MVP_top60_DNA_seed | MAPK1 | 0.713842 | BRCA | MCF7 | VX-11e | MVP_top60_DNA+RNA_seed | MAPK1 | 0.315106 | BRCA | CAL-148 | Ulixertinib |
| MVP_top60_DNA_seed | CDK2 | 0.715128 | BRCA | EFM-19 | MK-8776 | MVP_top60_DNA+RNA_seed | PIK3CA | 0.315274 | BRCA | DU-4475 | AZD8186 |
| MVP_top60_DNA_seed | ERBB2 | 0.715982 | BRCA | CAL-148 | Lapatinib | MVP_top60_DNA+RNA_seed | SRC | 0.315316 | BRCA | HCC1937 | Bosutinib |
| MVP_top60_DNA_seed | EGFR | 0.715982 | BRCA | CAL-148 | Lapatinib | MVP_top60_DNA+RNA_seed | SRC | 0.315473 | BRCA | T47D | Bosutinib |
| MVP_top60_DNA_seed | CDK2 | 0.716176 | BRCA | HCC1599 | Dinaciclib | MVP_top60_DNA+RNA_seed | EGFR | 0.316304 | BRCA | HCC1187 | Gefitinib |
| MVP_top60_DNA_seed | CDK1 | 0.716176 | BRCA | HCC1599 | Dinaciclib | MVP_top60_DNA+RNA_seed | ESR1 | 0.318846 | BRCA | CAL-148 | Tamoxifen |
| MVP_top60_DNA_seed | CDK1 | 0.71654 | BRCA | EFM-192A | RO-3306 | MVP_top60_DNA+RNA_seed | PIK3CA | 0.319258 | BRCA | HCC1395 | GNE-317 |
| MVP_top60_DNA_seed | AKT1 | 0.716764 | BRCA | JIMT-1 | GSK2110183B | MVP_top60_DNA+RNA_seed | EGFR | 0.319356 | BRCA | HCC1395 | Erlotinib |
| MVP_top60_DNA_seed | AKT1 | 0.727067 | BRCA | JIMT-1 | AT13148 | MVP_top60_DNA+RNA_seed | EGFR | 0.320019 | BRCA | MRK-nu-1 | Sapitinib |
| MVP_top60_DNA_seed | MDM2 | 0.730482 | BRCA | MDA-MB-330 | Serdemetan | MVP_top60_DNA+RNA_seed | ERBB2 | 0.320019 | BRCA | MRK-nu-1 | Sapitinib |
| MVP_top60_DNA_seed | AKT1 | 0.730861 | BRCA | MDA-MB-231 | MK-2206 | MVP_top60_DNA+RNA_seed | ERBB3 | 0.320019 | BRCA | MRK-nu-1 | Sapitinib |
| MVP_top60_DNA_seed | EGFR | 0.733586 | BRCA | HCC1500 | Osimertinib | MVP_top60_DNA+RNA_seed | AKT1 | 0.320294 | BRCA | MDA-MB-231 | Uprosertib |
| MVP_top60_DNA_seed | HDAC1 | 0.744191 | BRCA | JIMT-1 | PCI-34051 | MVP_top60_DNA+RNA_seed | EGFR | 0.320661 | BRCA | EVSA-T | Afatinib |
| MVP_top60_DNA_seed | EGFR | 0.744628 | BRCA | MFM-223 | AZD3759 | MVP_top60_DNA+RNA_seed | ERBB2 | 0.320661 | BRCA | EVSA-T | Afatinib |
| MVP_top60_DNA_seed | ERBB2 | 0.744935 | BRCA | MDA-MB-436 | Sapitinib | MVP_top60_DNA+RNA_seed | EP300 | 0.321388 | BRCA | MDA-MB-415 | SGC-CBP30 |
| MVP_top60_DNA_seed | EGFR | 0.744935 | BRCA | MDA-MB-436 | Sapitinib | MVP_top60_DNA+RNA_seed | CDK2 | 0.32236 | BRCA | HCC1569 | Dinaciclib |
| MVP_top60_DNA_seed | ERBB3 | 0.744935 | BRCA | MDA-MB-436 | Sapitinib | MVP_top60_DNA+RNA_seed | CDK1 | 0.32236 | BRCA | HCC1569 | Dinaciclib |
| MVP_top60_DNA_seed | EP300 | 0.745128 | BRCA | BT-20 | CPI-637 | MVP_top60_DNA+RNA_seed | EGFR | 0.323621 | BRCA | CAMA-1 | Gefitinib |
| MVP_top60_DNA_seed | MAPK3 | 0.745682 | BRCA | MDA-MB-436 | SCH772984 | MVP_top60_DNA+RNA_seed | MDM2 | 0.325432 | BRCA | HCC2218 | Nutlin-3a (-) |
| MVP_top60_DNA_seed | MAPK1 | 0.745682 | BRCA | MDA-MB-436 | SCH772984 | MVP_top60_DNA+RNA_seed | EP300 | 0.327685 | BRCA | HCC1428 | CPI-637 |
| MVP_top60_DNA_seed | MAPK1 | 0.745888 | BRCA | Hs-578-T | VX-11e | MVP_top60_DNA+RNA_seed | ESR1 | 0.328086 | BRCA | Hs-578-T | GDC0810 |
| MVP_top60_DNA_seed | SRC | 0.746056 | BRCA | HCC1954 | Bosutinib | MVP_top60_DNA+RNA_seed | MAPK3 | 0.329549 | BRCA | EFM-192A | Ulixertinib |
| MVP_top60_DNA_seed | EGFR | 0.747607 | BRCA | HCC1395 | Osimertinib | MVP_top60_DNA+RNA_seed | MAPK1 | 0.329549 | BRCA | EFM-192A | Ulixertinib |
| MVP_top60_DNA_seed | EGFR | 0.749401 | BRCA | CAMA-1 | Erlotinib | MVP_top60_DNA+RNA_seed | AKT1 | 0.330373 | BRCA | CAL-85-1 | Afuresertib |
| MVP_top60_DNA_seed | CDK1 | 0.749464 | BRCA | MDA-MB-330 | RO-3306 | MVP_top60_DNA+RNA_seed | HDAC1 | 0.332186 | BRCA | MRK-nu-1 | PCI-34051 |
| MVP_top60_DNA_seed | AKT1 | 0.749546 | BRCA | HCC38 | Uprosertib | MVP_top60_DNA+RNA_seed | MAPK1 | 0.332632 | BRCA | MFM-223 | VX-11e |
| MVP_top60_DNA_seed | HDAC1 | 0.750636 | BRCA | HCC1143 | PCI-34051 | MVP_top60_DNA+RNA_seed | ESR1 | 0.333839 | BRCA | MDA-MB-231 | Tamoxifen |
| MVP_top60_DNA_seed | SRC | 0.75065 | BRCA | HCC1187 | Bosutinib | MVP_top60_DNA+RNA_seed | MAPK1 | 0.337024 | BRCA | CAMA-1 | VX-11e |
| MVP_top60_DNA_seed | CDK2 | 0.75272 | BRCA | MFM-223 | MK-8776 | MVP_top60_DNA+RNA_seed | EP300 | 0.337381 | BRCA | ZR-75-30 | CPI-637 |
| MVP_top60_DNA_seed | AKT1 | 0.753729 | BRCA | MDA-MB-436 | Uprosertib | MVP_top60_DNA+RNA_seed | MAPK3 | 0.338626 | BRCA | HCC70 | Ulixertinib |
| MVP_top60_DNA_seed | ERBB2 | 0.754708 | BRCA | MFM-223 | Afatinib | MVP_top60_DNA+RNA_seed | MAPK1 | 0.338626 | BRCA | HCC70 | Ulixertinib |
| MVP_top60_DNA_seed | EGFR | 0.754708 | BRCA | MFM-223 | Afatinib | MVP_top60_DNA+RNA_seed | EGFR | 0.3401 | BRCA | MFM-223 | Lapatinib |
| MVP_top60_DNA_seed | MDM2 | 0.755925 | BRCA | HCC1937 | Serdemetan | MVP_top60_DNA+RNA_seed | ERBB2 | 0.3401 | BRCA | MFM-223 | Lapatinib |
| MVP_top60_DNA_seed | HDAC1 | 0.756079 | BRCA | BT-549 | Romidepsin | MVP_top60_DNA+RNA_seed | PLK1 | 0.341469 | BRCA | CAL-85-1 | BI-2536 |
| MVP_top60_DNA_seed | HDAC2 | 0.756079 | BRCA | BT-549 | Romidepsin | MVP_top60_DNA+RNA_seed | PIK3CA | 0.342549 | BRCA | HCC1599 | GNE-317 |
| MVP_top60_DNA_seed | TP53 | 0.756287 | BRCA | HCC38 | MIRA-1 | MVP_top60_DNA+RNA_seed | CDK2 | 0.343294 | BRCA | HCC1419 | AZD5438 |

| | | | | | | | | | | | |
|---|---|---|---|---|---|---|---|---|---|---|---|
| MVP_top60_DNA_seed | HDAC1 | 0.756574 | BRCA | HCC1187 | PCI-34051 | MVP_top60_DNA+RNA_seed | HSP90AA1 | 0.346003 | BRCA | EFM-192A | Luminespib |
| MVP_top60_DNA_seed | AKT1 | 0.758038 | BRCA | MDA-MB-157 | Ipatasertib | MVP_top60_DNA+RNA_seed | HSP90AB1 | 0.346003 | BRCA | EFM-192A | Luminespib |
| MVP_top60_DNA_seed | HSP90AA1 | 0.758999 | BRCA | HCC1187 | Tanespimycin | MVP_top60_DNA+RNA_seed | ATM | 0.346457 | BRCA | CAL-51 | KU-55933 |
| MVP_top60_DNA_seed | HSP90AB1 | 0.758999 | BRCA | HCC1187 | Tanespimycin | MVP_top60_DNA+RNA_seed | MAPK3 | 0.346819 | BRCA | COLO-824 | Ulixertinib |
| MVP_top60_DNA_seed | TP53 | 0.759119 | BRCA | HCC1395 | MIRA-1 | MVP_top60_DNA+RNA_seed | MAPK1 | 0.346819 | BRCA | COLO-824 | Ulixertinib |
| MVP_top60_DNA_seed | ESR1 | 0.759629 | BRCA | HCC1395 | GDC0810 | MVP_top60_DNA+RNA_seed | HSP90AA1 | 0.348532 | BRCA | BT-474 | Tanespimycin |
| MVP_top60_DNA_seed | MAPK3 | 0.760427 | BRCA | MDA-MB-157 | SCH772984 | MVP_top60_DNA+RNA_seed | HSP90AB1 | 0.348532 | BRCA | BT-474 | Tanespimycin |
| MVP_top60_DNA_seed | MAPK1 | 0.760427 | BRCA | MDA-MB-157 | SCH772984 | MVP_top60_DNA+RNA_seed | PIK3CA | 0.349377 | BRCA | HCC38 | GNE-317 |
| MVP_top60_DNA_seed | EGFR | 0.761561 | BRCA | MDA-MB-157 | AZD3759 | MVP_top60_DNA+RNA_seed | MAPK3 | 0.35042 | BRCA | MDA-MB-231 | Ulixertinib |
| MVP_top60_DNA_seed | MAPK3 | 0.762309 | BRCA | MDA-MB-157 | Ulixertinib | MVP_top60_DNA+RNA_seed | MAPK1 | 0.35042 | BRCA | MDA-MB-231 | Ulixertinib |
| MVP_top60_DNA_seed | MAPK1 | 0.762309 | BRCA | MDA-MB-157 | Ulixertinib | MVP_top60_DNA+RNA_seed | EGFR | 0.350665 | BRCA | EVSA-T | Osimertinib |
| MVP_top60_DNA_seed | EGFR | 0.763567 | BRCA | CAL-85-1 | Erlotinib | MVP_top60_DNA+RNA_seed | EGFR | 0.350742 | BRCA | Hs-578-T | Sapitinib |
| MVP_top60_DNA_seed | HSP90AA1 | 0.763884 | BRCA | HCC1395 | Tanespimycin | MVP_top60_DNA+RNA_seed | ERBB2 | 0.350742 | BRCA | Hs-578-T | Sapitinib |
| MVP_top60_DNA_seed | HSP90AB1 | 0.763884 | BRCA | HCC1395 | Tanespimycin | MVP_top60_DNA+RNA_seed | ERBB3 | 0.350742 | BRCA | Hs-578-T | Sapitinib |
| MVP_top60_DNA_seed | EGFR | 0.77343 | BRCA | BT-20 | Osimertinib | MVP_top60_DNA+RNA_seed | EGFR | 0.351015 | BRCA | HCC1187 | Afatinib |
| MVP_top60_DNA_seed | MDM2 | 0.775007 | BRCA | ZR-75-30 | Nutlin-3a (-) | MVP_top60_DNA+RNA_seed | ERBB2 | 0.351015 | BRCA | HCC1187 | Afatinib |
| MVP_top60_DNA_seed | ESR1 | 0.775338 | BRCA | BT-549 | GDC0810 | MVP_top60_DNA+RNA_seed | HDAC1 | 0.355513 | BRCA | HDQ-P1 | Entinostat |
| MVP_top60_DNA_seed | PIK3CA | 0.776656 | BRCA | HCC1937 | GNE-317 | MVP_top60_DNA+RNA_seed | EGFR | 0.35555 | BRCA | HCC1937 | AZD3759 |
| MVP_top60_DNA_seed | PIK3CA | 0.776945 | BRCA | HCC1187 | Alpelisib | MVP_top60_DNA+RNA_seed | EGFR | 0.355994 | BRCA | MDA-MB-231 | AZD3759 |
| MVP_top60_DNA_seed | MAPK3 | 0.779847 | BRCA | JIMT-1 | Ulixertinib | MVP_top60_DNA+RNA_seed | MDM2 | 0.356087 | BRCA | OCUB-M | Nutlin-3a (-) |
| MVP_top60_DNA_seed | MAPK1 | 0.779847 | BRCA | JIMT-1 | Ulixertinib | MVP_top60_DNA+RNA_seed | PIK3CA | 0.358545 | BRCA | HCC2218 | AZD8186 |
| MVP_top60_DNA_seed | HSP90AA1 | 0.780103 | BRCA | HCC1395 | Luminespib | MVP_top60_DNA+RNA_seed | MAPK3 | 0.361365 | BRCA | HCC1954 | Ulixertinib |
| MVP_top60_DNA_seed | HSP90AB1 | 0.780103 | BRCA | HCC1395 | Luminespib | MVP_top60_DNA+RNA_seed | MAPK1 | 0.361365 | BRCA | HCC1954 | Ulixertinib |
| MVP_top60_DNA_seed | AR | 0.780504 | BRCA | UACC-893 | Bicalutamide | MVP_top60_DNA+RNA_seed | MAPK3 | 0.364817 | BRCA | HCC1395 | SCH772984 |
| MVP_top60_DNA_seed | CDK2 | 0.781068 | BRCA | MDA-MB-415 | MK-8776 | MVP_top60_DNA+RNA_seed | MAPK1 | 0.364817 | BRCA | HCC1395 | SCH772984 |
| MVP_top60_DNA_seed | MAPK3 | 0.782028 | BRCA | HCC1599 | Ulixertinib | MVP_top60_DNA+RNA_seed | HDAC1 | 0.365853 | BRCA | AU565 | Romidepsin |
| MVP_top60_DNA_seed | MAPK1 | 0.782028 | BRCA | HCC1599 | Ulixertinib | MVP_top60_DNA+RNA_seed | HDAC2 | 0.365853 | BRCA | AU565 | Romidepsin |
| MVP_top60_DNA_seed | EGFR | 0.782638 | BRCA | MDA-MB-453 | AZD3759 | MVP_top60_DNA+RNA_seed | HSP90AA1 | 0.366832 | BRCA | BT-20 | Luminespib |
| MVP_top60_DNA_seed | EGFR | 0.785247 | BRCA | T47D | Gefitinib | MVP_top60_DNA+RNA_seed | HSP90AB1 | 0.366832 | BRCA | BT-20 | Luminespib |
| MVP_top60_DNA_seed | EGFR | 0.785567 | BRCA | MDA-MB-157 | Gefitinib | MVP_top60_DNA+RNA_seed | EGFR | 0.367445 | BRCA | CAMA-1 | Sapitinib |
| MVP_top60_DNA_seed | AKT1 | 0.786773 | BRCA | MDA-MB-231 | Afuresertib | MVP_top60_DNA+RNA_seed | ERBB2 | 0.367445 | BRCA | CAMA-1 | Sapitinib |
| MVP_top60_DNA_seed | AKT1 | 0.787601 | BRCA | MDA-MB-436 | AZD5363 | MVP_top60_DNA+RNA_seed | ERBB3 | 0.367445 | BRCA | CAMA-1 | Sapitinib |
| MVP_top60_DNA_seed | SRC | 0.787765 | BRCA | HCC1428 | Bosutinib | MVP_top60_DNA+RNA_seed | MAPK3 | 0.36984 | BRCA | HCC1395 | Ulixertinib |
| MVP_top60_DNA_seed | EGFR | 0.796342 | BRCA | Hs-578-T | AZD3759 | MVP_top60_DNA+RNA_seed | MAPK1 | 0.36984 | BRCA | HCC1395 | Ulixertinib |
| MVP_top60_DNA_seed | MDM2 | 0.79802 | BRCA | Hs-578-T | Serdemetan | MVP_top60_DNA+RNA_seed | AKT1 | 0.370756 | BRCA | HCC1937 | AZD5363 |
| MVP_top60_DNA_seed | EGFR | 0.799129 | BRCA | MCF7 | Osimertinib | MVP_top60_DNA+RNA_seed | CDK2 | 0.373398 | BRCA | T47D | AZD5438 |
| MVP_top60_DNA_seed | AKT1 | 0.803402 | BRCA | HCC38 | GSK2110183B | MVP_top60_DNA+RNA_seed | PIK3CA | 0.373971 | BRCA | BT-20 | AZD8186 |
| MVP_top60_DNA_seed | AKT1 | 0.804515 | BRCA | CAL-120 | Ipatasertib | MVP_top60_DNA+RNA_seed | MAPK1 | 0.374475 | BRCA | HCC70 | VX-11e |
| MVP_top60_DNA_seed | MAPK3 | 0.810001 | BRCA | MDA-MB-436 | Ulixertinib | MVP_top60_DNA+RNA_seed | MAPK3 | 0.377161 | BRCA | HCC1419 | SCH772984 |
| MVP_top60_DNA_seed | MAPK1 | 0.810001 | BRCA | MDA-MB-436 | Ulixertinib | MVP_top60_DNA+RNA_seed | MAPK1 | 0.377161 | BRCA | HCC1419 | SCH772984 |
| MVP_top60_DNA_seed | MAPK3 | 0.811564 | BRCA | OCUB-M | Ulixertinib | MVP_top60_DNA+RNA_seed | SRC | 0.379868 | BRCA | AU565 | Bosutinib |

| | | | | | | | | | | | |
|---|---|---|---|---|---|---|---|---|---|---|---|
| MVP_top60_DNA_seed | MAPK1 | 0.811564 | BRCA | OCUB-M | Ulixertinib | MVP_top60_DNA+RNA_seed | MDM2 | 0.381394 | BRCA | HCC202 | Nutlin-3a (-) |
| MVP_top60_DNA_seed | AKT1 | 0.811948 | BRCA | T47D | AT13148 | MVP_top60_DNA+RNA_seed | CDK2 | 0.38377 | BRCA | CAL-120 | Dinaciclib |
| MVP_top60_DNA_seed | ESR1 | 0.814043 | BRCA | MFM-223 | GDC0810 | MVP_top60_DNA+RNA_seed | CDK1 | 0.38377 | BRCA | CAL-120 | Dinaciclib |
| MVP_top60_DNA_seed | EP300 | 0.815785 | BRCA | HCC1954 | CPI-637 | MVP_top60_DNA+RNA_seed | CDK1 | 0.383838 | BRCA | CAL-85-1 | RO-3306 |
| MVP_top60_DNA_seed | EGFR | 0.817906 | BRCA | JIMT-1 | AZD3759 | MVP_top60_DNA+RNA_seed | EGFR | 0.383858 | BRCA | HCC70 | Afatinib |
| MVP_top60_DNA_seed | AKT1 | 0.818369 | BRCA | MDA-MB-157 | Uprosertib | MVP_top60_DNA+RNA_seed | ERBB2 | 0.383858 | BRCA | HCC70 | Afatinib |
| MVP_top60_DNA_seed | MAPK3 | 0.821243 | BRCA | BT-20 | Ulixertinib | MVP_top60_DNA+RNA_seed | MDM2 | 0.38466 | BRCA | HDQ-P1 | Serdemetan |
| MVP_top60_DNA_seed | MAPK1 | 0.821243 | BRCA | BT-20 | Ulixertinib | MVP_top60_DNA+RNA_seed | ESR1 | 0.386212 | BRCA | HCC2218 | Tamoxifen |
| MVP_top60_DNA_seed | EP300 | 0.821278 | BRCA | EFM-192A | CPI-637 | MVP_top60_DNA+RNA_seed | CDK2 | 0.388468 | BRCA | JIMT-1 | MK-8776 |
| MVP_top60_DNA_seed | AKT1 | 0.826809 | BRCA | HCC1500 | AZD5363 | MVP_top60_DNA+RNA_seed | ATM | 0.388842 | BRCA | HCC1599 | KU-55933 |
| MVP_top60_DNA_seed | HSP90AA1 | 0.827201 | BRCA | HCC1599 | Tanespimycin | MVP_top60_DNA+RNA_seed | TP53 | 0.389817 | BRCA | CAL-51 | MIRA-1 |
| MVP_top60_DNA_seed | HSP90AB1 | 0.827201 | BRCA | HCC1599 | Tanespimycin | MVP_top60_DNA+RNA_seed | CDK2 | 0.390245 | BRCA | MDA-MB-330 | AZD5438 |
| MVP_top60_DNA_seed | MDM2 | 0.827641 | BRCA | HCC1937 | Nutlin-3a (-) | MVP_top60_DNA+RNA_seed | HSP90AA1 | 0.390655 | BRCA | T47D | Luminespib |
| MVP_top60_DNA_seed | HSP90AA1 | 0.828761 | BRCA | CAMA-1 | Tanespimycin | MVP_top60_DNA+RNA_seed | HSP90AB1 | 0.390655 | BRCA | T47D | Luminespib |
| MVP_top60_DNA_seed | HSP90AB1 | 0.828761 | BRCA | CAMA-1 | Tanespimycin | MVP_top60_DNA+RNA_seed | EP300 | 0.393664 | BRCA | JIMT-1 | CPI-637 |
| MVP_top60_DNA_seed | EP300 | 0.829052 | BRCA | MFM-223 | CPI-637 | MVP_top60_DNA+RNA_seed | TP53 | 0.393963 | BRCA | YMB-1-E | MIRA-1 |
| MVP_top60_DNA_seed | MAPK3 | 0.831668 | BRCA | MRK-nu-1 | Ulixertinib | MVP_top60_DNA+RNA_seed | MAPK1 | 0.394005 | BRCA | MRK-nu-1 | VX-11e |
| MVP_top60_DNA_seed | MAPK1 | 0.831668 | BRCA | MRK-nu-1 | Ulixertinib | MVP_top60_DNA+RNA_seed | MAPK3 | 0.395886 | BRCA | COLO-824 | Ulixertinib |
| MVP_top60_DNA_seed | SRC | 0.832036 | BRCA | MDA-MB-453 | Bosutinib | MVP_top60_DNA+RNA_seed | MAPK1 | 0.395886 | BRCA | COLO-824 | Ulixertinib |
| MVP_top60_DNA_seed | ESR1 | 0.832082 | BRCA | CAL-120 | Tamoxifen | MVP_top60_DNA+RNA_seed | PIK3CA | 0.396797 | BRCA | OCUB-M | AZD8186 |
| MVP_top60_DNA_seed | AKT1 | 0.833884 | BRCA | MDA-MB-157 | Uprosertib | MVP_top60_DNA+RNA_seed | HSP90AA1 | 0.399075 | BRCA | CAMA-1 | Luminespib |
| MVP_top60_DNA_seed | ESR1 | 0.834192 | BRCA | HCC1599 | Tamoxifen | MVP_top60_DNA+RNA_seed | HSP90AB1 | 0.399075 | BRCA | CAMA-1 | Luminespib |
| MVP_top60_DNA_seed | SRC | 0.83669 | BRCA | CAMA-1 | Bosutinib | MVP_top60_DNA+RNA_seed | MAPK3 | 0.400136 | BRCA | T47D | SCH772984 |
| MVP_top60_DNA_seed | EGFR | 0.837306 | BRCA | Hs-578-T | Gefitinib | MVP_top60_DNA+RNA_seed | MAPK1 | 0.400136 | BRCA | T47D | SCH772984 |
| MVP_top60_DNA_seed | EP300 | 0.842184 | BRCA | HCC1187 | CPI-637 | MVP_top60_DNA+RNA_seed | AR | 0.401465 | BRCA | HCC1599 | Bicalutamide |
| MVP_top60_DNA_seed | ERBB2 | 0.843482 | BRCA | HCC1500 | Lapatinib | MVP_top60_DNA+RNA_seed | MAPK3 | 0.401974 | BRCA | Hs-578-T | Ulixertinib |
| MVP_top60_DNA_seed | EGFR | 0.843482 | BRCA | HCC1500 | Lapatinib | MVP_top60_DNA+RNA_seed | MAPK1 | 0.401974 | BRCA | Hs-578-T | Ulixertinib |
| MVP_top60_DNA_seed | MDM2 | 0.846014 | BRCA | HCC70 | Nutlin-3a (-) | MVP_top60_DNA+RNA_seed | HDAC1 | 0.402384 | BRCA | HDQ-P1 | PCI-34051 |
| MVP_top60_DNA_seed | AKT1 | 0.849992 | BRCA | JIMT-1 | Afuresertib | MVP_top60_DNA+RNA_seed | MDM2 | 0.402435 | BRCA | UACC-812 | Serdemetan |
| MVP_top60_DNA_seed | ESR1 | 0.850311 | BRCA | HCC1569 | GDC0810 | MVP_top60_DNA+RNA_seed | PIK3CA | 0.405435 | BRCA | UACC-812 | Buparlisib |
| MVP_top60_DNA_seed | MAPK3 | 0.851578 | BRCA | MDA-MB-361 | SCH772984 | MVP_top60_DNA+RNA_seed | CDK1 | 0.405889 | BRCA | ZR-75-30 | RO-3306 |
| MVP_top60_DNA_seed | MAPK1 | 0.851578 | BRCA | MDA-MB-361 | SCH772984 | MVP_top60_DNA+RNA_seed | MAPK3 | 0.40609 | BRCA | T47D | Ulixertinib |
| MVP_top60_DNA_seed | MAPK3 | 0.854308 | BRCA | HCC1937 | Ulixertinib | MVP_top60_DNA+RNA_seed | MAPK1 | 0.40609 | BRCA | T47D | Ulixertinib |
| MVP_top60_DNA_seed | MAPK1 | 0.854308 | BRCA | HCC1937 | Ulixertinib | MVP_top60_DNA+RNA_seed | MAPK3 | 0.406631 | BRCA | AU565 | Ulixertinib |
| MVP_top60_DNA_seed | MAPK1 | 0.857524 | BRCA | EFM-192A | VX-11e | MVP_top60_DNA+RNA_seed | MAPK1 | 0.406631 | BRCA | AU565 | Ulixertinib |
| MVP_top60_DNA_seed | PIK3CA | 0.857775 | BRCA | HCC2218 | Buparlisib | MVP_top60_DNA+RNA_seed | HSP90AA1 | 0.406809 | BRCA | HCC70 | Tanespimycin |
| MVP_top60_DNA_seed | MDM2 | 0.860345 | BRCA | HCC1500 | Serdemetan | MVP_top60_DNA+RNA_seed | HSP90AB1 | 0.406809 | BRCA | HCC70 | Tanespimycin |
| MVP_top60_DNA_seed | TP53 | 0.860765 | BRCA | MCF7 | MIRA-1 | MVP_top60_DNA+RNA_seed | CDK2 | 0.409051 | BRCA | BT-549 | AZD5438 |
| MVP_top60_DNA_seed | CDK1 | 0.861349 | BRCA | EFM-19 | RO-3306 | MVP_top60_DNA+RNA_seed | HSP90AA1 | 0.413036 | BRCA | UACC-893 | Elesclomol |
| MVP_top60_DNA_seed | CDK2 | 0.863469 | BRCA | BT-474 | AZD5438 | MVP_top60_DNA+RNA_seed | HSP90AB1 | 0.413036 | BRCA | UACC-893 | Elesclomol |
| MVP_top60_DNA_seed | ESR1 | 0.863828 | BRCA | YMB-1-E | GDC0810 | MVP_top60_DNA+RNA_seed | HDAC1 | 0.41397 | BRCA | HCC1187 | Entinostat |

| | | | | | | | | | | | |
|---|---|---|---|---|---|---|---|---|---|---|---|
| MVP_top60_DNA_seed | EP300 | 0.864272 | BRCA | T47D | SGC-CBP30 | MVP_top60_DNA+RNA_seed | HDAC1 | 0.414933 | BRCA | MDA-MB-231 | PCI-34051 |
| MVP_top60_DNA_seed | CREBBP | 0.864272 | BRCA | T47D | SGC-CBP30 | MVP_top60_DNA+RNA_seed | MDM2 | 0.415639 | BRCA | HCC1599 | Nutlin-3a (-) |
| MVP_top60_DNA_seed | EP300 | 0.869091 | BRCA | HCC1500 | CPI-637 | MVP_top60_DNA+RNA_seed | MDM2 | 0.416264 | BRCA | HCC1954 | Nutlin-3a (-) |
| MVP_top60_DNA_seed | EP300 | 0.869783 | BRCA | HCC38 | CPI-637 | MVP_top60_DNA+RNA_seed | AKT1 | 0.417909 | BRCA | HCC2218 | MK-2206 |
| MVP_top60_DNA_seed | PIK3CA | 0.8721 | BRCA | HCC1599 | AZD8186 | MVP_top60_DNA+RNA_seed | MDM2 | 0.419545 | BRCA | MCF7 | Serdemetan |
| MVP_top60_DNA_seed | MAPK1 | 0.873812 | BRCA | BT-549 | VX-11e | MVP_top60_DNA+RNA_seed | EGFR | 0.419947 | BRCA | MDA-MB-231 | Erlotinib |
| MVP_top60_DNA_seed | CDK2 | 0.875586 | BRCA | MFM-223 | AZD5438 | MVP_top60_DNA+RNA_seed | MAPK3 | 0.420019 | BRCA | HCC1954 | SCH772984 |
| MVP_top60_DNA_seed | MDM2 | 0.876346 | BRCA | CAL-85-1 | Nutlin-3a (-) | MVP_top60_DNA+RNA_seed | MAPK1 | 0.420019 | BRCA | HCC1954 | SCH772984 |
| MVP_top60_DNA_seed | MDM2 | 0.876849 | BRCA | EFM-19 | Serdemetan | MVP_top60_DNA+RNA_seed | EGFR | 0.420646 | BRCA | CAL-148 | Sapitinib |
| MVP_top60_DNA_seed | ESR1 | 0.876885 | BRCA | HCC1599 | GDC0810 | MVP_top60_DNA+RNA_seed | ERBB2 | 0.420646 | BRCA | CAL-148 | Sapitinib |
| MVP_top60_DNA_seed | AKT1 | 0.877952 | BRCA | HCC1143 | Uprosertib | MVP_top60_DNA+RNA_seed | ERBB3 | 0.420646 | BRCA | CAL-148 | Sapitinib |
| MVP_top60_DNA_seed | SRC | 0.878322 | BRCA | EFM-19 | Bosutinib | MVP_top60_DNA+RNA_seed | AKT1 | 0.420836 | BRCA | HCC1187 | Afuresertib |
| MVP_top60_DNA_seed | MAPK3 | 0.882717 | BRCA | UACC-812 | Ulixertinib | MVP_top60_DNA+RNA_seed | PIK3CA | 0.422393 | BRCA | HCC1419 | AZD8186 |
| MVP_top60_DNA_seed | MAPK1 | 0.882717 | BRCA | UACC-812 | Ulixertinib | MVP_top60_DNA+RNA_seed | MDM2 | 0.423598 | BRCA | CAL-51 | Serdemetan |
| MVP_top60_DNA_seed | MDM2 | 0.882745 | BRCA | HDQ-P1 | Nutlin-3a (-) | MVP_top60_DNA+RNA_seed | HDAC1 | 0.425495 | BRCA | HCC1419 | PCI-34051 |
| MVP_top60_DNA_seed | MAPK3 | 0.887218 | BRCA | MCF7 | Ulixertinib | MVP_top60_DNA+RNA_seed | MAPK1 | 0.427494 | BRCA | HCC2157 | VX-11e |
| MVP_top60_DNA_seed | MAPK1 | 0.887218 | BRCA | MCF7 | Ulixertinib | MVP_top60_DNA+RNA_seed | EP300 | 0.429734 | BRCA | EFM-19 | CPI-637 |
| MVP_top60_DNA_seed | ESR1 | 0.889689 | BRCA | YMB-1-E | Tamoxifen | MVP_top60_DNA+RNA_seed | HDAC1 | 0.432214 | BRCA | HCC70 | PCI-34051 |
| MVP_top60_DNA_seed | EP300 | 0.893926 | BRCA | MFM-223 | SGC-CBP30 | MVP_top60_DNA+RNA_seed | ESR1 | 0.432784 | BRCA | MDA-MB-157 | Tamoxifen |
| MVP_top60_DNA_seed | CREBBP | 0.893926 | BRCA | MFM-223 | SGC-CBP30 | MVP_top60_DNA+RNA_seed | MDM2 | 0.432936 | BRCA | MDA-MB-415 | Serdemetan |
| MVP_top60_DNA_seed | HSP90AA1 | 0.895976 | BRCA | CAL-120 | Tanespimycin | MVP_top60_DNA+RNA_seed | EGFR | 0.433847 | BRCA | HCC1187 | Sapitinib |
| MVP_top60_DNA_seed | HSP90AB1 | 0.895976 | BRCA | CAL-120 | Tanespimycin | MVP_top60_DNA+RNA_seed | ERBB2 | 0.433847 | BRCA | HCC1187 | Sapitinib |
| MVP_top60_DNA_seed | EGFR | 0.897531 | BRCA | MFM-223 | Erlotinib | MVP_top60_DNA+RNA_seed | ERBB3 | 0.433847 | BRCA | HCC1187 | Sapitinib |
| MVP_top60_DNA_seed | EP300 | 0.898403 | BRCA | CAL-120 | SGC-CBP30 | MVP_top60_DNA+RNA_seed | CDK2 | 0.434729 | BRCA | MDA-MB-453 | AZD5438 |
| MVP_top60_DNA_seed | CREBBP | 0.898403 | BRCA | CAL-120 | SGC-CBP30 | MVP_top60_DNA+RNA_seed | ESR1 | 0.436325 | BRCA | HCC1143 | GDC0810 |
| MVP_top60_DNA_seed | MAPK3 | 0.899405 | BRCA | BT-20 | Ulixertinib | MVP_top60_DNA+RNA_seed | EGFR | 0.437476 | BRCA | MDA-MB-415 | Afatinib |
| MVP_top60_DNA_seed | MAPK1 | 0.899405 | BRCA | BT-20 | Ulixertinib | MVP_top60_DNA+RNA_seed | ERBB2 | 0.437476 | BRCA | MDA-MB-415 | Afatinib |
| MVP_top60_DNA_seed | PIK3CA | 0.901775 | BRCA | BT-474 | GNE-317 | MVP_top60_DNA+RNA_seed | CDK1 | 0.438682 | BRCA | HCC38 | RO-3306 |
| MVP_top60_DNA_seed | MDM2 | 0.906853 | BRCA | MDA-MB-157 | Serdemetan | MVP_top60_DNA+RNA_seed | HDAC1 | 0.438968 | BRCA | BT-474 | Romidepsin |
| MVP_top60_DNA_seed | EP300 | 0.911587 | BRCA | CAL-51 | CPI-637 | MVP_top60_DNA+RNA_seed | HDAC2 | 0.438968 | BRCA | BT-474 | Romidepsin |
| MVP_top60_DNA_seed | PIK3CA | 0.911682 | BRCA | MRK-nu-1 | AZD8186 | MVP_top60_DNA+RNA_seed | EP300 | 0.439838 | BRCA | BT-474 | SGC-CBP30 |
| MVP_top60_DNA_seed | MAPK3 | 0.911947 | BRCA | MFM-223 | Ulixertinib | MVP_top60_DNA+RNA_seed | MDM2 | 0.43996 | BRCA | COLO-824 | Nutlin-3a (-) |
| MVP_top60_DNA_seed | MAPK1 | 0.911947 | BRCA | MFM-223 | Ulixertinib | MVP_top60_DNA+RNA_seed | ESR1 | 0.442264 | BRCA | MDA-MB-436 | GDC0810 |
| MVP_top60_DNA_seed | MAPK3 | 0.916835 | BRCA | EFM-19 | Ulixertinib | MVP_top60_DNA+RNA_seed | AKT1 | 0.443013 | BRCA | HDQ-P1 | MK-2206 |
| MVP_top60_DNA_seed | MAPK1 | 0.916835 | BRCA | EFM-19 | Ulixertinib | MVP_top60_DNA+RNA_seed | CDK2 | 0.445264 | BRCA | HCC38 | Dinaciclib |
| MVP_top60_DNA_seed | HDAC1 | 0.918275 | BRCA | BT-20 | Entinostat | MVP_top60_DNA+RNA_seed | CDK1 | 0.445264 | BRCA | HCC38 | Dinaciclib |
| MVP_top60_DNA_seed | PIK3CA | 0.919742 | BRCA | HCC1569 | Buparlisib | MVP_top60_DNA+RNA_seed | AKT1 | 0.447848 | BRCA | MFM-223 | AT13148 |
| MVP_top60_DNA_seed | PIK3CA | 0.9207 | BRCA | MFM-223 | AZD8186 | MVP_top60_DNA+RNA_seed | PLK1 | 0.448416 | BRCA | MDA-MB-361 | MK-1775 |
| MVP_top60_DNA_seed | EGFR | 0.924335 | BRCA | JIMT-1 | Osimertinib | MVP_top60_DNA+RNA_seed | EGFR | 0.44962 | BRCA | HCC1500 | Gefitinib |
| MVP_top60_DNA_seed | MAPK3 | 0.928352 | BRCA | CAL-148 | Ulixertinib | MVP_top60_DNA+RNA_seed | MDM2 | 0.450224 | BRCA | CAMA-1 | Serdemetan |
| MVP_top60_DNA_seed | MAPK1 | 0.928352 | BRCA | CAL-148 | Ulixertinib | MVP_top60_DNA+RNA_seed | CDK2 | 0.450349 | BRCA | UACC-893 | Dinaciclib |

| | | | | | | | | | | | |
|---|---|---|---|---|---|---|---|---|---|---|---|
| MVP_top60_DNA_seed | AR | 0.929318 | BRCA | MDA-MB-330 | Bicalutamide | MVP_top60_DNA+RNA_seed | CDK1 | 0.450349 | BRCA | UACC-893 | Dinaciclib |
| MVP_top60_DNA_seed | MAPK3 | 0.931203 | BRCA | EFM-19 | Ulixertinib | MVP_top60_DNA+RNA_seed | EP300 | 0.451761 | BRCA | HCC1395 | CPI-637 |
| MVP_top60_DNA_seed | MAPK1 | 0.931203 | BRCA | EFM-19 | Ulixertinib | MVP_top60_DNA+RNA_seed | AKT1 | 0.452331 | BRCA | HCC1143 | Afuresertib |
| MVP_top60_DNA_seed | MAPK3 | 0.938143 | BRCA | MDA-MB-330 | SCH772984 | MVP_top60_DNA+RNA_seed | EP300 | 0.453045 | BRCA | CAL-85-1 | CPI-637 |
| MVP_top60_DNA_seed | MAPK1 | 0.938143 | BRCA | MDA-MB-330 | SCH772984 | MVP_top60_DNA+RNA_seed | MAPK1 | 0.453514 | BRCA | HCC2218 | VX-11e |
| MVP_top60_DNA_seed | CDK2 | 0.939686 | BRCA | T47D | Dinaciclib | MVP_top60_DNA+RNA_seed | SRC | 0.453599 | BRCA | Hs-578-T | Bosutinib |
| MVP_top60_DNA_seed | CDK1 | 0.939686 | BRCA | T47D | Dinaciclib | MVP_top60_DNA+RNA_seed | TP53 | 0.453655 | BRCA | MDA-MB-468 | MIRA-1 |
| MVP_top60_DNA_seed | CDK2 | 0.939865 | BRCA | HCC1954 | MK-8776 | MVP_top60_DNA+RNA_seed | CDK2 | 0.454282 | BRCA | BT-20 | MK-8776 |
| MVP_top60_DNA_seed | MDM2 | 0.942024 | BRCA | JIMT-1 | Nutlin-3a (-) | MVP_top60_DNA+RNA_seed | CDK2 | 0.45477 | BRCA | MDA-MB-436 | MK-8776 |
| MVP_top60_DNA_seed | ESR1 | 0.942933 | BRCA | HCC1428 | GDC0810 | MVP_top60_DNA+RNA_seed | HDAC1 | 0.457268 | BRCA | HCC1419 | Entinostat |
| MVP_top60_DNA_seed | ERBB2 | 0.943402 | BRCA | HCC1500 | Sapitinib | MVP_top60_DNA+RNA_seed | AKT1 | 0.457967 | BRCA | MDA-MB-157 | AT13148 |
| MVP_top60_DNA_seed | EGFR | 0.943402 | BRCA | HCC1500 | Sapitinib | MVP_top60_DNA+RNA_seed | MAPK1 | 0.459014 | BRCA | HCC1187 | VX-11e |
| MVP_top60_DNA_seed | ERBB3 | 0.943402 | BRCA | HCC1500 | Sapitinib | MVP_top60_DNA+RNA_seed | PIK3CA | 0.459333 | BRCA | BT-549 | Buparlisib |
| MVP_top60_DNA_seed | SRC | 0.944668 | BRCA | EFM-192A | Bosutinib | MVP_top60_DNA+RNA_seed | EGFR | 0.459412 | BRCA | HCC70 | Sapitinib |
| MVP_top60_DNA_seed | AKT1 | 0.945834 | BRCA | HCC38 | Ipatasertib | MVP_top60_DNA+RNA_seed | ERBB2 | 0.459412 | BRCA | HCC70 | Sapitinib |
| MVP_top60_DNA_seed | SRC | 0.946258 | BRCA | UACC-812 | Bosutinib | MVP_top60_DNA+RNA_seed | ERBB3 | 0.459412 | BRCA | HCC70 | Sapitinib |
| MVP_top60_DNA_seed | EGFR | 0.949244 | BRCA | HCC1428 | Gefitinib | MVP_top60_DNA+RNA_seed | AKT1 | 0.462422 | BRCA | JIMT-1 | MK-2206 |
| MVP_top60_DNA_seed | AKT1 | 0.949387 | BRCA | HCC1419 | AT13148 | MVP_top60_DNA+RNA_seed | PLK1 | 0.462976 | BRCA | UACC-893 | BI-2536 |
| MVP_top60_DNA_seed | MAPK1 | 0.949954 | BRCA | MDA-MB-436 | VX-11e | MVP_top60_DNA+RNA_seed | MAPK1 | 0.464492 | BRCA | AU565 | VX-11e |
| MVP_top60_DNA_seed | PIK3CA | 0.951276 | BRCA | MDA-MB-436 | Alpelisib | MVP_top60_DNA+RNA_seed | MAPK1 | 0.465861 | BRCA | HCC1500 | VX-11e |
| MVP_top60_DNA_seed | HSP90AA1 | 0.953192 | BRCA | MDA-MB-436 | Tanespimycin | MVP_top60_DNA+RNA_seed | MDM2 | 0.466177 | BRCA | UACC-893 | Nutlin-3a (-) |
| MVP_top60_DNA_seed | HSP90AB1 | 0.953192 | BRCA | MDA-MB-436 | Tanespimycin | MVP_top60_DNA+RNA_seed | EGFR | 0.466315 | BRCA | EFM-192A | AZD3759 |
| MVP_top60_DNA_seed | MAPK1 | 0.963209 | BRCA | HCC1143 | VX-11e | MVP_top60_DNA+RNA_seed | MAPK1 | 0.466883 | BRCA | BT-20 | VX-11e |
| MVP_top60_DNA_seed | ESR1 | 0.963472 | BRCA | HCC38 | GDC0810 | MVP_top60_DNA+RNA_seed | CDK2 | 0.46745 | BRCA | MDA-MB-468 | MK-8776 |
| MVP_top60_DNA_seed | HDAC1 | 0.964269 | BRCA | BT-20 | Romidepsin | MVP_top60_DNA+RNA_seed | PIK3CA | 0.4689 | BRCA | MDA-MB-415 | Alpelisib |
| MVP_top60_DNA_seed | HDAC2 | 0.964269 | BRCA | BT-20 | Romidepsin | MVP_top60_DNA+RNA_seed | AKT1 | 0.470302 | BRCA | MDA-MB-468 | Uprosertib |
| MVP_top60_DNA_seed | MDM2 | 0.966999 | BRCA | HCC1143 | Nutlin-3a (-) | MVP_top60_DNA+RNA_seed | HSP90AA1 | 0.471624 | BRCA | MDA-MB-157 | Tanespimycin |
| MVP_top60_DNA_seed | EGFR | 0.968993 | BRCA | EFM-19 | Gefitinib | MVP_top60_DNA+RNA_seed | HSP90AB1 | 0.471624 | BRCA | MDA-MB-157 | Tanespimycin |
| MVP_top60_DNA_seed | ERBB2 | 0.971702 | BRCA | MDA-MB-436 | Lapatinib | MVP_top60_DNA+RNA_seed | AKT1 | 0.472093 | BRCA | HCC1143 | AZD5363 |
| MVP_top60_DNA_seed | EGFR | 0.971702 | BRCA | MDA-MB-436 | Lapatinib | MVP_top60_DNA+RNA_seed | MDM2 | 0.47277 | BRCA | BT-20 | Nutlin-3a (-) |
| MVP_top60_DNA_seed | AKT1 | 0.972069 | BRCA | CAL-120 | Uprosertib | MVP_top60_DNA+RNA_seed | ESR1 | 0.476116 | BRCA | BT-474 | GDC0810 |
| MVP_top60_DNA_seed | MAPK3 | 0.972174 | BRCA | MDA-MB-361 | Ulixertinib | MVP_top60_DNA+RNA_seed | MAPK3 | 0.476896 | BRCA | MFM-223 | SCH772984 |
| MVP_top60_DNA_seed | MAPK1 | 0.972174 | BRCA | MDA-MB-361 | Ulixertinib | MVP_top60_DNA+RNA_seed | MAPK1 | 0.476896 | BRCA | MFM-223 | SCH772984 |
| MVP_top60_DNA_seed | HDAC1 | 0.972261 | BRCA | MDA-MB-157 | PCI-34051 | MVP_top60_DNA+RNA_seed | AKT1 | 0.478651 | BRCA | MDA-MB-157 | Afuresertib |
| MVP_top60_DNA_seed | HDAC1 | 0.974598 | BRCA | MCF7 | Entinostat | MVP_top60_DNA+RNA_seed | CDK2 | 0.47911 | BRCA | YMB-1-E | Dinaciclib |
| MVP_top60_DNA_seed | MDM2 | 0.975374 | BRCA | BT-549 | Nutlin-3a (-) | MVP_top60_DNA+RNA_seed | CDK1 | 0.47911 | BRCA | YMB-1-E | Dinaciclib |
| MVP_top60_DNA_seed | CDK2 | 0.976235 | BRCA | MCF7 | Dinaciclib | MVP_top60_DNA+RNA_seed | MAPK3 | 0.479354 | BRCA | HCC1187 | Ulixertinib |
| MVP_top60_DNA_seed | CDK1 | 0.976235 | BRCA | MCF7 | Dinaciclib | MVP_top60_DNA+RNA_seed | MAPK1 | 0.479354 | BRCA | HCC1187 | Ulixertinib |
| MVP_top60_DNA_seed | CDK2 | 0.976615 | BRCA | COLO-824 | MK-8776 | MVP_top60_DNA+RNA_seed | EP300 | 0.480464 | BRCA | HCC1187 | SGC-CBP30 |
| MVP_top60_DNA_seed | ESR1 | 0.976971 | BRCA | MDA-MB-330 | Tamoxifen | MVP_top60_DNA+RNA_seed | EGFR | 0.481107 | BRCA | EFM-19 | AZD3759 |
| MVP_top60_DNA_seed | AKT1 | 0.978393 | BRCA | MRK-nu-1 | Afuresertib | MVP_top60_DNA+RNA_seed | AKT1 | 0.481963 | BRCA | HCC1143 | Uprosertib |

| | | | | | | | | | | | |
|---|---|---|---|---|---|---|---|---|---|---|---|
| MVP_top60_DNA_seed | HDAC1 | 0.979205 | BRCA | CAL-120 | Romidepsin | MVP_top60_DNA+RNA_seed | PLK1 | 0.482187 | BRCA | MDA-MB-453 | MK-1775 |
| MVP_top60_DNA_seed | HDAC2 | 0.979205 | BRCA | CAL-120 | Romidepsin | MVP_top60_DNA+RNA_seed | PLK1 | 0.483204 | BRCA | OCUB-M | MK-1775 |
| MVP_top60_DNA_seed | HDAC1 | 0.979627 | BRCA | HCC38 | Entinostat | MVP_top60_DNA+RNA_seed | ESR1 | 0.484817 | BRCA | MDA-MB-453 | GDC0810 |
| MVP_top60_DNA_seed | SRC | 0.980572 | BRCA | MDA-MB-157 | Bosutinib | MVP_top60_DNA+RNA_seed | ESR1 | 0.48503 | BRCA | MDA-MB-330 | GDC0810 |
| MVP_top60_DNA_seed | MAPK1 | 0.986755 | BRCA | EFM-19 | VX-11e | MVP_top60_DNA+RNA_seed | MAPK3 | 0.485736 | BRCA | MRK-nu-1 | SCH772984 |
| MVP_top60_DNA_seed | EGFR | 0.988119 | BRCA | CAL-85-1 | AZD3759 | MVP_top60_DNA+RNA_seed | MAPK1 | 0.485736 | BRCA | MRK-nu-1 | SCH772984 |
| MVP_top60_DNA_seed | CDK1 | 0.988345 | BRCA | HCC1187 | RO-3306 | MVP_top60_DNA+RNA_seed | PIK3CA | 0.487822 | BRCA | CAL-85-1 | AZD8186 |
| MVP_top60_DNA_seed | EP300 | 0.988393 | BRCA | ZR-75-30 | SGC-CBP30 | MVP_top60_DNA+RNA_seed | HSP90AA1 | 0.488262 | BRCA | HCC1500 | Luminespib |
| MVP_top60_DNA_seed | CREBBP | 0.988393 | BRCA | ZR-75-30 | SGC-CBP30 | MVP_top60_DNA+RNA_seed | HSP90AB1 | 0.488262 | BRCA | HCC1500 | Luminespib |
| MVP_top60_DNA_seed | CDK2 | 0.989393 | BRCA | HCC1187 | MK-8776 | MVP_top60_DNA+RNA_seed | EP300 | 0.488286 | BRCA | HCC1500 | SGC-CBP30 |
| MVP_top60_DNA_seed | CDK2 | 0.990504 | BRCA | HCC1143 | AZD5438 | MVP_top60_DNA+RNA_seed | SRC | 0.488445 | BRCA | JIMT-1 | Bosutinib |
| MVP_top60_DNA_seed | HSP90AA1 | 0.993092 | BRCA | HCC1954 | Luminespib | MVP_top60_DNA+RNA_seed | EGFR | 0.491244 | BRCA | UACC-812 | Gefitinib |
| MVP_top60_DNA_seed | HSP90AB1 | 0.993092 | BRCA | HCC1954 | Luminespib | MVP_top60_DNA+RNA_seed | EP300 | 0.492236 | BRCA | HCC38 | SGC-CBP30 |
| MVP_top60_DNA_seed | MAPK1 | 0.997509 | BRCA | ZR-75-30 | VX-11e | MVP_top60_DNA+RNA_seed | EGFR | 0.493629 | BRCA | HCC1599 | AZD3759 |
| MVP_top60_DNA_seed | HDAC1 | 1.000216 | BRCA | MDA-MB-330 | PCI-34051 | MVP_top60_DNA+RNA_seed | EP300 | 0.495564 | BRCA | HDQ-P1 | CPI-637 |
| MVP_top60_DNA_seed | MAPK3 | 1.002171 | BRCA | HCC2157 | SCH772984 | MVP_top60_DNA+RNA_seed | MAPK3 | 0.497144 | BRCA | HCC2157 | Ulixertinib |
| MVP_top60_DNA_seed | MAPK1 | 1.002171 | BRCA | HCC2157 | SCH772984 | MVP_top60_DNA+RNA_seed | MAPK1 | 0.497144 | BRCA | HCC2157 | Ulixertinib |
| MVP_top60_DNA_seed | MDM2 | 1.003158 | BRCA | MDA-MB-330 | Nutlin-3a (-) | MVP_top60_DNA+RNA_seed | AKT1 | 0.497206 | BRCA | T47D | Uprosertib |
| MVP_top60_DNA_seed | HSP90AA1 | 1.003725 | BRCA | HCC1428 | Tanespimycin | MVP_top60_DNA+RNA_seed | EP300 | 0.498955 | BRCA | HCC1954 | SGC-CBP30 |
| MVP_top60_DNA_seed | HSP90AB1 | 1.003725 | BRCA | HCC1428 | Tanespimycin | MVP_top60_DNA+RNA_seed | TP53 | 0.499196 | BRCA | T47D | MIRA-1 |
| MVP_top60_DNA_seed | EGFR | 1.003802 | BRCA | EFM-19 | Erlotinib | MVP_top60_DNA+RNA_seed | ATM | 0.499748 | BRCA | CAL-148 | KU-55933 |
| MVP_top60_DNA_seed | MAPK3 | 1.005207 | BRCA | HCC1187 | Ulixertinib | MVP_top60_DNA+RNA_seed | MAPK3 | 0.500147 | BRCA | JIMT-1 | Ulixertinib |
| MVP_top60_DNA_seed | MAPK1 | 1.005207 | BRCA | HCC1187 | Ulixertinib | MVP_top60_DNA+RNA_seed | MAPK1 | 0.500147 | BRCA | JIMT-1 | Ulixertinib |
| MVP_top60_DNA_seed | AKT1 | 1.005771 | BRCA | MDA-MB-231 | Ipatasertib | MVP_top60_DNA+RNA_seed | AKT1 | 0.50023 | BRCA | MDA-MB-231 | Uprosertib |
| MVP_top60_DNA_seed | MAPK1 | 1.007065 | BRCA | MDA-MB-175-VII | VX-11e | MVP_top60_DNA+RNA_seed | PIK3CA | 0.501662 | BRCA | HCC1937 | Alpelisib |
| MVP_top60_DNA_seed | MAPK1 | 1.009248 | BRCA | HCC1599 | VX-11e | MVP_top60_DNA+RNA_seed | HDAC1 | 0.504462 | BRCA | MDA-MB-157 | Entinostat |
| MVP_top60_DNA_seed | AKT1 | 1.011347 | BRCA | UACC-812 | Uprosertib | MVP_top60_DNA+RNA_seed | PLK1 | 0.506409 | BRCA | CAL-120 | MK-1775 |
| MVP_top60_DNA_seed | ATM | 1.011559 | BRCA | HCC1428 | KU-55933 | MVP_top60_DNA+RNA_seed | CDK2 | 0.507491 | BRCA | CAMA-1 | AZD5438 |
| MVP_top60_DNA_seed | ATM | 1.01437 | BRCA | JIMT-1 | KU-55933 | MVP_top60_DNA+RNA_seed | EGFR | 0.510696 | BRCA | MDA-MB-157 | Lapatinib |
| MVP_top60_DNA_seed | EP300 | 1.015238 | BRCA | MDA-MB-468 | CPI-637 | MVP_top60_DNA+RNA_seed | ERBB2 | 0.510696 | BRCA | MDA-MB-157 | Lapatinib |
| MVP_top60_DNA_seed | AKT1 | 1.016184 | BRCA | CAL-85-1 | Uprosertib | MVP_top60_DNA+RNA_seed | EGFR | 0.515466 | BRCA | CAMA-1 | Lapatinib |
| MVP_top60_DNA_seed | AKT1 | 1.019492 | BRCA | JIMT-1 | Uprosertib | MVP_top60_DNA+RNA_seed | ERBB2 | 0.515466 | BRCA | CAMA-1 | Lapatinib |
| MVP_top60_DNA_seed | MAPK3 | 1.019916 | BRCA | EFM-192A | SCH772984 | MVP_top60_DNA+RNA_seed | ESR1 | 0.516456 | BRCA | BT-474 | Tamoxifen |
| MVP_top60_DNA_seed | MAPK1 | 1.019916 | BRCA | EFM-192A | SCH772984 | MVP_top60_DNA+RNA_seed | CDK1 | 0.517717 | BRCA | HCC1419 | RO-3306 |
| MVP_top60_DNA_seed | EGFR | 1.021128 | BRCA | HCC38 | Osimertinib | MVP_top60_DNA+RNA_seed | EGFR | 0.518824 | BRCA | HCC1937 | Afatinib |
| MVP_top60_DNA_seed | EGFR | 1.031091 | BRCA | MDA-MB-436 | Erlotinib | MVP_top60_DNA+RNA_seed | ERBB2 | 0.518824 | BRCA | HCC1937 | Afatinib |
| MVP_top60_DNA_seed | AKT1 | 1.033067 | BRCA | MDA-MB-436 | Ipatasertib | MVP_top60_DNA+RNA_seed | SRC | 0.519776 | BRCA | HCC1500 | Bosutinib |
| MVP_top60_DNA_seed | EGFR | 1.035799 | BRCA | HCC1599 | Gefitinib | MVP_top60_DNA+RNA_seed | CDK2 | 0.520274 | BRCA | MCF7 | AZD5438 |
| MVP_top60_DNA_seed | TP53 | 1.036629 | BRCA | HCC1937 | MIRA-1 | MVP_top60_DNA+RNA_seed | EGFR | 0.52298 | BRCA | HCC1569 | Erlotinib |
| MVP_top60_DNA_seed | EP300 | 1.039723 | BRCA | MDA-MB-436 | CPI-637 | MVP_top60_DNA+RNA_seed | PIK3CA | 0.522996 | BRCA | MDA-MB-157 | GNE-317 |
| MVP_top60_DNA_seed | PIK3CA | 1.040492 | BRCA | HCC38 | Buparlisib | MVP_top60_DNA+RNA_seed | HDAC1 | 0.52851 | BRCA | HCC1143 | Romidepsin |

| | | | | | | | | | | | |
|---|---|---|---|---|---|---|---|---|---|---|---|
| MVP_top60_DNA_seed | PIK3CA | 1.045803 | BRCA | MDA-MB-231 | AZD8186 | MVP_top60_DNA+RNA_seed | HDAC2 | 0.52851 | BRCA | HCC1143 | Romidepsin |
| MVP_top60_DNA_seed | HSP90AA1 | 1.048811 | BRCA | ZR-75-30 | Luminespib | MVP_top60_DNA+RNA_seed | AKT1 | 0.529439 | BRCA | MDA-MB-436 | Afuresertib |
| MVP_top60_DNA_seed | HSP90AB1 | 1.048811 | BRCA | ZR-75-30 | Luminespib | MVP_top60_DNA+RNA_seed | PIK3CA | 0.530461 | BRCA | HCC38 | Alpelisib |
| MVP_top60_DNA_seed | ATM | 1.048889 | BRCA | UACC-893 | KU-55933 | MVP_top60_DNA+RNA_seed | AKT1 | 0.532261 | BRCA | HCC1187 | AZD5363 |
| MVP_top60_DNA_seed | HDAC1 | 1.050179 | BRCA | COLO-824 | Romidepsin | MVP_top60_DNA+RNA_seed | CDK2 | 0.532401 | BRCA | HCC2157 | MK-8776 |
| MVP_top60_DNA_seed | HDAC2 | 1.050179 | BRCA | COLO-824 | Romidepsin | MVP_top60_DNA+RNA_seed | PIK3CA | 0.533957 | BRCA | HCC1419 | Alpelisib |
| MVP_top60_DNA_seed | HDAC1 | 1.050224 | BRCA | MFM-223 | PCI-34051 | MVP_top60_DNA+RNA_seed | EP300 | 0.534131 | BRCA | EFM-19 | SGC-CBP30 |
| MVP_top60_DNA_seed | ATM | 1.056079 | BRCA | BT-474 | KU-55933 | MVP_top60_DNA+RNA_seed | PIK3CA | 0.535915 | BRCA | HCC38 | AZD8186 |
| MVP_top60_DNA_seed | EGFR | 1.058024 | BRCA | MDA-MB-157 | Osimertinib | MVP_top60_DNA+RNA_seed | MAPK3 | 0.535941 | BRCA | EFM-19 | SCH772984 |
| MVP_top60_DNA_seed | AKT1 | 1.063553 | BRCA | MDA-MB-436 | GSK2110183B | MVP_top60_DNA+RNA_seed | MAPK1 | 0.535941 | BRCA | EFM-19 | SCH772984 |
| MVP_top60_DNA_seed | HDAC1 | 1.071644 | BRCA | HCC38 | PCI-34051 | MVP_top60_DNA+RNA_seed | PLK1 | 0.53622 | BRCA | HCC70 | MK-1775 |
| MVP_top60_DNA_seed | PIK3CA | 1.071977 | BRCA | MDA-MB-436 | GNE-317 | MVP_top60_DNA+RNA_seed | ATM | 0.537588 | BRCA | MDA-MB-330 | KU-55933 |
| MVP_top60_DNA_seed | EP300 | 1.078404 | BRCA | BT-549 | SGC-CBP30 | MVP_top60_DNA+RNA_seed | HDAC1 | 0.539044 | BRCA | BT-20 | PCI-34051 |
| MVP_top60_DNA_seed | CREBBP | 1.078404 | BRCA | BT-549 | SGC-CBP30 | MVP_top60_DNA+RNA_seed | PIK3CA | 0.540628 | BRCA | JIMT-1 | AZD8186 |
| MVP_top60_DNA_seed | ESR1 | 1.079545 | BRCA | ZR-75-30 | GDC0810 | MVP_top60_DNA+RNA_seed | PIK3CA | 0.541288 | BRCA | BT-549 | Alpelisib |
| MVP_top60_DNA_seed | MAPK3 | 1.081281 | BRCA | HDQ-P1 | Ulixertinib | MVP_top60_DNA+RNA_seed | EGFR | 0.543644 | BRCA | MRK-nu-1 | Lapatinib |
| MVP_top60_DNA_seed | MAPK1 | 1.081281 | BRCA | HDQ-P1 | Ulixertinib | MVP_top60_DNA+RNA_seed | ERBB2 | 0.543644 | BRCA | MRK-nu-1 | Lapatinib |
| MVP_top60_DNA_seed | AKT1 | 1.081818 | BRCA | HCC38 | Uprosertib | MVP_top60_DNA+RNA_seed | HSP90AA1 | 0.545039 | BRCA | MFM-223 | Luminespib |
| MVP_top60_DNA_seed | HDAC1 | 1.084074 | BRCA | MCF7 | PCI-34051 | MVP_top60_DNA+RNA_seed | HSP90AB1 | 0.545039 | BRCA | MFM-223 | Luminespib |
| MVP_top60_DNA_seed | AKT1 | 1.084199 | BRCA | CAL-85-1 | Uprosertib | MVP_top60_DNA+RNA_seed | PLK1 | 0.545301 | BRCA | CAL-85-1 | MK-1775 |
| MVP_top60_DNA_seed | AKT1 | 1.084234 | BRCA | ZR-75-30 | Ipatasertib | MVP_top60_DNA+RNA_seed | CDK2 | 0.545852 | BRCA | MDA-MB-231 | MK-8776 |
| MVP_top60_DNA_seed | EGFR | 1.090574 | BRCA | MCF7 | Erlotinib | MVP_top60_DNA+RNA_seed | ESR1 | 0.547848 | BRCA | UACC-812 | GDC0810 |
| MVP_top60_DNA_seed | ERBB2 | 1.092291 | BRCA | CAMA-1 | Afatinib | MVP_top60_DNA+RNA_seed | EGFR | 0.548339 | BRCA | BT-549 | Osimertinib |
| MVP_top60_DNA_seed | EGFR | 1.092291 | BRCA | CAMA-1 | Afatinib | MVP_top60_DNA+RNA_seed | AKT1 | 0.549188 | BRCA | JIMT-1 | AZD5363 |
| MVP_top60_DNA_seed | MAPK3 | 1.093167 | BRCA | MCF7 | Ulixertinib | MVP_top60_DNA+RNA_seed | CDK2 | 0.550147 | BRCA | BT-549 | Dinaciclib |
| MVP_top60_DNA_seed | MAPK1 | 1.093167 | BRCA | MCF7 | Ulixertinib | MVP_top60_DNA+RNA_seed | CDK1 | 0.550147 | BRCA | BT-549 | Dinaciclib |
| MVP_top60_DNA_seed | HSP90AA1 | 1.094578 | BRCA | MDA-MB-231 | Tanespimycin | MVP_top60_DNA+RNA_seed | AKT1 | 0.552365 | BRCA | BT-20 | AT13148 |
| MVP_top60_DNA_seed | HSP90AB1 | 1.094578 | BRCA | MDA-MB-231 | Tanespimycin | MVP_top60_DNA+RNA_seed | HDAC1 | 0.553209 | BRCA | HCC1500 | Romidepsin |
| MVP_top60_DNA_seed | MAPK3 | 1.097862 | BRCA | BT-474 | Ulixertinib | MVP_top60_DNA+RNA_seed | HDAC2 | 0.553209 | BRCA | HCC1500 | Romidepsin |
| MVP_top60_DNA_seed | MAPK1 | 1.097862 | BRCA | BT-474 | Ulixertinib | MVP_top60_DNA+RNA_seed | SRC | 0.553834 | BRCA | HCC70 | Bosutinib |
| MVP_top60_DNA_seed | MAPK1 | 1.098267 | BRCA | MDA-MB-157 | VX-11e | MVP_top60_DNA+RNA_seed | AKT1 | 0.556612 | BRCA | MDA-MB-157 | AZD5363 |
| MVP_top60_DNA_seed | ERBB2 | 1.098533 | BRCA | BT-549 | Sapitinib | MVP_top60_DNA+RNA_seed | EGFR | 0.556828 | BRCA | EFM-19 | Lapatinib |
| MVP_top60_DNA_seed | EGFR | 1.098533 | BRCA | BT-549 | Sapitinib | MVP_top60_DNA+RNA_seed | ERBB2 | 0.556828 | BRCA | EFM-19 | Lapatinib |
| MVP_top60_DNA_seed | ERBB3 | 1.098533 | BRCA | BT-549 | Sapitinib | MVP_top60_DNA+RNA_seed | TP53 | 0.557608 | BRCA | ZR-75-30 | MIRA-1 |
| MVP_top60_DNA_seed | PIK3CA | 1.098639 | BRCA | HCC1419 | Buparlisib | MVP_top60_DNA+RNA_seed | EGFR | 0.559887 | BRCA | HCC1599 | Afatinib |
| MVP_top60_DNA_seed | TP53 | 1.099032 | BRCA | BT-549 | MIRA-1 | MVP_top60_DNA+RNA_seed | ERBB2 | 0.559887 | BRCA | HCC1599 | Afatinib |
| MVP_top60_DNA_seed | PIK3CA | 1.099167 | BRCA | BT-474 | Buparlisib | MVP_top60_DNA+RNA_seed | PIK3CA | 0.561574 | BRCA | MFM-223 | Buparlisib |
| MVP_top60_DNA_seed | PIK3CA | 1.100603 | BRCA | ZR-75-30 | GNE-317 | MVP_top60_DNA+RNA_seed | AKT1 | 0.561979 | BRCA | BT-549 | GSK2110183B |
| MVP_top60_DNA_seed | SRC | 1.101792 | BRCA | MDA-MB-436 | Bosutinib | MVP_top60_DNA+RNA_seed | MAPK3 | 0.562695 | BRCA | YMB-1-E | Ulixertinib |
| MVP_top60_DNA_seed | ERBB2 | 1.106644 | BRCA | MCF7 | Afatinib | MVP_top60_DNA+RNA_seed | MAPK1 | 0.562695 | BRCA | YMB-1-E | Ulixertinib |
| MVP_top60_DNA_seed | EGFR | 1.106644 | BRCA | MCF7 | Afatinib | MVP_top60_DNA+RNA_seed | PIK3CA | 0.56285 | BRCA | MDA-MB-231 | Buparlisib |

| | | | | | | | | | | | |
|---|---|---|---|---|---|---|---|---|---|---|---|
| MVP_top60_DNA_seed | MAPK3 | 1.112723 | BRCA | UACC-812 | Ulixertinib | MVP_top60_DNA+RNA_seed | EGFR | 0.565418 | BRCA | CAL-85-1 | Gefitinib |
| MVP_top60_DNA_seed | MAPK1 | 1.112723 | BRCA | UACC-812 | Ulixertinib | MVP_top60_DNA+RNA_seed | AKT1 | 0.565608 | BRCA | MRK-nu-1 | Ipatasertib |
| MVP_top60_DNA_seed | AKT1 | 1.117362 | BRCA | CAL-85-1 | MK-2206 | MVP_top60_DNA+RNA_seed | HDAC1 | 0.566626 | BRCA | HCC1937 | Entinostat |
| MVP_top60_DNA_seed | AKT1 | 1.117489 | BRCA | MCF7 | GSK2110183B | MVP_top60_DNA+RNA_seed | AKT1 | 0.568882 | BRCA | ZR-75-30 | AT13148 |
| MVP_top60_DNA_seed | HDAC1 | 1.119887 | BRCA | HCC1428 | Romidepsin | MVP_top60_DNA+RNA_seed | AKT1 | 0.570058 | BRCA | HCC1143 | MK-2206 |
| MVP_top60_DNA_seed | HDAC2 | 1.119887 | BRCA | HCC1428 | Romidepsin | MVP_top60_DNA+RNA_seed | HDAC1 | 0.570452 | BRCA | MFM-223 | Romidepsin |
| MVP_top60_DNA_seed | AKT1 | 1.120459 | BRCA | MDA-MB-157 | MK-2206 | MVP_top60_DNA+RNA_seed | HDAC2 | 0.570452 | BRCA | MFM-223 | Romidepsin |
| MVP_top60_DNA_seed | EGFR | 1.120563 | BRCA | MCF7 | AZD3759 | MVP_top60_DNA+RNA_seed | EGFR | 0.570613 | BRCA | HCC1395 | AZD3759 |
| MVP_top60_DNA_seed | CDK2 | 1.121775 | BRCA | HDQ-P1 | Dinaciclib | MVP_top60_DNA+RNA_seed | CDK2 | 0.572848 | BRCA | CAMA-1 | Dinaciclib |
| MVP_top60_DNA_seed | CDK1 | 1.121775 | BRCA | HDQ-P1 | Dinaciclib | MVP_top60_DNA+RNA_seed | CDK1 | 0.572848 | BRCA | CAMA-1 | Dinaciclib |
| MVP_top60_DNA_seed | MDM2 | 1.126843 | BRCA | HCC38 | Nutlin-3a (-) | MVP_top60_DNA+RNA_seed | HSP90AA1 | 0.573333 | BRCA | UACC-893 | CCT-018159 |
| MVP_top60_DNA_seed | MAPK1 | 1.138814 | BRCA | YMB-1-E | VX-11e | MVP_top60_DNA+RNA_seed | HSP90AB1 | 0.573333 | BRCA | UACC-893 | CCT-018159 |
| MVP_top60_DNA_seed | PIK3CA | 1.140993 | BRCA | HCC1937 | Buparlisib | MVP_top60_DNA+RNA_seed | HDAC1 | 0.573789 | BRCA | HCC1954 | PCI-34051 |
| MVP_top60_DNA_seed | TP53 | 1.142254 | BRCA | HCC1143 | MIRA-1 | MVP_top60_DNA+RNA_seed | EGFR | 0.574579 | BRCA | MDA-MB-231 | Gefitinib |
| MVP_top60_DNA_seed | HSP90AA1 | 1.143489 | BRCA | YMB-1-E | CCT-018159 | MVP_top60_DNA+RNA_seed | AKT1 | 0.575969 | BRCA | HCC1143 | AT13148 |
| MVP_top60_DNA_seed | HSP90AB1 | 1.143489 | BRCA | YMB-1-E | CCT-018159 | MVP_top60_DNA+RNA_seed | CDK1 | 0.579946 | BRCA | HCC1937 | RO-3306 |
| MVP_top60_DNA_seed | SRC | 1.145831 | BRCA | MCF7 | Bosutinib | MVP_top60_DNA+RNA_seed | AKT1 | 0.582031 | BRCA | BT-483 | Uprosertib |
| MVP_top60_DNA_seed | HDAC1 | 1.147347 | BRCA | EFM-192A | PCI-34051 | MVP_top60_DNA+RNA_seed | AKT1 | 0.58658 | BRCA | HDQ-P1 | GSK2110183B |
| MVP_top60_DNA_seed | AKT1 | 1.14755 | BRCA | HCC1187 | Uprosertib | MVP_top60_DNA+RNA_seed | HSP90AA1 | 0.587856 | BRCA | MDA-MB-436 | Luminespib |
| MVP_top60_DNA_seed | EGFR | 1.147568 | BRCA | BT-549 | AZD3759 | MVP_top60_DNA+RNA_seed | HSP90AB1 | 0.587856 | BRCA | MDA-MB-436 | Luminespib |
| MVP_top60_DNA_seed | PIK3CA | 1.149817 | BRCA | HCC1500 | Alpelisib | MVP_top60_DNA+RNA_seed | EGFR | 0.591309 | BRCA | CAL-51 | Afatinib |
| MVP_top60_DNA_seed | CDK2 | 1.151019 | BRCA | T47D | MK-8776 | MVP_top60_DNA+RNA_seed | ERBB2 | 0.591309 | BRCA | CAL-51 | Afatinib |
| MVP_top60_DNA_seed | MAPK1 | 1.151696 | BRCA | MDA-MB-361 | VX-11e | MVP_top60_DNA+RNA_seed | MDM2 | 0.593548 | BRCA | BT-549 | Serdemetan |
| MVP_top60_DNA_seed | ERBB2 | 1.153388 | BRCA | MCF7 | Sapitinib | MVP_top60_DNA+RNA_seed | EGFR | 0.596675 | BRCA | HCC1395 | Gefitinib |
| MVP_top60_DNA_seed | EGFR | 1.153388 | BRCA | MCF7 | Sapitinib | MVP_top60_DNA+RNA_seed | PLK1 | 0.596821 | BRCA | HCC2157 | MK-1775 |
| MVP_top60_DNA_seed | ERBB3 | 1.153388 | BRCA | MCF7 | Sapitinib | MVP_top60_DNA+RNA_seed | TP53 | 0.597125 | BRCA | COLO-824 | MIRA-1 |
| MVP_top60_DNA_seed | MDM2 | 1.153855 | BRCA | BT-474 | Nutlin-3a (-) | MVP_top60_DNA+RNA_seed | SRC | 0.597296 | BRCA | HCC1599 | Bosutinib |
| MVP_top60_DNA_seed | PIK3CA | 1.15996 | BRCA | MDA-MB-157 | AZD8186 | MVP_top60_DNA+RNA_seed | EGFR | 0.599271 | BRCA | HCC38 | AZD3759 |
| MVP_top60_DNA_seed | PIK3CA | 1.160204 | BRCA | MDA-MB-436 | Buparlisib | MVP_top60_DNA+RNA_seed | MAPK3 | 0.599324 | BRCA | Hs-578-T | Ulixertinib |
| MVP_top60_DNA_seed | AKT1 | 1.160429 | BRCA | HCC1500 | Afuresertib | MVP_top60_DNA+RNA_seed | MAPK1 | 0.599324 | BRCA | Hs-578-T | Ulixertinib |
| MVP_top60_DNA_seed | CDK2 | 1.161658 | BRCA | CAL-120 | MK-8776 | MVP_top60_DNA+RNA_seed | HDAC1 | 0.600131 | BRCA | MDA-MB-436 | Romidepsin |
| MVP_top60_DNA_seed | HDAC1 | 1.16298 | BRCA | MDA-MB-175-VII | Entinostat | MVP_top60_DNA+RNA_seed | HDAC2 | 0.600131 | BRCA | MDA-MB-436 | Romidepsin |
| MVP_top60_DNA_seed | TP53 | 1.167902 | BRCA | MFM-223 | MIRA-1 | MVP_top60_DNA+RNA_seed | EGFR | 0.600241 | BRCA | MDA-MB-231 | Osimertinib |
| MVP_top60_DNA_seed | MDM2 | 1.16826 | BRCA | BT-483 | Serdemetan | MVP_top60_DNA+RNA_seed | PIK3CA | 0.60026 | BRCA | ZR-75-30 | AZD8186 |
| MVP_top60_DNA_seed | MAPK3 | 1.168425 | BRCA | BT-474 | SCH772984 | MVP_top60_DNA+RNA_seed | PIK3CA | 0.600468 | BRCA | MCF7 | GNE-317 |
| MVP_top60_DNA_seed | MAPK1 | 1.168425 | BRCA | BT-474 | SCH772984 | MVP_top60_DNA+RNA_seed | HDAC1 | 0.600637 | BRCA | MDA-MB-415 | PCI-34051 |
| MVP_top60_DNA_seed | ERBB2 | 1.168961 | BRCA | BT-483 | Lapatinib | MVP_top60_DNA+RNA_seed | HDAC1 | 0.600773 | BRCA | JIMT-1 | Entinostat |
| MVP_top60_DNA_seed | EGFR | 1.168961 | BRCA | BT-483 | Lapatinib | MVP_top60_DNA+RNA_seed | HDAC1 | 0.60109 | BRCA | MDA-MB-415 | Entinostat |
| MVP_top60_DNA_seed | EGFR | 1.169755 | BRCA | BT-549 | Gefitinib | MVP_top60_DNA+RNA_seed | EGFR | 0.60238 | BRCA | HCC1428 | Afatinib |
| MVP_top60_DNA_seed | CDK1 | 1.17777 | BRCA | MDA-MB-157 | RO-3306 | MVP_top60_DNA+RNA_seed | ERBB2 | 0.60238 | BRCA | HCC1428 | Afatinib |
| MVP_top60_DNA_seed | MAPK1 | 1.178844 | BRCA | CAL-120 | VX-11e | MVP_top60_DNA+RNA_seed | MAPK3 | 0.603121 | BRCA | MDA-MB-175-VII | Ulixertinib |

| | | | | | | | | | | | |
|---|---|---|---|---|---|---|---|---|---|---|---|
| MVP_top60_DNA_seed | EP300 | 1.183683 | BRCA | MCF7 | SGC-CBP30 | MVP_top60_DNA+RNA_seed | MAPK1 | 0.603121 | BRCA | MDA-MB-175-VII | Ulixertinib |
| MVP_top60_DNA_seed | CREBBP | 1.183683 | BRCA | MCF7 | SGC-CBP30 | MVP_top60_DNA+RNA_seed | MAPK1 | 0.60327 | BRCA | HCC1937 | VX-11e |
| MVP_top60_DNA_seed | HDAC1 | 1.184385 | BRCA | MDA-MB-436 | PCI-34051 | MVP_top60_DNA+RNA_seed | AKT1 | 0.603419 | BRCA | HCC1500 | GSK2110183B |
| MVP_top60_DNA_seed | EGFR | 1.185089 | BRCA | MFM-223 | Osimertinib | MVP_top60_DNA+RNA_seed | PIK3CA | 0.604254 | BRCA | MDA-MB-157 | Alpelisib |
| MVP_top60_DNA_seed | AKT1 | 1.191457 | BRCA | HCC1500 | Uprosertib | MVP_top60_DNA+RNA_seed | MDM2 | 0.60526 | BRCA | HCC1143 | Serdemetan |
| MVP_top60_DNA_seed | ERBB2 | 1.194379 | BRCA | CAL-120 | Afatinib | MVP_top60_DNA+RNA_seed | AKT1 | 0.607114 | BRCA | MCF7 | AZD5363 |
| MVP_top60_DNA_seed | EGFR | 1.194379 | BRCA | CAL-120 | Afatinib | MVP_top60_DNA+RNA_seed | EGFR | 0.60732 | BRCA | MDA-MB-157 | Erlotinib |
| MVP_top60_DNA_seed | ESR1 | 1.197697 | BRCA | ZR-75-30 | Tamoxifen | MVP_top60_DNA+RNA_seed | MAPK3 | 0.610385 | BRCA | HCC1143 | Ulixertinib |
| MVP_top60_DNA_seed | EGFR | 1.198466 | BRCA | MCF7 | Gefitinib | MVP_top60_DNA+RNA_seed | MAPK1 | 0.610385 | BRCA | HCC1143 | Ulixertinib |
| MVP_top60_DNA_seed | SRC | 1.19926 | BRCA | MDA-MB-330 | Bosutinib | MVP_top60_DNA+RNA_seed | PIK3CA | 0.611107 | BRCA | CAMA-1 | Alpelisib |
| MVP_top60_DNA_seed | TP53 | 1.204967 | BRCA | MDA-MB-436 | MIRA-1 | MVP_top60_DNA+RNA_seed | EP300 | 0.612053 | BRCA | BT-483 | CPI-637 |
| MVP_top60_DNA_seed | CDK2 | 1.205544 | BRCA | CAL-120 | AZD5438 | MVP_top60_DNA+RNA_seed | TP53 | 0.612158 | BRCA | EFM-19 | MIRA-1 |
| MVP_top60_DNA_seed | ERBB2 | 1.207114 | BRCA | BT-483 | Sapitinib | MVP_top60_DNA+RNA_seed | HSP90AA1 | 0.613982 | BRCA | ZR-75-30 | Tanespimycin |
| MVP_top60_DNA_seed | EGFR | 1.207114 | BRCA | BT-483 | Sapitinib | MVP_top60_DNA+RNA_seed | HSP90AB1 | 0.613982 | BRCA | ZR-75-30 | Tanespimycin |
| MVP_top60_DNA_seed | ERBB3 | 1.207114 | BRCA | BT-483 | Sapitinib | MVP_top60_DNA+RNA_seed | EGFR | 0.615211 | BRCA | CAL-85-1 | Afatinib |
| MVP_top60_DNA_seed | MAPK3 | 1.212895 | BRCA | HCC1937 | Ulixertinib | MVP_top60_DNA+RNA_seed | ERBB2 | 0.615211 | BRCA | CAL-85-1 | Afatinib |
| MVP_top60_DNA_seed | MAPK1 | 1.212895 | BRCA | HCC1937 | Ulixertinib | MVP_top60_DNA+RNA_seed | CDK2 | 0.615699 | BRCA | CAMA-1 | MK-8776 |
| MVP_top60_DNA_seed | EGFR | 1.218625 | BRCA | HCC38 | Gefitinib | MVP_top60_DNA+RNA_seed | PLK1 | 0.61745 | BRCA | EFM-19 | BI-2536 |
| MVP_top60_DNA_seed | CDK2 | 1.223227 | BRCA | CAL-85-1 | MK-8776 | MVP_top60_DNA+RNA_seed | MAPK3 | 0.618559 | BRCA | MDA-MB-415 | Ulixertinib |
| MVP_top60_DNA_seed | TP53 | 1.22708 | BRCA | CAL-120 | MIRA-1 | MVP_top60_DNA+RNA_seed | MAPK1 | 0.618559 | BRCA | MDA-MB-415 | Ulixertinib |
| MVP_top60_DNA_seed | ERBB2 | 1.228839 | BRCA | MFM-223 | Sapitinib | MVP_top60_DNA+RNA_seed | PIK3CA | 0.618825 | BRCA | HCC1187 | Buparlisib |
| MVP_top60_DNA_seed | EGFR | 1.228839 | BRCA | MFM-223 | Sapitinib | MVP_top60_DNA+RNA_seed | AKT1 | 0.619 | BRCA | ZR-75-30 | Afuresertib |
| MVP_top60_DNA_seed | ERBB3 | 1.228839 | BRCA | MFM-223 | Sapitinib | MVP_top60_DNA+RNA_seed | AKT1 | 0.619932 | BRCA | MDA-MB-157 | GSK2110183B |
| MVP_top60_DNA_seed | CDK2 | 1.231689 | BRCA | HCC1937 | AZD5438 | MVP_top60_DNA+RNA_seed | MAPK3 | 0.621142 | BRCA | CAMA-1 | Ulixertinib |
| MVP_top60_DNA_seed | CDK2 | 1.235341 | BRCA | MDA-MB-330 | MK-8776 | MVP_top60_DNA+RNA_seed | MAPK1 | 0.621142 | BRCA | CAMA-1 | Ulixertinib |
| MVP_top60_DNA_seed | SRC | 1.237267 | BRCA | HCC38 | Bosutinib | MVP_top60_DNA+RNA_seed | HDAC1 | 0.62297 | BRCA | HCC38 | Romidepsin |
| MVP_top60_DNA_seed | ESR1 | 1.238623 | BRCA | CAL-120 | GDC0810 | MVP_top60_DNA+RNA_seed | HDAC2 | 0.62297 | BRCA | HCC38 | Romidepsin |
| MVP_top60_DNA_seed | PIK3CA | 1.246289 | BRCA | BT-483 | GNE-317 | MVP_top60_DNA+RNA_seed | MDM2 | 0.623686 | BRCA | EFM-19 | Nutlin-3a (-) |
| MVP_top60_DNA_seed | EGFR | 1.257101 | BRCA | MDA-MB-436 | Gefitinib | MVP_top60_DNA+RNA_seed | MAPK1 | 0.628375 | BRCA | MDA-MB-330 | VX-11e |
| MVP_top60_DNA_seed | ESR1 | 1.257127 | BRCA | HCC1428 | Tamoxifen | MVP_top60_DNA+RNA_seed | SRC | 0.628974 | BRCA | BT-549 | Bosutinib |
| MVP_top60_DNA_seed | MAPK1 | 1.257885 | BRCA | HCC1419 | VX-11e | MVP_top60_DNA+RNA_seed | AKT1 | 0.631516 | BRCA | ZR-75-30 | Uprosertib |
| MVP_top60_DNA_seed | HSP90AA1 | 1.25846 | BRCA | HCC1599 | CCT-018159 | MVP_top60_DNA+RNA_seed | AKT1 | 0.631719 | BRCA | MDA-MB-231 | GSK2110183B |
| MVP_top60_DNA_seed | HSP90AB1 | 1.25846 | BRCA | HCC1599 | CCT-018159 | MVP_top60_DNA+RNA_seed | EP300 | 0.634874 | BRCA | UACC-812 | CPI-637 |
| MVP_top60_DNA_seed | CDK1 | 1.258821 | BRCA | BT-474 | RO-3306 | MVP_top60_DNA+RNA_seed | HDAC1 | 0.644273 | BRCA | MCF7 | Romidepsin |
| MVP_top60_DNA_seed | CDK1 | 1.259151 | BRCA | BT-483 | RO-3306 | MVP_top60_DNA+RNA_seed | HDAC2 | 0.644273 | BRCA | MCF7 | Romidepsin |
| MVP_top60_DNA_seed | EGFR | 1.261003 | BRCA | Hs-578-T | Osimertinib | MVP_top60_DNA+RNA_seed | ATM | 0.645282 | BRCA | EFM-192A | KU-55933 |
| MVP_top60_DNA_seed | MDM2 | 1.264404 | BRCA | BT-474 | Serdemetan | MVP_top60_DNA+RNA_seed | PLK1 | 0.646539 | BRCA | MRK-nu-1 | MK-1775 |
| MVP_top60_DNA_seed | ERBB2 | 1.267354 | BRCA | HCC1187 | Lapatinib | MVP_top60_DNA+RNA_seed | MDM2 | 0.646593 | BRCA | MDA-MB-361 | Nutlin-3a (-) |
| MVP_top60_DNA_seed | EGFR | 1.267354 | BRCA | HCC1187 | Lapatinib | MVP_top60_DNA+RNA_seed | CDK2 | 0.647108 | BRCA | EFM-192A | AZD5438 |
| MVP_top60_DNA_seed | PIK3CA | 1.275595 | BRCA | CAL-120 | AZD8186 | MVP_top60_DNA+RNA_seed | MAPK3 | 0.647704 | BRCA | CAMA-1 | SCH772984 |
| MVP_top60_DNA_seed | EGFR | 1.284891 | BRCA | CAL-120 | Erlotinib | MVP_top60_DNA+RNA_seed | MAPK1 | 0.647704 | BRCA | CAMA-1 | SCH772984 |

| | | | | | | | | | | | |
|---|---|---|---|---|---|---|---|---|---|---|---|
| MVP_top60_DNA_seed | MDM2 | 1.288863 | BRCA | CAL-120 | Serdemetan | MVP_top60_DNA+RNA_seed | AKT1 | 0.651384 | BRCA | ZR-75-30 | AZD5363 |
| MVP_top60_DNA_seed | TP53 | 1.29189 | BRCA | BT-474 | MIRA-1 | MVP_top60_DNA+RNA_seed | EGFR | 0.651455 | BRCA | YMB-1-E | Gefitinib |
| MVP_top60_DNA_seed | AKT1 | 1.292071 | BRCA | MDA-MB-436 | AT13148 | MVP_top60_DNA+RNA_seed | MAPK3 | 0.652549 | BRCA | HCC1937 | SCH772984 |
| MVP_top60_DNA_seed | AKT1 | 1.295298 | BRCA | MDA-MB-231 | AZD5363 | MVP_top60_DNA+RNA_seed | MAPK1 | 0.652549 | BRCA | HCC1937 | SCH772984 |
| MVP_top60_DNA_seed | MAPK3 | 1.297188 | BRCA | HCC1500 | Ulixertinib | MVP_top60_DNA+RNA_seed | EP300 | 0.654191 | BRCA | HCC1143 | SGC-CBP30 |
| MVP_top60_DNA_seed | MAPK1 | 1.297188 | BRCA | HCC1500 | Ulixertinib | MVP_top60_DNA+RNA_seed | PLK1 | 0.654826 | BRCA | CAL-120 | BI-2536 |
| MVP_top60_DNA_seed | SRC | 1.298928 | BRCA | MFM-223 | Bosutinib | MVP_top60_DNA+RNA_seed | CDK1 | 0.655792 | BRCA | MFM-223 | RO-3306 |
| MVP_top60_DNA_seed | MDM2 | 1.300555 | BRCA | MDA-MB-436 | Serdemetan | MVP_top60_DNA+RNA_seed | MAPK3 | 0.656012 | BRCA | BT-549 | SCH772984 |
| MVP_top60_DNA_seed | ERBB2 | 1.300938 | BRCA | HCC1428 | Sapitinib | MVP_top60_DNA+RNA_seed | MAPK1 | 0.656012 | BRCA | BT-549 | SCH772984 |
| MVP_top60_DNA_seed | EGFR | 1.300938 | BRCA | HCC1428 | Sapitinib | MVP_top60_DNA+RNA_seed | EGFR | 0.656821 | BRCA | EFM-192A | Gefitinib |
| MVP_top60_DNA_seed | ERBB3 | 1.300938 | BRCA | HCC1428 | Sapitinib | MVP_top60_DNA+RNA_seed | CDK2 | 0.660289 | BRCA | EFM-19 | AZD5438 |
| MVP_top60_DNA_seed | ATM | 1.302381 | BRCA | MDA-MB-157 | KU-55933 | MVP_top60_DNA+RNA_seed | EGFR | 0.662206 | BRCA | YMB-1-E | Sapitinib |
| MVP_top60_DNA_seed | AKT1 | 1.305997 | BRCA | CAL-120 | AT13148 | MVP_top60_DNA+RNA_seed | ERBB2 | 0.662206 | BRCA | YMB-1-E | Sapitinib |
| MVP_top60_DNA_seed | HDAC1 | 1.307431 | BRCA | MDA-MB-436 | Entinostat | MVP_top60_DNA+RNA_seed | ERBB3 | 0.662206 | BRCA | YMB-1-E | Sapitinib |
| MVP_top60_DNA_seed | HDAC1 | 1.309236 | BRCA | HCC1937 | PCI-34051 | MVP_top60_DNA+RNA_seed | MDM2 | 0.664847 | BRCA | EFM-192A | Nutlin-3a (-) |
| MVP_top60_DNA_seed | EP300 | 1.312826 | BRCA | MDA-MB-436 | SGC-CBP30 | MVP_top60_DNA+RNA_seed | HSP90AA1 | 0.666444 | BRCA | EFM-19 | Tanespimycin |
| MVP_top60_DNA_seed | CREBBP | 1.312826 | BRCA | MDA-MB-436 | SGC-CBP30 | MVP_top60_DNA+RNA_seed | HSP90AB1 | 0.666444 | BRCA | EFM-19 | Tanespimycin |
| MVP_top60_DNA_seed | HSP90AA1 | 1.322485 | BRCA | HCC1428 | Luminespib | MVP_top60_DNA+RNA_seed | AKT1 | 0.667264 | BRCA | JIMT-1 | Uprosertib |
| MVP_top60_DNA_seed | HSP90AB1 | 1.322485 | BRCA | HCC1428 | Luminespib | MVP_top60_DNA+RNA_seed | TP53 | 0.671396 | BRCA | EFM-192A | MIRA-1 |
| MVP_top60_DNA_seed | MDM2 | 1.323683 | BRCA | BT-483 | Nutlin-3a (-) | MVP_top60_DNA+RNA_seed | HSP90AA1 | 0.673884 | BRCA | EFM-19 | Luminespib |
| MVP_top60_DNA_seed | AKT1 | 1.330708 | BRCA | MDA-MB-436 | MK-2206 | MVP_top60_DNA+RNA_seed | HSP90AB1 | 0.673884 | BRCA | EFM-19 | Luminespib |
| MVP_top60_DNA_seed | SRC | 1.332667 | BRCA | YMB-1-E | Bosutinib | MVP_top60_DNA+RNA_seed | EGFR | 0.67533 | BRCA | JIMT-1 | Gefitinib |
| MVP_top60_DNA_seed | EP300 | 1.340486 | BRCA | HCC1143 | CPI-637 | MVP_top60_DNA+RNA_seed | HDAC1 | 0.675533 | BRCA | HCC1428 | PCI-34051 |
| MVP_top60_DNA_seed | CDK2 | 1.343109 | BRCA | HDQ-P1 | MK-8776 | MVP_top60_DNA+RNA_seed | MAPK1 | 0.67569 | BRCA | JIMT-1 | VX-11e |
| MVP_top60_DNA_seed | EGFR | 1.352546 | BRCA | BT-549 | Erlotinib | MVP_top60_DNA+RNA_seed | AKT1 | 0.675987 | BRCA | HCC38 | Afuresertib |
| MVP_top60_DNA_seed | EP300 | 1.353012 | BRCA | CAL-120 | CPI-637 | MVP_top60_DNA+RNA_seed | PLK1 | 0.677287 | BRCA | HCC1395 | BI-2536 |
| MVP_top60_DNA_seed | EGFR | 1.353353 | BRCA | UACC-812 | AZD3759 | MVP_top60_DNA+RNA_seed | PIK3CA | 0.678447 | BRCA | MDA-MB-231 | Alpelisib |
| MVP_top60_DNA_seed | ERBB2 | 1.355541 | BRCA | MDA-MB-157 | Afatinib | MVP_top60_DNA+RNA_seed | EGFR | 0.679063 | BRCA | BT-20 | Afatinib |
| MVP_top60_DNA_seed | EGFR | 1.355541 | BRCA | MDA-MB-157 | Afatinib | MVP_top60_DNA+RNA_seed | ERBB2 | 0.679063 | BRCA | BT-20 | Afatinib |
| MVP_top60_DNA_seed | AKT1 | 1.355933 | BRCA | HCC38 | MK-2206 | MVP_top60_DNA+RNA_seed | EGFR | 0.67979 | BRCA | MRK-nu-1 | Osimertinib |
| MVP_top60_DNA_seed | EGFR | 1.359327 | BRCA | HCC38 | Erlotinib | MVP_top60_DNA+RNA_seed | MAPK1 | 0.680542 | BRCA | T47D | VX-11e |
| MVP_top60_DNA_seed | MAPK3 | 1.36602 | BRCA | CAL-148 | SCH772984 | MVP_top60_DNA+RNA_seed | PIK3CA | 0.685429 | BRCA | MRK-nu-1 | Alpelisib |
| MVP_top60_DNA_seed | MAPK1 | 1.36602 | BRCA | CAL-148 | SCH772984 | MVP_top60_DNA+RNA_seed | MAPK3 | 0.688152 | BRCA | T47D | Ulixertinib |
| MVP_top60_DNA_seed | CDK2 | 1.368153 | BRCA | HCC1428 | MK-8776 | MVP_top60_DNA+RNA_seed | MAPK1 | 0.688152 | BRCA | T47D | Ulixertinib |
| MVP_top60_DNA_seed | CDK2 | 1.368896 | BRCA | MCF7 | MK-8776 | MVP_top60_DNA+RNA_seed | PLK1 | 0.691406 | BRCA | HCC1954 | MK-1775 |
| MVP_top60_DNA_seed | AKT1 | 1.374834 | BRCA | BT-483 | Uprosertib | MVP_top60_DNA+RNA_seed | ESR1 | 0.69315 | BRCA | UACC-893 | GDC0810 |
| MVP_top60_DNA_seed | MDM2 | 1.378181 | BRCA | MFM-223 | Nutlin-3a (-) | MVP_top60_DNA+RNA_seed | AKT1 | 0.696434 | BRCA | HCC2218 | Afuresertib |
| MVP_top60_DNA_seed | MAPK3 | 1.378805 | BRCA | HCC1428 | Ulixertinib | MVP_top60_DNA+RNA_seed | AKT1 | 0.697626 | BRCA | MDA-MB-436 | Uprosertib |
| MVP_top60_DNA_seed | MAPK1 | 1.378805 | BRCA | HCC1428 | Ulixertinib | MVP_top60_DNA+RNA_seed | EGFR | 0.698003 | BRCA | MDA-MB-436 | Osimertinib |
| MVP_top60_DNA_seed | AKT1 | 1.381455 | BRCA | HCC1500 | Uprosertib | MVP_top60_DNA+RNA_seed | MDM2 | 0.699228 | BRCA | MDA-MB-231 | Nutlin-3a (-) |
| MVP_top60_DNA_seed | ESR1 | 1.381542 | BRCA | UACC-893 | Tamoxifen | MVP_top60_DNA+RNA_seed | MAPK3 | 0.699837 | BRCA | EVSA-T | SCH772984 |

| | | | | | | | | | | | |
|---|---|---|---|---|---|---|---|---|---|---|---|
| MVP_top60_DNA_seed | CDK2 | 1.385994 | BRCA | EFM-192A | MK-8776 | MVP_top60_DNA+RNA_seed | MAPK1 | 0.699837 | BRCA | EVSA-T | SCH772984 |
| MVP_top60_DNA_seed | MDM2 | 1.389214 | BRCA | MDA-MB-436 | Nutlin-3a (-) | MVP_top60_DNA+RNA_seed | SRC | 0.700187 | BRCA | COLO-824 | Bosutinib |
| MVP_top60_DNA_seed | HDAC1 | 1.391353 | BRCA | BT-549 | PCI-34051 | MVP_top60_DNA+RNA_seed | HSP90AA1 | 0.700301 | BRCA | HCC1143 | Tanespimycin |
| MVP_top60_DNA_seed | ERBB2 | 1.392792 | BRCA | YMB-1-E | Afatinib | MVP_top60_DNA+RNA_seed | HSP90AB1 | 0.700301 | BRCA | HCC1143 | Tanespimycin |
| MVP_top60_DNA_seed | EGFR | 1.392792 | BRCA | YMB-1-E | Afatinib | MVP_top60_DNA+RNA_seed | EGFR | 0.701297 | BRCA | Hs-578-T | Erlotinib |
| MVP_top60_DNA_seed | EP300 | 1.397503 | BRCA | MDA-MB-175-VII | CPI-637 | MVP_top60_DNA+RNA_seed | MAPK3 | 0.702898 | BRCA | HCC1954 | Ulixertinib |
| MVP_top60_DNA_seed | EGFR | 1.397503 | BRCA | CAL-85-1 | Osimertinib | MVP_top60_DNA+RNA_seed | MAPK1 | 0.702898 | BRCA | HCC1954 | Ulixertinib |
| MVP_top60_DNA_seed | MAPK3 | 1.401421 | BRCA | EFM-192A | Ulixertinib | MVP_top60_DNA+RNA_seed | HSP90AA1 | 0.703344 | BRCA | MDA-MB-468 | Tanespimycin |
| MVP_top60_DNA_seed | MAPK1 | 1.401421 | BRCA | EFM-192A | Ulixertinib | MVP_top60_DNA+RNA_seed | HSP90AB1 | 0.703344 | BRCA | MDA-MB-468 | Tanespimycin |
| MVP_top60_DNA_seed | HDAC1 | 1.404337 | BRCA | CAL-120 | Entinostat | MVP_top60_DNA+RNA_seed | MDM2 | 0.704055 | BRCA | MDA-MB-361 | Serdemetan |
| MVP_top60_DNA_seed | HDAC1 | 1.40823 | BRCA | HCC1187 | Romidepsin | MVP_top60_DNA+RNA_seed | MDM2 | 0.706376 | BRCA | MDA-MB-453 | Nutlin-3a (-) |
| MVP_top60_DNA_seed | HDAC2 | 1.40823 | BRCA | HCC1187 | Romidepsin | MVP_top60_DNA+RNA_seed | PLK1 | 0.70649 | BRCA | AU565 | MK-1775 |
| MVP_top60_DNA_seed | AKT1 | 1.408726 | BRCA | CAL-85-1 | AT13148 | MVP_top60_DNA+RNA_seed | EP300 | 0.706754 | BRCA | MDA-MB-361 | CPI-637 |
| MVP_top60_DNA_seed | SRC | 1.409147 | BRCA | BT-474 | Bosutinib | MVP_top60_DNA+RNA_seed | PIK3CA | 0.706951 | BRCA | MDA-MB-468 | Alpelisib |
| MVP_top60_DNA_seed | CDK2 | 1.417989 | BRCA | BT-474 | Dinaciclib | MVP_top60_DNA+RNA_seed | EP300 | 0.707106 | BRCA | COLO-824 | SGC-CBP30 |
| MVP_top60_DNA_seed | CDK1 | 1.417989 | BRCA | BT-474 | Dinaciclib | MVP_top60_DNA+RNA_seed | MDM2 | 0.709502 | BRCA | UACC-893 | Serdemetan |
| MVP_top60_DNA_seed | ERBB2 | 1.422056 | BRCA | HCC38 | Afatinib | MVP_top60_DNA+RNA_seed | PLK1 | 0.712753 | BRCA | MDA-MB-157 | BI-2536 |
| MVP_top60_DNA_seed | EGFR | 1.422056 | BRCA | HCC38 | Afatinib | MVP_top60_DNA+RNA_seed | MAPK1 | 0.713842 | BRCA | MCF7 | VX-11e |
| MVP_top60_DNA_seed | EGFR | 1.428389 | BRCA | ZR-75-30 | AZD3759 | MVP_top60_DNA+RNA_seed | CDK2 | 0.715128 | BRCA | EFM-19 | MK-8776 |
| MVP_top60_DNA_seed | MAPK3 | 1.434598 | BRCA | HCC1187 | SCH772984 | MVP_top60_DNA+RNA_seed | EGFR | 0.715982 | BRCA | CAL-148 | Lapatinib |
| MVP_top60_DNA_seed | MAPK1 | 1.434598 | BRCA | HCC1187 | SCH772984 | MVP_top60_DNA+RNA_seed | ERBB2 | 0.715982 | BRCA | CAL-148 | Lapatinib |
| MVP_top60_DNA_seed | MAPK3 | 1.43646 | BRCA | JIMT-1 | SCH772984 | MVP_top60_DNA+RNA_seed | CDK2 | 0.716176 | BRCA | HCC1599 | Dinaciclib |
| MVP_top60_DNA_seed | MAPK1 | 1.43646 | BRCA | JIMT-1 | SCH772984 | MVP_top60_DNA+RNA_seed | CDK1 | 0.716176 | BRCA | HCC1599 | Dinaciclib |
| MVP_top60_DNA_seed | MAPK3 | 1.441685 | BRCA | HCC38 | SCH772984 | MVP_top60_DNA+RNA_seed | CDK1 | 0.71654 | BRCA | EFM-192A | RO-3306 |
| MVP_top60_DNA_seed | MAPK1 | 1.441685 | BRCA | HCC38 | SCH772984 | MVP_top60_DNA+RNA_seed | AKT1 | 0.716764 | BRCA | JIMT-1 | GSK2110183B |
| MVP_top60_DNA_seed | AKT1 | 1.443381 | BRCA | HCC1187 | Uprosertib | MVP_top60_DNA+RNA_seed | AKT1 | 0.727067 | BRCA | JIMT-1 | AT13148 |
| MVP_top60_DNA_seed | TP53 | 1.444911 | BRCA | HCC1428 | MIRA-1 | MVP_top60_DNA+RNA_seed | MDM2 | 0.730482 | BRCA | MDA-MB-330 | Serdemetan |
| MVP_top60_DNA_seed | MAPK3 | 1.445339 | BRCA | HCC1599 | Ulixertinib | MVP_top60_DNA+RNA_seed | AKT1 | 0.730861 | BRCA | MDA-MB-231 | MK-2206 |
| MVP_top60_DNA_seed | MAPK1 | 1.445339 | BRCA | HCC1599 | Ulixertinib | MVP_top60_DNA+RNA_seed | PLK1 | 0.733164 | BRCA | HCC1187 | MK-1775 |
| MVP_top60_DNA_seed | CDK2 | 1.447728 | BRCA | JIMT-1 | Dinaciclib | MVP_top60_DNA+RNA_seed | EGFR | 0.733586 | BRCA | HCC1500 | Osimertinib |
| MVP_top60_DNA_seed | CDK1 | 1.447728 | BRCA | JIMT-1 | Dinaciclib | MVP_top60_DNA+RNA_seed | HDAC1 | 0.744191 | BRCA | JIMT-1 | PCI-34051 |
| MVP_top60_DNA_seed | AKT1 | 1.458537 | BRCA | ZR-75-30 | GSK2110183B | MVP_top60_DNA+RNA_seed | EGFR | 0.744628 | BRCA | MFM-223 | AZD3759 |
| MVP_top60_DNA_seed | HDAC1 | 1.458654 | BRCA | EFM-19 | Entinostat | MVP_top60_DNA+RNA_seed | EGFR | 0.744935 | BRCA | MDA-MB-436 | Sapitinib |
| MVP_top60_DNA_seed | EGFR | 1.458705 | BRCA | HCC1428 | Erlotinib | MVP_top60_DNA+RNA_seed | ERBB2 | 0.744935 | BRCA | MDA-MB-436 | Sapitinib |
| MVP_top60_DNA_seed | EGFR | 1.45958 | BRCA | HCC1599 | Erlotinib | MVP_top60_DNA+RNA_seed | ERBB3 | 0.744935 | BRCA | MDA-MB-436 | Sapitinib |
| MVP_top60_DNA_seed | HDAC1 | 1.459961 | BRCA | BT-474 | PCI-34051 | MVP_top60_DNA+RNA_seed | EP300 | 0.745128 | BRCA | BT-20 | CPI-637 |
| MVP_top60_DNA_seed | AKT1 | 1.466902 | BRCA | CAL-120 | GSK2110183B | MVP_top60_DNA+RNA_seed | MAPK3 | 0.745682 | BRCA | MDA-MB-436 | SCH772984 |
| MVP_top60_DNA_seed | EP300 | 1.467191 | BRCA | HCC1428 | SGC-CBP30 | MVP_top60_DNA+RNA_seed | MAPK1 | 0.745682 | BRCA | MDA-MB-436 | SCH772984 |
| MVP_top60_DNA_seed | CREBBP | 1.467191 | BRCA | HCC1428 | SGC-CBP30 | MVP_top60_DNA+RNA_seed | MAPK1 | 0.745888 | BRCA | Hs-578-T | VX-11e |
| MVP_top60_DNA_seed | CDK2 | 1.471313 | BRCA | MDA-MB-436 | AZD5438 | MVP_top60_DNA+RNA_seed | SRC | 0.746056 | BRCA | HCC1954 | Bosutinib |
| MVP_top60_DNA_seed | PIK3CA | 1.481992 | BRCA | HCC1500 | Buparlisib | MVP_top60_DNA+RNA_seed | EGFR | 0.747607 | BRCA | HCC1395 | Osimertinib |

| | | | | | | | | | | | |
|---|---|---|---|---|---|---|---|---|---|---|---|
| MVP_top60_DNA_seed | MDM2 | 1.5077 | BRCA | HCC38 | Serdemetan | MVP_top60_DNA+RNA_seed | EGFR | 0.749401 | BRCA | CAMA-1 | Erlotinib |
| MVP_top60_DNA_seed | HSP90AA1 | 1.509597 | BRCA | BT-483 | Tanespimycin | MVP_top60_DNA+RNA_seed | CDK1 | 0.749464 | BRCA | MDA-MB-330 | RO-3306 |
| MVP_top60_DNA_seed | HSP90AB1 | 1.509597 | BRCA | BT-483 | Tanespimycin | MVP_top60_DNA+RNA_seed | AKT1 | 0.749546 | BRCA | HCC38 | Uprosertib |
| MVP_top60_DNA_seed | MAPK3 | 1.509658 | BRCA | BT-474 | Ulixertinib | MVP_top60_DNA+RNA_seed | HDAC1 | 0.750636 | BRCA | HCC1143 | PCI-34051 |
| MVP_top60_DNA_seed | MAPK1 | 1.509658 | BRCA | BT-474 | Ulixertinib | MVP_top60_DNA+RNA_seed | SRC | 0.75065 | BRCA | HCC1187 | Bosutinib |
| MVP_top60_DNA_seed | AKT1 | 1.51075 | BRCA | HCC38 | AT13148 | MVP_top60_DNA+RNA_seed | CDK2 | 0.75272 | BRCA | MFM-223 | MK-8776 |
| MVP_top60_DNA_seed | CDK2 | 1.519447 | BRCA | UACC-812 | MK-8776 | MVP_top60_DNA+RNA_seed | AKT1 | 0.753729 | BRCA | MDA-MB-436 | Uprosertib |
| MVP_top60_DNA_seed | MAPK1 | 1.52302 | BRCA | UACC-812 | VX-11e | MVP_top60_DNA+RNA_seed | EGFR | 0.754708 | BRCA | MFM-223 | Afatinib |
| MVP_top60_DNA_seed | MAPK3 | 1.531234 | BRCA | HCC1428 | SCH772984 | MVP_top60_DNA+RNA_seed | ERBB2 | 0.754708 | BRCA | MFM-223 | Afatinib |
| MVP_top60_DNA_seed | MAPK1 | 1.531234 | BRCA | HCC1428 | SCH772984 | MVP_top60_DNA+RNA_seed | MDM2 | 0.755925 | BRCA | HCC1937 | Serdemetan |
| MVP_top60_DNA_seed | MAPK3 | 1.535184 | BRCA | MCF7 | SCH772984 | MVP_top60_DNA+RNA_seed | HDAC1 | 0.756079 | BRCA | BT-549 | Romidepsin |
| MVP_top60_DNA_seed | MAPK1 | 1.535184 | BRCA | MCF7 | SCH772984 | MVP_top60_DNA+RNA_seed | HDAC2 | 0.756079 | BRCA | BT-549 | Romidepsin |
| MVP_top60_DNA_seed | MAPK3 | 1.536895 | BRCA | HCC38 | Ulixertinib | MVP_top60_DNA+RNA_seed | TP53 | 0.756287 | BRCA | HCC38 | MIRA-1 |
| MVP_top60_DNA_seed | MAPK1 | 1.536895 | BRCA | HCC38 | Ulixertinib | MVP_top60_DNA+RNA_seed | HDAC1 | 0.756574 | BRCA | HCC1187 | PCI-34051 |
| MVP_top60_DNA_seed | EP300 | 1.540577 | BRCA | HCC1937 | CPI-637 | MVP_top60_DNA+RNA_seed | AKT1 | 0.758038 | BRCA | MDA-MB-157 | Ipatasertib |
| MVP_top60_DNA_seed | HDAC1 | 1.548354 | BRCA | HCC1937 | Romidepsin | MVP_top60_DNA+RNA_seed | HSP90AA1 | 0.758999 | BRCA | HCC1187 | Tanespimycin |
| MVP_top60_DNA_seed | HDAC2 | 1.548354 | BRCA | HCC1937 | Romidepsin | MVP_top60_DNA+RNA_seed | HSP90AB1 | 0.758999 | BRCA | HCC1187 | Tanespimycin |
| MVP_top60_DNA_seed | PIK3CA | 1.553813 | BRCA | HCC1187 | GNE-317 | MVP_top60_DNA+RNA_seed | TP53 | 0.759119 | BRCA | HCC1395 | MIRA-1 |
| MVP_top60_DNA_seed | CDK2 | 1.557598 | BRCA | BT-474 | MK-8776 | MVP_top60_DNA+RNA_seed | ESR1 | 0.759629 | BRCA | HCC1395 | GDC0810 |
| MVP_top60_DNA_seed | PIK3CA | 1.582609 | BRCA | HCC1419 | GNE-317 | MVP_top60_DNA+RNA_seed | MAPK3 | 0.760427 | BRCA | MDA-MB-157 | SCH772984 |
| MVP_top60_DNA_seed | AKT1 | 1.593483 | BRCA | HCC1428 | GSK2110183B | MVP_top60_DNA+RNA_seed | MAPK1 | 0.760427 | BRCA | MDA-MB-157 | SCH772984 |
| MVP_top60_DNA_seed | AKT1 | 1.594381 | BRCA | HCC1937 | AT13148 | MVP_top60_DNA+RNA_seed | EGFR | 0.761561 | BRCA | MDA-MB-157 | AZD3759 |
| MVP_top60_DNA_seed | HDAC1 | 1.595622 | BRCA | ZR-75-30 | PCI-34051 | MVP_top60_DNA+RNA_seed | MAPK3 | 0.762309 | BRCA | MDA-MB-157 | Ulixertinib |
| MVP_top60_DNA_seed | PIK3CA | 1.601177 | BRCA | HCC1500 | AZD8186 | MVP_top60_DNA+RNA_seed | MAPK1 | 0.762309 | BRCA | MDA-MB-157 | Ulixertinib |
| MVP_top60_DNA_seed | AKT1 | 1.610376 | BRCA | HCC1428 | Uprosertib | MVP_top60_DNA+RNA_seed | EGFR | 0.763567 | BRCA | CAL-85-1 | Erlotinib |
| MVP_top60_DNA_seed | MDM2 | 1.625789 | BRCA | CAL-120 | Nutlin-3a (-) | MVP_top60_DNA+RNA_seed | HSP90AA1 | 0.763884 | BRCA | HCC1395 | Tanespimycin |
| MVP_top60_DNA_seed | EGFR | 1.63989 | BRCA | MFM-223 | Gefitinib | MVP_top60_DNA+RNA_seed | HSP90AB1 | 0.763884 | BRCA | HCC1395 | Tanespimycin |
| MVP_top60_DNA_seed | CDK2 | 1.645599 | BRCA | MDA-MB-436 | Dinaciclib | MVP_top60_DNA+RNA_seed | EGFR | 0.77343 | BRCA | BT-20 | Osimertinib |
| MVP_top60_DNA_seed | CDK1 | 1.645599 | BRCA | MDA-MB-436 | Dinaciclib | MVP_top60_DNA+RNA_seed | MDM2 | 0.775007 | BRCA | ZR-75-30 | Nutlin-3a (-) |
| MVP_top60_DNA_seed | MDM2 | 1.675931 | BRCA | HCC1428 | Serdemetan | MVP_top60_DNA+RNA_seed | ESR1 | 0.775338 | BRCA | BT-549 | GDC0810 |
| MVP_top60_DNA_seed | CDK2 | 1.682691 | BRCA | ZR-75-30 | MK-8776 | MVP_top60_DNA+RNA_seed | PIK3CA | 0.776656 | BRCA | HCC1937 | GNE-317 |
| MVP_top60_DNA_seed | AKT1 | 1.687189 | BRCA | CAL-85-1 | AZD5363 | MVP_top60_DNA+RNA_seed | PIK3CA | 0.776945 | BRCA | HCC1187 | Alpelisib |
| MVP_top60_DNA_seed | PIK3CA | 1.70168 | BRCA | ZR-75-30 | Buparlisib | MVP_top60_DNA+RNA_seed | MAPK3 | 0.779847 | BRCA | JIMT-1 | Ulixertinib |
| MVP_top60_DNA_seed | MAPK3 | 1.705641 | BRCA | HCC1428 | Ulixertinib | MVP_top60_DNA+RNA_seed | MAPK1 | 0.779847 | BRCA | JIMT-1 | Ulixertinib |
| MVP_top60_DNA_seed | MAPK1 | 1.705641 | BRCA | HCC1428 | Ulixertinib | MVP_top60_DNA+RNA_seed | HSP90AA1 | 0.780103 | BRCA | HCC1395 | Luminespib |
| MVP_top60_DNA_seed | HDAC1 | 1.706539 | BRCA | ZR-75-30 | Entinostat | MVP_top60_DNA+RNA_seed | HSP90AB1 | 0.780103 | BRCA | HCC1395 | Luminespib |
| MVP_top60_DNA_seed | AKT1 | 1.709297 | BRCA | MRK-nu-1 | MK-2206 | MVP_top60_DNA+RNA_seed | AR | 0.780504 | BRCA | UACC-893 | Bicalutamide |
| MVP_top60_DNA_seed | MAPK1 | 1.714617 | BRCA | HCC1428 | VX-11e | MVP_top60_DNA+RNA_seed | CDK2 | 0.781068 | BRCA | MDA-MB-415 | MK-8776 |
| MVP_top60_DNA_seed | HSP90AA1 | 1.724402 | BRCA | T47D | Tanespimycin | MVP_top60_DNA+RNA_seed | MAPK3 | 0.782028 | BRCA | HCC1599 | Ulixertinib |
| MVP_top60_DNA_seed | HSP90AB1 | 1.724402 | BRCA | T47D | Tanespimycin | MVP_top60_DNA+RNA_seed | MAPK1 | 0.782028 | BRCA | HCC1599 | Ulixertinib |
| MVP_top60_DNA_seed | HSP90AA1 | 1.736226 | BRCA | UACC-812 | Tanespimycin | MVP_top60_DNA+RNA_seed | EGFR | 0.782638 | BRCA | MDA-MB-453 | AZD3759 |

| | | | | | | | | | | | |
|---|---|---|---|---|---|---|---|---|---|---|---|
| MVP_top60_DNA_seed | HSP90AB1 | 1.736226 | BRCA | UACC-812 | Tanespimycin | MVP_top60_DNA+RNA_seed | EGFR | 0.785247 | BRCA | T47D | Gefitinib |
| MVP_top60_DNA_seed | EGFR | 1.762253 | BRCA | ZR-75-30 | Gefitinib | MVP_top60_DNA+RNA_seed | EGFR | 0.785567 | BRCA | MDA-MB-157 | Gefitinib |
| MVP_top60_DNA_seed | MAPK3 | 1.76262 | BRCA | CAL-120 | Ulixertinib | MVP_top60_DNA+RNA_seed | AKT1 | 0.786773 | BRCA | MDA-MB-231 | Afuresertib |
| MVP_top60_DNA_seed | MAPK1 | 1.76262 | BRCA | CAL-120 | Ulixertinib | MVP_top60_DNA+RNA_seed | AKT1 | 0.787601 | BRCA | MDA-MB-436 | AZD5363 |
| MVP_top60_DNA_seed | AKT1 | 1.768638 | BRCA | HCC1500 | MK-2206 | MVP_top60_DNA+RNA_seed | SRC | 0.787765 | BRCA | HCC1428 | Bosutinib |
| MVP_top60_DNA_seed | MDM2 | 1.784761 | BRCA | HCC1428 | Nutlin-3a (-) | MVP_top60_DNA+RNA_seed | EGFR | 0.796342 | BRCA | Hs-578-T | AZD3759 |
| MVP_top60_DNA_seed | MAPK3 | 1.785123 | BRCA | CAL-120 | SCH772984 | MVP_top60_DNA+RNA_seed | MDM2 | 0.79802 | BRCA | Hs-578-T | Serdemetan |
| MVP_top60_DNA_seed | MAPK1 | 1.785123 | BRCA | CAL-120 | SCH772984 | MVP_top60_DNA+RNA_seed | EGFR | 0.799129 | BRCA | MCF7 | Osimertinib |
| MVP_top60_DNA_seed | MAPK3 | 1.788339 | BRCA | BT-549 | Ulixertinib | MVP_top60_DNA+RNA_seed | PLK1 | 0.801968 | BRCA | YMB-1-E | BI-2536 |
| MVP_top60_DNA_seed | MAPK1 | 1.788339 | BRCA | BT-549 | Ulixertinib | MVP_top60_DNA+RNA_seed | AKT1 | 0.803402 | BRCA | HCC38 | GSK2110183B |
| MVP_top60_DNA_seed | ESR1 | 1.790311 | BRCA | MDA-MB-175-VII | Tamoxifen | MVP_top60_DNA+RNA_seed | AKT1 | 0.804515 | BRCA | CAL-120 | Ipatasertib |
| MVP_top60_DNA_seed | MAPK3 | 1.806136 | BRCA | UACC-812 | SCH772984 | MVP_top60_DNA+RNA_seed | MAPK3 | 0.810001 | BRCA | MDA-MB-436 | Ulixertinib |
| MVP_top60_DNA_seed | MAPK1 | 1.806136 | BRCA | UACC-812 | SCH772984 | MVP_top60_DNA+RNA_seed | MAPK1 | 0.810001 | BRCA | MDA-MB-436 | Ulixertinib |
| MVP_top60_DNA_seed | CDK2 | 1.808025 | BRCA | HCC1500 | Dinaciclib | MVP_top60_DNA+RNA_seed | MAPK3 | 0.811564 | BRCA | OCUB-M | Ulixertinib |
| MVP_top60_DNA_seed | CDK1 | 1.808025 | BRCA | HCC1500 | Dinaciclib | MVP_top60_DNA+RNA_seed | MAPK1 | 0.811564 | BRCA | OCUB-M | Ulixertinib |
| MVP_top60_DNA_seed | EGFR | 1.808409 | BRCA | ZR-75-30 | Erlotinib | MVP_top60_DNA+RNA_seed | AKT1 | 0.811948 | BRCA | T47D | AT13148 |
| MVP_top60_DNA_seed | MAPK3 | 1.812945 | BRCA | BT-483 | Ulixertinib | MVP_top60_DNA+RNA_seed | ESR1 | 0.814043 | BRCA | MFM-223 | GDC0810 |
| MVP_top60_DNA_seed | MAPK1 | 1.812945 | BRCA | BT-483 | Ulixertinib | MVP_top60_DNA+RNA_seed | EP300 | 0.815785 | BRCA | HCC1954 | CPI-637 |
| MVP_top60_DNA_seed | EGFR | 1.813324 | BRCA | UACC-812 | Erlotinib | MVP_top60_DNA+RNA_seed | EGFR | 0.817906 | BRCA | JIMT-1 | AZD3759 |
| MVP_top60_DNA_seed | HDAC1 | 1.814019 | BRCA | MFM-223 | Entinostat | MVP_top60_DNA+RNA_seed | AKT1 | 0.818369 | BRCA | MDA-MB-157 | Uprosertib |
| MVP_top60_DNA_seed | TP53 | 1.829054 | BRCA | MDA-MB-175-VII | MIRA-1 | MVP_top60_DNA+RNA_seed | MAPK3 | 0.821243 | BRCA | BT-20 | Ulixertinib |
| MVP_top60_DNA_seed | MDM2 | 1.829666 | BRCA | HCC1599 | Serdemetan | MVP_top60_DNA+RNA_seed | MAPK1 | 0.821243 | BRCA | BT-20 | Ulixertinib |
| MVP_top60_DNA_seed | ERBB2 | 1.830561 | BRCA | CAL-120 | Sapitinib | MVP_top60_DNA+RNA_seed | EP300 | 0.821278 | BRCA | EFM-192A | CPI-637 |
| MVP_top60_DNA_seed | EGFR | 1.830561 | BRCA | CAL-120 | Sapitinib | MVP_top60_DNA+RNA_seed | AKT1 | 0.826809 | BRCA | HCC1500 | AZD5363 |
| MVP_top60_DNA_seed | ERBB3 | 1.830561 | BRCA | CAL-120 | Sapitinib | MVP_top60_DNA+RNA_seed | HSP90AA1 | 0.827201 | BRCA | HCC1599 | Tanespimycin |
| MVP_top60_DNA_seed | SRC | 1.831053 | BRCA | ZR-75-30 | Bosutinib | MVP_top60_DNA+RNA_seed | HSP90AB1 | 0.827201 | BRCA | HCC1599 | Tanespimycin |
| MVP_top60_DNA_seed | MAPK1 | 1.832258 | BRCA | BT-474 | VX-11e | MVP_top60_DNA+RNA_seed | MDM2 | 0.827641 | BRCA | HCC1937 | Nutlin-3a (-) |
| MVP_top60_DNA_seed | AKT1 | 1.837836 | BRCA | HCC1428 | Uprosertib | MVP_top60_DNA+RNA_seed | HSP90AA1 | 0.828761 | BRCA | CAMA-1 | Tanespimycin |
| MVP_top60_DNA_seed | MAPK3 | 1.859775 | BRCA | HCC1599 | SCH772984 | MVP_top60_DNA+RNA_seed | HSP90AB1 | 0.828761 | BRCA | CAMA-1 | Tanespimycin |
| MVP_top60_DNA_seed | MAPK1 | 1.859775 | BRCA | HCC1599 | SCH772984 | MVP_top60_DNA+RNA_seed | EP300 | 0.829052 | BRCA | MFM-223 | CPI-637 |
| MVP_top60_DNA_seed | ERBB2 | 1.860494 | BRCA | MDA-MB-436 | Afatinib | MVP_top60_DNA+RNA_seed | MAPK3 | 0.831668 | BRCA | MRK-nu-1 | Ulixertinib |
| MVP_top60_DNA_seed | EGFR | 1.860494 | BRCA | MDA-MB-436 | Afatinib | MVP_top60_DNA+RNA_seed | MAPK1 | 0.831668 | BRCA | MRK-nu-1 | Ulixertinib |
| MVP_top60_DNA_seed | AKT1 | 1.869833 | BRCA | HCC1428 | AT13148 | MVP_top60_DNA+RNA_seed | SRC | 0.832036 | BRCA | MDA-MB-453 | Bosutinib |
| MVP_top60_DNA_seed | AKT1 | 1.888303 | BRCA | BT-474 | AT13148 | MVP_top60_DNA+RNA_seed | ESR1 | 0.832082 | BRCA | CAL-120 | Tamoxifen |
| MVP_top60_DNA_seed | SRC | 1.895881 | BRCA | CAL-120 | Bosutinib | MVP_top60_DNA+RNA_seed | AKT1 | 0.833884 | BRCA | MDA-MB-157 | Uprosertib |
| MVP_top60_DNA_seed | TP53 | 1.908261 | BRCA | BT-483 | MIRA-1 | MVP_top60_DNA+RNA_seed | ESR1 | 0.834192 | BRCA | HCC1599 | Tamoxifen |
| MVP_top60_DNA_seed | AKT1 | 1.915116 | BRCA | BT-483 | MK-2206 | MVP_top60_DNA+RNA_seed | SRC | 0.83669 | BRCA | CAMA-1 | Bosutinib |
| MVP_top60_DNA_seed | PIK3CA | 1.92054 | BRCA | HCC1500 | GNE-317 | MVP_top60_DNA+RNA_seed | EGFR | 0.837306 | BRCA | Hs-578-T | Gefitinib |
| MVP_top60_DNA_seed | MAPK3 | 1.925076 | BRCA | HCC38 | Ulixertinib | MVP_top60_DNA+RNA_seed | EP300 | 0.842184 | BRCA | HCC1187 | CPI-637 |
| MVP_top60_DNA_seed | MAPK1 | 1.925076 | BRCA | HCC38 | Ulixertinib | MVP_top60_DNA+RNA_seed | EGFR | 0.843482 | BRCA | HCC1500 | Lapatinib |
| MVP_top60_DNA_seed | EGFR | 1.9396 | BRCA | BT-474 | AZD3759 | MVP_top60_DNA+RNA_seed | ERBB2 | 0.843482 | BRCA | HCC1500 | Lapatinib |

| | | | | | | | | | | | |
|---|---|---|---|---|---|---|---|---|---|---|---|
| MVP_top60_DNA_seed | CDK2 | 1.949143 | BRCA | HCC1419 | Dinaciclib | MVP_top60_DNA+RNA_seed | MDM2 | 0.846014 | BRCA | HCC70 | Nutlin-3a (-) |
| MVP_top60_DNA_seed | CDK1 | 1.949143 | BRCA | HCC1419 | Dinaciclib | MVP_top60_DNA+RNA_seed | AKT1 | 0.849992 | BRCA | JIMT-1 | Afuresertib |
| MVP_top60_DNA_seed | AKT1 | 1.964761 | BRCA | CAL-120 | MK-2206 | MVP_top60_DNA+RNA_seed | ESR1 | 0.850311 | BRCA | HCC1569 | GDC0810 |
| MVP_top60_DNA_seed | MDM2 | 1.969728 | BRCA | MFM-223 | Serdemetan | MVP_top60_DNA+RNA_seed | MAPK3 | 0.851578 | BRCA | MDA-MB-361 | SCH772984 |
| MVP_top60_DNA_seed | CDK2 | 1.972524 | BRCA | HCC38 | AZD5438 | MVP_top60_DNA+RNA_seed | MAPK1 | 0.851578 | BRCA | MDA-MB-361 | SCH772984 |
| MVP_top60_DNA_seed | SRC | 1.975869 | BRCA | MDA-MB-175-VII | Bosutinib | MVP_top60_DNA+RNA_seed | PLK1 | 0.852642 | BRCA | BT-20 | MK-1775 |
| MVP_top60_DNA_seed | EGFR | 1.985716 | BRCA | CAL-120 | Gefitinib | MVP_top60_DNA+RNA_seed | MAPK3 | 0.854308 | BRCA | HCC1937 | Ulixertinib |
| MVP_top60_DNA_seed | HDAC1 | 1.993809 | BRCA | CAL-120 | PCI-34051 | MVP_top60_DNA+RNA_seed | MAPK1 | 0.854308 | BRCA | HCC1937 | Ulixertinib |
| MVP_top60_DNA_seed | AKT1 | 2.007378 | BRCA | BT-483 | Ipatasertib | MVP_top60_DNA+RNA_seed | PLK1 | 0.854893 | BRCA | MDA-MB-361 | BI-2536 |
| MVP_top60_DNA_seed | ATM | 2.026632 | BRCA | MDA-MB-175-VII | KU-55933 | MVP_top60_DNA+RNA_seed | MAPK1 | 0.857524 | BRCA | EFM-192A | VX-11e |
| MVP_top60_DNA_seed | PIK3CA | 2.052789 | BRCA | CAL-120 | Alpelisib | MVP_top60_DNA+RNA_seed | PIK3CA | 0.857775 | BRCA | HCC2218 | Buparlisib |
| MVP_top60_DNA_seed | CDK2 | 2.057509 | BRCA | HCC1500 | AZD5438 | MVP_top60_DNA+RNA_seed | PLK1 | 0.857858 | BRCA | HCC1937 | BI-2536 |
| MVP_top60_DNA_seed | AKT1 | 2.05911 | BRCA | CAL-120 | Uprosertib | MVP_top60_DNA+RNA_seed | MDM2 | 0.860345 | BRCA | HCC1500 | Serdemetan |
| MVP_top60_DNA_seed | CDK2 | 2.072564 | BRCA | MDA-MB-175-VII | MK-8776 | MVP_top60_DNA+RNA_seed | TP53 | 0.860765 | BRCA | MCF7 | MIRA-1 |
| MVP_top60_DNA_seed | MAPK3 | 2.09055 | BRCA | ZR-75-30 | Ulixertinib | MVP_top60_DNA+RNA_seed | CDK1 | 0.861349 | BRCA | EFM-19 | RO-3306 |
| MVP_top60_DNA_seed | MAPK1 | 2.09055 | BRCA | ZR-75-30 | Ulixertinib | MVP_top60_DNA+RNA_seed | CDK2 | 0.863469 | BRCA | BT-474 | AZD5438 |
| MVP_top60_DNA_seed | HDAC1 | 2.097541 | BRCA | MRK-nu-1 | Romidepsin | MVP_top60_DNA+RNA_seed | ESR1 | 0.863828 | BRCA | YMB-1-E | GDC0810 |
| MVP_top60_DNA_seed | HDAC2 | 2.097541 | BRCA | MRK-nu-1 | Romidepsin | MVP_top60_DNA+RNA_seed | PLK1 | 0.863951 | BRCA | HCC1569 | BI-2536 |
| MVP_top60_DNA_seed | AKT1 | 2.115379 | BRCA | HCC1428 | MK-2206 | MVP_top60_DNA+RNA_seed | EP300 | 0.864272 | BRCA | T47D | SGC-CBP30 |
| MVP_top60_DNA_seed | ATM | 2.123038 | BRCA | BT-483 | KU-55933 | MVP_top60_DNA+RNA_seed | EP300 | 0.869091 | BRCA | HCC1500 | CPI-637 |
| MVP_top60_DNA_seed | PIK3CA | 2.131534 | BRCA | BT-483 | Buparlisib | MVP_top60_DNA+RNA_seed | EP300 | 0.869783 | BRCA | HCC38 | CPI-637 |
| MVP_top60_DNA_seed | ATM | 2.133594 | BRCA | CAL-120 | KU-55933 | MVP_top60_DNA+RNA_seed | PLK1 | 0.870907 | BRCA | HCC1428 | BI-2536 |
| MVP_top60_DNA_seed | EGFR | 2.14083 | BRCA | HCC1428 | Osimertinib | MVP_top60_DNA+RNA_seed | PIK3CA | 0.8721 | BRCA | HCC1599 | AZD8186 |
| MVP_top60_DNA_seed | EGFR | 2.146308 | BRCA | YMB-1-E | Erlotinib | MVP_top60_DNA+RNA_seed | MAPK1 | 0.873812 | BRCA | BT-549 | VX-11e |
| MVP_top60_DNA_seed | MDM2 | 2.16152 | BRCA | MDA-MB-175-VII | Serdemetan | MVP_top60_DNA+RNA_seed | CDK2 | 0.875586 | BRCA | MFM-223 | AZD5438 |
| MVP_top60_DNA_seed | HSP90AA1 | 2.166102 | BRCA | HCC1187 | Luminespib | MVP_top60_DNA+RNA_seed | MDM2 | 0.876346 | BRCA | CAL-85-1 | Nutlin-3a (-) |
| MVP_top60_DNA_seed | HSP90AB1 | 2.166102 | BRCA | HCC1187 | Luminespib | MVP_top60_DNA+RNA_seed | MDM2 | 0.876849 | BRCA | EFM-19 | Serdemetan |
| MVP_top60_DNA_seed | EGFR | 2.169547 | BRCA | CAL-120 | AZD3759 | MVP_top60_DNA+RNA_seed | ESR1 | 0.876885 | BRCA | HCC1599 | GDC0810 |
| MVP_top60_DNA_seed | ESR1 | 2.186943 | BRCA | MDA-MB-175-VII | GDC0810 | MVP_top60_DNA+RNA_seed | AKT1 | 0.877952 | BRCA | HCC1143 | Uprosertib |
| MVP_top60_DNA_seed | SRC | 2.212045 | BRCA | BT-483 | Bosutinib | MVP_top60_DNA+RNA_seed | SRC | 0.878322 | BRCA | EFM-19 | Bosutinib |
| MVP_top60_DNA_seed | EGFR | 2.219776 | BRCA | BT-483 | Erlotinib | MVP_top60_DNA+RNA_seed | MAPK3 | 0.882717 | BRCA | UACC-812 | Ulixertinib |
| MVP_top60_DNA_seed | AKT1 | 2.228904 | BRCA | BT-483 | Afuresertib | MVP_top60_DNA+RNA_seed | MAPK1 | 0.882717 | BRCA | UACC-812 | Ulixertinib |
| MVP_top60_DNA_seed | ERBB2 | 2.245622 | BRCA | BT-483 | Afatinib | MVP_top60_DNA+RNA_seed | MDM2 | 0.882745 | BRCA | HDQ-P1 | Nutlin-3a (-) |
| MVP_top60_DNA_seed | EGFR | 2.245622 | BRCA | BT-483 | Afatinib | MVP_top60_DNA+RNA_seed | MAPK3 | 0.887218 | BRCA | MCF7 | Ulixertinib |
| MVP_top60_DNA_seed | MAPK3 | 2.250265 | BRCA | MDA-MB-330 | Ulixertinib | MVP_top60_DNA+RNA_seed | MAPK1 | 0.887218 | BRCA | MCF7 | Ulixertinib |
| MVP_top60_DNA_seed | MAPK1 | 2.250265 | BRCA | MDA-MB-330 | Ulixertinib | MVP_top60_DNA+RNA_seed | ESR1 | 0.889689 | BRCA | YMB-1-E | Tamoxifen |
| MVP_top60_DNA_seed | EGFR | 2.282786 | BRCA | CAL-120 | Osimertinib | MVP_top60_DNA+RNA_seed | PLK1 | 0.891807 | BRCA | MDA-MB-330 | BI-2536 |
| MVP_top60_DNA_seed | ATM | 2.321949 | BRCA | ZR-75-30 | KU-55933 | MVP_top60_DNA+RNA_seed | EP300 | 0.893926 | BRCA | MFM-223 | SGC-CBP30 |
| MVP_top60_DNA_seed | PIK3CA | 2.322717 | BRCA | HCC1428 | GNE-317 | MVP_top60_DNA+RNA_seed | HSP90AA1 | 0.895976 | BRCA | CAL-120 | Tanespimycin |
| MVP_top60_DNA_seed | MAPK3 | 2.329971 | BRCA | BT-549 | Ulixertinib | MVP_top60_DNA+RNA_seed | HSP90AB1 | 0.895976 | BRCA | CAL-120 | Tanespimycin |
| MVP_top60_DNA_seed | MAPK1 | 2.329971 | BRCA | BT-549 | Ulixertinib | MVP_top60_DNA+RNA_seed | PLK1 | 0.897235 | BRCA | EFM-19 | MK-1775 |

| | | | | | | | | | | | |
|---|---|---|---|---|---|---|---|---|---|---|---|
| MVP_top60_DNA_seed | AKT1 | 2.340748 | BRCA | HCC1428 | Ipatasertib | MVP_top60_DNA+RNA_seed | EGFR | 0.897531 | BRCA | MFM-223 | Erlotinib |
| MVP_top60_DNA_seed | MAPK3 | 2.346661 | BRCA | BT-483 | SCH772984 | MVP_top60_DNA+RNA_seed | EP300 | 0.898403 | BRCA | CAL-120 | SGC-CBP30 |
| MVP_top60_DNA_seed | MAPK1 | 2.346661 | BRCA | BT-483 | SCH772984 | MVP_top60_DNA+RNA_seed | MAPK3 | 0.899405 | BRCA | BT-20 | Ulixertinib |
| MVP_top60_DNA_seed | MAPK1 | 2.353221 | BRCA | HCC38 | VX-11e | MVP_top60_DNA+RNA_seed | MAPK1 | 0.899405 | BRCA | BT-20 | Ulixertinib |
| MVP_top60_DNA_seed | EGFR | 2.361925 | BRCA | HCC1428 | AZD3759 | MVP_top60_DNA+RNA_seed | PIK3CA | 0.901775 | BRCA | BT-474 | GNE-317 |
| MVP_top60_DNA_seed | CDK2 | 2.362452 | BRCA | ZR-75-30 | Dinaciclib | MVP_top60_DNA+RNA_seed | MDM2 | 0.906853 | BRCA | MDA-MB-157 | Serdemetan |
| MVP_top60_DNA_seed | CDK1 | 2.362452 | BRCA | ZR-75-30 | Dinaciclib | MVP_top60_DNA+RNA_seed | EP300 | 0.911587 | BRCA | CAL-51 | CPI-637 |
| MVP_top60_DNA_seed | MAPK3 | 2.385763 | BRCA | ZR-75-30 | SCH772984 | MVP_top60_DNA+RNA_seed | PIK3CA | 0.911682 | BRCA | MRK-nu-1 | AZD8186 |
| MVP_top60_DNA_seed | MAPK1 | 2.385763 | BRCA | ZR-75-30 | SCH772984 | MVP_top60_DNA+RNA_seed | MAPK3 | 0.911947 | BRCA | MFM-223 | Ulixertinib |
| MVP_top60_DNA_seed | AKT1 | 2.406693 | BRCA | BT-483 | AZD5363 | MVP_top60_DNA+RNA_seed | MAPK1 | 0.911947 | BRCA | MFM-223 | Ulixertinib |
| MVP_top60_DNA_seed | ESR1 | 2.421825 | BRCA | BT-483 | Tamoxifen | MVP_top60_DNA+RNA_seed | PLK1 | 0.913904 | BRCA | HCC1500 | BI-2536 |
| MVP_top60_DNA_seed | ERBB2 | 2.43059 | BRCA | CAL-120 | Lapatinib | MVP_top60_DNA+RNA_seed | MAPK3 | 0.916835 | BRCA | EFM-19 | Ulixertinib |
| MVP_top60_DNA_seed | EGFR | 2.43059 | BRCA | CAL-120 | Lapatinib | MVP_top60_DNA+RNA_seed | MAPK1 | 0.916835 | BRCA | EFM-19 | Ulixertinib |
| MVP_top60_DNA_seed | EGFR | 2.493382 | BRCA | BT-483 | AZD3759 | MVP_top60_DNA+RNA_seed | HDAC1 | 0.918275 | BRCA | BT-20 | Entinostat |
| MVP_top60_DNA_seed | PIK3CA | 2.57169 | BRCA | HCC1428 | Buparlisib | MVP_top60_DNA+RNA_seed | PIK3CA | 0.919742 | BRCA | HCC1569 | Buparlisib |
| MVP_top60_DNA_seed | CDK2 | 2.579025 | BRCA | BT-483 | MK-8776 | MVP_top60_DNA+RNA_seed | PIK3CA | 0.9207 | BRCA | MFM-223 | AZD8186 |
| MVP_top60_DNA_seed | CDK2 | 2.612586 | BRCA | ZR-75-30 | AZD5438 | MVP_top60_DNA+RNA_seed | EGFR | 0.924335 | BRCA | JIMT-1 | Osimertinib |
| MVP_top60_DNA_seed | CDK2 | 2.634821 | BRCA | HCC1428 | Dinaciclib | MVP_top60_DNA+RNA_seed | MAPK3 | 0.928352 | BRCA | CAL-148 | Ulixertinib |
| MVP_top60_DNA_seed | CDK1 | 2.634821 | BRCA | HCC1428 | Dinaciclib | MVP_top60_DNA+RNA_seed | MAPK1 | 0.928352 | BRCA | CAL-148 | Ulixertinib |
| MVP_top60_DNA_seed | ESR1 | 2.666956 | BRCA | BT-483 | GDC0810 | MVP_top60_DNA+RNA_seed | AR | 0.929318 | BRCA | MDA-MB-330 | Bicalutamide |
| MVP_top60_DNA_seed | PIK3CA | 2.676332 | BRCA | HCC1428 | AZD8186 | MVP_top60_DNA+RNA_seed | MAPK3 | 0.931203 | BRCA | EFM-19 | Ulixertinib |
| MVP_top60_DNA_seed | EGFR | 2.679423 | BRCA | BT-483 | Osimertinib | MVP_top60_DNA+RNA_seed | MAPK1 | 0.931203 | BRCA | EFM-19 | Ulixertinib |
| MVP_top60_DNA_seed | CDK2 | 2.723613 | BRCA | HCC1937 | Dinaciclib | MVP_top60_DNA+RNA_seed | MAPK3 | 0.938143 | BRCA | MDA-MB-330 | SCH772984 |
| MVP_top60_DNA_seed | CDK1 | 2.723613 | BRCA | HCC1937 | Dinaciclib | MVP_top60_DNA+RNA_seed | MAPK1 | 0.938143 | BRCA | MDA-MB-330 | SCH772984 |
| MVP_top60_DNA_seed | EGFR | 2.737833 | BRCA | BT-483 | Gefitinib | MVP_top60_DNA+RNA_seed | CDK2 | 0.939686 | BRCA | T47D | Dinaciclib |
| MVP_top60_DNA_seed | HDAC1 | 2.7515 | BRCA | BT-483 | Entinostat | MVP_top60_DNA+RNA_seed | CDK1 | 0.939686 | BRCA | T47D | Dinaciclib |
| MVP_top60_DNA_seed | HDAC1 | 2.757894 | BRCA | ZR-75-30 | Romidepsin | MVP_top60_DNA+RNA_seed | CDK2 | 0.939865 | BRCA | HCC1954 | MK-8776 |
| MVP_top60_DNA_seed | HDAC2 | 2.757894 | BRCA | ZR-75-30 | Romidepsin | MVP_top60_DNA+RNA_seed | PLK1 | 0.941195 | BRCA | HCC70 | BI-2536 |
| MVP_top60_DNA_seed | HSP90AA1 | 2.775373 | BRCA | BT-483 | Luminespib | MVP_top60_DNA+RNA_seed | MDM2 | 0.942024 | BRCA | JIMT-1 | Nutlin-3a (-) |
| MVP_top60_DNA_seed | HSP90AB1 | 2.775373 | BRCA | BT-483 | Luminespib | MVP_top60_DNA+RNA_seed | ESR1 | 0.942933 | BRCA | HCC1428 | GDC0810 |
| MVP_top60_DNA_seed | AKT1 | 2.798344 | BRCA | HCC1428 | Afuresertib | MVP_top60_DNA+RNA_seed | EGFR | 0.943402 | BRCA | HCC1500 | Sapitinib |
| MVP_top60_DNA_seed | AKT1 | 2.839131 | BRCA | HCC1428 | AZD5363 | MVP_top60_DNA+RNA_seed | ERBB2 | 0.943402 | BRCA | HCC1500 | Sapitinib |
| MVP_top60_DNA_seed | PIK3CA | 2.896574 | BRCA | HCC1428 | Alpelisib | MVP_top60_DNA+RNA_seed | ERBB3 | 0.943402 | BRCA | HCC1500 | Sapitinib |
| MVP_top60_DNA_seed | PIK3CA | 2.896804 | BRCA | BT-483 | AZD8186 | MVP_top60_DNA+RNA_seed | SRC | 0.944668 | BRCA | EFM-192A | Bosutinib |
| MVP_top60_DNA_seed | MAPK3 | 2.939699 | BRCA | ZR-75-30 | Ulixertinib | MVP_top60_DNA+RNA_seed | AKT1 | 0.945834 | BRCA | HCC38 | Ipatasertib |
| MVP_top60_DNA_seed | MAPK1 | 2.939699 | BRCA | ZR-75-30 | Ulixertinib | MVP_top60_DNA+RNA_seed | SRC | 0.946258 | BRCA | UACC-812 | Bosutinib |
| MVP_top60_DNA_seed | AKT1 | 2.970367 | BRCA | BT-483 | AT13148 | MVP_top60_DNA+RNA_seed | EGFR | 0.949244 | BRCA | HCC1428 | Gefitinib |
| MVP_top60_DNA_seed | CDK2 | 3.028935 | BRCA | HCC1428 | AZD5438 | MVP_top60_DNA+RNA_seed | AKT1 | 0.949387 | BRCA | HCC1419 | AT13148 |
| MVP_top60_DNA_seed | MAPK3 | 3.151976 | BRCA | BT-483 | Ulixertinib | MVP_top60_DNA+RNA_seed | MAPK1 | 0.949954 | BRCA | MDA-MB-436 | VX-11e |
| MVP_top60_DNA_seed | MAPK1 | 3.151976 | BRCA | BT-483 | Ulixertinib | MVP_top60_DNA+RNA_seed | PIK3CA | 0.951276 | BRCA | MDA-MB-436 | Alpelisib |
| MVP_top60_DNA_seed | MAPK1 | 3.461689 | BRCA | BT-483 | VX-11e | MVP_top60_DNA+RNA_seed | HSP90AA1 | 0.953192 | BRCA | MDA-MB-436 | Tanespimycin |

| | | | | | | | | | | | |
|---|---|---|---|---|---|---|---|---|---|---|---|
| MVP_top60_DNA_seed | AR | -4.816697 | COADREAD | CaR-1 | Bicalutamide | MVP_top60_DNA+RNA_seed | HSP90AB1 | 0.953192 | BRCA | MDA-MB-436 | Tanespimycin |
| MVP_top60_DNA_seed | BRAF | -4.363396 | COADREAD | COLO-205 | SB590885 | MVP_top60_DNA+RNA_seed | MAPK1 | 0.963209 | BRCA | HCC1143 | VX-11e |
| MVP_top60_DNA_seed | BRAF | -4.227096 | COADREAD | COLO-205 | PLX-4720 | MVP_top60_DNA+RNA_seed | ESR1 | 0.963472 | BRCA | HCC38 | GDC0810 |
| MVP_top60_DNA_seed | AR | -4.097544 | COADREAD | SW948 | Bicalutamide | MVP_top60_DNA+RNA_seed | HDAC1 | 0.964269 | BRCA | BT-20 | Romidepsin |
| MVP_top60_DNA_seed | ERBB4 | -3.717325 | COADREAD | CCK-81 | PF-00299804 | MVP_top60_DNA+RNA_seed | HDAC2 | 0.964269 | BRCA | BT-20 | Romidepsin |
| MVP_top60_DNA_seed | ERBB3 | -3.706856 | COADREAD | DiFi | AZD8931 | MVP_top60_DNA+RNA_seed | MDM2 | 0.966999 | BRCA | HCC1143 | Nutlin-3a (-) |
| MVP_top60_DNA_seed | PIK3CB | -3.638272 | COADREAD | LoVo | AZD8186 | MVP_top60_DNA+RNA_seed | EGFR | 0.968993 | BRCA | EFM-19 | Gefitinib |
| MVP_top60_DNA_seed | PIK3CD | -3.638272 | COADREAD | LoVo | AZD8186 | MVP_top60_DNA+RNA_seed | EGFR | 0.971702 | BRCA | MDA-MB-436 | Lapatinib |
| MVP_top60_DNA_seed | IGF1R | -3.173323 | COADREAD | COLO-205 | BMS-754807 | MVP_top60_DNA+RNA_seed | ERBB2 | 0.971702 | BRCA | MDA-MB-436 | Lapatinib |
| MVP_top60_DNA_seed | IGF1R | -3.141067 | COADREAD | COLO-205 | Linsitinib | MVP_top60_DNA+RNA_seed | AKT1 | 0.972069 | BRCA | CAL-120 | Uprosertib |
| MVP_top60_DNA_seed | ERBB4 | -2.977733 | COADREAD | CCK-81 | CI-1033 | MVP_top60_DNA+RNA_seed | MAPK3 | 0.972174 | BRCA | MDA-MB-361 | Ulixertinib |
| MVP_top60_DNA_seed | ERBB3 | -2.973127 | COADREAD | SNU-407 | AZD8931 | MVP_top60_DNA+RNA_seed | MAPK1 | 0.972174 | BRCA | MDA-MB-361 | Ulixertinib |
| MVP_top60_DNA_seed | BRAF | -2.966581 | COADREAD | MDST8 | Dabrafenib | MVP_top60_DNA+RNA_seed | HDAC1 | 0.972261 | BRCA | MDA-MB-157 | PCI-34051 |
| MVP_top60_DNA_seed | PIK3CA | -2.885395 | COADREAD | SNU-407 | AZD8835 | MVP_top60_DNA+RNA_seed | HDAC1 | 0.974598 | BRCA | MCF7 | Entinostat |
| MVP_top60_DNA_seed | PIK3CD | -2.885395 | COADREAD | SNU-407 | AZD8835 | MVP_top60_DNA+RNA_seed | MDM2 | 0.975374 | BRCA | BT-549 | Nutlin-3a (-) |
| MVP_top60_DNA_seed | GSK3B | -2.627545 | COADREAD | RKO | KIN001-042 | MVP_top60_DNA+RNA_seed | CDK2 | 0.976235 | BRCA | MCF7 | Dinaciclib |
| MVP_top60_DNA_seed | IGF1R | -2.623082 | COADREAD | KM12 | BMS-536924 | MVP_top60_DNA+RNA_seed | CDK1 | 0.976235 | BRCA | MCF7 | Dinaciclib |
| MVP_top60_DNA_seed | MET | -2.601849 | COADREAD | KM12 | Crizotinib | MVP_top60_DNA+RNA_seed | CDK2 | 0.976615 | BRCA | COLO-824 | MK-8776 |
| MVP_top60_DNA_seed | AKT1 | -2.54632 | COADREAD | SNU-C1 | BAY AKT1 | MVP_top60_DNA+RNA_seed | ESR1 | 0.976971 | BRCA | MDA-MB-330 | Tamoxifen |
| MVP_top60_DNA_seed | PIK3CD | -2.510813 | COADREAD | SW1463 | Idelalisib | MVP_top60_DNA+RNA_seed | AKT1 | 0.978393 | BRCA | MRK-nu-1 | Afuresertib |
| MVP_top60_DNA_seed | IGF1R | -2.500607 | COADREAD | NCI-H716 | BMS-536924 | MVP_top60_DNA+RNA_seed | HDAC1 | 0.979205 | BRCA | CAL-120 | Romidepsin |
| MVP_top60_DNA_seed | BRAF | -2.486896 | COADREAD | RKO | AZ628 | MVP_top60_DNA+RNA_seed | HDAC2 | 0.979205 | BRCA | CAL-120 | Romidepsin |
| MVP_top60_DNA_seed | BRAF | -2.479845 | COADREAD | COLO-205 | PLX-4720 | MVP_top60_DNA+RNA_seed | HDAC1 | 0.979627 | BRCA | HCC38 | Entinostat |
| MVP_top60_DNA_seed | IGF1R | -2.459322 | COADREAD | LS-1034 | Linsitinib | MVP_top60_DNA+RNA_seed | SRC | 0.980572 | BRCA | MDA-MB-157 | Bosutinib |
| MVP_top60_DNA_seed | PTK2 | -2.454079 | COADREAD | COLO-205 | PF-562271 | MVP_top60_DNA+RNA_seed | MAPK1 | 0.986755 | BRCA | EFM-19 | VX-11e |
| MVP_top60_DNA_seed | MAPK1 | -2.436384 | COADREAD | HCT-116 | VX-11e | MVP_top60_DNA+RNA_seed | EGFR | 0.988119 | BRCA | CAL-85-1 | AZD3759 |
| MVP_top60_DNA_seed | PTK2 | -2.435368 | COADREAD | KM12 | PF-562271 | MVP_top60_DNA+RNA_seed | CDK1 | 0.988345 | BRCA | HCC1187 | RO-3306 |
| MVP_top60_DNA_seed | MAPK1 | -2.417666 | COADREAD | LS-513 | VX-11e | MVP_top60_DNA+RNA_seed | EP300 | 0.988393 | BRCA | ZR-75-30 | SGC-CBP30 |
| MVP_top60_DNA_seed | CDK2 | -2.311522 | COADREAD | CaR-1 | AZD5438 | MVP_top60_DNA+RNA_seed | CDK2 | 0.989393 | BRCA | HCC1187 | MK-8776 |
| MVP_top60_DNA_seed | CDK2 | -2.305651 | COADREAD | HCT-116 | PHA-793887 | MVP_top60_DNA+RNA_seed | CDK2 | 0.990504 | BRCA | HCC1143 | AZD5438 |
| MVP_top60_DNA_seed | MAPK1 | -2.288208 | COADREAD | LS-411N | VX-11e | MVP_top60_DNA+RNA_seed | HSP90AA1 | 0.993092 | BRCA | HCC1954 | Luminespib |
| MVP_top60_DNA_seed | BRAF | -2.286426 | COADREAD | HCT-116 | RAF_9304 | MVP_top60_DNA+RNA_seed | HSP90AB1 | 0.993092 | BRCA | HCC1954 | Luminespib |
| MVP_top60_DNA_seed | ATM | -2.26219 | COADREAD | LoVo | KU-55933 | MVP_top60_DNA+RNA_seed | MAPK1 | 0.997509 | BRCA | ZR-75-30 | VX-11e |
| MVP_top60_DNA_seed | IGF1R | -2.192941 | COADREAD | T84 | BMS-754807 | MVP_top60_DNA+RNA_seed | HDAC1 | 1.000216 | BRCA | MDA-MB-330 | PCI-34051 |
| MVP_top60_DNA_seed | IGF1R | -2.176422 | COADREAD | HT-29 | BMS-754807 | MVP_top60_DNA+RNA_seed | MAPK3 | 1.002171 | BRCA | HCC2157 | SCH772984 |
| MVP_top60_DNA_seed | BRAF | -2.16582 | COADREAD | CL-11 | RAF_9304 | MVP_top60_DNA+RNA_seed | MAPK1 | 1.002171 | BRCA | HCC2157 | SCH772984 |
| MVP_top60_DNA_seed | BRAF | -2.150984 | COADREAD | LS-411N | AZ628 | MVP_top60_DNA+RNA_seed | MDM2 | 1.003158 | BRCA | MDA-MB-330 | Nutlin-3a (-) |
| MVP_top60_DNA_seed | IGF1R | -2.144939 | COADREAD | SW48 | BMS-536924 | MVP_top60_DNA+RNA_seed | HSP90AA1 | 1.003725 | BRCA | HCC1428 | Tanespimycin |
| MVP_top60_DNA_seed | MET | -2.126138 | COADREAD | KM12 | SU11274 | MVP_top60_DNA+RNA_seed | HSP90AB1 | 1.003725 | BRCA | HCC1428 | Tanespimycin |
| MVP_top60_DNA_seed | GSK3B | -2.094343 | COADREAD | LoVo | SB216763 | MVP_top60_DNA+RNA_seed | EGFR | 1.003802 | BRCA | EFM-19 | Erlotinib |
| MVP_top60_DNA_seed | IGF1R | -2.083245 | COADREAD | HT-29 | Linsitinib | MVP_top60_DNA+RNA_seed | MAPK3 | 1.005207 | BRCA | HCC1187 | Ulixertinib |

| | | | | | | | | | | | |
|---|---|---|---|---|---|---|---|---|---|---|---|
| MVP_top60_DNA_seed | ERBB4 | -2.075892 | COADREAD | CCK-81 | AST-1306 | MVP_top60_DNA+RNA_seed | MAPK1 | 1.005207 | BRCA | HCC1187 | Ulixertinib |
| MVP_top60_DNA_seed | IGF1R | -2.074399 | COADREAD | LS-1034 | BMS-754807 | MVP_top60_DNA+RNA_seed | AKT1 | 1.005771 | BRCA | MDA-MB-231 | Ipatasertib |
| MVP_top60_DNA_seed | BRAF | -2.064105 | COADREAD | MDST8 | RAF_9304 | MVP_top60_DNA+RNA_seed | MAPK1 | 1.007065 | BRCA | MDA-MB-175-VII | VX-11e |
| MVP_top60_DNA_seed | AKT1 | -2.04894 | COADREAD | SNU-81 | GSK690693 | MVP_top60_DNA+RNA_seed | MAPK1 | 1.009248 | BRCA | HCC1599 | VX-11e |
| MVP_top60_DNA_seed | MAPK1 | -2.024269 | COADREAD | RKO | VX-11e | MVP_top60_DNA+RNA_seed | AKT1 | 1.011347 | BRCA | UACC-812 | Uprosertib |
| MVP_top60_DNA_seed | CREBBP | -2.019942 | COADREAD | NCI-H508 | I-CBP112 | MVP_top60_DNA+RNA_seed | ATM | 1.011559 | BRCA | HCC1428 | KU-55933 |
| MVP_top60_DNA_seed | IGF1R | -2.014989 | COADREAD | COLO-320-HSR | Linsitinib | MVP_top60_DNA+RNA_seed | ATM | 1.01437 | BRCA | JIMT-1 | KU-55933 |
| MVP_top60_DNA_seed | PIK3CB | -2.002653 | COADREAD | C2BBe1 | TGX221 | MVP_top60_DNA+RNA_seed | EP300 | 1.015238 | BRCA | MDA-MB-468 | CPI-637 |
| MVP_top60_DNA_seed | IGF1R | -1.984166 | COADREAD | KM12 | BMS-754807 | MVP_top60_DNA+RNA_seed | AKT1 | 1.016184 | BRCA | CAL-85-1 | Uprosertib |
| MVP_top60_DNA_seed | BRAF | -1.973443 | COADREAD | C2BBe1 | SB590885 | MVP_top60_DNA+RNA_seed | AKT1 | 1.019492 | BRCA | JIMT-1 | Uprosertib |
| MVP_top60_DNA_seed | GSK3B | -1.963705 | COADREAD | SW1417 | CHIR-99021 | MVP_top60_DNA+RNA_seed | MAPK3 | 1.019916 | BRCA | EFM-192A | SCH772984 |
| MVP_top60_DNA_seed | IGF1R | -1.955877 | COADREAD | HCC2998 | BMS-754807 | MVP_top60_DNA+RNA_seed | MAPK1 | 1.019916 | BRCA | EFM-192A | SCH772984 |
| MVP_top60_DNA_seed | AKT1 | -1.955554 | COADREAD | LS-411N | AKT inhibitor VIII | MVP_top60_DNA+RNA_seed | EGFR | 1.021128 | BRCA | HCC38 | Osimertinib |
| MVP_top60_DNA_seed | MAPK1 | -1.914302 | COADREAD | COLO-205 | VX-11e | MVP_top60_DNA+RNA_seed | EGFR | 1.031091 | BRCA | MDA-MB-436 | Erlotinib |
| MVP_top60_DNA_seed | ESR1 | -1.884625 | COADREAD | SW48 | Fulvestrant | MVP_top60_DNA+RNA_seed | AKT1 | 1.033067 | BRCA | MDA-MB-436 | Ipatasertib |
| MVP_top60_DNA_seed | IGF1R | -1.853022 | COADREAD | COLO-205 | BMS-536924 | MVP_top60_DNA+RNA_seed | EGFR | 1.035799 | BRCA | HCC1599 | Gefitinib |
| MVP_top60_DNA_seed | MAPK1 | -1.831489 | COADREAD | LS-1034 | VX-11e | MVP_top60_DNA+RNA_seed | TP53 | 1.036629 | BRCA | HCC1937 | MIRA-1 |
| MVP_top60_DNA_seed | MAPK1 | -1.830445 | COADREAD | HT-29 | VX-11e | MVP_top60_DNA+RNA_seed | EP300 | 1.039723 | BRCA | MDA-MB-436 | CPI-637 |
| MVP_top60_DNA_seed | AKT1 | -1.807413 | COADREAD | DiFi | BAY AKT1 | MVP_top60_DNA+RNA_seed | PIK3CA | 1.040492 | BRCA | HCC38 | Buparlisib |
| MVP_top60_DNA_seed | BRAF | -1.791122 | COADREAD | MDST8 | PLX-4720 | MVP_top60_DNA+RNA_seed | PIK3CA | 1.045803 | BRCA | MDA-MB-231 | AZD8186 |
| MVP_top60_DNA_seed | AR | -1.774714 | COADREAD | LS-180 | Bicalutamide | MVP_top60_DNA+RNA_seed | HSP90AA1 | 1.048811 | BRCA | ZR-75-30 | Luminespib |
| MVP_top60_DNA_seed | BRAF | -1.761566 | COADREAD | HT-29 | PLX-4720 | MVP_top60_DNA+RNA_seed | HSP90AB1 | 1.048811 | BRCA | ZR-75-30 | Luminespib |
| MVP_top60_DNA_seed | SRC | -1.743851 | COADREAD | DiFi | Bosutinib | MVP_top60_DNA+RNA_seed | ATM | 1.048889 | BRCA | UACC-893 | KU-55933 |
| MVP_top60_DNA_seed | MAPK14 | -1.741658 | COADREAD | CCK-81 | TAK-715 | MVP_top60_DNA+RNA_seed | HDAC1 | 1.050179 | BRCA | COLO-824 | Romidepsin |
| MVP_top60_DNA_seed | SRC | -1.734961 | COADREAD | SNU-407 | NVP-BHG712 | MVP_top60_DNA+RNA_seed | HDAC2 | 1.050179 | BRCA | COLO-824 | Romidepsin |
| MVP_top60_DNA_seed | MAPK1 | -1.722028 | COADREAD | SNU-175 | FR-180204 | MVP_top60_DNA+RNA_seed | HDAC1 | 1.050224 | BRCA | MFM-223 | PCI-34051 |
| MVP_top60_DNA_seed | AKT1 | -1.711913 | COADREAD | SNU-1040 | GSK690693 | MVP_top60_DNA+RNA_seed | PLK1 | 1.051407 | BRCA | HCC38 | BI-2536 |
| MVP_top60_DNA_seed | BRAF | -1.707595 | COADREAD | HT-29 | PLX-4720 | MVP_top60_DNA+RNA_seed | PLK1 | 1.052519 | BRCA | T47D | BI-2536 |
| MVP_top60_DNA_seed | BRAF | -1.697462 | COADREAD | RKO | RAF_9304 | MVP_top60_DNA+RNA_seed | ATM | 1.056079 | BRCA | BT-474 | KU-55933 |
| MVP_top60_DNA_seed | IGF1R | -1.693276 | COADREAD | T84 | Linsitinib | MVP_top60_DNA+RNA_seed | EGFR | 1.058024 | BRCA | MDA-MB-157 | Osimertinib |
| MVP_top60_DNA_seed | BRAF | -1.692775 | COADREAD | HT-29 | SB590885 | MVP_top60_DNA+RNA_seed | AKT1 | 1.063553 | BRCA | MDA-MB-436 | GSK2110183B |
| MVP_top60_DNA_seed | AKT1 | -1.679353 | COADREAD | COLO-205 | AKT inhibitor VIII | MVP_top60_DNA+RNA_seed | PLK1 | 1.063651 | BRCA | HCC1937 | MK-1775 |
| MVP_top60_DNA_seed | BRAF | -1.676996 | COADREAD | LS-411N | PLX-4720 | MVP_top60_DNA+RNA_seed | PLK1 | 1.065279 | BRCA | MDA-MB-436 | MK-1775 |
| MVP_top60_DNA_seed | BRAF | -1.67516 | COADREAD | HCC2998 | AZ628 | MVP_top60_DNA+RNA_seed | HDAC1 | 1.071644 | BRCA | HCC38 | PCI-34051 |
| MVP_top60_DNA_seed | PIK3CB | -1.654664 | COADREAD | LoVo | AZD6482 | MVP_top60_DNA+RNA_seed | PIK3CA | 1.071977 | BRCA | MDA-MB-436 | GNE-317 |
| MVP_top60_DNA_seed | AKT1 | -1.651386 | COADREAD | NCI-H508 | BAY AKT1 | MVP_top60_DNA+RNA_seed | PLK1 | 1.072475 | BRCA | HDQ-P1 | BI-2536 |
| MVP_top60_DNA_seed | AR | -1.63943 | COADREAD | HCT-116 | AZD3514 | MVP_top60_DNA+RNA_seed | PLK1 | 1.078397 | BRCA | HCC1500 | MK-1775 |
| MVP_top60_DNA_seed | MAPK1 | -1.636379 | COADREAD | NCI-H747 | VX-11e | MVP_top60_DNA+RNA_seed | EP300 | 1.078404 | BRCA | BT-549 | SGC-CBP30 |
| MVP_top60_DNA_seed | IGF1R | -1.625184 | COADREAD | GP5d | BMS-754807 | MVP_top60_DNA+RNA_seed | ESR1 | 1.079545 | BRCA | ZR-75-30 | GDC0810 |
| MVP_top60_DNA_seed | GSK3B | -1.599783 | COADREAD | SW48 | SB216763 | MVP_top60_DNA+RNA_seed | MAPK3 | 1.081281 | BRCA | HDQ-P1 | Ulixertinib |
| MVP_top60_DNA_seed | IGF1R | -1.599647 | COADREAD | HCC2998 | Linsitinib | MVP_top60_DNA+RNA_seed | MAPK1 | 1.081281 | BRCA | HDQ-P1 | Ulixertinib |

| | | | | | | | | | | | |
|---|---|---|---|---|---|---|---|---|---|---|---|
| MVP_top60_DNA_seed | IGF1R | -1.593094 | COADREAD | KM12 | BMS-536924 | MVP_top60_DNA+RNA_seed | AKT1 | 1.081818 | BRCA | HCC38 | Uprosertib |
| MVP_top60_DNA_seed | IGF1R | -1.591244 | COADREAD | LS-411N | GSK1904529A | MVP_top60_DNA+RNA_seed | HDAC1 | 1.084074 | BRCA | MCF7 | PCI-34051 |
| MVP_top60_DNA_seed | ERBB3 | -1.588256 | COADREAD | SW48 | AZD8931 | MVP_top60_DNA+RNA_seed | AKT1 | 1.084199 | BRCA | CAL-85-1 | Uprosertib |
| MVP_top60_DNA_seed | CREBBP | -1.587603 | COADREAD | RKO | I-CBP112 | MVP_top60_DNA+RNA_seed | AKT1 | 1.084234 | BRCA | ZR-75-30 | Ipatasertib |
| MVP_top60_DNA_seed | MAPK1 | -1.580473 | COADREAD | SNU-61 | VX-11e | MVP_top60_DNA+RNA_seed | EGFR | 1.090574 | BRCA | MCF7 | Erlotinib |
| MVP_top60_DNA_seed | IGF1R | -1.578694 | COADREAD | HCT-116 | BMS-536924 | MVP_top60_DNA+RNA_seed | EGFR | 1.092291 | BRCA | CAMA-1 | Afatinib |
| MVP_top60_DNA_seed | PIK3CA | -1.570969 | COADREAD | CaR-1 | PI3Ka_4409 | MVP_top60_DNA+RNA_seed | ERBB2 | 1.092291 | BRCA | CAMA-1 | Afatinib |
| MVP_top60_DNA_seed | PIK3CD | -1.570969 | COADREAD | CaR-1 | PI3Ka_4409 | MVP_top60_DNA+RNA_seed | MAPK3 | 1.093167 | BRCA | MCF7 | Ulixertinib |
| MVP_top60_DNA_seed | AKT1 | -1.564586 | COADREAD | NCI-H747 | AKT inhibitor VIII | MVP_top60_DNA+RNA_seed | MAPK1 | 1.093167 | BRCA | MCF7 | Ulixertinib |
| MVP_top60_DNA_seed | MAPK1 | -1.561288 | COADREAD | SNU-407 | FR-180204 | MVP_top60_DNA+RNA_seed | HSP90AA1 | 1.094578 | BRCA | MDA-MB-231 | Tanespimycin |
| MVP_top60_DNA_seed | GSK3B | -1.552821 | COADREAD | HCT-116 | AZD7969 | MVP_top60_DNA+RNA_seed | HSP90AB1 | 1.094578 | BRCA | MDA-MB-231 | Tanespimycin |
| MVP_top60_DNA_seed | HSP90AA1 | -1.540735 | COADREAD | HCT-15 | CCT-018159 | MVP_top60_DNA+RNA_seed | MAPK3 | 1.097862 | BRCA | BT-474 | Ulixertinib |
| MVP_top60_DNA_seed | HSP90AB1 | -1.540735 | COADREAD | HCT-15 | CCT-018159 | MVP_top60_DNA+RNA_seed | MAPK1 | 1.097862 | BRCA | BT-474 | Ulixertinib |
| MVP_top60_DNA_seed | MET | -1.540208 | COADREAD | CCK-81 | JNJ38877605 | MVP_top60_DNA+RNA_seed | MAPK1 | 1.098267 | BRCA | MDA-MB-157 | VX-11e |
| MVP_top60_DNA_seed | ATM | -1.540048 | COADREAD | LS-411N | KU-60019 | MVP_top60_DNA+RNA_seed | EGFR | 1.098533 | BRCA | BT-549 | Sapitinib |
| MVP_top60_DNA_seed | IGF1R | -1.529272 | COADREAD | COLO-320-HSR | BMS-754807 | MVP_top60_DNA+RNA_seed | ERBB2 | 1.098533 | BRCA | BT-549 | Sapitinib |
| MVP_top60_DNA_seed | PIK3CA | -1.527487 | COADREAD | T84 | AZD8835 | MVP_top60_DNA+RNA_seed | ERBB3 | 1.098533 | BRCA | BT-549 | Sapitinib |
| MVP_top60_DNA_seed | PIK3CD | -1.527487 | COADREAD | T84 | AZD8835 | MVP_top60_DNA+RNA_seed | PIK3CA | 1.098639 | BRCA | HCC1419 | Buparlisib |
| MVP_top60_DNA_seed | BRAF | -1.517703 | COADREAD | SK-CO-1 | SB590885 | MVP_top60_DNA+RNA_seed | TP53 | 1.099032 | BRCA | BT-549 | MIRA-1 |
| MVP_top60_DNA_seed | PIK3CB | -1.508336 | COADREAD | SNU-81 | AZD8186 | MVP_top60_DNA+RNA_seed | PIK3CA | 1.099167 | BRCA | BT-474 | Buparlisib |
| MVP_top60_DNA_seed | PIK3CD | -1.508336 | COADREAD | SNU-81 | AZD8186 | MVP_top60_DNA+RNA_seed | PIK3CA | 1.100603 | BRCA | ZR-75-30 | GNE-317 |
| MVP_top60_DNA_seed | HSP90AA1 | -1.49404 | COADREAD | SW48 | Tanespimycin | MVP_top60_DNA+RNA_seed | SRC | 1.101792 | BRCA | MDA-MB-436 | Bosutinib |
| MVP_top60_DNA_seed | HSP90AB1 | -1.49404 | COADREAD | SW48 | Tanespimycin | MVP_top60_DNA+RNA_seed | EGFR | 1.106644 | BRCA | MCF7 | Afatinib |
| MVP_top60_DNA_seed | MAPK1 | -1.487271 | COADREAD | SNU-407 | VX-11e | MVP_top60_DNA+RNA_seed | ERBB2 | 1.106644 | BRCA | MCF7 | Afatinib |
| MVP_top60_DNA_seed | BRAF | -1.481452 | COADREAD | HT-29 | RAF_9304 | MVP_top60_DNA+RNA_seed | MAPK3 | 1.112723 | BRCA | UACC-812 | Ulixertinib |
| MVP_top60_DNA_seed | PIK3CA | -1.479836 | COADREAD | SW48 | AZD8835 | MVP_top60_DNA+RNA_seed | MAPK1 | 1.112723 | BRCA | UACC-812 | Ulixertinib |
| MVP_top60_DNA_seed | PIK3CD | -1.479836 | COADREAD | SW48 | AZD8835 | MVP_top60_DNA+RNA_seed | AKT1 | 1.117362 | BRCA | CAL-85-1 | MK-2206 |
| MVP_top60_DNA_seed | HSP90AA1 | -1.475028 | COADREAD | COLO-205 | Tanespimycin | MVP_top60_DNA+RNA_seed | AKT1 | 1.117489 | BRCA | MCF7 | GSK2110183B |
| MVP_top60_DNA_seed | HSP90AB1 | -1.475028 | COADREAD | COLO-205 | Tanespimycin | MVP_top60_DNA+RNA_seed | HDAC1 | 1.119887 | BRCA | HCC1428 | Romidepsin |
| MVP_top60_DNA_seed | IGF1R | -1.474191 | COADREAD | LS-411N | BMS-754807 | MVP_top60_DNA+RNA_seed | HDAC2 | 1.119887 | BRCA | HCC1428 | Romidepsin |
| MVP_top60_DNA_seed | CDK2 | -1.465851 | COADREAD | SNU-175 | PHA-793887 | MVP_top60_DNA+RNA_seed | AKT1 | 1.120459 | BRCA | MDA-MB-157 | MK-2206 |
| MVP_top60_DNA_seed | AKT1 | -1.465165 | COADREAD | NCI-H508 | MK-2206 | MVP_top60_DNA+RNA_seed | EGFR | 1.120563 | BRCA | MCF7 | AZD3759 |
| MVP_top60_DNA_seed | PTK2 | -1.463438 | COADREAD | SW48 | PF-562271 | MVP_top60_DNA+RNA_seed | CDK2 | 1.121775 | BRCA | HDQ-P1 | Dinaciclib |
| MVP_top60_DNA_seed | IGF1R | -1.460412 | COADREAD | SNU-C1 | Linsitinib | MVP_top60_DNA+RNA_seed | CDK1 | 1.121775 | BRCA | HDQ-P1 | Dinaciclib |
| MVP_top60_DNA_seed | CDK2 | -1.458351 | COADREAD | SW48 | AZD5438 | MVP_top60_DNA+RNA_seed | MDM2 | 1.126843 | BRCA | HCC38 | Nutlin-3a (-) |
| MVP_top60_DNA_seed | MET | -1.456509 | COADREAD | SNU-175 | SU11274 | MVP_top60_DNA+RNA_seed | MAPK1 | 1.138814 | BRCA | YMB-1-E | VX-11e |
| MVP_top60_DNA_seed | MAPK1 | -1.44468 | COADREAD | HT-29 | FR-180204 | MVP_top60_DNA+RNA_seed | PIK3CA | 1.140993 | BRCA | HCC1937 | Buparlisib |
| MVP_top60_DNA_seed | PIK3CB | -1.436165 | COADREAD | LS-411N | TGX221 | MVP_top60_DNA+RNA_seed | TP53 | 1.142254 | BRCA | HCC1143 | MIRA-1 |
| MVP_top60_DNA_seed | GSK3B | -1.432673 | COADREAD | LoVo | CHIR-99021 | MVP_top60_DNA+RNA_seed | HSP90AA1 | 1.143489 | BRCA | YMB-1-E | CCT-018159 |
| MVP_top60_DNA_seed | ESR1 | -1.431237 | COADREAD | HCT-116 | Fulvestrant | MVP_top60_DNA+RNA_seed | HSP90AB1 | 1.143489 | BRCA | YMB-1-E | CCT-018159 |
| MVP_top60_DNA_seed | AR | -1.424971 | COADREAD | HT-115 | Bicalutamide | MVP_top60_DNA+RNA_seed | SRC | 1.145831 | BRCA | MCF7 | Bosutinib |

| | | | | | | | | | | | |
|---|---|---|---|---|---|---|---|---|---|---|---|
| MVP_top60_DNA_seed | IGF1R | -1.424352 | COADREAD | SNU-C1 | BMS-536924 | MVP_top60_DNA+RNA_seed | HDAC1 | 1.147347 | BRCA | EFM-192A | PCI-34051 |
| MVP_top60_DNA_seed | IGF1R | -1.420785 | COADREAD | NCI-H747 | Linsitinib | MVP_top60_DNA+RNA_seed | AKT1 | 1.14755 | BRCA | HCC1187 | Uprosertib |
| MVP_top60_DNA_seed | PIK3CD | -1.404039 | COADREAD | SNU-81 | Idelalisib | MVP_top60_DNA+RNA_seed | EGFR | 1.147568 | BRCA | BT-549 | AZD3759 |
| MVP_top60_DNA_seed | SRC | -1.403007 | COADREAD | LS-513 | NVP-BHG712 | MVP_top60_DNA+RNA_seed | PIK3CA | 1.149817 | BRCA | HCC1500 | Alpelisib |
| MVP_top60_DNA_seed | BRAF | -1.402812 | COADREAD | NCI-H747 | AZ628 | MVP_top60_DNA+RNA_seed | CDK2 | 1.151019 | BRCA | T47D | MK-8776 |
| MVP_top60_DNA_seed | AKT1 | -1.400421 | COADREAD | DiFi | MK-2206 | MVP_top60_DNA+RNA_seed | MAPK1 | 1.151696 | BRCA | MDA-MB-361 | VX-11e |
| MVP_top60_DNA_seed | BRAF | -1.385014 | COADREAD | RKO | Dabrafenib | MVP_top60_DNA+RNA_seed | EGFR | 1.153388 | BRCA | MCF7 | Sapitinib |
| MVP_top60_DNA_seed | AKT1 | -1.384673 | COADREAD | HCT-15 | AKT inhibitor VIII | MVP_top60_DNA+RNA_seed | ERBB2 | 1.153388 | BRCA | MCF7 | Sapitinib |
| MVP_top60_DNA_seed | HSP90AA1 | -1.383208 | COADREAD | LS-180 | Tanespimycin | MVP_top60_DNA+RNA_seed | ERBB3 | 1.153388 | BRCA | MCF7 | Sapitinib |
| MVP_top60_DNA_seed | HSP90AB1 | -1.383208 | COADREAD | LS-180 | Tanespimycin | MVP_top60_DNA+RNA_seed | MDM2 | 1.153855 | BRCA | BT-474 | Nutlin-3a (-) |
| MVP_top60_DNA_seed | MET | -1.372711 | COADREAD | KM12 | PHA-665752 | MVP_top60_DNA+RNA_seed | PLK1 | 1.158134 | BRCA | T47D | MK-1775 |
| MVP_top60_DNA_seed | IGF1R | -1.367994 | COADREAD | LS-411N | BMS-536924 | MVP_top60_DNA+RNA_seed | PIK3CA | 1.15996 | BRCA | MDA-MB-157 | AZD8186 |
| MVP_top60_DNA_seed | AKT1 | -1.365797 | COADREAD | LS-411N | A-443654 | MVP_top60_DNA+RNA_seed | PIK3CA | 1.160204 | BRCA | MDA-MB-436 | Buparlisib |
| MVP_top60_DNA_seed | MAPK14 | -1.364891 | COADREAD | SNU-175 | TAK-715 | MVP_top60_DNA+RNA_seed | AKT1 | 1.160429 | BRCA | HCC1500 | Afuresertib |
| MVP_top60_DNA_seed | PTK2 | -1.360297 | COADREAD | HCT-116 | PF-562271 | MVP_top60_DNA+RNA_seed | CDK2 | 1.161658 | BRCA | CAL-120 | MK-8776 |
| MVP_top60_DNA_seed | HSP90AA1 | -1.355717 | COADREAD | HT-29 | Tanespimycin | MVP_top60_DNA+RNA_seed | HDAC1 | 1.16298 | BRCA | MDA-MB-175-VII | Entinostat |
| MVP_top60_DNA_seed | HSP90AB1 | -1.355717 | COADREAD | HT-29 | Tanespimycin | MVP_top60_DNA+RNA_seed | TP53 | 1.167902 | BRCA | MFM-223 | MIRA-1 |
| MVP_top60_DNA_seed | PTK2 | -1.351991 | COADREAD | GP5d | PF-562271 | MVP_top60_DNA+RNA_seed | MDM2 | 1.16826 | BRCA | BT-483 | Serdemetan |
| MVP_top60_DNA_seed | MAPK1 | -1.345289 | COADREAD | GP5d | FR-180204 | MVP_top60_DNA+RNA_seed | MAPK3 | 1.168425 | BRCA | BT-474 | SCH772984 |
| MVP_top60_DNA_seed | IGF1R | -1.343888 | COADREAD | NCI-H747 | BMS-754807 | MVP_top60_DNA+RNA_seed | MAPK1 | 1.168425 | BRCA | BT-474 | SCH772984 |
| MVP_top60_DNA_seed | IGF1R | -1.341783 | COADREAD | SNU-C1 | BMS-754807 | MVP_top60_DNA+RNA_seed | EGFR | 1.168961 | BRCA | BT-483 | Lapatinib |
| MVP_top60_DNA_seed | BRAF | -1.331183 | COADREAD | LS-411N | PLX-4720 | MVP_top60_DNA+RNA_seed | ERBB2 | 1.168961 | BRCA | BT-483 | Lapatinib |
| MVP_top60_DNA_seed | GSK3B | -1.319173 | COADREAD | COLO-205 | CHIR-99021 | MVP_top60_DNA+RNA_seed | EGFR | 1.169755 | BRCA | BT-549 | Gefitinib |
| MVP_top60_DNA_seed | AKT1 | -1.318288 | COADREAD | HT-29 | AKT inhibitor VIII | MVP_top60_DNA+RNA_seed | CDK1 | 1.17777 | BRCA | MDA-MB-157 | RO-3306 |
| MVP_top60_DNA_seed | GSK3B | -1.314432 | COADREAD | SK-CO-1 | AZD7969 | MVP_top60_DNA+RNA_seed | MAPK1 | 1.178844 | BRCA | CAL-120 | VX-11e |
| MVP_top60_DNA_seed | ERBB3 | -1.305685 | COADREAD | HT-115 | AZD8931 | MVP_top60_DNA+RNA_seed | EP300 | 1.183683 | BRCA | MCF7 | SGC-CBP30 |
| MVP_top60_DNA_seed | GSK3B | -1.300526 | COADREAD | LS-513 | SB216763 | MVP_top60_DNA+RNA_seed | HDAC1 | 1.184385 | BRCA | MDA-MB-436 | PCI-34051 |
| MVP_top60_DNA_seed | MAPK14 | -1.300327 | COADREAD | NCI-H747 | TAK-715 | MVP_top60_DNA+RNA_seed | EGFR | 1.185089 | BRCA | MFM-223 | Osimertinib |
| MVP_top60_DNA_seed | PIK3CD | -1.298458 | COADREAD | NCI-H747 | Idelalisib | MVP_top60_DNA+RNA_seed | AKT1 | 1.191457 | BRCA | HCC1500 | Uprosertib |
| MVP_top60_DNA_seed | GSK3B | -1.293739 | COADREAD | KM12 | KIN001-042 | MVP_top60_DNA+RNA_seed | EGFR | 1.194379 | BRCA | CAL-120 | Afatinib |
| MVP_top60_DNA_seed | AKT1 | -1.289571 | COADREAD | HCT-116 | AKT inhibitor VIII | MVP_top60_DNA+RNA_seed | ERBB2 | 1.194379 | BRCA | CAL-120 | Afatinib |
| MVP_top60_DNA_seed | ATM | -1.289227 | COADREAD | HCT-116 | KU-60019 | MVP_top60_DNA+RNA_seed | PLK1 | 1.196486 | BRCA | CAMA-1 | MK-1775 |
| MVP_top60_DNA_seed | MAPK14 | -1.288425 | COADREAD | CW-2 | TAK-715 | MVP_top60_DNA+RNA_seed | ESR1 | 1.197697 | BRCA | ZR-75-30 | Tamoxifen |
| MVP_top60_DNA_seed | IGF1R | -1.287751 | COADREAD | COLO-320-HSR | BMS-536924 | MVP_top60_DNA+RNA_seed | EGFR | 1.198466 | BRCA | MCF7 | Gefitinib |
| MVP_top60_DNA_seed | HSP90AA1 | -1.278961 | COADREAD | RKO | SNX-2112 | MVP_top60_DNA+RNA_seed | SRC | 1.19926 | BRCA | MDA-MB-330 | Bosutinib |
| MVP_top60_DNA_seed | HSP90AB1 | -1.278961 | COADREAD | RKO | SNX-2112 | MVP_top60_DNA+RNA_seed | TP53 | 1.204967 | BRCA | MDA-MB-436 | MIRA-1 |
| MVP_top60_DNA_seed | IGF1R | -1.278891 | COADREAD | LS-1034 | BMS-536924 | MVP_top60_DNA+RNA_seed | CDK2 | 1.205544 | BRCA | CAL-120 | AZD5438 |
| MVP_top60_DNA_seed | MAPK1 | -1.273607 | COADREAD | SNU-1040 | FR-180204 | MVP_top60_DNA+RNA_seed | EGFR | 1.207114 | BRCA | BT-483 | Sapitinib |
| MVP_top60_DNA_seed | AR | -1.268741 | COADREAD | COLO-205 | Bicalutamide | MVP_top60_DNA+RNA_seed | ERBB2 | 1.207114 | BRCA | BT-483 | Sapitinib |
| MVP_top60_DNA_seed | IGF1R | -1.257458 | COADREAD | LS-411N | Linsitinib | MVP_top60_DNA+RNA_seed | ERBB3 | 1.207114 | BRCA | BT-483 | Sapitinib |
| MVP_top60_DNA_seed | BRAF | -1.25549 | COADREAD | RKO | SB590885 | MVP_top60_DNA+RNA_seed | MAPK3 | 1.212895 | BRCA | HCC1937 | Ulixertinib |

| | | | | | | | | | | | |
|---|---|---|---|---|---|---|---|---|---|---|---|
| MVP_top60_DNA_seed | CDK2 | -1.252024 | COADREAD | COLO-320-HSR | Seliciclib | MVP_top60_DNA+RNA_seed | MAPK1 | 1.212895 | BRCA | HCC1937 | Ulixertinib |
| MVP_top60_DNA_seed | BRAF | -1.246299 | COADREAD | COLO-205 | RAF_9304 | MVP_top60_DNA+RNA_seed | EGFR | 1.218625 | BRCA | HCC38 | Gefitinib |
| MVP_top60_DNA_seed | ESR1 | -1.245743 | COADREAD | LoVo | Fulvestrant | MVP_top60_DNA+RNA_seed | CDK2 | 1.223227 | BRCA | CAL-85-1 | MK-8776 |
| MVP_top60_DNA_seed | ERBB4 | -1.243392 | COADREAD | SNU-175 | PF-00299804 | MVP_top60_DNA+RNA_seed | TP53 | 1.22708 | BRCA | CAL-120 | MIRA-1 |
| MVP_top60_DNA_seed | AKT1 | -1.235951 | COADREAD | LS-411N | GSK690693 | MVP_top60_DNA+RNA_seed | EGFR | 1.228839 | BRCA | MFM-223 | Sapitinib |
| MVP_top60_DNA_seed | SRC | -1.231503 | COADREAD | SNU-C5 | Bosutinib | MVP_top60_DNA+RNA_seed | ERBB2 | 1.228839 | BRCA | MFM-223 | Sapitinib |
| MVP_top60_DNA_seed | HSP90AA1 | -1.22617 | COADREAD | CaR-1 | Tanespimycin | MVP_top60_DNA+RNA_seed | ERBB3 | 1.228839 | BRCA | MFM-223 | Sapitinib |
| MVP_top60_DNA_seed | HSP90AB1 | -1.22617 | COADREAD | CaR-1 | Tanespimycin | MVP_top60_DNA+RNA_seed | CDK2 | 1.231689 | BRCA | HCC1937 | AZD5438 |
| MVP_top60_DNA_seed | SRC | -1.224806 | COADREAD | C2BBe1 | Saracatinib | MVP_top60_DNA+RNA_seed | CDK2 | 1.235341 | BRCA | MDA-MB-330 | MK-8776 |
| MVP_top60_DNA_seed | SRC | -1.215717 | COADREAD | HCT-116 | NVP-BHG712 | MVP_top60_DNA+RNA_seed | SRC | 1.237267 | BRCA | HCC38 | Bosutinib |
| MVP_top60_DNA_seed | BRAF | -1.215661 | COADREAD | SNU-C5 | PLX-4720 | MVP_top60_DNA+RNA_seed | ESR1 | 1.238623 | BRCA | CAL-120 | GDC0810 |
| MVP_top60_DNA_seed | AKT1 | -1.214898 | COADREAD | LoVo | MK-2206 | MVP_top60_DNA+RNA_seed | PIK3CA | 1.246289 | BRCA | BT-483 | GNE-317 |
| MVP_top60_DNA_seed | MAPK14 | -1.214448 | COADREAD | RKO | TAK-715 | MVP_top60_DNA+RNA_seed | EGFR | 1.257101 | BRCA | MDA-MB-436 | Gefitinib |
| MVP_top60_DNA_seed | AKT1 | -1.211412 | COADREAD | NCI-H716 | BAY AKT1 | MVP_top60_DNA+RNA_seed | ESR1 | 1.257127 | BRCA | HCC1428 | Tamoxifen |
| MVP_top60_DNA_seed | GSK3B | -1.210092 | COADREAD | LoVo | KIN001-042 | MVP_top60_DNA+RNA_seed | MAPK1 | 1.257885 | BRCA | HCC1419 | VX-11e |
| MVP_top60_DNA_seed | BRAF | -1.207046 | COADREAD | CCK-81 | SB590885 | MVP_top60_DNA+RNA_seed | HSP90AA1 | 1.25846 | BRCA | HCC1599 | CCT-018159 |
| MVP_top60_DNA_seed | ERBB4 | -1.200828 | COADREAD | HT-115 | CI-1033 | MVP_top60_DNA+RNA_seed | HSP90AB1 | 1.25846 | BRCA | HCC1599 | CCT-018159 |
| MVP_top60_DNA_seed | HSP90AA1 | -1.199899 | COADREAD | HCT-116 | CCT-018159 | MVP_top60_DNA+RNA_seed | CDK1 | 1.258821 | BRCA | BT-474 | RO-3306 |
| MVP_top60_DNA_seed | HSP90AB1 | -1.199899 | COADREAD | HCT-116 | CCT-018159 | MVP_top60_DNA+RNA_seed | CDK1 | 1.259151 | BRCA | BT-483 | RO-3306 |
| MVP_top60_DNA_seed | HSP90AA1 | -1.193057 | COADREAD | SNU-175 | SNX-2112 | MVP_top60_DNA+RNA_seed | EGFR | 1.261003 | BRCA | Hs-578-T | Osimertinib |
| MVP_top60_DNA_seed | HSP90AB1 | -1.193057 | COADREAD | SNU-175 | SNX-2112 | MVP_top60_DNA+RNA_seed | MDM2 | 1.264404 | BRCA | BT-474 | Serdemetan |
| MVP_top60_DNA_seed | MAPK14 | -1.192349 | COADREAD | HT-115 | TAK-715 | MVP_top60_DNA+RNA_seed | EGFR | 1.267354 | BRCA | HCC1187 | Lapatinib |
| MVP_top60_DNA_seed | ATM | -1.191718 | COADREAD | MDST8 | torin2 | MVP_top60_DNA+RNA_seed | ERBB2 | 1.267354 | BRCA | HCC1187 | Lapatinib |
| MVP_top60_DNA_seed | AKT1 | -1.189434 | COADREAD | NCI-H747 | A-443654 | MVP_top60_DNA+RNA_seed | PIK3CA | 1.275595 | BRCA | CAL-120 | AZD8186 |
| MVP_top60_DNA_seed | CDK2 | -1.185414 | COADREAD | KM12 | Seliciclib | MVP_top60_DNA+RNA_seed | PLK1 | 1.276223 | BRCA | MDA-MB-175-VII | MK-1775 |
| MVP_top60_DNA_seed | HSP90AA1 | -1.182013 | COADREAD | SW48 | SNX-2112 | MVP_top60_DNA+RNA_seed | EGFR | 1.284891 | BRCA | CAL-120 | Erlotinib |
| MVP_top60_DNA_seed | HSP90AB1 | -1.182013 | COADREAD | SW48 | SNX-2112 | MVP_top60_DNA+RNA_seed | PLK1 | 1.287501 | BRCA | HCC1428 | MK-1775 |
| MVP_top60_DNA_seed | IGF1R | -1.179152 | COADREAD | KM12 | GSK1904529A | MVP_top60_DNA+RNA_seed | MDM2 | 1.288863 | BRCA | CAL-120 | Serdemetan |
| MVP_top60_DNA_seed | PIK3CA | -1.178649 | COADREAD | SK-CO-1 | AZD8835 | MVP_top60_DNA+RNA_seed | TP53 | 1.29189 | BRCA | BT-474 | MIRA-1 |
| MVP_top60_DNA_seed | PIK3CD | -1.178649 | COADREAD | SK-CO-1 | AZD8835 | MVP_top60_DNA+RNA_seed | AKT1 | 1.292071 | BRCA | MDA-MB-436 | AT13148 |
| MVP_top60_DNA_seed | IGF1R | -1.175946 | COADREAD | NCI-H747 | BMS-536924 | MVP_top60_DNA+RNA_seed | AKT1 | 1.295298 | BRCA | MDA-MB-231 | AZD5363 |
| MVP_top60_DNA_seed | AKT1 | -1.171344 | COADREAD | SNU-81 | MK-2206 | MVP_top60_DNA+RNA_seed | MAPK3 | 1.297188 | BRCA | HCC1500 | Ulixertinib |
| MVP_top60_DNA_seed | HSP90AA1 | -1.170093 | COADREAD | SW48 | CCT-018159 | MVP_top60_DNA+RNA_seed | MAPK1 | 1.297188 | BRCA | HCC1500 | Ulixertinib |
| MVP_top60_DNA_seed | HSP90AB1 | -1.170093 | COADREAD | SW48 | CCT-018159 | MVP_top60_DNA+RNA_seed | SRC | 1.298928 | BRCA | MFM-223 | Bosutinib |
| MVP_top60_DNA_seed | MAPK1 | -1.167058 | COADREAD | SW620 | VX-11e | MVP_top60_DNA+RNA_seed | MDM2 | 1.300555 | BRCA | MDA-MB-436 | Serdemetan |
| MVP_top60_DNA_seed | PIK3CD | -1.16705 | COADREAD | LoVo | IC-87114 | MVP_top60_DNA+RNA_seed | EGFR | 1.300938 | BRCA | HCC1428 | Sapitinib |
| MVP_top60_DNA_seed | HSP90AA1 | -1.165297 | COADREAD | SK-CO-1 | Tanespimycin | MVP_top60_DNA+RNA_seed | ERBB2 | 1.300938 | BRCA | HCC1428 | Sapitinib |
| MVP_top60_DNA_seed | HSP90AB1 | -1.165297 | COADREAD | SK-CO-1 | Tanespimycin | MVP_top60_DNA+RNA_seed | ERBB3 | 1.300938 | BRCA | HCC1428 | Sapitinib |
| MVP_top60_DNA_seed | GSK3B | -1.163151 | COADREAD | SW48 | AZD7969 | MVP_top60_DNA+RNA_seed | ATM | 1.302381 | BRCA | MDA-MB-157 | KU-55933 |
| MVP_top60_DNA_seed | HSP90AA1 | -1.161249 | COADREAD | KM12 | Tanespimycin | MVP_top60_DNA+RNA_seed | AKT1 | 1.305997 | BRCA | CAL-120 | AT13148 |
| MVP_top60_DNA_seed | HSP90AB1 | -1.161249 | COADREAD | KM12 | Tanespimycin | MVP_top60_DNA+RNA_seed | HDAC1 | 1.307431 | BRCA | MDA-MB-436 | Entinostat |

| | | | | | | | | | | | |
|---|---|---|---|---|---|---|---|---|---|---|---|
| MVP_top60_DNA_seed | HSP90AA1 | -1.155373 | COADREAD | LS-411N | SNX-2112 | MVP_top60_DNA+RNA_seed | PLK1 | 1.308867 | BRCA | MDA-MB-415 | BI-2536 |
| MVP_top60_DNA_seed | HSP90AB1 | -1.155373 | COADREAD | LS-411N | SNX-2112 | MVP_top60_DNA+RNA_seed | HDAC1 | 1.309236 | BRCA | HCC1937 | PCI-34051 |
| MVP_top60_DNA_seed | GSK3B | -1.147071 | COADREAD | RCM-1 | CHIR-99021 | MVP_top60_DNA+RNA_seed | EP300 | 1.312826 | BRCA | MDA-MB-436 | SGC-CBP30 |
| MVP_top60_DNA_seed | IGF1R | -1.145111 | COADREAD | LS-513 | Linsitinib | MVP_top60_DNA+RNA_seed | PLK1 | 1.315157 | BRCA | BT-474 | BI-2536 |
| MVP_top60_DNA_seed | SRC | -1.144012 | COADREAD | NCI-H747 | Saracatinib | MVP_top60_DNA+RNA_seed | HSP90AA1 | 1.322485 | BRCA | HCC1428 | Luminespib |
| MVP_top60_DNA_seed | ATM | -1.134419 | COADREAD | NCI-H716 | KU-60019 | MVP_top60_DNA+RNA_seed | HSP90AB1 | 1.322485 | BRCA | HCC1428 | Luminespib |
| MVP_top60_DNA_seed | BRAF | -1.132251 | COADREAD | LS-513 | AZ628 | MVP_top60_DNA+RNA_seed | MDM2 | 1.323683 | BRCA | BT-483 | Nutlin-3a (-) |
| MVP_top60_DNA_seed | ESR1 | -1.131395 | COADREAD | LS-180 | Tamoxifen | MVP_top60_DNA+RNA_seed | AKT1 | 1.330708 | BRCA | MDA-MB-436 | MK-2206 |
| MVP_top60_DNA_seed | ATM | -1.124209 | COADREAD | SNU-175 | CP466722 | MVP_top60_DNA+RNA_seed | SRC | 1.332667 | BRCA | YMB-1-E | Bosutinib |
| MVP_top60_DNA_seed | MAPK1 | -1.121201 | COADREAD | HCT-15 | VX-11e | MVP_top60_DNA+RNA_seed | EP300 | 1.340486 | BRCA | HCC1143 | CPI-637 |
| MVP_top60_DNA_seed | HSP90AA1 | -1.114351 | COADREAD | RKO | Luminespib | MVP_top60_DNA+RNA_seed | CDK2 | 1.343109 | BRCA | HDQ-P1 | MK-8776 |
| MVP_top60_DNA_seed | HSP90AB1 | -1.114351 | COADREAD | RKO | Luminespib | MVP_top60_DNA+RNA_seed | PLK1 | 1.348728 | BRCA | CAMA-1 | BI-2536 |
| MVP_top60_DNA_seed | MET | -1.11237 | COADREAD | LoVo | SU11274 | MVP_top60_DNA+RNA_seed | EGFR | 1.352546 | BRCA | BT-549 | Erlotinib |
| MVP_top60_DNA_seed | PIK3CB | -1.112036 | COADREAD | LS-180 | AZD6482 | MVP_top60_DNA+RNA_seed | EP300 | 1.353012 | BRCA | CAL-120 | CPI-637 |
| MVP_top60_DNA_seed | SRC | -1.111319 | COADREAD | LoVo | HG-6-71-01 | MVP_top60_DNA+RNA_seed | EGFR | 1.353353 | BRCA | UACC-812 | AZD3759 |
| MVP_top60_DNA_seed | AR | -1.110622 | COADREAD | SW48 | Bicalutamide | MVP_top60_DNA+RNA_seed | PLK1 | 1.354522 | BRCA | MDA-MB-330 | MK-1775 |
| MVP_top60_DNA_seed | MAPK14 | -1.109294 | COADREAD | KM12 | TAK-715 | MVP_top60_DNA+RNA_seed | EGFR | 1.355541 | BRCA | MDA-MB-157 | Afatinib |
| MVP_top60_DNA_seed | BRAF | -1.097259 | COADREAD | LoVo | HG6-64-1 | MVP_top60_DNA+RNA_seed | ERBB2 | 1.355541 | BRCA | MDA-MB-157 | Afatinib |
| MVP_top60_DNA_seed | IGF1R | -1.097252 | COADREAD | COLO-320-HSR | BMS-536924 | MVP_top60_DNA+RNA_seed | AKT1 | 1.355933 | BRCA | HCC38 | MK-2206 |
| MVP_top60_DNA_seed | AKT1 | -1.092442 | COADREAD | NCI-H747 | GSK690693 | MVP_top60_DNA+RNA_seed | EGFR | 1.359327 | BRCA | HCC38 | Erlotinib |
| MVP_top60_DNA_seed | SRC | -1.090901 | COADREAD | RKO | NVP-BHG712 | MVP_top60_DNA+RNA_seed | PLK1 | 1.359822 | BRCA | HCC2218 | BI-2536 |
| MVP_top60_DNA_seed | IGF1R | -1.089427 | COADREAD | CCK-81 | Linsitinib | MVP_top60_DNA+RNA_seed | MAPK3 | 1.36602 | BRCA | CAL-148 | SCH772984 |
| MVP_top60_DNA_seed | PIK3CA | -1.086516 | COADREAD | GP5d | AZD8835 | MVP_top60_DNA+RNA_seed | MAPK1 | 1.36602 | BRCA | CAL-148 | SCH772984 |
| MVP_top60_DNA_seed | PIK3CD | -1.086516 | COADREAD | GP5d | AZD8835 | MVP_top60_DNA+RNA_seed | CDK2 | 1.368153 | BRCA | HCC1428 | MK-8776 |
| MVP_top60_DNA_seed | PIK3CA | -1.084276 | COADREAD | DiFi | PI3Ka_4409 | MVP_top60_DNA+RNA_seed | CDK2 | 1.368896 | BRCA | MCF7 | MK-8776 |
| MVP_top60_DNA_seed | PIK3CD | -1.084276 | COADREAD | DiFi | PI3Ka_4409 | MVP_top60_DNA+RNA_seed | AKT1 | 1.374834 | BRCA | BT-483 | Uprosertib |
| MVP_top60_DNA_seed | PIK3CA | -1.081583 | COADREAD | SNU-407 | PI3Ka_4409 | MVP_top60_DNA+RNA_seed | MDM2 | 1.378181 | BRCA | MFM-223 | Nutlin-3a (-) |
| MVP_top60_DNA_seed | PIK3CD | -1.081583 | COADREAD | SNU-407 | PI3Ka_4409 | MVP_top60_DNA+RNA_seed | MAPK3 | 1.378805 | BRCA | HCC1428 | Ulixertinib |
| MVP_top60_DNA_seed | BRAF | -1.077068 | COADREAD | COLO-205 | HG6-64-1 | MVP_top60_DNA+RNA_seed | MAPK1 | 1.378805 | BRCA | HCC1428 | Ulixertinib |
| MVP_top60_DNA_seed | MAPK1 | -1.076615 | COADREAD | SW948 | VX-11e | MVP_top60_DNA+RNA_seed | AKT1 | 1.381455 | BRCA | HCC1500 | Uprosertib |
| MVP_top60_DNA_seed | MAPK1 | -1.075552 | COADREAD | SW48 | FR-180204 | MVP_top60_DNA+RNA_seed | ESR1 | 1.381542 | BRCA | UACC-893 | Tamoxifen |
| MVP_top60_DNA_seed | BRAF | -1.074232 | COADREAD | LS-513 | RAF_9304 | MVP_top60_DNA+RNA_seed | PLK1 | 1.382404 | BRCA | HCC1954 | BI-2536 |
| MVP_top60_DNA_seed | ERBB4 | -1.07416 | COADREAD | HT-115 | PF-00299804 | MVP_top60_DNA+RNA_seed | CDK2 | 1.385994 | BRCA | EFM-192A | MK-8776 |
| MVP_top60_DNA_seed | BRAF | -1.066656 | COADREAD | LS-411N | SB590885 | MVP_top60_DNA+RNA_seed | PLK1 | 1.38815 | BRCA | HCC1187 | BI-2536 |
| MVP_top60_DNA_seed | CDK2 | -1.064815 | COADREAD | HCT-116 | AZD5438 | MVP_top60_DNA+RNA_seed | MDM2 | 1.389214 | BRCA | MDA-MB-436 | Nutlin-3a (-) |
| MVP_top60_DNA_seed | AKT1 | -1.063981 | COADREAD | SNU-175 | AKT inhibitor VIII | MVP_top60_DNA+RNA_seed | HDAC1 | 1.391353 | BRCA | BT-549 | PCI-34051 |
| MVP_top60_DNA_seed | AKT1 | -1.061978 | COADREAD | LoVo | AKT inhibitor VIII | MVP_top60_DNA+RNA_seed | EGFR | 1.392792 | BRCA | YMB-1-E | Afatinib |
| MVP_top60_DNA_seed | PIK3CA | -1.060381 | COADREAD | NCI-H747 | AZD8835 | MVP_top60_DNA+RNA_seed | ERBB2 | 1.392792 | BRCA | YMB-1-E | Afatinib |
| MVP_top60_DNA_seed | PIK3CD | -1.060381 | COADREAD | NCI-H747 | AZD8835 | MVP_top60_DNA+RNA_seed | EP300 | 1.397503 | BRCA | MDA-MB-175-VII | CPI-637 |
| MVP_top60_DNA_seed | CDK2 | -1.059276 | COADREAD | HT-115 | AZD5438 | MVP_top60_DNA+RNA_seed | EGFR | 1.397503 | BRCA | CAL-85-1 | Osimertinib |
| MVP_top60_DNA_seed | AKT1 | -1.058893 | COADREAD | RKO | MK-2206 | MVP_top60_DNA+RNA_seed | MAPK3 | 1.401421 | BRCA | EFM-192A | Ulixertinib |

| | | | | | | | | | | | |
|---|---|---|---|---|---|---|---|---|---|---|---|
| MVP_top60_DNA_seed | CDK2 | -1.055658 | COADREAD | SW620 | PHA-793887 | MVP_top60_DNA+RNA_seed | MAPK1 | 1.401421 | BRCA | EFM-192A | Ulixertinib |
| MVP_top60_DNA_seed | PIK3CB | -1.053646 | COADREAD | RKO | AZD6482 | MVP_top60_DNA+RNA_seed | HDAC1 | 1.404337 | BRCA | CAL-120 | Entinostat |
| MVP_top60_DNA_seed | IGF1R | -1.052362 | COADREAD | RKO | BMS-536924 | MVP_top60_DNA+RNA_seed | HDAC1 | 1.40823 | BRCA | HCC1187 | Romidepsin |
| MVP_top60_DNA_seed | ESR1 | -1.051126 | COADREAD | HCT-116 | Tamoxifen | MVP_top60_DNA+RNA_seed | HDAC2 | 1.40823 | BRCA | HCC1187 | Romidepsin |
| MVP_top60_DNA_seed | BRAF | -1.048822 | COADREAD | LoVo | PLX-4720 | MVP_top60_DNA+RNA_seed | AKT1 | 1.408726 | BRCA | CAL-85-1 | AT13148 |
| MVP_top60_DNA_seed | ATM | -1.042063 | COADREAD | NCI-H716 | KU-55933 | MVP_top60_DNA+RNA_seed | SRC | 1.409147 | BRCA | BT-474 | Bosutinib |
| MVP_top60_DNA_seed | ESR1 | -1.041264 | COADREAD | MDST8 | Fulvestrant | MVP_top60_DNA+RNA_seed | CDK2 | 1.417989 | BRCA | BT-474 | Dinaciclib |
| MVP_top60_DNA_seed | MAPK1 | -1.037989 | COADREAD | SW837 | VX-11e | MVP_top60_DNA+RNA_seed | CDK1 | 1.417989 | BRCA | BT-474 | Dinaciclib |
| MVP_top60_DNA_seed | CREBBP | -1.036797 | COADREAD | HCC2998 | I-CBP112 | MVP_top60_DNA+RNA_seed | EGFR | 1.422056 | BRCA | HCC38 | Afatinib |
| MVP_top60_DNA_seed | AKT1 | -1.034944 | COADREAD | NCI-H716 | AKT inhibitor VIII | MVP_top60_DNA+RNA_seed | ERBB2 | 1.422056 | BRCA | HCC38 | Afatinib |
| MVP_top60_DNA_seed | PTK2 | -1.033563 | COADREAD | SW620 | PF-562271 | MVP_top60_DNA+RNA_seed | EGFR | 1.428389 | BRCA | ZR-75-30 | AZD3759 |
| MVP_top60_DNA_seed | CDK2 | -1.03127 | COADREAD | RKO | AZD5438 | MVP_top60_DNA+RNA_seed | MAPK3 | 1.434598 | BRCA | HCC1187 | SCH772984 |
| MVP_top60_DNA_seed | BRAF | -1.031048 | COADREAD | MDST8 | PLX-4720 | MVP_top60_DNA+RNA_seed | MAPK1 | 1.434598 | BRCA | HCC1187 | SCH772984 |
| MVP_top60_DNA_seed | IGF1R | -1.030512 | COADREAD | GP5d | Linsitinib | MVP_top60_DNA+RNA_seed | MAPK3 | 1.43646 | BRCA | JIMT-1 | SCH772984 |
| MVP_top60_DNA_seed | HSP90AA1 | -1.029666 | COADREAD | HCT-116 | SNX-2112 | MVP_top60_DNA+RNA_seed | MAPK1 | 1.43646 | BRCA | JIMT-1 | SCH772984 |
| MVP_top60_DNA_seed | HSP90AB1 | -1.029666 | COADREAD | HCT-116 | SNX-2112 | MVP_top60_DNA+RNA_seed | MAPK3 | 1.441685 | BRCA | HCC38 | SCH772984 |
| MVP_top60_DNA_seed | HSP90AA1 | -1.029424 | COADREAD | LS-513 | SNX-2112 | MVP_top60_DNA+RNA_seed | MAPK1 | 1.441685 | BRCA | HCC38 | SCH772984 |
| MVP_top60_DNA_seed | HSP90AB1 | -1.029424 | COADREAD | LS-513 | SNX-2112 | MVP_top60_DNA+RNA_seed | AKT1 | 1.443381 | BRCA | HCC1187 | Uprosertib |
| MVP_top60_DNA_seed | HSP90AA1 | -1.028081 | COADREAD | HT-29 | SNX-2112 | MVP_top60_DNA+RNA_seed | TP53 | 1.444911 | BRCA | HCC1428 | MIRA-1 |
| MVP_top60_DNA_seed | HSP90AB1 | -1.028081 | COADREAD | HT-29 | SNX-2112 | MVP_top60_DNA+RNA_seed | MAPK3 | 1.445339 | BRCA | HCC1599 | Ulixertinib |
| MVP_top60_DNA_seed | HSP90AA1 | -1.021061 | COADREAD | NCI-H508 | Tanespimycin | MVP_top60_DNA+RNA_seed | MAPK1 | 1.445339 | BRCA | HCC1599 | Ulixertinib |
| MVP_top60_DNA_seed | HSP90AB1 | -1.021061 | COADREAD | NCI-H508 | Tanespimycin | MVP_top60_DNA+RNA_seed | CDK2 | 1.447728 | BRCA | JIMT-1 | Dinaciclib |
| MVP_top60_DNA_seed | MAPK1 | -1.016098 | COADREAD | CCK-81 | FR-180204 | MVP_top60_DNA+RNA_seed | CDK1 | 1.447728 | BRCA | JIMT-1 | Dinaciclib |
| MVP_top60_DNA_seed | PIK3CA | -1.013241 | COADREAD | HCT-116 | PI3Ka_4409 | MVP_top60_DNA+RNA_seed | AKT1 | 1.458537 | BRCA | ZR-75-30 | GSK2110183B |
| MVP_top60_DNA_seed | PIK3CD | -1.013241 | COADREAD | HCT-116 | PI3Ka_4409 | MVP_top60_DNA+RNA_seed | HDAC1 | 1.458654 | BRCA | EFM-19 | Entinostat |
| MVP_top60_DNA_seed | ATM | -1.012839 | COADREAD | SNU-C5 | KU-55933 | MVP_top60_DNA+RNA_seed | EGFR | 1.458705 | BRCA | HCC1428 | Erlotinib |
| MVP_top60_DNA_seed | CDK2 | -1.007668 | COADREAD | MDST8 | AZD5438 | MVP_top60_DNA+RNA_seed | EGFR | 1.45958 | BRCA | HCC1599 | Erlotinib |
| MVP_top60_DNA_seed | MAPK1 | -1.006754 | COADREAD | LS-513 | FR-180204 | MVP_top60_DNA+RNA_seed | HDAC1 | 1.459961 | BRCA | BT-474 | PCI-34051 |
| MVP_top60_DNA_seed | HSP90AA1 | -1.006089 | COADREAD | SW620 | Luminespib | MVP_top60_DNA+RNA_seed | AKT1 | 1.466902 | BRCA | CAL-120 | GSK2110183B |
| MVP_top60_DNA_seed | HSP90AB1 | -1.006089 | COADREAD | SW620 | Luminespib | MVP_top60_DNA+RNA_seed | EP300 | 1.467191 | BRCA | HCC1428 | SGC-CBP30 |
| MVP_top60_DNA_seed | HSP90AA1 | -1.004215 | COADREAD | SW620 | CCT-018159 | MVP_top60_DNA+RNA_seed | CDK2 | 1.471313 | BRCA | MDA-MB-436 | AZD5438 |
| MVP_top60_DNA_seed | HSP90AB1 | -1.004215 | COADREAD | SW620 | CCT-018159 | MVP_top60_DNA+RNA_seed | PIK3CA | 1.481992 | BRCA | HCC1500 | Buparlisib |
| MVP_top60_DNA_seed | ERBB3 | -0.999809 | COADREAD | GP5d | AZD8931 | MVP_top60_DNA+RNA_seed | MDM2 | 1.5077 | BRCA | HCC38 | Serdemetan |
| MVP_top60_DNA_seed | BRAF | -0.99188 | COADREAD | LS-411N | Dabrafenib | MVP_top60_DNA+RNA_seed | HSP90AA1 | 1.509597 | BRCA | BT-483 | Tanespimycin |
| MVP_top60_DNA_seed | GSK3B | -0.990536 | COADREAD | MDST8 | AZD7969 | MVP_top60_DNA+RNA_seed | HSP90AB1 | 1.509597 | BRCA | BT-483 | Tanespimycin |
| MVP_top60_DNA_seed | BRAF | -0.990068 | COADREAD | HCT-116 | Dabrafenib | MVP_top60_DNA+RNA_seed | MAPK3 | 1.509658 | BRCA | BT-474 | Ulixertinib |
| MVP_top60_DNA_seed | MAPK1 | -0.989042 | COADREAD | CCK-81 | VX-11e | MVP_top60_DNA+RNA_seed | MAPK1 | 1.509658 | BRCA | BT-474 | Ulixertinib |
| MVP_top60_DNA_seed | ESR1 | -0.988402 | COADREAD | HT-115 | Tamoxifen | MVP_top60_DNA+RNA_seed | AKT1 | 1.51075 | BRCA | HCC38 | AT13148 |
| MVP_top60_DNA_seed | SRC | -0.987133 | COADREAD | HCT-116 | Bosutinib | MVP_top60_DNA+RNA_seed | CDK2 | 1.519447 | BRCA | UACC-812 | MK-8776 |
| MVP_top60_DNA_seed | PIK3CA | -0.984215 | COADREAD | GP5d | PI3Ka_4409 | MVP_top60_DNA+RNA_seed | MAPK1 | 1.52302 | BRCA | UACC-812 | VX-11e |
| MVP_top60_DNA_seed | PIK3CD | -0.984215 | COADREAD | GP5d | PI3Ka_4409 | MVP_top60_DNA+RNA_seed | MAPK3 | 1.531234 | BRCA | HCC1428 | SCH772984 |

| | | | | | | | | | | | |
|---|---|---|---|---|---|---|---|---|---|---|---|
| MVP_top60_DNA_seed | AKT1 | -0.983791 | COADREAD | COLO-205 | BAY AKT1 | MVP_top60_DNA+RNA_seed | MAPK1 | 1.531234 | BRCA | HCC1428 | SCH772984 |
| MVP_top60_DNA_seed | HSP90AA1 | -0.982615 | COADREAD | COLO-205 | SNX-2112 | MVP_top60_DNA+RNA_seed | MAPK3 | 1.535184 | BRCA | MCF7 | SCH772984 |
| MVP_top60_DNA_seed | HSP90AB1 | -0.982615 | COADREAD | COLO-205 | SNX-2112 | MVP_top60_DNA+RNA_seed | MAPK1 | 1.535184 | BRCA | MCF7 | SCH772984 |
| MVP_top60_DNA_seed | ATM | -0.98135 | COADREAD | KM12 | KU-60019 | MVP_top60_DNA+RNA_seed | MAPK3 | 1.536895 | BRCA | HCC38 | Ulixertinib |
| MVP_top60_DNA_seed | GSK3B | -0.978221 | COADREAD | HT-29 | SB216763 | MVP_top60_DNA+RNA_seed | MAPK1 | 1.536895 | BRCA | HCC38 | Ulixertinib |
| MVP_top60_DNA_seed | IGF1R | -0.976678 | COADREAD | SK-CO-1 | BMS-536924 | MVP_top60_DNA+RNA_seed | EP300 | 1.540577 | BRCA | HCC1937 | CPI-637 |
| MVP_top60_DNA_seed | GSK3B | -0.975218 | COADREAD | HCT-116 | CHIR-99021 | MVP_top60_DNA+RNA_seed | PLK1 | 1.546905 | BRCA | BT-483 | BI-2536 |
| MVP_top60_DNA_seed | ERBB3 | -0.973837 | COADREAD | LoVo | AZD8931 | MVP_top60_DNA+RNA_seed | HDAC1 | 1.548354 | BRCA | HCC1937 | Romidepsin |
| MVP_top60_DNA_seed | ERBB4 | -0.969238 | COADREAD | SNU-175 | CI-1033 | MVP_top60_DNA+RNA_seed | HDAC2 | 1.548354 | BRCA | HCC1937 | Romidepsin |
| MVP_top60_DNA_seed | IGF1R | -0.963142 | COADREAD | LS-1034 | BMS-536924 | MVP_top60_DNA+RNA_seed | PLK1 | 1.552509 | BRCA | UACC-812 | MK-1775 |
| MVP_top60_DNA_seed | HSP90AA1 | -0.96185 | COADREAD | SW620 | SNX-2112 | MVP_top60_DNA+RNA_seed | PIK3CA | 1.553813 | BRCA | HCC1187 | GNE-317 |
| MVP_top60_DNA_seed | HSP90AB1 | -0.96185 | COADREAD | SW620 | SNX-2112 | MVP_top60_DNA+RNA_seed | CDK2 | 1.557598 | BRCA | BT-474 | MK-8776 |
| MVP_top60_DNA_seed | MAPK1 | -0.959284 | COADREAD | SW48 | VX-11e | MVP_top60_DNA+RNA_seed | PLK1 | 1.569675 | BRCA | BT-474 | MK-1775 |
| MVP_top60_DNA_seed | ESR1 | -0.95816 | COADREAD | RKO | Fulvestrant | MVP_top60_DNA+RNA_seed | PIK3CA | 1.582609 | BRCA | HCC1419 | GNE-317 |
| MVP_top60_DNA_seed | GSK3B | -0.956867 | COADREAD | GP5d | KIN001-042 | MVP_top60_DNA+RNA_seed | PLK1 | 1.585866 | BRCA | UACC-893 | MK-1775 |
| MVP_top60_DNA_seed | BRAF | -0.953793 | COADREAD | LS-411N | RAF_9304 | MVP_top60_DNA+RNA_seed | AKT1 | 1.593483 | BRCA | HCC1428 | GSK2110183B |
| MVP_top60_DNA_seed | ERBB4 | -0.953289 | COADREAD | HT-115 | AST-1306 | MVP_top60_DNA+RNA_seed | AKT1 | 1.594381 | BRCA | HCC1937 | AT13148 |
| MVP_top60_DNA_seed | MAPK1 | -0.951821 | COADREAD | T84 | VX-11e | MVP_top60_DNA+RNA_seed | HDAC1 | 1.595622 | BRCA | ZR-75-30 | PCI-34051 |
| MVP_top60_DNA_seed | CDK2 | -0.95082 | COADREAD | RKO | Seliciclib | MVP_top60_DNA+RNA_seed | PLK1 | 1.598239 | BRCA | MDA-MB-436 | BI-2536 |
| MVP_top60_DNA_seed | GSK3B | -0.94936 | COADREAD | GP5d | CHIR-99021 | MVP_top60_DNA+RNA_seed | PLK1 | 1.600018 | BRCA | EFM-192A | BI-2536 |
| MVP_top60_DNA_seed | ERBB4 | -0.947921 | COADREAD | LoVo | PF-00299804 | MVP_top60_DNA+RNA_seed | PIK3CA | 1.601177 | BRCA | HCC1500 | AZD8186 |
| MVP_top60_DNA_seed | MAPK1 | -0.945212 | COADREAD | SNU-175 | VX-11e | MVP_top60_DNA+RNA_seed | AKT1 | 1.610376 | BRCA | HCC1428 | Uprosertib |
| MVP_top60_DNA_seed | MAPK14 | -0.938931 | COADREAD | LS-513 | TAK-715 | MVP_top60_DNA+RNA_seed | PLK1 | 1.61371 | BRCA | MDA-MB-175-VII | BI-2536 |
| MVP_top60_DNA_seed | SRC | -0.930433 | COADREAD | HT-29 | NVP-BHG712 | MVP_top60_DNA+RNA_seed | MDM2 | 1.625789 | BRCA | CAL-120 | Nutlin-3a (-) |
| MVP_top60_DNA_seed | PIK3CA | -0.928663 | COADREAD | HCT-116 | AZD8835 | MVP_top60_DNA+RNA_seed | EGFR | 1.63989 | BRCA | MFM-223 | Gefitinib |
| MVP_top60_DNA_seed | PIK3CD | -0.928663 | COADREAD | HCT-116 | AZD8835 | MVP_top60_DNA+RNA_seed | CDK2 | 1.645599 | BRCA | MDA-MB-436 | Dinaciclib |
| MVP_top60_DNA_seed | MAPK1 | -0.925878 | COADREAD | SNU-C2B | VX-11e | MVP_top60_DNA+RNA_seed | CDK1 | 1.645599 | BRCA | MDA-MB-436 | Dinaciclib |
| MVP_top60_DNA_seed | AKT1 | -0.923658 | COADREAD | LS-180 | AKT inhibitor VIII | MVP_top60_DNA+RNA_seed | PLK1 | 1.659986 | BRCA | HCC2218 | MK-1775 |
| MVP_top60_DNA_seed | SRC | -0.920611 | COADREAD | SNU-81 | Bosutinib | MVP_top60_DNA+RNA_seed | PLK1 | 1.672698 | BRCA | MCF7 | MK-1775 |
| MVP_top60_DNA_seed | ERBB4 | -0.917014 | COADREAD | LoVo | CI-1033 | MVP_top60_DNA+RNA_seed | MDM2 | 1.675931 | BRCA | HCC1428 | Serdemetan |
| MVP_top60_DNA_seed | AR | -0.91592 | COADREAD | LS-411N | AZD3514 | MVP_top60_DNA+RNA_seed | PLK1 | 1.682074 | BRCA | MDA-MB-415 | MK-1775 |
| MVP_top60_DNA_seed | CDK2 | -0.914337 | COADREAD | HT-29 | PHA-793887 | MVP_top60_DNA+RNA_seed | CDK2 | 1.682691 | BRCA | ZR-75-30 | MK-8776 |
| MVP_top60_DNA_seed | ERBB3 | -0.914211 | COADREAD | SW1463 | AZD8931 | MVP_top60_DNA+RNA_seed | AKT1 | 1.687189 | BRCA | CAL-85-1 | AZD5363 |
| MVP_top60_DNA_seed | SRC | -0.913402 | COADREAD | MDST8 | NVP-BHG712 | MVP_top60_DNA+RNA_seed | PIK3CA | 1.70168 | BRCA | ZR-75-30 | Buparlisib |
| MVP_top60_DNA_seed | BRAF | -0.905346 | COADREAD | SNU-C5 | Dabrafenib | MVP_top60_DNA+RNA_seed | MAPK3 | 1.705641 | BRCA | HCC1428 | Ulixertinib |
| MVP_top60_DNA_seed | HSP90AA1 | -0.901731 | COADREAD | HCT-15 | Elesclomol | MVP_top60_DNA+RNA_seed | MAPK1 | 1.705641 | BRCA | HCC1428 | Ulixertinib |
| MVP_top60_DNA_seed | HSP90AB1 | -0.901731 | COADREAD | HCT-15 | Elesclomol | MVP_top60_DNA+RNA_seed | HDAC1 | 1.706539 | BRCA | ZR-75-30 | Entinostat |
| MVP_top60_DNA_seed | ERBB3 | -0.896592 | COADREAD | HCT-116 | AZD8931 | MVP_top60_DNA+RNA_seed | AKT1 | 1.709297 | BRCA | MRK-nu-1 | MK-2206 |
| MVP_top60_DNA_seed | ERBB3 | -0.892663 | COADREAD | LS-513 | AZD8931 | MVP_top60_DNA+RNA_seed | MAPK1 | 1.714617 | BRCA | HCC1428 | VX-11e |
| MVP_top60_DNA_seed | MAPK1 | -0.892068 | COADREAD | MDST8 | VX-11e | MVP_top60_DNA+RNA_seed | HSP90AA1 | 1.724402 | BRCA | T47D | Tanespimycin |
| MVP_top60_DNA_seed | ATM | -0.890736 | COADREAD | LoVo | KU-60019 | MVP_top60_DNA+RNA_seed | HSP90AB1 | 1.724402 | BRCA | T47D | Tanespimycin |

| | | | | | | | | | | | |
|---|---|---|---|---|---|---|---|---|---|---|---|
| MVP_top60_DNA_seed | IGF1R | -0.886705 | COADREAD | HT-29 | BMS-536924 | MVP_top60_DNA+RNA_seed | HSP90AA1 | 1.736226 | BRCA | UACC-812 | Tanespimycin |
| MVP_top60_DNA_seed | MET | -0.885738 | COADREAD | LS-411N | Crizotinib | MVP_top60_DNA+RNA_seed | HSP90AB1 | 1.736226 | BRCA | UACC-812 | Tanespimycin |
| MVP_top60_DNA_seed | HSP90AA1 | -0.885297 | COADREAD | SW620 | Tanespimycin | MVP_top60_DNA+RNA_seed | EGFR | 1.762253 | BRCA | ZR-75-30 | Gefitinib |
| MVP_top60_DNA_seed | HSP90AB1 | -0.885297 | COADREAD | SW620 | Tanespimycin | MVP_top60_DNA+RNA_seed | MAPK3 | 1.76262 | BRCA | CAL-120 | Ulixertinib |
| MVP_top60_DNA_seed | AKT1 | -0.883882 | COADREAD | LS-513 | MK-2206 | MVP_top60_DNA+RNA_seed | MAPK1 | 1.76262 | BRCA | CAL-120 | Ulixertinib |
| MVP_top60_DNA_seed | GSK3B | -0.883203 | COADREAD | SNU-C5 | CHIR-99021 | MVP_top60_DNA+RNA_seed | AKT1 | 1.768638 | BRCA | HCC1500 | MK-2206 |
| MVP_top60_DNA_seed | SRC | -0.877365 | COADREAD | NCI-H747 | NVP-BHG712 | MVP_top60_DNA+RNA_seed | MDM2 | 1.784761 | BRCA | HCC1428 | Nutlin-3a (-) |
| MVP_top60_DNA_seed | PIK3CA | -0.869761 | COADREAD | RKO | AZD8835 | MVP_top60_DNA+RNA_seed | MAPK3 | 1.785123 | BRCA | CAL-120 | SCH772984 |
| MVP_top60_DNA_seed | PIK3CD | -0.869761 | COADREAD | RKO | AZD8835 | MVP_top60_DNA+RNA_seed | MAPK1 | 1.785123 | BRCA | CAL-120 | SCH772984 |
| MVP_top60_DNA_seed | BRAF | -0.866334 | COADREAD | HCC2998 | RAF_9304 | MVP_top60_DNA+RNA_seed | MAPK3 | 1.788339 | BRCA | BT-549 | Ulixertinib |
| MVP_top60_DNA_seed | GSK3B | -0.864362 | COADREAD | HT-115 | SB216763 | MVP_top60_DNA+RNA_seed | MAPK1 | 1.788339 | BRCA | BT-549 | Ulixertinib |
| MVP_top60_DNA_seed | PTK2 | -0.862029 | COADREAD | RKO | PF-562271 | MVP_top60_DNA+RNA_seed | ESR1 | 1.790311 | BRCA | MDA-MB-175-VII | Tamoxifen |
| MVP_top60_DNA_seed | BRAF | -0.858547 | COADREAD | SW948 | SB590885 | MVP_top60_DNA+RNA_seed | MAPK3 | 1.806136 | BRCA | UACC-812 | SCH772984 |
| MVP_top60_DNA_seed | GSK3B | -0.857376 | COADREAD | SW620 | AZD7969 | MVP_top60_DNA+RNA_seed | MAPK1 | 1.806136 | BRCA | UACC-812 | SCH772984 |
| MVP_top60_DNA_seed | GSK3B | -0.856485 | COADREAD | HT-115 | AZD7969 | MVP_top60_DNA+RNA_seed | CDK2 | 1.808025 | BRCA | HCC1500 | Dinaciclib |
| MVP_top60_DNA_seed | HSP90AA1 | -0.85375 | COADREAD | KM12 | CCT-018159 | MVP_top60_DNA+RNA_seed | CDK1 | 1.808025 | BRCA | HCC1500 | Dinaciclib |
| MVP_top60_DNA_seed | HSP90AB1 | -0.85375 | COADREAD | KM12 | CCT-018159 | MVP_top60_DNA+RNA_seed | EGFR | 1.808409 | BRCA | ZR-75-30 | Erlotinib |
| MVP_top60_DNA_seed | MAPK1 | -0.852633 | COADREAD | LS-411N | FR-180204 | MVP_top60_DNA+RNA_seed | MAPK3 | 1.812945 | BRCA | BT-483 | Ulixertinib |
| MVP_top60_DNA_seed | PIK3CB | -0.852167 | COADREAD | HCT-116 | AZD6482 | MVP_top60_DNA+RNA_seed | MAPK1 | 1.812945 | BRCA | BT-483 | Ulixertinib |
| MVP_top60_DNA_seed | MAPK14 | -0.851458 | COADREAD | SNU-1040 | TAK-715 | MVP_top60_DNA+RNA_seed | EGFR | 1.813324 | BRCA | UACC-812 | Erlotinib |
| MVP_top60_DNA_seed | AKT1 | -0.850234 | COADREAD | RKO | A-443654 | MVP_top60_DNA+RNA_seed | HDAC1 | 1.814019 | BRCA | MFM-223 | Entinostat |
| MVP_top60_DNA_seed | HSP90AA1 | -0.850035 | COADREAD | SNU-C2B | Luminespib | MVP_top60_DNA+RNA_seed | PLK1 | 1.820042 | BRCA | HCC1419 | MK-1775 |
| MVP_top60_DNA_seed | HSP90AB1 | -0.850035 | COADREAD | SNU-C2B | Luminespib | MVP_top60_DNA+RNA_seed | PLK1 | 1.826433 | BRCA | ZR-75-30 | BI-2536 |
| MVP_top60_DNA_seed | MET | -0.847099 | COADREAD | KM12 | JNJ38877605 | MVP_top60_DNA+RNA_seed | TP53 | 1.829054 | BRCA | MDA-MB-175-VII | MIRA-1 |
| MVP_top60_DNA_seed | GSK3B | -0.846871 | COADREAD | HCT-116 | SB216763 | MVP_top60_DNA+RNA_seed | MDM2 | 1.829666 | BRCA | HCC1599 | Serdemetan |
| MVP_top60_DNA_seed | HSP90AA1 | -0.845633 | COADREAD | MDST8 | SNX-2112 | MVP_top60_DNA+RNA_seed | EGFR | 1.830561 | BRCA | CAL-120 | Sapitinib |
| MVP_top60_DNA_seed | HSP90AB1 | -0.845633 | COADREAD | MDST8 | SNX-2112 | MVP_top60_DNA+RNA_seed | ERBB2 | 1.830561 | BRCA | CAL-120 | Sapitinib |
| MVP_top60_DNA_seed | PIK3CA | -0.844898 | COADREAD | SNU-C1 | AZD8835 | MVP_top60_DNA+RNA_seed | ERBB3 | 1.830561 | BRCA | CAL-120 | Sapitinib |
| MVP_top60_DNA_seed | PIK3CD | -0.844898 | COADREAD | SNU-C1 | AZD8835 | MVP_top60_DNA+RNA_seed | SRC | 1.831053 | BRCA | ZR-75-30 | Bosutinib |
| MVP_top60_DNA_seed | PIK3CA | -0.843751 | COADREAD | SW48 | PI3Ka_4409 | MVP_top60_DNA+RNA_seed | MAPK1 | 1.832258 | BRCA | BT-474 | VX-11e |
| MVP_top60_DNA_seed | PIK3CD | -0.843751 | COADREAD | SW48 | PI3Ka_4409 | MVP_top60_DNA+RNA_seed | AKT1 | 1.837836 | BRCA | HCC1428 | Uprosertib |
| MVP_top60_DNA_seed | AKT1 | -0.83972 | COADREAD | CCK-81 | AKT inhibitor VIII | MVP_top60_DNA+RNA_seed | PLK1 | 1.851474 | BRCA | HCC1419 | BI-2536 |
| MVP_top60_DNA_seed | MAPK1 | -0.838878 | COADREAD | SW1417 | VX-11e | MVP_top60_DNA+RNA_seed | MAPK3 | 1.859775 | BRCA | HCC1599 | SCH772984 |
| MVP_top60_DNA_seed | HSP90AA1 | -0.836348 | COADREAD | LS-513 | CCT-018159 | MVP_top60_DNA+RNA_seed | MAPK1 | 1.859775 | BRCA | HCC1599 | SCH772984 |
| MVP_top60_DNA_seed | HSP90AB1 | -0.836348 | COADREAD | LS-513 | CCT-018159 | MVP_top60_DNA+RNA_seed | EGFR | 1.860494 | BRCA | MDA-MB-436 | Afatinib |
| MVP_top60_DNA_seed | AKT1 | -0.833189 | COADREAD | SNU-81 | AKT inhibitor VIII | MVP_top60_DNA+RNA_seed | ERBB2 | 1.860494 | BRCA | MDA-MB-436 | Afatinib |
| MVP_top60_DNA_seed | MAPK1 | -0.832177 | COADREAD | COLO-205 | FR-180204 | MVP_top60_DNA+RNA_seed | AKT1 | 1.869833 | BRCA | HCC1428 | AT13148 |
| MVP_top60_DNA_seed | MET | -0.828905 | COADREAD | NCI-H716 | SU11274 | MVP_top60_DNA+RNA_seed | AKT1 | 1.888303 | BRCA | BT-474 | AT13148 |
| MVP_top60_DNA_seed | MET | -0.827957 | COADREAD | LoVo | JNJ38877605 | MVP_top60_DNA+RNA_seed | SRC | 1.895881 | BRCA | CAL-120 | Bosutinib |
| MVP_top60_DNA_seed | ATM | -0.825948 | COADREAD | CL-11 | KU-60019 | MVP_top60_DNA+RNA_seed | TP53 | 1.908261 | BRCA | BT-483 | MIRA-1 |
| MVP_top60_DNA_seed | MAPK1 | -0.819389 | COADREAD | HCC2998 | VX-11e | MVP_top60_DNA+RNA_seed | AKT1 | 1.915116 | BRCA | BT-483 | MK-2206 |

| | | | | | | | | | | | |
|---|---|---|---|---|---|---|---|---|---|---|---|
| MVP_top60_DNA_seed | HSP90AA1 | -0.815762 | COADREAD | HT55 | CCT-018159 | MVP_top60_DNA+RNA_seed | PIK3CA | 1.92054 | BRCA | HCC1500 | GNE-317 |
| MVP_top60_DNA_seed | HSP90AB1 | -0.815762 | COADREAD | HT55 | CCT-018159 | MVP_top60_DNA+RNA_seed | MAPK3 | 1.925076 | BRCA | HCC38 | Ulixertinib |
| MVP_top60_DNA_seed | HSP90AA1 | -0.814833 | COADREAD | COLO-678 | Tanespimycin | MVP_top60_DNA+RNA_seed | MAPK1 | 1.925076 | BRCA | HCC38 | Ulixertinib |
| MVP_top60_DNA_seed | HSP90AB1 | -0.814833 | COADREAD | COLO-678 | Tanespimycin | MVP_top60_DNA+RNA_seed | EGFR | 1.9396 | BRCA | BT-474 | AZD3759 |
| MVP_top60_DNA_seed | AR | -0.811173 | COADREAD | LS-180 | Bicalutamide | MVP_top60_DNA+RNA_seed | CDK2 | 1.949143 | BRCA | HCC1419 | Dinaciclib |
| MVP_top60_DNA_seed | IGF1R | -0.810778 | COADREAD | SW48 | BMS-754807 | MVP_top60_DNA+RNA_seed | CDK1 | 1.949143 | BRCA | HCC1419 | Dinaciclib |
| MVP_top60_DNA_seed | HSP90AA1 | -0.804421 | COADREAD | LoVo | CCT-018159 | MVP_top60_DNA+RNA_seed | AKT1 | 1.964761 | BRCA | CAL-120 | MK-2206 |
| MVP_top60_DNA_seed | HSP90AB1 | -0.804421 | COADREAD | LoVo | CCT-018159 | MVP_top60_DNA+RNA_seed | MDM2 | 1.969728 | BRCA | MFM-223 | Serdemetan |
| MVP_top60_DNA_seed | AR | -0.802513 | COADREAD | HCT-116 | Bicalutamide | MVP_top60_DNA+RNA_seed | CDK2 | 1.972524 | BRCA | HCC38 | AZD5438 |
| MVP_top60_DNA_seed | MET | -0.802238 | COADREAD | SNU-175 | JNJ38877605 | MVP_top60_DNA+RNA_seed | SRC | 1.975869 | BRCA | MDA-MB-175-VII | Bosutinib |
| MVP_top60_DNA_seed | BRAF | -0.801322 | COADREAD | NCI-H747 | RAF_9304 | MVP_top60_DNA+RNA_seed | EGFR | 1.985716 | BRCA | CAL-120 | Gefitinib |
| MVP_top60_DNA_seed | BRAF | -0.800854 | COADREAD | HCT-116 | HG6-64-1 | MVP_top60_DNA+RNA_seed | HDAC1 | 1.993809 | BRCA | CAL-120 | PCI-34051 |
| MVP_top60_DNA_seed | AR | -0.796608 | COADREAD | SW48 | AZD3514 | MVP_top60_DNA+RNA_seed | PLK1 | 2.00218 | BRCA | UACC-812 | BI-2536 |
| MVP_top60_DNA_seed | AR | -0.794147 | COADREAD | RKO | AZD3514 | MVP_top60_DNA+RNA_seed | AKT1 | 2.007378 | BRCA | BT-483 | Ipatasertib |
| MVP_top60_DNA_seed | AKT1 | -0.791737 | COADREAD | KM12 | GSK690693 | MVP_top60_DNA+RNA_seed | ATM | 2.026632 | BRCA | MDA-MB-175-VII | KU-55933 |
| MVP_top60_DNA_seed | AR | -0.791279 | COADREAD | T84 | Bicalutamide | MVP_top60_DNA+RNA_seed | PIK3CA | 2.052789 | BRCA | CAL-120 | Alpelisib |
| MVP_top60_DNA_seed | GSK3B | -0.790664 | COADREAD | LoVo | AZD7969 | MVP_top60_DNA+RNA_seed | CDK2 | 2.057509 | BRCA | HCC1500 | AZD5438 |
| MVP_top60_DNA_seed | ATM | -0.790348 | COADREAD | HCT-116 | torin2 | MVP_top60_DNA+RNA_seed | AKT1 | 2.05911 | BRCA | CAL-120 | Uprosertib |
| MVP_top60_DNA_seed | ATM | -0.790196 | COADREAD | RKO | KU-60019 | MVP_top60_DNA+RNA_seed | CDK2 | 2.072564 | BRCA | MDA-MB-175-VII | MK-8776 |
| MVP_top60_DNA_seed | HSP90AA1 | -0.786359 | COADREAD | LS-411N | Luminespib | MVP_top60_DNA+RNA_seed | MAPK3 | 2.09055 | BRCA | ZR-75-30 | Ulixertinib |
| MVP_top60_DNA_seed | HSP90AB1 | -0.786359 | COADREAD | LS-411N | Luminespib | MVP_top60_DNA+RNA_seed | MAPK1 | 2.09055 | BRCA | ZR-75-30 | Ulixertinib |
| MVP_top60_DNA_seed | HSP90AA1 | -0.782158 | COADREAD | RKO | Tanespimycin | MVP_top60_DNA+RNA_seed | HDAC1 | 2.097541 | BRCA | MRK-nu-1 | Romidepsin |
| MVP_top60_DNA_seed | HSP90AB1 | -0.782158 | COADREAD | RKO | Tanespimycin | MVP_top60_DNA+RNA_seed | HDAC2 | 2.097541 | BRCA | MRK-nu-1 | Romidepsin |
| MVP_top60_DNA_seed | GSK3B | -0.78029 | COADREAD | CaR-1 | SB216763 | MVP_top60_DNA+RNA_seed | AKT1 | 2.115379 | BRCA | HCC1428 | MK-2206 |
| MVP_top60_DNA_seed | AKT1 | -0.779899 | COADREAD | SNU-C1 | MK-2206 | MVP_top60_DNA+RNA_seed | ATM | 2.123038 | BRCA | BT-483 | KU-55933 |
| MVP_top60_DNA_seed | BRAF | -0.77774 | COADREAD | HCT-15 | HG6-64-1 | MVP_top60_DNA+RNA_seed | PIK3CA | 2.131534 | BRCA | BT-483 | Buparlisib |
| MVP_top60_DNA_seed | ATM | -0.776531 | COADREAD | SNU-81 | CP466722 | MVP_top60_DNA+RNA_seed | ATM | 2.133594 | BRCA | CAL-120 | KU-55933 |
| MVP_top60_DNA_seed | HSP90AA1 | -0.775295 | COADREAD | SNU-C2B | SNX-2112 | MVP_top60_DNA+RNA_seed | EGFR | 2.14083 | BRCA | HCC1428 | Osimertinib |
| MVP_top60_DNA_seed | HSP90AB1 | -0.775295 | COADREAD | SNU-C2B | SNX-2112 | MVP_top60_DNA+RNA_seed | EGFR | 2.146308 | BRCA | YMB-1-E | Erlotinib |
| MVP_top60_DNA_seed | HSP90AA1 | -0.77219 | COADREAD | DiFi | Tanespimycin | MVP_top60_DNA+RNA_seed | MDM2 | 2.16152 | BRCA | MDA-MB-175-VII | Serdemetan |
| MVP_top60_DNA_seed | HSP90AB1 | -0.77219 | COADREAD | DiFi | Tanespimycin | MVP_top60_DNA+RNA_seed | HSP90AA1 | 2.166102 | BRCA | HCC1187 | Luminespib |
| MVP_top60_DNA_seed | HSP90AA1 | -0.768092 | COADREAD | NCI-H747 | Luminespib | MVP_top60_DNA+RNA_seed | HSP90AB1 | 2.166102 | BRCA | HCC1187 | Luminespib |
| MVP_top60_DNA_seed | HSP90AB1 | -0.768092 | COADREAD | NCI-H747 | Luminespib | MVP_top60_DNA+RNA_seed | EGFR | 2.169547 | BRCA | CAL-120 | AZD3759 |
| MVP_top60_DNA_seed | MAPK14 | -0.765701 | COADREAD | HCT-116 | TAK-715 | MVP_top60_DNA+RNA_seed | ESR1 | 2.186943 | BRCA | MDA-MB-175-VII | GDC0810 |
| MVP_top60_DNA_seed | MAPK1 | -0.763799 | COADREAD | MDST8 | FR-180204 | MVP_top60_DNA+RNA_seed | SRC | 2.212045 | BRCA | BT-483 | Bosutinib |
| MVP_top60_DNA_seed | CDK2 | -0.758766 | COADREAD | SNU-61 | PHA-793887 | MVP_top60_DNA+RNA_seed | EGFR | 2.219776 | BRCA | BT-483 | Erlotinib |
| MVP_top60_DNA_seed | SRC | -0.754412 | COADREAD | SNU-C2B | Saracatinib | MVP_top60_DNA+RNA_seed | AKT1 | 2.228904 | BRCA | BT-483 | Afuresertib |
| MVP_top60_DNA_seed | GSK3B | -0.754014 | COADREAD | HCT-116 | CHIR-99021 | MVP_top60_DNA+RNA_seed | EGFR | 2.245622 | BRCA | BT-483 | Afatinib |
| MVP_top60_DNA_seed | IGF1R | -0.752856 | COADREAD | CL-11 | BMS-536924 | MVP_top60_DNA+RNA_seed | ERBB2 | 2.245622 | BRCA | BT-483 | Afatinib |
| MVP_top60_DNA_seed | GSK3B | -0.752029 | COADREAD | SNU-C5 | AZD7969 | MVP_top60_DNA+RNA_seed | MAPK3 | 2.250265 | BRCA | MDA-MB-330 | Ulixertinib |
| MVP_top60_DNA_seed | IGF1R | -0.747931 | COADREAD | LS-180 | BMS-754807 | MVP_top60_DNA+RNA_seed | MAPK1 | 2.250265 | BRCA | MDA-MB-330 | Ulixertinib |

| | | | | | | | | | | | |
|---|---|---|---|---|---|---|---|---|---|---|---|
| MVP_top60_DNA_seed | SRC | -0.746474 | COADREAD | LoVo | Bosutinib | MVP_top60_DNA+RNA_seed | EGFR | 2.282786 | BRCA | CAL-120 | Osimertinib |
| MVP_top60_DNA_seed | BRAF | -0.743223 | COADREAD | LoVo | Dabrafenib | MVP_top60_DNA+RNA_seed | ATM | 2.321949 | BRCA | ZR-75-30 | KU-55933 |
| MVP_top60_DNA_seed | BRAF | -0.740234 | COADREAD | GP5d | HG6-64-1 | MVP_top60_DNA+RNA_seed | PIK3CA | 2.322717 | BRCA | HCC1428 | GNE-317 |
| MVP_top60_DNA_seed | SRC | -0.739859 | COADREAD | MDST8 | Bosutinib | MVP_top60_DNA+RNA_seed | MAPK3 | 2.329971 | BRCA | BT-549 | Ulixertinib |
| MVP_top60_DNA_seed | AKT1 | -0.736683 | COADREAD | SNU-407 | GSK690693 | MVP_top60_DNA+RNA_seed | MAPK1 | 2.329971 | BRCA | BT-549 | Ulixertinib |
| MVP_top60_DNA_seed | FYN | -0.735427 | COADREAD | NCI-H747 | A-770041 | MVP_top60_DNA+RNA_seed | AKT1 | 2.340748 | BRCA | HCC1428 | Ipatasertib |
| MVP_top60_DNA_seed | SRC | -0.731384 | COADREAD | KM12 | Saracatinib | MVP_top60_DNA+RNA_seed | MAPK3 | 2.346661 | BRCA | BT-483 | SCH772984 |
| MVP_top60_DNA_seed | CDK2 | -0.730998 | COADREAD | SW620 | AZD5438 | MVP_top60_DNA+RNA_seed | MAPK1 | 2.346661 | BRCA | BT-483 | SCH772984 |
| MVP_top60_DNA_seed | CDK2 | -0.730341 | COADREAD | RKO | PHA-793887 | MVP_top60_DNA+RNA_seed | MAPK1 | 2.353221 | BRCA | HCC38 | VX-11e |
| MVP_top60_DNA_seed | MAPK1 | -0.727931 | COADREAD | RCM-1 | VX-11e | MVP_top60_DNA+RNA_seed | EGFR | 2.361925 | BRCA | HCC1428 | AZD3759 |
| MVP_top60_DNA_seed | MAPK1 | -0.726286 | COADREAD | SNU-C5 | VX-11e | MVP_top60_DNA+RNA_seed | CDK2 | 2.362452 | BRCA | ZR-75-30 | Dinaciclib |
| MVP_top60_DNA_seed | PIK3CA | -0.72489 | COADREAD | COLO-320-HSR | AZD8835 | MVP_top60_DNA+RNA_seed | CDK1 | 2.362452 | BRCA | ZR-75-30 | Dinaciclib |
| MVP_top60_DNA_seed | PIK3CD | -0.72489 | COADREAD | COLO-320-HSR | AZD8835 | MVP_top60_DNA+RNA_seed | MAPK3 | 2.385763 | BRCA | ZR-75-30 | SCH772984 |
| MVP_top60_DNA_seed | ERBB4 | -0.722778 | COADREAD | SW1116 | AST-1306 | MVP_top60_DNA+RNA_seed | MAPK1 | 2.385763 | BRCA | ZR-75-30 | SCH772984 |
| MVP_top60_DNA_seed | GSK3B | -0.721675 | COADREAD | SW48 | CHIR-99021 | MVP_top60_DNA+RNA_seed | PLK1 | 2.39159 | BRCA | EFM-192A | MK-1775 |
| MVP_top60_DNA_seed | CDK2 | -0.720069 | COADREAD | SNU-81 | PHA-793887 | MVP_top60_DNA+RNA_seed | AKT1 | 2.406693 | BRCA | BT-483 | AZD5363 |
| MVP_top60_DNA_seed | GSK3B | -0.716443 | COADREAD | LS-513 | CHIR-99021 | MVP_top60_DNA+RNA_seed | ESR1 | 2.421825 | BRCA | BT-483 | Tamoxifen |
| MVP_top60_DNA_seed | MET | -0.716231 | COADREAD | SW48 | SU11274 | MVP_top60_DNA+RNA_seed | EGFR | 2.43059 | BRCA | CAL-120 | Lapatinib |
| MVP_top60_DNA_seed | PIK3CA | -0.715688 | COADREAD | RKO | PI3Ka_4409 | MVP_top60_DNA+RNA_seed | ERBB2 | 2.43059 | BRCA | CAL-120 | Lapatinib |
| MVP_top60_DNA_seed | PIK3CD | -0.715688 | COADREAD | RKO | PI3Ka_4409 | MVP_top60_DNA+RNA_seed | PLK1 | 2.463964 | BRCA | YMB-1-E | MK-1775 |
| MVP_top60_DNA_seed | SRC | -0.713943 | COADREAD | SW48 | Bosutinib | MVP_top60_DNA+RNA_seed | EGFR | 2.493382 | BRCA | BT-483 | AZD3759 |
| MVP_top60_DNA_seed | AKT1 | -0.709351 | COADREAD | SNU-407 | AKT inhibitor VIII | MVP_top60_DNA+RNA_seed | PIK3CA | 2.57169 | BRCA | HCC1428 | Buparlisib |
| MVP_top60_DNA_seed | BRAF | -0.709088 | COADREAD | LoVo | RAF_9304 | MVP_top60_DNA+RNA_seed | CDK2 | 2.579025 | BRCA | BT-483 | MK-8776 |
| MVP_top60_DNA_seed | ATM | -0.709043 | COADREAD | HCT-116 | CP466722 | MVP_top60_DNA+RNA_seed | CDK2 | 2.612586 | BRCA | ZR-75-30 | AZD5438 |
| MVP_top60_DNA_seed | MET | -0.706053 | COADREAD | SNU-1040 | JNJ38877605 | MVP_top60_DNA+RNA_seed | CDK2 | 2.634821 | BRCA | HCC1428 | Dinaciclib |
| MVP_top60_DNA_seed | IGF1R | -0.705988 | COADREAD | HT55 | BMS-536924 | MVP_top60_DNA+RNA_seed | CDK1 | 2.634821 | BRCA | HCC1428 | Dinaciclib |
| MVP_top60_DNA_seed | MAPK14 | -0.704564 | COADREAD | COLO-320-HSR | TAK-715 | MVP_top60_DNA+RNA_seed | ESR1 | 2.666956 | BRCA | BT-483 | GDC0810 |
| MVP_top60_DNA_seed | ATM | -0.698908 | COADREAD | LS-180 | CP466722 | MVP_top60_DNA+RNA_seed | PIK3CA | 2.676332 | BRCA | HCC1428 | AZD8186 |
| MVP_top60_DNA_seed | MAPK14 | -0.698701 | COADREAD | SW620 | TAK-715 | MVP_top60_DNA+RNA_seed | EGFR | 2.679423 | BRCA | BT-483 | Osimertinib |
| MVP_top60_DNA_seed | MAPK1 | -0.697514 | COADREAD | C2BBe1 | FR-180204 | MVP_top60_DNA+RNA_seed | CDK2 | 2.723613 | BRCA | HCC1937 | Dinaciclib |
| MVP_top60_DNA_seed | PIK3CA | -0.694177 | COADREAD | COLO-205 | PI3Ka_4409 | MVP_top60_DNA+RNA_seed | CDK1 | 2.723613 | BRCA | HCC1937 | Dinaciclib |
| MVP_top60_DNA_seed | PIK3CD | -0.694177 | COADREAD | COLO-205 | PI3Ka_4409 | MVP_top60_DNA+RNA_seed | EGFR | 2.737833 | BRCA | BT-483 | Gefitinib |
| MVP_top60_DNA_seed | MAPK1 | -0.69316 | COADREAD | KM12 | VX-11e | MVP_top60_DNA+RNA_seed | HDAC1 | 2.7515 | BRCA | BT-483 | Entinostat |
| MVP_top60_DNA_seed | HSP90AA1 | -0.692597 | COADREAD | COLO-320-HSR | CCT-018159 | MVP_top60_DNA+RNA_seed | HDAC1 | 2.757894 | BRCA | ZR-75-30 | Romidepsin |
| MVP_top60_DNA_seed | HSP90AB1 | -0.692597 | COADREAD | COLO-320-HSR | CCT-018159 | MVP_top60_DNA+RNA_seed | HDAC2 | 2.757894 | BRCA | ZR-75-30 | Romidepsin |
| MVP_top60_DNA_seed | HSP90AA1 | -0.691748 | COADREAD | T84 | Luminespib | MVP_top60_DNA+RNA_seed | HSP90AA1 | 2.775373 | BRCA | BT-483 | Luminespib |
| MVP_top60_DNA_seed | HSP90AB1 | -0.691748 | COADREAD | T84 | Luminespib | MVP_top60_DNA+RNA_seed | HSP90AB1 | 2.775373 | BRCA | BT-483 | Luminespib |
| MVP_top60_DNA_seed | AKT1 | -0.689762 | COADREAD | GP5d | AKT inhibitor VIII | MVP_top60_DNA+RNA_seed | PLK1 | 2.788994 | BRCA | BT-483 | MK-1775 |
| MVP_top60_DNA_seed | HSP90AA1 | -0.685283 | COADREAD | MDST8 | CCT-018159 | MVP_top60_DNA+RNA_seed | AKT1 | 2.798344 | BRCA | HCC1428 | Afuresertib |
| MVP_top60_DNA_seed | HSP90AB1 | -0.685283 | COADREAD | MDST8 | CCT-018159 | MVP_top60_DNA+RNA_seed | AKT1 | 2.839131 | BRCA | HCC1428 | AZD5363 |
| MVP_top60_DNA_seed | BRAF | -0.684369 | COADREAD | KM12 | AZ628 | MVP_top60_DNA+RNA_seed | PIK3CA | 2.896574 | BRCA | HCC1428 | Alpelisib |

| | | | | | | | | | | | |
|---|---|---|---|---|---|---|---|---|---|---|---|
| MVP_top60_DNA_seed | IGF1R | -0.683448 | COADREAD | LS-513 | BMS-754807 | MVP_top60_DNA+RNA_seed | PIK3CA | 2.896804 | BRCA | BT-483 | AZD8186 |
| MVP_top60_DNA_seed | HSP90AA1 | -0.683062 | COADREAD | GP5d | Tanespimycin | MVP_top60_DNA+RNA_seed | MAPK3 | 2.939699 | BRCA | ZR-75-30 | Ulixertinib |
| MVP_top60_DNA_seed | HSP90AB1 | -0.683062 | COADREAD | GP5d | Tanespimycin | MVP_top60_DNA+RNA_seed | MAPK1 | 2.939699 | BRCA | ZR-75-30 | Ulixertinib |
| MVP_top60_DNA_seed | AKT1 | -0.679649 | COADREAD | GP5d | AKT inhibitor VIII | MVP_top60_DNA+RNA_seed | AKT1 | 2.970367 | BRCA | BT-483 | AT13148 |
| MVP_top60_DNA_seed | IGF1R | -0.676127 | COADREAD | SNU-C1 | BMS-536924 | MVP_top60_DNA+RNA_seed | PLK1 | 3.008957 | BRCA | ZR-75-30 | MK-1775 |
| MVP_top60_DNA_seed | BRAF | -0.675173 | COADREAD | CL-11 | AZ628 | MVP_top60_DNA+RNA_seed | CDK2 | 3.028935 | BRCA | HCC1428 | AZD5438 |
| MVP_top60_DNA_seed | ATM | -0.673183 | COADREAD | SNU-81 | KU-55933 | MVP_top60_DNA+RNA_seed | MAPK3 | 3.151976 | BRCA | BT-483 | Ulixertinib |
| MVP_top60_DNA_seed | AKT1 | -0.67277 | COADREAD | LS-513 | AKT inhibitor VIII | MVP_top60_DNA+RNA_seed | MAPK1 | 3.151976 | BRCA | BT-483 | Ulixertinib |
| MVP_top60_DNA_seed | CDK2 | -0.670997 | COADREAD | COLO-205 | PHA-793887 | MVP_top60_DNA+RNA_seed | MAPK1 | 3.461689 | BRCA | BT-483 | VX-11e |
| MVP_top60_DNA_seed | ERBB4 | -0.670845 | COADREAD | SNU-C2B | PF-00299804 | MVP_top60_DNA+RNA_seed | AR | -4.816697 | COADREAD | CaR-1 | Bicalutamide |
| MVP_top60_DNA_seed | BRAF | -0.662747 | COADREAD | SK-CO-1 | RAF_9304 | MVP_top60_DNA+RNA_seed | AR | -4.097544 | COADREAD | SW948 | Bicalutamide |
| MVP_top60_DNA_seed | CDK2 | -0.657704 | COADREAD | SNU-1040 | PHA-793887 | MVP_top60_DNA+RNA_seed | ERBB4 | -3.717325 | COADREAD | CCK-81 | PF-00299804 |
| MVP_top60_DNA_seed | SRC | -0.650585 | COADREAD | SW1417 | NVP-BHG712 | MVP_top60_DNA+RNA_seed | ERBB3 | -3.706856 | COADREAD | DiFi | AZD8931 |
| MVP_top60_DNA_seed | HSP90AA1 | -0.64811 | COADREAD | GP5d | SNX-2112 | MVP_top60_DNA+RNA_seed | PIK3CB | -3.638272 | COADREAD | LoVo | AZD8186 |
| MVP_top60_DNA_seed | HSP90AB1 | -0.64811 | COADREAD | GP5d | SNX-2112 | MVP_top60_DNA+RNA_seed | PIK3CD | -3.638272 | COADREAD | LoVo | AZD8186 |
| MVP_top60_DNA_seed | SRC | -0.646625 | COADREAD | NCI-H508 | Bosutinib | MVP_top60_DNA+RNA_seed | IGF1R | -3.173323 | COADREAD | COLO-205 | BMS-754807 |
| MVP_top60_DNA_seed | SRC | -0.642285 | COADREAD | GP5d | Bosutinib | MVP_top60_DNA+RNA_seed | IGF1R | -3.141067 | COADREAD | COLO-205 | Linsitinib |
| MVP_top60_DNA_seed | HSP90AA1 | -0.640286 | COADREAD | CCK-81 | SNX-2112 | MVP_top60_DNA+RNA_seed | ERBB4 | -2.977733 | COADREAD | CCK-81 | CI-1033 |
| MVP_top60_DNA_seed | HSP90AB1 | -0.640286 | COADREAD | CCK-81 | SNX-2112 | MVP_top60_DNA+RNA_seed | ERBB3 | -2.973127 | COADREAD | SNU-407 | AZD8931 |
| MVP_top60_DNA_seed | ERBB3 | -0.637258 | COADREAD | CCK-81 | AZD8931 | MVP_top60_DNA+RNA_seed | PIK3CA | -2.885395 | COADREAD | SNU-407 | AZD8835 |
| MVP_top60_DNA_seed | MAPK1 | -0.636414 | COADREAD | SNU-1040 | VX-11e | MVP_top60_DNA+RNA_seed | PIK3CD | -2.885395 | COADREAD | SNU-407 | AZD8835 |
| MVP_top60_DNA_seed | ATM | -0.635834 | COADREAD | CW-2 | KU-60019 | MVP_top60_DNA+RNA_seed | GSK3B | -2.627545 | COADREAD | RKO | KIN001-042 |
| MVP_top60_DNA_seed | FYN | -0.635272 | COADREAD | RKO | A-770041 | MVP_top60_DNA+RNA_seed | IGF1R | -2.623082 | COADREAD | KM12 | BMS-536924 |
| MVP_top60_DNA_seed | AR | -0.635145 | COADREAD | LS-180 | AZD3514 | MVP_top60_DNA+RNA_seed | MET | -2.601849 | COADREAD | KM12 | Crizotinib |
| MVP_top60_DNA_seed | SRC | -0.635092 | COADREAD | LS-1034 | NVP-BHG712 | MVP_top60_DNA+RNA_seed | AKT1 | -2.54632 | COADREAD | SNU-C1 | BAY AKT1 |
| MVP_top60_DNA_seed | SRC | -0.634814 | COADREAD | CCK-81 | Bosutinib | MVP_top60_DNA+RNA_seed | PIK3CD | -2.510813 | COADREAD | SW1463 | Idelalisib |
| MVP_top60_DNA_seed | AKT1 | -0.629589 | COADREAD | COLO-320-HSR | AKT inhibitor VIII | MVP_top60_DNA+RNA_seed | IGF1R | -2.500607 | COADREAD | NCI-H716 | BMS-536924 |
| MVP_top60_DNA_seed | CDK2 | -0.627736 | COADREAD | LS-513 | Seliciclib | MVP_top60_DNA+RNA_seed | IGF1R | -2.459322 | COADREAD | LS-1034 | Linsitinib |
| MVP_top60_DNA_seed | PIK3CA | -0.627517 | COADREAD | DiFi | AZD8835 | MVP_top60_DNA+RNA_seed | PTK2 | -2.454079 | COADREAD | COLO-205 | PF-562271 |
| MVP_top60_DNA_seed | PIK3CD | -0.627517 | COADREAD | DiFi | AZD8835 | MVP_top60_DNA+RNA_seed | PTK2 | -2.435368 | COADREAD | KM12 | PF-562271 |
| MVP_top60_DNA_seed | GSK3B | -0.626934 | COADREAD | GP5d | SB216763 | MVP_top60_DNA+RNA_seed | CDK2 | -2.311522 | COADREAD | CaR-1 | AZD5438 |
| MVP_top60_DNA_seed | PIK3CA | -0.624684 | COADREAD | SNU-C5 | AZD8835 | MVP_top60_DNA+RNA_seed | CDK2 | -2.305651 | COADREAD | HCT-116 | PHA-793887 |
| MVP_top60_DNA_seed | PIK3CD | -0.624684 | COADREAD | SNU-C5 | AZD8835 | MVP_top60_DNA+RNA_seed | ATM | -2.26219 | COADREAD | LoVo | KU-55933 |
| MVP_top60_DNA_seed | MAPK1 | -0.623281 | COADREAD | SNU-C5 | FR-180204 | MVP_top60_DNA+RNA_seed | IGF1R | -2.192941 | COADREAD | T84 | BMS-754807 |
| MVP_top60_DNA_seed | IGF1R | -0.622832 | COADREAD | SNU-C2B | BMS-536924 | MVP_top60_DNA+RNA_seed | IGF1R | -2.176422 | COADREAD | HT-29 | BMS-754807 |
| MVP_top60_DNA_seed | ESR1 | -0.622466 | COADREAD | SNU-C5 | Fulvestrant | MVP_top60_DNA+RNA_seed | IGF1R | -2.144939 | COADREAD | SW48 | BMS-536924 |
| MVP_top60_DNA_seed | ATM | -0.61809 | COADREAD | NCI-H747 | CP466722 | MVP_top60_DNA+RNA_seed | MET | -2.126138 | COADREAD | KM12 | SU11274 |
| MVP_top60_DNA_seed | AKT1 | -0.618028 | COADREAD | CCK-81 | MK-2206 | MVP_top60_DNA+RNA_seed | GSK3B | -2.094343 | COADREAD | LoVo | SB216763 |
| MVP_top60_DNA_seed | MAPK1 | -0.617848 | COADREAD | NCI-H747 | FR-180204 | MVP_top60_DNA+RNA_seed | IGF1R | -2.083245 | COADREAD | HT-29 | Linsitinib |
| MVP_top60_DNA_seed | ERBB4 | -0.617328 | COADREAD | SNU-175 | AST-1306 | MVP_top60_DNA+RNA_seed | ERBB4 | -2.075892 | COADREAD | CCK-81 | AST-1306 |
| MVP_top60_DNA_seed | BRAF | -0.614838 | COADREAD | NCI-H508 | PLX-4720 | MVP_top60_DNA+RNA_seed | IGF1R | -2.074399 | COADREAD | LS-1034 | BMS-754807 |

| | | | | | | | | | | | |
|---|---|---|---|---|---|---|---|---|---|---|---|
| MVP_top60_DNA_seed | CDK2 | -0.613517 | COADREAD | SNU-C5 | AZD5438 | MVP_top60_DNA+RNA_seed | AKT1 | -2.04894 | COADREAD | SNU-81 | GSK690693 |
| MVP_top60_DNA_seed | ERBB3 | -0.613457 | COADREAD | HT55 | AZD8931 | MVP_top60_DNA+RNA_seed | IGF1R | -2.014989 | COADREAD | COLO-320-HSR | Linsitinib |
| MVP_top60_DNA_seed | PIK3CA | -0.61323 | COADREAD | SK-CO-1 | PI3Ka_4409 | MVP_top60_DNA+RNA_seed | PIK3CB | -2.002653 | COADREAD | C2BBe1 | TGX221 |
| MVP_top60_DNA_seed | PIK3CD | -0.61323 | COADREAD | SK-CO-1 | PI3Ka_4409 | MVP_top60_DNA+RNA_seed | IGF1R | -1.984166 | COADREAD | KM12 | BMS-754807 |
| MVP_top60_DNA_seed | AKT1 | -0.613133 | COADREAD | KM12 | A-443654 | MVP_top60_DNA+RNA_seed | GSK3B | -1.963705 | COADREAD | SW1417 | CHIR-99021 |
| MVP_top60_DNA_seed | AKT1 | -0.612438 | COADREAD | HCT-15 | MK-2206 | MVP_top60_DNA+RNA_seed | IGF1R | -1.955877 | COADREAD | HCC2998 | BMS-754807 |
| MVP_top60_DNA_seed | HSP90AA1 | -0.611035 | COADREAD | NCI-H716 | SNX-2112 | MVP_top60_DNA+RNA_seed | AKT1 | -1.955554 | COADREAD | LS-411N | AKT inhibitor VIII |
| MVP_top60_DNA_seed | HSP90AB1 | -0.611035 | COADREAD | NCI-H716 | SNX-2112 | MVP_top60_DNA+RNA_seed | IGF1R | -1.853022 | COADREAD | COLO-205 | BMS-536924 |
| MVP_top60_DNA_seed | AKT1 | -0.606411 | COADREAD | NCI-H747 | MK-2206 | MVP_top60_DNA+RNA_seed | BCL2 | -1.836906 | COADREAD | SW620 | Obatoclax Mesylate |
| MVP_top60_DNA_seed | BRAF | -0.603733 | COADREAD | RKO | PLX-4720 | MVP_top60_DNA+RNA_seed | AKT1 | -1.807413 | COADREAD | DiFi | BAY AKT1 |
| MVP_top60_DNA_seed | SRC | -0.601542 | COADREAD | LS-411N | NVP-BHG712 | MVP_top60_DNA+RNA_seed | AR | -1.774714 | COADREAD | LS-180 | Bicalutamide |
| MVP_top60_DNA_seed | AKT1 | -0.598504 | COADREAD | LS-411N | AKT inhibitor VIII | MVP_top60_DNA+RNA_seed | SRC | -1.743851 | COADREAD | DiFi | Bosutinib |
| MVP_top60_DNA_seed | BRAF | -0.598427 | COADREAD | SNU-C5 | PLX-4720 | MVP_top60_DNA+RNA_seed | SRC | -1.734961 | COADREAD | SNU-407 | NVP-BHG712 |
| MVP_top60_DNA_seed | PIK3CD | -0.598091 | COADREAD | LS-123 | Idelalisib | MVP_top60_DNA+RNA_seed | AKT1 | -1.711913 | COADREAD | SNU-1040 | GSK690693 |
| MVP_top60_DNA_seed | MET | -0.597897 | COADREAD | GP5d | JNJ38877605 | MVP_top60_DNA+RNA_seed | IGF1R | -1.693276 | COADREAD | T84 | Linsitinib |
| MVP_top60_DNA_seed | HSP90AA1 | -0.596686 | COADREAD | SW48 | Luminespib | MVP_top60_DNA+RNA_seed | AKT1 | -1.679353 | COADREAD | COLO-205 | AKT inhibitor VIII |
| MVP_top60_DNA_seed | HSP90AB1 | -0.596686 | COADREAD | SW48 | Luminespib | MVP_top60_DNA+RNA_seed | PIK3CB | -1.654664 | COADREAD | LoVo | AZD6482 |
| MVP_top60_DNA_seed | SRC | -0.59592 | COADREAD | HCT-116 | HG-6-71-01 | MVP_top60_DNA+RNA_seed | AKT1 | -1.651386 | COADREAD | NCI-H508 | BAY AKT1 |
| MVP_top60_DNA_seed | SRC | -0.595253 | COADREAD | COLO-205 | NVP-BHG712 | MVP_top60_DNA+RNA_seed | AR | -1.63943 | COADREAD | HCT-116 | AZD3514 |
| MVP_top60_DNA_seed | ERBB4 | -0.593571 | COADREAD | NCI-H747 | AST-1306 | MVP_top60_DNA+RNA_seed | IGF1R | -1.625184 | COADREAD | GP5d | BMS-754807 |
| MVP_top60_DNA_seed | BRAF | -0.592205 | COADREAD | NCI-H716 | SB590885 | MVP_top60_DNA+RNA_seed | GSK3B | -1.599783 | COADREAD | SW48 | SB216763 |
| MVP_top60_DNA_seed | MAPK1 | -0.590452 | COADREAD | GP5d | VX-11e | MVP_top60_DNA+RNA_seed | IGF1R | -1.599647 | COADREAD | HCC2998 | Linsitinib |
| MVP_top60_DNA_seed | ATM | -0.587908 | COADREAD | CCK-81 | KU-60019 | MVP_top60_DNA+RNA_seed | IGF1R | -1.593094 | COADREAD | KM12 | BMS-536924 |
| MVP_top60_DNA_seed | PIK3CA | -0.586716 | COADREAD | SNU-C5 | PI3Ka_4409 | MVP_top60_DNA+RNA_seed | IGF1R | -1.591244 | COADREAD | LS-411N | GSK1904529A |
| MVP_top60_DNA_seed | PIK3CD | -0.586716 | COADREAD | SNU-C5 | PI3Ka_4409 | MVP_top60_DNA+RNA_seed | BCL2 | -1.591164 | COADREAD | MDST8 | Navitoclax |
| MVP_top60_DNA_seed | MET | -0.585958 | COADREAD | SNU-61 | JNJ38877605 | MVP_top60_DNA+RNA_seed | ERBB3 | -1.588256 | COADREAD | SW48 | AZD8931 |
| MVP_top60_DNA_seed | ATM | -0.584066 | COADREAD | COLO-320-HSR | KU-60019 | MVP_top60_DNA+RNA_seed | IGF1R | -1.578694 | COADREAD | HCT-116 | BMS-536924 |
| MVP_top60_DNA_seed | GSK3B | -0.582429 | COADREAD | NCI-H747 | CHIR-99021 | MVP_top60_DNA+RNA_seed | PIK3CA | -1.570969 | COADREAD | CaR-1 | PI3Ka_4409 |
| MVP_top60_DNA_seed | CREBBP | -0.581749 | COADREAD | SK-CO-1 | I-CBP112 | MVP_top60_DNA+RNA_seed | PIK3CD | -1.570969 | COADREAD | CaR-1 | PI3Ka_4409 |
| MVP_top60_DNA_seed | PIK3CB | -0.57936 | COADREAD | LS-123 | AZD8186 | MVP_top60_DNA+RNA_seed | AKT1 | -1.564586 | COADREAD | NCI-H747 | AKT inhibitor VIII |
| MVP_top60_DNA_seed | PIK3CD | -0.57936 | COADREAD | LS-123 | AZD8186 | MVP_top60_DNA+RNA_seed | GSK3B | -1.552821 | COADREAD | HCT-116 | AZD7969 |
| MVP_top60_DNA_seed | MET | -0.579084 | COADREAD | GP5d | SU11274 | MVP_top60_DNA+RNA_seed | HSP90AA1 | -1.540735 | COADREAD | HCT-15 | CCT-018159 |
| MVP_top60_DNA_seed | HSP90AA1 | -0.575255 | COADREAD | LS-1034 | SNX-2112 | MVP_top60_DNA+RNA_seed | HSP90AB1 | -1.540735 | COADREAD | HCT-15 | CCT-018159 |
| MVP_top60_DNA_seed | HSP90AB1 | -0.575255 | COADREAD | LS-1034 | SNX-2112 | MVP_top60_DNA+RNA_seed | MET | -1.540208 | COADREAD | CCK-81 | JNJ38877605 |
| MVP_top60_DNA_seed | PIK3CA | -0.572921 | COADREAD | SNU-C1 | PI3Ka_4409 | MVP_top60_DNA+RNA_seed | ATM | -1.540048 | COADREAD | LS-411N | KU-60019 |
| MVP_top60_DNA_seed | PIK3CD | -0.572921 | COADREAD | SNU-C1 | PI3Ka_4409 | MVP_top60_DNA+RNA_seed | IGF1R | -1.529272 | COADREAD | COLO-320-HSR | BMS-754807 |
| MVP_top60_DNA_seed | BRAF | -0.572903 | COADREAD | LS-411N | HG6-64-1 | MVP_top60_DNA+RNA_seed | PIK3CA | -1.527487 | COADREAD | T84 | AZD8835 |
| MVP_top60_DNA_seed | SRC | -0.572499 | COADREAD | C2BBe1 | Bosutinib | MVP_top60_DNA+RNA_seed | PIK3CD | -1.527487 | COADREAD | T84 | AZD8835 |
| MVP_top60_DNA_seed | PTK2 | -0.572351 | COADREAD | HCC2998 | PF-562271 | MVP_top60_DNA+RNA_seed | BCL2 | -1.508551 | COADREAD | HCT-116 | TW 37 |
| MVP_top60_DNA_seed | BRAF | -0.571538 | COADREAD | HCT-116 | SB590885 | MVP_top60_DNA+RNA_seed | PIK3CB | -1.508336 | COADREAD | SNU-81 | AZD8186 |
| MVP_top60_DNA_seed | IGF1R | -0.567744 | COADREAD | C2BBe1 | Linsitinib | MVP_top60_DNA+RNA_seed | PIK3CD | -1.508336 | COADREAD | SNU-81 | AZD8186 |

| | | | | | | | | | | | |
|---|---|---|---|---|---|---|---|---|---|---|---|
| MVP_top60_DNA_seed | AKT1 | -0.56542 | COADREAD | SNU-1040 | AKT inhibitor VIII | MVP_top60_DNA+RNA_seed | BCL2 | -1.503269 | COADREAD | HCC2998 | Obatoclax Mesylate |
| MVP_top60_DNA_seed | AKT1 | -0.56514 | COADREAD | SNU-C2B | A-443654 | MVP_top60_DNA+RNA_seed | HSP90AA1 | -1.49404 | COADREAD | SW48 | Tanespimycin |
| MVP_top60_DNA_seed | AKT1 | -0.564052 | COADREAD | SNU-C5 | MK-2206 | MVP_top60_DNA+RNA_seed | HSP90AB1 | -1.49404 | COADREAD | SW48 | Tanespimycin |
| MVP_top60_DNA_seed | SRC | -0.559757 | COADREAD | SW620 | NVP-BHG712 | MVP_top60_DNA+RNA_seed | PIK3CA | -1.479836 | COADREAD | SW48 | AZD8835 |
| MVP_top60_DNA_seed | ERBB3 | -0.558975 | COADREAD | NCI-H508 | AZD8931 | MVP_top60_DNA+RNA_seed | PIK3CD | -1.479836 | COADREAD | SW48 | AZD8835 |
| MVP_top60_DNA_seed | BRAF | -0.557702 | COADREAD | LS-180 | SB590885 | MVP_top60_DNA+RNA_seed | HSP90AA1 | -1.475028 | COADREAD | COLO-205 | Tanespimycin |
| MVP_top60_DNA_seed | CDK2 | -0.55705 | COADREAD | LS-513 | AZD5438 | MVP_top60_DNA+RNA_seed | HSP90AB1 | -1.475028 | COADREAD | COLO-205 | Tanespimycin |
| MVP_top60_DNA_seed | PIK3CD | -0.555741 | COADREAD | GP5d | IC-87114 | MVP_top60_DNA+RNA_seed | IGF1R | -1.474191 | COADREAD | LS-411N | BMS-754807 |
| MVP_top60_DNA_seed | MET | -0.548876 | COADREAD | NCI-H747 | JNJ38877605 | MVP_top60_DNA+RNA_seed | CDK2 | -1.465851 | COADREAD | SNU-175 | PHA-793887 |
| MVP_top60_DNA_seed | BRAF | -0.544497 | COADREAD | SW948 | RAF_9304 | MVP_top60_DNA+RNA_seed | AKT1 | -1.465165 | COADREAD | NCI-H508 | MK-2206 |
| MVP_top60_DNA_seed | PIK3CA | -0.542571 | COADREAD | LoVo | AZD8835 | MVP_top60_DNA+RNA_seed | PTK2 | -1.463438 | COADREAD | SW48 | PF-562271 |
| MVP_top60_DNA_seed | PIK3CD | -0.542571 | COADREAD | LoVo | AZD8835 | MVP_top60_DNA+RNA_seed | IGF1R | -1.460412 | COADREAD | SNU-C1 | Linsitinib |
| MVP_top60_DNA_seed | AR | -0.540983 | COADREAD | T84 | Bicalutamide | MVP_top60_DNA+RNA_seed | CDK2 | -1.458351 | COADREAD | SW48 | AZD5438 |
| MVP_top60_DNA_seed | PIK3CA | -0.539297 | COADREAD | NCI-H747 | PI3Ka_4409 | MVP_top60_DNA+RNA_seed | MET | -1.456509 | COADREAD | SNU-175 | SU11274 |
| MVP_top60_DNA_seed | PIK3CD | -0.539297 | COADREAD | NCI-H747 | PI3Ka_4409 | MVP_top60_DNA+RNA_seed | PIK3CB | -1.436165 | COADREAD | LS-411N | TGX221 |
| MVP_top60_DNA_seed | MET | -0.537811 | COADREAD | HCT-116 | SU11274 | MVP_top60_DNA+RNA_seed | GSK3B | -1.432673 | COADREAD | LoVo | CHIR-99021 |
| MVP_top60_DNA_seed | FYN | -0.537777 | COADREAD | C2BBe1 | A-770041 | MVP_top60_DNA+RNA_seed | AR | -1.424971 | COADREAD | HT-115 | Bicalutamide |
| MVP_top60_DNA_seed | MAPK1 | -0.537642 | COADREAD | LoVo | VX-11e | MVP_top60_DNA+RNA_seed | IGF1R | -1.424352 | COADREAD | SNU-C1 | BMS-536924 |
| MVP_top60_DNA_seed | PIK3CB | -0.536913 | COADREAD | HCT-116 | AZD8186 | MVP_top60_DNA+RNA_seed | IGF1R | -1.420785 | COADREAD | NCI-H747 | Linsitinib |
| MVP_top60_DNA_seed | PIK3CD | -0.536913 | COADREAD | HCT-116 | AZD8186 | MVP_top60_DNA+RNA_seed | PIK3CD | -1.404039 | COADREAD | SNU-81 | Idelalisib |
| MVP_top60_DNA_seed | PIK3CA | -0.534642 | COADREAD | HT-29 | AZD8835 | MVP_top60_DNA+RNA_seed | SRC | -1.403007 | COADREAD | LS-513 | NVP-BHG712 |
| MVP_top60_DNA_seed | PIK3CD | -0.534642 | COADREAD | HT-29 | AZD8835 | MVP_top60_DNA+RNA_seed | AKT1 | -1.400421 | COADREAD | DiFi | MK-2206 |
| MVP_top60_DNA_seed | HSP90AA1 | -0.534374 | COADREAD | HCC2998 | Luminespib | MVP_top60_DNA+RNA_seed | AKT1 | -1.384673 | COADREAD | HCT-15 | AKT inhibitor VIII |
| MVP_top60_DNA_seed | HSP90AB1 | -0.534374 | COADREAD | HCC2998 | Luminespib | MVP_top60_DNA+RNA_seed | HSP90AA1 | -1.383208 | COADREAD | LS-180 | Tanespimycin |
| MVP_top60_DNA_seed | HSP90AA1 | -0.534314 | COADREAD | SNU-61 | SNX-2112 | MVP_top60_DNA+RNA_seed | HSP90AB1 | -1.383208 | COADREAD | LS-180 | Tanespimycin |
| MVP_top60_DNA_seed | HSP90AB1 | -0.534314 | COADREAD | SNU-61 | SNX-2112 | MVP_top60_DNA+RNA_seed | MET | -1.372711 | COADREAD | KM12 | PHA-665752 |
| MVP_top60_DNA_seed | HSP90AA1 | -0.533815 | COADREAD | LoVo | Luminespib | MVP_top60_DNA+RNA_seed | IGF1R | -1.367994 | COADREAD | LS-411N | BMS-536924 |
| MVP_top60_DNA_seed | HSP90AB1 | -0.533815 | COADREAD | LoVo | Luminespib | MVP_top60_DNA+RNA_seed | AKT1 | -1.365797 | COADREAD | LS-411N | A-443654 |
| MVP_top60_DNA_seed | AR | -0.528166 | COADREAD | SNU-C2B | Bicalutamide | MVP_top60_DNA+RNA_seed | PTK2 | -1.360297 | COADREAD | HCT-116 | PF-562271 |
| MVP_top60_DNA_seed | HSP90AA1 | -0.525717 | COADREAD | SNU-407 | SNX-2112 | MVP_top60_DNA+RNA_seed | HSP90AA1 | -1.355717 | COADREAD | HT-29 | Tanespimycin |
| MVP_top60_DNA_seed | HSP90AB1 | -0.525717 | COADREAD | SNU-407 | SNX-2112 | MVP_top60_DNA+RNA_seed | HSP90AB1 | -1.355717 | COADREAD | HT-29 | Tanespimycin |
| MVP_top60_DNA_seed | PIK3CA | -0.521684 | COADREAD | NCI-H716 | PI3Ka_4409 | MVP_top60_DNA+RNA_seed | PTK2 | -1.351991 | COADREAD | GP5d | PF-562271 |
| MVP_top60_DNA_seed | PIK3CD | -0.521684 | COADREAD | NCI-H716 | PI3Ka_4409 | MVP_top60_DNA+RNA_seed | IGF1R | -1.343888 | COADREAD | NCI-H747 | BMS-754807 |
| MVP_top60_DNA_seed | HSP90AA1 | -0.520826 | COADREAD | RKO | CCT-018159 | MVP_top60_DNA+RNA_seed | IGF1R | -1.341783 | COADREAD | SNU-C1 | BMS-754807 |
| MVP_top60_DNA_seed | HSP90AB1 | -0.520826 | COADREAD | RKO | CCT-018159 | MVP_top60_DNA+RNA_seed | GSK3B | -1.319173 | COADREAD | COLO-205 | CHIR-99021 |
| MVP_top60_DNA_seed | ATM | -0.519922 | COADREAD | CL-11 | KU-55933 | MVP_top60_DNA+RNA_seed | AKT1 | -1.318288 | COADREAD | HT-29 | AKT inhibitor VIII |
| MVP_top60_DNA_seed | AR | -0.518724 | COADREAD | GP5d | Bicalutamide | MVP_top60_DNA+RNA_seed | GSK3B | -1.314432 | COADREAD | SK-CO-1 | AZD7969 |
| MVP_top60_DNA_seed | ATM | -0.518202 | COADREAD | SK-CO-1 | KU-55933 | MVP_top60_DNA+RNA_seed | ERBB3 | -1.305685 | COADREAD | HT-115 | AZD8931 |
| MVP_top60_DNA_seed | ERBB3 | -0.515605 | COADREAD | SW1116 | AZD8931 | MVP_top60_DNA+RNA_seed | GSK3B | -1.300526 | COADREAD | LS-513 | SB216763 |
| MVP_top60_DNA_seed | HSP90AA1 | -0.515085 | COADREAD | RKO | Elesclomol | MVP_top60_DNA+RNA_seed | PIK3CD | -1.298458 | COADREAD | NCI-H747 | Idelalisib |
| MVP_top60_DNA_seed | HSP90AB1 | -0.515085 | COADREAD | RKO | Elesclomol | MVP_top60_DNA+RNA_seed | GSK3B | -1.293739 | COADREAD | KM12 | KIN001-042 |

| | | | | | | | | | | | |
|---|---|---|---|---|---|---|---|---|---|---|---|
| MVP_top60_DNA_seed | BRAF | -0.513367 | COADREAD | CaR-1 | RAF_9304 | MVP_top60_DNA+RNA_seed | AKT1 | -1.289571 | COADREAD | HCT-116 | AKT inhibitor VIII |
| MVP_top60_DNA_seed | HSP90AA1 | -0.51296 | COADREAD | KM12 | Luminespib | MVP_top60_DNA+RNA_seed | ATM | -1.289227 | COADREAD | HCT-116 | KU-60019 |
| MVP_top60_DNA_seed | HSP90AB1 | -0.51296 | COADREAD | KM12 | Luminespib | MVP_top60_DNA+RNA_seed | IGF1R | -1.287751 | COADREAD | COLO-320-HSR | BMS-536924 |
| MVP_top60_DNA_seed | ATM | -0.511225 | COADREAD | T84 | CP466722 | MVP_top60_DNA+RNA_seed | BCL2 | -1.283707 | COADREAD | HT-29 | Obatoclax Mesylate |
| MVP_top60_DNA_seed | ATM | -0.510789 | COADREAD | SNU-1040 | CP466722 | MVP_top60_DNA+RNA_seed | BCL2 | -1.279003 | COADREAD | SK-CO-1 | Navitoclax |
| MVP_top60_DNA_seed | IGF1R | -0.507326 | COADREAD | SW620 | BMS-754807 | MVP_top60_DNA+RNA_seed | HSP90AA1 | -1.278961 | COADREAD | RKO | SNX-2112 |
| MVP_top60_DNA_seed | MET | -0.507046 | COADREAD | SW48 | JNJ38877605 | MVP_top60_DNA+RNA_seed | HSP90AB1 | -1.278961 | COADREAD | RKO | SNX-2112 |
| MVP_top60_DNA_seed | MAPK1 | -0.506187 | COADREAD | KM12 | FR-180204 | MVP_top60_DNA+RNA_seed | IGF1R | -1.278891 | COADREAD | LS-1034 | BMS-536924 |
| MVP_top60_DNA_seed | MET | -0.504784 | COADREAD | CCK-81 | SU11274 | MVP_top60_DNA+RNA_seed | BCL2 | -1.274032 | COADREAD | RKO | Obatoclax Mesylate |
| MVP_top60_DNA_seed | GSK3B | -0.502184 | COADREAD | CCK-81 | SB216763 | MVP_top60_DNA+RNA_seed | AR | -1.268741 | COADREAD | COLO-205 | Bicalutamide |
| MVP_top60_DNA_seed | CDK2 | -0.50212 | COADREAD | SNU-C2B | PHA-793887 | MVP_top60_DNA+RNA_seed | IGF1R | -1.257458 | COADREAD | LS-411N | Linsitinib |
| MVP_top60_DNA_seed | PIK3CB | -0.501409 | COADREAD | NCI-H716 | AZD8186 | MVP_top60_DNA+RNA_seed | CDK2 | -1.252024 | COADREAD | COLO-320-HSR | Seliciclib |
| MVP_top60_DNA_seed | PIK3CD | -0.501409 | COADREAD | NCI-H716 | AZD8186 | MVP_top60_DNA+RNA_seed | ERBB4 | -1.243392 | COADREAD | SNU-175 | PF-00299804 |
| MVP_top60_DNA_seed | BRAF | -0.495341 | COADREAD | MDST8 | AZ628 | MVP_top60_DNA+RNA_seed | AKT1 | -1.235951 | COADREAD | LS-411N | GSK690693 |
| MVP_top60_DNA_seed | CDK2 | -0.494179 | COADREAD | NCI-H747 | PHA-793887 | MVP_top60_DNA+RNA_seed | SRC | -1.231503 | COADREAD | SNU-C5 | Bosutinib |
| MVP_top60_DNA_seed | HSP90AA1 | -0.49181 | COADREAD | MDST8 | Elesclomol | MVP_top60_DNA+RNA_seed | HSP90AA1 | -1.22617 | COADREAD | CaR-1 | Tanespimycin |
| MVP_top60_DNA_seed | HSP90AB1 | -0.49181 | COADREAD | MDST8 | Elesclomol | MVP_top60_DNA+RNA_seed | HSP90AB1 | -1.22617 | COADREAD | CaR-1 | Tanespimycin |
| MVP_top60_DNA_seed | AKT1 | -0.491021 | COADREAD | RKO | AKT inhibitor VIII | MVP_top60_DNA+RNA_seed | SRC | -1.224806 | COADREAD | C2BBe1 | Saracatinib |
| MVP_top60_DNA_seed | GSK3B | -0.490984 | COADREAD | LoVo | CHIR-99021 | MVP_top60_DNA+RNA_seed | SRC | -1.215717 | COADREAD | HCT-116 | NVP-BHG712 |
| MVP_top60_DNA_seed | BRAF | -0.48826 | COADREAD | SNU-407 | RAF_9304 | MVP_top60_DNA+RNA_seed | AKT1 | -1.214898 | COADREAD | LoVo | MK-2206 |
| MVP_top60_DNA_seed | IGF1R | -0.487266 | COADREAD | C2BBe1 | BMS-754807 | MVP_top60_DNA+RNA_seed | AKT1 | -1.211412 | COADREAD | NCI-H716 | BAY AKT1 |
| MVP_top60_DNA_seed | GSK3B | -0.483272 | COADREAD | HT-115 | KIN001-042 | MVP_top60_DNA+RNA_seed | GSK3B | -1.210092 | COADREAD | LoVo | KIN001-042 |
| MVP_top60_DNA_seed | PTK2 | -0.478548 | COADREAD | HT-29 | PF-562271 | MVP_top60_DNA+RNA_seed | ERBB4 | -1.200828 | COADREAD | HT-115 | CI-1033 |
| MVP_top60_DNA_seed | CDK2 | -0.475131 | COADREAD | LoVo | AZD5438 | MVP_top60_DNA+RNA_seed | HSP90AA1 | -1.199899 | COADREAD | HCT-116 | CCT-018159 |
| MVP_top60_DNA_seed | BRAF | -0.474294 | COADREAD | NCI-H716 | RAF_9304 | MVP_top60_DNA+RNA_seed | HSP90AB1 | -1.199899 | COADREAD | HCT-116 | CCT-018159 |
| MVP_top60_DNA_seed | PIK3CA | -0.474138 | COADREAD | SW948 | PI3Ka_4409 | MVP_top60_DNA+RNA_seed | HSP90AA1 | -1.193057 | COADREAD | SNU-175 | SNX-2112 |
| MVP_top60_DNA_seed | PIK3CD | -0.474138 | COADREAD | SW948 | PI3Ka_4409 | MVP_top60_DNA+RNA_seed | HSP90AB1 | -1.193057 | COADREAD | SNU-175 | SNX-2112 |
| MVP_top60_DNA_seed | MET | -0.467249 | COADREAD | NCI-H716 | JNJ38877605 | MVP_top60_DNA+RNA_seed | ATM | -1.191718 | COADREAD | MDST8 | torin2 |
| MVP_top60_DNA_seed | AKT1 | -0.466331 | COADREAD | SNU-407 | MK-2206 | MVP_top60_DNA+RNA_seed | AKT1 | -1.189434 | COADREAD | NCI-H747 | A-443654 |
| MVP_top60_DNA_seed | HSP90AA1 | -0.465239 | COADREAD | SW620 | Elesclomol | MVP_top60_DNA+RNA_seed | CDK2 | -1.185414 | COADREAD | KM12 | Seliciclib |
| MVP_top60_DNA_seed | HSP90AB1 | -0.465239 | COADREAD | SW620 | Elesclomol | MVP_top60_DNA+RNA_seed | BCL2 | -1.184373 | COADREAD | HCT-15 | TW 37 |
| MVP_top60_DNA_seed | ATM | -0.463427 | COADREAD | COLO-320-HSR | CP466722 | MVP_top60_DNA+RNA_seed | HSP90AA1 | -1.182013 | COADREAD | SW48 | SNX-2112 |
| MVP_top60_DNA_seed | AKT1 | -0.463423 | COADREAD | COLO-205 | MK-2206 | MVP_top60_DNA+RNA_seed | HSP90AB1 | -1.182013 | COADREAD | SW48 | SNX-2112 |
| MVP_top60_DNA_seed | SRC | -0.462747 | COADREAD | SNU-175 | NVP-BHG712 | MVP_top60_DNA+RNA_seed | IGF1R | -1.179152 | COADREAD | KM12 | GSK1904529A |
| MVP_top60_DNA_seed | ATM | -0.461765 | COADREAD | CCK-81 | CP466722 | MVP_top60_DNA+RNA_seed | PIK3CA | -1.178649 | COADREAD | SK-CO-1 | AZD8835 |
| MVP_top60_DNA_seed | SRC | -0.461177 | COADREAD | HCT-15 | Bosutinib | MVP_top60_DNA+RNA_seed | PIK3CD | -1.178649 | COADREAD | SK-CO-1 | AZD8835 |
| MVP_top60_DNA_seed | PTK2 | -0.459939 | COADREAD | LS-411N | PF-562271 | MVP_top60_DNA+RNA_seed | IGF1R | -1.175946 | COADREAD | NCI-H747 | BMS-536924 |
| MVP_top60_DNA_seed | HSP90AA1 | -0.458332 | COADREAD | LS-123 | Luminespib | MVP_top60_DNA+RNA_seed | BCL2 | -1.17472 | COADREAD | COLO-205 | Obatoclax Mesylate |
| MVP_top60_DNA_seed | HSP90AB1 | -0.458332 | COADREAD | LS-123 | Luminespib | MVP_top60_DNA+RNA_seed | AKT1 | -1.171344 | COADREAD | SNU-81 | MK-2206 |
| MVP_top60_DNA_seed | IGF1R | -0.457392 | COADREAD | GP5d | BMS-536924 | MVP_top60_DNA+RNA_seed | HSP90AA1 | -1.170093 | COADREAD | SW48 | CCT-018159 |
| MVP_top60_DNA_seed | AR | -0.453972 | COADREAD | SK-CO-1 | Bicalutamide | MVP_top60_DNA+RNA_seed | HSP90AB1 | -1.170093 | COADREAD | SW48 | CCT-018159 |

| | | | | | | | | | | | |
|---|---|---|---|---|---|---|---|---|---|---|---|
| MVP_top60_DNA_seed | GSK3B | -0.452908 | COADREAD | KM12 | CHIR-99021 | MVP_top60_DNA+RNA_seed | PIK3CD | -1.16705 | COADREAD | LoVo | IC-87114 |
| MVP_top60_DNA_seed | GSK3B | -0.447552 | COADREAD | HCC2998 | AZD7969 | MVP_top60_DNA+RNA_seed | HSP90AA1 | -1.165297 | COADREAD | SK-CO-1 | Tanespimycin |
| MVP_top60_DNA_seed | AKT1 | -0.446491 | COADREAD | T84 | AKT inhibitor VIII | MVP_top60_DNA+RNA_seed | HSP90AB1 | -1.165297 | COADREAD | SK-CO-1 | Tanespimycin |
| MVP_top60_DNA_seed | HSP90AA1 | -0.442865 | COADREAD | SNU-C5 | Tanespimycin | MVP_top60_DNA+RNA_seed | GSK3B | -1.163151 | COADREAD | SW48 | AZD7969 |
| MVP_top60_DNA_seed | HSP90AB1 | -0.442865 | COADREAD | SNU-C5 | Tanespimycin | MVP_top60_DNA+RNA_seed | HSP90AA1 | -1.161249 | COADREAD | KM12 | Tanespimycin |
| MVP_top60_DNA_seed | GSK3B | -0.442639 | COADREAD | SW48 | CHIR-99021 | MVP_top60_DNA+RNA_seed | HSP90AB1 | -1.161249 | COADREAD | KM12 | Tanespimycin |
| MVP_top60_DNA_seed | SRC | -0.43469 | COADREAD | LS-123 | WH-4-023 | MVP_top60_DNA+RNA_seed | HSP90AA1 | -1.155373 | COADREAD | LS-411N | SNX-2112 |
| MVP_top60_DNA_seed | GSK3B | -0.433737 | COADREAD | HT-115 | CHIR-99021 | MVP_top60_DNA+RNA_seed | HSP90AB1 | -1.155373 | COADREAD | LS-411N | SNX-2112 |
| MVP_top60_DNA_seed | ERBB4 | -0.43373 | COADREAD | SW1116 | CI-1033 | MVP_top60_DNA+RNA_seed | GSK3B | -1.147071 | COADREAD | RCM-1 | CHIR-99021 |
| MVP_top60_DNA_seed | PIK3CA | -0.433547 | COADREAD | NCI-H508 | PI3Ka_4409 | MVP_top60_DNA+RNA_seed | IGF1R | -1.145111 | COADREAD | LS-513 | Linsitinib |
| MVP_top60_DNA_seed | PIK3CD | -0.433547 | COADREAD | NCI-H508 | PI3Ka_4409 | MVP_top60_DNA+RNA_seed | SRC | -1.144012 | COADREAD | NCI-H747 | Saracatinib |
| MVP_top60_DNA_seed | SRC | -0.433534 | COADREAD | SNU-407 | Bosutinib | MVP_top60_DNA+RNA_seed | ATM | -1.134419 | COADREAD | NCI-H716 | KU-60019 |
| MVP_top60_DNA_seed | ATM | -0.432508 | COADREAD | HCT-15 | CP466722 | MVP_top60_DNA+RNA_seed | ATM | -1.124209 | COADREAD | SNU-175 | CP466722 |
| MVP_top60_DNA_seed | ATM | -0.432033 | COADREAD | SNU-81 | KU-60019 | MVP_top60_DNA+RNA_seed | HSP90AA1 | -1.114351 | COADREAD | RKO | Luminespib |
| MVP_top60_DNA_seed | IGF1R | -0.431333 | COADREAD | SNU-407 | BMS-536924 | MVP_top60_DNA+RNA_seed | HSP90AB1 | -1.114351 | COADREAD | RKO | Luminespib |
| MVP_top60_DNA_seed | SRC | -0.430207 | COADREAD | NCI-H747 | WH-4-023 | MVP_top60_DNA+RNA_seed | MET | -1.11237 | COADREAD | LoVo | SU11274 |
| MVP_top60_DNA_seed | HSP90AA1 | -0.424943 | COADREAD | COLO-320-HSR | SNX-2112 | MVP_top60_DNA+RNA_seed | PIK3CB | -1.112036 | COADREAD | LS-180 | AZD6482 |
| MVP_top60_DNA_seed | HSP90AB1 | -0.424943 | COADREAD | COLO-320-HSR | SNX-2112 | MVP_top60_DNA+RNA_seed | SRC | -1.111319 | COADREAD | LoVo | HG-6-71-01 |
| MVP_top60_DNA_seed | MET | -0.422215 | COADREAD | HCT-116 | AZD6094 | MVP_top60_DNA+RNA_seed | AR | -1.110622 | COADREAD | SW48 | Bicalutamide |
| MVP_top60_DNA_seed | GSK3B | -0.420137 | COADREAD | SW620 | KIN001-042 | MVP_top60_DNA+RNA_seed | IGF1R | -1.097252 | COADREAD | COLO-320-HSR | BMS-536924 |
| MVP_top60_DNA_seed | HSP90AA1 | -0.419316 | COADREAD | CaR-1 | SNX-2112 | MVP_top60_DNA+RNA_seed | AKT1 | -1.092442 | COADREAD | NCI-H747 | GSK690693 |
| MVP_top60_DNA_seed | HSP90AB1 | -0.419316 | COADREAD | CaR-1 | SNX-2112 | MVP_top60_DNA+RNA_seed | SRC | -1.090901 | COADREAD | RKO | NVP-BHG712 |
| MVP_top60_DNA_seed | BRAF | -0.417936 | COADREAD | MDST8 | SB590885 | MVP_top60_DNA+RNA_seed | IGF1R | -1.089427 | COADREAD | CCK-81 | Linsitinib |
| MVP_top60_DNA_seed | GSK3B | -0.414456 | COADREAD | GP5d | AZD7969 | MVP_top60_DNA+RNA_seed | PIK3CA | -1.086516 | COADREAD | GP5d | AZD8835 |
| MVP_top60_DNA_seed | ATM | -0.414445 | COADREAD | KM12 | CP466722 | MVP_top60_DNA+RNA_seed | PIK3CD | -1.086516 | COADREAD | GP5d | AZD8835 |
| MVP_top60_DNA_seed | IGF1R | -0.413062 | COADREAD | T84 | BMS-536924 | MVP_top60_DNA+RNA_seed | PIK3CA | -1.084276 | COADREAD | DiFi | PI3Ka_4409 |
| MVP_top60_DNA_seed | GSK3B | -0.410285 | COADREAD | HCC2998 | CHIR-99021 | MVP_top60_DNA+RNA_seed | PIK3CD | -1.084276 | COADREAD | DiFi | PI3Ka_4409 |
| MVP_top60_DNA_seed | ERBB3 | -0.409705 | COADREAD | SK-CO-1 | AZD8931 | MVP_top60_DNA+RNA_seed | PIK3CA | -1.081583 | COADREAD | SNU-407 | PI3Ka_4409 |
| MVP_top60_DNA_seed | HSP90AA1 | -0.40962 | COADREAD | SNU-407 | Elesclomol | MVP_top60_DNA+RNA_seed | PIK3CD | -1.081583 | COADREAD | SNU-407 | PI3Ka_4409 |
| MVP_top60_DNA_seed | HSP90AB1 | -0.40962 | COADREAD | SNU-407 | Elesclomol | MVP_top60_DNA+RNA_seed | ERBB4 | -1.07416 | COADREAD | HT-115 | PF-00299804 |
| MVP_top60_DNA_seed | ESR1 | -0.408358 | COADREAD | LS-123 | Fulvestrant | MVP_top60_DNA+RNA_seed | CDK2 | -1.064815 | COADREAD | HCT-116 | AZD5438 |
| MVP_top60_DNA_seed | IGF1R | -0.405281 | COADREAD | LS-513 | BMS-536924 | MVP_top60_DNA+RNA_seed | AKT1 | -1.063981 | COADREAD | SNU-175 | AKT inhibitor VIII |
| MVP_top60_DNA_seed | GSK3B | -0.403294 | COADREAD | LS-1034 | AZD7969 | MVP_top60_DNA+RNA_seed | AKT1 | -1.061978 | COADREAD | LoVo | AKT inhibitor VIII |
| MVP_top60_DNA_seed | AKT1 | -0.401915 | COADREAD | HCC2998 | BAY AKT1 | MVP_top60_DNA+RNA_seed | PIK3CA | -1.060381 | COADREAD | NCI-H747 | AZD8835 |
| MVP_top60_DNA_seed | MAPK1 | -0.401676 | COADREAD | LS-1034 | FR-180204 | MVP_top60_DNA+RNA_seed | PIK3CD | -1.060381 | COADREAD | NCI-H747 | AZD8835 |
| MVP_top60_DNA_seed | CDK2 | -0.397909 | COADREAD | COLO-320-HSR | AZD5438 | MVP_top60_DNA+RNA_seed | CDK2 | -1.059276 | COADREAD | HT-115 | AZD5438 |
| MVP_top60_DNA_seed | HSP90AA1 | -0.396004 | COADREAD | C2BBe1 | Tanespimycin | MVP_top60_DNA+RNA_seed | AKT1 | -1.058893 | COADREAD | RKO | MK-2206 |
| MVP_top60_DNA_seed | HSP90AB1 | -0.396004 | COADREAD | C2BBe1 | Tanespimycin | MVP_top60_DNA+RNA_seed | CDK2 | -1.055658 | COADREAD | SW620 | PHA-793887 |
| MVP_top60_DNA_seed | CREBBP | -0.393195 | COADREAD | HT-29 | I-CBP112 | MVP_top60_DNA+RNA_seed | PIK3CB | -1.053646 | COADREAD | RKO | AZD6482 |
| MVP_top60_DNA_seed | MAPK14 | -0.391037 | COADREAD | LS-180 | TAK-715 | MVP_top60_DNA+RNA_seed | IGF1R | -1.052362 | COADREAD | RKO | BMS-536924 |
| MVP_top60_DNA_seed | ERBB4 | -0.390573 | COADREAD | NCI-H747 | PF-00299804 | MVP_top60_DNA+RNA_seed | ATM | -1.042063 | COADREAD | NCI-H716 | KU-55933 |

| | | | | | | | | | | | |
|---|---|---|---|---|---|---|---|---|---|---|---|
| MVP_top60_DNA_seed | FYN | -0.383507 | COADREAD | LS-123 | A-770041 | MVP_top60_DNA+RNA_seed | AKT1 | -1.034944 | COADREAD | NCI-H716 | AKT inhibitor VIII |
| MVP_top60_DNA_seed | BRAF | -0.383245 | COADREAD | HCT-15 | PLX-4720 | MVP_top60_DNA+RNA_seed | BCL2 | -1.034494 | COADREAD | SW1417 | Navitoclax |
| MVP_top60_DNA_seed | AKT1 | -0.382601 | COADREAD | KM12 | AKT inhibitor VIII | MVP_top60_DNA+RNA_seed | PTK2 | -1.033563 | COADREAD | SW620 | PF-562271 |
| MVP_top60_DNA_seed | ATM | -0.38166 | COADREAD | COLO-205 | CP466722 | MVP_top60_DNA+RNA_seed | CDK2 | -1.03127 | COADREAD | RKO | AZD5438 |
| MVP_top60_DNA_seed | ERBB3 | -0.38052 | COADREAD | NCI-H747 | AZD8931 | MVP_top60_DNA+RNA_seed | IGF1R | -1.030512 | COADREAD | GP5d | Linsitinib |
| MVP_top60_DNA_seed | HSP90AA1 | -0.376604 | COADREAD | T84 | SNX-2112 | MVP_top60_DNA+RNA_seed | HSP90AA1 | -1.029666 | COADREAD | HCT-116 | SNX-2112 |
| MVP_top60_DNA_seed | HSP90AB1 | -0.376604 | COADREAD | T84 | SNX-2112 | MVP_top60_DNA+RNA_seed | HSP90AB1 | -1.029666 | COADREAD | HCT-116 | SNX-2112 |
| MVP_top60_DNA_seed | GSK3B | -0.375297 | COADREAD | SW1116 | SB216763 | MVP_top60_DNA+RNA_seed | HSP90AA1 | -1.029424 | COADREAD | LS-513 | SNX-2112 |
| MVP_top60_DNA_seed | PIK3CB | -0.373521 | COADREAD | MDST8 | AZD8186 | MVP_top60_DNA+RNA_seed | HSP90AB1 | -1.029424 | COADREAD | LS-513 | SNX-2112 |
| MVP_top60_DNA_seed | PIK3CD | -0.373521 | COADREAD | MDST8 | AZD8186 | MVP_top60_DNA+RNA_seed | HSP90AA1 | -1.028081 | COADREAD | HT-29 | SNX-2112 |
| MVP_top60_DNA_seed | AR | -0.373086 | COADREAD | SNU-C2B | AZD3514 | MVP_top60_DNA+RNA_seed | HSP90AB1 | -1.028081 | COADREAD | HT-29 | SNX-2112 |
| MVP_top60_DNA_seed | PIK3CA | -0.36886 | COADREAD | LS-513 | PI3Ka_4409 | MVP_top60_DNA+RNA_seed | HSP90AA1 | -1.021061 | COADREAD | NCI-H508 | Tanespimycin |
| MVP_top60_DNA_seed | PIK3CD | -0.36886 | COADREAD | LS-513 | PI3Ka_4409 | MVP_top60_DNA+RNA_seed | HSP90AB1 | -1.021061 | COADREAD | NCI-H508 | Tanespimycin |
| MVP_top60_DNA_seed | IGF1R | -0.368033 | COADREAD | SW837 | Linsitinib | MVP_top60_DNA+RNA_seed | PIK3CA | -1.013241 | COADREAD | HCT-116 | PI3Ka_4409 |
| MVP_top60_DNA_seed | BRAF | -0.367373 | COADREAD | LS-180 | HG6-64-1 | MVP_top60_DNA+RNA_seed | PIK3CD | -1.013241 | COADREAD | HCT-116 | PI3Ka_4409 |
| MVP_top60_DNA_seed | PIK3CD | -0.366195 | COADREAD | SNU-81 | IC-87114 | MVP_top60_DNA+RNA_seed | ATM | -1.012839 | COADREAD | SNU-C5 | KU-55933 |
| MVP_top60_DNA_seed | HSP90AA1 | -0.365355 | COADREAD | HCT-15 | SNX-2112 | MVP_top60_DNA+RNA_seed | CDK2 | -1.007668 | COADREAD | MDST8 | AZD5438 |
| MVP_top60_DNA_seed | HSP90AB1 | -0.365355 | COADREAD | HCT-15 | SNX-2112 | MVP_top60_DNA+RNA_seed | HSP90AA1 | -1.006089 | COADREAD | SW620 | Luminespib |
| MVP_top60_DNA_seed | PIK3CB | -0.364882 | COADREAD | C2BBe1 | AZD6482 | MVP_top60_DNA+RNA_seed | HSP90AB1 | -1.006089 | COADREAD | SW620 | Luminespib |
| MVP_top60_DNA_seed | MAPK14 | -0.363481 | COADREAD | LS-411N | TAK-715 | MVP_top60_DNA+RNA_seed | HSP90AA1 | -1.004215 | COADREAD | SW620 | CCT-018159 |
| MVP_top60_DNA_seed | GSK3B | -0.363097 | COADREAD | CL-11 | SB216763 | MVP_top60_DNA+RNA_seed | HSP90AB1 | -1.004215 | COADREAD | SW620 | CCT-018159 |
| MVP_top60_DNA_seed | BRAF | -0.36118 | COADREAD | SW48 | RAF_9304 | MVP_top60_DNA+RNA_seed | ERBB3 | -0.999809 | COADREAD | GP5d | AZD8931 |
| MVP_top60_DNA_seed | ATM | -0.360689 | COADREAD | HCT-15 | KU-60019 | MVP_top60_DNA+RNA_seed | GSK3B | -0.990536 | COADREAD | MDST8 | AZD7969 |
| MVP_top60_DNA_seed | FYN | -0.35927 | COADREAD | SNU-C2B | A-770041 | MVP_top60_DNA+RNA_seed | SRC | -0.987133 | COADREAD | HCT-116 | Bosutinib |
| MVP_top60_DNA_seed | MET | -0.358717 | COADREAD | HT-115 | SU11274 | MVP_top60_DNA+RNA_seed | PIK3CA | -0.984215 | COADREAD | GP5d | PI3Ka_4409 |
| MVP_top60_DNA_seed | MAPK14 | -0.358244 | COADREAD | NCI-H716 | TAK-715 | MVP_top60_DNA+RNA_seed | PIK3CD | -0.984215 | COADREAD | GP5d | PI3Ka_4409 |
| MVP_top60_DNA_seed | AKT1 | -0.353981 | COADREAD | LS-513 | GSK690693 | MVP_top60_DNA+RNA_seed | AKT1 | -0.983791 | COADREAD | COLO-205 | BAY AKT1 |
| MVP_top60_DNA_seed | GSK3B | -0.353564 | COADREAD | NCI-H716 | SB216763 | MVP_top60_DNA+RNA_seed | HSP90AA1 | -0.982615 | COADREAD | COLO-205 | SNX-2112 |
| MVP_top60_DNA_seed | ATM | -0.353132 | COADREAD | RKO | CP466722 | MVP_top60_DNA+RNA_seed | HSP90AB1 | -0.982615 | COADREAD | COLO-205 | SNX-2112 |
| MVP_top60_DNA_seed | ATM | -0.353111 | COADREAD | SW620 | CP466722 | MVP_top60_DNA+RNA_seed | ATM | -0.98135 | COADREAD | KM12 | KU-60019 |
| MVP_top60_DNA_seed | ESR1 | -0.350112 | COADREAD | GP5d | Fulvestrant | MVP_top60_DNA+RNA_seed | GSK3B | -0.978221 | COADREAD | HT-29 | SB216763 |
| MVP_top60_DNA_seed | BRAF | -0.348715 | COADREAD | NCI-H747 | HG6-64-1 | MVP_top60_DNA+RNA_seed | IGF1R | -0.976678 | COADREAD | SK-CO-1 | BMS-536924 |
| MVP_top60_DNA_seed | AR | -0.348069 | COADREAD | SNU-407 | AZD3514 | MVP_top60_DNA+RNA_seed | GSK3B | -0.975218 | COADREAD | HCT-116 | CHIR-99021 |
| MVP_top60_DNA_seed | PIK3CA | -0.347499 | COADREAD | COLO-320-HSR | PI3Ka_4409 | MVP_top60_DNA+RNA_seed | ERBB3 | -0.973837 | COADREAD | LoVo | AZD8931 |
| MVP_top60_DNA_seed | PIK3CD | -0.347499 | COADREAD | COLO-320-HSR | PI3Ka_4409 | MVP_top60_DNA+RNA_seed | ERBB4 | -0.969238 | COADREAD | SNU-175 | CI-1033 |
| MVP_top60_DNA_seed | BRAF | -0.345525 | COADREAD | KM12 | HG6-64-1 | MVP_top60_DNA+RNA_seed | IGF1R | -0.963142 | COADREAD | LS-1034 | BMS-536924 |
| MVP_top60_DNA_seed | PIK3CA | -0.344309 | COADREAD | NCI-H716 | AZD8835 | MVP_top60_DNA+RNA_seed | HSP90AA1 | -0.96185 | COADREAD | SW620 | SNX-2112 |
| MVP_top60_DNA_seed | PIK3CD | -0.344309 | COADREAD | NCI-H716 | AZD8835 | MVP_top60_DNA+RNA_seed | HSP90AB1 | -0.96185 | COADREAD | SW620 | SNX-2112 |
| MVP_top60_DNA_seed | AR | -0.343672 | COADREAD | NCI-H747 | AZD3514 | MVP_top60_DNA+RNA_seed | GSK3B | -0.956867 | COADREAD | GP5d | KIN001-042 |
| MVP_top60_DNA_seed | ESR1 | -0.341378 | COADREAD | CW-2 | Fulvestrant | MVP_top60_DNA+RNA_seed | ERBB4 | -0.953289 | COADREAD | HT-115 | AST-1306 |
| MVP_top60_DNA_seed | GSK3B | -0.338959 | COADREAD | SK-CO-1 | CHIR-99021 | MVP_top60_DNA+RNA_seed | CDK2 | -0.95082 | COADREAD | RKO | Seliciclib |

| | | | | | | | | | | | |
|---|---|---|---|---|---|---|---|---|---|---|---|
| MVP_top60_DNA_seed | CDK2 | -0.338654 | COADREAD | LS-411N | Seliciclib | MVP_top60_DNA+RNA_seed | GSK3B | -0.94936 | COADREAD | GP5d | CHIR-99021 |
| MVP_top60_DNA_seed | MAPK1 | -0.337699 | COADREAD | HCT-116 | FR-180204 | MVP_top60_DNA+RNA_seed | ERBB4 | -0.947921 | COADREAD | LoVo | PF-00299804 |
| MVP_top60_DNA_seed | PIK3CA | -0.336371 | COADREAD | SW1463 | PI3Ka_4409 | MVP_top60_DNA+RNA_seed | SRC | -0.930433 | COADREAD | HT-29 | NVP-BHG712 |
| MVP_top60_DNA_seed | PIK3CD | -0.336371 | COADREAD | SW1463 | PI3Ka_4409 | MVP_top60_DNA+RNA_seed | PIK3CA | -0.928663 | COADREAD | HCT-116 | AZD8835 |
| MVP_top60_DNA_seed | GSK3B | -0.335235 | COADREAD | COLO-320-HSR | KIN001-042 | MVP_top60_DNA+RNA_seed | PIK3CD | -0.928663 | COADREAD | HCT-116 | AZD8835 |
| MVP_top60_DNA_seed | BRAF | -0.335172 | COADREAD | GP5d | RAF_9304 | MVP_top60_DNA+RNA_seed | AKT1 | -0.923658 | COADREAD | LS-180 | AKT inhibitor VIII |
| MVP_top60_DNA_seed | HSP90AA1 | -0.328917 | COADREAD | HCC2998 | Tanespimycin | MVP_top60_DNA+RNA_seed | SRC | -0.920611 | COADREAD | SNU-81 | Bosutinib |
| MVP_top60_DNA_seed | HSP90AB1 | -0.328917 | COADREAD | HCC2998 | Tanespimycin | MVP_top60_DNA+RNA_seed | BCL2 | -0.918475 | COADREAD | RKO | TW 37 |
| MVP_top60_DNA_seed | MET | -0.328176 | COADREAD | HT-29 | JNJ38877605 | MVP_top60_DNA+RNA_seed | ERBB4 | -0.917014 | COADREAD | LoVo | CI-1033 |
| MVP_top60_DNA_seed | HSP90AA1 | -0.324863 | COADREAD | NCI-H747 | SNX-2112 | MVP_top60_DNA+RNA_seed | AR | -0.91592 | COADREAD | LS-411N | AZD3514 |
| MVP_top60_DNA_seed | HSP90AB1 | -0.324863 | COADREAD | NCI-H747 | SNX-2112 | MVP_top60_DNA+RNA_seed | CDK2 | -0.914337 | COADREAD | HT-29 | PHA-793887 |
| MVP_top60_DNA_seed | GSK3B | -0.322784 | COADREAD | LS-513 | KIN001-042 | MVP_top60_DNA+RNA_seed | ERBB3 | -0.914211 | COADREAD | SW1463 | AZD8931 |
| MVP_top60_DNA_seed | ERBB4 | -0.320522 | COADREAD | NCI-H747 | CI-1033 | MVP_top60_DNA+RNA_seed | SRC | -0.913402 | COADREAD | MDST8 | NVP-BHG712 |
| MVP_top60_DNA_seed | PIK3CA | -0.319794 | COADREAD | COLO-205 | AZD8835 | MVP_top60_DNA+RNA_seed | HSP90AA1 | -0.901731 | COADREAD | HCT-15 | Elesclomol |
| MVP_top60_DNA_seed | PIK3CD | -0.319794 | COADREAD | COLO-205 | AZD8835 | MVP_top60_DNA+RNA_seed | HSP90AB1 | -0.901731 | COADREAD | HCT-15 | Elesclomol |
| MVP_top60_DNA_seed | MAPK1 | -0.319392 | COADREAD | SNU-61 | FR-180204 | MVP_top60_DNA+RNA_seed | ERBB3 | -0.896592 | COADREAD | HCT-116 | AZD8931 |
| MVP_top60_DNA_seed | SRC | -0.319177 | COADREAD | KM12 | NVP-BHG712 | MVP_top60_DNA+RNA_seed | ERBB3 | -0.892663 | COADREAD | LS-513 | AZD8931 |
| MVP_top60_DNA_seed | MAPK1 | -0.318629 | COADREAD | LS-180 | VX-11e | MVP_top60_DNA+RNA_seed | ATM | -0.890736 | COADREAD | LoVo | KU-60019 |
| MVP_top60_DNA_seed | PIK3CA | -0.318476 | COADREAD | MDST8 | AZD8835 | MVP_top60_DNA+RNA_seed | BCL2 | -0.890422 | COADREAD | LS-411N | Obatoclax Mesylate |
| MVP_top60_DNA_seed | PIK3CD | -0.318476 | COADREAD | MDST8 | AZD8835 | MVP_top60_DNA+RNA_seed | IGF1R | -0.886705 | COADREAD | HT-29 | BMS-536924 |
| MVP_top60_DNA_seed | HSP90AA1 | -0.317495 | COADREAD | GP5d | Luminespib | MVP_top60_DNA+RNA_seed | MET | -0.885738 | COADREAD | LS-411N | Crizotinib |
| MVP_top60_DNA_seed | HSP90AB1 | -0.317495 | COADREAD | GP5d | Luminespib | MVP_top60_DNA+RNA_seed | HSP90AA1 | -0.885297 | COADREAD | SW620 | Tanespimycin |
| MVP_top60_DNA_seed | AKT1 | -0.315839 | COADREAD | LS-180 | AKT inhibitor VIII | MVP_top60_DNA+RNA_seed | HSP90AB1 | -0.885297 | COADREAD | SW620 | Tanespimycin |
| MVP_top60_DNA_seed | PTK2 | -0.315161 | COADREAD | NCI-H747 | PF-562271 | MVP_top60_DNA+RNA_seed | AKT1 | -0.883882 | COADREAD | LS-513 | MK-2206 |
| MVP_top60_DNA_seed | AKT1 | -0.31205 | COADREAD | SW948 | GSK690693 | MVP_top60_DNA+RNA_seed | GSK3B | -0.883203 | COADREAD | SNU-C5 | CHIR-99021 |
| MVP_top60_DNA_seed | AR | -0.310652 | COADREAD | KM12 | Bicalutamide | MVP_top60_DNA+RNA_seed | BCL2 | -0.882171 | COADREAD | COLO-205 | Navitoclax |
| MVP_top60_DNA_seed | BRAF | -0.30831 | COADREAD | SW1417 | PLX-4720 | MVP_top60_DNA+RNA_seed | SRC | -0.877365 | COADREAD | NCI-H747 | NVP-BHG712 |
| MVP_top60_DNA_seed | IGF1R | -0.308205 | COADREAD | LoVo | BMS-536924 | MVP_top60_DNA+RNA_seed | PIK3CA | -0.869761 | COADREAD | RKO | AZD8835 |
| MVP_top60_DNA_seed | PIK3CD | -0.307003 | COADREAD | CCK-81 | Idelalisib | MVP_top60_DNA+RNA_seed | PIK3CD | -0.869761 | COADREAD | RKO | AZD8835 |
| MVP_top60_DNA_seed | ESR1 | -0.304683 | COADREAD | HT-115 | Fulvestrant | MVP_top60_DNA+RNA_seed | GSK3B | -0.864362 | COADREAD | HT-115 | SB216763 |
| MVP_top60_DNA_seed | IGF1R | -0.304123 | COADREAD | SNU-C2B | Linsitinib | MVP_top60_DNA+RNA_seed | PTK2 | -0.862029 | COADREAD | RKO | PF-562271 |
| MVP_top60_DNA_seed | BRAF | -0.303669 | COADREAD | SNU-C5 | RAF_9304 | MVP_top60_DNA+RNA_seed | GSK3B | -0.857376 | COADREAD | SW620 | AZD7969 |
| MVP_top60_DNA_seed | MET | -0.302398 | COADREAD | COLO-205 | AZD6094 | MVP_top60_DNA+RNA_seed | GSK3B | -0.856485 | COADREAD | HT-115 | AZD7969 |
| MVP_top60_DNA_seed | AR | -0.302285 | COADREAD | NCI-H716 | AZD3514 | MVP_top60_DNA+RNA_seed | HSP90AA1 | -0.85375 | COADREAD | KM12 | CCT-018159 |
| MVP_top60_DNA_seed | PTK2 | -0.302222 | COADREAD | LoVo | PF-562271 | MVP_top60_DNA+RNA_seed | HSP90AB1 | -0.85375 | COADREAD | KM12 | CCT-018159 |
| MVP_top60_DNA_seed | GSK3B | -0.300728 | COADREAD | LS-411N | CHIR-99021 | MVP_top60_DNA+RNA_seed | PIK3CB | -0.852167 | COADREAD | HCT-116 | AZD6482 |
| MVP_top60_DNA_seed | SRC | -0.298466 | COADREAD | MDST8 | HG-6-71-01 | MVP_top60_DNA+RNA_seed | AKT1 | -0.850234 | COADREAD | RKO | A-443654 |
| MVP_top60_DNA_seed | AKT1 | -0.297477 | COADREAD | GP5d | GSK690693 | MVP_top60_DNA+RNA_seed | HSP90AA1 | -0.850035 | COADREAD | SNU-C2B | Luminespib |
| MVP_top60_DNA_seed | GSK3B | -0.296121 | COADREAD | RKO | AZD7969 | MVP_top60_DNA+RNA_seed | HSP90AB1 | -0.850035 | COADREAD | SNU-C2B | Luminespib |
| MVP_top60_DNA_seed | MAPK1 | -0.292002 | COADREAD | C2BBe1 | VX-11e | MVP_top60_DNA+RNA_seed | MET | -0.847099 | COADREAD | KM12 | JNJ38877605 |
| MVP_top60_DNA_seed | GSK3B | -0.291615 | COADREAD | NCI-H747 | SB216763 | MVP_top60_DNA+RNA_seed | GSK3B | -0.846871 | COADREAD | HCT-116 | SB216763 |

| | | | | | | | | | | | |
|---|---|---|---|---|---|---|---|---|---|---|---|
| MVP_top60_DNA_seed | MET | -0.291471 | COADREAD | SW1417 | JNJ38877605 | MVP_top60_DNA+RNA_seed | HSP90AA1 | -0.845633 | COADREAD | MDST8 | SNX-2112 |
| MVP_top60_DNA_seed | SRC | -0.291213 | COADREAD | CCK-81 | NVP-BHG712 | MVP_top60_DNA+RNA_seed | HSP90AB1 | -0.845633 | COADREAD | MDST8 | SNX-2112 |
| MVP_top60_DNA_seed | IGF1R | -0.289911 | COADREAD | CaR-1 | BMS-536924 | MVP_top60_DNA+RNA_seed | PIK3CA | -0.844898 | COADREAD | SNU-C1 | AZD8835 |
| MVP_top60_DNA_seed | PIK3CD | -0.28945 | COADREAD | RKO | Idelalisib | MVP_top60_DNA+RNA_seed | PIK3CD | -0.844898 | COADREAD | SNU-C1 | AZD8835 |
| MVP_top60_DNA_seed | HSP90AA1 | -0.288218 | COADREAD | NCI-H747 | Tanespimycin | MVP_top60_DNA+RNA_seed | PIK3CA | -0.843751 | COADREAD | SW48 | PI3Ka_4409 |
| MVP_top60_DNA_seed | HSP90AB1 | -0.288218 | COADREAD | NCI-H747 | Tanespimycin | MVP_top60_DNA+RNA_seed | PIK3CD | -0.843751 | COADREAD | SW48 | PI3Ka_4409 |
| MVP_top60_DNA_seed | GSK3B | -0.285862 | COADREAD | MDST8 | CHIR-99021 | MVP_top60_DNA+RNA_seed | AKT1 | -0.83972 | COADREAD | CCK-81 | AKT inhibitor VIII |
| MVP_top60_DNA_seed | ATM | -0.283284 | COADREAD | SW48 | KU-55933 | MVP_top60_DNA+RNA_seed | HSP90AA1 | -0.836348 | COADREAD | LS-513 | CCT-018159 |
| MVP_top60_DNA_seed | GSK3B | -0.282916 | COADREAD | COLO-205 | SB216763 | MVP_top60_DNA+RNA_seed | HSP90AB1 | -0.836348 | COADREAD | LS-513 | CCT-018159 |
| MVP_top60_DNA_seed | GSK3B | -0.280222 | COADREAD | T84 | KIN001-042 | MVP_top60_DNA+RNA_seed | BCL2 | -0.834157 | COADREAD | LoVo | TW 37 |
| MVP_top60_DNA_seed | MAPK14 | -0.280024 | COADREAD | SNU-C2B | TAK-715 | MVP_top60_DNA+RNA_seed | AKT1 | -0.833189 | COADREAD | SNU-81 | AKT inhibitor VIII |
| MVP_top60_DNA_seed | IGF1R | -0.279373 | COADREAD | LS-513 | BMS-536924 | MVP_top60_DNA+RNA_seed | MET | -0.828905 | COADREAD | NCI-H716 | SU11274 |
| MVP_top60_DNA_seed | PIK3CB | -0.278332 | COADREAD | NCI-H747 | AZD8186 | MVP_top60_DNA+RNA_seed | MET | -0.827957 | COADREAD | LoVo | JNJ38877605 |
| MVP_top60_DNA_seed | PIK3CD | -0.278332 | COADREAD | NCI-H747 | AZD8186 | MVP_top60_DNA+RNA_seed | ATM | -0.825948 | COADREAD | CL-11 | KU-60019 |
| MVP_top60_DNA_seed | AKT1 | -0.277854 | COADREAD | COLO-320-HSR | BAY AKT1 | MVP_top60_DNA+RNA_seed | HSP90AA1 | -0.815762 | COADREAD | HT55 | CCT-018159 |
| MVP_top60_DNA_seed | ATM | -0.277802 | COADREAD | MDST8 | KU-60019 | MVP_top60_DNA+RNA_seed | HSP90AB1 | -0.815762 | COADREAD | HT55 | CCT-018159 |
| MVP_top60_DNA_seed | ATM | -0.273401 | COADREAD | SK-CO-1 | KU-60019 | MVP_top60_DNA+RNA_seed | HSP90AA1 | -0.814833 | COADREAD | COLO-678 | Tanespimycin |
| MVP_top60_DNA_seed | MAPK14 | -0.271996 | COADREAD | T84 | TAK-715 | MVP_top60_DNA+RNA_seed | HSP90AB1 | -0.814833 | COADREAD | COLO-678 | Tanespimycin |
| MVP_top60_DNA_seed | GSK3B | -0.269551 | COADREAD | RKO | CHIR-99021 | MVP_top60_DNA+RNA_seed | AR | -0.811173 | COADREAD | LS-180 | Bicalutamide |
| MVP_top60_DNA_seed | MET | -0.268429 | COADREAD | T84 | JNJ38877605 | MVP_top60_DNA+RNA_seed | IGF1R | -0.810778 | COADREAD | SW48 | BMS-754807 |
| MVP_top60_DNA_seed | PIK3CA | -0.2684 | COADREAD | CCK-81 | AZD8835 | MVP_top60_DNA+RNA_seed | BCL2 | -0.808263 | COADREAD | HCC2998 | Navitoclax |
| MVP_top60_DNA_seed | PIK3CD | -0.2684 | COADREAD | CCK-81 | AZD8835 | MVP_top60_DNA+RNA_seed | HSP90AA1 | -0.804421 | COADREAD | LoVo | CCT-018159 |
| MVP_top60_DNA_seed | IGF1R | -0.267056 | COADREAD | CCK-81 | GSK1904529A | MVP_top60_DNA+RNA_seed | HSP90AB1 | -0.804421 | COADREAD | LoVo | CCT-018159 |
| MVP_top60_DNA_seed | HSP90AA1 | -0.265654 | COADREAD | CW-2 | CCT-018159 | MVP_top60_DNA+RNA_seed | AR | -0.802513 | COADREAD | HCT-116 | Bicalutamide |
| MVP_top60_DNA_seed | HSP90AB1 | -0.265654 | COADREAD | CW-2 | CCT-018159 | MVP_top60_DNA+RNA_seed | MET | -0.802238 | COADREAD | SNU-175 | JNJ38877605 |
| MVP_top60_DNA_seed | BRAF | -0.265451 | COADREAD | SNU-407 | Dabrafenib | MVP_top60_DNA+RNA_seed | BCL2 | -0.797921 | COADREAD | SW1417 | Obatoclax Mesylate |
| MVP_top60_DNA_seed | BRAF | -0.265331 | COADREAD | HT-115 | RAF_9304 | MVP_top60_DNA+RNA_seed | AR | -0.796608 | COADREAD | SW48 | AZD3514 |
| MVP_top60_DNA_seed | ESR1 | -0.264947 | COADREAD | DiFi | Fulvestrant | MVP_top60_DNA+RNA_seed | AR | -0.794147 | COADREAD | RKO | AZD3514 |
| MVP_top60_DNA_seed | ERBB4 | -0.263149 | COADREAD | LS-513 | PF-00299804 | MVP_top60_DNA+RNA_seed | AKT1 | -0.791737 | COADREAD | KM12 | GSK690693 |
| MVP_top60_DNA_seed | AR | -0.261322 | COADREAD | COLO-320-HSR | Bicalutamide | MVP_top60_DNA+RNA_seed | AR | -0.791279 | COADREAD | T84 | Bicalutamide |
| MVP_top60_DNA_seed | IGF1R | -0.259315 | COADREAD | GP5d | GSK1904529A | MVP_top60_DNA+RNA_seed | GSK3B | -0.790664 | COADREAD | LoVo | AZD7969 |
| MVP_top60_DNA_seed | PIK3CD | -0.258431 | COADREAD | MDST8 | Idelalisib | MVP_top60_DNA+RNA_seed | ATM | -0.790348 | COADREAD | HCT-116 | torin2 |
| MVP_top60_DNA_seed | ERBB3 | -0.257199 | COADREAD | MDST8 | AZD8931 | MVP_top60_DNA+RNA_seed | ATM | -0.790196 | COADREAD | RKO | KU-60019 |
| MVP_top60_DNA_seed | CDK2 | -0.256615 | COADREAD | LS-1034 | Seliciclib | MVP_top60_DNA+RNA_seed | HSP90AA1 | -0.786359 | COADREAD | LS-411N | Luminespib |
| MVP_top60_DNA_seed | PIK3CD | -0.252629 | COADREAD | COLO-320-HSR | Idelalisib | MVP_top60_DNA+RNA_seed | HSP90AB1 | -0.786359 | COADREAD | LS-411N | Luminespib |
| MVP_top60_DNA_seed | HSP90AA1 | -0.251207 | COADREAD | HCT-116 | Tanespimycin | MVP_top60_DNA+RNA_seed | HSP90AA1 | -0.782158 | COADREAD | RKO | Tanespimycin |
| MVP_top60_DNA_seed | HSP90AB1 | -0.251207 | COADREAD | HCT-116 | Tanespimycin | MVP_top60_DNA+RNA_seed | HSP90AB1 | -0.782158 | COADREAD | RKO | Tanespimycin |
| MVP_top60_DNA_seed | PIK3CB | -0.248284 | COADREAD | SW48 | AZD8186 | MVP_top60_DNA+RNA_seed | GSK3B | -0.78029 | COADREAD | CaR-1 | SB216763 |
| MVP_top60_DNA_seed | PIK3CD | -0.248284 | COADREAD | SW48 | AZD8186 | MVP_top60_DNA+RNA_seed | AKT1 | -0.779899 | COADREAD | SNU-C1 | MK-2206 |
| MVP_top60_DNA_seed | MET | -0.24611 | COADREAD | NCI-H747 | SU11274 | MVP_top60_DNA+RNA_seed | ATM | -0.776531 | COADREAD | SNU-81 | CP466722 |
| MVP_top60_DNA_seed | MAPK1 | -0.245673 | COADREAD | NCI-H716 | VX-11e | MVP_top60_DNA+RNA_seed | HSP90AA1 | -0.775295 | COADREAD | SNU-C2B | SNX-2112 |

| | | | | | | | | | | | |
|---|---|---|---|---|---|---|---|---|---|---|---|
| MVP_top60_DNA_seed | GSK3B | -0.245573 | COADREAD | HCC2998 | KIN001-042 | MVP_top60_DNA+RNA_seed | HSP90AB1 | -0.775295 | COADREAD | SNU-C2B | SNX-2112 |
| MVP_top60_DNA_seed | ATM | -0.243581 | COADREAD | LS-411N | CP466722 | MVP_top60_DNA+RNA_seed | HSP90AA1 | -0.77219 | COADREAD | DiFi | Tanespimycin |
| MVP_top60_DNA_seed | BRAF | -0.242675 | COADREAD | SNU-C2B | RAF_9304 | MVP_top60_DNA+RNA_seed | HSP90AB1 | -0.77219 | COADREAD | DiFi | Tanespimycin |
| MVP_top60_DNA_seed | AR | -0.240472 | COADREAD | NCI-H716 | Bicalutamide | MVP_top60_DNA+RNA_seed | HSP90AA1 | -0.768092 | COADREAD | NCI-H747 | Luminespib |
| MVP_top60_DNA_seed | MET | -0.238431 | COADREAD | CW-2 | JNJ38877605 | MVP_top60_DNA+RNA_seed | HSP90AB1 | -0.768092 | COADREAD | NCI-H747 | Luminespib |
| MVP_top60_DNA_seed | BRAF | -0.238157 | COADREAD | NCI-H716 | PLX-4720 | MVP_top60_DNA+RNA_seed | CDK2 | -0.758766 | COADREAD | SNU-61 | PHA-793887 |
| MVP_top60_DNA_seed | IGF1R | -0.235945 | COADREAD | RKO | GSK1904529A | MVP_top60_DNA+RNA_seed | SRC | -0.754412 | COADREAD | SNU-C2B | Saracatinib |
| MVP_top60_DNA_seed | CDK2 | -0.235573 | COADREAD | LS-411N | PHA-793887 | MVP_top60_DNA+RNA_seed | GSK3B | -0.754014 | COADREAD | HCT-116 | CHIR-99021 |
| MVP_top60_DNA_seed | AR | -0.234824 | COADREAD | LoVo | AZD3514 | MVP_top60_DNA+RNA_seed | BCL2 | -0.753552 | COADREAD | LS-123 | Obatoclax Mesylate |
| MVP_top60_DNA_seed | CDK2 | -0.233403 | COADREAD | SNU-407 | AZD5438 | MVP_top60_DNA+RNA_seed | IGF1R | -0.752856 | COADREAD | CL-11 | BMS-536924 |
| MVP_top60_DNA_seed | IGF1R | -0.230477 | COADREAD | RKO | BMS-754807 | MVP_top60_DNA+RNA_seed | GSK3B | -0.752029 | COADREAD | SNU-C5 | AZD7969 |
| MVP_top60_DNA_seed | PTK2 | -0.229981 | COADREAD | COLO-320-HSR | PF-562271 | MVP_top60_DNA+RNA_seed | IGF1R | -0.747931 | COADREAD | LS-180 | BMS-754807 |
| MVP_top60_DNA_seed | PIK3CD | -0.228988 | COADREAD | CaR-1 | IC-87114 | MVP_top60_DNA+RNA_seed | SRC | -0.746474 | COADREAD | LoVo | Bosutinib |
| MVP_top60_DNA_seed | ERBB4 | -0.22807 | COADREAD | SNU-81 | AST-1306 | MVP_top60_DNA+RNA_seed | BCL2 | -0.743605 | COADREAD | HCT-116 | Obatoclax Mesylate |
| MVP_top60_DNA_seed | GSK3B | -0.22762 | COADREAD | NCI-H747 | KIN001-042 | MVP_top60_DNA+RNA_seed | SRC | -0.739859 | COADREAD | MDST8 | Bosutinib |
| MVP_top60_DNA_seed | GSK3B | -0.225722 | COADREAD | COLO-205 | KIN001-042 | MVP_top60_DNA+RNA_seed | AKT1 | -0.736683 | COADREAD | SNU-407 | GSK690693 |
| MVP_top60_DNA_seed | AKT1 | -0.222594 | COADREAD | HCT-116 | GSK690693 | MVP_top60_DNA+RNA_seed | FYN | -0.735427 | COADREAD | NCI-H747 | A-770041 |
| MVP_top60_DNA_seed | AR | -0.220921 | COADREAD | COLO-320-HSR | Bicalutamide | MVP_top60_DNA+RNA_seed | SRC | -0.731384 | COADREAD | KM12 | Saracatinib |
| MVP_top60_DNA_seed | BRAF | -0.220269 | COADREAD | SW837 | RAF_9304 | MVP_top60_DNA+RNA_seed | CDK2 | -0.730998 | COADREAD | SW620 | AZD5438 |
| MVP_top60_DNA_seed | MAPK14 | -0.219698 | COADREAD | GP5d | TAK-715 | MVP_top60_DNA+RNA_seed | CDK2 | -0.730341 | COADREAD | RKO | PHA-793887 |
| MVP_top60_DNA_seed | PIK3CD | -0.218565 | COADREAD | SNU-1040 | Idelalisib | MVP_top60_DNA+RNA_seed | PIK3CA | -0.72489 | COADREAD | COLO-320-HSR | AZD8835 |
| MVP_top60_DNA_seed | IGF1R | -0.216538 | COADREAD | CaR-1 | GSK1904529A | MVP_top60_DNA+RNA_seed | PIK3CD | -0.72489 | COADREAD | COLO-320-HSR | AZD8835 |
| MVP_top60_DNA_seed | AKT1 | -0.214771 | COADREAD | NCI-H747 | AKT inhibitor VIII | MVP_top60_DNA+RNA_seed | ERBB4 | -0.722778 | COADREAD | SW1116 | AST-1306 |
| MVP_top60_DNA_seed | ERBB4 | -0.214317 | COADREAD | SNU-61 | AST-1306 | MVP_top60_DNA+RNA_seed | GSK3B | -0.721675 | COADREAD | SW48 | CHIR-99021 |
| MVP_top60_DNA_seed | SRC | -0.210596 | COADREAD | GP5d | NVP-BHG712 | MVP_top60_DNA+RNA_seed | CDK2 | -0.720069 | COADREAD | SNU-81 | PHA-793887 |
| MVP_top60_DNA_seed | GSK3B | -0.206601 | COADREAD | LS-1034 | SB216763 | MVP_top60_DNA+RNA_seed | GSK3B | -0.716443 | COADREAD | LS-513 | CHIR-99021 |
| MVP_top60_DNA_seed | SRC | -0.206511 | COADREAD | LoVo | NVP-BHG712 | MVP_top60_DNA+RNA_seed | MET | -0.716231 | COADREAD | SW48 | SU11274 |
| MVP_top60_DNA_seed | ERBB3 | -0.205692 | COADREAD | SNU-81 | AZD8931 | MVP_top60_DNA+RNA_seed | PIK3CA | -0.715688 | COADREAD | RKO | PI3Ka_4409 |
| MVP_top60_DNA_seed | IGF1R | -0.203811 | COADREAD | LS-180 | Linsitinib | MVP_top60_DNA+RNA_seed | PIK3CD | -0.715688 | COADREAD | RKO | PI3Ka_4409 |
| MVP_top60_DNA_seed | AR | -0.201737 | COADREAD | RKO | Bicalutamide | MVP_top60_DNA+RNA_seed | SRC | -0.713943 | COADREAD | SW48 | Bosutinib |
| MVP_top60_DNA_seed | ERBB3 | -0.201522 | COADREAD | SNU-C2B | AZD8931 | MVP_top60_DNA+RNA_seed | BCL2 | -0.712483 | COADREAD | SNU-C2B | TW 37 |
| MVP_top60_DNA_seed | AKT1 | -0.20144 | COADREAD | CW-2 | AKT inhibitor VIII | MVP_top60_DNA+RNA_seed | AKT1 | -0.709351 | COADREAD | SNU-407 | AKT inhibitor VIII |
| MVP_top60_DNA_seed | IGF1R | -0.201062 | COADREAD | LS-411N | BMS-536924 | MVP_top60_DNA+RNA_seed | ATM | -0.709043 | COADREAD | HCT-116 | CP466722 |
| MVP_top60_DNA_seed | MAPK1 | -0.20072 | COADREAD | SW1417 | FR-180204 | MVP_top60_DNA+RNA_seed | MET | -0.706053 | COADREAD | SNU-1040 | JNJ38877605 |
| MVP_top60_DNA_seed | ERBB4 | -0.199301 | COADREAD | C2BBe1 | AST-1306 | MVP_top60_DNA+RNA_seed | IGF1R | -0.705988 | COADREAD | HT55 | BMS-536924 |
| MVP_top60_DNA_seed | SRC | -0.196719 | COADREAD | NCI-H716 | Bosutinib | MVP_top60_DNA+RNA_seed | ATM | -0.698908 | COADREAD | LS-180 | CP466722 |
| MVP_top60_DNA_seed | PIK3CD | -0.19634 | COADREAD | SNU-175 | IC-87114 | MVP_top60_DNA+RNA_seed | PIK3CA | -0.694177 | COADREAD | COLO-205 | PI3Ka_4409 |
| MVP_top60_DNA_seed | HSP90AA1 | -0.195888 | COADREAD | LS-513 | Tanespimycin | MVP_top60_DNA+RNA_seed | PIK3CD | -0.694177 | COADREAD | COLO-205 | PI3Ka_4409 |
| MVP_top60_DNA_seed | HSP90AB1 | -0.195888 | COADREAD | LS-513 | Tanespimycin | MVP_top60_DNA+RNA_seed | HSP90AA1 | -0.692597 | COADREAD | COLO-320-HSR | CCT-018159 |
| MVP_top60_DNA_seed | BRAF | -0.191986 | COADREAD | RCM-1 | Dabrafenib | MVP_top60_DNA+RNA_seed | HSP90AB1 | -0.692597 | COADREAD | COLO-320-HSR | CCT-018159 |
| MVP_top60_DNA_seed | AR | -0.191681 | COADREAD | CCK-81 | Bicalutamide | MVP_top60_DNA+RNA_seed | HSP90AA1 | -0.691748 | COADREAD | T84 | Luminespib |

| | | | | | | | | | | | |
|---|---|---|---|---|---|---|---|---|---|---|---|
| MVP_top60_DNA_seed | PIK3CA | -0.191277 | COADREAD | LS-180 | AZD8835 | MVP_top60_DNA+RNA_seed | HSP90AB1 | -0.691748 | COADREAD | T84 | Luminespib |
| MVP_top60_DNA_seed | PIK3CD | -0.191277 | COADREAD | LS-180 | AZD8835 | MVP_top60_DNA+RNA_seed | AKT1 | -0.689762 | COADREAD | GP5d | AKT inhibitor VIII |
| MVP_top60_DNA_seed | SRC | -0.190201 | COADREAD | LS-123 | Saracatinib | MVP_top60_DNA+RNA_seed | HSP90AA1 | -0.685283 | COADREAD | MDST8 | CCT-018159 |
| MVP_top60_DNA_seed | GSK3B | -0.190068 | COADREAD | HCT-15 | AZD7969 | MVP_top60_DNA+RNA_seed | HSP90AB1 | -0.685283 | COADREAD | MDST8 | CCT-018159 |
| MVP_top60_DNA_seed | ESR1 | -0.189055 | COADREAD | SK-CO-1 | Fulvestrant | MVP_top60_DNA+RNA_seed | IGF1R | -0.683448 | COADREAD | LS-513 | BMS-754807 |
| MVP_top60_DNA_seed | GSK3B | -0.187356 | COADREAD | NCI-H747 | CHIR-99021 | MVP_top60_DNA+RNA_seed | HSP90AA1 | -0.683062 | COADREAD | GP5d | Tanespimycin |
| MVP_top60_DNA_seed | HSP90AA1 | -0.186852 | COADREAD | COLO-320-HSR | Luminespib | MVP_top60_DNA+RNA_seed | HSP90AB1 | -0.683062 | COADREAD | GP5d | Tanespimycin |
| MVP_top60_DNA_seed | HSP90AB1 | -0.186852 | COADREAD | COLO-320-HSR | Luminespib | MVP_top60_DNA+RNA_seed | BCL2 | -0.682206 | COADREAD | MDST8 | TW 37 |
| MVP_top60_DNA_seed | MAPK14 | -0.184453 | COADREAD | SNU-407 | TAK-715 | MVP_top60_DNA+RNA_seed | AKT1 | -0.679649 | COADREAD | GP5d | AKT inhibitor VIII |
| MVP_top60_DNA_seed | HSP90AA1 | -0.183769 | COADREAD | LS-1034 | Luminespib | MVP_top60_DNA+RNA_seed | IGF1R | -0.676127 | COADREAD | SNU-C1 | BMS-536924 |
| MVP_top60_DNA_seed | HSP90AB1 | -0.183769 | COADREAD | LS-1034 | Luminespib | MVP_top60_DNA+RNA_seed | ATM | -0.673183 | COADREAD | SNU-81 | KU-55933 |
| MVP_top60_DNA_seed | SRC | -0.183704 | COADREAD | NCI-H716 | NVP-BHG712 | MVP_top60_DNA+RNA_seed | AKT1 | -0.67277 | COADREAD | LS-513 | AKT inhibitor VIII |
| MVP_top60_DNA_seed | PIK3CA | -0.182657 | COADREAD | LoVo | PI3Ka_4409 | MVP_top60_DNA+RNA_seed | CDK2 | -0.670997 | COADREAD | COLO-205 | PHA-793887 |
| MVP_top60_DNA_seed | PIK3CD | -0.182657 | COADREAD | LoVo | PI3Ka_4409 | MVP_top60_DNA+RNA_seed | ERBB4 | -0.670845 | COADREAD | SNU-C2B | PF-00299804 |
| MVP_top60_DNA_seed | AR | -0.182442 | COADREAD | CCK-81 | AZD3514 | MVP_top60_DNA+RNA_seed | BCL2 | -0.659074 | COADREAD | LoVo | Obatoclax Mesylate |
| MVP_top60_DNA_seed | ATM | -0.182176 | COADREAD | SW837 | CP466722 | MVP_top60_DNA+RNA_seed | CDK2 | -0.657704 | COADREAD | SNU-1040 | PHA-793887 |
| MVP_top60_DNA_seed | ERBB4 | -0.181255 | COADREAD | SNU-81 | CI-1033 | MVP_top60_DNA+RNA_seed | SRC | -0.650585 | COADREAD | SW1417 | NVP-BHG712 |
| MVP_top60_DNA_seed | AKT1 | -0.181097 | COADREAD | SNU-407 | BAY AKT1 | MVP_top60_DNA+RNA_seed | HSP90AA1 | -0.64811 | COADREAD | GP5d | SNX-2112 |
| MVP_top60_DNA_seed | SRC | -0.177613 | COADREAD | HCC2998 | NVP-BHG712 | MVP_top60_DNA+RNA_seed | HSP90AB1 | -0.64811 | COADREAD | GP5d | SNX-2112 |
| MVP_top60_DNA_seed | IGF1R | -0.176039 | COADREAD | HCC2998 | BMS-536924 | MVP_top60_DNA+RNA_seed | SRC | -0.646625 | COADREAD | NCI-H508 | Bosutinib |
| MVP_top60_DNA_seed | SRC | -0.175302 | COADREAD | C2BBe1 | NVP-BHG712 | MVP_top60_DNA+RNA_seed | SRC | -0.642285 | COADREAD | GP5d | Bosutinib |
| MVP_top60_DNA_seed | ESR1 | -0.173084 | COADREAD | HCT-15 | Fulvestrant | MVP_top60_DNA+RNA_seed | HSP90AA1 | -0.640286 | COADREAD | CCK-81 | SNX-2112 |
| MVP_top60_DNA_seed | ATM | -0.173003 | COADREAD | LoVo | torin2 | MVP_top60_DNA+RNA_seed | HSP90AB1 | -0.640286 | COADREAD | CCK-81 | SNX-2112 |
| MVP_top60_DNA_seed | PIK3CB | -0.171744 | COADREAD | KM12 | TGX221 | MVP_top60_DNA+RNA_seed | ERBB3 | -0.637258 | COADREAD | CCK-81 | AZD8931 |
| MVP_top60_DNA_seed | PIK3CA | -0.168674 | COADREAD | HCT-15 | AZD8835 | MVP_top60_DNA+RNA_seed | ATM | -0.635834 | COADREAD | CW-2 | KU-60019 |
| MVP_top60_DNA_seed | PIK3CD | -0.168674 | COADREAD | HCT-15 | AZD8835 | MVP_top60_DNA+RNA_seed | FYN | -0.635272 | COADREAD | RKO | A-770041 |
| MVP_top60_DNA_seed | AKT1 | -0.168247 | COADREAD | SW1463 | MK-2206 | MVP_top60_DNA+RNA_seed | AR | -0.635145 | COADREAD | LS-180 | AZD3514 |
| MVP_top60_DNA_seed | IGF1R | -0.164215 | COADREAD | COLO-678 | BMS-754807 | MVP_top60_DNA+RNA_seed | SRC | -0.635092 | COADREAD | LS-1034 | NVP-BHG712 |
| MVP_top60_DNA_seed | IGF1R | -0.164035 | COADREAD | LoVo | BMS-754807 | MVP_top60_DNA+RNA_seed | SRC | -0.634814 | COADREAD | CCK-81 | Bosutinib |
| MVP_top60_DNA_seed | MET | -0.162049 | COADREAD | SW837 | JNJ38877605 | MVP_top60_DNA+RNA_seed | AKT1 | -0.629589 | COADREAD | COLO-320-HSR | AKT inhibitor VIII |
| MVP_top60_DNA_seed | AKT1 | -0.160481 | COADREAD | NCI-H747 | BAY AKT1 | MVP_top60_DNA+RNA_seed | CDK2 | -0.627736 | COADREAD | LS-513 | Seliciclib |
| MVP_top60_DNA_seed | MET | -0.160139 | COADREAD | CaR-1 | JNJ38877605 | MVP_top60_DNA+RNA_seed | PIK3CA | -0.627517 | COADREAD | DiFi | AZD8835 |
| MVP_top60_DNA_seed | CDK2 | -0.155209 | COADREAD | LS-123 | PHA-793887 | MVP_top60_DNA+RNA_seed | PIK3CD | -0.627517 | COADREAD | DiFi | AZD8835 |
| MVP_top60_DNA_seed | BRAF | -0.154587 | COADREAD | LS-180 | RAF_9304 | MVP_top60_DNA+RNA_seed | GSK3B | -0.626934 | COADREAD | GP5d | SB216763 |
| MVP_top60_DNA_seed | MET | -0.153229 | COADREAD | RKO | Crizotinib | MVP_top60_DNA+RNA_seed | PIK3CA | -0.624684 | COADREAD | SNU-C5 | AZD8835 |
| MVP_top60_DNA_seed | MET | -0.152884 | COADREAD | C2BBe1 | PHA-665752 | MVP_top60_DNA+RNA_seed | PIK3CD | -0.624684 | COADREAD | SNU-C5 | AZD8835 |
| MVP_top60_DNA_seed | PIK3CD | -0.152481 | COADREAD | GP5d | Idelalisib | MVP_top60_DNA+RNA_seed | BCL2 | -0.624281 | COADREAD | HCT-15 | Obatoclax Mesylate |
| MVP_top60_DNA_seed | AR | -0.152241 | COADREAD | COLO-678 | AZD3514 | MVP_top60_DNA+RNA_seed | IGF1R | -0.622832 | COADREAD | SNU-C2B | BMS-536924 |
| MVP_top60_DNA_seed | ATM | -0.151994 | COADREAD | HCT-15 | KU-55933 | MVP_top60_DNA+RNA_seed | ATM | -0.61809 | COADREAD | NCI-H747 | CP466722 |
| MVP_top60_DNA_seed | AKT1 | -0.14922 | COADREAD | GP5d | MK-2206 | MVP_top60_DNA+RNA_seed | AKT1 | -0.618028 | COADREAD | CCK-81 | MK-2206 |
| MVP_top60_DNA_seed | BRAF | -0.147022 | COADREAD | SW1417 | RAF_9304 | MVP_top60_DNA+RNA_seed | ERBB4 | -0.617328 | COADREAD | SNU-175 | AST-1306 |

| | | | | | | | | | | | |
|---|---|---|---|---|---|---|---|---|---|---|---|
| MVP_top60_DNA_seed | ERBB4 | -0.144805 | COADREAD | LoVo | AST-1306 | MVP_top60_DNA+RNA_seed | CDK2 | -0.613517 | COADREAD | SNU-C5 | AZD5438 |
| MVP_top60_DNA_seed | AR | -0.143557 | COADREAD | COLO-205 | AZD3514 | MVP_top60_DNA+RNA_seed | ERBB3 | -0.613457 | COADREAD | HT55 | AZD8931 |
| MVP_top60_DNA_seed | ERBB4 | -0.142006 | COADREAD | SNU-61 | CI-1033 | MVP_top60_DNA+RNA_seed | PIK3CA | -0.61323 | COADREAD | SK-CO-1 | PI3Ka_4409 |
| MVP_top60_DNA_seed | CDK2 | -0.141569 | COADREAD | HCC2998 | AZD5438 | MVP_top60_DNA+RNA_seed | PIK3CD | -0.61323 | COADREAD | SK-CO-1 | PI3Ka_4409 |
| MVP_top60_DNA_seed | HSP90AA1 | -0.14001 | COADREAD | MDST8 | Tanespimycin | MVP_top60_DNA+RNA_seed | AKT1 | -0.613133 | COADREAD | KM12 | A-443654 |
| MVP_top60_DNA_seed | HSP90AB1 | -0.14001 | COADREAD | MDST8 | Tanespimycin | MVP_top60_DNA+RNA_seed | AKT1 | -0.612438 | COADREAD | HCT-15 | MK-2206 |
| MVP_top60_DNA_seed | ERBB4 | -0.138818 | COADREAD | SNU-1040 | PF-00299804 | MVP_top60_DNA+RNA_seed | HSP90AA1 | -0.611035 | COADREAD | NCI-H716 | SNX-2112 |
| MVP_top60_DNA_seed | GSK3B | -0.130824 | COADREAD | KM12 | SB216763 | MVP_top60_DNA+RNA_seed | HSP90AB1 | -0.611035 | COADREAD | NCI-H716 | SNX-2112 |
| MVP_top60_DNA_seed | PIK3CA | -0.130352 | COADREAD | T84 | PI3Ka_4409 | MVP_top60_DNA+RNA_seed | AKT1 | -0.606411 | COADREAD | NCI-H747 | MK-2206 |
| MVP_top60_DNA_seed | PIK3CD | -0.130352 | COADREAD | T84 | PI3Ka_4409 | MVP_top60_DNA+RNA_seed | SRC | -0.601542 | COADREAD | LS-411N | NVP-BHG712 |
| MVP_top60_DNA_seed | GSK3B | -0.126797 | COADREAD | HT-29 | CHIR-99021 | MVP_top60_DNA+RNA_seed | AKT1 | -0.598504 | COADREAD | LS-411N | AKT inhibitor VIII |
| MVP_top60_DNA_seed | ERBB4 | -0.125793 | COADREAD | GP5d | PF-00299804 | MVP_top60_DNA+RNA_seed | PIK3CD | -0.598091 | COADREAD | LS-123 | Idelalisib |
| MVP_top60_DNA_seed | BRAF | -0.125091 | COADREAD | LS-180 | PLX-4720 | MVP_top60_DNA+RNA_seed | MET | -0.597897 | COADREAD | GP5d | JNJ38877605 |
| MVP_top60_DNA_seed | ERBB4 | -0.122594 | COADREAD | SW1116 | PF-00299804 | MVP_top60_DNA+RNA_seed | HSP90AA1 | -0.596686 | COADREAD | SW48 | Luminespib |
| MVP_top60_DNA_seed | GSK3B | -0.122351 | COADREAD | SW48 | KIN001-042 | MVP_top60_DNA+RNA_seed | HSP90AB1 | -0.596686 | COADREAD | SW48 | Luminespib |
| MVP_top60_DNA_seed | PIK3CD | -0.121155 | COADREAD | SNU-407 | Idelalisib | MVP_top60_DNA+RNA_seed | SRC | -0.59592 | COADREAD | HCT-116 | HG-6-71-01 |
| MVP_top60_DNA_seed | AKT1 | -0.119968 | COADREAD | SW620 | AKT inhibitor VIII | MVP_top60_DNA+RNA_seed | SRC | -0.595253 | COADREAD | COLO-205 | NVP-BHG712 |
| MVP_top60_DNA_seed | CDK2 | -0.119071 | COADREAD | SNU-407 | PHA-793887 | MVP_top60_DNA+RNA_seed | ERBB4 | -0.593571 | COADREAD | NCI-H747 | AST-1306 |
| MVP_top60_DNA_seed | PIK3CA | -0.118668 | COADREAD | NCI-H508 | AZD8835 | MVP_top60_DNA+RNA_seed | ATM | -0.587908 | COADREAD | CCK-81 | KU-60019 |
| MVP_top60_DNA_seed | PIK3CD | -0.118668 | COADREAD | NCI-H508 | AZD8835 | MVP_top60_DNA+RNA_seed | PIK3CA | -0.586716 | COADREAD | SNU-C5 | PI3Ka_4409 |
| MVP_top60_DNA_seed | PIK3CA | -0.113655 | COADREAD | HT-115 | PI3Ka_4409 | MVP_top60_DNA+RNA_seed | PIK3CD | -0.586716 | COADREAD | SNU-C5 | PI3Ka_4409 |
| MVP_top60_DNA_seed | PIK3CD | -0.113655 | COADREAD | HT-115 | PI3Ka_4409 | MVP_top60_DNA+RNA_seed | MET | -0.585958 | COADREAD | SNU-61 | JNJ38877605 |
| MVP_top60_DNA_seed | BRAF | -0.109933 | COADREAD | SNU-C5 | SB590885 | MVP_top60_DNA+RNA_seed | ATM | -0.584066 | COADREAD | COLO-320-HSR | KU-60019 |
| MVP_top60_DNA_seed | SRC | -0.109291 | COADREAD | SW948 | NVP-BHG712 | MVP_top60_DNA+RNA_seed | GSK3B | -0.582429 | COADREAD | NCI-H747 | CHIR-99021 |
| MVP_top60_DNA_seed | ERBB4 | -0.108331 | COADREAD | SNU-C2B | AST-1306 | MVP_top60_DNA+RNA_seed | PIK3CB | -0.57936 | COADREAD | LS-123 | AZD8186 |
| MVP_top60_DNA_seed | PTK2 | -0.10814 | COADREAD | LS-180 | PF-562271 | MVP_top60_DNA+RNA_seed | PIK3CD | -0.57936 | COADREAD | LS-123 | AZD8186 |
| MVP_top60_DNA_seed | BRAF | -0.106926 | COADREAD | C2BBe1 | HG6-64-1 | MVP_top60_DNA+RNA_seed | MET | -0.579084 | COADREAD | GP5d | SU11274 |
| MVP_top60_DNA_seed | GSK3B | -0.105936 | COADREAD | LS-513 | AZD7969 | MVP_top60_DNA+RNA_seed | HSP90AA1 | -0.575255 | COADREAD | LS-1034 | SNX-2112 |
| MVP_top60_DNA_seed | HSP90AA1 | -0.103508 | COADREAD | SW837 | SNX-2112 | MVP_top60_DNA+RNA_seed | HSP90AB1 | -0.575255 | COADREAD | LS-1034 | SNX-2112 |
| MVP_top60_DNA_seed | HSP90AB1 | -0.103508 | COADREAD | SW837 | SNX-2112 | MVP_top60_DNA+RNA_seed | PIK3CA | -0.572921 | COADREAD | SNU-C1 | PI3Ka_4409 |
| MVP_top60_DNA_seed | GSK3B | -0.10253 | COADREAD | SNU-C2B | AZD7969 | MVP_top60_DNA+RNA_seed | PIK3CD | -0.572921 | COADREAD | SNU-C1 | PI3Ka_4409 |
| MVP_top60_DNA_seed | PIK3CA | -0.099566 | COADREAD | LS-513 | AZD8835 | MVP_top60_DNA+RNA_seed | SRC | -0.572499 | COADREAD | C2BBe1 | Bosutinib |
| MVP_top60_DNA_seed | PIK3CD | -0.099566 | COADREAD | LS-513 | AZD8835 | MVP_top60_DNA+RNA_seed | PTK2 | -0.572351 | COADREAD | HCC2998 | PF-562271 |
| MVP_top60_DNA_seed | ERBB4 | -0.098547 | COADREAD | RKO | PF-00299804 | MVP_top60_DNA+RNA_seed | IGF1R | -0.567744 | COADREAD | C2BBe1 | Linsitinib |
| MVP_top60_DNA_seed | IGF1R | -0.098332 | COADREAD | CCK-81 | BMS-536924 | MVP_top60_DNA+RNA_seed | AKT1 | -0.56542 | COADREAD | SNU-1040 | AKT inhibitor VIII |
| MVP_top60_DNA_seed | ERBB4 | -0.096883 | COADREAD | LS-180 | CI-1033 | MVP_top60_DNA+RNA_seed | AKT1 | -0.56514 | COADREAD | SNU-C2B | A-443654 |
| MVP_top60_DNA_seed | IGF1R | -0.094957 | COADREAD | HCT-116 | GSK1904529A | MVP_top60_DNA+RNA_seed | AKT1 | -0.564052 | COADREAD | SNU-C5 | MK-2206 |
| MVP_top60_DNA_seed | CDK2 | -0.094768 | COADREAD | SNU-C2B | AZD5438 | MVP_top60_DNA+RNA_seed | BCL2 | -0.560745 | COADREAD | SW48 | TW 37 |
| MVP_top60_DNA_seed | MET | -0.094336 | COADREAD | COLO-205 | JNJ38877605 | MVP_top60_DNA+RNA_seed | SRC | -0.559757 | COADREAD | SW620 | NVP-BHG712 |
| MVP_top60_DNA_seed | ESR1 | -0.093395 | COADREAD | LS-1034 | Fulvestrant | MVP_top60_DNA+RNA_seed | ERBB3 | -0.558975 | COADREAD | NCI-H508 | AZD8931 |
| MVP_top60_DNA_seed | BRAF | -0.092861 | COADREAD | CL-11 | Dabrafenib | MVP_top60_DNA+RNA_seed | CDK2 | -0.55705 | COADREAD | LS-513 | AZD5438 |

| | | | | | | | | | | | |
|---|---|---|---|---|---|---|---|---|---|---|---|
| MVP_top60_DNA_seed | PIK3CA | -0.092356 | COADREAD | HT-115 | AZD8835 | MVP_top60_DNA+RNA_seed | PIK3CD | -0.555741 | COADREAD | GP5d | IC-87114 |
| MVP_top60_DNA_seed | PIK3CD | -0.092356 | COADREAD | HT-115 | AZD8835 | MVP_top60_DNA+RNA_seed | MET | -0.548876 | COADREAD | NCI-H747 | JNJ38877605 |
| MVP_top60_DNA_seed | IGF1R | -0.091947 | COADREAD | SW948 | BMS-754807 | MVP_top60_DNA+RNA_seed | PIK3CA | -0.542571 | COADREAD | LoVo | AZD8835 |
| MVP_top60_DNA_seed | BRAF | -0.088897 | COADREAD | SK-CO-1 | Dabrafenib | MVP_top60_DNA+RNA_seed | PIK3CD | -0.542571 | COADREAD | LoVo | AZD8835 |
| MVP_top60_DNA_seed | ERBB4 | -0.088563 | COADREAD | SNU-1040 | AST-1306 | MVP_top60_DNA+RNA_seed | AR | -0.540983 | COADREAD | T84 | Bicalutamide |
| MVP_top60_DNA_seed | GSK3B | -0.086921 | COADREAD | HT55 | KIN001-042 | MVP_top60_DNA+RNA_seed | PIK3CA | -0.539297 | COADREAD | NCI-H747 | PI3Ka_4409 |
| MVP_top60_DNA_seed | PIK3CA | -0.086547 | COADREAD | LS-180 | PI3Ka_4409 | MVP_top60_DNA+RNA_seed | PIK3CD | -0.539297 | COADREAD | NCI-H747 | PI3Ka_4409 |
| MVP_top60_DNA_seed | PIK3CD | -0.086547 | COADREAD | LS-180 | PI3Ka_4409 | MVP_top60_DNA+RNA_seed | MET | -0.537811 | COADREAD | HCT-116 | SU11274 |
| MVP_top60_DNA_seed | AKT1 | -0.086122 | COADREAD | SNU-C1 | AKT inhibitor VIII | MVP_top60_DNA+RNA_seed | FYN | -0.537777 | COADREAD | C2BBe1 | A-770041 |
| MVP_top60_DNA_seed | IGF1R | -0.085648 | COADREAD | CCK-81 | BMS-754807 | MVP_top60_DNA+RNA_seed | PIK3CB | -0.536913 | COADREAD | HCT-116 | AZD8186 |
| MVP_top60_DNA_seed | GSK3B | -0.081325 | COADREAD | CCK-81 | CHIR-99021 | MVP_top60_DNA+RNA_seed | PIK3CD | -0.536913 | COADREAD | HCT-116 | AZD8186 |
| MVP_top60_DNA_seed | IGF1R | -0.080503 | COADREAD | SW620 | Linsitinib | MVP_top60_DNA+RNA_seed | PIK3CA | -0.534642 | COADREAD | HT-29 | AZD8835 |
| MVP_top60_DNA_seed | MET | -0.078778 | COADREAD | RKO | AZD6094 | MVP_top60_DNA+RNA_seed | PIK3CD | -0.534642 | COADREAD | HT-29 | AZD8835 |
| MVP_top60_DNA_seed | ERBB3 | -0.078348 | COADREAD | HCC2998 | AZD8931 | MVP_top60_DNA+RNA_seed | HSP90AA1 | -0.534374 | COADREAD | HCC2998 | Luminespib |
| MVP_top60_DNA_seed | MET | -0.077351 | COADREAD | CW-2 | SU11274 | MVP_top60_DNA+RNA_seed | HSP90AB1 | -0.534374 | COADREAD | HCC2998 | Luminespib |
| MVP_top60_DNA_seed | PIK3CA | -0.077205 | COADREAD | CCK-81 | PI3Ka_4409 | MVP_top60_DNA+RNA_seed | HSP90AA1 | -0.534314 | COADREAD | SNU-61 | SNX-2112 |
| MVP_top60_DNA_seed | PIK3CD | -0.077205 | COADREAD | CCK-81 | PI3Ka_4409 | MVP_top60_DNA+RNA_seed | HSP90AB1 | -0.534314 | COADREAD | SNU-61 | SNX-2112 |
| MVP_top60_DNA_seed | HSP90AA1 | -0.077139 | COADREAD | HCT-116 | Elesclomol | MVP_top60_DNA+RNA_seed | HSP90AA1 | -0.533815 | COADREAD | LoVo | Luminespib |
| MVP_top60_DNA_seed | HSP90AB1 | -0.077139 | COADREAD | HCT-116 | Elesclomol | MVP_top60_DNA+RNA_seed | HSP90AB1 | -0.533815 | COADREAD | LoVo | Luminespib |
| MVP_top60_DNA_seed | BRAF | -0.076556 | COADREAD | RKO | HG6-64-1 | MVP_top60_DNA+RNA_seed | AR | -0.528166 | COADREAD | SNU-C2B | Bicalutamide |
| MVP_top60_DNA_seed | HSP90AA1 | -0.072957 | COADREAD | LS-180 | Luminespib | MVP_top60_DNA+RNA_seed | HSP90AA1 | -0.525717 | COADREAD | SNU-407 | SNX-2112 |
| MVP_top60_DNA_seed | HSP90AB1 | -0.072957 | COADREAD | LS-180 | Luminespib | MVP_top60_DNA+RNA_seed | HSP90AB1 | -0.525717 | COADREAD | SNU-407 | SNX-2112 |
| MVP_top60_DNA_seed | AR | -0.071808 | COADREAD | HT55 | Bicalutamide | MVP_top60_DNA+RNA_seed | PIK3CA | -0.521684 | COADREAD | NCI-H716 | PI3Ka_4409 |
| MVP_top60_DNA_seed | MET | -0.071523 | COADREAD | RKO | SU11274 | MVP_top60_DNA+RNA_seed | PIK3CD | -0.521684 | COADREAD | NCI-H716 | PI3Ka_4409 |
| MVP_top60_DNA_seed | PIK3CB | -0.071368 | COADREAD | NCI-H747 | AZD6482 | MVP_top60_DNA+RNA_seed | HSP90AA1 | -0.520826 | COADREAD | RKO | CCT-018159 |
| MVP_top60_DNA_seed | IGF1R | -0.071209 | COADREAD | SNU-C2B | BMS-754807 | MVP_top60_DNA+RNA_seed | HSP90AB1 | -0.520826 | COADREAD | RKO | CCT-018159 |
| MVP_top60_DNA_seed | AKT1 | -0.069555 | COADREAD | HCC2998 | MK-2206 | MVP_top60_DNA+RNA_seed | ATM | -0.519922 | COADREAD | CL-11 | KU-55933 |
| MVP_top60_DNA_seed | HSP90AA1 | -0.069286 | COADREAD | LS-411N | Tanespimycin | MVP_top60_DNA+RNA_seed | AR | -0.518724 | COADREAD | GP5d | Bicalutamide |
| MVP_top60_DNA_seed | HSP90AB1 | -0.069286 | COADREAD | LS-411N | Tanespimycin | MVP_top60_DNA+RNA_seed | ATM | -0.518202 | COADREAD | SK-CO-1 | KU-55933 |
| MVP_top60_DNA_seed | HSP90AA1 | -0.068131 | COADREAD | HCT-15 | Tanespimycin | MVP_top60_DNA+RNA_seed | ERBB3 | -0.515605 | COADREAD | SW1116 | AZD8931 |
| MVP_top60_DNA_seed | HSP90AB1 | -0.068131 | COADREAD | HCT-15 | Tanespimycin | MVP_top60_DNA+RNA_seed | HSP90AA1 | -0.515085 | COADREAD | RKO | Elesclomol |
| MVP_top60_DNA_seed | AKT1 | -0.06744 | COADREAD | RKO | AKT inhibitor VIII | MVP_top60_DNA+RNA_seed | HSP90AB1 | -0.515085 | COADREAD | RKO | Elesclomol |
| MVP_top60_DNA_seed | ATM | -0.067091 | COADREAD | SNU-407 | CP466722 | MVP_top60_DNA+RNA_seed | HSP90AA1 | -0.51296 | COADREAD | KM12 | Luminespib |
| MVP_top60_DNA_seed | BRAF | -0.064654 | COADREAD | HT-29 | HG6-64-1 | MVP_top60_DNA+RNA_seed | HSP90AB1 | -0.51296 | COADREAD | KM12 | Luminespib |
| MVP_top60_DNA_seed | GSK3B | -0.063952 | COADREAD | CaR-1 | KIN001-042 | MVP_top60_DNA+RNA_seed | ATM | -0.511225 | COADREAD | T84 | CP466722 |
| MVP_top60_DNA_seed | BRAF | -0.063906 | COADREAD | HT55 | RAF_9304 | MVP_top60_DNA+RNA_seed | ATM | -0.510789 | COADREAD | SNU-1040 | CP466722 |
| MVP_top60_DNA_seed | GSK3B | -0.063397 | COADREAD | LS-123 | SB216763 | MVP_top60_DNA+RNA_seed | IGF1R | -0.507326 | COADREAD | SW620 | BMS-754807 |
| MVP_top60_DNA_seed | ERBB4 | -0.062005 | COADREAD | SNU-1040 | CI-1033 | MVP_top60_DNA+RNA_seed | MET | -0.507046 | COADREAD | SW48 | JNJ38877605 |
| MVP_top60_DNA_seed | ESR1 | -0.059254 | COADREAD | COLO-320-HSR | Tamoxifen | MVP_top60_DNA+RNA_seed | MET | -0.504784 | COADREAD | CCK-81 | SU11274 |
| MVP_top60_DNA_seed | ATM | -0.057984 | COADREAD | SW48 | torin2 | MVP_top60_DNA+RNA_seed | GSK3B | -0.502184 | COADREAD | CCK-81 | SB216763 |
| MVP_top60_DNA_seed | GSK3B | -0.057516 | COADREAD | DiFi | CHIR-99021 | MVP_top60_DNA+RNA_seed | CDK2 | -0.50212 | COADREAD | SNU-C2B | PHA-793887 |

| | | | | | | | | | | | |
|---|---|---|---|---|---|---|---|---|---|---|---|
| MVP_top60_DNA_seed | ERBB4 | -0.057181 | COADREAD | HCT-116 | PF-00299804 | MVP_top60_DNA+RNA_seed | PIK3CB | -0.501409 | COADREAD | NCI-H716 | AZD8186 |
| MVP_top60_DNA_seed | BRAF | -0.056508 | COADREAD | RCM-1 | SB590885 | MVP_top60_DNA+RNA_seed | PIK3CD | -0.501409 | COADREAD | NCI-H716 | AZD8186 |
| MVP_top60_DNA_seed | GSK3B | -0.055826 | COADREAD | COLO-678 | AZD7969 | MVP_top60_DNA+RNA_seed | BCL2 | -0.498464 | COADREAD | KM12 | Obatoclax Mesylate |
| MVP_top60_DNA_seed | AKT1 | -0.055389 | COADREAD | HCC2998 | GSK690693 | MVP_top60_DNA+RNA_seed | BCL2 | -0.494635 | COADREAD | C2BBe1 | TW 37 |
| MVP_top60_DNA_seed | SRC | -0.053608 | COADREAD | RKO | WH-4-023 | MVP_top60_DNA+RNA_seed | CDK2 | -0.494179 | COADREAD | NCI-H747 | PHA-793887 |
| MVP_top60_DNA_seed | IGF1R | -0.053112 | COADREAD | COLO-205 | GSK1904529A | MVP_top60_DNA+RNA_seed | HSP90AA1 | -0.49181 | COADREAD | MDST8 | Elesclomol |
| MVP_top60_DNA_seed | BRAF | -0.052428 | COADREAD | SW620 | HG6-64-1 | MVP_top60_DNA+RNA_seed | HSP90AB1 | -0.49181 | COADREAD | MDST8 | Elesclomol |
| MVP_top60_DNA_seed | BRAF | -0.052056 | COADREAD | NCI-H716 | HG6-64-1 | MVP_top60_DNA+RNA_seed | AKT1 | -0.491021 | COADREAD | RKO | AKT inhibitor VIII |
| MVP_top60_DNA_seed | BRAF | -0.051633 | COADREAD | LoVo | SB590885 | MVP_top60_DNA+RNA_seed | GSK3B | -0.490984 | COADREAD | LoVo | CHIR-99021 |
| MVP_top60_DNA_seed | GSK3B | -0.051287 | COADREAD | COLO-320-HSR | SB216763 | MVP_top60_DNA+RNA_seed | IGF1R | -0.487266 | COADREAD | C2BBe1 | BMS-754807 |
| MVP_top60_DNA_seed | MAPK14 | -0.050945 | COADREAD | SW48 | TAK-715 | MVP_top60_DNA+RNA_seed | GSK3B | -0.483272 | COADREAD | HT-115 | KIN001-042 |
| MVP_top60_DNA_seed | GSK3B | -0.047934 | COADREAD | HT-29 | KIN001-042 | MVP_top60_DNA+RNA_seed | BCL2 | -0.480042 | COADREAD | NCI-H747 | TW 37 |
| MVP_top60_DNA_seed | SRC | -0.047825 | COADREAD | SNU-C2B | NVP-BHG712 | MVP_top60_DNA+RNA_seed | PTK2 | -0.478548 | COADREAD | HT-29 | PF-562271 |
| MVP_top60_DNA_seed | IGF1R | -0.046129 | COADREAD | LS-123 | GSK1904529A | MVP_top60_DNA+RNA_seed | BCL2 | -0.475299 | COADREAD | NCI-H747 | Obatoclax Mesylate |
| MVP_top60_DNA_seed | BRAF | -0.046015 | COADREAD | SW620 | RAF_9304 | MVP_top60_DNA+RNA_seed | CDK2 | -0.475131 | COADREAD | LoVo | AZD5438 |
| MVP_top60_DNA_seed | AKT1 | -0.042683 | COADREAD | SK-CO-1 | MK-2206 | MVP_top60_DNA+RNA_seed | PIK3CA | -0.474138 | COADREAD | SW948 | PI3Ka_4409 |
| MVP_top60_DNA_seed | BRAF | -0.038668 | COADREAD | RCM-1 | RAF_9304 | MVP_top60_DNA+RNA_seed | PIK3CD | -0.474138 | COADREAD | SW948 | PI3Ka_4409 |
| MVP_top60_DNA_seed | ATM | -0.037897 | COADREAD | LS-180 | KU-55933 | MVP_top60_DNA+RNA_seed | BCL2 | -0.470225 | COADREAD | GP5d | Obatoclax Mesylate |
| MVP_top60_DNA_seed | MET | -0.037079 | COADREAD | LS-513 | JNJ38877605 | MVP_top60_DNA+RNA_seed | MET | -0.467249 | COADREAD | NCI-H716 | JNJ38877605 |
| MVP_top60_DNA_seed | ATM | -0.035626 | COADREAD | GP5d | CP466722 | MVP_top60_DNA+RNA_seed | AKT1 | -0.466331 | COADREAD | SNU-407 | MK-2206 |
| MVP_top60_DNA_seed | AKT1 | -0.035069 | COADREAD | COLO-205 | AKT inhibitor VIII | MVP_top60_DNA+RNA_seed | HSP90AA1 | -0.465239 | COADREAD | SW620 | Elesclomol |
| MVP_top60_DNA_seed | GSK3B | -0.034486 | COADREAD | RKO | SB216763 | MVP_top60_DNA+RNA_seed | HSP90AB1 | -0.465239 | COADREAD | SW620 | Elesclomol |
| MVP_top60_DNA_seed | CDK2 | -0.033882 | COADREAD | LS-1034 | AZD5438 | MVP_top60_DNA+RNA_seed | ATM | -0.463427 | COADREAD | COLO-320-HSR | CP466722 |
| MVP_top60_DNA_seed | MAPK14 | -0.03315 | COADREAD | HCC2998 | TAK-715 | MVP_top60_DNA+RNA_seed | AKT1 | -0.463423 | COADREAD | COLO-205 | MK-2206 |
| MVP_top60_DNA_seed | PIK3CD | -0.031025 | COADREAD | T84 | Idelalisib | MVP_top60_DNA+RNA_seed | SRC | -0.462747 | COADREAD | SNU-175 | NVP-BHG712 |
| MVP_top60_DNA_seed | CDK2 | -0.03029 | COADREAD | GP5d | PHA-793887 | MVP_top60_DNA+RNA_seed | ATM | -0.461765 | COADREAD | CCK-81 | CP466722 |
| MVP_top60_DNA_seed | SRC | -0.029833 | COADREAD | SNU-C2B | WH-4-023 | MVP_top60_DNA+RNA_seed | SRC | -0.461177 | COADREAD | HCT-15 | Bosutinib |
| MVP_top60_DNA_seed | CDK2 | -0.029594 | COADREAD | KM12 | PHA-793887 | MVP_top60_DNA+RNA_seed | PTK2 | -0.459939 | COADREAD | LS-411N | PF-562271 |
| MVP_top60_DNA_seed | AR | -0.025057 | COADREAD | CL-11 | AZD3514 | MVP_top60_DNA+RNA_seed | HSP90AA1 | -0.458332 | COADREAD | LS-123 | Luminespib |
| MVP_top60_DNA_seed | BRAF | -0.024449 | COADREAD | COLO-678 | HG6-64-1 | MVP_top60_DNA+RNA_seed | HSP90AB1 | -0.458332 | COADREAD | LS-123 | Luminespib |
| MVP_top60_DNA_seed | AR | -0.023246 | COADREAD | MDST8 | Bicalutamide | MVP_top60_DNA+RNA_seed | IGF1R | -0.457392 | COADREAD | GP5d | BMS-536924 |
| MVP_top60_DNA_seed | ERBB4 | -0.022913 | COADREAD | SW1417 | CI-1033 | MVP_top60_DNA+RNA_seed | AR | -0.453972 | COADREAD | SK-CO-1 | Bicalutamide |
| MVP_top60_DNA_seed | GSK3B | -0.017953 | COADREAD | NCI-H747 | AZD7969 | MVP_top60_DNA+RNA_seed | GSK3B | -0.452908 | COADREAD | KM12 | CHIR-99021 |
| MVP_top60_DNA_seed | MET | -0.016816 | COADREAD | HCC2998 | JNJ38877605 | MVP_top60_DNA+RNA_seed | GSK3B | -0.447552 | COADREAD | HCC2998 | AZD7969 |
| MVP_top60_DNA_seed | MAPK1 | -0.016381 | COADREAD | RKO | FR-180204 | MVP_top60_DNA+RNA_seed | AKT1 | -0.446491 | COADREAD | T84 | AKT inhibitor VIII |
| MVP_top60_DNA_seed | HSP90AA1 | -0.015843 | COADREAD | COLO-205 | Luminespib | MVP_top60_DNA+RNA_seed | HSP90AA1 | -0.442865 | COADREAD | SNU-C5 | Tanespimycin |
| MVP_top60_DNA_seed | HSP90AB1 | -0.015843 | COADREAD | COLO-205 | Luminespib | MVP_top60_DNA+RNA_seed | HSP90AB1 | -0.442865 | COADREAD | SNU-C5 | Tanespimycin |
| MVP_top60_DNA_seed | HSP90AA1 | -0.013194 | COADREAD | KM12 | SNX-2112 | MVP_top60_DNA+RNA_seed | GSK3B | -0.442639 | COADREAD | SW48 | CHIR-99021 |
| MVP_top60_DNA_seed | HSP90AB1 | -0.013194 | COADREAD | KM12 | SNX-2112 | MVP_top60_DNA+RNA_seed | BCL2 | -0.438734 | COADREAD | COLO-320-HSR | TW 37 |
| MVP_top60_DNA_seed | ATM | -0.012139 | COADREAD | RKO | KU-55933 | MVP_top60_DNA+RNA_seed | SRC | -0.43469 | COADREAD | LS-123 | WH-4-023 |
| MVP_top60_DNA_seed | AKT1 | -0.011248 | COADREAD | HCC2998 | AKT inhibitor VIII | MVP_top60_DNA+RNA_seed | GSK3B | -0.433737 | COADREAD | HT-115 | CHIR-99021 |

| | | | | | | | | | | | |
|---|---|---|---|---|---|---|---|---|---|---|---|
| MVP_top60_DNA_seed | ERBB3 | -0.010886 | COADREAD | LS-180 | AZD8931 | MVP_top60_DNA+RNA_seed | ERBB4 | -0.43373 | COADREAD | SW1116 | CI-1033 |
| MVP_top60_DNA_seed | AKT1 | -0.008845 | COADREAD | MDST8 | MK-2206 | MVP_top60_DNA+RNA_seed | PIK3CA | -0.433547 | COADREAD | NCI-H508 | PI3Ka_4409 |
| MVP_top60_DNA_seed | ERBB4 | -0.007733 | COADREAD | HT55 | AST-1306 | MVP_top60_DNA+RNA_seed | PIK3CD | -0.433547 | COADREAD | NCI-H508 | PI3Ka_4409 |
| MVP_top60_DNA_seed | PIK3CD | -0.007236 | COADREAD | LS-180 | Idelalisib | MVP_top60_DNA+RNA_seed | SRC | -0.433534 | COADREAD | SNU-407 | Bosutinib |
| MVP_top60_DNA_seed | MET | -0.006465 | COADREAD | NCI-H747 | AZD6094 | MVP_top60_DNA+RNA_seed | ATM | -0.432508 | COADREAD | HCT-15 | CP466722 |
| MVP_top60_DNA_seed | PTK2 | -0.005536 | COADREAD | LS-123 | PF-562271 | MVP_top60_DNA+RNA_seed | ATM | -0.432033 | COADREAD | SNU-81 | KU-60019 |
| MVP_top60_DNA_seed | AR | -0.00485 | COADREAD | HCC2998 | Bicalutamide | MVP_top60_DNA+RNA_seed | BCL2 | -0.432008 | COADREAD | NCI-H508 | Navitoclax |
| MVP_top60_DNA_seed | AR | -0.001807 | COADREAD | RKO | Bicalutamide | MVP_top60_DNA+RNA_seed | IGF1R | -0.431333 | COADREAD | SNU-407 | BMS-536924 |
| MVP_top60_DNA_seed | MAPK1 | -0.000707 | COADREAD | HCT-15 | FR-180204 | MVP_top60_DNA+RNA_seed | SRC | -0.430207 | COADREAD | NCI-H747 | WH-4-023 |
| MVP_top60_DNA_seed | HSP90AA1 | -0.000142 | COADREAD | C2BBe1 | Elesclomol | MVP_top60_DNA+RNA_seed | HSP90AA1 | -0.424943 | COADREAD | COLO-320-HSR | SNX-2112 |
| MVP_top60_DNA_seed | HSP90AB1 | -0.000142 | COADREAD | C2BBe1 | Elesclomol | MVP_top60_DNA+RNA_seed | HSP90AB1 | -0.424943 | COADREAD | COLO-320-HSR | SNX-2112 |
| MVP_top60_DNA_seed | AKT1 | 0.000203 | COADREAD | C2BBe1 | AKT inhibitor VIII | MVP_top60_DNA+RNA_seed | MET | -0.422215 | COADREAD | HCT-116 | AZD6094 |
| MVP_top60_DNA_seed | PIK3CB | 0.001164 | COADREAD | COLO-205 | AZD6482 | MVP_top60_DNA+RNA_seed | GSK3B | -0.420137 | COADREAD | SW620 | KIN001-042 |
| MVP_top60_DNA_seed | CDK2 | 0.001426 | COADREAD | HCC2998 | PHA-793887 | MVP_top60_DNA+RNA_seed | BCL2 | -0.420011 | COADREAD | DiFi | Navitoclax |
| MVP_top60_DNA_seed | BRAF | 0.005611 | COADREAD | LS-123 | RAF_9304 | MVP_top60_DNA+RNA_seed | HSP90AA1 | -0.419316 | COADREAD | CaR-1 | SNX-2112 |
| MVP_top60_DNA_seed | HSP90AA1 | 0.005744 | COADREAD | NCI-H716 | CCT-018159 | MVP_top60_DNA+RNA_seed | HSP90AB1 | -0.419316 | COADREAD | CaR-1 | SNX-2112 |
| MVP_top60_DNA_seed | HSP90AB1 | 0.005744 | COADREAD | NCI-H716 | CCT-018159 | MVP_top60_DNA+RNA_seed | GSK3B | -0.414456 | COADREAD | GP5d | AZD7969 |
| MVP_top60_DNA_seed | BRAF | 0.008408 | COADREAD | C2BBe1 | RAF_9304 | MVP_top60_DNA+RNA_seed | ATM | -0.414445 | COADREAD | KM12 | CP466722 |
| MVP_top60_DNA_seed | AKT1 | 0.009782 | COADREAD | RKO | GSK690693 | MVP_top60_DNA+RNA_seed | IGF1R | -0.413062 | COADREAD | T84 | BMS-536924 |
| MVP_top60_DNA_seed | ERBB4 | 0.01298 | COADREAD | CaR-1 | AST-1306 | MVP_top60_DNA+RNA_seed | GSK3B | -0.410285 | COADREAD | HCC2998 | CHIR-99021 |
| MVP_top60_DNA_seed | BRAF | 0.014266 | COADREAD | SW1417 | PLX-4720 | MVP_top60_DNA+RNA_seed | ERBB3 | -0.409705 | COADREAD | SK-CO-1 | AZD8931 |
| MVP_top60_DNA_seed | CDK2 | 0.016716 | COADREAD | NCI-H716 | AZD5438 | MVP_top60_DNA+RNA_seed | HSP90AA1 | -0.40962 | COADREAD | SNU-407 | Elesclomol |
| MVP_top60_DNA_seed | AKT1 | 0.018104 | COADREAD | LS-123 | A-443654 | MVP_top60_DNA+RNA_seed | HSP90AB1 | -0.40962 | COADREAD | SNU-407 | Elesclomol |
| MVP_top60_DNA_seed | HSP90AA1 | 0.018872 | COADREAD | MDST8 | Luminespib | MVP_top60_DNA+RNA_seed | IGF1R | -0.405281 | COADREAD | LS-513 | BMS-536924 |
| MVP_top60_DNA_seed | HSP90AB1 | 0.018872 | COADREAD | MDST8 | Luminespib | MVP_top60_DNA+RNA_seed | GSK3B | -0.403294 | COADREAD | LS-1034 | AZD7969 |
| MVP_top60_DNA_seed | HSP90AA1 | 0.020484 | COADREAD | COLO-678 | Elesclomol | MVP_top60_DNA+RNA_seed | AKT1 | -0.401915 | COADREAD | HCC2998 | BAY AKT1 |
| MVP_top60_DNA_seed | HSP90AB1 | 0.020484 | COADREAD | COLO-678 | Elesclomol | MVP_top60_DNA+RNA_seed | CDK2 | -0.397909 | COADREAD | COLO-320-HSR | AZD5438 |
| MVP_top60_DNA_seed | BRAF | 0.021297 | COADREAD | SW1116 | RAF_9304 | MVP_top60_DNA+RNA_seed | HSP90AA1 | -0.396004 | COADREAD | C2BBe1 | Tanespimycin |
| MVP_top60_DNA_seed | IGF1R | 0.021813 | COADREAD | C2BBe1 | BMS-536924 | MVP_top60_DNA+RNA_seed | HSP90AB1 | -0.396004 | COADREAD | C2BBe1 | Tanespimycin |
| MVP_top60_DNA_seed | HSP90AA1 | 0.021814 | COADREAD | LS-513 | Luminespib | MVP_top60_DNA+RNA_seed | ERBB4 | -0.390573 | COADREAD | NCI-H747 | PF-00299804 |
| MVP_top60_DNA_seed | HSP90AB1 | 0.021814 | COADREAD | LS-513 | Luminespib | MVP_top60_DNA+RNA_seed | BCL2 | -0.384425 | COADREAD | SW620 | Navitoclax |
| MVP_top60_DNA_seed | PIK3CA | 0.022367 | COADREAD | SNU-81 | AZD8835 | MVP_top60_DNA+RNA_seed | FYN | -0.383507 | COADREAD | LS-123 | A-770041 |
| MVP_top60_DNA_seed | PIK3CD | 0.022367 | COADREAD | SNU-81 | AZD8835 | MVP_top60_DNA+RNA_seed | BCL2 | -0.383268 | COADREAD | COLO-320-HSR | Obatoclax Mesylate |
| MVP_top60_DNA_seed | HSP90AA1 | 0.02393 | COADREAD | CCK-81 | CCT-018159 | MVP_top60_DNA+RNA_seed | AKT1 | -0.382601 | COADREAD | KM12 | AKT inhibitor VIII |
| MVP_top60_DNA_seed | HSP90AB1 | 0.02393 | COADREAD | CCK-81 | CCT-018159 | MVP_top60_DNA+RNA_seed | ATM | -0.38166 | COADREAD | COLO-205 | CP466722 |
| MVP_top60_DNA_seed | AR | 0.024116 | COADREAD | HT-29 | AZD3514 | MVP_top60_DNA+RNA_seed | ERBB3 | -0.38052 | COADREAD | NCI-H747 | AZD8931 |
| MVP_top60_DNA_seed | ERBB4 | 0.024327 | COADREAD | GP5d | AST-1306 | MVP_top60_DNA+RNA_seed | HSP90AA1 | -0.376604 | COADREAD | T84 | SNX-2112 |
| MVP_top60_DNA_seed | ERBB4 | 0.025599 | COADREAD | RKO | AST-1306 | MVP_top60_DNA+RNA_seed | HSP90AB1 | -0.376604 | COADREAD | T84 | SNX-2112 |
| MVP_top60_DNA_seed | CREBBP | 0.026964 | COADREAD | KM12 | I-CBP112 | MVP_top60_DNA+RNA_seed | GSK3B | -0.375297 | COADREAD | SW1116 | SB216763 |
| MVP_top60_DNA_seed | MAPK1 | 0.027922 | COADREAD | T84 | FR-180204 | MVP_top60_DNA+RNA_seed | PIK3CB | -0.373521 | COADREAD | MDST8 | AZD8186 |
| MVP_top60_DNA_seed | PIK3CA | 0.028046 | COADREAD | CL-11 | PI3Ka_4409 | MVP_top60_DNA+RNA_seed | PIK3CD | -0.373521 | COADREAD | MDST8 | AZD8186 |

| | | | | | | | | | | | |
|---|---|---|---|---|---|---|---|---|---|---|---|
| MVP_top60_DNA_seed | PIK3CD | 0.028046 | COADREAD | CL-11 | PI3Ka_4409 | MVP_top60_DNA+RNA_seed | AR | -0.373086 | COADREAD | SNU-C2B | AZD3514 |
| MVP_top60_DNA_seed | ESR1 | 0.028568 | COADREAD | HT-29 | Fulvestrant | MVP_top60_DNA+RNA_seed | PIK3CA | -0.36886 | COADREAD | LS-513 | PI3Ka_4409 |
| MVP_top60_DNA_seed | PIK3CB | 0.02924 | COADREAD | HCC2998 | TGX221 | MVP_top60_DNA+RNA_seed | PIK3CD | -0.36886 | COADREAD | LS-513 | PI3Ka_4409 |
| MVP_top60_DNA_seed | CDK2 | 0.029517 | COADREAD | HCC2998 | Seliciclib | MVP_top60_DNA+RNA_seed | IGF1R | -0.368033 | COADREAD | SW837 | Linsitinib |
| MVP_top60_DNA_seed | AKT1 | 0.030179 | COADREAD | SNU-C5 | BAY AKT1 | MVP_top60_DNA+RNA_seed | PIK3CD | -0.366195 | COADREAD | SNU-81 | IC-87114 |
| MVP_top60_DNA_seed | ERBB4 | 0.030699 | COADREAD | LS-180 | PF-00299804 | MVP_top60_DNA+RNA_seed | HSP90AA1 | -0.365355 | COADREAD | HCT-15 | SNX-2112 |
| MVP_top60_DNA_seed | HSP90AA1 | 0.031206 | COADREAD | SNU-C5 | Elesclomol | MVP_top60_DNA+RNA_seed | HSP90AB1 | -0.365355 | COADREAD | HCT-15 | SNX-2112 |
| MVP_top60_DNA_seed | HSP90AB1 | 0.031206 | COADREAD | SNU-C5 | Elesclomol | MVP_top60_DNA+RNA_seed | PIK3CB | -0.364882 | COADREAD | C2BBe1 | AZD6482 |
| MVP_top60_DNA_seed | GSK3B | 0.031994 | COADREAD | HCT-15 | CHIR-99021 | MVP_top60_DNA+RNA_seed | GSK3B | -0.363097 | COADREAD | CL-11 | SB216763 |
| MVP_top60_DNA_seed | BRAF | 0.036466 | COADREAD | SW1463 | SB590885 | MVP_top60_DNA+RNA_seed | ATM | -0.360689 | COADREAD | HCT-15 | KU-60019 |
| MVP_top60_DNA_seed | HSP90AA1 | 0.036958 | COADREAD | LS-123 | Elesclomol | MVP_top60_DNA+RNA_seed | FYN | -0.35927 | COADREAD | SNU-C2B | A-770041 |
| MVP_top60_DNA_seed | HSP90AB1 | 0.036958 | COADREAD | LS-123 | Elesclomol | MVP_top60_DNA+RNA_seed | MET | -0.358717 | COADREAD | HT-115 | SU11274 |
| MVP_top60_DNA_seed | PIK3CA | 0.037754 | COADREAD | LS-123 | PI3Ka_4409 | MVP_top60_DNA+RNA_seed | AKT1 | -0.353981 | COADREAD | LS-513 | GSK690693 |
| MVP_top60_DNA_seed | PIK3CD | 0.037754 | COADREAD | LS-123 | PI3Ka_4409 | MVP_top60_DNA+RNA_seed | GSK3B | -0.353564 | COADREAD | NCI-H716 | SB216763 |
| MVP_top60_DNA_seed | PIK3CA | 0.038105 | COADREAD | SW837 | PI3Ka_4409 | MVP_top60_DNA+RNA_seed | ATM | -0.353132 | COADREAD | RKO | CP466722 |
| MVP_top60_DNA_seed | PIK3CD | 0.038105 | COADREAD | SW837 | PI3Ka_4409 | MVP_top60_DNA+RNA_seed | ATM | -0.353111 | COADREAD | SW620 | CP466722 |
| MVP_top60_DNA_seed | FYN | 0.039886 | COADREAD | LS-1034 | A-770041 | MVP_top60_DNA+RNA_seed | AR | -0.348069 | COADREAD | SNU-407 | AZD3514 |
| MVP_top60_DNA_seed | PIK3CA | 0.039911 | COADREAD | SW948 | AZD8835 | MVP_top60_DNA+RNA_seed | PIK3CA | -0.347499 | COADREAD | COLO-320-HSR | PI3Ka_4409 |
| MVP_top60_DNA_seed | PIK3CD | 0.039911 | COADREAD | SW948 | AZD8835 | MVP_top60_DNA+RNA_seed | PIK3CD | -0.347499 | COADREAD | COLO-320-HSR | PI3Ka_4409 |
| MVP_top60_DNA_seed | AR | 0.041203 | COADREAD | NCI-H508 | AZD3514 | MVP_top60_DNA+RNA_seed | PIK3CA | -0.344309 | COADREAD | NCI-H716 | AZD8835 |
| MVP_top60_DNA_seed | SRC | 0.041774 | COADREAD | SW837 | Bosutinib | MVP_top60_DNA+RNA_seed | PIK3CD | -0.344309 | COADREAD | NCI-H716 | AZD8835 |
| MVP_top60_DNA_seed | IGF1R | 0.042076 | COADREAD | HT-115 | BMS-536924 | MVP_top60_DNA+RNA_seed | AR | -0.343672 | COADREAD | NCI-H747 | AZD3514 |
| MVP_top60_DNA_seed | AR | 0.044503 | COADREAD | HT-115 | Bicalutamide | MVP_top60_DNA+RNA_seed | GSK3B | -0.338959 | COADREAD | SK-CO-1 | CHIR-99021 |
| MVP_top60_DNA_seed | SRC | 0.04576 | COADREAD | SNU-1040 | NVP-BHG712 | MVP_top60_DNA+RNA_seed | CDK2 | -0.338654 | COADREAD | LS-411N | Seliciclib |
| MVP_top60_DNA_seed | AKT1 | 0.046622 | COADREAD | LS-180 | BAY AKT1 | MVP_top60_DNA+RNA_seed | BCL2 | -0.337918 | COADREAD | C2BBe1 | Navitoclax |
| MVP_top60_DNA_seed | HSP90AA1 | 0.046861 | COADREAD | SNU-C5 | SNX-2112 | MVP_top60_DNA+RNA_seed | PIK3CA | -0.336371 | COADREAD | SW1463 | PI3Ka_4409 |
| MVP_top60_DNA_seed | HSP90AB1 | 0.046861 | COADREAD | SNU-C5 | SNX-2112 | MVP_top60_DNA+RNA_seed | PIK3CD | -0.336371 | COADREAD | SW1463 | PI3Ka_4409 |
| MVP_top60_DNA_seed | BRAF | 0.046935 | COADREAD | CCK-81 | RAF_9304 | MVP_top60_DNA+RNA_seed | GSK3B | -0.335235 | COADREAD | COLO-320-HSR | KIN001-042 |
| MVP_top60_DNA_seed | AR | 0.047295 | COADREAD | HCC2998 | Bicalutamide | MVP_top60_DNA+RNA_seed | HSP90AA1 | -0.328917 | COADREAD | HCC2998 | Tanespimycin |
| MVP_top60_DNA_seed | ERBB3 | 0.047486 | COADREAD | SW620 | AZD8931 | MVP_top60_DNA+RNA_seed | HSP90AB1 | -0.328917 | COADREAD | HCC2998 | Tanespimycin |
| MVP_top60_DNA_seed | BRAF | 0.050947 | COADREAD | GP5d | PLX-4720 | MVP_top60_DNA+RNA_seed | MET | -0.328176 | COADREAD | HT-29 | JNJ38877605 |
| MVP_top60_DNA_seed | CREBBP | 0.052303 | COADREAD | COLO-205 | I-CBP112 | MVP_top60_DNA+RNA_seed | HSP90AA1 | -0.324863 | COADREAD | NCI-H747 | SNX-2112 |
| MVP_top60_DNA_seed | BRAF | 0.053849 | COADREAD | COLO-320-HSR | HG6-64-1 | MVP_top60_DNA+RNA_seed | HSP90AB1 | -0.324863 | COADREAD | NCI-H747 | SNX-2112 |
| MVP_top60_DNA_seed | HSP90AA1 | 0.0541 | COADREAD | SW948 | SNX-2112 | MVP_top60_DNA+RNA_seed | GSK3B | -0.322784 | COADREAD | LS-513 | KIN001-042 |
| MVP_top60_DNA_seed | HSP90AB1 | 0.0541 | COADREAD | SW948 | SNX-2112 | MVP_top60_DNA+RNA_seed | ERBB4 | -0.320522 | COADREAD | NCI-H747 | CI-1033 |
| MVP_top60_DNA_seed | AKT1 | 0.056641 | COADREAD | HCT-116 | AKT inhibitor VIII | MVP_top60_DNA+RNA_seed | PIK3CA | -0.319794 | COADREAD | COLO-205 | AZD8835 |
| MVP_top60_DNA_seed | PIK3CA | 0.057106 | COADREAD | HT-29 | PI3Ka_4409 | MVP_top60_DNA+RNA_seed | PIK3CD | -0.319794 | COADREAD | COLO-205 | AZD8835 |
| MVP_top60_DNA_seed | PIK3CD | 0.057106 | COADREAD | HT-29 | PI3Ka_4409 | MVP_top60_DNA+RNA_seed | SRC | -0.319177 | COADREAD | KM12 | NVP-BHG712 |
| MVP_top60_DNA_seed | IGF1R | 0.05728 | COADREAD | COLO-678 | GSK1904529A | MVP_top60_DNA+RNA_seed | PIK3CA | -0.318476 | COADREAD | MDST8 | AZD8835 |
| MVP_top60_DNA_seed | MET | 0.059495 | COADREAD | SW620 | JNJ38877605 | MVP_top60_DNA+RNA_seed | PIK3CD | -0.318476 | COADREAD | MDST8 | AZD8835 |
| MVP_top60_DNA_seed | ERBB4 | 0.06093 | COADREAD | COLO-320-HSR | PF-00299804 | MVP_top60_DNA+RNA_seed | HSP90AA1 | -0.317495 | COADREAD | GP5d | Luminespib |

| | | | | | | | | | | | |
|---|---|---|---|---|---|---|---|---|---|---|---|
| MVP_top60_DNA_seed | ATM | 0.062083 | COADREAD | SW620 | KU-60019 | MVP_top60_DNA+RNA_seed | HSP90AB1 | -0.317495 | COADREAD | GP5d | Luminespib |
| MVP_top60_DNA_seed | BRAF | 0.063099 | COADREAD | SK-CO-1 | PLX-4720 | MVP_top60_DNA+RNA_seed | AKT1 | -0.315839 | COADREAD | LS-180 | AKT inhibitor VIII |
| MVP_top60_DNA_seed | PIK3CA | 0.064424 | COADREAD | LS-1034 | PI3Ka_4409 | MVP_top60_DNA+RNA_seed | PTK2 | -0.315161 | COADREAD | NCI-H747 | PF-562271 |
| MVP_top60_DNA_seed | PIK3CD | 0.064424 | COADREAD | LS-1034 | PI3Ka_4409 | MVP_top60_DNA+RNA_seed | AKT1 | -0.31205 | COADREAD | SW948 | GSK690693 |
| MVP_top60_DNA_seed | BRAF | 0.06462 | COADREAD | SW1417 | Dabrafenib | MVP_top60_DNA+RNA_seed | AR | -0.310652 | COADREAD | KM12 | Bicalutamide |
| MVP_top60_DNA_seed | GSK3B | 0.066142 | COADREAD | SW620 | SB216763 | MVP_top60_DNA+RNA_seed | IGF1R | -0.308205 | COADREAD | LoVo | BMS-536924 |
| MVP_top60_DNA_seed | IGF1R | 0.066922 | COADREAD | SW620 | BMS-536924 | MVP_top60_DNA+RNA_seed | PIK3CD | -0.307003 | COADREAD | CCK-81 | Idelalisib |
| MVP_top60_DNA_seed | ERBB4 | 0.066961 | COADREAD | COLO-205 | PF-00299804 | MVP_top60_DNA+RNA_seed | IGF1R | -0.304123 | COADREAD | SNU-C2B | Linsitinib |
| MVP_top60_DNA_seed | BRAF | 0.067297 | COADREAD | NCI-H747 | Dabrafenib | MVP_top60_DNA+RNA_seed | MET | -0.302398 | COADREAD | COLO-205 | AZD6094 |
| MVP_top60_DNA_seed | CDK2 | 0.070221 | COADREAD | MDST8 | Seliciclib | MVP_top60_DNA+RNA_seed | BCL2 | -0.302391 | COADREAD | CW-2 | TW 37 |
| MVP_top60_DNA_seed | ESR1 | 0.07036 | COADREAD | COLO-205 | Tamoxifen | MVP_top60_DNA+RNA_seed | AR | -0.302285 | COADREAD | NCI-H716 | AZD3514 |
| MVP_top60_DNA_seed | CDK2 | 0.072017 | COADREAD | MDST8 | PHA-793887 | MVP_top60_DNA+RNA_seed | PTK2 | -0.302222 | COADREAD | LoVo | PF-562271 |
| MVP_top60_DNA_seed | AR | 0.073282 | COADREAD | HCT-116 | Bicalutamide | MVP_top60_DNA+RNA_seed | BCL2 | -0.301203 | COADREAD | T84 | Obatoclax Mesylate |
| MVP_top60_DNA_seed | BRAF | 0.074205 | COADREAD | SW48 | PLX-4720 | MVP_top60_DNA+RNA_seed | GSK3B | -0.300728 | COADREAD | LS-411N | CHIR-99021 |
| MVP_top60_DNA_seed | MAPK1 | 0.07461 | COADREAD | SNU-81 | VX-11e | MVP_top60_DNA+RNA_seed | SRC | -0.298466 | COADREAD | MDST8 | HG-6-71-01 |
| MVP_top60_DNA_seed | MET | 0.075353 | COADREAD | GP5d | AZD6094 | MVP_top60_DNA+RNA_seed | AKT1 | -0.297477 | COADREAD | GP5d | GSK690693 |
| MVP_top60_DNA_seed | MET | 0.075659 | COADREAD | CW-2 | PHA-665752 | MVP_top60_DNA+RNA_seed | GSK3B | -0.296121 | COADREAD | RKO | AZD7969 |
| MVP_top60_DNA_seed | ERBB3 | 0.07653 | COADREAD | HCT-15 | AZD8931 | MVP_top60_DNA+RNA_seed | GSK3B | -0.291615 | COADREAD | NCI-H747 | SB216763 |
| MVP_top60_DNA_seed | AR | 0.076685 | COADREAD | CL-11 | Bicalutamide | MVP_top60_DNA+RNA_seed | MET | -0.291471 | COADREAD | SW1417 | JNJ38877605 |
| MVP_top60_DNA_seed | GSK3B | 0.07669 | COADREAD | CCK-81 | AZD7969 | MVP_top60_DNA+RNA_seed | SRC | -0.291213 | COADREAD | CCK-81 | NVP-BHG712 |
| MVP_top60_DNA_seed | ERBB4 | 0.07703 | COADREAD | C2BBe1 | CI-1033 | MVP_top60_DNA+RNA_seed | IGF1R | -0.289911 | COADREAD | CaR-1 | BMS-536924 |
| MVP_top60_DNA_seed | PIK3CA | 0.078169 | COADREAD | SNU-1040 | PI3Ka_4409 | MVP_top60_DNA+RNA_seed | PIK3CD | -0.28945 | COADREAD | RKO | Idelalisib |
| MVP_top60_DNA_seed | PIK3CD | 0.078169 | COADREAD | SNU-1040 | PI3Ka_4409 | MVP_top60_DNA+RNA_seed | HSP90AA1 | -0.288218 | COADREAD | NCI-H747 | Tanespimycin |
| MVP_top60_DNA_seed | BRAF | 0.078758 | COADREAD | RKO | PLX-4720 | MVP_top60_DNA+RNA_seed | HSP90AB1 | -0.288218 | COADREAD | NCI-H747 | Tanespimycin |
| MVP_top60_DNA_seed | ATM | 0.079666 | COADREAD | HT-29 | CP466722 | MVP_top60_DNA+RNA_seed | BCL2 | -0.286388 | COADREAD | NCI-H716 | Navitoclax |
| MVP_top60_DNA_seed | GSK3B | 0.081653 | COADREAD | C2BBe1 | CHIR-99021 | MVP_top60_DNA+RNA_seed | GSK3B | -0.285862 | COADREAD | MDST8 | CHIR-99021 |
| MVP_top60_DNA_seed | GSK3B | 0.082098 | COADREAD | CCK-81 | KIN001-042 | MVP_top60_DNA+RNA_seed | ATM | -0.283284 | COADREAD | SW48 | KU-55933 |
| MVP_top60_DNA_seed | ATM | 0.083235 | COADREAD | C2BBe1 | KU-60019 | MVP_top60_DNA+RNA_seed | GSK3B | -0.282916 | COADREAD | COLO-205 | SB216763 |
| MVP_top60_DNA_seed | PIK3CB | 0.088006 | COADREAD | LS-123 | TGX221 | MVP_top60_DNA+RNA_seed | GSK3B | -0.280222 | COADREAD | T84 | KIN001-042 |
| MVP_top60_DNA_seed | ERBB4 | 0.089055 | COADREAD | CL-11 | AST-1306 | MVP_top60_DNA+RNA_seed | IGF1R | -0.279373 | COADREAD | LS-513 | BMS-536924 |
| MVP_top60_DNA_seed | CDK2 | 0.089275 | COADREAD | C2BBe1 | AZD5438 | MVP_top60_DNA+RNA_seed | PIK3CB | -0.278332 | COADREAD | NCI-H747 | AZD8186 |
| MVP_top60_DNA_seed | CREBBP | 0.089683 | COADREAD | SNU-C5 | I-CBP112 | MVP_top60_DNA+RNA_seed | PIK3CD | -0.278332 | COADREAD | NCI-H747 | AZD8186 |
| MVP_top60_DNA_seed | ATM | 0.090307 | COADREAD | CL-11 | torin2 | MVP_top60_DNA+RNA_seed | AKT1 | -0.277854 | COADREAD | COLO-320-HSR | BAY AKT1 |
| MVP_top60_DNA_seed | CDK2 | 0.09192 | COADREAD | SK-CO-1 | AZD5438 | MVP_top60_DNA+RNA_seed | ATM | -0.277802 | COADREAD | MDST8 | KU-60019 |
| MVP_top60_DNA_seed | SRC | 0.091963 | COADREAD | COLO-678 | Bosutinib | MVP_top60_DNA+RNA_seed | ATM | -0.273401 | COADREAD | SK-CO-1 | KU-60019 |
| MVP_top60_DNA_seed | BRAF | 0.094188 | COADREAD | HT-115 | Dabrafenib | MVP_top60_DNA+RNA_seed | GSK3B | -0.269551 | COADREAD | RKO | CHIR-99021 |
| MVP_top60_DNA_seed | ERBB4 | 0.095234 | COADREAD | LS-411N | PF-00299804 | MVP_top60_DNA+RNA_seed | MET | -0.268429 | COADREAD | T84 | JNJ38877605 |
| MVP_top60_DNA_seed | FYN | 0.095757 | COADREAD | COLO-320-HSR | A-770041 | MVP_top60_DNA+RNA_seed | PIK3CA | -0.2684 | COADREAD | CCK-81 | AZD8835 |
| MVP_top60_DNA_seed | PIK3CD | 0.096357 | COADREAD | RKO | IC-87114 | MVP_top60_DNA+RNA_seed | PIK3CD | -0.2684 | COADREAD | CCK-81 | AZD8835 |
| MVP_top60_DNA_seed | MET | 0.09665 | COADREAD | COLO-320-HSR | PHA-665752 | MVP_top60_DNA+RNA_seed | IGF1R | -0.267056 | COADREAD | CCK-81 | GSK1904529A |
| MVP_top60_DNA_seed | AKT1 | 0.101675 | COADREAD | SNU-C1 | AKT inhibitor VIII | MVP_top60_DNA+RNA_seed | HSP90AA1 | -0.265654 | COADREAD | CW-2 | CCT-018159 |

| | | | | | | | | | | | |
|---|---|---|---|---|---|---|---|---|---|---|---|
| MVP_top60_DNA_seed | BRAF | 0.103934 | COADREAD | HCT-15 | Dabrafenib | MVP_top60_DNA+RNA_seed | HSP90AB1 | -0.265654 | COADREAD | CW-2 | CCT-018159 |
| MVP_top60_DNA_seed | AKT1 | 0.105316 | COADREAD | MDST8 | GSK690693 | MVP_top60_DNA+RNA_seed | BCL2 | -0.264908 | COADREAD | GP5d | Navitoclax |
| MVP_top60_DNA_seed | AR | 0.105373 | COADREAD | LoVo | Bicalutamide | MVP_top60_DNA+RNA_seed | ERBB4 | -0.263149 | COADREAD | LS-513 | PF-00299804 |
| MVP_top60_DNA_seed | AKT1 | 0.105655 | COADREAD | CW-2 | GSK690693 | MVP_top60_DNA+RNA_seed | AR | -0.261322 | COADREAD | COLO-320-HSR | Bicalutamide |
| MVP_top60_DNA_seed | MAPK1 | 0.10581 | COADREAD | SW620 | FR-180204 | MVP_top60_DNA+RNA_seed | IGF1R | -0.259315 | COADREAD | GP5d | GSK1904529A |
| MVP_top60_DNA_seed | HSP90AA1 | 0.106424 | COADREAD | COLO-678 | CCT-018159 | MVP_top60_DNA+RNA_seed | PIK3CD | -0.258431 | COADREAD | MDST8 | Idelalisib |
| MVP_top60_DNA_seed | HSP90AB1 | 0.106424 | COADREAD | COLO-678 | CCT-018159 | MVP_top60_DNA+RNA_seed | ERBB3 | -0.257199 | COADREAD | MDST8 | AZD8931 |
| MVP_top60_DNA_seed | MET | 0.106542 | COADREAD | COLO-678 | JNJ38877605 | MVP_top60_DNA+RNA_seed | CDK2 | -0.256615 | COADREAD | LS-1034 | Seliciclib |
| MVP_top60_DNA_seed | BRAF | 0.109116 | COADREAD | C2BBe1 | PLX-4720 | MVP_top60_DNA+RNA_seed | PIK3CD | -0.252629 | COADREAD | COLO-320-HSR | Idelalisib |
| MVP_top60_DNA_seed | HSP90AA1 | 0.109734 | COADREAD | SK-CO-1 | Elesclomol | MVP_top60_DNA+RNA_seed | HSP90AA1 | -0.251207 | COADREAD | HCT-116 | Tanespimycin |
| MVP_top60_DNA_seed | HSP90AB1 | 0.109734 | COADREAD | SK-CO-1 | Elesclomol | MVP_top60_DNA+RNA_seed | HSP90AB1 | -0.251207 | COADREAD | HCT-116 | Tanespimycin |
| MVP_top60_DNA_seed | ERBB4 | 0.10991 | COADREAD | SW1417 | AST-1306 | MVP_top60_DNA+RNA_seed | PIK3CB | -0.248284 | COADREAD | SW48 | AZD8186 |
| MVP_top60_DNA_seed | MAPK1 | 0.109984 | COADREAD | CW-2 | VX-11e | MVP_top60_DNA+RNA_seed | PIK3CD | -0.248284 | COADREAD | SW48 | AZD8186 |
| MVP_top60_DNA_seed | IGF1R | 0.110374 | COADREAD | HCT-116 | BMS-754807 | MVP_top60_DNA+RNA_seed | MET | -0.24611 | COADREAD | NCI-H747 | SU11274 |
| MVP_top60_DNA_seed | PIK3CD | 0.110451 | COADREAD | HT-115 | IC-87114 | MVP_top60_DNA+RNA_seed | GSK3B | -0.245573 | COADREAD | HCC2998 | KIN001-042 |
| MVP_top60_DNA_seed | PIK3CB | 0.111096 | COADREAD | SW837 | AZD8186 | MVP_top60_DNA+RNA_seed | ATM | -0.243581 | COADREAD | LS-411N | CP466722 |
| MVP_top60_DNA_seed | PIK3CD | 0.111096 | COADREAD | SW837 | AZD8186 | MVP_top60_DNA+RNA_seed | AR | -0.240472 | COADREAD | NCI-H716 | Bicalutamide |
| MVP_top60_DNA_seed | PIK3CD | 0.111612 | COADREAD | HCT-116 | IC-87114 | MVP_top60_DNA+RNA_seed | MET | -0.238431 | COADREAD | CW-2 | JNJ38877605 |
| MVP_top60_DNA_seed | ATM | 0.114673 | COADREAD | GP5d | torin2 | MVP_top60_DNA+RNA_seed | IGF1R | -0.235945 | COADREAD | RKO | GSK1904529A |
| MVP_top60_DNA_seed | SRC | 0.114684 | COADREAD | CaR-1 | Bosutinib | MVP_top60_DNA+RNA_seed | CDK2 | -0.235573 | COADREAD | LS-411N | PHA-793887 |
| MVP_top60_DNA_seed | BRAF | 0.116283 | COADREAD | KM12 | Dabrafenib | MVP_top60_DNA+RNA_seed | AR | -0.234824 | COADREAD | LoVo | AZD3514 |
| MVP_top60_DNA_seed | SRC | 0.120193 | COADREAD | COLO-320-HSR | NVP-BHG712 | MVP_top60_DNA+RNA_seed | CDK2 | -0.233403 | COADREAD | SNU-407 | AZD5438 |
| MVP_top60_DNA_seed | CREBBP | 0.120425 | COADREAD | LS-411N | I-CBP112 | MVP_top60_DNA+RNA_seed | IGF1R | -0.230477 | COADREAD | RKO | BMS-754807 |
| MVP_top60_DNA_seed | AKT1 | 0.121509 | COADREAD | CL-11 | MK-2206 | MVP_top60_DNA+RNA_seed | PTK2 | -0.229981 | COADREAD | COLO-320-HSR | PF-562271 |
| MVP_top60_DNA_seed | ERBB3 | 0.123873 | COADREAD | C2BBe1 | AZD8931 | MVP_top60_DNA+RNA_seed | PIK3CD | -0.228988 | COADREAD | CaR-1 | IC-87114 |
| MVP_top60_DNA_seed | ATM | 0.125947 | COADREAD | HT-115 | torin2 | MVP_top60_DNA+RNA_seed | ERBB4 | -0.22807 | COADREAD | SNU-81 | AST-1306 |
| MVP_top60_DNA_seed | AKT1 | 0.127558 | COADREAD | SW620 | AKT inhibitor VIII | MVP_top60_DNA+RNA_seed | GSK3B | -0.22762 | COADREAD | NCI-H747 | KIN001-042 |
| MVP_top60_DNA_seed | CREBBP | 0.12826 | COADREAD | NCI-H747 | I-CBP112 | MVP_top60_DNA+RNA_seed | GSK3B | -0.225722 | COADREAD | COLO-205 | KIN001-042 |
| MVP_top60_DNA_seed | ERBB3 | 0.129399 | COADREAD | COLO-320-HSR | AZD8931 | MVP_top60_DNA+RNA_seed | AKT1 | -0.222594 | COADREAD | HCT-116 | GSK690693 |
| MVP_top60_DNA_seed | ERBB3 | 0.130766 | COADREAD | CW-2 | AZD8931 | MVP_top60_DNA+RNA_seed | AR | -0.220921 | COADREAD | COLO-320-HSR | Bicalutamide |
| MVP_top60_DNA_seed | PIK3CD | 0.133843 | COADREAD | LS-513 | Idelalisib | MVP_top60_DNA+RNA_seed | PIK3CD | -0.218565 | COADREAD | SNU-1040 | Idelalisib |
| MVP_top60_DNA_seed | SRC | 0.13453 | COADREAD | LS-411N | WH-4-023 | MVP_top60_DNA+RNA_seed | IGF1R | -0.216538 | COADREAD | CaR-1 | GSK1904529A |
| MVP_top60_DNA_seed | PIK3CD | 0.136648 | COADREAD | HT-29 | Idelalisib | MVP_top60_DNA+RNA_seed | AKT1 | -0.214771 | COADREAD | NCI-H747 | AKT inhibitor VIII |
| MVP_top60_DNA_seed | CDK2 | 0.137795 | COADREAD | NCI-H747 | AZD5438 | MVP_top60_DNA+RNA_seed | ERBB4 | -0.214317 | COADREAD | SNU-61 | AST-1306 |
| MVP_top60_DNA_seed | HSP90AA1 | 0.138381 | COADREAD | LS-180 | SNX-2112 | MVP_top60_DNA+RNA_seed | SRC | -0.210596 | COADREAD | GP5d | NVP-BHG712 |
| MVP_top60_DNA_seed | HSP90AB1 | 0.138381 | COADREAD | LS-180 | SNX-2112 | MVP_top60_DNA+RNA_seed | GSK3B | -0.206601 | COADREAD | LS-1034 | SB216763 |
| MVP_top60_DNA_seed | MET | 0.139109 | COADREAD | SNU-C5 | AZD6094 | MVP_top60_DNA+RNA_seed | SRC | -0.206511 | COADREAD | LoVo | NVP-BHG712 |
| MVP_top60_DNA_seed | ERBB3 | 0.139812 | COADREAD | SNU-C5 | AZD8931 | MVP_top60_DNA+RNA_seed | ERBB3 | -0.205692 | COADREAD | SNU-81 | AZD8931 |
| MVP_top60_DNA_seed | ERBB4 | 0.139863 | COADREAD | HT55 | PF-00299804 | MVP_top60_DNA+RNA_seed | IGF1R | -0.203811 | COADREAD | LS-180 | Linsitinib |
| MVP_top60_DNA_seed | AKT1 | 0.140191 | COADREAD | LS-1034 | GSK690693 | MVP_top60_DNA+RNA_seed | AR | -0.201737 | COADREAD | RKO | Bicalutamide |
| MVP_top60_DNA_seed | BRAF | 0.141169 | COADREAD | NCI-H716 | PLX-4720 | MVP_top60_DNA+RNA_seed | ERBB3 | -0.201522 | COADREAD | SNU-C2B | AZD8931 |

| | | | | | | | | | | | |
|---|---|---|---|---|---|---|---|---|---|---|---|
| MVP_top60_DNA_seed | HSP90AA1 | 0.141941 | COADREAD | HCC2998 | SNX-2112 | MVP_top60_DNA+RNA_seed | AKT1 | -0.20144 | COADREAD | CW-2 | AKT inhibitor VIII |
| MVP_top60_DNA_seed | HSP90AB1 | 0.141941 | COADREAD | HCC2998 | SNX-2112 | MVP_top60_DNA+RNA_seed | IGF1R | -0.201062 | COADREAD | LS-411N | BMS-536924 |
| MVP_top60_DNA_seed | IGF1R | 0.142495 | COADREAD | SW1417 | BMS-536924 | MVP_top60_DNA+RNA_seed | ERBB4 | -0.199301 | COADREAD | C2BBe1 | AST-1306 |
| MVP_top60_DNA_seed | GSK3B | 0.144005 | COADREAD | HCT-116 | KIN001-042 | MVP_top60_DNA+RNA_seed | SRC | -0.196719 | COADREAD | NCI-H716 | Bosutinib |
| MVP_top60_DNA_seed | AR | 0.144302 | COADREAD | SNU-407 | Bicalutamide | MVP_top60_DNA+RNA_seed | PIK3CD | -0.19634 | COADREAD | SNU-175 | IC-87114 |
| MVP_top60_DNA_seed | ERBB3 | 0.147705 | COADREAD | HT-29 | AZD8931 | MVP_top60_DNA+RNA_seed | HSP90AA1 | -0.195888 | COADREAD | LS-513 | Tanespimycin |
| MVP_top60_DNA_seed | PIK3CA | 0.148652 | COADREAD | LS-1034 | AZD8835 | MVP_top60_DNA+RNA_seed | HSP90AB1 | -0.195888 | COADREAD | LS-513 | Tanespimycin |
| MVP_top60_DNA_seed | PIK3CD | 0.148652 | COADREAD | LS-1034 | AZD8835 | MVP_top60_DNA+RNA_seed | AR | -0.191681 | COADREAD | CCK-81 | Bicalutamide |
| MVP_top60_DNA_seed | AKT1 | 0.149032 | COADREAD | NCI-H716 | GSK690693 | MVP_top60_DNA+RNA_seed | PIK3CA | -0.191277 | COADREAD | LS-180 | AZD8835 |
| MVP_top60_DNA_seed | AR | 0.151076 | COADREAD | COLO-205 | Bicalutamide | MVP_top60_DNA+RNA_seed | PIK3CD | -0.191277 | COADREAD | LS-180 | AZD8835 |
| MVP_top60_DNA_seed | PIK3CA | 0.151439 | COADREAD | HT55 | PI3Ka_4409 | MVP_top60_DNA+RNA_seed | SRC | -0.190201 | COADREAD | LS-123 | Saracatinib |
| MVP_top60_DNA_seed | PIK3CD | 0.151439 | COADREAD | HT55 | PI3Ka_4409 | MVP_top60_DNA+RNA_seed | GSK3B | -0.190068 | COADREAD | HCT-15 | AZD7969 |
| MVP_top60_DNA_seed | PIK3CB | 0.152207 | COADREAD | KM12 | AZD6482 | MVP_top60_DNA+RNA_seed | BCL2 | -0.189747 | COADREAD | SK-CO-1 | TW 37 |
| MVP_top60_DNA_seed | AR | 0.154311 | COADREAD | CCK-81 | Bicalutamide | MVP_top60_DNA+RNA_seed | GSK3B | -0.187356 | COADREAD | NCI-H747 | CHIR-99021 |
| MVP_top60_DNA_seed | ESR1 | 0.155172 | COADREAD | NCI-H747 | Fulvestrant | MVP_top60_DNA+RNA_seed | HSP90AA1 | -0.186852 | COADREAD | COLO-320-HSR | Luminespib |
| MVP_top60_DNA_seed | CREBBP | 0.156685 | COADREAD | NCI-H716 | I-CBP112 | MVP_top60_DNA+RNA_seed | HSP90AB1 | -0.186852 | COADREAD | COLO-320-HSR | Luminespib |
| MVP_top60_DNA_seed | IGF1R | 0.157714 | COADREAD | SW837 | BMS-754807 | MVP_top60_DNA+RNA_seed | HSP90AA1 | -0.183769 | COADREAD | LS-1034 | Luminespib |
| MVP_top60_DNA_seed | CDK2 | 0.157901 | COADREAD | C2BBe1 | PHA-793887 | MVP_top60_DNA+RNA_seed | HSP90AB1 | -0.183769 | COADREAD | LS-1034 | Luminespib |
| MVP_top60_DNA_seed | IGF1R | 0.158924 | COADREAD | SNU-C2B | BMS-536924 | MVP_top60_DNA+RNA_seed | SRC | -0.183704 | COADREAD | NCI-H716 | NVP-BHG712 |
| MVP_top60_DNA_seed | IGF1R | 0.159673 | COADREAD | MDST8 | BMS-536924 | MVP_top60_DNA+RNA_seed | PIK3CA | -0.182657 | COADREAD | LoVo | PI3Ka_4409 |
| MVP_top60_DNA_seed | SRC | 0.159973 | COADREAD | CaR-1 | HG-6-71-01 | MVP_top60_DNA+RNA_seed | PIK3CD | -0.182657 | COADREAD | LoVo | PI3Ka_4409 |
| MVP_top60_DNA_seed | ERBB4 | 0.160675 | COADREAD | SNU-C2B | CI-1033 | MVP_top60_DNA+RNA_seed | AR | -0.182442 | COADREAD | CCK-81 | AZD3514 |
| MVP_top60_DNA_seed | ESR1 | 0.161814 | COADREAD | RKO | Tamoxifen | MVP_top60_DNA+RNA_seed | ATM | -0.182176 | COADREAD | SW837 | CP466722 |
| MVP_top60_DNA_seed | IGF1R | 0.163869 | COADREAD | LS-123 | BMS-754807 | MVP_top60_DNA+RNA_seed | ERBB4 | -0.181255 | COADREAD | SNU-81 | CI-1033 |
| MVP_top60_DNA_seed | PIK3CA | 0.165612 | COADREAD | CL-11 | AZD8835 | MVP_top60_DNA+RNA_seed | AKT1 | -0.181097 | COADREAD | SNU-407 | BAY AKT1 |
| MVP_top60_DNA_seed | PIK3CD | 0.165612 | COADREAD | CL-11 | AZD8835 | MVP_top60_DNA+RNA_seed | SRC | -0.177613 | COADREAD | HCC2998 | NVP-BHG712 |
| MVP_top60_DNA_seed | ATM | 0.166745 | COADREAD | HCC2998 | CP466722 | MVP_top60_DNA+RNA_seed | IGF1R | -0.176039 | COADREAD | HCC2998 | BMS-536924 |
| MVP_top60_DNA_seed | BRAF | 0.167257 | COADREAD | GP5d | SB590885 | MVP_top60_DNA+RNA_seed | SRC | -0.175302 | COADREAD | C2BBe1 | NVP-BHG712 |
| MVP_top60_DNA_seed | SRC | 0.167936 | COADREAD | CL-11 | HG-6-71-01 | MVP_top60_DNA+RNA_seed | ATM | -0.173003 | COADREAD | LoVo | torin2 |
| MVP_top60_DNA_seed | PTK2 | 0.170637 | COADREAD | HT-115 | PF-562271 | MVP_top60_DNA+RNA_seed | PIK3CB | -0.171744 | COADREAD | KM12 | TGX221 |
| MVP_top60_DNA_seed | IGF1R | 0.172379 | COADREAD | DiFi | BMS-536924 | MVP_top60_DNA+RNA_seed | BCL2 | -0.170329 | COADREAD | RCM-1 | Navitoclax |
| MVP_top60_DNA_seed | AR | 0.173296 | COADREAD | SNU-C5 | AZD3514 | MVP_top60_DNA+RNA_seed | PIK3CA | -0.168674 | COADREAD | HCT-15 | AZD8835 |
| MVP_top60_DNA_seed | ERBB4 | 0.173582 | COADREAD | HCT-15 | PF-00299804 | MVP_top60_DNA+RNA_seed | PIK3CD | -0.168674 | COADREAD | HCT-15 | AZD8835 |
| MVP_top60_DNA_seed | HSP90AA1 | 0.174026 | COADREAD | T84 | CCT-018159 | MVP_top60_DNA+RNA_seed | AKT1 | -0.168247 | COADREAD | SW1463 | MK-2206 |
| MVP_top60_DNA_seed | HSP90AB1 | 0.174026 | COADREAD | T84 | CCT-018159 | MVP_top60_DNA+RNA_seed | IGF1R | -0.164215 | COADREAD | COLO-678 | BMS-754807 |
| MVP_top60_DNA_seed | CDK2 | 0.174582 | COADREAD | SNU-C5 | PHA-793887 | MVP_top60_DNA+RNA_seed | IGF1R | -0.164035 | COADREAD | LoVo | BMS-754807 |
| MVP_top60_DNA_seed | ERBB3 | 0.175298 | COADREAD | CaR-1 | AZD8931 | MVP_top60_DNA+RNA_seed | MET | -0.162049 | COADREAD | SW837 | JNJ38877605 |
| MVP_top60_DNA_seed | GSK3B | 0.175335 | COADREAD | MDST8 | KIN001-042 | MVP_top60_DNA+RNA_seed | AKT1 | -0.160481 | COADREAD | NCI-H747 | BAY AKT1 |
| MVP_top60_DNA_seed | SRC | 0.175812 | COADREAD | COLO-205 | Bosutinib | MVP_top60_DNA+RNA_seed | MET | -0.160139 | COADREAD | CaR-1 | JNJ38877605 |
| MVP_top60_DNA_seed | HSP90AA1 | 0.17594 | COADREAD | NCI-H747 | Elesclomol | MVP_top60_DNA+RNA_seed | CDK2 | -0.155209 | COADREAD | LS-123 | PHA-793887 |
| MVP_top60_DNA_seed | HSP90AB1 | 0.17594 | COADREAD | NCI-H747 | Elesclomol | MVP_top60_DNA+RNA_seed | MET | -0.153229 | COADREAD | RKO | Crizotinib |

| | | | | | | | | | | | |
|---|---|---|---|---|---|---|---|---|---|---|---|
| MVP_top60_DNA_seed | MET | 0.175963 | COADREAD | RKO | JNJ38877605 | MVP_top60_DNA+RNA_seed | MET | -0.152884 | COADREAD | C2BBe1 | PHA-665752 |
| MVP_top60_DNA_seed | PIK3CD | 0.176447 | COADREAD | HCT-116 | Idelalisib | MVP_top60_DNA+RNA_seed | PIK3CD | -0.152481 | COADREAD | GP5d | Idelalisib |
| MVP_top60_DNA_seed | MET | 0.178713 | COADREAD | SNU-C2B | JNJ38877605 | MVP_top60_DNA+RNA_seed | AR | -0.152241 | COADREAD | COLO-678 | AZD3514 |
| MVP_top60_DNA_seed | SRC | 0.179207 | COADREAD | HT-29 | Bosutinib | MVP_top60_DNA+RNA_seed | ATM | -0.151994 | COADREAD | HCT-15 | KU-55933 |
| MVP_top60_DNA_seed | AR | 0.179992 | COADREAD | GP5d | Bicalutamide | MVP_top60_DNA+RNA_seed | AKT1 | -0.14922 | COADREAD | GP5d | MK-2206 |
| MVP_top60_DNA_seed | PIK3CD | 0.180015 | COADREAD | COLO-205 | IC-87114 | MVP_top60_DNA+RNA_seed | ERBB4 | -0.144805 | COADREAD | LoVo | AST-1306 |
| MVP_top60_DNA_seed | MET | 0.180552 | COADREAD | LS-411N | PHA-665752 | MVP_top60_DNA+RNA_seed | AR | -0.143557 | COADREAD | COLO-205 | AZD3514 |
| MVP_top60_DNA_seed | MET | 0.182313 | COADREAD | SNU-C5 | SU11274 | MVP_top60_DNA+RNA_seed | ERBB4 | -0.142006 | COADREAD | SNU-61 | CI-1033 |
| MVP_top60_DNA_seed | BRAF | 0.182679 | COADREAD | SW48 | Dabrafenib | MVP_top60_DNA+RNA_seed | CDK2 | -0.141569 | COADREAD | HCC2998 | AZD5438 |
| MVP_top60_DNA_seed | GSK3B | 0.184741 | COADREAD | SNU-81 | AZD7969 | MVP_top60_DNA+RNA_seed | HSP90AA1 | -0.14001 | COADREAD | MDST8 | Tanespimycin |
| MVP_top60_DNA_seed | CREBBP | 0.184796 | COADREAD | COLO-320-HSR | I-CBP112 | MVP_top60_DNA+RNA_seed | HSP90AB1 | -0.14001 | COADREAD | MDST8 | Tanespimycin |
| MVP_top60_DNA_seed | PIK3CB | 0.186201 | COADREAD | CL-11 | AZD8186 | MVP_top60_DNA+RNA_seed | ERBB4 | -0.138818 | COADREAD | SNU-1040 | PF-00299804 |
| MVP_top60_DNA_seed | PIK3CD | 0.186201 | COADREAD | CL-11 | AZD8186 | MVP_top60_DNA+RNA_seed | GSK3B | -0.130824 | COADREAD | KM12 | SB216763 |
| MVP_top60_DNA_seed | ESR1 | 0.18731 | COADREAD | GP5d | Tamoxifen | MVP_top60_DNA+RNA_seed | PIK3CA | -0.130352 | COADREAD | T84 | PI3Ka_4409 |
| MVP_top60_DNA_seed | PIK3CB | 0.187846 | COADREAD | CaR-1 | AZD6482 | MVP_top60_DNA+RNA_seed | PIK3CD | -0.130352 | COADREAD | T84 | PI3Ka_4409 |
| MVP_top60_DNA_seed | MAPK14 | 0.188037 | COADREAD | SNU-C5 | TAK-715 | MVP_top60_DNA+RNA_seed | GSK3B | -0.126797 | COADREAD | HT-29 | CHIR-99021 |
| MVP_top60_DNA_seed | PIK3CB | 0.190635 | COADREAD | COLO-320-HSR | AZD6482 | MVP_top60_DNA+RNA_seed | ERBB4 | -0.125793 | COADREAD | GP5d | PF-00299804 |
| MVP_top60_DNA_seed | SRC | 0.191244 | COADREAD | T84 | NVP-BHG712 | MVP_top60_DNA+RNA_seed | ERBB4 | -0.122594 | COADREAD | SW1116 | PF-00299804 |
| MVP_top60_DNA_seed | ATM | 0.191918 | COADREAD | HCT-116 | KU-55933 | MVP_top60_DNA+RNA_seed | GSK3B | -0.122351 | COADREAD | SW48 | KIN001-042 |
| MVP_top60_DNA_seed | SRC | 0.193129 | COADREAD | RCM-1 | Bosutinib | MVP_top60_DNA+RNA_seed | PIK3CD | -0.121155 | COADREAD | SNU-407 | Idelalisib |
| MVP_top60_DNA_seed | HSP90AA1 | 0.194747 | COADREAD | NCI-H716 | Tanespimycin | MVP_top60_DNA+RNA_seed | AKT1 | -0.119968 | COADREAD | SW620 | AKT inhibitor VIII |
| MVP_top60_DNA_seed | HSP90AB1 | 0.194747 | COADREAD | NCI-H716 | Tanespimycin | MVP_top60_DNA+RNA_seed | CDK2 | -0.119071 | COADREAD | SNU-407 | PHA-793887 |
| MVP_top60_DNA_seed | BRAF | 0.195223 | COADREAD | SW48 | HG6-64-1 | MVP_top60_DNA+RNA_seed | PIK3CA | -0.118668 | COADREAD | NCI-H508 | AZD8835 |
| MVP_top60_DNA_seed | IGF1R | 0.195473 | COADREAD | LS-123 | Linsitinib | MVP_top60_DNA+RNA_seed | PIK3CD | -0.118668 | COADREAD | NCI-H508 | AZD8835 |
| MVP_top60_DNA_seed | AKT1 | 0.198189 | COADREAD | LS-123 | MK-2206 | MVP_top60_DNA+RNA_seed | PIK3CA | -0.113655 | COADREAD | HT-115 | PI3Ka_4409 |
| MVP_top60_DNA_seed | PIK3CA | 0.200779 | COADREAD | LS-123 | AZD8835 | MVP_top60_DNA+RNA_seed | PIK3CD | -0.113655 | COADREAD | HT-115 | PI3Ka_4409 |
| MVP_top60_DNA_seed | PIK3CD | 0.200779 | COADREAD | LS-123 | AZD8835 | MVP_top60_DNA+RNA_seed | SRC | -0.109291 | COADREAD | SW948 | NVP-BHG712 |
| MVP_top60_DNA_seed | MET | 0.200917 | COADREAD | SW620 | AZD6094 | MVP_top60_DNA+RNA_seed | ERBB4 | -0.108331 | COADREAD | SNU-C2B | AST-1306 |
| MVP_top60_DNA_seed | GSK3B | 0.204363 | COADREAD | SNU-C5 | KIN001-042 | MVP_top60_DNA+RNA_seed | PTK2 | -0.10814 | COADREAD | LS-180 | PF-562271 |
| MVP_top60_DNA_seed | HSP90AA1 | 0.204869 | COADREAD | HT-29 | Luminespib | MVP_top60_DNA+RNA_seed | GSK3B | -0.105936 | COADREAD | LS-513 | AZD7969 |
| MVP_top60_DNA_seed | HSP90AB1 | 0.204869 | COADREAD | HT-29 | Luminespib | MVP_top60_DNA+RNA_seed | HSP90AA1 | -0.103508 | COADREAD | SW837 | SNX-2112 |
| MVP_top60_DNA_seed | AKT1 | 0.207988 | COADREAD | LS-1034 | A-443654 | MVP_top60_DNA+RNA_seed | HSP90AB1 | -0.103508 | COADREAD | SW837 | SNX-2112 |
| MVP_top60_DNA_seed | PIK3CB | 0.20995 | COADREAD | RKO | AZD8186 | MVP_top60_DNA+RNA_seed | GSK3B | -0.10253 | COADREAD | SNU-C2B | AZD7969 |
| MVP_top60_DNA_seed | PIK3CD | 0.20995 | COADREAD | RKO | AZD8186 | MVP_top60_DNA+RNA_seed | PIK3CA | -0.099566 | COADREAD | LS-513 | AZD8835 |
| MVP_top60_DNA_seed | CDK2 | 0.21047 | COADREAD | CL-11 | AZD5438 | MVP_top60_DNA+RNA_seed | PIK3CD | -0.099566 | COADREAD | LS-513 | AZD8835 |
| MVP_top60_DNA_seed | MET | 0.212144 | COADREAD | NCI-H716 | AZD6094 | MVP_top60_DNA+RNA_seed | ERBB4 | -0.098547 | COADREAD | RKO | PF-00299804 |
| MVP_top60_DNA_seed | GSK3B | 0.215205 | COADREAD | COLO-320-HSR | AZD7969 | MVP_top60_DNA+RNA_seed | IGF1R | -0.098332 | COADREAD | CCK-81 | BMS-536924 |
| MVP_top60_DNA_seed | MET | 0.215728 | COADREAD | COLO-320-HSR | AZD6094 | MVP_top60_DNA+RNA_seed | ERBB4 | -0.096883 | COADREAD | LS-180 | CI-1033 |
| MVP_top60_DNA_seed | PIK3CD | 0.215953 | COADREAD | CCK-81 | IC-87114 | MVP_top60_DNA+RNA_seed | IGF1R | -0.094957 | COADREAD | HCT-116 | GSK1904529A |
| MVP_top60_DNA_seed | PIK3CD | 0.216039 | COADREAD | SW620 | Idelalisib | MVP_top60_DNA+RNA_seed | CDK2 | -0.094768 | COADREAD | SNU-C2B | AZD5438 |
| MVP_top60_DNA_seed | GSK3B | 0.216419 | COADREAD | SK-CO-1 | SB216763 | MVP_top60_DNA+RNA_seed | MET | -0.094336 | COADREAD | COLO-205 | JNJ38877605 |

| | | | | | | | | | | | |
|---|---|---|---|---|---|---|---|---|---|---|---|
| MVP_top60_DNA_seed | MET | 0.217132 | COADREAD | SK-CO-1 | AZD6094 | MVP_top60_DNA+RNA_seed | BCL2 | -0.09409 | COADREAD | SNU-C1 | Navitoclax |
| MVP_top60_DNA_seed | CDK2 | 0.217332 | COADREAD | SW48 | PHA-793887 | MVP_top60_DNA+RNA_seed | PIK3CA | -0.092356 | COADREAD | HT-115 | AZD8835 |
| MVP_top60_DNA_seed | IGF1R | 0.217401 | COADREAD | SW837 | BMS-536924 | MVP_top60_DNA+RNA_seed | PIK3CD | -0.092356 | COADREAD | HT-115 | AZD8835 |
| MVP_top60_DNA_seed | BRAF | 0.218369 | COADREAD | COLO-320-HSR | SB590885 | MVP_top60_DNA+RNA_seed | IGF1R | -0.091947 | COADREAD | SW948 | BMS-754807 |
| MVP_top60_DNA_seed | HSP90AA1 | 0.219143 | COADREAD | HT-115 | Tanespimycin | MVP_top60_DNA+RNA_seed | ERBB4 | -0.088563 | COADREAD | SNU-1040 | AST-1306 |
| MVP_top60_DNA_seed | HSP90AB1 | 0.219143 | COADREAD | HT-115 | Tanespimycin | MVP_top60_DNA+RNA_seed | GSK3B | -0.086921 | COADREAD | HT55 | KIN001-042 |
| MVP_top60_DNA_seed | AKT1 | 0.220574 | COADREAD | HT-29 | BAY AKT1 | MVP_top60_DNA+RNA_seed | PIK3CA | -0.086547 | COADREAD | LS-180 | PI3Ka_4409 |
| MVP_top60_DNA_seed | IGF1R | 0.220823 | COADREAD | NCI-H716 | BMS-536924 | MVP_top60_DNA+RNA_seed | PIK3CD | -0.086547 | COADREAD | LS-180 | PI3Ka_4409 |
| MVP_top60_DNA_seed | ATM | 0.220984 | COADREAD | CaR-1 | KU-55933 | MVP_top60_DNA+RNA_seed | AKT1 | -0.086122 | COADREAD | SNU-C1 | AKT inhibitor VIII |
| MVP_top60_DNA_seed | GSK3B | 0.222095 | COADREAD | CW-2 | KIN001-042 | MVP_top60_DNA+RNA_seed | IGF1R | -0.085648 | COADREAD | CCK-81 | BMS-754807 |
| MVP_top60_DNA_seed | IGF1R | 0.222953 | COADREAD | LS-123 | BMS-536924 | MVP_top60_DNA+RNA_seed | GSK3B | -0.081325 | COADREAD | CCK-81 | CHIR-99021 |
| MVP_top60_DNA_seed | PIK3CB | 0.223258 | COADREAD | COLO-205 | AZD8186 | MVP_top60_DNA+RNA_seed | IGF1R | -0.080503 | COADREAD | SW620 | Linsitinib |
| MVP_top60_DNA_seed | PIK3CD | 0.223258 | COADREAD | COLO-205 | AZD8186 | MVP_top60_DNA+RNA_seed | MET | -0.078778 | COADREAD | RKO | AZD6094 |
| MVP_top60_DNA_seed | PIK3CB | 0.223475 | COADREAD | SW48 | AZD6482 | MVP_top60_DNA+RNA_seed | ERBB3 | -0.078348 | COADREAD | HCC2998 | AZD8931 |
| MVP_top60_DNA_seed | AR | 0.223902 | COADREAD | GP5d | AZD3514 | MVP_top60_DNA+RNA_seed | MET | -0.077351 | COADREAD | CW-2 | SU11274 |
| MVP_top60_DNA_seed | IGF1R | 0.22413 | COADREAD | C2BBe1 | BMS-536924 | MVP_top60_DNA+RNA_seed | PIK3CA | -0.077205 | COADREAD | CCK-81 | PI3Ka_4409 |
| MVP_top60_DNA_seed | GSK3B | 0.224238 | COADREAD | HT-29 | AZD7969 | MVP_top60_DNA+RNA_seed | PIK3CD | -0.077205 | COADREAD | CCK-81 | PI3Ka_4409 |
| MVP_top60_DNA_seed | GSK3B | 0.226386 | COADREAD | SNU-C1 | KIN001-042 | MVP_top60_DNA+RNA_seed | HSP90AA1 | -0.077139 | COADREAD | HCT-116 | Elesclomol |
| MVP_top60_DNA_seed | GSK3B | 0.22647 | COADREAD | GP5d | CHIR-99021 | MVP_top60_DNA+RNA_seed | HSP90AB1 | -0.077139 | COADREAD | HCT-116 | Elesclomol |
| MVP_top60_DNA_seed | ERBB4 | 0.22687 | COADREAD | RKO | CI-1033 | MVP_top60_DNA+RNA_seed | HSP90AA1 | -0.072957 | COADREAD | LS-180 | Luminespib |
| MVP_top60_DNA_seed | ERBB4 | 0.229151 | COADREAD | COLO-205 | AST-1306 | MVP_top60_DNA+RNA_seed | HSP90AB1 | -0.072957 | COADREAD | LS-180 | Luminespib |
| MVP_top60_DNA_seed | MET | 0.230342 | COADREAD | SNU-C2B | SU11274 | MVP_top60_DNA+RNA_seed | AR | -0.071808 | COADREAD | HT55 | Bicalutamide |
| MVP_top60_DNA_seed | HSP90AA1 | 0.230513 | COADREAD | HCT-15 | Luminespib | MVP_top60_DNA+RNA_seed | MET | -0.071523 | COADREAD | RKO | SU11274 |
| MVP_top60_DNA_seed | HSP90AB1 | 0.230513 | COADREAD | HCT-15 | Luminespib | MVP_top60_DNA+RNA_seed | PIK3CB | -0.071368 | COADREAD | NCI-H747 | AZD6482 |
| MVP_top60_DNA_seed | ERBB4 | 0.231178 | COADREAD | SW837 | CI-1033 | MVP_top60_DNA+RNA_seed | IGF1R | -0.071209 | COADREAD | SNU-C2B | BMS-754807 |
| MVP_top60_DNA_seed | IGF1R | 0.231733 | COADREAD | NCI-H716 | BMS-754807 | MVP_top60_DNA+RNA_seed | AKT1 | -0.069555 | COADREAD | HCC2998 | MK-2206 |
| MVP_top60_DNA_seed | ATM | 0.232157 | COADREAD | LS-123 | CP466722 | MVP_top60_DNA+RNA_seed | HSP90AA1 | -0.069286 | COADREAD | LS-411N | Tanespimycin |
| MVP_top60_DNA_seed | HSP90AA1 | 0.234101 | COADREAD | LoVo | SNX-2112 | MVP_top60_DNA+RNA_seed | HSP90AB1 | -0.069286 | COADREAD | LS-411N | Tanespimycin |
| MVP_top60_DNA_seed | HSP90AB1 | 0.234101 | COADREAD | LoVo | SNX-2112 | MVP_top60_DNA+RNA_seed | HSP90AA1 | -0.068131 | COADREAD | HCT-15 | Tanespimycin |
| MVP_top60_DNA_seed | AKT1 | 0.23428 | COADREAD | RKO | BAY AKT1 | MVP_top60_DNA+RNA_seed | HSP90AB1 | -0.068131 | COADREAD | HCT-15 | Tanespimycin |
| MVP_top60_DNA_seed | AKT1 | 0.235043 | COADREAD | COLO-320-HSR | A-443654 | MVP_top60_DNA+RNA_seed | AKT1 | -0.06744 | COADREAD | RKO | AKT inhibitor VIII |
| MVP_top60_DNA_seed | IGF1R | 0.235689 | COADREAD | NCI-H716 | Linsitinib | MVP_top60_DNA+RNA_seed | ATM | -0.067091 | COADREAD | SNU-407 | CP466722 |
| MVP_top60_DNA_seed | PIK3CD | 0.241292 | COADREAD | LS-411N | IC-87114 | MVP_top60_DNA+RNA_seed | GSK3B | -0.063952 | COADREAD | CaR-1 | KIN001-042 |
| MVP_top60_DNA_seed | BRAF | 0.242776 | COADREAD | HCT-116 | PLX-4720 | MVP_top60_DNA+RNA_seed | GSK3B | -0.063397 | COADREAD | LS-123 | SB216763 |
| MVP_top60_DNA_seed | MAPK1 | 0.243459 | COADREAD | CL-11 | VX-11e | MVP_top60_DNA+RNA_seed | ERBB4 | -0.062005 | COADREAD | SNU-1040 | CI-1033 |
| MVP_top60_DNA_seed | MET | 0.247395 | COADREAD | RKO | PHA-665752 | MVP_top60_DNA+RNA_seed | ATM | -0.057984 | COADREAD | SW48 | torin2 |
| MVP_top60_DNA_seed | AKT1 | 0.247465 | COADREAD | HT-115 | MK-2206 | MVP_top60_DNA+RNA_seed | GSK3B | -0.057516 | COADREAD | DiFi | CHIR-99021 |
| MVP_top60_DNA_seed | IGF1R | 0.24781 | COADREAD | COLO-678 | BMS-536924 | MVP_top60_DNA+RNA_seed | ERBB4 | -0.057181 | COADREAD | HCT-116 | PF-00299804 |
| MVP_top60_DNA_seed | AKT1 | 0.249596 | COADREAD | SNU-C2B | AKT inhibitor VIII | MVP_top60_DNA+RNA_seed | GSK3B | -0.055826 | COADREAD | COLO-678 | AZD7969 |
| MVP_top60_DNA_seed | BRAF | 0.251086 | COADREAD | COLO-678 | RAF_9304 | MVP_top60_DNA+RNA_seed | AKT1 | -0.055389 | COADREAD | HCC2998 | GSK690693 |
| MVP_top60_DNA_seed | BRAF | 0.25152 | COADREAD | SW837 | PLX-4720 | MVP_top60_DNA+RNA_seed | SRC | -0.053608 | COADREAD | RKO | WH-4-023 |

| | | | | | | | | | | | |
|---|---|---|---|---|---|---|---|---|---|---|---|
| MVP_top60_DNA_seed | AR | 0.251656 | COADREAD | LS-513 | Bicalutamide | MVP_top60_DNA+RNA_seed | IGF1R | -0.053112 | COADREAD | COLO-205 | GSK1904529A |
| MVP_top60_DNA_seed | IGF1R | 0.252234 | COADREAD | SW48 | Linsitinib | MVP_top60_DNA+RNA_seed | GSK3B | -0.051287 | COADREAD | COLO-320-HSR | SB216763 |
| MVP_top60_DNA_seed | GSK3B | 0.255116 | COADREAD | SNU-175 | KIN001-042 | MVP_top60_DNA+RNA_seed | GSK3B | -0.047934 | COADREAD | HT-29 | KIN001-042 |
| MVP_top60_DNA_seed | ERBB4 | 0.256022 | COADREAD | HCC2998 | CI-1033 | MVP_top60_DNA+RNA_seed | SRC | -0.047825 | COADREAD | SNU-C2B | NVP-BHG712 |
| MVP_top60_DNA_seed | PIK3CB | 0.256785 | COADREAD | SK-CO-1 | AZD8186 | MVP_top60_DNA+RNA_seed | IGF1R | -0.046129 | COADREAD | LS-123 | GSK1904529A |
| MVP_top60_DNA_seed | PIK3CD | 0.256785 | COADREAD | SK-CO-1 | AZD8186 | MVP_top60_DNA+RNA_seed | AKT1 | -0.042683 | COADREAD | SK-CO-1 | MK-2206 |
| MVP_top60_DNA_seed | MAPK1 | 0.257187 | COADREAD | LS-123 | VX-11e | MVP_top60_DNA+RNA_seed | ATM | -0.037897 | COADREAD | LS-180 | KU-55933 |
| MVP_top60_DNA_seed | ESR1 | 0.257199 | COADREAD | MDST8 | Tamoxifen | MVP_top60_DNA+RNA_seed | MET | -0.037079 | COADREAD | LS-513 | JNJ38877605 |
| MVP_top60_DNA_seed | PIK3CB | 0.260368 | COADREAD | CCK-81 | AZD6482 | MVP_top60_DNA+RNA_seed | ATM | -0.035626 | COADREAD | GP5d | CP466722 |
| MVP_top60_DNA_seed | BRAF | 0.260894 | COADREAD | SW1417 | SB590885 | MVP_top60_DNA+RNA_seed | AKT1 | -0.035069 | COADREAD | COLO-205 | AKT inhibitor VIII |
| MVP_top60_DNA_seed | MET | 0.261201 | COADREAD | HCC2998 | AZD6094 | MVP_top60_DNA+RNA_seed | GSK3B | -0.034486 | COADREAD | RKO | SB216763 |
| MVP_top60_DNA_seed | CDK2 | 0.261956 | COADREAD | GP5d | AZD5438 | MVP_top60_DNA+RNA_seed | CDK2 | -0.033882 | COADREAD | LS-1034 | AZD5438 |
| MVP_top60_DNA_seed | AKT1 | 0.262406 | COADREAD | COLO-205 | GSK690693 | MVP_top60_DNA+RNA_seed | PIK3CD | -0.031025 | COADREAD | T84 | Idelalisib |
| MVP_top60_DNA_seed | HSP90AA1 | 0.263291 | COADREAD | SNU-C2B | Elesclomol | MVP_top60_DNA+RNA_seed | CDK2 | -0.03029 | COADREAD | GP5d | PHA-793887 |
| MVP_top60_DNA_seed | HSP90AB1 | 0.263291 | COADREAD | SNU-C2B | Elesclomol | MVP_top60_DNA+RNA_seed | SRC | -0.029833 | COADREAD | SNU-C2B | WH-4-023 |
| MVP_top60_DNA_seed | SRC | 0.263533 | COADREAD | SNU-C5 | NVP-BHG712 | MVP_top60_DNA+RNA_seed | CDK2 | -0.029594 | COADREAD | KM12 | PHA-793887 |
| MVP_top60_DNA_seed | ERBB3 | 0.264419 | COADREAD | RCM-1 | AZD8931 | MVP_top60_DNA+RNA_seed | AR | -0.025057 | COADREAD | CL-11 | AZD3514 |
| MVP_top60_DNA_seed | HSP90AA1 | 0.26456 | COADREAD | SW837 | Elesclomol | MVP_top60_DNA+RNA_seed | AR | -0.023246 | COADREAD | MDST8 | Bicalutamide |
| MVP_top60_DNA_seed | HSP90AB1 | 0.26456 | COADREAD | SW837 | Elesclomol | MVP_top60_DNA+RNA_seed | ERBB4 | -0.022913 | COADREAD | SW1417 | CI-1033 |
| MVP_top60_DNA_seed | SRC | 0.264585 | COADREAD | SW48 | HG-6-71-01 | MVP_top60_DNA+RNA_seed | GSK3B | -0.017953 | COADREAD | NCI-H747 | AZD7969 |
| MVP_top60_DNA_seed | MAPK14 | 0.265392 | COADREAD | SNU-61 | TAK-715 | MVP_top60_DNA+RNA_seed | MET | -0.016816 | COADREAD | HCC2998 | JNJ38877605 |
| MVP_top60_DNA_seed | PIK3CA | 0.265577 | COADREAD | HT55 | AZD8835 | MVP_top60_DNA+RNA_seed | HSP90AA1 | -0.015843 | COADREAD | COLO-205 | Luminespib |
| MVP_top60_DNA_seed | PIK3CD | 0.265577 | COADREAD | HT55 | AZD8835 | MVP_top60_DNA+RNA_seed | HSP90AB1 | -0.015843 | COADREAD | COLO-205 | Luminespib |
| MVP_top60_DNA_seed | ATM | 0.267102 | COADREAD | COLO-320-HSR | KU-55933 | MVP_top60_DNA+RNA_seed | HSP90AA1 | -0.013194 | COADREAD | KM12 | SNX-2112 |
| MVP_top60_DNA_seed | BRAF | 0.267861 | COADREAD | HCC2998 | PLX-4720 | MVP_top60_DNA+RNA_seed | HSP90AB1 | -0.013194 | COADREAD | KM12 | SNX-2112 |
| MVP_top60_DNA_seed | ESR1 | 0.267959 | COADREAD | LS-411N | Fulvestrant | MVP_top60_DNA+RNA_seed | ATM | -0.012139 | COADREAD | RKO | KU-55933 |
| MVP_top60_DNA_seed | BRAF | 0.26976 | COADREAD | SW1417 | HG6-64-1 | MVP_top60_DNA+RNA_seed | AKT1 | -0.011248 | COADREAD | HCC2998 | AKT inhibitor VIII |
| MVP_top60_DNA_seed | PIK3CA | 0.271829 | COADREAD | SW1463 | AZD8835 | MVP_top60_DNA+RNA_seed | ERBB3 | -0.010886 | COADREAD | LS-180 | AZD8931 |
| MVP_top60_DNA_seed | PIK3CD | 0.271829 | COADREAD | SW1463 | AZD8835 | MVP_top60_DNA+RNA_seed | AKT1 | -0.008845 | COADREAD | MDST8 | MK-2206 |
| MVP_top60_DNA_seed | ESR1 | 0.274396 | COADREAD | SW620 | Fulvestrant | MVP_top60_DNA+RNA_seed | ERBB4 | -0.007733 | COADREAD | HT55 | AST-1306 |
| MVP_top60_DNA_seed | ESR1 | 0.275404 | COADREAD | T84 | Fulvestrant | MVP_top60_DNA+RNA_seed | PIK3CD | -0.007236 | COADREAD | LS-180 | Idelalisib |
| MVP_top60_DNA_seed | ESR1 | 0.275613 | COADREAD | NCI-H716 | Tamoxifen | MVP_top60_DNA+RNA_seed | MET | -0.006465 | COADREAD | NCI-H747 | AZD6094 |
| MVP_top60_DNA_seed | CREBBP | 0.276505 | COADREAD | DiFi | I-CBP112 | MVP_top60_DNA+RNA_seed | PTK2 | -0.005536 | COADREAD | LS-123 | PF-562271 |
| MVP_top60_DNA_seed | IGF1R | 0.278901 | COADREAD | HT55 | BMS-754807 | MVP_top60_DNA+RNA_seed | AR | -0.00485 | COADREAD | HCC2998 | Bicalutamide |
| MVP_top60_DNA_seed | BRAF | 0.279003 | COADREAD | HCT-15 | PLX-4720 | MVP_top60_DNA+RNA_seed | AR | -0.001807 | COADREAD | RKO | Bicalutamide |
| MVP_top60_DNA_seed | ATM | 0.279929 | COADREAD | SNU-61 | CP466722 | MVP_top60_DNA+RNA_seed | HSP90AA1 | -0.000142 | COADREAD | C2BBe1 | Elesclomol |
| MVP_top60_DNA_seed | MAPK14 | 0.280273 | COADREAD | RCM-1 | TAK-715 | MVP_top60_DNA+RNA_seed | HSP90AB1 | -0.000142 | COADREAD | C2BBe1 | Elesclomol |
| MVP_top60_DNA_seed | ESR1 | 0.285545 | COADREAD | LS-411N | Tamoxifen | MVP_top60_DNA+RNA_seed | AKT1 | 0.000203 | COADREAD | C2BBe1 | AKT inhibitor VIII |
| MVP_top60_DNA_seed | IGF1R | 0.285579 | COADREAD | NCI-H508 | BMS-536924 | MVP_top60_DNA+RNA_seed | PIK3CB | 0.001164 | COADREAD | COLO-205 | AZD6482 |
| MVP_top60_DNA_seed | PIK3CA | 0.286264 | COADREAD | SNU-C2B | PI3Ka_4409 | MVP_top60_DNA+RNA_seed | CDK2 | 0.001426 | COADREAD | HCC2998 | PHA-793887 |
| MVP_top60_DNA_seed | PIK3CD | 0.286264 | COADREAD | SNU-C2B | PI3Ka_4409 | MVP_top60_DNA+RNA_seed | HSP90AA1 | 0.005744 | COADREAD | NCI-H716 | CCT-018159 |

| | | | | | | | | | | | |
|---|---|---|---|---|---|---|---|---|---|---|---|
| MVP_top60_DNA_seed | AKT1 | 0.286319 | COADREAD | COLO-320-HSR | MK-2206 | MVP_top60_DNA+RNA_seed | HSP90AB1 | 0.005744 | COADREAD | NCI-H716 | CCT-018159 |
| MVP_top60_DNA_seed | ERBB4 | 0.286685 | COADREAD | C2BBe1 | PF-00299804 | MVP_top60_DNA+RNA_seed | AKT1 | 0.009782 | COADREAD | RKO | GSK690693 |
| MVP_top60_DNA_seed | MET | 0.286702 | COADREAD | COLO-205 | SU11274 | MVP_top60_DNA+RNA_seed | ERBB4 | 0.01298 | COADREAD | CaR-1 | AST-1306 |
| MVP_top60_DNA_seed | ATM | 0.287877 | COADREAD | GP5d | KU-60019 | MVP_top60_DNA+RNA_seed | CDK2 | 0.016716 | COADREAD | NCI-H716 | AZD5438 |
| MVP_top60_DNA_seed | AKT1 | 0.287881 | COADREAD | HCT-116 | MK-2206 | MVP_top60_DNA+RNA_seed | AKT1 | 0.018104 | COADREAD | LS-123 | A-443654 |
| MVP_top60_DNA_seed | GSK3B | 0.289414 | COADREAD | NCI-H716 | KIN001-042 | MVP_top60_DNA+RNA_seed | HSP90AA1 | 0.018872 | COADREAD | MDST8 | Luminespib |
| MVP_top60_DNA_seed | ATM | 0.289545 | COADREAD | NCI-H508 | KU-55933 | MVP_top60_DNA+RNA_seed | HSP90AB1 | 0.018872 | COADREAD | MDST8 | Luminespib |
| MVP_top60_DNA_seed | BRAF | 0.289821 | COADREAD | KM12 | SB590885 | MVP_top60_DNA+RNA_seed | HSP90AA1 | 0.020484 | COADREAD | COLO-678 | Elesclomol |
| MVP_top60_DNA_seed | HSP90AA1 | 0.289851 | COADREAD | SK-CO-1 | CCT-018159 | MVP_top60_DNA+RNA_seed | HSP90AB1 | 0.020484 | COADREAD | COLO-678 | Elesclomol |
| MVP_top60_DNA_seed | HSP90AB1 | 0.289851 | COADREAD | SK-CO-1 | CCT-018159 | MVP_top60_DNA+RNA_seed | IGF1R | 0.021813 | COADREAD | C2BBe1 | BMS-536924 |
| MVP_top60_DNA_seed | BRAF | 0.294653 | COADREAD | LS-123 | AZ628 | MVP_top60_DNA+RNA_seed | HSP90AA1 | 0.021814 | COADREAD | LS-513 | Luminespib |
| MVP_top60_DNA_seed | PIK3CB | 0.296582 | COADREAD | SNU-C2B | AZD8186 | MVP_top60_DNA+RNA_seed | HSP90AB1 | 0.021814 | COADREAD | LS-513 | Luminespib |
| MVP_top60_DNA_seed | PIK3CD | 0.296582 | COADREAD | SNU-C2B | AZD8186 | MVP_top60_DNA+RNA_seed | PIK3CA | 0.022367 | COADREAD | SNU-81 | AZD8835 |
| MVP_top60_DNA_seed | SRC | 0.298139 | COADREAD | C2BBe1 | WH-4-023 | MVP_top60_DNA+RNA_seed | PIK3CD | 0.022367 | COADREAD | SNU-81 | AZD8835 |
| MVP_top60_DNA_seed | MET | 0.298202 | COADREAD | NCI-H747 | Crizotinib | MVP_top60_DNA+RNA_seed | HSP90AA1 | 0.02393 | COADREAD | CCK-81 | CCT-018159 |
| MVP_top60_DNA_seed | ATM | 0.299476 | COADREAD | LS-1034 | KU-60019 | MVP_top60_DNA+RNA_seed | HSP90AB1 | 0.02393 | COADREAD | CCK-81 | CCT-018159 |
| MVP_top60_DNA_seed | PIK3CD | 0.300009 | COADREAD | COLO-320-HSR | IC-87114 | MVP_top60_DNA+RNA_seed | AR | 0.024116 | COADREAD | HT-29 | AZD3514 |
| MVP_top60_DNA_seed | MAPK14 | 0.300438 | COADREAD | HCT-15 | TAK-715 | MVP_top60_DNA+RNA_seed | ERBB4 | 0.024327 | COADREAD | GP5d | AST-1306 |
| MVP_top60_DNA_seed | GSK3B | 0.301685 | COADREAD | LS-180 | SB216763 | MVP_top60_DNA+RNA_seed | ERBB4 | 0.025599 | COADREAD | RKO | AST-1306 |
| MVP_top60_DNA_seed | ESR1 | 0.302581 | COADREAD | DiFi | Tamoxifen | MVP_top60_DNA+RNA_seed | PIK3CA | 0.028046 | COADREAD | CL-11 | PI3Ka_4409 |
| MVP_top60_DNA_seed | ERBB3 | 0.302785 | COADREAD | RKO | AZD8931 | MVP_top60_DNA+RNA_seed | PIK3CD | 0.028046 | COADREAD | CL-11 | PI3Ka_4409 |
| MVP_top60_DNA_seed | BRAF | 0.30351 | COADREAD | T84 | RAF_9304 | MVP_top60_DNA+RNA_seed | PIK3CB | 0.02924 | COADREAD | HCC2998 | TGX221 |
| MVP_top60_DNA_seed | BRAF | 0.304312 | COADREAD | NCI-H716 | Dabrafenib | MVP_top60_DNA+RNA_seed | CDK2 | 0.029517 | COADREAD | HCC2998 | Seliciclib |
| MVP_top60_DNA_seed | BRAF | 0.31342 | COADREAD | HCC2998 | SB590885 | MVP_top60_DNA+RNA_seed | AKT1 | 0.030179 | COADREAD | SNU-C5 | BAY AKT1 |
| MVP_top60_DNA_seed | CDK2 | 0.313458 | COADREAD | SNU-81 | AZD5438 | MVP_top60_DNA+RNA_seed | ERBB4 | 0.030699 | COADREAD | LS-180 | PF-00299804 |
| MVP_top60_DNA_seed | ERBB4 | 0.314411 | COADREAD | SW837 | PF-00299804 | MVP_top60_DNA+RNA_seed | HSP90AA1 | 0.031206 | COADREAD | SNU-C5 | Elesclomol |
| MVP_top60_DNA_seed | AKT1 | 0.314682 | COADREAD | LS-1034 | AKT inhibitor VIII | MVP_top60_DNA+RNA_seed | HSP90AB1 | 0.031206 | COADREAD | SNU-C5 | Elesclomol |
| MVP_top60_DNA_seed | PIK3CA | 0.315191 | COADREAD | HCC2998 | PI3Ka_4409 | MVP_top60_DNA+RNA_seed | BCL2 | 0.031407 | COADREAD | HT55 | TW 37 |
| MVP_top60_DNA_seed | PIK3CD | 0.315191 | COADREAD | HCC2998 | PI3Ka_4409 | MVP_top60_DNA+RNA_seed | BCL2 | 0.031774 | COADREAD | LS-180 | Obatoclax Mesylate |
| MVP_top60_DNA_seed | AR | 0.316644 | COADREAD | SW837 | AZD3514 | MVP_top60_DNA+RNA_seed | GSK3B | 0.031994 | COADREAD | HCT-15 | CHIR-99021 |
| MVP_top60_DNA_seed | GSK3B | 0.317366 | COADREAD | HCT-15 | KIN001-042 | MVP_top60_DNA+RNA_seed | HSP90AA1 | 0.036958 | COADREAD | LS-123 | Elesclomol |
| MVP_top60_DNA_seed | BRAF | 0.318441 | COADREAD | SW620 | Dabrafenib | MVP_top60_DNA+RNA_seed | HSP90AB1 | 0.036958 | COADREAD | LS-123 | Elesclomol |
| MVP_top60_DNA_seed | MET | 0.318844 | COADREAD | NCI-H716 | PHA-665752 | MVP_top60_DNA+RNA_seed | PIK3CA | 0.037754 | COADREAD | LS-123 | PI3Ka_4409 |
| MVP_top60_DNA_seed | HSP90AA1 | 0.319765 | COADREAD | CW-2 | SNX-2112 | MVP_top60_DNA+RNA_seed | PIK3CD | 0.037754 | COADREAD | LS-123 | PI3Ka_4409 |
| MVP_top60_DNA_seed | HSP90AB1 | 0.319765 | COADREAD | CW-2 | SNX-2112 | MVP_top60_DNA+RNA_seed | PIK3CA | 0.038105 | COADREAD | SW837 | PI3Ka_4409 |
| MVP_top60_DNA_seed | PIK3CA | 0.320958 | COADREAD | HCT-15 | PI3Ka_4409 | MVP_top60_DNA+RNA_seed | PIK3CD | 0.038105 | COADREAD | SW837 | PI3Ka_4409 |
| MVP_top60_DNA_seed | PIK3CD | 0.320958 | COADREAD | HCT-15 | PI3Ka_4409 | MVP_top60_DNA+RNA_seed | FYN | 0.039886 | COADREAD | LS-1034 | A-770041 |
| MVP_top60_DNA_seed | HSP90AA1 | 0.321801 | COADREAD | C2BBe1 | Luminespib | MVP_top60_DNA+RNA_seed | PIK3CA | 0.039911 | COADREAD | SW948 | AZD8835 |
| MVP_top60_DNA_seed | HSP90AB1 | 0.321801 | COADREAD | C2BBe1 | Luminespib | MVP_top60_DNA+RNA_seed | PIK3CD | 0.039911 | COADREAD | SW948 | AZD8835 |
| MVP_top60_DNA_seed | GSK3B | 0.323895 | COADREAD | LS-180 | KIN001-042 | MVP_top60_DNA+RNA_seed | AR | 0.041203 | COADREAD | NCI-H508 | AZD3514 |
| MVP_top60_DNA_seed | HSP90AA1 | 0.325308 | COADREAD | LoVo | Elesclomol | MVP_top60_DNA+RNA_seed | SRC | 0.041774 | COADREAD | SW837 | Bosutinib |

| | | | | | | | | | | | |
|---|---|---|---|---|---|---|---|---|---|---|---|
| MVP_top60_DNA_seed | HSP90AB1 | 0.325308 | COADREAD | LoVo | Elesclomol | MVP_top60_DNA+RNA_seed | IGF1R | 0.042076 | COADREAD | HT-115 | BMS-536924 |
| MVP_top60_DNA_seed | ERBB4 | 0.325509 | COADREAD | GP5d | CI-1033 | MVP_top60_DNA+RNA_seed | AR | 0.044503 | COADREAD | HT-115 | Bicalutamide |
| MVP_top60_DNA_seed | ERBB4 | 0.326663 | COADREAD | SNU-81 | PF-00299804 | MVP_top60_DNA+RNA_seed | SRC | 0.04576 | COADREAD | SNU-1040 | NVP-BHG712 |
| MVP_top60_DNA_seed | HSP90AA1 | 0.327222 | COADREAD | HCC2998 | CCT-018159 | MVP_top60_DNA+RNA_seed | AKT1 | 0.046622 | COADREAD | LS-180 | BAY AKT1 |
| MVP_top60_DNA_seed | HSP90AB1 | 0.327222 | COADREAD | HCC2998 | CCT-018159 | MVP_top60_DNA+RNA_seed | HSP90AA1 | 0.046861 | COADREAD | SNU-C5 | SNX-2112 |
| MVP_top60_DNA_seed | GSK3B | 0.327658 | COADREAD | LS-180 | AZD7969 | MVP_top60_DNA+RNA_seed | HSP90AB1 | 0.046861 | COADREAD | SNU-C5 | SNX-2112 |
| MVP_top60_DNA_seed | MET | 0.329526 | COADREAD | MDST8 | AZD6094 | MVP_top60_DNA+RNA_seed | AR | 0.047295 | COADREAD | HCC2998 | Bicalutamide |
| MVP_top60_DNA_seed | PIK3CA | 0.331059 | COADREAD | SW1417 | PI3Ka_4409 | MVP_top60_DNA+RNA_seed | ERBB3 | 0.047486 | COADREAD | SW620 | AZD8931 |
| MVP_top60_DNA_seed | PIK3CD | 0.331059 | COADREAD | SW1417 | PI3Ka_4409 | MVP_top60_DNA+RNA_seed | HSP90AA1 | 0.0541 | COADREAD | SW948 | SNX-2112 |
| MVP_top60_DNA_seed | AKT1 | 0.331596 | COADREAD | LS-1034 | AKT inhibitor VIII | MVP_top60_DNA+RNA_seed | HSP90AB1 | 0.0541 | COADREAD | SW948 | SNX-2112 |
| MVP_top60_DNA_seed | BRAF | 0.332525 | COADREAD | GP5d | PLX-4720 | MVP_top60_DNA+RNA_seed | AKT1 | 0.056641 | COADREAD | HCT-116 | AKT inhibitor VIII |
| MVP_top60_DNA_seed | MET | 0.334734 | COADREAD | CaR-1 | SU11274 | MVP_top60_DNA+RNA_seed | PIK3CA | 0.057106 | COADREAD | HT-29 | PI3Ka_4409 |
| MVP_top60_DNA_seed | SRC | 0.337516 | COADREAD | RKO | Saracatinib | MVP_top60_DNA+RNA_seed | PIK3CD | 0.057106 | COADREAD | HT-29 | PI3Ka_4409 |
| MVP_top60_DNA_seed | AKT1 | 0.338185 | COADREAD | LS-123 | BAY AKT1 | MVP_top60_DNA+RNA_seed | IGF1R | 0.05728 | COADREAD | COLO-678 | GSK1904529A |
| MVP_top60_DNA_seed | SRC | 0.339198 | COADREAD | SNU-61 | NVP-BHG712 | MVP_top60_DNA+RNA_seed | MET | 0.059495 | COADREAD | SW620 | JNJ38877605 |
| MVP_top60_DNA_seed | BRAF | 0.340068 | COADREAD | NCI-H508 | RAF_9304 | MVP_top60_DNA+RNA_seed | ERBB4 | 0.06093 | COADREAD | COLO-320-HSR | PF-00299804 |
| MVP_top60_DNA_seed | ATM | 0.340794 | COADREAD | SW48 | CP466722 | MVP_top60_DNA+RNA_seed | ATM | 0.062083 | COADREAD | SW620 | KU-60019 |
| MVP_top60_DNA_seed | AKT1 | 0.341217 | COADREAD | LS-180 | MK-2206 | MVP_top60_DNA+RNA_seed | PIK3CA | 0.064424 | COADREAD | LS-1034 | PI3Ka_4409 |
| MVP_top60_DNA_seed | ERBB3 | 0.343092 | COADREAD | SW837 | AZD8931 | MVP_top60_DNA+RNA_seed | PIK3CD | 0.064424 | COADREAD | LS-1034 | PI3Ka_4409 |
| MVP_top60_DNA_seed | ESR1 | 0.344613 | COADREAD | NCI-H508 | Tamoxifen | MVP_top60_DNA+RNA_seed | BCL2 | 0.06521 | COADREAD | SW48 | Navitoclax |
| MVP_top60_DNA_seed | MET | 0.345193 | COADREAD | LoVo | AZD6094 | MVP_top60_DNA+RNA_seed | GSK3B | 0.066142 | COADREAD | SW620 | SB216763 |
| MVP_top60_DNA_seed | ATM | 0.345977 | COADREAD | NCI-H747 | KU-60019 | MVP_top60_DNA+RNA_seed | IGF1R | 0.066922 | COADREAD | SW620 | BMS-536924 |
| MVP_top60_DNA_seed | IGF1R | 0.346394 | COADREAD | NCI-H747 | BMS-536924 | MVP_top60_DNA+RNA_seed | ERBB4 | 0.066961 | COADREAD | COLO-205 | PF-00299804 |
| MVP_top60_DNA_seed | BRAF | 0.347518 | COADREAD | NCI-H508 | Dabrafenib | MVP_top60_DNA+RNA_seed | CDK2 | 0.070221 | COADREAD | MDST8 | Seliciclib |
| MVP_top60_DNA_seed | HSP90AA1 | 0.349892 | COADREAD | RCM-1 | Tanespimycin | MVP_top60_DNA+RNA_seed | CDK2 | 0.072017 | COADREAD | MDST8 | PHA-793887 |
| MVP_top60_DNA_seed | HSP90AB1 | 0.349892 | COADREAD | RCM-1 | Tanespimycin | MVP_top60_DNA+RNA_seed | AR | 0.073282 | COADREAD | HCT-116 | Bicalutamide |
| MVP_top60_DNA_seed | PIK3CD | 0.352255 | COADREAD | SNU-1040 | IC-87114 | MVP_top60_DNA+RNA_seed | MET | 0.075353 | COADREAD | GP5d | AZD6094 |
| MVP_top60_DNA_seed | GSK3B | 0.353586 | COADREAD | SNU-C2B | CHIR-99021 | MVP_top60_DNA+RNA_seed | MET | 0.075659 | COADREAD | CW-2 | PHA-665752 |
| MVP_top60_DNA_seed | ESR1 | 0.353633 | COADREAD | HCC2998 | Tamoxifen | MVP_top60_DNA+RNA_seed | ERBB3 | 0.07653 | COADREAD | HCT-15 | AZD8931 |
| MVP_top60_DNA_seed | PIK3CD | 0.35471 | COADREAD | LS-411N | Idelalisib | MVP_top60_DNA+RNA_seed | AR | 0.076685 | COADREAD | CL-11 | Bicalutamide |
| MVP_top60_DNA_seed | ATM | 0.356093 | COADREAD | HT-115 | KU-60019 | MVP_top60_DNA+RNA_seed | GSK3B | 0.07669 | COADREAD | CCK-81 | AZD7969 |
| MVP_top60_DNA_seed | HSP90AA1 | 0.356311 | COADREAD | HT55 | Tanespimycin | MVP_top60_DNA+RNA_seed | ERBB4 | 0.07703 | COADREAD | C2BBe1 | CI-1033 |
| MVP_top60_DNA_seed | HSP90AB1 | 0.356311 | COADREAD | HT55 | Tanespimycin | MVP_top60_DNA+RNA_seed | PIK3CA | 0.078169 | COADREAD | SNU-1040 | PI3Ka_4409 |
| MVP_top60_DNA_seed | SRC | 0.356529 | COADREAD | SW620 | Bosutinib | MVP_top60_DNA+RNA_seed | PIK3CD | 0.078169 | COADREAD | SNU-1040 | PI3Ka_4409 |
| MVP_top60_DNA_seed | ESR1 | 0.357815 | COADREAD | SW48 | Tamoxifen | MVP_top60_DNA+RNA_seed | ATM | 0.079666 | COADREAD | HT-29 | CP466722 |
| MVP_top60_DNA_seed | MET | 0.359117 | COADREAD | HT-115 | JNJ38877605 | MVP_top60_DNA+RNA_seed | GSK3B | 0.081653 | COADREAD | C2BBe1 | CHIR-99021 |
| MVP_top60_DNA_seed | AKT1 | 0.3598 | COADREAD | HT-115 | AKT inhibitor VIII | MVP_top60_DNA+RNA_seed | GSK3B | 0.082098 | COADREAD | CCK-81 | KIN001-042 |
| MVP_top60_DNA_seed | PIK3CB | 0.359917 | COADREAD | RKO | TGX221 | MVP_top60_DNA+RNA_seed | ATM | 0.083235 | COADREAD | C2BBe1 | KU-60019 |
| MVP_top60_DNA_seed | ESR1 | 0.361091 | COADREAD | SK-CO-1 | Tamoxifen | MVP_top60_DNA+RNA_seed | PIK3CB | 0.088006 | COADREAD | LS-123 | TGX221 |
| MVP_top60_DNA_seed | AKT1 | 0.364121 | COADREAD | CL-11 | BAY AKT1 | MVP_top60_DNA+RNA_seed | ERBB4 | 0.089055 | COADREAD | CL-11 | AST-1306 |
| MVP_top60_DNA_seed | ERBB4 | 0.364252 | COADREAD | HT-29 | PF-00299804 | MVP_top60_DNA+RNA_seed | CDK2 | 0.089275 | COADREAD | C2BBe1 | AZD5438 |

| | | | | | | | | | | | |
|---|---|---|---|---|---|---|---|---|---|---|---|
| MVP_top60_DNA_seed | AKT1 | 0.364678 | COADREAD | HT-29 | GSK690693 | MVP_top60_DNA+RNA_seed | ATM | 0.090307 | COADREAD | CL-11 | torin2 |
| MVP_top60_DNA_seed | ERBB4 | 0.364772 | COADREAD | LS-513 | CI-1033 | MVP_top60_DNA+RNA_seed | CDK2 | 0.09192 | COADREAD | SK-CO-1 | AZD5438 |
| MVP_top60_DNA_seed | ATM | 0.364793 | COADREAD | NCI-H716 | CP466722 | MVP_top60_DNA+RNA_seed | SRC | 0.091963 | COADREAD | COLO-678 | Bosutinib |
| MVP_top60_DNA_seed | BRAF | 0.36533 | COADREAD | DiFi | Dabrafenib | MVP_top60_DNA+RNA_seed | ERBB4 | 0.095234 | COADREAD | LS-411N | PF-00299804 |
| MVP_top60_DNA_seed | GSK3B | 0.366349 | COADREAD | MDST8 | SB216763 | MVP_top60_DNA+RNA_seed | FYN | 0.095757 | COADREAD | COLO-320-HSR | A-770041 |
| MVP_top60_DNA_seed | ATM | 0.366626 | COADREAD | LS-513 | CP466722 | MVP_top60_DNA+RNA_seed | PIK3CD | 0.096357 | COADREAD | RKO | IC-87114 |
| MVP_top60_DNA_seed | AKT1 | 0.366711 | COADREAD | SNU-61 | AKT inhibitor VIII | MVP_top60_DNA+RNA_seed | MET | 0.09665 | COADREAD | COLO-320-HSR | PHA-665752 |
| MVP_top60_DNA_seed | PIK3CD | 0.366829 | COADREAD | KM12 | Idelalisib | MVP_top60_DNA+RNA_seed | AKT1 | 0.101675 | COADREAD | SNU-C1 | AKT inhibitor VIII |
| MVP_top60_DNA_seed | GSK3B | 0.367105 | COADREAD | SNU-407 | AZD7969 | MVP_top60_DNA+RNA_seed | AKT1 | 0.105316 | COADREAD | MDST8 | GSK690693 |
| MVP_top60_DNA_seed | IGF1R | 0.367383 | COADREAD | HT55 | GSK1904529A | MVP_top60_DNA+RNA_seed | AR | 0.105373 | COADREAD | LoVo | Bicalutamide |
| MVP_top60_DNA_seed | MET | 0.367974 | COADREAD | HT-115 | AZD6094 | MVP_top60_DNA+RNA_seed | AKT1 | 0.105655 | COADREAD | CW-2 | GSK690693 |
| MVP_top60_DNA_seed | BRAF | 0.368817 | COADREAD | NCI-H747 | SB590885 | MVP_top60_DNA+RNA_seed | HSP90AA1 | 0.106424 | COADREAD | COLO-678 | CCT-018159 |
| MVP_top60_DNA_seed | PIK3CD | 0.369462 | COADREAD | SW620 | IC-87114 | MVP_top60_DNA+RNA_seed | HSP90AB1 | 0.106424 | COADREAD | COLO-678 | CCT-018159 |
| MVP_top60_DNA_seed | HSP90AA1 | 0.369557 | COADREAD | RCM-1 | CCT-018159 | MVP_top60_DNA+RNA_seed | MET | 0.106542 | COADREAD | COLO-678 | JNJ38877605 |
| MVP_top60_DNA_seed | HSP90AB1 | 0.369557 | COADREAD | RCM-1 | CCT-018159 | MVP_top60_DNA+RNA_seed | BCL2 | 0.1088 | COADREAD | LoVo | Navitoclax |
| MVP_top60_DNA_seed | BRAF | 0.371569 | COADREAD | CaR-1 | SB590885 | MVP_top60_DNA+RNA_seed | HSP90AA1 | 0.109734 | COADREAD | SK-CO-1 | Elesclomol |
| MVP_top60_DNA_seed | PIK3CB | 0.372144 | COADREAD | NCI-H747 | TGX221 | MVP_top60_DNA+RNA_seed | HSP90AB1 | 0.109734 | COADREAD | SK-CO-1 | Elesclomol |
| MVP_top60_DNA_seed | CDK2 | 0.372614 | COADREAD | C2BBe1 | Seliciclib | MVP_top60_DNA+RNA_seed | ERBB4 | 0.10991 | COADREAD | SW1417 | AST-1306 |
| MVP_top60_DNA_seed | ATM | 0.374839 | COADREAD | LS-1034 | CP466722 | MVP_top60_DNA+RNA_seed | IGF1R | 0.110374 | COADREAD | HCT-116 | BMS-754807 |
| MVP_top60_DNA_seed | AKT1 | 0.375724 | COADREAD | SW48 | AKT inhibitor VIII | MVP_top60_DNA+RNA_seed | PIK3CD | 0.110451 | COADREAD | HT-115 | IC-87114 |
| MVP_top60_DNA_seed | SRC | 0.377317 | COADREAD | HCT-15 | NVP-BHG712 | MVP_top60_DNA+RNA_seed | PIK3CB | 0.111096 | COADREAD | SW837 | AZD8186 |
| MVP_top60_DNA_seed | PIK3CB | 0.378235 | COADREAD | SNU-C1 | AZD8186 | MVP_top60_DNA+RNA_seed | PIK3CD | 0.111096 | COADREAD | SW837 | AZD8186 |
| MVP_top60_DNA_seed | PIK3CD | 0.378235 | COADREAD | SNU-C1 | AZD8186 | MVP_top60_DNA+RNA_seed | PIK3CD | 0.111612 | COADREAD | HCT-116 | IC-87114 |
| MVP_top60_DNA_seed | AKT1 | 0.379372 | COADREAD | LS-513 | BAY AKT1 | MVP_top60_DNA+RNA_seed | BCL2 | 0.114102 | COADREAD | SW620 | TW 37 |
| MVP_top60_DNA_seed | AR | 0.380128 | COADREAD | LoVo | Bicalutamide | MVP_top60_DNA+RNA_seed | ATM | 0.114673 | COADREAD | GP5d | torin2 |
| MVP_top60_DNA_seed | MET | 0.38041 | COADREAD | SNU-C2B | Crizotinib | MVP_top60_DNA+RNA_seed | SRC | 0.114684 | COADREAD | CaR-1 | Bosutinib |
| MVP_top60_DNA_seed | SRC | 0.38122 | COADREAD | SW48 | NVP-BHG712 | MVP_top60_DNA+RNA_seed | SRC | 0.120193 | COADREAD | COLO-320-HSR | NVP-BHG712 |
| MVP_top60_DNA_seed | ATM | 0.38181 | COADREAD | C2BBe1 | torin2 | MVP_top60_DNA+RNA_seed | AKT1 | 0.121509 | COADREAD | CL-11 | MK-2206 |
| MVP_top60_DNA_seed | ATM | 0.382052 | COADREAD | CaR-1 | torin2 | MVP_top60_DNA+RNA_seed | ERBB3 | 0.123873 | COADREAD | C2BBe1 | AZD8931 |
| MVP_top60_DNA_seed | HSP90AA1 | 0.382511 | COADREAD | C2BBe1 | SNX-2112 | MVP_top60_DNA+RNA_seed | ATM | 0.125947 | COADREAD | HT-115 | torin2 |
| MVP_top60_DNA_seed | HSP90AB1 | 0.382511 | COADREAD | C2BBe1 | SNX-2112 | MVP_top60_DNA+RNA_seed | AKT1 | 0.127558 | COADREAD | SW620 | AKT inhibitor VIII |
| MVP_top60_DNA_seed | PIK3CD | 0.385874 | COADREAD | SW48 | IC-87114 | MVP_top60_DNA+RNA_seed | ERBB3 | 0.129399 | COADREAD | COLO-320-HSR | AZD8931 |
| MVP_top60_DNA_seed | PIK3CA | 0.388994 | COADREAD | SNU-81 | PI3Ka_4409 | MVP_top60_DNA+RNA_seed | ERBB3 | 0.130766 | COADREAD | CW-2 | AZD8931 |
| MVP_top60_DNA_seed | PIK3CD | 0.388994 | COADREAD | SNU-81 | PI3Ka_4409 | MVP_top60_DNA+RNA_seed | PIK3CD | 0.133843 | COADREAD | LS-513 | Idelalisib |
| MVP_top60_DNA_seed | BRAF | 0.392273 | COADREAD | CCK-81 | HG6-64-1 | MVP_top60_DNA+RNA_seed | SRC | 0.13453 | COADREAD | LS-411N | WH-4-023 |
| MVP_top60_DNA_seed | BRAF | 0.392488 | COADREAD | SW620 | PLX-4720 | MVP_top60_DNA+RNA_seed | PIK3CD | 0.136648 | COADREAD | HT-29 | Idelalisib |
| MVP_top60_DNA_seed | BRAF | 0.392974 | COADREAD | C2BBe1 | AZ628 | MVP_top60_DNA+RNA_seed | CDK2 | 0.137795 | COADREAD | NCI-H747 | AZD5438 |
| MVP_top60_DNA_seed | MET | 0.393881 | COADREAD | RCM-1 | JNJ38877605 | MVP_top60_DNA+RNA_seed | HSP90AA1 | 0.138381 | COADREAD | LS-180 | SNX-2112 |
| MVP_top60_DNA_seed | PIK3CD | 0.39397 | COADREAD | LS-1034 | Idelalisib | MVP_top60_DNA+RNA_seed | HSP90AB1 | 0.138381 | COADREAD | LS-180 | SNX-2112 |
| MVP_top60_DNA_seed | MET | 0.394052 | COADREAD | SW48 | AZD6094 | MVP_top60_DNA+RNA_seed | MET | 0.139109 | COADREAD | SNU-C5 | AZD6094 |
| MVP_top60_DNA_seed | ATM | 0.395235 | COADREAD | HCT-15 | torin2 | MVP_top60_DNA+RNA_seed | ERBB3 | 0.139812 | COADREAD | SNU-C5 | AZD8931 |

| | | | | | | | | | | | |
|---|---|---|---|---|---|---|---|---|---|---|---|
| MVP_top60_DNA_seed | MET | 0.396228 | COADREAD | CCK-81 | AZD6094 | MVP_top60_DNA+RNA_seed | ERBB4 | 0.139863 | COADREAD | HT55 | PF-00299804 |
| MVP_top60_DNA_seed | AKT1 | 0.3965 | COADREAD | T84 | GSK690693 | MVP_top60_DNA+RNA_seed | AKT1 | 0.140191 | COADREAD | LS-1034 | GSK690693 |
| MVP_top60_DNA_seed | BRAF | 0.397356 | COADREAD | DiFi | RAF_9304 | MVP_top60_DNA+RNA_seed | HSP90AA1 | 0.141941 | COADREAD | HCC2998 | SNX-2112 |
| MVP_top60_DNA_seed | ATM | 0.397829 | COADREAD | HT-115 | KU-55933 | MVP_top60_DNA+RNA_seed | HSP90AB1 | 0.141941 | COADREAD | HCC2998 | SNX-2112 |
| MVP_top60_DNA_seed | MAPK1 | 0.39925 | COADREAD | HT-115 | VX-11e | MVP_top60_DNA+RNA_seed | IGF1R | 0.142495 | COADREAD | SW1417 | BMS-536924 |
| MVP_top60_DNA_seed | PIK3CB | 0.401272 | COADREAD | COLO-320-HSR | AZD8186 | MVP_top60_DNA+RNA_seed | GSK3B | 0.144005 | COADREAD | HCT-116 | KIN001-042 |
| MVP_top60_DNA_seed | PIK3CD | 0.401272 | COADREAD | COLO-320-HSR | AZD8186 | MVP_top60_DNA+RNA_seed | AR | 0.144302 | COADREAD | SNU-407 | Bicalutamide |
| MVP_top60_DNA_seed | AR | 0.402456 | COADREAD | SNU-81 | Bicalutamide | MVP_top60_DNA+RNA_seed | ERBB3 | 0.147705 | COADREAD | HT-29 | AZD8931 |
| MVP_top60_DNA_seed | GSK3B | 0.403251 | COADREAD | T84 | AZD7969 | MVP_top60_DNA+RNA_seed | BCL2 | 0.148433 | COADREAD | LS-513 | TW 37 |
| MVP_top60_DNA_seed | IGF1R | 0.403443 | COADREAD | LS-123 | BMS-536924 | MVP_top60_DNA+RNA_seed | PIK3CA | 0.148652 | COADREAD | LS-1034 | AZD8835 |
| MVP_top60_DNA_seed | PTK2 | 0.403786 | COADREAD | RCM-1 | PF-562271 | MVP_top60_DNA+RNA_seed | PIK3CD | 0.148652 | COADREAD | LS-1034 | AZD8835 |
| MVP_top60_DNA_seed | ATM | 0.403881 | COADREAD | C2BBe1 | KU-55933 | MVP_top60_DNA+RNA_seed | AKT1 | 0.149032 | COADREAD | NCI-H716 | GSK690693 |
| MVP_top60_DNA_seed | BRAF | 0.403922 | COADREAD | SW1463 | RAF_9304 | MVP_top60_DNA+RNA_seed | AR | 0.151076 | COADREAD | COLO-205 | Bicalutamide |
| MVP_top60_DNA_seed | SRC | 0.404198 | COADREAD | CL-11 | NVP-BHG712 | MVP_top60_DNA+RNA_seed | PIK3CA | 0.151439 | COADREAD | HT55 | PI3Ka_4409 |
| MVP_top60_DNA_seed | ATM | 0.404426 | COADREAD | MDST8 | KU-55933 | MVP_top60_DNA+RNA_seed | PIK3CD | 0.151439 | COADREAD | HT55 | PI3Ka_4409 |
| MVP_top60_DNA_seed | CDK2 | 0.406087 | COADREAD | CCK-81 | AZD5438 | MVP_top60_DNA+RNA_seed | PIK3CB | 0.152207 | COADREAD | KM12 | AZD6482 |
| MVP_top60_DNA_seed | IGF1R | 0.406436 | COADREAD | HCC2998 | GSK1904529A | MVP_top60_DNA+RNA_seed | AR | 0.154311 | COADREAD | CCK-81 | Bicalutamide |
| MVP_top60_DNA_seed | ERBB4 | 0.407448 | COADREAD | HT-29 | AST-1306 | MVP_top60_DNA+RNA_seed | IGF1R | 0.157714 | COADREAD | SW837 | BMS-754807 |
| MVP_top60_DNA_seed | MAPK14 | 0.409261 | COADREAD | SNU-81 | TAK-715 | MVP_top60_DNA+RNA_seed | CDK2 | 0.157901 | COADREAD | C2BBe1 | PHA-793887 |
| MVP_top60_DNA_seed | BRAF | 0.410079 | COADREAD | SW1116 | Dabrafenib | MVP_top60_DNA+RNA_seed | IGF1R | 0.158924 | COADREAD | SNU-C2B | BMS-536924 |
| MVP_top60_DNA_seed | ERBB4 | 0.410436 | COADREAD | LS-180 | AST-1306 | MVP_top60_DNA+RNA_seed | IGF1R | 0.159673 | COADREAD | MDST8 | BMS-536924 |
| MVP_top60_DNA_seed | GSK3B | 0.412051 | COADREAD | DiFi | AZD7969 | MVP_top60_DNA+RNA_seed | SRC | 0.159973 | COADREAD | CaR-1 | HG-6-71-01 |
| MVP_top60_DNA_seed | BRAF | 0.412899 | COADREAD | NCI-H716 | AZ628 | MVP_top60_DNA+RNA_seed | ERBB4 | 0.160675 | COADREAD | SNU-C2B | CI-1033 |
| MVP_top60_DNA_seed | PIK3CD | 0.412927 | COADREAD | C2BBe1 | Idelalisib | MVP_top60_DNA+RNA_seed | IGF1R | 0.163869 | COADREAD | LS-123 | BMS-754807 |
| MVP_top60_DNA_seed | CDK2 | 0.413889 | COADREAD | LS-180 | PHA-793887 | MVP_top60_DNA+RNA_seed | PIK3CA | 0.165612 | COADREAD | CL-11 | AZD8835 |
| MVP_top60_DNA_seed | PIK3CA | 0.416232 | COADREAD | MDST8 | PI3Ka_4409 | MVP_top60_DNA+RNA_seed | PIK3CD | 0.165612 | COADREAD | CL-11 | AZD8835 |
| MVP_top60_DNA_seed | PIK3CD | 0.416232 | COADREAD | MDST8 | PI3Ka_4409 | MVP_top60_DNA+RNA_seed | ATM | 0.166745 | COADREAD | HCC2998 | CP466722 |
| MVP_top60_DNA_seed | ERBB4 | 0.419277 | COADREAD | SNU-407 | CI-1033 | MVP_top60_DNA+RNA_seed | SRC | 0.167936 | COADREAD | CL-11 | HG-6-71-01 |
| MVP_top60_DNA_seed | CDK2 | 0.420278 | COADREAD | SW837 | PHA-793887 | MVP_top60_DNA+RNA_seed | PTK2 | 0.170637 | COADREAD | HT-115 | PF-562271 |
| MVP_top60_DNA_seed | HSP90AA1 | 0.422552 | COADREAD | SW837 | Tanespimycin | MVP_top60_DNA+RNA_seed | IGF1R | 0.172379 | COADREAD | DiFi | BMS-536924 |
| MVP_top60_DNA_seed | HSP90AB1 | 0.422552 | COADREAD | SW837 | Tanespimycin | MVP_top60_DNA+RNA_seed | AR | 0.173296 | COADREAD | SNU-C5 | AZD3514 |
| MVP_top60_DNA_seed | GSK3B | 0.42258 | COADREAD | MDST8 | CHIR-99021 | MVP_top60_DNA+RNA_seed | ERBB4 | 0.173582 | COADREAD | HCT-15 | PF-00299804 |
| MVP_top60_DNA_seed | AR | 0.425462 | COADREAD | HCT-15 | Bicalutamide | MVP_top60_DNA+RNA_seed | HSP90AA1 | 0.174026 | COADREAD | T84 | CCT-018159 |
| MVP_top60_DNA_seed | BRAF | 0.427269 | COADREAD | COLO-320-HSR | RAF_9304 | MVP_top60_DNA+RNA_seed | HSP90AB1 | 0.174026 | COADREAD | T84 | CCT-018159 |
| MVP_top60_DNA_seed | AKT1 | 0.429226 | COADREAD | SNU-175 | GSK690693 | MVP_top60_DNA+RNA_seed | CDK2 | 0.174582 | COADREAD | SNU-C5 | PHA-793887 |
| MVP_top60_DNA_seed | IGF1R | 0.429454 | COADREAD | LoVo | GSK1904529A | MVP_top60_DNA+RNA_seed | ERBB3 | 0.175298 | COADREAD | CaR-1 | AZD8931 |
| MVP_top60_DNA_seed | ESR1 | 0.430098 | COADREAD | CL-11 | Fulvestrant | MVP_top60_DNA+RNA_seed | GSK3B | 0.175335 | COADREAD | MDST8 | KIN001-042 |
| MVP_top60_DNA_seed | AKT1 | 0.430202 | COADREAD | SNU-C2B | AKT inhibitor VIII | MVP_top60_DNA+RNA_seed | SRC | 0.175812 | COADREAD | COLO-205 | Bosutinib |
| MVP_top60_DNA_seed | MET | 0.430599 | COADREAD | SNU-407 | SU11274 | MVP_top60_DNA+RNA_seed | HSP90AA1 | 0.17594 | COADREAD | NCI-H747 | Elesclomol |
| MVP_top60_DNA_seed | AR | 0.434984 | COADREAD | SW620 | Bicalutamide | MVP_top60_DNA+RNA_seed | HSP90AB1 | 0.17594 | COADREAD | NCI-H747 | Elesclomol |
| MVP_top60_DNA_seed | ERBB4 | 0.435346 | COADREAD | COLO-205 | CI-1033 | MVP_top60_DNA+RNA_seed | MET | 0.175963 | COADREAD | RKO | JNJ38877605 |

| | | | | | | | | | | | |
|---|---|---|---|---|---|---|---|---|---|---|---|
| MVP_top60_DNA_seed | GSK3B | 0.438428 | COADREAD | RCM-1 | CHIR-99021 | MVP_top60_DNA+RNA_seed | PIK3CD | 0.176447 | COADREAD | HCT-116 | Idelalisib |
| MVP_top60_DNA_seed | AKT1 | 0.438481 | COADREAD | SK-CO-1 | BAY AKT1 | MVP_top60_DNA+RNA_seed | MET | 0.178713 | COADREAD | SNU-C2B | JNJ38877605 |
| MVP_top60_DNA_seed | MAPK1 | 0.439334 | COADREAD | COLO-320-HSR | FR-180204 | MVP_top60_DNA+RNA_seed | SRC | 0.179207 | COADREAD | HT-29 | Bosutinib |
| MVP_top60_DNA_seed | AKT1 | 0.441706 | COADREAD | SNU-61 | MK-2206 | MVP_top60_DNA+RNA_seed | AR | 0.179992 | COADREAD | GP5d | Bicalutamide |
| MVP_top60_DNA_seed | IGF1R | 0.441836 | COADREAD | SNU-C2B | GSK1904529A | MVP_top60_DNA+RNA_seed | PIK3CD | 0.180015 | COADREAD | COLO-205 | IC-87114 |
| MVP_top60_DNA_seed | BRAF | 0.441962 | COADREAD | LS-123 | Dabrafenib | MVP_top60_DNA+RNA_seed | MET | 0.180552 | COADREAD | LS-411N | PHA-665752 |
| MVP_top60_DNA_seed | MET | 0.443924 | COADREAD | COLO-320-HSR | JNJ38877605 | MVP_top60_DNA+RNA_seed | MET | 0.182313 | COADREAD | SNU-C5 | SU11274 |
| MVP_top60_DNA_seed | BRAF | 0.444489 | COADREAD | CW-2 | RAF_9304 | MVP_top60_DNA+RNA_seed | GSK3B | 0.184741 | COADREAD | SNU-81 | AZD7969 |
| MVP_top60_DNA_seed | IGF1R | 0.446969 | COADREAD | HCT-15 | GSK1904529A | MVP_top60_DNA+RNA_seed | PIK3CB | 0.186201 | COADREAD | CL-11 | AZD8186 |
| MVP_top60_DNA_seed | PIK3CD | 0.447453 | COADREAD | SNU-407 | IC-87114 | MVP_top60_DNA+RNA_seed | PIK3CD | 0.186201 | COADREAD | CL-11 | AZD8186 |
| MVP_top60_DNA_seed | BRAF | 0.448024 | COADREAD | SK-CO-1 | PLX-4720 | MVP_top60_DNA+RNA_seed | PIK3CB | 0.187846 | COADREAD | CaR-1 | AZD6482 |
| MVP_top60_DNA_seed | BRAF | 0.450682 | COADREAD | RCM-1 | PLX-4720 | MVP_top60_DNA+RNA_seed | PIK3CB | 0.190635 | COADREAD | COLO-320-HSR | AZD6482 |
| MVP_top60_DNA_seed | ATM | 0.450843 | COADREAD | SW620 | torin2 | MVP_top60_DNA+RNA_seed | SRC | 0.191244 | COADREAD | T84 | NVP-BHG712 |
| MVP_top60_DNA_seed | GSK3B | 0.452079 | COADREAD | RCM-1 | SB216763 | MVP_top60_DNA+RNA_seed | ATM | 0.191918 | COADREAD | HCT-116 | KU-55933 |
| MVP_top60_DNA_seed | AKT1 | 0.454402 | COADREAD | SNU-C5 | GSK690693 | MVP_top60_DNA+RNA_seed | SRC | 0.193129 | COADREAD | RCM-1 | Bosutinib |
| MVP_top60_DNA_seed | PIK3CD | 0.45666 | COADREAD | MDST8 | IC-87114 | MVP_top60_DNA+RNA_seed | BCL2 | 0.193433 | COADREAD | NCI-H716 | TW 37 |
| MVP_top60_DNA_seed | ERBB4 | 0.457601 | COADREAD | CW-2 | CI-1033 | MVP_top60_DNA+RNA_seed | HSP90AA1 | 0.194747 | COADREAD | NCI-H716 | Tanespimycin |
| MVP_top60_DNA_seed | AKT1 | 0.458758 | COADREAD | HCT-15 | GSK690693 | MVP_top60_DNA+RNA_seed | HSP90AB1 | 0.194747 | COADREAD | NCI-H716 | Tanespimycin |
| MVP_top60_DNA_seed | PIK3CB | 0.458988 | COADREAD | MDST8 | AZD6482 | MVP_top60_DNA+RNA_seed | IGF1R | 0.195473 | COADREAD | LS-123 | Linsitinib |
| MVP_top60_DNA_seed | ATM | 0.459969 | COADREAD | SW837 | KU-60019 | MVP_top60_DNA+RNA_seed | AKT1 | 0.198189 | COADREAD | LS-123 | MK-2206 |
| MVP_top60_DNA_seed | AR | 0.460199 | COADREAD | LS-1034 | Bicalutamide | MVP_top60_DNA+RNA_seed | PIK3CA | 0.200779 | COADREAD | LS-123 | AZD8835 |
| MVP_top60_DNA_seed | GSK3B | 0.46273 | COADREAD | LS-513 | CHIR-99021 | MVP_top60_DNA+RNA_seed | PIK3CD | 0.200779 | COADREAD | LS-123 | AZD8835 |
| MVP_top60_DNA_seed | AR | 0.463933 | COADREAD | CL-11 | Bicalutamide | MVP_top60_DNA+RNA_seed | MET | 0.200917 | COADREAD | SW620 | AZD6094 |
| MVP_top60_DNA_seed | IGF1R | 0.464241 | COADREAD | RCM-1 | BMS-536924 | MVP_top60_DNA+RNA_seed | GSK3B | 0.204363 | COADREAD | SNU-C5 | KIN001-042 |
| MVP_top60_DNA_seed | IGF1R | 0.464875 | COADREAD | MDST8 | Linsitinib | MVP_top60_DNA+RNA_seed | HSP90AA1 | 0.204869 | COADREAD | HT-29 | Luminespib |
| MVP_top60_DNA_seed | PIK3CA | 0.465554 | COADREAD | SW837 | AZD8835 | MVP_top60_DNA+RNA_seed | HSP90AB1 | 0.204869 | COADREAD | HT-29 | Luminespib |
| MVP_top60_DNA_seed | PIK3CD | 0.465554 | COADREAD | SW837 | AZD8835 | MVP_top60_DNA+RNA_seed | AKT1 | 0.207988 | COADREAD | LS-1034 | A-443654 |
| MVP_top60_DNA_seed | CDK2 | 0.466013 | COADREAD | KM12 | AZD5438 | MVP_top60_DNA+RNA_seed | PIK3CB | 0.20995 | COADREAD | RKO | AZD8186 |
| MVP_top60_DNA_seed | SRC | 0.467377 | COADREAD | SNU-C2B | Bosutinib | MVP_top60_DNA+RNA_seed | PIK3CD | 0.20995 | COADREAD | RKO | AZD8186 |
| MVP_top60_DNA_seed | GSK3B | 0.46857 | COADREAD | CW-2 | AZD7969 | MVP_top60_DNA+RNA_seed | CDK2 | 0.21047 | COADREAD | CL-11 | AZD5438 |
| MVP_top60_DNA_seed | GSK3B | 0.469408 | COADREAD | CCK-81 | CHIR-99021 | MVP_top60_DNA+RNA_seed | MET | 0.212144 | COADREAD | NCI-H716 | AZD6094 |
| MVP_top60_DNA_seed | BRAF | 0.469949 | COADREAD | COLO-678 | Dabrafenib | MVP_top60_DNA+RNA_seed | BCL2 | 0.214948 | COADREAD | GP5d | TW 37 |
| MVP_top60_DNA_seed | ESR1 | 0.46998 | COADREAD | NCI-H747 | Tamoxifen | MVP_top60_DNA+RNA_seed | GSK3B | 0.215205 | COADREAD | COLO-320-HSR | AZD7969 |
| MVP_top60_DNA_seed | MAPK14 | 0.4705 | COADREAD | HT-29 | TAK-715 | MVP_top60_DNA+RNA_seed | MET | 0.215728 | COADREAD | COLO-320-HSR | AZD6094 |
| MVP_top60_DNA_seed | AKT1 | 0.471176 | COADREAD | SW620 | GSK690693 | MVP_top60_DNA+RNA_seed | PIK3CD | 0.215953 | COADREAD | CCK-81 | IC-87114 |
| MVP_top60_DNA_seed | ERBB4 | 0.471648 | COADREAD | CL-11 | CI-1033 | MVP_top60_DNA+RNA_seed | PIK3CD | 0.216039 | COADREAD | SW620 | Idelalisib |
| MVP_top60_DNA_seed | BRAF | 0.472184 | COADREAD | COLO-320-HSR | PLX-4720 | MVP_top60_DNA+RNA_seed | GSK3B | 0.216419 | COADREAD | SK-CO-1 | SB216763 |
| MVP_top60_DNA_seed | SRC | 0.472706 | COADREAD | COLO-320-HSR | Saracatinib | MVP_top60_DNA+RNA_seed | MET | 0.217132 | COADREAD | SK-CO-1 | AZD6094 |
| MVP_top60_DNA_seed | GSK3B | 0.472764 | COADREAD | SW948 | CHIR-99021 | MVP_top60_DNA+RNA_seed | CDK2 | 0.217332 | COADREAD | SW48 | PHA-793887 |
| MVP_top60_DNA_seed | ESR1 | 0.473311 | COADREAD | SNU-C2B | Fulvestrant | MVP_top60_DNA+RNA_seed | IGF1R | 0.217401 | COADREAD | SW837 | BMS-536924 |
| MVP_top60_DNA_seed | GSK3B | 0.473992 | COADREAD | LS-411N | AZD7969 | MVP_top60_DNA+RNA_seed | HSP90AA1 | 0.219143 | COADREAD | HT-115 | Tanespimycin |

| | | | | | | | | | | | |
|---|---|---|---|---|---|---|---|---|---|---|---|
| MVP_top60_DNA_seed | MAPK1 | 0.473994 | COADREAD | COLO-320-HSR | VX-11e | MVP_top60_DNA+RNA_seed | HSP90AB1 | 0.219143 | COADREAD | HT-115 | Tanespimycin |
| MVP_top60_DNA_seed | AKT1 | 0.474098 | COADREAD | SW48 | MK-2206 | MVP_top60_DNA+RNA_seed | AKT1 | 0.220574 | COADREAD | HT-29 | BAY AKT1 |
| MVP_top60_DNA_seed | PIK3CD | 0.474163 | COADREAD | HCC2998 | IC-87114 | MVP_top60_DNA+RNA_seed | IGF1R | 0.220823 | COADREAD | NCI-H716 | BMS-536924 |
| MVP_top60_DNA_seed | GSK3B | 0.474477 | COADREAD | LS-411N | CHIR-99021 | MVP_top60_DNA+RNA_seed | ATM | 0.220984 | COADREAD | CaR-1 | KU-55933 |
| MVP_top60_DNA_seed | HSP90AA1 | 0.476212 | COADREAD | LS-513 | Elesclomol | MVP_top60_DNA+RNA_seed | GSK3B | 0.222095 | COADREAD | CW-2 | KIN001-042 |
| MVP_top60_DNA_seed | HSP90AB1 | 0.476212 | COADREAD | LS-513 | Elesclomol | MVP_top60_DNA+RNA_seed | IGF1R | 0.222953 | COADREAD | LS-123 | BMS-536924 |
| MVP_top60_DNA_seed | MAPK1 | 0.476934 | COADREAD | SW1463 | VX-11e | MVP_top60_DNA+RNA_seed | PIK3CB | 0.223258 | COADREAD | COLO-205 | AZD8186 |
| MVP_top60_DNA_seed | AKT1 | 0.47805 | COADREAD | SNU-1040 | BAY AKT1 | MVP_top60_DNA+RNA_seed | PIK3CD | 0.223258 | COADREAD | COLO-205 | AZD8186 |
| MVP_top60_DNA_seed | GSK3B | 0.478233 | COADREAD | SW1116 | CHIR-99021 | MVP_top60_DNA+RNA_seed | PIK3CB | 0.223475 | COADREAD | SW48 | AZD6482 |
| MVP_top60_DNA_seed | SRC | 0.478334 | COADREAD | HT55 | NVP-BHG712 | MVP_top60_DNA+RNA_seed | AR | 0.223902 | COADREAD | GP5d | AZD3514 |
| MVP_top60_DNA_seed | SRC | 0.478455 | COADREAD | CaR-1 | NVP-BHG712 | MVP_top60_DNA+RNA_seed | IGF1R | 0.22413 | COADREAD | C2BBe1 | BMS-536924 |
| MVP_top60_DNA_seed | ERBB4 | 0.479359 | COADREAD | SW48 | AST-1306 | MVP_top60_DNA+RNA_seed | GSK3B | 0.224238 | COADREAD | HT-29 | AZD7969 |
| MVP_top60_DNA_seed | CREBBP | 0.479663 | COADREAD | LS-513 | I-CBP112 | MVP_top60_DNA+RNA_seed | GSK3B | 0.226386 | COADREAD | SNU-C1 | KIN001-042 |
| MVP_top60_DNA_seed | PIK3CD | 0.481279 | COADREAD | SNU-175 | Idelalisib | MVP_top60_DNA+RNA_seed | GSK3B | 0.22647 | COADREAD | GP5d | CHIR-99021 |
| MVP_top60_DNA_seed | AKT1 | 0.481529 | COADREAD | SW837 | AKT inhibitor VIII | MVP_top60_DNA+RNA_seed | ERBB4 | 0.22687 | COADREAD | RKO | CI-1033 |
| MVP_top60_DNA_seed | PIK3CD | 0.481735 | COADREAD | HT55 | IC-87114 | MVP_top60_DNA+RNA_seed | ERBB4 | 0.229151 | COADREAD | COLO-205 | AST-1306 |
| MVP_top60_DNA_seed | AKT1 | 0.485757 | COADREAD | HCT-15 | AKT inhibitor VIII | MVP_top60_DNA+RNA_seed | MET | 0.230342 | COADREAD | SNU-C2B | SU11274 |
| MVP_top60_DNA_seed | GSK3B | 0.487078 | COADREAD | KM12 | CHIR-99021 | MVP_top60_DNA+RNA_seed | HSP90AA1 | 0.230513 | COADREAD | HCT-15 | Luminespib |
| MVP_top60_DNA_seed | AKT1 | 0.487425 | COADREAD | CCK-81 | GSK690693 | MVP_top60_DNA+RNA_seed | HSP90AB1 | 0.230513 | COADREAD | HCT-15 | Luminespib |
| MVP_top60_DNA_seed | HSP90AA1 | 0.487872 | COADREAD | SNU-C2B | Tanespimycin | MVP_top60_DNA+RNA_seed | BCL2 | 0.230583 | COADREAD | CL-11 | Navitoclax |
| MVP_top60_DNA_seed | HSP90AB1 | 0.487872 | COADREAD | SNU-C2B | Tanespimycin | MVP_top60_DNA+RNA_seed | ERBB4 | 0.231178 | COADREAD | SW837 | CI-1033 |
| MVP_top60_DNA_seed | IGF1R | 0.487925 | COADREAD | SW48 | GSK1904529A | MVP_top60_DNA+RNA_seed | IGF1R | 0.231733 | COADREAD | NCI-H716 | BMS-754807 |
| MVP_top60_DNA_seed | MAPK14 | 0.48897 | COADREAD | LS-123 | TAK-715 | MVP_top60_DNA+RNA_seed | ATM | 0.232157 | COADREAD | LS-123 | CP466722 |
| MVP_top60_DNA_seed | GSK3B | 0.4899 | COADREAD | HCC2998 | SB216763 | MVP_top60_DNA+RNA_seed | HSP90AA1 | 0.234101 | COADREAD | LoVo | SNX-2112 |
| MVP_top60_DNA_seed | GSK3B | 0.490676 | COADREAD | C2BBe1 | KIN001-042 | MVP_top60_DNA+RNA_seed | HSP90AB1 | 0.234101 | COADREAD | LoVo | SNX-2112 |
| MVP_top60_DNA_seed | IGF1R | 0.492173 | COADREAD | HCT-116 | Linsitinib | MVP_top60_DNA+RNA_seed | AKT1 | 0.23428 | COADREAD | RKO | BAY AKT1 |
| MVP_top60_DNA_seed | BRAF | 0.493704 | COADREAD | HCC2998 | Dabrafenib | MVP_top60_DNA+RNA_seed | AKT1 | 0.235043 | COADREAD | COLO-320-HSR | A-443654 |
| MVP_top60_DNA_seed | PIK3CD | 0.494593 | COADREAD | NCI-H716 | Idelalisib | MVP_top60_DNA+RNA_seed | IGF1R | 0.235689 | COADREAD | NCI-H716 | Linsitinib |
| MVP_top60_DNA_seed | SRC | 0.495551 | COADREAD | NCI-H716 | WH-4-023 | MVP_top60_DNA+RNA_seed | PIK3CD | 0.241292 | COADREAD | LS-411N | IC-87114 |
| MVP_top60_DNA_seed | BRAF | 0.495644 | COADREAD | LS-1034 | AZ628 | MVP_top60_DNA+RNA_seed | BCL2 | 0.242021 | COADREAD | RCM-1 | Obatoclax Mesylate |
| MVP_top60_DNA_seed | SRC | 0.496924 | COADREAD | RKO | Bosutinib | MVP_top60_DNA+RNA_seed | MET | 0.247395 | COADREAD | RKO | PHA-665752 |
| MVP_top60_DNA_seed | FYN | 0.497943 | COADREAD | LS-411N | A-770041 | MVP_top60_DNA+RNA_seed | AKT1 | 0.247465 | COADREAD | HT-115 | MK-2206 |
| MVP_top60_DNA_seed | MET | 0.498615 | COADREAD | LS-513 | Crizotinib | MVP_top60_DNA+RNA_seed | IGF1R | 0.24781 | COADREAD | COLO-678 | BMS-536924 |
| MVP_top60_DNA_seed | CDK2 | 0.501652 | COADREAD | LS-1034 | PHA-793887 | MVP_top60_DNA+RNA_seed | AKT1 | 0.249596 | COADREAD | SNU-C2B | AKT inhibitor VIII |
| MVP_top60_DNA_seed | HSP90AA1 | 0.502148 | COADREAD | LS-123 | Tanespimycin | MVP_top60_DNA+RNA_seed | AR | 0.251656 | COADREAD | LS-513 | Bicalutamide |
| MVP_top60_DNA_seed | HSP90AB1 | 0.502148 | COADREAD | LS-123 | Tanespimycin | MVP_top60_DNA+RNA_seed | IGF1R | 0.252234 | COADREAD | SW48 | Linsitinib |
| MVP_top60_DNA_seed | AR | 0.503908 | COADREAD | SNU-1040 | Bicalutamide | MVP_top60_DNA+RNA_seed | GSK3B | 0.255116 | COADREAD | SNU-175 | KIN001-042 |
| MVP_top60_DNA_seed | PIK3CD | 0.505656 | COADREAD | RCM-1 | IC-87114 | MVP_top60_DNA+RNA_seed | ERBB4 | 0.256022 | COADREAD | HCC2998 | CI-1033 |
| MVP_top60_DNA_seed | GSK3B | 0.507042 | COADREAD | NCI-H508 | SB216763 | MVP_top60_DNA+RNA_seed | PIK3CB | 0.256785 | COADREAD | SK-CO-1 | AZD8186 |
| MVP_top60_DNA_seed | HSP90AA1 | 0.507691 | COADREAD | HCT-116 | Luminespib | MVP_top60_DNA+RNA_seed | PIK3CD | 0.256785 | COADREAD | SK-CO-1 | AZD8186 |
| MVP_top60_DNA_seed | HSP90AB1 | 0.507691 | COADREAD | HCT-116 | Luminespib | MVP_top60_DNA+RNA_seed | PIK3CB | 0.260368 | COADREAD | CCK-81 | AZD6482 |

| | | | | | | | | | | | |
|---|---|---|---|---|---|---|---|---|---|---|---|
| MVP_top60_DNA_seed | MET | 0.508738 | COADREAD | LS-180 | JNJ38877605 | MVP_top60_DNA+RNA_seed | MET | 0.261201 | COADREAD | HCC2998 | AZD6094 |
| MVP_top60_DNA_seed | MAPK1 | 0.510115 | COADREAD | SW837 | FR-180204 | MVP_top60_DNA+RNA_seed | CDK2 | 0.261956 | COADREAD | GP5d | AZD5438 |
| MVP_top60_DNA_seed | CDK2 | 0.510576 | COADREAD | NCI-H716 | PHA-793887 | MVP_top60_DNA+RNA_seed | AKT1 | 0.262406 | COADREAD | COLO-205 | GSK690693 |
| MVP_top60_DNA_seed | MAPK14 | 0.511293 | COADREAD | COLO-205 | TAK-715 | MVP_top60_DNA+RNA_seed | HSP90AA1 | 0.263291 | COADREAD | SNU-C2B | Elesclomol |
| MVP_top60_DNA_seed | MET | 0.512733 | COADREAD | SNU-1040 | SU11274 | MVP_top60_DNA+RNA_seed | HSP90AB1 | 0.263291 | COADREAD | SNU-C2B | Elesclomol |
| MVP_top60_DNA_seed | AR | 0.514015 | COADREAD | MDST8 | AZD3514 | MVP_top60_DNA+RNA_seed | SRC | 0.263533 | COADREAD | SNU-C5 | NVP-BHG712 |
| MVP_top60_DNA_seed | BRAF | 0.514964 | COADREAD | LoVo | PLX-4720 | MVP_top60_DNA+RNA_seed | ERBB3 | 0.264419 | COADREAD | RCM-1 | AZD8931 |
| MVP_top60_DNA_seed | SRC | 0.515083 | COADREAD | COLO-320-HSR | WH-4-023 | MVP_top60_DNA+RNA_seed | HSP90AA1 | 0.26456 | COADREAD | SW837 | Elesclomol |
| MVP_top60_DNA_seed | IGF1R | 0.515128 | COADREAD | MDST8 | BMS-536924 | MVP_top60_DNA+RNA_seed | HSP90AB1 | 0.26456 | COADREAD | SW837 | Elesclomol |
| MVP_top60_DNA_seed | HSP90AA1 | 0.515624 | COADREAD | SW837 | Luminespib | MVP_top60_DNA+RNA_seed | SRC | 0.264585 | COADREAD | SW48 | HG-6-71-01 |
| MVP_top60_DNA_seed | HSP90AB1 | 0.515624 | COADREAD | SW837 | Luminespib | MVP_top60_DNA+RNA_seed | PIK3CA | 0.265577 | COADREAD | HT55 | AZD8835 |
| MVP_top60_DNA_seed | AKT1 | 0.517484 | COADREAD | LS-513 | AKT inhibitor VIII | MVP_top60_DNA+RNA_seed | PIK3CD | 0.265577 | COADREAD | HT55 | AZD8835 |
| MVP_top60_DNA_seed | ESR1 | 0.518461 | COADREAD | COLO-678 | Fulvestrant | MVP_top60_DNA+RNA_seed | ATM | 0.267102 | COADREAD | COLO-320-HSR | KU-55933 |
| MVP_top60_DNA_seed | PIK3CD | 0.518923 | COADREAD | NCI-H747 | IC-87114 | MVP_top60_DNA+RNA_seed | PIK3CA | 0.271829 | COADREAD | SW1463 | AZD8835 |
| MVP_top60_DNA_seed | MET | 0.519471 | COADREAD | LS-1034 | AZD6094 | MVP_top60_DNA+RNA_seed | PIK3CD | 0.271829 | COADREAD | SW1463 | AZD8835 |
| MVP_top60_DNA_seed | PIK3CB | 0.519721 | COADREAD | MDST8 | TGX221 | MVP_top60_DNA+RNA_seed | IGF1R | 0.278901 | COADREAD | HT55 | BMS-754807 |
| MVP_top60_DNA_seed | AKT1 | 0.519895 | COADREAD | LS-411N | BAY AKT1 | MVP_top60_DNA+RNA_seed | ATM | 0.279929 | COADREAD | SNU-61 | CP466722 |
| MVP_top60_DNA_seed | PTK2 | 0.527031 | COADREAD | SNU-C1 | PF-562271 | MVP_top60_DNA+RNA_seed | IGF1R | 0.285579 | COADREAD | NCI-H508 | BMS-536924 |
| MVP_top60_DNA_seed | HSP90AA1 | 0.529101 | COADREAD | GP5d | Elesclomol | MVP_top60_DNA+RNA_seed | PIK3CA | 0.286264 | COADREAD | SNU-C2B | PI3Ka_4409 |
| MVP_top60_DNA_seed | HSP90AB1 | 0.529101 | COADREAD | GP5d | Elesclomol | MVP_top60_DNA+RNA_seed | PIK3CD | 0.286264 | COADREAD | SNU-C2B | PI3Ka_4409 |
| MVP_top60_DNA_seed | AKT1 | 0.530064 | COADREAD | CaR-1 | AKT inhibitor VIII | MVP_top60_DNA+RNA_seed | AKT1 | 0.286319 | COADREAD | COLO-320-HSR | MK-2206 |
| MVP_top60_DNA_seed | ERBB3 | 0.530317 | COADREAD | T84 | AZD8931 | MVP_top60_DNA+RNA_seed | ERBB4 | 0.286685 | COADREAD | C2BBe1 | PF-00299804 |
| MVP_top60_DNA_seed | MAPK1 | 0.531935 | COADREAD | COLO-678 | VX-11e | MVP_top60_DNA+RNA_seed | MET | 0.286702 | COADREAD | COLO-205 | SU11274 |
| MVP_top60_DNA_seed | ATM | 0.532861 | COADREAD | COLO-205 | KU-55933 | MVP_top60_DNA+RNA_seed | ATM | 0.287877 | COADREAD | GP5d | KU-60019 |
| MVP_top60_DNA_seed | BRAF | 0.533077 | COADREAD | HCC2998 | PLX-4720 | MVP_top60_DNA+RNA_seed | AKT1 | 0.287881 | COADREAD | HCT-116 | MK-2206 |
| MVP_top60_DNA_seed | PIK3CB | 0.534793 | COADREAD | HT-29 | AZD6482 | MVP_top60_DNA+RNA_seed | GSK3B | 0.289414 | COADREAD | NCI-H716 | KIN001-042 |
| MVP_top60_DNA_seed | GSK3B | 0.534964 | COADREAD | HT-29 | CHIR-99021 | MVP_top60_DNA+RNA_seed | ATM | 0.289545 | COADREAD | NCI-H508 | KU-55933 |
| MVP_top60_DNA_seed | ATM | 0.540154 | COADREAD | CaR-1 | CP466722 | MVP_top60_DNA+RNA_seed | HSP90AA1 | 0.289851 | COADREAD | SK-CO-1 | CCT-018159 |
| MVP_top60_DNA_seed | HSP90AA1 | 0.540994 | COADREAD | LS-411N | Elesclomol | MVP_top60_DNA+RNA_seed | HSP90AB1 | 0.289851 | COADREAD | SK-CO-1 | CCT-018159 |
| MVP_top60_DNA_seed | HSP90AB1 | 0.540994 | COADREAD | LS-411N | Elesclomol | MVP_top60_DNA+RNA_seed | BCL2 | 0.293029 | COADREAD | COLO-320-HSR | Navitoclax |
| MVP_top60_DNA_seed | ERBB4 | 0.541254 | COADREAD | HCT-116 | AST-1306 | MVP_top60_DNA+RNA_seed | PIK3CB | 0.296582 | COADREAD | SNU-C2B | AZD8186 |
| MVP_top60_DNA_seed | HSP90AA1 | 0.541643 | COADREAD | SNU-C2B | CCT-018159 | MVP_top60_DNA+RNA_seed | PIK3CD | 0.296582 | COADREAD | SNU-C2B | AZD8186 |
| MVP_top60_DNA_seed | HSP90AB1 | 0.541643 | COADREAD | SNU-C2B | CCT-018159 | MVP_top60_DNA+RNA_seed | SRC | 0.298139 | COADREAD | C2BBe1 | WH-4-023 |
| MVP_top60_DNA_seed | MET | 0.544746 | COADREAD | COLO-320-HSR | Crizotinib | MVP_top60_DNA+RNA_seed | MET | 0.298202 | COADREAD | NCI-H747 | Crizotinib |
| MVP_top60_DNA_seed | GSK3B | 0.54719 | COADREAD | SW620 | CHIR-99021 | MVP_top60_DNA+RNA_seed | ATM | 0.299476 | COADREAD | LS-1034 | KU-60019 |
| MVP_top60_DNA_seed | MET | 0.548115 | COADREAD | LS-513 | SU11274 | MVP_top60_DNA+RNA_seed | PIK3CD | 0.300009 | COADREAD | COLO-320-HSR | IC-87114 |
| MVP_top60_DNA_seed | CDK2 | 0.548416 | COADREAD | LS-180 | AZD5438 | MVP_top60_DNA+RNA_seed | GSK3B | 0.301685 | COADREAD | LS-180 | SB216763 |
| MVP_top60_DNA_seed | CDK2 | 0.549227 | COADREAD | NCI-H747 | Seliciclib | MVP_top60_DNA+RNA_seed | ERBB3 | 0.302785 | COADREAD | RKO | AZD8931 |
| MVP_top60_DNA_seed | MAPK14 | 0.550147 | COADREAD | SW948 | TAK-715 | MVP_top60_DNA+RNA_seed | CDK2 | 0.313458 | COADREAD | SNU-81 | AZD5438 |
| MVP_top60_DNA_seed | IGF1R | 0.553034 | COADREAD | RKO | BMS-536924 | MVP_top60_DNA+RNA_seed | ERBB4 | 0.314411 | COADREAD | SW837 | PF-00299804 |
| MVP_top60_DNA_seed | ERBB4 | 0.55471 | COADREAD | LS-123 | PF-00299804 | MVP_top60_DNA+RNA_seed | AKT1 | 0.314682 | COADREAD | LS-1034 | AKT inhibitor VIII |

| | | | | | | | | | | | |
|---|---|---|---|---|---|---|---|---|---|---|---|
| MVP_top60_DNA_seed | IGF1R | 0.554902 | COADREAD | SW1116 | BMS-536924 | MVP_top60_DNA+RNA_seed | PIK3CA | 0.315191 | COADREAD | HCC2998 | PI3Ka_4409 |
| MVP_top60_DNA_seed | CDK2 | 0.555807 | COADREAD | LS-123 | AZD5438 | MVP_top60_DNA+RNA_seed | PIK3CD | 0.315191 | COADREAD | HCC2998 | PI3Ka_4409 |
| MVP_top60_DNA_seed | ESR1 | 0.556211 | COADREAD | SNU-407 | Fulvestrant | MVP_top60_DNA+RNA_seed | AR | 0.316644 | COADREAD | SW837 | AZD3514 |
| MVP_top60_DNA_seed | ATM | 0.556719 | COADREAD | SW48 | KU-60019 | MVP_top60_DNA+RNA_seed | GSK3B | 0.317366 | COADREAD | HCT-15 | KIN001-042 |
| MVP_top60_DNA_seed | IGF1R | 0.561451 | COADREAD | LS-513 | GSK1904529A | MVP_top60_DNA+RNA_seed | MET | 0.318844 | COADREAD | NCI-H716 | PHA-665752 |
| MVP_top60_DNA_seed | SRC | 0.561701 | COADREAD | LS-1034 | WH-4-023 | MVP_top60_DNA+RNA_seed | HSP90AA1 | 0.319765 | COADREAD | CW-2 | SNX-2112 |
| MVP_top60_DNA_seed | PIK3CD | 0.561973 | COADREAD | CW-2 | Idelalisib | MVP_top60_DNA+RNA_seed | HSP90AB1 | 0.319765 | COADREAD | CW-2 | SNX-2112 |
| MVP_top60_DNA_seed | ERBB3 | 0.562343 | COADREAD | LS-411N | AZD8931 | MVP_top60_DNA+RNA_seed | PIK3CA | 0.320958 | COADREAD | HCT-15 | PI3Ka_4409 |
| MVP_top60_DNA_seed | MET | 0.56264 | COADREAD | HCC2998 | SU11274 | MVP_top60_DNA+RNA_seed | PIK3CD | 0.320958 | COADREAD | HCT-15 | PI3Ka_4409 |
| MVP_top60_DNA_seed | HSP90AA1 | 0.56365 | COADREAD | SNU-81 | CCT-018159 | MVP_top60_DNA+RNA_seed | HSP90AA1 | 0.321801 | COADREAD | C2BBe1 | Luminespib |
| MVP_top60_DNA_seed | HSP90AB1 | 0.56365 | COADREAD | SNU-81 | CCT-018159 | MVP_top60_DNA+RNA_seed | HSP90AB1 | 0.321801 | COADREAD | C2BBe1 | Luminespib |
| MVP_top60_DNA_seed | ATM | 0.564396 | COADREAD | SNU-C2B | torin2 | MVP_top60_DNA+RNA_seed | GSK3B | 0.323895 | COADREAD | LS-180 | KIN001-042 |
| MVP_top60_DNA_seed | PIK3CD | 0.565272 | COADREAD | SW48 | Idelalisib | MVP_top60_DNA+RNA_seed | HSP90AA1 | 0.325308 | COADREAD | LoVo | Elesclomol |
| MVP_top60_DNA_seed | AR | 0.566062 | COADREAD | SNU-C1 | AZD3514 | MVP_top60_DNA+RNA_seed | HSP90AB1 | 0.325308 | COADREAD | LoVo | Elesclomol |
| MVP_top60_DNA_seed | MET | 0.569503 | COADREAD | MDST8 | PHA-665752 | MVP_top60_DNA+RNA_seed | ERBB4 | 0.325509 | COADREAD | GP5d | CI-1033 |
| MVP_top60_DNA_seed | GSK3B | 0.570012 | COADREAD | SNU-C2B | CHIR-99021 | MVP_top60_DNA+RNA_seed | ERBB4 | 0.326663 | COADREAD | SNU-81 | PF-00299804 |
| MVP_top60_DNA_seed | PIK3CD | 0.570535 | COADREAD | SNU-C5 | Idelalisib | MVP_top60_DNA+RNA_seed | HSP90AA1 | 0.327222 | COADREAD | HCC2998 | CCT-018159 |
| MVP_top60_DNA_seed | GSK3B | 0.570749 | COADREAD | CL-11 | AZD7969 | MVP_top60_DNA+RNA_seed | HSP90AB1 | 0.327222 | COADREAD | HCC2998 | CCT-018159 |
| MVP_top60_DNA_seed | ESR1 | 0.573505 | COADREAD | CCK-81 | Fulvestrant | MVP_top60_DNA+RNA_seed | GSK3B | 0.327658 | COADREAD | LS-180 | AZD7969 |
| MVP_top60_DNA_seed | HSP90AA1 | 0.573523 | COADREAD | SW1417 | CCT-018159 | MVP_top60_DNA+RNA_seed | MET | 0.329526 | COADREAD | MDST8 | AZD6094 |
| MVP_top60_DNA_seed | HSP90AB1 | 0.573523 | COADREAD | SW1417 | CCT-018159 | MVP_top60_DNA+RNA_seed | PIK3CA | 0.331059 | COADREAD | SW1417 | PI3Ka_4409 |
| MVP_top60_DNA_seed | CDK2 | 0.57502 | COADREAD | CaR-1 | PHA-793887 | MVP_top60_DNA+RNA_seed | PIK3CD | 0.331059 | COADREAD | SW1417 | PI3Ka_4409 |
| MVP_top60_DNA_seed | IGF1R | 0.577806 | COADREAD | MDST8 | GSK1904529A | MVP_top60_DNA+RNA_seed | AKT1 | 0.331596 | COADREAD | LS-1034 | AKT inhibitor VIII |
| MVP_top60_DNA_seed | HSP90AA1 | 0.57793 | COADREAD | LS-411N | CCT-018159 | MVP_top60_DNA+RNA_seed | BCL2 | 0.331715 | COADREAD | SNU-407 | TW 37 |
| MVP_top60_DNA_seed | HSP90AB1 | 0.57793 | COADREAD | LS-411N | CCT-018159 | MVP_top60_DNA+RNA_seed | MET | 0.334734 | COADREAD | CaR-1 | SU11274 |
| MVP_top60_DNA_seed | BRAF | 0.57794 | COADREAD | HT-115 | PLX-4720 | MVP_top60_DNA+RNA_seed | SRC | 0.337516 | COADREAD | RKO | Saracatinib |
| MVP_top60_DNA_seed | AR | 0.578126 | COADREAD | MDST8 | Bicalutamide | MVP_top60_DNA+RNA_seed | AKT1 | 0.338185 | COADREAD | LS-123 | BAY AKT1 |
| MVP_top60_DNA_seed | MET | 0.578742 | COADREAD | HCT-15 | AZD6094 | MVP_top60_DNA+RNA_seed | SRC | 0.339198 | COADREAD | SNU-61 | NVP-BHG712 |
| MVP_top60_DNA_seed | BRAF | 0.581441 | COADREAD | LS-513 | Dabrafenib | MVP_top60_DNA+RNA_seed | ATM | 0.340794 | COADREAD | SW48 | CP466722 |
| MVP_top60_DNA_seed | ESR1 | 0.581494 | COADREAD | KM12 | Tamoxifen | MVP_top60_DNA+RNA_seed | AKT1 | 0.341217 | COADREAD | LS-180 | MK-2206 |
| MVP_top60_DNA_seed | AR | 0.58184 | COADREAD | NCI-H747 | Bicalutamide | MVP_top60_DNA+RNA_seed | ERBB3 | 0.343092 | COADREAD | SW837 | AZD8931 |
| MVP_top60_DNA_seed | PIK3CD | 0.584855 | COADREAD | CW-2 | IC-87114 | MVP_top60_DNA+RNA_seed | MET | 0.345193 | COADREAD | LoVo | AZD6094 |
| MVP_top60_DNA_seed | CDK2 | 0.587523 | COADREAD | COLO-678 | PHA-793887 | MVP_top60_DNA+RNA_seed | ATM | 0.345977 | COADREAD | NCI-H747 | KU-60019 |
| MVP_top60_DNA_seed | MAPK1 | 0.59011 | COADREAD | CW-2 | FR-180204 | MVP_top60_DNA+RNA_seed | IGF1R | 0.346394 | COADREAD | NCI-H747 | BMS-536924 |
| MVP_top60_DNA_seed | AKT1 | 0.590635 | COADREAD | SNU-81 | BAY AKT1 | MVP_top60_DNA+RNA_seed | HSP90AA1 | 0.349892 | COADREAD | RCM-1 | Tanespimycin |
| MVP_top60_DNA_seed | HSP90AA1 | 0.592651 | COADREAD | SW48 | Elesclomol | MVP_top60_DNA+RNA_seed | HSP90AB1 | 0.349892 | COADREAD | RCM-1 | Tanespimycin |
| MVP_top60_DNA_seed | HSP90AB1 | 0.592651 | COADREAD | SW48 | Elesclomol | MVP_top60_DNA+RNA_seed | PIK3CD | 0.352255 | COADREAD | SNU-1040 | IC-87114 |
| MVP_top60_DNA_seed | SRC | 0.592698 | COADREAD | C2BBe1 | HG-6-71-01 | MVP_top60_DNA+RNA_seed | GSK3B | 0.353586 | COADREAD | SNU-C2B | CHIR-99021 |
| MVP_top60_DNA_seed | MAPK1 | 0.593516 | COADREAD | SNU-C2B | FR-180204 | MVP_top60_DNA+RNA_seed | PIK3CD | 0.35471 | COADREAD | LS-411N | Idelalisib |
| MVP_top60_DNA_seed | CDK2 | 0.593583 | COADREAD | LS-513 | PHA-793887 | MVP_top60_DNA+RNA_seed | ATM | 0.356093 | COADREAD | HT-115 | KU-60019 |
| MVP_top60_DNA_seed | AKT1 | 0.594083 | COADREAD | CCK-81 | BAY AKT1 | MVP_top60_DNA+RNA_seed | HSP90AA1 | 0.356311 | COADREAD | HT55 | Tanespimycin |

| | | | | | | | | | | | |
|---|---|---|---|---|---|---|---|---|---|---|---|
| MVP_top60_DNA_seed | PIK3CA | 0.59458 | COADREAD | SNU-C2B | AZD8835 | MVP_top60_DNA+RNA_seed | HSP90AB1 | 0.356311 | COADREAD | HT55 | Tanespimycin |
| MVP_top60_DNA_seed | PIK3CD | 0.59458 | COADREAD | SNU-C2B | AZD8835 | MVP_top60_DNA+RNA_seed | SRC | 0.356529 | COADREAD | SW620 | Bosutinib |
| MVP_top60_DNA_seed | MAPK14 | 0.596473 | COADREAD | CL-11 | TAK-715 | MVP_top60_DNA+RNA_seed | MET | 0.359117 | COADREAD | HT-115 | JNJ38877605 |
| MVP_top60_DNA_seed | BRAF | 0.59687 | COADREAD | COLO-320-HSR | PLX-4720 | MVP_top60_DNA+RNA_seed | AKT1 | 0.3598 | COADREAD | HT-115 | AKT inhibitor VIII |
| MVP_top60_DNA_seed | AKT1 | 0.597049 | COADREAD | SNU-C5 | AKT inhibitor VIII | MVP_top60_DNA+RNA_seed | PIK3CB | 0.359917 | COADREAD | RKO | TGX221 |
| MVP_top60_DNA_seed | MET | 0.599906 | COADREAD | SW1417 | AZD6094 | MVP_top60_DNA+RNA_seed | AKT1 | 0.364121 | COADREAD | CL-11 | BAY AKT1 |
| MVP_top60_DNA_seed | GSK3B | 0.600258 | COADREAD | HT-115 | CHIR-99021 | MVP_top60_DNA+RNA_seed | ERBB4 | 0.364252 | COADREAD | HT-29 | PF-00299804 |
| MVP_top60_DNA_seed | HSP90AA1 | 0.601088 | COADREAD | HCC2998 | Elesclomol | MVP_top60_DNA+RNA_seed | AKT1 | 0.364678 | COADREAD | HT-29 | GSK690693 |
| MVP_top60_DNA_seed | HSP90AB1 | 0.601088 | COADREAD | HCC2998 | Elesclomol | MVP_top60_DNA+RNA_seed | ERBB4 | 0.364772 | COADREAD | LS-513 | CI-1033 |
| MVP_top60_DNA_seed | PIK3CB | 0.604291 | COADREAD | C2BBe1 | AZD8186 | MVP_top60_DNA+RNA_seed | ATM | 0.364793 | COADREAD | NCI-H716 | CP466722 |
| MVP_top60_DNA_seed | PIK3CD | 0.604291 | COADREAD | C2BBe1 | AZD8186 | MVP_top60_DNA+RNA_seed | GSK3B | 0.366349 | COADREAD | MDST8 | SB216763 |
| MVP_top60_DNA_seed | ERBB4 | 0.604541 | COADREAD | NCI-H716 | PF-00299804 | MVP_top60_DNA+RNA_seed | ATM | 0.366626 | COADREAD | LS-513 | CP466722 |
| MVP_top60_DNA_seed | HSP90AA1 | 0.607046 | COADREAD | HT-29 | Elesclomol | MVP_top60_DNA+RNA_seed | AKT1 | 0.366711 | COADREAD | SNU-61 | AKT inhibitor VIII |
| MVP_top60_DNA_seed | HSP90AB1 | 0.607046 | COADREAD | HT-29 | Elesclomol | MVP_top60_DNA+RNA_seed | PIK3CD | 0.366829 | COADREAD | KM12 | Idelalisib |
| MVP_top60_DNA_seed | ERBB4 | 0.609301 | COADREAD | LS-123 | CI-1033 | MVP_top60_DNA+RNA_seed | GSK3B | 0.367105 | COADREAD | SNU-407 | AZD7969 |
| MVP_top60_DNA_seed | AKT1 | 0.609895 | COADREAD | SW837 | BAY AKT1 | MVP_top60_DNA+RNA_seed | IGF1R | 0.367383 | COADREAD | HT55 | GSK1904529A |
| MVP_top60_DNA_seed | PIK3CD | 0.611504 | COADREAD | SW1417 | Idelalisib | MVP_top60_DNA+RNA_seed | MET | 0.367974 | COADREAD | HT-115 | AZD6094 |
| MVP_top60_DNA_seed | HSP90AA1 | 0.611952 | COADREAD | SNU-81 | Tanespimycin | MVP_top60_DNA+RNA_seed | PIK3CD | 0.369462 | COADREAD | SW620 | IC-87114 |
| MVP_top60_DNA_seed | HSP90AB1 | 0.611952 | COADREAD | SNU-81 | Tanespimycin | MVP_top60_DNA+RNA_seed | HSP90AA1 | 0.369557 | COADREAD | RCM-1 | CCT-018159 |
| MVP_top60_DNA_seed | AR | 0.612007 | COADREAD | SW837 | Bicalutamide | MVP_top60_DNA+RNA_seed | HSP90AB1 | 0.369557 | COADREAD | RCM-1 | CCT-018159 |
| MVP_top60_DNA_seed | MET | 0.613018 | COADREAD | LS-513 | AZD6094 | MVP_top60_DNA+RNA_seed | BCL2 | 0.371192 | COADREAD | LS-1034 | Obatoclax Mesylate |
| MVP_top60_DNA_seed | MAPK1 | 0.614419 | COADREAD | NCI-H716 | FR-180204 | MVP_top60_DNA+RNA_seed | PIK3CB | 0.372144 | COADREAD | NCI-H747 | TGX221 |
| MVP_top60_DNA_seed | HSP90AA1 | 0.616381 | COADREAD | LS-123 | SNX-2112 | MVP_top60_DNA+RNA_seed | CDK2 | 0.372614 | COADREAD | C2BBe1 | Seliciclib |
| MVP_top60_DNA_seed | HSP90AB1 | 0.616381 | COADREAD | LS-123 | SNX-2112 | MVP_top60_DNA+RNA_seed | ATM | 0.374839 | COADREAD | LS-1034 | CP466722 |
| MVP_top60_DNA_seed | CDK2 | 0.61762 | COADREAD | T84 | AZD5438 | MVP_top60_DNA+RNA_seed | AKT1 | 0.375724 | COADREAD | SW48 | AKT inhibitor VIII |
| MVP_top60_DNA_seed | PTK2 | 0.617731 | COADREAD | CaR-1 | PF-562271 | MVP_top60_DNA+RNA_seed | SRC | 0.377317 | COADREAD | HCT-15 | NVP-BHG712 |
| MVP_top60_DNA_seed | AKT1 | 0.617978 | COADREAD | SNU-C1 | GSK690693 | MVP_top60_DNA+RNA_seed | PIK3CB | 0.378235 | COADREAD | SNU-C1 | AZD8186 |
| MVP_top60_DNA_seed | IGF1R | 0.61896 | COADREAD | SW1417 | BMS-754807 | MVP_top60_DNA+RNA_seed | PIK3CD | 0.378235 | COADREAD | SNU-C1 | AZD8186 |
| MVP_top60_DNA_seed | ESR1 | 0.619263 | COADREAD | SNU-C1 | Tamoxifen | MVP_top60_DNA+RNA_seed | AKT1 | 0.379372 | COADREAD | LS-513 | BAY AKT1 |
| MVP_top60_DNA_seed | BRAF | 0.620732 | COADREAD | SW48 | SB590885 | MVP_top60_DNA+RNA_seed | AR | 0.380128 | COADREAD | LoVo | Bicalutamide |
| MVP_top60_DNA_seed | IGF1R | 0.622928 | COADREAD | RKO | Linsitinib | MVP_top60_DNA+RNA_seed | MET | 0.38041 | COADREAD | SNU-C2B | Crizotinib |
| MVP_top60_DNA_seed | BRAF | 0.623059 | COADREAD | HCT-116 | PLX-4720 | MVP_top60_DNA+RNA_seed | SRC | 0.38122 | COADREAD | SW48 | NVP-BHG712 |
| MVP_top60_DNA_seed | ERBB4 | 0.62344 | COADREAD | SW1417 | PF-00299804 | MVP_top60_DNA+RNA_seed | ATM | 0.38181 | COADREAD | C2BBe1 | torin2 |
| MVP_top60_DNA_seed | AKT1 | 0.624579 | COADREAD | SW1417 | MK-2206 | MVP_top60_DNA+RNA_seed | ATM | 0.382052 | COADREAD | CaR-1 | torin2 |
| MVP_top60_DNA_seed | SRC | 0.624669 | COADREAD | SNU-61 | Bosutinib | MVP_top60_DNA+RNA_seed | HSP90AA1 | 0.382511 | COADREAD | C2BBe1 | SNX-2112 |
| MVP_top60_DNA_seed | IGF1R | 0.624791 | COADREAD | COLO-320-HSR | GSK1904529A | MVP_top60_DNA+RNA_seed | HSP90AB1 | 0.382511 | COADREAD | C2BBe1 | SNX-2112 |
| MVP_top60_DNA_seed | MET | 0.625292 | COADREAD | KM12 | AZD6094 | MVP_top60_DNA+RNA_seed | PIK3CD | 0.385874 | COADREAD | SW48 | IC-87114 |
| MVP_top60_DNA_seed | BRAF | 0.62713 | COADREAD | LS-513 | SB590885 | MVP_top60_DNA+RNA_seed | PIK3CA | 0.388994 | COADREAD | SNU-81 | PI3Ka_4409 |
| MVP_top60_DNA_seed | HSP90AA1 | 0.627388 | COADREAD | CCK-81 | Luminespib | MVP_top60_DNA+RNA_seed | PIK3CD | 0.388994 | COADREAD | SNU-81 | PI3Ka_4409 |
| MVP_top60_DNA_seed | HSP90AB1 | 0.627388 | COADREAD | CCK-81 | Luminespib | MVP_top60_DNA+RNA_seed | MET | 0.393881 | COADREAD | RCM-1 | JNJ38877605 |
| MVP_top60_DNA_seed | BRAF | 0.628952 | COADREAD | SNU-1040 | Dabrafenib | MVP_top60_DNA+RNA_seed | PIK3CD | 0.39397 | COADREAD | LS-1034 | Idelalisib |

| | | | | | | | | | | | |
|---|---|---|---|---|---|---|---|---|---|---|---|
| MVP_top60_DNA_seed | BRAF | 0.631429 | COADREAD | GP5d | Dabrafenib | MVP_top60_DNA+RNA_seed | MET | 0.394052 | COADREAD | SW48 | AZD6094 |
| MVP_top60_DNA_seed | PIK3CD | 0.632522 | COADREAD | C2BBe1 | IC-87114 | MVP_top60_DNA+RNA_seed | ATM | 0.395235 | COADREAD | HCT-15 | torin2 |
| MVP_top60_DNA_seed | SRC | 0.633231 | COADREAD | GP5d | HG-6-71-01 | MVP_top60_DNA+RNA_seed | MET | 0.396228 | COADREAD | CCK-81 | AZD6094 |
| MVP_top60_DNA_seed | MAPK14 | 0.634239 | COADREAD | CaR-1 | TAK-715 | MVP_top60_DNA+RNA_seed | AKT1 | 0.3965 | COADREAD | T84 | GSK690693 |
| MVP_top60_DNA_seed | CREBBP | 0.637308 | COADREAD | SNU-81 | I-CBP112 | MVP_top60_DNA+RNA_seed | ATM | 0.397829 | COADREAD | HT-115 | KU-55933 |
| MVP_top60_DNA_seed | AKT1 | 0.637815 | COADREAD | HT55 | MK-2206 | MVP_top60_DNA+RNA_seed | BCL2 | 0.399354 | COADREAD | SW48 | Obatoclax Mesylate |
| MVP_top60_DNA_seed | GSK3B | 0.639382 | COADREAD | SNU-C2B | KIN001-042 | MVP_top60_DNA+RNA_seed | PIK3CB | 0.401272 | COADREAD | COLO-320-HSR | AZD8186 |
| MVP_top60_DNA_seed | CDK2 | 0.640485 | COADREAD | CW-2 | AZD5438 | MVP_top60_DNA+RNA_seed | PIK3CD | 0.401272 | COADREAD | COLO-320-HSR | AZD8186 |
| MVP_top60_DNA_seed | IGF1R | 0.641073 | COADREAD | HT-115 | GSK1904529A | MVP_top60_DNA+RNA_seed | AR | 0.402456 | COADREAD | SNU-81 | Bicalutamide |
| MVP_top60_DNA_seed | FYN | 0.641237 | COADREAD | KM12 | A-770041 | MVP_top60_DNA+RNA_seed | GSK3B | 0.403251 | COADREAD | T84 | AZD7969 |
| MVP_top60_DNA_seed | BRAF | 0.64165 | COADREAD | DiFi | PLX-4720 | MVP_top60_DNA+RNA_seed | IGF1R | 0.403443 | COADREAD | LS-123 | BMS-536924 |
| MVP_top60_DNA_seed | SRC | 0.644135 | COADREAD | HT-115 | Bosutinib | MVP_top60_DNA+RNA_seed | PTK2 | 0.403786 | COADREAD | RCM-1 | PF-562271 |
| MVP_top60_DNA_seed | ESR1 | 0.644588 | COADREAD | LoVo | Tamoxifen | MVP_top60_DNA+RNA_seed | ATM | 0.403881 | COADREAD | C2BBe1 | KU-55933 |
| MVP_top60_DNA_seed | BRAF | 0.645082 | COADREAD | HT-115 | SB590885 | MVP_top60_DNA+RNA_seed | SRC | 0.404198 | COADREAD | CL-11 | NVP-BHG712 |
| MVP_top60_DNA_seed | HSP90AA1 | 0.645617 | COADREAD | SW1417 | Tanespimycin | MVP_top60_DNA+RNA_seed | ATM | 0.404426 | COADREAD | MDST8 | KU-55933 |
| MVP_top60_DNA_seed | HSP90AB1 | 0.645617 | COADREAD | SW1417 | Tanespimycin | MVP_top60_DNA+RNA_seed | CDK2 | 0.406087 | COADREAD | CCK-81 | AZD5438 |
| MVP_top60_DNA_seed | AR | 0.645724 | COADREAD | LS-123 | AZD3514 | MVP_top60_DNA+RNA_seed | IGF1R | 0.406436 | COADREAD | HCC2998 | GSK1904529A |
| MVP_top60_DNA_seed | PIK3CD | 0.647094 | COADREAD | NCI-H716 | IC-87114 | MVP_top60_DNA+RNA_seed | ERBB4 | 0.407448 | COADREAD | HT-29 | AST-1306 |
| MVP_top60_DNA_seed | MAPK14 | 0.64805 | COADREAD | SW837 | TAK-715 | MVP_top60_DNA+RNA_seed | ERBB4 | 0.410436 | COADREAD | LS-180 | AST-1306 |
| MVP_top60_DNA_seed | AKT1 | 0.648404 | COADREAD | LS-180 | GSK690693 | MVP_top60_DNA+RNA_seed | GSK3B | 0.412051 | COADREAD | DiFi | AZD7969 |
| MVP_top60_DNA_seed | BRAF | 0.650184 | COADREAD | LS-513 | PLX-4720 | MVP_top60_DNA+RNA_seed | PIK3CD | 0.412927 | COADREAD | C2BBe1 | Idelalisib |
| MVP_top60_DNA_seed | ATM | 0.652341 | COADREAD | SNU-C2B | KU-60019 | MVP_top60_DNA+RNA_seed | CDK2 | 0.413889 | COADREAD | LS-180 | PHA-793887 |
| MVP_top60_DNA_seed | CDK2 | 0.653351 | COADREAD | RCM-1 | AZD5438 | MVP_top60_DNA+RNA_seed | PIK3CA | 0.416232 | COADREAD | MDST8 | PI3Ka_4409 |
| MVP_top60_DNA_seed | ATM | 0.654055 | COADREAD | SNU-407 | KU-55933 | MVP_top60_DNA+RNA_seed | PIK3CD | 0.416232 | COADREAD | MDST8 | PI3Ka_4409 |
| MVP_top60_DNA_seed | MAPK14 | 0.655036 | COADREAD | LS-1034 | TAK-715 | MVP_top60_DNA+RNA_seed | ERBB4 | 0.419277 | COADREAD | SNU-407 | CI-1033 |
| MVP_top60_DNA_seed | ESR1 | 0.65553 | COADREAD | RCM-1 | Fulvestrant | MVP_top60_DNA+RNA_seed | CDK2 | 0.420278 | COADREAD | SW837 | PHA-793887 |
| MVP_top60_DNA_seed | MAPK1 | 0.655668 | COADREAD | LS-123 | FR-180204 | MVP_top60_DNA+RNA_seed | HSP90AA1 | 0.422552 | COADREAD | SW837 | Tanespimycin |
| MVP_top60_DNA_seed | PIK3CB | 0.656293 | COADREAD | GP5d | AZD8186 | MVP_top60_DNA+RNA_seed | HSP90AB1 | 0.422552 | COADREAD | SW837 | Tanespimycin |
| MVP_top60_DNA_seed | PIK3CD | 0.656293 | COADREAD | GP5d | AZD8186 | MVP_top60_DNA+RNA_seed | GSK3B | 0.42258 | COADREAD | MDST8 | CHIR-99021 |
| MVP_top60_DNA_seed | PIK3CD | 0.657334 | COADREAD | SW1417 | IC-87114 | MVP_top60_DNA+RNA_seed | BCL2 | 0.423163 | COADREAD | KM12 | Navitoclax |
| MVP_top60_DNA_seed | ERBB4 | 0.657562 | COADREAD | NCI-H716 | AST-1306 | MVP_top60_DNA+RNA_seed | BCL2 | 0.425159 | COADREAD | KM12 | TW 37 |
| MVP_top60_DNA_seed | HSP90AA1 | 0.658876 | COADREAD | NCI-H716 | Elesclomol | MVP_top60_DNA+RNA_seed | AR | 0.425462 | COADREAD | HCT-15 | Bicalutamide |
| MVP_top60_DNA_seed | HSP90AB1 | 0.658876 | COADREAD | NCI-H716 | Elesclomol | MVP_top60_DNA+RNA_seed | AKT1 | 0.429226 | COADREAD | SNU-175 | GSK690693 |
| MVP_top60_DNA_seed | MET | 0.659345 | COADREAD | COLO-320-HSR | SU11274 | MVP_top60_DNA+RNA_seed | IGF1R | 0.429454 | COADREAD | LoVo | GSK1904529A |
| MVP_top60_DNA_seed | PTK2 | 0.660953 | COADREAD | C2BBe1 | PF-562271 | MVP_top60_DNA+RNA_seed | AKT1 | 0.430202 | COADREAD | SNU-C2B | AKT inhibitor VIII |
| MVP_top60_DNA_seed | CREBBP | 0.661153 | COADREAD | LS-1034 | I-CBP112 | MVP_top60_DNA+RNA_seed | MET | 0.430599 | COADREAD | SNU-407 | SU11274 |
| MVP_top60_DNA_seed | ERBB3 | 0.662739 | COADREAD | NCI-H716 | AZD8931 | MVP_top60_DNA+RNA_seed | AR | 0.434984 | COADREAD | SW620 | Bicalutamide |
| MVP_top60_DNA_seed | BRAF | 0.663698 | COADREAD | SNU-407 | PLX-4720 | MVP_top60_DNA+RNA_seed | ERBB4 | 0.435346 | COADREAD | COLO-205 | CI-1033 |
| MVP_top60_DNA_seed | MAPK1 | 0.663871 | COADREAD | HT-115 | FR-180204 | MVP_top60_DNA+RNA_seed | BCL2 | 0.436716 | COADREAD | SNU-C2B | Obatoclax Mesylate |
| MVP_top60_DNA_seed | BRAF | 0.664534 | COADREAD | SNU-C2B | PLX-4720 | MVP_top60_DNA+RNA_seed | GSK3B | 0.438428 | COADREAD | RCM-1 | CHIR-99021 |
| MVP_top60_DNA_seed | CDK2 | 0.668004 | COADREAD | HCT-15 | AZD5438 | MVP_top60_DNA+RNA_seed | AKT1 | 0.438481 | COADREAD | SK-CO-1 | BAY AKT1 |

| | | | | | | | | | | | |
|---|---|---|---|---|---|---|---|---|---|---|---|
| MVP_top60_DNA_seed | MET | 0.668154 | COADREAD | LS-513 | PHA-665752 | MVP_top60_DNA+RNA_seed | AKT1 | 0.441706 | COADREAD | SNU-61 | MK-2206 |
| MVP_top60_DNA_seed | ERBB4 | 0.668666 | COADREAD | LS-411N | AST-1306 | MVP_top60_DNA+RNA_seed | IGF1R | 0.441836 | COADREAD | SNU-C2B | GSK1904529A |
| MVP_top60_DNA_seed | PIK3CD | 0.670156 | COADREAD | HT-29 | IC-87114 | MVP_top60_DNA+RNA_seed | MET | 0.443924 | COADREAD | COLO-320-HSR | JNJ38877605 |
| MVP_top60_DNA_seed | GSK3B | 0.672734 | COADREAD | SW837 | CHIR-99021 | MVP_top60_DNA+RNA_seed | IGF1R | 0.446969 | COADREAD | HCT-15 | GSK1904529A |
| MVP_top60_DNA_seed | MET | 0.672841 | COADREAD | LS-411N | SU11274 | MVP_top60_DNA+RNA_seed | PIK3CD | 0.447453 | COADREAD | SNU-407 | IC-87114 |
| MVP_top60_DNA_seed | SRC | 0.676077 | COADREAD | KM12 | Bosutinib | MVP_top60_DNA+RNA_seed | ATM | 0.450843 | COADREAD | SW620 | torin2 |
| MVP_top60_DNA_seed | ATM | 0.677636 | COADREAD | SW1463 | KU-55933 | MVP_top60_DNA+RNA_seed | GSK3B | 0.452079 | COADREAD | RCM-1 | SB216763 |
| MVP_top60_DNA_seed | GSK3B | 0.679434 | COADREAD | NCI-H716 | AZD7969 | MVP_top60_DNA+RNA_seed | AKT1 | 0.454402 | COADREAD | SNU-C5 | GSK690693 |
| MVP_top60_DNA_seed | GSK3B | 0.68163 | COADREAD | CW-2 | CHIR-99021 | MVP_top60_DNA+RNA_seed | PIK3CD | 0.45666 | COADREAD | MDST8 | IC-87114 |
| MVP_top60_DNA_seed | PIK3CB | 0.681764 | COADREAD | SW1417 | AZD8186 | MVP_top60_DNA+RNA_seed | ERBB4 | 0.457601 | COADREAD | CW-2 | CI-1033 |
| MVP_top60_DNA_seed | PIK3CD | 0.681764 | COADREAD | SW1417 | AZD8186 | MVP_top60_DNA+RNA_seed | AKT1 | 0.458758 | COADREAD | HCT-15 | GSK690693 |
| MVP_top60_DNA_seed | HSP90AA1 | 0.682942 | COADREAD | SNU-407 | CCT-018159 | MVP_top60_DNA+RNA_seed | PIK3CB | 0.458988 | COADREAD | MDST8 | AZD6482 |
| MVP_top60_DNA_seed | HSP90AB1 | 0.682942 | COADREAD | SNU-407 | CCT-018159 | MVP_top60_DNA+RNA_seed | ATM | 0.459969 | COADREAD | SW837 | KU-60019 |
| MVP_top60_DNA_seed | AR | 0.684221 | COADREAD | SW48 | Bicalutamide | MVP_top60_DNA+RNA_seed | AR | 0.460199 | COADREAD | LS-1034 | Bicalutamide |
| MVP_top60_DNA_seed | MET | 0.686548 | COADREAD | T84 | AZD6094 | MVP_top60_DNA+RNA_seed | GSK3B | 0.46273 | COADREAD | LS-513 | CHIR-99021 |
| MVP_top60_DNA_seed | PIK3CA | 0.686876 | COADREAD | HCC2998 | AZD8835 | MVP_top60_DNA+RNA_seed | AR | 0.463933 | COADREAD | CL-11 | Bicalutamide |
| MVP_top60_DNA_seed | PIK3CD | 0.686876 | COADREAD | HCC2998 | AZD8835 | MVP_top60_DNA+RNA_seed | IGF1R | 0.464241 | COADREAD | RCM-1 | BMS-536924 |
| MVP_top60_DNA_seed | SRC | 0.687497 | COADREAD | LS-180 | NVP-BHG712 | MVP_top60_DNA+RNA_seed | IGF1R | 0.464875 | COADREAD | MDST8 | Linsitinib |
| MVP_top60_DNA_seed | MET | 0.687647 | COADREAD | MDST8 | Crizotinib | MVP_top60_DNA+RNA_seed | PIK3CA | 0.465554 | COADREAD | SW837 | AZD8835 |
| MVP_top60_DNA_seed | GSK3B | 0.69064 | COADREAD | COLO-678 | CHIR-99021 | MVP_top60_DNA+RNA_seed | PIK3CD | 0.465554 | COADREAD | SW837 | AZD8835 |
| MVP_top60_DNA_seed | CDK2 | 0.691077 | COADREAD | DiFi | AZD5438 | MVP_top60_DNA+RNA_seed | CDK2 | 0.466013 | COADREAD | KM12 | AZD5438 |
| MVP_top60_DNA_seed | ERBB3 | 0.693676 | COADREAD | COLO-678 | AZD8931 | MVP_top60_DNA+RNA_seed | BCL2 | 0.467368 | COADREAD | LS-180 | Navitoclax |
| MVP_top60_DNA_seed | HSP90AA1 | 0.694137 | COADREAD | COLO-678 | Luminespib | MVP_top60_DNA+RNA_seed | SRC | 0.467377 | COADREAD | SNU-C2B | Bosutinib |
| MVP_top60_DNA_seed | HSP90AB1 | 0.694137 | COADREAD | COLO-678 | Luminespib | MVP_top60_DNA+RNA_seed | GSK3B | 0.46857 | COADREAD | CW-2 | AZD7969 |
| MVP_top60_DNA_seed | GSK3B | 0.694624 | COADREAD | SW837 | CHIR-99021 | MVP_top60_DNA+RNA_seed | GSK3B | 0.469408 | COADREAD | CCK-81 | CHIR-99021 |
| MVP_top60_DNA_seed | BRAF | 0.695822 | COADREAD | SNU-C2B | HG6-64-1 | MVP_top60_DNA+RNA_seed | AKT1 | 0.471176 | COADREAD | SW620 | GSK690693 |
| MVP_top60_DNA_seed | CREBBP | 0.695838 | COADREAD | T84 | I-CBP112 | MVP_top60_DNA+RNA_seed | ERBB4 | 0.471648 | COADREAD | CL-11 | CI-1033 |
| MVP_top60_DNA_seed | BRAF | 0.698147 | COADREAD | COLO-320-HSR | Dabrafenib | MVP_top60_DNA+RNA_seed | SRC | 0.472706 | COADREAD | COLO-320-HSR | Saracatinib |
| MVP_top60_DNA_seed | GSK3B | 0.698274 | COADREAD | HCC2998 | CHIR-99021 | MVP_top60_DNA+RNA_seed | GSK3B | 0.472764 | COADREAD | SW948 | CHIR-99021 |
| MVP_top60_DNA_seed | MET | 0.700497 | COADREAD | SNU-407 | AZD6094 | MVP_top60_DNA+RNA_seed | GSK3B | 0.473992 | COADREAD | LS-411N | AZD7969 |
| MVP_top60_DNA_seed | CREBBP | 0.701969 | COADREAD | CL-11 | I-CBP112 | MVP_top60_DNA+RNA_seed | AKT1 | 0.474098 | COADREAD | SW48 | MK-2206 |
| MVP_top60_DNA_seed | AKT1 | 0.703024 | COADREAD | HT-29 | MK-2206 | MVP_top60_DNA+RNA_seed | PIK3CD | 0.474163 | COADREAD | HCC2998 | IC-87114 |
| MVP_top60_DNA_seed | HSP90AA1 | 0.703561 | COADREAD | CW-2 | Elesclomol | MVP_top60_DNA+RNA_seed | GSK3B | 0.474477 | COADREAD | LS-411N | CHIR-99021 |
| MVP_top60_DNA_seed | HSP90AB1 | 0.703561 | COADREAD | CW-2 | Elesclomol | MVP_top60_DNA+RNA_seed | HSP90AA1 | 0.476212 | COADREAD | LS-513 | Elesclomol |
| MVP_top60_DNA_seed | ATM | 0.705629 | COADREAD | SW1417 | CP466722 | MVP_top60_DNA+RNA_seed | HSP90AB1 | 0.476212 | COADREAD | LS-513 | Elesclomol |
| MVP_top60_DNA_seed | AR | 0.706151 | COADREAD | HT55 | AZD3514 | MVP_top60_DNA+RNA_seed | AKT1 | 0.47805 | COADREAD | SNU-1040 | BAY AKT1 |
| MVP_top60_DNA_seed | IGF1R | 0.707248 | COADREAD | MDST8 | BMS-754807 | MVP_top60_DNA+RNA_seed | GSK3B | 0.478233 | COADREAD | SW1116 | CHIR-99021 |
| MVP_top60_DNA_seed | PIK3CB | 0.707509 | COADREAD | CaR-1 | AZD8186 | MVP_top60_DNA+RNA_seed | SRC | 0.478334 | COADREAD | HT55 | NVP-BHG712 |
| MVP_top60_DNA_seed | PIK3CD | 0.707509 | COADREAD | CaR-1 | AZD8186 | MVP_top60_DNA+RNA_seed | SRC | 0.478455 | COADREAD | CaR-1 | NVP-BHG712 |
| MVP_top60_DNA_seed | ERBB4 | 0.707822 | COADREAD | KM12 | PF-00299804 | MVP_top60_DNA+RNA_seed | ERBB4 | 0.479359 | COADREAD | SW48 | AST-1306 |
| MVP_top60_DNA_seed | BRAF | 0.709396 | COADREAD | SNU-81 | SB590885 | MVP_top60_DNA+RNA_seed | BCL2 | 0.479844 | COADREAD | SW837 | Navitoclax |

| | | | | | | | | | | | |
|---|---|---|---|---|---|---|---|---|---|---|---|
| MVP_top60_DNA_seed | GSK3B | 0.710742 | COADREAD | LS-123 | KIN001-042 | MVP_top60_DNA+RNA_seed | PIK3CD | 0.481279 | COADREAD | SNU-175 | Idelalisib |
| MVP_top60_DNA_seed | GSK3B | 0.712087 | COADREAD | LS-411N | KIN001-042 | MVP_top60_DNA+RNA_seed | AKT1 | 0.481529 | COADREAD | SW837 | AKT inhibitor VIII |
| MVP_top60_DNA_seed | MET | 0.712138 | COADREAD | SNU-61 | AZD6094 | MVP_top60_DNA+RNA_seed | PIK3CD | 0.481735 | COADREAD | HT55 | IC-87114 |
| MVP_top60_DNA_seed | PIK3CB | 0.712668 | COADREAD | HT-115 | AZD6482 | MVP_top60_DNA+RNA_seed | AKT1 | 0.485757 | COADREAD | HCT-15 | AKT inhibitor VIII |
| MVP_top60_DNA_seed | GSK3B | 0.71276 | COADREAD | COLO-320-HSR | CHIR-99021 | MVP_top60_DNA+RNA_seed | GSK3B | 0.487078 | COADREAD | KM12 | CHIR-99021 |
| MVP_top60_DNA_seed | AKT1 | 0.71285 | COADREAD | C2BBe1 | GSK690693 | MVP_top60_DNA+RNA_seed | AKT1 | 0.487425 | COADREAD | CCK-81 | GSK690693 |
| MVP_top60_DNA_seed | MET | 0.713405 | COADREAD | LS-123 | JNJ38877605 | MVP_top60_DNA+RNA_seed | HSP90AA1 | 0.487872 | COADREAD | SNU-C2B | Tanespimycin |
| MVP_top60_DNA_seed | BRAF | 0.71386 | COADREAD | CCK-81 | PLX-4720 | MVP_top60_DNA+RNA_seed | HSP90AB1 | 0.487872 | COADREAD | SNU-C2B | Tanespimycin |
| MVP_top60_DNA_seed | ATM | 0.713882 | COADREAD | CaR-1 | KU-60019 | MVP_top60_DNA+RNA_seed | IGF1R | 0.487925 | COADREAD | SW48 | GSK1904529A |
| MVP_top60_DNA_seed | CDK2 | 0.714718 | COADREAD | SNU-C1 | AZD5438 | MVP_top60_DNA+RNA_seed | GSK3B | 0.4899 | COADREAD | HCC2998 | SB216763 |
| MVP_top60_DNA_seed | HSP90AA1 | 0.716149 | COADREAD | CCK-81 | Tanespimycin | MVP_top60_DNA+RNA_seed | GSK3B | 0.490676 | COADREAD | C2BBe1 | KIN001-042 |
| MVP_top60_DNA_seed | HSP90AB1 | 0.716149 | COADREAD | CCK-81 | Tanespimycin | MVP_top60_DNA+RNA_seed | IGF1R | 0.492173 | COADREAD | HCT-116 | Linsitinib |
| MVP_top60_DNA_seed | PTK2 | 0.716335 | COADREAD | LS-513 | PF-562271 | MVP_top60_DNA+RNA_seed | PIK3CD | 0.494593 | COADREAD | NCI-H716 | Idelalisib |
| MVP_top60_DNA_seed | HSP90AA1 | 0.717116 | COADREAD | SNU-1040 | SNX-2112 | MVP_top60_DNA+RNA_seed | SRC | 0.495551 | COADREAD | NCI-H716 | WH-4-023 |
| MVP_top60_DNA_seed | HSP90AB1 | 0.717116 | COADREAD | SNU-1040 | SNX-2112 | MVP_top60_DNA+RNA_seed | SRC | 0.496924 | COADREAD | RKO | Bosutinib |
| MVP_top60_DNA_seed | ERBB3 | 0.717602 | COADREAD | CL-11 | AZD8931 | MVP_top60_DNA+RNA_seed | FYN | 0.497943 | COADREAD | LS-411N | A-770041 |
| MVP_top60_DNA_seed | BRAF | 0.718315 | COADREAD | SNU-C1 | PLX-4720 | MVP_top60_DNA+RNA_seed | MET | 0.498615 | COADREAD | LS-513 | Crizotinib |
| MVP_top60_DNA_seed | GSK3B | 0.720694 | COADREAD | COLO-205 | CHIR-99021 | MVP_top60_DNA+RNA_seed | CDK2 | 0.501652 | COADREAD | LS-1034 | PHA-793887 |
| MVP_top60_DNA_seed | ESR1 | 0.723254 | COADREAD | CaR-1 | Tamoxifen | MVP_top60_DNA+RNA_seed | HSP90AA1 | 0.502148 | COADREAD | LS-123 | Tanespimycin |
| MVP_top60_DNA_seed | PIK3CB | 0.723515 | COADREAD | LS-411N | AZD6482 | MVP_top60_DNA+RNA_seed | HSP90AB1 | 0.502148 | COADREAD | LS-123 | Tanespimycin |
| MVP_top60_DNA_seed | AKT1 | 0.723742 | COADREAD | COLO-320-HSR | GSK690693 | MVP_top60_DNA+RNA_seed | AR | 0.503908 | COADREAD | SNU-1040 | Bicalutamide |
| MVP_top60_DNA_seed | AR | 0.724423 | COADREAD | LS-513 | Bicalutamide | MVP_top60_DNA+RNA_seed | PIK3CD | 0.505656 | COADREAD | RCM-1 | IC-87114 |
| MVP_top60_DNA_seed | MET | 0.72468 | COADREAD | LS-411N | AZD6094 | MVP_top60_DNA+RNA_seed | GSK3B | 0.507042 | COADREAD | NCI-H508 | SB216763 |
| MVP_top60_DNA_seed | PTK2 | 0.72578 | COADREAD | SNU-C2B | PF-562271 | MVP_top60_DNA+RNA_seed | HSP90AA1 | 0.507691 | COADREAD | HCT-116 | Luminespib |
| MVP_top60_DNA_seed | PIK3CD | 0.727068 | COADREAD | HCT-15 | IC-87114 | MVP_top60_DNA+RNA_seed | HSP90AB1 | 0.507691 | COADREAD | HCT-116 | Luminespib |
| MVP_top60_DNA_seed | ESR1 | 0.728028 | COADREAD | SW837 | Tamoxifen | MVP_top60_DNA+RNA_seed | BCL2 | 0.5081 | COADREAD | LS-1034 | TW 37 |
| MVP_top60_DNA_seed | HSP90AA1 | 0.728095 | COADREAD | SNU-61 | Tanespimycin | MVP_top60_DNA+RNA_seed | MET | 0.508738 | COADREAD | LS-180 | JNJ38877605 |
| MVP_top60_DNA_seed | HSP90AB1 | 0.728095 | COADREAD | SNU-61 | Tanespimycin | MVP_top60_DNA+RNA_seed | CDK2 | 0.510576 | COADREAD | NCI-H716 | PHA-793887 |
| MVP_top60_DNA_seed | AKT1 | 0.728839 | COADREAD | SNU-61 | GSK690693 | MVP_top60_DNA+RNA_seed | MET | 0.512733 | COADREAD | SNU-1040 | SU11274 |
| MVP_top60_DNA_seed | AKT1 | 0.729032 | COADREAD | SNU-C2B | MK-2206 | MVP_top60_DNA+RNA_seed | AR | 0.514015 | COADREAD | MDST8 | AZD3514 |
| MVP_top60_DNA_seed | BRAF | 0.729073 | COADREAD | DiFi | SB590885 | MVP_top60_DNA+RNA_seed | SRC | 0.515083 | COADREAD | COLO-320-HSR | WH-4-023 |
| MVP_top60_DNA_seed | SRC | 0.730608 | COADREAD | LS-513 | Saracatinib | MVP_top60_DNA+RNA_seed | IGF1R | 0.515128 | COADREAD | MDST8 | BMS-536924 |
| MVP_top60_DNA_seed | ESR1 | 0.730954 | COADREAD | CCK-81 | Tamoxifen | MVP_top60_DNA+RNA_seed | HSP90AA1 | 0.515624 | COADREAD | SW837 | Luminespib |
| MVP_top60_DNA_seed | MET | 0.73098 | COADREAD | LS-180 | SU11274 | MVP_top60_DNA+RNA_seed | HSP90AB1 | 0.515624 | COADREAD | SW837 | Luminespib |
| MVP_top60_DNA_seed | MET | 0.731449 | COADREAD | HT55 | AZD6094 | MVP_top60_DNA+RNA_seed | AKT1 | 0.517484 | COADREAD | LS-513 | AKT inhibitor VIII |
| MVP_top60_DNA_seed | ERBB3 | 0.732919 | COADREAD | LS-1034 | AZD8931 | MVP_top60_DNA+RNA_seed | PIK3CD | 0.518923 | COADREAD | NCI-H747 | IC-87114 |
| MVP_top60_DNA_seed | PIK3CD | 0.733169 | COADREAD | SNU-C5 | IC-87114 | MVP_top60_DNA+RNA_seed | MET | 0.519471 | COADREAD | LS-1034 | AZD6094 |
| MVP_top60_DNA_seed | GSK3B | 0.733918 | COADREAD | SW1463 | AZD7969 | MVP_top60_DNA+RNA_seed | PIK3CB | 0.519721 | COADREAD | MDST8 | TGX221 |
| MVP_top60_DNA_seed | MAPK14 | 0.734351 | COADREAD | C2BBe1 | TAK-715 | MVP_top60_DNA+RNA_seed | AKT1 | 0.519895 | COADREAD | LS-411N | BAY AKT1 |
| MVP_top60_DNA_seed | ERBB4 | 0.735259 | COADREAD | HT-29 | CI-1033 | MVP_top60_DNA+RNA_seed | PTK2 | 0.527031 | COADREAD | SNU-C1 | PF-562271 |
| MVP_top60_DNA_seed | HSP90AA1 | 0.735349 | COADREAD | SNU-C1 | Tanespimycin | MVP_top60_DNA+RNA_seed | HSP90AA1 | 0.529101 | COADREAD | GP5d | Elesclomol |

| | | | | | | | | | | | |
|---|---|---|---|---|---|---|---|---|---|---|---|
| MVP_top60_DNA_seed | HSP90AB1 | 0.735349 | COADREAD | SNU-C1 | Tanespimycin | MVP_top60_DNA+RNA_seed | HSP90AB1 | 0.529101 | COADREAD | GP5d | Elesclomol |
| MVP_top60_DNA_seed | PIK3CA | 0.737672 | COADREAD | CW-2 | AZD8835 | MVP_top60_DNA+RNA_seed | AKT1 | 0.530064 | COADREAD | CaR-1 | AKT inhibitor VIII |
| MVP_top60_DNA_seed | PIK3CD | 0.737672 | COADREAD | CW-2 | AZD8835 | MVP_top60_DNA+RNA_seed | ERBB3 | 0.530317 | COADREAD | T84 | AZD8931 |
| MVP_top60_DNA_seed | BRAF | 0.737854 | COADREAD | KM12 | PLX-4720 | MVP_top60_DNA+RNA_seed | ATM | 0.532861 | COADREAD | COLO-205 | KU-55933 |
| MVP_top60_DNA_seed | ERBB4 | 0.740807 | COADREAD | SNU-407 | AST-1306 | MVP_top60_DNA+RNA_seed | PIK3CB | 0.534793 | COADREAD | HT-29 | AZD6482 |
| MVP_top60_DNA_seed | MET | 0.741162 | COADREAD | SW620 | SU11274 | MVP_top60_DNA+RNA_seed | GSK3B | 0.534964 | COADREAD | HT-29 | CHIR-99021 |
| MVP_top60_DNA_seed | ERBB4 | 0.741743 | COADREAD | LS-123 | AST-1306 | MVP_top60_DNA+RNA_seed | ATM | 0.540154 | COADREAD | CaR-1 | CP466722 |
| MVP_top60_DNA_seed | ATM | 0.742282 | COADREAD | HCC2998 | KU-60019 | MVP_top60_DNA+RNA_seed | HSP90AA1 | 0.540994 | COADREAD | LS-411N | Elesclomol |
| MVP_top60_DNA_seed | AR | 0.742994 | COADREAD | SW1417 | Bicalutamide | MVP_top60_DNA+RNA_seed | HSP90AB1 | 0.540994 | COADREAD | LS-411N | Elesclomol |
| MVP_top60_DNA_seed | HSP90AA1 | 0.743353 | COADREAD | RCM-1 | SNX-2112 | MVP_top60_DNA+RNA_seed | ERBB4 | 0.541254 | COADREAD | HCT-116 | AST-1306 |
| MVP_top60_DNA_seed | HSP90AB1 | 0.743353 | COADREAD | RCM-1 | SNX-2112 | MVP_top60_DNA+RNA_seed | HSP90AA1 | 0.541643 | COADREAD | SNU-C2B | CCT-018159 |
| MVP_top60_DNA_seed | SRC | 0.744864 | COADREAD | SW837 | NVP-BHG712 | MVP_top60_DNA+RNA_seed | HSP90AB1 | 0.541643 | COADREAD | SNU-C2B | CCT-018159 |
| MVP_top60_DNA_seed | PIK3CA | 0.746074 | COADREAD | RCM-1 | AZD8835 | MVP_top60_DNA+RNA_seed | MET | 0.544746 | COADREAD | COLO-320-HSR | Crizotinib |
| MVP_top60_DNA_seed | PIK3CD | 0.746074 | COADREAD | RCM-1 | AZD8835 | MVP_top60_DNA+RNA_seed | GSK3B | 0.54719 | COADREAD | SW620 | CHIR-99021 |
| MVP_top60_DNA_seed | PIK3CB | 0.748571 | COADREAD | HT55 | AZD8186 | MVP_top60_DNA+RNA_seed | MET | 0.548115 | COADREAD | LS-513 | SU11274 |
| MVP_top60_DNA_seed | PIK3CD | 0.748571 | COADREAD | HT55 | AZD8186 | MVP_top60_DNA+RNA_seed | CDK2 | 0.548416 | COADREAD | LS-180 | AZD5438 |
| MVP_top60_DNA_seed | MAPK1 | 0.748726 | COADREAD | HCC2998 | FR-180204 | MVP_top60_DNA+RNA_seed | BCL2 | 0.548671 | COADREAD | SW837 | Obatoclax Mesylate |
| MVP_top60_DNA_seed | IGF1R | 0.749268 | COADREAD | T84 | GSK1904529A | MVP_top60_DNA+RNA_seed | CDK2 | 0.549227 | COADREAD | NCI-H747 | Seliciclib |
| MVP_top60_DNA_seed | PTK2 | 0.751637 | COADREAD | LS-1034 | PF-562271 | MVP_top60_DNA+RNA_seed | IGF1R | 0.553034 | COADREAD | RKO | BMS-536924 |
| MVP_top60_DNA_seed | ERBB4 | 0.753289 | COADREAD | RCM-1 | AST-1306 | MVP_top60_DNA+RNA_seed | ERBB4 | 0.55471 | COADREAD | LS-123 | PF-00299804 |
| MVP_top60_DNA_seed | AKT1 | 0.75329 | COADREAD | T84 | BAY AKT1 | MVP_top60_DNA+RNA_seed | IGF1R | 0.554902 | COADREAD | SW1116 | BMS-536924 |
| MVP_top60_DNA_seed | SRC | 0.754352 | COADREAD | MDST8 | Saracatinib | MVP_top60_DNA+RNA_seed | CDK2 | 0.555807 | COADREAD | LS-123 | AZD5438 |
| MVP_top60_DNA_seed | PTK2 | 0.755765 | COADREAD | CCK-81 | PF-562271 | MVP_top60_DNA+RNA_seed | ATM | 0.556719 | COADREAD | SW48 | KU-60019 |
| MVP_top60_DNA_seed | IGF1R | 0.756383 | COADREAD | LS-180 | BMS-536924 | MVP_top60_DNA+RNA_seed | IGF1R | 0.561451 | COADREAD | LS-513 | GSK1904529A |
| MVP_top60_DNA_seed | BRAF | 0.758533 | COADREAD | T84 | Dabrafenib | MVP_top60_DNA+RNA_seed | SRC | 0.561701 | COADREAD | LS-1034 | WH-4-023 |
| MVP_top60_DNA_seed | ERBB4 | 0.758946 | COADREAD | SNU-C5 | AST-1306 | MVP_top60_DNA+RNA_seed | PIK3CD | 0.561973 | COADREAD | CW-2 | Idelalisib |
| MVP_top60_DNA_seed | CDK2 | 0.759818 | COADREAD | CW-2 | Seliciclib | MVP_top60_DNA+RNA_seed | ERBB3 | 0.562343 | COADREAD | LS-411N | AZD8931 |
| MVP_top60_DNA_seed | AR | 0.760463 | COADREAD | NCI-H716 | Bicalutamide | MVP_top60_DNA+RNA_seed | MET | 0.56264 | COADREAD | HCC2998 | SU11274 |
| MVP_top60_DNA_seed | GSK3B | 0.762139 | COADREAD | LS-1034 | CHIR-99021 | MVP_top60_DNA+RNA_seed | HSP90AA1 | 0.56365 | COADREAD | SNU-81 | CCT-018159 |
| MVP_top60_DNA_seed | BRAF | 0.762312 | COADREAD | HCC2998 | HG6-64-1 | MVP_top60_DNA+RNA_seed | HSP90AB1 | 0.56365 | COADREAD | SNU-81 | CCT-018159 |
| MVP_top60_DNA_seed | AKT1 | 0.764144 | COADREAD | SW1463 | BAY AKT1 | MVP_top60_DNA+RNA_seed | ATM | 0.564396 | COADREAD | SNU-C2B | torin2 |
| MVP_top60_DNA_seed | ESR1 | 0.764235 | COADREAD | COLO-205 | Fulvestrant | MVP_top60_DNA+RNA_seed | PIK3CD | 0.565272 | COADREAD | SW48 | Idelalisib |
| MVP_top60_DNA_seed | PIK3CD | 0.764971 | COADREAD | KM12 | IC-87114 | MVP_top60_DNA+RNA_seed | AR | 0.566062 | COADREAD | SNU-C1 | AZD3514 |
| MVP_top60_DNA_seed | PTK2 | 0.765013 | COADREAD | MDST8 | PF-562271 | MVP_top60_DNA+RNA_seed | MET | 0.569503 | COADREAD | MDST8 | PHA-665752 |
| MVP_top60_DNA_seed | CDK2 | 0.766794 | COADREAD | CCK-81 | PHA-793887 | MVP_top60_DNA+RNA_seed | GSK3B | 0.570012 | COADREAD | SNU-C2B | CHIR-99021 |
| MVP_top60_DNA_seed | SRC | 0.767828 | COADREAD | HT-29 | HG-6-71-01 | MVP_top60_DNA+RNA_seed | PIK3CD | 0.570535 | COADREAD | SNU-C5 | Idelalisib |
| MVP_top60_DNA_seed | PIK3CD | 0.767996 | COADREAD | T84 | IC-87114 | MVP_top60_DNA+RNA_seed | GSK3B | 0.570749 | COADREAD | CL-11 | AZD7969 |
| MVP_top60_DNA_seed | AKT1 | 0.768115 | COADREAD | HCC2998 | AKT inhibitor VIII | MVP_top60_DNA+RNA_seed | HSP90AA1 | 0.573523 | COADREAD | SW1417 | CCT-018159 |
| MVP_top60_DNA_seed | AKT1 | 0.770802 | COADREAD | SW1417 | AKT inhibitor VIII | MVP_top60_DNA+RNA_seed | HSP90AB1 | 0.573523 | COADREAD | SW1417 | CCT-018159 |
| MVP_top60_DNA_seed | MET | 0.77132 | COADREAD | LS-123 | PHA-665752 | MVP_top60_DNA+RNA_seed | CDK2 | 0.57502 | COADREAD | CaR-1 | PHA-793887 |
| MVP_top60_DNA_seed | ERBB4 | 0.771923 | COADREAD | COLO-678 | AST-1306 | MVP_top60_DNA+RNA_seed | IGF1R | 0.577806 | COADREAD | MDST8 | GSK1904529A |

| | | | | | | | | | | | |
|---|---|---|---|---|---|---|---|---|---|---|---|
| MVP_top60_DNA_seed | ATM | 0.772567 | COADREAD | GP5d | KU-55933 | MVP_top60_DNA+RNA_seed | HSP90AA1 | 0.57793 | COADREAD | LS-411N | CCT-018159 |
| MVP_top60_DNA_seed | ERBB4 | 0.772891 | COADREAD | LS-513 | AST-1306 | MVP_top60_DNA+RNA_seed | HSP90AB1 | 0.57793 | COADREAD | LS-411N | CCT-018159 |
| MVP_top60_DNA_seed | BRAF | 0.774553 | COADREAD | SNU-C1 | Dabrafenib | MVP_top60_DNA+RNA_seed | AR | 0.578126 | COADREAD | MDST8 | Bicalutamide |
| MVP_top60_DNA_seed | ERBB4 | 0.775616 | COADREAD | SNU-407 | PF-00299804 | MVP_top60_DNA+RNA_seed | MET | 0.578742 | COADREAD | HCT-15 | AZD6094 |
| MVP_top60_DNA_seed | BRAF | 0.775816 | COADREAD | SNU-407 | SB590885 | MVP_top60_DNA+RNA_seed | AR | 0.58184 | COADREAD | NCI-H747 | Bicalutamide |
| MVP_top60_DNA_seed | ATM | 0.776763 | COADREAD | HCC2998 | KU-55933 | MVP_top60_DNA+RNA_seed | PIK3CD | 0.584855 | COADREAD | CW-2 | IC-87114 |
| MVP_top60_DNA_seed | AKT1 | 0.776865 | COADREAD | HT-115 | GSK690693 | MVP_top60_DNA+RNA_seed | CDK2 | 0.587523 | COADREAD | COLO-678 | PHA-793887 |
| MVP_top60_DNA_seed | AR | 0.776955 | COADREAD | RCM-1 | AZD3514 | MVP_top60_DNA+RNA_seed | AKT1 | 0.590635 | COADREAD | SNU-81 | BAY AKT1 |
| MVP_top60_DNA_seed | HSP90AA1 | 0.779328 | COADREAD | SNU-407 | Tanespimycin | MVP_top60_DNA+RNA_seed | HSP90AA1 | 0.592651 | COADREAD | SW48 | Elesclomol |
| MVP_top60_DNA_seed | HSP90AB1 | 0.779328 | COADREAD | SNU-407 | Tanespimycin | MVP_top60_DNA+RNA_seed | HSP90AB1 | 0.592651 | COADREAD | SW48 | Elesclomol |
| MVP_top60_DNA_seed | HSP90AA1 | 0.780158 | COADREAD | C2BBe1 | CCT-018159 | MVP_top60_DNA+RNA_seed | SRC | 0.592698 | COADREAD | C2BBe1 | HG-6-71-01 |
| MVP_top60_DNA_seed | HSP90AB1 | 0.780158 | COADREAD | C2BBe1 | CCT-018159 | MVP_top60_DNA+RNA_seed | CDK2 | 0.593583 | COADREAD | LS-513 | PHA-793887 |
| MVP_top60_DNA_seed | ESR1 | 0.781653 | COADREAD | HCC2998 | Fulvestrant | MVP_top60_DNA+RNA_seed | AKT1 | 0.594083 | COADREAD | CCK-81 | BAY AKT1 |
| MVP_top60_DNA_seed | PIK3CD | 0.781895 | COADREAD | CaR-1 | Idelalisib | MVP_top60_DNA+RNA_seed | PIK3CA | 0.59458 | COADREAD | SNU-C2B | AZD8835 |
| MVP_top60_DNA_seed | ATM | 0.782786 | COADREAD | LoVo | CP466722 | MVP_top60_DNA+RNA_seed | PIK3CD | 0.59458 | COADREAD | SNU-C2B | AZD8835 |
| MVP_top60_DNA_seed | FYN | 0.784415 | COADREAD | LS-513 | A-770041 | MVP_top60_DNA+RNA_seed | AKT1 | 0.597049 | COADREAD | SNU-C5 | AKT inhibitor VIII |
| MVP_top60_DNA_seed | PIK3CD | 0.785688 | COADREAD | HCC2998 | Idelalisib | MVP_top60_DNA+RNA_seed | MET | 0.599906 | COADREAD | SW1417 | AZD6094 |
| MVP_top60_DNA_seed | IGF1R | 0.787897 | COADREAD | HCT-15 | BMS-754807 | MVP_top60_DNA+RNA_seed | GSK3B | 0.600258 | COADREAD | HT-115 | CHIR-99021 |
| MVP_top60_DNA_seed | MET | 0.789716 | COADREAD | HCT-116 | JNJ38877605 | MVP_top60_DNA+RNA_seed | HSP90AA1 | 0.601088 | COADREAD | HCC2998 | Elesclomol |
| MVP_top60_DNA_seed | CDK2 | 0.790933 | COADREAD | LS-123 | Seliciclib | MVP_top60_DNA+RNA_seed | HSP90AB1 | 0.601088 | COADREAD | HCC2998 | Elesclomol |
| MVP_top60_DNA_seed | MET | 0.791202 | COADREAD | NCI-H508 | AZD6094 | MVP_top60_DNA+RNA_seed | PIK3CB | 0.604291 | COADREAD | C2BBe1 | AZD8186 |
| MVP_top60_DNA_seed | BRAF | 0.791578 | COADREAD | T84 | PLX-4720 | MVP_top60_DNA+RNA_seed | PIK3CD | 0.604291 | COADREAD | C2BBe1 | AZD8186 |
| MVP_top60_DNA_seed | GSK3B | 0.791862 | COADREAD | SW1417 | CHIR-99021 | MVP_top60_DNA+RNA_seed | ERBB4 | 0.604541 | COADREAD | NCI-H716 | PF-00299804 |
| MVP_top60_DNA_seed | ESR1 | 0.792319 | COADREAD | HCT-15 | Tamoxifen | MVP_top60_DNA+RNA_seed | HSP90AA1 | 0.607046 | COADREAD | HT-29 | Elesclomol |
| MVP_top60_DNA_seed | BRAF | 0.793588 | COADREAD | SNU-81 | RAF_9304 | MVP_top60_DNA+RNA_seed | HSP90AB1 | 0.607046 | COADREAD | HT-29 | Elesclomol |
| MVP_top60_DNA_seed | IGF1R | 0.795454 | COADREAD | HCT-15 | BMS-536924 | MVP_top60_DNA+RNA_seed | ERBB4 | 0.609301 | COADREAD | LS-123 | CI-1033 |
| MVP_top60_DNA_seed | ATM | 0.796034 | COADREAD | SNU-C2B | CP466722 | MVP_top60_DNA+RNA_seed | AKT1 | 0.609895 | COADREAD | SW837 | BAY AKT1 |
| MVP_top60_DNA_seed | PIK3CB | 0.796793 | COADREAD | HCC2998 | AZD8186 | MVP_top60_DNA+RNA_seed | PIK3CD | 0.611504 | COADREAD | SW1417 | Idelalisib |
| MVP_top60_DNA_seed | PIK3CD | 0.796793 | COADREAD | HCC2998 | AZD8186 | MVP_top60_DNA+RNA_seed | HSP90AA1 | 0.611952 | COADREAD | SNU-81 | Tanespimycin |
| MVP_top60_DNA_seed | HSP90AA1 | 0.797991 | COADREAD | SW1417 | SNX-2112 | MVP_top60_DNA+RNA_seed | HSP90AB1 | 0.611952 | COADREAD | SNU-81 | Tanespimycin |
| MVP_top60_DNA_seed | HSP90AB1 | 0.797991 | COADREAD | SW1417 | SNX-2112 | MVP_top60_DNA+RNA_seed | AR | 0.612007 | COADREAD | SW837 | Bicalutamide |
| MVP_top60_DNA_seed | AKT1 | 0.798423 | COADREAD | HT-29 | AKT inhibitor VIII | MVP_top60_DNA+RNA_seed | MET | 0.613018 | COADREAD | LS-513 | AZD6094 |
| MVP_top60_DNA_seed | AKT1 | 0.800284 | COADREAD | MDST8 | AKT inhibitor VIII | MVP_top60_DNA+RNA_seed | BCL2 | 0.615524 | COADREAD | HT-29 | TW 37 |
| MVP_top60_DNA_seed | PIK3CB | 0.802573 | COADREAD | SNU-C5 | AZD8186 | MVP_top60_DNA+RNA_seed | HSP90AA1 | 0.616381 | COADREAD | LS-123 | SNX-2112 |
| MVP_top60_DNA_seed | PIK3CD | 0.802573 | COADREAD | SNU-C5 | AZD8186 | MVP_top60_DNA+RNA_seed | HSP90AB1 | 0.616381 | COADREAD | LS-123 | SNX-2112 |
| MVP_top60_DNA_seed | AKT1 | 0.803413 | COADREAD | HT-115 | AKT inhibitor VIII | MVP_top60_DNA+RNA_seed | CDK2 | 0.61762 | COADREAD | T84 | AZD5438 |
| MVP_top60_DNA_seed | BRAF | 0.80362 | COADREAD | SW1463 | PLX-4720 | MVP_top60_DNA+RNA_seed | PTK2 | 0.617731 | COADREAD | CaR-1 | PF-562271 |
| MVP_top60_DNA_seed | MET | 0.80415 | COADREAD | C2BBe1 | SU11274 | MVP_top60_DNA+RNA_seed | AKT1 | 0.617978 | COADREAD | SNU-C1 | GSK690693 |
| MVP_top60_DNA_seed | AKT1 | 0.805855 | COADREAD | LoVo | GSK690693 | MVP_top60_DNA+RNA_seed | IGF1R | 0.61896 | COADREAD | SW1417 | BMS-754807 |
| MVP_top60_DNA_seed | ATM | 0.806017 | COADREAD | SW1417 | KU-60019 | MVP_top60_DNA+RNA_seed | IGF1R | 0.622928 | COADREAD | RKO | Linsitinib |
| MVP_top60_DNA_seed | IGF1R | 0.806249 | COADREAD | SW620 | GSK1904529A | MVP_top60_DNA+RNA_seed | ERBB4 | 0.62344 | COADREAD | SW1417 | PF-00299804 |

| | | | | | | | | | | | |
|---|---|---|---|---|---|---|---|---|---|---|---|
| MVP_top60_DNA_seed | SRC | 0.806405 | COADREAD | NCI-H716 | Saracatinib | MVP_top60_DNA+RNA_seed | AKT1 | 0.624579 | COADREAD | SW1417 | MK-2206 |
| MVP_top60_DNA_seed | SRC | 0.806858 | COADREAD | SW620 | HG-6-71-01 | MVP_top60_DNA+RNA_seed | SRC | 0.624669 | COADREAD | SNU-61 | Bosutinib |
| MVP_top60_DNA_seed | BRAF | 0.807135 | COADREAD | T84 | SB590885 | MVP_top60_DNA+RNA_seed | IGF1R | 0.624791 | COADREAD | COLO-320-HSR | GSK1904529A |
| MVP_top60_DNA_seed | MET | 0.808043 | COADREAD | SNU-C2B | AZD6094 | MVP_top60_DNA+RNA_seed | MET | 0.625292 | COADREAD | KM12 | AZD6094 |
| MVP_top60_DNA_seed | PIK3CB | 0.80833 | COADREAD | COLO-320-HSR | TGX221 | MVP_top60_DNA+RNA_seed | HSP90AA1 | 0.627388 | COADREAD | CCK-81 | Luminespib |
| MVP_top60_DNA_seed | SRC | 0.808896 | COADREAD | HT-115 | NVP-BHG712 | MVP_top60_DNA+RNA_seed | HSP90AB1 | 0.627388 | COADREAD | CCK-81 | Luminespib |
| MVP_top60_DNA_seed | CDK2 | 0.80973 | COADREAD | SW948 | AZD5438 | MVP_top60_DNA+RNA_seed | PIK3CD | 0.632522 | COADREAD | C2BBe1 | IC-87114 |
| MVP_top60_DNA_seed | AKT1 | 0.809879 | COADREAD | SW48 | GSK690693 | MVP_top60_DNA+RNA_seed | SRC | 0.633231 | COADREAD | GP5d | HG-6-71-01 |
| MVP_top60_DNA_seed | ATM | 0.810344 | COADREAD | LS-123 | KU-60019 | MVP_top60_DNA+RNA_seed | AKT1 | 0.637815 | COADREAD | HT55 | MK-2206 |
| MVP_top60_DNA_seed | ATM | 0.811286 | COADREAD | LS-411N | KU-55933 | MVP_top60_DNA+RNA_seed | GSK3B | 0.639382 | COADREAD | SNU-C2B | KIN001-042 |
| MVP_top60_DNA_seed | BRAF | 0.811666 | COADREAD | SW948 | PLX-4720 | MVP_top60_DNA+RNA_seed | CDK2 | 0.640485 | COADREAD | CW-2 | AZD5438 |
| MVP_top60_DNA_seed | BRAF | 0.813107 | COADREAD | LS-1034 | PLX-4720 | MVP_top60_DNA+RNA_seed | IGF1R | 0.641073 | COADREAD | HT-115 | GSK1904529A |
| MVP_top60_DNA_seed | BRAF | 0.813415 | COADREAD | SW620 | SB590885 | MVP_top60_DNA+RNA_seed | FYN | 0.641237 | COADREAD | KM12 | A-770041 |
| MVP_top60_DNA_seed | AKT1 | 0.813653 | COADREAD | LS-123 | GSK690693 | MVP_top60_DNA+RNA_seed | SRC | 0.644135 | COADREAD | HT-115 | Bosutinib |
| MVP_top60_DNA_seed | AR | 0.815172 | COADREAD | HCC2998 | AZD3514 | MVP_top60_DNA+RNA_seed | HSP90AA1 | 0.645617 | COADREAD | SW1417 | Tanespimycin |
| MVP_top60_DNA_seed | ERBB4 | 0.815839 | COADREAD | CaR-1 | PF-00299804 | MVP_top60_DNA+RNA_seed | HSP90AB1 | 0.645617 | COADREAD | SW1417 | Tanespimycin |
| MVP_top60_DNA_seed | MET | 0.81775 | COADREAD | LS-1034 | JNJ38877605 | MVP_top60_DNA+RNA_seed | AR | 0.645724 | COADREAD | LS-123 | AZD3514 |
| MVP_top60_DNA_seed | GSK3B | 0.818765 | COADREAD | LS-180 | CHIR-99021 | MVP_top60_DNA+RNA_seed | PIK3CD | 0.647094 | COADREAD | NCI-H716 | IC-87114 |
| MVP_top60_DNA_seed | ESR1 | 0.819673 | COADREAD | LS-513 | Tamoxifen | MVP_top60_DNA+RNA_seed | AKT1 | 0.648404 | COADREAD | LS-180 | GSK690693 |
| MVP_top60_DNA_seed | AKT1 | 0.820179 | COADREAD | LoVo | AKT inhibitor VIII | MVP_top60_DNA+RNA_seed | ATM | 0.652341 | COADREAD | SNU-C2B | KU-60019 |
| MVP_top60_DNA_seed | SRC | 0.82113 | COADREAD | SNU-C2B | HG-6-71-01 | MVP_top60_DNA+RNA_seed | CDK2 | 0.653351 | COADREAD | RCM-1 | AZD5438 |
| MVP_top60_DNA_seed | HSP90AA1 | 0.821289 | COADREAD | LS-1034 | Tanespimycin | MVP_top60_DNA+RNA_seed | ATM | 0.654055 | COADREAD | SNU-407 | KU-55933 |
| MVP_top60_DNA_seed | HSP90AB1 | 0.821289 | COADREAD | LS-1034 | Tanespimycin | MVP_top60_DNA+RNA_seed | PIK3CB | 0.656293 | COADREAD | GP5d | AZD8186 |
| MVP_top60_DNA_seed | MAPK1 | 0.821359 | COADREAD | HT55 | FR-180204 | MVP_top60_DNA+RNA_seed | PIK3CD | 0.656293 | COADREAD | GP5d | AZD8186 |
| MVP_top60_DNA_seed | PIK3CD | 0.823391 | COADREAD | SW837 | Idelalisib | MVP_top60_DNA+RNA_seed | PIK3CD | 0.657334 | COADREAD | SW1417 | IC-87114 |
| MVP_top60_DNA_seed | AKT1 | 0.824216 | COADREAD | COLO-320-HSR | AKT inhibitor VIII | MVP_top60_DNA+RNA_seed | ERBB4 | 0.657562 | COADREAD | NCI-H716 | AST-1306 |
| MVP_top60_DNA_seed | ERBB4 | 0.82472 | COADREAD | CL-11 | PF-00299804 | MVP_top60_DNA+RNA_seed | HSP90AA1 | 0.658876 | COADREAD | NCI-H716 | Elesclomol |
| MVP_top60_DNA_seed | ESR1 | 0.825257 | COADREAD | NCI-H508 | Fulvestrant | MVP_top60_DNA+RNA_seed | HSP90AB1 | 0.658876 | COADREAD | NCI-H716 | Elesclomol |
| MVP_top60_DNA_seed | SRC | 0.827859 | COADREAD | COLO-320-HSR | Bosutinib | MVP_top60_DNA+RNA_seed | MET | 0.659345 | COADREAD | COLO-320-HSR | SU11274 |
| MVP_top60_DNA_seed | MET | 0.828397 | COADREAD | CW-2 | AZD6094 | MVP_top60_DNA+RNA_seed | PTK2 | 0.660953 | COADREAD | C2BBe1 | PF-562271 |
| MVP_top60_DNA_seed | PIK3CB | 0.828743 | COADREAD | DiFi | AZD8186 | MVP_top60_DNA+RNA_seed | ERBB3 | 0.662739 | COADREAD | NCI-H716 | AZD8931 |
| MVP_top60_DNA_seed | PIK3CD | 0.828743 | COADREAD | DiFi | AZD8186 | MVP_top60_DNA+RNA_seed | CDK2 | 0.668004 | COADREAD | HCT-15 | AZD5438 |
| MVP_top60_DNA_seed | GSK3B | 0.83156 | COADREAD | COLO-320-HSR | CHIR-99021 | MVP_top60_DNA+RNA_seed | MET | 0.668154 | COADREAD | LS-513 | PHA-665752 |
| MVP_top60_DNA_seed | BRAF | 0.832385 | COADREAD | HT55 | SB590885 | MVP_top60_DNA+RNA_seed | ERBB4 | 0.668666 | COADREAD | LS-411N | AST-1306 |
| MVP_top60_DNA_seed | GSK3B | 0.832621 | COADREAD | LS-1034 | CHIR-99021 | MVP_top60_DNA+RNA_seed | PIK3CD | 0.670156 | COADREAD | HT-29 | IC-87114 |
| MVP_top60_DNA_seed | BRAF | 0.836697 | COADREAD | CL-11 | SB590885 | MVP_top60_DNA+RNA_seed | GSK3B | 0.672734 | COADREAD | SW837 | CHIR-99021 |
| MVP_top60_DNA_seed | MET | 0.837364 | COADREAD | C2BBe1 | Crizotinib | MVP_top60_DNA+RNA_seed | MET | 0.672841 | COADREAD | LS-411N | SU11274 |
| MVP_top60_DNA_seed | GSK3B | 0.838729 | COADREAD | RCM-1 | AZD7969 | MVP_top60_DNA+RNA_seed | SRC | 0.676077 | COADREAD | KM12 | Bosutinib |
| MVP_top60_DNA_seed | GSK3B | 0.840255 | COADREAD | NCI-H508 | AZD7969 | MVP_top60_DNA+RNA_seed | ATM | 0.677636 | COADREAD | SW1463 | KU-55933 |
| MVP_top60_DNA_seed | ATM | 0.840684 | COADREAD | SNU-C5 | KU-60019 | MVP_top60_DNA+RNA_seed | GSK3B | 0.679434 | COADREAD | NCI-H716 | AZD7969 |
| MVP_top60_DNA_seed | ATM | 0.84323 | COADREAD | CCK-81 | KU-55933 | MVP_top60_DNA+RNA_seed | GSK3B | 0.68163 | COADREAD | CW-2 | CHIR-99021 |

| | | | | | | | | | | | |
|---|---|---|---|---|---|---|---|---|---|---|---|
| MVP_top60_DNA_seed | GSK3B | 0.844888 | COADREAD | SW620 | CHIR-99021 | MVP_top60_DNA+RNA_seed | PIK3CB | 0.681764 | COADREAD | SW1417 | AZD8186 |
| MVP_top60_DNA_seed | PIK3CB | 0.845218 | COADREAD | NCI-H508 | AZD8186 | MVP_top60_DNA+RNA_seed | PIK3CD | 0.681764 | COADREAD | SW1417 | AZD8186 |
| MVP_top60_DNA_seed | PIK3CD | 0.845218 | COADREAD | NCI-H508 | AZD8186 | MVP_top60_DNA+RNA_seed | BCL2 | 0.681863 | COADREAD | SNU-407 | Navitoclax |
| MVP_top60_DNA_seed | AKT1 | 0.845542 | COADREAD | LS-123 | AKT inhibitor VIII | MVP_top60_DNA+RNA_seed | HSP90AA1 | 0.682942 | COADREAD | SNU-407 | CCT-018159 |
| MVP_top60_DNA_seed | PIK3CB | 0.847949 | COADREAD | T84 | AZD8186 | MVP_top60_DNA+RNA_seed | HSP90AB1 | 0.682942 | COADREAD | SNU-407 | CCT-018159 |
| MVP_top60_DNA_seed | PIK3CD | 0.847949 | COADREAD | T84 | AZD8186 | MVP_top60_DNA+RNA_seed | AR | 0.684221 | COADREAD | SW48 | Bicalutamide |
| MVP_top60_DNA_seed | BRAF | 0.848974 | COADREAD | SNU-C2B | Dabrafenib | MVP_top60_DNA+RNA_seed | MET | 0.686548 | COADREAD | T84 | AZD6094 |
| MVP_top60_DNA_seed | ERBB4 | 0.850863 | COADREAD | HCT-116 | CI-1033 | MVP_top60_DNA+RNA_seed | PIK3CA | 0.686876 | COADREAD | HCC2998 | AZD8835 |
| MVP_top60_DNA_seed | AKT1 | 0.851556 | COADREAD | KM12 | MK-2206 | MVP_top60_DNA+RNA_seed | PIK3CD | 0.686876 | COADREAD | HCC2998 | AZD8835 |
| MVP_top60_DNA_seed | BRAF | 0.851612 | COADREAD | LS-1034 | SB590885 | MVP_top60_DNA+RNA_seed | SRC | 0.687497 | COADREAD | LS-180 | NVP-BHG712 |
| MVP_top60_DNA_seed | ERBB4 | 0.852693 | COADREAD | COLO-678 | PF-00299804 | MVP_top60_DNA+RNA_seed | MET | 0.687647 | COADREAD | MDST8 | Crizotinib |
| MVP_top60_DNA_seed | AR | 0.85275 | COADREAD | SNU-C2B | Bicalutamide | MVP_top60_DNA+RNA_seed | GSK3B | 0.69064 | COADREAD | COLO-678 | CHIR-99021 |
| MVP_top60_DNA_seed | BRAF | 0.852871 | COADREAD | LS-123 | HG6-64-1 | MVP_top60_DNA+RNA_seed | CDK2 | 0.691077 | COADREAD | DiFi | AZD5438 |
| MVP_top60_DNA_seed | ATM | 0.853584 | COADREAD | C2BBe1 | CP466722 | MVP_top60_DNA+RNA_seed | ERBB3 | 0.693676 | COADREAD | COLO-678 | AZD8931 |
| MVP_top60_DNA_seed | MET | 0.855049 | COADREAD | C2BBe1 | AZD6094 | MVP_top60_DNA+RNA_seed | HSP90AA1 | 0.694137 | COADREAD | COLO-678 | Luminespib |
| MVP_top60_DNA_seed | ERBB4 | 0.857216 | COADREAD | HT55 | CI-1033 | MVP_top60_DNA+RNA_seed | HSP90AB1 | 0.694137 | COADREAD | COLO-678 | Luminespib |
| MVP_top60_DNA_seed | BRAF | 0.85741 | COADREAD | CCK-81 | Dabrafenib | MVP_top60_DNA+RNA_seed | GSK3B | 0.694624 | COADREAD | SW837 | CHIR-99021 |
| MVP_top60_DNA_seed | MET | 0.858423 | COADREAD | DiFi | AZD6094 | MVP_top60_DNA+RNA_seed | GSK3B | 0.698274 | COADREAD | HCC2998 | CHIR-99021 |
| MVP_top60_DNA_seed | CREBBP | 0.860626 | COADREAD | SNU-1040 | I-CBP112 | MVP_top60_DNA+RNA_seed | MET | 0.700497 | COADREAD | SNU-407 | AZD6094 |
| MVP_top60_DNA_seed | MET | 0.8609 | COADREAD | SNU-407 | JNJ38877605 | MVP_top60_DNA+RNA_seed | AKT1 | 0.703024 | COADREAD | HT-29 | MK-2206 |
| MVP_top60_DNA_seed | PIK3CB | 0.86119 | COADREAD | GP5d | AZD6482 | MVP_top60_DNA+RNA_seed | HSP90AA1 | 0.703561 | COADREAD | CW-2 | Elesclomol |
| MVP_top60_DNA_seed | AKT1 | 0.861273 | COADREAD | KM12 | AKT inhibitor VIII | MVP_top60_DNA+RNA_seed | HSP90AB1 | 0.703561 | COADREAD | CW-2 | Elesclomol |
| MVP_top60_DNA_seed | HSP90AA1 | 0.861536 | COADREAD | RCM-1 | Luminespib | MVP_top60_DNA+RNA_seed | ATM | 0.705629 | COADREAD | SW1417 | CP466722 |
| MVP_top60_DNA_seed | HSP90AB1 | 0.861536 | COADREAD | RCM-1 | Luminespib | MVP_top60_DNA+RNA_seed | AR | 0.706151 | COADREAD | HT55 | AZD3514 |
| MVP_top60_DNA_seed | BRAF | 0.864044 | COADREAD | NCI-H508 | PLX-4720 | MVP_top60_DNA+RNA_seed | IGF1R | 0.707248 | COADREAD | MDST8 | BMS-754807 |
| MVP_top60_DNA_seed | SRC | 0.864705 | COADREAD | KM12 | WH-4-023 | MVP_top60_DNA+RNA_seed | PIK3CB | 0.707509 | COADREAD | CaR-1 | AZD8186 |
| MVP_top60_DNA_seed | AR | 0.866284 | COADREAD | COLO-320-HSR | AZD3514 | MVP_top60_DNA+RNA_seed | PIK3CD | 0.707509 | COADREAD | CaR-1 | AZD8186 |
| MVP_top60_DNA_seed | GSK3B | 0.869442 | COADREAD | COLO-205 | AZD7969 | MVP_top60_DNA+RNA_seed | ERBB4 | 0.707822 | COADREAD | KM12 | PF-00299804 |
| MVP_top60_DNA_seed | ESR1 | 0.869452 | COADREAD | COLO-320-HSR | Fulvestrant | MVP_top60_DNA+RNA_seed | GSK3B | 0.710742 | COADREAD | LS-123 | KIN001-042 |
| MVP_top60_DNA_seed | IGF1R | 0.869922 | COADREAD | CL-11 | BMS-536924 | MVP_top60_DNA+RNA_seed | BCL2 | 0.71139 | COADREAD | RKO | Navitoclax |
| MVP_top60_DNA_seed | PIK3CD | 0.87006 | COADREAD | COLO-205 | Idelalisib | MVP_top60_DNA+RNA_seed | GSK3B | 0.712087 | COADREAD | LS-411N | KIN001-042 |
| MVP_top60_DNA_seed | BRAF | 0.870694 | COADREAD | SNU-1040 | PLX-4720 | MVP_top60_DNA+RNA_seed | MET | 0.712138 | COADREAD | SNU-61 | AZD6094 |
| MVP_top60_DNA_seed | IGF1R | 0.870781 | COADREAD | SW1463 | Linsitinib | MVP_top60_DNA+RNA_seed | PIK3CB | 0.712668 | COADREAD | HT-115 | AZD6482 |
| MVP_top60_DNA_seed | ATM | 0.871659 | COADREAD | LS-1034 | KU-55933 | MVP_top60_DNA+RNA_seed | GSK3B | 0.71276 | COADREAD | COLO-320-HSR | CHIR-99021 |
| MVP_top60_DNA_seed | SRC | 0.874426 | COADREAD | CW-2 | NVP-BHG712 | MVP_top60_DNA+RNA_seed | AKT1 | 0.71285 | COADREAD | C2BBe1 | GSK690693 |
| MVP_top60_DNA_seed | SRC | 0.875488 | COADREAD | RCM-1 | HG-6-71-01 | MVP_top60_DNA+RNA_seed | MET | 0.713405 | COADREAD | LS-123 | JNJ38877605 |
| MVP_top60_DNA_seed | MET | 0.876386 | COADREAD | HT-29 | SU11274 | MVP_top60_DNA+RNA_seed | BCL2 | 0.713459 | COADREAD | SNU-C5 | Navitoclax |
| MVP_top60_DNA_seed | HSP90AA1 | 0.876529 | COADREAD | CaR-1 | Luminespib | MVP_top60_DNA+RNA_seed | ATM | 0.713882 | COADREAD | CaR-1 | KU-60019 |
| MVP_top60_DNA_seed | HSP90AB1 | 0.876529 | COADREAD | CaR-1 | Luminespib | MVP_top60_DNA+RNA_seed | CDK2 | 0.714718 | COADREAD | SNU-C1 | AZD5438 |
| MVP_top60_DNA_seed | SRC | 0.877186 | COADREAD | COLO-678 | NVP-BHG712 | MVP_top60_DNA+RNA_seed | HSP90AA1 | 0.716149 | COADREAD | CCK-81 | Tanespimycin |
| MVP_top60_DNA_seed | HSP90AA1 | 0.878829 | COADREAD | LS-180 | CCT-018159 | MVP_top60_DNA+RNA_seed | HSP90AB1 | 0.716149 | COADREAD | CCK-81 | Tanespimycin |

| | | | | | | | | | | | |
|---|---|---|---|---|---|---|---|---|---|---|---|
| MVP_top60_DNA_seed | HSP90AB1 | 0.878829 | COADREAD | LS-180 | CCT-018159 | MVP_top60_DNA+RNA_seed | PTK2 | 0.716335 | COADREAD | LS-513 | PF-562271 |
| MVP_top60_DNA_seed | IGF1R | 0.879153 | COADREAD | LoVo | Linsitinib | MVP_top60_DNA+RNA_seed | BCL2 | 0.717018 | COADREAD | T84 | Navitoclax |
| MVP_top60_DNA_seed | MAPK1 | 0.881706 | COADREAD | HT55 | VX-11e | MVP_top60_DNA+RNA_seed | HSP90AA1 | 0.717116 | COADREAD | SNU-1040 | SNX-2112 |
| MVP_top60_DNA_seed | IGF1R | 0.881867 | COADREAD | SW948 | BMS-536924 | MVP_top60_DNA+RNA_seed | HSP90AB1 | 0.717116 | COADREAD | SNU-1040 | SNX-2112 |
| MVP_top60_DNA_seed | ERBB4 | 0.882399 | COADREAD | SW837 | AST-1306 | MVP_top60_DNA+RNA_seed | ERBB3 | 0.717602 | COADREAD | CL-11 | AZD8931 |
| MVP_top60_DNA_seed | BRAF | 0.882413 | COADREAD | HT-115 | HG6-64-1 | MVP_top60_DNA+RNA_seed | GSK3B | 0.720694 | COADREAD | COLO-205 | CHIR-99021 |
| MVP_top60_DNA_seed | MET | 0.882944 | COADREAD | NCI-H716 | Crizotinib | MVP_top60_DNA+RNA_seed | PIK3CB | 0.723515 | COADREAD | LS-411N | AZD6482 |
| MVP_top60_DNA_seed | AKT1 | 0.8857 | COADREAD | T84 | AKT inhibitor VIII | MVP_top60_DNA+RNA_seed | AKT1 | 0.723742 | COADREAD | COLO-320-HSR | GSK690693 |
| MVP_top60_DNA_seed | IGF1R | 0.885714 | COADREAD | HT55 | Linsitinib | MVP_top60_DNA+RNA_seed | AR | 0.724423 | COADREAD | LS-513 | Bicalutamide |
| MVP_top60_DNA_seed | BRAF | 0.886147 | COADREAD | SNU-81 | PLX-4720 | MVP_top60_DNA+RNA_seed | MET | 0.72468 | COADREAD | LS-411N | AZD6094 |
| MVP_top60_DNA_seed | MAPK14 | 0.887764 | COADREAD | HT55 | TAK-715 | MVP_top60_DNA+RNA_seed | PTK2 | 0.72578 | COADREAD | SNU-C2B | PF-562271 |
| MVP_top60_DNA_seed | BRAF | 0.889052 | COADREAD | HCT-15 | RAF_9304 | MVP_top60_DNA+RNA_seed | PIK3CD | 0.727068 | COADREAD | HCT-15 | IC-87114 |
| MVP_top60_DNA_seed | HSP90AA1 | 0.889417 | COADREAD | LoVo | Tanespimycin | MVP_top60_DNA+RNA_seed | HSP90AA1 | 0.728095 | COADREAD | SNU-61 | Tanespimycin |
| MVP_top60_DNA_seed | HSP90AB1 | 0.889417 | COADREAD | LoVo | Tanespimycin | MVP_top60_DNA+RNA_seed | HSP90AB1 | 0.728095 | COADREAD | SNU-61 | Tanespimycin |
| MVP_top60_DNA_seed | MAPK1 | 0.891646 | COADREAD | LoVo | FR-180204 | MVP_top60_DNA+RNA_seed | AKT1 | 0.728839 | COADREAD | SNU-61 | GSK690693 |
| MVP_top60_DNA_seed | PIK3CB | 0.892551 | COADREAD | LS-513 | TGX221 | MVP_top60_DNA+RNA_seed | AKT1 | 0.729032 | COADREAD | SNU-C2B | MK-2206 |
| MVP_top60_DNA_seed | GSK3B | 0.892758 | COADREAD | HCT-15 | SB216763 | MVP_top60_DNA+RNA_seed | SRC | 0.730608 | COADREAD | LS-513 | Saracatinib |
| MVP_top60_DNA_seed | PIK3CB | 0.893277 | COADREAD | HT-29 | AZD8186 | MVP_top60_DNA+RNA_seed | MET | 0.73098 | COADREAD | LS-180 | SU11274 |
| MVP_top60_DNA_seed | PIK3CD | 0.893277 | COADREAD | HT-29 | AZD8186 | MVP_top60_DNA+RNA_seed | MET | 0.731449 | COADREAD | HT55 | AZD6094 |
| MVP_top60_DNA_seed | AR | 0.893463 | COADREAD | SW1417 | AZD3514 | MVP_top60_DNA+RNA_seed | ERBB3 | 0.732919 | COADREAD | LS-1034 | AZD8931 |
| MVP_top60_DNA_seed | AKT1 | 0.893927 | COADREAD | LS-411N | MK-2206 | MVP_top60_DNA+RNA_seed | PIK3CD | 0.733169 | COADREAD | SNU-C5 | IC-87114 |
| MVP_top60_DNA_seed | CDK2 | 0.897315 | COADREAD | HT55 | AZD5438 | MVP_top60_DNA+RNA_seed | GSK3B | 0.733918 | COADREAD | SW1463 | AZD7969 |
| MVP_top60_DNA_seed | SRC | 0.897502 | COADREAD | LS-513 | Bosutinib | MVP_top60_DNA+RNA_seed | ERBB4 | 0.735259 | COADREAD | HT-29 | CI-1033 |
| MVP_top60_DNA_seed | AKT1 | 0.898225 | COADREAD | C2BBe1 | AKT inhibitor VIII | MVP_top60_DNA+RNA_seed | HSP90AA1 | 0.735349 | COADREAD | SNU-C1 | Tanespimycin |
| MVP_top60_DNA_seed | PIK3CA | 0.898392 | COADREAD | SW620 | AZD8835 | MVP_top60_DNA+RNA_seed | HSP90AB1 | 0.735349 | COADREAD | SNU-C1 | Tanespimycin |
| MVP_top60_DNA_seed | PIK3CD | 0.898392 | COADREAD | SW620 | AZD8835 | MVP_top60_DNA+RNA_seed | PIK3CA | 0.737672 | COADREAD | CW-2 | AZD8835 |
| MVP_top60_DNA_seed | ERBB4 | 0.89957 | COADREAD | HCC2998 | PF-00299804 | MVP_top60_DNA+RNA_seed | PIK3CD | 0.737672 | COADREAD | CW-2 | AZD8835 |
| MVP_top60_DNA_seed | GSK3B | 0.900713 | COADREAD | T84 | CHIR-99021 | MVP_top60_DNA+RNA_seed | ERBB4 | 0.740807 | COADREAD | SNU-407 | AST-1306 |
| MVP_top60_DNA_seed | AKT1 | 0.900715 | COADREAD | COLO-678 | AKT inhibitor VIII | MVP_top60_DNA+RNA_seed | MET | 0.741162 | COADREAD | SW620 | SU11274 |
| MVP_top60_DNA_seed | PIK3CD | 0.904648 | COADREAD | LS-1034 | IC-87114 | MVP_top60_DNA+RNA_seed | ERBB4 | 0.741743 | COADREAD | LS-123 | AST-1306 |
| MVP_top60_DNA_seed | SRC | 0.906083 | COADREAD | MDST8 | WH-4-023 | MVP_top60_DNA+RNA_seed | ATM | 0.742282 | COADREAD | HCC2998 | KU-60019 |
| MVP_top60_DNA_seed | ATM | 0.90616 | COADREAD | KM12 | KU-55933 | MVP_top60_DNA+RNA_seed | AR | 0.742994 | COADREAD | SW1417 | Bicalutamide |
| MVP_top60_DNA_seed | BRAF | 0.906177 | COADREAD | SW48 | PLX-4720 | MVP_top60_DNA+RNA_seed | HSP90AA1 | 0.743353 | COADREAD | RCM-1 | SNX-2112 |
| MVP_top60_DNA_seed | GSK3B | 0.90619 | COADREAD | HT55 | CHIR-99021 | MVP_top60_DNA+RNA_seed | HSP90AB1 | 0.743353 | COADREAD | RCM-1 | SNX-2112 |
| MVP_top60_DNA_seed | GSK3B | 0.910105 | COADREAD | SW837 | SB216763 | MVP_top60_DNA+RNA_seed | SRC | 0.744864 | COADREAD | SW837 | NVP-BHG712 |
| MVP_top60_DNA_seed | CREBBP | 0.911158 | COADREAD | SW1417 | I-CBP112 | MVP_top60_DNA+RNA_seed | PIK3CA | 0.746074 | COADREAD | RCM-1 | AZD8835 |
| MVP_top60_DNA_seed | SRC | 0.912493 | COADREAD | RCM-1 | NVP-BHG712 | MVP_top60_DNA+RNA_seed | PIK3CD | 0.746074 | COADREAD | RCM-1 | AZD8835 |
| MVP_top60_DNA_seed | BRAF | 0.914824 | COADREAD | C2BBe1 | PLX-4720 | MVP_top60_DNA+RNA_seed | PIK3CB | 0.748571 | COADREAD | HT55 | AZD8186 |
| MVP_top60_DNA_seed | GSK3B | 0.915056 | COADREAD | COLO-678 | SB216763 | MVP_top60_DNA+RNA_seed | PIK3CD | 0.748571 | COADREAD | HT55 | AZD8186 |
| MVP_top60_DNA_seed | IGF1R | 0.915548 | COADREAD | SW1463 | BMS-754807 | MVP_top60_DNA+RNA_seed | IGF1R | 0.749268 | COADREAD | T84 | GSK1904529A |
| MVP_top60_DNA_seed | FYN | 0.915818 | COADREAD | MDST8 | A-770041 | MVP_top60_DNA+RNA_seed | PTK2 | 0.751637 | COADREAD | LS-1034 | PF-562271 |

| | | | | | | | | | | | |
|---|---|---|---|---|---|---|---|---|---|---|---|
| MVP_top60_DNA_seed | AR | 0.916959 | COADREAD | CW-2 | AZD3514 | MVP_top60_DNA+RNA_seed | ERBB4 | 0.753289 | COADREAD | RCM-1 | AST-1306 |
| MVP_top60_DNA_seed | GSK3B | 0.917032 | COADREAD | T84 | CHIR-99021 | MVP_top60_DNA+RNA_seed | AKT1 | 0.75329 | COADREAD | T84 | BAY AKT1 |
| MVP_top60_DNA_seed | PIK3CA | 0.918071 | COADREAD | SW1116 | PI3Ka_4409 | MVP_top60_DNA+RNA_seed | SRC | 0.754352 | COADREAD | MDST8 | Saracatinib |
| MVP_top60_DNA_seed | PIK3CD | 0.918071 | COADREAD | SW1116 | PI3Ka_4409 | MVP_top60_DNA+RNA_seed | PTK2 | 0.755765 | COADREAD | CCK-81 | PF-562271 |
| MVP_top60_DNA_seed | ESR1 | 0.9185 | COADREAD | T84 | Tamoxifen | MVP_top60_DNA+RNA_seed | IGF1R | 0.756383 | COADREAD | LS-180 | BMS-536924 |
| MVP_top60_DNA_seed | BRAF | 0.919444 | COADREAD | SW1463 | Dabrafenib | MVP_top60_DNA+RNA_seed | ERBB4 | 0.758946 | COADREAD | SNU-C5 | AST-1306 |
| MVP_top60_DNA_seed | GSK3B | 0.919686 | COADREAD | SNU-407 | KIN001-042 | MVP_top60_DNA+RNA_seed | CDK2 | 0.759818 | COADREAD | CW-2 | Seliciclib |
| MVP_top60_DNA_seed | ESR1 | 0.925442 | COADREAD | SW837 | Fulvestrant | MVP_top60_DNA+RNA_seed | BCL2 | 0.76034 | COADREAD | HT-115 | TW 37 |
| MVP_top60_DNA_seed | GSK3B | 0.926944 | COADREAD | SW948 | KIN001-042 | MVP_top60_DNA+RNA_seed | AR | 0.760463 | COADREAD | NCI-H716 | Bicalutamide |
| MVP_top60_DNA_seed | PIK3CD | 0.927021 | COADREAD | SNU-C2B | Idelalisib | MVP_top60_DNA+RNA_seed | GSK3B | 0.762139 | COADREAD | LS-1034 | CHIR-99021 |
| MVP_top60_DNA_seed | MET | 0.927952 | COADREAD | LS-1034 | PHA-665752 | MVP_top60_DNA+RNA_seed | AKT1 | 0.764144 | COADREAD | SW1463 | BAY AKT1 |
| MVP_top60_DNA_seed | PTK2 | 0.927991 | COADREAD | SW948 | PF-562271 | MVP_top60_DNA+RNA_seed | PIK3CD | 0.764971 | COADREAD | KM12 | IC-87114 |
| MVP_top60_DNA_seed | MET | 0.932731 | COADREAD | CL-11 | Crizotinib | MVP_top60_DNA+RNA_seed | PTK2 | 0.765013 | COADREAD | MDST8 | PF-562271 |
| MVP_top60_DNA_seed | SRC | 0.933456 | COADREAD | LS-411N | Bosutinib | MVP_top60_DNA+RNA_seed | CDK2 | 0.766794 | COADREAD | CCK-81 | PHA-793887 |
| MVP_top60_DNA_seed | HSP90AA1 | 0.934681 | COADREAD | NCI-H747 | CCT-018159 | MVP_top60_DNA+RNA_seed | SRC | 0.767828 | COADREAD | HT-29 | HG-6-71-01 |
| MVP_top60_DNA_seed | HSP90AB1 | 0.934681 | COADREAD | NCI-H747 | CCT-018159 | MVP_top60_DNA+RNA_seed | PIK3CD | 0.767996 | COADREAD | T84 | IC-87114 |
| MVP_top60_DNA_seed | SRC | 0.936914 | COADREAD | HCT-15 | HG-6-71-01 | MVP_top60_DNA+RNA_seed | AKT1 | 0.768115 | COADREAD | HCC2998 | AKT inhibitor VIII |
| MVP_top60_DNA_seed | PTK2 | 0.93716 | COADREAD | COLO-678 | PF-562271 | MVP_top60_DNA+RNA_seed | AKT1 | 0.770802 | COADREAD | SW1417 | AKT inhibitor VIII |
| MVP_top60_DNA_seed | ESR1 | 0.938653 | COADREAD | SNU-407 | Tamoxifen | MVP_top60_DNA+RNA_seed | MET | 0.77132 | COADREAD | LS-123 | PHA-665752 |
| MVP_top60_DNA_seed | PIK3CB | 0.938659 | COADREAD | COLO-678 | AZD6482 | MVP_top60_DNA+RNA_seed | ERBB4 | 0.771923 | COADREAD | COLO-678 | AST-1306 |
| MVP_top60_DNA_seed | CREBBP | 0.942339 | COADREAD | CCK-81 | I-CBP112 | MVP_top60_DNA+RNA_seed | ATM | 0.772567 | COADREAD | GP5d | KU-55933 |
| MVP_top60_DNA_seed | ESR1 | 0.944617 | COADREAD | SNU-1040 | Tamoxifen | MVP_top60_DNA+RNA_seed | ERBB4 | 0.772891 | COADREAD | LS-513 | AST-1306 |
| MVP_top60_DNA_seed | BRAF | 0.946948 | COADREAD | LS-1034 | Dabrafenib | MVP_top60_DNA+RNA_seed | ERBB4 | 0.775616 | COADREAD | SNU-407 | PF-00299804 |
| MVP_top60_DNA_seed | MET | 0.948402 | COADREAD | LS-180 | AZD6094 | MVP_top60_DNA+RNA_seed | ATM | 0.776763 | COADREAD | HCC2998 | KU-55933 |
| MVP_top60_DNA_seed | MET | 0.95032 | COADREAD | MDST8 | SU11274 | MVP_top60_DNA+RNA_seed | AKT1 | 0.776865 | COADREAD | HT-115 | GSK690693 |
| MVP_top60_DNA_seed | HSP90AA1 | 0.950743 | COADREAD | COLO-678 | SNX-2112 | MVP_top60_DNA+RNA_seed | AR | 0.776955 | COADREAD | RCM-1 | AZD3514 |
| MVP_top60_DNA_seed | HSP90AB1 | 0.950743 | COADREAD | COLO-678 | SNX-2112 | MVP_top60_DNA+RNA_seed | HSP90AA1 | 0.779328 | COADREAD | SNU-407 | Tanespimycin |
| MVP_top60_DNA_seed | AKT1 | 0.951401 | COADREAD | HT55 | GSK690693 | MVP_top60_DNA+RNA_seed | HSP90AB1 | 0.779328 | COADREAD | SNU-407 | Tanespimycin |
| MVP_top60_DNA_seed | GSK3B | 0.952383 | COADREAD | SW1463 | CHIR-99021 | MVP_top60_DNA+RNA_seed | HSP90AA1 | 0.780158 | COADREAD | C2BBe1 | CCT-018159 |
| MVP_top60_DNA_seed | AR | 0.952588 | COADREAD | LS-513 | AZD3514 | MVP_top60_DNA+RNA_seed | HSP90AB1 | 0.780158 | COADREAD | C2BBe1 | CCT-018159 |
| MVP_top60_DNA_seed | CDK2 | 0.953799 | COADREAD | SW837 | AZD5438 | MVP_top60_DNA+RNA_seed | PIK3CD | 0.781895 | COADREAD | CaR-1 | Idelalisib |
| MVP_top60_DNA_seed | ERBB4 | 0.95393 | COADREAD | LS-411N | CI-1033 | MVP_top60_DNA+RNA_seed | ATM | 0.782786 | COADREAD | LoVo | CP466722 |
| MVP_top60_DNA_seed | AR | 0.955799 | COADREAD | KM12 | Bicalutamide | MVP_top60_DNA+RNA_seed | FYN | 0.784415 | COADREAD | LS-513 | A-770041 |
| MVP_top60_DNA_seed | AKT1 | 0.956203 | COADREAD | SW1417 | GSK690693 | MVP_top60_DNA+RNA_seed | PIK3CD | 0.785688 | COADREAD | HCC2998 | Idelalisib |
| MVP_top60_DNA_seed | ERBB4 | 0.956951 | COADREAD | SNU-61 | PF-00299804 | MVP_top60_DNA+RNA_seed | IGF1R | 0.787897 | COADREAD | HCT-15 | BMS-754807 |
| MVP_top60_DNA_seed | MET | 0.957992 | COADREAD | CL-11 | AZD6094 | MVP_top60_DNA+RNA_seed | MET | 0.789716 | COADREAD | HCT-116 | JNJ38877605 |
| MVP_top60_DNA_seed | MET | 0.963094 | COADREAD | SNU-C5 | JNJ38877605 | MVP_top60_DNA+RNA_seed | CDK2 | 0.790933 | COADREAD | LS-123 | Seliciclib |
| MVP_top60_DNA_seed | BRAF | 0.965513 | COADREAD | CCK-81 | PLX-4720 | MVP_top60_DNA+RNA_seed | MET | 0.791202 | COADREAD | NCI-H508 | AZD6094 |
| MVP_top60_DNA_seed | ESR1 | 0.965655 | COADREAD | SNU-C5 | Tamoxifen | MVP_top60_DNA+RNA_seed | GSK3B | 0.791862 | COADREAD | SW1417 | CHIR-99021 |
| MVP_top60_DNA_seed | HSP90AA1 | 0.965896 | COADREAD | COLO-320-HSR | Elesclomol | MVP_top60_DNA+RNA_seed | BCL2 | 0.794308 | COADREAD | SNU-C5 | TW 37 |
| MVP_top60_DNA_seed | HSP90AB1 | 0.965896 | COADREAD | COLO-320-HSR | Elesclomol | MVP_top60_DNA+RNA_seed | IGF1R | 0.795454 | COADREAD | HCT-15 | BMS-536924 |

| | | | | | | | | | | | |
|---|---|---|---|---|---|---|---|---|---|---|---|
| MVP_top60_DNA_seed | BRAF | 0.966486 | COADREAD | CaR-1 | HG6-64-1 | MVP_top60_DNA+RNA_seed | ATM | 0.796034 | COADREAD | SNU-C2B | CP466722 |
| MVP_top60_DNA_seed | BRAF | 0.969409 | COADREAD | DiFi | PLX-4720 | MVP_top60_DNA+RNA_seed | PIK3CB | 0.796793 | COADREAD | HCC2998 | AZD8186 |
| MVP_top60_DNA_seed | MET | 0.970831 | COADREAD | RCM-1 | AZD6094 | MVP_top60_DNA+RNA_seed | PIK3CD | 0.796793 | COADREAD | HCC2998 | AZD8186 |
| MVP_top60_DNA_seed | GSK3B | 0.972836 | COADREAD | NCI-H716 | CHIR-99021 | MVP_top60_DNA+RNA_seed | HSP90AA1 | 0.797991 | COADREAD | SW1417 | SNX-2112 |
| MVP_top60_DNA_seed | ATM | 0.975841 | COADREAD | DiFi | KU-55933 | MVP_top60_DNA+RNA_seed | HSP90AB1 | 0.797991 | COADREAD | SW1417 | SNX-2112 |
| MVP_top60_DNA_seed | GSK3B | 0.97726 | COADREAD | CL-11 | CHIR-99021 | MVP_top60_DNA+RNA_seed | AKT1 | 0.798423 | COADREAD | HT-29 | AKT inhibitor VIII |
| MVP_top60_DNA_seed | ERBB3 | 0.977719 | COADREAD | SW948 | AZD8931 | MVP_top60_DNA+RNA_seed | AKT1 | 0.800284 | COADREAD | MDST8 | AKT inhibitor VIII |
| MVP_top60_DNA_seed | ESR1 | 0.977994 | COADREAD | HT55 | Tamoxifen | MVP_top60_DNA+RNA_seed | PIK3CB | 0.802573 | COADREAD | SNU-C5 | AZD8186 |
| MVP_top60_DNA_seed | ATM | 0.979241 | COADREAD | SNU-C1 | KU-55933 | MVP_top60_DNA+RNA_seed | PIK3CD | 0.802573 | COADREAD | SNU-C5 | AZD8186 |
| MVP_top60_DNA_seed | CDK2 | 0.979413 | COADREAD | HT-29 | AZD5438 | MVP_top60_DNA+RNA_seed | AKT1 | 0.803413 | COADREAD | HT-115 | AKT inhibitor VIII |
| MVP_top60_DNA_seed | ESR1 | 0.981129 | COADREAD | LS-1034 | Tamoxifen | MVP_top60_DNA+RNA_seed | MET | 0.80415 | COADREAD | C2BBe1 | SU11274 |
| MVP_top60_DNA_seed | PIK3CD | 0.981438 | COADREAD | SNU-C2B | IC-87114 | MVP_top60_DNA+RNA_seed | AKT1 | 0.805855 | COADREAD | LoVo | GSK690693 |
| MVP_top60_DNA_seed | HSP90AA1 | 0.984452 | COADREAD | HT-115 | Elesclomol | MVP_top60_DNA+RNA_seed | ATM | 0.806017 | COADREAD | SW1417 | KU-60019 |
| MVP_top60_DNA_seed | HSP90AB1 | 0.984452 | COADREAD | HT-115 | Elesclomol | MVP_top60_DNA+RNA_seed | IGF1R | 0.806249 | COADREAD | SW620 | GSK1904529A |
| MVP_top60_DNA_seed | PTK2 | 0.986102 | COADREAD | NCI-H716 | PF-562271 | MVP_top60_DNA+RNA_seed | SRC | 0.806405 | COADREAD | NCI-H716 | Saracatinib |
| MVP_top60_DNA_seed | MET | 0.986849 | COADREAD | T84 | SU11274 | MVP_top60_DNA+RNA_seed | SRC | 0.806858 | COADREAD | SW620 | HG-6-71-01 |
| MVP_top60_DNA_seed | HSP90AA1 | 0.987062 | COADREAD | SW1116 | CCT-018159 | MVP_top60_DNA+RNA_seed | MET | 0.808043 | COADREAD | SNU-C2B | AZD6094 |
| MVP_top60_DNA_seed | HSP90AB1 | 0.987062 | COADREAD | SW1116 | CCT-018159 | MVP_top60_DNA+RNA_seed | PIK3CB | 0.80833 | COADREAD | COLO-320-HSR | TGX221 |
| MVP_top60_DNA_seed | BRAF | 0.988457 | COADREAD | HCT-15 | SB590885 | MVP_top60_DNA+RNA_seed | SRC | 0.808896 | COADREAD | HT-115 | NVP-BHG712 |
| MVP_top60_DNA_seed | BRAF | 0.989371 | COADREAD | LS-513 | HG6-64-1 | MVP_top60_DNA+RNA_seed | CDK2 | 0.80973 | COADREAD | SW948 | AZD5438 |
| MVP_top60_DNA_seed | BRAF | 0.989568 | COADREAD | CL-11 | PLX-4720 | MVP_top60_DNA+RNA_seed | AKT1 | 0.809879 | COADREAD | SW48 | GSK690693 |
| MVP_top60_DNA_seed | ERBB4 | 0.991626 | COADREAD | RCM-1 | PF-00299804 | MVP_top60_DNA+RNA_seed | ATM | 0.810344 | COADREAD | LS-123 | KU-60019 |
| MVP_top60_DNA_seed | ERBB4 | 0.99204 | COADREAD | SW48 | CI-1033 | MVP_top60_DNA+RNA_seed | ATM | 0.811286 | COADREAD | LS-411N | KU-55933 |
| MVP_top60_DNA_seed | HSP90AA1 | 0.992759 | COADREAD | CL-11 | Elesclomol | MVP_top60_DNA+RNA_seed | BCL2 | 0.813534 | COADREAD | HT-115 | Navitoclax |
| MVP_top60_DNA_seed | HSP90AB1 | 0.992759 | COADREAD | CL-11 | Elesclomol | MVP_top60_DNA+RNA_seed | AKT1 | 0.813653 | COADREAD | LS-123 | GSK690693 |
| MVP_top60_DNA_seed | HSP90AA1 | 0.994546 | COADREAD | HT-29 | CCT-018159 | MVP_top60_DNA+RNA_seed | AR | 0.815172 | COADREAD | HCC2998 | AZD3514 |
| MVP_top60_DNA_seed | HSP90AB1 | 0.994546 | COADREAD | HT-29 | CCT-018159 | MVP_top60_DNA+RNA_seed | ERBB4 | 0.815839 | COADREAD | CaR-1 | PF-00299804 |
| MVP_top60_DNA_seed | HSP90AA1 | 0.994863 | COADREAD | HT-115 | CCT-018159 | MVP_top60_DNA+RNA_seed | MET | 0.81775 | COADREAD | LS-1034 | JNJ38877605 |
| MVP_top60_DNA_seed | HSP90AB1 | 0.994863 | COADREAD | HT-115 | CCT-018159 | MVP_top60_DNA+RNA_seed | GSK3B | 0.818765 | COADREAD | LS-180 | CHIR-99021 |
| MVP_top60_DNA_seed | ERBB4 | 0.996167 | COADREAD | COLO-320-HSR | AST-1306 | MVP_top60_DNA+RNA_seed | AKT1 | 0.820179 | COADREAD | LoVo | AKT inhibitor VIII |
| MVP_top60_DNA_seed | GSK3B | 0.997646 | COADREAD | LS-1034 | KIN001-042 | MVP_top60_DNA+RNA_seed | SRC | 0.82113 | COADREAD | SNU-C2B | HG-6-71-01 |
| MVP_top60_DNA_seed | CDK2 | 0.999329 | COADREAD | SNU-1040 | AZD5438 | MVP_top60_DNA+RNA_seed | HSP90AA1 | 0.821289 | COADREAD | LS-1034 | Tanespimycin |
| MVP_top60_DNA_seed | GSK3B | 0.999358 | COADREAD | SNU-1040 | KIN001-042 | MVP_top60_DNA+RNA_seed | HSP90AB1 | 0.821289 | COADREAD | LS-1034 | Tanespimycin |
| MVP_top60_DNA_seed | BRAF | 0.999727 | COADREAD | C2BBe1 | Dabrafenib | MVP_top60_DNA+RNA_seed | PIK3CD | 0.823391 | COADREAD | SW837 | Idelalisib |
| MVP_top60_DNA_seed | IGF1R | 1.000739 | COADREAD | SNU-C5 | BMS-536924 | MVP_top60_DNA+RNA_seed | AKT1 | 0.824216 | COADREAD | COLO-320-HSR | AKT inhibitor VIII |
| MVP_top60_DNA_seed | MET | 1.001109 | COADREAD | LS-1034 | Crizotinib | MVP_top60_DNA+RNA_seed | ERBB4 | 0.82472 | COADREAD | CL-11 | PF-00299804 |
| MVP_top60_DNA_seed | IGF1R | 1.001822 | COADREAD | LS-180 | GSK1904529A | MVP_top60_DNA+RNA_seed | SRC | 0.827859 | COADREAD | COLO-320-HSR | Bosutinib |
| MVP_top60_DNA_seed | AR | 1.002418 | COADREAD | C2BBe1 | Bicalutamide | MVP_top60_DNA+RNA_seed | MET | 0.828397 | COADREAD | CW-2 | AZD6094 |
| MVP_top60_DNA_seed | IGF1R | 1.002533 | COADREAD | HT-115 | Linsitinib | MVP_top60_DNA+RNA_seed | PIK3CB | 0.828743 | COADREAD | DiFi | AZD8186 |
| MVP_top60_DNA_seed | ATM | 1.00267 | COADREAD | HT-115 | CP466722 | MVP_top60_DNA+RNA_seed | PIK3CD | 0.828743 | COADREAD | DiFi | AZD8186 |
| MVP_top60_DNA_seed | PIK3CB | 1.003549 | COADREAD | LS-513 | AZD8186 | MVP_top60_DNA+RNA_seed | GSK3B | 0.83156 | COADREAD | COLO-320-HSR | CHIR-99021 |

| | | | | | | | | | | | |
|---|---|---|---|---|---|---|---|---|---|---|---|
| MVP_top60_DNA_seed | PIK3CD | 1.003549 | COADREAD | LS-513 | AZD8186 | MVP_top60_DNA+RNA_seed | GSK3B | 0.832621 | COADREAD | LS-1034 | CHIR-99021 |
| MVP_top60_DNA_seed | SRC | 1.004673 | COADREAD | LS-123 | NVP-BHG712 | MVP_top60_DNA+RNA_seed | MET | 0.837364 | COADREAD | C2BBe1 | Crizotinib |
| MVP_top60_DNA_seed | PIK3CD | 1.006626 | COADREAD | HT-115 | Idelalisib | MVP_top60_DNA+RNA_seed | GSK3B | 0.838729 | COADREAD | RCM-1 | AZD7969 |
| MVP_top60_DNA_seed | MET | 1.007629 | COADREAD | MDST8 | JNJ38877605 | MVP_top60_DNA+RNA_seed | GSK3B | 0.840255 | COADREAD | NCI-H508 | AZD7969 |
| MVP_top60_DNA_seed | AKT1 | 1.009536 | COADREAD | SW48 | AKT inhibitor VIII | MVP_top60_DNA+RNA_seed | ATM | 0.840684 | COADREAD | SNU-C5 | KU-60019 |
| MVP_top60_DNA_seed | GSK3B | 1.009541 | COADREAD | T84 | SB216763 | MVP_top60_DNA+RNA_seed | ATM | 0.84323 | COADREAD | CCK-81 | KU-55933 |
| MVP_top60_DNA_seed | IGF1R | 1.011357 | COADREAD | NCI-H747 | GSK1904529A | MVP_top60_DNA+RNA_seed | GSK3B | 0.844888 | COADREAD | SW620 | CHIR-99021 |
| MVP_top60_DNA_seed | IGF1R | 1.011613 | COADREAD | COLO-678 | Linsitinib | MVP_top60_DNA+RNA_seed | PIK3CB | 0.845218 | COADREAD | NCI-H508 | AZD8186 |
| MVP_top60_DNA_seed | IGF1R | 1.012019 | COADREAD | HT-29 | GSK1904529A | MVP_top60_DNA+RNA_seed | PIK3CD | 0.845218 | COADREAD | NCI-H508 | AZD8186 |
| MVP_top60_DNA_seed | PIK3CA | 1.012971 | COADREAD | COLO-678 | AZD8835 | MVP_top60_DNA+RNA_seed | AKT1 | 0.845542 | COADREAD | LS-123 | AKT inhibitor VIII |
| MVP_top60_DNA_seed | PIK3CD | 1.012971 | COADREAD | COLO-678 | AZD8835 | MVP_top60_DNA+RNA_seed | PIK3CB | 0.847949 | COADREAD | T84 | AZD8186 |
| MVP_top60_DNA_seed | AKT1 | 1.01589 | COADREAD | SNU-C2B | GSK690693 | MVP_top60_DNA+RNA_seed | PIK3CD | 0.847949 | COADREAD | T84 | AZD8186 |
| MVP_top60_DNA_seed | BRAF | 1.017936 | COADREAD | SW1463 | HG6-64-1 | MVP_top60_DNA+RNA_seed | BCL2 | 0.848138 | COADREAD | CCK-81 | TW 37 |
| MVP_top60_DNA_seed | ATM | 1.01836 | COADREAD | DiFi | KU-60019 | MVP_top60_DNA+RNA_seed | BCL2 | 0.848694 | COADREAD | SW837 | TW 37 |
| MVP_top60_DNA_seed | BRAF | 1.02012 | COADREAD | SNU-81 | PLX-4720 | MVP_top60_DNA+RNA_seed | ERBB4 | 0.850863 | COADREAD | HCT-116 | CI-1033 |
| MVP_top60_DNA_seed | CREBBP | 1.02138 | COADREAD | SW620 | I-CBP112 | MVP_top60_DNA+RNA_seed | AKT1 | 0.851556 | COADREAD | KM12 | MK-2206 |
| MVP_top60_DNA_seed | BRAF | 1.025394 | COADREAD | LS-123 | PLX-4720 | MVP_top60_DNA+RNA_seed | ERBB4 | 0.852693 | COADREAD | COLO-678 | PF-00299804 |
| MVP_top60_DNA_seed | ERBB3 | 1.025529 | COADREAD | LS-123 | AZD8931 | MVP_top60_DNA+RNA_seed | AR | 0.85275 | COADREAD | SNU-C2B | Bicalutamide |
| MVP_top60_DNA_seed | PIK3CA | 1.026844 | COADREAD | CaR-1 | AZD8835 | MVP_top60_DNA+RNA_seed | ATM | 0.853584 | COADREAD | C2BBe1 | CP466722 |
| MVP_top60_DNA_seed | PIK3CD | 1.026844 | COADREAD | CaR-1 | AZD8835 | MVP_top60_DNA+RNA_seed | MET | 0.855049 | COADREAD | C2BBe1 | AZD6094 |
| MVP_top60_DNA_seed | BRAF | 1.027412 | COADREAD | LS-1034 | RAF_9304 | MVP_top60_DNA+RNA_seed | ERBB4 | 0.857216 | COADREAD | HT55 | CI-1033 |
| MVP_top60_DNA_seed | BRAF | 1.02833 | COADREAD | KM12 | PLX-4720 | MVP_top60_DNA+RNA_seed | MET | 0.858423 | COADREAD | DiFi | AZD6094 |
| MVP_top60_DNA_seed | HSP90AA1 | 1.029465 | COADREAD | CW-2 | Tanespimycin | MVP_top60_DNA+RNA_seed | MET | 0.8609 | COADREAD | SNU-407 | JNJ38877605 |
| MVP_top60_DNA_seed | HSP90AB1 | 1.029465 | COADREAD | CW-2 | Tanespimycin | MVP_top60_DNA+RNA_seed | BCL2 | 0.861063 | COADREAD | LS-411N | TW 37 |
| MVP_top60_DNA_seed | PIK3CD | 1.031825 | COADREAD | HCT-15 | Idelalisib | MVP_top60_DNA+RNA_seed | PIK3CB | 0.86119 | COADREAD | GP5d | AZD6482 |
| MVP_top60_DNA_seed | GSK3B | 1.03513 | COADREAD | LS-411N | SB216763 | MVP_top60_DNA+RNA_seed | AKT1 | 0.861273 | COADREAD | KM12 | AKT inhibitor VIII |
| MVP_top60_DNA_seed | MET | 1.03695 | COADREAD | SNU-C1 | Crizotinib | MVP_top60_DNA+RNA_seed | HSP90AA1 | 0.861536 | COADREAD | RCM-1 | Luminespib |
| MVP_top60_DNA_seed | IGF1R | 1.037901 | COADREAD | LS-1034 | GSK1904529A | MVP_top60_DNA+RNA_seed | HSP90AB1 | 0.861536 | COADREAD | RCM-1 | Luminespib |
| MVP_top60_DNA_seed | GSK3B | 1.038853 | COADREAD | SNU-C1 | CHIR-99021 | MVP_top60_DNA+RNA_seed | SRC | 0.864705 | COADREAD | KM12 | WH-4-023 |
| MVP_top60_DNA_seed | GSK3B | 1.039278 | COADREAD | SNU-C2B | SB216763 | MVP_top60_DNA+RNA_seed | AR | 0.866284 | COADREAD | COLO-320-HSR | AZD3514 |
| MVP_top60_DNA_seed | BRAF | 1.042429 | COADREAD | RCM-1 | PLX-4720 | MVP_top60_DNA+RNA_seed | GSK3B | 0.869442 | COADREAD | COLO-205 | AZD7969 |
| MVP_top60_DNA_seed | AR | 1.042827 | COADREAD | LS-411N | Bicalutamide | MVP_top60_DNA+RNA_seed | IGF1R | 0.869922 | COADREAD | CL-11 | BMS-536924 |
| MVP_top60_DNA_seed | BRAF | 1.043 | COADREAD | SNU-C2B | PLX-4720 | MVP_top60_DNA+RNA_seed | PIK3CD | 0.87006 | COADREAD | COLO-205 | Idelalisib |
| MVP_top60_DNA_seed | BRAF | 1.045255 | COADREAD | SNU-C2B | SB590885 | MVP_top60_DNA+RNA_seed | IGF1R | 0.870781 | COADREAD | SW1463 | Linsitinib |
| MVP_top60_DNA_seed | MET | 1.04701 | COADREAD | HCT-15 | SU11274 | MVP_top60_DNA+RNA_seed | ATM | 0.871659 | COADREAD | LS-1034 | KU-55933 |
| MVP_top60_DNA_seed | MET | 1.047465 | COADREAD | SW837 | AZD6094 | MVP_top60_DNA+RNA_seed | SRC | 0.874426 | COADREAD | CW-2 | NVP-BHG712 |
| MVP_top60_DNA_seed | CDK2 | 1.050294 | COADREAD | COLO-320-HSR | PHA-793887 | MVP_top60_DNA+RNA_seed | SRC | 0.875488 | COADREAD | RCM-1 | HG-6-71-01 |
| MVP_top60_DNA_seed | AR | 1.05138 | COADREAD | SNU-C1 | Bicalutamide | MVP_top60_DNA+RNA_seed | MET | 0.876386 | COADREAD | HT-29 | SU11274 |
| MVP_top60_DNA_seed | HSP90AA1 | 1.05243 | COADREAD | NCI-H716 | Luminespib | MVP_top60_DNA+RNA_seed | HSP90AA1 | 0.876529 | COADREAD | CaR-1 | Luminespib |
| MVP_top60_DNA_seed | HSP90AB1 | 1.05243 | COADREAD | NCI-H716 | Luminespib | MVP_top60_DNA+RNA_seed | HSP90AB1 | 0.876529 | COADREAD | CaR-1 | Luminespib |
| MVP_top60_DNA_seed | MET | 1.052442 | COADREAD | LS-1034 | SU11274 | MVP_top60_DNA+RNA_seed | SRC | 0.877186 | COADREAD | COLO-678 | NVP-BHG712 |

| | | | | | | | | | | | |
|---|---|---|---|---|---|---|---|---|---|---|---|
| MVP_top60_DNA_seed | AKT1 | 1.053858 | COADREAD | CW-2 | MK-2206 | MVP_top60_DNA+RNA_seed | HSP90AA1 | 0.878829 | COADREAD | LS-180 | CCT-018159 |
| MVP_top60_DNA_seed | AR | 1.05546 | COADREAD | LS-1034 | AZD3514 | MVP_top60_DNA+RNA_seed | HSP90AB1 | 0.878829 | COADREAD | LS-180 | CCT-018159 |
| MVP_top60_DNA_seed | BRAF | 1.055755 | COADREAD | LS-180 | PLX-4720 | MVP_top60_DNA+RNA_seed | IGF1R | 0.879153 | COADREAD | LoVo | Linsitinib |
| MVP_top60_DNA_seed | MET | 1.057071 | COADREAD | SNU-81 | AZD6094 | MVP_top60_DNA+RNA_seed | IGF1R | 0.881867 | COADREAD | SW948 | BMS-536924 |
| MVP_top60_DNA_seed | PIK3CB | 1.057664 | COADREAD | HCT-15 | AZD6482 | MVP_top60_DNA+RNA_seed | ERBB4 | 0.882399 | COADREAD | SW837 | AST-1306 |
| MVP_top60_DNA_seed | MET | 1.057885 | COADREAD | NCI-H747 | PHA-665752 | MVP_top60_DNA+RNA_seed | MET | 0.882944 | COADREAD | NCI-H716 | Crizotinib |
| MVP_top60_DNA_seed | GSK3B | 1.060105 | COADREAD | SNU-1040 | AZD7969 | MVP_top60_DNA+RNA_seed | AKT1 | 0.8857 | COADREAD | T84 | AKT inhibitor VIII |
| MVP_top60_DNA_seed | ERBB4 | 1.060906 | COADREAD | CW-2 | PF-00299804 | MVP_top60_DNA+RNA_seed | IGF1R | 0.885714 | COADREAD | HT55 | Linsitinib |
| MVP_top60_DNA_seed | SRC | 1.063157 | COADREAD | HCC2998 | Bosutinib | MVP_top60_DNA+RNA_seed | HSP90AA1 | 0.889417 | COADREAD | LoVo | Tanespimycin |
| MVP_top60_DNA_seed | ESR1 | 1.064091 | COADREAD | HT55 | Fulvestrant | MVP_top60_DNA+RNA_seed | HSP90AB1 | 0.889417 | COADREAD | LoVo | Tanespimycin |
| MVP_top60_DNA_seed | SRC | 1.064476 | COADREAD | SK-CO-1 | Bosutinib | MVP_top60_DNA+RNA_seed | PIK3CB | 0.892551 | COADREAD | LS-513 | TGX221 |
| MVP_top60_DNA_seed | BRAF | 1.06569 | COADREAD | LS-180 | Dabrafenib | MVP_top60_DNA+RNA_seed | GSK3B | 0.892758 | COADREAD | HCT-15 | SB216763 |
| MVP_top60_DNA_seed | PIK3CB | 1.066955 | COADREAD | LS-1034 | AZD8186 | MVP_top60_DNA+RNA_seed | PIK3CB | 0.893277 | COADREAD | HT-29 | AZD8186 |
| MVP_top60_DNA_seed | PIK3CD | 1.066955 | COADREAD | LS-1034 | AZD8186 | MVP_top60_DNA+RNA_seed | PIK3CD | 0.893277 | COADREAD | HT-29 | AZD8186 |
| MVP_top60_DNA_seed | AKT1 | 1.067371 | COADREAD | LS-1034 | BAY AKT1 | MVP_top60_DNA+RNA_seed | AR | 0.893463 | COADREAD | SW1417 | AZD3514 |
| MVP_top60_DNA_seed | GSK3B | 1.067397 | COADREAD | C2BBe1 | SB216763 | MVP_top60_DNA+RNA_seed | BCL2 | 0.893785 | COADREAD | NCI-H716 | Obatoclax Mesylate |
| MVP_top60_DNA_seed | ERBB3 | 1.069043 | COADREAD | SW1417 | AZD8931 | MVP_top60_DNA+RNA_seed | AKT1 | 0.893927 | COADREAD | LS-411N | MK-2206 |
| MVP_top60_DNA_seed | HSP90AA1 | 1.070183 | COADREAD | CL-11 | Tanespimycin | MVP_top60_DNA+RNA_seed | BCL2 | 0.896015 | COADREAD | C2BBe1 | Obatoclax Mesylate |
| MVP_top60_DNA_seed | HSP90AB1 | 1.070183 | COADREAD | CL-11 | Tanespimycin | MVP_top60_DNA+RNA_seed | CDK2 | 0.897315 | COADREAD | HT55 | AZD5438 |
| MVP_top60_DNA_seed | SRC | 1.07317 | COADREAD | HT-115 | HG-6-71-01 | MVP_top60_DNA+RNA_seed | SRC | 0.897502 | COADREAD | LS-513 | Bosutinib |
| MVP_top60_DNA_seed | HSP90AA1 | 1.074699 | COADREAD | SNU-C1 | SNX-2112 | MVP_top60_DNA+RNA_seed | AKT1 | 0.898225 | COADREAD | C2BBe1 | AKT inhibitor VIII |
| MVP_top60_DNA_seed | HSP90AB1 | 1.074699 | COADREAD | SNU-C1 | SNX-2112 | MVP_top60_DNA+RNA_seed | PIK3CA | 0.898392 | COADREAD | SW620 | AZD8835 |
| MVP_top60_DNA_seed | HSP90AA1 | 1.074842 | COADREAD | DiFi | CCT-018159 | MVP_top60_DNA+RNA_seed | PIK3CD | 0.898392 | COADREAD | SW620 | AZD8835 |
| MVP_top60_DNA_seed | HSP90AB1 | 1.074842 | COADREAD | DiFi | CCT-018159 | MVP_top60_DNA+RNA_seed | ERBB4 | 0.89957 | COADREAD | HCC2998 | PF-00299804 |
| MVP_top60_DNA_seed | HSP90AA1 | 1.075325 | COADREAD | HT55 | SNX-2112 | MVP_top60_DNA+RNA_seed | BCL2 | 0.900028 | COADREAD | HCT-116 | Navitoclax |
| MVP_top60_DNA_seed | HSP90AB1 | 1.075325 | COADREAD | HT55 | SNX-2112 | MVP_top60_DNA+RNA_seed | GSK3B | 0.900713 | COADREAD | T84 | CHIR-99021 |
| MVP_top60_DNA_seed | SRC | 1.075788 | COADREAD | NCI-H747 | Bosutinib | MVP_top60_DNA+RNA_seed | AKT1 | 0.900715 | COADREAD | COLO-678 | AKT inhibitor VIII |
| MVP_top60_DNA_seed | AR | 1.078102 | COADREAD | SW1417 | Bicalutamide | MVP_top60_DNA+RNA_seed | PIK3CD | 0.904648 | COADREAD | LS-1034 | IC-87114 |
| MVP_top60_DNA_seed | ERBB3 | 1.078566 | COADREAD | KM12 | AZD8931 | MVP_top60_DNA+RNA_seed | SRC | 0.906083 | COADREAD | MDST8 | WH-4-023 |
| MVP_top60_DNA_seed | AR | 1.079543 | COADREAD | LS-1034 | Bicalutamide | MVP_top60_DNA+RNA_seed | ATM | 0.90616 | COADREAD | KM12 | KU-55933 |
| MVP_top60_DNA_seed | BRAF | 1.08096 | COADREAD | LS-123 | PLX-4720 | MVP_top60_DNA+RNA_seed | GSK3B | 0.90619 | COADREAD | HT55 | CHIR-99021 |
| MVP_top60_DNA_seed | BRAF | 1.083645 | COADREAD | T84 | PLX-4720 | MVP_top60_DNA+RNA_seed | GSK3B | 0.910105 | COADREAD | SW837 | SB216763 |
| MVP_top60_DNA_seed | BRAF | 1.083854 | COADREAD | SNU-81 | Dabrafenib | MVP_top60_DNA+RNA_seed | SRC | 0.912493 | COADREAD | RCM-1 | NVP-BHG712 |
| MVP_top60_DNA_seed | MET | 1.084589 | COADREAD | SNU-1040 | AZD6094 | MVP_top60_DNA+RNA_seed | GSK3B | 0.915056 | COADREAD | COLO-678 | SB216763 |
| MVP_top60_DNA_seed | GSK3B | 1.085503 | COADREAD | SW948 | AZD7969 | MVP_top60_DNA+RNA_seed | IGF1R | 0.915548 | COADREAD | SW1463 | BMS-754807 |
| MVP_top60_DNA_seed | IGF1R | 1.086477 | COADREAD | SW837 | GSK1904529A | MVP_top60_DNA+RNA_seed | FYN | 0.915818 | COADREAD | MDST8 | A-770041 |
| MVP_top60_DNA_seed | MET | 1.091485 | COADREAD | COLO-678 | AZD6094 | MVP_top60_DNA+RNA_seed | AR | 0.916959 | COADREAD | CW-2 | AZD3514 |
| MVP_top60_DNA_seed | BRAF | 1.09231 | COADREAD | T84 | HG6-64-1 | MVP_top60_DNA+RNA_seed | GSK3B | 0.917032 | COADREAD | T84 | CHIR-99021 |
| MVP_top60_DNA_seed | AR | 1.09262 | COADREAD | LS-411N | Bicalutamide | MVP_top60_DNA+RNA_seed | PIK3CA | 0.918071 | COADREAD | SW1116 | PI3Ka_4409 |
| MVP_top60_DNA_seed | HSP90AA1 | 1.094997 | COADREAD | CaR-1 | Elesclomol | MVP_top60_DNA+RNA_seed | PIK3CD | 0.918071 | COADREAD | SW1116 | PI3Ka_4409 |
| MVP_top60_DNA_seed | HSP90AB1 | 1.094997 | COADREAD | CaR-1 | Elesclomol | MVP_top60_DNA+RNA_seed | GSK3B | 0.919686 | COADREAD | SNU-407 | KIN001-042 |

| | | | | | | | | | | | |
|---|---|---|---|---|---|---|---|---|---|---|---|
| MVP_top60_DNA_seed | IGF1R | 1.09674 | COADREAD | HT-115 | BMS-754807 | MVP_top60_DNA+RNA_seed | GSK3B | 0.926944 | COADREAD | SW948 | KIN001-042 |
| MVP_top60_DNA_seed | BRAF | 1.102298 | COADREAD | HT55 | PLX-4720 | MVP_top60_DNA+RNA_seed | PIK3CD | 0.927021 | COADREAD | SNU-C2B | Idelalisib |
| MVP_top60_DNA_seed | PIK3CB | 1.104725 | COADREAD | LS-1034 | TGX221 | MVP_top60_DNA+RNA_seed | MET | 0.927952 | COADREAD | LS-1034 | PHA-665752 |
| MVP_top60_DNA_seed | SRC | 1.105651 | COADREAD | LS-513 | WH-4-023 | MVP_top60_DNA+RNA_seed | PTK2 | 0.927991 | COADREAD | SW948 | PF-562271 |
| MVP_top60_DNA_seed | AKT1 | 1.106423 | COADREAD | MDST8 | AKT inhibitor VIII | MVP_top60_DNA+RNA_seed | MET | 0.932731 | COADREAD | CL-11 | Crizotinib |
| MVP_top60_DNA_seed | MET | 1.107667 | COADREAD | HT55 | SU11274 | MVP_top60_DNA+RNA_seed | SRC | 0.933456 | COADREAD | LS-411N | Bosutinib |
| MVP_top60_DNA_seed | CDK2 | 1.108322 | COADREAD | LS-411N | AZD5438 | MVP_top60_DNA+RNA_seed | HSP90AA1 | 0.934681 | COADREAD | NCI-H747 | CCT-018159 |
| MVP_top60_DNA_seed | AR | 1.110041 | COADREAD | SNU-C5 | Bicalutamide | MVP_top60_DNA+RNA_seed | HSP90AB1 | 0.934681 | COADREAD | NCI-H747 | CCT-018159 |
| MVP_top60_DNA_seed | AR | 1.111908 | COADREAD | HT-115 | AZD3514 | MVP_top60_DNA+RNA_seed | SRC | 0.936914 | COADREAD | HCT-15 | HG-6-71-01 |
| MVP_top60_DNA_seed | ATM | 1.112672 | COADREAD | RCM-1 | torin2 | MVP_top60_DNA+RNA_seed | PTK2 | 0.93716 | COADREAD | COLO-678 | PF-562271 |
| MVP_top60_DNA_seed | IGF1R | 1.114452 | COADREAD | HCT-15 | Linsitinib | MVP_top60_DNA+RNA_seed | PIK3CB | 0.938659 | COADREAD | COLO-678 | AZD6482 |
| MVP_top60_DNA_seed | AKT1 | 1.11694 | COADREAD | SW837 | GSK690693 | MVP_top60_DNA+RNA_seed | BCL2 | 0.938832 | COADREAD | SW1463 | Obatoclax Mesylate |
| MVP_top60_DNA_seed | PIK3CD | 1.118249 | COADREAD | SW837 | IC-87114 | MVP_top60_DNA+RNA_seed | MET | 0.948402 | COADREAD | LS-180 | AZD6094 |
| MVP_top60_DNA_seed | PIK3CD | 1.118331 | COADREAD | LoVo | Idelalisib | MVP_top60_DNA+RNA_seed | MET | 0.95032 | COADREAD | MDST8 | SU11274 |
| MVP_top60_DNA_seed | IGF1R | 1.119059 | COADREAD | NCI-H716 | GSK1904529A | MVP_top60_DNA+RNA_seed | HSP90AA1 | 0.950743 | COADREAD | COLO-678 | SNX-2112 |
| MVP_top60_DNA_seed | SRC | 1.123856 | COADREAD | HT55 | Bosutinib | MVP_top60_DNA+RNA_seed | HSP90AB1 | 0.950743 | COADREAD | COLO-678 | SNX-2112 |
| MVP_top60_DNA_seed | ESR1 | 1.124404 | COADREAD | C2BBe1 | Tamoxifen | MVP_top60_DNA+RNA_seed | AKT1 | 0.951401 | COADREAD | HT55 | GSK690693 |
| MVP_top60_DNA_seed | MET | 1.125191 | COADREAD | LS-123 | SU11274 | MVP_top60_DNA+RNA_seed | GSK3B | 0.952383 | COADREAD | SW1463 | CHIR-99021 |
| MVP_top60_DNA_seed | BRAF | 1.125855 | COADREAD | SNU-61 | Dabrafenib | MVP_top60_DNA+RNA_seed | AR | 0.952588 | COADREAD | LS-513 | AZD3514 |
| MVP_top60_DNA_seed | GSK3B | 1.127733 | COADREAD | HCT-15 | CHIR-99021 | MVP_top60_DNA+RNA_seed | CDK2 | 0.953799 | COADREAD | SW837 | AZD5438 |
| MVP_top60_DNA_seed | AR | 1.128407 | COADREAD | NCI-H747 | Bicalutamide | MVP_top60_DNA+RNA_seed | ERBB4 | 0.95393 | COADREAD | LS-411N | CI-1033 |
| MVP_top60_DNA_seed | AKT1 | 1.129865 | COADREAD | CaR-1 | MK-2206 | MVP_top60_DNA+RNA_seed | BCL2 | 0.954888 | COADREAD | SW1417 | TW 37 |
| MVP_top60_DNA_seed | GSK3B | 1.129898 | COADREAD | SNU-81 | CHIR-99021 | MVP_top60_DNA+RNA_seed | AR | 0.955799 | COADREAD | KM12 | Bicalutamide |
| MVP_top60_DNA_seed | MAPK1 | 1.130514 | COADREAD | SW948 | FR-180204 | MVP_top60_DNA+RNA_seed | AKT1 | 0.956203 | COADREAD | SW1417 | GSK690693 |
| MVP_top60_DNA_seed | AKT1 | 1.133795 | COADREAD | RCM-1 | GSK690693 | MVP_top60_DNA+RNA_seed | BCL2 | 0.956291 | COADREAD | SW1463 | Navitoclax |
| MVP_top60_DNA_seed | ERBB4 | 1.134712 | COADREAD | CaR-1 | CI-1033 | MVP_top60_DNA+RNA_seed | ERBB4 | 0.956951 | COADREAD | SNU-61 | PF-00299804 |
| MVP_top60_DNA_seed | AKT1 | 1.136258 | COADREAD | COLO-678 | GSK690693 | MVP_top60_DNA+RNA_seed | MET | 0.957992 | COADREAD | CL-11 | AZD6094 |
| MVP_top60_DNA_seed | MAPK14 | 1.137095 | COADREAD | MDST8 | TAK-715 | MVP_top60_DNA+RNA_seed | MET | 0.963094 | COADREAD | SNU-C5 | JNJ38877605 |
| MVP_top60_DNA_seed | IGF1R | 1.139989 | COADREAD | RCM-1 | BMS-754807 | MVP_top60_DNA+RNA_seed | HSP90AA1 | 0.965896 | COADREAD | COLO-320-HSR | Elesclomol |
| MVP_top60_DNA_seed | PIK3CD | 1.140565 | COADREAD | HT55 | Idelalisib | MVP_top60_DNA+RNA_seed | HSP90AB1 | 0.965896 | COADREAD | COLO-320-HSR | Elesclomol |
| MVP_top60_DNA_seed | MET | 1.14497 | COADREAD | SNU-C1 | SU11274 | MVP_top60_DNA+RNA_seed | MET | 0.970831 | COADREAD | RCM-1 | AZD6094 |
| MVP_top60_DNA_seed | AKT1 | 1.145261 | COADREAD | SW948 | MK-2206 | MVP_top60_DNA+RNA_seed | BCL2 | 0.971946 | COADREAD | CCK-81 | Obatoclax Mesylate |
| MVP_top60_DNA_seed | BRAF | 1.145662 | COADREAD | COLO-678 | SB590885 | MVP_top60_DNA+RNA_seed | GSK3B | 0.972836 | COADREAD | NCI-H716 | CHIR-99021 |
| MVP_top60_DNA_seed | PIK3CD | 1.14605 | COADREAD | LS-180 | IC-87114 | MVP_top60_DNA+RNA_seed | ATM | 0.975841 | COADREAD | DiFi | KU-55933 |
| MVP_top60_DNA_seed | CDK2 | 1.147262 | COADREAD | NCI-H508 | AZD5438 | MVP_top60_DNA+RNA_seed | GSK3B | 0.97726 | COADREAD | CL-11 | CHIR-99021 |
| MVP_top60_DNA_seed | SRC | 1.147311 | COADREAD | LS-1034 | Saracatinib | MVP_top60_DNA+RNA_seed | ERBB3 | 0.977719 | COADREAD | SW948 | AZD8931 |
| MVP_top60_DNA_seed | SRC | 1.147866 | COADREAD | CW-2 | WH-4-023 | MVP_top60_DNA+RNA_seed | ATM | 0.979241 | COADREAD | SNU-C1 | KU-55933 |
| MVP_top60_DNA_seed | BRAF | 1.148894 | COADREAD | CW-2 | Dabrafenib | MVP_top60_DNA+RNA_seed | CDK2 | 0.979413 | COADREAD | HT-29 | AZD5438 |
| MVP_top60_DNA_seed | BRAF | 1.149149 | COADREAD | CaR-1 | Dabrafenib | MVP_top60_DNA+RNA_seed | PIK3CD | 0.981438 | COADREAD | SNU-C2B | IC-87114 |
| MVP_top60_DNA_seed | ATM | 1.149447 | COADREAD | HT-29 | KU-55933 | MVP_top60_DNA+RNA_seed | HSP90AA1 | 0.984452 | COADREAD | HT-115 | Elesclomol |
| MVP_top60_DNA_seed | IGF1R | 1.150067 | COADREAD | KM12 | Linsitinib | MVP_top60_DNA+RNA_seed | HSP90AB1 | 0.984452 | COADREAD | HT-115 | Elesclomol |

| | | | | | | | | | | | |
|---|---|---|---|---|---|---|---|---|---|---|---|
| MVP_top60_DNA_seed | PIK3CB | 1.150792 | COADREAD | HT-115 | AZD8186 | MVP_top60_DNA+RNA_seed | PTK2 | 0.986102 | COADREAD | NCI-H716 | PF-562271 |
| MVP_top60_DNA_seed | PIK3CD | 1.150792 | COADREAD | HT-115 | AZD8186 | MVP_top60_DNA+RNA_seed | MET | 0.986849 | COADREAD | T84 | SU11274 |
| MVP_top60_DNA_seed | CREBBP | 1.15154 | COADREAD | SW837 | I-CBP112 | MVP_top60_DNA+RNA_seed | HSP90AA1 | 0.987062 | COADREAD | SW1116 | CCT-018159 |
| MVP_top60_DNA_seed | BRAF | 1.15232 | COADREAD | LS-1034 | PLX-4720 | MVP_top60_DNA+RNA_seed | HSP90AB1 | 0.987062 | COADREAD | SW1116 | CCT-018159 |
| MVP_top60_DNA_seed | BRAF | 1.155179 | COADREAD | HT-115 | PLX-4720 | MVP_top60_DNA+RNA_seed | ERBB4 | 0.991626 | COADREAD | RCM-1 | PF-00299804 |
| MVP_top60_DNA_seed | PIK3CB | 1.158064 | COADREAD | NCI-H716 | TGX221 | MVP_top60_DNA+RNA_seed | ERBB4 | 0.99204 | COADREAD | SW48 | CI-1033 |
| MVP_top60_DNA_seed | AR | 1.160064 | COADREAD | SNU-1040 | AZD3514 | MVP_top60_DNA+RNA_seed | HSP90AA1 | 0.992759 | COADREAD | CL-11 | Elesclomol |
| MVP_top60_DNA_seed | SRC | 1.161135 | COADREAD | SW1417 | Bosutinib | MVP_top60_DNA+RNA_seed | HSP90AB1 | 0.992759 | COADREAD | CL-11 | Elesclomol |
| MVP_top60_DNA_seed | PIK3CB | 1.162021 | COADREAD | SW620 | AZD6482 | MVP_top60_DNA+RNA_seed | HSP90AA1 | 0.994546 | COADREAD | HT-29 | CCT-018159 |
| MVP_top60_DNA_seed | IGF1R | 1.162825 | COADREAD | CL-11 | Linsitinib | MVP_top60_DNA+RNA_seed | HSP90AB1 | 0.994546 | COADREAD | HT-29 | CCT-018159 |
| MVP_top60_DNA_seed | MET | 1.163243 | COADREAD | SNU-C2B | PHA-665752 | MVP_top60_DNA+RNA_seed | HSP90AA1 | 0.994863 | COADREAD | HT-115 | CCT-018159 |
| MVP_top60_DNA_seed | GSK3B | 1.16492 | COADREAD | RCM-1 | KIN001-042 | MVP_top60_DNA+RNA_seed | HSP90AB1 | 0.994863 | COADREAD | HT-115 | CCT-018159 |
| MVP_top60_DNA_seed | MET | 1.166586 | COADREAD | CaR-1 | AZD6094 | MVP_top60_DNA+RNA_seed | ERBB4 | 0.996167 | COADREAD | COLO-320-HSR | AST-1306 |
| MVP_top60_DNA_seed | ESR1 | 1.167907 | COADREAD | RCM-1 | Tamoxifen | MVP_top60_DNA+RNA_seed | GSK3B | 0.997646 | COADREAD | LS-1034 | KIN001-042 |
| MVP_top60_DNA_seed | SRC | 1.168962 | COADREAD | LS-411N | Saracatinib | MVP_top60_DNA+RNA_seed | CDK2 | 0.999329 | COADREAD | SNU-1040 | AZD5438 |
| MVP_top60_DNA_seed | BRAF | 1.169008 | COADREAD | CW-2 | PLX-4720 | MVP_top60_DNA+RNA_seed | GSK3B | 0.999358 | COADREAD | SNU-1040 | KIN001-042 |
| MVP_top60_DNA_seed | GSK3B | 1.170348 | COADREAD | HT55 | CHIR-99021 | MVP_top60_DNA+RNA_seed | BCL2 | 0.999548 | COADREAD | SNU-C2B | Navitoclax |
| MVP_top60_DNA_seed | ATM | 1.170588 | COADREAD | SW1417 | KU-55933 | MVP_top60_DNA+RNA_seed | IGF1R | 1.000739 | COADREAD | SNU-C5 | BMS-536924 |
| MVP_top60_DNA_seed | IGF1R | 1.170919 | COADREAD | SW948 | Linsitinib | MVP_top60_DNA+RNA_seed | MET | 1.001109 | COADREAD | LS-1034 | Crizotinib |
| MVP_top60_DNA_seed | GSK3B | 1.172102 | COADREAD | CL-11 | CHIR-99021 | MVP_top60_DNA+RNA_seed | IGF1R | 1.001822 | COADREAD | LS-180 | GSK1904529A |
| MVP_top60_DNA_seed | HSP90AA1 | 1.173232 | COADREAD | SW1463 | SNX-2112 | MVP_top60_DNA+RNA_seed | AR | 1.002418 | COADREAD | C2BBe1 | Bicalutamide |
| MVP_top60_DNA_seed | HSP90AB1 | 1.173232 | COADREAD | SW1463 | SNX-2112 | MVP_top60_DNA+RNA_seed | IGF1R | 1.002533 | COADREAD | HT-115 | Linsitinib |
| MVP_top60_DNA_seed | AKT1 | 1.174656 | COADREAD | MDST8 | A-443654 | MVP_top60_DNA+RNA_seed | ATM | 1.00267 | COADREAD | HT-115 | CP466722 |
| MVP_top60_DNA_seed | CDK2 | 1.177889 | COADREAD | SNU-C1 | Seliciclib | MVP_top60_DNA+RNA_seed | PIK3CB | 1.003549 | COADREAD | LS-513 | AZD8186 |
| MVP_top60_DNA_seed | ERBB4 | 1.179365 | COADREAD | KM12 | AST-1306 | MVP_top60_DNA+RNA_seed | PIK3CD | 1.003549 | COADREAD | LS-513 | AZD8186 |
| MVP_top60_DNA_seed | PIK3CB | 1.180194 | COADREAD | SNU-C2B | AZD6482 | MVP_top60_DNA+RNA_seed | SRC | 1.004673 | COADREAD | LS-123 | NVP-BHG712 |
| MVP_top60_DNA_seed | AR | 1.180664 | COADREAD | COLO-678 | Bicalutamide | MVP_top60_DNA+RNA_seed | PIK3CD | 1.006626 | COADREAD | HT-115 | Idelalisib |
| MVP_top60_DNA_seed | BRAF | 1.181593 | COADREAD | CW-2 | PLX-4720 | MVP_top60_DNA+RNA_seed | MET | 1.007629 | COADREAD | MDST8 | JNJ38877605 |
| MVP_top60_DNA_seed | BRAF | 1.184731 | COADREAD | SW837 | Dabrafenib | MVP_top60_DNA+RNA_seed | AKT1 | 1.009536 | COADREAD | SW48 | AKT inhibitor VIII |
| MVP_top60_DNA_seed | SRC | 1.185125 | COADREAD | SW1116 | Bosutinib | MVP_top60_DNA+RNA_seed | GSK3B | 1.009541 | COADREAD | T84 | SB216763 |
| MVP_top60_DNA_seed | PIK3CD | 1.185505 | COADREAD | SW948 | Idelalisib | MVP_top60_DNA+RNA_seed | BCL2 | 1.010974 | COADREAD | COLO-678 | Navitoclax |
| MVP_top60_DNA_seed | FYN | 1.186654 | COADREAD | NCI-H716 | A-770041 | MVP_top60_DNA+RNA_seed | IGF1R | 1.011357 | COADREAD | NCI-H747 | GSK1904529A |
| MVP_top60_DNA_seed | PIK3CB | 1.189332 | COADREAD | SNU-407 | AZD8186 | MVP_top60_DNA+RNA_seed | IGF1R | 1.011613 | COADREAD | COLO-678 | Linsitinib |
| MVP_top60_DNA_seed | PIK3CD | 1.189332 | COADREAD | SNU-407 | AZD8186 | MVP_top60_DNA+RNA_seed | IGF1R | 1.012019 | COADREAD | HT-29 | GSK1904529A |
| MVP_top60_DNA_seed | PTK2 | 1.190048 | COADREAD | HT55 | PF-562271 | MVP_top60_DNA+RNA_seed | PIK3CA | 1.012971 | COADREAD | COLO-678 | AZD8835 |
| MVP_top60_DNA_seed | PIK3CB | 1.191466 | COADREAD | HT55 | AZD6482 | MVP_top60_DNA+RNA_seed | PIK3CD | 1.012971 | COADREAD | COLO-678 | AZD8835 |
| MVP_top60_DNA_seed | BRAF | 1.192472 | COADREAD | MDST8 | HG6-64-1 | MVP_top60_DNA+RNA_seed | AKT1 | 1.01589 | COADREAD | SNU-C2B | GSK690693 |
| MVP_top60_DNA_seed | ATM | 1.192943 | COADREAD | COLO-678 | KU-60019 | MVP_top60_DNA+RNA_seed | ATM | 1.01836 | COADREAD | DiFi | KU-60019 |
| MVP_top60_DNA_seed | ERBB4 | 1.19357 | COADREAD | NCI-H716 | CI-1033 | MVP_top60_DNA+RNA_seed | BCL2 | 1.024956 | COADREAD | SNU-81 | TW 37 |
| MVP_top60_DNA_seed | PIK3CB | 1.195353 | COADREAD | CCK-81 | AZD8186 | MVP_top60_DNA+RNA_seed | ERBB3 | 1.025529 | COADREAD | LS-123 | AZD8931 |
| MVP_top60_DNA_seed | PIK3CD | 1.195353 | COADREAD | CCK-81 | AZD8186 | MVP_top60_DNA+RNA_seed | PIK3CA | 1.026844 | COADREAD | CaR-1 | AZD8835 |

| | | | | | | | | | | | |
|---|---|---|---|---|---|---|---|---|---|---|---|
| MVP_top60_DNA_seed | BRAF | 1.195714 | COADREAD | CW-2 | AZ628 | MVP_top60_DNA+RNA_seed | PIK3CD | 1.026844 | COADREAD | CaR-1 | AZD8835 |
| MVP_top60_DNA_seed | MET | 1.196749 | COADREAD | CL-11 | PHA-665752 | MVP_top60_DNA+RNA_seed | HSP90AA1 | 1.029465 | COADREAD | CW-2 | Tanespimycin |
| MVP_top60_DNA_seed | BRAF | 1.198266 | COADREAD | SNU-407 | PLX-4720 | MVP_top60_DNA+RNA_seed | HSP90AB1 | 1.029465 | COADREAD | CW-2 | Tanespimycin |
| MVP_top60_DNA_seed | ERBB4 | 1.198507 | COADREAD | KM12 | CI-1033 | MVP_top60_DNA+RNA_seed | PIK3CD | 1.031825 | COADREAD | HCT-15 | Idelalisib |
| MVP_top60_DNA_seed | ATM | 1.199291 | COADREAD | CL-11 | CP466722 | MVP_top60_DNA+RNA_seed | GSK3B | 1.03513 | COADREAD | LS-411N | SB216763 |
| MVP_top60_DNA_seed | MAPK1 | 1.199654 | COADREAD | LS-180 | FR-180204 | MVP_top60_DNA+RNA_seed | BCL2 | 1.036287 | COADREAD | SW1116 | Navitoclax |
| MVP_top60_DNA_seed | BRAF | 1.200588 | COADREAD | SNU-C1 | HG6-64-1 | MVP_top60_DNA+RNA_seed | MET | 1.03695 | COADREAD | SNU-C1 | Crizotinib |
| MVP_top60_DNA_seed | BRAF | 1.20171 | COADREAD | SW948 | HG6-64-1 | MVP_top60_DNA+RNA_seed | IGF1R | 1.037901 | COADREAD | LS-1034 | GSK1904529A |
| MVP_top60_DNA_seed | HSP90AA1 | 1.202078 | COADREAD | CL-11 | CCT-018159 | MVP_top60_DNA+RNA_seed | GSK3B | 1.038853 | COADREAD | SNU-C1 | CHIR-99021 |
| MVP_top60_DNA_seed | HSP90AB1 | 1.202078 | COADREAD | CL-11 | CCT-018159 | MVP_top60_DNA+RNA_seed | GSK3B | 1.039278 | COADREAD | SNU-C2B | SB216763 |
| MVP_top60_DNA_seed | ERBB4 | 1.205621 | COADREAD | SW1463 | AST-1306 | MVP_top60_DNA+RNA_seed | AR | 1.042827 | COADREAD | LS-411N | Bicalutamide |
| MVP_top60_DNA_seed | SRC | 1.205656 | COADREAD | CL-11 | WH-4-023 | MVP_top60_DNA+RNA_seed | MET | 1.04701 | COADREAD | HCT-15 | SU11274 |
| MVP_top60_DNA_seed | AR | 1.206208 | COADREAD | SW1116 | Bicalutamide | MVP_top60_DNA+RNA_seed | MET | 1.047465 | COADREAD | SW837 | AZD6094 |
| MVP_top60_DNA_seed | GSK3B | 1.20847 | COADREAD | SW1417 | SB216763 | MVP_top60_DNA+RNA_seed | CDK2 | 1.050294 | COADREAD | COLO-320-HSR | PHA-793887 |
| MVP_top60_DNA_seed | SRC | 1.210641 | COADREAD | CL-11 | Saracatinib | MVP_top60_DNA+RNA_seed | AR | 1.05138 | COADREAD | SNU-C1 | Bicalutamide |
| MVP_top60_DNA_seed | PIK3CD | 1.211399 | COADREAD | LS-513 | IC-87114 | MVP_top60_DNA+RNA_seed | HSP90AA1 | 1.05243 | COADREAD | NCI-H716 | Luminespib |
| MVP_top60_DNA_seed | PIK3CA | 1.211845 | COADREAD | C2BBe1 | AZD8835 | MVP_top60_DNA+RNA_seed | HSP90AB1 | 1.05243 | COADREAD | NCI-H716 | Luminespib |
| MVP_top60_DNA_seed | PIK3CD | 1.211845 | COADREAD | C2BBe1 | AZD8835 | MVP_top60_DNA+RNA_seed | MET | 1.052442 | COADREAD | LS-1034 | SU11274 |
| MVP_top60_DNA_seed | AR | 1.21234 | COADREAD | CW-2 | Bicalutamide | MVP_top60_DNA+RNA_seed | AKT1 | 1.053858 | COADREAD | CW-2 | MK-2206 |
| MVP_top60_DNA_seed | GSK3B | 1.214647 | COADREAD | NCI-H508 | CHIR-99021 | MVP_top60_DNA+RNA_seed | AR | 1.05546 | COADREAD | LS-1034 | AZD3514 |
| MVP_top60_DNA_seed | FYN | 1.216698 | COADREAD | CL-11 | A-770041 | MVP_top60_DNA+RNA_seed | MET | 1.057071 | COADREAD | SNU-81 | AZD6094 |
| MVP_top60_DNA_seed | CDK2 | 1.218529 | COADREAD | SW1463 | AZD5438 | MVP_top60_DNA+RNA_seed | PIK3CB | 1.057664 | COADREAD | HCT-15 | AZD6482 |
| MVP_top60_DNA_seed | AKT1 | 1.220721 | COADREAD | CCK-81 | AKT inhibitor VIII | MVP_top60_DNA+RNA_seed | MET | 1.057885 | COADREAD | NCI-H747 | PHA-665752 |
| MVP_top60_DNA_seed | PIK3CB | 1.222042 | COADREAD | CL-11 | TGX221 | MVP_top60_DNA+RNA_seed | GSK3B | 1.060105 | COADREAD | SNU-1040 | AZD7969 |
| MVP_top60_DNA_seed | BRAF | 1.222679 | COADREAD | CW-2 | SB590885 | MVP_top60_DNA+RNA_seed | ERBB4 | 1.060906 | COADREAD | CW-2 | PF-00299804 |
| MVP_top60_DNA_seed | MET | 1.223582 | COADREAD | HT55 | JNJ38877605 | MVP_top60_DNA+RNA_seed | SRC | 1.063157 | COADREAD | HCC2998 | Bosutinib |
| MVP_top60_DNA_seed | ATM | 1.224194 | COADREAD | SNU-C5 | CP466722 | MVP_top60_DNA+RNA_seed | SRC | 1.064476 | COADREAD | SK-CO-1 | Bosutinib |
| MVP_top60_DNA_seed | AR | 1.227698 | COADREAD | RCM-1 | Bicalutamide | MVP_top60_DNA+RNA_seed | BCL2 | 1.066542 | COADREAD | CCK-81 | Navitoclax |
| MVP_top60_DNA_seed | PIK3CA | 1.228973 | COADREAD | SW1417 | AZD8835 | MVP_top60_DNA+RNA_seed | PIK3CB | 1.066955 | COADREAD | LS-1034 | AZD8186 |
| MVP_top60_DNA_seed | PIK3CD | 1.228973 | COADREAD | SW1417 | AZD8835 | MVP_top60_DNA+RNA_seed | PIK3CD | 1.066955 | COADREAD | LS-1034 | AZD8186 |
| MVP_top60_DNA_seed | GSK3B | 1.230826 | COADREAD | SNU-407 | CHIR-99021 | MVP_top60_DNA+RNA_seed | AKT1 | 1.067371 | COADREAD | LS-1034 | BAY AKT1 |
| MVP_top60_DNA_seed | PIK3CB | 1.231771 | COADREAD | HCT-15 | AZD8186 | MVP_top60_DNA+RNA_seed | GSK3B | 1.067397 | COADREAD | C2BBe1 | SB216763 |
| MVP_top60_DNA_seed | PIK3CD | 1.231771 | COADREAD | HCT-15 | AZD8186 | MVP_top60_DNA+RNA_seed | ERBB3 | 1.069043 | COADREAD | SW1417 | AZD8931 |
| MVP_top60_DNA_seed | AKT1 | 1.231922 | COADREAD | CaR-1 | GSK690693 | MVP_top60_DNA+RNA_seed | HSP90AA1 | 1.070183 | COADREAD | CL-11 | Tanespimycin |
| MVP_top60_DNA_seed | ATM | 1.232944 | COADREAD | SNU-407 | KU-60019 | MVP_top60_DNA+RNA_seed | HSP90AB1 | 1.070183 | COADREAD | CL-11 | Tanespimycin |
| MVP_top60_DNA_seed | CDK2 | 1.235913 | COADREAD | HT-115 | PHA-793887 | MVP_top60_DNA+RNA_seed | SRC | 1.07317 | COADREAD | HT-115 | HG-6-71-01 |
| MVP_top60_DNA_seed | PIK3CD | 1.241548 | COADREAD | LS-123 | IC-87114 | MVP_top60_DNA+RNA_seed | HSP90AA1 | 1.074699 | COADREAD | SNU-C1 | SNX-2112 |
| MVP_top60_DNA_seed | ATM | 1.24555 | COADREAD | T84 | torin2 | MVP_top60_DNA+RNA_seed | HSP90AB1 | 1.074699 | COADREAD | SNU-C1 | SNX-2112 |
| MVP_top60_DNA_seed | ESR1 | 1.246299 | COADREAD | CL-11 | Tamoxifen | MVP_top60_DNA+RNA_seed | HSP90AA1 | 1.074842 | COADREAD | DiFi | CCT-018159 |
| MVP_top60_DNA_seed | ERBB4 | 1.251237 | COADREAD | HCT-15 | AST-1306 | MVP_top60_DNA+RNA_seed | HSP90AB1 | 1.074842 | COADREAD | DiFi | CCT-018159 |
| MVP_top60_DNA_seed | IGF1R | 1.251773 | COADREAD | RCM-1 | GSK1904529A | MVP_top60_DNA+RNA_seed | HSP90AA1 | 1.075325 | COADREAD | HT55 | SNX-2112 |

| | | | | | | | | | | | |
|---|---|---|---|---|---|---|---|---|---|---|---|
| MVP_top60_DNA_seed | AR | 1.257528 | COADREAD | RCM-1 | Bicalutamide | MVP_top60_DNA+RNA_seed | HSP90AB1 | 1.075325 | COADREAD | HT55 | SNX-2112 |
| MVP_top60_DNA_seed | HSP90AA1 | 1.258347 | COADREAD | LS-1034 | CCT-018159 | MVP_top60_DNA+RNA_seed | SRC | 1.075788 | COADREAD | NCI-H747 | Bosutinib |
| MVP_top60_DNA_seed | HSP90AB1 | 1.258347 | COADREAD | LS-1034 | CCT-018159 | MVP_top60_DNA+RNA_seed | AR | 1.078102 | COADREAD | SW1417 | Bicalutamide |
| MVP_top60_DNA_seed | SRC | 1.260519 | COADREAD | SNU-C1 | Bosutinib | MVP_top60_DNA+RNA_seed | ERBB3 | 1.078566 | COADREAD | KM12 | AZD8931 |
| MVP_top60_DNA_seed | CDK2 | 1.261709 | COADREAD | COLO-205 | AZD5438 | MVP_top60_DNA+RNA_seed | AR | 1.079543 | COADREAD | LS-1034 | Bicalutamide |
| MVP_top60_DNA_seed | GSK3B | 1.261775 | COADREAD | NCI-H716 | CHIR-99021 | MVP_top60_DNA+RNA_seed | MET | 1.084589 | COADREAD | SNU-1040 | AZD6094 |
| MVP_top60_DNA_seed | BRAF | 1.262356 | COADREAD | SW948 | Dabrafenib | MVP_top60_DNA+RNA_seed | GSK3B | 1.085503 | COADREAD | SW948 | AZD7969 |
| MVP_top60_DNA_seed | CREBBP | 1.264696 | COADREAD | RCM-1 | I-CBP112 | MVP_top60_DNA+RNA_seed | IGF1R | 1.086477 | COADREAD | SW837 | GSK1904529A |
| MVP_top60_DNA_seed | GSK3B | 1.266487 | COADREAD | CaR-1 | CHIR-99021 | MVP_top60_DNA+RNA_seed | MET | 1.091485 | COADREAD | COLO-678 | AZD6094 |
| MVP_top60_DNA_seed | PIK3CB | 1.268566 | COADREAD | SW1417 | AZD6482 | MVP_top60_DNA+RNA_seed | AR | 1.09262 | COADREAD | LS-411N | Bicalutamide |
| MVP_top60_DNA_seed | CDK2 | 1.269282 | COADREAD | LoVo | PHA-793887 | MVP_top60_DNA+RNA_seed | HSP90AA1 | 1.094997 | COADREAD | CaR-1 | Elesclomol |
| MVP_top60_DNA_seed | ESR1 | 1.270116 | COADREAD | KM12 | Fulvestrant | MVP_top60_DNA+RNA_seed | HSP90AB1 | 1.094997 | COADREAD | CaR-1 | Elesclomol |
| MVP_top60_DNA_seed | AKT1 | 1.274606 | COADREAD | CL-11 | GSK690693 | MVP_top60_DNA+RNA_seed | IGF1R | 1.09674 | COADREAD | HT-115 | BMS-754807 |
| MVP_top60_DNA_seed | MAPK1 | 1.276925 | COADREAD | SNU-C1 | VX-11e | MVP_top60_DNA+RNA_seed | BCL2 | 1.098496 | COADREAD | LS-513 | Navitoclax |
| MVP_top60_DNA_seed | BRAF | 1.280658 | COADREAD | LS-123 | SB590885 | MVP_top60_DNA+RNA_seed | PIK3CB | 1.104725 | COADREAD | LS-1034 | TGX221 |
| MVP_top60_DNA_seed | ATM | 1.280746 | COADREAD | SW948 | CP466722 | MVP_top60_DNA+RNA_seed | SRC | 1.105651 | COADREAD | LS-513 | WH-4-023 |
| MVP_top60_DNA_seed | CDK2 | 1.282022 | COADREAD | RCM-1 | PHA-793887 | MVP_top60_DNA+RNA_seed | AKT1 | 1.106423 | COADREAD | MDST8 | AKT inhibitor VIII |
| MVP_top60_DNA_seed | ATM | 1.283206 | COADREAD | T84 | KU-55933 | MVP_top60_DNA+RNA_seed | MET | 1.107667 | COADREAD | HT55 | SU11274 |
| MVP_top60_DNA_seed | AR | 1.283711 | COADREAD | C2BBe1 | Bicalutamide | MVP_top60_DNA+RNA_seed | CDK2 | 1.108322 | COADREAD | LS-411N | AZD5438 |
| MVP_top60_DNA_seed | IGF1R | 1.285927 | COADREAD | CaR-1 | Linsitinib | MVP_top60_DNA+RNA_seed | AR | 1.110041 | COADREAD | SNU-C5 | Bicalutamide |
| MVP_top60_DNA_seed | AKT1 | 1.287341 | COADREAD | LS-123 | AKT inhibitor VIII | MVP_top60_DNA+RNA_seed | BCL2 | 1.110782 | COADREAD | CW-2 | Navitoclax |
| MVP_top60_DNA_seed | MAPK14 | 1.28873 | COADREAD | LoVo | TAK-715 | MVP_top60_DNA+RNA_seed | AR | 1.111908 | COADREAD | HT-115 | AZD3514 |
| MVP_top60_DNA_seed | IGF1R | 1.28928 | COADREAD | RCM-1 | Linsitinib | MVP_top60_DNA+RNA_seed | ATM | 1.112672 | COADREAD | RCM-1 | torin2 |
| MVP_top60_DNA_seed | PIK3CB | 1.289383 | COADREAD | SNU-C1 | AZD6482 | MVP_top60_DNA+RNA_seed | IGF1R | 1.114452 | COADREAD | HCT-15 | Linsitinib |
| MVP_top60_DNA_seed | PIK3CA | 1.290498 | COADREAD | RCM-1 | PI3Ka_4409 | MVP_top60_DNA+RNA_seed | AKT1 | 1.11694 | COADREAD | SW837 | GSK690693 |
| MVP_top60_DNA_seed | PIK3CD | 1.290498 | COADREAD | RCM-1 | PI3Ka_4409 | MVP_top60_DNA+RNA_seed | PIK3CD | 1.118249 | COADREAD | SW837 | IC-87114 |
| MVP_top60_DNA_seed | AKT1 | 1.292188 | COADREAD | SW837 | AKT inhibitor VIII | MVP_top60_DNA+RNA_seed | PIK3CD | 1.118331 | COADREAD | LoVo | Idelalisib |
| MVP_top60_DNA_seed | BRAF | 1.294629 | COADREAD | LS-1034 | HG6-64-1 | MVP_top60_DNA+RNA_seed | IGF1R | 1.119059 | COADREAD | NCI-H716 | GSK1904529A |
| MVP_top60_DNA_seed | ESR1 | 1.295856 | COADREAD | SW1417 | Tamoxifen | MVP_top60_DNA+RNA_seed | SRC | 1.123856 | COADREAD | HT55 | Bosutinib |
| MVP_top60_DNA_seed | IGF1R | 1.296603 | COADREAD | SW1417 | Linsitinib | MVP_top60_DNA+RNA_seed | MET | 1.125191 | COADREAD | LS-123 | SU11274 |
| MVP_top60_DNA_seed | ATM | 1.300873 | COADREAD | COLO-205 | KU-60019 | MVP_top60_DNA+RNA_seed | BCL2 | 1.125586 | COADREAD | SW948 | Navitoclax |
| MVP_top60_DNA_seed | ERBB4 | 1.301085 | COADREAD | HCC2998 | AST-1306 | MVP_top60_DNA+RNA_seed | BCL2 | 1.126136 | COADREAD | CaR-1 | Navitoclax |
| MVP_top60_DNA_seed | ERBB4 | 1.301972 | COADREAD | SW620 | PF-00299804 | MVP_top60_DNA+RNA_seed | GSK3B | 1.127733 | COADREAD | HCT-15 | CHIR-99021 |
| MVP_top60_DNA_seed | BRAF | 1.303785 | COADREAD | NCI-H747 | PLX-4720 | MVP_top60_DNA+RNA_seed | AR | 1.128407 | COADREAD | NCI-H747 | Bicalutamide |
| MVP_top60_DNA_seed | MET | 1.303883 | COADREAD | RCM-1 | SU11274 | MVP_top60_DNA+RNA_seed | AKT1 | 1.129865 | COADREAD | CaR-1 | MK-2206 |
| MVP_top60_DNA_seed | ERBB4 | 1.305854 | COADREAD | COLO-320-HSR | CI-1033 | MVP_top60_DNA+RNA_seed | GSK3B | 1.129898 | COADREAD | SNU-81 | CHIR-99021 |
| MVP_top60_DNA_seed | SRC | 1.307016 | COADREAD | LS-180 | Bosutinib | MVP_top60_DNA+RNA_seed | AKT1 | 1.133795 | COADREAD | RCM-1 | GSK690693 |
| MVP_top60_DNA_seed | BRAF | 1.307196 | COADREAD | COLO-320-HSR | AZ628 | MVP_top60_DNA+RNA_seed | ERBB4 | 1.134712 | COADREAD | CaR-1 | CI-1033 |
| MVP_top60_DNA_seed | MET | 1.308327 | COADREAD | LS-123 | AZD6094 | MVP_top60_DNA+RNA_seed | AKT1 | 1.136258 | COADREAD | COLO-678 | GSK690693 |
| MVP_top60_DNA_seed | ESR1 | 1.30911 | COADREAD | SNU-C2B | Tamoxifen | MVP_top60_DNA+RNA_seed | IGF1R | 1.139989 | COADREAD | RCM-1 | BMS-754807 |
| MVP_top60_DNA_seed | MET | 1.31119 | COADREAD | SW1417 | SU11274 | MVP_top60_DNA+RNA_seed | PIK3CD | 1.140565 | COADREAD | HT55 | Idelalisib |

| | | | | | | | | | | | |
|---|---|---|---|---|---|---|---|---|---|---|---|
| MVP_top60_DNA_seed | MAPK1 | 1.315987 | COADREAD | RCM-1 | FR-180204 | MVP_top60_DNA+RNA_seed | MET | 1.14497 | COADREAD | SNU-C1 | SU11274 |
| MVP_top60_DNA_seed | PIK3CD | 1.317011 | COADREAD | SNU-61 | Idelalisib | MVP_top60_DNA+RNA_seed | AKT1 | 1.145261 | COADREAD | SW948 | MK-2206 |
| MVP_top60_DNA_seed | PIK3CB | 1.317713 | COADREAD | SW837 | AZD6482 | MVP_top60_DNA+RNA_seed | PIK3CD | 1.14605 | COADREAD | LS-180 | IC-87114 |
| MVP_top60_DNA_seed | HSP90AA1 | 1.317911 | COADREAD | COLO-205 | CCT-018159 | MVP_top60_DNA+RNA_seed | CDK2 | 1.147262 | COADREAD | NCI-H508 | AZD5438 |
| MVP_top60_DNA_seed | HSP90AB1 | 1.317911 | COADREAD | COLO-205 | CCT-018159 | MVP_top60_DNA+RNA_seed | SRC | 1.147311 | COADREAD | LS-1034 | Saracatinib |
| MVP_top60_DNA_seed | HSP90AA1 | 1.318282 | COADREAD | CaR-1 | CCT-018159 | MVP_top60_DNA+RNA_seed | SRC | 1.147866 | COADREAD | CW-2 | WH-4-023 |
| MVP_top60_DNA_seed | HSP90AB1 | 1.318282 | COADREAD | CaR-1 | CCT-018159 | MVP_top60_DNA+RNA_seed | ATM | 1.149447 | COADREAD | HT-29 | KU-55933 |
| MVP_top60_DNA_seed | CREBBP | 1.319187 | COADREAD | LS-180 | I-CBP112 | MVP_top60_DNA+RNA_seed | IGF1R | 1.150067 | COADREAD | KM12 | Linsitinib |
| MVP_top60_DNA_seed | CDK2 | 1.31929 | COADREAD | HCT-15 | PHA-793887 | MVP_top60_DNA+RNA_seed | PIK3CB | 1.150792 | COADREAD | HT-115 | AZD8186 |
| MVP_top60_DNA_seed | ESR1 | 1.321187 | COADREAD | LS-513 | Fulvestrant | MVP_top60_DNA+RNA_seed | PIK3CD | 1.150792 | COADREAD | HT-115 | AZD8186 |
| MVP_top60_DNA_seed | AKT1 | 1.321285 | COADREAD | COLO-678 | BAY AKT1 | MVP_top60_DNA+RNA_seed | PIK3CB | 1.158064 | COADREAD | NCI-H716 | TGX221 |
| MVP_top60_DNA_seed | GSK3B | 1.325969 | COADREAD | SW1417 | KIN001-042 | MVP_top60_DNA+RNA_seed | AR | 1.160064 | COADREAD | SNU-1040 | AZD3514 |
| MVP_top60_DNA_seed | AR | 1.329117 | COADREAD | LS-123 | Bicalutamide | MVP_top60_DNA+RNA_seed | SRC | 1.161135 | COADREAD | SW1417 | Bosutinib |
| MVP_top60_DNA_seed | ATM | 1.330659 | COADREAD | RCM-1 | CP466722 | MVP_top60_DNA+RNA_seed | PIK3CB | 1.162021 | COADREAD | SW620 | AZD6482 |
| MVP_top60_DNA_seed | SRC | 1.330812 | COADREAD | LS-1034 | Bosutinib | MVP_top60_DNA+RNA_seed | IGF1R | 1.162825 | COADREAD | CL-11 | Linsitinib |
| MVP_top60_DNA_seed | GSK3B | 1.331955 | COADREAD | COLO-678 | CHIR-99021 | MVP_top60_DNA+RNA_seed | MET | 1.163243 | COADREAD | SNU-C2B | PHA-665752 |
| MVP_top60_DNA_seed | GSK3B | 1.334623 | COADREAD | CaR-1 | CHIR-99021 | MVP_top60_DNA+RNA_seed | GSK3B | 1.16492 | COADREAD | RCM-1 | KIN001-042 |
| MVP_top60_DNA_seed | AKT1 | 1.335525 | COADREAD | CaR-1 | AKT inhibitor VIII | MVP_top60_DNA+RNA_seed | MET | 1.166586 | COADREAD | CaR-1 | AZD6094 |
| MVP_top60_DNA_seed | AR | 1.336737 | COADREAD | COLO-678 | Bicalutamide | MVP_top60_DNA+RNA_seed | SRC | 1.168962 | COADREAD | LS-411N | Saracatinib |
| MVP_top60_DNA_seed | PIK3CA | 1.336968 | COADREAD | LS-411N | AZD8835 | MVP_top60_DNA+RNA_seed | GSK3B | 1.170348 | COADREAD | HT55 | CHIR-99021 |
| MVP_top60_DNA_seed | PIK3CD | 1.336968 | COADREAD | LS-411N | AZD8835 | MVP_top60_DNA+RNA_seed | ATM | 1.170588 | COADREAD | SW1417 | KU-55933 |
| MVP_top60_DNA_seed | HSP90AA1 | 1.337706 | COADREAD | SW837 | CCT-018159 | MVP_top60_DNA+RNA_seed | IGF1R | 1.170919 | COADREAD | SW948 | Linsitinib |
| MVP_top60_DNA_seed | HSP90AB1 | 1.337706 | COADREAD | SW837 | CCT-018159 | MVP_top60_DNA+RNA_seed | GSK3B | 1.172102 | COADREAD | CL-11 | CHIR-99021 |
| MVP_top60_DNA_seed | CDK2 | 1.338841 | COADREAD | T84 | PHA-793887 | MVP_top60_DNA+RNA_seed | HSP90AA1 | 1.173232 | COADREAD | SW1463 | SNX-2112 |
| MVP_top60_DNA_seed | ERBB4 | 1.339891 | COADREAD | CW-2 | AST-1306 | MVP_top60_DNA+RNA_seed | HSP90AB1 | 1.173232 | COADREAD | SW1463 | SNX-2112 |
| MVP_top60_DNA_seed | MAPK1 | 1.342536 | COADREAD | COLO-678 | FR-180204 | MVP_top60_DNA+RNA_seed | AKT1 | 1.174656 | COADREAD | MDST8 | A-443654 |
| MVP_top60_DNA_seed | GSK3B | 1.34414 | COADREAD | SW837 | KIN001-042 | MVP_top60_DNA+RNA_seed | CDK2 | 1.177889 | COADREAD | SNU-C1 | Seliciclib |
| MVP_top60_DNA_seed | PIK3CD | 1.346651 | COADREAD | CL-11 | Idelalisib | MVP_top60_DNA+RNA_seed | ERBB4 | 1.179365 | COADREAD | KM12 | AST-1306 |
| MVP_top60_DNA_seed | HSP90AA1 | 1.346959 | COADREAD | T84 | Tanespimycin | MVP_top60_DNA+RNA_seed | PIK3CB | 1.180194 | COADREAD | SNU-C2B | AZD6482 |
| MVP_top60_DNA_seed | HSP90AB1 | 1.346959 | COADREAD | T84 | Tanespimycin | MVP_top60_DNA+RNA_seed | AR | 1.180664 | COADREAD | COLO-678 | Bicalutamide |
| MVP_top60_DNA_seed | SRC | 1.348071 | COADREAD | SW1463 | NVP-BHG712 | MVP_top60_DNA+RNA_seed | BCL2 | 1.184081 | COADREAD | HT55 | Navitoclax |
| MVP_top60_DNA_seed | AR | 1.348203 | COADREAD | SW620 | AZD3514 | MVP_top60_DNA+RNA_seed | SRC | 1.185125 | COADREAD | SW1116 | Bosutinib |
| MVP_top60_DNA_seed | ESR1 | 1.348233 | COADREAD | NCI-H716 | Fulvestrant | MVP_top60_DNA+RNA_seed | PIK3CD | 1.185505 | COADREAD | SW948 | Idelalisib |
| MVP_top60_DNA_seed | AKT1 | 1.348357 | COADREAD | LS-513 | A-443654 | MVP_top60_DNA+RNA_seed | FYN | 1.186654 | COADREAD | NCI-H716 | A-770041 |
| MVP_top60_DNA_seed | SRC | 1.349841 | COADREAD | CL-11 | Bosutinib | MVP_top60_DNA+RNA_seed | PIK3CB | 1.189332 | COADREAD | SNU-407 | AZD8186 |
| MVP_top60_DNA_seed | PIK3CB | 1.351541 | COADREAD | HCC2998 | AZD6482 | MVP_top60_DNA+RNA_seed | PIK3CD | 1.189332 | COADREAD | SNU-407 | AZD8186 |
| MVP_top60_DNA_seed | ERBB4 | 1.352313 | COADREAD | T84 | CI-1033 | MVP_top60_DNA+RNA_seed | BCL2 | 1.189475 | COADREAD | LS-1034 | Navitoclax |
| MVP_top60_DNA_seed | GSK3B | 1.352354 | COADREAD | SW837 | AZD7969 | MVP_top60_DNA+RNA_seed | PTK2 | 1.190048 | COADREAD | HT55 | PF-562271 |
| MVP_top60_DNA_seed | ESR1 | 1.35607 | COADREAD | COLO-678 | Tamoxifen | MVP_top60_DNA+RNA_seed | PIK3CB | 1.191466 | COADREAD | HT55 | AZD6482 |
| MVP_top60_DNA_seed | AR | 1.359402 | COADREAD | SK-CO-1 | AZD3514 | MVP_top60_DNA+RNA_seed | ATM | 1.192943 | COADREAD | COLO-678 | KU-60019 |
| MVP_top60_DNA_seed | ATM | 1.362202 | COADREAD | SNU-C1 | KU-60019 | MVP_top60_DNA+RNA_seed | ERBB4 | 1.19357 | COADREAD | NCI-H716 | CI-1033 |

| | | | | | | | | | | | |
|---|---|---|---|---|---|---|---|---|---|---|---|
| MVP_top60_DNA_seed | CDK2 | 1.363823 | COADREAD | HT55 | PHA-793887 | MVP_top60_DNA+RNA_seed | PIK3CB | 1.195353 | COADREAD | CCK-81 | AZD8186 |
| MVP_top60_DNA_seed | CDK2 | 1.364032 | COADREAD | SW1417 | PHA-793887 | MVP_top60_DNA+RNA_seed | PIK3CD | 1.195353 | COADREAD | CCK-81 | AZD8186 |
| MVP_top60_DNA_seed | AKT1 | 1.366387 | COADREAD | RCM-1 | BAY AKT1 | MVP_top60_DNA+RNA_seed | MET | 1.196749 | COADREAD | CL-11 | PHA-665752 |
| MVP_top60_DNA_seed | ESR1 | 1.367209 | COADREAD | SNU-81 | Tamoxifen | MVP_top60_DNA+RNA_seed | ERBB4 | 1.198507 | COADREAD | KM12 | CI-1033 |
| MVP_top60_DNA_seed | CREBBP | 1.367367 | COADREAD | COLO-678 | I-CBP112 | MVP_top60_DNA+RNA_seed | ATM | 1.199291 | COADREAD | CL-11 | CP466722 |
| MVP_top60_DNA_seed | SRC | 1.369027 | COADREAD | T84 | Bosutinib | MVP_top60_DNA+RNA_seed | HSP90AA1 | 1.202078 | COADREAD | CL-11 | CCT-018159 |
| MVP_top60_DNA_seed | AKT1 | 1.370853 | COADREAD | SW620 | MK-2206 | MVP_top60_DNA+RNA_seed | HSP90AB1 | 1.202078 | COADREAD | CL-11 | CCT-018159 |
| MVP_top60_DNA_seed | PIK3CA | 1.377467 | COADREAD | CW-2 | PI3Ka_4409 | MVP_top60_DNA+RNA_seed | ERBB4 | 1.205621 | COADREAD | SW1463 | AST-1306 |
| MVP_top60_DNA_seed | PIK3CD | 1.377467 | COADREAD | CW-2 | PI3Ka_4409 | MVP_top60_DNA+RNA_seed | SRC | 1.205656 | COADREAD | CL-11 | WH-4-023 |
| MVP_top60_DNA_seed | ERBB4 | 1.378075 | COADREAD | SW620 | AST-1306 | MVP_top60_DNA+RNA_seed | BCL2 | 1.205714 | COADREAD | HT55 | Obatoclax Mesylate |
| MVP_top60_DNA_seed | GSK3B | 1.381005 | COADREAD | SNU-81 | KIN001-042 | MVP_top60_DNA+RNA_seed | AR | 1.206208 | COADREAD | SW1116 | Bicalutamide |
| MVP_top60_DNA_seed | AR | 1.382947 | COADREAD | DiFi | Bicalutamide | MVP_top60_DNA+RNA_seed | GSK3B | 1.20847 | COADREAD | SW1417 | SB216763 |
| MVP_top60_DNA_seed | MAPK1 | 1.383129 | COADREAD | CL-11 | FR-180204 | MVP_top60_DNA+RNA_seed | SRC | 1.210641 | COADREAD | CL-11 | Saracatinib |
| MVP_top60_DNA_seed | ERBB4 | 1.383224 | COADREAD | RCM-1 | CI-1033 | MVP_top60_DNA+RNA_seed | PIK3CD | 1.211399 | COADREAD | LS-513 | IC-87114 |
| MVP_top60_DNA_seed | IGF1R | 1.383854 | COADREAD | SW1417 | GSK1904529A | MVP_top60_DNA+RNA_seed | PIK3CA | 1.211845 | COADREAD | C2BBe1 | AZD8835 |
| MVP_top60_DNA_seed | BRAF | 1.384328 | COADREAD | LS-513 | PLX-4720 | MVP_top60_DNA+RNA_seed | PIK3CD | 1.211845 | COADREAD | C2BBe1 | AZD8835 |
| MVP_top60_DNA_seed | PIK3CD | 1.3844 | COADREAD | CL-11 | IC-87114 | MVP_top60_DNA+RNA_seed | AR | 1.21234 | COADREAD | CW-2 | Bicalutamide |
| MVP_top60_DNA_seed | ATM | 1.385027 | COADREAD | NCI-H747 | KU-55933 | MVP_top60_DNA+RNA_seed | GSK3B | 1.214647 | COADREAD | NCI-H508 | CHIR-99021 |
| MVP_top60_DNA_seed | ERBB4 | 1.385506 | COADREAD | T84 | AST-1306 | MVP_top60_DNA+RNA_seed | FYN | 1.216698 | COADREAD | CL-11 | A-770041 |
| MVP_top60_DNA_seed | PIK3CB | 1.388789 | COADREAD | RCM-1 | AZD8186 | MVP_top60_DNA+RNA_seed | CDK2 | 1.218529 | COADREAD | SW1463 | AZD5438 |
| MVP_top60_DNA_seed | PIK3CD | 1.388789 | COADREAD | RCM-1 | AZD8186 | MVP_top60_DNA+RNA_seed | AKT1 | 1.220721 | COADREAD | CCK-81 | AKT inhibitor VIII |
| MVP_top60_DNA_seed | ATM | 1.389542 | COADREAD | SW1116 | CP466722 | MVP_top60_DNA+RNA_seed | PIK3CB | 1.222042 | COADREAD | CL-11 | TGX221 |
| MVP_top60_DNA_seed | HSP90AA1 | 1.393507 | COADREAD | HT55 | Elesclomol | MVP_top60_DNA+RNA_seed | MET | 1.223582 | COADREAD | HT55 | JNJ38877605 |
| MVP_top60_DNA_seed | HSP90AB1 | 1.393507 | COADREAD | HT55 | Elesclomol | MVP_top60_DNA+RNA_seed | ATM | 1.224194 | COADREAD | SNU-C5 | CP466722 |
| MVP_top60_DNA_seed | HSP90AA1 | 1.393924 | COADREAD | GP5d | CCT-018159 | MVP_top60_DNA+RNA_seed | AR | 1.227698 | COADREAD | RCM-1 | Bicalutamide |
| MVP_top60_DNA_seed | HSP90AB1 | 1.393924 | COADREAD | GP5d | CCT-018159 | MVP_top60_DNA+RNA_seed | PIK3CA | 1.228973 | COADREAD | SW1417 | AZD8835 |
| MVP_top60_DNA_seed | HSP90AA1 | 1.394065 | COADREAD | SNU-81 | Elesclomol | MVP_top60_DNA+RNA_seed | PIK3CD | 1.228973 | COADREAD | SW1417 | AZD8835 |
| MVP_top60_DNA_seed | HSP90AB1 | 1.394065 | COADREAD | SNU-81 | Elesclomol | MVP_top60_DNA+RNA_seed | GSK3B | 1.230826 | COADREAD | SNU-407 | CHIR-99021 |
| MVP_top60_DNA_seed | BRAF | 1.394983 | COADREAD | CaR-1 | PLX-4720 | MVP_top60_DNA+RNA_seed | BCL2 | 1.231362 | COADREAD | HCT-15 | Navitoclax |
| MVP_top60_DNA_seed | AR | 1.395052 | COADREAD | SW1463 | Bicalutamide | MVP_top60_DNA+RNA_seed | PIK3CB | 1.231771 | COADREAD | HCT-15 | AZD8186 |
| MVP_top60_DNA_seed | MAPK14 | 1.397484 | COADREAD | SNU-C1 | TAK-715 | MVP_top60_DNA+RNA_seed | PIK3CD | 1.231771 | COADREAD | HCT-15 | AZD8186 |
| MVP_top60_DNA_seed | HSP90AA1 | 1.397948 | COADREAD | SW1417 | Elesclomol | MVP_top60_DNA+RNA_seed | AKT1 | 1.231922 | COADREAD | CaR-1 | GSK690693 |
| MVP_top60_DNA_seed | HSP90AB1 | 1.397948 | COADREAD | SW1417 | Elesclomol | MVP_top60_DNA+RNA_seed | ATM | 1.232944 | COADREAD | SNU-407 | KU-60019 |
| MVP_top60_DNA_seed | ATM | 1.398281 | COADREAD | CW-2 | torin2 | MVP_top60_DNA+RNA_seed | BCL2 | 1.233034 | COADREAD | SNU-61 | Navitoclax |
| MVP_top60_DNA_seed | PTK2 | 1.398282 | COADREAD | CL-11 | PF-562271 | MVP_top60_DNA+RNA_seed | CDK2 | 1.235913 | COADREAD | HT-115 | PHA-793887 |
| MVP_top60_DNA_seed | ATM | 1.398285 | COADREAD | SNU-C1 | CP466722 | MVP_top60_DNA+RNA_seed | PIK3CD | 1.241548 | COADREAD | LS-123 | IC-87114 |
| MVP_top60_DNA_seed | PIK3CD | 1.402634 | COADREAD | SNU-C1 | Idelalisib | MVP_top60_DNA+RNA_seed | ATM | 1.24555 | COADREAD | T84 | torin2 |
| MVP_top60_DNA_seed | IGF1R | 1.402843 | COADREAD | CL-11 | BMS-754807 | MVP_top60_DNA+RNA_seed | ERBB4 | 1.251237 | COADREAD | HCT-15 | AST-1306 |
| MVP_top60_DNA_seed | GSK3B | 1.404924 | COADREAD | C2BBe1 | CHIR-99021 | MVP_top60_DNA+RNA_seed | IGF1R | 1.251773 | COADREAD | RCM-1 | GSK1904529A |
| MVP_top60_DNA_seed | IGF1R | 1.40573 | COADREAD | CL-11 | GSK1904529A | MVP_top60_DNA+RNA_seed | AR | 1.257528 | COADREAD | RCM-1 | Bicalutamide |
| MVP_top60_DNA_seed | MET | 1.40775 | COADREAD | SW948 | AZD6094 | MVP_top60_DNA+RNA_seed | HSP90AA1 | 1.258347 | COADREAD | LS-1034 | CCT-018159 |

| | | | | | | | | | | | |
|---|---|---|---|---|---|---|---|---|---|---|---|
| MVP_top60_DNA_seed | AR | 1.40974 | COADREAD | SW948 | Bicalutamide | MVP_top60_DNA+RNA_seed | HSP90AB1 | 1.258347 | COADREAD | LS-1034 | CCT-018159 |
| MVP_top60_DNA_seed | AKT1 | 1.411196 | COADREAD | SNU-C1 | A-443654 | MVP_top60_DNA+RNA_seed | SRC | 1.260519 | COADREAD | SNU-C1 | Bosutinib |
| MVP_top60_DNA_seed | BRAF | 1.412692 | COADREAD | SNU-61 | SB590885 | MVP_top60_DNA+RNA_seed | CDK2 | 1.261709 | COADREAD | COLO-205 | AZD5438 |
| MVP_top60_DNA_seed | HSP90AA1 | 1.414203 | COADREAD | CCK-81 | Elesclomol | MVP_top60_DNA+RNA_seed | GSK3B | 1.261775 | COADREAD | NCI-H716 | CHIR-99021 |
| MVP_top60_DNA_seed | HSP90AB1 | 1.414203 | COADREAD | CCK-81 | Elesclomol | MVP_top60_DNA+RNA_seed | GSK3B | 1.266487 | COADREAD | CaR-1 | CHIR-99021 |
| MVP_top60_DNA_seed | ERBB4 | 1.414935 | COADREAD | T84 | PF-00299804 | MVP_top60_DNA+RNA_seed | PIK3CB | 1.268566 | COADREAD | SW1417 | AZD6482 |
| MVP_top60_DNA_seed | ATM | 1.416189 | COADREAD | CW-2 | KU-55933 | MVP_top60_DNA+RNA_seed | BCL2 | 1.268616 | COADREAD | LS-123 | TW 37 |
| MVP_top60_DNA_seed | ATM | 1.417218 | COADREAD | HT55 | KU-55933 | MVP_top60_DNA+RNA_seed | CDK2 | 1.269282 | COADREAD | LoVo | PHA-793887 |
| MVP_top60_DNA_seed | AKT1 | 1.417436 | COADREAD | RCM-1 | AKT inhibitor VIII | MVP_top60_DNA+RNA_seed | AKT1 | 1.274606 | COADREAD | CL-11 | GSK690693 |
| MVP_top60_DNA_seed | GSK3B | 1.419708 | COADREAD | CW-2 | CHIR-99021 | MVP_top60_DNA+RNA_seed | ATM | 1.280746 | COADREAD | SW948 | CP466722 |
| MVP_top60_DNA_seed | PTK2 | 1.422965 | COADREAD | T84 | PF-562271 | MVP_top60_DNA+RNA_seed | CDK2 | 1.282022 | COADREAD | RCM-1 | PHA-793887 |
| MVP_top60_DNA_seed | BRAF | 1.423385 | COADREAD | SNU-C1 | AZ628 | MVP_top60_DNA+RNA_seed | ATM | 1.283206 | COADREAD | T84 | KU-55933 |
| MVP_top60_DNA_seed | HSP90AA1 | 1.42473 | COADREAD | CL-11 | SNX-2112 | MVP_top60_DNA+RNA_seed | AR | 1.283711 | COADREAD | C2BBe1 | Bicalutamide |
| MVP_top60_DNA_seed | HSP90AB1 | 1.42473 | COADREAD | CL-11 | SNX-2112 | MVP_top60_DNA+RNA_seed | IGF1R | 1.285927 | COADREAD | CaR-1 | Linsitinib |
| MVP_top60_DNA_seed | PIK3CB | 1.428655 | COADREAD | T84 | AZD6482 | MVP_top60_DNA+RNA_seed | AKT1 | 1.287341 | COADREAD | LS-123 | AKT inhibitor VIII |
| MVP_top60_DNA_seed | BRAF | 1.43449 | COADREAD | SNU-1040 | RAF_9304 | MVP_top60_DNA+RNA_seed | IGF1R | 1.28928 | COADREAD | RCM-1 | Linsitinib |
| MVP_top60_DNA_seed | BRAF | 1.435047 | COADREAD | SW948 | PLX-4720 | MVP_top60_DNA+RNA_seed | PIK3CB | 1.289383 | COADREAD | SNU-C1 | AZD6482 |
| MVP_top60_DNA_seed | GSK3B | 1.437436 | COADREAD | C2BBe1 | AZD7969 | MVP_top60_DNA+RNA_seed | PIK3CA | 1.290498 | COADREAD | RCM-1 | PI3Ka_4409 |
| MVP_top60_DNA_seed | GSK3B | 1.440799 | COADREAD | LS-123 | CHIR-99021 | MVP_top60_DNA+RNA_seed | PIK3CD | 1.290498 | COADREAD | RCM-1 | PI3Ka_4409 |
| MVP_top60_DNA_seed | ERBB4 | 1.44087 | COADREAD | SW948 | PF-00299804 | MVP_top60_DNA+RNA_seed | BCL2 | 1.290881 | COADREAD | COLO-678 | Obatoclax Mesylate |
| MVP_top60_DNA_seed | PIK3CB | 1.443038 | COADREAD | CW-2 | TGX221 | MVP_top60_DNA+RNA_seed | AKT1 | 1.292188 | COADREAD | SW837 | AKT inhibitor VIII |
| MVP_top60_DNA_seed | PIK3CB | 1.443899 | COADREAD | LS-513 | AZD6482 | MVP_top60_DNA+RNA_seed | IGF1R | 1.296603 | COADREAD | SW1417 | Linsitinib |
| MVP_top60_DNA_seed | GSK3B | 1.446459 | COADREAD | SNU-61 | KIN001-042 | MVP_top60_DNA+RNA_seed | ATM | 1.300873 | COADREAD | COLO-205 | KU-60019 |
| MVP_top60_DNA_seed | AR | 1.450002 | COADREAD | HT-29 | Bicalutamide | MVP_top60_DNA+RNA_seed | ERBB4 | 1.301085 | COADREAD | HCC2998 | AST-1306 |
| MVP_top60_DNA_seed | ATM | 1.45201 | COADREAD | SW620 | KU-55933 | MVP_top60_DNA+RNA_seed | ERBB4 | 1.301972 | COADREAD | SW620 | PF-00299804 |
| MVP_top60_DNA_seed | HSP90AA1 | 1.457103 | COADREAD | COLO-320-HSR | Tanespimycin | MVP_top60_DNA+RNA_seed | MET | 1.303883 | COADREAD | RCM-1 | SU11274 |
| MVP_top60_DNA_seed | HSP90AB1 | 1.457103 | COADREAD | COLO-320-HSR | Tanespimycin | MVP_top60_DNA+RNA_seed | ERBB4 | 1.305854 | COADREAD | COLO-320-HSR | CI-1033 |
| MVP_top60_DNA_seed | ERBB3 | 1.458012 | COADREAD | SNU-61 | AZD8931 | MVP_top60_DNA+RNA_seed | SRC | 1.307016 | COADREAD | LS-180 | Bosutinib |
| MVP_top60_DNA_seed | PTK2 | 1.460327 | COADREAD | SW1463 | PF-562271 | MVP_top60_DNA+RNA_seed | MET | 1.308327 | COADREAD | LS-123 | AZD6094 |
| MVP_top60_DNA_seed | MET | 1.463742 | COADREAD | CW-2 | Crizotinib | MVP_top60_DNA+RNA_seed | MET | 1.31119 | COADREAD | SW1417 | SU11274 |
| MVP_top60_DNA_seed | PTK2 | 1.466061 | COADREAD | SW1417 | PF-562271 | MVP_top60_DNA+RNA_seed | PIK3CD | 1.317011 | COADREAD | SNU-61 | Idelalisib |
| MVP_top60_DNA_seed | BRAF | 1.466158 | COADREAD | COLO-678 | PLX-4720 | MVP_top60_DNA+RNA_seed | PIK3CB | 1.317713 | COADREAD | SW837 | AZD6482 |
| MVP_top60_DNA_seed | MET | 1.477444 | COADREAD | SNU-C1 | PHA-665752 | MVP_top60_DNA+RNA_seed | HSP90AA1 | 1.317911 | COADREAD | COLO-205 | CCT-018159 |
| MVP_top60_DNA_seed | AKT1 | 1.479086 | COADREAD | C2BBe1 | MK-2206 | MVP_top60_DNA+RNA_seed | HSP90AB1 | 1.317911 | COADREAD | COLO-205 | CCT-018159 |
| MVP_top60_DNA_seed | GSK3B | 1.482156 | COADREAD | SW948 | SB216763 | MVP_top60_DNA+RNA_seed | HSP90AA1 | 1.318282 | COADREAD | CaR-1 | CCT-018159 |
| MVP_top60_DNA_seed | HSP90AA1 | 1.484549 | COADREAD | KM12 | Elesclomol | MVP_top60_DNA+RNA_seed | HSP90AB1 | 1.318282 | COADREAD | CaR-1 | CCT-018159 |
| MVP_top60_DNA_seed | HSP90AB1 | 1.484549 | COADREAD | KM12 | Elesclomol | MVP_top60_DNA+RNA_seed | CDK2 | 1.31929 | COADREAD | HCT-15 | PHA-793887 |
| MVP_top60_DNA_seed | ERBB4 | 1.484886 | COADREAD | SNU-C1 | AST-1306 | MVP_top60_DNA+RNA_seed | AKT1 | 1.321285 | COADREAD | COLO-678 | BAY AKT1 |
| MVP_top60_DNA_seed | MET | 1.494176 | COADREAD | HCT-15 | JNJ38877605 | MVP_top60_DNA+RNA_seed | GSK3B | 1.325969 | COADREAD | SW1417 | KIN001-042 |
| MVP_top60_DNA_seed | ERBB3 | 1.496494 | COADREAD | COLO-205 | AZD8931 | MVP_top60_DNA+RNA_seed | BCL2 | 1.326186 | COADREAD | LS-411N | Navitoclax |
| MVP_top60_DNA_seed | MET | 1.498316 | COADREAD | CL-11 | JNJ38877605 | MVP_top60_DNA+RNA_seed | AR | 1.329117 | COADREAD | LS-123 | Bicalutamide |

| | | | | | | | | | | | |
|---|---|---|---|---|---|---|---|---|---|---|---|
| MVP_top60_DNA_seed | SRC | 1.499481 | COADREAD | CW-2 | Saracatinib | MVP_top60_DNA+RNA_seed | ATM | 1.330659 | COADREAD | RCM-1 | CP466722 |
| MVP_top60_DNA_seed | AKT1 | 1.500991 | COADREAD | SW1417 | BAY AKT1 | MVP_top60_DNA+RNA_seed | SRC | 1.330812 | COADREAD | LS-1034 | Bosutinib |
| MVP_top60_DNA_seed | SRC | 1.501389 | COADREAD | CW-2 | HG-6-71-01 | MVP_top60_DNA+RNA_seed | GSK3B | 1.331955 | COADREAD | COLO-678 | CHIR-99021 |
| MVP_top60_DNA_seed | ATM | 1.501923 | COADREAD | HT55 | KU-60019 | MVP_top60_DNA+RNA_seed | GSK3B | 1.334623 | COADREAD | CaR-1 | CHIR-99021 |
| MVP_top60_DNA_seed | ATM | 1.50684 | COADREAD | RCM-1 | KU-60019 | MVP_top60_DNA+RNA_seed | AKT1 | 1.335525 | COADREAD | CaR-1 | AKT inhibitor VIII |
| MVP_top60_DNA_seed | PIK3CD | 1.508703 | COADREAD | COLO-678 | IC-87114 | MVP_top60_DNA+RNA_seed | AR | 1.336737 | COADREAD | COLO-678 | Bicalutamide |
| MVP_top60_DNA_seed | PIK3CB | 1.510995 | COADREAD | SW620 | AZD8186 | MVP_top60_DNA+RNA_seed | PIK3CA | 1.336968 | COADREAD | LS-411N | AZD8835 |
| MVP_top60_DNA_seed | PIK3CD | 1.510995 | COADREAD | SW620 | AZD8186 | MVP_top60_DNA+RNA_seed | PIK3CD | 1.336968 | COADREAD | LS-411N | AZD8835 |
| MVP_top60_DNA_seed | AKT1 | 1.514339 | COADREAD | SW1116 | GSK690693 | MVP_top60_DNA+RNA_seed | HSP90AA1 | 1.337706 | COADREAD | SW837 | CCT-018159 |
| MVP_top60_DNA_seed | PIK3CB | 1.514443 | COADREAD | KM12 | AZD8186 | MVP_top60_DNA+RNA_seed | HSP90AB1 | 1.337706 | COADREAD | SW837 | CCT-018159 |
| MVP_top60_DNA_seed | PIK3CD | 1.514443 | COADREAD | KM12 | AZD8186 | MVP_top60_DNA+RNA_seed | CDK2 | 1.338841 | COADREAD | T84 | PHA-793887 |
| MVP_top60_DNA_seed | IGF1R | 1.519006 | COADREAD | CW-2 | BMS-536924 | MVP_top60_DNA+RNA_seed | ERBB4 | 1.339891 | COADREAD | CW-2 | AST-1306 |
| MVP_top60_DNA_seed | IGF1R | 1.519259 | COADREAD | SNU-81 | BMS-536924 | MVP_top60_DNA+RNA_seed | GSK3B | 1.34414 | COADREAD | SW837 | KIN001-042 |
| MVP_top60_DNA_seed | CREBBP | 1.521094 | COADREAD | SNU-C1 | I-CBP112 | MVP_top60_DNA+RNA_seed | PIK3CD | 1.346651 | COADREAD | CL-11 | Idelalisib |
| MVP_top60_DNA_seed | BRAF | 1.525145 | COADREAD | SW837 | HG6-64-1 | MVP_top60_DNA+RNA_seed | HSP90AA1 | 1.346959 | COADREAD | T84 | Tanespimycin |
| MVP_top60_DNA_seed | MET | 1.525519 | COADREAD | LS-123 | Crizotinib | MVP_top60_DNA+RNA_seed | HSP90AB1 | 1.346959 | COADREAD | T84 | Tanespimycin |
| MVP_top60_DNA_seed | MET | 1.526215 | COADREAD | SW1116 | AZD6094 | MVP_top60_DNA+RNA_seed | SRC | 1.348071 | COADREAD | SW1463 | NVP-BHG712 |
| MVP_top60_DNA_seed | GSK3B | 1.527269 | COADREAD | CaR-1 | AZD7969 | MVP_top60_DNA+RNA_seed | AR | 1.348203 | COADREAD | SW620 | AZD3514 |
| MVP_top60_DNA_seed | MET | 1.528584 | COADREAD | LS-411N | JNJ38877605 | MVP_top60_DNA+RNA_seed | AKT1 | 1.348357 | COADREAD | LS-513 | A-443654 |
| MVP_top60_DNA_seed | BRAF | 1.531545 | COADREAD | NCI-H747 | PLX-4720 | MVP_top60_DNA+RNA_seed | SRC | 1.349841 | COADREAD | CL-11 | Bosutinib |
| MVP_top60_DNA_seed | AR | 1.533553 | COADREAD | CaR-1 | Bicalutamide | MVP_top60_DNA+RNA_seed | PIK3CB | 1.351541 | COADREAD | HCC2998 | AZD6482 |
| MVP_top60_DNA_seed | SRC | 1.535019 | COADREAD | SNU-81 | NVP-BHG712 | MVP_top60_DNA+RNA_seed | ERBB4 | 1.352313 | COADREAD | T84 | CI-1033 |
| MVP_top60_DNA_seed | ERBB4 | 1.536211 | COADREAD | SW1463 | CI-1033 | MVP_top60_DNA+RNA_seed | GSK3B | 1.352354 | COADREAD | SW837 | AZD7969 |
| MVP_top60_DNA_seed | IGF1R | 1.536502 | COADREAD | CW-2 | Linsitinib | MVP_top60_DNA+RNA_seed | AR | 1.359402 | COADREAD | SK-CO-1 | AZD3514 |
| MVP_top60_DNA_seed | AKT1 | 1.537519 | COADREAD | LS-1034 | MK-2206 | MVP_top60_DNA+RNA_seed | ATM | 1.362202 | COADREAD | SNU-C1 | KU-60019 |
| MVP_top60_DNA_seed | BRAF | 1.53931 | COADREAD | CL-11 | HG6-64-1 | MVP_top60_DNA+RNA_seed | CDK2 | 1.363823 | COADREAD | HT55 | PHA-793887 |
| MVP_top60_DNA_seed | SRC | 1.539479 | COADREAD | SW948 | Bosutinib | MVP_top60_DNA+RNA_seed | CDK2 | 1.364032 | COADREAD | SW1417 | PHA-793887 |
| MVP_top60_DNA_seed | ESR1 | 1.540995 | COADREAD | C2BBe1 | Fulvestrant | MVP_top60_DNA+RNA_seed | AKT1 | 1.366387 | COADREAD | RCM-1 | BAY AKT1 |
| MVP_top60_DNA_seed | AKT1 | 1.542465 | COADREAD | C2BBe1 | A-443654 | MVP_top60_DNA+RNA_seed | SRC | 1.369027 | COADREAD | T84 | Bosutinib |
| MVP_top60_DNA_seed | PIK3CB | 1.543077 | COADREAD | LS-1034 | AZD6482 | MVP_top60_DNA+RNA_seed | AKT1 | 1.370853 | COADREAD | SW620 | MK-2206 |
| MVP_top60_DNA_seed | GSK3B | 1.546208 | COADREAD | SNU-1040 | CHIR-99021 | MVP_top60_DNA+RNA_seed | PIK3CA | 1.377467 | COADREAD | CW-2 | PI3Ka_4409 |
| MVP_top60_DNA_seed | ESR1 | 1.547258 | COADREAD | SW1417 | Fulvestrant | MVP_top60_DNA+RNA_seed | PIK3CD | 1.377467 | COADREAD | CW-2 | PI3Ka_4409 |
| MVP_top60_DNA_seed | MAPK1 | 1.548434 | COADREAD | CaR-1 | VX-11e | MVP_top60_DNA+RNA_seed | ERBB4 | 1.378075 | COADREAD | SW620 | AST-1306 |
| MVP_top60_DNA_seed | PIK3CD | 1.548541 | COADREAD | SW1116 | IC-87114 | MVP_top60_DNA+RNA_seed | GSK3B | 1.381005 | COADREAD | SNU-81 | KIN001-042 |
| MVP_top60_DNA_seed | PIK3CD | 1.550271 | COADREAD | RCM-1 | Idelalisib | MVP_top60_DNA+RNA_seed | BCL2 | 1.381457 | COADREAD | SNU-81 | Navitoclax |
| MVP_top60_DNA_seed | ESR1 | 1.553715 | COADREAD | CW-2 | Tamoxifen | MVP_top60_DNA+RNA_seed | AR | 1.382947 | COADREAD | DiFi | Bicalutamide |
| MVP_top60_DNA_seed | PIK3CB | 1.555231 | COADREAD | COLO-678 | AZD8186 | MVP_top60_DNA+RNA_seed | ERBB4 | 1.383224 | COADREAD | RCM-1 | CI-1033 |
| MVP_top60_DNA_seed | PIK3CD | 1.555231 | COADREAD | COLO-678 | AZD8186 | MVP_top60_DNA+RNA_seed | BCL2 | 1.383501 | COADREAD | NCI-H747 | Navitoclax |
| MVP_top60_DNA_seed | PIK3CB | 1.555269 | COADREAD | NCI-H716 | AZD6482 | MVP_top60_DNA+RNA_seed | IGF1R | 1.383854 | COADREAD | SW1417 | GSK1904529A |
| MVP_top60_DNA_seed | CDK2 | 1.556112 | COADREAD | COLO-678 | AZD5438 | MVP_top60_DNA+RNA_seed | PIK3CD | 1.3844 | COADREAD | CL-11 | IC-87114 |
| MVP_top60_DNA_seed | MAPK1 | 1.557318 | COADREAD | CaR-1 | FR-180204 | MVP_top60_DNA+RNA_seed | ATM | 1.385027 | COADREAD | NCI-H747 | KU-55933 |

| | | | | | | | | | | | |
|---|---|---|---|---|---|---|---|---|---|---|---|
| MVP_top60_DNA_seed | IGF1R | 1.558292 | COADREAD | SW1463 | BMS-536924 | MVP_top60_DNA+RNA_seed | ERBB4 | 1.385506 | COADREAD | T84 | AST-1306 |
| MVP_top60_DNA_seed | CDK2 | 1.559282 | COADREAD | CW-2 | PHA-793887 | MVP_top60_DNA+RNA_seed | PIK3CB | 1.388789 | COADREAD | RCM-1 | AZD8186 |
| MVP_top60_DNA_seed | MAPK1 | 1.559358 | COADREAD | SNU-81 | FR-180204 | MVP_top60_DNA+RNA_seed | PIK3CD | 1.388789 | COADREAD | RCM-1 | AZD8186 |
| MVP_top60_DNA_seed | HSP90AA1 | 1.563028 | COADREAD | SW948 | Tanespimycin | MVP_top60_DNA+RNA_seed | ATM | 1.389542 | COADREAD | SW1116 | CP466722 |
| MVP_top60_DNA_seed | HSP90AB1 | 1.563028 | COADREAD | SW948 | Tanespimycin | MVP_top60_DNA+RNA_seed | HSP90AA1 | 1.393507 | COADREAD | HT55 | Elesclomol |
| MVP_top60_DNA_seed | PIK3CA | 1.564331 | COADREAD | SNU-61 | AZD8835 | MVP_top60_DNA+RNA_seed | HSP90AB1 | 1.393507 | COADREAD | HT55 | Elesclomol |
| MVP_top60_DNA_seed | PIK3CD | 1.564331 | COADREAD | SNU-61 | AZD8835 | MVP_top60_DNA+RNA_seed | HSP90AA1 | 1.393924 | COADREAD | GP5d | CCT-018159 |
| MVP_top60_DNA_seed | BRAF | 1.56492 | COADREAD | NCI-H508 | SB590885 | MVP_top60_DNA+RNA_seed | HSP90AB1 | 1.393924 | COADREAD | GP5d | CCT-018159 |
| MVP_top60_DNA_seed | MET | 1.569886 | COADREAD | SW837 | SU11274 | MVP_top60_DNA+RNA_seed | HSP90AA1 | 1.394065 | COADREAD | SNU-81 | Elesclomol |
| MVP_top60_DNA_seed | PIK3CA | 1.574111 | COADREAD | SW1116 | AZD8835 | MVP_top60_DNA+RNA_seed | HSP90AB1 | 1.394065 | COADREAD | SNU-81 | Elesclomol |
| MVP_top60_DNA_seed | PIK3CD | 1.574111 | COADREAD | SW1116 | AZD8835 | MVP_top60_DNA+RNA_seed | AR | 1.395052 | COADREAD | SW1463 | Bicalutamide |
| MVP_top60_DNA_seed | MET | 1.5801 | COADREAD | C2BBe1 | JNJ38877605 | MVP_top60_DNA+RNA_seed | HSP90AA1 | 1.397948 | COADREAD | SW1417 | Elesclomol |
| MVP_top60_DNA_seed | HSP90AA1 | 1.580656 | COADREAD | HT-115 | SNX-2112 | MVP_top60_DNA+RNA_seed | HSP90AB1 | 1.397948 | COADREAD | SW1417 | Elesclomol |
| MVP_top60_DNA_seed | HSP90AB1 | 1.580656 | COADREAD | HT-115 | SNX-2112 | MVP_top60_DNA+RNA_seed | ATM | 1.398281 | COADREAD | CW-2 | torin2 |
| MVP_top60_DNA_seed | AKT1 | 1.586257 | COADREAD | NCI-H716 | AKT inhibitor VIII | MVP_top60_DNA+RNA_seed | PTK2 | 1.398282 | COADREAD | CL-11 | PF-562271 |
| MVP_top60_DNA_seed | AKT1 | 1.587305 | COADREAD | COLO-678 | AKT inhibitor VIII | MVP_top60_DNA+RNA_seed | ATM | 1.398285 | COADREAD | SNU-C1 | CP466722 |
| MVP_top60_DNA_seed | HSP90AA1 | 1.587993 | COADREAD | T84 | Elesclomol | MVP_top60_DNA+RNA_seed | PIK3CD | 1.402634 | COADREAD | SNU-C1 | Idelalisib |
| MVP_top60_DNA_seed | HSP90AB1 | 1.587993 | COADREAD | T84 | Elesclomol | MVP_top60_DNA+RNA_seed | IGF1R | 1.402843 | COADREAD | CL-11 | BMS-754807 |
| MVP_top60_DNA_seed | PIK3CB | 1.588505 | COADREAD | LS-411N | AZD8186 | MVP_top60_DNA+RNA_seed | GSK3B | 1.404924 | COADREAD | C2BBe1 | CHIR-99021 |
| MVP_top60_DNA_seed | PIK3CD | 1.588505 | COADREAD | LS-411N | AZD8186 | MVP_top60_DNA+RNA_seed | IGF1R | 1.40573 | COADREAD | CL-11 | GSK1904529A |
| MVP_top60_DNA_seed | AR | 1.591735 | COADREAD | SNU-81 | AZD3514 | MVP_top60_DNA+RNA_seed | MET | 1.40775 | COADREAD | SW948 | AZD6094 |
| MVP_top60_DNA_seed | MET | 1.592839 | COADREAD | SNU-81 | SU11274 | MVP_top60_DNA+RNA_seed | AR | 1.40974 | COADREAD | SW948 | Bicalutamide |
| MVP_top60_DNA_seed | GSK3B | 1.593079 | COADREAD | LS-180 | CHIR-99021 | MVP_top60_DNA+RNA_seed | AKT1 | 1.411196 | COADREAD | SNU-C1 | A-443654 |
| MVP_top60_DNA_seed | HSP90AA1 | 1.594595 | COADREAD | HT55 | Luminespib | MVP_top60_DNA+RNA_seed | HSP90AA1 | 1.414203 | COADREAD | CCK-81 | Elesclomol |
| MVP_top60_DNA_seed | HSP90AB1 | 1.594595 | COADREAD | HT55 | Luminespib | MVP_top60_DNA+RNA_seed | HSP90AB1 | 1.414203 | COADREAD | CCK-81 | Elesclomol |
| MVP_top60_DNA_seed | HSP90AA1 | 1.597363 | COADREAD | DiFi | Elesclomol | MVP_top60_DNA+RNA_seed | ERBB4 | 1.414935 | COADREAD | T84 | PF-00299804 |
| MVP_top60_DNA_seed | HSP90AB1 | 1.597363 | COADREAD | DiFi | Elesclomol | MVP_top60_DNA+RNA_seed | ATM | 1.416189 | COADREAD | CW-2 | KU-55933 |
| MVP_top60_DNA_seed | ATM | 1.598014 | COADREAD | RCM-1 | KU-55933 | MVP_top60_DNA+RNA_seed | ATM | 1.417218 | COADREAD | HT55 | KU-55933 |
| MVP_top60_DNA_seed | BRAF | 1.599263 | COADREAD | CL-11 | PLX-4720 | MVP_top60_DNA+RNA_seed | AKT1 | 1.417436 | COADREAD | RCM-1 | AKT inhibitor VIII |
| MVP_top60_DNA_seed | PIK3CB | 1.599804 | COADREAD | SW1116 | AZD8186 | MVP_top60_DNA+RNA_seed | GSK3B | 1.419708 | COADREAD | CW-2 | CHIR-99021 |
| MVP_top60_DNA_seed | PIK3CD | 1.599804 | COADREAD | SW1116 | AZD8186 | MVP_top60_DNA+RNA_seed | BCL2 | 1.420735 | COADREAD | RCM-1 | TW 37 |
| MVP_top60_DNA_seed | GSK3B | 1.600104 | COADREAD | CW-2 | SB216763 | MVP_top60_DNA+RNA_seed | BCL2 | 1.421772 | COADREAD | CaR-1 | Obatoclax Mesylate |
| MVP_top60_DNA_seed | AR | 1.605791 | COADREAD | T84 | AZD3514 | MVP_top60_DNA+RNA_seed | PTK2 | 1.422965 | COADREAD | T84 | PF-562271 |
| MVP_top60_DNA_seed | ERBB4 | 1.607297 | COADREAD | SNU-C5 | PF-00299804 | MVP_top60_DNA+RNA_seed | BCL2 | 1.424323 | COADREAD | COLO-678 | TW 37 |
| MVP_top60_DNA_seed | GSK3B | 1.609012 | COADREAD | CL-11 | KIN001-042 | MVP_top60_DNA+RNA_seed | HSP90AA1 | 1.42473 | COADREAD | CL-11 | SNX-2112 |
| MVP_top60_DNA_seed | PIK3CA | 1.61439 | COADREAD | COLO-678 | PI3Ka_4409 | MVP_top60_DNA+RNA_seed | HSP90AB1 | 1.42473 | COADREAD | CL-11 | SNX-2112 |
| MVP_top60_DNA_seed | PIK3CD | 1.61439 | COADREAD | COLO-678 | PI3Ka_4409 | MVP_top60_DNA+RNA_seed | PIK3CB | 1.428655 | COADREAD | T84 | AZD6482 |
| MVP_top60_DNA_seed | PTK2 | 1.616088 | COADREAD | HCT-15 | PF-562271 | MVP_top60_DNA+RNA_seed | GSK3B | 1.437436 | COADREAD | C2BBe1 | AZD7969 |
| MVP_top60_DNA_seed | HSP90AA1 | 1.618984 | COADREAD | SNU-81 | SNX-2112 | MVP_top60_DNA+RNA_seed | GSK3B | 1.440799 | COADREAD | LS-123 | CHIR-99021 |
| MVP_top60_DNA_seed | HSP90AB1 | 1.618984 | COADREAD | SNU-81 | SNX-2112 | MVP_top60_DNA+RNA_seed | ERBB4 | 1.44087 | COADREAD | SW948 | PF-00299804 |
| MVP_top60_DNA_seed | FYN | 1.61959 | COADREAD | CW-2 | A-770041 | MVP_top60_DNA+RNA_seed | PIK3CB | 1.443038 | COADREAD | CW-2 | TGX221 |

| | | | | | | | | | | | |
|---|---|---|---|---|---|---|---|---|---|---|---|
| MVP_top60_DNA_seed | ESR1 | 1.625184 | COADREAD | SNU-C1 | Fulvestrant | MVP_top60_DNA+RNA_seed | PIK3CB | 1.443899 | COADREAD | LS-513 | AZD6482 |
| MVP_top60_DNA_seed | BRAF | 1.627943 | COADREAD | RCM-1 | HG6-64-1 | MVP_top60_DNA+RNA_seed | GSK3B | 1.446459 | COADREAD | SNU-61 | KIN001-042 |
| MVP_top60_DNA_seed | BRAF | 1.633544 | COADREAD | SW1116 | PLX-4720 | MVP_top60_DNA+RNA_seed | AR | 1.450002 | COADREAD | HT-29 | Bicalutamide |
| MVP_top60_DNA_seed | ATM | 1.635518 | COADREAD | CW-2 | CP466722 | MVP_top60_DNA+RNA_seed | ATM | 1.45201 | COADREAD | SW620 | KU-55933 |
| MVP_top60_DNA_seed | HSP90AA1 | 1.636602 | COADREAD | SW1417 | Luminespib | MVP_top60_DNA+RNA_seed | HSP90AA1 | 1.457103 | COADREAD | COLO-320-HSR | Tanespimycin |
| MVP_top60_DNA_seed | HSP90AB1 | 1.636602 | COADREAD | SW1417 | Luminespib | MVP_top60_DNA+RNA_seed | HSP90AB1 | 1.457103 | COADREAD | COLO-320-HSR | Tanespimycin |
| MVP_top60_DNA_seed | AKT1 | 1.640221 | COADREAD | KM12 | BAY AKT1 | MVP_top60_DNA+RNA_seed | ERBB3 | 1.458012 | COADREAD | SNU-61 | AZD8931 |
| MVP_top60_DNA_seed | AKT1 | 1.640421 | COADREAD | SW948 | AKT inhibitor VIII | MVP_top60_DNA+RNA_seed | PTK2 | 1.460327 | COADREAD | SW1463 | PF-562271 |
| MVP_top60_DNA_seed | ERBB4 | 1.640477 | COADREAD | SW48 | PF-00299804 | MVP_top60_DNA+RNA_seed | MET | 1.463742 | COADREAD | CW-2 | Crizotinib |
| MVP_top60_DNA_seed | FYN | 1.64413 | COADREAD | SNU-C1 | A-770041 | MVP_top60_DNA+RNA_seed | PTK2 | 1.466061 | COADREAD | SW1417 | PF-562271 |
| MVP_top60_DNA_seed | AR | 1.647872 | COADREAD | SW948 | AZD3514 | MVP_top60_DNA+RNA_seed | MET | 1.477444 | COADREAD | SNU-C1 | PHA-665752 |
| MVP_top60_DNA_seed | BRAF | 1.648702 | COADREAD | COLO-678 | PLX-4720 | MVP_top60_DNA+RNA_seed | AKT1 | 1.479086 | COADREAD | C2BBe1 | MK-2206 |
| MVP_top60_DNA_seed | BRAF | 1.65166 | COADREAD | KM12 | RAF_9304 | MVP_top60_DNA+RNA_seed | GSK3B | 1.482156 | COADREAD | SW948 | SB216763 |
| MVP_top60_DNA_seed | SRC | 1.654173 | COADREAD | SNU-C1 | NVP-BHG712 | MVP_top60_DNA+RNA_seed | HSP90AA1 | 1.484549 | COADREAD | KM12 | Elesclomol |
| MVP_top60_DNA_seed | CDK2 | 1.655671 | COADREAD | CL-11 | PHA-793887 | MVP_top60_DNA+RNA_seed | HSP90AB1 | 1.484549 | COADREAD | KM12 | Elesclomol |
| MVP_top60_DNA_seed | BRAF | 1.660652 | COADREAD | SW1116 | SB590885 | MVP_top60_DNA+RNA_seed | ERBB4 | 1.484886 | COADREAD | SNU-C1 | AST-1306 |
| MVP_top60_DNA_seed | MET | 1.665653 | COADREAD | CL-11 | SU11274 | MVP_top60_DNA+RNA_seed | BCL2 | 1.489491 | COADREAD | HT-29 | Navitoclax |
| MVP_top60_DNA_seed | BRAF | 1.666703 | COADREAD | SW620 | PLX-4720 | MVP_top60_DNA+RNA_seed | MET | 1.494176 | COADREAD | HCT-15 | JNJ38877605 |
| MVP_top60_DNA_seed | PIK3CB | 1.672276 | COADREAD | SW1463 | AZD6482 | MVP_top60_DNA+RNA_seed | BCL2 | 1.496035 | COADREAD | LS-123 | Navitoclax |
| MVP_top60_DNA_seed | PIK3CA | 1.673282 | COADREAD | SW620 | PI3Ka_4409 | MVP_top60_DNA+RNA_seed | ERBB3 | 1.496494 | COADREAD | COLO-205 | AZD8931 |
| MVP_top60_DNA_seed | PIK3CD | 1.673282 | COADREAD | SW620 | PI3Ka_4409 | MVP_top60_DNA+RNA_seed | MET | 1.498316 | COADREAD | CL-11 | JNJ38877605 |
| MVP_top60_DNA_seed | PIK3CD | 1.682366 | COADREAD | SW948 | IC-87114 | MVP_top60_DNA+RNA_seed | SRC | 1.499481 | COADREAD | CW-2 | Saracatinib |
| MVP_top60_DNA_seed | CDK2 | 1.684807 | COADREAD | SNU-C1 | PHA-793887 | MVP_top60_DNA+RNA_seed | AKT1 | 1.500991 | COADREAD | SW1417 | BAY AKT1 |
| MVP_top60_DNA_seed | HSP90AA1 | 1.686523 | COADREAD | SW1463 | Luminespib | MVP_top60_DNA+RNA_seed | SRC | 1.501389 | COADREAD | CW-2 | HG-6-71-01 |
| MVP_top60_DNA_seed | HSP90AB1 | 1.686523 | COADREAD | SW1463 | Luminespib | MVP_top60_DNA+RNA_seed | ATM | 1.501923 | COADREAD | HT55 | KU-60019 |
| MVP_top60_DNA_seed | AR | 1.688054 | COADREAD | HCT-15 | AZD3514 | MVP_top60_DNA+RNA_seed | ATM | 1.50684 | COADREAD | RCM-1 | KU-60019 |
| MVP_top60_DNA_seed | AKT1 | 1.688538 | COADREAD | RCM-1 | AKT inhibitor VIII | MVP_top60_DNA+RNA_seed | PIK3CD | 1.508703 | COADREAD | COLO-678 | IC-87114 |
| MVP_top60_DNA_seed | ERBB3 | 1.690652 | COADREAD | SNU-1040 | AZD8931 | MVP_top60_DNA+RNA_seed | PIK3CB | 1.510995 | COADREAD | SW620 | AZD8186 |
| MVP_top60_DNA_seed | MET | 1.692116 | COADREAD | SNU-61 | SU11274 | MVP_top60_DNA+RNA_seed | PIK3CD | 1.510995 | COADREAD | SW620 | AZD8186 |
| MVP_top60_DNA_seed | ATM | 1.693951 | COADREAD | SW1463 | CP466722 | MVP_top60_DNA+RNA_seed | AKT1 | 1.514339 | COADREAD | SW1116 | GSK690693 |
| MVP_top60_DNA_seed | ATM | 1.695339 | COADREAD | MDST8 | CP466722 | MVP_top60_DNA+RNA_seed | PIK3CB | 1.514443 | COADREAD | KM12 | AZD8186 |
| MVP_top60_DNA_seed | ESR1 | 1.697024 | COADREAD | LS-180 | Fulvestrant | MVP_top60_DNA+RNA_seed | PIK3CD | 1.514443 | COADREAD | KM12 | AZD8186 |
| MVP_top60_DNA_seed | CDK2 | 1.69865 | COADREAD | SW948 | PHA-793887 | MVP_top60_DNA+RNA_seed | IGF1R | 1.519006 | COADREAD | CW-2 | BMS-536924 |
| MVP_top60_DNA_seed | PIK3CB | 1.702397 | COADREAD | CL-11 | AZD6482 | MVP_top60_DNA+RNA_seed | IGF1R | 1.519259 | COADREAD | SNU-81 | BMS-536924 |
| MVP_top60_DNA_seed | MET | 1.703109 | COADREAD | COLO-678 | SU11274 | MVP_top60_DNA+RNA_seed | MET | 1.525519 | COADREAD | LS-123 | Crizotinib |
| MVP_top60_DNA_seed | PIK3CB | 1.703131 | COADREAD | SW948 | AZD8186 | MVP_top60_DNA+RNA_seed | MET | 1.526215 | COADREAD | SW1116 | AZD6094 |
| MVP_top60_DNA_seed | PIK3CD | 1.703131 | COADREAD | SW948 | AZD8186 | MVP_top60_DNA+RNA_seed | GSK3B | 1.527269 | COADREAD | CaR-1 | AZD7969 |
| MVP_top60_DNA_seed | BRAF | 1.706309 | COADREAD | SNU-C1 | SB590885 | MVP_top60_DNA+RNA_seed | MET | 1.528584 | COADREAD | LS-411N | JNJ38877605 |
| MVP_top60_DNA_seed | MET | 1.707711 | COADREAD | SW948 | SU11274 | MVP_top60_DNA+RNA_seed | AR | 1.533553 | COADREAD | CaR-1 | Bicalutamide |
| MVP_top60_DNA_seed | ERBB4 | 1.710712 | COADREAD | LS-1034 | PF-00299804 | MVP_top60_DNA+RNA_seed | SRC | 1.535019 | COADREAD | SNU-81 | NVP-BHG712 |
| MVP_top60_DNA_seed | AKT1 | 1.722905 | COADREAD | CL-11 | A-443654 | MVP_top60_DNA+RNA_seed | ERBB4 | 1.536211 | COADREAD | SW1463 | CI-1033 |

| | | | | | | | | | | | |
|---|---|---|---|---|---|---|---|---|---|---|---|
| MVP_top60_DNA_seed | PIK3CD | 1.723226 | COADREAD | COLO-678 | Idelalisib | MVP_top60_DNA+RNA_seed | IGF1R | 1.536502 | COADREAD | CW-2 | Linsitinib |
| MVP_top60_DNA_seed | ATM | 1.724064 | COADREAD | SNU-C2B | KU-55933 | MVP_top60_DNA+RNA_seed | AKT1 | 1.537519 | COADREAD | LS-1034 | MK-2206 |
| MVP_top60_DNA_seed | SRC | 1.724106 | COADREAD | SNU-C1 | WH-4-023 | MVP_top60_DNA+RNA_seed | SRC | 1.539479 | COADREAD | SW948 | Bosutinib |
| MVP_top60_DNA_seed | BRAF | 1.725127 | COADREAD | SNU-61 | PLX-4720 | MVP_top60_DNA+RNA_seed | AKT1 | 1.542465 | COADREAD | C2BBe1 | A-443654 |
| MVP_top60_DNA_seed | MET | 1.725143 | COADREAD | HT-29 | AZD6094 | MVP_top60_DNA+RNA_seed | PIK3CB | 1.543077 | COADREAD | LS-1034 | AZD6482 |
| MVP_top60_DNA_seed | IGF1R | 1.727114 | COADREAD | C2BBe1 | GSK1904529A | MVP_top60_DNA+RNA_seed | GSK3B | 1.546208 | COADREAD | SNU-1040 | CHIR-99021 |
| MVP_top60_DNA_seed | AR | 1.731939 | COADREAD | KM12 | AZD3514 | MVP_top60_DNA+RNA_seed | PIK3CD | 1.548541 | COADREAD | SW1116 | IC-87114 |
| MVP_top60_DNA_seed | GSK3B | 1.740381 | COADREAD | KM12 | AZD7969 | MVP_top60_DNA+RNA_seed | PIK3CD | 1.550271 | COADREAD | RCM-1 | Idelalisib |
| MVP_top60_DNA_seed | SRC | 1.742021 | COADREAD | SNU-C1 | Saracatinib | MVP_top60_DNA+RNA_seed | PIK3CB | 1.555231 | COADREAD | COLO-678 | AZD8186 |
| MVP_top60_DNA_seed | AKT1 | 1.743835 | COADREAD | SW1463 | GSK690693 | MVP_top60_DNA+RNA_seed | PIK3CD | 1.555231 | COADREAD | COLO-678 | AZD8186 |
| MVP_top60_DNA_seed | MAPK14 | 1.743847 | COADREAD | SW1417 | TAK-715 | MVP_top60_DNA+RNA_seed | PIK3CB | 1.555269 | COADREAD | NCI-H716 | AZD6482 |
| MVP_top60_DNA_seed | SRC | 1.752543 | COADREAD | CW-2 | Bosutinib | MVP_top60_DNA+RNA_seed | CDK2 | 1.556112 | COADREAD | COLO-678 | AZD5438 |
| MVP_top60_DNA_seed | ESR1 | 1.752687 | COADREAD | SNU-81 | Fulvestrant | MVP_top60_DNA+RNA_seed | IGF1R | 1.558292 | COADREAD | SW1463 | BMS-536924 |
| MVP_top60_DNA_seed | AKT1 | 1.76016 | COADREAD | CL-11 | AKT inhibitor VIII | MVP_top60_DNA+RNA_seed | CDK2 | 1.559282 | COADREAD | CW-2 | PHA-793887 |
| MVP_top60_DNA_seed | AR | 1.760441 | COADREAD | SW1463 | AZD3514 | MVP_top60_DNA+RNA_seed | HSP90AA1 | 1.563028 | COADREAD | SW948 | Tanespimycin |
| MVP_top60_DNA_seed | CDK2 | 1.760467 | COADREAD | CL-11 | Seliciclib | MVP_top60_DNA+RNA_seed | HSP90AB1 | 1.563028 | COADREAD | SW948 | Tanespimycin |
| MVP_top60_DNA_seed | PIK3CD | 1.761672 | COADREAD | SNU-C1 | IC-87114 | MVP_top60_DNA+RNA_seed | PIK3CA | 1.564331 | COADREAD | SNU-61 | AZD8835 |
| MVP_top60_DNA_seed | AKT1 | 1.762327 | COADREAD | NCI-H716 | MK-2206 | MVP_top60_DNA+RNA_seed | PIK3CD | 1.564331 | COADREAD | SNU-61 | AZD8835 |
| MVP_top60_DNA_seed | AKT1 | 1.76279 | COADREAD | SNU-61 | BAY AKT1 | MVP_top60_DNA+RNA_seed | MET | 1.569886 | COADREAD | SW837 | SU11274 |
| MVP_top60_DNA_seed | SRC | 1.764506 | COADREAD | KM12 | HG-6-71-01 | MVP_top60_DNA+RNA_seed | PIK3CA | 1.574111 | COADREAD | SW1116 | AZD8835 |
| MVP_top60_DNA_seed | ESR1 | 1.765473 | COADREAD | SNU-61 | Tamoxifen | MVP_top60_DNA+RNA_seed | PIK3CD | 1.574111 | COADREAD | SW1116 | AZD8835 |
| MVP_top60_DNA_seed | AKT1 | 1.768714 | COADREAD | SW1116 | AKT inhibitor VIII | MVP_top60_DNA+RNA_seed | MET | 1.5801 | COADREAD | C2BBe1 | JNJ38877605 |
| MVP_top60_DNA_seed | IGF1R | 1.770993 | COADREAD | CW-2 | GSK1904529A | MVP_top60_DNA+RNA_seed | HSP90AA1 | 1.580656 | COADREAD | HT-115 | SNX-2112 |
| MVP_top60_DNA_seed | PIK3CA | 1.773409 | COADREAD | LS-411N | PI3Ka_4409 | MVP_top60_DNA+RNA_seed | HSP90AB1 | 1.580656 | COADREAD | HT-115 | SNX-2112 |
| MVP_top60_DNA_seed | PIK3CD | 1.773409 | COADREAD | LS-411N | PI3Ka_4409 | MVP_top60_DNA+RNA_seed | AKT1 | 1.586257 | COADREAD | NCI-H716 | AKT inhibitor VIII |
| MVP_top60_DNA_seed | HSP90AA1 | 1.773781 | COADREAD | LS-123 | CCT-018159 | MVP_top60_DNA+RNA_seed | AKT1 | 1.587305 | COADREAD | COLO-678 | AKT inhibitor VIII |
| MVP_top60_DNA_seed | HSP90AB1 | 1.773781 | COADREAD | LS-123 | CCT-018159 | MVP_top60_DNA+RNA_seed | HSP90AA1 | 1.587993 | COADREAD | T84 | Elesclomol |
| MVP_top60_DNA_seed | ESR1 | 1.780082 | COADREAD | SW948 | Tamoxifen | MVP_top60_DNA+RNA_seed | HSP90AB1 | 1.587993 | COADREAD | T84 | Elesclomol |
| MVP_top60_DNA_seed | ATM | 1.782405 | COADREAD | LS-513 | KU-55933 | MVP_top60_DNA+RNA_seed | PIK3CB | 1.588505 | COADREAD | LS-411N | AZD8186 |
| MVP_top60_DNA_seed | CREBBP | 1.785602 | COADREAD | SNU-61 | I-CBP112 | MVP_top60_DNA+RNA_seed | PIK3CD | 1.588505 | COADREAD | LS-411N | AZD8186 |
| MVP_top60_DNA_seed | AR | 1.789356 | COADREAD | CW-2 | Bicalutamide | MVP_top60_DNA+RNA_seed | AR | 1.591735 | COADREAD | SNU-81 | AZD3514 |
| MVP_top60_DNA_seed | CDK2 | 1.793992 | COADREAD | SW1116 | AZD5438 | MVP_top60_DNA+RNA_seed | MET | 1.592839 | COADREAD | SNU-81 | SU11274 |
| MVP_top60_DNA_seed | GSK3B | 1.796136 | COADREAD | SW1463 | CHIR-99021 | MVP_top60_DNA+RNA_seed | GSK3B | 1.593079 | COADREAD | LS-180 | CHIR-99021 |
| MVP_top60_DNA_seed | ERBB3 | 1.796449 | COADREAD | SNU-C1 | AZD8931 | MVP_top60_DNA+RNA_seed | HSP90AA1 | 1.594595 | COADREAD | HT55 | Luminespib |
| MVP_top60_DNA_seed | ATM | 1.796726 | COADREAD | COLO-678 | KU-55933 | MVP_top60_DNA+RNA_seed | HSP90AB1 | 1.594595 | COADREAD | HT55 | Luminespib |
| MVP_top60_DNA_seed | BRAF | 1.79791 | COADREAD | SNU-61 | RAF_9304 | MVP_top60_DNA+RNA_seed | HSP90AA1 | 1.597363 | COADREAD | DiFi | Elesclomol |
| MVP_top60_DNA_seed | BRAF | 1.798255 | COADREAD | SW837 | PLX-4720 | MVP_top60_DNA+RNA_seed | HSP90AB1 | 1.597363 | COADREAD | DiFi | Elesclomol |
| MVP_top60_DNA_seed | IGF1R | 1.799542 | COADREAD | CaR-1 | BMS-754807 | MVP_top60_DNA+RNA_seed | ATM | 1.598014 | COADREAD | RCM-1 | KU-55933 |
| MVP_top60_DNA_seed | PIK3CA | 1.799843 | COADREAD | SNU-61 | PI3Ka_4409 | MVP_top60_DNA+RNA_seed | PIK3CB | 1.599804 | COADREAD | SW1116 | AZD8186 |
| MVP_top60_DNA_seed | PIK3CD | 1.799843 | COADREAD | SNU-61 | PI3Ka_4409 | MVP_top60_DNA+RNA_seed | PIK3CD | 1.599804 | COADREAD | SW1116 | AZD8186 |
| MVP_top60_DNA_seed | SRC | 1.800697 | COADREAD | SW1463 | Bosutinib | MVP_top60_DNA+RNA_seed | GSK3B | 1.600104 | COADREAD | CW-2 | SB216763 |

| | | | | | | | | | | | |
|---|---|---|---|---|---|---|---|---|---|---|---|
| MVP_top60_DNA_seed | BRAF | 1.809291 | COADREAD | SNU-61 | PLX-4720 | MVP_top60_DNA+RNA_seed | AR | 1.605791 | COADREAD | T84 | AZD3514 |
| MVP_top60_DNA_seed | GSK3B | 1.811202 | COADREAD | SW948 | CHIR-99021 | MVP_top60_DNA+RNA_seed | ERBB4 | 1.607297 | COADREAD | SNU-C5 | PF-00299804 |
| MVP_top60_DNA_seed | AKT1 | 1.81131 | COADREAD | SW1417 | AKT inhibitor VIII | MVP_top60_DNA+RNA_seed | GSK3B | 1.609012 | COADREAD | CL-11 | KIN001-042 |
| MVP_top60_DNA_seed | GSK3B | 1.823985 | COADREAD | COLO-678 | KIN001-042 | MVP_top60_DNA+RNA_seed | PIK3CA | 1.61439 | COADREAD | COLO-678 | PI3Ka_4409 |
| MVP_top60_DNA_seed | CDK2 | 1.824851 | COADREAD | NCI-H716 | Seliciclib | MVP_top60_DNA+RNA_seed | PIK3CD | 1.61439 | COADREAD | COLO-678 | PI3Ka_4409 |
| MVP_top60_DNA_seed | PIK3CA | 1.826066 | COADREAD | KM12 | AZD8835 | MVP_top60_DNA+RNA_seed | PTK2 | 1.616088 | COADREAD | HCT-15 | PF-562271 |
| MVP_top60_DNA_seed | PIK3CD | 1.826066 | COADREAD | KM12 | AZD8835 | MVP_top60_DNA+RNA_seed | HSP90AA1 | 1.618984 | COADREAD | SNU-81 | SNX-2112 |
| MVP_top60_DNA_seed | ESR1 | 1.826796 | COADREAD | HT-29 | Tamoxifen | MVP_top60_DNA+RNA_seed | HSP90AB1 | 1.618984 | COADREAD | SNU-81 | SNX-2112 |
| MVP_top60_DNA_seed | ERBB4 | 1.837485 | COADREAD | COLO-678 | CI-1033 | MVP_top60_DNA+RNA_seed | FYN | 1.61959 | COADREAD | CW-2 | A-770041 |
| MVP_top60_DNA_seed | ESR1 | 1.837752 | COADREAD | LS-123 | Tamoxifen | MVP_top60_DNA+RNA_seed | ATM | 1.635518 | COADREAD | CW-2 | CP466722 |
| MVP_top60_DNA_seed | SRC | 1.839582 | COADREAD | LS-123 | Bosutinib | MVP_top60_DNA+RNA_seed | HSP90AA1 | 1.636602 | COADREAD | SW1417 | Luminespib |
| MVP_top60_DNA_seed | AR | 1.840858 | COADREAD | HT-29 | Bicalutamide | MVP_top60_DNA+RNA_seed | HSP90AB1 | 1.636602 | COADREAD | SW1417 | Luminespib |
| MVP_top60_DNA_seed | ESR1 | 1.841854 | COADREAD | SW1463 | Fulvestrant | MVP_top60_DNA+RNA_seed | AKT1 | 1.640221 | COADREAD | KM12 | BAY AKT1 |
| MVP_top60_DNA_seed | AKT1 | 1.841974 | COADREAD | SW1116 | BAY AKT1 | MVP_top60_DNA+RNA_seed | AKT1 | 1.640421 | COADREAD | SW948 | AKT inhibitor VIII |
| MVP_top60_DNA_seed | PIK3CB | 1.84767 | COADREAD | RCM-1 | AZD6482 | MVP_top60_DNA+RNA_seed | ERBB4 | 1.640477 | COADREAD | SW48 | PF-00299804 |
| MVP_top60_DNA_seed | AR | 1.855999 | COADREAD | SNU-61 | Bicalutamide | MVP_top60_DNA+RNA_seed | FYN | 1.64413 | COADREAD | SNU-C1 | A-770041 |
| MVP_top60_DNA_seed | AKT1 | 1.8599 | COADREAD | SW1463 | AKT inhibitor VIII | MVP_top60_DNA+RNA_seed | AR | 1.647872 | COADREAD | SW948 | AZD3514 |
| MVP_top60_DNA_seed | GSK3B | 1.864381 | COADREAD | LS-123 | AZD7969 | MVP_top60_DNA+RNA_seed | SRC | 1.654173 | COADREAD | SNU-C1 | NVP-BHG712 |
| MVP_top60_DNA_seed | CDK2 | 1.867422 | COADREAD | SW1116 | PHA-793887 | MVP_top60_DNA+RNA_seed | CDK2 | 1.655671 | COADREAD | CL-11 | PHA-793887 |
| MVP_top60_DNA_seed | ESR1 | 1.867827 | COADREAD | SNU-1040 | Fulvestrant | MVP_top60_DNA+RNA_seed | MET | 1.665653 | COADREAD | CL-11 | SU11274 |
| MVP_top60_DNA_seed | PIK3CB | 1.868278 | COADREAD | SNU-C1 | TGX221 | MVP_top60_DNA+RNA_seed | PIK3CB | 1.672276 | COADREAD | SW1463 | AZD6482 |
| MVP_top60_DNA_seed | PIK3CA | 1.869052 | COADREAD | C2BBe1 | PI3Ka_4409 | MVP_top60_DNA+RNA_seed | PIK3CA | 1.673282 | COADREAD | SW620 | PI3Ka_4409 |
| MVP_top60_DNA_seed | PIK3CD | 1.869052 | COADREAD | C2BBe1 | PI3Ka_4409 | MVP_top60_DNA+RNA_seed | PIK3CD | 1.673282 | COADREAD | SW620 | PI3Ka_4409 |
| MVP_top60_DNA_seed | HSP90AA1 | 1.870765 | COADREAD | COLO-205 | Elesclomol | MVP_top60_DNA+RNA_seed | PIK3CD | 1.682366 | COADREAD | SW948 | IC-87114 |
| MVP_top60_DNA_seed | HSP90AB1 | 1.870765 | COADREAD | COLO-205 | Elesclomol | MVP_top60_DNA+RNA_seed | CDK2 | 1.684807 | COADREAD | SNU-C1 | PHA-793887 |
| MVP_top60_DNA_seed | HSP90AA1 | 1.879065 | COADREAD | SW1116 | Elesclomol | MVP_top60_DNA+RNA_seed | HSP90AA1 | 1.686523 | COADREAD | SW1463 | Luminespib |
| MVP_top60_DNA_seed | HSP90AB1 | 1.879065 | COADREAD | SW1116 | Elesclomol | MVP_top60_DNA+RNA_seed | HSP90AB1 | 1.686523 | COADREAD | SW1463 | Luminespib |
| MVP_top60_DNA_seed | AKT1 | 1.882993 | COADREAD | T84 | MK-2206 | MVP_top60_DNA+RNA_seed | AR | 1.688054 | COADREAD | HCT-15 | AZD3514 |
| MVP_top60_DNA_seed | AR | 1.885556 | COADREAD | C2BBe1 | AZD3514 | MVP_top60_DNA+RNA_seed | AKT1 | 1.688538 | COADREAD | RCM-1 | AKT inhibitor VIII |
| MVP_top60_DNA_seed | IGF1R | 1.887598 | COADREAD | SNU-1040 | BMS-536924 | MVP_top60_DNA+RNA_seed | ERBB3 | 1.690652 | COADREAD | SNU-1040 | AZD8931 |
| MVP_top60_DNA_seed | BRAF | 1.89441 | COADREAD | SNU-C1 | PLX-4720 | MVP_top60_DNA+RNA_seed | MET | 1.692116 | COADREAD | SNU-61 | SU11274 |
| MVP_top60_DNA_seed | ESR1 | 1.895048 | COADREAD | CaR-1 | Fulvestrant | MVP_top60_DNA+RNA_seed | ATM | 1.693951 | COADREAD | SW1463 | CP466722 |
| MVP_top60_DNA_seed | AKT1 | 1.896555 | COADREAD | HT55 | AKT inhibitor VIII | MVP_top60_DNA+RNA_seed | ATM | 1.695339 | COADREAD | MDST8 | CP466722 |
| MVP_top60_DNA_seed | MAPK14 | 1.898698 | COADREAD | SW1463 | TAK-715 | MVP_top60_DNA+RNA_seed | BCL2 | 1.695823 | COADREAD | SW948 | Obatoclax Mesylate |
| MVP_top60_DNA_seed | BRAF | 1.90181 | COADREAD | HT55 | HG6-64-1 | MVP_top60_DNA+RNA_seed | CDK2 | 1.69865 | COADREAD | SW948 | PHA-793887 |
| MVP_top60_DNA_seed | PIK3CB | 1.902959 | COADREAD | LS-123 | AZD6482 | MVP_top60_DNA+RNA_seed | PIK3CB | 1.702397 | COADREAD | CL-11 | AZD6482 |
| MVP_top60_DNA_seed | ERBB4 | 1.906833 | COADREAD | SW620 | CI-1033 | MVP_top60_DNA+RNA_seed | MET | 1.703109 | COADREAD | COLO-678 | SU11274 |
| MVP_top60_DNA_seed | PIK3CB | 1.907161 | COADREAD | SNU-1040 | AZD8186 | MVP_top60_DNA+RNA_seed | PIK3CB | 1.703131 | COADREAD | SW948 | AZD8186 |
| MVP_top60_DNA_seed | PIK3CD | 1.907161 | COADREAD | SNU-1040 | AZD8186 | MVP_top60_DNA+RNA_seed | PIK3CD | 1.703131 | COADREAD | SW948 | AZD8186 |
| MVP_top60_DNA_seed | HSP90AA1 | 1.908858 | COADREAD | NCI-H508 | Elesclomol | MVP_top60_DNA+RNA_seed | MET | 1.707711 | COADREAD | SW948 | SU11274 |
| MVP_top60_DNA_seed | HSP90AB1 | 1.908858 | COADREAD | NCI-H508 | Elesclomol | MVP_top60_DNA+RNA_seed | ERBB4 | 1.710712 | COADREAD | LS-1034 | PF-00299804 |

| | | | | | | | | | | | |
|---|---|---|---|---|---|---|---|---|---|---|---|
| MVP_top60_DNA_seed | HSP90AA1 | 1.910671 | COADREAD | CL-11 | Luminespib | MVP_top60_DNA+RNA_seed | AKT1 | 1.722905 | COADREAD | CL-11 | A-443654 |
| MVP_top60_DNA_seed | HSP90AB1 | 1.910671 | COADREAD | CL-11 | Luminespib | MVP_top60_DNA+RNA_seed | PIK3CD | 1.723226 | COADREAD | COLO-678 | Idelalisib |
| MVP_top60_DNA_seed | ESR1 | 1.916879 | COADREAD | SW620 | Tamoxifen | MVP_top60_DNA+RNA_seed | ATM | 1.724064 | COADREAD | SNU-C2B | KU-55933 |
| MVP_top60_DNA_seed | GSK3B | 1.916896 | COADREAD | HT55 | AZD7969 | MVP_top60_DNA+RNA_seed | SRC | 1.724106 | COADREAD | SNU-C1 | WH-4-023 |
| MVP_top60_DNA_seed | AR | 1.924926 | COADREAD | SW620 | Bicalutamide | MVP_top60_DNA+RNA_seed | MET | 1.725143 | COADREAD | HT-29 | AZD6094 |
| MVP_top60_DNA_seed | PIK3CA | 1.928324 | COADREAD | SNU-1040 | AZD8835 | MVP_top60_DNA+RNA_seed | IGF1R | 1.727114 | COADREAD | C2BBe1 | GSK1904529A |
| MVP_top60_DNA_seed | PIK3CD | 1.928324 | COADREAD | SNU-1040 | AZD8835 | MVP_top60_DNA+RNA_seed | AR | 1.731939 | COADREAD | KM12 | AZD3514 |
| MVP_top60_DNA_seed | ATM | 1.9308 | COADREAD | SW837 | KU-55933 | MVP_top60_DNA+RNA_seed | GSK3B | 1.740381 | COADREAD | KM12 | AZD7969 |
| MVP_top60_DNA_seed | MET | 1.934629 | COADREAD | SNU-81 | JNJ38877605 | MVP_top60_DNA+RNA_seed | SRC | 1.742021 | COADREAD | SNU-C1 | Saracatinib |
| MVP_top60_DNA_seed | GSK3B | 1.935474 | COADREAD | LS-123 | CHIR-99021 | MVP_top60_DNA+RNA_seed | AKT1 | 1.743835 | COADREAD | SW1463 | GSK690693 |
| MVP_top60_DNA_seed | GSK3B | 1.940117 | COADREAD | SNU-61 | CHIR-99021 | MVP_top60_DNA+RNA_seed | SRC | 1.752543 | COADREAD | CW-2 | Bosutinib |
| MVP_top60_DNA_seed | IGF1R | 1.943844 | COADREAD | CW-2 | BMS-536924 | MVP_top60_DNA+RNA_seed | BCL2 | 1.757097 | COADREAD | LS-180 | TW 37 |
| MVP_top60_DNA_seed | AKT1 | 1.944805 | COADREAD | SW1116 | MK-2206 | MVP_top60_DNA+RNA_seed | AKT1 | 1.76016 | COADREAD | CL-11 | AKT inhibitor VIII |
| MVP_top60_DNA_seed | PIK3CB | 1.949729 | COADREAD | LS-180 | AZD8186 | MVP_top60_DNA+RNA_seed | AR | 1.760441 | COADREAD | SW1463 | AZD3514 |
| MVP_top60_DNA_seed | PIK3CD | 1.949729 | COADREAD | LS-180 | AZD8186 | MVP_top60_DNA+RNA_seed | CDK2 | 1.760467 | COADREAD | CL-11 | Seliciclib |
| MVP_top60_DNA_seed | MAPK1 | 1.952301 | COADREAD | SW1116 | VX-11e | MVP_top60_DNA+RNA_seed | PIK3CD | 1.761672 | COADREAD | SNU-C1 | IC-87114 |
| MVP_top60_DNA_seed | PIK3CB | 1.957446 | COADREAD | CW-2 | AZD6482 | MVP_top60_DNA+RNA_seed | AKT1 | 1.762327 | COADREAD | NCI-H716 | MK-2206 |
| MVP_top60_DNA_seed | PTK2 | 1.958593 | COADREAD | SW837 | PF-562271 | MVP_top60_DNA+RNA_seed | AKT1 | 1.76279 | COADREAD | SNU-61 | BAY AKT1 |
| MVP_top60_DNA_seed | AKT1 | 1.960269 | COADREAD | COLO-678 | MK-2206 | MVP_top60_DNA+RNA_seed | SRC | 1.764506 | COADREAD | KM12 | HG-6-71-01 |
| MVP_top60_DNA_seed | PIK3CB | 1.961016 | COADREAD | CW-2 | AZD8186 | MVP_top60_DNA+RNA_seed | AKT1 | 1.768714 | COADREAD | SW1116 | AKT inhibitor VIII |
| MVP_top60_DNA_seed | PIK3CD | 1.961016 | COADREAD | CW-2 | AZD8186 | MVP_top60_DNA+RNA_seed | IGF1R | 1.770993 | COADREAD | CW-2 | GSK1904529A |
| MVP_top60_DNA_seed | AR | 1.974818 | COADREAD | SW1116 | AZD3514 | MVP_top60_DNA+RNA_seed | PIK3CA | 1.773409 | COADREAD | LS-411N | PI3Ka_4409 |
| MVP_top60_DNA_seed | AR | 1.978266 | COADREAD | DiFi | AZD3514 | MVP_top60_DNA+RNA_seed | PIK3CD | 1.773409 | COADREAD | LS-411N | PI3Ka_4409 |
| MVP_top60_DNA_seed | HSP90AA1 | 1.992247 | COADREAD | SW1116 | Tanespimycin | MVP_top60_DNA+RNA_seed | HSP90AA1 | 1.773781 | COADREAD | LS-123 | CCT-018159 |
| MVP_top60_DNA_seed | HSP90AB1 | 1.992247 | COADREAD | SW1116 | Tanespimycin | MVP_top60_DNA+RNA_seed | HSP90AB1 | 1.773781 | COADREAD | LS-123 | CCT-018159 |
| MVP_top60_DNA_seed | PIK3CD | 1.99544 | COADREAD | SNU-61 | IC-87114 | MVP_top60_DNA+RNA_seed | ATM | 1.782405 | COADREAD | LS-513 | KU-55933 |
| MVP_top60_DNA_seed | MET | 1.999142 | COADREAD | SNU-C1 | AZD6094 | MVP_top60_DNA+RNA_seed | AR | 1.789356 | COADREAD | CW-2 | Bicalutamide |
| MVP_top60_DNA_seed | HSP90AA1 | 2.007253 | COADREAD | SNU-C5 | CCT-018159 | MVP_top60_DNA+RNA_seed | CDK2 | 1.793992 | COADREAD | SW1116 | AZD5438 |
| MVP_top60_DNA_seed | HSP90AB1 | 2.007253 | COADREAD | SNU-C5 | CCT-018159 | MVP_top60_DNA+RNA_seed | GSK3B | 1.796136 | COADREAD | SW1463 | CHIR-99021 |
| MVP_top60_DNA_seed | AKT1 | 2.008281 | COADREAD | CW-2 | AKT inhibitor VIII | MVP_top60_DNA+RNA_seed | ERBB3 | 1.796449 | COADREAD | SNU-C1 | AZD8931 |
| MVP_top60_DNA_seed | ATM | 2.012473 | COADREAD | COLO-678 | CP466722 | MVP_top60_DNA+RNA_seed | ATM | 1.796726 | COADREAD | COLO-678 | KU-55933 |
| MVP_top60_DNA_seed | ATM | 2.01646 | COADREAD | T84 | KU-60019 | MVP_top60_DNA+RNA_seed | IGF1R | 1.799542 | COADREAD | CaR-1 | BMS-754807 |
| MVP_top60_DNA_seed | PIK3CD | 2.01691 | COADREAD | SW1116 | Idelalisib | MVP_top60_DNA+RNA_seed | PIK3CA | 1.799843 | COADREAD | SNU-61 | PI3Ka_4409 |
| MVP_top60_DNA_seed | ERBB4 | 2.042327 | COADREAD | SW1463 | PF-00299804 | MVP_top60_DNA+RNA_seed | PIK3CD | 1.799843 | COADREAD | SNU-61 | PI3Ka_4409 |
| MVP_top60_DNA_seed | SRC | 2.043946 | COADREAD | HT55 | HG-6-71-01 | MVP_top60_DNA+RNA_seed | SRC | 1.800697 | COADREAD | SW1463 | Bosutinib |
| MVP_top60_DNA_seed | AKT1 | 2.053788 | COADREAD | CL-11 | AKT inhibitor VIII | MVP_top60_DNA+RNA_seed | BCL2 | 1.805778 | COADREAD | MDST8 | Obatoclax Mesylate |
| MVP_top60_DNA_seed | SRC | 2.053909 | COADREAD | SW1116 | NVP-BHG712 | MVP_top60_DNA+RNA_seed | BCL2 | 1.809052 | COADREAD | DiFi | TW 37 |
| MVP_top60_DNA_seed | GSK3B | 2.054105 | COADREAD | SW1463 | SB216763 | MVP_top60_DNA+RNA_seed | GSK3B | 1.811202 | COADREAD | SW948 | CHIR-99021 |
| MVP_top60_DNA_seed | CDK2 | 2.056485 | COADREAD | SW1463 | PHA-793887 | MVP_top60_DNA+RNA_seed | AKT1 | 1.81131 | COADREAD | SW1417 | AKT inhibitor VIII |
| MVP_top60_DNA_seed | CDK2 | 2.061113 | COADREAD | SW1417 | AZD5438 | MVP_top60_DNA+RNA_seed | GSK3B | 1.823985 | COADREAD | COLO-678 | KIN001-042 |
| MVP_top60_DNA_seed | MET | 2.061636 | COADREAD | SW1463 | AZD6094 | MVP_top60_DNA+RNA_seed | CDK2 | 1.824851 | COADREAD | NCI-H716 | Seliciclib |

| | | | | | | | | | | | |
|---|---|---|---|---|---|---|---|---|---|---|---|
| MVP_top60_DNA_seed | HSP90AA1 | 2.062466 | COADREAD | SNU-61 | Elesclomol | MVP_top60_DNA+RNA_seed | PIK3CA | 1.826066 | COADREAD | KM12 | AZD8835 |
| MVP_top60_DNA_seed | HSP90AB1 | 2.062466 | COADREAD | SNU-61 | Elesclomol | MVP_top60_DNA+RNA_seed | PIK3CD | 1.826066 | COADREAD | KM12 | AZD8835 |
| MVP_top60_DNA_seed | MET | 2.063711 | COADREAD | SW1116 | JNJ38877605 | MVP_top60_DNA+RNA_seed | ERBB4 | 1.837485 | COADREAD | COLO-678 | CI-1033 |
| MVP_top60_DNA_seed | AKT1 | 2.064504 | COADREAD | RCM-1 | MK-2206 | MVP_top60_DNA+RNA_seed | SRC | 1.839582 | COADREAD | LS-123 | Bosutinib |
| MVP_top60_DNA_seed | PIK3CB | 2.066858 | COADREAD | SW948 | AZD6482 | MVP_top60_DNA+RNA_seed | AR | 1.840858 | COADREAD | HT-29 | Bicalutamide |
| MVP_top60_DNA_seed | IGF1R | 2.071002 | COADREAD | SW948 | GSK1904529A | MVP_top60_DNA+RNA_seed | BCL2 | 1.841754 | COADREAD | CL-11 | Obatoclax Mesylate |
| MVP_top60_DNA_seed | GSK3B | 2.075879 | COADREAD | SNU-61 | SB216763 | MVP_top60_DNA+RNA_seed | AKT1 | 1.841974 | COADREAD | SW1116 | BAY AKT1 |
| MVP_top60_DNA_seed | IGF1R | 2.08387 | COADREAD | CW-2 | BMS-754807 | MVP_top60_DNA+RNA_seed | PIK3CB | 1.84767 | COADREAD | RCM-1 | AZD6482 |
| MVP_top60_DNA_seed | HSP90AA1 | 2.087691 | COADREAD | RCM-1 | Elesclomol | MVP_top60_DNA+RNA_seed | AR | 1.855999 | COADREAD | SNU-61 | Bicalutamide |
| MVP_top60_DNA_seed | HSP90AB1 | 2.087691 | COADREAD | RCM-1 | Elesclomol | MVP_top60_DNA+RNA_seed | AKT1 | 1.8599 | COADREAD | SW1463 | AKT inhibitor VIII |
| MVP_top60_DNA_seed | GSK3B | 2.090263 | COADREAD | SW1116 | KIN001-042 | MVP_top60_DNA+RNA_seed | GSK3B | 1.864381 | COADREAD | LS-123 | AZD7969 |
| MVP_top60_DNA_seed | MAPK14 | 2.092086 | COADREAD | COLO-678 | TAK-715 | MVP_top60_DNA+RNA_seed | CDK2 | 1.867422 | COADREAD | SW1116 | PHA-793887 |
| MVP_top60_DNA_seed | PIK3CA | 2.09473 | COADREAD | KM12 | PI3Ka_4409 | MVP_top60_DNA+RNA_seed | PIK3CB | 1.868278 | COADREAD | SNU-C1 | TGX221 |
| MVP_top60_DNA_seed | PIK3CD | 2.09473 | COADREAD | KM12 | PI3Ka_4409 | MVP_top60_DNA+RNA_seed | PIK3CA | 1.869052 | COADREAD | C2BBe1 | PI3Ka_4409 |
| MVP_top60_DNA_seed | AKT1 | 2.109723 | COADREAD | SW620 | BAY AKT1 | MVP_top60_DNA+RNA_seed | PIK3CD | 1.869052 | COADREAD | C2BBe1 | PI3Ka_4409 |
| MVP_top60_DNA_seed | ERBB4 | 2.111112 | COADREAD | HCT-15 | CI-1033 | MVP_top60_DNA+RNA_seed | HSP90AA1 | 1.870765 | COADREAD | COLO-205 | Elesclomol |
| MVP_top60_DNA_seed | ATM | 2.112754 | COADREAD | LS-123 | KU-55933 | MVP_top60_DNA+RNA_seed | HSP90AB1 | 1.870765 | COADREAD | COLO-205 | Elesclomol |
| MVP_top60_DNA_seed | ATM | 2.116676 | COADREAD | HT55 | torin2 | MVP_top60_DNA+RNA_seed | HSP90AA1 | 1.879065 | COADREAD | SW1116 | Elesclomol |
| MVP_top60_DNA_seed | IGF1R | 2.11867 | COADREAD | SNU-61 | BMS-536924 | MVP_top60_DNA+RNA_seed | HSP90AB1 | 1.879065 | COADREAD | SW1116 | Elesclomol |
| MVP_top60_DNA_seed | ERBB4 | 2.128012 | COADREAD | SNU-C5 | CI-1033 | MVP_top60_DNA+RNA_seed | BCL2 | 1.882473 | COADREAD | CL-11 | TW 37 |
| MVP_top60_DNA_seed | ESR1 | 2.134421 | COADREAD | SW1116 | Fulvestrant | MVP_top60_DNA+RNA_seed | AKT1 | 1.882993 | COADREAD | T84 | MK-2206 |
| MVP_top60_DNA_seed | HSP90AA1 | 2.135319 | COADREAD | SW1463 | Elesclomol | MVP_top60_DNA+RNA_seed | AR | 1.885556 | COADREAD | C2BBe1 | AZD3514 |
| MVP_top60_DNA_seed | HSP90AB1 | 2.135319 | COADREAD | SW1463 | Elesclomol | MVP_top60_DNA+RNA_seed | IGF1R | 1.887598 | COADREAD | SNU-1040 | BMS-536924 |
| MVP_top60_DNA_seed | MET | 2.140357 | COADREAD | SW1463 | SU11274 | MVP_top60_DNA+RNA_seed | BCL2 | 1.895019 | COADREAD | HCC2998 | TW 37 |
| MVP_top60_DNA_seed | ERBB4 | 2.142317 | COADREAD | LS-1034 | AST-1306 | MVP_top60_DNA+RNA_seed | AKT1 | 1.896555 | COADREAD | HT55 | AKT inhibitor VIII |
| MVP_top60_DNA_seed | ATM | 2.148201 | COADREAD | HT-29 | KU-60019 | MVP_top60_DNA+RNA_seed | PIK3CB | 1.902959 | COADREAD | LS-123 | AZD6482 |
| MVP_top60_DNA_seed | MET | 2.150234 | COADREAD | SW1116 | SU11274 | MVP_top60_DNA+RNA_seed | ERBB4 | 1.906833 | COADREAD | SW620 | CI-1033 |
| MVP_top60_DNA_seed | ESR1 | 2.152401 | COADREAD | SW948 | Fulvestrant | MVP_top60_DNA+RNA_seed | PIK3CB | 1.907161 | COADREAD | SNU-1040 | AZD8186 |
| MVP_top60_DNA_seed | PIK3CB | 2.16025 | COADREAD | SNU-61 | AZD8186 | MVP_top60_DNA+RNA_seed | PIK3CD | 1.907161 | COADREAD | SNU-1040 | AZD8186 |
| MVP_top60_DNA_seed | PIK3CD | 2.16025 | COADREAD | SNU-61 | AZD8186 | MVP_top60_DNA+RNA_seed | HSP90AA1 | 1.908858 | COADREAD | NCI-H508 | Elesclomol |
| MVP_top60_DNA_seed | GSK3B | 2.161546 | COADREAD | SW1463 | KIN001-042 | MVP_top60_DNA+RNA_seed | HSP90AB1 | 1.908858 | COADREAD | NCI-H508 | Elesclomol |
| MVP_top60_DNA_seed | MAPK1 | 2.17182 | COADREAD | SNU-C1 | FR-180204 | MVP_top60_DNA+RNA_seed | HSP90AA1 | 1.910671 | COADREAD | CL-11 | Luminespib |
| MVP_top60_DNA_seed | MET | 2.200625 | COADREAD | SW1463 | JNJ38877605 | MVP_top60_DNA+RNA_seed | HSP90AB1 | 1.910671 | COADREAD | CL-11 | Luminespib |
| MVP_top60_DNA_seed | GSK3B | 2.222205 | COADREAD | SNU-C1 | SB216763 | MVP_top60_DNA+RNA_seed | GSK3B | 1.916896 | COADREAD | HT55 | AZD7969 |
| MVP_top60_DNA_seed | ATM | 2.230647 | COADREAD | SW948 | KU-55933 | MVP_top60_DNA+RNA_seed | AR | 1.924926 | COADREAD | SW620 | Bicalutamide |
| MVP_top60_DNA_seed | ATM | 2.230884 | COADREAD | LS-513 | KU-60019 | MVP_top60_DNA+RNA_seed | PIK3CA | 1.928324 | COADREAD | SNU-1040 | AZD8835 |
| MVP_top60_DNA_seed | CREBBP | 2.239728 | COADREAD | LS-123 | I-CBP112 | MVP_top60_DNA+RNA_seed | PIK3CD | 1.928324 | COADREAD | SNU-1040 | AZD8835 |
| MVP_top60_DNA_seed | IGF1R | 2.245758 | COADREAD | SNU-C1 | GSK1904529A | MVP_top60_DNA+RNA_seed | ATM | 1.9308 | COADREAD | SW837 | KU-55933 |
| MVP_top60_DNA_seed | AR | 2.252173 | COADREAD | SNU-61 | AZD3514 | MVP_top60_DNA+RNA_seed | MET | 1.934629 | COADREAD | SNU-81 | JNJ38877605 |
| MVP_top60_DNA_seed | PTK2 | 2.256602 | COADREAD | CW-2 | PF-562271 | MVP_top60_DNA+RNA_seed | GSK3B | 1.935474 | COADREAD | LS-123 | CHIR-99021 |
| MVP_top60_DNA_seed | AR | 2.261199 | COADREAD | SW837 | Bicalutamide | MVP_top60_DNA+RNA_seed | GSK3B | 1.940117 | COADREAD | SNU-61 | CHIR-99021 |

| | | | | | | | | | | | |
|---|---|---|---|---|---|---|---|---|---|---|---|
| MVP_top60_DNA_seed | AR | 2.262446 | COADREAD | LS-123 | Bicalutamide | MVP_top60_DNA+RNA_seed | IGF1R | 1.943844 | COADREAD | CW-2 | BMS-536924 |
| MVP_top60_DNA_seed | MET | 2.273844 | COADREAD | SW948 | JNJ38877605 | MVP_top60_DNA+RNA_seed | AKT1 | 1.944805 | COADREAD | SW1116 | MK-2206 |
| MVP_top60_DNA_seed | SRC | 2.287595 | COADREAD | T84 | HG-6-71-01 | MVP_top60_DNA+RNA_seed | PIK3CB | 1.949729 | COADREAD | LS-180 | AZD8186 |
| MVP_top60_DNA_seed | GSK3B | 2.288262 | COADREAD | SNU-C1 | AZD7969 | MVP_top60_DNA+RNA_seed | PIK3CD | 1.949729 | COADREAD | LS-180 | AZD8186 |
| MVP_top60_DNA_seed | HSP90AA1 | 2.291011 | COADREAD | LS-180 | Elesclomol | MVP_top60_DNA+RNA_seed | PIK3CB | 1.957446 | COADREAD | CW-2 | AZD6482 |
| MVP_top60_DNA_seed | HSP90AB1 | 2.291011 | COADREAD | LS-180 | Elesclomol | MVP_top60_DNA+RNA_seed | PTK2 | 1.958593 | COADREAD | SW837 | PF-562271 |
| MVP_top60_DNA_seed | GSK3B | 2.295778 | COADREAD | SNU-C1 | CHIR-99021 | MVP_top60_DNA+RNA_seed | AKT1 | 1.960269 | COADREAD | COLO-678 | MK-2206 |
| MVP_top60_DNA_seed | ATM | 2.296439 | COADREAD | KM12 | torin2 | MVP_top60_DNA+RNA_seed | PIK3CB | 1.961016 | COADREAD | CW-2 | AZD8186 |
| MVP_top60_DNA_seed | PIK3CD | 2.297824 | COADREAD | SW1463 | IC-87114 | MVP_top60_DNA+RNA_seed | PIK3CD | 1.961016 | COADREAD | CW-2 | AZD8186 |
| MVP_top60_DNA_seed | ATM | 2.299113 | COADREAD | SW948 | KU-60019 | MVP_top60_DNA+RNA_seed | AR | 1.974818 | COADREAD | SW1116 | AZD3514 |
| MVP_top60_DNA_seed | ESR1 | 2.301975 | COADREAD | SNU-61 | Fulvestrant | MVP_top60_DNA+RNA_seed | AR | 1.978266 | COADREAD | DiFi | AZD3514 |
| MVP_top60_DNA_seed | ERBB4 | 2.303344 | COADREAD | MDST8 | CI-1033 | MVP_top60_DNA+RNA_seed | HSP90AA1 | 1.992247 | COADREAD | SW1116 | Tanespimycin |
| MVP_top60_DNA_seed | PIK3CB | 2.308875 | COADREAD | SW1463 | AZD8186 | MVP_top60_DNA+RNA_seed | HSP90AB1 | 1.992247 | COADREAD | SW1116 | Tanespimycin |
| MVP_top60_DNA_seed | PIK3CD | 2.308875 | COADREAD | SW1463 | AZD8186 | MVP_top60_DNA+RNA_seed | PIK3CD | 1.99544 | COADREAD | SNU-61 | IC-87114 |
| MVP_top60_DNA_seed | BRAF | 2.320515 | COADREAD | SW1116 | PLX-4720 | MVP_top60_DNA+RNA_seed | MET | 1.999142 | COADREAD | SNU-C1 | AZD6094 |
| MVP_top60_DNA_seed | AKT1 | 2.323803 | COADREAD | SW948 | AKT inhibitor VIII | MVP_top60_DNA+RNA_seed | HSP90AA1 | 2.007253 | COADREAD | SNU-C5 | CCT-018159 |
| MVP_top60_DNA_seed | GSK3B | 2.324651 | COADREAD | SNU-61 | AZD7969 | MVP_top60_DNA+RNA_seed | HSP90AB1 | 2.007253 | COADREAD | SNU-C5 | CCT-018159 |
| MVP_top60_DNA_seed | HSP90AA1 | 2.342817 | COADREAD | NCI-H508 | CCT-018159 | MVP_top60_DNA+RNA_seed | AKT1 | 2.008281 | COADREAD | CW-2 | AKT inhibitor VIII |
| MVP_top60_DNA_seed | HSP90AB1 | 2.342817 | COADREAD | NCI-H508 | CCT-018159 | MVP_top60_DNA+RNA_seed | ATM | 2.012473 | COADREAD | COLO-678 | CP466722 |
| MVP_top60_DNA_seed | HSP90AA1 | 2.372512 | COADREAD | SW1463 | Tanespimycin | MVP_top60_DNA+RNA_seed | ATM | 2.01646 | COADREAD | T84 | KU-60019 |
| MVP_top60_DNA_seed | HSP90AB1 | 2.372512 | COADREAD | SW1463 | Tanespimycin | MVP_top60_DNA+RNA_seed | PIK3CD | 2.01691 | COADREAD | SW1116 | Idelalisib |
| MVP_top60_DNA_seed | MAPK1 | 2.38132 | COADREAD | SW1116 | FR-180204 | MVP_top60_DNA+RNA_seed | BCL2 | 2.020191 | COADREAD | CaR-1 | TW 37 |
| MVP_top60_DNA_seed | ESR1 | 2.382835 | COADREAD | SW1116 | Tamoxifen | MVP_top60_DNA+RNA_seed | ERBB4 | 2.042327 | COADREAD | SW1463 | PF-00299804 |
| MVP_top60_DNA_seed | ATM | 2.384769 | COADREAD | LS-180 | KU-60019 | MVP_top60_DNA+RNA_seed | SRC | 2.043946 | COADREAD | HT55 | HG-6-71-01 |
| MVP_top60_DNA_seed | ERBB4 | 2.392871 | COADREAD | SNU-C1 | PF-00299804 | MVP_top60_DNA+RNA_seed | AKT1 | 2.053788 | COADREAD | CL-11 | AKT inhibitor VIII |
| MVP_top60_DNA_seed | ERBB4 | 2.394581 | COADREAD | LS-1034 | CI-1033 | MVP_top60_DNA+RNA_seed | SRC | 2.053909 | COADREAD | SW1116 | NVP-BHG712 |
| MVP_top60_DNA_seed | CREBBP | 2.419771 | COADREAD | SW1116 | I-CBP112 | MVP_top60_DNA+RNA_seed | GSK3B | 2.054105 | COADREAD | SW1463 | SB216763 |
| MVP_top60_DNA_seed | MET | 2.430743 | COADREAD | SNU-C1 | JNJ38877605 | MVP_top60_DNA+RNA_seed | CDK2 | 2.056485 | COADREAD | SW1463 | PHA-793887 |
| MVP_top60_DNA_seed | GSK3B | 2.431403 | COADREAD | SW1417 | AZD7969 | MVP_top60_DNA+RNA_seed | CDK2 | 2.061113 | COADREAD | SW1417 | AZD5438 |
| MVP_top60_DNA_seed | BRAF | 2.439146 | COADREAD | CW-2 | HG6-64-1 | MVP_top60_DNA+RNA_seed | MET | 2.061636 | COADREAD | SW1463 | AZD6094 |
| MVP_top60_DNA_seed | ATM | 2.446635 | COADREAD | SNU-61 | KU-55933 | MVP_top60_DNA+RNA_seed | HSP90AA1 | 2.062466 | COADREAD | SNU-61 | Elesclomol |
| MVP_top60_DNA_seed | MAPK14 | 2.472138 | COADREAD | SW1116 | TAK-715 | MVP_top60_DNA+RNA_seed | HSP90AB1 | 2.062466 | COADREAD | SNU-61 | Elesclomol |
| MVP_top60_DNA_seed | MAPK1 | 2.512924 | COADREAD | SW1463 | FR-180204 | MVP_top60_DNA+RNA_seed | MET | 2.063711 | COADREAD | SW1116 | JNJ38877605 |
| MVP_top60_DNA_seed | HSP90AA1 | 2.523607 | COADREAD | LS-1034 | Elesclomol | MVP_top60_DNA+RNA_seed | AKT1 | 2.064504 | COADREAD | RCM-1 | MK-2206 |
| MVP_top60_DNA_seed | HSP90AB1 | 2.523607 | COADREAD | LS-1034 | Elesclomol | MVP_top60_DNA+RNA_seed | PIK3CB | 2.066858 | COADREAD | SW948 | AZD6482 |
| MVP_top60_DNA_seed | ATM | 2.534752 | COADREAD | LS-180 | torin2 | MVP_top60_DNA+RNA_seed | IGF1R | 2.071002 | COADREAD | SW948 | GSK1904529A |
| MVP_top60_DNA_seed | AR | 2.569492 | COADREAD | NCI-H508 | Bicalutamide | MVP_top60_DNA+RNA_seed | GSK3B | 2.075879 | COADREAD | SNU-61 | SB216763 |
| MVP_top60_DNA_seed | HSP90AA1 | 2.569554 | COADREAD | SW1116 | SNX-2112 | MVP_top60_DNA+RNA_seed | IGF1R | 2.08387 | COADREAD | CW-2 | BMS-754807 |
| MVP_top60_DNA_seed | HSP90AB1 | 2.569554 | COADREAD | SW1116 | SNX-2112 | MVP_top60_DNA+RNA_seed | HSP90AA1 | 2.087691 | COADREAD | RCM-1 | Elesclomol |
| MVP_top60_DNA_seed | HSP90AA1 | 2.570803 | COADREAD | HT-115 | Luminespib | MVP_top60_DNA+RNA_seed | HSP90AB1 | 2.087691 | COADREAD | RCM-1 | Elesclomol |
| MVP_top60_DNA_seed | HSP90AB1 | 2.570803 | COADREAD | HT-115 | Luminespib | MVP_top60_DNA+RNA_seed | GSK3B | 2.090263 | COADREAD | SW1116 | KIN001-042 |

| | | | | | | | | | | | |
|---|---|---|---|---|---|---|---|---|---|---|---|
| MVP_top60_DNA_seed | AKT1 | 2.57214 | COADREAD | NCI-H716 | A-443654 | MVP_top60_DNA+RNA_seed | PIK3CA | 2.09473 | COADREAD | KM12 | PI3Ka_4409 |
| MVP_top60_DNA_seed | BRAF | 2.574624 | COADREAD | SW1463 | PLX-4720 | MVP_top60_DNA+RNA_seed | PIK3CD | 2.09473 | COADREAD | KM12 | PI3Ka_4409 |
| MVP_top60_DNA_seed | ESR1 | 2.616591 | COADREAD | SW1463 | Tamoxifen | MVP_top60_DNA+RNA_seed | AKT1 | 2.109723 | COADREAD | SW620 | BAY AKT1 |
| MVP_top60_DNA_seed | HSP90AA1 | 2.622291 | COADREAD | SNU-61 | CCT-018159 | MVP_top60_DNA+RNA_seed | ERBB4 | 2.111112 | COADREAD | HCT-15 | CI-1033 |
| MVP_top60_DNA_seed | HSP90AB1 | 2.622291 | COADREAD | SNU-61 | CCT-018159 | MVP_top60_DNA+RNA_seed | ATM | 2.112754 | COADREAD | LS-123 | KU-55933 |
| MVP_top60_DNA_seed | HSP90AA1 | 2.623684 | COADREAD | SW948 | CCT-018159 | MVP_top60_DNA+RNA_seed | ATM | 2.116676 | COADREAD | HT55 | torin2 |
| MVP_top60_DNA_seed | HSP90AB1 | 2.623684 | COADREAD | SW948 | CCT-018159 | MVP_top60_DNA+RNA_seed | IGF1R | 2.11867 | COADREAD | SNU-61 | BMS-536924 |
| MVP_top60_DNA_seed | ATM | 2.624244 | COADREAD | HT-29 | torin2 | MVP_top60_DNA+RNA_seed | ERBB4 | 2.128012 | COADREAD | SNU-C5 | CI-1033 |
| MVP_top60_DNA_seed | ATM | 2.62843 | COADREAD | HT55 | CP466722 | MVP_top60_DNA+RNA_seed | HSP90AA1 | 2.135319 | COADREAD | SW1463 | Elesclomol |
| MVP_top60_DNA_seed | CDK2 | 2.634465 | COADREAD | SNU-61 | AZD5438 | MVP_top60_DNA+RNA_seed | HSP90AB1 | 2.135319 | COADREAD | SW1463 | Elesclomol |
| MVP_top60_DNA_seed | CREBBP | 2.660717 | COADREAD | SW1463 | I-CBP112 | MVP_top60_DNA+RNA_seed | MET | 2.140357 | COADREAD | SW1463 | SU11274 |
| MVP_top60_DNA_seed | ATM | 2.713114 | COADREAD | SNU-61 | KU-60019 | MVP_top60_DNA+RNA_seed | ERBB4 | 2.142317 | COADREAD | LS-1034 | AST-1306 |
| MVP_top60_DNA_seed | ATM | 2.716752 | COADREAD | SW1116 | KU-60019 | MVP_top60_DNA+RNA_seed | ATM | 2.148201 | COADREAD | HT-29 | KU-60019 |
| MVP_top60_DNA_seed | HSP90AA1 | 2.731918 | COADREAD | SW948 | Luminespib | MVP_top60_DNA+RNA_seed | MET | 2.150234 | COADREAD | SW1116 | SU11274 |
| MVP_top60_DNA_seed | HSP90AB1 | 2.731918 | COADREAD | SW948 | Luminespib | MVP_top60_DNA+RNA_seed | PIK3CB | 2.16025 | COADREAD | SNU-61 | AZD8186 |
| MVP_top60_DNA_seed | AR | 2.734709 | COADREAD | SNU-C1 | Bicalutamide | MVP_top60_DNA+RNA_seed | PIK3CD | 2.16025 | COADREAD | SNU-61 | AZD8186 |
| MVP_top60_DNA_seed | ATM | 2.752819 | COADREAD | SW1116 | KU-55933 | MVP_top60_DNA+RNA_seed | BCL2 | 2.160746 | COADREAD | T84 | TW 37 |
| MVP_top60_DNA_seed | ERBB4 | 2.779688 | COADREAD | MDST8 | AST-1306 | MVP_top60_DNA+RNA_seed | GSK3B | 2.161546 | COADREAD | SW1463 | KIN001-042 |
| MVP_top60_DNA_seed | ERBB4 | 2.81898 | COADREAD | MDST8 | PF-00299804 | MVP_top60_DNA+RNA_seed | MET | 2.200625 | COADREAD | SW1463 | JNJ38877605 |
| MVP_top60_DNA_seed | GSK3B | 2.837699 | COADREAD | SW1116 | AZD7969 | MVP_top60_DNA+RNA_seed | GSK3B | 2.222205 | COADREAD | SNU-C1 | SB216763 |
| MVP_top60_DNA_seed | IGF1R | 2.844592 | COADREAD | SW1463 | GSK1904529A | MVP_top60_DNA+RNA_seed | ATM | 2.230647 | COADREAD | SW948 | KU-55933 |
| MVP_top60_DNA_seed | AKT1 | 2.854649 | COADREAD | SW1463 | AKT inhibitor VIII | MVP_top60_DNA+RNA_seed | ATM | 2.230884 | COADREAD | LS-513 | KU-60019 |
| MVP_top60_DNA_seed | HSP90AA1 | 2.88444 | COADREAD | SW948 | Elesclomol | MVP_top60_DNA+RNA_seed | IGF1R | 2.245758 | COADREAD | SNU-C1 | GSK1904529A |
| MVP_top60_DNA_seed | HSP90AB1 | 2.88444 | COADREAD | SW948 | Elesclomol | MVP_top60_DNA+RNA_seed | BCL2 | 2.246389 | COADREAD | HT-115 | Obatoclax Mesylate |
| MVP_top60_DNA_seed | AKT1 | 2.933637 | COADREAD | CW-2 | A-443654 | MVP_top60_DNA+RNA_seed | AR | 2.252173 | COADREAD | SNU-61 | AZD3514 |
| MVP_top60_DNA_seed | HSP90AA1 | 2.958141 | COADREAD | SNU-C1 | Elesclomol | MVP_top60_DNA+RNA_seed | PTK2 | 2.256602 | COADREAD | CW-2 | PF-562271 |
| MVP_top60_DNA_seed | HSP90AB1 | 2.958141 | COADREAD | SNU-C1 | Elesclomol | MVP_top60_DNA+RNA_seed | AR | 2.261199 | COADREAD | SW837 | Bicalutamide |
| MVP_top60_DNA_seed | BRAF | 2.958328 | COADREAD | SNU-C1 | RAF_9304 | MVP_top60_DNA+RNA_seed | AR | 2.262446 | COADREAD | LS-123 | Bicalutamide |
| MVP_top60_DNA_seed | ERBB4 | 2.99927 | COADREAD | SW948 | CI-1033 | MVP_top60_DNA+RNA_seed | MET | 2.273844 | COADREAD | SW948 | JNJ38877605 |
| MVP_top60_DNA_seed | ATM | 3.090585 | COADREAD | SW1463 | KU-60019 | MVP_top60_DNA+RNA_seed | SRC | 2.287595 | COADREAD | T84 | HG-6-71-01 |
| MVP_top60_DNA_seed | HSP90AA1 | 3.098864 | COADREAD | CW-2 | Luminespib | MVP_top60_DNA+RNA_seed | GSK3B | 2.288262 | COADREAD | SNU-C1 | AZD7969 |
| MVP_top60_DNA_seed | HSP90AB1 | 3.098864 | COADREAD | CW-2 | Luminespib | MVP_top60_DNA+RNA_seed | HSP90AA1 | 2.291011 | COADREAD | LS-180 | Elesclomol |
| MVP_top60_DNA_seed | ATM | 3.099592 | COADREAD | SW948 | torin2 | MVP_top60_DNA+RNA_seed | HSP90AB1 | 2.291011 | COADREAD | LS-180 | Elesclomol |
| MVP_top60_DNA_seed | ATM | 3.150686 | COADREAD | NCI-H508 | KU-60019 | MVP_top60_DNA+RNA_seed | GSK3B | 2.295778 | COADREAD | SNU-C1 | CHIR-99021 |
| MVP_top60_DNA_seed | HSP90AA1 | 3.294103 | COADREAD | SW1463 | CCT-018159 | MVP_top60_DNA+RNA_seed | ATM | 2.296439 | COADREAD | KM12 | torin2 |
| MVP_top60_DNA_seed | HSP90AB1 | 3.294103 | COADREAD | SW1463 | CCT-018159 | MVP_top60_DNA+RNA_seed | PIK3CD | 2.297824 | COADREAD | SW1463 | IC-87114 |
| MVP_top60_DNA_seed | HSP90AA1 | 3.321052 | COADREAD | SNU-C1 | Luminespib | MVP_top60_DNA+RNA_seed | ATM | 2.299113 | COADREAD | SW948 | KU-60019 |
| MVP_top60_DNA_seed | HSP90AB1 | 3.321052 | COADREAD | SNU-C1 | Luminespib | MVP_top60_DNA+RNA_seed | ERBB4 | 2.303344 | COADREAD | MDST8 | CI-1033 |
| MVP_top60_DNA_seed | ERBB4 | 3.373915 | COADREAD | SNU-C1 | CI-1033 | MVP_top60_DNA+RNA_seed | PIK3CB | 2.308875 | COADREAD | SW1463 | AZD8186 |
| MVP_top60_DNA_seed | ERBB4 | 3.418571 | COADREAD | SW948 | AST-1306 | MVP_top60_DNA+RNA_seed | PIK3CD | 2.308875 | COADREAD | SW1463 | AZD8186 |
| MVP_top60_DNA_seed | SRC | 3.514521 | COADREAD | LS-180 | HG-6-71-01 | MVP_top60_DNA+RNA_seed | AKT1 | 2.323803 | COADREAD | SW948 | AKT inhibitor VIII |

| | | | | | | | | | | | |
|---|---|---|---|---|---|---|---|---|---|---|---|
| MVP_top60_DNA_seed | HSP90AA1 | 3.526931 | COADREAD | SNU-C1 | CCT-018159 | MVP_top60_DNA+RNA_seed | GSK3B | 2.324651 | COADREAD | SNU-61 | AZD7969 |
| MVP_top60_DNA_seed | HSP90AB1 | 3.526931 | COADREAD | SNU-C1 | CCT-018159 | MVP_top60_DNA+RNA_seed | BCL2 | 2.327641 | COADREAD | LS-513 | Obatoclax Mesylate |
| MVP_top60_DNA_seed | SRC | 3.723935 | COADREAD | SW948 | HG-6-71-01 | MVP_top60_DNA+RNA_seed | HSP90AA1 | 2.342817 | COADREAD | NCI-H508 | CCT-018159 |
| MVP_top60_DNA_seed | AR | 3.81982 | COADREAD | SW1463 | Bicalutamide | MVP_top60_DNA+RNA_seed | HSP90AB1 | 2.342817 | COADREAD | NCI-H508 | CCT-018159 |
| MVP_top60_DNA_seed | ERBB3 | -4.39312 | COADREAD | DiFi | Sapitinib | MVP_top60_DNA+RNA_seed | HSP90AA1 | 2.372512 | COADREAD | SW1463 | Tanespimycin |
| MVP_top60_DNA_seed | ESR1 | -4.310294 | COADREAD | COLO-320-HSR | GDC0810 | MVP_top60_DNA+RNA_seed | HSP90AB1 | 2.372512 | COADREAD | SW1463 | Tanespimycin |
| MVP_top60_DNA_seed | BRAF | -3.715734 | COADREAD | COLO-205 | SB590885 | MVP_top60_DNA+RNA_seed | ATM | 2.384769 | COADREAD | LS-180 | KU-60019 |
| MVP_top60_DNA_seed | IGF1R | -3.54392 | COADREAD | KM12 | BMS-536924 | MVP_top60_DNA+RNA_seed | ERBB4 | 2.392871 | COADREAD | SNU-C1 | PF-00299804 |
| MVP_top60_DNA_seed | MAPK1 | -3.04239 | COADREAD | LoVo | VX-11e | MVP_top60_DNA+RNA_seed | ERBB4 | 2.394581 | COADREAD | LS-1034 | CI-1033 |
| MVP_top60_DNA_seed | MAPK1 | -2.800628 | COADREAD | SK-CO-1 | SCH772984 | MVP_top60_DNA+RNA_seed | MET | 2.430743 | COADREAD | SNU-C1 | JNJ38877605 |
| MVP_top60_DNA_seed | IGF1R | -2.714608 | COADREAD | HT55 | BMS-754807 | MVP_top60_DNA+RNA_seed | GSK3B | 2.431403 | COADREAD | SW1417 | AZD7969 |
| MVP_top60_DNA_seed | MAPK1 | -2.652843 | COADREAD | COLO-205 | SCH772984 | MVP_top60_DNA+RNA_seed | ATM | 2.446635 | COADREAD | SNU-61 | KU-55933 |
| MVP_top60_DNA_seed | IGF1R | -2.600273 | COADREAD | COLO-320-HSR | BMS-536924 | MVP_top60_DNA+RNA_seed | HSP90AA1 | 2.523607 | COADREAD | LS-1034 | Elesclomol |
| MVP_top60_DNA_seed | IGF1R | -2.580636 | COADREAD | HT55 | Linsitinib | MVP_top60_DNA+RNA_seed | HSP90AB1 | 2.523607 | COADREAD | LS-1034 | Elesclomol |
| MVP_top60_DNA_seed | ERBB3 | -2.560911 | COADREAD | CCK-81 | Sapitinib | MVP_top60_DNA+RNA_seed | ATM | 2.534752 | COADREAD | LS-180 | torin2 |
| MVP_top60_DNA_seed | IGF1R | -2.538889 | COADREAD | SK-CO-1 | BMS-754807 | MVP_top60_DNA+RNA_seed | BCL2 | 2.561382 | COADREAD | SNU-C1 | Obatoclax Mesylate |
| MVP_top60_DNA_seed | MAPK1 | -2.474314 | COADREAD | HCC2998 | SCH772984 | MVP_top60_DNA+RNA_seed | AR | 2.569492 | COADREAD | NCI-H508 | Bicalutamide |
| MVP_top60_DNA_seed | BRAF | -2.42515 | COADREAD | CCK-81 | SB590885 | MVP_top60_DNA+RNA_seed | HSP90AA1 | 2.569554 | COADREAD | SW1116 | SNX-2112 |
| MVP_top60_DNA_seed | BRAF | -2.424628 | COADREAD | COLO-205 | PLX-4720 | MVP_top60_DNA+RNA_seed | HSP90AB1 | 2.569554 | COADREAD | SW1116 | SNX-2112 |
| MVP_top60_DNA_seed | ERBB3 | -2.400359 | COADREAD | NCI-H508 | Sapitinib | MVP_top60_DNA+RNA_seed | HSP90AA1 | 2.570803 | COADREAD | HT-115 | Luminespib |
| MVP_top60_DNA_seed | IGF1R | -2.399669 | COADREAD | COLO-205 | BMS-754807 | MVP_top60_DNA+RNA_seed | HSP90AB1 | 2.570803 | COADREAD | HT-115 | Luminespib |
| MVP_top60_DNA_seed | IGF1R | -2.387482 | COADREAD | COLO-320-HSR | NVP-ADW742 | MVP_top60_DNA+RNA_seed | AKT1 | 2.57214 | COADREAD | NCI-H716 | A-443654 |
| MVP_top60_DNA_seed | BRAF | -2.336353 | COADREAD | COLO-205 | Dabrafenib | MVP_top60_DNA+RNA_seed | HSP90AA1 | 2.622291 | COADREAD | SNU-61 | CCT-018159 |
| MVP_top60_DNA_seed | MAPK1 | -2.32705 | COADREAD | CL-11 | VX-11e | MVP_top60_DNA+RNA_seed | HSP90AB1 | 2.622291 | COADREAD | SNU-61 | CCT-018159 |
| MVP_top60_DNA_seed | PIK3CA | -2.306434 | COADREAD | C2BBe1 | AZD8186 | MVP_top60_DNA+RNA_seed | HSP90AA1 | 2.623684 | COADREAD | SW948 | CCT-018159 |
| MVP_top60_DNA_seed | PIK3CB | -2.306434 | COADREAD | C2BBe1 | AZD8186 | MVP_top60_DNA+RNA_seed | HSP90AB1 | 2.623684 | COADREAD | SW948 | CCT-018159 |
| MVP_top60_DNA_seed | AKT1 | -2.292727 | COADREAD | NCI-H508 | Ipatasertib | MVP_top60_DNA+RNA_seed | ATM | 2.624244 | COADREAD | HT-29 | torin2 |
| MVP_top60_DNA_seed | MAPK1 | -2.285555 | COADREAD | HCT-116 | Ulixertinib | MVP_top60_DNA+RNA_seed | ATM | 2.62843 | COADREAD | HT55 | CP466722 |
| MVP_top60_DNA_seed | IGF1R | -2.28298 | COADREAD | SK-CO-1 | Linsitinib | MVP_top60_DNA+RNA_seed | CDK2 | 2.634465 | COADREAD | SNU-61 | AZD5438 |
| MVP_top60_DNA_seed | AKT1 | -2.263737 | COADREAD | NCI-H508 | Uprosertib | MVP_top60_DNA+RNA_seed | BCL2 | 2.636136 | COADREAD | SW1116 | TW 37 |
| MVP_top60_DNA_seed | IGF1R | -2.253406 | COADREAD | NCI-H716 | BMS-536924 | MVP_top60_DNA+RNA_seed | ATM | 2.713114 | COADREAD | SNU-61 | KU-60019 |
| MVP_top60_DNA_seed | MAPK1 | -2.249274 | COADREAD | HCC2998 | VX-11e | MVP_top60_DNA+RNA_seed | ATM | 2.716752 | COADREAD | SW1116 | KU-60019 |
| MVP_top60_DNA_seed | IGF1R | -2.243466 | COADREAD | KM12 | BMS-754807 | MVP_top60_DNA+RNA_seed | HSP90AA1 | 2.731918 | COADREAD | SW948 | Luminespib |
| MVP_top60_DNA_seed | IGF1R | -2.242433 | COADREAD | NCI-H716 | BMS-754807 | MVP_top60_DNA+RNA_seed | HSP90AB1 | 2.731918 | COADREAD | SW948 | Luminespib |
| MVP_top60_DNA_seed | MAPK1 | -2.215852 | COADREAD | SNU-C1 | SCH772984 | MVP_top60_DNA+RNA_seed | AR | 2.734709 | COADREAD | SNU-C1 | Bicalutamide |
| MVP_top60_DNA_seed | BRAF | -2.208516 | COADREAD | HT-29 | Dabrafenib | MVP_top60_DNA+RNA_seed | ATM | 2.752819 | COADREAD | SW1116 | KU-55933 |
| MVP_top60_DNA_seed | ESR1 | -2.204537 | COADREAD | COLO-320-HSR | Tamoxifen | MVP_top60_DNA+RNA_seed | BCL2 | 2.773481 | COADREAD | COLO-205 | TW 37 |
| MVP_top60_DNA_seed | MAPK1 | -2.194581 | COADREAD | HT-29 | SCH772984 | MVP_top60_DNA+RNA_seed | ERBB4 | 2.779688 | COADREAD | MDST8 | AST-1306 |
| MVP_top60_DNA_seed | MAPK1 | -2.178984 | COADREAD | MDST8 | SCH772984 | MVP_top60_DNA+RNA_seed | ERBB4 | 2.81898 | COADREAD | MDST8 | PF-00299804 |
| MVP_top60_DNA_seed | IGF1R | -2.176908 | COADREAD | COLO-205 | Linsitinib | MVP_top60_DNA+RNA_seed | GSK3B | 2.837699 | COADREAD | SW1116 | AZD7969 |
| MVP_top60_DNA_seed | MET | -2.172774 | COADREAD | KM12 | Crizotinib | MVP_top60_DNA+RNA_seed | IGF1R | 2.844592 | COADREAD | SW1463 | GSK1904529A |

| | | | | | | | | | | | |
|---|---|---|---|---|---|---|---|---|---|---|---|
| MVP_top60_DNA_seed | IGF1R | -2.169504 | COADREAD | SW48 | BMS-754807 | MVP_top60_DNA+RNA_seed | AKT1 | 2.854649 | COADREAD | SW1463 | AKT inhibitor VIII |
| MVP_top60_DNA_seed | PIK3CA | -2.162493 | COADREAD | NCI-H508 | Alpelisib | MVP_top60_DNA+RNA_seed | HSP90AA1 | 2.88444 | COADREAD | SW948 | Elesclomol |
| MVP_top60_DNA_seed | MAPK1 | -2.144906 | COADREAD | SK-CO-1 | Ulixertinib | MVP_top60_DNA+RNA_seed | HSP90AB1 | 2.88444 | COADREAD | SW948 | Elesclomol |
| MVP_top60_DNA_seed | MAPK1 | -2.144653 | COADREAD | HCC2998 | Ulixertinib | MVP_top60_DNA+RNA_seed | AKT1 | 2.933637 | COADREAD | CW-2 | A-443654 |
| MVP_top60_DNA_seed | PIK3CA | -2.120441 | COADREAD | LoVo | AZD8186 | MVP_top60_DNA+RNA_seed | HSP90AA1 | 2.958141 | COADREAD | SNU-C1 | Elesclomol |
| MVP_top60_DNA_seed | PIK3CB | -2.120441 | COADREAD | LoVo | AZD8186 | MVP_top60_DNA+RNA_seed | HSP90AB1 | 2.958141 | COADREAD | SNU-C1 | Elesclomol |
| MVP_top60_DNA_seed | MAPK1 | -2.099375 | COADREAD | COLO-205 | Ulixertinib | MVP_top60_DNA+RNA_seed | ERBB4 | 2.99927 | COADREAD | SW948 | CI-1033 |
| MVP_top60_DNA_seed | IGF1R | -2.073516 | COADREAD | SW48 | Linsitinib | MVP_top60_DNA+RNA_seed | ATM | 3.090585 | COADREAD | SW1463 | KU-60019 |
| MVP_top60_DNA_seed | MAPK1 | -2.054591 | COADREAD | HT-29 | VX-11e | MVP_top60_DNA+RNA_seed | BCL2 | 3.098837 | COADREAD | SNU-61 | TW 37 |
| MVP_top60_DNA_seed | GSK3B | -2.052661 | COADREAD | LS-513 | SB216763 | MVP_top60_DNA+RNA_seed | HSP90AA1 | 3.098864 | COADREAD | CW-2 | Luminespib |
| MVP_top60_DNA_seed | MAPK1 | -2.030259 | COADREAD | HCT-116 | SCH772984 | MVP_top60_DNA+RNA_seed | HSP90AB1 | 3.098864 | COADREAD | CW-2 | Luminespib |
| MVP_top60_DNA_seed | MAPK1 | -2.027697 | COADREAD | LS-513 | VX-11e | MVP_top60_DNA+RNA_seed | ATM | 3.099592 | COADREAD | SW948 | torin2 |
| MVP_top60_DNA_seed | AKT1 | -2.019189 | COADREAD | LoVo | Uprosertib | MVP_top60_DNA+RNA_seed | ATM | 3.150686 | COADREAD | NCI-H508 | KU-60019 |
| MVP_top60_DNA_seed | MAPK1 | -2.005738 | COADREAD | SK-CO-1 | VX-11e | MVP_top60_DNA+RNA_seed | BCL2 | 3.277545 | COADREAD | SW948 | TW 37 |
| MVP_top60_DNA_seed | MAPK1 | -1.977919 | COADREAD | HCT-116 | VX-11e | MVP_top60_DNA+RNA_seed | HSP90AA1 | 3.294103 | COADREAD | SW1463 | CCT-018159 |
| MVP_top60_DNA_seed | MAPK1 | -1.95416 | COADREAD | HT-29 | Ulixertinib | MVP_top60_DNA+RNA_seed | HSP90AB1 | 3.294103 | COADREAD | SW1463 | CCT-018159 |
| MVP_top60_DNA_seed | HSP90AA1 | -1.944093 | COADREAD | SW48 | Tanespimycin | MVP_top60_DNA+RNA_seed | HSP90AA1 | 3.321052 | COADREAD | SNU-C1 | Luminespib |
| MVP_top60_DNA_seed | HSP90AB1 | -1.944093 | COADREAD | SW48 | Tanespimycin | MVP_top60_DNA+RNA_seed | HSP90AB1 | 3.321052 | COADREAD | SNU-C1 | Luminespib |
| MVP_top60_DNA_seed | MAPK1 | -1.943735 | COADREAD | HCT-116 | Ulixertinib | MVP_top60_DNA+RNA_seed | ERBB4 | 3.373915 | COADREAD | SNU-C1 | CI-1033 |
| MVP_top60_DNA_seed | GSK3B | -1.941929 | COADREAD | HT-115 | CHIR-99021 | MVP_top60_DNA+RNA_seed | ERBB4 | 3.418571 | COADREAD | SW948 | AST-1306 |
| MVP_top60_DNA_seed | IGF1R | -1.936546 | COADREAD | COLO-205 | BMS-536924 | MVP_top60_DNA+RNA_seed | SRC | 3.514521 | COADREAD | LS-180 | HG-6-71-01 |
| MVP_top60_DNA_seed | IGF1R | -1.933365 | COADREAD | SW48 | BMS-536924 | MVP_top60_DNA+RNA_seed | HSP90AA1 | 3.526931 | COADREAD | SNU-C1 | CCT-018159 |
| MVP_top60_DNA_seed | MAPK1 | -1.926096 | COADREAD | LS-411N | Ulixertinib | MVP_top60_DNA+RNA_seed | HSP90AB1 | 3.526931 | COADREAD | SNU-C1 | CCT-018159 |
| MVP_top60_DNA_seed | AKT1 | -1.923966 | COADREAD | KM12 | Afuresertib | MVP_top60_DNA+RNA_seed | BCL2 | 3.666407 | COADREAD | SW1463 | TW 37 |
| MVP_top60_DNA_seed | IGF1R | -1.910036 | COADREAD | SK-CO-1 | BMS-536924 | MVP_top60_DNA+RNA_seed | SRC | 3.723935 | COADREAD | SW948 | HG-6-71-01 |
| MVP_top60_DNA_seed | MAPK1 | -1.907672 | COADREAD | MDST8 | Ulixertinib | MVP_top60_DNA+RNA_seed | AR | 3.81982 | COADREAD | SW1463 | Bicalutamide |
| MVP_top60_DNA_seed | AKT1 | -1.902874 | COADREAD | NCI-H508 | Afuresertib | MVP_top60_DNA+RNA_seed | BCL2 | 3.852817 | COADREAD | NCI-H508 | TW 37 |
| MVP_top60_DNA_seed | MAPK1 | -1.879203 | COADREAD | HCC2998 | Ulixertinib | MVP_top60_DNA+RNA_seed | BCL2 | 4.110897 | COADREAD | CW-2 | Obatoclax Mesylate |
| MVP_top60_DNA_seed | AKT1 | -1.862369 | COADREAD | NCI-H508 | MK-2206 | MVP_top60_DNA+RNA_seed | BCL2 | 5.243342 | COADREAD | SNU-C1 | TW 37 |
| MVP_top60_DNA_seed | IGF1R | -1.856477 | COADREAD | COLO-320-HSR | Linsitinib | MVP_top60_DNA+RNA_seed | ERBB3 | -4.39312 | COADREAD | DiFi | Sapitinib |
| MVP_top60_DNA_seed | GSK3B | -1.852956 | COADREAD | NCI-H508 | SB216763 | MVP_top60_DNA+RNA_seed | IGF1R | -3.54392 | COADREAD | KM12 | BMS-536924 |
| MVP_top60_DNA_seed | MAPK1 | -1.84253 | COADREAD | SK-CO-1 | Ulixertinib | MVP_top60_DNA+RNA_seed | IGF1R | -2.714608 | COADREAD | HT55 | BMS-754807 |
| MVP_top60_DNA_seed | IGF1R | -1.841411 | COADREAD | NCI-H508 | BMS-754807 | MVP_top60_DNA+RNA_seed | IGF1R | -2.600273 | COADREAD | COLO-320-HSR | BMS-536924 |
| MVP_top60_DNA_seed | AKT1 | -1.832173 | COADREAD | KM12 | Uprosertib | MVP_top60_DNA+RNA_seed | IGF1R | -2.580636 | COADREAD | HT55 | Linsitinib |
| MVP_top60_DNA_seed | GSK3B | -1.82968 | COADREAD | HT-115 | SB216763 | MVP_top60_DNA+RNA_seed | ERBB3 | -2.560911 | COADREAD | CCK-81 | Sapitinib |
| MVP_top60_DNA_seed | IGF1R | -1.826463 | COADREAD | COLO-205 | NVP-ADW742 | MVP_top60_DNA+RNA_seed | IGF1R | -2.538889 | COADREAD | SK-CO-1 | BMS-754807 |
| MVP_top60_DNA_seed | IGF1R | -1.799875 | COADREAD | SK-CO-1 | NVP-ADW742 | MVP_top60_DNA+RNA_seed | ERBB3 | -2.400359 | COADREAD | NCI-H508 | Sapitinib |
| MVP_top60_DNA_seed | IGF1R | -1.798393 | COADREAD | SNU-C1 | BMS-754807 | MVP_top60_DNA+RNA_seed | IGF1R | -2.399669 | COADREAD | COLO-205 | BMS-754807 |
| MVP_top60_DNA_seed | MAPK1 | -1.795416 | COADREAD | NCI-H747 | SCH772984 | MVP_top60_DNA+RNA_seed | IGF1R | -2.387482 | COADREAD | COLO-320-HSR | NVP-ADW742 |
| MVP_top60_DNA_seed | IGF1R | -1.793471 | COADREAD | LS-513 | Linsitinib | MVP_top60_DNA+RNA_seed | PIK3CA | -2.306434 | COADREAD | C2BBe1 | AZD8186 |
| MVP_top60_DNA_seed | BRAF | -1.782926 | COADREAD | MDST8 | Dabrafenib | MVP_top60_DNA+RNA_seed | PIK3CB | -2.306434 | COADREAD | C2BBe1 | AZD8186 |

| | | | | | | | | | | | |
|---|---|---|---|---|---|---|---|---|---|---|---|
| MVP_top60_DNA_seed | AKT1 | -1.748626 | COADREAD | LoVo | MK-2206 | MVP_top60_DNA+RNA_seed | AKT1 | -2.292727 | COADREAD | NCI-H508 | Ipatasertib |
| MVP_top60_DNA_seed | ESR1 | -1.748574 | COADREAD | CCK-81 | GDC0810 | MVP_top60_DNA+RNA_seed | IGF1R | -2.28298 | COADREAD | SK-CO-1 | Linsitinib |
| MVP_top60_DNA_seed | MAPK1 | -1.733668 | COADREAD | COLO-205 | VX-11e | MVP_top60_DNA+RNA_seed | AKT1 | -2.263737 | COADREAD | NCI-H508 | Uprosertib |
| MVP_top60_DNA_seed | IGF1R | -1.729344 | COADREAD | SNU-175 | BMS-754807 | MVP_top60_DNA+RNA_seed | IGF1R | -2.253406 | COADREAD | NCI-H716 | BMS-536924 |
| MVP_top60_DNA_seed | GSK3B | -1.728412 | COADREAD | CCK-81 | SB216763 | MVP_top60_DNA+RNA_seed | IGF1R | -2.243466 | COADREAD | KM12 | BMS-754807 |
| MVP_top60_DNA_seed | MAPK1 | -1.719827 | COADREAD | LS-411N | SCH772984 | MVP_top60_DNA+RNA_seed | IGF1R | -2.242433 | COADREAD | NCI-H716 | BMS-754807 |
| MVP_top60_DNA_seed | GSK3B | -1.706066 | COADREAD | COLO-205 | SB216763 | MVP_top60_DNA+RNA_seed | IGF1R | -2.176908 | COADREAD | COLO-205 | Linsitinib |
| MVP_top60_DNA_seed | AKT1 | -1.70417 | COADREAD | LoVo | GSK2110183B | MVP_top60_DNA+RNA_seed | MET | -2.172774 | COADREAD | KM12 | Crizotinib |
| MVP_top60_DNA_seed | MAPK1 | -1.701206 | COADREAD | HT55 | SCH772984 | MVP_top60_DNA+RNA_seed | IGF1R | -2.169504 | COADREAD | SW48 | BMS-754807 |
| MVP_top60_DNA_seed | IGF1R | -1.694994 | COADREAD | LS-411N | NVP-ADW742 | MVP_top60_DNA+RNA_seed | PIK3CA | -2.162493 | COADREAD | NCI-H508 | Alpelisib |
| MVP_top60_DNA_seed | MAPK1 | -1.689861 | COADREAD | LS-411N | VX-11e | MVP_top60_DNA+RNA_seed | PIK3CA | -2.120441 | COADREAD | LoVo | AZD8186 |
| MVP_top60_DNA_seed | IGF1R | -1.681765 | COADREAD | SW948 | BMS-754807 | MVP_top60_DNA+RNA_seed | PIK3CB | -2.120441 | COADREAD | LoVo | AZD8186 |
| MVP_top60_DNA_seed | AKT1 | -1.671478 | COADREAD | LS-411N | Uprosertib | MVP_top60_DNA+RNA_seed | BCL2 | -2.096024 | COADREAD | HT-29 | Obatoclax Mesylate |
| MVP_top60_DNA_seed | IGF1R | -1.643508 | COADREAD | NCI-H716 | Linsitinib | MVP_top60_DNA+RNA_seed | IGF1R | -2.073516 | COADREAD | SW48 | Linsitinib |
| MVP_top60_DNA_seed | IGF1R | -1.640865 | COADREAD | LS-513 | BMS-754807 | MVP_top60_DNA+RNA_seed | GSK3B | -2.052661 | COADREAD | LS-513 | SB216763 |
| MVP_top60_DNA_seed | ERBB3 | -1.629848 | COADREAD | CW-2 | Sapitinib | MVP_top60_DNA+RNA_seed | AKT1 | -2.019189 | COADREAD | LoVo | Uprosertib |
| MVP_top60_DNA_seed | HSP90AA1 | -1.622209 | COADREAD | GP5d | Tanespimycin | MVP_top60_DNA+RNA_seed | HSP90AA1 | -1.944093 | COADREAD | SW48 | Tanespimycin |
| MVP_top60_DNA_seed | HSP90AB1 | -1.622209 | COADREAD | GP5d | Tanespimycin | MVP_top60_DNA+RNA_seed | HSP90AB1 | -1.944093 | COADREAD | SW48 | Tanespimycin |
| MVP_top60_DNA_seed | MAPK1 | -1.618179 | COADREAD | COLO-205 | Ulixertinib | MVP_top60_DNA+RNA_seed | GSK3B | -1.941929 | COADREAD | HT-115 | CHIR-99021 |
| MVP_top60_DNA_seed | AKT1 | -1.598056 | COADREAD | LoVo | AT13148 | MVP_top60_DNA+RNA_seed | IGF1R | -1.936546 | COADREAD | COLO-205 | BMS-536924 |
| MVP_top60_DNA_seed | AKT1 | -1.596983 | COADREAD | SNU-C1 | GSK2110183B | MVP_top60_DNA+RNA_seed | IGF1R | -1.933365 | COADREAD | SW48 | BMS-536924 |
| MVP_top60_DNA_seed | MAPK1 | -1.595756 | COADREAD | MDST8 | VX-11e | MVP_top60_DNA+RNA_seed | AKT1 | -1.923966 | COADREAD | KM12 | Afuresertib |
| MVP_top60_DNA_seed | HSP90AA1 | -1.594319 | COADREAD | HT-29 | Tanespimycin | MVP_top60_DNA+RNA_seed | IGF1R | -1.910036 | COADREAD | SK-CO-1 | BMS-536924 |
| MVP_top60_DNA_seed | HSP90AB1 | -1.594319 | COADREAD | HT-29 | Tanespimycin | MVP_top60_DNA+RNA_seed | AKT1 | -1.902874 | COADREAD | NCI-H508 | Afuresertib |
| MVP_top60_DNA_seed | AKT1 | -1.5913 | COADREAD | KM12 | AZD5363 | MVP_top60_DNA+RNA_seed | AKT1 | -1.862369 | COADREAD | NCI-H508 | MK-2206 |
| MVP_top60_DNA_seed | BRAF | -1.585375 | COADREAD | HT-29 | PLX-4720 | MVP_top60_DNA+RNA_seed | IGF1R | -1.856477 | COADREAD | COLO-320-HSR | Linsitinib |
| MVP_top60_DNA_seed | AKT1 | -1.564764 | COADREAD | NCI-H508 | AZD5363 | MVP_top60_DNA+RNA_seed | GSK3B | -1.852956 | COADREAD | NCI-H508 | SB216763 |
| MVP_top60_DNA_seed | IGF1R | -1.54775 | COADREAD | SW48 | NVP-ADW742 | MVP_top60_DNA+RNA_seed | IGF1R | -1.841411 | COADREAD | NCI-H508 | BMS-754807 |
| MVP_top60_DNA_seed | MAPK1 | -1.540209 | COADREAD | KM12 | Ulixertinib | MVP_top60_DNA+RNA_seed | AKT1 | -1.832173 | COADREAD | KM12 | Uprosertib |
| MVP_top60_DNA_seed | AKT1 | -1.53445 | COADREAD | DiFi | Uprosertib | MVP_top60_DNA+RNA_seed | GSK3B | -1.82968 | COADREAD | HT-115 | SB216763 |
| MVP_top60_DNA_seed | ATM | -1.525765 | COADREAD | SK-CO-1 | KU-55933 | MVP_top60_DNA+RNA_seed | IGF1R | -1.826463 | COADREAD | COLO-205 | NVP-ADW742 |
| MVP_top60_DNA_seed | AKT1 | -1.519347 | COADREAD | SNU-81 | Ipatasertib | MVP_top60_DNA+RNA_seed | IGF1R | -1.799875 | COADREAD | SK-CO-1 | NVP-ADW742 |
| MVP_top60_DNA_seed | BRAF | -1.518973 | COADREAD | RKO | Dabrafenib | MVP_top60_DNA+RNA_seed | IGF1R | -1.798393 | COADREAD | SNU-C1 | BMS-754807 |
| MVP_top60_DNA_seed | AKT1 | -1.517079 | COADREAD | KM12 | GSK2110183B | MVP_top60_DNA+RNA_seed | IGF1R | -1.793471 | COADREAD | LS-513 | Linsitinib |
| MVP_top60_DNA_seed | PIK3CA | -1.501377 | COADREAD | GP5d | Alpelisib | MVP_top60_DNA+RNA_seed | AKT1 | -1.748626 | COADREAD | LoVo | MK-2206 |
| MVP_top60_DNA_seed | IGF1R | -1.499877 | COADREAD | SNU-C1 | Linsitinib | MVP_top60_DNA+RNA_seed | IGF1R | -1.729344 | COADREAD | SNU-175 | BMS-754807 |
| MVP_top60_DNA_seed | AKT1 | -1.498848 | COADREAD | DiFi | GSK2110183B | MVP_top60_DNA+RNA_seed | GSK3B | -1.728412 | COADREAD | CCK-81 | SB216763 |
| MVP_top60_DNA_seed | ERBB3 | -1.493277 | COADREAD | SW48 | Sapitinib | MVP_top60_DNA+RNA_seed | GSK3B | -1.706066 | COADREAD | COLO-205 | SB216763 |
| MVP_top60_DNA_seed | MAPK1 | -1.491133 | COADREAD | LS-411N | Ulixertinib | MVP_top60_DNA+RNA_seed | AKT1 | -1.70417 | COADREAD | LoVo | GSK2110183B |
| MVP_top60_DNA_seed | ESR1 | -1.487007 | COADREAD | T84 | GDC0810 | MVP_top60_DNA+RNA_seed | IGF1R | -1.694994 | COADREAD | LS-411N | NVP-ADW742 |
| MVP_top60_DNA_seed | MAPK1 | -1.486483 | COADREAD | SW48 | SCH772984 | MVP_top60_DNA+RNA_seed | IGF1R | -1.681765 | COADREAD | SW948 | BMS-754807 |

| | | | | | | | | | | | |
|---|---|---|---|---|---|---|---|---|---|---|---|
| MVP_top60_DNA_seed | MAPK1 | -1.484094 | COADREAD | SW1463 | Ulixertinib | MVP_top60_DNA+RNA_seed | AKT1 | -1.671478 | COADREAD | LS-411N | Uprosertib |
| MVP_top60_DNA_seed | ATM | -1.481905 | COADREAD | RKO | KU-55933 | MVP_top60_DNA+RNA_seed | IGF1R | -1.643508 | COADREAD | NCI-H716 | Linsitinib |
| MVP_top60_DNA_seed | ERBB3 | -1.459895 | COADREAD | SNU-175 | Sapitinib | MVP_top60_DNA+RNA_seed | IGF1R | -1.640865 | COADREAD | LS-513 | BMS-754807 |
| MVP_top60_DNA_seed | IGF1R | -1.459885 | COADREAD | SNU-175 | GSK1904529A | MVP_top60_DNA+RNA_seed | ERBB3 | -1.629848 | COADREAD | CW-2 | Sapitinib |
| MVP_top60_DNA_seed | MET | -1.45958 | COADREAD | NCI-H716 | Savolitinib | MVP_top60_DNA+RNA_seed | HSP90AA1 | -1.622209 | COADREAD | GP5d | Tanespimycin |
| MVP_top60_DNA_seed | ERBB3 | -1.456916 | COADREAD | LoVo | Sapitinib | MVP_top60_DNA+RNA_seed | HSP90AB1 | -1.622209 | COADREAD | GP5d | Tanespimycin |
| MVP_top60_DNA_seed | AKT1 | -1.451713 | COADREAD | GP5d | GSK2110183B | MVP_top60_DNA+RNA_seed | AKT1 | -1.598056 | COADREAD | LoVo | AT13148 |
| MVP_top60_DNA_seed | IGF1R | -1.443249 | COADREAD | HT-29 | Linsitinib | MVP_top60_DNA+RNA_seed | AKT1 | -1.596983 | COADREAD | SNU-C1 | GSK2110183B |
| MVP_top60_DNA_seed | MAPK1 | -1.440379 | COADREAD | SW48 | Ulixertinib | MVP_top60_DNA+RNA_seed | HSP90AA1 | -1.594319 | COADREAD | HT-29 | Tanespimycin |
| MVP_top60_DNA_seed | AKT1 | -1.436764 | COADREAD | SW948 | Uprosertib | MVP_top60_DNA+RNA_seed | HSP90AB1 | -1.594319 | COADREAD | HT-29 | Tanespimycin |
| MVP_top60_DNA_seed | AKT1 | -1.435505 | COADREAD | COLO-320-HSR | Afuresertib | MVP_top60_DNA+RNA_seed | AKT1 | -1.5913 | COADREAD | KM12 | AZD5363 |
| MVP_top60_DNA_seed | MAPK1 | -1.434243 | COADREAD | NCI-H747 | Ulixertinib | MVP_top60_DNA+RNA_seed | BCL2 | -1.579417 | COADREAD | HCT-116 | Obatoclax Mesylate |
| MVP_top60_DNA_seed | AKT1 | -1.432425 | COADREAD | NCI-H747 | Uprosertib | MVP_top60_DNA+RNA_seed | AKT1 | -1.564764 | COADREAD | NCI-H508 | AZD5363 |
| MVP_top60_DNA_seed | BRAF | -1.427691 | COADREAD | LS-411N | PLX-4720 | MVP_top60_DNA+RNA_seed | IGF1R | -1.54775 | COADREAD | SW48 | NVP-ADW742 |
| MVP_top60_DNA_seed | KRAS | -1.426956 | COADREAD | KM12 | KRAS (G12C) Inhibitor-1: | MVP_top60_DNA+RNA_seed | AKT1 | -1.53445 | COADREAD | DiFi | Uprosertib |
| MVP_top60_DNA_seed | IGF1R | -1.425122 | COADREAD | COLO-320-HSR | BMS-754807 | MVP_top60_DNA+RNA_seed | ATM | -1.525765 | COADREAD | SK-CO-1 | KU-55933 |
| MVP_top60_DNA_seed | MAPK1 | -1.420504 | COADREAD | HT55 | VX-11e | MVP_top60_DNA+RNA_seed | AKT1 | -1.519347 | COADREAD | SNU-81 | Ipatasertib |
| MVP_top60_DNA_seed | ATM | -1.418079 | COADREAD | CCK-81 | KU-55933 | MVP_top60_DNA+RNA_seed | AKT1 | -1.517079 | COADREAD | KM12 | GSK2110183B |
| MVP_top60_DNA_seed | BRAF | -1.412943 | COADREAD | HT-29 | SB590885 | MVP_top60_DNA+RNA_seed | PIK3CA | -1.501377 | COADREAD | GP5d | Alpelisib |
| MVP_top60_DNA_seed | SRC | -1.3937 | COADREAD | KM12 | Bosutinib | MVP_top60_DNA+RNA_seed | IGF1R | -1.499877 | COADREAD | SNU-C1 | Linsitinib |
| MVP_top60_DNA_seed | MAPK1 | -1.393525 | COADREAD | SNU-C1 | Ulixertinib | MVP_top60_DNA+RNA_seed | AKT1 | -1.498848 | COADREAD | DiFi | GSK2110183B |
| MVP_top60_DNA_seed | MET | -1.389328 | COADREAD | SW948 | Savolitinib | MVP_top60_DNA+RNA_seed | ERBB3 | -1.493277 | COADREAD | SW48 | Sapitinib |
| MVP_top60_DNA_seed | MAPK1 | -1.388175 | COADREAD | SW1417 | SCH772984 | MVP_top60_DNA+RNA_seed | ATM | -1.481905 | COADREAD | RKO | KU-55933 |
| MVP_top60_DNA_seed | GSK3B | -1.387434 | COADREAD | NCI-H747 | SB216763 | MVP_top60_DNA+RNA_seed | ERBB3 | -1.459895 | COADREAD | SNU-175 | Sapitinib |
| MVP_top60_DNA_seed | BRAF | -1.3808 | COADREAD | RCM-1 | SB590885 | MVP_top60_DNA+RNA_seed | IGF1R | -1.459885 | COADREAD | SNU-175 | GSK1904529A |
| MVP_top60_DNA_seed | AKT1 | -1.377157 | COADREAD | HCC2998 | Afuresertib | MVP_top60_DNA+RNA_seed | MET | -1.45958 | COADREAD | NCI-H716 | Savolitinib |
| MVP_top60_DNA_seed | ATM | -1.3741 | COADREAD | LoVo | KU-55933 | MVP_top60_DNA+RNA_seed | ERBB3 | -1.456916 | COADREAD | LoVo | Sapitinib |
| MVP_top60_DNA_seed | BRAF | -1.369627 | COADREAD | MDST8 | PLX-4720 | MVP_top60_DNA+RNA_seed | AKT1 | -1.451713 | COADREAD | GP5d | GSK2110183B |
| MVP_top60_DNA_seed | HSP90AA1 | -1.365428 | COADREAD | COLO-205 | Tanespimycin | MVP_top60_DNA+RNA_seed | IGF1R | -1.443249 | COADREAD | HT-29 | Linsitinib |
| MVP_top60_DNA_seed | HSP90AB1 | -1.365428 | COADREAD | COLO-205 | Tanespimycin | MVP_top60_DNA+RNA_seed | AKT1 | -1.436764 | COADREAD | SW948 | Uprosertib |
| MVP_top60_DNA_seed | MAPK1 | -1.356264 | COADREAD | LoVo | Ulixertinib | MVP_top60_DNA+RNA_seed | AKT1 | -1.435505 | COADREAD | COLO-320-HSR | Afuresertib |
| MVP_top60_DNA_seed | MAPK1 | -1.347363 | COADREAD | LS-513 | SCH772984 | MVP_top60_DNA+RNA_seed | BCL2 | -1.43424 | COADREAD | SW1417 | Navitoclax |
| MVP_top60_DNA_seed | AKT1 | -1.34604 | COADREAD | KM12 | Ipatasertib | MVP_top60_DNA+RNA_seed | AKT1 | -1.432425 | COADREAD | NCI-H747 | Uprosertib |
| MVP_top60_DNA_seed | AKT1 | -1.340581 | COADREAD | HT-115 | AT13148 | MVP_top60_DNA+RNA_seed | KRAS | -1.426956 | COADREAD | KM12 | KRAS (G12C) Inhibitor-1 |
| MVP_top60_DNA_seed | AKT1 | -1.32684 | COADREAD | SNU-C1 | Uprosertib | MVP_top60_DNA+RNA_seed | IGF1R | -1.425122 | COADREAD | COLO-320-HSR | BMS-754807 |
| MVP_top60_DNA_seed | MAPK1 | -1.326069 | COADREAD | KM12 | VX-11e | MVP_top60_DNA+RNA_seed | ATM | -1.418079 | COADREAD | CCK-81 | KU-55933 |
| MVP_top60_DNA_seed | MAPK1 | -1.323256 | COADREAD | HT-29 | Ulixertinib | MVP_top60_DNA+RNA_seed | BCL2 | -1.394963 | COADREAD | COLO-320-HSR | TW 37 |
| MVP_top60_DNA_seed | BRAF | -1.313137 | COADREAD | MDST8 | SB590885 | MVP_top60_DNA+RNA_seed | SRC | -1.3937 | COADREAD | KM12 | Bosutinib |
| MVP_top60_DNA_seed | ATM | -1.312324 | COADREAD | T84 | KU-55933 | MVP_top60_DNA+RNA_seed | MET | -1.389328 | COADREAD | SW948 | Savolitinib |
| MVP_top60_DNA_seed | CDK2 | -1.308692 | COADREAD | COLO-320-HSR | MK-8776 | MVP_top60_DNA+RNA_seed | GSK3B | -1.387434 | COADREAD | NCI-H747 | SB216763 |
| MVP_top60_DNA_seed | ESR1 | -1.304184 | COADREAD | SK-CO-1 | Tamoxifen | MVP_top60_DNA+RNA_seed | BCL2 | -1.383923 | COADREAD | SK-CO-1 | Navitoclax |

| | | | | | | | | | | | |
|---|---|---|---|---|---|---|---|---|---|---|---|
| MVP_top60_DNA_seed | MET | -1.302795 | COADREAD | SW48 | Crizotinib | MVP_top60_DNA+RNA_seed | AKT1 | -1.377157 | COADREAD | HCC2998 | Afuresertib |
| MVP_top60_DNA_seed | AKT1 | -1.302398 | COADREAD | GP5d | Uprosertib | MVP_top60_DNA+RNA_seed | ATM | -1.3741 | COADREAD | LoVo | KU-55933 |
| MVP_top60_DNA_seed | KRAS | -1.301712 | COADREAD | HCT-15 | KRAS (G12C) Inhibitor-12 | MVP_top60_DNA+RNA_seed | HSP90AA1 | -1.365428 | COADREAD | COLO-205 | Tanespimycin |
| MVP_top60_DNA_seed | AKT1 | -1.291894 | COADREAD | DiFi | AT13148 | MVP_top60_DNA+RNA_seed | HSP90AB1 | -1.365428 | COADREAD | COLO-205 | Tanespimycin |
| MVP_top60_DNA_seed | HSP90AA1 | -1.288081 | COADREAD | SK-CO-1 | Tanespimycin | MVP_top60_DNA+RNA_seed | BCL2 | -1.358924 | COADREAD | CL-11 | Obatoclax Mesylate |
| MVP_top60_DNA_seed | HSP90AB1 | -1.288081 | COADREAD | SK-CO-1 | Tanespimycin | MVP_top60_DNA+RNA_seed | AKT1 | -1.34604 | COADREAD | KM12 | Ipatasertib |
| MVP_top60_DNA_seed | HSP90AA1 | -1.285761 | COADREAD | SW620 | Tanespimycin | MVP_top60_DNA+RNA_seed | BCL2 | -1.34281 | COADREAD | RKO | Obatoclax Mesylate |
| MVP_top60_DNA_seed | HSP90AB1 | -1.285761 | COADREAD | SW620 | Tanespimycin | MVP_top60_DNA+RNA_seed | AKT1 | -1.340581 | COADREAD | HT-115 | AT13148 |
| MVP_top60_DNA_seed | MET | -1.283924 | COADREAD | KM12 | Savolitinib | MVP_top60_DNA+RNA_seed | AKT1 | -1.32684 | COADREAD | SNU-C1 | Uprosertib |
| MVP_top60_DNA_seed | HSP90AA1 | -1.278251 | COADREAD | HCC2998 | Tanespimycin | MVP_top60_DNA+RNA_seed | BCL2 | -1.322188 | COADREAD | HCT-116 | Sabutoclax |
| MVP_top60_DNA_seed | HSP90AB1 | -1.278251 | COADREAD | HCC2998 | Tanespimycin | MVP_top60_DNA+RNA_seed | BCL2 | -1.319103 | COADREAD | RKO | TW 37 |
| MVP_top60_DNA_seed | AKT1 | -1.27755 | COADREAD | LS-513 | MK-2206 | MVP_top60_DNA+RNA_seed | ATM | -1.312324 | COADREAD | T84 | KU-55933 |
| MVP_top60_DNA_seed | IGF1R | -1.277325 | COADREAD | HT-29 | BMS-536924 | MVP_top60_DNA+RNA_seed | CDK2 | -1.308692 | COADREAD | COLO-320-HSR | MK-8776 |
| MVP_top60_DNA_seed | AKT1 | -1.27498 | COADREAD | HCC2998 | GSK2110183B | MVP_top60_DNA+RNA_seed | MET | -1.302795 | COADREAD | SW48 | Crizotinib |
| MVP_top60_DNA_seed | ESR1 | -1.271957 | COADREAD | SK-CO-1 | GDC0810 | MVP_top60_DNA+RNA_seed | AKT1 | -1.302398 | COADREAD | GP5d | Uprosertib |
| MVP_top60_DNA_seed | IGF1R | -1.270367 | COADREAD | HT-29 | NVP-ADW742 | MVP_top60_DNA+RNA_seed | KRAS | -1.301712 | COADREAD | HCT-15 | KRAS (G12C) Inhibitor-1 |
| MVP_top60_DNA_seed | MAPK1 | -1.263702 | COADREAD | SNU-175 | SCH772984 | MVP_top60_DNA+RNA_seed | AKT1 | -1.291894 | COADREAD | DiFi | AT13148 |
| MVP_top60_DNA_seed | MAPK1 | -1.263247 | COADREAD | RKO | Ulixertinib | MVP_top60_DNA+RNA_seed | HSP90AA1 | -1.288081 | COADREAD | SK-CO-1 | Tanespimycin |
| MVP_top60_DNA_seed | MAPK1 | -1.260461 | COADREAD | SW1463 | VX-11e | MVP_top60_DNA+RNA_seed | HSP90AB1 | -1.288081 | COADREAD | SK-CO-1 | Tanespimycin |
| MVP_top60_DNA_seed | HSP90AA1 | -1.259449 | COADREAD | CCK-81 | Tanespimycin | MVP_top60_DNA+RNA_seed | HSP90AA1 | -1.285761 | COADREAD | SW620 | Tanespimycin |
| MVP_top60_DNA_seed | HSP90AB1 | -1.259449 | COADREAD | CCK-81 | Tanespimycin | MVP_top60_DNA+RNA_seed | HSP90AB1 | -1.285761 | COADREAD | SW620 | Tanespimycin |
| MVP_top60_DNA_seed | MET | -1.25731 | COADREAD | SK-CO-1 | Crizotinib | MVP_top60_DNA+RNA_seed | MET | -1.283924 | COADREAD | KM12 | Savolitinib |
| MVP_top60_DNA_seed | IGF1R | -1.252954 | COADREAD | NCI-H508 | Linsitinib | MVP_top60_DNA+RNA_seed | HSP90AA1 | -1.278251 | COADREAD | HCC2998 | Tanespimycin |
| MVP_top60_DNA_seed | MAPK1 | -1.247209 | COADREAD | KM12 | SCH772984 | MVP_top60_DNA+RNA_seed | HSP90AB1 | -1.278251 | COADREAD | HCC2998 | Tanespimycin |
| MVP_top60_DNA_seed | AKT1 | -1.246593 | COADREAD | SNU-C1 | Afuresertib | MVP_top60_DNA+RNA_seed | AKT1 | -1.27755 | COADREAD | LS-513 | MK-2206 |
| MVP_top60_DNA_seed | MAPK1 | -1.240634 | COADREAD | CL-11 | Ulixertinib | MVP_top60_DNA+RNA_seed | IGF1R | -1.277325 | COADREAD | HT-29 | BMS-536924 |
| MVP_top60_DNA_seed | MAPK1 | -1.2387 | COADREAD | RCM-1 | VX-11e | MVP_top60_DNA+RNA_seed | AKT1 | -1.27498 | COADREAD | HCC2998 | GSK2110183B |
| MVP_top60_DNA_seed | IGF1R | -1.236001 | COADREAD | HT55 | BMS-536924 | MVP_top60_DNA+RNA_seed | IGF1R | -1.270367 | COADREAD | HT-29 | NVP-ADW742 |
| MVP_top60_DNA_seed | BRAF | -1.232931 | COADREAD | SW1417 | Dabrafenib | MVP_top60_DNA+RNA_seed | BCL2 | -1.265263 | COADREAD | HCT-15 | TW 37 |
| MVP_top60_DNA_seed | GSK3B | -1.226249 | COADREAD | DiFi | SB216763 | MVP_top60_DNA+RNA_seed | HSP90AA1 | -1.259449 | COADREAD | CCK-81 | Tanespimycin |
| MVP_top60_DNA_seed | IGF1R | -1.224254 | COADREAD | CCK-81 | Linsitinib | MVP_top60_DNA+RNA_seed | HSP90AB1 | -1.259449 | COADREAD | CCK-81 | Tanespimycin |
| MVP_top60_DNA_seed | IGF1R | -1.221772 | COADREAD | SNU-175 | Linsitinib | MVP_top60_DNA+RNA_seed | MET | -1.25731 | COADREAD | SK-CO-1 | Crizotinib |
| MVP_top60_DNA_seed | AKT1 | -1.216936 | COADREAD | LS-513 | GSK2110183B | MVP_top60_DNA+RNA_seed | IGF1R | -1.252954 | COADREAD | NCI-H508 | Linsitinib |
| MVP_top60_DNA_seed | CREBBP | -1.215684 | COADREAD | HT-115 | SGC-CBP30 | MVP_top60_DNA+RNA_seed | AKT1 | -1.246593 | COADREAD | SNU-C1 | Afuresertib |
| MVP_top60_DNA_seed | MAPK1 | -1.212363 | COADREAD | CL-11 | Ulixertinib | MVP_top60_DNA+RNA_seed | BCL2 | -1.237918 | COADREAD | HCT-15 | Sabutoclax |
| MVP_top60_DNA_seed | MAPK1 | -1.208517 | COADREAD | SW620 | SCH772984 | MVP_top60_DNA+RNA_seed | IGF1R | -1.236001 | COADREAD | HT55 | BMS-536924 |
| MVP_top60_DNA_seed | PIK3CA | -1.201292 | COADREAD | SW48 | AZD8186 | MVP_top60_DNA+RNA_seed | BCL2 | -1.229243 | COADREAD | HCT-116 | TW 37 |
| MVP_top60_DNA_seed | PIK3CB | -1.201292 | COADREAD | SW48 | AZD8186 | MVP_top60_DNA+RNA_seed | GSK3B | -1.226249 | COADREAD | DiFi | SB216763 |
| MVP_top60_DNA_seed | CDK2 | -1.198675 | COADREAD | T84 | MK-8776 | MVP_top60_DNA+RNA_seed | IGF1R | -1.224254 | COADREAD | CCK-81 | Linsitinib |
| MVP_top60_DNA_seed | PIK3CA | -1.197354 | COADREAD | HCT-15 | GNE-317 | MVP_top60_DNA+RNA_seed | IGF1R | -1.221772 | COADREAD | SNU-175 | Linsitinib |
| MVP_top60_DNA_seed | AKT1 | -1.196322 | COADREAD | HCC2998 | Uprosertib | MVP_top60_DNA+RNA_seed | AKT1 | -1.216936 | COADREAD | LS-513 | GSK2110183B |

| | | | | | | | | | | | |
|---|---|---|---|---|---|---|---|---|---|---|---|
| MVP_top60_DNA_seed | MAPK1 | -1.194788 | COADREAD | LoVo | SCH772984 | MVP_top60_DNA+RNA_seed | PIK3CA | -1.201292 | COADREAD | SW48 | AZD8186 |
| MVP_top60_DNA_seed | MAPK1 | -1.193629 | COADREAD | LS-1034 | VX-11e | MVP_top60_DNA+RNA_seed | PIK3CB | -1.201292 | COADREAD | SW48 | AZD8186 |
| MVP_top60_DNA_seed | AKT1 | -1.193175 | COADREAD | HCT-15 | Uprosertib | MVP_top60_DNA+RNA_seed | CDK2 | -1.198675 | COADREAD | T84 | MK-8776 |
| MVP_top60_DNA_seed | MAPK1 | -1.190838 | COADREAD | RCM-1 | Ulixertinib | MVP_top60_DNA+RNA_seed | PIK3CA | -1.197354 | COADREAD | HCT-15 | GNE-317 |
| MVP_top60_DNA_seed | MAPK1 | -1.182939 | COADREAD | DiFi | VX-11e | MVP_top60_DNA+RNA_seed | AKT1 | -1.196322 | COADREAD | HCC2998 | Uprosertib |
| MVP_top60_DNA_seed | AKT1 | -1.181745 | COADREAD | MDST8 | Uprosertib | MVP_top60_DNA+RNA_seed | AKT1 | -1.193175 | COADREAD | HCT-15 | Uprosertib |
| MVP_top60_DNA_seed | MAPK1 | -1.180582 | COADREAD | LoVo | Ulixertinib | MVP_top60_DNA+RNA_seed | AKT1 | -1.181745 | COADREAD | MDST8 | Uprosertib |
| MVP_top60_DNA_seed | AKT1 | -1.177633 | COADREAD | SW948 | GSK2110183B | MVP_top60_DNA+RNA_seed | AKT1 | -1.177633 | COADREAD | SW948 | GSK2110183B |
| MVP_top60_DNA_seed | MAPK1 | -1.177431 | COADREAD | NCI-H716 | Ulixertinib | MVP_top60_DNA+RNA_seed | ERBB3 | -1.176306 | COADREAD | LS-513 | Sapitinib |
| MVP_top60_DNA_seed | ERBB3 | -1.176306 | COADREAD | LS-513 | Sapitinib | MVP_top60_DNA+RNA_seed | AKT1 | -1.17521 | COADREAD | HT55 | MK-2206 |
| MVP_top60_DNA_seed | AKT1 | -1.17521 | COADREAD | HT55 | MK-2206 | MVP_top60_DNA+RNA_seed | ERBB3 | -1.164165 | COADREAD | SK-CO-1 | Sapitinib |
| MVP_top60_DNA_seed | ERBB3 | -1.164165 | COADREAD | SK-CO-1 | Sapitinib | MVP_top60_DNA+RNA_seed | AKT1 | -1.162377 | COADREAD | LS-411N | Uprosertib |
| MVP_top60_DNA_seed | AKT1 | -1.162377 | COADREAD | LS-411N | Uprosertib | MVP_top60_DNA+RNA_seed | PIK3CA | -1.158287 | COADREAD | NCI-H508 | AZD8186 |
| MVP_top60_DNA_seed | MAPK1 | -1.160529 | COADREAD | CL-11 | SCH772984 | MVP_top60_DNA+RNA_seed | PIK3CB | -1.158287 | COADREAD | NCI-H508 | AZD8186 |
| MVP_top60_DNA_seed | PIK3CA | -1.158287 | COADREAD | NCI-H508 | AZD8186 | MVP_top60_DNA+RNA_seed | HSP90AA1 | -1.143462 | COADREAD | CaR-1 | Tanespimycin |
| MVP_top60_DNA_seed | PIK3CB | -1.158287 | COADREAD | NCI-H508 | AZD8186 | MVP_top60_DNA+RNA_seed | HSP90AB1 | -1.143462 | COADREAD | CaR-1 | Tanespimycin |
| MVP_top60_DNA_seed | BRAF | -1.156491 | COADREAD | LS-411N | Dabrafenib | MVP_top60_DNA+RNA_seed | IGF1R | -1.142594 | COADREAD | LS-411N | BMS-536924 |
| MVP_top60_DNA_seed | MAPK1 | -1.152998 | COADREAD | DiFi | SCH772984 | MVP_top60_DNA+RNA_seed | CDK2 | -1.13873 | COADREAD | HCT-116 | AZD5438 |
| MVP_top60_DNA_seed | MAPK1 | -1.149959 | COADREAD | SW48 | Ulixertinib | MVP_top60_DNA+RNA_seed | HSP90AA1 | -1.137224 | COADREAD | LS-513 | Tanespimycin |
| MVP_top60_DNA_seed | ESR1 | -1.147121 | COADREAD | T84 | Tamoxifen | MVP_top60_DNA+RNA_seed | HSP90AB1 | -1.137224 | COADREAD | LS-513 | Tanespimycin |
| MVP_top60_DNA_seed | HSP90AA1 | -1.143462 | COADREAD | CaR-1 | Tanespimycin | MVP_top60_DNA+RNA_seed | IGF1R | -1.134674 | COADREAD | CaR-1 | BMS-536924 |
| MVP_top60_DNA_seed | HSP90AB1 | -1.143462 | COADREAD | CaR-1 | Tanespimycin | MVP_top60_DNA+RNA_seed | AKT1 | -1.129699 | COADREAD | HT-29 | Ipatasertib |
| MVP_top60_DNA_seed | IGF1R | -1.142594 | COADREAD | LS-411N | BMS-536924 | MVP_top60_DNA+RNA_seed | PIK3CA | -1.125492 | COADREAD | SNU-81 | AZD8186 |
| MVP_top60_DNA_seed | CDK2 | -1.13873 | COADREAD | HCT-116 | AZD5438 | MVP_top60_DNA+RNA_seed | PIK3CB | -1.125492 | COADREAD | SNU-81 | AZD8186 |
| MVP_top60_DNA_seed | HSP90AA1 | -1.137224 | COADREAD | LS-513 | Tanespimycin | MVP_top60_DNA+RNA_seed | AKT1 | -1.121157 | COADREAD | NCI-H716 | Uprosertib |
| MVP_top60_DNA_seed | HSP90AB1 | -1.137224 | COADREAD | LS-513 | Tanespimycin | MVP_top60_DNA+RNA_seed | PIK3CA | -1.117837 | COADREAD | SW48 | GNE-317 |
| MVP_top60_DNA_seed | IGF1R | -1.134674 | COADREAD | CaR-1 | BMS-536924 | MVP_top60_DNA+RNA_seed | HSP90AA1 | -1.117113 | COADREAD | HT-29 | Elesclomol |
| MVP_top60_DNA_seed | AKT1 | -1.129699 | COADREAD | HT-29 | Ipatasertib | MVP_top60_DNA+RNA_seed | HSP90AB1 | -1.117113 | COADREAD | HT-29 | Elesclomol |
| MVP_top60_DNA_seed | PIK3CA | -1.125492 | COADREAD | SNU-81 | AZD8186 | MVP_top60_DNA+RNA_seed | HSP90AA1 | -1.106764 | COADREAD | HT55 | Tanespimycin |
| MVP_top60_DNA_seed | PIK3CB | -1.125492 | COADREAD | SNU-81 | AZD8186 | MVP_top60_DNA+RNA_seed | HSP90AB1 | -1.106764 | COADREAD | HT55 | Tanespimycin |
| MVP_top60_DNA_seed | BRAF | -1.125398 | COADREAD | LoVo | Dabrafenib | MVP_top60_DNA+RNA_seed | PIK3CA | -1.104252 | COADREAD | C2BBe1 | GNE-317 |
| MVP_top60_DNA_seed | AKT1 | -1.121157 | COADREAD | NCI-H716 | Uprosertib | MVP_top60_DNA+RNA_seed | TP53 | -1.101466 | COADREAD | COLO-320-HSR | MIRA-1 |
| MVP_top60_DNA_seed | PIK3CA | -1.117837 | COADREAD | SW48 | GNE-317 | MVP_top60_DNA+RNA_seed | BCL2 | -1.101204 | COADREAD | RCM-1 | Obatoclax Mesylate |
| MVP_top60_DNA_seed | HSP90AA1 | -1.117113 | COADREAD | HT-29 | Elesclomol | MVP_top60_DNA+RNA_seed | GSK3B | -1.098156 | COADREAD | HCT-116 | SB216763 |
| MVP_top60_DNA_seed | HSP90AB1 | -1.117113 | COADREAD | HT-29 | Elesclomol | MVP_top60_DNA+RNA_seed | AKT1 | -1.094777 | COADREAD | SNU-81 | GSK2110183B |
| MVP_top60_DNA_seed | MAPK1 | -1.114206 | COADREAD | RCM-1 | SCH772984 | MVP_top60_DNA+RNA_seed | AKT1 | -1.093191 | COADREAD | SNU-175 | Uprosertib |
| MVP_top60_DNA_seed | HSP90AA1 | -1.106764 | COADREAD | HT55 | Tanespimycin | MVP_top60_DNA+RNA_seed | AKT1 | -1.092239 | COADREAD | GP5d | Uprosertib |
| MVP_top60_DNA_seed | HSP90AB1 | -1.106764 | COADREAD | HT55 | Tanespimycin | MVP_top60_DNA+RNA_seed | BCL2 | -1.087546 | COADREAD | SW620 | TW 37 |
| MVP_top60_DNA_seed | MAPK1 | -1.104328 | COADREAD | NCI-H747 | Ulixertinib | MVP_top60_DNA+RNA_seed | HSP90AA1 | -1.084284 | COADREAD | SNU-175 | Tanespimycin |
| MVP_top60_DNA_seed | PIK3CA | -1.104252 | COADREAD | C2BBe1 | GNE-317 | MVP_top60_DNA+RNA_seed | HSP90AB1 | -1.084284 | COADREAD | SNU-175 | Tanespimycin |
| MVP_top60_DNA_seed | MAPK1 | -1.102105 | COADREAD | RKO | VX-11e | MVP_top60_DNA+RNA_seed | AKT1 | -1.080392 | COADREAD | NCI-H747 | Ipatasertib |

| | | | | | | | | | | | |
|---|---|---|---|---|---|---|---|---|---|---|---|
| MVP_top60_DNA_seed | TP53 | -1.101466 | COADREAD | COLO-320-HSR | MIRA-1 | MVP_top60_DNA+RNA_seed | BCL2 | -1.075701 | COADREAD | KM12 | Sabutoclax |
| MVP_top60_DNA_seed | GSK3B | -1.098156 | COADREAD | HCT-116 | SB216763 | MVP_top60_DNA+RNA_seed | GSK3B | -1.075151 | COADREAD | SK-CO-1 | SB216763 |
| MVP_top60_DNA_seed | AKT1 | -1.094777 | COADREAD | SNU-81 | GSK2110183B | MVP_top60_DNA+RNA_seed | AKT1 | -1.068429 | COADREAD | HCT-15 | AT13148 |
| MVP_top60_DNA_seed | AKT1 | -1.093191 | COADREAD | SNU-175 | Uprosertib | MVP_top60_DNA+RNA_seed | BCL2 | -1.061792 | COADREAD | CW-2 | TW 37 |
| MVP_top60_DNA_seed | MAPK1 | -1.092256 | COADREAD | COLO-320-HSR | Ulixertinib | MVP_top60_DNA+RNA_seed | IGF1R | -1.057035 | COADREAD | HT-115 | BMS-754807 |
| MVP_top60_DNA_seed | AKT1 | -1.092239 | COADREAD | GP5d | Uprosertib | MVP_top60_DNA+RNA_seed | AKT1 | -1.056328 | COADREAD | SNU-1040 | GSK2110183B |
| MVP_top60_DNA_seed | HSP90AA1 | -1.084284 | COADREAD | SNU-175 | Tanespimycin | MVP_top60_DNA+RNA_seed | MET | -1.054146 | COADREAD | LoVo | Crizotinib |
| MVP_top60_DNA_seed | HSP90AB1 | -1.084284 | COADREAD | SNU-175 | Tanespimycin | MVP_top60_DNA+RNA_seed | AKT1 | -1.053659 | COADREAD | HT-29 | GSK2110183B |
| MVP_top60_DNA_seed | AKT1 | -1.080392 | COADREAD | NCI-H747 | Ipatasertib | MVP_top60_DNA+RNA_seed | HSP90AA1 | -1.053348 | COADREAD | SW48 | Luminespib |
| MVP_top60_DNA_seed | GSK3B | -1.075151 | COADREAD | SK-CO-1 | SB216763 | MVP_top60_DNA+RNA_seed | HSP90AB1 | -1.053348 | COADREAD | SW48 | Luminespib |
| MVP_top60_DNA_seed | MAPK1 | -1.071254 | COADREAD | RKO | Ulixertinib | MVP_top60_DNA+RNA_seed | PIK3CA | -1.052704 | COADREAD | SW48 | Alpelisib |
| MVP_top60_DNA_seed | AKT1 | -1.068429 | COADREAD | HCT-15 | AT13148 | MVP_top60_DNA+RNA_seed | PIK3CA | -1.051702 | COADREAD | COLO-320-HSR | Buparlisib |
| MVP_top60_DNA_seed | MAPK1 | -1.067297 | COADREAD | NCI-H716 | Ulixertinib | MVP_top60_DNA+RNA_seed | PIK3CB | -1.051702 | COADREAD | COLO-320-HSR | Buparlisib |
| MVP_top60_DNA_seed | BRAF | -1.062641 | COADREAD | LoVo | SB590885 | MVP_top60_DNA+RNA_seed | PIK3CD | -1.051702 | COADREAD | COLO-320-HSR | Buparlisib |
| MVP_top60_DNA_seed | MAPK1 | -1.061441 | COADREAD | SW620 | Ulixertinib | MVP_top60_DNA+RNA_seed | PIK3CA | -1.051678 | COADREAD | T84 | Alpelisib |
| MVP_top60_DNA_seed | IGF1R | -1.057035 | COADREAD | HT-115 | BMS-754807 | MVP_top60_DNA+RNA_seed | IGF1R | -1.050817 | COADREAD | GP5d | BMS-754807 |
| MVP_top60_DNA_seed | ESR1 | -1.056926 | COADREAD | CCK-81 | Tamoxifen | MVP_top60_DNA+RNA_seed | AKT1 | -1.049326 | COADREAD | GP5d | Ipatasertib |
| MVP_top60_DNA_seed | BRAF | -1.05669 | COADREAD | HCT-116 | Dabrafenib | MVP_top60_DNA+RNA_seed | IGF1R | -1.044202 | COADREAD | CCK-81 | GSK1904529A |
| MVP_top60_DNA_seed | AKT1 | -1.056328 | COADREAD | SNU-1040 | GSK2110183B | MVP_top60_DNA+RNA_seed | AKT1 | -1.04317 | COADREAD | LS-411N | Ipatasertib |
| MVP_top60_DNA_seed | MET | -1.054146 | COADREAD | LoVo | Crizotinib | MVP_top60_DNA+RNA_seed | HSP90AA1 | -1.038462 | COADREAD | MDST8 | Tanespimycin |
| MVP_top60_DNA_seed | AKT1 | -1.053659 | COADREAD | HT-29 | GSK2110183B | MVP_top60_DNA+RNA_seed | HSP90AB1 | -1.038462 | COADREAD | MDST8 | Tanespimycin |
| MVP_top60_DNA_seed | HSP90AA1 | -1.053348 | COADREAD | SW48 | Luminespib | MVP_top60_DNA+RNA_seed | AKT1 | -1.037639 | COADREAD | LS-411N | AZD5363 |
| MVP_top60_DNA_seed | HSP90AB1 | -1.053348 | COADREAD | SW48 | Luminespib | MVP_top60_DNA+RNA_seed | ERBB3 | -1.036593 | COADREAD | SW1463 | Sapitinib |
| MVP_top60_DNA_seed | PIK3CA | -1.052704 | COADREAD | SW48 | Alpelisib | MVP_top60_DNA+RNA_seed | AKT1 | -1.03649 | COADREAD | SNU-407 | Uprosertib |
| MVP_top60_DNA_seed | PIK3CA | -1.051702 | COADREAD | COLO-320-HSR | Buparlisib | MVP_top60_DNA+RNA_seed | BCL2 | -1.036046 | COADREAD | SNU-175 | Sabutoclax |
| MVP_top60_DNA_seed | PIK3CB | -1.051702 | COADREAD | COLO-320-HSR | Buparlisib | MVP_top60_DNA+RNA_seed | AKT1 | -1.033779 | COADREAD | LS-411N | Afuresertib |
| MVP_top60_DNA_seed | PIK3CD | -1.051702 | COADREAD | COLO-320-HSR | Buparlisib | MVP_top60_DNA+RNA_seed | PIK3CA | -1.026602 | COADREAD | HCT-15 | Alpelisib |
| MVP_top60_DNA_seed | PIK3CA | -1.051678 | COADREAD | T84 | Alpelisib | MVP_top60_DNA+RNA_seed | AKT1 | -1.023996 | COADREAD | LS-411N | AT13148 |
| MVP_top60_DNA_seed | IGF1R | -1.050817 | COADREAD | GP5d | BMS-754807 | MVP_top60_DNA+RNA_seed | AKT1 | -1.022738 | COADREAD | COLO-205 | MK-2206 |
| MVP_top60_DNA_seed | AKT1 | -1.049326 | COADREAD | GP5d | Ipatasertib | MVP_top60_DNA+RNA_seed | AKT1 | -1.022477 | COADREAD | NCI-H747 | AZD5363 |
| MVP_top60_DNA_seed | IGF1R | -1.044202 | COADREAD | CCK-81 | GSK1904529A | MVP_top60_DNA+RNA_seed | ATM | -1.019802 | COADREAD | HCC2998 | KU-55933 |
| MVP_top60_DNA_seed | AKT1 | -1.04317 | COADREAD | LS-411N | Ipatasertib | MVP_top60_DNA+RNA_seed | AKT1 | -1.018008 | COADREAD | DiFi | MK-2206 |
| MVP_top60_DNA_seed | MAPK1 | -1.038917 | COADREAD | LS-180 | Ulixertinib | MVP_top60_DNA+RNA_seed | ERBB3 | -1.013938 | COADREAD | HT55 | Sapitinib |
| MVP_top60_DNA_seed | HSP90AA1 | -1.038462 | COADREAD | MDST8 | Tanespimycin | MVP_top60_DNA+RNA_seed | MET | -1.013764 | COADREAD | SNU-175 | Crizotinib |
| MVP_top60_DNA_seed | HSP90AB1 | -1.038462 | COADREAD | MDST8 | Tanespimycin | MVP_top60_DNA+RNA_seed | PIK3CA | -1.008135 | COADREAD | DiFi | Alpelisib |
| MVP_top60_DNA_seed | AKT1 | -1.037639 | COADREAD | LS-411N | AZD5363 | MVP_top60_DNA+RNA_seed | IGF1R | -1.005472 | COADREAD | HT-29 | BMS-754807 |
| MVP_top60_DNA_seed | ERBB3 | -1.036593 | COADREAD | SW1463 | Sapitinib | MVP_top60_DNA+RNA_seed | PIK3CA | -1.004211 | COADREAD | SK-CO-1 | AZD8186 |
| MVP_top60_DNA_seed | AKT1 | -1.03649 | COADREAD | SNU-407 | Uprosertib | MVP_top60_DNA+RNA_seed | PIK3CB | -1.004211 | COADREAD | SK-CO-1 | AZD8186 |
| MVP_top60_DNA_seed | AKT1 | -1.033779 | COADREAD | LS-411N | Afuresertib | MVP_top60_DNA+RNA_seed | AKT1 | -1.003914 | COADREAD | SW948 | Uprosertib |
| MVP_top60_DNA_seed | PIK3CA | -1.026602 | COADREAD | HCT-15 | Alpelisib | MVP_top60_DNA+RNA_seed | AKT1 | -1.000973 | COADREAD | LS-513 | Uprosertib |
| MVP_top60_DNA_seed | AKT1 | -1.023996 | COADREAD | LS-411N | AT13148 | MVP_top60_DNA+RNA_seed | AKT1 | -0.999078 | COADREAD | SW48 | AZD5363 |

| | | | | | | | | | | | |
|---|---|---|---|---|---|---|---|---|---|---|---|
| MVP_top60_DNA_seed | AKT1 | -1.022738 | COADREAD | COLO-205 | MK-2206 | MVP_top60_DNA+RNA_seed | AKT1 | -0.996298 | COADREAD | LoVo | AZD5363 |
| MVP_top60_DNA_seed | AKT1 | -1.022477 | COADREAD | NCI-H747 | AZD5363 | MVP_top60_DNA+RNA_seed | AKT1 | -0.994942 | COADREAD | COLO-320-HSR | Ipatasertib |
| MVP_top60_DNA_seed | ATM | -1.019802 | COADREAD | HCC2998 | KU-55933 | MVP_top60_DNA+RNA_seed | AKT1 | -0.994086 | COADREAD | HCC2998 | Ipatasertib |
| MVP_top60_DNA_seed | MAPK1 | -1.018328 | COADREAD | CCK-81 | VX-11e | MVP_top60_DNA+RNA_seed | PIK3CA | -0.993072 | COADREAD | MDST8 | GNE-317 |
| MVP_top60_DNA_seed | AKT1 | -1.018008 | COADREAD | DiFi | MK-2206 | MVP_top60_DNA+RNA_seed | HSP90AA1 | -0.992101 | COADREAD | RKO | Tanespimycin |
| MVP_top60_DNA_seed | MAPK1 | -1.017314 | COADREAD | NCI-H508 | Ulixertinib | MVP_top60_DNA+RNA_seed | HSP90AB1 | -0.992101 | COADREAD | RKO | Tanespimycin |
| MVP_top60_DNA_seed | ERBB3 | -1.013938 | COADREAD | HT55 | Sapitinib | MVP_top60_DNA+RNA_seed | CDK2 | -0.991802 | COADREAD | CCK-81 | MK-8776 |
| MVP_top60_DNA_seed | MET | -1.013764 | COADREAD | SNU-175 | Crizotinib | MVP_top60_DNA+RNA_seed | BCL2 | -0.991734 | COADREAD | NCI-H716 | Sabutoclax |
| MVP_top60_DNA_seed | MAPK1 | -1.00866 | COADREAD | NCI-H508 | SCH772984 | MVP_top60_DNA+RNA_seed | AKT1 | -0.985815 | COADREAD | HT-29 | Uprosertib |
| MVP_top60_DNA_seed | PIK3CA | -1.008135 | COADREAD | DiFi | Alpelisib | MVP_top60_DNA+RNA_seed | BCL2 | -0.981042 | COADREAD | NCI-H716 | Navitoclax |
| MVP_top60_DNA_seed | IGF1R | -1.005472 | COADREAD | HT-29 | BMS-754807 | MVP_top60_DNA+RNA_seed | AKT1 | -0.98037 | COADREAD | LoVo | Afuresertib |
| MVP_top60_DNA_seed | PIK3CA | -1.004211 | COADREAD | SK-CO-1 | AZD8186 | MVP_top60_DNA+RNA_seed | CDK2 | -0.979902 | COADREAD | GP5d | AZD5438 |
| MVP_top60_DNA_seed | PIK3CB | -1.004211 | COADREAD | SK-CO-1 | AZD8186 | MVP_top60_DNA+RNA_seed | PIK3CA | -0.977944 | COADREAD | SK-CO-1 | Buparlisib |
| MVP_top60_DNA_seed | AKT1 | -1.003914 | COADREAD | SW948 | Uprosertib | MVP_top60_DNA+RNA_seed | PIK3CB | -0.977944 | COADREAD | SK-CO-1 | Buparlisib |
| MVP_top60_DNA_seed | AKT1 | -1.000973 | COADREAD | LS-513 | Uprosertib | MVP_top60_DNA+RNA_seed | PIK3CD | -0.977944 | COADREAD | SK-CO-1 | Buparlisib |
| MVP_top60_DNA_seed | AKT1 | -0.999078 | COADREAD | SW48 | AZD5363 | MVP_top60_DNA+RNA_seed | HSP90AA1 | -0.975378 | COADREAD | DiFi | Tanespimycin |
| MVP_top60_DNA_seed | AKT1 | -0.996298 | COADREAD | LoVo | AZD5363 | MVP_top60_DNA+RNA_seed | HSP90AB1 | -0.975378 | COADREAD | DiFi | Tanespimycin |
| MVP_top60_DNA_seed | AKT1 | -0.994942 | COADREAD | COLO-320-HSR | Ipatasertib | MVP_top60_DNA+RNA_seed | AKT1 | -0.973728 | COADREAD | HT-29 | Uprosertib |
| MVP_top60_DNA_seed | BRAF | -0.994267 | COADREAD | RKO | SB590885 | MVP_top60_DNA+RNA_seed | IGF1R | -0.973556 | COADREAD | GP5d | GSK1904529A |
| MVP_top60_DNA_seed | AKT1 | -0.994086 | COADREAD | HCC2998 | Ipatasertib | MVP_top60_DNA+RNA_seed | IGF1R | -0.973306 | COADREAD | RKO | GSK1904529A |
| MVP_top60_DNA_seed | PIK3CA | -0.993072 | COADREAD | MDST8 | GNE-317 | MVP_top60_DNA+RNA_seed | AKT1 | -0.971616 | COADREAD | GP5d | MK-2206 |
| MVP_top60_DNA_seed | HSP90AA1 | -0.992101 | COADREAD | RKO | Tanespimycin | MVP_top60_DNA+RNA_seed | BCL2 | -0.965614 | COADREAD | LoVo | Sabutoclax |
| MVP_top60_DNA_seed | HSP90AB1 | -0.992101 | COADREAD | RKO | Tanespimycin | MVP_top60_DNA+RNA_seed | AKT1 | -0.964404 | COADREAD | LS-411N | GSK2110183B |
| MVP_top60_DNA_seed | CDK2 | -0.991802 | COADREAD | CCK-81 | MK-8776 | MVP_top60_DNA+RNA_seed | IGF1R | -0.963168 | COADREAD | MDST8 | NVP-ADW742 |
| MVP_top60_DNA_seed | AKT1 | -0.985815 | COADREAD | HT-29 | Uprosertib | MVP_top60_DNA+RNA_seed | SRC | -0.961426 | COADREAD | CCK-81 | Bosutinib |
| MVP_top60_DNA_seed | AKT1 | -0.98037 | COADREAD | LoVo | Afuresertib | MVP_top60_DNA+RNA_seed | BCL2 | -0.961162 | COADREAD | SW948 | Navitoclax |
| MVP_top60_DNA_seed | CDK2 | -0.979902 | COADREAD | GP5d | AZD5438 | MVP_top60_DNA+RNA_seed | AKT1 | -0.959918 | COADREAD | HT-29 | Afuresertib |
| MVP_top60_DNA_seed | PIK3CA | -0.977944 | COADREAD | SK-CO-1 | Buparlisib | MVP_top60_DNA+RNA_seed | MET | -0.959249 | COADREAD | SNU-175 | Savolitinib |
| MVP_top60_DNA_seed | PIK3CB | -0.977944 | COADREAD | SK-CO-1 | Buparlisib | MVP_top60_DNA+RNA_seed | PIK3CA | -0.958464 | COADREAD | NCI-H716 | AZD8186 |
| MVP_top60_DNA_seed | PIK3CD | -0.977944 | COADREAD | SK-CO-1 | Buparlisib | MVP_top60_DNA+RNA_seed | PIK3CB | -0.958464 | COADREAD | NCI-H716 | AZD8186 |
| MVP_top60_DNA_seed | HSP90AA1 | -0.975378 | COADREAD | DiFi | Tanespimycin | MVP_top60_DNA+RNA_seed | KRAS | -0.948158 | COADREAD | HT55 | KRAS (G12C) Inhibitor-1 |
| MVP_top60_DNA_seed | HSP90AB1 | -0.975378 | COADREAD | DiFi | Tanespimycin | MVP_top60_DNA+RNA_seed | MET | -0.945671 | COADREAD | HCT-116 | Crizotinib |
| MVP_top60_DNA_seed | AKT1 | -0.973728 | COADREAD | HT-29 | Uprosertib | MVP_top60_DNA+RNA_seed | AKT1 | -0.943595 | COADREAD | SW948 | AT13148 |
| MVP_top60_DNA_seed | IGF1R | -0.973556 | COADREAD | GP5d | GSK1904529A | MVP_top60_DNA+RNA_seed | BCL2 | -0.939081 | COADREAD | SW620 | Obatoclax Mesylate |
| MVP_top60_DNA_seed | IGF1R | -0.973306 | COADREAD | RKO | GSK1904529A | MVP_top60_DNA+RNA_seed | AKT1 | -0.937607 | COADREAD | SNU-C1 | MK-2206 |
| MVP_top60_DNA_seed | AKT1 | -0.971616 | COADREAD | GP5d | MK-2206 | MVP_top60_DNA+RNA_seed | CDK2 | -0.935773 | COADREAD | LS-411N | MK-8776 |
| MVP_top60_DNA_seed | AKT1 | -0.964404 | COADREAD | LS-411N | GSK2110183B | MVP_top60_DNA+RNA_seed | PIK3CA | -0.935631 | COADREAD | NCI-H747 | Alpelisib |
| MVP_top60_DNA_seed | IGF1R | -0.963168 | COADREAD | MDST8 | NVP-ADW742 | MVP_top60_DNA+RNA_seed | BCL2 | -0.934703 | COADREAD | DiFi | Navitoclax |
| MVP_top60_DNA_seed | MAPK1 | -0.961896 | COADREAD | NCI-H716 | SCH772984 | MVP_top60_DNA+RNA_seed | HSP90AA1 | -0.933817 | COADREAD | KM12 | Tanespimycin |
| MVP_top60_DNA_seed | SRC | -0.961426 | COADREAD | CCK-81 | Bosutinib | MVP_top60_DNA+RNA_seed | HSP90AB1 | -0.933817 | COADREAD | KM12 | Tanespimycin |
| MVP_top60_DNA_seed | AKT1 | -0.959918 | COADREAD | HT-29 | Afuresertib | MVP_top60_DNA+RNA_seed | IGF1R | -0.925635 | COADREAD | HT-115 | Linsitinib |

| | | | | | | | | | | | |
|---|---|---|---|---|---|---|---|---|---|---|---|
| MVP_top60_DNA_seed | MET | -0.959249 | COADREAD | SNU-175 | Savolitinib | MVP_top60_DNA+RNA_seed | PIK3CA | -0.924875 | COADREAD | CaR-1 | GNE-317 |
| MVP_top60_DNA_seed | PIK3CA | -0.958464 | COADREAD | NCI-H716 | AZD8186 | MVP_top60_DNA+RNA_seed | AKT1 | -0.921447 | COADREAD | SNU-407 | AT13148 |
| MVP_top60_DNA_seed | PIK3CB | -0.958464 | COADREAD | NCI-H716 | AZD8186 | MVP_top60_DNA+RNA_seed | AKT1 | -0.920156 | COADREAD | DiFi | Ipatasertib |
| MVP_top60_DNA_seed | MAPK1 | -0.948649 | COADREAD | SW1463 | Ulixertinib | MVP_top60_DNA+RNA_seed | IGF1R | -0.918849 | COADREAD | HT55 | NVP-ADW742 |
| MVP_top60_DNA_seed | ESR1 | -0.948181 | COADREAD | KM12 | Tamoxifen | MVP_top60_DNA+RNA_seed | AKT1 | -0.91841 | COADREAD | MDST8 | Ipatasertib |
| MVP_top60_DNA_seed | KRAS | -0.948158 | COADREAD | HT55 | KRAS (G12C) Inhibitor-1: | MVP_top60_DNA+RNA_seed | IGF1R | -0.917725 | COADREAD | SNU-C1 | NVP-ADW742 |
| MVP_top60_DNA_seed | MET | -0.945671 | COADREAD | HCT-116 | Crizotinib | MVP_top60_DNA+RNA_seed | CDK2 | -0.917715 | COADREAD | HCT-15 | AZD5438 |
| MVP_top60_DNA_seed | MAPK1 | -0.945122 | COADREAD | NCI-H747 | VX-11e | MVP_top60_DNA+RNA_seed | AKT1 | -0.914769 | COADREAD | LoVo | Ipatasertib |
| MVP_top60_DNA_seed | MAPK1 | -0.944194 | COADREAD | HT55 | Ulixertinib | MVP_top60_DNA+RNA_seed | PIK3CA | -0.911292 | COADREAD | COLO-205 | GNE-317 |
| MVP_top60_DNA_seed | AKT1 | -0.943595 | COADREAD | SW948 | AT13148 | MVP_top60_DNA+RNA_seed | ERBB3 | -0.911175 | COADREAD | NCI-H747 | Sapitinib |
| MVP_top60_DNA_seed | AKT1 | -0.937607 | COADREAD | SNU-C1 | MK-2206 | MVP_top60_DNA+RNA_seed | TP53 | -0.910323 | COADREAD | COLO-205 | MIRA-1 |
| MVP_top60_DNA_seed | CDK2 | -0.935773 | COADREAD | LS-411N | MK-8776 | MVP_top60_DNA+RNA_seed | AKT1 | -0.909866 | COADREAD | COLO-205 | Ipatasertib |
| MVP_top60_DNA_seed | PIK3CA | -0.935631 | COADREAD | NCI-H747 | Alpelisib | MVP_top60_DNA+RNA_seed | CDK2 | -0.906863 | COADREAD | LS-513 | AZD5438 |
| MVP_top60_DNA_seed | HSP90AA1 | -0.933817 | COADREAD | KM12 | Tanespimycin | MVP_top60_DNA+RNA_seed | GSK3B | -0.906425 | COADREAD | SNU-175 | SB216763 |
| MVP_top60_DNA_seed | HSP90AB1 | -0.933817 | COADREAD | KM12 | Tanespimycin | MVP_top60_DNA+RNA_seed | AKT1 | -0.903888 | COADREAD | NCI-H747 | Uprosertib |
| MVP_top60_DNA_seed | BRAF | -0.93309 | COADREAD | NCI-H716 | SB590885 | MVP_top60_DNA+RNA_seed | AKT1 | -0.902551 | COADREAD | HCC2998 | Uprosertib |
| MVP_top60_DNA_seed | MAPK1 | -0.930687 | COADREAD | RKO | SCH772984 | MVP_top60_DNA+RNA_seed | MET | -0.899268 | COADREAD | NCI-H716 | Crizotinib |
| MVP_top60_DNA_seed | IGF1R | -0.925635 | COADREAD | HT-115 | Linsitinib | MVP_top60_DNA+RNA_seed | PIK3CA | -0.897857 | COADREAD | HT55 | Alpelisib |
| MVP_top60_DNA_seed | PIK3CA | -0.924875 | COADREAD | CaR-1 | GNE-317 | MVP_top60_DNA+RNA_seed | BCL2 | -0.889617 | COADREAD | COLO-205 | Obatoclax Mesylate |
| MVP_top60_DNA_seed | AKT1 | -0.921447 | COADREAD | SNU-407 | AT13148 | MVP_top60_DNA+RNA_seed | PIK3CA | -0.889066 | COADREAD | CCK-81 | Alpelisib |
| MVP_top60_DNA_seed | AKT1 | -0.920156 | COADREAD | DiFi | Ipatasertib | MVP_top60_DNA+RNA_seed | BCL2 | -0.888953 | COADREAD | SK-CO-1 | TW 37 |
| MVP_top60_DNA_seed | IGF1R | -0.918849 | COADREAD | HT55 | NVP-ADW742 | MVP_top60_DNA+RNA_seed | HSP90AA1 | -0.888569 | COADREAD | RKO | Luminespib |
| MVP_top60_DNA_seed | AKT1 | -0.91841 | COADREAD | MDST8 | Ipatasertib | MVP_top60_DNA+RNA_seed | HSP90AB1 | -0.888569 | COADREAD | RKO | Luminespib |
| MVP_top60_DNA_seed | IGF1R | -0.917725 | COADREAD | SNU-C1 | NVP-ADW742 | MVP_top60_DNA+RNA_seed | MET | -0.887425 | COADREAD | SW948 | Crizotinib |
| MVP_top60_DNA_seed | CDK2 | -0.917715 | COADREAD | HCT-15 | AZD5438 | MVP_top60_DNA+RNA_seed | MET | -0.882153 | COADREAD | HCC2998 | Savolitinib |
| MVP_top60_DNA_seed | AKT1 | -0.914769 | COADREAD | LoVo | Ipatasertib | MVP_top60_DNA+RNA_seed | HSP90AA1 | -0.881529 | COADREAD | LS-513 | Luminespib |
| MVP_top60_DNA_seed | PIK3CA | -0.911292 | COADREAD | COLO-205 | GNE-317 | MVP_top60_DNA+RNA_seed | HSP90AB1 | -0.881529 | COADREAD | LS-513 | Luminespib |
| MVP_top60_DNA_seed | ERBB3 | -0.911175 | COADREAD | NCI-H747 | Sapitinib | MVP_top60_DNA+RNA_seed | SRC | -0.879342 | COADREAD | GP5d | Bosutinib |
| MVP_top60_DNA_seed | CREBBP | -0.911015 | COADREAD | COLO-320-HSR | SGC-CBP30 | MVP_top60_DNA+RNA_seed | MET | -0.876095 | COADREAD | COLO-205 | Crizotinib |
| MVP_top60_DNA_seed | TP53 | -0.910323 | COADREAD | COLO-205 | MIRA-1 | MVP_top60_DNA+RNA_seed | AKT1 | -0.868673 | COADREAD | DiFi | Uprosertib |
| MVP_top60_DNA_seed | AKT1 | -0.909866 | COADREAD | COLO-205 | Ipatasertib | MVP_top60_DNA+RNA_seed | BCL2 | -0.868456 | COADREAD | LS-411N | Sabutoclax |
| MVP_top60_DNA_seed | CDK2 | -0.906863 | COADREAD | LS-513 | AZD5438 | MVP_top60_DNA+RNA_seed | AKT1 | -0.863691 | COADREAD | SW48 | Ipatasertib |
| MVP_top60_DNA_seed | GSK3B | -0.906425 | COADREAD | SNU-175 | SB216763 | MVP_top60_DNA+RNA_seed | BCL2 | -0.853759 | COADREAD | HCT-15 | Obatoclax Mesylate |
| MVP_top60_DNA_seed | AKT1 | -0.903888 | COADREAD | NCI-H747 | Uprosertib | MVP_top60_DNA+RNA_seed | MET | -0.851195 | COADREAD | GP5d | Crizotinib |
| MVP_top60_DNA_seed | AKT1 | -0.902551 | COADREAD | HCC2998 | Uprosertib | MVP_top60_DNA+RNA_seed | MET | -0.849313 | COADREAD | LS-513 | Crizotinib |
| MVP_top60_DNA_seed | MET | -0.899268 | COADREAD | NCI-H716 | Crizotinib | MVP_top60_DNA+RNA_seed | CDK2 | -0.845193 | COADREAD | SNU-175 | Dinaciclib |
| MVP_top60_DNA_seed | PIK3CA | -0.897857 | COADREAD | HT55 | Alpelisib | MVP_top60_DNA+RNA_seed | AKT1 | -0.844687 | COADREAD | T84 | Ipatasertib |
| MVP_top60_DNA_seed | PIK3CA | -0.889066 | COADREAD | CCK-81 | Alpelisib | MVP_top60_DNA+RNA_seed | AKT1 | -0.842205 | COADREAD | LS-513 | Uprosertib |
| MVP_top60_DNA_seed | HSP90AA1 | -0.888569 | COADREAD | RKO | Luminespib | MVP_top60_DNA+RNA_seed | PIK3CA | -0.83054 | COADREAD | HT-29 | Alpelisib |
| MVP_top60_DNA_seed | HSP90AB1 | -0.888569 | COADREAD | RKO | Luminespib | MVP_top60_DNA+RNA_seed | IGF1R | -0.829072 | COADREAD | LS-1034 | Linsitinib |
| MVP_top60_DNA_seed | MET | -0.887425 | COADREAD | SW948 | Crizotinib | MVP_top60_DNA+RNA_seed | PIK3CA | -0.827186 | COADREAD | HCT-15 | Buparlisib |

| | | | | | | | | | | | |
|---|---|---|---|---|---|---|---|---|---|---|---|
| MVP_top60_DNA_seed | MET | -0.882153 | COADREAD | HCC2998 | Savolitinib | MVP_top60_DNA+RNA_seed | PIK3CB | -0.827186 | COADREAD | HCT-15 | Buparlisib |
| MVP_top60_DNA_seed | HSP90AA1 | -0.881529 | COADREAD | LS-513 | Luminespib | MVP_top60_DNA+RNA_seed | PIK3CD | -0.827186 | COADREAD | HCT-15 | Buparlisib |
| MVP_top60_DNA_seed | HSP90AB1 | -0.881529 | COADREAD | LS-513 | Luminespib | MVP_top60_DNA+RNA_seed | PIK3CA | -0.826032 | COADREAD | SK-CO-1 | GNE-317 |
| MVP_top60_DNA_seed | MAPK1 | -0.880618 | COADREAD | SW1417 | Ulixertinib | MVP_top60_DNA+RNA_seed | CDK2 | -0.825499 | COADREAD | DiFi | Dinaciclib |
| MVP_top60_DNA_seed | SRC | -0.879342 | COADREAD | GP5d | Bosutinib | MVP_top60_DNA+RNA_seed | BCL2 | -0.824979 | COADREAD | COLO-320-HSR | Venetoclax |
| MVP_top60_DNA_seed | MET | -0.876095 | COADREAD | COLO-205 | Crizotinib | MVP_top60_DNA+RNA_seed | BCL2 | -0.819826 | COADREAD | SNU-81 | Navitoclax |
| MVP_top60_DNA_seed | ESR1 | -0.874091 | COADREAD | HCT-15 | Tamoxifen | MVP_top60_DNA+RNA_seed | AKT1 | -0.818434 | COADREAD | MDST8 | Afuresertib |
| MVP_top60_DNA_seed | AKT1 | -0.868673 | COADREAD | DiFi | Uprosertib | MVP_top60_DNA+RNA_seed | CDK2 | -0.815487 | COADREAD | SNU-C5 | AZD5438 |
| MVP_top60_DNA_seed | MAPK1 | -0.867746 | COADREAD | SW948 | Ulixertinib | MVP_top60_DNA+RNA_seed | AKT1 | -0.813587 | COADREAD | SW948 | Ipatasertib |
| MVP_top60_DNA_seed | MAPK1 | -0.866447 | COADREAD | SW948 | VX-11e | MVP_top60_DNA+RNA_seed | PIK3CA | -0.80982 | COADREAD | GP5d | Buparlisib |
| MVP_top60_DNA_seed | AKT1 | -0.863691 | COADREAD | SW48 | Ipatasertib | MVP_top60_DNA+RNA_seed | PIK3CB | -0.80982 | COADREAD | GP5d | Buparlisib |
| MVP_top60_DNA_seed | CREBBP | -0.86339 | COADREAD | HCT-15 | SGC-CBP30 | MVP_top60_DNA+RNA_seed | PIK3CD | -0.80982 | COADREAD | GP5d | Buparlisib |
| MVP_top60_DNA_seed | BRAF | -0.8602 | COADREAD | RKO | PLX-4720 | MVP_top60_DNA+RNA_seed | PIK3CA | -0.808737 | COADREAD | COLO-320-HSR | Alpelisib |
| MVP_top60_DNA_seed | MAPK1 | -0.854207 | COADREAD | HT55 | Ulixertinib | MVP_top60_DNA+RNA_seed | AKT1 | -0.80825 | COADREAD | SNU-C1 | Ipatasertib |
| MVP_top60_DNA_seed | MET | -0.851195 | COADREAD | GP5d | Crizotinib | MVP_top60_DNA+RNA_seed | AKT1 | -0.80823 | COADREAD | SNU-175 | Uprosertib |
| MVP_top60_DNA_seed | MET | -0.849313 | COADREAD | LS-513 | Crizotinib | MVP_top60_DNA+RNA_seed | PIK3CA | -0.807439 | COADREAD | MDST8 | Buparlisib |
| MVP_top60_DNA_seed | CDK2 | -0.845193 | COADREAD | SNU-175 | Dinaciclib | MVP_top60_DNA+RNA_seed | PIK3CB | -0.807439 | COADREAD | MDST8 | Buparlisib |
| MVP_top60_DNA_seed | AKT1 | -0.844687 | COADREAD | T84 | Ipatasertib | MVP_top60_DNA+RNA_seed | PIK3CD | -0.807439 | COADREAD | MDST8 | Buparlisib |
| MVP_top60_DNA_seed | AKT1 | -0.842205 | COADREAD | LS-513 | Uprosertib | MVP_top60_DNA+RNA_seed | MET | -0.806078 | COADREAD | SNU-407 | Savolitinib |
| MVP_top60_DNA_seed | ESR1 | -0.84176 | COADREAD | HCC2998 | Tamoxifen | MVP_top60_DNA+RNA_seed | BCL2 | -0.805045 | COADREAD | COLO-205 | Navitoclax |
| MVP_top60_DNA_seed | ESR1 | -0.839229 | COADREAD | SNU-407 | GDC0810 | MVP_top60_DNA+RNA_seed | IGF1R | -0.799961 | COADREAD | NCI-H747 | Linsitinib |
| MVP_top60_DNA_seed | BRAF | -0.839029 | COADREAD | KM12 | SB590885 | MVP_top60_DNA+RNA_seed | IGF1R | -0.79871 | COADREAD | NCI-H747 | BMS-536924 |
| MVP_top60_DNA_seed | ESR1 | -0.838032 | COADREAD | LS-411N | Tamoxifen | MVP_top60_DNA+RNA_seed | AKT1 | -0.798293 | COADREAD | SNU-175 | GSK2110183B |
| MVP_top60_DNA_seed | PIK3CA | -0.83054 | COADREAD | HT-29 | Alpelisib | MVP_top60_DNA+RNA_seed | PIK3CA | -0.797717 | COADREAD | NCI-H747 | Buparlisib |
| MVP_top60_DNA_seed | IGF1R | -0.829072 | COADREAD | LS-1034 | Linsitinib | MVP_top60_DNA+RNA_seed | PIK3CB | -0.797717 | COADREAD | NCI-H747 | Buparlisib |
| MVP_top60_DNA_seed | ESR1 | -0.827791 | COADREAD | SW837 | GDC0810 | MVP_top60_DNA+RNA_seed | PIK3CD | -0.797717 | COADREAD | NCI-H747 | Buparlisib |
| MVP_top60_DNA_seed | BRAF | -0.827317 | COADREAD | SNU-C5 | Dabrafenib | MVP_top60_DNA+RNA_seed | PIK3CA | -0.795901 | COADREAD | NCI-H747 | GNE-317 |
| MVP_top60_DNA_seed | PIK3CA | -0.827186 | COADREAD | HCT-15 | Buparlisib | MVP_top60_DNA+RNA_seed | IGF1R | -0.794964 | COADREAD | HCC2998 | BMS-536924 |
| MVP_top60_DNA_seed | PIK3CB | -0.827186 | COADREAD | HCT-15 | Buparlisib | MVP_top60_DNA+RNA_seed | BCL2 | -0.794277 | COADREAD | COLO-320-HSR | Obatoclax Mesylate |
| MVP_top60_DNA_seed | PIK3CD | -0.827186 | COADREAD | HCT-15 | Buparlisib | MVP_top60_DNA+RNA_seed | ERBB3 | -0.793093 | COADREAD | SW1116 | Sapitinib |
| MVP_top60_DNA_seed | PIK3CA | -0.826032 | COADREAD | SK-CO-1 | GNE-317 | MVP_top60_DNA+RNA_seed | AKT1 | -0.792429 | COADREAD | HCT-116 | AT13148 |
| MVP_top60_DNA_seed | CDK2 | -0.825499 | COADREAD | DiFi | Dinaciclib | MVP_top60_DNA+RNA_seed | ERBB3 | -0.789675 | COADREAD | C2BBe1 | Sapitinib |
| MVP_top60_DNA_seed | MAPK1 | -0.822589 | COADREAD | SNU-175 | VX-11e | MVP_top60_DNA+RNA_seed | SRC | -0.789499 | COADREAD | HCT-15 | Bosutinib |
| MVP_top60_DNA_seed | AKT1 | -0.818434 | COADREAD | MDST8 | Afuresertib | MVP_top60_DNA+RNA_seed | AKT1 | -0.787336 | COADREAD | GP5d | Afuresertib |
| MVP_top60_DNA_seed | MAPK1 | -0.81808 | COADREAD | SNU-175 | Ulixertinib | MVP_top60_DNA+RNA_seed | IGF1R | -0.784542 | COADREAD | SNU-407 | Linsitinib |
| MVP_top60_DNA_seed | CDK2 | -0.815487 | COADREAD | SNU-C5 | AZD5438 | MVP_top60_DNA+RNA_seed | PIK3CA | -0.783071 | COADREAD | SW948 | Alpelisib |
| MVP_top60_DNA_seed | AKT1 | -0.813587 | COADREAD | SW948 | Ipatasertib | MVP_top60_DNA+RNA_seed | BCL2 | -0.776926 | COADREAD | SNU-C5 | Sabutoclax |
| MVP_top60_DNA_seed | PIK3CA | -0.80982 | COADREAD | GP5d | Buparlisib | MVP_top60_DNA+RNA_seed | MET | -0.776768 | COADREAD | COLO-320-HSR | Crizotinib |
| MVP_top60_DNA_seed | PIK3CB | -0.80982 | COADREAD | GP5d | Buparlisib | MVP_top60_DNA+RNA_seed | CDK2 | -0.776667 | COADREAD | GP5d | Dinaciclib |
| MVP_top60_DNA_seed | PIK3CD | -0.80982 | COADREAD | GP5d | Buparlisib | MVP_top60_DNA+RNA_seed | AKT1 | -0.775208 | COADREAD | COLO-205 | Uprosertib |
| MVP_top60_DNA_seed | BRAF | -0.809263 | COADREAD | LoVo | PLX-4720 | MVP_top60_DNA+RNA_seed | AKT1 | -0.774506 | COADREAD | SW948 | Afuresertib |

| | | | | | | | | | | | |
|---|---|---|---|---|---|---|---|---|---|---|---|
| MVP_top60_DNA_seed | PIK3CA | -0.808737 | COADREAD | COLO-320-HSR | Alpelisib | MVP_top60_DNA+RNA_seed | MET | -0.773405 | COADREAD | HT-115 | Crizotinib |
| MVP_top60_DNA_seed | MAPK1 | -0.808394 | COADREAD | SNU-407 | SCH772984 | MVP_top60_DNA+RNA_seed | PIK3CA | -0.772557 | COADREAD | LoVo | Buparlisib |
| MVP_top60_DNA_seed | AKT1 | -0.80825 | COADREAD | SNU-C1 | Ipatasertib | MVP_top60_DNA+RNA_seed | PIK3CB | -0.772557 | COADREAD | LoVo | Buparlisib |
| MVP_top60_DNA_seed | AKT1 | -0.80823 | COADREAD | SNU-175 | Uprosertib | MVP_top60_DNA+RNA_seed | PIK3CD | -0.772557 | COADREAD | LoVo | Buparlisib |
| MVP_top60_DNA_seed | PIK3CA | -0.807439 | COADREAD | MDST8 | Buparlisib | MVP_top60_DNA+RNA_seed | AKT1 | -0.771777 | COADREAD | CCK-81 | GSK2110183B |
| MVP_top60_DNA_seed | PIK3CB | -0.807439 | COADREAD | MDST8 | Buparlisib | MVP_top60_DNA+RNA_seed | AKT1 | -0.767971 | COADREAD | HCT-15 | GSK2110183B |
| MVP_top60_DNA_seed | PIK3CD | -0.807439 | COADREAD | MDST8 | Buparlisib | MVP_top60_DNA+RNA_seed | PIK3CA | -0.765258 | COADREAD | COLO-205 | Buparlisib |
| MVP_top60_DNA_seed | MAPK1 | -0.806161 | COADREAD | NCI-H716 | VX-11e | MVP_top60_DNA+RNA_seed | PIK3CB | -0.765258 | COADREAD | COLO-205 | Buparlisib |
| MVP_top60_DNA_seed | MET | -0.806078 | COADREAD | SNU-407 | Savolitinib | MVP_top60_DNA+RNA_seed | PIK3CD | -0.765258 | COADREAD | COLO-205 | Buparlisib |
| MVP_top60_DNA_seed | ESR1 | -0.804156 | COADREAD | LoVo | Tamoxifen | MVP_top60_DNA+RNA_seed | HSP90AA1 | -0.765218 | COADREAD | LS-411N | Tanespimycin |
| MVP_top60_DNA_seed | ESR1 | -0.803681 | COADREAD | HT-29 | Tamoxifen | MVP_top60_DNA+RNA_seed | HSP90AB1 | -0.765218 | COADREAD | LS-411N | Tanespimycin |
| MVP_top60_DNA_seed | IGF1R | -0.799961 | COADREAD | NCI-H747 | Linsitinib | MVP_top60_DNA+RNA_seed | PIK3CA | -0.758564 | COADREAD | SNU-175 | Buparlisib |
| MVP_top60_DNA_seed | IGF1R | -0.79871 | COADREAD | NCI-H747 | BMS-536924 | MVP_top60_DNA+RNA_seed | PIK3CB | -0.758564 | COADREAD | SNU-175 | Buparlisib |
| MVP_top60_DNA_seed | AKT1 | -0.798293 | COADREAD | SNU-175 | GSK2110183B | MVP_top60_DNA+RNA_seed | PIK3CD | -0.758564 | COADREAD | SNU-175 | Buparlisib |
| MVP_top60_DNA_seed | MAPK1 | -0.798163 | COADREAD | SNU-C1 | Ulixertinib | MVP_top60_DNA+RNA_seed | PIK3CA | -0.755599 | COADREAD | SW48 | Buparlisib |
| MVP_top60_DNA_seed | PIK3CA | -0.797717 | COADREAD | NCI-H747 | Buparlisib | MVP_top60_DNA+RNA_seed | PIK3CB | -0.755599 | COADREAD | SW48 | Buparlisib |
| MVP_top60_DNA_seed | PIK3CB | -0.797717 | COADREAD | NCI-H747 | Buparlisib | MVP_top60_DNA+RNA_seed | PIK3CD | -0.755599 | COADREAD | SW48 | Buparlisib |
| MVP_top60_DNA_seed | PIK3CD | -0.797717 | COADREAD | NCI-H747 | Buparlisib | MVP_top60_DNA+RNA_seed | PIK3CA | -0.753669 | COADREAD | LoVo | GNE-317 |
| MVP_top60_DNA_seed | PIK3CA | -0.795901 | COADREAD | NCI-H747 | GNE-317 | MVP_top60_DNA+RNA_seed | TP53 | -0.749332 | COADREAD | LS-513 | MIRA-1 |
| MVP_top60_DNA_seed | IGF1R | -0.794964 | COADREAD | HCC2998 | BMS-536924 | MVP_top60_DNA+RNA_seed | BCL2 | -0.749255 | COADREAD | SNU-1040 | Navitoclax |
| MVP_top60_DNA_seed | ESR1 | -0.793865 | COADREAD | RKO | Tamoxifen | MVP_top60_DNA+RNA_seed | IGF1R | -0.747772 | COADREAD | LoVo | BMS-754807 |
| MVP_top60_DNA_seed | ERBB3 | -0.793093 | COADREAD | SW1116 | Sapitinib | MVP_top60_DNA+RNA_seed | HSP90AA1 | -0.743574 | COADREAD | MDST8 | Luminespib |
| MVP_top60_DNA_seed | AKT1 | -0.792429 | COADREAD | HCT-116 | AT13148 | MVP_top60_DNA+RNA_seed | HSP90AB1 | -0.743574 | COADREAD | MDST8 | Luminespib |
| MVP_top60_DNA_seed | ERBB3 | -0.789675 | COADREAD | C2BBe1 | Sapitinib | MVP_top60_DNA+RNA_seed | AKT1 | -0.737455 | COADREAD | HCT-15 | Uprosertib |
| MVP_top60_DNA_seed | SRC | -0.789499 | COADREAD | HCT-15 | Bosutinib | MVP_top60_DNA+RNA_seed | PIK3CA | -0.735045 | COADREAD | SW948 | GNE-317 |
| MVP_top60_DNA_seed | MAPK1 | -0.78856 | COADREAD | SNU-175 | Ulixertinib | MVP_top60_DNA+RNA_seed | AKT1 | -0.734879 | COADREAD | MDST8 | AT13148 |
| MVP_top60_DNA_seed | AKT1 | -0.787336 | COADREAD | GP5d | Afuresertib | MVP_top60_DNA+RNA_seed | AKT1 | -0.734266 | COADREAD | HCC2998 | AZD5363 |
| MVP_top60_DNA_seed | BRAF | -0.785509 | COADREAD | SNU-C5 | SB590885 | MVP_top60_DNA+RNA_seed | PIK3CA | -0.732431 | COADREAD | C2BBe1 | Buparlisib |
| MVP_top60_DNA_seed | IGF1R | -0.784542 | COADREAD | SNU-407 | Linsitinib | MVP_top60_DNA+RNA_seed | PIK3CB | -0.732431 | COADREAD | C2BBe1 | Buparlisib |
| MVP_top60_DNA_seed | PIK3CA | -0.783071 | COADREAD | SW948 | Alpelisib | MVP_top60_DNA+RNA_seed | PIK3CD | -0.732431 | COADREAD | C2BBe1 | Buparlisib |
| MVP_top60_DNA_seed | MET | -0.776768 | COADREAD | COLO-320-HSR | Crizotinib | MVP_top60_DNA+RNA_seed | AKT1 | -0.729926 | COADREAD | NCI-H747 | Afuresertib |
| MVP_top60_DNA_seed | CDK2 | -0.776667 | COADREAD | GP5d | Dinaciclib | MVP_top60_DNA+RNA_seed | PIK3CA | -0.728024 | COADREAD | KM12 | Buparlisib |
| MVP_top60_DNA_seed | AKT1 | -0.775208 | COADREAD | COLO-205 | Uprosertib | MVP_top60_DNA+RNA_seed | PIK3CB | -0.728024 | COADREAD | KM12 | Buparlisib |
| MVP_top60_DNA_seed | AKT1 | -0.774506 | COADREAD | SW948 | Afuresertib | MVP_top60_DNA+RNA_seed | PIK3CD | -0.728024 | COADREAD | KM12 | Buparlisib |
| MVP_top60_DNA_seed | MET | -0.773405 | COADREAD | HT-115 | Crizotinib | MVP_top60_DNA+RNA_seed | AKT1 | -0.727106 | COADREAD | SW48 | Afuresertib |
| MVP_top60_DNA_seed | PIK3CA | -0.772557 | COADREAD | LoVo | Buparlisib | MVP_top60_DNA+RNA_seed | IGF1R | -0.725799 | COADREAD | NCI-H508 | NVP-ADW742 |
| MVP_top60_DNA_seed | PIK3CB | -0.772557 | COADREAD | LoVo | Buparlisib | MVP_top60_DNA+RNA_seed | AKT1 | -0.722594 | COADREAD | GP5d | AZD5363 |
| MVP_top60_DNA_seed | PIK3CD | -0.772557 | COADREAD | LoVo | Buparlisib | MVP_top60_DNA+RNA_seed | BCL2 | -0.721927 | COADREAD | SW48 | Navitoclax |
| MVP_top60_DNA_seed | AKT1 | -0.771777 | COADREAD | CCK-81 | GSK2110183B | MVP_top60_DNA+RNA_seed | PIK3CB | -0.719391 | COADREAD | SNU-81 | AZD6482 |
| MVP_top60_DNA_seed | AKT1 | -0.767971 | COADREAD | HCT-15 | GSK2110183B | MVP_top60_DNA+RNA_seed | HSP90AA1 | -0.717754 | COADREAD | SW620 | CCT-018159 |
| MVP_top60_DNA_seed | PIK3CA | -0.765258 | COADREAD | COLO-205 | Buparlisib | MVP_top60_DNA+RNA_seed | HSP90AB1 | -0.717754 | COADREAD | SW620 | CCT-018159 |

| | | | | | | | | | | | |
|---|---|---|---|---|---|---|---|---|---|---|---|
| MVP_top60_DNA_seed | PIK3CB | -0.765258 | COADREAD | COLO-205 | Buparlisib | MVP_top60_DNA+RNA_seed | IGF1R | -0.717488 | COADREAD | MDST8 | BMS-754807 |
| MVP_top60_DNA_seed | PIK3CD | -0.765258 | COADREAD | COLO-205 | Buparlisib | MVP_top60_DNA+RNA_seed | BCL2 | -0.712263 | COADREAD | SW837 | TW 37 |
| MVP_top60_DNA_seed | HSP90AA1 | -0.765218 | COADREAD | LS-411N | Tanespimycin | MVP_top60_DNA+RNA_seed | AKT1 | -0.70739 | COADREAD | SNU-1040 | Uprosertib |
| MVP_top60_DNA_seed | HSP90AB1 | -0.765218 | COADREAD | LS-411N | Tanespimycin | MVP_top60_DNA+RNA_seed | CDK2 | -0.704881 | COADREAD | CCK-81 | AZD5438 |
| MVP_top60_DNA_seed | MAPK1 | -0.764973 | COADREAD | LS-180 | SCH772984 | MVP_top60_DNA+RNA_seed | IGF1R | -0.703827 | COADREAD | HCC2998 | NVP-ADW742 |
| MVP_top60_DNA_seed | BRAF | -0.763068 | COADREAD | LS-513 | SB590885 | MVP_top60_DNA+RNA_seed | IGF1R | -0.703566 | COADREAD | SW1463 | Linsitinib |
| MVP_top60_DNA_seed | MAPK1 | -0.760252 | COADREAD | SW948 | SCH772984 | MVP_top60_DNA+RNA_seed | HSP90AA1 | -0.70147 | COADREAD | HCT-116 | Luminespib |
| MVP_top60_DNA_seed | PIK3CA | -0.758564 | COADREAD | SNU-175 | Buparlisib | MVP_top60_DNA+RNA_seed | HSP90AB1 | -0.70147 | COADREAD | HCT-116 | Luminespib |
| MVP_top60_DNA_seed | PIK3CB | -0.758564 | COADREAD | SNU-175 | Buparlisib | MVP_top60_DNA+RNA_seed | AKT1 | -0.69941 | COADREAD | LoVo | Uprosertib |
| MVP_top60_DNA_seed | PIK3CD | -0.758564 | COADREAD | SNU-175 | Buparlisib | MVP_top60_DNA+RNA_seed | BCL2 | -0.699163 | COADREAD | T84 | Venetoclax |
| MVP_top60_DNA_seed | PIK3CA | -0.755599 | COADREAD | SW48 | Buparlisib | MVP_top60_DNA+RNA_seed | PIK3CA | -0.698622 | COADREAD | COLO-320-HSR | GNE-317 |
| MVP_top60_DNA_seed | PIK3CB | -0.755599 | COADREAD | SW48 | Buparlisib | MVP_top60_DNA+RNA_seed | AKT1 | -0.697584 | COADREAD | HT-29 | MK-2206 |
| MVP_top60_DNA_seed | PIK3CD | -0.755599 | COADREAD | SW48 | Buparlisib | MVP_top60_DNA+RNA_seed | ERBB3 | -0.697092 | COADREAD | CaR-1 | Sapitinib |
| MVP_top60_DNA_seed | PIK3CA | -0.753669 | COADREAD | LoVo | GNE-317 | MVP_top60_DNA+RNA_seed | CDK2 | -0.696668 | COADREAD | NCI-H508 | Dinaciclib |
| MVP_top60_DNA_seed | TP53 | -0.749332 | COADREAD | LS-513 | MIRA-1 | MVP_top60_DNA+RNA_seed | AKT1 | -0.696195 | COADREAD | COLO-205 | Afuresertib |
| MVP_top60_DNA_seed | MAPK1 | -0.749246 | COADREAD | CCK-81 | Ulixertinib | MVP_top60_DNA+RNA_seed | BCL2 | -0.693477 | COADREAD | CCK-81 | TW 37 |
| MVP_top60_DNA_seed | BRAF | -0.747864 | COADREAD | SNU-C5 | PLX-4720 | MVP_top60_DNA+RNA_seed | CDK2 | -0.691649 | COADREAD | RKO | Dinaciclib |
| MVP_top60_DNA_seed | IGF1R | -0.747772 | COADREAD | LoVo | BMS-754807 | MVP_top60_DNA+RNA_seed | PIK3CA | -0.691121 | COADREAD | DiFi | GNE-317 |
| MVP_top60_DNA_seed | HSP90AA1 | -0.743574 | COADREAD | MDST8 | Luminespib | MVP_top60_DNA+RNA_seed | CDK2 | -0.688265 | COADREAD | RKO | AZD5438 |
| MVP_top60_DNA_seed | HSP90AB1 | -0.743574 | COADREAD | MDST8 | Luminespib | MVP_top60_DNA+RNA_seed | AKT1 | -0.687765 | COADREAD | SNU-C5 | Uprosertib |
| MVP_top60_DNA_seed | BRAF | -0.739477 | COADREAD | SK-CO-1 | SB590885 | MVP_top60_DNA+RNA_seed | GSK3B | -0.687113 | COADREAD | SW48 | SB216763 |
| MVP_top60_DNA_seed | AKT1 | -0.737455 | COADREAD | HCT-15 | Uprosertib | MVP_top60_DNA+RNA_seed | GSK3B | -0.686661 | COADREAD | LoVo | SB216763 |
| MVP_top60_DNA_seed | MAPK1 | -0.736083 | COADREAD | HT-115 | VX-11e | MVP_top60_DNA+RNA_seed | BCL2 | -0.684013 | COADREAD | SW48 | TW 37 |
| MVP_top60_DNA_seed | PIK3CA | -0.735045 | COADREAD | SW948 | GNE-317 | MVP_top60_DNA+RNA_seed | ATM | -0.679532 | COADREAD | SNU-C5 | KU-55933 |
| MVP_top60_DNA_seed | AKT1 | -0.734879 | COADREAD | MDST8 | AT13148 | MVP_top60_DNA+RNA_seed | CDK2 | -0.676557 | COADREAD | HT55 | Dinaciclib |
| MVP_top60_DNA_seed | AKT1 | -0.734266 | COADREAD | HCC2998 | AZD5363 | MVP_top60_DNA+RNA_seed | HSP90AA1 | -0.675576 | COADREAD | SW837 | Tanespimycin |
| MVP_top60_DNA_seed | PIK3CA | -0.732431 | COADREAD | C2BBe1 | Buparlisib | MVP_top60_DNA+RNA_seed | HSP90AB1 | -0.675576 | COADREAD | SW837 | Tanespimycin |
| MVP_top60_DNA_seed | PIK3CB | -0.732431 | COADREAD | C2BBe1 | Buparlisib | MVP_top60_DNA+RNA_seed | IGF1R | -0.674908 | COADREAD | CCK-81 | BMS-536924 |
| MVP_top60_DNA_seed | PIK3CD | -0.732431 | COADREAD | C2BBe1 | Buparlisib | MVP_top60_DNA+RNA_seed | AKT1 | -0.674059 | COADREAD | SNU-81 | Afuresertib |
| MVP_top60_DNA_seed | AKT1 | -0.729926 | COADREAD | NCI-H747 | Afuresertib | MVP_top60_DNA+RNA_seed | MET | -0.672931 | COADREAD | NCI-H508 | Crizotinib |
| MVP_top60_DNA_seed | PIK3CA | -0.728024 | COADREAD | KM12 | Buparlisib | MVP_top60_DNA+RNA_seed | CDK2 | -0.668117 | COADREAD | HCC2998 | Dinaciclib |
| MVP_top60_DNA_seed | PIK3CB | -0.728024 | COADREAD | KM12 | Buparlisib | MVP_top60_DNA+RNA_seed | PIK3CA | -0.661258 | COADREAD | DiFi | Buparlisib |
| MVP_top60_DNA_seed | PIK3CD | -0.728024 | COADREAD | KM12 | Buparlisib | MVP_top60_DNA+RNA_seed | PIK3CB | -0.661258 | COADREAD | DiFi | Buparlisib |
| MVP_top60_DNA_seed | AKT1 | -0.727106 | COADREAD | SW48 | Afuresertib | MVP_top60_DNA+RNA_seed | PIK3CD | -0.661258 | COADREAD | DiFi | Buparlisib |
| MVP_top60_DNA_seed | IGF1R | -0.725799 | COADREAD | NCI-H508 | NVP-ADW742 | MVP_top60_DNA+RNA_seed | BCL2 | -0.661175 | COADREAD | HT-115 | Navitoclax |
| MVP_top60_DNA_seed | AKT1 | -0.722594 | COADREAD | GP5d | AZD5363 | MVP_top60_DNA+RNA_seed | ATM | -0.660589 | COADREAD | MDST8 | KU-55933 |
| MVP_top60_DNA_seed | PIK3CB | -0.719391 | COADREAD | SNU-81 | AZD6482 | MVP_top60_DNA+RNA_seed | IGF1R | -0.659398 | COADREAD | NCI-H747 | NVP-ADW742 |
| MVP_top60_DNA_seed | HSP90AA1 | -0.717754 | COADREAD | SW620 | CCT-018159 | MVP_top60_DNA+RNA_seed | BCL2 | -0.659178 | COADREAD | SNU-C5 | Obatoclax Mesylate |
| MVP_top60_DNA_seed | HSP90AB1 | -0.717754 | COADREAD | SW620 | CCT-018159 | MVP_top60_DNA+RNA_seed | GSK3B | -0.657382 | COADREAD | MDST8 | SB216763 |
| MVP_top60_DNA_seed | IGF1R | -0.717488 | COADREAD | MDST8 | BMS-754807 | MVP_top60_DNA+RNA_seed | CDK2 | -0.65616 | COADREAD | KM12 | Dinaciclib |
| MVP_top60_DNA_seed | AKT1 | -0.70739 | COADREAD | SNU-1040 | Uprosertib | MVP_top60_DNA+RNA_seed | MET | -0.653521 | COADREAD | MDST8 | Crizotinib |

| | | | | | | | | | | | |
|---|---|---|---|---|---|---|---|---|---|---|---|
| MVP_top60_DNA_seed | CDK2 | -0.704881 | COADREAD | CCK-81 | AZD5438 | MVP_top60_DNA+RNA_seed | HSP90AA1 | -0.652925 | COADREAD | NCI-H716 | Tanespimycin |
| MVP_top60_DNA_seed | IGF1R | -0.703827 | COADREAD | HCC2998 | NVP-ADW742 | MVP_top60_DNA+RNA_seed | HSP90AB1 | -0.652925 | COADREAD | NCI-H716 | Tanespimycin |
| MVP_top60_DNA_seed | IGF1R | -0.703566 | COADREAD | SW1463 | Linsitinib | MVP_top60_DNA+RNA_seed | KRAS | -0.652878 | COADREAD | HCC2998 | [illegible]RAS (G12C) Inhibitor-1 |
| MVP_top60_DNA_seed | HSP90AA1 | -0.70147 | COADREAD | HCT-116 | Luminespib | MVP_top60_DNA+RNA_seed | IGF1R | -0.651443 | COADREAD | NCI-H716 | NVP-ADW742 |
| MVP_top60_DNA_seed | HSP90AB1 | -0.70147 | COADREAD | HCT-116 | Luminespib | MVP_top60_DNA+RNA_seed | IGF1R | -0.650109 | COADREAD | CaR-1 | BMS-754807 |
| MVP_top60_DNA_seed | AKT1 | -0.69941 | COADREAD | LoVo | Uprosertib | MVP_top60_DNA+RNA_seed | IGF1R | -0.643261 | COADREAD | CL-11 | BMS-754807 |
| MVP_top60_DNA_seed | MAPK1 | -0.698966 | COADREAD | SNU-61 | SCH772984 | MVP_top60_DNA+RNA_seed | AKT1 | -0.641756 | COADREAD | SNU-C1 | Uprosertib |
| MVP_top60_DNA_seed | PIK3CA | -0.698622 | COADREAD | COLO-320-HSR | GNE-317 | MVP_top60_DNA+RNA_seed | AKT1 | -0.638859 | COADREAD | NCI-H716 | MK-2206 |
| MVP_top60_DNA_seed | AKT1 | -0.697584 | COADREAD | HT-29 | MK-2206 | MVP_top60_DNA+RNA_seed | CDK2 | -0.636811 | COADREAD | SNU-C1 | AZD5438 |
| MVP_top60_DNA_seed | ERBB3 | -0.697092 | COADREAD | CaR-1 | Sapitinib | MVP_top60_DNA+RNA_seed | BCL2 | -0.633238 | COADREAD | NCI-H747 | Obatoclax Mesylate |
| MVP_top60_DNA_seed | CDK2 | -0.696668 | COADREAD | NCI-H508 | Dinaciclib | MVP_top60_DNA+RNA_seed | SRC | -0.633083 | COADREAD | CaR-1 | Bosutinib |
| MVP_top60_DNA_seed | MAPK1 | -0.696436 | COADREAD | SW837 | SCH772984 | MVP_top60_DNA+RNA_seed | IGF1R | -0.631978 | COADREAD | SNU-407 | BMS-536924 |
| MVP_top60_DNA_seed | AKT1 | -0.696195 | COADREAD | COLO-205 | Afuresertib | MVP_top60_DNA+RNA_seed | AKT1 | -0.631883 | COADREAD | HT55 | Afuresertib |
| MVP_top60_DNA_seed | CDK2 | -0.691649 | COADREAD | RKO | Dinaciclib | MVP_top60_DNA+RNA_seed | IGF1R | -0.630867 | COADREAD | DiFi | Linsitinib |
| MVP_top60_DNA_seed | PIK3CA | -0.691121 | COADREAD | DiFi | GNE-317 | MVP_top60_DNA+RNA_seed | HSP90AA1 | -0.630842 | COADREAD | SW620 | Luminespib |
| MVP_top60_DNA_seed | CDK2 | -0.688265 | COADREAD | RKO | AZD5438 | MVP_top60_DNA+RNA_seed | HSP90AB1 | -0.630842 | COADREAD | SW620 | Luminespib |
| MVP_top60_DNA_seed | AKT1 | -0.687765 | COADREAD | SNU-C5 | Uprosertib | MVP_top60_DNA+RNA_seed | IGF1R | -0.628694 | COADREAD | DiFi | GSK1904529A |
| MVP_top60_DNA_seed | GSK3B | -0.687113 | COADREAD | SW48 | SB216763 | MVP_top60_DNA+RNA_seed | AKT1 | -0.627836 | COADREAD | HT55 | Uprosertib |
| MVP_top60_DNA_seed | GSK3B | -0.686661 | COADREAD | LoVo | SB216763 | MVP_top60_DNA+RNA_seed | ATM | -0.620347 | COADREAD | SW837 | KU-55933 |
| MVP_top60_DNA_seed | MAPK1 | -0.680742 | COADREAD | SW1463 | SCH772984 | MVP_top60_DNA+RNA_seed | SRC | -0.616533 | COADREAD | SNU-175 | Bosutinib |
| MVP_top60_DNA_seed | ATM | -0.679532 | COADREAD | SNU-C5 | KU-55933 | MVP_top60_DNA+RNA_seed | CDK2 | -0.613728 | COADREAD | SW620 | Dinaciclib |
| MVP_top60_DNA_seed | MAPK1 | -0.67898 | COADREAD | SW620 | VX-11e | MVP_top60_DNA+RNA_seed | CDK2 | -0.613388 | COADREAD | CCK-81 | Dinaciclib |
| MVP_top60_DNA_seed | CREBBP | -0.677702 | COADREAD | RKO | SGC-CBP30 | MVP_top60_DNA+RNA_seed | AKT1 | -0.612859 | COADREAD | SNU-C5 | GSK2110183B |
| MVP_top60_DNA_seed | CDK2 | -0.676557 | COADREAD | HT55 | Dinaciclib | MVP_top60_DNA+RNA_seed | BCL2 | -0.612827 | COADREAD | RCM-1 | Navitoclax |
| MVP_top60_DNA_seed | HSP90AA1 | -0.675576 | COADREAD | SW837 | Tanespimycin | MVP_top60_DNA+RNA_seed | BCL2 | -0.609612 | COADREAD | GP5d | Sabutoclax |
| MVP_top60_DNA_seed | HSP90AB1 | -0.675576 | COADREAD | SW837 | Tanespimycin | MVP_top60_DNA+RNA_seed | ERBB3 | -0.608721 | COADREAD | GP5d | Sapitinib |
| MVP_top60_DNA_seed | IGF1R | -0.674908 | COADREAD | CCK-81 | BMS-536924 | MVP_top60_DNA+RNA_seed | HSP90AA1 | -0.608495 | COADREAD | GP5d | Luminespib |
| MVP_top60_DNA_seed | AKT1 | -0.674059 | COADREAD | SNU-81 | Afuresertib | MVP_top60_DNA+RNA_seed | HSP90AB1 | -0.608495 | COADREAD | GP5d | Luminespib |
| MVP_top60_DNA_seed | MET | -0.672931 | COADREAD | NCI-H508 | Crizotinib | MVP_top60_DNA+RNA_seed | IGF1R | -0.606034 | COADREAD | HCT-116 | GSK1904529A |
| MVP_top60_DNA_seed | ESR1 | -0.670136 | COADREAD | SW48 | Tamoxifen | MVP_top60_DNA+RNA_seed | BCL2 | -0.60573 | COADREAD | HT55 | Obatoclax Mesylate |
| MVP_top60_DNA_seed | CDK2 | -0.668117 | COADREAD | HCC2998 | Dinaciclib | MVP_top60_DNA+RNA_seed | HSP90AA1 | -0.605249 | COADREAD | NCI-H508 | Tanespimycin |
| MVP_top60_DNA_seed | MAPK1 | -0.666961 | COADREAD | SNU-407 | VX-11e | MVP_top60_DNA+RNA_seed | HSP90AB1 | -0.605249 | COADREAD | NCI-H508 | Tanespimycin |
| MVP_top60_DNA_seed | BRAF | -0.66488 | COADREAD | KM12 | Dabrafenib | MVP_top60_DNA+RNA_seed | PIK3CA | -0.60258 | COADREAD | SK-CO-1 | Alpelisib |
| MVP_top60_DNA_seed | PIK3CA | -0.661258 | COADREAD | DiFi | Buparlisib | MVP_top60_DNA+RNA_seed | PIK3CA | -0.602487 | COADREAD | SNU-C5 | Buparlisib |
| MVP_top60_DNA_seed | PIK3CB | -0.661258 | COADREAD | DiFi | Buparlisib | MVP_top60_DNA+RNA_seed | PIK3CB | -0.602487 | COADREAD | SNU-C5 | Buparlisib |
| MVP_top60_DNA_seed | PIK3CD | -0.661258 | COADREAD | DiFi | Buparlisib | MVP_top60_DNA+RNA_seed | PIK3CD | -0.602487 | COADREAD | SNU-C5 | Buparlisib |
| MVP_top60_DNA_seed | ATM | -0.660589 | COADREAD | MDST8 | KU-55933 | MVP_top60_DNA+RNA_seed | HSP90AA1 | -0.602038 | COADREAD | LoVo | Tanespimycin |
| MVP_top60_DNA_seed | IGF1R | -0.659398 | COADREAD | NCI-H747 | NVP-ADW742 | MVP_top60_DNA+RNA_seed | HSP90AB1 | -0.602038 | COADREAD | LoVo | Tanespimycin |
| MVP_top60_DNA_seed | BRAF | -0.658788 | COADREAD | NCI-H508 | Dabrafenib | MVP_top60_DNA+RNA_seed | AKT1 | -0.601917 | COADREAD | MDST8 | AZD5363 |
| MVP_top60_DNA_seed | GSK3B | -0.657382 | COADREAD | MDST8 | SB216763 | MVP_top60_DNA+RNA_seed | IGF1R | -0.601517 | COADREAD | HCC2998 | Linsitinib |
| MVP_top60_DNA_seed | CDK2 | -0.65616 | COADREAD | KM12 | Dinaciclib | MVP_top60_DNA+RNA_seed | MET | -0.601381 | COADREAD | HT55 | Crizotinib |

| | | | | | | | | | | | |
|---|---|---|---|---|---|---|---|---|---|---|---|
| MVP_top60_DNA_seed | MET | -0.653521 | COADREAD | MDST8 | Crizotinib | MVP_top60_DNA+RNA_seed | AKT1 | -0.601238 | COADREAD | SW948 | MK-2206 |
| MVP_top60_DNA_seed | HSP90AA1 | -0.652925 | COADREAD | NCI-H716 | Tanespimycin | MVP_top60_DNA+RNA_seed | BCL2 | -0.599313 | COADREAD | SNU-C2B | TW 37 |
| MVP_top60_DNA_seed | HSP90AB1 | -0.652925 | COADREAD | NCI-H716 | Tanespimycin | MVP_top60_DNA+RNA_seed | AKT1 | -0.598392 | COADREAD | NCI-H716 | AZD5363 |
| MVP_top60_DNA_seed | KRAS | -0.652878 | COADREAD | HCC2998 | KRAS (G12C) Inhibitor-12 | MVP_top60_DNA+RNA_seed | AKT1 | -0.597886 | COADREAD | T84 | Uprosertib |
| MVP_top60_DNA_seed | IGF1R | -0.651443 | COADREAD | NCI-H716 | NVP-ADW742 | MVP_top60_DNA+RNA_seed | IGF1R | -0.597299 | COADREAD | HCT-15 | BMS-754807 |
| MVP_top60_DNA_seed | IGF1R | -0.650109 | COADREAD | CaR-1 | BMS-754807 | MVP_top60_DNA+RNA_seed | HSP90AA1 | -0.595727 | COADREAD | NCI-H747 | Tanespimycin |
| MVP_top60_DNA_seed | CREBBP | -0.649786 | COADREAD | GP5d | SGC-CBP30 | MVP_top60_DNA+RNA_seed | HSP90AB1 | -0.595727 | COADREAD | NCI-H747 | Tanespimycin |
| MVP_top60_DNA_seed | BRAF | -0.647974 | COADREAD | COLO-320-HSR | PLX-4720 | MVP_top60_DNA+RNA_seed | PIK3CA | -0.594705 | COADREAD | HCT-116 | Buparlisib |
| MVP_top60_DNA_seed | IGF1R | -0.643261 | COADREAD | CL-11 | BMS-754807 | MVP_top60_DNA+RNA_seed | PIK3CB | -0.594705 | COADREAD | HCT-116 | Buparlisib |
| MVP_top60_DNA_seed | AKT1 | -0.641756 | COADREAD | SNU-C1 | Uprosertib | MVP_top60_DNA+RNA_seed | PIK3CD | -0.594705 | COADREAD | HCT-116 | Buparlisib |
| MVP_top60_DNA_seed | AKT1 | -0.638859 | COADREAD | NCI-H716 | MK-2206 | MVP_top60_DNA+RNA_seed | BCL2 | -0.594637 | COADREAD | CaR-1 | TW 37 |
| MVP_top60_DNA_seed | CDK2 | -0.636811 | COADREAD | SNU-C1 | AZD5438 | MVP_top60_DNA+RNA_seed | HSP90AA1 | -0.594622 | COADREAD | LS-411N | Luminespib |
| MVP_top60_DNA_seed | SRC | -0.633083 | COADREAD | CaR-1 | Bosutinib | MVP_top60_DNA+RNA_seed | HSP90AB1 | -0.594622 | COADREAD | LS-411N | Luminespib |
| MVP_top60_DNA_seed | IGF1R | -0.631978 | COADREAD | SNU-407 | BMS-536924 | MVP_top60_DNA+RNA_seed | CDK2 | -0.591854 | COADREAD | NCI-H716 | MK-8776 |
| MVP_top60_DNA_seed | AKT1 | -0.631883 | COADREAD | HT55 | Afuresertib | MVP_top60_DNA+RNA_seed | AKT1 | -0.590804 | COADREAD | HT55 | Ipatasertib |
| MVP_top60_DNA_seed | IGF1R | -0.630867 | COADREAD | DiFi | Linsitinib | MVP_top60_DNA+RNA_seed | BCL2 | -0.587208 | COADREAD | MDST8 | Navitoclax |
| MVP_top60_DNA_seed | HSP90AA1 | -0.630842 | COADREAD | SW620 | Luminespib | MVP_top60_DNA+RNA_seed | IGF1R | -0.585155 | COADREAD | LS-1034 | NVP-ADW742 |
| MVP_top60_DNA_seed | HSP90AB1 | -0.630842 | COADREAD | SW620 | Luminespib | MVP_top60_DNA+RNA_seed | IGF1R | -0.58411 | COADREAD | DiFi | BMS-754807 |
| MVP_top60_DNA_seed | IGF1R | -0.628694 | COADREAD | DiFi | GSK1904529A | MVP_top60_DNA+RNA_seed | CDK2 | -0.578235 | COADREAD | HT-115 | AZD5438 |
| MVP_top60_DNA_seed | AKT1 | -0.627836 | COADREAD | HT55 | Uprosertib | MVP_top60_DNA+RNA_seed | IGF1R | -0.578195 | COADREAD | SNU-407 | BMS-754807 |
| MVP_top60_DNA_seed | ATM | -0.620347 | COADREAD | SW837 | KU-55933 | MVP_top60_DNA+RNA_seed | AKT1 | -0.578057 | COADREAD | T84 | GSK2110183B |
| MVP_top60_DNA_seed | SRC | -0.616533 | COADREAD | SNU-175 | Bosutinib | MVP_top60_DNA+RNA_seed | SRC | -0.576678 | COADREAD | HT-29 | Bosutinib |
| MVP_top60_DNA_seed | MAPK1 | -0.61619 | COADREAD | T84 | VX-11e | MVP_top60_DNA+RNA_seed | MET | -0.575195 | COADREAD | SNU-407 | Crizotinib |
| MVP_top60_DNA_seed | CDK2 | -0.613728 | COADREAD | SW620 | Dinaciclib | MVP_top60_DNA+RNA_seed | SRC | -0.574902 | COADREAD | SNU-407 | Bosutinib |
| MVP_top60_DNA_seed | CDK2 | -0.613388 | COADREAD | CCK-81 | Dinaciclib | MVP_top60_DNA+RNA_seed | AKT1 | -0.573916 | COADREAD | RKO | GSK2110183B |
| MVP_top60_DNA_seed | AKT1 | -0.612859 | COADREAD | SNU-C5 | GSK2110183B | MVP_top60_DNA+RNA_seed | GSK3B | -0.5731 | COADREAD | RCM-1 | SB216763 |
| MVP_top60_DNA_seed | MAPK1 | -0.609937 | COADREAD | KM12 | Ulixertinib | MVP_top60_DNA+RNA_seed | AKT1 | -0.559787 | COADREAD | SNU-175 | MK-2206 |
| MVP_top60_DNA_seed | ERBB3 | -0.608721 | COADREAD | GP5d | Sapitinib | MVP_top60_DNA+RNA_seed | BCL2 | -0.558752 | COADREAD | CL-11 | Navitoclax |
| MVP_top60_DNA_seed | HSP90AA1 | -0.608495 | COADREAD | GP5d | Luminespib | MVP_top60_DNA+RNA_seed | GSK3B | -0.558552 | COADREAD | HT55 | SB216763 |
| MVP_top60_DNA_seed | HSP90AB1 | -0.608495 | COADREAD | GP5d | Luminespib | MVP_top60_DNA+RNA_seed | IGF1R | -0.555588 | COADREAD | SNU-407 | GSK1904529A |
| MVP_top60_DNA_seed | IGF1R | -0.606034 | COADREAD | HCT-116 | GSK1904529A | MVP_top60_DNA+RNA_seed | BCL2 | -0.5546 | COADREAD | MDST8 | Obatoclax Mesylate |
| MVP_top60_DNA_seed | HSP90AA1 | -0.605249 | COADREAD | NCI-H508 | Tanespimycin | MVP_top60_DNA+RNA_seed | PIK3CA | -0.551992 | COADREAD | COLO-320-HSR | AZD8186 |
| MVP_top60_DNA_seed | HSP90AB1 | -0.605249 | COADREAD | NCI-H508 | Tanespimycin | MVP_top60_DNA+RNA_seed | PIK3CB | -0.551992 | COADREAD | COLO-320-HSR | AZD8186 |
| MVP_top60_DNA_seed | PIK3CA | -0.60258 | COADREAD | SK-CO-1 | Alpelisib | MVP_top60_DNA+RNA_seed | BCL2 | -0.551954 | COADREAD | LS-1034 | TW 37 |
| MVP_top60_DNA_seed | PIK3CA | -0.602487 | COADREAD | SNU-C5 | Buparlisib | MVP_top60_DNA+RNA_seed | CDK2 | -0.549071 | COADREAD | SW620 | AZD5438 |
| MVP_top60_DNA_seed | PIK3CB | -0.602487 | COADREAD | SNU-C5 | Buparlisib | MVP_top60_DNA+RNA_seed | IGF1R | -0.547465 | COADREAD | HCT-15 | GSK1904529A |
| MVP_top60_DNA_seed | PIK3CD | -0.602487 | COADREAD | SNU-C5 | Buparlisib | MVP_top60_DNA+RNA_seed | IGF1R | -0.546986 | COADREAD | CCK-81 | BMS-754807 |
| MVP_top60_DNA_seed | HSP90AA1 | -0.602038 | COADREAD | LoVo | Tanespimycin | MVP_top60_DNA+RNA_seed | AKT1 | -0.545935 | COADREAD | SNU-C5 | AZD5363 |
| MVP_top60_DNA_seed | HSP90AB1 | -0.602038 | COADREAD | LoVo | Tanespimycin | MVP_top60_DNA+RNA_seed | CDK2 | -0.545803 | COADREAD | COLO-320-HSR | AZD5438 |
| MVP_top60_DNA_seed | AKT1 | -0.601917 | COADREAD | MDST8 | AZD5363 | MVP_top60_DNA+RNA_seed | BCL2 | -0.543643 | COADREAD | LoVo | Obatoclax Mesylate |
| MVP_top60_DNA_seed | IGF1R | -0.601517 | COADREAD | HCC2998 | Linsitinib | MVP_top60_DNA+RNA_seed | IGF1R | -0.538329 | COADREAD | HT-115 | NVP-ADW742 |

| | | | | | | | | | | | |
|---|---|---|---|---|---|---|---|---|---|---|---|
| MVP_top60_DNA_seed | MET | -0.601381 | COADREAD | HT55 | Crizotinib | MVP_top60_DNA+RNA_seed | BCL2 | -0.536832 | COADREAD | DiFi | Obatoclax Mesylate |
| MVP_top60_DNA_seed | MAPK1 | -0.601311 | COADREAD | GP5d | Ulixertinib | MVP_top60_DNA+RNA_seed | HSP90AA1 | -0.535356 | COADREAD | HCT-15 | Luminespib |
| MVP_top60_DNA_seed | AKT1 | -0.601238 | COADREAD | SW948 | MK-2206 | MVP_top60_DNA+RNA_seed | HSP90AB1 | -0.535356 | COADREAD | HCT-15 | Luminespib |
| MVP_top60_DNA_seed | AKT1 | -0.598392 | COADREAD | NCI-H716 | AZD5363 | MVP_top60_DNA+RNA_seed | MET | -0.531331 | COADREAD | HT-29 | Crizotinib |
| MVP_top60_DNA_seed | AKT1 | -0.597886 | COADREAD | T84 | Uprosertib | MVP_top60_DNA+RNA_seed | PIK3CA | -0.527602 | COADREAD | HT-115 | Buparlisib |
| MVP_top60_DNA_seed | IGF1R | -0.597299 | COADREAD | HCT-15 | BMS-754807 | MVP_top60_DNA+RNA_seed | PIK3CB | -0.527602 | COADREAD | HT-115 | Buparlisib |
| MVP_top60_DNA_seed | HSP90AA1 | -0.595727 | COADREAD | NCI-H747 | Tanespimycin | MVP_top60_DNA+RNA_seed | PIK3CD | -0.527602 | COADREAD | HT-115 | Buparlisib |
| MVP_top60_DNA_seed | HSP90AB1 | -0.595727 | COADREAD | NCI-H747 | Tanespimycin | MVP_top60_DNA+RNA_seed | IGF1R | -0.527302 | COADREAD | HCT-15 | NVP-ADW742 |
| MVP_top60_DNA_seed | PIK3CA | -0.594705 | COADREAD | HCT-116 | Buparlisib | MVP_top60_DNA+RNA_seed | BCL2 | -0.526831 | COADREAD | SW1116 | Navitoclax |
| MVP_top60_DNA_seed | PIK3CB | -0.594705 | COADREAD | HCT-116 | Buparlisib | MVP_top60_DNA+RNA_seed | ERBB3 | -0.525584 | COADREAD | RCM-1 | Sapitinib |
| MVP_top60_DNA_seed | PIK3CD | -0.594705 | COADREAD | HCT-116 | Buparlisib | MVP_top60_DNA+RNA_seed | CDK2 | -0.524328 | COADREAD | DiFi | AZD5438 |
| MVP_top60_DNA_seed | HSP90AA1 | -0.594622 | COADREAD | LS-411N | Luminespib | MVP_top60_DNA+RNA_seed | AKT1 | -0.522051 | COADREAD | KM12 | AT13148 |
| MVP_top60_DNA_seed | HSP90AB1 | -0.594622 | COADREAD | LS-411N | Luminespib | MVP_top60_DNA+RNA_seed | ERBB3 | -0.519172 | COADREAD | SNU-407 | Sapitinib |
| MVP_top60_DNA_seed | CREBBP | -0.592449 | COADREAD | DiFi | SGC-CBP30 | MVP_top60_DNA+RNA_seed | AKT1 | -0.516088 | COADREAD | HT55 | AZD5363 |
| MVP_top60_DNA_seed | CDK2 | -0.591854 | COADREAD | NCI-H716 | MK-8776 | MVP_top60_DNA+RNA_seed | AKT1 | -0.511879 | COADREAD | RKO | Uprosertib |
| MVP_top60_DNA_seed | AKT1 | -0.590804 | COADREAD | HT55 | Ipatasertib | MVP_top60_DNA+RNA_seed | BCL2 | -0.511543 | COADREAD | LS-123 | Navitoclax |
| MVP_top60_DNA_seed | BRAF | -0.585591 | COADREAD | HT-115 | SB590885 | MVP_top60_DNA+RNA_seed | AKT1 | -0.510288 | COADREAD | DiFi | Afuresertib |
| MVP_top60_DNA_seed | IGF1R | -0.585155 | COADREAD | LS-1034 | NVP-ADW742 | MVP_top60_DNA+RNA_seed | PIK3CA | -0.508767 | COADREAD | HCT-116 | Alpelisib |
| MVP_top60_DNA_seed | IGF1R | -0.58411 | COADREAD | DiFi | BMS-754807 | MVP_top60_DNA+RNA_seed | IGF1R | -0.50831 | COADREAD | CCK-81 | NVP-ADW742 |
| MVP_top60_DNA_seed | CDK2 | -0.578235 | COADREAD | HT-115 | AZD5438 | MVP_top60_DNA+RNA_seed | IGF1R | -0.505544 | COADREAD | MDST8 | Linsitinib |
| MVP_top60_DNA_seed | IGF1R | -0.578195 | COADREAD | SNU-407 | BMS-754807 | MVP_top60_DNA+RNA_seed | TP53 | -0.502326 | COADREAD | GP5d | MIRA-1 |
| MVP_top60_DNA_seed | AKT1 | -0.578057 | COADREAD | T84 | GSK2110183B | MVP_top60_DNA+RNA_seed | AKT1 | -0.501385 | COADREAD | SNU-175 | Ipatasertib |
| MVP_top60_DNA_seed | BRAF | -0.577844 | COADREAD | HCT-15 | PLX-4720 | MVP_top60_DNA+RNA_seed | AKT1 | -0.498827 | COADREAD | SNU-C1 | AZD5363 |
| MVP_top60_DNA_seed | SRC | -0.576678 | COADREAD | HT-29 | Bosutinib | MVP_top60_DNA+RNA_seed | ERBB3 | -0.493606 | COADREAD | HT-115 | Sapitinib |
| MVP_top60_DNA_seed | MET | -0.575195 | COADREAD | SNU-407 | Crizotinib | MVP_top60_DNA+RNA_seed | MET | -0.485965 | COADREAD | SNU-C1 | Crizotinib |
| MVP_top60_DNA_seed | SRC | -0.574902 | COADREAD | SNU-407 | Bosutinib | MVP_top60_DNA+RNA_seed | GSK3B | -0.485663 | COADREAD | HCT-15 | SB216763 |
| MVP_top60_DNA_seed | AKT1 | -0.573916 | COADREAD | RKO | GSK2110183B | MVP_top60_DNA+RNA_seed | PIK3CA | -0.484905 | COADREAD | CCK-81 | Buparlisib |
| MVP_top60_DNA_seed | GSK3B | -0.5731 | COADREAD | RCM-1 | SB216763 | MVP_top60_DNA+RNA_seed | PIK3CB | -0.484905 | COADREAD | CCK-81 | Buparlisib |
| MVP_top60_DNA_seed | ESR1 | -0.568499 | COADREAD | SW48 | GDC0810 | MVP_top60_DNA+RNA_seed | PIK3CD | -0.484905 | COADREAD | CCK-81 | Buparlisib |
| MVP_top60_DNA_seed | AKT1 | -0.559787 | COADREAD | SNU-175 | MK-2206 | MVP_top60_DNA+RNA_seed | BCL2 | -0.482388 | COADREAD | GP5d | Navitoclax |
| MVP_top60_DNA_seed | GSK3B | -0.558552 | COADREAD | HT55 | SB216763 | MVP_top60_DNA+RNA_seed | PIK3CA | -0.48204 | COADREAD | CaR-1 | AZD8186 |
| MVP_top60_DNA_seed | IGF1R | -0.555588 | COADREAD | SNU-407 | GSK1904529A | MVP_top60_DNA+RNA_seed | PIK3CB | -0.48204 | COADREAD | CaR-1 | AZD8186 |
| MVP_top60_DNA_seed | MAPK1 | -0.553474 | COADREAD | DiFi | Ulixertinib | MVP_top60_DNA+RNA_seed | MET | -0.481129 | COADREAD | CCK-81 | Crizotinib |
| MVP_top60_DNA_seed | PIK3CA | -0.551992 | COADREAD | COLO-320-HSR | AZD8186 | MVP_top60_DNA+RNA_seed | PIK3CA | -0.478742 | COADREAD | NCI-H747 | AZD8186 |
| MVP_top60_DNA_seed | PIK3CB | -0.551992 | COADREAD | COLO-320-HSR | AZD8186 | MVP_top60_DNA+RNA_seed | PIK3CB | -0.478742 | COADREAD | NCI-H747 | AZD8186 |
| MVP_top60_DNA_seed | CDK2 | -0.549071 | COADREAD | SW620 | AZD5438 | MVP_top60_DNA+RNA_seed | MET | -0.478251 | COADREAD | GP5d | Savolitinib |
| MVP_top60_DNA_seed | ESR1 | -0.54861 | COADREAD | GP5d | GDC0810 | MVP_top60_DNA+RNA_seed | BCL2 | -0.471868 | COADREAD | SK-CO-1 | Venetoclax |
| MVP_top60_DNA_seed | IGF1R | -0.547465 | COADREAD | HCT-15 | GSK1904529A | MVP_top60_DNA+RNA_seed | CDK2 | -0.466064 | COADREAD | COLO-320-HSR | Dinaciclib |
| MVP_top60_DNA_seed | IGF1R | -0.546986 | COADREAD | CCK-81 | BMS-754807 | MVP_top60_DNA+RNA_seed | ERBB3 | -0.465185 | COADREAD | LS-180 | Sapitinib |
| MVP_top60_DNA_seed | ESR1 | -0.546783 | COADREAD | LS-180 | GDC0810 | MVP_top60_DNA+RNA_seed | CDK2 | -0.464046 | COADREAD | HCT-15 | Dinaciclib |
| MVP_top60_DNA_seed | AKT1 | -0.545935 | COADREAD | SNU-C5 | AZD5363 | MVP_top60_DNA+RNA_seed | KRAS | -0.460015 | COADREAD | GP5d | [illegible]RAS (G12C) Inhibitor-1 |

| | | | | | | | | | | | |
|---|---|---|---|---|---|---|---|---|---|---|---|
| MVP_top60_DNA_seed | CDK2 | -0.545803 | COADREAD | COLO-320-HSR | AZD5438 | MVP_top60_DNA+RNA_seed | IGF1R | -0.459995 | COADREAD | SNU-C1 | BMS-536924 |
| MVP_top60_DNA_seed | ESR1 | -0.542719 | COADREAD | SW1463 | GDC0810 | MVP_top60_DNA+RNA_seed | AKT1 | -0.458333 | COADREAD | SNU-C5 | Ipatasertib |
| MVP_top60_DNA_seed | BRAF | -0.54219 | COADREAD | GP5d | SB590885 | MVP_top60_DNA+RNA_seed | KRAS | -0.452548 | COADREAD | LoVo | RAS (G12C) Inhibitor-1 |
| MVP_top60_DNA_seed | IGF1R | -0.538329 | COADREAD | HT-115 | NVP-ADW742 | MVP_top60_DNA+RNA_seed | AKT1 | -0.448912 | COADREAD | RKO | Uprosertib |
| MVP_top60_DNA_seed | HSP90AA1 | -0.535356 | COADREAD | HCT-15 | Luminespib | MVP_top60_DNA+RNA_seed | CDK2 | -0.447205 | COADREAD | HT-29 | Dinaciclib |
| MVP_top60_DNA_seed | HSP90AB1 | -0.535356 | COADREAD | HCT-15 | Luminespib | MVP_top60_DNA+RNA_seed | AKT1 | -0.447164 | COADREAD | COLO-320-HSR | MK-2206 |
| MVP_top60_DNA_seed | MET | -0.531331 | COADREAD | HT-29 | Crizotinib | MVP_top60_DNA+RNA_seed | AKT1 | -0.444805 | COADREAD | HCT-15 | AZD5363 |
| MVP_top60_DNA_seed | ESR1 | -0.529008 | COADREAD | HCC2998 | GDC0810 | MVP_top60_DNA+RNA_seed | HSP90AA1 | -0.442582 | COADREAD | SW948 | Tanespimycin |
| MVP_top60_DNA_seed | PIK3CA | -0.527602 | COADREAD | HT-115 | Buparlisib | MVP_top60_DNA+RNA_seed | HSP90AB1 | -0.442582 | COADREAD | SW948 | Tanespimycin |
| MVP_top60_DNA_seed | PIK3CB | -0.527602 | COADREAD | HT-115 | Buparlisib | MVP_top60_DNA+RNA_seed | ATM | -0.441003 | COADREAD | CaR-1 | KU-55933 |
| MVP_top60_DNA_seed | PIK3CD | -0.527602 | COADREAD | HT-115 | Buparlisib | MVP_top60_DNA+RNA_seed | IGF1R | -0.439876 | COADREAD | SNU-175 | BMS-536924 |
| MVP_top60_DNA_seed | IGF1R | -0.527302 | COADREAD | HCT-15 | NVP-ADW742 | MVP_top60_DNA+RNA_seed | AKT1 | -0.436189 | COADREAD | T84 | Afuresertib |
| MVP_top60_DNA_seed | MAPK1 | -0.526681 | COADREAD | CCK-81 | SCH772984 | MVP_top60_DNA+RNA_seed | KRAS | -0.43604 | COADREAD | SW620 | RAS (G12C) Inhibitor-1 |
| MVP_top60_DNA_seed | ERBB3 | -0.525584 | COADREAD | RCM-1 | Sapitinib | MVP_top60_DNA+RNA_seed | AR | -0.434107 | COADREAD | NCI-H508 | Bicalutamide |
| MVP_top60_DNA_seed | CDK2 | -0.524328 | COADREAD | DiFi | AZD5438 | MVP_top60_DNA+RNA_seed | PIK3CA | -0.432524 | COADREAD | SNU-407 | Buparlisib |
| MVP_top60_DNA_seed | ESR1 | -0.522876 | COADREAD | SNU-C5 | Tamoxifen | MVP_top60_DNA+RNA_seed | PIK3CB | -0.432524 | COADREAD | SNU-407 | Buparlisib |
| MVP_top60_DNA_seed | MAPK1 | -0.522405 | COADREAD | SNU-C5 | Ulixertinib | MVP_top60_DNA+RNA_seed | PIK3CD | -0.432524 | COADREAD | SNU-407 | Buparlisib |
| MVP_top60_DNA_seed | AKT1 | -0.522051 | COADREAD | KM12 | AT13148 | MVP_top60_DNA+RNA_seed | ERBB3 | -0.431845 | COADREAD | SNU-61 | Sapitinib |
| MVP_top60_DNA_seed | ERBB3 | -0.519172 | COADREAD | SNU-407 | Sapitinib | MVP_top60_DNA+RNA_seed | MET | -0.4315 | COADREAD | HCT-15 | Crizotinib |
| MVP_top60_DNA_seed | AKT1 | -0.516088 | COADREAD | HT55 | AZD5363 | MVP_top60_DNA+RNA_seed | AKT1 | -0.431361 | COADREAD | GP5d | AT13148 |
| MVP_top60_DNA_seed | AKT1 | -0.511879 | COADREAD | RKO | Uprosertib | MVP_top60_DNA+RNA_seed | IGF1R | -0.430286 | COADREAD | KM12 | NVP-ADW742 |
| MVP_top60_DNA_seed | MAPK1 | -0.510483 | COADREAD | NCI-H508 | VX-11e | MVP_top60_DNA+RNA_seed | IGF1R | -0.430152 | COADREAD | SW620 | BMS-754807 |
| MVP_top60_DNA_seed | AKT1 | -0.510288 | COADREAD | DiFi | Afuresertib | MVP_top60_DNA+RNA_seed | CDK2 | -0.429991 | COADREAD | HCC2998 | AZD5438 |
| MVP_top60_DNA_seed | PIK3CA | -0.508767 | COADREAD | HCT-116 | Alpelisib | MVP_top60_DNA+RNA_seed | TP53 | -0.424729 | COADREAD | DiFi | MIRA-1 |
| MVP_top60_DNA_seed | IGF1R | -0.50831 | COADREAD | CCK-81 | NVP-ADW742 | MVP_top60_DNA+RNA_seed | GSK3B | -0.424728 | COADREAD | SNU-81 | CHIR-99021 |
| MVP_top60_DNA_seed | IGF1R | -0.505544 | COADREAD | MDST8 | Linsitinib | MVP_top60_DNA+RNA_seed | GSK3B | -0.42337 | COADREAD | SNU-C1 | SB216763 |
| MVP_top60_DNA_seed | ESR1 | -0.503447 | COADREAD | SNU-407 | Tamoxifen | MVP_top60_DNA+RNA_seed | PIK3CA | -0.42315 | COADREAD | RKO | Alpelisib |
| MVP_top60_DNA_seed | ESR1 | -0.503372 | COADREAD | SW1463 | Tamoxifen | MVP_top60_DNA+RNA_seed | GSK3B | -0.422516 | COADREAD | T84 | SB216763 |
| MVP_top60_DNA_seed | MAPK1 | -0.502473 | COADREAD | LS-1034 | SCH772984 | MVP_top60_DNA+RNA_seed | SRC | -0.422471 | COADREAD | NCI-H508 | Bosutinib |
| MVP_top60_DNA_seed | TP53 | -0.502326 | COADREAD | GP5d | MIRA-1 | MVP_top60_DNA+RNA_seed | AKT1 | -0.412525 | COADREAD | SNU-81 | AZD5363 |
| MVP_top60_DNA_seed | AKT1 | -0.501385 | COADREAD | SNU-175 | Ipatasertib | MVP_top60_DNA+RNA_seed | PIK3CA | -0.408004 | COADREAD | HCT-116 | GNE-317 |
| MVP_top60_DNA_seed | AKT1 | -0.498827 | COADREAD | SNU-C1 | AZD5363 | MVP_top60_DNA+RNA_seed | BCL2 | -0.407064 | COADREAD | LS-180 | Venetoclax |
| MVP_top60_DNA_seed | ERBB3 | -0.493606 | COADREAD | HT-115 | Sapitinib | MVP_top60_DNA+RNA_seed | SRC | -0.40634 | COADREAD | SK-CO-1 | Bosutinib |
| MVP_top60_DNA_seed | MET | -0.485965 | COADREAD | SNU-C1 | Crizotinib | MVP_top60_DNA+RNA_seed | AKT1 | -0.405602 | COADREAD | HT55 | Uprosertib |
| MVP_top60_DNA_seed | GSK3B | -0.485663 | COADREAD | HCT-15 | SB216763 | MVP_top60_DNA+RNA_seed | AKT1 | -0.40203 | COADREAD | HCT-15 | Afuresertib |
| MVP_top60_DNA_seed | PIK3CA | -0.484905 | COADREAD | CCK-81 | Buparlisib | MVP_top60_DNA+RNA_seed | CDK2 | -0.399142 | COADREAD | LS-411N | Dinaciclib |
| MVP_top60_DNA_seed | PIK3CB | -0.484905 | COADREAD | CCK-81 | Buparlisib | MVP_top60_DNA+RNA_seed | PIK3CA | -0.39723 | COADREAD | SNU-407 | Alpelisib |
| MVP_top60_DNA_seed | PIK3CD | -0.484905 | COADREAD | CCK-81 | Buparlisib | MVP_top60_DNA+RNA_seed | IGF1R | -0.394395 | COADREAD | HCT-15 | Linsitinib |
| MVP_top60_DNA_seed | MAPK1 | -0.482359 | COADREAD | SW1116 | SCH772984 | MVP_top60_DNA+RNA_seed | ATM | -0.391766 | COADREAD | C2BBe1 | KU-55933 |
| MVP_top60_DNA_seed | PIK3CA | -0.48204 | COADREAD | CaR-1 | AZD8186 | MVP_top60_DNA+RNA_seed | BCL2 | -0.391707 | COADREAD | C2BBe1 | Obatoclax Mesylate |
| MVP_top60_DNA_seed | PIK3CB | -0.48204 | COADREAD | CaR-1 | AZD8186 | MVP_top60_DNA+RNA_seed | ERBB3 | -0.390288 | COADREAD | T84 | Sapitinib |

| | | | | | | | | | | | |
|---|---|---|---|---|---|---|---|---|---|---|---|
| MVP_top60_DNA_seed | CREBBP | -0.481788 | COADREAD | HCT-116 | SGC-CBP30 | MVP_top60_DNA+RNA_seed | PIK3CA | -0.389519 | COADREAD | LS-180 | GNE-317 |
| MVP_top60_DNA_seed | MET | -0.481129 | COADREAD | CCK-81 | Crizotinib | MVP_top60_DNA+RNA_seed | AKT1 | -0.389323 | COADREAD | SK-CO-1 | AT13148 |
| MVP_top60_DNA_seed | PIK3CA | -0.478742 | COADREAD | NCI-H747 | AZD8186 | MVP_top60_DNA+RNA_seed | AKT1 | -0.388886 | COADREAD | NCI-H508 | AT13148 |
| MVP_top60_DNA_seed | PIK3CB | -0.478742 | COADREAD | NCI-H747 | AZD8186 | MVP_top60_DNA+RNA_seed | HSP90AA1 | -0.387798 | COADREAD | HCT-15 | Tanespimycin |
| MVP_top60_DNA_seed | MET | -0.478251 | COADREAD | GP5d | Savolitinib | MVP_top60_DNA+RNA_seed | HSP90AB1 | -0.387798 | COADREAD | HCT-15 | Tanespimycin |
| MVP_top60_DNA_seed | ESR1 | -0.478066 | COADREAD | SNU-175 | GDC0810 | MVP_top60_DNA+RNA_seed | IGF1R | -0.384882 | COADREAD | T84 | NVP-ADW742 |
| MVP_top60_DNA_seed | MAPK1 | -0.476988 | COADREAD | SNU-C1 | VX-11e | MVP_top60_DNA+RNA_seed | MET | -0.381325 | COADREAD | RKO | Crizotinib |
| MVP_top60_DNA_seed | MAPK1 | -0.474075 | COADREAD | GP5d | VX-11e | MVP_top60_DNA+RNA_seed | BCL2 | -0.380397 | COADREAD | HCC2998 | Navitoclax |
| MVP_top60_DNA_seed | CDK2 | -0.466064 | COADREAD | COLO-320-HSR | Dinaciclib | MVP_top60_DNA+RNA_seed | PIK3CA | -0.380122 | COADREAD | COLO-205 | AZD8186 |
| MVP_top60_DNA_seed | ERBB3 | -0.465185 | COADREAD | LS-180 | Sapitinib | MVP_top60_DNA+RNA_seed | PIK3CB | -0.380122 | COADREAD | COLO-205 | AZD8186 |
| MVP_top60_DNA_seed | CDK2 | -0.464046 | COADREAD | HCT-15 | Dinaciclib | MVP_top60_DNA+RNA_seed | KRAS | -0.379246 | COADREAD | CW-2 | KRAS (G12C) Inhibitor-1 |
| MVP_top60_DNA_seed | MAPK1 | -0.46278 | COADREAD | SNU-C5 | SCH772984 | MVP_top60_DNA+RNA_seed | ATM | -0.378328 | COADREAD | SNU-C1 | KU-55933 |
| MVP_top60_DNA_seed | KRAS | -0.460015 | COADREAD | GP5d | KRAS (G12C) Inhibitor-1 | MVP_top60_DNA+RNA_seed | AKT1 | -0.377747 | COADREAD | DiFi | AZD5363 |
| MVP_top60_DNA_seed | IGF1R | -0.459995 | COADREAD | SNU-C1 | BMS-536924 | MVP_top60_DNA+RNA_seed | MET | -0.376923 | COADREAD | CaR-1 | Crizotinib |
| MVP_top60_DNA_seed | AKT1 | -0.458333 | COADREAD | SNU-C5 | Ipatasertib | MVP_top60_DNA+RNA_seed | HSP90AA1 | -0.376908 | COADREAD | COLO-320-HSR | Luminespib |
| MVP_top60_DNA_seed | MAPK1 | -0.453334 | COADREAD | LS-123 | Ulixertinib | MVP_top60_DNA+RNA_seed | HSP90AB1 | -0.376908 | COADREAD | COLO-320-HSR | Luminespib |
| MVP_top60_DNA_seed | KRAS | -0.452548 | COADREAD | LoVo | KRAS (G12C) Inhibitor-1 | MVP_top60_DNA+RNA_seed | BCL2 | -0.376226 | COADREAD | CCK-81 | Venetoclax |
| MVP_top60_DNA_seed | AKT1 | -0.448912 | COADREAD | RKO | Uprosertib | MVP_top60_DNA+RNA_seed | CDK2 | -0.375776 | COADREAD | CW-2 | AZD5438 |
| MVP_top60_DNA_seed | CDK2 | -0.447205 | COADREAD | HT-29 | Dinaciclib | MVP_top60_DNA+RNA_seed | AKT1 | -0.37564 | COADREAD | NCI-H716 | GSK2110183B |
| MVP_top60_DNA_seed | AKT1 | -0.447164 | COADREAD | COLO-320-HSR | MK-2206 | MVP_top60_DNA+RNA_seed | IGF1R | -0.372461 | COADREAD | HCT-116 | BMS-536924 |
| MVP_top60_DNA_seed | MAPK1 | -0.446404 | COADREAD | SW620 | Ulixertinib | MVP_top60_DNA+RNA_seed | AKT1 | -0.37215 | COADREAD | MDST8 | MK-2206 |
| MVP_top60_DNA_seed | AKT1 | -0.444805 | COADREAD | HCT-15 | AZD5363 | MVP_top60_DNA+RNA_seed | IGF1R | -0.370472 | COADREAD | LS-1034 | BMS-754807 |
| MVP_top60_DNA_seed | HSP90AA1 | -0.442582 | COADREAD | SW948 | Tanespimycin | MVP_top60_DNA+RNA_seed | AKT1 | -0.367896 | COADREAD | SNU-C5 | Uprosertib |
| MVP_top60_DNA_seed | HSP90AB1 | -0.442582 | COADREAD | SW948 | Tanespimycin | MVP_top60_DNA+RNA_seed | GSK3B | -0.366362 | COADREAD | GP5d | SB216763 |
| MVP_top60_DNA_seed | ATM | -0.441003 | COADREAD | CaR-1 | KU-55933 | MVP_top60_DNA+RNA_seed | TP53 | -0.365414 | COADREAD | CW-2 | MIRA-1 |
| MVP_top60_DNA_seed | IGF1R | -0.439876 | COADREAD | SNU-175 | BMS-536924 | MVP_top60_DNA+RNA_seed | TP53 | -0.364305 | COADREAD | HT55 | MIRA-1 |
| MVP_top60_DNA_seed | MAPK1 | -0.438036 | COADREAD | SNU-61 | Ulixertinib | MVP_top60_DNA+RNA_seed | CDK2 | -0.36337 | COADREAD | SNU-C5 | MK-8776 |
| MVP_top60_DNA_seed | AKT1 | -0.436189 | COADREAD | T84 | Afuresertib | MVP_top60_DNA+RNA_seed | AKT1 | -0.363181 | COADREAD | HCT-15 | Ipatasertib |
| MVP_top60_DNA_seed | KRAS | -0.43604 | COADREAD | SW620 | KRAS (G12C) Inhibitor-1 | MVP_top60_DNA+RNA_seed | AKT1 | -0.362766 | COADREAD | SW48 | MK-2206 |
| MVP_top60_DNA_seed | MAPK1 | -0.434149 | COADREAD | GP5d | Ulixertinib | MVP_top60_DNA+RNA_seed | IGF1R | -0.362289 | COADREAD | CW-2 | Linsitinib |
| MVP_top60_DNA_seed | AR | -0.434107 | COADREAD | NCI-H508 | Bicalutamide | MVP_top60_DNA+RNA_seed | AKT1 | -0.36171 | COADREAD | SW948 | AZD5363 |
| MVP_top60_DNA_seed | ESR1 | -0.432748 | COADREAD | LS-1034 | Tamoxifen | MVP_top60_DNA+RNA_seed | MET | -0.357141 | COADREAD | HCC2998 | Crizotinib |
| MVP_top60_DNA_seed | PIK3CA | -0.432524 | COADREAD | SNU-407 | Buparlisib | MVP_top60_DNA+RNA_seed | MET | -0.353982 | COADREAD | MDST8 | Savolitinib |
| MVP_top60_DNA_seed | PIK3CB | -0.432524 | COADREAD | SNU-407 | Buparlisib | MVP_top60_DNA+RNA_seed | BCL2 | -0.351069 | COADREAD | RKO | Sabutoclax |
| MVP_top60_DNA_seed | PIK3CD | -0.432524 | COADREAD | SNU-407 | Buparlisib | MVP_top60_DNA+RNA_seed | AKT1 | -0.350355 | COADREAD | SNU-C5 | Afuresertib |
| MVP_top60_DNA_seed | ERBB3 | -0.431845 | COADREAD | SNU-61 | Sapitinib | MVP_top60_DNA+RNA_seed | CDK2 | -0.346904 | COADREAD | HT-29 | MK-8776 |
| MVP_top60_DNA_seed | MET | -0.4315 | COADREAD | HCT-15 | Crizotinib | MVP_top60_DNA+RNA_seed | AKT1 | -0.346368 | COADREAD | CCK-81 | Uprosertib |
| MVP_top60_DNA_seed | AKT1 | -0.431361 | COADREAD | GP5d | AT13148 | MVP_top60_DNA+RNA_seed | IGF1R | -0.345984 | COADREAD | CaR-1 | GSK1904529A |
| MVP_top60_DNA_seed | IGF1R | -0.430286 | COADREAD | KM12 | NVP-ADW742 | MVP_top60_DNA+RNA_seed | BCL2 | -0.341638 | COADREAD | KM12 | Venetoclax |
| MVP_top60_DNA_seed | IGF1R | -0.430152 | COADREAD | SW620 | BMS-754807 | MVP_top60_DNA+RNA_seed | IGF1R | -0.340409 | COADREAD | HCT-116 | NVP-ADW742 |
| MVP_top60_DNA_seed | CDK2 | -0.429991 | COADREAD | HCC2998 | AZD5438 | MVP_top60_DNA+RNA_seed | IGF1R | -0.338341 | COADREAD | SNU-407 | NVP-ADW742 |

| | | | | | | | | | | | |
|---|---|---|---|---|---|---|---|---|---|---|---|
| MVP_top60_DNA_seed | TP53 | -0.424729 | COADREAD | DiFi | MIRA-1 | MVP_top60_DNA+RNA_seed | TP53 | -0.337806 | COADREAD | CCK-81 | MIRA-1 |
| MVP_top60_DNA_seed | GSK3B | -0.424728 | COADREAD | SNU-81 | CHIR-99021 | MVP_top60_DNA+RNA_seed | PIK3CA | -0.337026 | COADREAD | COLO-205 | Alpelisib |
| MVP_top60_DNA_seed | MAPK1 | -0.424336 | COADREAD | SW1116 | VX-11e | MVP_top60_DNA+RNA_seed | SRC | -0.33694 | COADREAD | HCT-116 | Bosutinib |
| MVP_top60_DNA_seed | GSK3B | -0.42337 | COADREAD | SNU-C1 | SB216763 | MVP_top60_DNA+RNA_seed | IGF1R | -0.33612 | COADREAD | T84 | BMS-754807 |
| MVP_top60_DNA_seed | PIK3CA | -0.42315 | COADREAD | RKO | Alpelisib | MVP_top60_DNA+RNA_seed | KRAS | -0.334025 | COADREAD | RKO | (RAS (G12C) Inhibitor-1 |
| MVP_top60_DNA_seed | GSK3B | -0.422516 | COADREAD | T84 | SB216763 | MVP_top60_DNA+RNA_seed | BCL2 | -0.330702 | COADREAD | SNU-C1 | Navitoclax |
| MVP_top60_DNA_seed | SRC | -0.422471 | COADREAD | NCI-H508 | Bosutinib | MVP_top60_DNA+RNA_seed | IGF1R | -0.330626 | COADREAD | DiFi | NVP-ADW742 |
| MVP_top60_DNA_seed | ESR1 | -0.415527 | COADREAD | NCI-H747 | Tamoxifen | MVP_top60_DNA+RNA_seed | AKT1 | -0.329953 | COADREAD | HCT-15 | MK-2206 |
| MVP_top60_DNA_seed | MAPK1 | -0.413334 | COADREAD | LS-180 | VX-11e | MVP_top60_DNA+RNA_seed | BCL2 | -0.329482 | COADREAD | HT55 | Sabutoclax |
| MVP_top60_DNA_seed | AKT1 | -0.412525 | COADREAD | SNU-81 | AZD5363 | MVP_top60_DNA+RNA_seed | MET | -0.328967 | COADREAD | LS-411N | Crizotinib |
| MVP_top60_DNA_seed | MAPK1 | -0.411285 | COADREAD | COLO-678 | VX-11e | MVP_top60_DNA+RNA_seed | CDK2 | -0.323208 | COADREAD | HCT-116 | Dinaciclib |
| MVP_top60_DNA_seed | PIK3CA | -0.408004 | COADREAD | HCT-116 | GNE-317 | MVP_top60_DNA+RNA_seed | PIK3CA | -0.321428 | COADREAD | CaR-1 | Alpelisib |
| MVP_top60_DNA_seed | SRC | -0.40634 | COADREAD | SK-CO-1 | Bosutinib | MVP_top60_DNA+RNA_seed | AKT1 | -0.319251 | COADREAD | RKO | Ipatasertib |
| MVP_top60_DNA_seed | AKT1 | -0.405602 | COADREAD | HT55 | Uprosertib | MVP_top60_DNA+RNA_seed | BCL2 | -0.3173 | COADREAD | SW620 | Navitoclax |
| MVP_top60_DNA_seed | BRAF | -0.404416 | COADREAD | HT55 | PLX-4720 | MVP_top60_DNA+RNA_seed | CDK2 | -0.314599 | COADREAD | HT55 | AZD5438 |
| MVP_top60_DNA_seed | AKT1 | -0.40203 | COADREAD | HCT-15 | Afuresertib | MVP_top60_DNA+RNA_seed | BCL2 | -0.310143 | COADREAD | SW837 | Navitoclax |
| MVP_top60_DNA_seed | MAPK1 | -0.401057 | COADREAD | COLO-678 | SCH772984 | MVP_top60_DNA+RNA_seed | AKT1 | -0.308496 | COADREAD | CL-11 | AT13148 |
| MVP_top60_DNA_seed | CDK2 | -0.399142 | COADREAD | LS-411N | Dinaciclib | MVP_top60_DNA+RNA_seed | IGF1R | -0.304362 | COADREAD | HCT-116 | BMS-754807 |
| MVP_top60_DNA_seed | PIK3CA | -0.39723 | COADREAD | SNU-407 | Alpelisib | MVP_top60_DNA+RNA_seed | AKT1 | -0.303332 | COADREAD | SNU-407 | Ipatasertib |
| MVP_top60_DNA_seed | CREBBP | -0.397201 | COADREAD | SNU-C5 | SGC-CBP30 | MVP_top60_DNA+RNA_seed | AKT1 | -0.301119 | COADREAD | CaR-1 | Afuresertib |
| MVP_top60_DNA_seed | IGF1R | -0.394395 | COADREAD | HCT-15 | Linsitinib | MVP_top60_DNA+RNA_seed | ERBB3 | -0.30004 | COADREAD | SNU-C2B | Sapitinib |
| MVP_top60_DNA_seed | BRAF | -0.394189 | COADREAD | HCC2998 | SB590885 | MVP_top60_DNA+RNA_seed | KRAS | -0.297829 | COADREAD | SW48 | (RAS (G12C) Inhibitor-1 |
| MVP_top60_DNA_seed | ATM | -0.391766 | COADREAD | C2BBe1 | KU-55933 | MVP_top60_DNA+RNA_seed | HSP90AA1 | -0.296247 | COADREAD | LS-180 | Tanespimycin |
| MVP_top60_DNA_seed | ERBB3 | -0.390288 | COADREAD | T84 | Sapitinib | MVP_top60_DNA+RNA_seed | HSP90AB1 | -0.296247 | COADREAD | LS-180 | Tanespimycin |
| MVP_top60_DNA_seed | PIK3CA | -0.389519 | COADREAD | LS-180 | GNE-317 | MVP_top60_DNA+RNA_seed | PIK3CA | -0.290618 | COADREAD | LS-411N | GNE-317 |
| MVP_top60_DNA_seed | AKT1 | -0.389323 | COADREAD | SK-CO-1 | AT13148 | MVP_top60_DNA+RNA_seed | AKT1 | -0.282943 | COADREAD | SNU-C5 | MK-2206 |
| MVP_top60_DNA_seed | AKT1 | -0.388886 | COADREAD | NCI-H508 | AT13148 | MVP_top60_DNA+RNA_seed | MET | -0.28226 | COADREAD | LS-1034 | Crizotinib |
| MVP_top60_DNA_seed | HSP90AA1 | -0.387798 | COADREAD | HCT-15 | Tanespimycin | MVP_top60_DNA+RNA_seed | AKT1 | -0.279904 | COADREAD | SW48 | Uprosertib |
| MVP_top60_DNA_seed | HSP90AB1 | -0.387798 | COADREAD | HCT-15 | Tanespimycin | MVP_top60_DNA+RNA_seed | ATM | -0.278066 | COADREAD | CW-2 | KU-55933 |
| MVP_top60_DNA_seed | IGF1R | -0.384882 | COADREAD | T84 | NVP-ADW742 | MVP_top60_DNA+RNA_seed | IGF1R | -0.275636 | COADREAD | HCT-116 | Linsitinib |
| MVP_top60_DNA_seed | MAPK1 | -0.383541 | COADREAD | SW1116 | Ulixertinib | MVP_top60_DNA+RNA_seed | AKT1 | -0.273662 | COADREAD | SW48 | GSK2110183B |
| MVP_top60_DNA_seed | MET | -0.381325 | COADREAD | RKO | Crizotinib | MVP_top60_DNA+RNA_seed | BCL2 | -0.270446 | COADREAD | SNU-407 | Sabutoclax |
| MVP_top60_DNA_seed | BRAF | -0.380895 | COADREAD | HCT-15 | Dabrafenib | MVP_top60_DNA+RNA_seed | IGF1R | -0.270091 | COADREAD | RKO | BMS-754807 |
| MVP_top60_DNA_seed | PIK3CA | -0.380122 | COADREAD | COLO-205 | AZD8186 | MVP_top60_DNA+RNA_seed | HSP90AA1 | -0.269612 | COADREAD | LoVo | Luminespib |
| MVP_top60_DNA_seed | PIK3CB | -0.380122 | COADREAD | COLO-205 | AZD8186 | MVP_top60_DNA+RNA_seed | HSP90AB1 | -0.269612 | COADREAD | LoVo | Luminespib |
| MVP_top60_DNA_seed | KRAS | -0.379246 | COADREAD | CW-2 | KRAS (G12C) Inhibitor-1: | MVP_top60_DNA+RNA_seed | CDK2 | -0.266927 | COADREAD | DiFi | MK-8776 |
| MVP_top60_DNA_seed | ATM | -0.378328 | COADREAD | SNU-C1 | KU-55933 | MVP_top60_DNA+RNA_seed | IGF1R | -0.263218 | COADREAD | NCI-H747 | BMS-754807 |
| MVP_top60_DNA_seed | AKT1 | -0.377747 | COADREAD | DiFi | AZD5363 | MVP_top60_DNA+RNA_seed | BCL2 | -0.262981 | COADREAD | LS-180 | Obatoclax Mesylate |
| MVP_top60_DNA_seed | ESR1 | -0.37729 | COADREAD | COLO-205 | GDC0810 | MVP_top60_DNA+RNA_seed | KRAS | -0.259001 | COADREAD | NCI-H716 | (RAS (G12C) Inhibitor-1 |
| MVP_top60_DNA_seed | MET | -0.376923 | COADREAD | CaR-1 | Crizotinib | MVP_top60_DNA+RNA_seed | GSK3B | -0.257894 | COADREAD | SNU-C5 | SB216763 |
| MVP_top60_DNA_seed | HSP90AA1 | -0.376908 | COADREAD | COLO-320-HSR | Luminespib | MVP_top60_DNA+RNA_seed | BCL2 | -0.256015 | COADREAD | NCI-H508 | Navitoclax |

| | | | | | | | | | | | |
|---|---|---|---|---|---|---|---|---|---|---|---|
| MVP_top60_DNA_seed | HSP90AB1 | -0.376908 | COADREAD | COLO-320-HSR | Luminespib | MVP_top60_DNA+RNA_seed | BCL2 | -0.253282 | COADREAD | HT-29 | Sabutoclax |
| MVP_top60_DNA_seed | CDK2 | -0.375776 | COADREAD | CW-2 | AZD5438 | MVP_top60_DNA+RNA_seed | MET | -0.252253 | COADREAD | SNU-1040 | Crizotinib |
| MVP_top60_DNA_seed | AKT1 | -0.37564 | COADREAD | NCI-H716 | GSK2110183B | MVP_top60_DNA+RNA_seed | IGF1R | -0.2505 | COADREAD | SNU-175 | NVP-ADW742 |
| MVP_top60_DNA_seed | BRAF | -0.375272 | COADREAD | SNU-81 | PLX-4720 | MVP_top60_DNA+RNA_seed | AKT1 | -0.250481 | COADREAD | HT-29 | AT13148 |
| MVP_top60_DNA_seed | MAPK1 | -0.37294 | COADREAD | SW837 | Ulixertinib | MVP_top60_DNA+RNA_seed | KRAS | -0.249931 | COADREAD | SNU-C5 | (RAS (G12C) Inhibitor-1 |
| MVP_top60_DNA_seed | IGF1R | -0.372461 | COADREAD | HCT-116 | BMS-536924 | MVP_top60_DNA+RNA_seed | PIK3CA | -0.24948 | COADREAD | KM12 | GNE-317 |
| MVP_top60_DNA_seed | AKT1 | -0.37215 | COADREAD | MDST8 | MK-2206 | MVP_top60_DNA+RNA_seed | SRC | -0.248181 | COADREAD | COLO-320-HSR | Bosutinib |
| MVP_top60_DNA_seed | IGF1R | -0.370472 | COADREAD | LS-1034 | BMS-754807 | MVP_top60_DNA+RNA_seed | AKT1 | -0.248065 | COADREAD | C2BBe1 | AT13148 |
| MVP_top60_DNA_seed | AKT1 | -0.367896 | COADREAD | SNU-C5 | Uprosertib | MVP_top60_DNA+RNA_seed | IGF1R | -0.245488 | COADREAD | CW-2 | NVP-ADW742 |
| MVP_top60_DNA_seed | GSK3B | -0.366362 | COADREAD | GP5d | SB216763 | MVP_top60_DNA+RNA_seed | MET | -0.245418 | COADREAD | HCT-15 | Savolitinib |
| MVP_top60_DNA_seed | TP53 | -0.365414 | COADREAD | CW-2 | MIRA-1 | MVP_top60_DNA+RNA_seed | CDK2 | -0.243002 | COADREAD | RCM-1 | AZD5438 |
| MVP_top60_DNA_seed | TP53 | -0.364305 | COADREAD | HT55 | MIRA-1 | MVP_top60_DNA+RNA_seed | BCL2 | -0.242525 | COADREAD | HCT-116 | Venetoclax |
| MVP_top60_DNA_seed | CDK2 | -0.36337 | COADREAD | SNU-C5 | MK-8776 | MVP_top60_DNA+RNA_seed | MET | -0.242401 | COADREAD | CW-2 | Savolitinib |
| MVP_top60_DNA_seed | AKT1 | -0.363181 | COADREAD | HCT-15 | Ipatasertib | MVP_top60_DNA+RNA_seed | AKT1 | -0.24221 | COADREAD | HCC2998 | MK-2206 |
| MVP_top60_DNA_seed | AKT1 | -0.362766 | COADREAD | SW48 | MK-2206 | MVP_top60_DNA+RNA_seed | BCL2 | -0.236581 | COADREAD | GP5d | Obatoclax Mesylate |
| MVP_top60_DNA_seed | IGF1R | -0.362289 | COADREAD | CW-2 | Linsitinib | MVP_top60_DNA+RNA_seed | AKT1 | -0.234881 | COADREAD | LS-180 | Afuresertib |
| MVP_top60_DNA_seed | AKT1 | -0.36171 | COADREAD | SW948 | AZD5363 | MVP_top60_DNA+RNA_seed | PIK3CA | -0.233439 | COADREAD | CaR-1 | Buparlisib |
| MVP_top60_DNA_seed | MET | -0.357141 | COADREAD | HCC2998 | Crizotinib | MVP_top60_DNA+RNA_seed | PIK3CB | -0.233439 | COADREAD | CaR-1 | Buparlisib |
| MVP_top60_DNA_seed | MET | -0.353982 | COADREAD | MDST8 | Savolitinib | MVP_top60_DNA+RNA_seed | PIK3CD | -0.233439 | COADREAD | CaR-1 | Buparlisib |
| MVP_top60_DNA_seed | CREBBP | -0.353598 | COADREAD | CaR-1 | SGC-CBP30 | MVP_top60_DNA+RNA_seed | AKT1 | -0.232703 | COADREAD | HT-29 | AZD5363 |
| MVP_top60_DNA_seed | AKT1 | -0.350355 | COADREAD | SNU-C5 | Afuresertib | MVP_top60_DNA+RNA_seed | BCL2 | -0.2326 | COADREAD | C2BBe1 | Navitoclax |
| MVP_top60_DNA_seed | CDK2 | -0.346904 | COADREAD | HT-29 | MK-8776 | MVP_top60_DNA+RNA_seed | PIK3CA | -0.227941 | COADREAD | LS-123 | GNE-317 |
| MVP_top60_DNA_seed | AKT1 | -0.346368 | COADREAD | CCK-81 | Uprosertib | MVP_top60_DNA+RNA_seed | IGF1R | -0.222524 | COADREAD | SW948 | Linsitinib |
| MVP_top60_DNA_seed | IGF1R | -0.345984 | COADREAD | CaR-1 | GSK1904529A | MVP_top60_DNA+RNA_seed | GSK3B | -0.215842 | COADREAD | CW-2 | SB216763 |
| MVP_top60_DNA_seed | IGF1R | -0.340409 | COADREAD | HCT-116 | NVP-ADW742 | MVP_top60_DNA+RNA_seed | IGF1R | -0.214374 | COADREAD | GP5d | NVP-ADW742 |
| MVP_top60_DNA_seed | IGF1R | -0.338341 | COADREAD | SNU-407 | NVP-ADW742 | MVP_top60_DNA+RNA_seed | BCL2 | -0.212399 | COADREAD | SNU-407 | Obatoclax Mesylate |
| MVP_top60_DNA_seed | MAPK1 | -0.338213 | COADREAD | CW-2 | Ulixertinib | MVP_top60_DNA+RNA_seed | KRAS | -0.211738 | COADREAD | LS-513 | (RAS (G12C) Inhibitor-1 |
| MVP_top60_DNA_seed | TP53 | -0.337806 | COADREAD | CCK-81 | MIRA-1 | MVP_top60_DNA+RNA_seed | IGF1R | -0.210481 | COADREAD | GP5d | BMS-536924 |
| MVP_top60_DNA_seed | BRAF | -0.337173 | COADREAD | SW1417 | PLX-4720 | MVP_top60_DNA+RNA_seed | SRC | -0.206284 | COADREAD | MDST8 | Bosutinib |
| MVP_top60_DNA_seed | PIK3CA | -0.337026 | COADREAD | COLO-205 | Alpelisib | MVP_top60_DNA+RNA_seed | HSP90AA1 | -0.204889 | COADREAD | CW-2 | Luminespib |
| MVP_top60_DNA_seed | SRC | -0.33694 | COADREAD | HCT-116 | Bosutinib | MVP_top60_DNA+RNA_seed | HSP90AB1 | -0.204889 | COADREAD | CW-2 | Luminespib |
| MVP_top60_DNA_seed | IGF1R | -0.33612 | COADREAD | T84 | BMS-754807 | MVP_top60_DNA+RNA_seed | AKT1 | -0.201099 | COADREAD | RKO | Afuresertib |
| MVP_top60_DNA_seed | MAPK1 | -0.335927 | COADREAD | T84 | Ulixertinib | MVP_top60_DNA+RNA_seed | MET | -0.195368 | COADREAD | COLO-320-HSR | Savolitinib |
| MVP_top60_DNA_seed | KRAS | -0.334025 | COADREAD | RKO | (RAS (G12C) Inhibitor-1: | MVP_top60_DNA+RNA_seed | PIK3CA | -0.194916 | COADREAD | HT-29 | GNE-317 |
| MVP_top60_DNA_seed | IGF1R | -0.330626 | COADREAD | DiFi | NVP-ADW742 | MVP_top60_DNA+RNA_seed | CDK2 | -0.194144 | COADREAD | MDST8 | Dinaciclib |
| MVP_top60_DNA_seed | AKT1 | -0.329953 | COADREAD | HCT-15 | MK-2206 | MVP_top60_DNA+RNA_seed | CDK2 | -0.194101 | COADREAD | SNU-1040 | Dinaciclib |
| MVP_top60_DNA_seed | MET | -0.328967 | COADREAD | LS-411N | Crizotinib | MVP_top60_DNA+RNA_seed | AKT1 | -0.193287 | COADREAD | CW-2 | Uprosertib |
| MVP_top60_DNA_seed | MAPK1 | -0.328244 | COADREAD | HCT-15 | VX-11e | MVP_top60_DNA+RNA_seed | PIK3CA | -0.192095 | COADREAD | LS-513 | Alpelisib |
| MVP_top60_DNA_seed | ESR1 | -0.326111 | COADREAD | DiFi | Tamoxifen | MVP_top60_DNA+RNA_seed | TP53 | -0.191951 | COADREAD | MDST8 | MIRA-1 |
| MVP_top60_DNA_seed | CDK2 | -0.323208 | COADREAD | HCT-116 | Dinaciclib | MVP_top60_DNA+RNA_seed | CDK2 | -0.191147 | COADREAD | SNU-175 | AZD5438 |
| MVP_top60_DNA_seed | PIK3CA | -0.321428 | COADREAD | CaR-1 | Alpelisib | MVP_top60_DNA+RNA_seed | PIK3CA | -0.190287 | COADREAD | HCC2998 | Buparlisib |

| | | | | | | | | | | | |
|---|---|---|---|---|---|---|---|---|---|---|---|
| MVP_top60_DNA_seed | AKT1 | -0.319251 | COADREAD | RKO | Ipatasertib | MVP_top60_DNA+RNA_seed | PIK3CB | -0.190287 | COADREAD | HCC2998 | Buparlisib |
| MVP_top60_DNA_seed | CDK2 | -0.314599 | COADREAD | HT55 | AZD5438 | MVP_top60_DNA+RNA_seed | PIK3CD | -0.190287 | COADREAD | HCC2998 | Buparlisib |
| MVP_top60_DNA_seed | AKT1 | -0.308496 | COADREAD | CL-11 | AT13148 | MVP_top60_DNA+RNA_seed | ATM | -0.186846 | COADREAD | NCI-H716 | KU-55933 |
| MVP_top60_DNA_seed | IGF1R | -0.304362 | COADREAD | HCT-116 | BMS-754807 | MVP_top60_DNA+RNA_seed | MET | -0.186487 | COADREAD | RCM-1 | Crizotinib |
| MVP_top60_DNA_seed | AKT1 | -0.303332 | COADREAD | SNU-407 | Ipatasertib | MVP_top60_DNA+RNA_seed | AKT1 | -0.186464 | COADREAD | CaR-1 | Uprosertib |
| MVP_top60_DNA_seed | ESR1 | -0.301355 | COADREAD | CaR-1 | Tamoxifen | MVP_top60_DNA+RNA_seed | ATM | -0.184286 | COADREAD | HT55 | KU-55933 |
| MVP_top60_DNA_seed | AKT1 | -0.301119 | COADREAD | CaR-1 | Afuresertib | MVP_top60_DNA+RNA_seed | CDK2 | -0.181144 | COADREAD | NCI-H508 | MK-8776 |
| MVP_top60_DNA_seed | ERBB3 | -0.30004 | COADREAD | SNU-C2B | Sapitinib | MVP_top60_DNA+RNA_seed | IGF1R | -0.180748 | COADREAD | SW1463 | BMS-754807 |
| MVP_top60_DNA_seed | KRAS | -0.297829 | COADREAD | SW48 | KRAS (G12C) Inhibitor-12 | MVP_top60_DNA+RNA_seed | BCL2 | -0.178684 | COADREAD | HCC2998 | Obatoclax Mesylate |
| MVP_top60_DNA_seed | HSP90AA1 | -0.296247 | COADREAD | LS-180 | Tanespimycin | MVP_top60_DNA+RNA_seed | IGF1R | -0.177609 | COADREAD | LS-411N | Linsitinib |
| MVP_top60_DNA_seed | HSP90AB1 | -0.296247 | COADREAD | LS-180 | Tanespimycin | MVP_top60_DNA+RNA_seed | HSP90AA1 | -0.176512 | COADREAD | HCT-116 | Tanespimycin |
| MVP_top60_DNA_seed | BRAF | -0.294989 | COADREAD | SK-CO-1 | Dabrafenib | MVP_top60_DNA+RNA_seed | HSP90AB1 | -0.176512 | COADREAD | HCT-116 | Tanespimycin |
| MVP_top60_DNA_seed | ESR1 | -0.294515 | COADREAD | COLO-205 | Tamoxifen | MVP_top60_DNA+RNA_seed | PIK3CA | -0.172189 | COADREAD | LS-180 | Alpelisib |
| MVP_top60_DNA_seed | BRAF | -0.292846 | COADREAD | LS-1034 | Dabrafenib | MVP_top60_DNA+RNA_seed | AKT1 | -0.170917 | COADREAD | HCT-116 | Afuresertib |
| MVP_top60_DNA_seed | PIK3CA | -0.290618 | COADREAD | LS-411N | GNE-317 | MVP_top60_DNA+RNA_seed | PIK3CA | -0.170209 | COADREAD | SNU-407 | AZD8186 |
| MVP_top60_DNA_seed | AKT1 | -0.282943 | COADREAD | SNU-C5 | MK-2206 | MVP_top60_DNA+RNA_seed | PIK3CB | -0.170209 | COADREAD | SNU-407 | AZD8186 |
| MVP_top60_DNA_seed | ESR1 | -0.282862 | COADREAD | MDST8 | GDC0810 | MVP_top60_DNA+RNA_seed | AKT1 | -0.170196 | COADREAD | SNU-407 | MK-2206 |
| MVP_top60_DNA_seed | MET | -0.28226 | COADREAD | LS-1034 | Crizotinib | MVP_top60_DNA+RNA_seed | MET | -0.169706 | COADREAD | SW620 | Crizotinib |
| MVP_top60_DNA_seed | ESR1 | -0.279984 | COADREAD | SW837 | Tamoxifen | MVP_top60_DNA+RNA_seed | IGF1R | -0.168659 | COADREAD | HCT-15 | BMS-536924 |
| MVP_top60_DNA_seed | AKT1 | -0.279904 | COADREAD | SW48 | Uprosertib | MVP_top60_DNA+RNA_seed | BCL2 | -0.168392 | COADREAD | CCK-81 | Obatoclax Mesylate |
| MVP_top60_DNA_seed | ATM | -0.278066 | COADREAD | CW-2 | KU-55933 | MVP_top60_DNA+RNA_seed | MET | -0.166036 | COADREAD | SNU-61 | Savolitinib |
| MVP_top60_DNA_seed | IGF1R | -0.275636 | COADREAD | HCT-116 | Linsitinib | MVP_top60_DNA+RNA_seed | PIK3CA | -0.164333 | COADREAD | HT-29 | AZD8186 |
| MVP_top60_DNA_seed | AKT1 | -0.273662 | COADREAD | SW48 | GSK2110183B | MVP_top60_DNA+RNA_seed | PIK3CB | -0.164333 | COADREAD | HT-29 | AZD8186 |
| MVP_top60_DNA_seed | IGF1R | -0.270091 | COADREAD | RKO | BMS-754807 | MVP_top60_DNA+RNA_seed | AKT1 | -0.164102 | COADREAD | CaR-1 | AT13148 |
| MVP_top60_DNA_seed | HSP90AA1 | -0.269612 | COADREAD | LoVo | Luminespib | MVP_top60_DNA+RNA_seed | AKT1 | -0.160398 | COADREAD | SNU-407 | AZD5363 |
| MVP_top60_DNA_seed | HSP90AB1 | -0.269612 | COADREAD | LoVo | Luminespib | MVP_top60_DNA+RNA_seed | PIK3CA | -0.158809 | COADREAD | HT-115 | Alpelisib |
| MVP_top60_DNA_seed | CDK2 | -0.266927 | COADREAD | DiFi | MK-8776 | MVP_top60_DNA+RNA_seed | BCL2 | -0.157904 | COADREAD | LS-411N | Obatoclax Mesylate |
| MVP_top60_DNA_seed | ESR1 | -0.266809 | COADREAD | C2BBe1 | Tamoxifen | MVP_top60_DNA+RNA_seed | CDK2 | -0.157654 | COADREAD | LS-513 | Dinaciclib |
| MVP_top60_DNA_seed | IGF1R | -0.263218 | COADREAD | NCI-H747 | BMS-754807 | MVP_top60_DNA+RNA_seed | MET | -0.157152 | COADREAD | SNU-C5 | Crizotinib |
| MVP_top60_DNA_seed | KRAS | -0.259001 | COADREAD | NCI-H716 | KRAS (G12C) Inhibitor-12 | MVP_top60_DNA+RNA_seed | PIK3CA | -0.156407 | COADREAD | MDST8 | AZD8186 |
| MVP_top60_DNA_seed | GSK3B | -0.257894 | COADREAD | SNU-C5 | SB216763 | MVP_top60_DNA+RNA_seed | PIK3CB | -0.156407 | COADREAD | MDST8 | AZD8186 |
| MVP_top60_DNA_seed | MET | -0.252253 | COADREAD | SNU-1040 | Crizotinib | MVP_top60_DNA+RNA_seed | CDK2 | -0.156344 | COADREAD | RKO | MK-8776 |
| MVP_top60_DNA_seed | IGF1R | -0.2505 | COADREAD | SNU-175 | NVP-ADW742 | MVP_top60_DNA+RNA_seed | PIK3CA | -0.155478 | COADREAD | SW948 | Buparlisib |
| MVP_top60_DNA_seed | AKT1 | -0.250481 | COADREAD | HT-29 | AT13148 | MVP_top60_DNA+RNA_seed | PIK3CB | -0.155478 | COADREAD | SW948 | Buparlisib |
| MVP_top60_DNA_seed | KRAS | -0.249931 | COADREAD | SNU-C5 | KRAS (G12C) Inhibitor-12 | MVP_top60_DNA+RNA_seed | PIK3CD | -0.155478 | COADREAD | SW948 | Buparlisib |
| MVP_top60_DNA_seed | PIK3CA | -0.24948 | COADREAD | KM12 | GNE-317 | MVP_top60_DNA+RNA_seed | AKT1 | -0.155375 | COADREAD | COLO-205 | Uprosertib |
| MVP_top60_DNA_seed | SRC | -0.248181 | COADREAD | COLO-320-HSR | Bosutinib | MVP_top60_DNA+RNA_seed | AKT1 | -0.154932 | COADREAD | HCT-116 | AZD5363 |
| MVP_top60_DNA_seed | AKT1 | -0.248065 | COADREAD | C2BBe1 | AT13148 | MVP_top60_DNA+RNA_seed | AKT1 | -0.154742 | COADREAD | NCI-H747 | AT13148 |
| MVP_top60_DNA_seed | IGF1R | -0.245488 | COADREAD | CW-2 | NVP-ADW742 | MVP_top60_DNA+RNA_seed | AKT1 | -0.153909 | COADREAD | HCT-116 | Uprosertib |
| MVP_top60_DNA_seed | MET | -0.245418 | COADREAD | HCT-15 | Savolitinib | MVP_top60_DNA+RNA_seed | AKT1 | -0.153752 | COADREAD | RKO | AZD5363 |
| MVP_top60_DNA_seed | CDK2 | -0.243002 | COADREAD | RCM-1 | AZD5438 | MVP_top60_DNA+RNA_seed | CDK2 | -0.153624 | COADREAD | LS-411N | AZD5438 |

| | | | | | | | | | | | |
|---|---|---|---|---|---|---|---|---|---|---|---|
| MVP_top60_DNA_seed | MET | -0.242401 | COADREAD | CW-2 | Savolitinib | MVP_top60_DNA+RNA_seed | IGF1R | -0.15142 | COADREAD | LS-411N | BMS-754807 |
| MVP_top60_DNA_seed | AKT1 | -0.24221 | COADREAD | HCC2998 | MK-2206 | MVP_top60_DNA+RNA_seed | AKT1 | -0.151309 | COADREAD | CW-2 | Afuresertib |
| MVP_top60_DNA_seed | MAPK1 | -0.241381 | COADREAD | SNU-C5 | Ulixertinib | MVP_top60_DNA+RNA_seed | CDK2 | -0.150535 | COADREAD | HCT-116 | MK-8776 |
| MVP_top60_DNA_seed | ESR1 | -0.238254 | COADREAD | MDST8 | Tamoxifen | MVP_top60_DNA+RNA_seed | CDK2 | -0.147607 | COADREAD | SNU-C5 | Dinaciclib |
| MVP_top60_DNA_seed | BRAF | -0.237639 | COADREAD | HT55 | Dabrafenib | MVP_top60_DNA+RNA_seed | MET | -0.145439 | COADREAD | DiFi | Savolitinib |
| MVP_top60_DNA_seed | AKT1 | -0.234881 | COADREAD | LS-180 | Afuresertib | MVP_top60_DNA+RNA_seed | AKT1 | -0.142633 | COADREAD | CW-2 | GSK2110183B |
| MVP_top60_DNA_seed | PIK3CA | -0.233439 | COADREAD | CaR-1 | Buparlisib | MVP_top60_DNA+RNA_seed | BCL2 | -0.14206 | COADREAD | HT-115 | Obatoclax Mesylate |
| MVP_top60_DNA_seed | PIK3CB | -0.233439 | COADREAD | CaR-1 | Buparlisib | MVP_top60_DNA+RNA_seed | AKT1 | -0.139264 | COADREAD | CCK-81 | MK-2206 |
| MVP_top60_DNA_seed | PIK3CD | -0.233439 | COADREAD | CaR-1 | Buparlisib | MVP_top60_DNA+RNA_seed | PIK3CA | -0.13814 | COADREAD | CW-2 | Alpelisib |
| MVP_top60_DNA_seed | AKT1 | -0.232703 | COADREAD | HT-29 | AZD5363 | MVP_top60_DNA+RNA_seed | PIK3CA | -0.136585 | COADREAD | LS-123 | AZD8186 |
| MVP_top60_DNA_seed | PIK3CA | -0.227941 | COADREAD | LS-123 | GNE-317 | MVP_top60_DNA+RNA_seed | PIK3CB | -0.136585 | COADREAD | LS-123 | AZD8186 |
| MVP_top60_DNA_seed | MAPK1 | -0.22742 | COADREAD | LS-180 | Ulixertinib | MVP_top60_DNA+RNA_seed | AKT1 | -0.13602 | COADREAD | COLO-320-HSR | Uprosertib |
| MVP_top60_DNA_seed | MAPK1 | -0.22698 | COADREAD | HCT-15 | Ulixertinib | MVP_top60_DNA+RNA_seed | SRC | -0.135012 | COADREAD | LoVo | Bosutinib |
| MVP_top60_DNA_seed | MAPK1 | -0.224412 | COADREAD | C2BBe1 | Ulixertinib | MVP_top60_DNA+RNA_seed | CDK2 | -0.130337 | COADREAD | SNU-C2B | MK-8776 |
| MVP_top60_DNA_seed | IGF1R | -0.222524 | COADREAD | SW948 | Linsitinib | MVP_top60_DNA+RNA_seed | AKT1 | -0.125655 | COADREAD | CCK-81 | Afuresertib |
| MVP_top60_DNA_seed | ESR1 | -0.221797 | COADREAD | RKO | GDC0810 | MVP_top60_DNA+RNA_seed | ATM | -0.125136 | COADREAD | KM12 | KU-55933 |
| MVP_top60_DNA_seed | GSK3B | -0.215842 | COADREAD | CW-2 | SB216763 | MVP_top60_DNA+RNA_seed | AKT1 | -0.124502 | COADREAD | SNU-81 | Uprosertib |
| MVP_top60_DNA_seed | IGF1R | -0.214374 | COADREAD | GP5d | NVP-ADW742 | MVP_top60_DNA+RNA_seed | CDK2 | -0.124217 | COADREAD | GP5d | MK-8776 |
| MVP_top60_DNA_seed | BRAF | -0.213705 | COADREAD | HCT-116 | PLX-4720 | MVP_top60_DNA+RNA_seed | SRC | -0.12386 | COADREAD | SNU-C1 | Bosutinib |
| MVP_top60_DNA_seed | CREBBP | -0.212776 | COADREAD | CCK-81 | SGC-CBP30 | MVP_top60_DNA+RNA_seed | AKT1 | -0.122725 | COADREAD | COLO-205 | AZD5363 |
| MVP_top60_DNA_seed | KRAS | -0.211738 | COADREAD | LS-513 | KRAS (G12C) Inhibitor-1: | MVP_top60_DNA+RNA_seed | ERBB3 | -0.121206 | COADREAD | HCT-15 | Sapitinib |
| MVP_top60_DNA_seed | IGF1R | -0.210481 | COADREAD | GP5d | BMS-536924 | MVP_top60_DNA+RNA_seed | HSP90AA1 | -0.121187 | COADREAD | NCI-H747 | Luminespib |
| MVP_top60_DNA_seed | ESR1 | -0.206538 | COADREAD | LoVo | GDC0810 | MVP_top60_DNA+RNA_seed | HSP90AB1 | -0.121187 | COADREAD | NCI-H747 | Luminespib |
| MVP_top60_DNA_seed | SRC | -0.206284 | COADREAD | MDST8 | Bosutinib | MVP_top60_DNA+RNA_seed | AKT1 | -0.12003 | COADREAD | NCI-H716 | Afuresertib |
| MVP_top60_DNA_seed | BRAF | -0.204938 | COADREAD | DiFi | SB590885 | MVP_top60_DNA+RNA_seed | CDK2 | -0.119412 | COADREAD | CL-11 | MK-8776 |
| MVP_top60_DNA_seed | HSP90AA1 | -0.204889 | COADREAD | CW-2 | Luminespib | MVP_top60_DNA+RNA_seed | IGF1R | -0.115586 | COADREAD | SNU-1040 | BMS-754807 |
| MVP_top60_DNA_seed | HSP90AB1 | -0.204889 | COADREAD | CW-2 | Luminespib | MVP_top60_DNA+RNA_seed | PIK3CA | -0.115279 | COADREAD | SNU-C5 | Alpelisib |
| MVP_top60_DNA_seed | AKT1 | -0.201099 | COADREAD | RKO | Afuresertib | MVP_top60_DNA+RNA_seed | PIK3CA | -0.115211 | COADREAD | LoVo | Alpelisib |
| MVP_top60_DNA_seed | MAPK1 | -0.199475 | COADREAD | SNU-407 | Ulixertinib | MVP_top60_DNA+RNA_seed | BCL2 | -0.110842 | COADREAD | SNU-175 | Venetoclax |
| MVP_top60_DNA_seed | MET | -0.195368 | COADREAD | COLO-320-HSR | Savolitinib | MVP_top60_DNA+RNA_seed | MET | -0.11 | COADREAD | CL-11 | Crizotinib |
| MVP_top60_DNA_seed | PIK3CA | -0.194916 | COADREAD | HT-29 | GNE-317 | MVP_top60_DNA+RNA_seed | BCL2 | -0.106568 | COADREAD | LS-1034 | Venetoclax |
| MVP_top60_DNA_seed | CDK2 | -0.194144 | COADREAD | MDST8 | Dinaciclib | MVP_top60_DNA+RNA_seed | BCL2 | -0.104284 | COADREAD | LS-1034 | Navitoclax |
| MVP_top60_DNA_seed | CDK2 | -0.194101 | COADREAD | SNU-1040 | Dinaciclib | MVP_top60_DNA+RNA_seed | HSP90AA1 | -0.102919 | COADREAD | SW620 | Elesclomol |
| MVP_top60_DNA_seed | AKT1 | -0.193287 | COADREAD | CW-2 | Uprosertib | MVP_top60_DNA+RNA_seed | HSP90AB1 | -0.102919 | COADREAD | SW620 | Elesclomol |
| MVP_top60_DNA_seed | PIK3CA | -0.192095 | COADREAD | LS-513 | Alpelisib | MVP_top60_DNA+RNA_seed | PIK3CA | -0.09687 | COADREAD | HT55 | AZD8186 |
| MVP_top60_DNA_seed | TP53 | -0.191951 | COADREAD | MDST8 | MIRA-1 | MVP_top60_DNA+RNA_seed | PIK3CB | -0.09687 | COADREAD | HT55 | AZD8186 |
| MVP_top60_DNA_seed | ESR1 | -0.191825 | COADREAD | SNU-C1 | Tamoxifen | MVP_top60_DNA+RNA_seed | BCL2 | -0.096729 | COADREAD | CCK-81 | Navitoclax |
| MVP_top60_DNA_seed | CDK2 | -0.191147 | COADREAD | SNU-175 | AZD5438 | MVP_top60_DNA+RNA_seed | AKT1 | -0.092923 | COADREAD | NCI-H747 | MK-2206 |
| MVP_top60_DNA_seed | MAPK1 | -0.190511 | COADREAD | LS-123 | SCH772984 | MVP_top60_DNA+RNA_seed | CDK2 | -0.090892 | COADREAD | NCI-H716 | AZD5438 |
| MVP_top60_DNA_seed | PIK3CA | -0.190287 | COADREAD | HCC2998 | Buparlisib | MVP_top60_DNA+RNA_seed | HSP90AA1 | -0.090335 | COADREAD | HCC2998 | Luminespib |
| MVP_top60_DNA_seed | PIK3CB | -0.190287 | COADREAD | HCC2998 | Buparlisib | MVP_top60_DNA+RNA_seed | HSP90AB1 | -0.090335 | COADREAD | HCC2998 | Luminespib |

| | | | | | | | | | | | |
|---|---|---|---|---|---|---|---|---|---|---|---|
| MVP_top60_DNA_seed | PIK3CD | -0.190287 | COADREAD | HCC2998 | Buparlisib | MVP_top60_DNA+RNA_seed | BCL2 | -0.089015 | COADREAD | SNU-175 | Obatoclax Mesylate |
| MVP_top60_DNA_seed | ATM | -0.186846 | COADREAD | NCI-H716 | KU-55933 | MVP_top60_DNA+RNA_seed | AKT1 | -0.087401 | COADREAD | RKO | AT13148 |
| MVP_top60_DNA_seed | MET | -0.186487 | COADREAD | RCM-1 | Crizotinib | MVP_top60_DNA+RNA_seed | CDK2 | -0.087358 | COADREAD | SNU-C1 | Dinaciclib |
| MVP_top60_DNA_seed | AKT1 | -0.186464 | COADREAD | CaR-1 | Uprosertib | MVP_top60_DNA+RNA_seed | CDK2 | -0.085834 | COADREAD | SNU-175 | MK-8776 |
| MVP_top60_DNA_seed | ATM | -0.184286 | COADREAD | HT55 | KU-55933 | MVP_top60_DNA+RNA_seed | PIK3CA | -0.085083 | COADREAD | T84 | AZD8186 |
| MVP_top60_DNA_seed | MAPK1 | -0.184135 | COADREAD | LS-1034 | Ulixertinib | MVP_top60_DNA+RNA_seed | PIK3CB | -0.085083 | COADREAD | T84 | AZD8186 |
| MVP_top60_DNA_seed | CDK2 | -0.181144 | COADREAD | NCI-H508 | MK-8776 | MVP_top60_DNA+RNA_seed | IGF1R | -0.081158 | COADREAD | SW837 | BMS-754807 |
| MVP_top60_DNA_seed | IGF1R | -0.180748 | COADREAD | SW1463 | BMS-754807 | MVP_top60_DNA+RNA_seed | AKT1 | -0.08107 | COADREAD | SNU-407 | Afuresertib |
| MVP_top60_DNA_seed | MAPK1 | -0.180469 | COADREAD | T84 | SCH772984 | MVP_top60_DNA+RNA_seed | PIK3CA | -0.080345 | COADREAD | HCC2998 | GNE-317 |
| MVP_top60_DNA_seed | BRAF | -0.18041 | COADREAD | HCT-116 | SB590885 | MVP_top60_DNA+RNA_seed | HSP90AA1 | -0.080165 | COADREAD | HT55 | Luminespib |
| MVP_top60_DNA_seed | MAPK1 | -0.178639 | COADREAD | GP5d | SCH772984 | MVP_top60_DNA+RNA_seed | HSP90AB1 | -0.080165 | COADREAD | HT55 | Luminespib |
| MVP_top60_DNA_seed | MAPK1 | -0.178358 | COADREAD | HT-115 | SCH772984 | MVP_top60_DNA+RNA_seed | HSP90AA1 | -0.077036 | COADREAD | SNU-C5 | Tanespimycin |
| MVP_top60_DNA_seed | IGF1R | -0.177609 | COADREAD | LS-411N | Linsitinib | MVP_top60_DNA+RNA_seed | HSP90AB1 | -0.077036 | COADREAD | SNU-C5 | Tanespimycin |
| MVP_top60_DNA_seed | HSP90AA1 | -0.176512 | COADREAD | HCT-116 | Tanespimycin | MVP_top60_DNA+RNA_seed | PIK3CA | -0.072293 | COADREAD | RCM-1 | Alpelisib |
| MVP_top60_DNA_seed | HSP90AB1 | -0.176512 | COADREAD | HCT-116 | Tanespimycin | MVP_top60_DNA+RNA_seed | PIK3CA | -0.072201 | COADREAD | GP5d | GNE-317 |
| MVP_top60_DNA_seed | ESR1 | -0.173756 | COADREAD | LS-180 | Tamoxifen | MVP_top60_DNA+RNA_seed | KRAS | -0.072151 | COADREAD | NCI-H747 | KRAS (G12C) Inhibitor-1 |
| MVP_top60_DNA_seed | PIK3CA | -0.172189 | COADREAD | LS-180 | Alpelisib | MVP_top60_DNA+RNA_seed | IGF1R | -0.072036 | COADREAD | HT-115 | GSK1904529A |
| MVP_top60_DNA_seed | AKT1 | -0.170917 | COADREAD | HCT-116 | Afuresertib | MVP_top60_DNA+RNA_seed | MET | -0.070178 | COADREAD | HT-115 | Savolitinib |
| MVP_top60_DNA_seed | PIK3CA | -0.170209 | COADREAD | SNU-407 | AZD8186 | MVP_top60_DNA+RNA_seed | CDK2 | -0.069316 | COADREAD | COLO-205 | Dinaciclib |
| MVP_top60_DNA_seed | PIK3CB | -0.170209 | COADREAD | SNU-407 | AZD8186 | MVP_top60_DNA+RNA_seed | KRAS | -0.069158 | COADREAD | HT-29 | KRAS (G12C) Inhibitor-1 |
| MVP_top60_DNA_seed | AKT1 | -0.170196 | COADREAD | SNU-407 | MK-2206 | MVP_top60_DNA+RNA_seed | PIK3CA | -0.066991 | COADREAD | CW-2 | Buparlisib |
| MVP_top60_DNA_seed | MET | -0.169706 | COADREAD | SW620 | Crizotinib | MVP_top60_DNA+RNA_seed | PIK3CB | -0.066991 | COADREAD | CW-2 | Buparlisib |
| MVP_top60_DNA_seed | ESR1 | -0.168973 | COADREAD | CaR-1 | GDC0810 | MVP_top60_DNA+RNA_seed | PIK3CD | -0.066991 | COADREAD | CW-2 | Buparlisib |
| MVP_top60_DNA_seed | IGF1R | -0.168659 | COADREAD | HCT-15 | BMS-536924 | MVP_top60_DNA+RNA_seed | BCL2 | -0.066892 | COADREAD | T84 | Obatoclax Mesylate |
| MVP_top60_DNA_seed | MET | -0.166036 | COADREAD | SNU-61 | Savolitinib | MVP_top60_DNA+RNA_seed | BCL2 | -0.062977 | COADREAD | COLO-205 | Venetoclax |
| MVP_top60_DNA_seed | PIK3CA | -0.164333 | COADREAD | HT-29 | AZD8186 | MVP_top60_DNA+RNA_seed | PIK3CA | -0.060572 | COADREAD | SW1463 | GNE-317 |
| MVP_top60_DNA_seed | PIK3CB | -0.164333 | COADREAD | HT-29 | AZD8186 | MVP_top60_DNA+RNA_seed | ATM | -0.059558 | COADREAD | HT-115 | KU-55933 |
| MVP_top60_DNA_seed | AKT1 | -0.164102 | COADREAD | CaR-1 | AT13148 | MVP_top60_DNA+RNA_seed | SRC | -0.058319 | COADREAD | CW-2 | Bosutinib |
| MVP_top60_DNA_seed | BRAF | -0.163102 | COADREAD | CaR-1 | Dabrafenib | MVP_top60_DNA+RNA_seed | PIK3CA | -0.058033 | COADREAD | HCC2998 | AZD8186 |
| MVP_top60_DNA_seed | AKT1 | -0.160398 | COADREAD | SNU-407 | AZD5363 | MVP_top60_DNA+RNA_seed | PIK3CB | -0.058033 | COADREAD | HCC2998 | AZD8186 |
| MVP_top60_DNA_seed | PIK3CA | -0.158809 | COADREAD | HT-115 | Alpelisib | MVP_top60_DNA+RNA_seed | AKT1 | -0.056117 | COADREAD | SK-CO-1 | Ipatasertib |
| MVP_top60_DNA_seed | CDK2 | -0.157654 | COADREAD | LS-513 | Dinaciclib | MVP_top60_DNA+RNA_seed | IGF1R | -0.055693 | COADREAD | COLO-320-HSR | GSK1904529A |
| MVP_top60_DNA_seed | MET | -0.157152 | COADREAD | SNU-C5 | Crizotinib | MVP_top60_DNA+RNA_seed | BCL2 | -0.055291 | COADREAD | SK-CO-1 | Obatoclax Mesylate |
| MVP_top60_DNA_seed | PIK3CA | -0.156407 | COADREAD | MDST8 | AZD8186 | MVP_top60_DNA+RNA_seed | KRAS | -0.055151 | COADREAD | HCT-116 | KRAS (G12C) Inhibitor-1 |
| MVP_top60_DNA_seed | PIK3CB | -0.156407 | COADREAD | MDST8 | AZD8186 | MVP_top60_DNA+RNA_seed | TP53 | -0.052856 | COADREAD | HCT-116 | MIRA-1 |
| MVP_top60_DNA_seed | CDK2 | -0.156344 | COADREAD | RKO | MK-8776 | MVP_top60_DNA+RNA_seed | IGF1R | -0.052827 | COADREAD | SNU-61 | GSK1904529A |
| MVP_top60_DNA_seed | PIK3CA | -0.155478 | COADREAD | SW948 | Buparlisib | MVP_top60_DNA+RNA_seed | MET | -0.051901 | COADREAD | HCT-116 | Savolitinib |
| MVP_top60_DNA_seed | PIK3CB | -0.155478 | COADREAD | SW948 | Buparlisib | MVP_top60_DNA+RNA_seed | ATM | -0.0517 | COADREAD | GP5d | KU-55933 |
| MVP_top60_DNA_seed | PIK3CD | -0.155478 | COADREAD | SW948 | Buparlisib | MVP_top60_DNA+RNA_seed | AKT1 | -0.049034 | COADREAD | LS-180 | Ipatasertib |
| MVP_top60_DNA_seed | AKT1 | -0.155375 | COADREAD | COLO-205 | Uprosertib | MVP_top60_DNA+RNA_seed | ERBB3 | -0.048857 | COADREAD | SNU-1040 | Sapitinib |
| MVP_top60_DNA_seed | AKT1 | -0.154932 | COADREAD | HCT-116 | AZD5363 | MVP_top60_DNA+RNA_seed | IGF1R | -0.046163 | COADREAD | SNU-C5 | NVP-ADW742 |

| | | | | | | | | | | | |
|---|---|---|---|---|---|---|---|---|---|---|---|
| MVP_top60_DNA_seed | AKT1 | -0.154742 | COADREAD | NCI-H747 | AT13148 | MVP_top60_DNA+RNA_seed | AKT1 | -0.041346 | COADREAD | SNU-407 | Uprosertib |
| MVP_top60_DNA_seed | AKT1 | -0.153909 | COADREAD | HCT-116 | Uprosertib | MVP_top60_DNA+RNA_seed | IGF1R | -0.041183 | COADREAD | SW1463 | NVP-ADW742 |
| MVP_top60_DNA_seed | AKT1 | -0.153752 | COADREAD | RKO | AZD5363 | MVP_top60_DNA+RNA_seed | IGF1R | -0.039374 | COADREAD | C2BBe1 | Linsitinib |
| MVP_top60_DNA_seed | CDK2 | -0.153624 | COADREAD | LS-411N | AZD5438 | MVP_top60_DNA+RNA_seed | AKT1 | -0.039019 | COADREAD | CL-11 | Uprosertib |
| MVP_top60_DNA_seed | IGF1R | -0.15142 | COADREAD | LS-411N | BMS-754807 | MVP_top60_DNA+RNA_seed | AKT1 | -0.036049 | COADREAD | COLO-320-HSR | AZD5363 |
| MVP_top60_DNA_seed | AKT1 | -0.151309 | COADREAD | CW-2 | Afuresertib | MVP_top60_DNA+RNA_seed | AKT1 | -0.035518 | COADREAD | KM12 | MK-2206 |
| MVP_top60_DNA_seed | CDK2 | -0.150535 | COADREAD | HCT-116 | MK-8776 | MVP_top60_DNA+RNA_seed | BCL2 | -0.035496 | COADREAD | SW1116 | TW 37 |
| MVP_top60_DNA_seed | CDK2 | -0.147607 | COADREAD | SNU-C5 | Dinaciclib | MVP_top60_DNA+RNA_seed | AKT1 | -0.035301 | COADREAD | CW-2 | Uprosertib |
| MVP_top60_DNA_seed | MET | -0.145439 | COADREAD | DiFi | Savolitinib | MVP_top60_DNA+RNA_seed | IGF1R | -0.034823 | COADREAD | LS-1034 | BMS-536924 |
| MVP_top60_DNA_seed | AKT1 | -0.142633 | COADREAD | CW-2 | GSK2110183B | MVP_top60_DNA+RNA_seed | KRAS | -0.034769 | COADREAD | SNU-1040 | KRAS (G12C) Inhibitor-1 |
| MVP_top60_DNA_seed | AKT1 | -0.139264 | COADREAD | CCK-81 | MK-2206 | MVP_top60_DNA+RNA_seed | AKT1 | -0.034231 | COADREAD | SNU-175 | Afuresertib |
| MVP_top60_DNA_seed | PIK3CA | -0.13814 | COADREAD | CW-2 | Alpelisib | MVP_top60_DNA+RNA_seed | AKT1 | -0.028649 | COADREAD | HT-115 | MK-2206 |
| MVP_top60_DNA_seed | PIK3CA | -0.136585 | COADREAD | LS-123 | AZD8186 | MVP_top60_DNA+RNA_seed | BCL2 | -0.02736 | COADREAD | GP5d | Venetoclax |
| MVP_top60_DNA_seed | PIK3CB | -0.136585 | COADREAD | LS-123 | AZD8186 | MVP_top60_DNA+RNA_seed | AKT1 | -0.027301 | COADREAD | SNU-1040 | MK-2206 |
| MVP_top60_DNA_seed | AKT1 | -0.13602 | COADREAD | COLO-320-HSR | Uprosertib | MVP_top60_DNA+RNA_seed | PIK3CA | -0.027099 | COADREAD | RKO | Buparlisib |
| MVP_top60_DNA_seed | SRC | -0.135012 | COADREAD | LoVo | Bosutinib | MVP_top60_DNA+RNA_seed | PIK3CB | -0.027099 | COADREAD | RKO | Buparlisib |
| MVP_top60_DNA_seed | ESR1 | -0.131471 | COADREAD | HCT-15 | GDC0810 | MVP_top60_DNA+RNA_seed | PIK3CD | -0.027099 | COADREAD | RKO | Buparlisib |
| MVP_top60_DNA_seed | CDK2 | -0.130337 | COADREAD | SNU-C2B | MK-8776 | MVP_top60_DNA+RNA_seed | PIK3CA | -0.026078 | COADREAD | SNU-C5 | AZD8186 |
| MVP_top60_DNA_seed | AKT1 | -0.125655 | COADREAD | CCK-81 | Afuresertib | MVP_top60_DNA+RNA_seed | PIK3CB | -0.026078 | COADREAD | SNU-C5 | AZD8186 |
| MVP_top60_DNA_seed | ATM | -0.125136 | COADREAD | KM12 | KU-55933 | MVP_top60_DNA+RNA_seed | ERBB3 | -0.025835 | COADREAD | COLO-678 | Sapitinib |
| MVP_top60_DNA_seed | AKT1 | -0.124502 | COADREAD | SNU-81 | Uprosertib | MVP_top60_DNA+RNA_seed | GSK3B | -0.025553 | COADREAD | SW1116 | SB216763 |
| MVP_top60_DNA_seed | CDK2 | -0.124217 | COADREAD | GP5d | MK-8776 | MVP_top60_DNA+RNA_seed | BCL2 | -0.023068 | COADREAD | RCM-1 | Sabutoclax |
| MVP_top60_DNA_seed | SRC | -0.12386 | COADREAD | SNU-C1 | Bosutinib | MVP_top60_DNA+RNA_seed | BCL2 | -0.021663 | COADREAD | SW48 | Venetoclax |
| MVP_top60_DNA_seed | AKT1 | -0.122725 | COADREAD | COLO-205 | AZD5363 | MVP_top60_DNA+RNA_seed | PIK3CA | -0.019827 | COADREAD | NCI-H508 | Buparlisib |
| MVP_top60_DNA_seed | ERBB3 | -0.121206 | COADREAD | HCT-15 | Sapitinib | MVP_top60_DNA+RNA_seed | PIK3CB | -0.019827 | COADREAD | NCI-H508 | Buparlisib |
| MVP_top60_DNA_seed | HSP90AA1 | -0.121187 | COADREAD | NCI-H747 | Luminespib | MVP_top60_DNA+RNA_seed | PIK3CD | -0.019827 | COADREAD | NCI-H508 | Buparlisib |
| MVP_top60_DNA_seed | HSP90AB1 | -0.121187 | COADREAD | NCI-H747 | Luminespib | MVP_top60_DNA+RNA_seed | AKT1 | -0.017883 | COADREAD | T84 | AZD5363 |
| MVP_top60_DNA_seed | AKT1 | -0.12003 | COADREAD | NCI-H716 | Afuresertib | MVP_top60_DNA+RNA_seed | AKT1 | -0.017067 | COADREAD | T84 | AT13148 |
| MVP_top60_DNA_seed | CREBBP | -0.119782 | COADREAD | SNU-61 | SGC-CBP30 | MVP_top60_DNA+RNA_seed | IGF1R | -0.016462 | COADREAD | LoVo | NVP-ADW742 |
| MVP_top60_DNA_seed | CDK2 | -0.119412 | COADREAD | CL-11 | MK-8776 | MVP_top60_DNA+RNA_seed | CDK2 | -0.014069 | COADREAD | SW948 | AZD5438 |
| MVP_top60_DNA_seed | MAPK1 | -0.11941 | COADREAD | DiFi | Ulixertinib | MVP_top60_DNA+RNA_seed | AKT1 | -0.013617 | COADREAD | LS-180 | Uprosertib |
| MVP_top60_DNA_seed | IGF1R | -0.115586 | COADREAD | SNU-1040 | BMS-754807 | MVP_top60_DNA+RNA_seed | IGF1R | -0.009753 | COADREAD | SW1116 | BMS-754807 |
| MVP_top60_DNA_seed | PIK3CA | -0.115279 | COADREAD | SNU-C5 | Alpelisib | MVP_top60_DNA+RNA_seed | CDK2 | -0.009737 | COADREAD | SNU-407 | AZD5438 |
| MVP_top60_DNA_seed | PIK3CA | -0.115211 | COADREAD | LoVo | Alpelisib | MVP_top60_DNA+RNA_seed | BCL2 | -0.005601 | COADREAD | SNU-175 | Navitoclax |
| MVP_top60_DNA_seed | MET | -0.11 | COADREAD | CL-11 | Crizotinib | MVP_top60_DNA+RNA_seed | BCL2 | -0.001539 | COADREAD | HCT-15 | Venetoclax |
| MVP_top60_DNA_seed | HSP90AA1 | -0.102919 | COADREAD | SW620 | Elesclomol | MVP_top60_DNA+RNA_seed | PIK3CA | -0.000728 | COADREAD | SNU-407 | GNE-317 |
| MVP_top60_DNA_seed | HSP90AB1 | -0.102919 | COADREAD | SW620 | Elesclomol | MVP_top60_DNA+RNA_seed | BCL2 | -0.000202 | COADREAD | MDST8 | Sabutoclax |
| MVP_top60_DNA_seed | MAPK1 | -0.099919 | COADREAD | SNU-81 | VX-11e | MVP_top60_DNA+RNA_seed | AKT1 | 0.000962 | COADREAD | CCK-81 | AZD5363 |
| MVP_top60_DNA_seed | PIK3CA | -0.09687 | COADREAD | HT55 | AZD8186 | MVP_top60_DNA+RNA_seed | HSP90AA1 | 0.001137 | COADREAD | CaR-1 | Luminespib |
| MVP_top60_DNA_seed | PIK3CB | -0.09687 | COADREAD | HT55 | AZD8186 | MVP_top60_DNA+RNA_seed | HSP90AB1 | 0.001137 | COADREAD | CaR-1 | Luminespib |
| MVP_top60_DNA_seed | ESR1 | -0.094348 | COADREAD | HT-29 | GDC0810 | MVP_top60_DNA+RNA_seed | GSK3B | 0.002209 | COADREAD | RKO | SB216763 |

| | | | | | | | | | | | |
|---|---|---|---|---|---|---|---|---|---|---|---|
| MVP_top60_DNA_seed | AKT1 | -0.092923 | COADREAD | NCI-H747 | MK-2206 | MVP_top60_DNA+RNA_seed | PIK3CA | 0.002855 | COADREAD | SNU-175 | AZD8186 |
| MVP_top60_DNA_seed | CDK2 | -0.090892 | COADREAD | NCI-H716 | AZD5438 | MVP_top60_DNA+RNA_seed | PIK3CB | 0.002855 | COADREAD | SNU-175 | AZD8186 |
| MVP_top60_DNA_seed | HSP90AA1 | -0.090335 | COADREAD | HCC2998 | Luminespib | MVP_top60_DNA+RNA_seed | BCL2 | 0.005693 | COADREAD | MDST8 | Venetoclax |
| MVP_top60_DNA_seed | HSP90AB1 | -0.090335 | COADREAD | HCC2998 | Luminespib | MVP_top60_DNA+RNA_seed | BCL2 | 0.006115 | COADREAD | LS-513 | Obatoclax Mesylate |
| MVP_top60_DNA_seed | AKT1 | -0.087401 | COADREAD | RKO | AT13148 | MVP_top60_DNA+RNA_seed | PIK3CA | 0.006593 | COADREAD | SNU-175 | GNE-317 |
| MVP_top60_DNA_seed | CDK2 | -0.087358 | COADREAD | SNU-C1 | Dinaciclib | MVP_top60_DNA+RNA_seed | IGF1R | 0.008174 | COADREAD | RCM-1 | Linsitinib |
| MVP_top60_DNA_seed | CDK2 | -0.085834 | COADREAD | SNU-175 | MK-8776 | MVP_top60_DNA+RNA_seed | PIK3CA | 0.0087 | COADREAD | HT-29 | Buparlisib |
| MVP_top60_DNA_seed | PIK3CA | -0.085083 | COADREAD | T84 | AZD8186 | MVP_top60_DNA+RNA_seed | PIK3CB | 0.0087 | COADREAD | HT-29 | Buparlisib |
| MVP_top60_DNA_seed | PIK3CB | -0.085083 | COADREAD | T84 | AZD8186 | MVP_top60_DNA+RNA_seed | PIK3CD | 0.0087 | COADREAD | HT-29 | Buparlisib |
| MVP_top60_DNA_seed | IGF1R | -0.081158 | COADREAD | SW837 | BMS-754807 | MVP_top60_DNA+RNA_seed | AKT1 | 0.013892 | COADREAD | C2BBe1 | Uprosertib |
| MVP_top60_DNA_seed | AKT1 | -0.08107 | COADREAD | SNU-407 | Afuresertib | MVP_top60_DNA+RNA_seed | BCL2 | 0.014344 | COADREAD | KM12 | Obatoclax Mesylate |
| MVP_top60_DNA_seed | PIK3CA | -0.080345 | COADREAD | HCC2998 | GNE-317 | MVP_top60_DNA+RNA_seed | HSP90AA1 | 0.014707 | COADREAD | SNU-C2B | Luminespib |
| MVP_top60_DNA_seed | HSP90AA1 | -0.080165 | COADREAD | HT55 | Luminespib | MVP_top60_DNA+RNA_seed | HSP90AB1 | 0.014707 | COADREAD | SNU-C2B | Luminespib |
| MVP_top60_DNA_seed | HSP90AB1 | -0.080165 | COADREAD | HT55 | Luminespib | MVP_top60_DNA+RNA_seed | AKT1 | 0.016988 | COADREAD | LS-123 | Uprosertib |
| MVP_top60_DNA_seed | HSP90AA1 | -0.077036 | COADREAD | SNU-C5 | Tanespimycin | MVP_top60_DNA+RNA_seed | PIK3CA | 0.017624 | COADREAD | CCK-81 | AZD8186 |
| MVP_top60_DNA_seed | HSP90AB1 | -0.077036 | COADREAD | SNU-C5 | Tanespimycin | MVP_top60_DNA+RNA_seed | PIK3CB | 0.017624 | COADREAD | CCK-81 | AZD8186 |
| MVP_top60_DNA_seed | MAPK1 | -0.074064 | COADREAD | LS-123 | VX-11e | MVP_top60_DNA+RNA_seed | ATM | 0.018433 | COADREAD | COLO-320-HSR | KU-55933 |
| MVP_top60_DNA_seed | BRAF | -0.073861 | COADREAD | COLO-320-HSR | SB590885 | MVP_top60_DNA+RNA_seed | MET | 0.020309 | COADREAD | COLO-205 | Savolitinib |
| MVP_top60_DNA_seed | PIK3CA | -0.072293 | COADREAD | RCM-1 | Alpelisib | MVP_top60_DNA+RNA_seed | MET | 0.020793 | COADREAD | SNU-81 | Savolitinib |
| MVP_top60_DNA_seed | PIK3CA | -0.072201 | COADREAD | GP5d | GNE-317 | MVP_top60_DNA+RNA_seed | AKT1 | 0.026323 | COADREAD | SNU-175 | AT13148 |
| MVP_top60_DNA_seed | KRAS | -0.072151 | COADREAD | NCI-H747 | KRAS (G12C) Inhibitor-1: | MVP_top60_DNA+RNA_seed | HSP90AA1 | 0.026626 | COADREAD | SK-CO-1 | Luminespib |
| MVP_top60_DNA_seed | IGF1R | -0.072036 | COADREAD | HT-115 | GSK1904529A | MVP_top60_DNA+RNA_seed | HSP90AB1 | 0.026626 | COADREAD | SK-CO-1 | Luminespib |
| MVP_top60_DNA_seed | MET | -0.070178 | COADREAD | HT-115 | Savolitinib | MVP_top60_DNA+RNA_seed | BCL2 | 0.027916 | COADREAD | LS-123 | Obatoclax Mesylate |
| MVP_top60_DNA_seed | CDK2 | -0.069316 | COADREAD | COLO-205 | Dinaciclib | MVP_top60_DNA+RNA_seed | CDK2 | 0.028078 | COADREAD | HCC2998 | MK-8776 |
| MVP_top60_DNA_seed | KRAS | -0.069158 | COADREAD | HT-29 | KRAS (G12C) Inhibitor-1: | MVP_top60_DNA+RNA_seed | BCL2 | 0.028273 | COADREAD | NCI-H716 | Obatoclax Mesylate |
| MVP_top60_DNA_seed | PIK3CA | -0.066991 | COADREAD | CW-2 | Buparlisib | MVP_top60_DNA+RNA_seed | CDK2 | 0.02885 | COADREAD | LS-180 | AZD5438 |
| MVP_top60_DNA_seed | PIK3CB | -0.066991 | COADREAD | CW-2 | Buparlisib | MVP_top60_DNA+RNA_seed | SRC | 0.031142 | COADREAD | SNU-C5 | Bosutinib |
| MVP_top60_DNA_seed | PIK3CD | -0.066991 | COADREAD | CW-2 | Buparlisib | MVP_top60_DNA+RNA_seed | TP53 | 0.031989 | COADREAD | HCC2998 | MIRA-1 |
| MVP_top60_DNA_seed | BRAF | -0.06537 | COADREAD | CW-2 | SB590885 | MVP_top60_DNA+RNA_seed | IGF1R | 0.032126 | COADREAD | CL-11 | BMS-536924 |
| MVP_top60_DNA_seed | PIK3CA | -0.060572 | COADREAD | SW1463 | GNE-317 | MVP_top60_DNA+RNA_seed | AKT1 | 0.032775 | COADREAD | COLO-205 | GSK2110183B |
| MVP_top60_DNA_seed | ATM | -0.059558 | COADREAD | HT-115 | KU-55933 | MVP_top60_DNA+RNA_seed | IGF1R | 0.033151 | COADREAD | CaR-1 | NVP-ADW742 |
| MVP_top60_DNA_seed | SRC | -0.058319 | COADREAD | CW-2 | Bosutinib | MVP_top60_DNA+RNA_seed | BCL2 | 0.036621 | COADREAD | SW1463 | Navitoclax |
| MVP_top60_DNA_seed | PIK3CA | -0.058033 | COADREAD | HCC2998 | AZD8186 | MVP_top60_DNA+RNA_seed | CDK2 | 0.03717 | COADREAD | LS-1034 | AZD5438 |
| MVP_top60_DNA_seed | PIK3CB | -0.058033 | COADREAD | HCC2998 | AZD8186 | MVP_top60_DNA+RNA_seed | BCL2 | 0.037179 | COADREAD | LS-411N | Navitoclax |
| MVP_top60_DNA_seed | AKT1 | -0.056117 | COADREAD | SK-CO-1 | Ipatasertib | MVP_top60_DNA+RNA_seed | CDK2 | 0.039368 | COADREAD | SW1463 | AZD5438 |
| MVP_top60_DNA_seed | IGF1R | -0.055693 | COADREAD | COLO-320-HSR | GSK1904529A | MVP_top60_DNA+RNA_seed | IGF1R | 0.039373 | COADREAD | T84 | BMS-536924 |
| MVP_top60_DNA_seed | KRAS | -0.055151 | COADREAD | HCT-116 | KRAS (G12C) Inhibitor-1: | MVP_top60_DNA+RNA_seed | BCL2 | 0.046147 | COADREAD | LoVo | Navitoclax |
| MVP_top60_DNA_seed | MAPK1 | -0.053791 | COADREAD | CCK-81 | Ulixertinib | MVP_top60_DNA+RNA_seed | BCL2 | 0.047729 | COADREAD | LS-180 | TW 37 |
| MVP_top60_DNA_seed | TP53 | -0.052856 | COADREAD | HCT-116 | MIRA-1 | MVP_top60_DNA+RNA_seed | BCL2 | 0.048668 | COADREAD | HCC2998 | Venetoclax |
| MVP_top60_DNA_seed | IGF1R | -0.052827 | COADREAD | SNU-61 | GSK1904529A | MVP_top60_DNA+RNA_seed | IGF1R | 0.051912 | COADREAD | LS-513 | GSK1904529A |
| MVP_top60_DNA_seed | MET | -0.051901 | COADREAD | HCT-116 | Savolitinib | MVP_top60_DNA+RNA_seed | KRAS | 0.053306 | COADREAD | LS-411N | KRAS (G12C) Inhibitor-1 |

| | | | | | | | | | | | |
|---|---|---|---|---|---|---|---|---|---|---|---|
| MVP_top60_DNA_seed | ATM | -0.0517 | COADREAD | GP5d | KU-55933 | MVP_top60_DNA+RNA_seed | IGF1R | 0.061116 | COADREAD | CaR-1 | Linsitinib |
| MVP_top60_DNA_seed | AKT1 | -0.049034 | COADREAD | LS-180 | Ipatasertib | MVP_top60_DNA+RNA_seed | CDK2 | 0.061367 | COADREAD | SW48 | MK-8776 |
| MVP_top60_DNA_seed | ERBB3 | -0.048857 | COADREAD | SNU-1040 | Sapitinib | MVP_top60_DNA+RNA_seed | CDK2 | 0.063397 | COADREAD | SW48 | AZD5438 |
| MVP_top60_DNA_seed | ESR1 | -0.048827 | COADREAD | RCM-1 | Tamoxifen | MVP_top60_DNA+RNA_seed | BCL2 | 0.064582 | COADREAD | CW-2 | Obatoclax Mesylate |
| MVP_top60_DNA_seed | BRAF | -0.046993 | COADREAD | NCI-H716 | Dabrafenib | MVP_top60_DNA+RNA_seed | PIK3CA | 0.06485 | COADREAD | DiFi | AZD8186 |
| MVP_top60_DNA_seed | IGF1R | -0.046163 | COADREAD | SNU-C5 | NVP-ADW742 | MVP_top60_DNA+RNA_seed | PIK3CB | 0.06485 | COADREAD | DiFi | AZD8186 |
| MVP_top60_DNA_seed | AKT1 | -0.041346 | COADREAD | SNU-407 | Uprosertib | MVP_top60_DNA+RNA_seed | TP53 | 0.065601 | COADREAD | SK-CO-1 | MIRA-1 |
| MVP_top60_DNA_seed | IGF1R | -0.041183 | COADREAD | SW1463 | NVP-ADW742 | MVP_top60_DNA+RNA_seed | BCL2 | 0.067447 | COADREAD | T84 | Navitoclax |
| MVP_top60_DNA_seed | IGF1R | -0.039374 | COADREAD | C2BBe1 | Linsitinib | MVP_top60_DNA+RNA_seed | PIK3CA | 0.067512 | COADREAD | SNU-175 | Alpelisib |
| MVP_top60_DNA_seed | AKT1 | -0.039019 | COADREAD | CL-11 | Uprosertib | MVP_top60_DNA+RNA_seed | BCL2 | 0.072477 | COADREAD | CCK-81 | Sabutoclax |
| MVP_top60_DNA_seed | AKT1 | -0.036049 | COADREAD | COLO-320-HSR | AZD5363 | MVP_top60_DNA+RNA_seed | CDK2 | 0.07268 | COADREAD | SNU-C2B | Dinaciclib |
| MVP_top60_DNA_seed | AKT1 | -0.035518 | COADREAD | KM12 | MK-2206 | MVP_top60_DNA+RNA_seed | MET | 0.073968 | COADREAD | SW1463 | Crizotinib |
| MVP_top60_DNA_seed | AKT1 | -0.035301 | COADREAD | CW-2 | Uprosertib | MVP_top60_DNA+RNA_seed | BCL2 | 0.074954 | COADREAD | HT55 | Navitoclax |
| MVP_top60_DNA_seed | IGF1R | -0.034823 | COADREAD | LS-1034 | BMS-536924 | MVP_top60_DNA+RNA_seed | IGF1R | 0.076402 | COADREAD | MDST8 | GSK1904529A |
| MVP_top60_DNA_seed | KRAS | -0.034769 | COADREAD | SNU-1040 | KRAS (G12C) Inhibitor-12 | MVP_top60_DNA+RNA_seed | HSP90AA1 | 0.080794 | COADREAD | SW1463 | Tanespimycin |
| MVP_top60_DNA_seed | AKT1 | -0.034231 | COADREAD | SNU-175 | Afuresertib | MVP_top60_DNA+RNA_seed | HSP90AB1 | 0.080794 | COADREAD | SW1463 | Tanespimycin |
| MVP_top60_DNA_seed | BRAF | -0.032983 | COADREAD | SW948 | Dabrafenib | MVP_top60_DNA+RNA_seed | AKT1 | 0.081107 | COADREAD | RKO | MK-2206 |
| MVP_top60_DNA_seed | BRAF | -0.032273 | COADREAD | HT55 | SB590885 | MVP_top60_DNA+RNA_seed | AKT1 | 0.08304 | COADREAD | SK-CO-1 | AZD5363 |
| MVP_top60_DNA_seed | AKT1 | -0.028649 | COADREAD | HT-115 | MK-2206 | MVP_top60_DNA+RNA_seed | IGF1R | 0.083047 | COADREAD | SNU-C5 | BMS-754807 |
| MVP_top60_DNA_seed | ESR1 | -0.027623 | COADREAD | HT55 | Tamoxifen | MVP_top60_DNA+RNA_seed | AKT1 | 0.083147 | COADREAD | SNU-81 | MK-2206 |
| MVP_top60_DNA_seed | AKT1 | -0.027301 | COADREAD | SNU-1040 | MK-2206 | MVP_top60_DNA+RNA_seed | PIK3CA | 0.085727 | COADREAD | GP5d | AZD8186 |
| MVP_top60_DNA_seed | PIK3CA | -0.027099 | COADREAD | RKO | Buparlisib | MVP_top60_DNA+RNA_seed | PIK3CB | 0.085727 | COADREAD | GP5d | AZD8186 |
| MVP_top60_DNA_seed | PIK3CB | -0.027099 | COADREAD | RKO | Buparlisib | MVP_top60_DNA+RNA_seed | CDK2 | 0.087181 | COADREAD | T84 | AZD5438 |
| MVP_top60_DNA_seed | PIK3CD | -0.027099 | COADREAD | RKO | Buparlisib | MVP_top60_DNA+RNA_seed | PIK3CA | 0.087524 | COADREAD | SW837 | GNE-317 |
| MVP_top60_DNA_seed | PIK3CA | -0.026078 | COADREAD | SNU-C5 | AZD8186 | MVP_top60_DNA+RNA_seed | BCL2 | 0.087657 | COADREAD | RKO | Navitoclax |
| MVP_top60_DNA_seed | PIK3CB | -0.026078 | COADREAD | SNU-C5 | AZD8186 | MVP_top60_DNA+RNA_seed | CDK2 | 0.088332 | COADREAD | LoVo | MK-8776 |
| MVP_top60_DNA_seed | ERBB3 | -0.025835 | COADREAD | COLO-678 | Sapitinib | MVP_top60_DNA+RNA_seed | TP53 | 0.089289 | COADREAD | HT-115 | MIRA-1 |
| MVP_top60_DNA_seed | GSK3B | -0.025553 | COADREAD | SW1116 | SB216763 | MVP_top60_DNA+RNA_seed | CDK2 | 0.090208 | COADREAD | MDST8 | AZD5438 |
| MVP_top60_DNA_seed | PIK3CA | -0.019827 | COADREAD | NCI-H508 | Buparlisib | MVP_top60_DNA+RNA_seed | IGF1R | 0.091019 | COADREAD | HCC2998 | BMS-754807 |
| MVP_top60_DNA_seed | PIK3CB | -0.019827 | COADREAD | NCI-H508 | Buparlisib | MVP_top60_DNA+RNA_seed | HSP90AA1 | 0.091237 | COADREAD | SNU-C5 | Luminespib |
| MVP_top60_DNA_seed | PIK3CD | -0.019827 | COADREAD | NCI-H508 | Buparlisib | MVP_top60_DNA+RNA_seed | HSP90AB1 | 0.091237 | COADREAD | SNU-C5 | Luminespib |
| MVP_top60_DNA_seed | AKT1 | -0.017883 | COADREAD | T84 | AZD5363 | MVP_top60_DNA+RNA_seed | IGF1R | 0.091488 | COADREAD | LS-411N | GSK1904529A |
| MVP_top60_DNA_seed | AKT1 | -0.017067 | COADREAD | T84 | AT13148 | MVP_top60_DNA+RNA_seed | AKT1 | 0.092447 | COADREAD | CaR-1 | MK-2206 |
| MVP_top60_DNA_seed | IGF1R | -0.016462 | COADREAD | LoVo | NVP-ADW742 | MVP_top60_DNA+RNA_seed | AKT1 | 0.093526 | COADREAD | NCI-H747 | GSK2110183B |
| MVP_top60_DNA_seed | CDK2 | -0.014069 | COADREAD | SW948 | AZD5438 | MVP_top60_DNA+RNA_seed | BCL2 | 0.099104 | COADREAD | HCT-116 | Navitoclax |
| MVP_top60_DNA_seed | AKT1 | -0.013617 | COADREAD | LS-180 | Uprosertib | MVP_top60_DNA+RNA_seed | AKT1 | 0.104612 | COADREAD | SNU-407 | GSK2110183B |
| MVP_top60_DNA_seed | IGF1R | -0.009753 | COADREAD | SW1116 | BMS-754807 | MVP_top60_DNA+RNA_seed | KRAS | 0.104894 | COADREAD | SNU-175 | KRAS (G12C) Inhibitor-1 |
| MVP_top60_DNA_seed | CDK2 | -0.009737 | COADREAD | SNU-407 | AZD5438 | MVP_top60_DNA+RNA_seed | AKT1 | 0.105069 | COADREAD | CaR-1 | Ipatasertib |
| MVP_top60_DNA_seed | PIK3CA | -0.000728 | COADREAD | SNU-407 | GNE-317 | MVP_top60_DNA+RNA_seed | BCL2 | 0.106261 | COADREAD | SNU-C2B | Sabutoclax |
| MVP_top60_DNA_seed | AKT1 | 0.000962 | COADREAD | CCK-81 | AZD5363 | MVP_top60_DNA+RNA_seed | BCL2 | 0.11134 | COADREAD | SW1463 | TW 37 |
| MVP_top60_DNA_seed | HSP90AA1 | 0.001137 | COADREAD | CaR-1 | Luminespib | MVP_top60_DNA+RNA_seed | TP53 | 0.111455 | COADREAD | SNU-175 | MIRA-1 |

| | | | | | | | | | | | |
|---|---|---|---|---|---|---|---|---|---|---|---|
| MVP_top60_DNA_seed | HSP90AB1 | 0.001137 | COADREAD | CaR-1 | Luminespib | MVP_top60_DNA+RNA_seed | AKT1 | 0.113815 | COADREAD | MDST8 | Uprosertib |
| MVP_top60_DNA_seed | MAPK1 | 0.00203 | COADREAD | SW948 | Ulixertinib | MVP_top60_DNA+RNA_seed | ERBB3 | 0.1163 | COADREAD | SW837 | Sapitinib |
| MVP_top60_DNA_seed | GSK3B | 0.002209 | COADREAD | RKO | SB216763 | MVP_top60_DNA+RNA_seed | BCL2 | 0.116449 | COADREAD | LS-513 | Sabutoclax |
| MVP_top60_DNA_seed | PIK3CA | 0.002855 | COADREAD | SNU-175 | AZD8186 | MVP_top60_DNA+RNA_seed | PIK3CA | 0.117066 | COADREAD | HCC2998 | Alpelisib |
| MVP_top60_DNA_seed | PIK3CB | 0.002855 | COADREAD | SNU-175 | AZD8186 | MVP_top60_DNA+RNA_seed | BCL2 | 0.117159 | COADREAD | HT-29 | TW 37 |
| MVP_top60_DNA_seed | MAPK1 | 0.005414 | COADREAD | COLO-678 | Ulixertinib | MVP_top60_DNA+RNA_seed | BCL2 | 0.119151 | COADREAD | SNU-407 | Venetoclax |
| MVP_top60_DNA_seed | PIK3CA | 0.006593 | COADREAD | SNU-175 | GNE-317 | MVP_top60_DNA+RNA_seed | MET | 0.119556 | COADREAD | RKO | Savolitinib |
| MVP_top60_DNA_seed | IGF1R | 0.008174 | COADREAD | RCM-1 | Linsitinib | MVP_top60_DNA+RNA_seed | BCL2 | 0.121134 | COADREAD | CaR-1 | Navitoclax |
| MVP_top60_DNA_seed | PIK3CA | 0.0087 | COADREAD | HT-29 | Buparlisib | MVP_top60_DNA+RNA_seed | IGF1R | 0.121399 | COADREAD | SNU-C5 | BMS-536924 |
| MVP_top60_DNA_seed | PIK3CB | 0.0087 | COADREAD | HT-29 | Buparlisib | MVP_top60_DNA+RNA_seed | ERBB3 | 0.127041 | COADREAD | LS-123 | Sapitinib |
| MVP_top60_DNA_seed | PIK3CD | 0.0087 | COADREAD | HT-29 | Buparlisib | MVP_top60_DNA+RNA_seed | AKT1 | 0.131777 | COADREAD | T84 | Uprosertib |
| MVP_top60_DNA_seed | AKT1 | 0.013892 | COADREAD | C2BBe1 | Uprosertib | MVP_top60_DNA+RNA_seed | CDK2 | 0.133356 | COADREAD | SNU-C1 | MK-8776 |
| MVP_top60_DNA_seed | HSP90AA1 | 0.014707 | COADREAD | SNU-C2B | Luminespib | MVP_top60_DNA+RNA_seed | AKT1 | 0.133532 | COADREAD | CaR-1 | AZD5363 |
| MVP_top60_DNA_seed | HSP90AB1 | 0.014707 | COADREAD | SNU-C2B | Luminespib | MVP_top60_DNA+RNA_seed | HSP90AA1 | 0.133877 | COADREAD | CW-2 | Tanespimycin |
| MVP_top60_DNA_seed | AKT1 | 0.016988 | COADREAD | LS-123 | Uprosertib | MVP_top60_DNA+RNA_seed | HSP90AB1 | 0.133877 | COADREAD | CW-2 | Tanespimycin |
| MVP_top60_DNA_seed | PIK3CA | 0.017624 | COADREAD | CCK-81 | AZD8186 | MVP_top60_DNA+RNA_seed | CDK2 | 0.140132 | COADREAD | SNU-81 | Dinaciclib |
| MVP_top60_DNA_seed | PIK3CB | 0.017624 | COADREAD | CCK-81 | AZD8186 | MVP_top60_DNA+RNA_seed | PIK3CA | 0.140505 | COADREAD | T84 | Buparlisib |
| MVP_top60_DNA_seed | ATM | 0.018433 | COADREAD | COLO-320-HSR | KU-55933 | MVP_top60_DNA+RNA_seed | PIK3CB | 0.140505 | COADREAD | T84 | Buparlisib |
| MVP_top60_DNA_seed | MET | 0.020309 | COADREAD | COLO-205 | Savolitinib | MVP_top60_DNA+RNA_seed | PIK3CD | 0.140505 | COADREAD | T84 | Buparlisib |
| MVP_top60_DNA_seed | MET | 0.020793 | COADREAD | SNU-81 | Savolitinib | MVP_top60_DNA+RNA_seed | IGF1R | 0.143267 | COADREAD | LS-180 | BMS-754807 |
| MVP_top60_DNA_seed | AKT1 | 0.026323 | COADREAD | SNU-175 | AT13148 | MVP_top60_DNA+RNA_seed | IGF1R | 0.145407 | COADREAD | CL-11 | Linsitinib |
| MVP_top60_DNA_seed | HSP90AA1 | 0.026626 | COADREAD | SK-CO-1 | Luminespib | MVP_top60_DNA+RNA_seed | BCL2 | 0.147972 | COADREAD | HCC2998 | Sabutoclax |
| MVP_top60_DNA_seed | HSP90AB1 | 0.026626 | COADREAD | SK-CO-1 | Luminespib | MVP_top60_DNA+RNA_seed | GSK3B | 0.151996 | COADREAD | SW837 | SB216763 |
| MVP_top60_DNA_seed | CDK2 | 0.028078 | COADREAD | HCC2998 | MK-8776 | MVP_top60_DNA+RNA_seed | PIK3CA | 0.152698 | COADREAD | HT55 | Buparlisib |
| MVP_top60_DNA_seed | CDK2 | 0.02885 | COADREAD | LS-180 | AZD5438 | MVP_top60_DNA+RNA_seed | PIK3CB | 0.152698 | COADREAD | HT55 | Buparlisib |
| MVP_top60_DNA_seed | MAPK1 | 0.03067 | COADREAD | LS-1034 | Ulixertinib | MVP_top60_DNA+RNA_seed | PIK3CD | 0.152698 | COADREAD | HT55 | Buparlisib |
| MVP_top60_DNA_seed | SRC | 0.031142 | COADREAD | SNU-C5 | Bosutinib | MVP_top60_DNA+RNA_seed | PIK3CA | 0.152922 | COADREAD | SNU-C5 | GNE-317 |
| MVP_top60_DNA_seed | TP53 | 0.031989 | COADREAD | HCC2998 | MIRA-1 | MVP_top60_DNA+RNA_seed | PIK3CA | 0.153117 | COADREAD | NCI-H508 | GNE-317 |
| MVP_top60_DNA_seed | IGF1R | 0.032126 | COADREAD | CL-11 | BMS-536924 | MVP_top60_DNA+RNA_seed | MET | 0.156133 | COADREAD | NCI-H747 | Crizotinib |
| MVP_top60_DNA_seed | AKT1 | 0.032775 | COADREAD | COLO-205 | GSK2110183B | MVP_top60_DNA+RNA_seed | IGF1R | 0.156334 | COADREAD | SW1116 | GSK1904529A |
| MVP_top60_DNA_seed | BRAF | 0.032811 | COADREAD | LS-411N | SB590885 | MVP_top60_DNA+RNA_seed | HSP90AA1 | 0.157377 | COADREAD | C2BBe1 | Luminespib |
| MVP_top60_DNA_seed | IGF1R | 0.033151 | COADREAD | CaR-1 | NVP-ADW742 | MVP_top60_DNA+RNA_seed | HSP90AB1 | 0.157377 | COADREAD | C2BBe1 | Luminespib |
| MVP_top60_DNA_seed | CDK2 | 0.03717 | COADREAD | LS-1034 | AZD5438 | MVP_top60_DNA+RNA_seed | IGF1R | 0.158503 | COADREAD | NCI-H508 | BMS-536924 |
| MVP_top60_DNA_seed | CDK2 | 0.039368 | COADREAD | SW1463 | AZD5438 | MVP_top60_DNA+RNA_seed | IGF1R | 0.159382 | COADREAD | MDST8 | BMS-536924 |
| MVP_top60_DNA_seed | IGF1R | 0.039373 | COADREAD | T84 | BMS-536924 | MVP_top60_DNA+RNA_seed | IGF1R | 0.159978 | COADREAD | C2BBe1 | NVP-ADW742 |
| MVP_top60_DNA_seed | ESR1 | 0.039662 | COADREAD | SNU-C5 | GDC0810 | MVP_top60_DNA+RNA_seed | AKT1 | 0.160392 | COADREAD | C2BBe1 | MK-2206 |
| MVP_top60_DNA_seed | ESR1 | 0.039865 | COADREAD | NCI-H508 | GDC0810 | MVP_top60_DNA+RNA_seed | IGF1R | 0.160443 | COADREAD | C2BBe1 | GSK1904529A |
| MVP_top60_DNA_seed | MAPK1 | 0.041196 | COADREAD | C2BBe1 | SCH772984 | MVP_top60_DNA+RNA_seed | IGF1R | 0.162529 | COADREAD | GP5d | Linsitinib |
| MVP_top60_DNA_seed | MAPK1 | 0.048729 | COADREAD | SNU-1040 | SCH772984 | MVP_top60_DNA+RNA_seed | BCL2 | 0.166262 | COADREAD | HCT-15 | Navitoclax |
| MVP_top60_DNA_seed | IGF1R | 0.051912 | COADREAD | LS-513 | GSK1904529A | MVP_top60_DNA+RNA_seed | MET | 0.168153 | COADREAD | LS-123 | Crizotinib |
| MVP_top60_DNA_seed | KRAS | 0.053306 | COADREAD | LS-411N | KRAS (G12C) Inhibitor-12 | MVP_top60_DNA+RNA_seed | AKT1 | 0.168676 | COADREAD | CW-2 | Ipatasertib |

| | | | | | | | | | | | |
|---|---|---|---|---|---|---|---|---|---|---|---|
| MVP_top60_DNA_seed | MAPK1 | 0.057068 | COADREAD | T84 | Ulixertinib | MVP_top60_DNA+RNA_seed | AKT1 | 0.170995 | COADREAD | SK-CO-1 | GSK2110183B |
| MVP_top60_DNA_seed | BRAF | 0.057755 | COADREAD | SNU-C1 | Dabrafenib | MVP_top60_DNA+RNA_seed | AKT1 | 0.171019 | COADREAD | C2BBe1 | Afuresertib |
| MVP_top60_DNA_seed | BRAF | 0.058484 | COADREAD | CaR-1 | PLX-4720 | MVP_top60_DNA+RNA_seed | KRAS | 0.172869 | COADREAD | SNU-407 | KRAS (G12C) Inhibitor-1 |
| MVP_top60_DNA_seed | MAPK1 | 0.059527 | COADREAD | C2BBe1 | VX-11e | MVP_top60_DNA+RNA_seed | BCL2 | 0.172915 | COADREAD | SW1116 | Venetoclax |
| MVP_top60_DNA_seed | IGF1R | 0.061116 | COADREAD | CaR-1 | Linsitinib | MVP_top60_DNA+RNA_seed | HSP90AA1 | 0.173816 | COADREAD | NCI-H716 | Luminespib |
| MVP_top60_DNA_seed | CDK2 | 0.061367 | COADREAD | SW48 | MK-8776 | MVP_top60_DNA+RNA_seed | HSP90AB1 | 0.173816 | COADREAD | NCI-H716 | Luminespib |
| MVP_top60_DNA_seed | MAPK1 | 0.06159 | COADREAD | SNU-C2B | SCH772984 | MVP_top60_DNA+RNA_seed | PIK3CA | 0.178998 | COADREAD | LS-180 | AZD8186 |
| MVP_top60_DNA_seed | CDK2 | 0.063397 | COADREAD | SW48 | AZD5438 | MVP_top60_DNA+RNA_seed | PIK3CB | 0.178998 | COADREAD | LS-180 | AZD8186 |
| MVP_top60_DNA_seed | PIK3CA | 0.06485 | COADREAD | DiFi | AZD8186 | MVP_top60_DNA+RNA_seed | SRC | 0.181088 | COADREAD | SW1463 | Bosutinib |
| MVP_top60_DNA_seed | PIK3CB | 0.06485 | COADREAD | DiFi | AZD8186 | MVP_top60_DNA+RNA_seed | SRC | 0.18288 | COADREAD | HCC2998 | Bosutinib |
| MVP_top60_DNA_seed | TP53 | 0.065601 | COADREAD | SK-CO-1 | MIRA-1 | MVP_top60_DNA+RNA_seed | HSP90AA1 | 0.184134 | COADREAD | SNU-C1 | Tanespimycin |
| MVP_top60_DNA_seed | PIK3CA | 0.067512 | COADREAD | SNU-175 | Alpelisib | MVP_top60_DNA+RNA_seed | HSP90AB1 | 0.184134 | COADREAD | SNU-C1 | Tanespimycin |
| MVP_top60_DNA_seed | CDK2 | 0.07268 | COADREAD | SNU-C2B | Dinaciclib | MVP_top60_DNA+RNA_seed | BCL2 | 0.184382 | COADREAD | LS-513 | Navitoclax |
| MVP_top60_DNA_seed | ESR1 | 0.073533 | COADREAD | SNU-175 | Tamoxifen | MVP_top60_DNA+RNA_seed | BCL2 | 0.18981 | COADREAD | NCI-H747 | Navitoclax |
| MVP_top60_DNA_seed | MET | 0.073968 | COADREAD | SW1463 | Crizotinib | MVP_top60_DNA+RNA_seed | IGF1R | 0.190012 | COADREAD | SNU-C2B | BMS-754807 |
| MVP_top60_DNA_seed | IGF1R | 0.076402 | COADREAD | MDST8 | GSK1904529A | MVP_top60_DNA+RNA_seed | AKT1 | 0.190732 | COADREAD | CW-2 | MK-2206 |
| MVP_top60_DNA_seed | CREBBP | 0.078039 | COADREAD | HCC2998 | SGC-CBP30 | MVP_top60_DNA+RNA_seed | CDK2 | 0.192185 | COADREAD | SNU-C2B | AZD5438 |
| MVP_top60_DNA_seed | BRAF | 0.079847 | COADREAD | LS-513 | Dabrafenib | MVP_top60_DNA+RNA_seed | HSP90AA1 | 0.1943 | COADREAD | RCM-1 | Tanespimycin |
| MVP_top60_DNA_seed | CREBBP | 0.080223 | COADREAD | SNU-81 | SGC-CBP30 | MVP_top60_DNA+RNA_seed | HSP90AB1 | 0.1943 | COADREAD | RCM-1 | Tanespimycin |
| MVP_top60_DNA_seed | HSP90AA1 | 0.080794 | COADREAD | SW1463 | Tanespimycin | MVP_top60_DNA+RNA_seed | BCL2 | 0.195229 | COADREAD | NCI-H508 | Venetoclax |
| MVP_top60_DNA_seed | HSP90AB1 | 0.080794 | COADREAD | SW1463 | Tanespimycin | MVP_top60_DNA+RNA_seed | MET | 0.195783 | COADREAD | T84 | Crizotinib |
| MVP_top60_DNA_seed | AKT1 | 0.081107 | COADREAD | RKO | MK-2206 | MVP_top60_DNA+RNA_seed | ATM | 0.195901 | COADREAD | LS-1034 | KU-55933 |
| MVP_top60_DNA_seed | AKT1 | 0.08304 | COADREAD | SK-CO-1 | AZD5363 | MVP_top60_DNA+RNA_seed | HSP90AA1 | 0.200103 | COADREAD | SNU-175 | Luminespib |
| MVP_top60_DNA_seed | IGF1R | 0.083047 | COADREAD | SNU-C5 | BMS-754807 | MVP_top60_DNA+RNA_seed | HSP90AB1 | 0.200103 | COADREAD | SNU-175 | Luminespib |
| MVP_top60_DNA_seed | AKT1 | 0.083147 | COADREAD | SNU-81 | MK-2206 | MVP_top60_DNA+RNA_seed | MET | 0.202306 | COADREAD | LS-180 | Crizotinib |
| MVP_top60_DNA_seed | PIK3CA | 0.085727 | COADREAD | GP5d | AZD8186 | MVP_top60_DNA+RNA_seed | IGF1R | 0.20234 | COADREAD | SW837 | Linsitinib |
| MVP_top60_DNA_seed | PIK3CB | 0.085727 | COADREAD | GP5d | AZD8186 | MVP_top60_DNA+RNA_seed | MET | 0.203052 | COADREAD | LS-1034 | Savolitinib |
| MVP_top60_DNA_seed | MAPK1 | 0.085904 | COADREAD | SNU-81 | SCH772984 | MVP_top60_DNA+RNA_seed | TP53 | 0.205136 | COADREAD | T84 | MIRA-1 |
| MVP_top60_DNA_seed | CDK2 | 0.087181 | COADREAD | T84 | AZD5438 | MVP_top60_DNA+RNA_seed | AKT1 | 0.206609 | COADREAD | LS-180 | Uprosertib |
| MVP_top60_DNA_seed | PIK3CA | 0.087524 | COADREAD | SW837 | GNE-317 | MVP_top60_DNA+RNA_seed | KRAS | 0.207918 | COADREAD | SK-CO-1 | KRAS (G12C) Inhibitor-1 |
| MVP_top60_DNA_seed | CDK2 | 0.088332 | COADREAD | LoVo | MK-8776 | MVP_top60_DNA+RNA_seed | BCL2 | 0.208829 | COADREAD | HT-29 | Venetoclax |
| MVP_top60_DNA_seed | TP53 | 0.089289 | COADREAD | HT-115 | MIRA-1 | MVP_top60_DNA+RNA_seed | AKT1 | 0.209509 | COADREAD | SK-CO-1 | MK-2206 |
| MVP_top60_DNA_seed | CDK2 | 0.090208 | COADREAD | MDST8 | AZD5438 | MVP_top60_DNA+RNA_seed | BCL2 | 0.213834 | COADREAD | SNU-C5 | Venetoclax |
| MVP_top60_DNA_seed | IGF1R | 0.091019 | COADREAD | HCC2998 | BMS-754807 | MVP_top60_DNA+RNA_seed | IGF1R | 0.217886 | COADREAD | SW948 | NVP-ADW742 |
| MVP_top60_DNA_seed | HSP90AA1 | 0.091237 | COADREAD | SNU-C5 | Luminespib | MVP_top60_DNA+RNA_seed | PIK3CA | 0.220829 | COADREAD | SNU-C1 | GNE-317 |
| MVP_top60_DNA_seed | HSP90AB1 | 0.091237 | COADREAD | SNU-C5 | Luminespib | MVP_top60_DNA+RNA_seed | AKT1 | 0.221768 | COADREAD | HCT-116 | Uprosertib |
| MVP_top60_DNA_seed | IGF1R | 0.091488 | COADREAD | LS-411N | GSK1904529A | MVP_top60_DNA+RNA_seed | CDK2 | 0.222386 | COADREAD | HT-29 | AZD5438 |
| MVP_top60_DNA_seed | ESR1 | 0.091634 | COADREAD | KM12 | GDC0810 | MVP_top60_DNA+RNA_seed | PIK3CA | 0.22992 | COADREAD | HCT-15 | AZD8186 |
| MVP_top60_DNA_seed | AKT1 | 0.092447 | COADREAD | CaR-1 | MK-2206 | MVP_top60_DNA+RNA_seed | PIK3CB | 0.22992 | COADREAD | HCT-15 | AZD8186 |
| MVP_top60_DNA_seed | ESR1 | 0.093177 | COADREAD | HT-115 | GDC0810 | MVP_top60_DNA+RNA_seed | BCL2 | 0.231617 | COADREAD | SNU-81 | Venetoclax |
| MVP_top60_DNA_seed | AKT1 | 0.093526 | COADREAD | NCI-H747 | GSK2110183B | MVP_top60_DNA+RNA_seed | AKT1 | 0.231937 | COADREAD | C2BBe1 | Ipatasertib |

| | | | | | | | | | | | |
|---|---|---|---|---|---|---|---|---|---|---|---|
| MVP_top60_DNA_seed | MAPK1 | 0.094644 | COADREAD | SW1417 | Ulixertinib | MVP_top60_DNA+RNA_seed | HSP90AA1 | 0.233464 | COADREAD | HT-115 | Luminespib |
| MVP_top60_DNA_seed | BRAF | 0.095294 | COADREAD | SW948 | PLX-4720 | MVP_top60_DNA+RNA_seed | HSP90AB1 | 0.233464 | COADREAD | HT-115 | Luminespib |
| MVP_top60_DNA_seed | ESR1 | 0.096672 | COADREAD | HCT-116 | GDC0810 | MVP_top60_DNA+RNA_seed | ATM | 0.233929 | COADREAD | SW1463 | KU-55933 |
| MVP_top60_DNA_seed | BRAF | 0.101765 | COADREAD | DiFi | PLX-4720 | MVP_top60_DNA+RNA_seed | CDK2 | 0.239916 | COADREAD | SW1463 | Dinaciclib |
| MVP_top60_DNA_seed | AKT1 | 0.104612 | COADREAD | SNU-407 | GSK2110183B | MVP_top60_DNA+RNA_seed | PIK3CA | 0.24029 | COADREAD | SW620 | GNE-317 |
| MVP_top60_DNA_seed | KRAS | 0.104894 | COADREAD | SNU-175 | KRAS (G12C) Inhibitor-1 | MVP_top60_DNA+RNA_seed | TP53 | 0.240471 | COADREAD | LS-180 | MIRA-1 |
| MVP_top60_DNA_seed | AKT1 | 0.105069 | COADREAD | CaR-1 | Ipatasertib | MVP_top60_DNA+RNA_seed | TP53 | 0.240883 | COADREAD | SW48 | MIRA-1 |
| MVP_top60_DNA_seed | MAPK1 | 0.106585 | COADREAD | HCT-15 | SCH772984 | MVP_top60_DNA+RNA_seed | IGF1R | 0.240937 | COADREAD | SW620 | BMS-536924 |
| MVP_top60_DNA_seed | ESR1 | 0.109967 | COADREAD | HT55 | GDC0810 | MVP_top60_DNA+RNA_seed | MET | 0.241227 | COADREAD | CW-2 | Crizotinib |
| MVP_top60_DNA_seed | TP53 | 0.111455 | COADREAD | SNU-175 | MIRA-1 | MVP_top60_DNA+RNA_seed | AKT1 | 0.242502 | COADREAD | SNU-81 | AT13148 |
| MVP_top60_DNA_seed | AKT1 | 0.113815 | COADREAD | MDST8 | Uprosertib | MVP_top60_DNA+RNA_seed | PIK3CA | 0.246012 | COADREAD | HT55 | GNE-317 |
| MVP_top60_DNA_seed | ERBB3 | 0.1163 | COADREAD | SW837 | Sapitinib | MVP_top60_DNA+RNA_seed | CDK2 | 0.24695 | COADREAD | COLO-205 | MK-8776 |
| MVP_top60_DNA_seed | PIK3CA | 0.117066 | COADREAD | HCC2998 | Alpelisib | MVP_top60_DNA+RNA_seed | AKT1 | 0.250915 | COADREAD | CCK-81 | Ipatasertib |
| MVP_top60_DNA_seed | MAPK1 | 0.11827 | COADREAD | CW-2 | SCH772984 | MVP_top60_DNA+RNA_seed | IGF1R | 0.253056 | COADREAD | COLO-205 | GSK1904529A |
| MVP_top60_DNA_seed | MET | 0.119556 | COADREAD | RKO | Savolitinib | MVP_top60_DNA+RNA_seed | HSP90AA1 | 0.257589 | COADREAD | T84 | Tanespimycin |
| MVP_top60_DNA_seed | IGF1R | 0.121399 | COADREAD | SNU-C5 | BMS-536924 | MVP_top60_DNA+RNA_seed | HSP90AB1 | 0.257589 | COADREAD | T84 | Tanespimycin |
| MVP_top60_DNA_seed | ERBB3 | 0.127041 | COADREAD | LS-123 | Sapitinib | MVP_top60_DNA+RNA_seed | ATM | 0.257987 | COADREAD | HCT-116 | KU-55933 |
| MVP_top60_DNA_seed | AKT1 | 0.131777 | COADREAD | T84 | Uprosertib | MVP_top60_DNA+RNA_seed | AKT1 | 0.259517 | COADREAD | SNU-175 | AZD5363 |
| MVP_top60_DNA_seed | CDK2 | 0.133356 | COADREAD | SNU-C1 | MK-8776 | MVP_top60_DNA+RNA_seed | SRC | 0.260611 | COADREAD | NCI-H716 | Bosutinib |
| MVP_top60_DNA_seed | AKT1 | 0.133532 | COADREAD | CaR-1 | AZD5363 | MVP_top60_DNA+RNA_seed | PIK3CB | 0.261939 | COADREAD | HT-115 | AZD6482 |
| MVP_top60_DNA_seed | BRAF | 0.133763 | COADREAD | SK-CO-1 | PLX-4720 | MVP_top60_DNA+RNA_seed | AKT1 | 0.262573 | COADREAD | HT55 | AT13148 |
| MVP_top60_DNA_seed | HSP90AA1 | 0.133877 | COADREAD | CW-2 | Tanespimycin | MVP_top60_DNA+RNA_seed | AKT1 | 0.264502 | COADREAD | LS-1034 | MK-2206 |
| MVP_top60_DNA_seed | HSP90AB1 | 0.133877 | COADREAD | CW-2 | Tanespimycin | MVP_top60_DNA+RNA_seed | IGF1R | 0.265741 | COADREAD | LoVo | GSK1904529A |
| MVP_top60_DNA_seed | MAPK1 | 0.138174 | COADREAD | SNU-C2B | VX-11e | MVP_top60_DNA+RNA_seed | SRC | 0.265978 | COADREAD | SW1116 | Bosutinib |
| MVP_top60_DNA_seed | CDK2 | 0.140132 | COADREAD | SNU-81 | Dinaciclib | MVP_top60_DNA+RNA_seed | AKT1 | 0.266056 | COADREAD | SNU-C2B | AT13148 |
| MVP_top60_DNA_seed | PIK3CA | 0.140505 | COADREAD | T84 | Buparlisib | MVP_top60_DNA+RNA_seed | TP53 | 0.268349 | COADREAD | SNU-81 | MIRA-1 |
| MVP_top60_DNA_seed | PIK3CB | 0.140505 | COADREAD | T84 | Buparlisib | MVP_top60_DNA+RNA_seed | TP53 | 0.268778 | COADREAD | RKO | MIRA-1 |
| MVP_top60_DNA_seed | PIK3CD | 0.140505 | COADREAD | T84 | Buparlisib | MVP_top60_DNA+RNA_seed | ATM | 0.269069 | COADREAD | SNU-81 | KU-55933 |
| MVP_top60_DNA_seed | ESR1 | 0.14262 | COADREAD | SNU-81 | GDC0810 | MVP_top60_DNA+RNA_seed | IGF1R | 0.274249 | COADREAD | LS-123 | BMS-754807 |
| MVP_top60_DNA_seed | IGF1R | 0.143267 | COADREAD | LS-180 | BMS-754807 | MVP_top60_DNA+RNA_seed | IGF1R | 0.275641 | COADREAD | RCM-1 | GSK1904529A |
| MVP_top60_DNA_seed | MAPK1 | 0.143397 | COADREAD | COLO-320-HSR | Ulixertinib | MVP_top60_DNA+RNA_seed | IGF1R | 0.276509 | COADREAD | SW620 | Linsitinib |
| MVP_top60_DNA_seed | IGF1R | 0.145407 | COADREAD | CL-11 | Linsitinib | MVP_top60_DNA+RNA_seed | AR | 0.277011 | COADREAD | SNU-81 | Bicalutamide |
| MVP_top60_DNA_seed | ESR1 | 0.146832 | COADREAD | GP5d | Tamoxifen | MVP_top60_DNA+RNA_seed | MET | 0.278346 | COADREAD | CCK-81 | Savolitinib |
| MVP_top60_DNA_seed | GSK3B | 0.151996 | COADREAD | SW837 | SB216763 | MVP_top60_DNA+RNA_seed | IGF1R | 0.279788 | COADREAD | SNU-61 | Linsitinib |
| MVP_top60_DNA_seed | PIK3CA | 0.152698 | COADREAD | HT55 | Buparlisib | MVP_top60_DNA+RNA_seed | TP53 | 0.280323 | COADREAD | HCT-15 | MIRA-1 |
| MVP_top60_DNA_seed | PIK3CB | 0.152698 | COADREAD | HT55 | Buparlisib | MVP_top60_DNA+RNA_seed | CDK2 | 0.281513 | COADREAD | COLO-205 | AZD5438 |
| MVP_top60_DNA_seed | PIK3CD | 0.152698 | COADREAD | HT55 | Buparlisib | MVP_top60_DNA+RNA_seed | AKT1 | 0.287187 | COADREAD | HCT-116 | MK-2206 |
| MVP_top60_DNA_seed | PIK3CA | 0.152922 | COADREAD | SNU-C5 | GNE-317 | MVP_top60_DNA+RNA_seed | BCL2 | 0.288041 | COADREAD | KM12 | Navitoclax |
| MVP_top60_DNA_seed | PIK3CA | 0.153117 | COADREAD | NCI-H508 | GNE-317 | MVP_top60_DNA+RNA_seed | AKT1 | 0.288509 | COADREAD | LS-180 | MK-2206 |
| MVP_top60_DNA_seed | MET | 0.156133 | COADREAD | NCI-H747 | Crizotinib | MVP_top60_DNA+RNA_seed | IGF1R | 0.289791 | COADREAD | CW-2 | BMS-536924 |
| MVP_top60_DNA_seed | IGF1R | 0.156334 | COADREAD | SW1116 | GSK1904529A | MVP_top60_DNA+RNA_seed | MET | 0.301048 | COADREAD | SW1116 | Crizotinib |

| | | | | | | | | | | | |
|---|---|---|---|---|---|---|---|---|---|---|---|
| MVP_top60_DNA_seed | HSP90AA1 | 0.157377 | COADREAD | C2BBe1 | Luminespib | MVP_top60_DNA+RNA_seed | AKT1 | 0.306525 | COADREAD | LS-180 | AZD5363 |
| MVP_top60_DNA_seed | HSP90AB1 | 0.157377 | COADREAD | C2BBe1 | Luminespib | MVP_top60_DNA+RNA_seed | CDK2 | 0.310858 | COADREAD | CW-2 | Dinaciclib |
| MVP_top60_DNA_seed | IGF1R | 0.158503 | COADREAD | NCI-H508 | BMS-536924 | MVP_top60_DNA+RNA_seed | PIK3CA | 0.310936 | COADREAD | T84 | GNE-317 |
| MVP_top60_DNA_seed | IGF1R | 0.159382 | COADREAD | MDST8 | BMS-536924 | MVP_top60_DNA+RNA_seed | BCL2 | 0.312103 | COADREAD | NCI-H508 | Sabutoclax |
| MVP_top60_DNA_seed | IGF1R | 0.159978 | COADREAD | C2BBe1 | NVP-ADW742 | MVP_top60_DNA+RNA_seed | HSP90AA1 | 0.31541 | COADREAD | RCM-1 | Luminespib |
| MVP_top60_DNA_seed | AKT1 | 0.160392 | COADREAD | C2BBe1 | MK-2206 | MVP_top60_DNA+RNA_seed | HSP90AB1 | 0.31541 | COADREAD | RCM-1 | Luminespib |
| MVP_top60_DNA_seed | IGF1R | 0.160443 | COADREAD | C2BBe1 | GSK1904529A | MVP_top60_DNA+RNA_seed | BCL2 | 0.320145 | COADREAD | T84 | TW 37 |
| MVP_top60_DNA_seed | CREBBP | 0.16168 | COADREAD | SNU-175 | SGC-CBP30 | MVP_top60_DNA+RNA_seed | CDK2 | 0.325177 | COADREAD | T84 | Dinaciclib |
| MVP_top60_DNA_seed | IGF1R | 0.162529 | COADREAD | GP5d | Linsitinib | MVP_top60_DNA+RNA_seed | CDK2 | 0.333163 | COADREAD | LoVo | Dinaciclib |
| MVP_top60_DNA_seed | MET | 0.168153 | COADREAD | LS-123 | Crizotinib | MVP_top60_DNA+RNA_seed | CDK2 | 0.334538 | COADREAD | LS-180 | Dinaciclib |
| MVP_top60_DNA_seed | AKT1 | 0.168676 | COADREAD | CW-2 | Ipatasertib | MVP_top60_DNA+RNA_seed | ATM | 0.339776 | COADREAD | SNU-175 | KU-55933 |
| MVP_top60_DNA_seed | MAPK1 | 0.169831 | COADREAD | CaR-1 | VX-11e | MVP_top60_DNA+RNA_seed | GSK3B | 0.339982 | COADREAD | SNU-407 | SB216763 |
| MVP_top60_DNA_seed | AKT1 | 0.170995 | COADREAD | SK-CO-1 | GSK2110183B | MVP_top60_DNA+RNA_seed | IGF1R | 0.34113 | COADREAD | RCM-1 | BMS-754807 |
| MVP_top60_DNA_seed | AKT1 | 0.171019 | COADREAD | C2BBe1 | Afuresertib | MVP_top60_DNA+RNA_seed | HSP90AA1 | 0.342956 | COADREAD | SNU-61 | Tanespimycin |
| MVP_top60_DNA_seed | KRAS | 0.172869 | COADREAD | SNU-407 | KRAS (G12C) Inhibitor-1[illegible] | MVP_top60_DNA+RNA_seed | HSP90AB1 | 0.342956 | COADREAD | SNU-61 | Tanespimycin |
| MVP_top60_DNA_seed | HSP90AA1 | 0.173816 | COADREAD | NCI-H716 | Luminespib | MVP_top60_DNA+RNA_seed | BCL2 | 0.344004 | COADREAD | DiFi | Venetoclax |
| MVP_top60_DNA_seed | HSP90AB1 | 0.173816 | COADREAD | NCI-H716 | Luminespib | MVP_top60_DNA+RNA_seed | SRC | 0.344414 | COADREAD | SNU-81 | Bosutinib |
| MVP_top60_DNA_seed | PIK3CA | 0.178998 | COADREAD | LS-180 | AZD8186 | MVP_top60_DNA+RNA_seed | AKT1 | 0.345796 | COADREAD | HCC2998 | AT13148 |
| MVP_top60_DNA_seed | PIK3CB | 0.178998 | COADREAD | LS-180 | AZD8186 | MVP_top60_DNA+RNA_seed | AKT1 | 0.346405 | COADREAD | RCM-1 | Uprosertib |
| MVP_top60_DNA_seed | SRC | 0.181088 | COADREAD | SW1463 | Bosutinib | MVP_top60_DNA+RNA_seed | HSP90AA1 | 0.346482 | COADREAD | LS-1034 | Tanespimycin |
| MVP_top60_DNA_seed | SRC | 0.18288 | COADREAD | HCC2998 | Bosutinib | MVP_top60_DNA+RNA_seed | HSP90AB1 | 0.346482 | COADREAD | LS-1034 | Tanespimycin |
| MVP_top60_DNA_seed | HSP90AA1 | 0.184134 | COADREAD | SNU-C1 | Tanespimycin | MVP_top60_DNA+RNA_seed | HSP90AA1 | 0.346893 | COADREAD | HT-115 | Tanespimycin |
| MVP_top60_DNA_seed | HSP90AB1 | 0.184134 | COADREAD | SNU-C1 | Tanespimycin | MVP_top60_DNA+RNA_seed | HSP90AB1 | 0.346893 | COADREAD | HT-115 | Tanespimycin |
| MVP_top60_DNA_seed | BRAF | 0.188235 | COADREAD | T84 | PLX-4720 | MVP_top60_DNA+RNA_seed | CDK2 | 0.347127 | COADREAD | HCT-15 | MK-8776 |
| MVP_top60_DNA_seed | IGF1R | 0.190012 | COADREAD | SNU-C2B | BMS-754807 | MVP_top60_DNA+RNA_seed | AKT1 | 0.347505 | COADREAD | HCT-116 | Ipatasertib |
| MVP_top60_DNA_seed | CREBBP | 0.190718 | COADREAD | KM12 | SGC-CBP30 | MVP_top60_DNA+RNA_seed | PIK3CA | 0.348843 | COADREAD | SNU-81 | GNE-317 |
| MVP_top60_DNA_seed | AKT1 | 0.190732 | COADREAD | CW-2 | MK-2206 | MVP_top60_DNA+RNA_seed | BCL2 | 0.350696 | COADREAD | SW948 | Venetoclax |
| MVP_top60_DNA_seed | CDK2 | 0.192185 | COADREAD | SNU-C2B | AZD5438 | MVP_top60_DNA+RNA_seed | CDK2 | 0.351148 | COADREAD | SNU-81 | AZD5438 |
| MVP_top60_DNA_seed | BRAF | 0.192778 | COADREAD | HCC2998 | Dabrafenib | MVP_top60_DNA+RNA_seed | IGF1R | 0.354776 | COADREAD | RKO | Linsitinib |
| MVP_top60_DNA_seed | HSP90AA1 | 0.1943 | COADREAD | RCM-1 | Tanespimycin | MVP_top60_DNA+RNA_seed | AKT1 | 0.355918 | COADREAD | COLO-320-HSR | GSK2110183B |
| MVP_top60_DNA_seed | HSP90AB1 | 0.1943 | COADREAD | RCM-1 | Tanespimycin | MVP_top60_DNA+RNA_seed | BCL2 | 0.356429 | COADREAD | RCM-1 | TW 37 |
| MVP_top60_DNA_seed | MET | 0.195783 | COADREAD | T84 | Crizotinib | MVP_top60_DNA+RNA_seed | AKT1 | 0.359065 | COADREAD | SNU-C2B | Uprosertib |
| MVP_top60_DNA_seed | ATM | 0.195901 | COADREAD | LS-1034 | KU-55933 | MVP_top60_DNA+RNA_seed | MET | 0.360053 | COADREAD | HT55 | Savolitinib |
| MVP_top60_DNA_seed | HSP90AA1 | 0.200103 | COADREAD | SNU-175 | Luminespib | MVP_top60_DNA+RNA_seed | MET | 0.360843 | COADREAD | SNU-C5 | Savolitinib |
| MVP_top60_DNA_seed | HSP90AB1 | 0.200103 | COADREAD | SNU-175 | Luminespib | MVP_top60_DNA+RNA_seed | PIK3CA | 0.364046 | COADREAD | LS-123 | Buparlisib |
| MVP_top60_DNA_seed | MET | 0.202306 | COADREAD | LS-180 | Crizotinib | MVP_top60_DNA+RNA_seed | PIK3CB | 0.364046 | COADREAD | LS-123 | Buparlisib |
| MVP_top60_DNA_seed | IGF1R | 0.20234 | COADREAD | SW837 | Linsitinib | MVP_top60_DNA+RNA_seed | PIK3CD | 0.364046 | COADREAD | LS-123 | Buparlisib |
| MVP_top60_DNA_seed | MET | 0.203052 | COADREAD | LS-1034 | Savolitinib | MVP_top60_DNA+RNA_seed | CDK2 | 0.367209 | COADREAD | SNU-407 | Dinaciclib |
| MVP_top60_DNA_seed | TP53 | 0.205136 | COADREAD | T84 | MIRA-1 | MVP_top60_DNA+RNA_seed | IGF1R | 0.371435 | COADREAD | SNU-C5 | Linsitinib |
| MVP_top60_DNA_seed | AKT1 | 0.206609 | COADREAD | LS-180 | Uprosertib | MVP_top60_DNA+RNA_seed | AKT1 | 0.373851 | COADREAD | LS-180 | AT13148 |
| MVP_top60_DNA_seed | MAPK1 | 0.207909 | COADREAD | SNU-81 | Ulixertinib | MVP_top60_DNA+RNA_seed | AKT1 | 0.377633 | COADREAD | CW-2 | AT13148 |

| | | | | | | | | | | | |
|---|---|---|---|---|---|---|---|---|---|---|---|
| MVP_top60_DNA_seed | KRAS | 0.207918 | COADREAD | SK-CO-1 | KRAS (G12C) Inhibitor-12 | MVP_top60_DNA+RNA_seed | MET | 0.380239 | COADREAD | SW837 | Crizotinib |
| MVP_top60_DNA_seed | CREBBP | 0.209413 | COADREAD | SW620 | SGC-CBP30 | MVP_top60_DNA+RNA_seed | ATM | 0.380863 | COADREAD | CL-11 | KU-55933 |
| MVP_top60_DNA_seed | AKT1 | 0.209509 | COADREAD | SK-CO-1 | MK-2206 | MVP_top60_DNA+RNA_seed | PIK3CA | 0.381011 | COADREAD | SW620 | Buparlisib |
| MVP_top60_DNA_seed | ESR1 | 0.216763 | COADREAD | CL-11 | Tamoxifen | MVP_top60_DNA+RNA_seed | PIK3CB | 0.381011 | COADREAD | SW620 | Buparlisib |
| MVP_top60_DNA_seed | ESR1 | 0.217498 | COADREAD | HT-115 | Tamoxifen | MVP_top60_DNA+RNA_seed | PIK3CD | 0.381011 | COADREAD | SW620 | Buparlisib |
| MVP_top60_DNA_seed | IGF1R | 0.217886 | COADREAD | SW948 | NVP-ADW742 | MVP_top60_DNA+RNA_seed | SRC | 0.381465 | COADREAD | COLO-678 | Bosutinib |
| MVP_top60_DNA_seed | PIK3CA | 0.220829 | COADREAD | SNU-C1 | GNE-317 | MVP_top60_DNA+RNA_seed | TP53 | 0.382935 | COADREAD | CaR-1 | MIRA-1 |
| MVP_top60_DNA_seed | AKT1 | 0.221768 | COADREAD | HCT-116 | Uprosertib | MVP_top60_DNA+RNA_seed | AKT1 | 0.386093 | COADREAD | C2BBe1 | AZD5363 |
| MVP_top60_DNA_seed | CDK2 | 0.222386 | COADREAD | HT-29 | AZD5438 | MVP_top60_DNA+RNA_seed | PIK3CA | 0.389427 | COADREAD | NCI-H716 | Buparlisib |
| MVP_top60_DNA_seed | PIK3CA | 0.22992 | COADREAD | HCT-15 | AZD8186 | MVP_top60_DNA+RNA_seed | PIK3CB | 0.389427 | COADREAD | NCI-H716 | Buparlisib |
| MVP_top60_DNA_seed | PIK3CB | 0.22992 | COADREAD | HCT-15 | AZD8186 | MVP_top60_DNA+RNA_seed | PIK3CD | 0.389427 | COADREAD | NCI-H716 | Buparlisib |
| MVP_top60_DNA_seed | AKT1 | 0.231937 | COADREAD | C2BBe1 | Ipatasertib | MVP_top60_DNA+RNA_seed | BCL2 | 0.39055 | COADREAD | LS-123 | TW 37 |
| MVP_top60_DNA_seed | HSP90AA1 | 0.233464 | COADREAD | HT-115 | Luminespib | MVP_top60_DNA+RNA_seed | HSP90AA1 | 0.394059 | COADREAD | SW1417 | Tanespimycin |
| MVP_top60_DNA_seed | HSP90AB1 | 0.233464 | COADREAD | HT-115 | Luminespib | MVP_top60_DNA+RNA_seed | HSP90AB1 | 0.394059 | COADREAD | SW1417 | Tanespimycin |
| MVP_top60_DNA_seed | ATM | 0.233929 | COADREAD | SW1463 | KU-55933 | MVP_top60_DNA+RNA_seed | KRAS | 0.394184 | COADREAD | DiFi | KRAS (G12C) Inhibitor-1 |
| MVP_top60_DNA_seed | CDK2 | 0.239916 | COADREAD | SW1463 | Dinaciclib | MVP_top60_DNA+RNA_seed | TP53 | 0.396854 | COADREAD | NCI-H508 | MIRA-1 |
| MVP_top60_DNA_seed | PIK3CA | 0.24029 | COADREAD | SW620 | GNE-317 | MVP_top60_DNA+RNA_seed | HSP90AA1 | 0.402172 | COADREAD | CCK-81 | Luminespib |
| MVP_top60_DNA_seed | TP53 | 0.240471 | COADREAD | LS-180 | MIRA-1 | MVP_top60_DNA+RNA_seed | HSP90AB1 | 0.402172 | COADREAD | CCK-81 | Luminespib |
| MVP_top60_DNA_seed | TP53 | 0.240883 | COADREAD | SW48 | MIRA-1 | MVP_top60_DNA+RNA_seed | CDK2 | 0.402514 | COADREAD | SK-CO-1 | MK-8776 |
| MVP_top60_DNA_seed | IGF1R | 0.240937 | COADREAD | SW620 | BMS-536924 | MVP_top60_DNA+RNA_seed | HSP90AA1 | 0.403948 | COADREAD | NCI-H508 | Elesclomol |
| MVP_top60_DNA_seed | CREBBP | 0.241042 | COADREAD | HT55 | SGC-CBP30 | MVP_top60_DNA+RNA_seed | HSP90AB1 | 0.403948 | COADREAD | NCI-H508 | Elesclomol |
| MVP_top60_DNA_seed | MET | 0.241227 | COADREAD | CW-2 | Crizotinib | MVP_top60_DNA+RNA_seed | IGF1R | 0.408569 | COADREAD | SW1116 | BMS-536924 |
| MVP_top60_DNA_seed | AKT1 | 0.242502 | COADREAD | SNU-81 | AT13148 | MVP_top60_DNA+RNA_seed | IGF1R | 0.408777 | COADREAD | LS-1034 | GSK1904529A |
| MVP_top60_DNA_seed | CREBBP | 0.24371 | COADREAD | SW948 | SGC-CBP30 | MVP_top60_DNA+RNA_seed | PIK3CA | 0.409103 | COADREAD | SW837 | AZD8186 |
| MVP_top60_DNA_seed | PIK3CA | 0.246012 | COADREAD | HT55 | GNE-317 | MVP_top60_DNA+RNA_seed | PIK3CB | 0.409103 | COADREAD | SW837 | AZD8186 |
| MVP_top60_DNA_seed | CDK2 | 0.24695 | COADREAD | COLO-205 | MK-8776 | MVP_top60_DNA+RNA_seed | MET | 0.409787 | COADREAD | C2BBe1 | Savolitinib |
| MVP_top60_DNA_seed | BRAF | 0.248667 | COADREAD | COLO-320-HSR | Dabrafenib | MVP_top60_DNA+RNA_seed | PIK3CA | 0.411115 | COADREAD | HT-115 | GNE-317 |
| MVP_top60_DNA_seed | AKT1 | 0.250915 | COADREAD | CCK-81 | Ipatasertib | MVP_top60_DNA+RNA_seed | PIK3CA | 0.414408 | COADREAD | CCK-81 | GNE-317 |
| MVP_top60_DNA_seed | IGF1R | 0.253056 | COADREAD | COLO-205 | GSK1904529A | MVP_top60_DNA+RNA_seed | SRC | 0.417419 | COADREAD | DiFi | Bosutinib |
| MVP_top60_DNA_seed | HSP90AA1 | 0.257589 | COADREAD | T84 | Tanespimycin | MVP_top60_DNA+RNA_seed | PIK3CA | 0.425256 | COADREAD | SW1463 | AZD8186 |
| MVP_top60_DNA_seed | HSP90AB1 | 0.257589 | COADREAD | T84 | Tanespimycin | MVP_top60_DNA+RNA_seed | PIK3CB | 0.425256 | COADREAD | SW1463 | AZD8186 |
| MVP_top60_DNA_seed | ATM | 0.257987 | COADREAD | HCT-116 | KU-55933 | MVP_top60_DNA+RNA_seed | PIK3CA | 0.426361 | COADREAD | HCT-116 | AZD8186 |
| MVP_top60_DNA_seed | CREBBP | 0.258034 | COADREAD | LoVo | SGC-CBP30 | MVP_top60_DNA+RNA_seed | PIK3CB | 0.426361 | COADREAD | HCT-116 | AZD8186 |
| MVP_top60_DNA_seed | AKT1 | 0.259517 | COADREAD | SNU-175 | AZD5363 | MVP_top60_DNA+RNA_seed | IGF1R | 0.426749 | COADREAD | HT55 | GSK1904529A |
| MVP_top60_DNA_seed | SRC | 0.260611 | COADREAD | NCI-H716 | Bosutinib | MVP_top60_DNA+RNA_seed | IGF1R | 0.426915 | COADREAD | SW1417 | BMS-754807 |
| MVP_top60_DNA_seed | BRAF | 0.26104 | COADREAD | SNU-407 | Dabrafenib | MVP_top60_DNA+RNA_seed | IGF1R | 0.427135 | COADREAD | SNU-81 | GSK1904529A |
| MVP_top60_DNA_seed | PIK3CB | 0.261939 | COADREAD | HT-115 | AZD6482 | MVP_top60_DNA+RNA_seed | MET | 0.427246 | COADREAD | T84 | Savolitinib |
| MVP_top60_DNA_seed | BRAF | 0.262301 | COADREAD | LS-1034 | SB590885 | MVP_top60_DNA+RNA_seed | HSP90AA1 | 0.42778 | COADREAD | HT-115 | CCT-018159 |
| MVP_top60_DNA_seed | AKT1 | 0.262573 | COADREAD | HT55 | AT13148 | MVP_top60_DNA+RNA_seed | HSP90AB1 | 0.42778 | COADREAD | HT-115 | CCT-018159 |
| MVP_top60_DNA_seed | AKT1 | 0.264502 | COADREAD | LS-1034 | MK-2206 | MVP_top60_DNA+RNA_seed | ERBB3 | 0.427965 | COADREAD | RKO | Sapitinib |
| MVP_top60_DNA_seed | MAPK1 | 0.264682 | COADREAD | SW1417 | VX-11e | MVP_top60_DNA+RNA_seed | SRC | 0.430052 | COADREAD | RKO | Bosutinib |

| | | | | | | | | | | | |
|---|---|---|---|---|---|---|---|---|---|---|---|
| MVP_top60_DNA_seed | IGF1R | 0.265741 | COADREAD | LoVo | GSK1904529A | MVP_top60_DNA+RNA_seed | GSK3B | 0.430959 | COADREAD | NCI-H508 | CHIR-99021 |
| MVP_top60_DNA_seed | SRC | 0.265978 | COADREAD | SW1116 | Bosutinib | MVP_top60_DNA+RNA_seed | ATM | 0.433564 | COADREAD | HCT-15 | KU-55933 |
| MVP_top60_DNA_seed | AKT1 | 0.266056 | COADREAD | SNU-C2B | AT13148 | MVP_top60_DNA+RNA_seed | CDK2 | 0.434983 | COADREAD | SW48 | Dinaciclib |
| MVP_top60_DNA_seed | MAPK1 | 0.266657 | COADREAD | SW837 | VX-11e | MVP_top60_DNA+RNA_seed | TP53 | 0.436215 | COADREAD | KM12 | MIRA-1 |
| MVP_top60_DNA_seed | TP53 | 0.268349 | COADREAD | SNU-81 | MIRA-1 | MVP_top60_DNA+RNA_seed | CDK2 | 0.439335 | COADREAD | SW1116 | AZD5438 |
| MVP_top60_DNA_seed | TP53 | 0.268778 | COADREAD | RKO | MIRA-1 | MVP_top60_DNA+RNA_seed | AKT1 | 0.43987 | COADREAD | LS-1034 | Uprosertib |
| MVP_top60_DNA_seed | ATM | 0.269069 | COADREAD | SNU-81 | KU-55933 | MVP_top60_DNA+RNA_seed | CDK2 | 0.441192 | COADREAD | NCI-H716 | Dinaciclib |
| MVP_top60_DNA_seed | BRAF | 0.272616 | COADREAD | NCI-H716 | PLX-4720 | MVP_top60_DNA+RNA_seed | BCL2 | 0.442636 | COADREAD | LS-180 | Navitoclax |
| MVP_top60_DNA_seed | IGF1R | 0.274249 | COADREAD | LS-123 | BMS-754807 | MVP_top60_DNA+RNA_seed | CDK2 | 0.443945 | COADREAD | C2BBe1 | Dinaciclib |
| MVP_top60_DNA_seed | IGF1R | 0.275641 | COADREAD | RCM-1 | GSK1904529A | MVP_top60_DNA+RNA_seed | AKT1 | 0.447395 | COADREAD | RCM-1 | Uprosertib |
| MVP_top60_DNA_seed | IGF1R | 0.276509 | COADREAD | SW620 | Linsitinib | MVP_top60_DNA+RNA_seed | PIK3CA | 0.447664 | COADREAD | LS-180 | Buparlisib |
| MVP_top60_DNA_seed | AR | 0.277011 | COADREAD | SNU-81 | Bicalutamide | MVP_top60_DNA+RNA_seed | PIK3CB | 0.447664 | COADREAD | LS-180 | Buparlisib |
| MVP_top60_DNA_seed | MET | 0.278346 | COADREAD | CCK-81 | Savolitinib | MVP_top60_DNA+RNA_seed | PIK3CD | 0.447664 | COADREAD | LS-180 | Buparlisib |
| MVP_top60_DNA_seed | IGF1R | 0.279788 | COADREAD | SNU-61 | Linsitinib | MVP_top60_DNA+RNA_seed | IGF1R | 0.448033 | COADREAD | RKO | BMS-536924 |
| MVP_top60_DNA_seed | TP53 | 0.280323 | COADREAD | HCT-15 | MIRA-1 | MVP_top60_DNA+RNA_seed | IGF1R | 0.449633 | COADREAD | CW-2 | GSK1904529A |
| MVP_top60_DNA_seed | CDK2 | 0.281513 | COADREAD | COLO-205 | AZD5438 | MVP_top60_DNA+RNA_seed | HSP90AA1 | 0.452968 | COADREAD | LS-180 | Luminespib |
| MVP_top60_DNA_seed | BRAF | 0.286585 | COADREAD | SNU-175 | SB590885 | MVP_top60_DNA+RNA_seed | HSP90AB1 | 0.452968 | COADREAD | LS-180 | Luminespib |
| MVP_top60_DNA_seed | AKT1 | 0.287187 | COADREAD | HCT-116 | MK-2206 | MVP_top60_DNA+RNA_seed | AKT1 | 0.455982 | COADREAD | HCT-116 | GSK2110183B |
| MVP_top60_DNA_seed | AKT1 | 0.288509 | COADREAD | LS-180 | MK-2206 | MVP_top60_DNA+RNA_seed | PIK3CA | 0.45689 | COADREAD | LS-411N | Buparlisib |
| MVP_top60_DNA_seed | IGF1R | 0.289791 | COADREAD | CW-2 | BMS-536924 | MVP_top60_DNA+RNA_seed | PIK3CB | 0.45689 | COADREAD | LS-411N | Buparlisib |
| MVP_top60_DNA_seed | BRAF | 0.29153 | COADREAD | GP5d | PLX-4720 | MVP_top60_DNA+RNA_seed | PIK3CD | 0.45689 | COADREAD | LS-411N | Buparlisib |
| MVP_top60_DNA_seed | BRAF | 0.300419 | COADREAD | RCM-1 | Dabrafenib | MVP_top60_DNA+RNA_seed | IGF1R | 0.458215 | COADREAD | SW1463 | BMS-536924 |
| MVP_top60_DNA_seed | MAPK1 | 0.300668 | COADREAD | RCM-1 | Ulixertinib | MVP_top60_DNA+RNA_seed | PIK3CA | 0.460474 | COADREAD | SNU-C1 | Alpelisib |
| MVP_top60_DNA_seed | MET | 0.301048 | COADREAD | SW1116 | Crizotinib | MVP_top60_DNA+RNA_seed | SRC | 0.463796 | COADREAD | LS-513 | Bosutinib |
| MVP_top60_DNA_seed | ESR1 | 0.301196 | COADREAD | CW-2 | GDC0810 | MVP_top60_DNA+RNA_seed | BCL2 | 0.468663 | COADREAD | CaR-1 | Venetoclax |
| MVP_top60_DNA_seed | ESR1 | 0.30313 | COADREAD | CW-2 | Tamoxifen | MVP_top60_DNA+RNA_seed | BCL2 | 0.469528 | COADREAD | SNU-1040 | Sabutoclax |
| MVP_top60_DNA_seed | AKT1 | 0.306525 | COADREAD | LS-180 | AZD5363 | MVP_top60_DNA+RNA_seed | IGF1R | 0.469798 | COADREAD | LS-180 | GSK1904529A |
| MVP_top60_DNA_seed | CDK2 | 0.310858 | COADREAD | CW-2 | Dinaciclib | MVP_top60_DNA+RNA_seed | AKT1 | 0.474809 | COADREAD | LS-123 | MK-2206 |
| MVP_top60_DNA_seed | PIK3CA | 0.310936 | COADREAD | T84 | GNE-317 | MVP_top60_DNA+RNA_seed | AKT1 | 0.478817 | COADREAD | LS-1034 | Afuresertib |
| MVP_top60_DNA_seed | BRAF | 0.314626 | COADREAD | HT-115 | PLX-4720 | MVP_top60_DNA+RNA_seed | IGF1R | 0.47973 | COADREAD | T84 | Linsitinib |
| MVP_top60_DNA_seed | HSP90AA1 | 0.31541 | COADREAD | RCM-1 | Luminespib | MVP_top60_DNA+RNA_seed | ERBB3 | 0.480324 | COADREAD | SW1417 | Sapitinib |
| MVP_top60_DNA_seed | HSP90AB1 | 0.31541 | COADREAD | RCM-1 | Luminespib | MVP_top60_DNA+RNA_seed | IGF1R | 0.481295 | COADREAD | DiFi | BMS-536924 |
| MVP_top60_DNA_seed | MAPK1 | 0.318223 | COADREAD | SNU-C2B | Ulixertinib | MVP_top60_DNA+RNA_seed | AKT1 | 0.482012 | COADREAD | CL-11 | Uprosertib |
| MVP_top60_DNA_seed | CDK2 | 0.325177 | COADREAD | T84 | Dinaciclib | MVP_top60_DNA+RNA_seed | TP53 | 0.484948 | COADREAD | LS-411N | MIRA-1 |
| MVP_top60_DNA_seed | BRAF | 0.328334 | COADREAD | HCT-15 | SB590885 | MVP_top60_DNA+RNA_seed | HSP90AA1 | 0.485452 | COADREAD | SNU-407 | Luminespib |
| MVP_top60_DNA_seed | MAPK1 | 0.329492 | COADREAD | CaR-1 | SCH772984 | MVP_top60_DNA+RNA_seed | HSP90AB1 | 0.485452 | COADREAD | SNU-407 | Luminespib |
| MVP_top60_DNA_seed | CDK2 | 0.333163 | COADREAD | LoVo | Dinaciclib | MVP_top60_DNA+RNA_seed | BCL2 | 0.488418 | COADREAD | LS-1034 | Obatoclax Mesylate |
| MVP_top60_DNA_seed | CDK2 | 0.334538 | COADREAD | LS-180 | Dinaciclib | MVP_top60_DNA+RNA_seed | AKT1 | 0.48941 | COADREAD | T84 | MK-2206 |
| MVP_top60_DNA_seed | MAPK1 | 0.334547 | COADREAD | HT-115 | Ulixertinib | MVP_top60_DNA+RNA_seed | IGF1R | 0.490018 | COADREAD | T84 | GSK1904529A |
| MVP_top60_DNA_seed | ATM | 0.339776 | COADREAD | SNU-175 | KU-55933 | MVP_top60_DNA+RNA_seed | SRC | 0.495182 | COADREAD | LS-180 | Bosutinib |
| MVP_top60_DNA_seed | GSK3B | 0.339982 | COADREAD | SNU-407 | SB216763 | MVP_top60_DNA+RNA_seed | BCL2 | 0.497536 | COADREAD | NCI-H747 | Venetoclax |

| | | | | | | | | | | | |
|---|---|---|---|---|---|---|---|---|---|---|---|
| MVP_top60_DNA_seed | IGF1R | 0.34113 | COADREAD | RCM-1 | BMS-754807 | MVP_top60_DNA+RNA_seed | IGF1R | 0.497754 | COADREAD | SNU-1040 | Linsitinib |
| MVP_top60_DNA_seed | HSP90AA1 | 0.342956 | COADREAD | SNU-61 | Tanespimycin | MVP_top60_DNA+RNA_seed | IGF1R | 0.498675 | COADREAD | KM12 | Linsitinib |
| MVP_top60_DNA_seed | HSP90AB1 | 0.342956 | COADREAD | SNU-61 | Tanespimycin | MVP_top60_DNA+RNA_seed | BCL2 | 0.503234 | COADREAD | SK-CO-1 | Sabutoclax |
| MVP_top60_DNA_seed | MAPK1 | 0.343915 | COADREAD | CW-2 | VX-11e | MVP_top60_DNA+RNA_seed | IGF1R | 0.503699 | COADREAD | SW837 | GSK1904529A |
| MVP_top60_DNA_seed | SRC | 0.344414 | COADREAD | SNU-81 | Bosutinib | MVP_top60_DNA+RNA_seed | PIK3CA | 0.504079 | COADREAD | SNU-81 | Buparlisib |
| MVP_top60_DNA_seed | AKT1 | 0.345796 | COADREAD | HCC2998 | AT13148 | MVP_top60_DNA+RNA_seed | PIK3CB | 0.504079 | COADREAD | SNU-81 | Buparlisib |
| MVP_top60_DNA_seed | BRAF | 0.346097 | COADREAD | SNU-61 | SB590885 | MVP_top60_DNA+RNA_seed | PIK3CD | 0.504079 | COADREAD | SNU-81 | Buparlisib |
| MVP_top60_DNA_seed | AKT1 | 0.346405 | COADREAD | RCM-1 | Uprosertib | MVP_top60_DNA+RNA_seed | PIK3CB | 0.504248 | COADREAD | NCI-H508 | AZD6482 |
| MVP_top60_DNA_seed | HSP90AA1 | 0.346482 | COADREAD | LS-1034 | Tanespimycin | MVP_top60_DNA+RNA_seed | MET | 0.506067 | COADREAD | SK-CO-1 | Savolitinib |
| MVP_top60_DNA_seed | HSP90AB1 | 0.346482 | COADREAD | LS-1034 | Tanespimycin | MVP_top60_DNA+RNA_seed | IGF1R | 0.508267 | COADREAD | SW1417 | BMS-536924 |
| MVP_top60_DNA_seed | HSP90AA1 | 0.346893 | COADREAD | HT-115 | Tanespimycin | MVP_top60_DNA+RNA_seed | IGF1R | 0.510458 | COADREAD | LS-180 | NVP-ADW742 |
| MVP_top60_DNA_seed | HSP90AB1 | 0.346893 | COADREAD | HT-115 | Tanespimycin | MVP_top60_DNA+RNA_seed | MET | 0.513714 | COADREAD | CL-11 | Savolitinib |
| MVP_top60_DNA_seed | CDK2 | 0.347127 | COADREAD | HCT-15 | MK-8776 | MVP_top60_DNA+RNA_seed | BCL2 | 0.513922 | COADREAD | CW-2 | Navitoclax |
| MVP_top60_DNA_seed | AKT1 | 0.347505 | COADREAD | HCT-116 | Ipatasertib | MVP_top60_DNA+RNA_seed | IGF1R | 0.515822 | COADREAD | SK-CO-1 | GSK1904529A |
| MVP_top60_DNA_seed | PIK3CA | 0.348843 | COADREAD | SNU-81 | GNE-317 | MVP_top60_DNA+RNA_seed | IGF1R | 0.515918 | COADREAD | SW837 | NVP-ADW742 |
| MVP_top60_DNA_seed | CDK2 | 0.351148 | COADREAD | SNU-81 | AZD5438 | MVP_top60_DNA+RNA_seed | AKT1 | 0.518349 | COADREAD | COLO-320-HSR | AT13148 |
| MVP_top60_DNA_seed | IGF1R | 0.354776 | COADREAD | RKO | Linsitinib | MVP_top60_DNA+RNA_seed | TP53 | 0.519311 | COADREAD | LoVo | MIRA-1 |
| MVP_top60_DNA_seed | AKT1 | 0.355918 | COADREAD | COLO-320-HSR | GSK2110183B | MVP_top60_DNA+RNA_seed | SRC | 0.520276 | COADREAD | SW837 | Bosutinib |
| MVP_top60_DNA_seed | BRAF | 0.356624 | COADREAD | SW1463 | SB590885 | MVP_top60_DNA+RNA_seed | IGF1R | 0.521335 | COADREAD | CW-2 | BMS-754807 |
| MVP_top60_DNA_seed | BRAF | 0.358405 | COADREAD | SW837 | SB590885 | MVP_top60_DNA+RNA_seed | IGF1R | 0.521623 | COADREAD | HCC2998 | GSK1904529A |
| MVP_top60_DNA_seed | AKT1 | 0.359065 | COADREAD | SNU-C2B | Uprosertib | MVP_top60_DNA+RNA_seed | PIK3CA | 0.522049 | COADREAD | C2BBe1 | Alpelisib |
| MVP_top60_DNA_seed | MET | 0.360053 | COADREAD | HT55 | Savolitinib | MVP_top60_DNA+RNA_seed | ATM | 0.522708 | COADREAD | DiFi | KU-55933 |
| MVP_top60_DNA_seed | MET | 0.360843 | COADREAD | SNU-C5 | Savolitinib | MVP_top60_DNA+RNA_seed | HSP90AA1 | 0.523204 | COADREAD | COLO-320-HSR | Tanespimycin |
| MVP_top60_DNA_seed | PIK3CA | 0.364046 | COADREAD | LS-123 | Buparlisib | MVP_top60_DNA+RNA_seed | HSP90AB1 | 0.523204 | COADREAD | COLO-320-HSR | Tanespimycin |
| MVP_top60_DNA_seed | PIK3CB | 0.364046 | COADREAD | LS-123 | Buparlisib | MVP_top60_DNA+RNA_seed | PIK3CA | 0.523605 | COADREAD | SW1417 | AZD8186 |
| MVP_top60_DNA_seed | PIK3CD | 0.364046 | COADREAD | LS-123 | Buparlisib | MVP_top60_DNA+RNA_seed | PIK3CB | 0.523605 | COADREAD | SW1417 | AZD8186 |
| MVP_top60_DNA_seed | CDK2 | 0.367209 | COADREAD | SNU-407 | Dinaciclib | MVP_top60_DNA+RNA_seed | PIK3CA | 0.523866 | COADREAD | SNU-C2B | GNE-317 |
| MVP_top60_DNA_seed | IGF1R | 0.371435 | COADREAD | SNU-C5 | Linsitinib | MVP_top60_DNA+RNA_seed | SRC | 0.526936 | COADREAD | NCI-H747 | Bosutinib |
| MVP_top60_DNA_seed | AKT1 | 0.373851 | COADREAD | LS-180 | AT13148 | MVP_top60_DNA+RNA_seed | SRC | 0.531947 | COADREAD | LS-123 | Bosutinib |
| MVP_top60_DNA_seed | AKT1 | 0.377633 | COADREAD | CW-2 | AT13148 | MVP_top60_DNA+RNA_seed | CDK2 | 0.532351 | COADREAD | KM12 | MK-8776 |
| MVP_top60_DNA_seed | BRAF | 0.379977 | COADREAD | CW-2 | Dabrafenib | MVP_top60_DNA+RNA_seed | IGF1R | 0.535322 | COADREAD | HT-29 | GSK1904529A |
| MVP_top60_DNA_seed | MET | 0.380239 | COADREAD | SW837 | Crizotinib | MVP_top60_DNA+RNA_seed | HSP90AA1 | 0.540233 | COADREAD | KM12 | Luminespib |
| MVP_top60_DNA_seed | ATM | 0.380863 | COADREAD | CL-11 | KU-55933 | MVP_top60_DNA+RNA_seed | HSP90AB1 | 0.540233 | COADREAD | KM12 | Luminespib |
| MVP_top60_DNA_seed | PIK3CA | 0.381011 | COADREAD | SW620 | Buparlisib | MVP_top60_DNA+RNA_seed | IGF1R | 0.540851 | COADREAD | SW1417 | Linsitinib |
| MVP_top60_DNA_seed | PIK3CB | 0.381011 | COADREAD | SW620 | Buparlisib | MVP_top60_DNA+RNA_seed | PIK3CA | 0.546419 | COADREAD | SW1116 | Alpelisib |
| MVP_top60_DNA_seed | PIK3CD | 0.381011 | COADREAD | SW620 | Buparlisib | MVP_top60_DNA+RNA_seed | CDK2 | 0.548117 | COADREAD | LS-180 | MK-8776 |
| MVP_top60_DNA_seed | SRC | 0.381465 | COADREAD | COLO-678 | Bosutinib | MVP_top60_DNA+RNA_seed | IGF1R | 0.54904 | COADREAD | SW1116 | Linsitinib |
| MVP_top60_DNA_seed | TP53 | 0.382935 | COADREAD | CaR-1 | MIRA-1 | MVP_top60_DNA+RNA_seed | GSK3B | 0.550503 | COADREAD | SW1463 | SB216763 |
| MVP_top60_DNA_seed | AKT1 | 0.386093 | COADREAD | C2BBe1 | AZD5363 | MVP_top60_DNA+RNA_seed | AKT1 | 0.550628 | COADREAD | SNU-C1 | AT13148 |
| MVP_top60_DNA_seed | PIK3CA | 0.389427 | COADREAD | NCI-H716 | Buparlisib | MVP_top60_DNA+RNA_seed | KRAS | 0.556824 | COADREAD | CaR-1 | KRAS (G12C) Inhibitor-1 |
| MVP_top60_DNA_seed | PIK3CB | 0.389427 | COADREAD | NCI-H716 | Buparlisib | MVP_top60_DNA+RNA_seed | CDK2 | 0.557318 | COADREAD | HT-115 | Dinaciclib |

| | | | | | | | | | | | |
|---|---|---|---|---|---|---|---|---|---|---|---|
| MVP_top60_DNA_seed | PIK3CD | 0.389427 | COADREAD | NCI-H716 | Buparlisib | MVP_top60_DNA+RNA_seed | GSK3B | 0.557342 | COADREAD | SNU-81 | SB216763 |
| MVP_top60_DNA_seed | HSP90AA1 | 0.394059 | COADREAD | SW1417 | Tanespimycin | MVP_top60_DNA+RNA_seed | AKT1 | 0.560673 | COADREAD | LS-513 | AT13148 |
| MVP_top60_DNA_seed | HSP90AB1 | 0.394059 | COADREAD | SW1417 | Tanespimycin | MVP_top60_DNA+RNA_seed | ERBB3 | 0.562308 | COADREAD | HT-29 | Sapitinib |
| MVP_top60_DNA_seed | KRAS | 0.394184 | COADREAD | DiFi | KRAS (G12C) Inhibitor-12 | MVP_top60_DNA+RNA_seed | MET | 0.568018 | COADREAD | NCI-H747 | Savolitinib |
| MVP_top60_DNA_seed | TP53 | 0.396854 | COADREAD | NCI-H508 | MIRA-1 | MVP_top60_DNA+RNA_seed | MET | 0.56804 | COADREAD | SW48 | Savolitinib |
| MVP_top60_DNA_seed | CREBBP | 0.400822 | COADREAD | CW-2 | SGC-CBP30 | MVP_top60_DNA+RNA_seed | SRC | 0.572456 | COADREAD | COLO-205 | Bosutinib |
| MVP_top60_DNA_seed | HSP90AA1 | 0.402172 | COADREAD | CCK-81 | Luminespib | MVP_top60_DNA+RNA_seed | PIK3CA | 0.57669 | COADREAD | LS-123 | Alpelisib |
| MVP_top60_DNA_seed | HSP90AB1 | 0.402172 | COADREAD | CCK-81 | Luminespib | MVP_top60_DNA+RNA_seed | BCL2 | 0.583665 | COADREAD | C2BBe1 | Sabutoclax |
| MVP_top60_DNA_seed | CDK2 | 0.402514 | COADREAD | SK-CO-1 | MK-8776 | MVP_top60_DNA+RNA_seed | IGF1R | 0.584069 | COADREAD | SNU-C2B | Linsitinib |
| MVP_top60_DNA_seed | HSP90AA1 | 0.403948 | COADREAD | NCI-H508 | Elesclomol | MVP_top60_DNA+RNA_seed | BCL2 | 0.586675 | COADREAD | DiFi | Sabutoclax |
| MVP_top60_DNA_seed | HSP90AB1 | 0.403948 | COADREAD | NCI-H508 | Elesclomol | MVP_top60_DNA+RNA_seed | GSK3B | 0.588418 | COADREAD | HCC2998 | SB216763 |
| MVP_top60_DNA_seed | IGF1R | 0.408569 | COADREAD | SW1116 | BMS-536924 | MVP_top60_DNA+RNA_seed | IGF1R | 0.590916 | COADREAD | RCM-1 | NVP-ADW742 |
| MVP_top60_DNA_seed | IGF1R | 0.408777 | COADREAD | LS-1034 | GSK1904529A | MVP_top60_DNA+RNA_seed | IGF1R | 0.592348 | COADREAD | NCI-H716 | GSK1904529A |
| MVP_top60_DNA_seed | PIK3CA | 0.409103 | COADREAD | SW837 | AZD8186 | MVP_top60_DNA+RNA_seed | ATM | 0.59262 | COADREAD | NCI-H747 | KU-55933 |
| MVP_top60_DNA_seed | PIK3CB | 0.409103 | COADREAD | SW837 | AZD8186 | MVP_top60_DNA+RNA_seed | CDK2 | 0.592711 | COADREAD | LS-1034 | MK-8776 |
| MVP_top60_DNA_seed | MET | 0.409787 | COADREAD | C2BBe1 | Savolitinib | MVP_top60_DNA+RNA_seed | BCL2 | 0.595644 | COADREAD | CW-2 | Sabutoclax |
| MVP_top60_DNA_seed | PIK3CA | 0.411115 | COADREAD | HT-115 | GNE-317 | MVP_top60_DNA+RNA_seed | ERBB3 | 0.596864 | COADREAD | HCT-116 | Sapitinib |
| MVP_top60_DNA_seed | PIK3CA | 0.414408 | COADREAD | CCK-81 | GNE-317 | MVP_top60_DNA+RNA_seed | ATM | 0.596974 | COADREAD | SNU-407 | KU-55933 |
| MVP_top60_DNA_seed | SRC | 0.417419 | COADREAD | DiFi | Bosutinib | MVP_top60_DNA+RNA_seed | BCL2 | 0.600024 | COADREAD | SW1417 | TW 37 |
| MVP_top60_DNA_seed | MAPK1 | 0.419059 | COADREAD | COLO-320-HSR | VX-11e | MVP_top60_DNA+RNA_seed | IGF1R | 0.602145 | COADREAD | LS-180 | Linsitinib |
| MVP_top60_DNA_seed | BRAF | 0.421528 | COADREAD | SW837 | PLX-4720 | MVP_top60_DNA+RNA_seed | HSP90AA1 | 0.603249 | COADREAD | SNU-407 | Tanespimycin |
| MVP_top60_DNA_seed | PIK3CA | 0.425256 | COADREAD | SW1463 | AZD8186 | MVP_top60_DNA+RNA_seed | HSP90AB1 | 0.603249 | COADREAD | SNU-407 | Tanespimycin |
| MVP_top60_DNA_seed | PIK3CB | 0.425256 | COADREAD | SW1463 | AZD8186 | MVP_top60_DNA+RNA_seed | PIK3CA | 0.603429 | COADREAD | NCI-H716 | Alpelisib |
| MVP_top60_DNA_seed | PIK3CA | 0.426361 | COADREAD | HCT-116 | AZD8186 | MVP_top60_DNA+RNA_seed | AKT1 | 0.605266 | COADREAD | MDST8 | GSK2110183B |
| MVP_top60_DNA_seed | PIK3CB | 0.426361 | COADREAD | HCT-116 | AZD8186 | MVP_top60_DNA+RNA_seed | AKT1 | 0.607749 | COADREAD | CCK-81 | AT13148 |
| MVP_top60_DNA_seed | IGF1R | 0.426749 | COADREAD | HT55 | GSK1904529A | MVP_top60_DNA+RNA_seed | MET | 0.607996 | COADREAD | SNU-C1 | Savolitinib |
| MVP_top60_DNA_seed | IGF1R | 0.426915 | COADREAD | SW1417 | BMS-754807 | MVP_top60_DNA+RNA_seed | BCL2 | 0.608581 | COADREAD | COLO-320-HSR | Navitoclax |
| MVP_top60_DNA_seed | IGF1R | 0.427135 | COADREAD | SNU-81 | GSK1904529A | MVP_top60_DNA+RNA_seed | MET | 0.610292 | COADREAD | NCI-H508 | Savolitinib |
| MVP_top60_DNA_seed | MET | 0.427246 | COADREAD | T84 | Savolitinib | MVP_top60_DNA+RNA_seed | TP53 | 0.611766 | COADREAD | LS-1034 | MIRA-1 |
| MVP_top60_DNA_seed | BRAF | 0.427606 | COADREAD | CCK-81 | PLX-4720 | MVP_top60_DNA+RNA_seed | BCL2 | 0.612557 | COADREAD | SNU-61 | Venetoclax |
| MVP_top60_DNA_seed | HSP90AA1 | 0.42778 | COADREAD | HT-115 | CCT-018159 | MVP_top60_DNA+RNA_seed | BCL2 | 0.613219 | COADREAD | NCI-H747 | Sabutoclax |
| MVP_top60_DNA_seed | HSP90AB1 | 0.42778 | COADREAD | HT-115 | CCT-018159 | MVP_top60_DNA+RNA_seed | HSP90AA1 | 0.61425 | COADREAD | T84 | Luminespib |
| MVP_top60_DNA_seed | ERBB3 | 0.427965 | COADREAD | RKO | Sapitinib | MVP_top60_DNA+RNA_seed | HSP90AB1 | 0.61425 | COADREAD | T84 | Luminespib |
| MVP_top60_DNA_seed | SRC | 0.430052 | COADREAD | RKO | Bosutinib | MVP_top60_DNA+RNA_seed | GSK3B | 0.616759 | COADREAD | COLO-320-HSR | SB216763 |
| MVP_top60_DNA_seed | ESR1 | 0.430837 | COADREAD | NCI-H747 | GDC0810 | MVP_top60_DNA+RNA_seed | IGF1R | 0.622143 | COADREAD | COLO-678 | NVP-ADW742 |
| MVP_top60_DNA_seed | GSK3B | 0.430959 | COADREAD | NCI-H508 | CHIR-99021 | MVP_top60_DNA+RNA_seed | SRC | 0.623608 | COADREAD | HT-115 | Bosutinib |
| MVP_top60_DNA_seed | BRAF | 0.432668 | COADREAD | SW948 | SB590885 | MVP_top60_DNA+RNA_seed | CDK2 | 0.623834 | COADREAD | RCM-1 | Dinaciclib |
| MVP_top60_DNA_seed | ATM | 0.433564 | COADREAD | HCT-15 | KU-55933 | MVP_top60_DNA+RNA_seed | AKT1 | 0.623846 | COADREAD | SK-CO-1 | Afuresertib |
| MVP_top60_DNA_seed | MAPK1 | 0.434153 | COADREAD | SW837 | Ulixertinib | MVP_top60_DNA+RNA_seed | ERBB3 | 0.623875 | COADREAD | SNU-81 | Sapitinib |
| MVP_top60_DNA_seed | BRAF | 0.434516 | COADREAD | CaR-1 | SB590885 | MVP_top60_DNA+RNA_seed | IGF1R | 0.627355 | COADREAD | NCI-H747 | GSK1904529A |
| MVP_top60_DNA_seed | CDK2 | 0.434983 | COADREAD | SW48 | Dinaciclib | MVP_top60_DNA+RNA_seed | AKT1 | 0.627417 | COADREAD | NCI-H716 | AT13148 |

| | | | | | | | | | | | |
|---|---|---|---|---|---|---|---|---|---|---|---|
| MVP_top60_DNA_seed | TP53 | 0.436215 | COADREAD | KM12 | MIRA-1 | MVP_top60_DNA+RNA_seed | ATM | 0.629547 | COADREAD | NCI-H508 | KU-55933 |
| MVP_top60_DNA_seed | CDK2 | 0.439335 | COADREAD | SW1116 | AZD5438 | MVP_top60_DNA+RNA_seed | GSK3B | 0.630959 | COADREAD | LS-180 | SB216763 |
| MVP_top60_DNA_seed | MAPK1 | 0.439605 | COADREAD | CaR-1 | Ulixertinib | MVP_top60_DNA+RNA_seed | MET | 0.632827 | COADREAD | CaR-1 | Savolitinib |
| MVP_top60_DNA_seed | AKT1 | 0.43987 | COADREAD | LS-1034 | Uprosertib | MVP_top60_DNA+RNA_seed | BCL2 | 0.634429 | COADREAD | SW837 | Venetoclax |
| MVP_top60_DNA_seed | CDK2 | 0.441192 | COADREAD | NCI-H716 | Dinaciclib | MVP_top60_DNA+RNA_seed | SRC | 0.63506 | COADREAD | SW948 | Bosutinib |
| MVP_top60_DNA_seed | BRAF | 0.441368 | COADREAD | SNU-81 | Dabrafenib | MVP_top60_DNA+RNA_seed | MET | 0.635349 | COADREAD | SNU-C2B | Crizotinib |
| MVP_top60_DNA_seed | CDK2 | 0.443945 | COADREAD | C2BBe1 | Dinaciclib | MVP_top60_DNA+RNA_seed | BCL2 | 0.63647 | COADREAD | SW620 | Sabutoclax |
| MVP_top60_DNA_seed | AKT1 | 0.447395 | COADREAD | RCM-1 | Uprosertib | MVP_top60_DNA+RNA_seed | CDK2 | 0.637685 | COADREAD | SNU-81 | MK-8776 |
| MVP_top60_DNA_seed | PIK3CA | 0.447664 | COADREAD | LS-180 | Buparlisib | MVP_top60_DNA+RNA_seed | AR | 0.6402 | COADREAD | SW620 | Bicalutamide |
| MVP_top60_DNA_seed | PIK3CB | 0.447664 | COADREAD | LS-180 | Buparlisib | MVP_top60_DNA+RNA_seed | AKT1 | 0.640662 | COADREAD | SW620 | AT13148 |
| MVP_top60_DNA_seed | PIK3CD | 0.447664 | COADREAD | LS-180 | Buparlisib | MVP_top60_DNA+RNA_seed | PIK3CA | 0.645643 | COADREAD | CL-11 | Alpelisib |
| MVP_top60_DNA_seed | IGF1R | 0.448033 | COADREAD | RKO | BMS-536924 | MVP_top60_DNA+RNA_seed | TP53 | 0.646022 | COADREAD | SW1463 | MIRA-1 |
| MVP_top60_DNA_seed | BRAF | 0.448205 | COADREAD | LS-180 | SB590885 | MVP_top60_DNA+RNA_seed | IGF1R | 0.651094 | COADREAD | SW620 | NVP-ADW742 |
| MVP_top60_DNA_seed | IGF1R | 0.449633 | COADREAD | CW-2 | GSK1904529A | MVP_top60_DNA+RNA_seed | ATM | 0.651683 | COADREAD | LS-513 | KU-55933 |
| MVP_top60_DNA_seed | BRAF | 0.450449 | COADREAD | HT-115 | Dabrafenib | MVP_top60_DNA+RNA_seed | PIK3CA | 0.651935 | COADREAD | HT-115 | AZD8186 |
| MVP_top60_DNA_seed | HSP90AA1 | 0.452968 | COADREAD | LS-180 | Luminespib | MVP_top60_DNA+RNA_seed | PIK3CB | 0.651935 | COADREAD | HT-115 | AZD8186 |
| MVP_top60_DNA_seed | HSP90AB1 | 0.452968 | COADREAD | LS-180 | Luminespib | MVP_top60_DNA+RNA_seed | BCL2 | 0.656542 | COADREAD | C2BBe1 | Venetoclax |
| MVP_top60_DNA_seed | MAPK1 | 0.454737 | COADREAD | CW-2 | Ulixertinib | MVP_top60_DNA+RNA_seed | BCL2 | 0.661719 | COADREAD | CL-11 | Sabutoclax |
| MVP_top60_DNA_seed | AKT1 | 0.455982 | COADREAD | HCT-116 | GSK2110183B | MVP_top60_DNA+RNA_seed | ATM | 0.668588 | COADREAD | COLO-205 | KU-55933 |
| MVP_top60_DNA_seed | PIK3CA | 0.45689 | COADREAD | LS-411N | Buparlisib | MVP_top60_DNA+RNA_seed | BCL2 | 0.670628 | COADREAD | SW48 | Sabutoclax |
| MVP_top60_DNA_seed | PIK3CB | 0.45689 | COADREAD | LS-411N | Buparlisib | MVP_top60_DNA+RNA_seed | AKT1 | 0.672407 | COADREAD | SW1463 | GSK2110183B |
| MVP_top60_DNA_seed | PIK3CD | 0.45689 | COADREAD | LS-411N | Buparlisib | MVP_top60_DNA+RNA_seed | BCL2 | 0.672801 | COADREAD | SW837 | Obatoclax Mesylate |
| MVP_top60_DNA_seed | IGF1R | 0.458215 | COADREAD | SW1463 | BMS-536924 | MVP_top60_DNA+RNA_seed | IGF1R | 0.673319 | COADREAD | SNU-C5 | GSK1904529A |
| MVP_top60_DNA_seed | PIK3CA | 0.460474 | COADREAD | SNU-C1 | Alpelisib | MVP_top60_DNA+RNA_seed | IGF1R | 0.677836 | COADREAD | RCM-1 | BMS-536924 |
| MVP_top60_DNA_seed | SRC | 0.463796 | COADREAD | LS-513 | Bosutinib | MVP_top60_DNA+RNA_seed | KRAS | 0.679288 | COADREAD | SW948 | (RAS (G12C) Inhibitor-1 |
| MVP_top60_DNA_seed | IGF1R | 0.469798 | COADREAD | LS-180 | GSK1904529A | MVP_top60_DNA+RNA_seed | AKT1 | 0.679479 | COADREAD | SW1463 | Ipatasertib |
| MVP_top60_DNA_seed | BRAF | 0.472625 | COADREAD | CL-11 | Dabrafenib | MVP_top60_DNA+RNA_seed | BCL2 | 0.681511 | COADREAD | SW48 | Obatoclax Mesylate |
| MVP_top60_DNA_seed | MAPK1 | 0.473382 | COADREAD | LS-123 | Ulixertinib | MVP_top60_DNA+RNA_seed | IGF1R | 0.682794 | COADREAD | SNU-C1 | GSK1904529A |
| MVP_top60_DNA_seed | BRAF | 0.473968 | COADREAD | SW620 | Dabrafenib | MVP_top60_DNA+RNA_seed | IGF1R | 0.682813 | COADREAD | LoVo | Linsitinib |
| MVP_top60_DNA_seed | AKT1 | 0.474809 | COADREAD | LS-123 | MK-2206 | MVP_top60_DNA+RNA_seed | PIK3CA | 0.683489 | COADREAD | CL-11 | AZD8186 |
| MVP_top60_DNA_seed | AKT1 | 0.478817 | COADREAD | LS-1034 | Afuresertib | MVP_top60_DNA+RNA_seed | PIK3CB | 0.683489 | COADREAD | CL-11 | AZD8186 |
| MVP_top60_DNA_seed | IGF1R | 0.47973 | COADREAD | T84 | Linsitinib | MVP_top60_DNA+RNA_seed | CDK2 | 0.683873 | COADREAD | SK-CO-1 | AZD5438 |
| MVP_top60_DNA_seed | ERBB3 | 0.480324 | COADREAD | SW1417 | Sapitinib | MVP_top60_DNA+RNA_seed | BCL2 | 0.686717 | COADREAD | COLO-205 | Sabutoclax |
| MVP_top60_DNA_seed | IGF1R | 0.481295 | COADREAD | DiFi | BMS-536924 | MVP_top60_DNA+RNA_seed | KRAS | 0.687276 | COADREAD | HT-115 | (RAS (G12C) Inhibitor-1 |
| MVP_top60_DNA_seed | AKT1 | 0.482012 | COADREAD | CL-11 | Uprosertib | MVP_top60_DNA+RNA_seed | BCL2 | 0.689168 | COADREAD | RKO | Venetoclax |
| MVP_top60_DNA_seed | TP53 | 0.484948 | COADREAD | LS-411N | MIRA-1 | MVP_top60_DNA+RNA_seed | IGF1R | 0.689312 | COADREAD | SNU-C2B | NVP-ADW742 |
| MVP_top60_DNA_seed | HSP90AA1 | 0.485452 | COADREAD | SNU-407 | Luminespib | MVP_top60_DNA+RNA_seed | ATM | 0.694096 | COADREAD | SW948 | KU-55933 |
| MVP_top60_DNA_seed | HSP90AB1 | 0.485452 | COADREAD | SNU-407 | Luminespib | MVP_top60_DNA+RNA_seed | AKT1 | 0.696135 | COADREAD | CCK-81 | Uprosertib |
| MVP_top60_DNA_seed | AKT1 | 0.48941 | COADREAD | T84 | MK-2206 | MVP_top60_DNA+RNA_seed | MET | 0.697373 | COADREAD | DiFi | Crizotinib |
| MVP_top60_DNA_seed | IGF1R | 0.490018 | COADREAD | T84 | GSK1904529A | MVP_top60_DNA+RNA_seed | BCL2 | 0.70103 | COADREAD | SNU-C5 | Navitoclax |
| MVP_top60_DNA_seed | SRC | 0.495182 | COADREAD | LS-180 | Bosutinib | MVP_top60_DNA+RNA_seed | HSP90AA1 | 0.703988 | COADREAD | SNU-81 | Tanespimycin |

| | | | | | | | | | | | |
|---|---|---|---|---|---|---|---|---|---|---|---|
| MVP_top60_DNA_seed | BRAF | 0.495745 | COADREAD | SNU-1040 | Dabrafenib | MVP_top60_DNA+RNA_seed | HSP90AB1 | 0.703988 | COADREAD | SNU-81 | Tanespimycin |
| MVP_top60_DNA_seed | ESR1 | 0.497389 | COADREAD | SNU-C2B | Tamoxifen | MVP_top60_DNA+RNA_seed | ATM | 0.704125 | COADREAD | HT-29 | KU-55933 |
| MVP_top60_DNA_seed | IGF1R | 0.497754 | COADREAD | SNU-1040 | Linsitinib | MVP_top60_DNA+RNA_seed | AKT1 | 0.70472 | COADREAD | NCI-H716 | Ipatasertib |
| MVP_top60_DNA_seed | IGF1R | 0.498675 | COADREAD | KM12 | Linsitinib | MVP_top60_DNA+RNA_seed | AKT1 | 0.707649 | COADREAD | LS-1034 | Ipatasertib |
| MVP_top60_DNA_seed | IGF1R | 0.503699 | COADREAD | SW837 | GSK1904529A | MVP_top60_DNA+RNA_seed | HSP90AA1 | 0.71371 | COADREAD | HT-29 | Luminespib |
| MVP_top60_DNA_seed | PIK3CA | 0.504079 | COADREAD | SNU-81 | Buparlisib | MVP_top60_DNA+RNA_seed | HSP90AB1 | 0.71371 | COADREAD | HT-29 | Luminespib |
| MVP_top60_DNA_seed | PIK3CB | 0.504079 | COADREAD | SNU-81 | Buparlisib | MVP_top60_DNA+RNA_seed | CDK2 | 0.713724 | COADREAD | CaR-1 | MK-8776 |
| MVP_top60_DNA_seed | PIK3CD | 0.504079 | COADREAD | SNU-81 | Buparlisib | MVP_top60_DNA+RNA_seed | AKT1 | 0.716489 | COADREAD | HT-115 | Uprosertib |
| MVP_top60_DNA_seed | PIK3CB | 0.504248 | COADREAD | NCI-H508 | AZD6482 | MVP_top60_DNA+RNA_seed | PIK3CA | 0.716953 | COADREAD | RCM-1 | AZD8186 |
| MVP_top60_DNA_seed | MET | 0.506067 | COADREAD | SK-CO-1 | Savolitinib | MVP_top60_DNA+RNA_seed | PIK3CB | 0.716953 | COADREAD | RCM-1 | AZD8186 |
| MVP_top60_DNA_seed | IGF1R | 0.508267 | COADREAD | SW1417 | BMS-536924 | MVP_top60_DNA+RNA_seed | AR | 0.719069 | COADREAD | HT-115 | Bicalutamide |
| MVP_top60_DNA_seed | BRAF | 0.50889 | COADREAD | C2BBe1 | SB590885 | MVP_top60_DNA+RNA_seed | SRC | 0.720088 | COADREAD | SNU-61 | Bosutinib |
| MVP_top60_DNA_seed | ESR1 | 0.509846 | COADREAD | NCI-H508 | Tamoxifen | MVP_top60_DNA+RNA_seed | IGF1R | 0.720252 | COADREAD | SNU-61 | NVP-ADW742 |
| MVP_top60_DNA_seed | IGF1R | 0.510458 | COADREAD | LS-180 | NVP-ADW742 | MVP_top60_DNA+RNA_seed | BCL2 | 0.722743 | COADREAD | CL-11 | Venetoclax |
| MVP_top60_DNA_seed | MET | 0.513714 | COADREAD | CL-11 | Savolitinib | MVP_top60_DNA+RNA_seed | BCL2 | 0.722825 | COADREAD | RCM-1 | Venetoclax |
| MVP_top60_DNA_seed | BRAF | 0.515803 | COADREAD | SNU-175 | Dabrafenib | MVP_top60_DNA+RNA_seed | SRC | 0.722874 | COADREAD | RCM-1 | Bosutinib |
| MVP_top60_DNA_seed | IGF1R | 0.515822 | COADREAD | SK-CO-1 | GSK1904529A | MVP_top60_DNA+RNA_seed | IGF1R | 0.725128 | COADREAD | COLO-678 | BMS-754807 |
| MVP_top60_DNA_seed | IGF1R | 0.515918 | COADREAD | SW837 | NVP-ADW742 | MVP_top60_DNA+RNA_seed | BCL2 | 0.731864 | COADREAD | SNU-C1 | Sabutoclax |
| MVP_top60_DNA_seed | AKT1 | 0.518349 | COADREAD | COLO-320-HSR | AT13148 | MVP_top60_DNA+RNA_seed | PIK3CA | 0.732518 | COADREAD | CL-11 | Buparlisib |
| MVP_top60_DNA_seed | TP53 | 0.519311 | COADREAD | LoVo | MIRA-1 | MVP_top60_DNA+RNA_seed | PIK3CB | 0.732518 | COADREAD | CL-11 | Buparlisib |
| MVP_top60_DNA_seed | SRC | 0.520276 | COADREAD | SW837 | Bosutinib | MVP_top60_DNA+RNA_seed | PIK3CD | 0.732518 | COADREAD | CL-11 | Buparlisib |
| MVP_top60_DNA_seed | IGF1R | 0.521335 | COADREAD | CW-2 | BMS-754807 | MVP_top60_DNA+RNA_seed | CDK2 | 0.734773 | COADREAD | LoVo | AZD5438 |
| MVP_top60_DNA_seed | IGF1R | 0.521623 | COADREAD | HCC2998 | GSK1904529A | MVP_top60_DNA+RNA_seed | ERBB3 | 0.735416 | COADREAD | NCI-H716 | Sapitinib |
| MVP_top60_DNA_seed | PIK3CA | 0.522049 | COADREAD | C2BBe1 | Alpelisib | MVP_top60_DNA+RNA_seed | AKT1 | 0.737225 | COADREAD | LS-180 | GSK2110183B |
| MVP_top60_DNA_seed | ATM | 0.522708 | COADREAD | DiFi | KU-55933 | MVP_top60_DNA+RNA_seed | HSP90AA1 | 0.737419 | COADREAD | LS-123 | Tanespimycin |
| MVP_top60_DNA_seed | HSP90AA1 | 0.523204 | COADREAD | COLO-320-HSR | Tanespimycin | MVP_top60_DNA+RNA_seed | HSP90AB1 | 0.737419 | COADREAD | LS-123 | Tanespimycin |
| MVP_top60_DNA_seed | HSP90AB1 | 0.523204 | COADREAD | COLO-320-HSR | Tanespimycin | MVP_top60_DNA+RNA_seed | SRC | 0.737466 | COADREAD | HT55 | Bosutinib |
| MVP_top60_DNA_seed | PIK3CA | 0.523605 | COADREAD | SW1417 | AZD8186 | MVP_top60_DNA+RNA_seed | ATM | 0.740156 | COADREAD | SW48 | KU-55933 |
| MVP_top60_DNA_seed | PIK3CB | 0.523605 | COADREAD | SW1417 | AZD8186 | MVP_top60_DNA+RNA_seed | PIK3CA | 0.743821 | COADREAD | RKO | AZD8186 |
| MVP_top60_DNA_seed | PIK3CA | 0.523866 | COADREAD | SNU-C2B | GNE-317 | MVP_top60_DNA+RNA_seed | PIK3CB | 0.743821 | COADREAD | RKO | AZD8186 |
| MVP_top60_DNA_seed | SRC | 0.526936 | COADREAD | NCI-H747 | Bosutinib | MVP_top60_DNA+RNA_seed | IGF1R | 0.745857 | COADREAD | SNU-1040 | GSK1904529A |
| MVP_top60_DNA_seed | CREBBP | 0.527851 | COADREAD | MDST8 | SGC-CBP30 | MVP_top60_DNA+RNA_seed | SRC | 0.749238 | COADREAD | SW1417 | Bosutinib |
| MVP_top60_DNA_seed | SRC | 0.531947 | COADREAD | LS-123 | Bosutinib | MVP_top60_DNA+RNA_seed | HSP90AA1 | 0.750969 | COADREAD | SW1116 | Luminespib |
| MVP_top60_DNA_seed | CDK2 | 0.532351 | COADREAD | KM12 | MK-8776 | MVP_top60_DNA+RNA_seed | HSP90AB1 | 0.750969 | COADREAD | SW1116 | Luminespib |
| MVP_top60_DNA_seed | ESR1 | 0.533077 | COADREAD | HCT-116 | Tamoxifen | MVP_top60_DNA+RNA_seed | IGF1R | 0.750997 | COADREAD | SNU-81 | Linsitinib |
| MVP_top60_DNA_seed | IGF1R | 0.535322 | COADREAD | HT-29 | GSK1904529A | MVP_top60_DNA+RNA_seed | TP53 | 0.752339 | COADREAD | CL-11 | MIRA-1 |
| MVP_top60_DNA_seed | HSP90AA1 | 0.540233 | COADREAD | KM12 | Luminespib | MVP_top60_DNA+RNA_seed | AKT1 | 0.755021 | COADREAD | SNU-61 | GSK2110183B |
| MVP_top60_DNA_seed | HSP90AB1 | 0.540233 | COADREAD | KM12 | Luminespib | MVP_top60_DNA+RNA_seed | IGF1R | 0.756837 | COADREAD | HT-115 | BMS-536924 |
| MVP_top60_DNA_seed | IGF1R | 0.540851 | COADREAD | SW1417 | Linsitinib | MVP_top60_DNA+RNA_seed | MET | 0.757627 | COADREAD | SW837 | Savolitinib |
| MVP_top60_DNA_seed | BRAF | 0.542197 | COADREAD | SNU-81 | SB590885 | MVP_top60_DNA+RNA_seed | BCL2 | 0.759172 | COADREAD | SNU-C2B | Venetoclax |
| MVP_top60_DNA_seed | CREBBP | 0.544873 | COADREAD | LS-513 | SGC-CBP30 | MVP_top60_DNA+RNA_seed | MET | 0.759862 | COADREAD | SW1463 | Savolitinib |

| | | | | | | | | | | | |
|---|---|---|---|---|---|---|---|---|---|---|---|
| MVP_top60_DNA_seed | BRAF | 0.544978 | COADREAD | NCI-H508 | SB590885 | MVP_top60_DNA+RNA_seed | IGF1R | 0.765411 | COADREAD | NCI-H508 | GSK1904529A |
| MVP_top60_DNA_seed | PIK3CA | 0.546419 | COADREAD | SW1116 | Alpelisib | MVP_top60_DNA+RNA_seed | AKT1 | 0.765427 | COADREAD | SW1463 | MK-2206 |
| MVP_top60_DNA_seed | MAPK1 | 0.546547 | COADREAD | COLO-678 | Ulixertinib | MVP_top60_DNA+RNA_seed | MET | 0.767108 | COADREAD | LS-180 | Savolitinib |
| MVP_top60_DNA_seed | CDK2 | 0.548117 | COADREAD | LS-180 | MK-8776 | MVP_top60_DNA+RNA_seed | PIK3CA | 0.769725 | COADREAD | SW620 | Alpelisib |
| MVP_top60_DNA_seed | IGF1R | 0.54904 | COADREAD | SW1116 | Linsitinib | MVP_top60_DNA+RNA_seed | IGF1R | 0.770165 | COADREAD | SW620 | GSK1904529A |
| MVP_top60_DNA_seed | GSK3B | 0.550503 | COADREAD | SW1463 | SB216763 | MVP_top60_DNA+RNA_seed | AKT1 | 0.77104 | COADREAD | SNU-C2B | Uprosertib |
| MVP_top60_DNA_seed | AKT1 | 0.550628 | COADREAD | SNU-C1 | AT13148 | MVP_top60_DNA+RNA_seed | IGF1R | 0.771369 | COADREAD | LoVo | BMS-536924 |
| MVP_top60_DNA_seed | ESR1 | 0.55357 | COADREAD | LS-1034 | GDC0810 | MVP_top60_DNA+RNA_seed | AKT1 | 0.77257 | COADREAD | LS-1034 | GSK2110183B |
| MVP_top60_DNA_seed | MAPK1 | 0.554591 | COADREAD | SNU-C5 | VX-11e | MVP_top60_DNA+RNA_seed | MET | 0.773781 | COADREAD | LoVo | Savolitinib |
| MVP_top60_DNA_seed | CREBBP | 0.554804 | COADREAD | SW1116 | SGC-CBP30 | MVP_top60_DNA+RNA_seed | IGF1R | 0.776209 | COADREAD | C2BBe1 | BMS-754807 |
| MVP_top60_DNA_seed | KRAS | 0.556824 | COADREAD | CaR-1 | KRAS (G12C) Inhibitor-1[illegible] | MVP_top60_DNA+RNA_seed | CDK2 | 0.779508 | COADREAD | NCI-H747 | Dinaciclib |
| MVP_top60_DNA_seed | CDK2 | 0.557318 | COADREAD | HT-115 | Dinaciclib | MVP_top60_DNA+RNA_seed | AKT1 | 0.780483 | COADREAD | LS-411N | MK-2206 |
| MVP_top60_DNA_seed | GSK3B | 0.557342 | COADREAD | SNU-81 | SB216763 | MVP_top60_DNA+RNA_seed | AKT1 | 0.782482 | COADREAD | SW1417 | Uprosertib |
| MVP_top60_DNA_seed | AKT1 | 0.560673 | COADREAD | LS-513 | AT13148 | MVP_top60_DNA+RNA_seed | IGF1R | 0.784403 | COADREAD | CL-11 | NVP-ADW742 |
| MVP_top60_DNA_seed | ERBB3 | 0.562308 | COADREAD | HT-29 | Sapitinib | MVP_top60_DNA+RNA_seed | AKT1 | 0.788046 | COADREAD | SNU-61 | Uprosertib |
| MVP_top60_DNA_seed | MET | 0.568018 | COADREAD | NCI-H747 | Savolitinib | MVP_top60_DNA+RNA_seed | BCL2 | 0.788552 | COADREAD | HT-115 | Venetoclax |
| MVP_top60_DNA_seed | MET | 0.56804 | COADREAD | SW48 | Savolitinib | MVP_top60_DNA+RNA_seed | BCL2 | 0.790361 | COADREAD | SW1463 | Venetoclax |
| MVP_top60_DNA_seed | ESR1 | 0.56981 | COADREAD | SW1417 | GDC0810 | MVP_top60_DNA+RNA_seed | AKT1 | 0.791063 | COADREAD | HT-115 | GSK2110183B |
| MVP_top60_DNA_seed | SRC | 0.572456 | COADREAD | COLO-205 | Bosutinib | MVP_top60_DNA+RNA_seed | BCL2 | 0.793526 | COADREAD | SNU-61 | Navitoclax |
| MVP_top60_DNA_seed | BRAF | 0.574168 | COADREAD | LS-513 | PLX-4720 | MVP_top60_DNA+RNA_seed | PIK3CA | 0.794784 | COADREAD | SNU-C2B | Buparlisib |
| MVP_top60_DNA_seed | BRAF | 0.574511 | COADREAD | T84 | SB590885 | MVP_top60_DNA+RNA_seed | PIK3CB | 0.794784 | COADREAD | SNU-C2B | Buparlisib |
| MVP_top60_DNA_seed | PIK3CA | 0.57669 | COADREAD | LS-123 | Alpelisib | MVP_top60_DNA+RNA_seed | PIK3CD | 0.794784 | COADREAD | SNU-C2B | Buparlisib |
| MVP_top60_DNA_seed | BRAF | 0.582661 | COADREAD | SW1417 | SB590885 | MVP_top60_DNA+RNA_seed | GSK3B | 0.796314 | COADREAD | LS-123 | SB216763 |
| MVP_top60_DNA_seed | IGF1R | 0.584069 | COADREAD | SNU-C2B | Linsitinib | MVP_top60_DNA+RNA_seed | GSK3B | 0.797398 | COADREAD | LS-1034 | SB216763 |
| MVP_top60_DNA_seed | BRAF | 0.585585 | COADREAD | SW48 | PLX-4720 | MVP_top60_DNA+RNA_seed | BCL2 | 0.802487 | COADREAD | SNU-61 | Obatoclax Mesylate |
| MVP_top60_DNA_seed | GSK3B | 0.588418 | COADREAD | HCC2998 | SB216763 | MVP_top60_DNA+RNA_seed | ATM | 0.804114 | COADREAD | RCM-1 | KU-55933 |
| MVP_top60_DNA_seed | IGF1R | 0.590916 | COADREAD | RCM-1 | NVP-ADW742 | MVP_top60_DNA+RNA_seed | ATM | 0.808598 | COADREAD | SW1116 | KU-55933 |
| MVP_top60_DNA_seed | IGF1R | 0.592348 | COADREAD | NCI-H716 | GSK1904529A | MVP_top60_DNA+RNA_seed | PIK3CA | 0.811062 | COADREAD | SW1417 | GNE-317 |
| MVP_top60_DNA_seed | ATM | 0.59262 | COADREAD | NCI-H747 | KU-55933 | MVP_top60_DNA+RNA_seed | ERBB3 | 0.813782 | COADREAD | COLO-320-HSR | Sapitinib |
| MVP_top60_DNA_seed | CDK2 | 0.592711 | COADREAD | LS-1034 | MK-8776 | MVP_top60_DNA+RNA_seed | GSK3B | 0.815496 | COADREAD | C2BBe1 | SB216763 |
| MVP_top60_DNA_seed | ERBB3 | 0.596864 | COADREAD | HCT-116 | Sapitinib | MVP_top60_DNA+RNA_seed | BCL2 | 0.816095 | COADREAD | LS-411N | Venetoclax |
| MVP_top60_DNA_seed | ATM | 0.596974 | COADREAD | SNU-407 | KU-55933 | MVP_top60_DNA+RNA_seed | IGF1R | 0.816213 | COADREAD | KM12 | GSK1904529A |
| MVP_top60_DNA_seed | ESR1 | 0.600066 | COADREAD | DiFi | GDC0810 | MVP_top60_DNA+RNA_seed | GSK3B | 0.816802 | COADREAD | SNU-61 | SB216763 |
| MVP_top60_DNA_seed | IGF1R | 0.602145 | COADREAD | LS-180 | Linsitinib | MVP_top60_DNA+RNA_seed | HSP90AA1 | 0.818692 | COADREAD | LS-1034 | Luminespib |
| MVP_top60_DNA_seed | HSP90AA1 | 0.603249 | COADREAD | SNU-407 | Tanespimycin | MVP_top60_DNA+RNA_seed | HSP90AB1 | 0.818692 | COADREAD | LS-1034 | Luminespib |
| MVP_top60_DNA_seed | HSP90AB1 | 0.603249 | COADREAD | SNU-407 | Tanespimycin | MVP_top60_DNA+RNA_seed | CDK2 | 0.8187 | COADREAD | HT-115 | MK-8776 |
| MVP_top60_DNA_seed | PIK3CA | 0.603429 | COADREAD | NCI-H716 | Alpelisib | MVP_top60_DNA+RNA_seed | CDK2 | 0.82011 | COADREAD | SNU-61 | AZD5438 |
| MVP_top60_DNA_seed | AKT1 | 0.605266 | COADREAD | MDST8 | GSK2110183B | MVP_top60_DNA+RNA_seed | SRC | 0.820985 | COADREAD | LS-1034 | Bosutinib |
| MVP_top60_DNA_seed | MAPK1 | 0.60569 | COADREAD | HCT-15 | Ulixertinib | MVP_top60_DNA+RNA_seed | CDK2 | 0.822609 | COADREAD | CaR-1 | Dinaciclib |
| MVP_top60_DNA_seed | BRAF | 0.60628 | COADREAD | SW1463 | Dabrafenib | MVP_top60_DNA+RNA_seed | AKT1 | 0.822871 | COADREAD | SW837 | Ipatasertib |
| MVP_top60_DNA_seed | AKT1 | 0.607749 | COADREAD | CCK-81 | AT13148 | MVP_top60_DNA+RNA_seed | MET | 0.823566 | COADREAD | SW1417 | Crizotinib |

| | | | | | | | | | | | |
|---|---|---|---|---|---|---|---|---|---|---|---|
| MVP_top60_DNA_seed | MET | 0.607996 | COADREAD | SNU-C1 | Savolitinib | MVP_top60_DNA+RNA_seed | AKT1 | 0.823825 | COADREAD | CW-2 | AZD5363 |
| MVP_top60_DNA_seed | MET | 0.610292 | COADREAD | NCI-H508 | Savolitinib | MVP_top60_DNA+RNA_seed | AKT1 | 0.826853 | COADREAD | C2BBe1 | Uprosertib |
| MVP_top60_DNA_seed | BRAF | 0.610297 | COADREAD | SNU-407 | SB590885 | MVP_top60_DNA+RNA_seed | IGF1R | 0.828099 | COADREAD | SNU-81 | BMS-754807 |
| MVP_top60_DNA_seed | TP53 | 0.611766 | COADREAD | LS-1034 | MIRA-1 | MVP_top60_DNA+RNA_seed | AKT1 | 0.828707 | COADREAD | NCI-H716 | Uprosertib |
| MVP_top60_DNA_seed | CREBBP | 0.61181 | COADREAD | SK-CO-1 | SGC-CBP30 | MVP_top60_DNA+RNA_seed | SRC | 0.837197 | COADREAD | SW48 | Bosutinib |
| MVP_top60_DNA_seed | HSP90AA1 | 0.61425 | COADREAD | T84 | Luminespib | MVP_top60_DNA+RNA_seed | CDK2 | 0.840863 | COADREAD | C2BBe1 | AZD5438 |
| MVP_top60_DNA_seed | HSP90AB1 | 0.61425 | COADREAD | T84 | Luminespib | MVP_top60_DNA+RNA_seed | BCL2 | 0.843018 | COADREAD | SNU-407 | Navitoclax |
| MVP_top60_DNA_seed | ESR1 | 0.614401 | COADREAD | SNU-C2B | GDC0810 | MVP_top60_DNA+RNA_seed | PIK3CA | 0.843262 | COADREAD | SNU-61 | Alpelisib |
| MVP_top60_DNA_seed | GSK3B | 0.616759 | COADREAD | COLO-320-HSR | SB216763 | MVP_top60_DNA+RNA_seed | HSP90AA1 | 0.843682 | COADREAD | COLO-678 | Tanespimycin |
| MVP_top60_DNA_seed | IGF1R | 0.622143 | COADREAD | COLO-678 | NVP-ADW742 | MVP_top60_DNA+RNA_seed | HSP90AB1 | 0.843682 | COADREAD | COLO-678 | Tanespimycin |
| MVP_top60_DNA_seed | SRC | 0.623608 | COADREAD | HT-115 | Bosutinib | MVP_top60_DNA+RNA_seed | BCL2 | 0.843791 | COADREAD | LoVo | Venetoclax |
| MVP_top60_DNA_seed | CDK2 | 0.623834 | COADREAD | RCM-1 | Dinaciclib | MVP_top60_DNA+RNA_seed | BCL2 | 0.844824 | COADREAD | HT55 | Venetoclax |
| MVP_top60_DNA_seed | AKT1 | 0.623846 | COADREAD | SK-CO-1 | Afuresertib | MVP_top60_DNA+RNA_seed | AKT1 | 0.84644 | COADREAD | RCM-1 | Afuresertib |
| MVP_top60_DNA_seed | ERBB3 | 0.623875 | COADREAD | SNU-81 | Sapitinib | MVP_top60_DNA+RNA_seed | PIK3CA | 0.846979 | COADREAD | SNU-C1 | Buparlisib |
| MVP_top60_DNA_seed | IGF1R | 0.627355 | COADREAD | NCI-H747 | GSK1904529A | MVP_top60_DNA+RNA_seed | PIK3CB | 0.846979 | COADREAD | SNU-C1 | Buparlisib |
| MVP_top60_DNA_seed | AKT1 | 0.627417 | COADREAD | NCI-H716 | AT13148 | MVP_top60_DNA+RNA_seed | PIK3CD | 0.846979 | COADREAD | SNU-C1 | Buparlisib |
| MVP_top60_DNA_seed | ATM | 0.629547 | COADREAD | NCI-H508 | KU-55933 | MVP_top60_DNA+RNA_seed | BCL2 | 0.848141 | COADREAD | HT-29 | Navitoclax |
| MVP_top60_DNA_seed | GSK3B | 0.630959 | COADREAD | LS-180 | SB216763 | MVP_top60_DNA+RNA_seed | PIK3CA | 0.849273 | COADREAD | SW837 | Buparlisib |
| MVP_top60_DNA_seed | MET | 0.632827 | COADREAD | CaR-1 | Savolitinib | MVP_top60_DNA+RNA_seed | PIK3CB | 0.849273 | COADREAD | SW837 | Buparlisib |
| MVP_top60_DNA_seed | SRC | 0.63506 | COADREAD | SW948 | Bosutinib | MVP_top60_DNA+RNA_seed | PIK3CD | 0.849273 | COADREAD | SW837 | Buparlisib |
| MVP_top60_DNA_seed | MET | 0.635349 | COADREAD | SNU-C2B | Crizotinib | MVP_top60_DNA+RNA_seed | AKT1 | 0.850724 | COADREAD | SNU-C2B | MK-2206 |
| MVP_top60_DNA_seed | CDK2 | 0.637685 | COADREAD | SNU-81 | MK-8776 | MVP_top60_DNA+RNA_seed | KRAS | 0.850839 | COADREAD | COLO-320-HSR | RAS (G12C) Inhibitor-1 |
| MVP_top60_DNA_seed | CREBBP | 0.63996 | COADREAD | LS-1034 | SGC-CBP30 | MVP_top60_DNA+RNA_seed | BCL2 | 0.851359 | COADREAD | HT-115 | Sabutoclax |
| MVP_top60_DNA_seed | AR | 0.6402 | COADREAD | SW620 | Bicalutamide | MVP_top60_DNA+RNA_seed | BCL2 | 0.852221 | COADREAD | CW-2 | Venetoclax |
| MVP_top60_DNA_seed | AKT1 | 0.640662 | COADREAD | SW620 | AT13148 | MVP_top60_DNA+RNA_seed | AKT1 | 0.855748 | COADREAD | CaR-1 | Uprosertib |
| MVP_top60_DNA_seed | BRAF | 0.641139 | COADREAD | NCI-H747 | SB590885 | MVP_top60_DNA+RNA_seed | HSP90AA1 | 0.859349 | COADREAD | LS-123 | Luminespib |
| MVP_top60_DNA_seed | BRAF | 0.643356 | COADREAD | T84 | Dabrafenib | MVP_top60_DNA+RNA_seed | HSP90AB1 | 0.859349 | COADREAD | LS-123 | Luminespib |
| MVP_top60_DNA_seed | PIK3CA | 0.645643 | COADREAD | CL-11 | Alpelisib | MVP_top60_DNA+RNA_seed | AKT1 | 0.865128 | COADREAD | SW1417 | AT13148 |
| MVP_top60_DNA_seed | TP53 | 0.646022 | COADREAD | SW1463 | MIRA-1 | MVP_top60_DNA+RNA_seed | IGF1R | 0.868323 | COADREAD | SW948 | BMS-536924 |
| MVP_top60_DNA_seed | IGF1R | 0.651094 | COADREAD | SW620 | NVP-ADW742 | MVP_top60_DNA+RNA_seed | PIK3CA | 0.87246 | COADREAD | SW948 | AZD8186 |
| MVP_top60_DNA_seed | ATM | 0.651683 | COADREAD | LS-513 | KU-55933 | MVP_top60_DNA+RNA_seed | PIK3CB | 0.87246 | COADREAD | SW948 | AZD8186 |
| MVP_top60_DNA_seed | PIK3CA | 0.651935 | COADREAD | HT-115 | AZD8186 | MVP_top60_DNA+RNA_seed | TP53 | 0.872614 | COADREAD | HT-29 | MIRA-1 |
| MVP_top60_DNA_seed | PIK3CB | 0.651935 | COADREAD | HT-115 | AZD8186 | MVP_top60_DNA+RNA_seed | BCL2 | 0.874488 | COADREAD | SW620 | Venetoclax |
| MVP_top60_DNA_seed | ESR1 | 0.654687 | COADREAD | LS-411N | GDC0810 | MVP_top60_DNA+RNA_seed | PIK3CA | 0.87469 | COADREAD | SW1463 | Alpelisib |
| MVP_top60_DNA_seed | ESR1 | 0.655454 | COADREAD | SW948 | GDC0810 | MVP_top60_DNA+RNA_seed | AKT1 | 0.876086 | COADREAD | SK-CO-1 | Uprosertib |
| MVP_top60_DNA_seed | CREBBP | 0.660766 | COADREAD | COLO-205 | SGC-CBP30 | MVP_top60_DNA+RNA_seed | TP53 | 0.877092 | COADREAD | SW948 | MIRA-1 |
| MVP_top60_DNA_seed | ATM | 0.668588 | COADREAD | COLO-205 | KU-55933 | MVP_top60_DNA+RNA_seed | BCL2 | 0.878836 | COADREAD | SW1417 | Obatoclax Mesylate |
| MVP_top60_DNA_seed | AKT1 | 0.672407 | COADREAD | SW1463 | GSK2110183B | MVP_top60_DNA+RNA_seed | MET | 0.880076 | COADREAD | LS-411N | Savolitinib |
| MVP_top60_DNA_seed | IGF1R | 0.673319 | COADREAD | SNU-C5 | GSK1904529A | MVP_top60_DNA+RNA_seed | AKT1 | 0.880895 | COADREAD | C2BBe1 | GSK2110183B |
| MVP_top60_DNA_seed | IGF1R | 0.677836 | COADREAD | RCM-1 | BMS-536924 | MVP_top60_DNA+RNA_seed | ERBB3 | 0.881579 | COADREAD | COLO-205 | Sapitinib |
| MVP_top60_DNA_seed | KRAS | 0.679288 | COADREAD | SW948 | KRAS (G12C) Inhibitor-1 | MVP_top60_DNA+RNA_seed | AKT1 | 0.883365 | COADREAD | SW1116 | AZD5363 |

| | | | | | | | | | | | |
|---|---|---|---|---|---|---|---|---|---|---|---|
| MVP_top60_DNA_seed | AKT1 | 0.679479 | COADREAD | SW1463 | Ipatasertib | MVP_top60_DNA+RNA_seed | AKT1 | 0.885174 | COADREAD | CaR-1 | GSK2110183B |
| MVP_top60_DNA_seed | ESR1 | 0.68111 | COADREAD | SW620 | Tamoxifen | MVP_top60_DNA+RNA_seed | IGF1R | 0.889294 | COADREAD | COLO-678 | BMS-536924 |
| MVP_top60_DNA_seed | IGF1R | 0.682794 | COADREAD | SNU-C1 | GSK1904529A | MVP_top60_DNA+RNA_seed | TP53 | 0.892991 | COADREAD | SNU-407 | MIRA-1 |
| MVP_top60_DNA_seed | IGF1R | 0.682813 | COADREAD | LoVo | Linsitinib | MVP_top60_DNA+RNA_seed | PIK3CA | 0.893689 | COADREAD | LS-1034 | Alpelisib |
| MVP_top60_DNA_seed | PIK3CA | 0.683489 | COADREAD | CL-11 | AZD8186 | MVP_top60_DNA+RNA_seed | BCL2 | 0.893881 | COADREAD | SNU-C2B | Obatoclax Mesylate |
| MVP_top60_DNA_seed | PIK3CB | 0.683489 | COADREAD | CL-11 | AZD8186 | MVP_top60_DNA+RNA_seed | MET | 0.897007 | COADREAD | C2BBe1 | Crizotinib |
| MVP_top60_DNA_seed | CDK2 | 0.683873 | COADREAD | SK-CO-1 | AZD5438 | MVP_top60_DNA+RNA_seed | BCL2 | 0.898638 | COADREAD | SW1116 | Obatoclax Mesylate |
| MVP_top60_DNA_seed | BRAF | 0.685516 | COADREAD | SW48 | SB590885 | MVP_top60_DNA+RNA_seed | ERBB3 | 0.902886 | COADREAD | MDST8 | Sapitinib |
| MVP_top60_DNA_seed | KRAS | 0.687276 | COADREAD | HT-115 | KRAS (G12C) Inhibitor-12 | MVP_top60_DNA+RNA_seed | AKT1 | 0.904056 | COADREAD | RCM-1 | GSK2110183B |
| MVP_top60_DNA_seed | IGF1R | 0.689312 | COADREAD | SNU-C2B | NVP-ADW742 | MVP_top60_DNA+RNA_seed | MET | 0.907059 | COADREAD | SNU-61 | Crizotinib |
| MVP_top60_DNA_seed | BRAF | 0.693735 | COADREAD | CW-2 | PLX-4720 | MVP_top60_DNA+RNA_seed | MET | 0.907575 | COADREAD | SNU-81 | Crizotinib |
| MVP_top60_DNA_seed | ATM | 0.694096 | COADREAD | SW948 | KU-55933 | MVP_top60_DNA+RNA_seed | SRC | 0.908465 | COADREAD | C2BBe1 | Bosutinib |
| MVP_top60_DNA_seed | AKT1 | 0.696135 | COADREAD | CCK-81 | Uprosertib | MVP_top60_DNA+RNA_seed | AKT1 | 0.912681 | COADREAD | LS-123 | Afuresertib |
| MVP_top60_DNA_seed | CREBBP | 0.697322 | COADREAD | T84 | SGC-CBP30 | MVP_top60_DNA+RNA_seed | TP53 | 0.920579 | COADREAD | SNU-61 | MIRA-1 |
| MVP_top60_DNA_seed | MET | 0.697373 | COADREAD | DiFi | Crizotinib | MVP_top60_DNA+RNA_seed | SRC | 0.924767 | COADREAD | T84 | Bosutinib |
| MVP_top60_DNA_seed | MAPK1 | 0.702223 | COADREAD | SNU-C2B | Ulixertinib | MVP_top60_DNA+RNA_seed | BCL2 | 0.926272 | COADREAD | COLO-320-HSR | Sabutoclax |
| MVP_top60_DNA_seed | HSP90AA1 | 0.703988 | COADREAD | SNU-81 | Tanespimycin | MVP_top60_DNA+RNA_seed | IGF1R | 0.93072 | COADREAD | SW837 | BMS-536924 |
| MVP_top60_DNA_seed | HSP90AB1 | 0.703988 | COADREAD | SNU-81 | Tanespimycin | MVP_top60_DNA+RNA_seed | GSK3B | 0.930787 | COADREAD | KM12 | SB216763 |
| MVP_top60_DNA_seed | ATM | 0.704125 | COADREAD | HT-29 | KU-55933 | MVP_top60_DNA+RNA_seed | CDK2 | 0.936504 | COADREAD | SW1116 | MK-8776 |
| MVP_top60_DNA_seed | AKT1 | 0.70472 | COADREAD | NCI-H716 | Ipatasertib | MVP_top60_DNA+RNA_seed | AKT1 | 0.938897 | COADREAD | SNU-61 | Ipatasertib |
| MVP_top60_DNA_seed | AKT1 | 0.707649 | COADREAD | LS-1034 | Ipatasertib | MVP_top60_DNA+RNA_seed | BCL2 | 0.940441 | COADREAD | COLO-678 | Navitoclax |
| MVP_top60_DNA_seed | ESR1 | 0.711646 | COADREAD | SW948 | Tamoxifen | MVP_top60_DNA+RNA_seed | AKT1 | 0.941062 | COADREAD | SW837 | MK-2206 |
| MVP_top60_DNA_seed | HSP90AA1 | 0.71371 | COADREAD | HT-29 | Luminespib | MVP_top60_DNA+RNA_seed | AKT1 | 0.946644 | COADREAD | HT55 | GSK2110183B |
| MVP_top60_DNA_seed | HSP90AB1 | 0.71371 | COADREAD | HT-29 | Luminespib | MVP_top60_DNA+RNA_seed | AKT1 | 0.952554 | COADREAD | SW1417 | Uprosertib |
| MVP_top60_DNA_seed | CDK2 | 0.713724 | COADREAD | CaR-1 | MK-8776 | MVP_top60_DNA+RNA_seed | BCL2 | 0.956171 | COADREAD | CaR-1 | Obatoclax Mesylate |
| MVP_top60_DNA_seed | AKT1 | 0.716489 | COADREAD | HT-115 | Uprosertib | MVP_top60_DNA+RNA_seed | BCL2 | 0.958396 | COADREAD | SW1463 | Obatoclax Mesylate |
| MVP_top60_DNA_seed | PIK3CA | 0.716953 | COADREAD | RCM-1 | AZD8186 | MVP_top60_DNA+RNA_seed | AKT1 | 0.963189 | COADREAD | CL-11 | Afuresertib |
| MVP_top60_DNA_seed | PIK3CB | 0.716953 | COADREAD | RCM-1 | AZD8186 | MVP_top60_DNA+RNA_seed | ATM | 0.966502 | COADREAD | SNU-61 | KU-55933 |
| MVP_top60_DNA_seed | AR | 0.719069 | COADREAD | HT-115 | Bicalutamide | MVP_top60_DNA+RNA_seed | AKT1 | 0.966865 | COADREAD | SNU-61 | AZD5363 |
| MVP_top60_DNA_seed | SRC | 0.720088 | COADREAD | SNU-61 | Bosutinib | MVP_top60_DNA+RNA_seed | PIK3CA | 0.97764 | COADREAD | SW1417 | Alpelisib |
| MVP_top60_DNA_seed | IGF1R | 0.720252 | COADREAD | SNU-61 | NVP-ADW742 | MVP_top60_DNA+RNA_seed | HSP90AA1 | 0.980103 | COADREAD | NCI-H508 | Luminespib |
| MVP_top60_DNA_seed | SRC | 0.722874 | COADREAD | RCM-1 | Bosutinib | MVP_top60_DNA+RNA_seed | HSP90AB1 | 0.980103 | COADREAD | NCI-H508 | Luminespib |
| MVP_top60_DNA_seed | IGF1R | 0.725128 | COADREAD | COLO-678 | BMS-754807 | MVP_top60_DNA+RNA_seed | AKT1 | 0.983078 | COADREAD | HT-115 | Ipatasertib |
| MVP_top60_DNA_seed | ESR1 | 0.726687 | COADREAD | CL-11 | GDC0810 | MVP_top60_DNA+RNA_seed | PIK3CA | 0.98394 | COADREAD | SNU-61 | Buparlisib |
| MVP_top60_DNA_seed | CREBBP | 0.72941 | COADREAD | SW837 | SGC-CBP30 | MVP_top60_DNA+RNA_seed | PIK3CB | 0.98394 | COADREAD | SNU-61 | Buparlisib |
| MVP_top60_DNA_seed | BRAF | 0.729667 | COADREAD | C2BBe1 | PLX-4720 | MVP_top60_DNA+RNA_seed | PIK3CD | 0.98394 | COADREAD | SNU-61 | Buparlisib |
| MVP_top60_DNA_seed | MAPK1 | 0.732135 | COADREAD | CaR-1 | Ulixertinib | MVP_top60_DNA+RNA_seed | AKT1 | 0.986802 | COADREAD | SNU-C5 | AT13148 |
| MVP_top60_DNA_seed | PIK3CA | 0.732518 | COADREAD | CL-11 | Buparlisib | MVP_top60_DNA+RNA_seed | AKT1 | 0.987999 | COADREAD | SW1116 | GSK2110183B |
| MVP_top60_DNA_seed | PIK3CB | 0.732518 | COADREAD | CL-11 | Buparlisib | MVP_top60_DNA+RNA_seed | KRAS | 0.989445 | COADREAD | CL-11 | KRAS (G12C) Inhibitor-1 |
| MVP_top60_DNA_seed | PIK3CD | 0.732518 | COADREAD | CL-11 | Buparlisib | MVP_top60_DNA+RNA_seed | AKT1 | 0.990615 | COADREAD | LS-1034 | AT13148 |
| MVP_top60_DNA_seed | CDK2 | 0.734773 | COADREAD | LoVo | AZD5438 | MVP_top60_DNA+RNA_seed | IGF1R | 0.9931 | COADREAD | LS-123 | Linsitinib |

| | | | | | | | | | | | |
|---|---|---|---|---|---|---|---|---|---|---|---|
| MVP_top60_DNA_seed | ERBB3 | 0.735416 | COADREAD | NCI-H716 | Sapitinib | MVP_top60_DNA+RNA_seed | BCL2 | 0.994049 | COADREAD | SNU-C1 | Venetoclax |
| MVP_top60_DNA_seed | AKT1 | 0.737225 | COADREAD | LS-180 | GSK2110183B | MVP_top60_DNA+RNA_seed | CDK2 | 0.998336 | COADREAD | SNU-1040 | AZD5438 |
| MVP_top60_DNA_seed | HSP90AA1 | 0.737419 | COADREAD | LS-123 | Tanespimycin | MVP_top60_DNA+RNA_seed | HSP90AA1 | 1.016775 | COADREAD | CL-11 | Luminespib |
| MVP_top60_DNA_seed | HSP90AB1 | 0.737419 | COADREAD | LS-123 | Tanespimycin | MVP_top60_DNA+RNA_seed | HSP90AB1 | 1.016775 | COADREAD | CL-11 | Luminespib |
| MVP_top60_DNA_seed | SRC | 0.737466 | COADREAD | HT55 | Bosutinib | MVP_top60_DNA+RNA_seed | ATM | 1.018111 | COADREAD | LS-411N | KU-55933 |
| MVP_top60_DNA_seed | ATM | 0.740156 | COADREAD | SW48 | KU-55933 | MVP_top60_DNA+RNA_seed | CDK2 | 1.018869 | COADREAD | SW948 | MK-8776 |
| MVP_top60_DNA_seed | PIK3CA | 0.743821 | COADREAD | RKO | AZD8186 | MVP_top60_DNA+RNA_seed | AKT1 | 1.019547 | COADREAD | SNU-61 | Afuresertib |
| MVP_top60_DNA_seed | PIK3CB | 0.743821 | COADREAD | RKO | AZD8186 | MVP_top60_DNA+RNA_seed | MET | 1.021199 | COADREAD | HT-29 | Savolitinib |
| MVP_top60_DNA_seed | IGF1R | 0.745857 | COADREAD | SNU-1040 | GSK1904529A | MVP_top60_DNA+RNA_seed | CDK2 | 1.024019 | COADREAD | CW-2 | MK-8776 |
| MVP_top60_DNA_seed | SRC | 0.749238 | COADREAD | SW1417 | Bosutinib | MVP_top60_DNA+RNA_seed | AKT1 | 1.025358 | COADREAD | SW1463 | Uprosertib |
| MVP_top60_DNA_seed | ESR1 | 0.750267 | COADREAD | SNU-61 | Tamoxifen | MVP_top60_DNA+RNA_seed | ATM | 1.028655 | COADREAD | LS-180 | KU-55933 |
| MVP_top60_DNA_seed | HSP90AA1 | 0.750969 | COADREAD | SW1116 | Luminespib | MVP_top60_DNA+RNA_seed | PIK3CA | 1.029105 | COADREAD | MDST8 | Alpelisib |
| MVP_top60_DNA_seed | HSP90AB1 | 0.750969 | COADREAD | SW1116 | Luminespib | MVP_top60_DNA+RNA_seed | AKT1 | 1.030377 | COADREAD | SNU-C2B | Ipatasertib |
| MVP_top60_DNA_seed | IGF1R | 0.750997 | COADREAD | SNU-81 | Linsitinib | MVP_top60_DNA+RNA_seed | HSP90AA1 | 1.033457 | COADREAD | SW1417 | Luminespib |
| MVP_top60_DNA_seed | TP53 | 0.752339 | COADREAD | CL-11 | MIRA-1 | MVP_top60_DNA+RNA_seed | HSP90AB1 | 1.033457 | COADREAD | SW1417 | Luminespib |
| MVP_top60_DNA_seed | AKT1 | 0.755021 | COADREAD | SNU-61 | GSK2110183B | MVP_top60_DNA+RNA_seed | PIK3CB | 1.042915 | COADREAD | HT-29 | AZD6482 |
| MVP_top60_DNA_seed | IGF1R | 0.756837 | COADREAD | HT-115 | BMS-536924 | MVP_top60_DNA+RNA_seed | TP53 | 1.043725 | COADREAD | SNU-C5 | MIRA-1 |
| MVP_top60_DNA_seed | MET | 0.757627 | COADREAD | SW837 | Savolitinib | MVP_top60_DNA+RNA_seed | BCL2 | 1.043946 | COADREAD | NCI-H716 | Venetoclax |
| MVP_top60_DNA_seed | MET | 0.759862 | COADREAD | SW1463 | Savolitinib | MVP_top60_DNA+RNA_seed | TP53 | 1.048788 | COADREAD | SNU-C1 | MIRA-1 |
| MVP_top60_DNA_seed | BRAF | 0.76091 | COADREAD | CCK-81 | Dabrafenib | MVP_top60_DNA+RNA_seed | KRAS | 1.051334 | COADREAD | CCK-81 | KRAS (G12C) Inhibitor-1 |
| MVP_top60_DNA_seed | IGF1R | 0.765411 | COADREAD | NCI-H508 | GSK1904529A | MVP_top60_DNA+RNA_seed | KRAS | 1.053421 | COADREAD | MDST8 | KRAS (G12C) Inhibitor-1 |
| MVP_top60_DNA_seed | AKT1 | 0.765427 | COADREAD | SW1463 | MK-2206 | MVP_top60_DNA+RNA_seed | PIK3CA | 1.054854 | COADREAD | SNU-C2B | Alpelisib |
| MVP_top60_DNA_seed | MET | 0.767108 | COADREAD | LS-180 | Savolitinib | MVP_top60_DNA+RNA_seed | ERBB3 | 1.057069 | COADREAD | SW948 | Sapitinib |
| MVP_top60_DNA_seed | PIK3CA | 0.769725 | COADREAD | SW620 | Alpelisib | MVP_top60_DNA+RNA_seed | AKT1 | 1.065601 | COADREAD | SW837 | GSK2110183B |
| MVP_top60_DNA_seed | IGF1R | 0.770165 | COADREAD | SW620 | GSK1904529A | MVP_top60_DNA+RNA_seed | HSP90AA1 | 1.066249 | COADREAD | C2BBe1 | Tanespimycin |
| MVP_top60_DNA_seed | AKT1 | 0.77104 | COADREAD | SNU-C2B | Uprosertib | MVP_top60_DNA+RNA_seed | HSP90AB1 | 1.066249 | COADREAD | C2BBe1 | Tanespimycin |
| MVP_top60_DNA_seed | IGF1R | 0.771369 | COADREAD | LoVo | BMS-536924 | MVP_top60_DNA+RNA_seed | PIK3CA | 1.068569 | COADREAD | SNU-C2B | AZD8186 |
| MVP_top60_DNA_seed | AKT1 | 0.77257 | COADREAD | LS-1034 | GSK2110183B | MVP_top60_DNA+RNA_seed | PIK3CB | 1.068569 | COADREAD | SNU-C2B | AZD8186 |
| MVP_top60_DNA_seed | MET | 0.773781 | COADREAD | LoVo | Savolitinib | MVP_top60_DNA+RNA_seed | IGF1R | 1.071973 | COADREAD | LS-180 | BMS-536924 |
| MVP_top60_DNA_seed | IGF1R | 0.776209 | COADREAD | C2BBe1 | BMS-754807 | MVP_top60_DNA+RNA_seed | TP53 | 1.072246 | COADREAD | SW837 | MIRA-1 |
| MVP_top60_DNA_seed | CDK2 | 0.779508 | COADREAD | NCI-H747 | Dinaciclib | MVP_top60_DNA+RNA_seed | AKT1 | 1.073691 | COADREAD | SNU-C2B | AZD5363 |
| MVP_top60_DNA_seed | AKT1 | 0.780483 | COADREAD | LS-411N | MK-2206 | MVP_top60_DNA+RNA_seed | CDK2 | 1.074076 | COADREAD | SW837 | MK-8776 |
| MVP_top60_DNA_seed | AKT1 | 0.782482 | COADREAD | SW1417 | Uprosertib | MVP_top60_DNA+RNA_seed | GSK3B | 1.078365 | COADREAD | CaR-1 | SB216763 |
| MVP_top60_DNA_seed | IGF1R | 0.784403 | COADREAD | CL-11 | NVP-ADW742 | MVP_top60_DNA+RNA_seed | TP53 | 1.083472 | COADREAD | NCI-H716 | MIRA-1 |
| MVP_top60_DNA_seed | BRAF | 0.785674 | COADREAD | CL-11 | SB590885 | MVP_top60_DNA+RNA_seed | TP53 | 1.084915 | COADREAD | SW620 | MIRA-1 |
| MVP_top60_DNA_seed | ESR1 | 0.786607 | COADREAD | SW1116 | GDC0810 | MVP_top60_DNA+RNA_seed | ERBB3 | 1.08603 | COADREAD | HCC2998 | Sapitinib |
| MVP_top60_DNA_seed | AKT1 | 0.788046 | COADREAD | SNU-61 | Uprosertib | MVP_top60_DNA+RNA_seed | BCL2 | 1.092495 | COADREAD | CaR-1 | Sabutoclax |
| MVP_top60_DNA_seed | BRAF | 0.788804 | COADREAD | LS-180 | Dabrafenib | MVP_top60_DNA+RNA_seed | CDK2 | 1.095237 | COADREAD | HT55 | MK-8776 |
| MVP_top60_DNA_seed | AKT1 | 0.791063 | COADREAD | HT-115 | GSK2110183B | MVP_top60_DNA+RNA_seed | ERBB3 | 1.108986 | COADREAD | SNU-C5 | Sapitinib |
| MVP_top60_DNA_seed | PIK3CA | 0.794784 | COADREAD | SNU-C2B | Buparlisib | MVP_top60_DNA+RNA_seed | KRAS | 1.118692 | COADREAD | SW837 | KRAS (G12C) Inhibitor-1 |
| MVP_top60_DNA_seed | PIK3CB | 0.794784 | COADREAD | SNU-C2B | Buparlisib | MVP_top60_DNA+RNA_seed | KRAS | 1.119802 | COADREAD | LS-1034 | KRAS (G12C) Inhibitor-1 |

| | | | | | | | | | | | |
|---|---|---|---|---|---|---|---|---|---|---|---|
| MVP_top60_DNA_seed | PIK3CD | 0.794784 | COADREAD | SNU-C2B | Buparlisib | MVP_top60_DNA+RNA_seed | KRAS | 1.127681 | COADREAD | SNU-61 | KRAS (G12C) Inhibitor-1 |
| MVP_top60_DNA_seed | GSK3B | 0.796314 | COADREAD | LS-123 | SB216763 | MVP_top60_DNA+RNA_seed | AKT1 | 1.128005 | COADREAD | SK-CO-1 | Uprosertib |
| MVP_top60_DNA_seed | GSK3B | 0.797398 | COADREAD | LS-1034 | SB216763 | MVP_top60_DNA+RNA_seed | IGF1R | 1.128152 | COADREAD | RKO | NVP-ADW742 |
| MVP_top60_DNA_seed | BRAF | 0.803115 | COADREAD | RCM-1 | PLX-4720 | MVP_top60_DNA+RNA_seed | AKT1 | 1.13044 | COADREAD | SNU-61 | MK-2206 |
| MVP_top60_DNA_seed | ATM | 0.804114 | COADREAD | RCM-1 | KU-55933 | MVP_top60_DNA+RNA_seed | KRAS | 1.130554 | COADREAD | NCI-H508 | KRAS (G12C) Inhibitor-1 |
| MVP_top60_DNA_seed | ATM | 0.808598 | COADREAD | SW1116 | KU-55933 | MVP_top60_DNA+RNA_seed | SRC | 1.137335 | COADREAD | LS-411N | Bosutinib |
| MVP_top60_DNA_seed | PIK3CA | 0.811062 | COADREAD | SW1417 | GNE-317 | MVP_top60_DNA+RNA_seed | AKT1 | 1.138447 | COADREAD | SW1417 | Afuresertib |
| MVP_top60_DNA_seed | BRAF | 0.812777 | COADREAD | SW620 | PLX-4720 | MVP_top60_DNA+RNA_seed | ERBB3 | 1.139987 | COADREAD | KM12 | Sapitinib |
| MVP_top60_DNA_seed | ERBB3 | 0.813782 | COADREAD | COLO-320-HSR | Sapitinib | MVP_top60_DNA+RNA_seed | AKT1 | 1.141984 | COADREAD | CL-11 | Ipatasertib |
| MVP_top60_DNA_seed | GSK3B | 0.815496 | COADREAD | C2BBe1 | SB216763 | MVP_top60_DNA+RNA_seed | IGF1R | 1.142881 | COADREAD | SNU-81 | BMS-536924 |
| MVP_top60_DNA_seed | IGF1R | 0.816213 | COADREAD | KM12 | GSK1904529A | MVP_top60_DNA+RNA_seed | PIK3CA | 1.142895 | COADREAD | LS-1034 | GNE-317 |
| MVP_top60_DNA_seed | ESR1 | 0.816365 | COADREAD | RCM-1 | GDC0810 | MVP_top60_DNA+RNA_seed | ERBB3 | 1.148021 | COADREAD | LS-1034 | Sapitinib |
| MVP_top60_DNA_seed | GSK3B | 0.816802 | COADREAD | SNU-61 | SB216763 | MVP_top60_DNA+RNA_seed | ERBB3 | 1.151692 | COADREAD | SW620 | Sapitinib |
| MVP_top60_DNA_seed | HSP90AA1 | 0.818692 | COADREAD | LS-1034 | Luminespib | MVP_top60_DNA+RNA_seed | CDK2 | 1.159001 | COADREAD | SW1463 | MK-8776 |
| MVP_top60_DNA_seed | HSP90AB1 | 0.818692 | COADREAD | LS-1034 | Luminespib | MVP_top60_DNA+RNA_seed | PIK3CA | 1.160921 | COADREAD | SNU-1040 | Alpelisib |
| MVP_top60_DNA_seed | CDK2 | 0.8187 | COADREAD | HT-115 | MK-8776 | MVP_top60_DNA+RNA_seed | IGF1R | 1.169089 | COADREAD | SNU-C2B | GSK1904529A |
| MVP_top60_DNA_seed | CDK2 | 0.82011 | COADREAD | SNU-61 | AZD5438 | MVP_top60_DNA+RNA_seed | IGF1R | 1.170627 | COADREAD | CL-11 | GSK1904529A |
| MVP_top60_DNA_seed | SRC | 0.820985 | COADREAD | LS-1034 | Bosutinib | MVP_top60_DNA+RNA_seed | AKT1 | 1.178653 | COADREAD | COLO-205 | AT13148 |
| MVP_top60_DNA_seed | CDK2 | 0.822609 | COADREAD | CaR-1 | Dinaciclib | MVP_top60_DNA+RNA_seed | BCL2 | 1.179634 | COADREAD | SW837 | Sabutoclax |
| MVP_top60_DNA_seed | AKT1 | 0.822871 | COADREAD | SW837 | Ipatasertib | MVP_top60_DNA+RNA_seed | IGF1R | 1.182053 | COADREAD | SW948 | GSK1904529A |
| MVP_top60_DNA_seed | MET | 0.823566 | COADREAD | SW1417 | Crizotinib | MVP_top60_DNA+RNA_seed | GSK3B | 1.190387 | COADREAD | SW948 | SB216763 |
| MVP_top60_DNA_seed | AKT1 | 0.823825 | COADREAD | CW-2 | AZD5363 | MVP_top60_DNA+RNA_seed | AKT1 | 1.191813 | COADREAD | SW837 | Afuresertib |
| MVP_top60_DNA_seed | AKT1 | 0.826853 | COADREAD | C2BBe1 | Uprosertib | MVP_top60_DNA+RNA_seed | AKT1 | 1.192618 | COADREAD | SW1463 | AZD5363 |
| MVP_top60_DNA_seed | IGF1R | 0.828099 | COADREAD | SNU-81 | BMS-754807 | MVP_top60_DNA+RNA_seed | HSP90AA1 | 1.192722 | COADREAD | COLO-205 | Luminespib |
| MVP_top60_DNA_seed | AKT1 | 0.828707 | COADREAD | NCI-H716 | Uprosertib | MVP_top60_DNA+RNA_seed | HSP90AB1 | 1.192722 | COADREAD | COLO-205 | Luminespib |
| MVP_top60_DNA_seed | ESR1 | 0.828887 | COADREAD | SW1116 | Tamoxifen | MVP_top60_DNA+RNA_seed | AKT1 | 1.193481 | COADREAD | SNU-C2B | Afuresertib |
| MVP_top60_DNA_seed | CREBBP | 0.833515 | COADREAD | SNU-C1 | SGC-CBP30 | MVP_top60_DNA+RNA_seed | IGF1R | 1.198687 | COADREAD | SW1417 | GSK1904529A |
| MVP_top60_DNA_seed | BRAF | 0.83446 | COADREAD | SNU-C1 | SB590885 | MVP_top60_DNA+RNA_seed | KRAS | 1.198722 | COADREAD | C2BBe1 | KRAS (G12C) Inhibitor-1 |
| MVP_top60_DNA_seed | SRC | 0.837197 | COADREAD | SW48 | Bosutinib | MVP_top60_DNA+RNA_seed | KRAS | 1.199411 | COADREAD | COLO-205 | KRAS (G12C) Inhibitor-1 |
| MVP_top60_DNA_seed | CREBBP | 0.837422 | COADREAD | SNU-407 | SGC-CBP30 | MVP_top60_DNA+RNA_seed | KRAS | 1.200548 | COADREAD | T84 | KRAS (G12C) Inhibitor-1 |
| MVP_top60_DNA_seed | BRAF | 0.838636 | COADREAD | KM12 | PLX-4720 | MVP_top60_DNA+RNA_seed | IGF1R | 1.211296 | COADREAD | SNU-C2B | BMS-536924 |
| MVP_top60_DNA_seed | BRAF | 0.839014 | COADREAD | GP5d | Dabrafenib | MVP_top60_DNA+RNA_seed | PIK3CA | 1.217937 | COADREAD | NCI-H716 | GNE-317 |
| MVP_top60_DNA_seed | CDK2 | 0.840863 | COADREAD | C2BBe1 | AZD5438 | MVP_top60_DNA+RNA_seed | AKT1 | 1.218117 | COADREAD | COLO-320-HSR | Uprosertib |
| MVP_top60_DNA_seed | PIK3CA | 0.843262 | COADREAD | SNU-61 | Alpelisib | MVP_top60_DNA+RNA_seed | BCL2 | 1.220206 | COADREAD | SNU-C1 | TW 37 |
| MVP_top60_DNA_seed | HSP90AA1 | 0.843682 | COADREAD | COLO-678 | Tanespimycin | MVP_top60_DNA+RNA_seed | CDK2 | 1.225324 | COADREAD | CaR-1 | AZD5438 |
| MVP_top60_DNA_seed | HSP90AB1 | 0.843682 | COADREAD | COLO-678 | Tanespimycin | MVP_top60_DNA+RNA_seed | KRAS | 1.230805 | COADREAD | SNU-81 | KRAS (G12C) Inhibitor-1 |
| MVP_top60_DNA_seed | AKT1 | 0.84644 | COADREAD | RCM-1 | Afuresertib | MVP_top60_DNA+RNA_seed | AKT1 | 1.232889 | COADREAD | CL-11 | GSK2110183B |
| MVP_top60_DNA_seed | PIK3CA | 0.846979 | COADREAD | SNU-C1 | Buparlisib | MVP_top60_DNA+RNA_seed | TP53 | 1.236705 | COADREAD | LS-123 | MIRA-1 |
| MVP_top60_DNA_seed | PIK3CB | 0.846979 | COADREAD | SNU-C1 | Buparlisib | MVP_top60_DNA+RNA_seed | AKT1 | 1.238996 | COADREAD | SW1417 | AZD5363 |
| MVP_top60_DNA_seed | PIK3CD | 0.846979 | COADREAD | SNU-C1 | Buparlisib | MVP_top60_DNA+RNA_seed | KRAS | 1.240603 | COADREAD | SNU-C1 | KRAS (G12C) Inhibitor-1 |
| MVP_top60_DNA_seed | PIK3CA | 0.849273 | COADREAD | SW837 | Buparlisib | MVP_top60_DNA+RNA_seed | TP53 | 1.244492 | COADREAD | SW1116 | MIRA-1 |

| | | | | | | | | | | | |
|---|---|---|---|---|---|---|---|---|---|---|---|
| MVP_top60_DNA_seed | PIK3CB | 0.849273 | COADREAD | SW837 | Buparlisib | MVP_top60_DNA+RNA_seed | PIK3CA | 1.245301 | COADREAD | CW-2 | AZD8186 |
| MVP_top60_DNA_seed | PIK3CD | 0.849273 | COADREAD | SW837 | Buparlisib | MVP_top60_DNA+RNA_seed | PIK3CB | 1.245301 | COADREAD | CW-2 | AZD8186 |
| MVP_top60_DNA_seed | AKT1 | 0.850724 | COADREAD | SNU-C2B | MK-2206 | MVP_top60_DNA+RNA_seed | HSP90AA1 | 1.250543 | COADREAD | SNU-81 | CCT-018159 |
| MVP_top60_DNA_seed | KRAS | 0.850839 | COADREAD | COLO-320-HSR | KRAS (G12C) Inhibitor-1 | MVP_top60_DNA+RNA_seed | HSP90AB1 | 1.250543 | COADREAD | SNU-81 | CCT-018159 |
| MVP_top60_DNA_seed | BRAF | 0.855065 | COADREAD | SNU-175 | PLX-4720 | MVP_top60_DNA+RNA_seed | BCL2 | 1.25087 | COADREAD | LS-123 | Venetoclax |
| MVP_top60_DNA_seed | AKT1 | 0.855748 | COADREAD | CaR-1 | Uprosertib | MVP_top60_DNA+RNA_seed | GSK3B | 1.25174 | COADREAD | CL-11 | SB216763 |
| MVP_top60_DNA_seed | HSP90AA1 | 0.859349 | COADREAD | LS-123 | Luminespib | MVP_top60_DNA+RNA_seed | PIK3CA | 1.253307 | COADREAD | RCM-1 | GNE-317 |
| MVP_top60_DNA_seed | HSP90AB1 | 0.859349 | COADREAD | LS-123 | Luminespib | MVP_top60_DNA+RNA_seed | CDK2 | 1.255163 | COADREAD | SW837 | AZD5438 |
| MVP_top60_DNA_seed | AKT1 | 0.865128 | COADREAD | SW1417 | AT13148 | MVP_top60_DNA+RNA_seed | BCL2 | 1.255489 | COADREAD | SW948 | TW 37 |
| MVP_top60_DNA_seed | IGF1R | 0.868323 | COADREAD | SW948 | BMS-536924 | MVP_top60_DNA+RNA_seed | PIK3CA | 1.257325 | COADREAD | SNU-61 | AZD8186 |
| MVP_top60_DNA_seed | PIK3CA | 0.87246 | COADREAD | SW948 | AZD8186 | MVP_top60_DNA+RNA_seed | PIK3CB | 1.257325 | COADREAD | SNU-61 | AZD8186 |
| MVP_top60_DNA_seed | PIK3CB | 0.87246 | COADREAD | SW948 | AZD8186 | MVP_top60_DNA+RNA_seed | GSK3B | 1.258602 | COADREAD | LS-411N | SB216763 |
| MVP_top60_DNA_seed | TP53 | 0.872614 | COADREAD | HT-29 | MIRA-1 | MVP_top60_DNA+RNA_seed | HSP90AA1 | 1.259702 | COADREAD | SNU-1040 | Luminespib |
| MVP_top60_DNA_seed | PIK3CA | 0.87469 | COADREAD | SW1463 | Alpelisib | MVP_top60_DNA+RNA_seed | HSP90AB1 | 1.259702 | COADREAD | SNU-1040 | Luminespib |
| MVP_top60_DNA_seed | AKT1 | 0.876086 | COADREAD | SK-CO-1 | Uprosertib | MVP_top60_DNA+RNA_seed | AKT1 | 1.261687 | COADREAD | HT-115 | Afuresertib |
| MVP_top60_DNA_seed | TP53 | 0.877092 | COADREAD | SW948 | MIRA-1 | MVP_top60_DNA+RNA_seed | MET | 1.264003 | COADREAD | SW1116 | Savolitinib |
| MVP_top60_DNA_seed | MET | 0.880076 | COADREAD | LS-411N | Savolitinib | MVP_top60_DNA+RNA_seed | PIK3CA | 1.270247 | COADREAD | SW1463 | Buparlisib |
| MVP_top60_DNA_seed | AKT1 | 0.880895 | COADREAD | C2BBe1 | GSK2110183B | MVP_top60_DNA+RNA_seed | PIK3CB | 1.270247 | COADREAD | SW1463 | Buparlisib |
| MVP_top60_DNA_seed | ERBB3 | 0.881579 | COADREAD | COLO-205 | Sapitinib | MVP_top60_DNA+RNA_seed | PIK3CD | 1.270247 | COADREAD | SW1463 | Buparlisib |
| MVP_top60_DNA_seed | AKT1 | 0.883365 | COADREAD | SW1116 | AZD5363 | MVP_top60_DNA+RNA_seed | BCL2 | 1.276438 | COADREAD | LoVo | TW 37 |
| MVP_top60_DNA_seed | AKT1 | 0.885174 | COADREAD | CaR-1 | GSK2110183B | MVP_top60_DNA+RNA_seed | KRAS | 1.278234 | COADREAD | RCM-1 | KRAS (G12C) Inhibitor-1 |
| MVP_top60_DNA_seed | IGF1R | 0.889294 | COADREAD | COLO-678 | BMS-536924 | MVP_top60_DNA+RNA_seed | HSP90AA1 | 1.28332 | COADREAD | NCI-H508 | CCT-018159 |
| MVP_top60_DNA_seed | TP53 | 0.892991 | COADREAD | SNU-407 | MIRA-1 | MVP_top60_DNA+RNA_seed | HSP90AB1 | 1.28332 | COADREAD | NCI-H508 | CCT-018159 |
| MVP_top60_DNA_seed | PIK3CA | 0.893689 | COADREAD | LS-1034 | Alpelisib | MVP_top60_DNA+RNA_seed | AKT1 | 1.283844 | COADREAD | SW837 | AZD5363 |
| MVP_top60_DNA_seed | MET | 0.897007 | COADREAD | C2BBe1 | Crizotinib | MVP_top60_DNA+RNA_seed | CDK2 | 1.285963 | COADREAD | NCI-H747 | MK-8776 |
| MVP_top60_DNA_seed | ERBB3 | 0.902886 | COADREAD | MDST8 | Sapitinib | MVP_top60_DNA+RNA_seed | BCL2 | 1.288961 | COADREAD | COLO-678 | Venetoclax |
| MVP_top60_DNA_seed | AKT1 | 0.904056 | COADREAD | RCM-1 | GSK2110183B | MVP_top60_DNA+RNA_seed | AKT1 | 1.290754 | COADREAD | SW1417 | Ipatasertib |
| MVP_top60_DNA_seed | MET | 0.907059 | COADREAD | SNU-61 | Crizotinib | MVP_top60_DNA+RNA_seed | BCL2 | 1.291165 | COADREAD | SNU-C1 | Obatoclax Mesylate |
| MVP_top60_DNA_seed | MET | 0.907575 | COADREAD | SNU-81 | Crizotinib | MVP_top60_DNA+RNA_seed | BCL2 | 1.297244 | COADREAD | SNU-C2B | Navitoclax |
| MVP_top60_DNA_seed | SRC | 0.908465 | COADREAD | C2BBe1 | Bosutinib | MVP_top60_DNA+RNA_seed | CDK2 | 1.298364 | COADREAD | SNU-61 | Dinaciclib |
| MVP_top60_DNA_seed | AKT1 | 0.912681 | COADREAD | LS-123 | Afuresertib | MVP_top60_DNA+RNA_seed | AKT1 | 1.298439 | COADREAD | CL-11 | MK-2206 |
| MVP_top60_DNA_seed | TP53 | 0.920579 | COADREAD | SNU-61 | MIRA-1 | MVP_top60_DNA+RNA_seed | BCL2 | 1.30063 | COADREAD | LS-123 | Sabutoclax |
| MVP_top60_DNA_seed | BRAF | 0.922635 | COADREAD | SW1116 | SB590885 | MVP_top60_DNA+RNA_seed | KRAS | 1.302596 | COADREAD | LS-180 | KRAS (G12C) Inhibitor-1 |
| MVP_top60_DNA_seed | SRC | 0.924767 | COADREAD | T84 | Bosutinib | MVP_top60_DNA+RNA_seed | AKT1 | 1.309442 | COADREAD | RCM-1 | MK-2206 |
| MVP_top60_DNA_seed | ESR1 | 0.930229 | COADREAD | SNU-61 | GDC0810 | MVP_top60_DNA+RNA_seed | IGF1R | 1.318148 | COADREAD | SNU-61 | BMS-754807 |
| MVP_top60_DNA_seed | IGF1R | 0.93072 | COADREAD | SW837 | BMS-536924 | MVP_top60_DNA+RNA_seed | IGF1R | 1.318668 | COADREAD | LS-123 | BMS-536924 |
| MVP_top60_DNA_seed | GSK3B | 0.930787 | COADREAD | KM12 | SB216763 | MVP_top60_DNA+RNA_seed | MET | 1.329244 | COADREAD | RCM-1 | Savolitinib |
| MVP_top60_DNA_seed | CDK2 | 0.936504 | COADREAD | SW1116 | MK-8776 | MVP_top60_DNA+RNA_seed | AKT1 | 1.332579 | COADREAD | SW1463 | Afuresertib |
| MVP_top60_DNA_seed | AKT1 | 0.938897 | COADREAD | SNU-61 | Ipatasertib | MVP_top60_DNA+RNA_seed | CDK2 | 1.340203 | COADREAD | SW948 | Dinaciclib |
| MVP_top60_DNA_seed | AKT1 | 0.941062 | COADREAD | SW837 | MK-2206 | MVP_top60_DNA+RNA_seed | CDK2 | 1.342173 | COADREAD | LS-1034 | Dinaciclib |
| MVP_top60_DNA_seed | AKT1 | 0.946644 | COADREAD | HT55 | GSK2110183B | MVP_top60_DNA+RNA_seed | HSP90AA1 | 1.348454 | COADREAD | SNU-81 | Elesclomol |

| | | | | | | | | | | | |
|---|---|---|---|---|---|---|---|---|---|---|---|
| MVP_top60_DNA_seed | BRAF | 0.949576 | COADREAD | SW48 | Dabrafenib | MVP_top60_DNA+RNA_seed | HSP90AB1 | 1.348454 | COADREAD | SNU-81 | Elesclomol |
| MVP_top60_DNA_seed | AKT1 | 0.952554 | COADREAD | SW1417 | Uprosertib | MVP_top60_DNA+RNA_seed | ATM | 1.357096 | COADREAD | SW1417 | KU-55933 |
| MVP_top60_DNA_seed | BRAF | 0.958502 | COADREAD | SW1463 | PLX-4720 | MVP_top60_DNA+RNA_seed | AKT1 | 1.358021 | COADREAD | RCM-1 | Ipatasertib |
| MVP_top60_DNA_seed | AKT1 | 0.963189 | COADREAD | CL-11 | Afuresertib | MVP_top60_DNA+RNA_seed | PIK3CA | 1.368682 | COADREAD | CL-11 | GNE-317 |
| MVP_top60_DNA_seed | ATM | 0.966502 | COADREAD | SNU-61 | KU-55933 | MVP_top60_DNA+RNA_seed | TP53 | 1.369924 | COADREAD | RCM-1 | MIRA-1 |
| MVP_top60_DNA_seed | AKT1 | 0.966865 | COADREAD | SNU-61 | AZD5363 | MVP_top60_DNA+RNA_seed | PIK3CA | 1.369951 | COADREAD | LS-411N | Alpelisib |
| MVP_top60_DNA_seed | PIK3CA | 0.97764 | COADREAD | SW1417 | Alpelisib | MVP_top60_DNA+RNA_seed | CDK2 | 1.375303 | COADREAD | LS-123 | AZD5438 |
| MVP_top60_DNA_seed | HSP90AA1 | 0.980103 | COADREAD | NCI-H508 | Luminespib | MVP_top60_DNA+RNA_seed | AKT1 | 1.377814 | COADREAD | SW1116 | MK-2206 |
| MVP_top60_DNA_seed | HSP90AB1 | 0.980103 | COADREAD | NCI-H508 | Luminespib | MVP_top60_DNA+RNA_seed | IGF1R | 1.385151 | COADREAD | SNU-81 | NVP-ADW742 |
| MVP_top60_DNA_seed | AKT1 | 0.983078 | COADREAD | HT-115 | Ipatasertib | MVP_top60_DNA+RNA_seed | PIK3CA | 1.385943 | COADREAD | SW620 | AZD8186 |
| MVP_top60_DNA_seed | PIK3CA | 0.98394 | COADREAD | SNU-61 | Buparlisib | MVP_top60_DNA+RNA_seed | PIK3CB | 1.385943 | COADREAD | SW620 | AZD8186 |
| MVP_top60_DNA_seed | PIK3CB | 0.98394 | COADREAD | SNU-61 | Buparlisib | MVP_top60_DNA+RNA_seed | AKT1 | 1.386481 | COADREAD | COLO-678 | Uprosertib |
| MVP_top60_DNA_seed | PIK3CD | 0.98394 | COADREAD | SNU-61 | Buparlisib | MVP_top60_DNA+RNA_seed | BCL2 | 1.386985 | COADREAD | SW1417 | Venetoclax |
| MVP_top60_DNA_seed | AKT1 | 0.986802 | COADREAD | SNU-C5 | AT13148 | MVP_top60_DNA+RNA_seed | PIK3CA | 1.393056 | COADREAD | SW1417 | Buparlisib |
| MVP_top60_DNA_seed | AKT1 | 0.987999 | COADREAD | SW1116 | GSK2110183B | MVP_top60_DNA+RNA_seed | PIK3CB | 1.393056 | COADREAD | SW1417 | Buparlisib |
| MVP_top60_DNA_seed | BRAF | 0.988845 | COADREAD | HCC2998 | PLX-4720 | MVP_top60_DNA+RNA_seed | PIK3CD | 1.393056 | COADREAD | SW1417 | Buparlisib |
| MVP_top60_DNA_seed | KRAS | 0.989445 | COADREAD | CL-11 | KRAS (G12C) Inhibitor-1 | MVP_top60_DNA+RNA_seed | AKT1 | 1.400861 | COADREAD | RCM-1 | AT13148 |
| MVP_top60_DNA_seed | AKT1 | 0.990615 | COADREAD | LS-1034 | AT13148 | MVP_top60_DNA+RNA_seed | HSP90AA1 | 1.401067 | COADREAD | HT-29 | CCT-018159 |
| MVP_top60_DNA_seed | IGF1R | 0.9931 | COADREAD | LS-123 | Linsitinib | MVP_top60_DNA+RNA_seed | HSP90AB1 | 1.401067 | COADREAD | HT-29 | CCT-018159 |
| MVP_top60_DNA_seed | CDK2 | 0.998336 | COADREAD | SNU-1040 | AZD5438 | MVP_top60_DNA+RNA_seed | AKT1 | 1.402894 | COADREAD | LS-123 | AT13148 |
| MVP_top60_DNA_seed | CREBBP | 1.009512 | COADREAD | SW48 | SGC-CBP30 | MVP_top60_DNA+RNA_seed | CDK2 | 1.407263 | COADREAD | SW837 | Dinaciclib |
| MVP_top60_DNA_seed | BRAF | 1.013611 | COADREAD | DiFi | Dabrafenib | MVP_top60_DNA+RNA_seed | PIK3CA | 1.412744 | COADREAD | SNU-81 | Alpelisib |
| MVP_top60_DNA_seed | HSP90AA1 | 1.016775 | COADREAD | CL-11 | Luminespib | MVP_top60_DNA+RNA_seed | IGF1R | 1.418309 | COADREAD | LS-123 | NVP-ADW742 |
| MVP_top60_DNA_seed | HSP90AB1 | 1.016775 | COADREAD | CL-11 | Luminespib | MVP_top60_DNA+RNA_seed | BCL2 | 1.419445 | COADREAD | SW1116 | Sabutoclax |
| MVP_top60_DNA_seed | ATM | 1.018111 | COADREAD | LS-411N | KU-55933 | MVP_top60_DNA+RNA_seed | IGF1R | 1.422658 | COADREAD | SW48 | GSK1904529A |
| MVP_top60_DNA_seed | CDK2 | 1.018869 | COADREAD | SW948 | MK-8776 | MVP_top60_DNA+RNA_seed | IGF1R | 1.42769 | COADREAD | SW1463 | GSK1904529A |
| MVP_top60_DNA_seed | AKT1 | 1.019547 | COADREAD | SNU-61 | Afuresertib | MVP_top60_DNA+RNA_seed | AKT1 | 1.429917 | COADREAD | CL-11 | AZD5363 |
| MVP_top60_DNA_seed | MET | 1.021199 | COADREAD | HT-29 | Savolitinib | MVP_top60_DNA+RNA_seed | AKT1 | 1.431469 | COADREAD | SW1116 | Uprosertib |
| MVP_top60_DNA_seed | CDK2 | 1.024019 | COADREAD | CW-2 | MK-8776 | MVP_top60_DNA+RNA_seed | BCL2 | 1.433531 | COADREAD | SNU-81 | Sabutoclax |
| MVP_top60_DNA_seed | AKT1 | 1.025358 | COADREAD | SW1463 | Uprosertib | MVP_top60_DNA+RNA_seed | MET | 1.43807 | COADREAD | SNU-C2B | Savolitinib |
| MVP_top60_DNA_seed | BRAF | 1.02829 | COADREAD | NCI-H508 | PLX-4720 | MVP_top60_DNA+RNA_seed | KRAS | 1.442541 | COADREAD | SW1463 | KRAS (G12C) Inhibitor-1 |
| MVP_top60_DNA_seed | ATM | 1.028655 | COADREAD | LS-180 | KU-55933 | MVP_top60_DNA+RNA_seed | HSP90AA1 | 1.443852 | COADREAD | HT-115 | Elesclomol |
| MVP_top60_DNA_seed | PIK3CA | 1.029105 | COADREAD | MDST8 | Alpelisib | MVP_top60_DNA+RNA_seed | HSP90AB1 | 1.443852 | COADREAD | HT-115 | Elesclomol |
| MVP_top60_DNA_seed | AKT1 | 1.030377 | COADREAD | SNU-C2B | Ipatasertib | MVP_top60_DNA+RNA_seed | BCL2 | 1.444813 | COADREAD | COLO-678 | Obatoclax Mesylate |
| MVP_top60_DNA_seed | HSP90AA1 | 1.033457 | COADREAD | SW1417 | Luminespib | MVP_top60_DNA+RNA_seed | AKT1 | 1.448984 | COADREAD | LS-123 | Ipatasertib |
| MVP_top60_DNA_seed | HSP90AB1 | 1.033457 | COADREAD | SW1417 | Luminespib | MVP_top60_DNA+RNA_seed | KRAS | 1.460753 | COADREAD | SNU-C2B | KRAS (G12C) Inhibitor-1 |
| MVP_top60_DNA_seed | PIK3CB | 1.042915 | COADREAD | HT-29 | AZD6482 | MVP_top60_DNA+RNA_seed | IGF1R | 1.462317 | COADREAD | SNU-61 | BMS-536924 |
| MVP_top60_DNA_seed | TP53 | 1.043725 | COADREAD | SNU-C5 | MIRA-1 | MVP_top60_DNA+RNA_seed | MET | 1.462578 | COADREAD | LS-123 | Savolitinib |
| MVP_top60_DNA_seed | TP53 | 1.048788 | COADREAD | SNU-C1 | MIRA-1 | MVP_top60_DNA+RNA_seed | AKT1 | 1.467641 | COADREAD | SW837 | Uprosertib |
| MVP_top60_DNA_seed | KRAS | 1.051334 | COADREAD | CCK-81 | KRAS (G12C) Inhibitor-1 | MVP_top60_DNA+RNA_seed | PIK3CA | 1.474853 | COADREAD | SW837 | Alpelisib |
| MVP_top60_DNA_seed | KRAS | 1.053421 | COADREAD | MDST8 | KRAS (G12C) Inhibitor-1 | MVP_top60_DNA+RNA_seed | HSP90AA1 | 1.483148 | COADREAD | SW1116 | Tanespimycin |

| | | | | | | | | | | | |
|---|---|---|---|---|---|---|---|---|---|---|---|
| MVP_top60_DNA_seed | PIK3CA | 1.054854 | COADREAD | SNU-C2B | Alpelisib | MVP_top60_DNA+RNA_seed | HSP90AB1 | 1.483148 | COADREAD | SW1116 | Tanespimycin |
| MVP_top60_DNA_seed | ERBB3 | 1.057069 | COADREAD | SW948 | Sapitinib | MVP_top60_DNA+RNA_seed | SRC | 1.484129 | COADREAD | CL-11 | Bosutinib |
| MVP_top60_DNA_seed | CREBBP | 1.058306 | COADREAD | SW1463 | SGC-CBP30 | MVP_top60_DNA+RNA_seed | BCL2 | 1.48434 | COADREAD | LS-180 | Sabutoclax |
| MVP_top60_DNA_seed | AKT1 | 1.065601 | COADREAD | SW837 | GSK2110183B | MVP_top60_DNA+RNA_seed | PIK3CA | 1.489188 | COADREAD | KM12 | Alpelisib |
| MVP_top60_DNA_seed | HSP90AA1 | 1.066249 | COADREAD | C2BBe1 | Tanespimycin | MVP_top60_DNA+RNA_seed | PIK3CA | 1.493804 | COADREAD | RKO | GNE-317 |
| MVP_top60_DNA_seed | HSP90AB1 | 1.066249 | COADREAD | C2BBe1 | Tanespimycin | MVP_top60_DNA+RNA_seed | AKT1 | 1.498516 | COADREAD | LS-123 | GSK2110183B |
| MVP_top60_DNA_seed | PIK3CA | 1.068569 | COADREAD | SNU-C2B | AZD8186 | MVP_top60_DNA+RNA_seed | PIK3CA | 1.499855 | COADREAD | LS-1034 | Buparlisib |
| MVP_top60_DNA_seed | PIK3CB | 1.068569 | COADREAD | SNU-C2B | AZD8186 | MVP_top60_DNA+RNA_seed | PIK3CB | 1.499855 | COADREAD | LS-1034 | Buparlisib |
| MVP_top60_DNA_seed | IGF1R | 1.071973 | COADREAD | LS-180 | BMS-536924 | MVP_top60_DNA+RNA_seed | PIK3CD | 1.499855 | COADREAD | LS-1034 | Buparlisib |
| MVP_top60_DNA_seed | TP53 | 1.072246 | COADREAD | SW837 | MIRA-1 | MVP_top60_DNA+RNA_seed | BCL2 | 1.503152 | COADREAD | SW948 | Obatoclax Mesylate |
| MVP_top60_DNA_seed | AKT1 | 1.073691 | COADREAD | SNU-C2B | AZD5363 | MVP_top60_DNA+RNA_seed | AKT1 | 1.507258 | COADREAD | SW837 | Uprosertib |
| MVP_top60_DNA_seed | CDK2 | 1.074076 | COADREAD | SW837 | MK-8776 | MVP_top60_DNA+RNA_seed | AKT1 | 1.507316 | COADREAD | KM12 | Uprosertib |
| MVP_top60_DNA_seed | ESR1 | 1.076816 | COADREAD | NCI-H716 | GDC0810 | MVP_top60_DNA+RNA_seed | SRC | 1.508488 | COADREAD | SW620 | Bosutinib |
| MVP_top60_DNA_seed | GSK3B | 1.078365 | COADREAD | CaR-1 | SB216763 | MVP_top60_DNA+RNA_seed | AKT1 | 1.510797 | COADREAD | SW1463 | AT13148 |
| MVP_top60_DNA_seed | CREBBP | 1.083281 | COADREAD | SNU-1040 | SGC-CBP30 | MVP_top60_DNA+RNA_seed | GSK3B | 1.511319 | COADREAD | SNU-1040 | SB216763 |
| MVP_top60_DNA_seed | TP53 | 1.083472 | COADREAD | NCI-H716 | MIRA-1 | MVP_top60_DNA+RNA_seed | BCL2 | 1.511484 | COADREAD | T84 | Sabutoclax |
| MVP_top60_DNA_seed | TP53 | 1.084915 | COADREAD | SW620 | MIRA-1 | MVP_top60_DNA+RNA_seed | AKT1 | 1.512685 | COADREAD | RCM-1 | AZD5363 |
| MVP_top60_DNA_seed | ERBB3 | 1.08603 | COADREAD | HCC2998 | Sapitinib | MVP_top60_DNA+RNA_seed | MET | 1.526885 | COADREAD | COLO-678 | Crizotinib |
| MVP_top60_DNA_seed | BRAF | 1.089549 | COADREAD | C2BBe1 | Dabrafenib | MVP_top60_DNA+RNA_seed | CDK2 | 1.537395 | COADREAD | CL-11 | AZD5438 |
| MVP_top60_DNA_seed | ESR1 | 1.090528 | COADREAD | SNU-C1 | GDC0810 | MVP_top60_DNA+RNA_seed | PIK3CA | 1.548009 | COADREAD | LS-411N | AZD8186 |
| MVP_top60_DNA_seed | BRAF | 1.094671 | COADREAD | SNU-61 | PLX-4720 | MVP_top60_DNA+RNA_seed | PIK3CB | 1.548009 | COADREAD | LS-411N | AZD8186 |
| MVP_top60_DNA_seed | CDK2 | 1.095237 | COADREAD | HT55 | MK-8776 | MVP_top60_DNA+RNA_seed | AKT1 | 1.5539 | COADREAD | SW620 | AZD5363 |
| MVP_top60_DNA_seed | BRAF | 1.104791 | COADREAD | SNU-1040 | PLX-4720 | MVP_top60_DNA+RNA_seed | AKT1 | 1.568909 | COADREAD | SW1116 | Ipatasertib |
| MVP_top60_DNA_seed | BRAF | 1.108184 | COADREAD | SNU-407 | PLX-4720 | MVP_top60_DNA+RNA_seed | AKT1 | 1.580196 | COADREAD | SW837 | AT13148 |
| MVP_top60_DNA_seed | ERBB3 | 1.108986 | COADREAD | SNU-C5 | Sapitinib | MVP_top60_DNA+RNA_seed | HSP90AA1 | 1.584491 | COADREAD | SW1463 | Luminespib |
| MVP_top60_DNA_seed | KRAS | 1.118692 | COADREAD | SW837 | KRAS (G12C) Inhibitor-12 | MVP_top60_DNA+RNA_seed | HSP90AB1 | 1.584491 | COADREAD | SW1463 | Luminespib |
| MVP_top60_DNA_seed | KRAS | 1.119802 | COADREAD | LS-1034 | KRAS (G12C) Inhibitor-12 | MVP_top60_DNA+RNA_seed | AKT1 | 1.593113 | COADREAD | SW620 | Ipatasertib |
| MVP_top60_DNA_seed | KRAS | 1.127681 | COADREAD | SNU-61 | KRAS (G12C) Inhibitor-12 | MVP_top60_DNA+RNA_seed | AKT1 | 1.593433 | COADREAD | SNU-C2B | GSK2110183B |
| MVP_top60_DNA_seed | AKT1 | 1.128005 | COADREAD | SK-CO-1 | Uprosertib | MVP_top60_DNA+RNA_seed | PIK3CA | 1.603978 | COADREAD | SNU-C1 | AZD8186 |
| MVP_top60_DNA_seed | IGF1R | 1.128152 | COADREAD | RKO | NVP-ADW742 | MVP_top60_DNA+RNA_seed | PIK3CB | 1.603978 | COADREAD | SNU-C1 | AZD8186 |
| MVP_top60_DNA_seed | AKT1 | 1.13044 | COADREAD | SNU-61 | MK-2206 | MVP_top60_DNA+RNA_seed | KRAS | 1.604202 | COADREAD | SW1116 | KRAS (G12C) Inhibitor-1 |
| MVP_top60_DNA_seed | KRAS | 1.130554 | COADREAD | NCI-H508 | KRAS (G12C) Inhibitor-12 | MVP_top60_DNA+RNA_seed | CDK2 | 1.627002 | COADREAD | CL-11 | Dinaciclib |
| MVP_top60_DNA_seed | SRC | 1.137335 | COADREAD | LS-411N | Bosutinib | MVP_top60_DNA+RNA_seed | HSP90AA1 | 1.630206 | COADREAD | SW837 | Luminespib |
| MVP_top60_DNA_seed | AKT1 | 1.138447 | COADREAD | SW1417 | Afuresertib | MVP_top60_DNA+RNA_seed | HSP90AB1 | 1.630206 | COADREAD | SW837 | Luminespib |
| MVP_top60_DNA_seed | ERBB3 | 1.139987 | COADREAD | KM12 | Sapitinib | MVP_top60_DNA+RNA_seed | ATM | 1.637887 | COADREAD | LS-123 | KU-55933 |
| MVP_top60_DNA_seed | AKT1 | 1.141984 | COADREAD | CL-11 | Ipatasertib | MVP_top60_DNA+RNA_seed | ERBB3 | 1.651232 | COADREAD | LS-411N | Sapitinib |
| MVP_top60_DNA_seed | IGF1R | 1.142881 | COADREAD | SNU-81 | BMS-536924 | MVP_top60_DNA+RNA_seed | PIK3CA | 1.652514 | COADREAD | LS-1034 | AZD8186 |
| MVP_top60_DNA_seed | PIK3CA | 1.142895 | COADREAD | LS-1034 | GNE-317 | MVP_top60_DNA+RNA_seed | PIK3CB | 1.652514 | COADREAD | LS-1034 | AZD8186 |
| MVP_top60_DNA_seed | CREBBP | 1.143201 | COADREAD | LS-180 | SGC-CBP30 | MVP_top60_DNA+RNA_seed | PIK3CB | 1.65465 | COADREAD | SW620 | AZD6482 |
| MVP_top60_DNA_seed | ERBB3 | 1.148021 | COADREAD | LS-1034 | Sapitinib | MVP_top60_DNA+RNA_seed | AKT1 | 1.659385 | COADREAD | SW1463 | Uprosertib |
| MVP_top60_DNA_seed | ERBB3 | 1.151692 | COADREAD | SW620 | Sapitinib | MVP_top60_DNA+RNA_seed | BCL2 | 1.661613 | COADREAD | SW1417 | Sabutoclax |

| | | | | | | | | | | | |
|---|---|---|---|---|---|---|---|---|---|---|---|
| MVP_top60_DNA_seed | ESR1 | 1.152723 | COADREAD | SW620 | GDC0810 | MVP_top60_DNA+RNA_seed | TP53 | 1.663562 | COADREAD | C2BBe1 | MIRA-1 |
| MVP_top60_DNA_seed | CDK2 | 1.159001 | COADREAD | SW1463 | MK-8776 | MVP_top60_DNA+RNA_seed | CDK2 | 1.669117 | COADREAD | RCM-1 | MK-8776 |
| MVP_top60_DNA_seed | PIK3CA | 1.160921 | COADREAD | SNU-1040 | Alpelisib | MVP_top60_DNA+RNA_seed | IGF1R | 1.670378 | COADREAD | SW1116 | NVP-ADW742 |
| MVP_top60_DNA_seed | BRAF | 1.162674 | COADREAD | LS-1034 | PLX-4720 | MVP_top60_DNA+RNA_seed | PIK3CA | 1.670726 | COADREAD | KM12 | AZD8186 |
| MVP_top60_DNA_seed | IGF1R | 1.169089 | COADREAD | SNU-C2B | GSK1904529A | MVP_top60_DNA+RNA_seed | PIK3CB | 1.670726 | COADREAD | KM12 | AZD8186 |
| MVP_top60_DNA_seed | IGF1R | 1.170627 | COADREAD | CL-11 | GSK1904529A | MVP_top60_DNA+RNA_seed | AKT1 | 1.690126 | COADREAD | SW1116 | AT13148 |
| MVP_top60_DNA_seed | AKT1 | 1.178653 | COADREAD | COLO-205 | AT13148 | MVP_top60_DNA+RNA_seed | IGF1R | 1.698916 | COADREAD | SW1417 | NVP-ADW742 |
| MVP_top60_DNA_seed | IGF1R | 1.182053 | COADREAD | SW948 | GSK1904529A | MVP_top60_DNA+RNA_seed | AKT1 | 1.717056 | COADREAD | LS-1034 | AZD5363 |
| MVP_top60_DNA_seed | GSK3B | 1.190387 | COADREAD | SW948 | SB216763 | MVP_top60_DNA+RNA_seed | TP53 | 1.727207 | COADREAD | NCI-H747 | MIRA-1 |
| MVP_top60_DNA_seed | AKT1 | 1.191813 | COADREAD | SW837 | Afuresertib | MVP_top60_DNA+RNA_seed | CDK2 | 1.733373 | COADREAD | NCI-H747 | AZD5438 |
| MVP_top60_DNA_seed | AKT1 | 1.192618 | COADREAD | SW1463 | AZD5363 | MVP_top60_DNA+RNA_seed | MET | 1.734326 | COADREAD | COLO-678 | Savolitinib |
| MVP_top60_DNA_seed | HSP90AA1 | 1.192722 | COADREAD | COLO-205 | Luminespib | MVP_top60_DNA+RNA_seed | AKT1 | 1.73635 | COADREAD | LS-1034 | Uprosertib |
| MVP_top60_DNA_seed | HSP90AB1 | 1.192722 | COADREAD | COLO-205 | Luminespib | MVP_top60_DNA+RNA_seed | BCL2 | 1.759343 | COADREAD | SNU-61 | Sabutoclax |
| MVP_top60_DNA_seed | AKT1 | 1.193481 | COADREAD | SNU-C2B | Afuresertib | MVP_top60_DNA+RNA_seed | HSP90AA1 | 1.759718 | COADREAD | CL-11 | Tanespimycin |
| MVP_top60_DNA_seed | IGF1R | 1.198687 | COADREAD | SW1417 | GSK1904529A | MVP_top60_DNA+RNA_seed | HSP90AB1 | 1.759718 | COADREAD | CL-11 | Tanespimycin |
| MVP_top60_DNA_seed | KRAS | 1.198722 | COADREAD | C2BBe1 | KRAS (G12C) Inhibitor-12 | MVP_top60_DNA+RNA_seed | ATM | 1.762167 | COADREAD | SNU-C2B | KU-55933 |
| MVP_top60_DNA_seed | KRAS | 1.199411 | COADREAD | COLO-205 | KRAS (G12C) Inhibitor-12 | MVP_top60_DNA+RNA_seed | ERBB3 | 1.768894 | COADREAD | SNU-C1 | Sapitinib |
| MVP_top60_DNA_seed | KRAS | 1.200548 | COADREAD | T84 | KRAS (G12C) Inhibitor-12 | MVP_top60_DNA+RNA_seed | AKT1 | 1.769358 | COADREAD | SW1116 | Uprosertib |
| MVP_top60_DNA_seed | BRAF | 1.202437 | COADREAD | SW837 | Dabrafenib | MVP_top60_DNA+RNA_seed | PIK3CA | 1.771889 | COADREAD | SW1116 | Buparlisib |
| MVP_top60_DNA_seed | IGF1R | 1.211296 | COADREAD | SNU-C2B | BMS-536924 | MVP_top60_DNA+RNA_seed | PIK3CB | 1.771889 | COADREAD | SW1116 | Buparlisib |
| MVP_top60_DNA_seed | MAPK1 | 1.215319 | COADREAD | COLO-320-HSR | SCH772984 | MVP_top60_DNA+RNA_seed | PIK3CD | 1.771889 | COADREAD | SW1116 | Buparlisib |
| MVP_top60_DNA_seed | PIK3CA | 1.217937 | COADREAD | NCI-H716 | GNE-317 | MVP_top60_DNA+RNA_seed | AKT1 | 1.774513 | COADREAD | SW620 | Uprosertib |
| MVP_top60_DNA_seed | AKT1 | 1.218117 | COADREAD | COLO-320-HSR | Uprosertib | MVP_top60_DNA+RNA_seed | GSK3B | 1.774967 | COADREAD | SNU-C2B | SB216763 |
| MVP_top60_DNA_seed | CDK2 | 1.225324 | COADREAD | CaR-1 | AZD5438 | MVP_top60_DNA+RNA_seed | AKT1 | 1.775962 | COADREAD | HT-115 | AZD5363 |
| MVP_top60_DNA_seed | KRAS | 1.230805 | COADREAD | SNU-81 | KRAS (G12C) Inhibitor-12 | MVP_top60_DNA+RNA_seed | BCL2 | 1.777162 | COADREAD | LS-1034 | Sabutoclax |
| MVP_top60_DNA_seed | AKT1 | 1.232889 | COADREAD | CL-11 | GSK2110183B | MVP_top60_DNA+RNA_seed | HSP90AA1 | 1.784675 | COADREAD | SNU-C2B | Tanespimycin |
| MVP_top60_DNA_seed | TP53 | 1.236705 | COADREAD | LS-123 | MIRA-1 | MVP_top60_DNA+RNA_seed | HSP90AB1 | 1.784675 | COADREAD | SNU-C2B | Tanespimycin |
| MVP_top60_DNA_seed | AKT1 | 1.238996 | COADREAD | SW1417 | AZD5363 | MVP_top60_DNA+RNA_seed | AKT1 | 1.799962 | COADREAD | LS-123 | Uprosertib |
| MVP_top60_DNA_seed | KRAS | 1.240603 | COADREAD | SNU-C1 | KRAS (G12C) Inhibitor-12 | MVP_top60_DNA+RNA_seed | GSK3B | 1.806404 | COADREAD | NCI-H716 | SB216763 |
| MVP_top60_DNA_seed | BRAF | 1.244345 | COADREAD | LS-123 | Dabrafenib | MVP_top60_DNA+RNA_seed | GSK3B | 1.817415 | COADREAD | HT-29 | CHIR-99021 |
| MVP_top60_DNA_seed | TP53 | 1.244492 | COADREAD | SW1116 | MIRA-1 | MVP_top60_DNA+RNA_seed | AKT1 | 1.830566 | COADREAD | SW1417 | GSK2110183B |
| MVP_top60_DNA_seed | PIK3CA | 1.245301 | COADREAD | CW-2 | AZD8186 | MVP_top60_DNA+RNA_seed | MET | 1.834107 | COADREAD | SW620 | Savolitinib |
| MVP_top60_DNA_seed | PIK3CB | 1.245301 | COADREAD | CW-2 | AZD8186 | MVP_top60_DNA+RNA_seed | AKT1 | 1.834918 | COADREAD | LS-123 | AZD5363 |
| MVP_top60_DNA_seed | MAPK1 | 1.250228 | COADREAD | SW1116 | Ulixertinib | MVP_top60_DNA+RNA_seed | PIK3CA | 1.863039 | COADREAD | CW-2 | GNE-317 |
| MVP_top60_DNA_seed | HSP90AA1 | 1.250543 | COADREAD | SNU-81 | CCT-018159 | MVP_top60_DNA+RNA_seed | AR | 1.872618 | COADREAD | HT-29 | Bicalutamide |
| MVP_top60_DNA_seed | HSP90AB1 | 1.250543 | COADREAD | SNU-81 | CCT-018159 | MVP_top60_DNA+RNA_seed | HSP90AA1 | 1.876363 | COADREAD | SNU-61 | Luminespib |
| MVP_top60_DNA_seed | GSK3B | 1.25174 | COADREAD | CL-11 | SB216763 | MVP_top60_DNA+RNA_seed | HSP90AB1 | 1.876363 | COADREAD | SNU-61 | Luminespib |
| MVP_top60_DNA_seed | PIK3CA | 1.253307 | COADREAD | RCM-1 | GNE-317 | MVP_top60_DNA+RNA_seed | PIK3CA | 1.876554 | COADREAD | SW1116 | GNE-317 |
| MVP_top60_DNA_seed | CDK2 | 1.255163 | COADREAD | SW837 | AZD5438 | MVP_top60_DNA+RNA_seed | CDK2 | 1.878235 | COADREAD | C2BBe1 | MK-8776 |
| MVP_top60_DNA_seed | PIK3CA | 1.257325 | COADREAD | SNU-61 | AZD8186 | MVP_top60_DNA+RNA_seed | CDK2 | 1.88685 | COADREAD | SW1116 | Dinaciclib |
| MVP_top60_DNA_seed | PIK3CB | 1.257325 | COADREAD | SNU-61 | AZD8186 | MVP_top60_DNA+RNA_seed | BCL2 | 1.901099 | COADREAD | COLO-678 | Sabutoclax |

| | | | | | | | | | | | |
|---|---|---|---|---|---|---|---|---|---|---|---|
| MVP_top60_DNA_seed | GSK3B | 1.258602 | COADREAD | LS-411N | SB216763 | MVP_top60_DNA+RNA_seed | CDK2 | 1.907372 | COADREAD | LS-123 | MK-8776 |
| MVP_top60_DNA_seed | HSP90AA1 | 1.259702 | COADREAD | SNU-1040 | Luminespib | MVP_top60_DNA+RNA_seed | PIK3CA | 1.907945 | COADREAD | COLO-678 | AZD8186 |
| MVP_top60_DNA_seed | HSP90AB1 | 1.259702 | COADREAD | SNU-1040 | Luminespib | MVP_top60_DNA+RNA_seed | PIK3CB | 1.907945 | COADREAD | COLO-678 | AZD8186 |
| MVP_top60_DNA_seed | AKT1 | 1.261687 | COADREAD | HT-115 | Afuresertib | MVP_top60_DNA+RNA_seed | IGF1R | 1.908475 | COADREAD | COLO-678 | Linsitinib |
| MVP_top60_DNA_seed | MET | 1.264003 | COADREAD | SW1116 | Savolitinib | MVP_top60_DNA+RNA_seed | AKT1 | 1.910749 | COADREAD | SW620 | Afuresertib |
| MVP_top60_DNA_seed | PIK3CA | 1.270247 | COADREAD | SW1463 | Buparlisib | MVP_top60_DNA+RNA_seed | AKT1 | 1.913541 | COADREAD | SW1417 | MK-2206 |
| MVP_top60_DNA_seed | PIK3CB | 1.270247 | COADREAD | SW1463 | Buparlisib | MVP_top60_DNA+RNA_seed | CDK2 | 1.949386 | COADREAD | SW620 | MK-8776 |
| MVP_top60_DNA_seed | PIK3CD | 1.270247 | COADREAD | SW1463 | Buparlisib | MVP_top60_DNA+RNA_seed | HSP90AA1 | 1.957433 | COADREAD | SW948 | Luminespib |
| MVP_top60_DNA_seed | CREBBP | 1.278118 | COADREAD | LS-411N | SGC-CBP30 | MVP_top60_DNA+RNA_seed | HSP90AB1 | 1.957433 | COADREAD | SW948 | Luminespib |
| MVP_top60_DNA_seed | KRAS | 1.278234 | COADREAD | RCM-1 | KRAS (G12C) Inhibitor-12 | MVP_top60_DNA+RNA_seed | AKT1 | 1.960755 | COADREAD | SW1116 | Afuresertib |
| MVP_top60_DNA_seed | HSP90AA1 | 1.28332 | COADREAD | NCI-H508 | CCT-018159 | MVP_top60_DNA+RNA_seed | IGF1R | 1.961416 | COADREAD | LS-123 | GSK1904529A |
| MVP_top60_DNA_seed | HSP90AB1 | 1.28332 | COADREAD | NCI-H508 | CCT-018159 | MVP_top60_DNA+RNA_seed | PIK3CA | 1.98122 | COADREAD | SW1116 | AZD8186 |
| MVP_top60_DNA_seed | AKT1 | 1.283844 | COADREAD | SW837 | AZD5363 | MVP_top60_DNA+RNA_seed | PIK3CB | 1.98122 | COADREAD | SW1116 | AZD8186 |
| MVP_top60_DNA_seed | CDK2 | 1.285963 | COADREAD | NCI-H747 | MK-8776 | MVP_top60_DNA+RNA_seed | HSP90AA1 | 1.982823 | COADREAD | COLO-678 | Luminespib |
| MVP_top60_DNA_seed | BRAF | 1.289872 | COADREAD | SNU-C2B | SB590885 | MVP_top60_DNA+RNA_seed | HSP90AB1 | 1.982823 | COADREAD | COLO-678 | Luminespib |
| MVP_top60_DNA_seed | AKT1 | 1.290754 | COADREAD | SW1417 | Ipatasertib | MVP_top60_DNA+RNA_seed | ERBB3 | 1.989226 | COADREAD | CL-11 | Sapitinib |
| MVP_top60_DNA_seed | CDK2 | 1.298364 | COADREAD | SNU-61 | Dinaciclib | MVP_top60_DNA+RNA_seed | SRC | 1.996035 | COADREAD | SNU-C2B | Bosutinib |
| MVP_top60_DNA_seed | AKT1 | 1.298439 | COADREAD | CL-11 | MK-2206 | MVP_top60_DNA+RNA_seed | BCL2 | 1.999939 | COADREAD | SW948 | Sabutoclax |
| MVP_top60_DNA_seed | KRAS | 1.302596 | COADREAD | LS-180 | KRAS (G12C) Inhibitor-12 | MVP_top60_DNA+RNA_seed | KRAS | 2.035954 | COADREAD | LS-123 | KRAS (G12C) Inhibitor-1 |
| MVP_top60_DNA_seed | AKT1 | 1.309442 | COADREAD | RCM-1 | MK-2206 | MVP_top60_DNA+RNA_seed | TP53 | 2.051736 | COADREAD | SNU-C2B | MIRA-1 |
| MVP_top60_DNA_seed | IGF1R | 1.318148 | COADREAD | SNU-61 | BMS-754807 | MVP_top60_DNA+RNA_seed | CDK2 | 2.099169 | COADREAD | COLO-678 | Dinaciclib |
| MVP_top60_DNA_seed | IGF1R | 1.318668 | COADREAD | LS-123 | BMS-536924 | MVP_top60_DNA+RNA_seed | BCL2 | 2.1292 | COADREAD | SW1463 | Sabutoclax |
| MVP_top60_DNA_seed | BRAF | 1.329114 | COADREAD | SW1116 | Dabrafenib | MVP_top60_DNA+RNA_seed | BCL2 | 2.13624 | COADREAD | SNU-1040 | Obatoclax Mesylate |
| MVP_top60_DNA_seed | MET | 1.329244 | COADREAD | RCM-1 | Savolitinib | MVP_top60_DNA+RNA_seed | AKT1 | 2.147336 | COADREAD | SW620 | MK-2206 |
| MVP_top60_DNA_seed | AKT1 | 1.332579 | COADREAD | SW1463 | Afuresertib | MVP_top60_DNA+RNA_seed | HSP90AA1 | 2.148666 | COADREAD | DiFi | Luminespib |
| MVP_top60_DNA_seed | CREBBP | 1.340021 | COADREAD | RCM-1 | SGC-CBP30 | MVP_top60_DNA+RNA_seed | HSP90AB1 | 2.148666 | COADREAD | DiFi | Luminespib |
| MVP_top60_DNA_seed | CDK2 | 1.340203 | COADREAD | SW948 | Dinaciclib | MVP_top60_DNA+RNA_seed | CDK2 | 2.170457 | COADREAD | LS-123 | Dinaciclib |
| MVP_top60_DNA_seed | CDK2 | 1.342173 | COADREAD | LS-1034 | Dinaciclib | MVP_top60_DNA+RNA_seed | IGF1R | 2.193558 | COADREAD | C2BBe1 | BMS-536924 |
| MVP_top60_DNA_seed | BRAF | 1.347972 | COADREAD | LS-180 | PLX-4720 | MVP_top60_DNA+RNA_seed | AKT1 | 2.212207 | COADREAD | SW620 | GSK2110183B |
| MVP_top60_DNA_seed | HSP90AA1 | 1.348454 | COADREAD | SNU-81 | Elesclomol | MVP_top60_DNA+RNA_seed | GSK3B | 2.214252 | COADREAD | SW620 | CHIR-99021 |
| MVP_top60_DNA_seed | HSP90AB1 | 1.348454 | COADREAD | SNU-81 | Elesclomol | MVP_top60_DNA+RNA_seed | CDK2 | 2.22528 | COADREAD | KM12 | AZD5438 |
| MVP_top60_DNA_seed | ATM | 1.357096 | COADREAD | SW1417 | KU-55933 | MVP_top60_DNA+RNA_seed | GSK3B | 2.225873 | COADREAD | SW620 | SB216763 |
| MVP_top60_DNA_seed | AKT1 | 1.358021 | COADREAD | RCM-1 | Ipatasertib | MVP_top60_DNA+RNA_seed | BCL2 | 2.238148 | COADREAD | LS-513 | TW 37 |
| MVP_top60_DNA_seed | CREBBP | 1.3653 | COADREAD | NCI-H716 | SGC-CBP30 | MVP_top60_DNA+RNA_seed | CDK2 | 2.242149 | COADREAD | SW1417 | MK-8776 |
| MVP_top60_DNA_seed | PIK3CA | 1.368682 | COADREAD | CL-11 | GNE-317 | MVP_top60_DNA+RNA_seed | AKT1 | 2.296319 | COADREAD | COLO-678 | GSK2110183B |
| MVP_top60_DNA_seed | TP53 | 1.369924 | COADREAD | RCM-1 | MIRA-1 | MVP_top60_DNA+RNA_seed | PIK3CA | 2.296675 | COADREAD | RCM-1 | Buparlisib |
| MVP_top60_DNA_seed | PIK3CA | 1.369951 | COADREAD | LS-411N | Alpelisib | MVP_top60_DNA+RNA_seed | PIK3CB | 2.296675 | COADREAD | RCM-1 | Buparlisib |
| MVP_top60_DNA_seed | CDK2 | 1.375303 | COADREAD | LS-123 | AZD5438 | MVP_top60_DNA+RNA_seed | PIK3CD | 2.296675 | COADREAD | RCM-1 | Buparlisib |
| MVP_top60_DNA_seed | AKT1 | 1.377814 | COADREAD | SW1116 | MK-2206 | MVP_top60_DNA+RNA_seed | BCL2 | 2.306806 | COADREAD | SNU-81 | Obatoclax Mesylate |
| MVP_top60_DNA_seed | ESR1 | 1.378446 | COADREAD | SNU-81 | Tamoxifen | MVP_top60_DNA+RNA_seed | HSP90AA1 | 2.312686 | COADREAD | SNU-81 | Luminespib |
| MVP_top60_DNA_seed | IGF1R | 1.385151 | COADREAD | SNU-81 | NVP-ADW742 | MVP_top60_DNA+RNA_seed | HSP90AB1 | 2.312686 | COADREAD | SNU-81 | Luminespib |

| | | | | | | | | | | | |
|---|---|---|---|---|---|---|---|---|---|---|---|
| MVP_top60_DNA_seed | PIK3CA | 1.385943 | COADREAD | SW620 | AZD8186 | MVP_top60_DNA+RNA_seed | CDK2 | 2.369786 | COADREAD | COLO-678 | AZD5438 |
| MVP_top60_DNA_seed | PIK3CB | 1.385943 | COADREAD | SW620 | AZD8186 | MVP_top60_DNA+RNA_seed | ATM | 2.380035 | COADREAD | SW620 | KU-55933 |
| MVP_top60_DNA_seed | AKT1 | 1.386481 | COADREAD | COLO-678 | Uprosertib | MVP_top60_DNA+RNA_seed | AKT1 | 2.399471 | COADREAD | COLO-678 | Uprosertib |
| MVP_top60_DNA_seed | PIK3CA | 1.393056 | COADREAD | SW1417 | Buparlisib | MVP_top60_DNA+RNA_seed | KRAS | 2.416192 | COADREAD | SW1417 | KRAS (G12C) Inhibitor-1 |
| MVP_top60_DNA_seed | PIK3CB | 1.393056 | COADREAD | SW1417 | Buparlisib | MVP_top60_DNA+RNA_seed | MET | 2.443539 | COADREAD | SW1417 | Savolitinib |
| MVP_top60_DNA_seed | PIK3CD | 1.393056 | COADREAD | SW1417 | Buparlisib | MVP_top60_DNA+RNA_seed | AKT1 | 2.456908 | COADREAD | SW620 | Uprosertib |
| MVP_top60_DNA_seed | CREBBP | 1.395471 | COADREAD | SNU-C2B | SGC-CBP30 | MVP_top60_DNA+RNA_seed | TP53 | 2.502555 | COADREAD | SW1417 | MIRA-1 |
| MVP_top60_DNA_seed | AKT1 | 1.400861 | COADREAD | RCM-1 | AT13148 | MVP_top60_DNA+RNA_seed | PIK3CA | 2.581429 | COADREAD | COLO-678 | Alpelisib |
| MVP_top60_DNA_seed | HSP90AA1 | 1.401067 | COADREAD | HT-29 | CCT-018159 | MVP_top60_DNA+RNA_seed | CDK2 | 2.585337 | COADREAD | COLO-678 | MK-8776 |
| MVP_top60_DNA_seed | HSP90AB1 | 1.401067 | COADREAD | HT-29 | CCT-018159 | MVP_top60_DNA+RNA_seed | GSK3B | 2.613515 | COADREAD | HT-29 | SB216763 |
| MVP_top60_DNA_seed | AKT1 | 1.402894 | COADREAD | LS-123 | AT13148 | MVP_top60_DNA+RNA_seed | AKT1 | 2.679064 | COADREAD | COLO-678 | Ipatasertib |
| MVP_top60_DNA_seed | CDK2 | 1.407263 | COADREAD | SW837 | Dinaciclib | MVP_top60_DNA+RNA_seed | KRAS | 2.680078 | COADREAD | COLO-678 | KRAS (G12C) Inhibitor-1 |
| MVP_top60_DNA_seed | PIK3CA | 1.412744 | COADREAD | SNU-81 | Alpelisib | MVP_top60_DNA+RNA_seed | CDK2 | 2.758567 | COADREAD | SW1417 | Dinaciclib |
| MVP_top60_DNA_seed | ESR1 | 1.41587 | COADREAD | LS-123 | Tamoxifen | MVP_top60_DNA+RNA_seed | CDK2 | 2.785664 | COADREAD | SW1417 | AZD5438 |
| MVP_top60_DNA_seed | IGF1R | 1.418309 | COADREAD | LS-123 | NVP-ADW742 | MVP_top60_DNA+RNA_seed | AKT1 | 2.852562 | COADREAD | COLO-678 | AT13148 |
| MVP_top60_DNA_seed | IGF1R | 1.422658 | COADREAD | SW48 | GSK1904529A | MVP_top60_DNA+RNA_seed | PIK3CA | 2.855048 | COADREAD | SNU-61 | GNE-317 |
| MVP_top60_DNA_seed | IGF1R | 1.42769 | COADREAD | SW1463 | GSK1904529A | MVP_top60_DNA+RNA_seed | ATM | 2.874474 | COADREAD | COLO-678 | KU-55933 |
| MVP_top60_DNA_seed | AKT1 | 1.429917 | COADREAD | CL-11 | AZD5363 | MVP_top60_DNA+RNA_seed | AKT1 | 3.074071 | COADREAD | COLO-678 | MK-2206 |
| MVP_top60_DNA_seed | AKT1 | 1.431469 | COADREAD | SW1116 | Uprosertib | MVP_top60_DNA+RNA_seed | AKT1 | 3.093275 | COADREAD | COLO-678 | Afuresertib |
| MVP_top60_DNA_seed | MET | 1.43807 | COADREAD | SNU-C2B | Savolitinib | MVP_top60_DNA+RNA_seed | HSP90AA1 | 3.149368 | COADREAD | SNU-C1 | Luminespib |
| MVP_top60_DNA_seed | KRAS | 1.442541 | COADREAD | SW1463 | KRAS (G12C) Inhibitor-1 | MVP_top60_DNA+RNA_seed | HSP90AB1 | 3.149368 | COADREAD | SNU-C1 | Luminespib |
| MVP_top60_DNA_seed | HSP90AA1 | 1.443852 | COADREAD | HT-115 | Elesclomol | MVP_top60_DNA+RNA_seed | CDK2 | 3.18751 | COADREAD | SK-CO-1 | Dinaciclib |
| MVP_top60_DNA_seed | HSP90AB1 | 1.443852 | COADREAD | HT-115 | Elesclomol | MVP_top60_DNA+RNA_seed | TP53 | 3.209116 | COADREAD | COLO-678 | MIRA-1 |
| MVP_top60_DNA_seed | AKT1 | 1.448984 | COADREAD | LS-123 | Ipatasertib | MVP_top60_DNA+RNA_seed | AKT1 | 3.325058 | COADREAD | COLO-678 | AZD5363 |
| MVP_top60_DNA_seed | ESR1 | 1.453873 | COADREAD | C2BBe1 | GDC0810 | MVP_top60_DNA+RNA_seed | GSK3B | 3.462526 | COADREAD | COLO-678 | SB216763 |
| MVP_top60_DNA_seed | BRAF | 1.455466 | COADREAD | NCI-H747 | Dabrafenib | MVP_top60_DNA+RNA_seed | PIK3CA | 3.578788 | COADREAD | COLO-678 | Buparlisib |
| MVP_top60_DNA_seed | ESR1 | 1.456377 | COADREAD | SW1417 | Tamoxifen | MVP_top60_DNA+RNA_seed | PIK3CB | 3.578788 | COADREAD | COLO-678 | Buparlisib |
| MVP_top60_DNA_seed | BRAF | 1.457877 | COADREAD | SNU-C1 | PLX-4720 | MVP_top60_DNA+RNA_seed | PIK3CD | 3.578788 | COADREAD | COLO-678 | Buparlisib |
| MVP_top60_DNA_seed | KRAS | 1.460753 | COADREAD | SNU-C2B | KRAS (G12C) Inhibitor-1 | MVP_top60_DNA+RNA_seed | GSK3B | 3.787917 | COADREAD | SW1417 | SB216763 |
| MVP_top60_DNA_seed | IGF1R | 1.462317 | COADREAD | SNU-61 | BMS-536924 | MVP_top60_DNA+RNA_seed | PIK3CA | 3.798474 | COADREAD | COLO-678 | GNE-317 |
| MVP_top60_DNA_seed | MET | 1.462578 | COADREAD | LS-123 | Savolitinib | MVP_top60_DNA+RNA_seed | IGF1R | 3.813965 | COADREAD | COLO-678 | GSK1904529A |
| MVP_top60_DNA_seed | AKT1 | 1.467641 | COADREAD | SW837 | Uprosertib | MVP_top60_DNA+RNA_seed | PIK3CB | -4.113669 | GBM | Becker | TGX221 |
| MVP_top60_DNA_seed | PIK3CA | 1.474853 | COADREAD | SW837 | Alpelisib | MVP_top60_DNA+RNA_seed | MET | -3.74326 | GBM | DK-MG | AZD6094 |
| MVP_top60_DNA_seed | HSP90AA1 | 1.483148 | COADREAD | SW1116 | Tanespimycin | MVP_top60_DNA+RNA_seed | IGF1R | -3.591653 | GBM | KS-1 | Linsitinib |
| MVP_top60_DNA_seed | HSP90AB1 | 1.483148 | COADREAD | SW1116 | Tanespimycin | MVP_top60_DNA+RNA_seed | MET | -3.14451 | GBM | DK-MG | JNJ38877605 |
| MVP_top60_DNA_seed | SRC | 1.484129 | COADREAD | CL-11 | Bosutinib | MVP_top60_DNA+RNA_seed | PIK3CB | -3.09566 | GBM | D-542MG | TGX221 |
| MVP_top60_DNA_seed | PIK3CA | 1.489188 | COADREAD | KM12 | Alpelisib | MVP_top60_DNA+RNA_seed | PDGFRA | -3.050319 | GBM | D-247MG | Amuvatinib |
| MVP_top60_DNA_seed | PIK3CA | 1.493804 | COADREAD | RKO | GNE-317 | MVP_top60_DNA+RNA_seed | KIT | -3.050319 | GBM | D-247MG | Amuvatinib |
| MVP_top60_DNA_seed | AKT1 | 1.498516 | COADREAD | LS-123 | GSK2110183B | MVP_top60_DNA+RNA_seed | PDGFRA | -2.850486 | GBM | D-247MG | Masitinib |
| MVP_top60_DNA_seed | PIK3CA | 1.499855 | COADREAD | LS-1034 | Buparlisib | MVP_top60_DNA+RNA_seed | PDGFRB | -2.850486 | GBM | D-247MG | Masitinib |
| MVP_top60_DNA_seed | PIK3CB | 1.499855 | COADREAD | LS-1034 | Buparlisib | MVP_top60_DNA+RNA_seed | KIT | -2.850486 | GBM | D-247MG | Masitinib |

| | | | | | | | | | | | |
|---|---|---|---|---|---|---|---|---|---|---|---|
| MVP_top60_DNA_seed | PIK3CD | 1.499855 | COADREAD | LS-1034 | Buparlisib | MVP_top60_DNA+RNA_seed | PDGFRB | -2.660137 | GBM | 42-MG-BA | Pazopanib |
| MVP_top60_DNA_seed | AKT1 | 1.507258 | COADREAD | SW837 | Uprosertib | MVP_top60_DNA+RNA_seed | KIT | -2.660137 | GBM | 42-MG-BA | Pazopanib |
| MVP_top60_DNA_seed | AKT1 | 1.507316 | COADREAD | KM12 | Uprosertib | MVP_top60_DNA+RNA_seed | MET | -2.594024 | GBM | Becker | AZD6094 |
| MVP_top60_DNA_seed | SRC | 1.508488 | COADREAD | SW620 | Bosutinib | MVP_top60_DNA+RNA_seed | IGF1R | -2.585227 | GBM | KS-1 | BMS-754807 |
| MVP_top60_DNA_seed | AKT1 | 1.510797 | COADREAD | SW1463 | AT13148 | MVP_top60_DNA+RNA_seed | PIK3CB | -2.539864 | GBM | CAS-1 | TGX221 |
| MVP_top60_DNA_seed | GSK3B | 1.511319 | COADREAD | SNU-1040 | SB216763 | MVP_top60_DNA+RNA_seed | PIK3CB | -2.443509 | GBM | Becker | AZD6482 |
| MVP_top60_DNA_seed | AKT1 | 1.512685 | COADREAD | RCM-1 | AZD5363 | MVP_top60_DNA+RNA_seed | JAK2 | -2.386879 | GBM | DK-MG | Fedratinib |
| MVP_top60_DNA_seed | BRAF | 1.516738 | COADREAD | LS-123 | SB590885 | MVP_top60_DNA+RNA_seed | PDGFRA | -2.358508 | GBM | KS-1 | Amuvatinib |
| MVP_top60_DNA_seed | ESR1 | 1.525581 | COADREAD | NCI-H716 | Tamoxifen | MVP_top60_DNA+RNA_seed | KIT | -2.358508 | GBM | KS-1 | Amuvatinib |
| MVP_top60_DNA_seed | MET | 1.526885 | COADREAD | COLO-678 | Crizotinib | MVP_top60_DNA+RNA_seed | IGF1R | -2.311177 | GBM | KS-1 | BMS-536924 |
| MVP_top60_DNA_seed | CDK2 | 1.537395 | COADREAD | CL-11 | AZD5438 | MVP_top60_DNA+RNA_seed | KIT | -2.272548 | GBM | YH-13 | Motesanib |
| MVP_top60_DNA_seed | PIK3CA | 1.548009 | COADREAD | LS-411N | AZD8186 | MVP_top60_DNA+RNA_seed | PIK3CB | -2.191283 | GBM | D-247MG | TGX221 |
| MVP_top60_DNA_seed | PIK3CB | 1.548009 | COADREAD | LS-411N | AZD8186 | MVP_top60_DNA+RNA_seed | MET | -2.107339 | GBM | YH-13 | JNJ38877605 |
| MVP_top60_DNA_seed | AKT1 | 1.5539 | COADREAD | SW620 | AZD5363 | MVP_top60_DNA+RNA_seed | KIT | -2.085568 | GBM | DK-MG | OSI-930 |
| MVP_top60_DNA_seed | AKT1 | 1.568909 | COADREAD | SW1116 | Ipatasertib | MVP_top60_DNA+RNA_seed | MET | -2.004192 | GBM | SW1088 | AZD6094 |
| MVP_top60_DNA_seed | AKT1 | 1.580196 | COADREAD | SW837 | AT13148 | MVP_top60_DNA+RNA_seed | KIT | -1.79081 | GBM | 42-MG-BA | Motesanib |
| MVP_top60_DNA_seed | HSP90AA1 | 1.584491 | COADREAD | SW1463 | Luminespib | MVP_top60_DNA+RNA_seed | MET | -1.781652 | GBM | SF295 | AZD6094 |
| MVP_top60_DNA_seed | HSP90AB1 | 1.584491 | COADREAD | SW1463 | Luminespib | MVP_top60_DNA+RNA_seed | MET | -1.780441 | GBM | Becker | PHA-665752 |
| MVP_top60_DNA_seed | AKT1 | 1.593113 | COADREAD | SW620 | Ipatasertib | MVP_top60_DNA+RNA_seed | AKT1 | -1.767868 | GBM | YKG-1 | AKT inhibitor VIII |
| MVP_top60_DNA_seed | AKT1 | 1.593433 | COADREAD | SNU-C2B | GSK2110183B | MVP_top60_DNA+RNA_seed | MET | -1.754126 | GBM | YH-13 | AZD6094 |
| MVP_top60_DNA_seed | CREBBP | 1.593796 | COADREAD | HT-29 | SGC-CBP30 | MVP_top60_DNA+RNA_seed | PIK3CB | -1.745856 | GBM | SW1088 | AZD8186 |
| MVP_top60_DNA_seed | BRAF | 1.602098 | COADREAD | CL-11 | PLX-4720 | MVP_top60_DNA+RNA_seed | PIK3CD | -1.745856 | GBM | SW1088 | AZD8186 |
| MVP_top60_DNA_seed | PIK3CA | 1.603978 | COADREAD | SNU-C1 | AZD8186 | MVP_top60_DNA+RNA_seed | KIT | -1.727975 | GBM | SW1088 | Motesanib |
| MVP_top60_DNA_seed | PIK3CB | 1.603978 | COADREAD | SNU-C1 | AZD8186 | MVP_top60_DNA+RNA_seed | PIK3CB | -1.70907 | GBM | LN-18 | AZD8186 |
| MVP_top60_DNA_seed | KRAS | 1.604202 | COADREAD | SW1116 | KRAS (G12C) Inhibitor-1[illegible] | MVP_top60_DNA+RNA_seed | PIK3CD | -1.70907 | GBM | LN-18 | AZD8186 |
| MVP_top60_DNA_seed | MAPK1 | 1.625305 | COADREAD | C2BBe1 | Ulixertinib | MVP_top60_DNA+RNA_seed | PDGFRA | -1.687812 | GBM | AM-38 | Amuvatinib |
| MVP_top60_DNA_seed | CDK2 | 1.627002 | COADREAD | CL-11 | Dinaciclib | MVP_top60_DNA+RNA_seed | KIT | -1.687812 | GBM | AM-38 | Amuvatinib |
| MVP_top60_DNA_seed | HSP90AA1 | 1.630206 | COADREAD | SW837 | Luminespib | MVP_top60_DNA+RNA_seed | PIK3CD | -1.677454 | GBM | D-423MG | Idelalisib |
| MVP_top60_DNA_seed | HSP90AB1 | 1.630206 | COADREAD | SW837 | Luminespib | MVP_top60_DNA+RNA_seed | MET | -1.675041 | GBM | D-247MG | JNJ38877605 |
| MVP_top60_DNA_seed | ATM | 1.637887 | COADREAD | LS-123 | KU-55933 | MVP_top60_DNA+RNA_seed | PDGFRB | -1.65303 | GBM | SNB75 | Pazopanib |
| MVP_top60_DNA_seed | ERBB3 | 1.651232 | COADREAD | LS-411N | Sapitinib | MVP_top60_DNA+RNA_seed | KIT | -1.65303 | GBM | SNB75 | Pazopanib |
| MVP_top60_DNA_seed | PIK3CA | 1.652514 | COADREAD | LS-1034 | AZD8186 | MVP_top60_DNA+RNA_seed | CDK4 | -1.652208 | GBM | D-263MG | Palbociclib |
| MVP_top60_DNA_seed | PIK3CB | 1.652514 | COADREAD | LS-1034 | AZD8186 | MVP_top60_DNA+RNA_seed | PIK3CB | -1.637693 | GBM | LN-18 | AZD6482 |
| MVP_top60_DNA_seed | PIK3CB | 1.65465 | COADREAD | SW620 | AZD6482 | MVP_top60_DNA+RNA_seed | PDGFRB | -1.571127 | GBM | SF126 | Pazopanib |
| MVP_top60_DNA_seed | BRAF | 1.658252 | COADREAD | SW1116 | PLX-4720 | MVP_top60_DNA+RNA_seed | KIT | -1.571127 | GBM | SF126 | Pazopanib |
| MVP_top60_DNA_seed | AKT1 | 1.659385 | COADREAD | SW1463 | Uprosertib | MVP_top60_DNA+RNA_seed | KIT | -1.559146 | GBM | D-247MG | OSI-930 |
| MVP_top60_DNA_seed | TP53 | 1.663562 | COADREAD | C2BBe1 | MIRA-1 | MVP_top60_DNA+RNA_seed | PIK3CB | -1.555074 | GBM | D-542MG | AZD6482 |
| MVP_top60_DNA_seed | CDK2 | 1.669117 | COADREAD | RCM-1 | MK-8776 | MVP_top60_DNA+RNA_seed | EGFR | -1.546145 | GBM | KS-1 | Erlotinib |
| MVP_top60_DNA_seed | IGF1R | 1.670378 | COADREAD | SW1116 | NVP-ADW742 | MVP_top60_DNA+RNA_seed | PDGFRB | -1.524173 | GBM | M059J | Pazopanib |
| MVP_top60_DNA_seed | PIK3CA | 1.670726 | COADREAD | KM12 | AZD8186 | MVP_top60_DNA+RNA_seed | KIT | -1.524173 | GBM | M059J | Pazopanib |
| MVP_top60_DNA_seed | PIK3CB | 1.670726 | COADREAD | KM12 | AZD8186 | MVP_top60_DNA+RNA_seed | JAK3 | -1.48412 | GBM | Becker | JAK3_7406 |

| | | | | | | | | | | | |
|---|---|---|---|---|---|---|---|---|---|---|---|
| MVP_top60_DNA_seed | CREBBP | 1.689366 | COADREAD | NCI-H747 | SGC-CBP30 | MVP_top60_DNA+RNA_seed | AKT1 | -1.481662 | GBM | KS-1 | A-443654 |
| MVP_top60_DNA_seed | AKT1 | 1.690126 | COADREAD | SW1116 | AT13148 | MVP_top60_DNA+RNA_seed | KIT | -1.471504 | GBM | D-263MG | Motesanib |
| MVP_top60_DNA_seed | BRAF | 1.692513 | COADREAD | SNU-C2B | PLX-4720 | MVP_top60_DNA+RNA_seed | JAK1 | -1.462585 | GBM | D-263MG | JAK1_3715 |
| MVP_top60_DNA_seed | IGF1R | 1.698916 | COADREAD | SW1417 | NVP-ADW742 | MVP_top60_DNA+RNA_seed | CDK4 | -1.453012 | GBM | U251 | Palbociclib |
| MVP_top60_DNA_seed | AKT1 | 1.717056 | COADREAD | LS-1034 | AZD5363 | MVP_top60_DNA+RNA_seed | PIK3CB | -1.451875 | GBM | D-392MG | TGX221 |
| MVP_top60_DNA_seed | CREBBP | 1.726824 | COADREAD | SW1417 | SGC-CBP30 | MVP_top60_DNA+RNA_seed | PIK3CB | -1.417765 | GBM | SW1088 | AZD6482 |
| MVP_top60_DNA_seed | TP53 | 1.727207 | COADREAD | NCI-H747 | MIRA-1 | MVP_top60_DNA+RNA_seed | PDGFRB | -1.412645 | GBM | YH-13 | Pazopanib |
| MVP_top60_DNA_seed | BRAF | 1.729052 | COADREAD | LS-123 | PLX-4720 | MVP_top60_DNA+RNA_seed | KIT | -1.412645 | GBM | YH-13 | Pazopanib |
| MVP_top60_DNA_seed | CDK2 | 1.733373 | COADREAD | NCI-H747 | AZD5438 | MVP_top60_DNA+RNA_seed | PIK3CB | -1.398947 | GBM | SNB75 | AZD6482 |
| MVP_top60_DNA_seed | MET | 1.734326 | COADREAD | COLO-678 | Savolitinib | MVP_top60_DNA+RNA_seed | PIK3CB | -1.390615 | GBM | KS-1 | TGX221 |
| MVP_top60_DNA_seed | AKT1 | 1.73635 | COADREAD | LS-1034 | Uprosertib | MVP_top60_DNA+RNA_seed | MET | -1.390184 | GBM | Becker | Crizotinib |
| MVP_top60_DNA_seed | BRAF | 1.746145 | COADREAD | NCI-H747 | PLX-4720 | MVP_top60_DNA+RNA_seed | MET | -1.38624 | GBM | KS-1 | JNJ38877605 |
| MVP_top60_DNA_seed | HSP90AA1 | 1.759718 | COADREAD | CL-11 | Tanespimycin | MVP_top60_DNA+RNA_seed | PDGFRB | -1.383928 | GBM | A172 | Pazopanib |
| MVP_top60_DNA_seed | HSP90AB1 | 1.759718 | COADREAD | CL-11 | Tanespimycin | MVP_top60_DNA+RNA_seed | KIT | -1.383928 | GBM | A172 | Pazopanib |
| MVP_top60_DNA_seed | ATM | 1.762167 | COADREAD | SNU-C2B | KU-55933 | MVP_top60_DNA+RNA_seed | PIK3CB | -1.38113 | GBM | SNB75 | TGX221 |
| MVP_top60_DNA_seed | ERBB3 | 1.768894 | COADREAD | SNU-C1 | Sapitinib | MVP_top60_DNA+RNA_seed | IGF1R | -1.379357 | GBM | GB-1 | Linsitinib |
| MVP_top60_DNA_seed | AKT1 | 1.769358 | COADREAD | SW1116 | Uprosertib | MVP_top60_DNA+RNA_seed | IGF1R | -1.368132 | GBM | GB-1 | BMS-536924 |
| MVP_top60_DNA_seed | PIK3CA | 1.771889 | COADREAD | SW1116 | Buparlisib | MVP_top60_DNA+RNA_seed | JAK3 | -1.365446 | GBM | D-566MG | JAK3_7406 |
| MVP_top60_DNA_seed | PIK3CB | 1.771889 | COADREAD | SW1116 | Buparlisib | MVP_top60_DNA+RNA_seed | PIK3CB | -1.364576 | GBM | SF126 | TGX221 |
| MVP_top60_DNA_seed | PIK3CD | 1.771889 | COADREAD | SW1116 | Buparlisib | MVP_top60_DNA+RNA_seed | KIT | -1.356714 | GBM | YH-13 | Axitinib |
| MVP_top60_DNA_seed | AKT1 | 1.774513 | COADREAD | SW620 | Uprosertib | MVP_top60_DNA+RNA_seed | KIT | -1.349951 | GBM | D-566MG | Axitinib |
| MVP_top60_DNA_seed | GSK3B | 1.774967 | COADREAD | SNU-C2B | SB216763 | MVP_top60_DNA+RNA_seed | KIT | -1.348018 | GBM | LN-18 | Motesanib |
| MVP_top60_DNA_seed | AKT1 | 1.775962 | COADREAD | HT-115 | AZD5363 | MVP_top60_DNA+RNA_seed | MET | -1.326086 | GBM | Becker | JNJ38877605 |
| MVP_top60_DNA_seed | HSP90AA1 | 1.784675 | COADREAD | SNU-C2B | Tanespimycin | MVP_top60_DNA+RNA_seed | PIK3CD | -1.317009 | GBM | CAS-1 | IC-87114 |
| MVP_top60_DNA_seed | HSP90AB1 | 1.784675 | COADREAD | SNU-C2B | Tanespimycin | MVP_top60_DNA+RNA_seed | MET | -1.309462 | GBM | KS-1 | Crizotinib |
| MVP_top60_DNA_seed | AKT1 | 1.799962 | COADREAD | LS-123 | Uprosertib | MVP_top60_DNA+RNA_seed | MET | -1.299862 | GBM | U-87-MG | JNJ38877605 |
| MVP_top60_DNA_seed | GSK3B | 1.806404 | COADREAD | NCI-H716 | SB216763 | MVP_top60_DNA+RNA_seed | IGF1R | -1.284005 | GBM | KS-1 | BMS-536924 |
| MVP_top60_DNA_seed | GSK3B | 1.817415 | COADREAD | HT-29 | CHIR-99021 | MVP_top60_DNA+RNA_seed | MET | -1.257587 | GBM | AM-38 | SU11274 |
| MVP_top60_DNA_seed | AKT1 | 1.830566 | COADREAD | SW1417 | GSK2110183B | MVP_top60_DNA+RNA_seed | JAK3 | -1.248949 | GBM | 42-MG-BA | JAK3_7406 |
| MVP_top60_DNA_seed | MET | 1.834107 | COADREAD | SW620 | Savolitinib | MVP_top60_DNA+RNA_seed | PIK3CD | -1.24375 | GBM | D-566MG | Idelalisib |
| MVP_top60_DNA_seed | AKT1 | 1.834918 | COADREAD | LS-123 | AZD5363 | MVP_top60_DNA+RNA_seed | MET | -1.243092 | GBM | D-392MG | AZD6094 |
| MVP_top60_DNA_seed | PIK3CA | 1.863039 | COADREAD | CW-2 | GNE-317 | MVP_top60_DNA+RNA_seed | PDGFRB | -1.240521 | GBM | D-392MG | Pazopanib |
| MVP_top60_DNA_seed | AR | 1.872618 | COADREAD | HT-29 | Bicalutamide | MVP_top60_DNA+RNA_seed | KIT | -1.240521 | GBM | D-392MG | Pazopanib |
| MVP_top60_DNA_seed | HSP90AA1 | 1.876363 | COADREAD | SNU-61 | Luminespib | MVP_top60_DNA+RNA_seed | JAK2 | -1.238767 | GBM | KS-1 | Fedratinib |
| MVP_top60_DNA_seed | HSP90AB1 | 1.876363 | COADREAD | SNU-61 | Luminespib | MVP_top60_DNA+RNA_seed | PIK3CB | -1.236397 | GBM | YH-13 | AZD8186 |
| MVP_top60_DNA_seed | PIK3CA | 1.876554 | COADREAD | SW1116 | GNE-317 | MVP_top60_DNA+RNA_seed | PIK3CD | -1.236397 | GBM | YH-13 | AZD8186 |
| MVP_top60_DNA_seed | CDK2 | 1.878235 | COADREAD | C2BBe1 | MK-8776 | MVP_top60_DNA+RNA_seed | PDGFRB | -1.216698 | GBM | DK-MG | Pazopanib |
| MVP_top60_DNA_seed | ESR1 | 1.879763 | COADREAD | COLO-678 | Tamoxifen | MVP_top60_DNA+RNA_seed | KIT | -1.216698 | GBM | DK-MG | Pazopanib |
| MVP_top60_DNA_seed | BRAF | 1.88081 | COADREAD | SNU-C2B | Dabrafenib | MVP_top60_DNA+RNA_seed | PIK3CB | -1.205508 | GBM | SF295 | AZD6482 |
| MVP_top60_DNA_seed | CDK2 | 1.88685 | COADREAD | SW1116 | Dinaciclib | MVP_top60_DNA+RNA_seed | AKT1 | -1.202663 | GBM | Becker | AKT inhibitor VIII |
| MVP_top60_DNA_seed | CDK2 | 1.907372 | COADREAD | LS-123 | MK-8776 | MVP_top60_DNA+RNA_seed | EGFR | -1.189085 | GBM | U251 | CUDC-101 |

| | | | | | | | | | | | |
|---|---|---|---|---|---|---|---|---|---|---|---|
| MVP_top60_DNA_seed | PIK3CA | 1.907945 | COADREAD | COLO-678 | AZD8186 | MVP_top60_DNA+RNA_seed | MET | -1.179916 | GBM | U-118-MG | AZD6094 |
| MVP_top60_DNA_seed | PIK3CB | 1.907945 | COADREAD | COLO-678 | AZD8186 | MVP_top60_DNA+RNA_seed | KIT | -1.170538 | GBM | 8-MG-BA | Sorafenib |
| MVP_top60_DNA_seed | IGF1R | 1.908475 | COADREAD | COLO-678 | Linsitinib | MVP_top60_DNA+RNA_seed | IGF1R | -1.16911 | GBM | GB-1 | BMS-754807 |
| MVP_top60_DNA_seed | AKT1 | 1.910749 | COADREAD | SW620 | Afuresertib | MVP_top60_DNA+RNA_seed | EGFR | -1.16436 | GBM | DK-MG | Pelitinib |
| MVP_top60_DNA_seed | AKT1 | 1.913541 | COADREAD | SW1417 | MK-2206 | MVP_top60_DNA+RNA_seed | AKT1 | -1.162183 | GBM | SW1088 | MK-2206 |
| MVP_top60_DNA_seed | CDK2 | 1.949386 | COADREAD | SW620 | MK-8776 | MVP_top60_DNA+RNA_seed | PDGFRB | -1.161978 | GBM | GB-1 | Pazopanib |
| MVP_top60_DNA_seed | HSP90AA1 | 1.957433 | COADREAD | SW948 | Luminespib | MVP_top60_DNA+RNA_seed | KIT | -1.161978 | GBM | GB-1 | Pazopanib |
| MVP_top60_DNA_seed | HSP90AB1 | 1.957433 | COADREAD | SW948 | Luminespib | MVP_top60_DNA+RNA_seed | AKT1 | -1.16184 | GBM | SK-MG-1 | AKT inhibitor VIII |
| MVP_top60_DNA_seed | AKT1 | 1.960755 | COADREAD | SW1116 | Afuresertib | MVP_top60_DNA+RNA_seed | KIT | -1.154051 | GBM | AM-38 | Motesanib |
| MVP_top60_DNA_seed | IGF1R | 1.961416 | COADREAD | LS-123 | GSK1904529A | MVP_top60_DNA+RNA_seed | CDK4 | -1.153023 | GBM | YH-13 | Palbociclib |
| MVP_top60_DNA_seed | PIK3CA | 1.98122 | COADREAD | SW1116 | AZD8186 | MVP_top60_DNA+RNA_seed | PIK3CB | -1.139554 | GBM | LN-405 | AZD6482 |
| MVP_top60_DNA_seed | PIK3CB | 1.98122 | COADREAD | SW1116 | AZD8186 | MVP_top60_DNA+RNA_seed | MET | -1.126494 | GBM | D-263MG | JNJ38877605 |
| MVP_top60_DNA_seed | HSP90AA1 | 1.982823 | COADREAD | COLO-678 | Luminespib | MVP_top60_DNA+RNA_seed | CDK4 | -1.122478 | GBM | KS-1 | CGP-082996 |
| MVP_top60_DNA_seed | HSP90AB1 | 1.982823 | COADREAD | COLO-678 | Luminespib | MVP_top60_DNA+RNA_seed | JAK3 | -1.116311 | GBM | 8-MG-BA | JAK3_7406 |
| MVP_top60_DNA_seed | CREBBP | 1.984159 | COADREAD | LS-123 | SGC-CBP30 | MVP_top60_DNA+RNA_seed | PDGFRA | -1.114523 | GBM | DK-MG | Masitinib |
| MVP_top60_DNA_seed | ERBB3 | 1.989226 | COADREAD | CL-11 | Sapitinib | MVP_top60_DNA+RNA_seed | PDGFRB | -1.114523 | GBM | DK-MG | Masitinib |
| MVP_top60_DNA_seed | SRC | 1.996035 | COADREAD | SNU-C2B | Bosutinib | MVP_top60_DNA+RNA_seed | KIT | -1.114523 | GBM | DK-MG | Masitinib |
| MVP_top60_DNA_seed | KRAS | 2.035954 | COADREAD | LS-123 | KRAS (G12C) Inhibitor-1[illegible] | MVP_top60_DNA+RNA_seed | JAK2 | -1.113785 | GBM | 8-MG-BA | AZD1480 |
| MVP_top60_DNA_seed | TP53 | 2.051736 | COADREAD | SNU-C2B | MIRA-1 | MVP_top60_DNA+RNA_seed | JAK1 | -1.113785 | GBM | 8-MG-BA | AZD1480 |
| MVP_top60_DNA_seed | CDK2 | 2.099169 | COADREAD | COLO-678 | Dinaciclib | MVP_top60_DNA+RNA_seed | PIK3CA | -1.113586 | GBM | SK-MG-1 | AZD8835 |
| MVP_top60_DNA_seed | ESR1 | 2.103383 | COADREAD | LS-123 | GDC0810 | MVP_top60_DNA+RNA_seed | PIK3CD | -1.113586 | GBM | SK-MG-1 | AZD8835 |
| MVP_top60_DNA_seed | AKT1 | 2.147336 | COADREAD | SW620 | MK-2206 | MVP_top60_DNA+RNA_seed | PIK3CB | -1.108965 | GBM | D-566MG | AZD6482 |
| MVP_top60_DNA_seed | HSP90AA1 | 2.148666 | COADREAD | DiFi | Luminespib | MVP_top60_DNA+RNA_seed | CDK4 | -1.097164 | GBM | GB-1 | CGP-082996 |
| MVP_top60_DNA_seed | HSP90AB1 | 2.148666 | COADREAD | DiFi | Luminespib | MVP_top60_DNA+RNA_seed | MET | -1.08852 | GBM | LN-229 | JNJ38877605 |
| MVP_top60_DNA_seed | BRAF | 2.158822 | COADREAD | COLO-678 | Dabrafenib | MVP_top60_DNA+RNA_seed | JAK2 | -1.087596 | GBM | D-247MG | Fedratinib |
| MVP_top60_DNA_seed | CDK2 | 2.170457 | COADREAD | LS-123 | Dinaciclib | MVP_top60_DNA+RNA_seed | KIT | -1.085279 | GBM | 8-MG-BA | OSI-930 |
| MVP_top60_DNA_seed | CREBBP | 2.172285 | COADREAD | CL-11 | SGC-CBP30 | MVP_top60_DNA+RNA_seed | PIK3CB | -1.082312 | GBM | D-263MG | TGX221 |
| MVP_top60_DNA_seed | IGF1R | 2.193558 | COADREAD | C2BBe1 | BMS-536924 | MVP_top60_DNA+RNA_seed | JAK2 | -1.082204 | GBM | T98G | AZD1480 |
| MVP_top60_DNA_seed | AKT1 | 2.212207 | COADREAD | SW620 | GSK2110183B | MVP_top60_DNA+RNA_seed | JAK1 | -1.082204 | GBM | T98G | AZD1480 |
| MVP_top60_DNA_seed | GSK3B | 2.214252 | COADREAD | SW620 | CHIR-99021 | MVP_top60_DNA+RNA_seed | IGF1R | -1.07684 | GBM | D-542MG | GSK1904529A |
| MVP_top60_DNA_seed | CDK2 | 2.22528 | COADREAD | KM12 | AZD5438 | MVP_top60_DNA+RNA_seed | EP300 | -1.073287 | GBM | A172 | I-CBP112 |
| MVP_top60_DNA_seed | GSK3B | 2.225873 | COADREAD | SW620 | SB216763 | MVP_top60_DNA+RNA_seed | CREBBP | -1.073287 | GBM | A172 | I-CBP112 |
| MVP_top60_DNA_seed | CDK2 | 2.242149 | COADREAD | SW1417 | MK-8776 | MVP_top60_DNA+RNA_seed | JAK3 | -1.059803 | GBM | YH-13 | WHI-P97 |
| MVP_top60_DNA_seed | AKT1 | 2.296319 | COADREAD | COLO-678 | GSK2110183B | MVP_top60_DNA+RNA_seed | KIT | -1.051823 | GBM | D-392MG | Sorafenib |
| MVP_top60_DNA_seed | CREBBP | 2.296349 | COADREAD | C2BBe1 | SGC-CBP30 | MVP_top60_DNA+RNA_seed | PDGFRB | -1.044783 | GBM | CAS-1 | Pazopanib |
| MVP_top60_DNA_seed | PIK3CA | 2.296675 | COADREAD | RCM-1 | Buparlisib | MVP_top60_DNA+RNA_seed | KIT | -1.044783 | GBM | CAS-1 | Pazopanib |
| MVP_top60_DNA_seed | PIK3CB | 2.296675 | COADREAD | RCM-1 | Buparlisib | MVP_top60_DNA+RNA_seed | KIT | -1.044413 | GBM | A172 | Axitinib |
| MVP_top60_DNA_seed | PIK3CD | 2.296675 | COADREAD | RCM-1 | Buparlisib | MVP_top60_DNA+RNA_seed | EGFR | -1.028082 | GBM | Becker | Erlotinib |
| MVP_top60_DNA_seed | HSP90AA1 | 2.312686 | COADREAD | SNU-81 | Luminespib | MVP_top60_DNA+RNA_seed | PIK3CB | -1.027586 | GBM | 8-MG-BA | AZD6482 |
| MVP_top60_DNA_seed | HSP90AB1 | 2.312686 | COADREAD | SNU-81 | Luminespib | MVP_top60_DNA+RNA_seed | AKT1 | -1.026708 | GBM | SNB75 | A-443654 |
| MVP_top60_DNA_seed | CDK2 | 2.369786 | COADREAD | COLO-678 | AZD5438 | MVP_top60_DNA+RNA_seed | PIK3CD | -1.02601 | GBM | YH-13 | IC-87114 |

| | | | | | | | | | | | |
|---|---|---|---|---|---|---|---|---|---|---|---|
| MVP_top60_DNA_seed | BRAF | 2.373089 | COADREAD | SW620 | SB590885 | MVP_top60_DNA+RNA_seed | JAK3 | -1.022166 | GBM | U251 | JAK3_7406 |
| MVP_top60_DNA_seed | ATM | 2.380035 | COADREAD | SW620 | KU-55933 | MVP_top60_DNA+RNA_seed | IGF1R | -1.018088 | GBM | 42-MG-BA | GSK1904529A |
| MVP_top60_DNA_seed | AKT1 | 2.399471 | COADREAD | COLO-678 | Uprosertib | MVP_top60_DNA+RNA_seed | PIK3CB | -1.014967 | GBM | SF126 | AZD6482 |
| MVP_top60_DNA_seed | BRAF | 2.411923 | COADREAD | COLO-678 | SB590885 | MVP_top60_DNA+RNA_seed | MET | -1.012152 | GBM | 8-MG-BA | SU11274 |
| MVP_top60_DNA_seed | KRAS | 2.416192 | COADREAD | SW1417 | KRAS (G12C) Inhibitor-12 | MVP_top60_DNA+RNA_seed | CDK4 | -1.011535 | GBM | SF268 | Palbociclib |
| MVP_top60_DNA_seed | MET | 2.443539 | COADREAD | SW1417 | Savolitinib | MVP_top60_DNA+RNA_seed | KIT | -1.008269 | GBM | D-247MG | Sorafenib |
| MVP_top60_DNA_seed | AKT1 | 2.456908 | COADREAD | SW620 | Uprosertib | MVP_top60_DNA+RNA_seed | KIT | -1.000371 | GBM | U251 | OSI-930 |
| MVP_top60_DNA_seed | TP53 | 2.502555 | COADREAD | SW1417 | MIRA-1 | MVP_top60_DNA+RNA_seed | PIK3CB | -0.987533 | GBM | T98G | AZD8186 |
| MVP_top60_DNA_seed | PIK3CA | 2.581429 | COADREAD | COLO-678 | Alpelisib | MVP_top60_DNA+RNA_seed | PIK3CD | -0.987533 | GBM | T98G | AZD8186 |
| MVP_top60_DNA_seed | CDK2 | 2.585337 | COADREAD | COLO-678 | MK-8776 | MVP_top60_DNA+RNA_seed | KIT | -0.979923 | GBM | DK-MG | Axitinib |
| MVP_top60_DNA_seed | GSK3B | 2.613515 | COADREAD | HT-29 | SB216763 | MVP_top60_DNA+RNA_seed | IGF1R | -0.974694 | GBM | LN-18 | GSK1904529A |
| MVP_top60_DNA_seed | AKT1 | 2.679064 | COADREAD | COLO-678 | Ipatasertib | MVP_top60_DNA+RNA_seed | AKT1 | -0.961593 | GBM | YH-13 | MK-2206 |
| MVP_top60_DNA_seed | KRAS | 2.680078 | COADREAD | COLO-678 | KRAS (G12C) Inhibitor-12 | MVP_top60_DNA+RNA_seed | PIK3CB | -0.9608 | GBM | 42-MG-BA | AZD6482 |
| MVP_top60_DNA_seed | CDK2 | 2.758567 | COADREAD | SW1417 | Dinaciclib | MVP_top60_DNA+RNA_seed | EGFR | -0.954878 | GBM | U-87-MG | Afatinib |
| MVP_top60_DNA_seed | CDK2 | 2.785664 | COADREAD | SW1417 | AZD5438 | MVP_top60_DNA+RNA_seed | JAK3 | -0.952038 | GBM | LNZTA3WT4 | JAK3_7406 |
| MVP_top60_DNA_seed | AKT1 | 2.852562 | COADREAD | COLO-678 | AT13148 | MVP_top60_DNA+RNA_seed | KIT | -0.949293 | GBM | SF126 | Axitinib |
| MVP_top60_DNA_seed | PIK3CA | 2.855048 | COADREAD | SNU-61 | GNE-317 | MVP_top60_DNA+RNA_seed | PIK3CB | -0.949007 | GBM | D-263MG | AZD8186 |
| MVP_top60_DNA_seed | ATM | 2.874474 | COADREAD | COLO-678 | KU-55933 | MVP_top60_DNA+RNA_seed | PIK3CD | -0.949007 | GBM | D-263MG | AZD8186 |
| MVP_top60_DNA_seed | CREBBP | 2.934182 | COADREAD | COLO-678 | SGC-CBP30 | MVP_top60_DNA+RNA_seed | KIT | -0.948321 | GBM | D-392MG | Axitinib |
| MVP_top60_DNA_seed | AKT1 | 3.074071 | COADREAD | COLO-678 | MK-2206 | MVP_top60_DNA+RNA_seed | EP300 | -0.945062 | GBM | YH-13 | I-CBP112 |
| MVP_top60_DNA_seed | AKT1 | 3.093275 | COADREAD | COLO-678 | Afuresertib | MVP_top60_DNA+RNA_seed | CREBBP | -0.945062 | GBM | YH-13 | I-CBP112 |
| MVP_top60_DNA_seed | HSP90AA1 | 3.149368 | COADREAD | SNU-C1 | Luminespib | MVP_top60_DNA+RNA_seed | PDGFRA | -0.941443 | GBM | D-423MG | Amuvatinib |
| MVP_top60_DNA_seed | HSP90AB1 | 3.149368 | COADREAD | SNU-C1 | Luminespib | MVP_top60_DNA+RNA_seed | KIT | -0.941443 | GBM | D-423MG | Amuvatinib |
| MVP_top60_DNA_seed | CDK2 | 3.18751 | COADREAD | SK-CO-1 | Dinaciclib | MVP_top60_DNA+RNA_seed | EGFR | -0.941235 | GBM | KS-1 | Lapatinib |
| MVP_top60_DNA_seed | TP53 | 3.209116 | COADREAD | COLO-678 | MIRA-1 | MVP_top60_DNA+RNA_seed | JAK1 | -0.939957 | GBM | SF295 | JAK1_3715 |
| MVP_top60_DNA_seed | BRAF | 3.295513 | COADREAD | COLO-678 | PLX-4720 | MVP_top60_DNA+RNA_seed | JAK2 | -0.934605 | GBM | 42-MG-BA | AZD1480 |
| MVP_top60_DNA_seed | AKT1 | 3.325058 | COADREAD | COLO-678 | AZD5363 | MVP_top60_DNA+RNA_seed | JAK1 | -0.934605 | GBM | 42-MG-BA | AZD1480 |
| MVP_top60_DNA_seed | GSK3B | 3.462526 | COADREAD | COLO-678 | SB216763 | MVP_top60_DNA+RNA_seed | PIK3CD | -0.933098 | GBM | Becker | Idelalisib |
| MVP_top60_DNA_seed | PIK3CA | 3.578788 | COADREAD | COLO-678 | Buparlisib | MVP_top60_DNA+RNA_seed | PIK3CA | -0.932259 | GBM | D-566MG | PI3Ka_4409 |
| MVP_top60_DNA_seed | PIK3CB | 3.578788 | COADREAD | COLO-678 | Buparlisib | MVP_top60_DNA+RNA_seed | PIK3CD | -0.932259 | GBM | D-566MG | PI3Ka_4409 |
| MVP_top60_DNA_seed | PIK3CD | 3.578788 | COADREAD | COLO-678 | Buparlisib | MVP_top60_DNA+RNA_seed | EP300 | -0.930668 | GBM | Becker | I-CBP112 |
| MVP_top60_DNA_seed | GSK3B | 3.787917 | COADREAD | SW1417 | SB216763 | MVP_top60_DNA+RNA_seed | CREBBP | -0.930668 | GBM | Becker | I-CBP112 |
| MVP_top60_DNA_seed | ESR1 | 3.78917 | COADREAD | COLO-678 | GDC0810 | MVP_top60_DNA+RNA_seed | CDK4 | -0.93056 | GBM | YKG-1 | Palbociclib |
| MVP_top60_DNA_seed | PIK3CA | 3.798474 | COADREAD | COLO-678 | GNE-317 | MVP_top60_DNA+RNA_seed | JAK3 | -0.924884 | GBM | YH-13 | JAK3_7406 |
| MVP_top60_DNA_seed | IGF1R | 3.813965 | COADREAD | COLO-678 | GSK1904529A | MVP_top60_DNA+RNA_seed | EGFR | -0.924194 | GBM | SF268 | CUDC-101 |
| MVP_top60_DNA_seed | PIK3CB | -4.113669 | GBM | Becker | TGX221 | MVP_top60_DNA+RNA_seed | EGFR | -0.919244 | GBM | A172 | Cetuximab |
| MVP_top60_DNA_seed | PIK3CB | -3.09566 | GBM | D-542MG | TGX221 | MVP_top60_DNA+RNA_seed | PDGFRB | -0.914208 | GBM | D-566MG | Pazopanib |
| MVP_top60_DNA_seed | PDGFRA | -3.050319 | GBM | D-247MG | Amuvatinib | MVP_top60_DNA+RNA_seed | KIT | -0.914208 | GBM | D-566MG | Pazopanib |
| MVP_top60_DNA_seed | PDGFRA | -2.850486 | GBM | D-247MG | Masitinib | MVP_top60_DNA+RNA_seed | MET | -0.914114 | GBM | D-566MG | AZD6094 |
| MVP_top60_DNA_seed | PDGFRB | -2.850486 | GBM | D-247MG | Masitinib | MVP_top60_DNA+RNA_seed | PIK3CD | -0.914057 | GBM | U-87-MG | IC-87114 |
| MVP_top60_DNA_seed | PDGFRB | -2.660137 | GBM | 42-MG-BA | Pazopanib | MVP_top60_DNA+RNA_seed | JAK1 | -0.910931 | GBM | 8-MG-BA | JAK1_3715 |

| | | | | | | | | | | | |
|---|---|---|---|---|---|---|---|---|---|---|---|
| MVP_top60_DNA_seed | PIK3CB | -2.539864 | GBM | CAS-1 | TGX221 | MVP_top60_DNA+RNA_seed | CDK4 | -0.9049 | GBM | Becker | Palbociclib |
| MVP_top60_DNA_seed | PIK3CB | -2.443509 | GBM | Becker | AZD6482 | MVP_top60_DNA+RNA_seed | PDGFRB | -0.904738 | GBM | T98G | Pazopanib |
| MVP_top60_DNA_seed | JAK2 | -2.386879 | GBM | DK-MG | Fedratinib | MVP_top60_DNA+RNA_seed | KIT | -0.904738 | GBM | T98G | Pazopanib |
| MVP_top60_DNA_seed | PDGFRA | -2.358508 | GBM | KS-1 | Amuvatinib | MVP_top60_DNA+RNA_seed | PDGFRB | -0.900213 | GBM | SW1088 | Pazopanib |
| MVP_top60_DNA_seed | PIK3CB | -2.191283 | GBM | D-247MG | TGX221 | MVP_top60_DNA+RNA_seed | KIT | -0.900213 | GBM | SW1088 | Pazopanib |
| MVP_top60_DNA_seed | PIK3CB | -1.745856 | GBM | SW1088 | AZD8186 | MVP_top60_DNA+RNA_seed | AKT1 | -0.895138 | GBM | SW1088 | BAY AKT1 |
| MVP_top60_DNA_seed | PIK3CD | -1.745856 | GBM | SW1088 | AZD8186 | MVP_top60_DNA+RNA_seed | JAK3 | -0.89306 | GBM | SK-MG-1 | JAK3_7406 |
| MVP_top60_DNA_seed | PIK3CB | -1.70907 | GBM | LN-18 | AZD8186 | MVP_top60_DNA+RNA_seed | PIK3CB | -0.889132 | GBM | GB-1 | TGX221 |
| MVP_top60_DNA_seed | PIK3CD | -1.70907 | GBM | LN-18 | AZD8186 | MVP_top60_DNA+RNA_seed | PIK3CB | -0.885152 | GBM | D-566MG | AZD8186 |
| MVP_top60_DNA_seed | PDGFRA | -1.687812 | GBM | AM-38 | Amuvatinib | MVP_top60_DNA+RNA_seed | PIK3CD | -0.885152 | GBM | D-566MG | AZD8186 |
| MVP_top60_DNA_seed | PIK3CD | -1.677454 | GBM | D-423MG | Idelalisib | MVP_top60_DNA+RNA_seed | CDK4 | -0.881495 | GBM | T98G | Palbociclib |
| MVP_top60_DNA_seed | PDGFRB | -1.65303 | GBM | SNB75 | Pazopanib | MVP_top60_DNA+RNA_seed | PIK3CB | -0.878634 | GBM | CAS-1 | AZD6482 |
| MVP_top60_DNA_seed | CDK4 | -1.652208 | GBM | D-263MG | Palbociclib | MVP_top60_DNA+RNA_seed | MET | -0.876415 | GBM | KS-1 | PHA-665752 |
| MVP_top60_DNA_seed | PIK3CB | -1.637693 | GBM | LN-18 | AZD6482 | MVP_top60_DNA+RNA_seed | CDK4 | -0.874117 | GBM | SW1088 | Palbociclib |
| MVP_top60_DNA_seed | PDGFRB | -1.571127 | GBM | SF126 | Pazopanib | MVP_top60_DNA+RNA_seed | JAK2 | -0.867376 | GBM | D-263MG | AZD1480 |
| MVP_top60_DNA_seed | PIK3CB | -1.555074 | GBM | D-542MG | AZD6482 | MVP_top60_DNA+RNA_seed | JAK1 | -0.867376 | GBM | D-263MG | AZD1480 |
| MVP_top60_DNA_seed | EGFR | -1.546145 | GBM | KS-1 | Erlotinib | MVP_top60_DNA+RNA_seed | PIK3CB | -0.859912 | GBM | T98G | AZD6482 |
| MVP_top60_DNA_seed | PDGFRB | -1.524173 | GBM | M059J | Pazopanib | MVP_top60_DNA+RNA_seed | EGFR | -0.859529 | GBM | GB-1 | CUDC-101 |
| MVP_top60_DNA_seed | JAK3 | -1.48412 | GBM | Becker | JAK3_7406 | MVP_top60_DNA+RNA_seed | JAK3 | -0.854999 | GBM | D-392MG | JAK3_7406 |
| MVP_top60_DNA_seed | JAK1 | -1.462585 | GBM | D-263MG | JAK1_3715 | MVP_top60_DNA+RNA_seed | AKT1 | -0.850062 | GBM | 8-MG-BA | A-443654 |
| MVP_top60_DNA_seed | CDK4 | -1.453012 | GBM | U251 | Palbociclib | MVP_top60_DNA+RNA_seed | KIT | -0.843251 | GBM | DK-MG | Motesanib |
| MVP_top60_DNA_seed | PIK3CB | -1.451875 | GBM | D-392MG | TGX221 | MVP_top60_DNA+RNA_seed | JAK3 | -0.835926 | GBM | D-263MG | JAK3_7406 |
| MVP_top60_DNA_seed | PIK3CB | -1.417765 | GBM | SW1088 | AZD6482 | MVP_top60_DNA+RNA_seed | PDGFRA | -0.835034 | GBM | U251 | Amuvatinib |
| MVP_top60_DNA_seed | PDGFRB | -1.412645 | GBM | YH-13 | Pazopanib | MVP_top60_DNA+RNA_seed | KIT | -0.835034 | GBM | U251 | Amuvatinib |
| MVP_top60_DNA_seed | PIK3CB | -1.398947 | GBM | SNB75 | AZD6482 | MVP_top60_DNA+RNA_seed | PIK3CB | -0.827562 | GBM | D-247MG | AZD6482 |
| MVP_top60_DNA_seed | PIK3CB | -1.390615 | GBM | KS-1 | TGX221 | MVP_top60_DNA+RNA_seed | AKT1 | -0.822761 | GBM | D-423MG | AKT inhibitor VIII |
| MVP_top60_DNA_seed | PDGFRB | -1.383928 | GBM | A172 | Pazopanib | MVP_top60_DNA+RNA_seed | JAK2 | -0.817538 | GBM | KS-1 | AZD1480 |
| MVP_top60_DNA_seed | PIK3CB | -1.38113 | GBM | SNB75 | TGX221 | MVP_top60_DNA+RNA_seed | JAK1 | -0.817538 | GBM | KS-1 | AZD1480 |
| MVP_top60_DNA_seed | JAK3 | -1.365446 | GBM | D-566MG | JAK3_7406 | MVP_top60_DNA+RNA_seed | IGF1R | -0.810703 | GBM | T98G | BMS-536924 |
| MVP_top60_DNA_seed | PIK3CB | -1.364576 | GBM | SF126 | TGX221 | MVP_top60_DNA+RNA_seed | PDGFRB | -0.810695 | GBM | KS-1 | Pazopanib |
| MVP_top60_DNA_seed | PIK3CD | -1.317009 | GBM | CAS-1 | IC-87114 | MVP_top60_DNA+RNA_seed | KIT | -0.810695 | GBM | KS-1 | Pazopanib |
| MVP_top60_DNA_seed | JAK3 | -1.248949 | GBM | 42-MG-BA | JAK3_7406 | MVP_top60_DNA+RNA_seed | PIK3CB | -0.802736 | GBM | LN-405 | AZD8186 |
| MVP_top60_DNA_seed | PIK3CD | -1.24375 | GBM | D-566MG | Idelalisib | MVP_top60_DNA+RNA_seed | PIK3CD | -0.802736 | GBM | LN-405 | AZD8186 |
| MVP_top60_DNA_seed | PDGFRB | -1.240521 | GBM | D-392MG | Pazopanib | MVP_top60_DNA+RNA_seed | KIT | -0.802047 | GBM | SF126 | Motesanib |
| MVP_top60_DNA_seed | JAK2 | -1.238767 | GBM | KS-1 | Fedratinib | MVP_top60_DNA+RNA_seed | JAK2 | -0.801648 | GBM | D-392MG | Fedratinib |
| MVP_top60_DNA_seed | PIK3CB | -1.236397 | GBM | YH-13 | AZD8186 | MVP_top60_DNA+RNA_seed | PIK3CA | -0.799841 | GBM | SK-MG-1 | PI3Ka_4409 |
| MVP_top60_DNA_seed | PIK3CD | -1.236397 | GBM | YH-13 | AZD8186 | MVP_top60_DNA+RNA_seed | PIK3CD | -0.799841 | GBM | SK-MG-1 | PI3Ka_4409 |
| MVP_top60_DNA_seed | PDGFRB | -1.216698 | GBM | DK-MG | Pazopanib | MVP_top60_DNA+RNA_seed | AKT1 | -0.794791 | GBM | D-566MG | AKT inhibitor VIII |
| MVP_top60_DNA_seed | PIK3CB | -1.205508 | GBM | SF295 | AZD6482 | MVP_top60_DNA+RNA_seed | MET | -0.782686 | GBM | U-118-MG | JNJ38877605 |
| MVP_top60_DNA_seed | EGFR | -1.189085 | GBM | U251 | CUDC-101 | MVP_top60_DNA+RNA_seed | IGF1R | -0.780471 | GBM | D-423MG | GSK1904529A |
| MVP_top60_DNA_seed | EGFR | -1.16436 | GBM | DK-MG | Pelitinib | MVP_top60_DNA+RNA_seed | PDGFRB | -0.779205 | GBM | LN-405 | Pazopanib |

| | | | | | | | | | | | |
|---|---|---|---|---|---|---|---|---|---|---|---|
| MVP_top60_DNA_seed | PDGFRB | -1.161978 | GBM | GB-1 | Pazopanib | MVP_top60_DNA+RNA_seed | KIT | -0.779205 | GBM | LN-405 | Pazopanib |
| MVP_top60_DNA_seed | CDK4 | -1.153023 | GBM | YH-13 | Palbociclib | MVP_top60_DNA+RNA_seed | KIT | -0.777983 | GBM | SF126 | Sorafenib |
| MVP_top60_DNA_seed | PIK3CB | -1.139554 | GBM | LN-405 | AZD6482 | MVP_top60_DNA+RNA_seed | PIK3CB | -0.77574 | GBM | D-423MG | AZD6482 |
| MVP_top60_DNA_seed | CDK4 | -1.122478 | GBM | KS-1 | CGP-082996 | MVP_top60_DNA+RNA_seed | PIK3CA | -0.775331 | GBM | T98G | AZD8835 |
| MVP_top60_DNA_seed | JAK3 | -1.116311 | GBM | 8-MG-BA | JAK3_7406 | MVP_top60_DNA+RNA_seed | PIK3CD | -0.775331 | GBM | T98G | AZD8835 |
| MVP_top60_DNA_seed | PDGFRA | -1.114523 | GBM | DK-MG | Masitinib | MVP_top60_DNA+RNA_seed | PIK3CA | -0.767765 | GBM | 42-MG-BA | AZD8835 |
| MVP_top60_DNA_seed | PDGFRB | -1.114523 | GBM | DK-MG | Masitinib | MVP_top60_DNA+RNA_seed | PIK3CD | -0.767765 | GBM | 42-MG-BA | AZD8835 |
| MVP_top60_DNA_seed | JAK2 | -1.113785 | GBM | 8-MG-BA | AZD1480 | MVP_top60_DNA+RNA_seed | CDK4 | -0.764434 | GBM | SF295 | Palbociclib |
| MVP_top60_DNA_seed | JAK1 | -1.113785 | GBM | 8-MG-BA | AZD1480 | MVP_top60_DNA+RNA_seed | IGF1R | -0.763665 | GBM | DK-MG | BMS-754807 |
| MVP_top60_DNA_seed | PIK3CA | -1.113586 | GBM | SK-MG-1 | AZD8835 | MVP_top60_DNA+RNA_seed | AKT1 | -0.754725 | GBM | SF126 | AKT inhibitor VIII |
| MVP_top60_DNA_seed | PIK3CD | -1.113586 | GBM | SK-MG-1 | AZD8835 | MVP_top60_DNA+RNA_seed | EGFR | -0.747323 | GBM | Becker | AZD8931 |
| MVP_top60_DNA_seed | PIK3CB | -1.108965 | GBM | D-566MG | AZD6482 | MVP_top60_DNA+RNA_seed | ERBB3 | -0.747323 | GBM | Becker | AZD8931 |
| MVP_top60_DNA_seed | CDK4 | -1.097164 | GBM | GB-1 | CGP-082996 | MVP_top60_DNA+RNA_seed | JAK1 | -0.743094 | GBM | SF268 | JAK1_3715 |
| MVP_top60_DNA_seed | JAK2 | -1.087596 | GBM | D-247MG | Fedratinib | MVP_top60_DNA+RNA_seed | AKT1 | -0.735313 | GBM | D-263MG | BAY AKT1 |
| MVP_top60_DNA_seed | PIK3CB | -1.082312 | GBM | D-263MG | TGX221 | MVP_top60_DNA+RNA_seed | AKT1 | -0.733042 | GBM | KS-1 | MK-2206 |
| MVP_top60_DNA_seed | JAK2 | -1.082204 | GBM | T98G | AZD1480 | MVP_top60_DNA+RNA_seed | CDK4 | -0.730447 | GBM | D-423MG | Palbociclib |
| MVP_top60_DNA_seed | JAK1 | -1.082204 | GBM | T98G | AZD1480 | MVP_top60_DNA+RNA_seed | EGFR | -0.730374 | GBM | D-542MG | Cetuximab |
| MVP_top60_DNA_seed | EP300 | -1.073287 | GBM | A172 | I-CBP112 | MVP_top60_DNA+RNA_seed | EGFR | -0.724267 | GBM | KS-1 | CI-1033 |
| MVP_top60_DNA_seed | JAK3 | -1.059803 | GBM | YH-13 | WHI-P97 | MVP_top60_DNA+RNA_seed | PIK3CB | -0.724031 | GBM | Becker | AZD8186 |
| MVP_top60_DNA_seed | PDGFRB | -1.044783 | GBM | CAS-1 | Pazopanib | MVP_top60_DNA+RNA_seed | PIK3CD | -0.724031 | GBM | Becker | AZD8186 |
| MVP_top60_DNA_seed | EGFR | -1.028082 | GBM | Becker | Erlotinib | MVP_top60_DNA+RNA_seed | CDK4 | -0.720499 | GBM | D-247MG | Palbociclib |
| MVP_top60_DNA_seed | PIK3CB | -1.027586 | GBM | 8-MG-BA | AZD6482 | MVP_top60_DNA+RNA_seed | AKT1 | -0.720135 | GBM | SK-MG-1 | MK-2206 |
| MVP_top60_DNA_seed | PIK3CD | -1.02601 | GBM | YH-13 | IC-87114 | MVP_top60_DNA+RNA_seed | PDGFRA | -0.712378 | GBM | U251 | Masitinib |
| MVP_top60_DNA_seed | JAK3 | -1.022166 | GBM | U251 | JAK3_7406 | MVP_top60_DNA+RNA_seed | PDGFRB | -0.712378 | GBM | U251 | Masitinib |
| MVP_top60_DNA_seed | PIK3CB | -1.014967 | GBM | SF126 | AZD6482 | MVP_top60_DNA+RNA_seed | KIT | -0.712378 | GBM | U251 | Masitinib |
| MVP_top60_DNA_seed | CDK4 | -1.011535 | GBM | SF268 | Palbociclib | MVP_top60_DNA+RNA_seed | PIK3CB | -0.710063 | GBM | DBTRG-05MG | AZD6482 |
| MVP_top60_DNA_seed | PIK3CB | -0.987533 | GBM | T98G | AZD8186 | MVP_top60_DNA+RNA_seed | CDK4 | -0.709831 | GBM | DK-MG | Palbociclib |
| MVP_top60_DNA_seed | PIK3CD | -0.987533 | GBM | T98G | AZD8186 | MVP_top60_DNA+RNA_seed | PIK3CB | -0.707295 | GBM | U251 | AZD6482 |
| MVP_top60_DNA_seed | PIK3CB | -0.9608 | GBM | 42-MG-BA | AZD6482 | MVP_top60_DNA+RNA_seed | MET | -0.704954 | GBM | SNB75 | JNJ38877605 |
| MVP_top60_DNA_seed | EGFR | -0.954878 | GBM | U-87-MG | Afatinib | MVP_top60_DNA+RNA_seed | PIK3CB | -0.704705 | GBM | D-423MG | AZD8186 |
| MVP_top60_DNA_seed | JAK3 | -0.952038 | GBM | LNZTA3WT4 | JAK3_7406 | MVP_top60_DNA+RNA_seed | PIK3CD | -0.704705 | GBM | D-423MG | AZD8186 |
| MVP_top60_DNA_seed | PIK3CB | -0.949007 | GBM | D-263MG | AZD8186 | MVP_top60_DNA+RNA_seed | EGFR | -0.704283 | GBM | YH-13 | AZD8931 |
| MVP_top60_DNA_seed | PIK3CD | -0.949007 | GBM | D-263MG | AZD8186 | MVP_top60_DNA+RNA_seed | ERBB3 | -0.704283 | GBM | YH-13 | AZD8931 |
| MVP_top60_DNA_seed | EP300 | -0.945062 | GBM | YH-13 | I-CBP112 | MVP_top60_DNA+RNA_seed | JAK2 | -0.699706 | GBM | D-566MG | AZD1480 |
| MVP_top60_DNA_seed | PDGFRA | -0.941443 | GBM | D-423MG | Amuvatinib | MVP_top60_DNA+RNA_seed | JAK1 | -0.699706 | GBM | D-566MG | AZD1480 |
| MVP_top60_DNA_seed | EGFR | -0.941235 | GBM | KS-1 | Lapatinib | MVP_top60_DNA+RNA_seed | IGF1R | -0.697732 | GBM | A172 | Linsitinib |
| MVP_top60_DNA_seed | JAK1 | -0.939957 | GBM | SF295 | JAK1_3715 | MVP_top60_DNA+RNA_seed | AKT1 | -0.692603 | GBM | D-566MG | AKT inhibitor VIII |
| MVP_top60_DNA_seed | JAK2 | -0.934605 | GBM | 42-MG-BA | AZD1480 | MVP_top60_DNA+RNA_seed | AKT1 | -0.686814 | GBM | SF295 | AKT inhibitor VIII |
| MVP_top60_DNA_seed | JAK1 | -0.934605 | GBM | 42-MG-BA | AZD1480 | MVP_top60_DNA+RNA_seed | JAK3 | -0.682131 | GBM | D-247MG | WHI-P97 |
| MVP_top60_DNA_seed | PIK3CD | -0.933098 | GBM | Becker | Idelalisib | MVP_top60_DNA+RNA_seed | CDK4 | -0.680791 | GBM | SNB75 | CGP-082996 |
| MVP_top60_DNA_seed | PIK3CA | -0.932259 | GBM | D-566MG | PI3Ka_4409 | MVP_top60_DNA+RNA_seed | MET | -0.679463 | GBM | D-542MG | JNJ38877605 |

| | | | | | | | | | | | |
|---|---|---|---|---|---|---|---|---|---|---|---|
| MVP_top60_DNA_seed | PIK3CD | -0.932259 | GBM | D-566MG | PI3Ka_4409 | MVP_top60_DNA+RNA_seed | AKT1 | -0.679222 | GBM | 8-MG-BA | GSK690693 |
| MVP_top60_DNA_seed | EP300 | -0.930668 | GBM | Becker | I-CBP112 | MVP_top60_DNA+RNA_seed | MET | -0.678098 | GBM | YH-13 | SU11274 |
| MVP_top60_DNA_seed | CDK4 | -0.93056 | GBM | YKG-1 | Palbociclib | MVP_top60_DNA+RNA_seed | AKT1 | -0.673093 | GBM | 42-MG-BA | BAY AKT1 |
| MVP_top60_DNA_seed | JAK3 | -0.924884 | GBM | YH-13 | JAK3_7406 | MVP_top60_DNA+RNA_seed | EGFR | -0.671917 | GBM | U-87-MG | Cetuximab |
| MVP_top60_DNA_seed | EGFR | -0.924194 | GBM | SF268 | CUDC-101 | MVP_top60_DNA+RNA_seed | PDGFRB | -0.671466 | GBM | U-118-MG | Pazopanib |
| MVP_top60_DNA_seed | EGFR | -0.919244 | GBM | A172 | Cetuximab | MVP_top60_DNA+RNA_seed | KIT | -0.671466 | GBM | U-118-MG | Pazopanib |
| MVP_top60_DNA_seed | PDGFRB | -0.914208 | GBM | D-566MG | Pazopanib | MVP_top60_DNA+RNA_seed | AKT1 | -0.666883 | GBM | D-423MG | GSK690693 |
| MVP_top60_DNA_seed | PIK3CD | -0.914057 | GBM | U-87-MG | IC-87114 | MVP_top60_DNA+RNA_seed | PIK3CB | -0.665021 | GBM | DK-MG | AZD8186 |
| MVP_top60_DNA_seed | JAK1 | -0.910931 | GBM | 8-MG-BA | JAK1_3715 | MVP_top60_DNA+RNA_seed | PIK3CD | -0.665021 | GBM | DK-MG | AZD8186 |
| MVP_top60_DNA_seed | CDK4 | -0.9049 | GBM | Becker | Palbociclib | MVP_top60_DNA+RNA_seed | JAK2 | -0.664566 | GBM | A172 | AZD1480 |
| MVP_top60_DNA_seed | PDGFRB | -0.904738 | GBM | T98G | Pazopanib | MVP_top60_DNA+RNA_seed | JAK1 | -0.664566 | GBM | A172 | AZD1480 |
| MVP_top60_DNA_seed | PDGFRB | -0.900213 | GBM | SW1088 | Pazopanib | MVP_top60_DNA+RNA_seed | PIK3CB | -0.66251 | GBM | SK-MG-1 | AZD6482 |
| MVP_top60_DNA_seed | JAK3 | -0.89306 | GBM | SK-MG-1 | JAK3_7406 | MVP_top60_DNA+RNA_seed | AKT1 | -0.656272 | GBM | 8-MG-BA | AKT inhibitor VIII |
| MVP_top60_DNA_seed | PIK3CB | -0.889132 | GBM | GB-1 | TGX221 | MVP_top60_DNA+RNA_seed | AKT1 | -0.655994 | GBM | GB-1 | A-443654 |
| MVP_top60_DNA_seed | PIK3CB | -0.885152 | GBM | D-566MG | AZD8186 | MVP_top60_DNA+RNA_seed | JAK2 | -0.655213 | GBM | D-566MG | Fedratinib |
| MVP_top60_DNA_seed | PIK3CD | -0.885152 | GBM | D-566MG | AZD8186 | MVP_top60_DNA+RNA_seed | EGFR | -0.652341 | GBM | SW1088 | Gefitinib |
| MVP_top60_DNA_seed | CDK4 | -0.881495 | GBM | T98G | Palbociclib | MVP_top60_DNA+RNA_seed | JAK1 | -0.651118 | GBM | SF126 | JAK1_3715 |
| MVP_top60_DNA_seed | PIK3CB | -0.878634 | GBM | CAS-1 | AZD6482 | MVP_top60_DNA+RNA_seed | JAK3 | -0.648632 | GBM | KS-1 | JAK3_7406 |
| MVP_top60_DNA_seed | CDK4 | -0.874117 | GBM | SW1088 | Palbociclib | MVP_top60_DNA+RNA_seed | PDGFRA | -0.646932 | GBM | SK-MG-1 | Masitinib |
| MVP_top60_DNA_seed | JAK2 | -0.867376 | GBM | D-263MG | AZD1480 | MVP_top60_DNA+RNA_seed | PDGFRB | -0.646932 | GBM | SK-MG-1 | Masitinib |
| MVP_top60_DNA_seed | JAK1 | -0.867376 | GBM | D-263MG | AZD1480 | MVP_top60_DNA+RNA_seed | KIT | -0.646932 | GBM | SK-MG-1 | Masitinib |
| MVP_top60_DNA_seed | PIK3CB | -0.859912 | GBM | T98G | AZD6482 | MVP_top60_DNA+RNA_seed | EGFR | -0.645346 | GBM | 8-MG-BA | Pelitinib |
| MVP_top60_DNA_seed | EGFR | -0.859529 | GBM | GB-1 | CUDC-101 | MVP_top60_DNA+RNA_seed | PIK3CD | -0.637272 | GBM | YH-13 | Idelalisib |
| MVP_top60_DNA_seed | JAK3 | -0.854999 | GBM | D-392MG | JAK3_7406 | MVP_top60_DNA+RNA_seed | PDGFRB | -0.637122 | GBM | SF268 | Pazopanib |
| MVP_top60_DNA_seed | JAK3 | -0.835926 | GBM | D-263MG | JAK3_7406 | MVP_top60_DNA+RNA_seed | KIT | -0.637122 | GBM | SF268 | Pazopanib |
| MVP_top60_DNA_seed | PDGFRA | -0.835034 | GBM | U251 | Amuvatinib | MVP_top60_DNA+RNA_seed | KIT | -0.633538 | GBM | 8-MG-BA | Axitinib |
| MVP_top60_DNA_seed | PIK3CB | -0.827562 | GBM | D-247MG | AZD6482 | MVP_top60_DNA+RNA_seed | EGFR | -0.630226 | GBM | D-566MG | CUDC-101 |
| MVP_top60_DNA_seed | JAK2 | -0.817538 | GBM | KS-1 | AZD1480 | MVP_top60_DNA+RNA_seed | IGF1R | -0.628055 | GBM | DK-MG | Linsitinib |
| MVP_top60_DNA_seed | JAK1 | -0.817538 | GBM | KS-1 | AZD1480 | MVP_top60_DNA+RNA_seed | PDGFRA | -0.624878 | GBM | 8-MG-BA | Masitinib |
| MVP_top60_DNA_seed | PDGFRB | -0.810695 | GBM | KS-1 | Pazopanib | MVP_top60_DNA+RNA_seed | PDGFRB | -0.624878 | GBM | 8-MG-BA | Masitinib |
| MVP_top60_DNA_seed | PIK3CB | -0.802736 | GBM | LN-405 | AZD8186 | MVP_top60_DNA+RNA_seed | KIT | -0.624878 | GBM | 8-MG-BA | Masitinib |
| MVP_top60_DNA_seed | PIK3CD | -0.802736 | GBM | LN-405 | AZD8186 | MVP_top60_DNA+RNA_seed | KIT | -0.622387 | GBM | SF295 | Axitinib |
| MVP_top60_DNA_seed | JAK2 | -0.801648 | GBM | D-392MG | Fedratinib | MVP_top60_DNA+RNA_seed | EGFR | -0.622044 | GBM | YKG-1 | Afatinib |
| MVP_top60_DNA_seed | PIK3CA | -0.799841 | GBM | SK-MG-1 | PI3Ka_4409 | MVP_top60_DNA+RNA_seed | EGFR | -0.621896 | GBM | D-423MG | CUDC-101 |
| MVP_top60_DNA_seed | PIK3CD | -0.799841 | GBM | SK-MG-1 | PI3Ka_4409 | MVP_top60_DNA+RNA_seed | JAK2 | -0.618495 | GBM | CAS-1 | Ruxolitinib |
| MVP_top60_DNA_seed | PDGFRB | -0.779205 | GBM | LN-405 | Pazopanib | MVP_top60_DNA+RNA_seed | JAK1 | -0.618495 | GBM | CAS-1 | Ruxolitinib |
| MVP_top60_DNA_seed | PIK3CB | -0.77574 | GBM | D-423MG | AZD6482 | MVP_top60_DNA+RNA_seed | JAK2 | -0.615902 | GBM | SF268 | AZD1480 |
| MVP_top60_DNA_seed | PIK3CA | -0.775331 | GBM | T98G | AZD8835 | MVP_top60_DNA+RNA_seed | JAK1 | -0.615902 | GBM | SF268 | AZD1480 |
| MVP_top60_DNA_seed | PIK3CD | -0.775331 | GBM | T98G | AZD8835 | MVP_top60_DNA+RNA_seed | IGF1R | -0.615499 | GBM | SNB75 | BMS-536924 |
| MVP_top60_DNA_seed | PIK3CA | -0.767765 | GBM | 42-MG-BA | AZD8835 | MVP_top60_DNA+RNA_seed | PIK3CB | -0.603426 | GBM | SNB75 | AZD8186 |
| MVP_top60_DNA_seed | PIK3CD | -0.767765 | GBM | 42-MG-BA | AZD8835 | MVP_top60_DNA+RNA_seed | PIK3CD | -0.603426 | GBM | SNB75 | AZD8186 |

| | | | | | | | | | | | |
|---|---|---|---|---|---|---|---|---|---|---|---|
| MVP_top60_DNA_seed | CDK4 | -0.764434 | GBM | SF295 | Palbociclib | MVP_top60_DNA+RNA_seed | AKT1 | -0.601736 | GBM | SNB75 | AKT inhibitor VIII |
| MVP_top60_DNA_seed | EGFR | -0.747323 | GBM | Becker | AZD8931 | MVP_top60_DNA+RNA_seed | JAK2 | -0.600826 | GBM | YH-13 | AZD1480 |
| MVP_top60_DNA_seed | JAK1 | -0.743094 | GBM | SF268 | JAK1_3715 | MVP_top60_DNA+RNA_seed | JAK1 | -0.600826 | GBM | YH-13 | AZD1480 |
| MVP_top60_DNA_seed | CDK4 | -0.730447 | GBM | D-423MG | Palbociclib | MVP_top60_DNA+RNA_seed | JAK3 | -0.598046 | GBM | AM-38 | WHI-P97 |
| MVP_top60_DNA_seed | EGFR | -0.730374 | GBM | D-542MG | Cetuximab | MVP_top60_DNA+RNA_seed | JAK2 | -0.594365 | GBM | M059J | AZD1480 |
| MVP_top60_DNA_seed | EGFR | -0.724267 | GBM | KS-1 | CI-1033 | MVP_top60_DNA+RNA_seed | JAK1 | -0.594365 | GBM | M059J | AZD1480 |
| MVP_top60_DNA_seed | PIK3CB | -0.724031 | GBM | Becker | AZD8186 | MVP_top60_DNA+RNA_seed | PIK3CD | -0.591109 | GBM | U251 | Idelalisib |
| MVP_top60_DNA_seed | PIK3CD | -0.724031 | GBM | Becker | AZD8186 | MVP_top60_DNA+RNA_seed | CDK4 | -0.589324 | GBM | GB-1 | Palbociclib |
| MVP_top60_DNA_seed | CDK4 | -0.720499 | GBM | D-247MG | Palbociclib | MVP_top60_DNA+RNA_seed | PIK3CD | -0.587312 | GBM | D-542MG | IC-87114 |
| MVP_top60_DNA_seed | PDGFRA | -0.712378 | GBM | U251 | Masitinib | MVP_top60_DNA+RNA_seed | MET | -0.582692 | GBM | T98G | AZD6094 |
| MVP_top60_DNA_seed | PDGFRB | -0.712378 | GBM | U251 | Masitinib | MVP_top60_DNA+RNA_seed | PIK3CB | -0.581513 | GBM | KS-1 | AZD8186 |
| MVP_top60_DNA_seed | PIK3CB | -0.710063 | GBM | DBTRG-05MG | AZD6482 | MVP_top60_DNA+RNA_seed | PIK3CD | -0.581513 | GBM | KS-1 | AZD8186 |
| MVP_top60_DNA_seed | CDK4 | -0.709831 | GBM | DK-MG | Palbociclib | MVP_top60_DNA+RNA_seed | PIK3CB | -0.570406 | GBM | D-392MG | AZD6482 |
| MVP_top60_DNA_seed | PIK3CB | -0.707295 | GBM | U251 | AZD6482 | MVP_top60_DNA+RNA_seed | AKT1 | -0.568093 | GBM | D-423MG | MK-2206 |
| MVP_top60_DNA_seed | PIK3CB | -0.704705 | GBM | D-423MG | AZD8186 | MVP_top60_DNA+RNA_seed | JAK2 | -0.565376 | GBM | LN-18 | Ruxolitinib |
| MVP_top60_DNA_seed | PIK3CD | -0.704705 | GBM | D-423MG | AZD8186 | MVP_top60_DNA+RNA_seed | JAK1 | -0.565376 | GBM | LN-18 | Ruxolitinib |
| MVP_top60_DNA_seed | EGFR | -0.704283 | GBM | YH-13 | AZD8931 | MVP_top60_DNA+RNA_seed | MET | -0.559366 | GBM | 42-MG-BA | JNJ38877605 |
| MVP_top60_DNA_seed | JAK2 | -0.699706 | GBM | D-566MG | AZD1480 | MVP_top60_DNA+RNA_seed | EGFR | -0.55904 | GBM | SF295 | Afatinib |
| MVP_top60_DNA_seed | JAK1 | -0.699706 | GBM | D-566MG | AZD1480 | MVP_top60_DNA+RNA_seed | PIK3CA | -0.558372 | GBM | YH-13 | PI3Ka_4409 |
| MVP_top60_DNA_seed | JAK3 | -0.682131 | GBM | D-247MG | WHI-P97 | MVP_top60_DNA+RNA_seed | PIK3CD | -0.558372 | GBM | YH-13 | PI3Ka_4409 |
| MVP_top60_DNA_seed | CDK4 | -0.680791 | GBM | SNB75 | CGP-082996 | MVP_top60_DNA+RNA_seed | EGFR | -0.556108 | GBM | KS-1 | Pelitinib |
| MVP_top60_DNA_seed | EGFR | -0.671917 | GBM | U-87-MG | Cetuximab | MVP_top60_DNA+RNA_seed | PIK3CB | -0.553003 | GBM | 42-MG-BA | AZD8186 |
| MVP_top60_DNA_seed | PDGFRB | -0.671466 | GBM | U-118-MG | Pazopanib | MVP_top60_DNA+RNA_seed | PIK3CD | -0.553003 | GBM | 42-MG-BA | AZD8186 |
| MVP_top60_DNA_seed | PIK3CB | -0.665021 | GBM | DK-MG | AZD8186 | MVP_top60_DNA+RNA_seed | PIK3CB | -0.55188 | GBM | SF268 | AZD6482 |
| MVP_top60_DNA_seed | PIK3CD | -0.665021 | GBM | DK-MG | AZD8186 | MVP_top60_DNA+RNA_seed | PDGFRA | -0.548783 | GBM | D-566MG | Masitinib |
| MVP_top60_DNA_seed | JAK2 | -0.664566 | GBM | A172 | AZD1480 | MVP_top60_DNA+RNA_seed | PDGFRB | -0.548783 | GBM | D-566MG | Masitinib |
| MVP_top60_DNA_seed | JAK1 | -0.664566 | GBM | A172 | AZD1480 | MVP_top60_DNA+RNA_seed | KIT | -0.548783 | GBM | D-566MG | Masitinib |
| MVP_top60_DNA_seed | PIK3CB | -0.66251 | GBM | SK-MG-1 | AZD6482 | MVP_top60_DNA+RNA_seed | PIK3CA | -0.543971 | GBM | 8-MG-BA | AZD8835 |
| MVP_top60_DNA_seed | JAK2 | -0.655213 | GBM | D-566MG | Fedratinib | MVP_top60_DNA+RNA_seed | PIK3CD | -0.543971 | GBM | 8-MG-BA | AZD8835 |
| MVP_top60_DNA_seed | EGFR | -0.652341 | GBM | SW1088 | Gefitinib | MVP_top60_DNA+RNA_seed | AKT1 | -0.542697 | GBM | LN-18 | AKT inhibitor VIII |
| MVP_top60_DNA_seed | JAK1 | -0.651118 | GBM | SF126 | JAK1_3715 | MVP_top60_DNA+RNA_seed | IGF1R | -0.539905 | GBM | 8-MG-BA | BMS-536924 |
| MVP_top60_DNA_seed | JAK3 | -0.648632 | GBM | KS-1 | JAK3_7406 | MVP_top60_DNA+RNA_seed | EGFR | -0.539459 | GBM | D-542MG | Erlotinib |
| MVP_top60_DNA_seed | PDGFRA | -0.646932 | GBM | SK-MG-1 | Masitinib | MVP_top60_DNA+RNA_seed | KIT | -0.537768 | GBM | LN-405 | Motesanib |
| MVP_top60_DNA_seed | PDGFRB | -0.646932 | GBM | SK-MG-1 | Masitinib | MVP_top60_DNA+RNA_seed | IGF1R | -0.535072 | GBM | SF126 | BMS-754807 |
| MVP_top60_DNA_seed | EGFR | -0.645346 | GBM | 8-MG-BA | Pelitinib | MVP_top60_DNA+RNA_seed | JAK2 | -0.533473 | GBM | Becker | AZD1480 |
| MVP_top60_DNA_seed | PIK3CD | -0.637272 | GBM | YH-13 | Idelalisib | MVP_top60_DNA+RNA_seed | JAK1 | -0.533473 | GBM | Becker | AZD1480 |
| MVP_top60_DNA_seed | PDGFRB | -0.637122 | GBM | SF268 | Pazopanib | MVP_top60_DNA+RNA_seed | EGFR | -0.53299 | GBM | CAS-1 | Pelitinib |
| MVP_top60_DNA_seed | EGFR | -0.630226 | GBM | D-566MG | CUDC-101 | MVP_top60_DNA+RNA_seed | MET | -0.53145 | GBM | SF126 | AZD6094 |
| MVP_top60_DNA_seed | PDGFRA | -0.624878 | GBM | 8-MG-BA | Masitinib | MVP_top60_DNA+RNA_seed | JAK2 | -0.531129 | GBM | SK-MG-1 | AZD1480 |
| MVP_top60_DNA_seed | PDGFRB | -0.624878 | GBM | 8-MG-BA | Masitinib | MVP_top60_DNA+RNA_seed | JAK1 | -0.531129 | GBM | SK-MG-1 | AZD1480 |
| MVP_top60_DNA_seed | EGFR | -0.622044 | GBM | YKG-1 | Afatinib | MVP_top60_DNA+RNA_seed | KIT | -0.529625 | GBM | D-247MG | Motesanib |

| | | | | | | | | | | | |
|---|---|---|---|---|---|---|---|---|---|---|---|
| MVP_top60_DNA_seed | EGFR | -0.621896 | GBM | D-423MG | CUDC-101 | MVP_top60_DNA+RNA_seed | IGF1R | -0.528733 | GBM | SNB75 | Linsitinib |
| MVP_top60_DNA_seed | JAK2 | -0.618495 | GBM | CAS-1 | Ruxolitinib | MVP_top60_DNA+RNA_seed | EGFR | -0.528168 | GBM | YH-13 | AST-1306 |
| MVP_top60_DNA_seed | JAK1 | -0.618495 | GBM | CAS-1 | Ruxolitinib | MVP_top60_DNA+RNA_seed | PIK3CA | -0.522935 | GBM | SF126 | AZD8835 |
| MVP_top60_DNA_seed | JAK2 | -0.615902 | GBM | SF268 | AZD1480 | MVP_top60_DNA+RNA_seed | PIK3CD | -0.522935 | GBM | SF126 | AZD8835 |
| MVP_top60_DNA_seed | JAK1 | -0.615902 | GBM | SF268 | AZD1480 | MVP_top60_DNA+RNA_seed | CDK4 | -0.522497 | GBM | KS-1 | Palbociclib |
| MVP_top60_DNA_seed | PIK3CB | -0.603426 | GBM | SNB75 | AZD8186 | MVP_top60_DNA+RNA_seed | IGF1R | -0.521672 | GBM | YKG-1 | BMS-754807 |
| MVP_top60_DNA_seed | PIK3CD | -0.603426 | GBM | SNB75 | AZD8186 | MVP_top60_DNA+RNA_seed | EGFR | -0.521341 | GBM | D-392MG | Lapatinib |
| MVP_top60_DNA_seed | JAK2 | -0.600826 | GBM | YH-13 | AZD1480 | MVP_top60_DNA+RNA_seed | KIT | -0.518883 | GBM | LNZTA3WT4 | Axitinib |
| MVP_top60_DNA_seed | JAK1 | -0.600826 | GBM | YH-13 | AZD1480 | MVP_top60_DNA+RNA_seed | PIK3CB | -0.518628 | GBM | SK-MG-1 | AZD8186 |
| MVP_top60_DNA_seed | JAK3 | -0.598046 | GBM | AM-38 | WHI-P97 | MVP_top60_DNA+RNA_seed | PIK3CD | -0.518628 | GBM | SK-MG-1 | AZD8186 |
| MVP_top60_DNA_seed | JAK2 | -0.594365 | GBM | M059J | AZD1480 | MVP_top60_DNA+RNA_seed | PDGFRA | -0.516043 | GBM | YKG-1 | Masitinib |
| MVP_top60_DNA_seed | JAK1 | -0.594365 | GBM | M059J | AZD1480 | MVP_top60_DNA+RNA_seed | PDGFRB | -0.516043 | GBM | YKG-1 | Masitinib |
| MVP_top60_DNA_seed | PIK3CD | -0.591109 | GBM | U251 | Idelalisib | MVP_top60_DNA+RNA_seed | KIT | -0.516043 | GBM | YKG-1 | Masitinib |
| MVP_top60_DNA_seed | CDK4 | -0.589324 | GBM | GB-1 | Palbociclib | MVP_top60_DNA+RNA_seed | PIK3CB | -0.510226 | GBM | YH-13 | AZD6482 |
| MVP_top60_DNA_seed | PIK3CD | -0.587312 | GBM | D-542MG | IC-87114 | MVP_top60_DNA+RNA_seed | AKT1 | -0.502234 | GBM | Becker | GSK690693 |
| MVP_top60_DNA_seed | PIK3CB | -0.581513 | GBM | KS-1 | AZD8186 | MVP_top60_DNA+RNA_seed | EGFR | -0.501998 | GBM | A172 | Gefitinib |
| MVP_top60_DNA_seed | PIK3CD | -0.581513 | GBM | KS-1 | AZD8186 | MVP_top60_DNA+RNA_seed | PIK3CB | -0.499687 | GBM | SF295 | AZD8186 |
| MVP_top60_DNA_seed | PIK3CB | -0.570406 | GBM | D-392MG | AZD6482 | MVP_top60_DNA+RNA_seed | PIK3CD | -0.499687 | GBM | SF295 | AZD8186 |
| MVP_top60_DNA_seed | JAK2 | -0.565376 | GBM | LN-18 | Ruxolitinib | MVP_top60_DNA+RNA_seed | EGFR | -0.498351 | GBM | GB-1 | Cetuximab |
| MVP_top60_DNA_seed | JAK1 | -0.565376 | GBM | LN-18 | Ruxolitinib | MVP_top60_DNA+RNA_seed | EP300 | -0.49642 | GBM | LN-18 | I-CBP112 |
| MVP_top60_DNA_seed | EGFR | -0.55904 | GBM | SF295 | Afatinib | MVP_top60_DNA+RNA_seed | CREBBP | -0.49642 | GBM | LN-18 | I-CBP112 |
| MVP_top60_DNA_seed | PIK3CA | -0.558372 | GBM | YH-13 | PI3Ka_4409 | MVP_top60_DNA+RNA_seed | JAK3 | -0.490977 | GBM | SF268 | JAK3_7406 |
| MVP_top60_DNA_seed | PIK3CD | -0.558372 | GBM | YH-13 | PI3Ka_4409 | MVP_top60_DNA+RNA_seed | JAK2 | -0.490785 | GBM | CAS-1 | Fedratinib |
| MVP_top60_DNA_seed | EGFR | -0.556108 | GBM | KS-1 | Pelitinib | MVP_top60_DNA+RNA_seed | CDK4 | -0.488739 | GBM | SK-MG-1 | Palbociclib |
| MVP_top60_DNA_seed | PIK3CB | -0.553003 | GBM | 42-MG-BA | AZD8186 | MVP_top60_DNA+RNA_seed | AKT1 | -0.484228 | GBM | 42-MG-BA | MK-2206 |
| MVP_top60_DNA_seed | PIK3CD | -0.553003 | GBM | 42-MG-BA | AZD8186 | MVP_top60_DNA+RNA_seed | PIK3CB | -0.480287 | GBM | KS-1 | AZD6482 |
| MVP_top60_DNA_seed | PIK3CB | -0.55188 | GBM | SF268 | AZD6482 | MVP_top60_DNA+RNA_seed | KIT | -0.476714 | GBM | D-542MG | Motesanib |
| MVP_top60_DNA_seed | PDGFRA | -0.548783 | GBM | D-566MG | Masitinib | MVP_top60_DNA+RNA_seed | PIK3CD | -0.475201 | GBM | D-423MG | IC-87114 |
| MVP_top60_DNA_seed | PDGFRB | -0.548783 | GBM | D-566MG | Masitinib | MVP_top60_DNA+RNA_seed | AKT1 | -0.474865 | GBM | SF126 | MK-2206 |
| MVP_top60_DNA_seed | PIK3CA | -0.543971 | GBM | 8-MG-BA | AZD8835 | MVP_top60_DNA+RNA_seed | MET | -0.469059 | GBM | U-87-MG | PHA-665752 |
| MVP_top60_DNA_seed | PIK3CD | -0.543971 | GBM | 8-MG-BA | AZD8835 | MVP_top60_DNA+RNA_seed | AKT1 | -0.468963 | GBM | YKG-1 | GSK690693 |
| MVP_top60_DNA_seed | EGFR | -0.539459 | GBM | D-542MG | Erlotinib | MVP_top60_DNA+RNA_seed | KIT | -0.467782 | GBM | D-423MG | Axitinib |
| MVP_top60_DNA_seed | JAK2 | -0.533473 | GBM | Becker | AZD1480 | MVP_top60_DNA+RNA_seed | PIK3CA | -0.465958 | GBM | 8-MG-BA | PI3Ka_4409 |
| MVP_top60_DNA_seed | JAK1 | -0.533473 | GBM | Becker | AZD1480 | MVP_top60_DNA+RNA_seed | PIK3CD | -0.465958 | GBM | 8-MG-BA | PI3Ka_4409 |
| MVP_top60_DNA_seed | EGFR | -0.53299 | GBM | CAS-1 | Pelitinib | MVP_top60_DNA+RNA_seed | KIT | -0.464692 | GBM | U251 | Motesanib |
| MVP_top60_DNA_seed | JAK2 | -0.531129 | GBM | SK-MG-1 | AZD1480 | MVP_top60_DNA+RNA_seed | EGFR | -0.464147 | GBM | DK-MG | CI-1033 |
| MVP_top60_DNA_seed | JAK1 | -0.531129 | GBM | SK-MG-1 | AZD1480 | MVP_top60_DNA+RNA_seed | EGFR | -0.463366 | GBM | SF268 | Cetuximab |
| MVP_top60_DNA_seed | EGFR | -0.528168 | GBM | YH-13 | AST-1306 | MVP_top60_DNA+RNA_seed | JAK3 | -0.463359 | GBM | A172 | JAK3_7406 |
| MVP_top60_DNA_seed | PIK3CA | -0.522935 | GBM | SF126 | AZD8835 | MVP_top60_DNA+RNA_seed | PDGFRA | -0.456348 | GBM | A172 | Masitinib |
| MVP_top60_DNA_seed | PIK3CD | -0.522935 | GBM | SF126 | AZD8835 | MVP_top60_DNA+RNA_seed | PDGFRB | -0.456348 | GBM | A172 | Masitinib |
| MVP_top60_DNA_seed | CDK4 | -0.522497 | GBM | KS-1 | Palbociclib | MVP_top60_DNA+RNA_seed | KIT | -0.456348 | GBM | A172 | Masitinib |

| | | | | | | | | | | | |
|---|---|---|---|---|---|---|---|---|---|---|---|
| MVP_top60_DNA_seed | EGFR | -0.521341 | GBM | D-392MG | Lapatinib | MVP_top60_DNA+RNA_seed | KIT | -0.450407 | GBM | SF268 | Axitinib |
| MVP_top60_DNA_seed | PIK3CB | -0.518628 | GBM | SK-MG-1 | AZD8186 | MVP_top60_DNA+RNA_seed | PIK3CA | -0.449847 | GBM | 42-MG-BA | PI3Ka_4409 |
| MVP_top60_DNA_seed | PIK3CD | -0.518628 | GBM | SK-MG-1 | AZD8186 | MVP_top60_DNA+RNA_seed | PIK3CD | -0.449847 | GBM | 42-MG-BA | PI3Ka_4409 |
| MVP_top60_DNA_seed | PDGFRA | -0.516043 | GBM | YKG-1 | Masitinib | MVP_top60_DNA+RNA_seed | MET | -0.448495 | GBM | GB-1 | SU11274 |
| MVP_top60_DNA_seed | PDGFRB | -0.516043 | GBM | YKG-1 | Masitinib | MVP_top60_DNA+RNA_seed | AKT1 | -0.437838 | GBM | KS-1 | GSK690693 |
| MVP_top60_DNA_seed | PIK3CB | -0.510226 | GBM | YH-13 | AZD6482 | MVP_top60_DNA+RNA_seed | KIT | -0.433233 | GBM | T98G | Axitinib |
| MVP_top60_DNA_seed | EGFR | -0.501998 | GBM | A172 | Gefitinib | MVP_top60_DNA+RNA_seed | AKT1 | -0.431926 | GBM | T98G | MK-2206 |
| MVP_top60_DNA_seed | PIK3CB | -0.499687 | GBM | SF295 | AZD8186 | MVP_top60_DNA+RNA_seed | AKT1 | -0.431813 | GBM | 8-MG-BA | BAY AKT1 |
| MVP_top60_DNA_seed | PIK3CD | -0.499687 | GBM | SF295 | AZD8186 | MVP_top60_DNA+RNA_seed | JAK2 | -0.425989 | GBM | LNZTA3WT4 | AZD1480 |
| MVP_top60_DNA_seed | EGFR | -0.498351 | GBM | GB-1 | Cetuximab | MVP_top60_DNA+RNA_seed | JAK1 | -0.425989 | GBM | LNZTA3WT4 | AZD1480 |
| MVP_top60_DNA_seed | EP300 | -0.49642 | GBM | LN-18 | I-CBP112 | MVP_top60_DNA+RNA_seed | EGFR | -0.42457 | GBM | Becker | Lapatinib |
| MVP_top60_DNA_seed | JAK3 | -0.490977 | GBM | SF268 | JAK3_7406 | MVP_top60_DNA+RNA_seed | MET | -0.424115 | GBM | LN-405 | JNJ38877605 |
| MVP_top60_DNA_seed | JAK2 | -0.490785 | GBM | CAS-1 | Fedratinib | MVP_top60_DNA+RNA_seed | EGFR | -0.42128 | GBM | D-247MG | Lapatinib |
| MVP_top60_DNA_seed | CDK4 | -0.488739 | GBM | SK-MG-1 | Palbociclib | MVP_top60_DNA+RNA_seed | PIK3CA | -0.419597 | GBM | KS-1 | PI3Ka_4409 |
| MVP_top60_DNA_seed | PIK3CB | -0.480287 | GBM | KS-1 | AZD6482 | MVP_top60_DNA+RNA_seed | PIK3CD | -0.419597 | GBM | KS-1 | PI3Ka_4409 |
| MVP_top60_DNA_seed | PIK3CD | -0.475201 | GBM | D-423MG | IC-87114 | MVP_top60_DNA+RNA_seed | JAK3 | -0.418466 | GBM | GB-1 | WHI-P97 |
| MVP_top60_DNA_seed | PIK3CA | -0.465958 | GBM | 8-MG-BA | PI3Ka_4409 | MVP_top60_DNA+RNA_seed | PIK3CA | -0.415715 | GBM | YH-13 | AZD8835 |
| MVP_top60_DNA_seed | PIK3CD | -0.465958 | GBM | 8-MG-BA | PI3Ka_4409 | MVP_top60_DNA+RNA_seed | PIK3CD | -0.415715 | GBM | YH-13 | AZD8835 |
| MVP_top60_DNA_seed | EGFR | -0.464147 | GBM | DK-MG | CI-1033 | MVP_top60_DNA+RNA_seed | JAK3 | -0.415412 | GBM | D-542MG | JAK3_7406 |
| MVP_top60_DNA_seed | EGFR | -0.463366 | GBM | SF268 | Cetuximab | MVP_top60_DNA+RNA_seed | KIT | -0.414221 | GBM | D-542MG | Sorafenib |
| MVP_top60_DNA_seed | JAK3 | -0.463359 | GBM | A172 | JAK3_7406 | MVP_top60_DNA+RNA_seed | AKT1 | -0.411399 | GBM | U251 | MK-2206 |
| MVP_top60_DNA_seed | PDGFRA | -0.456348 | GBM | A172 | Masitinib | MVP_top60_DNA+RNA_seed | KIT | -0.409304 | GBM | D-392MG | Motesanib |
| MVP_top60_DNA_seed | PDGFRB | -0.456348 | GBM | A172 | Masitinib | MVP_top60_DNA+RNA_seed | KIT | -0.408316 | GBM | 42-MG-BA | Axitinib |
| MVP_top60_DNA_seed | PIK3CA | -0.449847 | GBM | 42-MG-BA | PI3Ka_4409 | MVP_top60_DNA+RNA_seed | EP300 | -0.406708 | GBM | SF268 | I-CBP112 |
| MVP_top60_DNA_seed | PIK3CD | -0.449847 | GBM | 42-MG-BA | PI3Ka_4409 | MVP_top60_DNA+RNA_seed | CREBBP | -0.406708 | GBM | SF268 | I-CBP112 |
| MVP_top60_DNA_seed | JAK2 | -0.425989 | GBM | LNZTA3WT4 | AZD1480 | MVP_top60_DNA+RNA_seed | PIK3CD | -0.406369 | GBM | AM-38 | Idelalisib |
| MVP_top60_DNA_seed | JAK1 | -0.425989 | GBM | LNZTA3WT4 | AZD1480 | MVP_top60_DNA+RNA_seed | IGF1R | -0.404695 | GBM | SF126 | BMS-536924 |
| MVP_top60_DNA_seed | EGFR | -0.42457 | GBM | Becker | Lapatinib | MVP_top60_DNA+RNA_seed | MET | -0.404401 | GBM | U251 | JNJ38877605 |
| MVP_top60_DNA_seed | EGFR | -0.42128 | GBM | D-247MG | Lapatinib | MVP_top60_DNA+RNA_seed | EGFR | -0.40314 | GBM | YH-13 | Afatinib |
| MVP_top60_DNA_seed | PIK3CA | -0.419597 | GBM | KS-1 | PI3Ka_4409 | MVP_top60_DNA+RNA_seed | KIT | -0.402132 | GBM | SK-MG-1 | Motesanib |
| MVP_top60_DNA_seed | PIK3CD | -0.419597 | GBM | KS-1 | PI3Ka_4409 | MVP_top60_DNA+RNA_seed | AKT1 | -0.401086 | GBM | U251 | AKT inhibitor VIII |
| MVP_top60_DNA_seed | JAK3 | -0.418466 | GBM | GB-1 | WHI-P97 | MVP_top60_DNA+RNA_seed | IGF1R | -0.398564 | GBM | GB-1 | BMS-536924 |
| MVP_top60_DNA_seed | PIK3CA | -0.415715 | GBM | YH-13 | AZD8835 | MVP_top60_DNA+RNA_seed | EGFR | -0.398414 | GBM | SF126 | Cetuximab |
| MVP_top60_DNA_seed | PIK3CD | -0.415715 | GBM | YH-13 | AZD8835 | MVP_top60_DNA+RNA_seed | PIK3CB | -0.397953 | GBM | CAS-1 | AZD8186 |
| MVP_top60_DNA_seed | JAK3 | -0.415412 | GBM | D-542MG | JAK3_7406 | MVP_top60_DNA+RNA_seed | PIK3CD | -0.397953 | GBM | CAS-1 | AZD8186 |
| MVP_top60_DNA_seed | EP300 | -0.406708 | GBM | SF268 | I-CBP112 | MVP_top60_DNA+RNA_seed | EGFR | -0.397437 | GBM | D-247MG | CUDC-101 |
| MVP_top60_DNA_seed | PIK3CD | -0.406369 | GBM | AM-38 | Idelalisib | MVP_top60_DNA+RNA_seed | CDK4 | -0.396022 | GBM | LN-18 | Palbociclib |
| MVP_top60_DNA_seed | EGFR | -0.40314 | GBM | YH-13 | Afatinib | MVP_top60_DNA+RNA_seed | KIT | -0.393166 | GBM | D-263MG | Axitinib |
| MVP_top60_DNA_seed | EGFR | -0.398414 | GBM | SF126 | Cetuximab | MVP_top60_DNA+RNA_seed | AKT1 | -0.393001 | GBM | YH-13 | GSK690693 |
| MVP_top60_DNA_seed | PIK3CB | -0.397953 | GBM | CAS-1 | AZD8186 | MVP_top60_DNA+RNA_seed | EGFR | -0.392801 | GBM | U251 | AZD8931 |
| MVP_top60_DNA_seed | PIK3CD | -0.397953 | GBM | CAS-1 | AZD8186 | MVP_top60_DNA+RNA_seed | ERBB3 | -0.392801 | GBM | U251 | AZD8931 |

| | | | | | | | | | | | |
|---|---|---|---|---|---|---|---|---|---|---|---|
| MVP_top60_DNA_seed | EGFR | -0.397437 | GBM | D-247MG | CUDC-101 | MVP_top60_DNA+RNA_seed | IGF1R | -0.386565 | GBM | 42-MG-BA | BMS-536924 |
| MVP_top60_DNA_seed | CDK4 | -0.396022 | GBM | LN-18 | Palbociclib | MVP_top60_DNA+RNA_seed | CDK4 | -0.384323 | GBM | 8-MG-BA | CGP-082996 |
| MVP_top60_DNA_seed | EGFR | -0.392801 | GBM | U251 | AZD8931 | MVP_top60_DNA+RNA_seed | EGFR | -0.383351 | GBM | SF295 | Gefitinib |
| MVP_top60_DNA_seed | CDK4 | -0.384323 | GBM | 8-MG-BA | CGP-082996 | MVP_top60_DNA+RNA_seed | EGFR | -0.378426 | GBM | KS-1 | CUDC-101 |
| MVP_top60_DNA_seed | EGFR | -0.383351 | GBM | SF295 | Gefitinib | MVP_top60_DNA+RNA_seed | JAK2 | -0.377517 | GBM | 42-MG-BA | Fedratinib |
| MVP_top60_DNA_seed | EGFR | -0.378426 | GBM | KS-1 | CUDC-101 | MVP_top60_DNA+RNA_seed | KIT | -0.377473 | GBM | SF295 | Motesanib |
| MVP_top60_DNA_seed | JAK2 | -0.377517 | GBM | 42-MG-BA | Fedratinib | MVP_top60_DNA+RNA_seed | PIK3CB | -0.37565 | GBM | SF268 | TGX221 |
| MVP_top60_DNA_seed | PIK3CB | -0.37565 | GBM | SF268 | TGX221 | MVP_top60_DNA+RNA_seed | KIT | -0.374604 | GBM | D-542MG | OSI-930 |
| MVP_top60_DNA_seed | EGFR | -0.373229 | GBM | D-566MG | Gefitinib | MVP_top60_DNA+RNA_seed | EGFR | -0.373229 | GBM | D-566MG | Gefitinib |
| MVP_top60_DNA_seed | PIK3CD | -0.373199 | GBM | D-263MG | IC-87114 | MVP_top60_DNA+RNA_seed | PIK3CD | -0.373199 | GBM | D-263MG | IC-87114 |
| MVP_top60_DNA_seed | JAK2 | -0.36797 | GBM | YKG-1 | AZD1480 | MVP_top60_DNA+RNA_seed | KIT | -0.371311 | GBM | YKG-1 | OSI-930 |
| MVP_top60_DNA_seed | JAK1 | -0.36797 | GBM | YKG-1 | AZD1480 | MVP_top60_DNA+RNA_seed | JAK2 | -0.36797 | GBM | YKG-1 | AZD1480 |
| MVP_top60_DNA_seed | EGFR | -0.366672 | GBM | SK-MG-1 | AZD8931 | MVP_top60_DNA+RNA_seed | JAK1 | -0.36797 | GBM | YKG-1 | AZD1480 |
| MVP_top60_DNA_seed | EGFR | -0.36401 | GBM | T98G | Cetuximab | MVP_top60_DNA+RNA_seed | EGFR | -0.366672 | GBM | SK-MG-1 | AZD8931 |
| MVP_top60_DNA_seed | PDGFRB | -0.359892 | GBM | Becker | Pazopanib | MVP_top60_DNA+RNA_seed | ERBB3 | -0.366672 | GBM | SK-MG-1 | AZD8931 |
| MVP_top60_DNA_seed | EGFR | -0.357658 | GBM | SF268 | AZD8931 | MVP_top60_DNA+RNA_seed | EGFR | -0.36401 | GBM | T98G | Cetuximab |
| MVP_top60_DNA_seed | PDGFRB | -0.356805 | GBM | D-423MG | Pazopanib | MVP_top60_DNA+RNA_seed | PDGFRB | -0.359892 | GBM | Becker | Pazopanib |
| MVP_top60_DNA_seed | PIK3CD | -0.35571 | GBM | SK-MG-1 | Idelalisib | MVP_top60_DNA+RNA_seed | KIT | -0.359892 | GBM | Becker | Pazopanib |
| MVP_top60_DNA_seed | PIK3CD | -0.354333 | GBM | U-118-MG | IC-87114 | MVP_top60_DNA+RNA_seed | EGFR | -0.357658 | GBM | SF268 | AZD8931 |
| MVP_top60_DNA_seed | EP300 | -0.352271 | GBM | 42-MG-BA | I-CBP112 | MVP_top60_DNA+RNA_seed | ERBB3 | -0.357658 | GBM | SF268 | AZD8931 |
| MVP_top60_DNA_seed | JAK2 | -0.346322 | GBM | CAS-1 | AZD1480 | MVP_top60_DNA+RNA_seed | AKT1 | -0.356901 | GBM | SF268 | AKT inhibitor VIII |
| MVP_top60_DNA_seed | JAK1 | -0.346322 | GBM | CAS-1 | AZD1480 | MVP_top60_DNA+RNA_seed | PDGFRB | -0.356805 | GBM | D-423MG | Pazopanib |
| MVP_top60_DNA_seed | PDGFRA | -0.345152 | GBM | SF268 | Masitinib | MVP_top60_DNA+RNA_seed | KIT | -0.356805 | GBM | D-423MG | Pazopanib |
| MVP_top60_DNA_seed | PDGFRB | -0.345152 | GBM | SF268 | Masitinib | MVP_top60_DNA+RNA_seed | PIK3CD | -0.35571 | GBM | SK-MG-1 | Idelalisib |
| MVP_top60_DNA_seed | JAK2 | -0.343199 | GBM | SW1088 | AZD1480 | MVP_top60_DNA+RNA_seed | PIK3CD | -0.354333 | GBM | U-118-MG | IC-87114 |
| MVP_top60_DNA_seed | JAK1 | -0.343199 | GBM | SW1088 | AZD1480 | MVP_top60_DNA+RNA_seed | EP300 | -0.352271 | GBM | 42-MG-BA | I-CBP112 |
| MVP_top60_DNA_seed | PIK3CD | -0.341783 | GBM | KS-1 | Idelalisib | MVP_top60_DNA+RNA_seed | CREBBP | -0.352271 | GBM | 42-MG-BA | I-CBP112 |
| MVP_top60_DNA_seed | EGFR | -0.341445 | GBM | D-542MG | AZD8931 | MVP_top60_DNA+RNA_seed | IGF1R | -0.348416 | GBM | LN-18 | BMS-754807 |
| MVP_top60_DNA_seed | JAK3 | -0.339821 | GBM | U-87-MG | JAK3_7406 | MVP_top60_DNA+RNA_seed | KIT | -0.347847 | GBM | T98G | Motesanib |
| MVP_top60_DNA_seed | JAK1 | -0.338013 | GBM | LN-18 | JAK1_3715 | MVP_top60_DNA+RNA_seed | JAK2 | -0.346322 | GBM | CAS-1 | AZD1480 |
| MVP_top60_DNA_seed | EGFR | -0.335944 | GBM | D-566MG | Afatinib | MVP_top60_DNA+RNA_seed | JAK1 | -0.346322 | GBM | CAS-1 | AZD1480 |
| MVP_top60_DNA_seed | PIK3CD | -0.335535 | GBM | LN-18 | IC-87114 | MVP_top60_DNA+RNA_seed | PDGFRA | -0.345152 | GBM | SF268 | Masitinib |
| MVP_top60_DNA_seed | EGFR | -0.331832 | GBM | AM-38 | Pelitinib | MVP_top60_DNA+RNA_seed | PDGFRB | -0.345152 | GBM | SF268 | Masitinib |
| MVP_top60_DNA_seed | PIK3CA | -0.327198 | GBM | KS-1 | AZD8835 | MVP_top60_DNA+RNA_seed | KIT | -0.345152 | GBM | SF268 | Masitinib |
| MVP_top60_DNA_seed | PIK3CD | -0.327198 | GBM | KS-1 | AZD8835 | MVP_top60_DNA+RNA_seed | AKT1 | -0.343938 | GBM | Becker | A-443654 |
| MVP_top60_DNA_seed | PIK3CB | -0.319046 | GBM | YKG-1 | AZD6482 | MVP_top60_DNA+RNA_seed | IGF1R | -0.343736 | GBM | DBTRG-05MG | BMS-536924 |
| MVP_top60_DNA_seed | PDGFRA | -0.317828 | GBM | D-423MG | Masitinib | MVP_top60_DNA+RNA_seed | JAK2 | -0.343199 | GBM | SW1088 | AZD1480 |
| MVP_top60_DNA_seed | PDGFRB | -0.317828 | GBM | D-423MG | Masitinib | MVP_top60_DNA+RNA_seed | JAK1 | -0.343199 | GBM | SW1088 | AZD1480 |
| MVP_top60_DNA_seed | JAK3 | -0.312285 | GBM | SF126 | JAK3_7406 | MVP_top60_DNA+RNA_seed | MET | -0.341919 | GBM | YKG-1 | SU11274 |
| MVP_top60_DNA_seed | PDGFRA | -0.311239 | GBM | AM-38 | Masitinib | MVP_top60_DNA+RNA_seed | PIK3CD | -0.341783 | GBM | KS-1 | Idelalisib |
| MVP_top60_DNA_seed | PDGFRB | -0.311239 | GBM | AM-38 | Masitinib | MVP_top60_DNA+RNA_seed | EGFR | -0.341445 | GBM | D-542MG | AZD8931 |

| | | | | | | | | | | | |
|---|---|---|---|---|---|---|---|---|---|---|---|
| MVP_top60_DNA_seed | JAK1 | -0.311032 | GBM | YH-13 | JAK1_3715 | MVP_top60_DNA+RNA_seed | ERBB3 | -0.341445 | GBM | D-542MG | AZD8931 |
| MVP_top60_DNA_seed | EGFR | -0.309137 | GBM | 8-MG-BA | Gefitinib | MVP_top60_DNA+RNA_seed | AKT1 | -0.340705 | GBM | 42-MG-BA | AKT inhibitor VIII |
| MVP_top60_DNA_seed | JAK2 | -0.305045 | GBM | D-423MG | Fedratinib | MVP_top60_DNA+RNA_seed | IGF1R | -0.34066 | GBM | U251 | GSK1904529A |
| MVP_top60_DNA_seed | PIK3CB | -0.30388 | GBM | 8-MG-BA | AZD8186 | MVP_top60_DNA+RNA_seed | KIT | -0.339833 | GBM | SW1088 | Axitinib |
| MVP_top60_DNA_seed | PIK3CD | -0.30388 | GBM | 8-MG-BA | AZD8186 | MVP_top60_DNA+RNA_seed | JAK3 | -0.339821 | GBM | U-87-MG | JAK3_7406 |
| MVP_top60_DNA_seed | CDK4 | -0.302098 | GBM | D-566MG | Palbociclib | MVP_top60_DNA+RNA_seed | JAK1 | -0.338013 | GBM | LN-18 | JAK1_3715 |
| MVP_top60_DNA_seed | PDGFRA | -0.299402 | GBM | YH-13 | Masitinib | MVP_top60_DNA+RNA_seed | EGFR | -0.335944 | GBM | D-566MG | Afatinib |
| MVP_top60_DNA_seed | PDGFRB | -0.299402 | GBM | YH-13 | Masitinib | MVP_top60_DNA+RNA_seed | PIK3CD | -0.335535 | GBM | LN-18 | IC-87114 |
| MVP_top60_DNA_seed | EGFR | -0.297176 | GBM | D-392MG | Erlotinib | MVP_top60_DNA+RNA_seed | MET | -0.333492 | GBM | GB-1 | PHA-665752 |
| MVP_top60_DNA_seed | PDGFRA | -0.296604 | GBM | SF295 | Masitinib | MVP_top60_DNA+RNA_seed | KIT | -0.333278 | GBM | A172 | OSI-930 |
| MVP_top60_DNA_seed | PDGFRB | -0.296604 | GBM | SF295 | Masitinib | MVP_top60_DNA+RNA_seed | EGFR | -0.331832 | GBM | AM-38 | Pelitinib |
| MVP_top60_DNA_seed | PIK3CD | -0.296182 | GBM | D-392MG | Idelalisib | MVP_top60_DNA+RNA_seed | PIK3CA | -0.327198 | GBM | KS-1 | AZD8835 |
| MVP_top60_DNA_seed | PDGFRA | -0.294946 | GBM | LNZTA3WT4 | Amuvatinib | MVP_top60_DNA+RNA_seed | PIK3CD | -0.327198 | GBM | KS-1 | AZD8835 |
| MVP_top60_DNA_seed | EGFR | -0.290201 | GBM | SF268 | Lapatinib | MVP_top60_DNA+RNA_seed | PIK3CB | -0.319046 | GBM | YKG-1 | AZD6482 |
| MVP_top60_DNA_seed | PIK3CD | -0.284676 | GBM | D-566MG | IC-87114 | MVP_top60_DNA+RNA_seed | PDGFRA | -0.317828 | GBM | D-423MG | Masitinib |
| MVP_top60_DNA_seed | EGFR | -0.283792 | GBM | U-87-MG | Afatinib | MVP_top60_DNA+RNA_seed | PDGFRB | -0.317828 | GBM | D-423MG | Masitinib |
| MVP_top60_DNA_seed | CDK4 | -0.283675 | GBM | SF268 | CGP-082996 | MVP_top60_DNA+RNA_seed | KIT | -0.317828 | GBM | D-423MG | Masitinib |
| MVP_top60_DNA_seed | JAK3 | -0.279524 | GBM | 8-MG-BA | WHI-P97 | MVP_top60_DNA+RNA_seed | AKT1 | -0.317762 | GBM | SF126 | A-443654 |
| MVP_top60_DNA_seed | EGFR | -0.277359 | GBM | D-247MG | Erlotinib | MVP_top60_DNA+RNA_seed | IGF1R | -0.317466 | GBM | SNB75 | BMS-754807 |
| MVP_top60_DNA_seed | EGFR | -0.27486 | GBM | KS-1 | AZD8931 | MVP_top60_DNA+RNA_seed | JAK3 | -0.312285 | GBM | SF126 | JAK3_7406 |
| MVP_top60_DNA_seed | PDGFRA | -0.27437 | GBM | SF126 | Masitinib | MVP_top60_DNA+RNA_seed | MET | -0.311856 | GBM | LN-18 | JNJ38877605 |
| MVP_top60_DNA_seed | PDGFRB | -0.27437 | GBM | SF126 | Masitinib | MVP_top60_DNA+RNA_seed | PDGFRA | -0.311239 | GBM | AM-38 | Masitinib |
| MVP_top60_DNA_seed | JAK3 | -0.274145 | GBM | SW1088 | JAK3_7406 | MVP_top60_DNA+RNA_seed | PDGFRB | -0.311239 | GBM | AM-38 | Masitinib |
| MVP_top60_DNA_seed | PIK3CA | -0.271978 | GBM | T98G | PI3Ka_4409 | MVP_top60_DNA+RNA_seed | KIT | -0.311239 | GBM | AM-38 | Masitinib |
| MVP_top60_DNA_seed | PIK3CD | -0.271978 | GBM | T98G | PI3Ka_4409 | MVP_top60_DNA+RNA_seed | JAK1 | -0.311032 | GBM | YH-13 | JAK1_3715 |
| MVP_top60_DNA_seed | EGFR | -0.270597 | GBM | KS-1 | PF-00299804 | MVP_top60_DNA+RNA_seed | EGFR | -0.309137 | GBM | 8-MG-BA | Gefitinib |
| MVP_top60_DNA_seed | EGFR | -0.269198 | GBM | SF126 | AZD8931 | MVP_top60_DNA+RNA_seed | IGF1R | -0.308932 | GBM | Becker | BMS-536924 |
| MVP_top60_DNA_seed | EGFR | -0.259102 | GBM | D-542MG | Lapatinib | MVP_top60_DNA+RNA_seed | IGF1R | -0.306488 | GBM | D-542MG | BMS-754807 |
| MVP_top60_DNA_seed | EGFR | -0.252138 | GBM | SNB75 | AZD8931 | MVP_top60_DNA+RNA_seed | JAK2 | -0.305045 | GBM | D-423MG | Fedratinib |
| MVP_top60_DNA_seed | PDGFRA | -0.250136 | GBM | GB-1 | Amuvatinib | MVP_top60_DNA+RNA_seed | MET | -0.304978 | GBM | CAS-1 | JNJ38877605 |
| MVP_top60_DNA_seed | EGFR | -0.249357 | GBM | 8-MG-BA | CUDC-101 | MVP_top60_DNA+RNA_seed | PIK3CB | -0.30388 | GBM | 8-MG-BA | AZD8186 |
| MVP_top60_DNA_seed | PIK3CB | -0.241235 | GBM | LNZTA3WT4 | AZD6482 | MVP_top60_DNA+RNA_seed | PIK3CD | -0.30388 | GBM | 8-MG-BA | AZD8186 |
| MVP_top60_DNA_seed | JAK1 | -0.240181 | GBM | A172 | JAK1_3715 | MVP_top60_DNA+RNA_seed | MET | -0.303078 | GBM | GB-1 | Crizotinib |
| MVP_top60_DNA_seed | EP300 | -0.235579 | GBM | 8-MG-BA | I-CBP112 | MVP_top60_DNA+RNA_seed | CDK4 | -0.302098 | GBM | D-566MG | Palbociclib |
| MVP_top60_DNA_seed | PIK3CA | -0.233947 | GBM | Becker | AZD8835 | MVP_top60_DNA+RNA_seed | IGF1R | -0.300954 | GBM | YH-13 | GSK1904529A |
| MVP_top60_DNA_seed | PIK3CD | -0.233947 | GBM | Becker | AZD8835 | MVP_top60_DNA+RNA_seed | PDGFRA | -0.299402 | GBM | YH-13 | Masitinib |
| MVP_top60_DNA_seed | EGFR | -0.233569 | GBM | DBTRG-05MG | Pelitinib | MVP_top60_DNA+RNA_seed | PDGFRB | -0.299402 | GBM | YH-13 | Masitinib |
| MVP_top60_DNA_seed | PIK3CD | -0.231003 | GBM | 8-MG-BA | Idelalisib | MVP_top60_DNA+RNA_seed | KIT | -0.299402 | GBM | YH-13 | Masitinib |
| MVP_top60_DNA_seed | PDGFRB | -0.226997 | GBM | 8-MG-BA | Pazopanib | MVP_top60_DNA+RNA_seed | EGFR | -0.297176 | GBM | D-392MG | Erlotinib |
| MVP_top60_DNA_seed | PDGFRA | -0.226488 | GBM | KS-1 | Masitinib | MVP_top60_DNA+RNA_seed | PDGFRA | -0.296604 | GBM | SF295 | Masitinib |
| MVP_top60_DNA_seed | PDGFRB | -0.226488 | GBM | KS-1 | Masitinib | MVP_top60_DNA+RNA_seed | PDGFRB | -0.296604 | GBM | SF295 | Masitinib |

| | | | | | | | | | | | |
|---|---|---|---|---|---|---|---|---|---|---|---|
| MVP_top60_DNA_seed | PIK3CA | -0.225268 | GBM | SNB75 | AZD8835 | MVP_top60_DNA+RNA_seed | KIT | -0.296604 | GBM | SF295 | Masitinib |
| MVP_top60_DNA_seed | PIK3CD | -0.225268 | GBM | SNB75 | AZD8835 | MVP_top60_DNA+RNA_seed | PIK3CD | -0.296182 | GBM | D-392MG | Idelalisib |
| MVP_top60_DNA_seed | JAK3 | -0.223313 | GBM | D-566MG | WHI-P97 | MVP_top60_DNA+RNA_seed | PDGFRA | -0.294946 | GBM | LNZTA3WT4 | Amuvatinib |
| MVP_top60_DNA_seed | CDK4 | -0.222332 | GBM | U-118-MG | Palbociclib | MVP_top60_DNA+RNA_seed | KIT | -0.294946 | GBM | LNZTA3WT4 | Amuvatinib |
| MVP_top60_DNA_seed | PIK3CA | -0.221465 | GBM | D-423MG | AZD8835 | MVP_top60_DNA+RNA_seed | EGFR | -0.290201 | GBM | SF268 | Lapatinib |
| MVP_top60_DNA_seed | PIK3CD | -0.221465 | GBM | D-423MG | AZD8835 | MVP_top60_DNA+RNA_seed | IGF1R | -0.289231 | GBM | SF268 | BMS-536924 |
| MVP_top60_DNA_seed | EGFR | -0.218535 | GBM | D-423MG | Pelitinib | MVP_top60_DNA+RNA_seed | MET | -0.28793 | GBM | U251 | SU11274 |
| MVP_top60_DNA_seed | EGFR | -0.217538 | GBM | SK-MG-1 | Cetuximab | MVP_top60_DNA+RNA_seed | AKT1 | -0.287139 | GBM | LN-18 | MK-2206 |
| MVP_top60_DNA_seed | EGFR | -0.208843 | GBM | LN-229 | Afatinib | MVP_top60_DNA+RNA_seed | PIK3CD | -0.284676 | GBM | D-566MG | IC-87114 |
| MVP_top60_DNA_seed | EGFR | -0.207617 | GBM | LN-405 | Gefitinib | MVP_top60_DNA+RNA_seed | MET | -0.283913 | GBM | LN-18 | AZD6094 |
| MVP_top60_DNA_seed | JAK3 | -0.196411 | GBM | DK-MG | JAK3_7406 | MVP_top60_DNA+RNA_seed | EGFR | -0.283792 | GBM | U-87-MG | Afatinib |
| MVP_top60_DNA_seed | EGFR | -0.192046 | GBM | GB-1 | AST-1306 | MVP_top60_DNA+RNA_seed | CDK4 | -0.283675 | GBM | SF268 | CGP-082996 |
| MVP_top60_DNA_seed | EGFR | -0.192031 | GBM | U-87-MG | AZD8931 | MVP_top60_DNA+RNA_seed | AKT1 | -0.282183 | GBM | 42-MG-BA | GSK690693 |
| MVP_top60_DNA_seed | EGFR | -0.190368 | GBM | KS-1 | Afatinib | MVP_top60_DNA+RNA_seed | JAK3 | -0.279524 | GBM | 8-MG-BA | WHI-P97 |
| MVP_top60_DNA_seed | EGFR | -0.186879 | GBM | YKG-1 | CI-1033 | MVP_top60_DNA+RNA_seed | EGFR | -0.277359 | GBM | D-247MG | Erlotinib |
| MVP_top60_DNA_seed | JAK2 | -0.182608 | GBM | SF268 | Fedratinib | MVP_top60_DNA+RNA_seed | IGF1R | -0.276754 | GBM | SNB75 | GSK1904529A |
| MVP_top60_DNA_seed | JAK2 | -0.182107 | GBM | 8-MG-BA | Ruxolitinib | MVP_top60_DNA+RNA_seed | EGFR | -0.27486 | GBM | KS-1 | AZD8931 |
| MVP_top60_DNA_seed | JAK1 | -0.182107 | GBM | 8-MG-BA | Ruxolitinib | MVP_top60_DNA+RNA_seed | ERBB3 | -0.27486 | GBM | KS-1 | AZD8931 |
| MVP_top60_DNA_seed | EGFR | -0.180533 | GBM | KS-1 | Afatinib | MVP_top60_DNA+RNA_seed | PDGFRA | -0.27437 | GBM | SF126 | Masitinib |
| MVP_top60_DNA_seed | EGFR | -0.179381 | GBM | DBTRG-05MG | Gefitinib | MVP_top60_DNA+RNA_seed | PDGFRB | -0.27437 | GBM | SF126 | Masitinib |
| MVP_top60_DNA_seed | JAK3 | -0.179199 | GBM | SNB75 | JAK3_7406 | MVP_top60_DNA+RNA_seed | KIT | -0.27437 | GBM | SF126 | Masitinib |
| MVP_top60_DNA_seed | JAK2 | -0.173952 | GBM | 8-MG-BA | Fedratinib | MVP_top60_DNA+RNA_seed | JAK3 | -0.274145 | GBM | SW1088 | JAK3_7406 |
| MVP_top60_DNA_seed | EGFR | -0.171645 | GBM | U251 | Pelitinib | MVP_top60_DNA+RNA_seed | PIK3CA | -0.271978 | GBM | T98G | PI3Ka_4409 |
| MVP_top60_DNA_seed | PDGFRB | -0.169749 | GBM | U251 | Pazopanib | MVP_top60_DNA+RNA_seed | PIK3CD | -0.271978 | GBM | T98G | PI3Ka_4409 |
| MVP_top60_DNA_seed | EGFR | -0.158119 | GBM | D-542MG | CUDC-101 | MVP_top60_DNA+RNA_seed | EGFR | -0.270597 | GBM | KS-1 | PF-00299804 |
| MVP_top60_DNA_seed | EGFR | -0.157381 | GBM | 8-MG-BA | Cetuximab | MVP_top60_DNA+RNA_seed | EGFR | -0.269198 | GBM | SF126 | AZD8931 |
| MVP_top60_DNA_seed | PIK3CD | -0.152861 | GBM | LN-405 | Idelalisib | MVP_top60_DNA+RNA_seed | ERBB3 | -0.269198 | GBM | SF126 | AZD8931 |
| MVP_top60_DNA_seed | PDGFRB | -0.152528 | GBM | LNZTA3WT4 | Pazopanib | MVP_top60_DNA+RNA_seed | IGF1R | -0.266409 | GBM | SF295 | BMS-754807 |
| MVP_top60_DNA_seed | JAK1 | -0.151369 | GBM | SK-MG-1 | JAK1_3715 | MVP_top60_DNA+RNA_seed | AKT1 | -0.26342 | GBM | D-263MG | MK-2206 |
| MVP_top60_DNA_seed | EGFR | -0.146351 | GBM | A172 | CUDC-101 | MVP_top60_DNA+RNA_seed | AKT1 | -0.263352 | GBM | SF295 | MK-2206 |
| MVP_top60_DNA_seed | PIK3CA | -0.143762 | GBM | SW1088 | PI3Ka_4409 | MVP_top60_DNA+RNA_seed | MET | -0.261263 | GBM | KS-1 | AZD6094 |
| MVP_top60_DNA_seed | PIK3CD | -0.143762 | GBM | SW1088 | PI3Ka_4409 | MVP_top60_DNA+RNA_seed | EGFR | -0.259102 | GBM | D-542MG | Lapatinib |
| MVP_top60_DNA_seed | EGFR | -0.143505 | GBM | GB-1 | CI-1033 | MVP_top60_DNA+RNA_seed | AKT1 | -0.256089 | GBM | SF295 | BAY AKT1 |
| MVP_top60_DNA_seed | EGFR | -0.142824 | GBM | YKG-1 | Cetuximab | MVP_top60_DNA+RNA_seed | EGFR | -0.252138 | GBM | SNB75 | AZD8931 |
| MVP_top60_DNA_seed | EGFR | -0.13723 | GBM | DK-MG | PF-00299804 | MVP_top60_DNA+RNA_seed | ERBB3 | -0.252138 | GBM | SNB75 | AZD8931 |
| MVP_top60_DNA_seed | JAK2 | -0.136964 | GBM | D-247MG | Ruxolitinib | MVP_top60_DNA+RNA_seed | PDGFRA | -0.250136 | GBM | GB-1 | Amuvatinib |
| MVP_top60_DNA_seed | JAK1 | -0.136964 | GBM | D-247MG | Ruxolitinib | MVP_top60_DNA+RNA_seed | KIT | -0.250136 | GBM | GB-1 | Amuvatinib |
| MVP_top60_DNA_seed | PIK3CB | -0.136687 | GBM | LN-229 | AZD6482 | MVP_top60_DNA+RNA_seed | EGFR | -0.249357 | GBM | 8-MG-BA | CUDC-101 |
| MVP_top60_DNA_seed | PIK3CB | -0.13328 | GBM | U-87-MG | TGX221 | MVP_top60_DNA+RNA_seed | IGF1R | -0.242647 | GBM | DBTRG-05MG | BMS-754807 |
| MVP_top60_DNA_seed | PIK3CD | -0.13142 | GBM | T98G | Idelalisib | MVP_top60_DNA+RNA_seed | PIK3CB | -0.241235 | GBM | LNZTA3WT4 | AZD6482 |
| MVP_top60_DNA_seed | CDK4 | -0.130802 | GBM | AM-38 | Palbociclib | MVP_top60_DNA+RNA_seed | JAK1 | -0.240181 | GBM | A172 | JAK1_3715 |

| | | | | | | | | | | | |
|---|---|---|---|---|---|---|---|---|---|---|---|
| MVP_top60_DNA_seed | PIK3CA | -0.12983 | GBM | D-263MG | PI3Ka_4409 | MVP_top60_DNA+RNA_seed | AKT1 | -0.238595 | GBM | U-118-MG | AKT inhibitor VIII |
| MVP_top60_DNA_seed | PIK3CD | -0.12983 | GBM | D-263MG | PI3Ka_4409 | MVP_top60_DNA+RNA_seed | EP300 | -0.235579 | GBM | 8-MG-BA | I-CBP112 |
| MVP_top60_DNA_seed | JAK2 | -0.128126 | GBM | A172 | Fedratinib | MVP_top60_DNA+RNA_seed | CREBBP | -0.235579 | GBM | 8-MG-BA | I-CBP112 |
| MVP_top60_DNA_seed | PDGFRA | -0.12509 | GBM | U-87-MG | Masitinib | MVP_top60_DNA+RNA_seed | IGF1R | -0.234516 | GBM | 8-MG-BA | BMS-536924 |
| MVP_top60_DNA_seed | PDGFRB | -0.12509 | GBM | U-87-MG | Masitinib | MVP_top60_DNA+RNA_seed | MET | -0.234333 | GBM | YKG-1 | JNJ38877605 |
| MVP_top60_DNA_seed | JAK2 | -0.124539 | GBM | SK-MG-1 | Fedratinib | MVP_top60_DNA+RNA_seed | PIK3CA | -0.233947 | GBM | Becker | AZD8835 |
| MVP_top60_DNA_seed | EGFR | -0.119082 | GBM | T98G | AZD8931 | MVP_top60_DNA+RNA_seed | PIK3CD | -0.233947 | GBM | Becker | AZD8835 |
| MVP_top60_DNA_seed | PDGFRB | -0.118261 | GBM | LN-18 | Pazopanib | MVP_top60_DNA+RNA_seed | EGFR | -0.233569 | GBM | DBTRG-05MG | Pelitinib |
| MVP_top60_DNA_seed | EGFR | -0.116551 | GBM | SF126 | CI-1033 | MVP_top60_DNA+RNA_seed | IGF1R | -0.232859 | GBM | D-263MG | BMS-536924 |
| MVP_top60_DNA_seed | JAK1 | -0.11532 | GBM | D-566MG | JAK1_3715 | MVP_top60_DNA+RNA_seed | PIK3CD | -0.231003 | GBM | 8-MG-BA | Idelalisib |
| MVP_top60_DNA_seed | PDGFRA | -0.110574 | GBM | CAS-1 | Amuvatinib | MVP_top60_DNA+RNA_seed | KIT | -0.229462 | GBM | SF268 | Motesanib |
| MVP_top60_DNA_seed | PIK3CA | -0.109203 | GBM | D-263MG | AZD8835 | MVP_top60_DNA+RNA_seed | IGF1R | -0.228332 | GBM | SF126 | GSK1904529A |
| MVP_top60_DNA_seed | PIK3CD | -0.109203 | GBM | D-263MG | AZD8835 | MVP_top60_DNA+RNA_seed | PDGFRB | -0.226997 | GBM | 8-MG-BA | Pazopanib |
| MVP_top60_DNA_seed | EGFR | -0.108835 | GBM | M059J | AZD8931 | MVP_top60_DNA+RNA_seed | KIT | -0.226997 | GBM | 8-MG-BA | Pazopanib |
| MVP_top60_DNA_seed | PDGFRA | -0.107572 | GBM | SF268 | Amuvatinib | MVP_top60_DNA+RNA_seed | PDGFRA | -0.226488 | GBM | KS-1 | Masitinib |
| MVP_top60_DNA_seed | JAK3 | -0.102274 | GBM | T98G | WHI-P97 | MVP_top60_DNA+RNA_seed | PDGFRB | -0.226488 | GBM | KS-1 | Masitinib |
| MVP_top60_DNA_seed | PIK3CB | -0.099929 | GBM | D-542MG | AZD8186 | MVP_top60_DNA+RNA_seed | KIT | -0.226488 | GBM | KS-1 | Masitinib |
| MVP_top60_DNA_seed | PIK3CD | -0.099929 | GBM | D-542MG | AZD8186 | MVP_top60_DNA+RNA_seed | AKT1 | -0.226397 | GBM | D-542MG | MK-2206 |
| MVP_top60_DNA_seed | JAK2 | -0.097213 | GBM | DK-MG | AZD1480 | MVP_top60_DNA+RNA_seed | PIK3CA | -0.225268 | GBM | SNB75 | AZD8835 |
| MVP_top60_DNA_seed | JAK1 | -0.097213 | GBM | DK-MG | AZD1480 | MVP_top60_DNA+RNA_seed | PIK3CD | -0.225268 | GBM | SNB75 | AZD8835 |
| MVP_top60_DNA_seed | EGFR | -0.095753 | GBM | CAS-1 | AST-1306 | MVP_top60_DNA+RNA_seed | JAK3 | -0.223313 | GBM | D-566MG | WHI-P97 |
| MVP_top60_DNA_seed | JAK2 | -0.094521 | GBM | A172 | Ruxolitinib | MVP_top60_DNA+RNA_seed | CDK4 | -0.222332 | GBM | U-118-MG | Palbociclib |
| MVP_top60_DNA_seed | JAK1 | -0.094521 | GBM | A172 | Ruxolitinib | MVP_top60_DNA+RNA_seed | PIK3CA | -0.221465 | GBM | D-423MG | AZD8835 |
| MVP_top60_DNA_seed | JAK2 | -0.09292 | GBM | D-566MG | Ruxolitinib | MVP_top60_DNA+RNA_seed | PIK3CD | -0.221465 | GBM | D-423MG | AZD8835 |
| MVP_top60_DNA_seed | JAK1 | -0.09292 | GBM | D-566MG | Ruxolitinib | MVP_top60_DNA+RNA_seed | EGFR | -0.218535 | GBM | D-423MG | Pelitinib |
| MVP_top60_DNA_seed | JAK3 | -0.09253 | GBM | DK-MG | WHI-P97 | MVP_top60_DNA+RNA_seed | KIT | -0.218461 | GBM | Becker | Axitinib |
| MVP_top60_DNA_seed | JAK2 | -0.092216 | GBM | T98G | Ruxolitinib | MVP_top60_DNA+RNA_seed | EGFR | -0.217538 | GBM | SK-MG-1 | Cetuximab |
| MVP_top60_DNA_seed | JAK1 | -0.092216 | GBM | T98G | Ruxolitinib | MVP_top60_DNA+RNA_seed | KIT | -0.217188 | GBM | D-392MG | OSI-930 |
| MVP_top60_DNA_seed | JAK2 | -0.088868 | GBM | YH-13 | Fedratinib | MVP_top60_DNA+RNA_seed | KIT | -0.216375 | GBM | GB-1 | Sorafenib |
| MVP_top60_DNA_seed | JAK1 | -0.086346 | GBM | T98G | JAK1_3715 | MVP_top60_DNA+RNA_seed | EGFR | -0.208843 | GBM | LN-229 | Afatinib |
| MVP_top60_DNA_seed | EGFR | -0.085668 | GBM | AM-38 | Gefitinib | MVP_top60_DNA+RNA_seed | EGFR | -0.207617 | GBM | LN-405 | Gefitinib |
| MVP_top60_DNA_seed | PIK3CB | -0.083218 | GBM | D-263MG | AZD6482 | MVP_top60_DNA+RNA_seed | IGF1R | -0.207582 | GBM | SW1088 | BMS-754807 |
| MVP_top60_DNA_seed | PIK3CD | -0.078886 | GBM | D-247MG | IC-87114 | MVP_top60_DNA+RNA_seed | IGF1R | -0.204489 | GBM | SW1088 | BMS-536924 |
| MVP_top60_DNA_seed | EGFR | -0.078794 | GBM | M059J | CUDC-101 | MVP_top60_DNA+RNA_seed | MET | -0.200692 | GBM | Becker | SU11274 |
| MVP_top60_DNA_seed | JAK2 | -0.07739 | GBM | D-247MG | AZD1480 | MVP_top60_DNA+RNA_seed | KIT | -0.198885 | GBM | LNZTA3WT4 | OSI-930 |
| MVP_top60_DNA_seed | JAK1 | -0.07739 | GBM | D-247MG | AZD1480 | MVP_top60_DNA+RNA_seed | AKT1 | -0.196799 | GBM | Becker | MK-2206 |
| MVP_top60_DNA_seed | PIK3CD | -0.076874 | GBM | SF295 | Idelalisib | MVP_top60_DNA+RNA_seed | JAK3 | -0.196411 | GBM | DK-MG | JAK3_7406 |
| MVP_top60_DNA_seed | PIK3CB | -0.076718 | GBM | GB-1 | AZD6482 | MVP_top60_DNA+RNA_seed | AKT1 | -0.196151 | GBM | U-118-MG | GSK690693 |
| MVP_top60_DNA_seed | JAK1 | -0.074043 | GBM | D-423MG | JAK1_3715 | MVP_top60_DNA+RNA_seed | EGFR | -0.192046 | GBM | GB-1 | AST-1306 |
| MVP_top60_DNA_seed | CDK4 | -0.07251 | GBM | D-392MG | Palbociclib | MVP_top60_DNA+RNA_seed | EGFR | -0.192031 | GBM | U-87-MG | AZD8931 |
| MVP_top60_DNA_seed | JAK2 | -0.071016 | GBM | SW1088 | Fedratinib | MVP_top60_DNA+RNA_seed | ERBB3 | -0.192031 | GBM | U-87-MG | AZD8931 |

| | | | | | | | | | | | |
|---|---|---|---|---|---|---|---|---|---|---|---|
| MVP_top60_DNA_seed | PIK3CD | -0.06567 | GBM | GB-1 | IC-87114 | MVP_top60_DNA+RNA_seed | EGFR | -0.190368 | GBM | KS-1 | Afatinib |
| MVP_top60_DNA_seed | JAK2 | -0.063483 | GBM | U251 | Ruxolitinib | MVP_top60_DNA+RNA_seed | EGFR | -0.186879 | GBM | YKG-1 | CI-1033 |
| MVP_top60_DNA_seed | JAK1 | -0.063483 | GBM | U251 | Ruxolitinib | MVP_top60_DNA+RNA_seed | MET | -0.183454 | GBM | D-263MG | AZD6094 |
| MVP_top60_DNA_seed | JAK2 | -0.060648 | GBM | AM-38 | Ruxolitinib | MVP_top60_DNA+RNA_seed | JAK2 | -0.182608 | GBM | SF268 | Fedratinib |
| MVP_top60_DNA_seed | JAK1 | -0.060648 | GBM | AM-38 | Ruxolitinib | MVP_top60_DNA+RNA_seed | JAK2 | -0.182107 | GBM | 8-MG-BA | Ruxolitinib |
| MVP_top60_DNA_seed | JAK3 | -0.057614 | GBM | Becker | WHI-P97 | MVP_top60_DNA+RNA_seed | JAK1 | -0.182107 | GBM | 8-MG-BA | Ruxolitinib |
| MVP_top60_DNA_seed | PIK3CD | -0.057539 | GBM | 8-MG-BA | IC-87114 | MVP_top60_DNA+RNA_seed | EGFR | -0.180533 | GBM | KS-1 | Afatinib |
| MVP_top60_DNA_seed | EGFR | -0.057445 | GBM | AM-38 | AST-1306 | MVP_top60_DNA+RNA_seed | AKT1 | -0.180512 | GBM | SK-MG-1 | BAY AKT1 |
| MVP_top60_DNA_seed | PIK3CD | -0.053876 | GBM | U-87-MG | Idelalisib | MVP_top60_DNA+RNA_seed | EGFR | -0.179381 | GBM | DBTRG-05MG | Gefitinib |
| MVP_top60_DNA_seed | JAK2 | -0.046987 | GBM | AM-38 | Fedratinib | MVP_top60_DNA+RNA_seed | JAK3 | -0.179199 | GBM | SNB75 | JAK3_7406 |
| MVP_top60_DNA_seed | EGFR | -0.046977 | GBM | AM-38 | CUDC-101 | MVP_top60_DNA+RNA_seed | JAK2 | -0.173952 | GBM | 8-MG-BA | Fedratinib |
| MVP_top60_DNA_seed | EGFR | -0.038738 | GBM | D-423MG | Cetuximab | MVP_top60_DNA+RNA_seed | KIT | -0.173465 | GBM | LN-18 | Axitinib |
| MVP_top60_DNA_seed | JAK1 | -0.037805 | GBM | U251 | JAK1_3715 | MVP_top60_DNA+RNA_seed | EGFR | -0.171645 | GBM | U251 | Pelitinib |
| MVP_top60_DNA_seed | JAK1 | -0.037624 | GBM | 42-MG-BA | JAK1_3715 | MVP_top60_DNA+RNA_seed | PDGFRB | -0.169749 | GBM | U251 | Pazopanib |
| MVP_top60_DNA_seed | JAK3 | -0.03534 | GBM | LN-229 | WHI-P97 | MVP_top60_DNA+RNA_seed | KIT | -0.169749 | GBM | U251 | Pazopanib |
| MVP_top60_DNA_seed | EGFR | -0.033848 | GBM | YH-13 | CI-1033 | MVP_top60_DNA+RNA_seed | AKT1 | -0.168991 | GBM | SNB75 | BAY AKT1 |
| MVP_top60_DNA_seed | PIK3CD | -0.033518 | GBM | D-247MG | Idelalisib | MVP_top60_DNA+RNA_seed | AKT1 | -0.166577 | GBM | GB-1 | MK-2206 |
| MVP_top60_DNA_seed | PIK3CB | -0.030548 | GBM | M059J | AZD6482 | MVP_top60_DNA+RNA_seed | MET | -0.161147 | GBM | 8-MG-BA | AZD6094 |
| MVP_top60_DNA_seed | JAK2 | -0.03013 | GBM | SF126 | AZD1480 | MVP_top60_DNA+RNA_seed | EGFR | -0.158119 | GBM | D-542MG | CUDC-101 |
| MVP_top60_DNA_seed | JAK1 | -0.03013 | GBM | SF126 | AZD1480 | MVP_top60_DNA+RNA_seed | AKT1 | -0.157911 | GBM | U251 | GSK690693 |
| MVP_top60_DNA_seed | EGFR | -0.029853 | GBM | 42-MG-BA | CUDC-101 | MVP_top60_DNA+RNA_seed | EGFR | -0.157381 | GBM | 8-MG-BA | Cetuximab |
| MVP_top60_DNA_seed | EGFR | -0.024533 | GBM | KS-1 | AST-1306 | MVP_top60_DNA+RNA_seed | PIK3CD | -0.152861 | GBM | LN-405 | Idelalisib |
| MVP_top60_DNA_seed | EGFR | -0.021607 | GBM | 8-MG-BA | Erlotinib | MVP_top60_DNA+RNA_seed | PDGFRB | -0.152528 | GBM | LNZTA3WT4 | Pazopanib |
| MVP_top60_DNA_seed | EP300 | -0.018012 | GBM | T98G | I-CBP112 | MVP_top60_DNA+RNA_seed | KIT | -0.152528 | GBM | LNZTA3WT4 | Pazopanib |
| MVP_top60_DNA_seed | JAK2 | -0.014916 | GBM | U251 | AZD1480 | MVP_top60_DNA+RNA_seed | JAK1 | -0.151369 | GBM | SK-MG-1 | JAK1_3715 |
| MVP_top60_DNA_seed | JAK1 | -0.014916 | GBM | U251 | AZD1480 | MVP_top60_DNA+RNA_seed | KIT | -0.151015 | GBM | SK-MG-1 | Axitinib |
| MVP_top60_DNA_seed | EGFR | -0.013575 | GBM | SF126 | AST-1306 | MVP_top60_DNA+RNA_seed | AKT1 | -0.14893 | GBM | LN-405 | MK-2206 |
| MVP_top60_DNA_seed | EGFR | -0.011909 | GBM | YKG-1 | PF-00299804 | MVP_top60_DNA+RNA_seed | EGFR | -0.146351 | GBM | A172 | CUDC-101 |
| MVP_top60_DNA_seed | PIK3CB | -0.011033 | GBM | U-87-MG | AZD6482 | MVP_top60_DNA+RNA_seed | PIK3CA | -0.143762 | GBM | SW1088 | PI3Ka_4409 |
| MVP_top60_DNA_seed | EGFR | -0.010827 | GBM | LN-229 | AST-1306 | MVP_top60_DNA+RNA_seed | PIK3CD | -0.143762 | GBM | SW1088 | PI3Ka_4409 |
| MVP_top60_DNA_seed | PDGFRA | -0.009646 | GBM | LN-18 | Masitinib | MVP_top60_DNA+RNA_seed | EGFR | -0.143505 | GBM | GB-1 | CI-1033 |
| MVP_top60_DNA_seed | PDGFRB | -0.009646 | GBM | LN-18 | Masitinib | MVP_top60_DNA+RNA_seed | EGFR | -0.142824 | GBM | YKG-1 | Cetuximab |
| MVP_top60_DNA_seed | PIK3CB | -0.009252 | GBM | M059J | AZD8186 | MVP_top60_DNA+RNA_seed | EGFR | -0.13723 | GBM | DK-MG | PF-00299804 |
| MVP_top60_DNA_seed | PIK3CD | -0.009252 | GBM | M059J | AZD8186 | MVP_top60_DNA+RNA_seed | JAK2 | -0.136964 | GBM | D-247MG | Ruxolitinib |
| MVP_top60_DNA_seed | JAK1 | -0.007254 | GBM | D-247MG | JAK1_3715 | MVP_top60_DNA+RNA_seed | JAK1 | -0.136964 | GBM | D-247MG | Ruxolitinib |
| MVP_top60_DNA_seed | EP300 | -0.006046 | GBM | U251 | I-CBP112 | MVP_top60_DNA+RNA_seed | PIK3CB | -0.136687 | GBM | LN-229 | AZD6482 |
| MVP_top60_DNA_seed | CDK4 | -0.000924 | GBM | LNZTA3WT4 | Palbociclib | MVP_top60_DNA+RNA_seed | AKT1 | -0.133288 | GBM | SNB75 | MK-2206 |
| MVP_top60_DNA_seed | PDGFRA | -0.000135 | GBM | 8-MG-BA | Amuvatinib | MVP_top60_DNA+RNA_seed | PIK3CB | -0.13328 | GBM | U-87-MG | TGX221 |
| MVP_top60_DNA_seed | PDGFRA | -0.000123 | GBM | D-542MG | Amuvatinib | MVP_top60_DNA+RNA_seed | PIK3CD | -0.13142 | GBM | T98G | Idelalisib |
| MVP_top60_DNA_seed | JAK2 | -6.20E-05 | GBM | T98G | Fedratinib | MVP_top60_DNA+RNA_seed | CDK4 | -0.130802 | GBM | AM-38 | Palbociclib |
| MVP_top60_DNA_seed | JAK2 | 0.004519 | GBM | M059J | Fedratinib | MVP_top60_DNA+RNA_seed | PIK3CA | -0.12983 | GBM | D-263MG | PI3Ka_4409 |

| | | | | | | | | | | | |
|---|---|---|---|---|---|---|---|---|---|---|---|
| MVP_top60_DNA_seed | PIK3CA | 0.005288 | GBM | D-423MG | PI3Ka_4409 | MVP_top60_DNA+RNA_seed | PIK3CD | -0.12983 | GBM | D-263MG | PI3Ka_4409 |
| MVP_top60_DNA_seed | PIK3CD | 0.005288 | GBM | D-423MG | PI3Ka_4409 | MVP_top60_DNA+RNA_seed | JAK2 | -0.128126 | GBM | A172 | Fedratinib |
| MVP_top60_DNA_seed | JAK2 | 0.005289 | GBM | SF126 | Fedratinib | MVP_top60_DNA+RNA_seed | IGF1R | -0.125677 | GBM | CAS-1 | Linsitinib |
| MVP_top60_DNA_seed | EGFR | 0.006771 | GBM | SNB75 | Lapatinib | MVP_top60_DNA+RNA_seed | PDGFRA | -0.12509 | GBM | U-87-MG | Masitinib |
| MVP_top60_DNA_seed | EGFR | 0.007005 | GBM | YKG-1 | Pelitinib | MVP_top60_DNA+RNA_seed | PDGFRB | -0.12509 | GBM | U-87-MG | Masitinib |
| MVP_top60_DNA_seed | PDGFRA | 0.009774 | GBM | LN-405 | Masitinib | MVP_top60_DNA+RNA_seed | KIT | -0.12509 | GBM | U-87-MG | Masitinib |
| MVP_top60_DNA_seed | PDGFRB | 0.009774 | GBM | LN-405 | Masitinib | MVP_top60_DNA+RNA_seed | JAK2 | -0.124539 | GBM | SK-MG-1 | Fedratinib |
| MVP_top60_DNA_seed | EGFR | 0.012495 | GBM | U251 | Cetuximab | MVP_top60_DNA+RNA_seed | EGFR | -0.119082 | GBM | T98G | AZD8931 |
| MVP_top60_DNA_seed | EGFR | 0.01419 | GBM | U251 | PF-00299804 | MVP_top60_DNA+RNA_seed | ERBB3 | -0.119082 | GBM | T98G | AZD8931 |
| MVP_top60_DNA_seed | JAK2 | 0.014493 | GBM | SF268 | Ruxolitinib | MVP_top60_DNA+RNA_seed | PDGFRB | -0.118261 | GBM | LN-18 | Pazopanib |
| MVP_top60_DNA_seed | JAK1 | 0.014493 | GBM | SF268 | Ruxolitinib | MVP_top60_DNA+RNA_seed | KIT | -0.118261 | GBM | LN-18 | Pazopanib |
| MVP_top60_DNA_seed | EGFR | 0.01544 | GBM | 8-MG-BA | CI-1033 | MVP_top60_DNA+RNA_seed | EGFR | -0.116551 | GBM | SF126 | CI-1033 |
| MVP_top60_DNA_seed | PIK3CD | 0.017774 | GBM | T98G | IC-87114 | MVP_top60_DNA+RNA_seed | JAK1 | -0.11532 | GBM | D-566MG | JAK1_3715 |
| MVP_top60_DNA_seed | EGFR | 0.020324 | GBM | D-423MG | Afatinib | MVP_top60_DNA+RNA_seed | IGF1R | -0.113169 | GBM | T98G | GSK1904529A |
| MVP_top60_DNA_seed | JAK3 | 0.024142 | GBM | D-423MG | WHI-P97 | MVP_top60_DNA+RNA_seed | MET | -0.112525 | GBM | AM-38 | JNJ38877605 |
| MVP_top60_DNA_seed | PIK3CD | 0.024665 | GBM | SF126 | Idelalisib | MVP_top60_DNA+RNA_seed | PDGFRA | -0.110574 | GBM | CAS-1 | Amuvatinib |
| MVP_top60_DNA_seed | EGFR | 0.029027 | GBM | 8-MG-BA | Lapatinib | MVP_top60_DNA+RNA_seed | KIT | -0.110574 | GBM | CAS-1 | Amuvatinib |
| MVP_top60_DNA_seed | PIK3CB | 0.030563 | GBM | D-247MG | AZD8186 | MVP_top60_DNA+RNA_seed | PIK3CA | -0.109203 | GBM | D-263MG | AZD8835 |
| MVP_top60_DNA_seed | PIK3CD | 0.030563 | GBM | D-247MG | AZD8186 | MVP_top60_DNA+RNA_seed | PIK3CD | -0.109203 | GBM | D-263MG | AZD8835 |
| MVP_top60_DNA_seed | EGFR | 0.032605 | GBM | LN-18 | AST-1306 | MVP_top60_DNA+RNA_seed | EGFR | -0.108835 | GBM | M059J | AZD8931 |
| MVP_top60_DNA_seed | EGFR | 0.037946 | GBM | LNZTA3WT4 | AST-1306 | MVP_top60_DNA+RNA_seed | ERBB3 | -0.108835 | GBM | M059J | AZD8931 |
| MVP_top60_DNA_seed | PIK3CD | 0.039439 | GBM | 42-MG-BA | IC-87114 | MVP_top60_DNA+RNA_seed | MET | -0.108304 | GBM | DBTRG-05MG | JNJ38877605 |
| MVP_top60_DNA_seed | PIK3CB | 0.042089 | GBM | U-118-MG | AZD8186 | MVP_top60_DNA+RNA_seed | PDGFRA | -0.107572 | GBM | SF268 | Amuvatinib |
| MVP_top60_DNA_seed | PIK3CD | 0.042089 | GBM | U-118-MG | AZD8186 | MVP_top60_DNA+RNA_seed | KIT | -0.107572 | GBM | SF268 | Amuvatinib |
| MVP_top60_DNA_seed | EGFR | 0.04348 | GBM | D-423MG | PF-00299804 | MVP_top60_DNA+RNA_seed | JAK3 | -0.102274 | GBM | T98G | WHI-P97 |
| MVP_top60_DNA_seed | EGFR | 0.044373 | GBM | U251 | AST-1306 | MVP_top60_DNA+RNA_seed | AKT1 | -0.10009 | GBM | A172 | AKT inhibitor VIII |
| MVP_top60_DNA_seed | EGFR | 0.044568 | GBM | D-263MG | AST-1306 | MVP_top60_DNA+RNA_seed | PIK3CB | -0.099929 | GBM | D-542MG | AZD8186 |
| MVP_top60_DNA_seed | EGFR | 0.045027 | GBM | LN-229 | CI-1033 | MVP_top60_DNA+RNA_seed | PIK3CD | -0.099929 | GBM | D-542MG | AZD8186 |
| MVP_top60_DNA_seed | PDGFRA | 0.047682 | GBM | A172 | Amuvatinib | MVP_top60_DNA+RNA_seed | MET | -0.097435 | GBM | DK-MG | SU11274 |
| MVP_top60_DNA_seed | EGFR | 0.04783 | GBM | SK-MG-1 | CUDC-101 | MVP_top60_DNA+RNA_seed | JAK2 | -0.097213 | GBM | DK-MG | AZD1480 |
| MVP_top60_DNA_seed | PIK3CA | 0.052866 | GBM | GB-1 | AZD8835 | MVP_top60_DNA+RNA_seed | JAK1 | -0.097213 | GBM | DK-MG | AZD1480 |
| MVP_top60_DNA_seed | PIK3CD | 0.052866 | GBM | GB-1 | AZD8835 | MVP_top60_DNA+RNA_seed | EGFR | -0.095753 | GBM | CAS-1 | AST-1306 |
| MVP_top60_DNA_seed | EGFR | 0.053342 | GBM | LN-18 | Cetuximab | MVP_top60_DNA+RNA_seed | JAK2 | -0.094521 | GBM | A172 | Ruxolitinib |
| MVP_top60_DNA_seed | JAK2 | 0.05367 | GBM | SNB75 | Fedratinib | MVP_top60_DNA+RNA_seed | JAK1 | -0.094521 | GBM | A172 | Ruxolitinib |
| MVP_top60_DNA_seed | EGFR | 0.055296 | GBM | LN-18 | AZD8931 | MVP_top60_DNA+RNA_seed | AKT1 | -0.093756 | GBM | U-87-MG | GSK690693 |
| MVP_top60_DNA_seed | PIK3CD | 0.055351 | GBM | A172 | IC-87114 | MVP_top60_DNA+RNA_seed | JAK2 | -0.09292 | GBM | D-566MG | Ruxolitinib |
| MVP_top60_DNA_seed | EGFR | 0.066144 | GBM | SF295 | AZD8931 | MVP_top60_DNA+RNA_seed | JAK1 | -0.09292 | GBM | D-566MG | Ruxolitinib |
| MVP_top60_DNA_seed | EGFR | 0.067477 | GBM | LN-18 | PF-00299804 | MVP_top60_DNA+RNA_seed | JAK3 | -0.09253 | GBM | DK-MG | WHI-P97 |
| MVP_top60_DNA_seed | PIK3CA | 0.068246 | GBM | D-566MG | AZD8835 | MVP_top60_DNA+RNA_seed | JAK2 | -0.092216 | GBM | T98G | Ruxolitinib |
| MVP_top60_DNA_seed | PIK3CD | 0.068246 | GBM | D-566MG | AZD8835 | MVP_top60_DNA+RNA_seed | JAK1 | -0.092216 | GBM | T98G | Ruxolitinib |
| MVP_top60_DNA_seed | EGFR | 0.078088 | GBM | 8-MG-BA | Afatinib | MVP_top60_DNA+RNA_seed | JAK2 | -0.088868 | GBM | YH-13 | Fedratinib |

| | | | | | | | | | | | |
|---|---|---|---|---|---|---|---|---|---|---|---|
| MVP_top60_DNA_seed | EGFR | 0.083535 | GBM | U251 | CI-1033 | MVP_top60_DNA+RNA_seed | IGF1R | -0.086558 | GBM | 8-MG-BA | Linsitinib |
| MVP_top60_DNA_seed | PIK3CD | 0.083699 | GBM | YKG-1 | Idelalisib | MVP_top60_DNA+RNA_seed | JAK1 | -0.086346 | GBM | T98G | JAK1_3715 |
| MVP_top60_DNA_seed | PIK3CA | 0.084452 | GBM | DBTRG-05MG | AZD8835 | MVP_top60_DNA+RNA_seed | EGFR | -0.085668 | GBM | AM-38 | Gefitinib |
| MVP_top60_DNA_seed | PIK3CD | 0.084452 | GBM | DBTRG-05MG | AZD8835 | MVP_top60_DNA+RNA_seed | AKT1 | -0.083939 | GBM | AM-38 | AKT inhibitor VIII |
| MVP_top60_DNA_seed | PDGFRA | 0.085676 | GBM | YKG-1 | Amuvatinib | MVP_top60_DNA+RNA_seed | PIK3CB | -0.083218 | GBM | D-263MG | AZD6482 |
| MVP_top60_DNA_seed | PIK3CD | 0.086688 | GBM | CAS-1 | Idelalisib | MVP_top60_DNA+RNA_seed | KIT | -0.079933 | GBM | D-566MG | OSI-930 |
| MVP_top60_DNA_seed | PIK3CD | 0.086806 | GBM | DBTRG-05MG | Idelalisib | MVP_top60_DNA+RNA_seed | PIK3CD | -0.078886 | GBM | D-247MG | IC-87114 |
| MVP_top60_DNA_seed | EGFR | 0.087284 | GBM | 8-MG-BA | AST-1306 | MVP_top60_DNA+RNA_seed | EGFR | -0.078794 | GBM | M059J | CUDC-101 |
| MVP_top60_DNA_seed | PIK3CB | 0.089032 | GBM | DBTRG-05MG | AZD8186 | MVP_top60_DNA+RNA_seed | JAK2 | -0.07739 | GBM | D-247MG | AZD1480 |
| MVP_top60_DNA_seed | PIK3CD | 0.089032 | GBM | DBTRG-05MG | AZD8186 | MVP_top60_DNA+RNA_seed | JAK1 | -0.07739 | GBM | D-247MG | AZD1480 |
| MVP_top60_DNA_seed | JAK3 | 0.09014 | GBM | D-423MG | JAK3_7406 | MVP_top60_DNA+RNA_seed | PIK3CD | -0.076874 | GBM | SF295 | Idelalisib |
| MVP_top60_DNA_seed | PIK3CB | 0.092072 | GBM | YKG-1 | AZD8186 | MVP_top60_DNA+RNA_seed | PIK3CB | -0.076718 | GBM | GB-1 | AZD6482 |
| MVP_top60_DNA_seed | PIK3CD | 0.092072 | GBM | YKG-1 | AZD8186 | MVP_top60_DNA+RNA_seed | JAK1 | -0.074043 | GBM | D-423MG | JAK1_3715 |
| MVP_top60_DNA_seed | CDK4 | 0.09599 | GBM | DBTRG-05MG | Palbociclib | MVP_top60_DNA+RNA_seed | CDK4 | -0.07251 | GBM | D-392MG | Palbociclib |
| MVP_top60_DNA_seed | EGFR | 0.100039 | GBM | SK-MG-1 | PF-00299804 | MVP_top60_DNA+RNA_seed | JAK2 | -0.071016 | GBM | SW1088 | Fedratinib |
| MVP_top60_DNA_seed | JAK3 | 0.100345 | GBM | LN-229 | JAK3_7406 | MVP_top60_DNA+RNA_seed | AKT1 | -0.070053 | GBM | DK-MG | AKT inhibitor VIII |
| MVP_top60_DNA_seed | EGFR | 0.100493 | GBM | D-566MG | CI-1033 | MVP_top60_DNA+RNA_seed | PIK3CD | -0.06567 | GBM | GB-1 | IC-87114 |
| MVP_top60_DNA_seed | EGFR | 0.101348 | GBM | YKG-1 | AZD8931 | MVP_top60_DNA+RNA_seed | JAK2 | -0.063483 | GBM | U251 | Ruxolitinib |
| MVP_top60_DNA_seed | EGFR | 0.101937 | GBM | LN-229 | AZD8931 | MVP_top60_DNA+RNA_seed | JAK1 | -0.063483 | GBM | U251 | Ruxolitinib |
| MVP_top60_DNA_seed | PDGFRB | 0.1034 | GBM | SK-MG-1 | Pazopanib | MVP_top60_DNA+RNA_seed | JAK2 | -0.060648 | GBM | AM-38 | Ruxolitinib |
| MVP_top60_DNA_seed | JAK3 | 0.106687 | GBM | SF126 | WHI-P97 | MVP_top60_DNA+RNA_seed | JAK1 | -0.060648 | GBM | AM-38 | Ruxolitinib |
| MVP_top60_DNA_seed | EGFR | 0.106911 | GBM | D-542MG | AST-1306 | MVP_top60_DNA+RNA_seed | AKT1 | -0.058609 | GBM | LN-18 | BAY AKT1 |
| MVP_top60_DNA_seed | EGFR | 0.109626 | GBM | LN-229 | PF-00299804 | MVP_top60_DNA+RNA_seed | JAK3 | -0.057614 | GBM | Becker | WHI-P97 |
| MVP_top60_DNA_seed | EGFR | 0.120889 | GBM | AM-38 | CI-1033 | MVP_top60_DNA+RNA_seed | PIK3CD | -0.057539 | GBM | 8-MG-BA | IC-87114 |
| MVP_top60_DNA_seed | EGFR | 0.122608 | GBM | 42-MG-BA | CI-1033 | MVP_top60_DNA+RNA_seed | EGFR | -0.057445 | GBM | AM-38 | AST-1306 |
| MVP_top60_DNA_seed | EGFR | 0.122964 | GBM | YH-13 | Pelitinib | MVP_top60_DNA+RNA_seed | PIK3CD | -0.053876 | GBM | U-87-MG | Idelalisib |
| MVP_top60_DNA_seed | EGFR | 0.126075 | GBM | D-247MG | AZD8931 | MVP_top60_DNA+RNA_seed | KIT | -0.052292 | GBM | Becker | Sorafenib |
| MVP_top60_DNA_seed | PIK3CD | 0.129297 | GBM | U251 | IC-87114 | MVP_top60_DNA+RNA_seed | AKT1 | -0.050544 | GBM | T98G | AKT inhibitor VIII |
| MVP_top60_DNA_seed | EP300 | 0.129432 | GBM | SF126 | I-CBP112 | MVP_top60_DNA+RNA_seed | AKT1 | -0.049952 | GBM | SF268 | A-443654 |
| MVP_top60_DNA_seed | EGFR | 0.129469 | GBM | LN-229 | Afatinib | MVP_top60_DNA+RNA_seed | JAK2 | -0.046987 | GBM | AM-38 | Fedratinib |
| MVP_top60_DNA_seed | PDGFRA | 0.132873 | GBM | Becker | Amuvatinib | MVP_top60_DNA+RNA_seed | EGFR | -0.046977 | GBM | AM-38 | CUDC-101 |
| MVP_top60_DNA_seed | EGFR | 0.133898 | GBM | 8-MG-BA | PF-00299804 | MVP_top60_DNA+RNA_seed | AKT1 | -0.041296 | GBM | SNB75 | GSK690693 |
| MVP_top60_DNA_seed | EGFR | 0.134567 | GBM | D-423MG | AZD8931 | MVP_top60_DNA+RNA_seed | MET | -0.040372 | GBM | D-566MG | JNJ38877605 |
| MVP_top60_DNA_seed | EGFR | 0.140854 | GBM | D-566MG | Pelitinib | MVP_top60_DNA+RNA_seed | AKT1 | -0.040037 | GBM | D-392MG | AKT inhibitor VIII |
| MVP_top60_DNA_seed | EGFR | 0.156294 | GBM | D-566MG | AST-1306 | MVP_top60_DNA+RNA_seed | AKT1 | -0.039726 | GBM | CAS-1 | GSK690693 |
| MVP_top60_DNA_seed | PIK3CA | 0.158394 | GBM | LNZTA3WT4 | AZD8835 | MVP_top60_DNA+RNA_seed | AKT1 | -0.038967 | GBM | SK-MG-1 | GSK690693 |
| MVP_top60_DNA_seed | PIK3CD | 0.158394 | GBM | LNZTA3WT4 | AZD8835 | MVP_top60_DNA+RNA_seed | EGFR | -0.038738 | GBM | D-423MG | Cetuximab |
| MVP_top60_DNA_seed | EP300 | 0.158996 | GBM | D-263MG | I-CBP112 | MVP_top60_DNA+RNA_seed | JAK1 | -0.037805 | GBM | U251 | JAK1_3715 |
| MVP_top60_DNA_seed | EGFR | 0.160378 | GBM | AM-38 | PF-00299804 | MVP_top60_DNA+RNA_seed | JAK1 | -0.037624 | GBM | 42-MG-BA | JAK1_3715 |
| MVP_top60_DNA_seed | EGFR | 0.160936 | GBM | Becker | Pelitinib | MVP_top60_DNA+RNA_seed | KIT | -0.035361 | GBM | LN-18 | OSI-930 |
| MVP_top60_DNA_seed | PIK3CA | 0.163953 | GBM | GB-1 | PI3Ka_4409 | MVP_top60_DNA+RNA_seed | JAK3 | -0.03534 | GBM | LN-229 | WHI-P97 |

| | | | | | | | | | | | |
|---|---|---|---|---|---|---|---|---|---|---|---|
| MVP_top60_DNA_seed | PIK3CD | 0.163953 | GBM | GB-1 | PI3Ka_4409 | MVP_top60_DNA+RNA_seed | EGFR | -0.033848 | GBM | YH-13 | CI-1033 |
| MVP_top60_DNA_seed | EGFR | 0.165266 | GBM | U-87-MG | AST-1306 | MVP_top60_DNA+RNA_seed | AKT1 | -0.033618 | GBM | SK-MG-1 | AKT inhibitor VIII |
| MVP_top60_DNA_seed | EGFR | 0.167431 | GBM | SNB75 | AST-1306 | MVP_top60_DNA+RNA_seed | PIK3CD | -0.033518 | GBM | D-247MG | Idelalisib |
| MVP_top60_DNA_seed | EGFR | 0.167731 | GBM | DBTRG-05MG | CUDC-101 | MVP_top60_DNA+RNA_seed | PIK3CB | -0.030548 | GBM | M059J | AZD6482 |
| MVP_top60_DNA_seed | PIK3CD | 0.170874 | GBM | YKG-1 | IC-87114 | MVP_top60_DNA+RNA_seed | IGF1R | -0.03037 | GBM | D-566MG | BMS-536924 |
| MVP_top60_DNA_seed | JAK3 | 0.17125 | GBM | YKG-1 | WHI-P97 | MVP_top60_DNA+RNA_seed | JAK2 | -0.03013 | GBM | SF126 | AZD1480 |
| MVP_top60_DNA_seed | EGFR | 0.171606 | GBM | D-392MG | Cetuximab | MVP_top60_DNA+RNA_seed | JAK1 | -0.03013 | GBM | SF126 | AZD1480 |
| MVP_top60_DNA_seed | PDGFRA | 0.176476 | GBM | DBTRG-05MG | Masitinib | MVP_top60_DNA+RNA_seed | EGFR | -0.029853 | GBM | 42-MG-BA | CUDC-101 |
| MVP_top60_DNA_seed | PDGFRB | 0.176476 | GBM | DBTRG-05MG | Masitinib | MVP_top60_DNA+RNA_seed | IGF1R | -0.029149 | GBM | A172 | BMS-754807 |
| MVP_top60_DNA_seed | EGFR | 0.176952 | GBM | SF126 | Erlotinib | MVP_top60_DNA+RNA_seed | IGF1R | -0.028948 | GBM | SF295 | GSK1904529A |
| MVP_top60_DNA_seed | EP300 | 0.177199 | GBM | SF295 | I-CBP112 | MVP_top60_DNA+RNA_seed | EGFR | -0.024533 | GBM | KS-1 | AST-1306 |
| MVP_top60_DNA_seed | PIK3CD | 0.177961 | GBM | GB-1 | Idelalisib | MVP_top60_DNA+RNA_seed | KIT | -0.021645 | GBM | SF295 | OSI-930 |
| MVP_top60_DNA_seed | JAK2 | 0.181685 | GBM | 42-MG-BA | Ruxolitinib | MVP_top60_DNA+RNA_seed | EGFR | -0.021607 | GBM | 8-MG-BA | Erlotinib |
| MVP_top60_DNA_seed | JAK1 | 0.181685 | GBM | 42-MG-BA | Ruxolitinib | MVP_top60_DNA+RNA_seed | IGF1R | -0.020663 | GBM | 8-MG-BA | BMS-754807 |
| MVP_top60_DNA_seed | EGFR | 0.185456 | GBM | A172 | AZD8931 | MVP_top60_DNA+RNA_seed | EP300 | -0.018012 | GBM | T98G | I-CBP112 |
| MVP_top60_DNA_seed | EGFR | 0.188649 | GBM | SF295 | CUDC-101 | MVP_top60_DNA+RNA_seed | CREBBP | -0.018012 | GBM | T98G | I-CBP112 |
| MVP_top60_DNA_seed | PIK3CD | 0.192243 | GBM | Becker | IC-87114 | MVP_top60_DNA+RNA_seed | KIT | -0.016623 | GBM | U-118-MG | OSI-930 |
| MVP_top60_DNA_seed | EGFR | 0.193134 | GBM | YH-13 | CUDC-101 | MVP_top60_DNA+RNA_seed | IGF1R | -0.015568 | GBM | D-423MG | Linsitinib |
| MVP_top60_DNA_seed | PIK3CD | 0.195409 | GBM | SW1088 | IC-87114 | MVP_top60_DNA+RNA_seed | JAK2 | -0.014916 | GBM | U251 | AZD1480 |
| MVP_top60_DNA_seed | PIK3CA | 0.197067 | GBM | SF295 | AZD8835 | MVP_top60_DNA+RNA_seed | JAK1 | -0.014916 | GBM | U251 | AZD1480 |
| MVP_top60_DNA_seed | PIK3CD | 0.197067 | GBM | SF295 | AZD8835 | MVP_top60_DNA+RNA_seed | EGFR | -0.013575 | GBM | SF126 | AST-1306 |
| MVP_top60_DNA_seed | PIK3CD | 0.19915 | GBM | M059J | Idelalisib | MVP_top60_DNA+RNA_seed | EGFR | -0.011909 | GBM | YKG-1 | PF-00299804 |
| MVP_top60_DNA_seed | EGFR | 0.203313 | GBM | LN-229 | CUDC-101 | MVP_top60_DNA+RNA_seed | PIK3CB | -0.011033 | GBM | U-87-MG | AZD6482 |
| MVP_top60_DNA_seed | EGFR | 0.207045 | GBM | GB-1 | Lapatinib | MVP_top60_DNA+RNA_seed | EGFR | -0.010827 | GBM | LN-229 | AST-1306 |
| MVP_top60_DNA_seed | PIK3CA | 0.21589 | GBM | SF295 | PI3Ka_4409 | MVP_top60_DNA+RNA_seed | PDGFRA | -0.009646 | GBM | LN-18 | Masitinib |
| MVP_top60_DNA_seed | PIK3CD | 0.21589 | GBM | SF295 | PI3Ka_4409 | MVP_top60_DNA+RNA_seed | PDGFRB | -0.009646 | GBM | LN-18 | Masitinib |
| MVP_top60_DNA_seed | EGFR | 0.216187 | GBM | SK-MG-1 | AST-1306 | MVP_top60_DNA+RNA_seed | KIT | -0.009646 | GBM | LN-18 | Masitinib |
| MVP_top60_DNA_seed | PIK3CA | 0.217007 | GBM | SNB75 | PI3Ka_4409 | MVP_top60_DNA+RNA_seed | PIK3CB | -0.009252 | GBM | M059J | AZD8186 |
| MVP_top60_DNA_seed | PIK3CD | 0.217007 | GBM | SNB75 | PI3Ka_4409 | MVP_top60_DNA+RNA_seed | PIK3CD | -0.009252 | GBM | M059J | AZD8186 |
| MVP_top60_DNA_seed | PIK3CD | 0.217184 | GBM | KS-1 | IC-87114 | MVP_top60_DNA+RNA_seed | JAK1 | -0.007254 | GBM | D-247MG | JAK1_3715 |
| MVP_top60_DNA_seed | EGFR | 0.22013 | GBM | YKG-1 | AST-1306 | MVP_top60_DNA+RNA_seed | EP300 | -0.006046 | GBM | U251 | I-CBP112 |
| MVP_top60_DNA_seed | EGFR | 0.220175 | GBM | LN-229 | Gefitinib | MVP_top60_DNA+RNA_seed | CREBBP | -0.006046 | GBM | U251 | I-CBP112 |
| MVP_top60_DNA_seed | EP300 | 0.224737 | GBM | U-87-MG | I-CBP112 | MVP_top60_DNA+RNA_seed | CDK4 | -0.000924 | GBM | LNZTA3WT4 | Palbociclib |
| MVP_top60_DNA_seed | PDGFRA | 0.225253 | GBM | M059J | Masitinib | MVP_top60_DNA+RNA_seed | PDGFRA | -0.000135 | GBM | 8-MG-BA | Amuvatinib |
| MVP_top60_DNA_seed | PDGFRB | 0.225253 | GBM | M059J | Masitinib | MVP_top60_DNA+RNA_seed | KIT | -0.000135 | GBM | 8-MG-BA | Amuvatinib |
| MVP_top60_DNA_seed | JAK3 | 0.225464 | GBM | A172 | WHI-P97 | MVP_top60_DNA+RNA_seed | PDGFRA | -0.000123 | GBM | D-542MG | Amuvatinib |
| MVP_top60_DNA_seed | PIK3CD | 0.229603 | GBM | SF268 | Idelalisib | MVP_top60_DNA+RNA_seed | KIT | -0.000123 | GBM | D-542MG | Amuvatinib |
| MVP_top60_DNA_seed | EGFR | 0.231418 | GBM | GB-1 | Afatinib | MVP_top60_DNA+RNA_seed | JAK2 | -6.20E-05 | GBM | T98G | Fedratinib |
| MVP_top60_DNA_seed | PIK3CB | 0.231575 | GBM | A172 | AZD6482 | MVP_top60_DNA+RNA_seed | IGF1R | 0.002559 | GBM | LNZTA3WT4 | GSK1904529A |
| MVP_top60_DNA_seed | PIK3CD | 0.237523 | GBM | LNZTA3WT4 | IC-87114 | MVP_top60_DNA+RNA_seed | AKT1 | 0.004168 | GBM | SF126 | BAY AKT1 |
| MVP_top60_DNA_seed | JAK3 | 0.239205 | GBM | LN-18 | WHI-P97 | MVP_top60_DNA+RNA_seed | JAK2 | 0.004519 | GBM | M059J | Fedratinib |

| | | | | | | | | | | | |
|---|---|---|---|---|---|---|---|---|---|---|---|
| MVP_top60_DNA_seed | EGFR | 0.239334 | GBM | D-263MG | Erlotinib | MVP_top60_DNA+RNA_seed | PIK3CA | 0.005288 | GBM | D-423MG | PI3Ka_4409 |
| MVP_top60_DNA_seed | PDGFRB | 0.240315 | GBM | DBTRG-05MG | Pazopanib | MVP_top60_DNA+RNA_seed | PIK3CD | 0.005288 | GBM | D-423MG | PI3Ka_4409 |
| MVP_top60_DNA_seed | PIK3CD | 0.240982 | GBM | M059J | IC-87114 | MVP_top60_DNA+RNA_seed | JAK2 | 0.005289 | GBM | SF126 | Fedratinib |
| MVP_top60_DNA_seed | EGFR | 0.241185 | GBM | YH-13 | Gefitinib | MVP_top60_DNA+RNA_seed | KIT | 0.006691 | GBM | CAS-1 | Axitinib |
| MVP_top60_DNA_seed | JAK2 | 0.246647 | GBM | LNZTA3WT4 | Ruxolitinib | MVP_top60_DNA+RNA_seed | EGFR | 0.006771 | GBM | SNB75 | Lapatinib |
| MVP_top60_DNA_seed | JAK1 | 0.246647 | GBM | LNZTA3WT4 | Ruxolitinib | MVP_top60_DNA+RNA_seed | EGFR | 0.007005 | GBM | YKG-1 | Pelitinib |
| MVP_top60_DNA_seed | EGFR | 0.248324 | GBM | LN-18 | CI-1033 | MVP_top60_DNA+RNA_seed | AKT1 | 0.00705 | GBM | GB-1 | AKT inhibitor VIII |
| MVP_top60_DNA_seed | CDK4 | 0.248666 | GBM | SF126 | CGP-082996 | MVP_top60_DNA+RNA_seed | PDGFRA | 0.009774 | GBM | LN-405 | Masitinib |
| MVP_top60_DNA_seed | JAK2 | 0.250642 | GBM | YKG-1 | Fedratinib | MVP_top60_DNA+RNA_seed | PDGFRB | 0.009774 | GBM | LN-405 | Masitinib |
| MVP_top60_DNA_seed | EP300 | 0.25214 | GBM | LNZTA3WT4 | I-CBP112 | MVP_top60_DNA+RNA_seed | KIT | 0.009774 | GBM | LN-405 | Masitinib |
| MVP_top60_DNA_seed | EGFR | 0.255388 | GBM | Becker | Cetuximab | MVP_top60_DNA+RNA_seed | EGFR | 0.012495 | GBM | U251 | Cetuximab |
| MVP_top60_DNA_seed | EGFR | 0.25594 | GBM | SF295 | Cetuximab | MVP_top60_DNA+RNA_seed | EGFR | 0.01419 | GBM | U251 | PF-00299804 |
| MVP_top60_DNA_seed | EGFR | 0.261073 | GBM | KS-1 | Gefitinib | MVP_top60_DNA+RNA_seed | JAK2 | 0.014493 | GBM | SF268 | Ruxolitinib |
| MVP_top60_DNA_seed | CDK4 | 0.267169 | GBM | D-247MG | CGP-082996 | MVP_top60_DNA+RNA_seed | JAK1 | 0.014493 | GBM | SF268 | Ruxolitinib |
| MVP_top60_DNA_seed | JAK1 | 0.268802 | GBM | Becker | JAK1_3715 | MVP_top60_DNA+RNA_seed | EGFR | 0.01544 | GBM | 8-MG-BA | CI-1033 |
| MVP_top60_DNA_seed | EGFR | 0.269698 | GBM | LN-18 | Afatinib | MVP_top60_DNA+RNA_seed | PIK3CD | 0.017774 | GBM | T98G | IC-87114 |
| MVP_top60_DNA_seed | EGFR | 0.274048 | GBM | SF126 | CUDC-101 | MVP_top60_DNA+RNA_seed | EGFR | 0.020324 | GBM | D-423MG | Afatinib |
| MVP_top60_DNA_seed | JAK2 | 0.274497 | GBM | YH-13 | Ruxolitinib | MVP_top60_DNA+RNA_seed | KIT | 0.022968 | GBM | Becker | OSI-930 |
| MVP_top60_DNA_seed | JAK1 | 0.274497 | GBM | YH-13 | Ruxolitinib | MVP_top60_DNA+RNA_seed | MET | 0.023052 | GBM | U251 | AZD6094 |
| MVP_top60_DNA_seed | JAK2 | 0.27474 | GBM | SF295 | Ruxolitinib | MVP_top60_DNA+RNA_seed | JAK3 | 0.024142 | GBM | D-423MG | WHI-P97 |
| MVP_top60_DNA_seed | JAK1 | 0.27474 | GBM | SF295 | Ruxolitinib | MVP_top60_DNA+RNA_seed | PIK3CD | 0.024665 | GBM | SF126 | Idelalisib |
| MVP_top60_DNA_seed | EGFR | 0.276723 | GBM | DBTRG-05MG | AZD8931 | MVP_top60_DNA+RNA_seed | EGFR | 0.029027 | GBM | 8-MG-BA | Lapatinib |
| MVP_top60_DNA_seed | EGFR | 0.276843 | GBM | SW1088 | Afatinib | MVP_top60_DNA+RNA_seed | PIK3CB | 0.030563 | GBM | D-247MG | AZD8186 |
| MVP_top60_DNA_seed | EGFR | 0.279396 | GBM | 8-MG-BA | AZD8931 | MVP_top60_DNA+RNA_seed | PIK3CD | 0.030563 | GBM | D-247MG | AZD8186 |
| MVP_top60_DNA_seed | EGFR | 0.282952 | GBM | CAS-1 | PF-00299804 | MVP_top60_DNA+RNA_seed | EGFR | 0.032605 | GBM | LN-18 | AST-1306 |
| MVP_top60_DNA_seed | EGFR | 0.286063 | GBM | SNB75 | Afatinib | MVP_top60_DNA+RNA_seed | AKT1 | 0.034792 | GBM | D-566MG | GSK690693 |
| MVP_top60_DNA_seed | EGFR | 0.287584 | GBM | D-263MG | AZD8931 | MVP_top60_DNA+RNA_seed | MET | 0.036508 | GBM | M059J | JNJ38877605 |
| MVP_top60_DNA_seed | JAK2 | 0.288193 | GBM | M059J | Ruxolitinib | MVP_top60_DNA+RNA_seed | IGF1R | 0.037822 | GBM | SF126 | Linsitinib |
| MVP_top60_DNA_seed | JAK1 | 0.288193 | GBM | M059J | Ruxolitinib | MVP_top60_DNA+RNA_seed | EGFR | 0.037946 | GBM | LNZTA3WT4 | AST-1306 |
| MVP_top60_DNA_seed | EGFR | 0.290069 | GBM | U251 | Gefitinib | MVP_top60_DNA+RNA_seed | PIK3CD | 0.039439 | GBM | 42-MG-BA | IC-87114 |
| MVP_top60_DNA_seed | PIK3CB | 0.290127 | GBM | A172 | AZD8186 | MVP_top60_DNA+RNA_seed | KIT | 0.039707 | GBM | A172 | Motesanib |
| MVP_top60_DNA_seed | PIK3CD | 0.290127 | GBM | A172 | AZD8186 | MVP_top60_DNA+RNA_seed | PIK3CB | 0.042089 | GBM | U-118-MG | AZD8186 |
| MVP_top60_DNA_seed | EGFR | 0.292605 | GBM | D-247MG | AST-1306 | MVP_top60_DNA+RNA_seed | PIK3CD | 0.042089 | GBM | U-118-MG | AZD8186 |
| MVP_top60_DNA_seed | EP300 | 0.293796 | GBM | SK-MG-1 | I-CBP112 | MVP_top60_DNA+RNA_seed | AKT1 | 0.042359 | GBM | SF295 | AKT inhibitor VIII |
| MVP_top60_DNA_seed | EGFR | 0.294226 | GBM | GB-1 | Pelitinib | MVP_top60_DNA+RNA_seed | EGFR | 0.04348 | GBM | D-423MG | PF-00299804 |
| MVP_top60_DNA_seed | JAK3 | 0.29455 | GBM | M059J | WHI-P97 | MVP_top60_DNA+RNA_seed | EGFR | 0.044373 | GBM | U251 | AST-1306 |
| MVP_top60_DNA_seed | JAK2 | 0.297464 | GBM | DBTRG-05MG | Fedratinib | MVP_top60_DNA+RNA_seed | EGFR | 0.044568 | GBM | D-263MG | AST-1306 |
| MVP_top60_DNA_seed | JAK3 | 0.299226 | GBM | YKG-1 | JAK3_7406 | MVP_top60_DNA+RNA_seed | EGFR | 0.045027 | GBM | LN-229 | CI-1033 |
| MVP_top60_DNA_seed | EGFR | 0.3009 | GBM | Becker | Afatinib | MVP_top60_DNA+RNA_seed | AKT1 | 0.047258 | GBM | LNZTA3WT4 | AKT inhibitor VIII |
| MVP_top60_DNA_seed | EGFR | 0.306633 | GBM | LN-18 | Gefitinib | MVP_top60_DNA+RNA_seed | PDGFRA | 0.047682 | GBM | A172 | Amuvatinib |
| MVP_top60_DNA_seed | JAK1 | 0.30713 | GBM | M059J | JAK1_3715 | MVP_top60_DNA+RNA_seed | KIT | 0.047682 | GBM | A172 | Amuvatinib |

| | | | | | | | | | | | |
|---|---|---|---|---|---|---|---|---|---|---|---|
| MVP_top60_DNA_seed | EGFR | 0.309742 | GBM | D-392MG | CUDC-101 | MVP_top60_DNA+RNA_seed | EGFR | 0.04783 | GBM | SK-MG-1 | CUDC-101 |
| MVP_top60_DNA_seed | EGFR | 0.310051 | GBM | LN-18 | Afatinib | MVP_top60_DNA+RNA_seed | IGF1R | 0.051388 | GBM | AM-38 | BMS-754807 |
| MVP_top60_DNA_seed | PIK3CD | 0.3104 | GBM | DBTRG-05MG | IC-87114 | MVP_top60_DNA+RNA_seed | PIK3CA | 0.052866 | GBM | GB-1 | AZD8835 |
| MVP_top60_DNA_seed | EGFR | 0.311172 | GBM | M059J | CI-1033 | MVP_top60_DNA+RNA_seed | PIK3CD | 0.052866 | GBM | GB-1 | AZD8835 |
| MVP_top60_DNA_seed | PIK3CB | 0.311403 | GBM | 8-MG-BA | TGX221 | MVP_top60_DNA+RNA_seed | EGFR | 0.053342 | GBM | LN-18 | Cetuximab |
| MVP_top60_DNA_seed | EGFR | 0.313652 | GBM | CAS-1 | CI-1033 | MVP_top60_DNA+RNA_seed | JAK2 | 0.05367 | GBM | SNB75 | Fedratinib |
| MVP_top60_DNA_seed | EGFR | 0.315478 | GBM | KS-1 | Cetuximab | MVP_top60_DNA+RNA_seed | EGFR | 0.055296 | GBM | LN-18 | AZD8931 |
| MVP_top60_DNA_seed | EGFR | 0.317128 | GBM | GB-1 | PF-00299804 | MVP_top60_DNA+RNA_seed | ERBB3 | 0.055296 | GBM | LN-18 | AZD8931 |
| MVP_top60_DNA_seed | PIK3CB | 0.319059 | GBM | SF126 | AZD8186 | MVP_top60_DNA+RNA_seed | PIK3CD | 0.055351 | GBM | A172 | IC-87114 |
| MVP_top60_DNA_seed | PIK3CD | 0.319059 | GBM | SF126 | AZD8186 | MVP_top60_DNA+RNA_seed | AKT1 | 0.056668 | GBM | SF126 | GSK690693 |
| MVP_top60_DNA_seed | JAK2 | 0.321281 | GBM | Becker | Fedratinib | MVP_top60_DNA+RNA_seed | KIT | 0.058221 | GBM | D-566MG | Motesanib |
| MVP_top60_DNA_seed | EGFR | 0.324796 | GBM | U251 | Afatinib | MVP_top60_DNA+RNA_seed | KIT | 0.059381 | GBM | LNZTA3WT4 | Motesanib |
| MVP_top60_DNA_seed | EGFR | 0.32485 | GBM | D-566MG | Afatinib | MVP_top60_DNA+RNA_seed | EGFR | 0.066144 | GBM | SF295 | AZD8931 |
| MVP_top60_DNA_seed | EGFR | 0.325862 | GBM | D-263MG | Lapatinib | MVP_top60_DNA+RNA_seed | ERBB3 | 0.066144 | GBM | SF295 | AZD8931 |
| MVP_top60_DNA_seed | JAK1 | 0.326462 | GBM | YKG-1 | JAK1_3715 | MVP_top60_DNA+RNA_seed | KIT | 0.066404 | GBM | LN-405 | OSI-930 |
| MVP_top60_DNA_seed | EP300 | 0.328635 | GBM | LN-405 | I-CBP112 | MVP_top60_DNA+RNA_seed | IGF1R | 0.067015 | GBM | D-566MG | BMS-754807 |
| MVP_top60_DNA_seed | EGFR | 0.328818 | GBM | D-423MG | CI-1033 | MVP_top60_DNA+RNA_seed | EGFR | 0.067477 | GBM | LN-18 | PF-00299804 |
| MVP_top60_DNA_seed | PIK3CA | 0.329017 | GBM | U-87-MG | PI3Ka_4409 | MVP_top60_DNA+RNA_seed | PIK3CA | 0.068246 | GBM | D-566MG | AZD8835 |
| MVP_top60_DNA_seed | PIK3CD | 0.329017 | GBM | U-87-MG | PI3Ka_4409 | MVP_top60_DNA+RNA_seed | PIK3CD | 0.068246 | GBM | D-566MG | AZD8835 |
| MVP_top60_DNA_seed | PDGFRA | 0.331779 | GBM | T98G | Amuvatinib | MVP_top60_DNA+RNA_seed | MET | 0.068545 | GBM | LN-229 | AZD6094 |
| MVP_top60_DNA_seed | JAK2 | 0.334619 | GBM | LN-405 | AZD1480 | MVP_top60_DNA+RNA_seed | IGF1R | 0.069649 | GBM | SK-MG-1 | BMS-754807 |
| MVP_top60_DNA_seed | JAK1 | 0.334619 | GBM | LN-405 | AZD1480 | MVP_top60_DNA+RNA_seed | IGF1R | 0.070181 | GBM | AM-38 | Linsitinib |
| MVP_top60_DNA_seed | PIK3CA | 0.338585 | GBM | D-542MG | PI3Ka_4409 | MVP_top60_DNA+RNA_seed | IGF1R | 0.071379 | GBM | A172 | GSK1904529A |
| MVP_top60_DNA_seed | PIK3CD | 0.338585 | GBM | D-542MG | PI3Ka_4409 | MVP_top60_DNA+RNA_seed | KIT | 0.072607 | GBM | CAS-1 | OSI-930 |
| MVP_top60_DNA_seed | EGFR | 0.34101 | GBM | DK-MG | AST-1306 | MVP_top60_DNA+RNA_seed | IGF1R | 0.076222 | GBM | 8-MG-BA | GSK1904529A |
| MVP_top60_DNA_seed | PIK3CA | 0.341573 | GBM | YKG-1 | AZD8835 | MVP_top60_DNA+RNA_seed | EGFR | 0.078088 | GBM | 8-MG-BA | Afatinib |
| MVP_top60_DNA_seed | PIK3CD | 0.341573 | GBM | YKG-1 | AZD8835 | MVP_top60_DNA+RNA_seed | IGF1R | 0.079256 | GBM | Becker | BMS-754807 |
| MVP_top60_DNA_seed | PIK3CB | 0.342544 | GBM | SF268 | AZD8186 | MVP_top60_DNA+RNA_seed | AKT1 | 0.081256 | GBM | SF295 | GSK690693 |
| MVP_top60_DNA_seed | PIK3CD | 0.342544 | GBM | SF268 | AZD8186 | MVP_top60_DNA+RNA_seed | KIT | 0.083062 | GBM | T98G | OSI-930 |
| MVP_top60_DNA_seed | JAK2 | 0.343693 | GBM | DBTRG-05MG | AZD1480 | MVP_top60_DNA+RNA_seed | EGFR | 0.083535 | GBM | U251 | CI-1033 |
| MVP_top60_DNA_seed | JAK1 | 0.343693 | GBM | DBTRG-05MG | AZD1480 | MVP_top60_DNA+RNA_seed | PIK3CD | 0.083699 | GBM | YKG-1 | Idelalisib |
| MVP_top60_DNA_seed | JAK2 | 0.34528 | GBM | D-423MG | Ruxolitinib | MVP_top60_DNA+RNA_seed | PIK3CA | 0.084452 | GBM | DBTRG-05MG | AZD8835 |
| MVP_top60_DNA_seed | JAK1 | 0.34528 | GBM | D-423MG | Ruxolitinib | MVP_top60_DNA+RNA_seed | PIK3CD | 0.084452 | GBM | DBTRG-05MG | AZD8835 |
| MVP_top60_DNA_seed | JAK2 | 0.347541 | GBM | SF126 | Ruxolitinib | MVP_top60_DNA+RNA_seed | PDGFRA | 0.085676 | GBM | YKG-1 | Amuvatinib |
| MVP_top60_DNA_seed | JAK1 | 0.347541 | GBM | SF126 | Ruxolitinib | MVP_top60_DNA+RNA_seed | KIT | 0.085676 | GBM | YKG-1 | Amuvatinib |
| MVP_top60_DNA_seed | EGFR | 0.349775 | GBM | T98G | Afatinib | MVP_top60_DNA+RNA_seed | PIK3CD | 0.086688 | GBM | CAS-1 | Idelalisib |
| MVP_top60_DNA_seed | PIK3CB | 0.352562 | GBM | U-87-MG | AZD8186 | MVP_top60_DNA+RNA_seed | PIK3CD | 0.086806 | GBM | DBTRG-05MG | Idelalisib |
| MVP_top60_DNA_seed | PIK3CD | 0.352562 | GBM | U-87-MG | AZD8186 | MVP_top60_DNA+RNA_seed | EGFR | 0.087284 | GBM | 8-MG-BA | AST-1306 |
| MVP_top60_DNA_seed | JAK3 | 0.353054 | GBM | U-118-MG | WHI-P97 | MVP_top60_DNA+RNA_seed | PIK3CB | 0.089032 | GBM | DBTRG-05MG | AZD8186 |
| MVP_top60_DNA_seed | PDGFRA | 0.353126 | GBM | D-392MG | Masitinib | MVP_top60_DNA+RNA_seed | PIK3CD | 0.089032 | GBM | DBTRG-05MG | AZD8186 |
| MVP_top60_DNA_seed | PDGFRB | 0.353126 | GBM | D-392MG | Masitinib | MVP_top60_DNA+RNA_seed | JAK3 | 0.09014 | GBM | D-423MG | JAK3_7406 |

| | | | | | | | | | | | |
|---|---|---|---|---|---|---|---|---|---|---|---|
| MVP_top60_DNA_seed | JAK3 | 0.355561 | GBM | LN-405 | JAK3_7406 | MVP_top60_DNA+RNA_seed | AKT1 | 0.091325 | GBM | T98G | BAY AKT1 |
| MVP_top60_DNA_seed | JAK2 | 0.35741 | GBM | KS-1 | Ruxolitinib | MVP_top60_DNA+RNA_seed | PIK3CB | 0.092072 | GBM | YKG-1 | AZD8186 |
| MVP_top60_DNA_seed | JAK1 | 0.35741 | GBM | KS-1 | Ruxolitinib | MVP_top60_DNA+RNA_seed | PIK3CD | 0.092072 | GBM | YKG-1 | AZD8186 |
| MVP_top60_DNA_seed | PIK3CD | 0.357926 | GBM | DK-MG | IC-87114 | MVP_top60_DNA+RNA_seed | KIT | 0.09315 | GBM | 8-MG-BA | Imatinib |
| MVP_top60_DNA_seed | EGFR | 0.359435 | GBM | T98G | Pelitinib | MVP_top60_DNA+RNA_seed | CDK4 | 0.09599 | GBM | DBTRG-05MG | Palbociclib |
| MVP_top60_DNA_seed | JAK2 | 0.361371 | GBM | SNB75 | Ruxolitinib | MVP_top60_DNA+RNA_seed | EGFR | 0.100039 | GBM | SK-MG-1 | PF-00299804 |
| MVP_top60_DNA_seed | JAK1 | 0.361371 | GBM | SNB75 | Ruxolitinib | MVP_top60_DNA+RNA_seed | MET | 0.100221 | GBM | D-566MG | SU11274 |
| MVP_top60_DNA_seed | PDGFRA | 0.361447 | GBM | T98G | Masitinib | MVP_top60_DNA+RNA_seed | JAK3 | 0.100345 | GBM | LN-229 | JAK3_7406 |
| MVP_top60_DNA_seed | PDGFRB | 0.361447 | GBM | T98G | Masitinib | MVP_top60_DNA+RNA_seed | EGFR | 0.100493 | GBM | D-566MG | CI-1033 |
| MVP_top60_DNA_seed | EGFR | 0.361453 | GBM | D-392MG | AST-1306 | MVP_top60_DNA+RNA_seed | EGFR | 0.101348 | GBM | YKG-1 | AZD8931 |
| MVP_top60_DNA_seed | EGFR | 0.362003 | GBM | U251 | Afatinib | MVP_top60_DNA+RNA_seed | ERBB3 | 0.101348 | GBM | YKG-1 | AZD8931 |
| MVP_top60_DNA_seed | EGFR | 0.365374 | GBM | M059J | Cetuximab | MVP_top60_DNA+RNA_seed | EGFR | 0.101937 | GBM | LN-229 | AZD8931 |
| MVP_top60_DNA_seed | EGFR | 0.36579 | GBM | SF126 | PF-00299804 | MVP_top60_DNA+RNA_seed | ERBB3 | 0.101937 | GBM | LN-229 | AZD8931 |
| MVP_top60_DNA_seed | JAK2 | 0.367046 | GBM | LN-405 | Ruxolitinib | MVP_top60_DNA+RNA_seed | PDGFRB | 0.1034 | GBM | SK-MG-1 | Pazopanib |
| MVP_top60_DNA_seed | JAK1 | 0.367046 | GBM | LN-405 | Ruxolitinib | MVP_top60_DNA+RNA_seed | KIT | 0.1034 | GBM | SK-MG-1 | Pazopanib |
| MVP_top60_DNA_seed | PIK3CD | 0.367282 | GBM | 42-MG-BA | Idelalisib | MVP_top60_DNA+RNA_seed | KIT | 0.103442 | GBM | U251 | Axitinib |
| MVP_top60_DNA_seed | EGFR | 0.368807 | GBM | D-566MG | AZD8931 | MVP_top60_DNA+RNA_seed | KIT | 0.1047 | GBM | YKG-1 | Axitinib |
| MVP_top60_DNA_seed | PIK3CA | 0.369082 | GBM | D-247MG | PI3Ka_4409 | MVP_top60_DNA+RNA_seed | JAK3 | 0.106687 | GBM | SF126 | WHI-P97 |
| MVP_top60_DNA_seed | PIK3CD | 0.369082 | GBM | D-247MG | PI3Ka_4409 | MVP_top60_DNA+RNA_seed | EGFR | 0.106911 | GBM | D-542MG | AST-1306 |
| MVP_top60_DNA_seed | EGFR | 0.374295 | GBM | YH-13 | PF-00299804 | MVP_top60_DNA+RNA_seed | AKT1 | 0.107879 | GBM | M059J | AKT inhibitor VIII |
| MVP_top60_DNA_seed | JAK3 | 0.37634 | GBM | T98G | JAK3_7406 | MVP_top60_DNA+RNA_seed | EGFR | 0.109626 | GBM | LN-229 | PF-00299804 |
| MVP_top60_DNA_seed | EGFR | 0.376762 | GBM | U-118-MG | CUDC-101 | MVP_top60_DNA+RNA_seed | KIT | 0.115877 | GBM | KS-1 | Sorafenib |
| MVP_top60_DNA_seed | EGFR | 0.376843 | GBM | M059J | Gefitinib | MVP_top60_DNA+RNA_seed | IGF1R | 0.116595 | GBM | U-118-MG | GSK1904529A |
| MVP_top60_DNA_seed | EGFR | 0.379354 | GBM | 42-MG-BA | Cetuximab | MVP_top60_DNA+RNA_seed | KIT | 0.117192 | GBM | SW1088 | OSI-930 |
| MVP_top60_DNA_seed | PIK3CA | 0.387437 | GBM | Becker | PI3Ka_4409 | MVP_top60_DNA+RNA_seed | EGFR | 0.120889 | GBM | AM-38 | CI-1033 |
| MVP_top60_DNA_seed | PIK3CD | 0.387437 | GBM | Becker | PI3Ka_4409 | MVP_top60_DNA+RNA_seed | MET | 0.122155 | GBM | U-87-MG | AZD6094 |
| MVP_top60_DNA_seed | JAK3 | 0.388814 | GBM | LNZTA3WT4 | WHI-P97 | MVP_top60_DNA+RNA_seed | EGFR | 0.122608 | GBM | 42-MG-BA | CI-1033 |
| MVP_top60_DNA_seed | EGFR | 0.39038 | GBM | A172 | Pelitinib | MVP_top60_DNA+RNA_seed | EGFR | 0.122964 | GBM | YH-13 | Pelitinib |
| MVP_top60_DNA_seed | EGFR | 0.390957 | GBM | D-392MG | Gefitinib | MVP_top60_DNA+RNA_seed | EGFR | 0.126075 | GBM | D-247MG | AZD8931 |
| MVP_top60_DNA_seed | JAK1 | 0.398139 | GBM | SW1088 | JAK1_3715 | MVP_top60_DNA+RNA_seed | ERBB3 | 0.126075 | GBM | D-247MG | AZD8931 |
| MVP_top60_DNA_seed | CDK4 | 0.398742 | GBM | U-87-MG | Palbociclib | MVP_top60_DNA+RNA_seed | PIK3CD | 0.129297 | GBM | U251 | IC-87114 |
| MVP_top60_DNA_seed | EGFR | 0.3991 | GBM | D-392MG | AZD8931 | MVP_top60_DNA+RNA_seed | EP300 | 0.129432 | GBM | SF126 | I-CBP112 |
| MVP_top60_DNA_seed | EGFR | 0.399554 | GBM | M059J | AST-1306 | MVP_top60_DNA+RNA_seed | CREBBP | 0.129432 | GBM | SF126 | I-CBP112 |
| MVP_top60_DNA_seed | EGFR | 0.403308 | GBM | SF268 | Pelitinib | MVP_top60_DNA+RNA_seed | EGFR | 0.129469 | GBM | LN-229 | Afatinib |
| MVP_top60_DNA_seed | PDGFRA | 0.408351 | GBM | D-392MG | Amuvatinib | MVP_top60_DNA+RNA_seed | AKT1 | 0.131431 | GBM | DK-MG | AKT inhibitor VIII |
| MVP_top60_DNA_seed | PIK3CB | 0.409753 | GBM | U-118-MG | AZD6482 | MVP_top60_DNA+RNA_seed | PDGFRA | 0.132873 | GBM | Becker | Amuvatinib |
| MVP_top60_DNA_seed | EGFR | 0.41024 | GBM | 8-MG-BA | Afatinib | MVP_top60_DNA+RNA_seed | KIT | 0.132873 | GBM | Becker | Amuvatinib |
| MVP_top60_DNA_seed | JAK3 | 0.411957 | GBM | SW1088 | WHI-P97 | MVP_top60_DNA+RNA_seed | EGFR | 0.133898 | GBM | 8-MG-BA | PF-00299804 |
| MVP_top60_DNA_seed | EP300 | 0.414046 | GBM | SNB75 | I-CBP112 | MVP_top60_DNA+RNA_seed | EGFR | 0.134567 | GBM | D-423MG | AZD8931 |
| MVP_top60_DNA_seed | PIK3CD | 0.414412 | GBM | LNZTA3WT4 | Idelalisib | MVP_top60_DNA+RNA_seed | ERBB3 | 0.134567 | GBM | D-423MG | AZD8931 |
| MVP_top60_DNA_seed | PIK3CA | 0.416305 | GBM | SF268 | PI3Ka_4409 | MVP_top60_DNA+RNA_seed | AKT1 | 0.138068 | GBM | D-566MG | MK-2206 |

| | | | | | | | | | | | |
|---|---|---|---|---|---|---|---|---|---|---|---|
| MVP_top60_DNA_seed | PIK3CD | 0.416305 | GBM | SF268 | PI3Ka_4409 | MVP_top60_DNA+RNA_seed | KIT | 0.138209 | GBM | 8-MG-BA | Motesanib |
| MVP_top60_DNA_seed | JAK3 | 0.417274 | GBM | SF295 | JAK3_7406 | MVP_top60_DNA+RNA_seed | KIT | 0.139832 | GBM | SF126 | Imatinib |
| MVP_top60_DNA_seed | PDGFRB | 0.422007 | GBM | LN-229 | Pazopanib | MVP_top60_DNA+RNA_seed | EGFR | 0.140854 | GBM | D-566MG | Pelitinib |
| MVP_top60_DNA_seed | EP300 | 0.424253 | GBM | YKG-1 | I-CBP112 | MVP_top60_DNA+RNA_seed | KIT | 0.141263 | GBM | D-423MG | OSI-930 |
| MVP_top60_DNA_seed | JAK2 | 0.425922 | GBM | Becker | Ruxolitinib | MVP_top60_DNA+RNA_seed | MET | 0.143615 | GBM | SF268 | SU11274 |
| MVP_top60_DNA_seed | JAK1 | 0.425922 | GBM | Becker | Ruxolitinib | MVP_top60_DNA+RNA_seed | MET | 0.151754 | GBM | M059J | AZD6094 |
| MVP_top60_DNA_seed | EGFR | 0.427033 | GBM | SK-MG-1 | Afatinib | MVP_top60_DNA+RNA_seed | EGFR | 0.156294 | GBM | D-566MG | AST-1306 |
| MVP_top60_DNA_seed | EGFR | 0.433777 | GBM | SK-MG-1 | Afatinib | MVP_top60_DNA+RNA_seed | PIK3CA | 0.158394 | GBM | LNZTA3WT4 | AZD8835 |
| MVP_top60_DNA_seed | EGFR | 0.436719 | GBM | SF268 | Afatinib | MVP_top60_DNA+RNA_seed | PIK3CD | 0.158394 | GBM | LNZTA3WT4 | AZD8835 |
| MVP_top60_DNA_seed | PDGFRA | 0.440862 | GBM | SW1088 | Masitinib | MVP_top60_DNA+RNA_seed | EP300 | 0.158996 | GBM | D-263MG | I-CBP112 |
| MVP_top60_DNA_seed | PDGFRB | 0.440862 | GBM | SW1088 | Masitinib | MVP_top60_DNA+RNA_seed | CREBBP | 0.158996 | GBM | D-263MG | I-CBP112 |
| MVP_top60_DNA_seed | JAK2 | 0.442271 | GBM | YKG-1 | Ruxolitinib | MVP_top60_DNA+RNA_seed | MET | 0.159503 | GBM | U-87-MG | Crizotinib |
| MVP_top60_DNA_seed | JAK1 | 0.442271 | GBM | YKG-1 | Ruxolitinib | MVP_top60_DNA+RNA_seed | EGFR | 0.160378 | GBM | AM-38 | PF-00299804 |
| MVP_top60_DNA_seed | JAK2 | 0.442399 | GBM | U251 | Fedratinib | MVP_top60_DNA+RNA_seed | EGFR | 0.160936 | GBM | Becker | Pelitinib |
| MVP_top60_DNA_seed | JAK2 | 0.449394 | GBM | D-542MG | AZD1480 | MVP_top60_DNA+RNA_seed | PIK3CA | 0.163953 | GBM | GB-1 | PI3Ka_4409 |
| MVP_top60_DNA_seed | JAK1 | 0.449394 | GBM | D-542MG | AZD1480 | MVP_top60_DNA+RNA_seed | PIK3CD | 0.163953 | GBM | GB-1 | PI3Ka_4409 |
| MVP_top60_DNA_seed | PIK3CD | 0.45067 | GBM | SF295 | IC-87114 | MVP_top60_DNA+RNA_seed | EGFR | 0.165266 | GBM | U-87-MG | AST-1306 |
| MVP_top60_DNA_seed | JAK2 | 0.451114 | GBM | GB-1 | Fedratinib | MVP_top60_DNA+RNA_seed | AKT1 | 0.166531 | GBM | 8-MG-BA | AKT inhibitor VIII |
| MVP_top60_DNA_seed | JAK2 | 0.452717 | GBM | LN-229 | Ruxolitinib | MVP_top60_DNA+RNA_seed | EGFR | 0.167431 | GBM | SNB75 | AST-1306 |
| MVP_top60_DNA_seed | JAK1 | 0.452717 | GBM | LN-229 | Ruxolitinib | MVP_top60_DNA+RNA_seed | EGFR | 0.167731 | GBM | DBTRG-05MG | CUDC-101 |
| MVP_top60_DNA_seed | EGFR | 0.45681 | GBM | LNZTA3WT4 | CI-1033 | MVP_top60_DNA+RNA_seed | PIK3CD | 0.170874 | GBM | YKG-1 | IC-87114 |
| MVP_top60_DNA_seed | EGFR | 0.462306 | GBM | D-423MG | AST-1306 | MVP_top60_DNA+RNA_seed | JAK3 | 0.17125 | GBM | YKG-1 | WHI-P97 |
| MVP_top60_DNA_seed | JAK3 | 0.462981 | GBM | D-392MG | WHI-P97 | MVP_top60_DNA+RNA_seed | EGFR | 0.171606 | GBM | D-392MG | Cetuximab |
| MVP_top60_DNA_seed | PDGFRA | 0.466775 | GBM | SF295 | Amuvatinib | MVP_top60_DNA+RNA_seed | PDGFRA | 0.176476 | GBM | DBTRG-05MG | Masitinib |
| MVP_top60_DNA_seed | EGFR | 0.468353 | GBM | DK-MG | CUDC-101 | MVP_top60_DNA+RNA_seed | PDGFRB | 0.176476 | GBM | DBTRG-05MG | Masitinib |
| MVP_top60_DNA_seed | EGFR | 0.46971 | GBM | DK-MG | Cetuximab | MVP_top60_DNA+RNA_seed | KIT | 0.176476 | GBM | DBTRG-05MG | Masitinib |
| MVP_top60_DNA_seed | EGFR | 0.474493 | GBM | 42-MG-BA | PF-00299804 | MVP_top60_DNA+RNA_seed | EGFR | 0.176952 | GBM | SF126 | Erlotinib |
| MVP_top60_DNA_seed | EGFR | 0.476711 | GBM | SNB75 | CUDC-101 | MVP_top60_DNA+RNA_seed | EP300 | 0.177199 | GBM | SF295 | I-CBP112 |
| MVP_top60_DNA_seed | EGFR | 0.480983 | GBM | LN-229 | Cetuximab | MVP_top60_DNA+RNA_seed | CREBBP | 0.177199 | GBM | SF295 | I-CBP112 |
| MVP_top60_DNA_seed | PIK3CA | 0.481802 | GBM | LN-405 | AZD8835 | MVP_top60_DNA+RNA_seed | PIK3CD | 0.177961 | GBM | GB-1 | Idelalisib |
| MVP_top60_DNA_seed | PIK3CD | 0.481802 | GBM | LN-405 | AZD8835 | MVP_top60_DNA+RNA_seed | JAK2 | 0.181685 | GBM | 42-MG-BA | Ruxolitinib |
| MVP_top60_DNA_seed | EGFR | 0.481954 | GBM | LN-405 | AST-1306 | MVP_top60_DNA+RNA_seed | JAK1 | 0.181685 | GBM | 42-MG-BA | Ruxolitinib |
| MVP_top60_DNA_seed | EGFR | 0.485131 | GBM | SW1088 | AST-1306 | MVP_top60_DNA+RNA_seed | IGF1R | 0.183734 | GBM | D-263MG | BMS-536924 |
| MVP_top60_DNA_seed | PIK3CA | 0.488392 | GBM | SW1088 | AZD8835 | MVP_top60_DNA+RNA_seed | EGFR | 0.185456 | GBM | A172 | AZD8931 |
| MVP_top60_DNA_seed | PIK3CD | 0.488392 | GBM | SW1088 | AZD8835 | MVP_top60_DNA+RNA_seed | ERBB3 | 0.185456 | GBM | A172 | AZD8931 |
| MVP_top60_DNA_seed | JAK3 | 0.489965 | GBM | U251 | WHI-P97 | MVP_top60_DNA+RNA_seed | EGFR | 0.188649 | GBM | SF295 | CUDC-101 |
| MVP_top60_DNA_seed | PIK3CD | 0.497239 | GBM | SW1088 | Idelalisib | MVP_top60_DNA+RNA_seed | MET | 0.189552 | GBM | 8-MG-BA | PHA-665752 |
| MVP_top60_DNA_seed | EGFR | 0.498474 | GBM | D-542MG | PF-00299804 | MVP_top60_DNA+RNA_seed | AKT1 | 0.191065 | GBM | CAS-1 | BAY AKT1 |
| MVP_top60_DNA_seed | EGFR | 0.500836 | GBM | LN-229 | Pelitinib | MVP_top60_DNA+RNA_seed | PIK3CD | 0.192243 | GBM | Becker | IC-87114 |
| MVP_top60_DNA_seed | PIK3CA | 0.50228 | GBM | M059J | PI3Ka_4409 | MVP_top60_DNA+RNA_seed | EGFR | 0.193134 | GBM | YH-13 | CUDC-101 |
| MVP_top60_DNA_seed | PIK3CD | 0.50228 | GBM | M059J | PI3Ka_4409 | MVP_top60_DNA+RNA_seed | KIT | 0.19325 | GBM | Becker | Imatinib |

| | | | | | | | | | | | |
|---|---|---|---|---|---|---|---|---|---|---|---|
| MVP_top60_DNA_seed | JAK2 | 0.502598 | GBM | GB-1 | Ruxolitinib | MVP_top60_DNA+RNA_seed | AKT1 | 0.193463 | GBM | D-542MG | AKT inhibitor VIII |
| MVP_top60_DNA_seed | JAK1 | 0.502598 | GBM | GB-1 | Ruxolitinib | MVP_top60_DNA+RNA_seed | PIK3CD | 0.195409 | GBM | SW1088 | IC-87114 |
| MVP_top60_DNA_seed | PIK3CD | 0.503721 | GBM | LN-229 | IC-87114 | MVP_top60_DNA+RNA_seed | PIK3CA | 0.197067 | GBM | SF295 | AZD8835 |
| MVP_top60_DNA_seed | PIK3CD | 0.508152 | GBM | SNB75 | Idelalisib | MVP_top60_DNA+RNA_seed | PIK3CD | 0.197067 | GBM | SF295 | AZD8835 |
| MVP_top60_DNA_seed | PIK3CD | 0.512967 | GBM | LN-18 | Idelalisib | MVP_top60_DNA+RNA_seed | PIK3CD | 0.19915 | GBM | M059J | Idelalisib |
| MVP_top60_DNA_seed | JAK3 | 0.517839 | GBM | CAS-1 | JAK3_7406 | MVP_top60_DNA+RNA_seed | KIT | 0.201687 | GBM | D-263MG | Sorafenib |
| MVP_top60_DNA_seed | EGFR | 0.51882 | GBM | U-118-MG | Gefitinib | MVP_top60_DNA+RNA_seed | EGFR | 0.203313 | GBM | LN-229 | CUDC-101 |
| MVP_top60_DNA_seed | PDGFRB | 0.520885 | GBM | D-263MG | Pazopanib | MVP_top60_DNA+RNA_seed | KIT | 0.203491 | GBM | U-87-MG | Imatinib |
| MVP_top60_DNA_seed | PIK3CD | 0.522128 | GBM | SF268 | IC-87114 | MVP_top60_DNA+RNA_seed | EGFR | 0.207045 | GBM | GB-1 | Lapatinib |
| MVP_top60_DNA_seed | JAK1 | 0.523465 | GBM | LNZTA3WT4 | JAK1_3715 | MVP_top60_DNA+RNA_seed | AKT1 | 0.213223 | GBM | T98G | GSK690693 |
| MVP_top60_DNA_seed | EGFR | 0.525061 | GBM | YKG-1 | Gefitinib | MVP_top60_DNA+RNA_seed | KIT | 0.213831 | GBM | DBTRG-05MG | Axitinib |
| MVP_top60_DNA_seed | PDGFRA | 0.525591 | GBM | D-566MG | Amuvatinib | MVP_top60_DNA+RNA_seed | PIK3CA | 0.21589 | GBM | SF295 | PI3Ka_4409 |
| MVP_top60_DNA_seed | EGFR | 0.528014 | GBM | D-263MG | Afatinib | MVP_top60_DNA+RNA_seed | PIK3CD | 0.21589 | GBM | SF295 | PI3Ka_4409 |
| MVP_top60_DNA_seed | PIK3CD | 0.528328 | GBM | LN-405 | IC-87114 | MVP_top60_DNA+RNA_seed | MET | 0.216128 | GBM | LN-18 | SU11274 |
| MVP_top60_DNA_seed | JAK2 | 0.530674 | GBM | D-423MG | AZD1480 | MVP_top60_DNA+RNA_seed | EGFR | 0.216187 | GBM | SK-MG-1 | AST-1306 |
| MVP_top60_DNA_seed | JAK1 | 0.530674 | GBM | D-423MG | AZD1480 | MVP_top60_DNA+RNA_seed | PIK3CA | 0.217007 | GBM | SNB75 | PI3Ka_4409 |
| MVP_top60_DNA_seed | EGFR | 0.530853 | GBM | D-423MG | Gefitinib | MVP_top60_DNA+RNA_seed | PIK3CD | 0.217007 | GBM | SNB75 | PI3Ka_4409 |
| MVP_top60_DNA_seed | EGFR | 0.534881 | GBM | SF295 | Afatinib | MVP_top60_DNA+RNA_seed | PIK3CD | 0.217184 | GBM | KS-1 | IC-87114 |
| MVP_top60_DNA_seed | EGFR | 0.535548 | GBM | A172 | Afatinib | MVP_top60_DNA+RNA_seed | AKT1 | 0.217966 | GBM | LN-405 | BAY AKT1 |
| MVP_top60_DNA_seed | CDK4 | 0.537366 | GBM | SF126 | Palbociclib | MVP_top60_DNA+RNA_seed | EGFR | 0.22013 | GBM | YKG-1 | AST-1306 |
| MVP_top60_DNA_seed | PIK3CA | 0.543295 | GBM | LN-229 | AZD8835 | MVP_top60_DNA+RNA_seed | EGFR | 0.220175 | GBM | LN-229 | Gefitinib |
| MVP_top60_DNA_seed | PIK3CD | 0.543295 | GBM | LN-229 | AZD8835 | MVP_top60_DNA+RNA_seed | AKT1 | 0.222344 | GBM | SW1088 | GSK690693 |
| MVP_top60_DNA_seed | EGFR | 0.545301 | GBM | LN-18 | CUDC-101 | MVP_top60_DNA+RNA_seed | EP300 | 0.224737 | GBM | U-87-MG | I-CBP112 |
| MVP_top60_DNA_seed | PDGFRA | 0.545993 | GBM | LN-18 | Amuvatinib | MVP_top60_DNA+RNA_seed | CREBBP | 0.224737 | GBM | U-87-MG | I-CBP112 |
| MVP_top60_DNA_seed | EGFR | 0.546202 | GBM | Becker | Afatinib | MVP_top60_DNA+RNA_seed | IGF1R | 0.225029 | GBM | SW1088 | Linsitinib |
| MVP_top60_DNA_seed | PIK3CA | 0.546258 | GBM | LN-405 | PI3Ka_4409 | MVP_top60_DNA+RNA_seed | AKT1 | 0.225153 | GBM | YH-13 | AKT inhibitor VIII |
| MVP_top60_DNA_seed | PIK3CD | 0.546258 | GBM | LN-405 | PI3Ka_4409 | MVP_top60_DNA+RNA_seed | PDGFRA | 0.225253 | GBM | M059J | Masitinib |
| MVP_top60_DNA_seed | EP300 | 0.547231 | GBM | GB-1 | I-CBP112 | MVP_top60_DNA+RNA_seed | PDGFRB | 0.225253 | GBM | M059J | Masitinib |
| MVP_top60_DNA_seed | EGFR | 0.548579 | GBM | SW1088 | AZD8931 | MVP_top60_DNA+RNA_seed | KIT | 0.225253 | GBM | M059J | Masitinib |
| MVP_top60_DNA_seed | PDGFRA | 0.551922 | GBM | CAS-1 | Masitinib | MVP_top60_DNA+RNA_seed | JAK3 | 0.225464 | GBM | A172 | WHI-P97 |
| MVP_top60_DNA_seed | PDGFRB | 0.551922 | GBM | CAS-1 | Masitinib | MVP_top60_DNA+RNA_seed | AKT1 | 0.227969 | GBM | U251 | AKT inhibitor VIII |
| MVP_top60_DNA_seed | PIK3CB | 0.554465 | GBM | LN-229 | AZD8186 | MVP_top60_DNA+RNA_seed | PIK3CD | 0.229603 | GBM | SF268 | Idelalisib |
| MVP_top60_DNA_seed | PIK3CD | 0.554465 | GBM | LN-229 | AZD8186 | MVP_top60_DNA+RNA_seed | KIT | 0.229827 | GBM | D-247MG | Imatinib |
| MVP_top60_DNA_seed | PIK3CA | 0.554576 | GBM | U-87-MG | AZD8835 | MVP_top60_DNA+RNA_seed | EGFR | 0.231418 | GBM | GB-1 | Afatinib |
| MVP_top60_DNA_seed | PIK3CD | 0.554576 | GBM | U-87-MG | AZD8835 | MVP_top60_DNA+RNA_seed | PIK3CB | 0.231575 | GBM | A172 | AZD6482 |
| MVP_top60_DNA_seed | JAK3 | 0.556436 | GBM | SF268 | WHI-P97 | MVP_top60_DNA+RNA_seed | IGF1R | 0.232091 | GBM | D-423MG | BMS-754807 |
| MVP_top60_DNA_seed | PDGFRA | 0.556644 | GBM | U-118-MG | Masitinib | MVP_top60_DNA+RNA_seed | PIK3CD | 0.237523 | GBM | LNZTA3WT4 | IC-87114 |
| MVP_top60_DNA_seed | PDGFRB | 0.556644 | GBM | U-118-MG | Masitinib | MVP_top60_DNA+RNA_seed | JAK3 | 0.239205 | GBM | LN-18 | WHI-P97 |
| MVP_top60_DNA_seed | EGFR | 0.557431 | GBM | SF126 | Pelitinib | MVP_top60_DNA+RNA_seed | EGFR | 0.239334 | GBM | D-263MG | Erlotinib |
| MVP_top60_DNA_seed | JAK3 | 0.558137 | GBM | M059J | JAK3_7406 | MVP_top60_DNA+RNA_seed | PDGFRB | 0.240315 | GBM | DBTRG-05MG | Pazopanib |
| MVP_top60_DNA_seed | JAK2 | 0.559856 | GBM | SNB75 | AZD1480 | MVP_top60_DNA+RNA_seed | KIT | 0.240315 | GBM | DBTRG-05MG | Pazopanib |

| | | | | | | | | | | | |
|---|---|---|---|---|---|---|---|---|---|---|---|
| MVP_top60_DNA_seed | JAK1 | 0.559856 | GBM | SNB75 | AZD1480 | MVP_top60_DNA+RNA_seed | PIK3CD | 0.240982 | GBM | M059J | IC-87114 |
| MVP_top60_DNA_seed | EGFR | 0.562046 | GBM | DBTRG-05MG | AST-1306 | MVP_top60_DNA+RNA_seed | EGFR | 0.241185 | GBM | YH-13 | Gefitinib |
| MVP_top60_DNA_seed | PDGFRA | 0.562627 | GBM | YH-13 | Amuvatinib | MVP_top60_DNA+RNA_seed | KIT | 0.243139 | GBM | D-542MG | Axitinib |
| MVP_top60_DNA_seed | PIK3CA | 0.569583 | GBM | U251 | AZD8835 | MVP_top60_DNA+RNA_seed | KIT | 0.243608 | GBM | 42-MG-BA | OSI-930 |
| MVP_top60_DNA_seed | PIK3CD | 0.569583 | GBM | U251 | AZD8835 | MVP_top60_DNA+RNA_seed | KIT | 0.245897 | GBM | U-87-MG | Motesanib |
| MVP_top60_DNA_seed | JAK2 | 0.575505 | GBM | LN-18 | AZD1480 | MVP_top60_DNA+RNA_seed | JAK2 | 0.246647 | GBM | LNZTA3WT4 | Ruxolitinib |
| MVP_top60_DNA_seed | JAK1 | 0.575505 | GBM | LN-18 | AZD1480 | MVP_top60_DNA+RNA_seed | JAK1 | 0.246647 | GBM | LNZTA3WT4 | Ruxolitinib |
| MVP_top60_DNA_seed | EGFR | 0.57615 | GBM | D-263MG | CI-1033 | MVP_top60_DNA+RNA_seed | EGFR | 0.248324 | GBM | LN-18 | CI-1033 |
| MVP_top60_DNA_seed | PIK3CB | 0.576309 | GBM | DK-MG | AZD6482 | MVP_top60_DNA+RNA_seed | CDK4 | 0.248666 | GBM | SF126 | CGP-082996 |
| MVP_top60_DNA_seed | EGFR | 0.578175 | GBM | 42-MG-BA | Afatinib | MVP_top60_DNA+RNA_seed | JAK2 | 0.250642 | GBM | YKG-1 | Fedratinib |
| MVP_top60_DNA_seed | JAK3 | 0.580369 | GBM | 42-MG-BA | WHI-P97 | MVP_top60_DNA+RNA_seed | AKT1 | 0.251932 | GBM | SF268 | AKT inhibitor VIII |
| MVP_top60_DNA_seed | EGFR | 0.584384 | GBM | D-566MG | PF-00299804 | MVP_top60_DNA+RNA_seed | EP300 | 0.25214 | GBM | LNZTA3WT4 | I-CBP112 |
| MVP_top60_DNA_seed | JAK2 | 0.587011 | GBM | SK-MG-1 | Ruxolitinib | MVP_top60_DNA+RNA_seed | CREBBP | 0.25214 | GBM | LNZTA3WT4 | I-CBP112 |
| MVP_top60_DNA_seed | JAK1 | 0.587011 | GBM | SK-MG-1 | Ruxolitinib | MVP_top60_DNA+RNA_seed | IGF1R | 0.254918 | GBM | KS-1 | GSK1904529A |
| MVP_top60_DNA_seed | EGFR | 0.588596 | GBM | T98G | Afatinib | MVP_top60_DNA+RNA_seed | EGFR | 0.255388 | GBM | Becker | Cetuximab |
| MVP_top60_DNA_seed | EGFR | 0.59153 | GBM | SK-MG-1 | CI-1033 | MVP_top60_DNA+RNA_seed | EGFR | 0.25594 | GBM | SF295 | Cetuximab |
| MVP_top60_DNA_seed | EGFR | 0.592515 | GBM | D-263MG | PF-00299804 | MVP_top60_DNA+RNA_seed | IGF1R | 0.259696 | GBM | SF268 | BMS-754807 |
| MVP_top60_DNA_seed | EGFR | 0.593466 | GBM | GB-1 | Afatinib | MVP_top60_DNA+RNA_seed | EGFR | 0.261073 | GBM | KS-1 | Gefitinib |
| MVP_top60_DNA_seed | PIK3CA | 0.594421 | GBM | LN-229 | PI3Ka_4409 | MVP_top60_DNA+RNA_seed | IGF1R | 0.261896 | GBM | M059J | BMS-754807 |
| MVP_top60_DNA_seed | PIK3CD | 0.594421 | GBM | LN-229 | PI3Ka_4409 | MVP_top60_DNA+RNA_seed | MET | 0.263791 | GBM | D-247MG | Crizotinib |
| MVP_top60_DNA_seed | EGFR | 0.596205 | GBM | LN-405 | Cetuximab | MVP_top60_DNA+RNA_seed | CDK4 | 0.267169 | GBM | D-247MG | CGP-082996 |
| MVP_top60_DNA_seed | EGFR | 0.599575 | GBM | D-263MG | Cetuximab | MVP_top60_DNA+RNA_seed | JAK1 | 0.268802 | GBM | Becker | JAK1_3715 |
| MVP_top60_DNA_seed | EGFR | 0.603259 | GBM | Becker | AST-1306 | MVP_top60_DNA+RNA_seed | EGFR | 0.269698 | GBM | LN-18 | Afatinib |
| MVP_top60_DNA_seed | PDGFRA | 0.604991 | GBM | 42-MG-BA | Masitinib | MVP_top60_DNA+RNA_seed | IGF1R | 0.270066 | GBM | SK-MG-1 | Linsitinib |
| MVP_top60_DNA_seed | PDGFRB | 0.604991 | GBM | 42-MG-BA | Masitinib | MVP_top60_DNA+RNA_seed | AKT1 | 0.271082 | GBM | D-263MG | A-443654 |
| MVP_top60_DNA_seed | PDGFRA | 0.608412 | GBM | U-118-MG | Amuvatinib | MVP_top60_DNA+RNA_seed | AKT1 | 0.272942 | GBM | DBTRG-05MG | GSK690693 |
| MVP_top60_DNA_seed | EGFR | 0.611763 | GBM | DK-MG | Afatinib | MVP_top60_DNA+RNA_seed | EGFR | 0.274048 | GBM | SF126 | CUDC-101 |
| MVP_top60_DNA_seed | EGFR | 0.613775 | GBM | T98G | Gefitinib | MVP_top60_DNA+RNA_seed | JAK2 | 0.274497 | GBM | YH-13 | Ruxolitinib |
| MVP_top60_DNA_seed | JAK2 | 0.616488 | GBM | D-263MG | Ruxolitinib | MVP_top60_DNA+RNA_seed | JAK1 | 0.274497 | GBM | YH-13 | Ruxolitinib |
| MVP_top60_DNA_seed | JAK1 | 0.616488 | GBM | D-263MG | Ruxolitinib | MVP_top60_DNA+RNA_seed | JAK2 | 0.27474 | GBM | SF295 | Ruxolitinib |
| MVP_top60_DNA_seed | JAK2 | 0.617868 | GBM | D-392MG | Ruxolitinib | MVP_top60_DNA+RNA_seed | JAK1 | 0.27474 | GBM | SF295 | Ruxolitinib |
| MVP_top60_DNA_seed | JAK1 | 0.617868 | GBM | D-392MG | Ruxolitinib | MVP_top60_DNA+RNA_seed | EGFR | 0.276723 | GBM | DBTRG-05MG | AZD8931 |
| MVP_top60_DNA_seed | PDGFRA | 0.619007 | GBM | SF126 | Amuvatinib | MVP_top60_DNA+RNA_seed | ERBB3 | 0.276723 | GBM | DBTRG-05MG | AZD8931 |
| MVP_top60_DNA_seed | EGFR | 0.620391 | GBM | 42-MG-BA | AST-1306 | MVP_top60_DNA+RNA_seed | EGFR | 0.276843 | GBM | SW1088 | Afatinib |
| MVP_top60_DNA_seed | PIK3CD | 0.620537 | GBM | AM-38 | IC-87114 | MVP_top60_DNA+RNA_seed | MET | 0.27843 | GBM | SF268 | JNJ38877605 |
| MVP_top60_DNA_seed | EGFR | 0.62599 | GBM | SF268 | PF-00299804 | MVP_top60_DNA+RNA_seed | AKT1 | 0.279139 | GBM | Becker | BAY AKT1 |
| MVP_top60_DNA_seed | EGFR | 0.627451 | GBM | A172 | Afatinib | MVP_top60_DNA+RNA_seed | AKT1 | 0.27928 | GBM | LNZTA3WT4 | MK-2206 |
| MVP_top60_DNA_seed | JAK2 | 0.630573 | GBM | D-542MG | Ruxolitinib | MVP_top60_DNA+RNA_seed | EGFR | 0.279396 | GBM | 8-MG-BA | AZD8931 |
| MVP_top60_DNA_seed | JAK1 | 0.630573 | GBM | D-542MG | Ruxolitinib | MVP_top60_DNA+RNA_seed | ERBB3 | 0.279396 | GBM | 8-MG-BA | AZD8931 |
| MVP_top60_DNA_seed | EP300 | 0.631881 | GBM | LN-229 | I-CBP112 | MVP_top60_DNA+RNA_seed | AKT1 | 0.281106 | GBM | AM-38 | MK-2206 |
| MVP_top60_DNA_seed | EP300 | 0.632516 | GBM | D-542MG | I-CBP112 | MVP_top60_DNA+RNA_seed | EGFR | 0.282952 | GBM | CAS-1 | PF-00299804 |

| | | | | | | | | | | | |
|---|---|---|---|---|---|---|---|---|---|---|---|
| MVP_top60_DNA_seed | EGFR | 0.636266 | GBM | LNZTA3WT4 | Cetuximab | MVP_top60_DNA+RNA_seed | MET | 0.284318 | GBM | D-263MG | SU11274 |
| MVP_top60_DNA_seed | EGFR | 0.639196 | GBM | D-392MG | PF-00299804 | MVP_top60_DNA+RNA_seed | KIT | 0.285659 | GBM | KS-1 | Axitinib |
| MVP_top60_DNA_seed | EGFR | 0.643621 | GBM | Becker | CI-1033 | MVP_top60_DNA+RNA_seed | EGFR | 0.286063 | GBM | SNB75 | Afatinib |
| MVP_top60_DNA_seed | EGFR | 0.645694 | GBM | CAS-1 | CUDC-101 | MVP_top60_DNA+RNA_seed | EGFR | 0.287584 | GBM | D-263MG | AZD8931 |
| MVP_top60_DNA_seed | EGFR | 0.648066 | GBM | U-118-MG | PF-00299804 | MVP_top60_DNA+RNA_seed | ERBB3 | 0.287584 | GBM | D-263MG | AZD8931 |
| MVP_top60_DNA_seed | EGFR | 0.648765 | GBM | D-263MG | Gefitinib | MVP_top60_DNA+RNA_seed | JAK2 | 0.288193 | GBM | M059J | Ruxolitinib |
| MVP_top60_DNA_seed | EGFR | 0.651567 | GBM | D-247MG | Cetuximab | MVP_top60_DNA+RNA_seed | JAK1 | 0.288193 | GBM | M059J | Ruxolitinib |
| MVP_top60_DNA_seed | EGFR | 0.654301 | GBM | SF295 | PF-00299804 | MVP_top60_DNA+RNA_seed | EGFR | 0.290069 | GBM | U251 | Gefitinib |
| MVP_top60_DNA_seed | PIK3CA | 0.659763 | GBM | SF268 | AZD8835 | MVP_top60_DNA+RNA_seed | PIK3CB | 0.290127 | GBM | A172 | AZD8186 |
| MVP_top60_DNA_seed | PIK3CD | 0.659763 | GBM | SF268 | AZD8835 | MVP_top60_DNA+RNA_seed | PIK3CD | 0.290127 | GBM | A172 | AZD8186 |
| MVP_top60_DNA_seed | PIK3CA | 0.660103 | GBM | D-247MG | AZD8835 | MVP_top60_DNA+RNA_seed | EGFR | 0.292605 | GBM | D-247MG | AST-1306 |
| MVP_top60_DNA_seed | PIK3CD | 0.660103 | GBM | D-247MG | AZD8835 | MVP_top60_DNA+RNA_seed | EP300 | 0.293796 | GBM | SK-MG-1 | I-CBP112 |
| MVP_top60_DNA_seed | EGFR | 0.660387 | GBM | U-87-MG | PF-00299804 | MVP_top60_DNA+RNA_seed | CREBBP | 0.293796 | GBM | SK-MG-1 | I-CBP112 |
| MVP_top60_DNA_seed | EGFR | 0.661416 | GBM | 42-MG-BA | Gefitinib | MVP_top60_DNA+RNA_seed | EGFR | 0.294226 | GBM | GB-1 | Pelitinib |
| MVP_top60_DNA_seed | EGFR | 0.666765 | GBM | CAS-1 | Lapatinib | MVP_top60_DNA+RNA_seed | JAK3 | 0.29455 | GBM | M059J | WHI-P97 |
| MVP_top60_DNA_seed | JAK2 | 0.672527 | GBM | GB-1 | AZD1480 | MVP_top60_DNA+RNA_seed | MET | 0.295714 | GBM | 42-MG-BA | AZD6094 |
| MVP_top60_DNA_seed | JAK1 | 0.672527 | GBM | GB-1 | AZD1480 | MVP_top60_DNA+RNA_seed | MET | 0.29742 | GBM | LN-405 | AZD6094 |
| MVP_top60_DNA_seed | EGFR | 0.674141 | GBM | A172 | AST-1306 | MVP_top60_DNA+RNA_seed | JAK2 | 0.297464 | GBM | DBTRG-05MG | Fedratinib |
| MVP_top60_DNA_seed | EGFR | 0.675234 | GBM | M059J | PF-00299804 | MVP_top60_DNA+RNA_seed | JAK3 | 0.299226 | GBM | YKG-1 | JAK3_7406 |
| MVP_top60_DNA_seed | EGFR | 0.677006 | GBM | D-263MG | Afatinib | MVP_top60_DNA+RNA_seed | KIT | 0.300276 | GBM | AM-38 | Imatinib |
| MVP_top60_DNA_seed | EGFR | 0.6825 | GBM | D-392MG | Afatinib | MVP_top60_DNA+RNA_seed | AKT1 | 0.300557 | GBM | YKG-1 | MK-2206 |
| MVP_top60_DNA_seed | EGFR | 0.686034 | GBM | U-87-MG | CUDC-101 | MVP_top60_DNA+RNA_seed | EGFR | 0.3009 | GBM | Becker | Afatinib |
| MVP_top60_DNA_seed | EGFR | 0.689103 | GBM | M059J | Afatinib | MVP_top60_DNA+RNA_seed | IGF1R | 0.304318 | GBM | D-423MG | BMS-536924 |
| MVP_top60_DNA_seed | PIK3CA | 0.689415 | GBM | CAS-1 | PI3Ka_4409 | MVP_top60_DNA+RNA_seed | EGFR | 0.306633 | GBM | LN-18 | Gefitinib |
| MVP_top60_DNA_seed | PIK3CD | 0.689415 | GBM | CAS-1 | PI3Ka_4409 | MVP_top60_DNA+RNA_seed | JAK1 | 0.30713 | GBM | M059J | JAK1_3715 |
| MVP_top60_DNA_seed | JAK2 | 0.691275 | GBM | SF295 | AZD1480 | MVP_top60_DNA+RNA_seed | EGFR | 0.309742 | GBM | D-392MG | CUDC-101 |
| MVP_top60_DNA_seed | JAK1 | 0.691275 | GBM | SF295 | AZD1480 | MVP_top60_DNA+RNA_seed | EGFR | 0.310051 | GBM | LN-18 | Afatinib |
| MVP_top60_DNA_seed | PIK3CB | 0.693304 | GBM | GB-1 | AZD8186 | MVP_top60_DNA+RNA_seed | PIK3CD | 0.3104 | GBM | DBTRG-05MG | IC-87114 |
| MVP_top60_DNA_seed | PIK3CD | 0.693304 | GBM | GB-1 | AZD8186 | MVP_top60_DNA+RNA_seed | EGFR | 0.311172 | GBM | M059J | CI-1033 |
| MVP_top60_DNA_seed | EGFR | 0.695419 | GBM | U-118-MG | AZD8931 | MVP_top60_DNA+RNA_seed | PIK3CB | 0.311403 | GBM | 8-MG-BA | TGX221 |
| MVP_top60_DNA_seed | EGFR | 0.695518 | GBM | Becker | Gefitinib | MVP_top60_DNA+RNA_seed | MET | 0.312563 | GBM | SK-MG-1 | JNJ38877605 |
| MVP_top60_DNA_seed | EGFR | 0.698169 | GBM | 42-MG-BA | Afatinib | MVP_top60_DNA+RNA_seed | KIT | 0.313197 | GBM | D-247MG | Axitinib |
| MVP_top60_DNA_seed | EGFR | 0.698915 | GBM | 42-MG-BA | AZD8931 | MVP_top60_DNA+RNA_seed | EGFR | 0.313652 | GBM | CAS-1 | CI-1033 |
| MVP_top60_DNA_seed | EGFR | 0.699406 | GBM | YKG-1 | Afatinib | MVP_top60_DNA+RNA_seed | MET | 0.314453 | GBM | LNZTA3WT4 | AZD6094 |
| MVP_top60_DNA_seed | EGFR | 0.699493 | GBM | CAS-1 | Cetuximab | MVP_top60_DNA+RNA_seed | EGFR | 0.315478 | GBM | KS-1 | Cetuximab |
| MVP_top60_DNA_seed | EGFR | 0.699883 | GBM | SK-MG-1 | Pelitinib | MVP_top60_DNA+RNA_seed | EGFR | 0.317128 | GBM | GB-1 | PF-00299804 |
| MVP_top60_DNA_seed | PDGFRA | 0.701246 | GBM | GB-1 | Masitinib | MVP_top60_DNA+RNA_seed | PIK3CB | 0.319059 | GBM | SF126 | AZD8186 |
| MVP_top60_DNA_seed | PDGFRB | 0.701246 | GBM | GB-1 | Masitinib | MVP_top60_DNA+RNA_seed | PIK3CD | 0.319059 | GBM | SF126 | AZD8186 |
| MVP_top60_DNA_seed | PIK3CD | 0.702257 | GBM | SF126 | IC-87114 | MVP_top60_DNA+RNA_seed | MET | 0.321128 | GBM | AM-38 | PHA-665752 |
| MVP_top60_DNA_seed | PDGFRA | 0.706838 | GBM | D-263MG | Amuvatinib | MVP_top60_DNA+RNA_seed | MET | 0.321203 | GBM | M059J | SU11274 |
| MVP_top60_DNA_seed | EGFR | 0.708314 | GBM | T98G | AST-1306 | MVP_top60_DNA+RNA_seed | JAK2 | 0.321281 | GBM | Becker | Fedratinib |

| | | | | | | | | | | | |
|---|---|---|---|---|---|---|---|---|---|---|---|
| MVP_top60_DNA_seed | EGFR | 0.709652 | GBM | SW1088 | Pelitinib | MVP_top60_DNA+RNA_seed | MET | 0.321414 | GBM | 8-MG-BA | JNJ38877605 |
| MVP_top60_DNA_seed | PIK3CD | 0.714056 | GBM | SNB75 | IC-87114 | MVP_top60_DNA+RNA_seed | EGFR | 0.324796 | GBM | U251 | Afatinib |
| MVP_top60_DNA_seed | EGFR | 0.715007 | GBM | U-87-MG | CI-1033 | MVP_top60_DNA+RNA_seed | EGFR | 0.32485 | GBM | D-566MG | Afatinib |
| MVP_top60_DNA_seed | EGFR | 0.716797 | GBM | YH-13 | Afatinib | MVP_top60_DNA+RNA_seed | EGFR | 0.325862 | GBM | D-263MG | Lapatinib |
| MVP_top60_DNA_seed | JAK1 | 0.717719 | GBM | GB-1 | JAK1_3715 | MVP_top60_DNA+RNA_seed | JAK1 | 0.326462 | GBM | YKG-1 | JAK1_3715 |
| MVP_top60_DNA_seed | PIK3CD | 0.72555 | GBM | SK-MG-1 | IC-87114 | MVP_top60_DNA+RNA_seed | EP300 | 0.328635 | GBM | LN-405 | I-CBP112 |
| MVP_top60_DNA_seed | PIK3CD | 0.726805 | GBM | A172 | Idelalisib | MVP_top60_DNA+RNA_seed | CREBBP | 0.328635 | GBM | LN-405 | I-CBP112 |
| MVP_top60_DNA_seed | JAK3 | 0.730156 | GBM | LN-18 | JAK3_7406 | MVP_top60_DNA+RNA_seed | EGFR | 0.328818 | GBM | D-423MG | CI-1033 |
| MVP_top60_DNA_seed | EGFR | 0.730349 | GBM | 42-MG-BA | Pelitinib | MVP_top60_DNA+RNA_seed | PIK3CA | 0.329017 | GBM | U-87-MG | PI3Ka_4409 |
| MVP_top60_DNA_seed | EGFR | 0.732461 | GBM | SF126 | Gefitinib | MVP_top60_DNA+RNA_seed | PIK3CD | 0.329017 | GBM | U-87-MG | PI3Ka_4409 |
| MVP_top60_DNA_seed | JAK2 | 0.735079 | GBM | U-118-MG | AZD1480 | MVP_top60_DNA+RNA_seed | MET | 0.329194 | GBM | A172 | AZD6094 |
| MVP_top60_DNA_seed | JAK1 | 0.735079 | GBM | U-118-MG | AZD1480 | MVP_top60_DNA+RNA_seed | PDGFRA | 0.331779 | GBM | T98G | Amuvatinib |
| MVP_top60_DNA_seed | PIK3CA | 0.735691 | GBM | U-118-MG | PI3Ka_4409 | MVP_top60_DNA+RNA_seed | KIT | 0.331779 | GBM | T98G | Amuvatinib |
| MVP_top60_DNA_seed | PIK3CD | 0.735691 | GBM | U-118-MG | PI3Ka_4409 | MVP_top60_DNA+RNA_seed | MET | 0.333479 | GBM | SF126 | Crizotinib |
| MVP_top60_DNA_seed | PIK3CA | 0.737695 | GBM | LN-18 | AZD8835 | MVP_top60_DNA+RNA_seed | JAK2 | 0.334619 | GBM | LN-405 | AZD1480 |
| MVP_top60_DNA_seed | PIK3CD | 0.737695 | GBM | LN-18 | AZD8835 | MVP_top60_DNA+RNA_seed | JAK1 | 0.334619 | GBM | LN-405 | AZD1480 |
| MVP_top60_DNA_seed | PIK3CB | 0.739064 | GBM | LNZTA3WT4 | AZD8186 | MVP_top60_DNA+RNA_seed | KIT | 0.336955 | GBM | SF268 | Sorafenib |
| MVP_top60_DNA_seed | PIK3CD | 0.739064 | GBM | LNZTA3WT4 | AZD8186 | MVP_top60_DNA+RNA_seed | PIK3CA | 0.338585 | GBM | D-542MG | PI3Ka_4409 |
| MVP_top60_DNA_seed | PIK3CA | 0.739802 | GBM | LN-18 | PI3Ka_4409 | MVP_top60_DNA+RNA_seed | PIK3CD | 0.338585 | GBM | D-542MG | PI3Ka_4409 |
| MVP_top60_DNA_seed | PIK3CD | 0.739802 | GBM | LN-18 | PI3Ka_4409 | MVP_top60_DNA+RNA_seed | MET | 0.339852 | GBM | SK-MG-1 | SU11274 |
| MVP_top60_DNA_seed | EGFR | 0.741936 | GBM | D-392MG | CI-1033 | MVP_top60_DNA+RNA_seed | EGFR | 0.34101 | GBM | DK-MG | AST-1306 |
| MVP_top60_DNA_seed | PDGFRA | 0.742057 | GBM | DBTRG-05MG | Amuvatinib | MVP_top60_DNA+RNA_seed | PIK3CA | 0.341573 | GBM | YKG-1 | AZD8835 |
| MVP_top60_DNA_seed | PIK3CB | 0.745823 | GBM | AM-38 | AZD6482 | MVP_top60_DNA+RNA_seed | PIK3CD | 0.341573 | GBM | YKG-1 | AZD8835 |
| MVP_top60_DNA_seed | EGFR | 0.748635 | GBM | AM-38 | Lapatinib | MVP_top60_DNA+RNA_seed | PIK3CB | 0.342544 | GBM | SF268 | AZD8186 |
| MVP_top60_DNA_seed | EGFR | 0.753517 | GBM | SF126 | Afatinib | MVP_top60_DNA+RNA_seed | PIK3CD | 0.342544 | GBM | SF268 | AZD8186 |
| MVP_top60_DNA_seed | PIK3CB | 0.753534 | GBM | AM-38 | TGX221 | MVP_top60_DNA+RNA_seed | JAK2 | 0.343693 | GBM | DBTRG-05MG | AZD1480 |
| MVP_top60_DNA_seed | JAK1 | 0.754363 | GBM | LN-229 | JAK1_3715 | MVP_top60_DNA+RNA_seed | JAK1 | 0.343693 | GBM | DBTRG-05MG | AZD1480 |
| MVP_top60_DNA_seed | JAK2 | 0.754412 | GBM | D-542MG | Fedratinib | MVP_top60_DNA+RNA_seed | AKT1 | 0.344506 | GBM | DK-MG | BAY AKT1 |
| MVP_top60_DNA_seed | EGFR | 0.755547 | GBM | U-87-MG | Lapatinib | MVP_top60_DNA+RNA_seed | JAK2 | 0.34528 | GBM | D-423MG | Ruxolitinib |
| MVP_top60_DNA_seed | JAK1 | 0.762739 | GBM | U-87-MG | JAK1_3715 | MVP_top60_DNA+RNA_seed | JAK1 | 0.34528 | GBM | D-423MG | Ruxolitinib |
| MVP_top60_DNA_seed | EGFR | 0.774173 | GBM | SF295 | Pelitinib | MVP_top60_DNA+RNA_seed | JAK2 | 0.347541 | GBM | SF126 | Ruxolitinib |
| MVP_top60_DNA_seed | EGFR | 0.775129 | GBM | AM-38 | Afatinib | MVP_top60_DNA+RNA_seed | JAK1 | 0.347541 | GBM | SF126 | Ruxolitinib |
| MVP_top60_DNA_seed | EGFR | 0.778509 | GBM | D-392MG | Pelitinib | MVP_top60_DNA+RNA_seed | KIT | 0.348106 | GBM | SNB75 | Motesanib |
| MVP_top60_DNA_seed | EGFR | 0.780786 | GBM | AM-38 | Erlotinib | MVP_top60_DNA+RNA_seed | MET | 0.349628 | GBM | D-423MG | AZD6094 |
| MVP_top60_DNA_seed | JAK2 | 0.781469 | GBM | DK-MG | Ruxolitinib | MVP_top60_DNA+RNA_seed | EGFR | 0.349775 | GBM | T98G | Afatinib |
| MVP_top60_DNA_seed | JAK1 | 0.781469 | GBM | DK-MG | Ruxolitinib | MVP_top60_DNA+RNA_seed | MET | 0.350701 | GBM | AM-38 | AZD6094 |
| MVP_top60_DNA_seed | EGFR | 0.782544 | GBM | LN-405 | CUDC-101 | MVP_top60_DNA+RNA_seed | PIK3CB | 0.352562 | GBM | U-87-MG | AZD8186 |
| MVP_top60_DNA_seed | EGFR | 0.782733 | GBM | SNB75 | Cetuximab | MVP_top60_DNA+RNA_seed | PIK3CD | 0.352562 | GBM | U-87-MG | AZD8186 |
| MVP_top60_DNA_seed | PIK3CA | 0.785392 | GBM | YKG-1 | PI3Ka_4409 | MVP_top60_DNA+RNA_seed | JAK3 | 0.353054 | GBM | U-118-MG | WHI-P97 |
| MVP_top60_DNA_seed | PIK3CD | 0.785392 | GBM | YKG-1 | PI3Ka_4409 | MVP_top60_DNA+RNA_seed | PDGFRA | 0.353126 | GBM | D-392MG | Masitinib |
| MVP_top60_DNA_seed | EGFR | 0.785877 | GBM | SF126 | Lapatinib | MVP_top60_DNA+RNA_seed | PDGFRB | 0.353126 | GBM | D-392MG | Masitinib |

| | | | | | | | | | | | |
|---|---|---|---|---|---|---|---|---|---|---|---|
| MVP_top60_DNA_seed | EGFR | 0.786171 | GBM | LN-405 | AZD8931 | MVP_top60_DNA+RNA_seed | KIT | 0.353126 | GBM | D-392MG | Masitinib |
| MVP_top60_DNA_seed | JAK3 | 0.789399 | GBM | KS-1 | WHI-P97 | MVP_top60_DNA+RNA_seed | IGF1R | 0.354013 | GBM | M059J | GSK1904529A |
| MVP_top60_DNA_seed | EGFR | 0.792512 | GBM | LN-405 | Afatinib | MVP_top60_DNA+RNA_seed | AKT1 | 0.354471 | GBM | LN-18 | GSK690693 |
| MVP_top60_DNA_seed | EGFR | 0.798119 | GBM | SF126 | Afatinib | MVP_top60_DNA+RNA_seed | JAK3 | 0.355561 | GBM | LN-405 | JAK3_7406 |
| MVP_top60_DNA_seed | EGFR | 0.799122 | GBM | U-87-MG | Gefitinib | MVP_top60_DNA+RNA_seed | JAK2 | 0.35741 | GBM | KS-1 | Ruxolitinib |
| MVP_top60_DNA_seed | EGFR | 0.803099 | GBM | GB-1 | AZD8931 | MVP_top60_DNA+RNA_seed | JAK1 | 0.35741 | GBM | KS-1 | Ruxolitinib |
| MVP_top60_DNA_seed | JAK1 | 0.809169 | GBM | D-392MG | JAK1_3715 | MVP_top60_DNA+RNA_seed | PIK3CD | 0.357926 | GBM | DK-MG | IC-87114 |
| MVP_top60_DNA_seed | EGFR | 0.81115 | GBM | SF295 | AST-1306 | MVP_top60_DNA+RNA_seed | KIT | 0.358007 | GBM | U-118-MG | Axitinib |
| MVP_top60_DNA_seed | EGFR | 0.813821 | GBM | LNZTA3WT4 | PF-00299804 | MVP_top60_DNA+RNA_seed | EGFR | 0.359435 | GBM | T98G | Pelitinib |
| MVP_top60_DNA_seed | PIK3CD | 0.824271 | GBM | LN-229 | Idelalisib | MVP_top60_DNA+RNA_seed | JAK2 | 0.361371 | GBM | SNB75 | Ruxolitinib |
| MVP_top60_DNA_seed | EGFR | 0.824329 | GBM | SF268 | Erlotinib | MVP_top60_DNA+RNA_seed | JAK1 | 0.361371 | GBM | SNB75 | Ruxolitinib |
| MVP_top60_DNA_seed | JAK2 | 0.830263 | GBM | U-118-MG | Ruxolitinib | MVP_top60_DNA+RNA_seed | PDGFRA | 0.361447 | GBM | T98G | Masitinib |
| MVP_top60_DNA_seed | JAK1 | 0.830263 | GBM | U-118-MG | Ruxolitinib | MVP_top60_DNA+RNA_seed | PDGFRB | 0.361447 | GBM | T98G | Masitinib |
| MVP_top60_DNA_seed | PIK3CA | 0.841522 | GBM | CAS-1 | AZD8835 | MVP_top60_DNA+RNA_seed | KIT | 0.361447 | GBM | T98G | Masitinib |
| MVP_top60_DNA_seed | PIK3CD | 0.841522 | GBM | CAS-1 | AZD8835 | MVP_top60_DNA+RNA_seed | EGFR | 0.361453 | GBM | D-392MG | AST-1306 |
| MVP_top60_DNA_seed | EGFR | 0.848985 | GBM | T98G | PF-00299804 | MVP_top60_DNA+RNA_seed | IGF1R | 0.361468 | GBM | SF268 | GSK1904529A |
| MVP_top60_DNA_seed | EGFR | 0.85078 | GBM | D-423MG | Afatinib | MVP_top60_DNA+RNA_seed | KIT | 0.362003 | GBM | D-263MG | OSI-930 |
| MVP_top60_DNA_seed | EGFR | 0.852691 | GBM | LNZTA3WT4 | Afatinib | MVP_top60_DNA+RNA_seed | EGFR | 0.362003 | GBM | U251 | Afatinib |
| MVP_top60_DNA_seed | EGFR | 0.854566 | GBM | GB-1 | Gefitinib | MVP_top60_DNA+RNA_seed | KIT | 0.362144 | GBM | DBTRG-05MG | Motesanib |
| MVP_top60_DNA_seed | PDGFRA | 0.861017 | GBM | SNB75 | Masitinib | MVP_top60_DNA+RNA_seed | KIT | 0.363008 | GBM | KS-1 | Motesanib |
| MVP_top60_DNA_seed | PDGFRB | 0.861017 | GBM | SNB75 | Masitinib | MVP_top60_DNA+RNA_seed | EGFR | 0.365374 | GBM | M059J | Cetuximab |
| MVP_top60_DNA_seed | JAK1 | 0.861634 | GBM | LN-405 | JAK1_3715 | MVP_top60_DNA+RNA_seed | MET | 0.36562 | GBM | 42-MG-BA | SU11274 |
| MVP_top60_DNA_seed | PIK3CA | 0.866778 | GBM | SF126 | PI3Ka_4409 | MVP_top60_DNA+RNA_seed | EGFR | 0.36579 | GBM | SF126 | PF-00299804 |
| MVP_top60_DNA_seed | PIK3CD | 0.866778 | GBM | SF126 | PI3Ka_4409 | MVP_top60_DNA+RNA_seed | JAK2 | 0.367046 | GBM | LN-405 | Ruxolitinib |
| MVP_top60_DNA_seed | EGFR | 0.868661 | GBM | M059J | Afatinib | MVP_top60_DNA+RNA_seed | JAK1 | 0.367046 | GBM | LN-405 | Ruxolitinib |
| MVP_top60_DNA_seed | PDGFRA | 0.869844 | GBM | LNZTA3WT4 | Masitinib | MVP_top60_DNA+RNA_seed | PIK3CD | 0.367282 | GBM | 42-MG-BA | Idelalisib |
| MVP_top60_DNA_seed | PDGFRB | 0.869844 | GBM | LNZTA3WT4 | Masitinib | MVP_top60_DNA+RNA_seed | AKT1 | 0.368043 | GBM | SF268 | MK-2206 |
| MVP_top60_DNA_seed | EGFR | 0.871061 | GBM | SK-MG-1 | Gefitinib | MVP_top60_DNA+RNA_seed | EGFR | 0.368807 | GBM | D-566MG | AZD8931 |
| MVP_top60_DNA_seed | PIK3CA | 0.873925 | GBM | U251 | PI3Ka_4409 | MVP_top60_DNA+RNA_seed | ERBB3 | 0.368807 | GBM | D-566MG | AZD8931 |
| MVP_top60_DNA_seed | PIK3CD | 0.873925 | GBM | U251 | PI3Ka_4409 | MVP_top60_DNA+RNA_seed | PIK3CA | 0.369082 | GBM | D-247MG | PI3Ka_4409 |
| MVP_top60_DNA_seed | EGFR | 0.875322 | GBM | LNZTA3WT4 | Afatinib | MVP_top60_DNA+RNA_seed | PIK3CD | 0.369082 | GBM | D-247MG | PI3Ka_4409 |
| MVP_top60_DNA_seed | JAK3 | 0.879609 | GBM | SF295 | WHI-P97 | MVP_top60_DNA+RNA_seed | MET | 0.373602 | GBM | SK-MG-1 | AZD6094 |
| MVP_top60_DNA_seed | JAK2 | 0.888191 | GBM | SW1088 | Ruxolitinib | MVP_top60_DNA+RNA_seed | EGFR | 0.374295 | GBM | YH-13 | PF-00299804 |
| MVP_top60_DNA_seed | JAK1 | 0.888191 | GBM | SW1088 | Ruxolitinib | MVP_top60_DNA+RNA_seed | KIT | 0.375078 | GBM | U-118-MG | Motesanib |
| MVP_top60_DNA_seed | EGFR | 0.893863 | GBM | SF268 | AST-1306 | MVP_top60_DNA+RNA_seed | MET | 0.375264 | GBM | YKG-1 | AZD6094 |
| MVP_top60_DNA_seed | JAK2 | 0.89522 | GBM | SF295 | Fedratinib | MVP_top60_DNA+RNA_seed | AKT1 | 0.37633 | GBM | 8-MG-BA | MK-2206 |
| MVP_top60_DNA_seed | PDGFRA | 0.896973 | GBM | 42-MG-BA | Amuvatinib | MVP_top60_DNA+RNA_seed | JAK3 | 0.37634 | GBM | T98G | JAK3_7406 |
| MVP_top60_DNA_seed | PIK3CD | 0.904813 | GBM | D-392MG | IC-87114 | MVP_top60_DNA+RNA_seed | IGF1R | 0.376543 | GBM | 42-MG-BA | BMS-754807 |
| MVP_top60_DNA_seed | EGFR | 0.905008 | GBM | CAS-1 | AZD8931 | MVP_top60_DNA+RNA_seed | EGFR | 0.376762 | GBM | U-118-MG | CUDC-101 |
| MVP_top60_DNA_seed | EGFR | 0.905612 | GBM | LNZTA3WT4 | Pelitinib | MVP_top60_DNA+RNA_seed | EGFR | 0.376843 | GBM | M059J | Gefitinib |
| MVP_top60_DNA_seed | EGFR | 0.907562 | GBM | U-118-MG | AST-1306 | MVP_top60_DNA+RNA_seed | AKT1 | 0.379339 | GBM | D-423MG | AKT inhibitor VIII |

| | | | | | | | | | | | |
|---|---|---|---|---|---|---|---|---|---|---|---|
| MVP_top60_DNA_seed | PDGFRA | 0.907839 | GBM | D-542MG | Masitinib | MVP_top60_DNA+RNA_seed | EGFR | 0.379354 | GBM | 42-MG-BA | Cetuximab |
| MVP_top60_DNA_seed | PDGFRB | 0.907839 | GBM | D-542MG | Masitinib | MVP_top60_DNA+RNA_seed | AKT1 | 0.381136 | GBM | SF268 | GSK690693 |
| MVP_top60_DNA_seed | EGFR | 0.911172 | GBM | Becker | CUDC-101 | MVP_top60_DNA+RNA_seed | IGF1R | 0.381948 | GBM | U-118-MG | BMS-754807 |
| MVP_top60_DNA_seed | JAK3 | 0.917445 | GBM | SK-MG-1 | WHI-P97 | MVP_top60_DNA+RNA_seed | PIK3CA | 0.387437 | GBM | Becker | PI3Ka_4409 |
| MVP_top60_DNA_seed | EGFR | 0.918206 | GBM | LN-405 | PF-00299804 | MVP_top60_DNA+RNA_seed | PIK3CD | 0.387437 | GBM | Becker | PI3Ka_4409 |
| MVP_top60_DNA_seed | EGFR | 0.922185 | GBM | A172 | PF-00299804 | MVP_top60_DNA+RNA_seed | JAK3 | 0.388814 | GBM | LNZTA3WT4 | WHI-P97 |
| MVP_top60_DNA_seed | EGFR | 0.922255 | GBM | SNB75 | PF-00299804 | MVP_top60_DNA+RNA_seed | IGF1R | 0.38978 | GBM | U251 | BMS-754807 |
| MVP_top60_DNA_seed | EGFR | 0.928215 | GBM | SW1088 | PF-00299804 | MVP_top60_DNA+RNA_seed | AKT1 | 0.390295 | GBM | 42-MG-BA | AKT inhibitor VIII |
| MVP_top60_DNA_seed | EGFR | 0.930239 | GBM | SF268 | Afatinib | MVP_top60_DNA+RNA_seed | EGFR | 0.39038 | GBM | A172 | Pelitinib |
| MVP_top60_DNA_seed | EGFR | 0.930559 | GBM | DBTRG-05MG | Cetuximab | MVP_top60_DNA+RNA_seed | AKT1 | 0.390565 | GBM | LN-405 | GSK690693 |
| MVP_top60_DNA_seed | EGFR | 0.934676 | GBM | D-247MG | Gefitinib | MVP_top60_DNA+RNA_seed | IGF1R | 0.390761 | GBM | YKG-1 | BMS-536924 |
| MVP_top60_DNA_seed | JAK1 | 0.935662 | GBM | D-542MG | JAK1_3715 | MVP_top60_DNA+RNA_seed | EGFR | 0.390957 | GBM | D-392MG | Gefitinib |
| MVP_top60_DNA_seed | JAK2 | 0.945487 | GBM | U-87-MG | Ruxolitinib | MVP_top60_DNA+RNA_seed | KIT | 0.39282 | GBM | M059J | Axitinib |
| MVP_top60_DNA_seed | JAK1 | 0.945487 | GBM | U-87-MG | Ruxolitinib | MVP_top60_DNA+RNA_seed | IGF1R | 0.39468 | GBM | CAS-1 | BMS-536924 |
| MVP_top60_DNA_seed | PIK3CB | 0.946816 | GBM | D-392MG | AZD8186 | MVP_top60_DNA+RNA_seed | IGF1R | 0.395168 | GBM | YKG-1 | GSK1904529A |
| MVP_top60_DNA_seed | PIK3CD | 0.946816 | GBM | D-392MG | AZD8186 | MVP_top60_DNA+RNA_seed | IGF1R | 0.397126 | GBM | LN-18 | BMS-536924 |
| MVP_top60_DNA_seed | EGFR | 0.948982 | GBM | DBTRG-05MG | PF-00299804 | MVP_top60_DNA+RNA_seed | JAK1 | 0.398139 | GBM | SW1088 | JAK1_3715 |
| MVP_top60_DNA_seed | EGFR | 0.950111 | GBM | LN-18 | Pelitinib | MVP_top60_DNA+RNA_seed | CDK4 | 0.398742 | GBM | U-87-MG | Palbociclib |
| MVP_top60_DNA_seed | EGFR | 0.950739 | GBM | D-263MG | CUDC-101 | MVP_top60_DNA+RNA_seed | EGFR | 0.3991 | GBM | D-392MG | AZD8931 |
| MVP_top60_DNA_seed | JAK3 | 0.951276 | GBM | D-542MG | WHI-P97 | MVP_top60_DNA+RNA_seed | ERBB3 | 0.3991 | GBM | D-392MG | AZD8931 |
| MVP_top60_DNA_seed | PIK3CB | 0.955373 | GBM | U251 | AZD8186 | MVP_top60_DNA+RNA_seed | EGFR | 0.399554 | GBM | M059J | AST-1306 |
| MVP_top60_DNA_seed | PIK3CD | 0.955373 | GBM | U251 | AZD8186 | MVP_top60_DNA+RNA_seed | EGFR | 0.403308 | GBM | SF268 | Pelitinib |
| MVP_top60_DNA_seed | EP300 | 0.956974 | GBM | AM-38 | I-CBP112 | MVP_top60_DNA+RNA_seed | MET | 0.404379 | GBM | A172 | JNJ38877605 |
| MVP_top60_DNA_seed | EP300 | 0.960338 | GBM | D-392MG | I-CBP112 | MVP_top60_DNA+RNA_seed | AKT1 | 0.406902 | GBM | D-247MG | MK-2206 |
| MVP_top60_DNA_seed | EGFR | 0.963154 | GBM | D-247MG | PF-00299804 | MVP_top60_DNA+RNA_seed | IGF1R | 0.40741 | GBM | Becker | GSK1904529A |
| MVP_top60_DNA_seed | EGFR | 0.966872 | GBM | LN-405 | Afatinib | MVP_top60_DNA+RNA_seed | PDGFRA | 0.408351 | GBM | D-392MG | Amuvatinib |
| MVP_top60_DNA_seed | EGFR | 0.973124 | GBM | GB-1 | Erlotinib | MVP_top60_DNA+RNA_seed | KIT | 0.408351 | GBM | D-392MG | Amuvatinib |
| MVP_top60_DNA_seed | CDK4 | 0.97739 | GBM | 42-MG-BA | Palbociclib | MVP_top60_DNA+RNA_seed | PIK3CB | 0.409753 | GBM | U-118-MG | AZD6482 |
| MVP_top60_DNA_seed | JAK2 | 0.990506 | GBM | U-87-MG | Fedratinib | MVP_top60_DNA+RNA_seed | MET | 0.409838 | GBM | SF268 | Crizotinib |
| MVP_top60_DNA_seed | PDGFRA | 0.9916 | GBM | M059J | Amuvatinib | MVP_top60_DNA+RNA_seed | EGFR | 0.41024 | GBM | 8-MG-BA | Afatinib |
| MVP_top60_DNA_seed | JAK3 | 0.993898 | GBM | GB-1 | JAK3_7406 | MVP_top60_DNA+RNA_seed | JAK3 | 0.411957 | GBM | SW1088 | WHI-P97 |
| MVP_top60_DNA_seed | EGFR | 0.997767 | GBM | U-118-MG | Cetuximab | MVP_top60_DNA+RNA_seed | EP300 | 0.414046 | GBM | SNB75 | I-CBP112 |
| MVP_top60_DNA_seed | JAK2 | 0.998788 | GBM | LN-405 | Fedratinib | MVP_top60_DNA+RNA_seed | CREBBP | 0.414046 | GBM | SNB75 | I-CBP112 |
| MVP_top60_DNA_seed | PDGFRB | 1.001311 | GBM | SF295 | Pazopanib | MVP_top60_DNA+RNA_seed | PIK3CD | 0.414412 | GBM | LNZTA3WT4 | Idelalisib |
| MVP_top60_DNA_seed | PIK3CA | 1.004612 | GBM | M059J | AZD8835 | MVP_top60_DNA+RNA_seed | PIK3CA | 0.416305 | GBM | SF268 | PI3Ka_4409 |
| MVP_top60_DNA_seed | PIK3CD | 1.004612 | GBM | M059J | AZD8835 | MVP_top60_DNA+RNA_seed | PIK3CD | 0.416305 | GBM | SF268 | PI3Ka_4409 |
| MVP_top60_DNA_seed | EGFR | 1.007573 | GBM | DK-MG | Gefitinib | MVP_top60_DNA+RNA_seed | JAK3 | 0.417274 | GBM | SF295 | JAK3_7406 |
| MVP_top60_DNA_seed | JAK2 | 1.011303 | GBM | DBTRG-05MG | Ruxolitinib | MVP_top60_DNA+RNA_seed | AKT1 | 0.417977 | GBM | LNZTA3WT4 | BAY AKT1 |
| MVP_top60_DNA_seed | JAK1 | 1.011303 | GBM | DBTRG-05MG | Ruxolitinib | MVP_top60_DNA+RNA_seed | IGF1R | 0.419067 | GBM | LN-229 | BMS-754807 |
| MVP_top60_DNA_seed | JAK1 | 1.01297 | GBM | U-118-MG | JAK1_3715 | MVP_top60_DNA+RNA_seed | MET | 0.421369 | GBM | 8-MG-BA | Crizotinib |
| MVP_top60_DNA_seed | EGFR | 1.013694 | GBM | U-118-MG | Afatinib | MVP_top60_DNA+RNA_seed | AKT1 | 0.421806 | GBM | KS-1 | BAY AKT1 |

| | | | | | | | | | | | |
|---|---|---|---|---|---|---|---|---|---|---|---|
| MVP_top60_DNA_seed | JAK3 | 1.015945 | GBM | LN-405 | WHI-P97 | MVP_top60_DNA+RNA_seed | PDGFRB | 0.422007 | GBM | LN-229 | Pazopanib |
| MVP_top60_DNA_seed | JAK2 | 1.017056 | GBM | LN-229 | AZD1480 | MVP_top60_DNA+RNA_seed | KIT | 0.422007 | GBM | LN-229 | Pazopanib |
| MVP_top60_DNA_seed | JAK1 | 1.017056 | GBM | LN-229 | AZD1480 | MVP_top60_DNA+RNA_seed | EP300 | 0.424253 | GBM | YKG-1 | I-CBP112 |
| MVP_top60_DNA_seed | EGFR | 1.023486 | GBM | DBTRG-05MG | Afatinib | MVP_top60_DNA+RNA_seed | CREBBP | 0.424253 | GBM | YKG-1 | I-CBP112 |
| MVP_top60_DNA_seed | EGFR | 1.028719 | GBM | D-392MG | Afatinib | MVP_top60_DNA+RNA_seed | JAK2 | 0.425922 | GBM | Becker | Ruxolitinib |
| MVP_top60_DNA_seed | EGFR | 1.030523 | GBM | SW1088 | Afatinib | MVP_top60_DNA+RNA_seed | JAK1 | 0.425922 | GBM | Becker | Ruxolitinib |
| MVP_top60_DNA_seed | EGFR | 1.034087 | GBM | Becker | PF-00299804 | MVP_top60_DNA+RNA_seed | IGF1R | 0.425988 | GBM | LN-18 | Linsitinib |
| MVP_top60_DNA_seed | EP300 | 1.040931 | GBM | SW1088 | I-CBP112 | MVP_top60_DNA+RNA_seed | EGFR | 0.427033 | GBM | SK-MG-1 | Afatinib |
| MVP_top60_DNA_seed | EGFR | 1.041416 | GBM | DBTRG-05MG | Afatinib | MVP_top60_DNA+RNA_seed | AKT1 | 0.429491 | GBM | GB-1 | BAY AKT1 |
| MVP_top60_DNA_seed | EGFR | 1.041725 | GBM | SF268 | Gefitinib | MVP_top60_DNA+RNA_seed | MET | 0.429697 | GBM | D-392MG | JNJ38877605 |
| MVP_top60_DNA_seed | PIK3CA | 1.044933 | GBM | DBTRG-05MG | PI3Ka_4409 | MVP_top60_DNA+RNA_seed | IGF1R | 0.430807 | GBM | D-263MG | BMS-754807 |
| MVP_top60_DNA_seed | PIK3CD | 1.044933 | GBM | DBTRG-05MG | PI3Ka_4409 | MVP_top60_DNA+RNA_seed | EGFR | 0.433777 | GBM | SK-MG-1 | Afatinib |
| MVP_top60_DNA_seed | JAK1 | 1.046499 | GBM | AM-38 | JAK1_3715 | MVP_top60_DNA+RNA_seed | IGF1R | 0.43442 | GBM | SF295 | BMS-536924 |
| MVP_top60_DNA_seed | EGFR | 1.047505 | GBM | CAS-1 | Erlotinib | MVP_top60_DNA+RNA_seed | MET | 0.434855 | GBM | A172 | SU11274 |
| MVP_top60_DNA_seed | CDK4 | 1.050188 | GBM | 8-MG-BA | Palbociclib | MVP_top60_DNA+RNA_seed | EGFR | 0.436719 | GBM | SF268 | Afatinib |
| MVP_top60_DNA_seed | EGFR | 1.051861 | GBM | D-542MG | Afatinib | MVP_top60_DNA+RNA_seed | PDGFRA | 0.440862 | GBM | SW1088 | Masitinib |
| MVP_top60_DNA_seed | JAK2 | 1.053148 | GBM | D-392MG | AZD1480 | MVP_top60_DNA+RNA_seed | PDGFRB | 0.440862 | GBM | SW1088 | Masitinib |
| MVP_top60_DNA_seed | JAK1 | 1.053148 | GBM | D-392MG | AZD1480 | MVP_top60_DNA+RNA_seed | KIT | 0.440862 | GBM | SW1088 | Masitinib |
| MVP_top60_DNA_seed | PDGFRA | 1.054111 | GBM | LN-405 | Amuvatinib | MVP_top60_DNA+RNA_seed | JAK2 | 0.442271 | GBM | YKG-1 | Ruxolitinib |
| MVP_top60_DNA_seed | EGFR | 1.060722 | GBM | DK-MG | Afatinib | MVP_top60_DNA+RNA_seed | JAK1 | 0.442271 | GBM | YKG-1 | Ruxolitinib |
| MVP_top60_DNA_seed | EGFR | 1.06613 | GBM | D-542MG | Afatinib | MVP_top60_DNA+RNA_seed | JAK2 | 0.442399 | GBM | U251 | Fedratinib |
| MVP_top60_DNA_seed | EGFR | 1.074244 | GBM | AM-38 | Cetuximab | MVP_top60_DNA+RNA_seed | MET | 0.44292 | GBM | SF268 | AZD6094 |
| MVP_top60_DNA_seed | CDK4 | 1.080758 | GBM | CAS-1 | Palbociclib | MVP_top60_DNA+RNA_seed | JAK2 | 0.449394 | GBM | D-542MG | AZD1480 |
| MVP_top60_DNA_seed | PIK3CA | 1.081435 | GBM | A172 | PI3Ka_4409 | MVP_top60_DNA+RNA_seed | JAK1 | 0.449394 | GBM | D-542MG | AZD1480 |
| MVP_top60_DNA_seed | PIK3CD | 1.081435 | GBM | A172 | PI3Ka_4409 | MVP_top60_DNA+RNA_seed | PIK3CD | 0.45067 | GBM | SF295 | IC-87114 |
| MVP_top60_DNA_seed | EGFR | 1.084594 | GBM | CAS-1 | Afatinib | MVP_top60_DNA+RNA_seed | JAK2 | 0.451114 | GBM | GB-1 | Fedratinib |
| MVP_top60_DNA_seed | EGFR | 1.088397 | GBM | SNB75 | Pelitinib | MVP_top60_DNA+RNA_seed | JAK2 | 0.452717 | GBM | LN-229 | Ruxolitinib |
| MVP_top60_DNA_seed | EGFR | 1.091108 | GBM | M059J | Pelitinib | MVP_top60_DNA+RNA_seed | JAK1 | 0.452717 | GBM | LN-229 | Ruxolitinib |
| MVP_top60_DNA_seed | EGFR | 1.092916 | GBM | T98G | CUDC-101 | MVP_top60_DNA+RNA_seed | EGFR | 0.45681 | GBM | LNZTA3WT4 | CI-1033 |
| MVP_top60_DNA_seed | EGFR | 1.093055 | GBM | SW1088 | Cetuximab | MVP_top60_DNA+RNA_seed | KIT | 0.458569 | GBM | M059J | Motesanib |
| MVP_top60_DNA_seed | EGFR | 1.093282 | GBM | D-542MG | CI-1033 | MVP_top60_DNA+RNA_seed | AKT1 | 0.459727 | GBM | KS-1 | AKT inhibitor VIII |
| MVP_top60_DNA_seed | JAK1 | 1.098248 | GBM | SNB75 | JAK1_3715 | MVP_top60_DNA+RNA_seed | IGF1R | 0.460638 | GBM | SK-MG-1 | GSK1904529A |
| MVP_top60_DNA_seed | EP300 | 1.104118 | GBM | KS-1 | I-CBP112 | MVP_top60_DNA+RNA_seed | EGFR | 0.462306 | GBM | D-423MG | AST-1306 |
| MVP_top60_DNA_seed | EGFR | 1.114354 | GBM | SNB75 | CI-1033 | MVP_top60_DNA+RNA_seed | KIT | 0.462812 | GBM | CAS-1 | Imatinib |
| MVP_top60_DNA_seed | EGFR | 1.11661 | GBM | LNZTA3WT4 | Gefitinib | MVP_top60_DNA+RNA_seed | JAK3 | 0.462981 | GBM | D-392MG | WHI-P97 |
| MVP_top60_DNA_seed | PIK3CA | 1.117505 | GBM | D-392MG | AZD8835 | MVP_top60_DNA+RNA_seed | PDGFRA | 0.466775 | GBM | SF295 | Amuvatinib |
| MVP_top60_DNA_seed | PIK3CD | 1.117505 | GBM | D-392MG | AZD8835 | MVP_top60_DNA+RNA_seed | KIT | 0.466775 | GBM | SF295 | Amuvatinib |
| MVP_top60_DNA_seed | EGFR | 1.123878 | GBM | AM-38 | Afatinib | MVP_top60_DNA+RNA_seed | EGFR | 0.468353 | GBM | DK-MG | CUDC-101 |
| MVP_top60_DNA_seed | CDK4 | 1.124655 | GBM | U-87-MG | CGP-082996 | MVP_top60_DNA+RNA_seed | EGFR | 0.46971 | GBM | DK-MG | Cetuximab |
| MVP_top60_DNA_seed | EGFR | 1.128897 | GBM | D-247MG | Afatinib | MVP_top60_DNA+RNA_seed | EGFR | 0.474493 | GBM | 42-MG-BA | PF-00299804 |
| MVP_top60_DNA_seed | PDGFRB | 1.129922 | GBM | AM-38 | Pazopanib | MVP_top60_DNA+RNA_seed | EGFR | 0.476711 | GBM | SNB75 | CUDC-101 |

| | | | | | | | | | | | |
|---|---|---|---|---|---|---|---|---|---|---|---|
| MVP_top60_DNA_seed | PIK3CA | 1.153257 | GBM | D-392MG | PI3Ka_4409 | MVP_top60_DNA+RNA_seed | AKT1 | 0.478053 | GBM | D-392MG | AKT inhibitor VIII |
| MVP_top60_DNA_seed | PIK3CD | 1.153257 | GBM | D-392MG | PI3Ka_4409 | MVP_top60_DNA+RNA_seed | EGFR | 0.480983 | GBM | LN-229 | Cetuximab |
| MVP_top60_DNA_seed | EGFR | 1.164147 | GBM | LN-405 | Pelitinib | MVP_top60_DNA+RNA_seed | PIK3CA | 0.481802 | GBM | LN-405 | AZD8835 |
| MVP_top60_DNA_seed | PIK3CA | 1.164432 | GBM | DK-MG | PI3Ka_4409 | MVP_top60_DNA+RNA_seed | PIK3CD | 0.481802 | GBM | LN-405 | AZD8835 |
| MVP_top60_DNA_seed | PIK3CD | 1.164432 | GBM | DK-MG | PI3Ka_4409 | MVP_top60_DNA+RNA_seed | EGFR | 0.481954 | GBM | LN-405 | AST-1306 |
| MVP_top60_DNA_seed | EGFR | 1.169363 | GBM | D-247MG | Afatinib | MVP_top60_DNA+RNA_seed | EGFR | 0.485131 | GBM | SW1088 | AST-1306 |
| MVP_top60_DNA_seed | PIK3CD | 1.171712 | GBM | D-542MG | Idelalisib | MVP_top60_DNA+RNA_seed | MET | 0.486467 | GBM | GB-1 | AZD6094 |
| MVP_top60_DNA_seed | JAK1 | 1.172708 | GBM | DBTRG-05MG | JAK1_3715 | MVP_top60_DNA+RNA_seed | AKT1 | 0.488055 | GBM | YKG-1 | AKT inhibitor VIII |
| MVP_top60_DNA_seed | PIK3CA | 1.178644 | GBM | U-118-MG | AZD8835 | MVP_top60_DNA+RNA_seed | PIK3CA | 0.488392 | GBM | SW1088 | AZD8835 |
| MVP_top60_DNA_seed | PIK3CD | 1.178644 | GBM | U-118-MG | AZD8835 | MVP_top60_DNA+RNA_seed | PIK3CD | 0.488392 | GBM | SW1088 | AZD8835 |
| MVP_top60_DNA_seed | JAK3 | 1.179073 | GBM | D-263MG | WHI-P97 | MVP_top60_DNA+RNA_seed | MET | 0.489502 | GBM | D-423MG | JNJ38877605 |
| MVP_top60_DNA_seed | EGFR | 1.183565 | GBM | D-542MG | Gefitinib | MVP_top60_DNA+RNA_seed | JAK3 | 0.489965 | GBM | U251 | WHI-P97 |
| MVP_top60_DNA_seed | JAK3 | 1.1858 | GBM | DBTRG-05MG | WHI-P97 | MVP_top60_DNA+RNA_seed | IGF1R | 0.492257 | GBM | DBTRG-05MG | Linsitinib |
| MVP_top60_DNA_seed | PDGFRB | 1.192739 | GBM | U-87-MG | Pazopanib | MVP_top60_DNA+RNA_seed | KIT | 0.497011 | GBM | DBTRG-05MG | OSI-930 |
| MVP_top60_DNA_seed | PIK3CD | 1.197102 | GBM | DK-MG | Idelalisib | MVP_top60_DNA+RNA_seed | PIK3CD | 0.497239 | GBM | SW1088 | Idelalisib |
| MVP_top60_DNA_seed | EGFR | 1.217418 | GBM | U-87-MG | Erlotinib | MVP_top60_DNA+RNA_seed | AKT1 | 0.497786 | GBM | LN-229 | AKT inhibitor VIII |
| MVP_top60_DNA_seed | JAK2 | 1.219291 | GBM | LN-229 | Fedratinib | MVP_top60_DNA+RNA_seed | EGFR | 0.498474 | GBM | D-542MG | PF-00299804 |
| MVP_top60_DNA_seed | JAK1 | 1.220882 | GBM | CAS-1 | JAK1_3715 | MVP_top60_DNA+RNA_seed | KIT | 0.499232 | GBM | U-87-MG | Sorafenib |
| MVP_top60_DNA_seed | CDK4 | 1.232426 | GBM | D-263MG | CGP-082996 | MVP_top60_DNA+RNA_seed | EGFR | 0.500836 | GBM | LN-229 | Pelitinib |
| MVP_top60_DNA_seed | EGFR | 1.234474 | GBM | CAS-1 | Afatinib | MVP_top60_DNA+RNA_seed | PIK3CA | 0.50228 | GBM | M059J | PI3Ka_4409 |
| MVP_top60_DNA_seed | JAK2 | 1.23889 | GBM | LN-18 | Fedratinib | MVP_top60_DNA+RNA_seed | PIK3CD | 0.50228 | GBM | M059J | PI3Ka_4409 |
| MVP_top60_DNA_seed | JAK2 | 1.239607 | GBM | LNZTA3WT4 | Fedratinib | MVP_top60_DNA+RNA_seed | JAK2 | 0.502598 | GBM | GB-1 | Ruxolitinib |
| MVP_top60_DNA_seed | EGFR | 1.248653 | GBM | SNB75 | Afatinib | MVP_top60_DNA+RNA_seed | JAK1 | 0.502598 | GBM | GB-1 | Ruxolitinib |
| MVP_top60_DNA_seed | CDK4 | 1.249184 | GBM | AM-38 | CGP-082996 | MVP_top60_DNA+RNA_seed | AKT1 | 0.503154 | GBM | U-118-MG | BAY AKT1 |
| MVP_top60_DNA_seed | PIK3CD | 1.252617 | GBM | D-263MG | Idelalisib | MVP_top60_DNA+RNA_seed | PIK3CD | 0.503721 | GBM | LN-229 | IC-87114 |
| MVP_top60_DNA_seed | JAK3 | 1.253604 | GBM | AM-38 | JAK3_7406 | MVP_top60_DNA+RNA_seed | AKT1 | 0.505319 | GBM | YH-13 | BAY AKT1 |
| MVP_top60_DNA_seed | EGFR | 1.261068 | GBM | U-118-MG | Afatinib | MVP_top60_DNA+RNA_seed | AKT1 | 0.508087 | GBM | LNZTA3WT4 | GSK690693 |
| MVP_top60_DNA_seed | EGFR | 1.280577 | GBM | CAS-1 | Gefitinib | MVP_top60_DNA+RNA_seed | PIK3CD | 0.508152 | GBM | SNB75 | Idelalisib |
| MVP_top60_DNA_seed | PDGFRA | 1.280743 | GBM | LN-229 | Amuvatinib | MVP_top60_DNA+RNA_seed | IGF1R | 0.509871 | GBM | LN-229 | GSK1904529A |
| MVP_top60_DNA_seed | PIK3CD | 1.28076 | GBM | U-118-MG | Idelalisib | MVP_top60_DNA+RNA_seed | PIK3CD | 0.512967 | GBM | LN-18 | Idelalisib |
| MVP_top60_DNA_seed | PIK3CB | 1.286163 | GBM | AM-38 | AZD8186 | MVP_top60_DNA+RNA_seed | JAK3 | 0.517839 | GBM | CAS-1 | JAK3_7406 |
| MVP_top60_DNA_seed | PIK3CD | 1.286163 | GBM | AM-38 | AZD8186 | MVP_top60_DNA+RNA_seed | EGFR | 0.51882 | GBM | U-118-MG | Gefitinib |
| MVP_top60_DNA_seed | JAK2 | 1.289316 | GBM | AM-38 | AZD1480 | MVP_top60_DNA+RNA_seed | PDGFRB | 0.520885 | GBM | D-263MG | Pazopanib |
| MVP_top60_DNA_seed | JAK1 | 1.289316 | GBM | AM-38 | AZD1480 | MVP_top60_DNA+RNA_seed | KIT | 0.520885 | GBM | D-263MG | Pazopanib |
| MVP_top60_DNA_seed | PIK3CA | 1.298605 | GBM | A172 | AZD8835 | MVP_top60_DNA+RNA_seed | PIK3CD | 0.522128 | GBM | SF268 | IC-87114 |
| MVP_top60_DNA_seed | PIK3CD | 1.298605 | GBM | A172 | AZD8835 | MVP_top60_DNA+RNA_seed | JAK1 | 0.523465 | GBM | LNZTA3WT4 | JAK1_3715 |
| MVP_top60_DNA_seed | PDGFRA | 1.317396 | GBM | Becker | Masitinib | MVP_top60_DNA+RNA_seed | EGFR | 0.525061 | GBM | YKG-1 | Gefitinib |
| MVP_top60_DNA_seed | PDGFRB | 1.317396 | GBM | Becker | Masitinib | MVP_top60_DNA+RNA_seed | PDGFRA | 0.525591 | GBM | D-566MG | Amuvatinib |
| MVP_top60_DNA_seed | EGFR | 1.318872 | GBM | DK-MG | AZD8931 | MVP_top60_DNA+RNA_seed | KIT | 0.525591 | GBM | D-566MG | Amuvatinib |
| MVP_top60_DNA_seed | EGFR | 1.321979 | GBM | SNB75 | Erlotinib | MVP_top60_DNA+RNA_seed | AKT1 | 0.52596 | GBM | YKG-1 | BAY AKT1 |
| MVP_top60_DNA_seed | PDGFRA | 1.330843 | GBM | SW1088 | Amuvatinib | MVP_top60_DNA+RNA_seed | EGFR | 0.528014 | GBM | D-263MG | Afatinib |

| | | | | | | | | | | | |
|---|---|---|---|---|---|---|---|---|---|---|---|
| MVP_top60_DNA_seed | JAK3 | 1.338655 | GBM | U-87-MG | WHI-P97 | MVP_top60_DNA+RNA_seed | PIK3CD | 0.528328 | GBM | LN-405 | IC-87114 |
| MVP_top60_DNA_seed | PDGFRA | 1.340726 | GBM | SNB75 | Amuvatinib | MVP_top60_DNA+RNA_seed | JAK2 | 0.530674 | GBM | D-423MG | AZD1480 |
| MVP_top60_DNA_seed | PDGFRA | 1.356297 | GBM | DK-MG | Amuvatinib | MVP_top60_DNA+RNA_seed | JAK1 | 0.530674 | GBM | D-423MG | AZD1480 |
| MVP_top60_DNA_seed | CDK4 | 1.357365 | GBM | LN-405 | Palbociclib | MVP_top60_DNA+RNA_seed | EGFR | 0.530853 | GBM | D-423MG | Gefitinib |
| MVP_top60_DNA_seed | EGFR | 1.364496 | GBM | U-118-MG | Pelitinib | MVP_top60_DNA+RNA_seed | AKT1 | 0.530999 | GBM | AM-38 | GSK690693 |
| MVP_top60_DNA_seed | EP300 | 1.376447 | GBM | CAS-1 | I-CBP112 | MVP_top60_DNA+RNA_seed | MET | 0.532933 | GBM | D-247MG | SU11274 |
| MVP_top60_DNA_seed | CDK4 | 1.396558 | GBM | Becker | CGP-082996 | MVP_top60_DNA+RNA_seed | MET | 0.533329 | GBM | LNZTA3WT4 | SU11274 |
| MVP_top60_DNA_seed | EGFR | 1.39991 | GBM | U-87-MG | Pelitinib | MVP_top60_DNA+RNA_seed | KIT | 0.534042 | GBM | SF268 | Imatinib |
| MVP_top60_DNA_seed | JAK1 | 1.413885 | GBM | KS-1 | JAK1_3715 | MVP_top60_DNA+RNA_seed | EGFR | 0.534881 | GBM | SF295 | Afatinib |
| MVP_top60_DNA_seed | EGFR | 1.433478 | GBM | SF268 | CI-1033 | MVP_top60_DNA+RNA_seed | EGFR | 0.535548 | GBM | A172 | Afatinib |
| MVP_top60_DNA_seed | EGFR | 1.439725 | GBM | D-263MG | Pelitinib | MVP_top60_DNA+RNA_seed | KIT | 0.53667 | GBM | LN-229 | Motesanib |
| MVP_top60_DNA_seed | JAK1 | 1.49302 | GBM | DK-MG | JAK1_3715 | MVP_top60_DNA+RNA_seed | CDK4 | 0.537366 | GBM | SF126 | Palbociclib |
| MVP_top60_DNA_seed | PDGFRA | 1.501964 | GBM | U-87-MG | Amuvatinib | MVP_top60_DNA+RNA_seed | PIK3CA | 0.543295 | GBM | LN-229 | AZD8835 |
| MVP_top60_DNA_seed | EGFR | 1.516762 | GBM | D-247MG | Pelitinib | MVP_top60_DNA+RNA_seed | PIK3CD | 0.543295 | GBM | LN-229 | AZD8835 |
| MVP_top60_DNA_seed | PIK3CA | 1.518444 | GBM | DK-MG | AZD8835 | MVP_top60_DNA+RNA_seed | MET | 0.545051 | GBM | SNB75 | SU11274 |
| MVP_top60_DNA_seed | PIK3CD | 1.518444 | GBM | DK-MG | AZD8835 | MVP_top60_DNA+RNA_seed | EGFR | 0.545301 | GBM | LN-18 | CUDC-101 |
| MVP_top60_DNA_seed | EGFR | 1.520523 | GBM | SNB75 | Gefitinib | MVP_top60_DNA+RNA_seed | PDGFRA | 0.545993 | GBM | LN-18 | Amuvatinib |
| MVP_top60_DNA_seed | PDGFRA | 1.533048 | GBM | LN-229 | Masitinib | MVP_top60_DNA+RNA_seed | KIT | 0.545993 | GBM | LN-18 | Amuvatinib |
| MVP_top60_DNA_seed | PDGFRB | 1.533048 | GBM | LN-229 | Masitinib | MVP_top60_DNA+RNA_seed | EGFR | 0.546202 | GBM | Becker | Afatinib |
| MVP_top60_DNA_seed | EGFR | 1.541826 | GBM | SW1088 | CI-1033 | MVP_top60_DNA+RNA_seed | PIK3CA | 0.546258 | GBM | LN-405 | PI3Ka_4409 |
| MVP_top60_DNA_seed | JAK3 | 1.543824 | GBM | CAS-1 | WHI-P97 | MVP_top60_DNA+RNA_seed | PIK3CD | 0.546258 | GBM | LN-405 | PI3Ka_4409 |
| MVP_top60_DNA_seed | EGFR | 1.544437 | GBM | U-118-MG | CI-1033 | MVP_top60_DNA+RNA_seed | IGF1R | 0.546632 | GBM | D-247MG | BMS-536924 |
| MVP_top60_DNA_seed | EGFR | 1.553352 | GBM | D-542MG | Pelitinib | MVP_top60_DNA+RNA_seed | EP300 | 0.547231 | GBM | GB-1 | I-CBP112 |
| MVP_top60_DNA_seed | CDK4 | 1.554434 | GBM | M059J | Palbociclib | MVP_top60_DNA+RNA_seed | CREBBP | 0.547231 | GBM | GB-1 | I-CBP112 |
| MVP_top60_DNA_seed | EGFR | 1.571145 | GBM | D-247MG | CI-1033 | MVP_top60_DNA+RNA_seed | MET | 0.5477 | GBM | AM-38 | Crizotinib |
| MVP_top60_DNA_seed | JAK3 | 1.634691 | GBM | SNB75 | WHI-P97 | MVP_top60_DNA+RNA_seed | IGF1R | 0.548533 | GBM | U-87-MG | BMS-536924 |
| MVP_top60_DNA_seed | PIK3CA | 1.655623 | GBM | LNZTA3WT4 | PI3Ka_4409 | MVP_top60_DNA+RNA_seed | EGFR | 0.548579 | GBM | SW1088 | AZD8931 |
| MVP_top60_DNA_seed | PIK3CD | 1.655623 | GBM | LNZTA3WT4 | PI3Ka_4409 | MVP_top60_DNA+RNA_seed | ERBB3 | 0.548579 | GBM | SW1088 | AZD8931 |
| MVP_top60_DNA_seed | EP300 | 1.655741 | GBM | U-118-MG | I-CBP112 | MVP_top60_DNA+RNA_seed | IGF1R | 0.550277 | GBM | LN-229 | Linsitinib |
| MVP_top60_DNA_seed | JAK2 | 1.668018 | GBM | D-263MG | Fedratinib | MVP_top60_DNA+RNA_seed | PDGFRA | 0.551922 | GBM | CAS-1 | Masitinib |
| MVP_top60_DNA_seed | EGFR | 1.672108 | GBM | AM-38 | AZD8931 | MVP_top60_DNA+RNA_seed | PDGFRB | 0.551922 | GBM | CAS-1 | Masitinib |
| MVP_top60_DNA_seed | CDK4 | 1.678389 | GBM | D-542MG | Palbociclib | MVP_top60_DNA+RNA_seed | KIT | 0.551922 | GBM | CAS-1 | Masitinib |
| MVP_top60_DNA_seed | CDK4 | 1.67935 | GBM | CAS-1 | CGP-082996 | MVP_top60_DNA+RNA_seed | PIK3CB | 0.554465 | GBM | LN-229 | AZD8186 |
| MVP_top60_DNA_seed | JAK2 | 1.697654 | GBM | U-87-MG | AZD1480 | MVP_top60_DNA+RNA_seed | PIK3CD | 0.554465 | GBM | LN-229 | AZD8186 |
| MVP_top60_DNA_seed | JAK1 | 1.697654 | GBM | U-87-MG | AZD1480 | MVP_top60_DNA+RNA_seed | PIK3CA | 0.554576 | GBM | U-87-MG | AZD8835 |
| MVP_top60_DNA_seed | JAK3 | 1.701603 | GBM | D-247MG | JAK3_7406 | MVP_top60_DNA+RNA_seed | PIK3CD | 0.554576 | GBM | U-87-MG | AZD8835 |
| MVP_top60_DNA_seed | JAK3 | 1.722374 | GBM | U-118-MG | JAK3_7406 | MVP_top60_DNA+RNA_seed | JAK3 | 0.556436 | GBM | SF268 | WHI-P97 |
| MVP_top60_DNA_seed | JAK3 | 1.728853 | GBM | DBTRG-05MG | JAK3_7406 | MVP_top60_DNA+RNA_seed | PDGFRA | 0.556644 | GBM | U-118-MG | Masitinib |
| MVP_top60_DNA_seed | PDGFRA | 1.747289 | GBM | D-263MG | Masitinib | MVP_top60_DNA+RNA_seed | PDGFRB | 0.556644 | GBM | U-118-MG | Masitinib |
| MVP_top60_DNA_seed | PDGFRB | 1.747289 | GBM | D-263MG | Masitinib | MVP_top60_DNA+RNA_seed | KIT | 0.556644 | GBM | U-118-MG | Masitinib |
| MVP_top60_DNA_seed | EGFR | 1.79721 | GBM | SF295 | CI-1033 | MVP_top60_DNA+RNA_seed | EGFR | 0.557431 | GBM | SF126 | Pelitinib |

| | | | | | | | | | | | |
|---|---|---|---|---|---|---|---|---|---|---|---|
| MVP_top60_DNA_seed | PIK3CA | 1.818212 | GBM | AM-38 | AZD8835 | MVP_top60_DNA+RNA_seed | JAK3 | 0.558137 | GBM | M059J | JAK3_7406 |
| MVP_top60_DNA_seed | PIK3CD | 1.818212 | GBM | AM-38 | AZD8835 | MVP_top60_DNA+RNA_seed | KIT | 0.55828 | GBM | GB-1 | Axitinib |
| MVP_top60_DNA_seed | EGFR | 1.824225 | GBM | A172 | CI-1033 | MVP_top60_DNA+RNA_seed | MET | 0.559708 | GBM | D-247MG | AZD6094 |
| MVP_top60_DNA_seed | PIK3CA | 1.830186 | GBM | AM-38 | PI3Ka_4409 | MVP_top60_DNA+RNA_seed | JAK2 | 0.559856 | GBM | SNB75 | AZD1480 |
| MVP_top60_DNA_seed | PIK3CD | 1.830186 | GBM | AM-38 | PI3Ka_4409 | MVP_top60_DNA+RNA_seed | JAK1 | 0.559856 | GBM | SNB75 | AZD1480 |
| MVP_top60_DNA_seed | EGFR | 1.873966 | GBM | T98G | CI-1033 | MVP_top60_DNA+RNA_seed | AKT1 | 0.561983 | GBM | LN-18 | AKT inhibitor VIII |
| MVP_top60_DNA_seed | EGFR | 1.886643 | GBM | LNZTA3WT4 | AZD8931 | MVP_top60_DNA+RNA_seed | EGFR | 0.562046 | GBM | DBTRG-05MG | AST-1306 |
| MVP_top60_DNA_seed | EGFR | 1.894947 | GBM | DBTRG-05MG | CI-1033 | MVP_top60_DNA+RNA_seed | PDGFRA | 0.562627 | GBM | YH-13 | Amuvatinib |
| MVP_top60_DNA_seed | EGFR | 1.929988 | GBM | LN-405 | CI-1033 | MVP_top60_DNA+RNA_seed | KIT | 0.562627 | GBM | YH-13 | Amuvatinib |
| MVP_top60_DNA_seed | EGFR | 1.939787 | GBM | SW1088 | CUDC-101 | MVP_top60_DNA+RNA_seed | AKT1 | 0.567066 | GBM | SNB75 | AKT inhibitor VIII |
| MVP_top60_DNA_seed | CDK4 | 1.965836 | GBM | SNB75 | Palbociclib | MVP_top60_DNA+RNA_seed | PIK3CA | 0.569583 | GBM | U251 | AZD8835 |
| MVP_top60_DNA_seed | EP300 | 1.999887 | GBM | DK-MG | I-CBP112 | MVP_top60_DNA+RNA_seed | PIK3CD | 0.569583 | GBM | U251 | AZD8835 |
| MVP_top60_DNA_seed | JAK2 | 2.010632 | GBM | U-118-MG | Fedratinib | MVP_top60_DNA+RNA_seed | MET | 0.57095 | GBM | SF295 | SU11274 |
| MVP_top60_DNA_seed | EGFR | 2.206699 | GBM | LNZTA3WT4 | CUDC-101 | MVP_top60_DNA+RNA_seed | AKT1 | 0.572794 | GBM | SF126 | AKT inhibitor VIII |
| MVP_top60_DNA_seed | PIK3CA | -1.942775 | GBM | YH-13 | AZD8186 | MVP_top60_DNA+RNA_seed | JAK2 | 0.575505 | GBM | LN-18 | AZD1480 |
| MVP_top60_DNA_seed | PIK3CB | -1.942775 | GBM | YH-13 | AZD8186 | MVP_top60_DNA+RNA_seed | JAK1 | 0.575505 | GBM | LN-18 | AZD1480 |
| MVP_top60_DNA_seed | PIK3CA | -1.437528 | GBM | SW1088 | AZD8186 | MVP_top60_DNA+RNA_seed | AKT1 | 0.575856 | GBM | A172 | AKT inhibitor VIII |
| MVP_top60_DNA_seed | PIK3CB | -1.437528 | GBM | SW1088 | AZD8186 | MVP_top60_DNA+RNA_seed | EGFR | 0.57615 | GBM | D-263MG | CI-1033 |
| MVP_top60_DNA_seed | EP300 | -1.43111 | GBM | AM-38 | SGC-CBP30 | MVP_top60_DNA+RNA_seed | PIK3CB | 0.576309 | GBM | DK-MG | AZD6482 |
| MVP_top60_DNA_seed | JAK2 | -1.419946 | GBM | AM-38 | Ruxolitinib | MVP_top60_DNA+RNA_seed | AKT1 | 0.576443 | GBM | U-87-MG | BAY AKT1 |
| MVP_top60_DNA_seed | JAK1 | -1.419946 | GBM | AM-38 | Ruxolitinib | MVP_top60_DNA+RNA_seed | EGFR | 0.578175 | GBM | 42-MG-BA | Afatinib |
| MVP_top60_DNA_seed | CDK4 | -1.409729 | GBM | D-245MG | Ribociclib | MVP_top60_DNA+RNA_seed | JAK3 | 0.580369 | GBM | 42-MG-BA | WHI-P97 |
| MVP_top60_DNA_seed | PIK3CA | -1.38032 | GBM | YH-13 | GNE-317 | MVP_top60_DNA+RNA_seed | EGFR | 0.584384 | GBM | D-566MG | PF-00299804 |
| MVP_top60_DNA_seed | JAK2 | -1.352688 | GBM | A172 | JAK_8517 | MVP_top60_DNA+RNA_seed | MET | 0.585509 | GBM | DBTRG-05MG | SU11274 |
| MVP_top60_DNA_seed | JAK1 | -1.352688 | GBM | A172 | JAK_8517 | MVP_top60_DNA+RNA_seed | JAK2 | 0.587011 | GBM | SK-MG-1 | Ruxolitinib |
| MVP_top60_DNA_seed | JAK2 | -1.33318 | GBM | KS-1 | JAK_8517 | MVP_top60_DNA+RNA_seed | JAK1 | 0.587011 | GBM | SK-MG-1 | Ruxolitinib |
| MVP_top60_DNA_seed | JAK1 | -1.33318 | GBM | KS-1 | JAK_8517 | MVP_top60_DNA+RNA_seed | EGFR | 0.588596 | GBM | T98G | Afatinib |
| MVP_top60_DNA_seed | PIK3CA | -1.301092 | GBM | D-247MG | Alpelisib | MVP_top60_DNA+RNA_seed | EGFR | 0.59153 | GBM | SK-MG-1 | CI-1033 |
| MVP_top60_DNA_seed | PIK3CA | -1.29419 | GBM | Becker | AZD8186 | MVP_top60_DNA+RNA_seed | EGFR | 0.592515 | GBM | D-263MG | PF-00299804 |
| MVP_top60_DNA_seed | PIK3CB | -1.29419 | GBM | Becker | AZD8186 | MVP_top60_DNA+RNA_seed | EGFR | 0.593466 | GBM | GB-1 | Afatinib |
| MVP_top60_DNA_seed | PIK3CA | -1.278532 | GBM | D-423MG | AZD8186 | MVP_top60_DNA+RNA_seed | MET | 0.593909 | GBM | SF126 | PHA-665752 |
| MVP_top60_DNA_seed | PIK3CB | -1.278532 | GBM | D-423MG | AZD8186 | MVP_top60_DNA+RNA_seed | PIK3CA | 0.594421 | GBM | LN-229 | PI3Ka_4409 |
| MVP_top60_DNA_seed | PIK3CA | -1.198811 | GBM | SW1088 | GNE-317 | MVP_top60_DNA+RNA_seed | PIK3CD | 0.594421 | GBM | LN-229 | PI3Ka_4409 |
| MVP_top60_DNA_seed | PIK3CA | -1.193069 | GBM | 8-MG-BA | Buparlisib | MVP_top60_DNA+RNA_seed | EGFR | 0.596205 | GBM | LN-405 | Cetuximab |
| MVP_top60_DNA_seed | PIK3CB | -1.193069 | GBM | 8-MG-BA | Buparlisib | MVP_top60_DNA+RNA_seed | AKT1 | 0.596268 | GBM | LN-229 | AKT inhibitor VIII |
| MVP_top60_DNA_seed | PIK3CD | -1.193069 | GBM | 8-MG-BA | Buparlisib | MVP_top60_DNA+RNA_seed | KIT | 0.59876 | GBM | AM-38 | OSI-930 |
| MVP_top60_DNA_seed | JAK2 | -1.171273 | GBM | CAS-1 | JAK_8517 | MVP_top60_DNA+RNA_seed | EGFR | 0.599575 | GBM | D-263MG | Cetuximab |
| MVP_top60_DNA_seed | JAK1 | -1.171273 | GBM | CAS-1 | JAK_8517 | MVP_top60_DNA+RNA_seed | MET | 0.600264 | GBM | GB-1 | JNJ38877605 |
| MVP_top60_DNA_seed | TP53 | -1.090076 | GBM | LNZTA3WT4 | MIRA-1 | MVP_top60_DNA+RNA_seed | EGFR | 0.603259 | GBM | Becker | AST-1306 |
| MVP_top60_DNA_seed | JAK1 | -1.089505 | GBM | 8-MG-BA | JAK1_8709 | MVP_top60_DNA+RNA_seed | PDGFRA | 0.604991 | GBM | 42-MG-BA | Masitinib |
| MVP_top60_DNA_seed | EP300 | -1.083943 | GBM | SF268 | SGC-CBP30 | MVP_top60_DNA+RNA_seed | PDGFRB | 0.604991 | GBM | 42-MG-BA | Masitinib |

| | | | | | | | | | | | |
|---|---|---|---|---|---|---|---|---|---|---|---|
| MVP_top60_DNA_seed | PIK3CA | -1.074354 | GBM | 42-MG-BA | GNE-317 | MVP_top60_DNA+RNA_seed | KIT | 0.604991 | GBM | 42-MG-BA | Masitinib |
| MVP_top60_DNA_seed | EGFR | -1.068129 | GBM | D-423MG | AZD3759 | MVP_top60_DNA+RNA_seed | PDGFRA | 0.608412 | GBM | U-118-MG | Amuvatinib |
| MVP_top60_DNA_seed | JAK2 | -1.054067 | GBM | D-423MG | Ruxolitinib | MVP_top60_DNA+RNA_seed | KIT | 0.608412 | GBM | U-118-MG | Amuvatinib |
| MVP_top60_DNA_seed | JAK1 | -1.054067 | GBM | D-423MG | Ruxolitinib | MVP_top60_DNA+RNA_seed | EGFR | 0.611763 | GBM | DK-MG | Afatinib |
| MVP_top60_DNA_seed | PIK3CA | -1.03274 | GBM | SK-MG-1 | AZD8186 | MVP_top60_DNA+RNA_seed | EGFR | 0.613775 | GBM | T98G | Gefitinib |
| MVP_top60_DNA_seed | PIK3CB | -1.03274 | GBM | SK-MG-1 | AZD8186 | MVP_top60_DNA+RNA_seed | IGF1R | 0.614338 | GBM | A172 | BMS-536924 |
| MVP_top60_DNA_seed | CDK4 | -1.023605 | GBM | YH-13 | Ribociclib | MVP_top60_DNA+RNA_seed | JAK2 | 0.616488 | GBM | D-263MG | Ruxolitinib |
| MVP_top60_DNA_seed | CDK4 | -1.003435 | GBM | A172 | Ribociclib | MVP_top60_DNA+RNA_seed | JAK1 | 0.616488 | GBM | D-263MG | Ruxolitinib |
| MVP_top60_DNA_seed | PIK3CA | -0.987237 | GBM | D-542MG | AZD8186 | MVP_top60_DNA+RNA_seed | JAK2 | 0.617868 | GBM | D-392MG | Ruxolitinib |
| MVP_top60_DNA_seed | PIK3CB | -0.987237 | GBM | D-542MG | AZD8186 | MVP_top60_DNA+RNA_seed | JAK1 | 0.617868 | GBM | D-392MG | Ruxolitinib |
| MVP_top60_DNA_seed | PIK3CA | -0.98207 | GBM | D-423MG | GNE-317 | MVP_top60_DNA+RNA_seed | PDGFRA | 0.619007 | GBM | SF126 | Amuvatinib |
| MVP_top60_DNA_seed | EP300 | -0.978372 | GBM | 42-MG-BA | SGC-CBP30 | MVP_top60_DNA+RNA_seed | KIT | 0.619007 | GBM | SF126 | Amuvatinib |
| MVP_top60_DNA_seed | PIK3CA | -0.967452 | GBM | A172 | Buparlisib | MVP_top60_DNA+RNA_seed | IGF1R | 0.61938 | GBM | U251 | Linsitinib |
| MVP_top60_DNA_seed | PIK3CB | -0.967452 | GBM | A172 | Buparlisib | MVP_top60_DNA+RNA_seed | EGFR | 0.620391 | GBM | 42-MG-BA | AST-1306 |
| MVP_top60_DNA_seed | PIK3CD | -0.967452 | GBM | A172 | Buparlisib | MVP_top60_DNA+RNA_seed | PIK3CD | 0.620537 | GBM | AM-38 | IC-87114 |
| MVP_top60_DNA_seed | TP53 | -0.962093 | GBM | D-566MG | MIRA-1 | MVP_top60_DNA+RNA_seed | EGFR | 0.62599 | GBM | SF268 | PF-00299804 |
| MVP_top60_DNA_seed | JAK2 | -0.961467 | GBM | T98G | GSK2276186C | MVP_top60_DNA+RNA_seed | KIT | 0.626929 | GBM | GB-1 | Imatinib |
| MVP_top60_DNA_seed | JAK1 | -0.961467 | GBM | T98G | GSK2276186C | MVP_top60_DNA+RNA_seed | EGFR | 0.627451 | GBM | A172 | Afatinib |
| MVP_top60_DNA_seed | JAK3 | -0.961467 | GBM | T98G | GSK2276186C | MVP_top60_DNA+RNA_seed | AKT1 | 0.629206 | GBM | D-392MG | GSK690693 |
| MVP_top60_DNA_seed | JAK2 | -0.953326 | GBM | D-245MG | AZ960 | MVP_top60_DNA+RNA_seed | MET | 0.630173 | GBM | LN-229 | SU11274 |
| MVP_top60_DNA_seed | JAK3 | -0.953326 | GBM | D-245MG | AZ960 | MVP_top60_DNA+RNA_seed | JAK2 | 0.630573 | GBM | D-542MG | Ruxolitinib |
| MVP_top60_DNA_seed | JAK2 | -0.942085 | GBM | D-566MG | AZ960 | MVP_top60_DNA+RNA_seed | JAK1 | 0.630573 | GBM | D-542MG | Ruxolitinib |
| MVP_top60_DNA_seed | JAK3 | -0.942085 | GBM | D-566MG | AZ960 | MVP_top60_DNA+RNA_seed | AKT1 | 0.630709 | GBM | SW1088 | AKT inhibitor VIII |
| MVP_top60_DNA_seed | JAK2 | -0.939447 | GBM | T98G | AZ960 | MVP_top60_DNA+RNA_seed | EP300 | 0.631881 | GBM | LN-229 | I-CBP112 |
| MVP_top60_DNA_seed | JAK3 | -0.939447 | GBM | T98G | AZ960 | MVP_top60_DNA+RNA_seed | CREBBP | 0.631881 | GBM | LN-229 | I-CBP112 |
| MVP_top60_DNA_seed | JAK2 | -0.92737 | GBM | YH-13 | JAK_8517 | MVP_top60_DNA+RNA_seed | EP300 | 0.632516 | GBM | D-542MG | I-CBP112 |
| MVP_top60_DNA_seed | JAK1 | -0.92737 | GBM | YH-13 | JAK_8517 | MVP_top60_DNA+RNA_seed | CREBBP | 0.632516 | GBM | D-542MG | I-CBP112 |
| MVP_top60_DNA_seed | PIK3CA | -0.927 | GBM | A172 | GNE-317 | MVP_top60_DNA+RNA_seed | KIT | 0.633945 | GBM | SNB75 | Axitinib |
| MVP_top60_DNA_seed | PIK3CA | -0.90808 | GBM | SK-MG-1 | Buparlisib | MVP_top60_DNA+RNA_seed | EGFR | 0.636266 | GBM | LNZTA3WT4 | Cetuximab |
| MVP_top60_DNA_seed | PIK3CB | -0.90808 | GBM | SK-MG-1 | Buparlisib | MVP_top60_DNA+RNA_seed | AKT1 | 0.637893 | GBM | M059J | GSK690693 |
| MVP_top60_DNA_seed | PIK3CD | -0.90808 | GBM | SK-MG-1 | Buparlisib | MVP_top60_DNA+RNA_seed | EGFR | 0.639196 | GBM | D-392MG | PF-00299804 |
| MVP_top60_DNA_seed | JAK2 | -0.904073 | GBM | 42-MG-BA | GSK2276186C | MVP_top60_DNA+RNA_seed | EGFR | 0.643621 | GBM | Becker | CI-1033 |
| MVP_top60_DNA_seed | JAK1 | -0.904073 | GBM | 42-MG-BA | GSK2276186C | MVP_top60_DNA+RNA_seed | AKT1 | 0.644279 | GBM | M059J | MK-2206 |
| MVP_top60_DNA_seed | JAK3 | -0.904073 | GBM | 42-MG-BA | GSK2276186C | MVP_top60_DNA+RNA_seed | EGFR | 0.645694 | GBM | CAS-1 | CUDC-101 |
| MVP_top60_DNA_seed | PIK3CA | -0.874547 | GBM | 8-MG-BA | GNE-317 | MVP_top60_DNA+RNA_seed | MET | 0.646874 | GBM | T98G | SU11274 |
| MVP_top60_DNA_seed | PIK3CA | -0.871281 | GBM | 42-MG-BA | AZD8186 | MVP_top60_DNA+RNA_seed | EGFR | 0.648066 | GBM | U-118-MG | PF-00299804 |
| MVP_top60_DNA_seed | PIK3CB | -0.871281 | GBM | 42-MG-BA | AZD8186 | MVP_top60_DNA+RNA_seed | EGFR | 0.648765 | GBM | D-263MG | Gefitinib |
| MVP_top60_DNA_seed | JAK2 | -0.865471 | GBM | A172 | AZ960 | MVP_top60_DNA+RNA_seed | MET | 0.648865 | GBM | SW1088 | SU11274 |
| MVP_top60_DNA_seed | JAK3 | -0.865471 | GBM | A172 | AZ960 | MVP_top60_DNA+RNA_seed | EGFR | 0.651567 | GBM | D-247MG | Cetuximab |
| MVP_top60_DNA_seed | PIK3CB | -0.862436 | GBM | D-566MG | AZD6482 | MVP_top60_DNA+RNA_seed | AKT1 | 0.654029 | GBM | KS-1 | AKT inhibitor VIII |
| MVP_top60_DNA_seed | PIK3CA | -0.856878 | GBM | SF295 | AZD8186 | MVP_top60_DNA+RNA_seed | EGFR | 0.654301 | GBM | SF295 | PF-00299804 |

| | | | | | | | | | | | |
|---|---|---|---|---|---|---|---|---|---|---|---|
| MVP_top60_DNA_seed | PIK3CB | -0.856878 | GBM | SF295 | AZD8186 | MVP_top60_DNA+RNA_seed | AKT1 | 0.655202 | GBM | M059J | AKT inhibitor VIII |
| MVP_top60_DNA_seed | EP300 | -0.853833 | GBM | YH-13 | CPI-637 | MVP_top60_DNA+RNA_seed | PIK3CA | 0.659763 | GBM | SF268 | AZD8835 |
| MVP_top60_DNA_seed | EGFR | -0.844443 | GBM | LN-229 | Erlotinib | MVP_top60_DNA+RNA_seed | PIK3CD | 0.659763 | GBM | SF268 | AZD8835 |
| MVP_top60_DNA_seed | PIK3CA | -0.832556 | GBM | YH-13 | Buparlisib | MVP_top60_DNA+RNA_seed | PIK3CA | 0.660103 | GBM | D-247MG | AZD8835 |
| MVP_top60_DNA_seed | PIK3CB | -0.832556 | GBM | YH-13 | Buparlisib | MVP_top60_DNA+RNA_seed | PIK3CD | 0.660103 | GBM | D-247MG | AZD8835 |
| MVP_top60_DNA_seed | PIK3CD | -0.832556 | GBM | YH-13 | Buparlisib | MVP_top60_DNA+RNA_seed | EGFR | 0.660387 | GBM | U-87-MG | PF-00299804 |
| MVP_top60_DNA_seed | JAK2 | -0.831074 | GBM | 42-MG-BA | JAK_8517 | MVP_top60_DNA+RNA_seed | KIT | 0.660892 | GBM | SK-MG-1 | OSI-930 |
| MVP_top60_DNA_seed | JAK1 | -0.831074 | GBM | 42-MG-BA | JAK_8517 | MVP_top60_DNA+RNA_seed | EGFR | 0.661416 | GBM | 42-MG-BA | Gefitinib |
| MVP_top60_DNA_seed | JAK2 | -0.826606 | GBM | YH-13 | AZ960 | MVP_top60_DNA+RNA_seed | IGF1R | 0.66233 | GBM | GB-1 | GSK1904529A |
| MVP_top60_DNA_seed | JAK3 | -0.826606 | GBM | YH-13 | AZ960 | MVP_top60_DNA+RNA_seed | IGF1R | 0.662961 | GBM | T98G | BMS-754807 |
| MVP_top60_DNA_seed | PIK3CB | -0.805151 | GBM | D-423MG | AZD6482 | MVP_top60_DNA+RNA_seed | MET | 0.66648 | GBM | KS-1 | SU11274 |
| MVP_top60_DNA_seed | EP300 | -0.804523 | GBM | KS-1 | CPI-637 | MVP_top60_DNA+RNA_seed | EGFR | 0.666765 | GBM | CAS-1 | Lapatinib |
| MVP_top60_DNA_seed | EP300 | -0.799497 | GBM | U-87-MG | SGC-CBP30 | MVP_top60_DNA+RNA_seed | AKT1 | 0.668449 | GBM | YH-13 | AKT inhibitor VIII |
| MVP_top60_DNA_seed | JAK2 | -0.794066 | GBM | SF126 | AZ960 | MVP_top60_DNA+RNA_seed | JAK2 | 0.672527 | GBM | GB-1 | AZD1480 |
| MVP_top60_DNA_seed | JAK3 | -0.794066 | GBM | SF126 | AZ960 | MVP_top60_DNA+RNA_seed | JAK1 | 0.672527 | GBM | GB-1 | AZD1480 |
| MVP_top60_DNA_seed | EGFR | -0.783918 | GBM | 8-MG-BA | Lapatinib | MVP_top60_DNA+RNA_seed | EGFR | 0.674141 | GBM | A172 | AST-1306 |
| MVP_top60_DNA_seed | JAK2 | -0.780905 | GBM | LN-229 | Ruxolitinib | MVP_top60_DNA+RNA_seed | MET | 0.674804 | GBM | SW1088 | JNJ38877605 |
| MVP_top60_DNA_seed | JAK1 | -0.780905 | GBM | LN-229 | Ruxolitinib | MVP_top60_DNA+RNA_seed | EGFR | 0.675234 | GBM | M059J | PF-00299804 |
| MVP_top60_DNA_seed | JAK2 | -0.774509 | GBM | SW1088 | JAK_8517 | MVP_top60_DNA+RNA_seed | EGFR | 0.677006 | GBM | D-263MG | Afatinib |
| MVP_top60_DNA_seed | JAK1 | -0.774509 | GBM | SW1088 | JAK_8517 | MVP_top60_DNA+RNA_seed | MET | 0.677019 | GBM | SF268 | PHA-665752 |
| MVP_top60_DNA_seed | JAK2 | -0.763715 | GBM | T98G | JAK_8517 | MVP_top60_DNA+RNA_seed | IGF1R | 0.677721 | GBM | Becker | BMS-536924 |
| MVP_top60_DNA_seed | JAK1 | -0.763715 | GBM | T98G | JAK_8517 | MVP_top60_DNA+RNA_seed | AKT1 | 0.679784 | GBM | CAS-1 | MK-2206 |
| MVP_top60_DNA_seed | EP300 | -0.763421 | GBM | U-87-MG | CPI-637 | MVP_top60_DNA+RNA_seed | IGF1R | 0.681557 | GBM | LNZTA3WT4 | Linsitinib |
| MVP_top60_DNA_seed | JAK2 | -0.760054 | GBM | 8-MG-BA | GSK2276186C | MVP_top60_DNA+RNA_seed | EGFR | 0.6825 | GBM | D-392MG | Afatinib |
| MVP_top60_DNA_seed | JAK1 | -0.760054 | GBM | 8-MG-BA | GSK2276186C | MVP_top60_DNA+RNA_seed | AKT1 | 0.683453 | GBM | U251 | BAY AKT1 |
| MVP_top60_DNA_seed | JAK3 | -0.760054 | GBM | 8-MG-BA | GSK2276186C | MVP_top60_DNA+RNA_seed | EGFR | 0.686034 | GBM | U-87-MG | CUDC-101 |
| MVP_top60_DNA_seed | PIK3CA | -0.750057 | GBM | YH-13 | Alpelisib | MVP_top60_DNA+RNA_seed | EGFR | 0.689103 | GBM | M059J | Afatinib |
| MVP_top60_DNA_seed | EGFR | -0.74154 | GBM | D-566MG | Lapatinib | MVP_top60_DNA+RNA_seed | PIK3CA | 0.689415 | GBM | CAS-1 | PI3Ka_4409 |
| MVP_top60_DNA_seed | JAK2 | -0.737726 | GBM | KS-1 | AZ960 | MVP_top60_DNA+RNA_seed | PIK3CD | 0.689415 | GBM | CAS-1 | PI3Ka_4409 |
| MVP_top60_DNA_seed | JAK3 | -0.737726 | GBM | KS-1 | AZ960 | MVP_top60_DNA+RNA_seed | JAK2 | 0.691275 | GBM | SF295 | AZD1480 |
| MVP_top60_DNA_seed | PIK3CA | -0.725529 | GBM | SF295 | GNE-317 | MVP_top60_DNA+RNA_seed | JAK1 | 0.691275 | GBM | SF295 | AZD1480 |
| MVP_top60_DNA_seed | PIK3CA | -0.719633 | GBM | SF126 | Alpelisib | MVP_top60_DNA+RNA_seed | AKT1 | 0.692256 | GBM | U-87-MG | AKT inhibitor VIII |
| MVP_top60_DNA_seed | EP300 | -0.716364 | GBM | KS-1 | SGC-CBP30 | MVP_top60_DNA+RNA_seed | PIK3CB | 0.693304 | GBM | GB-1 | AZD8186 |
| MVP_top60_DNA_seed | PIK3CA | -0.715394 | GBM | D-247MG | AZD8186 | MVP_top60_DNA+RNA_seed | PIK3CD | 0.693304 | GBM | GB-1 | AZD8186 |
| MVP_top60_DNA_seed | PIK3CB | -0.715394 | GBM | D-247MG | AZD8186 | MVP_top60_DNA+RNA_seed | EGFR | 0.695419 | GBM | U-118-MG | AZD8931 |
| MVP_top60_DNA_seed | PIK3CA | -0.702953 | GBM | LN-18 | AZD8186 | MVP_top60_DNA+RNA_seed | ERBB3 | 0.695419 | GBM | U-118-MG | AZD8931 |
| MVP_top60_DNA_seed | PIK3CB | -0.702953 | GBM | LN-18 | AZD8186 | MVP_top60_DNA+RNA_seed | EGFR | 0.695518 | GBM | Becker | Gefitinib |
| MVP_top60_DNA_seed | PIK3CA | -0.696871 | GBM | D-245MG | Alpelisib | MVP_top60_DNA+RNA_seed | EGFR | 0.698169 | GBM | 42-MG-BA | Afatinib |
| MVP_top60_DNA_seed | EGFR | -0.69609 | GBM | LN-18 | Lapatinib | MVP_top60_DNA+RNA_seed | EGFR | 0.698915 | GBM | 42-MG-BA | AZD8931 |
| MVP_top60_DNA_seed | JAK2 | -0.695968 | GBM | 42-MG-BA | Ruxolitinib | MVP_top60_DNA+RNA_seed | ERBB3 | 0.698915 | GBM | 42-MG-BA | AZD8931 |
| MVP_top60_DNA_seed | JAK1 | -0.695968 | GBM | 42-MG-BA | Ruxolitinib | MVP_top60_DNA+RNA_seed | EGFR | 0.699406 | GBM | YKG-1 | Afatinib |

| | | | | | | | | | | | |
|---|---|---|---|---|---|---|---|---|---|---|---|
| MVP_top60_DNA_seed | EGFR | -0.690795 | GBM | SF126 | Osimertinib | MVP_top60_DNA+RNA_seed | EGFR | 0.699493 | GBM | CAS-1 | Cetuximab |
| MVP_top60_DNA_seed | PIK3CB | -0.680806 | GBM | D-247MG | AZD6482 | MVP_top60_DNA+RNA_seed | EGFR | 0.699883 | GBM | SK-MG-1 | Pelitinib |
| MVP_top60_DNA_seed | CDK4 | -0.676501 | GBM | LNZTA3WT4 | Palbociclib | MVP_top60_DNA+RNA_seed | PDGFRA | 0.701246 | GBM | GB-1 | Masitinib |
| MVP_top60_DNA_seed | JAK2 | -0.671886 | GBM | DK-MG | JAK_8517 | MVP_top60_DNA+RNA_seed | PDGFRB | 0.701246 | GBM | GB-1 | Masitinib |
| MVP_top60_DNA_seed | JAK1 | -0.671886 | GBM | DK-MG | JAK_8517 | MVP_top60_DNA+RNA_seed | KIT | 0.701246 | GBM | GB-1 | Masitinib |
| MVP_top60_DNA_seed | PIK3CA | -0.667689 | GBM | SNB75 | AZD8186 | MVP_top60_DNA+RNA_seed | PIK3CD | 0.702257 | GBM | SF126 | IC-87114 |
| MVP_top60_DNA_seed | PIK3CB | -0.667689 | GBM | SNB75 | AZD8186 | MVP_top60_DNA+RNA_seed | IGF1R | 0.703257 | GBM | AM-38 | GSK1904529A |
| MVP_top60_DNA_seed | PIK3CA | -0.66427 | GBM | 42-MG-BA | Buparlisib | MVP_top60_DNA+RNA_seed | PDGFRA | 0.706838 | GBM | D-263MG | Amuvatinib |
| MVP_top60_DNA_seed | PIK3CB | -0.66427 | GBM | 42-MG-BA | Buparlisib | MVP_top60_DNA+RNA_seed | KIT | 0.706838 | GBM | D-263MG | Amuvatinib |
| MVP_top60_DNA_seed | PIK3CD | -0.66427 | GBM | 42-MG-BA | Buparlisib | MVP_top60_DNA+RNA_seed | IGF1R | 0.707753 | GBM | M059J | BMS-536924 |
| MVP_top60_DNA_seed | CDK4 | -0.663907 | GBM | DK-MG | Ribociclib | MVP_top60_DNA+RNA_seed | EGFR | 0.708314 | GBM | T98G | AST-1306 |
| MVP_top60_DNA_seed | PIK3CB | -0.657151 | GBM | D-542MG | AZD6482 | MVP_top60_DNA+RNA_seed | EGFR | 0.709652 | GBM | SW1088 | Pelitinib |
| MVP_top60_DNA_seed | PIK3CA | -0.654801 | GBM | D-566MG | AZD8186 | MVP_top60_DNA+RNA_seed | PIK3CD | 0.714056 | GBM | SNB75 | IC-87114 |
| MVP_top60_DNA_seed | PIK3CB | -0.654801 | GBM | D-566MG | AZD8186 | MVP_top60_DNA+RNA_seed | EGFR | 0.715007 | GBM | U-87-MG | CI-1033 |
| MVP_top60_DNA_seed | JAK2 | -0.653326 | GBM | M059J | JAK_8517 | MVP_top60_DNA+RNA_seed | KIT | 0.715535 | GBM | YKG-1 | Motesanib |
| MVP_top60_DNA_seed | JAK1 | -0.653326 | GBM | M059J | JAK_8517 | MVP_top60_DNA+RNA_seed | MET | 0.715563 | GBM | D-423MG | SU11274 |
| MVP_top60_DNA_seed | EGFR | -0.641273 | GBM | D-423MG | Lapatinib | MVP_top60_DNA+RNA_seed | KIT | 0.715967 | GBM | SF126 | OSI-930 |
| MVP_top60_DNA_seed | JAK2 | -0.634959 | GBM | D-263MG | JAK_8517 | MVP_top60_DNA+RNA_seed | KIT | 0.716577 | GBM | GB-1 | OSI-930 |
| MVP_top60_DNA_seed | JAK1 | -0.634959 | GBM | D-263MG | JAK_8517 | MVP_top60_DNA+RNA_seed | EGFR | 0.716797 | GBM | YH-13 | Afatinib |
| MVP_top60_DNA_seed | PIK3CA | -0.634461 | GBM | SK-MG-1 | GNE-317 | MVP_top60_DNA+RNA_seed | MET | 0.717172 | GBM | D-542MG | AZD6094 |
| MVP_top60_DNA_seed | JAK2 | -0.629177 | GBM | SNB75 | AZ960 | MVP_top60_DNA+RNA_seed | JAK1 | 0.717719 | GBM | GB-1 | JAK1_3715 |
| MVP_top60_DNA_seed | JAK3 | -0.629177 | GBM | SNB75 | AZ960 | MVP_top60_DNA+RNA_seed | PIK3CD | 0.72555 | GBM | SK-MG-1 | IC-87114 |
| MVP_top60_DNA_seed | EP300 | -0.628276 | GBM | LN-18 | SGC-CBP30 | MVP_top60_DNA+RNA_seed | PIK3CD | 0.726805 | GBM | A172 | Idelalisib |
| MVP_top60_DNA_seed | JAK2 | -0.624481 | GBM | U-87-MG | Ruxolitinib | MVP_top60_DNA+RNA_seed | JAK3 | 0.730156 | GBM | LN-18 | JAK3_7406 |
| MVP_top60_DNA_seed | JAK1 | -0.624481 | GBM | U-87-MG | Ruxolitinib | MVP_top60_DNA+RNA_seed | EGFR | 0.730349 | GBM | 42-MG-BA | Pelitinib |
| MVP_top60_DNA_seed | PIK3CA | -0.608372 | GBM | LN-18 | Buparlisib | MVP_top60_DNA+RNA_seed | EGFR | 0.732461 | GBM | SF126 | Gefitinib |
| MVP_top60_DNA_seed | PIK3CB | -0.608372 | GBM | LN-18 | Buparlisib | MVP_top60_DNA+RNA_seed | MET | 0.733456 | GBM | LNZTA3WT4 | JNJ38877605 |
| MVP_top60_DNA_seed | PIK3CD | -0.608372 | GBM | LN-18 | Buparlisib | MVP_top60_DNA+RNA_seed | MET | 0.734537 | GBM | SF126 | SU11274 |
| MVP_top60_DNA_seed | PIK3CA | -0.605729 | GBM | SK-MG-1 | Alpelisib | MVP_top60_DNA+RNA_seed | JAK2 | 0.735079 | GBM | U-118-MG | AZD1480 |
| MVP_top60_DNA_seed | PIK3CA | -0.603704 | GBM | U-118-MG | GNE-317 | MVP_top60_DNA+RNA_seed | JAK1 | 0.735079 | GBM | U-118-MG | AZD1480 |
| MVP_top60_DNA_seed | PIK3CA | -0.598305 | GBM | Becker | Buparlisib | MVP_top60_DNA+RNA_seed | PIK3CA | 0.735691 | GBM | U-118-MG | PI3Ka_4409 |
| MVP_top60_DNA_seed | PIK3CB | -0.598305 | GBM | Becker | Buparlisib | MVP_top60_DNA+RNA_seed | PIK3CD | 0.735691 | GBM | U-118-MG | PI3Ka_4409 |
| MVP_top60_DNA_seed | PIK3CD | -0.598305 | GBM | Becker | Buparlisib | MVP_top60_DNA+RNA_seed | AKT1 | 0.736383 | GBM | D-247MG | A-443654 |
| MVP_top60_DNA_seed | JAK2 | -0.572111 | GBM | D-542MG | AZ960 | MVP_top60_DNA+RNA_seed | PIK3CA | 0.737695 | GBM | LN-18 | AZD8835 |
| MVP_top60_DNA_seed | JAK3 | -0.572111 | GBM | D-542MG | AZ960 | MVP_top60_DNA+RNA_seed | PIK3CD | 0.737695 | GBM | LN-18 | AZD8835 |
| MVP_top60_DNA_seed | PIK3CA | -0.564623 | GBM | D-566MG | GNE-317 | MVP_top60_DNA+RNA_seed | AKT1 | 0.738989 | GBM | D-392MG | BAY AKT1 |
| MVP_top60_DNA_seed | TP53 | -0.55726 | GBM | SF295 | MIRA-1 | MVP_top60_DNA+RNA_seed | PIK3CB | 0.739064 | GBM | LNZTA3WT4 | AZD8186 |
| MVP_top60_DNA_seed | PIK3CA | -0.547656 | GBM | M059J | GNE-317 | MVP_top60_DNA+RNA_seed | PIK3CD | 0.739064 | GBM | LNZTA3WT4 | AZD8186 |
| MVP_top60_DNA_seed | PIK3CA | -0.537568 | GBM | SNB75 | Alpelisib | MVP_top60_DNA+RNA_seed | PIK3CA | 0.739802 | GBM | LN-18 | PI3Ka_4409 |
| MVP_top60_DNA_seed | PIK3CA | -0.530889 | GBM | U-118-MG | AZD8186 | MVP_top60_DNA+RNA_seed | PIK3CD | 0.739802 | GBM | LN-18 | PI3Ka_4409 |
| MVP_top60_DNA_seed | PIK3CB | -0.530889 | GBM | U-118-MG | AZD8186 | MVP_top60_DNA+RNA_seed | MET | 0.741021 | GBM | T98G | JNJ38877605 |

| | | | | | | | | | | | |
|---|---|---|---|---|---|---|---|---|---|---|---|
| MVP_top60_DNA_seed | JAK2 | -0.525712 | GBM | D-566MG | Ruxolitinib | MVP_top60_DNA+RNA_seed | EGFR | 0.741936 | GBM | D-392MG | CI-1033 |
| MVP_top60_DNA_seed | JAK1 | -0.525712 | GBM | D-566MG | Ruxolitinib | MVP_top60_DNA+RNA_seed | PDGFRA | 0.742057 | GBM | DBTRG-05MG | Amuvatinib |
| MVP_top60_DNA_seed | PIK3CA | -0.524473 | GBM | SF126 | AZD8186 | MVP_top60_DNA+RNA_seed | KIT | 0.742057 | GBM | DBTRG-05MG | Amuvatinib |
| MVP_top60_DNA_seed | PIK3CB | -0.524473 | GBM | SF126 | AZD8186 | MVP_top60_DNA+RNA_seed | AKT1 | 0.744301 | GBM | A172 | GSK690693 |
| MVP_top60_DNA_seed | JAK2 | -0.522878 | GBM | KS-1 | GSK2276186C | MVP_top60_DNA+RNA_seed | IGF1R | 0.74566 | GBM | D-247MG | BMS-536924 |
| MVP_top60_DNA_seed | JAK1 | -0.522878 | GBM | KS-1 | GSK2276186C | MVP_top60_DNA+RNA_seed | PIK3CB | 0.745823 | GBM | AM-38 | AZD6482 |
| MVP_top60_DNA_seed | JAK3 | -0.522878 | GBM | KS-1 | GSK2276186C | MVP_top60_DNA+RNA_seed | EGFR | 0.748635 | GBM | AM-38 | Lapatinib |
| MVP_top60_DNA_seed | PIK3CA | -0.518104 | GBM | Becker | GNE-317 | MVP_top60_DNA+RNA_seed | IGF1R | 0.748675 | GBM | D-566MG | Linsitinib |
| MVP_top60_DNA_seed | EGFR | -0.514164 | GBM | SF126 | Lapatinib | MVP_top60_DNA+RNA_seed | IGF1R | 0.753342 | GBM | T98G | Linsitinib |
| MVP_top60_DNA_seed | EP300 | -0.512248 | GBM | A172 | CPI-637 | MVP_top60_DNA+RNA_seed | EGFR | 0.753517 | GBM | SF126 | Afatinib |
| MVP_top60_DNA_seed | CDK4 | -0.506462 | GBM | D-263MG | Ribociclib | MVP_top60_DNA+RNA_seed | PIK3CB | 0.753534 | GBM | AM-38 | TGX221 |
| MVP_top60_DNA_seed | JAK2 | -0.501097 | GBM | LN-18 | Ruxolitinib | MVP_top60_DNA+RNA_seed | JAK1 | 0.754363 | GBM | LN-229 | JAK1_3715 |
| MVP_top60_DNA_seed | JAK1 | -0.501097 | GBM | LN-18 | Ruxolitinib | MVP_top60_DNA+RNA_seed | JAK2 | 0.754412 | GBM | D-542MG | Fedratinib |
| MVP_top60_DNA_seed | CDK4 | -0.498988 | GBM | A172 | Palbociclib | MVP_top60_DNA+RNA_seed | AKT1 | 0.755492 | GBM | SF268 | BAY AKT1 |
| MVP_top60_DNA_seed | JAK2 | -0.492503 | GBM | D-245MG | Ruxolitinib | MVP_top60_DNA+RNA_seed | EGFR | 0.755547 | GBM | U-87-MG | Lapatinib |
| MVP_top60_DNA_seed | JAK1 | -0.492503 | GBM | D-245MG | Ruxolitinib | MVP_top60_DNA+RNA_seed | IGF1R | 0.758306 | GBM | SF126 | BMS-536924 |
| MVP_top60_DNA_seed | TP53 | -0.487817 | GBM | GB-1 | MIRA-1 | MVP_top60_DNA+RNA_seed | IGF1R | 0.761501 | GBM | SF295 | Linsitinib |
| MVP_top60_DNA_seed | PIK3CA | -0.482165 | GBM | LN-18 | GNE-317 | MVP_top60_DNA+RNA_seed | JAK1 | 0.762739 | GBM | U-87-MG | JAK1_3715 |
| MVP_top60_DNA_seed | JAK2 | -0.479701 | GBM | 42-MG-BA | AZ960 | MVP_top60_DNA+RNA_seed | KIT | 0.766075 | GBM | Becker | Motesanib |
| MVP_top60_DNA_seed | JAK3 | -0.479701 | GBM | 42-MG-BA | AZ960 | MVP_top60_DNA+RNA_seed | AKT1 | 0.772329 | GBM | Becker | AKT inhibitor VIII |
| MVP_top60_DNA_seed | EGFR | -0.478769 | GBM | SF295 | AZD3759 | MVP_top60_DNA+RNA_seed | EGFR | 0.774173 | GBM | SF295 | Pelitinib |
| MVP_top60_DNA_seed | JAK2 | -0.472731 | GBM | CAS-1 | AZ960 | MVP_top60_DNA+RNA_seed | EGFR | 0.775129 | GBM | AM-38 | Afatinib |
| MVP_top60_DNA_seed | JAK3 | -0.472731 | GBM | CAS-1 | AZ960 | MVP_top60_DNA+RNA_seed | IGF1R | 0.777536 | GBM | LNZTA3WT4 | BMS-536924 |
| MVP_top60_DNA_seed | EGFR | -0.470561 | GBM | LN-18 | Erlotinib | MVP_top60_DNA+RNA_seed | EGFR | 0.778509 | GBM | D-392MG | Pelitinib |
| MVP_top60_DNA_seed | JAK2 | -0.46123 | GBM | D-263MG | AZ960 | MVP_top60_DNA+RNA_seed | EGFR | 0.780786 | GBM | AM-38 | Erlotinib |
| MVP_top60_DNA_seed | JAK3 | -0.46123 | GBM | D-263MG | AZ960 | MVP_top60_DNA+RNA_seed | JAK2 | 0.781469 | GBM | DK-MG | Ruxolitinib |
| MVP_top60_DNA_seed | JAK2 | -0.45918 | GBM | D-247MG | AZ960 | MVP_top60_DNA+RNA_seed | JAK1 | 0.781469 | GBM | DK-MG | Ruxolitinib |
| MVP_top60_DNA_seed | JAK3 | -0.45918 | GBM | D-247MG | AZ960 | MVP_top60_DNA+RNA_seed | EGFR | 0.782544 | GBM | LN-405 | CUDC-101 |
| MVP_top60_DNA_seed | PIK3CB | -0.447853 | GBM | SNB75 | AZD6482 | MVP_top60_DNA+RNA_seed | EGFR | 0.782733 | GBM | SNB75 | Cetuximab |
| MVP_top60_DNA_seed | EGFR | -0.439711 | GBM | 8-MG-BA | Gefitinib | MVP_top60_DNA+RNA_seed | PIK3CA | 0.785392 | GBM | YKG-1 | PI3Ka_4409 |
| MVP_top60_DNA_seed | JAK2 | -0.41508 | GBM | DK-MG | AZ960 | MVP_top60_DNA+RNA_seed | PIK3CD | 0.785392 | GBM | YKG-1 | PI3Ka_4409 |
| MVP_top60_DNA_seed | JAK3 | -0.41508 | GBM | DK-MG | AZ960 | MVP_top60_DNA+RNA_seed | EGFR | 0.785877 | GBM | SF126 | Lapatinib |
| MVP_top60_DNA_seed | JAK2 | -0.412733 | GBM | D-392MG | JAK_8517 | MVP_top60_DNA+RNA_seed | EGFR | 0.786171 | GBM | LN-405 | AZD8931 |
| MVP_top60_DNA_seed | JAK1 | -0.412733 | GBM | D-392MG | JAK_8517 | MVP_top60_DNA+RNA_seed | ERBB3 | 0.786171 | GBM | LN-405 | AZD8931 |
| MVP_top60_DNA_seed | EGFR | -0.409178 | GBM | YKG-1 | Sapitinib | MVP_top60_DNA+RNA_seed | KIT | 0.786733 | GBM | D-263MG | Imatinib |
| MVP_top60_DNA_seed | JAK2 | -0.40139 | GBM | YKG-1 | AZ960 | MVP_top60_DNA+RNA_seed | JAK3 | 0.789399 | GBM | KS-1 | WHI-P97 |
| MVP_top60_DNA_seed | JAK3 | -0.40139 | GBM | YKG-1 | AZ960 | MVP_top60_DNA+RNA_seed | EGFR | 0.792512 | GBM | LN-405 | Afatinib |
| MVP_top60_DNA_seed | PIK3CA | -0.400543 | GBM | D-247MG | GNE-317 | MVP_top60_DNA+RNA_seed | AKT1 | 0.79335 | GBM | GB-1 | GSK690693 |
| MVP_top60_DNA_seed | PIK3CA | -0.397583 | GBM | 8-MG-BA | Alpelisib | MVP_top60_DNA+RNA_seed | KIT | 0.795167 | GBM | YH-13 | OSI-930 |
| MVP_top60_DNA_seed | TP53 | -0.396213 | GBM | D-423MG | MIRA-1 | MVP_top60_DNA+RNA_seed | KIT | 0.795643 | GBM | LN-229 | OSI-930 |
| MVP_top60_DNA_seed | EGFR | -0.393865 | GBM | GB-1 | Lapatinib | MVP_top60_DNA+RNA_seed | EGFR | 0.798119 | GBM | SF126 | Afatinib |

| | | | | | | | | | | | |
|---|---|---|---|---|---|---|---|---|---|---|---|
| MVP_top60_DNA_seed | PIK3CA | -0.390477 | GBM | SW1088 | Buparlisib | MVP_top60_DNA+RNA_seed | EGFR | 0.799122 | GBM | U-87-MG | Gefitinib |
| MVP_top60_DNA_seed | PIK3CB | -0.390477 | GBM | SW1088 | Buparlisib | MVP_top60_DNA+RNA_seed | EGFR | 0.803099 | GBM | GB-1 | AZD8931 |
| MVP_top60_DNA_seed | PIK3CD | -0.390477 | GBM | SW1088 | Buparlisib | MVP_top60_DNA+RNA_seed | ERBB3 | 0.803099 | GBM | GB-1 | AZD8931 |
| MVP_top60_DNA_seed | PIK3CA | -0.384948 | GBM | U-87-MG | AZD8186 | MVP_top60_DNA+RNA_seed | MET | 0.805781 | GBM | CAS-1 | AZD6094 |
| MVP_top60_DNA_seed | PIK3CB | -0.384948 | GBM | U-87-MG | AZD8186 | MVP_top60_DNA+RNA_seed | JAK1 | 0.809169 | GBM | D-392MG | JAK1_3715 |
| MVP_top60_DNA_seed | EGFR | -0.377618 | GBM | YKG-1 | Lapatinib | MVP_top60_DNA+RNA_seed | IGF1R | 0.809893 | GBM | LN-405 | BMS-536924 |
| MVP_top60_DNA_seed | JAK1 | -0.375317 | GBM | LN-18 | JAK1_8709 | MVP_top60_DNA+RNA_seed | EGFR | 0.81115 | GBM | SF295 | AST-1306 |
| MVP_top60_DNA_seed | CDK4 | -0.373544 | GBM | LNZTA3WT4 | Ribociclib | MVP_top60_DNA+RNA_seed | EGFR | 0.813821 | GBM | LNZTA3WT4 | PF-00299804 |
| MVP_top60_DNA_seed | JAK2 | -0.370197 | GBM | 8-MG-BA | AZ960 | MVP_top60_DNA+RNA_seed | KIT | 0.813889 | GBM | D-423MG | Motesanib |
| MVP_top60_DNA_seed | JAK3 | -0.370197 | GBM | 8-MG-BA | AZ960 | MVP_top60_DNA+RNA_seed | IGF1R | 0.815647 | GBM | SK-MG-1 | BMS-536924 |
| MVP_top60_DNA_seed | CDK4 | -0.369037 | GBM | Becker | Palbociclib | MVP_top60_DNA+RNA_seed | MET | 0.815817 | GBM | DBTRG-05MG | AZD6094 |
| MVP_top60_DNA_seed | EP300 | -0.359928 | GBM | SW1088 | CPI-637 | MVP_top60_DNA+RNA_seed | KIT | 0.820614 | GBM | AM-38 | Axitinib |
| MVP_top60_DNA_seed | EGFR | -0.358775 | GBM | SNB75 | AZD3759 | MVP_top60_DNA+RNA_seed | MET | 0.82169 | GBM | CAS-1 | Crizotinib |
| MVP_top60_DNA_seed | EGFR | -0.356905 | GBM | LN-229 | Sapitinib | MVP_top60_DNA+RNA_seed | AKT1 | 0.822446 | GBM | CAS-1 | A-443654 |
| MVP_top60_DNA_seed | CDK4 | -0.355037 | GBM | YH-13 | Palbociclib | MVP_top60_DNA+RNA_seed | PIK3CD | 0.824271 | GBM | LN-229 | Idelalisib |
| MVP_top60_DNA_seed | PIK3CA | -0.350083 | GBM | 8-MG-BA | AZD8186 | MVP_top60_DNA+RNA_seed | EGFR | 0.824329 | GBM | SF268 | Erlotinib |
| MVP_top60_DNA_seed | PIK3CB | -0.350083 | GBM | 8-MG-BA | AZD8186 | MVP_top60_DNA+RNA_seed | IGF1R | 0.827645 | GBM | YH-13 | BMS-536924 |
| MVP_top60_DNA_seed | PIK3CA | -0.343418 | GBM | DK-MG | AZD8186 | MVP_top60_DNA+RNA_seed | IGF1R | 0.829285 | GBM | YH-13 | BMS-754807 |
| MVP_top60_DNA_seed | PIK3CB | -0.343418 | GBM | DK-MG | AZD8186 | MVP_top60_DNA+RNA_seed | KIT | 0.829619 | GBM | M059J | OSI-930 |
| MVP_top60_DNA_seed | EGFR | -0.339388 | GBM | GB-1 | Erlotinib | MVP_top60_DNA+RNA_seed | JAK2 | 0.830263 | GBM | U-118-MG | Ruxolitinib |
| MVP_top60_DNA_seed | PIK3CA | -0.335624 | GBM | 42-MG-BA | Alpelisib | MVP_top60_DNA+RNA_seed | JAK1 | 0.830263 | GBM | U-118-MG | Ruxolitinib |
| MVP_top60_DNA_seed | EGFR | -0.334036 | GBM | YKG-1 | AZD3759 | MVP_top60_DNA+RNA_seed | AKT1 | 0.830819 | GBM | T98G | AKT inhibitor VIII |
| MVP_top60_DNA_seed | PIK3CA | -0.330178 | GBM | GB-1 | GNE-317 | MVP_top60_DNA+RNA_seed | PIK3CA | 0.841522 | GBM | CAS-1 | AZD8835 |
| MVP_top60_DNA_seed | PIK3CA | -0.322309 | GBM | T98G | GNE-317 | MVP_top60_DNA+RNA_seed | PIK3CD | 0.841522 | GBM | CAS-1 | AZD8835 |
| MVP_top60_DNA_seed | TP53 | -0.319202 | GBM | KS-1 | MIRA-1 | MVP_top60_DNA+RNA_seed | AKT1 | 0.841655 | GBM | LN-405 | AKT inhibitor VIII |
| MVP_top60_DNA_seed | EGFR | -0.317899 | GBM | D-423MG | Osimertinib | MVP_top60_DNA+RNA_seed | IGF1R | 0.842502 | GBM | CAS-1 | BMS-754807 |
| MVP_top60_DNA_seed | JAK2 | -0.311776 | GBM | Becker | JAK_8517 | MVP_top60_DNA+RNA_seed | EGFR | 0.848985 | GBM | T98G | PF-00299804 |
| MVP_top60_DNA_seed | JAK1 | -0.311776 | GBM | Becker | JAK_8517 | MVP_top60_DNA+RNA_seed | EGFR | 0.85078 | GBM | D-423MG | Afatinib |
| MVP_top60_DNA_seed | TP53 | -0.309237 | GBM | LN-18 | MIRA-1 | MVP_top60_DNA+RNA_seed | EGFR | 0.852691 | GBM | LNZTA3WT4 | Afatinib |
| MVP_top60_DNA_seed | JAK2 | -0.307654 | GBM | SW1088 | AZ960 | MVP_top60_DNA+RNA_seed | EGFR | 0.854566 | GBM | GB-1 | Gefitinib |
| MVP_top60_DNA_seed | JAK3 | -0.307654 | GBM | SW1088 | AZ960 | MVP_top60_DNA+RNA_seed | MET | 0.855538 | GBM | D-542MG | SU11274 |
| MVP_top60_DNA_seed | PIK3CA | -0.292366 | GBM | D-392MG | Buparlisib | MVP_top60_DNA+RNA_seed | KIT | 0.856831 | GBM | KS-1 | Imatinib |
| MVP_top60_DNA_seed | PIK3CB | -0.292366 | GBM | D-392MG | Buparlisib | MVP_top60_DNA+RNA_seed | IGF1R | 0.858487 | GBM | SF268 | Linsitinib |
| MVP_top60_DNA_seed | PIK3CD | -0.292366 | GBM | D-392MG | Buparlisib | MVP_top60_DNA+RNA_seed | IGF1R | 0.858671 | GBM | 42-MG-BA | Linsitinib |
| MVP_top60_DNA_seed | TP53 | -0.290196 | GBM | D-247MG | MIRA-1 | MVP_top60_DNA+RNA_seed | PDGFRA | 0.861017 | GBM | SNB75 | Masitinib |
| MVP_top60_DNA_seed | PIK3CA | -0.286346 | GBM | D-542MG | GNE-317 | MVP_top60_DNA+RNA_seed | PDGFRB | 0.861017 | GBM | SNB75 | Masitinib |
| MVP_top60_DNA_seed | EP300 | -0.285626 | GBM | Becker | CPI-637 | MVP_top60_DNA+RNA_seed | KIT | 0.861017 | GBM | SNB75 | Masitinib |
| MVP_top60_DNA_seed | CDK4 | -0.271687 | GBM | 42-MG-BA | Ribociclib | MVP_top60_DNA+RNA_seed | JAK1 | 0.861634 | GBM | LN-405 | JAK1_3715 |
| MVP_top60_DNA_seed | EGFR | -0.26647 | GBM | DK-MG | Osimertinib | MVP_top60_DNA+RNA_seed | IGF1R | 0.862723 | GBM | LN-405 | GSK1904529A |
| MVP_top60_DNA_seed | CDK4 | -0.262916 | GBM | D-263MG | Palbociclib | MVP_top60_DNA+RNA_seed | AKT1 | 0.863748 | GBM | D-247MG | GSK690693 |
| MVP_top60_DNA_seed | PIK3CA | -0.260705 | GBM | AM-38 | Buparlisib | MVP_top60_DNA+RNA_seed | PIK3CA | 0.866778 | GBM | SF126 | PI3Ka_4409 |

| | | | | | | | | | | | |
|---|---|---|---|---|---|---|---|---|---|---|---|
| MVP_top60_DNA_seed | PIK3CB | -0.260705 | GBM | AM-38 | Buparlisib | MVP_top60_DNA+RNA_seed | PIK3CD | 0.866778 | GBM | SF126 | PI3Ka_4409 |
| MVP_top60_DNA_seed | PIK3CD | -0.260705 | GBM | AM-38 | Buparlisib | MVP_top60_DNA+RNA_seed | EGFR | 0.868661 | GBM | M059J | Afatinib |
| MVP_top60_DNA_seed | EGFR | -0.258488 | GBM | SF126 | Gefitinib | MVP_top60_DNA+RNA_seed | PDGFRA | 0.869844 | GBM | LNZTA3WT4 | Masitinib |
| MVP_top60_DNA_seed | PIK3CB | -0.257592 | GBM | SF295 | AZD6482 | MVP_top60_DNA+RNA_seed | PDGFRB | 0.869844 | GBM | LNZTA3WT4 | Masitinib |
| MVP_top60_DNA_seed | EGFR | -0.252165 | GBM | U-87-MG | Erlotinib | MVP_top60_DNA+RNA_seed | KIT | 0.869844 | GBM | LNZTA3WT4 | Masitinib |
| MVP_top60_DNA_seed | TP53 | -0.246912 | GBM | A172 | MIRA-1 | MVP_top60_DNA+RNA_seed | EGFR | 0.871061 | GBM | SK-MG-1 | Gefitinib |
| MVP_top60_DNA_seed | TP53 | -0.241901 | GBM | YH-13 | MIRA-1 | MVP_top60_DNA+RNA_seed | PIK3CA | 0.873925 | GBM | U251 | PI3Ka_4409 |
| MVP_top60_DNA_seed | JAK2 | -0.241079 | GBM | GB-1 | Ruxolitinib | MVP_top60_DNA+RNA_seed | PIK3CD | 0.873925 | GBM | U251 | PI3Ka_4409 |
| MVP_top60_DNA_seed | JAK1 | -0.241079 | GBM | GB-1 | Ruxolitinib | MVP_top60_DNA+RNA_seed | EGFR | 0.875322 | GBM | LNZTA3WT4 | Afatinib |
| MVP_top60_DNA_seed | PIK3CA | -0.235878 | GBM | SNB75 | GNE-317 | MVP_top60_DNA+RNA_seed | MET | 0.87565 | GBM | LN-405 | SU11274 |
| MVP_top60_DNA_seed | EGFR | -0.235615 | GBM | LN-229 | Osimertinib | MVP_top60_DNA+RNA_seed | JAK3 | 0.879609 | GBM | SF295 | WHI-P97 |
| MVP_top60_DNA_seed | JAK2 | -0.235033 | GBM | D-423MG | AZ960 | MVP_top60_DNA+RNA_seed | AKT1 | 0.884385 | GBM | CAS-1 | AKT inhibitor VIII |
| MVP_top60_DNA_seed | JAK3 | -0.235033 | GBM | D-423MG | AZ960 | MVP_top60_DNA+RNA_seed | KIT | 0.886004 | GBM | AM-38 | Sorafenib |
| MVP_top60_DNA_seed | CDK4 | -0.231495 | GBM | D-566MG | Palbociclib | MVP_top60_DNA+RNA_seed | JAK2 | 0.888191 | GBM | SW1088 | Ruxolitinib |
| MVP_top60_DNA_seed | EGFR | -0.23003 | GBM | AM-38 | Erlotinib | MVP_top60_DNA+RNA_seed | JAK1 | 0.888191 | GBM | SW1088 | Ruxolitinib |
| MVP_top60_DNA_seed | EGFR | -0.225836 | GBM | 42-MG-BA | Lapatinib | MVP_top60_DNA+RNA_seed | EGFR | 0.893863 | GBM | SF268 | AST-1306 |
| MVP_top60_DNA_seed | PIK3CA | -0.219946 | GBM | D-245MG | AZD8186 | MVP_top60_DNA+RNA_seed | JAK2 | 0.89522 | GBM | SF295 | Fedratinib |
| MVP_top60_DNA_seed | PIK3CB | -0.219946 | GBM | D-245MG | AZD8186 | MVP_top60_DNA+RNA_seed | PDGFRA | 0.896973 | GBM | 42-MG-BA | Amuvatinib |
| MVP_top60_DNA_seed | PIK3CA | -0.212897 | GBM | D-566MG | Buparlisib | MVP_top60_DNA+RNA_seed | KIT | 0.896973 | GBM | 42-MG-BA | Amuvatinib |
| MVP_top60_DNA_seed | PIK3CB | -0.212897 | GBM | D-566MG | Buparlisib | MVP_top60_DNA+RNA_seed | AKT1 | 0.898074 | GBM | DK-MG | MK-2206 |
| MVP_top60_DNA_seed | PIK3CD | -0.212897 | GBM | D-566MG | Buparlisib | MVP_top60_DNA+RNA_seed | IGF1R | 0.903375 | GBM | U-118-MG | Linsitinib |
| MVP_top60_DNA_seed | CDK4 | -0.208943 | GBM | SF295 | Palbociclib | MVP_top60_DNA+RNA_seed | PIK3CD | 0.904813 | GBM | D-392MG | IC-87114 |
| MVP_top60_DNA_seed | TP53 | -0.206846 | GBM | 8-MG-BA | MIRA-1 | MVP_top60_DNA+RNA_seed | EGFR | 0.905008 | GBM | CAS-1 | AZD8931 |
| MVP_top60_DNA_seed | PIK3CA | -0.202898 | GBM | CAS-1 | AZD8186 | MVP_top60_DNA+RNA_seed | ERBB3 | 0.905008 | GBM | CAS-1 | AZD8931 |
| MVP_top60_DNA_seed | PIK3CB | -0.202898 | GBM | CAS-1 | AZD8186 | MVP_top60_DNA+RNA_seed | EGFR | 0.905612 | GBM | LNZTA3WT4 | Pelitinib |
| MVP_top60_DNA_seed | EGFR | -0.19335 | GBM | D-245MG | Sapitinib | MVP_top60_DNA+RNA_seed | KIT | 0.907527 | GBM | GB-1 | Motesanib |
| MVP_top60_DNA_seed | PIK3CB | -0.190454 | GBM | D-245MG | AZD6482 | MVP_top60_DNA+RNA_seed | EGFR | 0.907562 | GBM | U-118-MG | AST-1306 |
| MVP_top60_DNA_seed | PIK3CA | -0.186516 | GBM | U-118-MG | Buparlisib | MVP_top60_DNA+RNA_seed | PDGFRA | 0.907839 | GBM | D-542MG | Masitinib |
| MVP_top60_DNA_seed | PIK3CB | -0.186516 | GBM | U-118-MG | Buparlisib | MVP_top60_DNA+RNA_seed | PDGFRB | 0.907839 | GBM | D-542MG | Masitinib |
| MVP_top60_DNA_seed | PIK3CD | -0.186516 | GBM | U-118-MG | Buparlisib | MVP_top60_DNA+RNA_seed | KIT | 0.907839 | GBM | D-542MG | Masitinib |
| MVP_top60_DNA_seed | JAK1 | -0.184306 | GBM | YH-13 | JAK1_8709 | MVP_top60_DNA+RNA_seed | IGF1R | 0.908247 | GBM | LN-405 | BMS-754807 |
| MVP_top60_DNA_seed | CDK4 | -0.181636 | GBM | YKG-1 | Palbociclib | MVP_top60_DNA+RNA_seed | EGFR | 0.911172 | GBM | Becker | CUDC-101 |
| MVP_top60_DNA_seed | EGFR | -0.17646 | GBM | SW1088 | Sapitinib | MVP_top60_DNA+RNA_seed | AKT1 | 0.913756 | GBM | LN-229 | GSK690693 |
| MVP_top60_DNA_seed | EP300 | -0.171448 | GBM | AM-38 | CPI-637 | MVP_top60_DNA+RNA_seed | JAK3 | 0.917445 | GBM | SK-MG-1 | WHI-P97 |
| MVP_top60_DNA_seed | TP53 | -0.170488 | GBM | 42-MG-BA | MIRA-1 | MVP_top60_DNA+RNA_seed | EGFR | 0.918206 | GBM | LN-405 | PF-00299804 |
| MVP_top60_DNA_seed | JAK2 | -0.169915 | GBM | Becker | AZ960 | MVP_top60_DNA+RNA_seed | IGF1R | 0.919957 | GBM | D-392MG | BMS-536924 |
| MVP_top60_DNA_seed | JAK3 | -0.169915 | GBM | Becker | AZ960 | MVP_top60_DNA+RNA_seed | MET | 0.920329 | GBM | D-247MG | PHA-665752 |
| MVP_top60_DNA_seed | JAK2 | -0.162817 | GBM | U-118-MG | JAK_8517 | MVP_top60_DNA+RNA_seed | EGFR | 0.922185 | GBM | A172 | PF-00299804 |
| MVP_top60_DNA_seed | JAK1 | -0.162817 | GBM | U-118-MG | JAK_8517 | MVP_top60_DNA+RNA_seed | EGFR | 0.922255 | GBM | SNB75 | PF-00299804 |
| MVP_top60_DNA_seed | TP53 | -0.162285 | GBM | D-263MG | MIRA-1 | MVP_top60_DNA+RNA_seed | MET | 0.923461 | GBM | U-87-MG | SU11274 |
| MVP_top60_DNA_seed | JAK2 | -0.144879 | GBM | M059J | AZ960 | MVP_top60_DNA+RNA_seed | MET | 0.92679 | GBM | D-263MG | PHA-665752 |

| | | | | | | | | | | | |
|---|---|---|---|---|---|---|---|---|---|---|---|
| MVP_top60_DNA_seed | JAK3 | -0.144879 | GBM | M059J | AZ960 | MVP_top60_DNA+RNA_seed | IGF1R | 0.927427 | GBM | SF268 | BMS-536924 |
| MVP_top60_DNA_seed | JAK1 | -0.143102 | GBM | AM-38 | JAK1_8709 | MVP_top60_DNA+RNA_seed | EGFR | 0.928215 | GBM | SW1088 | PF-00299804 |
| MVP_top60_DNA_seed | PIK3CA | -0.139669 | GBM | D-542MG | Alpelisib | MVP_top60_DNA+RNA_seed | EGFR | 0.930239 | GBM | SF268 | Afatinib |
| MVP_top60_DNA_seed | EGFR | -0.138111 | GBM | 42-MG-BA | Osimertinib | MVP_top60_DNA+RNA_seed | EGFR | 0.930559 | GBM | DBTRG-05MG | Cetuximab |
| MVP_top60_DNA_seed | CDK4 | -0.137955 | GBM | AM-38 | Palbociclib | MVP_top60_DNA+RNA_seed | AKT1 | 0.93239 | GBM | CAS-1 | AKT inhibitor VIII |
| MVP_top60_DNA_seed | JAK2 | -0.128926 | GBM | M059J | Ruxolitinib | MVP_top60_DNA+RNA_seed | EGFR | 0.934676 | GBM | D-247MG | Gefitinib |
| MVP_top60_DNA_seed | JAK1 | -0.128926 | GBM | M059J | Ruxolitinib | MVP_top60_DNA+RNA_seed | JAK1 | 0.935662 | GBM | D-542MG | JAK1_3715 |
| MVP_top60_DNA_seed | PIK3CA | -0.126408 | GBM | T98G | AZD8186 | MVP_top60_DNA+RNA_seed | AKT1 | 0.939254 | GBM | D-263MG | GSK690693 |
| MVP_top60_DNA_seed | PIK3CB | -0.126408 | GBM | T98G | AZD8186 | MVP_top60_DNA+RNA_seed | IGF1R | 0.943781 | GBM | LNZTA3WT4 | BMS-754807 |
| MVP_top60_DNA_seed | EGFR | -0.12271 | GBM | 8-MG-BA | Erlotinib | MVP_top60_DNA+RNA_seed | JAK2 | 0.945487 | GBM | U-87-MG | Ruxolitinib |
| MVP_top60_DNA_seed | CDK4 | -0.12219 | GBM | D-566MG | Ribociclib | MVP_top60_DNA+RNA_seed | JAK1 | 0.945487 | GBM | U-87-MG | Ruxolitinib |
| MVP_top60_DNA_seed | PIK3CA | -0.121559 | GBM | D-263MG | GNE-317 | MVP_top60_DNA+RNA_seed | KIT | 0.94659 | GBM | SF268 | OSI-930 |
| MVP_top60_DNA_seed | JAK1 | -0.119898 | GBM | 42-MG-BA | JAK1_8709 | MVP_top60_DNA+RNA_seed | PIK3CB | 0.946816 | GBM | D-392MG | AZD8186 |
| MVP_top60_DNA_seed | PIK3CB | -0.118698 | GBM | SF126 | AZD6482 | MVP_top60_DNA+RNA_seed | PIK3CD | 0.946816 | GBM | D-392MG | AZD8186 |
| MVP_top60_DNA_seed | EGFR | -0.116719 | GBM | A172 | Afatinib | MVP_top60_DNA+RNA_seed | EGFR | 0.948982 | GBM | DBTRG-05MG | PF-00299804 |
| MVP_top60_DNA_seed | PIK3CA | -0.110897 | GBM | M059J | AZD8186 | MVP_top60_DNA+RNA_seed | EGFR | 0.950111 | GBM | LN-18 | Pelitinib |
| MVP_top60_DNA_seed | PIK3CB | -0.110897 | GBM | M059J | AZD8186 | MVP_top60_DNA+RNA_seed | IGF1R | 0.950547 | GBM | YH-13 | Linsitinib |
| MVP_top60_DNA_seed | JAK1 | -0.109389 | GBM | DK-MG | JAK1_8709 | MVP_top60_DNA+RNA_seed | MET | 0.950641 | GBM | SF295 | JNJ38877605 |
| MVP_top60_DNA_seed | PIK3CA | -0.105879 | GBM | U-87-MG | Buparlisib | MVP_top60_DNA+RNA_seed | EGFR | 0.950739 | GBM | D-263MG | CUDC-101 |
| MVP_top60_DNA_seed | PIK3CB | -0.105879 | GBM | U-87-MG | Buparlisib | MVP_top60_DNA+RNA_seed | JAK3 | 0.951276 | GBM | D-542MG | WHI-P97 |
| MVP_top60_DNA_seed | PIK3CD | -0.105879 | GBM | U-87-MG | Buparlisib | MVP_top60_DNA+RNA_seed | PIK3CB | 0.955373 | GBM | U251 | AZD8186 |
| MVP_top60_DNA_seed | CDK4 | -0.102853 | GBM | D-247MG | Palbociclib | MVP_top60_DNA+RNA_seed | PIK3CD | 0.955373 | GBM | U251 | AZD8186 |
| MVP_top60_DNA_seed | EGFR | -0.0997 | GBM | CAS-1 | Lapatinib | MVP_top60_DNA+RNA_seed | EP300 | 0.956974 | GBM | AM-38 | I-CBP112 |
| MVP_top60_DNA_seed | JAK2 | -0.099094 | GBM | SK-MG-1 | AZ960 | MVP_top60_DNA+RNA_seed | CREBBP | 0.956974 | GBM | AM-38 | I-CBP112 |
| MVP_top60_DNA_seed | JAK3 | -0.099094 | GBM | SK-MG-1 | AZ960 | MVP_top60_DNA+RNA_seed | IGF1R | 0.957772 | GBM | D-263MG | GSK1904529A |
| MVP_top60_DNA_seed | JAK1 | -0.098785 | GBM | U-87-MG | JAK1_8709 | MVP_top60_DNA+RNA_seed | EP300 | 0.960338 | GBM | D-392MG | I-CBP112 |
| MVP_top60_DNA_seed | JAK2 | -0.092393 | GBM | Becker | GSK2276186C | MVP_top60_DNA+RNA_seed | CREBBP | 0.960338 | GBM | D-392MG | I-CBP112 |
| MVP_top60_DNA_seed | JAK1 | -0.092393 | GBM | Becker | GSK2276186C | MVP_top60_DNA+RNA_seed | EGFR | 0.963154 | GBM | D-247MG | PF-00299804 |
| MVP_top60_DNA_seed | JAK3 | -0.092393 | GBM | Becker | GSK2276186C | MVP_top60_DNA+RNA_seed | IGF1R | 0.965513 | GBM | SW1088 | GSK1904529A |
| MVP_top60_DNA_seed | CDK4 | -0.086184 | GBM | KS-1 | Ribociclib | MVP_top60_DNA+RNA_seed | EGFR | 0.966872 | GBM | LN-405 | Afatinib |
| MVP_top60_DNA_seed | EGFR | -0.082638 | GBM | SNB75 | Osimertinib | MVP_top60_DNA+RNA_seed | EGFR | 0.973124 | GBM | GB-1 | Erlotinib |
| MVP_top60_DNA_seed | PIK3CA | -0.081116 | GBM | A172 | AZD8186 | MVP_top60_DNA+RNA_seed | AKT1 | 0.97335 | GBM | SW1088 | AKT inhibitor VIII |
| MVP_top60_DNA_seed | PIK3CB | -0.081116 | GBM | A172 | AZD8186 | MVP_top60_DNA+RNA_seed | IGF1R | 0.97566 | GBM | D-542MG | Linsitinib |
| MVP_top60_DNA_seed | PIK3CA | -0.074798 | GBM | D-423MG | Buparlisib | MVP_top60_DNA+RNA_seed | CDK4 | 0.97739 | GBM | 42-MG-BA | Palbociclib |
| MVP_top60_DNA_seed | PIK3CB | -0.074798 | GBM | D-423MG | Buparlisib | MVP_top60_DNA+RNA_seed | IGF1R | 0.986778 | GBM | U251 | BMS-536924 |
| MVP_top60_DNA_seed | PIK3CD | -0.074798 | GBM | D-423MG | Buparlisib | MVP_top60_DNA+RNA_seed | JAK2 | 0.990506 | GBM | U-87-MG | Fedratinib |
| MVP_top60_DNA_seed | JAK2 | -0.066034 | GBM | GB-1 | AZ960 | MVP_top60_DNA+RNA_seed | PDGFRA | 0.9916 | GBM | M059J | Amuvatinib |
| MVP_top60_DNA_seed | JAK3 | -0.066034 | GBM | GB-1 | AZ960 | MVP_top60_DNA+RNA_seed | KIT | 0.9916 | GBM | M059J | Amuvatinib |
| MVP_top60_DNA_seed | EGFR | -0.06101 | GBM | LN-229 | Gefitinib | MVP_top60_DNA+RNA_seed | IGF1R | 0.992275 | GBM | CAS-1 | BMS-536924 |
| MVP_top60_DNA_seed | EP300 | -0.05961 | GBM | SK-MG-1 | CPI-637 | MVP_top60_DNA+RNA_seed | JAK3 | 0.993898 | GBM | GB-1 | JAK3_7406 |
| MVP_top60_DNA_seed | EGFR | -0.059066 | GBM | 8-MG-BA | Sapitinib | MVP_top60_DNA+RNA_seed | EGFR | 0.997767 | GBM | U-118-MG | Cetuximab |

| | | | | | | | | | | | |
|---|---|---|---|---|---|---|---|---|---|---|---|
| MVP_top60_DNA_seed | EGFR | -0.057422 | GBM | A172 | Gefitinib | MVP_top60_DNA+RNA_seed | JAK2 | 0.998788 | GBM | LN-405 | Fedratinib |
| MVP_top60_DNA_seed | EGFR | -0.055395 | GBM | D-542MG | Lapatinib | MVP_top60_DNA+RNA_seed | PDGFRB | 1.001311 | GBM | SF295 | Pazopanib |
| MVP_top60_DNA_seed | CDK4 | -0.053038 | GBM | U251 | Palbociclib | MVP_top60_DNA+RNA_seed | KIT | 1.001311 | GBM | SF295 | Pazopanib |
| MVP_top60_DNA_seed | EGFR | -0.052 | GBM | 42-MG-BA | Afatinib | MVP_top60_DNA+RNA_seed | PIK3CA | 1.004612 | GBM | M059J | AZD8835 |
| MVP_top60_DNA_seed | EGFR | -0.050785 | GBM | A172 | Osimertinib | MVP_top60_DNA+RNA_seed | PIK3CD | 1.004612 | GBM | M059J | AZD8835 |
| MVP_top60_DNA_seed | PIK3CA | -0.048167 | GBM | GB-1 | Buparlisib | MVP_top60_DNA+RNA_seed | EGFR | 1.007573 | GBM | DK-MG | Gefitinib |
| MVP_top60_DNA_seed | PIK3CB | -0.048167 | GBM | GB-1 | Buparlisib | MVP_top60_DNA+RNA_seed | AKT1 | 1.008843 | GBM | LN-405 | AKT inhibitor VIII |
| MVP_top60_DNA_seed | PIK3CD | -0.048167 | GBM | GB-1 | Buparlisib | MVP_top60_DNA+RNA_seed | MET | 1.011092 | GBM | D-263MG | Crizotinib |
| MVP_top60_DNA_seed | EGFR | -0.042264 | GBM | D-566MG | Erlotinib | MVP_top60_DNA+RNA_seed | JAK2 | 1.011303 | GBM | DBTRG-05MG | Ruxolitinib |
| MVP_top60_DNA_seed | PIK3CA | -0.034133 | GBM | AM-38 | GNE-317 | MVP_top60_DNA+RNA_seed | JAK1 | 1.011303 | GBM | DBTRG-05MG | Ruxolitinib |
| MVP_top60_DNA_seed | TP53 | -0.03173 | GBM | M059J | MIRA-1 | MVP_top60_DNA+RNA_seed | JAK1 | 1.01297 | GBM | U-118-MG | JAK1_3715 |
| MVP_top60_DNA_seed | EGFR | -0.0309 | GBM | SF295 | Osimertinib | MVP_top60_DNA+RNA_seed | EGFR | 1.013694 | GBM | U-118-MG | Afatinib |
| MVP_top60_DNA_seed | CDK4 | -0.030002 | GBM | U-87-MG | Palbociclib | MVP_top60_DNA+RNA_seed | JAK3 | 1.015945 | GBM | LN-405 | WHI-P97 |
| MVP_top60_DNA_seed | CDK4 | -0.021426 | GBM | 8-MG-BA | Palbociclib | MVP_top60_DNA+RNA_seed | JAK2 | 1.017056 | GBM | LN-229 | AZD1480 |
| MVP_top60_DNA_seed | JAK2 | -0.018538 | GBM | LN-18 | JAK_8517 | MVP_top60_DNA+RNA_seed | JAK1 | 1.017056 | GBM | LN-229 | AZD1480 |
| MVP_top60_DNA_seed | JAK1 | -0.018538 | GBM | LN-18 | JAK_8517 | MVP_top60_DNA+RNA_seed | KIT | 1.017944 | GBM | SNB75 | Imatinib |
| MVP_top60_DNA_seed | PIK3CA | -0.017547 | GBM | CAS-1 | Buparlisib | MVP_top60_DNA+RNA_seed | AKT1 | 1.021154 | GBM | D-542MG | GSK690693 |
| MVP_top60_DNA_seed | PIK3CB | -0.017547 | GBM | CAS-1 | Buparlisib | MVP_top60_DNA+RNA_seed | EGFR | 1.023486 | GBM | DBTRG-05MG | Afatinib |
| MVP_top60_DNA_seed | PIK3CD | -0.017547 | GBM | CAS-1 | Buparlisib | MVP_top60_DNA+RNA_seed | AKT1 | 1.025237 | GBM | DBTRG-05MG | AKT inhibitor VIII |
| MVP_top60_DNA_seed | JAK2 | -0.016788 | GBM | D-392MG | AZ960 | MVP_top60_DNA+RNA_seed | EGFR | 1.028719 | GBM | D-392MG | Afatinib |
| MVP_top60_DNA_seed | JAK3 | -0.016788 | GBM | D-392MG | AZ960 | MVP_top60_DNA+RNA_seed | EGFR | 1.030523 | GBM | SW1088 | Afatinib |
| MVP_top60_DNA_seed | EGFR | -0.015156 | GBM | U-87-MG | AZD3759 | MVP_top60_DNA+RNA_seed | EGFR | 1.034087 | GBM | Becker | PF-00299804 |
| MVP_top60_DNA_seed | EGFR | -0.014364 | GBM | YH-13 | Erlotinib | MVP_top60_DNA+RNA_seed | IGF1R | 1.035985 | GBM | Becker | Linsitinib |
| MVP_top60_DNA_seed | CDK4 | -0.012447 | GBM | 8-MG-BA | Ribociclib | MVP_top60_DNA+RNA_seed | KIT | 1.040731 | GBM | KS-1 | OSI-930 |
| MVP_top60_DNA_seed | JAK2 | -0.011785 | GBM | U251 | JAK_8517 | MVP_top60_DNA+RNA_seed | EP300 | 1.040931 | GBM | SW1088 | I-CBP112 |
| MVP_top60_DNA_seed | JAK1 | -0.011785 | GBM | U251 | JAK_8517 | MVP_top60_DNA+RNA_seed | CREBBP | 1.040931 | GBM | SW1088 | I-CBP112 |
| MVP_top60_DNA_seed | EGFR | -0.011375 | GBM | YKG-1 | Osimertinib | MVP_top60_DNA+RNA_seed | EGFR | 1.041416 | GBM | DBTRG-05MG | Afatinib |
| MVP_top60_DNA_seed | EP300 | -0.010627 | GBM | M059J | SGC-CBP30 | MVP_top60_DNA+RNA_seed | EGFR | 1.041725 | GBM | SF268 | Gefitinib |
| MVP_top60_DNA_seed | EP300 | -0.008614 | GBM | U251 | CPI-637 | MVP_top60_DNA+RNA_seed | PIK3CA | 1.044933 | GBM | DBTRG-05MG | PI3Ka_4409 |
| MVP_top60_DNA_seed | EGFR | -0.007482 | GBM | YH-13 | Osimertinib | MVP_top60_DNA+RNA_seed | PIK3CD | 1.044933 | GBM | DBTRG-05MG | PI3Ka_4409 |
| MVP_top60_DNA_seed | JAK2 | -0.005964 | GBM | SF295 | AZ960 | MVP_top60_DNA+RNA_seed | JAK1 | 1.046499 | GBM | AM-38 | JAK1_3715 |
| MVP_top60_DNA_seed | JAK3 | -0.005964 | GBM | SF295 | AZ960 | MVP_top60_DNA+RNA_seed | EGFR | 1.047505 | GBM | CAS-1 | Erlotinib |
| MVP_top60_DNA_seed | EGFR | -0.005381 | GBM | SF126 | Sapitinib | MVP_top60_DNA+RNA_seed | MET | 1.050095 | GBM | D-392MG | SU11274 |
| MVP_top60_DNA_seed | JAK2 | -0.001596 | GBM | D-392MG | GSK2276186C | MVP_top60_DNA+RNA_seed | CDK4 | 1.050188 | GBM | 8-MG-BA | Palbociclib |
| MVP_top60_DNA_seed | JAK1 | -0.001596 | GBM | D-392MG | GSK2276186C | MVP_top60_DNA+RNA_seed | EGFR | 1.051861 | GBM | D-542MG | Afatinib |
| MVP_top60_DNA_seed | JAK3 | -0.001596 | GBM | D-392MG | GSK2276186C | MVP_top60_DNA+RNA_seed | JAK2 | 1.053148 | GBM | D-392MG | AZD1480 |
| MVP_top60_DNA_seed | EGFR | 0.002614 | GBM | M059J | Erlotinib | MVP_top60_DNA+RNA_seed | JAK1 | 1.053148 | GBM | D-392MG | AZD1480 |
| MVP_top60_DNA_seed | EGFR | 0.004394 | GBM | U251 | Erlotinib | MVP_top60_DNA+RNA_seed | PDGFRA | 1.054111 | GBM | LN-405 | Amuvatinib |
| MVP_top60_DNA_seed | EP300 | 0.00589 | GBM | 8-MG-BA | SGC-CBP30 | MVP_top60_DNA+RNA_seed | KIT | 1.054111 | GBM | LN-405 | Amuvatinib |
| MVP_top60_DNA_seed | TP53 | 0.007679 | GBM | T98G | MIRA-1 | MVP_top60_DNA+RNA_seed | MET | 1.05898 | GBM | SNB75 | PHA-665752 |
| MVP_top60_DNA_seed | JAK2 | 0.00891 | GBM | T98G | Ruxolitinib | MVP_top60_DNA+RNA_seed | EGFR | 1.060722 | GBM | DK-MG | Afatinib |

| | | | | | | | | | | | |
|---|---|---|---|---|---|---|---|---|---|---|---|
| MVP_top60_DNA_seed | JAK1 | 0.00891 | GBM | T98G | Ruxolitinib | MVP_top60_DNA+RNA_seed | EGFR | 1.06613 | GBM | D-542MG | Afatinib |
| MVP_top60_DNA_seed | EGFR | 0.008988 | GBM | KS-1 | Sapitinib | MVP_top60_DNA+RNA_seed | MET | 1.067721 | GBM | SF126 | JNJ38877605 |
| MVP_top60_DNA_seed | EGFR | 0.010215 | GBM | SF268 | Erlotinib | MVP_top60_DNA+RNA_seed | IGF1R | 1.069011 | GBM | M059J | Linsitinib |
| MVP_top60_DNA_seed | PIK3CA | 0.015188 | GBM | D-263MG | Buparlisib | MVP_top60_DNA+RNA_seed | MET | 1.072362 | GBM | SNB75 | AZD6094 |
| MVP_top60_DNA_seed | PIK3CB | 0.015188 | GBM | D-263MG | Buparlisib | MVP_top60_DNA+RNA_seed | EGFR | 1.074244 | GBM | AM-38 | Cetuximab |
| MVP_top60_DNA_seed | PIK3CD | 0.015188 | GBM | D-263MG | Buparlisib | MVP_top60_DNA+RNA_seed | IGF1R | 1.078397 | GBM | U-118-MG | BMS-536924 |
| MVP_top60_DNA_seed | PIK3CA | 0.019902 | GBM | KS-1 | GNE-317 | MVP_top60_DNA+RNA_seed | CDK4 | 1.080758 | GBM | CAS-1 | Palbociclib |
| MVP_top60_DNA_seed | PIK3CA | 0.022618 | GBM | U-87-MG | GNE-317 | MVP_top60_DNA+RNA_seed | PIK3CA | 1.081435 | GBM | A172 | PI3Ka_4409 |
| MVP_top60_DNA_seed | EGFR | 0.027993 | GBM | A172 | Lapatinib | MVP_top60_DNA+RNA_seed | PIK3CD | 1.081435 | GBM | A172 | PI3Ka_4409 |
| MVP_top60_DNA_seed | JAK2 | 0.028488 | GBM | U251 | GSK2276186C | MVP_top60_DNA+RNA_seed | AKT1 | 1.082133 | GBM | GB-1 | AKT inhibitor VIII |
| MVP_top60_DNA_seed | JAK1 | 0.028488 | GBM | U251 | GSK2276186C | MVP_top60_DNA+RNA_seed | EGFR | 1.084594 | GBM | CAS-1 | Afatinib |
| MVP_top60_DNA_seed | JAK3 | 0.028488 | GBM | U251 | GSK2276186C | MVP_top60_DNA+RNA_seed | EGFR | 1.088397 | GBM | SNB75 | Pelitinib |
| MVP_top60_DNA_seed | EGFR | 0.029904 | GBM | YKG-1 | Gefitinib | MVP_top60_DNA+RNA_seed | EGFR | 1.091108 | GBM | M059J | Pelitinib |
| MVP_top60_DNA_seed | EGFR | 0.030789 | GBM | 8-MG-BA | Afatinib | MVP_top60_DNA+RNA_seed | EGFR | 1.092916 | GBM | T98G | CUDC-101 |
| MVP_top60_DNA_seed | CDK4 | 0.031313 | GBM | SW1088 | Palbociclib | MVP_top60_DNA+RNA_seed | EGFR | 1.093055 | GBM | SW1088 | Cetuximab |
| MVP_top60_DNA_seed | EGFR | 0.035055 | GBM | LN-18 | Sapitinib | MVP_top60_DNA+RNA_seed | EGFR | 1.093282 | GBM | D-542MG | CI-1033 |
| MVP_top60_DNA_seed | EGFR | 0.036472 | GBM | DK-MG | Erlotinib | MVP_top60_DNA+RNA_seed | JAK1 | 1.098248 | GBM | SNB75 | JAK1_3715 |
| MVP_top60_DNA_seed | EP300 | 0.043633 | GBM | M059J | CPI-637 | MVP_top60_DNA+RNA_seed | EP300 | 1.104118 | GBM | KS-1 | I-CBP112 |
| MVP_top60_DNA_seed | PIK3CA | 0.053743 | GBM | D-245MG | GNE-317 | MVP_top60_DNA+RNA_seed | CREBBP | 1.104118 | GBM | KS-1 | I-CBP112 |
| MVP_top60_DNA_seed | JAK2 | 0.055117 | GBM | CAS-1 | GSK2276186C | MVP_top60_DNA+RNA_seed | IGF1R | 1.104686 | GBM | D-392MG | BMS-754807 |
| MVP_top60_DNA_seed | JAK1 | 0.055117 | GBM | CAS-1 | GSK2276186C | MVP_top60_DNA+RNA_seed | EGFR | 1.114354 | GBM | SNB75 | CI-1033 |
| MVP_top60_DNA_seed | JAK3 | 0.055117 | GBM | CAS-1 | GSK2276186C | MVP_top60_DNA+RNA_seed | EGFR | 1.11661 | GBM | LNZTA3WT4 | Gefitinib |
| MVP_top60_DNA_seed | TP53 | 0.063766 | GBM | SNB75 | MIRA-1 | MVP_top60_DNA+RNA_seed | IGF1R | 1.116789 | GBM | DK-MG | BMS-536924 |
| MVP_top60_DNA_seed | EGFR | 0.065361 | GBM | D-566MG | Gefitinib | MVP_top60_DNA+RNA_seed | PIK3CA | 1.117505 | GBM | D-392MG | AZD8835 |
| MVP_top60_DNA_seed | EGFR | 0.067748 | GBM | LNZTA3WT4 | Afatinib | MVP_top60_DNA+RNA_seed | PIK3CD | 1.117505 | GBM | D-392MG | AZD8835 |
| MVP_top60_DNA_seed | PIK3CA | 0.070068 | GBM | D-247MG | Buparlisib | MVP_top60_DNA+RNA_seed | IGF1R | 1.123498 | GBM | AM-38 | BMS-536924 |
| MVP_top60_DNA_seed | PIK3CB | 0.070068 | GBM | D-247MG | Buparlisib | MVP_top60_DNA+RNA_seed | EGFR | 1.123878 | GBM | AM-38 | Afatinib |
| MVP_top60_DNA_seed | PIK3CD | 0.070068 | GBM | D-247MG | Buparlisib | MVP_top60_DNA+RNA_seed | CDK4 | 1.124655 | GBM | U-87-MG | CGP-082996 |
| MVP_top60_DNA_seed | EP300 | 0.073544 | GBM | GB-1 | SGC-CBP30 | MVP_top60_DNA+RNA_seed | EGFR | 1.128897 | GBM | D-247MG | Afatinib |
| MVP_top60_DNA_seed | CDK4 | 0.076872 | GBM | D-423MG | Palbociclib | MVP_top60_DNA+RNA_seed | PDGFRB | 1.129922 | GBM | AM-38 | Pazopanib |
| MVP_top60_DNA_seed | EP300 | 0.077697 | GBM | LN-229 | CPI-637 | MVP_top60_DNA+RNA_seed | KIT | 1.129922 | GBM | AM-38 | Pazopanib |
| MVP_top60_DNA_seed | CDK4 | 0.080189 | GBM | D-245MG | Palbociclib | MVP_top60_DNA+RNA_seed | IGF1R | 1.133144 | GBM | LN-229 | BMS-536924 |
| MVP_top60_DNA_seed | PIK3CA | 0.08024 | GBM | CAS-1 | GNE-317 | MVP_top60_DNA+RNA_seed | KIT | 1.13374 | GBM | U-87-MG | Axitinib |
| MVP_top60_DNA_seed | TP53 | 0.090294 | GBM | SF126 | MIRA-1 | MVP_top60_DNA+RNA_seed | PIK3CA | 1.153257 | GBM | D-392MG | PI3Ka_4409 |
| MVP_top60_DNA_seed | JAK2 | 0.091665 | GBM | 8-MG-BA | JAK_8517 | MVP_top60_DNA+RNA_seed | PIK3CD | 1.153257 | GBM | D-392MG | PI3Ka_4409 |
| MVP_top60_DNA_seed | JAK1 | 0.091665 | GBM | 8-MG-BA | JAK_8517 | MVP_top60_DNA+RNA_seed | AKT1 | 1.159887 | GBM | LNZTA3WT4 | AKT inhibitor VIII |
| MVP_top60_DNA_seed | EGFR | 0.091768 | GBM | U-118-MG | AZD3759 | MVP_top60_DNA+RNA_seed | AKT1 | 1.163744 | GBM | AM-38 | AKT inhibitor VIII |
| MVP_top60_DNA_seed | PIK3CA | 0.098691 | GBM | YKG-1 | GNE-317 | MVP_top60_DNA+RNA_seed | EGFR | 1.164147 | GBM | LN-405 | Pelitinib |
| MVP_top60_DNA_seed | EGFR | 0.100732 | GBM | SF295 | Lapatinib | MVP_top60_DNA+RNA_seed | PIK3CA | 1.164432 | GBM | DK-MG | PI3Ka_4409 |
| MVP_top60_DNA_seed | PIK3CA | 0.106176 | GBM | SF126 | GNE-317 | MVP_top60_DNA+RNA_seed | PIK3CD | 1.164432 | GBM | DK-MG | PI3Ka_4409 |
| MVP_top60_DNA_seed | EGFR | 0.110759 | GBM | U-118-MG | Osimertinib | MVP_top60_DNA+RNA_seed | EGFR | 1.169363 | GBM | D-247MG | Afatinib |

| | | | | | | | | | | | |
|---|---|---|---|---|---|---|---|---|---|---|---|
| MVP_top60_DNA_seed | EGFR | 0.114846 | GBM | SK-MG-1 | Afatinib | MVP_top60_DNA+RNA_seed | PIK3CD | 1.171712 | GBM | D-542MG | Idelalisib |
| MVP_top60_DNA_seed | EGFR | 0.115143 | GBM | DK-MG | Sapitinib | MVP_top60_DNA+RNA_seed | JAK1 | 1.172708 | GBM | DBTRG-05MG | JAK1_3715 |
| MVP_top60_DNA_seed | PIK3CA | 0.115924 | GBM | D-542MG | Buparlisib | MVP_top60_DNA+RNA_seed | PIK3CA | 1.178644 | GBM | U-118-MG | AZD8835 |
| MVP_top60_DNA_seed | PIK3CB | 0.115924 | GBM | D-542MG | Buparlisib | MVP_top60_DNA+RNA_seed | PIK3CD | 1.178644 | GBM | U-118-MG | AZD8835 |
| MVP_top60_DNA_seed | PIK3CD | 0.115924 | GBM | D-542MG | Buparlisib | MVP_top60_DNA+RNA_seed | JAK3 | 1.179073 | GBM | D-263MG | WHI-P97 |
| MVP_top60_DNA_seed | EGFR | 0.118834 | GBM | D-542MG | Osimertinib | MVP_top60_DNA+RNA_seed | IGF1R | 1.179493 | GBM | YKG-1 | Linsitinib |
| MVP_top60_DNA_seed | EGFR | 0.124166 | GBM | SF126 | AZD3759 | MVP_top60_DNA+RNA_seed | IGF1R | 1.182859 | GBM | SNB75 | BMS-536924 |
| MVP_top60_DNA_seed | EGFR | 0.12491 | GBM | AM-38 | AZD3759 | MVP_top60_DNA+RNA_seed | KIT | 1.183016 | GBM | LN-229 | Axitinib |
| MVP_top60_DNA_seed | CDK4 | 0.128484 | GBM | SF268 | Ribociclib | MVP_top60_DNA+RNA_seed | EGFR | 1.183565 | GBM | D-542MG | Gefitinib |
| MVP_top60_DNA_seed | EGFR | 0.132447 | GBM | D-566MG | AZD3759 | MVP_top60_DNA+RNA_seed | JAK3 | 1.1858 | GBM | DBTRG-05MG | WHI-P97 |
| MVP_top60_DNA_seed | EGFR | 0.134693 | GBM | LNZTA3WT4 | Lapatinib | MVP_top60_DNA+RNA_seed | IGF1R | 1.186496 | GBM | D-263MG | Linsitinib |
| MVP_top60_DNA_seed | PIK3CA | 0.135088 | GBM | KS-1 | Alpelisib | MVP_top60_DNA+RNA_seed | PDGFRB | 1.192739 | GBM | U-87-MG | Pazopanib |
| MVP_top60_DNA_seed | EGFR | 0.135413 | GBM | T98G | Gefitinib | MVP_top60_DNA+RNA_seed | KIT | 1.192739 | GBM | U-87-MG | Pazopanib |
| MVP_top60_DNA_seed | PIK3CA | 0.136491 | GBM | T98G | Buparlisib | MVP_top60_DNA+RNA_seed | PIK3CD | 1.197102 | GBM | DK-MG | Idelalisib |
| MVP_top60_DNA_seed | PIK3CB | 0.136491 | GBM | T98G | Buparlisib | MVP_top60_DNA+RNA_seed | AKT1 | 1.207541 | GBM | DK-MG | GSK690693 |
| MVP_top60_DNA_seed | PIK3CD | 0.136491 | GBM | T98G | Buparlisib | MVP_top60_DNA+RNA_seed | EGFR | 1.217418 | GBM | U-87-MG | Erlotinib |
| MVP_top60_DNA_seed | EGFR | 0.148283 | GBM | DK-MG | Afatinib | MVP_top60_DNA+RNA_seed | JAK2 | 1.219291 | GBM | LN-229 | Fedratinib |
| MVP_top60_DNA_seed | EGFR | 0.149432 | GBM | D-263MG | AZD3759 | MVP_top60_DNA+RNA_seed | MET | 1.219599 | GBM | SNB75 | Crizotinib |
| MVP_top60_DNA_seed | CDK4 | 0.152862 | GBM | SF126 | Ribociclib | MVP_top60_DNA+RNA_seed | JAK1 | 1.220882 | GBM | CAS-1 | JAK1_3715 |
| MVP_top60_DNA_seed | JAK2 | 0.158329 | GBM | LN-18 | AZ960 | MVP_top60_DNA+RNA_seed | CDK4 | 1.232426 | GBM | D-263MG | CGP-082996 |
| MVP_top60_DNA_seed | JAK3 | 0.158329 | GBM | LN-18 | AZ960 | MVP_top60_DNA+RNA_seed | EGFR | 1.234474 | GBM | CAS-1 | Afatinib |
| MVP_top60_DNA_seed | CDK4 | 0.169151 | GBM | D-423MG | Ribociclib | MVP_top60_DNA+RNA_seed | JAK2 | 1.23889 | GBM | LN-18 | Fedratinib |
| MVP_top60_DNA_seed | CDK4 | 0.169831 | GBM | LN-18 | Ribociclib | MVP_top60_DNA+RNA_seed | JAK2 | 1.239607 | GBM | LNZTA3WT4 | Fedratinib |
| MVP_top60_DNA_seed | EGFR | 0.17414 | GBM | U-118-MG | Erlotinib | MVP_top60_DNA+RNA_seed | AKT1 | 1.246588 | GBM | D-263MG | AKT inhibitor VIII |
| MVP_top60_DNA_seed | EGFR | 0.175123 | GBM | D-423MG | Gefitinib | MVP_top60_DNA+RNA_seed | EGFR | 1.248653 | GBM | SNB75 | Afatinib |
| MVP_top60_DNA_seed | EGFR | 0.177191 | GBM | SF268 | Afatinib | MVP_top60_DNA+RNA_seed | CDK4 | 1.249184 | GBM | AM-38 | CGP-082996 |
| MVP_top60_DNA_seed | EGFR | 0.180797 | GBM | A172 | AZD3759 | MVP_top60_DNA+RNA_seed | PIK3CD | 1.252617 | GBM | D-263MG | Idelalisib |
| MVP_top60_DNA_seed | EGFR | 0.181391 | GBM | A172 | Erlotinib | MVP_top60_DNA+RNA_seed | AKT1 | 1.253262 | GBM | A172 | BAY AKT1 |
| MVP_top60_DNA_seed | EGFR | 0.183624 | GBM | D-245MG | Afatinib | MVP_top60_DNA+RNA_seed | JAK3 | 1.253604 | GBM | AM-38 | JAK3_7406 |
| MVP_top60_DNA_seed | CDK4 | 0.18412 | GBM | DK-MG | Palbociclib | MVP_top60_DNA+RNA_seed | IGF1R | 1.257264 | GBM | AM-38 | BMS-536924 |
| MVP_top60_DNA_seed | EGFR | 0.192616 | GBM | D-245MG | Osimertinib | MVP_top60_DNA+RNA_seed | MET | 1.25765 | GBM | U-118-MG | SU11274 |
| MVP_top60_DNA_seed | EGFR | 0.192717 | GBM | SF268 | Gefitinib | MVP_top60_DNA+RNA_seed | EGFR | 1.261068 | GBM | U-118-MG | Afatinib |
| MVP_top60_DNA_seed | CDK4 | 0.195795 | GBM | U-118-MG | Palbociclib | MVP_top60_DNA+RNA_seed | EGFR | 1.280577 | GBM | CAS-1 | Gefitinib |
| MVP_top60_DNA_seed | EGFR | 0.197386 | GBM | D-247MG | Osimertinib | MVP_top60_DNA+RNA_seed | PDGFRA | 1.280743 | GBM | LN-229 | Amuvatinib |
| MVP_top60_DNA_seed | EGFR | 0.200155 | GBM | AM-38 | Lapatinib | MVP_top60_DNA+RNA_seed | KIT | 1.280743 | GBM | LN-229 | Amuvatinib |
| MVP_top60_DNA_seed | PIK3CA | 0.209158 | GBM | U-87-MG | Alpelisib | MVP_top60_DNA+RNA_seed | PIK3CD | 1.28076 | GBM | U-118-MG | Idelalisib |
| MVP_top60_DNA_seed | EGFR | 0.209204 | GBM | U251 | AZD3759 | MVP_top60_DNA+RNA_seed | PIK3CB | 1.286163 | GBM | AM-38 | AZD8186 |
| MVP_top60_DNA_seed | EGFR | 0.20971 | GBM | YH-13 | Afatinib | MVP_top60_DNA+RNA_seed | PIK3CD | 1.286163 | GBM | AM-38 | AZD8186 |
| MVP_top60_DNA_seed | PIK3CA | 0.216864 | GBM | D-392MG | GNE-317 | MVP_top60_DNA+RNA_seed | KIT | 1.286309 | GBM | CAS-1 | Motesanib |
| MVP_top60_DNA_seed | JAK2 | 0.219615 | GBM | A172 | GSK2276186C | MVP_top60_DNA+RNA_seed | JAK2 | 1.289316 | GBM | AM-38 | AZD1480 |
| MVP_top60_DNA_seed | JAK1 | 0.219615 | GBM | A172 | GSK2276186C | MVP_top60_DNA+RNA_seed | JAK1 | 1.289316 | GBM | AM-38 | AZD1480 |

| | | | | | | | | | | | |
|---|---|---|---|---|---|---|---|---|---|---|---|
| MVP_top60_DNA_seed | JAK3 | 0.219615 | GBM | A172 | GSK2276186C | MVP_top60_DNA+RNA_seed | KIT | 1.294529 | GBM | U-87-MG | OSI-930 |
| MVP_top60_DNA_seed | JAK2 | 0.222005 | GBM | SF268 | Ruxolitinib | MVP_top60_DNA+RNA_seed | PIK3CA | 1.298605 | GBM | A172 | AZD8835 |
| MVP_top60_DNA_seed | JAK1 | 0.222005 | GBM | SF268 | Ruxolitinib | MVP_top60_DNA+RNA_seed | PIK3CD | 1.298605 | GBM | A172 | AZD8835 |
| MVP_top60_DNA_seed | PIK3CA | 0.22435 | GBM | KS-1 | Buparlisib | MVP_top60_DNA+RNA_seed | IGF1R | 1.303786 | GBM | CAS-1 | GSK1904529A |
| MVP_top60_DNA_seed | PIK3CB | 0.22435 | GBM | KS-1 | Buparlisib | MVP_top60_DNA+RNA_seed | PDGFRA | 1.317396 | GBM | Becker | Masitinib |
| MVP_top60_DNA_seed | PIK3CD | 0.22435 | GBM | KS-1 | Buparlisib | MVP_top60_DNA+RNA_seed | PDGFRB | 1.317396 | GBM | Becker | Masitinib |
| MVP_top60_DNA_seed | EGFR | 0.225471 | GBM | M059J | Afatinib | MVP_top60_DNA+RNA_seed | KIT | 1.317396 | GBM | Becker | Masitinib |
| MVP_top60_DNA_seed | PIK3CA | 0.226173 | GBM | D-566MG | Alpelisib | MVP_top60_DNA+RNA_seed | EGFR | 1.318872 | GBM | DK-MG | AZD8931 |
| MVP_top60_DNA_seed | EGFR | 0.232195 | GBM | M059J | Lapatinib | MVP_top60_DNA+RNA_seed | ERBB3 | 1.318872 | GBM | DK-MG | AZD8931 |
| MVP_top60_DNA_seed | PIK3CA | 0.232417 | GBM | D-392MG | AZD8186 | MVP_top60_DNA+RNA_seed | EGFR | 1.321979 | GBM | SNB75 | Erlotinib |
| MVP_top60_DNA_seed | PIK3CB | 0.232417 | GBM | D-392MG | AZD8186 | MVP_top60_DNA+RNA_seed | PDGFRA | 1.330843 | GBM | SW1088 | Amuvatinib |
| MVP_top60_DNA_seed | EGFR | 0.233071 | GBM | DK-MG | AZD3759 | MVP_top60_DNA+RNA_seed | KIT | 1.330843 | GBM | SW1088 | Amuvatinib |
| MVP_top60_DNA_seed | JAK2 | 0.233557 | GBM | GB-1 | JAK_8517 | MVP_top60_DNA+RNA_seed | AKT1 | 1.33193 | GBM | U-118-MG | MK-2206 |
| MVP_top60_DNA_seed | JAK1 | 0.233557 | GBM | GB-1 | JAK_8517 | MVP_top60_DNA+RNA_seed | MET | 1.33723 | GBM | CAS-1 | PHA-665752 |
| MVP_top60_DNA_seed | JAK2 | 0.235612 | GBM | A172 | Ruxolitinib | MVP_top60_DNA+RNA_seed | JAK3 | 1.338655 | GBM | U-87-MG | WHI-P97 |
| MVP_top60_DNA_seed | JAK1 | 0.235612 | GBM | A172 | Ruxolitinib | MVP_top60_DNA+RNA_seed | PDGFRA | 1.340726 | GBM | SNB75 | Amuvatinib |
| MVP_top60_DNA_seed | EGFR | 0.237068 | GBM | U-87-MG | Gefitinib | MVP_top60_DNA+RNA_seed | KIT | 1.340726 | GBM | SNB75 | Amuvatinib |
| MVP_top60_DNA_seed | EGFR | 0.241009 | GBM | U-87-MG | Sapitinib | MVP_top60_DNA+RNA_seed | IGF1R | 1.349924 | GBM | LN-405 | Linsitinib |
| MVP_top60_DNA_seed | JAK2 | 0.241593 | GBM | AM-38 | AZ960 | MVP_top60_DNA+RNA_seed | PDGFRA | 1.356297 | GBM | DK-MG | Amuvatinib |
| MVP_top60_DNA_seed | JAK3 | 0.241593 | GBM | AM-38 | AZ960 | MVP_top60_DNA+RNA_seed | KIT | 1.356297 | GBM | DK-MG | Amuvatinib |
| MVP_top60_DNA_seed | JAK1 | 0.243108 | GBM | U-118-MG | JAK1_8709 | MVP_top60_DNA+RNA_seed | CDK4 | 1.357365 | GBM | LN-405 | Palbociclib |
| MVP_top60_DNA_seed | EGFR | 0.255549 | GBM | LN-229 | AZD3759 | MVP_top60_DNA+RNA_seed | EGFR | 1.364496 | GBM | U-118-MG | Pelitinib |
| MVP_top60_DNA_seed | JAK2 | 0.260931 | GBM | DK-MG | Ruxolitinib | MVP_top60_DNA+RNA_seed | EP300 | 1.376447 | GBM | CAS-1 | I-CBP112 |
| MVP_top60_DNA_seed | JAK1 | 0.260931 | GBM | DK-MG | Ruxolitinib | MVP_top60_DNA+RNA_seed | CREBBP | 1.376447 | GBM | CAS-1 | I-CBP112 |
| MVP_top60_DNA_seed | CDK4 | 0.261538 | GBM | SF295 | Ribociclib | MVP_top60_DNA+RNA_seed | CDK4 | 1.396558 | GBM | Becker | CGP-082996 |
| MVP_top60_DNA_seed | TP53 | 0.264647 | GBM | YKG-1 | MIRA-1 | MVP_top60_DNA+RNA_seed | EGFR | 1.39991 | GBM | U-87-MG | Pelitinib |
| MVP_top60_DNA_seed | JAK2 | 0.265974 | GBM | SF268 | AZ960 | MVP_top60_DNA+RNA_seed | IGF1R | 1.405104 | GBM | D-392MG | Linsitinib |
| MVP_top60_DNA_seed | JAK3 | 0.265974 | GBM | SF268 | AZ960 | MVP_top60_DNA+RNA_seed | IGF1R | 1.405494 | GBM | U-87-MG | Linsitinib |
| MVP_top60_DNA_seed | JAK2 | 0.267109 | GBM | SF295 | Ruxolitinib | MVP_top60_DNA+RNA_seed | JAK1 | 1.413885 | GBM | KS-1 | JAK1_3715 |
| MVP_top60_DNA_seed | JAK1 | 0.267109 | GBM | SF295 | Ruxolitinib | MVP_top60_DNA+RNA_seed | EGFR | 1.433478 | GBM | SF268 | CI-1033 |
| MVP_top60_DNA_seed | EGFR | 0.269102 | GBM | 42-MG-BA | Sapitinib | MVP_top60_DNA+RNA_seed | EGFR | 1.439725 | GBM | D-263MG | Pelitinib |
| MVP_top60_DNA_seed | EGFR | 0.27457 | GBM | SW1088 | Erlotinib | MVP_top60_DNA+RNA_seed | IGF1R | 1.451009 | GBM | U-87-MG | BMS-754807 |
| MVP_top60_DNA_seed | EGFR | 0.275207 | GBM | D-542MG | Gefitinib | MVP_top60_DNA+RNA_seed | MET | 1.456393 | GBM | CAS-1 | SU11274 |
| MVP_top60_DNA_seed | JAK2 | 0.279236 | GBM | D-392MG | Ruxolitinib | MVP_top60_DNA+RNA_seed | KIT | 1.475745 | GBM | SNB75 | OSI-930 |
| MVP_top60_DNA_seed | JAK1 | 0.279236 | GBM | D-392MG | Ruxolitinib | MVP_top60_DNA+RNA_seed | IGF1R | 1.487304 | GBM | U-87-MG | GSK1904529A |
| MVP_top60_DNA_seed | EGFR | 0.280485 | GBM | Becker | AZD3759 | MVP_top60_DNA+RNA_seed | JAK1 | 1.49302 | GBM | DK-MG | JAK1_3715 |
| MVP_top60_DNA_seed | JAK2 | 0.286986 | GBM | DK-MG | GSK2276186C | MVP_top60_DNA+RNA_seed | IGF1R | 1.494617 | GBM | D-392MG | GSK1904529A |
| MVP_top60_DNA_seed | JAK1 | 0.286986 | GBM | DK-MG | GSK2276186C | MVP_top60_DNA+RNA_seed | PDGFRA | 1.501964 | GBM | U-87-MG | Amuvatinib |
| MVP_top60_DNA_seed | JAK3 | 0.286986 | GBM | DK-MG | GSK2276186C | MVP_top60_DNA+RNA_seed | KIT | 1.501964 | GBM | U-87-MG | Amuvatinib |
| MVP_top60_DNA_seed | CDK4 | 0.290798 | GBM | U-87-MG | Ribociclib | MVP_top60_DNA+RNA_seed | AKT1 | 1.503678 | GBM | DBTRG-05MG | AKT inhibitor VIII |
| MVP_top60_DNA_seed | JAK2 | 0.29155 | GBM | D-263MG | Ruxolitinib | MVP_top60_DNA+RNA_seed | AKT1 | 1.507165 | GBM | U-87-MG | AKT inhibitor VIII |

| | | | | | | | | | | | |
|---|---|---|---|---|---|---|---|---|---|---|---|
| MVP_top60_DNA_seed | JAK1 | 0.29155 | GBM | D-263MG | Ruxolitinib | MVP_top60_DNA+RNA_seed | EGFR | 1.516762 | GBM | D-247MG | Pelitinib |
| MVP_top60_DNA_seed | EGFR | 0.293897 | GBM | SW1088 | Osimertinib | MVP_top60_DNA+RNA_seed | PIK3CA | 1.518444 | GBM | DK-MG | AZD8835 |
| MVP_top60_DNA_seed | EGFR | 0.294884 | GBM | T98G | Lapatinib | MVP_top60_DNA+RNA_seed | PIK3CD | 1.518444 | GBM | DK-MG | AZD8835 |
| MVP_top60_DNA_seed | EGFR | 0.296246 | GBM | CAS-1 | Sapitinib | MVP_top60_DNA+RNA_seed | EGFR | 1.520523 | GBM | SNB75 | Gefitinib |
| MVP_top60_DNA_seed | TP53 | 0.299196 | GBM | AM-38 | MIRA-1 | MVP_top60_DNA+RNA_seed | PDGFRA | 1.533048 | GBM | LN-229 | Masitinib |
| MVP_top60_DNA_seed | CDK4 | 0.305782 | GBM | CAS-1 | Palbociclib | MVP_top60_DNA+RNA_seed | PDGFRB | 1.533048 | GBM | LN-229 | Masitinib |
| MVP_top60_DNA_seed | TP53 | 0.3112 | GBM | D-542MG | MIRA-1 | MVP_top60_DNA+RNA_seed | KIT | 1.533048 | GBM | LN-229 | Masitinib |
| MVP_top60_DNA_seed | PIK3CA | 0.311367 | GBM | SF126 | Buparlisib | MVP_top60_DNA+RNA_seed | EGFR | 1.541826 | GBM | SW1088 | CI-1033 |
| MVP_top60_DNA_seed | PIK3CB | 0.311367 | GBM | SF126 | Buparlisib | MVP_top60_DNA+RNA_seed | IGF1R | 1.542405 | GBM | DBTRG-05MG | GSK1904529A |
| MVP_top60_DNA_seed | PIK3CD | 0.311367 | GBM | SF126 | Buparlisib | MVP_top60_DNA+RNA_seed | JAK3 | 1.543824 | GBM | CAS-1 | WHI-P97 |
| MVP_top60_DNA_seed | TP53 | 0.313675 | GBM | Becker | MIRA-1 | MVP_top60_DNA+RNA_seed | EGFR | 1.544437 | GBM | U-118-MG | CI-1033 |
| MVP_top60_DNA_seed | EP300 | 0.31425 | GBM | Becker | SGC-CBP30 | MVP_top60_DNA+RNA_seed | EGFR | 1.553352 | GBM | D-542MG | Pelitinib |
| MVP_top60_DNA_seed | JAK2 | 0.315852 | GBM | 8-MG-BA | Ruxolitinib | MVP_top60_DNA+RNA_seed | CDK4 | 1.554434 | GBM | M059J | Palbociclib |
| MVP_top60_DNA_seed | JAK1 | 0.315852 | GBM | 8-MG-BA | Ruxolitinib | MVP_top60_DNA+RNA_seed | EGFR | 1.571145 | GBM | D-247MG | CI-1033 |
| MVP_top60_DNA_seed | PIK3CA | 0.319196 | GBM | D-423MG | Alpelisib | MVP_top60_DNA+RNA_seed | AKT1 | 1.58128 | GBM | LN-229 | BAY AKT1 |
| MVP_top60_DNA_seed | EGFR | 0.327969 | GBM | D-423MG | Erlotinib | MVP_top60_DNA+RNA_seed | KIT | 1.599698 | GBM | LN-405 | Axitinib |
| MVP_top60_DNA_seed | PIK3CA | 0.328473 | GBM | SF295 | Alpelisib | MVP_top60_DNA+RNA_seed | IGF1R | 1.612647 | GBM | D-542MG | BMS-536924 |
| MVP_top60_DNA_seed | EP300 | 0.33234 | GBM | D-392MG | SGC-CBP30 | MVP_top60_DNA+RNA_seed | AKT1 | 1.617832 | GBM | D-542MG | AKT inhibitor VIII |
| MVP_top60_DNA_seed | EGFR | 0.334174 | GBM | 8-MG-BA | AZD3759 | MVP_top60_DNA+RNA_seed | JAK3 | 1.634691 | GBM | SNB75 | WHI-P97 |
| MVP_top60_DNA_seed | EGFR | 0.334651 | GBM | D-423MG | Afatinib | MVP_top60_DNA+RNA_seed | PIK3CA | 1.655623 | GBM | LNZTA3WT4 | PI3Ka_4409 |
| MVP_top60_DNA_seed | EGFR | 0.337738 | GBM | D-566MG | Osimertinib | MVP_top60_DNA+RNA_seed | PIK3CD | 1.655623 | GBM | LNZTA3WT4 | PI3Ka_4409 |
| MVP_top60_DNA_seed | PIK3CA | 0.340646 | GBM | M059J | Buparlisib | MVP_top60_DNA+RNA_seed | EP300 | 1.655741 | GBM | U-118-MG | I-CBP112 |
| MVP_top60_DNA_seed | PIK3CB | 0.340646 | GBM | M059J | Buparlisib | MVP_top60_DNA+RNA_seed | CREBBP | 1.655741 | GBM | U-118-MG | I-CBP112 |
| MVP_top60_DNA_seed | PIK3CD | 0.340646 | GBM | M059J | Buparlisib | MVP_top60_DNA+RNA_seed | IGF1R | 1.662753 | GBM | U-87-MG | BMS-536924 |
| MVP_top60_DNA_seed | JAK1 | 0.350672 | GBM | KS-1 | JAK1_8709 | MVP_top60_DNA+RNA_seed | AKT1 | 1.666308 | GBM | DBTRG-05MG | MK-2206 |
| MVP_top60_DNA_seed | PIK3CA | 0.351358 | GBM | LN-405 | AZD8186 | MVP_top60_DNA+RNA_seed | JAK2 | 1.668018 | GBM | D-263MG | Fedratinib |
| MVP_top60_DNA_seed | PIK3CB | 0.351358 | GBM | LN-405 | AZD8186 | MVP_top60_DNA+RNA_seed | KIT | 1.670904 | GBM | CAS-1 | Sorafenib |
| MVP_top60_DNA_seed | JAK1 | 0.353868 | GBM | SF268 | JAK1_8709 | MVP_top60_DNA+RNA_seed | EGFR | 1.672108 | GBM | AM-38 | AZD8931 |
| MVP_top60_DNA_seed | EGFR | 0.364198 | GBM | SF268 | Sapitinib | MVP_top60_DNA+RNA_seed | ERBB3 | 1.672108 | GBM | AM-38 | AZD8931 |
| MVP_top60_DNA_seed | EGFR | 0.364904 | GBM | D-247MG | Sapitinib | MVP_top60_DNA+RNA_seed | CDK4 | 1.678389 | GBM | D-542MG | Palbociclib |
| MVP_top60_DNA_seed | EGFR | 0.366709 | GBM | AM-38 | Gefitinib | MVP_top60_DNA+RNA_seed | CDK4 | 1.67935 | GBM | CAS-1 | CGP-082996 |
| MVP_top60_DNA_seed | PIK3CA | 0.370219 | GBM | SF268 | Buparlisib | MVP_top60_DNA+RNA_seed | AKT1 | 1.682823 | GBM | D-392MG | MK-2206 |
| MVP_top60_DNA_seed | PIK3CB | 0.370219 | GBM | SF268 | Buparlisib | MVP_top60_DNA+RNA_seed | JAK2 | 1.697654 | GBM | U-87-MG | AZD1480 |
| MVP_top60_DNA_seed | PIK3CD | 0.370219 | GBM | SF268 | Buparlisib | MVP_top60_DNA+RNA_seed | JAK1 | 1.697654 | GBM | U-87-MG | AZD1480 |
| MVP_top60_DNA_seed | CDK4 | 0.375127 | GBM | D-392MG | Ribociclib | MVP_top60_DNA+RNA_seed | JAK3 | 1.701603 | GBM | D-247MG | JAK3_7406 |
| MVP_top60_DNA_seed | JAK1 | 0.377577 | GBM | A172 | JAK1_8709 | MVP_top60_DNA+RNA_seed | JAK3 | 1.722374 | GBM | U-118-MG | JAK3_7406 |
| MVP_top60_DNA_seed | EP300 | 0.377944 | GBM | 42-MG-BA | CPI-637 | MVP_top60_DNA+RNA_seed | AKT1 | 1.726836 | GBM | U-87-MG | MK-2206 |
| MVP_top60_DNA_seed | EGFR | 0.37903 | GBM | SF295 | Gefitinib | MVP_top60_DNA+RNA_seed | JAK3 | 1.728853 | GBM | DBTRG-05MG | JAK3_7406 |
| MVP_top60_DNA_seed | EGFR | 0.379162 | GBM | YKG-1 | Erlotinib | MVP_top60_DNA+RNA_seed | AKT1 | 1.73069 | GBM | D-542MG | BAY AKT1 |
| MVP_top60_DNA_seed | EGFR | 0.382277 | GBM | D-247MG | Lapatinib | MVP_top60_DNA+RNA_seed | AKT1 | 1.745475 | GBM | D-247MG | AKT inhibitor VIII |
| MVP_top60_DNA_seed | JAK2 | 0.384084 | GBM | U-118-MG | AZ960 | MVP_top60_DNA+RNA_seed | IGF1R | 1.745932 | GBM | DK-MG | GSK1904529A |

| | | | | | | | | | | | |
|---|---|---|---|---|---|---|---|---|---|---|---|
| MVP_top60_DNA_seed | JAK3 | 0.384084 | GBM | U-118-MG | AZ960 | MVP_top60_DNA+RNA_seed | PDGFRA | 1.747289 | GBM | D-263MG | Masitinib |
| MVP_top60_DNA_seed | PIK3CA | 0.384515 | GBM | DK-MG | GNE-317 | MVP_top60_DNA+RNA_seed | PDGFRB | 1.747289 | GBM | D-263MG | Masitinib |
| MVP_top60_DNA_seed | EGFR | 0.384935 | GBM | 42-MG-BA | AZD3759 | MVP_top60_DNA+RNA_seed | KIT | 1.747289 | GBM | D-263MG | Masitinib |
| MVP_top60_DNA_seed | CDK4 | 0.385871 | GBM | SF126 | Palbociclib | MVP_top60_DNA+RNA_seed | AKT1 | 1.761224 | GBM | AM-38 | BAY AKT1 |
| MVP_top60_DNA_seed | EGFR | 0.3876 | GBM | KS-1 | Afatinib | MVP_top60_DNA+RNA_seed | AKT1 | 1.766713 | GBM | D-247MG | AKT inhibitor VIII |
| MVP_top60_DNA_seed | EGFR | 0.393013 | GBM | YH-13 | Sapitinib | MVP_top60_DNA+RNA_seed | EGFR | 1.79721 | GBM | SF295 | CI-1033 |
| MVP_top60_DNA_seed | EGFR | 0.393487 | GBM | SK-MG-1 | Osimertinib | MVP_top60_DNA+RNA_seed | PIK3CA | 1.818212 | GBM | AM-38 | AZD8835 |
| MVP_top60_DNA_seed | CDK4 | 0.394151 | GBM | D-542MG | Ribociclib | MVP_top60_DNA+RNA_seed | PIK3CD | 1.818212 | GBM | AM-38 | AZD8835 |
| MVP_top60_DNA_seed | EP300 | 0.398977 | GBM | SK-MG-1 | SGC-CBP30 | MVP_top60_DNA+RNA_seed | EGFR | 1.824225 | GBM | A172 | CI-1033 |
| MVP_top60_DNA_seed | EGFR | 0.400711 | GBM | D-245MG | Gefitinib | MVP_top60_DNA+RNA_seed | PIK3CA | 1.830186 | GBM | AM-38 | PI3Ka_4409 |
| MVP_top60_DNA_seed | EGFR | 0.402606 | GBM | SK-MG-1 | Erlotinib | MVP_top60_DNA+RNA_seed | PIK3CD | 1.830186 | GBM | AM-38 | PI3Ka_4409 |
| MVP_top60_DNA_seed | PIK3CA | 0.404616 | GBM | DK-MG | Buparlisib | MVP_top60_DNA+RNA_seed | EGFR | 1.873966 | GBM | T98G | CI-1033 |
| MVP_top60_DNA_seed | PIK3CB | 0.404616 | GBM | DK-MG | Buparlisib | MVP_top60_DNA+RNA_seed | AKT1 | 1.878771 | GBM | D-263MG | AKT inhibitor VIII |
| MVP_top60_DNA_seed | PIK3CD | 0.404616 | GBM | DK-MG | Buparlisib | MVP_top60_DNA+RNA_seed | EGFR | 1.886643 | GBM | LNZTA3WT4 | AZD8931 |
| MVP_top60_DNA_seed | CDK4 | 0.410973 | GBM | Becker | Ribociclib | MVP_top60_DNA+RNA_seed | ERBB3 | 1.886643 | GBM | LNZTA3WT4 | AZD8931 |
| MVP_top60_DNA_seed | EGFR | 0.412974 | GBM | D-566MG | Sapitinib | MVP_top60_DNA+RNA_seed | EGFR | 1.894947 | GBM | DBTRG-05MG | CI-1033 |
| MVP_top60_DNA_seed | EGFR | 0.417341 | GBM | U-118-MG | Gefitinib | MVP_top60_DNA+RNA_seed | EGFR | 1.929988 | GBM | LN-405 | CI-1033 |
| MVP_top60_DNA_seed | PIK3CA | 0.417712 | GBM | YKG-1 | AZD8186 | MVP_top60_DNA+RNA_seed | EGFR | 1.939787 | GBM | SW1088 | CUDC-101 |
| MVP_top60_DNA_seed | PIK3CB | 0.417712 | GBM | YKG-1 | AZD8186 | MVP_top60_DNA+RNA_seed | CDK4 | 1.965836 | GBM | SNB75 | Palbociclib |
| MVP_top60_DNA_seed | EP300 | 0.420427 | GBM | D-263MG | CPI-637 | MVP_top60_DNA+RNA_seed | AKT1 | 1.99002 | GBM | AM-38 | A-443654 |
| MVP_top60_DNA_seed | JAK2 | 0.422478 | GBM | YH-13 | GSK2276186C | MVP_top60_DNA+RNA_seed | EP300 | 1.999887 | GBM | DK-MG | I-CBP112 |
| MVP_top60_DNA_seed | JAK1 | 0.422478 | GBM | YH-13 | GSK2276186C | MVP_top60_DNA+RNA_seed | CREBBP | 1.999887 | GBM | DK-MG | I-CBP112 |
| MVP_top60_DNA_seed | JAK3 | 0.422478 | GBM | YH-13 | GSK2276186C | MVP_top60_DNA+RNA_seed | JAK2 | 2.010632 | GBM | U-118-MG | Fedratinib |
| MVP_top60_DNA_seed | JAK1 | 0.4246 | GBM | LN-229 | JAK1_8709 | MVP_top60_DNA+RNA_seed | AKT1 | 2.052036 | GBM | U-118-MG | AKT inhibitor VIII |
| MVP_top60_DNA_seed | EGFR | 0.431461 | GBM | LNZTA3WT4 | AZD3759 | MVP_top60_DNA+RNA_seed | EGFR | 2.206699 | GBM | LNZTA3WT4 | CUDC-101 |
| MVP_top60_DNA_seed | PIK3CA | 0.437291 | GBM | DBTRG-05MG | AZD8186 | MVP_top60_DNA+RNA_seed | KIT | 2.210286 | GBM | SNB75 | Sorafenib |
| MVP_top60_DNA_seed | PIK3CB | 0.437291 | GBM | DBTRG-05MG | AZD8186 | MVP_top60_DNA+RNA_seed | AKT1 | 2.402846 | GBM | U-87-MG | A-443654 |
| MVP_top60_DNA_seed | PIK3CA | 0.443899 | GBM | SF295 | Buparlisib | MVP_top60_DNA+RNA_seed | KIT | -2.444714 | GBM | 42-MG-BA | Motesanib |
| MVP_top60_DNA_seed | PIK3CB | 0.443899 | GBM | SF295 | Buparlisib | MVP_top60_DNA+RNA_seed | IGF1R | -2.37215 | GBM | D-423MG | GSK1904529A |
| MVP_top60_DNA_seed | PIK3CD | 0.443899 | GBM | SF295 | Buparlisib | MVP_top60_DNA+RNA_seed | PIK3CA | -1.942775 | GBM | YH-13 | AZD8186 |
| MVP_top60_DNA_seed | EGFR | 0.447714 | GBM | D-423MG | Sapitinib | MVP_top60_DNA+RNA_seed | PIK3CB | -1.942775 | GBM | YH-13 | AZD8186 |
| MVP_top60_DNA_seed | JAK1 | 0.455118 | GBM | GB-1 | JAK1_8709 | MVP_top60_DNA+RNA_seed | AKT1 | -1.89534 | GBM | D-245MG | AT13148 |
| MVP_top60_DNA_seed | EGFR | 0.455249 | GBM | M059J | AZD3759 | MVP_top60_DNA+RNA_seed | MET | -1.830462 | GBM | DK-MG | Savolitinib |
| MVP_top60_DNA_seed | TP53 | 0.47018 | GBM | D-245MG | MIRA-1 | MVP_top60_DNA+RNA_seed | KRAS | -1.755479 | GBM | SF126 | :RAS (G12C) Inhibitor-1 |
| MVP_top60_DNA_seed | PIK3CA | 0.47052 | GBM | U251 | Buparlisib | MVP_top60_DNA+RNA_seed | KRAS | -1.45397 | GBM | D-423MG | :RAS (G12C) Inhibitor-1 |
| MVP_top60_DNA_seed | PIK3CB | 0.47052 | GBM | U251 | Buparlisib | MVP_top60_DNA+RNA_seed | PIK3CA | -1.437528 | GBM | SW1088 | AZD8186 |
| MVP_top60_DNA_seed | PIK3CD | 0.47052 | GBM | U251 | Buparlisib | MVP_top60_DNA+RNA_seed | PIK3CB | -1.437528 | GBM | SW1088 | AZD8186 |
| MVP_top60_DNA_seed | JAK2 | 0.471794 | GBM | KS-1 | Ruxolitinib | MVP_top60_DNA+RNA_seed | EP300 | -1.43111 | GBM | AM-38 | SGC-CBP30 |
| MVP_top60_DNA_seed | JAK1 | 0.471794 | GBM | KS-1 | Ruxolitinib | MVP_top60_DNA+RNA_seed | CREBBP | -1.43111 | GBM | AM-38 | SGC-CBP30 |
| MVP_top60_DNA_seed | CDK4 | 0.472011 | GBM | D-392MG | Palbociclib | MVP_top60_DNA+RNA_seed | KIT | -1.42003 | GBM | D-245MG | Axitinib |
| MVP_top60_DNA_seed | EGFR | 0.475655 | GBM | A172 | Sapitinib | MVP_top60_DNA+RNA_seed | JAK2 | -1.419946 | GBM | AM-38 | Ruxolitinib |

| | | | | | | | | | | | |
|---|---|---|---|---|---|---|---|---|---|---|---|
| MVP_top60_DNA_seed | CDK4 | 0.480546 | GBM | SK-MG-1 | Palbociclib | MVP_top60_DNA+RNA_seed | JAK1 | -1.419946 | GBM | AM-38 | Ruxolitinib |
| MVP_top60_DNA_seed | EGFR | 0.4854 | GBM | KS-1 | Erlotinib | MVP_top60_DNA+RNA_seed | AKT1 | -1.4173 | GBM | D-566MG | AT13148 |
| MVP_top60_DNA_seed | JAK2 | 0.490187 | GBM | SF126 | Ruxolitinib | MVP_top60_DNA+RNA_seed | CDK4 | -1.409729 | GBM | D-245MG | Ribociclib |
| MVP_top60_DNA_seed | JAK1 | 0.490187 | GBM | SF126 | Ruxolitinib | MVP_top60_DNA+RNA_seed | AKT1 | -1.405284 | GBM | D-423MG | AT13148 |
| MVP_top60_DNA_seed | TP53 | 0.49485 | GBM | U251 | MIRA-1 | MVP_top60_DNA+RNA_seed | KIT | -1.403938 | GBM | DK-MG | Motesanib |
| MVP_top60_DNA_seed | JAK2 | 0.502099 | GBM | SK-MG-1 | Ruxolitinib | MVP_top60_DNA+RNA_seed | PIK3CA | -1.38032 | GBM | YH-13 | GNE-317 |
| MVP_top60_DNA_seed | JAK1 | 0.502099 | GBM | SK-MG-1 | Ruxolitinib | MVP_top60_DNA+RNA_seed | JAK2 | -1.352688 | GBM | A172 | JAK_8517 |
| MVP_top60_DNA_seed | EGFR | 0.503098 | GBM | D-263MG | Gefitinib | MVP_top60_DNA+RNA_seed | JAK1 | -1.352688 | GBM | A172 | JAK_8517 |
| MVP_top60_DNA_seed | CDK4 | 0.505729 | GBM | KS-1 | Palbociclib | MVP_top60_DNA+RNA_seed | JAK2 | -1.33318 | GBM | KS-1 | JAK_8517 |
| MVP_top60_DNA_seed | EGFR | 0.506116 | GBM | LN-229 | Afatinib | MVP_top60_DNA+RNA_seed | JAK1 | -1.33318 | GBM | KS-1 | JAK_8517 |
| MVP_top60_DNA_seed | EGFR | 0.509287 | GBM | U-87-MG | Osimertinib | MVP_top60_DNA+RNA_seed | PIK3CA | -1.301092 | GBM | D-247MG | Alpelisib |
| MVP_top60_DNA_seed | EGFR | 0.510367 | GBM | SNB75 | Gefitinib | MVP_top60_DNA+RNA_seed | IGF1R | -1.299163 | GBM | D-566MG | GSK1904529A |
| MVP_top60_DNA_seed | PIK3CA | 0.512704 | GBM | T98G | Alpelisib | MVP_top60_DNA+RNA_seed | PIK3CA | -1.29419 | GBM | Becker | AZD8186 |
| MVP_top60_DNA_seed | EGFR | 0.514688 | GBM | SW1088 | Afatinib | MVP_top60_DNA+RNA_seed | PIK3CB | -1.29419 | GBM | Becker | AZD8186 |
| MVP_top60_DNA_seed | CDK4 | 0.515567 | GBM | T98G | Palbociclib | MVP_top60_DNA+RNA_seed | AKT1 | -1.290316 | GBM | 42-MG-BA | GSK2110183B |
| MVP_top60_DNA_seed | EGFR | 0.520009 | GBM | LN-405 | Gefitinib | MVP_top60_DNA+RNA_seed | AKT1 | -1.285764 | GBM | CAS-1 | AT13148 |
| MVP_top60_DNA_seed | EGFR | 0.526224 | GBM | D-542MG | Afatinib | MVP_top60_DNA+RNA_seed | PIK3CA | -1.278532 | GBM | D-423MG | AZD8186 |
| MVP_top60_DNA_seed | CDK4 | 0.527481 | GBM | AM-38 | Ribociclib | MVP_top60_DNA+RNA_seed | PIK3CB | -1.278532 | GBM | D-423MG | AZD8186 |
| MVP_top60_DNA_seed | EP300 | 0.528751 | GBM | D-392MG | CPI-637 | MVP_top60_DNA+RNA_seed | AKT1 | -1.252036 | GBM | 42-MG-BA | Afuresertib |
| MVP_top60_DNA_seed | EGFR | 0.530148 | GBM | D-542MG | Sapitinib | MVP_top60_DNA+RNA_seed | PIK3CA | -1.198811 | GBM | SW1088 | GNE-317 |
| MVP_top60_DNA_seed | CDK4 | 0.533868 | GBM | LN-18 | Palbociclib | MVP_top60_DNA+RNA_seed | PIK3CA | -1.193069 | GBM | 8-MG-BA | Buparlisib |
| MVP_top60_DNA_seed | JAK2 | 0.536203 | GBM | SF268 | JAK_8517 | MVP_top60_DNA+RNA_seed | PIK3CB | -1.193069 | GBM | 8-MG-BA | Buparlisib |
| MVP_top60_DNA_seed | JAK1 | 0.536203 | GBM | SF268 | JAK_8517 | MVP_top60_DNA+RNA_seed | PIK3CD | -1.193069 | GBM | 8-MG-BA | Buparlisib |
| MVP_top60_DNA_seed | EGFR | 0.537031 | GBM | 42-MG-BA | Erlotinib | MVP_top60_DNA+RNA_seed | KIT | -1.177564 | GBM | LN-405 | Motesanib |
| MVP_top60_DNA_seed | CDK4 | 0.537833 | GBM | D-247MG | Ribociclib | MVP_top60_DNA+RNA_seed | JAK2 | -1.171273 | GBM | CAS-1 | JAK_8517 |
| MVP_top60_DNA_seed | JAK1 | 0.546057 | GBM | CAS-1 | JAK1_8709 | MVP_top60_DNA+RNA_seed | JAK1 | -1.171273 | GBM | CAS-1 | JAK_8517 |
| MVP_top60_DNA_seed | PIK3CA | 0.546515 | GBM | YKG-1 | Buparlisib | MVP_top60_DNA+RNA_seed | AKT1 | -1.149746 | GBM | SF295 | Afuresertib |
| MVP_top60_DNA_seed | PIK3CB | 0.546515 | GBM | YKG-1 | Buparlisib | MVP_top60_DNA+RNA_seed | MET | -1.12781 | GBM | 8-MG-BA | Savolitinib |
| MVP_top60_DNA_seed | PIK3CD | 0.546515 | GBM | YKG-1 | Buparlisib | MVP_top60_DNA+RNA_seed | AKT1 | -1.118439 | GBM | LN-18 | AT13148 |
| MVP_top60_DNA_seed | EGFR | 0.546951 | GBM | DK-MG | Lapatinib | MVP_top60_DNA+RNA_seed | KIT | -1.097832 | GBM | D-423MG | Sorafenib |
| MVP_top60_DNA_seed | EGFR | 0.547239 | GBM | SF268 | Lapatinib | MVP_top60_DNA+RNA_seed | TP53 | -1.090076 | GBM | LNZTA3WT4 | MIRA-1 |
| MVP_top60_DNA_seed | EP300 | 0.549773 | GBM | SF268 | CPI-637 | MVP_top60_DNA+RNA_seed | JAK1 | -1.089505 | GBM | 8-MG-BA | JAK1_8709 |
| MVP_top60_DNA_seed | CDK4 | 0.557772 | GBM | SK-MG-1 | Ribociclib | MVP_top60_DNA+RNA_seed | EP300 | -1.083943 | GBM | SF268 | SGC-CBP30 |
| MVP_top60_DNA_seed | JAK2 | 0.558113 | GBM | Becker | Ruxolitinib | MVP_top60_DNA+RNA_seed | CREBBP | -1.083943 | GBM | SF268 | SGC-CBP30 |
| MVP_top60_DNA_seed | JAK1 | 0.558113 | GBM | Becker | Ruxolitinib | MVP_top60_DNA+RNA_seed | PIK3CA | -1.074354 | GBM | 42-MG-BA | GNE-317 |
| MVP_top60_DNA_seed | EGFR | 0.56236 | GBM | 8-MG-BA | Osimertinib | MVP_top60_DNA+RNA_seed | AKT1 | -1.073083 | GBM | SW1088 | AZD5363 |
| MVP_top60_DNA_seed | EP300 | 0.570706 | GBM | LN-405 | SGC-CBP30 | MVP_top60_DNA+RNA_seed | MET | -1.071618 | GBM | U-118-MG | Savolitinib |
| MVP_top60_DNA_seed | EGFR | 0.575883 | GBM | D-542MG | AZD3759 | MVP_top60_DNA+RNA_seed | EGFR | -1.068129 | GBM | D-423MG | AZD3759 |
| MVP_top60_DNA_seed | EGFR | 0.579175 | GBM | YH-13 | Lapatinib | MVP_top60_DNA+RNA_seed | JAK2 | -1.054067 | GBM | D-423MG | Ruxolitinib |
| MVP_top60_DNA_seed | JAK2 | 0.579378 | GBM | U251 | Ruxolitinib | MVP_top60_DNA+RNA_seed | JAK1 | -1.054067 | GBM | D-423MG | Ruxolitinib |
| MVP_top60_DNA_seed | JAK1 | 0.579378 | GBM | U251 | Ruxolitinib | MVP_top60_DNA+RNA_seed | AKT1 | -1.047481 | GBM | SW1088 | AT13148 |

| | | | | | | | | | | | |
|---|---|---|---|---|---|---|---|---|---|---|---|
| MVP_top60_DNA_seed | EGFR | 0.587528 | GBM | 42-MG-BA | Gefitinib | MVP_top60_DNA+RNA_seed | AKT1 | -1.036275 | GBM | SF126 | AZD5363 |
| MVP_top60_DNA_seed | JAK1 | 0.588174 | GBM | D-263MG | JAK1_8709 | MVP_top60_DNA+RNA_seed | KRAS | -1.033108 | GBM | SW1088 | KRAS (G12C) Inhibitor-1 |
| MVP_top60_DNA_seed | JAK2 | 0.590174 | GBM | DBTRG-05MG | JAK_8517 | MVP_top60_DNA+RNA_seed | PIK3CA | -1.03274 | GBM | SK-MG-1 | AZD8186 |
| MVP_top60_DNA_seed | JAK1 | 0.590174 | GBM | DBTRG-05MG | JAK_8517 | MVP_top60_DNA+RNA_seed | PIK3CB | -1.03274 | GBM | SK-MG-1 | AZD8186 |
| MVP_top60_DNA_seed | EGFR | 0.590257 | GBM | M059J | Sapitinib | MVP_top60_DNA+RNA_seed | AKT1 | -1.025874 | GBM | 42-MG-BA | Ipatasertib |
| MVP_top60_DNA_seed | EGFR | 0.595524 | GBM | SK-MG-1 | AZD3759 | MVP_top60_DNA+RNA_seed | CDK4 | -1.023605 | GBM | YH-13 | Ribociclib |
| MVP_top60_DNA_seed | EGFR | 0.595763 | GBM | D-542MG | Erlotinib | MVP_top60_DNA+RNA_seed | CDK4 | -1.003435 | GBM | A172 | Ribociclib |
| MVP_top60_DNA_seed | EGFR | 0.595809 | GBM | D-245MG | Lapatinib | MVP_top60_DNA+RNA_seed | AKT1 | -1.00225 | GBM | 42-MG-BA | AZD5363 |
| MVP_top60_DNA_seed | JAK2 | 0.601885 | GBM | U-87-MG | AZ960 | MVP_top60_DNA+RNA_seed | AKT1 | -0.995681 | GBM | 8-MG-BA | Afuresertib |
| MVP_top60_DNA_seed | JAK3 | 0.601885 | GBM | U-87-MG | AZ960 | MVP_top60_DNA+RNA_seed | PIK3CA | -0.987237 | GBM | D-542MG | AZD8186 |
| MVP_top60_DNA_seed | JAK2 | 0.603406 | GBM | SF268 | GSK2276186C | MVP_top60_DNA+RNA_seed | PIK3CB | -0.987237 | GBM | D-542MG | AZD8186 |
| MVP_top60_DNA_seed | JAK1 | 0.603406 | GBM | SF268 | GSK2276186C | MVP_top60_DNA+RNA_seed | PIK3CA | -0.98207 | GBM | D-423MG | GNE-317 |
| MVP_top60_DNA_seed | JAK3 | 0.603406 | GBM | SF268 | GSK2276186C | MVP_top60_DNA+RNA_seed | EP300 | -0.978372 | GBM | 42-MG-BA | SGC-CBP30 |
| MVP_top60_DNA_seed | CDK4 | 0.605292 | GBM | GB-1 | Palbociclib | MVP_top60_DNA+RNA_seed | CREBBP | -0.978372 | GBM | 42-MG-BA | SGC-CBP30 |
| MVP_top60_DNA_seed | JAK2 | 0.611251 | GBM | LN-18 | GSK2276186C | MVP_top60_DNA+RNA_seed | IGF1R | -0.977713 | GBM | 8-MG-BA | NVP-ADW742 |
| MVP_top60_DNA_seed | JAK1 | 0.611251 | GBM | LN-18 | GSK2276186C | MVP_top60_DNA+RNA_seed | PIK3CA | -0.967452 | GBM | A172 | Buparlisib |
| MVP_top60_DNA_seed | JAK3 | 0.611251 | GBM | LN-18 | GSK2276186C | MVP_top60_DNA+RNA_seed | PIK3CB | -0.967452 | GBM | A172 | Buparlisib |
| MVP_top60_DNA_seed | CDK4 | 0.61874 | GBM | YKG-1 | Ribociclib | MVP_top60_DNA+RNA_seed | PIK3CD | -0.967452 | GBM | A172 | Buparlisib |
| MVP_top60_DNA_seed | EGFR | 0.621027 | GBM | LN-18 | Gefitinib | MVP_top60_DNA+RNA_seed | KRAS | -0.963113 | GBM | 8-MG-BA | KRAS (G12C) Inhibitor-1 |
| MVP_top60_DNA_seed | EP300 | 0.62222 | GBM | LN-18 | CPI-637 | MVP_top60_DNA+RNA_seed | TP53 | -0.962093 | GBM | D-566MG | MIRA-1 |
| MVP_top60_DNA_seed | EGFR | 0.627379 | GBM | Becker | Osimertinib | MVP_top60_DNA+RNA_seed | JAK2 | -0.961467 | GBM | T98G | GSK2276186C |
| MVP_top60_DNA_seed | EGFR | 0.628974 | GBM | GB-1 | Sapitinib | MVP_top60_DNA+RNA_seed | JAK1 | -0.961467 | GBM | T98G | GSK2276186C |
| MVP_top60_DNA_seed | EGFR | 0.629459 | GBM | Becker | Afatinib | MVP_top60_DNA+RNA_seed | JAK3 | -0.961467 | GBM | T98G | GSK2276186C |
| MVP_top60_DNA_seed | JAK1 | 0.631924 | GBM | T98G | JAK1_8709 | MVP_top60_DNA+RNA_seed | JAK2 | -0.953326 | GBM | D-245MG | AZ960 |
| MVP_top60_DNA_seed | EGFR | 0.635474 | GBM | D-247MG | Afatinib | MVP_top60_DNA+RNA_seed | JAK3 | -0.953326 | GBM | D-245MG | AZ960 |
| MVP_top60_DNA_seed | EP300 | 0.638921 | GBM | U-118-MG | SGC-CBP30 | MVP_top60_DNA+RNA_seed | JAK2 | -0.942085 | GBM | D-566MG | AZ960 |
| MVP_top60_DNA_seed | EGFR | 0.639898 | GBM | T98G | AZD3759 | MVP_top60_DNA+RNA_seed | JAK3 | -0.942085 | GBM | D-566MG | AZ960 |
| MVP_top60_DNA_seed | TP53 | 0.645376 | GBM | LN-229 | MIRA-1 | MVP_top60_DNA+RNA_seed | JAK2 | -0.939447 | GBM | T98G | AZ960 |
| MVP_top60_DNA_seed | TP53 | 0.648846 | GBM | U-118-MG | MIRA-1 | MVP_top60_DNA+RNA_seed | JAK3 | -0.939447 | GBM | T98G | AZ960 |
| MVP_top60_DNA_seed | EP300 | 0.651391 | GBM | T98G | SGC-CBP30 | MVP_top60_DNA+RNA_seed | MET | -0.932644 | GBM | Becker | Savolitinib |
| MVP_top60_DNA_seed | EGFR | 0.662326 | GBM | YH-13 | AZD3759 | MVP_top60_DNA+RNA_seed | JAK2 | -0.92737 | GBM | YH-13 | JAK_8517 |
| MVP_top60_DNA_seed | EP300 | 0.662332 | GBM | 8-MG-BA | CPI-637 | MVP_top60_DNA+RNA_seed | JAK1 | -0.92737 | GBM | YH-13 | JAK_8517 |
| MVP_top60_DNA_seed | CDK4 | 0.663048 | GBM | SW1088 | Ribociclib | MVP_top60_DNA+RNA_seed | PIK3CA | -0.927 | GBM | A172 | GNE-317 |
| MVP_top60_DNA_seed | JAK2 | 0.664893 | GBM | AM-38 | GSK2276186C | MVP_top60_DNA+RNA_seed | PIK3CA | -0.90808 | GBM | SK-MG-1 | Buparlisib |
| MVP_top60_DNA_seed | JAK1 | 0.664893 | GBM | AM-38 | GSK2276186C | MVP_top60_DNA+RNA_seed | PIK3CB | -0.90808 | GBM | SK-MG-1 | Buparlisib |
| MVP_top60_DNA_seed | JAK3 | 0.664893 | GBM | AM-38 | GSK2276186C | MVP_top60_DNA+RNA_seed | PIK3CD | -0.90808 | GBM | SK-MG-1 | Buparlisib |
| MVP_top60_DNA_seed | EGFR | 0.666117 | GBM | DBTRG-05MG | Erlotinib | MVP_top60_DNA+RNA_seed | JAK2 | -0.904073 | GBM | 42-MG-BA | GSK2276186C |
| MVP_top60_DNA_seed | EGFR | 0.667379 | GBM | AM-38 | Osimertinib | MVP_top60_DNA+RNA_seed | JAK1 | -0.904073 | GBM | 42-MG-BA | GSK2276186C |
| MVP_top60_DNA_seed | TP53 | 0.673706 | GBM | D-392MG | MIRA-1 | MVP_top60_DNA+RNA_seed | JAK3 | -0.904073 | GBM | 42-MG-BA | GSK2276186C |
| MVP_top60_DNA_seed | EGFR | 0.686248 | GBM | AM-38 | Afatinib | MVP_top60_DNA+RNA_seed | AKT1 | -0.889507 | GBM | SF295 | AZD5363 |
| MVP_top60_DNA_seed | EGFR | 0.688348 | GBM | CAS-1 | Osimertinib | MVP_top60_DNA+RNA_seed | AKT1 | -0.880307 | GBM | SF126 | Afuresertib |

| | | | | | | | | | | | |
|---|---|---|---|---|---|---|---|---|---|---|---|
| MVP_top60_DNA_seed | PIK3CA | 0.689663 | GBM | SF268 | GNE-317 | MVP_top60_DNA+RNA_seed | AKT1 | -0.878073 | GBM | U-118-MG | AZD5363 |
| MVP_top60_DNA_seed | CDK4 | 0.691742 | GBM | M059J | Ribociclib | MVP_top60_DNA+RNA_seed | PIK3CA | -0.874547 | GBM | 8-MG-BA | GNE-317 |
| MVP_top60_DNA_seed | EGFR | 0.702379 | GBM | D-392MG | Osimertinib | MVP_top60_DNA+RNA_seed | PIK3CA | -0.871281 | GBM | 42-MG-BA | AZD8186 |
| MVP_top60_DNA_seed | JAK2 | 0.70349 | GBM | DBTRG-05MG | AZ960 | MVP_top60_DNA+RNA_seed | PIK3CB | -0.871281 | GBM | 42-MG-BA | AZD8186 |
| MVP_top60_DNA_seed | JAK3 | 0.70349 | GBM | DBTRG-05MG | AZ960 | MVP_top60_DNA+RNA_seed | AKT1 | -0.87011 | GBM | SK-MG-1 | AZD5363 |
| MVP_top60_DNA_seed | CDK4 | 0.703697 | GBM | U251 | Ribociclib | MVP_top60_DNA+RNA_seed | JAK2 | -0.865471 | GBM | A172 | AZ960 |
| MVP_top60_DNA_seed | CDK4 | 0.704074 | GBM | SNB75 | Ribociclib | MVP_top60_DNA+RNA_seed | JAK3 | -0.865471 | GBM | A172 | AZ960 |
| MVP_top60_DNA_seed | EGFR | 0.706663 | GBM | M059J | Osimertinib | MVP_top60_DNA+RNA_seed | PIK3CB | -0.862436 | GBM | D-566MG | AZD6482 |
| MVP_top60_DNA_seed | EP300 | 0.71015 | GBM | D-263MG | SGC-CBP30 | MVP_top60_DNA+RNA_seed | PIK3CA | -0.856878 | GBM | SF295 | AZD8186 |
| MVP_top60_DNA_seed | TP53 | 0.714854 | GBM | U-87-MG | MIRA-1 | MVP_top60_DNA+RNA_seed | PIK3CB | -0.856878 | GBM | SF295 | AZD8186 |
| MVP_top60_DNA_seed | EGFR | 0.71768 | GBM | T98G | Afatinib | MVP_top60_DNA+RNA_seed | EP300 | -0.853833 | GBM | YH-13 | CPI-637 |
| MVP_top60_DNA_seed | JAK2 | 0.720744 | GBM | SK-MG-1 | GSK2276186C | MVP_top60_DNA+RNA_seed | EGFR | -0.844443 | GBM | LN-229 | Erlotinib |
| MVP_top60_DNA_seed | JAK1 | 0.720744 | GBM | SK-MG-1 | GSK2276186C | MVP_top60_DNA+RNA_seed | AKT1 | -0.842198 | GBM | D-263MG | AT13148 |
| MVP_top60_DNA_seed | JAK3 | 0.720744 | GBM | SK-MG-1 | GSK2276186C | MVP_top60_DNA+RNA_seed | AKT1 | -0.84149 | GBM | 8-MG-BA | AT13148 |
| MVP_top60_DNA_seed | PIK3CA | 0.722615 | GBM | SF268 | Alpelisib | MVP_top60_DNA+RNA_seed | AKT1 | -0.840815 | GBM | SW1088 | Ipatasertib |
| MVP_top60_DNA_seed | EGFR | 0.722859 | GBM | SW1088 | Lapatinib | MVP_top60_DNA+RNA_seed | AKT1 | -0.833797 | GBM | 42-MG-BA | MK-2206 |
| MVP_top60_DNA_seed | PIK3CA | 0.724599 | GBM | DBTRG-05MG | Buparlisib | MVP_top60_DNA+RNA_seed | AKT1 | -0.832999 | GBM | SK-MG-1 | Afuresertib |
| MVP_top60_DNA_seed | PIK3CB | 0.724599 | GBM | DBTRG-05MG | Buparlisib | MVP_top60_DNA+RNA_seed | PIK3CA | -0.832556 | GBM | YH-13 | Buparlisib |
| MVP_top60_DNA_seed | PIK3CD | 0.724599 | GBM | DBTRG-05MG | Buparlisib | MVP_top60_DNA+RNA_seed | PIK3CB | -0.832556 | GBM | YH-13 | Buparlisib |
| MVP_top60_DNA_seed | PIK3CA | 0.725636 | GBM | KS-1 | AZD8186 | MVP_top60_DNA+RNA_seed | PIK3CD | -0.832556 | GBM | YH-13 | Buparlisib |
| MVP_top60_DNA_seed | PIK3CB | 0.725636 | GBM | KS-1 | AZD8186 | MVP_top60_DNA+RNA_seed | JAK2 | -0.831074 | GBM | 42-MG-BA | JAK_8517 |
| MVP_top60_DNA_seed | EGFR | 0.7261 | GBM | Becker | Gefitinib | MVP_top60_DNA+RNA_seed | JAK1 | -0.831074 | GBM | 42-MG-BA | JAK_8517 |
| MVP_top60_DNA_seed | PIK3CA | 0.731918 | GBM | LN-229 | Buparlisib | MVP_top60_DNA+RNA_seed | JAK2 | -0.826606 | GBM | YH-13 | AZ960 |
| MVP_top60_DNA_seed | PIK3CB | 0.731918 | GBM | LN-229 | Buparlisib | MVP_top60_DNA+RNA_seed | JAK3 | -0.826606 | GBM | YH-13 | AZ960 |
| MVP_top60_DNA_seed | PIK3CD | 0.731918 | GBM | LN-229 | Buparlisib | MVP_top60_DNA+RNA_seed | IGF1R | -0.807963 | GBM | LN-18 | BMS-536924 |
| MVP_top60_DNA_seed | EP300 | 0.733056 | GBM | T98G | CPI-637 | MVP_top60_DNA+RNA_seed | KIT | -0.806644 | GBM | 42-MG-BA | Axitinib |
| MVP_top60_DNA_seed | JAK2 | 0.733416 | GBM | LN-405 | GSK2276186C | MVP_top60_DNA+RNA_seed | KRAS | -0.805941 | GBM | D-566MG | KRAS (G12C) Inhibitor-1 |
| MVP_top60_DNA_seed | JAK1 | 0.733416 | GBM | LN-405 | GSK2276186C | MVP_top60_DNA+RNA_seed | PIK3CB | -0.805151 | GBM | D-423MG | AZD6482 |
| MVP_top60_DNA_seed | JAK3 | 0.733416 | GBM | LN-405 | GSK2276186C | MVP_top60_DNA+RNA_seed | EP300 | -0.804523 | GBM | KS-1 | CPI-637 |
| MVP_top60_DNA_seed | EGFR | 0.738527 | GBM | LN-229 | Lapatinib | MVP_top60_DNA+RNA_seed | EP300 | -0.799497 | GBM | U-87-MG | SGC-CBP30 |
| MVP_top60_DNA_seed | CDK4 | 0.744679 | GBM | 42-MG-BA | Palbociclib | MVP_top60_DNA+RNA_seed | CREBBP | -0.799497 | GBM | U-87-MG | SGC-CBP30 |
| MVP_top60_DNA_seed | PIK3CA | 0.746592 | GBM | AM-38 | Alpelisib | MVP_top60_DNA+RNA_seed | JAK2 | -0.794066 | GBM | SF126 | AZ960 |
| MVP_top60_DNA_seed | EP300 | 0.746626 | GBM | YH-13 | SGC-CBP30 | MVP_top60_DNA+RNA_seed | JAK3 | -0.794066 | GBM | SF126 | AZ960 |
| MVP_top60_DNA_seed | EGFR | 0.746715 | GBM | CAS-1 | Gefitinib | MVP_top60_DNA+RNA_seed | AKT1 | -0.785484 | GBM | SW1088 | MK-2206 |
| MVP_top60_DNA_seed | JAK2 | 0.756556 | GBM | U-118-MG | Ruxolitinib | MVP_top60_DNA+RNA_seed | EGFR | -0.783918 | GBM | 8-MG-BA | Lapatinib |
| MVP_top60_DNA_seed | JAK1 | 0.756556 | GBM | U-118-MG | Ruxolitinib | MVP_top60_DNA+RNA_seed | JAK2 | -0.780905 | GBM | LN-229 | Ruxolitinib |
| MVP_top60_DNA_seed | JAK2 | 0.758348 | GBM | D-263MG | GSK2276186C | MVP_top60_DNA+RNA_seed | JAK1 | -0.780905 | GBM | LN-229 | Ruxolitinib |
| MVP_top60_DNA_seed | JAK1 | 0.758348 | GBM | D-263MG | GSK2276186C | MVP_top60_DNA+RNA_seed | IGF1R | -0.776813 | GBM | LN-18 | GSK1904529A |
| MVP_top60_DNA_seed | JAK3 | 0.758348 | GBM | D-263MG | GSK2276186C | MVP_top60_DNA+RNA_seed | JAK2 | -0.774509 | GBM | SW1088 | JAK_8517 |
| MVP_top60_DNA_seed | EGFR | 0.758851 | GBM | SK-MG-1 | Lapatinib | MVP_top60_DNA+RNA_seed | JAK1 | -0.774509 | GBM | SW1088 | JAK_8517 |
| MVP_top60_DNA_seed | EGFR | 0.760432 | GBM | D-392MG | AZD3759 | MVP_top60_DNA+RNA_seed | AKT1 | -0.773699 | GBM | SF126 | AT13148 |

| | | | | | | | | | | | |
|---|---|---|---|---|---|---|---|---|---|---|---|
| MVP_top60_DNA_seed | PIK3CA | 0.762769 | GBM | LN-405 | Alpelisib | MVP_top60_DNA+RNA_seed | KIT | -0.768842 | GBM | M059J | Axitinib |
| MVP_top60_DNA_seed | EP300 | 0.777915 | GBM | LN-405 | CPI-637 | MVP_top60_DNA+RNA_seed | JAK2 | -0.763715 | GBM | T98G | JAK_8517 |
| MVP_top60_DNA_seed | EGFR | 0.780566 | GBM | D-392MG | Afatinib | MVP_top60_DNA+RNA_seed | JAK1 | -0.763715 | GBM | T98G | JAK_8517 |
| MVP_top60_DNA_seed | EGFR | 0.787238 | GBM | SW1088 | AZD3759 | MVP_top60_DNA+RNA_seed | EP300 | -0.763421 | GBM | U-87-MG | CPI-637 |
| MVP_top60_DNA_seed | CDK4 | 0.792326 | GBM | SNB75 | Palbociclib | MVP_top60_DNA+RNA_seed | JAK2 | -0.760054 | GBM | 8-MG-BA | GSK2276186C |
| MVP_top60_DNA_seed | JAK1 | 0.793498 | GBM | U251 | JAK1_8709 | MVP_top60_DNA+RNA_seed | JAK1 | -0.760054 | GBM | 8-MG-BA | GSK2276186C |
| MVP_top60_DNA_seed | EGFR | 0.803738 | GBM | CAS-1 | Afatinib | MVP_top60_DNA+RNA_seed | JAK3 | -0.760054 | GBM | 8-MG-BA | GSK2276186C |
| MVP_top60_DNA_seed | EGFR | 0.809603 | GBM | LN-405 | Erlotinib | MVP_top60_DNA+RNA_seed | KIT | -0.756471 | GBM | A172 | Axitinib |
| MVP_top60_DNA_seed | EGFR | 0.810487 | GBM | U251 | Gefitinib | MVP_top60_DNA+RNA_seed | AKT1 | -0.750838 | GBM | SW1088 | Uprosertib |
| MVP_top60_DNA_seed | JAK1 | 0.811211 | GBM | SK-MG-1 | JAK1_8709 | MVP_top60_DNA+RNA_seed | PIK3CA | -0.750057 | GBM | YH-13 | Alpelisib |
| MVP_top60_DNA_seed | EGFR | 0.811865 | GBM | SNB75 | Sapitinib | MVP_top60_DNA+RNA_seed | AKT1 | -0.747736 | GBM | 8-MG-BA | GSK2110183B |
| MVP_top60_DNA_seed | EGFR | 0.820737 | GBM | D-245MG | AZD3759 | MVP_top60_DNA+RNA_seed | EGFR | -0.74154 | GBM | D-566MG | Lapatinib |
| MVP_top60_DNA_seed | EGFR | 0.822151 | GBM | D-247MG | Gefitinib | MVP_top60_DNA+RNA_seed | JAK2 | -0.737726 | GBM | KS-1 | AZ960 |
| MVP_top60_DNA_seed | JAK2 | 0.823085 | GBM | YH-13 | Ruxolitinib | MVP_top60_DNA+RNA_seed | JAK3 | -0.737726 | GBM | KS-1 | AZ960 |
| MVP_top60_DNA_seed | JAK1 | 0.823085 | GBM | YH-13 | Ruxolitinib | MVP_top60_DNA+RNA_seed | AKT1 | -0.727618 | GBM | 42-MG-BA | Uprosertib |
| MVP_top60_DNA_seed | EGFR | 0.823413 | GBM | KS-1 | Osimertinib | MVP_top60_DNA+RNA_seed | PIK3CA | -0.725529 | GBM | SF295 | GNE-317 |
| MVP_top60_DNA_seed | CDK4 | 0.825781 | GBM | T98G | Ribociclib | MVP_top60_DNA+RNA_seed | PIK3CA | -0.719633 | GBM | SF126 | Alpelisib |
| MVP_top60_DNA_seed | EGFR | 0.835318 | GBM | YH-13 | Gefitinib | MVP_top60_DNA+RNA_seed | AKT1 | -0.717833 | GBM | SF268 | AT13148 |
| MVP_top60_DNA_seed | PIK3CA | 0.837061 | GBM | LN-229 | Alpelisib | MVP_top60_DNA+RNA_seed | EP300 | -0.716364 | GBM | KS-1 | SGC-CBP30 |
| MVP_top60_DNA_seed | JAK2 | 0.843237 | GBM | U-118-MG | GSK2276186C | MVP_top60_DNA+RNA_seed | CREBBP | -0.716364 | GBM | KS-1 | SGC-CBP30 |
| MVP_top60_DNA_seed | JAK1 | 0.843237 | GBM | U-118-MG | GSK2276186C | MVP_top60_DNA+RNA_seed | PIK3CA | -0.715394 | GBM | D-247MG | AZD8186 |
| MVP_top60_DNA_seed | JAK3 | 0.843237 | GBM | U-118-MG | GSK2276186C | MVP_top60_DNA+RNA_seed | PIK3CB | -0.715394 | GBM | D-247MG | AZD8186 |
| MVP_top60_DNA_seed | PIK3CA | 0.846997 | GBM | M059J | Alpelisib | MVP_top60_DNA+RNA_seed | KIT | -0.71489 | GBM | SW1088 | Axitinib |
| MVP_top60_DNA_seed | EGFR | 0.849635 | GBM | T98G | Sapitinib | MVP_top60_DNA+RNA_seed | AKT1 | -0.709578 | GBM | U-118-MG | Afuresertib |
| MVP_top60_DNA_seed | EP300 | 0.850144 | GBM | A172 | SGC-CBP30 | MVP_top60_DNA+RNA_seed | MET | -0.707835 | GBM | 42-MG-BA | Savolitinib |
| MVP_top60_DNA_seed | TP53 | 0.851192 | GBM | SK-MG-1 | MIRA-1 | MVP_top60_DNA+RNA_seed | PIK3CA | -0.702953 | GBM | LN-18 | AZD8186 |
| MVP_top60_DNA_seed | EP300 | 0.854304 | GBM | LNZTA3WT4 | CPI-637 | MVP_top60_DNA+RNA_seed | PIK3CB | -0.702953 | GBM | LN-18 | AZD8186 |
| MVP_top60_DNA_seed | EGFR | 0.860964 | GBM | SF295 | Sapitinib | MVP_top60_DNA+RNA_seed | KIT | -0.702457 | GBM | D-566MG | Sorafenib |
| MVP_top60_DNA_seed | EGFR | 0.871233 | GBM | LN-405 | AZD3759 | MVP_top60_DNA+RNA_seed | PIK3CA | -0.696871 | GBM | D-245MG | Alpelisib |
| MVP_top60_DNA_seed | TP53 | 0.871374 | GBM | SF268 | MIRA-1 | MVP_top60_DNA+RNA_seed | EGFR | -0.69609 | GBM | LN-18 | Lapatinib |
| MVP_top60_DNA_seed | EGFR | 0.87195 | GBM | D-263MG | Erlotinib | MVP_top60_DNA+RNA_seed | JAK2 | -0.695968 | GBM | 42-MG-BA | Ruxolitinib |
| MVP_top60_DNA_seed | JAK2 | 0.874697 | GBM | M059J | GSK2276186C | MVP_top60_DNA+RNA_seed | JAK1 | -0.695968 | GBM | 42-MG-BA | Ruxolitinib |
| MVP_top60_DNA_seed | JAK1 | 0.874697 | GBM | M059J | GSK2276186C | MVP_top60_DNA+RNA_seed | AKT1 | -0.694469 | GBM | LN-405 | GSK2110183B |
| MVP_top60_DNA_seed | JAK3 | 0.874697 | GBM | M059J | GSK2276186C | MVP_top60_DNA+RNA_seed | EGFR | -0.690795 | GBM | SF126 | Osimertinib |
| MVP_top60_DNA_seed | PIK3CA | 0.879163 | GBM | CAS-1 | Alpelisib | MVP_top60_DNA+RNA_seed | PIK3CB | -0.680806 | GBM | D-247MG | AZD6482 |
| MVP_top60_DNA_seed | PIK3CA | 0.879601 | GBM | LN-229 | AZD8186 | MVP_top60_DNA+RNA_seed | AKT1 | -0.679377 | GBM | SF126 | MK-2206 |
| MVP_top60_DNA_seed | PIK3CB | 0.879601 | GBM | LN-229 | AZD8186 | MVP_top60_DNA+RNA_seed | IGF1R | -0.678936 | GBM | 42-MG-BA | NVP-ADW742 |
| MVP_top60_DNA_seed | PIK3CA | 0.879981 | GBM | D-263MG | AZD8186 | MVP_top60_DNA+RNA_seed | KIT | -0.678608 | GBM | DK-MG | Axitinib |
| MVP_top60_DNA_seed | PIK3CB | 0.879981 | GBM | D-263MG | AZD8186 | MVP_top60_DNA+RNA_seed | CDK4 | -0.676501 | GBM | LNZTA3WT4 | Palbociclib |
| MVP_top60_DNA_seed | PIK3CA | 0.883991 | GBM | D-392MG | Alpelisib | MVP_top60_DNA+RNA_seed | KIT | -0.674652 | GBM | 8-MG-BA | Sorafenib |
| MVP_top60_DNA_seed | EGFR | 0.884814 | GBM | D-392MG | Gefitinib | MVP_top60_DNA+RNA_seed | IGF1R | -0.672119 | GBM | GB-1 | BMS-536924 |

| | | | | | | | | | | | |
|---|---|---|---|---|---|---|---|---|---|---|---|
| MVP_top60_DNA_seed | EGFR | 0.884914 | GBM | KS-1 | Gefitinib | MVP_top60_DNA+RNA_seed | JAK2 | -0.671886 | GBM | DK-MG | JAK_8517 |
| MVP_top60_DNA_seed | PIK3CA | 0.885395 | GBM | SNB75 | Buparlisib | MVP_top60_DNA+RNA_seed | JAK1 | -0.671886 | GBM | DK-MG | JAK_8517 |
| MVP_top60_DNA_seed | PIK3CB | 0.885395 | GBM | SNB75 | Buparlisib | MVP_top60_DNA+RNA_seed | PIK3CA | -0.667689 | GBM | SNB75 | AZD8186 |
| MVP_top60_DNA_seed | PIK3CD | 0.885395 | GBM | SNB75 | Buparlisib | MVP_top60_DNA+RNA_seed | PIK3CB | -0.667689 | GBM | SNB75 | AZD8186 |
| MVP_top60_DNA_seed | EGFR | 0.888304 | GBM | SK-MG-1 | Sapitinib | MVP_top60_DNA+RNA_seed | PIK3CA | -0.66427 | GBM | 42-MG-BA | Buparlisib |
| MVP_top60_DNA_seed | JAK2 | 0.895027 | GBM | YKG-1 | Ruxolitinib | MVP_top60_DNA+RNA_seed | PIK3CB | -0.66427 | GBM | 42-MG-BA | Buparlisib |
| MVP_top60_DNA_seed | JAK1 | 0.895027 | GBM | YKG-1 | Ruxolitinib | MVP_top60_DNA+RNA_seed | PIK3CD | -0.66427 | GBM | 42-MG-BA | Buparlisib |
| MVP_top60_DNA_seed | EGFR | 0.895519 | GBM | SF126 | Erlotinib | MVP_top60_DNA+RNA_seed | CDK4 | -0.663907 | GBM | DK-MG | Ribociclib |
| MVP_top60_DNA_seed | CDK4 | 0.899147 | GBM | SF268 | Palbociclib | MVP_top60_DNA+RNA_seed | MET | -0.663646 | GBM | SK-MG-1 | Savolitinib |
| MVP_top60_DNA_seed | JAK2 | 0.907261 | GBM | GB-1 | GSK2276186C | MVP_top60_DNA+RNA_seed | KIT | -0.660411 | GBM | LN-18 | Sorafenib |
| MVP_top60_DNA_seed | JAK1 | 0.907261 | GBM | GB-1 | GSK2276186C | MVP_top60_DNA+RNA_seed | AKT1 | -0.657316 | GBM | SK-MG-1 | AT13148 |
| MVP_top60_DNA_seed | JAK3 | 0.907261 | GBM | GB-1 | GSK2276186C | MVP_top60_DNA+RNA_seed | PIK3CB | -0.657151 | GBM | D-542MG | AZD6482 |
| MVP_top60_DNA_seed | EGFR | 0.908304 | GBM | SF268 | AZD3759 | MVP_top60_DNA+RNA_seed | PIK3CA | -0.654801 | GBM | D-566MG | AZD8186 |
| MVP_top60_DNA_seed | EGFR | 0.910745 | GBM | LNZTA3WT4 | Sapitinib | MVP_top60_DNA+RNA_seed | PIK3CB | -0.654801 | GBM | D-566MG | AZD8186 |
| MVP_top60_DNA_seed | CDK4 | 0.917941 | GBM | LN-229 | Ribociclib | MVP_top60_DNA+RNA_seed | JAK2 | -0.653326 | GBM | M059J | JAK_8517 |
| MVP_top60_DNA_seed | PIK3CA | 0.926442 | GBM | LN-18 | Alpelisib | MVP_top60_DNA+RNA_seed | JAK1 | -0.653326 | GBM | M059J | JAK_8517 |
| MVP_top60_DNA_seed | JAK2 | 0.926962 | GBM | D-542MG | Ruxolitinib | MVP_top60_DNA+RNA_seed | AKT1 | -0.650979 | GBM | 42-MG-BA | Uprosertib |
| MVP_top60_DNA_seed | JAK1 | 0.926962 | GBM | D-542MG | Ruxolitinib | MVP_top60_DNA+RNA_seed | EGFR | -0.641273 | GBM | D-423MG | Lapatinib |
| MVP_top60_DNA_seed | EGFR | 0.931245 | GBM | U251 | Osimertinib | MVP_top60_DNA+RNA_seed | AKT1 | -0.636361 | GBM | YH-13 | Ipatasertib |
| MVP_top60_DNA_seed | EGFR | 0.931524 | GBM | SF126 | Afatinib | MVP_top60_DNA+RNA_seed | AKT1 | -0.636361 | GBM | SW1088 | Uprosertib |
| MVP_top60_DNA_seed | EGFR | 0.932418 | GBM | CAS-1 | Erlotinib | MVP_top60_DNA+RNA_seed | JAK2 | -0.634959 | GBM | D-263MG | JAK_8517 |
| MVP_top60_DNA_seed | EGFR | 0.938563 | GBM | T98G | Erlotinib | MVP_top60_DNA+RNA_seed | JAK1 | -0.634959 | GBM | D-263MG | JAK_8517 |
| MVP_top60_DNA_seed | PIK3CA | 0.938589 | GBM | DK-MG | Alpelisib | MVP_top60_DNA+RNA_seed | KIT | -0.634731 | GBM | D-392MG | Axitinib |
| MVP_top60_DNA_seed | EGFR | 0.941274 | GBM | D-566MG | Afatinib | MVP_top60_DNA+RNA_seed | AKT1 | -0.634659 | GBM | SK-MG-1 | MK-2206 |
| MVP_top60_DNA_seed | EGFR | 0.945968 | GBM | D-247MG | Erlotinib | MVP_top60_DNA+RNA_seed | PIK3CA | -0.634461 | GBM | SK-MG-1 | GNE-317 |
| MVP_top60_DNA_seed | EGFR | 0.949568 | GBM | U-118-MG | Afatinib | MVP_top60_DNA+RNA_seed | AKT1 | -0.630079 | GBM | YH-13 | MK-2206 |
| MVP_top60_DNA_seed | JAK2 | 0.954454 | GBM | LN-229 | GSK2276186C | MVP_top60_DNA+RNA_seed | JAK2 | -0.629177 | GBM | SNB75 | AZ960 |
| MVP_top60_DNA_seed | JAK1 | 0.954454 | GBM | LN-229 | GSK2276186C | MVP_top60_DNA+RNA_seed | JAK3 | -0.629177 | GBM | SNB75 | AZ960 |
| MVP_top60_DNA_seed | JAK3 | 0.954454 | GBM | LN-229 | GSK2276186C | MVP_top60_DNA+RNA_seed | EP300 | -0.628276 | GBM | LN-18 | SGC-CBP30 |
| MVP_top60_DNA_seed | EGFR | 0.955973 | GBM | T98G | Osimertinib | MVP_top60_DNA+RNA_seed | CREBBP | -0.628276 | GBM | LN-18 | SGC-CBP30 |
| MVP_top60_DNA_seed | EGFR | 0.956074 | GBM | D-263MG | Sapitinib | MVP_top60_DNA+RNA_seed | JAK2 | -0.624481 | GBM | U-87-MG | Ruxolitinib |
| MVP_top60_DNA_seed | JAK2 | 0.957853 | GBM | U-87-MG | GSK2276186C | MVP_top60_DNA+RNA_seed | JAK1 | -0.624481 | GBM | U-87-MG | Ruxolitinib |
| MVP_top60_DNA_seed | JAK1 | 0.957853 | GBM | U-87-MG | GSK2276186C | MVP_top60_DNA+RNA_seed | IGF1R | -0.624332 | GBM | KS-1 | BMS-536924 |
| MVP_top60_DNA_seed | JAK3 | 0.957853 | GBM | U-87-MG | GSK2276186C | MVP_top60_DNA+RNA_seed | IGF1R | -0.622476 | GBM | D-566MG | BMS-754807 |
| MVP_top60_DNA_seed | EGFR | 0.965185 | GBM | SNB75 | Lapatinib | MVP_top60_DNA+RNA_seed | KIT | -0.615117 | GBM | SF126 | Axitinib |
| MVP_top60_DNA_seed | JAK2 | 0.965438 | GBM | LN-405 | AZ960 | MVP_top60_DNA+RNA_seed | PIK3CA | -0.608372 | GBM | LN-18 | Buparlisib |
| MVP_top60_DNA_seed | JAK3 | 0.965438 | GBM | LN-405 | AZ960 | MVP_top60_DNA+RNA_seed | PIK3CB | -0.608372 | GBM | LN-18 | Buparlisib |
| MVP_top60_DNA_seed | EGFR | 0.972104 | GBM | LN-18 | AZD3759 | MVP_top60_DNA+RNA_seed | PIK3CD | -0.608372 | GBM | LN-18 | Buparlisib |
| MVP_top60_DNA_seed | EGFR | 0.973538 | GBM | SF295 | Afatinib | MVP_top60_DNA+RNA_seed | PIK3CA | -0.605729 | GBM | SK-MG-1 | Alpelisib |
| MVP_top60_DNA_seed | TP53 | 0.975512 | GBM | DK-MG | MIRA-1 | MVP_top60_DNA+RNA_seed | PIK3CA | -0.603704 | GBM | U-118-MG | GNE-317 |
| MVP_top60_DNA_seed | EGFR | 0.979684 | GBM | SK-MG-1 | Gefitinib | MVP_top60_DNA+RNA_seed | KIT | -0.600814 | GBM | 42-MG-BA | Sorafenib |

| | | | | | | | | | | | |
|---|---|---|---|---|---|---|---|---|---|---|---|
| MVP_top60_DNA_seed | EGFR | 0.980727 | GBM | LN-405 | Sapitinib | MVP_top60_DNA+RNA_seed | AKT1 | -0.599176 | GBM | YH-13 | AZD5363 |
| MVP_top60_DNA_seed | JAK2 | 0.985713 | GBM | LN-405 | Ruxolitinib | MVP_top60_DNA+RNA_seed | PIK3CA | -0.598305 | GBM | Becker | Buparlisib |
| MVP_top60_DNA_seed | JAK1 | 0.985713 | GBM | LN-405 | Ruxolitinib | MVP_top60_DNA+RNA_seed | PIK3CB | -0.598305 | GBM | Becker | Buparlisib |
| MVP_top60_DNA_seed | EGFR | 0.986548 | GBM | D-263MG | Osimertinib | MVP_top60_DNA+RNA_seed | PIK3CD | -0.598305 | GBM | Becker | Buparlisib |
| MVP_top60_DNA_seed | EGFR | 0.9873 | GBM | D-263MG | Afatinib | MVP_top60_DNA+RNA_seed | IGF1R | -0.595477 | GBM | A172 | BMS-754807 |
| MVP_top60_DNA_seed | EGFR | 0.991396 | GBM | M059J | Gefitinib | MVP_top60_DNA+RNA_seed | KIT | -0.588784 | GBM | D-263MG | Axitinib |
| MVP_top60_DNA_seed | EGFR | 1.000318 | GBM | YKG-1 | Afatinib | MVP_top60_DNA+RNA_seed | AKT1 | -0.580436 | GBM | YH-13 | AT13148 |
| MVP_top60_DNA_seed | CDK4 | 1.00091 | GBM | LN-405 | Ribociclib | MVP_top60_DNA+RNA_seed | KIT | -0.579761 | GBM | SF295 | Sorafenib |
| MVP_top60_DNA_seed | PIK3CA | 1.021972 | GBM | GB-1 | Alpelisib | MVP_top60_DNA+RNA_seed | JAK2 | -0.572111 | GBM | D-542MG | AZ960 |
| MVP_top60_DNA_seed | EGFR | 1.023988 | GBM | Becker | Sapitinib | MVP_top60_DNA+RNA_seed | JAK3 | -0.572111 | GBM | D-542MG | AZ960 |
| MVP_top60_DNA_seed | EP300 | 1.024816 | GBM | U251 | SGC-CBP30 | MVP_top60_DNA+RNA_seed | AKT1 | -0.571166 | GBM | 8-MG-BA | Uprosertib |
| MVP_top60_DNA_seed | EGFR | 1.027722 | GBM | U251 | Afatinib | MVP_top60_DNA+RNA_seed | AKT1 | -0.570988 | GBM | SF295 | Uprosertib |
| MVP_top60_DNA_seed | PIK3CA | 1.030939 | GBM | Becker | Alpelisib | MVP_top60_DNA+RNA_seed | AKT1 | -0.570131 | GBM | 8-MG-BA | AZD5363 |
| MVP_top60_DNA_seed | JAK1 | 1.03481 | GBM | SW1088 | JAK1_8709 | MVP_top60_DNA+RNA_seed | MET | -0.566054 | GBM | A172 | Savolitinib |
| MVP_top60_DNA_seed | EGFR | 1.039273 | GBM | LN-18 | Afatinib | MVP_top60_DNA+RNA_seed | AKT1 | -0.565336 | GBM | Becker | AZD5363 |
| MVP_top60_DNA_seed | EGFR | 1.039408 | GBM | KS-1 | AZD3759 | MVP_top60_DNA+RNA_seed | PIK3CA | -0.564623 | GBM | D-566MG | GNE-317 |
| MVP_top60_DNA_seed | EGFR | 1.04507 | GBM | LN-18 | Osimertinib | MVP_top60_DNA+RNA_seed | IGF1R | -0.56132 | GBM | LN-18 | NVP-ADW742 |
| MVP_top60_DNA_seed | EGFR | 1.049049 | GBM | D-247MG | AZD3759 | MVP_top60_DNA+RNA_seed | TP53 | -0.55726 | GBM | SF295 | MIRA-1 |
| MVP_top60_DNA_seed | PIK3CA | 1.04963 | GBM | SW1088 | Alpelisib | MVP_top60_DNA+RNA_seed | IGF1R | -0.55627 | GBM | GB-1 | NVP-ADW742 |
| MVP_top60_DNA_seed | JAK2 | 1.052645 | GBM | SW1088 | GSK2276186C | MVP_top60_DNA+RNA_seed | AKT1 | -0.552834 | GBM | M059J | AT13148 |
| MVP_top60_DNA_seed | JAK1 | 1.052645 | GBM | SW1088 | GSK2276186C | MVP_top60_DNA+RNA_seed | PIK3CA | -0.547656 | GBM | M059J | GNE-317 |
| MVP_top60_DNA_seed | JAK3 | 1.052645 | GBM | SW1088 | GSK2276186C | MVP_top60_DNA+RNA_seed | IGF1R | -0.546722 | GBM | SW1088 | GSK1904529A |
| MVP_top60_DNA_seed | EGFR | 1.056207 | GBM | SW1088 | Gefitinib | MVP_top60_DNA+RNA_seed | KIT | -0.545149 | GBM | D-423MG | Axitinib |
| MVP_top60_DNA_seed | EGFR | 1.063211 | GBM | GB-1 | Gefitinib | MVP_top60_DNA+RNA_seed | PIK3CA | -0.537568 | GBM | SNB75 | Alpelisib |
| MVP_top60_DNA_seed | JAK2 | 1.073445 | GBM | LN-229 | AZ960 | MVP_top60_DNA+RNA_seed | AKT1 | -0.536185 | GBM | SF295 | Ipatasertib |
| MVP_top60_DNA_seed | JAK3 | 1.073445 | GBM | LN-229 | AZ960 | MVP_top60_DNA+RNA_seed | PIK3CA | -0.530889 | GBM | U-118-MG | AZD8186 |
| MVP_top60_DNA_seed | EGFR | 1.086889 | GBM | GB-1 | Afatinib | MVP_top60_DNA+RNA_seed | PIK3CB | -0.530889 | GBM | U-118-MG | AZD8186 |
| MVP_top60_DNA_seed | EGFR | 1.088469 | GBM | Becker | Erlotinib | MVP_top60_DNA+RNA_seed | JAK2 | -0.525712 | GBM | D-566MG | Ruxolitinib |
| MVP_top60_DNA_seed | JAK1 | 1.100288 | GBM | D-392MG | JAK1_8709 | MVP_top60_DNA+RNA_seed | JAK1 | -0.525712 | GBM | D-566MG | Ruxolitinib |
| MVP_top60_DNA_seed | TP53 | 1.104613 | GBM | LN-405 | MIRA-1 | MVP_top60_DNA+RNA_seed | PIK3CA | -0.524473 | GBM | SF126 | AZD8186 |
| MVP_top60_DNA_seed | JAK2 | 1.112096 | GBM | CAS-1 | Ruxolitinib | MVP_top60_DNA+RNA_seed | PIK3CB | -0.524473 | GBM | SF126 | AZD8186 |
| MVP_top60_DNA_seed | JAK1 | 1.112096 | GBM | CAS-1 | Ruxolitinib | MVP_top60_DNA+RNA_seed | MET | -0.523242 | GBM | DK-MG | Crizotinib |
| MVP_top60_DNA_seed | EGFR | 1.128677 | GBM | AM-38 | Sapitinib | MVP_top60_DNA+RNA_seed | JAK2 | -0.522878 | GBM | KS-1 | GSK2276186C |
| MVP_top60_DNA_seed | EGFR | 1.130392 | GBM | D-392MG | Erlotinib | MVP_top60_DNA+RNA_seed | JAK1 | -0.522878 | GBM | KS-1 | GSK2276186C |
| MVP_top60_DNA_seed | PIK3CB | 1.139373 | GBM | YKG-1 | AZD6482 | MVP_top60_DNA+RNA_seed | JAK3 | -0.522878 | GBM | KS-1 | GSK2276186C |
| MVP_top60_DNA_seed | EGFR | 1.14068 | GBM | D-392MG | Sapitinib | MVP_top60_DNA+RNA_seed | PIK3CA | -0.518104 | GBM | Becker | GNE-317 |
| MVP_top60_DNA_seed | CDK4 | 1.157249 | GBM | U-118-MG | Ribociclib | MVP_top60_DNA+RNA_seed | EGFR | -0.514164 | GBM | SF126 | Lapatinib |
| MVP_top60_DNA_seed | PIK3CA | 1.171426 | GBM | U251 | AZD8186 | MVP_top60_DNA+RNA_seed | EP300 | -0.512248 | GBM | A172 | CPI-637 |
| MVP_top60_DNA_seed | PIK3CB | 1.171426 | GBM | U251 | AZD8186 | MVP_top60_DNA+RNA_seed | CDK4 | -0.506462 | GBM | D-263MG | Ribociclib |
| MVP_top60_DNA_seed | CDK4 | 1.178543 | GBM | M059J | Palbociclib | MVP_top60_DNA+RNA_seed | AKT1 | -0.502023 | GBM | SW1088 | GSK2110183B |
| MVP_top60_DNA_seed | PIK3CA | 1.180057 | GBM | A172 | Alpelisib | MVP_top60_DNA+RNA_seed | JAK2 | -0.501097 | GBM | LN-18 | Ruxolitinib |

| | | | | | | | | | | | |
|---|---|---|---|---|---|---|---|---|---|---|---|
| MVP_top60_DNA_seed | CDK4 | 1.180443 | GBM | CAS-1 | Ribociclib | MVP_top60_DNA+RNA_seed | JAK1 | -0.501097 | GBM | LN-18 | Ruxolitinib |
| MVP_top60_DNA_seed | EGFR | 1.180972 | GBM | U251 | Lapatinib | MVP_top60_DNA+RNA_seed | CDK4 | -0.498988 | GBM | A172 | Palbociclib |
| MVP_top60_DNA_seed | JAK2 | 1.181315 | GBM | SNB75 | Ruxolitinib | MVP_top60_DNA+RNA_seed | AKT1 | -0.494239 | GBM | CAS-1 | AZD5363 |
| MVP_top60_DNA_seed | JAK1 | 1.181315 | GBM | SNB75 | Ruxolitinib | MVP_top60_DNA+RNA_seed | JAK2 | -0.492503 | GBM | D-245MG | Ruxolitinib |
| MVP_top60_DNA_seed | TP53 | 1.185029 | GBM | SW1088 | MIRA-1 | MVP_top60_DNA+RNA_seed | JAK1 | -0.492503 | GBM | D-245MG | Ruxolitinib |
| MVP_top60_DNA_seed | EGFR | 1.186611 | GBM | SF268 | Osimertinib | MVP_top60_DNA+RNA_seed | TP53 | -0.487817 | GBM | GB-1 | MIRA-1 |
| MVP_top60_DNA_seed | EGFR | 1.187698 | GBM | DBTRG-05MG | Osimertinib | MVP_top60_DNA+RNA_seed | KIT | -0.485328 | GBM | CAS-1 | Axitinib |
| MVP_top60_DNA_seed | EGFR | 1.201006 | GBM | LN-405 | Osimertinib | MVP_top60_DNA+RNA_seed | AKT1 | -0.484583 | GBM | D-423MG | Afuresertib |
| MVP_top60_DNA_seed | PIK3CA | 1.219196 | GBM | SF268 | AZD8186 | MVP_top60_DNA+RNA_seed | AKT1 | -0.482181 | GBM | SF126 | Ipatasertib |
| MVP_top60_DNA_seed | PIK3CB | 1.219196 | GBM | SF268 | AZD8186 | MVP_top60_DNA+RNA_seed | PIK3CA | -0.482165 | GBM | LN-18 | GNE-317 |
| MVP_top60_DNA_seed | PIK3CA | 1.220952 | GBM | LN-229 | GNE-317 | MVP_top60_DNA+RNA_seed | JAK2 | -0.479701 | GBM | 42-MG-BA | AZ960 |
| MVP_top60_DNA_seed | JAK2 | 1.222014 | GBM | LN-405 | JAK_8517 | MVP_top60_DNA+RNA_seed | JAK3 | -0.479701 | GBM | 42-MG-BA | AZ960 |
| MVP_top60_DNA_seed | JAK1 | 1.222014 | GBM | LN-405 | JAK_8517 | MVP_top60_DNA+RNA_seed | EGFR | -0.478769 | GBM | SF295 | AZD3759 |
| MVP_top60_DNA_seed | EGFR | 1.222906 | GBM | U-118-MG | Lapatinib | MVP_top60_DNA+RNA_seed | JAK2 | -0.472731 | GBM | CAS-1 | AZ960 |
| MVP_top60_DNA_seed | JAK2 | 1.231012 | GBM | D-247MG | Ruxolitinib | MVP_top60_DNA+RNA_seed | JAK3 | -0.472731 | GBM | CAS-1 | AZ960 |
| MVP_top60_DNA_seed | JAK1 | 1.231012 | GBM | D-247MG | Ruxolitinib | MVP_top60_DNA+RNA_seed | IGF1R | -0.471944 | GBM | D-423MG | BMS-536924 |
| MVP_top60_DNA_seed | EGFR | 1.234905 | GBM | DK-MG | Gefitinib | MVP_top60_DNA+RNA_seed | EGFR | -0.470561 | GBM | LN-18 | Erlotinib |
| MVP_top60_DNA_seed | PIK3CA | 1.23908 | GBM | U251 | Alpelisib | MVP_top60_DNA+RNA_seed | IGF1R | -0.470181 | GBM | D-423MG | NVP-ADW742 |
| MVP_top60_DNA_seed | EGFR | 1.247878 | GBM | CAS-1 | AZD3759 | MVP_top60_DNA+RNA_seed | JAK2 | -0.46123 | GBM | D-263MG | AZ960 |
| MVP_top60_DNA_seed | TP53 | 1.249454 | GBM | CAS-1 | MIRA-1 | MVP_top60_DNA+RNA_seed | JAK3 | -0.46123 | GBM | D-263MG | AZ960 |
| MVP_top60_DNA_seed | EGFR | 1.260517 | GBM | SNB75 | Erlotinib | MVP_top60_DNA+RNA_seed | JAK2 | -0.45918 | GBM | D-247MG | AZ960 |
| MVP_top60_DNA_seed | CDK4 | 1.263292 | GBM | LN-405 | Palbociclib | MVP_top60_DNA+RNA_seed | JAK3 | -0.45918 | GBM | D-247MG | AZ960 |
| MVP_top60_DNA_seed | JAK2 | 1.273339 | GBM | SK-MG-1 | JAK_8517 | MVP_top60_DNA+RNA_seed | KIT | -0.456512 | GBM | LNZTA3WT4 | Axitinib |
| MVP_top60_DNA_seed | JAK1 | 1.273339 | GBM | SK-MG-1 | JAK_8517 | MVP_top60_DNA+RNA_seed | AKT1 | -0.448679 | GBM | SK-MG-1 | Ipatasertib |
| MVP_top60_DNA_seed | JAK2 | 1.280343 | GBM | AM-38 | JAK_8517 | MVP_top60_DNA+RNA_seed | PIK3CB | -0.447853 | GBM | SNB75 | AZD6482 |
| MVP_top60_DNA_seed | JAK1 | 1.280343 | GBM | AM-38 | JAK_8517 | MVP_top60_DNA+RNA_seed | IGF1R | -0.440978 | GBM | A172 | NVP-ADW742 |
| MVP_top60_DNA_seed | EGFR | 1.283699 | GBM | D-263MG | Lapatinib | MVP_top60_DNA+RNA_seed | IGF1R | -0.440327 | GBM | AM-38 | NVP-ADW742 |
| MVP_top60_DNA_seed | EGFR | 1.288037 | GBM | D-245MG | Erlotinib | MVP_top60_DNA+RNA_seed | EGFR | -0.439711 | GBM | 8-MG-BA | Gefitinib |
| MVP_top60_DNA_seed | JAK1 | 1.289234 | GBM | M059J | JAK1_8709 | MVP_top60_DNA+RNA_seed | AKT1 | -0.438613 | GBM | D-542MG | Afuresertib |
| MVP_top60_DNA_seed | CDK4 | 1.299051 | GBM | D-542MG | Palbociclib | MVP_top60_DNA+RNA_seed | IGF1R | -0.436665 | GBM | GB-1 | BMS-754807 |
| MVP_top60_DNA_seed | JAK2 | 1.303162 | GBM | U251 | AZ960 | MVP_top60_DNA+RNA_seed | IGF1R | -0.43542 | GBM | 8-MG-BA | BMS-536924 |
| MVP_top60_DNA_seed | JAK3 | 1.303162 | GBM | U251 | AZ960 | MVP_top60_DNA+RNA_seed | IGF1R | -0.431678 | GBM | SF126 | BMS-754807 |
| MVP_top60_DNA_seed | PIK3CA | 1.306821 | GBM | DBTRG-05MG | GNE-317 | MVP_top60_DNA+RNA_seed | MET | -0.424542 | GBM | D-247MG | Crizotinib |
| MVP_top60_DNA_seed | CDK4 | 1.331047 | GBM | GB-1 | Ribociclib | MVP_top60_DNA+RNA_seed | KIT | -0.422334 | GBM | AM-38 | Sorafenib |
| MVP_top60_DNA_seed | EP300 | 1.335778 | GBM | LN-229 | SGC-CBP30 | MVP_top60_DNA+RNA_seed | IGF1R | -0.418449 | GBM | D-247MG | BMS-754807 |
| MVP_top60_DNA_seed | JAK2 | 1.352287 | GBM | SW1088 | Ruxolitinib | MVP_top60_DNA+RNA_seed | IGF1R | -0.418446 | GBM | SK-MG-1 | NVP-ADW742 |
| MVP_top60_DNA_seed | JAK1 | 1.352287 | GBM | SW1088 | Ruxolitinib | MVP_top60_DNA+RNA_seed | MET | -0.416426 | GBM | LN-18 | Savolitinib |
| MVP_top60_DNA_seed | PIK3CA | 1.361822 | GBM | LN-405 | Buparlisib | MVP_top60_DNA+RNA_seed | JAK2 | -0.41508 | GBM | DK-MG | AZ960 |
| MVP_top60_DNA_seed | PIK3CB | 1.361822 | GBM | LN-405 | Buparlisib | MVP_top60_DNA+RNA_seed | JAK3 | -0.41508 | GBM | DK-MG | AZ960 |
| MVP_top60_DNA_seed | PIK3CD | 1.361822 | GBM | LN-405 | Buparlisib | MVP_top60_DNA+RNA_seed | IGF1R | -0.415053 | GBM | U-87-MG | NVP-ADW742 |
| MVP_top60_DNA_seed | EGFR | 1.375176 | GBM | SNB75 | Afatinib | MVP_top60_DNA+RNA_seed | JAK2 | -0.412733 | GBM | D-392MG | JAK_8517 |

| | | | | | | | | | | | |
|---|---|---|---|---|---|---|---|---|---|---|---|
| MVP_top60_DNA_seed | EGFR | 1.378183 | GBM | DBTRG-05MG | AZD3759 | MVP_top60_DNA+RNA_seed | JAK1 | -0.412733 | GBM | D-392MG | JAK_8517 |
| MVP_top60_DNA_seed | EP300 | 1.379405 | GBM | CAS-1 | CPI-637 | MVP_top60_DNA+RNA_seed | EGFR | -0.409178 | GBM | YKG-1 | Sapitinib |
| MVP_top60_DNA_seed | EGFR | 1.39833 | GBM | GB-1 | AZD3759 | MVP_top60_DNA+RNA_seed | ERBB3 | -0.409178 | GBM | YKG-1 | Sapitinib |
| MVP_top60_DNA_seed | EGFR | 1.401945 | GBM | U251 | Sapitinib | MVP_top60_DNA+RNA_seed | KRAS | -0.405085 | GBM | T98G | RAS (G12C) Inhibitor-1 |
| MVP_top60_DNA_seed | JAK1 | 1.403396 | GBM | LN-405 | JAK1_8709 | MVP_top60_DNA+RNA_seed | AKT1 | -0.403725 | GBM | SW1088 | Afuresertib |
| MVP_top60_DNA_seed | EP300 | 1.403904 | GBM | GB-1 | CPI-637 | MVP_top60_DNA+RNA_seed | JAK2 | -0.40139 | GBM | YKG-1 | AZ960 |
| MVP_top60_DNA_seed | EP300 | 1.41132 | GBM | DK-MG | SGC-CBP30 | MVP_top60_DNA+RNA_seed | JAK3 | -0.40139 | GBM | YKG-1 | AZ960 |
| MVP_top60_DNA_seed | EP300 | 1.419186 | GBM | SW1088 | SGC-CBP30 | MVP_top60_DNA+RNA_seed | PIK3CA | -0.400543 | GBM | D-247MG | GNE-317 |
| MVP_top60_DNA_seed | EGFR | 1.425539 | GBM | U-87-MG | Afatinib | MVP_top60_DNA+RNA_seed | KRAS | -0.39874 | GBM | 42-MG-BA | RAS (G12C) Inhibitor-1 |
| MVP_top60_DNA_seed | EGFR | 1.430242 | GBM | U-87-MG | Lapatinib | MVP_top60_DNA+RNA_seed | PIK3CA | -0.397583 | GBM | 8-MG-BA | Alpelisib |
| MVP_top60_DNA_seed | EGFR | 1.446664 | GBM | DBTRG-05MG | Afatinib | MVP_top60_DNA+RNA_seed | TP53 | -0.396213 | GBM | D-423MG | MIRA-1 |
| MVP_top60_DNA_seed | PIK3CA | 1.45936 | GBM | YKG-1 | Alpelisib | MVP_top60_DNA+RNA_seed | EGFR | -0.393865 | GBM | GB-1 | Lapatinib |
| MVP_top60_DNA_seed | EGFR | 1.4674 | GBM | Becker | Lapatinib | MVP_top60_DNA+RNA_seed | MET | -0.392921 | GBM | KS-1 | Crizotinib |
| MVP_top60_DNA_seed | JAK1 | 1.481197 | GBM | Becker | JAK1_8709 | MVP_top60_DNA+RNA_seed | PIK3CA | -0.390477 | GBM | SW1088 | Buparlisib |
| MVP_top60_DNA_seed | CDK4 | 1.489772 | GBM | LN-229 | Palbociclib | MVP_top60_DNA+RNA_seed | PIK3CB | -0.390477 | GBM | SW1088 | Buparlisib |
| MVP_top60_DNA_seed | EP300 | 1.492985 | GBM | U-118-MG | CPI-637 | MVP_top60_DNA+RNA_seed | PIK3CD | -0.390477 | GBM | SW1088 | Buparlisib |
| MVP_top60_DNA_seed | PIK3CA | 1.520659 | GBM | GB-1 | AZD8186 | MVP_top60_DNA+RNA_seed | AKT1 | -0.386898 | GBM | KS-1 | GSK2110183B |
| MVP_top60_DNA_seed | PIK3CB | 1.520659 | GBM | GB-1 | AZD8186 | MVP_top60_DNA+RNA_seed | KIT | -0.385714 | GBM | D-566MG | Motesanib |
| MVP_top60_DNA_seed | EGFR | 1.52125 | GBM | LN-405 | Afatinib | MVP_top60_DNA+RNA_seed | PIK3CA | -0.384948 | GBM | U-87-MG | AZD8186 |
| MVP_top60_DNA_seed | JAK2 | 1.540296 | GBM | U-87-MG | JAK_8517 | MVP_top60_DNA+RNA_seed | PIK3CB | -0.384948 | GBM | U-87-MG | AZD8186 |
| MVP_top60_DNA_seed | JAK1 | 1.540296 | GBM | U-87-MG | JAK_8517 | MVP_top60_DNA+RNA_seed | EGFR | -0.377618 | GBM | YKG-1 | Lapatinib |
| MVP_top60_DNA_seed | EP300 | 1.544784 | GBM | DK-MG | CPI-637 | MVP_top60_DNA+RNA_seed | KIT | -0.37585 | GBM | A172 | Sorafenib |
| MVP_top60_DNA_seed | PIK3CA | 1.552749 | GBM | D-245MG | Buparlisib | MVP_top60_DNA+RNA_seed | JAK1 | -0.375317 | GBM | LN-18 | JAK1_8709 |
| MVP_top60_DNA_seed | PIK3CB | 1.552749 | GBM | D-245MG | Buparlisib | MVP_top60_DNA+RNA_seed | AKT1 | -0.373823 | GBM | LN-18 | Uprosertib |
| MVP_top60_DNA_seed | PIK3CD | 1.552749 | GBM | D-245MG | Buparlisib | MVP_top60_DNA+RNA_seed | CDK4 | -0.373544 | GBM | LNZTA3WT4 | Ribociclib |
| MVP_top60_DNA_seed | EGFR | 1.562033 | GBM | KS-1 | Lapatinib | MVP_top60_DNA+RNA_seed | JAK2 | -0.370197 | GBM | 8-MG-BA | AZ960 |
| MVP_top60_DNA_seed | EP300 | 1.591641 | GBM | DBTRG-05MG | CPI-637 | MVP_top60_DNA+RNA_seed | JAK3 | -0.370197 | GBM | 8-MG-BA | AZ960 |
| MVP_top60_DNA_seed | PIK3CA | 1.637238 | GBM | D-263MG | Alpelisib | MVP_top60_DNA+RNA_seed | IGF1R | -0.369624 | GBM | D-245MG | BMS-536924 |
| MVP_top60_DNA_seed | JAK2 | 1.644075 | GBM | DBTRG-05MG | Ruxolitinib | MVP_top60_DNA+RNA_seed | CDK4 | -0.369037 | GBM | Becker | Palbociclib |
| MVP_top60_DNA_seed | JAK1 | 1.644075 | GBM | DBTRG-05MG | Ruxolitinib | MVP_top60_DNA+RNA_seed | EP300 | -0.359928 | GBM | SW1088 | CPI-637 |
| MVP_top60_DNA_seed | PIK3CA | 1.682586 | GBM | AM-38 | AZD8186 | MVP_top60_DNA+RNA_seed | EGFR | -0.358775 | GBM | SNB75 | AZD3759 |
| MVP_top60_DNA_seed | PIK3CB | 1.682586 | GBM | AM-38 | AZD8186 | MVP_top60_DNA+RNA_seed | AKT1 | -0.357906 | GBM | M059J | AZD5363 |
| MVP_top60_DNA_seed | CDK4 | 1.691322 | GBM | DBTRG-05MG | Ribociclib | MVP_top60_DNA+RNA_seed | KRAS | -0.357651 | GBM | KS-1 | RAS (G12C) Inhibitor-1 |
| MVP_top60_DNA_seed | EGFR | 1.716475 | GBM | SF295 | Erlotinib | MVP_top60_DNA+RNA_seed | EGFR | -0.356905 | GBM | LN-229 | Sapitinib |
| MVP_top60_DNA_seed | JAK1 | 1.744043 | GBM | DBTRG-05MG | JAK1_8709 | MVP_top60_DNA+RNA_seed | ERBB3 | -0.356905 | GBM | LN-229 | Sapitinib |
| MVP_top60_DNA_seed | EGFR | 1.747985 | GBM | LN-405 | Lapatinib | MVP_top60_DNA+RNA_seed | AKT1 | -0.356283 | GBM | YH-13 | Uprosertib |
| MVP_top60_DNA_seed | EGFR | 1.750427 | GBM | U-118-MG | Sapitinib | MVP_top60_DNA+RNA_seed | AKT1 | -0.355475 | GBM | LNZTA3WT4 | AT13148 |
| MVP_top60_DNA_seed | EGFR | 1.767129 | GBM | DBTRG-05MG | Sapitinib | MVP_top60_DNA+RNA_seed | CDK4 | -0.355037 | GBM | YH-13 | Palbociclib |
| MVP_top60_DNA_seed | EP300 | 1.777659 | GBM | CAS-1 | SGC-CBP30 | MVP_top60_DNA+RNA_seed | AKT1 | -0.352326 | GBM | 8-MG-BA | Ipatasertib |
| MVP_top60_DNA_seed | EGFR | 1.805704 | GBM | GB-1 | Osimertinib | MVP_top60_DNA+RNA_seed | AKT1 | -0.350279 | GBM | T98G | GSK2110183B |
| MVP_top60_DNA_seed | JAK2 | 1.824469 | GBM | LN-229 | JAK_8517 | MVP_top60_DNA+RNA_seed | PIK3CA | -0.350083 | GBM | 8-MG-BA | AZD8186 |

| | | | | | | | | | | | |
|---|---|---|---|---|---|---|---|---|---|---|---|
| MVP_top60_DNA_seed | JAK1 | 1.824469 | GBM | LN-229 | JAK_8517 | MVP_top60_DNA+RNA_seed | PIK3CB | -0.350083 | GBM | 8-MG-BA | AZD8186 |
| MVP_top60_DNA_seed | PIK3CA | 1.925213 | GBM | U-118-MG | Alpelisib | MVP_top60_DNA+RNA_seed | MET | -0.349904 | GBM | U-87-MG | Savolitinib |
| MVP_top60_DNA_seed | PIK3CA | 1.941553 | GBM | LN-405 | GNE-317 | MVP_top60_DNA+RNA_seed | PIK3CA | -0.343418 | GBM | DK-MG | AZD8186 |
| MVP_top60_DNA_seed | JAK2 | 2.016043 | GBM | DBTRG-05MG | GSK2276186C | MVP_top60_DNA+RNA_seed | PIK3CB | -0.343418 | GBM | DK-MG | AZD8186 |
| MVP_top60_DNA_seed | JAK1 | 2.016043 | GBM | DBTRG-05MG | GSK2276186C | MVP_top60_DNA+RNA_seed | AKT1 | -0.340469 | GBM | CAS-1 | Ipatasertib |
| MVP_top60_DNA_seed | JAK3 | 2.016043 | GBM | DBTRG-05MG | GSK2276186C | MVP_top60_DNA+RNA_seed | AKT1 | -0.339909 | GBM | SF295 | MK-2206 |
| MVP_top60_DNA_seed | CDK4 | 2.065252 | GBM | DBTRG-05MG | Palbociclib | MVP_top60_DNA+RNA_seed | EGFR | -0.339388 | GBM | GB-1 | Erlotinib |
| MVP_top60_DNA_seed | EGFR | 2.079235 | GBM | D-392MG | Lapatinib | MVP_top60_DNA+RNA_seed | AKT1 | -0.336008 | GBM | 42-MG-BA | AT13148 |
| MVP_top60_DNA_seed | EGFR | 2.094157 | GBM | DBTRG-05MG | Gefitinib | MVP_top60_DNA+RNA_seed | PIK3CA | -0.335624 | GBM | 42-MG-BA | Alpelisib |
| MVP_top60_DNA_seed | TP53 | 2.125063 | GBM | DBTRG-05MG | MIRA-1 | MVP_top60_DNA+RNA_seed | KIT | -0.334249 | GBM | SF126 | Sorafenib |
| MVP_top60_DNA_seed | PIK3CA | 2.300414 | GBM | U251 | GNE-317 | MVP_top60_DNA+RNA_seed | EGFR | -0.334036 | GBM | YKG-1 | AZD3759 |
| MVP_top60_DNA_seed | PIK3CA | 2.542779 | GBM | DBTRG-05MG | Alpelisib | MVP_top60_DNA+RNA_seed | AKT1 | -0.333941 | GBM | T98G | AT13148 |
| MVP_top60_DNA_seed | EGFR | 2.576791 | GBM | DBTRG-05MG | Lapatinib | MVP_top60_DNA+RNA_seed | KIT | -0.333345 | GBM | D-247MG | Motesanib |
| MVP_top60_DNA_seed | EP300 | 2.840979 | GBM | DBTRG-05MG | SGC-CBP30 | MVP_top60_DNA+RNA_seed | AKT1 | -0.33315 | GBM | U-87-MG | Uprosertib |
| MVP_top60_DNA_seed | EGFR | -4.644851 | HNSC | JHU-029 | Cetuximab | MVP_top60_DNA+RNA_seed | IGF1R | -0.331958 | GBM | SF126 | GSK1904529A |
| MVP_top60_DNA_seed | EGFR | -3.919324 | HNSC | BB30-HNC | Erlotinib | MVP_top60_DNA+RNA_seed | PIK3CA | -0.330178 | GBM | GB-1 | GNE-317 |
| MVP_top60_DNA_seed | EGFR | -3.734406 | HNSC | BB30-HNC | Lapatinib | MVP_top60_DNA+RNA_seed | AKT1 | -0.327494 | GBM | DK-MG | AT13148 |
| MVP_top60_DNA_seed | EGFR | -3.616271 | HNSC | BICR78 | Gefitinib | MVP_top60_DNA+RNA_seed | IGF1R | -0.326896 | GBM | 42-MG-BA | BMS-536924 |
| MVP_top60_DNA_seed | EGFR | -3.537032 | HNSC | PCI-4B | Gefitinib | MVP_top60_DNA+RNA_seed | MET | -0.324195 | GBM | Becker | Crizotinib |
| MVP_top60_DNA_seed | EGFR | -3.484307 | HNSC | DOK | Gefitinib | MVP_top60_DNA+RNA_seed | PIK3CA | -0.322309 | GBM | T98G | GNE-317 |
| MVP_top60_DNA_seed | AR | -3.398851 | HNSC | HSC-3 | Bicalutamide | MVP_top60_DNA+RNA_seed | IGF1R | -0.319696 | GBM | CAS-1 | BMS-754807 |
| MVP_top60_DNA_seed | EGFR | -3.368979 | HNSC | PCI-6A | Gefitinib | MVP_top60_DNA+RNA_seed | AKT1 | -0.319287 | GBM | KS-1 | AT13148 |
| MVP_top60_DNA_seed | EGFR | -3.357327 | HNSC | HSC-3 | Afatinib | MVP_top60_DNA+RNA_seed | TP53 | -0.319202 | GBM | KS-1 | MIRA-1 |
| MVP_top60_DNA_seed | EGFR | -3.320364 | HNSC | CAL-27 | Afatinib | MVP_top60_DNA+RNA_seed | EGFR | -0.317899 | GBM | D-423MG | Osimertinib |
| MVP_top60_DNA_seed | EGFR | -3.297692 | HNSC | JHU-029 | AZD8931 | MVP_top60_DNA+RNA_seed | AKT1 | -0.313085 | GBM | LNZTA3WT4 | Uprosertib |
| MVP_top60_DNA_seed | EGFR | -3.291909 | HNSC | HSC-4 | Gefitinib | MVP_top60_DNA+RNA_seed | JAK2 | -0.311776 | GBM | Becker | JAK_8517 |
| MVP_top60_DNA_seed | EGFR | -3.283504 | HNSC | HSC-4 | Cetuximab | MVP_top60_DNA+RNA_seed | JAK1 | -0.311776 | GBM | Becker | JAK_8517 |
| MVP_top60_DNA_seed | EGFR | -3.283305 | HNSC | SAT | AZD8931 | MVP_top60_DNA+RNA_seed | TP53 | -0.309237 | GBM | LN-18 | MIRA-1 |
| MVP_top60_DNA_seed | AR | -3.223444 | HNSC | SCC-25 | Bicalutamide | MVP_top60_DNA+RNA_seed | JAK2 | -0.307654 | GBM | SW1088 | AZ960 |
| MVP_top60_DNA_seed | EGFR | -3.21519 | HNSC | Ca9-22 | AZD8931 | MVP_top60_DNA+RNA_seed | JAK3 | -0.307654 | GBM | SW1088 | AZ960 |
| MVP_top60_DNA_seed | EGFR | -3.204473 | HNSC | HSC-4 | Afatinib | MVP_top60_DNA+RNA_seed | AKT1 | -0.307646 | GBM | D-566MG | Ipatasertib |
| MVP_top60_DNA_seed | EGFR | -3.141427 | HNSC | PCI-4B | Afatinib | MVP_top60_DNA+RNA_seed | AKT1 | -0.302204 | GBM | SNB75 | Ipatasertib |
| MVP_top60_DNA_seed | EGFR | -3.135639 | HNSC | SCC-9 | Afatinib | MVP_top60_DNA+RNA_seed | AKT1 | -0.301254 | GBM | YH-13 | Uprosertib |
| MVP_top60_DNA_seed | EGFR | -3.109147 | HNSC | Ca9-22 | Afatinib | MVP_top60_DNA+RNA_seed | IGF1R | -0.296519 | GBM | SK-MG-1 | GSK1904529A |
| MVP_top60_DNA_seed | EGFR | -3.107691 | HNSC | CAL-27 | AZD8931 | MVP_top60_DNA+RNA_seed | AKT1 | -0.296471 | GBM | YKG-1 | AZD5363 |
| MVP_top60_DNA_seed | EGFR | -3.097346 | HNSC | JHU-029 | Afatinib | MVP_top60_DNA+RNA_seed | MET | -0.293524 | GBM | SW1088 | Crizotinib |
| MVP_top60_DNA_seed | EGFR | -3.074312 | HNSC | DOK | Afatinib | MVP_top60_DNA+RNA_seed | PIK3CA | -0.292366 | GBM | D-392MG | Buparlisib |
| MVP_top60_DNA_seed | EGFR | -3.068971 | HNSC | BICR78 | Afatinib | MVP_top60_DNA+RNA_seed | PIK3CB | -0.292366 | GBM | D-392MG | Buparlisib |
| MVP_top60_DNA_seed | EGFR | -3.017944 | HNSC | SCC-15 | Afatinib | MVP_top60_DNA+RNA_seed | PIK3CD | -0.292366 | GBM | D-392MG | Buparlisib |
| MVP_top60_DNA_seed | EGFR | -2.90094 | HNSC | SAT | PF-00299804 | MVP_top60_DNA+RNA_seed | MET | -0.290745 | GBM | D-566MG | Crizotinib |
| MVP_top60_DNA_seed | EGFR | -2.878281 | HNSC | SAT | Afatinib | MVP_top60_DNA+RNA_seed | TP53 | -0.290196 | GBM | D-247MG | MIRA-1 |

| | | | | | | | | | | | |
|---|---|---|---|---|---|---|---|---|---|---|---|
| MVP_top60_DNA_seed | EGFR | -2.861255 | HNSC | PCI-6A | Afatinib | MVP_top60_DNA+RNA_seed | IGF1R | -0.28646 | GBM | AM-38 | GSK1904529A |
| MVP_top60_DNA_seed | EGFR | -2.849463 | HNSC | SCC-9 | AST-1306 | MVP_top60_DNA+RNA_seed | PIK3CA | -0.286346 | GBM | D-542MG | GNE-317 |
| MVP_top60_DNA_seed | EGFR | -2.835836 | HNSC | JHU-011 | Afatinib | MVP_top60_DNA+RNA_seed | EP300 | -0.285626 | GBM | Becker | CPI-637 |
| MVP_top60_DNA_seed | EGFR | -2.815631 | HNSC | SCC-15 | PF-00299804 | MVP_top60_DNA+RNA_seed | AKT1 | -0.283079 | GBM | DK-MG | Uprosertib |
| MVP_top60_DNA_seed | EGFR | -2.769954 | HNSC | SAT | Afatinib | MVP_top60_DNA+RNA_seed | KIT | -0.281704 | GBM | D-247MG | Axitinib |
| MVP_top60_DNA_seed | EGFR | -2.729879 | HNSC | DOK | AZD8931 | MVP_top60_DNA+RNA_seed | IGF1R | -0.280373 | GBM | A172 | GSK1904529A |
| MVP_top60_DNA_seed | EGFR | -2.685176 | HNSC | SCC-9 | Cetuximab | MVP_top60_DNA+RNA_seed | AKT1 | -0.277986 | GBM | KS-1 | Uprosertib |
| MVP_top60_DNA_seed | EGFR | -2.665682 | HNSC | SCC-15 | AZD8931 | MVP_top60_DNA+RNA_seed | AKT1 | -0.277813 | GBM | SK-MG-1 | Uprosertib |
| MVP_top60_DNA_seed | AR | -2.616887 | HNSC | SAS | Bicalutamide | MVP_top60_DNA+RNA_seed | IGF1R | -0.277233 | GBM | D-566MG | BMS-536924 |
| MVP_top60_DNA_seed | PIK3CA | -2.592722 | HNSC | CAL-33 | AZD8835 | MVP_top60_DNA+RNA_seed | AKT1 | -0.275387 | GBM | D-423MG | MK-2206 |
| MVP_top60_DNA_seed | SRC | -2.583369 | HNSC | HSC-4 | Bosutinib | MVP_top60_DNA+RNA_seed | CDK4 | -0.271687 | GBM | 42-MG-BA | Ribociclib |
| MVP_top60_DNA_seed | EGFR | -2.577328 | HNSC | SAT | Gefitinib | MVP_top60_DNA+RNA_seed | EGFR | -0.26647 | GBM | DK-MG | Osimertinib |
| MVP_top60_DNA_seed | EGFR | -2.572577 | HNSC | SAT | Cetuximab | MVP_top60_DNA+RNA_seed | AKT1 | -0.266232 | GBM | M059J | Ipatasertib |
| MVP_top60_DNA_seed | EGFR | -2.56328 | HNSC | SCC-15 | AST-1306 | MVP_top60_DNA+RNA_seed | CDK4 | -0.262916 | GBM | D-263MG | Palbociclib |
| MVP_top60_DNA_seed | EGFR | -2.558771 | HNSC | DOK | Afatinib | MVP_top60_DNA+RNA_seed | AKT1 | -0.260874 | GBM | M059J | Uprosertib |
| MVP_top60_DNA_seed | EGFR | -2.544152 | HNSC | CAL-33 | AZD8931 | MVP_top60_DNA+RNA_seed | PIK3CA | -0.260705 | GBM | AM-38 | Buparlisib |
| MVP_top60_DNA_seed | EGFR | -2.524839 | HNSC | HSC-4 | AZD8931 | MVP_top60_DNA+RNA_seed | PIK3CB | -0.260705 | GBM | AM-38 | Buparlisib |
| MVP_top60_DNA_seed | EGFR | -2.517314 | HNSC | HSC-3 | AZD8931 | MVP_top60_DNA+RNA_seed | PIK3CD | -0.260705 | GBM | AM-38 | Buparlisib |
| MVP_top60_DNA_seed | EGFR | -2.507355 | HNSC | CAL-27 | Gefitinib | MVP_top60_DNA+RNA_seed | IGF1R | -0.259973 | GBM | D-263MG | BMS-536924 |
| MVP_top60_DNA_seed | EGFR | -2.505071 | HNSC | SCC-15 | Cetuximab | MVP_top60_DNA+RNA_seed | KIT | -0.259552 | GBM | 8-MG-BA | Motesanib |
| MVP_top60_DNA_seed | EGFR | -2.498475 | HNSC | HSC-2 | Gefitinib | MVP_top60_DNA+RNA_seed | IGF1R | -0.258701 | GBM | D-263MG | GSK1904529A |
| MVP_top60_DNA_seed | EGFR | -2.461381 | HNSC | JHU-022 | AZD8931 | MVP_top60_DNA+RNA_seed | EGFR | -0.258488 | GBM | SF126 | Gefitinib |
| MVP_top60_DNA_seed | EGFR | -2.452059 | HNSC | JHU-011 | Gefitinib | MVP_top60_DNA+RNA_seed | PIK3CB | -0.257592 | GBM | SF295 | AZD6482 |
| MVP_top60_DNA_seed | EGFR | -2.420302 | HNSC | HSC-4 | Afatinib | MVP_top60_DNA+RNA_seed | IGF1R | -0.252193 | GBM | DK-MG | BMS-754807 |
| MVP_top60_DNA_seed | AR | -2.418248 | HNSC | PE-CA-PJ15 | Bicalutamide | MVP_top60_DNA+RNA_seed | EGFR | -0.252165 | GBM | U-87-MG | Erlotinib |
| MVP_top60_DNA_seed | EGFR | -2.417639 | HNSC | JHU-029 | Afatinib | MVP_top60_DNA+RNA_seed | AKT1 | -0.249747 | GBM | LNZTA3WT4 | MK-2206 |
| MVP_top60_DNA_seed | EGFR | -2.407403 | HNSC | CAL-27 | Afatinib | MVP_top60_DNA+RNA_seed | AKT1 | -0.247947 | GBM | SK-MG-1 | Uprosertib |
| MVP_top60_DNA_seed | EGFR | -2.404439 | HNSC | Ca9-22 | Afatinib | MVP_top60_DNA+RNA_seed | TP53 | -0.246912 | GBM | A172 | MIRA-1 |
| MVP_top60_DNA_seed | SRC | -2.362284 | HNSC | BICR78 | Bosutinib | MVP_top60_DNA+RNA_seed | TP53 | -0.241901 | GBM | YH-13 | MIRA-1 |
| MVP_top60_DNA_seed | SMARCA4 | -2.36046 | HNSC | CAL-27 | PFI-3 | MVP_top60_DNA+RNA_seed | JAK2 | -0.241079 | GBM | GB-1 | Ruxolitinib |
| MVP_top60_DNA_seed | PIK3CA | -2.355176 | HNSC | JHU-029 | PI3Ka_4409 | MVP_top60_DNA+RNA_seed | JAK1 | -0.241079 | GBM | GB-1 | Ruxolitinib |
| MVP_top60_DNA_seed | EGFR | -2.35515 | HNSC | SCC-9 | PF-00299804 | MVP_top60_DNA+RNA_seed | PIK3CA | -0.235878 | GBM | SNB75 | GNE-317 |
| MVP_top60_DNA_seed | AR | -2.354815 | HNSC | KOSC-2 | Bicalutamide | MVP_top60_DNA+RNA_seed | EGFR | -0.235615 | GBM | LN-229 | Osimertinib |
| MVP_top60_DNA_seed | EGFR | -2.328828 | HNSC | SAS | Cetuximab | MVP_top60_DNA+RNA_seed | JAK2 | -0.235033 | GBM | D-423MG | AZ960 |
| MVP_top60_DNA_seed | AR | -2.322207 | HNSC | HSC-3 | AZD3514 | MVP_top60_DNA+RNA_seed | JAK3 | -0.235033 | GBM | D-423MG | AZ960 |
| MVP_top60_DNA_seed | EGFR | -2.318591 | HNSC | JHU-022 | Gefitinib | MVP_top60_DNA+RNA_seed | MET | -0.232003 | GBM | D-423MG | Crizotinib |
| MVP_top60_DNA_seed | SRC | -2.292887 | HNSC | PCI-6A | Bosutinib | MVP_top60_DNA+RNA_seed | CDK4 | -0.231495 | GBM | D-566MG | Palbociclib |
| MVP_top60_DNA_seed | NTRK1 | -2.286338 | HNSC | HSC-4 | GW441756 | MVP_top60_DNA+RNA_seed | EGFR | -0.23003 | GBM | AM-38 | Erlotinib |
| MVP_top60_DNA_seed | EGFR | -2.26906 | HNSC | HSC-3 | Cetuximab | MVP_top60_DNA+RNA_seed | AKT1 | -0.226733 | GBM | 8-MG-BA | MK-2206 |
| MVP_top60_DNA_seed | EGFR | -2.261333 | HNSC | FADU | Afatinib | MVP_top60_DNA+RNA_seed | EGFR | -0.225836 | GBM | 42-MG-BA | Lapatinib |
| MVP_top60_DNA_seed | MDM2 | -2.254735 | HNSC | KOSC-2 | Serdemetan | MVP_top60_DNA+RNA_seed | IGF1R | -0.224167 | GBM | KS-1 | BMS-754807 |

| | | | | | | | | | | | |
|---|---|---|---|---|---|---|---|---|---|---|---|
| MVP_top60_DNA_seed | EGFR | -2.209351 | HNSC | PCI-4B | Afatinib | MVP_top60_DNA+RNA_seed | AKT1 | -0.22194 | GBM | CAS-1 | Afuresertib |
| MVP_top60_DNA_seed | EGFR | -2.209088 | HNSC | CAL-33 | Gefitinib | MVP_top60_DNA+RNA_seed | PIK3CA | -0.219946 | GBM | D-245MG | AZD8186 |
| MVP_top60_DNA_seed | SMARCA4 | -2.203612 | HNSC | BICR22 | PFI-3 | MVP_top60_DNA+RNA_seed | PIK3CB | -0.219946 | GBM | D-245MG | AZD8186 |
| MVP_top60_DNA_seed | SRC | -2.20092 | HNSC | JHU-011 | Bosutinib | MVP_top60_DNA+RNA_seed | AKT1 | -0.217717 | GBM | SF126 | Uprosertib |
| MVP_top60_DNA_seed | EGFR | -2.195564 | HNSC | OSC-19 | Afatinib | MVP_top60_DNA+RNA_seed | PIK3CA | -0.212897 | GBM | D-566MG | Buparlisib |
| MVP_top60_DNA_seed | ESR1 | -2.188832 | HNSC | BB49-HNC | Tamoxifen | MVP_top60_DNA+RNA_seed | PIK3CB | -0.212897 | GBM | D-566MG | Buparlisib |
| MVP_top60_DNA_seed | EGFR | -2.187051 | HNSC | CAL-27 | AST-1306 | MVP_top60_DNA+RNA_seed | PIK3CD | -0.212897 | GBM | D-566MG | Buparlisib |
| MVP_top60_DNA_seed | EGFR | -2.149642 | HNSC | SCC-9 | Pelitinib | MVP_top60_DNA+RNA_seed | KIT | -0.212426 | GBM | YKG-1 | Sorafenib |
| MVP_top60_DNA_seed | SRC | -2.146544 | HNSC | DOK | Bosutinib | MVP_top60_DNA+RNA_seed | AKT1 | -0.211411 | GBM | D-542MG | AZD5363 |
| MVP_top60_DNA_seed | EGFR | -2.146392 | HNSC | HSC-2 | Cetuximab | MVP_top60_DNA+RNA_seed | CDK4 | -0.208943 | GBM | SF295 | Palbociclib |
| MVP_top60_DNA_seed | HSP90AA1 | -2.142554 | HNSC | HSC-3 | Tanespimycin | MVP_top60_DNA+RNA_seed | KIT | -0.208197 | GBM | D-245MG | Motesanib |
| MVP_top60_DNA_seed | EGFR | -2.135437 | HNSC | CAL-33 | Afatinib | MVP_top60_DNA+RNA_seed | TP53 | -0.206846 | GBM | 8-MG-BA | MIRA-1 |
| MVP_top60_DNA_seed | EGFR | -2.128159 | HNSC | BB30-HNC | Afatinib | MVP_top60_DNA+RNA_seed | AKT1 | -0.206453 | GBM | YH-13 | GSK2110183B |
| MVP_top60_DNA_seed | EGFR | -2.101722 | HNSC | SCC-9 | CI-1033 | MVP_top60_DNA+RNA_seed | PIK3CA | -0.202898 | GBM | CAS-1 | AZD8186 |
| MVP_top60_DNA_seed | SRC | -2.088613 | HNSC | JHU-022 | Bosutinib | MVP_top60_DNA+RNA_seed | PIK3CB | -0.202898 | GBM | CAS-1 | AZD8186 |
| MVP_top60_DNA_seed | EGFR | -2.073805 | HNSC | JHU-029 | Gefitinib | MVP_top60_DNA+RNA_seed | MET | -0.202124 | GBM | AM-38 | Savolitinib |
| MVP_top60_DNA_seed | SRC | -2.072769 | HNSC | JHU-029 | Bosutinib | MVP_top60_DNA+RNA_seed | MET | -0.202031 | GBM | SF268 | Savolitinib |
| MVP_top60_DNA_seed | HSP90AA1 | -2.069089 | HNSC | FADU | Tanespimycin | MVP_top60_DNA+RNA_seed | AKT1 | -0.200535 | GBM | D-423MG | AZD5363 |
| MVP_top60_DNA_seed | EGFR | -2.0645 | HNSC | PCI-4B | AZD8931 | MVP_top60_DNA+RNA_seed | MET | -0.199337 | GBM | A172 | Crizotinib |
| MVP_top60_DNA_seed | AKT1 | -2.061699 | HNSC | BICR10 | MK-2206 | MVP_top60_DNA+RNA_seed | IGF1R | -0.199204 | GBM | T98G | BMS-536924 |
| MVP_top60_DNA_seed | EGFR | -2.044606 | HNSC | HO-1-u-1 | Afatinib | MVP_top60_DNA+RNA_seed | EGFR | -0.19335 | GBM | D-245MG | Sapitinib |
| MVP_top60_DNA_seed | CDK2 | -2.033806 | HNSC | BB49-HNC | AZD5438 | MVP_top60_DNA+RNA_seed | ERBB3 | -0.19335 | GBM | D-245MG | Sapitinib |
| MVP_top60_DNA_seed | SMARCA4 | -2.011572 | HNSC | JHU-029 | PFI-3 | MVP_top60_DNA+RNA_seed | AKT1 | -0.192325 | GBM | D-247MG | AT13148 |
| MVP_top60_DNA_seed | CDK2 | -2.007055 | HNSC | BICR22 | AZD5438 | MVP_top60_DNA+RNA_seed | PIK3CB | -0.190454 | GBM | D-245MG | AZD6482 |
| MVP_top60_DNA_seed | CDK2 | -2.006513 | HNSC | HSC-2 | AZD5438 | MVP_top60_DNA+RNA_seed | PIK3CA | -0.186516 | GBM | U-118-MG | Buparlisib |
| MVP_top60_DNA_seed | EGFR | -2.005404 | HNSC | BB30-HNC | Afatinib | MVP_top60_DNA+RNA_seed | PIK3CB | -0.186516 | GBM | U-118-MG | Buparlisib |
| MVP_top60_DNA_seed | EGFR | -2.002407 | HNSC | FADU | AZD8931 | MVP_top60_DNA+RNA_seed | PIK3CD | -0.186516 | GBM | U-118-MG | Buparlisib |
| MVP_top60_DNA_seed | EGFR | -1.996737 | HNSC | BHY | AST-1306 | MVP_top60_DNA+RNA_seed | JAK1 | -0.184306 | GBM | YH-13 | JAK1_8709 |
| MVP_top60_DNA_seed | SRC | -1.99661 | HNSC | PCI-4B | Bosutinib | MVP_top60_DNA+RNA_seed | IGF1R | -0.182896 | GBM | SW1088 | BMS-536924 |
| MVP_top60_DNA_seed | EGFR | -1.988704 | HNSC | SCC-9 | AZD8931 | MVP_top60_DNA+RNA_seed | AKT1 | -0.18213 | GBM | D-247MG | Ipatasertib |
| MVP_top60_DNA_seed | EGFR | -1.985278 | HNSC | BB30-HNC | AST-1306 | MVP_top60_DNA+RNA_seed | CDK4 | -0.181636 | GBM | YKG-1 | Palbociclib |
| MVP_top60_DNA_seed | AR | -1.969553 | HNSC | BICR78 | AZD3514 | MVP_top60_DNA+RNA_seed | AKT1 | -0.180479 | GBM | LN-18 | AZD5363 |
| MVP_top60_DNA_seed | EGFR | -1.966485 | HNSC | SCC90 | Cetuximab | MVP_top60_DNA+RNA_seed | EGFR | -0.17646 | GBM | SW1088 | Sapitinib |
| MVP_top60_DNA_seed | EGFR | -1.955558 | HNSC | HSC-2 | Afatinib | MVP_top60_DNA+RNA_seed | ERBB3 | -0.17646 | GBM | SW1088 | Sapitinib |
| MVP_top60_DNA_seed | EGFR | -1.949914 | HNSC | JHU-022 | Afatinib | MVP_top60_DNA+RNA_seed | AKT1 | -0.172119 | GBM | SF295 | AT13148 |
| MVP_top60_DNA_seed | NTRK1 | -1.936917 | HNSC | SAT | GW441756 | MVP_top60_DNA+RNA_seed | EP300 | -0.171448 | GBM | AM-38 | CPI-637 |
| MVP_top60_DNA_seed | ESR1 | -1.933342 | HNSC | KON | Tamoxifen | MVP_top60_DNA+RNA_seed | TP53 | -0.170488 | GBM | 42-MG-BA | MIRA-1 |
| MVP_top60_DNA_seed | EGFR | -1.926396 | HNSC | HSC-3 | Afatinib | MVP_top60_DNA+RNA_seed | KIT | -0.17028 | GBM | D-566MG | Axitinib |
| MVP_top60_DNA_seed | HSP90AA1 | -1.915538 | HNSC | KON | Tanespimycin | MVP_top60_DNA+RNA_seed | JAK2 | -0.169915 | GBM | Becker | AZ960 |
| MVP_top60_DNA_seed | NTRK1 | -1.910623 | HNSC | BICR78 | GW441756 | MVP_top60_DNA+RNA_seed | JAK3 | -0.169915 | GBM | Becker | AZ960 |
| MVP_top60_DNA_seed | PIK3CA | -1.907966 | HNSC | BB49-HNC | AZD8835 | MVP_top60_DNA+RNA_seed | AKT1 | -0.168918 | GBM | DK-MG | Ipatasertib |

| | | | | | | | | | | | |
|---|---|---|---|---|---|---|---|---|---|---|---|
| MVP_top60_DNA_seed | ESR1 | -1.892157 | HNSC | KOSC-2 | Fulvestrant | MVP_top60_DNA+RNA_seed | JAK2 | -0.162817 | GBM | U-118-MG | JAK_8517 |
| MVP_top60_DNA_seed | RAC1 | -1.886122 | HNSC | PCI-38 | EHT-1864 | MVP_top60_DNA+RNA_seed | JAK1 | -0.162817 | GBM | U-118-MG | JAK_8517 |
| MVP_top60_DNA_seed | AR | -1.884322 | HNSC | CAL-27 | AZD3514 | MVP_top60_DNA+RNA_seed | TP53 | -0.162285 | GBM | D-263MG | MIRA-1 |
| MVP_top60_DNA_seed | EGFR | -1.881316 | HNSC | SAT | AST-1306 | MVP_top60_DNA+RNA_seed | KRAS | -0.159613 | GBM | SF295 | KRAS (G12C) Inhibitor-1 |
| MVP_top60_DNA_seed | EGFR | -1.88097 | HNSC | HO-1-N-1 | Afatinib | MVP_top60_DNA+RNA_seed | IGF1R | -0.154917 | GBM | D-263MG | BMS-754807 |
| MVP_top60_DNA_seed | ESR1 | -1.878531 | HNSC | Detroit562 | Tamoxifen | MVP_top60_DNA+RNA_seed | IGF1R | -0.154382 | GBM | CAS-1 | NVP-ADW742 |
| MVP_top60_DNA_seed | EGFR | -1.876535 | HNSC | Ca9-22 | AST-1306 | MVP_top60_DNA+RNA_seed | IGF1R | -0.152942 | GBM | 8-MG-BA | Linsitinib |
| MVP_top60_DNA_seed | EGFR | -1.873166 | HNSC | FADU | Gefitinib | MVP_top60_DNA+RNA_seed | AKT1 | -0.149859 | GBM | LN-18 | Uprosertib |
| MVP_top60_DNA_seed | HSP90AA1 | -1.868938 | HNSC | FADU | CCT-018159 | MVP_top60_DNA+RNA_seed | KIT | -0.149323 | GBM | D-423MG | Motesanib |
| MVP_top60_DNA_seed | ESR1 | -1.86727 | HNSC | Ca9-22 | Fulvestrant | MVP_top60_DNA+RNA_seed | JAK2 | -0.144879 | GBM | M059J | AZ960 |
| MVP_top60_DNA_seed | HDAC1 | -1.863047 | HNSC | Ca9-22 | JQ12 | MVP_top60_DNA+RNA_seed | JAK3 | -0.144879 | GBM | M059J | AZ960 |
| MVP_top60_DNA_seed | EGFR | -1.861327 | HNSC | SCC-15 | CI-1033 | MVP_top60_DNA+RNA_seed | JAK1 | -0.143102 | GBM | AM-38 | JAK1_8709 |
| MVP_top60_DNA_seed | AKT1 | -1.86076 | HNSC | SAT | BAY AKT1 | MVP_top60_DNA+RNA_seed | AKT1 | -0.139981 | GBM | Becker | Uprosertib |
| MVP_top60_DNA_seed | SRC | -1.858924 | HNSC | SAT | Bosutinib | MVP_top60_DNA+RNA_seed | PIK3CA | -0.139669 | GBM | D-542MG | Alpelisib |
| MVP_top60_DNA_seed | EGFR | -1.853894 | HNSC | HSC-2 | Afatinib | MVP_top60_DNA+RNA_seed | IGF1R | -0.138627 | GBM | YH-13 | BMS-754807 |
| MVP_top60_DNA_seed | PIK3CA | -1.848721 | HNSC | JHU-029 | AZD8835 | MVP_top60_DNA+RNA_seed | EGFR | -0.138111 | GBM | 42-MG-BA | Osimertinib |
| MVP_top60_DNA_seed | EGFR | -1.847063 | HNSC | HO-1-u-1 | Gefitinib | MVP_top60_DNA+RNA_seed | CDK4 | -0.137955 | GBM | AM-38 | Palbociclib |
| MVP_top60_DNA_seed | EGFR | -1.84096 | HNSC | JHU-022 | Afatinib | MVP_top60_DNA+RNA_seed | AKT1 | -0.134692 | GBM | U-118-MG | Ipatasertib |
| MVP_top60_DNA_seed | EGFR | -1.838059 | HNSC | HO-1-N-1 | Afatinib | MVP_top60_DNA+RNA_seed | IGF1R | -0.134525 | GBM | D-392MG | BMS-536924 |
| MVP_top60_DNA_seed | PIK3CA | -1.837295 | HNSC | HSC-2 | PI3Ka_4409 | MVP_top60_DNA+RNA_seed | IGF1R | -0.134354 | GBM | SK-MG-1 | Linsitinib |
| MVP_top60_DNA_seed | AR | -1.833733 | HNSC | CAL-27 | Bicalutamide | MVP_top60_DNA+RNA_seed | JAK2 | -0.128926 | GBM | M059J | Ruxolitinib |
| MVP_top60_DNA_seed | NTRK1 | -1.825294 | HNSC | BB49-HNC | AZD1332 | MVP_top60_DNA+RNA_seed | JAK1 | -0.128926 | GBM | M059J | Ruxolitinib |
| MVP_top60_DNA_seed | HSP90AA1 | -1.821952 | HNSC | JHU-011 | Tanespimycin | MVP_top60_DNA+RNA_seed | KIT | -0.128427 | GBM | SF295 | Axitinib |
| MVP_top60_DNA_seed | ESR1 | -1.818345 | HNSC | HSC-3 | Fulvestrant | MVP_top60_DNA+RNA_seed | PIK3CA | -0.126408 | GBM | T98G | AZD8186 |
| MVP_top60_DNA_seed | SRC | -1.816354 | HNSC | BB30-HNC | Saracatinib | MVP_top60_DNA+RNA_seed | PIK3CB | -0.126408 | GBM | T98G | AZD8186 |
| MVP_top60_DNA_seed | PIK3CA | -1.796513 | HNSC | CAL-33 | PI3Ka_4409 | MVP_top60_DNA+RNA_seed | AKT1 | -0.124344 | GBM | SK-MG-1 | GSK2110183B |
| MVP_top60_DNA_seed | ESR1 | -1.794197 | HNSC | CAL-27 | Tamoxifen | MVP_top60_DNA+RNA_seed | IGF1R | -0.124087 | GBM | GB-1 | Linsitinib |
| MVP_top60_DNA_seed | SRC | -1.791462 | HNSC | KON | Bosutinib | MVP_top60_DNA+RNA_seed | EGFR | -0.12271 | GBM | 8-MG-BA | Erlotinib |
| MVP_top60_DNA_seed | AKT1 | -1.786302 | HNSC | JHU-029 | AKT inhibitor VIII | MVP_top60_DNA+RNA_seed | CDK4 | -0.12219 | GBM | D-566MG | Ribociclib |
| MVP_top60_DNA_seed | PIK3CA | -1.785323 | HNSC | DOK | AZD8835 | MVP_top60_DNA+RNA_seed | PIK3CA | -0.121559 | GBM | D-263MG | GNE-317 |
| MVP_top60_DNA_seed | PIK3CA | -1.782516 | HNSC | HSC-3 | PI3Ka_4409 | MVP_top60_DNA+RNA_seed | JAK1 | -0.119898 | GBM | 42-MG-BA | JAK1_8709 |
| MVP_top60_DNA_seed | ESR1 | -1.762484 | HNSC | PCI-38 | Fulvestrant | MVP_top60_DNA+RNA_seed | AKT1 | -0.119848 | GBM | D-542MG | AT13148 |
| MVP_top60_DNA_seed | EGFR | -1.755606 | HNSC | BICR78 | AZD8931 | MVP_top60_DNA+RNA_seed | MET | -0.119519 | GBM | LN-229 | Savolitinib |
| MVP_top60_DNA_seed | EGFR | -1.754129 | HNSC | PCI-38 | Gefitinib | MVP_top60_DNA+RNA_seed | PIK3CB | -0.118698 | GBM | SF126 | AZD6482 |
| MVP_top60_DNA_seed | NTRK1 | -1.74913 | HNSC | PCI-15A | GW441756 | MVP_top60_DNA+RNA_seed | EGFR | -0.116719 | GBM | A172 | Afatinib |
| MVP_top60_DNA_seed | EGFR | -1.739624 | HNSC | BB30-HNC | AZD8931 | MVP_top60_DNA+RNA_seed | AKT1 | -0.115227 | GBM | D-542MG | MK-2206 |
| MVP_top60_DNA_seed | EGFR | -1.737444 | HNSC | PCI-4B | CI-1033 | MVP_top60_DNA+RNA_seed | AKT1 | -0.111483 | GBM | GB-1 | Afuresertib |
| MVP_top60_DNA_seed | EGFR | -1.714523 | HNSC | BHY | Afatinib | MVP_top60_DNA+RNA_seed | AKT1 | -0.11139 | GBM | D-247MG | MK-2206 |
| MVP_top60_DNA_seed | EGFR | -1.711667 | HNSC | BHY | Cetuximab | MVP_top60_DNA+RNA_seed | PIK3CA | -0.110897 | GBM | M059J | AZD8186 |
| MVP_top60_DNA_seed | EGFR | -1.699656 | HNSC | SCC90 | Afatinib | MVP_top60_DNA+RNA_seed | PIK3CB | -0.110897 | GBM | M059J | AZD8186 |
| MVP_top60_DNA_seed | SRC | -1.693545 | HNSC | PCI-38 | Bosutinib | MVP_top60_DNA+RNA_seed | JAK1 | -0.109389 | GBM | DK-MG | JAK1_8709 |

| | | | | | | | | | | | |
|---|---|---|---|---|---|---|---|---|---|---|---|
| MVP_top60_DNA_seed | ESR1 | -1.69107 | HNSC | HSC-3 | Tamoxifen | MVP_top60_DNA+RNA_seed | AKT1 | -0.107964 | GBM | SNB75 | MK-2206 |
| MVP_top60_DNA_seed | PIK3CA | -1.678015 | HNSC | SCC-9 | PI3Ka_4409 | MVP_top60_DNA+RNA_seed | PIK3CA | -0.105879 | GBM | U-87-MG | Buparlisib |
| MVP_top60_DNA_seed | EGFR | -1.673231 | HNSC | PCI-15A | Gefitinib | MVP_top60_DNA+RNA_seed | PIK3CB | -0.105879 | GBM | U-87-MG | Buparlisib |
| MVP_top60_DNA_seed | SRC | -1.670905 | HNSC | LB771-HNC | Bosutinib | MVP_top60_DNA+RNA_seed | PIK3CD | -0.105879 | GBM | U-87-MG | Buparlisib |
| MVP_top60_DNA_seed | SRC | -1.670409 | HNSC | CAL-33 | Bosutinib | MVP_top60_DNA+RNA_seed | AKT1 | -0.104101 | GBM | YKG-1 | AT13148 |
| MVP_top60_DNA_seed | EGFR | -1.670006 | HNSC | PCI-6A | Afatinib | MVP_top60_DNA+RNA_seed | CDK4 | -0.102853 | GBM | D-247MG | Palbociclib |
| MVP_top60_DNA_seed | SMARCA4 | -1.666284 | HNSC | HSC-3 | PFI-3 | MVP_top60_DNA+RNA_seed | KIT | -0.09975 | GBM | SF126 | Motesanib |
| MVP_top60_DNA_seed | SRC | -1.657499 | HNSC | BB30-HNC | Bosutinib | MVP_top60_DNA+RNA_seed | EGFR | -0.0997 | GBM | CAS-1 | Lapatinib |
| MVP_top60_DNA_seed | CDK2 | -1.650056 | HNSC | HO-1-N-1 | AZD5438 | MVP_top60_DNA+RNA_seed | AKT1 | -0.099349 | GBM | T98G | Uprosertib |
| MVP_top60_DNA_seed | EGFR | -1.649695 | HNSC | BHY | AZD8931 | MVP_top60_DNA+RNA_seed | JAK2 | -0.099094 | GBM | SK-MG-1 | AZ960 |
| MVP_top60_DNA_seed | HSP90AA1 | -1.647891 | HNSC | BB30-HNC | CCT-018159 | MVP_top60_DNA+RNA_seed | JAK3 | -0.099094 | GBM | SK-MG-1 | AZ960 |
| MVP_top60_DNA_seed | HSP90AA1 | -1.644215 | HNSC | JHU-022 | Elesclomol | MVP_top60_DNA+RNA_seed | JAK1 | -0.098785 | GBM | U-87-MG | JAK1_8709 |
| MVP_top60_DNA_seed | HSP90AA1 | -1.643061 | HNSC | HSC-2 | CCT-018159 | MVP_top60_DNA+RNA_seed | JAK2 | -0.092393 | GBM | Becker | GSK2276186C |
| MVP_top60_DNA_seed | EGFR | -1.641906 | HNSC | PCI-15A | AST-1306 | MVP_top60_DNA+RNA_seed | JAK1 | -0.092393 | GBM | Becker | GSK2276186C |
| MVP_top60_DNA_seed | EGFR | -1.637374 | HNSC | CAL-33 | Afatinib | MVP_top60_DNA+RNA_seed | JAK3 | -0.092393 | GBM | Becker | GSK2276186C |
| MVP_top60_DNA_seed | EGFR | -1.635801 | HNSC | PCI-4B | PF-00299804 | MVP_top60_DNA+RNA_seed | IGF1R | -0.090118 | GBM | LN-18 | BMS-754807 |
| MVP_top60_DNA_seed | CDK2 | -1.634374 | HNSC | Detroit562 | AZD5438 | MVP_top60_DNA+RNA_seed | IGF1R | -0.088571 | GBM | A172 | BMS-536924 |
| MVP_top60_DNA_seed | EGFR | -1.61617 | HNSC | BHY | Afatinib | MVP_top60_DNA+RNA_seed | CDK4 | -0.086184 | GBM | KS-1 | Ribociclib |
| MVP_top60_DNA_seed | EP300 | -1.615592 | HNSC | LB771-HNC | I-CBP112 | MVP_top60_DNA+RNA_seed | AKT1 | -0.083082 | GBM | D-247MG | Uprosertib |
| MVP_top60_DNA_seed | CREBBP | -1.615592 | HNSC | LB771-HNC | I-CBP112 | MVP_top60_DNA+RNA_seed | EGFR | -0.082638 | GBM | SNB75 | Osimertinib |
| MVP_top60_DNA_seed | AKT1 | -1.603401 | HNSC | SCC90 | AKT inhibitor VIII | MVP_top60_DNA+RNA_seed | PIK3CA | -0.081116 | GBM | A172 | AZD8186 |
| MVP_top60_DNA_seed | EGFR | -1.595951 | HNSC | PE-CA-PJ15 | AST-1306 | MVP_top60_DNA+RNA_seed | PIK3CB | -0.081116 | GBM | A172 | AZD8186 |
| MVP_top60_DNA_seed | AR | -1.591649 | HNSC | OSC-20 | Bicalutamide | MVP_top60_DNA+RNA_seed | AKT1 | -0.076703 | GBM | CAS-1 | Uprosertib |
| MVP_top60_DNA_seed | HDAC1 | -1.581489 | HNSC | JHU-029 | Dacinostat | MVP_top60_DNA+RNA_seed | KIT | -0.076167 | GBM | D-542MG | Sorafenib |
| MVP_top60_DNA_seed | EGFR | -1.578351 | HNSC | PCI-4B | AST-1306 | MVP_top60_DNA+RNA_seed | PIK3CA | -0.074798 | GBM | D-423MG | Buparlisib |
| MVP_top60_DNA_seed | NTRK1 | -1.570224 | HNSC | SCC90 | GW441756 | MVP_top60_DNA+RNA_seed | PIK3CB | -0.074798 | GBM | D-423MG | Buparlisib |
| MVP_top60_DNA_seed | HSP90AA1 | -1.567286 | HNSC | PCI-38 | Tanespimycin | MVP_top60_DNA+RNA_seed | PIK3CD | -0.074798 | GBM | D-423MG | Buparlisib |
| MVP_top60_DNA_seed | HDAC1 | -1.560998 | HNSC | PCI-6A | Dacinostat | MVP_top60_DNA+RNA_seed | AKT1 | -0.071645 | GBM | D-566MG | AZD5363 |
| MVP_top60_DNA_seed | SRC | -1.560412 | HNSC | SKN-3 | Bosutinib | MVP_top60_DNA+RNA_seed | IGF1R | -0.06858 | GBM | 8-MG-BA | GSK1904529A |
| MVP_top60_DNA_seed | EGFR | -1.560063 | HNSC | SCC90 | AZD8931 | MVP_top60_DNA+RNA_seed | IGF1R | -0.067668 | GBM | 42-MG-BA | BMS-754807 |
| MVP_top60_DNA_seed | AKT1 | -1.554152 | HNSC | BB30-HNC | AKT inhibitor VIII | MVP_top60_DNA+RNA_seed | AKT1 | -0.066044 | GBM | 8-MG-BA | Uprosertib |
| MVP_top60_DNA_seed | HDAC1 | -1.552383 | HNSC | JHU-011 | Dacinostat | MVP_top60_DNA+RNA_seed | JAK2 | -0.066034 | GBM | GB-1 | AZ960 |
| MVP_top60_DNA_seed | EGFR | -1.55195 | HNSC | SKN-3 | Gefitinib | MVP_top60_DNA+RNA_seed | JAK3 | -0.066034 | GBM | GB-1 | AZ960 |
| MVP_top60_DNA_seed | HSP90AA1 | -1.544523 | HNSC | HO-1-u-1 | Tanespimycin | MVP_top60_DNA+RNA_seed | EGFR | -0.06101 | GBM | LN-229 | Gefitinib |
| MVP_top60_DNA_seed | HSP90AA1 | -1.534773 | HNSC | PCI-30 | Tanespimycin | MVP_top60_DNA+RNA_seed | EP300 | -0.05961 | GBM | SK-MG-1 | CPI-637 |
| MVP_top60_DNA_seed | EGFR | -1.53223 | HNSC | FADU | Afatinib | MVP_top60_DNA+RNA_seed | AKT1 | -0.059306 | GBM | DK-MG | Uprosertib |
| MVP_top60_DNA_seed | AR | -1.531699 | HNSC | JHU-011 | AZD3514 | MVP_top60_DNA+RNA_seed | EGFR | -0.059066 | GBM | 8-MG-BA | Sapitinib |
| MVP_top60_DNA_seed | SRC | -1.529339 | HNSC | CAL-27 | Bosutinib | MVP_top60_DNA+RNA_seed | ERBB3 | -0.059066 | GBM | 8-MG-BA | Sapitinib |
| MVP_top60_DNA_seed | EGFR | -1.525474 | HNSC | OSC-19 | Afatinib | MVP_top60_DNA+RNA_seed | IGF1R | -0.058826 | GBM | U-118-MG | NVP-ADW742 |
| MVP_top60_DNA_seed | EGFR | -1.514295 | HNSC | BB30-HNC | Cetuximab | MVP_top60_DNA+RNA_seed | AKT1 | -0.05759 | GBM | SNB75 | AZD5363 |
| MVP_top60_DNA_seed | AKT1 | -1.510364 | HNSC | BICR10 | AKT inhibitor VIII | MVP_top60_DNA+RNA_seed | EGFR | -0.057422 | GBM | A172 | Gefitinib |

| | | | | | | | | | | | |
|---|---|---|---|---|---|---|---|---|---|---|---|
| MVP_top60_DNA_seed | EGFR | -1.505685 | HNSC | Ca9-22 | Gefitinib | MVP_top60_DNA+RNA_seed | AKT1 | -0.056498 | GBM | U-87-MG | Ipatasertib |
| MVP_top60_DNA_seed | CDK2 | -1.501826 | HNSC | FADU | AZD5438 | MVP_top60_DNA+RNA_seed | EGFR | -0.055395 | GBM | D-542MG | Lapatinib |
| MVP_top60_DNA_seed | AR | -1.496793 | HNSC | HO-1-N-1 | AZD3514 | MVP_top60_DNA+RNA_seed | CDK4 | -0.053038 | GBM | U251 | Palbociclib |
| MVP_top60_DNA_seed | EGFR | -1.494677 | HNSC | HSC-2 | AZD8931 | MVP_top60_DNA+RNA_seed | EGFR | -0.052 | GBM | 42-MG-BA | Afatinib |
| MVP_top60_DNA_seed | HSP90AA1 | -1.494105 | HNSC | DOK | Tanespimycin | MVP_top60_DNA+RNA_seed | EGFR | -0.050785 | GBM | A172 | Osimertinib |
| MVP_top60_DNA_seed | PIK3CA | -1.492015 | HNSC | DOK | PI3Ka_4409 | MVP_top60_DNA+RNA_seed | AKT1 | -0.049438 | GBM | SNB75 | AT13148 |
| MVP_top60_DNA_seed | ESR1 | -1.488211 | HNSC | JHU-029 | Tamoxifen | MVP_top60_DNA+RNA_seed | PIK3CA | -0.048167 | GBM | GB-1 | Buparlisib |
| MVP_top60_DNA_seed | PIK3CA | -1.487289 | HNSC | BICR10 | AZD8835 | MVP_top60_DNA+RNA_seed | PIK3CB | -0.048167 | GBM | GB-1 | Buparlisib |
| MVP_top60_DNA_seed | MDM2 | -1.487182 | HNSC | BICR22 | Serdemetan | MVP_top60_DNA+RNA_seed | PIK3CD | -0.048167 | GBM | GB-1 | Buparlisib |
| MVP_top60_DNA_seed | EGFR | -1.482341 | HNSC | HO-1-N-1 | Gefitinib | MVP_top60_DNA+RNA_seed | IGF1R | -0.042407 | GBM | SW1088 | BMS-754807 |
| MVP_top60_DNA_seed | EGFR | -1.47772 | HNSC | PCI-6A | PF-00299804 | MVP_top60_DNA+RNA_seed | EGFR | -0.042264 | GBM | D-566MG | Erlotinib |
| MVP_top60_DNA_seed | AR | -1.47237 | HNSC | BB49-HNC | Bicalutamide | MVP_top60_DNA+RNA_seed | IGF1R | -0.040794 | GBM | D-566MG | Linsitinib |
| MVP_top60_DNA_seed | AR | -1.466825 | HNSC | JHU-029 | AZD3514 | MVP_top60_DNA+RNA_seed | MET | -0.039125 | GBM | D-566MG | Savolitinib |
| MVP_top60_DNA_seed | PIK3CA | -1.463342 | HNSC | BICR22 | PI3Ka_4409 | MVP_top60_DNA+RNA_seed | PIK3CA | -0.034133 | GBM | AM-38 | GNE-317 |
| MVP_top60_DNA_seed | ESR1 | -1.462735 | HNSC | SAT | Fulvestrant | MVP_top60_DNA+RNA_seed | TP53 | -0.03173 | GBM | M059J | MIRA-1 |
| MVP_top60_DNA_seed | SMARCA4 | -1.460472 | HNSC | SCC-9 | PFI-3 | MVP_top60_DNA+RNA_seed | EGFR | -0.0309 | GBM | SF295 | Osimertinib |
| MVP_top60_DNA_seed | HDAC1 | -1.459815 | HNSC | Ca9-22 | Dacinostat | MVP_top60_DNA+RNA_seed | MET | -0.030863 | GBM | 42-MG-BA | Crizotinib |
| MVP_top60_DNA_seed | HSP90AA1 | -1.458934 | HNSC | Ca9-22 | CCT-018159 | MVP_top60_DNA+RNA_seed | CDK4 | -0.030002 | GBM | U-87-MG | Palbociclib |
| MVP_top60_DNA_seed | AR | -1.444556 | HNSC | SCC-15 | AZD3514 | MVP_top60_DNA+RNA_seed | CDK4 | -0.021426 | GBM | 8-MG-BA | Palbociclib |
| MVP_top60_DNA_seed | SRC | -1.437832 | HNSC | HO-1-N-1 | Bosutinib | MVP_top60_DNA+RNA_seed | JAK2 | -0.018538 | GBM | LN-18 | JAK_8517 |
| MVP_top60_DNA_seed | EGFR | -1.437435 | HNSC | CAL-27 | Pelitinib | MVP_top60_DNA+RNA_seed | JAK1 | -0.018538 | GBM | LN-18 | JAK_8517 |
| MVP_top60_DNA_seed | SMARCA4 | -1.43625 | HNSC | HO-1-u-1 | PFI-3 | MVP_top60_DNA+RNA_seed | PIK3CA | -0.017547 | GBM | CAS-1 | Buparlisib |
| MVP_top60_DNA_seed | NTRK1 | -1.433588 | HNSC | KON | GW441756 | MVP_top60_DNA+RNA_seed | PIK3CB | -0.017547 | GBM | CAS-1 | Buparlisib |
| MVP_top60_DNA_seed | EGFR | -1.432868 | HNSC | BB30-HNC | Pelitinib | MVP_top60_DNA+RNA_seed | PIK3CD | -0.017547 | GBM | CAS-1 | Buparlisib |
| MVP_top60_DNA_seed | EGFR | -1.430248 | HNSC | SAS | Afatinib | MVP_top60_DNA+RNA_seed | JAK2 | -0.016788 | GBM | D-392MG | AZ960 |
| MVP_top60_DNA_seed | SMARCA4 | -1.428647 | HNSC | SCC-15 | PFI-3 | MVP_top60_DNA+RNA_seed | JAK3 | -0.016788 | GBM | D-392MG | AZ960 |
| MVP_top60_DNA_seed | AKT1 | -1.426163 | HNSC | PCI-15A | MK-2206 | MVP_top60_DNA+RNA_seed | EGFR | -0.015156 | GBM | U-87-MG | AZD3759 |
| MVP_top60_DNA_seed | PIK3CA | -1.423077 | HNSC | HSC-4 | PI3Ka_4409 | MVP_top60_DNA+RNA_seed | EGFR | -0.014364 | GBM | YH-13 | Erlotinib |
| MVP_top60_DNA_seed | AR | -1.420247 | HNSC | HO-1-N-1 | Bicalutamide | MVP_top60_DNA+RNA_seed | AKT1 | -0.013279 | GBM | D-245MG | AZD5363 |
| MVP_top60_DNA_seed | EGFR | -1.418758 | HNSC | BICR78 | Afatinib | MVP_top60_DNA+RNA_seed | CDK4 | -0.012447 | GBM | 8-MG-BA | Ribociclib |
| MVP_top60_DNA_seed | ESR1 | -1.416282 | HNSC | CAL-33 | Fulvestrant | MVP_top60_DNA+RNA_seed | AKT1 | -0.012302 | GBM | D-542MG | Uprosertib |
| MVP_top60_DNA_seed | HSP90AA1 | -1.412108 | HNSC | CAL-33 | CCT-018159 | MVP_top60_DNA+RNA_seed | JAK2 | -0.011785 | GBM | U251 | JAK_8517 |
| MVP_top60_DNA_seed | HSP90AA1 | -1.411594 | HNSC | JHU-022 | Tanespimycin | MVP_top60_DNA+RNA_seed | JAK1 | -0.011785 | GBM | U251 | JAK_8517 |
| MVP_top60_DNA_seed | AKT1 | -1.40823 | HNSC | JHU-011 | AKT inhibitor VIII | MVP_top60_DNA+RNA_seed | EGFR | -0.011375 | GBM | YKG-1 | Osimertinib |
| MVP_top60_DNA_seed | EGFR | -1.407675 | HNSC | CAL-27 | PF-00299804 | MVP_top60_DNA+RNA_seed | EP300 | -0.010627 | GBM | M059J | SGC-CBP30 |
| MVP_top60_DNA_seed | HSP90AA1 | -1.40592 | HNSC | Ca9-22 | Elesclomol | MVP_top60_DNA+RNA_seed | CREBBP | -0.010627 | GBM | M059J | SGC-CBP30 |
| MVP_top60_DNA_seed | NTRK1 | -1.40514 | HNSC | PCI-38 | AZD1332 | MVP_top60_DNA+RNA_seed | EP300 | -0.008614 | GBM | U251 | CPI-637 |
| MVP_top60_DNA_seed | EGFR | -1.397728 | HNSC | BB30-HNC | Gefitinib | MVP_top60_DNA+RNA_seed | MET | -0.008463 | GBM | U251 | Savolitinib |
| MVP_top60_DNA_seed | EGFR | -1.395582 | HNSC | PCI-15A | AZD8931 | MVP_top60_DNA+RNA_seed | EGFR | -0.007482 | GBM | YH-13 | Osimertinib |
| MVP_top60_DNA_seed | NTRK1 | -1.38965 | HNSC | PCI-38 | GW441756 | MVP_top60_DNA+RNA_seed | JAK2 | -0.005964 | GBM | SF295 | AZ960 |
| MVP_top60_DNA_seed | SRC | -1.388673 | HNSC | HSC-3 | Bosutinib | MVP_top60_DNA+RNA_seed | JAK3 | -0.005964 | GBM | SF295 | AZ960 |

| | | | | | | | | | | | |
|---|---|---|---|---|---|---|---|---|---|---|---|
| MVP_top60_DNA_seed | PIK3CA | -1.387447 | HNSC | BB49-HNC | PI3Ka_4409 | MVP_top60_DNA+RNA_seed | EGFR | -0.005381 | GBM | SF126 | Sapitinib |
| MVP_top60_DNA_seed | AKT1 | -1.38067 | HNSC | PCI-38 | AKT inhibitor VIII | MVP_top60_DNA+RNA_seed | ERBB3 | -0.005381 | GBM | SF126 | Sapitinib |
| MVP_top60_DNA_seed | HSP90AA1 | -1.367406 | HNSC | HSC-3 | Elesclomol | MVP_top60_DNA+RNA_seed | JAK2 | -0.001596 | GBM | D-392MG | GSK2276186C |
| MVP_top60_DNA_seed | PIK3CA | -1.367361 | HNSC | SCC-25 | PI3Ka_4409 | MVP_top60_DNA+RNA_seed | JAK1 | -0.001596 | GBM | D-392MG | GSK2276186C |
| MVP_top60_DNA_seed | ESR1 | -1.364695 | HNSC | CAL-27 | Fulvestrant | MVP_top60_DNA+RNA_seed | JAK3 | -0.001596 | GBM | D-392MG | GSK2276186C |
| MVP_top60_DNA_seed | EGFR | -1.356298 | HNSC | SCC-15 | Gefitinib | MVP_top60_DNA+RNA_seed | KIT | -0.001062 | GBM | U-118-MG | Sorafenib |
| MVP_top60_DNA_seed | NTRK1 | -1.356152 | HNSC | CAL-33 | GW441756 | MVP_top60_DNA+RNA_seed | AKT1 | 0.002269 | GBM | YKG-1 | Afuresertib |
| MVP_top60_DNA_seed | AKT1 | -1.347827 | HNSC | DOK | MK-2206 | MVP_top60_DNA+RNA_seed | EGFR | 0.002614 | GBM | M059J | Erlotinib |
| MVP_top60_DNA_seed | HSP90AA1 | -1.34219 | HNSC | HSC-4 | Elesclomol | MVP_top60_DNA+RNA_seed | EGFR | 0.004394 | GBM | U251 | Erlotinib |
| MVP_top60_DNA_seed | HSP90AA1 | -1.341298 | HNSC | HO-1-N-1 | Tanespimycin | MVP_top60_DNA+RNA_seed | IGF1R | 0.00468 | GBM | 42-MG-BA | GSK1904529A |
| MVP_top60_DNA_seed | HSP90AA1 | -1.340844 | HNSC | HSC-3 | Luminespib | MVP_top60_DNA+RNA_seed | EP300 | 0.00589 | GBM | 8-MG-BA | SGC-CBP30 |
| MVP_top60_DNA_seed | HSP90AA1 | -1.340209 | HNSC | SAS | CCT-018159 | MVP_top60_DNA+RNA_seed | CREBBP | 0.00589 | GBM | 8-MG-BA | SGC-CBP30 |
| MVP_top60_DNA_seed | MDM2 | -1.337219 | HNSC | PCI-4B | Serdemetan | MVP_top60_DNA+RNA_seed | TP53 | 0.007679 | GBM | T98G | MIRA-1 |
| MVP_top60_DNA_seed | EGFR | -1.336818 | HNSC | PCI-6A | CUDC-101 | MVP_top60_DNA+RNA_seed | JAK2 | 0.00891 | GBM | T98G | Ruxolitinib |
| MVP_top60_DNA_seed | EGFR | -1.329261 | HNSC | SCC-25 | Afatinib | MVP_top60_DNA+RNA_seed | JAK1 | 0.00891 | GBM | T98G | Ruxolitinib |
| MVP_top60_DNA_seed | EGFR | -1.328331 | HNSC | OSC-20 | Gefitinib | MVP_top60_DNA+RNA_seed | EGFR | 0.008988 | GBM | KS-1 | Sapitinib |
| MVP_top60_DNA_seed | HSP90AA1 | -1.327116 | HNSC | BB49-HNC | Tanespimycin | MVP_top60_DNA+RNA_seed | ERBB3 | 0.008988 | GBM | KS-1 | Sapitinib |
| MVP_top60_DNA_seed | SRC | -1.32466 | HNSC | FADU | Bosutinib | MVP_top60_DNA+RNA_seed | AKT1 | 0.009022 | GBM | M059J | Uprosertib |
| MVP_top60_DNA_seed | HSP90AA1 | -1.320702 | HNSC | Detroit562 | Tanespimycin | MVP_top60_DNA+RNA_seed | EGFR | 0.010215 | GBM | SF268 | Erlotinib |
| MVP_top60_DNA_seed | EGFR | -1.320389 | HNSC | PCI-38 | AZD8931 | MVP_top60_DNA+RNA_seed | AKT1 | 0.010466 | GBM | SNB75 | Afuresertib |
| MVP_top60_DNA_seed | EGFR | -1.320279 | HNSC | HSC-3 | Gefitinib | MVP_top60_DNA+RNA_seed | KIT | 0.010475 | GBM | YH-13 | Motesanib |
| MVP_top60_DNA_seed | HSP90AA1 | -1.31805 | HNSC | HSC-2 | Luminespib | MVP_top60_DNA+RNA_seed | PIK3CA | 0.015188 | GBM | D-263MG | Buparlisib |
| MVP_top60_DNA_seed | AKT1 | -1.311764 | HNSC | KON | MK-2206 | MVP_top60_DNA+RNA_seed | PIK3CB | 0.015188 | GBM | D-263MG | Buparlisib |
| MVP_top60_DNA_seed | MAPK1 | -1.309958 | HNSC | SAS | FR-180204 | MVP_top60_DNA+RNA_seed | PIK3CD | 0.015188 | GBM | D-263MG | Buparlisib |
| MVP_top60_DNA_seed | MAPK3 | -1.309958 | HNSC | SAS | FR-180204 | MVP_top60_DNA+RNA_seed | AKT1 | 0.016882 | GBM | U-118-MG | Uprosertib |
| MVP_top60_DNA_seed | EGFR | -1.308128 | HNSC | PCI-6A | AZD8931 | MVP_top60_DNA+RNA_seed | IGF1R | 0.019398 | GBM | SF126 | BMS-536924 |
| MVP_top60_DNA_seed | ESR1 | -1.306826 | HNSC | HO-1-N-1 | Tamoxifen | MVP_top60_DNA+RNA_seed | PIK3CA | 0.019902 | GBM | KS-1 | GNE-317 |
| MVP_top60_DNA_seed | MDM2 | -1.306716 | HNSC | HSC-3 | Serdemetan | MVP_top60_DNA+RNA_seed | PIK3CA | 0.022618 | GBM | U-87-MG | GNE-317 |
| MVP_top60_DNA_seed | AKT1 | -1.300432 | HNSC | KON | AKT inhibitor VIII | MVP_top60_DNA+RNA_seed | EGFR | 0.027993 | GBM | A172 | Lapatinib |
| MVP_top60_DNA_seed | PIK3CA | -1.298999 | HNSC | HSC-2 | AZD8835 | MVP_top60_DNA+RNA_seed | JAK2 | 0.028488 | GBM | U251 | GSK2276186C |
| MVP_top60_DNA_seed | NTRK1 | -1.295705 | HNSC | JHU-022 | GW441756 | MVP_top60_DNA+RNA_seed | JAK1 | 0.028488 | GBM | U251 | GSK2276186C |
| MVP_top60_DNA_seed | HSP90AA1 | -1.293905 | HNSC | FADU | Elesclomol | MVP_top60_DNA+RNA_seed | JAK3 | 0.028488 | GBM | U251 | GSK2276186C |
| MVP_top60_DNA_seed | CDK2 | -1.293311 | HNSC | CAL-27 | AZD5438 | MVP_top60_DNA+RNA_seed | EGFR | 0.029904 | GBM | YKG-1 | Gefitinib |
| MVP_top60_DNA_seed | HSP90AA1 | -1.27943 | HNSC | Ca9-22 | Luminespib | MVP_top60_DNA+RNA_seed | EGFR | 0.030789 | GBM | 8-MG-BA | Afatinib |
| MVP_top60_DNA_seed | CDK2 | -1.273665 | HNSC | PCI-38 | AZD5438 | MVP_top60_DNA+RNA_seed | CDK4 | 0.031313 | GBM | SW1088 | Palbociclib |
| MVP_top60_DNA_seed | SRC | -1.269328 | HNSC | BHY | Bosutinib | MVP_top60_DNA+RNA_seed | EGFR | 0.035055 | GBM | LN-18 | Sapitinib |
| MVP_top60_DNA_seed | HSP90AA1 | -1.269239 | HNSC | PCI-4B | Tanespimycin | MVP_top60_DNA+RNA_seed | ERBB3 | 0.035055 | GBM | LN-18 | Sapitinib |
| MVP_top60_DNA_seed | ESR1 | -1.266869 | HNSC | HO-1-N-1 | Fulvestrant | MVP_top60_DNA+RNA_seed | MET | 0.03628 | GBM | 8-MG-BA | Crizotinib |
| MVP_top60_DNA_seed | EGFR | -1.261107 | HNSC | DOK | AST-1306 | MVP_top60_DNA+RNA_seed | EGFR | 0.036472 | GBM | DK-MG | Erlotinib |
| MVP_top60_DNA_seed | EGFR | -1.256667 | HNSC | PCI-15A | Cetuximab | MVP_top60_DNA+RNA_seed | IGF1R | 0.037052 | GBM | Becker | BMS-754807 |
| MVP_top60_DNA_seed | HSP90AA1 | -1.249909 | HNSC | PCI-4B | CCT-018159 | MVP_top60_DNA+RNA_seed | AKT1 | 0.038208 | GBM | LN-405 | Uprosertib |

| | | | | | | | | | | | |
|---|---|---|---|---|---|---|---|---|---|---|---|
| MVP_top60_DNA_seed | ESR1 | -1.248514 | HNSC | SCC-25 | Tamoxifen | MVP_top60_DNA+RNA_seed | IGF1R | 0.038525 | GBM | SF295 | BMS-754807 |
| MVP_top60_DNA_seed | MDM2 | -1.24635 | HNSC | PCI-15A | Serdemetan | MVP_top60_DNA+RNA_seed | AKT1 | 0.041703 | GBM | A172 | Uprosertib |
| MVP_top60_DNA_seed | EGFR | -1.243164 | HNSC | DOK | PF-00299804 | MVP_top60_DNA+RNA_seed | EP300 | 0.043633 | GBM | M059J | CPI-637 |
| MVP_top60_DNA_seed | EGFR | -1.242007 | HNSC | SCC90 | Afatinib | MVP_top60_DNA+RNA_seed | AKT1 | 0.049579 | GBM | GB-1 | AT13148 |
| MVP_top60_DNA_seed | EGFR | -1.241919 | HNSC | BHY | Pelitinib | MVP_top60_DNA+RNA_seed | AKT1 | 0.051755 | GBM | D-566MG | Afuresertib |
| MVP_top60_DNA_seed | EGFR | -1.232298 | HNSC | SCC-25 | Pelitinib | MVP_top60_DNA+RNA_seed | PIK3CA | 0.053743 | GBM | D-245MG | GNE-317 |
| MVP_top60_DNA_seed | EGFR | -1.230817 | HNSC | PE-CA-PJ15 | Afatinib | MVP_top60_DNA+RNA_seed | JAK2 | 0.055117 | GBM | CAS-1 | GSK2276186C |
| MVP_top60_DNA_seed | AKT1 | -1.228785 | HNSC | SCC-9 | AKT inhibitor VIII | MVP_top60_DNA+RNA_seed | JAK1 | 0.055117 | GBM | CAS-1 | GSK2276186C |
| MVP_top60_DNA_seed | MDM2 | -1.225689 | HNSC | FADU | Serdemetan | MVP_top60_DNA+RNA_seed | JAK3 | 0.055117 | GBM | CAS-1 | GSK2276186C |
| MVP_top60_DNA_seed | EGFR | -1.225354 | HNSC | BHY | Gefitinib | MVP_top60_DNA+RNA_seed | AKT1 | 0.055236 | GBM | M059J | Afuresertib |
| MVP_top60_DNA_seed | AKT1 | -1.224708 | HNSC | JHU-011 | MK-2206 | MVP_top60_DNA+RNA_seed | AKT1 | 0.057558 | GBM | DK-MG | MK-2206 |
| MVP_top60_DNA_seed | ESR1 | -1.224568 | HNSC | FADU | Tamoxifen | MVP_top60_DNA+RNA_seed | IGF1R | 0.05865 | GBM | M059J | BMS-754807 |
| MVP_top60_DNA_seed | SRC | -1.223875 | HNSC | BICR22 | Bosutinib | MVP_top60_DNA+RNA_seed | AKT1 | 0.059325 | GBM | CAS-1 | Uprosertib |
| MVP_top60_DNA_seed | HSP90AA1 | -1.21769 | HNSC | PCI-6A | Tanespimycin | MVP_top60_DNA+RNA_seed | KIT | 0.060232 | GBM | U251 | Sorafenib |
| MVP_top60_DNA_seed | MAPK1 | -1.214048 | HNSC | Ca9-22 | FR-180204 | MVP_top60_DNA+RNA_seed | KIT | 0.060283 | GBM | LN-18 | Axitinib |
| MVP_top60_DNA_seed | MAPK3 | -1.214048 | HNSC | Ca9-22 | FR-180204 | MVP_top60_DNA+RNA_seed | AKT1 | 0.061964 | GBM | LN-229 | GSK2110183B |
| MVP_top60_DNA_seed | AKT1 | -1.210741 | HNSC | CAL-33 | AKT inhibitor VIII | MVP_top60_DNA+RNA_seed | TP53 | 0.063766 | GBM | SNB75 | MIRA-1 |
| MVP_top60_DNA_seed | HSP90AA1 | -1.208872 | HNSC | PE-CA-PJ15 | Tanespimycin | MVP_top60_DNA+RNA_seed | AKT1 | 0.064787 | GBM | U-87-MG | Afuresertib |
| MVP_top60_DNA_seed | AKT1 | -1.201096 | HNSC | BB49-HNC | BAY AKT1 | MVP_top60_DNA+RNA_seed | EGFR | 0.065361 | GBM | D-566MG | Gefitinib |
| MVP_top60_DNA_seed | PIK3CA | -1.199689 | HNSC | SAT | AZD8835 | MVP_top60_DNA+RNA_seed | EGFR | 0.067748 | GBM | LNZTA3WT4 | Afatinib |
| MVP_top60_DNA_seed | RAC1 | -1.19828 | HNSC | LB771-HNC | EHT-1864 | MVP_top60_DNA+RNA_seed | MET | 0.067944 | GBM | CAS-1 | Savolitinib |
| MVP_top60_DNA_seed | HSP90AA1 | -1.195921 | HNSC | KOSC-2 | Elesclomol | MVP_top60_DNA+RNA_seed | PIK3CA | 0.070068 | GBM | D-247MG | Buparlisib |
| MVP_top60_DNA_seed | RAC1 | -1.194694 | HNSC | PCI-4B | EHT-1864 | MVP_top60_DNA+RNA_seed | PIK3CB | 0.070068 | GBM | D-247MG | Buparlisib |
| MVP_top60_DNA_seed | EGFR | -1.194669 | HNSC | SCC-25 | Afatinib | MVP_top60_DNA+RNA_seed | PIK3CD | 0.070068 | GBM | D-247MG | Buparlisib |
| MVP_top60_DNA_seed | EGFR | -1.194208 | HNSC | HO-1-N-1 | AZD8931 | MVP_top60_DNA+RNA_seed | KRAS | 0.073121 | GBM | YH-13 | KRAS (G12C) Inhibitor-1 |
| MVP_top60_DNA_seed | AKT1 | -1.192205 | HNSC | JHU-029 | BAY AKT1 | MVP_top60_DNA+RNA_seed | EP300 | 0.073544 | GBM | GB-1 | SGC-CBP30 |
| MVP_top60_DNA_seed | HSP90AA1 | -1.192054 | HNSC | HSC-4 | CCT-018159 | MVP_top60_DNA+RNA_seed | CREBBP | 0.073544 | GBM | GB-1 | SGC-CBP30 |
| MVP_top60_DNA_seed | HSP90AA1 | -1.191721 | HNSC | JHU-029 | Luminespib | MVP_top60_DNA+RNA_seed | CDK4 | 0.076872 | GBM | D-423MG | Palbociclib |
| MVP_top60_DNA_seed | AR | -1.189547 | HNSC | HSC-2 | Bicalutamide | MVP_top60_DNA+RNA_seed | IGF1R | 0.077048 | GBM | SF268 | NVP-ADW742 |
| MVP_top60_DNA_seed | HSP90AA1 | -1.189518 | HNSC | BB30-HNC | Luminespib | MVP_top60_DNA+RNA_seed | EP300 | 0.077697 | GBM | LN-229 | CPI-637 |
| MVP_top60_DNA_seed | HSP90AA1 | -1.18678 | HNSC | PE-CA-PJ15 | CCT-018159 | MVP_top60_DNA+RNA_seed | CDK4 | 0.080189 | GBM | D-245MG | Palbociclib |
| MVP_top60_DNA_seed | HDAC1 | -1.176525 | HNSC | Ca9-22 | AR-42 | MVP_top60_DNA+RNA_seed | PIK3CA | 0.08024 | GBM | CAS-1 | GNE-317 |
| MVP_top60_DNA_seed | PIK3CA | -1.172892 | HNSC | SCC-9 | AZD8835 | MVP_top60_DNA+RNA_seed | KRAS | 0.082869 | GBM | SF268 | KRAS (G12C) Inhibitor-1 |
| MVP_top60_DNA_seed | AR | -1.168383 | HNSC | PCI-4B | AZD3514 | MVP_top60_DNA+RNA_seed | IGF1R | 0.083708 | GBM | YKG-1 | BMS-754807 |
| MVP_top60_DNA_seed | MAPK1 | -1.168145 | HNSC | SCC-25 | VX-11e | MVP_top60_DNA+RNA_seed | IGF1R | 0.084282 | GBM | AM-38 | BMS-754807 |
| MVP_top60_DNA_seed | PIK3CA | -1.164822 | HNSC | CAL-27 | PI3Ka_4409 | MVP_top60_DNA+RNA_seed | MET | 0.085066 | GBM | SF295 | Crizotinib |
| MVP_top60_DNA_seed | EGFR | -1.160657 | HNSC | SAS | Afatinib | MVP_top60_DNA+RNA_seed | IGF1R | 0.085717 | GBM | CAS-1 | BMS-536924 |
| MVP_top60_DNA_seed | HSP90AA1 | -1.160125 | HNSC | OSC-19 | CCT-018159 | MVP_top60_DNA+RNA_seed | AKT1 | 0.089598 | GBM | D-423MG | Ipatasertib |
| MVP_top60_DNA_seed | PIK3CA | -1.158928 | HNSC | CAL-27 | AZD8835 | MVP_top60_DNA+RNA_seed | TP53 | 0.090294 | GBM | SF126 | MIRA-1 |
| MVP_top60_DNA_seed | HSP90AA1 | -1.15652 | HNSC | PCI-4B | Luminespib | MVP_top60_DNA+RNA_seed | JAK2 | 0.091665 | GBM | 8-MG-BA | JAK_8517 |
| MVP_top60_DNA_seed | EGFR | -1.15583 | HNSC | PE-CA-PJ15 | Gefitinib | MVP_top60_DNA+RNA_seed | JAK1 | 0.091665 | GBM | 8-MG-BA | JAK_8517 |

| | | | | | | | | | | | |
|---|---|---|---|---|---|---|---|---|---|---|---|
| MVP_top60_DNA_seed | PIK3CA | -1.155628 | HNSC | PE-CA-PJ15 | PI3Ka_4409 | MVP_top60_DNA+RNA_seed | EGFR | 0.091768 | GBM | U-118-MG | AZD3759 |
| MVP_top60_DNA_seed | EGFR | -1.153564 | HNSC | PE-CA-PJ15 | CI-1033 | MVP_top60_DNA+RNA_seed | AKT1 | 0.095678 | GBM | D-245MG | Ipatasertib |
| MVP_top60_DNA_seed | EGFR | -1.144676 | HNSC | LB771-HNC | Lapatinib | MVP_top60_DNA+RNA_seed | PIK3CA | 0.098691 | GBM | YKG-1 | GNE-317 |
| MVP_top60_DNA_seed | AR | -1.143425 | HNSC | BB30-HNC | AZD3514 | MVP_top60_DNA+RNA_seed | IGF1R | 0.098986 | GBM | LNZTA3WT4 | BMS-754807 |
| MVP_top60_DNA_seed | RAC1 | -1.142579 | HNSC | SAS | EHT-1864 | MVP_top60_DNA+RNA_seed | AKT1 | 0.099142 | GBM | U251 | Afuresertib |
| MVP_top60_DNA_seed | HSP90AA1 | -1.141789 | HNSC | BICR22 | Tanespimycin | MVP_top60_DNA+RNA_seed | EGFR | 0.100732 | GBM | SF295 | Lapatinib |
| MVP_top60_DNA_seed | EP300 | -1.141697 | HNSC | Ca9-22 | I-CBP112 | MVP_top60_DNA+RNA_seed | PIK3CA | 0.106176 | GBM | SF126 | GNE-317 |
| MVP_top60_DNA_seed | CREBBP | -1.141697 | HNSC | Ca9-22 | I-CBP112 | MVP_top60_DNA+RNA_seed | AKT1 | 0.109301 | GBM | A172 | Afuresertib |
| MVP_top60_DNA_seed | HSP90AA1 | -1.1352 | HNSC | KON | CCT-018159 | MVP_top60_DNA+RNA_seed | EGFR | 0.110759 | GBM | U-118-MG | Osimertinib |
| MVP_top60_DNA_seed | EGFR | -1.132485 | HNSC | Ca9-22 | Cetuximab | MVP_top60_DNA+RNA_seed | AKT1 | 0.112449 | GBM | Becker | Uprosertib |
| MVP_top60_DNA_seed | EGFR | -1.131126 | HNSC | SAS | Gefitinib | MVP_top60_DNA+RNA_seed | MET | 0.113882 | GBM | AM-38 | Crizotinib |
| MVP_top60_DNA_seed | MDM2 | -1.130911 | HNSC | PCI-6A | Serdemetan | MVP_top60_DNA+RNA_seed | EGFR | 0.114846 | GBM | SK-MG-1 | Afatinib |
| MVP_top60_DNA_seed | ESR1 | -1.13087 | HNSC | BB30-HNC | Fulvestrant | MVP_top60_DNA+RNA_seed | EGFR | 0.115143 | GBM | DK-MG | Sapitinib |
| MVP_top60_DNA_seed | HDAC1 | -1.130786 | HNSC | JHU-029 | JQ12 | MVP_top60_DNA+RNA_seed | ERBB3 | 0.115143 | GBM | DK-MG | Sapitinib |
| MVP_top60_DNA_seed | EGFR | -1.127785 | HNSC | SKN-3 | Afatinib | MVP_top60_DNA+RNA_seed | PIK3CA | 0.115924 | GBM | D-542MG | Buparlisib |
| MVP_top60_DNA_seed | HSP90AA1 | -1.127717 | HNSC | KON | Luminespib | MVP_top60_DNA+RNA_seed | PIK3CB | 0.115924 | GBM | D-542MG | Buparlisib |
| MVP_top60_DNA_seed | PIK3CA | -1.127244 | HNSC | HSC-4 | AZD8835 | MVP_top60_DNA+RNA_seed | PIK3CD | 0.115924 | GBM | D-542MG | Buparlisib |
| MVP_top60_DNA_seed | EGFR | -1.126541 | HNSC | CAL-33 | Pelitinib | MVP_top60_DNA+RNA_seed | EGFR | 0.118834 | GBM | D-542MG | Osimertinib |
| MVP_top60_DNA_seed | MDM2 | -1.126137 | HNSC | BB30-HNC | Serdemetan | MVP_top60_DNA+RNA_seed | IGF1R | 0.118875 | GBM | U-118-MG | GSK1904529A |
| MVP_top60_DNA_seed | HSP90AA1 | -1.12509 | HNSC | JHU-011 | Luminespib | MVP_top60_DNA+RNA_seed | IGF1R | 0.121465 | GBM | DBTRG-05MG | BMS-536924 |
| MVP_top60_DNA_seed | EGFR | -1.12419 | HNSC | Ca9-22 | CUDC-101 | MVP_top60_DNA+RNA_seed | KIT | 0.122009 | GBM | LN-18 | Motesanib |
| MVP_top60_DNA_seed | AR | -1.119908 | HNSC | HSC-4 | AZD3514 | MVP_top60_DNA+RNA_seed | KIT | 0.122163 | GBM | GB-1 | Sorafenib |
| MVP_top60_DNA_seed | ESR1 | -1.119098 | HNSC | BICR22 | Fulvestrant | MVP_top60_DNA+RNA_seed | EGFR | 0.124166 | GBM | SF126 | AZD3759 |
| MVP_top60_DNA_seed | SMARCA4 | -1.118597 | HNSC | PCI-4B | PFI-3 | MVP_top60_DNA+RNA_seed | EGFR | 0.12491 | GBM | AM-38 | AZD3759 |
| MVP_top60_DNA_seed | HDAC1 | -1.116984 | HNSC | KOSC-2 | JQ12 | MVP_top60_DNA+RNA_seed | AKT1 | 0.126616 | GBM | D-263MG | Afuresertib |
| MVP_top60_DNA_seed | AKT1 | -1.114976 | HNSC | SCC-25 | AKT inhibitor VIII | MVP_top60_DNA+RNA_seed | CDK4 | 0.128484 | GBM | SF268 | Ribociclib |
| MVP_top60_DNA_seed | HSP90AA1 | -1.11277 | HNSC | BICR22 | Elesclomol | MVP_top60_DNA+RNA_seed | KIT | 0.129903 | GBM | LNZTA3WT4 | Motesanib |
| MVP_top60_DNA_seed | ESR1 | -1.106481 | HNSC | SCC-15 | Tamoxifen | MVP_top60_DNA+RNA_seed | EGFR | 0.132447 | GBM | D-566MG | AZD3759 |
| MVP_top60_DNA_seed | AR | -1.102502 | HNSC | BB49-HNC | AZD3514 | MVP_top60_DNA+RNA_seed | EGFR | 0.134693 | GBM | LNZTA3WT4 | Lapatinib |
| MVP_top60_DNA_seed | HSP90AA1 | -1.100712 | HNSC | HO-1-u-1 | Elesclomol | MVP_top60_DNA+RNA_seed | PIK3CA | 0.135088 | GBM | KS-1 | Alpelisib |
| MVP_top60_DNA_seed | HSP90AA1 | -1.100074 | HNSC | Detroit562 | CCT-018159 | MVP_top60_DNA+RNA_seed | EGFR | 0.135413 | GBM | T98G | Gefitinib |
| MVP_top60_DNA_seed | HDAC1 | -1.097368 | HNSC | PCI-6A | CAY10603 | MVP_top60_DNA+RNA_seed | MET | 0.136027 | GBM | YKG-1 | Crizotinib |
| MVP_top60_DNA_seed | SRC | -1.09439 | HNSC | KOSC-2 | Bosutinib | MVP_top60_DNA+RNA_seed | IGF1R | 0.136327 | GBM | D-542MG | BMS-754807 |
| MVP_top60_DNA_seed | HSP90AA1 | -1.092209 | HNSC | BHY | Luminespib | MVP_top60_DNA+RNA_seed | PIK3CA | 0.136491 | GBM | T98G | Buparlisib |
| MVP_top60_DNA_seed | HSP90AA1 | -1.087166 | HNSC | PCI-4B | SNX-2112 | MVP_top60_DNA+RNA_seed | PIK3CB | 0.136491 | GBM | T98G | Buparlisib |
| MVP_top60_DNA_seed | HDAC1 | -1.087024 | HNSC | BB30-HNC | Parthenolide | MVP_top60_DNA+RNA_seed | PIK3CD | 0.136491 | GBM | T98G | Buparlisib |
| MVP_top60_DNA_seed | AR | -1.086856 | HNSC | Ca9-22 | AZD3514 | MVP_top60_DNA+RNA_seed | KIT | 0.139652 | GBM | T98G | Motesanib |
| MVP_top60_DNA_seed | EGFR | -1.081415 | HNSC | BB30-HNC | CI-1033 | MVP_top60_DNA+RNA_seed | KIT | 0.140043 | GBM | D-245MG | Sorafenib |
| MVP_top60_DNA_seed | EGFR | -1.079556 | HNSC | PCI-4B | Cetuximab | MVP_top60_DNA+RNA_seed | IGF1R | 0.143144 | GBM | U-118-MG | BMS-754807 |
| MVP_top60_DNA_seed | HDAC1 | -1.073451 | HNSC | PCI-6A | Belinostat | MVP_top60_DNA+RNA_seed | AKT1 | 0.143791 | GBM | Becker | AT13148 |
| MVP_top60_DNA_seed | EGFR | -1.071859 | HNSC | PCI-6A | AST-1306 | MVP_top60_DNA+RNA_seed | MET | 0.145675 | GBM | D-245MG | Crizotinib |

| | | | | | | | | | | | |
|---|---|---|---|---|---|---|---|---|---|---|---|
| MVP_top60_DNA_seed | EGFR | -1.071845 | HNSC | BICR22 | Afatinib | MVP_top60_DNA+RNA_seed | KIT | 0.146053 | GBM | A172 | Motesanib |
| MVP_top60_DNA_seed | EGFR | -1.069631 | HNSC | SAT | CI-1033 | MVP_top60_DNA+RNA_seed | AKT1 | 0.147993 | GBM | YKG-1 | Ipatasertib |
| MVP_top60_DNA_seed | NTRK1 | -1.06624 | HNSC | LB771-HNC | GW441756 | MVP_top60_DNA+RNA_seed | EGFR | 0.148283 | GBM | DK-MG | Afatinib |
| MVP_top60_DNA_seed | HSP90AA1 | -1.064945 | HNSC | BICR31 | Luminespib | MVP_top60_DNA+RNA_seed | IGF1R | 0.148485 | GBM | KS-1 | Linsitinib |
| MVP_top60_DNA_seed | EGFR | -1.060927 | HNSC | BICR22 | Afatinib | MVP_top60_DNA+RNA_seed | EGFR | 0.149432 | GBM | D-263MG | AZD3759 |
| MVP_top60_DNA_seed | SRC | -1.057408 | HNSC | DOK | HG-6-71-01 | MVP_top60_DNA+RNA_seed | MET | 0.152406 | GBM | SNB75 | Crizotinib |
| MVP_top60_DNA_seed | HSP90AA1 | -1.057063 | HNSC | CAL-27 | CCT-018159 | MVP_top60_DNA+RNA_seed | CDK4 | 0.152862 | GBM | SF126 | Ribociclib |
| MVP_top60_DNA_seed | HDAC1 | -1.055762 | HNSC | PCI-6A | AR-42 | MVP_top60_DNA+RNA_seed | AKT1 | 0.154672 | GBM | U-87-MG | AZD5363 |
| MVP_top60_DNA_seed | SMARCA4 | -1.046805 | HNSC | HSC-4 | PFI-3 | MVP_top60_DNA+RNA_seed | IGF1R | 0.155079 | GBM | LNZTA3WT4 | Linsitinib |
| MVP_top60_DNA_seed | EGFR | -1.044366 | HNSC | BHY | PF-00299804 | MVP_top60_DNA+RNA_seed | AKT1 | 0.155615 | GBM | U251 | GSK2110183B |
| MVP_top60_DNA_seed | AKT1 | -1.037493 | HNSC | Detroit562 | AKT inhibitor VIII | MVP_top60_DNA+RNA_seed | JAK2 | 0.158329 | GBM | LN-18 | AZ960 |
| MVP_top60_DNA_seed | EGFR | -1.03016 | HNSC | LB771-HNC | Erlotinib | MVP_top60_DNA+RNA_seed | JAK3 | 0.158329 | GBM | LN-18 | AZ960 |
| MVP_top60_DNA_seed | CDK2 | -1.027376 | HNSC | KOSC-2 | AZD5438 | MVP_top60_DNA+RNA_seed | AKT1 | 0.160586 | GBM | YH-13 | Afuresertib |
| MVP_top60_DNA_seed | HSP90AA1 | -1.026625 | HNSC | PCI-38 | CCT-018159 | MVP_top60_DNA+RNA_seed | AKT1 | 0.16675 | GBM | AM-38 | AT13148 |
| MVP_top60_DNA_seed | EGFR | -1.024361 | HNSC | CAL-27 | Cetuximab | MVP_top60_DNA+RNA_seed | CDK4 | 0.169151 | GBM | D-423MG | Ribociclib |
| MVP_top60_DNA_seed | EGFR | -1.023667 | HNSC | PE-CA-PJ15 | AZD8931 | MVP_top60_DNA+RNA_seed | CDK4 | 0.169831 | GBM | LN-18 | Ribociclib |
| MVP_top60_DNA_seed | SRC | -1.02351 | HNSC | Ca9-22 | Bosutinib | MVP_top60_DNA+RNA_seed | IGF1R | 0.173431 | GBM | LN-229 | GSK1904529A |
| MVP_top60_DNA_seed | EGFR | -1.020661 | HNSC | BICR22 | Gefitinib | MVP_top60_DNA+RNA_seed | EGFR | 0.17414 | GBM | U-118-MG | Erlotinib |
| MVP_top60_DNA_seed | EGFR | -1.019949 | HNSC | SKN-3 | AST-1306 | MVP_top60_DNA+RNA_seed | EGFR | 0.175123 | GBM | D-423MG | Gefitinib |
| MVP_top60_DNA_seed | AR | -1.019105 | HNSC | SCC-25 | Bicalutamide | MVP_top60_DNA+RNA_seed | IGF1R | 0.175822 | GBM | DK-MG | NVP-ADW742 |
| MVP_top60_DNA_seed | HSP90AA1 | -1.018032 | HNSC | PCI-6A | Elesclomol | MVP_top60_DNA+RNA_seed | MET | 0.17696 | GBM | LN-18 | Crizotinib |
| MVP_top60_DNA_seed | EGFR | -1.017153 | HNSC | BHY | CI-1033 | MVP_top60_DNA+RNA_seed | EGFR | 0.177191 | GBM | SF268 | Afatinib |
| MVP_top60_DNA_seed | MDM2 | -1.017003 | HNSC | BB49-HNC | Nutlin-3a (-) | MVP_top60_DNA+RNA_seed | EGFR | 0.180797 | GBM | A172 | AZD3759 |
| MVP_top60_DNA_seed | HSP90AA1 | -1.01556 | HNSC | PCI-6A | SNX-2112 | MVP_top60_DNA+RNA_seed | EGFR | 0.181391 | GBM | A172 | Erlotinib |
| MVP_top60_DNA_seed | EGFR | -1.013786 | HNSC | KOSC-2 | Afatinib | MVP_top60_DNA+RNA_seed | EGFR | 0.183624 | GBM | D-245MG | Afatinib |
| MVP_top60_DNA_seed | AR | -1.013666 | HNSC | OSC-19 | AZD3514 | MVP_top60_DNA+RNA_seed | CDK4 | 0.18412 | GBM | DK-MG | Palbociclib |
| MVP_top60_DNA_seed | EGFR | -1.011551 | HNSC | KOSC-2 | AZD8931 | MVP_top60_DNA+RNA_seed | MET | 0.186555 | GBM | T98G | Savolitinib |
| MVP_top60_DNA_seed | EGFR | -1.011262 | HNSC | SCC-4 | Gefitinib | MVP_top60_DNA+RNA_seed | KIT | 0.188521 | GBM | AM-38 | Axitinib |
| MVP_top60_DNA_seed | MDM2 | -1.010345 | HNSC | HO-1-u-1 | Serdemetan | MVP_top60_DNA+RNA_seed | AKT1 | 0.188828 | GBM | A172 | AT13148 |
| MVP_top60_DNA_seed | AKT1 | -1.009041 | HNSC | PCI-6A | AKT inhibitor VIII | MVP_top60_DNA+RNA_seed | AKT1 | 0.190767 | GBM | DK-MG | Afuresertib |
| MVP_top60_DNA_seed | PIK3CA | -1.008638 | HNSC | BB30-HNC | PI3Ka_4409 | MVP_top60_DNA+RNA_seed | EGFR | 0.192616 | GBM | D-245MG | Osimertinib |
| MVP_top60_DNA_seed | EGFR | -1.006504 | HNSC | SCC-25 | Cetuximab | MVP_top60_DNA+RNA_seed | EGFR | 0.192717 | GBM | SF268 | Gefitinib |
| MVP_top60_DNA_seed | HSP90AA1 | -1.005418 | HNSC | SCC-9 | SNX-2112 | MVP_top60_DNA+RNA_seed | MET | 0.193744 | GBM | GB-1 | Savolitinib |
| MVP_top60_DNA_seed | HSP90AA1 | -1.00181 | HNSC | BB30-HNC | SNX-2112 | MVP_top60_DNA+RNA_seed | KRAS | 0.194769 | GBM | GB-1 | ⁝RAS (G12C) Inhibitor-1 |
| MVP_top60_DNA_seed | PIK3CA | -1.001532 | HNSC | BICR10 | PI3Ka_4409 | MVP_top60_DNA+RNA_seed | CDK4 | 0.195795 | GBM | U-118-MG | Palbociclib |
| MVP_top60_DNA_seed | SRC | -1.001193 | HNSC | BICR10 | Bosutinib | MVP_top60_DNA+RNA_seed | KRAS | 0.196854 | GBM | D-263MG | ⁝RAS (G12C) Inhibitor-1 |
| MVP_top60_DNA_seed | PIK3CA | -1.000452 | HNSC | JHU-011 | PI3Ka_4409 | MVP_top60_DNA+RNA_seed | IGF1R | 0.19725 | GBM | D-423MG | BMS-754807 |
| MVP_top60_DNA_seed | EGFR | -0.99551 | HNSC | OSC-19 | AST-1306 | MVP_top60_DNA+RNA_seed | EGFR | 0.197386 | GBM | D-247MG | Osimertinib |
| MVP_top60_DNA_seed | PIK3CA | -0.994369 | HNSC | SAT | PI3Ka_4409 | MVP_top60_DNA+RNA_seed | KIT | 0.197784 | GBM | AM-38 | Motesanib |
| MVP_top60_DNA_seed | NTRK1 | -0.990624 | HNSC | HSC-3 | AZD1332 | MVP_top60_DNA+RNA_seed | AKT1 | 0.198443 | GBM | U-118-MG | AT13148 |
| MVP_top60_DNA_seed | NTRK1 | -0.990553 | HNSC | JHU-029 | GW441756 | MVP_top60_DNA+RNA_seed | EGFR | 0.200155 | GBM | AM-38 | Lapatinib |

| | | | | | | | | | | | |
|---|---|---|---|---|---|---|---|---|---|---|---|
| MVP_top60_DNA_seed | RAC1 | -0.989888 | HNSC | Detroit562 | EHT-1864 | MVP_top60_DNA+RNA_seed | AKT1 | 0.203041 | GBM | SF268 | Uprosertib |
| MVP_top60_DNA_seed | MDM2 | -0.98868 | HNSC | PCI-38 | Serdemetan | MVP_top60_DNA+RNA_seed | MET | 0.208345 | GBM | YH-13 | Savolitinib |
| MVP_top60_DNA_seed | CDK2 | -0.986882 | HNSC | JHU-011 | PHA-793887 | MVP_top60_DNA+RNA_seed | PIK3CA | 0.209158 | GBM | U-87-MG | Alpelisib |
| MVP_top60_DNA_seed | CDK2 | -0.986437 | HNSC | PCI-4B | PHA-793887 | MVP_top60_DNA+RNA_seed | EGFR | 0.209204 | GBM | U251 | AZD3759 |
| MVP_top60_DNA_seed | AR | -0.981907 | HNSC | SAT | AZD3514 | MVP_top60_DNA+RNA_seed | EGFR | 0.20971 | GBM | YH-13 | Afatinib |
| MVP_top60_DNA_seed | EGFR | -0.981126 | HNSC | DOK | CI-1033 | MVP_top60_DNA+RNA_seed | KIT | 0.211268 | GBM | T98G | Axitinib |
| MVP_top60_DNA_seed | NTRK1 | -0.973394 | HNSC | FADU | GW441756 | MVP_top60_DNA+RNA_seed | IGF1R | 0.212021 | GBM | M059J | BMS-536924 |
| MVP_top60_DNA_seed | PIK3CA | -0.973117 | HNSC | SCC-15 | PI3Ka_4409 | MVP_top60_DNA+RNA_seed | KIT | 0.21285 | GBM | YH-13 | Axitinib |
| MVP_top60_DNA_seed | EGFR | -0.970885 | HNSC | SCC-25 | Gefitinib | MVP_top60_DNA+RNA_seed | PIK3CA | 0.216864 | GBM | D-392MG | GNE-317 |
| MVP_top60_DNA_seed | EGFR | -0.969156 | HNSC | SCC-4 | Afatinib | MVP_top60_DNA+RNA_seed | JAK2 | 0.219615 | GBM | A172 | GSK2276186C |
| MVP_top60_DNA_seed | HSP90AA1 | -0.968131 | HNSC | BB30-HNC | Tanespimycin | MVP_top60_DNA+RNA_seed | JAK1 | 0.219615 | GBM | A172 | GSK2276186C |
| MVP_top60_DNA_seed | EGFR | -0.966081 | HNSC | PCI-38 | Cetuximab | MVP_top60_DNA+RNA_seed | JAK3 | 0.219615 | GBM | A172 | GSK2276186C |
| MVP_top60_DNA_seed | NTRK1 | -0.962889 | HNSC | Ca9-22 | AZD1332 | MVP_top60_DNA+RNA_seed | AKT1 | 0.220143 | GBM | LN-405 | AZD5363 |
| MVP_top60_DNA_seed | PIK3CA | -0.955939 | HNSC | Detroit562 | PI3Ka_4409 | MVP_top60_DNA+RNA_seed | MET | 0.22032 | GBM | SF126 | Crizotinib |
| MVP_top60_DNA_seed | CDK2 | -0.951727 | HNSC | HO-1-u-1 | AZD5438 | MVP_top60_DNA+RNA_seed | JAK2 | 0.222005 | GBM | SF268 | Ruxolitinib |
| MVP_top60_DNA_seed | MDM2 | -0.950079 | HNSC | KON | Serdemetan | MVP_top60_DNA+RNA_seed | JAK1 | 0.222005 | GBM | SF268 | Ruxolitinib |
| MVP_top60_DNA_seed | AR | -0.949349 | HNSC | SAS | AZD3514 | MVP_top60_DNA+RNA_seed | PIK3CA | 0.22435 | GBM | KS-1 | Buparlisib |
| MVP_top60_DNA_seed | AR | -0.944751 | HNSC | HSC-3 | Bicalutamide | MVP_top60_DNA+RNA_seed | PIK3CB | 0.22435 | GBM | KS-1 | Buparlisib |
| MVP_top60_DNA_seed | HDAC1 | -0.944182 | HNSC | KOSC-2 | Dacinostat | MVP_top60_DNA+RNA_seed | PIK3CD | 0.22435 | GBM | KS-1 | Buparlisib |
| MVP_top60_DNA_seed | AR | -0.942769 | HNSC | KON | Bicalutamide | MVP_top60_DNA+RNA_seed | EGFR | 0.225471 | GBM | M059J | Afatinib |
| MVP_top60_DNA_seed | MAPK1 | -0.938368 | HNSC | PCI-6A | FR-180204 | MVP_top60_DNA+RNA_seed | PIK3CA | 0.226173 | GBM | D-566MG | Alpelisib |
| MVP_top60_DNA_seed | MAPK3 | -0.938368 | HNSC | PCI-6A | FR-180204 | MVP_top60_DNA+RNA_seed | EGFR | 0.232195 | GBM | M059J | Lapatinib |
| MVP_top60_DNA_seed | SMARCA4 | -0.937996 | HNSC | BICR78 | PFI-3 | MVP_top60_DNA+RNA_seed | PIK3CA | 0.232417 | GBM | D-392MG | AZD8186 |
| MVP_top60_DNA_seed | CDK2 | -0.93622 | HNSC | SAS | AZD5438 | MVP_top60_DNA+RNA_seed | PIK3CB | 0.232417 | GBM | D-392MG | AZD8186 |
| MVP_top60_DNA_seed | HSP90AA1 | -0.933882 | HNSC | JHU-011 | CCT-018159 | MVP_top60_DNA+RNA_seed | EGFR | 0.233071 | GBM | DK-MG | AZD3759 |
| MVP_top60_DNA_seed | EGFR | -0.932541 | HNSC | SCC-25 | CI-1033 | MVP_top60_DNA+RNA_seed | KIT | 0.233351 | GBM | SF268 | Axitinib |
| MVP_top60_DNA_seed | CDK2 | -0.931342 | HNSC | SCC-9 | PHA-793887 | MVP_top60_DNA+RNA_seed | JAK2 | 0.233557 | GBM | GB-1 | JAK_8517 |
| MVP_top60_DNA_seed | MDM2 | -0.929094 | HNSC | DOK | Serdemetan | MVP_top60_DNA+RNA_seed | JAK1 | 0.233557 | GBM | GB-1 | JAK_8517 |
| MVP_top60_DNA_seed | EGFR | -0.929002 | HNSC | SCC-15 | Pelitinib | MVP_top60_DNA+RNA_seed | AKT1 | 0.234744 | GBM | M059J | GSK2110183B |
| MVP_top60_DNA_seed | HDAC1 | -0.928022 | HNSC | Ca9-22 | CAY10603 | MVP_top60_DNA+RNA_seed | KIT | 0.235526 | GBM | SK-MG-1 | Axitinib |
| MVP_top60_DNA_seed | EP300 | -0.923946 | HNSC | PCI-4B | I-CBP112 | MVP_top60_DNA+RNA_seed | JAK2 | 0.235612 | GBM | A172 | Ruxolitinib |
| MVP_top60_DNA_seed | CREBBP | -0.923946 | HNSC | PCI-4B | I-CBP112 | MVP_top60_DNA+RNA_seed | JAK1 | 0.235612 | GBM | A172 | Ruxolitinib |
| MVP_top60_DNA_seed | SRC | -0.923 | HNSC | BB30-HNC | WH-4-023 | MVP_top60_DNA+RNA_seed | EGFR | 0.237068 | GBM | U-87-MG | Gefitinib |
| MVP_top60_DNA_seed | PIK3CA | -0.916753 | HNSC | JHU-022 | PI3Ka_4409 | MVP_top60_DNA+RNA_seed | IGF1R | 0.238684 | GBM | LN-405 | BMS-754807 |
| MVP_top60_DNA_seed | ESR1 | -0.910123 | HNSC | BICR78 | Tamoxifen | MVP_top60_DNA+RNA_seed | AKT1 | 0.240253 | GBM | U251 | Ipatasertib |
| MVP_top60_DNA_seed | SRC | -0.909871 | HNSC | HSC-4 | HG-6-71-01 | MVP_top60_DNA+RNA_seed | EGFR | 0.241009 | GBM | U-87-MG | Sapitinib |
| MVP_top60_DNA_seed | HDAC1 | -0.909442 | HNSC | OSC-20 | JQ12 | MVP_top60_DNA+RNA_seed | ERBB3 | 0.241009 | GBM | U-87-MG | Sapitinib |
| MVP_top60_DNA_seed | EGFR | -0.908857 | HNSC | SCC-15 | Afatinib | MVP_top60_DNA+RNA_seed | JAK2 | 0.241593 | GBM | AM-38 | AZ960 |
| MVP_top60_DNA_seed | HSP90AA1 | -0.908666 | HNSC | BICR78 | Tanespimycin | MVP_top60_DNA+RNA_seed | JAK3 | 0.241593 | GBM | AM-38 | AZ960 |
| MVP_top60_DNA_seed | PIK3CA | -0.908083 | HNSC | SCC-15 | AZD8835 | MVP_top60_DNA+RNA_seed | MET | 0.242769 | GBM | D-423MG | Savolitinib |
| MVP_top60_DNA_seed | AR | -0.899002 | HNSC | PCI-38 | AZD3514 | MVP_top60_DNA+RNA_seed | JAK1 | 0.243108 | GBM | U-118-MG | JAK1_8709 |

| | | | | | | | | | | | |
|---|---|---|---|---|---|---|---|---|---|---|---|
| MVP_top60_DNA_seed | EGFR | -0.898467 | HNSC | OSC-20 | Cetuximab | MVP_top60_DNA+RNA_seed | KIT | 0.243335 | GBM | SF268 | Sorafenib |
| MVP_top60_DNA_seed | MDM2 | -0.898452 | HNSC | JHU-011 | Serdemetan | MVP_top60_DNA+RNA_seed | KIT | 0.248351 | GBM | U251 | Axitinib |
| MVP_top60_DNA_seed | PIK3CA | -0.897997 | HNSC | PE-CA-PJ15 | AZD8835 | MVP_top60_DNA+RNA_seed | AKT1 | 0.248502 | GBM | Becker | Afuresertib |
| MVP_top60_DNA_seed | EGFR | -0.896753 | HNSC | LB771-HNC | Gefitinib | MVP_top60_DNA+RNA_seed | IGF1R | 0.248785 | GBM | SK-MG-1 | BMS-536924 |
| MVP_top60_DNA_seed | EP300 | -0.891471 | HNSC | JHU-022 | I-CBP112 | MVP_top60_DNA+RNA_seed | AKT1 | 0.249589 | GBM | A172 | AZD5363 |
| MVP_top60_DNA_seed | CREBBP | -0.891471 | HNSC | JHU-022 | I-CBP112 | MVP_top60_DNA+RNA_seed | IGF1R | 0.250307 | GBM | U251 | GSK1904529A |
| MVP_top60_DNA_seed | HSP90AA1 | -0.890848 | HNSC | SCC-4 | CCT-018159 | MVP_top60_DNA+RNA_seed | AKT1 | 0.251046 | GBM | U251 | AZD5363 |
| MVP_top60_DNA_seed | EGFR | -0.889973 | HNSC | CAL-27 | CI-1033 | MVP_top60_DNA+RNA_seed | AKT1 | 0.25396 | GBM | D-245MG | Afuresertib |
| MVP_top60_DNA_seed | PIK3CA | -0.889663 | HNSC | KOSC-2 | PI3Ka_4409 | MVP_top60_DNA+RNA_seed | EGFR | 0.255549 | GBM | LN-229 | AZD3759 |
| MVP_top60_DNA_seed | SRC | -0.888673 | HNSC | CAL-33 | HG-6-71-01 | MVP_top60_DNA+RNA_seed | MET | 0.258192 | GBM | YH-13 | Crizotinib |
| MVP_top60_DNA_seed | HSP90AA1 | -0.884189 | HNSC | JHU-011 | SNX-2112 | MVP_top60_DNA+RNA_seed | JAK2 | 0.260931 | GBM | DK-MG | Ruxolitinib |
| MVP_top60_DNA_seed | RAC1 | -0.883201 | HNSC | HO-1-u-1 | EHT-1864 | MVP_top60_DNA+RNA_seed | JAK1 | 0.260931 | GBM | DK-MG | Ruxolitinib |
| MVP_top60_DNA_seed | PIK3CA | -0.881566 | HNSC | BHY | PI3Ka_4409 | MVP_top60_DNA+RNA_seed | CDK4 | 0.261538 | GBM | SF295 | Ribociclib |
| MVP_top60_DNA_seed | HSP90AA1 | -0.879137 | HNSC | PCI-4B | Elesclomol | MVP_top60_DNA+RNA_seed | KIT | 0.262318 | GBM | CAS-1 | Motesanib |
| MVP_top60_DNA_seed | HSP90AA1 | -0.878333 | HNSC | OSC-20 | Tanespimycin | MVP_top60_DNA+RNA_seed | IGF1R | 0.263783 | GBM | AM-38 | Linsitinib |
| MVP_top60_DNA_seed | CDK2 | -0.876739 | HNSC | JHU-011 | AZD5438 | MVP_top60_DNA+RNA_seed | TP53 | 0.264647 | GBM | YKG-1 | MIRA-1 |
| MVP_top60_DNA_seed | EGFR | -0.875657 | HNSC | JHU-011 | Pelitinib | MVP_top60_DNA+RNA_seed | KIT | 0.264698 | GBM | Becker | Axitinib |
| MVP_top60_DNA_seed | AKT1 | -0.87438 | HNSC | BICR22 | BAY AKT1 | MVP_top60_DNA+RNA_seed | IGF1R | 0.265222 | GBM | D-247MG | NVP-ADW742 |
| MVP_top60_DNA_seed | NTRK1 | -0.874336 | HNSC | HO-1-N-1 | GW441756 | MVP_top60_DNA+RNA_seed | JAK2 | 0.265974 | GBM | SF268 | AZ960 |
| MVP_top60_DNA_seed | HSP90AA1 | -0.873876 | HNSC | PCI-30 | Elesclomol | MVP_top60_DNA+RNA_seed | JAK3 | 0.265974 | GBM | SF268 | AZ960 |
| MVP_top60_DNA_seed | HDAC1 | -0.870819 | HNSC | PCI-4B | JQ12 | MVP_top60_DNA+RNA_seed | IGF1R | 0.266157 | GBM | LN-18 | Linsitinib |
| MVP_top60_DNA_seed | AKT1 | -0.869501 | HNSC | BICR78 | AKT inhibitor VIII | MVP_top60_DNA+RNA_seed | JAK2 | 0.267109 | GBM | SF295 | Ruxolitinib |
| MVP_top60_DNA_seed | EGFR | -0.868355 | HNSC | PCI-6A | CI-1033 | MVP_top60_DNA+RNA_seed | JAK1 | 0.267109 | GBM | SF295 | Ruxolitinib |
| MVP_top60_DNA_seed | NTRK1 | -0.867661 | HNSC | JHU-011 | GW441756 | MVP_top60_DNA+RNA_seed | EGFR | 0.269102 | GBM | 42-MG-BA | Sapitinib |
| MVP_top60_DNA_seed | HSP90AA1 | -0.866439 | HNSC | BICR78 | Elesclomol | MVP_top60_DNA+RNA_seed | ERBB3 | 0.269102 | GBM | 42-MG-BA | Sapitinib |
| MVP_top60_DNA_seed | PIK3CA | -0.863537 | HNSC | BICR78 | AZD8835 | MVP_top60_DNA+RNA_seed | MET | 0.27313 | GBM | SF126 | Savolitinib |
| MVP_top60_DNA_seed | NTRK1 | -0.86299 | HNSC | BICR10 | AZD1332 | MVP_top60_DNA+RNA_seed | MET | 0.274326 | GBM | SK-MG-1 | Crizotinib |
| MVP_top60_DNA_seed | RAC1 | -0.862894 | HNSC | JHU-011 | EHT-1864 | MVP_top60_DNA+RNA_seed | EGFR | 0.27457 | GBM | SW1088 | Erlotinib |
| MVP_top60_DNA_seed | AR | -0.861028 | HNSC | BICR22 | AZD3514 | MVP_top60_DNA+RNA_seed | EGFR | 0.275207 | GBM | D-542MG | Gefitinib |
| MVP_top60_DNA_seed | CDK2 | -0.860013 | HNSC | HSC-3 | AZD5438 | MVP_top60_DNA+RNA_seed | AKT1 | 0.27842 | GBM | LN-18 | Afuresertib |
| MVP_top60_DNA_seed | MDM2 | -0.859359 | HNSC | JHU-029 | Serdemetan | MVP_top60_DNA+RNA_seed | JAK2 | 0.279236 | GBM | D-392MG | Ruxolitinib |
| MVP_top60_DNA_seed | EGFR | -0.858079 | HNSC | SCC-25 | CUDC-101 | MVP_top60_DNA+RNA_seed | JAK1 | 0.279236 | GBM | D-392MG | Ruxolitinib |
| MVP_top60_DNA_seed | AKT1 | -0.855566 | HNSC | SCC-9 | AKT inhibitor VIII | MVP_top60_DNA+RNA_seed | EGFR | 0.280485 | GBM | Becker | AZD3759 |
| MVP_top60_DNA_seed | HSP90AA1 | -0.84635 | HNSC | CAL-33 | Elesclomol | MVP_top60_DNA+RNA_seed | IGF1R | 0.283076 | GBM | U-118-MG | BMS-536924 |
| MVP_top60_DNA_seed | HSP90AA1 | -0.846304 | HNSC | HSC-3 | CCT-018159 | MVP_top60_DNA+RNA_seed | AKT1 | 0.285533 | GBM | KS-1 | Uprosertib |
| MVP_top60_DNA_seed | ESR1 | -0.844597 | HNSC | FADU | Fulvestrant | MVP_top60_DNA+RNA_seed | JAK2 | 0.286986 | GBM | DK-MG | GSK2276186C |
| MVP_top60_DNA_seed | CDK2 | -0.843906 | HNSC | KON | AZD5438 | MVP_top60_DNA+RNA_seed | JAK1 | 0.286986 | GBM | DK-MG | GSK2276186C |
| MVP_top60_DNA_seed | EGFR | -0.840608 | HNSC | Ca9-22 | CI-1033 | MVP_top60_DNA+RNA_seed | JAK3 | 0.286986 | GBM | DK-MG | GSK2276186C |
| MVP_top60_DNA_seed | AR | -0.837603 | HNSC | HSC-2 | AZD3514 | MVP_top60_DNA+RNA_seed | IGF1R | 0.287026 | GBM | SNB75 | BMS-536924 |
| MVP_top60_DNA_seed | HDAC1 | -0.837045 | HNSC | Ca9-22 | Belinostat | MVP_top60_DNA+RNA_seed | CDK4 | 0.290798 | GBM | U-87-MG | Ribociclib |
| MVP_top60_DNA_seed | SRC | -0.836426 | HNSC | OSC-20 | Bosutinib | MVP_top60_DNA+RNA_seed | JAK2 | 0.29155 | GBM | D-263MG | Ruxolitinib |

| | | | | | | | | | | | |
|---|---|---|---|---|---|---|---|---|---|---|---|
| MVP_top60_DNA_seed | HDAC1 | -0.832136 | HNSC | SCC-25 | Belinostat | MVP_top60_DNA+RNA_seed | JAK1 | 0.29155 | GBM | D-263MG | Ruxolitinib |
| MVP_top60_DNA_seed | PIK3CA | -0.830883 | HNSC | BB30-HNC | AZD8835 | MVP_top60_DNA+RNA_seed | IGF1R | 0.293691 | GBM | D-245MG | BMS-754807 |
| MVP_top60_DNA_seed | AKT1 | -0.830541 | HNSC | Ca9-22 | AKT inhibitor VIII | MVP_top60_DNA+RNA_seed | EGFR | 0.293897 | GBM | SW1088 | Osimertinib |
| MVP_top60_DNA_seed | EGFR | -0.828051 | HNSC | BHY | CUDC-101 | MVP_top60_DNA+RNA_seed | EGFR | 0.294884 | GBM | T98G | Lapatinib |
| MVP_top60_DNA_seed | HSP90AA1 | -0.826251 | HNSC | KOSC-2 | Tanespimycin | MVP_top60_DNA+RNA_seed | AKT1 | 0.29497 | GBM | LN-18 | MK-2206 |
| MVP_top60_DNA_seed | EGFR | -0.822188 | HNSC | BICR78 | Cetuximab | MVP_top60_DNA+RNA_seed | EGFR | 0.296246 | GBM | CAS-1 | Sapitinib |
| MVP_top60_DNA_seed | HSP90AA1 | -0.822036 | HNSC | OSC-20 | Luminespib | MVP_top60_DNA+RNA_seed | ERBB3 | 0.296246 | GBM | CAS-1 | Sapitinib |
| MVP_top60_DNA_seed | HSP90AA1 | -0.817667 | HNSC | SAT | Tanespimycin | MVP_top60_DNA+RNA_seed | KIT | 0.296414 | GBM | KS-1 | Axitinib |
| MVP_top60_DNA_seed | PIK3CA | -0.815559 | HNSC | HO-1-N-1 | PI3Ka_4409 | MVP_top60_DNA+RNA_seed | AKT1 | 0.296595 | GBM | U-118-MG | Uprosertib |
| MVP_top60_DNA_seed | PIK3CA | -0.815453 | HNSC | Detroit562 | AZD8835 | MVP_top60_DNA+RNA_seed | IGF1R | 0.297202 | GBM | 8-MG-BA | BMS-754807 |
| MVP_top60_DNA_seed | MAPK1 | -0.812022 | HNSC | SCC-9 | VX-11e | MVP_top60_DNA+RNA_seed | MET | 0.299004 | GBM | T98G | Crizotinib |
| MVP_top60_DNA_seed | EGFR | -0.811788 | HNSC | BB30-HNC | PF-00299804 | MVP_top60_DNA+RNA_seed | TP53 | 0.299196 | GBM | AM-38 | MIRA-1 |
| MVP_top60_DNA_seed | EGFR | -0.806252 | HNSC | SCC-9 | CUDC-101 | MVP_top60_DNA+RNA_seed | CDK4 | 0.305782 | GBM | CAS-1 | Palbociclib |
| MVP_top60_DNA_seed | HDAC1 | -0.801662 | HNSC | PCI-4B | Tubastatin A | MVP_top60_DNA+RNA_seed | IGF1R | 0.308899 | GBM | SF268 | BMS-754807 |
| MVP_top60_DNA_seed | PIK3CA | -0.80159 | HNSC | OSC-20 | PI3Ka_4409 | MVP_top60_DNA+RNA_seed | KIT | 0.310924 | GBM | YKG-1 | Motesanib |
| MVP_top60_DNA_seed | HDAC1 | -0.80099 | HNSC | SCC-25 | Tubastatin A | MVP_top60_DNA+RNA_seed | TP53 | 0.3112 | GBM | D-542MG | MIRA-1 |
| MVP_top60_DNA_seed | EGFR | -0.80078 | HNSC | SAS | CUDC-101 | MVP_top60_DNA+RNA_seed | PIK3CA | 0.311367 | GBM | SF126 | Buparlisib |
| MVP_top60_DNA_seed | EGFR | -0.798053 | HNSC | JHU-022 | Cetuximab | MVP_top60_DNA+RNA_seed | PIK3CB | 0.311367 | GBM | SF126 | Buparlisib |
| MVP_top60_DNA_seed | AKT1 | -0.794895 | HNSC | JHU-029 | MK-2206 | MVP_top60_DNA+RNA_seed | PIK3CD | 0.311367 | GBM | SF126 | Buparlisib |
| MVP_top60_DNA_seed | EGFR | -0.794791 | HNSC | CAL-27 | CUDC-101 | MVP_top60_DNA+RNA_seed | MET | 0.31239 | GBM | LNZTA3WT4 | Crizotinib |
| MVP_top60_DNA_seed | HSP90AA1 | -0.793791 | HNSC | JHU-029 | Elesclomol | MVP_top60_DNA+RNA_seed | TP53 | 0.313675 | GBM | Becker | MIRA-1 |
| MVP_top60_DNA_seed | HSP90AA1 | -0.792846 | HNSC | SAS | Tanespimycin | MVP_top60_DNA+RNA_seed | EP300 | 0.31425 | GBM | Becker | SGC-CBP30 |
| MVP_top60_DNA_seed | HDAC1 | -0.791023 | HNSC | SCC-4 | JQ12 | MVP_top60_DNA+RNA_seed | CREBBP | 0.31425 | GBM | Becker | SGC-CBP30 |
| MVP_top60_DNA_seed | RAC1 | -0.791008 | HNSC | DOK | EHT-1864 | MVP_top60_DNA+RNA_seed | JAK2 | 0.315852 | GBM | 8-MG-BA | Ruxolitinib |
| MVP_top60_DNA_seed | EGFR | -0.786671 | HNSC | SKN-3 | CI-1033 | MVP_top60_DNA+RNA_seed | JAK1 | 0.315852 | GBM | 8-MG-BA | Ruxolitinib |
| MVP_top60_DNA_seed | AR | -0.786007 | HNSC | SKN-3 | AZD3514 | MVP_top60_DNA+RNA_seed | PIK3CA | 0.319196 | GBM | D-423MG | Alpelisib |
| MVP_top60_DNA_seed | CDK2 | -0.780628 | HNSC | SAS | PHA-793887 | MVP_top60_DNA+RNA_seed | AKT1 | 0.320756 | GBM | DK-MG | AZD5363 |
| MVP_top60_DNA_seed | MDM2 | -0.778165 | HNSC | SAT | Serdemetan | MVP_top60_DNA+RNA_seed | IGF1R | 0.321301 | GBM | SNB75 | BMS-754807 |
| MVP_top60_DNA_seed | HSP90AA1 | -0.775957 | HNSC | KOSC-2 | Luminespib | MVP_top60_DNA+RNA_seed | KIT | 0.321616 | GBM | SF295 | Motesanib |
| MVP_top60_DNA_seed | EGFR | -0.775938 | HNSC | Detroit562 | AZD8931 | MVP_top60_DNA+RNA_seed | MET | 0.324084 | GBM | SF295 | Savolitinib |
| MVP_top60_DNA_seed | HSP90AA1 | -0.775761 | HNSC | OSC-19 | Tanespimycin | MVP_top60_DNA+RNA_seed | AKT1 | 0.325138 | GBM | T98G | Uprosertib |
| MVP_top60_DNA_seed | HSP90AA1 | -0.775257 | HNSC | SAS | SNX-2112 | MVP_top60_DNA+RNA_seed | AKT1 | 0.326187 | GBM | KS-1 | Afuresertib |
| MVP_top60_DNA_seed | HSP90AA1 | -0.773815 | HNSC | OSC-19 | Luminespib | MVP_top60_DNA+RNA_seed | KIT | 0.326282 | GBM | DK-MG | Sorafenib |
| MVP_top60_DNA_seed | EGFR | -0.769713 | HNSC | SAS | Pelitinib | MVP_top60_DNA+RNA_seed | AKT1 | 0.327828 | GBM | GB-1 | AZD5363 |
| MVP_top60_DNA_seed | AKT1 | -0.76907 | HNSC | HSC-3 | BAY AKT1 | MVP_top60_DNA+RNA_seed | IGF1R | 0.327958 | GBM | U-87-MG | GSK1904529A |
| MVP_top60_DNA_seed | ESR1 | -0.768939 | HNSC | LB771-HNC | Fulvestrant | MVP_top60_DNA+RNA_seed | EGFR | 0.327969 | GBM | D-423MG | Erlotinib |
| MVP_top60_DNA_seed | HSP90AA1 | -0.76891 | HNSC | SCC-9 | Luminespib | MVP_top60_DNA+RNA_seed | PIK3CA | 0.328473 | GBM | SF295 | Alpelisib |
| MVP_top60_DNA_seed | SRC | -0.768287 | HNSC | OSC-19 | Bosutinib | MVP_top60_DNA+RNA_seed | IGF1R | 0.331179 | GBM | DK-MG | GSK1904529A |
| MVP_top60_DNA_seed | HDAC1 | -0.765809 | HNSC | HO-1-N-1 | JQ12 | MVP_top60_DNA+RNA_seed | EP300 | 0.33234 | GBM | D-392MG | SGC-CBP30 |
| MVP_top60_DNA_seed | CDK2 | -0.763583 | HNSC | JHU-029 | AZD5438 | MVP_top60_DNA+RNA_seed | CREBBP | 0.33234 | GBM | D-392MG | SGC-CBP30 |
| MVP_top60_DNA_seed | AR | -0.763438 | HNSC | Detroit562 | AZD3514 | MVP_top60_DNA+RNA_seed | EGFR | 0.334174 | GBM | 8-MG-BA | AZD3759 |

| | | | | | | | | | | | |
|---|---|---|---|---|---|---|---|---|---|---|---|
| MVP_top60_DNA_seed | EGFR | -0.7612 | HNSC | HO-1-N-1 | AST-1306 | MVP_top60_DNA+RNA_seed | EGFR | 0.334651 | GBM | D-423MG | Afatinib |
| MVP_top60_DNA_seed | AKT1 | -0.759857 | HNSC | SCC-9 | GSK690693 | MVP_top60_DNA+RNA_seed | EGFR | 0.337738 | GBM | D-566MG | Osimertinib |
| MVP_top60_DNA_seed | MDM2 | -0.759246 | HNSC | PE-CA-PJ15 | Serdemetan | MVP_top60_DNA+RNA_seed | AKT1 | 0.338928 | GBM | U-87-MG | MK-2206 |
| MVP_top60_DNA_seed | SMARCA4 | -0.758926 | HNSC | HO-1-N-1 | PFI-3 | MVP_top60_DNA+RNA_seed | PIK3CA | 0.340646 | GBM | M059J | Buparlisib |
| MVP_top60_DNA_seed | HSP90AA1 | -0.758748 | HNSC | PCI-38 | Elesclomol | MVP_top60_DNA+RNA_seed | PIK3CB | 0.340646 | GBM | M059J | Buparlisib |
| MVP_top60_DNA_seed | EGFR | -0.75252 | HNSC | LB771-HNC | Cetuximab | MVP_top60_DNA+RNA_seed | PIK3CD | 0.340646 | GBM | M059J | Buparlisib |
| MVP_top60_DNA_seed | EGFR | -0.750142 | HNSC | SAT | Pelitinib | MVP_top60_DNA+RNA_seed | AKT1 | 0.341412 | GBM | CAS-1 | GSK2110183B |
| MVP_top60_DNA_seed | HDAC1 | -0.748248 | HNSC | PCI-6A | Tubastatin A | MVP_top60_DNA+RNA_seed | AKT1 | 0.34229 | GBM | U251 | Uprosertib |
| MVP_top60_DNA_seed | SRC | -0.7448 | HNSC | BB49-HNC | Saracatinib | MVP_top60_DNA+RNA_seed | IGF1R | 0.343612 | GBM | D-392MG | BMS-754807 |
| MVP_top60_DNA_seed | ESR1 | -0.744397 | HNSC | BB49-HNC | Fulvestrant | MVP_top60_DNA+RNA_seed | AKT1 | 0.34584 | GBM | A172 | Ipatasertib |
| MVP_top60_DNA_seed | ESR1 | -0.742778 | HNSC | JHU-022 | Fulvestrant | MVP_top60_DNA+RNA_seed | KRAS | 0.347473 | GBM | D-247MG | KRAS (G12C) Inhibitor-1 |
| MVP_top60_DNA_seed | EGFR | -0.742571 | HNSC | PCI-6A | Pelitinib | MVP_top60_DNA+RNA_seed | MET | 0.349385 | GBM | D-263MG | Crizotinib |
| MVP_top60_DNA_seed | NTRK1 | -0.742258 | HNSC | HO-1-N-1 | AZD1332 | MVP_top60_DNA+RNA_seed | JAK1 | 0.350672 | GBM | KS-1 | JAK1_8709 |
| MVP_top60_DNA_seed | AKT1 | -0.741752 | HNSC | SAT | MK-2206 | MVP_top60_DNA+RNA_seed | PIK3CA | 0.351358 | GBM | LN-405 | AZD8186 |
| MVP_top60_DNA_seed | EGFR | -0.739858 | HNSC | PE-CA-PJ15 | PF-00299804 | MVP_top60_DNA+RNA_seed | PIK3CB | 0.351358 | GBM | LN-405 | AZD8186 |
| MVP_top60_DNA_seed | EGFR | -0.738501 | HNSC | SKN-3 | AZD8931 | MVP_top60_DNA+RNA_seed | IGF1R | 0.352866 | GBM | YKG-1 | NVP-ADW742 |
| MVP_top60_DNA_seed | EGFR | -0.734379 | HNSC | CAL-33 | AST-1306 | MVP_top60_DNA+RNA_seed | JAK1 | 0.353868 | GBM | SF268 | JAK1_8709 |
| MVP_top60_DNA_seed | HDAC1 | -0.730857 | HNSC | KON | JQ12 | MVP_top60_DNA+RNA_seed | AKT1 | 0.358083 | GBM | SNB75 | Uprosertib |
| MVP_top60_DNA_seed | HSP90AA1 | -0.729702 | HNSC | CAL-27 | SNX-2112 | MVP_top60_DNA+RNA_seed | EGFR | 0.364198 | GBM | SF268 | Sapitinib |
| MVP_top60_DNA_seed | HDAC1 | -0.727542 | HNSC | HSC-3 | Dacinostat | MVP_top60_DNA+RNA_seed | ERBB3 | 0.364198 | GBM | SF268 | Sapitinib |
| MVP_top60_DNA_seed | NTRK1 | -0.72666 | HNSC | BICR10 | GW441756 | MVP_top60_DNA+RNA_seed | AKT1 | 0.36434 | GBM | LN-405 | Afuresertib |
| MVP_top60_DNA_seed | AKT1 | -0.726067 | HNSC | HO-1-u-1 | AKT inhibitor VIII | MVP_top60_DNA+RNA_seed | AKT1 | 0.364442 | GBM | LN-229 | Uprosertib |
| MVP_top60_DNA_seed | HSP90AA1 | -0.722007 | HNSC | PCI-15A | CCT-018159 | MVP_top60_DNA+RNA_seed | EGFR | 0.364904 | GBM | D-247MG | Sapitinib |
| MVP_top60_DNA_seed | EGFR | -0.721788 | HNSC | SKN-3 | CUDC-101 | MVP_top60_DNA+RNA_seed | ERBB3 | 0.364904 | GBM | D-247MG | Sapitinib |
| MVP_top60_DNA_seed | EGFR | -0.720897 | HNSC | KON | AZD8931 | MVP_top60_DNA+RNA_seed | EGFR | 0.366709 | GBM | AM-38 | Gefitinib |
| MVP_top60_DNA_seed | NTRK1 | -0.720545 | HNSC | HSC-2 | AZD1332 | MVP_top60_DNA+RNA_seed | MET | 0.366936 | GBM | D-392MG | Crizotinib |
| MVP_top60_DNA_seed | EGFR | -0.717468 | HNSC | PCI-15A | Afatinib | MVP_top60_DNA+RNA_seed | IGF1R | 0.367295 | GBM | YH-13 | GSK1904529A |
| MVP_top60_DNA_seed | HDAC1 | -0.714958 | HNSC | BB30-HNC | Tubastatin A | MVP_top60_DNA+RNA_seed | IGF1R | 0.369178 | GBM | AM-38 | BMS-536924 |
| MVP_top60_DNA_seed | ESR1 | -0.71476 | HNSC | HSC-4 | Tamoxifen | MVP_top60_DNA+RNA_seed | AKT1 | 0.369408 | GBM | T98G | Ipatasertib |
| MVP_top60_DNA_seed | CDK2 | -0.713399 | HNSC | OSC-20 | AZD5438 | MVP_top60_DNA+RNA_seed | PIK3CA | 0.370219 | GBM | SF268 | Buparlisib |
| MVP_top60_DNA_seed | AR | -0.712986 | HNSC | HO-1-u-1 | Bicalutamide | MVP_top60_DNA+RNA_seed | PIK3CB | 0.370219 | GBM | SF268 | Buparlisib |
| MVP_top60_DNA_seed | PIK3CA | -0.712964 | HNSC | LB771-HNC | AZD8835 | MVP_top60_DNA+RNA_seed | PIK3CD | 0.370219 | GBM | SF268 | Buparlisib |
| MVP_top60_DNA_seed | ESR1 | -0.712219 | HNSC | BICR22 | Tamoxifen | MVP_top60_DNA+RNA_seed | AKT1 | 0.371167 | GBM | U-118-MG | GSK2110183B |
| MVP_top60_DNA_seed | AKT1 | -0.705155 | HNSC | HSC-4 | BAY AKT1 | MVP_top60_DNA+RNA_seed | IGF1R | 0.37177 | GBM | LN-229 | Linsitinib |
| MVP_top60_DNA_seed | MDM2 | -0.700869 | HNSC | PCI-38 | Nutlin-3a (-) | MVP_top60_DNA+RNA_seed | IGF1R | 0.371804 | GBM | D-566MG | NVP-ADW742 |
| MVP_top60_DNA_seed | MDM2 | -0.698114 | HNSC | CAL-33 | Serdemetan | MVP_top60_DNA+RNA_seed | MET | 0.373531 | GBM | LN-405 | Savolitinib |
| MVP_top60_DNA_seed | HDAC1 | -0.694784 | HNSC | PCI-6A | JQ12 | MVP_top60_DNA+RNA_seed | AKT1 | 0.373541 | GBM | AM-38 | AZD5363 |
| MVP_top60_DNA_seed | HSP90AA1 | -0.693865 | HNSC | OSC-20 | SNX-2112 | MVP_top60_DNA+RNA_seed | CDK4 | 0.375127 | GBM | D-392MG | Ribociclib |
| MVP_top60_DNA_seed | ESR1 | -0.690622 | HNSC | PCI-4B | Fulvestrant | MVP_top60_DNA+RNA_seed | KIT | 0.376224 | GBM | T98G | Sorafenib |
| MVP_top60_DNA_seed | AKT1 | -0.6884 | HNSC | CAL-33 | MK-2206 | MVP_top60_DNA+RNA_seed | MET | 0.376545 | GBM | D-392MG | Savolitinib |
| MVP_top60_DNA_seed | HDAC1 | -0.687978 | HNSC | PCI-15A | JQ12 | MVP_top60_DNA+RNA_seed | JAK1 | 0.377577 | GBM | A172 | JAK1_8709 |

| | | | | | | | | | | | |
|---|---|---|---|---|---|---|---|---|---|---|---|
| MVP_top60_DNA_seed | NTRK1 | -0.687088 | HNSC | HO-1-u-1 | AZD1332 | MVP_top60_DNA+RNA_seed | EP300 | 0.377944 | GBM | 42-MG-BA | CPI-637 |
| MVP_top60_DNA_seed | EP300 | -0.682664 | HNSC | HSC-4 | I-CBP112 | MVP_top60_DNA+RNA_seed | EGFR | 0.37903 | GBM | SF295 | Gefitinib |
| MVP_top60_DNA_seed | CREBBP | -0.682664 | HNSC | HSC-4 | I-CBP112 | MVP_top60_DNA+RNA_seed | EGFR | 0.379162 | GBM | YKG-1 | Erlotinib |
| MVP_top60_DNA_seed | SMARCA4 | -0.681699 | HNSC | BB49-HNC | PFI-3 | MVP_top60_DNA+RNA_seed | EGFR | 0.382277 | GBM | D-247MG | Lapatinib |
| MVP_top60_DNA_seed | EP300 | -0.680555 | HNSC | HSC-2 | I-CBP112 | MVP_top60_DNA+RNA_seed | JAK2 | 0.384084 | GBM | U-118-MG | AZ960 |
| MVP_top60_DNA_seed | CREBBP | -0.680555 | HNSC | HSC-2 | I-CBP112 | MVP_top60_DNA+RNA_seed | JAK3 | 0.384084 | GBM | U-118-MG | AZ960 |
| MVP_top60_DNA_seed | HSP90AA1 | -0.679678 | HNSC | SCC-25 | CCT-018159 | MVP_top60_DNA+RNA_seed | PIK3CA | 0.384515 | GBM | DK-MG | GNE-317 |
| MVP_top60_DNA_seed | AKT1 | -0.679345 | HNSC | HSC-2 | AKT inhibitor VIII | MVP_top60_DNA+RNA_seed | EGFR | 0.384935 | GBM | 42-MG-BA | AZD3759 |
| MVP_top60_DNA_seed | EGFR | -0.677712 | HNSC | Detroit562 | Gefitinib | MVP_top60_DNA+RNA_seed | CDK4 | 0.385871 | GBM | SF126 | Palbociclib |
| MVP_top60_DNA_seed | PIK3CA | -0.676735 | HNSC | SCC-25 | AZD8835 | MVP_top60_DNA+RNA_seed | EGFR | 0.3876 | GBM | KS-1 | Afatinib |
| MVP_top60_DNA_seed | SRC | -0.676713 | HNSC | SCC-15 | Bosutinib | MVP_top60_DNA+RNA_seed | IGF1R | 0.389603 | GBM | D-542MG | GSK1904529A |
| MVP_top60_DNA_seed | AR | -0.676403 | HNSC | KON | AZD3514 | MVP_top60_DNA+RNA_seed | AKT1 | 0.3915 | GBM | D-263MG | Uprosertib |
| MVP_top60_DNA_seed | HSP90AA1 | -0.675872 | HNSC | BICR78 | CCT-018159 | MVP_top60_DNA+RNA_seed | AKT1 | 0.391925 | GBM | U251 | Uprosertib |
| MVP_top60_DNA_seed | CDK2 | -0.674685 | HNSC | PCI-4B | AZD5438 | MVP_top60_DNA+RNA_seed | EGFR | 0.393013 | GBM | YH-13 | Sapitinib |
| MVP_top60_DNA_seed | ESR1 | -0.674558 | HNSC | Ca9-22 | Tamoxifen | MVP_top60_DNA+RNA_seed | ERBB3 | 0.393013 | GBM | YH-13 | Sapitinib |
| MVP_top60_DNA_seed | PIK3CA | -0.673846 | HNSC | KON | PI3Ka_4409 | MVP_top60_DNA+RNA_seed | EGFR | 0.393487 | GBM | SK-MG-1 | Osimertinib |
| MVP_top60_DNA_seed | EGFR | -0.672143 | HNSC | PCI-6A | Cetuximab | MVP_top60_DNA+RNA_seed | CDK4 | 0.394151 | GBM | D-542MG | Ribociclib |
| MVP_top60_DNA_seed | HSP90AA1 | -0.670157 | HNSC | SAT | CCT-018159 | MVP_top60_DNA+RNA_seed | AKT1 | 0.395109 | GBM | D-566MG | Uprosertib |
| MVP_top60_DNA_seed | AKT1 | -0.669537 | HNSC | BICR78 | MK-2206 | MVP_top60_DNA+RNA_seed | KIT | 0.395221 | GBM | YKG-1 | Axitinib |
| MVP_top60_DNA_seed | HSP90AA1 | -0.666713 | HNSC | HO-1-N-1 | CCT-018159 | MVP_top60_DNA+RNA_seed | KIT | 0.396427 | GBM | U251 | Motesanib |
| MVP_top60_DNA_seed | HDAC1 | -0.665479 | HNSC | BB49-HNC | JQ12 | MVP_top60_DNA+RNA_seed | EP300 | 0.398977 | GBM | SK-MG-1 | SGC-CBP30 |
| MVP_top60_DNA_seed | EGFR | -0.665289 | HNSC | Ca9-22 | Pelitinib | MVP_top60_DNA+RNA_seed | CREBBP | 0.398977 | GBM | SK-MG-1 | SGC-CBP30 |
| MVP_top60_DNA_seed | HSP90AA1 | -0.663736 | HNSC | BB30-HNC | Elesclomol | MVP_top60_DNA+RNA_seed | EGFR | 0.400711 | GBM | D-245MG | Gefitinib |
| MVP_top60_DNA_seed | HDAC1 | -0.663438 | HNSC | BICR78 | JQ12 | MVP_top60_DNA+RNA_seed | IGF1R | 0.402573 | GBM | KS-1 | NVP-ADW742 |
| MVP_top60_DNA_seed | HSP90AA1 | -0.662712 | HNSC | KON | Elesclomol | MVP_top60_DNA+RNA_seed | EGFR | 0.402606 | GBM | SK-MG-1 | Erlotinib |
| MVP_top60_DNA_seed | HSP90AA1 | -0.661944 | HNSC | KOSC-2 | CCT-018159 | MVP_top60_DNA+RNA_seed | PIK3CA | 0.404616 | GBM | DK-MG | Buparlisib |
| MVP_top60_DNA_seed | HDAC1 | -0.660543 | HNSC | PCI-4B | Dacinostat | MVP_top60_DNA+RNA_seed | PIK3CB | 0.404616 | GBM | DK-MG | Buparlisib |
| MVP_top60_DNA_seed | EGFR | -0.658841 | HNSC | PCI-15A | Pelitinib | MVP_top60_DNA+RNA_seed | PIK3CD | 0.404616 | GBM | DK-MG | Buparlisib |
| MVP_top60_DNA_seed | MAPK1 | -0.656434 | HNSC | BB30-HNC | VX-11e | MVP_top60_DNA+RNA_seed | KIT | 0.406433 | GBM | D-263MG | Motesanib |
| MVP_top60_DNA_seed | RAC1 | -0.655489 | HNSC | HSC-2 | EHT-1864 | MVP_top60_DNA+RNA_seed | MET | 0.406956 | GBM | SF268 | Crizotinib |
| MVP_top60_DNA_seed | EGFR | -0.654594 | HNSC | Detroit562 | Afatinib | MVP_top60_DNA+RNA_seed | CDK4 | 0.410973 | GBM | Becker | Ribociclib |
| MVP_top60_DNA_seed | NTRK1 | -0.65397 | HNSC | PCI-4B | AZD1332 | MVP_top60_DNA+RNA_seed | EGFR | 0.412974 | GBM | D-566MG | Sapitinib |
| MVP_top60_DNA_seed | RAC1 | -0.652965 | HNSC | SCC-4 | EHT-1864 | MVP_top60_DNA+RNA_seed | ERBB3 | 0.412974 | GBM | D-566MG | Sapitinib |
| MVP_top60_DNA_seed | PIK3CA | -0.65167 | HNSC | HSC-3 | AZD8835 | MVP_top60_DNA+RNA_seed | EGFR | 0.417341 | GBM | U-118-MG | Gefitinib |
| MVP_top60_DNA_seed | HSP90AA1 | -0.651539 | HNSC | SCC-4 | SNX-2112 | MVP_top60_DNA+RNA_seed | PIK3CA | 0.417712 | GBM | YKG-1 | AZD8186 |
| MVP_top60_DNA_seed | MDM2 | -0.648756 | HNSC | Detroit562 | Serdemetan | MVP_top60_DNA+RNA_seed | PIK3CB | 0.417712 | GBM | YKG-1 | AZD8186 |
| MVP_top60_DNA_seed | EGFR | -0.64787 | HNSC | PCI-4B | CUDC-101 | MVP_top60_DNA+RNA_seed | EP300 | 0.420427 | GBM | D-263MG | CPI-637 |
| MVP_top60_DNA_seed | EGFR | -0.646649 | HNSC | KOSC-2 | CUDC-101 | MVP_top60_DNA+RNA_seed | JAK2 | 0.422478 | GBM | YH-13 | GSK2276186C |
| MVP_top60_DNA_seed | AKT1 | -0.645599 | HNSC | PCI-4B | AKT inhibitor VIII | MVP_top60_DNA+RNA_seed | JAK1 | 0.422478 | GBM | YH-13 | GSK2276186C |
| MVP_top60_DNA_seed | EGFR | -0.644513 | HNSC | LB771-HNC | AZD8931 | MVP_top60_DNA+RNA_seed | JAK3 | 0.422478 | GBM | YH-13 | GSK2276186C |
| MVP_top60_DNA_seed | EGFR | -0.644027 | HNSC | SCC-4 | AST-1306 | MVP_top60_DNA+RNA_seed | JAK1 | 0.4246 | GBM | LN-229 | JAK1_8709 |

| | | | | | | | | | | | |
|---|---|---|---|---|---|---|---|---|---|---|---|
| MVP_top60_DNA_seed | AKT1 | -0.643688 | HNSC | BICR22 | MK-2206 | MVP_top60_DNA+RNA_seed | AKT1 | 0.428083 | GBM | CAS-1 | MK-2206 |
| MVP_top60_DNA_seed | CDK2 | -0.64341 | HNSC | BICR78 | AZD5438 | MVP_top60_DNA+RNA_seed | IGF1R | 0.428728 | GBM | YKG-1 | BMS-536924 |
| MVP_top60_DNA_seed | EGFR | -0.641026 | HNSC | OSC-19 | CUDC-101 | MVP_top60_DNA+RNA_seed | EGFR | 0.431461 | GBM | LNZTA3WT4 | AZD3759 |
| MVP_top60_DNA_seed | HSP90AA1 | -0.638462 | HNSC | SCC-4 | Tanespimycin | MVP_top60_DNA+RNA_seed | AKT1 | 0.431873 | GBM | YKG-1 | MK-2206 |
| MVP_top60_DNA_seed | MAPK1 | -0.635832 | HNSC | SAS | VX-11e | MVP_top60_DNA+RNA_seed | KIT | 0.433319 | GBM | SF268 | Motesanib |
| MVP_top60_DNA_seed | HDAC1 | -0.631317 | HNSC | BHY | Dacinostat | MVP_top60_DNA+RNA_seed | IGF1R | 0.435173 | GBM | SF268 | GSK1904529A |
| MVP_top60_DNA_seed | HSP90AA1 | -0.627431 | HNSC | BB49-HNC | Luminespib | MVP_top60_DNA+RNA_seed | MET | 0.436031 | GBM | U-87-MG | Crizotinib |
| MVP_top60_DNA_seed | HSP90AA1 | -0.627229 | HNSC | HSC-3 | SNX-2112 | MVP_top60_DNA+RNA_seed | PIK3CA | 0.437291 | GBM | DBTRG-05MG | AZD8186 |
| MVP_top60_DNA_seed | EGFR | -0.626668 | HNSC | KON | Afatinib | MVP_top60_DNA+RNA_seed | PIK3CB | 0.437291 | GBM | DBTRG-05MG | AZD8186 |
| MVP_top60_DNA_seed | EGFR | -0.624682 | HNSC | HO-1-N-1 | CI-1033 | MVP_top60_DNA+RNA_seed | AKT1 | 0.437756 | GBM | LN-405 | Uprosertib |
| MVP_top60_DNA_seed | EGFR | -0.621652 | HNSC | PCI-15A | Afatinib | MVP_top60_DNA+RNA_seed | IGF1R | 0.440595 | GBM | M059J | GSK1904529A |
| MVP_top60_DNA_seed | HSP90AA1 | -0.620488 | HNSC | JHU-022 | CCT-018159 | MVP_top60_DNA+RNA_seed | KIT | 0.442759 | GBM | SK-MG-1 | Motesanib |
| MVP_top60_DNA_seed | AKT1 | -0.619914 | HNSC | CAL-33 | AKT inhibitor VIII | MVP_top60_DNA+RNA_seed | PIK3CA | 0.443899 | GBM | SF295 | Buparlisib |
| MVP_top60_DNA_seed | MDM2 | -0.61772 | HNSC | BICR10 | Serdemetan | MVP_top60_DNA+RNA_seed | PIK3CB | 0.443899 | GBM | SF295 | Buparlisib |
| MVP_top60_DNA_seed | HSP90AA1 | -0.617508 | HNSC | SCC-25 | SNX-2112 | MVP_top60_DNA+RNA_seed | PIK3CD | 0.443899 | GBM | SF295 | Buparlisib |
| MVP_top60_DNA_seed | HSP90AA1 | -0.615037 | HNSC | JHU-011 | Elesclomol | MVP_top60_DNA+RNA_seed | EGFR | 0.447714 | GBM | D-423MG | Sapitinib |
| MVP_top60_DNA_seed | PIK3CA | -0.612883 | HNSC | Ca9-22 | AZD8835 | MVP_top60_DNA+RNA_seed | ERBB3 | 0.447714 | GBM | D-423MG | Sapitinib |
| MVP_top60_DNA_seed | AR | -0.612569 | HNSC | HO-1-u-1 | Bicalutamide | MVP_top60_DNA+RNA_seed | KIT | 0.452088 | GBM | Becker | Motesanib |
| MVP_top60_DNA_seed | HDAC1 | -0.611718 | HNSC | OSC-19 | Belinostat | MVP_top60_DNA+RNA_seed | JAK1 | 0.455118 | GBM | GB-1 | JAK1_8709 |
| MVP_top60_DNA_seed | AKT1 | -0.611083 | HNSC | HO-1-u-1 | GSK690693 | MVP_top60_DNA+RNA_seed | EGFR | 0.455249 | GBM | M059J | AZD3759 |
| MVP_top60_DNA_seed | EGFR | -0.610816 | HNSC | HO-1-N-1 | Cetuximab | MVP_top60_DNA+RNA_seed | KRAS | 0.459518 | GBM | YKG-1 | KRAS (G12C) Inhibitor-1 |
| MVP_top60_DNA_seed | HSP90AA1 | -0.607766 | HNSC | BB49-HNC | CCT-018159 | MVP_top60_DNA+RNA_seed | IGF1R | 0.464404 | GBM | SF295 | GSK1904529A |
| MVP_top60_DNA_seed | AR | -0.607372 | HNSC | PCI-6A | Bicalutamide | MVP_top60_DNA+RNA_seed | AKT1 | 0.469395 | GBM | D-566MG | MK-2206 |
| MVP_top60_DNA_seed | EGFR | -0.606523 | HNSC | BICR78 | CUDC-101 | MVP_top60_DNA+RNA_seed | TP53 | 0.47018 | GBM | D-245MG | MIRA-1 |
| MVP_top60_DNA_seed | HDAC1 | -0.604948 | HNSC | CAL-33 | JQ12 | MVP_top60_DNA+RNA_seed | PIK3CA | 0.47052 | GBM | U251 | Buparlisib |
| MVP_top60_DNA_seed | HSP90AA1 | -0.604575 | HNSC | CAL-33 | SNX-2112 | MVP_top60_DNA+RNA_seed | PIK3CB | 0.47052 | GBM | U251 | Buparlisib |
| MVP_top60_DNA_seed | PIK3CA | -0.604195 | HNSC | HO-1-N-1 | AZD8835 | MVP_top60_DNA+RNA_seed | PIK3CD | 0.47052 | GBM | U251 | Buparlisib |
| MVP_top60_DNA_seed | EGFR | -0.603608 | HNSC | BICR31 | CUDC-101 | MVP_top60_DNA+RNA_seed | KIT | 0.47121 | GBM | D-263MG | Sorafenib |
| MVP_top60_DNA_seed | HSP90AA1 | -0.603361 | HNSC | HO-1-u-1 | CCT-018159 | MVP_top60_DNA+RNA_seed | JAK2 | 0.471794 | GBM | KS-1 | Ruxolitinib |
| MVP_top60_DNA_seed | PIK3CA | -0.602938 | HNSC | SKN-3 | PI3Ka_4409 | MVP_top60_DNA+RNA_seed | JAK1 | 0.471794 | GBM | KS-1 | Ruxolitinib |
| MVP_top60_DNA_seed | NTRK1 | -0.601424 | HNSC | CAL-27 | AZD1332 | MVP_top60_DNA+RNA_seed | CDK4 | 0.472011 | GBM | D-392MG | Palbociclib |
| MVP_top60_DNA_seed | HSP90AA1 | -0.601317 | HNSC | SCC-9 | Tanespimycin | MVP_top60_DNA+RNA_seed | AKT1 | 0.475099 | GBM | D-392MG | AZD5363 |
| MVP_top60_DNA_seed | CDK2 | -0.600095 | HNSC | HSC-4 | AZD5438 | MVP_top60_DNA+RNA_seed | EGFR | 0.475655 | GBM | A172 | Sapitinib |
| MVP_top60_DNA_seed | AR | -0.597817 | HNSC | HSC-4 | Bicalutamide | MVP_top60_DNA+RNA_seed | ERBB3 | 0.475655 | GBM | A172 | Sapitinib |
| MVP_top60_DNA_seed | RAC1 | -0.594635 | HNSC | Ca9-22 | EHT-1864 | MVP_top60_DNA+RNA_seed | KRAS | 0.480169 | GBM | D-392MG | KRAS (G12C) Inhibitor-1 |
| MVP_top60_DNA_seed | HSP90AA1 | -0.594579 | HNSC | BICR10 | Tanespimycin | MVP_top60_DNA+RNA_seed | CDK4 | 0.480546 | GBM | SK-MG-1 | Palbociclib |
| MVP_top60_DNA_seed | MAPK1 | -0.588621 | HNSC | BHY | VX-11e | MVP_top60_DNA+RNA_seed | IGF1R | 0.480923 | GBM | DK-MG | BMS-536924 |
| MVP_top60_DNA_seed | NTRK1 | -0.586788 | HNSC | Detroit562 | AZD1332 | MVP_top60_DNA+RNA_seed | EGFR | 0.4854 | GBM | KS-1 | Erlotinib |
| MVP_top60_DNA_seed | RAC1 | -0.582885 | HNSC | PCI-15A | EHT-1864 | MVP_top60_DNA+RNA_seed | AKT1 | 0.486479 | GBM | D-423MG | Uprosertib |
| MVP_top60_DNA_seed | MDM2 | -0.582663 | HNSC | PCI-4B | Nutlin-3a (-) | MVP_top60_DNA+RNA_seed | MET | 0.486935 | GBM | KS-1 | Savolitinib |
| MVP_top60_DNA_seed | EGFR | -0.58251 | HNSC | SAS | AZD8931 | MVP_top60_DNA+RNA_seed | IGF1R | 0.487016 | GBM | YH-13 | BMS-536924 |

| | | | | | | | | | | | |
|---|---|---|---|---|---|---|---|---|---|---|---|
| MVP_top60_DNA_seed | NTRK1 | -0.580811 | HNSC | DOK | AZD1332 | MVP_top60_DNA+RNA_seed | KRAS | 0.487506 | GBM | U251 | KRAS (G12C) Inhibitor-1 |
| MVP_top60_DNA_seed | MDM2 | -0.578823 | HNSC | OSC-20 | Serdemetan | MVP_top60_DNA+RNA_seed | AKT1 | 0.489138 | GBM | M059J | MK-2206 |
| MVP_top60_DNA_seed | HDAC1 | -0.577903 | HNSC | PCI-4B | Belinostat | MVP_top60_DNA+RNA_seed | JAK2 | 0.490187 | GBM | SF126 | Ruxolitinib |
| MVP_top60_DNA_seed | CDK2 | -0.576438 | HNSC | PCI-6A | PHA-793887 | MVP_top60_DNA+RNA_seed | JAK1 | 0.490187 | GBM | SF126 | Ruxolitinib |
| MVP_top60_DNA_seed | EGFR | -0.573356 | HNSC | OSC-20 | CI-1033 | MVP_top60_DNA+RNA_seed | TP53 | 0.49485 | GBM | U251 | MIRA-1 |
| MVP_top60_DNA_seed | RAC1 | -0.572322 | HNSC | BICR22 | EHT-1864 | MVP_top60_DNA+RNA_seed | AKT1 | 0.495212 | GBM | AM-38 | Uprosertib |
| MVP_top60_DNA_seed | SRC | -0.567134 | HNSC | SAS | Bosutinib | MVP_top60_DNA+RNA_seed | AKT1 | 0.496564 | GBM | LN-405 | MK-2206 |
| MVP_top60_DNA_seed | AKT1 | -0.566754 | HNSC | LB771-HNC | AKT inhibitor VIII | MVP_top60_DNA+RNA_seed | JAK2 | 0.502099 | GBM | SK-MG-1 | Ruxolitinib |
| MVP_top60_DNA_seed | EGFR | -0.566552 | HNSC | SCC-4 | AZD8931 | MVP_top60_DNA+RNA_seed | JAK1 | 0.502099 | GBM | SK-MG-1 | Ruxolitinib |
| MVP_top60_DNA_seed | HSP90AA1 | -0.564194 | HNSC | CAL-33 | Luminespib | MVP_top60_DNA+RNA_seed | AKT1 | 0.502519 | GBM | GB-1 | Uprosertib |
| MVP_top60_DNA_seed | HDAC1 | -0.563613 | HNSC | JHU-011 | JQ12 | MVP_top60_DNA+RNA_seed | EGFR | 0.503098 | GBM | D-263MG | Gefitinib |
| MVP_top60_DNA_seed | EGFR | -0.559901 | HNSC | BICR31 | Pelitinib | MVP_top60_DNA+RNA_seed | CDK4 | 0.505729 | GBM | KS-1 | Palbociclib |
| MVP_top60_DNA_seed | SRC | -0.557209 | HNSC | HO-1-u-1 | Bosutinib | MVP_top60_DNA+RNA_seed | EGFR | 0.506116 | GBM | LN-229 | Afatinib |
| MVP_top60_DNA_seed | SRC | -0.556927 | HNSC | HSC-2 | Bosutinib | MVP_top60_DNA+RNA_seed | IGF1R | 0.506791 | GBM | YKG-1 | Linsitinib |
| MVP_top60_DNA_seed | HDAC1 | -0.555545 | HNSC | JHU-011 | Belinostat | MVP_top60_DNA+RNA_seed | MET | 0.507071 | GBM | D-263MG | Savolitinib |
| MVP_top60_DNA_seed | AKT1 | -0.55363 | HNSC | BHY | AKT inhibitor VIII | MVP_top60_DNA+RNA_seed | EGFR | 0.509287 | GBM | U-87-MG | Osimertinib |
| MVP_top60_DNA_seed | AKT1 | -0.551154 | HNSC | SCC-25 | BAY AKT1 | MVP_top60_DNA+RNA_seed | EGFR | 0.510367 | GBM | SNB75 | Gefitinib |
| MVP_top60_DNA_seed | AKT1 | -0.548206 | HNSC | LB771-HNC | A-443654 | MVP_top60_DNA+RNA_seed | PIK3CA | 0.512704 | GBM | T98G | Alpelisib |
| MVP_top60_DNA_seed | MAPK1 | -0.54759 | HNSC | BB30-HNC | FR-180204 | MVP_top60_DNA+RNA_seed | EGFR | 0.514688 | GBM | SW1088 | Afatinib |
| MVP_top60_DNA_seed | MAPK3 | -0.54759 | HNSC | BB30-HNC | FR-180204 | MVP_top60_DNA+RNA_seed | CDK4 | 0.515567 | GBM | T98G | Palbociclib |
| MVP_top60_DNA_seed | ESR1 | -0.547424 | HNSC | OSC-19 | Fulvestrant | MVP_top60_DNA+RNA_seed | IGF1R | 0.518199 | GBM | Becker | GSK1904529A |
| MVP_top60_DNA_seed | HSP90AA1 | -0.547191 | HNSC | SCC90 | Elesclomol | MVP_top60_DNA+RNA_seed | KIT | 0.518554 | GBM | KS-1 | Motesanib |
| MVP_top60_DNA_seed | EGFR | -0.546777 | HNSC | SCC-15 | CUDC-101 | MVP_top60_DNA+RNA_seed | EGFR | 0.520009 | GBM | LN-405 | Gefitinib |
| MVP_top60_DNA_seed | HDAC1 | -0.545472 | HNSC | HSC-3 | JQ12 | MVP_top60_DNA+RNA_seed | IGF1R | 0.521538 | GBM | U251 | NVP-ADW742 |
| MVP_top60_DNA_seed | PIK3CA | -0.54341 | HNSC | KOSC-2 | AZD8835 | MVP_top60_DNA+RNA_seed | IGF1R | 0.524796 | GBM | D-542MG | NVP-ADW742 |
| MVP_top60_DNA_seed | AR | -0.541137 | HNSC | PCI-38 | Bicalutamide | MVP_top60_DNA+RNA_seed | EGFR | 0.526224 | GBM | D-542MG | Afatinib |
| MVP_top60_DNA_seed | MDM2 | -0.540537 | HNSC | HO-1-N-1 | Serdemetan | MVP_top60_DNA+RNA_seed | CDK4 | 0.527481 | GBM | AM-38 | Ribociclib |
| MVP_top60_DNA_seed | EGFR | -0.539933 | HNSC | JHU-022 | AST-1306 | MVP_top60_DNA+RNA_seed | IGF1R | 0.527661 | GBM | D-542MG | BMS-536924 |
| MVP_top60_DNA_seed | AR | -0.53852 | HNSC | JHU-029 | Bicalutamide | MVP_top60_DNA+RNA_seed | EP300 | 0.528751 | GBM | D-392MG | CPI-637 |
| MVP_top60_DNA_seed | NTRK1 | -0.537617 | HNSC | KOSC-2 | AZD1332 | MVP_top60_DNA+RNA_seed | EGFR | 0.530148 | GBM | D-542MG | Sapitinib |
| MVP_top60_DNA_seed | NTRK1 | -0.537453 | HNSC | BICR22 | GW441756 | MVP_top60_DNA+RNA_seed | ERBB3 | 0.530148 | GBM | D-542MG | Sapitinib |
| MVP_top60_DNA_seed | EGFR | -0.536595 | HNSC | OSC-20 | Afatinib | MVP_top60_DNA+RNA_seed | KIT | 0.530999 | GBM | SNB75 | Sorafenib |
| MVP_top60_DNA_seed | AKT1 | -0.536181 | HNSC | BB30-HNC | A-443654 | MVP_top60_DNA+RNA_seed | CDK4 | 0.533868 | GBM | LN-18 | Palbociclib |
| MVP_top60_DNA_seed | HSP90AA1 | -0.536003 | HNSC | DOK | Elesclomol | MVP_top60_DNA+RNA_seed | AKT1 | 0.53396 | GBM | D-245MG | MK-2206 |
| MVP_top60_DNA_seed | HSP90AA1 | -0.534664 | HNSC | Ca9-22 | SNX-2112 | MVP_top60_DNA+RNA_seed | IGF1R | 0.535296 | GBM | SF268 | BMS-536924 |
| MVP_top60_DNA_seed | MDM2 | -0.534456 | HNSC | SAS | Serdemetan | MVP_top60_DNA+RNA_seed | JAK2 | 0.536203 | GBM | SF268 | JAK_8517 |
| MVP_top60_DNA_seed | CDK2 | -0.53392 | HNSC | LB771-HNC | Seliciclib | MVP_top60_DNA+RNA_seed | JAK1 | 0.536203 | GBM | SF268 | JAK_8517 |
| MVP_top60_DNA_seed | HDAC1 | -0.533403 | HNSC | JHU-011 | CAY10603 | MVP_top60_DNA+RNA_seed | EGFR | 0.537031 | GBM | 42-MG-BA | Erlotinib |
| MVP_top60_DNA_seed | HSP90AA1 | -0.531192 | HNSC | BHY | SNX-2112 | MVP_top60_DNA+RNA_seed | CDK4 | 0.537833 | GBM | D-247MG | Ribociclib |
| MVP_top60_DNA_seed | EGFR | -0.526347 | HNSC | PE-CA-PJ15 | Pelitinib | MVP_top60_DNA+RNA_seed | AKT1 | 0.54568 | GBM | D-542MG | Ipatasertib |
| MVP_top60_DNA_seed | EGFR | -0.523403 | HNSC | SCC-25 | AST-1306 | MVP_top60_DNA+RNA_seed | JAK1 | 0.546057 | GBM | CAS-1 | JAK1_8709 |

| | | | | | | | | | | | |
|---|---|---|---|---|---|---|---|---|---|---|---|
| MVP_top60_DNA_seed | MAPK1 | -0.520127 | HNSC | BHY | FR-180204 | MVP_top60_DNA+RNA_seed | PIK3CA | 0.546515 | GBM | YKG-1 | Buparlisib |
| MVP_top60_DNA_seed | MAPK3 | -0.520127 | HNSC | BHY | FR-180204 | MVP_top60_DNA+RNA_seed | PIK3CB | 0.546515 | GBM | YKG-1 | Buparlisib |
| MVP_top60_DNA_seed | HDAC1 | -0.519978 | HNSC | PCI-4B | CAY10603 | MVP_top60_DNA+RNA_seed | PIK3CD | 0.546515 | GBM | YKG-1 | Buparlisib |
| MVP_top60_DNA_seed | HDAC1 | -0.519659 | HNSC | SCC-25 | CAY10603 | MVP_top60_DNA+RNA_seed | EGFR | 0.546951 | GBM | DK-MG | Lapatinib |
| MVP_top60_DNA_seed | MDM2 | -0.518754 | HNSC | BICR78 | Serdemetan | MVP_top60_DNA+RNA_seed | EGFR | 0.547239 | GBM | SF268 | Lapatinib |
| MVP_top60_DNA_seed | AKT1 | -0.516964 | HNSC | BICR10 | BAY AKT1 | MVP_top60_DNA+RNA_seed | AKT1 | 0.547524 | GBM | T98G | AZD5363 |
| MVP_top60_DNA_seed | PIK3CA | -0.516948 | HNSC | OSC-19 | PI3Ka_4409 | MVP_top60_DNA+RNA_seed | EP300 | 0.549773 | GBM | SF268 | CPI-637 |
| MVP_top60_DNA_seed | EGFR | -0.515787 | HNSC | DOK | Pelitinib | MVP_top60_DNA+RNA_seed | AKT1 | 0.552849 | GBM | Becker | GSK2110183B |
| MVP_top60_DNA_seed | PIK3CA | -0.515743 | HNSC | BICR22 | AZD8835 | MVP_top60_DNA+RNA_seed | CDK4 | 0.557772 | GBM | SK-MG-1 | Ribociclib |
| MVP_top60_DNA_seed | HSP90AA1 | -0.515173 | HNSC | BB49-HNC | SNX-2112 | MVP_top60_DNA+RNA_seed | JAK2 | 0.558113 | GBM | Becker | Ruxolitinib |
| MVP_top60_DNA_seed | HDAC1 | -0.51315 | HNSC | HO-1-u-1 | Tubastatin A | MVP_top60_DNA+RNA_seed | JAK1 | 0.558113 | GBM | Becker | Ruxolitinib |
| MVP_top60_DNA_seed | AR | -0.511771 | HNSC | KON | Bicalutamide | MVP_top60_DNA+RNA_seed | KIT | 0.562065 | GBM | M059J | Motesanib |
| MVP_top60_DNA_seed | AKT1 | -0.510087 | HNSC | OSC-19 | BAY AKT1 | MVP_top60_DNA+RNA_seed | EGFR | 0.56236 | GBM | 8-MG-BA | Osimertinib |
| MVP_top60_DNA_seed | AKT1 | -0.506852 | HNSC | SCC-9 | BAY AKT1 | MVP_top60_DNA+RNA_seed | MET | 0.564768 | GBM | DBTRG-05MG | Savolitinib |
| MVP_top60_DNA_seed | AKT1 | -0.506573 | HNSC | HO-1-u-1 | MK-2206 | MVP_top60_DNA+RNA_seed | AKT1 | 0.570444 | GBM | D-263MG | AZD5363 |
| MVP_top60_DNA_seed | HSP90AA1 | -0.505614 | HNSC | PCI-30 | CCT-018159 | MVP_top60_DNA+RNA_seed | EP300 | 0.570706 | GBM | LN-405 | SGC-CBP30 |
| MVP_top60_DNA_seed | EGFR | -0.504491 | HNSC | BICR22 | AZD8931 | MVP_top60_DNA+RNA_seed | CREBBP | 0.570706 | GBM | LN-405 | SGC-CBP30 |
| MVP_top60_DNA_seed | CDK2 | -0.503602 | HNSC | BB30-HNC | Seliciclib | MVP_top60_DNA+RNA_seed | MET | 0.571369 | GBM | SW1088 | Savolitinib |
| MVP_top60_DNA_seed | CDK2 | -0.502724 | HNSC | SKN-3 | AZD5438 | MVP_top60_DNA+RNA_seed | EGFR | 0.575883 | GBM | D-542MG | AZD3759 |
| MVP_top60_DNA_seed | AR | -0.500061 | HNSC | HO-1-N-1 | Bicalutamide | MVP_top60_DNA+RNA_seed | EGFR | 0.579175 | GBM | YH-13 | Lapatinib |
| MVP_top60_DNA_seed | HDAC1 | -0.499976 | HNSC | KON | Dacinostat | MVP_top60_DNA+RNA_seed | JAK2 | 0.579378 | GBM | U251 | Ruxolitinib |
| MVP_top60_DNA_seed | NTRK1 | -0.49988 | HNSC | HSC-3 | GW441756 | MVP_top60_DNA+RNA_seed | JAK1 | 0.579378 | GBM | U251 | Ruxolitinib |
| MVP_top60_DNA_seed | HSP90AA1 | -0.498145 | HNSC | CAL-27 | Elesclomol | MVP_top60_DNA+RNA_seed | AKT1 | 0.580762 | GBM | LN-229 | AT13148 |
| MVP_top60_DNA_seed | ESR1 | -0.496803 | HNSC | BICR10 | Tamoxifen | MVP_top60_DNA+RNA_seed | AKT1 | 0.58515 | GBM | D-245MG | Uprosertib |
| MVP_top60_DNA_seed | HDAC1 | -0.492305 | HNSC | SCC-25 | AR-42 | MVP_top60_DNA+RNA_seed | EGFR | 0.587528 | GBM | 42-MG-BA | Gefitinib |
| MVP_top60_DNA_seed | HSP90AA1 | -0.491011 | HNSC | CAL-27 | Luminespib | MVP_top60_DNA+RNA_seed | AKT1 | 0.587909 | GBM | YKG-1 | Uprosertib |
| MVP_top60_DNA_seed | HSP90AA1 | -0.489692 | HNSC | SCC-15 | CCT-018159 | MVP_top60_DNA+RNA_seed | JAK1 | 0.588174 | GBM | D-263MG | JAK1_8709 |
| MVP_top60_DNA_seed | HDAC1 | -0.489415 | HNSC | PCI-4B | AR-42 | MVP_top60_DNA+RNA_seed | KRAS | 0.588581 | GBM | SNB75 | [illegible]RAS (G12C) Inhibitor-1 |
| MVP_top60_DNA_seed | AR | -0.483224 | HNSC | HO-1-u-1 | AZD3514 | MVP_top60_DNA+RNA_seed | IGF1R | 0.5901 | GBM | T98G | BMS-754807 |
| MVP_top60_DNA_seed | HSP90AA1 | -0.481152 | HNSC | PCI-6A | Luminespib | MVP_top60_DNA+RNA_seed | JAK2 | 0.590174 | GBM | DBTRG-05MG | JAK_8517 |
| MVP_top60_DNA_seed | PIK3CA | -0.480643 | HNSC | OSC-20 | AZD8835 | MVP_top60_DNA+RNA_seed | JAK1 | 0.590174 | GBM | DBTRG-05MG | JAK_8517 |
| MVP_top60_DNA_seed | MDM2 | -0.479401 | HNSC | SKN-3 | Serdemetan | MVP_top60_DNA+RNA_seed | EGFR | 0.590257 | GBM | M059J | Sapitinib |
| MVP_top60_DNA_seed | EGFR | -0.476078 | HNSC | OSC-20 | AST-1306 | MVP_top60_DNA+RNA_seed | ERBB3 | 0.590257 | GBM | M059J | Sapitinib |
| MVP_top60_DNA_seed | AR | -0.475565 | HNSC | HSC-2 | Bicalutamide | MVP_top60_DNA+RNA_seed | AKT1 | 0.595379 | GBM | A172 | MK-2206 |
| MVP_top60_DNA_seed | HDAC1 | -0.474759 | HNSC | SCC-4 | Dacinostat | MVP_top60_DNA+RNA_seed | EGFR | 0.595524 | GBM | SK-MG-1 | AZD3759 |
| MVP_top60_DNA_seed | EGFR | -0.473945 | HNSC | KOSC-2 | Gefitinib | MVP_top60_DNA+RNA_seed | EGFR | 0.595763 | GBM | D-542MG | Erlotinib |
| MVP_top60_DNA_seed | HSP90AA1 | -0.473022 | HNSC | BICR10 | Elesclomol | MVP_top60_DNA+RNA_seed | EGFR | 0.595809 | GBM | D-245MG | Lapatinib |
| MVP_top60_DNA_seed | MDM2 | -0.470005 | HNSC | KON | Nutlin-3a (-) | MVP_top60_DNA+RNA_seed | KIT | 0.600619 | GBM | U-87-MG | Sorafenib |
| MVP_top60_DNA_seed | HSP90AA1 | -0.46882 | HNSC | HSC-2 | Tanespimycin | MVP_top60_DNA+RNA_seed | JAK2 | 0.601885 | GBM | U-87-MG | AZ960 |
| MVP_top60_DNA_seed | ESR1 | -0.468659 | HNSC | PCI-4B | Tamoxifen | MVP_top60_DNA+RNA_seed | JAK3 | 0.601885 | GBM | U-87-MG | AZ960 |
| MVP_top60_DNA_seed | HDAC1 | -0.46823 | HNSC | BICR78 | AR-42 | MVP_top60_DNA+RNA_seed | JAK2 | 0.603406 | GBM | SF268 | GSK2276186C |

| | | | | | | | | | | | |
|---|---|---|---|---|---|---|---|---|---|---|---|
| MVP_top60_DNA_seed | AR | -0.465139 | HNSC | KOSC-2 | AZD3514 | MVP_top60_DNA+RNA_seed | JAK1 | 0.603406 | GBM | SF268 | GSK2276186C |
| MVP_top60_DNA_seed | HDAC1 | -0.464385 | HNSC | BICR78 | Belinostat | MVP_top60_DNA+RNA_seed | JAK3 | 0.603406 | GBM | SF268 | GSK2276186C |
| MVP_top60_DNA_seed | HSP90AA1 | -0.461681 | HNSC | BICR31 | SNX-2112 | MVP_top60_DNA+RNA_seed | CDK4 | 0.605292 | GBM | GB-1 | Palbociclib |
| MVP_top60_DNA_seed | HSP90AA1 | -0.459316 | HNSC | CAL-27 | Tanespimycin | MVP_top60_DNA+RNA_seed | KRAS | 0.608359 | GBM | Becker | KRAS (G12C) Inhibitor-1 |
| MVP_top60_DNA_seed | EGFR | -0.459251 | HNSC | JHU-011 | AST-1306 | MVP_top60_DNA+RNA_seed | KIT | 0.611181 | GBM | SNB75 | Motesanib |
| MVP_top60_DNA_seed | CDK2 | -0.459234 | HNSC | PE-CA-PJ15 | AZD5438 | MVP_top60_DNA+RNA_seed | JAK2 | 0.611251 | GBM | LN-18 | GSK2276186C |
| MVP_top60_DNA_seed | HDAC1 | -0.454363 | HNSC | CAL-27 | Belinostat | MVP_top60_DNA+RNA_seed | JAK1 | 0.611251 | GBM | LN-18 | GSK2276186C |
| MVP_top60_DNA_seed | NTRK1 | -0.45356 | HNSC | KON | AZD1332 | MVP_top60_DNA+RNA_seed | JAK3 | 0.611251 | GBM | LN-18 | GSK2276186C |
| MVP_top60_DNA_seed | AR | -0.45318 | HNSC | FADU | AZD3514 | MVP_top60_DNA+RNA_seed | IGF1R | 0.612347 | GBM | D-423MG | Linsitinib |
| MVP_top60_DNA_seed | HDAC1 | -0.452197 | HNSC | BHY | Belinostat | MVP_top60_DNA+RNA_seed | CDK4 | 0.61874 | GBM | YKG-1 | Ribociclib |
| MVP_top60_DNA_seed | EGFR | -0.45137 | HNSC | BB49-HNC | AZD8931 | MVP_top60_DNA+RNA_seed | EGFR | 0.621027 | GBM | LN-18 | Gefitinib |
| MVP_top60_DNA_seed | EGFR | -0.447716 | HNSC | HSC-4 | CUDC-101 | MVP_top60_DNA+RNA_seed | KIT | 0.621581 | GBM | 8-MG-BA | Axitinib |
| MVP_top60_DNA_seed | SMARCA4 | -0.446803 | HNSC | KOSC-2 | PFI-3 | MVP_top60_DNA+RNA_seed | EP300 | 0.62222 | GBM | LN-18 | CPI-637 |
| MVP_top60_DNA_seed | EGFR | -0.446738 | HNSC | KON | Afatinib | MVP_top60_DNA+RNA_seed | KIT | 0.62534 | GBM | Becker | Sorafenib |
| MVP_top60_DNA_seed | HDAC1 | -0.444413 | HNSC | SAS | Belinostat | MVP_top60_DNA+RNA_seed | EGFR | 0.627379 | GBM | Becker | Osimertinib |
| MVP_top60_DNA_seed | PIK3CA | -0.444205 | HNSC | Ca9-22 | PI3Ka_4409 | MVP_top60_DNA+RNA_seed | EGFR | 0.628974 | GBM | GB-1 | Sapitinib |
| MVP_top60_DNA_seed | AKT1 | -0.443214 | HNSC | JHU-011 | GSK690693 | MVP_top60_DNA+RNA_seed | ERBB3 | 0.628974 | GBM | GB-1 | Sapitinib |
| MVP_top60_DNA_seed | HSP90AA1 | -0.441948 | HNSC | SCC-15 | Luminespib | MVP_top60_DNA+RNA_seed | EGFR | 0.629459 | GBM | Becker | Afatinib |
| MVP_top60_DNA_seed | PIK3CA | -0.44101 | HNSC | KON | AZD8835 | MVP_top60_DNA+RNA_seed | IGF1R | 0.630114 | GBM | 42-MG-BA | Linsitinib |
| MVP_top60_DNA_seed | HSP90AA1 | -0.440879 | HNSC | SCC-25 | Tanespimycin | MVP_top60_DNA+RNA_seed | JAK1 | 0.631924 | GBM | T98G | JAK1_8709 |
| MVP_top60_DNA_seed | AKT1 | -0.44023 | HNSC | CAL-27 | AKT inhibitor VIII | MVP_top60_DNA+RNA_seed | AKT1 | 0.634715 | GBM | A172 | Uprosertib |
| MVP_top60_DNA_seed | EP300 | -0.437778 | HNSC | BB49-HNC | I-CBP112 | MVP_top60_DNA+RNA_seed | EGFR | 0.635474 | GBM | D-247MG | Afatinib |
| MVP_top60_DNA_seed | CREBBP | -0.437778 | HNSC | BB49-HNC | I-CBP112 | MVP_top60_DNA+RNA_seed | MET | 0.635615 | GBM | M059J | Crizotinib |
| MVP_top60_DNA_seed | AR | -0.437325 | HNSC | JHU-022 | AZD3514 | MVP_top60_DNA+RNA_seed | EP300 | 0.638921 | GBM | U-118-MG | SGC-CBP30 |
| MVP_top60_DNA_seed | EP300 | -0.434773 | HNSC | BICR78 | I-CBP112 | MVP_top60_DNA+RNA_seed | CREBBP | 0.638921 | GBM | U-118-MG | SGC-CBP30 |
| MVP_top60_DNA_seed | CREBBP | -0.434773 | HNSC | BICR78 | I-CBP112 | MVP_top60_DNA+RNA_seed | EGFR | 0.639898 | GBM | T98G | AZD3759 |
| MVP_top60_DNA_seed | AR | -0.433685 | HNSC | JHU-011 | Bicalutamide | MVP_top60_DNA+RNA_seed | KIT | 0.643224 | GBM | U-87-MG | Axitinib |
| MVP_top60_DNA_seed | HSP90AA1 | -0.429992 | HNSC | SCC-9 | CCT-018159 | MVP_top60_DNA+RNA_seed | KIT | 0.643827 | GBM | YH-13 | Sorafenib |
| MVP_top60_DNA_seed | EGFR | -0.428998 | HNSC | KOSC-2 | Afatinib | MVP_top60_DNA+RNA_seed | TP53 | 0.645376 | GBM | LN-229 | MIRA-1 |
| MVP_top60_DNA_seed | NTRK1 | -0.426982 | HNSC | BICR22 | AZD1332 | MVP_top60_DNA+RNA_seed | KIT | 0.647223 | GBM | D-542MG | Motesanib |
| MVP_top60_DNA_seed | MDM2 | -0.426451 | HNSC | BB49-HNC | Serdemetan | MVP_top60_DNA+RNA_seed | TP53 | 0.648846 | GBM | U-118-MG | MIRA-1 |
| MVP_top60_DNA_seed | CDK2 | -0.424363 | HNSC | CAL-27 | PHA-793887 | MVP_top60_DNA+RNA_seed | EP300 | 0.651391 | GBM | T98G | SGC-CBP30 |
| MVP_top60_DNA_seed | HDAC1 | -0.421759 | HNSC | HSC-2 | JQ12 | MVP_top60_DNA+RNA_seed | CREBBP | 0.651391 | GBM | T98G | SGC-CBP30 |
| MVP_top60_DNA_seed | ESR1 | -0.419222 | HNSC | BICR10 | Fulvestrant | MVP_top60_DNA+RNA_seed | AKT1 | 0.661783 | GBM | DBTRG-05MG | AZD5363 |
| MVP_top60_DNA_seed | AR | -0.417242 | HNSC | BICR22 | Bicalutamide | MVP_top60_DNA+RNA_seed | IGF1R | 0.661933 | GBM | D-392MG | Linsitinib |
| MVP_top60_DNA_seed | HDAC1 | -0.416552 | HNSC | OSC-19 | CAY10603 | MVP_top60_DNA+RNA_seed | EGFR | 0.662326 | GBM | YH-13 | AZD3759 |
| MVP_top60_DNA_seed | HSP90AA1 | -0.414548 | HNSC | JHU-029 | CCT-018159 | MVP_top60_DNA+RNA_seed | EP300 | 0.662332 | GBM | 8-MG-BA | CPI-637 |
| MVP_top60_DNA_seed | SRC | -0.413096 | HNSC | SCC-15 | WH-4-023 | MVP_top60_DNA+RNA_seed | CDK4 | 0.663048 | GBM | SW1088 | Ribociclib |
| MVP_top60_DNA_seed | MDM2 | -0.411407 | HNSC | HSC-4 | Serdemetan | MVP_top60_DNA+RNA_seed | JAK2 | 0.664893 | GBM | AM-38 | GSK2276186C |
| MVP_top60_DNA_seed | HSP90AA1 | -0.411134 | HNSC | HO-1-N-1 | Elesclomol | MVP_top60_DNA+RNA_seed | JAK1 | 0.664893 | GBM | AM-38 | GSK2276186C |
| MVP_top60_DNA_seed | MAPK1 | -0.408128 | HNSC | Ca9-22 | VX-11e | MVP_top60_DNA+RNA_seed | JAK3 | 0.664893 | GBM | AM-38 | GSK2276186C |

| | | | | | | | | | | | |
|---|---|---|---|---|---|---|---|---|---|---|---|
| MVP_top60_DNA_seed | HDAC1 | -0.407465 | HNSC | SCC-9 | Belinostat | MVP_top60_DNA+RNA_seed | MET | 0.665786 | GBM | YKG-1 | Savolitinib |
| MVP_top60_DNA_seed | EGFR | -0.405624 | HNSC | BICR31 | CI-1033 | MVP_top60_DNA+RNA_seed | IGF1R | 0.665968 | GBM | LN-229 | BMS-536924 |
| MVP_top60_DNA_seed | EGFR | -0.405519 | HNSC | BB30-HNC | CUDC-101 | MVP_top60_DNA+RNA_seed | EGFR | 0.666117 | GBM | DBTRG-05MG | Erlotinib |
| MVP_top60_DNA_seed | AKT1 | -0.405268 | HNSC | SCC-25 | AKT inhibitor VIII | MVP_top60_DNA+RNA_seed | EGFR | 0.667379 | GBM | AM-38 | Osimertinib |
| MVP_top60_DNA_seed | AKT1 | -0.402367 | HNSC | HO-1-u-1 | AKT inhibitor VIII | MVP_top60_DNA+RNA_seed | TP53 | 0.673706 | GBM | D-392MG | MIRA-1 |
| MVP_top60_DNA_seed | HSP90AA1 | -0.399186 | HNSC | PCI-15A | Luminespib | MVP_top60_DNA+RNA_seed | AKT1 | 0.678384 | GBM | SF268 | MK-2206 |
| MVP_top60_DNA_seed | HSP90AA1 | -0.398779 | HNSC | HO-1-N-1 | Luminespib | MVP_top60_DNA+RNA_seed | AKT1 | 0.683076 | GBM | D-263MG | GSK2110183B |
| MVP_top60_DNA_seed | HDAC1 | -0.396259 | HNSC | BB30-HNC | Dacinostat | MVP_top60_DNA+RNA_seed | EGFR | 0.686248 | GBM | AM-38 | Afatinib |
| MVP_top60_DNA_seed | AKT1 | -0.395948 | HNSC | PCI-6A | MK-2206 | MVP_top60_DNA+RNA_seed | EGFR | 0.688348 | GBM | CAS-1 | Osimertinib |
| MVP_top60_DNA_seed | HSP90AA1 | -0.395352 | HNSC | SCC-25 | Elesclomol | MVP_top60_DNA+RNA_seed | AKT1 | 0.688466 | GBM | LN-18 | GSK2110183B |
| MVP_top60_DNA_seed | EGFR | -0.395348 | HNSC | OSC-19 | Pelitinib | MVP_top60_DNA+RNA_seed | AKT1 | 0.689467 | GBM | Becker | MK-2206 |
| MVP_top60_DNA_seed | AKT1 | -0.39483 | HNSC | BB30-HNC | MK-2206 | MVP_top60_DNA+RNA_seed | PIK3CA | 0.689663 | GBM | SF268 | GNE-317 |
| MVP_top60_DNA_seed | HDAC1 | -0.393769 | HNSC | KOSC-2 | AR-42 | MVP_top60_DNA+RNA_seed | IGF1R | 0.691106 | GBM | DBTRG-05MG | BMS-754807 |
| MVP_top60_DNA_seed | EP300 | -0.393161 | HNSC | HSC-3 | I-CBP112 | MVP_top60_DNA+RNA_seed | CDK4 | 0.691742 | GBM | M059J | Ribociclib |
| MVP_top60_DNA_seed | CREBBP | -0.393161 | HNSC | HSC-3 | I-CBP112 | MVP_top60_DNA+RNA_seed | KRAS | 0.693397 | GBM | D-245MG | :RAS (G12C) Inhibitor-1 |
| MVP_top60_DNA_seed | EGFR | -0.393133 | HNSC | JHU-011 | CUDC-101 | MVP_top60_DNA+RNA_seed | IGF1R | 0.700711 | GBM | Becker | BMS-536924 |
| MVP_top60_DNA_seed | EGFR | -0.39127 | HNSC | SAT | CUDC-101 | MVP_top60_DNA+RNA_seed | EGFR | 0.702379 | GBM | D-392MG | Osimertinib |
| MVP_top60_DNA_seed | AKT1 | -0.387096 | HNSC | SCC-25 | MK-2206 | MVP_top60_DNA+RNA_seed | JAK2 | 0.70349 | GBM | DBTRG-05MG | AZ960 |
| MVP_top60_DNA_seed | MAPK1 | -0.387087 | HNSC | PCI-4B | VX-11e | MVP_top60_DNA+RNA_seed | JAK3 | 0.70349 | GBM | DBTRG-05MG | AZ960 |
| MVP_top60_DNA_seed | MAPK1 | -0.381963 | HNSC | PCI-6A | VX-11e | MVP_top60_DNA+RNA_seed | CDK4 | 0.703697 | GBM | U251 | Ribociclib |
| MVP_top60_DNA_seed | HSP90AA1 | -0.374988 | HNSC | SAS | Elesclomol | MVP_top60_DNA+RNA_seed | CDK4 | 0.704074 | GBM | SNB75 | Ribociclib |
| MVP_top60_DNA_seed | HDAC1 | -0.374425 | HNSC | OSC-19 | AR-42 | MVP_top60_DNA+RNA_seed | AKT1 | 0.705739 | GBM | Becker | Ipatasertib |
| MVP_top60_DNA_seed | HSP90AA1 | -0.372124 | HNSC | BHY | Tanespimycin | MVP_top60_DNA+RNA_seed | EGFR | 0.706663 | GBM | M059J | Osimertinib |
| MVP_top60_DNA_seed | EGFR | -0.370604 | HNSC | PCI-38 | Afatinib | MVP_top60_DNA+RNA_seed | EP300 | 0.71015 | GBM | D-263MG | SGC-CBP30 |
| MVP_top60_DNA_seed | ESR1 | -0.366048 | HNSC | HO-1-u-1 | Fulvestrant | MVP_top60_DNA+RNA_seed | CREBBP | 0.71015 | GBM | D-263MG | SGC-CBP30 |
| MVP_top60_DNA_seed | MAPK1 | -0.361688 | HNSC | JHU-011 | FR-180204 | MVP_top60_DNA+RNA_seed | AKT1 | 0.711969 | GBM | KS-1 | AZD5363 |
| MVP_top60_DNA_seed | MAPK3 | -0.361688 | HNSC | JHU-011 | FR-180204 | MVP_top60_DNA+RNA_seed | TP53 | 0.714854 | GBM | U-87-MG | MIRA-1 |
| MVP_top60_DNA_seed | HSP90AA1 | -0.360249 | HNSC | SKN-3 | SNX-2112 | MVP_top60_DNA+RNA_seed | KIT | 0.716923 | GBM | D-392MG | Motesanib |
| MVP_top60_DNA_seed | HDAC1 | -0.359021 | HNSC | HO-1-u-1 | Belinostat | MVP_top60_DNA+RNA_seed | EGFR | 0.71768 | GBM | T98G | Afatinib |
| MVP_top60_DNA_seed | HDAC1 | -0.358854 | HNSC | SCC-9 | JQ12 | MVP_top60_DNA+RNA_seed | AKT1 | 0.719955 | GBM | D-392MG | Ipatasertib |
| MVP_top60_DNA_seed | MAPK1 | -0.357791 | HNSC | CAL-27 | FR-180204 | MVP_top60_DNA+RNA_seed | JAK2 | 0.720744 | GBM | SK-MG-1 | GSK2276186C |
| MVP_top60_DNA_seed | MAPK3 | -0.357791 | HNSC | CAL-27 | FR-180204 | MVP_top60_DNA+RNA_seed | JAK1 | 0.720744 | GBM | SK-MG-1 | GSK2276186C |
| MVP_top60_DNA_seed | EGFR | -0.357256 | HNSC | HSC-3 | AST-1306 | MVP_top60_DNA+RNA_seed | JAK3 | 0.720744 | GBM | SK-MG-1 | GSK2276186C |
| MVP_top60_DNA_seed | SMARCA4 | -0.357076 | HNSC | Detroit562 | PFI-3 | MVP_top60_DNA+RNA_seed | IGF1R | 0.722157 | GBM | GB-1 | GSK1904529A |
| MVP_top60_DNA_seed | CDK2 | -0.356069 | HNSC | BB30-HNC | AZD5438 | MVP_top60_DNA+RNA_seed | PIK3CA | 0.722615 | GBM | SF268 | Alpelisib |
| MVP_top60_DNA_seed | HDAC1 | -0.355733 | HNSC | BHY | AR-42 | MVP_top60_DNA+RNA_seed | EGFR | 0.722859 | GBM | SW1088 | Lapatinib |
| MVP_top60_DNA_seed | HSP90AA1 | -0.350979 | HNSC | PCI-38 | SNX-2112 | MVP_top60_DNA+RNA_seed | PIK3CA | 0.724599 | GBM | DBTRG-05MG | Buparlisib |
| MVP_top60_DNA_seed | PIK3CA | -0.346995 | HNSC | LB771-HNC | PI3Ka_4409 | MVP_top60_DNA+RNA_seed | PIK3CB | 0.724599 | GBM | DBTRG-05MG | Buparlisib |
| MVP_top60_DNA_seed | SRC | -0.343377 | HNSC | BHY | NVP-BHG712 | MVP_top60_DNA+RNA_seed | PIK3CD | 0.724599 | GBM | DBTRG-05MG | Buparlisib |
| MVP_top60_DNA_seed | CDK2 | -0.343102 | HNSC | HO-1-u-1 | PHA-793887 | MVP_top60_DNA+RNA_seed | PIK3CA | 0.725636 | GBM | KS-1 | AZD8186 |
| MVP_top60_DNA_seed | HDAC1 | -0.342119 | HNSC | CAL-27 | AR-42 | MVP_top60_DNA+RNA_seed | PIK3CB | 0.725636 | GBM | KS-1 | AZD8186 |

| | | | | | | | | | | | |
|---|---|---|---|---|---|---|---|---|---|---|---|
| MVP_top60_DNA_seed | HSP90AA1 | -0.340233 | HNSC | DOK | CCT-018159 | MVP_top60_DNA+RNA_seed | EGFR | 0.7261 | GBM | Becker | Gefitinib |
| MVP_top60_DNA_seed | PIK3CA | -0.339834 | HNSC | JHU-011 | AZD8835 | MVP_top60_DNA+RNA_seed | KIT | 0.726755 | GBM | LN-405 | Axitinib |
| MVP_top60_DNA_seed | HDAC1 | -0.337983 | HNSC | OSC-19 | JQ12 | MVP_top60_DNA+RNA_seed | IGF1R | 0.730823 | GBM | D-247MG | BMS-536924 |
| MVP_top60_DNA_seed | MAPK1 | -0.336858 | HNSC | JHU-011 | VX-11e | MVP_top60_DNA+RNA_seed | PIK3CA | 0.731918 | GBM | LN-229 | Buparlisib |
| MVP_top60_DNA_seed | MAPK1 | -0.333052 | HNSC | SCC-9 | FR-180204 | MVP_top60_DNA+RNA_seed | PIK3CB | 0.731918 | GBM | LN-229 | Buparlisib |
| MVP_top60_DNA_seed | MAPK3 | -0.333052 | HNSC | SCC-9 | FR-180204 | MVP_top60_DNA+RNA_seed | PIK3CD | 0.731918 | GBM | LN-229 | Buparlisib |
| MVP_top60_DNA_seed | MDM2 | -0.330542 | HNSC | JHU-022 | Serdemetan | MVP_top60_DNA+RNA_seed | EP300 | 0.733056 | GBM | T98G | CPI-637 |
| MVP_top60_DNA_seed | AKT1 | -0.325633 | HNSC | PCI-15A | GSK690693 | MVP_top60_DNA+RNA_seed | JAK2 | 0.733416 | GBM | LN-405 | GSK2276186C |
| MVP_top60_DNA_seed | AKT1 | -0.323604 | HNSC | OSC-20 | AKT inhibitor VIII | MVP_top60_DNA+RNA_seed | JAK1 | 0.733416 | GBM | LN-405 | GSK2276186C |
| MVP_top60_DNA_seed | AR | -0.323427 | HNSC | SAT | Bicalutamide | MVP_top60_DNA+RNA_seed | JAK3 | 0.733416 | GBM | LN-405 | GSK2276186C |
| MVP_top60_DNA_seed | CDK2 | -0.322873 | HNSC | SAT | AZD5438 | MVP_top60_DNA+RNA_seed | IGF1R | 0.737837 | GBM | D-263MG | Linsitinib |
| MVP_top60_DNA_seed | EGFR | -0.322 | HNSC | SCC-25 | AZD8931 | MVP_top60_DNA+RNA_seed | EGFR | 0.738527 | GBM | LN-229 | Lapatinib |
| MVP_top60_DNA_seed | PIK3CA | -0.31855 | HNSC | BICR78 | PI3Ka_4409 | MVP_top60_DNA+RNA_seed | CDK4 | 0.744679 | GBM | 42-MG-BA | Palbociclib |
| MVP_top60_DNA_seed | EGFR | -0.317332 | HNSC | FADU | Pelitinib | MVP_top60_DNA+RNA_seed | PIK3CA | 0.746592 | GBM | AM-38 | Alpelisib |
| MVP_top60_DNA_seed | AKT1 | -0.317186 | HNSC | BB30-HNC | GSK690693 | MVP_top60_DNA+RNA_seed | EP300 | 0.746626 | GBM | YH-13 | SGC-CBP30 |
| MVP_top60_DNA_seed | CDK2 | -0.315916 | HNSC | BHY | PHA-793887 | MVP_top60_DNA+RNA_seed | CREBBP | 0.746626 | GBM | YH-13 | SGC-CBP30 |
| MVP_top60_DNA_seed | NTRK1 | -0.315368 | HNSC | SAT | AZD1332 | MVP_top60_DNA+RNA_seed | EGFR | 0.746715 | GBM | CAS-1 | Gefitinib |
| MVP_top60_DNA_seed | HSP90AA1 | -0.314659 | HNSC | PCI-38 | Luminespib | MVP_top60_DNA+RNA_seed | AKT1 | 0.74905 | GBM | GB-1 | Uprosertib |
| MVP_top60_DNA_seed | ESR1 | -0.313095 | HNSC | HSC-4 | Fulvestrant | MVP_top60_DNA+RNA_seed | JAK2 | 0.756556 | GBM | U-118-MG | Ruxolitinib |
| MVP_top60_DNA_seed | EGFR | -0.310731 | HNSC | OSC-20 | Pelitinib | MVP_top60_DNA+RNA_seed | JAK1 | 0.756556 | GBM | U-118-MG | Ruxolitinib |
| MVP_top60_DNA_seed | HSP90AA1 | -0.309452 | HNSC | SCC-15 | Elesclomol | MVP_top60_DNA+RNA_seed | JAK2 | 0.758348 | GBM | D-263MG | GSK2276186C |
| MVP_top60_DNA_seed | NTRK1 | -0.309294 | HNSC | PE-CA-PJ15 | AZD1332 | MVP_top60_DNA+RNA_seed | JAK1 | 0.758348 | GBM | D-263MG | GSK2276186C |
| MVP_top60_DNA_seed | EGFR | -0.308459 | HNSC | PCI-15A | CI-1033 | MVP_top60_DNA+RNA_seed | JAK3 | 0.758348 | GBM | D-263MG | GSK2276186C |
| MVP_top60_DNA_seed | PIK3CA | -0.308458 | HNSC | OSC-19 | AZD8835 | MVP_top60_DNA+RNA_seed | EGFR | 0.758851 | GBM | SK-MG-1 | Lapatinib |
| MVP_top60_DNA_seed | PIK3CA | -0.308287 | HNSC | PCI-4B | AZD8835 | MVP_top60_DNA+RNA_seed | KIT | 0.759812 | GBM | LN-229 | Sorafenib |
| MVP_top60_DNA_seed | AR | -0.308276 | HNSC | PCI-15A | Bicalutamide | MVP_top60_DNA+RNA_seed | EGFR | 0.760432 | GBM | D-392MG | AZD3759 |
| MVP_top60_DNA_seed | HSP90AA1 | -0.307936 | HNSC | Detroit562 | Luminespib | MVP_top60_DNA+RNA_seed | AKT1 | 0.762703 | GBM | LN-405 | Ipatasertib |
| MVP_top60_DNA_seed | HDAC1 | -0.307422 | HNSC | JHU-011 | AR-42 | MVP_top60_DNA+RNA_seed | PIK3CA | 0.762769 | GBM | LN-405 | Alpelisib |
| MVP_top60_DNA_seed | HSP90AA1 | -0.306775 | HNSC | SCC-4 | Luminespib | MVP_top60_DNA+RNA_seed | AKT1 | 0.76375 | GBM | T98G | Afuresertib |
| MVP_top60_DNA_seed | CDK2 | -0.306357 | HNSC | PCI-15A | AZD5438 | MVP_top60_DNA+RNA_seed | KRAS | 0.766321 | GBM | U-87-MG | KRAS (G12C) Inhibitor-1 |
| MVP_top60_DNA_seed | RAC1 | -0.305792 | HNSC | SCC-25 | EHT-1864 | MVP_top60_DNA+RNA_seed | AKT1 | 0.770565 | GBM | T98G | MK-2206 |
| MVP_top60_DNA_seed | AKT1 | -0.305791 | HNSC | HO-1-N-1 | MK-2206 | MVP_top60_DNA+RNA_seed | EP300 | 0.777915 | GBM | LN-405 | CPI-637 |
| MVP_top60_DNA_seed | HDAC1 | -0.305619 | HNSC | BB30-HNC | Entinostat | MVP_top60_DNA+RNA_seed | EGFR | 0.780566 | GBM | D-392MG | Afatinib |
| MVP_top60_DNA_seed | HDAC3 | -0.305619 | HNSC | BB30-HNC | Entinostat | MVP_top60_DNA+RNA_seed | AKT1 | 0.782174 | GBM | U-87-MG | Uprosertib |
| MVP_top60_DNA_seed | ESR1 | -0.305386 | HNSC | SKN-3 | Fulvestrant | MVP_top60_DNA+RNA_seed | AKT1 | 0.783045 | GBM | AM-38 | GSK2110183B |
| MVP_top60_DNA_seed | HDAC1 | -0.305361 | HNSC | SAS | CAY10603 | MVP_top60_DNA+RNA_seed | EGFR | 0.787238 | GBM | SW1088 | AZD3759 |
| MVP_top60_DNA_seed | HDAC1 | -0.305145 | HNSC | CAL-27 | CAY10603 | MVP_top60_DNA+RNA_seed | CDK4 | 0.792326 | GBM | SNB75 | Palbociclib |
| MVP_top60_DNA_seed | AKT1 | -0.304426 | HNSC | PCI-6A | AKT inhibitor VIII | MVP_top60_DNA+RNA_seed | AKT1 | 0.792665 | GBM | DBTRG-05MG | Afuresertib |
| MVP_top60_DNA_seed | HDAC1 | -0.299183 | HNSC | SCC-4 | Tubastatin A | MVP_top60_DNA+RNA_seed | JAK1 | 0.793498 | GBM | U251 | JAK1_8709 |
| MVP_top60_DNA_seed | SRC | -0.298665 | HNSC | SCC-9 | NVP-BHG712 | MVP_top60_DNA+RNA_seed | IGF1R | 0.794822 | GBM | YKG-1 | GSK1904529A |
| MVP_top60_DNA_seed | CDK2 | -0.2968 | HNSC | HO-1-N-1 | PHA-793887 | MVP_top60_DNA+RNA_seed | AKT1 | 0.801914 | GBM | D-392MG | Afuresertib |

| | | | | | | | | | | | |
|---|---|---|---|---|---|---|---|---|---|---|---|
| MVP_top60_DNA_seed | RAC1 | -0.296218 | HNSC | CAL-33 | EHT-1864 | MVP_top60_DNA+RNA_seed | AKT1 | 0.80302 | GBM | SF268 | GSK2110183B |
| MVP_top60_DNA_seed | EGFR | -0.295869 | HNSC | JHU-011 | CI-1033 | MVP_top60_DNA+RNA_seed | EGFR | 0.803738 | GBM | CAS-1 | Afatinib |
| MVP_top60_DNA_seed | AR | -0.291491 | HNSC | DOK | AZD3514 | MVP_top60_DNA+RNA_seed | KIT | 0.807835 | GBM | M059J | Sorafenib |
| MVP_top60_DNA_seed | AKT1 | -0.291199 | HNSC | OSC-20 | AKT inhibitor VIII | MVP_top60_DNA+RNA_seed | EGFR | 0.809603 | GBM | LN-405 | Erlotinib |
| MVP_top60_DNA_seed | HDAC1 | -0.288865 | HNSC | BB30-HNC | JQ12 | MVP_top60_DNA+RNA_seed | EGFR | 0.810487 | GBM | U251 | Gefitinib |
| MVP_top60_DNA_seed | HDAC1 | -0.288084 | HNSC | OSC-20 | Belinostat | MVP_top60_DNA+RNA_seed | JAK1 | 0.811211 | GBM | SK-MG-1 | JAK1_8709 |
| MVP_top60_DNA_seed | MDM2 | -0.287176 | HNSC | CAL-33 | Nutlin-3a (-) | MVP_top60_DNA+RNA_seed | EGFR | 0.811865 | GBM | SNB75 | Sapitinib |
| MVP_top60_DNA_seed | ESR1 | -0.287024 | HNSC | PCI-15A | Fulvestrant | MVP_top60_DNA+RNA_seed | ERBB3 | 0.811865 | GBM | SNB75 | Sapitinib |
| MVP_top60_DNA_seed | PIK3CA | -0.286122 | HNSC | BHY | AZD8835 | MVP_top60_DNA+RNA_seed | KRAS | 0.816596 | GBM | LN-18 | :RAS (G12C) Inhibitor-1 |
| MVP_top60_DNA_seed | CDK2 | -0.285559 | HNSC | BB49-HNC | Seliciclib | MVP_top60_DNA+RNA_seed | IGF1R | 0.819267 | GBM | LN-405 | GSK1904529A |
| MVP_top60_DNA_seed | HDAC1 | -0.284499 | HNSC | BHY | Tubastatin A | MVP_top60_DNA+RNA_seed | EGFR | 0.820737 | GBM | D-245MG | AZD3759 |
| MVP_top60_DNA_seed | HDAC1 | -0.284493 | HNSC | KOSC-2 | Belinostat | MVP_top60_DNA+RNA_seed | IGF1R | 0.821598 | GBM | T98G | Linsitinib |
| MVP_top60_DNA_seed | HDAC1 | -0.282111 | HNSC | BHY | CAY10603 | MVP_top60_DNA+RNA_seed | EGFR | 0.822151 | GBM | D-247MG | Gefitinib |
| MVP_top60_DNA_seed | EGFR | -0.281983 | HNSC | CAL-33 | CUDC-101 | MVP_top60_DNA+RNA_seed | JAK2 | 0.823085 | GBM | YH-13 | Ruxolitinib |
| MVP_top60_DNA_seed | AKT1 | -0.275397 | HNSC | SCC-15 | BAY AKT1 | MVP_top60_DNA+RNA_seed | JAK1 | 0.823085 | GBM | YH-13 | Ruxolitinib |
| MVP_top60_DNA_seed | EP300 | -0.275109 | HNSC | SAS | I-CBP112 | MVP_top60_DNA+RNA_seed | EGFR | 0.823413 | GBM | KS-1 | Osimertinib |
| MVP_top60_DNA_seed | CREBBP | -0.275109 | HNSC | SAS | I-CBP112 | MVP_top60_DNA+RNA_seed | AKT1 | 0.825342 | GBM | D-263MG | Ipatasertib |
| MVP_top60_DNA_seed | AR | -0.274248 | HNSC | BB30-HNC | Bicalutamide | MVP_top60_DNA+RNA_seed | KIT | 0.825736 | GBM | SW1088 | Motesanib |
| MVP_top60_DNA_seed | HSP90AA1 | -0.273629 | HNSC | FADU | Luminespib | MVP_top60_DNA+RNA_seed | CDK4 | 0.825781 | GBM | T98G | Ribociclib |
| MVP_top60_DNA_seed | MAPK1 | -0.272386 | HNSC | HO-1-N-1 | FR-180204 | MVP_top60_DNA+RNA_seed | IGF1R | 0.828195 | GBM | U-87-MG | BMS-536924 |
| MVP_top60_DNA_seed | MAPK3 | -0.272386 | HNSC | HO-1-N-1 | FR-180204 | MVP_top60_DNA+RNA_seed | AKT1 | 0.833892 | GBM | DK-MG | GSK2110183B |
| MVP_top60_DNA_seed | EGFR | -0.271233 | HNSC | SKN-3 | PF-00299804 | MVP_top60_DNA+RNA_seed | KIT | 0.835293 | GBM | U-118-MG | Axitinib |
| MVP_top60_DNA_seed | EGFR | -0.271065 | HNSC | JHU-022 | CI-1033 | MVP_top60_DNA+RNA_seed | EGFR | 0.835318 | GBM | YH-13 | Gefitinib |
| MVP_top60_DNA_seed | HSP90AA1 | -0.269596 | HNSC | HO-1-u-1 | SNX-2112 | MVP_top60_DNA+RNA_seed | AKT1 | 0.836194 | GBM | D-392MG | AT13148 |
| MVP_top60_DNA_seed | PIK3CA | -0.269128 | HNSC | HO-1-u-1 | AZD8835 | MVP_top60_DNA+RNA_seed | PIK3CA | 0.837061 | GBM | LN-229 | Alpelisib |
| MVP_top60_DNA_seed | NTRK1 | -0.268439 | HNSC | HSC-4 | AZD1332 | MVP_top60_DNA+RNA_seed | JAK2 | 0.843237 | GBM | U-118-MG | GSK2276186C |
| MVP_top60_DNA_seed | EGFR | -0.264464 | HNSC | JHU-011 | AZD8931 | MVP_top60_DNA+RNA_seed | JAK1 | 0.843237 | GBM | U-118-MG | GSK2276186C |
| MVP_top60_DNA_seed | EGFR | -0.2638 | HNSC | OSC-20 | Afatinib | MVP_top60_DNA+RNA_seed | JAK3 | 0.843237 | GBM | U-118-MG | GSK2276186C |
| MVP_top60_DNA_seed | ESR1 | -0.26256 | HNSC | BHY | Tamoxifen | MVP_top60_DNA+RNA_seed | KRAS | 0.844269 | GBM | LN-229 | :RAS (G12C) Inhibitor-1 |
| MVP_top60_DNA_seed | AR | -0.261366 | HNSC | CAL-33 | Bicalutamide | MVP_top60_DNA+RNA_seed | PIK3CA | 0.846997 | GBM | M059J | Alpelisib |
| MVP_top60_DNA_seed | ESR1 | -0.261326 | HNSC | Detroit562 | Fulvestrant | MVP_top60_DNA+RNA_seed | EGFR | 0.849635 | GBM | T98G | Sapitinib |
| MVP_top60_DNA_seed | HDAC1 | -0.259282 | HNSC | PCI-15A | Dacinostat | MVP_top60_DNA+RNA_seed | ERBB3 | 0.849635 | GBM | T98G | Sapitinib |
| MVP_top60_DNA_seed | HDAC1 | -0.256715 | HNSC | HO-1-N-1 | Dacinostat | MVP_top60_DNA+RNA_seed | EP300 | 0.850144 | GBM | A172 | SGC-CBP30 |
| MVP_top60_DNA_seed | MAPK1 | -0.25532 | HNSC | OSC-19 | FR-180204 | MVP_top60_DNA+RNA_seed | CREBBP | 0.850144 | GBM | A172 | SGC-CBP30 |
| MVP_top60_DNA_seed | MAPK3 | -0.25532 | HNSC | OSC-19 | FR-180204 | MVP_top60_DNA+RNA_seed | TP53 | 0.851192 | GBM | SK-MG-1 | MIRA-1 |
| MVP_top60_DNA_seed | RAC1 | -0.252686 | HNSC | HSC-4 | EHT-1864 | MVP_top60_DNA+RNA_seed | EP300 | 0.854304 | GBM | LNZTA3WT4 | CPI-637 |
| MVP_top60_DNA_seed | CDK2 | -0.252618 | HNSC | PCI-38 | PHA-793887 | MVP_top60_DNA+RNA_seed | KRAS | 0.860011 | GBM | D-542MG | :RAS (G12C) Inhibitor-1 |
| MVP_top60_DNA_seed | HDAC1 | -0.252479 | HNSC | OSC-20 | AR-42 | MVP_top60_DNA+RNA_seed | EGFR | 0.860964 | GBM | SF295 | Sapitinib |
| MVP_top60_DNA_seed | RAC1 | -0.251157 | HNSC | BB30-HNC | EHT-1864 | MVP_top60_DNA+RNA_seed | ERBB3 | 0.860964 | GBM | SF295 | Sapitinib |
| MVP_top60_DNA_seed | EGFR | -0.248849 | HNSC | PCI-38 | Afatinib | MVP_top60_DNA+RNA_seed | IGF1R | 0.867189 | GBM | D-392MG | GSK1904529A |
| MVP_top60_DNA_seed | EGFR | -0.248735 | HNSC | OSC-20 | CUDC-101 | MVP_top60_DNA+RNA_seed | MET | 0.869562 | GBM | SNB75 | Savolitinib |

| | | | | | | | | | | | |
|---|---|---|---|---|---|---|---|---|---|---|---|
| MVP_top60_DNA_seed | ESR1 | -0.248235 | HNSC | SKN-3 | Tamoxifen | MVP_top60_DNA+RNA_seed | EGFR | 0.871233 | GBM | LN-405 | AZD3759 |
| MVP_top60_DNA_seed | AR | -0.246779 | HNSC | CAL-27 | Bicalutamide | MVP_top60_DNA+RNA_seed | TP53 | 0.871374 | GBM | SF268 | MIRA-1 |
| MVP_top60_DNA_seed | HDAC1 | -0.246343 | HNSC | HSC-2 | Dacinostat | MVP_top60_DNA+RNA_seed | EGFR | 0.87195 | GBM | D-263MG | Erlotinib |
| MVP_top60_DNA_seed | EGFR | -0.245514 | HNSC | SCC90 | Gefitinib | MVP_top60_DNA+RNA_seed | IGF1R | 0.873569 | GBM | U251 | BMS-754807 |
| MVP_top60_DNA_seed | AR | -0.245483 | HNSC | JHU-022 | Bicalutamide | MVP_top60_DNA+RNA_seed | JAK2 | 0.874697 | GBM | M059J | GSK2276186C |
| MVP_top60_DNA_seed | HSP90AA1 | -0.243522 | HNSC | OSC-20 | CCT-018159 | MVP_top60_DNA+RNA_seed | JAK1 | 0.874697 | GBM | M059J | GSK2276186C |
| MVP_top60_DNA_seed | CDK2 | -0.24294 | HNSC | PCI-15A | PHA-793887 | MVP_top60_DNA+RNA_seed | JAK3 | 0.874697 | GBM | M059J | GSK2276186C |
| MVP_top60_DNA_seed | AKT1 | -0.241237 | HNSC | BB30-HNC | BAY AKT1 | MVP_top60_DNA+RNA_seed | IGF1R | 0.876092 | GBM | SF126 | Linsitinib |
| MVP_top60_DNA_seed | ESR1 | -0.240019 | HNSC | JHU-022 | Tamoxifen | MVP_top60_DNA+RNA_seed | PIK3CA | 0.879163 | GBM | CAS-1 | Alpelisib |
| MVP_top60_DNA_seed | PIK3CA | -0.237089 | HNSC | FADU | PI3Ka_4409 | MVP_top60_DNA+RNA_seed | PIK3CA | 0.879601 | GBM | LN-229 | AZD8186 |
| MVP_top60_DNA_seed | ESR1 | -0.234808 | HNSC | PE-CA-PJ15 | Fulvestrant | MVP_top60_DNA+RNA_seed | PIK3CB | 0.879601 | GBM | LN-229 | AZD8186 |
| MVP_top60_DNA_seed | AR | -0.229476 | HNSC | PCI-4B | Bicalutamide | MVP_top60_DNA+RNA_seed | AKT1 | 0.879952 | GBM | U-118-MG | MK-2206 |
| MVP_top60_DNA_seed | HDAC1 | -0.229282 | HNSC | SCC-9 | CAY10603 | MVP_top60_DNA+RNA_seed | PIK3CA | 0.879981 | GBM | D-263MG | AZD8186 |
| MVP_top60_DNA_seed | EGFR | -0.227981 | HNSC | OSC-19 | AZD8931 | MVP_top60_DNA+RNA_seed | PIK3CB | 0.879981 | GBM | D-263MG | AZD8186 |
| MVP_top60_DNA_seed | AKT1 | -0.226893 | HNSC | CAL-33 | GSK690693 | MVP_top60_DNA+RNA_seed | KIT | 0.880972 | GBM | SK-MG-1 | Sorafenib |
| MVP_top60_DNA_seed | HDAC1 | -0.226587 | HNSC | SKN-3 | JQ12 | MVP_top60_DNA+RNA_seed | PIK3CA | 0.883991 | GBM | D-392MG | Alpelisib |
| MVP_top60_DNA_seed | NTRK1 | -0.225698 | HNSC | CAL-33 | AZD1332 | MVP_top60_DNA+RNA_seed | EGFR | 0.884814 | GBM | D-392MG | Gefitinib |
| MVP_top60_DNA_seed | EGFR | -0.222778 | HNSC | SCC-4 | Cetuximab | MVP_top60_DNA+RNA_seed | EGFR | 0.884914 | GBM | KS-1 | Gefitinib |
| MVP_top60_DNA_seed | HDAC1 | -0.220227 | HNSC | HO-1-u-1 | CAY10603 | MVP_top60_DNA+RNA_seed | PIK3CA | 0.885395 | GBM | SNB75 | Buparlisib |
| MVP_top60_DNA_seed | MDM2 | -0.219531 | HNSC | HSC-3 | Nutlin-3a (-) | MVP_top60_DNA+RNA_seed | PIK3CB | 0.885395 | GBM | SNB75 | Buparlisib |
| MVP_top60_DNA_seed | SRC | -0.219398 | HNSC | PCI-15A | Bosutinib | MVP_top60_DNA+RNA_seed | PIK3CD | 0.885395 | GBM | SNB75 | Buparlisib |
| MVP_top60_DNA_seed | SRC | -0.218367 | HNSC | PCI-30 | Bosutinib | MVP_top60_DNA+RNA_seed | KIT | 0.886867 | GBM | GB-1 | Motesanib |
| MVP_top60_DNA_seed | EGFR | -0.215609 | HNSC | HO-1-u-1 | Afatinib | MVP_top60_DNA+RNA_seed | EGFR | 0.888304 | GBM | SK-MG-1 | Sapitinib |
| MVP_top60_DNA_seed | AKT1 | -0.213027 | HNSC | BICR31 | AKT inhibitor VIII | MVP_top60_DNA+RNA_seed | ERBB3 | 0.888304 | GBM | SK-MG-1 | Sapitinib |
| MVP_top60_DNA_seed | HDAC1 | -0.212583 | HNSC | SCC-9 | AR-42 | MVP_top60_DNA+RNA_seed | KIT | 0.894385 | GBM | LN-229 | Motesanib |
| MVP_top60_DNA_seed | HDAC1 | -0.20974 | HNSC | OSC-20 | CAY10603 | MVP_top60_DNA+RNA_seed | JAK2 | 0.895027 | GBM | YKG-1 | Ruxolitinib |
| MVP_top60_DNA_seed | EGFR | -0.208986 | HNSC | OSC-19 | Gefitinib | MVP_top60_DNA+RNA_seed | JAK1 | 0.895027 | GBM | YKG-1 | Ruxolitinib |
| MVP_top60_DNA_seed | EGFR | -0.208858 | HNSC | SCC-25 | PF-00299804 | MVP_top60_DNA+RNA_seed | EGFR | 0.895519 | GBM | SF126 | Erlotinib |
| MVP_top60_DNA_seed | AKT1 | -0.208296 | HNSC | SCC-25 | GSK690693 | MVP_top60_DNA+RNA_seed | IGF1R | 0.897273 | GBM | LN-229 | NVP-ADW742 |
| MVP_top60_DNA_seed | AR | -0.207764 | HNSC | PCI-6A | AZD3514 | MVP_top60_DNA+RNA_seed | IGF1R | 0.897559 | GBM | SK-MG-1 | BMS-754807 |
| MVP_top60_DNA_seed | EGFR | -0.206489 | HNSC | SCC-4 | PF-00299804 | MVP_top60_DNA+RNA_seed | CDK4 | 0.899147 | GBM | SF268 | Palbociclib |
| MVP_top60_DNA_seed | AKT1 | -0.204697 | HNSC | HO-1-u-1 | BAY AKT1 | MVP_top60_DNA+RNA_seed | IGF1R | 0.899404 | GBM | D-392MG | NVP-ADW742 |
| MVP_top60_DNA_seed | SRC | -0.201497 | HNSC | Detroit562 | Bosutinib | MVP_top60_DNA+RNA_seed | MET | 0.899932 | GBM | M059J | Savolitinib |
| MVP_top60_DNA_seed | CDK2 | -0.199515 | HNSC | SCC-25 | PHA-793887 | MVP_top60_DNA+RNA_seed | JAK2 | 0.907261 | GBM | GB-1 | GSK2276186C |
| MVP_top60_DNA_seed | AR | -0.199421 | HNSC | BHY | AZD3514 | MVP_top60_DNA+RNA_seed | JAK1 | 0.907261 | GBM | GB-1 | GSK2276186C |
| MVP_top60_DNA_seed | PIK3CA | -0.197253 | HNSC | PCI-4B | PI3Ka_4409 | MVP_top60_DNA+RNA_seed | JAK3 | 0.907261 | GBM | GB-1 | GSK2276186C |
| MVP_top60_DNA_seed | HDAC1 | -0.196486 | HNSC | BICR78 | CAY10603 | MVP_top60_DNA+RNA_seed | EGFR | 0.908304 | GBM | SF268 | AZD3759 |
| MVP_top60_DNA_seed | RAC1 | -0.194656 | HNSC | PE-CA-PJ15 | EHT-1864 | MVP_top60_DNA+RNA_seed | EGFR | 0.910745 | GBM | LNZTA3WT4 | Sapitinib |
| MVP_top60_DNA_seed | RAC1 | -0.19381 | HNSC | BHY | EHT-1864 | MVP_top60_DNA+RNA_seed | ERBB3 | 0.910745 | GBM | LNZTA3WT4 | Sapitinib |
| MVP_top60_DNA_seed | HDAC1 | -0.192664 | HNSC | CAL-27 | Tubastatin A | MVP_top60_DNA+RNA_seed | KIT | 0.914464 | GBM | SW1088 | Sorafenib |
| MVP_top60_DNA_seed | EGFR | -0.191744 | HNSC | PE-CA-PJ15 | Cetuximab | MVP_top60_DNA+RNA_seed | AKT1 | 0.915512 | GBM | GB-1 | MK-2206 |

| | | | | | | | | | | | |
|---|---|---|---|---|---|---|---|---|---|---|---|
| MVP_top60_DNA_seed | EGFR | -0.189631 | HNSC | OSC-20 | AZD8931 | MVP_top60_DNA+RNA_seed | AKT1 | 0.916615 | GBM | LN-229 | AZD5363 |
| MVP_top60_DNA_seed | PIK3CA | -0.188754 | HNSC | PCI-15A | PI3Ka_4409 | MVP_top60_DNA+RNA_seed | AKT1 | 0.916835 | GBM | DBTRG-05MG | Ipatasertib |
| MVP_top60_DNA_seed | PIK3CA | -0.18721 | HNSC | PCI-6A | AZD8835 | MVP_top60_DNA+RNA_seed | CDK4 | 0.917941 | GBM | LN-229 | Ribociclib |
| MVP_top60_DNA_seed | HDAC1 | -0.186798 | HNSC | HO-1-N-1 | Belinostat | MVP_top60_DNA+RNA_seed | PIK3CA | 0.926442 | GBM | LN-18 | Alpelisib |
| MVP_top60_DNA_seed | AKT1 | -0.186632 | HNSC | LB771-HNC | BAY AKT1 | MVP_top60_DNA+RNA_seed | JAK2 | 0.926962 | GBM | D-542MG | Ruxolitinib |
| MVP_top60_DNA_seed | SRC | -0.184638 | HNSC | LB771-HNC | Saracatinib | MVP_top60_DNA+RNA_seed | JAK1 | 0.926962 | GBM | D-542MG | Ruxolitinib |
| MVP_top60_DNA_seed | SRC | -0.184352 | HNSC | SCC-4 | Bosutinib | MVP_top60_DNA+RNA_seed | EGFR | 0.931245 | GBM | U251 | Osimertinib |
| MVP_top60_DNA_seed | AKT1 | -0.183637 | HNSC | PCI-4B | GSK690693 | MVP_top60_DNA+RNA_seed | EGFR | 0.931524 | GBM | SF126 | Afatinib |
| MVP_top60_DNA_seed | HDAC1 | -0.183535 | HNSC | KOSC-2 | CAY10603 | MVP_top60_DNA+RNA_seed | IGF1R | 0.931814 | GBM | M059J | Linsitinib |
| MVP_top60_DNA_seed | NTRK1 | -0.182733 | HNSC | JHU-029 | AZD1332 | MVP_top60_DNA+RNA_seed | EGFR | 0.932418 | GBM | CAS-1 | Erlotinib |
| MVP_top60_DNA_seed | PIK3CA | -0.179975 | HNSC | PCI-15A | AZD8835 | MVP_top60_DNA+RNA_seed | EGFR | 0.938563 | GBM | T98G | Erlotinib |
| MVP_top60_DNA_seed | MDM2 | -0.179714 | HNSC | PCI-6A | Nutlin-3a (-) | MVP_top60_DNA+RNA_seed | PIK3CA | 0.938589 | GBM | DK-MG | Alpelisib |
| MVP_top60_DNA_seed | EGFR | -0.174681 | HNSC | PE-CA-PJ15 | CUDC-101 | MVP_top60_DNA+RNA_seed | IGF1R | 0.939065 | GBM | SW1088 | NVP-ADW742 |
| MVP_top60_DNA_seed | HSP90AA1 | -0.173954 | HNSC | BICR10 | CCT-018159 | MVP_top60_DNA+RNA_seed | EGFR | 0.941274 | GBM | D-566MG | Afatinib |
| MVP_top60_DNA_seed | NTRK1 | -0.173046 | HNSC | SCC-25 | AZD1332 | MVP_top60_DNA+RNA_seed | KIT | 0.942809 | GBM | U-118-MG | Motesanib |
| MVP_top60_DNA_seed | SRC | -0.171439 | HNSC | SCC-25 | NVP-BHG712 | MVP_top60_DNA+RNA_seed | EGFR | 0.945968 | GBM | D-247MG | Erlotinib |
| MVP_top60_DNA_seed | EGFR | -0.170332 | HNSC | BICR22 | CUDC-101 | MVP_top60_DNA+RNA_seed | KRAS | 0.945971 | GBM | DK-MG | (RAS (G12C) Inhibitor-1 |
| MVP_top60_DNA_seed | MDM2 | -0.16996 | HNSC | JHU-011 | Nutlin-3a (-) | MVP_top60_DNA+RNA_seed | AKT1 | 0.946305 | GBM | D-263MG | MK-2206 |
| MVP_top60_DNA_seed | RAC1 | -0.169444 | HNSC | SCC-9 | EHT-1864 | MVP_top60_DNA+RNA_seed | EGFR | 0.949568 | GBM | U-118-MG | Afatinib |
| MVP_top60_DNA_seed | PIK3CA | -0.168298 | HNSC | PCI-38 | PI3Ka_4409 | MVP_top60_DNA+RNA_seed | AKT1 | 0.950444 | GBM | GB-1 | Ipatasertib |
| MVP_top60_DNA_seed | AKT1 | -0.16751 | HNSC | DOK | AKT inhibitor VIII | MVP_top60_DNA+RNA_seed | JAK2 | 0.954454 | GBM | LN-229 | GSK2276186C |
| MVP_top60_DNA_seed | NTRK1 | -0.167122 | HNSC | PCI-4B | GW441756 | MVP_top60_DNA+RNA_seed | JAK1 | 0.954454 | GBM | LN-229 | GSK2276186C |
| MVP_top60_DNA_seed | MDM2 | -0.165435 | HNSC | HO-1-N-1 | Nutlin-3a (-) | MVP_top60_DNA+RNA_seed | JAK3 | 0.954454 | GBM | LN-229 | GSK2276186C |
| MVP_top60_DNA_seed | MAPK1 | -0.165424 | HNSC | SCC-25 | FR-180204 | MVP_top60_DNA+RNA_seed | AKT1 | 0.954752 | GBM | SF268 | Ipatasertib |
| MVP_top60_DNA_seed | MAPK3 | -0.165424 | HNSC | SCC-25 | FR-180204 | MVP_top60_DNA+RNA_seed | EGFR | 0.955973 | GBM | T98G | Osimertinib |
| MVP_top60_DNA_seed | AKT1 | -0.163931 | HNSC | OSC-20 | BAY AKT1 | MVP_top60_DNA+RNA_seed | EGFR | 0.956074 | GBM | D-263MG | Sapitinib |
| MVP_top60_DNA_seed | SMARCA4 | -0.162626 | HNSC | Ca9-22 | PFI-3 | MVP_top60_DNA+RNA_seed | ERBB3 | 0.956074 | GBM | D-263MG | Sapitinib |
| MVP_top60_DNA_seed | NTRK1 | -0.159662 | HNSC | BB30-HNC | AZD1332 | MVP_top60_DNA+RNA_seed | IGF1R | 0.957849 | GBM | YH-13 | NVP-ADW742 |
| MVP_top60_DNA_seed | SMARCA4 | -0.156831 | HNSC | HSC-2 | PFI-3 | MVP_top60_DNA+RNA_seed | JAK2 | 0.957853 | GBM | U-87-MG | GSK2276186C |
| MVP_top60_DNA_seed | MAPK1 | -0.154963 | HNSC | PCI-38 | VX-11e | MVP_top60_DNA+RNA_seed | JAK1 | 0.957853 | GBM | U-87-MG | GSK2276186C |
| MVP_top60_DNA_seed | HDAC1 | -0.151443 | HNSC | SAS | AR-42 | MVP_top60_DNA+RNA_seed | JAK3 | 0.957853 | GBM | U-87-MG | GSK2276186C |
| MVP_top60_DNA_seed | CDK2 | -0.150916 | HNSC | Ca9-22 | AZD5438 | MVP_top60_DNA+RNA_seed | KIT | 0.958155 | GBM | D-542MG | Axitinib |
| MVP_top60_DNA_seed | HDAC1 | -0.150213 | HNSC | OSC-19 | Dacinostat | MVP_top60_DNA+RNA_seed | KIT | 0.958709 | GBM | LN-229 | Axitinib |
| MVP_top60_DNA_seed | NTRK1 | -0.147081 | HNSC | PCI-15A | AZD1332 | MVP_top60_DNA+RNA_seed | KIT | 0.959321 | GBM | D-247MG | Sorafenib |
| MVP_top60_DNA_seed | MDM2 | -0.14698 | HNSC | BB30-HNC | Nutlin-3a (-) | MVP_top60_DNA+RNA_seed | IGF1R | 0.959782 | GBM | D-245MG | Linsitinib |
| MVP_top60_DNA_seed | SRC | -0.143459 | HNSC | SAS | NVP-BHG712 | MVP_top60_DNA+RNA_seed | IGF1R | 0.96089 | GBM | SF268 | Linsitinib |
| MVP_top60_DNA_seed | RAC1 | -0.143103 | HNSC | SKN-3 | EHT-1864 | MVP_top60_DNA+RNA_seed | IGF1R | 0.961558 | GBM | SF295 | NVP-ADW742 |
| MVP_top60_DNA_seed | SRC | -0.140856 | HNSC | PCI-6A | NVP-BHG712 | MVP_top60_DNA+RNA_seed | AKT1 | 0.963615 | GBM | SF268 | Uprosertib |
| MVP_top60_DNA_seed | RAC1 | -0.139818 | HNSC | SAT | EHT-1864 | MVP_top60_DNA+RNA_seed | EGFR | 0.965185 | GBM | SNB75 | Lapatinib |
| MVP_top60_DNA_seed | MAPK1 | -0.137472 | HNSC | SCC-4 | VX-11e | MVP_top60_DNA+RNA_seed | JAK2 | 0.965438 | GBM | LN-405 | AZ960 |
| MVP_top60_DNA_seed | EP300 | -0.13624 | HNSC | PCI-38 | I-CBP112 | MVP_top60_DNA+RNA_seed | JAK3 | 0.965438 | GBM | LN-405 | AZ960 |

| | | | | | | | | | | | |
|---|---|---|---|---|---|---|---|---|---|---|---|
| MVP_top60_DNA_seed | CREBBP | -0.13624 | HNSC | PCI-38 | I-CBP112 | MVP_top60_DNA+RNA_seed | IGF1R | 0.967218 | GBM | LN-405 | Linsitinib |
| MVP_top60_DNA_seed | AR | -0.135124 | HNSC | BB49-HNC | Bicalutamide | MVP_top60_DNA+RNA_seed | EGFR | 0.972104 | GBM | LN-18 | AZD3759 |
| MVP_top60_DNA_seed | CDK2 | -0.134306 | HNSC | JHU-022 | AZD5438 | MVP_top60_DNA+RNA_seed | MET | 0.972825 | GBM | D-245MG | Savolitinib |
| MVP_top60_DNA_seed | NTRK1 | -0.132468 | HNSC | SCC-15 | AZD1332 | MVP_top60_DNA+RNA_seed | EGFR | 0.973538 | GBM | SF295 | Afatinib |
| MVP_top60_DNA_seed | NTRK1 | -0.128158 | HNSC | SCC-15 | GW441756 | MVP_top60_DNA+RNA_seed | TP53 | 0.975512 | GBM | DK-MG | MIRA-1 |
| MVP_top60_DNA_seed | SMARCA4 | -0.127594 | HNSC | PCI-6A | PFI-3 | MVP_top60_DNA+RNA_seed | EGFR | 0.979684 | GBM | SK-MG-1 | Gefitinib |
| MVP_top60_DNA_seed | MAPK1 | -0.127067 | HNSC | HO-1-u-1 | FR-180204 | MVP_top60_DNA+RNA_seed | EGFR | 0.980727 | GBM | LN-405 | Sapitinib |
| MVP_top60_DNA_seed | MAPK3 | -0.127067 | HNSC | HO-1-u-1 | FR-180204 | MVP_top60_DNA+RNA_seed | ERBB3 | 0.980727 | GBM | LN-405 | Sapitinib |
| MVP_top60_DNA_seed | PIK3CA | -0.126852 | HNSC | PCI-6A | PI3Ka_4409 | MVP_top60_DNA+RNA_seed | JAK2 | 0.985713 | GBM | LN-405 | Ruxolitinib |
| MVP_top60_DNA_seed | EGFR | -0.126282 | HNSC | PE-CA-PJ15 | Afatinib | MVP_top60_DNA+RNA_seed | JAK1 | 0.985713 | GBM | LN-405 | Ruxolitinib |
| MVP_top60_DNA_seed | ESR1 | -0.125149 | HNSC | HSC-2 | Tamoxifen | MVP_top60_DNA+RNA_seed | EGFR | 0.986548 | GBM | D-263MG | Osimertinib |
| MVP_top60_DNA_seed | EGFR | -0.124493 | HNSC | BICR10 | AZD8931 | MVP_top60_DNA+RNA_seed | EGFR | 0.9873 | GBM | D-263MG | Afatinib |
| MVP_top60_DNA_seed | AKT1 | -0.124261 | HNSC | BB30-HNC | AKT inhibitor VIII | MVP_top60_DNA+RNA_seed | EGFR | 0.991396 | GBM | M059J | Gefitinib |
| MVP_top60_DNA_seed | EGFR | -0.123245 | HNSC | SCC-4 | CI-1033 | MVP_top60_DNA+RNA_seed | MET | 0.995912 | GBM | D-247MG | Savolitinib |
| MVP_top60_DNA_seed | EGFR | -0.11955 | HNSC | KOSC-2 | CI-1033 | MVP_top60_DNA+RNA_seed | AKT1 | 0.999886 | GBM | AM-38 | Uprosertib |
| MVP_top60_DNA_seed | EGFR | -0.117539 | HNSC | JHU-011 | PF-00299804 | MVP_top60_DNA+RNA_seed | EGFR | 1.000318 | GBM | YKG-1 | Afatinib |
| MVP_top60_DNA_seed | MAPK1 | -0.117309 | HNSC | CAL-33 | FR-180204 | MVP_top60_DNA+RNA_seed | AKT1 | 1.000548 | GBM | AM-38 | MK-2206 |
| MVP_top60_DNA_seed | MAPK3 | -0.117309 | HNSC | CAL-33 | FR-180204 | MVP_top60_DNA+RNA_seed | CDK4 | 1.00091 | GBM | LN-405 | Ribociclib |
| MVP_top60_DNA_seed | EP300 | -0.117288 | HNSC | PCI-30 | I-CBP112 | MVP_top60_DNA+RNA_seed | PIK3CA | 1.021972 | GBM | GB-1 | Alpelisib |
| MVP_top60_DNA_seed | CREBBP | -0.117288 | HNSC | PCI-30 | I-CBP112 | MVP_top60_DNA+RNA_seed | EGFR | 1.023988 | GBM | Becker | Sapitinib |
| MVP_top60_DNA_seed | AKT1 | -0.115725 | HNSC | KON | BAY AKT1 | MVP_top60_DNA+RNA_seed | ERBB3 | 1.023988 | GBM | Becker | Sapitinib |
| MVP_top60_DNA_seed | HSP90AA1 | -0.114597 | HNSC | HSC-2 | SNX-2112 | MVP_top60_DNA+RNA_seed | EP300 | 1.024816 | GBM | U251 | SGC-CBP30 |
| MVP_top60_DNA_seed | SRC | -0.113509 | HNSC | Ca9-22 | NVP-BHG712 | MVP_top60_DNA+RNA_seed | CREBBP | 1.024816 | GBM | U251 | SGC-CBP30 |
| MVP_top60_DNA_seed | NTRK1 | -0.113312 | HNSC | FADU | AZD1332 | MVP_top60_DNA+RNA_seed | AKT1 | 1.027237 | GBM | D-247MG | AZD5363 |
| MVP_top60_DNA_seed | AR | -0.113035 | HNSC | FADU | Bicalutamide | MVP_top60_DNA+RNA_seed | EGFR | 1.027722 | GBM | U251 | Afatinib |
| MVP_top60_DNA_seed | AKT1 | -0.110252 | HNSC | JHU-011 | AKT inhibitor VIII | MVP_top60_DNA+RNA_seed | AKT1 | 1.03031 | GBM | U-87-MG | GSK2110183B |
| MVP_top60_DNA_seed | SRC | -0.110119 | HNSC | BB49-HNC | WH-4-023 | MVP_top60_DNA+RNA_seed | PIK3CA | 1.030939 | GBM | Becker | Alpelisib |
| MVP_top60_DNA_seed | HDAC1 | -0.109786 | HNSC | HO-1-u-1 | AR-42 | MVP_top60_DNA+RNA_seed | AKT1 | 1.03098 | GBM | KS-1 | MK-2206 |
| MVP_top60_DNA_seed | HSP90AA1 | -0.108892 | HNSC | PCI-6A | CCT-018159 | MVP_top60_DNA+RNA_seed | AKT1 | 1.032456 | GBM | D-392MG | Uprosertib |
| MVP_top60_DNA_seed | EGFR | -0.105528 | HNSC | OSC-19 | CI-1033 | MVP_top60_DNA+RNA_seed | JAK1 | 1.03481 | GBM | SW1088 | JAK1_8709 |
| MVP_top60_DNA_seed | AR | -0.103497 | HNSC | PCI-15A | Bicalutamide | MVP_top60_DNA+RNA_seed | EGFR | 1.039273 | GBM | LN-18 | Afatinib |
| MVP_top60_DNA_seed | MAPK1 | -0.101061 | HNSC | SKN-3 | FR-180204 | MVP_top60_DNA+RNA_seed | EGFR | 1.039408 | GBM | KS-1 | AZD3759 |
| MVP_top60_DNA_seed | MAPK3 | -0.101061 | HNSC | SKN-3 | FR-180204 | MVP_top60_DNA+RNA_seed | EGFR | 1.04507 | GBM | LN-18 | Osimertinib |
| MVP_top60_DNA_seed | AKT1 | -0.101059 | HNSC | HSC-4 | GSK690693 | MVP_top60_DNA+RNA_seed | MET | 1.047005 | GBM | D-542MG | Savolitinib |
| MVP_top60_DNA_seed | HDAC1 | -0.098764 | HNSC | SAT | Belinostat | MVP_top60_DNA+RNA_seed | EGFR | 1.049049 | GBM | D-247MG | AZD3759 |
| MVP_top60_DNA_seed | PIK3CA | -0.094636 | HNSC | SKN-3 | AZD8835 | MVP_top60_DNA+RNA_seed | PIK3CA | 1.04963 | GBM | SW1088 | Alpelisib |
| MVP_top60_DNA_seed | EGFR | -0.094629 | HNSC | HSC-2 | CUDC-101 | MVP_top60_DNA+RNA_seed | JAK2 | 1.052645 | GBM | SW1088 | GSK2276186C |
| MVP_top60_DNA_seed | SRC | -0.093481 | HNSC | OSC-19 | NVP-BHG712 | MVP_top60_DNA+RNA_seed | JAK1 | 1.052645 | GBM | SW1088 | GSK2276186C |
| MVP_top60_DNA_seed | AR | -0.092385 | HNSC | Ca9-22 | Bicalutamide | MVP_top60_DNA+RNA_seed | JAK3 | 1.052645 | GBM | SW1088 | GSK2276186C |
| MVP_top60_DNA_seed | CDK2 | -0.090763 | HNSC | BB30-HNC | PHA-793887 | MVP_top60_DNA+RNA_seed | KIT | 1.05376 | GBM | GB-1 | Axitinib |
| MVP_top60_DNA_seed | EGFR | -0.086477 | HNSC | HSC-4 | Pelitinib | MVP_top60_DNA+RNA_seed | EGFR | 1.056207 | GBM | SW1088 | Gefitinib |

| | | | | | | | | | | | |
|---|---|---|---|---|---|---|---|---|---|---|---|
| MVP_top60_DNA_seed | MAPK1 | -0.082804 | HNSC | BICR31 | FR-180204 | MVP_top60_DNA+RNA_seed | IGF1R | 1.060155 | GBM | Becker | NVP-ADW742 |
| MVP_top60_DNA_seed | MAPK3 | -0.082804 | HNSC | BICR31 | FR-180204 | MVP_top60_DNA+RNA_seed | EGFR | 1.063211 | GBM | GB-1 | Gefitinib |
| MVP_top60_DNA_seed | MDM2 | -0.081573 | HNSC | Ca9-22 | Serdemetan | MVP_top60_DNA+RNA_seed | KIT | 1.065315 | GBM | LN-405 | Sorafenib |
| MVP_top60_DNA_seed | EGFR | -0.079897 | HNSC | HSC-3 | Pelitinib | MVP_top60_DNA+RNA_seed | AKT1 | 1.071557 | GBM | D-247MG | Afuresertib |
| MVP_top60_DNA_seed | AKT1 | -0.079756 | HNSC | OSC-20 | GSK690693 | MVP_top60_DNA+RNA_seed | KRAS | 1.072045 | GBM | A172 | KRAS (G12C) Inhibitor-1 |
| MVP_top60_DNA_seed | HDAC1 | -0.078075 | HNSC | BHY | JQ12 | MVP_top60_DNA+RNA_seed | JAK2 | 1.073445 | GBM | LN-229 | AZ960 |
| MVP_top60_DNA_seed | AKT1 | -0.075423 | HNSC | OSC-19 | MK-2206 | MVP_top60_DNA+RNA_seed | JAK3 | 1.073445 | GBM | LN-229 | AZ960 |
| MVP_top60_DNA_seed | AKT1 | -0.073999 | HNSC | PCI-30 | MK-2206 | MVP_top60_DNA+RNA_seed | AKT1 | 1.077059 | GBM | D-392MG | GSK2110183B |
| MVP_top60_DNA_seed | SMARCA4 | -0.073923 | HNSC | BHY | PFI-3 | MVP_top60_DNA+RNA_seed | IGF1R | 1.07706 | GBM | U251 | Linsitinib |
| MVP_top60_DNA_seed | PIK3CA | -0.073796 | HNSC | PCI-38 | AZD8835 | MVP_top60_DNA+RNA_seed | AKT1 | 1.078303 | GBM | SF268 | AZD5363 |
| MVP_top60_DNA_seed | AKT1 | -0.072313 | HNSC | SCC90 | MK-2206 | MVP_top60_DNA+RNA_seed | KIT | 1.080951 | GBM | SNB75 | Axitinib |
| MVP_top60_DNA_seed | CDK2 | -0.068767 | HNSC | OSC-19 | AZD5438 | MVP_top60_DNA+RNA_seed | EGFR | 1.086889 | GBM | GB-1 | Afatinib |
| MVP_top60_DNA_seed | AKT1 | -0.066323 | HNSC | SCC-4 | MK-2206 | MVP_top60_DNA+RNA_seed | EGFR | 1.088469 | GBM | Becker | Erlotinib |
| MVP_top60_DNA_seed | HSP90AA1 | -0.066234 | HNSC | Detroit562 | Elesclomol | MVP_top60_DNA+RNA_seed | IGF1R | 1.089655 | GBM | A172 | Linsitinib |
| MVP_top60_DNA_seed | ESR1 | -0.066142 | HNSC | SCC-4 | Fulvestrant | MVP_top60_DNA+RNA_seed | MET | 1.094619 | GBM | CAS-1 | Crizotinib |
| MVP_top60_DNA_seed | SRC | -0.065876 | HNSC | SCC90 | Bosutinib | MVP_top60_DNA+RNA_seed | KIT | 1.095995 | GBM | CAS-1 | Sorafenib |
| MVP_top60_DNA_seed | EGFR | -0.065761 | HNSC | PCI-38 | CUDC-101 | MVP_top60_DNA+RNA_seed | JAK1 | 1.100288 | GBM | D-392MG | JAK1_8709 |
| MVP_top60_DNA_seed | EP300 | -0.064188 | HNSC | KOSC-2 | I-CBP112 | MVP_top60_DNA+RNA_seed | MET | 1.102133 | GBM | GB-1 | Crizotinib |
| MVP_top60_DNA_seed | CREBBP | -0.064188 | HNSC | KOSC-2 | I-CBP112 | MVP_top60_DNA+RNA_seed | IGF1R | 1.102519 | GBM | YH-13 | Linsitinib |
| MVP_top60_DNA_seed | PIK3CA | -0.063626 | HNSC | SCC-4 | PI3Ka_4409 | MVP_top60_DNA+RNA_seed | IGF1R | 1.103677 | GBM | U251 | BMS-536924 |
| MVP_top60_DNA_seed | EGFR | -0.06153 | HNSC | HO-1-u-1 | Cetuximab | MVP_top60_DNA+RNA_seed | TP53 | 1.104613 | GBM | LN-405 | MIRA-1 |
| MVP_top60_DNA_seed | NTRK1 | -0.060861 | HNSC | PCI-30 | AZD1332 | MVP_top60_DNA+RNA_seed | IGF1R | 1.106072 | GBM | D-247MG | GSK1904529A |
| MVP_top60_DNA_seed | EGFR | -0.058944 | HNSC | SCC-4 | CUDC-101 | MVP_top60_DNA+RNA_seed | AKT1 | 1.108785 | GBM | A172 | GSK2110183B |
| MVP_top60_DNA_seed | AR | -0.057192 | HNSC | Ca9-22 | Bicalutamide | MVP_top60_DNA+RNA_seed | JAK2 | 1.112096 | GBM | CAS-1 | Ruxolitinib |
| MVP_top60_DNA_seed | AKT1 | -0.050811 | HNSC | CAL-27 | BAY AKT1 | MVP_top60_DNA+RNA_seed | JAK1 | 1.112096 | GBM | CAS-1 | Ruxolitinib |
| MVP_top60_DNA_seed | SMARCA4 | -0.050073 | HNSC | JHU-011 | PFI-3 | MVP_top60_DNA+RNA_seed | IGF1R | 1.127513 | GBM | SF126 | NVP-ADW742 |
| MVP_top60_DNA_seed | MDM2 | -0.049964 | HNSC | SCC-4 | Serdemetan | MVP_top60_DNA+RNA_seed | EGFR | 1.128677 | GBM | AM-38 | Sapitinib |
| MVP_top60_DNA_seed | RAC1 | -0.048658 | HNSC | BICR78 | EHT-1864 | MVP_top60_DNA+RNA_seed | ERBB3 | 1.128677 | GBM | AM-38 | Sapitinib |
| MVP_top60_DNA_seed | MAPK1 | -0.046823 | HNSC | SAT | VX-11e | MVP_top60_DNA+RNA_seed | EGFR | 1.130392 | GBM | D-392MG | Erlotinib |
| MVP_top60_DNA_seed | HSP90AA1 | -0.046788 | HNSC | SCC-15 | SNX-2112 | MVP_top60_DNA+RNA_seed | PIK3CB | 1.139373 | GBM | YKG-1 | AZD6482 |
| MVP_top60_DNA_seed | RAC1 | -0.045772 | HNSC | CAL-27 | EHT-1864 | MVP_top60_DNA+RNA_seed | EGFR | 1.14068 | GBM | D-392MG | Sapitinib |
| MVP_top60_DNA_seed | MAPK1 | -0.04464 | HNSC | HSC-2 | FR-180204 | MVP_top60_DNA+RNA_seed | ERBB3 | 1.14068 | GBM | D-392MG | Sapitinib |
| MVP_top60_DNA_seed | MAPK3 | -0.04464 | HNSC | HSC-2 | FR-180204 | MVP_top60_DNA+RNA_seed | AKT1 | 1.145723 | GBM | KS-1 | Ipatasertib |
| MVP_top60_DNA_seed | HSP90AA1 | -0.039599 | HNSC | BICR10 | Luminespib | MVP_top60_DNA+RNA_seed | KRAS | 1.146305 | GBM | AM-38 | KRAS (G12C) Inhibitor-1 |
| MVP_top60_DNA_seed | AR | -0.037754 | HNSC | OSC-20 | Bicalutamide | MVP_top60_DNA+RNA_seed | IGF1R | 1.154929 | GBM | SNB75 | NVP-ADW742 |
| MVP_top60_DNA_seed | MDM2 | -0.035261 | HNSC | OSC-19 | Nutlin-3a (-) | MVP_top60_DNA+RNA_seed | IGF1R | 1.155049 | GBM | SNB75 | GSK1904529A |
| MVP_top60_DNA_seed | HDAC1 | -0.034493 | HNSC | PCI-38 | Belinostat | MVP_top60_DNA+RNA_seed | CDK4 | 1.157249 | GBM | U-118-MG | Ribociclib |
| MVP_top60_DNA_seed | HDAC1 | -0.034468 | HNSC | CAL-27 | Dacinostat | MVP_top60_DNA+RNA_seed | IGF1R | 1.164171 | GBM | D-245MG | NVP-ADW742 |
| MVP_top60_DNA_seed | SRC | -0.033995 | HNSC | SCC-25 | Bosutinib | MVP_top60_DNA+RNA_seed | IGF1R | 1.167469 | GBM | LN-229 | BMS-754807 |
| MVP_top60_DNA_seed | ESR1 | -0.033906 | HNSC | SCC-25 | Fulvestrant | MVP_top60_DNA+RNA_seed | PIK3CA | 1.171426 | GBM | U251 | AZD8186 |
| MVP_top60_DNA_seed | HDAC1 | -0.033637 | HNSC | BICR31 | Dacinostat | MVP_top60_DNA+RNA_seed | PIK3CB | 1.171426 | GBM | U251 | AZD8186 |

| | | | | | | | | | | | |
|---|---|---|---|---|---|---|---|---|---|---|---|
| MVP_top60_DNA_seed | PIK3CA | -0.030052 | HNSC | HO-1-u-1 | PI3Ka_4409 | MVP_top60_DNA+RNA_seed | CDK4 | 1.178543 | GBM | M059J | Palbociclib |
| MVP_top60_DNA_seed | MDM2 | -0.028914 | HNSC | Ca9-22 | Nutlin-3a (-) | MVP_top60_DNA+RNA_seed | PIK3CA | 1.180057 | GBM | A172 | Alpelisib |
| MVP_top60_DNA_seed | AR | -0.028875 | HNSC | BICR10 | Bicalutamide | MVP_top60_DNA+RNA_seed | CDK4 | 1.180443 | GBM | CAS-1 | Ribociclib |
| MVP_top60_DNA_seed | MDM2 | -0.026415 | HNSC | SAT | Nutlin-3a (-) | MVP_top60_DNA+RNA_seed | EGFR | 1.180972 | GBM | U251 | Lapatinib |
| MVP_top60_DNA_seed | ESR1 | -0.025456 | HNSC | SAS | Fulvestrant | MVP_top60_DNA+RNA_seed | JAK2 | 1.181315 | GBM | SNB75 | Ruxolitinib |
| MVP_top60_DNA_seed | RAC1 | -0.024507 | HNSC | KOSC-2 | EHT-1864 | MVP_top60_DNA+RNA_seed | JAK1 | 1.181315 | GBM | SNB75 | Ruxolitinib |
| MVP_top60_DNA_seed | AKT1 | -0.024157 | HNSC | KOSC-2 | AKT inhibitor VIII | MVP_top60_DNA+RNA_seed | TP53 | 1.185029 | GBM | SW1088 | MIRA-1 |
| MVP_top60_DNA_seed | EP300 | -0.016155 | HNSC | HO-1-u-1 | I-CBP112 | MVP_top60_DNA+RNA_seed | EGFR | 1.186611 | GBM | SF268 | Osimertinib |
| MVP_top60_DNA_seed | CREBBP | -0.016155 | HNSC | HO-1-u-1 | I-CBP112 | MVP_top60_DNA+RNA_seed | EGFR | 1.187698 | GBM | DBTRG-05MG | Osimertinib |
| MVP_top60_DNA_seed | NTRK1 | -0.011703 | HNSC | BICR78 | AZD1332 | MVP_top60_DNA+RNA_seed | IGF1R | 1.191329 | GBM | CAS-1 | GSK1904529A |
| MVP_top60_DNA_seed | HDAC1 | -0.011469 | HNSC | SCC-4 | Belinostat | MVP_top60_DNA+RNA_seed | IGF1R | 1.195532 | GBM | T98G | GSK1904529A |
| MVP_top60_DNA_seed | AR | -0.008732 | HNSC | LB771-HNC | Bicalutamide | MVP_top60_DNA+RNA_seed | EGFR | 1.201006 | GBM | LN-405 | Osimertinib |
| MVP_top60_DNA_seed | HSP90AA1 | -0.007565 | HNSC | BICR22 | SNX-2112 | MVP_top60_DNA+RNA_seed | MET | 1.216838 | GBM | D-542MG | Crizotinib |
| MVP_top60_DNA_seed | RAC1 | -0.007335 | HNSC | PCI-6A | EHT-1864 | MVP_top60_DNA+RNA_seed | PIK3CA | 1.219196 | GBM | SF268 | AZD8186 |
| MVP_top60_DNA_seed | HDAC1 | -0.007307 | HNSC | SCC-25 | Dacinostat | MVP_top60_DNA+RNA_seed | PIK3CB | 1.219196 | GBM | SF268 | AZD8186 |
| MVP_top60_DNA_seed | EP300 | -0.006743 | HNSC | CAL-27 | I-CBP112 | MVP_top60_DNA+RNA_seed | AKT1 | 1.219213 | GBM | SF268 | Afuresertib |
| MVP_top60_DNA_seed | CREBBP | -0.006743 | HNSC | CAL-27 | I-CBP112 | MVP_top60_DNA+RNA_seed | PIK3CA | 1.220952 | GBM | LN-229 | GNE-317 |
| MVP_top60_DNA_seed | EGFR | -0.005869 | HNSC | JHU-022 | CUDC-101 | MVP_top60_DNA+RNA_seed | JAK2 | 1.222014 | GBM | LN-405 | JAK_8517 |
| MVP_top60_DNA_seed | PIK3CA | -0.005308 | HNSC | FADU | AZD8835 | MVP_top60_DNA+RNA_seed | JAK1 | 1.222014 | GBM | LN-405 | JAK_8517 |
| MVP_top60_DNA_seed | MDM2 | -0.002807 | HNSC | SAS | Nutlin-3a (-) | MVP_top60_DNA+RNA_seed | EGFR | 1.222906 | GBM | U-118-MG | Lapatinib |
| MVP_top60_DNA_seed | NTRK1 | -0.001013 | HNSC | HO-1-u-1 | GW441756 | MVP_top60_DNA+RNA_seed | IGF1R | 1.226106 | GBM | D-245MG | GSK1904529A |
| MVP_top60_DNA_seed | CDK2 | 0.001242 | HNSC | OSC-20 | PHA-793887 | MVP_top60_DNA+RNA_seed | AKT1 | 1.226209 | GBM | DBTRG-05MG | Uprosertib |
| MVP_top60_DNA_seed | MDM2 | 0.001418 | HNSC | DOK | Nutlin-3a (-) | MVP_top60_DNA+RNA_seed | IGF1R | 1.226709 | GBM | Becker | Linsitinib |
| MVP_top60_DNA_seed | HDAC1 | 0.004016 | HNSC | BB49-HNC | Entinostat | MVP_top60_DNA+RNA_seed | JAK2 | 1.231012 | GBM | D-247MG | Ruxolitinib |
| MVP_top60_DNA_seed | HDAC3 | 0.004016 | HNSC | BB49-HNC | Entinostat | MVP_top60_DNA+RNA_seed | JAK1 | 1.231012 | GBM | D-247MG | Ruxolitinib |
| MVP_top60_DNA_seed | AKT1 | 0.004838 | HNSC | SCC90 | BAY AKT1 | MVP_top60_DNA+RNA_seed | EGFR | 1.234905 | GBM | DK-MG | Gefitinib |
| MVP_top60_DNA_seed | EP300 | 0.007036 | HNSC | HO-1-N-1 | I-CBP112 | MVP_top60_DNA+RNA_seed | PIK3CA | 1.23908 | GBM | U251 | Alpelisib |
| MVP_top60_DNA_seed | CREBBP | 0.007036 | HNSC | HO-1-N-1 | I-CBP112 | MVP_top60_DNA+RNA_seed | AKT1 | 1.244011 | GBM | LN-18 | Ipatasertib |
| MVP_top60_DNA_seed | HDAC1 | 0.008615 | HNSC | BB30-HNC | Belinostat | MVP_top60_DNA+RNA_seed | EGFR | 1.247878 | GBM | CAS-1 | AZD3759 |
| MVP_top60_DNA_seed | MAPK1 | 0.008868 | HNSC | CAL-27 | VX-11e | MVP_top60_DNA+RNA_seed | KRAS | 1.249052 | GBM | LN-405 | KRAS (G12C) Inhibitor-1 |
| MVP_top60_DNA_seed | NTRK1 | 0.009099 | HNSC | JHU-022 | AZD1332 | MVP_top60_DNA+RNA_seed | TP53 | 1.249454 | GBM | CAS-1 | MIRA-1 |
| MVP_top60_DNA_seed | EP300 | 0.00981 | HNSC | JHU-011 | I-CBP112 | MVP_top60_DNA+RNA_seed | KRAS | 1.253882 | GBM | M059J | KRAS (G12C) Inhibitor-1 |
| MVP_top60_DNA_seed | CREBBP | 0.00981 | HNSC | JHU-011 | I-CBP112 | MVP_top60_DNA+RNA_seed | IGF1R | 1.258393 | GBM | SW1088 | Linsitinib |
| MVP_top60_DNA_seed | HDAC1 | 0.014089 | HNSC | Ca9-22 | Tubastatin A | MVP_top60_DNA+RNA_seed | EGFR | 1.260517 | GBM | SNB75 | Erlotinib |
| MVP_top60_DNA_seed | SMARCA4 | 0.014594 | HNSC | KON | PFI-3 | MVP_top60_DNA+RNA_seed | CDK4 | 1.263292 | GBM | LN-405 | Palbociclib |
| MVP_top60_DNA_seed | RAC1 | 0.017771 | HNSC | HSC-3 | EHT-1864 | MVP_top60_DNA+RNA_seed | JAK2 | 1.273339 | GBM | SK-MG-1 | JAK_8517 |
| MVP_top60_DNA_seed | HDAC1 | 0.018968 | HNSC | SKN-3 | Belinostat | MVP_top60_DNA+RNA_seed | JAK1 | 1.273339 | GBM | SK-MG-1 | JAK_8517 |
| MVP_top60_DNA_seed | EGFR | 0.019358 | HNSC | KON | Cetuximab | MVP_top60_DNA+RNA_seed | JAK2 | 1.280343 | GBM | AM-38 | JAK_8517 |
| MVP_top60_DNA_seed | HDAC1 | 0.021623 | HNSC | SAT | AR-42 | MVP_top60_DNA+RNA_seed | JAK1 | 1.280343 | GBM | AM-38 | JAK_8517 |
| MVP_top60_DNA_seed | AR | 0.022112 | HNSC | SCC-15 | Bicalutamide | MVP_top60_DNA+RNA_seed | EGFR | 1.283699 | GBM | D-263MG | Lapatinib |
| MVP_top60_DNA_seed | AKT1 | 0.022854 | HNSC | PCI-38 | MK-2206 | MVP_top60_DNA+RNA_seed | EGFR | 1.288037 | GBM | D-245MG | Erlotinib |

| | | | | | | | | | | | |
|---|---|---|---|---|---|---|---|---|---|---|---|
| MVP_top60_DNA_seed | HDAC1 | 0.022921 | HNSC | PCI-38 | CAY10603 | MVP_top60_DNA+RNA_seed | JAK1 | 1.289234 | GBM | M059J | JAK1_8709 |
| MVP_top60_DNA_seed | HSP90AA1 | 0.02501 | HNSC | SCC90 | Tanespimycin | MVP_top60_DNA+RNA_seed | CDK4 | 1.299051 | GBM | D-542MG | Palbociclib |
| MVP_top60_DNA_seed | HDAC1 | 0.026279 | HNSC | PCI-38 | Tubastatin A | MVP_top60_DNA+RNA_seed | JAK2 | 1.303162 | GBM | U251 | AZ960 |
| MVP_top60_DNA_seed | EGFR | 0.027558 | HNSC | HO-1-u-1 | AZD8931 | MVP_top60_DNA+RNA_seed | JAK3 | 1.303162 | GBM | U251 | AZ960 |
| MVP_top60_DNA_seed | MDM2 | 0.027614 | HNSC | PE-CA-PJ15 | Nutlin-3a (-) | MVP_top60_DNA+RNA_seed | PIK3CA | 1.306821 | GBM | DBTRG-05MG | GNE-317 |
| MVP_top60_DNA_seed | AR | 0.028063 | HNSC | JHU-022 | Bicalutamide | MVP_top60_DNA+RNA_seed | MET | 1.313941 | GBM | U-118-MG | Crizotinib |
| MVP_top60_DNA_seed | HSP90AA1 | 0.028317 | HNSC | BICR22 | Luminespib | MVP_top60_DNA+RNA_seed | IGF1R | 1.328038 | GBM | SF295 | BMS-536924 |
| MVP_top60_DNA_seed | SRC | 0.030168 | HNSC | LB771-HNC | WH-4-023 | MVP_top60_DNA+RNA_seed | CDK4 | 1.331047 | GBM | GB-1 | Ribociclib |
| MVP_top60_DNA_seed | EP300 | 0.030847 | HNSC | PCI-6A | I-CBP112 | MVP_top60_DNA+RNA_seed | EP300 | 1.335778 | GBM | LN-229 | SGC-CBP30 |
| MVP_top60_DNA_seed | CREBBP | 0.030847 | HNSC | PCI-6A | I-CBP112 | MVP_top60_DNA+RNA_seed | CREBBP | 1.335778 | GBM | LN-229 | SGC-CBP30 |
| MVP_top60_DNA_seed | AR | 0.033381 | HNSC | PCI-38 | Bicalutamide | MVP_top60_DNA+RNA_seed | AKT1 | 1.338705 | GBM | D-263MG | Uprosertib |
| MVP_top60_DNA_seed | AR | 0.035926 | HNSC | BICR10 | AZD3514 | MVP_top60_DNA+RNA_seed | IGF1R | 1.344317 | GBM | D-263MG | NVP-ADW742 |
| MVP_top60_DNA_seed | HDAC1 | 0.037184 | HNSC | HO-1-N-1 | AR-42 | MVP_top60_DNA+RNA_seed | AKT1 | 1.351829 | GBM | U251 | MK-2206 |
| MVP_top60_DNA_seed | AKT1 | 0.037833 | HNSC | HSC-3 | AKT inhibitor VIII | MVP_top60_DNA+RNA_seed | JAK2 | 1.352287 | GBM | SW1088 | Ruxolitinib |
| MVP_top60_DNA_seed | NTRK1 | 0.038298 | HNSC | BHY | AZD1332 | MVP_top60_DNA+RNA_seed | JAK1 | 1.352287 | GBM | SW1088 | Ruxolitinib |
| MVP_top60_DNA_seed | AR | 0.038919 | HNSC | PE-CA-PJ15 | AZD3514 | MVP_top60_DNA+RNA_seed | KRAS | 1.353915 | GBM | SK-MG-1 | KRAS (G12C) Inhibitor-1 |
| MVP_top60_DNA_seed | SRC | 0.041682 | HNSC | BB30-HNC | NVP-BHG712 | MVP_top60_DNA+RNA_seed | PIK3CA | 1.361822 | GBM | LN-405 | Buparlisib |
| MVP_top60_DNA_seed | AKT1 | 0.043972 | HNSC | BHY | AKT inhibitor VIII | MVP_top60_DNA+RNA_seed | PIK3CB | 1.361822 | GBM | LN-405 | Buparlisib |
| MVP_top60_DNA_seed | CDK2 | 0.044214 | HNSC | Ca9-22 | PHA-793887 | MVP_top60_DNA+RNA_seed | PIK3CD | 1.361822 | GBM | LN-405 | Buparlisib |
| MVP_top60_DNA_seed | HSP90AA1 | 0.0455 | HNSC | JHU-022 | Luminespib | MVP_top60_DNA+RNA_seed | IGF1R | 1.374268 | GBM | LN-405 | BMS-536924 |
| MVP_top60_DNA_seed | HDAC1 | 0.046674 | HNSC | BB30-HNC | CAY10603 | MVP_top60_DNA+RNA_seed | IGF1R | 1.374383 | GBM | SF295 | Linsitinib |
| MVP_top60_DNA_seed | EGFR | 0.049135 | HNSC | HO-1-u-1 | CUDC-101 | MVP_top60_DNA+RNA_seed | EGFR | 1.375176 | GBM | SNB75 | Afatinib |
| MVP_top60_DNA_seed | HDAC1 | 0.049967 | HNSC | PCI-38 | AR-42 | MVP_top60_DNA+RNA_seed | EGFR | 1.378183 | GBM | DBTRG-05MG | AZD3759 |
| MVP_top60_DNA_seed | RAC1 | 0.051773 | HNSC | FADU | EHT-1864 | MVP_top60_DNA+RNA_seed | AKT1 | 1.37841 | GBM | GB-1 | GSK2110183B |
| MVP_top60_DNA_seed | NTRK1 | 0.052064 | HNSC | LB771-HNC | AZD1332 | MVP_top60_DNA+RNA_seed | AKT1 | 1.378761 | GBM | U251 | AT13148 |
| MVP_top60_DNA_seed | AKT1 | 0.052664 | HNSC | PCI-38 | GSK690693 | MVP_top60_DNA+RNA_seed | EP300 | 1.379405 | GBM | CAS-1 | CPI-637 |
| MVP_top60_DNA_seed | AR | 0.059507 | HNSC | PCI-6A | Bicalutamide | MVP_top60_DNA+RNA_seed | MET | 1.381276 | GBM | LN-405 | Crizotinib |
| MVP_top60_DNA_seed | EGFR | 0.060277 | HNSC | KOSC-2 | Cetuximab | MVP_top60_DNA+RNA_seed | IGF1R | 1.394123 | GBM | SNB75 | Linsitinib |
| MVP_top60_DNA_seed | AR | 0.06039 | HNSC | BB30-HNC | Bicalutamide | MVP_top60_DNA+RNA_seed | EGFR | 1.39833 | GBM | GB-1 | AZD3759 |
| MVP_top60_DNA_seed | ESR1 | 0.062206 | HNSC | CAL-33 | Tamoxifen | MVP_top60_DNA+RNA_seed | EGFR | 1.401945 | GBM | U251 | Sapitinib |
| MVP_top60_DNA_seed | RAC1 | 0.062618 | HNSC | HO-1-N-1 | EHT-1864 | MVP_top60_DNA+RNA_seed | ERBB3 | 1.401945 | GBM | U251 | Sapitinib |
| MVP_top60_DNA_seed | NTRK1 | 0.065074 | HNSC | PCI-6A | GW441756 | MVP_top60_DNA+RNA_seed | JAK1 | 1.403396 | GBM | LN-405 | JAK1_8709 |
| MVP_top60_DNA_seed | HSP90AA1 | 0.067518 | HNSC | PCI-15A | Tanespimycin | MVP_top60_DNA+RNA_seed | EP300 | 1.403904 | GBM | GB-1 | CPI-637 |
| MVP_top60_DNA_seed | AKT1 | 0.067654 | HNSC | PCI-6A | GSK690693 | MVP_top60_DNA+RNA_seed | EP300 | 1.41132 | GBM | DK-MG | SGC-CBP30 |
| MVP_top60_DNA_seed | HDAC1 | 0.068508 | HNSC | SKN-3 | CAY10603 | MVP_top60_DNA+RNA_seed | CREBBP | 1.41132 | GBM | DK-MG | SGC-CBP30 |
| MVP_top60_DNA_seed | HSP90AA1 | 0.068736 | HNSC | OSC-19 | SNX-2112 | MVP_top60_DNA+RNA_seed | IGF1R | 1.411941 | GBM | DK-MG | Linsitinib |
| MVP_top60_DNA_seed | AKT1 | 0.069534 | HNSC | PE-CA-PJ15 | BAY AKT1 | MVP_top60_DNA+RNA_seed | EP300 | 1.419186 | GBM | SW1088 | SGC-CBP30 |
| MVP_top60_DNA_seed | EGFR | 0.073295 | HNSC | OSC-19 | Cetuximab | MVP_top60_DNA+RNA_seed | CREBBP | 1.419186 | GBM | SW1088 | SGC-CBP30 |
| MVP_top60_DNA_seed | MDM2 | 0.079862 | HNSC | BICR10 | Nutlin-3a (-) | MVP_top60_DNA+RNA_seed | AKT1 | 1.420985 | GBM | LN-405 | AT13148 |
| MVP_top60_DNA_seed | AR | 0.080858 | HNSC | OSC-19 | Bicalutamide | MVP_top60_DNA+RNA_seed | EGFR | 1.425539 | GBM | U-87-MG | Afatinib |
| MVP_top60_DNA_seed | EGFR | 0.082892 | HNSC | HO-1-N-1 | CUDC-101 | MVP_top60_DNA+RNA_seed | EGFR | 1.430242 | GBM | U-87-MG | Lapatinib |

| | | | | | | | | | | | |
|---|---|---|---|---|---|---|---|---|---|---|---|
| MVP_top60_DNA_seed | SRC | 0.085109 | HNSC | PCI-38 | NVP-BHG712 | MVP_top60_DNA+RNA_seed | IGF1R | 1.431986 | GBM | D-542MG | Linsitinib |
| MVP_top60_DNA_seed | EP300 | 0.085716 | HNSC | KON | I-CBP112 | MVP_top60_DNA+RNA_seed | EGFR | 1.446664 | GBM | DBTRG-05MG | Afatinib |
| MVP_top60_DNA_seed | CREBBP | 0.085716 | HNSC | KON | I-CBP112 | MVP_top60_DNA+RNA_seed | PIK3CA | 1.45936 | GBM | YKG-1 | Alpelisib |
| MVP_top60_DNA_seed | ESR1 | 0.08619 | HNSC | OSC-19 | Tamoxifen | MVP_top60_DNA+RNA_seed | AKT1 | 1.461602 | GBM | LN-229 | Ipatasertib |
| MVP_top60_DNA_seed | HDAC1 | 0.086242 | HNSC | SCC-15 | Belinostat | MVP_top60_DNA+RNA_seed | AKT1 | 1.461672 | GBM | D-392MG | Uprosertib |
| MVP_top60_DNA_seed | AKT1 | 0.090839 | HNSC | JHU-022 | MK-2206 | MVP_top60_DNA+RNA_seed | MET | 1.466188 | GBM | U251 | Crizotinib |
| MVP_top60_DNA_seed | EGFR | 0.091691 | HNSC | KOSC-2 | AST-1306 | MVP_top60_DNA+RNA_seed | EGFR | 1.4674 | GBM | Becker | Lapatinib |
| MVP_top60_DNA_seed | HDAC1 | 0.093128 | HNSC | HO-1-N-1 | CAY10603 | MVP_top60_DNA+RNA_seed | JAK1 | 1.481197 | GBM | Becker | JAK1_8709 |
| MVP_top60_DNA_seed | CDK2 | 0.097821 | HNSC | BHY | AZD5438 | MVP_top60_DNA+RNA_seed | IGF1R | 1.485479 | GBM | CAS-1 | Linsitinib |
| MVP_top60_DNA_seed | EGFR | 0.099217 | HNSC | LB771-HNC | Pelitinib | MVP_top60_DNA+RNA_seed | CDK4 | 1.489772 | GBM | LN-229 | Palbociclib |
| MVP_top60_DNA_seed | EGFR | 0.099634 | HNSC | PCI-30 | Cetuximab | MVP_top60_DNA+RNA_seed | EP300 | 1.492985 | GBM | U-118-MG | CPI-637 |
| MVP_top60_DNA_seed | EP300 | 0.100455 | HNSC | SKN-3 | I-CBP112 | MVP_top60_DNA+RNA_seed | AKT1 | 1.497497 | GBM | D-392MG | MK-2206 |
| MVP_top60_DNA_seed | CREBBP | 0.100455 | HNSC | SKN-3 | I-CBP112 | MVP_top60_DNA+RNA_seed | IGF1R | 1.506456 | GBM | T98G | NVP-ADW742 |
| MVP_top60_DNA_seed | HDAC1 | 0.100843 | HNSC | SCC-4 | CAY10603 | MVP_top60_DNA+RNA_seed | PIK3CA | 1.520659 | GBM | GB-1 | AZD8186 |
| MVP_top60_DNA_seed | AKT1 | 0.100932 | HNSC | SCC-4 | AKT inhibitor VIII | MVP_top60_DNA+RNA_seed | PIK3CB | 1.520659 | GBM | GB-1 | AZD8186 |
| MVP_top60_DNA_seed | AR | 0.102086 | HNSC | SCC-25 | AZD3514 | MVP_top60_DNA+RNA_seed | EGFR | 1.52125 | GBM | LN-405 | Afatinib |
| MVP_top60_DNA_seed | EGFR | 0.102262 | HNSC | HSC-2 | Pelitinib | MVP_top60_DNA+RNA_seed | JAK2 | 1.540296 | GBM | U-87-MG | JAK_8517 |
| MVP_top60_DNA_seed | EGFR | 0.10446 | HNSC | Ca9-22 | PF-00299804 | MVP_top60_DNA+RNA_seed | JAK1 | 1.540296 | GBM | U-87-MG | JAK_8517 |
| MVP_top60_DNA_seed | HDAC1 | 0.10645 | HNSC | SAT | CAY10603 | MVP_top60_DNA+RNA_seed | EP300 | 1.544784 | GBM | DK-MG | CPI-637 |
| MVP_top60_DNA_seed | EGFR | 0.109355 | HNSC | LB771-HNC | Afatinib | MVP_top60_DNA+RNA_seed | PIK3CA | 1.552749 | GBM | D-245MG | Buparlisib |
| MVP_top60_DNA_seed | HDAC1 | 0.117103 | HNSC | SCC-15 | CAY10603 | MVP_top60_DNA+RNA_seed | PIK3CB | 1.552749 | GBM | D-245MG | Buparlisib |
| MVP_top60_DNA_seed | EGFR | 0.117211 | HNSC | FADU | CUDC-101 | MVP_top60_DNA+RNA_seed | PIK3CD | 1.552749 | GBM | D-245MG | Buparlisib |
| MVP_top60_DNA_seed | EGFR | 0.118243 | HNSC | HSC-3 | CI-1033 | MVP_top60_DNA+RNA_seed | AKT1 | 1.559274 | GBM | U-87-MG | AT13148 |
| MVP_top60_DNA_seed | AKT1 | 0.119125 | HNSC | JHU-022 | BAY AKT1 | MVP_top60_DNA+RNA_seed | IGF1R | 1.560514 | GBM | U-87-MG | Linsitinib |
| MVP_top60_DNA_seed | NTRK1 | 0.123694 | HNSC | PE-CA-PJ15 | GW441756 | MVP_top60_DNA+RNA_seed | EGFR | 1.562033 | GBM | KS-1 | Lapatinib |
| MVP_top60_DNA_seed | PIK3CA | 0.127621 | HNSC | SAS | AZD8835 | MVP_top60_DNA+RNA_seed | IGF1R | 1.562932 | GBM | M059J | NVP-ADW742 |
| MVP_top60_DNA_seed | EGFR | 0.128153 | HNSC | BICR31 | PF-00299804 | MVP_top60_DNA+RNA_seed | IGF1R | 1.576059 | GBM | KS-1 | GSK1904529A |
| MVP_top60_DNA_seed | SRC | 0.129418 | HNSC | CAL-27 | HG-6-71-01 | MVP_top60_DNA+RNA_seed | EP300 | 1.591641 | GBM | DBTRG-05MG | CPI-637 |
| MVP_top60_DNA_seed | ESR1 | 0.130031 | HNSC | PCI-6A | Fulvestrant | MVP_top60_DNA+RNA_seed | AKT1 | 1.594318 | GBM | AM-38 | Ipatasertib |
| MVP_top60_DNA_seed | EGFR | 0.130726 | HNSC | HO-1-N-1 | Pelitinib | MVP_top60_DNA+RNA_seed | AKT1 | 1.603431 | GBM | DBTRG-05MG | Uprosertib |
| MVP_top60_DNA_seed | EGFR | 0.138009 | HNSC | SCC-4 | Afatinib | MVP_top60_DNA+RNA_seed | KRAS | 1.607531 | GBM | U-118-MG | KRAS (G12C) Inhibitor-1 |
| MVP_top60_DNA_seed | AKT1 | 0.140438 | HNSC | SCC90 | AKT inhibitor VIII | MVP_top60_DNA+RNA_seed | AKT1 | 1.633293 | GBM | AM-38 | Afuresertib |
| MVP_top60_DNA_seed | MAPK1 | 0.140711 | HNSC | HSC-3 | FR-180204 | MVP_top60_DNA+RNA_seed | PIK3CA | 1.637238 | GBM | D-263MG | Alpelisib |
| MVP_top60_DNA_seed | MAPK3 | 0.140711 | HNSC | HSC-3 | FR-180204 | MVP_top60_DNA+RNA_seed | JAK2 | 1.644075 | GBM | DBTRG-05MG | Ruxolitinib |
| MVP_top60_DNA_seed | HDAC1 | 0.142216 | HNSC | OSC-20 | Dacinostat | MVP_top60_DNA+RNA_seed | JAK1 | 1.644075 | GBM | DBTRG-05MG | Ruxolitinib |
| MVP_top60_DNA_seed | HSP90AA1 | 0.14344 | HNSC | HO-1-N-1 | SNX-2112 | MVP_top60_DNA+RNA_seed | AKT1 | 1.644691 | GBM | LN-229 | Afuresertib |
| MVP_top60_DNA_seed | AKT1 | 0.14549 | HNSC | HO-1-N-1 | BAY AKT1 | MVP_top60_DNA+RNA_seed | MET | 1.668273 | GBM | LN-229 | Crizotinib |
| MVP_top60_DNA_seed | EGFR | 0.147506 | HNSC | CAL-33 | CI-1033 | MVP_top60_DNA+RNA_seed | IGF1R | 1.672385 | GBM | D-247MG | Linsitinib |
| MVP_top60_DNA_seed | EGFR | 0.149242 | HNSC | BICR10 | Cetuximab | MVP_top60_DNA+RNA_seed | PIK3CA | 1.682586 | GBM | AM-38 | AZD8186 |
| MVP_top60_DNA_seed | HSP90AA1 | 0.150865 | HNSC | BICR22 | CCT-018159 | MVP_top60_DNA+RNA_seed | PIK3CB | 1.682586 | GBM | AM-38 | AZD8186 |
| MVP_top60_DNA_seed | HSP90AA1 | 0.153454 | HNSC | SAS | Luminespib | MVP_top60_DNA+RNA_seed | CDK4 | 1.691322 | GBM | DBTRG-05MG | Ribociclib |

| | | | | | | | | | | | |
|---|---|---|---|---|---|---|---|---|---|---|---|
| MVP_top60_DNA_seed | NTRK1 | 0.153842 | HNSC | CAL-27 | GW441756 | MVP_top60_DNA+RNA_seed | EGFR | 1.716475 | GBM | SF295 | Erlotinib |
| MVP_top60_DNA_seed | EGFR | 0.154134 | HNSC | HSC-3 | CUDC-101 | MVP_top60_DNA+RNA_seed | JAK1 | 1.744043 | GBM | DBTRG-05MG | JAK1_8709 |
| MVP_top60_DNA_seed | HDAC1 | 0.157995 | HNSC | JHU-011 | Tubastatin A | MVP_top60_DNA+RNA_seed | AKT1 | 1.745721 | GBM | LN-229 | Uprosertib |
| MVP_top60_DNA_seed | HSP90AA1 | 0.158745 | HNSC | SCC90 | Luminespib | MVP_top60_DNA+RNA_seed | AKT1 | 1.745815 | GBM | DBTRG-05MG | GSK2110183B |
| MVP_top60_DNA_seed | ESR1 | 0.161516 | HNSC | HO-1-u-1 | Tamoxifen | MVP_top60_DNA+RNA_seed | EGFR | 1.747985 | GBM | LN-405 | Lapatinib |
| MVP_top60_DNA_seed | CDK2 | 0.162866 | HNSC | CAL-33 | AZD5438 | MVP_top60_DNA+RNA_seed | EGFR | 1.750427 | GBM | U-118-MG | Sapitinib |
| MVP_top60_DNA_seed | HDAC1 | 0.163136 | HNSC | CAL-33 | Belinostat | MVP_top60_DNA+RNA_seed | ERBB3 | 1.750427 | GBM | U-118-MG | Sapitinib |
| MVP_top60_DNA_seed | RAC1 | 0.163528 | HNSC | KON | EHT-1864 | MVP_top60_DNA+RNA_seed | KIT | 1.751138 | GBM | DBTRG-05MG | Sorafenib |
| MVP_top60_DNA_seed | MAPK1 | 0.164276 | HNSC | SKN-3 | VX-11e | MVP_top60_DNA+RNA_seed | IGF1R | 1.764852 | GBM | LN-405 | NVP-ADW742 |
| MVP_top60_DNA_seed | AKT1 | 0.165134 | HNSC | LB771-HNC | GSK690693 | MVP_top60_DNA+RNA_seed | EGFR | 1.767129 | GBM | DBTRG-05MG | Sapitinib |
| MVP_top60_DNA_seed | AR | 0.165395 | HNSC | PCI-4B | Bicalutamide | MVP_top60_DNA+RNA_seed | ERBB3 | 1.767129 | GBM | DBTRG-05MG | Sapitinib |
| MVP_top60_DNA_seed | HDAC1 | 0.165407 | HNSC | BICR78 | Dacinostat | MVP_top60_DNA+RNA_seed | EP300 | 1.777659 | GBM | CAS-1 | SGC-CBP30 |
| MVP_top60_DNA_seed | HDAC1 | 0.167671 | HNSC | BB30-HNC | AR-42 | MVP_top60_DNA+RNA_seed | CREBBP | 1.777659 | GBM | CAS-1 | SGC-CBP30 |
| MVP_top60_DNA_seed | HDAC1 | 0.1685 | HNSC | CAL-33 | CAY10603 | MVP_top60_DNA+RNA_seed | EGFR | 1.805704 | GBM | GB-1 | Osimertinib |
| MVP_top60_DNA_seed | AKT1 | 0.170963 | HNSC | BHY | GSK690693 | MVP_top60_DNA+RNA_seed | KIT | 1.822663 | GBM | KS-1 | Sorafenib |
| MVP_top60_DNA_seed | EP300 | 0.172735 | HNSC | Detroit562 | I-CBP112 | MVP_top60_DNA+RNA_seed | JAK2 | 1.824469 | GBM | LN-229 | JAK_8517 |
| MVP_top60_DNA_seed | CREBBP | 0.172735 | HNSC | Detroit562 | I-CBP112 | MVP_top60_DNA+RNA_seed | JAK1 | 1.824469 | GBM | LN-229 | JAK_8517 |
| MVP_top60_DNA_seed | SRC | 0.173335 | HNSC | JHU-011 | NVP-BHG712 | MVP_top60_DNA+RNA_seed | KIT | 1.831669 | GBM | DBTRG-05MG | Motesanib |
| MVP_top60_DNA_seed | MDM2 | 0.173679 | HNSC | BHY | Serdemetan | MVP_top60_DNA+RNA_seed | IGF1R | 1.840908 | GBM | U-118-MG | Linsitinib |
| MVP_top60_DNA_seed | HDAC1 | 0.174078 | HNSC | PE-CA-PJ15 | Dacinostat | MVP_top60_DNA+RNA_seed | AKT1 | 1.844739 | GBM | DBTRG-05MG | AT13148 |
| MVP_top60_DNA_seed | HDAC1 | 0.174478 | HNSC | PE-CA-PJ15 | Belinostat | MVP_top60_DNA+RNA_seed | KRAS | 1.875351 | GBM | CAS-1 | RAS (G12C) Inhibitor-1 |
| MVP_top60_DNA_seed | AR | 0.175932 | HNSC | BICR31 | Bicalutamide | MVP_top60_DNA+RNA_seed | AKT1 | 1.876375 | GBM | LN-229 | MK-2206 |
| MVP_top60_DNA_seed | AR | 0.177094 | HNSC | PE-CA-PJ15 | Bicalutamide | MVP_top60_DNA+RNA_seed | PIK3CA | 1.925213 | GBM | U-118-MG | Alpelisib |
| MVP_top60_DNA_seed | AR | 0.177706 | HNSC | BHY | Bicalutamide | MVP_top60_DNA+RNA_seed | PIK3CA | 1.941553 | GBM | LN-405 | GNE-317 |
| MVP_top60_DNA_seed | HDAC1 | 0.177833 | HNSC | SKN-3 | AR-42 | MVP_top60_DNA+RNA_seed | IGF1R | 2.013736 | GBM | DBTRG-05MG | NVP-ADW742 |
| MVP_top60_DNA_seed | MDM2 | 0.180583 | HNSC | SCC-25 | Serdemetan | MVP_top60_DNA+RNA_seed | JAK2 | 2.016043 | GBM | DBTRG-05MG | GSK2276186C |
| MVP_top60_DNA_seed | EP300 | 0.181608 | HNSC | BHY | I-CBP112 | MVP_top60_DNA+RNA_seed | JAK1 | 2.016043 | GBM | DBTRG-05MG | GSK2276186C |
| MVP_top60_DNA_seed | CREBBP | 0.181608 | HNSC | BHY | I-CBP112 | MVP_top60_DNA+RNA_seed | JAK3 | 2.016043 | GBM | DBTRG-05MG | GSK2276186C |
| MVP_top60_DNA_seed | HSP90AA1 | 0.184746 | HNSC | PE-CA-PJ15 | Elesclomol | MVP_top60_DNA+RNA_seed | CDK4 | 2.065252 | GBM | DBTRG-05MG | Palbociclib |
| MVP_top60_DNA_seed | AKT1 | 0.188854 | HNSC | SAS | GSK690693 | MVP_top60_DNA+RNA_seed | EGFR | 2.079235 | GBM | D-392MG | Lapatinib |
| MVP_top60_DNA_seed | EGFR | 0.18944 | HNSC | PCI-30 | AZD8931 | MVP_top60_DNA+RNA_seed | EGFR | 2.094157 | GBM | DBTRG-05MG | Gefitinib |
| MVP_top60_DNA_seed | HSP90AA1 | 0.189534 | HNSC | Ca9-22 | Tanespimycin | MVP_top60_DNA+RNA_seed | TP53 | 2.125063 | GBM | DBTRG-05MG | MIRA-1 |
| MVP_top60_DNA_seed | HDAC1 | 0.191997 | HNSC | SCC-4 | AR-42 | MVP_top60_DNA+RNA_seed | IGF1R | 2.176697 | GBM | DBTRG-05MG | GSK1904529A |
| MVP_top60_DNA_seed | HDAC1 | 0.193396 | HNSC | BICR22 | AR-42 | MVP_top60_DNA+RNA_seed | IGF1R | 2.194829 | GBM | DBTRG-05MG | Linsitinib |
| MVP_top60_DNA_seed | HDAC1 | 0.193549 | HNSC | DOK | CAY10603 | MVP_top60_DNA+RNA_seed | KIT | 2.201168 | GBM | D-392MG | Sorafenib |
| MVP_top60_DNA_seed | MAPK1 | 0.194067 | HNSC | BICR78 | FR-180204 | MVP_top60_DNA+RNA_seed | KIT | 2.251505 | GBM | DBTRG-05MG | Axitinib |
| MVP_top60_DNA_seed | MAPK3 | 0.194067 | HNSC | BICR78 | FR-180204 | MVP_top60_DNA+RNA_seed | AKT1 | 2.260821 | GBM | DBTRG-05MG | MK-2206 |
| MVP_top60_DNA_seed | HSP90AA1 | 0.198415 | HNSC | BICR10 | SNX-2112 | MVP_top60_DNA+RNA_seed | PIK3CA | 2.300414 | GBM | U251 | GNE-317 |
| MVP_top60_DNA_seed | NTRK1 | 0.200323 | HNSC | Detroit562 | GW441756 | MVP_top60_DNA+RNA_seed | KRAS | 2.310501 | GBM | DBTRG-05MG | RAS (G12C) Inhibitor-1 |
| MVP_top60_DNA_seed | CDK2 | 0.200488 | HNSC | SKN-3 | PHA-793887 | MVP_top60_DNA+RNA_seed | PIK3CA | 2.542779 | GBM | DBTRG-05MG | Alpelisib |
| MVP_top60_DNA_seed | HDAC1 | 0.204924 | HNSC | LB771-HNC | Belinostat | MVP_top60_DNA+RNA_seed | EGFR | 2.576791 | GBM | DBTRG-05MG | Lapatinib |

| | | | | | | | | | | | |
|---|---|---|---|---|---|---|---|---|---|---|---|
| MVP_top60_DNA_seed | EP300 | 0.205309 | HNSC | PCI-15A | I-CBP112 | MVP_top60_DNA+RNA_seed | EP300 | 2.840979 | GBM | DBTRG-05MG | SGC-CBP30 |
| MVP_top60_DNA_seed | CREBBP | 0.205309 | HNSC | PCI-15A | I-CBP112 | MVP_top60_DNA+RNA_seed | CREBBP | 2.840979 | GBM | DBTRG-05MG | SGC-CBP30 |
| MVP_top60_DNA_seed | NTRK1 | 0.209305 | HNSC | OSC-19 | AZD1332 | MVP_top60_DNA+RNA_seed | MET | 2.885711 | GBM | DBTRG-05MG | Crizotinib |
| MVP_top60_DNA_seed | EGFR | 0.211812 | HNSC | KOSC-2 | PF-00299804 | MVP_top60_DNA+RNA_seed | MTOR | -3.86697 | KIRC | 786-0 | CRT0105446 |
| MVP_top60_DNA_seed | ESR1 | 0.211866 | HNSC | JHU-011 | Tamoxifen | MVP_top60_DNA+RNA_seed | PIK3CD | -3.688259 | KIRC | RCC-FG2 | AZD8186 |
| MVP_top60_DNA_seed | HSP90AA1 | 0.212484 | HNSC | HO-1-u-1 | Luminespib | MVP_top60_DNA+RNA_seed | PIK3CD | -3.016708 | KIRC | A498 | AZD8186 |
| MVP_top60_DNA_seed | AKT1 | 0.212785 | HNSC | KOSC-2 | MK-2206 | MVP_top60_DNA+RNA_seed | AR | -2.858498 | KIRC | U031 | AZD3514 |
| MVP_top60_DNA_seed | AKT1 | 0.2141 | HNSC | KOSC-2 | BAY AKT1 | MVP_top60_DNA+RNA_seed | PIK3CD | -2.644204 | KIRC | RCC-JF | AZD8186 |
| MVP_top60_DNA_seed | NTRK1 | 0.215562 | HNSC | BHY | GW441756 | MVP_top60_DNA+RNA_seed | MTOR | -2.526586 | KIRC | RCC-JW | Temsirolimus |
| MVP_top60_DNA_seed | AKT1 | 0.218379 | HNSC | BICR22 | AKT inhibitor VIII | MVP_top60_DNA+RNA_seed | PIK3CD | -2.464178 | KIRC | U031 | AZD8186 |
| MVP_top60_DNA_seed | HSP90AA1 | 0.222127 | HNSC | JHU-022 | SNX-2112 | MVP_top60_DNA+RNA_seed | FGFR4 | -2.4591 | KIRC | U031 | FGFR_0939 |
| MVP_top60_DNA_seed | EGFR | 0.22688 | HNSC | KON | Gefitinib | MVP_top60_DNA+RNA_seed | SMARCA4 | -2.283803 | KIRC | A498 | PFI-3 |
| MVP_top60_DNA_seed | HSP90AA1 | 0.228292 | HNSC | SAT | SNX-2112 | MVP_top60_DNA+RNA_seed | SMARCA2 | -2.283803 | KIRC | A498 | PFI-3 |
| MVP_top60_DNA_seed | RAC1 | 0.229469 | HNSC | JHU-022 | EHT-1864 | MVP_top60_DNA+RNA_seed | ATM | -2.249208 | KIRC | SW156 | KU-60019 |
| MVP_top60_DNA_seed | CDK2 | 0.230105 | HNSC | HSC-3 | PHA-793887 | MVP_top60_DNA+RNA_seed | PIK3CD | -2.249095 | KIRC | CAL-54 | AZD8186 |
| MVP_top60_DNA_seed | AKT1 | 0.233098 | HNSC | PE-CA-PJ15 | MK-2206 | MVP_top60_DNA+RNA_seed | MTOR | -2.133136 | KIRC | LB996-RCC | CRT0105446 |
| MVP_top60_DNA_seed | HSP90AA1 | 0.233575 | HNSC | LB771-HNC | CCT-018159 | MVP_top60_DNA+RNA_seed | IGF1R | -2.071902 | KIRC | CAL-54 | GSK1904529A |
| MVP_top60_DNA_seed | HDAC1 | 0.233851 | HNSC | Detroit562 | Dacinostat | MVP_top60_DNA+RNA_seed | EP300 | -1.956195 | KIRC | U031 | I-CBP112 |
| MVP_top60_DNA_seed | PIK3CA | 0.236136 | HNSC | SCC-4 | AZD8835 | MVP_top60_DNA+RNA_seed | PIK3CD | -1.946539 | KIRC | OS-RC-2 | AZD8186 |
| MVP_top60_DNA_seed | MAPK1 | 0.238208 | HNSC | PE-CA-PJ15 | VX-11e | MVP_top60_DNA+RNA_seed | HDAC1 | -1.93648 | KIRC | OS-RC-2 | Tubastatin A |
| MVP_top60_DNA_seed | AR | 0.239112 | HNSC | Detroit562 | Bicalutamide | MVP_top60_DNA+RNA_seed | ATM | -1.906513 | KIRC | LB2241-RCC | KU-55933 |
| MVP_top60_DNA_seed | AKT1 | 0.239553 | HNSC | PCI-6A | BAY AKT1 | MVP_top60_DNA+RNA_seed | AR | -1.844146 | KIRC | RCC-AB | Bicalutamide |
| MVP_top60_DNA_seed | HDAC1 | 0.239965 | HNSC | JHU-022 | Belinostat | MVP_top60_DNA+RNA_seed | IGF1R | -1.843809 | KIRC | 786-0 | GSK1904529A |
| MVP_top60_DNA_seed | CDK2 | 0.240616 | HNSC | OSC-19 | PHA-793887 | MVP_top60_DNA+RNA_seed | PIK3CD | -1.839539 | KIRC | LB996-RCC | AZD8186 |
| MVP_top60_DNA_seed | HDAC1 | 0.241698 | HNSC | SCC-15 | AR-42 | MVP_top60_DNA+RNA_seed | PIK3CD | -1.804286 | KIRC | CAKI-1 | AZD8186 |
| MVP_top60_DNA_seed | AKT1 | 0.242523 | HNSC | CAL-27 | MK-2206 | MVP_top60_DNA+RNA_seed | PIK3CD | -1.785491 | KIRC | HA7-RCC | AZD8186 |
| MVP_top60_DNA_seed | AKT1 | 0.244209 | HNSC | Detroit562 | AKT inhibitor VIII | MVP_top60_DNA+RNA_seed | MTOR | -1.780475 | KIRC | BB65-RCC | Temsirolimus |
| MVP_top60_DNA_seed | AKT1 | 0.245841 | HNSC | PCI-4B | MK-2206 | MVP_top60_DNA+RNA_seed | ATM | -1.768834 | KIRC | VMRC-RCW | KU-55933 |
| MVP_top60_DNA_seed | AR | 0.24701 | HNSC | LB771-HNC | AZD3514 | MVP_top60_DNA+RNA_seed | MTOR | -1.768524 | KIRC | BFTC-909 | CRT0105446 |
| MVP_top60_DNA_seed | HDAC1 | 0.247221 | HNSC | LB771-HNC | AR-42 | MVP_top60_DNA+RNA_seed | HSP90AA1 | -1.765769 | KIRC | 786-0 | CCT-018159 |
| MVP_top60_DNA_seed | EGFR | 0.247403 | HNSC | BICR10 | CI-1033 | MVP_top60_DNA+RNA_seed | HSP90AB1 | -1.765769 | KIRC | 786-0 | CCT-018159 |
| MVP_top60_DNA_seed | SMARCA4 | 0.248882 | HNSC | PCI-15A | PFI-3 | MVP_top60_DNA+RNA_seed | PIK3CD | -1.761642 | KIRC | LB2241-RCC | AZD8186 |
| MVP_top60_DNA_seed | MDM2 | 0.2493 | HNSC | PCI-30 | Nutlin-3a (-) | MVP_top60_DNA+RNA_seed | ATM | -1.754731 | KIRC | RCC-FG2 | KU-55933 |
| MVP_top60_DNA_seed | AKT1 | 0.25133 | HNSC | Ca9-22 | BAY AKT1 | MVP_top60_DNA+RNA_seed | MTOR | -1.744366 | KIRC | RCC-MF | Temsirolimus |
| MVP_top60_DNA_seed | AKT1 | 0.252389 | HNSC | BICR22 | GSK690693 | MVP_top60_DNA+RNA_seed | HSP90AA1 | -1.728269 | KIRC | RCC-JF | Elesclomol |
| MVP_top60_DNA_seed | EGFR | 0.252427 | HNSC | JHU-022 | Pelitinib | MVP_top60_DNA+RNA_seed | HSP90AB1 | -1.728269 | KIRC | RCC-JF | Elesclomol |
| MVP_top60_DNA_seed | AKT1 | 0.254528 | HNSC | BICR78 | GSK690693 | MVP_top60_DNA+RNA_seed | PIK3CD | -1.718753 | KIRC | RCC-FG2 | IC-87114 |
| MVP_top60_DNA_seed | EGFR | 0.255118 | HNSC | HO-1-u-1 | Pelitinib | MVP_top60_DNA+RNA_seed | PIK3CD | -1.662756 | KIRC | ACHN | AZD8186 |
| MVP_top60_DNA_seed | HSP90AA1 | 0.256084 | HNSC | DOK | SNX-2112 | MVP_top60_DNA+RNA_seed | FGFR4 | -1.657996 | KIRC | RCC-JW | FGFR_0939 |
| MVP_top60_DNA_seed | AKT1 | 0.256346 | HNSC | LB771-HNC | MK-2206 | MVP_top60_DNA+RNA_seed | PIK3CD | -1.652401 | KIRC | RCC-AB | Idelalisib |
| MVP_top60_DNA_seed | MDM2 | 0.256719 | HNSC | JHU-022 | Nutlin-3a (-) | MVP_top60_DNA+RNA_seed | ATM | -1.652006 | KIRC | LB1047-RCC | KU-60019 |

| | | | | | | | | | | | |
|---|---|---|---|---|---|---|---|---|---|---|---|
| MVP_top60_DNA_seed | NTRK1 | 0.257061 | HNSC | JHU-011 | AZD1332 | MVP_top60_DNA+RNA_seed | PIK3CD | -1.642083 | KIRC | RXF393 | AZD8186 |
| MVP_top60_DNA_seed | HDAC1 | 0.25917 | HNSC | HO-1-N-1 | Tubastatin A | MVP_top60_DNA+RNA_seed | ESR1 | -1.62741 | KIRC | 769-P | Fulvestrant |
| MVP_top60_DNA_seed | MDM2 | 0.261017 | HNSC | CAL-27 | Nutlin-3a (-) | MVP_top60_DNA+RNA_seed | IGF1R | -1.588944 | KIRC | LB996-RCC | GSK1904529A |
| MVP_top60_DNA_seed | SRC | 0.263241 | HNSC | PE-CA-PJ15 | Bosutinib | MVP_top60_DNA+RNA_seed | HSP90AA1 | -1.587905 | KIRC | 786-0 | Luminespib |
| MVP_top60_DNA_seed | CDK2 | 0.267713 | HNSC | BICR22 | PHA-793887 | MVP_top60_DNA+RNA_seed | HSP90AB1 | -1.587905 | KIRC | 786-0 | Luminespib |
| MVP_top60_DNA_seed | HDAC1 | 0.2683 | HNSC | CAL-33 | AR-42 | MVP_top60_DNA+RNA_seed | ATM | -1.586484 | KIRC | U031 | KU-60019 |
| MVP_top60_DNA_seed | EGFR | 0.270589 | HNSC | BICR78 | Pelitinib | MVP_top60_DNA+RNA_seed | PIK3CD | -1.578398 | KIRC | RCC-JW | AZD8186 |
| MVP_top60_DNA_seed | CDK2 | 0.271675 | HNSC | CAL-33 | PHA-793887 | MVP_top60_DNA+RNA_seed | MTOR | -1.571169 | KIRC | 769-P | CRT0105446 |
| MVP_top60_DNA_seed | NTRK1 | 0.271944 | HNSC | SAS | AZD1332 | MVP_top60_DNA+RNA_seed | PIK3CD | -1.531889 | KIRC | CAL-54 | IC-87114 |
| MVP_top60_DNA_seed | AKT1 | 0.272004 | HNSC | LB771-HNC | AKT inhibitor VIII | MVP_top60_DNA+RNA_seed | MTOR | -1.523669 | KIRC | ACHN | CRT0105446 |
| MVP_top60_DNA_seed | HDAC1 | 0.274096 | HNSC | SAS | JQ12 | MVP_top60_DNA+RNA_seed | MTOR | -1.510239 | KIRC | LB996-RCC | JW-7-52-1 |
| MVP_top60_DNA_seed | HDAC1 | 0.274934 | HNSC | SKN-3 | Dacinostat | MVP_top60_DNA+RNA_seed | EP300 | -1.472768 | KIRC | RCC-JW | I-CBP112 |
| MVP_top60_DNA_seed | NTRK1 | 0.275942 | HNSC | SKN-3 | GW441756 | MVP_top60_DNA+RNA_seed | MTOR | -1.469224 | KIRC | HA7-RCC | JW-7-52-1 |
| MVP_top60_DNA_seed | SRC | 0.276312 | HNSC | PCI-4B | NVP-BHG712 | MVP_top60_DNA+RNA_seed | SMARCA4 | -1.468826 | KIRC | U031 | PFI-3 |
| MVP_top60_DNA_seed | MDM2 | 0.27838 | HNSC | OSC-20 | Nutlin-3a (-) | MVP_top60_DNA+RNA_seed | SMARCA2 | -1.468826 | KIRC | U031 | PFI-3 |
| MVP_top60_DNA_seed | NTRK1 | 0.279338 | HNSC | SAS | GW441756 | MVP_top60_DNA+RNA_seed | PIK3CA | -1.467061 | KIRC | LB1047-RCC | AZD8835 |
| MVP_top60_DNA_seed | ESR1 | 0.282281 | HNSC | KON | Fulvestrant | MVP_top60_DNA+RNA_seed | PIK3CD | -1.467061 | KIRC | LB1047-RCC | AZD8835 |
| MVP_top60_DNA_seed | ESR1 | 0.283474 | HNSC | BICR78 | Fulvestrant | MVP_top60_DNA+RNA_seed | MTOR | -1.466339 | KIRC | CAL-54 | CRT0105446 |
| MVP_top60_DNA_seed | EGFR | 0.285051 | HNSC | BICR31 | AST-1306 | MVP_top60_DNA+RNA_seed | MTOR | -1.463518 | KIRC | RCC-AB | Voxtalisib |
| MVP_top60_DNA_seed | AR | 0.285959 | HNSC | BHY | Bicalutamide | MVP_top60_DNA+RNA_seed | MTOR | -1.440285 | KIRC | OS-RC-2 | JW-7-52-1 |
| MVP_top60_DNA_seed | MAPK1 | 0.289206 | HNSC | OSC-19 | VX-11e | MVP_top60_DNA+RNA_seed | MTOR | -1.43107 | KIRC | LB1047-RCC | JW-7-52-1 |
| MVP_top60_DNA_seed | ESR1 | 0.295163 | HNSC | PE-CA-PJ15 | Tamoxifen | MVP_top60_DNA+RNA_seed | AR | -1.426413 | KIRC | LB2241-RCC | AZD3514 |
| MVP_top60_DNA_seed | MDM2 | 0.295168 | HNSC | SCC-15 | Nutlin-3a (-) | MVP_top60_DNA+RNA_seed | PIK3CD | -1.415123 | KIRC | OS-RC-2 | Idelalisib |
| MVP_top60_DNA_seed | AR | 0.295268 | HNSC | SCC-4 | AZD3514 | MVP_top60_DNA+RNA_seed | HDAC1 | -1.396808 | KIRC | 786-0 | JQ12 |
| MVP_top60_DNA_seed | AKT1 | 0.295479 | HNSC | BB49-HNC | MK-2206 | MVP_top60_DNA+RNA_seed | HDAC1 | -1.394774 | KIRC | 786-0 | Dacinostat |
| MVP_top60_DNA_seed | AKT1 | 0.296951 | HNSC | PCI-30 | AKT inhibitor VIII | MVP_top60_DNA+RNA_seed | PIK3CD | -1.383624 | KIRC | KMRC-1 | AZD8186 |
| MVP_top60_DNA_seed | SRC | 0.301505 | HNSC | SKN-3 | NVP-BHG712 | MVP_top60_DNA+RNA_seed | MTOR | -1.377506 | KIRC | RCC-ER | CRT0105446 |
| MVP_top60_DNA_seed | ESR1 | 0.305715 | HNSC | PCI-6A | Tamoxifen | MVP_top60_DNA+RNA_seed | HDAC1 | -1.370527 | KIRC | BB65-RCC | Parthenolide |
| MVP_top60_DNA_seed | HDAC1 | 0.309456 | HNSC | PE-CA-PJ15 | JQ12 | MVP_top60_DNA+RNA_seed | HDAC1 | -1.370414 | KIRC | BB65-RCC | JQ12 |
| MVP_top60_DNA_seed | HDAC1 | 0.3098 | HNSC | OSC-20 | Tubastatin A | MVP_top60_DNA+RNA_seed | HSP90AA1 | -1.368965 | KIRC | RXF393 | CCT-018159 |
| MVP_top60_DNA_seed | NTRK1 | 0.311803 | HNSC | SKN-3 | AZD1332 | MVP_top60_DNA+RNA_seed | HSP90AB1 | -1.368965 | KIRC | RXF393 | CCT-018159 |
| MVP_top60_DNA_seed | SMARCA4 | 0.312848 | HNSC | OSC-19 | PFI-3 | MVP_top60_DNA+RNA_seed | IGF1R | -1.365836 | KIRC | LB1047-RCC | GSK1904529A |
| MVP_top60_DNA_seed | AKT1 | 0.312963 | HNSC | SCC-15 | MK-2206 | MVP_top60_DNA+RNA_seed | MTOR | -1.364338 | KIRC | RXF393 | Temsirolimus |
| MVP_top60_DNA_seed | EGFR | 0.314093 | HNSC | HO-1-N-1 | PF-00299804 | MVP_top60_DNA+RNA_seed | PIK3CD | -1.357428 | KIRC | 769-P | AZD8186 |
| MVP_top60_DNA_seed | NTRK1 | 0.314614 | HNSC | OSC-20 | AZD1332 | MVP_top60_DNA+RNA_seed | AR | -1.327946 | KIRC | CAKI-1 | AZD3514 |
| MVP_top60_DNA_seed | SRC | 0.320398 | HNSC | CAL-27 | NVP-BHG712 | MVP_top60_DNA+RNA_seed | ESR1 | -1.322515 | KIRC | LB2241-RCC | Fulvestrant |
| MVP_top60_DNA_seed | MAPK1 | 0.322778 | HNSC | BICR78 | VX-11e | MVP_top60_DNA+RNA_seed | IGF1R | -1.297166 | KIRC | LB1047-RCC | BMS-536924 |
| MVP_top60_DNA_seed | AR | 0.325984 | HNSC | Detroit562 | Bicalutamide | MVP_top60_DNA+RNA_seed | IGF1R | -1.294234 | KIRC | LB1047-RCC | BMS-536924 |
| MVP_top60_DNA_seed | AKT1 | 0.327363 | HNSC | HO-1-N-1 | AKT inhibitor VIII | MVP_top60_DNA+RNA_seed | HSP90AA1 | -1.293755 | KIRC | RXF393 | Luminespib |
| MVP_top60_DNA_seed | CDK2 | 0.327902 | HNSC | FADU | PHA-793887 | MVP_top60_DNA+RNA_seed | HSP90AB1 | -1.293755 | KIRC | RXF393 | Luminespib |
| MVP_top60_DNA_seed | HDAC1 | 0.327933 | HNSC | HSC-2 | CAY10603 | MVP_top60_DNA+RNA_seed | MTOR | -1.291163 | KIRC | OS-RC-2 | Temsirolimus |

| | | | | | | | | | | | |
|---|---|---|---|---|---|---|---|---|---|---|---|
| MVP_top60_DNA_seed | HDAC1 | 0.32866 | HNSC | BICR10 | JQ12 | MVP_top60_DNA+RNA_seed | ATM | -1.284397 | KIRC | BFTC-909 | KU-55933 |
| MVP_top60_DNA_seed | AR | 0.330896 | HNSC | OSC-20 | AZD3514 | MVP_top60_DNA+RNA_seed | HSP90AA1 | -1.282383 | KIRC | U031 | CCT-018159 |
| MVP_top60_DNA_seed | PIK3CA | 0.333339 | HNSC | SCC90 | AZD8835 | MVP_top60_DNA+RNA_seed | HSP90AB1 | -1.282383 | KIRC | U031 | CCT-018159 |
| MVP_top60_DNA_seed | MDM2 | 0.335878 | HNSC | BICR78 | Nutlin-3a (-) | MVP_top60_DNA+RNA_seed | FGFR4 | -1.281831 | KIRC | HA7-RCC | FGFR_0939 |
| MVP_top60_DNA_seed | EGFR | 0.338073 | HNSC | PCI-38 | Pelitinib | MVP_top60_DNA+RNA_seed | ESR1 | -1.261861 | KIRC | HA7-RCC | Tamoxifen |
| MVP_top60_DNA_seed | AKT1 | 0.338279 | HNSC | PCI-30 | BAY AKT1 | MVP_top60_DNA+RNA_seed | HDAC1 | -1.25374 | KIRC | 769-P | JQ12 |
| MVP_top60_DNA_seed | HDAC1 | 0.338612 | HNSC | BICR31 | JQ12 | MVP_top60_DNA+RNA_seed | HSP90AA1 | -1.238835 | KIRC | RCC-JF | Tanespimycin |
| MVP_top60_DNA_seed | NTRK1 | 0.339011 | HNSC | SCC-4 | GW441756 | MVP_top60_DNA+RNA_seed | HSP90AB1 | -1.238835 | KIRC | RCC-JF | Tanespimycin |
| MVP_top60_DNA_seed | EGFR | 0.340194 | HNSC | Detroit562 | Pelitinib | MVP_top60_DNA+RNA_seed | PIK3CD | -1.237243 | KIRC | RCC-JF | Idelalisib |
| MVP_top60_DNA_seed | MAPK1 | 0.340754 | HNSC | LB771-HNC | FR-180204 | MVP_top60_DNA+RNA_seed | MTOR | -1.236075 | KIRC | RCC-FG2 | Temsirolimus |
| MVP_top60_DNA_seed | MAPK3 | 0.340754 | HNSC | LB771-HNC | FR-180204 | MVP_top60_DNA+RNA_seed | HSP90AA1 | -1.235766 | KIRC | HA7-RCC | Elesclomol |
| MVP_top60_DNA_seed | MAPK1 | 0.341157 | HNSC | PCI-4B | FR-180204 | MVP_top60_DNA+RNA_seed | HSP90AB1 | -1.235766 | KIRC | HA7-RCC | Elesclomol |
| MVP_top60_DNA_seed | MAPK3 | 0.341157 | HNSC | PCI-4B | FR-180204 | MVP_top60_DNA+RNA_seed | PIK3CD | -1.233329 | KIRC | VMRC-RCZ | Idelalisib |
| MVP_top60_DNA_seed | CDK2 | 0.342009 | HNSC | SCC-15 | AZD5438 | MVP_top60_DNA+RNA_seed | HSP90AA1 | -1.232476 | KIRC | LB1047-RCC | Tanespimycin |
| MVP_top60_DNA_seed | SMARCA4 | 0.345183 | HNSC | PCI-38 | PFI-3 | MVP_top60_DNA+RNA_seed | HSP90AB1 | -1.232476 | KIRC | LB1047-RCC | Tanespimycin |
| MVP_top60_DNA_seed | CDK2 | 0.345317 | HNSC | PCI-30 | AZD5438 | MVP_top60_DNA+RNA_seed | HSP90AA1 | -1.228895 | KIRC | RCC-MF | Elesclomol |
| MVP_top60_DNA_seed | HDAC1 | 0.345565 | HNSC | SCC-15 | Tubastatin A | MVP_top60_DNA+RNA_seed | HSP90AB1 | -1.228895 | KIRC | RCC-MF | Elesclomol |
| MVP_top60_DNA_seed | AKT1 | 0.346144 | HNSC | SAS | AKT inhibitor VIII | MVP_top60_DNA+RNA_seed | MTOR | -1.226459 | KIRC | RCC-JW | QL-VIII-58 |
| MVP_top60_DNA_seed | HDAC1 | 0.346734 | HNSC | SCC-25 | JQ12 | MVP_top60_DNA+RNA_seed | ESR1 | -1.215325 | KIRC | BFTC-909 | Tamoxifen |
| MVP_top60_DNA_seed | HDAC1 | 0.3472 | HNSC | HSC-2 | Belinostat | MVP_top60_DNA+RNA_seed | MTOR | -1.211035 | KIRC | LB2241-RCC | CRT0105446 |
| MVP_top60_DNA_seed | HSP90AA1 | 0.3472 | HNSC | BHY | Elesclomol | MVP_top60_DNA+RNA_seed | MTOR | -1.209743 | KIRC | BB65-RCC | JW-7-52-1 |
| MVP_top60_DNA_seed | RAC1 | 0.34734 | HNSC | SCC-15 | EHT-1864 | MVP_top60_DNA+RNA_seed | MTOR | -1.209592 | KIRC | RCC-JF | Temsirolimus |
| MVP_top60_DNA_seed | MAPK1 | 0.349764 | HNSC | HO-1-N-1 | VX-11e | MVP_top60_DNA+RNA_seed | IGF1R | -1.203689 | KIRC | KMRC-20 | GSK1904529A |
| MVP_top60_DNA_seed | HDAC1 | 0.353132 | HNSC | Detroit562 | JQ12 | MVP_top60_DNA+RNA_seed | HDAC1 | -1.192423 | KIRC | RCC-MF | Tubastatin A |
| MVP_top60_DNA_seed | EGFR | 0.35358 | HNSC | SKN-3 | Afatinib | MVP_top60_DNA+RNA_seed | HDAC1 | -1.190828 | KIRC | VMRC-RCW | JQ12 |
| MVP_top60_DNA_seed | HDAC1 | 0.355026 | HNSC | CAL-33 | Dacinostat | MVP_top60_DNA+RNA_seed | ATM | -1.190312 | KIRC | CAL-54 | KU-55933 |
| MVP_top60_DNA_seed | MDM2 | 0.359806 | HNSC | SCC-4 | Nutlin-3a (-) | MVP_top60_DNA+RNA_seed | HSP90AA1 | -1.186982 | KIRC | BFTC-909 | Tanespimycin |
| MVP_top60_DNA_seed | AR | 0.360813 | HNSC | BICR22 | Bicalutamide | MVP_top60_DNA+RNA_seed | HSP90AB1 | -1.186982 | KIRC | BFTC-909 | Tanespimycin |
| MVP_top60_DNA_seed | AKT1 | 0.360968 | HNSC | SCC-15 | AKT inhibitor VIII | MVP_top60_DNA+RNA_seed | HSP90AA1 | -1.18548 | KIRC | KMRC-20 | Elesclomol |
| MVP_top60_DNA_seed | AKT1 | 0.369358 | HNSC | Detroit562 | GSK690693 | MVP_top60_DNA+RNA_seed | HSP90AB1 | -1.18548 | KIRC | KMRC-20 | Elesclomol |
| MVP_top60_DNA_seed | MDM2 | 0.370962 | HNSC | BHY | Nutlin-3a (-) | MVP_top60_DNA+RNA_seed | IGF1R | -1.176876 | KIRC | LB1047-RCC | Linsitinib |
| MVP_top60_DNA_seed | HDAC1 | 0.371942 | HNSC | BICR31 | CAY10603 | MVP_top60_DNA+RNA_seed | IGF1R | -1.170012 | KIRC | LB2241-RCC | BMS-536924 |
| MVP_top60_DNA_seed | AKT1 | 0.371978 | HNSC | Detroit562 | MK-2206 | MVP_top60_DNA+RNA_seed | ATM | -1.168808 | KIRC | LB1047-RCC | CP466722 |
| MVP_top60_DNA_seed | MAPK1 | 0.373049 | HNSC | HSC-2 | VX-11e | MVP_top60_DNA+RNA_seed | ATM | -1.168706 | KIRC | A704 | KU-55933 |
| MVP_top60_DNA_seed | AKT1 | 0.37652 | HNSC | BB49-HNC | AKT inhibitor VIII | MVP_top60_DNA+RNA_seed | ATM | -1.156213 | KIRC | LB1047-RCC | torin2 |
| MVP_top60_DNA_seed | MDM2 | 0.377034 | HNSC | BICR22 | Nutlin-3a (-) | MVP_top60_DNA+RNA_seed | MTOR | -1.156213 | KIRC | LB1047-RCC | torin2 |
| MVP_top60_DNA_seed | HDAC1 | 0.377432 | HNSC | HSC-3 | Belinostat | MVP_top60_DNA+RNA_seed | HSP90AA1 | -1.143307 | KIRC | BFTC-909 | CCT-018159 |
| MVP_top60_DNA_seed | HDAC1 | 0.38317 | HNSC | PE-CA-PJ15 | CAY10603 | MVP_top60_DNA+RNA_seed | HSP90AB1 | -1.143307 | KIRC | BFTC-909 | CCT-018159 |
| MVP_top60_DNA_seed | AKT1 | 0.386636 | HNSC | PCI-15A | AKT inhibitor VIII | MVP_top60_DNA+RNA_seed | IGF1R | -1.137449 | KIRC | LB996-RCC | BMS-536924 |
| MVP_top60_DNA_seed | AKT1 | 0.388784 | HNSC | PCI-4B | AKT inhibitor VIII | MVP_top60_DNA+RNA_seed | ATM | -1.127976 | KIRC | CAKI-1 | CP466722 |
| MVP_top60_DNA_seed | HDAC1 | 0.391133 | HNSC | JHU-022 | AR-42 | MVP_top60_DNA+RNA_seed | HDAC1 | -1.116641 | KIRC | LB2241-RCC | Parthenolide |

| | | | | | | | | | | | |
|---|---|---|---|---|---|---|---|---|---|---|---|
| MVP_top60_DNA_seed | HSP90AA1 | 0.392976 | HNSC | FADU | SNX-2112 | MVP_top60_DNA+RNA_seed | ATM | -1.112846 | KIRC | U031 | KU-55933 |
| MVP_top60_DNA_seed | MAPK1 | 0.392994 | HNSC | LB771-HNC | VX-11e | MVP_top60_DNA+RNA_seed | PIK3CD | -1.11088 | KIRC | VMRC-RCZ | AZD8186 |
| MVP_top60_DNA_seed | EGFR | 0.396502 | HNSC | PCI-15A | PF-00299804 | MVP_top60_DNA+RNA_seed | EP300 | -1.106521 | KIRC | 786-0 | I-CBP112 |
| MVP_top60_DNA_seed | MAPK1 | 0.397661 | HNSC | CAL-33 | VX-11e | MVP_top60_DNA+RNA_seed | PIK3CA | -1.104616 | KIRC | LB2241-RCC | AZD8835 |
| MVP_top60_DNA_seed | EGFR | 0.401214 | HNSC | JHU-022 | PF-00299804 | MVP_top60_DNA+RNA_seed | PIK3CD | -1.104616 | KIRC | LB2241-RCC | AZD8835 |
| MVP_top60_DNA_seed | SMARCA4 | 0.402645 | HNSC | SCC-25 | PFI-3 | MVP_top60_DNA+RNA_seed | IGF1R | -1.090061 | KIRC | HA7-RCC | Linsitinib |
| MVP_top60_DNA_seed | AR | 0.40441 | HNSC | OSC-19 | Bicalutamide | MVP_top60_DNA+RNA_seed | MTOR | -1.079655 | KIRC | SW156 | CRT0105446 |
| MVP_top60_DNA_seed | SRC | 0.407572 | HNSC | HSC-3 | NVP-BHG712 | MVP_top60_DNA+RNA_seed | PIK3CA | -1.073661 | KIRC | BFTC-909 | AZD8835 |
| MVP_top60_DNA_seed | EGFR | 0.408531 | HNSC | OSC-20 | PF-00299804 | MVP_top60_DNA+RNA_seed | PIK3CD | -1.073661 | KIRC | BFTC-909 | AZD8835 |
| MVP_top60_DNA_seed | HDAC1 | 0.410763 | HNSC | HO-1-u-1 | Dacinostat | MVP_top60_DNA+RNA_seed | PIK3CD | -1.067321 | KIRC | SN12C | IC-87114 |
| MVP_top60_DNA_seed | MAPK1 | 0.411234 | HNSC | Detroit562 | FR-180204 | MVP_top60_DNA+RNA_seed | PIK3CD | -1.058185 | KIRC | RCC-MF | AZD8186 |
| MVP_top60_DNA_seed | MAPK3 | 0.411234 | HNSC | Detroit562 | FR-180204 | MVP_top60_DNA+RNA_seed | MTOR | -1.057194 | KIRC | RCC-MF | CRT0105446 |
| MVP_top60_DNA_seed | ESR1 | 0.412301 | HNSC | SCC90 | Fulvestrant | MVP_top60_DNA+RNA_seed | HSP90AA1 | -1.057067 | KIRC | RCC-JF | SNX-2112 |
| MVP_top60_DNA_seed | HDAC1 | 0.412624 | HNSC | BICR22 | Tubastatin A | MVP_top60_DNA+RNA_seed | HSP90AB1 | -1.057067 | KIRC | RCC-JF | SNX-2112 |
| MVP_top60_DNA_seed | HDAC1 | 0.412854 | HNSC | BICR31 | Belinostat | MVP_top60_DNA+RNA_seed | PIK3CD | -1.055819 | KIRC | SN12C | AZD8186 |
| MVP_top60_DNA_seed | CDK2 | 0.412974 | HNSC | PCI-6A | AZD5438 | MVP_top60_DNA+RNA_seed | HDAC1 | -1.053452 | KIRC | LB1047-RCC | JQ12 |
| MVP_top60_DNA_seed | AKT1 | 0.413138 | HNSC | Ca9-22 | GSK690693 | MVP_top60_DNA+RNA_seed | PIK3CD | -1.048886 | KIRC | NCC021 | Idelalisib |
| MVP_top60_DNA_seed | HSP90AA1 | 0.413321 | HNSC | LB771-HNC | SNX-2112 | MVP_top60_DNA+RNA_seed | PIK3CA | -1.042838 | KIRC | LB2241-RCC | PI3Ka_4409 |
| MVP_top60_DNA_seed | HDAC1 | 0.414359 | HNSC | SAS | Dacinostat | MVP_top60_DNA+RNA_seed | PIK3CD | -1.042838 | KIRC | LB2241-RCC | PI3Ka_4409 |
| MVP_top60_DNA_seed | HDAC1 | 0.415596 | HNSC | FADU | Belinostat | MVP_top60_DNA+RNA_seed | MTOR | -1.041572 | KIRC | A704 | CRT0105446 |
| MVP_top60_DNA_seed | EP300 | 0.417527 | HNSC | OSC-19 | I-CBP112 | MVP_top60_DNA+RNA_seed | IGF1R | -1.041409 | KIRC | A498 | BMS-536924 |
| MVP_top60_DNA_seed | CREBBP | 0.417527 | HNSC | OSC-19 | I-CBP112 | MVP_top60_DNA+RNA_seed | HDAC1 | -1.040398 | KIRC | RCC-AB | Belinostat |
| MVP_top60_DNA_seed | HSP90AA1 | 0.418231 | HNSC | SAT | Luminespib | MVP_top60_DNA+RNA_seed | AR | -1.037407 | KIRC | NCC010 | Bicalutamide |
| MVP_top60_DNA_seed | RAC1 | 0.419426 | HNSC | OSC-20 | EHT-1864 | MVP_top60_DNA+RNA_seed | HDAC1 | -1.034082 | KIRC | CAKI-1 | JQ12 |
| MVP_top60_DNA_seed | EGFR | 0.42368 | HNSC | Detroit562 | CUDC-101 | MVP_top60_DNA+RNA_seed | PIK3CD | -1.032167 | KIRC | OS-RC-2 | IC-87114 |
| MVP_top60_DNA_seed | HDAC1 | 0.425733 | HNSC | HO-1-u-1 | JQ12 | MVP_top60_DNA+RNA_seed | MTOR | -1.029474 | KIRC | CAKI-1 | CRT0105446 |
| MVP_top60_DNA_seed | AKT1 | 0.427489 | HNSC | PCI-38 | BAY AKT1 | MVP_top60_DNA+RNA_seed | AR | -1.028907 | KIRC | LB1047-RCC | Bicalutamide |
| MVP_top60_DNA_seed | SRC | 0.429131 | HNSC | SCC-4 | NVP-BHG712 | MVP_top60_DNA+RNA_seed | PIK3CA | -1.017018 | KIRC | LB1047-RCC | PI3Ka_4409 |
| MVP_top60_DNA_seed | CDK2 | 0.430675 | HNSC | SCC-15 | PHA-793887 | MVP_top60_DNA+RNA_seed | PIK3CD | -1.017018 | KIRC | LB1047-RCC | PI3Ka_4409 |
| MVP_top60_DNA_seed | SMARCA4 | 0.430925 | HNSC | JHU-022 | PFI-3 | MVP_top60_DNA+RNA_seed | PIK3CD | -1.001522 | KIRC | 786-0 | Idelalisib |
| MVP_top60_DNA_seed | AKT1 | 0.436366 | HNSC | DOK | AKT inhibitor VIII | MVP_top60_DNA+RNA_seed | HDAC1 | -0.994458 | KIRC | RCC-AB | CAY10603 |
| MVP_top60_DNA_seed | MAPK1 | 0.438575 | HNSC | HO-1-u-1 | VX-11e | MVP_top60_DNA+RNA_seed | HSP90AA1 | -0.990287 | KIRC | BB65-RCC | Tanespimycin |
| MVP_top60_DNA_seed | ESR1 | 0.439655 | HNSC | LB771-HNC | Tamoxifen | MVP_top60_DNA+RNA_seed | HSP90AB1 | -0.990287 | KIRC | BB65-RCC | Tanespimycin |
| MVP_top60_DNA_seed | AR | 0.441794 | HNSC | BICR78 | Bicalutamide | MVP_top60_DNA+RNA_seed | PTK2 | -0.990199 | KIRC | 786-0 | PF-562271 |
| MVP_top60_DNA_seed | HSP90AA1 | 0.442062 | HNSC | SCC-15 | Tanespimycin | MVP_top60_DNA+RNA_seed | PIK3CD | -0.986495 | KIRC | NCC021 | IC-87114 |
| MVP_top60_DNA_seed | ESR1 | 0.442082 | HNSC | PCI-30 | Fulvestrant | MVP_top60_DNA+RNA_seed | MTOR | -0.980109 | KIRC | RCC-JW | CRT0105446 |
| MVP_top60_DNA_seed | NTRK1 | 0.445766 | HNSC | Ca9-22 | GW441756 | MVP_top60_DNA+RNA_seed | IGF1R | -0.979638 | KIRC | BB65-RCC | Linsitinib |
| MVP_top60_DNA_seed | HDAC1 | 0.448329 | HNSC | SCC-9 | Tubastatin A | MVP_top60_DNA+RNA_seed | MTOR | -0.975483 | KIRC | BB65-RCC | CRT0105446 |
| MVP_top60_DNA_seed | HSP90AA1 | 0.448735 | HNSC | HSC-2 | Elesclomol | MVP_top60_DNA+RNA_seed | MTOR | -0.974448 | KIRC | KMRC-20 | CRT0105446 |
| MVP_top60_DNA_seed | ESR1 | 0.450207 | HNSC | PCI-15A | Tamoxifen | MVP_top60_DNA+RNA_seed | ATM | -0.972057 | KIRC | LB1047-RCC | KU-55933 |
| MVP_top60_DNA_seed | HDAC1 | 0.453616 | HNSC | OSC-19 | Tubastatin A | MVP_top60_DNA+RNA_seed | MTOR | -0.971259 | KIRC | 769-P | Torin 2 |

| | | | | | | | | | | | |
|---|---|---|---|---|---|---|---|---|---|---|---|
| MVP_top60_DNA_seed | HDAC1 | 0.45571 | HNSC | HSC-3 | CAY10603 | MVP_top60_DNA+RNA_seed | ATM | -0.970238 | KIRC | NCC021 | KU-55933 |
| MVP_top60_DNA_seed | AKT1 | 0.460701 | HNSC | HSC-3 | GSK690693 | MVP_top60_DNA+RNA_seed | SMARCA4 | -0.967205 | KIRC | HA7-RCC | PFI-3 |
| MVP_top60_DNA_seed | AR | 0.464387 | HNSC | SCC-4 | Bicalutamide | MVP_top60_DNA+RNA_seed | SMARCA2 | -0.967205 | KIRC | HA7-RCC | PFI-3 |
| MVP_top60_DNA_seed | ESR1 | 0.464862 | HNSC | BHY | Fulvestrant | MVP_top60_DNA+RNA_seed | MTOR | -0.966708 | KIRC | TK10 | CRT0105446 |
| MVP_top60_DNA_seed | MAPK1 | 0.465582 | HNSC | PE-CA-PJ15 | FR-180204 | MVP_top60_DNA+RNA_seed | SMARCA4 | -0.964557 | KIRC | SN12C | PFI-3 |
| MVP_top60_DNA_seed | MAPK3 | 0.465582 | HNSC | PE-CA-PJ15 | FR-180204 | MVP_top60_DNA+RNA_seed | SMARCA2 | -0.964557 | KIRC | SN12C | PFI-3 |
| MVP_top60_DNA_seed | MDM2 | 0.467935 | HNSC | SKN-3 | Nutlin-3a (-) | MVP_top60_DNA+RNA_seed | IGF1R | -0.964275 | KIRC | LB996-RCC | BMS-536924 |
| MVP_top60_DNA_seed | AKT1 | 0.471588 | HNSC | SKN-3 | MK-2206 | MVP_top60_DNA+RNA_seed | PIK3CD | -0.962027 | KIRC | U031 | IC-87114 |
| MVP_top60_DNA_seed | MAPK1 | 0.472357 | HNSC | HSC-4 | FR-180204 | MVP_top60_DNA+RNA_seed | PIK3CD | -0.959565 | KIRC | NCC010 | AZD8186 |
| MVP_top60_DNA_seed | MAPK3 | 0.472357 | HNSC | HSC-4 | FR-180204 | MVP_top60_DNA+RNA_seed | ATM | -0.957689 | KIRC | RCC-FG2 | KU-60019 |
| MVP_top60_DNA_seed | NTRK1 | 0.472405 | HNSC | HSC-2 | GW441756 | MVP_top60_DNA+RNA_seed | ATM | -0.956909 | KIRC | 769-P | KU-60019 |
| MVP_top60_DNA_seed | HDAC1 | 0.472545 | HNSC | HSC-4 | Belinostat | MVP_top60_DNA+RNA_seed | FGFR4 | -0.9546 | KIRC | LB1047-RCC | FGFR_0939 |
| MVP_top60_DNA_seed | HSP90AA1 | 0.473562 | HNSC | HSC-4 | Luminespib | MVP_top60_DNA+RNA_seed | HSP90AA1 | -0.951836 | KIRC | 769-P | Luminespib |
| MVP_top60_DNA_seed | HDAC1 | 0.473874 | HNSC | SAT | Tubastatin A | MVP_top60_DNA+RNA_seed | HSP90AB1 | -0.951836 | KIRC | 769-P | Luminespib |
| MVP_top60_DNA_seed | EGFR | 0.473966 | HNSC | Detroit562 | Afatinib | MVP_top60_DNA+RNA_seed | HSP90AA1 | -0.943485 | KIRC | RCC-FG2 | Elesclomol |
| MVP_top60_DNA_seed | MAPK1 | 0.474022 | HNSC | OSC-20 | VX-11e | MVP_top60_DNA+RNA_seed | HSP90AB1 | -0.943485 | KIRC | RCC-FG2 | Elesclomol |
| MVP_top60_DNA_seed | MAPK1 | 0.474244 | HNSC | OSC-20 | FR-180204 | MVP_top60_DNA+RNA_seed | PTK2 | -0.940364 | KIRC | CAKI-1 | PF-562271 |
| MVP_top60_DNA_seed | MAPK3 | 0.474244 | HNSC | OSC-20 | FR-180204 | MVP_top60_DNA+RNA_seed | HSP90AA1 | -0.933335 | KIRC | RCC-JW | Elesclomol |
| MVP_top60_DNA_seed | HSP90AA1 | 0.475059 | HNSC | SKN-3 | CCT-018159 | MVP_top60_DNA+RNA_seed | HSP90AB1 | -0.933335 | KIRC | RCC-JW | Elesclomol |
| MVP_top60_DNA_seed | EP300 | 0.475526 | HNSC | PE-CA-PJ15 | I-CBP112 | MVP_top60_DNA+RNA_seed | PIK3CA | -0.932926 | KIRC | U031 | AZD8835 |
| MVP_top60_DNA_seed | CREBBP | 0.475526 | HNSC | PE-CA-PJ15 | I-CBP112 | MVP_top60_DNA+RNA_seed | PIK3CD | -0.932926 | KIRC | U031 | AZD8835 |
| MVP_top60_DNA_seed | SRC | 0.479404 | HNSC | LB771-HNC | HG-6-71-01 | MVP_top60_DNA+RNA_seed | PIK3CD | -0.930707 | KIRC | RXF393 | Idelalisib |
| MVP_top60_DNA_seed | HSP90AA1 | 0.481377 | HNSC | OSC-20 | Elesclomol | MVP_top60_DNA+RNA_seed | HSP90AA1 | -0.930451 | KIRC | SN12C | Tanespimycin |
| MVP_top60_DNA_seed | EGFR | 0.484314 | HNSC | OSC-19 | PF-00299804 | MVP_top60_DNA+RNA_seed | HSP90AB1 | -0.930451 | KIRC | SN12C | Tanespimycin |
| MVP_top60_DNA_seed | MAPK1 | 0.485402 | HNSC | FADU | FR-180204 | MVP_top60_DNA+RNA_seed | IGF1R | -0.928467 | KIRC | RCC10RGB | Linsitinib |
| MVP_top60_DNA_seed | MAPK3 | 0.485402 | HNSC | FADU | FR-180204 | MVP_top60_DNA+RNA_seed | HSP90AA1 | -0.921568 | KIRC | LB1047-RCC | Elesclomol |
| MVP_top60_DNA_seed | AR | 0.48683 | HNSC | BICR10 | Bicalutamide | MVP_top60_DNA+RNA_seed | HSP90AB1 | -0.921568 | KIRC | LB1047-RCC | Elesclomol |
| MVP_top60_DNA_seed | AKT1 | 0.487295 | HNSC | BHY | BAY AKT1 | MVP_top60_DNA+RNA_seed | ESR1 | -0.921313 | KIRC | KMRC-1 | Tamoxifen |
| MVP_top60_DNA_seed | HDAC1 | 0.487509 | HNSC | HSC-3 | Tubastatin A | MVP_top60_DNA+RNA_seed | FGFR4 | -0.919515 | KIRC | RCC-AB | FGFR_0939 |
| MVP_top60_DNA_seed | HSP90AA1 | 0.488027 | HNSC | SKN-3 | Elesclomol | MVP_top60_DNA+RNA_seed | ESR1 | -0.90322 | KIRC | CAKI-1 | Tamoxifen |
| MVP_top60_DNA_seed | EP300 | 0.488521 | HNSC | SCC-25 | I-CBP112 | MVP_top60_DNA+RNA_seed | FGFR4 | -0.892812 | KIRC | 786-0 | FGFR_0939 |
| MVP_top60_DNA_seed | CREBBP | 0.488521 | HNSC | SCC-25 | I-CBP112 | MVP_top60_DNA+RNA_seed | PIK3CD | -0.892059 | KIRC | U031 | Idelalisib |
| MVP_top60_DNA_seed | HSP90AA1 | 0.488698 | HNSC | PE-CA-PJ15 | Luminespib | MVP_top60_DNA+RNA_seed | HSP90AA1 | -0.891472 | KIRC | 769-P | Elesclomol |
| MVP_top60_DNA_seed | HDAC1 | 0.489609 | HNSC | BICR22 | Belinostat | MVP_top60_DNA+RNA_seed | HSP90AB1 | -0.891472 | KIRC | 769-P | Elesclomol |
| MVP_top60_DNA_seed | SMARCA4 | 0.492434 | HNSC | PE-CA-PJ15 | PFI-3 | MVP_top60_DNA+RNA_seed | AR | -0.889988 | KIRC | A704 | Bicalutamide |
| MVP_top60_DNA_seed | HDAC1 | 0.494282 | HNSC | SAT | Dacinostat | MVP_top60_DNA+RNA_seed | ESR1 | -0.886543 | KIRC | BFTC-909 | Fulvestrant |
| MVP_top60_DNA_seed | HSP90AA1 | 0.502566 | HNSC | HSC-4 | SNX-2112 | MVP_top60_DNA+RNA_seed | MTOR | -0.881007 | KIRC | RXF393 | JW-7-52-1 |
| MVP_top60_DNA_seed | MAPK1 | 0.50351 | HNSC | SCC-4 | FR-180204 | MVP_top60_DNA+RNA_seed | IGF1R | -0.87944 | KIRC | A704 | BMS-536924 |
| MVP_top60_DNA_seed | MAPK3 | 0.50351 | HNSC | SCC-4 | FR-180204 | MVP_top60_DNA+RNA_seed | ATM | -0.876664 | KIRC | RCC-JW | CP466722 |
| MVP_top60_DNA_seed | MDM2 | 0.507673 | HNSC | PCI-30 | Serdemetan | MVP_top60_DNA+RNA_seed | MTOR | -0.87654 | KIRC | RCC-JF | WYE-125132 |
| MVP_top60_DNA_seed | MAPK1 | 0.508595 | HNSC | KON | FR-180204 | MVP_top60_DNA+RNA_seed | FGFR4 | -0.875187 | KIRC | A704 | FGFR_0939 |

| | | | | | | | | | | | |
|---|---|---|---|---|---|---|---|---|---|---|---|
| MVP_top60_DNA_seed | MAPK3 | 0.508595 | HNSC | KON | FR-180204 | MVP_top60_DNA+RNA_seed | IGF1R | -0.874576 | KIRC | ACHN | GSK1904529A |
| MVP_top60_DNA_seed | HDAC1 | 0.510655 | HNSC | PE-CA-PJ15 | AR-42 | MVP_top60_DNA+RNA_seed | HSP90AA1 | -0.873516 | KIRC | RXF393 | Tanespimycin |
| MVP_top60_DNA_seed | HSP90AA1 | 0.518023 | HNSC | PE-CA-PJ15 | SNX-2112 | MVP_top60_DNA+RNA_seed | HSP90AB1 | -0.873516 | KIRC | RXF393 | Tanespimycin |
| MVP_top60_DNA_seed | MAPK1 | 0.522897 | HNSC | JHU-022 | VX-11e | MVP_top60_DNA+RNA_seed | HSP90AA1 | -0.871823 | KIRC | VMRC-RCZ | Luminespib |
| MVP_top60_DNA_seed | NTRK1 | 0.523003 | HNSC | SCC-4 | AZD1332 | MVP_top60_DNA+RNA_seed | HSP90AB1 | -0.871823 | KIRC | VMRC-RCZ | Luminespib |
| MVP_top60_DNA_seed | MDM2 | 0.523603 | HNSC | CAL-27 | Serdemetan | MVP_top60_DNA+RNA_seed | HDAC1 | -0.870557 | KIRC | TK10 | Parthenolide |
| MVP_top60_DNA_seed | MAPK1 | 0.525892 | HNSC | HSC-3 | VX-11e | MVP_top60_DNA+RNA_seed | HSP90AA1 | -0.86888 | KIRC | LB2241-RCC | Tanespimycin |
| MVP_top60_DNA_seed | AKT1 | 0.527184 | HNSC | CAL-27 | AKT inhibitor VIII | MVP_top60_DNA+RNA_seed | HSP90AB1 | -0.86888 | KIRC | LB2241-RCC | Tanespimycin |
| MVP_top60_DNA_seed | HDAC1 | 0.527502 | HNSC | HSC-4 | CAY10603 | MVP_top60_DNA+RNA_seed | ATM | -0.867323 | KIRC | RCC-MF | torin2 |
| MVP_top60_DNA_seed | EGFR | 0.527682 | HNSC | Detroit562 | CI-1033 | MVP_top60_DNA+RNA_seed | MTOR | -0.867323 | KIRC | RCC-MF | torin2 |
| MVP_top60_DNA_seed | ESR1 | 0.528035 | HNSC | PCI-38 | Tamoxifen | MVP_top60_DNA+RNA_seed | HDAC1 | -0.861044 | KIRC | RCC-JF | CAY10603 |
| MVP_top60_DNA_seed | SRC | 0.528273 | HNSC | HSC-2 | NVP-BHG712 | MVP_top60_DNA+RNA_seed | MTOR | -0.857557 | KIRC | LB2241-RCC | Temsirolimus |
| MVP_top60_DNA_seed | HDAC1 | 0.529051 | HNSC | BICR31 | AR-42 | MVP_top60_DNA+RNA_seed | ESR1 | -0.855608 | KIRC | 786-0 | Fulvestrant |
| MVP_top60_DNA_seed | AKT1 | 0.530778 | HNSC | HSC-2 | GSK690693 | MVP_top60_DNA+RNA_seed | PIK3CD | -0.852974 | KIRC | LB2241-RCC | IC-87114 |
| MVP_top60_DNA_seed | EGFR | 0.531159 | HNSC | Detroit562 | Cetuximab | MVP_top60_DNA+RNA_seed | HSP90AA1 | -0.852251 | KIRC | TK10 | Luminespib |
| MVP_top60_DNA_seed | CDK2 | 0.536905 | HNSC | SCC-4 | AZD5438 | MVP_top60_DNA+RNA_seed | HSP90AB1 | -0.852251 | KIRC | TK10 | Luminespib |
| MVP_top60_DNA_seed | AKT1 | 0.540341 | HNSC | SCC-15 | GSK690693 | MVP_top60_DNA+RNA_seed | PIK3CD | -0.846935 | KIRC | RCC-JW | Idelalisib |
| MVP_top60_DNA_seed | AR | 0.542335 | HNSC | SAS | Bicalutamide | MVP_top60_DNA+RNA_seed | ATM | -0.843736 | KIRC | RCC-JF | torin2 |
| MVP_top60_DNA_seed | NTRK1 | 0.548108 | HNSC | BB30-HNC | GW441756 | MVP_top60_DNA+RNA_seed | MTOR | -0.843736 | KIRC | RCC-JF | torin2 |
| MVP_top60_DNA_seed | HSP90AA1 | 0.549907 | HNSC | SCC-9 | Elesclomol | MVP_top60_DNA+RNA_seed | MTOR | -0.842286 | KIRC | 769-P | WYE-125132 |
| MVP_top60_DNA_seed | HSP90AA1 | 0.551442 | HNSC | BHY | CCT-018159 | MVP_top60_DNA+RNA_seed | ESR1 | -0.841428 | KIRC | 769-P | Tamoxifen |
| MVP_top60_DNA_seed | AKT1 | 0.551549 | HNSC | PCI-38 | AKT inhibitor VIII | MVP_top60_DNA+RNA_seed | IGF1R | -0.837284 | KIRC | OS-RC-2 | BMS-536924 |
| MVP_top60_DNA_seed | MAPK1 | 0.553548 | HNSC | HSC-4 | VX-11e | MVP_top60_DNA+RNA_seed | ATM | -0.836316 | KIRC | LB2241-RCC | KU-60019 |
| MVP_top60_DNA_seed | AKT1 | 0.555939 | HNSC | HO-1-N-1 | GSK690693 | MVP_top60_DNA+RNA_seed | IGF1R | -0.834593 | KIRC | CAKI-1 | GSK1904529A |
| MVP_top60_DNA_seed | SRC | 0.556337 | HNSC | BB49-HNC | NVP-BHG712 | MVP_top60_DNA+RNA_seed | MTOR | -0.833686 | KIRC | HA7-RCC | CRT0105446 |
| MVP_top60_DNA_seed | ESR1 | 0.557131 | HNSC | JHU-029 | Fulvestrant | MVP_top60_DNA+RNA_seed | ATM | -0.832563 | KIRC | RCC-ER | KU-60019 |
| MVP_top60_DNA_seed | EGFR | 0.559733 | HNSC | BICR22 | PF-00299804 | MVP_top60_DNA+RNA_seed | PTK2 | -0.831707 | KIRC | RCC10RGB | PF-562271 |
| MVP_top60_DNA_seed | EGFR | 0.560161 | HNSC | BICR10 | Pelitinib | MVP_top60_DNA+RNA_seed | HSP90AA1 | -0.828999 | KIRC | TK10 | Tanespimycin |
| MVP_top60_DNA_seed | CDK2 | 0.565534 | HNSC | LB771-HNC | PHA-793887 | MVP_top60_DNA+RNA_seed | HSP90AB1 | -0.828999 | KIRC | TK10 | Tanespimycin |
| MVP_top60_DNA_seed | EGFR | 0.568115 | HNSC | PCI-30 | Afatinib | MVP_top60_DNA+RNA_seed | ATM | -0.828326 | KIRC | 786-0 | CP466722 |
| MVP_top60_DNA_seed | ESR1 | 0.56852 | HNSC | DOK | Fulvestrant | MVP_top60_DNA+RNA_seed | HSP90AA1 | -0.828051 | KIRC | VMRC-RCW | Luminespib |
| MVP_top60_DNA_seed | ESR1 | 0.57055 | HNSC | JHU-011 | Fulvestrant | MVP_top60_DNA+RNA_seed | HSP90AB1 | -0.828051 | KIRC | VMRC-RCW | Luminespib |
| MVP_top60_DNA_seed | SRC | 0.572144 | HNSC | PE-CA-PJ15 | NVP-BHG712 | MVP_top60_DNA+RNA_seed | HSP90AA1 | -0.824943 | KIRC | 786-0 | SNX-2112 |
| MVP_top60_DNA_seed | AKT1 | 0.572664 | HNSC | CAL-27 | GSK690693 | MVP_top60_DNA+RNA_seed | HSP90AB1 | -0.824943 | KIRC | 786-0 | SNX-2112 |
| MVP_top60_DNA_seed | HDAC1 | 0.572982 | HNSC | SCC-15 | Dacinostat | MVP_top60_DNA+RNA_seed | IGF1R | -0.822088 | KIRC | U031 | GSK1904529A |
| MVP_top60_DNA_seed | NTRK1 | 0.573586 | HNSC | PCI-6A | AZD1332 | MVP_top60_DNA+RNA_seed | PIK3CD | -0.821972 | KIRC | SW156 | Idelalisib |
| MVP_top60_DNA_seed | CDK2 | 0.574394 | HNSC | KOSC-2 | PHA-793887 | MVP_top60_DNA+RNA_seed | HDAC1 | -0.821809 | KIRC | HA7-RCC | Dacinostat |
| MVP_top60_DNA_seed | AKT1 | 0.574736 | HNSC | FADU | AKT inhibitor VIII | MVP_top60_DNA+RNA_seed | MTOR | -0.821255 | KIRC | RCC-FG2 | CRT0105446 |
| MVP_top60_DNA_seed | AR | 0.575575 | HNSC | LB771-HNC | Bicalutamide | MVP_top60_DNA+RNA_seed | PIK3CA | -0.820282 | KIRC | LB996-RCC | AZD8835 |
| MVP_top60_DNA_seed | EGFR | 0.577624 | HNSC | BICR10 | Afatinib | MVP_top60_DNA+RNA_seed | PIK3CD | -0.820282 | KIRC | LB996-RCC | AZD8835 |
| MVP_top60_DNA_seed | PIK3CA | 0.58502 | HNSC | PCI-30 | AZD8835 | MVP_top60_DNA+RNA_seed | AR | -0.811472 | KIRC | 769-P | AZD3514 |

| | | | | | | | | | | | |
|---|---|---|---|---|---|---|---|---|---|---|---|
| MVP_top60_DNA_seed | ESR1 | 0.585909 | HNSC | HSC-2 | Fulvestrant | MVP_top60_DNA+RNA_seed | HSP90AA1 | -0.808879 | KIRC | BB65-RCC | Luminespib |
| MVP_top60_DNA_seed | SRC | 0.589626 | HNSC | LB771-HNC | NVP-BHG712 | MVP_top60_DNA+RNA_seed | HSP90AB1 | -0.808879 | KIRC | BB65-RCC | Luminespib |
| MVP_top60_DNA_seed | AKT1 | 0.589787 | HNSC | OSC-19 | AKT inhibitor VIII | MVP_top60_DNA+RNA_seed | HSP90AA1 | -0.803576 | KIRC | 786-0 | Tanespimycin |
| MVP_top60_DNA_seed | CDK2 | 0.590862 | HNSC | DOK | PHA-793887 | MVP_top60_DNA+RNA_seed | HSP90AB1 | -0.803576 | KIRC | 786-0 | Tanespimycin |
| MVP_top60_DNA_seed | NTRK1 | 0.592003 | HNSC | KOSC-2 | GW441756 | MVP_top60_DNA+RNA_seed | MTOR | -0.803544 | KIRC | RCC-JF | Apitolisib |
| MVP_top60_DNA_seed | MDM2 | 0.592815 | HNSC | SCC-9 | Serdemetan | MVP_top60_DNA+RNA_seed | MTOR | -0.802529 | KIRC | NCC021 | Apitolisib |
| MVP_top60_DNA_seed | AKT1 | 0.595697 | HNSC | SAS | AKT inhibitor VIII | MVP_top60_DNA+RNA_seed | MTOR | -0.801307 | KIRC | TK10 | JW-7-52-1 |
| MVP_top60_DNA_seed | AR | 0.599051 | HNSC | JHU-029 | Bicalutamide | MVP_top60_DNA+RNA_seed | IGF1R | -0.799903 | KIRC | LB1047-RCC | BMS-754807 |
| MVP_top60_DNA_seed | HSP90AA1 | 0.599888 | HNSC | BICR78 | Luminespib | MVP_top60_DNA+RNA_seed | HDAC1 | -0.798647 | KIRC | RCC-MF | CAY10603 |
| MVP_top60_DNA_seed | HDAC1 | 0.60058 | HNSC | HSC-3 | AR-42 | MVP_top60_DNA+RNA_seed | HSP90AA1 | -0.796719 | KIRC | A498 | Tanespimycin |
| MVP_top60_DNA_seed | AKT1 | 0.606019 | HNSC | Detroit562 | BAY AKT1 | MVP_top60_DNA+RNA_seed | HSP90AB1 | -0.796719 | KIRC | A498 | Tanespimycin |
| MVP_top60_DNA_seed | EP300 | 0.607355 | HNSC | BICR22 | I-CBP112 | MVP_top60_DNA+RNA_seed | SMARCA4 | -0.79587 | KIRC | 769-P | PFI-3 |
| MVP_top60_DNA_seed | CREBBP | 0.607355 | HNSC | BICR22 | I-CBP112 | MVP_top60_DNA+RNA_seed | SMARCA2 | -0.79587 | KIRC | 769-P | PFI-3 |
| MVP_top60_DNA_seed | HSP90AA1 | 0.607646 | HNSC | LB771-HNC | Elesclomol | MVP_top60_DNA+RNA_seed | EP300 | -0.795086 | KIRC | CAKI-1 | I-CBP112 |
| MVP_top60_DNA_seed | CDK2 | 0.610283 | HNSC | PE-CA-PJ15 | PHA-793887 | MVP_top60_DNA+RNA_seed | MTOR | -0.792318 | KIRC | RCC-JF | Torin 2 |
| MVP_top60_DNA_seed | HDAC1 | 0.612491 | HNSC | LB771-HNC | Tubastatin A | MVP_top60_DNA+RNA_seed | PIK3CD | -0.789079 | KIRC | RCC-FG2 | Idelalisib |
| MVP_top60_DNA_seed | CDK2 | 0.61392 | HNSC | HSC-4 | PHA-793887 | MVP_top60_DNA+RNA_seed | ATM | -0.78383 | KIRC | RCC-JF | KU-55933 |
| MVP_top60_DNA_seed | AKT1 | 0.61415 | HNSC | OSC-19 | GSK690693 | MVP_top60_DNA+RNA_seed | HSP90AA1 | -0.782752 | KIRC | 786-0 | Elesclomol |
| MVP_top60_DNA_seed | HSP90AA1 | 0.614548 | HNSC | SCC-25 | Luminespib | MVP_top60_DNA+RNA_seed | HSP90AB1 | -0.782752 | KIRC | 786-0 | Elesclomol |
| MVP_top60_DNA_seed | CDK2 | 0.616367 | HNSC | HSC-2 | PHA-793887 | MVP_top60_DNA+RNA_seed | HDAC1 | -0.781576 | KIRC | RCC-AB | AR-42 |
| MVP_top60_DNA_seed | MDM2 | 0.617761 | HNSC | HSC-2 | Nutlin-3a (-) | MVP_top60_DNA+RNA_seed | MTOR | -0.780837 | KIRC | A498 | QL-VIII-58 |
| MVP_top60_DNA_seed | AR | 0.618524 | HNSC | SKN-3 | Bicalutamide | MVP_top60_DNA+RNA_seed | HSP90AA1 | -0.779809 | KIRC | RXF393 | Elesclomol |
| MVP_top60_DNA_seed | AKT1 | 0.618563 | HNSC | KON | GSK690693 | MVP_top60_DNA+RNA_seed | HSP90AB1 | -0.779809 | KIRC | RXF393 | Elesclomol |
| MVP_top60_DNA_seed | AKT1 | 0.619215 | HNSC | OSC-20 | MK-2206 | MVP_top60_DNA+RNA_seed | MTOR | -0.777761 | KIRC | 786-0 | Temsirolimus |
| MVP_top60_DNA_seed | AKT1 | 0.623177 | HNSC | HSC-2 | MK-2206 | MVP_top60_DNA+RNA_seed | FGFR4 | -0.775996 | KIRC | NCC010 | FGFR_3831 |
| MVP_top60_DNA_seed | ESR1 | 0.629282 | HNSC | KOSC-2 | Tamoxifen | MVP_top60_DNA+RNA_seed | AR | -0.775798 | KIRC | ACHN | Bicalutamide |
| MVP_top60_DNA_seed | EGFR | 0.630217 | HNSC | BB49-HNC | Afatinib | MVP_top60_DNA+RNA_seed | PIK3CD | -0.769772 | KIRC | LB1047-RCC | AZD8186 |
| MVP_top60_DNA_seed | HDAC1 | 0.63376 | HNSC | KOSC-2 | Tubastatin A | MVP_top60_DNA+RNA_seed | MTOR | -0.759369 | KIRC | RCC-AB | WYE-125132 |
| MVP_top60_DNA_seed | HDAC1 | 0.634034 | HNSC | HSC-2 | Tubastatin A | MVP_top60_DNA+RNA_seed | ATM | -0.756578 | KIRC | CAKI-1 | KU-60019 |
| MVP_top60_DNA_seed | HDAC1 | 0.636258 | HNSC | SCC-9 | Dacinostat | MVP_top60_DNA+RNA_seed | ATM | -0.752802 | KIRC | RCC-MF | KU-60019 |
| MVP_top60_DNA_seed | MAPK1 | 0.640768 | HNSC | SCC-15 | VX-11e | MVP_top60_DNA+RNA_seed | ESR1 | -0.751521 | KIRC | RXF393 | Fulvestrant |
| MVP_top60_DNA_seed | MAPK1 | 0.643407 | HNSC | PCI-15A | FR-180204 | MVP_top60_DNA+RNA_seed | PIK3CD | -0.751263 | KIRC | CAKI-1 | IC-87114 |
| MVP_top60_DNA_seed | MAPK3 | 0.643407 | HNSC | PCI-15A | FR-180204 | MVP_top60_DNA+RNA_seed | FGFR4 | -0.749345 | KIRC | BFTC-909 | FGFR_0939 |
| MVP_top60_DNA_seed | EGFR | 0.644343 | HNSC | BICR10 | Afatinib | MVP_top60_DNA+RNA_seed | IGF1R | -0.74885 | KIRC | RXF393 | BMS-536924 |
| MVP_top60_DNA_seed | AKT1 | 0.646234 | HNSC | SKN-3 | BAY AKT1 | MVP_top60_DNA+RNA_seed | MTOR | -0.747246 | KIRC | CAL-54 | WYE-125132 |
| MVP_top60_DNA_seed | HDAC1 | 0.653125 | HNSC | LB771-HNC | JQ12 | MVP_top60_DNA+RNA_seed | ESR1 | -0.738511 | KIRC | LB1047-RCC | Fulvestrant |
| MVP_top60_DNA_seed | HDAC1 | 0.653596 | HNSC | FADU | Dacinostat | MVP_top60_DNA+RNA_seed | FGFR4 | -0.73585 | KIRC | RCC10RGB | FGFR_0939 |
| MVP_top60_DNA_seed | AR | 0.654142 | HNSC | SCC-9 | Bicalutamide | MVP_top60_DNA+RNA_seed | ATM | -0.735699 | KIRC | RCC-JF | CP466722 |
| MVP_top60_DNA_seed | AKT1 | 0.655031 | HNSC | HSC-2 | BAY AKT1 | MVP_top60_DNA+RNA_seed | IGF1R | -0.734846 | KIRC | LB996-RCC | BMS-754807 |
| MVP_top60_DNA_seed | AKT1 | 0.659337 | HNSC | BB49-HNC | AKT inhibitor VIII | MVP_top60_DNA+RNA_seed | HSP90AA1 | -0.727153 | KIRC | RCC-JW | Tanespimycin |
| MVP_top60_DNA_seed | AKT1 | 0.660384 | HNSC | SCC-4 | GSK690693 | MVP_top60_DNA+RNA_seed | HSP90AB1 | -0.727153 | KIRC | RCC-JW | Tanespimycin |

| | | | | | | | | | | | |
|---|---|---|---|---|---|---|---|---|---|---|---|
| MVP_top60_DNA_seed | EGFR | 0.660686 | HNSC | BB49-HNC | Lapatinib | MVP_top60_DNA+RNA_seed | AR | -0.725522 | KIRC | 786-0 | Bicalutamide |
| MVP_top60_DNA_seed | SMARCA4 | 0.66114 | HNSC | BICR10 | PFI-3 | MVP_top60_DNA+RNA_seed | HDAC1 | -0.725432 | KIRC | RCC-MF | Belinostat |
| MVP_top60_DNA_seed | SRC | 0.66798 | HNSC | Detroit562 | NVP-BHG712 | MVP_top60_DNA+RNA_seed | HDAC1 | -0.724644 | KIRC | OS-RC-2 | Parthenolide |
| MVP_top60_DNA_seed | EGFR | 0.66925 | HNSC | BICR78 | CI-1033 | MVP_top60_DNA+RNA_seed | AR | -0.723897 | KIRC | BFTC-909 | Bicalutamide |
| MVP_top60_DNA_seed | MDM2 | 0.669659 | HNSC | HO-1-u-1 | Nutlin-3a (-) | MVP_top60_DNA+RNA_seed | PTK2 | -0.715194 | KIRC | LB1047-RCC | PF-562271 |
| MVP_top60_DNA_seed | HDAC1 | 0.670165 | HNSC | HSC-2 | AR-42 | MVP_top60_DNA+RNA_seed | FGFR4 | -0.713748 | KIRC | A704 | FGFR_3831 |
| MVP_top60_DNA_seed | EGFR | 0.671439 | HNSC | FADU | CI-1033 | MVP_top60_DNA+RNA_seed | MTOR | -0.713535 | KIRC | LB996-RCC | Voxtalisib |
| MVP_top60_DNA_seed | CDK2 | 0.672741 | HNSC | LB771-HNC | AZD5438 | MVP_top60_DNA+RNA_seed | PIK3CD | -0.707945 | KIRC | HA7-RCC | Idelalisib |
| MVP_top60_DNA_seed | MAPK1 | 0.672748 | HNSC | KON | VX-11e | MVP_top60_DNA+RNA_seed | ESR1 | -0.6988 | KIRC | 786-0 | Tamoxifen |
| MVP_top60_DNA_seed | EGFR | 0.673509 | HNSC | CAL-33 | PF-00299804 | MVP_top60_DNA+RNA_seed | MTOR | -0.698256 | KIRC | SW156 | Temsirolimus |
| MVP_top60_DNA_seed | NTRK1 | 0.675322 | HNSC | SCC-9 | AZD1332 | MVP_top60_DNA+RNA_seed | HSP90AA1 | -0.696204 | KIRC | RCC-ER | Elesclomol |
| MVP_top60_DNA_seed | EP300 | 0.675778 | HNSC | BICR10 | I-CBP112 | MVP_top60_DNA+RNA_seed | HSP90AB1 | -0.696204 | KIRC | RCC-ER | Elesclomol |
| MVP_top60_DNA_seed | CREBBP | 0.675778 | HNSC | BICR10 | I-CBP112 | MVP_top60_DNA+RNA_seed | MTOR | -0.695836 | KIRC | 769-P | Apitolisib |
| MVP_top60_DNA_seed | EGFR | 0.676881 | HNSC | PCI-30 | Gefitinib | MVP_top60_DNA+RNA_seed | PIK3CD | -0.695272 | KIRC | A704 | AZD8186 |
| MVP_top60_DNA_seed | AKT1 | 0.678496 | HNSC | BICR78 | BAY AKT1 | MVP_top60_DNA+RNA_seed | ESR1 | -0.694744 | KIRC | CAL-54 | Tamoxifen |
| MVP_top60_DNA_seed | SMARCA4 | 0.680159 | HNSC | OSC-20 | PFI-3 | MVP_top60_DNA+RNA_seed | IGF1R | -0.694288 | KIRC | SW156 | GSK1904529A |
| MVP_top60_DNA_seed | AKT1 | 0.684338 | HNSC | DOK | GSK690693 | MVP_top60_DNA+RNA_seed | HDAC1 | -0.690676 | KIRC | VMRC-RCW | Dacinostat |
| MVP_top60_DNA_seed | MAPK1 | 0.687608 | HNSC | BICR31 | VX-11e | MVP_top60_DNA+RNA_seed | HDAC1 | -0.689737 | KIRC | RCC-JF | AR-42 |
| MVP_top60_DNA_seed | AKT1 | 0.68826 | HNSC | PCI-4B | BAY AKT1 | MVP_top60_DNA+RNA_seed | HDAC1 | -0.689617 | KIRC | RCC-MF | AR-42 |
| MVP_top60_DNA_seed | SMARCA4 | 0.689036 | HNSC | SAS | PFI-3 | MVP_top60_DNA+RNA_seed | HSP90AA1 | -0.6824 | KIRC | ACHN | Luminespib |
| MVP_top60_DNA_seed | CDK2 | 0.694082 | HNSC | Detroit562 | PHA-793887 | MVP_top60_DNA+RNA_seed | HSP90AB1 | -0.6824 | KIRC | ACHN | Luminespib |
| MVP_top60_DNA_seed | HDAC1 | 0.695798 | HNSC | FADU | AR-42 | MVP_top60_DNA+RNA_seed | PIK3CD | -0.67555 | KIRC | RCC-ER | AZD8186 |
| MVP_top60_DNA_seed | EGFR | 0.697799 | HNSC | SKN-3 | Pelitinib | MVP_top60_DNA+RNA_seed | MTOR | -0.673839 | KIRC | HA7-RCC | WYE-125132 |
| MVP_top60_DNA_seed | HDAC1 | 0.698619 | HNSC | SCC90 | Dacinostat | MVP_top60_DNA+RNA_seed | MTOR | -0.673735 | KIRC | CAL-54 | Apitolisib |
| MVP_top60_DNA_seed | EGFR | 0.698906 | HNSC | SKN-3 | Cetuximab | MVP_top60_DNA+RNA_seed | ATM | -0.672621 | KIRC | RCC-ER | KU-55933 |
| MVP_top60_DNA_seed | AR | 0.700497 | HNSC | SCC90 | Bicalutamide | MVP_top60_DNA+RNA_seed | PIK3CD | -0.670612 | KIRC | TK10 | AZD8186 |
| MVP_top60_DNA_seed | MAPK1 | 0.70088 | HNSC | BICR22 | VX-11e | MVP_top60_DNA+RNA_seed | ATM | -0.668881 | KIRC | RCC-MF | CP466722 |
| MVP_top60_DNA_seed | EGFR | 0.703171 | HNSC | BICR22 | Cetuximab | MVP_top60_DNA+RNA_seed | ATM | -0.668878 | KIRC | SW156 | CP466722 |
| MVP_top60_DNA_seed | RAC1 | 0.704009 | HNSC | PCI-30 | EHT-1864 | MVP_top60_DNA+RNA_seed | HDAC1 | -0.666335 | KIRC | 769-P | AR-42 |
| MVP_top60_DNA_seed | HDAC1 | 0.704891 | HNSC | BICR22 | CAY10603 | MVP_top60_DNA+RNA_seed | HDAC1 | -0.663409 | KIRC | OS-RC-2 | JQ12 |
| MVP_top60_DNA_seed | EGFR | 0.70509 | HNSC | PCI-30 | Afatinib | MVP_top60_DNA+RNA_seed | MTOR | -0.657279 | KIRC | RCC-FG2 | Voxtalisib |
| MVP_top60_DNA_seed | SRC | 0.705121 | HNSC | HO-1-u-1 | NVP-BHG712 | MVP_top60_DNA+RNA_seed | FGFR4 | -0.655322 | KIRC | RCC-FG2 | FGFR_0939 |
| MVP_top60_DNA_seed | HDAC1 | 0.70882 | HNSC | KON | Tubastatin A | MVP_top60_DNA+RNA_seed | ATM | -0.655116 | KIRC | CAL-54 | CP466722 |
| MVP_top60_DNA_seed | AKT1 | 0.710162 | HNSC | HSC-3 | AKT inhibitor VIII | MVP_top60_DNA+RNA_seed | FGFR4 | -0.65324 | KIRC | RXF393 | FGFR_3831 |
| MVP_top60_DNA_seed | MAPK1 | 0.710611 | HNSC | Detroit562 | VX-11e | MVP_top60_DNA+RNA_seed | FGFR4 | -0.652829 | KIRC | LB996-RCC | FGFR_0939 |
| MVP_top60_DNA_seed | SRC | 0.710772 | HNSC | CAL-33 | NVP-BHG712 | MVP_top60_DNA+RNA_seed | PTK2 | -0.651546 | KIRC | KMRC-1 | PF-562271 |
| MVP_top60_DNA_seed | AKT1 | 0.712353 | HNSC | FADU | MK-2206 | MVP_top60_DNA+RNA_seed | HDAC1 | -0.651089 | KIRC | LB1047-RCC | Dacinostat |
| MVP_top60_DNA_seed | SRC | 0.714308 | HNSC | SCC-9 | Bosutinib | MVP_top60_DNA+RNA_seed | ESR1 | -0.649392 | KIRC | A704 | Tamoxifen |
| MVP_top60_DNA_seed | HDAC1 | 0.716151 | HNSC | Detroit562 | Tubastatin A | MVP_top60_DNA+RNA_seed | AR | -0.648355 | KIRC | LB996-RCC | AZD3514 |
| MVP_top60_DNA_seed | HDAC1 | 0.716493 | HNSC | DOK | AR-42 | MVP_top60_DNA+RNA_seed | MTOR | -0.646077 | KIRC | RXF393 | QL-VIII-58 |
| MVP_top60_DNA_seed | CDK2 | 0.717834 | HNSC | JHU-022 | PHA-793887 | MVP_top60_DNA+RNA_seed | HSP90AA1 | -0.644704 | KIRC | U031 | Elesclomol |

| | | | | | | | | | | | |
|---|---|---|---|---|---|---|---|---|---|---|---|
| MVP_top60_DNA_seed | AKT1 | 0.718392 | HNSC | JHU-022 | AKT inhibitor VIII | MVP_top60_DNA+RNA_seed | HSP90AB1 | -0.644704 | KIRC | U031 | Elesclomol |
| MVP_top60_DNA_seed | HDAC1 | 0.718736 | HNSC | LB771-HNC | Dacinostat | MVP_top60_DNA+RNA_seed | ATM | -0.638226 | KIRC | 769-P | CP466722 |
| MVP_top60_DNA_seed | AKT1 | 0.724061 | HNSC | FADU | GSK690693 | MVP_top60_DNA+RNA_seed | ESR1 | -0.638038 | KIRC | LB2241-RCC | Tamoxifen |
| MVP_top60_DNA_seed | HSP90AA1 | 0.728261 | HNSC | CAL-33 | Tanespimycin | MVP_top60_DNA+RNA_seed | ATM | -0.637547 | KIRC | ACHN | CP466722 |
| MVP_top60_DNA_seed | ESR1 | 0.729962 | HNSC | OSC-20 | Fulvestrant | MVP_top60_DNA+RNA_seed | HSP90AA1 | -0.63567 | KIRC | RXF393 | SNX-2112 |
| MVP_top60_DNA_seed | AR | 0.73006 | HNSC | SCC-4 | Bicalutamide | MVP_top60_DNA+RNA_seed | HSP90AB1 | -0.63567 | KIRC | RXF393 | SNX-2112 |
| MVP_top60_DNA_seed | HSP90AA1 | 0.730512 | HNSC | LB771-HNC | Tanespimycin | MVP_top60_DNA+RNA_seed | MTOR | -0.635272 | KIRC | RCC-AB | CRT0105446 |
| MVP_top60_DNA_seed | HSP90AA1 | 0.731027 | HNSC | BICR78 | SNX-2112 | MVP_top60_DNA+RNA_seed | HDAC1 | -0.631559 | KIRC | RCC-JF | Belinostat |
| MVP_top60_DNA_seed | SMARCA4 | 0.731564 | HNSC | SKN-3 | PFI-3 | MVP_top60_DNA+RNA_seed | MTOR | -0.628922 | KIRC | LB1047-RCC | Temsirolimus |
| MVP_top60_DNA_seed | MAPK1 | 0.73288 | HNSC | KOSC-2 | VX-11e | MVP_top60_DNA+RNA_seed | IGF1R | -0.61667 | KIRC | TK10 | BMS-536924 |
| MVP_top60_DNA_seed | EP300 | 0.732923 | HNSC | SCC-4 | I-CBP112 | MVP_top60_DNA+RNA_seed | ATM | -0.616643 | KIRC | VMRC-RCZ | CP466722 |
| MVP_top60_DNA_seed | CREBBP | 0.732923 | HNSC | SCC-4 | I-CBP112 | MVP_top60_DNA+RNA_seed | MTOR | -0.615934 | KIRC | U031 | WYE-125132 |
| MVP_top60_DNA_seed | SRC | 0.733287 | HNSC | SCC-15 | NVP-BHG712 | MVP_top60_DNA+RNA_seed | ESR1 | -0.615863 | KIRC | U031 | Fulvestrant |
| MVP_top60_DNA_seed | CDK2 | 0.733636 | HNSC | BICR78 | PHA-793887 | MVP_top60_DNA+RNA_seed | ATM | -0.61374 | KIRC | RCC-AB | CP466722 |
| MVP_top60_DNA_seed | ESR1 | 0.734174 | HNSC | SCC-4 | Tamoxifen | MVP_top60_DNA+RNA_seed | MTOR | -0.610969 | KIRC | CAL-54 | Voxtalisib |
| MVP_top60_DNA_seed | EGFR | 0.734671 | HNSC | PCI-30 | Pelitinib | MVP_top60_DNA+RNA_seed | FGFR4 | -0.609385 | KIRC | RCC-JF | FGFR_0939 |
| MVP_top60_DNA_seed | EP300 | 0.735199 | HNSC | OSC-20 | I-CBP112 | MVP_top60_DNA+RNA_seed | MTOR | -0.606272 | KIRC | VMRC-RCZ | Torin 2 |
| MVP_top60_DNA_seed | CREBBP | 0.735199 | HNSC | OSC-20 | I-CBP112 | MVP_top60_DNA+RNA_seed | MTOR | -0.605071 | KIRC | A704 | Voxtalisib |
| MVP_top60_DNA_seed | AR | 0.73679 | HNSC | BICR78 | Bicalutamide | MVP_top60_DNA+RNA_seed | PIK3CD | -0.604726 | KIRC | BB65-RCC | Idelalisib |
| MVP_top60_DNA_seed | HDAC1 | 0.738394 | HNSC | FADU | CAY10603 | MVP_top60_DNA+RNA_seed | HDAC1 | -0.601798 | KIRC | 769-P | Dacinostat |
| MVP_top60_DNA_seed | EP300 | 0.739014 | HNSC | JHU-029 | I-CBP112 | MVP_top60_DNA+RNA_seed | HSP90AA1 | -0.601193 | KIRC | VMRC-RCZ | Tanespimycin |
| MVP_top60_DNA_seed | CREBBP | 0.739014 | HNSC | JHU-029 | I-CBP112 | MVP_top60_DNA+RNA_seed | HSP90AB1 | -0.601193 | KIRC | VMRC-RCZ | Tanespimycin |
| MVP_top60_DNA_seed | AKT1 | 0.741573 | HNSC | PE-CA-PJ15 | GSK690693 | MVP_top60_DNA+RNA_seed | MTOR | -0.600449 | KIRC | 786-0 | WYE-125132 |
| MVP_top60_DNA_seed | MDM2 | 0.743198 | HNSC | LB771-HNC | Serdemetan | MVP_top60_DNA+RNA_seed | MTOR | -0.599065 | KIRC | A704 | JW-7-52-1 |
| MVP_top60_DNA_seed | MAPK1 | 0.743655 | HNSC | PCI-38 | FR-180204 | MVP_top60_DNA+RNA_seed | MTOR | -0.599004 | KIRC | RCC-AB | Torin 2 |
| MVP_top60_DNA_seed | MAPK3 | 0.743655 | HNSC | PCI-38 | FR-180204 | MVP_top60_DNA+RNA_seed | HDAC1 | -0.597087 | KIRC | OS-RC-2 | Entinostat |
| MVP_top60_DNA_seed | AKT1 | 0.745301 | HNSC | SCC-15 | A-443654 | MVP_top60_DNA+RNA_seed | HDAC1 | -0.595429 | KIRC | 769-P | CAY10603 |
| MVP_top60_DNA_seed | NTRK1 | 0.74652 | HNSC | OSC-19 | GW441756 | MVP_top60_DNA+RNA_seed | PIK3CD | -0.59431 | KIRC | LB1047-RCC | Idelalisib |
| MVP_top60_DNA_seed | EGFR | 0.749982 | HNSC | SCC-15 | Lapatinib | MVP_top60_DNA+RNA_seed | PIK3CD | -0.591978 | KIRC | 786-0 | IC-87114 |
| MVP_top60_DNA_seed | EGFR | 0.760552 | HNSC | SCC-4 | Pelitinib | MVP_top60_DNA+RNA_seed | AR | -0.591923 | KIRC | HA7-RCC | AZD3514 |
| MVP_top60_DNA_seed | MAPK1 | 0.761112 | HNSC | FADU | VX-11e | MVP_top60_DNA+RNA_seed | HDAC1 | -0.583668 | KIRC | 769-P | Tubastatin A |
| MVP_top60_DNA_seed | HDAC1 | 0.762836 | HNSC | FADU | Tubastatin A | MVP_top60_DNA+RNA_seed | MTOR | -0.57318 | KIRC | RCC-AB | Apitolisib |
| MVP_top60_DNA_seed | CDK2 | 0.763781 | HNSC | BICR10 | AZD5438 | MVP_top60_DNA+RNA_seed | ESR1 | -0.568046 | KIRC | BB65-RCC | Tamoxifen |
| MVP_top60_DNA_seed | HSP90AA1 | 0.770493 | HNSC | BB49-HNC | Elesclomol | MVP_top60_DNA+RNA_seed | HDAC1 | -0.566739 | KIRC | RXF393 | Belinostat |
| MVP_top60_DNA_seed | SMARCA4 | 0.770944 | HNSC | LB771-HNC | PFI-3 | MVP_top60_DNA+RNA_seed | PTK2 | -0.564299 | KIRC | RXF393 | PF-562271 |
| MVP_top60_DNA_seed | HDAC1 | 0.772902 | HNSC | SCC-15 | JQ12 | MVP_top60_DNA+RNA_seed | PIK3CD | -0.561983 | KIRC | RCC-ER | IC-87114 |
| MVP_top60_DNA_seed | HSP90AA1 | 0.773874 | HNSC | PCI-30 | Luminespib | MVP_top60_DNA+RNA_seed | MTOR | -0.561634 | KIRC | RCC-FG2 | Apitolisib |
| MVP_top60_DNA_seed | CDK2 | 0.77627 | HNSC | SCC-9 | AZD5438 | MVP_top60_DNA+RNA_seed | IGF1R | -0.558891 | KIRC | RCC-ER | BMS-536924 |
| MVP_top60_DNA_seed | AKT1 | 0.777565 | HNSC | HSC-4 | AKT inhibitor VIII | MVP_top60_DNA+RNA_seed | AR | -0.556415 | KIRC | CAL-54 | AZD3514 |
| MVP_top60_DNA_seed | HDAC1 | 0.778143 | HNSC | CAL-33 | Tubastatin A | MVP_top60_DNA+RNA_seed | HSP90AA1 | -0.553513 | KIRC | KMRC-20 | Tanespimycin |
| MVP_top60_DNA_seed | EGFR | 0.785271 | HNSC | BICR10 | AST-1306 | MVP_top60_DNA+RNA_seed | HSP90AB1 | -0.553513 | KIRC | KMRC-20 | Tanespimycin |

| | | | | | | | | | | | |
|---|---|---|---|---|---|---|---|---|---|---|---|
| MVP_top60_DNA_seed | HSP90AA1 | 0.78784 | HNSC | SCC90 | CCT-018159 | MVP_top60_DNA+RNA_seed | HDAC1 | -0.550534 | KIRC | 769-P | Belinostat |
| MVP_top60_DNA_seed | AKT1 | 0.790616 | HNSC | BICR31 | GSK690693 | MVP_top60_DNA+RNA_seed | AR | -0.550534 | KIRC | RCC-AB | AZD3514 |
| MVP_top60_DNA_seed | EGFR | 0.790688 | HNSC | HSC-3 | PF-00299804 | MVP_top60_DNA+RNA_seed | IGF1R | -0.550316 | KIRC | A704 | GSK1904529A |
| MVP_top60_DNA_seed | CDK2 | 0.793025 | HNSC | SCC-25 | AZD5438 | MVP_top60_DNA+RNA_seed | PIK3CA | -0.549567 | KIRC | RCC-MF | AZD8835 |
| MVP_top60_DNA_seed | HDAC1 | 0.794715 | HNSC | Detroit562 | AR-42 | MVP_top60_DNA+RNA_seed | PIK3CD | -0.549567 | KIRC | RCC-MF | AZD8835 |
| MVP_top60_DNA_seed | RAC1 | 0.796955 | HNSC | OSC-19 | EHT-1864 | MVP_top60_DNA+RNA_seed | ATM | -0.547212 | KIRC | VMRC-RCZ | torin2 |
| MVP_top60_DNA_seed | NTRK1 | 0.798526 | HNSC | SCC-25 | GW441756 | MVP_top60_DNA+RNA_seed | MTOR | -0.547212 | KIRC | VMRC-RCZ | torin2 |
| MVP_top60_DNA_seed | SRC | 0.801594 | HNSC | HO-1-N-1 | NVP-BHG712 | MVP_top60_DNA+RNA_seed | ESR1 | -0.546801 | KIRC | RCC-JF | Fulvestrant |
| MVP_top60_DNA_seed | HDAC1 | 0.80286 | HNSC | SAS | Tubastatin A | MVP_top60_DNA+RNA_seed | PIK3CD | -0.546633 | KIRC | SW156 | IC-87114 |
| MVP_top60_DNA_seed | HDAC1 | 0.804958 | HNSC | BB49-HNC | Belinostat | MVP_top60_DNA+RNA_seed | HDAC1 | -0.544592 | KIRC | NCC021 | Belinostat |
| MVP_top60_DNA_seed | MDM2 | 0.805626 | HNSC | Detroit562 | Nutlin-3a (-) | MVP_top60_DNA+RNA_seed | ATM | -0.540539 | KIRC | BFTC-909 | KU-60019 |
| MVP_top60_DNA_seed | HDAC1 | 0.80673 | HNSC | PCI-15A | Tubastatin A | MVP_top60_DNA+RNA_seed | HDAC1 | -0.538626 | KIRC | LB1047-RCC | AR-42 |
| MVP_top60_DNA_seed | ESR1 | 0.809425 | HNSC | SAS | Tamoxifen | MVP_top60_DNA+RNA_seed | HDAC1 | -0.536125 | KIRC | NCC021 | CAY10603 |
| MVP_top60_DNA_seed | CDK2 | 0.814611 | HNSC | BICR31 | PHA-793887 | MVP_top60_DNA+RNA_seed | EP300 | -0.532319 | KIRC | LB2241-RCC | I-CBP112 |
| MVP_top60_DNA_seed | HDAC1 | 0.816963 | HNSC | LB771-HNC | CAY10603 | MVP_top60_DNA+RNA_seed | MTOR | -0.530932 | KIRC | RCC-FG2 | WYE-125132 |
| MVP_top60_DNA_seed | HDAC1 | 0.820585 | HNSC | HSC-4 | AR-42 | MVP_top60_DNA+RNA_seed | HDAC1 | -0.526125 | KIRC | LB1047-RCC | CAY10603 |
| MVP_top60_DNA_seed | AKT1 | 0.822236 | HNSC | HSC-3 | MK-2206 | MVP_top60_DNA+RNA_seed | PIK3CD | -0.526093 | KIRC | A704 | Idelalisib |
| MVP_top60_DNA_seed | HDAC1 | 0.824822 | HNSC | HSC-4 | Dacinostat | MVP_top60_DNA+RNA_seed | EP300 | -0.522934 | KIRC | RCC-ER | I-CBP112 |
| MVP_top60_DNA_seed | EGFR | 0.824914 | HNSC | HSC-2 | PF-00299804 | MVP_top60_DNA+RNA_seed | HDAC1 | -0.517879 | KIRC | LB1047-RCC | Belinostat |
| MVP_top60_DNA_seed | ESR1 | 0.830469 | HNSC | OSC-20 | Tamoxifen | MVP_top60_DNA+RNA_seed | PTK2 | -0.515469 | KIRC | 769-P | PF-562271 |
| MVP_top60_DNA_seed | EGFR | 0.842231 | HNSC | Detroit562 | PF-00299804 | MVP_top60_DNA+RNA_seed | ATM | -0.513805 | KIRC | NCC021 | KU-60019 |
| MVP_top60_DNA_seed | AKT1 | 0.84503 | HNSC | Ca9-22 | AKT inhibitor VIII | MVP_top60_DNA+RNA_seed | MTOR | -0.513638 | KIRC | KMRC-1 | Temsirolimus |
| MVP_top60_DNA_seed | CDK2 | 0.845117 | HNSC | DOK | AZD5438 | MVP_top60_DNA+RNA_seed | MTOR | -0.51299 | KIRC | ACHN | Apitolisib |
| MVP_top60_DNA_seed | AKT1 | 0.845321 | HNSC | KOSC-2 | GSK690693 | MVP_top60_DNA+RNA_seed | HDAC1 | -0.512198 | KIRC | NCC021 | AR-42 |
| MVP_top60_DNA_seed | MDM2 | 0.851928 | HNSC | FADU | Nutlin-3a (-) | MVP_top60_DNA+RNA_seed | MTOR | -0.510538 | KIRC | HA7-RCC | Apitolisib |
| MVP_top60_DNA_seed | MDM2 | 0.852393 | HNSC | PCI-15A | Nutlin-3a (-) | MVP_top60_DNA+RNA_seed | HDAC1 | -0.507052 | KIRC | RXF393 | CAY10603 |
| MVP_top60_DNA_seed | PIK3CA | 0.854607 | HNSC | PCI-30 | PI3Ka_4409 | MVP_top60_DNA+RNA_seed | HDAC1 | -0.503431 | KIRC | 786-0 | CAY10603 |
| MVP_top60_DNA_seed | AKT1 | 0.855409 | HNSC | SKN-3 | GSK690693 | MVP_top60_DNA+RNA_seed | HSP90AA1 | -0.502193 | KIRC | U031 | Tanespimycin |
| MVP_top60_DNA_seed | NTRK1 | 0.856254 | HNSC | OSC-20 | GW441756 | MVP_top60_DNA+RNA_seed | HSP90AB1 | -0.502193 | KIRC | U031 | Tanespimycin |
| MVP_top60_DNA_seed | AKT1 | 0.857457 | HNSC | BHY | MK-2206 | MVP_top60_DNA+RNA_seed | ATM | -0.500712 | KIRC | U031 | CP466722 |
| MVP_top60_DNA_seed | HDAC1 | 0.859514 | HNSC | BICR10 | CAY10603 | MVP_top60_DNA+RNA_seed | ATM | -0.497329 | KIRC | RCC-MF | KU-55933 |
| MVP_top60_DNA_seed | CDK2 | 0.860021 | HNSC | SAT | PHA-793887 | MVP_top60_DNA+RNA_seed | PIK3CD | -0.496752 | KIRC | TK10 | IC-87114 |
| MVP_top60_DNA_seed | AKT1 | 0.862678 | HNSC | BICR10 | GSK690693 | MVP_top60_DNA+RNA_seed | FGFR4 | -0.494367 | KIRC | 769-P | FGFR_0939 |
| MVP_top60_DNA_seed | AR | 0.864336 | HNSC | PCI-30 | AZD3514 | MVP_top60_DNA+RNA_seed | HDAC1 | -0.493316 | KIRC | RCC-AB | Tubastatin A |
| MVP_top60_DNA_seed | SMARCA4 | 0.865323 | HNSC | SCC-4 | PFI-3 | MVP_top60_DNA+RNA_seed | ATM | -0.491558 | KIRC | RCC-JW | KU-55933 |
| MVP_top60_DNA_seed | EGFR | 0.878324 | HNSC | SCC-9 | Gefitinib | MVP_top60_DNA+RNA_seed | IGF1R | -0.491347 | KIRC | A704 | BMS-536924 |
| MVP_top60_DNA_seed | AKT1 | 0.880408 | HNSC | SCC-4 | BAY AKT1 | MVP_top60_DNA+RNA_seed | HSP90AA1 | -0.489348 | KIRC | ACHN | CCT-018159 |
| MVP_top60_DNA_seed | EGFR | 0.887553 | HNSC | BB49-HNC | Cetuximab | MVP_top60_DNA+RNA_seed | HSP90AB1 | -0.489348 | KIRC | ACHN | CCT-018159 |
| MVP_top60_DNA_seed | MAPK1 | 0.889631 | HNSC | PCI-15A | VX-11e | MVP_top60_DNA+RNA_seed | HDAC1 | -0.489213 | KIRC | OS-RC-2 | Dacinostat |
| MVP_top60_DNA_seed | HDAC1 | 0.892657 | HNSC | DOK | Dacinostat | MVP_top60_DNA+RNA_seed | HSP90AA1 | -0.487504 | KIRC | LB996-RCC | Luminespib |
| MVP_top60_DNA_seed | HDAC1 | 0.896203 | HNSC | LB771-HNC | Entinostat | MVP_top60_DNA+RNA_seed | HSP90AB1 | -0.487504 | KIRC | LB996-RCC | Luminespib |

| | | | | | | | | | | | |
|---|---|---|---|---|---|---|---|---|---|---|---|
| MVP_top60_DNA_seed | HDAC3 | 0.896203 | HNSC | LB771-HNC | Entinostat | MVP_top60_DNA+RNA_seed | FGFR4 | -0.486649 | KIRC | NCC021 | FGFR_0939 |
| MVP_top60_DNA_seed | CDK2 | 0.896678 | HNSC | SCC-4 | PHA-793887 | MVP_top60_DNA+RNA_seed | MTOR | -0.485078 | KIRC | NCC010 | CRT0105446 |
| MVP_top60_DNA_seed | AKT1 | 0.897716 | HNSC | FADU | AKT inhibitor VIII | MVP_top60_DNA+RNA_seed | ATM | -0.484012 | KIRC | LB996-RCC | torin2 |
| MVP_top60_DNA_seed | EP300 | 0.899385 | HNSC | SCC-9 | I-CBP112 | MVP_top60_DNA+RNA_seed | MTOR | -0.484012 | KIRC | LB996-RCC | torin2 |
| MVP_top60_DNA_seed | CREBBP | 0.899385 | HNSC | SCC-9 | I-CBP112 | MVP_top60_DNA+RNA_seed | PIK3CA | -0.480993 | KIRC | U031 | PI3Ka_4409 |
| MVP_top60_DNA_seed | MDM2 | 0.899899 | HNSC | HSC-2 | Serdemetan | MVP_top60_DNA+RNA_seed | PIK3CD | -0.480993 | KIRC | U031 | PI3Ka_4409 |
| MVP_top60_DNA_seed | AKT1 | 0.900216 | HNSC | JHU-022 | GSK690693 | MVP_top60_DNA+RNA_seed | MTOR | -0.478844 | KIRC | RCC-ER | Temsirolimus |
| MVP_top60_DNA_seed | HDAC1 | 0.900565 | HNSC | PE-CA-PJ15 | Tubastatin A | MVP_top60_DNA+RNA_seed | ATM | -0.477268 | KIRC | A498 | KU-60019 |
| MVP_top60_DNA_seed | SRC | 0.901642 | HNSC | SAT | NVP-BHG712 | MVP_top60_DNA+RNA_seed | HDAC1 | -0.475168 | KIRC | VMRC-RCZ | JQ12 |
| MVP_top60_DNA_seed | AR | 0.910886 | HNSC | PCI-30 | Bicalutamide | MVP_top60_DNA+RNA_seed | ATM | -0.473667 | KIRC | HA7-RCC | KU-55933 |
| MVP_top60_DNA_seed | HDAC1 | 0.926173 | HNSC | SKN-3 | Tubastatin A | MVP_top60_DNA+RNA_seed | MTOR | -0.473038 | KIRC | CAKI-1 | Temsirolimus |
| MVP_top60_DNA_seed | EGFR | 0.932089 | HNSC | BB49-HNC | Afatinib | MVP_top60_DNA+RNA_seed | HSP90AA1 | -0.472837 | KIRC | 769-P | Tanespimycin |
| MVP_top60_DNA_seed | AKT1 | 0.932576 | HNSC | HO-1-N-1 | AKT inhibitor VIII | MVP_top60_DNA+RNA_seed | HSP90AB1 | -0.472837 | KIRC | 769-P | Tanespimycin |
| MVP_top60_DNA_seed | EGFR | 0.933939 | HNSC | KON | Pelitinib | MVP_top60_DNA+RNA_seed | HSP90AA1 | -0.466489 | KIRC | LB2241-RCC | CCT-018159 |
| MVP_top60_DNA_seed | AKT1 | 0.937033 | HNSC | BICR31 | AKT inhibitor VIII | MVP_top60_DNA+RNA_seed | HSP90AB1 | -0.466489 | KIRC | LB2241-RCC | CCT-018159 |
| MVP_top60_DNA_seed | AR | 0.938101 | HNSC | SCC-15 | Bicalutamide | MVP_top60_DNA+RNA_seed | MTOR | -0.465714 | KIRC | NCC021 | CRT0105446 |
| MVP_top60_DNA_seed | MDM2 | 0.938479 | HNSC | SCC-25 | Nutlin-3a (-) | MVP_top60_DNA+RNA_seed | FGFR4 | -0.461804 | KIRC | SN12C | FGFR_0939 |
| MVP_top60_DNA_seed | HDAC1 | 0.940197 | HNSC | JHU-022 | Dacinostat | MVP_top60_DNA+RNA_seed | PIK3CD | -0.460103 | KIRC | RCC-JF | IC-87114 |
| MVP_top60_DNA_seed | HDAC1 | 0.941307 | HNSC | JHU-022 | CAY10603 | MVP_top60_DNA+RNA_seed | HSP90AA1 | -0.459791 | KIRC | SW156 | Elesclomol |
| MVP_top60_DNA_seed | SRC | 0.943234 | HNSC | FADU | NVP-BHG712 | MVP_top60_DNA+RNA_seed | HSP90AB1 | -0.459791 | KIRC | SW156 | Elesclomol |
| MVP_top60_DNA_seed | MAPK1 | 0.944338 | HNSC | BICR22 | FR-180204 | MVP_top60_DNA+RNA_seed | MTOR | -0.456863 | KIRC | RCC-ER | Voxtalisib |
| MVP_top60_DNA_seed | MAPK3 | 0.944338 | HNSC | BICR22 | FR-180204 | MVP_top60_DNA+RNA_seed | FGFR4 | -0.453204 | KIRC | LB2241-RCC | FGFR_3831 |
| MVP_top60_DNA_seed | HDAC1 | 0.949845 | HNSC | JHU-022 | Tubastatin A | MVP_top60_DNA+RNA_seed | PIK3CD | -0.452815 | KIRC | RCC-MF | Idelalisib |
| MVP_top60_DNA_seed | HDAC1 | 0.952773 | HNSC | BICR31 | Tubastatin A | MVP_top60_DNA+RNA_seed | ESR1 | -0.449796 | KIRC | LB1047-RCC | Tamoxifen |
| MVP_top60_DNA_seed | SRC | 0.957223 | HNSC | PCI-15A | NVP-BHG712 | MVP_top60_DNA+RNA_seed | PTK2 | -0.448709 | KIRC | ACHN | PF-562271 |
| MVP_top60_DNA_seed | AR | 0.958284 | HNSC | SKN-3 | Bicalutamide | MVP_top60_DNA+RNA_seed | HDAC1 | -0.448511 | KIRC | CAKI-1 | AR-42 |
| MVP_top60_DNA_seed | HDAC1 | 0.959075 | HNSC | PCI-30 | Dacinostat | MVP_top60_DNA+RNA_seed | SMARCA4 | -0.445229 | KIRC | BFTC-909 | PFI-3 |
| MVP_top60_DNA_seed | EGFR | 0.95966 | HNSC | BICR10 | Gefitinib | MVP_top60_DNA+RNA_seed | SMARCA2 | -0.445229 | KIRC | BFTC-909 | PFI-3 |
| MVP_top60_DNA_seed | AR | 0.964184 | HNSC | HSC-4 | Bicalutamide | MVP_top60_DNA+RNA_seed | PIK3CA | -0.444494 | KIRC | RCC-JW | AZD8835 |
| MVP_top60_DNA_seed | HDAC1 | 0.972444 | HNSC | BB49-HNC | Tubastatin A | MVP_top60_DNA+RNA_seed | PIK3CD | -0.444494 | KIRC | RCC-JW | AZD8835 |
| MVP_top60_DNA_seed | EGFR | 0.972515 | HNSC | SCC-15 | Erlotinib | MVP_top60_DNA+RNA_seed | HDAC1 | -0.442551 | KIRC | TK10 | Dacinostat |
| MVP_top60_DNA_seed | NTRK1 | 0.973461 | HNSC | DOK | GW441756 | MVP_top60_DNA+RNA_seed | PTK2 | -0.441759 | KIRC | LB2241-RCC | PF-562271 |
| MVP_top60_DNA_seed | CDK2 | 0.974804 | HNSC | PCI-30 | PHA-793887 | MVP_top60_DNA+RNA_seed | HDAC1 | -0.441566 | KIRC | CAL-54 | Dacinostat |
| MVP_top60_DNA_seed | EGFR | 0.975696 | HNSC | HO-1-u-1 | PF-00299804 | MVP_top60_DNA+RNA_seed | IGF1R | -0.439168 | KIRC | 769-P | BMS-754807 |
| MVP_top60_DNA_seed | HDAC1 | 0.979429 | HNSC | CAL-27 | JQ12 | MVP_top60_DNA+RNA_seed | HDAC1 | -0.438616 | KIRC | BB65-RCC | Entinostat |
| MVP_top60_DNA_seed | AKT1 | 0.979726 | HNSC | OSC-19 | AKT inhibitor VIII | MVP_top60_DNA+RNA_seed | ATM | -0.43689 | KIRC | SW156 | KU-55933 |
| MVP_top60_DNA_seed | AKT1 | 0.982681 | HNSC | SAS | BAY AKT1 | MVP_top60_DNA+RNA_seed | MTOR | -0.436864 | KIRC | RCC-JF | Voxtalisib |
| MVP_top60_DNA_seed | HSP90AA1 | 0.984339 | HNSC | Detroit562 | SNX-2112 | MVP_top60_DNA+RNA_seed | HDAC1 | -0.435283 | KIRC | U031 | JQ12 |
| MVP_top60_DNA_seed | MDM2 | 0.985936 | HNSC | LB771-HNC | Nutlin-3a (-) | MVP_top60_DNA+RNA_seed | HSP90AA1 | -0.434772 | KIRC | A704 | Elesclomol |
| MVP_top60_DNA_seed | CDK2 | 0.986554 | HNSC | SCC-15 | Seliciclib | MVP_top60_DNA+RNA_seed | HSP90AB1 | -0.434772 | KIRC | A704 | Elesclomol |
| MVP_top60_DNA_seed | EGFR | 0.987194 | HNSC | LB771-HNC | Afatinib | MVP_top60_DNA+RNA_seed | HSP90AA1 | -0.433688 | KIRC | BFTC-909 | Luminespib |

| | | | | | | | | | | | |
|---|---|---|---|---|---|---|---|---|---|---|---|
| MVP_top60_DNA_seed | EGFR | 0.988739 | HNSC | SCC-9 | Afatinib | MVP_top60_DNA+RNA_seed | HSP90AB1 | -0.433688 | KIRC | BFTC-909 | Luminespib |
| MVP_top60_DNA_seed | CDK2 | 0.990719 | HNSC | KON | PHA-793887 | MVP_top60_DNA+RNA_seed | IGF1R | -0.433101 | KIRC | VMRC-RCW | GSK1904529A |
| MVP_top60_DNA_seed | AKT1 | 0.999395 | HNSC | PE-CA-PJ15 | AKT inhibitor VIII | MVP_top60_DNA+RNA_seed | PIK3CD | -0.429563 | KIRC | BB65-RCC | AZD8186 |
| MVP_top60_DNA_seed | AKT1 | 1.000111 | HNSC | PCI-30 | GSK690693 | MVP_top60_DNA+RNA_seed | PIK3CD | -0.428048 | KIRC | LB996-RCC | IC-87114 |
| MVP_top60_DNA_seed | HDAC1 | 1.001622 | HNSC | Detroit562 | Belinostat | MVP_top60_DNA+RNA_seed | HSP90AA1 | -0.420771 | KIRC | RCC-JF | CCT-018159 |
| MVP_top60_DNA_seed | MDM2 | 1.001747 | HNSC | JHU-029 | Nutlin-3a (-) | MVP_top60_DNA+RNA_seed | HSP90AB1 | -0.420771 | KIRC | RCC-JF | CCT-018159 |
| MVP_top60_DNA_seed | SRC | 1.001765 | HNSC | HSC-4 | NVP-BHG712 | MVP_top60_DNA+RNA_seed | ESR1 | -0.418613 | KIRC | RCC-ER | Tamoxifen |
| MVP_top60_DNA_seed | EGFR | 1.007329 | HNSC | BICR10 | PF-00299804 | MVP_top60_DNA+RNA_seed | ATM | -0.417627 | KIRC | RCC-ER | torin2 |
| MVP_top60_DNA_seed | MAPK1 | 1.008782 | HNSC | SAT | FR-180204 | MVP_top60_DNA+RNA_seed | MTOR | -0.417627 | KIRC | RCC-ER | torin2 |
| MVP_top60_DNA_seed | MAPK3 | 1.008782 | HNSC | SAT | FR-180204 | MVP_top60_DNA+RNA_seed | HSP90AA1 | -0.416513 | KIRC | LB1047-RCC | SNX-2112 |
| MVP_top60_DNA_seed | EP300 | 1.013716 | HNSC | SCC-15 | I-CBP112 | MVP_top60_DNA+RNA_seed | HSP90AB1 | -0.416513 | KIRC | LB1047-RCC | SNX-2112 |
| MVP_top60_DNA_seed | CREBBP | 1.013716 | HNSC | SCC-15 | I-CBP112 | MVP_top60_DNA+RNA_seed | ATM | -0.415485 | KIRC | TK10 | torin2 |
| MVP_top60_DNA_seed | HDAC1 | 1.014278 | HNSC | BICR10 | Belinostat | MVP_top60_DNA+RNA_seed | MTOR | -0.415485 | KIRC | TK10 | torin2 |
| MVP_top60_DNA_seed | SRC | 1.018269 | HNSC | KON | NVP-BHG712 | MVP_top60_DNA+RNA_seed | MTOR | -0.415406 | KIRC | RCC-ER | Apitolisib |
| MVP_top60_DNA_seed | HSP90AA1 | 1.021414 | HNSC | SAT | Elesclomol | MVP_top60_DNA+RNA_seed | HDAC1 | -0.414639 | KIRC | LB996-RCC | Entinostat |
| MVP_top60_DNA_seed | SRC | 1.027801 | HNSC | OSC-20 | NVP-BHG712 | MVP_top60_DNA+RNA_seed | MTOR | -0.413125 | KIRC | NCC021 | WYE-125132 |
| MVP_top60_DNA_seed | MAPK1 | 1.028634 | HNSC | BICR10 | FR-180204 | MVP_top60_DNA+RNA_seed | HSP90AA1 | -0.412263 | KIRC | CAKI-1 | Elesclomol |
| MVP_top60_DNA_seed | MAPK3 | 1.028634 | HNSC | BICR10 | FR-180204 | MVP_top60_DNA+RNA_seed | HSP90AB1 | -0.412263 | KIRC | CAKI-1 | Elesclomol |
| MVP_top60_DNA_seed | SRC | 1.031101 | HNSC | DOK | NVP-BHG712 | MVP_top60_DNA+RNA_seed | ESR1 | -0.405903 | KIRC | A704 | Fulvestrant |
| MVP_top60_DNA_seed | MAPK1 | 1.031453 | HNSC | KOSC-2 | FR-180204 | MVP_top60_DNA+RNA_seed | FGFR4 | -0.405782 | KIRC | BFTC-909 | FGFR_3831 |
| MVP_top60_DNA_seed | MAPK3 | 1.031453 | HNSC | KOSC-2 | FR-180204 | MVP_top60_DNA+RNA_seed | ATM | -0.401212 | KIRC | 786-0 | KU-60019 |
| MVP_top60_DNA_seed | RAC1 | 1.031535 | HNSC | JHU-029 | EHT-1864 | MVP_top60_DNA+RNA_seed | MTOR | -0.401189 | KIRC | ACHN | WYE-125132 |
| MVP_top60_DNA_seed | EGFR | 1.046491 | HNSC | LB771-HNC | CI-1033 | MVP_top60_DNA+RNA_seed | ATM | -0.400257 | KIRC | 786-0 | KU-55933 |
| MVP_top60_DNA_seed | HDAC1 | 1.049265 | HNSC | PCI-38 | JQ12 | MVP_top60_DNA+RNA_seed | PIK3CD | -0.399945 | KIRC | ACHN | Idelalisib |
| MVP_top60_DNA_seed | HDAC1 | 1.052733 | HNSC | BICR10 | Tubastatin A | MVP_top60_DNA+RNA_seed | MTOR | -0.399932 | KIRC | KMRC-20 | Temsirolimus |
| MVP_top60_DNA_seed | HDAC1 | 1.053879 | HNSC | Detroit562 | CAY10603 | MVP_top60_DNA+RNA_seed | IGF1R | -0.388749 | KIRC | LB2241-RCC | GSK1904529A |
| MVP_top60_DNA_seed | AKT1 | 1.055281 | HNSC | PCI-30 | AKT inhibitor VIII | MVP_top60_DNA+RNA_seed | MTOR | -0.387022 | KIRC | RCC-JW | Apitolisib |
| MVP_top60_DNA_seed | EGFR | 1.059552 | HNSC | BB49-HNC | CUDC-101 | MVP_top60_DNA+RNA_seed | HSP90AA1 | -0.386267 | KIRC | BB65-RCC | Elesclomol |
| MVP_top60_DNA_seed | AKT1 | 1.065534 | HNSC | BB49-HNC | GSK690693 | MVP_top60_DNA+RNA_seed | HSP90AB1 | -0.386267 | KIRC | BB65-RCC | Elesclomol |
| MVP_top60_DNA_seed | EGFR | 1.0669 | HNSC | BICR78 | PF-00299804 | MVP_top60_DNA+RNA_seed | HDAC1 | -0.38576 | KIRC | LB1047-RCC | Entinostat |
| MVP_top60_DNA_seed | ESR1 | 1.072844 | HNSC | SCC-9 | Tamoxifen | MVP_top60_DNA+RNA_seed | MTOR | -0.385203 | KIRC | RCC-JF | CRT0105446 |
| MVP_top60_DNA_seed | MDM2 | 1.076399 | HNSC | KOSC-2 | Nutlin-3a (-) | MVP_top60_DNA+RNA_seed | SMARCA4 | -0.385199 | KIRC | LB1047-RCC | PFI-3 |
| MVP_top60_DNA_seed | HDAC1 | 1.078025 | HNSC | BICR78 | Tubastatin A | MVP_top60_DNA+RNA_seed | SMARCA2 | -0.385199 | KIRC | LB1047-RCC | PFI-3 |
| MVP_top60_DNA_seed | HDAC1 | 1.08006 | HNSC | BB49-HNC | CAY10603 | MVP_top60_DNA+RNA_seed | PIK3CD | -0.383928 | KIRC | RCC10RGB | AZD8186 |
| MVP_top60_DNA_seed | MDM2 | 1.081002 | HNSC | SCC-9 | Nutlin-3a (-) | MVP_top60_DNA+RNA_seed | PIK3CA | -0.383181 | KIRC | CAL-54 | AZD8835 |
| MVP_top60_DNA_seed | EGFR | 1.083037 | HNSC | BB49-HNC | Gefitinib | MVP_top60_DNA+RNA_seed | PIK3CD | -0.383181 | KIRC | CAL-54 | AZD8835 |
| MVP_top60_DNA_seed | MAPK1 | 1.085853 | HNSC | SCC-15 | FR-180204 | MVP_top60_DNA+RNA_seed | PIK3CA | -0.382826 | KIRC | NCC010 | PI3Ka_4409 |
| MVP_top60_DNA_seed | MAPK3 | 1.085853 | HNSC | SCC-15 | FR-180204 | MVP_top60_DNA+RNA_seed | PIK3CD | -0.382826 | KIRC | NCC010 | PI3Ka_4409 |
| MVP_top60_DNA_seed | EGFR | 1.086696 | HNSC | Detroit562 | AST-1306 | MVP_top60_DNA+RNA_seed | HDAC1 | -0.378905 | KIRC | LB2241-RCC | Entinostat |
| MVP_top60_DNA_seed | SRC | 1.088993 | HNSC | BICR31 | NVP-BHG712 | MVP_top60_DNA+RNA_seed | HSP90AA1 | -0.375789 | KIRC | 769-P | SNX-2112 |
| MVP_top60_DNA_seed | MAPK1 | 1.089863 | HNSC | DOK | VX-11e | MVP_top60_DNA+RNA_seed | HSP90AB1 | -0.375789 | KIRC | 769-P | SNX-2112 |

| | | | | | | | | | | | |
|---|---|---|---|---|---|---|---|---|---|---|---|
| MVP_top60_DNA_seed | MAPK1 | 1.097409 | HNSC | BB49-HNC | FR-180204 | MVP_top60_DNA+RNA_seed | ATM | -0.375658 | KIRC | RCC-FG2 | torin2 |
| MVP_top60_DNA_seed | MAPK3 | 1.097409 | HNSC | BB49-HNC | FR-180204 | MVP_top60_DNA+RNA_seed | MTOR | -0.375658 | KIRC | RCC-FG2 | torin2 |
| MVP_top60_DNA_seed | AKT1 | 1.097496 | HNSC | PCI-15A | AKT inhibitor VIII | MVP_top60_DNA+RNA_seed | FGFR4 | -0.374645 | KIRC | HA7-RCC | FGFR_3831 |
| MVP_top60_DNA_seed | AKT1 | 1.098242 | HNSC | BB49-HNC | A-443654 | MVP_top60_DNA+RNA_seed | MTOR | -0.3736 | KIRC | LB2241-RCC | Apitolisib |
| MVP_top60_DNA_seed | AKT1 | 1.104906 | HNSC | Ca9-22 | MK-2206 | MVP_top60_DNA+RNA_seed | PIK3CA | -0.37179 | KIRC | NCC010 | AZD8835 |
| MVP_top60_DNA_seed | EGFR | 1.119183 | HNSC | BB49-HNC | Erlotinib | MVP_top60_DNA+RNA_seed | PIK3CD | -0.37179 | KIRC | NCC010 | AZD8835 |
| MVP_top60_DNA_seed | EGFR | 1.126308 | HNSC | PCI-15A | CUDC-101 | MVP_top60_DNA+RNA_seed | HSP90AA1 | -0.370234 | KIRC | RCC-AB | SNX-2112 |
| MVP_top60_DNA_seed | HSP90AA1 | 1.127396 | HNSC | SCC-4 | Elesclomol | MVP_top60_DNA+RNA_seed | HSP90AB1 | -0.370234 | KIRC | RCC-AB | SNX-2112 |
| MVP_top60_DNA_seed | AKT1 | 1.131152 | HNSC | SAS | MK-2206 | MVP_top60_DNA+RNA_seed | MTOR | -0.368045 | KIRC | TK10 | Apitolisib |
| MVP_top60_DNA_seed | ESR1 | 1.13633 | HNSC | SCC-15 | Fulvestrant | MVP_top60_DNA+RNA_seed | HSP90AA1 | -0.368018 | KIRC | OS-RC-2 | Tanespimycin |
| MVP_top60_DNA_seed | AKT1 | 1.145327 | HNSC | BICR22 | AKT inhibitor VIII | MVP_top60_DNA+RNA_seed | HSP90AB1 | -0.368018 | KIRC | OS-RC-2 | Tanespimycin |
| MVP_top60_DNA_seed | AR | 1.145937 | HNSC | KOSC-2 | Bicalutamide | MVP_top60_DNA+RNA_seed | AR | -0.364709 | KIRC | ACHN | AZD3514 |
| MVP_top60_DNA_seed | EGFR | 1.147293 | HNSC | HO-1-u-1 | CI-1033 | MVP_top60_DNA+RNA_seed | HDAC1 | -0.362238 | KIRC | TK10 | JQ12 |
| MVP_top60_DNA_seed | MAPK1 | 1.148234 | HNSC | JHU-022 | FR-180204 | MVP_top60_DNA+RNA_seed | PIK3CA | -0.356994 | KIRC | TK10 | AZD8835 |
| MVP_top60_DNA_seed | MAPK3 | 1.148234 | HNSC | JHU-022 | FR-180204 | MVP_top60_DNA+RNA_seed | PIK3CD | -0.356994 | KIRC | TK10 | AZD8835 |
| MVP_top60_DNA_seed | MAPK1 | 1.153329 | HNSC | BICR10 | VX-11e | MVP_top60_DNA+RNA_seed | FGFR4 | -0.3568 | KIRC | LB1047-RCC | FGFR_3831 |
| MVP_top60_DNA_seed | HDAC1 | 1.170607 | HNSC | KON | Belinostat | MVP_top60_DNA+RNA_seed | HSP90AA1 | -0.355801 | KIRC | BB65-RCC | SNX-2112 |
| MVP_top60_DNA_seed | EGFR | 1.172833 | HNSC | BICR10 | CUDC-101 | MVP_top60_DNA+RNA_seed | HSP90AB1 | -0.355801 | KIRC | BB65-RCC | SNX-2112 |
| MVP_top60_DNA_seed | NTRK1 | 1.173897 | HNSC | SCC-9 | GW441756 | MVP_top60_DNA+RNA_seed | PIK3CD | -0.353447 | KIRC | VMRC-RCZ | IC-87114 |
| MVP_top60_DNA_seed | SRC | 1.173995 | HNSC | KOSC-2 | NVP-BHG712 | MVP_top60_DNA+RNA_seed | ATM | -0.353272 | KIRC | RXF393 | CP466722 |
| MVP_top60_DNA_seed | SRC | 1.178905 | HNSC | SCC-15 | Saracatinib | MVP_top60_DNA+RNA_seed | PIK3CD | -0.352659 | KIRC | RCC-ER | Idelalisib |
| MVP_top60_DNA_seed | AKT1 | 1.183497 | HNSC | SAT | GSK690693 | MVP_top60_DNA+RNA_seed | MTOR | -0.351612 | KIRC | RCC-ER | Torin 2 |
| MVP_top60_DNA_seed | SRC | 1.185763 | HNSC | BICR78 | NVP-BHG712 | MVP_top60_DNA+RNA_seed | HDAC1 | -0.348724 | KIRC | ACHN | JQ12 |
| MVP_top60_DNA_seed | SMARCA4 | 1.191441 | HNSC | SCC90 | PFI-3 | MVP_top60_DNA+RNA_seed | ATM | -0.348576 | KIRC | KMRC-20 | KU-55933 |
| MVP_top60_DNA_seed | AR | 1.195561 | HNSC | CAL-33 | AZD3514 | MVP_top60_DNA+RNA_seed | HDAC1 | -0.346426 | KIRC | CAKI-1 | Belinostat |
| MVP_top60_DNA_seed | HDAC1 | 1.202499 | HNSC | BICR22 | JQ12 | MVP_top60_DNA+RNA_seed | HDAC1 | -0.345943 | KIRC | 786-0 | Belinostat |
| MVP_top60_DNA_seed | HDAC1 | 1.212364 | HNSC | BB49-HNC | Dacinostat | MVP_top60_DNA+RNA_seed | MTOR | -0.344726 | KIRC | RCC-JF | QL-VIII-58 |
| MVP_top60_DNA_seed | AKT1 | 1.212834 | HNSC | KON | AKT inhibitor VIII | MVP_top60_DNA+RNA_seed | IGF1R | -0.344405 | KIRC | CAL-54 | BMS-536924 |
| MVP_top60_DNA_seed | EGFR | 1.214272 | HNSC | KON | CUDC-101 | MVP_top60_DNA+RNA_seed | IGF1R | -0.344079 | KIRC | 786-0 | BMS-754807 |
| MVP_top60_DNA_seed | HDAC1 | 1.232295 | HNSC | PCI-30 | Belinostat | MVP_top60_DNA+RNA_seed | HSP90AA1 | -0.342318 | KIRC | LB2241-RCC | Luminespib |
| MVP_top60_DNA_seed | SRC | 1.237585 | HNSC | BICR22 | NVP-BHG712 | MVP_top60_DNA+RNA_seed | HSP90AB1 | -0.342318 | KIRC | LB2241-RCC | Luminespib |
| MVP_top60_DNA_seed | HDAC1 | 1.239928 | HNSC | HSC-4 | Tubastatin A | MVP_top60_DNA+RNA_seed | PIK3CA | -0.339491 | KIRC | RCC-FG2 | PI3Ka_4409 |
| MVP_top60_DNA_seed | AKT1 | 1.24201 | HNSC | BICR78 | AKT inhibitor VIII | MVP_top60_DNA+RNA_seed | PIK3CD | -0.339491 | KIRC | RCC-FG2 | PI3Ka_4409 |
| MVP_top60_DNA_seed | HDAC1 | 1.243458 | HNSC | BICR10 | Dacinostat | MVP_top60_DNA+RNA_seed | HSP90AA1 | -0.338799 | KIRC | LB996-RCC | SNX-2112 |
| MVP_top60_DNA_seed | HDAC1 | 1.244693 | HNSC | PCI-30 | Tubastatin A | MVP_top60_DNA+RNA_seed | HSP90AB1 | -0.338799 | KIRC | LB996-RCC | SNX-2112 |
| MVP_top60_DNA_seed | EGFR | 1.246443 | HNSC | BICR22 | Pelitinib | MVP_top60_DNA+RNA_seed | MTOR | -0.338631 | KIRC | LB2241-RCC | Torin 2 |
| MVP_top60_DNA_seed | NTRK1 | 1.248387 | HNSC | PCI-30 | GW441756 | MVP_top60_DNA+RNA_seed | PIK3CA | -0.33443 | KIRC | LB996-RCC | PI3Ka_4409 |
| MVP_top60_DNA_seed | HDAC1 | 1.263017 | HNSC | PCI-38 | Dacinostat | MVP_top60_DNA+RNA_seed | PIK3CD | -0.33443 | KIRC | LB996-RCC | PI3Ka_4409 |
| MVP_top60_DNA_seed | HSP90AA1 | 1.264022 | HNSC | PCI-30 | SNX-2112 | MVP_top60_DNA+RNA_seed | IGF1R | -0.332812 | KIRC | CAL-54 | Linsitinib |
| MVP_top60_DNA_seed | ESR1 | 1.264635 | HNSC | DOK | Tamoxifen | MVP_top60_DNA+RNA_seed | ATM | -0.332384 | KIRC | RCC-JF | KU-60019 |
| MVP_top60_DNA_seed | EGFR | 1.270692 | HNSC | HSC-4 | PF-00299804 | MVP_top60_DNA+RNA_seed | PIK3CD | -0.331329 | KIRC | CAL-54 | Idelalisib |

| | | | | | | | | | | | |
|---|---|---|---|---|---|---|---|---|---|---|---|
| MVP_top60_DNA_seed | HSP90AA1 | 1.275884 | HNSC | SKN-3 | Luminespib | MVP_top60_DNA+RNA_seed | HDAC1 | -0.32918 | KIRC | BFTC-909 | Dacinostat |
| MVP_top60_DNA_seed | HSP90AA1 | 1.276988 | HNSC | KOSC-2 | SNX-2112 | MVP_top60_DNA+RNA_seed | IGF1R | -0.32822 | KIRC | HA7-RCC | BMS-754807 |
| MVP_top60_DNA_seed | HDAC1 | 1.291347 | HNSC | LB771-HNC | Parthenolide | MVP_top60_DNA+RNA_seed | MTOR | -0.327969 | KIRC | RCC-MF | Voxtalisib |
| MVP_top60_DNA_seed | CDK2 | 1.294219 | HNSC | BICR10 | PHA-793887 | MVP_top60_DNA+RNA_seed | MTOR | -0.327316 | KIRC | LB2241-RCC | WYE-125132 |
| MVP_top60_DNA_seed | EGFR | 1.294778 | HNSC | BB49-HNC | Pelitinib | MVP_top60_DNA+RNA_seed | ESR1 | -0.32296 | KIRC | VMRC-RCZ | Fulvestrant |
| MVP_top60_DNA_seed | SRC | 1.299155 | HNSC | JHU-022 | NVP-BHG712 | MVP_top60_DNA+RNA_seed | ATM | -0.322313 | KIRC | CAL-54 | KU-60019 |
| MVP_top60_DNA_seed | HDAC1 | 1.299344 | HNSC | HSC-4 | JQ12 | MVP_top60_DNA+RNA_seed | EP300 | -0.320314 | KIRC | BB65-RCC | I-CBP112 |
| MVP_top60_DNA_seed | SRC | 1.301046 | HNSC | BICR10 | NVP-BHG712 | MVP_top60_DNA+RNA_seed | PIK3CA | -0.318491 | KIRC | SN12C | AZD8835 |
| MVP_top60_DNA_seed | ESR1 | 1.303993 | HNSC | PCI-30 | Tamoxifen | MVP_top60_DNA+RNA_seed | PIK3CD | -0.318491 | KIRC | SN12C | AZD8835 |
| MVP_top60_DNA_seed | SMARCA4 | 1.305308 | HNSC | PCI-30 | PFI-3 | MVP_top60_DNA+RNA_seed | AR | -0.317659 | KIRC | A704 | Bicalutamide |
| MVP_top60_DNA_seed | AR | 1.312474 | HNSC | SAT | Bicalutamide | MVP_top60_DNA+RNA_seed | ESR1 | -0.317198 | KIRC | RXF393 | Tamoxifen |
| MVP_top60_DNA_seed | NTRK1 | 1.31579 | HNSC | BB49-HNC | GW441756 | MVP_top60_DNA+RNA_seed | PIK3CA | -0.316568 | KIRC | RCC-JF | AZD8835 |
| MVP_top60_DNA_seed | AKT1 | 1.323411 | HNSC | PE-CA-PJ15 | AKT inhibitor VIII | MVP_top60_DNA+RNA_seed | PIK3CD | -0.316568 | KIRC | RCC-JF | AZD8835 |
| MVP_top60_DNA_seed | HDAC1 | 1.344355 | HNSC | BICR22 | Dacinostat | MVP_top60_DNA+RNA_seed | MTOR | -0.312126 | KIRC | VMRC-RCZ | Apitolisib |
| MVP_top60_DNA_seed | EGFR | 1.344461 | HNSC | KOSC-2 | Pelitinib | MVP_top60_DNA+RNA_seed | MTOR | -0.311661 | KIRC | RCC-FG2 | Torin 2 |
| MVP_top60_DNA_seed | MAPK1 | 1.356474 | HNSC | PCI-30 | FR-180204 | MVP_top60_DNA+RNA_seed | EP300 | -0.31123 | KIRC | LB1047-RCC | I-CBP112 |
| MVP_top60_DNA_seed | MAPK3 | 1.356474 | HNSC | PCI-30 | FR-180204 | MVP_top60_DNA+RNA_seed | HDAC1 | -0.305504 | KIRC | 786-0 | AR-42 |
| MVP_top60_DNA_seed | HDAC1 | 1.364789 | HNSC | DOK | Tubastatin A | MVP_top60_DNA+RNA_seed | FGFR4 | -0.302501 | KIRC | NCC010 | FGFR_0939 |
| MVP_top60_DNA_seed | AKT1 | 1.371184 | HNSC | HSC-4 | AKT inhibitor VIII | MVP_top60_DNA+RNA_seed | HDAC1 | -0.302237 | KIRC | BB65-RCC | CAY10603 |
| MVP_top60_DNA_seed | MDM2 | 1.373264 | HNSC | OSC-19 | Serdemetan | MVP_top60_DNA+RNA_seed | SMARCA4 | -0.300719 | KIRC | LB2241-RCC | PFI-3 |
| MVP_top60_DNA_seed | AKT1 | 1.386059 | HNSC | HSC-2 | AKT inhibitor VIII | MVP_top60_DNA+RNA_seed | SMARCA2 | -0.300719 | KIRC | LB2241-RCC | PFI-3 |
| MVP_top60_DNA_seed | MDM2 | 1.390795 | HNSC | SCC90 | Serdemetan | MVP_top60_DNA+RNA_seed | ESR1 | -0.300407 | KIRC | NCC010 | Tamoxifen |
| MVP_top60_DNA_seed | MAPK1 | 1.392803 | HNSC | BB49-HNC | VX-11e | MVP_top60_DNA+RNA_seed | FGFR4 | -0.296539 | KIRC | LB2241-RCC | FGFR_0939 |
| MVP_top60_DNA_seed | HSP90AA1 | 1.400958 | HNSC | KON | SNX-2112 | MVP_top60_DNA+RNA_seed | EP300 | -0.296421 | KIRC | RCC-MF | I-CBP112 |
| MVP_top60_DNA_seed | RAC1 | 1.405096 | HNSC | BICR10 | EHT-1864 | MVP_top60_DNA+RNA_seed | ATM | -0.295025 | KIRC | BB65-RCC | CP466722 |
| MVP_top60_DNA_seed | AR | 1.4197 | HNSC | PCI-15A | AZD3514 | MVP_top60_DNA+RNA_seed | HSP90AA1 | -0.294717 | KIRC | NCC010 | Elesclomol |
| MVP_top60_DNA_seed | HDAC1 | 1.430576 | HNSC | BB49-HNC | Parthenolide | MVP_top60_DNA+RNA_seed | HSP90AB1 | -0.294717 | KIRC | NCC010 | Elesclomol |
| MVP_top60_DNA_seed | ESR1 | 1.431673 | HNSC | SCC90 | Tamoxifen | MVP_top60_DNA+RNA_seed | MTOR | -0.292081 | KIRC | ACHN | Voxtalisib |
| MVP_top60_DNA_seed | RAC1 | 1.435435 | HNSC | BB49-HNC | EHT-1864 | MVP_top60_DNA+RNA_seed | PTK2 | -0.290598 | KIRC | SN12C | PF-562271 |
| MVP_top60_DNA_seed | AKT1 | 1.437458 | HNSC | PCI-15A | BAY AKT1 | MVP_top60_DNA+RNA_seed | PIK3CD | -0.28864 | KIRC | 786-0 | AZD8186 |
| MVP_top60_DNA_seed | AKT1 | 1.445724 | HNSC | SKN-3 | AKT inhibitor VIII | MVP_top60_DNA+RNA_seed | IGF1R | -0.283457 | KIRC | NCC010 | BMS-536924 |
| MVP_top60_DNA_seed | EGFR | 1.446486 | HNSC | HO-1-u-1 | AST-1306 | MVP_top60_DNA+RNA_seed | IGF1R | -0.282708 | KIRC | CAKI-1 | Linsitinib |
| MVP_top60_DNA_seed | CDK2 | 1.451112 | HNSC | BB49-HNC | PHA-793887 | MVP_top60_DNA+RNA_seed | IGF1R | -0.281105 | KIRC | 786-0 | Linsitinib |
| MVP_top60_DNA_seed | AR | 1.47573 | HNSC | PCI-30 | Bicalutamide | MVP_top60_DNA+RNA_seed | ATM | -0.280281 | KIRC | KMRC-20 | KU-60019 |
| MVP_top60_DNA_seed | AKT1 | 1.476536 | HNSC | SCC-15 | AKT inhibitor VIII | MVP_top60_DNA+RNA_seed | HSP90AA1 | -0.27881 | KIRC | SN12C | Elesclomol |
| MVP_top60_DNA_seed | HDAC1 | 1.488293 | HNSC | SAT | JQ12 | MVP_top60_DNA+RNA_seed | HSP90AB1 | -0.27881 | KIRC | SN12C | Elesclomol |
| MVP_top60_DNA_seed | AKT1 | 1.502319 | HNSC | SAT | AKT inhibitor VIII | MVP_top60_DNA+RNA_seed | IGF1R | -0.27567 | KIRC | HA7-RCC | BMS-536924 |
| MVP_top60_DNA_seed | AR | 1.503456 | HNSC | SCC90 | AZD3514 | MVP_top60_DNA+RNA_seed | MTOR | -0.269349 | KIRC | 786-0 | Torin 2 |
| MVP_top60_DNA_seed | EGFR | 1.504344 | HNSC | BICR22 | CI-1033 | MVP_top60_DNA+RNA_seed | ESR1 | -0.268063 | KIRC | NCC010 | Fulvestrant |
| MVP_top60_DNA_seed | MAPK1 | 1.505091 | HNSC | PCI-30 | VX-11e | MVP_top60_DNA+RNA_seed | FGFR4 | -0.262488 | KIRC | 769-P | FGFR_3831 |
| MVP_top60_DNA_seed | SRC | 1.508236 | HNSC | BB49-HNC | Bosutinib | MVP_top60_DNA+RNA_seed | PIK3CA | -0.26233 | KIRC | RCC-ER | AZD8835 |

| | | | | | | | | | | | |
|---|---|---|---|---|---|---|---|---|---|---|---|
| MVP_top60_DNA_seed | HSP90AA1 | 1.517451 | HNSC | JHU-029 | Tanespimycin | MVP_top60_DNA+RNA_seed | PIK3CD | -0.26233 | KIRC | RCC-ER | AZD8835 |
| MVP_top60_DNA_seed | EGFR | 1.523539 | HNSC | BICR22 | AST-1306 | MVP_top60_DNA+RNA_seed | HSP90AA1 | -0.262043 | KIRC | CAL-54 | CCT-018159 |
| MVP_top60_DNA_seed | PIK3CA | 1.534807 | HNSC | SAS | PI3Ka_4409 | MVP_top60_DNA+RNA_seed | HSP90AB1 | -0.262043 | KIRC | CAL-54 | CCT-018159 |
| MVP_top60_DNA_seed | AKT1 | 1.536421 | HNSC | SCC-4 | AKT inhibitor VIII | MVP_top60_DNA+RNA_seed | FGFR4 | -0.261901 | KIRC | KMRC-20 | FGFR_0939 |
| MVP_top60_DNA_seed | HSP90AA1 | 1.55453 | HNSC | PCI-15A | SNX-2112 | MVP_top60_DNA+RNA_seed | IGF1R | -0.261281 | KIRC | LB2241-RCC | Linsitinib |
| MVP_top60_DNA_seed | HDAC1 | 1.554851 | HNSC | PCI-30 | JQ12 | MVP_top60_DNA+RNA_seed | PIK3CA | -0.260492 | KIRC | HA7-RCC | AZD8835 |
| MVP_top60_DNA_seed | HDAC1 | 1.566303 | HNSC | KON | AR-42 | MVP_top60_DNA+RNA_seed | PIK3CD | -0.260492 | KIRC | HA7-RCC | AZD8835 |
| MVP_top60_DNA_seed | EGFR | 1.566477 | HNSC | PCI-30 | CUDC-101 | MVP_top60_DNA+RNA_seed | FGFR4 | -0.258943 | KIRC | RCC-JW | FGFR_3831 |
| MVP_top60_DNA_seed | EP300 | 1.575352 | HNSC | SCC90 | I-CBP112 | MVP_top60_DNA+RNA_seed | HDAC1 | -0.256534 | KIRC | 786-0 | Tubastatin A |
| MVP_top60_DNA_seed | CREBBP | 1.575352 | HNSC | SCC90 | I-CBP112 | MVP_top60_DNA+RNA_seed | HDAC1 | -0.255494 | KIRC | RXF393 | AR-42 |
| MVP_top60_DNA_seed | EGFR | 1.594524 | HNSC | LB771-HNC | PF-00299804 | MVP_top60_DNA+RNA_seed | HSP90AA1 | -0.255008 | KIRC | BB65-RCC | CCT-018159 |
| MVP_top60_DNA_seed | EGFR | 1.605547 | HNSC | LB771-HNC | AST-1306 | MVP_top60_DNA+RNA_seed | HSP90AB1 | -0.255008 | KIRC | BB65-RCC | CCT-018159 |
| MVP_top60_DNA_seed | MDM2 | 1.618252 | HNSC | SCC90 | Nutlin-3a (-) | MVP_top60_DNA+RNA_seed | MTOR | -0.253649 | KIRC | LB1047-RCC | Apitolisib |
| MVP_top60_DNA_seed | ESR1 | 1.619394 | HNSC | SCC-9 | Fulvestrant | MVP_top60_DNA+RNA_seed | FGFR4 | -0.248438 | KIRC | VMRC-RCZ | FGFR_0939 |
| MVP_top60_DNA_seed | HDAC1 | 1.634765 | HNSC | BICR10 | AR-42 | MVP_top60_DNA+RNA_seed | IGF1R | -0.246271 | KIRC | BB65-RCC | BMS-754807 |
| MVP_top60_DNA_seed | AKT1 | 1.634907 | HNSC | SCC-9 | MK-2206 | MVP_top60_DNA+RNA_seed | PIK3CA | -0.243047 | KIRC | RCC-MF | PI3Ka_4409 |
| MVP_top60_DNA_seed | HDAC1 | 1.646745 | HNSC | BB49-HNC | AR-42 | MVP_top60_DNA+RNA_seed | PIK3CD | -0.243047 | KIRC | RCC-MF | PI3Ka_4409 |
| MVP_top60_DNA_seed | MAPK1 | 1.652858 | HNSC | DOK | FR-180204 | MVP_top60_DNA+RNA_seed | FGFR4 | -0.241968 | KIRC | RCC-MF | FGFR_0939 |
| MVP_top60_DNA_seed | MAPK3 | 1.652858 | HNSC | DOK | FR-180204 | MVP_top60_DNA+RNA_seed | MTOR | -0.241817 | KIRC | 786-0 | Apitolisib |
| MVP_top60_DNA_seed | CDK2 | 1.654937 | HNSC | SCC90 | AZD5438 | MVP_top60_DNA+RNA_seed | PIK3CD | -0.239066 | KIRC | KMRC-1 | Idelalisib |
| MVP_top60_DNA_seed | AR | 1.674905 | HNSC | SCC-9 | AZD3514 | MVP_top60_DNA+RNA_seed | MTOR | -0.238001 | KIRC | LB1047-RCC | Voxtalisib |
| MVP_top60_DNA_seed | HDAC1 | 1.680245 | HNSC | DOK | JQ12 | MVP_top60_DNA+RNA_seed | MTOR | -0.237854 | KIRC | U031 | Apitolisib |
| MVP_top60_DNA_seed | HDAC1 | 1.699525 | HNSC | FADU | JQ12 | MVP_top60_DNA+RNA_seed | MTOR | -0.237017 | KIRC | LB2241-RCC | QL-VIII-58 |
| MVP_top60_DNA_seed | AR | 1.702281 | HNSC | SCC-9 | Bicalutamide | MVP_top60_DNA+RNA_seed | ATM | -0.233803 | KIRC | OS-RC-2 | CP466722 |
| MVP_top60_DNA_seed | AKT1 | 1.706161 | HNSC | BICR10 | AKT inhibitor VIII | MVP_top60_DNA+RNA_seed | HDAC1 | -0.232907 | KIRC | VMRC-RCZ | Belinostat |
| MVP_top60_DNA_seed | AKT1 | 1.717603 | HNSC | JHU-022 | AKT inhibitor VIII | MVP_top60_DNA+RNA_seed | MTOR | -0.230743 | KIRC | RCC-JW | Torin 2 |
| MVP_top60_DNA_seed | SRC | 1.739651 | HNSC | PCI-30 | NVP-BHG712 | MVP_top60_DNA+RNA_seed | MTOR | -0.229284 | KIRC | U031 | Voxtalisib |
| MVP_top60_DNA_seed | PIK3CA | 1.761379 | HNSC | SCC90 | PI3Ka_4409 | MVP_top60_DNA+RNA_seed | ATM | -0.226613 | KIRC | LB996-RCC | KU-60019 |
| MVP_top60_DNA_seed | HDAC1 | 1.773274 | HNSC | PCI-30 | AR-42 | MVP_top60_DNA+RNA_seed | MTOR | -0.226178 | KIRC | RCC-MF | Apitolisib |
| MVP_top60_DNA_seed | HSP90AA1 | 1.801136 | HNSC | SKN-3 | Tanespimycin | MVP_top60_DNA+RNA_seed | HDAC1 | -0.226001 | KIRC | BB65-RCC | Belinostat |
| MVP_top60_DNA_seed | HDAC1 | 1.839277 | HNSC | KON | CAY10603 | MVP_top60_DNA+RNA_seed | ESR1 | -0.223061 | KIRC | HA7-RCC | Fulvestrant |
| MVP_top60_DNA_seed | AKT1 | 1.850309 | HNSC | SKN-3 | AKT inhibitor VIII | MVP_top60_DNA+RNA_seed | SMARCA4 | -0.2226 | KIRC | 786-0 | PFI-3 |
| MVP_top60_DNA_seed | EGFR | 1.857349 | HNSC | BB49-HNC | PF-00299804 | MVP_top60_DNA+RNA_seed | SMARCA2 | -0.2226 | KIRC | 786-0 | PFI-3 |
| MVP_top60_DNA_seed | NTRK1 | 1.857399 | HNSC | SCC90 | AZD1332 | MVP_top60_DNA+RNA_seed | HDAC1 | -0.221206 | KIRC | CAL-54 | AR-42 |
| MVP_top60_DNA_seed | EGFR | 1.883831 | HNSC | HSC-2 | AST-1306 | MVP_top60_DNA+RNA_seed | HSP90AA1 | -0.220771 | KIRC | RCC-JW | CCT-018159 |
| MVP_top60_DNA_seed | AR | 1.902923 | HNSC | SCC90 | Bicalutamide | MVP_top60_DNA+RNA_seed | HSP90AB1 | -0.220771 | KIRC | RCC-JW | CCT-018159 |
| MVP_top60_DNA_seed | AR | 2.107969 | HNSC | DOK | Bicalutamide | MVP_top60_DNA+RNA_seed | IGF1R | -0.220006 | KIRC | CAKI-1 | BMS-754807 |
| MVP_top60_DNA_seed | EGFR | 2.114542 | HNSC | HSC-2 | CI-1033 | MVP_top60_DNA+RNA_seed | FGFR4 | -0.219742 | KIRC | RXF393 | FGFR_0939 |
| MVP_top60_DNA_seed | RAC1 | 2.124311 | HNSC | SCC90 | EHT-1864 | MVP_top60_DNA+RNA_seed | HSP90AA1 | -0.219293 | KIRC | OS-RC-2 | Luminespib |
| MVP_top60_DNA_seed | AKT1 | 2.131141 | HNSC | SAT | AKT inhibitor VIII | MVP_top60_DNA+RNA_seed | HSP90AB1 | -0.219293 | KIRC | OS-RC-2 | Luminespib |
| MVP_top60_DNA_seed | HDAC1 | 2.132668 | HNSC | JHU-022 | JQ12 | MVP_top60_DNA+RNA_seed | AR | -0.218286 | KIRC | LB996-RCC | Bicalutamide |

| | | | | | | | | | | | |
|---|---|---|---|---|---|---|---|---|---|---|---|
| MVP_top60_DNA_seed | EGFR | 2.159594 | HNSC | KON | CI-1033 | MVP_top60_DNA+RNA_seed | MTOR | -0.216544 | KIRC | OS-RC-2 | QL-VIII-58 |
| MVP_top60_DNA_seed | HSP90AA1 | 2.267654 | HNSC | PCI-15A | Elesclomol | MVP_top60_DNA+RNA_seed | MTOR | -0.21581 | KIRC | SN12C | Apitolisib |
| MVP_top60_DNA_seed | EGFR | 2.292391 | HNSC | LB771-HNC | CUDC-101 | MVP_top60_DNA+RNA_seed | PTK2 | -0.215323 | KIRC | VMRC-RCZ | PF-562271 |
| MVP_top60_DNA_seed | EGFR | 2.318979 | HNSC | PCI-38 | CI-1033 | MVP_top60_DNA+RNA_seed | IGF1R | -0.215234 | KIRC | OS-RC-2 | BMS-754807 |
| MVP_top60_DNA_seed | HSP90AA1 | 2.356172 | HNSC | OSC-19 | Elesclomol | MVP_top60_DNA+RNA_seed | HSP90AA1 | -0.214167 | KIRC | HA7-RCC | Tanespimycin |
| MVP_top60_DNA_seed | EGFR | 2.363263 | HNSC | SAS | CI-1033 | MVP_top60_DNA+RNA_seed | HSP90AB1 | -0.214167 | KIRC | HA7-RCC | Tanespimycin |
| MVP_top60_DNA_seed | HDAC1 | 2.369562 | HNSC | PCI-15A | CAY10603 | MVP_top60_DNA+RNA_seed | AR | -0.211203 | KIRC | 769-P | Bicalutamide |
| MVP_top60_DNA_seed | HDAC1 | 2.381934 | HNSC | SCC90 | JQ12 | MVP_top60_DNA+RNA_seed | PIK3CD | -0.209011 | KIRC | RCC-MF | IC-87114 |
| MVP_top60_DNA_seed | EGFR | 2.55609 | HNSC | KON | PF-00299804 | MVP_top60_DNA+RNA_seed | PIK3CA | -0.207929 | KIRC | BFTC-909 | PI3Ka_4409 |
| MVP_top60_DNA_seed | HDAC1 | 2.599563 | HNSC | PCI-15A | AR-42 | MVP_top60_DNA+RNA_seed | PIK3CD | -0.207929 | KIRC | BFTC-909 | PI3Ka_4409 |
| MVP_top60_DNA_seed | HDAC1 | 2.653739 | HNSC | PCI-15A | Belinostat | MVP_top60_DNA+RNA_seed | PTK2 | -0.207899 | KIRC | OS-RC-2 | PF-562271 |
| MVP_top60_DNA_seed | EGFR | 2.65391 | HNSC | HSC-4 | CI-1033 | MVP_top60_DNA+RNA_seed | IGF1R | -0.203158 | KIRC | NCC021 | BMS-536924 |
| MVP_top60_DNA_seed | MDM2 | 2.687119 | HNSC | SCC-15 | Serdemetan | MVP_top60_DNA+RNA_seed | MTOR | -0.19988 | KIRC | HA7-RCC | Temsirolimus |
| MVP_top60_DNA_seed | EGFR | 2.700139 | HNSC | SAS | PF-00299804 | MVP_top60_DNA+RNA_seed | FGFR4 | -0.199676 | KIRC | SW156 | FGFR_0939 |
| MVP_top60_DNA_seed | EGFR | 2.718759 | HNSC | BB49-HNC | CI-1033 | MVP_top60_DNA+RNA_seed | MTOR | -0.19637 | KIRC | BB65-RCC | Voxtalisib |
| MVP_top60_DNA_seed | HSP90AA1 | 2.730801 | HNSC | LB771-HNC | Luminespib | MVP_top60_DNA+RNA_seed | ATM | -0.196073 | KIRC | HA7-RCC | KU-60019 |
| MVP_top60_DNA_seed | EGFR | 2.840812 | HNSC | KON | AST-1306 | MVP_top60_DNA+RNA_seed | ATM | -0.195957 | KIRC | A704 | CP466722 |
| MVP_top60_DNA_seed | EGFR | 2.960294 | HNSC | FADU | AST-1306 | MVP_top60_DNA+RNA_seed | HDAC1 | -0.195199 | KIRC | BB65-RCC | AR-42 |
| MVP_top60_DNA_seed | EGFR | 2.982329 | HNSC | FADU | PF-00299804 | MVP_top60_DNA+RNA_seed | MTOR | -0.194274 | KIRC | CAL-54 | Temsirolimus |
| MVP_top60_DNA_seed | EGFR | 2.985193 | HNSC | SAS | AST-1306 | MVP_top60_DNA+RNA_seed | MTOR | -0.193932 | KIRC | NCC021 | Torin 2 |
| MVP_top60_DNA_seed | EGFR | 3.017311 | HNSC | PCI-38 | AST-1306 | MVP_top60_DNA+RNA_seed | IGF1R | -0.19389 | KIRC | VMRC-RCW | Linsitinib |
| MVP_top60_DNA_seed | EGFR | 3.045156 | HNSC | PCI-38 | PF-00299804 | MVP_top60_DNA+RNA_seed | PIK3CD | -0.19283 | KIRC | RCC-AB | IC-87114 |
| MVP_top60_DNA_seed | HDAC1 | 3.06302 | HNSC | PCI-30 | CAY10603 | MVP_top60_DNA+RNA_seed | PIK3CA | -0.189765 | KIRC | A704 | AZD8835 |
| MVP_top60_DNA_seed | EGFR | 3.244078 | HNSC | BICR78 | AST-1306 | MVP_top60_DNA+RNA_seed | PIK3CD | -0.189765 | KIRC | A704 | AZD8835 |
| MVP_top60_DNA_seed | EGFR | 3.390525 | HNSC | HSC-4 | AST-1306 | MVP_top60_DNA+RNA_seed | ATM | -0.189101 | KIRC | RCC-AB | KU-55933 |
| MVP_top60_DNA_seed | EGFR | 3.456292 | HNSC | BB49-HNC | AST-1306 | MVP_top60_DNA+RNA_seed | HSP90AA1 | -0.188808 | KIRC | LB1047-RCC | CCT-018159 |
| MVP_top60_DNA_seed | EGFR | -3.566915 | HNSC | DOK | Gefitinib | MVP_top60_DNA+RNA_seed | HSP90AB1 | -0.188808 | KIRC | LB1047-RCC | CCT-018159 |
| MVP_top60_DNA_seed | EGFR | -3.221494 | HNSC | JHU-022 | Sapitinib | MVP_top60_DNA+RNA_seed | MTOR | -0.187079 | KIRC | HA7-RCC | Voxtalisib |
| MVP_top60_DNA_seed | EGFR | -3.049336 | HNSC | JHU-022 | Afatinib | MVP_top60_DNA+RNA_seed | PTK2 | -0.185545 | KIRC | TK10 | PF-562271 |
| MVP_top60_DNA_seed | EGFR | -2.97915 | HNSC | DOK | Erlotinib | MVP_top60_DNA+RNA_seed | IGF1R | -0.184294 | KIRC | RXF393 | BMS-754807 |
| MVP_top60_DNA_seed | EGFR | -2.947957 | HNSC | BB30-HNC | Sapitinib | MVP_top60_DNA+RNA_seed | IGF1R | -0.184019 | KIRC | VMRC-RCZ | Linsitinib |
| MVP_top60_DNA_seed | EGFR | -2.90684 | HNSC | JHU-022 | Gefitinib | MVP_top60_DNA+RNA_seed | HDAC1 | -0.181892 | KIRC | RCC-JW | Belinostat |
| MVP_top60_DNA_seed | EGFR | -2.860554 | HNSC | PCI-15A | Sapitinib | MVP_top60_DNA+RNA_seed | PIK3CD | -0.17599 | KIRC | HA7-RCC | IC-87114 |
| MVP_top60_DNA_seed | EGFR | -2.773562 | HNSC | CAL-27 | Sapitinib | MVP_top60_DNA+RNA_seed | HSP90AA1 | -0.175101 | KIRC | SN12C | CCT-018159 |
| MVP_top60_DNA_seed | EGFR | -2.722374 | HNSC | SCC90 | Sapitinib | MVP_top60_DNA+RNA_seed | HSP90AB1 | -0.175101 | KIRC | SN12C | CCT-018159 |
| MVP_top60_DNA_seed | EGFR | -2.705987 | HNSC | CAL-27 | Afatinib | MVP_top60_DNA+RNA_seed | MTOR | -0.172219 | KIRC | VMRC-RCZ | Temsirolimus |
| MVP_top60_DNA_seed | AKT1 | -2.704943 | HNSC | HO-1-u-1 | Afuresertib | MVP_top60_DNA+RNA_seed | MTOR | -0.165496 | KIRC | LB996-RCC | Apitolisib |
| MVP_top60_DNA_seed | EGFR | -2.645496 | HNSC | JHU-029 | AZD3759 | MVP_top60_DNA+RNA_seed | HSP90AA1 | -0.164817 | KIRC | LB1047-RCC | Luminespib |
| MVP_top60_DNA_seed | EGFR | -2.644366 | HNSC | Ca9-22 | Gefitinib | MVP_top60_DNA+RNA_seed | HSP90AB1 | -0.164817 | KIRC | LB1047-RCC | Luminespib |
| MVP_top60_DNA_seed | EGFR | -2.620066 | HNSC | DOK | Sapitinib | MVP_top60_DNA+RNA_seed | HSP90AA1 | -0.164124 | KIRC | CAKI-1 | Tanespimycin |
| MVP_top60_DNA_seed | AKT1 | -2.582768 | HNSC | HO-1-u-1 | Ipatasertib | MVP_top60_DNA+RNA_seed | HSP90AB1 | -0.164124 | KIRC | CAKI-1 | Tanespimycin |

| | | | | | | | | | | | |
|---|---|---|---|---|---|---|---|---|---|---|---|
| MVP_top60_DNA_seed | EGFR | -2.539374 | HNSC | JHU-029 | Afatinib | MVP_top60_DNA+RNA_seed | HDAC1 | -0.162284 | KIRC | BFTC-909 | JQ12 |
| MVP_top60_DNA_seed | EGFR | -2.492557 | HNSC | DOK | AZD3759 | MVP_top60_DNA+RNA_seed | PIK3CD | -0.160427 | KIRC | CAKI-1 | Idelalisib |
| MVP_top60_DNA_seed | AKT1 | -2.481373 | HNSC | HO-1-u-1 | Uprosertib | MVP_top60_DNA+RNA_seed | MTOR | -0.158671 | KIRC | TK10 | Torin 2 |
| MVP_top60_DNA_seed | EGFR | -2.471929 | HNSC | JHU-022 | AZD3759 | MVP_top60_DNA+RNA_seed | HDAC1 | -0.157209 | KIRC | TK10 | Entinostat |
| MVP_top60_DNA_seed | EGFR | -2.445423 | HNSC | PCI-6A | AZD3759 | MVP_top60_DNA+RNA_seed | IGF1R | -0.154537 | KIRC | VMRC-RCW | BMS-754807 |
| MVP_top60_DNA_seed | AKT1 | -2.431201 | HNSC | HO-1-u-1 | AZD5363 | MVP_top60_DNA+RNA_seed | HDAC1 | -0.152026 | KIRC | RCC-JW | CAY10603 |
| MVP_top60_DNA_seed | EGFR | -2.422715 | HNSC | PCI-15A | AZD3759 | MVP_top60_DNA+RNA_seed | IGF1R | -0.145662 | KIRC | TK10 | BMS-754807 |
| MVP_top60_DNA_seed | AKT1 | -2.406612 | HNSC | SCC90 | Uprosertib | MVP_top60_DNA+RNA_seed | HDAC1 | -0.145571 | KIRC | U031 | Dacinostat |
| MVP_top60_DNA_seed | EGFR | -2.394435 | HNSC | BB30-HNC | Afatinib | MVP_top60_DNA+RNA_seed | HDAC1 | -0.142646 | KIRC | VMRC-RCZ | CAY10603 |
| MVP_top60_DNA_seed | EGFR | -2.354548 | HNSC | BB30-HNC | Gefitinib | MVP_top60_DNA+RNA_seed | IGF1R | -0.142552 | KIRC | RCC10RGB | BMS-536924 |
| MVP_top60_DNA_seed | EGFR | -2.353248 | HNSC | SAT | Gefitinib | MVP_top60_DNA+RNA_seed | HDAC1 | -0.137204 | KIRC | VMRC-RCZ | AR-42 |
| MVP_top60_DNA_seed | EGFR | -2.340962 | HNSC | BB30-HNC | Erlotinib | MVP_top60_DNA+RNA_seed | PIK3CD | -0.135731 | KIRC | RCC-AB | AZD8186 |
| MVP_top60_DNA_seed | RAC1 | -2.33522 | HNSC | HSC-4 | EHT-1864 | MVP_top60_DNA+RNA_seed | HSP90AA1 | -0.135203 | KIRC | LB996-RCC | Tanespimycin |
| MVP_top60_DNA_seed | EGFR | -2.324371 | HNSC | JHU-029 | Sapitinib | MVP_top60_DNA+RNA_seed | HSP90AB1 | -0.135203 | KIRC | LB996-RCC | Tanespimycin |
| MVP_top60_DNA_seed | EGFR | -2.275323 | HNSC | SAT | Sapitinib | MVP_top60_DNA+RNA_seed | IGF1R | -0.134765 | KIRC | ACHN | BMS-536924 |
| MVP_top60_DNA_seed | EGFR | -2.263832 | HNSC | Ca9-22 | Sapitinib | MVP_top60_DNA+RNA_seed | HDAC1 | -0.12785 | KIRC | KMRC-1 | Belinostat |
| MVP_top60_DNA_seed | EGFR | -2.25111 | HNSC | SCC-15 | Sapitinib | MVP_top60_DNA+RNA_seed | MTOR | -0.12488 | KIRC | TK10 | WYE-125132 |
| MVP_top60_DNA_seed | PIK3CA | -2.249485 | HNSC | DOK | Alpelisib | MVP_top60_DNA+RNA_seed | AR | -0.124031 | KIRC | RCC-JW | AZD3514 |
| MVP_top60_DNA_seed | EGFR | -2.244848 | HNSC | HO-1-u-1 | Gefitinib | MVP_top60_DNA+RNA_seed | MTOR | -0.117217 | KIRC | U031 | CRT0105446 |
| MVP_top60_DNA_seed | EGFR | -2.229231 | HNSC | SAT | Afatinib | MVP_top60_DNA+RNA_seed | FGFR4 | -0.116518 | KIRC | A498 | FGFR_3831 |
| MVP_top60_DNA_seed | EGFR | -2.214032 | HNSC | DOK | Osimertinib | MVP_top60_DNA+RNA_seed | AR | -0.114329 | KIRC | 786-0 | Bicalutamide |
| MVP_top60_DNA_seed | EGFR | -2.201715 | HNSC | PCI-15A | Gefitinib | MVP_top60_DNA+RNA_seed | PIK3CA | -0.113309 | KIRC | SN12C | PI3Ka_4409 |
| MVP_top60_DNA_seed | EGFR | -2.197485 | HNSC | HO-1-u-1 | Sapitinib | MVP_top60_DNA+RNA_seed | PIK3CD | -0.113309 | KIRC | SN12C | PI3Ka_4409 |
| MVP_top60_DNA_seed | EGFR | -2.181597 | HNSC | DOK | Afatinib | MVP_top60_DNA+RNA_seed | FGFR4 | -0.112227 | KIRC | 786-0 | FGFR_3831 |
| MVP_top60_DNA_seed | EGFR | -2.154724 | HNSC | BB30-HNC | AZD3759 | MVP_top60_DNA+RNA_seed | IGF1R | -0.112111 | KIRC | 769-P | GSK1904529A |
| MVP_top60_DNA_seed | EGFR | -2.113556 | HNSC | CAL-27 | Gefitinib | MVP_top60_DNA+RNA_seed | HDAC1 | -0.1111 | KIRC | RCC-FG2 | Belinostat |
| MVP_top60_DNA_seed | EGFR | -2.110055 | HNSC | JHU-022 | Osimertinib | MVP_top60_DNA+RNA_seed | IGF1R | -0.109975 | KIRC | U031 | Linsitinib |
| MVP_top60_DNA_seed | EGFR | -2.096094 | HNSC | Ca9-22 | Erlotinib | MVP_top60_DNA+RNA_seed | FGFR4 | -0.109773 | KIRC | BB65-RCC | FGFR_0939 |
| MVP_top60_DNA_seed | EGFR | -2.086745 | HNSC | FADU | Gefitinib | MVP_top60_DNA+RNA_seed | PIK3CD | -0.108683 | KIRC | RCC-JW | IC-87114 |
| MVP_top60_DNA_seed | EGFR | -2.085032 | HNSC | JHU-029 | Gefitinib | MVP_top60_DNA+RNA_seed | MTOR | -0.107345 | KIRC | RCC-JW | WYE-125132 |
| MVP_top60_DNA_seed | EGFR | -2.06152 | HNSC | SCC-15 | Gefitinib | MVP_top60_DNA+RNA_seed | MTOR | -0.107326 | KIRC | HA7-RCC | Torin 2 |
| MVP_top60_DNA_seed | PIK3CA | -2.022601 | HNSC | BB49-HNC | AZD8186 | MVP_top60_DNA+RNA_seed | ATM | -0.10658 | KIRC | ACHN | KU-60019 |
| MVP_top60_DNA_seed | EGFR | -2.02171 | HNSC | JHU-011 | AZD3759 | MVP_top60_DNA+RNA_seed | HDAC1 | -0.100476 | KIRC | U031 | Tubastatin A |
| MVP_top60_DNA_seed | PIK3CA | -2.021446 | HNSC | BB49-HNC | Alpelisib | MVP_top60_DNA+RNA_seed | PIK3CA | -0.095383 | KIRC | ACHN | AZD8835 |
| MVP_top60_DNA_seed | EGFR | -2.012184 | HNSC | JHU-029 | Erlotinib | MVP_top60_DNA+RNA_seed | PIK3CD | -0.095383 | KIRC | ACHN | AZD8835 |
| MVP_top60_DNA_seed | EGFR | -2.00951 | HNSC | CAL-27 | AZD3759 | MVP_top60_DNA+RNA_seed | ESR1 | -0.094579 | KIRC | ACHN | Fulvestrant |
| MVP_top60_DNA_seed | EGFR | -2.004977 | HNSC | SCC90 | Afatinib | MVP_top60_DNA+RNA_seed | HDAC1 | -0.09437 | KIRC | BB65-RCC | Dacinostat |
| MVP_top60_DNA_seed | EGFR | -1.996162 | HNSC | SAT | Osimertinib | MVP_top60_DNA+RNA_seed | MTOR | -0.089895 | KIRC | LB1047-RCC | Torin 2 |
| MVP_top60_DNA_seed | PIK3CA | -1.98395 | HNSC | BB49-HNC | Buparlisib | MVP_top60_DNA+RNA_seed | MTOR | -0.087372 | KIRC | SN12C | WYE-125132 |
| MVP_top60_DNA_seed | AKT1 | -1.967462 | HNSC | SCC90 | GSK2110183B | MVP_top60_DNA+RNA_seed | AR | -0.086792 | KIRC | U031 | Bicalutamide |
| MVP_top60_DNA_seed | EGFR | -1.9495 | HNSC | Ca9-22 | AZD3759 | MVP_top60_DNA+RNA_seed | PIK3CA | -0.082612 | KIRC | NCC021 | PI3Ka_4409 |

| | | | | | | | | | | | |
|---|---|---|---|---|---|---|---|---|---|---|---|
| MVP_top60_DNA_seed | AKT1 | -1.941813 | HNSC | DOK | Afuresertib | MVP_top60_DNA+RNA_seed | PIK3CD | -0.082612 | KIRC | NCC021 | PI3Ka_4409 |
| MVP_top60_DNA_seed | EGFR | -1.938119 | HNSC | HSC-4 | Gefitinib | MVP_top60_DNA+RNA_seed | MTOR | -0.08219 | KIRC | RCC-MF | Torin 2 |
| MVP_top60_DNA_seed | ESR1 | -1.930756 | HNSC | HO-1-u-1 | GDC0810 | MVP_top60_DNA+RNA_seed | FGFR4 | -0.080865 | KIRC | SN12C | FGFR_3831 |
| MVP_top60_DNA_seed | EGFR | -1.928244 | HNSC | FADU | Afatinib | MVP_top60_DNA+RNA_seed | IGF1R | -0.076025 | KIRC | RXF393 | Linsitinib |
| MVP_top60_DNA_seed | EGFR | -1.924365 | HNSC | FADU | Sapitinib | MVP_top60_DNA+RNA_seed | ATM | -0.07542 | KIRC | NCC010 | KU-55933 |
| MVP_top60_DNA_seed | EGFR | -1.918719 | HNSC | BB30-HNC | Osimertinib | MVP_top60_DNA+RNA_seed | EP300 | -0.073864 | KIRC | LB996-RCC | I-CBP112 |
| MVP_top60_DNA_seed | MAPK1 | -1.912366 | HNSC | JHU-029 | SCH772984 | MVP_top60_DNA+RNA_seed | HSP90AA1 | -0.06895 | KIRC | NCC010 | CCT-018159 |
| MVP_top60_DNA_seed | MAPK3 | -1.912366 | HNSC | JHU-029 | SCH772984 | MVP_top60_DNA+RNA_seed | HSP90AB1 | -0.06895 | KIRC | NCC010 | CCT-018159 |
| MVP_top60_DNA_seed | EGFR | -1.891622 | HNSC | PCI-15A | Lapatinib | MVP_top60_DNA+RNA_seed | IGF1R | -0.068784 | KIRC | TK10 | Linsitinib |
| MVP_top60_DNA_seed | HSP90AA1 | -1.890437 | HNSC | DOK | Tanespimycin | MVP_top60_DNA+RNA_seed | EP300 | -0.065939 | KIRC | TK10 | I-CBP112 |
| MVP_top60_DNA_seed | MAPK1 | -1.888301 | HNSC | SCC90 | SCH772984 | MVP_top60_DNA+RNA_seed | MTOR | -0.065909 | KIRC | SN12C | Temsirolimus |
| MVP_top60_DNA_seed | MAPK3 | -1.888301 | HNSC | SCC90 | SCH772984 | MVP_top60_DNA+RNA_seed | PIK3CA | -0.065568 | KIRC | VMRC-RCZ | AZD8835 |
| MVP_top60_DNA_seed | AKT1 | -1.88144 | HNSC | DOK | AZD5363 | MVP_top60_DNA+RNA_seed | PIK3CD | -0.065568 | KIRC | VMRC-RCZ | AZD8835 |
| MVP_top60_DNA_seed | EGFR | -1.873626 | HNSC | BB30-HNC | Lapatinib | MVP_top60_DNA+RNA_seed | MTOR | -0.065247 | KIRC | CAL-54 | Torin 2 |
| MVP_top60_DNA_seed | EGFR | -1.850098 | HNSC | PCI-15A | Osimertinib | MVP_top60_DNA+RNA_seed | AR | -0.064388 | KIRC | HA7-RCC | Bicalutamide |
| MVP_top60_DNA_seed | EGFR | -1.845065 | HNSC | JHU-029 | Lapatinib | MVP_top60_DNA+RNA_seed | PIK3CD | -0.061552 | KIRC | ACHN | IC-87114 |
| MVP_top60_DNA_seed | EGFR | -1.838298 | HNSC | JHU-022 | Lapatinib | MVP_top60_DNA+RNA_seed | HDAC1 | -0.059196 | KIRC | RCC-FG2 | CAY10603 |
| MVP_top60_DNA_seed | EGFR | -1.834029 | HNSC | SAT | AZD3759 | MVP_top60_DNA+RNA_seed | HDAC1 | -0.057742 | KIRC | CAKI-1 | CAY10603 |
| MVP_top60_DNA_seed | CDK2 | -1.82642 | HNSC | HO-1-u-1 | MK-8776 | MVP_top60_DNA+RNA_seed | HDAC1 | -0.056974 | KIRC | RCC-JW | AR-42 |
| MVP_top60_DNA_seed | AKT1 | -1.823186 | HNSC | BB49-HNC | AT13148 | MVP_top60_DNA+RNA_seed | EP300 | -0.0555 | KIRC | NCC021 | I-CBP112 |
| MVP_top60_DNA_seed | HSP90AA1 | -1.822219 | HNSC | FADU | Tanespimycin | MVP_top60_DNA+RNA_seed | HDAC1 | -0.054711 | KIRC | LB1047-RCC | Tubastatin A |
| MVP_top60_DNA_seed | EGFR | -1.818565 | HNSC | HSC-3 | Gefitinib | MVP_top60_DNA+RNA_seed | HSP90AA1 | -0.054437 | KIRC | BFTC-909 | Elesclomol |
| MVP_top60_DNA_seed | ESR1 | -1.769275 | HNSC | HO-1-u-1 | Tamoxifen | MVP_top60_DNA+RNA_seed | HSP90AB1 | -0.054437 | KIRC | BFTC-909 | Elesclomol |
| MVP_top60_DNA_seed | PIK3CA | -1.76856 | HNSC | HSC-2 | Alpelisib | MVP_top60_DNA+RNA_seed | FGFR4 | -0.052158 | KIRC | KMRC-1 | FGFR_0939 |
| MVP_top60_DNA_seed | EGFR | -1.768406 | HNSC | Ca9-22 | Lapatinib | MVP_top60_DNA+RNA_seed | MTOR | -0.049196 | KIRC | BFTC-909 | Temsirolimus |
| MVP_top60_DNA_seed | PIK3CA | -1.763759 | HNSC | JHU-011 | Alpelisib | MVP_top60_DNA+RNA_seed | AR | -0.045334 | KIRC | BFTC-909 | AZD3514 |
| MVP_top60_DNA_seed | EGFR | -1.74209 | HNSC | KOSC-2 | AZD3759 | MVP_top60_DNA+RNA_seed | FGFR4 | -0.044434 | KIRC | LB996-RCC | FGFR_3831 |
| MVP_top60_DNA_seed | EGFR | -1.738885 | HNSC | SCC-9 | Sapitinib | MVP_top60_DNA+RNA_seed | PIK3CD | -0.044269 | KIRC | 769-P | Idelalisib |
| MVP_top60_DNA_seed | EGFR | -1.735728 | HNSC | HSC-3 | Erlotinib | MVP_top60_DNA+RNA_seed | ESR1 | -0.042171 | KIRC | RCC-AB | Tamoxifen |
| MVP_top60_DNA_seed | EGFR | -1.724549 | HNSC | HO-1-u-1 | Erlotinib | MVP_top60_DNA+RNA_seed | IGF1R | -0.039219 | KIRC | A498 | BMS-536924 |
| MVP_top60_DNA_seed | EGFR | -1.722712 | HNSC | JHU-011 | Gefitinib | MVP_top60_DNA+RNA_seed | IGF1R | -0.038349 | KIRC | LB996-RCC | Linsitinib |
| MVP_top60_DNA_seed | PIK3CA | -1.722505 | HNSC | CAL-33 | Alpelisib | MVP_top60_DNA+RNA_seed | HDAC1 | -0.037955 | KIRC | KMRC-1 | CAY10603 |
| MVP_top60_DNA_seed | EGFR | -1.716556 | HNSC | Detroit562 | AZD3759 | MVP_top60_DNA+RNA_seed | HSP90AA1 | -0.037007 | KIRC | OS-RC-2 | Elesclomol |
| MVP_top60_DNA_seed | EGFR | -1.711139 | HNSC | PCI-15A | Erlotinib | MVP_top60_DNA+RNA_seed | HSP90AB1 | -0.037007 | KIRC | OS-RC-2 | Elesclomol |
| MVP_top60_DNA_seed | EGFR | -1.708645 | HNSC | SCC90 | Gefitinib | MVP_top60_DNA+RNA_seed | HDAC1 | -0.035003 | KIRC | RCC-JF | Tubastatin A |
| MVP_top60_DNA_seed | EGFR | -1.688246 | HNSC | FADU | Osimertinib | MVP_top60_DNA+RNA_seed | HDAC1 | -0.034271 | KIRC | KMRC-1 | AR-42 |
| MVP_top60_DNA_seed | TP53 | -1.685645 | HNSC | HO-1-N-1 | MIRA-1 | MVP_top60_DNA+RNA_seed | IGF1R | -0.030642 | KIRC | OS-RC-2 | BMS-536924 |
| MVP_top60_DNA_seed | EGFR | -1.676853 | HNSC | HSC-4 | Erlotinib | MVP_top60_DNA+RNA_seed | IGF1R | -0.028584 | KIRC | RCC-MF | BMS-536924 |
| MVP_top60_DNA_seed | EGFR | -1.676852 | HNSC | HO-1-u-1 | Osimertinib | MVP_top60_DNA+RNA_seed | MTOR | -0.02818 | KIRC | RCC-JW | Voxtalisib |
| MVP_top60_DNA_seed | EGFR | -1.674509 | HNSC | SCC90 | Lapatinib | MVP_top60_DNA+RNA_seed | MTOR | -0.027266 | KIRC | RCC-ER | WYE-125132 |
| MVP_top60_DNA_seed | EGFR | -1.667578 | HNSC | PCI-4B | Sapitinib | MVP_top60_DNA+RNA_seed | HDAC1 | -0.023813 | KIRC | LB2241-RCC | Dacinostat |

| | | | | | | | | | | | |
|---|---|---|---|---|---|---|---|---|---|---|---|
| MVP_top60_DNA_seed | AKT1 | -1.667141 | HNSC | DOK | Ipatasertib | MVP_top60_DNA+RNA_seed | IGF1R | -0.019203 | KIRC | RCC-FG2 | BMS-536924 |
| MVP_top60_DNA_seed | HSP90AA1 | -1.660769 | HNSC | PCI-15A | Tanespimycin | MVP_top60_DNA+RNA_seed | HSP90AA1 | -0.016759 | KIRC | SN12C | Luminespib |
| MVP_top60_DNA_seed | EGFR | -1.658695 | HNSC | SCC90 | Erlotinib | MVP_top60_DNA+RNA_seed | HSP90AB1 | -0.016759 | KIRC | SN12C | Luminespib |
| MVP_top60_DNA_seed | MAPK1 | -1.648933 | HNSC | SAS | SCH772984 | MVP_top60_DNA+RNA_seed | HSP90AA1 | -0.014276 | KIRC | TK10 | Elesclomol |
| MVP_top60_DNA_seed | MAPK3 | -1.648933 | HNSC | SAS | SCH772984 | MVP_top60_DNA+RNA_seed | HSP90AB1 | -0.014276 | KIRC | TK10 | Elesclomol |
| MVP_top60_DNA_seed | EGFR | -1.625996 | HNSC | HSC-2 | Sapitinib | MVP_top60_DNA+RNA_seed | HSP90AA1 | -0.012909 | KIRC | LB2241-RCC | Elesclomol |
| MVP_top60_DNA_seed | NTRK1 | -1.622622 | HNSC | BB49-HNC | AZD1332 | MVP_top60_DNA+RNA_seed | HSP90AB1 | -0.012909 | KIRC | LB2241-RCC | Elesclomol |
| MVP_top60_DNA_seed | EGFR | -1.622364 | HNSC | BHY | Gefitinib | MVP_top60_DNA+RNA_seed | PIK3CA | -0.008739 | KIRC | BB65-RCC | AZD8835 |
| MVP_top60_DNA_seed | EGFR | -1.621704 | HNSC | JHU-011 | Sapitinib | MVP_top60_DNA+RNA_seed | PIK3CD | -0.008739 | KIRC | BB65-RCC | AZD8835 |
| MVP_top60_DNA_seed | EGFR | -1.620646 | HNSC | FADU | AZD3759 | MVP_top60_DNA+RNA_seed | HSP90AA1 | -0.008217 | KIRC | U031 | Luminespib |
| MVP_top60_DNA_seed | NTRK1 | -1.620469 | HNSC | Detroit562 | GW441756 | MVP_top60_DNA+RNA_seed | HSP90AB1 | -0.008217 | KIRC | U031 | Luminespib |
| MVP_top60_DNA_seed | AKT1 | -1.588291 | HNSC | HO-1-u-1 | AT13148 | MVP_top60_DNA+RNA_seed | MTOR | -0.002093 | KIRC | LB1047-RCC | WYE-125132 |
| MVP_top60_DNA_seed | HSP90AA1 | -1.585949 | HNSC | BB30-HNC | Tanespimycin | MVP_top60_DNA+RNA_seed | AR | 0.002661 | KIRC | TK10 | AZD3514 |
| MVP_top60_DNA_seed | EGFR | -1.580619 | HNSC | SAS | Erlotinib | MVP_top60_DNA+RNA_seed | IGF1R | 0.003094 | KIRC | VMRC-RCZ | BMS-754807 |
| MVP_top60_DNA_seed | HSP90AA1 | -1.576869 | HNSC | HO-1-u-1 | Tanespimycin | MVP_top60_DNA+RNA_seed | SMARCA4 | 0.006007 | KIRC | RCC-JW | PFI-3 |
| MVP_top60_DNA_seed | HSP90AA1 | -1.576837 | HNSC | JHU-011 | Tanespimycin | MVP_top60_DNA+RNA_seed | SMARCA2 | 0.006007 | KIRC | RCC-JW | PFI-3 |
| MVP_top60_DNA_seed | EGFR | -1.576254 | HNSC | OSC-19 | AZD3759 | MVP_top60_DNA+RNA_seed | HDAC1 | 0.008618 | KIRC | ACHN | Dacinostat |
| MVP_top60_DNA_seed | PIK3CA | -1.575222 | HNSC | BB49-HNC | GNE-317 | MVP_top60_DNA+RNA_seed | HSP90AA1 | 0.009337 | KIRC | RCC-FG2 | Tanespimycin |
| MVP_top60_DNA_seed | AKT1 | -1.564704 | HNSC | Detroit562 | AT13148 | MVP_top60_DNA+RNA_seed | HSP90AB1 | 0.009337 | KIRC | RCC-FG2 | Tanespimycin |
| MVP_top60_DNA_seed | EGFR | -1.563303 | HNSC | PCI-15A | Afatinib | MVP_top60_DNA+RNA_seed | HSP90AA1 | 0.010175 | KIRC | KMRC-1 | Tanespimycin |
| MVP_top60_DNA_seed | EGFR | -1.54836 | HNSC | CAL-27 | Lapatinib | MVP_top60_DNA+RNA_seed | HSP90AB1 | 0.010175 | KIRC | KMRC-1 | Tanespimycin |
| MVP_top60_DNA_seed | EGFR | -1.539547 | HNSC | JHU-022 | Erlotinib | MVP_top60_DNA+RNA_seed | ESR1 | 0.016265 | KIRC | RCC-JW | Tamoxifen |
| MVP_top60_DNA_seed | AKT1 | -1.538778 | HNSC | HSC-3 | AT13148 | MVP_top60_DNA+RNA_seed | HDAC1 | 0.020514 | KIRC | CAL-54 | CAY10603 |
| MVP_top60_DNA_seed | EGFR | -1.536124 | HNSC | CAL-27 | Erlotinib | MVP_top60_DNA+RNA_seed | MTOR | 0.020699 | KIRC | BB65-RCC | QL-VIII-58 |
| MVP_top60_DNA_seed | EGFR | -1.532678 | HNSC | DOK | Lapatinib | MVP_top60_DNA+RNA_seed | ATM | 0.021403 | KIRC | HA7-RCC | CP466722 |
| MVP_top60_DNA_seed | EGFR | -1.52903 | HNSC | FADU | Lapatinib | MVP_top60_DNA+RNA_seed | FGFR4 | 0.022847 | KIRC | RCC-AB | FGFR_3831 |
| MVP_top60_DNA_seed | RAC1 | -1.523188 | HNSC | SAS | EHT-1864 | MVP_top60_DNA+RNA_seed | MTOR | 0.024029 | KIRC | BB65-RCC | WYE-125132 |
| MVP_top60_DNA_seed | EGFR | -1.522742 | HNSC | HSC-2 | Erlotinib | MVP_top60_DNA+RNA_seed | HDAC1 | 0.026564 | KIRC | RCC10RGB | Dacinostat |
| MVP_top60_DNA_seed | EGFR | -1.522426 | HNSC | PCI-6A | Sapitinib | MVP_top60_DNA+RNA_seed | AR | 0.030987 | KIRC | VMRC-RCW | Bicalutamide |
| MVP_top60_DNA_seed | EGFR | -1.51585 | HNSC | KOSC-2 | Sapitinib | MVP_top60_DNA+RNA_seed | HDAC1 | 0.031744 | KIRC | RCC-FG2 | AR-42 |
| MVP_top60_DNA_seed | PIK3CA | -1.51221 | HNSC | HO-1-u-1 | GNE-317 | MVP_top60_DNA+RNA_seed | MTOR | 0.032636 | KIRC | LB1047-RCC | CRT0105446 |
| MVP_top60_DNA_seed | EGFR | -1.512014 | HNSC | SAT | Lapatinib | MVP_top60_DNA+RNA_seed | PIK3CA | 0.033875 | KIRC | RCC-JW | PI3Ka_4409 |
| MVP_top60_DNA_seed | EGFR | -1.504427 | HNSC | PCI-6A | Lapatinib | MVP_top60_DNA+RNA_seed | PIK3CD | 0.033875 | KIRC | RCC-JW | PI3Ka_4409 |
| MVP_top60_DNA_seed | EGFR | -1.500202 | HNSC | HSC-2 | Gefitinib | MVP_top60_DNA+RNA_seed | PIK3CA | 0.034229 | KIRC | RCC-FG2 | AZD8835 |
| MVP_top60_DNA_seed | SRC | -1.493221 | HNSC | DOK | Bosutinib | MVP_top60_DNA+RNA_seed | PIK3CD | 0.034229 | KIRC | RCC-FG2 | AZD8835 |
| MVP_top60_DNA_seed | HSP90AA1 | -1.485925 | HNSC | HSC-2 | Tanespimycin | MVP_top60_DNA+RNA_seed | FGFR4 | 0.035524 | KIRC | CAKI-1 | FGFR_0939 |
| MVP_top60_DNA_seed | EGFR | -1.481878 | HNSC | FADU | Erlotinib | MVP_top60_DNA+RNA_seed | ESR1 | 0.047327 | KIRC | SW156 | Fulvestrant |
| MVP_top60_DNA_seed | MAPK1 | -1.480324 | HNSC | SAS | VX-11e | MVP_top60_DNA+RNA_seed | ATM | 0.050515 | KIRC | SN12C | KU-55933 |
| MVP_top60_DNA_seed | CDK2 | -1.479895 | HNSC | PCI-15A | Dinaciclib | MVP_top60_DNA+RNA_seed | AR | 0.054523 | KIRC | BFTC-909 | Bicalutamide |
| MVP_top60_DNA_seed | MAPK1 | -1.477812 | HNSC | DOK | Ulixertinib | MVP_top60_DNA+RNA_seed | AR | 0.05492 | KIRC | RCC-FG2 | Bicalutamide |
| MVP_top60_DNA_seed | MAPK3 | -1.477812 | HNSC | DOK | Ulixertinib | MVP_top60_DNA+RNA_seed | ATM | 0.056401 | KIRC | TK10 | KU-55933 |

| | | | | | | | | | | | |
|---|---|---|---|---|---|---|---|---|---|---|---|
| MVP_top60_DNA_seed | ESR1 | -1.476128 | HNSC | Detroit562 | Tamoxifen | MVP_top60_DNA+RNA_seed | MTOR | 0.059497 | KIRC | VMRC-RCZ | Voxtalisib |
| MVP_top60_DNA_seed | EGFR | -1.473384 | HNSC | SCC90 | AZD3759 | MVP_top60_DNA+RNA_seed | PIK3CD | 0.059697 | KIRC | A704 | IC-87114 |
| MVP_top60_DNA_seed | EGFR | -1.466893 | HNSC | CAL-33 | Gefitinib | MVP_top60_DNA+RNA_seed | IGF1R | 0.061786 | KIRC | 786-0 | BMS-536924 |
| MVP_top60_DNA_seed | CDK2 | -1.462514 | HNSC | BICR22 | Dinaciclib | MVP_top60_DNA+RNA_seed | EP300 | 0.063953 | KIRC | CAL-54 | I-CBP112 |
| MVP_top60_DNA_seed | EGFR | -1.459199 | HNSC | Ca9-22 | Afatinib | MVP_top60_DNA+RNA_seed | MTOR | 0.064233 | KIRC | CAKI-1 | Apitolisib |
| MVP_top60_DNA_seed | MAPK1 | -1.454562 | HNSC | SAS | Ulixertinib | MVP_top60_DNA+RNA_seed | FGFR4 | 0.06514 | KIRC | ACHN | FGFR_3831 |
| MVP_top60_DNA_seed | MAPK3 | -1.454562 | HNSC | SAS | Ulixertinib | MVP_top60_DNA+RNA_seed | HSP90AA1 | 0.06533 | KIRC | TK10 | CCT-018159 |
| MVP_top60_DNA_seed | AKT1 | -1.454299 | HNSC | JHU-011 | Ipatasertib | MVP_top60_DNA+RNA_seed | HSP90AB1 | 0.06533 | KIRC | TK10 | CCT-018159 |
| MVP_top60_DNA_seed | AKT1 | -1.453223 | HNSC | BB49-HNC | AZD5363 | MVP_top60_DNA+RNA_seed | MTOR | 0.065656 | KIRC | SN12C | Voxtalisib |
| MVP_top60_DNA_seed | CDK2 | -1.452132 | HNSC | BICR10 | MK-8776 | MVP_top60_DNA+RNA_seed | HSP90AA1 | 0.067311 | KIRC | HA7-RCC | Luminespib |
| MVP_top60_DNA_seed | EGFR | -1.449294 | HNSC | HO-1-u-1 | Lapatinib | MVP_top60_DNA+RNA_seed | HSP90AB1 | 0.067311 | KIRC | HA7-RCC | Luminespib |
| MVP_top60_DNA_seed | HSP90AA1 | -1.447799 | HNSC | KON | Tanespimycin | MVP_top60_DNA+RNA_seed | HSP90AA1 | 0.07142 | KIRC | A498 | Elesclomol |
| MVP_top60_DNA_seed | EGFR | -1.446331 | HNSC | SCC-15 | Afatinib | MVP_top60_DNA+RNA_seed | HSP90AB1 | 0.07142 | KIRC | A498 | Elesclomol |
| MVP_top60_DNA_seed | HSP90AA1 | -1.445119 | HNSC | HSC-3 | Tanespimycin | MVP_top60_DNA+RNA_seed | PIK3CA | 0.076929 | KIRC | ACHN | PI3Ka_4409 |
| MVP_top60_DNA_seed | MAPK1 | -1.443779 | HNSC | SAS | Ulixertinib | MVP_top60_DNA+RNA_seed | PIK3CD | 0.076929 | KIRC | ACHN | PI3Ka_4409 |
| MVP_top60_DNA_seed | MAPK3 | -1.443779 | HNSC | SAS | Ulixertinib | MVP_top60_DNA+RNA_seed | IGF1R | 0.077851 | KIRC | BB65-RCC | BMS-536924 |
| MVP_top60_DNA_seed | EGFR | -1.431127 | HNSC | KON | AZD3759 | MVP_top60_DNA+RNA_seed | MTOR | 0.079732 | KIRC | RCC-MF | WYE-125132 |
| MVP_top60_DNA_seed | EGFR | -1.428412 | HNSC | SAT | Erlotinib | MVP_top60_DNA+RNA_seed | HSP90AA1 | 0.084589 | KIRC | RCC-FG2 | SNX-2112 |
| MVP_top60_DNA_seed | EGFR | -1.425397 | HNSC | CAL-33 | Sapitinib | MVP_top60_DNA+RNA_seed | HSP90AB1 | 0.084589 | KIRC | RCC-FG2 | SNX-2112 |
| MVP_top60_DNA_seed | EGFR | -1.420914 | HNSC | PCI-6A | Afatinib | MVP_top60_DNA+RNA_seed | PIK3CA | 0.085709 | KIRC | CAL-54 | PI3Ka_4409 |
| MVP_top60_DNA_seed | CDK2 | -1.419259 | HNSC | Detroit562 | AZD5438 | MVP_top60_DNA+RNA_seed | PIK3CD | 0.085709 | KIRC | CAL-54 | PI3Ka_4409 |
| MVP_top60_DNA_seed | HSP90AA1 | -1.418773 | HNSC | HO-1-N-1 | Tanespimycin | MVP_top60_DNA+RNA_seed | FGFR4 | 0.088982 | KIRC | RCC-ER | FGFR_0939 |
| MVP_top60_DNA_seed | HSP90AA1 | -1.417106 | HNSC | BB30-HNC | CCT-018159 | MVP_top60_DNA+RNA_seed | MTOR | 0.090156 | KIRC | OS-RC-2 | Apitolisib |
| MVP_top60_DNA_seed | EGFR | -1.414253 | HNSC | SCC-9 | Afatinib | MVP_top60_DNA+RNA_seed | ESR1 | 0.090517 | KIRC | SN12C | Fulvestrant |
| MVP_top60_DNA_seed | MAPK1 | -1.412862 | HNSC | DOK | SCH772984 | MVP_top60_DNA+RNA_seed | AR | 0.093295 | KIRC | RCC-FG2 | AZD3514 |
| MVP_top60_DNA_seed | MAPK3 | -1.412862 | HNSC | DOK | SCH772984 | MVP_top60_DNA+RNA_seed | PIK3CD | 0.093615 | KIRC | BFTC-909 | Idelalisib |
| MVP_top60_DNA_seed | EGFR | -1.407549 | HNSC | PCI-6A | Gefitinib | MVP_top60_DNA+RNA_seed | ATM | 0.093799 | KIRC | TK10 | CP466722 |
| MVP_top60_DNA_seed | EGFR | -1.40703 | HNSC | SCC-15 | Erlotinib | MVP_top60_DNA+RNA_seed | HDAC1 | 0.098215 | KIRC | VMRC-RCZ | Dacinostat |
| MVP_top60_DNA_seed | HDAC1 | -1.405076 | HNSC | SCC-9 | Romidepsin | MVP_top60_DNA+RNA_seed | ESR1 | 0.100674 | KIRC | LB996-RCC | Fulvestrant |
| MVP_top60_DNA_seed | HDAC3 | -1.405076 | HNSC | SCC-9 | Romidepsin | MVP_top60_DNA+RNA_seed | HDAC1 | 0.101049 | KIRC | RXF393 | JQ12 |
| MVP_top60_DNA_seed | AKT1 | -1.404149 | HNSC | JHU-011 | AT13148 | MVP_top60_DNA+RNA_seed | ATM | 0.1012 | KIRC | RCC-ER | CP466722 |
| MVP_top60_DNA_seed | EGFR | -1.396957 | HNSC | HSC-3 | Osimertinib | MVP_top60_DNA+RNA_seed | ATM | 0.109715 | KIRC | RCC-AB | KU-60019 |
| MVP_top60_DNA_seed | MAPK1 | -1.394683 | HNSC | JHU-011 | VX-11e | MVP_top60_DNA+RNA_seed | HDAC1 | 0.114896 | KIRC | ACHN | CAY10603 |
| MVP_top60_DNA_seed | EGFR | -1.391369 | HNSC | HSC-2 | Afatinib | MVP_top60_DNA+RNA_seed | ATM | 0.122948 | KIRC | NCC021 | CP466722 |
| MVP_top60_DNA_seed | EGFR | -1.390252 | HNSC | JHU-011 | Lapatinib | MVP_top60_DNA+RNA_seed | PIK3CD | 0.128322 | KIRC | BFTC-909 | AZD8186 |
| MVP_top60_DNA_seed | AKT1 | -1.389811 | HNSC | BB49-HNC | Uprosertib | MVP_top60_DNA+RNA_seed | PIK3CD | 0.128476 | KIRC | BB65-RCC | IC-87114 |
| MVP_top60_DNA_seed | EP300 | -1.388387 | HNSC | SAS | CPI-637 | MVP_top60_DNA+RNA_seed | MTOR | 0.128534 | KIRC | TK10 | QL-VIII-58 |
| MVP_top60_DNA_seed | MDM2 | -1.3779 | HNSC | PCI-15A | Serdemetan | MVP_top60_DNA+RNA_seed | ATM | 0.128879 | KIRC | OS-RC-2 | KU-60019 |
| MVP_top60_DNA_seed | EP300 | -1.374951 | HNSC | HO-1-N-1 | SGC-CBP30 | MVP_top60_DNA+RNA_seed | IGF1R | 0.130416 | KIRC | HA7-RCC | BMS-536924 |
| MVP_top60_DNA_seed | CREBBP | -1.374951 | HNSC | HO-1-N-1 | SGC-CBP30 | MVP_top60_DNA+RNA_seed | ESR1 | 0.13196 | KIRC | KMRC-20 | Fulvestrant |
| MVP_top60_DNA_seed | EGFR | -1.373266 | HNSC | CAL-33 | Erlotinib | MVP_top60_DNA+RNA_seed | AR | 0.132516 | KIRC | LB2241-RCC | Bicalutamide |

| | | | | | | | | | | | |
|---|---|---|---|---|---|---|---|---|---|---|---|
| MVP_top60_DNA_seed | MAPK1 | -1.371382 | HNSC | PCI-15A | SCH772984 | MVP_top60_DNA+RNA_seed | MTOR | 0.135197 | KIRC | CAKI-1 | WYE-125132 |
| MVP_top60_DNA_seed | MAPK3 | -1.371382 | HNSC | PCI-15A | SCH772984 | MVP_top60_DNA+RNA_seed | IGF1R | 0.136863 | KIRC | RCC-JF | BMS-536924 |
| MVP_top60_DNA_seed | PIK3CA | -1.367034 | HNSC | BB30-HNC | Alpelisib | MVP_top60_DNA+RNA_seed | HDAC1 | 0.1372 | KIRC | ACHN | Belinostat |
| MVP_top60_DNA_seed | HDAC1 | -1.366658 | HNSC | PCI-15A | Entinostat | MVP_top60_DNA+RNA_seed | PIK3CA | 0.140781 | KIRC | A704 | PI3Ka_4409 |
| MVP_top60_DNA_seed | HDAC3 | -1.366658 | HNSC | PCI-15A | Entinostat | MVP_top60_DNA+RNA_seed | PIK3CD | 0.140781 | KIRC | A704 | PI3Ka_4409 |
| MVP_top60_DNA_seed | AKT1 | -1.361407 | HNSC | HO-1-u-1 | MK-2206 | MVP_top60_DNA+RNA_seed | MTOR | 0.140894 | KIRC | LB996-RCC | QL-VIII-58 |
| MVP_top60_DNA_seed | AKT1 | -1.360925 | HNSC | DOK | Uprosertib | MVP_top60_DNA+RNA_seed | IGF1R | 0.141853 | KIRC | OS-RC-2 | GSK1904529A |
| MVP_top60_DNA_seed | ESR1 | -1.359012 | HNSC | HSC-3 | Tamoxifen | MVP_top60_DNA+RNA_seed | HDAC1 | 0.143764 | KIRC | OS-RC-2 | CAY10603 |
| MVP_top60_DNA_seed | EGFR | -1.357321 | HNSC | HO-1-u-1 | Afatinib | MVP_top60_DNA+RNA_seed | ATM | 0.146853 | KIRC | NCC010 | KU-60019 |
| MVP_top60_DNA_seed | MAPK1 | -1.351359 | HNSC | PCI-15A | Ulixertinib | MVP_top60_DNA+RNA_seed | HSP90AA1 | 0.146882 | KIRC | VMRC-RCW | Elesclomol |
| MVP_top60_DNA_seed | MAPK3 | -1.351359 | HNSC | PCI-15A | Ulixertinib | MVP_top60_DNA+RNA_seed | HSP90AB1 | 0.146882 | KIRC | VMRC-RCW | Elesclomol |
| MVP_top60_DNA_seed | PIK3CA | -1.341142 | HNSC | DOK | AZD8186 | MVP_top60_DNA+RNA_seed | AR | 0.151006 | KIRC | U031 | Bicalutamide |
| MVP_top60_DNA_seed | EGFR | -1.340987 | HNSC | PCI-6A | Erlotinib | MVP_top60_DNA+RNA_seed | SMARCA4 | 0.151841 | KIRC | A704 | PFI-3 |
| MVP_top60_DNA_seed | MAPK1 | -1.340668 | HNSC | JHU-011 | Ulixertinib | MVP_top60_DNA+RNA_seed | SMARCA2 | 0.151841 | KIRC | A704 | PFI-3 |
| MVP_top60_DNA_seed | MAPK3 | -1.340668 | HNSC | JHU-011 | Ulixertinib | MVP_top60_DNA+RNA_seed | FGFR4 | 0.152314 | KIRC | RCC-JF | FGFR_3831 |
| MVP_top60_DNA_seed | MDM2 | -1.338593 | HNSC | PCI-4B | Serdemetan | MVP_top60_DNA+RNA_seed | HDAC1 | 0.153297 | KIRC | A704 | Dacinostat |
| MVP_top60_DNA_seed | EGFR | -1.331883 | HNSC | HSC-3 | Lapatinib | MVP_top60_DNA+RNA_seed | IGF1R | 0.155366 | KIRC | ACHN | BMS-754807 |
| MVP_top60_DNA_seed | HSP90AA1 | -1.329409 | HNSC | DOK | Luminespib | MVP_top60_DNA+RNA_seed | HDAC1 | 0.158543 | KIRC | RCC10RGB | Parthenolide |
| MVP_top60_DNA_seed | MAPK1 | -1.32687 | HNSC | BB30-HNC | SCH772984 | MVP_top60_DNA+RNA_seed | PIK3CA | 0.159012 | KIRC | RCC-AB | PI3Ka_4409 |
| MVP_top60_DNA_seed | MAPK3 | -1.32687 | HNSC | BB30-HNC | SCH772984 | MVP_top60_DNA+RNA_seed | PIK3CD | 0.159012 | KIRC | RCC-AB | PI3Ka_4409 |
| MVP_top60_DNA_seed | EGFR | -1.322439 | HNSC | KON | Lapatinib | MVP_top60_DNA+RNA_seed | ESR1 | 0.16386 | KIRC | RCC-FG2 | Tamoxifen |
| MVP_top60_DNA_seed | AKT1 | -1.320117 | HNSC | PCI-15A | AZD5363 | MVP_top60_DNA+RNA_seed | MTOR | 0.165214 | KIRC | NCC021 | Temsirolimus |
| MVP_top60_DNA_seed | EGFR | -1.319664 | HNSC | BHY | Sapitinib | MVP_top60_DNA+RNA_seed | IGF1R | 0.165313 | KIRC | RXF393 | BMS-536924 |
| MVP_top60_DNA_seed | MAPK1 | -1.31937 | HNSC | BB30-HNC | VX-11e | MVP_top60_DNA+RNA_seed | AR | 0.168559 | KIRC | NCC010 | AZD3514 |
| MVP_top60_DNA_seed | AKT1 | -1.317264 | HNSC | PCI-15A | Ipatasertib | MVP_top60_DNA+RNA_seed | PIK3CA | 0.171023 | KIRC | NCC021 | AZD8835 |
| MVP_top60_DNA_seed | EGFR | -1.316737 | HNSC | Ca9-22 | Osimertinib | MVP_top60_DNA+RNA_seed | PIK3CD | 0.171023 | KIRC | NCC021 | AZD8835 |
| MVP_top60_DNA_seed | EGFR | -1.315206 | HNSC | SAS | AZD3759 | MVP_top60_DNA+RNA_seed | HDAC1 | 0.171671 | KIRC | VMRC-RCW | AR-42 |
| MVP_top60_DNA_seed | PIK3CA | -1.314742 | HNSC | HO-1-u-1 | Alpelisib | MVP_top60_DNA+RNA_seed | MTOR | 0.173076 | KIRC | 769-P | Voxtalisib |
| MVP_top60_DNA_seed | EGFR | -1.314119 | HNSC | Detroit562 | Lapatinib | MVP_top60_DNA+RNA_seed | HSP90AA1 | 0.173693 | KIRC | ACHN | Tanespimycin |
| MVP_top60_DNA_seed | EGFR | -1.307325 | HNSC | KOSC-2 | Gefitinib | MVP_top60_DNA+RNA_seed | HSP90AB1 | 0.173693 | KIRC | ACHN | Tanespimycin |
| MVP_top60_DNA_seed | EGFR | -1.297427 | HNSC | Detroit562 | Erlotinib | MVP_top60_DNA+RNA_seed | HDAC1 | 0.174118 | KIRC | OS-RC-2 | Belinostat |
| MVP_top60_DNA_seed | HSP90AA1 | -1.297143 | HNSC | PCI-30 | Tanespimycin | MVP_top60_DNA+RNA_seed | FGFR4 | 0.175746 | KIRC | NCC021 | FGFR_3831 |
| MVP_top60_DNA_seed | EGFR | -1.291146 | HNSC | JHU-011 | Erlotinib | MVP_top60_DNA+RNA_seed | IGF1R | 0.176696 | KIRC | A704 | BMS-754807 |
| MVP_top60_DNA_seed | PIK3CA | -1.29089 | HNSC | DOK | Buparlisib | MVP_top60_DNA+RNA_seed | AR | 0.176825 | KIRC | RCC-MF | AZD3514 |
| MVP_top60_DNA_seed | SRC | -1.279975 | HNSC | HSC-4 | Bosutinib | MVP_top60_DNA+RNA_seed | HSP90AA1 | 0.176982 | KIRC | RCC-MF | SNX-2112 |
| MVP_top60_DNA_seed | NTRK1 | -1.279097 | HNSC | DOK | AZD1332 | MVP_top60_DNA+RNA_seed | HSP90AB1 | 0.176982 | KIRC | RCC-MF | SNX-2112 |
| MVP_top60_DNA_seed | MAPK1 | -1.27804 | HNSC | KON | VX-11e | MVP_top60_DNA+RNA_seed | IGF1R | 0.178395 | KIRC | LB2241-RCC | BMS-754807 |
| MVP_top60_DNA_seed | EGFR | -1.275631 | HNSC | HO-1-N-1 | Sapitinib | MVP_top60_DNA+RNA_seed | MTOR | 0.183035 | KIRC | VMRC-RCZ | QL-VIII-58 |
| MVP_top60_DNA_seed | EGFR | -1.27329 | HNSC | CAL-33 | AZD3759 | MVP_top60_DNA+RNA_seed | EP300 | 0.184209 | KIRC | NCC010 | I-CBP112 |
| MVP_top60_DNA_seed | EGFR | -1.270592 | HNSC | KOSC-2 | Lapatinib | MVP_top60_DNA+RNA_seed | HSP90AA1 | 0.184505 | KIRC | KMRC-1 | Luminespib |
| MVP_top60_DNA_seed | EGFR | -1.262116 | HNSC | PCI-6A | Osimertinib | MVP_top60_DNA+RNA_seed | HSP90AB1 | 0.184505 | KIRC | KMRC-1 | Luminespib |

| | | | | | | | | | | | |
|---|---|---|---|---|---|---|---|---|---|---|---|
| MVP_top60_DNA_seed | SMARCA4 | -1.260373 | HNSC | SAS | PFI3 | MVP_top60_DNA+RNA_seed | ATM | 0.186303 | KIRC | RCC-FG2 | CP466722 |
| MVP_top60_DNA_seed | EGFR | -1.256541 | HNSC | HSC-3 | Sapitinib | MVP_top60_DNA+RNA_seed | ESR1 | 0.188831 | KIRC | A498 | Fulvestrant |
| MVP_top60_DNA_seed | SRC | -1.255965 | HNSC | BB30-HNC | Bosutinib | MVP_top60_DNA+RNA_seed | HDAC1 | 0.19005 | KIRC | RXF393 | Dacinostat |
| MVP_top60_DNA_seed | ESR1 | -1.243868 | HNSC | LB771-HNC | GDC0810 | MVP_top60_DNA+RNA_seed | PTK2 | 0.190574 | KIRC | U031 | PF-562271 |
| MVP_top60_DNA_seed | AKT1 | -1.2377 | HNSC | BB49-HNC | MK-2206 | MVP_top60_DNA+RNA_seed | HDAC1 | 0.191654 | KIRC | SW156 | Dacinostat |
| MVP_top60_DNA_seed | EGFR | -1.237474 | HNSC | HO-1-N-1 | Afatinib | MVP_top60_DNA+RNA_seed | HSP90AA1 | 0.191987 | KIRC | KMRC-20 | CCT-018159 |
| MVP_top60_DNA_seed | NTRK1 | -1.235891 | HNSC | HSC-2 | AZD1332 | MVP_top60_DNA+RNA_seed | HSP90AB1 | 0.191987 | KIRC | KMRC-20 | CCT-018159 |
| MVP_top60_DNA_seed | EGFR | -1.234589 | HNSC | HSC-2 | Lapatinib | MVP_top60_DNA+RNA_seed | MTOR | 0.19565 | KIRC | A704 | Temsirolimus |
| MVP_top60_DNA_seed | AKT1 | -1.234202 | HNSC | Detroit562 | AZD5363 | MVP_top60_DNA+RNA_seed | SMARCA4 | 0.196914 | KIRC | BB65-RCC | PFI-3 |
| MVP_top60_DNA_seed | EGFR | -1.233816 | HNSC | CAL-33 | Lapatinib | MVP_top60_DNA+RNA_seed | SMARCA2 | 0.196914 | KIRC | BB65-RCC | PFI-3 |
| MVP_top60_DNA_seed | EGFR | -1.231545 | HNSC | HO-1-N-1 | Erlotinib | MVP_top60_DNA+RNA_seed | ATM | 0.198167 | KIRC | VMRC-RCZ | KU-60019 |
| MVP_top60_DNA_seed | EGFR | -1.230585 | HNSC | CAL-27 | Osimertinib | MVP_top60_DNA+RNA_seed | IGF1R | 0.202634 | KIRC | 769-P | Linsitinib |
| MVP_top60_DNA_seed | AKT1 | -1.224287 | HNSC | JHU-029 | AT13148 | MVP_top60_DNA+RNA_seed | PIK3CD | 0.204294 | KIRC | VMRC-RCW | IC-87114 |
| MVP_top60_DNA_seed | AKT1 | -1.221817 | HNSC | PCI-15A | Uprosertib | MVP_top60_DNA+RNA_seed | HSP90AA1 | 0.207005 | KIRC | OS-RC-2 | SNX-2112 |
| MVP_top60_DNA_seed | HSP90AA1 | -1.21894 | HNSC | PCI-15A | CCT-018159 | MVP_top60_DNA+RNA_seed | HSP90AB1 | 0.207005 | KIRC | OS-RC-2 | SNX-2112 |
| MVP_top60_DNA_seed | EGFR | -1.214262 | HNSC | OSC-19 | Osimertinib | MVP_top60_DNA+RNA_seed | HDAC1 | 0.213002 | KIRC | ACHN | AR-42 |
| MVP_top60_DNA_seed | HSP90AA1 | -1.212298 | HNSC | SCC-9 | Tanespimycin | MVP_top60_DNA+RNA_seed | EP300 | 0.21393 | KIRC | A704 | I-CBP112 |
| MVP_top60_DNA_seed | EGFR | -1.212064 | HNSC | PCI-4B | Afatinib | MVP_top60_DNA+RNA_seed | IGF1R | 0.219001 | KIRC | U031 | BMS-536924 |
| MVP_top60_DNA_seed | ESR1 | -1.211795 | HNSC | HSC-2 | Tamoxifen | MVP_top60_DNA+RNA_seed | AR | 0.219165 | KIRC | LB1047-RCC | AZD3514 |
| MVP_top60_DNA_seed | PIK3CA | -1.206075 | HNSC | PCI-15A | Alpelisib | MVP_top60_DNA+RNA_seed | IGF1R | 0.219513 | KIRC | BB65-RCC | BMS-536924 |
| MVP_top60_DNA_seed | AKT1 | -1.199652 | HNSC | JHU-011 | Afuresertib | MVP_top60_DNA+RNA_seed | PIK3CD | 0.219955 | KIRC | TK10 | Idelalisib |
| MVP_top60_DNA_seed | EGFR | -1.197485 | HNSC | HO-1-N-1 | Lapatinib | MVP_top60_DNA+RNA_seed | MTOR | 0.220647 | KIRC | NCC010 | Temsirolimus |
| MVP_top60_DNA_seed | AKT1 | -1.19452 | HNSC | BB49-HNC | Uprosertib | MVP_top60_DNA+RNA_seed | PIK3CD | 0.225197 | KIRC | KMRC-20 | AZD8186 |
| MVP_top60_DNA_seed | PIK3CA | -1.194457 | HNSC | JHU-029 | Alpelisib | MVP_top60_DNA+RNA_seed | IGF1R | 0.225708 | KIRC | SN12C | Linsitinib |
| MVP_top60_DNA_seed | AKT1 | -1.194338 | HNSC | DOK | MK-2206 | MVP_top60_DNA+RNA_seed | IGF1R | 0.228172 | KIRC | TK10 | BMS-536924 |
| MVP_top60_DNA_seed | AKT1 | -1.193791 | HNSC | Detroit562 | Uprosertib | MVP_top60_DNA+RNA_seed | PIK3CD | 0.23385 | KIRC | SW156 | AZD8186 |
| MVP_top60_DNA_seed | EGFR | -1.19356 | HNSC | Detroit562 | Sapitinib | MVP_top60_DNA+RNA_seed | PIK3CA | 0.236688 | KIRC | RCC-JF | PI3Ka_4409 |
| MVP_top60_DNA_seed | PIK3CA | -1.186851 | HNSC | BICR10 | Buparlisib | MVP_top60_DNA+RNA_seed | PIK3CD | 0.236688 | KIRC | RCC-JF | PI3Ka_4409 |
| MVP_top60_DNA_seed | EGFR | -1.186033 | HNSC | KON | Osimertinib | MVP_top60_DNA+RNA_seed | IGF1R | 0.239396 | KIRC | U031 | BMS-754807 |
| MVP_top60_DNA_seed | CDK2 | -1.168483 | HNSC | BICR22 | AZD5438 | MVP_top60_DNA+RNA_seed | FGFR4 | 0.242805 | KIRC | CAKI-1 | FGFR_3831 |
| MVP_top60_DNA_seed | HSP90AA1 | -1.164639 | HNSC | OSC-19 | Tanespimycin | MVP_top60_DNA+RNA_seed | ATM | 0.250514 | KIRC | CAKI-1 | KU-55933 |
| MVP_top60_DNA_seed | AR | -1.156615 | HNSC | BB30-HNC | Bicalutamide | MVP_top60_DNA+RNA_seed | HDAC1 | 0.253088 | KIRC | NCC021 | Tubastatin A |
| MVP_top60_DNA_seed | HDAC1 | -1.154789 | HNSC | HSC-2 | Romidepsin | MVP_top60_DNA+RNA_seed | AR | 0.25331 | KIRC | SN12C | AZD3514 |
| MVP_top60_DNA_seed | HDAC3 | -1.154789 | HNSC | HSC-2 | Romidepsin | MVP_top60_DNA+RNA_seed | FGFR4 | 0.253419 | KIRC | TK10 | FGFR_0939 |
| MVP_top60_DNA_seed | MDM2 | -1.152922 | HNSC | JHU-011 | Serdemetan | MVP_top60_DNA+RNA_seed | MTOR | 0.253633 | KIRC | 786-0 | Voxtalisib |
| MVP_top60_DNA_seed | PIK3CA | -1.147633 | HNSC | BICR22 | Alpelisib | MVP_top60_DNA+RNA_seed | HDAC1 | 0.255939 | KIRC | KMRC-1 | JQ12 |
| MVP_top60_DNA_seed | SRC | -1.146949 | HNSC | SCC-9 | Bosutinib | MVP_top60_DNA+RNA_seed | PIK3CA | 0.25607 | KIRC | 769-P | PI3Ka_4409 |
| MVP_top60_DNA_seed | PIK3CA | -1.144714 | HNSC | PCI-15A | GNE-317 | MVP_top60_DNA+RNA_seed | PIK3CD | 0.25607 | KIRC | 769-P | PI3Ka_4409 |
| MVP_top60_DNA_seed | AKT1 | -1.14314 | HNSC | Detroit562 | Ipatasertib | MVP_top60_DNA+RNA_seed | HDAC1 | 0.259922 | KIRC | OS-RC-2 | AR-42 |
| MVP_top60_DNA_seed | HSP90AA1 | -1.137755 | HNSC | HSC-2 | Luminespib | MVP_top60_DNA+RNA_seed | MTOR | 0.260014 | KIRC | OS-RC-2 | WYE-125132 |
| MVP_top60_DNA_seed | ESR1 | -1.136273 | HNSC | Detroit562 | GDC0810 | MVP_top60_DNA+RNA_seed | HDAC1 | 0.260916 | KIRC | LB996-RCC | Dacinostat |

| | | | | | | | | | | | |
|---|---|---|---|---|---|---|---|---|---|---|---|
| MVP_top60_DNA_seed | EP300 | -1.136168 | HNSC | SAS | SGC-CBP30 | MVP_top60_DNA+RNA_seed | MTOR | 0.267406 | KIRC | SW156 | Torin 2 |
| MVP_top60_DNA_seed | CREBBP | -1.136168 | HNSC | SAS | SGC-CBP30 | MVP_top60_DNA+RNA_seed | HDAC1 | 0.26879 | KIRC | RXF393 | Tubastatin A |
| MVP_top60_DNA_seed | RAC1 | -1.136128 | HNSC | HSC-2 | EHT-1864 | MVP_top60_DNA+RNA_seed | HDAC1 | 0.277604 | KIRC | U031 | Belinostat |
| MVP_top60_DNA_seed | EGFR | -1.129991 | HNSC | SCC-9 | Lapatinib | MVP_top60_DNA+RNA_seed | AR | 0.279041 | KIRC | LB1047-RCC | Bicalutamide |
| MVP_top60_DNA_seed | HSP90AA1 | -1.124919 | HNSC | SAS | Tanespimycin | MVP_top60_DNA+RNA_seed | IGF1R | 0.279672 | KIRC | SW156 | Linsitinib |
| MVP_top60_DNA_seed | ESR1 | -1.124043 | HNSC | HSC-3 | GDC0810 | MVP_top60_DNA+RNA_seed | HSP90AA1 | 0.282268 | KIRC | KMRC-1 | Elesclomol |
| MVP_top60_DNA_seed | MDM2 | -1.117938 | HNSC | BB49-HNC | Nutlin-3a (-) | MVP_top60_DNA+RNA_seed | HSP90AB1 | 0.282268 | KIRC | KMRC-1 | Elesclomol |
| MVP_top60_DNA_seed | EGFR | -1.1169 | HNSC | PE-CA-PJ15 | Gefitinib | MVP_top60_DNA+RNA_seed | MTOR | 0.282364 | KIRC | U031 | Torin 2 |
| MVP_top60_DNA_seed | PIK3CA | -1.115337 | HNSC | Detroit562 | GNE-317 | MVP_top60_DNA+RNA_seed | FGFR4 | 0.284177 | KIRC | KMRC-20 | FGFR_3831 |
| MVP_top60_DNA_seed | AR | -1.113085 | HNSC | SCC-25 | Bicalutamide | MVP_top60_DNA+RNA_seed | IGF1R | 0.285501 | KIRC | A704 | Linsitinib |
| MVP_top60_DNA_seed | SRC | -1.112621 | HNSC | SAS | Bosutinib | MVP_top60_DNA+RNA_seed | IGF1R | 0.285858 | KIRC | BFTC-909 | BMS-754807 |
| MVP_top60_DNA_seed | HDAC1 | -1.112268 | HNSC | BICR22 | Romidepsin | MVP_top60_DNA+RNA_seed | PIK3CD | 0.287095 | KIRC | KMRC-1 | IC-87114 |
| MVP_top60_DNA_seed | HDAC3 | -1.112268 | HNSC | BICR22 | Romidepsin | MVP_top60_DNA+RNA_seed | PIK3CD | 0.28924 | KIRC | VMRC-RCW | Idelalisib |
| MVP_top60_DNA_seed | EGFR | -1.110228 | HNSC | BHY | AZD3759 | MVP_top60_DNA+RNA_seed | HDAC1 | 0.293258 | KIRC | CAL-54 | JQ12 |
| MVP_top60_DNA_seed | ESR1 | -1.109124 | HNSC | FADU | GDC0810 | MVP_top60_DNA+RNA_seed | AR | 0.295495 | KIRC | 769-P | Bicalutamide |
| MVP_top60_DNA_seed | MAPK1 | -1.108667 | HNSC | SCC-9 | Ulixertinib | MVP_top60_DNA+RNA_seed | MTOR | 0.29829 | KIRC | LB996-RCC | Torin 2 |
| MVP_top60_DNA_seed | MAPK3 | -1.108667 | HNSC | SCC-9 | Ulixertinib | MVP_top60_DNA+RNA_seed | MTOR | 0.302319 | KIRC | BB65-RCC | Apitolisib |
| MVP_top60_DNA_seed | NTRK1 | -1.10841 | HNSC | BICR31 | GW441756 | MVP_top60_DNA+RNA_seed | AR | 0.304577 | KIRC | LB2241-RCC | Bicalutamide |
| MVP_top60_DNA_seed | PIK3CA | -1.10622 | HNSC | SAS | GNE-317 | MVP_top60_DNA+RNA_seed | HSP90AA1 | 0.313541 | KIRC | 769-P | CCT-018159 |
| MVP_top60_DNA_seed | EP300 | -1.102607 | HNSC | HSC-2 | SGC-CBP30 | MVP_top60_DNA+RNA_seed | HSP90AB1 | 0.313541 | KIRC | 769-P | CCT-018159 |
| MVP_top60_DNA_seed | CREBBP | -1.102607 | HNSC | HSC-2 | SGC-CBP30 | MVP_top60_DNA+RNA_seed | MTOR | 0.313632 | KIRC | 769-P | Temsirolimus |
| MVP_top60_DNA_seed | HSP90AA1 | -1.101738 | HNSC | PCI-6A | Tanespimycin | MVP_top60_DNA+RNA_seed | PIK3CA | 0.32219 | KIRC | RCC10RGB | AZD8835 |
| MVP_top60_DNA_seed | AKT1 | -1.10133 | HNSC | JHU-029 | Ipatasertib | MVP_top60_DNA+RNA_seed | PIK3CD | 0.32219 | KIRC | RCC10RGB | AZD8835 |
| MVP_top60_DNA_seed | AKT1 | -1.101119 | HNSC | PCI-15A | Afuresertib | MVP_top60_DNA+RNA_seed | MTOR | 0.322768 | KIRC | BB65-RCC | Torin 2 |
| MVP_top60_DNA_seed | AKT1 | -1.098849 | HNSC | DOK | Uprosertib | MVP_top60_DNA+RNA_seed | IGF1R | 0.323292 | KIRC | BFTC-909 | Linsitinib |
| MVP_top60_DNA_seed | NTRK1 | -1.084335 | HNSC | Ca9-22 | AZD1332 | MVP_top60_DNA+RNA_seed | ESR1 | 0.327371 | KIRC | BB65-RCC | Fulvestrant |
| MVP_top60_DNA_seed | EP300 | -1.083089 | HNSC | BICR22 | SGC-CBP30 | MVP_top60_DNA+RNA_seed | AR | 0.328079 | KIRC | HA7-RCC | Bicalutamide |
| MVP_top60_DNA_seed | CREBBP | -1.083089 | HNSC | BICR22 | SGC-CBP30 | MVP_top60_DNA+RNA_seed | AR | 0.32844 | KIRC | CAKI-1 | Bicalutamide |
| MVP_top60_DNA_seed | SRC | -1.080277 | HNSC | HO-1-N-1 | Bosutinib | MVP_top60_DNA+RNA_seed | AR | 0.329403 | KIRC | LB996-RCC | Bicalutamide |
| MVP_top60_DNA_seed | ESR1 | -1.076048 | HNSC | HSC-2 | GDC0810 | MVP_top60_DNA+RNA_seed | HDAC1 | 0.331657 | KIRC | HA7-RCC | JQ12 |
| MVP_top60_DNA_seed | AKT1 | -1.07143 | HNSC | BB49-HNC | Ipatasertib | MVP_top60_DNA+RNA_seed | HDAC1 | 0.335832 | KIRC | HA7-RCC | Entinostat |
| MVP_top60_DNA_seed | AKT1 | -1.069208 | HNSC | HSC-2 | GSK2110183B | MVP_top60_DNA+RNA_seed | EP300 | 0.336442 | KIRC | RCC-AB | I-CBP112 |
| MVP_top60_DNA_seed | RAC1 | -1.068947 | HNSC | BHY | EHT-1864 | MVP_top60_DNA+RNA_seed | HDAC1 | 0.341634 | KIRC | BFTC-909 | CAY10603 |
| MVP_top60_DNA_seed | SRC | -1.06876 | HNSC | JHU-011 | Bosutinib | MVP_top60_DNA+RNA_seed | MTOR | 0.342756 | KIRC | ACHN | Torin 2 |
| MVP_top60_DNA_seed | EGFR | -1.064843 | HNSC | SAS | Sapitinib | MVP_top60_DNA+RNA_seed | HDAC1 | 0.343958 | KIRC | U031 | CAY10603 |
| MVP_top60_DNA_seed | HSP90AA1 | -1.063471 | HNSC | PCI-15A | Luminespib | MVP_top60_DNA+RNA_seed | MTOR | 0.346814 | KIRC | LB2241-RCC | Voxtalisib |
| MVP_top60_DNA_seed | SRC | -1.061763 | HNSC | PCI-15A | Bosutinib | MVP_top60_DNA+RNA_seed | AR | 0.347497 | KIRC | RCC-JF | Bicalutamide |
| MVP_top60_DNA_seed | HSP90AA1 | -1.060273 | HNSC | HSC-3 | Luminespib | MVP_top60_DNA+RNA_seed | HDAC1 | 0.349436 | KIRC | SW156 | JQ12 |
| MVP_top60_DNA_seed | HSP90AA1 | -1.059632 | HNSC | PCI-4B | CCT-018159 | MVP_top60_DNA+RNA_seed | EP300 | 0.350056 | KIRC | 769-P | I-CBP112 |
| MVP_top60_DNA_seed | EGFR | -1.057981 | HNSC | HSC-3 | AZD3759 | MVP_top60_DNA+RNA_seed | MTOR | 0.354215 | KIRC | KMRC-1 | WYE-125132 |
| MVP_top60_DNA_seed | EGFR | -1.052744 | HNSC | JHU-029 | Osimertinib | MVP_top60_DNA+RNA_seed | HDAC1 | 0.355072 | KIRC | VMRC-RCW | CAY10603 |

| | | | | | | | | | | | |
|---|---|---|---|---|---|---|---|---|---|---|---|
| MVP_top60_DNA_seed | RAC1 | -1.052294 | HNSC | SCC-25 | EHT-1864 | MVP_top60_DNA+RNA_seed | HDAC1 | 0.357702 | KIRC | VMRC-RCW | Belinostat |
| MVP_top60_DNA_seed | MDM2 | -1.052129 | HNSC | HO-1-u-1 | Serdemetan | MVP_top60_DNA+RNA_seed | HDAC1 | 0.35877 | KIRC | CAL-54 | Tubastatin A |
| MVP_top60_DNA_seed | CDK2 | -1.048801 | HNSC | HSC-2 | Dinaciclib | MVP_top60_DNA+RNA_seed | IGF1R | 0.358879 | KIRC | BFTC-909 | BMS-536924 |
| MVP_top60_DNA_seed | SRC | -1.048417 | HNSC | FADU | Bosutinib | MVP_top60_DNA+RNA_seed | MTOR | 0.359669 | KIRC | A498 | Temsirolimus |
| MVP_top60_DNA_seed | HSP90AA1 | -1.047687 | HNSC | BICR22 | Tanespimycin | MVP_top60_DNA+RNA_seed | IGF1R | 0.362483 | KIRC | A498 | Linsitinib |
| MVP_top60_DNA_seed | PIK3CA | -1.044558 | HNSC | HO-1-N-1 | Buparlisib | MVP_top60_DNA+RNA_seed | MTOR | 0.363125 | KIRC | NCC021 | Voxtalisib |
| MVP_top60_DNA_seed | EGFR | -1.0434 | HNSC | PCI-4B | AZD3759 | MVP_top60_DNA+RNA_seed | PIK3CA | 0.371572 | KIRC | RCC-ER | PI3Ka_4409 |
| MVP_top60_DNA_seed | HSP90AA1 | -1.040802 | HNSC | JHU-011 | Luminespib | MVP_top60_DNA+RNA_seed | PIK3CD | 0.371572 | KIRC | RCC-ER | PI3Ka_4409 |
| MVP_top60_DNA_seed | SRC | -1.040663 | HNSC | JHU-022 | Bosutinib | MVP_top60_DNA+RNA_seed | MTOR | 0.381037 | KIRC | TK10 | Temsirolimus |
| MVP_top60_DNA_seed | MAPK1 | -1.033641 | HNSC | PCI-15A | VX-11e | MVP_top60_DNA+RNA_seed | FGFR4 | 0.386611 | KIRC | KMRC-1 | FGFR_3831 |
| MVP_top60_DNA_seed | MAPK1 | -1.032869 | HNSC | KON | Ulixertinib | MVP_top60_DNA+RNA_seed | MTOR | 0.387309 | KIRC | ACHN | Temsirolimus |
| MVP_top60_DNA_seed | MAPK3 | -1.032869 | HNSC | KON | Ulixertinib | MVP_top60_DNA+RNA_seed | IGF1R | 0.387543 | KIRC | KMRC-20 | BMS-536924 |
| MVP_top60_DNA_seed | HSP90AA1 | -1.031237 | HNSC | JHU-029 | Luminespib | MVP_top60_DNA+RNA_seed | MTOR | 0.390748 | KIRC | OS-RC-2 | Torin 2 |
| MVP_top60_DNA_seed | MAPK1 | -1.027429 | HNSC | JHU-029 | Ulixertinib | MVP_top60_DNA+RNA_seed | HDAC1 | 0.393168 | KIRC | KMRC-1 | Tubastatin A |
| MVP_top60_DNA_seed | MAPK3 | -1.027429 | HNSC | JHU-029 | Ulixertinib | MVP_top60_DNA+RNA_seed | HSP90AA1 | 0.397485 | KIRC | RCC-AB | Tanespimycin |
| MVP_top60_DNA_seed | EGFR | -1.025666 | HNSC | OSC-19 | Lapatinib | MVP_top60_DNA+RNA_seed | HSP90AB1 | 0.397485 | KIRC | RCC-AB | Tanespimycin |
| MVP_top60_DNA_seed | HSP90AA1 | -1.020833 | HNSC | JHU-022 | Tanespimycin | MVP_top60_DNA+RNA_seed | PIK3CA | 0.398346 | KIRC | TK10 | PI3Ka_4409 |
| MVP_top60_DNA_seed | MDM2 | -1.01965 | HNSC | Detroit562 | Serdemetan | MVP_top60_DNA+RNA_seed | PIK3CD | 0.398346 | KIRC | TK10 | PI3Ka_4409 |
| MVP_top60_DNA_seed | MAPK1 | -1.013543 | HNSC | JHU-011 | SCH772984 | MVP_top60_DNA+RNA_seed | AR | 0.398827 | KIRC | BB65-RCC | Bicalutamide |
| MVP_top60_DNA_seed | MAPK3 | -1.013543 | HNSC | JHU-011 | SCH772984 | MVP_top60_DNA+RNA_seed | MTOR | 0.398925 | KIRC | BFTC-909 | Voxtalisib |
| MVP_top60_DNA_seed | AKT1 | -1.011976 | HNSC | KON | AT13148 | MVP_top60_DNA+RNA_seed | MTOR | 0.400861 | KIRC | RCC-AB | QL-VIII-58 |
| MVP_top60_DNA_seed | SRC | -1.010523 | HNSC | SAT | Bosutinib | MVP_top60_DNA+RNA_seed | ATM | 0.402397 | KIRC | LB996-RCC | CP466722 |
| MVP_top60_DNA_seed | EGFR | -1.010163 | HNSC | SAS | Osimertinib | MVP_top60_DNA+RNA_seed | HSP90AA1 | 0.404428 | KIRC | A704 | SNX-2112 |
| MVP_top60_DNA_seed | CDK2 | -1.009923 | HNSC | BB30-HNC | Dinaciclib | MVP_top60_DNA+RNA_seed | HSP90AB1 | 0.404428 | KIRC | A704 | SNX-2112 |
| MVP_top60_DNA_seed | EGFR | -1.00795 | HNSC | JHU-011 | Osimertinib | MVP_top60_DNA+RNA_seed | AR | 0.406671 | KIRC | KMRC-1 | Bicalutamide |
| MVP_top60_DNA_seed | AKT1 | -1.006933 | HNSC | JHU-011 | AZD5363 | MVP_top60_DNA+RNA_seed | SMARCA4 | 0.407255 | KIRC | RCC-JF | PFI-3 |
| MVP_top60_DNA_seed | NTRK1 | -1.004657 | HNSC | PCI-15A | GW441756 | MVP_top60_DNA+RNA_seed | SMARCA2 | 0.407255 | KIRC | RCC-JF | PFI-3 |
| MVP_top60_DNA_seed | AKT1 | -1.004088 | HNSC | BB49-HNC | GSK2110183B | MVP_top60_DNA+RNA_seed | FGFR4 | 0.407777 | KIRC | CAL-54 | FGFR_3831 |
| MVP_top60_DNA_seed | HSP90AA1 | -1.00062 | HNSC | BB30-HNC | Luminespib | MVP_top60_DNA+RNA_seed | HDAC1 | 0.413154 | KIRC | RCC10RGB | JQ12 |
| MVP_top60_DNA_seed | RAC1 | -0.995265 | HNSC | CAL-33 | EHT-1864 | MVP_top60_DNA+RNA_seed | MTOR | 0.415762 | KIRC | LB1047-RCC | QL-VIII-58 |
| MVP_top60_DNA_seed | NTRK1 | -0.994767 | HNSC | PCI-4B | AZD1332 | MVP_top60_DNA+RNA_seed | ESR1 | 0.416579 | KIRC | U031 | Tamoxifen |
| MVP_top60_DNA_seed | EGFR | -0.993096 | HNSC | HSC-2 | AZD3759 | MVP_top60_DNA+RNA_seed | AR | 0.421284 | KIRC | CAL-54 | Bicalutamide |
| MVP_top60_DNA_seed | EGFR | -0.991019 | HNSC | PCI-4B | Lapatinib | MVP_top60_DNA+RNA_seed | AR | 0.421926 | KIRC | SW156 | Bicalutamide |
| MVP_top60_DNA_seed | EGFR | -0.988638 | HNSC | PE-CA-PJ15 | AZD3759 | MVP_top60_DNA+RNA_seed | PIK3CA | 0.423394 | KIRC | HA7-RCC | PI3Ka_4409 |
| MVP_top60_DNA_seed | ESR1 | -0.987898 | HNSC | FADU | Tamoxifen | MVP_top60_DNA+RNA_seed | PIK3CD | 0.423394 | KIRC | HA7-RCC | PI3Ka_4409 |
| MVP_top60_DNA_seed | EGFR | -0.987349 | HNSC | OSC-19 | Sapitinib | MVP_top60_DNA+RNA_seed | HDAC1 | 0.424162 | KIRC | CAL-54 | Belinostat |
| MVP_top60_DNA_seed | CDK2 | -0.984799 | HNSC | SCC90 | Dinaciclib | MVP_top60_DNA+RNA_seed | ESR1 | 0.424468 | KIRC | RCC-AB | Fulvestrant |
| MVP_top60_DNA_seed | PIK3CA | -0.983473 | HNSC | HO-1-u-1 | Buparlisib | MVP_top60_DNA+RNA_seed | ATM | 0.42595 | KIRC | SN12C | KU-60019 |
| MVP_top60_DNA_seed | PIK3CA | -0.980902 | HNSC | HSC-3 | GNE-317 | MVP_top60_DNA+RNA_seed | HSP90AA1 | 0.42759 | KIRC | RCC-MF | Tanespimycin |
| MVP_top60_DNA_seed | PIK3CA | -0.973561 | HNSC | KON | GNE-317 | MVP_top60_DNA+RNA_seed | HSP90AB1 | 0.42759 | KIRC | RCC-MF | Tanespimycin |
| MVP_top60_DNA_seed | MAPK1 | -0.973409 | HNSC | HO-1-N-1 | Ulixertinib | MVP_top60_DNA+RNA_seed | MTOR | 0.431119 | KIRC | SN12C | Torin 2 |

| | | | | | | | | | | | |
|---|---|---|---|---|---|---|---|---|---|---|---|
| MVP_top60_DNA_seed | MAPK3 | -0.973409 | HNSC | HO-1-N-1 | Ulixertinib | MVP_top60_DNA+RNA_seed | HDAC1 | 0.433588 | KIRC | BFTC-909 | AR-42 |
| MVP_top60_DNA_seed | HSP90AA1 | -0.972068 | HNSC | KOSC-2 | Luminespib | MVP_top60_DNA+RNA_seed | ESR1 | 0.433774 | KIRC | LB996-RCC | Tamoxifen |
| MVP_top60_DNA_seed | EGFR | -0.971919 | HNSC | SAS | Afatinib | MVP_top60_DNA+RNA_seed | AR | 0.434903 | KIRC | A498 | Bicalutamide |
| MVP_top60_DNA_seed | PIK3CA | -0.970934 | HNSC | JHU-011 | GNE-317 | MVP_top60_DNA+RNA_seed | HSP90AA1 | 0.435157 | KIRC | RCC-AB | Elesclomol |
| MVP_top60_DNA_seed | MAPK1 | -0.970914 | HNSC | JHU-029 | VX-11e | MVP_top60_DNA+RNA_seed | HSP90AB1 | 0.435157 | KIRC | RCC-AB | Elesclomol |
| MVP_top60_DNA_seed | MAPK1 | -0.969762 | HNSC | SCC90 | Ulixertinib | MVP_top60_DNA+RNA_seed | PTK2 | 0.438164 | KIRC | BB65-RCC | PF-562271 |
| MVP_top60_DNA_seed | MAPK3 | -0.969762 | HNSC | SCC90 | Ulixertinib | MVP_top60_DNA+RNA_seed | MTOR | 0.438687 | KIRC | SN12C | CRT0105446 |
| MVP_top60_DNA_seed | EGFR | -0.966735 | HNSC | SCC-15 | Osimertinib | MVP_top60_DNA+RNA_seed | ATM | 0.439653 | KIRC | BFTC-909 | CP466722 |
| MVP_top60_DNA_seed | CDK2 | -0.962159 | HNSC | HSC-2 | AZD5438 | MVP_top60_DNA+RNA_seed | PIK3CA | 0.440249 | KIRC | RCC-AB | AZD8835 |
| MVP_top60_DNA_seed | SRC | -0.961011 | HNSC | JHU-029 | Bosutinib | MVP_top60_DNA+RNA_seed | PIK3CD | 0.440249 | KIRC | RCC-AB | AZD8835 |
| MVP_top60_DNA_seed | MAPK1 | -0.943122 | HNSC | Ca9-22 | VX-11e | MVP_top60_DNA+RNA_seed | IGF1R | 0.445384 | KIRC | TK10 | GSK1904529A |
| MVP_top60_DNA_seed | MAPK1 | -0.941141 | HNSC | BICR78 | Ulixertinib | MVP_top60_DNA+RNA_seed | EP300 | 0.445612 | KIRC | RCC-JF | I-CBP112 |
| MVP_top60_DNA_seed | MAPK3 | -0.941141 | HNSC | BICR78 | Ulixertinib | MVP_top60_DNA+RNA_seed | MTOR | 0.446464 | KIRC | SW156 | Apitolisib |
| MVP_top60_DNA_seed | AKT1 | -0.941033 | HNSC | PCI-6A | Ipatasertib | MVP_top60_DNA+RNA_seed | IGF1R | 0.447103 | KIRC | BB65-RCC | GSK1904529A |
| MVP_top60_DNA_seed | MDM2 | -0.937468 | HNSC | BB30-HNC | Serdemetan | MVP_top60_DNA+RNA_seed | ESR1 | 0.448765 | KIRC | TK10 | Tamoxifen |
| MVP_top60_DNA_seed | AKT1 | -0.936856 | HNSC | BB49-HNC | Afuresertib | MVP_top60_DNA+RNA_seed | HDAC1 | 0.45089 | KIRC | A704 | JQ12 |
| MVP_top60_DNA_seed | CDK2 | -0.935403 | HNSC | HO-1-N-1 | AZD5438 | MVP_top60_DNA+RNA_seed | SMARCA4 | 0.452822 | KIRC | CAL-54 | PFI-3 |
| MVP_top60_DNA_seed | HDAC1 | -0.934552 | HNSC | BB49-HNC | Romidepsin | MVP_top60_DNA+RNA_seed | SMARCA2 | 0.452822 | KIRC | CAL-54 | PFI-3 |
| MVP_top60_DNA_seed | HDAC3 | -0.934552 | HNSC | BB49-HNC | Romidepsin | MVP_top60_DNA+RNA_seed | ATM | 0.453773 | KIRC | VMRC-RCZ | KU-55933 |
| MVP_top60_DNA_seed | PIK3CA | -0.934206 | HNSC | CAL-27 | Buparlisib | MVP_top60_DNA+RNA_seed | AR | 0.455729 | KIRC | KMRC-1 | AZD3514 |
| MVP_top60_DNA_seed | PIK3CA | -0.934102 | HNSC | KON | Alpelisib | MVP_top60_DNA+RNA_seed | HDAC1 | 0.458187 | KIRC | BFTC-909 | Belinostat |
| MVP_top60_DNA_seed | HSP90AA1 | -0.931606 | HNSC | BICR31 | CCT-018159 | MVP_top60_DNA+RNA_seed | PIK3CD | 0.459552 | KIRC | 769-P | IC-87114 |
| MVP_top60_DNA_seed | HSP90AA1 | -0.929941 | HNSC | SAS | Luminespib | MVP_top60_DNA+RNA_seed | HDAC1 | 0.459754 | KIRC | LB2241-RCC | Belinostat |
| MVP_top60_DNA_seed | HSP90AA1 | -0.92894 | HNSC | KOSC-2 | Tanespimycin | MVP_top60_DNA+RNA_seed | ATM | 0.459825 | KIRC | RXF393 | KU-60019 |
| MVP_top60_DNA_seed | HDAC1 | -0.924238 | HNSC | SAS | Romidepsin | MVP_top60_DNA+RNA_seed | AR | 0.45992 | KIRC | RCC-JW | Bicalutamide |
| MVP_top60_DNA_seed | HDAC3 | -0.924238 | HNSC | SAS | Romidepsin | MVP_top60_DNA+RNA_seed | ATM | 0.460985 | KIRC | SN12C | CP466722 |
| MVP_top60_DNA_seed | CDK2 | -0.924031 | HNSC | SAS | Dinaciclib | MVP_top60_DNA+RNA_seed | PIK3CA | 0.461901 | KIRC | RCC10RGB | PI3Ka_4409 |
| MVP_top60_DNA_seed | AKT1 | -0.918677 | HNSC | JHU-011 | MK-2206 | MVP_top60_DNA+RNA_seed | PIK3CD | 0.461901 | KIRC | RCC10RGB | PI3Ka_4409 |
| MVP_top60_DNA_seed | EGFR | -0.917916 | HNSC | CAL-33 | Osimertinib | MVP_top60_DNA+RNA_seed | MTOR | 0.467187 | KIRC | KMRC-1 | Apitolisib |
| MVP_top60_DNA_seed | HSP90AA1 | -0.913148 | HNSC | PE-CA-PJ15 | Tanespimycin | MVP_top60_DNA+RNA_seed | PTK2 | 0.469594 | KIRC | CAL-54 | PF-562271 |
| MVP_top60_DNA_seed | PIK3CA | -0.911513 | HNSC | BICR10 | GNE-317 | MVP_top60_DNA+RNA_seed | FGFR4 | 0.476178 | KIRC | BB65-RCC | FGFR_3831 |
| MVP_top60_DNA_seed | AKT1 | -0.909078 | HNSC | HSC-4 | GSK2110183B | MVP_top60_DNA+RNA_seed | PIK3CD | 0.477235 | KIRC | BFTC-909 | IC-87114 |
| MVP_top60_DNA_seed | MAPK1 | -0.908715 | HNSC | PE-CA-PJ15 | SCH772984 | MVP_top60_DNA+RNA_seed | SMARCA4 | 0.481007 | KIRC | OS-RC-2 | PFI-3 |
| MVP_top60_DNA_seed | MAPK3 | -0.908715 | HNSC | PE-CA-PJ15 | SCH772984 | MVP_top60_DNA+RNA_seed | SMARCA2 | 0.481007 | KIRC | OS-RC-2 | PFI-3 |
| MVP_top60_DNA_seed | RAC1 | -0.905649 | HNSC | BICR31 | EHT-1864 | MVP_top60_DNA+RNA_seed | IGF1R | 0.482713 | KIRC | KMRC-1 | GSK1904529A |
| MVP_top60_DNA_seed | PIK3CA | -0.902106 | HNSC | KON | AZD8186 | MVP_top60_DNA+RNA_seed | EP300 | 0.485906 | KIRC | SW156 | I-CBP112 |
| MVP_top60_DNA_seed | CDK2 | -0.900336 | HNSC | KON | Dinaciclib | MVP_top60_DNA+RNA_seed | HDAC1 | 0.487359 | KIRC | ACHN | Tubastatin A |
| MVP_top60_DNA_seed | HDAC1 | -0.899176 | HNSC | Ca9-22 | Romidepsin | MVP_top60_DNA+RNA_seed | MTOR | 0.489913 | KIRC | KMRC-1 | Voxtalisib |
| MVP_top60_DNA_seed | HDAC3 | -0.899176 | HNSC | Ca9-22 | Romidepsin | MVP_top60_DNA+RNA_seed | HSP90AA1 | 0.492077 | KIRC | BFTC-909 | SNX-2112 |
| MVP_top60_DNA_seed | EGFR | -0.896622 | HNSC | KON | Gefitinib | MVP_top60_DNA+RNA_seed | HSP90AB1 | 0.492077 | KIRC | BFTC-909 | SNX-2112 |
| MVP_top60_DNA_seed | CDK2 | -0.896226 | HNSC | JHU-011 | Dinaciclib | MVP_top60_DNA+RNA_seed | SMARCA4 | 0.494497 | KIRC | ACHN | PFI-3 |

| | | | | | | | | | | | |
|---|---|---|---|---|---|---|---|---|---|---|---|
| MVP_top60_DNA_seed | HSP90AA1 | -0.893876 | HNSC | FADU | Luminespib | MVP_top60_DNA+RNA_seed | SMARCA2 | 0.494497 | KIRC | ACHN | PFI-3 |
| MVP_top60_DNA_seed | HSP90AA1 | -0.8929 | HNSC | PCI-15A | Elesclomol | MVP_top60_DNA+RNA_seed | HDAC1 | 0.494739 | KIRC | SN12C | AR-42 |
| MVP_top60_DNA_seed | HSP90AA1 | -0.891373 | HNSC | PE-CA-PJ15 | CCT-018159 | MVP_top60_DNA+RNA_seed | MTOR | 0.495631 | KIRC | CAKI-1 | Voxtalisib |
| MVP_top60_DNA_seed | EGFR | -0.890074 | HNSC | HSC-4 | Osimertinib | MVP_top60_DNA+RNA_seed | ATM | 0.496336 | KIRC | KMRC-1 | CP466722 |
| MVP_top60_DNA_seed | MAPK1 | -0.887439 | HNSC | KOSC-2 | SCH772984 | MVP_top60_DNA+RNA_seed | SMARCA4 | 0.497987 | KIRC | NCC010 | PFI-3 |
| MVP_top60_DNA_seed | MAPK3 | -0.887439 | HNSC | KOSC-2 | SCH772984 | MVP_top60_DNA+RNA_seed | SMARCA2 | 0.497987 | KIRC | NCC010 | PFI-3 |
| MVP_top60_DNA_seed | AR | -0.887134 | HNSC | PE-CA-PJ15 | Bicalutamide | MVP_top60_DNA+RNA_seed | HDAC1 | 0.499504 | KIRC | LB996-RCC | AR-42 |
| MVP_top60_DNA_seed | NTRK1 | -0.886993 | HNSC | HSC-3 | AZD1332 | MVP_top60_DNA+RNA_seed | SMARCA4 | 0.502387 | KIRC | CAKI-1 | PFI-3 |
| MVP_top60_DNA_seed | HDAC1 | -0.884265 | HNSC | PCI-15A | PCI-34051 | MVP_top60_DNA+RNA_seed | SMARCA2 | 0.502387 | KIRC | CAKI-1 | PFI-3 |
| MVP_top60_DNA_seed | PIK3CA | -0.884112 | HNSC | Detroit562 | Alpelisib | MVP_top60_DNA+RNA_seed | HDAC1 | 0.505199 | KIRC | LB2241-RCC | CAY10603 |
| MVP_top60_DNA_seed | SRC | -0.883149 | HNSC | HSC-2 | Bosutinib | MVP_top60_DNA+RNA_seed | ESR1 | 0.507974 | KIRC | SN12C | Tamoxifen |
| MVP_top60_DNA_seed | EGFR | -0.879514 | HNSC | HSC-2 | Osimertinib | MVP_top60_DNA+RNA_seed | AR | 0.510566 | KIRC | TK10 | Bicalutamide |
| MVP_top60_DNA_seed | AKT1 | -0.878703 | HNSC | OSC-19 | Afuresertib | MVP_top60_DNA+RNA_seed | ATM | 0.515395 | KIRC | RXF393 | KU-55933 |
| MVP_top60_DNA_seed | EGFR | -0.877786 | HNSC | HO-1-N-1 | Gefitinib | MVP_top60_DNA+RNA_seed | AR | 0.51613 | KIRC | 786-0 | AZD3514 |
| MVP_top60_DNA_seed | EGFR | -0.877422 | HNSC | HSC-4 | AZD3759 | MVP_top60_DNA+RNA_seed | AR | 0.51877 | KIRC | SW156 | AZD3514 |
| MVP_top60_DNA_seed | AKT1 | -0.87492 | HNSC | JHU-029 | MK-2206 | MVP_top60_DNA+RNA_seed | ESR1 | 0.521158 | KIRC | RCC-MF | Fulvestrant |
| MVP_top60_DNA_seed | AKT1 | -0.874855 | HNSC | BICR10 | Uprosertib | MVP_top60_DNA+RNA_seed | FGFR4 | 0.522441 | KIRC | A498 | FGFR_0939 |
| MVP_top60_DNA_seed | SRC | -0.873353 | HNSC | PCI-6A | Bosutinib | MVP_top60_DNA+RNA_seed | HDAC1 | 0.526035 | KIRC | SN12C | CAY10603 |
| MVP_top60_DNA_seed | PIK3CA | -0.866948 | HNSC | PCI-6A | Alpelisib | MVP_top60_DNA+RNA_seed | IGF1R | 0.531131 | KIRC | KMRC-1 | BMS-754807 |
| MVP_top60_DNA_seed | EGFR | -0.86584 | HNSC | SAS | Lapatinib | MVP_top60_DNA+RNA_seed | PIK3CD | 0.53466 | KIRC | SN12C | Idelalisib |
| MVP_top60_DNA_seed | EGFR | -0.860572 | HNSC | HO-1-N-1 | AZD3759 | MVP_top60_DNA+RNA_seed | ESR1 | 0.534732 | KIRC | RCC-MF | Tamoxifen |
| MVP_top60_DNA_seed | EGFR | -0.857898 | HNSC | Detroit562 | Osimertinib | MVP_top60_DNA+RNA_seed | PIK3CA | 0.537162 | KIRC | 769-P | AZD8835 |
| MVP_top60_DNA_seed | EGFR | -0.853133 | HNSC | JHU-011 | Afatinib | MVP_top60_DNA+RNA_seed | PIK3CD | 0.537162 | KIRC | 769-P | AZD8835 |
| MVP_top60_DNA_seed | SRC | -0.850158 | HNSC | HSC-3 | Bosutinib | MVP_top60_DNA+RNA_seed | HSP90AA1 | 0.537846 | KIRC | NCC021 | Elesclomol |
| MVP_top60_DNA_seed | AKT1 | -0.845656 | HNSC | KOSC-2 | Uprosertib | MVP_top60_DNA+RNA_seed | HSP90AB1 | 0.537846 | KIRC | NCC021 | Elesclomol |
| MVP_top60_DNA_seed | EGFR | -0.843431 | HNSC | PCI-4B | Gefitinib | MVP_top60_DNA+RNA_seed | EP300 | 0.537992 | KIRC | HA7-RCC | I-CBP112 |
| MVP_top60_DNA_seed | AKT1 | -0.841031 | HNSC | OSC-19 | Uprosertib | MVP_top60_DNA+RNA_seed | AR | 0.539733 | KIRC | ACHN | Bicalutamide |
| MVP_top60_DNA_seed | MAPK1 | -0.839115 | HNSC | BB30-HNC | Ulixertinib | MVP_top60_DNA+RNA_seed | IGF1R | 0.549114 | KIRC | CAL-54 | BMS-754807 |
| MVP_top60_DNA_seed | MAPK3 | -0.839115 | HNSC | BB30-HNC | Ulixertinib | MVP_top60_DNA+RNA_seed | PIK3CD | 0.550738 | KIRC | LB1047-RCC | IC-87114 |
| MVP_top60_DNA_seed | SRC | -0.837996 | HNSC | KOSC-2 | Bosutinib | MVP_top60_DNA+RNA_seed | ATM | 0.555257 | KIRC | 769-P | KU-55933 |
| MVP_top60_DNA_seed | NTRK1 | -0.833171 | HNSC | HSC-4 | GW441756 | MVP_top60_DNA+RNA_seed | FGFR4 | 0.558079 | KIRC | U031 | FGFR_3831 |
| MVP_top60_DNA_seed | EGFR | -0.832533 | HNSC | Detroit562 | Afatinib | MVP_top60_DNA+RNA_seed | ESR1 | 0.559316 | KIRC | RCC-JF | Tamoxifen |
| MVP_top60_DNA_seed | EGFR | -0.831568 | HNSC | BICR78 | Erlotinib | MVP_top60_DNA+RNA_seed | HDAC1 | 0.56114 | KIRC | RCC-JW | Tubastatin A |
| MVP_top60_DNA_seed | HSP90AA1 | -0.831514 | HNSC | Detroit562 | Tanespimycin | MVP_top60_DNA+RNA_seed | HSP90AA1 | 0.56261 | KIRC | RCC-ER | Tanespimycin |
| MVP_top60_DNA_seed | EGFR | -0.831166 | HNSC | HSC-4 | Sapitinib | MVP_top60_DNA+RNA_seed | HSP90AB1 | 0.56261 | KIRC | RCC-ER | Tanespimycin |
| MVP_top60_DNA_seed | MAPK1 | -0.829531 | HNSC | KOSC-2 | VX-11e | MVP_top60_DNA+RNA_seed | AR | 0.56316 | KIRC | BB65-RCC | Bicalutamide |
| MVP_top60_DNA_seed | NTRK1 | -0.827797 | HNSC | HO-1-N-1 | AZD1332 | MVP_top60_DNA+RNA_seed | IGF1R | 0.563864 | KIRC | CAKI-1 | BMS-536924 |
| MVP_top60_DNA_seed | HDAC1 | -0.827294 | HNSC | KOSC-2 | PCI-34051 | MVP_top60_DNA+RNA_seed | SMARCA4 | 0.569708 | KIRC | RCC-ER | PFI-3 |
| MVP_top60_DNA_seed | HSP90AA1 | -0.82514 | HNSC | BICR10 | Luminespib | MVP_top60_DNA+RNA_seed | SMARCA2 | 0.569708 | KIRC | RCC-ER | PFI-3 |
| MVP_top60_DNA_seed | HSP90AA1 | -0.824165 | HNSC | OSC-19 | CCT-018159 | MVP_top60_DNA+RNA_seed | HDAC1 | 0.570152 | KIRC | RCC-ER | CAY10603 |
| MVP_top60_DNA_seed | EP300 | -0.823827 | HNSC | HO-1-N-1 | CPI-637 | MVP_top60_DNA+RNA_seed | PIK3CD | 0.571295 | KIRC | RXF393 | IC-87114 |

| | | | | | | | | | | | |
|---|---|---|---|---|---|---|---|---|---|---|---|
| MVP_top60_DNA_seed | AKT1 | -0.823577 | HNSC | HSC-3 | AZD5363 | MVP_top60_DNA+RNA_seed | AR | 0.572224 | KIRC | A704 | AZD3514 |
| MVP_top60_DNA_seed | NTRK1 | -0.820962 | HNSC | BICR10 | AZD1332 | MVP_top60_DNA+RNA_seed | HSP90AA1 | 0.572348 | KIRC | SW156 | Tanespimycin |
| MVP_top60_DNA_seed | AKT1 | -0.820252 | HNSC | HSC-4 | Ipatasertib | MVP_top60_DNA+RNA_seed | HSP90AB1 | 0.572348 | KIRC | SW156 | Tanespimycin |
| MVP_top60_DNA_seed | MAPK1 | -0.819685 | HNSC | PCI-4B | SCH772984 | MVP_top60_DNA+RNA_seed | AR | 0.572749 | KIRC | KMRC-20 | Bicalutamide |
| MVP_top60_DNA_seed | MAPK3 | -0.819685 | HNSC | PCI-4B | SCH772984 | MVP_top60_DNA+RNA_seed | HSP90AA1 | 0.572821 | KIRC | KMRC-1 | SNX-2112 |
| MVP_top60_DNA_seed | HSP90AA1 | -0.814784 | HNSC | HO-1-u-1 | Luminespib | MVP_top60_DNA+RNA_seed | HSP90AB1 | 0.572821 | KIRC | KMRC-1 | SNX-2112 |
| MVP_top60_DNA_seed | CDK2 | -0.813877 | HNSC | CAL-27 | AZD5438 | MVP_top60_DNA+RNA_seed | IGF1R | 0.57292 | KIRC | KMRC-20 | BMS-754807 |
| MVP_top60_DNA_seed | HSP90AA1 | -0.813198 | HNSC | SCC-4 | Tanespimycin | MVP_top60_DNA+RNA_seed | PTK2 | 0.573075 | KIRC | BFTC-909 | PF-562271 |
| MVP_top60_DNA_seed | HSP90AA1 | -0.812063 | HNSC | OSC-20 | Tanespimycin | MVP_top60_DNA+RNA_seed | HDAC1 | 0.575334 | KIRC | VMRC-RCW | Tubastatin A |
| MVP_top60_DNA_seed | ESR1 | -0.811135 | HNSC | OSC-20 | GDC0810 | MVP_top60_DNA+RNA_seed | HDAC1 | 0.577764 | KIRC | LB996-RCC | JQ12 |
| MVP_top60_DNA_seed | MAPK1 | -0.809782 | HNSC | JHU-022 | VX-11e | MVP_top60_DNA+RNA_seed | AR | 0.58066 | KIRC | SW156 | Bicalutamide |
| MVP_top60_DNA_seed | PIK3CA | -0.807013 | HNSC | SAT | Alpelisib | MVP_top60_DNA+RNA_seed | IGF1R | 0.580986 | KIRC | RCC10RGB | BMS-754807 |
| MVP_top60_DNA_seed | AKT1 | -0.804835 | HNSC | HSC-2 | Uprosertib | MVP_top60_DNA+RNA_seed | MTOR | 0.582225 | KIRC | VMRC-RCW | Voxtalisib |
| MVP_top60_DNA_seed | NTRK1 | -0.80395 | HNSC | BICR22 | AZD1332 | MVP_top60_DNA+RNA_seed | HSP90AA1 | 0.583768 | KIRC | VMRC-RCZ | Elesclomol |
| MVP_top60_DNA_seed | ESR1 | -0.803937 | HNSC | CAL-33 | GDC0810 | MVP_top60_DNA+RNA_seed | HSP90AB1 | 0.583768 | KIRC | VMRC-RCZ | Elesclomol |
| MVP_top60_DNA_seed | HSP90AA1 | -0.803563 | HNSC | JHU-029 | CCT-018159 | MVP_top60_DNA+RNA_seed | AR | 0.583842 | KIRC | VMRC-RCZ | Bicalutamide |
| MVP_top60_DNA_seed | EGFR | -0.79933 | HNSC | KON | Sapitinib | MVP_top60_DNA+RNA_seed | IGF1R | 0.58521 | KIRC | SN12C | BMS-754807 |
| MVP_top60_DNA_seed | CDK2 | -0.797025 | HNSC | KON | MK-8776 | MVP_top60_DNA+RNA_seed | AR | 0.587555 | KIRC | SN12C | Bicalutamide |
| MVP_top60_DNA_seed | CDK2 | -0.794034 | HNSC | PCI-15A | MK-8776 | MVP_top60_DNA+RNA_seed | ATM | 0.589285 | KIRC | TK10 | KU-60019 |
| MVP_top60_DNA_seed | AKT1 | -0.793916 | HNSC | BICR10 | AT13148 | MVP_top60_DNA+RNA_seed | HSP90AA1 | 0.590802 | KIRC | SW156 | CCT-018159 |
| MVP_top60_DNA_seed | NTRK1 | -0.793297 | HNSC | Detroit562 | AZD1332 | MVP_top60_DNA+RNA_seed | HSP90AB1 | 0.590802 | KIRC | SW156 | CCT-018159 |
| MVP_top60_DNA_seed | HSP90AA1 | -0.792777 | HNSC | BB49-HNC | Luminespib | MVP_top60_DNA+RNA_seed | PIK3CA | 0.592257 | KIRC | CAKI-1 | PI3Ka_4409 |
| MVP_top60_DNA_seed | ESR1 | -0.790133 | HNSC | PCI-15A | GDC0810 | MVP_top60_DNA+RNA_seed | PIK3CD | 0.592257 | KIRC | CAKI-1 | PI3Ka_4409 |
| MVP_top60_DNA_seed | TP53 | -0.78828 | HNSC | OSC-19 | MIRA-1 | MVP_top60_DNA+RNA_seed | HDAC1 | 0.596687 | KIRC | CAKI-1 | Dacinostat |
| MVP_top60_DNA_seed | HDAC1 | -0.78748 | HNSC | HO-1-N-1 | Romidepsin | MVP_top60_DNA+RNA_seed | SMARCA4 | 0.597513 | KIRC | KMRC-20 | PFI-3 |
| MVP_top60_DNA_seed | HDAC3 | -0.78748 | HNSC | HO-1-N-1 | Romidepsin | MVP_top60_DNA+RNA_seed | SMARCA2 | 0.597513 | KIRC | KMRC-20 | PFI-3 |
| MVP_top60_DNA_seed | CDK2 | -0.784967 | HNSC | HSC-3 | AZD5438 | MVP_top60_DNA+RNA_seed | ESR1 | 0.599873 | KIRC | TK10 | Fulvestrant |
| MVP_top60_DNA_seed | CDK2 | -0.783632 | HNSC | SCC-9 | Dinaciclib | MVP_top60_DNA+RNA_seed | PIK3CD | 0.604406 | KIRC | KMRC-20 | IC-87114 |
| MVP_top60_DNA_seed | AKT1 | -0.782949 | HNSC | HO-1-N-1 | AZD5363 | MVP_top60_DNA+RNA_seed | IGF1R | 0.60459 | KIRC | BFTC-909 | GSK1904529A |
| MVP_top60_DNA_seed | TP53 | -0.78149 | HNSC | PCI-15A | MIRA-1 | MVP_top60_DNA+RNA_seed | ESR1 | 0.610087 | KIRC | CAKI-1 | Fulvestrant |
| MVP_top60_DNA_seed | SMARCA4 | -0.78103 | HNSC | Detroit562 | PFI3 | MVP_top60_DNA+RNA_seed | SMARCA4 | 0.610283 | KIRC | LB996-RCC | PFI-3 |
| MVP_top60_DNA_seed | PIK3CA | -0.780878 | HNSC | Detroit562 | Buparlisib | MVP_top60_DNA+RNA_seed | SMARCA2 | 0.610283 | KIRC | LB996-RCC | PFI-3 |
| MVP_top60_DNA_seed | AKT1 | -0.780586 | HNSC | HSC-3 | Afuresertib | MVP_top60_DNA+RNA_seed | ESR1 | 0.61112 | KIRC | NCC021 | Tamoxifen |
| MVP_top60_DNA_seed | EGFR | -0.779405 | HNSC | BICR78 | Gefitinib | MVP_top60_DNA+RNA_seed | AR | 0.611362 | KIRC | SN12C | Bicalutamide |
| MVP_top60_DNA_seed | NTRK1 | -0.777906 | HNSC | CAL-33 | AZD1332 | MVP_top60_DNA+RNA_seed | MTOR | 0.612345 | KIRC | RCC-AB | Temsirolimus |
| MVP_top60_DNA_seed | EGFR | -0.777502 | HNSC | PCI-4B | Erlotinib | MVP_top60_DNA+RNA_seed | ATM | 0.612393 | KIRC | RCC10RGB | KU-60019 |
| MVP_top60_DNA_seed | AKT1 | -0.776642 | HNSC | JHU-011 | Uprosertib | MVP_top60_DNA+RNA_seed | ATM | 0.620442 | KIRC | ACHN | KU-55933 |
| MVP_top60_DNA_seed | MDM2 | -0.77291 | HNSC | FADU | Serdemetan | MVP_top60_DNA+RNA_seed | FGFR4 | 0.621739 | KIRC | VMRC-RCZ | FGFR_3831 |
| MVP_top60_DNA_seed | AKT1 | -0.772201 | HNSC | BICR10 | MK-2206 | MVP_top60_DNA+RNA_seed | HSP90AA1 | 0.623264 | KIRC | HA7-RCC | SNX-2112 |
| MVP_top60_DNA_seed | MAPK1 | -0.772007 | HNSC | OSC-19 | VX-11e | MVP_top60_DNA+RNA_seed | HSP90AB1 | 0.623264 | KIRC | HA7-RCC | SNX-2112 |
| MVP_top60_DNA_seed | AKT1 | -0.771813 | HNSC | KON | Ipatasertib | MVP_top60_DNA+RNA_seed | ATM | 0.639135 | KIRC | BB65-RCC | KU-60019 |

| | | | | | | | | | | | |
|---|---|---|---|---|---|---|---|---|---|---|---|
| MVP_top60_DNA_seed | HSP90AA1 | -0.771701 | HNSC | BICR22 | Luminespib | MVP_top60_DNA+RNA_seed | IGF1R | 0.641284 | KIRC | OS-RC-2 | Linsitinib |
| MVP_top60_DNA_seed | CDK2 | -0.768874 | HNSC | JHU-029 | Dinaciclib | MVP_top60_DNA+RNA_seed | HSP90AA1 | 0.642705 | KIRC | KMRC-1 | CCT-018159 |
| MVP_top60_DNA_seed | HSP90AA1 | -0.76828 | HNSC | PCI-4B | Luminespib | MVP_top60_DNA+RNA_seed | HSP90AB1 | 0.642705 | KIRC | KMRC-1 | CCT-018159 |
| MVP_top60_DNA_seed | CDK2 | -0.761983 | HNSC | CAL-27 | Dinaciclib | MVP_top60_DNA+RNA_seed | EP300 | 0.647817 | KIRC | KMRC-1 | I-CBP112 |
| MVP_top60_DNA_seed | HDAC1 | -0.761056 | HNSC | SCC-4 | Romidepsin | MVP_top60_DNA+RNA_seed | HSP90AA1 | 0.650698 | KIRC | VMRC-RCW | Tanespimycin |
| MVP_top60_DNA_seed | HDAC3 | -0.761056 | HNSC | SCC-4 | Romidepsin | MVP_top60_DNA+RNA_seed | HSP90AB1 | 0.650698 | KIRC | VMRC-RCW | Tanespimycin |
| MVP_top60_DNA_seed | PIK3CA | -0.760683 | HNSC | JHU-029 | AZD8186 | MVP_top60_DNA+RNA_seed | IGF1R | 0.650788 | KIRC | HA7-RCC | GSK1904529A |
| MVP_top60_DNA_seed | MAPK1 | -0.756893 | HNSC | BB49-HNC | Ulixertinib | MVP_top60_DNA+RNA_seed | HDAC1 | 0.652301 | KIRC | VMRC-RCZ | Tubastatin A |
| MVP_top60_DNA_seed | MAPK3 | -0.756893 | HNSC | BB49-HNC | Ulixertinib | MVP_top60_DNA+RNA_seed | HDAC1 | 0.655598 | KIRC | RCC10RGB | CAY10603 |
| MVP_top60_DNA_seed | PIK3CA | -0.755008 | HNSC | JHU-011 | Buparlisib | MVP_top60_DNA+RNA_seed | HDAC1 | 0.656522 | KIRC | KMRC-20 | Belinostat |
| MVP_top60_DNA_seed | PIK3CA | -0.753698 | HNSC | HO-1-N-1 | Alpelisib | MVP_top60_DNA+RNA_seed | ESR1 | 0.65667 | KIRC | KMRC-20 | Tamoxifen |
| MVP_top60_DNA_seed | AKT1 | -0.744519 | HNSC | BB30-HNC | AT13148 | MVP_top60_DNA+RNA_seed | PIK3CA | 0.657681 | KIRC | KMRC-20 | AZD8835 |
| MVP_top60_DNA_seed | EGFR | -0.743462 | HNSC | HO-1-N-1 | Osimertinib | MVP_top60_DNA+RNA_seed | PIK3CD | 0.657681 | KIRC | KMRC-20 | AZD8835 |
| MVP_top60_DNA_seed | AKT1 | -0.741362 | HNSC | Detroit562 | GSK2110183B | MVP_top60_DNA+RNA_seed | MTOR | 0.657814 | KIRC | BFTC-909 | Apitolisib |
| MVP_top60_DNA_seed | EGFR | -0.740882 | HNSC | SCC-9 | AZD3759 | MVP_top60_DNA+RNA_seed | SMARCA4 | 0.658517 | KIRC | RCC-AB | PFI-3 |
| MVP_top60_DNA_seed | CDK2 | -0.740742 | HNSC | BB49-HNC | MK-8776 | MVP_top60_DNA+RNA_seed | SMARCA2 | 0.658517 | KIRC | RCC-AB | PFI-3 |
| MVP_top60_DNA_seed | HSP90AA1 | -0.740052 | HNSC | BICR31 | Elesclomol | MVP_top60_DNA+RNA_seed | HDAC1 | 0.66256 | KIRC | LB2241-RCC | AR-42 |
| MVP_top60_DNA_seed | AKT1 | -0.736535 | HNSC | DOK | GSK2110183B | MVP_top60_DNA+RNA_seed | HSP90AA1 | 0.662564 | KIRC | CAKI-1 | CCT-018159 |
| MVP_top60_DNA_seed | AKT1 | -0.733552 | HNSC | KOSC-2 | Afuresertib | MVP_top60_DNA+RNA_seed | HSP90AB1 | 0.662564 | KIRC | CAKI-1 | CCT-018159 |
| MVP_top60_DNA_seed | AKT1 | -0.730215 | HNSC | HSC-4 | AZD5363 | MVP_top60_DNA+RNA_seed | FGFR4 | 0.668249 | KIRC | OS-RC-2 | FGFR_0939 |
| MVP_top60_DNA_seed | AKT1 | -0.729099 | HNSC | BICR31 | MK-2206 | MVP_top60_DNA+RNA_seed | IGF1R | 0.668936 | KIRC | KMRC-1 | BMS-536924 |
| MVP_top60_DNA_seed | CDK2 | -0.727675 | HNSC | PCI-4B | Dinaciclib | MVP_top60_DNA+RNA_seed | IGF1R | 0.670099 | KIRC | VMRC-RCZ | BMS-536924 |
| MVP_top60_DNA_seed | AKT1 | -0.727111 | HNSC | OSC-19 | Ipatasertib | MVP_top60_DNA+RNA_seed | HSP90AA1 | 0.670993 | KIRC | A704 | Tanespimycin |
| MVP_top60_DNA_seed | ESR1 | -0.72368 | HNSC | BICR10 | GDC0810 | MVP_top60_DNA+RNA_seed | HSP90AB1 | 0.670993 | KIRC | A704 | Tanespimycin |
| MVP_top60_DNA_seed | HSP90AA1 | -0.723386 | HNSC | KON | Luminespib | MVP_top60_DNA+RNA_seed | PTK2 | 0.686751 | KIRC | SW156 | PF-562271 |
| MVP_top60_DNA_seed | MDM2 | -0.71991 | HNSC | PCI-6A | Serdemetan | MVP_top60_DNA+RNA_seed | ESR1 | 0.690444 | KIRC | RCC-FG2 | Fulvestrant |
| MVP_top60_DNA_seed | MDM2 | -0.717551 | HNSC | KON | Serdemetan | MVP_top60_DNA+RNA_seed | ATM | 0.693701 | KIRC | KMRC-20 | CP466722 |
| MVP_top60_DNA_seed | EGFR | -0.716552 | HNSC | PCI-4B | Osimertinib | MVP_top60_DNA+RNA_seed | PIK3CA | 0.695183 | KIRC | SW156 | AZD8835 |
| MVP_top60_DNA_seed | CDK2 | -0.715915 | HNSC | HSC-4 | MK-8776 | MVP_top60_DNA+RNA_seed | PIK3CD | 0.695183 | KIRC | SW156 | AZD8835 |
| MVP_top60_DNA_seed | AKT1 | -0.714185 | HNSC | SAT | Uprosertib | MVP_top60_DNA+RNA_seed | SMARCA4 | 0.695263 | KIRC | KMRC-1 | PFI-3 |
| MVP_top60_DNA_seed | PIK3CA | -0.713635 | HNSC | BB30-HNC | AZD8186 | MVP_top60_DNA+RNA_seed | SMARCA2 | 0.695263 | KIRC | KMRC-1 | PFI-3 |
| MVP_top60_DNA_seed | AKT1 | -0.711837 | HNSC | SCC90 | AZD5363 | MVP_top60_DNA+RNA_seed | HSP90AA1 | 0.704135 | KIRC | VMRC-RCZ | SNX-2112 |
| MVP_top60_DNA_seed | AR | -0.710343 | HNSC | BICR31 | Bicalutamide | MVP_top60_DNA+RNA_seed | HSP90AB1 | 0.704135 | KIRC | VMRC-RCZ | SNX-2112 |
| MVP_top60_DNA_seed | MAPK1 | -0.710156 | HNSC | KON | SCH772984 | MVP_top60_DNA+RNA_seed | MTOR | 0.70542 | KIRC | OS-RC-2 | Voxtalisib |
| MVP_top60_DNA_seed | MAPK3 | -0.710156 | HNSC | KON | SCH772984 | MVP_top60_DNA+RNA_seed | AR | 0.711723 | KIRC | OS-RC-2 | Bicalutamide |
| MVP_top60_DNA_seed | EGFR | -0.709695 | HNSC | KON | Erlotinib | MVP_top60_DNA+RNA_seed | SMARCA4 | 0.712956 | KIRC | NCC021 | PFI-3 |
| MVP_top60_DNA_seed | AKT1 | -0.709467 | HNSC | HSC-2 | Uprosertib | MVP_top60_DNA+RNA_seed | SMARCA2 | 0.712956 | KIRC | NCC021 | PFI-3 |
| MVP_top60_DNA_seed | MAPK1 | -0.707642 | HNSC | SCC90 | Ulixertinib | MVP_top60_DNA+RNA_seed | ATM | 0.714991 | KIRC | VMRC-RCW | CP466722 |
| MVP_top60_DNA_seed | MAPK3 | -0.707642 | HNSC | SCC90 | Ulixertinib | MVP_top60_DNA+RNA_seed | MTOR | 0.720702 | KIRC | KMRC-20 | WYE-125132 |
| MVP_top60_DNA_seed | RAC1 | -0.7073 | HNSC | CAL-27 | EHT-1864 | MVP_top60_DNA+RNA_seed | EP300 | 0.720728 | KIRC | ACHN | I-CBP112 |
| MVP_top60_DNA_seed | AKT1 | -0.706884 | HNSC | PCI-15A | AT13148 | MVP_top60_DNA+RNA_seed | ESR1 | 0.726762 | KIRC | ACHN | Tamoxifen |

| | | | | | | | | | | | |
|---|---|---|---|---|---|---|---|---|---|---|---|
| MVP_top60_DNA_seed | HSP90AA1 | -0.705869 | HNSC | SCC-9 | Luminespib | MVP_top60_DNA+RNA_seed | ESR1 | 0.728505 | KIRC | VMRC-RCZ | Tamoxifen |
| MVP_top60_DNA_seed | HSP90AA1 | -0.697875 | HNSC | PCI-30 | Luminespib | MVP_top60_DNA+RNA_seed | IGF1R | 0.730686 | KIRC | SW156 | BMS-754807 |
| MVP_top60_DNA_seed | MAPK1 | -0.696681 | HNSC | Detroit562 | SCH772984 | MVP_top60_DNA+RNA_seed | HSP90AA1 | 0.734993 | KIRC | CAKI-1 | SNX-2112 |
| MVP_top60_DNA_seed | MAPK3 | -0.696681 | HNSC | Detroit562 | SCH772984 | MVP_top60_DNA+RNA_seed | HSP90AB1 | 0.734993 | KIRC | CAKI-1 | SNX-2112 |
| MVP_top60_DNA_seed | EGFR | -0.695647 | HNSC | KOSC-2 | Erlotinib | MVP_top60_DNA+RNA_seed | MTOR | 0.735312 | KIRC | KMRC-1 | CRT0105446 |
| MVP_top60_DNA_seed | PIK3CA | -0.693506 | HNSC | SCC-9 | Alpelisib | MVP_top60_DNA+RNA_seed | FGFR4 | 0.736576 | KIRC | ACHN | FGFR_0939 |
| MVP_top60_DNA_seed | CDK2 | -0.693199 | HNSC | HO-1-N-1 | Dinaciclib | MVP_top60_DNA+RNA_seed | ATM | 0.738205 | KIRC | BB65-RCC | KU-55933 |
| MVP_top60_DNA_seed | NTRK1 | -0.692934 | HNSC | BB30-HNC | GW441756 | MVP_top60_DNA+RNA_seed | HSP90AA1 | 0.743747 | KIRC | ACHN | SNX-2112 |
| MVP_top60_DNA_seed | EP300 | -0.691513 | HNSC | DOK | SGC-CBP30 | MVP_top60_DNA+RNA_seed | HSP90AB1 | 0.743747 | KIRC | ACHN | SNX-2112 |
| MVP_top60_DNA_seed | CREBBP | -0.691513 | HNSC | DOK | SGC-CBP30 | MVP_top60_DNA+RNA_seed | EP300 | 0.757749 | KIRC | KMRC-20 | I-CBP112 |
| MVP_top60_DNA_seed | MAPK1 | -0.689372 | HNSC | HO-1-N-1 | SCH772984 | MVP_top60_DNA+RNA_seed | IGF1R | 0.758986 | KIRC | KMRC-1 | Linsitinib |
| MVP_top60_DNA_seed | MAPK3 | -0.689372 | HNSC | HO-1-N-1 | SCH772984 | MVP_top60_DNA+RNA_seed | HSP90AA1 | 0.765625 | KIRC | VMRC-RCW | SNX-2112 |
| MVP_top60_DNA_seed | SRC | -0.68925 | HNSC | HO-1-u-1 | Bosutinib | MVP_top60_DNA+RNA_seed | HSP90AB1 | 0.765625 | KIRC | VMRC-RCW | SNX-2112 |
| MVP_top60_DNA_seed | AKT1 | -0.689134 | HNSC | HSC-4 | Afuresertib | MVP_top60_DNA+RNA_seed | MTOR | 0.769037 | KIRC | KMRC-20 | Voxtalisib |
| MVP_top60_DNA_seed | EGFR | -0.686369 | HNSC | KOSC-2 | Osimertinib | MVP_top60_DNA+RNA_seed | HDAC1 | 0.77053 | KIRC | LB996-RCC | Belinostat |
| MVP_top60_DNA_seed | EGFR | -0.68068 | HNSC | HSC-3 | Afatinib | MVP_top60_DNA+RNA_seed | IGF1R | 0.77547 | KIRC | SN12C | GSK1904529A |
| MVP_top60_DNA_seed | CDK2 | -0.68048 | HNSC | HSC-3 | Dinaciclib | MVP_top60_DNA+RNA_seed | MTOR | 0.775958 | KIRC | RCC10RGB | WYE-125132 |
| MVP_top60_DNA_seed | AKT1 | -0.679618 | HNSC | PCI-15A | MK-2206 | MVP_top60_DNA+RNA_seed | HSP90AA1 | 0.781887 | KIRC | NCC021 | SNX-2112 |
| MVP_top60_DNA_seed | ESR1 | -0.679339 | HNSC | BB49-HNC | Tamoxifen | MVP_top60_DNA+RNA_seed | HSP90AB1 | 0.781887 | KIRC | NCC021 | SNX-2112 |
| MVP_top60_DNA_seed | EGFR | -0.678077 | HNSC | BHY | Osimertinib | MVP_top60_DNA+RNA_seed | MTOR | 0.782525 | KIRC | RCC-MF | QL-VIII-58 |
| MVP_top60_DNA_seed | CDK2 | -0.676075 | HNSC | SAT | Dinaciclib | MVP_top60_DNA+RNA_seed | IGF1R | 0.78354 | KIRC | A498 | BMS-754807 |
| MVP_top60_DNA_seed | AKT1 | -0.675114 | HNSC | SAT | Uprosertib | MVP_top60_DNA+RNA_seed | MTOR | 0.790226 | KIRC | KMRC-20 | QL-VIII-58 |
| MVP_top60_DNA_seed | CDK2 | -0.671341 | HNSC | PCI-6A | Dinaciclib | MVP_top60_DNA+RNA_seed | ATM | 0.791789 | KIRC | KMRC-1 | KU-60019 |
| MVP_top60_DNA_seed | PIK3CA | -0.669504 | HNSC | BICR10 | Alpelisib | MVP_top60_DNA+RNA_seed | HSP90AA1 | 0.791947 | KIRC | RCC-JW | SNX-2112 |
| MVP_top60_DNA_seed | HSP90AA1 | -0.669168 | HNSC | CAL-33 | Luminespib | MVP_top60_DNA+RNA_seed | HSP90AB1 | 0.791947 | KIRC | RCC-JW | SNX-2112 |
| MVP_top60_DNA_seed | HSP90AA1 | -0.668226 | HNSC | OSC-19 | Luminespib | MVP_top60_DNA+RNA_seed | HDAC1 | 0.794676 | KIRC | CAKI-1 | Tubastatin A |
| MVP_top60_DNA_seed | RAC1 | -0.667587 | HNSC | BICR22 | EHT-1864 | MVP_top60_DNA+RNA_seed | IGF1R | 0.800728 | KIRC | SN12C | BMS-536924 |
| MVP_top60_DNA_seed | EP300 | -0.667086 | HNSC | DOK | CPI-637 | MVP_top60_DNA+RNA_seed | PTK2 | 0.80618 | KIRC | LB996-RCC | PF-562271 |
| MVP_top60_DNA_seed | EGFR | -0.665577 | HNSC | LB771-HNC | Erlotinib | MVP_top60_DNA+RNA_seed | MTOR | 0.808347 | KIRC | LB996-RCC | WYE-125132 |
| MVP_top60_DNA_seed | PIK3CA | -0.66092 | HNSC | HO-1-N-1 | AZD8186 | MVP_top60_DNA+RNA_seed | SMARCA4 | 0.808487 | KIRC | TK10 | PFI-3 |
| MVP_top60_DNA_seed | SMARCA4 | -0.657658 | HNSC | BB30-HNC | PFI3 | MVP_top60_DNA+RNA_seed | SMARCA2 | 0.808487 | KIRC | TK10 | PFI-3 |
| MVP_top60_DNA_seed | ESR1 | -0.65464 | HNSC | JHU-011 | GDC0810 | MVP_top60_DNA+RNA_seed | AR | 0.808629 | KIRC | CAKI-1 | Bicalutamide |
| MVP_top60_DNA_seed | ESR1 | -0.65052 | HNSC | SCC-9 | GDC0810 | MVP_top60_DNA+RNA_seed | SMARCA4 | 0.813777 | KIRC | RCC-FG2 | PFI-3 |
| MVP_top60_DNA_seed | EGFR | -0.649704 | HNSC | BB49-HNC | Lapatinib | MVP_top60_DNA+RNA_seed | SMARCA2 | 0.813777 | KIRC | RCC-FG2 | PFI-3 |
| MVP_top60_DNA_seed | MAPK1 | -0.649247 | HNSC | BICR22 | SCH772984 | MVP_top60_DNA+RNA_seed | ESR1 | 0.815815 | KIRC | KMRC-1 | Fulvestrant |
| MVP_top60_DNA_seed | MAPK3 | -0.649247 | HNSC | BICR22 | SCH772984 | MVP_top60_DNA+RNA_seed | PIK3CD | 0.817737 | KIRC | RCC10RGB | Idelalisib |
| MVP_top60_DNA_seed | HSP90AA1 | -0.647543 | HNSC | HO-1-N-1 | Luminespib | MVP_top60_DNA+RNA_seed | FGFR4 | 0.824612 | KIRC | TK10 | FGFR_3831 |
| MVP_top60_DNA_seed | MAPK1 | -0.644349 | HNSC | BICR31 | VX-11e | MVP_top60_DNA+RNA_seed | MTOR | 0.825966 | KIRC | VMRC-RCW | Torin 2 |
| MVP_top60_DNA_seed | CDK2 | -0.643866 | HNSC | PCI-4B | AZD5438 | MVP_top60_DNA+RNA_seed | HDAC1 | 0.82818 | KIRC | SN12C | Belinostat |
| MVP_top60_DNA_seed | ESR1 | -0.642122 | HNSC | HO-1-N-1 | Tamoxifen | MVP_top60_DNA+RNA_seed | PIK3CA | 0.835264 | KIRC | RXF393 | PI3Ka_4409 |
| MVP_top60_DNA_seed | MDM2 | -0.641919 | HNSC | BICR31 | Nutlin-3a (-) | MVP_top60_DNA+RNA_seed | PIK3CD | 0.835264 | KIRC | RXF393 | PI3Ka_4409 |

| | | | | | | | | | | | |
|---|---|---|---|---|---|---|---|---|---|---|---|
| MVP_top60_DNA_seed | HDAC1 | -0.639105 | HNSC | KON | PCI-34051 | MVP_top60_DNA+RNA_seed | EP300 | 0.843744 | KIRC | VMRC-RCZ | I-CBP112 |
| MVP_top60_DNA_seed | HSP90AA1 | -0.635525 | HNSC | CAL-27 | Luminespib | MVP_top60_DNA+RNA_seed | HDAC1 | 0.850195 | KIRC | TK10 | Tubastatin A |
| MVP_top60_DNA_seed | PIK3CA | -0.633116 | HNSC | HSC-2 | Buparlisib | MVP_top60_DNA+RNA_seed | PIK3CA | 0.852257 | KIRC | KMRC-20 | PI3Ka_4409 |
| MVP_top60_DNA_seed | PIK3CA | -0.632378 | HNSC | HSC-3 | Alpelisib | MVP_top60_DNA+RNA_seed | PIK3CD | 0.852257 | KIRC | KMRC-20 | PI3Ka_4409 |
| MVP_top60_DNA_seed | PIK3CA | -0.629658 | HNSC | Detroit562 | AZD8186 | MVP_top60_DNA+RNA_seed | HSP90AA1 | 0.853442 | KIRC | CAL-54 | Tanespimycin |
| MVP_top60_DNA_seed | EP300 | -0.625395 | HNSC | CAL-27 | SGC-CBP30 | MVP_top60_DNA+RNA_seed | HSP90AB1 | 0.853442 | KIRC | CAL-54 | Tanespimycin |
| MVP_top60_DNA_seed | CREBBP | -0.625395 | HNSC | CAL-27 | SGC-CBP30 | MVP_top60_DNA+RNA_seed | HDAC1 | 0.860175 | KIRC | RXF393 | Parthenolide |
| MVP_top60_DNA_seed | EP300 | -0.623916 | HNSC | BICR78 | CPI-637 | MVP_top60_DNA+RNA_seed | HDAC1 | 0.86166 | KIRC | LB996-RCC | Parthenolide |
| MVP_top60_DNA_seed | EP300 | -0.623598 | HNSC | LB771-HNC | SGC-CBP30 | MVP_top60_DNA+RNA_seed | SMARCA4 | 0.862322 | KIRC | VMRC-RCZ | PFI-3 |
| MVP_top60_DNA_seed | CREBBP | -0.623598 | HNSC | LB771-HNC | SGC-CBP30 | MVP_top60_DNA+RNA_seed | SMARCA2 | 0.862322 | KIRC | VMRC-RCZ | PFI-3 |
| MVP_top60_DNA_seed | MDM2 | -0.621345 | HNSC | HSC-2 | Serdemetan | MVP_top60_DNA+RNA_seed | PIK3CA | 0.863319 | KIRC | RXF393 | AZD8835 |
| MVP_top60_DNA_seed | AKT1 | -0.620879 | HNSC | HSC-2 | MK-2206 | MVP_top60_DNA+RNA_seed | PIK3CD | 0.863319 | KIRC | RXF393 | AZD8835 |
| MVP_top60_DNA_seed | MAPK1 | -0.619273 | HNSC | HO-1-u-1 | VX-11e | MVP_top60_DNA+RNA_seed | EP300 | 0.864749 | KIRC | BFTC-909 | I-CBP112 |
| MVP_top60_DNA_seed | EGFR | -0.618647 | HNSC | SCC-9 | Erlotinib | MVP_top60_DNA+RNA_seed | IGF1R | 0.865133 | KIRC | KMRC-20 | Linsitinib |
| MVP_top60_DNA_seed | EGFR | -0.6174 | HNSC | BB49-HNC | Gefitinib | MVP_top60_DNA+RNA_seed | HDAC1 | 0.866579 | KIRC | KMRC-20 | Dacinostat |
| MVP_top60_DNA_seed | SMARCA4 | -0.611582 | HNSC | HO-1-N-1 | PFI3 | MVP_top60_DNA+RNA_seed | HSP90AA1 | 0.86672 | KIRC | CAKI-1 | Luminespib |
| MVP_top60_DNA_seed | HDAC1 | -0.611306 | HNSC | CAL-27 | Romidepsin | MVP_top60_DNA+RNA_seed | HSP90AB1 | 0.86672 | KIRC | CAKI-1 | Luminespib |
| MVP_top60_DNA_seed | HDAC3 | -0.611306 | HNSC | CAL-27 | Romidepsin | MVP_top60_DNA+RNA_seed | HDAC1 | 0.867107 | KIRC | KMRC-20 | JQ12 |
| MVP_top60_DNA_seed | PIK3CA | -0.609705 | HNSC | SAS | Alpelisib | MVP_top60_DNA+RNA_seed | HDAC1 | 0.867711 | KIRC | KMRC-20 | CAY10603 |
| MVP_top60_DNA_seed | EGFR | -0.6087 | HNSC | SCC-9 | Gefitinib | MVP_top60_DNA+RNA_seed | MTOR | 0.868267 | KIRC | A498 | CRT0105446 |
| MVP_top60_DNA_seed | HDAC1 | -0.608274 | HNSC | JHU-029 | Entinostat | MVP_top60_DNA+RNA_seed | AR | 0.873634 | KIRC | RCC-MF | Bicalutamide |
| MVP_top60_DNA_seed | HDAC3 | -0.608274 | HNSC | JHU-029 | Entinostat | MVP_top60_DNA+RNA_seed | AR | 0.874335 | KIRC | CAL-54 | Bicalutamide |
| MVP_top60_DNA_seed | AKT1 | -0.607963 | HNSC | JHU-029 | Afuresertib | MVP_top60_DNA+RNA_seed | HSP90AA1 | 0.874677 | KIRC | VMRC-RCZ | CCT-018159 |
| MVP_top60_DNA_seed | HSP90AA1 | -0.607124 | HNSC | PCI-6A | Elesclomol | MVP_top60_DNA+RNA_seed | HSP90AB1 | 0.874677 | KIRC | VMRC-RCZ | CCT-018159 |
| MVP_top60_DNA_seed | AKT1 | -0.606544 | HNSC | SAT | GSK2110183B | MVP_top60_DNA+RNA_seed | MTOR | 0.876488 | KIRC | CAKI-1 | Torin 2 |
| MVP_top60_DNA_seed | CDK2 | -0.604789 | HNSC | Detroit562 | Dinaciclib | MVP_top60_DNA+RNA_seed | PIK3CD | 0.88096 | KIRC | LB996-RCC | Idelalisib |
| MVP_top60_DNA_seed | EP300 | -0.60384 | HNSC | PCI-4B | SGC-CBP30 | MVP_top60_DNA+RNA_seed | HSP90AA1 | 0.89066 | KIRC | NCC010 | Tanespimycin |
| MVP_top60_DNA_seed | CREBBP | -0.60384 | HNSC | PCI-4B | SGC-CBP30 | MVP_top60_DNA+RNA_seed | HSP90AB1 | 0.89066 | KIRC | NCC010 | Tanespimycin |
| MVP_top60_DNA_seed | CDK2 | -0.6037 | HNSC | SAS | MK-8776 | MVP_top60_DNA+RNA_seed | MTOR | 0.899723 | KIRC | BFTC-909 | WYE-125132 |
| MVP_top60_DNA_seed | NTRK1 | -0.601872 | HNSC | CAL-33 | GW441756 | MVP_top60_DNA+RNA_seed | HDAC1 | 0.901104 | KIRC | BFTC-909 | Tubastatin A |
| MVP_top60_DNA_seed | AKT1 | -0.597463 | HNSC | Detroit562 | Uprosertib | MVP_top60_DNA+RNA_seed | MTOR | 0.907342 | KIRC | KMRC-20 | Apitolisib |
| MVP_top60_DNA_seed | PIK3CA | -0.596823 | HNSC | BB30-HNC | Buparlisib | MVP_top60_DNA+RNA_seed | HDAC1 | 0.907583 | KIRC | KMRC-20 | Tubastatin A |
| MVP_top60_DNA_seed | EP300 | -0.596788 | HNSC | FADU | SGC-CBP30 | MVP_top60_DNA+RNA_seed | PTK2 | 0.921292 | KIRC | A498 | PF-562271 |
| MVP_top60_DNA_seed | CREBBP | -0.596788 | HNSC | FADU | SGC-CBP30 | MVP_top60_DNA+RNA_seed | ESR1 | 0.923461 | KIRC | CAL-54 | Fulvestrant |
| MVP_top60_DNA_seed | AKT1 | -0.595844 | HNSC | Detroit562 | Afuresertib | MVP_top60_DNA+RNA_seed | MTOR | 0.932015 | KIRC | VMRC-RCZ | CRT0105446 |
| MVP_top60_DNA_seed | AKT1 | -0.594992 | HNSC | HO-1-N-1 | GSK2110183B | MVP_top60_DNA+RNA_seed | PIK3CA | 0.932058 | KIRC | A498 | PI3Ka_4409 |
| MVP_top60_DNA_seed | CDK2 | -0.594746 | HNSC | SAS | AZD5438 | MVP_top60_DNA+RNA_seed | PIK3CD | 0.932058 | KIRC | A498 | PI3Ka_4409 |
| MVP_top60_DNA_seed | AKT1 | -0.591027 | HNSC | JHU-022 | Afuresertib | MVP_top60_DNA+RNA_seed | FGFR4 | 0.932604 | KIRC | CAL-54 | FGFR_0939 |
| MVP_top60_DNA_seed | HDAC1 | -0.590412 | HNSC | BICR78 | Romidepsin | MVP_top60_DNA+RNA_seed | AR | 0.937033 | KIRC | TK10 | Bicalutamide |
| MVP_top60_DNA_seed | HDAC3 | -0.590412 | HNSC | BICR78 | Romidepsin | MVP_top60_DNA+RNA_seed | EP300 | 0.937501 | KIRC | SN12C | I-CBP112 |
| MVP_top60_DNA_seed | MAPK1 | -0.58739 | HNSC | SCC-9 | SCH772984 | MVP_top60_DNA+RNA_seed | AR | 0.937686 | KIRC | NCC021 | Bicalutamide |

| | | | | | | | | | | | |
|---|---|---|---|---|---|---|---|---|---|---|---|
| MVP_top60_DNA_seed | MAPK3 | -0.58739 | HNSC | SCC-9 | SCH772984 | MVP_top60_DNA+RNA_seed | EP300 | 0.940137 | KIRC | RCC-FG2 | I-CBP112 |
| MVP_top60_DNA_seed | HSP90AA1 | -0.587259 | HNSC | PE-CA-PJ15 | Luminespib | MVP_top60_DNA+RNA_seed | HDAC1 | 0.940443 | KIRC | U031 | AR-42 |
| MVP_top60_DNA_seed | PIK3CA | -0.58603 | HNSC | SAS | AZD8186 | MVP_top60_DNA+RNA_seed | SMARCA4 | 0.940658 | KIRC | RCC-MF | PFI-3 |
| MVP_top60_DNA_seed | ESR1 | -0.583537 | HNSC | CAL-27 | GDC0810 | MVP_top60_DNA+RNA_seed | SMARCA2 | 0.940658 | KIRC | RCC-MF | PFI-3 |
| MVP_top60_DNA_seed | AKT1 | -0.583445 | HNSC | JHU-029 | AZD5363 | MVP_top60_DNA+RNA_seed | PIK3CA | 0.944562 | KIRC | KMRC-1 | PI3Ka_4409 |
| MVP_top60_DNA_seed | SRC | -0.577833 | HNSC | Detroit562 | Bosutinib | MVP_top60_DNA+RNA_seed | PIK3CD | 0.944562 | KIRC | KMRC-1 | PI3Ka_4409 |
| MVP_top60_DNA_seed | PIK3CA | -0.577704 | HNSC | HSC-3 | Buparlisib | MVP_top60_DNA+RNA_seed | HDAC1 | 0.946225 | KIRC | RCC-FG2 | Tubastatin A |
| MVP_top60_DNA_seed | EGFR | -0.576154 | HNSC | SCC-15 | AZD3759 | MVP_top60_DNA+RNA_seed | HDAC1 | 0.946908 | KIRC | SW156 | Tubastatin A |
| MVP_top60_DNA_seed | EGFR | -0.576086 | HNSC | SCC90 | Osimertinib | MVP_top60_DNA+RNA_seed | SMARCA4 | 0.950296 | KIRC | SW156 | PFI-3 |
| MVP_top60_DNA_seed | EGFR | -0.574644 | HNSC | Detroit562 | Gefitinib | MVP_top60_DNA+RNA_seed | SMARCA2 | 0.950296 | KIRC | SW156 | PFI-3 |
| MVP_top60_DNA_seed | HSP90AA1 | -0.574041 | HNSC | KOSC-2 | Elesclomol | MVP_top60_DNA+RNA_seed | HDAC1 | 0.95699 | KIRC | LB996-RCC | Tubastatin A |
| MVP_top60_DNA_seed | EGFR | -0.57293 | HNSC | SAS | Gefitinib | MVP_top60_DNA+RNA_seed | ESR1 | 0.957567 | KIRC | RCC-JW | Fulvestrant |
| MVP_top60_DNA_seed | SRC | -0.572634 | HNSC | Ca9-22 | Bosutinib | MVP_top60_DNA+RNA_seed | HDAC1 | 0.958161 | KIRC | RCC-ER | Tubastatin A |
| MVP_top60_DNA_seed | EGFR | -0.570607 | HNSC | BHY | Erlotinib | MVP_top60_DNA+RNA_seed | MTOR | 0.960748 | KIRC | RCC10RGB | Voxtalisib |
| MVP_top60_DNA_seed | HSP90AA1 | -0.567928 | HNSC | BICR31 | Luminespib | MVP_top60_DNA+RNA_seed | ESR1 | 0.965101 | KIRC | RCC-ER | Fulvestrant |
| MVP_top60_DNA_seed | RAC1 | -0.565654 | HNSC | LB771-HNC | EHT-1864 | MVP_top60_DNA+RNA_seed | PIK3CA | 0.975273 | KIRC | CAKI-1 | AZD8835 |
| MVP_top60_DNA_seed | CDK2 | -0.564791 | HNSC | DOK | Dinaciclib | MVP_top60_DNA+RNA_seed | PIK3CD | 0.975273 | KIRC | CAKI-1 | AZD8835 |
| MVP_top60_DNA_seed | EGFR | -0.561826 | HNSC | PE-CA-PJ15 | Osimertinib | MVP_top60_DNA+RNA_seed | HSP90AA1 | 0.979962 | KIRC | RCC-AB | CCT-018159 |
| MVP_top60_DNA_seed | TP53 | -0.558305 | HNSC | BHY | MIRA-1 | MVP_top60_DNA+RNA_seed | HSP90AB1 | 0.979962 | KIRC | RCC-AB | CCT-018159 |
| MVP_top60_DNA_seed | AKT1 | -0.557605 | HNSC | PCI-4B | Uprosertib | MVP_top60_DNA+RNA_seed | HDAC1 | 0.991384 | KIRC | SW156 | Belinostat |
| MVP_top60_DNA_seed | MAPK1 | -0.557563 | HNSC | Ca9-22 | Ulixertinib | MVP_top60_DNA+RNA_seed | HDAC1 | 0.994399 | KIRC | RCC-ER | AR-42 |
| MVP_top60_DNA_seed | MAPK3 | -0.557563 | HNSC | Ca9-22 | Ulixertinib | MVP_top60_DNA+RNA_seed | PIK3CD | 0.999026 | KIRC | RCC10RGB | IC-87114 |
| MVP_top60_DNA_seed | RAC1 | -0.555967 | HNSC | KON | EHT-1864 | MVP_top60_DNA+RNA_seed | HDAC1 | 1.000032 | KIRC | LB2241-RCC | Tubastatin A |
| MVP_top60_DNA_seed | CDK2 | -0.554689 | HNSC | PCI-6A | MK-8776 | MVP_top60_DNA+RNA_seed | AR | 1.002883 | KIRC | RCC-JF | AZD3514 |
| MVP_top60_DNA_seed | CDK2 | -0.554402 | HNSC | SCC-9 | AZD5438 | MVP_top60_DNA+RNA_seed | HSP90AA1 | 1.013877 | KIRC | RCC-FG2 | CCT-018159 |
| MVP_top60_DNA_seed | MDM2 | -0.554306 | HNSC | BICR31 | Serdemetan | MVP_top60_DNA+RNA_seed | HSP90AB1 | 1.013877 | KIRC | RCC-FG2 | CCT-018159 |
| MVP_top60_DNA_seed | EGFR | -0.553369 | HNSC | BICR78 | Sapitinib | MVP_top60_DNA+RNA_seed | HDAC1 | 1.020949 | KIRC | BB65-RCC | Tubastatin A |
| MVP_top60_DNA_seed | AKT1 | -0.553316 | HNSC | KOSC-2 | Ipatasertib | MVP_top60_DNA+RNA_seed | ESR1 | 1.02107 | KIRC | SW156 | Tamoxifen |
| MVP_top60_DNA_seed | HSP90AA1 | -0.553306 | HNSC | SCC90 | Tanespimycin | MVP_top60_DNA+RNA_seed | HDAC1 | 1.022491 | KIRC | SN12C | Dacinostat |
| MVP_top60_DNA_seed | MAPK1 | -0.549833 | HNSC | HSC-3 | Ulixertinib | MVP_top60_DNA+RNA_seed | HDAC1 | 1.039307 | KIRC | HA7-RCC | Tubastatin A |
| MVP_top60_DNA_seed | MAPK3 | -0.549833 | HNSC | HSC-3 | Ulixertinib | MVP_top60_DNA+RNA_seed | HDAC1 | 1.046399 | KIRC | SW156 | CAY10603 |
| MVP_top60_DNA_seed | PIK3CA | -0.549105 | HNSC | BICR22 | Buparlisib | MVP_top60_DNA+RNA_seed | ATM | 1.071908 | KIRC | LB996-RCC | KU-55933 |
| MVP_top60_DNA_seed | RAC1 | -0.544203 | HNSC | Detroit562 | EHT-1864 | MVP_top60_DNA+RNA_seed | PIK3CD | 1.074467 | KIRC | KMRC-20 | Idelalisib |
| MVP_top60_DNA_seed | AKT1 | -0.541002 | HNSC | Detroit562 | MK-2206 | MVP_top60_DNA+RNA_seed | HDAC1 | 1.075032 | KIRC | HA7-RCC | Parthenolide |
| MVP_top60_DNA_seed | HDAC1 | -0.54093 | HNSC | FADU | Romidepsin | MVP_top60_DNA+RNA_seed | MTOR | 1.078733 | KIRC | RCC-FG2 | QL-VIII-58 |
| MVP_top60_DNA_seed | HDAC3 | -0.54093 | HNSC | FADU | Romidepsin | MVP_top60_DNA+RNA_seed | PTK2 | 1.079902 | KIRC | HA7-RCC | PF-562271 |
| MVP_top60_DNA_seed | AKT1 | -0.539243 | HNSC | BICR10 | Uprosertib | MVP_top60_DNA+RNA_seed | HDAC1 | 1.083219 | KIRC | LB2241-RCC | JQ12 |
| MVP_top60_DNA_seed | AKT1 | -0.537585 | HNSC | BICR31 | AT13148 | MVP_top60_DNA+RNA_seed | MTOR | 1.084701 | KIRC | OS-RC-2 | CRT0105446 |
| MVP_top60_DNA_seed | AKT1 | -0.537052 | HNSC | KOSC-2 | AZD5363 | MVP_top60_DNA+RNA_seed | MTOR | 1.086138 | KIRC | RCC-ER | QL-VIII-58 |
| MVP_top60_DNA_seed | PIK3CA | -0.536458 | HNSC | Ca9-22 | Alpelisib | MVP_top60_DNA+RNA_seed | HDAC1 | 1.090848 | KIRC | KMRC-20 | AR-42 |
| MVP_top60_DNA_seed | EP300 | -0.534575 | HNSC | BICR78 | SGC-CBP30 | MVP_top60_DNA+RNA_seed | HDAC1 | 1.107478 | KIRC | LB1047-RCC | Parthenolide |

| | | | | | | | | | | | |
|---|---|---|---|---|---|---|---|---|---|---|---|
| MVP_top60_DNA_seed | CREBBP | -0.534575 | HNSC | BICR78 | SGC-CBP30 | MVP_top60_DNA+RNA_seed | MTOR | 1.1109 | KIRC | VMRC-RCW | Temsirolimus |
| MVP_top60_DNA_seed | EGFR | -0.534235 | HNSC | OSC-20 | Sapitinib | MVP_top60_DNA+RNA_seed | PIK3CA | 1.117208 | KIRC | 786-0 | AZD8835 |
| MVP_top60_DNA_seed | MAPK1 | -0.531401 | HNSC | SCC-4 | Ulixertinib | MVP_top60_DNA+RNA_seed | PIK3CD | 1.117208 | KIRC | 786-0 | AZD8835 |
| MVP_top60_DNA_seed | MAPK3 | -0.531401 | HNSC | SCC-4 | Ulixertinib | MVP_top60_DNA+RNA_seed | HDAC1 | 1.126561 | KIRC | SW156 | AR-42 |
| MVP_top60_DNA_seed | AKT1 | -0.530389 | HNSC | BICR22 | GSK2110183B | MVP_top60_DNA+RNA_seed | HDAC1 | 1.129408 | KIRC | RCC10RGB | AR-42 |
| MVP_top60_DNA_seed | EGFR | -0.529966 | HNSC | BICR22 | Lapatinib | MVP_top60_DNA+RNA_seed | MTOR | 1.131446 | KIRC | VMRC-RCW | WYE-125132 |
| MVP_top60_DNA_seed | AKT1 | -0.528816 | HNSC | CAL-33 | GSK2110183B | MVP_top60_DNA+RNA_seed | HSP90AA1 | 1.13204 | KIRC | NCC021 | Tanespimycin |
| MVP_top60_DNA_seed | RAC1 | -0.528493 | HNSC | Ca9-22 | EHT-1864 | MVP_top60_DNA+RNA_seed | HSP90AB1 | 1.13204 | KIRC | NCC021 | Tanespimycin |
| MVP_top60_DNA_seed | AKT1 | -0.525599 | HNSC | BICR10 | Afuresertib | MVP_top60_DNA+RNA_seed | AR | 1.132228 | KIRC | VMRC-RCZ | Bicalutamide |
| MVP_top60_DNA_seed | AKT1 | -0.523197 | HNSC | CAL-33 | AT13148 | MVP_top60_DNA+RNA_seed | HDAC1 | 1.134844 | KIRC | A498 | JQ12 |
| MVP_top60_DNA_seed | SMARCA4 | -0.520351 | HNSC | PCI-15A | PFI3 | MVP_top60_DNA+RNA_seed | MTOR | 1.13807 | KIRC | U031 | Temsirolimus |
| MVP_top60_DNA_seed | MDM2 | -0.51965 | HNSC | PE-CA-PJ15 | Serdemetan | MVP_top60_DNA+RNA_seed | HSP90AA1 | 1.142809 | KIRC | SN12C | SNX-2112 |
| MVP_top60_DNA_seed | MAPK1 | -0.518316 | HNSC | DOK | Ulixertinib | MVP_top60_DNA+RNA_seed | HSP90AB1 | 1.142809 | KIRC | SN12C | SNX-2112 |
| MVP_top60_DNA_seed | MAPK3 | -0.518316 | HNSC | DOK | Ulixertinib | MVP_top60_DNA+RNA_seed | MTOR | 1.150638 | KIRC | SW156 | Voxtalisib |
| MVP_top60_DNA_seed | HSP90AA1 | -0.514583 | HNSC | SAT | Tanespimycin | MVP_top60_DNA+RNA_seed | FGFR4 | 1.151913 | KIRC | SW156 | FGFR_3831 |
| MVP_top60_DNA_seed | EP300 | -0.514444 | HNSC | PCI-30 | CPI-637 | MVP_top60_DNA+RNA_seed | IGF1R | 1.153982 | KIRC | RCC-JW | BMS-536924 |
| MVP_top60_DNA_seed | HSP90AA1 | -0.513158 | HNSC | KON | CCT-018159 | MVP_top60_DNA+RNA_seed | FGFR4 | 1.15443 | KIRC | OS-RC-2 | FGFR_3831 |
| MVP_top60_DNA_seed | EGFR | -0.512212 | HNSC | BHY | Lapatinib | MVP_top60_DNA+RNA_seed | IGF1R | 1.167513 | KIRC | LB2241-RCC | BMS-536924 |
| MVP_top60_DNA_seed | MDM2 | -0.511454 | HNSC | JHU-029 | Serdemetan | MVP_top60_DNA+RNA_seed | PIK3CA | 1.170483 | KIRC | VMRC-RCZ | PI3Ka_4409 |
| MVP_top60_DNA_seed | AKT1 | -0.510336 | HNSC | BICR10 | AZD5363 | MVP_top60_DNA+RNA_seed | PIK3CD | 1.170483 | KIRC | VMRC-RCZ | PI3Ka_4409 |
| MVP_top60_DNA_seed | PIK3CA | -0.508338 | HNSC | BB30-HNC | GNE-317 | MVP_top60_DNA+RNA_seed | IGF1R | 1.176388 | KIRC | ACHN | Linsitinib |
| MVP_top60_DNA_seed | HSP90AA1 | -0.507311 | HNSC | PCI-4B | Tanespimycin | MVP_top60_DNA+RNA_seed | AR | 1.190661 | KIRC | RCC-ER | Bicalutamide |
| MVP_top60_DNA_seed | AKT1 | -0.502137 | HNSC | SCC-9 | Uprosertib | MVP_top60_DNA+RNA_seed | FGFR4 | 1.190802 | KIRC | RCC-FG2 | FGFR_3831 |
| MVP_top60_DNA_seed | ESR1 | -0.500976 | HNSC | BICR22 | GDC0810 | MVP_top60_DNA+RNA_seed | PIK3CA | 1.194837 | KIRC | BB65-RCC | PI3Ka_4409 |
| MVP_top60_DNA_seed | MAPK1 | -0.50084 | HNSC | PCI-6A | VX-11e | MVP_top60_DNA+RNA_seed | PIK3CD | 1.194837 | KIRC | BB65-RCC | PI3Ka_4409 |
| MVP_top60_DNA_seed | AKT1 | -0.50046 | HNSC | KON | MK-2206 | MVP_top60_DNA+RNA_seed | IGF1R | 1.209531 | KIRC | A498 | GSK1904529A |
| MVP_top60_DNA_seed | SMARCA4 | -0.496825 | HNSC | SKN-3 | PFI3 | MVP_top60_DNA+RNA_seed | MTOR | 1.210289 | KIRC | A704 | Torin 2 |
| MVP_top60_DNA_seed | EGFR | -0.495506 | HNSC | SCC-25 | AZD3759 | MVP_top60_DNA+RNA_seed | PIK3CD | 1.216171 | KIRC | LB2241-RCC | Idelalisib |
| MVP_top60_DNA_seed | HSP90AA1 | -0.494648 | HNSC | KOSC-2 | CCT-018159 | MVP_top60_DNA+RNA_seed | HSP90AA1 | 1.222358 | KIRC | CAL-54 | SNX-2112 |
| MVP_top60_DNA_seed | MAPK1 | -0.493771 | HNSC | PCI-4B | VX-11e | MVP_top60_DNA+RNA_seed | HSP90AB1 | 1.222358 | KIRC | CAL-54 | SNX-2112 |
| MVP_top60_DNA_seed | AKT1 | -0.493619 | HNSC | PCI-4B | GSK2110183B | MVP_top60_DNA+RNA_seed | AR | 1.231843 | KIRC | OS-RC-2 | AZD3514 |
| MVP_top60_DNA_seed | AKT1 | -0.491447 | HNSC | OSC-19 | AT13148 | MVP_top60_DNA+RNA_seed | AR | 1.231861 | KIRC | VMRC-RCZ | AZD3514 |
| MVP_top60_DNA_seed | EGFR | -0.49046 | HNSC | SCC-15 | Lapatinib | MVP_top60_DNA+RNA_seed | AR | 1.238458 | KIRC | RXF393 | Bicalutamide |
| MVP_top60_DNA_seed | AKT1 | -0.489386 | HNSC | SAT | AT13148 | MVP_top60_DNA+RNA_seed | ATM | 1.242675 | KIRC | KMRC-1 | KU-55933 |
| MVP_top60_DNA_seed | EP300 | -0.488947 | HNSC | Detroit562 | CPI-637 | MVP_top60_DNA+RNA_seed | IGF1R | 1.251603 | KIRC | VMRC-RCZ | GSK1904529A |
| MVP_top60_DNA_seed | MAPK1 | -0.488611 | HNSC | PE-CA-PJ15 | VX-11e | MVP_top60_DNA+RNA_seed | HDAC1 | 1.26503 | KIRC | RXF393 | Entinostat |
| MVP_top60_DNA_seed | MAPK1 | -0.488097 | HNSC | DOK | VX-11e | MVP_top60_DNA+RNA_seed | MTOR | 1.269411 | KIRC | KMRC-1 | Torin 2 |
| MVP_top60_DNA_seed | HDAC1 | -0.487848 | HNSC | BB49-HNC | PCI-34051 | MVP_top60_DNA+RNA_seed | ESR1 | 1.280738 | KIRC | NCC021 | Fulvestrant |
| MVP_top60_DNA_seed | HDAC1 | -0.486651 | HNSC | HSC-3 | Romidepsin | MVP_top60_DNA+RNA_seed | HSP90AA1 | 1.285857 | KIRC | SW156 | SNX-2112 |
| MVP_top60_DNA_seed | HDAC3 | -0.486651 | HNSC | HSC-3 | Romidepsin | MVP_top60_DNA+RNA_seed | HSP90AB1 | 1.285857 | KIRC | SW156 | SNX-2112 |
| MVP_top60_DNA_seed | AKT1 | -0.486504 | HNSC | BICR22 | Uprosertib | MVP_top60_DNA+RNA_seed | HSP90AA1 | 1.293296 | KIRC | CAL-54 | Elesclomol |

| | | | | | | | | | | | |
|---|---|---|---|---|---|---|---|---|---|---|---|
| MVP_top60_DNA_seed | HDAC1 | -0.486152 | HNSC | DOK | Romidepsin | MVP_top60_DNA+RNA_seed | HSP90AB1 | 1.293296 | KIRC | CAL-54 | Elesclomol |
| MVP_top60_DNA_seed | HDAC3 | -0.486152 | HNSC | DOK | Romidepsin | MVP_top60_DNA+RNA_seed | AR | 1.295094 | KIRC | KMRC-20 | AZD3514 |
| MVP_top60_DNA_seed | NTRK1 | -0.485782 | HNSC | SKN-3 | GW441756 | MVP_top60_DNA+RNA_seed | MTOR | 1.297411 | KIRC | RXF393 | Apitolisib |
| MVP_top60_DNA_seed | PIK3CA | -0.485251 | HNSC | CAL-27 | AZD8186 | MVP_top60_DNA+RNA_seed | AR | 1.311373 | KIRC | NCC021 | AZD3514 |
| MVP_top60_DNA_seed | AKT1 | -0.48512 | HNSC | SCC-25 | AT13148 | MVP_top60_DNA+RNA_seed | PTK2 | 1.318476 | KIRC | VMRC-RCW | PF-562271 |
| MVP_top60_DNA_seed | ESR1 | -0.483734 | HNSC | OSC-20 | Tamoxifen | MVP_top60_DNA+RNA_seed | MTOR | 1.319536 | KIRC | NCC010 | QL-VIII-58 |
| MVP_top60_DNA_seed | MAPK1 | -0.483678 | HNSC | BB49-HNC | SCH772984 | MVP_top60_DNA+RNA_seed | ESR1 | 1.338676 | KIRC | OS-RC-2 | Tamoxifen |
| MVP_top60_DNA_seed | MAPK3 | -0.483678 | HNSC | BB49-HNC | SCH772984 | MVP_top60_DNA+RNA_seed | IGF1R | 1.339708 | KIRC | RCC10RGB | GSK1904529A |
| MVP_top60_DNA_seed | SRC | -0.483352 | HNSC | PCI-4B | Bosutinib | MVP_top60_DNA+RNA_seed | IGF1R | 1.346113 | KIRC | RCC-AB | BMS-536924 |
| MVP_top60_DNA_seed | SRC | -0.483206 | HNSC | KON | Bosutinib | MVP_top60_DNA+RNA_seed | EP300 | 1.352305 | KIRC | OS-RC-2 | I-CBP112 |
| MVP_top60_DNA_seed | MAPK1 | -0.478877 | HNSC | HO-1-N-1 | VX-11e | MVP_top60_DNA+RNA_seed | EP300 | 1.354702 | KIRC | RXF393 | I-CBP112 |
| MVP_top60_DNA_seed | EGFR | -0.478473 | HNSC | BICR22 | Gefitinib | MVP_top60_DNA+RNA_seed | MTOR | 1.37493 | KIRC | VMRC-RCW | Apitolisib |
| MVP_top60_DNA_seed | AKT1 | -0.476654 | HNSC | BB30-HNC | Ipatasertib | MVP_top60_DNA+RNA_seed | AR | 1.375245 | KIRC | BB65-RCC | AZD3514 |
| MVP_top60_DNA_seed | RAC1 | -0.47463 | HNSC | PCI-4B | EHT-1864 | MVP_top60_DNA+RNA_seed | IGF1R | 1.394086 | KIRC | 769-P | BMS-536924 |
| MVP_top60_DNA_seed | TP53 | -0.473581 | HNSC | BB30-HNC | MIRA-1 | MVP_top60_DNA+RNA_seed | MTOR | 1.395824 | KIRC | KMRC-20 | Torin 2 |
| MVP_top60_DNA_seed | EGFR | -0.471159 | HNSC | BB49-HNC | Sapitinib | MVP_top60_DNA+RNA_seed | MTOR | 1.40829 | KIRC | RCC10RGB | Apitolisib |
| MVP_top60_DNA_seed | MAPK1 | -0.468153 | HNSC | Detroit562 | Ulixertinib | MVP_top60_DNA+RNA_seed | HSP90AA1 | 1.423532 | KIRC | U031 | SNX-2112 |
| MVP_top60_DNA_seed | MAPK3 | -0.468153 | HNSC | Detroit562 | Ulixertinib | MVP_top60_DNA+RNA_seed | HSP90AB1 | 1.423532 | KIRC | U031 | SNX-2112 |
| MVP_top60_DNA_seed | ESR1 | -0.467243 | HNSC | SAS | Tamoxifen | MVP_top60_DNA+RNA_seed | SMARCA4 | 1.436441 | KIRC | RXF393 | PFI-3 |
| MVP_top60_DNA_seed | PIK3CA | -0.465323 | HNSC | JHU-022 | GNE-317 | MVP_top60_DNA+RNA_seed | SMARCA2 | 1.436441 | KIRC | RXF393 | PFI-3 |
| MVP_top60_DNA_seed | PIK3CA | -0.46297 | HNSC | SCC-4 | Alpelisib | MVP_top60_DNA+RNA_seed | PIK3CA | 1.438003 | KIRC | SW156 | PI3Ka_4409 |
| MVP_top60_DNA_seed | AKT1 | -0.460967 | HNSC | HSC-3 | Uprosertib | MVP_top60_DNA+RNA_seed | PIK3CD | 1.438003 | KIRC | SW156 | PI3Ka_4409 |
| MVP_top60_DNA_seed | HSP90AA1 | -0.458565 | HNSC | BB30-HNC | Elesclomol | MVP_top60_DNA+RNA_seed | ESR1 | 1.442765 | KIRC | RCC10RGB | Fulvestrant |
| MVP_top60_DNA_seed | MAPK1 | -0.457582 | HNSC | JHU-022 | Ulixertinib | MVP_top60_DNA+RNA_seed | MTOR | 1.453888 | KIRC | RXF393 | Torin 2 |
| MVP_top60_DNA_seed | MAPK3 | -0.457582 | HNSC | JHU-022 | Ulixertinib | MVP_top60_DNA+RNA_seed | MTOR | 1.47322 | KIRC | TK10 | Voxtalisib |
| MVP_top60_DNA_seed | AKT1 | -0.457204 | HNSC | HSC-3 | MK-2206 | MVP_top60_DNA+RNA_seed | IGF1R | 1.4808 | KIRC | SW156 | BMS-536924 |
| MVP_top60_DNA_seed | CDK2 | -0.456519 | HNSC | OSC-19 | Dinaciclib | MVP_top60_DNA+RNA_seed | HSP90AA1 | 1.487972 | KIRC | HA7-RCC | CCT-018159 |
| MVP_top60_DNA_seed | SRC | -0.455521 | HNSC | CAL-33 | Bosutinib | MVP_top60_DNA+RNA_seed | HSP90AB1 | 1.487972 | KIRC | HA7-RCC | CCT-018159 |
| MVP_top60_DNA_seed | ESR1 | -0.455322 | HNSC | CAL-33 | Tamoxifen | MVP_top60_DNA+RNA_seed | HSP90AA1 | 1.488387 | KIRC | RCC10RGB | Luminespib |
| MVP_top60_DNA_seed | AKT1 | -0.453204 | HNSC | HSC-3 | Ipatasertib | MVP_top60_DNA+RNA_seed | HSP90AB1 | 1.488387 | KIRC | RCC10RGB | Luminespib |
| MVP_top60_DNA_seed | EGFR | -0.451922 | HNSC | PE-CA-PJ15 | Afatinib | MVP_top60_DNA+RNA_seed | HDAC1 | 1.492434 | KIRC | SN12C | Tubastatin A |
| MVP_top60_DNA_seed | HDAC1 | -0.449528 | HNSC | JHU-011 | PCI-34051 | MVP_top60_DNA+RNA_seed | PIK3CA | 1.498648 | KIRC | OS-RC-2 | PI3Ka_4409 |
| MVP_top60_DNA_seed | EGFR | -0.447915 | HNSC | PE-CA-PJ15 | Lapatinib | MVP_top60_DNA+RNA_seed | PIK3CD | 1.498648 | KIRC | OS-RC-2 | PI3Ka_4409 |
| MVP_top60_DNA_seed | HSP90AA1 | -0.447536 | HNSC | SKN-3 | Luminespib | MVP_top60_DNA+RNA_seed | HDAC1 | 1.50906 | KIRC | SN12C | JQ12 |
| MVP_top60_DNA_seed | ESR1 | -0.4462 | HNSC | Ca9-22 | Tamoxifen | MVP_top60_DNA+RNA_seed | ATM | 1.517968 | KIRC | OS-RC-2 | KU-55933 |
| MVP_top60_DNA_seed | ESR1 | -0.44055 | HNSC | BICR78 | Tamoxifen | MVP_top60_DNA+RNA_seed | MTOR | 1.531906 | KIRC | RXF393 | WYE-125132 |
| MVP_top60_DNA_seed | MAPK1 | -0.439799 | HNSC | Ca9-22 | Ulixertinib | MVP_top60_DNA+RNA_seed | PIK3CA | 1.532589 | KIRC | OS-RC-2 | AZD8835 |
| MVP_top60_DNA_seed | MAPK3 | -0.439799 | HNSC | Ca9-22 | Ulixertinib | MVP_top60_DNA+RNA_seed | PIK3CD | 1.532589 | KIRC | OS-RC-2 | AZD8835 |
| MVP_top60_DNA_seed | AKT1 | -0.439305 | HNSC | HSC-4 | Uprosertib | MVP_top60_DNA+RNA_seed | HDAC1 | 1.537922 | KIRC | A498 | CAY10603 |
| MVP_top60_DNA_seed | AKT1 | -0.438107 | HNSC | BICR10 | Ipatasertib | MVP_top60_DNA+RNA_seed | HSP90AA1 | 1.542734 | KIRC | SW156 | Luminespib |
| MVP_top60_DNA_seed | PIK3CA | -0.437463 | HNSC | OSC-19 | GNE-317 | MVP_top60_DNA+RNA_seed | HSP90AB1 | 1.542734 | KIRC | SW156 | Luminespib |

| | | | | | | | | | | | |
|---|---|---|---|---|---|---|---|---|---|---|---|
| MVP_top60_DNA_seed | PIK3CA | -0.432018 | HNSC | OSC-19 | AZD8186 | MVP_top60_DNA+RNA_seed | AR | 1.546026 | KIRC | RXF393 | Bicalutamide |
| MVP_top60_DNA_seed | MAPK1 | -0.431043 | HNSC | PCI-6A | Ulixertinib | MVP_top60_DNA+RNA_seed | MTOR | 1.551482 | KIRC | RCC10RGB | JW-7-52-1 |
| MVP_top60_DNA_seed | MAPK3 | -0.431043 | HNSC | PCI-6A | Ulixertinib | MVP_top60_DNA+RNA_seed | FGFR4 | 1.559144 | KIRC | RCC-MF | FGFR_3831 |
| MVP_top60_DNA_seed | MAPK1 | -0.427567 | HNSC | SCC-9 | VX-11e | MVP_top60_DNA+RNA_seed | HSP90AA1 | 1.564544 | KIRC | RCC10RGB | SNX-2112 |
| MVP_top60_DNA_seed | HDAC1 | -0.424687 | HNSC | PCI-4B | PCI-34051 | MVP_top60_DNA+RNA_seed | HSP90AB1 | 1.564544 | KIRC | RCC10RGB | SNX-2112 |
| MVP_top60_DNA_seed | CDK2 | -0.419442 | HNSC | HSC-4 | Dinaciclib | MVP_top60_DNA+RNA_seed | ATM | 1.568488 | KIRC | A498 | KU-55933 |
| MVP_top60_DNA_seed | PIK3CA | -0.41911 | HNSC | HO-1-N-1 | GNE-317 | MVP_top60_DNA+RNA_seed | PIK3CA | 1.571109 | KIRC | 786-0 | PI3Ka_4409 |
| MVP_top60_DNA_seed | NTRK1 | -0.416722 | HNSC | HO-1-N-1 | GW441756 | MVP_top60_DNA+RNA_seed | PIK3CD | 1.571109 | KIRC | 786-0 | PI3Ka_4409 |
| MVP_top60_DNA_seed | SMARCA4 | -0.416047 | HNSC | BICR31 | PFI3 | MVP_top60_DNA+RNA_seed | ATM | 1.571332 | KIRC | RCC-JW | KU-60019 |
| MVP_top60_DNA_seed | EP300 | -0.415567 | HNSC | Ca9-22 | SGC-CBP30 | MVP_top60_DNA+RNA_seed | MTOR | 1.582794 | KIRC | NCC021 | QL-VIII-58 |
| MVP_top60_DNA_seed | CREBBP | -0.415567 | HNSC | Ca9-22 | SGC-CBP30 | MVP_top60_DNA+RNA_seed | HSP90AA1 | 1.583639 | KIRC | A498 | CCT-018159 |
| MVP_top60_DNA_seed | PIK3CA | -0.415452 | HNSC | KOSC-2 | Alpelisib | MVP_top60_DNA+RNA_seed | HSP90AB1 | 1.583639 | KIRC | A498 | CCT-018159 |
| MVP_top60_DNA_seed | MAPK1 | -0.415055 | HNSC | Detroit562 | VX-11e | MVP_top60_DNA+RNA_seed | AR | 1.583993 | KIRC | RCC-ER | AZD3514 |
| MVP_top60_DNA_seed | MAPK1 | -0.413076 | HNSC | HO-1-u-1 | SCH772984 | MVP_top60_DNA+RNA_seed | EP300 | 1.605165 | KIRC | A498 | I-CBP112 |
| MVP_top60_DNA_seed | MAPK3 | -0.413076 | HNSC | HO-1-u-1 | SCH772984 | MVP_top60_DNA+RNA_seed | AR | 1.619653 | KIRC | OS-RC-2 | Bicalutamide |
| MVP_top60_DNA_seed | RAC1 | -0.412183 | HNSC | SAT | EHT-1864 | MVP_top60_DNA+RNA_seed | HSP90AA1 | 1.632913 | KIRC | KMRC-20 | SNX-2112 |
| MVP_top60_DNA_seed | MAPK1 | -0.410179 | HNSC | KOSC-2 | Ulixertinib | MVP_top60_DNA+RNA_seed | HSP90AB1 | 1.632913 | KIRC | KMRC-20 | SNX-2112 |
| MVP_top60_DNA_seed | MAPK3 | -0.410179 | HNSC | KOSC-2 | Ulixertinib | MVP_top60_DNA+RNA_seed | AR | 1.639079 | KIRC | RCC10RGB | AZD3514 |
| MVP_top60_DNA_seed | PIK3CA | -0.409117 | HNSC | PCI-4B | Alpelisib | MVP_top60_DNA+RNA_seed | MTOR | 1.653092 | KIRC | A704 | QL-VIII-58 |
| MVP_top60_DNA_seed | MDM2 | -0.408745 | HNSC | KOSC-2 | Serdemetan | MVP_top60_DNA+RNA_seed | IGF1R | 1.657419 | KIRC | RXF393 | GSK1904529A |
| MVP_top60_DNA_seed | CDK2 | -0.40863 | HNSC | KOSC-2 | Dinaciclib | MVP_top60_DNA+RNA_seed | HDAC1 | 1.682652 | KIRC | A498 | Tubastatin A |
| MVP_top60_DNA_seed | HSP90AA1 | -0.404555 | HNSC | JHU-011 | CCT-018159 | MVP_top60_DNA+RNA_seed | HSP90AA1 | 1.690657 | KIRC | CAL-54 | Luminespib |
| MVP_top60_DNA_seed | HDAC1 | -0.404122 | HNSC | PCI-6A | Entinostat | MVP_top60_DNA+RNA_seed | HSP90AB1 | 1.690657 | KIRC | CAL-54 | Luminespib |
| MVP_top60_DNA_seed | HDAC3 | -0.404122 | HNSC | PCI-6A | Entinostat | MVP_top60_DNA+RNA_seed | MTOR | 1.696703 | KIRC | A704 | WYE-125132 |
| MVP_top60_DNA_seed | RAC1 | -0.403295 | HNSC | OSC-20 | EHT-1864 | MVP_top60_DNA+RNA_seed | PIK3CD | 1.699136 | KIRC | A498 | Idelalisib |
| MVP_top60_DNA_seed | EGFR | -0.402325 | HNSC | SKN-3 | Gefitinib | MVP_top60_DNA+RNA_seed | HDAC1 | 1.711416 | KIRC | KMRC-1 | Dacinostat |
| MVP_top60_DNA_seed | AKT1 | -0.399211 | HNSC | BB30-HNC | Uprosertib | MVP_top60_DNA+RNA_seed | MTOR | 1.722473 | KIRC | RXF393 | CRT0105446 |
| MVP_top60_DNA_seed | CDK2 | -0.39906 | HNSC | SCC-25 | Dinaciclib | MVP_top60_DNA+RNA_seed | FGFR4 | 1.7371 | KIRC | RCC-ER | FGFR_3831 |
| MVP_top60_DNA_seed | MAPK1 | -0.398342 | HNSC | BB49-HNC | Ulixertinib | MVP_top60_DNA+RNA_seed | PTK2 | 1.756325 | KIRC | KMRC-20 | PF-562271 |
| MVP_top60_DNA_seed | MAPK3 | -0.398342 | HNSC | BB49-HNC | Ulixertinib | MVP_top60_DNA+RNA_seed | AR | 1.758796 | KIRC | A498 | AZD3514 |
| MVP_top60_DNA_seed | TP53 | -0.395709 | HNSC | BICR31 | MIRA-1 | MVP_top60_DNA+RNA_seed | PIK3CA | 1.768022 | KIRC | KMRC-1 | AZD8835 |
| MVP_top60_DNA_seed | MDM2 | -0.395313 | HNSC | HSC-3 | Serdemetan | MVP_top60_DNA+RNA_seed | PIK3CD | 1.768022 | KIRC | KMRC-1 | AZD8835 |
| MVP_top60_DNA_seed | ESR1 | -0.394931 | HNSC | DOK | Tamoxifen | MVP_top60_DNA+RNA_seed | AR | 1.774656 | KIRC | RCC10RGB | Bicalutamide |
| MVP_top60_DNA_seed | EGFR | -0.393985 | HNSC | OSC-19 | Gefitinib | MVP_top60_DNA+RNA_seed | HSP90AA1 | 1.79002 | KIRC | OS-RC-2 | CCT-018159 |
| MVP_top60_DNA_seed | AKT1 | -0.391947 | HNSC | BICR31 | Afuresertib | MVP_top60_DNA+RNA_seed | HSP90AB1 | 1.79002 | KIRC | OS-RC-2 | CCT-018159 |
| MVP_top60_DNA_seed | PIK3CA | -0.391237 | HNSC | LB771-HNC | Alpelisib | MVP_top60_DNA+RNA_seed | HSP90AA1 | 1.793778 | KIRC | LB2241-RCC | SNX-2112 |
| MVP_top60_DNA_seed | PIK3CA | -0.388996 | HNSC | JHU-029 | Buparlisib | MVP_top60_DNA+RNA_seed | HSP90AB1 | 1.793778 | KIRC | LB2241-RCC | SNX-2112 |
| MVP_top60_DNA_seed | CDK2 | -0.387344 | HNSC | PE-CA-PJ15 | Dinaciclib | MVP_top60_DNA+RNA_seed | HDAC1 | 1.810855 | KIRC | RCC-ER | Belinostat |
| MVP_top60_DNA_seed | HDAC1 | -0.386443 | HNSC | BICR10 | Romidepsin | MVP_top60_DNA+RNA_seed | ESR1 | 1.814996 | KIRC | OS-RC-2 | Fulvestrant |
| MVP_top60_DNA_seed | HDAC3 | -0.386443 | HNSC | BICR10 | Romidepsin | MVP_top60_DNA+RNA_seed | ATM | 1.827843 | KIRC | A498 | CP466722 |
| MVP_top60_DNA_seed | RAC1 | -0.385646 | HNSC | DOK | EHT-1864 | MVP_top60_DNA+RNA_seed | HSP90AA1 | 1.829653 | KIRC | ACHN | Elesclomol |

| | | | | | | | | | | | |
|---|---|---|---|---|---|---|---|---|---|---|---|
| MVP_top60_DNA_seed | HDAC1 | -0.385235 | HNSC | JHU-011 | Entinostat | MVP_top60_DNA+RNA_seed | HSP90AB1 | 1.829653 | KIRC | ACHN | Elesclomol |
| MVP_top60_DNA_seed | HDAC3 | -0.385235 | HNSC | JHU-011 | Entinostat | MVP_top60_DNA+RNA_seed | HDAC1 | 1.835936 | KIRC | A498 | AR-42 |
| MVP_top60_DNA_seed | PIK3CA | -0.385124 | HNSC | BICR22 | GNE-317 | MVP_top60_DNA+RNA_seed | HSP90AA1 | 1.862574 | KIRC | LB996-RCC | CCT-018159 |
| MVP_top60_DNA_seed | PIK3CA | -0.383097 | HNSC | BICR31 | Alpelisib | MVP_top60_DNA+RNA_seed | HSP90AB1 | 1.862574 | KIRC | LB996-RCC | CCT-018159 |
| MVP_top60_DNA_seed | EGFR | -0.381151 | HNSC | BICR78 | Lapatinib | MVP_top60_DNA+RNA_seed | HDAC1 | 1.872589 | KIRC | A498 | Parthenolide |
| MVP_top60_DNA_seed | AKT1 | -0.379766 | HNSC | KON | AZD5363 | MVP_top60_DNA+RNA_seed | MTOR | 1.875297 | KIRC | RXF393 | Voxtalisib |
| MVP_top60_DNA_seed | CDK2 | -0.378383 | HNSC | SCC90 | AZD5438 | MVP_top60_DNA+RNA_seed | HDAC1 | 1.892178 | KIRC | RCC10RGB | Tubastatin A |
| MVP_top60_DNA_seed | MAPK1 | -0.376215 | HNSC | SKN-3 | SCH772984 | MVP_top60_DNA+RNA_seed | MTOR | 1.908838 | KIRC | RCC10RGB | CRT0105446 |
| MVP_top60_DNA_seed | MAPK3 | -0.376215 | HNSC | SKN-3 | SCH772984 | MVP_top60_DNA+RNA_seed | MTOR | 1.91931 | KIRC | A704 | Apitolisib |
| MVP_top60_DNA_seed | TP53 | -0.372726 | HNSC | JHU-011 | MIRA-1 | MVP_top60_DNA+RNA_seed | HSP90AA1 | 1.970553 | KIRC | RCC-ER | CCT-018159 |
| MVP_top60_DNA_seed | SRC | -0.372525 | HNSC | SCC-4 | Bosutinib | MVP_top60_DNA+RNA_seed | HSP90AB1 | 1.970553 | KIRC | RCC-ER | CCT-018159 |
| MVP_top60_DNA_seed | AR | -0.37193 | HNSC | BHY | Bicalutamide | MVP_top60_DNA+RNA_seed | ESR1 | 2.010775 | KIRC | A498 | Tamoxifen |
| MVP_top60_DNA_seed | HDAC1 | -0.371564 | HNSC | PE-CA-PJ15 | PCI-34051 | MVP_top60_DNA+RNA_seed | ATM | 2.043105 | KIRC | RCC10RGB | CP466722 |
| MVP_top60_DNA_seed | MAPK1 | -0.367597 | HNSC | JHU-022 | SCH772984 | MVP_top60_DNA+RNA_seed | FGFR4 | 2.045145 | KIRC | RCC10RGB | FGFR_3831 |
| MVP_top60_DNA_seed | MAPK3 | -0.367597 | HNSC | JHU-022 | SCH772984 | MVP_top60_DNA+RNA_seed | AR | 2.050606 | KIRC | RXF393 | AZD3514 |
| MVP_top60_DNA_seed | HDAC1 | -0.367444 | HNSC | KOSC-2 | Entinostat | MVP_top60_DNA+RNA_seed | AR | 2.05533 | KIRC | A498 | Bicalutamide |
| MVP_top60_DNA_seed | HDAC3 | -0.367444 | HNSC | KOSC-2 | Entinostat | MVP_top60_DNA+RNA_seed | MTOR | 2.059937 | KIRC | RCC10RGB | Torin 2 |
| MVP_top60_DNA_seed | EGFR | -0.364893 | HNSC | CAL-33 | Afatinib | MVP_top60_DNA+RNA_seed | ATM | 2.072145 | KIRC | RCC10RGB | KU-55933 |
| MVP_top60_DNA_seed | AKT1 | -0.363532 | HNSC | PCI-4B | Afuresertib | MVP_top60_DNA+RNA_seed | HSP90AA1 | 2.072658 | KIRC | RCC-MF | CCT-018159 |
| MVP_top60_DNA_seed | MDM2 | -0.362902 | HNSC | SAT | Serdemetan | MVP_top60_DNA+RNA_seed | HSP90AB1 | 2.072658 | KIRC | RCC-MF | CCT-018159 |
| MVP_top60_DNA_seed | SRC | -0.362849 | HNSC | SKN-3 | Bosutinib | MVP_top60_DNA+RNA_seed | HSP90AA1 | 2.073883 | KIRC | A498 | SNX-2112 |
| MVP_top60_DNA_seed | EGFR | -0.362382 | HNSC | KOSC-2 | Afatinib | MVP_top60_DNA+RNA_seed | HSP90AB1 | 2.073883 | KIRC | A498 | SNX-2112 |
| MVP_top60_DNA_seed | AKT1 | -0.360204 | HNSC | BB30-HNC | MK-2206 | MVP_top60_DNA+RNA_seed | HSP90AA1 | 2.085682 | KIRC | RCC-ER | SNX-2112 |
| MVP_top60_DNA_seed | ESR1 | -0.359623 | HNSC | BB30-HNC | GDC0810 | MVP_top60_DNA+RNA_seed | HSP90AB1 | 2.085682 | KIRC | RCC-ER | SNX-2112 |
| MVP_top60_DNA_seed | AKT1 | -0.358757 | HNSC | PCI-4B | Ipatasertib | MVP_top60_DNA+RNA_seed | HSP90AA1 | 2.090851 | KIRC | RCC10RGB | Tanespimycin |
| MVP_top60_DNA_seed | HSP90AA1 | -0.35561 | HNSC | SCC-15 | Luminespib | MVP_top60_DNA+RNA_seed | HSP90AB1 | 2.090851 | KIRC | RCC10RGB | Tanespimycin |
| MVP_top60_DNA_seed | CDK2 | -0.354096 | HNSC | Ca9-22 | Dinaciclib | MVP_top60_DNA+RNA_seed | MTOR | 2.111663 | KIRC | RCC10RGB | Temsirolimus |
| MVP_top60_DNA_seed | SMARCA4 | -0.353835 | HNSC | KON | PFI3 | MVP_top60_DNA+RNA_seed | PIK3CA | 2.144494 | KIRC | A498 | AZD8835 |
| MVP_top60_DNA_seed | SMARCA4 | -0.353257 | HNSC | CAL-27 | PFI3 | MVP_top60_DNA+RNA_seed | PIK3CD | 2.144494 | KIRC | A498 | AZD8835 |
| MVP_top60_DNA_seed | PIK3CA | -0.352846 | HNSC | PCI-15A | Buparlisib | MVP_top60_DNA+RNA_seed | AR | 2.148962 | KIRC | RCC10RGB | Bicalutamide |
| MVP_top60_DNA_seed | PIK3CA | -0.352723 | HNSC | HSC-2 | AZD8186 | MVP_top60_DNA+RNA_seed | ATM | 2.163411 | KIRC | LB2241-RCC | CP466722 |
| MVP_top60_DNA_seed | EGFR | -0.346459 | HNSC | SCC-9 | Osimertinib | MVP_top60_DNA+RNA_seed | HSP90AA1 | 2.235188 | KIRC | RCC10RGB | CCT-018159 |
| MVP_top60_DNA_seed | PIK3CA | -0.345583 | HNSC | SCC-25 | Alpelisib | MVP_top60_DNA+RNA_seed | HSP90AB1 | 2.235188 | KIRC | RCC10RGB | CCT-018159 |
| MVP_top60_DNA_seed | EGFR | -0.344186 | HNSC | BICR78 | Osimertinib | MVP_top60_DNA+RNA_seed | PIK3CD | 2.298621 | KIRC | A498 | IC-87114 |
| MVP_top60_DNA_seed | AKT1 | -0.34332 | HNSC | Ca9-22 | AT13148 | MVP_top60_DNA+RNA_seed | MTOR | 2.356098 | KIRC | LB2241-RCC | JW-7-52-1 |
| MVP_top60_DNA_seed | EGFR | -0.34138 | HNSC | BICR31 | AZD3759 | MVP_top60_DNA+RNA_seed | HDAC1 | 2.410439 | KIRC | TK10 | AR-42 |
| MVP_top60_DNA_seed | HDAC1 | -0.34099 | HNSC | SAS | Entinostat | MVP_top60_DNA+RNA_seed | HDAC1 | 2.455744 | KIRC | TK10 | Belinostat |
| MVP_top60_DNA_seed | HDAC3 | -0.34099 | HNSC | SAS | Entinostat | MVP_top60_DNA+RNA_seed | HSP90AA1 | 2.498394 | KIRC | KMRC-20 | Luminespib |
| MVP_top60_DNA_seed | PIK3CA | -0.337761 | HNSC | JHU-022 | Alpelisib | MVP_top60_DNA+RNA_seed | HSP90AB1 | 2.498394 | KIRC | KMRC-20 | Luminespib |
| MVP_top60_DNA_seed | EGFR | -0.336172 | HNSC | SCC-4 | Afatinib | MVP_top60_DNA+RNA_seed | HDAC1 | 2.524926 | KIRC | HA7-RCC | AR-42 |
| MVP_top60_DNA_seed | CDK2 | -0.335989 | HNSC | CAL-33 | Dinaciclib | MVP_top60_DNA+RNA_seed | HDAC1 | 2.600514 | KIRC | RCC10RGB | Belinostat |

| | | | | | | | | | | | |
|---|---|---|---|---|---|---|---|---|---|---|---|
| MVP_top60_DNA_seed | ESR1 | -0.335021 | HNSC | BICR22 | Tamoxifen | MVP_top60_DNA+RNA_seed | HDAC1 | 2.634638 | KIRC | RCC10RGB | Entinostat |
| MVP_top60_DNA_seed | HDAC1 | -0.334877 | HNSC | SCC-15 | Romidepsin | MVP_top60_DNA+RNA_seed | HDAC1 | 2.708613 | KIRC | HA7-RCC | CAY10603 |
| MVP_top60_DNA_seed | HDAC3 | -0.334877 | HNSC | SCC-15 | Romidepsin | MVP_top60_DNA+RNA_seed | HDAC1 | 2.763816 | KIRC | TK10 | CAY10603 |
| MVP_top60_DNA_seed | EGFR | -0.33389 | HNSC | PE-CA-PJ15 | Erlotinib | MVP_top60_DNA+RNA_seed | HDAC1 | 2.788661 | KIRC | LB996-RCC | CAY10603 |
| MVP_top60_DNA_seed | EGFR | -0.332914 | HNSC | HSC-4 | Lapatinib | MVP_top60_DNA+RNA_seed | HDAC1 | 2.828967 | KIRC | A498 | Belinostat |
| MVP_top60_DNA_seed | MAPK1 | -0.329279 | HNSC | SAT | VX-11e | MVP_top60_DNA+RNA_seed | MTOR | 2.846558 | KIRC | A498 | WYE-125132 |
| MVP_top60_DNA_seed | HSP90AA1 | -0.324407 | HNSC | OSC-20 | Luminespib | MVP_top60_DNA+RNA_seed | HDAC1 | 2.950869 | KIRC | HA7-RCC | Belinostat |
| MVP_top60_DNA_seed | PIK3CA | -0.32346 | HNSC | JHU-011 | AZD8186 | MVP_top60_DNA+RNA_seed | MTOR | 3.018643 | KIRC | A498 | JW-7-52-1 |
| MVP_top60_DNA_seed | AKT1 | -0.323162 | HNSC | BICR10 | GSK2110183B | MVP_top60_DNA+RNA_seed | MTOR | 3.086888 | KIRC | A498 | Voxtalisib |
| MVP_top60_DNA_seed | SMARCA4 | -0.321748 | HNSC | JHU-011 | PFI3 | MVP_top60_DNA+RNA_seed | HDAC1 | 3.100667 | KIRC | A498 | Dacinostat |
| MVP_top60_DNA_seed | MAPK1 | -0.317753 | HNSC | Ca9-22 | SCH772984 | MVP_top60_DNA+RNA_seed | HSP90AA1 | 3.151748 | KIRC | RCC10RGB | Elesclomol |
| MVP_top60_DNA_seed | MAPK3 | -0.317753 | HNSC | Ca9-22 | SCH772984 | MVP_top60_DNA+RNA_seed | HSP90AB1 | 3.151748 | KIRC | RCC10RGB | Elesclomol |
| MVP_top60_DNA_seed | EP300 | -0.31703 | HNSC | Ca9-22 | CPI-637 | MVP_top60_DNA+RNA_seed | HSP90AA1 | 3.177217 | KIRC | A498 | Luminespib |
| MVP_top60_DNA_seed | EGFR | -0.316069 | HNSC | PE-CA-PJ15 | Sapitinib | MVP_top60_DNA+RNA_seed | HSP90AB1 | 3.177217 | KIRC | A498 | Luminespib |
| MVP_top60_DNA_seed | PIK3CA | -0.310817 | HNSC | BICR10 | AZD8186 | MVP_top60_DNA+RNA_seed | MTOR | 3.421029 | KIRC | A498 | Torin 2 |
| MVP_top60_DNA_seed | EGFR | -0.310743 | HNSC | HO-1-u-1 | AZD3759 | MVP_top60_DNA+RNA_seed | MTOR | 3.659331 | KIRC | A498 | Apitolisib |
| MVP_top60_DNA_seed | MAPK1 | -0.308868 | HNSC | CAL-27 | SCH772984 | MVP_top60_DNA+RNA_seed | PIK3CA | -3.218483 | KIRC | RCC-FG2 | AZD8186 |
| MVP_top60_DNA_seed | MAPK3 | -0.308868 | HNSC | CAL-27 | SCH772984 | MVP_top60_DNA+RNA_seed | PIK3CA | -2.68479 | KIRC | CAL-54 | AZD8186 |
| MVP_top60_DNA_seed | EP300 | -0.307139 | HNSC | HSC-2 | CPI-637 | MVP_top60_DNA+RNA_seed | PIK3CA | -2.420553 | KIRC | A498 | AZD8186 |
| MVP_top60_DNA_seed | HSP90AA1 | -0.298015 | HNSC | Detroit562 | Luminespib | MVP_top60_DNA+RNA_seed | PIK3CA | -2.259398 | KIRC | RCC-JF | AZD8186 |
| MVP_top60_DNA_seed | MAPK1 | -0.297997 | HNSC | SKN-3 | Ulixertinib | MVP_top60_DNA+RNA_seed | PIK3CA | -2.125832 | KIRC | KMRC-1 | AZD8186 |
| MVP_top60_DNA_seed | MAPK3 | -0.297997 | HNSC | SKN-3 | Ulixertinib | MVP_top60_DNA+RNA_seed | ATM | -2.07997 | KIRC | RCC-ER | KU-55933 |
| MVP_top60_DNA_seed | CDK2 | -0.297767 | HNSC | BB49-HNC | AZD5438 | MVP_top60_DNA+RNA_seed | PIK3CA | -1.724921 | KIRC | RXF393 | AZD8186 |
| MVP_top60_DNA_seed | PIK3CA | -0.297602 | HNSC | SCC-15 | AZD8186 | MVP_top60_DNA+RNA_seed | PIK3CA | -1.572785 | KIRC | U031 | AZD8186 |
| MVP_top60_DNA_seed | AKT1 | -0.297355 | HNSC | KON | Afuresertib | MVP_top60_DNA+RNA_seed | PIK3CA | -1.560281 | KIRC | BFTC-909 | Buparlisib |
| MVP_top60_DNA_seed | MDM2 | -0.296701 | HNSC | BB30-HNC | Nutlin-3a (-) | MVP_top60_DNA+RNA_seed | PIK3CD | -1.560281 | KIRC | BFTC-909 | Buparlisib |
| MVP_top60_DNA_seed | CDK2 | -0.295698 | HNSC | HSC-3 | MK-8776 | MVP_top60_DNA+RNA_seed | MTOR | -1.510518 | KIRC | RCC-FG2 | Temsirolimus |
| MVP_top60_DNA_seed | HDAC1 | -0.294584 | HNSC | CAL-33 | Romidepsin | MVP_top60_DNA+RNA_seed | HSP90AA1 | -1.502983 | KIRC | RCC-JF | Luminespib |
| MVP_top60_DNA_seed | HDAC3 | -0.294584 | HNSC | CAL-33 | Romidepsin | MVP_top60_DNA+RNA_seed | HSP90AB1 | -1.502983 | KIRC | RCC-JF | Luminespib |
| MVP_top60_DNA_seed | AKT1 | -0.291238 | HNSC | SAS | Ipatasertib | MVP_top60_DNA+RNA_seed | HSP90AA1 | -1.458065 | KIRC | RCC-JF | Elesclomol |
| MVP_top60_DNA_seed | TP53 | -0.291126 | HNSC | HO-1-u-1 | MIRA-1 | MVP_top60_DNA+RNA_seed | HSP90AB1 | -1.458065 | KIRC | RCC-JF | Elesclomol |
| MVP_top60_DNA_seed | SMARCA4 | -0.290927 | HNSC | HSC-2 | PFI3 | MVP_top60_DNA+RNA_seed | HSP90AA1 | -1.427487 | KIRC | RCC-JF | Tanespimycin |
| MVP_top60_DNA_seed | PIK3CA | -0.290049 | HNSC | PCI-15A | AZD8186 | MVP_top60_DNA+RNA_seed | HSP90AB1 | -1.427487 | KIRC | RCC-JF | Tanespimycin |
| MVP_top60_DNA_seed | MDM2 | -0.284967 | HNSC | PCI-15A | Nutlin-3a (-) | MVP_top60_DNA+RNA_seed | TP53 | -1.367967 | KIRC | RCC-ER | MIRA-1 |
| MVP_top60_DNA_seed | EGFR | -0.283019 | HNSC | BICR78 | AZD3759 | MVP_top60_DNA+RNA_seed | ESR1 | -1.356483 | KIRC | A498 | Tamoxifen |
| MVP_top60_DNA_seed | CDK2 | -0.28136 | HNSC | SCC-4 | Dinaciclib | MVP_top60_DNA+RNA_seed | PIK3CA | -1.355301 | KIRC | OS-RC-2 | AZD8186 |
| MVP_top60_DNA_seed | PIK3CA | -0.280666 | HNSC | KOSC-2 | Buparlisib | MVP_top60_DNA+RNA_seed | PIK3CA | -1.348011 | KIRC | NCC021 | AZD8186 |
| MVP_top60_DNA_seed | ESR1 | -0.279959 | HNSC | CAL-27 | Tamoxifen | MVP_top60_DNA+RNA_seed | PIK3CA | -1.337051 | KIRC | RCC-ER | AZD8186 |
| MVP_top60_DNA_seed | EP300 | -0.278086 | HNSC | Detroit562 | SGC-CBP30 | MVP_top60_DNA+RNA_seed | PIK3CA | -1.332343 | KIRC | CAL-54 | GNE-317 |
| MVP_top60_DNA_seed | CREBBP | -0.278086 | HNSC | Detroit562 | SGC-CBP30 | MVP_top60_DNA+RNA_seed | PIK3CA | -1.291194 | KIRC | BFTC-909 | GNE-317 |
| MVP_top60_DNA_seed | HDAC1 | -0.277318 | HNSC | PCI-6A | PCI-34051 | MVP_top60_DNA+RNA_seed | IGF1R | -1.272041 | KIRC | RCC-MF | GSK1904529A |

| | | | | | | | | | | | |
|---|---|---|---|---|---|---|---|---|---|---|---|
| MVP_top60_DNA_seed | AKT1 | -0.277219 | HNSC | CAL-33 | Uprosertib | MVP_top60_DNA+RNA_seed | HSP90AA1 | -1.266588 | KIRC | BFTC-909 | Tanespimycin |
| MVP_top60_DNA_seed | EGFR | -0.274885 | HNSC | OSC-20 | Gefitinib | MVP_top60_DNA+RNA_seed | HSP90AB1 | -1.266588 | KIRC | BFTC-909 | Tanespimycin |
| MVP_top60_DNA_seed | ESR1 | -0.274452 | HNSC | PCI-4B | GDC0810 | MVP_top60_DNA+RNA_seed | ESR1 | -1.241312 | KIRC | BFTC-909 | Tamoxifen |
| MVP_top60_DNA_seed | PIK3CA | -0.270935 | HNSC | Ca9-22 | Buparlisib | MVP_top60_DNA+RNA_seed | ESR1 | -1.224183 | KIRC | CAL-54 | Tamoxifen |
| MVP_top60_DNA_seed | AKT1 | -0.268177 | HNSC | BICR22 | Afuresertib | MVP_top60_DNA+RNA_seed | PIK3CA | -1.203648 | KIRC | CAL-54 | Buparlisib |
| MVP_top60_DNA_seed | EP300 | -0.265323 | HNSC | FADU | CPI-637 | MVP_top60_DNA+RNA_seed | PIK3CD | -1.203648 | KIRC | CAL-54 | Buparlisib |
| MVP_top60_DNA_seed | CDK2 | -0.264372 | HNSC | LB771-HNC | Dinaciclib | MVP_top60_DNA+RNA_seed | ATM | -1.184273 | KIRC | A704 | KU-55933 |
| MVP_top60_DNA_seed | PIK3CA | -0.26389 | HNSC | CAL-33 | GNE-317 | MVP_top60_DNA+RNA_seed | PIK3CA | -1.184112 | KIRC | A704 | AZD8186 |
| MVP_top60_DNA_seed | MAPK1 | -0.263802 | HNSC | CAL-27 | VX-11e | MVP_top60_DNA+RNA_seed | PIK3CA | -1.176332 | KIRC | BFTC-909 | Alpelisib |
| MVP_top60_DNA_seed | PIK3CA | -0.26291 | HNSC | OSC-19 | Alpelisib | MVP_top60_DNA+RNA_seed | MTOR | -1.159736 | KIRC | RCC-JF | Temsirolimus |
| MVP_top60_DNA_seed | EGFR | -0.262649 | HNSC | BICR31 | Lapatinib | MVP_top60_DNA+RNA_seed | IGF1R | -1.152004 | KIRC | CAL-54 | BMS-536924 |
| MVP_top60_DNA_seed | SMARCA4 | -0.261047 | HNSC | HSC-3 | PFI3 | MVP_top60_DNA+RNA_seed | HDAC1 | -1.140414 | KIRC | RCC-FG2 | PCI-34051 |
| MVP_top60_DNA_seed | PIK3CA | -0.260552 | HNSC | SCC-15 | Buparlisib | MVP_top60_DNA+RNA_seed | ATM | -1.134385 | KIRC | NCC010 | KU-55933 |
| MVP_top60_DNA_seed | AKT1 | -0.254743 | HNSC | SCC90 | Uprosertib | MVP_top60_DNA+RNA_seed | PIK3CA | -1.12485 | KIRC | RCC-JF | GNE-317 |
| MVP_top60_DNA_seed | MAPK1 | -0.254277 | HNSC | BICR31 | SCH772984 | MVP_top60_DNA+RNA_seed | IGF1R | -1.097138 | KIRC | LB996-RCC | BMS-536924 |
| MVP_top60_DNA_seed | MAPK3 | -0.254277 | HNSC | BICR31 | SCH772984 | MVP_top60_DNA+RNA_seed | PIK3CA | -1.065702 | KIRC | 786-0 | AZD8186 |
| MVP_top60_DNA_seed | PIK3CA | -0.251431 | HNSC | KON | Buparlisib | MVP_top60_DNA+RNA_seed | PIK3CA | -1.056549 | KIRC | LB996-RCC | AZD8186 |
| MVP_top60_DNA_seed | EP300 | -0.251026 | HNSC | SCC-9 | SGC-CBP30 | MVP_top60_DNA+RNA_seed | PIK3CA | -1.053786 | KIRC | CAKI-1 | AZD8186 |
| MVP_top60_DNA_seed | CREBBP | -0.251026 | HNSC | SCC-9 | SGC-CBP30 | MVP_top60_DNA+RNA_seed | HSP90AA1 | -1.016732 | KIRC | BB65-RCC | Tanespimycin |
| MVP_top60_DNA_seed | HSP90AA1 | -0.249848 | HNSC | SCC-4 | Luminespib | MVP_top60_DNA+RNA_seed | HSP90AB1 | -1.016732 | KIRC | BB65-RCC | Tanespimycin |
| MVP_top60_DNA_seed | NTRK1 | -0.24916 | HNSC | SCC-15 | AZD1332 | MVP_top60_DNA+RNA_seed | PIK3CA | -1.016439 | KIRC | LB996-RCC | GNE-317 |
| MVP_top60_DNA_seed | CDK2 | -0.248604 | HNSC | DOK | AZD5438 | MVP_top60_DNA+RNA_seed | HSP90AA1 | -1.008454 | KIRC | RCC-FG2 | Luminespib |
| MVP_top60_DNA_seed | HDAC1 | -0.248442 | HNSC | Detroit562 | Entinostat | MVP_top60_DNA+RNA_seed | HSP90AB1 | -1.008454 | KIRC | RCC-FG2 | Luminespib |
| MVP_top60_DNA_seed | HDAC3 | -0.248442 | HNSC | Detroit562 | Entinostat | MVP_top60_DNA+RNA_seed | EP300 | -1.003073 | KIRC | 769-P | CPI-637 |
| MVP_top60_DNA_seed | SMARCA4 | -0.248208 | HNSC | KOSC-2 | PFI3 | MVP_top60_DNA+RNA_seed | HSP90AA1 | -0.997573 | KIRC | RCC-JF | CCT-018159 |
| MVP_top60_DNA_seed | PIK3CA | -0.245717 | HNSC | PE-CA-PJ15 | Alpelisib | MVP_top60_DNA+RNA_seed | HSP90AB1 | -0.997573 | KIRC | RCC-JF | CCT-018159 |
| MVP_top60_DNA_seed | PIK3CA | -0.24413 | HNSC | CAL-33 | AZD8186 | MVP_top60_DNA+RNA_seed | PIK3CA | -0.993458 | KIRC | RCC-FG2 | GNE-317 |
| MVP_top60_DNA_seed | AKT1 | -0.237971 | HNSC | JHU-022 | Uprosertib | MVP_top60_DNA+RNA_seed | PIK3CA | -0.985424 | KIRC | LB996-RCC | Buparlisib |
| MVP_top60_DNA_seed | EP300 | -0.237897 | HNSC | HSC-3 | CPI-637 | MVP_top60_DNA+RNA_seed | PIK3CD | -0.985424 | KIRC | LB996-RCC | Buparlisib |
| MVP_top60_DNA_seed | AKT1 | -0.237559 | HNSC | HSC-2 | AZD5363 | MVP_top60_DNA+RNA_seed | HSP90AA1 | -0.962962 | KIRC | LB996-RCC | Luminespib |
| MVP_top60_DNA_seed | CDK2 | -0.236212 | HNSC | JHU-011 | MK-8776 | MVP_top60_DNA+RNA_seed | HSP90AB1 | -0.962962 | KIRC | LB996-RCC | Luminespib |
| MVP_top60_DNA_seed | PIK3CA | -0.234926 | HNSC | HSC-4 | AZD8186 | MVP_top60_DNA+RNA_seed | PIK3CA | -0.946664 | KIRC | RCC-ER | Alpelisib |
| MVP_top60_DNA_seed | CDK2 | -0.232088 | HNSC | SCC-15 | Dinaciclib | MVP_top60_DNA+RNA_seed | IGF1R | -0.941836 | KIRC | RCC-ER | BMS-536924 |
| MVP_top60_DNA_seed | HSP90AA1 | -0.229695 | HNSC | SAT | Luminespib | MVP_top60_DNA+RNA_seed | HSP90AA1 | -0.940046 | KIRC | LB996-RCC | Tanespimycin |
| MVP_top60_DNA_seed | HDAC1 | -0.228364 | HNSC | HO-1-N-1 | Entinostat | MVP_top60_DNA+RNA_seed | HSP90AB1 | -0.940046 | KIRC | LB996-RCC | Tanespimycin |
| MVP_top60_DNA_seed | HDAC3 | -0.228364 | HNSC | HO-1-N-1 | Entinostat | MVP_top60_DNA+RNA_seed | PIK3CA | -0.911076 | KIRC | LB1047-RCC | Alpelisib |
| MVP_top60_DNA_seed | SMARCA4 | -0.227483 | HNSC | FADU | PFI3 | MVP_top60_DNA+RNA_seed | IGF1R | -0.890138 | KIRC | RCC-ER | BMS-754807 |
| MVP_top60_DNA_seed | PIK3CA | -0.226396 | HNSC | CAL-33 | Buparlisib | MVP_top60_DNA+RNA_seed | ATM | -0.888756 | KIRC | RCC-JW | KU-55933 |
| MVP_top60_DNA_seed | NTRK1 | -0.226396 | HNSC | HSC-2 | GW441756 | MVP_top60_DNA+RNA_seed | PIK3CA | -0.870576 | KIRC | HA7-RCC | AZD8186 |
| MVP_top60_DNA_seed | RAC1 | -0.225399 | HNSC | PCI-6A | EHT-1864 | MVP_top60_DNA+RNA_seed | IGF1R | -0.863271 | KIRC | U031 | GSK1904529A |
| MVP_top60_DNA_seed | TP53 | -0.225344 | HNSC | HSC-2 | MIRA-1 | MVP_top60_DNA+RNA_seed | IGF1R | -0.8624 | KIRC | A704 | BMS-536924 |

| | | | | | | | | | | | |
|---|---|---|---|---|---|---|---|---|---|---|---|
| MVP_top60_DNA_seed | AKT1 | -0.225279 | HNSC | BB30-HNC | Afuresertib | MVP_top60_DNA+RNA_seed | PIK3CA | -0.854873 | KIRC | 769-P | GNE-317 |
| MVP_top60_DNA_seed | HDAC1 | -0.223855 | HNSC | OSC-20 | PCI-34051 | MVP_top60_DNA+RNA_seed | PIK3CA | -0.852015 | KIRC | LB2241-RCC | GNE-317 |
| MVP_top60_DNA_seed | PIK3CA | -0.223246 | HNSC | SAS | Buparlisib | MVP_top60_DNA+RNA_seed | HSP90AA1 | -0.849202 | KIRC | U031 | CCT-018159 |
| MVP_top60_DNA_seed | SMARCA4 | -0.223226 | HNSC | DOK | PFI3 | MVP_top60_DNA+RNA_seed | HSP90AB1 | -0.849202 | KIRC | U031 | CCT-018159 |
| MVP_top60_DNA_seed | MAPK1 | -0.221889 | HNSC | SCC-15 | Ulixertinib | MVP_top60_DNA+RNA_seed | HSP90AA1 | -0.847173 | KIRC | RCC-FG2 | CCT-018159 |
| MVP_top60_DNA_seed | MAPK3 | -0.221889 | HNSC | SCC-15 | Ulixertinib | MVP_top60_DNA+RNA_seed | HSP90AB1 | -0.847173 | KIRC | RCC-FG2 | CCT-018159 |
| MVP_top60_DNA_seed | EGFR | -0.221134 | HNSC | SKN-3 | Sapitinib | MVP_top60_DNA+RNA_seed | IGF1R | -0.816724 | KIRC | NCC021 | BMS-536924 |
| MVP_top60_DNA_seed | MAPK1 | -0.219833 | HNSC | SAT | SCH772984 | MVP_top60_DNA+RNA_seed | HSP90AA1 | -0.805852 | KIRC | RXF393 | Luminespib |
| MVP_top60_DNA_seed | MAPK3 | -0.219833 | HNSC | SAT | SCH772984 | MVP_top60_DNA+RNA_seed | HSP90AB1 | -0.805852 | KIRC | RXF393 | Luminespib |
| MVP_top60_DNA_seed | TP53 | -0.219795 | HNSC | PCI-4B | MIRA-1 | MVP_top60_DNA+RNA_seed | PIK3CA | -0.803391 | KIRC | RCC10RGB | AZD8186 |
| MVP_top60_DNA_seed | HSP90AA1 | -0.218658 | HNSC | Ca9-22 | Luminespib | MVP_top60_DNA+RNA_seed | PIK3CA | -0.803262 | KIRC | ACHN | GNE-317 |
| MVP_top60_DNA_seed | ESR1 | -0.217722 | HNSC | SAS | GDC0810 | MVP_top60_DNA+RNA_seed | PIK3CA | -0.797416 | KIRC | ACHN | AZD8186 |
| MVP_top60_DNA_seed | MDM2 | -0.216828 | HNSC | KOSC-2 | Nutlin-3a (-) | MVP_top60_DNA+RNA_seed | HDAC1 | -0.795837 | KIRC | RCC-JF | PCI-34051 |
| MVP_top60_DNA_seed | ESR1 | -0.215398 | HNSC | LB771-HNC | Tamoxifen | MVP_top60_DNA+RNA_seed | HSP90AA1 | -0.789065 | KIRC | RXF393 | CCT-018159 |
| MVP_top60_DNA_seed | MAPK1 | -0.21232 | HNSC | PCI-4B | Ulixertinib | MVP_top60_DNA+RNA_seed | HSP90AB1 | -0.789065 | KIRC | RXF393 | CCT-018159 |
| MVP_top60_DNA_seed | MAPK3 | -0.21232 | HNSC | PCI-4B | Ulixertinib | MVP_top60_DNA+RNA_seed | AR | -0.785124 | KIRC | U031 | Bicalutamide |
| MVP_top60_DNA_seed | AKT1 | -0.209974 | HNSC | HO-1-N-1 | Ipatasertib | MVP_top60_DNA+RNA_seed | ATM | -0.783711 | KIRC | RCC10RGB | KU-55933 |
| MVP_top60_DNA_seed | SRC | -0.209083 | HNSC | LB771-HNC | Bosutinib | MVP_top60_DNA+RNA_seed | IGF1R | -0.774009 | KIRC | LB996-RCC | GSK1904529A |
| MVP_top60_DNA_seed | MDM2 | -0.206936 | HNSC | DOK | Serdemetan | MVP_top60_DNA+RNA_seed | AR | -0.765839 | KIRC | RCC-JF | Bicalutamide |
| MVP_top60_DNA_seed | EGFR | -0.204147 | HNSC | PCI-30 | Osimertinib | MVP_top60_DNA+RNA_seed | IGF1R | -0.763919 | KIRC | U031 | BMS-536924 |
| MVP_top60_DNA_seed | HSP90AA1 | -0.203658 | HNSC | BICR31 | Tanespimycin | MVP_top60_DNA+RNA_seed | MTOR | -0.755409 | KIRC | A498 | Temsirolimus |
| MVP_top60_DNA_seed | EGFR | -0.202864 | HNSC | SKN-3 | Erlotinib | MVP_top60_DNA+RNA_seed | PIK3CA | -0.736097 | KIRC | BFTC-909 | AZD8186 |
| MVP_top60_DNA_seed | EP300 | -0.200462 | HNSC | SCC-9 | CPI-637 | MVP_top60_DNA+RNA_seed | ATM | -0.729976 | KIRC | RCC-FG2 | KU-55933 |
| MVP_top60_DNA_seed | EGFR | -0.199622 | HNSC | SCC-25 | Osimertinib | MVP_top60_DNA+RNA_seed | PIK3CA | -0.721635 | KIRC | U031 | GNE-317 |
| MVP_top60_DNA_seed | HDAC1 | -0.198828 | HNSC | OSC-19 | Entinostat | MVP_top60_DNA+RNA_seed | HDAC1 | -0.715943 | KIRC | NCC010 | PCI-34051 |
| MVP_top60_DNA_seed | HDAC3 | -0.198828 | HNSC | OSC-19 | Entinostat | MVP_top60_DNA+RNA_seed | IGF1R | -0.708279 | KIRC | NCC010 | BMS-754807 |
| MVP_top60_DNA_seed | ESR1 | -0.197468 | HNSC | HO-1-N-1 | GDC0810 | MVP_top60_DNA+RNA_seed | ATM | -0.702862 | KIRC | LB996-RCC | KU-55933 |
| MVP_top60_DNA_seed | HSP90AA1 | -0.193768 | HNSC | CAL-27 | Tanespimycin | MVP_top60_DNA+RNA_seed | EP300 | -0.699633 | KIRC | CAL-54 | CPI-637 |
| MVP_top60_DNA_seed | EGFR | -0.190175 | HNSC | OSC-20 | Osimertinib | MVP_top60_DNA+RNA_seed | PIK3CA | -0.699149 | KIRC | RCC-ER | Buparlisib |
| MVP_top60_DNA_seed | MDM2 | -0.188456 | HNSC | HO-1-N-1 | Serdemetan | MVP_top60_DNA+RNA_seed | PIK3CD | -0.699149 | KIRC | RCC-ER | Buparlisib |
| MVP_top60_DNA_seed | PIK3CA | -0.188249 | HNSC | FADU | Alpelisib | MVP_top60_DNA+RNA_seed | HSP90AA1 | -0.697923 | KIRC | BB65-RCC | Luminespib |
| MVP_top60_DNA_seed | AKT1 | -0.187781 | HNSC | PCI-6A | AZD5363 | MVP_top60_DNA+RNA_seed | HSP90AB1 | -0.697923 | KIRC | BB65-RCC | Luminespib |
| MVP_top60_DNA_seed | AKT1 | -0.18751 | HNSC | KON | Uprosertib | MVP_top60_DNA+RNA_seed | IGF1R | -0.695017 | KIRC | A498 | NVP-ADW742 |
| MVP_top60_DNA_seed | HSP90AA1 | -0.187008 | HNSC | JHU-029 | Tanespimycin | MVP_top60_DNA+RNA_seed | ESR1 | -0.678958 | KIRC | 786-0 | GDC0810 |
| MVP_top60_DNA_seed | EGFR | -0.186371 | HNSC | OSC-20 | Erlotinib | MVP_top60_DNA+RNA_seed | HDAC1 | -0.670549 | KIRC | CAKI-1 | PCI-34051 |
| MVP_top60_DNA_seed | TP53 | -0.184433 | HNSC | SCC-25 | MIRA-1 | MVP_top60_DNA+RNA_seed | HSP90AA1 | -0.668507 | KIRC | RCC-JW | Tanespimycin |
| MVP_top60_DNA_seed | SRC | -0.182647 | HNSC | OSC-19 | Bosutinib | MVP_top60_DNA+RNA_seed | HSP90AB1 | -0.668507 | KIRC | RCC-JW | Tanespimycin |
| MVP_top60_DNA_seed | AKT1 | -0.179677 | HNSC | KOSC-2 | AT13148 | MVP_top60_DNA+RNA_seed | ATM | -0.659454 | KIRC | CAL-54 | KU-55933 |
| MVP_top60_DNA_seed | MAPK1 | -0.177088 | HNSC | OSC-19 | SCH772984 | MVP_top60_DNA+RNA_seed | MTOR | -0.655868 | KIRC | RCC-ER | Temsirolimus |
| MVP_top60_DNA_seed | MAPK3 | -0.177088 | HNSC | OSC-19 | SCH772984 | MVP_top60_DNA+RNA_seed | IGF1R | -0.647047 | KIRC | LB996-RCC | NVP-ADW742 |
| MVP_top60_DNA_seed | AKT1 | -0.176665 | HNSC | CAL-33 | Uprosertib | MVP_top60_DNA+RNA_seed | IGF1R | -0.646483 | KIRC | A704 | NVP-ADW742 |

| | | | | | | | | | | | |
|---|---|---|---|---|---|---|---|---|---|---|---|
| MVP_top60_DNA_seed | RAC1 | -0.173999 | HNSC | BICR78 | EHT-1864 | MVP_top60_DNA+RNA_seed | HSP90AA1 | -0.644857 | KIRC | BFTC-909 | Luminespib |
| MVP_top60_DNA_seed | AKT1 | -0.171089 | HNSC | SAS | AZD5363 | MVP_top60_DNA+RNA_seed | HSP90AB1 | -0.644857 | KIRC | BFTC-909 | Luminespib |
| MVP_top60_DNA_seed | AKT1 | -0.17105 | HNSC | JHU-029 | Uprosertib | MVP_top60_DNA+RNA_seed | PIK3CA | -0.641962 | KIRC | RCC-JW | AZD8186 |
| MVP_top60_DNA_seed | HSP90AA1 | -0.169945 | HNSC | JHU-022 | Luminespib | MVP_top60_DNA+RNA_seed | TP53 | -0.640654 | KIRC | ACHN | MIRA-1 |
| MVP_top60_DNA_seed | MAPK1 | -0.169746 | HNSC | CAL-27 | Ulixertinib | MVP_top60_DNA+RNA_seed | EP300 | -0.628687 | KIRC | A704 | SGC-CBP30 |
| MVP_top60_DNA_seed | MAPK3 | -0.169746 | HNSC | CAL-27 | Ulixertinib | MVP_top60_DNA+RNA_seed | IGF1R | -0.621436 | KIRC | BB65-RCC | NVP-ADW742 |
| MVP_top60_DNA_seed | EGFR | -0.167285 | HNSC | SCC-4 | Sapitinib | MVP_top60_DNA+RNA_seed | PIK3CA | -0.606246 | KIRC | LB1047-RCC | GNE-317 |
| MVP_top60_DNA_seed | PIK3CA | -0.16585 | HNSC | SCC-9 | AZD8186 | MVP_top60_DNA+RNA_seed | IGF1R | -0.604646 | KIRC | A498 | BMS-536924 |
| MVP_top60_DNA_seed | AKT1 | -0.165796 | HNSC | JHU-022 | Ipatasertib | MVP_top60_DNA+RNA_seed | IGF1R | -0.604169 | KIRC | RCC-ER | GSK1904529A |
| MVP_top60_DNA_seed | ESR1 | -0.16568 | HNSC | BICR10 | Tamoxifen | MVP_top60_DNA+RNA_seed | PIK3CA | -0.592024 | KIRC | RCC-AB | GNE-317 |
| MVP_top60_DNA_seed | HDAC1 | -0.159885 | HNSC | BB30-HNC | Entinostat | MVP_top60_DNA+RNA_seed | HSP90AA1 | -0.586516 | KIRC | NCC010 | CCT-018159 |
| MVP_top60_DNA_seed | HDAC3 | -0.159885 | HNSC | BB30-HNC | Entinostat | MVP_top60_DNA+RNA_seed | HSP90AB1 | -0.586516 | KIRC | NCC010 | CCT-018159 |
| MVP_top60_DNA_seed | TP53 | -0.158759 | HNSC | DOK | MIRA-1 | MVP_top60_DNA+RNA_seed | PIK3CA | -0.585482 | KIRC | LB2241-RCC | AZD8186 |
| MVP_top60_DNA_seed | AKT1 | -0.157281 | HNSC | CAL-27 | AZD5363 | MVP_top60_DNA+RNA_seed | PIK3CA | -0.579237 | KIRC | CAKI-1 | GNE-317 |
| MVP_top60_DNA_seed | PIK3CA | -0.155686 | HNSC | JHU-022 | AZD8186 | MVP_top60_DNA+RNA_seed | HSP90AA1 | -0.578011 | KIRC | RCC-FG2 | Tanespimycin |
| MVP_top60_DNA_seed | HDAC1 | -0.153391 | HNSC | Detroit562 | Romidepsin | MVP_top60_DNA+RNA_seed | HSP90AB1 | -0.578011 | KIRC | RCC-FG2 | Tanespimycin |
| MVP_top60_DNA_seed | HDAC3 | -0.153391 | HNSC | Detroit562 | Romidepsin | MVP_top60_DNA+RNA_seed | HDAC1 | -0.554545 | KIRC | BFTC-909 | Romidepsin |
| MVP_top60_DNA_seed | NTRK1 | -0.152886 | HNSC | SAT | GW441756 | MVP_top60_DNA+RNA_seed | EP300 | -0.553506 | KIRC | NCC010 | SGC-CBP30 |
| MVP_top60_DNA_seed | CDK2 | -0.152158 | HNSC | CAL-33 | AZD5438 | MVP_top60_DNA+RNA_seed | TP53 | -0.5454 | KIRC | CAKI-1 | MIRA-1 |
| MVP_top60_DNA_seed | ESR1 | -0.150732 | HNSC | BB49-HNC | GDC0810 | MVP_top60_DNA+RNA_seed | IGF1R | -0.541972 | KIRC | RCC-JF | NVP-ADW742 |
| MVP_top60_DNA_seed | AKT1 | -0.147298 | HNSC | Ca9-22 | Uprosertib | MVP_top60_DNA+RNA_seed | ESR1 | -0.541189 | KIRC | U031 | GDC0810 |
| MVP_top60_DNA_seed | NTRK1 | -0.145888 | HNSC | SAS | GW441756 | MVP_top60_DNA+RNA_seed | HSP90AA1 | -0.534673 | KIRC | 769-P | Tanespimycin |
| MVP_top60_DNA_seed | HSP90AA1 | -0.138679 | HNSC | PCI-4B | Elesclomol | MVP_top60_DNA+RNA_seed | HSP90AB1 | -0.534673 | KIRC | 769-P | Tanespimycin |
| MVP_top60_DNA_seed | AKT1 | -0.137032 | HNSC | Ca9-22 | AZD5363 | MVP_top60_DNA+RNA_seed | IGF1R | -0.529765 | KIRC | RXF393 | BMS-536924 |
| MVP_top60_DNA_seed | RAC1 | -0.135565 | HNSC | KOSC-2 | EHT-1864 | MVP_top60_DNA+RNA_seed | MTOR | -0.527579 | KIRC | RCC-MF | Temsirolimus |
| MVP_top60_DNA_seed | HDAC1 | -0.135181 | HNSC | HO-1-N-1 | PCI-34051 | MVP_top60_DNA+RNA_seed | PIK3CA | -0.527564 | KIRC | 786-0 | GNE-317 |
| MVP_top60_DNA_seed | MDM2 | -0.133878 | HNSC | PCI-4B | Nutlin-3a (-) | MVP_top60_DNA+RNA_seed | IGF1R | -0.526387 | KIRC | NCC010 | GSK1904529A |
| MVP_top60_DNA_seed | AR | -0.133093 | HNSC | PCI-15A | Bicalutamide | MVP_top60_DNA+RNA_seed | HSP90AA1 | -0.520353 | KIRC | 786-0 | Tanespimycin |
| MVP_top60_DNA_seed | EP300 | -0.131582 | HNSC | BICR22 | CPI-637 | MVP_top60_DNA+RNA_seed | HSP90AB1 | -0.520353 | KIRC | 786-0 | Tanespimycin |
| MVP_top60_DNA_seed | AKT1 | -0.130046 | HNSC | HSC-2 | Afuresertib | MVP_top60_DNA+RNA_seed | IGF1R | -0.515093 | KIRC | RCC-JF | BMS-536924 |
| MVP_top60_DNA_seed | TP53 | -0.128942 | HNSC | KON | MIRA-1 | MVP_top60_DNA+RNA_seed | IGF1R | -0.509704 | KIRC | RCC-FG2 | BMS-536924 |
| MVP_top60_DNA_seed | RAC1 | -0.128441 | HNSC | SCC-15 | EHT-1864 | MVP_top60_DNA+RNA_seed | ESR1 | -0.501132 | KIRC | 786-0 | Tamoxifen |
| MVP_top60_DNA_seed | MAPK1 | -0.126861 | HNSC | FADU | SCH772984 | MVP_top60_DNA+RNA_seed | IGF1R | -0.500808 | KIRC | BFTC-909 | NVP-ADW742 |
| MVP_top60_DNA_seed | MAPK3 | -0.126861 | HNSC | FADU | SCH772984 | MVP_top60_DNA+RNA_seed | IGF1R | -0.489531 | KIRC | 786-0 | NVP-ADW742 |
| MVP_top60_DNA_seed | NTRK1 | -0.123622 | HNSC | HSC-3 | GW441756 | MVP_top60_DNA+RNA_seed | HDAC1 | -0.47977 | KIRC | RCC-ER | PCI-34051 |
| MVP_top60_DNA_seed | EGFR | -0.121489 | HNSC | BHY | Afatinib | MVP_top60_DNA+RNA_seed | HSP90AA1 | -0.472305 | KIRC | RXF393 | Tanespimycin |
| MVP_top60_DNA_seed | AKT1 | -0.118582 | HNSC | SAS | Afuresertib | MVP_top60_DNA+RNA_seed | HSP90AB1 | -0.472305 | KIRC | RXF393 | Tanespimycin |
| MVP_top60_DNA_seed | PIK3CA | -0.113985 | HNSC | SAT | GNE-317 | MVP_top60_DNA+RNA_seed | SMARCA4 | -0.472138 | KIRC | A704 | PFI3 |
| MVP_top60_DNA_seed | HSP90AA1 | -0.109438 | HNSC | BHY | CCT-018159 | MVP_top60_DNA+RNA_seed | SMARCA2 | -0.472138 | KIRC | A704 | PFI3 |
| MVP_top60_DNA_seed | NTRK1 | -0.104141 | HNSC | BICR78 | GW441756 | MVP_top60_DNA+RNA_seed | ESR1 | -0.468891 | KIRC | LB996-RCC | Tamoxifen |
| MVP_top60_DNA_seed | EGFR | -0.100642 | HNSC | SCC-4 | Lapatinib | MVP_top60_DNA+RNA_seed | MTOR | -0.460266 | KIRC | RXF393 | Temsirolimus |

| | | | | | | | | | | | |
|---|---|---|---|---|---|---|---|---|---|---|---|
| MVP_top60_DNA_seed | HDAC1 | -0.098316 | HNSC | BB30-HNC | PCI-34051 | MVP_top60_DNA+RNA_seed | PIK3CA | -0.456333 | KIRC | RCC-MF | GNE-317 |
| MVP_top60_DNA_seed | MAPK1 | -0.09453 | HNSC | HSC-3 | SCH772984 | MVP_top60_DNA+RNA_seed | EP300 | -0.456099 | KIRC | LB996-RCC | CPI-637 |
| MVP_top60_DNA_seed | MAPK3 | -0.09453 | HNSC | HSC-3 | SCH772984 | MVP_top60_DNA+RNA_seed | ESR1 | -0.448495 | KIRC | RCC-ER | Tamoxifen |
| MVP_top60_DNA_seed | HDAC1 | -0.09412 | HNSC | SAS | PCI-34051 | MVP_top60_DNA+RNA_seed | TP53 | -0.444953 | KIRC | BFTC-909 | MIRA-1 |
| MVP_top60_DNA_seed | SMARCA4 | -0.093109 | HNSC | BICR22 | PFI3 | MVP_top60_DNA+RNA_seed | PIK3CA | -0.435472 | KIRC | RCC-JF | Alpelisib |
| MVP_top60_DNA_seed | AKT1 | -0.091409 | HNSC | JHU-022 | GSK2110183B | MVP_top60_DNA+RNA_seed | HDAC1 | -0.432309 | KIRC | RCC-MF | PCI-34051 |
| MVP_top60_DNA_seed | TP53 | -0.09071 | HNSC | JHU-029 | MIRA-1 | MVP_top60_DNA+RNA_seed | SMARCA4 | -0.427139 | KIRC | NCC010 | PFI3 |
| MVP_top60_DNA_seed | ESR1 | -0.089786 | HNSC | HSC-4 | Tamoxifen | MVP_top60_DNA+RNA_seed | SMARCA2 | -0.427139 | KIRC | NCC010 | PFI3 |
| MVP_top60_DNA_seed | ESR1 | -0.088438 | HNSC | Ca9-22 | GDC0810 | MVP_top60_DNA+RNA_seed | IGF1R | -0.416927 | KIRC | RCC-ER | NVP-ADW742 |
| MVP_top60_DNA_seed | MAPK1 | -0.084804 | HNSC | BB49-HNC | VX-11e | MVP_top60_DNA+RNA_seed | IGF1R | -0.410611 | KIRC | LB1047-RCC | BMS-536924 |
| MVP_top60_DNA_seed | EGFR | -0.08302 | HNSC | BICR31 | Osimertinib | MVP_top60_DNA+RNA_seed | HDAC1 | -0.409844 | KIRC | 786-0 | PCI-34051 |
| MVP_top60_DNA_seed | AKT1 | -0.082628 | HNSC | HO-1-N-1 | Uprosertib | MVP_top60_DNA+RNA_seed | PIK3CA | -0.40875 | KIRC | LB1047-RCC | AZD8186 |
| MVP_top60_DNA_seed | MDM2 | -0.082518 | HNSC | Ca9-22 | Serdemetan | MVP_top60_DNA+RNA_seed | IGF1R | -0.406177 | KIRC | RCC-ER | Linsitinib |
| MVP_top60_DNA_seed | MDM2 | -0.081356 | HNSC | SCC-4 | Serdemetan | MVP_top60_DNA+RNA_seed | EP300 | -0.404447 | KIRC | RCC-JW | CPI-637 |
| MVP_top60_DNA_seed | TP53 | -0.081288 | HNSC | SAS | MIRA-1 | MVP_top60_DNA+RNA_seed | IGF1R | -0.397706 | KIRC | U031 | BMS-754807 |
| MVP_top60_DNA_seed | AKT1 | -0.080641 | HNSC | OSC-19 | MK-2206 | MVP_top60_DNA+RNA_seed | MTOR | -0.394806 | KIRC | CAL-54 | Temsirolimus |
| MVP_top60_DNA_seed | HDAC1 | -0.079903 | HNSC | JHU-029 | PCI-34051 | MVP_top60_DNA+RNA_seed | IGF1R | -0.388324 | KIRC | KMRC-20 | NVP-ADW742 |
| MVP_top60_DNA_seed | HDAC1 | -0.078352 | HNSC | SAT | Romidepsin | MVP_top60_DNA+RNA_seed | ESR1 | -0.378197 | KIRC | RCC-ER | GDC0810 |
| MVP_top60_DNA_seed | HDAC3 | -0.078352 | HNSC | SAT | Romidepsin | MVP_top60_DNA+RNA_seed | MTOR | -0.372529 | KIRC | SW156 | Temsirolimus |
| MVP_top60_DNA_seed | EGFR | -0.077713 | HNSC | SCC-4 | Osimertinib | MVP_top60_DNA+RNA_seed | TP53 | -0.370749 | KIRC | RCC-MF | MIRA-1 |
| MVP_top60_DNA_seed | TP53 | -0.077083 | HNSC | PCI-6A | MIRA-1 | MVP_top60_DNA+RNA_seed | IGF1R | -0.364509 | KIRC | LB996-RCC | BMS-754807 |
| MVP_top60_DNA_seed | AKT1 | -0.074836 | HNSC | CAL-33 | Afuresertib | MVP_top60_DNA+RNA_seed | HDAC1 | -0.364396 | KIRC | RCC-ER | Entinostat |
| MVP_top60_DNA_seed | MAPK1 | -0.072264 | HNSC | CAL-33 | SCH772984 | MVP_top60_DNA+RNA_seed | IGF1R | -0.361563 | KIRC | NCC010 | BMS-536924 |
| MVP_top60_DNA_seed | MAPK3 | -0.072264 | HNSC | CAL-33 | SCH772984 | MVP_top60_DNA+RNA_seed | MTOR | -0.357142 | KIRC | OS-RC-2 | Temsirolimus |
| MVP_top60_DNA_seed | PIK3CA | -0.070107 | HNSC | PCI-30 | GNE-317 | MVP_top60_DNA+RNA_seed | PIK3CA | -0.341345 | KIRC | RCC-FG2 | Alpelisib |
| MVP_top60_DNA_seed | HSP90AA1 | -0.069527 | HNSC | SCC-25 | Tanespimycin | MVP_top60_DNA+RNA_seed | HSP90AA1 | -0.331886 | KIRC | KMRC-20 | Tanespimycin |
| MVP_top60_DNA_seed | MAPK1 | -0.064901 | HNSC | FADU | Ulixertinib | MVP_top60_DNA+RNA_seed | HSP90AB1 | -0.331886 | KIRC | KMRC-20 | Tanespimycin |
| MVP_top60_DNA_seed | MAPK3 | -0.064901 | HNSC | FADU | Ulixertinib | MVP_top60_DNA+RNA_seed | IGF1R | -0.329485 | KIRC | RCC-MF | BMS-536924 |
| MVP_top60_DNA_seed | ESR1 | -0.061123 | HNSC | SAT | Tamoxifen | MVP_top60_DNA+RNA_seed | HDAC1 | -0.32946 | KIRC | RXF393 | Romidepsin |
| MVP_top60_DNA_seed | AR | -0.060329 | HNSC | KON | Bicalutamide | MVP_top60_DNA+RNA_seed | MTOR | -0.316733 | KIRC | BFTC-909 | Temsirolimus |
| MVP_top60_DNA_seed | HDAC1 | -0.060244 | HNSC | JHU-022 | Romidepsin | MVP_top60_DNA+RNA_seed | MTOR | -0.316635 | KIRC | RCC-JW | Temsirolimus |
| MVP_top60_DNA_seed | HDAC3 | -0.060244 | HNSC | JHU-022 | Romidepsin | MVP_top60_DNA+RNA_seed | HSP90AA1 | -0.314135 | KIRC | RCC-ER | CCT-018159 |
| MVP_top60_DNA_seed | AKT1 | -0.060076 | HNSC | SCC90 | MK-2206 | MVP_top60_DNA+RNA_seed | HSP90AB1 | -0.314135 | KIRC | RCC-ER | CCT-018159 |
| MVP_top60_DNA_seed | EGFR | -0.059399 | HNSC | BICR10 | Sapitinib | MVP_top60_DNA+RNA_seed | HSP90AA1 | -0.311872 | KIRC | CAKI-1 | Elesclomol |
| MVP_top60_DNA_seed | AKT1 | -0.057118 | HNSC | KOSC-2 | MK-2206 | MVP_top60_DNA+RNA_seed | HSP90AB1 | -0.311872 | KIRC | CAKI-1 | Elesclomol |
| MVP_top60_DNA_seed | ESR1 | -0.056738 | HNSC | PCI-30 | Tamoxifen | MVP_top60_DNA+RNA_seed | ESR1 | -0.303033 | KIRC | ACHN | Tamoxifen |
| MVP_top60_DNA_seed | AKT1 | -0.055612 | HNSC | HO-1-N-1 | AT13148 | MVP_top60_DNA+RNA_seed | HSP90AA1 | -0.302433 | KIRC | RCC-FG2 | Elesclomol |
| MVP_top60_DNA_seed | MAPK1 | -0.052071 | HNSC | JHU-022 | Ulixertinib | MVP_top60_DNA+RNA_seed | HSP90AB1 | -0.302433 | KIRC | RCC-FG2 | Elesclomol |
| MVP_top60_DNA_seed | MAPK3 | -0.052071 | HNSC | JHU-022 | Ulixertinib | MVP_top60_DNA+RNA_seed | HDAC1 | -0.301455 | KIRC | RCC-JF | Entinostat |
| MVP_top60_DNA_seed | EP300 | -0.050997 | HNSC | LB771-HNC | CPI-637 | MVP_top60_DNA+RNA_seed | ATM | -0.300438 | KIRC | KMRC-20 | KU-55933 |
| MVP_top60_DNA_seed | PIK3CA | -0.048748 | HNSC | CAL-27 | Alpelisib | MVP_top60_DNA+RNA_seed | MTOR | -0.29049 | KIRC | U031 | Temsirolimus |

| | | | | | | | | | | | |
|---|---|---|---|---|---|---|---|---|---|---|---|
| MVP_top60_DNA_seed | TP53 | -0.048623 | HNSC | FADU | MIRA-1 | MVP_top60_DNA+RNA_seed | IGF1R | -0.289694 | KIRC | 769-P | NVP-ADW742 |
| MVP_top60_DNA_seed | EP300 | -0.048338 | HNSC | SCC-4 | SGC-CBP30 | MVP_top60_DNA+RNA_seed | HDAC1 | -0.287285 | KIRC | BB65-RCC | PCI-34051 |
| MVP_top60_DNA_seed | CREBBP | -0.048338 | HNSC | SCC-4 | SGC-CBP30 | MVP_top60_DNA+RNA_seed | IGF1R | -0.286215 | KIRC | KMRC-20 | BMS-536924 |
| MVP_top60_DNA_seed | EGFR | -0.047268 | HNSC | SCC-25 | Sapitinib | MVP_top60_DNA+RNA_seed | HDAC1 | -0.275823 | KIRC | U031 | PCI-34051 |
| MVP_top60_DNA_seed | AKT1 | -0.045856 | HNSC | HSC-2 | Ipatasertib | MVP_top60_DNA+RNA_seed | TP53 | -0.275164 | KIRC | 769-P | MIRA-1 |
| MVP_top60_DNA_seed | MDM2 | -0.045783 | HNSC | PCI-6A | Nutlin-3a (-) | MVP_top60_DNA+RNA_seed | HSP90AA1 | -0.273761 | KIRC | NCC010 | Luminespib |
| MVP_top60_DNA_seed | MAPK1 | -0.043159 | HNSC | CAL-27 | Ulixertinib | MVP_top60_DNA+RNA_seed | HSP90AB1 | -0.273761 | KIRC | NCC010 | Luminespib |
| MVP_top60_DNA_seed | MAPK3 | -0.043159 | HNSC | CAL-27 | Ulixertinib | MVP_top60_DNA+RNA_seed | IGF1R | -0.271285 | KIRC | NCC021 | BMS-754807 |
| MVP_top60_DNA_seed | MDM2 | -0.043091 | HNSC | SAS | Serdemetan | MVP_top60_DNA+RNA_seed | IGF1R | -0.266696 | KIRC | A704 | BMS-754807 |
| MVP_top60_DNA_seed | ESR1 | -0.042469 | HNSC | BICR31 | GDC0810 | MVP_top60_DNA+RNA_seed | EP300 | -0.258428 | KIRC | 769-P | SGC-CBP30 |
| MVP_top60_DNA_seed | AKT1 | -0.041637 | HNSC | HO-1-N-1 | MK-2206 | MVP_top60_DNA+RNA_seed | PIK3CA | -0.253051 | KIRC | U031 | Alpelisib |
| MVP_top60_DNA_seed | AKT1 | -0.040978 | HNSC | LB771-HNC | Uprosertib | MVP_top60_DNA+RNA_seed | HDAC1 | -0.25068 | KIRC | RCC-JW | Romidepsin |
| MVP_top60_DNA_seed | AKT1 | -0.039831 | HNSC | LB771-HNC | MK-2206 | MVP_top60_DNA+RNA_seed | HDAC1 | -0.25019 | KIRC | RXF393 | PCI-34051 |
| MVP_top60_DNA_seed | TP53 | -0.0398 | HNSC | KOSC-2 | MIRA-1 | MVP_top60_DNA+RNA_seed | PIK3CA | -0.250163 | KIRC | CAKI-1 | Alpelisib |
| MVP_top60_DNA_seed | RAC1 | -0.039384 | HNSC | HO-1-N-1 | EHT-1864 | MVP_top60_DNA+RNA_seed | HDAC1 | -0.249587 | KIRC | CAKI-1 | Entinostat |
| MVP_top60_DNA_seed | MDM2 | -0.037962 | HNSC | CAL-33 | Serdemetan | MVP_top60_DNA+RNA_seed | HSP90AA1 | -0.240579 | KIRC | RXF393 | Elesclomol |
| MVP_top60_DNA_seed | MAPK1 | -0.034227 | HNSC | HO-1-N-1 | Ulixertinib | MVP_top60_DNA+RNA_seed | HSP90AB1 | -0.240579 | KIRC | RXF393 | Elesclomol |
| MVP_top60_DNA_seed | MAPK3 | -0.034227 | HNSC | HO-1-N-1 | Ulixertinib | MVP_top60_DNA+RNA_seed | PIK3CA | -0.237404 | KIRC | A704 | Alpelisib |
| MVP_top60_DNA_seed | PIK3CA | -0.032409 | HNSC | SCC-15 | GNE-317 | MVP_top60_DNA+RNA_seed | IGF1R | -0.233819 | KIRC | BFTC-909 | Linsitinib |
| MVP_top60_DNA_seed | EGFR | -0.028932 | HNSC | BICR31 | Gefitinib | MVP_top60_DNA+RNA_seed | SMARCA4 | -0.230155 | KIRC | SN12C | PFI3 |
| MVP_top60_DNA_seed | PIK3CA | -0.028222 | HNSC | OSC-19 | Buparlisib | MVP_top60_DNA+RNA_seed | SMARCA2 | -0.230155 | KIRC | SN12C | PFI3 |
| MVP_top60_DNA_seed | NTRK1 | -0.027374 | HNSC | KON | GW441756 | MVP_top60_DNA+RNA_seed | IGF1R | -0.230135 | KIRC | CAKI-1 | BMS-536924 |
| MVP_top60_DNA_seed | HDAC1 | -0.026659 | HNSC | KON | Entinostat | MVP_top60_DNA+RNA_seed | IGF1R | -0.223204 | KIRC | CAL-54 | NVP-ADW742 |
| MVP_top60_DNA_seed | HDAC3 | -0.026659 | HNSC | KON | Entinostat | MVP_top60_DNA+RNA_seed | ESR1 | -0.22078 | KIRC | KMRC-1 | Tamoxifen |
| MVP_top60_DNA_seed | MAPK1 | -0.026455 | HNSC | Detroit562 | Ulixertinib | MVP_top60_DNA+RNA_seed | IGF1R | -0.214301 | KIRC | LB2241-RCC | BMS-536924 |
| MVP_top60_DNA_seed | MAPK3 | -0.026455 | HNSC | Detroit562 | Ulixertinib | MVP_top60_DNA+RNA_seed | PIK3CA | -0.210648 | KIRC | NCC021 | Alpelisib |
| MVP_top60_DNA_seed | MAPK1 | -0.025676 | HNSC | HSC-3 | VX-11e | MVP_top60_DNA+RNA_seed | PIK3CA | -0.210197 | KIRC | RCC-MF | Buparlisib |
| MVP_top60_DNA_seed | MDM2 | -0.02339 | HNSC | SCC-25 | Serdemetan | MVP_top60_DNA+RNA_seed | PIK3CD | -0.210197 | KIRC | RCC-MF | Buparlisib |
| MVP_top60_DNA_seed | SMARCA4 | -0.022609 | HNSC | PCI-6A | PFI3 | MVP_top60_DNA+RNA_seed | HDAC1 | -0.20468 | KIRC | CAL-54 | Romidepsin |
| MVP_top60_DNA_seed | PIK3CA | -0.021695 | HNSC | FADU | Buparlisib | MVP_top60_DNA+RNA_seed | EP300 | -0.201961 | KIRC | SN12C | SGC-CBP30 |
| MVP_top60_DNA_seed | CDK2 | -0.018692 | HNSC | SCC-4 | AZD5438 | MVP_top60_DNA+RNA_seed | IGF1R | -0.200347 | KIRC | BB65-RCC | GSK1904529A |
| MVP_top60_DNA_seed | EGFR | -0.018177 | HNSC | KON | Afatinib | MVP_top60_DNA+RNA_seed | ESR1 | -0.198563 | KIRC | KMRC-20 | Tamoxifen |
| MVP_top60_DNA_seed | HSP90AA1 | -0.014816 | HNSC | SCC90 | Luminespib | MVP_top60_DNA+RNA_seed | ATM | -0.189255 | KIRC | RCC-JF | KU-55933 |
| MVP_top60_DNA_seed | HDAC1 | -0.013841 | HNSC | CAL-27 | Entinostat | MVP_top60_DNA+RNA_seed | PIK3CA | -0.188336 | KIRC | 786-0 | Buparlisib |
| MVP_top60_DNA_seed | HDAC3 | -0.013841 | HNSC | CAL-27 | Entinostat | MVP_top60_DNA+RNA_seed | PIK3CD | -0.188336 | KIRC | 786-0 | Buparlisib |
| MVP_top60_DNA_seed | MAPK1 | -0.013703 | HNSC | FADU | Ulixertinib | MVP_top60_DNA+RNA_seed | MTOR | -0.185475 | KIRC | TK10 | Temsirolimus |
| MVP_top60_DNA_seed | MAPK3 | -0.013703 | HNSC | FADU | Ulixertinib | MVP_top60_DNA+RNA_seed | PIK3CA | -0.173889 | KIRC | RCC-JF | Buparlisib |
| MVP_top60_DNA_seed | MAPK1 | -0.013156 | HNSC | BICR78 | Ulixertinib | MVP_top60_DNA+RNA_seed | PIK3CD | -0.173889 | KIRC | RCC-JF | Buparlisib |
| MVP_top60_DNA_seed | MAPK3 | -0.013156 | HNSC | BICR78 | Ulixertinib | MVP_top60_DNA+RNA_seed | IGF1R | -0.168394 | KIRC | CAKI-1 | BMS-754807 |
| MVP_top60_DNA_seed | PIK3CA | -0.011501 | HNSC | PE-CA-PJ15 | AZD8186 | MVP_top60_DNA+RNA_seed | PIK3CA | -0.159832 | KIRC | NCC010 | AZD8186 |
| MVP_top60_DNA_seed | HDAC1 | -0.011107 | HNSC | PCI-4B | Entinostat | MVP_top60_DNA+RNA_seed | PIK3CA | -0.158216 | KIRC | HA7-RCC | GNE-317 |

| | | | | | | | | | | | |
|---|---|---|---|---|---|---|---|---|---|---|---|
| MVP_top60_DNA_seed | HDAC3 | -0.011107 | HNSC | PCI-4B | Entinostat | MVP_top60_DNA+RNA_seed | TP53 | -0.157811 | KIRC | RCC-JW | MIRA-1 |
| MVP_top60_DNA_seed | AKT1 | -0.009218 | HNSC | HO-1-N-1 | Uprosertib | MVP_top60_DNA+RNA_seed | ESR1 | -0.156406 | KIRC | A498 | GDC0810 |
| MVP_top60_DNA_seed | AKT1 | -0.007431 | HNSC | SCC-4 | Uprosertib | MVP_top60_DNA+RNA_seed | EP300 | -0.156034 | KIRC | HA7-RCC | SGC-CBP30 |
| MVP_top60_DNA_seed | CDK2 | -0.005326 | HNSC | FADU | AZD5438 | MVP_top60_DNA+RNA_seed | IGF1R | -0.153255 | KIRC | ACHN | Linsitinib |
| MVP_top60_DNA_seed | AKT1 | -0.002598 | HNSC | Ca9-22 | Uprosertib | MVP_top60_DNA+RNA_seed | HDAC1 | -0.153153 | KIRC | U031 | Entinostat |
| MVP_top60_DNA_seed | EGFR | 0.002625 | HNSC | BICR22 | AZD3759 | MVP_top60_DNA+RNA_seed | HDAC1 | -0.152531 | KIRC | LB1047-RCC | Entinostat |
| MVP_top60_DNA_seed | PIK3CA | 0.00364 | HNSC | HO-1-u-1 | AZD8186 | MVP_top60_DNA+RNA_seed | PIK3CA | -0.146273 | KIRC | RCC-MF | Alpelisib |
| MVP_top60_DNA_seed | CDK2 | 0.004586 | HNSC | SAT | AZD5438 | MVP_top60_DNA+RNA_seed | IGF1R | -0.141101 | KIRC | LB2241-RCC | BMS-754807 |
| MVP_top60_DNA_seed | AKT1 | 0.006771 | HNSC | CAL-33 | AZD5363 | MVP_top60_DNA+RNA_seed | PIK3CA | -0.139308 | KIRC | KMRC-1 | GNE-317 |
| MVP_top60_DNA_seed | ESR1 | 0.007562 | HNSC | KON | GDC0810 | MVP_top60_DNA+RNA_seed | ATM | -0.13407 | KIRC | HA7-RCC | KU-55933 |
| MVP_top60_DNA_seed | AKT1 | 0.00819 | HNSC | PCI-6A | Afuresertib | MVP_top60_DNA+RNA_seed | HSP90AA1 | -0.131923 | KIRC | SW156 | Luminespib |
| MVP_top60_DNA_seed | SMARCA4 | 0.008959 | HNSC | SCC-15 | PFI3 | MVP_top60_DNA+RNA_seed | HSP90AB1 | -0.131923 | KIRC | SW156 | Luminespib |
| MVP_top60_DNA_seed | MDM2 | 0.011078 | HNSC | DOK | Nutlin-3a (-) | MVP_top60_DNA+RNA_seed | PIK3CA | -0.129105 | KIRC | HA7-RCC | Alpelisib |
| MVP_top60_DNA_seed | MAPK1 | 0.012137 | HNSC | SCC90 | VX-11e | MVP_top60_DNA+RNA_seed | TP53 | -0.118043 | KIRC | RCC-FG2 | MIRA-1 |
| MVP_top60_DNA_seed | MAPK1 | 0.01234 | HNSC | HO-1-u-1 | Ulixertinib | MVP_top60_DNA+RNA_seed | IGF1R | -0.108973 | KIRC | SW156 | GSK1904529A |
| MVP_top60_DNA_seed | MAPK3 | 0.01234 | HNSC | HO-1-u-1 | Ulixertinib | MVP_top60_DNA+RNA_seed | ATM | -0.108044 | KIRC | SW156 | KU-55933 |
| MVP_top60_DNA_seed | AKT1 | 0.01375 | HNSC | PCI-6A | Uprosertib | MVP_top60_DNA+RNA_seed | PIK3CA | -0.105575 | KIRC | LB2241-RCC | Alpelisib |
| MVP_top60_DNA_seed | HDAC1 | 0.013833 | HNSC | SKN-3 | Entinostat | MVP_top60_DNA+RNA_seed | PIK3CA | -0.10051 | KIRC | RCC-FG2 | Buparlisib |
| MVP_top60_DNA_seed | HDAC3 | 0.013833 | HNSC | SKN-3 | Entinostat | MVP_top60_DNA+RNA_seed | PIK3CD | -0.10051 | KIRC | RCC-FG2 | Buparlisib |
| MVP_top60_DNA_seed | MAPK1 | 0.014203 | HNSC | BICR22 | Ulixertinib | MVP_top60_DNA+RNA_seed | AR | -0.092604 | KIRC | RXF393 | Bicalutamide |
| MVP_top60_DNA_seed | MAPK3 | 0.014203 | HNSC | BICR22 | Ulixertinib | MVP_top60_DNA+RNA_seed | IGF1R | -0.09092 | KIRC | NCC010 | NVP-ADW742 |
| MVP_top60_DNA_seed | AKT1 | 0.01432 | HNSC | JHU-022 | Uprosertib | MVP_top60_DNA+RNA_seed | HDAC1 | -0.089297 | KIRC | 769-P | PCI-34051 |
| MVP_top60_DNA_seed | ESR1 | 0.01479 | HNSC | SCC-4 | GDC0810 | MVP_top60_DNA+RNA_seed | IGF1R | -0.086369 | KIRC | CAL-54 | Linsitinib |
| MVP_top60_DNA_seed | PIK3CA | 0.017505 | HNSC | BHY | AZD8186 | MVP_top60_DNA+RNA_seed | PIK3CA | -0.085348 | KIRC | A498 | Buparlisib |
| MVP_top60_DNA_seed | PIK3CA | 0.019557 | HNSC | LB771-HNC | Buparlisib | MVP_top60_DNA+RNA_seed | PIK3CD | -0.085348 | KIRC | A498 | Buparlisib |
| MVP_top60_DNA_seed | AKT1 | 0.022684 | HNSC | HSC-4 | Uprosertib | MVP_top60_DNA+RNA_seed | HSP90AA1 | -0.083914 | KIRC | ACHN | Tanespimycin |
| MVP_top60_DNA_seed | SMARCA4 | 0.022878 | HNSC | SCC-4 | PFI3 | MVP_top60_DNA+RNA_seed | HSP90AB1 | -0.083914 | KIRC | ACHN | Tanespimycin |
| MVP_top60_DNA_seed | MDM2 | 0.023264 | HNSC | JHU-029 | Nutlin-3a (-) | MVP_top60_DNA+RNA_seed | MTOR | -0.081958 | KIRC | 769-P | Temsirolimus |
| MVP_top60_DNA_seed | EP300 | 0.023715 | HNSC | HSC-4 | CPI-637 | MVP_top60_DNA+RNA_seed | PIK3CA | -0.081313 | KIRC | ACHN | Buparlisib |
| MVP_top60_DNA_seed | TP53 | 0.023865 | HNSC | BICR22 | MIRA-1 | MVP_top60_DNA+RNA_seed | PIK3CD | -0.081313 | KIRC | ACHN | Buparlisib |
| MVP_top60_DNA_seed | MAPK1 | 0.026715 | HNSC | BICR78 | VX-11e | MVP_top60_DNA+RNA_seed | PIK3CA | -0.079578 | KIRC | U031 | Buparlisib |
| MVP_top60_DNA_seed | HSP90AA1 | 0.028165 | HNSC | BHY | Luminespib | MVP_top60_DNA+RNA_seed | PIK3CD | -0.079578 | KIRC | U031 | Buparlisib |
| MVP_top60_DNA_seed | TP53 | 0.031459 | HNSC | CAL-33 | MIRA-1 | MVP_top60_DNA+RNA_seed | IGF1R | -0.079044 | KIRC | HA7-RCC | BMS-536924 |
| MVP_top60_DNA_seed | ESR1 | 0.034625 | HNSC | KOSC-2 | GDC0810 | MVP_top60_DNA+RNA_seed | HSP90AA1 | -0.074655 | KIRC | NCC021 | Luminespib |
| MVP_top60_DNA_seed | PIK3CA | 0.035033 | HNSC | PE-CA-PJ15 | GNE-317 | MVP_top60_DNA+RNA_seed | HSP90AB1 | -0.074655 | KIRC | NCC021 | Luminespib |
| MVP_top60_DNA_seed | MAPK1 | 0.035272 | HNSC | SCC-25 | SCH772984 | MVP_top60_DNA+RNA_seed | PIK3CA | -0.072205 | KIRC | A498 | GNE-317 |
| MVP_top60_DNA_seed | MAPK3 | 0.035272 | HNSC | SCC-25 | SCH772984 | MVP_top60_DNA+RNA_seed | HSP90AA1 | -0.070257 | KIRC | SW156 | Elesclomol |
| MVP_top60_DNA_seed | SMARCA4 | 0.035778 | HNSC | BICR78 | PFI3 | MVP_top60_DNA+RNA_seed | HSP90AB1 | -0.070257 | KIRC | SW156 | Elesclomol |
| MVP_top60_DNA_seed | MAPK1 | 0.036784 | HNSC | SAT | Ulixertinib | MVP_top60_DNA+RNA_seed | PIK3CA | -0.069469 | KIRC | CAL-54 | Alpelisib |
| MVP_top60_DNA_seed | MAPK3 | 0.036784 | HNSC | SAT | Ulixertinib | MVP_top60_DNA+RNA_seed | SMARCA4 | -0.068164 | KIRC | RCC-ER | PFI3 |
| MVP_top60_DNA_seed | AKT1 | 0.036786 | HNSC | JHU-022 | AZD5363 | MVP_top60_DNA+RNA_seed | SMARCA2 | -0.068164 | KIRC | RCC-ER | PFI3 |

| | | | | | | | | | | | |
|---|---|---|---|---|---|---|---|---|---|---|---|
| MVP_top60_DNA_seed | SRC | 0.037466 | HNSC | CAL-27 | Bosutinib | MVP_top60_DNA+RNA_seed | HSP90AA1 | -0.066749 | KIRC | RCC-ER | Luminespib |
| MVP_top60_DNA_seed | RAC1 | 0.040961 | HNSC | PE-CA-PJ15 | EHT-1864 | MVP_top60_DNA+RNA_seed | HSP90AB1 | -0.066749 | KIRC | RCC-ER | Luminespib |
| MVP_top60_DNA_seed | RAC1 | 0.041149 | HNSC | OSC-19 | EHT-1864 | MVP_top60_DNA+RNA_seed | HSP90AA1 | -0.065315 | KIRC | A498 | Tanespimycin |
| MVP_top60_DNA_seed | AKT1 | 0.042139 | HNSC | SCC-9 | Uprosertib | MVP_top60_DNA+RNA_seed | HSP90AB1 | -0.065315 | KIRC | A498 | Tanespimycin |
| MVP_top60_DNA_seed | ESR1 | 0.043736 | HNSC | BB30-HNC | Tamoxifen | MVP_top60_DNA+RNA_seed | EP300 | -0.065191 | KIRC | RCC-JW | SGC-CBP30 |
| MVP_top60_DNA_seed | TP53 | 0.04735 | HNSC | PE-CA-PJ15 | MIRA-1 | MVP_top60_DNA+RNA_seed | IGF1R | -0.062528 | KIRC | A704 | GSK1904529A |
| MVP_top60_DNA_seed | MAPK1 | 0.048483 | HNSC | SCC-9 | Ulixertinib | MVP_top60_DNA+RNA_seed | IGF1R | -0.060254 | KIRC | RCC-JF | BMS-754807 |
| MVP_top60_DNA_seed | MAPK3 | 0.048483 | HNSC | SCC-9 | Ulixertinib | MVP_top60_DNA+RNA_seed | PIK3CA | -0.055843 | KIRC | RCC-ER | GNE-317 |
| MVP_top60_DNA_seed | NTRK1 | 0.050179 | HNSC | DOK | GW441756 | MVP_top60_DNA+RNA_seed | IGF1R | -0.049247 | KIRC | BFTC-909 | BMS-536924 |
| MVP_top60_DNA_seed | TP53 | 0.050948 | HNSC | HSC-3 | MIRA-1 | MVP_top60_DNA+RNA_seed | ESR1 | -0.045743 | KIRC | BFTC-909 | GDC0810 |
| MVP_top60_DNA_seed | RAC1 | 0.052339 | HNSC | BB30-HNC | EHT-1864 | MVP_top60_DNA+RNA_seed | IGF1R | -0.045352 | KIRC | SN12C | BMS-536924 |
| MVP_top60_DNA_seed | MAPK1 | 0.053729 | HNSC | BICR78 | SCH772984 | MVP_top60_DNA+RNA_seed | HSP90AA1 | -0.040101 | KIRC | RCC-ER | Elesclomol |
| MVP_top60_DNA_seed | MAPK3 | 0.053729 | HNSC | BICR78 | SCH772984 | MVP_top60_DNA+RNA_seed | HSP90AB1 | -0.040101 | KIRC | RCC-ER | Elesclomol |
| MVP_top60_DNA_seed | MAPK1 | 0.05471 | HNSC | SCC-25 | VX-11e | MVP_top60_DNA+RNA_seed | IGF1R | -0.031113 | KIRC | ACHN | BMS-536924 |
| MVP_top60_DNA_seed | ESR1 | 0.055183 | HNSC | JHU-022 | Tamoxifen | MVP_top60_DNA+RNA_seed | ATM | -0.023913 | KIRC | NCC021 | KU-55933 |
| MVP_top60_DNA_seed | NTRK1 | 0.056407 | HNSC | KOSC-2 | GW441756 | MVP_top60_DNA+RNA_seed | ESR1 | -0.019002 | KIRC | RCC10RGB | Tamoxifen |
| MVP_top60_DNA_seed | EGFR | 0.056562 | HNSC | LB771-HNC | Osimertinib | MVP_top60_DNA+RNA_seed | HSP90AA1 | -0.015529 | KIRC | ACHN | Luminespib |
| MVP_top60_DNA_seed | AKT1 | 0.056746 | HNSC | BICR22 | Uprosertib | MVP_top60_DNA+RNA_seed | HSP90AB1 | -0.015529 | KIRC | ACHN | Luminespib |
| MVP_top60_DNA_seed | EGFR | 0.057949 | HNSC | SCC-25 | Gefitinib | MVP_top60_DNA+RNA_seed | HSP90AA1 | -0.010222 | KIRC | HA7-RCC | Tanespimycin |
| MVP_top60_DNA_seed | HDAC1 | 0.059736 | HNSC | FADU | PCI-34051 | MVP_top60_DNA+RNA_seed | HSP90AB1 | -0.010222 | KIRC | HA7-RCC | Tanespimycin |
| MVP_top60_DNA_seed | AKT1 | 0.061078 | HNSC | SKN-3 | Afuresertib | MVP_top60_DNA+RNA_seed | IGF1R | -0.006827 | KIRC | A704 | Linsitinib |
| MVP_top60_DNA_seed | SMARCA4 | 0.065394 | HNSC | SCC90 | PFI3 | MVP_top60_DNA+RNA_seed | SMARCA4 | 0.002601 | KIRC | BFTC-909 | PFI3 |
| MVP_top60_DNA_seed | NTRK1 | 0.067295 | HNSC | SKN-3 | AZD1332 | MVP_top60_DNA+RNA_seed | SMARCA2 | 0.002601 | KIRC | BFTC-909 | PFI3 |
| MVP_top60_DNA_seed | PIK3CA | 0.067968 | HNSC | SAT | AZD8186 | MVP_top60_DNA+RNA_seed | PIK3CA | 0.007027 | KIRC | RCC10RGB | GNE-317 |
| MVP_top60_DNA_seed | AKT1 | 0.06848 | HNSC | BICR22 | Ipatasertib | MVP_top60_DNA+RNA_seed | PIK3CA | 0.009887 | KIRC | A704 | Buparlisib |
| MVP_top60_DNA_seed | EGFR | 0.068589 | HNSC | BB49-HNC | Osimertinib | MVP_top60_DNA+RNA_seed | PIK3CD | 0.009887 | KIRC | A704 | Buparlisib |
| MVP_top60_DNA_seed | NTRK1 | 0.06938 | HNSC | HO-1-u-1 | GW441756 | MVP_top60_DNA+RNA_seed | EP300 | 0.013002 | KIRC | RCC10RGB | SGC-CBP30 |
| MVP_top60_DNA_seed | EGFR | 0.072645 | HNSC | SCC-25 | Lapatinib | MVP_top60_DNA+RNA_seed | HSP90AA1 | 0.013567 | KIRC | CAL-54 | Luminespib |
| MVP_top60_DNA_seed | MAPK1 | 0.074138 | HNSC | HSC-2 | VX-11e | MVP_top60_DNA+RNA_seed | HSP90AB1 | 0.013567 | KIRC | CAL-54 | Luminespib |
| MVP_top60_DNA_seed | HDAC1 | 0.075657 | HNSC | PCI-4B | Romidepsin | MVP_top60_DNA+RNA_seed | TP53 | 0.014332 | KIRC | RCC-JF | MIRA-1 |
| MVP_top60_DNA_seed | HDAC3 | 0.075657 | HNSC | PCI-4B | Romidepsin | MVP_top60_DNA+RNA_seed | HSP90AA1 | 0.023327 | KIRC | NCC010 | Tanespimycin |
| MVP_top60_DNA_seed | MDM2 | 0.077546 | HNSC | JHU-011 | Nutlin-3a (-) | MVP_top60_DNA+RNA_seed | HSP90AB1 | 0.023327 | KIRC | NCC010 | Tanespimycin |
| MVP_top60_DNA_seed | NTRK1 | 0.077548 | HNSC | CAL-27 | AZD1332 | MVP_top60_DNA+RNA_seed | PIK3CA | 0.026851 | KIRC | NCC010 | Alpelisib |
| MVP_top60_DNA_seed | CDK2 | 0.07938 | HNSC | BB30-HNC | MK-8776 | MVP_top60_DNA+RNA_seed | HSP90AA1 | 0.030298 | KIRC | CAKI-1 | Luminespib |
| MVP_top60_DNA_seed | CDK2 | 0.079738 | HNSC | SKN-3 | MK-8776 | MVP_top60_DNA+RNA_seed | HSP90AB1 | 0.030298 | KIRC | CAKI-1 | Luminespib |
| MVP_top60_DNA_seed | NTRK1 | 0.079907 | HNSC | SCC-9 | GW441756 | MVP_top60_DNA+RNA_seed | EP300 | 0.030997 | KIRC | ACHN | CPI-637 |
| MVP_top60_DNA_seed | PIK3CA | 0.08018 | HNSC | JHU-029 | GNE-317 | MVP_top60_DNA+RNA_seed | TP53 | 0.036065 | KIRC | NCC021 | MIRA-1 |
| MVP_top60_DNA_seed | RAC1 | 0.081459 | HNSC | SCC-4 | EHT-1864 | MVP_top60_DNA+RNA_seed | MTOR | 0.036414 | KIRC | LB996-RCC | Temsirolimus |
| MVP_top60_DNA_seed | MAPK1 | 0.082192 | HNSC | CAL-33 | Ulixertinib | MVP_top60_DNA+RNA_seed | HDAC1 | 0.037521 | KIRC | A498 | PCI-34051 |
| MVP_top60_DNA_seed | MAPK3 | 0.082192 | HNSC | CAL-33 | Ulixertinib | MVP_top60_DNA+RNA_seed | HDAC1 | 0.041232 | KIRC | SW156 | Entinostat |
| MVP_top60_DNA_seed | HDAC1 | 0.084345 | HNSC | SKN-3 | Romidepsin | MVP_top60_DNA+RNA_seed | TP53 | 0.041486 | KIRC | LB2241-RCC | MIRA-1 |

| | | | | | | | | | | | |
|---|---|---|---|---|---|---|---|---|---|---|---|
| MVP_top60_DNA_seed | HDAC3 | 0.084345 | HNSC | SKN-3 | Romidepsin | MVP_top60_DNA+RNA_seed | SMARCA4 | 0.046874 | KIRC | RCC-JF | PFI3 |
| MVP_top60_DNA_seed | SMARCA4 | 0.086456 | HNSC | BB49-HNC | PFI3 | MVP_top60_DNA+RNA_seed | SMARCA2 | 0.046874 | KIRC | RCC-JF | PFI3 |
| MVP_top60_DNA_seed | AKT1 | 0.100108 | HNSC | BB30-HNC | AZD5363 | MVP_top60_DNA+RNA_seed | EP300 | 0.051717 | KIRC | 786-0 | CPI-637 |
| MVP_top60_DNA_seed | CDK2 | 0.100711 | HNSC | LB771-HNC | AZD5438 | MVP_top60_DNA+RNA_seed | IGF1R | 0.053463 | KIRC | 786-0 | Linsitinib |
| MVP_top60_DNA_seed | MAPK1 | 0.100733 | HNSC | PCI-6A | SCH772984 | MVP_top60_DNA+RNA_seed | ESR1 | 0.054421 | KIRC | A704 | Tamoxifen |
| MVP_top60_DNA_seed | MAPK3 | 0.100733 | HNSC | PCI-6A | SCH772984 | MVP_top60_DNA+RNA_seed | HDAC1 | 0.0588 | KIRC | RCC-AB | Entinostat |
| MVP_top60_DNA_seed | SMARCA4 | 0.101108 | HNSC | JHU-029 | PFI3 | MVP_top60_DNA+RNA_seed | IGF1R | 0.064071 | KIRC | A498 | Linsitinib |
| MVP_top60_DNA_seed | SMARCA4 | 0.101499 | HNSC | OSC-19 | PFI3 | MVP_top60_DNA+RNA_seed | PIK3CA | 0.06458 | KIRC | SW156 | AZD8186 |
| MVP_top60_DNA_seed | CDK2 | 0.106393 | HNSC | LB771-HNC | MK-8776 | MVP_top60_DNA+RNA_seed | PIK3CA | 0.065939 | KIRC | RCC-AB | AZD8186 |
| MVP_top60_DNA_seed | PIK3CA | 0.107142 | HNSC | PCI-6A | Buparlisib | MVP_top60_DNA+RNA_seed | IGF1R | 0.06825 | KIRC | RXF393 | Linsitinib |
| MVP_top60_DNA_seed | ESR1 | 0.110018 | HNSC | SCC-4 | Tamoxifen | MVP_top60_DNA+RNA_seed | HDAC1 | 0.070282 | KIRC | KMRC-20 | Entinostat |
| MVP_top60_DNA_seed | RAC1 | 0.112929 | HNSC | HSC-3 | EHT-1864 | MVP_top60_DNA+RNA_seed | EP300 | 0.070755 | KIRC | BFTC-909 | CPI-637 |
| MVP_top60_DNA_seed | ESR1 | 0.113685 | HNSC | SCC-15 | Tamoxifen | MVP_top60_DNA+RNA_seed | SMARCA4 | 0.078614 | KIRC | RCC-JW | PFI3 |
| MVP_top60_DNA_seed | MDM2 | 0.115006 | HNSC | CAL-33 | Nutlin-3a (-) | MVP_top60_DNA+RNA_seed | SMARCA2 | 0.078614 | KIRC | RCC-JW | PFI3 |
| MVP_top60_DNA_seed | SRC | 0.116307 | HNSC | PE-CA-PJ15 | Bosutinib | MVP_top60_DNA+RNA_seed | PIK3CA | 0.081977 | KIRC | NCC021 | GNE-317 |
| MVP_top60_DNA_seed | EGFR | 0.116622 | HNSC | BICR22 | Osimertinib | MVP_top60_DNA+RNA_seed | TP53 | 0.082333 | KIRC | NCC010 | MIRA-1 |
| MVP_top60_DNA_seed | MDM2 | 0.118617 | HNSC | PE-CA-PJ15 | Nutlin-3a (-) | MVP_top60_DNA+RNA_seed | SMARCA4 | 0.084828 | KIRC | 786-0 | PFI3 |
| MVP_top60_DNA_seed | EGFR | 0.118732 | HNSC | SKN-3 | Lapatinib | MVP_top60_DNA+RNA_seed | SMARCA2 | 0.084828 | KIRC | 786-0 | PFI3 |
| MVP_top60_DNA_seed | CDK2 | 0.120823 | HNSC | BHY | Dinaciclib | MVP_top60_DNA+RNA_seed | ESR1 | 0.087509 | KIRC | RCC-JF | Tamoxifen |
| MVP_top60_DNA_seed | MDM2 | 0.124582 | HNSC | SCC-25 | Nutlin-3a (-) | MVP_top60_DNA+RNA_seed | SMARCA4 | 0.093692 | KIRC | 769-P | PFI3 |
| MVP_top60_DNA_seed | HDAC1 | 0.126105 | HNSC | Detroit562 | PCI-34051 | MVP_top60_DNA+RNA_seed | SMARCA2 | 0.093692 | KIRC | 769-P | PFI3 |
| MVP_top60_DNA_seed | EGFR | 0.126785 | HNSC | SKN-3 | Afatinib | MVP_top60_DNA+RNA_seed | HDAC1 | 0.094458 | KIRC | RCC-MF | Entinostat |
| MVP_top60_DNA_seed | AKT1 | 0.12688 | HNSC | HSC-3 | Uprosertib | MVP_top60_DNA+RNA_seed | PIK3CA | 0.096705 | KIRC | RXF393 | GNE-317 |
| MVP_top60_DNA_seed | MDM2 | 0.127435 | HNSC | KON | Nutlin-3a (-) | MVP_top60_DNA+RNA_seed | IGF1R | 0.097983 | KIRC | CAL-54 | BMS-754807 |
| MVP_top60_DNA_seed | SMARCA4 | 0.128163 | HNSC | CAL-33 | PFI3 | MVP_top60_DNA+RNA_seed | PIK3CA | 0.103392 | KIRC | RCC-MF | AZD8186 |
| MVP_top60_DNA_seed | AKT1 | 0.129076 | HNSC | BICR22 | AZD5363 | MVP_top60_DNA+RNA_seed | TP53 | 0.113903 | KIRC | U031 | MIRA-1 |
| MVP_top60_DNA_seed | HSP90AA1 | 0.133097 | HNSC | Ca9-22 | Tanespimycin | MVP_top60_DNA+RNA_seed | ATM | 0.116633 | KIRC | BB65-RCC | KU-55933 |
| MVP_top60_DNA_seed | MAPK1 | 0.134404 | HNSC | CAL-33 | Ulixertinib | MVP_top60_DNA+RNA_seed | IGF1R | 0.118136 | KIRC | LB1047-RCC | BMS-754807 |
| MVP_top60_DNA_seed | MAPK3 | 0.134404 | HNSC | CAL-33 | Ulixertinib | MVP_top60_DNA+RNA_seed | HSP90AA1 | 0.127148 | KIRC | CAL-54 | Tanespimycin |
| MVP_top60_DNA_seed | EGFR | 0.134749 | HNSC | PCI-30 | AZD3759 | MVP_top60_DNA+RNA_seed | HSP90AB1 | 0.127148 | KIRC | CAL-54 | Tanespimycin |
| MVP_top60_DNA_seed | AKT1 | 0.140923 | HNSC | PCI-30 | Ipatasertib | MVP_top60_DNA+RNA_seed | HSP90AA1 | 0.127691 | KIRC | 786-0 | Luminespib |
| MVP_top60_DNA_seed | AKT1 | 0.141465 | HNSC | PCI-4B | AT13148 | MVP_top60_DNA+RNA_seed | HSP90AB1 | 0.127691 | KIRC | 786-0 | Luminespib |
| MVP_top60_DNA_seed | MDM2 | 0.143945 | HNSC | HO-1-N-1 | Nutlin-3a (-) | MVP_top60_DNA+RNA_seed | EP300 | 0.129753 | KIRC | ACHN | SGC-CBP30 |
| MVP_top60_DNA_seed | EP300 | 0.145181 | HNSC | HSC-3 | SGC-CBP30 | MVP_top60_DNA+RNA_seed | HDAC1 | 0.142093 | KIRC | A704 | PCI-34051 |
| MVP_top60_DNA_seed | CREBBP | 0.145181 | HNSC | HSC-3 | SGC-CBP30 | MVP_top60_DNA+RNA_seed | SMARCA4 | 0.142904 | KIRC | LB996-RCC | PFI3 |
| MVP_top60_DNA_seed | EGFR | 0.146254 | HNSC | PCI-30 | Afatinib | MVP_top60_DNA+RNA_seed | SMARCA2 | 0.142904 | KIRC | LB996-RCC | PFI3 |
| MVP_top60_DNA_seed | AKT1 | 0.14729 | HNSC | SAS | GSK2110183B | MVP_top60_DNA+RNA_seed | PIK3CA | 0.143317 | KIRC | 769-P | AZD8186 |
| MVP_top60_DNA_seed | HDAC1 | 0.147343 | HNSC | Ca9-22 | Entinostat | MVP_top60_DNA+RNA_seed | HSP90AA1 | 0.143976 | KIRC | LB2241-RCC | Tanespimycin |
| MVP_top60_DNA_seed | HDAC3 | 0.147343 | HNSC | Ca9-22 | Entinostat | MVP_top60_DNA+RNA_seed | HSP90AB1 | 0.143976 | KIRC | LB2241-RCC | Tanespimycin |
| MVP_top60_DNA_seed | SRC | 0.147457 | HNSC | SCC90 | Bosutinib | MVP_top60_DNA+RNA_seed | IGF1R | 0.144922 | KIRC | RCC-FG2 | NVP-ADW742 |
| MVP_top60_DNA_seed | AKT1 | 0.15093 | HNSC | PCI-30 | AZD5363 | MVP_top60_DNA+RNA_seed | AR | 0.146122 | KIRC | LB2241-RCC | Bicalutamide |

| | | | | | | | | | | | |
|---|---|---|---|---|---|---|---|---|---|---|---|
| MVP_top60_DNA_seed | AR | 0.158824 | HNSC | KOSC-2 | Bicalutamide | MVP_top60_DNA+RNA_seed | IGF1R | 0.151229 | KIRC | BB65-RCC | BMS-536924 |
| MVP_top60_DNA_seed | AKT1 | 0.159402 | HNSC | PCI-4B | MK-2206 | MVP_top60_DNA+RNA_seed | HDAC1 | 0.151447 | KIRC | RCC-FG2 | Entinostat |
| MVP_top60_DNA_seed | NTRK1 | 0.164984 | HNSC | SCC-15 | GW441756 | MVP_top60_DNA+RNA_seed | ATM | 0.151675 | KIRC | RCC-MF | KU-55933 |
| MVP_top60_DNA_seed | NTRK1 | 0.167001 | HNSC | JHU-029 | GW441756 | MVP_top60_DNA+RNA_seed | HDAC1 | 0.156351 | KIRC | A498 | Romidepsin |
| MVP_top60_DNA_seed | AKT1 | 0.167132 | HNSC | BICR31 | Uprosertib | MVP_top60_DNA+RNA_seed | ESR1 | 0.157262 | KIRC | NCC010 | GDC0810 |
| MVP_top60_DNA_seed | NTRK1 | 0.167522 | HNSC | LB771-HNC | GW441756 | MVP_top60_DNA+RNA_seed | SMARCA4 | 0.15789 | KIRC | BB65-RCC | PFI3 |
| MVP_top60_DNA_seed | AKT1 | 0.168166 | HNSC | BICR22 | AT13148 | MVP_top60_DNA+RNA_seed | SMARCA2 | 0.15789 | KIRC | BB65-RCC | PFI3 |
| MVP_top60_DNA_seed | MAPK1 | 0.168649 | HNSC | HSC-4 | Ulixertinib | MVP_top60_DNA+RNA_seed | ESR1 | 0.157911 | KIRC | SW156 | GDC0810 |
| MVP_top60_DNA_seed | MAPK3 | 0.168649 | HNSC | HSC-4 | Ulixertinib | MVP_top60_DNA+RNA_seed | HSP90AA1 | 0.157954 | KIRC | KMRC-1 | Tanespimycin |
| MVP_top60_DNA_seed | MAPK1 | 0.169146 | HNSC | SCC-15 | SCH772984 | MVP_top60_DNA+RNA_seed | HSP90AB1 | 0.157954 | KIRC | KMRC-1 | Tanespimycin |
| MVP_top60_DNA_seed | MAPK3 | 0.169146 | HNSC | SCC-15 | SCH772984 | MVP_top60_DNA+RNA_seed | IGF1R | 0.159967 | KIRC | RCC-MF | Linsitinib |
| MVP_top60_DNA_seed | CDK2 | 0.17031 | HNSC | SCC-15 | AZD5438 | MVP_top60_DNA+RNA_seed | PIK3CA | 0.161126 | KIRC | KMRC-20 | AZD8186 |
| MVP_top60_DNA_seed | CDK2 | 0.170805 | HNSC | PCI-30 | MK-8776 | MVP_top60_DNA+RNA_seed | IGF1R | 0.166927 | KIRC | 769-P | BMS-754807 |
| MVP_top60_DNA_seed | AKT1 | 0.171088 | HNSC | SAT | MK-2206 | MVP_top60_DNA+RNA_seed | IGF1R | 0.167139 | KIRC | U031 | NVP-ADW742 |
| MVP_top60_DNA_seed | EGFR | 0.175318 | HNSC | OSC-19 | Afatinib | MVP_top60_DNA+RNA_seed | IGF1R | 0.168683 | KIRC | BFTC-909 | BMS-754807 |
| MVP_top60_DNA_seed | CDK2 | 0.175326 | HNSC | OSC-19 | MK-8776 | MVP_top60_DNA+RNA_seed | PIK3CA | 0.171731 | KIRC | 769-P | Buparlisib |
| MVP_top60_DNA_seed | EGFR | 0.175848 | HNSC | BICR22 | Afatinib | MVP_top60_DNA+RNA_seed | PIK3CD | 0.171731 | KIRC | 769-P | Buparlisib |
| MVP_top60_DNA_seed | MAPK1 | 0.176386 | HNSC | SCC-4 | SCH772984 | MVP_top60_DNA+RNA_seed | ESR1 | 0.1727 | KIRC | 769-P | Tamoxifen |
| MVP_top60_DNA_seed | MAPK3 | 0.176386 | HNSC | SCC-4 | SCH772984 | MVP_top60_DNA+RNA_seed | TP53 | 0.175628 | KIRC | HA7-RCC | MIRA-1 |
| MVP_top60_DNA_seed | HDAC1 | 0.180278 | HNSC | BICR22 | Entinostat | MVP_top60_DNA+RNA_seed | PIK3CA | 0.17665 | KIRC | VMRC-RCW | AZD8186 |
| MVP_top60_DNA_seed | HDAC3 | 0.180278 | HNSC | BICR22 | Entinostat | MVP_top60_DNA+RNA_seed | ESR1 | 0.177494 | KIRC | SN12C | Tamoxifen |
| MVP_top60_DNA_seed | HSP90AA1 | 0.183904 | HNSC | OSC-19 | Elesclomol | MVP_top60_DNA+RNA_seed | IGF1R | 0.185537 | KIRC | ACHN | GSK1904529A |
| MVP_top60_DNA_seed | AKT1 | 0.185022 | HNSC | LB771-HNC | Uprosertib | MVP_top60_DNA+RNA_seed | IGF1R | 0.185626 | KIRC | NCC021 | NVP-ADW742 |
| MVP_top60_DNA_seed | PIK3CA | 0.189775 | HNSC | SCC-15 | Alpelisib | MVP_top60_DNA+RNA_seed | PIK3CA | 0.18568 | KIRC | SN12C | AZD8186 |
| MVP_top60_DNA_seed | MAPK1 | 0.190661 | HNSC | BICR22 | VX-11e | MVP_top60_DNA+RNA_seed | PIK3CA | 0.188458 | KIRC | NCC010 | Buparlisib |
| MVP_top60_DNA_seed | MAPK1 | 0.192091 | HNSC | SKN-3 | VX-11e | MVP_top60_DNA+RNA_seed | PIK3CD | 0.188458 | KIRC | NCC010 | Buparlisib |
| MVP_top60_DNA_seed | HDAC1 | 0.193651 | HNSC | HSC-2 | PCI-34051 | MVP_top60_DNA+RNA_seed | HDAC1 | 0.189757 | KIRC | SN12C | Romidepsin |
| MVP_top60_DNA_seed | CDK2 | 0.194097 | HNSC | SKN-3 | AZD5438 | MVP_top60_DNA+RNA_seed | PIK3CA | 0.190648 | KIRC | SN12C | GNE-317 |
| MVP_top60_DNA_seed | ESR1 | 0.194915 | HNSC | SKN-3 | Tamoxifen | MVP_top60_DNA+RNA_seed | PIK3CA | 0.192211 | KIRC | NCC021 | Buparlisib |
| MVP_top60_DNA_seed | HDAC1 | 0.195883 | HNSC | SAT | Entinostat | MVP_top60_DNA+RNA_seed | PIK3CD | 0.192211 | KIRC | NCC021 | Buparlisib |
| MVP_top60_DNA_seed | HDAC3 | 0.195883 | HNSC | SAT | Entinostat | MVP_top60_DNA+RNA_seed | PIK3CA | 0.193138 | KIRC | KMRC-20 | Buparlisib |
| MVP_top60_DNA_seed | HSP90AA1 | 0.196499 | HNSC | KON | Elesclomol | MVP_top60_DNA+RNA_seed | PIK3CD | 0.193138 | KIRC | KMRC-20 | Buparlisib |
| MVP_top60_DNA_seed | NTRK1 | 0.197022 | HNSC | JHU-011 | GW441756 | MVP_top60_DNA+RNA_seed | SMARCA4 | 0.197023 | KIRC | CAL-54 | PFI3 |
| MVP_top60_DNA_seed | EGFR | 0.205221 | HNSC | BB49-HNC | Afatinib | MVP_top60_DNA+RNA_seed | SMARCA2 | 0.197023 | KIRC | CAL-54 | PFI3 |
| MVP_top60_DNA_seed | PIK3CA | 0.205448 | HNSC | BICR31 | AZD8186 | MVP_top60_DNA+RNA_seed | SMARCA4 | 0.197154 | KIRC | RCC-MF | PFI3 |
| MVP_top60_DNA_seed | NTRK1 | 0.205539 | HNSC | JHU-022 | GW441756 | MVP_top60_DNA+RNA_seed | SMARCA2 | 0.197154 | KIRC | RCC-MF | PFI3 |
| MVP_top60_DNA_seed | NTRK1 | 0.211684 | HNSC | FADU | AZD1332 | MVP_top60_DNA+RNA_seed | EP300 | 0.202254 | KIRC | SN12C | CPI-637 |
| MVP_top60_DNA_seed | AKT1 | 0.211901 | HNSC | OSC-19 | AZD5363 | MVP_top60_DNA+RNA_seed | IGF1R | 0.205418 | KIRC | 786-0 | BMS-754807 |
| MVP_top60_DNA_seed | AKT1 | 0.213817 | HNSC | LB771-HNC | Ipatasertib | MVP_top60_DNA+RNA_seed | IGF1R | 0.207612 | KIRC | RCC-AB | GSK1904529A |
| MVP_top60_DNA_seed | AKT1 | 0.219929 | HNSC | SCC90 | Afuresertib | MVP_top60_DNA+RNA_seed | IGF1R | 0.209522 | KIRC | KMRC-20 | GSK1904529A |
| MVP_top60_DNA_seed | AKT1 | 0.223764 | HNSC | SAS | Uprosertib | MVP_top60_DNA+RNA_seed | IGF1R | 0.210616 | KIRC | RCC10RGB | BMS-536924 |

| | | | | | | | | | | | |
|---|---|---|---|---|---|---|---|---|---|---|---|
| MVP_top60_DNA_seed | SRC | 0.224024 | HNSC | SCC-15 | Bosutinib | MVP_top60_DNA+RNA_seed | IGF1R | 0.210986 | KIRC | BFTC-909 | GSK1904529A |
| MVP_top60_DNA_seed | TP53 | 0.225644 | HNSC | Ca9-22 | MIRA-1 | MVP_top60_DNA+RNA_seed | AR | 0.211074 | KIRC | NCC010 | Bicalutamide |
| MVP_top60_DNA_seed | MAPK1 | 0.228542 | HNSC | BICR31 | Ulixertinib | MVP_top60_DNA+RNA_seed | ESR1 | 0.216767 | KIRC | RCC-JW | GDC0810 |
| MVP_top60_DNA_seed | MAPK3 | 0.228542 | HNSC | BICR31 | Ulixertinib | MVP_top60_DNA+RNA_seed | HDAC1 | 0.217962 | KIRC | OS-RC-2 | Entinostat |
| MVP_top60_DNA_seed | AKT1 | 0.230862 | HNSC | BICR31 | Ipatasertib | MVP_top60_DNA+RNA_seed | IGF1R | 0.222724 | KIRC | U031 | Linsitinib |
| MVP_top60_DNA_seed | SMARCA4 | 0.231458 | HNSC | HSC-4 | PFI3 | MVP_top60_DNA+RNA_seed | ESR1 | 0.225203 | KIRC | RCC-JF | GDC0810 |
| MVP_top60_DNA_seed | MDM2 | 0.23342 | HNSC | FADU | Nutlin-3a (-) | MVP_top60_DNA+RNA_seed | TP53 | 0.22885 | KIRC | TK10 | MIRA-1 |
| MVP_top60_DNA_seed | CDK2 | 0.235588 | HNSC | FADU | Dinaciclib | MVP_top60_DNA+RNA_seed | ATM | 0.229116 | KIRC | RXF393 | KU-55933 |
| MVP_top60_DNA_seed | SMARCA4 | 0.236253 | HNSC | OSC-20 | PFI3 | MVP_top60_DNA+RNA_seed | IGF1R | 0.230284 | KIRC | LB996-RCC | Linsitinib |
| MVP_top60_DNA_seed | EP300 | 0.238164 | HNSC | JHU-022 | CPI-637 | MVP_top60_DNA+RNA_seed | AR | 0.236802 | KIRC | LB1047-RCC | Bicalutamide |
| MVP_top60_DNA_seed | HSP90AA1 | 0.241721 | HNSC | JHU-029 | Elesclomol | MVP_top60_DNA+RNA_seed | MTOR | 0.238514 | KIRC | ACHN | Temsirolimus |
| MVP_top60_DNA_seed | PIK3CA | 0.246362 | HNSC | JHU-022 | Buparlisib | MVP_top60_DNA+RNA_seed | HSP90AA1 | 0.240452 | KIRC | NCC021 | CCT-018159 |
| MVP_top60_DNA_seed | EP300 | 0.258044 | HNSC | SKN-3 | CPI-637 | MVP_top60_DNA+RNA_seed | HSP90AB1 | 0.240452 | KIRC | NCC021 | CCT-018159 |
| MVP_top60_DNA_seed | HDAC1 | 0.260042 | HNSC | BICR78 | Entinostat | MVP_top60_DNA+RNA_seed | HSP90AA1 | 0.242824 | KIRC | OS-RC-2 | Tanespimycin |
| MVP_top60_DNA_seed | HDAC3 | 0.260042 | HNSC | BICR78 | Entinostat | MVP_top60_DNA+RNA_seed | HSP90AB1 | 0.242824 | KIRC | OS-RC-2 | Tanespimycin |
| MVP_top60_DNA_seed | MDM2 | 0.26047 | HNSC | HO-1-u-1 | Nutlin-3a (-) | MVP_top60_DNA+RNA_seed | EP300 | 0.24472 | KIRC | A498 | SGC-CBP30 |
| MVP_top60_DNA_seed | MDM2 | 0.261391 | HNSC | BHY | Serdemetan | MVP_top60_DNA+RNA_seed | HSP90AA1 | 0.245388 | KIRC | U031 | Luminespib |
| MVP_top60_DNA_seed | PIK3CA | 0.266267 | HNSC | HSC-3 | AZD8186 | MVP_top60_DNA+RNA_seed | HSP90AB1 | 0.245388 | KIRC | U031 | Luminespib |
| MVP_top60_DNA_seed | HSP90AA1 | 0.266882 | HNSC | BHY | Tanespimycin | MVP_top60_DNA+RNA_seed | IGF1R | 0.258351 | KIRC | RCC-JF | GSK1904529A |
| MVP_top60_DNA_seed | AKT1 | 0.269873 | HNSC | PE-CA-PJ15 | AT13148 | MVP_top60_DNA+RNA_seed | TP53 | 0.258536 | KIRC | BB65-RCC | MIRA-1 |
| MVP_top60_DNA_seed | EP300 | 0.270755 | HNSC | CAL-27 | CPI-637 | MVP_top60_DNA+RNA_seed | PIK3CA | 0.259897 | KIRC | RXF393 | Alpelisib |
| MVP_top60_DNA_seed | MDM2 | 0.271056 | HNSC | HSC-4 | Nutlin-3a (-) | MVP_top60_DNA+RNA_seed | AR | 0.262855 | KIRC | SW156 | Bicalutamide |
| MVP_top60_DNA_seed | PIK3CA | 0.27247 | HNSC | BHY | Alpelisib | MVP_top60_DNA+RNA_seed | HDAC1 | 0.263668 | KIRC | LB1047-RCC | PCI-34051 |
| MVP_top60_DNA_seed | PIK3CA | 0.27338 | HNSC | PCI-30 | AZD8186 | MVP_top60_DNA+RNA_seed | IGF1R | 0.271211 | KIRC | 786-0 | BMS-536924 |
| MVP_top60_DNA_seed | EP300 | 0.273557 | HNSC | BICR10 | CPI-637 | MVP_top60_DNA+RNA_seed | IGF1R | 0.271785 | KIRC | A498 | BMS-754807 |
| MVP_top60_DNA_seed | PIK3CA | 0.274282 | HNSC | PCI-6A | AZD8186 | MVP_top60_DNA+RNA_seed | MTOR | 0.276441 | KIRC | RCC-AB | Temsirolimus |
| MVP_top60_DNA_seed | AKT1 | 0.274657 | HNSC | HO-1-N-1 | Afuresertib | MVP_top60_DNA+RNA_seed | PIK3CA | 0.285639 | KIRC | KMRC-20 | GNE-317 |
| MVP_top60_DNA_seed | PIK3CA | 0.275438 | HNSC | CAL-27 | GNE-317 | MVP_top60_DNA+RNA_seed | IGF1R | 0.296533 | KIRC | RCC-MF | BMS-754807 |
| MVP_top60_DNA_seed | HSP90AA1 | 0.275764 | HNSC | JHU-011 | Elesclomol | MVP_top60_DNA+RNA_seed | IGF1R | 0.298273 | KIRC | LB1047-RCC | GSK1904529A |
| MVP_top60_DNA_seed | PIK3CA | 0.279707 | HNSC | FADU | GNE-317 | MVP_top60_DNA+RNA_seed | HDAC1 | 0.299265 | KIRC | NCC010 | Entinostat |
| MVP_top60_DNA_seed | SMARCA4 | 0.282343 | HNSC | SCC-25 | PFI3 | MVP_top60_DNA+RNA_seed | EP300 | 0.3007 | KIRC | KMRC-20 | CPI-637 |
| MVP_top60_DNA_seed | TP53 | 0.283242 | HNSC | Detroit562 | MIRA-1 | MVP_top60_DNA+RNA_seed | HDAC1 | 0.301587 | KIRC | LB996-RCC | Entinostat |
| MVP_top60_DNA_seed | AKT1 | 0.284005 | HNSC | PE-CA-PJ15 | Afuresertib | MVP_top60_DNA+RNA_seed | TP53 | 0.304421 | KIRC | CAL-54 | MIRA-1 |
| MVP_top60_DNA_seed | ESR1 | 0.285434 | HNSC | BICR78 | GDC0810 | MVP_top60_DNA+RNA_seed | TP53 | 0.307187 | KIRC | 786-0 | MIRA-1 |
| MVP_top60_DNA_seed | EP300 | 0.288131 | HNSC | SCC-4 | CPI-637 | MVP_top60_DNA+RNA_seed | PIK3CA | 0.30787 | KIRC | 769-P | Alpelisib |
| MVP_top60_DNA_seed | EGFR | 0.288412 | HNSC | SCC-25 | Erlotinib | MVP_top60_DNA+RNA_seed | IGF1R | 0.311208 | KIRC | KMRC-20 | BMS-754807 |
| MVP_top60_DNA_seed | MDM2 | 0.291201 | HNSC | SAS | Nutlin-3a (-) | MVP_top60_DNA+RNA_seed | HDAC1 | 0.31588 | KIRC | BFTC-909 | Entinostat |
| MVP_top60_DNA_seed | AKT1 | 0.29201 | HNSC | SAS | MK-2206 | MVP_top60_DNA+RNA_seed | MTOR | 0.316734 | KIRC | KMRC-1 | Temsirolimus |
| MVP_top60_DNA_seed | HSP90AA1 | 0.293424 | HNSC | HSC-4 | Luminespib | MVP_top60_DNA+RNA_seed | PIK3CA | 0.322928 | KIRC | NCC010 | GNE-317 |
| MVP_top60_DNA_seed | EGFR | 0.293669 | HNSC | BICR22 | Sapitinib | MVP_top60_DNA+RNA_seed | ATM | 0.325635 | KIRC | ACHN | KU-55933 |
| MVP_top60_DNA_seed | CDK2 | 0.297603 | HNSC | JHU-022 | MK-8776 | MVP_top60_DNA+RNA_seed | IGF1R | 0.32936 | KIRC | RCC-MF | NVP-ADW742 |

| | | | | | | | | | | | |
|---|---|---|---|---|---|---|---|---|---|---|---|
| MVP_top60_DNA_seed | AKT1 | 0.299091 | HNSC | CAL-33 | MK-2206 | MVP_top60_DNA+RNA_seed | IGF1R | 0.333165 | KIRC | RCC10RGB | NVP-ADW742 |
| MVP_top60_DNA_seed | NTRK1 | 0.300273 | HNSC | SAT | AZD1332 | MVP_top60_DNA+RNA_seed | IGF1R | 0.339529 | KIRC | RCC-JW | BMS-536924 |
| MVP_top60_DNA_seed | NTRK1 | 0.301375 | HNSC | PCI-6A | GW441756 | MVP_top60_DNA+RNA_seed | MTOR | 0.342166 | KIRC | NCC010 | Temsirolimus |
| MVP_top60_DNA_seed | AKT1 | 0.302331 | HNSC | PE-CA-PJ15 | MK-2206 | MVP_top60_DNA+RNA_seed | HSP90AA1 | 0.345824 | KIRC | 769-P | Luminespib |
| MVP_top60_DNA_seed | AKT1 | 0.303573 | HNSC | PCI-6A | MK-2206 | MVP_top60_DNA+RNA_seed | HSP90AB1 | 0.345824 | KIRC | 769-P | Luminespib |
| MVP_top60_DNA_seed | CDK2 | 0.303901 | HNSC | BICR10 | Dinaciclib | MVP_top60_DNA+RNA_seed | HSP90AA1 | 0.348258 | KIRC | LB2241-RCC | Luminespib |
| MVP_top60_DNA_seed | EGFR | 0.307039 | HNSC | OSC-19 | Erlotinib | MVP_top60_DNA+RNA_seed | HSP90AB1 | 0.348258 | KIRC | LB2241-RCC | Luminespib |
| MVP_top60_DNA_seed | ESR1 | 0.307556 | HNSC | PCI-6A | GDC0810 | MVP_top60_DNA+RNA_seed | HSP90AA1 | 0.349726 | KIRC | RCC-AB | Luminespib |
| MVP_top60_DNA_seed | MAPK1 | 0.307784 | HNSC | SKN-3 | Ulixertinib | MVP_top60_DNA+RNA_seed | HSP90AB1 | 0.349726 | KIRC | RCC-AB | Luminespib |
| MVP_top60_DNA_seed | MAPK3 | 0.307784 | HNSC | SKN-3 | Ulixertinib | MVP_top60_DNA+RNA_seed | HSP90AA1 | 0.351779 | KIRC | SN12C | Luminespib |
| MVP_top60_DNA_seed | MDM2 | 0.311357 | HNSC | HSC-3 | Nutlin-3a (-) | MVP_top60_DNA+RNA_seed | HSP90AB1 | 0.351779 | KIRC | SN12C | Luminespib |
| MVP_top60_DNA_seed | AKT1 | 0.312126 | HNSC | BICR78 | Afuresertib | MVP_top60_DNA+RNA_seed | ATM | 0.352015 | KIRC | BFTC-909 | KU-55933 |
| MVP_top60_DNA_seed | CDK2 | 0.315639 | HNSC | BB49-HNC | Dinaciclib | MVP_top60_DNA+RNA_seed | HDAC1 | 0.353183 | KIRC | RXF393 | Entinostat |
| MVP_top60_DNA_seed | TP53 | 0.317842 | HNSC | CAL-27 | MIRA-1 | MVP_top60_DNA+RNA_seed | HSP90AA1 | 0.355254 | KIRC | RCC-ER | Tanespimycin |
| MVP_top60_DNA_seed | MDM2 | 0.319317 | HNSC | CAL-27 | Serdemetan | MVP_top60_DNA+RNA_seed | HSP90AB1 | 0.355254 | KIRC | RCC-ER | Tanespimycin |
| MVP_top60_DNA_seed | CDK2 | 0.320903 | HNSC | Detroit562 | MK-8776 | MVP_top60_DNA+RNA_seed | HDAC1 | 0.355568 | KIRC | RCC10RGB | PCI-34051 |
| MVP_top60_DNA_seed | EP300 | 0.323972 | HNSC | HO-1-u-1 | CPI-637 | MVP_top60_DNA+RNA_seed | PIK3CA | 0.359376 | KIRC | OS-RC-2 | GNE-317 |
| MVP_top60_DNA_seed | MAPK1 | 0.325161 | HNSC | PE-CA-PJ15 | Ulixertinib | MVP_top60_DNA+RNA_seed | MTOR | 0.363112 | KIRC | CAKI-1 | Temsirolimus |
| MVP_top60_DNA_seed | MAPK3 | 0.325161 | HNSC | PE-CA-PJ15 | Ulixertinib | MVP_top60_DNA+RNA_seed | HSP90AA1 | 0.364493 | KIRC | RCC-MF | Elesclomol |
| MVP_top60_DNA_seed | HDAC1 | 0.328084 | HNSC | BICR31 | PCI-34051 | MVP_top60_DNA+RNA_seed | HSP90AB1 | 0.364493 | KIRC | RCC-MF | Elesclomol |
| MVP_top60_DNA_seed | CDK2 | 0.332192 | HNSC | SKN-3 | Dinaciclib | MVP_top60_DNA+RNA_seed | IGF1R | 0.364926 | KIRC | RXF393 | GSK1904529A |
| MVP_top60_DNA_seed | CDK2 | 0.333608 | HNSC | Ca9-22 | MK-8776 | MVP_top60_DNA+RNA_seed | HDAC1 | 0.366339 | KIRC | ACHN | PCI-34051 |
| MVP_top60_DNA_seed | NTRK1 | 0.337551 | HNSC | SCC-25 | GW441756 | MVP_top60_DNA+RNA_seed | HDAC1 | 0.366415 | KIRC | NCC021 | Entinostat |
| MVP_top60_DNA_seed | PIK3CA | 0.338833 | HNSC | PCI-6A | GNE-317 | MVP_top60_DNA+RNA_seed | ESR1 | 0.367631 | KIRC | CAL-54 | GDC0810 |
| MVP_top60_DNA_seed | MAPK1 | 0.338928 | HNSC | HO-1-u-1 | Ulixertinib | MVP_top60_DNA+RNA_seed | HSP90AA1 | 0.377435 | KIRC | NCC021 | Elesclomol |
| MVP_top60_DNA_seed | MAPK3 | 0.338928 | HNSC | HO-1-u-1 | Ulixertinib | MVP_top60_DNA+RNA_seed | HSP90AB1 | 0.377435 | KIRC | NCC021 | Elesclomol |
| MVP_top60_DNA_seed | MAPK1 | 0.340516 | HNSC | HSC-4 | Ulixertinib | MVP_top60_DNA+RNA_seed | ESR1 | 0.38195 | KIRC | RXF393 | GDC0810 |
| MVP_top60_DNA_seed | MAPK3 | 0.340516 | HNSC | HSC-4 | Ulixertinib | MVP_top60_DNA+RNA_seed | SMARCA4 | 0.387165 | KIRC | KMRC-20 | PFI3 |
| MVP_top60_DNA_seed | MAPK1 | 0.344101 | HNSC | HSC-2 | Ulixertinib | MVP_top60_DNA+RNA_seed | SMARCA2 | 0.387165 | KIRC | KMRC-20 | PFI3 |
| MVP_top60_DNA_seed | MAPK3 | 0.344101 | HNSC | HSC-2 | Ulixertinib | MVP_top60_DNA+RNA_seed | EP300 | 0.388821 | KIRC | BFTC-909 | SGC-CBP30 |
| MVP_top60_DNA_seed | AKT1 | 0.346917 | HNSC | PCI-4B | AZD5363 | MVP_top60_DNA+RNA_seed | IGF1R | 0.39041 | KIRC | KMRC-1 | BMS-536924 |
| MVP_top60_DNA_seed | AKT1 | 0.347425 | HNSC | SCC-15 | Afuresertib | MVP_top60_DNA+RNA_seed | HDAC1 | 0.40033 | KIRC | BB65-RCC | Entinostat |
| MVP_top60_DNA_seed | AKT1 | 0.34986 | HNSC | PE-CA-PJ15 | Ipatasertib | MVP_top60_DNA+RNA_seed | ESR1 | 0.401056 | KIRC | OS-RC-2 | GDC0810 |
| MVP_top60_DNA_seed | ESR1 | 0.351352 | HNSC | SCC-15 | GDC0810 | MVP_top60_DNA+RNA_seed | TP53 | 0.403485 | KIRC | SW156 | MIRA-1 |
| MVP_top60_DNA_seed | NTRK1 | 0.351835 | HNSC | BICR22 | GW441756 | MVP_top60_DNA+RNA_seed | ESR1 | 0.403867 | KIRC | ACHN | GDC0810 |
| MVP_top60_DNA_seed | RAC1 | 0.352332 | HNSC | PCI-15A | EHT-1864 | MVP_top60_DNA+RNA_seed | HSP90AA1 | 0.403962 | KIRC | LB1047-RCC | Tanespimycin |
| MVP_top60_DNA_seed | PIK3CA | 0.35721 | HNSC | LB771-HNC | GNE-317 | MVP_top60_DNA+RNA_seed | HSP90AB1 | 0.403962 | KIRC | LB1047-RCC | Tanespimycin |
| MVP_top60_DNA_seed | EGFR | 0.358587 | HNSC | HSC-4 | Afatinib | MVP_top60_DNA+RNA_seed | SMARCA4 | 0.40752 | KIRC | RCC-FG2 | PFI3 |
| MVP_top60_DNA_seed | MAPK1 | 0.359045 | HNSC | HSC-2 | SCH772984 | MVP_top60_DNA+RNA_seed | SMARCA2 | 0.40752 | KIRC | RCC-FG2 | PFI3 |
| MVP_top60_DNA_seed | MAPK3 | 0.359045 | HNSC | HSC-2 | SCH772984 | MVP_top60_DNA+RNA_seed | MTOR | 0.415946 | KIRC | RCC10RGB | Temsirolimus |
| MVP_top60_DNA_seed | EGFR | 0.359154 | HNSC | BICR78 | Afatinib | MVP_top60_DNA+RNA_seed | IGF1R | 0.419324 | KIRC | HA7-RCC | BMS-754807 |

| | | | | | | | | | | | |
|---|---|---|---|---|---|---|---|---|---|---|---|
| MVP_top60_DNA_seed | NTRK1 | 0.359226 | HNSC | Ca9-22 | GW441756 | MVP_top60_DNA+RNA_seed | ATM | 0.427488 | KIRC | U031 | KU-55933 |
| MVP_top60_DNA_seed | AKT1 | 0.359367 | HNSC | SCC-9 | MK-2206 | MVP_top60_DNA+RNA_seed | PIK3CA | 0.427528 | KIRC | SN12C | Buparlisib |
| MVP_top60_DNA_seed | HDAC1 | 0.364315 | HNSC | LB771-HNC | Romidepsin | MVP_top60_DNA+RNA_seed | PIK3CD | 0.427528 | KIRC | SN12C | Buparlisib |
| MVP_top60_DNA_seed | HDAC3 | 0.364315 | HNSC | LB771-HNC | Romidepsin | MVP_top60_DNA+RNA_seed | IGF1R | 0.433283 | KIRC | HA7-RCC | NVP-ADW742 |
| MVP_top60_DNA_seed | MAPK1 | 0.36466 | HNSC | BICR10 | Ulixertinib | MVP_top60_DNA+RNA_seed | HSP90AA1 | 0.438647 | KIRC | KMRC-1 | Luminespib |
| MVP_top60_DNA_seed | MAPK3 | 0.36466 | HNSC | BICR10 | Ulixertinib | MVP_top60_DNA+RNA_seed | HSP90AB1 | 0.438647 | KIRC | KMRC-1 | Luminespib |
| MVP_top60_DNA_seed | MAPK1 | 0.364839 | HNSC | CAL-33 | VX-11e | MVP_top60_DNA+RNA_seed | PIK3CA | 0.439711 | KIRC | BB65-RCC | AZD8186 |
| MVP_top60_DNA_seed | PIK3CA | 0.365457 | HNSC | Ca9-22 | GNE-317 | MVP_top60_DNA+RNA_seed | IGF1R | 0.441179 | KIRC | SW156 | BMS-754807 |
| MVP_top60_DNA_seed | NTRK1 | 0.367598 | HNSC | FADU | GW441756 | MVP_top60_DNA+RNA_seed | HSP90AA1 | 0.446857 | KIRC | RCC-JW | Luminespib |
| MVP_top60_DNA_seed | NTRK1 | 0.370521 | HNSC | SCC-4 | AZD1332 | MVP_top60_DNA+RNA_seed | HSP90AB1 | 0.446857 | KIRC | RCC-JW | Luminespib |
| MVP_top60_DNA_seed | AKT1 | 0.370889 | HNSC | SCC90 | Ipatasertib | MVP_top60_DNA+RNA_seed | ESR1 | 0.447783 | KIRC | RCC-JW | Tamoxifen |
| MVP_top60_DNA_seed | TP53 | 0.371511 | HNSC | PCI-30 | MIRA-1 | MVP_top60_DNA+RNA_seed | IGF1R | 0.448653 | KIRC | RCC10RGB | Linsitinib |
| MVP_top60_DNA_seed | NTRK1 | 0.372239 | HNSC | JHU-022 | AZD1332 | MVP_top60_DNA+RNA_seed | HDAC1 | 0.465005 | KIRC | KMRC-20 | PCI-34051 |
| MVP_top60_DNA_seed | SRC | 0.373603 | HNSC | BICR78 | Bosutinib | MVP_top60_DNA+RNA_seed | HDAC1 | 0.467398 | KIRC | A704 | Entinostat |
| MVP_top60_DNA_seed | MDM2 | 0.374993 | HNSC | Detroit562 | Nutlin-3a (-) | MVP_top60_DNA+RNA_seed | PIK3CA | 0.469685 | KIRC | RCC-JW | Alpelisib |
| MVP_top60_DNA_seed | HDAC1 | 0.382075 | HNSC | BICR31 | Entinostat | MVP_top60_DNA+RNA_seed | ESR1 | 0.4738 | KIRC | RCC-MF | GDC0810 |
| MVP_top60_DNA_seed | HDAC3 | 0.382075 | HNSC | BICR31 | Entinostat | MVP_top60_DNA+RNA_seed | EP300 | 0.480629 | KIRC | LB996-RCC | SGC-CBP30 |
| MVP_top60_DNA_seed | SMARCA4 | 0.383314 | HNSC | PE-CA-PJ15 | PFI3 | MVP_top60_DNA+RNA_seed | PIK3CA | 0.481959 | KIRC | LB2241-RCC | Buparlisib |
| MVP_top60_DNA_seed | NTRK1 | 0.385665 | HNSC | CAL-27 | GW441756 | MVP_top60_DNA+RNA_seed | PIK3CD | 0.481959 | KIRC | LB2241-RCC | Buparlisib |
| MVP_top60_DNA_seed | AKT1 | 0.388687 | HNSC | OSC-20 | Uprosertib | MVP_top60_DNA+RNA_seed | ESR1 | 0.48518 | KIRC | KMRC-20 | GDC0810 |
| MVP_top60_DNA_seed | PIK3CA | 0.388751 | HNSC | PCI-4B | Buparlisib | MVP_top60_DNA+RNA_seed | ESR1 | 0.490899 | KIRC | BB65-RCC | GDC0810 |
| MVP_top60_DNA_seed | MAPK1 | 0.38925 | HNSC | SCC-15 | VX-11e | MVP_top60_DNA+RNA_seed | ESR1 | 0.49772 | KIRC | KMRC-1 | GDC0810 |
| MVP_top60_DNA_seed | PIK3CA | 0.389747 | HNSC | PCI-30 | Alpelisib | MVP_top60_DNA+RNA_seed | PIK3CA | 0.498048 | KIRC | SW156 | Alpelisib |
| MVP_top60_DNA_seed | MDM2 | 0.394123 | HNSC | HSC-2 | Nutlin-3a (-) | MVP_top60_DNA+RNA_seed | EP300 | 0.498388 | KIRC | KMRC-1 | CPI-637 |
| MVP_top60_DNA_seed | MDM2 | 0.397231 | HNSC | BHY | Nutlin-3a (-) | MVP_top60_DNA+RNA_seed | TP53 | 0.50176 | KIRC | OS-RC-2 | MIRA-1 |
| MVP_top60_DNA_seed | PIK3CA | 0.39866 | HNSC | BICR78 | Alpelisib | MVP_top60_DNA+RNA_seed | EP300 | 0.511925 | KIRC | 786-0 | SGC-CBP30 |
| MVP_top60_DNA_seed | AKT1 | 0.400849 | HNSC | LB771-HNC | Afuresertib | MVP_top60_DNA+RNA_seed | EP300 | 0.514848 | KIRC | RXF393 | SGC-CBP30 |
| MVP_top60_DNA_seed | PIK3CA | 0.402136 | HNSC | BICR22 | AZD8186 | MVP_top60_DNA+RNA_seed | HSP90AA1 | 0.51719 | KIRC | A704 | Luminespib |
| MVP_top60_DNA_seed | EP300 | 0.410584 | HNSC | BICR10 | SGC-CBP30 | MVP_top60_DNA+RNA_seed | HSP90AB1 | 0.51719 | KIRC | A704 | Luminespib |
| MVP_top60_DNA_seed | CREBBP | 0.410584 | HNSC | BICR10 | SGC-CBP30 | MVP_top60_DNA+RNA_seed | HDAC1 | 0.520097 | KIRC | 786-0 | Romidepsin |
| MVP_top60_DNA_seed | TP53 | 0.411201 | HNSC | SCC-9 | MIRA-1 | MVP_top60_DNA+RNA_seed | HSP90AA1 | 0.520841 | KIRC | RCC-MF | CCT-018159 |
| MVP_top60_DNA_seed | ESR1 | 0.411897 | HNSC | SKN-3 | GDC0810 | MVP_top60_DNA+RNA_seed | HSP90AB1 | 0.520841 | KIRC | RCC-MF | CCT-018159 |
| MVP_top60_DNA_seed | PIK3CA | 0.412275 | HNSC | BICR31 | Buparlisib | MVP_top60_DNA+RNA_seed | PIK3CA | 0.521165 | KIRC | LB1047-RCC | Buparlisib |
| MVP_top60_DNA_seed | HSP90AA1 | 0.416118 | HNSC | HSC-4 | Tanespimycin | MVP_top60_DNA+RNA_seed | PIK3CD | 0.521165 | KIRC | LB1047-RCC | Buparlisib |
| MVP_top60_DNA_seed | EGFR | 0.417643 | HNSC | BICR31 | Sapitinib | MVP_top60_DNA+RNA_seed | IGF1R | 0.521406 | KIRC | VMRC-RCW | NVP-ADW742 |
| MVP_top60_DNA_seed | MAPK1 | 0.418088 | HNSC | BICR10 | SCH772984 | MVP_top60_DNA+RNA_seed | PIK3CA | 0.526806 | KIRC | A704 | GNE-317 |
| MVP_top60_DNA_seed | MAPK3 | 0.418088 | HNSC | BICR10 | SCH772984 | MVP_top60_DNA+RNA_seed | IGF1R | 0.528916 | KIRC | VMRC-RCW | BMS-536924 |
| MVP_top60_DNA_seed | EGFR | 0.419483 | HNSC | BICR10 | Erlotinib | MVP_top60_DNA+RNA_seed | IGF1R | 0.536162 | KIRC | OS-RC-2 | GSK1904529A |
| MVP_top60_DNA_seed | AKT1 | 0.420459 | HNSC | LB771-HNC | AZD5363 | MVP_top60_DNA+RNA_seed | IGF1R | 0.537402 | KIRC | 769-P | Linsitinib |
| MVP_top60_DNA_seed | EGFR | 0.420559 | HNSC | BICR10 | Afatinib | MVP_top60_DNA+RNA_seed | PIK3CA | 0.548043 | KIRC | LB996-RCC | Alpelisib |
| MVP_top60_DNA_seed | MAPK1 | 0.421642 | HNSC | BICR22 | Ulixertinib | MVP_top60_DNA+RNA_seed | IGF1R | 0.550424 | KIRC | RXF393 | BMS-754807 |

| | | | | | | | | | | | |
|---|---|---|---|---|---|---|---|---|---|---|---|
| MVP_top60_DNA_seed | MAPK3 | 0.421642 | HNSC | BICR22 | Ulixertinib | MVP_top60_DNA+RNA_seed | HSP90AA1 | 0.552355 | KIRC | BB65-RCC | Elesclomol |
| MVP_top60_DNA_seed | MDM2 | 0.422419 | HNSC | Ca9-22 | Nutlin-3a (-) | MVP_top60_DNA+RNA_seed | HSP90AB1 | 0.552355 | KIRC | BB65-RCC | Elesclomol |
| MVP_top60_DNA_seed | AKT1 | 0.423037 | HNSC | SCC-15 | Uprosertib | MVP_top60_DNA+RNA_seed | TP53 | 0.553529 | KIRC | RXF393 | MIRA-1 |
| MVP_top60_DNA_seed | NTRK1 | 0.431927 | HNSC | SAS | AZD1332 | MVP_top60_DNA+RNA_seed | IGF1R | 0.556082 | KIRC | NCC021 | GSK1904529A |
| MVP_top60_DNA_seed | AKT1 | 0.435461 | HNSC | CAL-27 | AT13148 | MVP_top60_DNA+RNA_seed | IGF1R | 0.557377 | KIRC | A498 | GSK1904529A |
| MVP_top60_DNA_seed | EP300 | 0.435589 | HNSC | HSC-4 | SGC-CBP30 | MVP_top60_DNA+RNA_seed | MTOR | 0.560461 | KIRC | 786-0 | Temsirolimus |
| MVP_top60_DNA_seed | CREBBP | 0.435589 | HNSC | HSC-4 | SGC-CBP30 | MVP_top60_DNA+RNA_seed | IGF1R | 0.565225 | KIRC | HA7-RCC | Linsitinib |
| MVP_top60_DNA_seed | AR | 0.43845 | HNSC | JHU-029 | Bicalutamide | MVP_top60_DNA+RNA_seed | IGF1R | 0.565473 | KIRC | OS-RC-2 | Linsitinib |
| MVP_top60_DNA_seed | PIK3CA | 0.438892 | HNSC | KOSC-2 | GNE-317 | MVP_top60_DNA+RNA_seed | PIK3CA | 0.565548 | KIRC | RCC-AB | Buparlisib |
| MVP_top60_DNA_seed | AKT1 | 0.439082 | HNSC | HSC-3 | GSK2110183B | MVP_top60_DNA+RNA_seed | PIK3CD | 0.565548 | KIRC | RCC-AB | Buparlisib |
| MVP_top60_DNA_seed | AKT1 | 0.440009 | HNSC | PCI-30 | Afuresertib | MVP_top60_DNA+RNA_seed | PIK3CA | 0.574285 | KIRC | TK10 | GNE-317 |
| MVP_top60_DNA_seed | CDK2 | 0.441219 | HNSC | KOSC-2 | MK-8776 | MVP_top60_DNA+RNA_seed | AR | 0.575036 | KIRC | RCC-FG2 | Bicalutamide |
| MVP_top60_DNA_seed | ESR1 | 0.441309 | HNSC | OSC-19 | GDC0810 | MVP_top60_DNA+RNA_seed | HDAC1 | 0.578521 | KIRC | HA7-RCC | Entinostat |
| MVP_top60_DNA_seed | MDM2 | 0.442806 | HNSC | SKN-3 | Nutlin-3a (-) | MVP_top60_DNA+RNA_seed | MTOR | 0.579023 | KIRC | LB1047-RCC | Temsirolimus |
| MVP_top60_DNA_seed | MAPK1 | 0.44463 | HNSC | LB771-HNC | VX-11e | MVP_top60_DNA+RNA_seed | PIK3CA | 0.580145 | KIRC | RCC-JW | Buparlisib |
| MVP_top60_DNA_seed | ESR1 | 0.44568 | HNSC | SCC-9 | Tamoxifen | MVP_top60_DNA+RNA_seed | PIK3CD | 0.580145 | KIRC | RCC-JW | Buparlisib |
| MVP_top60_DNA_seed | SMARCA4 | 0.445691 | HNSC | JHU-022 | PFI3 | MVP_top60_DNA+RNA_seed | IGF1R | 0.584229 | KIRC | RCC-FG2 | BMS-754807 |
| MVP_top60_DNA_seed | ESR1 | 0.446435 | HNSC | KOSC-2 | Tamoxifen | MVP_top60_DNA+RNA_seed | PIK3CA | 0.587686 | KIRC | OS-RC-2 | Alpelisib |
| MVP_top60_DNA_seed | HSP90AA1 | 0.447444 | HNSC | CAL-33 | Tanespimycin | MVP_top60_DNA+RNA_seed | PIK3CA | 0.590619 | KIRC | RCC-JW | GNE-317 |
| MVP_top60_DNA_seed | AKT1 | 0.44987 | HNSC | HSC-2 | AT13148 | MVP_top60_DNA+RNA_seed | MTOR | 0.59116 | KIRC | KMRC-20 | Temsirolimus |
| MVP_top60_DNA_seed | NTRK1 | 0.451112 | HNSC | PE-CA-PJ15 | GW441756 | MVP_top60_DNA+RNA_seed | ATM | 0.59152 | KIRC | 786-0 | KU-55933 |
| MVP_top60_DNA_seed | AKT1 | 0.451214 | HNSC | BHY | AT13148 | MVP_top60_DNA+RNA_seed | PIK3CA | 0.592076 | KIRC | CAKI-1 | Buparlisib |
| MVP_top60_DNA_seed | SMARCA4 | 0.452913 | HNSC | SCC-9 | PFI3 | MVP_top60_DNA+RNA_seed | PIK3CD | 0.592076 | KIRC | CAKI-1 | Buparlisib |
| MVP_top60_DNA_seed | MDM2 | 0.457752 | HNSC | SCC-4 | Nutlin-3a (-) | MVP_top60_DNA+RNA_seed | HDAC1 | 0.595867 | KIRC | LB996-RCC | Romidepsin |
| MVP_top60_DNA_seed | PIK3CA | 0.461071 | HNSC | SCC-4 | Buparlisib | MVP_top60_DNA+RNA_seed | HDAC1 | 0.596364 | KIRC | LB2241-RCC | Entinostat |
| MVP_top60_DNA_seed | EP300 | 0.461192 | HNSC | SKN-3 | SGC-CBP30 | MVP_top60_DNA+RNA_seed | PIK3CA | 0.59711 | KIRC | KMRC-1 | Buparlisib |
| MVP_top60_DNA_seed | CREBBP | 0.461192 | HNSC | SKN-3 | SGC-CBP30 | MVP_top60_DNA+RNA_seed | PIK3CD | 0.59711 | KIRC | KMRC-1 | Buparlisib |
| MVP_top60_DNA_seed | EGFR | 0.462138 | HNSC | SCC-4 | Gefitinib | MVP_top60_DNA+RNA_seed | MTOR | 0.599933 | KIRC | A704 | Temsirolimus |
| MVP_top60_DNA_seed | EP300 | 0.462448 | HNSC | OSC-20 | SGC-CBP30 | MVP_top60_DNA+RNA_seed | MTOR | 0.600924 | KIRC | NCC021 | Temsirolimus |
| MVP_top60_DNA_seed | CREBBP | 0.462448 | HNSC | OSC-20 | SGC-CBP30 | MVP_top60_DNA+RNA_seed | HDAC1 | 0.603041 | KIRC | LB996-RCC | PCI-34051 |
| MVP_top60_DNA_seed | TP53 | 0.463248 | HNSC | OSC-20 | MIRA-1 | MVP_top60_DNA+RNA_seed | IGF1R | 0.609547 | KIRC | BB65-RCC | Linsitinib |
| MVP_top60_DNA_seed | MAPK1 | 0.464205 | HNSC | FADU | VX-11e | MVP_top60_DNA+RNA_seed | IGF1R | 0.609641 | KIRC | ACHN | BMS-754807 |
| MVP_top60_DNA_seed | MAPK1 | 0.466475 | HNSC | SCC-4 | VX-11e | MVP_top60_DNA+RNA_seed | IGF1R | 0.609994 | KIRC | CAL-54 | GSK1904529A |
| MVP_top60_DNA_seed | MAPK1 | 0.466541 | HNSC | HSC-3 | Ulixertinib | MVP_top60_DNA+RNA_seed | SMARCA4 | 0.612045 | KIRC | U031 | PFI3 |
| MVP_top60_DNA_seed | MAPK3 | 0.466541 | HNSC | HSC-3 | Ulixertinib | MVP_top60_DNA+RNA_seed | SMARCA2 | 0.612045 | KIRC | U031 | PFI3 |
| MVP_top60_DNA_seed | MDM2 | 0.46892 | HNSC | CAL-27 | Nutlin-3a (-) | MVP_top60_DNA+RNA_seed | HSP90AA1 | 0.621203 | KIRC | NCC010 | Elesclomol |
| MVP_top60_DNA_seed | AKT1 | 0.469549 | HNSC | FADU | GSK2110183B | MVP_top60_DNA+RNA_seed | HSP90AB1 | 0.621203 | KIRC | NCC010 | Elesclomol |
| MVP_top60_DNA_seed | EGFR | 0.47045 | HNSC | PCI-30 | Erlotinib | MVP_top60_DNA+RNA_seed | HDAC1 | 0.63027 | KIRC | LB2241-RCC | PCI-34051 |
| MVP_top60_DNA_seed | AKT1 | 0.476357 | HNSC | CAL-27 | Uprosertib | MVP_top60_DNA+RNA_seed | IGF1R | 0.630389 | KIRC | 769-P | BMS-536924 |
| MVP_top60_DNA_seed | EP300 | 0.476849 | HNSC | SCC-15 | SGC-CBP30 | MVP_top60_DNA+RNA_seed | HDAC1 | 0.63155 | KIRC | CAL-54 | PCI-34051 |
| MVP_top60_DNA_seed | CREBBP | 0.476849 | HNSC | SCC-15 | SGC-CBP30 | MVP_top60_DNA+RNA_seed | PIK3CA | 0.631708 | KIRC | HA7-RCC | Buparlisib |

| | | | | | | | | | | | |
|---|---|---|---|---|---|---|---|---|---|---|---|
| MVP_top60_DNA_seed | AKT1 | 0.482261 | HNSC | BICR78 | AZD5363 | MVP_top60_DNA+RNA_seed | PIK3CD | 0.631708 | KIRC | HA7-RCC | Buparlisib |
| MVP_top60_DNA_seed | TP53 | 0.483098 | HNSC | SCC-15 | MIRA-1 | MVP_top60_DNA+RNA_seed | TP53 | 0.632368 | KIRC | A704 | MIRA-1 |
| MVP_top60_DNA_seed | PIK3CA | 0.483358 | HNSC | OSC-20 | Buparlisib | MVP_top60_DNA+RNA_seed | HDAC1 | 0.632386 | KIRC | HA7-RCC | Romidepsin |
| MVP_top60_DNA_seed | CDK2 | 0.487713 | HNSC | JHU-022 | AZD5438 | MVP_top60_DNA+RNA_seed | ESR1 | 0.634383 | KIRC | BB65-RCC | Tamoxifen |
| MVP_top60_DNA_seed | NTRK1 | 0.488906 | HNSC | OSC-19 | GW441756 | MVP_top60_DNA+RNA_seed | EP300 | 0.636231 | KIRC | A704 | CPI-637 |
| MVP_top60_DNA_seed | HDAC1 | 0.491511 | HNSC | CAL-27 | PCI-34051 | MVP_top60_DNA+RNA_seed | HSP90AA1 | 0.636838 | KIRC | SN12C | Tanespimycin |
| MVP_top60_DNA_seed | EGFR | 0.493056 | HNSC | BICR22 | Erlotinib | MVP_top60_DNA+RNA_seed | HSP90AB1 | 0.636838 | KIRC | SN12C | Tanespimycin |
| MVP_top60_DNA_seed | ESR1 | 0.495594 | HNSC | PE-CA-PJ15 | GDC0810 | MVP_top60_DNA+RNA_seed | IGF1R | 0.637966 | KIRC | CAKI-1 | GSK1904529A |
| MVP_top60_DNA_seed | NTRK1 | 0.495898 | HNSC | BHY | GW441756 | MVP_top60_DNA+RNA_seed | AR | 0.638532 | KIRC | HA7-RCC | Bicalutamide |
| MVP_top60_DNA_seed | MAPK1 | 0.499831 | HNSC | HSC-2 | Ulixertinib | MVP_top60_DNA+RNA_seed | HDAC1 | 0.646978 | KIRC | KMRC-20 | Romidepsin |
| MVP_top60_DNA_seed | MAPK3 | 0.499831 | HNSC | HSC-2 | Ulixertinib | MVP_top60_DNA+RNA_seed | TP53 | 0.653787 | KIRC | LB1047-RCC | MIRA-1 |
| MVP_top60_DNA_seed | AR | 0.502126 | HNSC | PCI-6A | Bicalutamide | MVP_top60_DNA+RNA_seed | ESR1 | 0.656359 | KIRC | 769-P | GDC0810 |
| MVP_top60_DNA_seed | ESR1 | 0.502428 | HNSC | BICR31 | Tamoxifen | MVP_top60_DNA+RNA_seed | ATM | 0.663352 | KIRC | LB1047-RCC | KU-55933 |
| MVP_top60_DNA_seed | CDK2 | 0.503952 | HNSC | HSC-2 | MK-8776 | MVP_top60_DNA+RNA_seed | AR | 0.664752 | KIRC | RCC-ER | Bicalutamide |
| MVP_top60_DNA_seed | AKT1 | 0.506028 | HNSC | BICR31 | AZD5363 | MVP_top60_DNA+RNA_seed | HDAC1 | 0.665242 | KIRC | A498 | Entinostat |
| MVP_top60_DNA_seed | CDK2 | 0.509731 | HNSC | PE-CA-PJ15 | MK-8776 | MVP_top60_DNA+RNA_seed | ATM | 0.675747 | KIRC | A498 | KU-55933 |
| MVP_top60_DNA_seed | MAPK1 | 0.511926 | HNSC | OSC-20 | VX-11e | MVP_top60_DNA+RNA_seed | TP53 | 0.679959 | KIRC | KMRC-1 | MIRA-1 |
| MVP_top60_DNA_seed | AKT1 | 0.512472 | HNSC | SCC-4 | GSK2110183B | MVP_top60_DNA+RNA_seed | ESR1 | 0.682241 | KIRC | RCC-FG2 | GDC0810 |
| MVP_top60_DNA_seed | EGFR | 0.515893 | HNSC | OSC-20 | Afatinib | MVP_top60_DNA+RNA_seed | IGF1R | 0.693713 | KIRC | ACHN | NVP-ADW742 |
| MVP_top60_DNA_seed | ESR1 | 0.516194 | HNSC | SCC-25 | GDC0810 | MVP_top60_DNA+RNA_seed | ATM | 0.694802 | KIRC | 769-P | KU-55933 |
| MVP_top60_DNA_seed | TP53 | 0.51814 | HNSC | BB49-HNC | MIRA-1 | MVP_top60_DNA+RNA_seed | SMARCA4 | 0.695265 | KIRC | LB2241-RCC | PFI3 |
| MVP_top60_DNA_seed | NTRK1 | 0.520246 | HNSC | PCI-4B | GW441756 | MVP_top60_DNA+RNA_seed | SMARCA2 | 0.695265 | KIRC | LB2241-RCC | PFI3 |
| MVP_top60_DNA_seed | HSP90AA1 | 0.520466 | HNSC | PCI-6A | Luminespib | MVP_top60_DNA+RNA_seed | TP53 | 0.706908 | KIRC | SN12C | MIRA-1 |
| MVP_top60_DNA_seed | HDAC1 | 0.525527 | HNSC | PE-CA-PJ15 | Entinostat | MVP_top60_DNA+RNA_seed | ESR1 | 0.709393 | KIRC | A704 | GDC0810 |
| MVP_top60_DNA_seed | HDAC3 | 0.525527 | HNSC | PE-CA-PJ15 | Entinostat | MVP_top60_DNA+RNA_seed | IGF1R | 0.711614 | KIRC | RCC-JW | NVP-ADW742 |
| MVP_top60_DNA_seed | HDAC1 | 0.525781 | HNSC | BICR22 | PCI-34051 | MVP_top60_DNA+RNA_seed | HDAC1 | 0.717405 | KIRC | NCC021 | PCI-34051 |
| MVP_top60_DNA_seed | SRC | 0.52666 | HNSC | BICR31 | Bosutinib | MVP_top60_DNA+RNA_seed | PIK3CA | 0.720841 | KIRC | RCC-AB | Alpelisib |
| MVP_top60_DNA_seed | EGFR | 0.530243 | HNSC | SKN-3 | AZD3759 | MVP_top60_DNA+RNA_seed | IGF1R | 0.721112 | KIRC | LB2241-RCC | NVP-ADW742 |
| MVP_top60_DNA_seed | PIK3CA | 0.531023 | HNSC | Ca9-22 | AZD8186 | MVP_top60_DNA+RNA_seed | HDAC1 | 0.726804 | KIRC | RCC-JW | Entinostat |
| MVP_top60_DNA_seed | AKT1 | 0.541608 | HNSC | SKN-3 | AZD5363 | MVP_top60_DNA+RNA_seed | HDAC1 | 0.728364 | KIRC | ACHN | Entinostat |
| MVP_top60_DNA_seed | HDAC1 | 0.545579 | HNSC | DOK | PCI-34051 | MVP_top60_DNA+RNA_seed | PIK3CA | 0.730589 | KIRC | OS-RC-2 | Buparlisib |
| MVP_top60_DNA_seed | AKT1 | 0.547381 | HNSC | CAL-27 | Uprosertib | MVP_top60_DNA+RNA_seed | PIK3CD | 0.730589 | KIRC | OS-RC-2 | Buparlisib |
| MVP_top60_DNA_seed | AKT1 | 0.55138 | HNSC | SCC-4 | Uprosertib | MVP_top60_DNA+RNA_seed | HDAC1 | 0.737516 | KIRC | 769-P | Entinostat |
| MVP_top60_DNA_seed | AKT1 | 0.554126 | HNSC | SAT | AZD5363 | MVP_top60_DNA+RNA_seed | SMARCA4 | 0.738045 | KIRC | OS-RC-2 | PFI3 |
| MVP_top60_DNA_seed | PIK3CA | 0.555687 | HNSC | HSC-4 | GNE-317 | MVP_top60_DNA+RNA_seed | SMARCA2 | 0.738045 | KIRC | OS-RC-2 | PFI3 |
| MVP_top60_DNA_seed | ESR1 | 0.55864 | HNSC | JHU-029 | GDC0810 | MVP_top60_DNA+RNA_seed | ESR1 | 0.743112 | KIRC | LB996-RCC | GDC0810 |
| MVP_top60_DNA_seed | AKT1 | 0.558876 | HNSC | SCC-25 | Afuresertib | MVP_top60_DNA+RNA_seed | IGF1R | 0.743566 | KIRC | SW156 | BMS-536924 |
| MVP_top60_DNA_seed | AKT1 | 0.560424 | HNSC | Ca9-22 | GSK2110183B | MVP_top60_DNA+RNA_seed | EP300 | 0.745621 | KIRC | A498 | CPI-637 |
| MVP_top60_DNA_seed | SMARCA4 | 0.564503 | HNSC | BICR10 | PFI3 | MVP_top60_DNA+RNA_seed | ATM | 0.753981 | KIRC | KMRC-1 | KU-55933 |
| MVP_top60_DNA_seed | ESR1 | 0.566344 | HNSC | BHY | GDC0810 | MVP_top60_DNA+RNA_seed | HSP90AA1 | 0.754848 | KIRC | RCC-AB | Tanespimycin |
| MVP_top60_DNA_seed | PIK3CA | 0.569282 | HNSC | SCC-4 | AZD8186 | MVP_top60_DNA+RNA_seed | HSP90AB1 | 0.754848 | KIRC | RCC-AB | Tanespimycin |

| | | | | | | | | | | | |
|---|---|---|---|---|---|---|---|---|---|---|---|
| MVP_top60_DNA_seed | CDK2 | 0.571194 | HNSC | CAL-33 | MK-8776 | MVP_top60_DNA+RNA_seed | HSP90AA1 | 0.756761 | KIRC | RCC-AB | Elesclomol |
| MVP_top60_DNA_seed | AKT1 | 0.571818 | HNSC | JHU-022 | MK-2206 | MVP_top60_DNA+RNA_seed | HSP90AB1 | 0.756761 | KIRC | RCC-AB | Elesclomol |
| MVP_top60_DNA_seed | EGFR | 0.57184 | HNSC | BICR10 | Gefitinib | MVP_top60_DNA+RNA_seed | ESR1 | 0.758063 | KIRC | SN12C | GDC0810 |
| MVP_top60_DNA_seed | AKT1 | 0.573462 | HNSC | SCC-9 | GSK2110183B | MVP_top60_DNA+RNA_seed | SMARCA4 | 0.758393 | KIRC | ACHN | PFI3 |
| MVP_top60_DNA_seed | MAPK1 | 0.573514 | HNSC | SCC-15 | Ulixertinib | MVP_top60_DNA+RNA_seed | SMARCA2 | 0.758393 | KIRC | ACHN | PFI3 |
| MVP_top60_DNA_seed | MAPK3 | 0.573514 | HNSC | SCC-15 | Ulixertinib | MVP_top60_DNA+RNA_seed | HSP90AA1 | 0.762159 | KIRC | U031 | Tanespimycin |
| MVP_top60_DNA_seed | AKT1 | 0.574995 | HNSC | BICR78 | Ipatasertib | MVP_top60_DNA+RNA_seed | HSP90AB1 | 0.762159 | KIRC | U031 | Tanespimycin |
| MVP_top60_DNA_seed | MAPK1 | 0.577772 | HNSC | BHY | VX-11e | MVP_top60_DNA+RNA_seed | IGF1R | 0.771331 | KIRC | RCC-AB | Linsitinib |
| MVP_top60_DNA_seed | AKT1 | 0.581508 | HNSC | BICR78 | AT13148 | MVP_top60_DNA+RNA_seed | HDAC1 | 0.773843 | KIRC | BFTC-909 | PCI-34051 |
| MVP_top60_DNA_seed | MAPK1 | 0.584053 | HNSC | SCC-25 | Ulixertinib | MVP_top60_DNA+RNA_seed | IGF1R | 0.776197 | KIRC | HA7-RCC | GSK1904529A |
| MVP_top60_DNA_seed | MAPK3 | 0.584053 | HNSC | SCC-25 | Ulixertinib | MVP_top60_DNA+RNA_seed | HSP90AA1 | 0.796496 | KIRC | TK10 | Tanespimycin |
| MVP_top60_DNA_seed | PIK3CA | 0.587562 | HNSC | FADU | AZD8186 | MVP_top60_DNA+RNA_seed | HSP90AB1 | 0.796496 | KIRC | TK10 | Tanespimycin |
| MVP_top60_DNA_seed | MDM2 | 0.588566 | HNSC | OSC-19 | Nutlin-3a (-) | MVP_top60_DNA+RNA_seed | TP53 | 0.80106 | KIRC | KMRC-20 | MIRA-1 |
| MVP_top60_DNA_seed | MAPK1 | 0.594275 | HNSC | SAT | Ulixertinib | MVP_top60_DNA+RNA_seed | SMARCA4 | 0.809915 | KIRC | RXF393 | PFI3 |
| MVP_top60_DNA_seed | MAPK3 | 0.594275 | HNSC | SAT | Ulixertinib | MVP_top60_DNA+RNA_seed | SMARCA2 | 0.809915 | KIRC | RXF393 | PFI3 |
| MVP_top60_DNA_seed | ESR1 | 0.597193 | HNSC | HSC-4 | GDC0810 | MVP_top60_DNA+RNA_seed | ATM | 0.811201 | KIRC | CAKI-1 | KU-55933 |
| MVP_top60_DNA_seed | ESR1 | 0.60062 | HNSC | OSC-19 | Tamoxifen | MVP_top60_DNA+RNA_seed | ATM | 0.818165 | KIRC | RCC-AB | KU-55933 |
| MVP_top60_DNA_seed | CDK2 | 0.603903 | HNSC | BICR31 | Dinaciclib | MVP_top60_DNA+RNA_seed | TP53 | 0.82094 | KIRC | LB996-RCC | MIRA-1 |
| MVP_top60_DNA_seed | SRC | 0.611592 | HNSC | BHY | Bosutinib | MVP_top60_DNA+RNA_seed | SMARCA4 | 0.829859 | KIRC | A498 | PFI3 |
| MVP_top60_DNA_seed | EGFR | 0.611984 | HNSC | BICR10 | Lapatinib | MVP_top60_DNA+RNA_seed | SMARCA2 | 0.829859 | KIRC | A498 | PFI3 |
| MVP_top60_DNA_seed | EGFR | 0.612375 | HNSC | PCI-30 | Gefitinib | MVP_top60_DNA+RNA_seed | ATM | 0.830618 | KIRC | LB2241-RCC | KU-55933 |
| MVP_top60_DNA_seed | HSP90AA1 | 0.612983 | HNSC | SKN-3 | Tanespimycin | MVP_top60_DNA+RNA_seed | HSP90AA1 | 0.831608 | KIRC | SW156 | Tanespimycin |
| MVP_top60_DNA_seed | HDAC1 | 0.613657 | HNSC | HSC-4 | Romidepsin | MVP_top60_DNA+RNA_seed | HSP90AB1 | 0.831608 | KIRC | SW156 | Tanespimycin |
| MVP_top60_DNA_seed | HDAC3 | 0.613657 | HNSC | HSC-4 | Romidepsin | MVP_top60_DNA+RNA_seed | SMARCA4 | 0.833466 | KIRC | SW156 | PFI3 |
| MVP_top60_DNA_seed | HDAC1 | 0.613692 | HNSC | HSC-2 | Entinostat | MVP_top60_DNA+RNA_seed | SMARCA2 | 0.833466 | KIRC | SW156 | PFI3 |
| MVP_top60_DNA_seed | HDAC3 | 0.613692 | HNSC | HSC-2 | Entinostat | MVP_top60_DNA+RNA_seed | IGF1R | 0.834671 | KIRC | RCC-JF | Linsitinib |
| MVP_top60_DNA_seed | SMARCA4 | 0.617023 | HNSC | PCI-4B | PFI3 | MVP_top60_DNA+RNA_seed | HDAC1 | 0.839066 | KIRC | RCC-AB | PCI-34051 |
| MVP_top60_DNA_seed | TP53 | 0.619898 | HNSC | HSC-4 | MIRA-1 | MVP_top60_DNA+RNA_seed | IGF1R | 0.843463 | KIRC | RXF393 | NVP-ADW742 |
| MVP_top60_DNA_seed | MAPK1 | 0.620034 | HNSC | OSC-20 | SCH772984 | MVP_top60_DNA+RNA_seed | IGF1R | 0.84891 | KIRC | NCC021 | Linsitinib |
| MVP_top60_DNA_seed | MAPK3 | 0.620034 | HNSC | OSC-20 | SCH772984 | MVP_top60_DNA+RNA_seed | EP300 | 0.850651 | KIRC | KMRC-20 | SGC-CBP30 |
| MVP_top60_DNA_seed | MAPK1 | 0.621499 | HNSC | PCI-30 | SCH772984 | MVP_top60_DNA+RNA_seed | ESR1 | 0.866638 | KIRC | OS-RC-2 | Tamoxifen |
| MVP_top60_DNA_seed | MAPK3 | 0.621499 | HNSC | PCI-30 | SCH772984 | MVP_top60_DNA+RNA_seed | HDAC1 | 0.883983 | KIRC | 786-0 | Entinostat |
| MVP_top60_DNA_seed | SMARCA4 | 0.625828 | HNSC | Ca9-22 | PFI3 | MVP_top60_DNA+RNA_seed | IGF1R | 0.89416 | KIRC | KMRC-20 | Linsitinib |
| MVP_top60_DNA_seed | NTRK1 | 0.629658 | HNSC | PCI-30 | GW441756 | MVP_top60_DNA+RNA_seed | AR | 0.895 | KIRC | BB65-RCC | Bicalutamide |
| MVP_top60_DNA_seed | AKT1 | 0.630153 | HNSC | Ca9-22 | Afuresertib | MVP_top60_DNA+RNA_seed | IGF1R | 0.895887 | KIRC | SN12C | Linsitinib |
| MVP_top60_DNA_seed | CDK2 | 0.632657 | HNSC | BICR31 | MK-8776 | MVP_top60_DNA+RNA_seed | PIK3CA | 0.9 | KIRC | BB65-RCC | Buparlisib |
| MVP_top60_DNA_seed | CDK2 | 0.63326 | HNSC | FADU | MK-8776 | MVP_top60_DNA+RNA_seed | PIK3CD | 0.9 | KIRC | BB65-RCC | Buparlisib |
| MVP_top60_DNA_seed | PIK3CA | 0.634428 | HNSC | SCC-9 | Buparlisib | MVP_top60_DNA+RNA_seed | ESR1 | 0.901768 | KIRC | LB2241-RCC | GDC0810 |
| MVP_top60_DNA_seed | HDAC1 | 0.639701 | HNSC | HSC-3 | PCI-34051 | MVP_top60_DNA+RNA_seed | IGF1R | 0.902805 | KIRC | LB2241-RCC | GSK1904529A |
| MVP_top60_DNA_seed | AKT1 | 0.640086 | HNSC | BHY | AZD5363 | MVP_top60_DNA+RNA_seed | IGF1R | 0.90767 | KIRC | OS-RC-2 | NVP-ADW742 |
| MVP_top60_DNA_seed | MAPK1 | 0.640247 | HNSC | HSC-4 | SCH772984 | MVP_top60_DNA+RNA_seed | SMARCA4 | 0.914798 | KIRC | NCC021 | PFI3 |

| | | | | | | | | | | | |
|---|---|---|---|---|---|---|---|---|---|---|---|
| MVP_top60_DNA_seed | MAPK3 | 0.640247 | HNSC | HSC-4 | SCH772984 | MVP_top60_DNA+RNA_seed | SMARCA2 | 0.914798 | KIRC | NCC021 | PFI3 |
| MVP_top60_DNA_seed | HDAC1 | 0.641536 | HNSC | SCC-25 | PCI-34051 | MVP_top60_DNA+RNA_seed | HDAC1 | 0.916293 | KIRC | SW156 | PCI-34051 |
| MVP_top60_DNA_seed | ESR1 | 0.643439 | HNSC | DOK | GDC0810 | MVP_top60_DNA+RNA_seed | ESR1 | 0.922205 | KIRC | RCC10RGB | GDC0810 |
| MVP_top60_DNA_seed | EP300 | 0.646766 | HNSC | SCC-15 | CPI-637 | MVP_top60_DNA+RNA_seed | HSP90AA1 | 0.924473 | KIRC | SW156 | CCT-018159 |
| MVP_top60_DNA_seed | HDAC1 | 0.647267 | HNSC | FADU | Entinostat | MVP_top60_DNA+RNA_seed | HSP90AB1 | 0.924473 | KIRC | SW156 | CCT-018159 |
| MVP_top60_DNA_seed | HDAC3 | 0.647267 | HNSC | FADU | Entinostat | MVP_top60_DNA+RNA_seed | TP53 | 0.925183 | KIRC | RCC-AB | MIRA-1 |
| MVP_top60_DNA_seed | CDK2 | 0.647957 | HNSC | BHY | MK-8776 | MVP_top60_DNA+RNA_seed | HSP90AA1 | 0.932291 | KIRC | CAKI-1 | Tanespimycin |
| MVP_top60_DNA_seed | CDK2 | 0.648118 | HNSC | SCC-25 | MK-8776 | MVP_top60_DNA+RNA_seed | HSP90AB1 | 0.932291 | KIRC | CAKI-1 | Tanespimycin |
| MVP_top60_DNA_seed | AKT1 | 0.654175 | HNSC | PCI-6A | AT13148 | MVP_top60_DNA+RNA_seed | ATM | 0.933906 | KIRC | OS-RC-2 | KU-55933 |
| MVP_top60_DNA_seed | PIK3CA | 0.655852 | HNSC | SCC-4 | GNE-317 | MVP_top60_DNA+RNA_seed | HDAC1 | 0.937418 | KIRC | ACHN | Romidepsin |
| MVP_top60_DNA_seed | MDM2 | 0.658888 | HNSC | BICR22 | Nutlin-3a (-) | MVP_top60_DNA+RNA_seed | TP53 | 0.939831 | KIRC | VMRC-RCW | MIRA-1 |
| MVP_top60_DNA_seed | MDM2 | 0.659993 | HNSC | OSC-20 | Serdemetan | MVP_top60_DNA+RNA_seed | HSP90AA1 | 0.942826 | KIRC | RCC-AB | CCT-018159 |
| MVP_top60_DNA_seed | HDAC1 | 0.663203 | HNSC | BB49-HNC | Entinostat | MVP_top60_DNA+RNA_seed | HSP90AB1 | 0.942826 | KIRC | RCC-AB | CCT-018159 |
| MVP_top60_DNA_seed | HDAC3 | 0.663203 | HNSC | BB49-HNC | Entinostat | MVP_top60_DNA+RNA_seed | ESR1 | 0.945693 | KIRC | NCC021 | GDC0810 |
| MVP_top60_DNA_seed | AR | 0.664435 | HNSC | JHU-011 | Bicalutamide | MVP_top60_DNA+RNA_seed | TP53 | 0.954891 | KIRC | A498 | MIRA-1 |
| MVP_top60_DNA_seed | SRC | 0.664538 | HNSC | SCC-25 | Bosutinib | MVP_top60_DNA+RNA_seed | HDAC1 | 0.955875 | KIRC | VMRC-RCZ | Romidepsin |
| MVP_top60_DNA_seed | MDM2 | 0.667172 | HNSC | HSC-4 | Serdemetan | MVP_top60_DNA+RNA_seed | SMARCA4 | 0.961015 | KIRC | LB1047-RCC | PFI3 |
| MVP_top60_DNA_seed | CDK2 | 0.669958 | HNSC | JHU-029 | MK-8776 | MVP_top60_DNA+RNA_seed | SMARCA2 | 0.961015 | KIRC | LB1047-RCC | PFI3 |
| MVP_top60_DNA_seed | MDM2 | 0.672807 | HNSC | PCI-30 | Nutlin-3a (-) | MVP_top60_DNA+RNA_seed | HDAC1 | 0.961034 | KIRC | CAL-54 | Entinostat |
| MVP_top60_DNA_seed | MAPK1 | 0.675233 | HNSC | BHY | SCH772984 | MVP_top60_DNA+RNA_seed | AR | 0.966853 | KIRC | RCC-MF | Bicalutamide |
| MVP_top60_DNA_seed | MAPK3 | 0.675233 | HNSC | BHY | SCH772984 | MVP_top60_DNA+RNA_seed | IGF1R | 0.968792 | KIRC | 786-0 | GSK1904529A |
| MVP_top60_DNA_seed | PIK3CA | 0.675844 | HNSC | KOSC-2 | AZD8186 | MVP_top60_DNA+RNA_seed | EP300 | 0.969571 | KIRC | VMRC-RCZ | SGC-CBP30 |
| MVP_top60_DNA_seed | MAPK1 | 0.675901 | HNSC | BICR10 | VX-11e | MVP_top60_DNA+RNA_seed | ESR1 | 0.972005 | KIRC | HA7-RCC | GDC0810 |
| MVP_top60_DNA_seed | EP300 | 0.678557 | HNSC | CAL-33 | SGC-CBP30 | MVP_top60_DNA+RNA_seed | ATM | 0.972123 | KIRC | SN12C | KU-55933 |
| MVP_top60_DNA_seed | CREBBP | 0.678557 | HNSC | CAL-33 | SGC-CBP30 | MVP_top60_DNA+RNA_seed | IGF1R | 0.979498 | KIRC | LB1047-RCC | Linsitinib |
| MVP_top60_DNA_seed | AKT1 | 0.679805 | HNSC | SCC-9 | AZD5363 | MVP_top60_DNA+RNA_seed | HSP90AA1 | 0.997302 | KIRC | TK10 | Luminespib |
| MVP_top60_DNA_seed | HDAC1 | 0.68052 | HNSC | Ca9-22 | PCI-34051 | MVP_top60_DNA+RNA_seed | HSP90AB1 | 0.997302 | KIRC | TK10 | Luminespib |
| MVP_top60_DNA_seed | HDAC1 | 0.682346 | HNSC | OSC-20 | Romidepsin | MVP_top60_DNA+RNA_seed | HSP90AA1 | 0.998496 | KIRC | LB1047-RCC | Elesclomol |
| MVP_top60_DNA_seed | HDAC3 | 0.682346 | HNSC | OSC-20 | Romidepsin | MVP_top60_DNA+RNA_seed | HSP90AB1 | 0.998496 | KIRC | LB1047-RCC | Elesclomol |
| MVP_top60_DNA_seed | AKT1 | 0.683795 | HNSC | Ca9-22 | MK-2206 | MVP_top60_DNA+RNA_seed | EP300 | 0.999916 | KIRC | VMRC-RCZ | CPI-637 |
| MVP_top60_DNA_seed | PIK3CA | 0.685895 | HNSC | PE-CA-PJ15 | Buparlisib | MVP_top60_DNA+RNA_seed | IGF1R | 1.000174 | KIRC | RCC-JW | GSK1904529A |
| MVP_top60_DNA_seed | AR | 0.687031 | HNSC | PCI-4B | Bicalutamide | MVP_top60_DNA+RNA_seed | IGF1R | 1.00236 | KIRC | OS-RC-2 | BMS-536924 |
| MVP_top60_DNA_seed | TP53 | 0.692683 | HNSC | BICR10 | MIRA-1 | MVP_top60_DNA+RNA_seed | IGF1R | 1.005405 | KIRC | CAKI-1 | NVP-ADW742 |
| MVP_top60_DNA_seed | EGFR | 0.693043 | HNSC | SKN-3 | Osimertinib | MVP_top60_DNA+RNA_seed | IGF1R | 1.008275 | KIRC | VMRC-RCW | GSK1904529A |
| MVP_top60_DNA_seed | EGFR | 0.695242 | HNSC | SCC-4 | AZD3759 | MVP_top60_DNA+RNA_seed | ESR1 | 1.011666 | KIRC | RCC-MF | Tamoxifen |
| MVP_top60_DNA_seed | TP53 | 0.696963 | HNSC | BICR78 | MIRA-1 | MVP_top60_DNA+RNA_seed | IGF1R | 1.020508 | KIRC | NCC010 | Linsitinib |
| MVP_top60_DNA_seed | AKT1 | 0.697399 | HNSC | SCC-15 | MK-2206 | MVP_top60_DNA+RNA_seed | PIK3CA | 1.021881 | KIRC | BB65-RCC | Alpelisib |
| MVP_top60_DNA_seed | CDK2 | 0.698203 | HNSC | Ca9-22 | AZD5438 | MVP_top60_DNA+RNA_seed | ESR1 | 1.059703 | KIRC | HA7-RCC | Tamoxifen |
| MVP_top60_DNA_seed | HSP90AA1 | 0.699838 | HNSC | PE-CA-PJ15 | Elesclomol | MVP_top60_DNA+RNA_seed | MTOR | 1.063152 | KIRC | BB65-RCC | Temsirolimus |
| MVP_top60_DNA_seed | EGFR | 0.700744 | HNSC | PCI-30 | Lapatinib | MVP_top60_DNA+RNA_seed | HSP90AA1 | 1.069167 | KIRC | A704 | Tanespimycin |
| MVP_top60_DNA_seed | HDAC1 | 0.703144 | HNSC | HSC-3 | Entinostat | MVP_top60_DNA+RNA_seed | HSP90AB1 | 1.069167 | KIRC | A704 | Tanespimycin |

| | | | | | | | | | | | |
|---|---|---|---|---|---|---|---|---|---|---|---|
| MVP_top60_DNA_seed | HDAC3 | 0.703144 | HNSC | HSC-3 | Entinostat | MVP_top60_DNA+RNA_seed | PIK3CA | 1.072953 | KIRC | KMRC-20 | Alpelisib |
| MVP_top60_DNA_seed | MDM2 | 0.705927 | HNSC | LB771-HNC | Nutlin-3a (-) | MVP_top60_DNA+RNA_seed | IGF1R | 1.076541 | KIRC | 769-P | GSK1904529A |
| MVP_top60_DNA_seed | AKT1 | 0.706098 | HNSC | CAL-33 | Ipatasertib | MVP_top60_DNA+RNA_seed | MTOR | 1.097443 | KIRC | HA7-RCC | Temsirolimus |
| MVP_top60_DNA_seed | HDAC1 | 0.711231 | HNSC | LB771-HNC | PCI-34051 | MVP_top60_DNA+RNA_seed | ESR1 | 1.101727 | KIRC | NCC010 | Tamoxifen |
| MVP_top60_DNA_seed | PIK3CA | 0.713261 | HNSC | OSC-20 | AZD8186 | MVP_top60_DNA+RNA_seed | ESR1 | 1.10184 | KIRC | VMRC-RCZ | Tamoxifen |
| MVP_top60_DNA_seed | MDM2 | 0.715355 | HNSC | SCC-15 | Nutlin-3a (-) | MVP_top60_DNA+RNA_seed | PIK3CA | 1.104382 | KIRC | VMRC-RCW | Alpelisib |
| MVP_top60_DNA_seed | AKT1 | 0.715604 | HNSC | HSC-4 | MK-2206 | MVP_top60_DNA+RNA_seed | HSP90AA1 | 1.108952 | KIRC | HA7-RCC | Elesclomol |
| MVP_top60_DNA_seed | MDM2 | 0.720901 | HNSC | PCI-30 | Serdemetan | MVP_top60_DNA+RNA_seed | HSP90AB1 | 1.108952 | KIRC | HA7-RCC | Elesclomol |
| MVP_top60_DNA_seed | HDAC1 | 0.721318 | HNSC | SKN-3 | PCI-34051 | MVP_top60_DNA+RNA_seed | ESR1 | 1.11942 | KIRC | LB1047-RCC | GDC0810 |
| MVP_top60_DNA_seed | MDM2 | 0.725433 | HNSC | SCC-9 | Nutlin-3a (-) | MVP_top60_DNA+RNA_seed | PIK3CA | 1.121707 | KIRC | BB65-RCC | GNE-317 |
| MVP_top60_DNA_seed | ESR1 | 0.730609 | HNSC | PCI-15A | Tamoxifen | MVP_top60_DNA+RNA_seed | AR | 1.127689 | KIRC | RCC-AB | Bicalutamide |
| MVP_top60_DNA_seed | MAPK1 | 0.732869 | HNSC | SCC-4 | Ulixertinib | MVP_top60_DNA+RNA_seed | ESR1 | 1.129094 | KIRC | LB2241-RCC | Tamoxifen |
| MVP_top60_DNA_seed | MAPK3 | 0.732869 | HNSC | SCC-4 | Ulixertinib | MVP_top60_DNA+RNA_seed | PIK3CA | 1.131475 | KIRC | RCC10RGB | Buparlisib |
| MVP_top60_DNA_seed | CDK2 | 0.733044 | HNSC | JHU-022 | Dinaciclib | MVP_top60_DNA+RNA_seed | PIK3CD | 1.131475 | KIRC | RCC10RGB | Buparlisib |
| MVP_top60_DNA_seed | EP300 | 0.734617 | HNSC | BB49-HNC | SGC-CBP30 | MVP_top60_DNA+RNA_seed | IGF1R | 1.133849 | KIRC | RCC-FG2 | GSK1904529A |
| MVP_top60_DNA_seed | CREBBP | 0.734617 | HNSC | BB49-HNC | SGC-CBP30 | MVP_top60_DNA+RNA_seed | TP53 | 1.134189 | KIRC | RCC10RGB | MIRA-1 |
| MVP_top60_DNA_seed | HSP90AA1 | 0.735298 | HNSC | SCC-15 | Tanespimycin | MVP_top60_DNA+RNA_seed | SMARCA4 | 1.168113 | KIRC | HA7-RCC | PFI3 |
| MVP_top60_DNA_seed | TP53 | 0.737087 | HNSC | SKN-3 | MIRA-1 | MVP_top60_DNA+RNA_seed | SMARCA2 | 1.168113 | KIRC | HA7-RCC | PFI3 |
| MVP_top60_DNA_seed | CDK2 | 0.73788 | HNSC | CAL-27 | MK-8776 | MVP_top60_DNA+RNA_seed | PIK3CA | 1.168539 | KIRC | ACHN | Alpelisib |
| MVP_top60_DNA_seed | HDAC1 | 0.739052 | HNSC | SCC-9 | Entinostat | MVP_top60_DNA+RNA_seed | HDAC1 | 1.169682 | KIRC | HA7-RCC | PCI-34051 |
| MVP_top60_DNA_seed | HDAC3 | 0.739052 | HNSC | SCC-9 | Entinostat | MVP_top60_DNA+RNA_seed | SMARCA4 | 1.170472 | KIRC | RCC-AB | PFI3 |
| MVP_top60_DNA_seed | SMARCA4 | 0.743748 | HNSC | BHY | PFI3 | MVP_top60_DNA+RNA_seed | SMARCA2 | 1.170472 | KIRC | RCC-AB | PFI3 |
| MVP_top60_DNA_seed | AKT1 | 0.74802 | HNSC | DOK | AT13148 | MVP_top60_DNA+RNA_seed | HDAC1 | 1.175065 | KIRC | RCC-JW | PCI-34051 |
| MVP_top60_DNA_seed | AKT1 | 0.748466 | HNSC | SCC-4 | MK-2206 | MVP_top60_DNA+RNA_seed | HDAC1 | 1.175395 | KIRC | A704 | Romidepsin |
| MVP_top60_DNA_seed | HDAC1 | 0.7495 | HNSC | SAT | PCI-34051 | MVP_top60_DNA+RNA_seed | IGF1R | 1.175795 | KIRC | OS-RC-2 | BMS-754807 |
| MVP_top60_DNA_seed | SRC | 0.75054 | HNSC | BICR22 | Bosutinib | MVP_top60_DNA+RNA_seed | IGF1R | 1.183257 | KIRC | SN12C | NVP-ADW742 |
| MVP_top60_DNA_seed | HSP90AA1 | 0.760832 | HNSC | BICR78 | Tanespimycin | MVP_top60_DNA+RNA_seed | PIK3CA | 1.195073 | KIRC | A498 | Alpelisib |
| MVP_top60_DNA_seed | PIK3CA | 0.76117 | HNSC | HSC-4 | Buparlisib | MVP_top60_DNA+RNA_seed | IGF1R | 1.196698 | KIRC | LB2241-RCC | Linsitinib |
| MVP_top60_DNA_seed | AKT1 | 0.763581 | HNSC | FADU | AZD5363 | MVP_top60_DNA+RNA_seed | PIK3CA | 1.218478 | KIRC | SW156 | GNE-317 |
| MVP_top60_DNA_seed | AKT1 | 0.763974 | HNSC | FADU | MK-2206 | MVP_top60_DNA+RNA_seed | IGF1R | 1.225311 | KIRC | CAKI-1 | Linsitinib |
| MVP_top60_DNA_seed | AKT1 | 0.766565 | HNSC | CAL-27 | Ipatasertib | MVP_top60_DNA+RNA_seed | IGF1R | 1.226438 | KIRC | SN12C | BMS-754807 |
| MVP_top60_DNA_seed | HDAC1 | 0.767635 | HNSC | JHU-022 | PCI-34051 | MVP_top60_DNA+RNA_seed | IGF1R | 1.22687 | KIRC | RCC-AB | BMS-754807 |
| MVP_top60_DNA_seed | PIK3CA | 0.770498 | HNSC | BHY | GNE-317 | MVP_top60_DNA+RNA_seed | HSP90AA1 | 1.234003 | KIRC | LB2241-RCC | CCT-018159 |
| MVP_top60_DNA_seed | PIK3CA | 0.77162 | HNSC | OSC-20 | Alpelisib | MVP_top60_DNA+RNA_seed | HSP90AB1 | 1.234003 | KIRC | LB2241-RCC | CCT-018159 |
| MVP_top60_DNA_seed | AKT1 | 0.773405 | HNSC | FADU | AT13148 | MVP_top60_DNA+RNA_seed | ESR1 | 1.23426 | KIRC | U031 | Tamoxifen |
| MVP_top60_DNA_seed | EP300 | 0.773758 | HNSC | CAL-33 | CPI-637 | MVP_top60_DNA+RNA_seed | HSP90AA1 | 1.234402 | KIRC | RCC-MF | Tanespimycin |
| MVP_top60_DNA_seed | RAC1 | 0.784407 | HNSC | JHU-022 | EHT-1864 | MVP_top60_DNA+RNA_seed | HSP90AB1 | 1.234402 | KIRC | RCC-MF | Tanespimycin |
| MVP_top60_DNA_seed | CDK2 | 0.784533 | HNSC | OSC-20 | AZD5438 | MVP_top60_DNA+RNA_seed | HSP90AA1 | 1.236386 | KIRC | VMRC-RCW | CCT-018159 |
| MVP_top60_DNA_seed | EGFR | 0.787115 | HNSC | OSC-20 | Lapatinib | MVP_top60_DNA+RNA_seed | HSP90AB1 | 1.236386 | KIRC | VMRC-RCW | CCT-018159 |
| MVP_top60_DNA_seed | SMARCA4 | 0.791809 | HNSC | SAT | PFI3 | MVP_top60_DNA+RNA_seed | IGF1R | 1.237731 | KIRC | RCC-AB | BMS-536924 |
| MVP_top60_DNA_seed | TP53 | 0.79274 | HNSC | SCC-4 | MIRA-1 | MVP_top60_DNA+RNA_seed | PIK3CA | 1.239671 | KIRC | RCC10RGB | Alpelisib |

| | | | | | | | | | | | |
|---|---|---|---|---|---|---|---|---|---|---|---|
| MVP_top60_DNA_seed | EGFR | 0.796499 | HNSC | OSC-20 | AZD3759 | MVP_top60_DNA+RNA_seed | ATM | 1.247355 | KIRC | VMRC-RCZ | KU-55933 |
| MVP_top60_DNA_seed | RAC1 | 0.797373 | HNSC | SCC-9 | EHT-1864 | MVP_top60_DNA+RNA_seed | PIK3CA | 1.24889 | KIRC | 786-0 | Alpelisib |
| MVP_top60_DNA_seed | AKT1 | 0.79788 | HNSC | CAL-27 | GSK2110183B | MVP_top60_DNA+RNA_seed | HDAC1 | 1.251491 | KIRC | VMRC-RCW | PCI-34051 |
| MVP_top60_DNA_seed | EGFR | 0.803149 | HNSC | BICR10 | AZD3759 | MVP_top60_DNA+RNA_seed | AR | 1.252801 | KIRC | VMRC-RCW | Bicalutamide |
| MVP_top60_DNA_seed | EGFR | 0.805253 | HNSC | PCI-30 | Sapitinib | MVP_top60_DNA+RNA_seed | IGF1R | 1.260442 | KIRC | RCC-JW | Linsitinib |
| MVP_top60_DNA_seed | NTRK1 | 0.806157 | HNSC | OSC-20 | GW441756 | MVP_top60_DNA+RNA_seed | EP300 | 1.269614 | KIRC | CAL-54 | SGC-CBP30 |
| MVP_top60_DNA_seed | MDM2 | 0.807297 | HNSC | SCC-9 | Serdemetan | MVP_top60_DNA+RNA_seed | MTOR | 1.274526 | KIRC | LB2241-RCC | Temsirolimus |
| MVP_top60_DNA_seed | EGFR | 0.810132 | HNSC | LB771-HNC | Lapatinib | MVP_top60_DNA+RNA_seed | HSP90AA1 | 1.277113 | KIRC | A498 | Luminespib |
| MVP_top60_DNA_seed | NTRK1 | 0.810363 | HNSC | SCC-4 | GW441756 | MVP_top60_DNA+RNA_seed | HSP90AB1 | 1.277113 | KIRC | A498 | Luminespib |
| MVP_top60_DNA_seed | SRC | 0.812195 | HNSC | OSC-20 | Bosutinib | MVP_top60_DNA+RNA_seed | HSP90AA1 | 1.277347 | KIRC | U031 | Elesclomol |
| MVP_top60_DNA_seed | AKT1 | 0.817874 | HNSC | SCC-25 | Uprosertib | MVP_top60_DNA+RNA_seed | HSP90AB1 | 1.277347 | KIRC | U031 | Elesclomol |
| MVP_top60_DNA_seed | CDK2 | 0.821217 | HNSC | OSC-20 | Dinaciclib | MVP_top60_DNA+RNA_seed | HDAC1 | 1.281265 | KIRC | SN12C | PCI-34051 |
| MVP_top60_DNA_seed | RAC1 | 0.824371 | HNSC | JHU-029 | EHT-1864 | MVP_top60_DNA+RNA_seed | EP300 | 1.282249 | KIRC | OS-RC-2 | CPI-637 |
| MVP_top60_DNA_seed | AKT1 | 0.826387 | HNSC | SKN-3 | Uprosertib | MVP_top60_DNA+RNA_seed | IGF1R | 1.283073 | KIRC | VMRC-RCW | BMS-754807 |
| MVP_top60_DNA_seed | MDM2 | 0.827042 | HNSC | SCC90 | Nutlin-3a (-) | MVP_top60_DNA+RNA_seed | MTOR | 1.297446 | KIRC | SN12C | Temsirolimus |
| MVP_top60_DNA_seed | HDAC1 | 0.830529 | HNSC | SCC-4 | PCI-34051 | MVP_top60_DNA+RNA_seed | IGF1R | 1.310829 | KIRC | LB1047-RCC | NVP-ADW742 |
| MVP_top60_DNA_seed | MAPK1 | 0.830606 | HNSC | LB771-HNC | SCH772984 | MVP_top60_DNA+RNA_seed | IGF1R | 1.318535 | KIRC | KMRC-1 | NVP-ADW742 |
| MVP_top60_DNA_seed | MAPK3 | 0.830606 | HNSC | LB771-HNC | SCH772984 | MVP_top60_DNA+RNA_seed | HDAC1 | 1.318589 | KIRC | VMRC-RCW | Entinostat |
| MVP_top60_DNA_seed | CDK2 | 0.830911 | HNSC | HO-1-N-1 | MK-8776 | MVP_top60_DNA+RNA_seed | PIK3CA | 1.320015 | KIRC | KMRC-1 | Alpelisib |
| MVP_top60_DNA_seed | AKT1 | 0.831882 | HNSC | SAT | Ipatasertib | MVP_top60_DNA+RNA_seed | IGF1R | 1.327066 | KIRC | SW156 | Linsitinib |
| MVP_top60_DNA_seed | PIK3CA | 0.833152 | HNSC | SKN-3 | AZD8186 | MVP_top60_DNA+RNA_seed | ESR1 | 1.333047 | KIRC | RCC-AB | GDC0810 |
| MVP_top60_DNA_seed | AKT1 | 0.834869 | HNSC | FADU | Uprosertib | MVP_top60_DNA+RNA_seed | IGF1R | 1.336044 | KIRC | TK10 | BMS-536924 |
| MVP_top60_DNA_seed | AKT1 | 0.837222 | HNSC | PE-CA-PJ15 | AZD5363 | MVP_top60_DNA+RNA_seed | IGF1R | 1.338909 | KIRC | RCC-FG2 | Linsitinib |
| MVP_top60_DNA_seed | HDAC1 | 0.84284 | HNSC | OSC-19 | PCI-34051 | MVP_top60_DNA+RNA_seed | AR | 1.34485 | KIRC | NCC021 | Bicalutamide |
| MVP_top60_DNA_seed | EGFR | 0.844159 | HNSC | BB49-HNC | Erlotinib | MVP_top60_DNA+RNA_seed | EP300 | 1.346693 | KIRC | OS-RC-2 | SGC-CBP30 |
| MVP_top60_DNA_seed | HDAC1 | 0.844777 | HNSC | BHY | PCI-34051 | MVP_top60_DNA+RNA_seed | HSP90AA1 | 1.347491 | KIRC | BB65-RCC | CCT-018159 |
| MVP_top60_DNA_seed | AKT1 | 0.845572 | HNSC | FADU | Ipatasertib | MVP_top60_DNA+RNA_seed | HSP90AB1 | 1.347491 | KIRC | BB65-RCC | CCT-018159 |
| MVP_top60_DNA_seed | CDK2 | 0.845796 | HNSC | BICR22 | MK-8776 | MVP_top60_DNA+RNA_seed | PIK3CA | 1.351246 | KIRC | VMRC-RCZ | AZD8186 |
| MVP_top60_DNA_seed | CDK2 | 0.851828 | HNSC | SCC-9 | MK-8776 | MVP_top60_DNA+RNA_seed | HSP90AA1 | 1.385036 | KIRC | OS-RC-2 | Luminespib |
| MVP_top60_DNA_seed | CDK2 | 0.855629 | HNSC | OSC-20 | MK-8776 | MVP_top60_DNA+RNA_seed | HSP90AB1 | 1.385036 | KIRC | OS-RC-2 | Luminespib |
| MVP_top60_DNA_seed | PIK3CA | 0.85678 | HNSC | PCI-4B | GNE-317 | MVP_top60_DNA+RNA_seed | ESR1 | 1.389136 | KIRC | RCC-FG2 | Tamoxifen |
| MVP_top60_DNA_seed | AKT1 | 0.862579 | HNSC | HSC-4 | AT13148 | MVP_top60_DNA+RNA_seed | HDAC1 | 1.397825 | KIRC | TK10 | Entinostat |
| MVP_top60_DNA_seed | HSP90AA1 | 0.864379 | HNSC | BICR78 | Luminespib | MVP_top60_DNA+RNA_seed | HSP90AA1 | 1.419594 | KIRC | CAKI-1 | CCT-018159 |
| MVP_top60_DNA_seed | ESR1 | 0.866929 | HNSC | SCC90 | Tamoxifen | MVP_top60_DNA+RNA_seed | HSP90AB1 | 1.419594 | KIRC | CAKI-1 | CCT-018159 |
| MVP_top60_DNA_seed | ESR1 | 0.868382 | HNSC | PCI-6A | Tamoxifen | MVP_top60_DNA+RNA_seed | ATM | 1.423753 | KIRC | VMRC-RCW | KU-55933 |
| MVP_top60_DNA_seed | CDK2 | 0.86921 | HNSC | BICR78 | MK-8776 | MVP_top60_DNA+RNA_seed | IGF1R | 1.427648 | KIRC | KMRC-1 | BMS-754807 |
| MVP_top60_DNA_seed | HDAC1 | 0.872801 | HNSC | CAL-33 | PCI-34051 | MVP_top60_DNA+RNA_seed | SMARCA4 | 1.431023 | KIRC | CAKI-1 | PFI3 |
| MVP_top60_DNA_seed | HDAC1 | 0.873084 | HNSC | DOK | Entinostat | MVP_top60_DNA+RNA_seed | SMARCA2 | 1.431023 | KIRC | CAKI-1 | PFI3 |
| MVP_top60_DNA_seed | HDAC3 | 0.873084 | HNSC | DOK | Entinostat | MVP_top60_DNA+RNA_seed | HDAC1 | 1.44453 | KIRC | TK10 | Romidepsin |
| MVP_top60_DNA_seed | AKT1 | 0.874726 | HNSC | SKN-3 | GSK2110183B | MVP_top60_DNA+RNA_seed | PIK3CA | 1.454868 | KIRC | TK10 | AZD8186 |
| MVP_top60_DNA_seed | SMARCA4 | 0.879654 | HNSC | LB771-HNC | PFI3 | MVP_top60_DNA+RNA_seed | PIK3CA | 1.45568 | KIRC | RXF393 | Buparlisib |

| | | | | | | | | | | | |
|---|---|---|---|---|---|---|---|---|---|---|---|
| MVP_top60_DNA_seed | AKT1 | 0.882982 | HNSC | BHY | Afuresertib | MVP_top60_DNA+RNA_seed | PIK3CD | 1.45568 | KIRC | RXF393 | Buparlisib |
| MVP_top60_DNA_seed | RAC1 | 0.886912 | HNSC | JHU-011 | EHT-1864 | MVP_top60_DNA+RNA_seed | HSP90AA1 | 1.481733 | KIRC | HA7-RCC | Luminespib |
| MVP_top60_DNA_seed | AKT1 | 0.896763 | HNSC | BICR78 | MK-2206 | MVP_top60_DNA+RNA_seed | HSP90AB1 | 1.481733 | KIRC | HA7-RCC | Luminespib |
| MVP_top60_DNA_seed | MDM2 | 0.89704 | HNSC | JHU-022 | Serdemetan | MVP_top60_DNA+RNA_seed | PIK3CA | 1.495973 | KIRC | SN12C | Alpelisib |
| MVP_top60_DNA_seed | PIK3CA | 0.898326 | HNSC | PCI-30 | Buparlisib | MVP_top60_DNA+RNA_seed | HSP90AA1 | 1.506671 | KIRC | KMRC-20 | Luminespib |
| MVP_top60_DNA_seed | AKT1 | 0.902163 | HNSC | SCC-15 | GSK2110183B | MVP_top60_DNA+RNA_seed | HSP90AB1 | 1.506671 | KIRC | KMRC-20 | Luminespib |
| MVP_top60_DNA_seed | AKT1 | 0.906913 | HNSC | CAL-27 | MK-2206 | MVP_top60_DNA+RNA_seed | HSP90AA1 | 1.507212 | KIRC | LB2241-RCC | Elesclomol |
| MVP_top60_DNA_seed | MDM2 | 0.911241 | HNSC | BICR10 | Nutlin-3a (-) | MVP_top60_DNA+RNA_seed | HSP90AB1 | 1.507212 | KIRC | LB2241-RCC | Elesclomol |
| MVP_top60_DNA_seed | MDM2 | 0.912133 | HNSC | OSC-20 | Nutlin-3a (-) | MVP_top60_DNA+RNA_seed | ESR1 | 1.509048 | KIRC | NCC021 | Tamoxifen |
| MVP_top60_DNA_seed | AR | 0.914436 | HNSC | OSC-19 | Bicalutamide | MVP_top60_DNA+RNA_seed | PIK3CA | 1.520458 | KIRC | TK10 | Buparlisib |
| MVP_top60_DNA_seed | EGFR | 0.916453 | HNSC | BICR31 | Erlotinib | MVP_top60_DNA+RNA_seed | PIK3CD | 1.520458 | KIRC | TK10 | Buparlisib |
| MVP_top60_DNA_seed | PIK3CA | 0.917451 | HNSC | BICR31 | GNE-317 | MVP_top60_DNA+RNA_seed | HDAC1 | 1.520889 | KIRC | OS-RC-2 | PCI-34051 |
| MVP_top60_DNA_seed | ESR1 | 0.917589 | HNSC | KON | Tamoxifen | MVP_top60_DNA+RNA_seed | HDAC1 | 1.521877 | KIRC | 769-P | Romidepsin |
| MVP_top60_DNA_seed | AKT1 | 0.919006 | HNSC | SCC-25 | AZD5363 | MVP_top60_DNA+RNA_seed | IGF1R | 1.549533 | KIRC | BB65-RCC | BMS-754807 |
| MVP_top60_DNA_seed | AKT1 | 0.923638 | HNSC | SAT | Afuresertib | MVP_top60_DNA+RNA_seed | IGF1R | 1.553473 | KIRC | SN12C | GSK1904529A |
| MVP_top60_DNA_seed | ESR1 | 0.925274 | HNSC | JHU-011 | Tamoxifen | MVP_top60_DNA+RNA_seed | HSP90AA1 | 1.559635 | KIRC | LB1047-RCC | Luminespib |
| MVP_top60_DNA_seed | HDAC1 | 0.92528 | HNSC | BICR78 | PCI-34051 | MVP_top60_DNA+RNA_seed | HSP90AB1 | 1.559635 | KIRC | LB1047-RCC | Luminespib |
| MVP_top60_DNA_seed | MAPK1 | 0.925914 | HNSC | BICR10 | Ulixertinib | MVP_top60_DNA+RNA_seed | AR | 1.572884 | KIRC | CAKI-1 | Bicalutamide |
| MVP_top60_DNA_seed | MAPK3 | 0.925914 | HNSC | BICR10 | Ulixertinib | MVP_top60_DNA+RNA_seed | MTOR | 1.576607 | KIRC | VMRC-RCZ | Temsirolimus |
| MVP_top60_DNA_seed | MAPK1 | 0.92664 | HNSC | LB771-HNC | Ulixertinib | MVP_top60_DNA+RNA_seed | IGF1R | 1.586322 | KIRC | TK10 | Linsitinib |
| MVP_top60_DNA_seed | MAPK3 | 0.92664 | HNSC | LB771-HNC | Ulixertinib | MVP_top60_DNA+RNA_seed | HDAC1 | 1.596102 | KIRC | NCC010 | Romidepsin |
| MVP_top60_DNA_seed | HDAC1 | 0.936625 | HNSC | HSC-4 | PCI-34051 | MVP_top60_DNA+RNA_seed | ESR1 | 1.600789 | KIRC | CAKI-1 | GDC0810 |
| MVP_top60_DNA_seed | CDK2 | 0.944462 | HNSC | DOK | MK-8776 | MVP_top60_DNA+RNA_seed | ESR1 | 1.601588 | KIRC | LB1047-RCC | Tamoxifen |
| MVP_top60_DNA_seed | AKT1 | 0.95102 | HNSC | Ca9-22 | Ipatasertib | MVP_top60_DNA+RNA_seed | ESR1 | 1.602068 | KIRC | RXF393 | Tamoxifen |
| MVP_top60_DNA_seed | HDAC1 | 0.952598 | HNSC | CAL-33 | Entinostat | MVP_top60_DNA+RNA_seed | ATM | 1.624105 | KIRC | TK10 | KU-55933 |
| MVP_top60_DNA_seed | HDAC3 | 0.952598 | HNSC | CAL-33 | Entinostat | MVP_top60_DNA+RNA_seed | HSP90AA1 | 1.637244 | KIRC | LB1047-RCC | CCT-018159 |
| MVP_top60_DNA_seed | AKT1 | 0.964992 | HNSC | SCC-9 | Afuresertib | MVP_top60_DNA+RNA_seed | HSP90AB1 | 1.637244 | KIRC | LB1047-RCC | CCT-018159 |
| MVP_top60_DNA_seed | PIK3CA | 0.968149 | HNSC | SKN-3 | GNE-317 | MVP_top60_DNA+RNA_seed | PIK3CA | 1.648698 | KIRC | VMRC-RCW | GNE-317 |
| MVP_top60_DNA_seed | EGFR | 0.974133 | HNSC | SCC-4 | Erlotinib | MVP_top60_DNA+RNA_seed | IGF1R | 1.653349 | KIRC | KMRC-1 | Linsitinib |
| MVP_top60_DNA_seed | EGFR | 0.977702 | HNSC | LB771-HNC | AZD3759 | MVP_top60_DNA+RNA_seed | HSP90AA1 | 1.679947 | KIRC | NCC021 | Tanespimycin |
| MVP_top60_DNA_seed | PIK3CA | 0.980543 | HNSC | LB771-HNC | AZD8186 | MVP_top60_DNA+RNA_seed | HSP90AB1 | 1.679947 | KIRC | NCC021 | Tanespimycin |
| MVP_top60_DNA_seed | AKT1 | 0.981333 | HNSC | PE-CA-PJ15 | Uprosertib | MVP_top60_DNA+RNA_seed | TP53 | 1.681904 | KIRC | VMRC-RCZ | MIRA-1 |
| MVP_top60_DNA_seed | AKT1 | 0.984442 | HNSC | SCC-4 | Ipatasertib | MVP_top60_DNA+RNA_seed | HDAC1 | 1.684918 | KIRC | OS-RC-2 | Romidepsin |
| MVP_top60_DNA_seed | AKT1 | 0.986695 | HNSC | PCI-30 | AT13148 | MVP_top60_DNA+RNA_seed | EP300 | 1.693076 | KIRC | TK10 | CPI-637 |
| MVP_top60_DNA_seed | AKT1 | 0.987301 | HNSC | SKN-3 | MK-2206 | MVP_top60_DNA+RNA_seed | ESR1 | 1.699914 | KIRC | SW156 | Tamoxifen |
| MVP_top60_DNA_seed | PIK3CA | 0.993353 | HNSC | PCI-4B | AZD8186 | MVP_top60_DNA+RNA_seed | IGF1R | 1.711974 | KIRC | KMRC-1 | GSK1904529A |
| MVP_top60_DNA_seed | AKT1 | 0.999321 | HNSC | BICR78 | Uprosertib | MVP_top60_DNA+RNA_seed | EP300 | 1.715111 | KIRC | RCC10RGB | CPI-637 |
| MVP_top60_DNA_seed | AKT1 | 1.005076 | HNSC | SAS | Uprosertib | MVP_top60_DNA+RNA_seed | PIK3CA | 1.741052 | KIRC | VMRC-RCZ | GNE-317 |
| MVP_top60_DNA_seed | AKT1 | 1.009659 | HNSC | SKN-3 | Ipatasertib | MVP_top60_DNA+RNA_seed | HSP90AA1 | 1.757652 | KIRC | HA7-RCC | CCT-018159 |
| MVP_top60_DNA_seed | RAC1 | 1.010736 | HNSC | FADU | EHT-1864 | MVP_top60_DNA+RNA_seed | HSP90AB1 | 1.757652 | KIRC | HA7-RCC | CCT-018159 |
| MVP_top60_DNA_seed | EGFR | 1.011777 | HNSC | BB49-HNC | AZD3759 | MVP_top60_DNA+RNA_seed | PIK3CA | 1.764756 | KIRC | SW156 | Buparlisib |

| | | | | | | | | | | | |
|---|---|---|---|---|---|---|---|---|---|---|---|
| MVP_top60_DNA_seed | CDK2 | 1.014634 | HNSC | SCC-4 | MK-8776 | MVP_top60_DNA+RNA_seed | PIK3CD | 1.764756 | KIRC | SW156 | Buparlisib |
| MVP_top60_DNA_seed | ESR1 | 1.017718 | HNSC | JHU-029 | Tamoxifen | MVP_top60_DNA+RNA_seed | HSP90AA1 | 1.774957 | KIRC | VMRC-RCZ | Tanespimycin |
| MVP_top60_DNA_seed | CDK2 | 1.018509 | HNSC | SAT | MK-8776 | MVP_top60_DNA+RNA_seed | HSP90AB1 | 1.774957 | KIRC | VMRC-RCZ | Tanespimycin |
| MVP_top60_DNA_seed | ESR1 | 1.021474 | HNSC | BHY | Tamoxifen | MVP_top60_DNA+RNA_seed | IGF1R | 1.782101 | KIRC | SW156 | NVP-ADW742 |
| MVP_top60_DNA_seed | EP300 | 1.021511 | HNSC | SAT | CPI-637 | MVP_top60_DNA+RNA_seed | HDAC1 | 1.832891 | KIRC | RCC10RGB | Entinostat |
| MVP_top60_DNA_seed | AKT1 | 1.021735 | HNSC | BHY | Ipatasertib | MVP_top60_DNA+RNA_seed | IGF1R | 1.835023 | KIRC | RCC-JW | BMS-754807 |
| MVP_top60_DNA_seed | AKT1 | 1.025658 | HNSC | CAL-27 | Afuresertib | MVP_top60_DNA+RNA_seed | HSP90AA1 | 1.844277 | KIRC | VMRC-RCW | Luminespib |
| MVP_top60_DNA_seed | TP53 | 1.029272 | HNSC | SAT | MIRA-1 | MVP_top60_DNA+RNA_seed | HSP90AB1 | 1.844277 | KIRC | VMRC-RCW | Luminespib |
| MVP_top60_DNA_seed | NTRK1 | 1.03143 | HNSC | HSC-4 | AZD1332 | MVP_top60_DNA+RNA_seed | ESR1 | 1.869394 | KIRC | RCC-AB | Tamoxifen |
| MVP_top60_DNA_seed | AKT1 | 1.04076 | HNSC | PCI-30 | Uprosertib | MVP_top60_DNA+RNA_seed | IGF1R | 1.884492 | KIRC | RCC10RGB | BMS-754807 |
| MVP_top60_DNA_seed | EGFR | 1.041565 | HNSC | LB771-HNC | Gefitinib | MVP_top60_DNA+RNA_seed | IGF1R | 1.917243 | KIRC | RCC-AB | NVP-ADW742 |
| MVP_top60_DNA_seed | HDAC1 | 1.04205 | HNSC | JHU-022 | Entinostat | MVP_top60_DNA+RNA_seed | IGF1R | 1.923712 | KIRC | TK10 | BMS-754807 |
| MVP_top60_DNA_seed | HDAC3 | 1.04205 | HNSC | JHU-022 | Entinostat | MVP_top60_DNA+RNA_seed | ESR1 | 1.931838 | KIRC | VMRC-RCW | GDC0810 |
| MVP_top60_DNA_seed | MDM2 | 1.043345 | HNSC | SAT | Nutlin-3a (-) | MVP_top60_DNA+RNA_seed | SMARCA4 | 1.986562 | KIRC | TK10 | PFI3 |
| MVP_top60_DNA_seed | PIK3CA | 1.047838 | HNSC | BICR78 | Buparlisib | MVP_top60_DNA+RNA_seed | SMARCA2 | 1.986562 | KIRC | TK10 | PFI3 |
| MVP_top60_DNA_seed | PIK3CA | 1.048236 | HNSC | SKN-3 | Buparlisib | MVP_top60_DNA+RNA_seed | SMARCA4 | 1.989225 | KIRC | RCC10RGB | PFI3 |
| MVP_top60_DNA_seed | EGFR | 1.051072 | HNSC | LB771-HNC | Sapitinib | MVP_top60_DNA+RNA_seed | SMARCA2 | 1.989225 | KIRC | RCC10RGB | PFI3 |
| MVP_top60_DNA_seed | PIK3CA | 1.05214 | HNSC | SCC90 | AZD8186 | MVP_top60_DNA+RNA_seed | HDAC1 | 1.995917 | KIRC | RCC10RGB | Romidepsin |
| MVP_top60_DNA_seed | AKT1 | 1.052608 | HNSC | BICR78 | Uprosertib | MVP_top60_DNA+RNA_seed | ESR1 | 2.093508 | KIRC | VMRC-RCW | Tamoxifen |
| MVP_top60_DNA_seed | AKT1 | 1.05274 | HNSC | OSC-20 | AZD5363 | MVP_top60_DNA+RNA_seed | IGF1R | 2.101741 | KIRC | RCC10RGB | GSK1904529A |
| MVP_top60_DNA_seed | ESR1 | 1.053722 | HNSC | PCI-4B | Tamoxifen | MVP_top60_DNA+RNA_seed | ESR1 | 2.143903 | KIRC | CAKI-1 | Tamoxifen |
| MVP_top60_DNA_seed | SRC | 1.059588 | HNSC | PCI-30 | Bosutinib | MVP_top60_DNA+RNA_seed | HSP90AA1 | 2.148358 | KIRC | VMRC-RCW | Elesclomol |
| MVP_top60_DNA_seed | EP300 | 1.066557 | HNSC | SAT | SGC-CBP30 | MVP_top60_DNA+RNA_seed | HSP90AB1 | 2.148358 | KIRC | VMRC-RCW | Elesclomol |
| MVP_top60_DNA_seed | CREBBP | 1.066557 | HNSC | SAT | SGC-CBP30 | MVP_top60_DNA+RNA_seed | PIK3CA | 2.185823 | KIRC | TK10 | Alpelisib |
| MVP_top60_DNA_seed | MDM2 | 1.068479 | HNSC | BICR22 | Serdemetan | MVP_top60_DNA+RNA_seed | ESR1 | 2.191646 | KIRC | VMRC-RCZ | GDC0810 |
| MVP_top60_DNA_seed | CDK2 | 1.071971 | HNSC | SCC-15 | MK-8776 | MVP_top60_DNA+RNA_seed | SMARCA4 | 2.203211 | KIRC | VMRC-RCZ | PFI3 |
| MVP_top60_DNA_seed | AKT1 | 1.077354 | HNSC | SCC-9 | Ipatasertib | MVP_top60_DNA+RNA_seed | SMARCA2 | 2.203211 | KIRC | VMRC-RCZ | PFI3 |
| MVP_top60_DNA_seed | ESR1 | 1.078528 | HNSC | SCC90 | GDC0810 | MVP_top60_DNA+RNA_seed | IGF1R | 2.212351 | KIRC | VMRC-RCZ | GSK1904529A |
| MVP_top60_DNA_seed | PIK3CA | 1.080515 | HNSC | BICR78 | AZD8186 | MVP_top60_DNA+RNA_seed | PIK3CA | 2.237278 | KIRC | VMRC-RCZ | Buparlisib |
| MVP_top60_DNA_seed | CDK2 | 1.084335 | HNSC | PCI-4B | MK-8776 | MVP_top60_DNA+RNA_seed | PIK3CD | 2.237278 | KIRC | VMRC-RCZ | Buparlisib |
| MVP_top60_DNA_seed | HDAC1 | 1.084437 | HNSC | SCC-9 | PCI-34051 | MVP_top60_DNA+RNA_seed | HDAC1 | 2.238131 | KIRC | VMRC-RCZ | PCI-34051 |
| MVP_top60_DNA_seed | PIK3CA | 1.084815 | HNSC | SAT | Buparlisib | MVP_top60_DNA+RNA_seed | HSP90AA1 | 2.299331 | KIRC | RCC-MF | Luminespib |
| MVP_top60_DNA_seed | AKT1 | 1.085947 | HNSC | SCC-15 | Uprosertib | MVP_top60_DNA+RNA_seed | HSP90AB1 | 2.299331 | KIRC | RCC-MF | Luminespib |
| MVP_top60_DNA_seed | PIK3CA | 1.086612 | HNSC | OSC-20 | GNE-317 | MVP_top60_DNA+RNA_seed | HDAC1 | 2.301832 | KIRC | VMRC-RCZ | Entinostat |
| MVP_top60_DNA_seed | MDM2 | 1.091434 | HNSC | BICR78 | Nutlin-3a (-) | MVP_top60_DNA+RNA_seed | MTOR | 2.324809 | KIRC | VMRC-RCW | Temsirolimus |
| MVP_top60_DNA_seed | TP53 | 1.097963 | HNSC | SCC90 | MIRA-1 | MVP_top60_DNA+RNA_seed | HDAC1 | 2.335545 | KIRC | TK10 | PCI-34051 |
| MVP_top60_DNA_seed | EGFR | 1.101466 | HNSC | BICR10 | Osimertinib | MVP_top60_DNA+RNA_seed | IGF1R | 2.425755 | KIRC | VMRC-RCZ | BMS-536924 |
| MVP_top60_DNA_seed | PIK3CA | 1.103864 | HNSC | SCC-9 | GNE-317 | MVP_top60_DNA+RNA_seed | SMARCA4 | 2.435711 | KIRC | VMRC-RCW | PFI3 |
| MVP_top60_DNA_seed | MAPK1 | 1.10537 | HNSC | PCI-30 | VX-11e | MVP_top60_DNA+RNA_seed | SMARCA2 | 2.435711 | KIRC | VMRC-RCW | PFI3 |
| MVP_top60_DNA_seed | AKT1 | 1.105817 | HNSC | SCC-4 | Afuresertib | MVP_top60_DNA+RNA_seed | ESR1 | 2.437983 | KIRC | TK10 | GDC0810 |
| MVP_top60_DNA_seed | PIK3CA | 1.113835 | HNSC | HSC-4 | Alpelisib | MVP_top60_DNA+RNA_seed | IGF1R | 2.442234 | KIRC | VMRC-RCZ | Linsitinib |

| | | | | | | | | | | | |
|---|---|---|---|---|---|---|---|---|---|---|---|
| MVP_top60_DNA_seed | AKT1 | 1.119829 | HNSC | OSC-20 | GSK2110183B | MVP_top60_DNA+RNA_seed | IGF1R | 2.446206 | KIRC | VMRC-RCW | Linsitinib |
| MVP_top60_DNA_seed | AKT1 | 1.120921 | HNSC | OSC-20 | AT13148 | MVP_top60_DNA+RNA_seed | HDAC1 | 2.471239 | KIRC | SN12C | Entinostat |
| MVP_top60_DNA_seed | AKT1 | 1.128635 | HNSC | SCC-25 | MK-2206 | MVP_top60_DNA+RNA_seed | IGF1R | 2.629637 | KIRC | TK10 | NVP-ADW742 |
| MVP_top60_DNA_seed | AKT1 | 1.13029 | HNSC | LB771-HNC | GSK2110183B | MVP_top60_DNA+RNA_seed | EP300 | 2.747749 | KIRC | TK10 | SGC-CBP30 |
| MVP_top60_DNA_seed | HDAC1 | 1.141261 | HNSC | BICR10 | Entinostat | MVP_top60_DNA+RNA_seed | IGF1R | 2.754441 | KIRC | VMRC-RCZ | BMS-754807 |
| MVP_top60_DNA_seed | HDAC3 | 1.141261 | HNSC | BICR10 | Entinostat | MVP_top60_DNA+RNA_seed | HSP90AA1 | 2.801677 | KIRC | VMRC-RCW | Tanespimycin |
| MVP_top60_DNA_seed | HDAC1 | 1.151543 | HNSC | SCC-4 | Entinostat | MVP_top60_DNA+RNA_seed | HSP90AB1 | 2.801677 | KIRC | VMRC-RCW | Tanespimycin |
| MVP_top60_DNA_seed | HDAC3 | 1.151543 | HNSC | SCC-4 | Entinostat | MVP_top60_DNA+RNA_seed | HSP90AA1 | 2.810492 | KIRC | RCC10RGB | Tanespimycin |
| MVP_top60_DNA_seed | RAC1 | 1.158652 | HNSC | SCC90 | EHT-1864 | MVP_top60_DNA+RNA_seed | HSP90AB1 | 2.810492 | KIRC | RCC10RGB | Tanespimycin |
| MVP_top60_DNA_seed | TP53 | 1.159736 | HNSC | JHU-022 | MIRA-1 | MVP_top60_DNA+RNA_seed | IGF1R | 2.866762 | KIRC | TK10 | GSK1904529A |
| MVP_top60_DNA_seed | TP53 | 1.159898 | HNSC | LB771-HNC | MIRA-1 | MVP_top60_DNA+RNA_seed | PIK3CA | 2.888068 | KIRC | VMRC-RCZ | Alpelisib |
| MVP_top60_DNA_seed | AKT1 | 1.165636 | HNSC | BICR78 | GSK2110183B | MVP_top60_DNA+RNA_seed | ESR1 | 2.932502 | KIRC | TK10 | Tamoxifen |
| MVP_top60_DNA_seed | ESR1 | 1.166551 | HNSC | SAT | GDC0810 | MVP_top60_DNA+RNA_seed | IGF1R | 2.964638 | KIRC | VMRC-RCZ | NVP-ADW742 |
| MVP_top60_DNA_seed | PIK3CA | 1.171475 | HNSC | SKN-3 | Alpelisib | MVP_top60_DNA+RNA_seed | HSP90AA1 | 3.168001 | KIRC | VMRC-RCZ | Luminespib |
| MVP_top60_DNA_seed | HDAC1 | 1.17437 | HNSC | BHY | Entinostat | MVP_top60_DNA+RNA_seed | HSP90AB1 | 3.168001 | KIRC | VMRC-RCZ | Luminespib |
| MVP_top60_DNA_seed | HDAC3 | 1.17437 | HNSC | BHY | Entinostat | MVP_top60_DNA+RNA_seed | PIK3CA | 3.212868 | KIRC | VMRC-RCW | Buparlisib |
| MVP_top60_DNA_seed | MDM2 | 1.174936 | HNSC | SCC-15 | Serdemetan | MVP_top60_DNA+RNA_seed | PIK3CD | 3.212868 | KIRC | VMRC-RCW | Buparlisib |
| MVP_top60_DNA_seed | MAPK1 | 1.177578 | HNSC | PCI-30 | Ulixertinib | MVP_top60_DNA+RNA_seed | HSP90AA1 | 3.345674 | KIRC | RCC10RGB | Luminespib |
| MVP_top60_DNA_seed | MAPK3 | 1.177578 | HNSC | PCI-30 | Ulixertinib | MVP_top60_DNA+RNA_seed | HSP90AB1 | 3.345674 | KIRC | RCC10RGB | Luminespib |
| MVP_top60_DNA_seed | AKT1 | 1.178177 | HNSC | OSC-20 | MK-2206 | MVP_top60_DNA+RNA_seed | EGFR | -5.4375 | LUAD | NCI-H1648 | Lapatinib |
| MVP_top60_DNA_seed | ESR1 | 1.17935 | HNSC | PCI-30 | GDC0810 | MVP_top60_DNA+RNA_seed | EGFR | -5.341957 | LUAD | HCC-827 | Afatinib |
| MVP_top60_DNA_seed | AKT1 | 1.184001 | HNSC | BICR22 | MK-2206 | MVP_top60_DNA+RNA_seed | MET | -5.097923 | LUAD | NCI-H1993 | AZD6094 |
| MVP_top60_DNA_seed | ESR1 | 1.193992 | HNSC | PE-CA-PJ15 | Tamoxifen | MVP_top60_DNA+RNA_seed | EGFR | -5.064987 | LUAD | NCI-H1648 | Erlotinib |
| MVP_top60_DNA_seed | HSP90AA1 | 1.19482 | HNSC | PCI-6A | CCT-018159 | MVP_top60_DNA+RNA_seed | EGFR | -4.873695 | LUAD | H3255 | Gefitinib |
| MVP_top60_DNA_seed | PIK3CA | 1.205565 | HNSC | BICR78 | GNE-317 | MVP_top60_DNA+RNA_seed | EGFR | -4.396938 | LUAD | HCC-827 | Gefitinib |
| MVP_top60_DNA_seed | AKT1 | 1.206387 | HNSC | PCI-30 | MK-2206 | MVP_top60_DNA+RNA_seed | EGFR | -4.210085 | LUAD | H3255 | Afatinib |
| MVP_top60_DNA_seed | CDK2 | 1.2093 | HNSC | BICR78 | Dinaciclib | MVP_top60_DNA+RNA_seed | EGFR | -4.088424 | LUAD | HCC-827 | Afatinib |
| MVP_top60_DNA_seed | MAPK1 | 1.210057 | HNSC | OSC-19 | Ulixertinib | MVP_top60_DNA+RNA_seed | EGFR | -4.081692 | LUAD | H3255 | Afatinib |
| MVP_top60_DNA_seed | MAPK3 | 1.210057 | HNSC | OSC-19 | Ulixertinib | MVP_top60_DNA+RNA_seed | EGFR | -3.986983 | LUAD | H3255 | Cetuximab |
| MVP_top60_DNA_seed | NTRK1 | 1.213696 | HNSC | SCC-9 | AZD1332 | MVP_top60_DNA+RNA_seed | SMARCA4 | -3.886817 | LUAD | NCI-H1650 | PFI-3 |
| MVP_top60_DNA_seed | ESR1 | 1.21992 | HNSC | SCC-25 | Tamoxifen | MVP_top60_DNA+RNA_seed | EGFR | -3.635979 | LUAD | PC-14 | Afatinib |
| MVP_top60_DNA_seed | RAC1 | 1.230965 | HNSC | HO-1-u-1 | EHT-1864 | MVP_top60_DNA+RNA_seed | MET | -3.344635 | LUAD | RERF-LC-MS | AZD6094 |
| MVP_top60_DNA_seed | ESR1 | 1.231004 | HNSC | JHU-022 | GDC0810 | MVP_top60_DNA+RNA_seed | SRC | -3.312942 | LUAD | HCC-827 | Bosutinib |
| MVP_top60_DNA_seed | HDAC1 | 1.231978 | HNSC | SCC-15 | Entinostat | MVP_top60_DNA+RNA_seed | EGFR | -3.30175 | LUAD | EKVX | Lapatinib |
| MVP_top60_DNA_seed | HDAC3 | 1.231978 | HNSC | SCC-15 | Entinostat | MVP_top60_DNA+RNA_seed | EGFR | -3.29092 | LUAD | PC-3_[JPC-3] | Afatinib |
| MVP_top60_DNA_seed | EP300 | 1.24698 | HNSC | SCC90 | SGC-CBP30 | MVP_top60_DNA+RNA_seed | EGFR | -3.289851 | LUAD | NCI-H1568 | Cetuximab |
| MVP_top60_DNA_seed | CREBBP | 1.24698 | HNSC | SCC90 | SGC-CBP30 | MVP_top60_DNA+RNA_seed | EGFR | -3.250339 | LUAD | EMC-BAC-1 | Cetuximab |
| MVP_top60_DNA_seed | MAPK1 | 1.277441 | HNSC | PCI-30 | Ulixertinib | MVP_top60_DNA+RNA_seed | EGFR | -3.247407 | LUAD | Calu-3 | PF-00299804 |
| MVP_top60_DNA_seed | MAPK3 | 1.277441 | HNSC | PCI-30 | Ulixertinib | MVP_top60_DNA+RNA_seed | EGFR | -3.241959 | LUAD | EKVX | Erlotinib |
| MVP_top60_DNA_seed | PIK3CA | 1.278254 | HNSC | SCC90 | Alpelisib | MVP_top60_DNA+RNA_seed | EGFR | -3.137964 | LUAD | H3255 | AZD8931 |
| MVP_top60_DNA_seed | HDAC1 | 1.280992 | HNSC | BICR10 | PCI-34051 | MVP_top60_DNA+RNA_seed | EGFR | -3.128579 | LUAD | COR-L105 | Pelitinib |

| | | | | | | | | | | | |
|---|---|---|---|---|---|---|---|---|---|---|---|
| MVP_top60_DNA_seed | NTRK1 | 1.283032 | HNSC | BICR78 | AZD1332 | MVP_top60_DNA+RNA_seed | EGFR | -3.121584 | LUAD | Calu-3 | AST-1306 |
| MVP_top60_DNA_seed | AKT1 | 1.285299 | HNSC | BHY | Uprosertib | MVP_top60_DNA+RNA_seed | EGFR | -3.094491 | LUAD | HCC-827 | PF-00299804 |
| MVP_top60_DNA_seed | HDAC1 | 1.297564 | HNSC | SCC-25 | Entinostat | MVP_top60_DNA+RNA_seed | EGFR | -3.061442 | LUAD | PC-3_[JPC-3] | Afatinib |
| MVP_top60_DNA_seed | HDAC3 | 1.297564 | HNSC | SCC-25 | Entinostat | MVP_top60_DNA+RNA_seed | EGFR | -3.048994 | LUAD | PC-14 | Afatinib |
| MVP_top60_DNA_seed | HDAC1 | 1.298562 | HNSC | SCC-15 | PCI-34051 | MVP_top60_DNA+RNA_seed | EGFR | -3.006622 | LUAD | HCC-827 | AZD8931 |
| MVP_top60_DNA_seed | AKT1 | 1.300263 | HNSC | SAS | AT13148 | MVP_top60_DNA+RNA_seed | HSP90AA1 | -2.986095 | LUAD | A549 | Tanespimycin |
| MVP_top60_DNA_seed | AKT1 | 1.303854 | HNSC | OSC-20 | Ipatasertib | MVP_top60_DNA+RNA_seed | EGFR | -2.962385 | LUAD | PC-14 | AZD8931 |
| MVP_top60_DNA_seed | MAPK1 | 1.308048 | HNSC | LB771-HNC | Ulixertinib | MVP_top60_DNA+RNA_seed | EGFR | -2.961245 | LUAD | NCI-H1623 | PF-00299804 |
| MVP_top60_DNA_seed | MAPK3 | 1.308048 | HNSC | LB771-HNC | Ulixertinib | MVP_top60_DNA+RNA_seed | EGFR | -2.959705 | LUAD | NCI-H1666 | Cetuximab |
| MVP_top60_DNA_seed | AKT1 | 1.315351 | HNSC | SCC90 | AT13148 | MVP_top60_DNA+RNA_seed | EGFR | -2.947302 | LUAD | EMC-BAC-1 | Afatinib |
| MVP_top60_DNA_seed | CDK2 | 1.316967 | HNSC | SCC90 | MK-8776 | MVP_top60_DNA+RNA_seed | EGFR | -2.942942 | LUAD | COR-L105 | Afatinib |
| MVP_top60_DNA_seed | AKT1 | 1.317668 | HNSC | OSC-20 | Uprosertib | MVP_top60_DNA+RNA_seed | MDM2 | -2.854602 | LUAD | HCC-78 | Serdemetan |
| MVP_top60_DNA_seed | AKT1 | 1.318083 | HNSC | SCC-4 | AZD5363 | MVP_top60_DNA+RNA_seed | IGF1R | -2.847594 | LUAD | NCI-H2342 | GSK1904529A |
| MVP_top60_DNA_seed | AKT1 | 1.325702 | HNSC | SKN-3 | AT13148 | MVP_top60_DNA+RNA_seed | EGFR | -2.79094 | LUAD | Calu-3 | CI-1033 |
| MVP_top60_DNA_seed | HDAC1 | 1.330086 | HNSC | HSC-4 | Entinostat | MVP_top60_DNA+RNA_seed | SRC | -2.787059 | LUAD | NCI-H1648 | Saracatinib |
| MVP_top60_DNA_seed | HDAC3 | 1.330086 | HNSC | HSC-4 | Entinostat | MVP_top60_DNA+RNA_seed | AKT1 | -2.773876 | LUAD | NCI-H3122 | AKT inhibitor VIII |
| MVP_top60_DNA_seed | PIK3CA | 1.349365 | HNSC | SCC-25 | AZD8186 | MVP_top60_DNA+RNA_seed | IGF1R | -2.773783 | LUAD | NCI-H2122 | BMS-754807 |
| MVP_top60_DNA_seed | NTRK1 | 1.359636 | HNSC | SCC90 | GW441756 | MVP_top60_DNA+RNA_seed | MAPK14 | -2.753005 | LUAD | PC-3_[JPC-3] | TAK-715 |
| MVP_top60_DNA_seed | EP300 | 1.368441 | HNSC | JHU-022 | SGC-CBP30 | MVP_top60_DNA+RNA_seed | EGFR | -2.730621 | LUAD | EMC-BAC-1 | Afatinib |
| MVP_top60_DNA_seed | CREBBP | 1.368441 | HNSC | JHU-022 | SGC-CBP30 | MVP_top60_DNA+RNA_seed | EGFR | -2.717338 | LUAD | EMC-BAC-1 | AZD8931 |
| MVP_top60_DNA_seed | MDM2 | 1.371424 | HNSC | OSC-19 | Serdemetan | MVP_top60_DNA+RNA_seed | EGFR | -2.707261 | LUAD | NCI-H1781 | PF-00299804 |
| MVP_top60_DNA_seed | AKT1 | 1.380644 | HNSC | FADU | Uprosertib | MVP_top60_DNA+RNA_seed | EGFR | -2.697602 | LUAD | H3255 | PF-00299804 |
| MVP_top60_DNA_seed | NTRK1 | 1.381135 | HNSC | LB771-HNC | AZD1332 | MVP_top60_DNA+RNA_seed | EGFR | -2.690145 | LUAD | NCI-H1623 | CI-1033 |
| MVP_top60_DNA_seed | RAC1 | 1.389903 | HNSC | SKN-3 | EHT-1864 | MVP_top60_DNA+RNA_seed | EGFR | -2.651437 | LUAD | NCI-H1568 | AZD8931 |
| MVP_top60_DNA_seed | AKT1 | 1.39623 | HNSC | SCC-25 | Ipatasertib | MVP_top60_DNA+RNA_seed | EGFR | -2.64536 | LUAD | NCI-H1623 | AST-1306 |
| MVP_top60_DNA_seed | AKT1 | 1.397851 | HNSC | FADU | Afuresertib | MVP_top60_DNA+RNA_seed | EGFR | -2.630997 | LUAD | HCC-827 | CI-1033 |
| MVP_top60_DNA_seed | AKT1 | 1.409904 | HNSC | SCC-15 | Ipatasertib | MVP_top60_DNA+RNA_seed | EGFR | -2.624419 | LUAD | PC-3_[JPC-3] | Gefitinib |
| MVP_top60_DNA_seed | EGFR | 1.410529 | HNSC | LB771-HNC | Afatinib | MVP_top60_DNA+RNA_seed | SRC | -2.604488 | LUAD | LC-2-ad | WH-4-023 |
| MVP_top60_DNA_seed | EP300 | 1.413502 | HNSC | OSC-20 | CPI-637 | MVP_top60_DNA+RNA_seed | BRAF | -2.600217 | LUAD | Calu-6 | AZ628 |
| MVP_top60_DNA_seed | AKT1 | 1.426154 | HNSC | SCC-15 | AT13148 | MVP_top60_DNA+RNA_seed | EGFR | -2.576548 | LUAD | NCI-H1623 | Afatinib |
| MVP_top60_DNA_seed | NTRK1 | 1.440895 | HNSC | OSC-20 | AZD1332 | MVP_top60_DNA+RNA_seed | AKT1 | -2.525469 | LUAD | NCI-H2342 | AKT inhibitor VIII |
| MVP_top60_DNA_seed | MDM2 | 1.447244 | HNSC | JHU-022 | Nutlin-3a (-) | MVP_top60_DNA+RNA_seed | IGF1R | -2.520279 | LUAD | NCI-H2122 | Linsitinib |
| MVP_top60_DNA_seed | HSP90AA1 | 1.450847 | HNSC | SCC-25 | CCT-018159 | MVP_top60_DNA+RNA_seed | MET | -2.48919 | LUAD | NCI-H2023 | AZD6094 |
| MVP_top60_DNA_seed | CDK2 | 1.457415 | HNSC | HSC-4 | AZD5438 | MVP_top60_DNA+RNA_seed | IGF1R | -2.475094 | LUAD | NCI-H1355 | BMS-754807 |
| MVP_top60_DNA_seed | MDM2 | 1.458693 | HNSC | SKN-3 | Serdemetan | MVP_top60_DNA+RNA_seed | IGF1R | -2.469308 | LUAD | NCI-H1355 | Linsitinib |
| MVP_top60_DNA_seed | MAPK1 | 1.467661 | HNSC | HSC-4 | VX-11e | MVP_top60_DNA+RNA_seed | SRC | -2.458888 | LUAD | NCI-H1755 | NVP-BHG712 |
| MVP_top60_DNA_seed | EP300 | 1.47024 | HNSC | SCC90 | CPI-637 | MVP_top60_DNA+RNA_seed | EGFR | -2.449449 | LUAD | NCI-H1666 | AZD8931 |
| MVP_top60_DNA_seed | AKT1 | 1.47457 | HNSC | SCC-4 | AT13148 | MVP_top60_DNA+RNA_seed | EGFR | -2.431793 | LUAD | H3255 | Pelitinib |
| MVP_top60_DNA_seed | AKT1 | 1.476381 | HNSC | SCC-15 | AZD5363 | MVP_top60_DNA+RNA_seed | EGFR | -2.403001 | LUAD | NCI-H1623 | Afatinib |
| MVP_top60_DNA_seed | AKT1 | 1.486621 | HNSC | SCC-9 | AT13148 | MVP_top60_DNA+RNA_seed | EGFR | -2.400405 | LUAD | NCI-H1568 | Afatinib |
| MVP_top60_DNA_seed | CDK2 | 1.498273 | HNSC | BICR78 | AZD5438 | MVP_top60_DNA+RNA_seed | EGFR | -2.390909 | LUAD | NCI-H1666 | Afatinib |

| | | | | | | | | | | | |
|---|---|---|---|---|---|---|---|---|---|---|---|
| MVP_top60_DNA_seed | AKT1 | 1.498828 | HNSC | SKN-3 | Uprosertib | MVP_top60_DNA+RNA_seed | EGFR | -2.390045 | LUAD | PC-14 | Gefitinib |
| MVP_top60_DNA_seed | HDAC1 | 1.500016 | HNSC | SCC90 | PCI-34051 | MVP_top60_DNA+RNA_seed | EGFR | -2.375309 | LUAD | NCI-H1975 | Pelitinib |
| MVP_top60_DNA_seed | MDM2 | 1.5071 | HNSC | LB771-HNC | Serdemetan | MVP_top60_DNA+RNA_seed | EGFR | -2.373737 | LUAD | HCC-827 | Pelitinib |
| MVP_top60_DNA_seed | MAPK1 | 1.511524 | HNSC | OSC-20 | Ulixertinib | MVP_top60_DNA+RNA_seed | HSP90AA1 | -2.371967 | LUAD | ABC-1 | Elesclomol |
| MVP_top60_DNA_seed | MAPK3 | 1.511524 | HNSC | OSC-20 | Ulixertinib | MVP_top60_DNA+RNA_seed | EGFR | -2.36446 | LUAD | NCI-H1781 | Afatinib |
| MVP_top60_DNA_seed | EGFR | 1.529307 | HNSC | SCC-25 | Afatinib | MVP_top60_DNA+RNA_seed | AKT1 | -2.358798 | LUAD | NCI-H1755 | AKT inhibitor VIII |
| MVP_top60_DNA_seed | MDM2 | 1.596871 | HNSC | SCC90 | Serdemetan | MVP_top60_DNA+RNA_seed | AKT1 | -2.351603 | LUAD | NCI-H1563 | AKT inhibitor VIII |
| MVP_top60_DNA_seed | NTRK1 | 1.597971 | HNSC | SCC90 | AZD1332 | MVP_top60_DNA+RNA_seed | EGFR | -2.333781 | LUAD | H3255 | CI-1033 |
| MVP_top60_DNA_seed | AKT1 | 1.606337 | HNSC | LB771-HNC | AT13148 | MVP_top60_DNA+RNA_seed | BRAF | -2.314637 | LUAD | LC-2-ad | HG6-64-1 |
| MVP_top60_DNA_seed | HSP90AA1 | 1.648471 | HNSC | BHY | Elesclomol | MVP_top60_DNA+RNA_seed | EGFR | -2.301404 | LUAD | H3255 | AST-1306 |
| MVP_top60_DNA_seed | EGFR | 1.656269 | HNSC | BICR31 | Afatinib | MVP_top60_DNA+RNA_seed | EGFR | -2.300195 | LUAD | Calu-3 | Afatinib |
| MVP_top60_DNA_seed | HSP90AA1 | 1.671851 | HNSC | SCC-25 | Elesclomol | MVP_top60_DNA+RNA_seed | IGF1R | -2.275211 | LUAD | LC-2-ad | BMS-536924 |
| MVP_top60_DNA_seed | PIK3CA | 1.68388 | HNSC | BHY | Buparlisib | MVP_top60_DNA+RNA_seed | HSP90AA1 | -2.268609 | LUAD | NCI-H3122 | CCT-018159 |
| MVP_top60_DNA_seed | PIK3CA | 1.696895 | HNSC | SCC90 | GNE-317 | MVP_top60_DNA+RNA_seed | EGFR | -2.236769 | LUAD | NCI-H1666 | Afatinib |
| MVP_top60_DNA_seed | CDK2 | 1.708453 | HNSC | BICR10 | AZD5438 | MVP_top60_DNA+RNA_seed | BRAF | -2.163684 | LUAD | EMC-BAC-2 | Dabrafenib |
| MVP_top60_DNA_seed | MDM2 | 1.72561 | HNSC | BICR78 | Serdemetan | MVP_top60_DNA+RNA_seed | AKT1 | -2.125675 | LUAD | NCI-H1437 | AKT inhibitor VIII |
| MVP_top60_DNA_seed | AKT1 | 1.7268 | HNSC | OSC-20 | Afuresertib | MVP_top60_DNA+RNA_seed | EGFR | -2.11529 | LUAD | Calu-3 | AZD8931 |
| MVP_top60_DNA_seed | MAPK1 | 1.73732 | HNSC | OSC-20 | Ulixertinib | MVP_top60_DNA+RNA_seed | AKT1 | -2.112531 | LUAD | NCI-H1975 | BAY AKT1 |
| MVP_top60_DNA_seed | MAPK3 | 1.73732 | HNSC | OSC-20 | Ulixertinib | MVP_top60_DNA+RNA_seed | EGFR | -2.102305 | LUAD | PC-3_[JPC-3] | Pelitinib |
| MVP_top60_DNA_seed | HDAC1 | 1.796846 | HNSC | OSC-20 | Entinostat | MVP_top60_DNA+RNA_seed | EGFR | -2.096039 | LUAD | PC-3_[JPC-3] | AZD8931 |
| MVP_top60_DNA_seed | HDAC3 | 1.796846 | HNSC | OSC-20 | Entinostat | MVP_top60_DNA+RNA_seed | AKT1 | -2.080775 | LUAD | LC-2-ad | A-443654 |
| MVP_top60_DNA_seed | PIK3CA | 1.801314 | HNSC | SCC90 | Buparlisib | MVP_top60_DNA+RNA_seed | HSP90AA1 | -2.045994 | LUAD | HCC-44 | Tanespimycin |
| MVP_top60_DNA_seed | PIK3CA | 1.832461 | HNSC | SCC-25 | GNE-317 | MVP_top60_DNA+RNA_seed | EGFR | -2.022879 | LUAD | NCI-H1666 | Gefitinib |
| MVP_top60_DNA_seed | HSP90AA1 | 1.998461 | HNSC | LB771-HNC | Tanespimycin | MVP_top60_DNA+RNA_seed | EGFR | -2.022512 | LUAD | NCI-H1781 | Pelitinib |
| MVP_top60_DNA_seed | AKT1 | 2.035544 | HNSC | JHU-022 | AT13148 | MVP_top60_DNA+RNA_seed | AURKA | -2.021489 | LUAD | LC-2-ad | Tozasertib |
| MVP_top60_DNA_seed | HDAC1 | 2.063865 | HNSC | LB771-HNC | Entinostat | MVP_top60_DNA+RNA_seed | EGFR | -1.990933 | LUAD | EMC-BAC-1 | Gefitinib |
| MVP_top60_DNA_seed | HDAC3 | 2.063865 | HNSC | LB771-HNC | Entinostat | MVP_top60_DNA+RNA_seed | EGFR | -1.990647 | LUAD | NCI-H1568 | PF-00299804 |
| MVP_top60_DNA_seed | AKT1 | 2.100703 | HNSC | BHY | MK-2206 | MVP_top60_DNA+RNA_seed | EGFR | -1.961863 | LUAD | PC-14 | Pelitinib |
| MVP_top60_DNA_seed | RAC1 | 2.142936 | HNSC | PCI-30 | EHT-1864 | MVP_top60_DNA+RNA_seed | IGF1R | -1.958465 | LUAD | NCI-H1648 | BMS-536924 |
| MVP_top60_DNA_seed | MAPK1 | 2.263031 | HNSC | BHY | Ulixertinib | MVP_top60_DNA+RNA_seed | EGFR | -1.944651 | LUAD | NCI-H1781 | AST-1306 |
| MVP_top60_DNA_seed | MAPK3 | 2.263031 | HNSC | BHY | Ulixertinib | MVP_top60_DNA+RNA_seed | EGFR | -1.928812 | LUAD | COR-L105 | Afatinib |
| MVP_top60_DNA_seed | HDAC1 | 2.270484 | HNSC | SCC90 | Entinostat | MVP_top60_DNA+RNA_seed | EGFR | -1.913401 | LUAD | NCI-H1623 | AZD8931 |
| MVP_top60_DNA_seed | HDAC3 | 2.270484 | HNSC | SCC90 | Entinostat | MVP_top60_DNA+RNA_seed | ATM | -1.905401 | LUAD | HCC-44 | KU-60019 |
| MVP_top60_DNA_seed | HDAC1 | 2.436313 | HNSC | SCC90 | Romidepsin | MVP_top60_DNA+RNA_seed | AKT1 | -1.900726 | LUAD | NCI-H1563 | MK-2206 |
| MVP_top60_DNA_seed | HDAC3 | 2.436313 | HNSC | SCC90 | Romidepsin | MVP_top60_DNA+RNA_seed | IGF1R | -1.89741 | LUAD | NCI-H358 | BMS-754807 |
| MVP_top60_DNA_seed | HSP90AA1 | 2.499784 | HNSC | SCC-25 | Luminespib | MVP_top60_DNA+RNA_seed | MDM2 | -1.894813 | LUAD | EMC-BAC-2 | Nutlin-3a (-) |
| MVP_top60_DNA_seed | HSP90AA1 | 2.986528 | HNSC | LB771-HNC | Luminespib | MVP_top60_DNA+RNA_seed | BRAF | -1.88687 | LUAD | NCI-H2122 | PLX-4720 |
| MVP_top60_DNA_seed | PIK3CA | 3.472935 | HNSC | SCC-25 | Buparlisib | MVP_top60_DNA+RNA_seed | MET | -1.882533 | LUAD | NCI-H2291 | JNJ38877605 |
| MVP_top60_DNA_seed | MTOR | -3.86697 | KIRC | 786-0 | CRT0105446 | MVP_top60_DNA+RNA_seed | EGFR | -1.877617 | LUAD | COR-L105 | AZD8931 |
| MVP_top60_DNA_seed | PIK3CD | -3.688259 | KIRC | RCC-FG2 | AZD8186 | MVP_top60_DNA+RNA_seed | IGF1R | -1.874303 | LUAD | HCC-78 | BMS-536924 |
| MVP_top60_DNA_seed | PARP1 | -3.082848 | KIRC | RXF393 | Rucaparib | MVP_top60_DNA+RNA_seed | SRC | -1.8739 | LUAD | H3255 | Bosutinib |

| | | | | | | | | | | | |
|---|---|---|---|---|---|---|---|---|---|---|---|
| MVP_top60_DNA_seed | PIK3CD | -3.016708 | KIRC | A498 | AZD8186 | MVP_top60_DNA+RNA_seed | AURKA | -1.865741 | LUAD | PC-14 | Genentech Cpd 10 |
| MVP_top60_DNA_seed | PARP1 | -2.91434 | KIRC | OS-RC-2 | Veliparib | MVP_top60_DNA+RNA_seed | EGFR | -1.861887 | LUAD | NCI-H1781 | CI-1033 |
| MVP_top60_DNA_seed | AR | -2.858498 | KIRC | U031 | AZD3514 | MVP_top60_DNA+RNA_seed | AKT1 | -1.851751 | LUAD | NCI-H1755 | AKT inhibitor VIII |
| MVP_top60_DNA_seed | PIK3CD | -2.644204 | KIRC | RCC-JF | AZD8186 | MVP_top60_DNA+RNA_seed | EGFR | -1.846051 | LUAD | NCI-H358 | Pelitinib |
| MVP_top60_DNA_seed | MTOR | -2.526586 | KIRC | RCC-JW | Temsirolimus | MVP_top60_DNA+RNA_seed | HDAC1 | -1.831845 | LUAD | Calu-3 | JQ12 |
| MVP_top60_DNA_seed | PARP1 | -2.508858 | KIRC | U031 | Olaparib | MVP_top60_DNA+RNA_seed | IGF1R | -1.82253 | LUAD | NCI-H358 | Linsitinib |
| MVP_top60_DNA_seed | PIK3CD | -2.464178 | KIRC | U031 | AZD8186 | MVP_top60_DNA+RNA_seed | HDAC1 | -1.806475 | LUAD | NCI-H1975 | Entinostat |
| MVP_top60_DNA_seed | FGFR4 | -2.4591 | KIRC | U031 | FGFR_0939 | MVP_top60_DNA+RNA_seed | HSP90AA1 | -1.799429 | LUAD | LXF-289 | Tanespimycin |
| MVP_top60_DNA_seed | PARP1 | -2.420123 | KIRC | 769-P | Rucaparib | MVP_top60_DNA+RNA_seed | EGFR | -1.797091 | LUAD | COR-L105 | PF-00299804 |
| MVP_top60_DNA_seed | SMARCA4 | -2.283803 | KIRC | A498 | PFI-3 | MVP_top60_DNA+RNA_seed | MAPK14 | -1.78644 | LUAD | NCI-H2087 | TAK-715 |
| MVP_top60_DNA_seed | ATM | -2.249208 | KIRC | SW156 | KU-60019 | MVP_top60_DNA+RNA_seed | EGFR | -1.777737 | LUAD | NCI-H1944 | Cetuximab |
| MVP_top60_DNA_seed | PIK3CD | -2.249095 | KIRC | CAL-54 | AZD8186 | MVP_top60_DNA+RNA_seed | MET | -1.763636 | LUAD | NCI-H2347 | JNJ38877605 |
| MVP_top60_DNA_seed | MTOR | -2.133136 | KIRC | LB996-RCC | CRT0105446 | MVP_top60_DNA+RNA_seed | HDAC1 | -1.759001 | LUAD | LC-2-ad | JQ12 |
| MVP_top60_DNA_seed | IGF1R | -2.071902 | KIRC | CAL-54 | GSK1904529A | MVP_top60_DNA+RNA_seed | EGFR | -1.758171 | LUAD | NCI-H1568 | CI-1033 |
| MVP_top60_DNA_seed | EP300 | -1.956195 | KIRC | U031 | I-CBP112 | MVP_top60_DNA+RNA_seed | SRC | -1.758019 | LUAD | PC-14 | Bosutinib |
| MVP_top60_DNA_seed | PIK3CD | -1.946539 | KIRC | OS-RC-2 | AZD8186 | MVP_top60_DNA+RNA_seed | MET | -1.752393 | LUAD | RERF-LC-MS | JNJ38877605 |
| MVP_top60_DNA_seed | HDAC1 | -1.93648 | KIRC | OS-RC-2 | Tubastatin A | MVP_top60_DNA+RNA_seed | AKT1 | -1.750753 | LUAD | NCI-H2228 | AKT inhibitor VIII |
| MVP_top60_DNA_seed | ATM | -1.906513 | KIRC | LB2241-RCC | KU-55933 | MVP_top60_DNA+RNA_seed | IGF1R | -1.730043 | LUAD | NCI-H1944 | GSK1904529A |
| MVP_top60_DNA_seed | AR | -1.844146 | KIRC | RCC-AB | Bicalutamide | MVP_top60_DNA+RNA_seed | EGFR | -1.724238 | LUAD | PC-3_[JPC-3] | Cetuximab |
| MVP_top60_DNA_seed | IGF1R | -1.843809 | KIRC | 786-0 | GSK1904529A | MVP_top60_DNA+RNA_seed | HSP90AA1 | -1.717851 | LUAD | NCI-H3122 | Tanespimycin |
| MVP_top60_DNA_seed | PIK3CD | -1.839539 | KIRC | LB996-RCC | AZD8186 | MVP_top60_DNA+RNA_seed | IGF1R | -1.715622 | LUAD | HCC-78 | BMS-754807 |
| MVP_top60_DNA_seed | PIK3CD | -1.804286 | KIRC | CAKI-1 | AZD8186 | MVP_top60_DNA+RNA_seed | AKT1 | -1.71489 | LUAD | HCC-44 | MK-2206 |
| MVP_top60_DNA_seed | PIK3CD | -1.785491 | KIRC | HA7-RCC | AZD8186 | MVP_top60_DNA+RNA_seed | EGFR | -1.706973 | LUAD | NCI-H1975 | Cetuximab |
| MVP_top60_DNA_seed | MTOR | -1.780475 | KIRC | BB65-RCC | Temsirolimus | MVP_top60_DNA+RNA_seed | AKT1 | -1.706946 | LUAD | LXF-289 | AKT inhibitor VIII |
| MVP_top60_DNA_seed | ATM | -1.768834 | KIRC | VMRC-RCW | KU-55933 | MVP_top60_DNA+RNA_seed | SRC | -1.705187 | LUAD | Calu-6 | NVP-BHG712 |
| MVP_top60_DNA_seed | MTOR | -1.768524 | KIRC | BFTC-909 | CRT0105446 | MVP_top60_DNA+RNA_seed | ATM | -1.694277 | LUAD | NCI-H1838 | KU-55933 |
| MVP_top60_DNA_seed | HSP90AA1 | -1.765769 | KIRC | 786-0 | CCT-018159 | MVP_top60_DNA+RNA_seed | PIK3CA | -1.691383 | LUAD | NCI-H1944 | AZD8835 |
| MVP_top60_DNA_seed | HSP90AB1 | -1.765769 | KIRC | 786-0 | CCT-018159 | MVP_top60_DNA+RNA_seed | BRAF | -1.690759 | LUAD | EMC-BAC-2 | PLX-4720 |
| MVP_top60_DNA_seed | PIK3CD | -1.761642 | KIRC | LB2241-RCC | AZD8186 | MVP_top60_DNA+RNA_seed | EGFR | -1.690751 | LUAD | NCI-H1648 | Afatinib |
| MVP_top60_DNA_seed | PARP1 | -1.754909 | KIRC | RCC-JW | PARP_9495 | MVP_top60_DNA+RNA_seed | AURKA | -1.679723 | LUAD | NCI-H358 | Genentech Cpd 10 |
| MVP_top60_DNA_seed | ATM | -1.754731 | KIRC | RCC-FG2 | KU-55933 | MVP_top60_DNA+RNA_seed | IGF1R | -1.670658 | LUAD | LC-2-ad | Linsitinib |
| MVP_top60_DNA_seed | MTOR | -1.744366 | KIRC | RCC-MF | Temsirolimus | MVP_top60_DNA+RNA_seed | EGFR | -1.666768 | LUAD | COR-L105 | AST-1306 |
| MVP_top60_DNA_seed | HSP90AA1 | -1.728269 | KIRC | RCC-JF | Elesclomol | MVP_top60_DNA+RNA_seed | IGF1R | -1.655968 | LUAD | NCI-H2009 | GSK1904529A |
| MVP_top60_DNA_seed | HSP90AB1 | -1.728269 | KIRC | RCC-JF | Elesclomol | MVP_top60_DNA+RNA_seed | IGF1R | -1.651727 | LUAD | NCI-H1437 | BMS-754807 |
| MVP_top60_DNA_seed | PIK3CD | -1.718753 | KIRC | RCC-FG2 | IC-87114 | MVP_top60_DNA+RNA_seed | EGFR | -1.649373 | LUAD | NCI-H1755 | Pelitinib |
| MVP_top60_DNA_seed | PIK3CD | -1.662756 | KIRC | ACHN | AZD8186 | MVP_top60_DNA+RNA_seed | EGFR | -1.648424 | LUAD | NCI-H1623 | Gefitinib |
| MVP_top60_DNA_seed | FGFR4 | -1.657996 | KIRC | RCC-JW | FGFR_0939 | MVP_top60_DNA+RNA_seed | IGF1R | -1.646996 | LUAD | NCI-H1975 | BMS-536924 |
| MVP_top60_DNA_seed | PIK3CD | -1.652401 | KIRC | RCC-AB | Idelalisib | MVP_top60_DNA+RNA_seed | HSP90AA1 | -1.640404 | LUAD | NCI-H1975 | CCT-018159 |
| MVP_top60_DNA_seed | ATM | -1.652006 | KIRC | LB1047-RCC | KU-60019 | MVP_top60_DNA+RNA_seed | MET | -1.638251 | LUAD | LC-2-ad | Crizotinib |
| MVP_top60_DNA_seed | PARP1 | -1.649342 | KIRC | LB996-RCC | Rucaparib | MVP_top60_DNA+RNA_seed | MDM2 | -1.638181 | LUAD | NCI-H1650 | Nutlin-3a (-) |
| MVP_top60_DNA_seed | PIK3CD | -1.642083 | KIRC | RXF393 | AZD8186 | MVP_top60_DNA+RNA_seed | EGFR | -1.63695 | LUAD | EMC-BAC-1 | Pelitinib |

| | | | | | | | | | | | |
|---|---|---|---|---|---|---|---|---|---|---|---|
| MVP_top60_DNA_seed | ESR1 | -1.62741 | KIRC | 769-P | Fulvestrant | MVP_top60_DNA+RNA_seed | HSP90AA1 | -1.632857 | LUAD | A549 | CCT-018159 |
| MVP_top60_DNA_seed | PARP1 | -1.607998 | KIRC | RCC-JF | Olaparib | MVP_top60_DNA+RNA_seed | IGF1R | -1.62212 | LUAD | ABC-1 | Linsitinib |
| MVP_top60_DNA_seed | IGF1R | -1.588944 | KIRC | LB996-RCC | GSK1904529A | MVP_top60_DNA+RNA_seed | EGFR | -1.619433 | LUAD | NCI-H322M | Gefitinib |
| MVP_top60_DNA_seed | HSP90AA1 | -1.587905 | KIRC | 786-0 | Luminespib | MVP_top60_DNA+RNA_seed | IGF1R | -1.618078 | LUAD | A549 | BMS-536924 |
| MVP_top60_DNA_seed | HSP90AB1 | -1.587905 | KIRC | 786-0 | Luminespib | MVP_top60_DNA+RNA_seed | EGFR | -1.61604 | LUAD | HCC-827 | CUDC-101 |
| MVP_top60_DNA_seed | PARP1 | -1.587068 | KIRC | BFTC-909 | Olaparib | MVP_top60_DNA+RNA_seed | PIK3CA | -1.614816 | LUAD | NCI-H1648 | PI3Ka_4409 |
| MVP_top60_DNA_seed | ATM | -1.586484 | KIRC | U031 | KU-60019 | MVP_top60_DNA+RNA_seed | IGF1R | -1.614691 | LUAD | NCI-H2228 | BMS-754807 |
| MVP_top60_DNA_seed | PIK3CD | -1.578398 | KIRC | RCC-JW | AZD8186 | MVP_top60_DNA+RNA_seed | IGF1R | -1.605288 | LUAD | NCI-H1355 | BMS-536924 |
| MVP_top60_DNA_seed | MTOR | -1.571169 | KIRC | 769-P | CRT0105446 | MVP_top60_DNA+RNA_seed | HSP90AA1 | -1.595321 | LUAD | NCI-H2030 | CCT-018159 |
| MVP_top60_DNA_seed | PARP1 | -1.570261 | KIRC | LB2241-RCC | Rucaparib | MVP_top60_DNA+RNA_seed | MAPK14 | -1.59469 | LUAD | HCC-827 | TAK-715 |
| MVP_top60_DNA_seed | PIK3CD | -1.531889 | KIRC | CAL-54 | IC-87114 | MVP_top60_DNA+RNA_seed | EGFR | -1.591446 | LUAD | PC-3_[JPC-3] | AST-1306 |
| MVP_top60_DNA_seed | MTOR | -1.523669 | KIRC | ACHN | CRT0105446 | MVP_top60_DNA+RNA_seed | IGF1R | -1.582989 | LUAD | ABC-1 | BMS-754807 |
| MVP_top60_DNA_seed | MTOR | -1.510239 | KIRC | LB996-RCC | JW-7-52-1 | MVP_top60_DNA+RNA_seed | HSP90AA1 | -1.579497 | LUAD | NCI-H1755 | CCT-018159 |
| MVP_top60_DNA_seed | EP300 | -1.472768 | KIRC | RCC-JW | I-CBP112 | MVP_top60_DNA+RNA_seed | EGFR | -1.578569 | LUAD | NCI-H1975 | Afatinib |
| MVP_top60_DNA_seed | PARP1 | -1.47192 | KIRC | CAL-54 | Veliparib | MVP_top60_DNA+RNA_seed | ATM | -1.569982 | LUAD | LXF-289 | KU-60019 |
| MVP_top60_DNA_seed | MTOR | -1.469224 | KIRC | HA7-RCC | JW-7-52-1 | MVP_top60_DNA+RNA_seed | EGFR | -1.56282 | LUAD | NCI-H1648 | PF-00299804 |
| MVP_top60_DNA_seed | SMARCA4 | -1.468826 | KIRC | U031 | PFI-3 | MVP_top60_DNA+RNA_seed | IGF1R | -1.55855 | LUAD | LC-2-ad | BMS-536924 |
| MVP_top60_DNA_seed | PIK3CA | -1.467061 | KIRC | LB1047-RCC | AZD8835 | MVP_top60_DNA+RNA_seed | AKT1 | -1.552066 | LUAD | NCI-H1437 | AKT inhibitor VIII |
| MVP_top60_DNA_seed | PIK3CD | -1.467061 | KIRC | LB1047-RCC | AZD8835 | MVP_top60_DNA+RNA_seed | EGFR | -1.549928 | LUAD | COR-L105 | CI-1033 |
| MVP_top60_DNA_seed | MTOR | -1.466339 | KIRC | CAL-54 | CRT0105446 | MVP_top60_DNA+RNA_seed | ATM | -1.546048 | LUAD | A549 | KU-55933 |
| MVP_top60_DNA_seed | MTOR | -1.463518 | KIRC | RCC-AB | Voxtalisib | MVP_top60_DNA+RNA_seed | AKT1 | -1.544812 | LUAD | NCI-H2030 | AKT inhibitor VIII |
| MVP_top60_DNA_seed | MTOR | -1.440285 | KIRC | OS-RC-2 | JW-7-52-1 | MVP_top60_DNA+RNA_seed | IGF1R | -1.536601 | LUAD | NCI-H1437 | GSK1904529A |
| MVP_top60_DNA_seed | MTOR | -1.43107 | KIRC | LB1047-RCC | JW-7-52-1 | MVP_top60_DNA+RNA_seed | MDM2 | -1.536493 | LUAD | A549 | Nutlin-3a (-) |
| MVP_top60_DNA_seed | AR | -1.426413 | KIRC | LB2241-RCC | AZD3514 | MVP_top60_DNA+RNA_seed | HSP90AA1 | -1.528909 | LUAD | NCI-H3122 | Luminespib |
| MVP_top60_DNA_seed | PARP1 | -1.423276 | KIRC | RCC-JF | PARP_9495 | MVP_top60_DNA+RNA_seed | HDAC1 | -1.525115 | LUAD | NCI-H1944 | Dacinostat |
| MVP_top60_DNA_seed | PARP1 | -1.415431 | KIRC | U031 | PARP_9482 | MVP_top60_DNA+RNA_seed | MAPK14 | -1.52012 | LUAD | PC-14 | TAK-715 |
| MVP_top60_DNA_seed | PIK3CD | -1.415123 | KIRC | OS-RC-2 | Idelalisib | MVP_top60_DNA+RNA_seed | AKT1 | -1.514808 | LUAD | NCI-H1975 | MK-2206 |
| MVP_top60_DNA_seed | PARP1 | -1.403806 | KIRC | KMRC-1 | Veliparib | MVP_top60_DNA+RNA_seed | EGFR | -1.505636 | LUAD | NCI-H1648 | CI-1033 |
| MVP_top60_DNA_seed | PARP1 | -1.397083 | KIRC | LB1047-RCC | PARP_9495 | MVP_top60_DNA+RNA_seed | SRC | -1.503924 | LUAD | COR-L105 | Bosutinib |
| MVP_top60_DNA_seed | HDAC1 | -1.396808 | KIRC | 786-0 | JQ12 | MVP_top60_DNA+RNA_seed | HSP90AA1 | -1.500664 | LUAD | NCI-H23 | Tanespimycin |
| MVP_top60_DNA_seed | HDAC1 | -1.394774 | KIRC | 786-0 | Dacinostat | MVP_top60_DNA+RNA_seed | IGF1R | -1.495104 | LUAD | NCI-H1437 | BMS-536924 |
| MVP_top60_DNA_seed | PARP1 | -1.389095 | KIRC | 769-P | Olaparib | MVP_top60_DNA+RNA_seed | ATM | -1.491805 | LUAD | NCI-H1755 | CP466722 |
| MVP_top60_DNA_seed | PIK3CD | -1.383624 | KIRC | KMRC-1 | AZD8186 | MVP_top60_DNA+RNA_seed | SMARCA4 | -1.475455 | LUAD | NCI-H1666 | PFI-3 |
| MVP_top60_DNA_seed | MTOR | -1.377506 | KIRC | RCC-ER | CRT0105446 | MVP_top60_DNA+RNA_seed | MET | -1.469508 | LUAD | NCI-H1781 | JNJ38877605 |
| MVP_top60_DNA_seed | HDAC1 | -1.370527 | KIRC | BB65-RCC | Parthenolide | MVP_top60_DNA+RNA_seed | IGF1R | -1.46746 | LUAD | NCI-H650 | Linsitinib |
| MVP_top60_DNA_seed | HDAC1 | -1.370414 | KIRC | BB65-RCC | JQ12 | MVP_top60_DNA+RNA_seed | ATM | -1.465962 | LUAD | NCI-H2030 | KU-55933 |
| MVP_top60_DNA_seed | HSP90AA1 | -1.368965 | KIRC | RXF393 | CCT-018159 | MVP_top60_DNA+RNA_seed | IGF1R | -1.46417 | LUAD | NCI-H1666 | BMS-536924 |
| MVP_top60_DNA_seed | HSP90AB1 | -1.368965 | KIRC | RXF393 | CCT-018159 | MVP_top60_DNA+RNA_seed | AURKA | -1.460825 | LUAD | LXF-289 | HG-5-113-01 |
| MVP_top60_DNA_seed | IGF1R | -1.365836 | KIRC | LB1047-RCC | GSK1904529A | MVP_top60_DNA+RNA_seed | HDAC1 | -1.455237 | LUAD | HOP-62 | JQ12 |
| MVP_top60_DNA_seed | MTOR | -1.364338 | KIRC | RXF393 | Temsirolimus | MVP_top60_DNA+RNA_seed | AKT1 | -1.454531 | LUAD | LXF-289 | MK-2206 |
| MVP_top60_DNA_seed | PIK3CD | -1.357428 | KIRC | 769-P | AZD8186 | MVP_top60_DNA+RNA_seed | ATM | -1.448614 | LUAD | LXF-289 | KU-55933 |

| | | | | | | | | | | | |
|---|---|---|---|---|---|---|---|---|---|---|---|
| MVP_top60_DNA_seed | AR | -1.327946 | KIRC | CAKI-1 | AZD3514 | MVP_top60_DNA+RNA_seed | IGF1R | -1.448517 | LUAD | NCI-H838 | BMS-536924 |
| MVP_top60_DNA_seed | ESR1 | -1.322515 | KIRC | LB2241-RCC | Fulvestrant | MVP_top60_DNA+RNA_seed | EGFR | -1.447333 | LUAD | EKVX | Afatinib |
| MVP_top60_DNA_seed | PARP1 | -1.301369 | KIRC | U031 | PARP_0108 | MVP_top60_DNA+RNA_seed | HSP90AA1 | -1.446474 | LUAD | PC-14 | CCT-018159 |
| MVP_top60_DNA_seed | IGF1R | -1.297166 | KIRC | LB1047-RCC | BMS-536924 | MVP_top60_DNA+RNA_seed | HDAC1 | -1.445597 | LUAD | LC-2-ad | Dacinostat |
| MVP_top60_DNA_seed | IGF1R | -1.294234 | KIRC | LB1047-RCC | BMS-536924 | MVP_top60_DNA+RNA_seed | HSP90AA1 | -1.428277 | LUAD | NCI-H1563 | Elesclomol |
| MVP_top60_DNA_seed | HSP90AA1 | -1.293755 | KIRC | RXF393 | Luminespib | MVP_top60_DNA+RNA_seed | FGFR1 | -1.426335 | LUAD | LC-2-ad | FGFR_3831 |
| MVP_top60_DNA_seed | HSP90AB1 | -1.293755 | KIRC | RXF393 | Luminespib | MVP_top60_DNA+RNA_seed | BRAF | -1.415592 | LUAD | NCI-H1781 | PLX-4720 |
| MVP_top60_DNA_seed | PARP1 | -1.292036 | KIRC | 786-0 | Veliparib | MVP_top60_DNA+RNA_seed | HSP90AA1 | -1.413799 | LUAD | NCI-H1975 | Tanespimycin |
| MVP_top60_DNA_seed | MTOR | -1.291163 | KIRC | OS-RC-2 | Temsirolimus | MVP_top60_DNA+RNA_seed | AKT1 | -1.408717 | LUAD | NCI-H1944 | AKT inhibitor VIII |
| MVP_top60_DNA_seed | ATM | -1.284397 | KIRC | BFTC-909 | KU-55933 | MVP_top60_DNA+RNA_seed | HDAC1 | -1.408523 | LUAD | HCC-44 | Tubastatin A |
| MVP_top60_DNA_seed | HSP90AA1 | -1.282383 | KIRC | U031 | CCT-018159 | MVP_top60_DNA+RNA_seed | IGF1R | -1.402368 | LUAD | LC-2-ad | BMS-754807 |
| MVP_top60_DNA_seed | HSP90AB1 | -1.282383 | KIRC | U031 | CCT-018159 | MVP_top60_DNA+RNA_seed | EGFR | -1.401813 | LUAD | NCI-H1355 | Erlotinib |
| MVP_top60_DNA_seed | FGFR4 | -1.281831 | KIRC | HA7-RCC | FGFR_0939 | MVP_top60_DNA+RNA_seed | IGF1R | -1.398164 | LUAD | ABC-1 | BMS-536924 |
| MVP_top60_DNA_seed | ESR1 | -1.261861 | KIRC | HA7-RCC | Tamoxifen | MVP_top60_DNA+RNA_seed | IGF1R | -1.395073 | LUAD | NCI-H2228 | BMS-536924 |
| MVP_top60_DNA_seed | HDAC1 | -1.25374 | KIRC | 769-P | JQ12 | MVP_top60_DNA+RNA_seed | HDAC1 | -1.392221 | LUAD | NCI-H522 | Dacinostat |
| MVP_top60_DNA_seed | HSP90AA1 | -1.238835 | KIRC | RCC-JF | Tanespimycin | MVP_top60_DNA+RNA_seed | ATM | -1.392128 | LUAD | NCI-H2030 | KU-60019 |
| MVP_top60_DNA_seed | HSP90AB1 | -1.238835 | KIRC | RCC-JF | Tanespimycin | MVP_top60_DNA+RNA_seed | AURKA | -1.387894 | LUAD | LXF-289 | Tozasertib |
| MVP_top60_DNA_seed | PIK3CD | -1.237243 | KIRC | RCC-JF | Idelalisib | MVP_top60_DNA+RNA_seed | EGFR | -1.387586 | LUAD | RERF-LC-KJ | Cetuximab |
| MVP_top60_DNA_seed | MTOR | -1.236075 | KIRC | RCC-FG2 | Temsirolimus | MVP_top60_DNA+RNA_seed | AKT1 | -1.384174 | LUAD | COR-L105 | MK-2206 |
| MVP_top60_DNA_seed | HSP90AA1 | -1.235766 | KIRC | HA7-RCC | Elesclomol | MVP_top60_DNA+RNA_seed | EGFR | -1.381905 | LUAD | NCI-H1648 | AZD8931 |
| MVP_top60_DNA_seed | HSP90AB1 | -1.235766 | KIRC | HA7-RCC | Elesclomol | MVP_top60_DNA+RNA_seed | IGF1R | -1.381452 | LUAD | SK-LU-1 | BMS-754807 |
| MVP_top60_DNA_seed | PIK3CD | -1.233329 | KIRC | VMRC-RCZ | Idelalisib | MVP_top60_DNA+RNA_seed | MDM2 | -1.378541 | LUAD | NCI-H2122 | Nutlin-3a (-) |
| MVP_top60_DNA_seed | HSP90AA1 | -1.232476 | KIRC | LB1047-RCC | Tanespimycin | MVP_top60_DNA+RNA_seed | AURKA | -1.375024 | LUAD | LXF-289 | ZM447439 |
| MVP_top60_DNA_seed | HSP90AB1 | -1.232476 | KIRC | LB1047-RCC | Tanespimycin | MVP_top60_DNA+RNA_seed | MDM2 | -1.374781 | LUAD | NCI-H1651 | Serdemetan |
| MVP_top60_DNA_seed | HSP90AA1 | -1.228895 | KIRC | RCC-MF | Elesclomol | MVP_top60_DNA+RNA_seed | HDAC1 | -1.374094 | LUAD | HCC-44 | JQ12 |
| MVP_top60_DNA_seed | HSP90AB1 | -1.228895 | KIRC | RCC-MF | Elesclomol | MVP_top60_DNA+RNA_seed | HSP90AA1 | -1.374089 | LUAD | PC-14 | Tanespimycin |
| MVP_top60_DNA_seed | MTOR | -1.226459 | KIRC | RCC-JW | QL-VIII-58 | MVP_top60_DNA+RNA_seed | EGFR | -1.372142 | LUAD | NCI-H1648 | AST-1306 |
| MVP_top60_DNA_seed | ESR1 | -1.215325 | KIRC | BFTC-909 | Tamoxifen | MVP_top60_DNA+RNA_seed | SRC | -1.36834 | LUAD | NCI-H1648 | WH-4-023 |
| MVP_top60_DNA_seed | MTOR | -1.211035 | KIRC | LB2241-RCC | CRT0105446 | MVP_top60_DNA+RNA_seed | HSP90AA1 | -1.366877 | LUAD | SK-LU-1 | Tanespimycin |
| MVP_top60_DNA_seed | MTOR | -1.209743 | KIRC | BB65-RCC | JW-7-52-1 | MVP_top60_DNA+RNA_seed | MET | -1.36463 | LUAD | HOP-62 | JNJ38877605 |
| MVP_top60_DNA_seed | MTOR | -1.209592 | KIRC | RCC-JF | Temsirolimus | MVP_top60_DNA+RNA_seed | MET | -1.36347 | LUAD | NCI-H2342 | JNJ38877605 |
| MVP_top60_DNA_seed | IGF1R | -1.203689 | KIRC | KMRC-20 | GSK1904529A | MVP_top60_DNA+RNA_seed | HSP90AA1 | -1.35179 | LUAD | NCI-H838 | Tanespimycin |
| MVP_top60_DNA_seed | HDAC1 | -1.192423 | KIRC | RCC-MF | Tubastatin A | MVP_top60_DNA+RNA_seed | AKT1 | -1.346078 | LUAD | Calu-3 | AKT inhibitor VIII |
| MVP_top60_DNA_seed | HDAC1 | -1.190828 | KIRC | VMRC-RCW | JQ12 | MVP_top60_DNA+RNA_seed | IGF1R | -1.343221 | LUAD | NCI-H2085 | GSK1904529A |
| MVP_top60_DNA_seed | ATM | -1.190312 | KIRC | CAL-54 | KU-55933 | MVP_top60_DNA+RNA_seed | HDAC1 | -1.332298 | LUAD | NCI-H1975 | Dacinostat |
| MVP_top60_DNA_seed | HSP90AA1 | -1.186982 | KIRC | BFTC-909 | Tanespimycin | MVP_top60_DNA+RNA_seed | EGFR | -1.331915 | LUAD | NCI-H1568 | AST-1306 |
| MVP_top60_DNA_seed | HSP90AB1 | -1.186982 | KIRC | BFTC-909 | Tanespimycin | MVP_top60_DNA+RNA_seed | AKT1 | -1.318789 | LUAD | NCI-H838 | MK-2206 |
| MVP_top60_DNA_seed | HSP90AA1 | -1.18548 | KIRC | KMRC-20 | Elesclomol | MVP_top60_DNA+RNA_seed | IGF1R | -1.318446 | LUAD | HCC-44 | GSK1904529A |
| MVP_top60_DNA_seed | HSP90AB1 | -1.18548 | KIRC | KMRC-20 | Elesclomol | MVP_top60_DNA+RNA_seed | HSP90AA1 | -1.317377 | LUAD | COR-L105 | Elesclomol |
| MVP_top60_DNA_seed | PARP1 | -1.182638 | KIRC | HA7-RCC | Olaparib | MVP_top60_DNA+RNA_seed | AURKA | -1.315361 | LUAD | NCI-H1944 | Genentech Cpd 10 |
| MVP_top60_DNA_seed | IGF1R | -1.176876 | KIRC | LB1047-RCC | Linsitinib | MVP_top60_DNA+RNA_seed | HDAC1 | -1.311584 | LUAD | NCI-H2122 | JQ12 |

| | | | | | | | | | | | |
|---|---|---|---|---|---|---|---|---|---|---|---|
| MVP_top60_DNA_seed | IGF1R | -1.170012 | KIRC | LB2241-RCC | BMS-536924 | MVP_top60_DNA+RNA_seed | PIK3CA | -1.311541 | LUAD | NCI-H1648 | AZD8835 |
| MVP_top60_DNA_seed | ATM | -1.168808 | KIRC | LB1047-RCC | CP466722 | MVP_top60_DNA+RNA_seed | IGF1R | -1.309301 | LUAD | NCI-H1975 | Linsitinib |
| MVP_top60_DNA_seed | ATM | -1.168706 | KIRC | A704 | KU-55933 | MVP_top60_DNA+RNA_seed | IGF1R | -1.305055 | LUAD | NCI-H1355 | BMS-536924 |
| MVP_top60_DNA_seed | PARP1 | -1.162839 | KIRC | CAL-54 | Rucaparib | MVP_top60_DNA+RNA_seed | EGFR | -1.300028 | LUAD | EKVX | Afatinib |
| MVP_top60_DNA_seed | ATM | -1.156213 | KIRC | LB1047-RCC | torin2 | MVP_top60_DNA+RNA_seed | HSP90AA1 | -1.299512 | LUAD | NCI-H2030 | Tanespimycin |
| MVP_top60_DNA_seed | MTOR | -1.156213 | KIRC | LB1047-RCC | torin2 | MVP_top60_DNA+RNA_seed | SRC | -1.297763 | LUAD | HOP-62 | WH-4-023 |
| MVP_top60_DNA_seed | PARP1 | -1.156087 | KIRC | SN12C | Rucaparib | MVP_top60_DNA+RNA_seed | IGF1R | -1.296557 | LUAD | NCI-H1944 | BMS-536924 |
| MVP_top60_DNA_seed | HSP90AA1 | -1.143307 | KIRC | BFTC-909 | CCT-018159 | MVP_top60_DNA+RNA_seed | EGFR | -1.296158 | LUAD | PC-3_[JPC-3] | PF-00299804 |
| MVP_top60_DNA_seed | HSP90AB1 | -1.143307 | KIRC | BFTC-909 | CCT-018159 | MVP_top60_DNA+RNA_seed | SRC | -1.291483 | LUAD | HOP-62 | Saracatinib |
| MVP_top60_DNA_seed | PARP1 | -1.140059 | KIRC | RCC-JW | Talazoparib | MVP_top60_DNA+RNA_seed | EGFR | -1.285613 | LUAD | NCI-H2030 | AZD8931 |
| MVP_top60_DNA_seed | IGF1R | -1.137449 | KIRC | LB996-RCC | BMS-536924 | MVP_top60_DNA+RNA_seed | SMARCA4 | -1.280394 | LUAD | VMRC-LCD | PFI-3 |
| MVP_top60_DNA_seed | ATM | -1.127976 | KIRC | CAKI-1 | CP466722 | MVP_top60_DNA+RNA_seed | EGFR | -1.275361 | LUAD | NCI-H322M | CI-1033 |
| MVP_top60_DNA_seed | PARP1 | -1.124406 | KIRC | RCC-JF | Olaparib | MVP_top60_DNA+RNA_seed | EGFR | -1.272352 | LUAD | NCI-H322M | PF-00299804 |
| MVP_top60_DNA_seed | HDAC1 | -1.116641 | KIRC | LB2241-RCC | Parthenolide | MVP_top60_DNA+RNA_seed | SRC | -1.272267 | LUAD | PC-3_[JPC-3] | Bosutinib |
| MVP_top60_DNA_seed | ATM | -1.112846 | KIRC | U031 | KU-55933 | MVP_top60_DNA+RNA_seed | AURKA | -1.270062 | LUAD | ABC-1 | ZM447439 |
| MVP_top60_DNA_seed | PIK3CD | -1.11088 | KIRC | VMRC-RCZ | AZD8186 | MVP_top60_DNA+RNA_seed | BRAF | -1.26912 | LUAD | Calu-6 | Dabrafenib |
| MVP_top60_DNA_seed | EP300 | -1.106521 | KIRC | 786-0 | I-CBP112 | MVP_top60_DNA+RNA_seed | ATM | -1.266769 | LUAD | NCI-H1437 | KU-55933 |
| MVP_top60_DNA_seed | PIK3CA | -1.104616 | KIRC | LB2241-RCC | AZD8835 | MVP_top60_DNA+RNA_seed | IGF1R | -1.252872 | LUAD | NCI-H1792 | BMS-536924 |
| MVP_top60_DNA_seed | PIK3CD | -1.104616 | KIRC | LB2241-RCC | AZD8835 | MVP_top60_DNA+RNA_seed | SRC | -1.250732 | LUAD | NCI-H1693 | Bosutinib |
| MVP_top60_DNA_seed | IGF1R | -1.090061 | KIRC | HA7-RCC | Linsitinib | MVP_top60_DNA+RNA_seed | AKT1 | -1.246958 | LUAD | RERF-LC-KJ | AKT inhibitor VIII |
| MVP_top60_DNA_seed | MTOR | -1.079655 | KIRC | SW156 | CRT0105446 | MVP_top60_DNA+RNA_seed | EGFR | -1.246743 | LUAD | PC-3_[JPC-3] | CI-1033 |
| MVP_top60_DNA_seed | PIK3CA | -1.073661 | KIRC | BFTC-909 | AZD8835 | MVP_top60_DNA+RNA_seed | EGFR | -1.244831 | LUAD | NCI-H358 | Afatinib |
| MVP_top60_DNA_seed | PIK3CD | -1.073661 | KIRC | BFTC-909 | AZD8835 | MVP_top60_DNA+RNA_seed | FGFR1 | -1.244259 | LUAD | HCC-44 | FGFR_3831 |
| MVP_top60_DNA_seed | PIK3CD | -1.067321 | KIRC | SN12C | IC-87114 | MVP_top60_DNA+RNA_seed | SRC | -1.243449 | LUAD | NCI-H1975 | WH-4-023 |
| MVP_top60_DNA_seed | PARP1 | -1.060765 | KIRC | LB1047-RCC | Talazoparib | MVP_top60_DNA+RNA_seed | EGFR | -1.241932 | LUAD | NCI-H1693 | Afatinib |
| MVP_top60_DNA_seed | PIK3CD | -1.058185 | KIRC | RCC-MF | AZD8186 | MVP_top60_DNA+RNA_seed | IGF1R | -1.24086 | LUAD | NCI-H3122 | GSK1904529A |
| MVP_top60_DNA_seed | MTOR | -1.057194 | KIRC | RCC-MF | CRT0105446 | MVP_top60_DNA+RNA_seed | EGFR | -1.240674 | LUAD | NCI-H322M | Erlotinib |
| MVP_top60_DNA_seed | HSP90AA1 | -1.057067 | KIRC | RCC-JF | SNX-2112 | MVP_top60_DNA+RNA_seed | IGF1R | -1.237844 | LUAD | NCI-H650 | BMS-754807 |
| MVP_top60_DNA_seed | HSP90AB1 | -1.057067 | KIRC | RCC-JF | SNX-2112 | MVP_top60_DNA+RNA_seed | EGFR | -1.237654 | LUAD | NCI-H1568 | Afatinib |
| MVP_top60_DNA_seed | PIK3CD | -1.055819 | KIRC | SN12C | AZD8186 | MVP_top60_DNA+RNA_seed | PIK3CA | -1.237086 | LUAD | HCC-78 | AZD8835 |
| MVP_top60_DNA_seed | PARP1 | -1.055208 | KIRC | LB1047-RCC | PARP_9482 | MVP_top60_DNA+RNA_seed | FGFR1 | -1.236905 | LUAD | NCI-H2122 | FGFR_3831 |
| MVP_top60_DNA_seed | HDAC1 | -1.053452 | KIRC | LB1047-RCC | JQ12 | MVP_top60_DNA+RNA_seed | MDM2 | -1.236763 | LUAD | NCI-H358 | Nutlin-3a (-) |
| MVP_top60_DNA_seed | PIK3CD | -1.048886 | KIRC | NCC021 | Idelalisib | MVP_top60_DNA+RNA_seed | IGF1R | -1.229092 | LUAD | COR-L105 | BMS-754807 |
| MVP_top60_DNA_seed | PIK3CA | -1.042838 | KIRC | LB2241-RCC | PI3Ka_4409 | MVP_top60_DNA+RNA_seed | IGF1R | -1.227584 | LUAD | RERF-LC-KJ | Linsitinib |
| MVP_top60_DNA_seed | PIK3CD | -1.042838 | KIRC | LB2241-RCC | PI3Ka_4409 | MVP_top60_DNA+RNA_seed | MAPK14 | -1.223952 | LUAD | NCI-H358 | TAK-715 |
| MVP_top60_DNA_seed | PARP1 | -1.042395 | KIRC | BFTC-909 | Talazoparib | MVP_top60_DNA+RNA_seed | MET | -1.222975 | LUAD | HOP-62 | SU11274 |
| MVP_top60_DNA_seed | MTOR | -1.041572 | KIRC | A704 | CRT0105446 | MVP_top60_DNA+RNA_seed | HSP90AA1 | -1.222171 | LUAD | NCI-H3122 | SNX-2112 |
| MVP_top60_DNA_seed | IGF1R | -1.041409 | KIRC | A498 | BMS-536924 | MVP_top60_DNA+RNA_seed | MDM2 | -1.221706 | LUAD | NCI-H1693 | Serdemetan |
| MVP_top60_DNA_seed | HDAC1 | -1.040398 | KIRC | RCC-AB | Belinostat | MVP_top60_DNA+RNA_seed | MET | -1.217885 | LUAD | EMC-BAC-2 | AZD6094 |
| MVP_top60_DNA_seed | AR | -1.037407 | KIRC | NCC010 | Bicalutamide | MVP_top60_DNA+RNA_seed | IGF1R | -1.217236 | LUAD | NCI-H441 | BMS-754807 |
| MVP_top60_DNA_seed | PARP1 | -1.03458 | KIRC | BFTC-909 | PARP_9482 | MVP_top60_DNA+RNA_seed | IGF1R | -1.215199 | LUAD | NCI-H1648 | BMS-754807 |

| | | | | | | | | | | | |
|---|---|---|---|---|---|---|---|---|---|---|---|
| MVP_top60_DNA_seed | HDAC1 | -1.034082 | KIRC | CAKI-1 | JQ12 | MVP_top60_DNA+RNA_seed | HDAC1 | -1.215023 | LUAD | NCI-H2122 | Dacinostat |
| MVP_top60_DNA_seed | PARP1 | -1.033413 | KIRC | LB2241-RCC | PARP_9482 | MVP_top60_DNA+RNA_seed | IGF1R | -1.214805 | LUAD | NCI-H441 | Linsitinib |
| MVP_top60_DNA_seed | PIK3CD | -1.032167 | KIRC | OS-RC-2 | IC-87114 | MVP_top60_DNA+RNA_seed | MAPK14 | -1.214686 | LUAD | NCI-H1944 | TAK-715 |
| MVP_top60_DNA_seed | MTOR | -1.029474 | KIRC | CAKI-1 | CRT0105446 | MVP_top60_DNA+RNA_seed | EGFR | -1.210845 | LUAD | LC-2-ad | Erlotinib |
| MVP_top60_DNA_seed | AR | -1.028907 | KIRC | LB1047-RCC | Bicalutamide | MVP_top60_DNA+RNA_seed | BRAF | -1.206378 | LUAD | HCC-827 | PLX-4720 |
| MVP_top60_DNA_seed | PARP1 | -1.027826 | KIRC | LB996-RCC | PARP_9495 | MVP_top60_DNA+RNA_seed | AKT1 | -1.206372 | LUAD | NCI-H1993 | AKT inhibitor VIII |
| MVP_top60_DNA_seed | PARP1 | -1.025977 | KIRC | RCC-JF | PARP_9482 | MVP_top60_DNA+RNA_seed | HDAC1 | -1.206119 | LUAD | NCI-H1944 | JQ12 |
| MVP_top60_DNA_seed | PIK3CA | -1.017018 | KIRC | LB1047-RCC | PI3Ka_4409 | MVP_top60_DNA+RNA_seed | EGFR | -1.205867 | LUAD | COR-L105 | Gefitinib |
| MVP_top60_DNA_seed | PIK3CD | -1.017018 | KIRC | LB1047-RCC | PI3Ka_4409 | MVP_top60_DNA+RNA_seed | HSP90AA1 | -1.203443 | LUAD | HCC-44 | Luminespib |
| MVP_top60_DNA_seed | PARP1 | -1.009496 | KIRC | BB65-RCC | Veliparib | MVP_top60_DNA+RNA_seed | AURKA | -1.198895 | LUAD | LC-2-ad | ZM447439 |
| MVP_top60_DNA_seed | PARP1 | -1.009087 | KIRC | BFTC-909 | PARP_0108 | MVP_top60_DNA+RNA_seed | AKT1 | -1.197261 | LUAD | NCI-H1355 | A-443654 |
| MVP_top60_DNA_seed | PARP1 | -1.006494 | KIRC | RCC-JF | Talazoparib | MVP_top60_DNA+RNA_seed | HDAC1 | -1.196574 | LUAD | LC-2-ad | Tubastatin A |
| MVP_top60_DNA_seed | PARP1 | -1.002635 | KIRC | 769-P | Olaparib | MVP_top60_DNA+RNA_seed | SRC | -1.191638 | LUAD | LXF-289 | Saracatinib |
| MVP_top60_DNA_seed | PIK3CD | -1.001522 | KIRC | 786-0 | Idelalisib | MVP_top60_DNA+RNA_seed | AURKA | -1.187684 | LUAD | EMC-BAC-2 | HG-5-113-01 |
| MVP_top60_DNA_seed | HDAC1 | -0.994458 | KIRC | RCC-AB | CAY10603 | MVP_top60_DNA+RNA_seed | EGFR | -1.186951 | LUAD | NCI-H1975 | Afatinib |
| MVP_top60_DNA_seed | HSP90AA1 | -0.990287 | KIRC | BB65-RCC | Tanespimycin | MVP_top60_DNA+RNA_seed | MET | -1.185481 | LUAD | H3255 | JNJ38877605 |
| MVP_top60_DNA_seed | HSP90AB1 | -0.990287 | KIRC | BB65-RCC | Tanespimycin | MVP_top60_DNA+RNA_seed | HSP90AA1 | -1.184703 | LUAD | LC-2-ad | Luminespib |
| MVP_top60_DNA_seed | PTK2 | -0.990199 | KIRC | 786-0 | PF-562271 | MVP_top60_DNA+RNA_seed | BRAF | -1.18469 | LUAD | NCI-H1648 | PLX-4720 |
| MVP_top60_DNA_seed | PIK3CD | -0.986495 | KIRC | NCC021 | IC-87114 | MVP_top60_DNA+RNA_seed | BRAF | -1.184439 | LUAD | NCI-H1648 | AZ628 |
| MVP_top60_DNA_seed | MTOR | -0.980109 | KIRC | RCC-JW | CRT0105446 | MVP_top60_DNA+RNA_seed | HSP90AA1 | -1.183128 | LUAD | NCI-H1975 | Elesclomol |
| MVP_top60_DNA_seed | IGF1R | -0.979638 | KIRC | BB65-RCC | Linsitinib | MVP_top60_DNA+RNA_seed | IGF1R | -1.181323 | LUAD | SK-LU-1 | Linsitinib |
| MVP_top60_DNA_seed | MTOR | -0.975483 | KIRC | BB65-RCC | CRT0105446 | MVP_top60_DNA+RNA_seed | HSP90AA1 | -1.181132 | LUAD | HCC-827 | CCT-018159 |
| MVP_top60_DNA_seed | MTOR | -0.974448 | KIRC | KMRC-20 | CRT0105446 | MVP_top60_DNA+RNA_seed | AKT1 | -1.179511 | LUAD | ABC-1 | MK-2206 |
| MVP_top60_DNA_seed | ATM | -0.972057 | KIRC | LB1047-RCC | KU-55933 | MVP_top60_DNA+RNA_seed | HDAC1 | -1.178192 | LUAD | NCI-H1755 | CAY10603 |
| MVP_top60_DNA_seed | MTOR | -0.971259 | KIRC | 769-P | Torin 2 | MVP_top60_DNA+RNA_seed | BRAF | -1.177737 | LUAD | NCI-H1975 | HG6-64-1 |
| MVP_top60_DNA_seed | ATM | -0.970238 | KIRC | NCC021 | KU-55933 | MVP_top60_DNA+RNA_seed | EGFR | -1.175376 | LUAD | NCI-H1355 | Cetuximab |
| MVP_top60_DNA_seed | SMARCA4 | -0.967205 | KIRC | HA7-RCC | PFI-3 | MVP_top60_DNA+RNA_seed | EGFR | -1.174421 | LUAD | NCI-H2122 | AZD8931 |
| MVP_top60_DNA_seed | MTOR | -0.966708 | KIRC | TK10 | CRT0105446 | MVP_top60_DNA+RNA_seed | ATM | -1.170695 | LUAD | HCC-78 | KU-60019 |
| MVP_top60_DNA_seed | PARP1 | -0.965866 | KIRC | CAKI-1 | Olaparib | MVP_top60_DNA+RNA_seed | EGFR | -1.164955 | LUAD | NCI-H1648 | Afatinib |
| MVP_top60_DNA_seed | SMARCA4 | -0.964557 | KIRC | SN12C | PFI-3 | MVP_top60_DNA+RNA_seed | IGF1R | -1.164914 | LUAD | EKVX | BMS-754807 |
| MVP_top60_DNA_seed | IGF1R | -0.964275 | KIRC | LB996-RCC | BMS-536924 | MVP_top60_DNA+RNA_seed | HSP90AA1 | -1.164778 | LUAD | NCI-H358 | SNX-2112 |
| MVP_top60_DNA_seed | PIK3CD | -0.962027 | KIRC | U031 | IC-87114 | MVP_top60_DNA+RNA_seed | AKT1 | -1.16414 | LUAD | NCI-H1975 | AKT inhibitor VIII |
| MVP_top60_DNA_seed | PIK3CD | -0.959565 | KIRC | NCC010 | AZD8186 | MVP_top60_DNA+RNA_seed | HSP90AA1 | -1.16255 | LUAD | NCI-H1944 | Tanespimycin |
| MVP_top60_DNA_seed | ATM | -0.957689 | KIRC | RCC-FG2 | KU-60019 | MVP_top60_DNA+RNA_seed | AURKA | -1.160347 | LUAD | NCI-H2122 | HG-5-113-01 |
| MVP_top60_DNA_seed | ATM | -0.956909 | KIRC | 769-P | KU-60019 | MVP_top60_DNA+RNA_seed | ATM | -1.151655 | LUAD | NCI-H358 | CP466722 |
| MVP_top60_DNA_seed | FGFR4 | -0.9546 | KIRC | LB1047-RCC | FGFR_0939 | MVP_top60_DNA+RNA_seed | HSP90AA1 | -1.151099 | LUAD | NCI-H1975 | Luminespib |
| MVP_top60_DNA_seed | HSP90AA1 | -0.951836 | KIRC | 769-P | Luminespib | MVP_top60_DNA+RNA_seed | AURKA | -1.150993 | LUAD | ABC-1 | HG-5-113-01 |
| MVP_top60_DNA_seed | HSP90AB1 | -0.951836 | KIRC | 769-P | Luminespib | MVP_top60_DNA+RNA_seed | SMARCA4 | -1.147041 | LUAD | HCC-78 | PFI-3 |
| MVP_top60_DNA_seed | PARP1 | -0.951476 | KIRC | U031 | PARP_9495 | MVP_top60_DNA+RNA_seed | HDAC1 | -1.146965 | LUAD | NCI-H1755 | Belinostat |
| MVP_top60_DNA_seed | PARP1 | -0.949877 | KIRC | 769-P | Talazoparib | MVP_top60_DNA+RNA_seed | HDAC1 | -1.144915 | LUAD | NCI-H1975 | JQ12 |
| MVP_top60_DNA_seed | PARP1 | -0.94781 | KIRC | RCC-JW | PARP_9482 | MVP_top60_DNA+RNA_seed | ATM | -1.144747 | LUAD | NCI-H1648 | KU-55933 |

| | | | | | | | | | | | |
|---|---|---|---|---|---|---|---|---|---|---|---|
| MVP_top60_DNA_seed | HSP90AA1 | -0.943485 | KIRC | RCC-FG2 | Elesclomol | MVP_top60_DNA+RNA_seed | IGF1R | -1.142682 | LUAD | HCC-44 | Linsitinib |
| MVP_top60_DNA_seed | HSP90AB1 | -0.943485 | KIRC | RCC-FG2 | Elesclomol | MVP_top60_DNA+RNA_seed | EGFR | -1.142091 | LUAD | Calu-3 | Pelitinib |
| MVP_top60_DNA_seed | PARP1 | -0.940848 | KIRC | LB1047-RCC | PARP_0108 | MVP_top60_DNA+RNA_seed | IGF1R | -1.141099 | LUAD | EKVX | Linsitinib |
| MVP_top60_DNA_seed | PTK2 | -0.940364 | KIRC | CAKI-1 | PF-562271 | MVP_top60_DNA+RNA_seed | AURKA | -1.136609 | LUAD | NCI-H1437 | Genentech Cpd 10 |
| MVP_top60_DNA_seed | HSP90AA1 | -0.933335 | KIRC | RCC-JW | Elesclomol | MVP_top60_DNA+RNA_seed | BRAF | -1.134449 | LUAD | NCI-H2405 | HG6-64-1 |
| MVP_top60_DNA_seed | HSP90AB1 | -0.933335 | KIRC | RCC-JW | Elesclomol | MVP_top60_DNA+RNA_seed | IGF1R | -1.134186 | LUAD | HCC-827 | GSK1904529A |
| MVP_top60_DNA_seed | PIK3CA | -0.932926 | KIRC | U031 | AZD8835 | MVP_top60_DNA+RNA_seed | AURKA | -1.131602 | LUAD | HCC-78 | Alisertib |
| MVP_top60_DNA_seed | PIK3CD | -0.932926 | KIRC | U031 | AZD8835 | MVP_top60_DNA+RNA_seed | EGFR | -1.131497 | LUAD | NCI-H1975 | PF-00299804 |
| MVP_top60_DNA_seed | PIK3CD | -0.930707 | KIRC | RXF393 | Idelalisib | MVP_top60_DNA+RNA_seed | BRAF | -1.131076 | LUAD | NCI-H522 | HG6-64-1 |
| MVP_top60_DNA_seed | HSP90AA1 | -0.930451 | KIRC | SN12C | Tanespimycin | MVP_top60_DNA+RNA_seed | PIK3CA | -1.130296 | LUAD | NCI-H2122 | PI3Ka_4409 |
| MVP_top60_DNA_seed | HSP90AB1 | -0.930451 | KIRC | SN12C | Tanespimycin | MVP_top60_DNA+RNA_seed | HDAC1 | -1.1295 | LUAD | LC-2-ad | Belinostat |
| MVP_top60_DNA_seed | IGF1R | -0.928467 | KIRC | RCC10RGB | Linsitinib | MVP_top60_DNA+RNA_seed | IGF1R | -1.128832 | LUAD | HOP-62 | BMS-536924 |
| MVP_top60_DNA_seed | HSP90AA1 | -0.921568 | KIRC | LB1047-RCC | Elesclomol | MVP_top60_DNA+RNA_seed | ATM | -1.127841 | LUAD | RERF-LC-KJ | KU-60019 |
| MVP_top60_DNA_seed | HSP90AB1 | -0.921568 | KIRC | LB1047-RCC | Elesclomol | MVP_top60_DNA+RNA_seed | ATM | -1.124553 | LUAD | EMC-BAC-2 | KU-60019 |
| MVP_top60_DNA_seed | ESR1 | -0.921313 | KIRC | KMRC-1 | Tamoxifen | MVP_top60_DNA+RNA_seed | EGFR | -1.123297 | LUAD | NCI-H1573 | Afatinib |
| MVP_top60_DNA_seed | FGFR4 | -0.919515 | KIRC | RCC-AB | FGFR_0939 | MVP_top60_DNA+RNA_seed | HSP90AA1 | -1.122215 | LUAD | NCI-H1993 | CCT-018159 |
| MVP_top60_DNA_seed | PARP1 | -0.906638 | KIRC | CAKI-1 | PARP_9482 | MVP_top60_DNA+RNA_seed | HDAC1 | -1.114996 | LUAD | NCI-H1437 | JQ12 |
| MVP_top60_DNA_seed | ESR1 | -0.90322 | KIRC | CAKI-1 | Tamoxifen | MVP_top60_DNA+RNA_seed | SRC | -1.113928 | LUAD | SK-LU-1 | Bosutinib |
| MVP_top60_DNA_seed | PARP1 | -0.90199 | KIRC | KMRC-1 | Rucaparib | MVP_top60_DNA+RNA_seed | PIK3CA | -1.113559 | LUAD | NCI-H1975 | AZD8835 |
| MVP_top60_DNA_seed | PARP1 | -0.893332 | KIRC | RCC-JF | PARP_0108 | MVP_top60_DNA+RNA_seed | MDM2 | -1.111722 | LUAD | NCI-H2087 | Nutlin-3a (-) |
| MVP_top60_DNA_seed | FGFR4 | -0.892812 | KIRC | 786-0 | FGFR_0939 | MVP_top60_DNA+RNA_seed | ATM | -1.111413 | LUAD | A549 | torin2 |
| MVP_top60_DNA_seed | PIK3CD | -0.892059 | KIRC | U031 | Idelalisib | MVP_top60_DNA+RNA_seed | BRAF | -1.109823 | LUAD | HOP-62 | HG6-64-1 |
| MVP_top60_DNA_seed | HSP90AA1 | -0.891472 | KIRC | 769-P | Elesclomol | MVP_top60_DNA+RNA_seed | AURKA | -1.104659 | LUAD | VMRC-LCD | HG-5-113-01 |
| MVP_top60_DNA_seed | HSP90AB1 | -0.891472 | KIRC | 769-P | Elesclomol | MVP_top60_DNA+RNA_seed | EGFR | -1.102787 | LUAD | NCI-H1975 | AST-1306 |
| MVP_top60_DNA_seed | PARP1 | -0.890006 | KIRC | HA7-RCC | PARP_9482 | MVP_top60_DNA+RNA_seed | MET | -1.101569 | LUAD | NCI-H522 | JNJ38877605 |
| MVP_top60_DNA_seed | AR | -0.889988 | KIRC | A704 | Bicalutamide | MVP_top60_DNA+RNA_seed | PTK2 | -1.099467 | LUAD | SK-LU-1 | PF-562271 |
| MVP_top60_DNA_seed | ESR1 | -0.886543 | KIRC | BFTC-909 | Fulvestrant | MVP_top60_DNA+RNA_seed | SRC | -1.09296 | LUAD | PC-14 | HG-6-71-01 |
| MVP_top60_DNA_seed | MTOR | -0.881007 | KIRC | RXF393 | JW-7-52-1 | MVP_top60_DNA+RNA_seed | PIK3CA | -1.087568 | LUAD | NCI-H1944 | PI3Ka_4409 |
| MVP_top60_DNA_seed | IGF1R | -0.87944 | KIRC | A704 | BMS-536924 | MVP_top60_DNA+RNA_seed | PIK3CA | -1.086683 | LUAD | NCI-H441 | PI3Ka_4409 |
| MVP_top60_DNA_seed | ATM | -0.876664 | KIRC | RCC-JW | CP466722 | MVP_top60_DNA+RNA_seed | SRC | -1.085245 | LUAD | NCI-H1975 | Saracatinib |
| MVP_top60_DNA_seed | PARP1 | -0.876581 | KIRC | RCC-MF | Talazoparib | MVP_top60_DNA+RNA_seed | AKT1 | -1.084907 | LUAD | EMC-BAC-2 | MK-2206 |
| MVP_top60_DNA_seed | MTOR | -0.87654 | KIRC | RCC-JF | WYE-125132 | MVP_top60_DNA+RNA_seed | EGFR | -1.084571 | LUAD | EKVX | Cetuximab |
| MVP_top60_DNA_seed | FGFR4 | -0.875187 | KIRC | A704 | FGFR_0939 | MVP_top60_DNA+RNA_seed | AKT1 | -1.082286 | LUAD | HCC-44 | AKT inhibitor VIII |
| MVP_top60_DNA_seed | IGF1R | -0.874576 | KIRC | ACHN | GSK1904529A | MVP_top60_DNA+RNA_seed | MET | -1.081373 | LUAD | NCI-H23 | AZD6094 |
| MVP_top60_DNA_seed | HSP90AA1 | -0.873516 | KIRC | RXF393 | Tanespimycin | MVP_top60_DNA+RNA_seed | PTK2 | -1.079412 | LUAD | LC-2-ad | PF-562271 |
| MVP_top60_DNA_seed | HSP90AB1 | -0.873516 | KIRC | RXF393 | Tanespimycin | MVP_top60_DNA+RNA_seed | HSP90AA1 | -1.079375 | LUAD | NCI-H2085 | Elesclomol |
| MVP_top60_DNA_seed | HSP90AA1 | -0.871823 | KIRC | VMRC-RCZ | Luminespib | MVP_top60_DNA+RNA_seed | HSP90AA1 | -1.078961 | LUAD | NCI-H1792 | CCT-018159 |
| MVP_top60_DNA_seed | HSP90AB1 | -0.871823 | KIRC | VMRC-RCZ | Luminespib | MVP_top60_DNA+RNA_seed | ATM | -1.077634 | LUAD | NCI-H2122 | torin2 |
| MVP_top60_DNA_seed | PARP1 | -0.870574 | KIRC | RCC-JW | PARP_0108 | MVP_top60_DNA+RNA_seed | SRC | -1.075479 | LUAD | LXF-289 | WH-4-023 |
| MVP_top60_DNA_seed | HDAC1 | -0.870557 | KIRC | TK10 | Parthenolide | MVP_top60_DNA+RNA_seed | IGF1R | -1.07359 | LUAD | NCI-H1755 | BMS-536924 |
| MVP_top60_DNA_seed | HSP90AA1 | -0.86888 | KIRC | LB2241-RCC | Tanespimycin | MVP_top60_DNA+RNA_seed | MET | -1.072892 | LUAD | HCC-44 | AZD6094 |

| | | | | | | | | | | | |
|---|---|---|---|---|---|---|---|---|---|---|---|
| MVP_top60_DNA_seed | HSP90AB1 | -0.86888 | KIRC | LB2241-RCC | Tanespimycin | MVP_top60_DNA+RNA_seed | HSP90AA1 | -1.070715 | LUAD | NCI-H2030 | Luminespib |
| MVP_top60_DNA_seed | ATM | -0.867323 | KIRC | RCC-MF | torin2 | MVP_top60_DNA+RNA_seed | AURKA | -1.069444 | LUAD | HCC-44 | Genentech Cpd 10 |
| MVP_top60_DNA_seed | MTOR | -0.867323 | KIRC | RCC-MF | torin2 | MVP_top60_DNA+RNA_seed | HSP90AA1 | -1.069428 | LUAD | NCI-H1755 | Tanespimycin |
| MVP_top60_DNA_seed | PARP1 | -0.86263 | KIRC | HA7-RCC | PARP_0108 | MVP_top60_DNA+RNA_seed | ATM | -1.06612 | LUAD | NCI-H1648 | KU-60019 |
| MVP_top60_DNA_seed | HDAC1 | -0.861044 | KIRC | RCC-JF | CAY10603 | MVP_top60_DNA+RNA_seed | AKT1 | -1.060228 | LUAD | HCC-78 | AKT inhibitor VIII |
| MVP_top60_DNA_seed | PARP1 | -0.858156 | KIRC | NCC010 | PARP_9495 | MVP_top60_DNA+RNA_seed | HDAC1 | -1.058888 | LUAD | NCI-H2342 | JQ12 |
| MVP_top60_DNA_seed | MTOR | -0.857557 | KIRC | LB2241-RCC | Temsirolimus | MVP_top60_DNA+RNA_seed | AKT1 | -1.055067 | LUAD | SW1573 | BAY AKT1 |
| MVP_top60_DNA_seed | ESR1 | -0.855608 | KIRC | 786-0 | Fulvestrant | MVP_top60_DNA+RNA_seed | FGFR1 | -1.054026 | LUAD | NCI-H838 | FGFR_3831 |
| MVP_top60_DNA_seed | PARP1 | -0.855511 | KIRC | A704 | PARP_9495 | MVP_top60_DNA+RNA_seed | SRC | -1.051514 | LUAD | ABC-1 | Bosutinib |
| MVP_top60_DNA_seed | PARP1 | -0.853839 | KIRC | LB2241-RCC | Talazoparib | MVP_top60_DNA+RNA_seed | HSP90AA1 | -1.05091 | LUAD | NCI-H838 | Elesclomol |
| MVP_top60_DNA_seed | PARP1 | -0.85343 | KIRC | 769-P | PARP_9482 | MVP_top60_DNA+RNA_seed | HSP90AA1 | -1.050058 | LUAD | RERF-LC-MS | CCT-018159 |
| MVP_top60_DNA_seed | PIK3CD | -0.852974 | KIRC | LB2241-RCC | IC-87114 | MVP_top60_DNA+RNA_seed | HDAC1 | -1.043651 | LUAD | NCI-H358 | Tubastatin A |
| MVP_top60_DNA_seed | HSP90AA1 | -0.852251 | KIRC | TK10 | Luminespib | MVP_top60_DNA+RNA_seed | BRAF | -1.043425 | LUAD | NCI-H2030 | HG6-64-1 |
| MVP_top60_DNA_seed | HSP90AB1 | -0.852251 | KIRC | TK10 | Luminespib | MVP_top60_DNA+RNA_seed | HDAC1 | -1.043009 | LUAD | NCI-H1975 | Tubastatin A |
| MVP_top60_DNA_seed | PARP1 | -0.850841 | KIRC | RCC-JF | Rucaparib | MVP_top60_DNA+RNA_seed | IGF1R | -1.04002 | LUAD | EMC-BAC-1 | BMS-536924 |
| MVP_top60_DNA_seed | PIK3CD | -0.846935 | KIRC | RCC-JW | Idelalisib | MVP_top60_DNA+RNA_seed | AKT1 | -1.039806 | LUAD | NCI-H2228 | MK-2206 |
| MVP_top60_DNA_seed | ATM | -0.843736 | KIRC | RCC-JF | torin2 | MVP_top60_DNA+RNA_seed | IGF1R | -1.037951 | LUAD | LXF-289 | BMS-536924 |
| MVP_top60_DNA_seed | MTOR | -0.843736 | KIRC | RCC-JF | torin2 | MVP_top60_DNA+RNA_seed | HDAC1 | -1.036798 | LUAD | NCI-H3122 | JQ12 |
| MVP_top60_DNA_seed | MTOR | -0.842286 | KIRC | 769-P | WYE-125132 | MVP_top60_DNA+RNA_seed | HSP90AA1 | -1.036185 | LUAD | NCI-H2122 | Luminespib |
| MVP_top60_DNA_seed | ESR1 | -0.841428 | KIRC | 769-P | Tamoxifen | MVP_top60_DNA+RNA_seed | AKT1 | -1.035957 | LUAD | NCI-H1944 | MK-2206 |
| MVP_top60_DNA_seed | IGF1R | -0.837284 | KIRC | OS-RC-2 | BMS-536924 | MVP_top60_DNA+RNA_seed | HSP90AA1 | -1.03557 | LUAD | NCI-H1792 | Tanespimycin |
| MVP_top60_DNA_seed | ATM | -0.836316 | KIRC | LB2241-RCC | KU-60019 | MVP_top60_DNA+RNA_seed | EGFR | -1.035253 | LUAD | EMC-BAC-2 | Pelitinib |
| MVP_top60_DNA_seed | IGF1R | -0.834593 | KIRC | CAKI-1 | GSK1904529A | MVP_top60_DNA+RNA_seed | ATM | -1.033447 | LUAD | NCI-H1437 | CP466722 |
| MVP_top60_DNA_seed | MTOR | -0.833686 | KIRC | HA7-RCC | CRT0105446 | MVP_top60_DNA+RNA_seed | MDM2 | -1.028881 | LUAD | NCI-H2342 | Serdemetan |
| MVP_top60_DNA_seed | ATM | -0.832563 | KIRC | RCC-ER | KU-60019 | MVP_top60_DNA+RNA_seed | HSP90AA1 | -1.028481 | LUAD | HOP-62 | Luminespib |
| MVP_top60_DNA_seed | PTK2 | -0.831707 | KIRC | RCC10RGB | PF-562271 | MVP_top60_DNA+RNA_seed | AKT1 | -1.028287 | LUAD | NCI-H2009 | AKT inhibitor VIII |
| MVP_top60_DNA_seed | PARP1 | -0.830522 | KIRC | LB2241-RCC | PARP_0108 | MVP_top60_DNA+RNA_seed | AURKA | -1.027453 | LUAD | NCI-H2030 | Genentech Cpd 10 |
| MVP_top60_DNA_seed | HSP90AA1 | -0.828999 | KIRC | TK10 | Tanespimycin | MVP_top60_DNA+RNA_seed | HSP90AA1 | -1.026593 | LUAD | NCI-H1944 | Elesclomol |
| MVP_top60_DNA_seed | HSP90AB1 | -0.828999 | KIRC | TK10 | Tanespimycin | MVP_top60_DNA+RNA_seed | SRC | -1.024607 | LUAD | EMC-BAC-1 | Bosutinib |
| MVP_top60_DNA_seed | ATM | -0.828326 | KIRC | 786-0 | CP466722 | MVP_top60_DNA+RNA_seed | HSP90AA1 | -1.021354 | LUAD | PC-14 | SNX-2112 |
| MVP_top60_DNA_seed | HSP90AA1 | -0.828051 | KIRC | VMRC-RCW | Luminespib | MVP_top60_DNA+RNA_seed | PIK3CA | -1.020125 | LUAD | NCI-H1355 | PI3Ka_4409 |
| MVP_top60_DNA_seed | HSP90AB1 | -0.828051 | KIRC | VMRC-RCW | Luminespib | MVP_top60_DNA+RNA_seed | HSP90AA1 | -1.010517 | LUAD | HOP-62 | SNX-2112 |
| MVP_top60_DNA_seed | HSP90AA1 | -0.824943 | KIRC | 786-0 | SNX-2112 | MVP_top60_DNA+RNA_seed | ATM | -1.008731 | LUAD | NCI-H2085 | KU-55933 |
| MVP_top60_DNA_seed | HSP90AB1 | -0.824943 | KIRC | 786-0 | SNX-2112 | MVP_top60_DNA+RNA_seed | HSP90AA1 | -1.008368 | LUAD | HCC-44 | SNX-2112 |
| MVP_top60_DNA_seed | IGF1R | -0.822088 | KIRC | U031 | GSK1904529A | MVP_top60_DNA+RNA_seed | AURKA | -1.006379 | LUAD | NCI-H2087 | Genentech Cpd 10 |
| MVP_top60_DNA_seed | PIK3CD | -0.821972 | KIRC | SW156 | Idelalisib | MVP_top60_DNA+RNA_seed | HSP90AA1 | -1.004941 | LUAD | NCI-H1944 | CCT-018159 |
| MVP_top60_DNA_seed | HDAC1 | -0.821809 | KIRC | HA7-RCC | Dacinostat | MVP_top60_DNA+RNA_seed | AKT1 | -1.004335 | LUAD | NCI-H2342 | BAY AKT1 |
| MVP_top60_DNA_seed | MTOR | -0.821255 | KIRC | RCC-FG2 | CRT0105446 | MVP_top60_DNA+RNA_seed | BRAF | -1.001934 | LUAD | NCI-H1666 | PLX-4720 |
| MVP_top60_DNA_seed | PIK3CA | -0.820282 | KIRC | LB996-RCC | AZD8835 | MVP_top60_DNA+RNA_seed | BRAF | -1.001378 | LUAD | NCI-H2342 | HG6-64-1 |
| MVP_top60_DNA_seed | PIK3CD | -0.820282 | KIRC | LB996-RCC | AZD8835 | MVP_top60_DNA+RNA_seed | EGFR | -1.001148 | LUAD | NCI-H322M | AST-1306 |
| MVP_top60_DNA_seed | AR | -0.811472 | KIRC | 769-P | AZD3514 | MVP_top60_DNA+RNA_seed | IGF1R | -1.000676 | LUAD | NCI-H838 | Linsitinib |

| | | | | | | | | | | | |
|---|---|---|---|---|---|---|---|---|---|---|---|
| MVP_top60_DNA_seed | HSP90AA1 | -0.808879 | KIRC | BB65-RCC | Luminespib | MVP_top60_DNA+RNA_seed | HDAC1 | -0.999116 | LUAD | HCC-44 | Dacinostat |
| MVP_top60_DNA_seed | HSP90AB1 | -0.808879 | KIRC | BB65-RCC | Luminespib | MVP_top60_DNA+RNA_seed | SRC | -0.998977 | LUAD | A549 | HG-6-71-01 |
| MVP_top60_DNA_seed | HSP90AA1 | -0.803576 | KIRC | 786-0 | Tanespimycin | MVP_top60_DNA+RNA_seed | MET | -0.998882 | LUAD | NCI-H1648 | JNJ38877605 |
| MVP_top60_DNA_seed | HSP90AB1 | -0.803576 | KIRC | 786-0 | Tanespimycin | MVP_top60_DNA+RNA_seed | HSP90AA1 | -0.998357 | LUAD | NCI-H1437 | CCT-018159 |
| MVP_top60_DNA_seed | MTOR | -0.803544 | KIRC | RCC-JF | Apitolisib | MVP_top60_DNA+RNA_seed | SRC | -0.996903 | LUAD | NCI-H2030 | Bosutinib |
| MVP_top60_DNA_seed | MTOR | -0.802529 | KIRC | NCC021 | Apitolisib | MVP_top60_DNA+RNA_seed | EGFR | -0.992871 | LUAD | NCI-H1648 | Gefitinib |
| MVP_top60_DNA_seed | MTOR | -0.801307 | KIRC | TK10 | JW-7-52-1 | MVP_top60_DNA+RNA_seed | AURKA | -0.992257 | LUAD | HCC-44 | ZM447439 |
| MVP_top60_DNA_seed | IGF1R | -0.799903 | KIRC | LB1047-RCC | BMS-754807 | MVP_top60_DNA+RNA_seed | ATM | -0.992236 | LUAD | NCI-H1648 | torin2 |
| MVP_top60_DNA_seed | PARP1 | -0.799298 | KIRC | RCC-MF | PARP_9482 | MVP_top60_DNA+RNA_seed | FGFR1 | -0.991899 | LUAD | NCI-H838 | AZD4547 |
| MVP_top60_DNA_seed | HDAC1 | -0.798647 | KIRC | RCC-MF | CAY10603 | MVP_top60_DNA+RNA_seed | PIK3CA | -0.989945 | LUAD | SW1573 | AZD8835 |
| MVP_top60_DNA_seed | HSP90AA1 | -0.796719 | KIRC | A498 | Tanespimycin | MVP_top60_DNA+RNA_seed | MET | -0.987813 | LUAD | LC-2-ad | JNJ38877605 |
| MVP_top60_DNA_seed | HSP90AB1 | -0.796719 | KIRC | A498 | Tanespimycin | MVP_top60_DNA+RNA_seed | PIK3CA | -0.983477 | LUAD | NCI-H1623 | PI3Ka_4409 |
| MVP_top60_DNA_seed | SMARCA4 | -0.79587 | KIRC | 769-P | PFI-3 | MVP_top60_DNA+RNA_seed | PIK3CA | -0.978632 | LUAD | HCC-44 | PI3Ka_4409 |
| MVP_top60_DNA_seed | EP300 | -0.795086 | KIRC | CAKI-1 | I-CBP112 | MVP_top60_DNA+RNA_seed | AURKA | -0.978187 | LUAD | NCI-H1648 | Genentech Cpd 10 |
| MVP_top60_DNA_seed | MTOR | -0.792318 | KIRC | RCC-JF | Torin 2 | MVP_top60_DNA+RNA_seed | EGFR | -0.974108 | LUAD | NCI-H1975 | CI-1033 |
| MVP_top60_DNA_seed | PIK3CD | -0.789079 | KIRC | RCC-FG2 | Idelalisib | MVP_top60_DNA+RNA_seed | IGF1R | -0.973175 | LUAD | NCI-H2030 | BMS-536924 |
| MVP_top60_DNA_seed | ATM | -0.78383 | KIRC | RCC-JF | KU-55933 | MVP_top60_DNA+RNA_seed | AKT1 | -0.971271 | LUAD | HCC-44 | BAY AKT1 |
| MVP_top60_DNA_seed | HSP90AA1 | -0.782752 | KIRC | 786-0 | Elesclomol | MVP_top60_DNA+RNA_seed | ATM | -0.970509 | LUAD | NCI-H1734 | KU-60019 |
| MVP_top60_DNA_seed | HSP90AB1 | -0.782752 | KIRC | 786-0 | Elesclomol | MVP_top60_DNA+RNA_seed | EGFR | -0.968627 | LUAD | NCI-H1944 | AZD8931 |
| MVP_top60_DNA_seed | HDAC1 | -0.781576 | KIRC | RCC-AB | AR-42 | MVP_top60_DNA+RNA_seed | HDAC1 | -0.96772 | LUAD | HOP-62 | Tubastatin A |
| MVP_top60_DNA_seed | MTOR | -0.780837 | KIRC | A498 | QL-VIII-58 | MVP_top60_DNA+RNA_seed | ATM | -0.967298 | LUAD | LXF-289 | torin2 |
| MVP_top60_DNA_seed | HSP90AA1 | -0.779809 | KIRC | RXF393 | Elesclomol | MVP_top60_DNA+RNA_seed | EGFR | -0.966415 | LUAD | EKVX | Gefitinib |
| MVP_top60_DNA_seed | HSP90AB1 | -0.779809 | KIRC | RXF393 | Elesclomol | MVP_top60_DNA+RNA_seed | HSP90AA1 | -0.962868 | LUAD | NCI-H2122 | Tanespimycin |
| MVP_top60_DNA_seed | MTOR | -0.777761 | KIRC | 786-0 | Temsirolimus | MVP_top60_DNA+RNA_seed | IGF1R | -0.961389 | LUAD | NCI-H2122 | BMS-536924 |
| MVP_top60_DNA_seed | FGFR4 | -0.775996 | KIRC | NCC010 | FGFR_3831 | MVP_top60_DNA+RNA_seed | AURKA | -0.959286 | LUAD | NCI-H1755 | Genentech Cpd 10 |
| MVP_top60_DNA_seed | AR | -0.775798 | KIRC | ACHN | Bicalutamide | MVP_top60_DNA+RNA_seed | FGFR1 | -0.959115 | LUAD | EMC-BAC-2 | FGFR_3831 |
| MVP_top60_DNA_seed | PARP1 | -0.775609 | KIRC | CAKI-1 | PARP_9495 | MVP_top60_DNA+RNA_seed | HSP90AA1 | -0.958201 | LUAD | NCI-H2342 | Luminespib |
| MVP_top60_DNA_seed | PARP1 | -0.775195 | KIRC | VMRC-RCZ | Rucaparib | MVP_top60_DNA+RNA_seed | SRC | -0.956777 | LUAD | LC-2-ad | Saracatinib |
| MVP_top60_DNA_seed | PIK3CD | -0.769772 | KIRC | LB1047-RCC | AZD8186 | MVP_top60_DNA+RNA_seed | HSP90AA1 | -0.955631 | LUAD | EMC-BAC-2 | Elesclomol |
| MVP_top60_DNA_seed | PARP1 | -0.761981 | KIRC | HA7-RCC | Talazoparib | MVP_top60_DNA+RNA_seed | BRAF | -0.95384 | LUAD | NCI-H358 | HG6-64-1 |
| MVP_top60_DNA_seed | MTOR | -0.759369 | KIRC | RCC-AB | WYE-125132 | MVP_top60_DNA+RNA_seed | MET | -0.953581 | LUAD | NCI-H1975 | AZD6094 |
| MVP_top60_DNA_seed | ATM | -0.756578 | KIRC | CAKI-1 | KU-60019 | MVP_top60_DNA+RNA_seed | EGFR | -0.951567 | LUAD | Calu-3 | Afatinib |
| MVP_top60_DNA_seed | ATM | -0.752802 | KIRC | RCC-MF | KU-60019 | MVP_top60_DNA+RNA_seed | HSP90AA1 | -0.947397 | LUAD | RERF-LC-MS | Elesclomol |
| MVP_top60_DNA_seed | ESR1 | -0.751521 | KIRC | RXF393 | Fulvestrant | MVP_top60_DNA+RNA_seed | IGF1R | -0.945771 | LUAD | EKVX | BMS-536924 |
| MVP_top60_DNA_seed | PIK3CD | -0.751263 | KIRC | CAKI-1 | IC-87114 | MVP_top60_DNA+RNA_seed | PIK3CA | -0.94476 | LUAD | NCI-H2228 | PI3Ka_4409 |
| MVP_top60_DNA_seed | FGFR4 | -0.749345 | KIRC | BFTC-909 | FGFR_0939 | MVP_top60_DNA+RNA_seed | PTK2 | -0.936408 | LUAD | NCI-H2122 | PF-562271 |
| MVP_top60_DNA_seed | IGF1R | -0.74885 | KIRC | RXF393 | BMS-536924 | MVP_top60_DNA+RNA_seed | HSP90AA1 | -0.933845 | LUAD | EMC-BAC-2 | Tanespimycin |
| MVP_top60_DNA_seed | MTOR | -0.747246 | KIRC | CAL-54 | WYE-125132 | MVP_top60_DNA+RNA_seed | MDM2 | -0.933725 | LUAD | NCI-H1568 | Serdemetan |
| MVP_top60_DNA_seed | PARP1 | -0.741032 | KIRC | U031 | Rucaparib | MVP_top60_DNA+RNA_seed | HSP90AA1 | -0.932636 | LUAD | SK-LU-1 | Luminespib |
| MVP_top60_DNA_seed | ESR1 | -0.738511 | KIRC | LB1047-RCC | Fulvestrant | MVP_top60_DNA+RNA_seed | EGFR | -0.931851 | LUAD | NCI-H1650 | Gefitinib |
| MVP_top60_DNA_seed | PARP1 | -0.738299 | KIRC | KMRC-20 | Olaparib | MVP_top60_DNA+RNA_seed | HDAC1 | -0.931749 | LUAD | LC-2-ad | Entinostat |

| | | | | | | | | | | | |
|---|---|---|---|---|---|---|---|---|---|---|---|
| MVP_top60_DNA_seed | FGFR4 | -0.73585 | KIRC | RCC10RGB | FGFR_0939 | MVP_top60_DNA+RNA_seed | AKT1 | -0.931429 | LUAD | NCI-H358 | AKT inhibitor VIII |
| MVP_top60_DNA_seed | ATM | -0.735699 | KIRC | RCC-JF | CP466722 | MVP_top60_DNA+RNA_seed | PTK2 | -0.930421 | LUAD | NCI-H650 | PF-562271 |
| MVP_top60_DNA_seed | IGF1R | -0.734846 | KIRC | LB996-RCC | BMS-754807 | MVP_top60_DNA+RNA_seed | MDM2 | -0.929405 | LUAD | RERF-LC-MS | Serdemetan |
| MVP_top60_DNA_seed | HSP90AA1 | -0.727153 | KIRC | RCC-JW | Tanespimycin | MVP_top60_DNA+RNA_seed | HSP90AA1 | -0.929309 | LUAD | NCI-H650 | SNX-2112 |
| MVP_top60_DNA_seed | HSP90AB1 | -0.727153 | KIRC | RCC-JW | Tanespimycin | MVP_top60_DNA+RNA_seed | EGFR | -0.926824 | LUAD | A549 | Afatinib |
| MVP_top60_DNA_seed | AR | -0.725522 | KIRC | 786-0 | Bicalutamide | MVP_top60_DNA+RNA_seed | SRC | -0.926307 | LUAD | COR-L105 | NVP-BHG712 |
| MVP_top60_DNA_seed | HDAC1 | -0.725432 | KIRC | RCC-MF | Belinostat | MVP_top60_DNA+RNA_seed | PTK2 | -0.92623 | LUAD | HCC-44 | PF-562271 |
| MVP_top60_DNA_seed | HDAC1 | -0.724644 | KIRC | OS-RC-2 | Parthenolide | MVP_top60_DNA+RNA_seed | MDM2 | -0.925226 | LUAD | NCI-H2087 | Serdemetan |
| MVP_top60_DNA_seed | AR | -0.723897 | KIRC | BFTC-909 | Bicalutamide | MVP_top60_DNA+RNA_seed | PTK2 | -0.925092 | LUAD | NCI-H358 | PF-562271 |
| MVP_top60_DNA_seed | PTK2 | -0.715194 | KIRC | LB1047-RCC | PF-562271 | MVP_top60_DNA+RNA_seed | HSP90AA1 | -0.923668 | LUAD | NCI-H1437 | Tanespimycin |
| MVP_top60_DNA_seed | FGFR4 | -0.713748 | KIRC | A704 | FGFR_3831 | MVP_top60_DNA+RNA_seed | MET | -0.923329 | LUAD | NCI-H1993 | JNJ38877605 |
| MVP_top60_DNA_seed | MTOR | -0.713535 | KIRC | LB996-RCC | Voxtalisib | MVP_top60_DNA+RNA_seed | MET | -0.923318 | LUAD | LXF-289 | PHA-665752 |
| MVP_top60_DNA_seed | PIK3CD | -0.707945 | KIRC | HA7-RCC | Idelalisib | MVP_top60_DNA+RNA_seed | EGFR | -0.920786 | LUAD | NCI-H650 | Cetuximab |
| MVP_top60_DNA_seed | ESR1 | -0.6988 | KIRC | 786-0 | Tamoxifen | MVP_top60_DNA+RNA_seed | IGF1R | -0.919482 | LUAD | RERF-LC-KJ | GSK1904529A |
| MVP_top60_DNA_seed | MTOR | -0.698256 | KIRC | SW156 | Temsirolimus | MVP_top60_DNA+RNA_seed | HDAC1 | -0.917954 | LUAD | NCI-H1437 | Dacinostat |
| MVP_top60_DNA_seed | HSP90AA1 | -0.696204 | KIRC | RCC-ER | Elesclomol | MVP_top60_DNA+RNA_seed | AKT1 | -0.916742 | LUAD | NCI-H2085 | MK-2206 |
| MVP_top60_DNA_seed | HSP90AB1 | -0.696204 | KIRC | RCC-ER | Elesclomol | MVP_top60_DNA+RNA_seed | SMARCA4 | -0.916037 | LUAD | EKVX | PFI-3 |
| MVP_top60_DNA_seed | MTOR | -0.695836 | KIRC | 769-P | Apitolisib | MVP_top60_DNA+RNA_seed | AURKA | -0.914346 | LUAD | A549 | HG-5-113-01 |
| MVP_top60_DNA_seed | PARP1 | -0.695658 | KIRC | LB2241-RCC | PARP_9495 | MVP_top60_DNA+RNA_seed | IGF1R | -0.913895 | LUAD | HCC-44 | BMS-754807 |
| MVP_top60_DNA_seed | PIK3CD | -0.695272 | KIRC | A704 | AZD8186 | MVP_top60_DNA+RNA_seed | BRAF | -0.913088 | LUAD | COR-L105 | HG6-64-1 |
| MVP_top60_DNA_seed | ESR1 | -0.694744 | KIRC | CAL-54 | Tamoxifen | MVP_top60_DNA+RNA_seed | ATM | -0.909217 | LUAD | LC-2-ad | CP466722 |
| MVP_top60_DNA_seed | IGF1R | -0.694288 | KIRC | SW156 | GSK1904529A | MVP_top60_DNA+RNA_seed | AURKA | -0.907529 | LUAD | EMC-BAC-1 | HG-5-113-01 |
| MVP_top60_DNA_seed | PARP1 | -0.69344 | KIRC | SW156 | Veliparib | MVP_top60_DNA+RNA_seed | HDAC1 | -0.906758 | LUAD | HOP-62 | Dacinostat |
| MVP_top60_DNA_seed | HDAC1 | -0.690676 | KIRC | VMRC-RCW | Dacinostat | MVP_top60_DNA+RNA_seed | FGFR1 | -0.906192 | LUAD | NCI-H1355 | PD173074 |
| MVP_top60_DNA_seed | PARP1 | -0.690128 | KIRC | RCC-JF | Veliparib | MVP_top60_DNA+RNA_seed | EGFR | -0.904502 | LUAD | NCI-H1781 | AZD8931 |
| MVP_top60_DNA_seed | HDAC1 | -0.689737 | KIRC | RCC-JF | AR-42 | MVP_top60_DNA+RNA_seed | HSP90AA1 | -0.901973 | LUAD | NCI-H1693 | Elesclomol |
| MVP_top60_DNA_seed | HDAC1 | -0.689617 | KIRC | RCC-MF | AR-42 | MVP_top60_DNA+RNA_seed | EGFR | -0.901723 | LUAD | NCI-H1755 | CUDC-101 |
| MVP_top60_DNA_seed | HSP90AA1 | -0.6824 | KIRC | ACHN | Luminespib | MVP_top60_DNA+RNA_seed | ATM | -0.899538 | LUAD | NCI-H838 | KU-55933 |
| MVP_top60_DNA_seed | HSP90AB1 | -0.6824 | KIRC | ACHN | Luminespib | MVP_top60_DNA+RNA_seed | HSP90AA1 | -0.895562 | LUAD | NCI-H1355 | Elesclomol |
| MVP_top60_DNA_seed | PARP1 | -0.680961 | KIRC | 769-P | PARP_9495 | MVP_top60_DNA+RNA_seed | BRAF | -0.893033 | LUAD | NCI-H1792 | HG6-64-1 |
| MVP_top60_DNA_seed | PIK3CD | -0.67555 | KIRC | RCC-ER | AZD8186 | MVP_top60_DNA+RNA_seed | PIK3CA | -0.892877 | LUAD | SW1573 | PI3Ka_4409 |
| MVP_top60_DNA_seed | MTOR | -0.673839 | KIRC | HA7-RCC | WYE-125132 | MVP_top60_DNA+RNA_seed | HSP90AA1 | -0.892459 | LUAD | EMC-BAC-1 | Tanespimycin |
| MVP_top60_DNA_seed | MTOR | -0.673735 | KIRC | CAL-54 | Apitolisib | MVP_top60_DNA+RNA_seed | AKT1 | -0.891721 | LUAD | NCI-H1648 | AKT inhibitor VIII |
| MVP_top60_DNA_seed | ATM | -0.672621 | KIRC | RCC-ER | KU-55933 | MVP_top60_DNA+RNA_seed | EGFR | -0.889269 | LUAD | COR-L105 | CUDC-101 |
| MVP_top60_DNA_seed | PIK3CD | -0.670612 | KIRC | TK10 | AZD8186 | MVP_top60_DNA+RNA_seed | HSP90AA1 | -0.88717 | LUAD | NCI-H2023 | CCT-018159 |
| MVP_top60_DNA_seed | ATM | -0.668881 | KIRC | RCC-MF | CP466722 | MVP_top60_DNA+RNA_seed | BRAF | -0.884074 | LUAD | Calu-6 | PLX-4720 |
| MVP_top60_DNA_seed | ATM | -0.668878 | KIRC | SW156 | CP466722 | MVP_top60_DNA+RNA_seed | MAPK14 | -0.88285 | LUAD | EMC-BAC-1 | TAK-715 |
| MVP_top60_DNA_seed | HDAC1 | -0.666335 | KIRC | 769-P | AR-42 | MVP_top60_DNA+RNA_seed | HDAC1 | -0.882527 | LUAD | LC-2-ad | CAY10603 |
| MVP_top60_DNA_seed | HDAC1 | -0.663409 | KIRC | OS-RC-2 | JQ12 | MVP_top60_DNA+RNA_seed | AURKA | -0.88191 | LUAD | NCI-H1648 | HG-5-113-01 |
| MVP_top60_DNA_seed | PARP1 | -0.658612 | KIRC | CAKI-1 | Rucaparib | MVP_top60_DNA+RNA_seed | AKT1 | -0.88095 | LUAD | NCI-H2009 | MK-2206 |
| MVP_top60_DNA_seed | MTOR | -0.657279 | KIRC | RCC-FG2 | Voxtalisib | MVP_top60_DNA+RNA_seed | HSP90AA1 | -0.880453 | LUAD | NCI-H650 | Luminespib |

| | | | | | | | | | | | |
|---|---|---|---|---|---|---|---|---|---|---|---|
| MVP_top60_DNA_seed | FGFR4 | -0.655322 | KIRC | RCC-FG2 | FGFR_0939 | MVP_top60_DNA+RNA_seed | MAPK14 | -0.878941 | LUAD | HOP-62 | TAK-715 |
| MVP_top60_DNA_seed | ATM | -0.655116 | KIRC | CAL-54 | CP466722 | MVP_top60_DNA+RNA_seed | MDM2 | -0.877497 | LUAD | LXF-289 | Serdemetan |
| MVP_top60_DNA_seed | FGFR4 | -0.65324 | KIRC | RXF393 | FGFR_3831 | MVP_top60_DNA+RNA_seed | HSP90AA1 | -0.87501 | LUAD | NCI-H2009 | Luminespib |
| MVP_top60_DNA_seed | FGFR4 | -0.652829 | KIRC | LB996-RCC | FGFR_0939 | MVP_top60_DNA+RNA_seed | HSP90AA1 | -0.874584 | LUAD | NCI-H2228 | Luminespib |
| MVP_top60_DNA_seed | PARP1 | -0.652758 | KIRC | LB996-RCC | Talazoparib | MVP_top60_DNA+RNA_seed | PIK3CA | -0.874333 | LUAD | NCI-H1975 | PI3Ka_4409 |
| MVP_top60_DNA_seed | PTK2 | -0.651546 | KIRC | KMRC-1 | PF-562271 | MVP_top60_DNA+RNA_seed | HSP90AA1 | -0.874079 | LUAD | RERF-LC-KJ | Elesclomol |
| MVP_top60_DNA_seed | HDAC1 | -0.651089 | KIRC | LB1047-RCC | Dacinostat | MVP_top60_DNA+RNA_seed | PIK3CA | -0.872118 | LUAD | RERF-LC-MS | AZD8835 |
| MVP_top60_DNA_seed | ESR1 | -0.649392 | KIRC | A704 | Tamoxifen | MVP_top60_DNA+RNA_seed | AURKA | -0.871966 | LUAD | NCI-H1651 | ZM447439 |
| MVP_top60_DNA_seed | AR | -0.648355 | KIRC | LB996-RCC | AZD3514 | MVP_top60_DNA+RNA_seed | AKT1 | -0.870995 | LUAD | NCI-H1993 | BAY AKT1 |
| MVP_top60_DNA_seed | MTOR | -0.646077 | KIRC | RXF393 | QL-VIII-58 | MVP_top60_DNA+RNA_seed | AKT1 | -0.869636 | LUAD | HOP-62 | A-443654 |
| MVP_top60_DNA_seed | PARP1 | -0.645493 | KIRC | CAKI-1 | PARP_0108 | MVP_top60_DNA+RNA_seed | EGFR | -0.868239 | LUAD | HCC-44 | Afatinib |
| MVP_top60_DNA_seed | HSP90AA1 | -0.644704 | KIRC | U031 | Elesclomol | MVP_top60_DNA+RNA_seed | MET | -0.86671 | LUAD | LXF-289 | JNJ38877605 |
| MVP_top60_DNA_seed | HSP90AB1 | -0.644704 | KIRC | U031 | Elesclomol | MVP_top60_DNA+RNA_seed | HDAC1 | -0.865955 | LUAD | NCI-H1755 | AR-42 |
| MVP_top60_DNA_seed | PARP1 | -0.643163 | KIRC | HA7-RCC | PARP_9495 | MVP_top60_DNA+RNA_seed | AKT1 | -0.86437 | LUAD | ABC-1 | GSK690693 |
| MVP_top60_DNA_seed | ATM | -0.638226 | KIRC | 769-P | CP466722 | MVP_top60_DNA+RNA_seed | IGF1R | -0.864116 | LUAD | NCI-H1648 | BMS-536924 |
| MVP_top60_DNA_seed | ESR1 | -0.638038 | KIRC | LB2241-RCC | Tamoxifen | MVP_top60_DNA+RNA_seed | EGFR | -0.861462 | LUAD | H3255 | CUDC-101 |
| MVP_top60_DNA_seed | ATM | -0.637547 | KIRC | ACHN | CP466722 | MVP_top60_DNA+RNA_seed | PIK3CA | -0.860426 | LUAD | Calu-3 | PI3Ka_4409 |
| MVP_top60_DNA_seed | PARP1 | -0.636099 | KIRC | ACHN | PARP_0108 | MVP_top60_DNA+RNA_seed | EGFR | -0.858535 | LUAD | PC-14 | CUDC-101 |
| MVP_top60_DNA_seed | HSP90AA1 | -0.63567 | KIRC | RXF393 | SNX-2112 | MVP_top60_DNA+RNA_seed | AURKA | -0.855929 | LUAD | HOP-62 | Genentech Cpd 10 |
| MVP_top60_DNA_seed | HSP90AB1 | -0.63567 | KIRC | RXF393 | SNX-2112 | MVP_top60_DNA+RNA_seed | IGF1R | -0.855027 | LUAD | NCI-H1944 | Linsitinib |
| MVP_top60_DNA_seed | MTOR | -0.635272 | KIRC | RCC-AB | CRT0105446 | MVP_top60_DNA+RNA_seed | SRC | -0.853721 | LUAD | NCI-H1975 | Bosutinib |
| MVP_top60_DNA_seed | HDAC1 | -0.631559 | KIRC | RCC-JF | Belinostat | MVP_top60_DNA+RNA_seed | BRAF | -0.850744 | LUAD | PC-14 | HG6-64-1 |
| MVP_top60_DNA_seed | MTOR | -0.628922 | KIRC | LB1047-RCC | Temsirolimus | MVP_top60_DNA+RNA_seed | IGF1R | -0.850236 | LUAD | NCI-H1975 | BMS-754807 |
| MVP_top60_DNA_seed | PARP1 | -0.623073 | KIRC | SN12C | Veliparib | MVP_top60_DNA+RNA_seed | EGFR | -0.849954 | LUAD | LC-2-ad | Lapatinib |
| MVP_top60_DNA_seed | PARP1 | -0.61823 | KIRC | BB65-RCC | Talazoparib | MVP_top60_DNA+RNA_seed | EGFR | -0.849942 | LUAD | NCI-H2122 | Afatinib |
| MVP_top60_DNA_seed | IGF1R | -0.61667 | KIRC | TK10 | BMS-536924 | MVP_top60_DNA+RNA_seed | EGFR | -0.84962 | LUAD | LC-2-ad | AZD8931 |
| MVP_top60_DNA_seed | ATM | -0.616643 | KIRC | VMRC-RCZ | CP466722 | MVP_top60_DNA+RNA_seed | BRAF | -0.846585 | LUAD | NCI-H358 | PLX-4720 |
| MVP_top60_DNA_seed | MTOR | -0.615934 | KIRC | U031 | WYE-125132 | MVP_top60_DNA+RNA_seed | AKT1 | -0.845301 | LUAD | NCI-H1563 | AKT inhibitor VIII |
| MVP_top60_DNA_seed | ESR1 | -0.615863 | KIRC | U031 | Fulvestrant | MVP_top60_DNA+RNA_seed | HSP90AA1 | -0.842669 | LUAD | HOP-62 | Tanespimycin |
| MVP_top60_DNA_seed | ATM | -0.61374 | KIRC | RCC-AB | CP466722 | MVP_top60_DNA+RNA_seed | AKT1 | -0.839841 | LUAD | LXF-289 | A-443654 |
| MVP_top60_DNA_seed | MTOR | -0.610969 | KIRC | CAL-54 | Voxtalisib | MVP_top60_DNA+RNA_seed | ATM | -0.839102 | LUAD | SK-LU-1 | CP466722 |
| MVP_top60_DNA_seed | FGFR4 | -0.609385 | KIRC | RCC-JF | FGFR_0939 | MVP_top60_DNA+RNA_seed | PIK3CA | -0.839079 | LUAD | HCC-827 | AZD8835 |
| MVP_top60_DNA_seed | MTOR | -0.606272 | KIRC | VMRC-RCZ | Torin 2 | MVP_top60_DNA+RNA_seed | EGFR | -0.837233 | LUAD | NCI-H2030 | Pelitinib |
| MVP_top60_DNA_seed | MTOR | -0.605071 | KIRC | A704 | Voxtalisib | MVP_top60_DNA+RNA_seed | FGFR1 | -0.836622 | LUAD | HCC-44 | PD173074 |
| MVP_top60_DNA_seed | PIK3CD | -0.604726 | KIRC | BB65-RCC | Idelalisib | MVP_top60_DNA+RNA_seed | HSP90AA1 | -0.836021 | LUAD | NCI-H23 | Elesclomol |
| MVP_top60_DNA_seed | HDAC1 | -0.601798 | KIRC | 769-P | Dacinostat | MVP_top60_DNA+RNA_seed | AKT1 | -0.833823 | LUAD | RERF-LC-KJ | AKT inhibitor VIII |
| MVP_top60_DNA_seed | HSP90AA1 | -0.601193 | KIRC | VMRC-RCZ | Tanespimycin | MVP_top60_DNA+RNA_seed | PTK2 | -0.833327 | LUAD | HOP-62 | PF-562271 |
| MVP_top60_DNA_seed | HSP90AB1 | -0.601193 | KIRC | VMRC-RCZ | Tanespimycin | MVP_top60_DNA+RNA_seed | BRAF | -0.833131 | LUAD | NCI-H3122 | SB590885 |
| MVP_top60_DNA_seed | MTOR | -0.600449 | KIRC | 786-0 | WYE-125132 | MVP_top60_DNA+RNA_seed | BRAF | -0.832747 | LUAD | NCI-H358 | PLX-4720 |
| MVP_top60_DNA_seed | MTOR | -0.599065 | KIRC | A704 | JW-7-52-1 | MVP_top60_DNA+RNA_seed | AKT1 | -0.832216 | LUAD | NCI-H2030 | AKT inhibitor VIII |
| MVP_top60_DNA_seed | MTOR | -0.599004 | KIRC | RCC-AB | Torin 2 | MVP_top60_DNA+RNA_seed | MET | -0.828569 | LUAD | NCI-H1734 | JNJ38877605 |

| | | | | | | | | | | | |
|---|---|---|---|---|---|---|---|---|---|---|---|
| MVP_top60_DNA_seed | PARP1 | -0.597426 | KIRC | SN12C | PARP_9482 | MVP_top60_DNA+RNA_seed | IGF1R | -0.82853 | LUAD | NCI-H650 | GSK1904529A |
| MVP_top60_DNA_seed | HDAC1 | -0.597087 | KIRC | OS-RC-2 | Entinostat | MVP_top60_DNA+RNA_seed | HSP90AA1 | -0.82841 | LUAD | SK-LU-1 | Elesclomol |
| MVP_top60_DNA_seed | HDAC1 | -0.595429 | KIRC | 769-P | CAY10603 | MVP_top60_DNA+RNA_seed | AKT1 | -0.828284 | LUAD | LXF-289 | BAY AKT1 |
| MVP_top60_DNA_seed | PIK3CD | -0.59431 | KIRC | LB1047-RCC | Idelalisib | MVP_top60_DNA+RNA_seed | HSP90AA1 | -0.827909 | LUAD | NCI-H2405 | Luminespib |
| MVP_top60_DNA_seed | PIK3CD | -0.591978 | KIRC | 786-0 | IC-87114 | MVP_top60_DNA+RNA_seed | MET | -0.826485 | LUAD | NCI-H1623 | JNJ38877605 |
| MVP_top60_DNA_seed | AR | -0.591923 | KIRC | HA7-RCC | AZD3514 | MVP_top60_DNA+RNA_seed | AKT1 | -0.826474 | LUAD | NCI-H1975 | A-443654 |
| MVP_top60_DNA_seed | PARP1 | -0.586431 | KIRC | RCC-ER | PARP_9495 | MVP_top60_DNA+RNA_seed | AURKA | -0.825324 | LUAD | HOP-62 | Tozasertib |
| MVP_top60_DNA_seed | HDAC1 | -0.583668 | KIRC | 769-P | Tubastatin A | MVP_top60_DNA+RNA_seed | HSP90AA1 | -0.821418 | LUAD | A549 | Elesclomol |
| MVP_top60_DNA_seed | MTOR | -0.57318 | KIRC | RCC-AB | Apitolisib | MVP_top60_DNA+RNA_seed | BRAF | -0.821024 | LUAD | SK-LU-1 | HG6-64-1 |
| MVP_top60_DNA_seed | ESR1 | -0.568046 | KIRC | BB65-RCC | Tamoxifen | MVP_top60_DNA+RNA_seed | MET | -0.820218 | LUAD | EMC-BAC-1 | JNJ38877605 |
| MVP_top60_DNA_seed | HDAC1 | -0.566739 | KIRC | RXF393 | Belinostat | MVP_top60_DNA+RNA_seed | EGFR | -0.819897 | LUAD | NCI-H358 | CI-1033 |
| MVP_top60_DNA_seed | PARP1 | -0.56525 | KIRC | RXF393 | PARP_9495 | MVP_top60_DNA+RNA_seed | BRAF | -0.81973 | LUAD | NCI-H1792 | RAF_9304 |
| MVP_top60_DNA_seed | PTK2 | -0.564299 | KIRC | RXF393 | PF-562271 | MVP_top60_DNA+RNA_seed | PTK2 | -0.817794 | LUAD | NCI-H3122 | PF-562271 |
| MVP_top60_DNA_seed | PARP1 | -0.563971 | KIRC | RCC-MF | PARP_0108 | MVP_top60_DNA+RNA_seed | AURKA | -0.81424 | LUAD | HCC-78 | GSK1070916 |
| MVP_top60_DNA_seed | PIK3CD | -0.561983 | KIRC | RCC-ER | IC-87114 | MVP_top60_DNA+RNA_seed | ATM | -0.813508 | LUAD | NCI-H2023 | KU-60019 |
| MVP_top60_DNA_seed | MTOR | -0.561634 | KIRC | RCC-FG2 | Apitolisib | MVP_top60_DNA+RNA_seed | AURKA | -0.812365 | LUAD | HCC-78 | Genentech Cpd 10 |
| MVP_top60_DNA_seed | IGF1R | -0.558891 | KIRC | RCC-ER | BMS-536924 | MVP_top60_DNA+RNA_seed | BRAF | -0.811809 | LUAD | NCI-H2405 | RAF_9304 |
| MVP_top60_DNA_seed | PARP1 | -0.558551 | KIRC | RCC-AB | PARP_9482 | MVP_top60_DNA+RNA_seed | AKT1 | -0.810689 | LUAD | NCI-H1648 | A-443654 |
| MVP_top60_DNA_seed | AR | -0.556415 | KIRC | CAL-54 | AZD3514 | MVP_top60_DNA+RNA_seed | PIK3CA | -0.809024 | LUAD | NCI-H1623 | AZD8835 |
| MVP_top60_DNA_seed | HSP90AA1 | -0.553513 | KIRC | KMRC-20 | Tanespimycin | MVP_top60_DNA+RNA_seed | AKT1 | -0.809019 | LUAD | NCI-H522 | AKT inhibitor VIII |
| MVP_top60_DNA_seed | HSP90AB1 | -0.553513 | KIRC | KMRC-20 | Tanespimycin | MVP_top60_DNA+RNA_seed | HSP90AA1 | -0.808824 | LUAD | NCI-H1648 | Luminespib |
| MVP_top60_DNA_seed | HDAC1 | -0.550534 | KIRC | 769-P | Belinostat | MVP_top60_DNA+RNA_seed | ATM | -0.807586 | LUAD | NCI-H322M | KU-55933 |
| MVP_top60_DNA_seed | AR | -0.550534 | KIRC | RCC-AB | AZD3514 | MVP_top60_DNA+RNA_seed | HSP90AA1 | -0.807239 | LUAD | SW1573 | Elesclomol |
| MVP_top60_DNA_seed | IGF1R | -0.550316 | KIRC | A704 | GSK1904529A | MVP_top60_DNA+RNA_seed | HSP90AA1 | -0.806797 | LUAD | NCI-H1755 | Elesclomol |
| MVP_top60_DNA_seed | PIK3CA | -0.549567 | KIRC | RCC-MF | AZD8835 | MVP_top60_DNA+RNA_seed | ATM | -0.806019 | LUAD | COR-L105 | CP466722 |
| MVP_top60_DNA_seed | PIK3CD | -0.549567 | KIRC | RCC-MF | AZD8835 | MVP_top60_DNA+RNA_seed | HDAC1 | -0.804132 | LUAD | NCI-H522 | JQ12 |
| MVP_top60_DNA_seed | PARP1 | -0.548625 | KIRC | RXF393 | Talazoparib | MVP_top60_DNA+RNA_seed | AKT1 | -0.803284 | LUAD | Calu-6 | A-443654 |
| MVP_top60_DNA_seed | PARP1 | -0.548431 | KIRC | RCC-AB | PARP_9495 | MVP_top60_DNA+RNA_seed | AKT1 | -0.803149 | LUAD | LXF-289 | AKT inhibitor VIII |
| MVP_top60_DNA_seed | ATM | -0.547212 | KIRC | VMRC-RCZ | torin2 | MVP_top60_DNA+RNA_seed | HSP90AA1 | -0.801552 | LUAD | NCI-H1755 | SNX-2112 |
| MVP_top60_DNA_seed | MTOR | -0.547212 | KIRC | VMRC-RCZ | torin2 | MVP_top60_DNA+RNA_seed | ATM | -0.801269 | LUAD | RERF-LC-KJ | KU-55933 |
| MVP_top60_DNA_seed | ESR1 | -0.546801 | KIRC | RCC-JF | Fulvestrant | MVP_top60_DNA+RNA_seed | PTK2 | -0.80083 | LUAD | NCI-H2087 | PF-562271 |
| MVP_top60_DNA_seed | PIK3CD | -0.546633 | KIRC | SW156 | IC-87114 | MVP_top60_DNA+RNA_seed | MET | -0.800421 | LUAD | NCI-H1651 | JNJ38877605 |
| MVP_top60_DNA_seed | PARP1 | -0.545845 | KIRC | SN12C | PARP_0108 | MVP_top60_DNA+RNA_seed | HSP90AA1 | -0.800327 | LUAD | COR-L105 | Luminespib |
| MVP_top60_DNA_seed | PARP1 | -0.545411 | KIRC | 786-0 | Olaparib | MVP_top60_DNA+RNA_seed | HDAC1 | -0.798706 | LUAD | ABC-1 | JQ12 |
| MVP_top60_DNA_seed | HDAC1 | -0.544592 | KIRC | NCC021 | Belinostat | MVP_top60_DNA+RNA_seed | HDAC1 | -0.798075 | LUAD | LC-2-ad | AR-42 |
| MVP_top60_DNA_seed | ATM | -0.540539 | KIRC | BFTC-909 | KU-60019 | MVP_top60_DNA+RNA_seed | MDM2 | -0.797776 | LUAD | NCI-H1755 | Serdemetan |
| MVP_top60_DNA_seed | HDAC1 | -0.538626 | KIRC | LB1047-RCC | AR-42 | MVP_top60_DNA+RNA_seed | MDM2 | -0.797588 | LUAD | EMC-BAC-1 | Serdemetan |
| MVP_top60_DNA_seed | HDAC1 | -0.536125 | KIRC | NCC021 | CAY10603 | MVP_top60_DNA+RNA_seed | EGFR | -0.797095 | LUAD | NCI-H322M | Afatinib |
| MVP_top60_DNA_seed | EP300 | -0.532319 | KIRC | LB2241-RCC | I-CBP112 | MVP_top60_DNA+RNA_seed | BRAF | -0.795748 | LUAD | NCI-H1975 | Dabrafenib |
| MVP_top60_DNA_seed | MTOR | -0.530932 | KIRC | RCC-FG2 | WYE-125132 | MVP_top60_DNA+RNA_seed | HSP90AA1 | -0.794547 | LUAD | NCI-H1651 | CCT-018159 |
| MVP_top60_DNA_seed | HDAC1 | -0.526125 | KIRC | LB1047-RCC | CAY10603 | MVP_top60_DNA+RNA_seed | BRAF | -0.793222 | LUAD | LC-2-ad | PLX-4720 |

| | | | | | | | | | | | |
|---|---|---|---|---|---|---|---|---|---|---|---|
| MVP_top60_DNA_seed | PIK3CD | -0.526093 | KIRC | A704 | Idelalisib | MVP_top60_DNA+RNA_seed | HDAC1 | -0.792252 | LUAD | HCC-44 | CAY10603 |
| MVP_top60_DNA_seed | EP300 | -0.522934 | KIRC | RCC-ER | I-CBP112 | MVP_top60_DNA+RNA_seed | HDAC1 | -0.790653 | LUAD | NCI-H2228 | Dacinostat |
| MVP_top60_DNA_seed | PARP1 | -0.522633 | KIRC | BB65-RCC | PARP_9482 | MVP_top60_DNA+RNA_seed | IGF1R | -0.789349 | LUAD | NCI-H358 | BMS-536924 |
| MVP_top60_DNA_seed | HDAC1 | -0.517879 | KIRC | LB1047-RCC | Belinostat | MVP_top60_DNA+RNA_seed | EGFR | -0.788934 | LUAD | COR-L105 | Cetuximab |
| MVP_top60_DNA_seed | PTK2 | -0.515469 | KIRC | 769-P | PF-562271 | MVP_top60_DNA+RNA_seed | BRAF | -0.788367 | LUAD | A549 | HG6-64-1 |
| MVP_top60_DNA_seed | PARP1 | -0.513963 | KIRC | NCC010 | Talazoparib | MVP_top60_DNA+RNA_seed | EGFR | -0.788279 | LUAD | HCC-78 | Afatinib |
| MVP_top60_DNA_seed | ATM | -0.513805 | KIRC | NCC021 | KU-60019 | MVP_top60_DNA+RNA_seed | MET | -0.788167 | LUAD | SK-LU-1 | JNJ38877605 |
| MVP_top60_DNA_seed | MTOR | -0.513638 | KIRC | KMRC-1 | Temsirolimus | MVP_top60_DNA+RNA_seed | SMARCA4 | -0.784166 | LUAD | NCI-H1792 | PFI-3 |
| MVP_top60_DNA_seed | MTOR | -0.51299 | KIRC | ACHN | Apitolisib | MVP_top60_DNA+RNA_seed | PIK3CA | -0.784165 | LUAD | Calu-3 | AZD8835 |
| MVP_top60_DNA_seed | HDAC1 | -0.512198 | KIRC | NCC021 | AR-42 | MVP_top60_DNA+RNA_seed | EGFR | -0.782968 | LUAD | NCI-H1693 | CUDC-101 |
| MVP_top60_DNA_seed | MTOR | -0.510538 | KIRC | HA7-RCC | Apitolisib | MVP_top60_DNA+RNA_seed | AKT1 | -0.782192 | LUAD | NCI-H3122 | BAY AKT1 |
| MVP_top60_DNA_seed | HDAC1 | -0.507052 | KIRC | RXF393 | CAY10603 | MVP_top60_DNA+RNA_seed | ATM | -0.78189 | LUAD | LXF-289 | CP466722 |
| MVP_top60_DNA_seed | HDAC1 | -0.503431 | KIRC | 786-0 | CAY10603 | MVP_top60_DNA+RNA_seed | IGF1R | -0.780734 | LUAD | NCI-H1563 | GSK1904529A |
| MVP_top60_DNA_seed | HSP90AA1 | -0.502193 | KIRC | U031 | Tanespimycin | MVP_top60_DNA+RNA_seed | ATM | -0.779539 | LUAD | NCI-H23 | CP466722 |
| MVP_top60_DNA_seed | HSP90AB1 | -0.502193 | KIRC | U031 | Tanespimycin | MVP_top60_DNA+RNA_seed | EGFR | -0.778734 | LUAD | HCC-78 | CUDC-101 |
| MVP_top60_DNA_seed | PARP1 | -0.501648 | KIRC | RCC-JW | Olaparib | MVP_top60_DNA+RNA_seed | BRAF | -0.77557 | LUAD | LXF-289 | HG6-64-1 |
| MVP_top60_DNA_seed | ATM | -0.500712 | KIRC | U031 | CP466722 | MVP_top60_DNA+RNA_seed | HSP90AA1 | -0.772694 | LUAD | NCI-H23 | CCT-018159 |
| MVP_top60_DNA_seed | PARP1 | -0.500294 | KIRC | SW156 | Rucaparib | MVP_top60_DNA+RNA_seed | AURKA | -0.770918 | LUAD | VMRC-LCD | ZM447439 |
| MVP_top60_DNA_seed | PARP1 | -0.497935 | KIRC | LB996-RCC | PARP_9482 | MVP_top60_DNA+RNA_seed | EGFR | -0.770762 | LUAD | PC-14 | PF-00299804 |
| MVP_top60_DNA_seed | ATM | -0.497329 | KIRC | RCC-MF | KU-55933 | MVP_top60_DNA+RNA_seed | HSP90AA1 | -0.770752 | LUAD | NCI-H1944 | SNX-2112 |
| MVP_top60_DNA_seed | PIK3CD | -0.496752 | KIRC | TK10 | IC-87114 | MVP_top60_DNA+RNA_seed | HDAC1 | -0.766864 | LUAD | PC-3_[JPC-3] | JQ12 |
| MVP_top60_DNA_seed | FGFR4 | -0.494367 | KIRC | 769-P | FGFR_0939 | MVP_top60_DNA+RNA_seed | MET | -0.76305 | LUAD | LXF-289 | AZD6094 |
| MVP_top60_DNA_seed | HDAC1 | -0.493316 | KIRC | RCC-AB | Tubastatin A | MVP_top60_DNA+RNA_seed | SRC | -0.762393 | LUAD | NCI-H2087 | NVP-BHG712 |
| MVP_top60_DNA_seed | ATM | -0.491558 | KIRC | RCC-JW | KU-55933 | MVP_top60_DNA+RNA_seed | AKT1 | -0.762254 | LUAD | NCI-H2228 | AKT inhibitor VIII |
| MVP_top60_DNA_seed | IGF1R | -0.491347 | KIRC | A704 | BMS-536924 | MVP_top60_DNA+RNA_seed | AKT1 | -0.762151 | LUAD | NCI-H1650 | AKT inhibitor VIII |
| MVP_top60_DNA_seed | HSP90AA1 | -0.489348 | KIRC | ACHN | CCT-018159 | MVP_top60_DNA+RNA_seed | HSP90AA1 | -0.762084 | LUAD | NCI-H1650 | Luminespib |
| MVP_top60_DNA_seed | HSP90AB1 | -0.489348 | KIRC | ACHN | CCT-018159 | MVP_top60_DNA+RNA_seed | HSP90AA1 | -0.762009 | LUAD | NCI-H2087 | Tanespimycin |
| MVP_top60_DNA_seed | HDAC1 | -0.489213 | KIRC | OS-RC-2 | Dacinostat | MVP_top60_DNA+RNA_seed | MAPK14 | -0.76174 | LUAD | NCI-H23 | TAK-715 |
| MVP_top60_DNA_seed | HSP90AA1 | -0.487504 | KIRC | LB996-RCC | Luminespib | MVP_top60_DNA+RNA_seed | BRAF | -0.7615 | LUAD | NCI-H2087 | Dabrafenib |
| MVP_top60_DNA_seed | HSP90AB1 | -0.487504 | KIRC | LB996-RCC | Luminespib | MVP_top60_DNA+RNA_seed | PIK3CA | -0.760682 | LUAD | HCC-78 | PI3Ka_4409 |
| MVP_top60_DNA_seed | FGFR4 | -0.486649 | KIRC | NCC021 | FGFR_0939 | MVP_top60_DNA+RNA_seed | BRAF | -0.755734 | LUAD | EMC-BAC-2 | PLX-4720 |
| MVP_top60_DNA_seed | MTOR | -0.485078 | KIRC | NCC010 | CRT0105446 | MVP_top60_DNA+RNA_seed | FGFR1 | -0.755444 | LUAD | LXF-289 | PD173074 |
| MVP_top60_DNA_seed | ATM | -0.484012 | KIRC | LB996-RCC | torin2 | MVP_top60_DNA+RNA_seed | EGFR | -0.755404 | LUAD | NCI-H2405 | Cetuximab |
| MVP_top60_DNA_seed | MTOR | -0.484012 | KIRC | LB996-RCC | torin2 | MVP_top60_DNA+RNA_seed | ATM | -0.754715 | LUAD | A549 | KU-60019 |
| MVP_top60_DNA_seed | PIK3CA | -0.480993 | KIRC | U031 | PI3Ka_4409 | MVP_top60_DNA+RNA_seed | ATM | -0.754364 | LUAD | NCI-H1755 | KU-60019 |
| MVP_top60_DNA_seed | PIK3CD | -0.480993 | KIRC | U031 | PI3Ka_4409 | MVP_top60_DNA+RNA_seed | PTK2 | -0.753482 | LUAD | LXF-289 | PF-562271 |
| MVP_top60_DNA_seed | MTOR | -0.478844 | KIRC | RCC-ER | Temsirolimus | MVP_top60_DNA+RNA_seed | EGFR | -0.753402 | LUAD | LC-2-ad | CUDC-101 |
| MVP_top60_DNA_seed | ATM | -0.477268 | KIRC | A498 | KU-60019 | MVP_top60_DNA+RNA_seed | MAPK14 | -0.752196 | LUAD | NCI-H2085 | TAK-715 |
| MVP_top60_DNA_seed | HDAC1 | -0.475168 | KIRC | VMRC-RCZ | JQ12 | MVP_top60_DNA+RNA_seed | HSP90AA1 | -0.749954 | LUAD | NCI-H1568 | Luminespib |
| MVP_top60_DNA_seed | ATM | -0.473667 | KIRC | HA7-RCC | KU-55933 | MVP_top60_DNA+RNA_seed | MET | -0.749332 | LUAD | HCC-78 | JNJ38877605 |
| MVP_top60_DNA_seed | MTOR | -0.473038 | KIRC | CAKI-1 | Temsirolimus | MVP_top60_DNA+RNA_seed | FGFR1 | -0.746649 | LUAD | NCI-H1781 | FGFR_3831 |

| | | | | | | | | | | | |
|---|---|---|---|---|---|---|---|---|---|---|---|
| MVP_top60_DNA_seed | HSP90AA1 | -0.472837 | KIRC | 769-P | Tanespimycin | MVP_top60_DNA+RNA_seed | HDAC1 | -0.74659 | LUAD | NCI-H1568 | JQ12 |
| MVP_top60_DNA_seed | HSP90AB1 | -0.472837 | KIRC | 769-P | Tanespimycin | MVP_top60_DNA+RNA_seed | HDAC1 | -0.743949 | LUAD | HCC-44 | AR-42 |
| MVP_top60_DNA_seed | PARP1 | -0.469973 | KIRC | BFTC-909 | Rucaparib | MVP_top60_DNA+RNA_seed | BRAF | -0.742221 | LUAD | H3255 | RAF_9304 |
| MVP_top60_DNA_seed | HSP90AA1 | -0.466489 | KIRC | LB2241-RCC | CCT-018159 | MVP_top60_DNA+RNA_seed | MET | -0.74079 | LUAD | NCI-H2087 | JNJ38877605 |
| MVP_top60_DNA_seed | HSP90AB1 | -0.466489 | KIRC | LB2241-RCC | CCT-018159 | MVP_top60_DNA+RNA_seed | HDAC1 | -0.740112 | LUAD | NCI-H1568 | Dacinostat |
| MVP_top60_DNA_seed | MTOR | -0.465714 | KIRC | NCC021 | CRT0105446 | MVP_top60_DNA+RNA_seed | ATM | -0.73971 | LUAD | NCI-H1648 | CP466722 |
| MVP_top60_DNA_seed | FGFR4 | -0.461804 | KIRC | SN12C | FGFR_0939 | MVP_top60_DNA+RNA_seed | MDM2 | -0.739485 | LUAD | NCI-H1666 | Nutlin-3a (-) |
| MVP_top60_DNA_seed | PARP1 | -0.460508 | KIRC | 786-0 | PARP_0108 | MVP_top60_DNA+RNA_seed | BRAF | -0.739242 | LUAD | A549 | Dabrafenib |
| MVP_top60_DNA_seed | PARP1 | -0.46024 | KIRC | LB2241-RCC | Olaparib | MVP_top60_DNA+RNA_seed | FGFR1 | -0.737829 | LUAD | SK-LU-1 | PD173074 |
| MVP_top60_DNA_seed | PIK3CD | -0.460103 | KIRC | RCC-JF | IC-87114 | MVP_top60_DNA+RNA_seed | ATM | -0.737179 | LUAD | HCC-44 | KU-55933 |
| MVP_top60_DNA_seed | HSP90AA1 | -0.459791 | KIRC | SW156 | Elesclomol | MVP_top60_DNA+RNA_seed | MDM2 | -0.736887 | LUAD | NCI-H1563 | Nutlin-3a (-) |
| MVP_top60_DNA_seed | HSP90AB1 | -0.459791 | KIRC | SW156 | Elesclomol | MVP_top60_DNA+RNA_seed | EGFR | -0.735907 | LUAD | NCI-H1693 | Afatinib |
| MVP_top60_DNA_seed | MTOR | -0.456863 | KIRC | RCC-ER | Voxtalisib | MVP_top60_DNA+RNA_seed | MDM2 | -0.735497 | LUAD | ABC-1 | Nutlin-3a (-) |
| MVP_top60_DNA_seed | FGFR4 | -0.453204 | KIRC | LB2241-RCC | FGFR_3831 | MVP_top60_DNA+RNA_seed | HSP90AA1 | -0.732356 | LUAD | NCI-H1666 | Tanespimycin |
| MVP_top60_DNA_seed | PIK3CD | -0.452815 | KIRC | RCC-MF | Idelalisib | MVP_top60_DNA+RNA_seed | EGFR | -0.732134 | LUAD | NCI-H1838 | Afatinib |
| MVP_top60_DNA_seed | ESR1 | -0.449796 | KIRC | LB1047-RCC | Tamoxifen | MVP_top60_DNA+RNA_seed | AURKA | -0.727087 | LUAD | COR-L105 | Genentech Cpd 10 |
| MVP_top60_DNA_seed | PTK2 | -0.448709 | KIRC | ACHN | PF-562271 | MVP_top60_DNA+RNA_seed | AURKA | -0.726525 | LUAD | NCI-H522 | CD532 |
| MVP_top60_DNA_seed | HDAC1 | -0.448511 | KIRC | CAKI-1 | AR-42 | MVP_top60_DNA+RNA_seed | ATM | -0.725217 | LUAD | NCI-H2030 | CP466722 |
| MVP_top60_DNA_seed | SMARCA4 | -0.445229 | KIRC | BFTC-909 | PFI-3 | MVP_top60_DNA+RNA_seed | EGFR | -0.724683 | LUAD | NCI-H1781 | Afatinib |
| MVP_top60_DNA_seed | PIK3CA | -0.444494 | KIRC | RCC-JW | AZD8835 | MVP_top60_DNA+RNA_seed | HSP90AA1 | -0.724433 | LUAD | NCI-H1568 | CCT-018159 |
| MVP_top60_DNA_seed | PIK3CD | -0.444494 | KIRC | RCC-JW | AZD8835 | MVP_top60_DNA+RNA_seed | BRAF | -0.723066 | LUAD | NCI-H2228 | SB590885 |
| MVP_top60_DNA_seed | HDAC1 | -0.442551 | KIRC | TK10 | Dacinostat | MVP_top60_DNA+RNA_seed | EGFR | -0.722475 | LUAD | EKVX | AST-1306 |
| MVP_top60_DNA_seed | PTK2 | -0.441759 | KIRC | LB2241-RCC | PF-562271 | MVP_top60_DNA+RNA_seed | AKT1 | -0.722092 | LUAD | NCI-H2085 | AKT inhibitor VIII |
| MVP_top60_DNA_seed | HDAC1 | -0.441566 | KIRC | CAL-54 | Dacinostat | MVP_top60_DNA+RNA_seed | PIK3CA | -0.722031 | LUAD | RERF-LC-KJ | PI3Ka_4409 |
| MVP_top60_DNA_seed | IGF1R | -0.439168 | KIRC | 769-P | BMS-754807 | MVP_top60_DNA+RNA_seed | HSP90AA1 | -0.721398 | LUAD | NCI-H2023 | Tanespimycin |
| MVP_top60_DNA_seed | HDAC1 | -0.438616 | KIRC | BB65-RCC | Entinostat | MVP_top60_DNA+RNA_seed | HSP90AA1 | -0.718576 | LUAD | EMC-BAC-1 | Elesclomol |
| MVP_top60_DNA_seed | ATM | -0.43689 | KIRC | SW156 | KU-55933 | MVP_top60_DNA+RNA_seed | AURKA | -0.716165 | LUAD | Calu-6 | ZM447439 |
| MVP_top60_DNA_seed | MTOR | -0.436864 | KIRC | RCC-JF | Voxtalisib | MVP_top60_DNA+RNA_seed | ATM | -0.714051 | LUAD | NCI-H1563 | CP466722 |
| MVP_top60_DNA_seed | HDAC1 | -0.435283 | KIRC | U031 | JQ12 | MVP_top60_DNA+RNA_seed | HSP90AA1 | -0.713767 | LUAD | NCI-H522 | Luminespib |
| MVP_top60_DNA_seed | PARP1 | -0.434847 | KIRC | SN12C | Olaparib | MVP_top60_DNA+RNA_seed | BRAF | -0.7137 | LUAD | LXF-289 | PLX-4720 |
| MVP_top60_DNA_seed | PARP1 | -0.434811 | KIRC | TK10 | PARP_9495 | MVP_top60_DNA+RNA_seed | PTK2 | -0.712438 | LUAD | NCI-H1975 | PF-562271 |
| MVP_top60_DNA_seed | HSP90AA1 | -0.434772 | KIRC | A704 | Elesclomol | MVP_top60_DNA+RNA_seed | EGFR | -0.710761 | LUAD | LXF-289 | Cetuximab |
| MVP_top60_DNA_seed | HSP90AB1 | -0.434772 | KIRC | A704 | Elesclomol | MVP_top60_DNA+RNA_seed | HSP90AA1 | -0.708337 | LUAD | NCI-H1355 | Tanespimycin |
| MVP_top60_DNA_seed | HSP90AA1 | -0.433688 | KIRC | BFTC-909 | Luminespib | MVP_top60_DNA+RNA_seed | AKT1 | -0.707251 | LUAD | NCI-H2023 | MK-2206 |
| MVP_top60_DNA_seed | HSP90AB1 | -0.433688 | KIRC | BFTC-909 | Luminespib | MVP_top60_DNA+RNA_seed | SRC | -0.706545 | LUAD | NCI-H23 | Bosutinib |
| MVP_top60_DNA_seed | IGF1R | -0.433101 | KIRC | VMRC-RCW | GSK1904529A | MVP_top60_DNA+RNA_seed | BRAF | -0.701023 | LUAD | NCI-H2291 | RAF_9304 |
| MVP_top60_DNA_seed | PIK3CD | -0.429563 | KIRC | BB65-RCC | AZD8186 | MVP_top60_DNA+RNA_seed | AKT1 | -0.700709 | LUAD | HCC-78 | BAY AKT1 |
| MVP_top60_DNA_seed | PIK3CD | -0.428048 | KIRC | LB996-RCC | IC-87114 | MVP_top60_DNA+RNA_seed | ATM | -0.700102 | LUAD | PC-14 | CP466722 |
| MVP_top60_DNA_seed | PARP1 | -0.422507 | KIRC | ACHN | Rucaparib | MVP_top60_DNA+RNA_seed | AURKA | -0.698983 | LUAD | NCI-H1781 | ZM447439 |
| MVP_top60_DNA_seed | HSP90AA1 | -0.420771 | KIRC | RCC-JF | CCT-018159 | MVP_top60_DNA+RNA_seed | AURKA | -0.697135 | LUAD | NCI-H1355 | Genentech Cpd 10 |
| MVP_top60_DNA_seed | HSP90AB1 | -0.420771 | KIRC | RCC-JF | CCT-018159 | MVP_top60_DNA+RNA_seed | EGFR | -0.696327 | LUAD | ABC-1 | CI-1033 |

| | | | | | | | | | | | |
|---|---|---|---|---|---|---|---|---|---|---|---|
| MVP_top60_DNA_seed | ESR1 | -0.418613 | KIRC | RCC-ER | Tamoxifen | MVP_top60_DNA+RNA_seed | IGF1R | -0.695728 | LUAD | NCI-H1975 | BMS-536924 |
| MVP_top60_DNA_seed | ATM | -0.417627 | KIRC | RCC-ER | torin2 | MVP_top60_DNA+RNA_seed | BRAF | -0.694328 | LUAD | SK-LU-1 | PLX-4720 |
| MVP_top60_DNA_seed | MTOR | -0.417627 | KIRC | RCC-ER | torin2 | MVP_top60_DNA+RNA_seed | HDAC1 | -0.691868 | LUAD | HCC-44 | Belinostat |
| MVP_top60_DNA_seed | HSP90AA1 | -0.416513 | KIRC | LB1047-RCC | SNX-2112 | MVP_top60_DNA+RNA_seed | BRAF | -0.690777 | LUAD | NCI-H2228 | PLX-4720 |
| MVP_top60_DNA_seed | HSP90AB1 | -0.416513 | KIRC | LB1047-RCC | SNX-2112 | MVP_top60_DNA+RNA_seed | IGF1R | -0.689977 | LUAD | SW1573 | BMS-536924 |
| MVP_top60_DNA_seed | ATM | -0.415485 | KIRC | TK10 | torin2 | MVP_top60_DNA+RNA_seed | BRAF | -0.689639 | LUAD | VMRC-LCD | PLX-4720 |
| MVP_top60_DNA_seed | MTOR | -0.415485 | KIRC | TK10 | torin2 | MVP_top60_DNA+RNA_seed | HSP90AA1 | -0.688588 | LUAD | NCI-H2030 | SNX-2112 |
| MVP_top60_DNA_seed | MTOR | -0.415406 | KIRC | RCC-ER | Apitolisib | MVP_top60_DNA+RNA_seed | ATM | -0.688418 | LUAD | NCI-H2009 | KU-60019 |
| MVP_top60_DNA_seed | HDAC1 | -0.414639 | KIRC | LB996-RCC | Entinostat | MVP_top60_DNA+RNA_seed | AURKA | -0.686987 | LUAD | HCC-78 | HG-5-113-01 |
| MVP_top60_DNA_seed | MTOR | -0.413125 | KIRC | NCC021 | WYE-125132 | MVP_top60_DNA+RNA_seed | BRAF | -0.686418 | LUAD | NCI-H358 | RAF_9304 |
| MVP_top60_DNA_seed | HSP90AA1 | -0.412263 | KIRC | CAKI-1 | Elesclomol | MVP_top60_DNA+RNA_seed | MAPK14 | -0.683061 | LUAD | H3255 | TAK-715 |
| MVP_top60_DNA_seed | HSP90AB1 | -0.412263 | KIRC | CAKI-1 | Elesclomol | MVP_top60_DNA+RNA_seed | AURKA | -0.68265 | LUAD | H3255 | Genentech Cpd 10 |
| MVP_top60_DNA_seed | PARP1 | -0.411863 | KIRC | NCC010 | PARP_9482 | MVP_top60_DNA+RNA_seed | PIK3CA | -0.681276 | LUAD | NCI-H1568 | PI3Ka_4409 |
| MVP_top60_DNA_seed | PARP1 | -0.411475 | KIRC | BB65-RCC | PARP_0108 | MVP_top60_DNA+RNA_seed | HSP90AA1 | -0.681092 | LUAD | NCI-H2347 | CCT-018159 |
| MVP_top60_DNA_seed | ESR1 | -0.405903 | KIRC | A704 | Fulvestrant | MVP_top60_DNA+RNA_seed | EGFR | -0.678975 | LUAD | NCI-H2342 | CI-1033 |
| MVP_top60_DNA_seed | FGFR4 | -0.405782 | KIRC | BFTC-909 | FGFR_3831 | MVP_top60_DNA+RNA_seed | HSP90AA1 | -0.678869 | LUAD | LXF-289 | Luminespib |
| MVP_top60_DNA_seed | ATM | -0.401212 | KIRC | 786-0 | KU-60019 | MVP_top60_DNA+RNA_seed | AURKA | -0.677445 | LUAD | ABC-1 | Alisertib |
| MVP_top60_DNA_seed | MTOR | -0.401189 | KIRC | ACHN | WYE-125132 | MVP_top60_DNA+RNA_seed | HSP90AA1 | -0.676301 | LUAD | EMC-BAC-2 | SNX-2112 |
| MVP_top60_DNA_seed | PARP1 | -0.400676 | KIRC | BB65-RCC | PARP_9495 | MVP_top60_DNA+RNA_seed | AURKA | -0.676034 | LUAD | SK-LU-1 | HG-5-113-01 |
| MVP_top60_DNA_seed | ATM | -0.400257 | KIRC | 786-0 | KU-55933 | MVP_top60_DNA+RNA_seed | ATM | -0.67364 | LUAD | EMC-BAC-2 | CP466722 |
| MVP_top60_DNA_seed | PIK3CD | -0.399945 | KIRC | ACHN | Idelalisib | MVP_top60_DNA+RNA_seed | MET | -0.673055 | LUAD | NCI-H2405 | JNJ38877605 |
| MVP_top60_DNA_seed | MTOR | -0.399932 | KIRC | KMRC-20 | Temsirolimus | MVP_top60_DNA+RNA_seed | HSP90AA1 | -0.672728 | LUAD | RERF-LC-KJ | CCT-018159 |
| MVP_top60_DNA_seed | PARP1 | -0.393944 | KIRC | RCC-AB | PARP_0108 | MVP_top60_DNA+RNA_seed | SRC | -0.672019 | LUAD | EMC-BAC-2 | Bosutinib |
| MVP_top60_DNA_seed | PARP1 | -0.393388 | KIRC | BFTC-909 | PARP_9495 | MVP_top60_DNA+RNA_seed | IGF1R | -0.668918 | LUAD | NCI-H1648 | Linsitinib |
| MVP_top60_DNA_seed | IGF1R | -0.388749 | KIRC | LB2241-RCC | GSK1904529A | MVP_top60_DNA+RNA_seed | HSP90AA1 | -0.668779 | LUAD | COR-L105 | SNX-2112 |
| MVP_top60_DNA_seed | MTOR | -0.387022 | KIRC | RCC-JW | Apitolisib | MVP_top60_DNA+RNA_seed | MET | -0.668265 | LUAD | NCI-H441 | AZD6094 |
| MVP_top60_DNA_seed | HSP90AA1 | -0.386267 | KIRC | BB65-RCC | Elesclomol | MVP_top60_DNA+RNA_seed | HDAC1 | -0.667757 | LUAD | HOP-62 | CAY10603 |
| MVP_top60_DNA_seed | HSP90AB1 | -0.386267 | KIRC | BB65-RCC | Elesclomol | MVP_top60_DNA+RNA_seed | ATM | -0.667739 | LUAD | NCI-H358 | KU-55933 |
| MVP_top60_DNA_seed | HDAC1 | -0.38576 | KIRC | LB1047-RCC | Entinostat | MVP_top60_DNA+RNA_seed | AKT1 | -0.666628 | LUAD | EMC-BAC-1 | AKT inhibitor VIII |
| MVP_top60_DNA_seed | MTOR | -0.385203 | KIRC | RCC-JF | CRT0105446 | MVP_top60_DNA+RNA_seed | EGFR | -0.666492 | LUAD | NCI-H1792 | AZD8931 |
| MVP_top60_DNA_seed | SMARCA4 | -0.385199 | KIRC | LB1047-RCC | PFI-3 | MVP_top60_DNA+RNA_seed | MDM2 | -0.666324 | LUAD | NCI-H1944 | Nutlin-3a (-) |
| MVP_top60_DNA_seed | PIK3CD | -0.383928 | KIRC | RCC10RGB | AZD8186 | MVP_top60_DNA+RNA_seed | EGFR | -0.664554 | LUAD | NCI-H2030 | Afatinib |
| MVP_top60_DNA_seed | PIK3CA | -0.383181 | KIRC | CAL-54 | AZD8835 | MVP_top60_DNA+RNA_seed | IGF1R | -0.664243 | LUAD | NCI-H1734 | BMS-754807 |
| MVP_top60_DNA_seed | PIK3CD | -0.383181 | KIRC | CAL-54 | AZD8835 | MVP_top60_DNA+RNA_seed | HDAC1 | -0.66281 | LUAD | Calu-6 | Parthenolide |
| MVP_top60_DNA_seed | PIK3CA | -0.382826 | KIRC | NCC010 | PI3Ka_4409 | MVP_top60_DNA+RNA_seed | HDAC1 | -0.662196 | LUAD | LXF-289 | Entinostat |
| MVP_top60_DNA_seed | PIK3CD | -0.382826 | KIRC | NCC010 | PI3Ka_4409 | MVP_top60_DNA+RNA_seed | PTK2 | -0.662071 | LUAD | Calu-6 | PF-562271 |
| MVP_top60_DNA_seed | HDAC1 | -0.378905 | KIRC | LB2241-RCC | Entinostat | MVP_top60_DNA+RNA_seed | HSP90AA1 | -0.660037 | LUAD | LXF-289 | Elesclomol |
| MVP_top60_DNA_seed | HSP90AA1 | -0.375789 | KIRC | 769-P | SNX-2112 | MVP_top60_DNA+RNA_seed | EGFR | -0.659382 | LUAD | PC-14 | AST-1306 |
| MVP_top60_DNA_seed | HSP90AB1 | -0.375789 | KIRC | 769-P | SNX-2112 | MVP_top60_DNA+RNA_seed | EGFR | -0.657954 | LUAD | NCI-H2085 | Gefitinib |
| MVP_top60_DNA_seed | ATM | -0.375658 | KIRC | RCC-FG2 | torin2 | MVP_top60_DNA+RNA_seed | BRAF | -0.657135 | LUAD | SK-LU-1 | PLX-4720 |
| MVP_top60_DNA_seed | MTOR | -0.375658 | KIRC | RCC-FG2 | torin2 | MVP_top60_DNA+RNA_seed | EGFR | -0.656322 | LUAD | NCI-H522 | Afatinib |

| | | | | | | | | | | | |
|---|---|---|---|---|---|---|---|---|---|---|---|
| MVP_top60_DNA_seed | FGFR4 | -0.374645 | KIRC | HA7-RCC | FGFR_3831 | MVP_top60_DNA+RNA_seed | PIK3CA | -0.6562 | LUAD | EMC-BAC-2 | PI3Ka_4409 |
| MVP_top60_DNA_seed | MTOR | -0.3736 | KIRC | LB2241-RCC | Apitolisib | MVP_top60_DNA+RNA_seed | HSP90AA1 | -0.655859 | LUAD | NCI-H2347 | Tanespimycin |
| MVP_top60_DNA_seed | PIK3CA | -0.37179 | KIRC | NCC010 | AZD8835 | MVP_top60_DNA+RNA_seed | BRAF | -0.655503 | LUAD | PC-14 | RAF_9304 |
| MVP_top60_DNA_seed | PIK3CD | -0.37179 | KIRC | NCC010 | AZD8835 | MVP_top60_DNA+RNA_seed | MDM2 | -0.654886 | LUAD | NCI-H23 | Serdemetan |
| MVP_top60_DNA_seed | HSP90AA1 | -0.370234 | KIRC | RCC-AB | SNX-2112 | MVP_top60_DNA+RNA_seed | HDAC1 | -0.653899 | LUAD | NCI-H358 | Belinostat |
| MVP_top60_DNA_seed | HSP90AB1 | -0.370234 | KIRC | RCC-AB | SNX-2112 | MVP_top60_DNA+RNA_seed | IGF1R | -0.653337 | LUAD | NCI-H1651 | Linsitinib |
| MVP_top60_DNA_seed | MTOR | -0.368045 | KIRC | TK10 | Apitolisib | MVP_top60_DNA+RNA_seed | IGF1R | -0.65133 | LUAD | NCI-H522 | Linsitinib |
| MVP_top60_DNA_seed | HSP90AA1 | -0.368018 | KIRC | OS-RC-2 | Tanespimycin | MVP_top60_DNA+RNA_seed | ATM | -0.651048 | LUAD | LC-2-ad | KU-55933 |
| MVP_top60_DNA_seed | HSP90AB1 | -0.368018 | KIRC | OS-RC-2 | Tanespimycin | MVP_top60_DNA+RNA_seed | AURKA | -0.649323 | LUAD | NCI-H23 | Genentech Cpd 10 |
| MVP_top60_DNA_seed | AR | -0.364709 | KIRC | ACHN | AZD3514 | MVP_top60_DNA+RNA_seed | MET | -0.648208 | LUAD | NCI-H23 | JNJ38877605 |
| MVP_top60_DNA_seed | HDAC1 | -0.362238 | KIRC | TK10 | JQ12 | MVP_top60_DNA+RNA_seed | AKT1 | -0.6474 | LUAD | NCI-H1648 | AKT inhibitor VIII |
| MVP_top60_DNA_seed | PIK3CA | -0.356994 | KIRC | TK10 | AZD8835 | MVP_top60_DNA+RNA_seed | PIK3CA | -0.646999 | LUAD | NCI-H1755 | AZD8835 |
| MVP_top60_DNA_seed | PIK3CD | -0.356994 | KIRC | TK10 | AZD8835 | MVP_top60_DNA+RNA_seed | EGFR | -0.646658 | LUAD | LXF-289 | Erlotinib |
| MVP_top60_DNA_seed | PARP1 | -0.356894 | KIRC | RCC-MF | PARP_9495 | MVP_top60_DNA+RNA_seed | HDAC1 | -0.646524 | LUAD | NCI-H1944 | CAY10603 |
| MVP_top60_DNA_seed | PARP1 | -0.356832 | KIRC | LB996-RCC | Olaparib | MVP_top60_DNA+RNA_seed | BRAF | -0.645122 | LUAD | NCI-H1792 | PLX-4720 |
| MVP_top60_DNA_seed | FGFR4 | -0.3568 | KIRC | LB1047-RCC | FGFR_3831 | MVP_top60_DNA+RNA_seed | BRAF | -0.644732 | LUAD | LXF-289 | SB590885 |
| MVP_top60_DNA_seed | HSP90AA1 | -0.355801 | KIRC | BB65-RCC | SNX-2112 | MVP_top60_DNA+RNA_seed | SRC | -0.643993 | LUAD | NCI-H1792 | Bosutinib |
| MVP_top60_DNA_seed | HSP90AB1 | -0.355801 | KIRC | BB65-RCC | SNX-2112 | MVP_top60_DNA+RNA_seed | SRC | -0.642236 | LUAD | NCI-H1355 | WH-4-023 |
| MVP_top60_DNA_seed | PIK3CD | -0.353447 | KIRC | VMRC-RCZ | IC-87114 | MVP_top60_DNA+RNA_seed | MET | -0.641834 | LUAD | PC-14 | SU11274 |
| MVP_top60_DNA_seed | ATM | -0.353272 | KIRC | RXF393 | CP466722 | MVP_top60_DNA+RNA_seed | BRAF | -0.641561 | LUAD | NCI-H1650 | PLX-4720 |
| MVP_top60_DNA_seed | PIK3CD | -0.352659 | KIRC | RCC-ER | Idelalisib | MVP_top60_DNA+RNA_seed | SRC | -0.641479 | LUAD | HCC-44 | NVP-BHG712 |
| MVP_top60_DNA_seed | MTOR | -0.351612 | KIRC | RCC-ER | Torin 2 | MVP_top60_DNA+RNA_seed | ATM | -0.639384 | LUAD | NCI-H1793 | KU-60019 |
| MVP_top60_DNA_seed | HDAC1 | -0.348724 | KIRC | ACHN | JQ12 | MVP_top60_DNA+RNA_seed | HDAC1 | -0.638633 | LUAD | NCI-H358 | CAY10603 |
| MVP_top60_DNA_seed | ATM | -0.348576 | KIRC | KMRC-20 | KU-55933 | MVP_top60_DNA+RNA_seed | AURKA | -0.638101 | LUAD | NCI-H2085 | ZM447439 |
| MVP_top60_DNA_seed | HDAC1 | -0.346426 | KIRC | CAKI-1 | Belinostat | MVP_top60_DNA+RNA_seed | MET | -0.634907 | LUAD | NCI-H1792 | JNJ38877605 |
| MVP_top60_DNA_seed | HDAC1 | -0.345943 | KIRC | 786-0 | Belinostat | MVP_top60_DNA+RNA_seed | PIK3CA | -0.633208 | LUAD | NCI-H2342 | PI3Ka_4409 |
| MVP_top60_DNA_seed | MTOR | -0.344726 | KIRC | RCC-JF | QL-VIII-58 | MVP_top60_DNA+RNA_seed | AURKA | -0.632865 | LUAD | NCI-H358 | ZM447439 |
| MVP_top60_DNA_seed | IGF1R | -0.344405 | KIRC | CAL-54 | BMS-536924 | MVP_top60_DNA+RNA_seed | HSP90AA1 | -0.632264 | LUAD | NCI-H1355 | Luminespib |
| MVP_top60_DNA_seed | IGF1R | -0.344079 | KIRC | 786-0 | BMS-754807 | MVP_top60_DNA+RNA_seed | EGFR | -0.632175 | LUAD | NCI-H358 | PF-00299804 |
| MVP_top60_DNA_seed | PARP1 | -0.343566 | KIRC | NCC021 | PARP_9495 | MVP_top60_DNA+RNA_seed | BRAF | -0.631858 | LUAD | NCI-H1755 | HG6-64-1 |
| MVP_top60_DNA_seed | HSP90AA1 | -0.342318 | KIRC | LB2241-RCC | Luminespib | MVP_top60_DNA+RNA_seed | AURKA | -0.630396 | LUAD | LC-2-ad | Genentech Cpd 10 |
| MVP_top60_DNA_seed | HSP90AB1 | -0.342318 | KIRC | LB2241-RCC | Luminespib | MVP_top60_DNA+RNA_seed | HDAC1 | -0.625641 | LUAD | HCC-78 | Belinostat |
| MVP_top60_DNA_seed | PIK3CA | -0.339491 | KIRC | RCC-FG2 | PI3Ka_4409 | MVP_top60_DNA+RNA_seed | HDAC1 | -0.62523 | LUAD | NCI-H1648 | Belinostat |
| MVP_top60_DNA_seed | PIK3CD | -0.339491 | KIRC | RCC-FG2 | PI3Ka_4409 | MVP_top60_DNA+RNA_seed | HSP90AA1 | -0.623574 | LUAD | H3255 | Tanespimycin |
| MVP_top60_DNA_seed | HSP90AA1 | -0.338799 | KIRC | LB996-RCC | SNX-2112 | MVP_top60_DNA+RNA_seed | IGF1R | -0.623259 | LUAD | NCI-H1651 | BMS-754807 |
| MVP_top60_DNA_seed | HSP90AB1 | -0.338799 | KIRC | LB996-RCC | SNX-2112 | MVP_top60_DNA+RNA_seed | EGFR | -0.622534 | LUAD | NCI-H1563 | Pelitinib |
| MVP_top60_DNA_seed | MTOR | -0.338631 | KIRC | LB2241-RCC | Torin 2 | MVP_top60_DNA+RNA_seed | MDM2 | -0.620165 | LUAD | NCI-H1650 | Serdemetan |
| MVP_top60_DNA_seed | PIK3CA | -0.33443 | KIRC | LB996-RCC | PI3Ka_4409 | MVP_top60_DNA+RNA_seed | SRC | -0.618727 | LUAD | NCI-H358 | NVP-BHG712 |
| MVP_top60_DNA_seed | PIK3CD | -0.33443 | KIRC | LB996-RCC | PI3Ka_4409 | MVP_top60_DNA+RNA_seed | EGFR | -0.618378 | LUAD | NCI-H358 | Afatinib |
| MVP_top60_DNA_seed | IGF1R | -0.332812 | KIRC | CAL-54 | Linsitinib | MVP_top60_DNA+RNA_seed | HDAC1 | -0.618053 | LUAD | Calu-6 | Dacinostat |
| MVP_top60_DNA_seed | ATM | -0.332384 | KIRC | RCC-JF | KU-60019 | MVP_top60_DNA+RNA_seed | SMARCA4 | -0.617358 | LUAD | NCI-H1944 | PFI-3 |

| | | | | | | | | | | | |
|---|---|---|---|---|---|---|---|---|---|---|---|
| MVP_top60_DNA_seed | PIK3CD | -0.331329 | KIRC | CAL-54 | Idelalisib | MVP_top60_DNA+RNA_seed | EGFR | -0.617067 | LUAD | NCI-H1944 | Pelitinib |
| MVP_top60_DNA_seed | HDAC1 | -0.32918 | KIRC | BFTC-909 | Dacinostat | MVP_top60_DNA+RNA_seed | BRAF | -0.61523 | LUAD | SW1573 | HG6-64-1 |
| MVP_top60_DNA_seed | IGF1R | -0.32822 | KIRC | HA7-RCC | BMS-754807 | MVP_top60_DNA+RNA_seed | HDAC1 | -0.615097 | LUAD | NCI-H1693 | AR-42 |
| MVP_top60_DNA_seed | MTOR | -0.327969 | KIRC | RCC-MF | Voxtalisib | MVP_top60_DNA+RNA_seed | BRAF | -0.61272 | LUAD | NCI-H2030 | PLX-4720 |
| MVP_top60_DNA_seed | MTOR | -0.327316 | KIRC | LB2241-RCC | WYE-125132 | MVP_top60_DNA+RNA_seed | PIK3CA | -0.612568 | LUAD | PC-14 | AZD8835 |
| MVP_top60_DNA_seed | PARP1 | -0.325859 | KIRC | BFTC-909 | Veliparib | MVP_top60_DNA+RNA_seed | PIK3CA | -0.611118 | LUAD | NCI-H1792 | AZD8835 |
| MVP_top60_DNA_seed | ESR1 | -0.32296 | KIRC | VMRC-RCZ | Fulvestrant | MVP_top60_DNA+RNA_seed | MET | -0.610997 | LUAD | RERF-LC-MS | SU11274 |
| MVP_top60_DNA_seed | ATM | -0.322313 | KIRC | CAL-54 | KU-60019 | MVP_top60_DNA+RNA_seed | HSP90AA1 | -0.609813 | LUAD | NCI-H1648 | Tanespimycin |
| MVP_top60_DNA_seed | EP300 | -0.320314 | KIRC | BB65-RCC | I-CBP112 | MVP_top60_DNA+RNA_seed | HSP90AA1 | -0.608125 | LUAD | NCI-H650 | Tanespimycin |
| MVP_top60_DNA_seed | PIK3CA | -0.318491 | KIRC | SN12C | AZD8835 | MVP_top60_DNA+RNA_seed | BRAF | -0.60789 | LUAD | NCI-H1975 | PLX-4720 |
| MVP_top60_DNA_seed | PIK3CD | -0.318491 | KIRC | SN12C | AZD8835 | MVP_top60_DNA+RNA_seed | MAPK14 | -0.607833 | LUAD | NCI-H1437 | TAK-715 |
| MVP_top60_DNA_seed | AR | -0.317659 | KIRC | A704 | Bicalutamide | MVP_top60_DNA+RNA_seed | EGFR | -0.607518 | LUAD | LC-2-ad | Pelitinib |
| MVP_top60_DNA_seed | ESR1 | -0.317198 | KIRC | RXF393 | Tamoxifen | MVP_top60_DNA+RNA_seed | IGF1R | -0.605174 | LUAD | NCI-H522 | GSK1904529A |
| MVP_top60_DNA_seed | PIK3CA | -0.316568 | KIRC | RCC-JF | AZD8835 | MVP_top60_DNA+RNA_seed | HSP90AA1 | -0.60477 | LUAD | LC-2-ad | Tanespimycin |
| MVP_top60_DNA_seed | PIK3CD | -0.316568 | KIRC | RCC-JF | AZD8835 | MVP_top60_DNA+RNA_seed | HDAC1 | -0.604037 | LUAD | PC-14 | Tubastatin A |
| MVP_top60_DNA_seed | PARP1 | -0.313652 | KIRC | ACHN | PARP_9482 | MVP_top60_DNA+RNA_seed | IGF1R | -0.603982 | LUAD | NCI-H2228 | Linsitinib |
| MVP_top60_DNA_seed | MTOR | -0.312126 | KIRC | VMRC-RCZ | Apitolisib | MVP_top60_DNA+RNA_seed | HDAC1 | -0.603575 | LUAD | HCC-78 | Dacinostat |
| MVP_top60_DNA_seed | MTOR | -0.311661 | KIRC | RCC-FG2 | Torin 2 | MVP_top60_DNA+RNA_seed | HDAC1 | -0.602551 | LUAD | Calu-6 | Belinostat |
| MVP_top60_DNA_seed | EP300 | -0.31123 | KIRC | LB1047-RCC | I-CBP112 | MVP_top60_DNA+RNA_seed | IGF1R | -0.599947 | LUAD | NCI-H1623 | Linsitinib |
| MVP_top60_DNA_seed | HDAC1 | -0.305504 | KIRC | 786-0 | AR-42 | MVP_top60_DNA+RNA_seed | BRAF | -0.599793 | LUAD | NCI-H1666 | RAF_9304 |
| MVP_top60_DNA_seed | FGFR4 | -0.302501 | KIRC | NCC010 | FGFR_0939 | MVP_top60_DNA+RNA_seed | EGFR | -0.598844 | LUAD | ABC-1 | CUDC-101 |
| MVP_top60_DNA_seed | HDAC1 | -0.302237 | KIRC | BB65-RCC | CAY10603 | MVP_top60_DNA+RNA_seed | MDM2 | -0.598743 | LUAD | NCI-H1734 | Serdemetan |
| MVP_top60_DNA_seed | SMARCA4 | -0.300719 | KIRC | LB2241-RCC | PFI-3 | MVP_top60_DNA+RNA_seed | EGFR | -0.597975 | LUAD | NCI-H522 | CUDC-101 |
| MVP_top60_DNA_seed | ESR1 | -0.300407 | KIRC | NCC010 | Tamoxifen | MVP_top60_DNA+RNA_seed | HDAC1 | -0.597717 | LUAD | NCI-H1944 | AR-42 |
| MVP_top60_DNA_seed | FGFR4 | -0.296539 | KIRC | LB2241-RCC | FGFR_0939 | MVP_top60_DNA+RNA_seed | HSP90AA1 | -0.597047 | LUAD | NCI-H2405 | CCT-018159 |
| MVP_top60_DNA_seed | EP300 | -0.296421 | KIRC | RCC-MF | I-CBP112 | MVP_top60_DNA+RNA_seed | ATM | -0.595568 | LUAD | NCI-H2122 | CP466722 |
| MVP_top60_DNA_seed | ATM | -0.295025 | KIRC | BB65-RCC | CP466722 | MVP_top60_DNA+RNA_seed | AURKA | -0.595024 | LUAD | EMC-BAC-1 | GSK1070916 |
| MVP_top60_DNA_seed | HSP90AA1 | -0.294717 | KIRC | NCC010 | Elesclomol | MVP_top60_DNA+RNA_seed | SRC | -0.594104 | LUAD | EMC-BAC-1 | NVP-BHG712 |
| MVP_top60_DNA_seed | HSP90AB1 | -0.294717 | KIRC | NCC010 | Elesclomol | MVP_top60_DNA+RNA_seed | HSP90AA1 | -0.592571 | LUAD | NCI-H1944 | Luminespib |
| MVP_top60_DNA_seed | MTOR | -0.292081 | KIRC | ACHN | Voxtalisib | MVP_top60_DNA+RNA_seed | HDAC1 | -0.590629 | LUAD | NCI-H1355 | JQ12 |
| MVP_top60_DNA_seed | PTK2 | -0.290598 | KIRC | SN12C | PF-562271 | MVP_top60_DNA+RNA_seed | AKT1 | -0.590362 | LUAD | NCI-H2291 | BAY AKT1 |
| MVP_top60_DNA_seed | PIK3CD | -0.28864 | KIRC | 786-0 | AZD8186 | MVP_top60_DNA+RNA_seed | HDAC1 | -0.589611 | LUAD | PC-14 | Belinostat |
| MVP_top60_DNA_seed | IGF1R | -0.283457 | KIRC | NCC010 | BMS-536924 | MVP_top60_DNA+RNA_seed | AKT1 | -0.589545 | LUAD | NCI-H2023 | BAY AKT1 |
| MVP_top60_DNA_seed | IGF1R | -0.282708 | KIRC | CAKI-1 | Linsitinib | MVP_top60_DNA+RNA_seed | EGFR | -0.588143 | LUAD | NCI-H1648 | CUDC-101 |
| MVP_top60_DNA_seed | PARP1 | -0.281856 | KIRC | LB996-RCC | PARP_0108 | MVP_top60_DNA+RNA_seed | IGF1R | -0.587678 | LUAD | NCI-H522 | BMS-754807 |
| MVP_top60_DNA_seed | IGF1R | -0.281105 | KIRC | 786-0 | Linsitinib | MVP_top60_DNA+RNA_seed | MDM2 | -0.587523 | LUAD | NCI-H2347 | Serdemetan |
| MVP_top60_DNA_seed | ATM | -0.280281 | KIRC | KMRC-20 | KU-60019 | MVP_top60_DNA+RNA_seed | BRAF | -0.586351 | LUAD | A549 | RAF_9304 |
| MVP_top60_DNA_seed | HSP90AA1 | -0.27881 | KIRC | SN12C | Elesclomol | MVP_top60_DNA+RNA_seed | SRC | -0.585827 | LUAD | RERF-LC-KJ | Bosutinib |
| MVP_top60_DNA_seed | HSP90AB1 | -0.27881 | KIRC | SN12C | Elesclomol | MVP_top60_DNA+RNA_seed | AURKA | -0.584803 | LUAD | PC-3_[JPC-3] | Genentech Cpd 10 |
| MVP_top60_DNA_seed | IGF1R | -0.27567 | KIRC | HA7-RCC | BMS-536924 | MVP_top60_DNA+RNA_seed | AURKA | -0.58403 | LUAD | NCI-H358 | GSK1070916 |
| MVP_top60_DNA_seed | MTOR | -0.269349 | KIRC | 786-0 | Torin 2 | MVP_top60_DNA+RNA_seed | HDAC1 | -0.583787 | LUAD | NCI-H1648 | AR-42 |

| | | | | | | | | | | | |
|---|---|---|---|---|---|---|---|---|---|---|---|
| MVP_top60_DNA_seed | ESR1 | -0.268063 | KIRC | NCC010 | Fulvestrant | MVP_top60_DNA+RNA_seed | PTK2 | -0.583637 | LUAD | NCI-H838 | PF-562271 |
| MVP_top60_DNA_seed | PARP1 | -0.265606 | KIRC | RCC-AB | Olaparib | MVP_top60_DNA+RNA_seed | HSP90AA1 | -0.582992 | LUAD | NCI-H2009 | Tanespimycin |
| MVP_top60_DNA_seed | PARP1 | -0.26254 | KIRC | 769-P | PARP_0108 | MVP_top60_DNA+RNA_seed | MDM2 | -0.582278 | LUAD | NCI-H2085 | Nutlin-3a (-) |
| MVP_top60_DNA_seed | FGFR4 | -0.262488 | KIRC | 769-P | FGFR_3831 | MVP_top60_DNA+RNA_seed | HSP90AA1 | -0.582159 | LUAD | NCI-H2085 | Luminespib |
| MVP_top60_DNA_seed | PIK3CA | -0.26233 | KIRC | RCC-ER | AZD8835 | MVP_top60_DNA+RNA_seed | SRC | -0.581158 | LUAD | NCI-H1650 | Bosutinib |
| MVP_top60_DNA_seed | PIK3CD | -0.26233 | KIRC | RCC-ER | AZD8835 | MVP_top60_DNA+RNA_seed | AKT1 | -0.580852 | LUAD | NCI-H2085 | AKT inhibitor VIII |
| MVP_top60_DNA_seed | HSP90AA1 | -0.262043 | KIRC | CAL-54 | CCT-018159 | MVP_top60_DNA+RNA_seed | HDAC1 | -0.579974 | LUAD | PC-14 | AR-42 |
| MVP_top60_DNA_seed | HSP90AB1 | -0.262043 | KIRC | CAL-54 | CCT-018159 | MVP_top60_DNA+RNA_seed | HSP90AA1 | -0.578678 | LUAD | NCI-H650 | CCT-018159 |
| MVP_top60_DNA_seed | FGFR4 | -0.261901 | KIRC | KMRC-20 | FGFR_0939 | MVP_top60_DNA+RNA_seed | EGFR | -0.578358 | LUAD | NCI-H1355 | AZD8931 |
| MVP_top60_DNA_seed | IGF1R | -0.261281 | KIRC | LB2241-RCC | Linsitinib | MVP_top60_DNA+RNA_seed | ATM | -0.578029 | LUAD | NCI-H1435 | KU-60019 |
| MVP_top60_DNA_seed | PIK3CA | -0.260492 | KIRC | HA7-RCC | AZD8835 | MVP_top60_DNA+RNA_seed | EGFR | -0.577925 | LUAD | NCI-H1666 | AST-1306 |
| MVP_top60_DNA_seed | PIK3CD | -0.260492 | KIRC | HA7-RCC | AZD8835 | MVP_top60_DNA+RNA_seed | BRAF | -0.577473 | LUAD | NCI-H1648 | RAF_9304 |
| MVP_top60_DNA_seed | PARP1 | -0.259802 | KIRC | RCC-AB | Talazoparib | MVP_top60_DNA+RNA_seed | EGFR | -0.575972 | LUAD | NCI-H2342 | AST-1306 |
| MVP_top60_DNA_seed | PARP1 | -0.259323 | KIRC | RXF393 | PARP_9482 | MVP_top60_DNA+RNA_seed | SRC | -0.575802 | LUAD | NCI-H2009 | Bosutinib |
| MVP_top60_DNA_seed | FGFR4 | -0.258943 | KIRC | RCC-JW | FGFR_3831 | MVP_top60_DNA+RNA_seed | AURKA | -0.574982 | LUAD | NCI-H2009 | ZM447439 |
| MVP_top60_DNA_seed | HDAC1 | -0.256534 | KIRC | 786-0 | Tubastatin A | MVP_top60_DNA+RNA_seed | MDM2 | -0.574738 | LUAD | PC-14 | Serdemetan |
| MVP_top60_DNA_seed | HDAC1 | -0.255494 | KIRC | RXF393 | AR-42 | MVP_top60_DNA+RNA_seed | BRAF | -0.570587 | LUAD | NCI-H1755 | PLX-4720 |
| MVP_top60_DNA_seed | HSP90AA1 | -0.255008 | KIRC | BB65-RCC | CCT-018159 | MVP_top60_DNA+RNA_seed | MET | -0.569336 | LUAD | LC-2-ad | PHA-665752 |
| MVP_top60_DNA_seed | HSP90AB1 | -0.255008 | KIRC | BB65-RCC | CCT-018159 | MVP_top60_DNA+RNA_seed | PTK2 | -0.568282 | LUAD | NCI-H1650 | PF-562271 |
| MVP_top60_DNA_seed | MTOR | -0.253649 | KIRC | LB1047-RCC | Apitolisib | MVP_top60_DNA+RNA_seed | EGFR | -0.566939 | LUAD | NCI-H358 | AST-1306 |
| MVP_top60_DNA_seed | PARP1 | -0.253603 | KIRC | RCC-JW | Olaparib | MVP_top60_DNA+RNA_seed | HSP90AA1 | -0.565504 | LUAD | PC-14 | Luminespib |
| MVP_top60_DNA_seed | FGFR4 | -0.248438 | KIRC | VMRC-RCZ | FGFR_0939 | MVP_top60_DNA+RNA_seed | ATM | -0.565224 | LUAD | NCI-H1563 | KU-60019 |
| MVP_top60_DNA_seed | IGF1R | -0.246271 | KIRC | BB65-RCC | BMS-754807 | MVP_top60_DNA+RNA_seed | AURKA | -0.564473 | LUAD | NCI-H650 | Genentech Cpd 10 |
| MVP_top60_DNA_seed | PARP1 | -0.244866 | KIRC | NCC010 | PARP_0108 | MVP_top60_DNA+RNA_seed | SMARCA4 | -0.564417 | LUAD | A549 | PFI-3 |
| MVP_top60_DNA_seed | PIK3CA | -0.243047 | KIRC | RCC-MF | PI3Ka_4409 | MVP_top60_DNA+RNA_seed | MDM2 | -0.564314 | LUAD | HOP-62 | Serdemetan |
| MVP_top60_DNA_seed | PIK3CD | -0.243047 | KIRC | RCC-MF | PI3Ka_4409 | MVP_top60_DNA+RNA_seed | ATM | -0.563834 | LUAD | NCI-H1435 | KU-55933 |
| MVP_top60_DNA_seed | FGFR4 | -0.241968 | KIRC | RCC-MF | FGFR_0939 | MVP_top60_DNA+RNA_seed | AURKA | -0.56132 | LUAD | NCI-H1755 | HG-5-113-01 |
| MVP_top60_DNA_seed | MTOR | -0.241817 | KIRC | 786-0 | Apitolisib | MVP_top60_DNA+RNA_seed | EGFR | -0.56006 | LUAD | A549 | AZD8931 |
| MVP_top60_DNA_seed | PIK3CD | -0.239066 | KIRC | KMRC-1 | Idelalisib | MVP_top60_DNA+RNA_seed | IGF1R | -0.559957 | LUAD | NCI-H1734 | Linsitinib |
| MVP_top60_DNA_seed | MTOR | -0.238001 | KIRC | LB1047-RCC | Voxtalisib | MVP_top60_DNA+RNA_seed | BRAF | -0.559891 | LUAD | HCC-44 | HG6-64-1 |
| MVP_top60_DNA_seed | PARP1 | -0.237976 | KIRC | SN12C | Talazoparib | MVP_top60_DNA+RNA_seed | IGF1R | -0.559808 | LUAD | EMC-BAC-2 | BMS-536924 |
| MVP_top60_DNA_seed | MTOR | -0.237854 | KIRC | U031 | Apitolisib | MVP_top60_DNA+RNA_seed | PIK3CA | -0.557845 | LUAD | NCI-H838 | AZD8835 |
| MVP_top60_DNA_seed | MTOR | -0.237017 | KIRC | LB2241-RCC | QL-VIII-58 | MVP_top60_DNA+RNA_seed | EGFR | -0.557688 | LUAD | LXF-289 | Lapatinib |
| MVP_top60_DNA_seed | ATM | -0.233803 | KIRC | OS-RC-2 | CP466722 | MVP_top60_DNA+RNA_seed | ATM | -0.557659 | LUAD | NCI-H1568 | KU-60019 |
| MVP_top60_DNA_seed | HDAC1 | -0.232907 | KIRC | VMRC-RCZ | Belinostat | MVP_top60_DNA+RNA_seed | AKT1 | -0.556928 | LUAD | Calu-3 | BAY AKT1 |
| MVP_top60_DNA_seed | MTOR | -0.230743 | KIRC | RCC-JW | Torin 2 | MVP_top60_DNA+RNA_seed | AURKA | -0.556336 | LUAD | NCI-H2087 | GSK1070916 |
| MVP_top60_DNA_seed | MTOR | -0.229284 | KIRC | U031 | Voxtalisib | MVP_top60_DNA+RNA_seed | HSP90AA1 | -0.555759 | LUAD | NCI-H2009 | Elesclomol |
| MVP_top60_DNA_seed | ATM | -0.226613 | KIRC | LB996-RCC | KU-60019 | MVP_top60_DNA+RNA_seed | PIK3CA | -0.55444 | LUAD | NCI-H2122 | AZD8835 |
| MVP_top60_DNA_seed | MTOR | -0.226178 | KIRC | RCC-MF | Apitolisib | MVP_top60_DNA+RNA_seed | IGF1R | -0.551359 | LUAD | NCI-H2342 | BMS-754807 |
| MVP_top60_DNA_seed | HDAC1 | -0.226001 | KIRC | BB65-RCC | Belinostat | MVP_top60_DNA+RNA_seed | HSP90AA1 | -0.550425 | LUAD | NCI-H1650 | Tanespimycin |
| MVP_top60_DNA_seed | ESR1 | -0.223061 | KIRC | HA7-RCC | Fulvestrant | MVP_top60_DNA+RNA_seed | BRAF | -0.549916 | LUAD | NCI-H441 | PLX-4720 |

| | | | | | | | | | | | |
|---|---|---|---|---|---|---|---|---|---|---|---|
| MVP_top60_DNA_seed | SMARCA4 | -0.2226 | KIRC | 786-0 | PFI-3 | MVP_top60_DNA+RNA_seed | EGFR | -0.548656 | LUAD | RERF-LC-KJ | AZD8931 |
| MVP_top60_DNA_seed | HDAC1 | -0.221206 | KIRC | CAL-54 | AR-42 | MVP_top60_DNA+RNA_seed | EGFR | -0.548372 | LUAD | NCI-H2085 | Afatinib |
| MVP_top60_DNA_seed | HSP90AA1 | -0.220771 | KIRC | RCC-JW | CCT-018159 | MVP_top60_DNA+RNA_seed | MAPK14 | -0.547337 | LUAD | ABC-1 | TAK-715 |
| MVP_top60_DNA_seed | HSP90AB1 | -0.220771 | KIRC | RCC-JW | CCT-018159 | MVP_top60_DNA+RNA_seed | SRC | -0.547096 | LUAD | NCI-H322M | Bosutinib |
| MVP_top60_DNA_seed | IGF1R | -0.220006 | KIRC | CAKI-1 | BMS-754807 | MVP_top60_DNA+RNA_seed | FGFR1 | -0.546775 | LUAD | A549 | FGFR_3831 |
| MVP_top60_DNA_seed | FGFR4 | -0.219742 | KIRC | RXF393 | FGFR_0939 | MVP_top60_DNA+RNA_seed | ATM | -0.544287 | LUAD | NCI-H1355 | CP466722 |
| MVP_top60_DNA_seed | HSP90AA1 | -0.219293 | KIRC | OS-RC-2 | Luminespib | MVP_top60_DNA+RNA_seed | MDM2 | -0.543896 | LUAD | NCI-H2023 | Serdemetan |
| MVP_top60_DNA_seed | HSP90AB1 | -0.219293 | KIRC | OS-RC-2 | Luminespib | MVP_top60_DNA+RNA_seed | ATM | -0.543291 | LUAD | PC-3_[JPC-3] | KU-60019 |
| MVP_top60_DNA_seed | AR | -0.218286 | KIRC | LB996-RCC | Bicalutamide | MVP_top60_DNA+RNA_seed | AURKA | -0.541447 | LUAD | NCI-H1648 | Tozasertib |
| MVP_top60_DNA_seed | MTOR | -0.216544 | KIRC | OS-RC-2 | QL-VIII-58 | MVP_top60_DNA+RNA_seed | EGFR | -0.540564 | LUAD | HCC-78 | AST-1306 |
| MVP_top60_DNA_seed | MTOR | -0.21581 | KIRC | SN12C | Apitolisib | MVP_top60_DNA+RNA_seed | HSP90AA1 | -0.540459 | LUAD | NCI-H1651 | Luminespib |
| MVP_top60_DNA_seed | PTK2 | -0.215323 | KIRC | VMRC-RCZ | PF-562271 | MVP_top60_DNA+RNA_seed | ATM | -0.539294 | LUAD | HCC-827 | KU-60019 |
| MVP_top60_DNA_seed | IGF1R | -0.215234 | KIRC | OS-RC-2 | BMS-754807 | MVP_top60_DNA+RNA_seed | AURKA | -0.538243 | LUAD | NCI-H2228 | Genentech Cpd 10 |
| MVP_top60_DNA_seed | HSP90AA1 | -0.214167 | KIRC | HA7-RCC | Tanespimycin | MVP_top60_DNA+RNA_seed | FGFR1 | -0.538199 | LUAD | H3255 | FGFR_3831 |
| MVP_top60_DNA_seed | HSP90AB1 | -0.214167 | KIRC | HA7-RCC | Tanespimycin | MVP_top60_DNA+RNA_seed | EGFR | -0.536147 | LUAD | NCI-H1435 | AST-1306 |
| MVP_top60_DNA_seed | AR | -0.211203 | KIRC | 769-P | Bicalutamide | MVP_top60_DNA+RNA_seed | FGFR1 | -0.536039 | LUAD | NCI-H1573 | AZD4547 |
| MVP_top60_DNA_seed | PIK3CD | -0.209011 | KIRC | RCC-MF | IC-87114 | MVP_top60_DNA+RNA_seed | HSP90AA1 | -0.53594 | LUAD | HOP-62 | Elesclomol |
| MVP_top60_DNA_seed | PIK3CA | -0.207929 | KIRC | BFTC-909 | PI3Ka_4409 | MVP_top60_DNA+RNA_seed | AKT1 | -0.535642 | LUAD | NCI-H1435 | AKT inhibitor VIII |
| MVP_top60_DNA_seed | PIK3CD | -0.207929 | KIRC | BFTC-909 | PI3Ka_4409 | MVP_top60_DNA+RNA_seed | EGFR | -0.533329 | LUAD | NCI-H1975 | AZD8931 |
| MVP_top60_DNA_seed | PTK2 | -0.207899 | KIRC | OS-RC-2 | PF-562271 | MVP_top60_DNA+RNA_seed | BRAF | -0.533062 | LUAD | LC-2-ad | RAF_9304 |
| MVP_top60_DNA_seed | IGF1R | -0.203158 | KIRC | NCC021 | BMS-536924 | MVP_top60_DNA+RNA_seed | PTK2 | -0.532298 | LUAD | A549 | PF-562271 |
| MVP_top60_DNA_seed | PARP1 | -0.203005 | KIRC | LB1047-RCC | Rucaparib | MVP_top60_DNA+RNA_seed | ATM | -0.528237 | LUAD | PC-3_[JPC-3] | CP466722 |
| MVP_top60_DNA_seed | MTOR | -0.19988 | KIRC | HA7-RCC | Temsirolimus | MVP_top60_DNA+RNA_seed | HDAC1 | -0.520482 | LUAD | NCI-H1944 | Belinostat |
| MVP_top60_DNA_seed | FGFR4 | -0.199676 | KIRC | SW156 | FGFR_0939 | MVP_top60_DNA+RNA_seed | BRAF | -0.51982 | LUAD | PC-3_[JPC-3] | HG6-64-1 |
| MVP_top60_DNA_seed | PARP1 | -0.19965 | KIRC | RXF393 | Olaparib | MVP_top60_DNA+RNA_seed | EGFR | -0.519382 | LUAD | NCI-H358 | CUDC-101 |
| MVP_top60_DNA_seed | PARP1 | -0.198292 | KIRC | CAKI-1 | Veliparib | MVP_top60_DNA+RNA_seed | MET | -0.51645 | LUAD | NCI-H1563 | JNJ38877605 |
| MVP_top60_DNA_seed | MTOR | -0.19637 | KIRC | BB65-RCC | Voxtalisib | MVP_top60_DNA+RNA_seed | PIK3CA | -0.516128 | LUAD | NCI-H1568 | AZD8835 |
| MVP_top60_DNA_seed | ATM | -0.196073 | KIRC | HA7-RCC | KU-60019 | MVP_top60_DNA+RNA_seed | IGF1R | -0.515823 | LUAD | NCI-H2087 | BMS-536924 |
| MVP_top60_DNA_seed | ATM | -0.195957 | KIRC | A704 | CP466722 | MVP_top60_DNA+RNA_seed | MAPK14 | -0.514844 | LUAD | NCI-H3122 | TAK-715 |
| MVP_top60_DNA_seed | HDAC1 | -0.195199 | KIRC | BB65-RCC | AR-42 | MVP_top60_DNA+RNA_seed | HSP90AA1 | -0.514611 | LUAD | PC-14 | Elesclomol |
| MVP_top60_DNA_seed | MTOR | -0.194274 | KIRC | CAL-54 | Temsirolimus | MVP_top60_DNA+RNA_seed | HDAC1 | -0.51439 | LUAD | COR-L105 | Tubastatin A |
| MVP_top60_DNA_seed | MTOR | -0.193932 | KIRC | NCC021 | Torin 2 | MVP_top60_DNA+RNA_seed | HSP90AA1 | -0.512257 | LUAD | Calu-6 | CCT-018159 |
| MVP_top60_DNA_seed | IGF1R | -0.19389 | KIRC | VMRC-RCW | Linsitinib | MVP_top60_DNA+RNA_seed | HSP90AA1 | -0.512241 | LUAD | NCI-H358 | Luminespib |
| MVP_top60_DNA_seed | PIK3CD | -0.19283 | KIRC | RCC-AB | IC-87114 | MVP_top60_DNA+RNA_seed | BRAF | -0.512051 | LUAD | RERF-LC-MS | PLX-4720 |
| MVP_top60_DNA_seed | PIK3CA | -0.189765 | KIRC | A704 | AZD8835 | MVP_top60_DNA+RNA_seed | ATM | -0.511908 | LUAD | NCI-H1975 | KU-60019 |
| MVP_top60_DNA_seed | PIK3CD | -0.189765 | KIRC | A704 | AZD8835 | MVP_top60_DNA+RNA_seed | ATM | -0.511369 | LUAD | NCI-H2023 | KU-55933 |
| MVP_top60_DNA_seed | ATM | -0.189101 | KIRC | RCC-AB | KU-55933 | MVP_top60_DNA+RNA_seed | ATM | -0.510955 | LUAD | NCI-H1437 | KU-60019 |
| MVP_top60_DNA_seed | HSP90AA1 | -0.188808 | KIRC | LB1047-RCC | CCT-018159 | MVP_top60_DNA+RNA_seed | FGFR1 | -0.508405 | LUAD | HOP-62 | FGFR_3831 |
| MVP_top60_DNA_seed | HSP90AB1 | -0.188808 | KIRC | LB1047-RCC | CCT-018159 | MVP_top60_DNA+RNA_seed | AURKA | -0.508019 | LUAD | LXF-289 | Genentech Cpd 10 |
| MVP_top60_DNA_seed | MTOR | -0.187079 | KIRC | HA7-RCC | Voxtalisib | MVP_top60_DNA+RNA_seed | HDAC1 | -0.507902 | LUAD | COR-L105 | CAY10603 |
| MVP_top60_DNA_seed | PARP1 | -0.186745 | KIRC | KMRC-1 | PARP_9495 | MVP_top60_DNA+RNA_seed | HDAC1 | -0.50664 | LUAD | NCI-H650 | Dacinostat |

| | | | | | | | | | | | |
|---|---|---|---|---|---|---|---|---|---|---|---|
| MVP_top60_DNA_seed | PTK2 | -0.185545 | KIRC | TK10 | PF-562271 | MVP_top60_DNA+RNA_seed | BRAF | -0.50553 | LUAD | HCC-44 | RAF_9304 |
| MVP_top60_DNA_seed | IGF1R | -0.184294 | KIRC | RXF393 | BMS-754807 | MVP_top60_DNA+RNA_seed | FGFR1 | -0.505281 | LUAD | NCI-H1755 | FGFR_3831 |
| MVP_top60_DNA_seed | IGF1R | -0.184019 | KIRC | VMRC-RCZ | Linsitinib | MVP_top60_DNA+RNA_seed | MDM2 | -0.503327 | LUAD | NCI-H1437 | Serdemetan |
| MVP_top60_DNA_seed | HDAC1 | -0.181892 | KIRC | RCC-JW | Belinostat | MVP_top60_DNA+RNA_seed | ATM | -0.503248 | LUAD | NCI-H838 | KU-60019 |
| MVP_top60_DNA_seed | PIK3CD | -0.17599 | KIRC | HA7-RCC | IC-87114 | MVP_top60_DNA+RNA_seed | HDAC1 | -0.501606 | LUAD | Calu-6 | AR-42 |
| MVP_top60_DNA_seed | HSP90AA1 | -0.175101 | KIRC | SN12C | CCT-018159 | MVP_top60_DNA+RNA_seed | ATM | -0.50156 | LUAD | SK-LU-1 | KU-55933 |
| MVP_top60_DNA_seed | HSP90AB1 | -0.175101 | KIRC | SN12C | CCT-018159 | MVP_top60_DNA+RNA_seed | FGFR1 | -0.499779 | LUAD | NCI-H1944 | FGFR_3831 |
| MVP_top60_DNA_seed | MTOR | -0.172219 | KIRC | VMRC-RCZ | Temsirolimus | MVP_top60_DNA+RNA_seed | MAPK14 | -0.498906 | LUAD | LXF-289 | TAK-715 |
| MVP_top60_DNA_seed | MTOR | -0.165496 | KIRC | LB996-RCC | Apitolisib | MVP_top60_DNA+RNA_seed | AKT1 | -0.498901 | LUAD | NCI-H1734 | AKT inhibitor VIII |
| MVP_top60_DNA_seed | HSP90AA1 | -0.164817 | KIRC | LB1047-RCC | Luminespib | MVP_top60_DNA+RNA_seed | AKT1 | -0.496511 | LUAD | A549 | MK-2206 |
| MVP_top60_DNA_seed | HSP90AB1 | -0.164817 | KIRC | LB1047-RCC | Luminespib | MVP_top60_DNA+RNA_seed | PIK3CA | -0.496282 | LUAD | NCI-H1563 | AZD8835 |
| MVP_top60_DNA_seed | HSP90AA1 | -0.164124 | KIRC | CAKI-1 | Tanespimycin | MVP_top60_DNA+RNA_seed | MAPK14 | -0.496197 | LUAD | COR-L105 | TAK-715 |
| MVP_top60_DNA_seed | HSP90AB1 | -0.164124 | KIRC | CAKI-1 | Tanespimycin | MVP_top60_DNA+RNA_seed | BRAF | -0.496184 | LUAD | HCC-827 | RAF_9304 |
| MVP_top60_DNA_seed | HDAC1 | -0.162284 | KIRC | BFTC-909 | JQ12 | MVP_top60_DNA+RNA_seed | MDM2 | -0.496133 | LUAD | NCI-H1944 | Serdemetan |
| MVP_top60_DNA_seed | PIK3CD | -0.160427 | KIRC | CAKI-1 | Idelalisib | MVP_top60_DNA+RNA_seed | IGF1R | -0.496074 | LUAD | COR-L105 | GSK1904529A |
| MVP_top60_DNA_seed | PARP1 | -0.15956 | KIRC | KMRC-1 | PARP_0108 | MVP_top60_DNA+RNA_seed | AKT1 | -0.495576 | LUAD | PC-14 | AKT inhibitor VIII |
| MVP_top60_DNA_seed | MTOR | -0.158671 | KIRC | TK10 | Torin 2 | MVP_top60_DNA+RNA_seed | EGFR | -0.493927 | LUAD | EMC-BAC-1 | CUDC-101 |
| MVP_top60_DNA_seed | HDAC1 | -0.157209 | KIRC | TK10 | Entinostat | MVP_top60_DNA+RNA_seed | SRC | -0.493685 | LUAD | NCI-H2405 | NVP-BHG712 |
| MVP_top60_DNA_seed | IGF1R | -0.154537 | KIRC | VMRC-RCW | BMS-754807 | MVP_top60_DNA+RNA_seed | AURKA | -0.492988 | LUAD | NCI-H1355 | HG-5-113-01 |
| MVP_top60_DNA_seed | HDAC1 | -0.152026 | KIRC | RCC-JW | CAY10603 | MVP_top60_DNA+RNA_seed | HDAC1 | -0.492319 | LUAD | ABC-1 | Dacinostat |
| MVP_top60_DNA_seed | IGF1R | -0.145662 | KIRC | TK10 | BMS-754807 | MVP_top60_DNA+RNA_seed | EGFR | -0.490492 | LUAD | NCI-H2085 | Pelitinib |
| MVP_top60_DNA_seed | HDAC1 | -0.145571 | KIRC | U031 | Dacinostat | MVP_top60_DNA+RNA_seed | FGFR1 | -0.489928 | LUAD | VMRC-LCD | FGFR_3831 |
| MVP_top60_DNA_seed | HDAC1 | -0.142646 | KIRC | VMRC-RCZ | CAY10603 | MVP_top60_DNA+RNA_seed | HDAC1 | -0.489708 | LUAD | LXF-289 | JQ12 |
| MVP_top60_DNA_seed | IGF1R | -0.142552 | KIRC | RCC10RGB | BMS-536924 | MVP_top60_DNA+RNA_seed | BRAF | -0.488453 | LUAD | Calu-6 | RAF_9304 |
| MVP_top60_DNA_seed | HDAC1 | -0.137204 | KIRC | VMRC-RCZ | AR-42 | MVP_top60_DNA+RNA_seed | ATM | -0.48727 | LUAD | RERF-LC-MS | KU-60019 |
| MVP_top60_DNA_seed | PIK3CD | -0.135731 | KIRC | RCC-AB | AZD8186 | MVP_top60_DNA+RNA_seed | BRAF | -0.485888 | LUAD | NCI-H2030 | RAF_9304 |
| MVP_top60_DNA_seed | HSP90AA1 | -0.135203 | KIRC | LB996-RCC | Tanespimycin | MVP_top60_DNA+RNA_seed | AKT1 | -0.485555 | LUAD | EKVX | BAY AKT1 |
| MVP_top60_DNA_seed | HSP90AB1 | -0.135203 | KIRC | LB996-RCC | Tanespimycin | MVP_top60_DNA+RNA_seed | ATM | -0.485232 | LUAD | RERF-LC-KJ | CP466722 |
| MVP_top60_DNA_seed | IGF1R | -0.134765 | KIRC | ACHN | BMS-536924 | MVP_top60_DNA+RNA_seed | AKT1 | -0.485027 | LUAD | HCC-78 | GSK690693 |
| MVP_top60_DNA_seed | PARP1 | -0.132789 | KIRC | 786-0 | Rucaparib | MVP_top60_DNA+RNA_seed | HDAC1 | -0.484117 | LUAD | NCI-H3122 | CAY10603 |
| MVP_top60_DNA_seed | HDAC1 | -0.12785 | KIRC | KMRC-1 | Belinostat | MVP_top60_DNA+RNA_seed | BRAF | -0.482571 | LUAD | NCI-H2087 | RAF_9304 |
| MVP_top60_DNA_seed | MTOR | -0.12488 | KIRC | TK10 | WYE-125132 | MVP_top60_DNA+RNA_seed | EGFR | -0.481605 | LUAD | NCI-H322M | Lapatinib |
| MVP_top60_DNA_seed | AR | -0.124031 | KIRC | RCC-JW | AZD3514 | MVP_top60_DNA+RNA_seed | AKT1 | -0.480728 | LUAD | ABC-1 | AKT inhibitor VIII |
| MVP_top60_DNA_seed | MTOR | -0.117217 | KIRC | U031 | CRT0105446 | MVP_top60_DNA+RNA_seed | BRAF | -0.480584 | LUAD | NCI-H2087 | HG6-64-1 |
| MVP_top60_DNA_seed | FGFR4 | -0.116518 | KIRC | A498 | FGFR_3831 | MVP_top60_DNA+RNA_seed | EGFR | -0.480015 | LUAD | NCI-H1648 | Pelitinib |
| MVP_top60_DNA_seed | AR | -0.114329 | KIRC | 786-0 | Bicalutamide | MVP_top60_DNA+RNA_seed | MET | -0.479848 | LUAD | NCI-H2030 | JNJ38877605 |
| MVP_top60_DNA_seed | PIK3CA | -0.113309 | KIRC | SN12C | PI3Ka_4409 | MVP_top60_DNA+RNA_seed | EGFR | -0.479488 | LUAD | EMC-BAC-1 | PF-00299804 |
| MVP_top60_DNA_seed | PIK3CD | -0.113309 | KIRC | SN12C | PI3Ka_4409 | MVP_top60_DNA+RNA_seed | EGFR | -0.479127 | LUAD | HCC-44 | Gefitinib |
| MVP_top60_DNA_seed | FGFR4 | -0.112227 | KIRC | 786-0 | FGFR_3831 | MVP_top60_DNA+RNA_seed | MET | -0.478375 | LUAD | PC-14 | AZD6094 |
| MVP_top60_DNA_seed | IGF1R | -0.112111 | KIRC | 769-P | GSK1904529A | MVP_top60_DNA+RNA_seed | HDAC1 | -0.474231 | LUAD | NCI-H3122 | Belinostat |
| MVP_top60_DNA_seed | PARP1 | -0.111427 | KIRC | VMRC-RCW | Veliparib | MVP_top60_DNA+RNA_seed | HDAC1 | -0.47385 | LUAD | NCI-H1648 | CAY10603 |

| | | | | | | | | | | | |
|---|---|---|---|---|---|---|---|---|---|---|---|
| MVP_top60_DNA_seed | HDAC1 | -0.1111 | KIRC | RCC-FG2 | Belinostat | MVP_top60_DNA+RNA_seed | ATM | -0.473727 | LUAD | NCI-H2087 | CP466722 |
| MVP_top60_DNA_seed | IGF1R | -0.109975 | KIRC | U031 | Linsitinib | MVP_top60_DNA+RNA_seed | MET | -0.47316 | LUAD | LC-2-ad | SU11274 |
| MVP_top60_DNA_seed | FGFR4 | -0.109773 | KIRC | BB65-RCC | FGFR_0939 | MVP_top60_DNA+RNA_seed | EGFR | -0.472183 | LUAD | HCC-44 | Cetuximab |
| MVP_top60_DNA_seed | PIK3CD | -0.108683 | KIRC | RCC-JW | IC-87114 | MVP_top60_DNA+RNA_seed | HDAC1 | -0.472101 | LUAD | NCI-H2087 | Tubastatin A |
| MVP_top60_DNA_seed | PARP1 | -0.107548 | KIRC | RCC-FG2 | PARP_9495 | MVP_top60_DNA+RNA_seed | EGFR | -0.471048 | LUAD | EMC-BAC-2 | Afatinib |
| MVP_top60_DNA_seed | MTOR | -0.107345 | KIRC | RCC-JW | WYE-125132 | MVP_top60_DNA+RNA_seed | EGFR | -0.470369 | LUAD | PC-14 | Cetuximab |
| MVP_top60_DNA_seed | MTOR | -0.107326 | KIRC | HA7-RCC | Torin 2 | MVP_top60_DNA+RNA_seed | ATM | -0.469854 | LUAD | Calu-6 | KU-60019 |
| MVP_top60_DNA_seed | ATM | -0.10658 | KIRC | ACHN | KU-60019 | MVP_top60_DNA+RNA_seed | IGF1R | -0.469304 | LUAD | NCI-H1437 | Linsitinib |
| MVP_top60_DNA_seed | PARP1 | -0.104656 | KIRC | RCC-ER | Talazoparib | MVP_top60_DNA+RNA_seed | HDAC1 | -0.469182 | LUAD | NCI-H3122 | Tubastatin A |
| MVP_top60_DNA_seed | HDAC1 | -0.100476 | KIRC | U031 | Tubastatin A | MVP_top60_DNA+RNA_seed | AURKA | -0.468556 | LUAD | H3255 | GSK1070916 |
| MVP_top60_DNA_seed | PIK3CA | -0.095383 | KIRC | ACHN | AZD8835 | MVP_top60_DNA+RNA_seed | EGFR | -0.466621 | LUAD | NCI-H1563 | Cetuximab |
| MVP_top60_DNA_seed | PIK3CD | -0.095383 | KIRC | ACHN | AZD8835 | MVP_top60_DNA+RNA_seed | SRC | -0.465303 | LUAD | LXF-289 | HG-6-71-01 |
| MVP_top60_DNA_seed | ESR1 | -0.094579 | KIRC | ACHN | Fulvestrant | MVP_top60_DNA+RNA_seed | EGFR | -0.465094 | LUAD | ABC-1 | Pelitinib |
| MVP_top60_DNA_seed | HDAC1 | -0.09437 | KIRC | BB65-RCC | Dacinostat | MVP_top60_DNA+RNA_seed | MET | -0.46465 | LUAD | NCI-H23 | SU11274 |
| MVP_top60_DNA_seed | MTOR | -0.089895 | KIRC | LB1047-RCC | Torin 2 | MVP_top60_DNA+RNA_seed | PIK3CA | -0.46407 | LUAD | HCC-44 | AZD8835 |
| MVP_top60_DNA_seed | MTOR | -0.087372 | KIRC | SN12C | WYE-125132 | MVP_top60_DNA+RNA_seed | FGFR1 | -0.463435 | LUAD | NCI-H838 | PD173074 |
| MVP_top60_DNA_seed | AR | -0.086792 | KIRC | U031 | Bicalutamide | MVP_top60_DNA+RNA_seed | HSP90AA1 | -0.461996 | LUAD | HCC-44 | CCT-018159 |
| MVP_top60_DNA_seed | PIK3CA | -0.082612 | KIRC | NCC021 | PI3Ka_4409 | MVP_top60_DNA+RNA_seed | AKT1 | -0.461306 | LUAD | LC-2-ad | AKT inhibitor VIII |
| MVP_top60_DNA_seed | PIK3CD | -0.082612 | KIRC | NCC021 | PI3Ka_4409 | MVP_top60_DNA+RNA_seed | EGFR | -0.459764 | LUAD | NCI-H1755 | Cetuximab |
| MVP_top60_DNA_seed | MTOR | -0.08219 | KIRC | RCC-MF | Torin 2 | MVP_top60_DNA+RNA_seed | AURKA | -0.45933 | LUAD | PC-14 | HG-5-113-01 |
| MVP_top60_DNA_seed | FGFR4 | -0.080865 | KIRC | SN12C | FGFR_3831 | MVP_top60_DNA+RNA_seed | ATM | -0.459248 | LUAD | NCI-H1944 | CP466722 |
| MVP_top60_DNA_seed | IGF1R | -0.076025 | KIRC | RXF393 | Linsitinib | MVP_top60_DNA+RNA_seed | AKT1 | -0.458409 | LUAD | H3255 | AKT inhibitor VIII |
| MVP_top60_DNA_seed | ATM | -0.07542 | KIRC | NCC010 | KU-55933 | MVP_top60_DNA+RNA_seed | AKT1 | -0.457943 | LUAD | PC-14 | GSK690693 |
| MVP_top60_DNA_seed | PARP1 | -0.074112 | KIRC | ACHN | Olaparib | MVP_top60_DNA+RNA_seed | MAPK14 | -0.457369 | LUAD | NCI-H1648 | TAK-715 |
| MVP_top60_DNA_seed | EP300 | -0.073864 | KIRC | LB996-RCC | I-CBP112 | MVP_top60_DNA+RNA_seed | HSP90AA1 | -0.457181 | LUAD | VMRC-LCD | CCT-018159 |
| MVP_top60_DNA_seed | PARP1 | -0.070991 | KIRC | BFTC-909 | Olaparib | MVP_top60_DNA+RNA_seed | AKT1 | -0.457181 | LUAD | EMC-BAC-1 | GSK690693 |
| MVP_top60_DNA_seed | HSP90AA1 | -0.06895 | KIRC | NCC010 | CCT-018159 | MVP_top60_DNA+RNA_seed | IGF1R | -0.45607 | LUAD | NCI-H1944 | BMS-754807 |
| MVP_top60_DNA_seed | HSP90AB1 | -0.06895 | KIRC | NCC010 | CCT-018159 | MVP_top60_DNA+RNA_seed | AKT1 | -0.454312 | LUAD | RERF-LC-KJ | MK-2206 |
| MVP_top60_DNA_seed | IGF1R | -0.068784 | KIRC | TK10 | Linsitinib | MVP_top60_DNA+RNA_seed | EGFR | -0.452975 | LUAD | NCI-H1944 | Afatinib |
| MVP_top60_DNA_seed | EP300 | -0.065939 | KIRC | TK10 | I-CBP112 | MVP_top60_DNA+RNA_seed | HSP90AA1 | -0.449618 | LUAD | HCC-78 | Tanespimycin |
| MVP_top60_DNA_seed | MTOR | -0.065909 | KIRC | SN12C | Temsirolimus | MVP_top60_DNA+RNA_seed | BRAF | -0.449512 | LUAD | HCC-44 | PLX-4720 |
| MVP_top60_DNA_seed | PIK3CA | -0.065568 | KIRC | VMRC-RCZ | AZD8835 | MVP_top60_DNA+RNA_seed | AKT1 | -0.449272 | LUAD | HCC-44 | AKT inhibitor VIII |
| MVP_top60_DNA_seed | PIK3CD | -0.065568 | KIRC | VMRC-RCZ | AZD8835 | MVP_top60_DNA+RNA_seed | HDAC1 | -0.448433 | LUAD | NCI-H2228 | JQ12 |
| MVP_top60_DNA_seed | MTOR | -0.065247 | KIRC | CAL-54 | Torin 2 | MVP_top60_DNA+RNA_seed | ATM | -0.448269 | LUAD | NCI-H2291 | KU-60019 |
| MVP_top60_DNA_seed | AR | -0.064388 | KIRC | HA7-RCC | Bicalutamide | MVP_top60_DNA+RNA_seed | MDM2 | -0.448117 | LUAD | SK-LU-1 | Nutlin-3a (-) |
| MVP_top60_DNA_seed | PIK3CD | -0.061552 | KIRC | ACHN | IC-87114 | MVP_top60_DNA+RNA_seed | PIK3CA | -0.44744 | LUAD | LC-2-ad | AZD8835 |
| MVP_top60_DNA_seed | HDAC1 | -0.059196 | KIRC | RCC-FG2 | CAY10603 | MVP_top60_DNA+RNA_seed | EGFR | -0.447421 | LUAD | HCC-78 | Cetuximab |
| MVP_top60_DNA_seed | HDAC1 | -0.057742 | KIRC | CAKI-1 | CAY10603 | MVP_top60_DNA+RNA_seed | PIK3CA | -0.447255 | LUAD | EMC-BAC-2 | AZD8835 |
| MVP_top60_DNA_seed | HDAC1 | -0.056974 | KIRC | RCC-JW | AR-42 | MVP_top60_DNA+RNA_seed | EGFR | -0.447188 | LUAD | NCI-H1944 | CUDC-101 |
| MVP_top60_DNA_seed | EP300 | -0.0555 | KIRC | NCC021 | I-CBP112 | MVP_top60_DNA+RNA_seed | MET | -0.447078 | LUAD | ABC-1 | SU11274 |
| MVP_top60_DNA_seed | HDAC1 | -0.054711 | KIRC | LB1047-RCC | Tubastatin A | MVP_top60_DNA+RNA_seed | EGFR | -0.446944 | LUAD | NCI-H1568 | Pelitinib |

| | | | | | | | | | | | |
|---|---|---|---|---|---|---|---|---|---|---|---|
| MVP_top60_DNA_seed | HSP90AA1 | -0.054437 | KIRC | BFTC-909 | Elesclomol | MVP_top60_DNA+RNA_seed | IGF1R | -0.446699 | LUAD | LXF-289 | BMS-754807 |
| MVP_top60_DNA_seed | HSP90AB1 | -0.054437 | KIRC | BFTC-909 | Elesclomol | MVP_top60_DNA+RNA_seed | AURKA | -0.445561 | LUAD | NCI-H1666 | ZM447439 |
| MVP_top60_DNA_seed | PARP1 | -0.05405 | KIRC | KMRC-1 | PARP_9482 | MVP_top60_DNA+RNA_seed | HDAC1 | -0.443942 | LUAD | HOP-62 | Entinostat |
| MVP_top60_DNA_seed | FGFR4 | -0.052158 | KIRC | KMRC-1 | FGFR_0939 | MVP_top60_DNA+RNA_seed | HDAC1 | -0.443765 | LUAD | HCC-78 | CAY10603 |
| MVP_top60_DNA_seed | MTOR | -0.049196 | KIRC | BFTC-909 | Temsirolimus | MVP_top60_DNA+RNA_seed | HSP90AA1 | -0.44359 | LUAD | NCI-H2087 | Luminespib |
| MVP_top60_DNA_seed | AR | -0.045334 | KIRC | BFTC-909 | AZD3514 | MVP_top60_DNA+RNA_seed | IGF1R | -0.443565 | LUAD | RERF-LC-MS | BMS-536924 |
| MVP_top60_DNA_seed | FGFR4 | -0.044434 | KIRC | LB996-RCC | FGFR_3831 | MVP_top60_DNA+RNA_seed | AURKA | -0.44272 | LUAD | EMC-BAC-1 | ZM447439 |
| MVP_top60_DNA_seed | PIK3CD | -0.044269 | KIRC | 769-P | Idelalisib | MVP_top60_DNA+RNA_seed | FGFR1 | -0.440694 | LUAD | NCI-H2228 | AZD4547 |
| MVP_top60_DNA_seed | ESR1 | -0.042171 | KIRC | RCC-AB | Tamoxifen | MVP_top60_DNA+RNA_seed | EGFR | -0.440563 | LUAD | NCI-H358 | AZD8931 |
| MVP_top60_DNA_seed | IGF1R | -0.039219 | KIRC | A498 | BMS-536924 | MVP_top60_DNA+RNA_seed | IGF1R | -0.440173 | LUAD | EKVX | BMS-536924 |
| MVP_top60_DNA_seed | IGF1R | -0.038349 | KIRC | LB996-RCC | Linsitinib | MVP_top60_DNA+RNA_seed | PTK2 | -0.439991 | LUAD | Calu-3 | PF-562271 |
| MVP_top60_DNA_seed | HDAC1 | -0.037955 | KIRC | KMRC-1 | CAY10603 | MVP_top60_DNA+RNA_seed | AKT1 | -0.43862 | LUAD | COR-L105 | AKT inhibitor VIII |
| MVP_top60_DNA_seed | HSP90AA1 | -0.037007 | KIRC | OS-RC-2 | Elesclomol | MVP_top60_DNA+RNA_seed | MET | -0.438141 | LUAD | HCC-44 | JNJ38877605 |
| MVP_top60_DNA_seed | HSP90AB1 | -0.037007 | KIRC | OS-RC-2 | Elesclomol | MVP_top60_DNA+RNA_seed | HDAC1 | -0.436736 | LUAD | NCI-H358 | AR-42 |
| MVP_top60_DNA_seed | HDAC1 | -0.035003 | KIRC | RCC-JF | Tubastatin A | MVP_top60_DNA+RNA_seed | EGFR | -0.436441 | LUAD | NCI-H2087 | AST-1306 |
| MVP_top60_DNA_seed | HDAC1 | -0.034271 | KIRC | KMRC-1 | AR-42 | MVP_top60_DNA+RNA_seed | HDAC1 | -0.435535 | LUAD | HOP-62 | Belinostat |
| MVP_top60_DNA_seed | PARP1 | -0.03249 | KIRC | ACHN | Olaparib | MVP_top60_DNA+RNA_seed | FGFR1 | -0.434147 | LUAD | HCC-78 | FGFR_3831 |
| MVP_top60_DNA_seed | PARP1 | -0.031227 | KIRC | A704 | Veliparib | MVP_top60_DNA+RNA_seed | AURKA | -0.43412 | LUAD | ABC-1 | Genentech Cpd 10 |
| MVP_top60_DNA_seed | IGF1R | -0.030642 | KIRC | OS-RC-2 | BMS-536924 | MVP_top60_DNA+RNA_seed | HDAC1 | -0.432696 | LUAD | Calu-3 | Dacinostat |
| MVP_top60_DNA_seed | IGF1R | -0.028584 | KIRC | RCC-MF | BMS-536924 | MVP_top60_DNA+RNA_seed | BRAF | -0.432684 | LUAD | NCI-H1666 | Dabrafenib |
| MVP_top60_DNA_seed | MTOR | -0.02818 | KIRC | RCC-JW | Voxtalisib | MVP_top60_DNA+RNA_seed | PIK3CA | -0.43188 | LUAD | NCI-H1666 | AZD8835 |
| MVP_top60_DNA_seed | MTOR | -0.027266 | KIRC | RCC-ER | WYE-125132 | MVP_top60_DNA+RNA_seed | AKT1 | -0.431308 | LUAD | NCI-H1563 | BAY AKT1 |
| MVP_top60_DNA_seed | HDAC1 | -0.023813 | KIRC | LB2241-RCC | Dacinostat | MVP_top60_DNA+RNA_seed | PIK3CA | -0.430608 | LUAD | LXF-289 | PI3Ka_4409 |
| MVP_top60_DNA_seed | IGF1R | -0.019203 | KIRC | RCC-FG2 | BMS-536924 | MVP_top60_DNA+RNA_seed | HSP90AA1 | -0.42824 | LUAD | LC-2-ad | SNX-2112 |
| MVP_top60_DNA_seed | HSP90AA1 | -0.016759 | KIRC | SN12C | Luminespib | MVP_top60_DNA+RNA_seed | HSP90AA1 | -0.427461 | LUAD | SW1573 | Tanespimycin |
| MVP_top60_DNA_seed | HSP90AB1 | -0.016759 | KIRC | SN12C | Luminespib | MVP_top60_DNA+RNA_seed | HSP90AA1 | -0.426312 | LUAD | NCI-H1437 | Luminespib |
| MVP_top60_DNA_seed | PARP1 | -0.015749 | KIRC | VMRC-RCZ | PARP_9495 | MVP_top60_DNA+RNA_seed | HDAC1 | -0.421876 | LUAD | Calu-6 | JQ12 |
| MVP_top60_DNA_seed | HSP90AA1 | -0.014276 | KIRC | TK10 | Elesclomol | MVP_top60_DNA+RNA_seed | BRAF | -0.420679 | LUAD | NCI-H2291 | Dabrafenib |
| MVP_top60_DNA_seed | HSP90AB1 | -0.014276 | KIRC | TK10 | Elesclomol | MVP_top60_DNA+RNA_seed | EGFR | -0.419999 | LUAD | NCI-H2087 | Cetuximab |
| MVP_top60_DNA_seed | HSP90AA1 | -0.012909 | KIRC | LB2241-RCC | Elesclomol | MVP_top60_DNA+RNA_seed | IGF1R | -0.4196 | LUAD | NCI-H838 | BMS-754807 |
| MVP_top60_DNA_seed | HSP90AB1 | -0.012909 | KIRC | LB2241-RCC | Elesclomol | MVP_top60_DNA+RNA_seed | ATM | -0.417586 | LUAD | HCC-44 | CP466722 |
| MVP_top60_DNA_seed | PARP1 | -0.011457 | KIRC | NCC021 | Olaparib | MVP_top60_DNA+RNA_seed | EGFR | -0.416219 | LUAD | RERF-LC-KJ | Afatinib |
| MVP_top60_DNA_seed | PIK3CA | -0.008739 | KIRC | BB65-RCC | AZD8835 | MVP_top60_DNA+RNA_seed | EGFR | -0.416023 | LUAD | NCI-H1355 | Afatinib |
| MVP_top60_DNA_seed | PIK3CD | -0.008739 | KIRC | BB65-RCC | AZD8835 | MVP_top60_DNA+RNA_seed | HSP90AA1 | -0.415991 | LUAD | HCC-44 | Elesclomol |
| MVP_top60_DNA_seed | HSP90AA1 | -0.008217 | KIRC | U031 | Luminespib | MVP_top60_DNA+RNA_seed | AKT1 | -0.414767 | LUAD | NCI-H1781 | AKT inhibitor VIII |
| MVP_top60_DNA_seed | HSP90AB1 | -0.008217 | KIRC | U031 | Luminespib | MVP_top60_DNA+RNA_seed | HSP90AA1 | -0.414557 | LUAD | PC-3_[JPC-3] | Luminespib |
| MVP_top60_DNA_seed | MTOR | -0.002093 | KIRC | LB1047-RCC | WYE-125132 | MVP_top60_DNA+RNA_seed | BRAF | -0.414052 | LUAD | PC-14 | PLX-4720 |
| MVP_top60_DNA_seed | AR | 0.002661 | KIRC | TK10 | AZD3514 | MVP_top60_DNA+RNA_seed | MAPK14 | -0.41338 | LUAD | HCC-78 | TAK-715 |
| MVP_top60_DNA_seed | IGF1R | 0.003094 | KIRC | VMRC-RCZ | BMS-754807 | MVP_top60_DNA+RNA_seed | AURKA | -0.411687 | LUAD | RERF-LC-KJ | Genentech Cpd 10 |
| MVP_top60_DNA_seed | SMARCA4 | 0.006007 | KIRC | RCC-JW | PFI-3 | MVP_top60_DNA+RNA_seed | AURKA | -0.408729 | LUAD | LC-2-ad | Alisertib |
| MVP_top60_DNA_seed | HDAC1 | 0.008618 | KIRC | ACHN | Dacinostat | MVP_top60_DNA+RNA_seed | PIK3CA | -0.408583 | LUAD | A549 | AZD8835 |

| | | | | | | | | | | | |
|---|---|---|---|---|---|---|---|---|---|---|---|
| MVP_top60_DNA_seed | HSP90AA1 | 0.009337 | KIRC | RCC-FG2 | Tanespimycin | MVP_top60_DNA+RNA_seed | HSP90AA1 | -0.407265 | LUAD | NCI-H2030 | Elesclomol |
| MVP_top60_DNA_seed | HSP90AB1 | 0.009337 | KIRC | RCC-FG2 | Tanespimycin | MVP_top60_DNA+RNA_seed | FGFR1 | -0.40521 | LUAD | NCI-H2228 | PD173074 |
| MVP_top60_DNA_seed | HSP90AA1 | 0.010175 | KIRC | KMRC-1 | Tanespimycin | MVP_top60_DNA+RNA_seed | BRAF | -0.405073 | LUAD | HCC-827 | HG6-64-1 |
| MVP_top60_DNA_seed | HSP90AB1 | 0.010175 | KIRC | KMRC-1 | Tanespimycin | MVP_top60_DNA+RNA_seed | ATM | -0.403417 | LUAD | SW1573 | torin2 |
| MVP_top60_DNA_seed | PARP1 | 0.013974 | KIRC | CAL-54 | PARP_9495 | MVP_top60_DNA+RNA_seed | AURKA | -0.402924 | LUAD | NCI-H1437 | HG-5-113-01 |
| MVP_top60_DNA_seed | ESR1 | 0.016265 | KIRC | RCC-JW | Tamoxifen | MVP_top60_DNA+RNA_seed | BRAF | -0.40254 | LUAD | NCI-H1734 | HG6-64-1 |
| MVP_top60_DNA_seed | HDAC1 | 0.020514 | KIRC | CAL-54 | CAY10603 | MVP_top60_DNA+RNA_seed | AKT1 | -0.402022 | LUAD | NCI-H2122 | AKT inhibitor VIII |
| MVP_top60_DNA_seed | MTOR | 0.020699 | KIRC | BB65-RCC | QL-VIII-58 | MVP_top60_DNA+RNA_seed | IGF1R | -0.401103 | LUAD | HCC-44 | BMS-536924 |
| MVP_top60_DNA_seed | ATM | 0.021403 | KIRC | HA7-RCC | CP466722 | MVP_top60_DNA+RNA_seed | ATM | -0.401095 | LUAD | NCI-H2291 | KU-55933 |
| MVP_top60_DNA_seed | FGFR4 | 0.022847 | KIRC | RCC-AB | FGFR_3831 | MVP_top60_DNA+RNA_seed | HSP90AA1 | -0.40005 | LUAD | NCI-H2087 | SNX-2112 |
| MVP_top60_DNA_seed | MTOR | 0.024029 | KIRC | BB65-RCC | WYE-125132 | MVP_top60_DNA+RNA_seed | AKT1 | -0.399703 | LUAD | NCI-H2122 | AKT inhibitor VIII |
| MVP_top60_DNA_seed | HDAC1 | 0.026564 | KIRC | RCC10RGB | Dacinostat | MVP_top60_DNA+RNA_seed | EGFR | -0.399537 | LUAD | EKVX | CI-1033 |
| MVP_top60_DNA_seed | AR | 0.030987 | KIRC | VMRC-RCW | Bicalutamide | MVP_top60_DNA+RNA_seed | ATM | -0.398691 | LUAD | NCI-H1650 | KU-60019 |
| MVP_top60_DNA_seed | HDAC1 | 0.031744 | KIRC | RCC-FG2 | AR-42 | MVP_top60_DNA+RNA_seed | PIK3CA | -0.398438 | LUAD | NCI-H23 | PI3Ka_4409 |
| MVP_top60_DNA_seed | MTOR | 0.032636 | KIRC | LB1047-RCC | CRT0105446 | MVP_top60_DNA+RNA_seed | AURKA | -0.39743 | LUAD | NCI-H1568 | Genentech Cpd 10 |
| MVP_top60_DNA_seed | PIK3CA | 0.033875 | KIRC | RCC-JW | PI3Ka_4409 | MVP_top60_DNA+RNA_seed | PIK3CA | -0.397218 | LUAD | NCI-H1355 | AZD8835 |
| MVP_top60_DNA_seed | PIK3CD | 0.033875 | KIRC | RCC-JW | PI3Ka_4409 | MVP_top60_DNA+RNA_seed | HDAC1 | -0.396289 | LUAD | HOP-62 | AR-42 |
| MVP_top60_DNA_seed | PIK3CA | 0.034229 | KIRC | RCC-FG2 | AZD8835 | MVP_top60_DNA+RNA_seed | AURKA | -0.392518 | LUAD | NCI-H2087 | ZM447439 |
| MVP_top60_DNA_seed | PIK3CD | 0.034229 | KIRC | RCC-FG2 | AZD8835 | MVP_top60_DNA+RNA_seed | HDAC1 | -0.391694 | LUAD | NCI-H1975 | Belinostat |
| MVP_top60_DNA_seed | FGFR4 | 0.035524 | KIRC | CAKI-1 | FGFR_0939 | MVP_top60_DNA+RNA_seed | ATM | -0.391139 | LUAD | NCI-H1666 | KU-55933 |
| MVP_top60_DNA_seed | PARP1 | 0.041083 | KIRC | A498 | PARP_9482 | MVP_top60_DNA+RNA_seed | MET | -0.390652 | LUAD | NCI-H1435 | JNJ38877605 |
| MVP_top60_DNA_seed | ESR1 | 0.047327 | KIRC | SW156 | Fulvestrant | MVP_top60_DNA+RNA_seed | BRAF | -0.388667 | LUAD | NCI-H1793 | HG6-64-1 |
| MVP_top60_DNA_seed | PARP1 | 0.048492 | KIRC | SN12C | Olaparib | MVP_top60_DNA+RNA_seed | AKT1 | -0.386641 | LUAD | NCI-H1651 | AKT inhibitor VIII |
| MVP_top60_DNA_seed | ATM | 0.050515 | KIRC | SN12C | KU-55933 | MVP_top60_DNA+RNA_seed | HDAC1 | -0.386465 | LUAD | HCC-78 | AR-42 |
| MVP_top60_DNA_seed | AR | 0.054523 | KIRC | BFTC-909 | Bicalutamide | MVP_top60_DNA+RNA_seed | BRAF | -0.385887 | LUAD | NCI-H23 | RAF_9304 |
| MVP_top60_DNA_seed | AR | 0.05492 | KIRC | RCC-FG2 | Bicalutamide | MVP_top60_DNA+RNA_seed | AKT1 | -0.384716 | LUAD | NCI-H1944 | AKT inhibitor VIII |
| MVP_top60_DNA_seed | ATM | 0.056401 | KIRC | TK10 | KU-55933 | MVP_top60_DNA+RNA_seed | IGF1R | -0.384488 | LUAD | RERF-LC-KJ | BMS-754807 |
| MVP_top60_DNA_seed | MTOR | 0.059497 | KIRC | VMRC-RCZ | Voxtalisib | MVP_top60_DNA+RNA_seed | AURKA | -0.383222 | LUAD | ABC-1 | GSK1070916 |
| MVP_top60_DNA_seed | PIK3CD | 0.059697 | KIRC | A704 | IC-87114 | MVP_top60_DNA+RNA_seed | MET | -0.382173 | LUAD | NCI-H1563 | AZD6094 |
| MVP_top60_DNA_seed | IGF1R | 0.061786 | KIRC | 786-0 | BMS-536924 | MVP_top60_DNA+RNA_seed | BRAF | -0.381501 | LUAD | NCI-H2030 | SB590885 |
| MVP_top60_DNA_seed | EP300 | 0.063953 | KIRC | CAL-54 | I-CBP112 | MVP_top60_DNA+RNA_seed | IGF1R | -0.380848 | LUAD | SK-LU-1 | BMS-536924 |
| MVP_top60_DNA_seed | PARP1 | 0.064193 | KIRC | CAKI-1 | Talazoparib | MVP_top60_DNA+RNA_seed | ATM | -0.380128 | LUAD | NCI-H2228 | KU-55933 |
| MVP_top60_DNA_seed | MTOR | 0.064233 | KIRC | CAKI-1 | Apitolisib | MVP_top60_DNA+RNA_seed | ATM | -0.38011 | LUAD | NCI-H1993 | KU-60019 |
| MVP_top60_DNA_seed | FGFR4 | 0.06514 | KIRC | ACHN | FGFR_3831 | MVP_top60_DNA+RNA_seed | PIK3CA | -0.379714 | LUAD | NCI-H2030 | PI3Ka_4409 |
| MVP_top60_DNA_seed | HSP90AA1 | 0.06533 | KIRC | TK10 | CCT-018159 | MVP_top60_DNA+RNA_seed | ATM | -0.378262 | LUAD | NCI-H2405 | CP466722 |
| MVP_top60_DNA_seed | HSP90AB1 | 0.06533 | KIRC | TK10 | CCT-018159 | MVP_top60_DNA+RNA_seed | IGF1R | -0.376376 | LUAD | NCI-H2030 | Linsitinib |
| MVP_top60_DNA_seed | MTOR | 0.065656 | KIRC | SN12C | Voxtalisib | MVP_top60_DNA+RNA_seed | HDAC1 | -0.376176 | LUAD | NCI-H522 | AR-42 |
| MVP_top60_DNA_seed | HSP90AA1 | 0.067311 | KIRC | HA7-RCC | Luminespib | MVP_top60_DNA+RNA_seed | IGF1R | -0.376119 | LUAD | NCI-H1573 | BMS-536924 |
| MVP_top60_DNA_seed | HSP90AB1 | 0.067311 | KIRC | HA7-RCC | Luminespib | MVP_top60_DNA+RNA_seed | BRAF | -0.371361 | LUAD | PC-14 | PLX-4720 |
| MVP_top60_DNA_seed | HSP90AA1 | 0.07142 | KIRC | A498 | Elesclomol | MVP_top60_DNA+RNA_seed | BRAF | -0.369763 | LUAD | LXF-289 | PLX-4720 |
| MVP_top60_DNA_seed | HSP90AB1 | 0.07142 | KIRC | A498 | Elesclomol | MVP_top60_DNA+RNA_seed | IGF1R | -0.368771 | LUAD | NCI-H838 | GSK1904529A |

| | | | | | | | | | | | |
|---|---|---|---|---|---|---|---|---|---|---|---|
| MVP_top60_DNA_seed | PIK3CA | 0.076929 | KIRC | ACHN | PI3Ka_4409 | MVP_top60_DNA+RNA_seed | BRAF | -0.367398 | LUAD | NCI-H650 | Dabrafenib |
| MVP_top60_DNA_seed | PIK3CD | 0.076929 | KIRC | ACHN | PI3Ka_4409 | MVP_top60_DNA+RNA_seed | HSP90AA1 | -0.364479 | LUAD | NCI-H2122 | CCT-018159 |
| MVP_top60_DNA_seed | IGF1R | 0.077851 | KIRC | BB65-RCC | BMS-536924 | MVP_top60_DNA+RNA_seed | EGFR | -0.362356 | LUAD | NCI-H2342 | Afatinib |
| MVP_top60_DNA_seed | MTOR | 0.079732 | KIRC | RCC-MF | WYE-125132 | MVP_top60_DNA+RNA_seed | IGF1R | -0.362298 | LUAD | COR-L105 | Linsitinib |
| MVP_top60_DNA_seed | HSP90AA1 | 0.084589 | KIRC | RCC-FG2 | SNX-2112 | MVP_top60_DNA+RNA_seed | IGF1R | -0.361303 | LUAD | H3255 | BMS-536924 |
| MVP_top60_DNA_seed | HSP90AB1 | 0.084589 | KIRC | RCC-FG2 | SNX-2112 | MVP_top60_DNA+RNA_seed | HSP90AA1 | -0.361204 | LUAD | NCI-H1650 | Elesclomol |
| MVP_top60_DNA_seed | PIK3CA | 0.085709 | KIRC | CAL-54 | PI3Ka_4409 | MVP_top60_DNA+RNA_seed | EGFR | -0.361079 | LUAD | HOP-62 | CUDC-101 |
| MVP_top60_DNA_seed | PIK3CD | 0.085709 | KIRC | CAL-54 | PI3Ka_4409 | MVP_top60_DNA+RNA_seed | PIK3CA | -0.360276 | LUAD | NCI-H2023 | AZD8835 |
| MVP_top60_DNA_seed | PARP1 | 0.086233 | KIRC | RCC-MF | Veliparib | MVP_top60_DNA+RNA_seed | EGFR | -0.359683 | LUAD | LXF-289 | AZD8931 |
| MVP_top60_DNA_seed | PARP1 | 0.088477 | KIRC | 786-0 | PARP_9482 | MVP_top60_DNA+RNA_seed | BRAF | -0.358977 | LUAD | LC-2-ad | AZ628 |
| MVP_top60_DNA_seed | FGFR4 | 0.088982 | KIRC | RCC-ER | FGFR_0939 | MVP_top60_DNA+RNA_seed | ATM | -0.358288 | LUAD | NCI-H2347 | KU-60019 |
| MVP_top60_DNA_seed | MTOR | 0.090156 | KIRC | OS-RC-2 | Apitolisib | MVP_top60_DNA+RNA_seed | AURKA | -0.357924 | LUAD | NCI-H1437 | GSK1070916 |
| MVP_top60_DNA_seed | ESR1 | 0.090517 | KIRC | SN12C | Fulvestrant | MVP_top60_DNA+RNA_seed | MDM2 | -0.357765 | LUAD | NCI-H2009 | Serdemetan |
| MVP_top60_DNA_seed | AR | 0.093295 | KIRC | RCC-FG2 | AZD3514 | MVP_top60_DNA+RNA_seed | MDM2 | -0.357401 | LUAD | HCC-827 | Nutlin-3a (-) |
| MVP_top60_DNA_seed | PIK3CD | 0.093615 | KIRC | BFTC-909 | Idelalisib | MVP_top60_DNA+RNA_seed | HDAC1 | -0.357358 | LUAD | NCI-H3122 | Dacinostat |
| MVP_top60_DNA_seed | ATM | 0.093799 | KIRC | TK10 | CP466722 | MVP_top60_DNA+RNA_seed | HSP90AA1 | -0.356973 | LUAD | LC-2-ad | Elesclomol |
| MVP_top60_DNA_seed | PARP1 | 0.096314 | KIRC | VMRC-RCW | Olaparib | MVP_top60_DNA+RNA_seed | FGFR1 | -0.356116 | LUAD | LC-2-ad | PD173074 |
| MVP_top60_DNA_seed | HDAC1 | 0.098215 | KIRC | VMRC-RCZ | Dacinostat | MVP_top60_DNA+RNA_seed | PTK2 | -0.355751 | LUAD | NCI-H2009 | PF-562271 |
| MVP_top60_DNA_seed | PARP1 | 0.099853 | KIRC | NCC021 | Talazoparib | MVP_top60_DNA+RNA_seed | BRAF | -0.35569 | LUAD | NCI-H1650 | HG6-64-1 |
| MVP_top60_DNA_seed | ESR1 | 0.100674 | KIRC | LB996-RCC | Fulvestrant | MVP_top60_DNA+RNA_seed | AKT1 | -0.354109 | LUAD | NCI-H2030 | BAY AKT1 |
| MVP_top60_DNA_seed | PARP1 | 0.100965 | KIRC | ACHN | PARP_9495 | MVP_top60_DNA+RNA_seed | IGF1R | -0.354083 | LUAD | NCI-H2405 | GSK1904529A |
| MVP_top60_DNA_seed | HDAC1 | 0.101049 | KIRC | RXF393 | JQ12 | MVP_top60_DNA+RNA_seed | EGFR | -0.354053 | LUAD | EMC-BAC-2 | AZD8931 |
| MVP_top60_DNA_seed | ATM | 0.1012 | KIRC | RCC-ER | CP466722 | MVP_top60_DNA+RNA_seed | EGFR | -0.353626 | LUAD | EKVX | PF-00299804 |
| MVP_top60_DNA_seed | PARP1 | 0.108533 | KIRC | CAL-54 | PARP_0108 | MVP_top60_DNA+RNA_seed | EGFR | -0.353107 | LUAD | NCI-H1792 | Gefitinib |
| MVP_top60_DNA_seed | ATM | 0.109715 | KIRC | RCC-AB | KU-60019 | MVP_top60_DNA+RNA_seed | HDAC1 | -0.351502 | LUAD | EMC-BAC-2 | CAY10603 |
| MVP_top60_DNA_seed | HDAC1 | 0.114896 | KIRC | ACHN | CAY10603 | MVP_top60_DNA+RNA_seed | SRC | -0.349839 | LUAD | PC-14 | NVP-BHG712 |
| MVP_top60_DNA_seed | ATM | 0.122948 | KIRC | NCC021 | CP466722 | MVP_top60_DNA+RNA_seed | HSP90AA1 | -0.349315 | LUAD | ABC-1 | Luminespib |
| MVP_top60_DNA_seed | PARP1 | 0.123502 | KIRC | BB65-RCC | Rucaparib | MVP_top60_DNA+RNA_seed | IGF1R | -0.348873 | LUAD | PC-14 | BMS-536924 |
| MVP_top60_DNA_seed | PIK3CD | 0.128322 | KIRC | BFTC-909 | AZD8186 | MVP_top60_DNA+RNA_seed | HDAC1 | -0.348675 | LUAD | EMC-BAC-2 | Belinostat |
| MVP_top60_DNA_seed | PIK3CD | 0.128476 | KIRC | BB65-RCC | IC-87114 | MVP_top60_DNA+RNA_seed | EGFR | -0.34688 | LUAD | NCI-H23 | Lapatinib |
| MVP_top60_DNA_seed | MTOR | 0.128534 | KIRC | TK10 | QL-VIII-58 | MVP_top60_DNA+RNA_seed | HSP90AA1 | -0.346649 | LUAD | SK-LU-1 | CCT-018159 |
| MVP_top60_DNA_seed | ATM | 0.128879 | KIRC | OS-RC-2 | KU-60019 | MVP_top60_DNA+RNA_seed | PIK3CA | -0.346425 | LUAD | NCI-H1666 | PI3Ka_4409 |
| MVP_top60_DNA_seed | IGF1R | 0.130416 | KIRC | HA7-RCC | BMS-536924 | MVP_top60_DNA+RNA_seed | AURKA | -0.344724 | LUAD | EMC-BAC-1 | Genentech Cpd 10 |
| MVP_top60_DNA_seed | ESR1 | 0.13196 | KIRC | KMRC-20 | Fulvestrant | MVP_top60_DNA+RNA_seed | ATM | -0.343404 | LUAD | PC-14 | KU-60019 |
| MVP_top60_DNA_seed | AR | 0.132516 | KIRC | LB2241-RCC | Bicalutamide | MVP_top60_DNA+RNA_seed | EGFR | -0.342656 | LUAD | NCI-H1792 | Pelitinib |
| MVP_top60_DNA_seed | MTOR | 0.135197 | KIRC | CAKI-1 | WYE-125132 | MVP_top60_DNA+RNA_seed | MET | -0.342002 | LUAD | NCI-H1975 | JNJ38877605 |
| MVP_top60_DNA_seed | IGF1R | 0.136863 | KIRC | RCC-JF | BMS-536924 | MVP_top60_DNA+RNA_seed | HDAC1 | -0.341973 | LUAD | NCI-H1355 | Parthenolide |
| MVP_top60_DNA_seed | HDAC1 | 0.1372 | KIRC | ACHN | Belinostat | MVP_top60_DNA+RNA_seed | ATM | -0.341272 | LUAD | NCI-H1793 | KU-55933 |
| MVP_top60_DNA_seed | PARP1 | 0.139504 | KIRC | CAL-54 | PARP_9482 | MVP_top60_DNA+RNA_seed | EGFR | -0.338522 | LUAD | NCI-H1734 | Afatinib |
| MVP_top60_DNA_seed | PIK3CA | 0.140781 | KIRC | A704 | PI3Ka_4409 | MVP_top60_DNA+RNA_seed | MET | -0.337777 | LUAD | LXF-289 | Crizotinib |
| MVP_top60_DNA_seed | PIK3CD | 0.140781 | KIRC | A704 | PI3Ka_4409 | MVP_top60_DNA+RNA_seed | BRAF | -0.337194 | LUAD | NCI-H1568 | RAF_9304 |

| | | | | | | | | | | | |
|---|---|---|---|---|---|---|---|---|---|---|---|
| MVP_top60_DNA_seed | MTOR | 0.140894 | KIRC | LB996-RCC | QL-VIII-58 | MVP_top60_DNA+RNA_seed | PTK2 | -0.335939 | LUAD | COR-L105 | PF-562271 |
| MVP_top60_DNA_seed | IGF1R | 0.141853 | KIRC | OS-RC-2 | GSK1904529A | MVP_top60_DNA+RNA_seed | MET | -0.335149 | LUAD | NCI-H1944 | AZD6094 |
| MVP_top60_DNA_seed | HDAC1 | 0.143764 | KIRC | OS-RC-2 | CAY10603 | MVP_top60_DNA+RNA_seed | SMARCA4 | -0.334159 | LUAD | NCI-H2030 | PFI-3 |
| MVP_top60_DNA_seed | ATM | 0.146853 | KIRC | NCC010 | KU-60019 | MVP_top60_DNA+RNA_seed | EGFR | -0.332907 | LUAD | PC-14 | CI-1033 |
| MVP_top60_DNA_seed | HSP90AA1 | 0.146882 | KIRC | VMRC-RCW | Elesclomol | MVP_top60_DNA+RNA_seed | IGF1R | -0.332222 | LUAD | NCI-H2405 | Linsitinib |
| MVP_top60_DNA_seed | HSP90AB1 | 0.146882 | KIRC | VMRC-RCW | Elesclomol | MVP_top60_DNA+RNA_seed | BRAF | -0.332218 | LUAD | LC-2-ad | PLX-4720 |
| MVP_top60_DNA_seed | AR | 0.151006 | KIRC | U031 | Bicalutamide | MVP_top60_DNA+RNA_seed | HSP90AA1 | -0.331667 | LUAD | HCC-78 | Luminespib |
| MVP_top60_DNA_seed | SMARCA4 | 0.151841 | KIRC | A704 | PFI-3 | MVP_top60_DNA+RNA_seed | HDAC1 | -0.330251 | LUAD | NCI-H1975 | Parthenolide |
| MVP_top60_DNA_seed | FGFR4 | 0.152314 | KIRC | RCC-JF | FGFR_3831 | MVP_top60_DNA+RNA_seed | BRAF | -0.330048 | LUAD | NCI-H2009 | HG6-64-1 |
| MVP_top60_DNA_seed | HDAC1 | 0.153297 | KIRC | A704 | Dacinostat | MVP_top60_DNA+RNA_seed | HDAC1 | -0.329632 | LUAD | EMC-BAC-2 | AR-42 |
| MVP_top60_DNA_seed | IGF1R | 0.155366 | KIRC | ACHN | BMS-754807 | MVP_top60_DNA+RNA_seed | EGFR | -0.329395 | LUAD | NCI-H2342 | Cetuximab |
| MVP_top60_DNA_seed | HDAC1 | 0.158543 | KIRC | RCC10RGB | Parthenolide | MVP_top60_DNA+RNA_seed | MDM2 | -0.329016 | LUAD | NCI-H1648 | Nutlin-3a (-) |
| MVP_top60_DNA_seed | PIK3CA | 0.159012 | KIRC | RCC-AB | PI3Ka_4409 | MVP_top60_DNA+RNA_seed | ATM | -0.32872 | LUAD | H3255 | KU-60019 |
| MVP_top60_DNA_seed | PIK3CD | 0.159012 | KIRC | RCC-AB | PI3Ka_4409 | MVP_top60_DNA+RNA_seed | AURKA | -0.328572 | LUAD | COR-L105 | ZM447439 |
| MVP_top60_DNA_seed | PARP1 | 0.161056 | KIRC | KMRC-20 | PARP_0108 | MVP_top60_DNA+RNA_seed | MET | -0.327084 | LUAD | NCI-H322M | JNJ38877605 |
| MVP_top60_DNA_seed | PARP1 | 0.161631 | KIRC | CAKI-1 | Olaparib | MVP_top60_DNA+RNA_seed | HSP90AA1 | -0.326522 | LUAD | NCI-H838 | CCT-018159 |
| MVP_top60_DNA_seed | ESR1 | 0.16386 | KIRC | RCC-FG2 | Tamoxifen | MVP_top60_DNA+RNA_seed | AURKA | -0.324213 | LUAD | PC-14 | GSK1070916 |
| MVP_top60_DNA_seed | MTOR | 0.165214 | KIRC | NCC021 | Temsirolimus | MVP_top60_DNA+RNA_seed | SRC | -0.323515 | LUAD | NCI-H1792 | NVP-BHG712 |
| MVP_top60_DNA_seed | IGF1R | 0.165313 | KIRC | RXF393 | BMS-536924 | MVP_top60_DNA+RNA_seed | HDAC1 | -0.323348 | LUAD | NCI-H1355 | Dacinostat |
| MVP_top60_DNA_seed | AR | 0.168559 | KIRC | NCC010 | AZD3514 | MVP_top60_DNA+RNA_seed | AURKA | -0.322665 | LUAD | NCI-H2122 | ZM447439 |
| MVP_top60_DNA_seed | PIK3CA | 0.171023 | KIRC | NCC021 | AZD8835 | MVP_top60_DNA+RNA_seed | EGFR | -0.321502 | LUAD | VMRC-LCD | AZD8931 |
| MVP_top60_DNA_seed | PIK3CD | 0.171023 | KIRC | NCC021 | AZD8835 | MVP_top60_DNA+RNA_seed | IGF1R | -0.320594 | LUAD | NCI-H1623 | BMS-754807 |
| MVP_top60_DNA_seed | HDAC1 | 0.171671 | KIRC | VMRC-RCW | AR-42 | MVP_top60_DNA+RNA_seed | AKT1 | -0.320444 | LUAD | NCI-H2347 | AKT inhibitor VIII |
| MVP_top60_DNA_seed | MTOR | 0.173076 | KIRC | 769-P | Voxtalisib | MVP_top60_DNA+RNA_seed | IGF1R | -0.319484 | LUAD | LC-2-ad | GSK1904529A |
| MVP_top60_DNA_seed | HSP90AA1 | 0.173693 | KIRC | ACHN | Tanespimycin | MVP_top60_DNA+RNA_seed | MDM2 | -0.319427 | LUAD | SW1573 | Nutlin-3a (-) |
| MVP_top60_DNA_seed | HSP90AB1 | 0.173693 | KIRC | ACHN | Tanespimycin | MVP_top60_DNA+RNA_seed | IGF1R | -0.318502 | LUAD | NCI-H1792 | BMS-754807 |
| MVP_top60_DNA_seed | HDAC1 | 0.174118 | KIRC | OS-RC-2 | Belinostat | MVP_top60_DNA+RNA_seed | IGF1R | -0.318294 | LUAD | NCI-H1993 | Linsitinib |
| MVP_top60_DNA_seed | PARP1 | 0.175011 | KIRC | RXF393 | PARP_0108 | MVP_top60_DNA+RNA_seed | AURKA | -0.31814 | LUAD | NCI-H2228 | GSK1070916 |
| MVP_top60_DNA_seed | FGFR4 | 0.175746 | KIRC | NCC021 | FGFR_3831 | MVP_top60_DNA+RNA_seed | AKT1 | -0.317699 | LUAD | NCI-H1623 | MK-2206 |
| MVP_top60_DNA_seed | IGF1R | 0.176696 | KIRC | A704 | BMS-754807 | MVP_top60_DNA+RNA_seed | PIK3CA | -0.317112 | LUAD | NCI-H358 | AZD8835 |
| MVP_top60_DNA_seed | AR | 0.176825 | KIRC | RCC-MF | AZD3514 | MVP_top60_DNA+RNA_seed | IGF1R | -0.316427 | LUAD | NCI-H1568 | Linsitinib |
| MVP_top60_DNA_seed | HSP90AA1 | 0.176982 | KIRC | RCC-MF | SNX-2112 | MVP_top60_DNA+RNA_seed | EGFR | -0.315379 | LUAD | NCI-H441 | Afatinib |
| MVP_top60_DNA_seed | HSP90AB1 | 0.176982 | KIRC | RCC-MF | SNX-2112 | MVP_top60_DNA+RNA_seed | EGFR | -0.315364 | LUAD | NCI-H23 | Cetuximab |
| MVP_top60_DNA_seed | IGF1R | 0.178395 | KIRC | LB2241-RCC | BMS-754807 | MVP_top60_DNA+RNA_seed | AURKA | -0.314849 | LUAD | SK-LU-1 | Genentech Cpd 10 |
| MVP_top60_DNA_seed | MTOR | 0.183035 | KIRC | VMRC-RCZ | QL-VIII-58 | MVP_top60_DNA+RNA_seed | MET | -0.314713 | LUAD | NCI-H1781 | AZD6094 |
| MVP_top60_DNA_seed | EP300 | 0.184209 | KIRC | NCC010 | I-CBP112 | MVP_top60_DNA+RNA_seed | SRC | -0.313683 | LUAD | NCI-H2030 | NVP-BHG712 |
| MVP_top60_DNA_seed | HSP90AA1 | 0.184505 | KIRC | KMRC-1 | Luminespib | MVP_top60_DNA+RNA_seed | MET | -0.312782 | LUAD | NCI-H1648 | Crizotinib |
| MVP_top60_DNA_seed | HSP90AB1 | 0.184505 | KIRC | KMRC-1 | Luminespib | MVP_top60_DNA+RNA_seed | ATM | -0.312659 | LUAD | NCI-H1781 | KU-60019 |
| MVP_top60_DNA_seed | ATM | 0.186303 | KIRC | RCC-FG2 | CP466722 | MVP_top60_DNA+RNA_seed | MET | -0.31155 | LUAD | NCI-H1944 | JNJ38877605 |
| MVP_top60_DNA_seed | ESR1 | 0.188831 | KIRC | A498 | Fulvestrant | MVP_top60_DNA+RNA_seed | SRC | -0.311253 | LUAD | NCI-H1755 | Bosutinib |
| MVP_top60_DNA_seed | HDAC1 | 0.19005 | KIRC | RXF393 | Dacinostat | MVP_top60_DNA+RNA_seed | EGFR | -0.309056 | LUAD | EMC-BAC-2 | CUDC-101 |

| | | | | | | | | | | | |
|---|---|---|---|---|---|---|---|---|---|---|---|
| MVP_top60_DNA_seed | PTK2 | 0.190574 | KIRC | U031 | PF-562271 | MVP_top60_DNA+RNA_seed | FGFR1 | -0.305574 | LUAD | NCI-H2291 | PD173074 |
| MVP_top60_DNA_seed | HDAC1 | 0.191654 | KIRC | SW156 | Dacinostat | MVP_top60_DNA+RNA_seed | IGF1R | -0.305561 | LUAD | HOP-62 | BMS-754807 |
| MVP_top60_DNA_seed | HSP90AA1 | 0.191987 | KIRC | KMRC-20 | CCT-018159 | MVP_top60_DNA+RNA_seed | AKT1 | -0.305088 | LUAD | LC-2-ad | GSK690693 |
| MVP_top60_DNA_seed | HSP90AB1 | 0.191987 | KIRC | KMRC-20 | CCT-018159 | MVP_top60_DNA+RNA_seed | ATM | -0.304998 | LUAD | NCI-H1355 | KU-55933 |
| MVP_top60_DNA_seed | MTOR | 0.19565 | KIRC | A704 | Temsirolimus | MVP_top60_DNA+RNA_seed | EGFR | -0.30368 | LUAD | HCC-78 | Pelitinib |
| MVP_top60_DNA_seed | SMARCA4 | 0.196914 | KIRC | BB65-RCC | PFI-3 | MVP_top60_DNA+RNA_seed | HDAC1 | -0.303521 | LUAD | ABC-1 | AR-42 |
| MVP_top60_DNA_seed | ATM | 0.198167 | KIRC | VMRC-RCZ | KU-60019 | MVP_top60_DNA+RNA_seed | HDAC1 | -0.302572 | LUAD | NCI-H1651 | Dacinostat |
| MVP_top60_DNA_seed | PARP1 | 0.201973 | KIRC | LB2241-RCC | Veliparib | MVP_top60_DNA+RNA_seed | HDAC1 | -0.302102 | LUAD | NCI-H3122 | AR-42 |
| MVP_top60_DNA_seed | IGF1R | 0.202634 | KIRC | 769-P | Linsitinib | MVP_top60_DNA+RNA_seed | MDM2 | -0.301973 | LUAD | NCI-H1648 | Serdemetan |
| MVP_top60_DNA_seed | PIK3CD | 0.204294 | KIRC | VMRC-RCW | IC-87114 | MVP_top60_DNA+RNA_seed | IGF1R | -0.301722 | LUAD | NCI-H3122 | BMS-754807 |
| MVP_top60_DNA_seed | PARP1 | 0.20496 | KIRC | NCC021 | PARP_9482 | MVP_top60_DNA+RNA_seed | AKT1 | -0.301337 | LUAD | NCI-H1355 | BAY AKT1 |
| MVP_top60_DNA_seed | PARP1 | 0.206252 | KIRC | A498 | PARP_0108 | MVP_top60_DNA+RNA_seed | PTK2 | -0.300498 | LUAD | NCI-H1792 | PF-562271 |
| MVP_top60_DNA_seed | HSP90AA1 | 0.207005 | KIRC | OS-RC-2 | SNX-2112 | MVP_top60_DNA+RNA_seed | MET | -0.299869 | LUAD | EMC-BAC-1 | SU11274 |
| MVP_top60_DNA_seed | HSP90AB1 | 0.207005 | KIRC | OS-RC-2 | SNX-2112 | MVP_top60_DNA+RNA_seed | EGFR | -0.299839 | LUAD | NCI-H1734 | CUDC-101 |
| MVP_top60_DNA_seed | HDAC1 | 0.213002 | KIRC | ACHN | AR-42 | MVP_top60_DNA+RNA_seed | HSP90AA1 | -0.299531 | LUAD | HOP-62 | CCT-018159 |
| MVP_top60_DNA_seed | EP300 | 0.21393 | KIRC | A704 | I-CBP112 | MVP_top60_DNA+RNA_seed | AURKA | -0.299099 | LUAD | SW1573 | HG-5-113-01 |
| MVP_top60_DNA_seed | PARP1 | 0.216055 | KIRC | CAL-54 | Talazoparib | MVP_top60_DNA+RNA_seed | AKT1 | -0.297716 | LUAD | H3255 | AKT inhibitor VIII |
| MVP_top60_DNA_seed | IGF1R | 0.219001 | KIRC | U031 | BMS-536924 | MVP_top60_DNA+RNA_seed | HDAC1 | -0.296217 | LUAD | ABC-1 | CAY10603 |
| MVP_top60_DNA_seed | AR | 0.219165 | KIRC | LB1047-RCC | AZD3514 | MVP_top60_DNA+RNA_seed | MET | -0.295283 | LUAD | NCI-H2030 | SU11274 |
| MVP_top60_DNA_seed | IGF1R | 0.219513 | KIRC | BB65-RCC | BMS-536924 | MVP_top60_DNA+RNA_seed | HDAC1 | -0.293775 | LUAD | NCI-H522 | Belinostat |
| MVP_top60_DNA_seed | PIK3CD | 0.219955 | KIRC | TK10 | Idelalisib | MVP_top60_DNA+RNA_seed | HSP90AA1 | -0.293298 | LUAD | RERF-LC-KJ | Tanespimycin |
| MVP_top60_DNA_seed | MTOR | 0.220647 | KIRC | NCC010 | Temsirolimus | MVP_top60_DNA+RNA_seed | EGFR | -0.292033 | LUAD | NCI-H2342 | AZD8931 |
| MVP_top60_DNA_seed | PARP1 | 0.221047 | KIRC | 786-0 | Talazoparib | MVP_top60_DNA+RNA_seed | AKT1 | -0.291412 | LUAD | NCI-H1693 | MK-2206 |
| MVP_top60_DNA_seed | PARP1 | 0.223247 | KIRC | 786-0 | Olaparib | MVP_top60_DNA+RNA_seed | HSP90AA1 | -0.29078 | LUAD | NCI-H1781 | Elesclomol |
| MVP_top60_DNA_seed | PIK3CD | 0.225197 | KIRC | KMRC-20 | AZD8186 | MVP_top60_DNA+RNA_seed | BRAF | -0.290296 | LUAD | NCI-H650 | PLX-4720 |
| MVP_top60_DNA_seed | PARP1 | 0.225533 | KIRC | KMRC-20 | PARP_9482 | MVP_top60_DNA+RNA_seed | MET | -0.289833 | LUAD | NCI-H1792 | SU11274 |
| MVP_top60_DNA_seed | IGF1R | 0.225708 | KIRC | SN12C | Linsitinib | MVP_top60_DNA+RNA_seed | EGFR | -0.289521 | LUAD | NCI-H2291 | Afatinib |
| MVP_top60_DNA_seed | PARP1 | 0.225987 | KIRC | U031 | Veliparib | MVP_top60_DNA+RNA_seed | BRAF | -0.289088 | LUAD | NCI-H1573 | RAF_9304 |
| MVP_top60_DNA_seed | IGF1R | 0.228172 | KIRC | TK10 | BMS-536924 | MVP_top60_DNA+RNA_seed | HSP90AA1 | -0.286845 | LUAD | NCI-H1792 | Luminespib |
| MVP_top60_DNA_seed | PIK3CD | 0.23385 | KIRC | SW156 | AZD8186 | MVP_top60_DNA+RNA_seed | HDAC1 | -0.286826 | LUAD | NCI-H2347 | AR-42 |
| MVP_top60_DNA_seed | PIK3CA | 0.236688 | KIRC | RCC-JF | PI3Ka_4409 | MVP_top60_DNA+RNA_seed | HSP90AA1 | -0.286614 | LUAD | NCI-H2087 | CCT-018159 |
| MVP_top60_DNA_seed | PIK3CD | 0.236688 | KIRC | RCC-JF | PI3Ka_4409 | MVP_top60_DNA+RNA_seed | PTK2 | -0.284743 | LUAD | NCI-H1648 | PF-562271 |
| MVP_top60_DNA_seed | IGF1R | 0.239396 | KIRC | U031 | BMS-754807 | MVP_top60_DNA+RNA_seed | PIK3CA | -0.28472 | LUAD | NCI-H2023 | PI3Ka_4409 |
| MVP_top60_DNA_seed | FGFR4 | 0.242805 | KIRC | CAKI-1 | FGFR_3831 | MVP_top60_DNA+RNA_seed | AKT1 | -0.283923 | LUAD | LC-2-ad | AKT inhibitor VIII |
| MVP_top60_DNA_seed | ATM | 0.250514 | KIRC | CAKI-1 | KU-55933 | MVP_top60_DNA+RNA_seed | MAPK14 | -0.28285 | LUAD | HCC-44 | TAK-715 |
| MVP_top60_DNA_seed | HDAC1 | 0.253088 | KIRC | NCC021 | Tubastatin A | MVP_top60_DNA+RNA_seed | IGF1R | -0.281819 | LUAD | NCI-H1650 | BMS-536924 |
| MVP_top60_DNA_seed | AR | 0.25331 | KIRC | SN12C | AZD3514 | MVP_top60_DNA+RNA_seed | AURKA | -0.281556 | LUAD | NCI-H1355 | ZM447439 |
| MVP_top60_DNA_seed | FGFR4 | 0.253419 | KIRC | TK10 | FGFR_0939 | MVP_top60_DNA+RNA_seed | ATM | -0.281519 | LUAD | NCI-H2347 | CP466722 |
| MVP_top60_DNA_seed | MTOR | 0.253633 | KIRC | 786-0 | Voxtalisib | MVP_top60_DNA+RNA_seed | SRC | -0.279777 | LUAD | NCI-H1648 | NVP-BHG712 |
| MVP_top60_DNA_seed | HDAC1 | 0.255939 | KIRC | KMRC-1 | JQ12 | MVP_top60_DNA+RNA_seed | FGFR1 | -0.279715 | LUAD | NCI-H23 | AZD4547 |
| MVP_top60_DNA_seed | PIK3CA | 0.25607 | KIRC | 769-P | PI3Ka_4409 | MVP_top60_DNA+RNA_seed | EGFR | -0.279504 | LUAD | NCI-H322M | CUDC-101 |

| | | | | | | | | | | | |
|---|---|---|---|---|---|---|---|---|---|---|---|
| MVP_top60_DNA_seed | PIK3CD | 0.25607 | KIRC | 769-P | PI3Ka_4409 | MVP_top60_DNA+RNA_seed | HDAC1 | -0.279173 | LUAD | RERF-LC-MS | JQ12 |
| MVP_top60_DNA_seed | HDAC1 | 0.259922 | KIRC | OS-RC-2 | AR-42 | MVP_top60_DNA+RNA_seed | AURKA | -0.279101 | LUAD | NCI-H1975 | Genentech Cpd 10 |
| MVP_top60_DNA_seed | MTOR | 0.260014 | KIRC | OS-RC-2 | WYE-125132 | MVP_top60_DNA+RNA_seed | EGFR | -0.278131 | LUAD | NCI-H1993 | Pelitinib |
| MVP_top60_DNA_seed | HDAC1 | 0.260916 | KIRC | LB996-RCC | Dacinostat | MVP_top60_DNA+RNA_seed | HDAC1 | -0.276324 | LUAD | NCI-H1975 | CAY10603 |
| MVP_top60_DNA_seed | MTOR | 0.267406 | KIRC | SW156 | Torin 2 | MVP_top60_DNA+RNA_seed | AURKA | -0.276235 | LUAD | NCI-H1944 | GSK1070916 |
| MVP_top60_DNA_seed | HDAC1 | 0.26879 | KIRC | RXF393 | Tubastatin A | MVP_top60_DNA+RNA_seed | IGF1R | -0.273746 | LUAD | NCI-H1568 | BMS-754807 |
| MVP_top60_DNA_seed | HDAC1 | 0.277604 | KIRC | U031 | Belinostat | MVP_top60_DNA+RNA_seed | AURKA | -0.273146 | LUAD | NCI-H650 | CD532 |
| MVP_top60_DNA_seed | AR | 0.279041 | KIRC | LB1047-RCC | Bicalutamide | MVP_top60_DNA+RNA_seed | HSP90AA1 | -0.270646 | LUAD | NCI-H1793 | Tanespimycin |
| MVP_top60_DNA_seed | IGF1R | 0.279672 | KIRC | SW156 | Linsitinib | MVP_top60_DNA+RNA_seed | ATM | -0.269359 | LUAD | NCI-H1437 | torin2 |
| MVP_top60_DNA_seed | HSP90AA1 | 0.282268 | KIRC | KMRC-1 | Elesclomol | MVP_top60_DNA+RNA_seed | EGFR | -0.267463 | LUAD | RERF-LC-KJ | Gefitinib |
| MVP_top60_DNA_seed | HSP90AB1 | 0.282268 | KIRC | KMRC-1 | Elesclomol | MVP_top60_DNA+RNA_seed | AURKA | -0.266977 | LUAD | A549 | ZM447439 |
| MVP_top60_DNA_seed | MTOR | 0.282364 | KIRC | U031 | Torin 2 | MVP_top60_DNA+RNA_seed | AURKA | -0.266601 | LUAD | NCI-H1755 | GSK1070916 |
| MVP_top60_DNA_seed | FGFR4 | 0.284177 | KIRC | KMRC-20 | FGFR_3831 | MVP_top60_DNA+RNA_seed | FGFR1 | -0.266485 | LUAD | NCI-H322M | PD173074 |
| MVP_top60_DNA_seed | IGF1R | 0.285501 | KIRC | A704 | Linsitinib | MVP_top60_DNA+RNA_seed | IGF1R | -0.26601 | LUAD | NCI-H2291 | BMS-536924 |
| MVP_top60_DNA_seed | IGF1R | 0.285858 | KIRC | BFTC-909 | BMS-754807 | MVP_top60_DNA+RNA_seed | HDAC1 | -0.265555 | LUAD | NCI-H1623 | Belinostat |
| MVP_top60_DNA_seed | PIK3CD | 0.287095 | KIRC | KMRC-1 | IC-87114 | MVP_top60_DNA+RNA_seed | BRAF | -0.264939 | LUAD | NCI-H2009 | PLX-4720 |
| MVP_top60_DNA_seed | PIK3CD | 0.28924 | KIRC | VMRC-RCW | Idelalisib | MVP_top60_DNA+RNA_seed | HDAC1 | -0.263539 | LUAD | COR-L105 | AR-42 |
| MVP_top60_DNA_seed | HDAC1 | 0.293258 | KIRC | CAL-54 | JQ12 | MVP_top60_DNA+RNA_seed | IGF1R | -0.262586 | LUAD | NCI-H441 | BMS-536924 |
| MVP_top60_DNA_seed | AR | 0.295495 | KIRC | 769-P | Bicalutamide | MVP_top60_DNA+RNA_seed | EGFR | -0.259758 | LUAD | NCI-H358 | Gefitinib |
| MVP_top60_DNA_seed | MTOR | 0.29829 | KIRC | LB996-RCC | Torin 2 | MVP_top60_DNA+RNA_seed | ATM | -0.257904 | LUAD | NCI-H1650 | KU-55933 |
| MVP_top60_DNA_seed | MTOR | 0.302319 | KIRC | BB65-RCC | Apitolisib | MVP_top60_DNA+RNA_seed | HSP90AA1 | -0.257085 | LUAD | NCI-H650 | Elesclomol |
| MVP_top60_DNA_seed | PARP1 | 0.304519 | KIRC | SW156 | Olaparib | MVP_top60_DNA+RNA_seed | HDAC1 | -0.256181 | LUAD | NCI-H1975 | AR-42 |
| MVP_top60_DNA_seed | AR | 0.304577 | KIRC | LB2241-RCC | Bicalutamide | MVP_top60_DNA+RNA_seed | ATM | -0.25578 | LUAD | NCI-H522 | CP466722 |
| MVP_top60_DNA_seed | PARP1 | 0.304886 | KIRC | OS-RC-2 | PARP_9482 | MVP_top60_DNA+RNA_seed | EGFR | -0.255374 | LUAD | HCC-44 | AZD8931 |
| MVP_top60_DNA_seed | HSP90AA1 | 0.313541 | KIRC | 769-P | CCT-018159 | MVP_top60_DNA+RNA_seed | AKT1 | -0.251692 | LUAD | NCI-H358 | AKT inhibitor VIII |
| MVP_top60_DNA_seed | HSP90AB1 | 0.313541 | KIRC | 769-P | CCT-018159 | MVP_top60_DNA+RNA_seed | BRAF | -0.251225 | LUAD | NCI-H2087 | PLX-4720 |
| MVP_top60_DNA_seed | MTOR | 0.313632 | KIRC | 769-P | Temsirolimus | MVP_top60_DNA+RNA_seed | PIK3CA | -0.250988 | LUAD | NCI-H2009 | AZD8835 |
| MVP_top60_DNA_seed | PARP1 | 0.322082 | KIRC | OS-RC-2 | PARP_0108 | MVP_top60_DNA+RNA_seed | MDM2 | -0.250532 | LUAD | NCI-H1573 | Nutlin-3a (-) |
| MVP_top60_DNA_seed | PIK3CA | 0.32219 | KIRC | RCC10RGB | AZD8835 | MVP_top60_DNA+RNA_seed | FGFR1 | -0.250275 | LUAD | NCI-H2009 | FGFR_3831 |
| MVP_top60_DNA_seed | PIK3CD | 0.32219 | KIRC | RCC10RGB | AZD8835 | MVP_top60_DNA+RNA_seed | AKT1 | -0.250192 | LUAD | NCI-H1437 | MK-2206 |
| MVP_top60_DNA_seed | MTOR | 0.322768 | KIRC | BB65-RCC | Torin 2 | MVP_top60_DNA+RNA_seed | SRC | -0.248791 | LUAD | NCI-H358 | Bosutinib |
| MVP_top60_DNA_seed | IGF1R | 0.323292 | KIRC | BFTC-909 | Linsitinib | MVP_top60_DNA+RNA_seed | PIK3CA | -0.248382 | LUAD | NCI-H2030 | AZD8835 |
| MVP_top60_DNA_seed | PARP1 | 0.324638 | KIRC | A498 | Olaparib | MVP_top60_DNA+RNA_seed | HDAC1 | -0.247494 | LUAD | NCI-H358 | Dacinostat |
| MVP_top60_DNA_seed | ESR1 | 0.327371 | KIRC | BB65-RCC | Fulvestrant | MVP_top60_DNA+RNA_seed | FGFR1 | -0.247401 | LUAD | LC-2-ad | AZD4547 |
| MVP_top60_DNA_seed | AR | 0.328079 | KIRC | HA7-RCC | Bicalutamide | MVP_top60_DNA+RNA_seed | HDAC1 | -0.245823 | LUAD | LXF-289 | Dacinostat |
| MVP_top60_DNA_seed | AR | 0.32844 | KIRC | CAKI-1 | Bicalutamide | MVP_top60_DNA+RNA_seed | PIK3CA | -0.244965 | LUAD | NCI-H1437 | PI3Ka_4409 |
| MVP_top60_DNA_seed | AR | 0.329403 | KIRC | LB996-RCC | Bicalutamide | MVP_top60_DNA+RNA_seed | BRAF | -0.244779 | LUAD | NCI-H1993 | Dabrafenib |
| MVP_top60_DNA_seed | HDAC1 | 0.331657 | KIRC | HA7-RCC | JQ12 | MVP_top60_DNA+RNA_seed | AURKA | -0.244557 | LUAD | NCI-H2291 | HG-5-113-01 |
| MVP_top60_DNA_seed | PARP1 | 0.335587 | KIRC | RCC-ER | PARP_9482 | MVP_top60_DNA+RNA_seed | AURKA | -0.244257 | LUAD | NCI-H2085 | Genentech Cpd 10 |
| MVP_top60_DNA_seed | HDAC1 | 0.335832 | KIRC | HA7-RCC | Entinostat | MVP_top60_DNA+RNA_seed | HSP90AA1 | -0.24408 | LUAD | NCI-H838 | SNX-2112 |
| MVP_top60_DNA_seed | EP300 | 0.336442 | KIRC | RCC-AB | I-CBP112 | MVP_top60_DNA+RNA_seed | MET | -0.24308 | LUAD | NCI-H2291 | AZD6094 |

| | | | | | | | | | | | |
|---|---|---|---|---|---|---|---|---|---|---|---|
| MVP_top60_DNA_seed | PARP1 | 0.339036 | KIRC | RCC-FG2 | PARP_9482 | MVP_top60_DNA+RNA_seed | PIK3CA | -0.241774 | LUAD | NCI-H2228 | AZD8835 |
| MVP_top60_DNA_seed | HDAC1 | 0.341634 | KIRC | BFTC-909 | CAY10603 | MVP_top60_DNA+RNA_seed | HDAC1 | -0.241482 | LUAD | PC-14 | CAY10603 |
| MVP_top60_DNA_seed | PARP1 | 0.342488 | KIRC | RCC-FG2 | Olaparib | MVP_top60_DNA+RNA_seed | AKT1 | -0.241002 | LUAD | NCI-H1792 | AKT inhibitor VIII |
| MVP_top60_DNA_seed | MTOR | 0.342756 | KIRC | ACHN | Torin 2 | MVP_top60_DNA+RNA_seed | AKT1 | -0.240609 | LUAD | NCI-H2347 | AKT inhibitor VIII |
| MVP_top60_DNA_seed | HDAC1 | 0.343958 | KIRC | U031 | CAY10603 | MVP_top60_DNA+RNA_seed | MAPK14 | -0.239574 | LUAD | NCI-H2122 | TAK-715 |
| MVP_top60_DNA_seed | MTOR | 0.346814 | KIRC | LB2241-RCC | Voxtalisib | MVP_top60_DNA+RNA_seed | HSP90AA1 | -0.239561 | LUAD | RERF-LC-MS | SNX-2112 |
| MVP_top60_DNA_seed | AR | 0.347497 | KIRC | RCC-JF | Bicalutamide | MVP_top60_DNA+RNA_seed | HDAC1 | -0.239207 | LUAD | NCI-H1623 | AR-42 |
| MVP_top60_DNA_seed | HDAC1 | 0.349436 | KIRC | SW156 | JQ12 | MVP_top60_DNA+RNA_seed | MDM2 | -0.235361 | LUAD | LXF-289 | Nutlin-3a (-) |
| MVP_top60_DNA_seed | EP300 | 0.350056 | KIRC | 769-P | I-CBP112 | MVP_top60_DNA+RNA_seed | AKT1 | -0.235291 | LUAD | NCI-H838 | AKT inhibitor VIII |
| MVP_top60_DNA_seed | PARP1 | 0.351424 | KIRC | TK10 | Talazoparib | MVP_top60_DNA+RNA_seed | EGFR | -0.235027 | LUAD | NCI-H23 | CI-1033 |
| MVP_top60_DNA_seed | MTOR | 0.354215 | KIRC | KMRC-1 | WYE-125132 | MVP_top60_DNA+RNA_seed | EGFR | -0.234702 | LUAD | NCI-H522 | PF-00299804 |
| MVP_top60_DNA_seed | HDAC1 | 0.355072 | KIRC | VMRC-RCW | CAY10603 | MVP_top60_DNA+RNA_seed | ATM | -0.233806 | LUAD | Calu-6 | CP466722 |
| MVP_top60_DNA_seed | HDAC1 | 0.357702 | KIRC | VMRC-RCW | Belinostat | MVP_top60_DNA+RNA_seed | HDAC1 | -0.233588 | LUAD | NCI-H2030 | JQ12 |
| MVP_top60_DNA_seed | HDAC1 | 0.35877 | KIRC | CAL-54 | Tubastatin A | MVP_top60_DNA+RNA_seed | AKT1 | -0.233523 | LUAD | NCI-H1755 | BAY AKT1 |
| MVP_top60_DNA_seed | IGF1R | 0.358879 | KIRC | BFTC-909 | BMS-536924 | MVP_top60_DNA+RNA_seed | HDAC1 | -0.232895 | LUAD | NCI-H1734 | Belinostat |
| MVP_top60_DNA_seed | MTOR | 0.359669 | KIRC | A498 | Temsirolimus | MVP_top60_DNA+RNA_seed | MDM2 | -0.229979 | LUAD | NCI-H2023 | Nutlin-3a (-) |
| MVP_top60_DNA_seed | IGF1R | 0.362483 | KIRC | A498 | Linsitinib | MVP_top60_DNA+RNA_seed | MDM2 | -0.229418 | LUAD | COR-L105 | Nutlin-3a (-) |
| MVP_top60_DNA_seed | MTOR | 0.363125 | KIRC | NCC021 | Voxtalisib | MVP_top60_DNA+RNA_seed | HDAC1 | -0.22941 | LUAD | Calu-6 | CAY10603 |
| MVP_top60_DNA_seed | PARP1 | 0.366599 | KIRC | OS-RC-2 | Talazoparib | MVP_top60_DNA+RNA_seed | AURKA | -0.227752 | LUAD | NCI-H1792 | Genentech Cpd 10 |
| MVP_top60_DNA_seed | PIK3CA | 0.371572 | KIRC | RCC-ER | PI3Ka_4409 | MVP_top60_DNA+RNA_seed | ATM | -0.226244 | LUAD | HCC-78 | KU-55933 |
| MVP_top60_DNA_seed | PIK3CD | 0.371572 | KIRC | RCC-ER | PI3Ka_4409 | MVP_top60_DNA+RNA_seed | BRAF | -0.225985 | LUAD | NCI-H2023 | RAF_9304 |
| MVP_top60_DNA_seed | PARP1 | 0.373357 | KIRC | RXF393 | Olaparib | MVP_top60_DNA+RNA_seed | BRAF | -0.225945 | LUAD | NCI-H2405 | Dabrafenib |
| MVP_top60_DNA_seed | MTOR | 0.381037 | KIRC | TK10 | Temsirolimus | MVP_top60_DNA+RNA_seed | AKT1 | -0.22507 | LUAD | NCI-H2087 | AKT inhibitor VIII |
| MVP_top60_DNA_seed | FGFR4 | 0.386611 | KIRC | KMRC-1 | FGFR_3831 | MVP_top60_DNA+RNA_seed | ATM | -0.224889 | LUAD | NCI-H2085 | CP466722 |
| MVP_top60_DNA_seed | MTOR | 0.387309 | KIRC | ACHN | Temsirolimus | MVP_top60_DNA+RNA_seed | IGF1R | -0.224233 | LUAD | HOP-62 | Linsitinib |
| MVP_top60_DNA_seed | IGF1R | 0.387543 | KIRC | KMRC-20 | BMS-536924 | MVP_top60_DNA+RNA_seed | AURKA | -0.222816 | LUAD | NCI-H1437 | ZM447439 |
| MVP_top60_DNA_seed | MTOR | 0.390748 | KIRC | OS-RC-2 | Torin 2 | MVP_top60_DNA+RNA_seed | SRC | -0.222191 | LUAD | NCI-H1793 | Bosutinib |
| MVP_top60_DNA_seed | HDAC1 | 0.393168 | KIRC | KMRC-1 | Tubastatin A | MVP_top60_DNA+RNA_seed | EGFR | -0.221968 | LUAD | NCI-H358 | Cetuximab |
| MVP_top60_DNA_seed | PARP1 | 0.393671 | KIRC | RCC-JW | Veliparib | MVP_top60_DNA+RNA_seed | PIK3CA | -0.220916 | LUAD | NCI-H838 | PI3Ka_4409 |
| MVP_top60_DNA_seed | HSP90AA1 | 0.397485 | KIRC | RCC-AB | Tanespimycin | MVP_top60_DNA+RNA_seed | IGF1R | -0.219407 | LUAD | LXF-289 | Linsitinib |
| MVP_top60_DNA_seed | HSP90AB1 | 0.397485 | KIRC | RCC-AB | Tanespimycin | MVP_top60_DNA+RNA_seed | HDAC1 | -0.219281 | LUAD | NCI-H2342 | Dacinostat |
| MVP_top60_DNA_seed | PIK3CA | 0.398346 | KIRC | TK10 | PI3Ka_4409 | MVP_top60_DNA+RNA_seed | BRAF | -0.219197 | LUAD | SK-LU-1 | Dabrafenib |
| MVP_top60_DNA_seed | PIK3CD | 0.398346 | KIRC | TK10 | PI3Ka_4409 | MVP_top60_DNA+RNA_seed | HSP90AA1 | -0.218363 | LUAD | PC-3_[JPC-3] | CCT-018159 |
| MVP_top60_DNA_seed | AR | 0.398827 | KIRC | BB65-RCC | Bicalutamide | MVP_top60_DNA+RNA_seed | AKT1 | -0.217588 | LUAD | NCI-H2228 | GSK690693 |
| MVP_top60_DNA_seed | MTOR | 0.398925 | KIRC | BFTC-909 | Voxtalisib | MVP_top60_DNA+RNA_seed | AKT1 | -0.216813 | LUAD | NCI-H1975 | GSK690693 |
| MVP_top60_DNA_seed | MTOR | 0.400861 | KIRC | RCC-AB | QL-VIII-58 | MVP_top60_DNA+RNA_seed | EGFR | -0.215806 | LUAD | Calu-6 | AST-1306 |
| MVP_top60_DNA_seed | PARP1 | 0.401021 | KIRC | A704 | PARP_9482 | MVP_top60_DNA+RNA_seed | BRAF | -0.215525 | LUAD | HCC-44 | SB590885 |
| MVP_top60_DNA_seed | ATM | 0.402397 | KIRC | LB996-RCC | CP466722 | MVP_top60_DNA+RNA_seed | EGFR | -0.215374 | LUAD | NCI-H2023 | Cetuximab |
| MVP_top60_DNA_seed | HSP90AA1 | 0.404428 | KIRC | A704 | SNX-2112 | MVP_top60_DNA+RNA_seed | HSP90AA1 | -0.215281 | LUAD | VMRC-LCD | Elesclomol |
| MVP_top60_DNA_seed | HSP90AB1 | 0.404428 | KIRC | A704 | SNX-2112 | MVP_top60_DNA+RNA_seed | IGF1R | -0.215138 | LUAD | A549 | BMS-754807 |
| MVP_top60_DNA_seed | PARP1 | 0.405736 | KIRC | RCC-FG2 | Talazoparib | MVP_top60_DNA+RNA_seed | BRAF | -0.214903 | LUAD | HOP-62 | AZ628 |

| | | | | | | | | | | | |
|---|---|---|---|---|---|---|---|---|---|---|---|
| MVP_top60_DNA_seed | AR | 0.406671 | KIRC | KMRC-1 | Bicalutamide | MVP_top60_DNA+RNA_seed | BRAF | -0.214636 | LUAD | RERF-LC-KJ | RAF_9304 |
| MVP_top60_DNA_seed | SMARCA4 | 0.407255 | KIRC | RCC-JF | PFI-3 | MVP_top60_DNA+RNA_seed | EGFR | -0.213125 | LUAD | ABC-1 | Cetuximab |
| MVP_top60_DNA_seed | FGFR4 | 0.407777 | KIRC | CAL-54 | FGFR_3831 | MVP_top60_DNA+RNA_seed | ATM | -0.213007 | LUAD | ABC-1 | CP466722 |
| MVP_top60_DNA_seed | PARP1 | 0.412274 | KIRC | A704 | Talazoparib | MVP_top60_DNA+RNA_seed | IGF1R | -0.212976 | LUAD | NCI-H1563 | BMS-536924 |
| MVP_top60_DNA_seed | HDAC1 | 0.413154 | KIRC | RCC10RGB | JQ12 | MVP_top60_DNA+RNA_seed | IGF1R | -0.212563 | LUAD | NCI-H1792 | Linsitinib |
| MVP_top60_DNA_seed | MTOR | 0.415762 | KIRC | LB1047-RCC | QL-VIII-58 | MVP_top60_DNA+RNA_seed | AURKA | -0.211863 | LUAD | NCI-H1755 | ZM447439 |
| MVP_top60_DNA_seed | ESR1 | 0.416579 | KIRC | U031 | Tamoxifen | MVP_top60_DNA+RNA_seed | EGFR | -0.211454 | LUAD | NCI-H2342 | PF-00299804 |
| MVP_top60_DNA_seed | AR | 0.421284 | KIRC | CAL-54 | Bicalutamide | MVP_top60_DNA+RNA_seed | EGFR | -0.211322 | LUAD | NCI-H1666 | PF-00299804 |
| MVP_top60_DNA_seed | AR | 0.421926 | KIRC | SW156 | Bicalutamide | MVP_top60_DNA+RNA_seed | HDAC1 | -0.211311 | LUAD | PC-3_[JPC-3] | Dacinostat |
| MVP_top60_DNA_seed | PIK3CA | 0.423394 | KIRC | HA7-RCC | PI3Ka_4409 | MVP_top60_DNA+RNA_seed | BRAF | -0.210228 | LUAD | A549 | PLX-4720 |
| MVP_top60_DNA_seed | PIK3CD | 0.423394 | KIRC | HA7-RCC | PI3Ka_4409 | MVP_top60_DNA+RNA_seed | BRAF | -0.208467 | LUAD | NCI-H1755 | Dabrafenib |
| MVP_top60_DNA_seed | HDAC1 | 0.424162 | KIRC | CAL-54 | Belinostat | MVP_top60_DNA+RNA_seed | HDAC1 | -0.208076 | LUAD | NCI-H1651 | AR-42 |
| MVP_top60_DNA_seed | ESR1 | 0.424468 | KIRC | RCC-AB | Fulvestrant | MVP_top60_DNA+RNA_seed | AURKA | -0.207546 | LUAD | NCI-H2347 | Alisertib |
| MVP_top60_DNA_seed | ATM | 0.42595 | KIRC | SN12C | KU-60019 | MVP_top60_DNA+RNA_seed | BRAF | -0.205165 | LUAD | NCI-H1755 | PLX-4720 |
| MVP_top60_DNA_seed | HSP90AA1 | 0.42759 | KIRC | RCC-MF | Tanespimycin | MVP_top60_DNA+RNA_seed | HDAC1 | -0.205146 | LUAD | NCI-H1734 | CAY10603 |
| MVP_top60_DNA_seed | HSP90AB1 | 0.42759 | KIRC | RCC-MF | Tanespimycin | MVP_top60_DNA+RNA_seed | AKT1 | -0.205144 | LUAD | Calu-3 | AKT inhibitor VIII |
| MVP_top60_DNA_seed | MTOR | 0.431119 | KIRC | SN12C | Torin 2 | MVP_top60_DNA+RNA_seed | FGFR1 | -0.204864 | LUAD | SW1573 | FGFR_3831 |
| MVP_top60_DNA_seed | HDAC1 | 0.433588 | KIRC | BFTC-909 | AR-42 | MVP_top60_DNA+RNA_seed | FGFR1 | -0.203856 | LUAD | EMC-BAC-1 | FGFR_3831 |
| MVP_top60_DNA_seed | ESR1 | 0.433774 | KIRC | LB996-RCC | Tamoxifen | MVP_top60_DNA+RNA_seed | EGFR | -0.20341 | LUAD | ABC-1 | PF-00299804 |
| MVP_top60_DNA_seed | AR | 0.434903 | KIRC | A498 | Bicalutamide | MVP_top60_DNA+RNA_seed | BRAF | -0.202006 | LUAD | EMC-BAC-1 | RAF_9304 |
| MVP_top60_DNA_seed | HSP90AA1 | 0.435157 | KIRC | RCC-AB | Elesclomol | MVP_top60_DNA+RNA_seed | SRC | -0.201811 | LUAD | NCI-H1355 | Saracatinib |
| MVP_top60_DNA_seed | HSP90AB1 | 0.435157 | KIRC | RCC-AB | Elesclomol | MVP_top60_DNA+RNA_seed | HSP90AA1 | -0.20147 | LUAD | NCI-H358 | Tanespimycin |
| MVP_top60_DNA_seed | PARP1 | 0.436805 | KIRC | NCC021 | Veliparib | MVP_top60_DNA+RNA_seed | MDM2 | -0.20101 | LUAD | NCI-H1792 | Serdemetan |
| MVP_top60_DNA_seed | PARP1 | 0.437984 | KIRC | BB65-RCC | Olaparib | MVP_top60_DNA+RNA_seed | AKT1 | -0.198877 | LUAD | NCI-H1792 | AKT inhibitor VIII |
| MVP_top60_DNA_seed | PTK2 | 0.438164 | KIRC | BB65-RCC | PF-562271 | MVP_top60_DNA+RNA_seed | HSP90AA1 | -0.197683 | LUAD | NCI-H2228 | Tanespimycin |
| MVP_top60_DNA_seed | MTOR | 0.438687 | KIRC | SN12C | CRT0105446 | MVP_top60_DNA+RNA_seed | EGFR | -0.197594 | LUAD | HCC-78 | CI-1033 |
| MVP_top60_DNA_seed | ATM | 0.439653 | KIRC | BFTC-909 | CP466722 | MVP_top60_DNA+RNA_seed | HDAC1 | -0.197306 | LUAD | COR-L105 | Belinostat |
| MVP_top60_DNA_seed | PIK3CA | 0.440249 | KIRC | RCC-AB | AZD8835 | MVP_top60_DNA+RNA_seed | AURKA | -0.197242 | LUAD | A549 | Genentech Cpd 10 |
| MVP_top60_DNA_seed | PIK3CD | 0.440249 | KIRC | RCC-AB | AZD8835 | MVP_top60_DNA+RNA_seed | HDAC1 | -0.197104 | LUAD | NCI-H1650 | JQ12 |
| MVP_top60_DNA_seed | PARP1 | 0.440396 | KIRC | RCC-ER | PARP_0108 | MVP_top60_DNA+RNA_seed | EGFR | -0.196598 | LUAD | NCI-H1693 | Gefitinib |
| MVP_top60_DNA_seed | PARP1 | 0.44176 | KIRC | HA7-RCC | Veliparib | MVP_top60_DNA+RNA_seed | AKT1 | -0.196325 | LUAD | NCI-H1781 | BAY AKT1 |
| MVP_top60_DNA_seed | PARP1 | 0.441961 | KIRC | CAL-54 | Olaparib | MVP_top60_DNA+RNA_seed | SRC | -0.196224 | LUAD | A549 | Bosutinib |
| MVP_top60_DNA_seed | IGF1R | 0.445384 | KIRC | TK10 | GSK1904529A | MVP_top60_DNA+RNA_seed | BRAF | -0.196063 | LUAD | NCI-H2347 | Dabrafenib |
| MVP_top60_DNA_seed | EP300 | 0.445612 | KIRC | RCC-JF | I-CBP112 | MVP_top60_DNA+RNA_seed | ATM | -0.195846 | LUAD | NCI-H2342 | KU-60019 |
| MVP_top60_DNA_seed | PARP1 | 0.445906 | KIRC | 769-P | Veliparib | MVP_top60_DNA+RNA_seed | SMARCA4 | -0.194096 | LUAD | NCI-H1651 | PFI-3 |
| MVP_top60_DNA_seed | MTOR | 0.446464 | KIRC | SW156 | Apitolisib | MVP_top60_DNA+RNA_seed | HDAC1 | -0.193826 | LUAD | NCI-H2347 | JQ12 |
| MVP_top60_DNA_seed | IGF1R | 0.447103 | KIRC | BB65-RCC | GSK1904529A | MVP_top60_DNA+RNA_seed | EGFR | -0.193496 | LUAD | NCI-H2087 | Pelitinib |
| MVP_top60_DNA_seed | ESR1 | 0.448765 | KIRC | TK10 | Tamoxifen | MVP_top60_DNA+RNA_seed | HSP90AA1 | -0.192795 | LUAD | NCI-H2347 | Elesclomol |
| MVP_top60_DNA_seed | HDAC1 | 0.45089 | KIRC | A704 | JQ12 | MVP_top60_DNA+RNA_seed | FGFR1 | -0.191277 | LUAD | NCI-H2030 | PD173074 |
| MVP_top60_DNA_seed | SMARCA4 | 0.452822 | KIRC | CAL-54 | PFI-3 | MVP_top60_DNA+RNA_seed | EGFR | -0.191133 | LUAD | NCI-H1975 | Erlotinib |
| MVP_top60_DNA_seed | ATM | 0.453773 | KIRC | VMRC-RCZ | KU-55933 | MVP_top60_DNA+RNA_seed | EGFR | -0.190391 | LUAD | Calu-6 | CUDC-101 |

| | | | | | | | | | | | |
|---|---|---|---|---|---|---|---|---|---|---|---|
| MVP_top60_DNA_seed | PARP1 | 0.453829 | KIRC | RCC-MF | Olaparib | MVP_top60_DNA+RNA_seed | BRAF | -0.189353 | LUAD | NCI-H23 | AZ628 |
| MVP_top60_DNA_seed | AR | 0.455729 | KIRC | KMRC-1 | AZD3514 | MVP_top60_DNA+RNA_seed | PIK3CA | -0.188454 | LUAD | H3255 | AZD8835 |
| MVP_top60_DNA_seed | HDAC1 | 0.458187 | KIRC | BFTC-909 | Belinostat | MVP_top60_DNA+RNA_seed | EGFR | -0.187417 | LUAD | NCI-H1435 | CI-1033 |
| MVP_top60_DNA_seed | PIK3CD | 0.459552 | KIRC | 769-P | IC-87114 | MVP_top60_DNA+RNA_seed | SRC | -0.186276 | LUAD | NCI-H2122 | HG-6-71-01 |
| MVP_top60_DNA_seed | HDAC1 | 0.459754 | KIRC | LB2241-RCC | Belinostat | MVP_top60_DNA+RNA_seed | ATM | -0.185191 | LUAD | NCI-H1693 | KU-60019 |
| MVP_top60_DNA_seed | ATM | 0.459825 | KIRC | RXF393 | KU-60019 | MVP_top60_DNA+RNA_seed | HSP90AA1 | -0.185056 | LUAD | NCI-H358 | CCT-018159 |
| MVP_top60_DNA_seed | AR | 0.45992 | KIRC | RCC-JW | Bicalutamide | MVP_top60_DNA+RNA_seed | IGF1R | -0.184774 | LUAD | COR-L105 | BMS-536924 |
| MVP_top60_DNA_seed | ATM | 0.460985 | KIRC | SN12C | CP466722 | MVP_top60_DNA+RNA_seed | MET | -0.183608 | LUAD | NCI-H1838 | JNJ38877605 |
| MVP_top60_DNA_seed | PIK3CA | 0.461901 | KIRC | RCC10RGB | PI3Ka_4409 | MVP_top60_DNA+RNA_seed | HDAC1 | -0.183601 | LUAD | NCI-H650 | CAY10603 |
| MVP_top60_DNA_seed | PIK3CD | 0.461901 | KIRC | RCC10RGB | PI3Ka_4409 | MVP_top60_DNA+RNA_seed | AKT1 | -0.183386 | LUAD | Calu-3 | GSK690693 |
| MVP_top60_DNA_seed | PARP1 | 0.466126 | KIRC | A704 | Olaparib | MVP_top60_DNA+RNA_seed | PTK2 | -0.182473 | LUAD | NCI-H2347 | PF-562271 |
| MVP_top60_DNA_seed | MTOR | 0.467187 | KIRC | KMRC-1 | Apitolisib | MVP_top60_DNA+RNA_seed | FGFR1 | -0.182266 | LUAD | NCI-H1755 | AZD4547 |
| MVP_top60_DNA_seed | PTK2 | 0.469594 | KIRC | CAL-54 | PF-562271 | MVP_top60_DNA+RNA_seed | HSP90AA1 | -0.181576 | LUAD | NCI-H1734 | Luminespib |
| MVP_top60_DNA_seed | FGFR4 | 0.476178 | KIRC | BB65-RCC | FGFR_3831 | MVP_top60_DNA+RNA_seed | HDAC1 | -0.180821 | LUAD | NCI-H522 | CAY10603 |
| MVP_top60_DNA_seed | PIK3CD | 0.477235 | KIRC | BFTC-909 | IC-87114 | MVP_top60_DNA+RNA_seed | PIK3CA | -0.179248 | LUAD | LXF-289 | AZD8835 |
| MVP_top60_DNA_seed | PARP1 | 0.48034 | KIRC | RCC10RGB | PARP_9495 | MVP_top60_DNA+RNA_seed | EGFR | -0.178781 | LUAD | NCI-H1666 | CI-1033 |
| MVP_top60_DNA_seed | SMARCA4 | 0.481007 | KIRC | OS-RC-2 | PFI-3 | MVP_top60_DNA+RNA_seed | EGFR | -0.178584 | LUAD | ABC-1 | Afatinib |
| MVP_top60_DNA_seed | IGF1R | 0.482713 | KIRC | KMRC-1 | GSK1904529A | MVP_top60_DNA+RNA_seed | SRC | -0.177753 | LUAD | NCI-H1568 | Bosutinib |
| MVP_top60_DNA_seed | EP300 | 0.485906 | KIRC | SW156 | I-CBP112 | MVP_top60_DNA+RNA_seed | SRC | -0.176779 | LUAD | PC-3_[JPC-3] | NVP-BHG712 |
| MVP_top60_DNA_seed | HDAC1 | 0.487359 | KIRC | ACHN | Tubastatin A | MVP_top60_DNA+RNA_seed | HDAC1 | -0.176288 | LUAD | PC-3_[JPC-3] | AR-42 |
| MVP_top60_DNA_seed | MTOR | 0.489913 | KIRC | KMRC-1 | Voxtalisib | MVP_top60_DNA+RNA_seed | HDAC1 | -0.175075 | LUAD | EMC-BAC-2 | Tubastatin A |
| MVP_top60_DNA_seed | HSP90AA1 | 0.492077 | KIRC | BFTC-909 | SNX-2112 | MVP_top60_DNA+RNA_seed | EGFR | -0.17504 | LUAD | NCI-H1838 | Erlotinib |
| MVP_top60_DNA_seed | HSP90AB1 | 0.492077 | KIRC | BFTC-909 | SNX-2112 | MVP_top60_DNA+RNA_seed | HDAC1 | -0.174994 | LUAD | NCI-H2030 | Dacinostat |
| MVP_top60_DNA_seed | SMARCA4 | 0.494497 | KIRC | ACHN | PFI-3 | MVP_top60_DNA+RNA_seed | EGFR | -0.17427 | LUAD | SK-LU-1 | Gefitinib |
| MVP_top60_DNA_seed | HDAC1 | 0.494739 | KIRC | SN12C | AR-42 | MVP_top60_DNA+RNA_seed | SRC | -0.174075 | LUAD | NCI-H1666 | Bosutinib |
| MVP_top60_DNA_seed | MTOR | 0.495631 | KIRC | CAKI-1 | Voxtalisib | MVP_top60_DNA+RNA_seed | EGFR | -0.173758 | LUAD | LXF-289 | Pelitinib |
| MVP_top60_DNA_seed | ATM | 0.496336 | KIRC | KMRC-1 | CP466722 | MVP_top60_DNA+RNA_seed | HDAC1 | -0.173691 | LUAD | LXF-289 | CAY10603 |
| MVP_top60_DNA_seed | SMARCA4 | 0.497987 | KIRC | NCC010 | PFI-3 | MVP_top60_DNA+RNA_seed | EGFR | -0.173503 | LUAD | HOP-62 | AZD8931 |
| MVP_top60_DNA_seed | HDAC1 | 0.499504 | KIRC | LB996-RCC | AR-42 | MVP_top60_DNA+RNA_seed | BRAF | -0.171539 | LUAD | NCI-H2228 | HG6-64-1 |
| MVP_top60_DNA_seed | SMARCA4 | 0.502387 | KIRC | CAKI-1 | PFI-3 | MVP_top60_DNA+RNA_seed | EGFR | -0.171272 | LUAD | NCI-H1355 | Lapatinib |
| MVP_top60_DNA_seed | HDAC1 | 0.505199 | KIRC | LB2241-RCC | CAY10603 | MVP_top60_DNA+RNA_seed | PIK3CA | -0.171097 | LUAD | NCI-H2291 | PI3Ka_4409 |
| MVP_top60_DNA_seed | PARP1 | 0.505559 | KIRC | KMRC-20 | Rucaparib | MVP_top60_DNA+RNA_seed | EGFR | -0.170644 | LUAD | NCI-H3122 | Pelitinib |
| MVP_top60_DNA_seed | ESR1 | 0.507974 | KIRC | SN12C | Tamoxifen | MVP_top60_DNA+RNA_seed | FGFR1 | -0.169764 | LUAD | LXF-289 | AZD4547 |
| MVP_top60_DNA_seed | AR | 0.510566 | KIRC | TK10 | Bicalutamide | MVP_top60_DNA+RNA_seed | HDAC1 | -0.168968 | LUAD | RERF-LC-MS | Belinostat |
| MVP_top60_DNA_seed | PARP1 | 0.515264 | KIRC | RCC-FG2 | Veliparib | MVP_top60_DNA+RNA_seed | EGFR | -0.16888 | LUAD | HCC-44 | CUDC-101 |
| MVP_top60_DNA_seed | ATM | 0.515395 | KIRC | RXF393 | KU-55933 | MVP_top60_DNA+RNA_seed | MDM2 | -0.16704 | LUAD | Calu-6 | Nutlin-3a (-) |
| MVP_top60_DNA_seed | AR | 0.51613 | KIRC | 786-0 | AZD3514 | MVP_top60_DNA+RNA_seed | FGFR1 | -0.166974 | LUAD | RERF-LC-MS | FGFR_3831 |
| MVP_top60_DNA_seed | AR | 0.51877 | KIRC | SW156 | AZD3514 | MVP_top60_DNA+RNA_seed | BRAF | -0.166512 | LUAD | NCI-H322M | PLX-4720 |
| MVP_top60_DNA_seed | ESR1 | 0.521158 | KIRC | RCC-MF | Fulvestrant | MVP_top60_DNA+RNA_seed | EGFR | -0.166289 | LUAD | EMC-BAC-1 | CI-1033 |
| MVP_top60_DNA_seed | FGFR4 | 0.522441 | KIRC | A498 | FGFR_0939 | MVP_top60_DNA+RNA_seed | AURKA | -0.165527 | LUAD | NCI-H1944 | CD532 |
| MVP_top60_DNA_seed | PARP1 | 0.523577 | KIRC | A704 | PARP_0108 | MVP_top60_DNA+RNA_seed | IGF1R | -0.165383 | LUAD | NCI-H2009 | BMS-536924 |

| | | | | | | | | | | | |
|---|---|---|---|---|---|---|---|---|---|---|---|
| MVP_top60_DNA_seed | HDAC1 | 0.526035 | KIRC | SN12C | CAY10603 | MVP_top60_DNA+RNA_seed | AKT1 | -0.165204 | LUAD | COR-L105 | AKT inhibitor VIII |
| MVP_top60_DNA_seed | IGF1R | 0.531131 | KIRC | KMRC-1 | BMS-754807 | MVP_top60_DNA+RNA_seed | PIK3CA | -0.16513 | LUAD | NCI-H1755 | PI3Ka_4409 |
| MVP_top60_DNA_seed | PARP1 | 0.531881 | KIRC | ACHN | Veliparib | MVP_top60_DNA+RNA_seed | PTK2 | -0.162876 | LUAD | NCI-H1568 | PF-562271 |
| MVP_top60_DNA_seed | PARP1 | 0.533033 | KIRC | LB2241-RCC | Olaparib | MVP_top60_DNA+RNA_seed | HSP90AA1 | -0.160389 | LUAD | NCI-H2405 | Tanespimycin |
| MVP_top60_DNA_seed | PIK3CD | 0.53466 | KIRC | SN12C | Idelalisib | MVP_top60_DNA+RNA_seed | HSP90AA1 | -0.160351 | LUAD | NCI-H2291 | Tanespimycin |
| MVP_top60_DNA_seed | ESR1 | 0.534732 | KIRC | RCC-MF | Tamoxifen | MVP_top60_DNA+RNA_seed | BRAF | -0.160091 | LUAD | H3255 | HG6-64-1 |
| MVP_top60_DNA_seed | PIK3CA | 0.537162 | KIRC | 769-P | AZD8835 | MVP_top60_DNA+RNA_seed | EGFR | -0.159555 | LUAD | NCI-H2030 | Gefitinib |
| MVP_top60_DNA_seed | PIK3CD | 0.537162 | KIRC | 769-P | AZD8835 | MVP_top60_DNA+RNA_seed | HDAC1 | -0.158975 | LUAD | NCI-H2023 | JQ12 |
| MVP_top60_DNA_seed | HSP90AA1 | 0.537846 | KIRC | NCC021 | Elesclomol | MVP_top60_DNA+RNA_seed | EGFR | -0.158541 | LUAD | HOP-62 | Pelitinib |
| MVP_top60_DNA_seed | HSP90AB1 | 0.537846 | KIRC | NCC021 | Elesclomol | MVP_top60_DNA+RNA_seed | AKT1 | -0.15815 | LUAD | HOP-62 | AKT inhibitor VIII |
| MVP_top60_DNA_seed | EP300 | 0.537992 | KIRC | HA7-RCC | I-CBP112 | MVP_top60_DNA+RNA_seed | IGF1R | -0.157391 | LUAD | NCI-H2087 | BMS-754807 |
| MVP_top60_DNA_seed | AR | 0.539733 | KIRC | ACHN | Bicalutamide | MVP_top60_DNA+RNA_seed | SRC | -0.156846 | LUAD | HOP-62 | NVP-BHG712 |
| MVP_top60_DNA_seed | IGF1R | 0.549114 | KIRC | CAL-54 | BMS-754807 | MVP_top60_DNA+RNA_seed | AKT1 | -0.156247 | LUAD | NCI-H1355 | MK-2206 |
| MVP_top60_DNA_seed | PIK3CD | 0.550738 | KIRC | LB1047-RCC | IC-87114 | MVP_top60_DNA+RNA_seed | EGFR | -0.156022 | LUAD | NCI-H1435 | PF-00299804 |
| MVP_top60_DNA_seed | ATM | 0.555257 | KIRC | 769-P | KU-55933 | MVP_top60_DNA+RNA_seed | MET | -0.155097 | LUAD | VMRC-LCD | AZD6094 |
| MVP_top60_DNA_seed | FGFR4 | 0.558079 | KIRC | U031 | FGFR_3831 | MVP_top60_DNA+RNA_seed | EGFR | -0.15483 | LUAD | HOP-62 | Lapatinib |
| MVP_top60_DNA_seed | ESR1 | 0.559316 | KIRC | RCC-JF | Tamoxifen | MVP_top60_DNA+RNA_seed | MAPK14 | -0.154526 | LUAD | NCI-H1355 | TAK-715 |
| MVP_top60_DNA_seed | HDAC1 | 0.56114 | KIRC | RCC-JW | Tubastatin A | MVP_top60_DNA+RNA_seed | IGF1R | -0.152819 | LUAD | SW1573 | BMS-754807 |
| MVP_top60_DNA_seed | HSP90AA1 | 0.56261 | KIRC | RCC-ER | Tanespimycin | MVP_top60_DNA+RNA_seed | EGFR | -0.151868 | LUAD | NCI-H1563 | Afatinib |
| MVP_top60_DNA_seed | HSP90AB1 | 0.56261 | KIRC | RCC-ER | Tanespimycin | MVP_top60_DNA+RNA_seed | FGFR1 | -0.149514 | LUAD | PC-14 | PD173074 |
| MVP_top60_DNA_seed | AR | 0.56316 | KIRC | BB65-RCC | Bicalutamide | MVP_top60_DNA+RNA_seed | EGFR | -0.149422 | LUAD | NCI-H1944 | Gefitinib |
| MVP_top60_DNA_seed | IGF1R | 0.563864 | KIRC | CAKI-1 | BMS-536924 | MVP_top60_DNA+RNA_seed | IGF1R | -0.14871 | LUAD | NCI-H1648 | GSK1904529A |
| MVP_top60_DNA_seed | SMARCA4 | 0.569708 | KIRC | RCC-ER | PFI-3 | MVP_top60_DNA+RNA_seed | ATM | -0.148524 | LUAD | NCI-H1944 | KU-60019 |
| MVP_top60_DNA_seed | HDAC1 | 0.570152 | KIRC | RCC-ER | CAY10603 | MVP_top60_DNA+RNA_seed | HDAC1 | -0.148386 | LUAD | LXF-289 | Belinostat |
| MVP_top60_DNA_seed | PIK3CD | 0.571295 | KIRC | RXF393 | IC-87114 | MVP_top60_DNA+RNA_seed | ATM | -0.148223 | LUAD | EMC-BAC-1 | CP466722 |
| MVP_top60_DNA_seed | AR | 0.572224 | KIRC | A704 | AZD3514 | MVP_top60_DNA+RNA_seed | BRAF | -0.14743 | LUAD | NCI-H23 | SB590885 |
| MVP_top60_DNA_seed | HSP90AA1 | 0.572348 | KIRC | SW156 | Tanespimycin | MVP_top60_DNA+RNA_seed | HDAC1 | -0.14683 | LUAD | A549 | JQ12 |
| MVP_top60_DNA_seed | HSP90AB1 | 0.572348 | KIRC | SW156 | Tanespimycin | MVP_top60_DNA+RNA_seed | FGFR1 | -0.146012 | LUAD | NCI-H2228 | FGFR_3831 |
| MVP_top60_DNA_seed | AR | 0.572749 | KIRC | KMRC-20 | Bicalutamide | MVP_top60_DNA+RNA_seed | EGFR | -0.145112 | LUAD | NCI-H1623 | CUDC-101 |
| MVP_top60_DNA_seed | HSP90AA1 | 0.572821 | KIRC | KMRC-1 | SNX-2112 | MVP_top60_DNA+RNA_seed | BRAF | -0.144388 | LUAD | NCI-H2347 | RAF_9304 |
| MVP_top60_DNA_seed | HSP90AB1 | 0.572821 | KIRC | KMRC-1 | SNX-2112 | MVP_top60_DNA+RNA_seed | HDAC1 | -0.143669 | LUAD | EKVX | JQ12 |
| MVP_top60_DNA_seed | IGF1R | 0.57292 | KIRC | KMRC-20 | BMS-754807 | MVP_top60_DNA+RNA_seed | BRAF | -0.143143 | LUAD | LXF-289 | RAF_9304 |
| MVP_top60_DNA_seed | PTK2 | 0.573075 | KIRC | BFTC-909 | PF-562271 | MVP_top60_DNA+RNA_seed | BRAF | -0.142131 | LUAD | NCI-H1568 | PLX-4720 |
| MVP_top60_DNA_seed | PARP1 | 0.57486 | KIRC | SW156 | PARP_9495 | MVP_top60_DNA+RNA_seed | AKT1 | -0.141885 | LUAD | NCI-H1944 | GSK690693 |
| MVP_top60_DNA_seed | HDAC1 | 0.575334 | KIRC | VMRC-RCW | Tubastatin A | MVP_top60_DNA+RNA_seed | EGFR | -0.141755 | LUAD | HCC-78 | Afatinib |
| MVP_top60_DNA_seed | PARP1 | 0.577121 | KIRC | TK10 | PARP_9482 | MVP_top60_DNA+RNA_seed | FGFR1 | -0.141636 | LUAD | PC-14 | FGFR_3831 |
| MVP_top60_DNA_seed | HDAC1 | 0.577764 | KIRC | LB996-RCC | JQ12 | MVP_top60_DNA+RNA_seed | MDM2 | -0.141423 | LUAD | NCI-H23 | Nutlin-3a (-) |
| MVP_top60_DNA_seed | AR | 0.58066 | KIRC | SW156 | Bicalutamide | MVP_top60_DNA+RNA_seed | ATM | -0.141313 | LUAD | PC-14 | torin2 |
| MVP_top60_DNA_seed | IGF1R | 0.580986 | KIRC | RCC10RGB | BMS-754807 | MVP_top60_DNA+RNA_seed | HDAC1 | -0.140213 | LUAD | NCI-H23 | Belinostat |
| MVP_top60_DNA_seed | PARP1 | 0.581805 | KIRC | TK10 | PARP_0108 | MVP_top60_DNA+RNA_seed | EGFR | -0.139195 | LUAD | NCI-H1944 | PF-00299804 |
| MVP_top60_DNA_seed | MTOR | 0.582225 | KIRC | VMRC-RCW | Voxtalisib | MVP_top60_DNA+RNA_seed | MAPK14 | -0.138875 | LUAD | NCI-H2030 | TAK-715 |

| | | | | | | | | | | | |
|---|---|---|---|---|---|---|---|---|---|---|---|
| MVP_top60_DNA_seed | HSP90AA1 | 0.583768 | KIRC | VMRC-RCZ | Elesclomol | MVP_top60_DNA+RNA_seed | EGFR | -0.138089 | LUAD | NCI-H1573 | AZD8931 |
| MVP_top60_DNA_seed | HSP90AB1 | 0.583768 | KIRC | VMRC-RCZ | Elesclomol | MVP_top60_DNA+RNA_seed | EGFR | -0.138058 | LUAD | Calu-6 | PF-00299804 |
| MVP_top60_DNA_seed | AR | 0.583842 | KIRC | VMRC-RCZ | Bicalutamide | MVP_top60_DNA+RNA_seed | SRC | -0.138032 | LUAD | NCI-H2009 | NVP-BHG712 |
| MVP_top60_DNA_seed | IGF1R | 0.58521 | KIRC | SN12C | BMS-754807 | MVP_top60_DNA+RNA_seed | EGFR | -0.137569 | LUAD | NCI-H650 | CUDC-101 |
| MVP_top60_DNA_seed | PARP1 | 0.586286 | KIRC | NCC021 | PARP_0108 | MVP_top60_DNA+RNA_seed | SRC | -0.136877 | LUAD | NCI-H2023 | Bosutinib |
| MVP_top60_DNA_seed | AR | 0.587555 | KIRC | SN12C | Bicalutamide | MVP_top60_DNA+RNA_seed | AKT1 | -0.13635 | LUAD | NCI-H2405 | AKT inhibitor VIII |
| MVP_top60_DNA_seed | ATM | 0.589285 | KIRC | TK10 | KU-60019 | MVP_top60_DNA+RNA_seed | EGFR | -0.135806 | LUAD | NCI-H1944 | Afatinib |
| MVP_top60_DNA_seed | PARP1 | 0.58935 | KIRC | U031 | Olaparib | MVP_top60_DNA+RNA_seed | ATM | -0.135725 | LUAD | EMC-BAC-1 | KU-60019 |
| MVP_top60_DNA_seed | HSP90AA1 | 0.590802 | KIRC | SW156 | CCT-018159 | MVP_top60_DNA+RNA_seed | AKT1 | -0.1354 | LUAD | EKVX | MK-2206 |
| MVP_top60_DNA_seed | HSP90AB1 | 0.590802 | KIRC | SW156 | CCT-018159 | MVP_top60_DNA+RNA_seed | AKT1 | -0.134984 | LUAD | RERF-LC-MS | AKT inhibitor VIII |
| MVP_top60_DNA_seed | PIK3CA | 0.592257 | KIRC | CAKI-1 | PI3Ka_4409 | MVP_top60_DNA+RNA_seed | HDAC1 | -0.131321 | LUAD | NCI-H838 | CAY10603 |
| MVP_top60_DNA_seed | PIK3CD | 0.592257 | KIRC | CAKI-1 | PI3Ka_4409 | MVP_top60_DNA+RNA_seed | HDAC1 | -0.129792 | LUAD | NCI-H2405 | Dacinostat |
| MVP_top60_DNA_seed | HDAC1 | 0.596687 | KIRC | CAKI-1 | Dacinostat | MVP_top60_DNA+RNA_seed | HSP90AA1 | -0.12927 | LUAD | NCI-H1693 | Luminespib |
| MVP_top60_DNA_seed | SMARCA4 | 0.597513 | KIRC | KMRC-20 | PFI-3 | MVP_top60_DNA+RNA_seed | EGFR | -0.128712 | LUAD | NCI-H2087 | PF-00299804 |
| MVP_top60_DNA_seed | PARP1 | 0.598769 | KIRC | VMRC-RCZ | PARP_9482 | MVP_top60_DNA+RNA_seed | EGFR | -0.128246 | LUAD | LXF-289 | Afatinib |
| MVP_top60_DNA_seed | ESR1 | 0.599873 | KIRC | TK10 | Fulvestrant | MVP_top60_DNA+RNA_seed | AURKA | -0.126652 | LUAD | EMC-BAC-2 | Alisertib |
| MVP_top60_DNA_seed | PIK3CD | 0.604406 | KIRC | KMRC-20 | IC-87114 | MVP_top60_DNA+RNA_seed | MET | -0.125826 | LUAD | NCI-H2023 | JNJ38877605 |
| MVP_top60_DNA_seed | IGF1R | 0.60459 | KIRC | BFTC-909 | GSK1904529A | MVP_top60_DNA+RNA_seed | AKT1 | -0.125757 | LUAD | NCI-H2085 | GSK690693 |
| MVP_top60_DNA_seed | ESR1 | 0.610087 | KIRC | CAKI-1 | Fulvestrant | MVP_top60_DNA+RNA_seed | SMARCA4 | -0.124364 | LUAD | HCC-44 | PFI-3 |
| MVP_top60_DNA_seed | SMARCA4 | 0.610283 | KIRC | LB996-RCC | PFI-3 | MVP_top60_DNA+RNA_seed | BRAF | -0.122172 | LUAD | NCI-H1355 | HG6-64-1 |
| MVP_top60_DNA_seed | ESR1 | 0.61112 | KIRC | NCC021 | Tamoxifen | MVP_top60_DNA+RNA_seed | BRAF | -0.122092 | LUAD | H3255 | SB590885 |
| MVP_top60_DNA_seed | AR | 0.611362 | KIRC | SN12C | Bicalutamide | MVP_top60_DNA+RNA_seed | HSP90AA1 | -0.12174 | LUAD | LXF-289 | SNX-2112 |
| MVP_top60_DNA_seed | MTOR | 0.612345 | KIRC | RCC-AB | Temsirolimus | MVP_top60_DNA+RNA_seed | AURKA | -0.121274 | LUAD | HCC-78 | ZM447439 |
| MVP_top60_DNA_seed | ATM | 0.612393 | KIRC | RCC10RGB | KU-60019 | MVP_top60_DNA+RNA_seed | EGFR | -0.12069 | LUAD | NCI-H1355 | Gefitinib |
| MVP_top60_DNA_seed | PARP1 | 0.61396 | KIRC | SN12C | PARP_9495 | MVP_top60_DNA+RNA_seed | AURKA | -0.119206 | LUAD | HCC-44 | GSK1070916 |
| MVP_top60_DNA_seed | PARP1 | 0.61495 | KIRC | RCC-ER | Veliparib | MVP_top60_DNA+RNA_seed | EGFR | -0.118356 | LUAD | NCI-H1944 | AST-1306 |
| MVP_top60_DNA_seed | ATM | 0.620442 | KIRC | ACHN | KU-55933 | MVP_top60_DNA+RNA_seed | HSP90AA1 | -0.11782 | LUAD | Calu-3 | Luminespib |
| MVP_top60_DNA_seed | FGFR4 | 0.621739 | KIRC | VMRC-RCZ | FGFR_3831 | MVP_top60_DNA+RNA_seed | ATM | -0.11774 | LUAD | EMC-BAC-1 | KU-55933 |
| MVP_top60_DNA_seed | HSP90AA1 | 0.623264 | KIRC | HA7-RCC | SNX-2112 | MVP_top60_DNA+RNA_seed | AKT1 | -0.11683 | LUAD | Calu-3 | MK-2206 |
| MVP_top60_DNA_seed | HSP90AB1 | 0.623264 | KIRC | HA7-RCC | SNX-2112 | MVP_top60_DNA+RNA_seed | HDAC1 | -0.116341 | LUAD | NCI-H1437 | CAY10603 |
| MVP_top60_DNA_seed | PARP1 | 0.629751 | KIRC | RXF393 | Veliparib | MVP_top60_DNA+RNA_seed | SRC | -0.115851 | LUAD | NCI-H1563 | HG-6-71-01 |
| MVP_top60_DNA_seed | PARP1 | 0.634342 | KIRC | SW156 | PARP_0108 | MVP_top60_DNA+RNA_seed | ATM | -0.115737 | LUAD | HOP-62 | KU-60019 |
| MVP_top60_DNA_seed | PARP1 | 0.63637 | KIRC | 786-0 | PARP_9495 | MVP_top60_DNA+RNA_seed | EGFR | -0.114885 | LUAD | NCI-H1623 | Cetuximab |
| MVP_top60_DNA_seed | ATM | 0.639135 | KIRC | BB65-RCC | KU-60019 | MVP_top60_DNA+RNA_seed | HDAC1 | -0.113477 | LUAD | NCI-H1573 | AR-42 |
| MVP_top60_DNA_seed | PARP1 | 0.640424 | KIRC | RCC-JW | Rucaparib | MVP_top60_DNA+RNA_seed | AKT1 | -0.113025 | LUAD | NCI-H1648 | GSK690693 |
| MVP_top60_DNA_seed | IGF1R | 0.641284 | KIRC | OS-RC-2 | Linsitinib | MVP_top60_DNA+RNA_seed | IGF1R | -0.112736 | LUAD | A549 | Linsitinib |
| MVP_top60_DNA_seed | HSP90AA1 | 0.642705 | KIRC | KMRC-1 | CCT-018159 | MVP_top60_DNA+RNA_seed | EGFR | -0.11196 | LUAD | NCI-H522 | AZD8931 |
| MVP_top60_DNA_seed | HSP90AB1 | 0.642705 | KIRC | KMRC-1 | CCT-018159 | MVP_top60_DNA+RNA_seed | EGFR | -0.111926 | LUAD | NCI-H1792 | Cetuximab |
| MVP_top60_DNA_seed | EP300 | 0.647817 | KIRC | KMRC-1 | I-CBP112 | MVP_top60_DNA+RNA_seed | SRC | -0.111829 | LUAD | LXF-289 | Bosutinib |
| MVP_top60_DNA_seed | PARP1 | 0.649936 | KIRC | VMRC-RCZ | Veliparib | MVP_top60_DNA+RNA_seed | AKT1 | -0.111196 | LUAD | HCC-44 | GSK690693 |
| MVP_top60_DNA_seed | HSP90AA1 | 0.650698 | KIRC | VMRC-RCW | Tanespimycin | MVP_top60_DNA+RNA_seed | HSP90AA1 | -0.11116 | LUAD | NCI-H1975 | SNX-2112 |

| | | | | | | | | | | | |
|---|---|---|---|---|---|---|---|---|---|---|---|
| MVP_top60_DNA_seed | HSP90AB1 | 0.650698 | KIRC | VMRC-RCW | Tanespimycin | MVP_top60_DNA+RNA_seed | HDAC1 | -0.110692 | LUAD | COR-L105 | JQ12 |
| MVP_top60_DNA_seed | IGF1R | 0.650788 | KIRC | HA7-RCC | GSK1904529A | MVP_top60_DNA+RNA_seed | EGFR | -0.110083 | LUAD | NCI-H1792 | AST-1306 |
| MVP_top60_DNA_seed | PARP1 | 0.651679 | KIRC | RCC-ER | Olaparib | MVP_top60_DNA+RNA_seed | PTK2 | -0.109519 | LUAD | RERF-LC-MS | PF-562271 |
| MVP_top60_DNA_seed | HDAC1 | 0.652301 | KIRC | VMRC-RCZ | Tubastatin A | MVP_top60_DNA+RNA_seed | BRAF | -0.109253 | LUAD | Calu-6 | HG6-64-1 |
| MVP_top60_DNA_seed | HDAC1 | 0.655598 | KIRC | RCC10RGB | CAY10603 | MVP_top60_DNA+RNA_seed | EGFR | -0.108825 | LUAD | LC-2-ad | AST-1306 |
| MVP_top60_DNA_seed | HDAC1 | 0.656522 | KIRC | KMRC-20 | Belinostat | MVP_top60_DNA+RNA_seed | AKT1 | -0.108126 | LUAD | NCI-H2023 | AKT inhibitor VIII |
| MVP_top60_DNA_seed | ESR1 | 0.65667 | KIRC | KMRC-20 | Tamoxifen | MVP_top60_DNA+RNA_seed | IGF1R | -0.107424 | LUAD | NCI-H2023 | BMS-536924 |
| MVP_top60_DNA_seed | PARP1 | 0.657019 | KIRC | NCC010 | Rucaparib | MVP_top60_DNA+RNA_seed | PTK2 | -0.1074 | LUAD | HCC-78 | PF-562271 |
| MVP_top60_DNA_seed | PIK3CA | 0.657681 | KIRC | KMRC-20 | AZD8835 | MVP_top60_DNA+RNA_seed | MET | -0.107012 | LUAD | NCI-H1568 | AZD6094 |
| MVP_top60_DNA_seed | PIK3CD | 0.657681 | KIRC | KMRC-20 | AZD8835 | MVP_top60_DNA+RNA_seed | BRAF | -0.106883 | LUAD | NCI-H1944 | RAF_9304 |
| MVP_top60_DNA_seed | MTOR | 0.657814 | KIRC | BFTC-909 | Apitolisib | MVP_top60_DNA+RNA_seed | PIK3CA | -0.106684 | LUAD | A549 | PI3Ka_4409 |
| MVP_top60_DNA_seed | SMARCA4 | 0.658517 | KIRC | RCC-AB | PFI-3 | MVP_top60_DNA+RNA_seed | HDAC1 | -0.106235 | LUAD | HOP-62 | Parthenolide |
| MVP_top60_DNA_seed | PARP1 | 0.659664 | KIRC | KMRC-20 | Talazoparib | MVP_top60_DNA+RNA_seed | MET | -0.105608 | LUAD | NCI-H23 | PHA-665752 |
| MVP_top60_DNA_seed | HDAC1 | 0.66256 | KIRC | LB2241-RCC | AR-42 | MVP_top60_DNA+RNA_seed | BRAF | -0.105565 | LUAD | NCI-H2085 | HG6-64-1 |
| MVP_top60_DNA_seed | HSP90AA1 | 0.662564 | KIRC | CAKI-1 | CCT-018159 | MVP_top60_DNA+RNA_seed | HDAC1 | -0.105466 | LUAD | NCI-H1734 | AR-42 |
| MVP_top60_DNA_seed | HSP90AB1 | 0.662564 | KIRC | CAKI-1 | CCT-018159 | MVP_top60_DNA+RNA_seed | AURKA | -0.105292 | LUAD | NCI-H1650 | ZM447439 |
| MVP_top60_DNA_seed | PARP1 | 0.667423 | KIRC | RCC-FG2 | PARP_0108 | MVP_top60_DNA+RNA_seed | SRC | -0.10519 | LUAD | NCI-H1944 | NVP-BHG712 |
| MVP_top60_DNA_seed | FGFR4 | 0.668249 | KIRC | OS-RC-2 | FGFR_0939 | MVP_top60_DNA+RNA_seed | BRAF | -0.104639 | LUAD | NCI-H1651 | HG6-64-1 |
| MVP_top60_DNA_seed | IGF1R | 0.668936 | KIRC | KMRC-1 | BMS-536924 | MVP_top60_DNA+RNA_seed | ATM | -0.104426 | LUAD | HCC-78 | CP466722 |
| MVP_top60_DNA_seed | IGF1R | 0.670099 | KIRC | VMRC-RCZ | BMS-536924 | MVP_top60_DNA+RNA_seed | EGFR | -0.104168 | LUAD | HCC-78 | PF-00299804 |
| MVP_top60_DNA_seed | HSP90AA1 | 0.670993 | KIRC | A704 | Tanespimycin | MVP_top60_DNA+RNA_seed | SRC | -0.103125 | LUAD | NCI-H23 | NVP-BHG712 |
| MVP_top60_DNA_seed | HSP90AB1 | 0.670993 | KIRC | A704 | Tanespimycin | MVP_top60_DNA+RNA_seed | SMARCA4 | -0.101686 | LUAD | Calu-3 | PFI-3 |
| MVP_top60_DNA_seed | PTK2 | 0.686751 | KIRC | SW156 | PF-562271 | MVP_top60_DNA+RNA_seed | IGF1R | -0.101395 | LUAD | NCI-H1792 | GSK1904529A |
| MVP_top60_DNA_seed | PARP1 | 0.688024 | KIRC | BB65-RCC | Olaparib | MVP_top60_DNA+RNA_seed | ATM | -0.100832 | LUAD | HOP-62 | CP466722 |
| MVP_top60_DNA_seed | ESR1 | 0.690444 | KIRC | RCC-FG2 | Fulvestrant | MVP_top60_DNA+RNA_seed | HDAC1 | -0.100681 | LUAD | NCI-H838 | Belinostat |
| MVP_top60_DNA_seed | ATM | 0.693701 | KIRC | KMRC-20 | CP466722 | MVP_top60_DNA+RNA_seed | EGFR | -0.09912 | LUAD | LC-2-ad | Cetuximab |
| MVP_top60_DNA_seed | PIK3CA | 0.695183 | KIRC | SW156 | AZD8835 | MVP_top60_DNA+RNA_seed | PTK2 | -0.097558 | LUAD | NCI-H2085 | PF-562271 |
| MVP_top60_DNA_seed | PIK3CD | 0.695183 | KIRC | SW156 | AZD8835 | MVP_top60_DNA+RNA_seed | EGFR | -0.097104 | LUAD | NCI-H2347 | PF-00299804 |
| MVP_top60_DNA_seed | SMARCA4 | 0.695263 | KIRC | KMRC-1 | PFI-3 | MVP_top60_DNA+RNA_seed | EGFR | -0.095243 | LUAD | HOP-62 | CI-1033 |
| MVP_top60_DNA_seed | PARP1 | 0.699191 | KIRC | LB1047-RCC | Veliparib | MVP_top60_DNA+RNA_seed | FGFR1 | -0.094244 | LUAD | HCC-44 | AZD4547 |
| MVP_top60_DNA_seed | PARP1 | 0.700463 | KIRC | TK10 | Rucaparib | MVP_top60_DNA+RNA_seed | SRC | -0.093479 | LUAD | LC-2-ad | NVP-BHG712 |
| MVP_top60_DNA_seed | HSP90AA1 | 0.704135 | KIRC | VMRC-RCZ | SNX-2112 | MVP_top60_DNA+RNA_seed | MDM2 | -0.093083 | LUAD | NCI-H2030 | Nutlin-3a (-) |
| MVP_top60_DNA_seed | HSP90AB1 | 0.704135 | KIRC | VMRC-RCZ | SNX-2112 | MVP_top60_DNA+RNA_seed | MET | -0.092929 | LUAD | HCC-78 | AZD6094 |
| MVP_top60_DNA_seed | MTOR | 0.70542 | KIRC | OS-RC-2 | Voxtalisib | MVP_top60_DNA+RNA_seed | AURKA | -0.090412 | LUAD | NCI-H1792 | GSK1070916 |
| MVP_top60_DNA_seed | PARP1 | 0.70841 | KIRC | OS-RC-2 | Olaparib | MVP_top60_DNA+RNA_seed | EGFR | -0.08911 | LUAD | SK-LU-1 | Pelitinib |
| MVP_top60_DNA_seed | PARP1 | 0.710111 | KIRC | NCC010 | Veliparib | MVP_top60_DNA+RNA_seed | FGFR1 | -0.088425 | LUAD | NCI-H1944 | AZD4547 |
| MVP_top60_DNA_seed | AR | 0.711723 | KIRC | OS-RC-2 | Bicalutamide | MVP_top60_DNA+RNA_seed | AURKA | -0.088163 | LUAD | NCI-H1944 | HG-5-113-01 |
| MVP_top60_DNA_seed | SMARCA4 | 0.712956 | KIRC | NCC021 | PFI-3 | MVP_top60_DNA+RNA_seed | MDM2 | -0.088098 | LUAD | VMRC-LCD | Serdemetan |
| MVP_top60_DNA_seed | ATM | 0.714991 | KIRC | VMRC-RCW | CP466722 | MVP_top60_DNA+RNA_seed | AKT1 | -0.086511 | LUAD | RERF-LC-MS | BAY AKT1 |
| MVP_top60_DNA_seed | MTOR | 0.720702 | KIRC | KMRC-20 | WYE-125132 | MVP_top60_DNA+RNA_seed | AKT1 | -0.085574 | LUAD | NCI-H650 | AKT inhibitor VIII |
| MVP_top60_DNA_seed | EP300 | 0.720728 | KIRC | ACHN | I-CBP112 | MVP_top60_DNA+RNA_seed | SRC | -0.084794 | LUAD | SW1573 | HG-6-71-01 |

| | | | | | | | | | | | |
|---|---|---|---|---|---|---|---|---|---|---|---|
| MVP_top60_DNA_seed | ESR1 | 0.726762 | KIRC | ACHN | Tamoxifen | MVP_top60_DNA+RNA_seed | HDAC1 | -0.084097 | LUAD | NCI-H1437 | AR-42 |
| MVP_top60_DNA_seed | PARP1 | 0.727267 | KIRC | SW156 | PARP_9482 | MVP_top60_DNA+RNA_seed | FGFR1 | -0.082982 | LUAD | Calu-6 | FGFR_3831 |
| MVP_top60_DNA_seed | ESR1 | 0.728505 | KIRC | VMRC-RCZ | Tamoxifen | MVP_top60_DNA+RNA_seed | HDAC1 | -0.082448 | LUAD | NCI-H2087 | Belinostat |
| MVP_top60_DNA_seed | IGF1R | 0.730686 | KIRC | SW156 | BMS-754807 | MVP_top60_DNA+RNA_seed | HDAC1 | -0.080712 | LUAD | NCI-H1648 | Dacinostat |
| MVP_top60_DNA_seed | HSP90AA1 | 0.734993 | KIRC | CAKI-1 | SNX-2112 | MVP_top60_DNA+RNA_seed | SRC | -0.080599 | LUAD | NCI-H2085 | Bosutinib |
| MVP_top60_DNA_seed | HSP90AB1 | 0.734993 | KIRC | CAKI-1 | SNX-2112 | MVP_top60_DNA+RNA_seed | HDAC1 | -0.080162 | LUAD | LXF-289 | AR-42 |
| MVP_top60_DNA_seed | MTOR | 0.735312 | KIRC | KMRC-1 | CRT0105446 | MVP_top60_DNA+RNA_seed | AURKA | -0.079302 | LUAD | NCI-H838 | GSK1070916 |
| MVP_top60_DNA_seed | FGFR4 | 0.736576 | KIRC | ACHN | FGFR_0939 | MVP_top60_DNA+RNA_seed | MET | -0.079123 | LUAD | EMC-BAC-2 | JNJ38877605 |
| MVP_top60_DNA_seed | ATM | 0.738205 | KIRC | BB65-RCC | KU-55933 | MVP_top60_DNA+RNA_seed | FGFR1 | -0.078476 | LUAD | NCI-H1666 | PD173074 |
| MVP_top60_DNA_seed | HSP90AA1 | 0.743747 | KIRC | ACHN | SNX-2112 | MVP_top60_DNA+RNA_seed | EGFR | -0.078303 | LUAD | NCI-H1792 | CI-1033 |
| MVP_top60_DNA_seed | HSP90AB1 | 0.743747 | KIRC | ACHN | SNX-2112 | MVP_top60_DNA+RNA_seed | MET | -0.077802 | LUAD | PC-14 | JNJ38877605 |
| MVP_top60_DNA_seed | PARP1 | 0.754222 | KIRC | VMRC-RCZ | PARP_0108 | MVP_top60_DNA+RNA_seed | ATM | -0.075855 | LUAD | A549 | CP466722 |
| MVP_top60_DNA_seed | EP300 | 0.757749 | KIRC | KMRC-20 | I-CBP112 | MVP_top60_DNA+RNA_seed | AURKA | -0.075733 | LUAD | SK-LU-1 | GSK1070916 |
| MVP_top60_DNA_seed | PARP1 | 0.758588 | KIRC | LB1047-RCC | Olaparib | MVP_top60_DNA+RNA_seed | FGFR1 | -0.075472 | LUAD | NCI-H1693 | AZD4547 |
| MVP_top60_DNA_seed | IGF1R | 0.758986 | KIRC | KMRC-1 | Linsitinib | MVP_top60_DNA+RNA_seed | PTK2 | -0.075042 | LUAD | NCI-H1781 | PF-562271 |
| MVP_top60_DNA_seed | HSP90AA1 | 0.765625 | KIRC | VMRC-RCW | SNX-2112 | MVP_top60_DNA+RNA_seed | AURKA | -0.073935 | LUAD | NCI-H1648 | ZM447439 |
| MVP_top60_DNA_seed | HSP90AB1 | 0.765625 | KIRC | VMRC-RCW | SNX-2112 | MVP_top60_DNA+RNA_seed | HSP90AA1 | -0.072311 | LUAD | NCI-H2405 | Elesclomol |
| MVP_top60_DNA_seed | PARP1 | 0.766862 | KIRC | KMRC-1 | Olaparib | MVP_top60_DNA+RNA_seed | AURKA | -0.072279 | LUAD | NCI-H23 | ZM447439 |
| MVP_top60_DNA_seed | MTOR | 0.769037 | KIRC | KMRC-20 | Voxtalisib | MVP_top60_DNA+RNA_seed | IGF1R | -0.071287 | LUAD | NCI-H23 | BMS-536924 |
| MVP_top60_DNA_seed | HDAC1 | 0.77053 | KIRC | LB996-RCC | Belinostat | MVP_top60_DNA+RNA_seed | AKT1 | -0.069534 | LUAD | NCI-H358 | GSK690693 |
| MVP_top60_DNA_seed | IGF1R | 0.77547 | KIRC | SN12C | GSK1904529A | MVP_top60_DNA+RNA_seed | HDAC1 | -0.068977 | LUAD | NCI-H2087 | JQ12 |
| MVP_top60_DNA_seed | PARP1 | 0.77576 | KIRC | OS-RC-2 | Rucaparib | MVP_top60_DNA+RNA_seed | MAPK14 | -0.068747 | LUAD | LC-2-ad | TAK-715 |
| MVP_top60_DNA_seed | MTOR | 0.775958 | KIRC | RCC10RGB | WYE-125132 | MVP_top60_DNA+RNA_seed | MAPK14 | -0.068287 | LUAD | NCI-H2342 | TAK-715 |
| MVP_top60_DNA_seed | HSP90AA1 | 0.781887 | KIRC | NCC021 | SNX-2112 | MVP_top60_DNA+RNA_seed | ATM | -0.068203 | LUAD | NCI-H1755 | KU-55933 |
| MVP_top60_DNA_seed | HSP90AB1 | 0.781887 | KIRC | NCC021 | SNX-2112 | MVP_top60_DNA+RNA_seed | HSP90AA1 | -0.068129 | LUAD | HCC-827 | Elesclomol |
| MVP_top60_DNA_seed | MTOR | 0.782525 | KIRC | RCC-MF | QL-VIII-58 | MVP_top60_DNA+RNA_seed | HSP90AA1 | -0.068085 | LUAD | EKVX | Elesclomol |
| MVP_top60_DNA_seed | IGF1R | 0.78354 | KIRC | A498 | BMS-754807 | MVP_top60_DNA+RNA_seed | HDAC1 | -0.067442 | LUAD | NCI-H1666 | Belinostat |
| MVP_top60_DNA_seed | MTOR | 0.790226 | KIRC | KMRC-20 | QL-VIII-58 | MVP_top60_DNA+RNA_seed | HDAC1 | -0.066834 | LUAD | RERF-LC-MS | AR-42 |
| MVP_top60_DNA_seed | ATM | 0.791789 | KIRC | KMRC-1 | KU-60019 | MVP_top60_DNA+RNA_seed | EGFR | -0.065994 | LUAD | NCI-H2291 | AST-1306 |
| MVP_top60_DNA_seed | HSP90AA1 | 0.791947 | KIRC | RCC-JW | SNX-2112 | MVP_top60_DNA+RNA_seed | BRAF | -0.064811 | LUAD | VMRC-LCD | SB590885 |
| MVP_top60_DNA_seed | HSP90AB1 | 0.791947 | KIRC | RCC-JW | SNX-2112 | MVP_top60_DNA+RNA_seed | FGFR1 | -0.06452 | LUAD | NCI-H1793 | AZD4547 |
| MVP_top60_DNA_seed | PARP1 | 0.792841 | KIRC | TK10 | Veliparib | MVP_top60_DNA+RNA_seed | SMARCA4 | -0.064269 | LUAD | LXF-289 | PFI-3 |
| MVP_top60_DNA_seed | HDAC1 | 0.794676 | KIRC | CAKI-1 | Tubastatin A | MVP_top60_DNA+RNA_seed | EGFR | -0.063764 | LUAD | LXF-289 | CUDC-101 |
| MVP_top60_DNA_seed | IGF1R | 0.800728 | KIRC | SN12C | BMS-536924 | MVP_top60_DNA+RNA_seed | AKT1 | -0.063253 | LUAD | NCI-H23 | GSK690693 |
| MVP_top60_DNA_seed | PTK2 | 0.80618 | KIRC | LB996-RCC | PF-562271 | MVP_top60_DNA+RNA_seed | EGFR | -0.062741 | LUAD | SW1573 | Cetuximab |
| MVP_top60_DNA_seed | MTOR | 0.808347 | KIRC | LB996-RCC | WYE-125132 | MVP_top60_DNA+RNA_seed | AKT1 | -0.062579 | LUAD | HCC-78 | MK-2206 |
| MVP_top60_DNA_seed | SMARCA4 | 0.808487 | KIRC | TK10 | PFI-3 | MVP_top60_DNA+RNA_seed | MET | -0.06207 | LUAD | NCI-H522 | AZD6094 |
| MVP_top60_DNA_seed | AR | 0.808629 | KIRC | CAKI-1 | Bicalutamide | MVP_top60_DNA+RNA_seed | PIK3CA | -0.061745 | LUAD | SK-LU-1 | AZD8835 |
| MVP_top60_DNA_seed | SMARCA4 | 0.813777 | KIRC | RCC-FG2 | PFI-3 | MVP_top60_DNA+RNA_seed | AURKA | -0.060406 | LUAD | NCI-H1648 | CD532 |
| MVP_top60_DNA_seed | ESR1 | 0.815815 | KIRC | KMRC-1 | Fulvestrant | MVP_top60_DNA+RNA_seed | BRAF | -0.058737 | LUAD | NCI-H1944 | HG6-64-1 |
| MVP_top60_DNA_seed | PIK3CD | 0.817737 | KIRC | RCC10RGB | Idelalisib | MVP_top60_DNA+RNA_seed | FGFR1 | -0.058704 | LUAD | NCI-H1563 | AZD4547 |

| | | | | | | | | | | | |
|---|---|---|---|---|---|---|---|---|---|---|---|
| MVP_top60_DNA_seed | FGFR4 | 0.824612 | KIRC | TK10 | FGFR_3831 | MVP_top60_DNA+RNA_seed | AURKA | -0.058351 | LUAD | PC-14 | Alisertib |
| MVP_top60_DNA_seed | MTOR | 0.825966 | KIRC | VMRC-RCW | Torin 2 | MVP_top60_DNA+RNA_seed | MET | -0.058205 | LUAD | NCI-H1435 | AZD6094 |
| MVP_top60_DNA_seed | HDAC1 | 0.82818 | KIRC | SN12C | Belinostat | MVP_top60_DNA+RNA_seed | AKT1 | -0.058031 | LUAD | NCI-H1623 | AKT inhibitor VIII |
| MVP_top60_DNA_seed | PARP1 | 0.833361 | KIRC | NCC010 | Olaparib | MVP_top60_DNA+RNA_seed | BRAF | -0.05781 | LUAD | NCI-H2023 | PLX-4720 |
| MVP_top60_DNA_seed | PIK3CA | 0.835264 | KIRC | RXF393 | PI3Ka_4409 | MVP_top60_DNA+RNA_seed | EGFR | -0.05778 | LUAD | HCC-44 | Pelitinib |
| MVP_top60_DNA_seed | PIK3CD | 0.835264 | KIRC | RXF393 | PI3Ka_4409 | MVP_top60_DNA+RNA_seed | HSP90AA1 | -0.056342 | LUAD | NCI-H2228 | Elesclomol |
| MVP_top60_DNA_seed | PARP1 | 0.842629 | KIRC | NCC021 | Rucaparib | MVP_top60_DNA+RNA_seed | BRAF | -0.055686 | LUAD | NCI-H1573 | Dabrafenib |
| MVP_top60_DNA_seed | EP300 | 0.843744 | KIRC | VMRC-RCZ | I-CBP112 | MVP_top60_DNA+RNA_seed | HSP90AA1 | -0.055172 | LUAD | NCI-H1666 | CCT-018159 |
| MVP_top60_DNA_seed | PARP1 | 0.844513 | KIRC | VMRC-RCZ | Talazoparib | MVP_top60_DNA+RNA_seed | PIK3CA | -0.053497 | LUAD | LC-2-ad | PI3Ka_4409 |
| MVP_top60_DNA_seed | HDAC1 | 0.850195 | KIRC | TK10 | Tubastatin A | MVP_top60_DNA+RNA_seed | MAPK14 | -0.053013 | LUAD | NCI-H522 | TAK-715 |
| MVP_top60_DNA_seed | PARP1 | 0.850591 | KIRC | RCC-FG2 | Olaparib | MVP_top60_DNA+RNA_seed | AKT1 | -0.051924 | LUAD | H3255 | BAY AKT1 |
| MVP_top60_DNA_seed | PIK3CA | 0.852257 | KIRC | KMRC-20 | PI3Ka_4409 | MVP_top60_DNA+RNA_seed | HDAC1 | -0.05048 | LUAD | HCC-78 | Tubastatin A |
| MVP_top60_DNA_seed | PIK3CD | 0.852257 | KIRC | KMRC-20 | PI3Ka_4409 | MVP_top60_DNA+RNA_seed | BRAF | -0.050473 | LUAD | PC-3_[JPC-3] | RAF_9304 |
| MVP_top60_DNA_seed | HSP90AA1 | 0.853442 | KIRC | CAL-54 | Tanespimycin | MVP_top60_DNA+RNA_seed | AURKA | -0.050425 | LUAD | NCI-H1563 | HG-5-113-01 |
| MVP_top60_DNA_seed | HSP90AB1 | 0.853442 | KIRC | CAL-54 | Tanespimycin | MVP_top60_DNA+RNA_seed | HDAC1 | -0.050074 | LUAD | NCI-H23 | AR-42 |
| MVP_top60_DNA_seed | PARP1 | 0.854163 | KIRC | LB1047-RCC | Olaparib | MVP_top60_DNA+RNA_seed | IGF1R | -0.04824 | LUAD | NCI-H2405 | BMS-754807 |
| MVP_top60_DNA_seed | PARP1 | 0.856694 | KIRC | KMRC-20 | Veliparib | MVP_top60_DNA+RNA_seed | PIK3CA | -0.048167 | LUAD | EMC-BAC-1 | PI3Ka_4409 |
| MVP_top60_DNA_seed | PARP1 | 0.857424 | KIRC | RCC-MF | Olaparib | MVP_top60_DNA+RNA_seed | AURKA | -0.047776 | LUAD | RERF-LC-KJ | ZM447439 |
| MVP_top60_DNA_seed | HDAC1 | 0.860175 | KIRC | RXF393 | Parthenolide | MVP_top60_DNA+RNA_seed | HDAC1 | -0.047074 | LUAD | NCI-H1437 | Belinostat |
| MVP_top60_DNA_seed | HDAC1 | 0.86166 | KIRC | LB996-RCC | Parthenolide | MVP_top60_DNA+RNA_seed | AURKA | -0.046747 | LUAD | NCI-H2030 | GSK1070916 |
| MVP_top60_DNA_seed | SMARCA4 | 0.862322 | KIRC | VMRC-RCZ | PFI-3 | MVP_top60_DNA+RNA_seed | AURKA | -0.04667 | LUAD | COR-L105 | CD532 |
| MVP_top60_DNA_seed | PIK3CA | 0.863319 | KIRC | RXF393 | AZD8835 | MVP_top60_DNA+RNA_seed | ATM | -0.045788 | LUAD | NCI-H650 | CP466722 |
| MVP_top60_DNA_seed | PIK3CD | 0.863319 | KIRC | RXF393 | AZD8835 | MVP_top60_DNA+RNA_seed | AURKA | -0.044808 | LUAD | COR-L105 | GSK1070916 |
| MVP_top60_DNA_seed | EP300 | 0.864749 | KIRC | BFTC-909 | I-CBP112 | MVP_top60_DNA+RNA_seed | MET | -0.0448 | LUAD | NCI-H358 | JNJ38877605 |
| MVP_top60_DNA_seed | IGF1R | 0.865133 | KIRC | KMRC-20 | Linsitinib | MVP_top60_DNA+RNA_seed | BRAF | -0.043217 | LUAD | NCI-H2122 | PLX-4720 |
| MVP_top60_DNA_seed | HDAC1 | 0.866579 | KIRC | KMRC-20 | Dacinostat | MVP_top60_DNA+RNA_seed | AURKA | -0.041592 | LUAD | NCI-H2087 | HG-5-113-01 |
| MVP_top60_DNA_seed | HSP90AA1 | 0.86672 | KIRC | CAKI-1 | Luminespib | MVP_top60_DNA+RNA_seed | AKT1 | -0.040953 | LUAD | NCI-H2342 | MK-2206 |
| MVP_top60_DNA_seed | HSP90AB1 | 0.86672 | KIRC | CAKI-1 | Luminespib | MVP_top60_DNA+RNA_seed | BRAF | -0.04087 | LUAD | NCI-H650 | HG6-64-1 |
| MVP_top60_DNA_seed | PARP1 | 0.866894 | KIRC | VMRC-RCZ | Olaparib | MVP_top60_DNA+RNA_seed | AURKA | -0.038891 | LUAD | NCI-H1734 | GSK1070916 |
| MVP_top60_DNA_seed | PARP1 | 0.867003 | KIRC | NCC010 | Olaparib | MVP_top60_DNA+RNA_seed | MET | -0.038381 | LUAD | NCI-H1792 | AZD6094 |
| MVP_top60_DNA_seed | HDAC1 | 0.867107 | KIRC | KMRC-20 | JQ12 | MVP_top60_DNA+RNA_seed | AKT1 | -0.038326 | LUAD | NCI-H1666 | BAY AKT1 |
| MVP_top60_DNA_seed | HDAC1 | 0.867711 | KIRC | KMRC-20 | CAY10603 | MVP_top60_DNA+RNA_seed | PIK3CA | -0.037592 | LUAD | NCI-H2342 | AZD8835 |
| MVP_top60_DNA_seed | MTOR | 0.868267 | KIRC | A498 | CRT0105446 | MVP_top60_DNA+RNA_seed | MAPK14 | -0.034942 | LUAD | EMC-BAC-2 | TAK-715 |
| MVP_top60_DNA_seed | AR | 0.873634 | KIRC | RCC-MF | Bicalutamide | MVP_top60_DNA+RNA_seed | MDM2 | -0.034711 | LUAD | NCI-H1793 | Nutlin-3a (-) |
| MVP_top60_DNA_seed | AR | 0.874335 | KIRC | CAL-54 | Bicalutamide | MVP_top60_DNA+RNA_seed | FGFR1 | -0.034512 | LUAD | NCI-H1648 | FGFR_3831 |
| MVP_top60_DNA_seed | HSP90AA1 | 0.874677 | KIRC | VMRC-RCZ | CCT-018159 | MVP_top60_DNA+RNA_seed | EGFR | -0.03403 | LUAD | NCI-H1838 | AST-1306 |
| MVP_top60_DNA_seed | HSP90AB1 | 0.874677 | KIRC | VMRC-RCZ | CCT-018159 | MVP_top60_DNA+RNA_seed | EGFR | -0.033332 | LUAD | A549 | Cetuximab |
| MVP_top60_DNA_seed | MTOR | 0.876488 | KIRC | CAKI-1 | Torin 2 | MVP_top60_DNA+RNA_seed | BRAF | -0.032685 | LUAD | NCI-H1975 | AZ628 |
| MVP_top60_DNA_seed | PIK3CD | 0.88096 | KIRC | LB996-RCC | Idelalisib | MVP_top60_DNA+RNA_seed | HSP90AA1 | -0.032393 | LUAD | NCI-H1355 | CCT-018159 |
| MVP_top60_DNA_seed | HSP90AA1 | 0.89066 | KIRC | NCC010 | Tanespimycin | MVP_top60_DNA+RNA_seed | EGFR | -0.03211 | LUAD | NCI-H23 | PF-00299804 |
| MVP_top60_DNA_seed | HSP90AB1 | 0.89066 | KIRC | NCC010 | Tanespimycin | MVP_top60_DNA+RNA_seed | HDAC1 | -0.031983 | LUAD | NCI-H1435 | JQ12 |

| | | | | | | | | | | | |
|---|---|---|---|---|---|---|---|---|---|---|---|
| MVP_top60_DNA_seed | MTOR | 0.899723 | KIRC | BFTC-909 | WYE-125132 | MVP_top60_DNA+RNA_seed | EGFR | -0.030434 | LUAD | NCI-H2030 | CI-1033 |
| MVP_top60_DNA_seed | HDAC1 | 0.901104 | KIRC | BFTC-909 | Tubastatin A | MVP_top60_DNA+RNA_seed | IGF1R | -0.029919 | LUAD | NCI-H2009 | BMS-754807 |
| MVP_top60_DNA_seed | PARP1 | 0.902039 | KIRC | HA7-RCC | Olaparib | MVP_top60_DNA+RNA_seed | IGF1R | -0.029791 | LUAD | NCI-H2030 | BMS-754807 |
| MVP_top60_DNA_seed | MTOR | 0.907342 | KIRC | KMRC-20 | Apitolisib | MVP_top60_DNA+RNA_seed | SRC | -0.029533 | LUAD | NCI-H2085 | NVP-BHG712 |
| MVP_top60_DNA_seed | HDAC1 | 0.907583 | KIRC | KMRC-20 | Tubastatin A | MVP_top60_DNA+RNA_seed | AURKA | -0.029091 | LUAD | EMC-BAC-2 | ZM447439 |
| MVP_top60_DNA_seed | PTK2 | 0.921292 | KIRC | A498 | PF-562271 | MVP_top60_DNA+RNA_seed | HDAC1 | -0.029058 | LUAD | NCI-H2087 | AR-42 |
| MVP_top60_DNA_seed | PARP1 | 0.923003 | KIRC | VMRC-RCZ | Olaparib | MVP_top60_DNA+RNA_seed | BRAF | -0.027466 | LUAD | EMC-BAC-1 | SB590885 |
| MVP_top60_DNA_seed | ESR1 | 0.923461 | KIRC | CAL-54 | Fulvestrant | MVP_top60_DNA+RNA_seed | HDAC1 | -0.027446 | LUAD | NCI-H2085 | Dacinostat |
| MVP_top60_DNA_seed | PARP1 | 0.930681 | KIRC | A498 | Talazoparib | MVP_top60_DNA+RNA_seed | EGFR | -0.026287 | LUAD | NCI-H522 | CI-1033 |
| MVP_top60_DNA_seed | PARP1 | 0.931761 | KIRC | RCC-AB | Veliparib | MVP_top60_DNA+RNA_seed | EGFR | -0.02625 | LUAD | EMC-BAC-2 | Cetuximab |
| MVP_top60_DNA_seed | MTOR | 0.932015 | KIRC | VMRC-RCZ | CRT0105446 | MVP_top60_DNA+RNA_seed | AURKA | -0.025173 | LUAD | H3255 | ZM447439 |
| MVP_top60_DNA_seed | PIK3CA | 0.932058 | KIRC | A498 | PI3Ka_4409 | MVP_top60_DNA+RNA_seed | AURKA | -0.025067 | LUAD | NCI-H1648 | Alisertib |
| MVP_top60_DNA_seed | PIK3CD | 0.932058 | KIRC | A498 | PI3Ka_4409 | MVP_top60_DNA+RNA_seed | AURKA | -0.024617 | LUAD | EMC-BAC-2 | GSK1070916 |
| MVP_top60_DNA_seed | FGFR4 | 0.932604 | KIRC | CAL-54 | FGFR_0939 | MVP_top60_DNA+RNA_seed | EGFR | -0.024421 | LUAD | NCI-H2087 | CUDC-101 |
| MVP_top60_DNA_seed | AR | 0.937033 | KIRC | TK10 | Bicalutamide | MVP_top60_DNA+RNA_seed | EGFR | -0.024176 | LUAD | NCI-H650 | CI-1033 |
| MVP_top60_DNA_seed | EP300 | 0.937501 | KIRC | SN12C | I-CBP112 | MVP_top60_DNA+RNA_seed | PIK3CA | -0.023811 | LUAD | NCI-H2009 | PI3Ka_4409 |
| MVP_top60_DNA_seed | AR | 0.937686 | KIRC | NCC021 | Bicalutamide | MVP_top60_DNA+RNA_seed | HSP90AA1 | -0.023167 | LUAD | SW1573 | Luminespib |
| MVP_top60_DNA_seed | EP300 | 0.940137 | KIRC | RCC-FG2 | I-CBP112 | MVP_top60_DNA+RNA_seed | HDAC1 | -0.022385 | LUAD | NCI-H650 | Belinostat |
| MVP_top60_DNA_seed | HDAC1 | 0.940443 | KIRC | U031 | AR-42 | MVP_top60_DNA+RNA_seed | EGFR | -0.021439 | LUAD | NCI-H1838 | CI-1033 |
| MVP_top60_DNA_seed | SMARCA4 | 0.940658 | KIRC | RCC-MF | PFI-3 | MVP_top60_DNA+RNA_seed | HDAC1 | -0.020088 | LUAD | NCI-H23 | CAY10603 |
| MVP_top60_DNA_seed | PARP1 | 0.943328 | KIRC | RCC-AB | Olaparib | MVP_top60_DNA+RNA_seed | HDAC1 | -0.019902 | LUAD | EMC-BAC-1 | CAY10603 |
| MVP_top60_DNA_seed | PIK3CA | 0.944562 | KIRC | KMRC-1 | PI3Ka_4409 | MVP_top60_DNA+RNA_seed | AURKA | -0.018517 | LUAD | RERF-LC-KJ | GSK1070916 |
| MVP_top60_DNA_seed | PIK3CD | 0.944562 | KIRC | KMRC-1 | PI3Ka_4409 | MVP_top60_DNA+RNA_seed | HSP90AA1 | -0.018315 | LUAD | NCI-H1781 | Tanespimycin |
| MVP_top60_DNA_seed | HDAC1 | 0.946225 | KIRC | RCC-FG2 | Tubastatin A | MVP_top60_DNA+RNA_seed | HDAC1 | -0.018079 | LUAD | NCI-H23 | Tubastatin A |
| MVP_top60_DNA_seed | HDAC1 | 0.946908 | KIRC | SW156 | Tubastatin A | MVP_top60_DNA+RNA_seed | EGFR | -0.01726 | LUAD | NCI-H522 | AST-1306 |
| MVP_top60_DNA_seed | SMARCA4 | 0.950296 | KIRC | SW156 | PFI-3 | MVP_top60_DNA+RNA_seed | ATM | -0.016552 | LUAD | RERF-LC-MS | CP466722 |
| MVP_top60_DNA_seed | HDAC1 | 0.95699 | KIRC | LB996-RCC | Tubastatin A | MVP_top60_DNA+RNA_seed | EGFR | -0.015355 | LUAD | NCI-H3122 | CUDC-101 |
| MVP_top60_DNA_seed | ESR1 | 0.957567 | KIRC | RCC-JW | Fulvestrant | MVP_top60_DNA+RNA_seed | HSP90AA1 | -0.014435 | LUAD | NCI-H2023 | Luminespib |
| MVP_top60_DNA_seed | HDAC1 | 0.958161 | KIRC | RCC-ER | Tubastatin A | MVP_top60_DNA+RNA_seed | EGFR | -0.013815 | LUAD | NCI-H2023 | Gefitinib |
| MVP_top60_DNA_seed | MTOR | 0.960748 | KIRC | RCC10RGB | Voxtalisib | MVP_top60_DNA+RNA_seed | AKT1 | -0.012952 | LUAD | NCI-H1734 | GSK690693 |
| MVP_top60_DNA_seed | ESR1 | 0.965101 | KIRC | RCC-ER | Fulvestrant | MVP_top60_DNA+RNA_seed | SRC | -0.012772 | LUAD | SK-LU-1 | NVP-BHG712 |
| MVP_top60_DNA_seed | PARP1 | 0.967099 | KIRC | SW156 | Talazoparib | MVP_top60_DNA+RNA_seed | AURKA | -0.012542 | LUAD | NCI-H838 | ZM447439 |
| MVP_top60_DNA_seed | PARP1 | 0.972792 | KIRC | A704 | Olaparib | MVP_top60_DNA+RNA_seed | IGF1R | -0.012454 | LUAD | HCC-827 | Linsitinib |
| MVP_top60_DNA_seed | PIK3CA | 0.975273 | KIRC | CAKI-1 | AZD8835 | MVP_top60_DNA+RNA_seed | HDAC1 | -0.011317 | LUAD | NCI-H1623 | CAY10603 |
| MVP_top60_DNA_seed | PIK3CD | 0.975273 | KIRC | CAKI-1 | AZD8835 | MVP_top60_DNA+RNA_seed | EGFR | -0.011308 | LUAD | PC-3_[JPC-3] | CUDC-101 |
| MVP_top60_DNA_seed | PARP1 | 0.9793 | KIRC | RCC-FG2 | Rucaparib | MVP_top60_DNA+RNA_seed | EGFR | -0.010331 | LUAD | NCI-H2030 | AST-1306 |
| MVP_top60_DNA_seed | PARP1 | 0.979474 | KIRC | RCC10RGB | PARP_9482 | MVP_top60_DNA+RNA_seed | EGFR | -0.009533 | LUAD | HOP-62 | AST-1306 |
| MVP_top60_DNA_seed | HSP90AA1 | 0.979962 | KIRC | RCC-AB | CCT-018159 | MVP_top60_DNA+RNA_seed | SRC | -0.008841 | LUAD | NCI-H1993 | NVP-BHG712 |
| MVP_top60_DNA_seed | HSP90AB1 | 0.979962 | KIRC | RCC-AB | CCT-018159 | MVP_top60_DNA+RNA_seed | BRAF | -0.008536 | LUAD | NCI-H2009 | PLX-4720 |
| MVP_top60_DNA_seed | HDAC1 | 0.991384 | KIRC | SW156 | Belinostat | MVP_top60_DNA+RNA_seed | EGFR | -0.008081 | LUAD | LC-2-ad | PF-00299804 |
| MVP_top60_DNA_seed | HDAC1 | 0.994399 | KIRC | RCC-ER | AR-42 | MVP_top60_DNA+RNA_seed | AURKA | -0.006705 | LUAD | NCI-H650 | HG-5-113-01 |

| | | | | | | | | | | | |
|---|---|---|---|---|---|---|---|---|---|---|---|
| MVP_top60_DNA_seed | PIK3CD | 0.999026 | KIRC | RCC10RGB | IC-87114 | MVP_top60_DNA+RNA_seed | HDAC1 | -0.006091 | LUAD | NCI-H2087 | CAY10603 |
| MVP_top60_DNA_seed | HDAC1 | 1.000032 | KIRC | LB2241-RCC | Tubastatin A | MVP_top60_DNA+RNA_seed | HDAC1 | -0.005837 | LUAD | NCI-H1693 | CAY10603 |
| MVP_top60_DNA_seed | PARP1 | 1.000805 | KIRC | HA7-RCC | Rucaparib | MVP_top60_DNA+RNA_seed | PIK3CA | -0.005836 | LUAD | NCI-H1563 | PI3Ka_4409 |
| MVP_top60_DNA_seed | AR | 1.002883 | KIRC | RCC-JF | AZD3514 | MVP_top60_DNA+RNA_seed | HDAC1 | -0.005655 | LUAD | EMC-BAC-1 | Tubastatin A |
| MVP_top60_DNA_seed | HSP90AA1 | 1.013877 | KIRC | RCC-FG2 | CCT-018159 | MVP_top60_DNA+RNA_seed | EGFR | -0.005511 | LUAD | NCI-H23 | AST-1306 |
| MVP_top60_DNA_seed | HSP90AB1 | 1.013877 | KIRC | RCC-FG2 | CCT-018159 | MVP_top60_DNA+RNA_seed | AKT1 | -0.005169 | LUAD | NCI-H1975 | AKT inhibitor VIII |
| MVP_top60_DNA_seed | PARP1 | 1.014851 | KIRC | A498 | Veliparib | MVP_top60_DNA+RNA_seed | BRAF | -0.004932 | LUAD | ABC-1 | HG6-64-1 |
| MVP_top60_DNA_seed | HDAC1 | 1.020949 | KIRC | BB65-RCC | Tubastatin A | MVP_top60_DNA+RNA_seed | IGF1R | -0.003819 | LUAD | NCI-H2347 | Linsitinib |
| MVP_top60_DNA_seed | ESR1 | 1.02107 | KIRC | SW156 | Tamoxifen | MVP_top60_DNA+RNA_seed | EGFR | -0.003693 | LUAD | NCI-H322M | Pelitinib |
| MVP_top60_DNA_seed | PARP1 | 1.021523 | KIRC | SW156 | Olaparib | MVP_top60_DNA+RNA_seed | AURKA | -0.00367 | LUAD | Calu-3 | Alisertib |
| MVP_top60_DNA_seed | HDAC1 | 1.022491 | KIRC | SN12C | Dacinostat | MVP_top60_DNA+RNA_seed | MET | -0.003056 | LUAD | NCI-H522 | SU11274 |
| MVP_top60_DNA_seed | PARP1 | 1.034094 | KIRC | KMRC-20 | Olaparib | MVP_top60_DNA+RNA_seed | AURKA | -0.002764 | LUAD | HOP-62 | GSK1070916 |
| MVP_top60_DNA_seed | HDAC1 | 1.039307 | KIRC | HA7-RCC | Tubastatin A | MVP_top60_DNA+RNA_seed | EGFR | -0.002066 | LUAD | NCI-H1792 | Afatinib |
| MVP_top60_DNA_seed | PARP1 | 1.044735 | KIRC | RCC-AB | Rucaparib | MVP_top60_DNA+RNA_seed | HDAC1 | -0.001696 | LUAD | NCI-H2347 | Belinostat |
| MVP_top60_DNA_seed | HDAC1 | 1.046399 | KIRC | SW156 | CAY10603 | MVP_top60_DNA+RNA_seed | ATM | -0.001618 | LUAD | Calu-3 | CP466722 |
| MVP_top60_DNA_seed | PARP1 | 1.059595 | KIRC | TK10 | Olaparib | MVP_top60_DNA+RNA_seed | SRC | -0.001381 | LUAD | SW1573 | Bosutinib |
| MVP_top60_DNA_seed | ATM | 1.071908 | KIRC | LB996-RCC | KU-55933 | MVP_top60_DNA+RNA_seed | EGFR | -0.000333 | LUAD | EMC-BAC-1 | AST-1306 |
| MVP_top60_DNA_seed | PIK3CD | 1.074467 | KIRC | KMRC-20 | Idelalisib | MVP_top60_DNA+RNA_seed | ATM | 0.000401 | LUAD | PC-14 | KU-55933 |
| MVP_top60_DNA_seed | HDAC1 | 1.075032 | KIRC | HA7-RCC | Parthenolide | MVP_top60_DNA+RNA_seed | AKT1 | 0.000643 | LUAD | NCI-H1944 | BAY AKT1 |
| MVP_top60_DNA_seed | MTOR | 1.078733 | KIRC | RCC-FG2 | QL-VIII-58 | MVP_top60_DNA+RNA_seed | IGF1R | 0.001576 | LUAD | SK-LU-1 | GSK1904529A |
| MVP_top60_DNA_seed | PTK2 | 1.079902 | KIRC | HA7-RCC | PF-562271 | MVP_top60_DNA+RNA_seed | FGFR1 | 0.002597 | LUAD | NCI-H1792 | FGFR_3831 |
| MVP_top60_DNA_seed | HDAC1 | 1.083219 | KIRC | LB2241-RCC | JQ12 | MVP_top60_DNA+RNA_seed | HDAC1 | 0.002935 | LUAD | NCI-H2009 | Dacinostat |
| MVP_top60_DNA_seed | MTOR | 1.084701 | KIRC | OS-RC-2 | CRT0105446 | MVP_top60_DNA+RNA_seed | SRC | 0.003275 | LUAD | NCI-H3122 | NVP-BHG712 |
| MVP_top60_DNA_seed | MTOR | 1.086138 | KIRC | RCC-ER | QL-VIII-58 | MVP_top60_DNA+RNA_seed | HDAC1 | 0.004402 | LUAD | NCI-H1648 | Tubastatin A |
| MVP_top60_DNA_seed | PARP1 | 1.090683 | KIRC | NCC021 | Olaparib | MVP_top60_DNA+RNA_seed | ATM | 0.004544 | LUAD | NCI-H1792 | KU-55933 |
| MVP_top60_DNA_seed | HDAC1 | 1.090848 | KIRC | KMRC-20 | AR-42 | MVP_top60_DNA+RNA_seed | FGFR1 | 0.004972 | LUAD | NCI-H3122 | FGFR_3831 |
| MVP_top60_DNA_seed | HDAC1 | 1.107478 | KIRC | LB1047-RCC | Parthenolide | MVP_top60_DNA+RNA_seed | MET | 0.0058 | LUAD | RERF-LC-KJ | AZD6094 |
| MVP_top60_DNA_seed | MTOR | 1.1109 | KIRC | VMRC-RCW | Temsirolimus | MVP_top60_DNA+RNA_seed | IGF1R | 0.006282 | LUAD | SW1573 | Linsitinib |
| MVP_top60_DNA_seed | PIK3CA | 1.117208 | KIRC | 786-0 | AZD8835 | MVP_top60_DNA+RNA_seed | AKT1 | 0.006813 | LUAD | HOP-62 | AKT inhibitor VIII |
| MVP_top60_DNA_seed | PIK3CD | 1.117208 | KIRC | 786-0 | AZD8835 | MVP_top60_DNA+RNA_seed | HDAC1 | 0.007332 | LUAD | NCI-H650 | AR-42 |
| MVP_top60_DNA_seed | HDAC1 | 1.126561 | KIRC | SW156 | AR-42 | MVP_top60_DNA+RNA_seed | EGFR | 0.007981 | LUAD | NCI-H1573 | Gefitinib |
| MVP_top60_DNA_seed | HDAC1 | 1.129408 | KIRC | RCC10RGB | AR-42 | MVP_top60_DNA+RNA_seed | HDAC1 | 0.008029 | LUAD | NCI-H1651 | Belinostat |
| MVP_top60_DNA_seed | MTOR | 1.131446 | KIRC | VMRC-RCW | WYE-125132 | MVP_top60_DNA+RNA_seed | ATM | 0.008962 | LUAD | VMRC-LCD | KU-60019 |
| MVP_top60_DNA_seed | HSP90AA1 | 1.13204 | KIRC | NCC021 | Tanespimycin | MVP_top60_DNA+RNA_seed | AURKA | 0.010628 | LUAD | NCI-H1568 | GSK1070916 |
| MVP_top60_DNA_seed | HSP90AB1 | 1.13204 | KIRC | NCC021 | Tanespimycin | MVP_top60_DNA+RNA_seed | HDAC1 | 0.011024 | LUAD | COR-L105 | Dacinostat |
| MVP_top60_DNA_seed | AR | 1.132228 | KIRC | VMRC-RCZ | Bicalutamide | MVP_top60_DNA+RNA_seed | AKT1 | 0.011316 | LUAD | NCI-H2347 | BAY AKT1 |
| MVP_top60_DNA_seed | HDAC1 | 1.134844 | KIRC | A498 | JQ12 | MVP_top60_DNA+RNA_seed | HDAC1 | 0.0115 | LUAD | ABC-1 | Belinostat |
| MVP_top60_DNA_seed | PARP1 | 1.135199 | KIRC | RCC10RGB | PARP_0108 | MVP_top60_DNA+RNA_seed | AKT1 | 0.012084 | LUAD | NCI-H1573 | AKT inhibitor VIII |
| MVP_top60_DNA_seed | MTOR | 1.13807 | KIRC | U031 | Temsirolimus | MVP_top60_DNA+RNA_seed | EGFR | 0.012487 | LUAD | Calu-6 | Afatinib |
| MVP_top60_DNA_seed | HSP90AA1 | 1.142809 | KIRC | SN12C | SNX-2112 | MVP_top60_DNA+RNA_seed | EGFR | 0.01318 | LUAD | NCI-H1792 | PF-00299804 |
| MVP_top60_DNA_seed | HSP90AB1 | 1.142809 | KIRC | SN12C | SNX-2112 | MVP_top60_DNA+RNA_seed | FGFR1 | 0.014596 | LUAD | NCI-H1568 | AZD4547 |

| | | | | | | | | | | | |
|---|---|---|---|---|---|---|---|---|---|---|---|
| MVP_top60_DNA_seed | MTOR | 1.150638 | KIRC | SW156 | Voxtalisib | MVP_top60_DNA+RNA_seed | HSP90AA1 | 0.015087 | LUAD | NCI-H2342 | Elesclomol |
| MVP_top60_DNA_seed | FGFR4 | 1.151913 | KIRC | SW156 | FGFR_3831 | MVP_top60_DNA+RNA_seed | ATM | 0.015879 | LUAD | NCI-H358 | KU-60019 |
| MVP_top60_DNA_seed | IGF1R | 1.153982 | KIRC | RCC-JW | BMS-536924 | MVP_top60_DNA+RNA_seed | MDM2 | 0.020238 | LUAD | NCI-H1437 | Nutlin-3a (-) |
| MVP_top60_DNA_seed | FGFR4 | 1.15443 | KIRC | OS-RC-2 | FGFR_3831 | MVP_top60_DNA+RNA_seed | PIK3CA | 0.021332 | LUAD | NCI-H2085 | AZD8835 |
| MVP_top60_DNA_seed | PARP1 | 1.160321 | KIRC | RCC10RGB | Rucaparib | MVP_top60_DNA+RNA_seed | PIK3CA | 0.022361 | LUAD | RERF-LC-MS | PI3Ka_4409 |
| MVP_top60_DNA_seed | IGF1R | 1.167513 | KIRC | LB2241-RCC | BMS-536924 | MVP_top60_DNA+RNA_seed | EGFR | 0.022476 | LUAD | NCI-H2347 | Cetuximab |
| MVP_top60_DNA_seed | PIK3CA | 1.170483 | KIRC | VMRC-RCZ | PI3Ka_4409 | MVP_top60_DNA+RNA_seed | HSP90AA1 | 0.022896 | LUAD | EMC-BAC-1 | SNX-2112 |
| MVP_top60_DNA_seed | PIK3CD | 1.170483 | KIRC | VMRC-RCZ | PI3Ka_4409 | MVP_top60_DNA+RNA_seed | EGFR | 0.023214 | LUAD | A549 | Afatinib |
| MVP_top60_DNA_seed | IGF1R | 1.176388 | KIRC | ACHN | Linsitinib | MVP_top60_DNA+RNA_seed | AURKA | 0.023434 | LUAD | SK-LU-1 | ZM447439 |
| MVP_top60_DNA_seed | PARP1 | 1.178479 | KIRC | RCC10RGB | Talazoparib | MVP_top60_DNA+RNA_seed | HSP90AA1 | 0.02429 | LUAD | NCI-H1623 | CCT-018159 |
| MVP_top60_DNA_seed | PARP1 | 1.18405 | KIRC | OS-RC-2 | Olaparib | MVP_top60_DNA+RNA_seed | EGFR | 0.024668 | LUAD | NCI-H650 | Pelitinib |
| MVP_top60_DNA_seed | PARP1 | 1.188081 | KIRC | RCC-MF | Rucaparib | MVP_top60_DNA+RNA_seed | HSP90AA1 | 0.024737 | LUAD | NCI-H1734 | Elesclomol |
| MVP_top60_DNA_seed | AR | 1.190661 | KIRC | RCC-ER | Bicalutamide | MVP_top60_DNA+RNA_seed | EGFR | 0.026267 | LUAD | LC-2-ad | CI-1033 |
| MVP_top60_DNA_seed | FGFR4 | 1.190802 | KIRC | RCC-FG2 | FGFR_3831 | MVP_top60_DNA+RNA_seed | MET | 0.028147 | LUAD | NCI-H3122 | JNJ38877605 |
| MVP_top60_DNA_seed | PIK3CA | 1.194837 | KIRC | BB65-RCC | PI3Ka_4409 | MVP_top60_DNA+RNA_seed | AURKA | 0.030073 | LUAD | NCI-H1975 | HG-5-113-01 |
| MVP_top60_DNA_seed | PIK3CD | 1.194837 | KIRC | BB65-RCC | PI3Ka_4409 | MVP_top60_DNA+RNA_seed | BRAF | 0.030821 | LUAD | SW1573 | Dabrafenib |
| MVP_top60_DNA_seed | PARP1 | 1.195855 | KIRC | CAL-54 | Olaparib | MVP_top60_DNA+RNA_seed | HSP90AA1 | 0.031034 | LUAD | NCI-H2023 | Elesclomol |
| MVP_top60_DNA_seed | PARP1 | 1.209415 | KIRC | A498 | Olaparib | MVP_top60_DNA+RNA_seed | BRAF | 0.031066 | LUAD | EKVX | PLX-4720 |
| MVP_top60_DNA_seed | IGF1R | 1.209531 | KIRC | A498 | GSK1904529A | MVP_top60_DNA+RNA_seed | IGF1R | 0.031382 | LUAD | PC-14 | GSK1904529A |
| MVP_top60_DNA_seed | MTOR | 1.210289 | KIRC | A704 | Torin 2 | MVP_top60_DNA+RNA_seed | AKT1 | 0.031511 | LUAD | NCI-H1781 | AKT inhibitor VIII |
| MVP_top60_DNA_seed | PARP1 | 1.213527 | KIRC | RCC-ER | Rucaparib | MVP_top60_DNA+RNA_seed | HDAC1 | 0.032313 | LUAD | NCI-H2030 | CAY10603 |
| MVP_top60_DNA_seed | PARP1 | 1.213859 | KIRC | KMRC-20 | PARP_9495 | MVP_top60_DNA+RNA_seed | ATM | 0.03253 | LUAD | NCI-H1975 | CP466722 |
| MVP_top60_DNA_seed | PIK3CD | 1.216171 | KIRC | LB2241-RCC | Idelalisib | MVP_top60_DNA+RNA_seed | AKT1 | 0.032543 | LUAD | NCI-H838 | BAY AKT1 |
| MVP_top60_DNA_seed | HSP90AA1 | 1.222358 | KIRC | CAL-54 | SNX-2112 | MVP_top60_DNA+RNA_seed | EGFR | 0.033544 | LUAD | NCI-H2405 | Pelitinib |
| MVP_top60_DNA_seed | HSP90AB1 | 1.222358 | KIRC | CAL-54 | SNX-2112 | MVP_top60_DNA+RNA_seed | BRAF | 0.033563 | LUAD | LC-2-ad | SB590885 |
| MVP_top60_DNA_seed | AR | 1.231843 | KIRC | OS-RC-2 | AZD3514 | MVP_top60_DNA+RNA_seed | BRAF | 0.034241 | LUAD | HOP-62 | PLX-4720 |
| MVP_top60_DNA_seed | AR | 1.231861 | KIRC | VMRC-RCZ | AZD3514 | MVP_top60_DNA+RNA_seed | EGFR | 0.034841 | LUAD | NCI-H2347 | AST-1306 |
| MVP_top60_DNA_seed | AR | 1.238458 | KIRC | RXF393 | Bicalutamide | MVP_top60_DNA+RNA_seed | BRAF | 0.035379 | LUAD | NCI-H838 | HG6-64-1 |
| MVP_top60_DNA_seed | ATM | 1.242675 | KIRC | KMRC-1 | KU-55933 | MVP_top60_DNA+RNA_seed | HDAC1 | 0.037451 | LUAD | HCC-827 | JQ12 |
| MVP_top60_DNA_seed | IGF1R | 1.251603 | KIRC | VMRC-RCZ | GSK1904529A | MVP_top60_DNA+RNA_seed | HSP90AA1 | 0.037625 | LUAD | NCI-H1573 | CCT-018159 |
| MVP_top60_DNA_seed | HDAC1 | 1.26503 | KIRC | RXF393 | Entinostat | MVP_top60_DNA+RNA_seed | BRAF | 0.038252 | LUAD | NCI-H441 | RAF_9304 |
| MVP_top60_DNA_seed | MTOR | 1.269411 | KIRC | KMRC-1 | Torin 2 | MVP_top60_DNA+RNA_seed | HSP90AA1 | 0.039488 | LUAD | NCI-H1573 | Tanespimycin |
| MVP_top60_DNA_seed | ESR1 | 1.280738 | KIRC | NCC021 | Fulvestrant | MVP_top60_DNA+RNA_seed | ATM | 0.039839 | LUAD | NCI-H2347 | KU-55933 |
| MVP_top60_DNA_seed | HSP90AA1 | 1.285857 | KIRC | SW156 | SNX-2112 | MVP_top60_DNA+RNA_seed | HDAC1 | 0.039956 | LUAD | NCI-H2030 | Tubastatin A |
| MVP_top60_DNA_seed | HSP90AB1 | 1.285857 | KIRC | SW156 | SNX-2112 | MVP_top60_DNA+RNA_seed | EGFR | 0.040419 | LUAD | NCI-H23 | AZD8931 |
| MVP_top60_DNA_seed | HSP90AA1 | 1.293296 | KIRC | CAL-54 | Elesclomol | MVP_top60_DNA+RNA_seed | AKT1 | 0.042646 | LUAD | NCI-H2085 | BAY AKT1 |
| MVP_top60_DNA_seed | HSP90AB1 | 1.293296 | KIRC | CAL-54 | Elesclomol | MVP_top60_DNA+RNA_seed | IGF1R | 0.042749 | LUAD | LXF-289 | GSK1904529A |
| MVP_top60_DNA_seed | AR | 1.295094 | KIRC | KMRC-20 | AZD3514 | MVP_top60_DNA+RNA_seed | SRC | 0.043018 | LUAD | NCI-H1563 | NVP-BHG712 |
| MVP_top60_DNA_seed | MTOR | 1.297411 | KIRC | RXF393 | Apitolisib | MVP_top60_DNA+RNA_seed | FGFR1 | 0.043029 | LUAD | NCI-H1792 | AZD4547 |
| MVP_top60_DNA_seed | AR | 1.311373 | KIRC | NCC021 | AZD3514 | MVP_top60_DNA+RNA_seed | BRAF | 0.043922 | LUAD | COR-L105 | RAF_9304 |
| MVP_top60_DNA_seed | PTK2 | 1.318476 | KIRC | VMRC-RCW | PF-562271 | MVP_top60_DNA+RNA_seed | HDAC1 | 0.044605 | LUAD | NCI-H1355 | Entinostat |

| | | | | | | | | | | | |
|---|---|---|---|---|---|---|---|---|---|---|---|
| MVP_top60_DNA_seed | MTOR | 1.319536 | KIRC | NCC010 | QL-VIII-58 | MVP_top60_DNA+RNA_seed | ATM | 0.045103 | LUAD | NCI-H3122 | KU-60019 |
| MVP_top60_DNA_seed | ESR1 | 1.338676 | KIRC | OS-RC-2 | Tamoxifen | MVP_top60_DNA+RNA_seed | EGFR | 0.04546 | LUAD | NCI-H2347 | CUDC-101 |
| MVP_top60_DNA_seed | IGF1R | 1.339708 | KIRC | RCC10RGB | GSK1904529A | MVP_top60_DNA+RNA_seed | EGFR | 0.046171 | LUAD | Calu-3 | CUDC-101 |
| MVP_top60_DNA_seed | PARP1 | 1.340199 | KIRC | A498 | PARP_9495 | MVP_top60_DNA+RNA_seed | HSP90AA1 | 0.04695 | LUAD | NCI-H522 | Tanespimycin |
| MVP_top60_DNA_seed | IGF1R | 1.346113 | KIRC | RCC-AB | BMS-536924 | MVP_top60_DNA+RNA_seed | BRAF | 0.047228 | LUAD | NCI-H2030 | Dabrafenib |
| MVP_top60_DNA_seed | EP300 | 1.352305 | KIRC | OS-RC-2 | I-CBP112 | MVP_top60_DNA+RNA_seed | MAPK14 | 0.049288 | LUAD | RERF-LC-KJ | TAK-715 |
| MVP_top60_DNA_seed | EP300 | 1.354702 | KIRC | RXF393 | I-CBP112 | MVP_top60_DNA+RNA_seed | BRAF | 0.050009 | LUAD | PC-14 | SB590885 |
| MVP_top60_DNA_seed | MTOR | 1.37493 | KIRC | VMRC-RCW | Apitolisib | MVP_top60_DNA+RNA_seed | ATM | 0.050298 | LUAD | H3255 | CP466722 |
| MVP_top60_DNA_seed | AR | 1.375245 | KIRC | BB65-RCC | AZD3514 | MVP_top60_DNA+RNA_seed | BRAF | 0.052559 | LUAD | EMC-BAC-1 | PLX-4720 |
| MVP_top60_DNA_seed | PARP1 | 1.38241 | KIRC | TK10 | Olaparib | MVP_top60_DNA+RNA_seed | HDAC1 | 0.052975 | LUAD | NCI-H441 | Dacinostat |
| MVP_top60_DNA_seed | IGF1R | 1.394086 | KIRC | 769-P | BMS-536924 | MVP_top60_DNA+RNA_seed | IGF1R | 0.053062 | LUAD | HCC-827 | BMS-536924 |
| MVP_top60_DNA_seed | MTOR | 1.395824 | KIRC | KMRC-20 | Torin 2 | MVP_top60_DNA+RNA_seed | MAPK14 | 0.054073 | LUAD | NCI-H1975 | TAK-715 |
| MVP_top60_DNA_seed | MTOR | 1.40829 | KIRC | RCC10RGB | Apitolisib | MVP_top60_DNA+RNA_seed | MET | 0.054235 | LUAD | NCI-H1666 | PHA-665752 |
| MVP_top60_DNA_seed | HSP90AA1 | 1.423532 | KIRC | U031 | SNX-2112 | MVP_top60_DNA+RNA_seed | SRC | 0.055716 | LUAD | NCI-H838 | Bosutinib |
| MVP_top60_DNA_seed | HSP90AB1 | 1.423532 | KIRC | U031 | SNX-2112 | MVP_top60_DNA+RNA_seed | FGFR1 | 0.056761 | LUAD | NCI-H1651 | AZD4547 |
| MVP_top60_DNA_seed | SMARCA4 | 1.436441 | KIRC | RXF393 | PFI-3 | MVP_top60_DNA+RNA_seed | AKT1 | 0.057221 | LUAD | NCI-H1650 | AKT inhibitor VIII |
| MVP_top60_DNA_seed | PIK3CA | 1.438003 | KIRC | SW156 | PI3Ka_4409 | MVP_top60_DNA+RNA_seed | MET | 0.057489 | LUAD | RERF-LC-KJ | JNJ38877605 |
| MVP_top60_DNA_seed | PIK3CD | 1.438003 | KIRC | SW156 | PI3Ka_4409 | MVP_top60_DNA+RNA_seed | BRAF | 0.058446 | LUAD | NCI-H23 | Dabrafenib |
| MVP_top60_DNA_seed | ESR1 | 1.442765 | KIRC | RCC10RGB | Fulvestrant | MVP_top60_DNA+RNA_seed | PIK3CA | 0.058583 | LUAD | NCI-H1993 | PI3Ka_4409 |
| MVP_top60_DNA_seed | MTOR | 1.453888 | KIRC | RXF393 | Torin 2 | MVP_top60_DNA+RNA_seed | AKT1 | 0.059093 | LUAD | NCI-H322M | AKT inhibitor VIII |
| MVP_top60_DNA_seed | PARP1 | 1.466399 | KIRC | RCC-ER | Olaparib | MVP_top60_DNA+RNA_seed | HSP90AA1 | 0.059208 | LUAD | NCI-H1563 | Tanespimycin |
| MVP_top60_DNA_seed | MTOR | 1.47322 | KIRC | TK10 | Voxtalisib | MVP_top60_DNA+RNA_seed | ATM | 0.060047 | LUAD | NCI-H3122 | CP466722 |
| MVP_top60_DNA_seed | IGF1R | 1.4808 | KIRC | SW156 | BMS-536924 | MVP_top60_DNA+RNA_seed | FGFR1 | 0.060867 | LUAD | NCI-H2085 | PD173074 |
| MVP_top60_DNA_seed | HSP90AA1 | 1.487972 | KIRC | HA7-RCC | CCT-018159 | MVP_top60_DNA+RNA_seed | FGFR1 | 0.061006 | LUAD | VMRC-LCD | AZD4547 |
| MVP_top60_DNA_seed | HSP90AB1 | 1.487972 | KIRC | HA7-RCC | CCT-018159 | MVP_top60_DNA+RNA_seed | IGF1R | 0.061168 | LUAD | NCI-H1755 | BMS-754807 |
| MVP_top60_DNA_seed | HSP90AA1 | 1.488387 | KIRC | RCC10RGB | Luminespib | MVP_top60_DNA+RNA_seed | MET | 0.062196 | LUAD | NCI-H1651 | SU11274 |
| MVP_top60_DNA_seed | HSP90AB1 | 1.488387 | KIRC | RCC10RGB | Luminespib | MVP_top60_DNA+RNA_seed | MDM2 | 0.062939 | LUAD | HCC-827 | Serdemetan |
| MVP_top60_DNA_seed | PARP1 | 1.492138 | KIRC | ACHN | Talazoparib | MVP_top60_DNA+RNA_seed | EGFR | 0.064074 | LUAD | Calu-6 | Pelitinib |
| MVP_top60_DNA_seed | HDAC1 | 1.492434 | KIRC | SN12C | Tubastatin A | MVP_top60_DNA+RNA_seed | AURKA | 0.065749 | LUAD | Calu-6 | CD532 |
| MVP_top60_DNA_seed | PIK3CA | 1.498648 | KIRC | OS-RC-2 | PI3Ka_4409 | MVP_top60_DNA+RNA_seed | MET | 0.0661 | LUAD | NCI-H2009 | AZD6094 |
| MVP_top60_DNA_seed | PIK3CD | 1.498648 | KIRC | OS-RC-2 | PI3Ka_4409 | MVP_top60_DNA+RNA_seed | AURKA | 0.066318 | LUAD | NCI-H1975 | Tozasertib |
| MVP_top60_DNA_seed | HDAC1 | 1.50906 | KIRC | SN12C | JQ12 | MVP_top60_DNA+RNA_seed | MDM2 | 0.067187 | LUAD | NCI-H1975 | Serdemetan |
| MVP_top60_DNA_seed | ATM | 1.517968 | KIRC | OS-RC-2 | KU-55933 | MVP_top60_DNA+RNA_seed | AKT1 | 0.068159 | LUAD | LC-2-ad | BAY AKT1 |
| MVP_top60_DNA_seed | PARP1 | 1.525245 | KIRC | OS-RC-2 | PARP_9495 | MVP_top60_DNA+RNA_seed | SRC | 0.068545 | LUAD | NCI-H2342 | Bosutinib |
| MVP_top60_DNA_seed | MTOR | 1.531906 | KIRC | RXF393 | WYE-125132 | MVP_top60_DNA+RNA_seed | MDM2 | 0.068805 | LUAD | HCC-44 | Nutlin-3a (-) |
| MVP_top60_DNA_seed | PIK3CA | 1.532589 | KIRC | OS-RC-2 | AZD8835 | MVP_top60_DNA+RNA_seed | EGFR | 0.068918 | LUAD | Calu-3 | Gefitinib |
| MVP_top60_DNA_seed | PIK3CD | 1.532589 | KIRC | OS-RC-2 | AZD8835 | MVP_top60_DNA+RNA_seed | BRAF | 0.068992 | LUAD | NCI-H2009 | RAF_9304 |
| MVP_top60_DNA_seed | HDAC1 | 1.537922 | KIRC | A498 | CAY10603 | MVP_top60_DNA+RNA_seed | HDAC1 | 0.069424 | LUAD | RERF-LC-MS | CAY10603 |
| MVP_top60_DNA_seed | HSP90AA1 | 1.542734 | KIRC | SW156 | Luminespib | MVP_top60_DNA+RNA_seed | AKT1 | 0.070384 | LUAD | EMC-BAC-1 | MK-2206 |
| MVP_top60_DNA_seed | HSP90AB1 | 1.542734 | KIRC | SW156 | Luminespib | MVP_top60_DNA+RNA_seed | FGFR1 | 0.070647 | LUAD | NCI-H2405 | AZD4547 |
| MVP_top60_DNA_seed | AR | 1.546026 | KIRC | RXF393 | Bicalutamide | MVP_top60_DNA+RNA_seed | MET | 0.070683 | LUAD | SW1573 | JNJ38877605 |

| | | | | | | | | | | | |
|---|---|---|---|---|---|---|---|---|---|---|---|
| MVP_top60_DNA_seed | MTOR | 1.551482 | KIRC | RCC10RGB | JW-7-52-1 | MVP_top60_DNA+RNA_seed | ATM | 0.07131 | LUAD | NCI-H1693 | CP466722 |
| MVP_top60_DNA_seed | FGFR4 | 1.559144 | KIRC | RCC-MF | FGFR_3831 | MVP_top60_DNA+RNA_seed | EGFR | 0.071393 | LUAD | EMC-BAC-2 | Afatinib |
| MVP_top60_DNA_seed | HSP90AA1 | 1.564544 | KIRC | RCC10RGB | SNX-2112 | MVP_top60_DNA+RNA_seed | AURKA | 0.071721 | LUAD | NCI-H1975 | GSK1070916 |
| MVP_top60_DNA_seed | HSP90AB1 | 1.564544 | KIRC | RCC10RGB | SNX-2112 | MVP_top60_DNA+RNA_seed | SMARCA4 | 0.071758 | LUAD | NCI-H1435 | PFI-3 |
| MVP_top60_DNA_seed | ATM | 1.568488 | KIRC | A498 | KU-55933 | MVP_top60_DNA+RNA_seed | AKT1 | 0.07226 | LUAD | EKVX | AKT inhibitor VIII |
| MVP_top60_DNA_seed | PIK3CA | 1.571109 | KIRC | 786-0 | PI3Ka_4409 | MVP_top60_DNA+RNA_seed | ATM | 0.073309 | LUAD | NCI-H1944 | KU-55933 |
| MVP_top60_DNA_seed | PIK3CD | 1.571109 | KIRC | 786-0 | PI3Ka_4409 | MVP_top60_DNA+RNA_seed | AURKA | 0.07333 | LUAD | NCI-H23 | GSK1070916 |
| MVP_top60_DNA_seed | ATM | 1.571332 | KIRC | RCC-JW | KU-60019 | MVP_top60_DNA+RNA_seed | MET | 0.073767 | LUAD | HOP-62 | AZD6094 |
| MVP_top60_DNA_seed | PARP1 | 1.573232 | KIRC | A498 | Rucaparib | MVP_top60_DNA+RNA_seed | EGFR | 0.074843 | LUAD | NCI-H1650 | Afatinib |
| MVP_top60_DNA_seed | MTOR | 1.582794 | KIRC | NCC021 | QL-VIII-58 | MVP_top60_DNA+RNA_seed | ATM | 0.075005 | LUAD | NCI-H1793 | CP466722 |
| MVP_top60_DNA_seed | HSP90AA1 | 1.583639 | KIRC | A498 | CCT-018159 | MVP_top60_DNA+RNA_seed | HDAC1 | 0.075354 | LUAD | NCI-H2030 | Belinostat |
| MVP_top60_DNA_seed | HSP90AB1 | 1.583639 | KIRC | A498 | CCT-018159 | MVP_top60_DNA+RNA_seed | IGF1R | 0.07543 | LUAD | NCI-H1651 | GSK1904529A |
| MVP_top60_DNA_seed | AR | 1.583993 | KIRC | RCC-ER | AZD3514 | MVP_top60_DNA+RNA_seed | HSP90AA1 | 0.076717 | LUAD | RERF-LC-KJ | Luminespib |
| MVP_top60_DNA_seed | EP300 | 1.605165 | KIRC | A498 | I-CBP112 | MVP_top60_DNA+RNA_seed | FGFR1 | 0.077484 | LUAD | NCI-H358 | PD173074 |
| MVP_top60_DNA_seed | AR | 1.619653 | KIRC | OS-RC-2 | Bicalutamide | MVP_top60_DNA+RNA_seed | HDAC1 | 0.078824 | LUAD | NCI-H838 | AR-42 |
| MVP_top60_DNA_seed | HSP90AA1 | 1.632913 | KIRC | KMRC-20 | SNX-2112 | MVP_top60_DNA+RNA_seed | ATM | 0.079111 | LUAD | NCI-H23 | KU-60019 |
| MVP_top60_DNA_seed | HSP90AB1 | 1.632913 | KIRC | KMRC-20 | SNX-2112 | MVP_top60_DNA+RNA_seed | EGFR | 0.079141 | LUAD | NCI-H3122 | Afatinib |
| MVP_top60_DNA_seed | AR | 1.639079 | KIRC | RCC10RGB | AZD3514 | MVP_top60_DNA+RNA_seed | AURKA | 0.079601 | LUAD | NCI-H2347 | HG-5-113-01 |
| MVP_top60_DNA_seed | MTOR | 1.653092 | KIRC | A704 | QL-VIII-58 | MVP_top60_DNA+RNA_seed | HDAC1 | 0.080527 | LUAD | NCI-H1651 | CAY10603 |
| MVP_top60_DNA_seed | IGF1R | 1.657419 | KIRC | RXF393 | GSK1904529A | MVP_top60_DNA+RNA_seed | PTK2 | 0.080757 | LUAD | NCI-H1437 | PF-562271 |
| MVP_top60_DNA_seed | PARP1 | 1.681237 | KIRC | RCC10RGB | Olaparib | MVP_top60_DNA+RNA_seed | AKT1 | 0.081894 | LUAD | NCI-H1355 | AKT inhibitor VIII |
| MVP_top60_DNA_seed | HDAC1 | 1.682652 | KIRC | A498 | Tubastatin A | MVP_top60_DNA+RNA_seed | AKT1 | 0.083347 | LUAD | HCC-827 | AKT inhibitor VIII |
| MVP_top60_DNA_seed | HSP90AA1 | 1.690657 | KIRC | CAL-54 | Luminespib | MVP_top60_DNA+RNA_seed | BRAF | 0.084937 | LUAD | HCC-44 | PLX-4720 |
| MVP_top60_DNA_seed | HSP90AB1 | 1.690657 | KIRC | CAL-54 | Luminespib | MVP_top60_DNA+RNA_seed | EGFR | 0.084963 | LUAD | NCI-H1435 | Afatinib |
| MVP_top60_DNA_seed | MTOR | 1.696703 | KIRC | A704 | WYE-125132 | MVP_top60_DNA+RNA_seed | EGFR | 0.085954 | LUAD | NCI-H1573 | AST-1306 |
| MVP_top60_DNA_seed | PIK3CD | 1.699136 | KIRC | A498 | Idelalisib | MVP_top60_DNA+RNA_seed | EGFR | 0.086348 | LUAD | NCI-H1793 | Gefitinib |
| MVP_top60_DNA_seed | HDAC1 | 1.711416 | KIRC | KMRC-1 | Dacinostat | MVP_top60_DNA+RNA_seed | HSP90AA1 | 0.086513 | LUAD | NCI-H2347 | Luminespib |
| MVP_top60_DNA_seed | MTOR | 1.722473 | KIRC | RXF393 | CRT0105446 | MVP_top60_DNA+RNA_seed | IGF1R | 0.086624 | LUAD | NCI-H2009 | Linsitinib |
| MVP_top60_DNA_seed | FGFR4 | 1.7371 | KIRC | RCC-ER | FGFR_3831 | MVP_top60_DNA+RNA_seed | AURKA | 0.08672 | LUAD | NCI-H522 | Genentech Cpd 10 |
| MVP_top60_DNA_seed | PTK2 | 1.756325 | KIRC | KMRC-20 | PF-562271 | MVP_top60_DNA+RNA_seed | HDAC1 | 0.088673 | LUAD | NCI-H2347 | Tubastatin A |
| MVP_top60_DNA_seed | AR | 1.758796 | KIRC | A498 | AZD3514 | MVP_top60_DNA+RNA_seed | IGF1R | 0.090597 | LUAD | NCI-H2023 | BMS-754807 |
| MVP_top60_DNA_seed | PIK3CA | 1.768022 | KIRC | KMRC-1 | AZD8835 | MVP_top60_DNA+RNA_seed | AURKA | 0.092787 | LUAD | NCI-H1792 | HG-5-113-01 |
| MVP_top60_DNA_seed | PIK3CD | 1.768022 | KIRC | KMRC-1 | AZD8835 | MVP_top60_DNA+RNA_seed | FGFR1 | 0.094044 | LUAD | NCI-H2030 | FGFR_3831 |
| MVP_top60_DNA_seed | AR | 1.774656 | KIRC | RCC10RGB | Bicalutamide | MVP_top60_DNA+RNA_seed | BRAF | 0.094367 | LUAD | NCI-H1651 | Dabrafenib |
| MVP_top60_DNA_seed | HSP90AA1 | 1.79002 | KIRC | OS-RC-2 | CCT-018159 | MVP_top60_DNA+RNA_seed | EGFR | 0.094966 | LUAD | RERF-LC-MS | AZD8931 |
| MVP_top60_DNA_seed | HSP90AB1 | 1.79002 | KIRC | OS-RC-2 | CCT-018159 | MVP_top60_DNA+RNA_seed | HDAC1 | 0.096712 | LUAD | NCI-H1666 | AR-42 |
| MVP_top60_DNA_seed | HSP90AA1 | 1.793778 | KIRC | LB2241-RCC | SNX-2112 | MVP_top60_DNA+RNA_seed | AKT1 | 0.097889 | LUAD | NCI-H3122 | MK-2206 |
| MVP_top60_DNA_seed | HSP90AB1 | 1.793778 | KIRC | LB2241-RCC | SNX-2112 | MVP_top60_DNA+RNA_seed | MDM2 | 0.099277 | LUAD | NCI-H2122 | Serdemetan |
| MVP_top60_DNA_seed | HDAC1 | 1.810855 | KIRC | RCC-ER | Belinostat | MVP_top60_DNA+RNA_seed | MDM2 | 0.099306 | LUAD | NCI-H358 | Serdemetan |
| MVP_top60_DNA_seed | ESR1 | 1.814996 | KIRC | OS-RC-2 | Fulvestrant | MVP_top60_DNA+RNA_seed | SMARCA4 | 0.099896 | LUAD | NCI-H1975 | PFI-3 |
| MVP_top60_DNA_seed | ATM | 1.827843 | KIRC | A498 | CP466722 | MVP_top60_DNA+RNA_seed | PIK3CA | 0.100394 | LUAD | NCI-H1792 | PI3Ka_4409 |

| | | | | | | | | | | | |
|---|---|---|---|---|---|---|---|---|---|---|---|
| MVP_top60_DNA_seed | HSP90AA1 | 1.829653 | KIRC | ACHN | Elesclomol | MVP_top60_DNA+RNA_seed | BRAF | 0.101724 | LUAD | LXF-289 | AZ628 |
| MVP_top60_DNA_seed | HSP90AB1 | 1.829653 | KIRC | ACHN | Elesclomol | MVP_top60_DNA+RNA_seed | MET | 0.102188 | LUAD | NCI-H3122 | SU11274 |
| MVP_top60_DNA_seed | HDAC1 | 1.835936 | KIRC | A498 | AR-42 | MVP_top60_DNA+RNA_seed | MAPK14 | 0.102869 | LUAD | NCI-H2405 | TAK-715 |
| MVP_top60_DNA_seed | HSP90AA1 | 1.862574 | KIRC | LB996-RCC | CCT-018159 | MVP_top60_DNA+RNA_seed | HDAC1 | 0.10294 | LUAD | LXF-289 | Parthenolide |
| MVP_top60_DNA_seed | HSP90AB1 | 1.862574 | KIRC | LB996-RCC | CCT-018159 | MVP_top60_DNA+RNA_seed | AURKA | 0.103542 | LUAD | NCI-H1781 | HG-5-113-01 |
| MVP_top60_DNA_seed | HDAC1 | 1.872589 | KIRC | A498 | Parthenolide | MVP_top60_DNA+RNA_seed | FGFR1 | 0.10405 | LUAD | HCC-78 | PD173074 |
| MVP_top60_DNA_seed | MTOR | 1.875297 | KIRC | RXF393 | Voxtalisib | MVP_top60_DNA+RNA_seed | AURKA | 0.105207 | LUAD | PC-3_[JPC-3] | Alisertib |
| MVP_top60_DNA_seed | HDAC1 | 1.892178 | KIRC | RCC10RGB | Tubastatin A | MVP_top60_DNA+RNA_seed | EGFR | 0.105223 | LUAD | NCI-H2030 | Cetuximab |
| MVP_top60_DNA_seed | MTOR | 1.908838 | KIRC | RCC10RGB | CRT0105446 | MVP_top60_DNA+RNA_seed | HDAC1 | 0.10561 | LUAD | NCI-H2087 | Dacinostat |
| MVP_top60_DNA_seed | MTOR | 1.91931 | KIRC | A704 | Apitolisib | MVP_top60_DNA+RNA_seed | BRAF | 0.106132 | LUAD | NCI-H2122 | RAF_9304 |
| MVP_top60_DNA_seed | HSP90AA1 | 1.970553 | KIRC | RCC-ER | CCT-018159 | MVP_top60_DNA+RNA_seed | EGFR | 0.106379 | LUAD | NCI-H2023 | Afatinib |
| MVP_top60_DNA_seed | HSP90AB1 | 1.970553 | KIRC | RCC-ER | CCT-018159 | MVP_top60_DNA+RNA_seed | FGFR1 | 0.107135 | LUAD | NCI-H1648 | PD173074 |
| MVP_top60_DNA_seed | PARP1 | 1.974776 | KIRC | RCC10RGB | Veliparib | MVP_top60_DNA+RNA_seed | MDM2 | 0.107372 | LUAD | Calu-3 | Serdemetan |
| MVP_top60_DNA_seed | ESR1 | 2.010775 | KIRC | A498 | Tamoxifen | MVP_top60_DNA+RNA_seed | BRAF | 0.107619 | LUAD | ABC-1 | Dabrafenib |
| MVP_top60_DNA_seed | ATM | 2.043105 | KIRC | RCC10RGB | CP466722 | MVP_top60_DNA+RNA_seed | MAPK14 | 0.108043 | LUAD | NCI-H1755 | TAK-715 |
| MVP_top60_DNA_seed | FGFR4 | 2.045145 | KIRC | RCC10RGB | FGFR_3831 | MVP_top60_DNA+RNA_seed | EGFR | 0.108768 | LUAD | LXF-289 | Afatinib |
| MVP_top60_DNA_seed | AR | 2.050606 | KIRC | RXF393 | AZD3514 | MVP_top60_DNA+RNA_seed | AKT1 | 0.109341 | LUAD | NCI-H1650 | MK-2206 |
| MVP_top60_DNA_seed | AR | 2.05533 | KIRC | A498 | Bicalutamide | MVP_top60_DNA+RNA_seed | FGFR1 | 0.110362 | LUAD | SK-LU-1 | AZD4547 |
| MVP_top60_DNA_seed | MTOR | 2.059937 | KIRC | RCC10RGB | Torin 2 | MVP_top60_DNA+RNA_seed | MDM2 | 0.111032 | LUAD | NCI-H1838 | Serdemetan |
| MVP_top60_DNA_seed | ATM | 2.072145 | KIRC | RCC10RGB | KU-55933 | MVP_top60_DNA+RNA_seed | HDAC1 | 0.11186 | LUAD | HCC-827 | Belinostat |
| MVP_top60_DNA_seed | HSP90AA1 | 2.072658 | KIRC | RCC-MF | CCT-018159 | MVP_top60_DNA+RNA_seed | AKT1 | 0.112574 | LUAD | COR-L105 | GSK690693 |
| MVP_top60_DNA_seed | HSP90AB1 | 2.072658 | KIRC | RCC-MF | CCT-018159 | MVP_top60_DNA+RNA_seed | EGFR | 0.112963 | LUAD | NCI-H1793 | AZD8931 |
| MVP_top60_DNA_seed | HSP90AA1 | 2.073883 | KIRC | A498 | SNX-2112 | MVP_top60_DNA+RNA_seed | EGFR | 0.113173 | LUAD | NCI-H650 | AST-1306 |
| MVP_top60_DNA_seed | HSP90AB1 | 2.073883 | KIRC | A498 | SNX-2112 | MVP_top60_DNA+RNA_seed | PTK2 | 0.113588 | LUAD | H3255 | PF-562271 |
| MVP_top60_DNA_seed | HSP90AA1 | 2.085682 | KIRC | RCC-ER | SNX-2112 | MVP_top60_DNA+RNA_seed | EGFR | 0.114856 | LUAD | NCI-H1563 | CI-1033 |
| MVP_top60_DNA_seed | HSP90AB1 | 2.085682 | KIRC | RCC-ER | SNX-2112 | MVP_top60_DNA+RNA_seed | FGFR1 | 0.115337 | LUAD | EMC-BAC-2 | AZD4547 |
| MVP_top60_DNA_seed | HSP90AA1 | 2.090851 | KIRC | RCC10RGB | Tanespimycin | MVP_top60_DNA+RNA_seed | BRAF | 0.11597 | LUAD | NCI-H2228 | RAF_9304 |
| MVP_top60_DNA_seed | HSP90AB1 | 2.090851 | KIRC | RCC10RGB | Tanespimycin | MVP_top60_DNA+RNA_seed | EGFR | 0.118719 | LUAD | VMRC-LCD | Cetuximab |
| MVP_top60_DNA_seed | MTOR | 2.111663 | KIRC | RCC10RGB | Temsirolimus | MVP_top60_DNA+RNA_seed | EGFR | 0.119269 | LUAD | HOP-62 | Cetuximab |
| MVP_top60_DNA_seed | PARP1 | 2.136773 | KIRC | RCC10RGB | Olaparib | MVP_top60_DNA+RNA_seed | HSP90AA1 | 0.119605 | LUAD | NCI-H522 | SNX-2112 |
| MVP_top60_DNA_seed | PIK3CA | 2.144494 | KIRC | A498 | AZD8835 | MVP_top60_DNA+RNA_seed | AURKA | 0.119938 | LUAD | NCI-H3122 | GSK1070916 |
| MVP_top60_DNA_seed | PIK3CD | 2.144494 | KIRC | A498 | AZD8835 | MVP_top60_DNA+RNA_seed | EGFR | 0.120587 | LUAD | NCI-H2405 | Gefitinib |
| MVP_top60_DNA_seed | AR | 2.148962 | KIRC | RCC10RGB | Bicalutamide | MVP_top60_DNA+RNA_seed | EGFR | 0.120819 | LUAD | RERF-LC-MS | Afatinib |
| MVP_top60_DNA_seed | ATM | 2.163411 | KIRC | LB2241-RCC | CP466722 | MVP_top60_DNA+RNA_seed | EGFR | 0.120835 | LUAD | NCI-H2347 | CI-1033 |
| MVP_top60_DNA_seed | HSP90AA1 | 2.235188 | KIRC | RCC10RGB | CCT-018159 | MVP_top60_DNA+RNA_seed | AURKA | 0.122198 | LUAD | PC-14 | ZM447439 |
| MVP_top60_DNA_seed | HSP90AB1 | 2.235188 | KIRC | RCC10RGB | CCT-018159 | MVP_top60_DNA+RNA_seed | EGFR | 0.122922 | LUAD | NCI-H1755 | AZD8931 |
| MVP_top60_DNA_seed | PIK3CD | 2.298621 | KIRC | A498 | IC-87114 | MVP_top60_DNA+RNA_seed | HDAC1 | 0.1232 | LUAD | HCC-78 | JQ12 |
| MVP_top60_DNA_seed | MTOR | 2.356098 | KIRC | LB2241-RCC | JW-7-52-1 | MVP_top60_DNA+RNA_seed | SRC | 0.123697 | LUAD | HCC-78 | NVP-BHG712 |
| MVP_top60_DNA_seed | HDAC1 | 2.410439 | KIRC | TK10 | AR-42 | MVP_top60_DNA+RNA_seed | HDAC1 | 0.125726 | LUAD | NCI-H1648 | JQ12 |
| MVP_top60_DNA_seed | HDAC1 | 2.455744 | KIRC | TK10 | Belinostat | MVP_top60_DNA+RNA_seed | EGFR | 0.126278 | LUAD | NCI-H1781 | CUDC-101 |
| MVP_top60_DNA_seed | HSP90AA1 | 2.498394 | KIRC | KMRC-20 | Luminespib | MVP_top60_DNA+RNA_seed | EGFR | 0.127341 | LUAD | HOP-62 | PF-00299804 |

| | | | | | | | | | | | |
|---|---|---|---|---|---|---|---|---|---|---|---|
| MVP_top60_DNA_seed | HSP90AB1 | 2.498394 | KIRC | KMRC-20 | Luminespib | MVP_top60_DNA+RNA_seed | AKT1 | 0.127859 | LUAD | NCI-H23 | A-443654 |
| MVP_top60_DNA_seed | HDAC1 | 2.524926 | KIRC | HA7-RCC | AR-42 | MVP_top60_DNA+RNA_seed | HSP90AA1 | 0.129047 | LUAD | NCI-H522 | CCT-018159 |
| MVP_top60_DNA_seed | HDAC1 | 2.600514 | KIRC | RCC10RGB | Belinostat | MVP_top60_DNA+RNA_seed | HSP90AA1 | 0.129654 | LUAD | RERF-LC-MS | Tanespimycin |
| MVP_top60_DNA_seed | HDAC1 | 2.634638 | KIRC | RCC10RGB | Entinostat | MVP_top60_DNA+RNA_seed | PIK3CA | 0.13005 | LUAD | PC-14 | PI3Ka_4409 |
| MVP_top60_DNA_seed | HDAC1 | 2.708613 | KIRC | HA7-RCC | CAY10603 | MVP_top60_DNA+RNA_seed | IGF1R | 0.13059 | LUAD | NCI-H2347 | BMS-536924 |
| MVP_top60_DNA_seed | HDAC1 | 2.763816 | KIRC | TK10 | CAY10603 | MVP_top60_DNA+RNA_seed | EGFR | 0.132407 | LUAD | NCI-H2347 | Afatinib |
| MVP_top60_DNA_seed | HDAC1 | 2.788661 | KIRC | LB996-RCC | CAY10603 | MVP_top60_DNA+RNA_seed | HSP90AA1 | 0.132426 | LUAD | NCI-H1651 | Tanespimycin |
| MVP_top60_DNA_seed | HDAC1 | 2.828967 | KIRC | A498 | Belinostat | MVP_top60_DNA+RNA_seed | HSP90AA1 | 0.133602 | LUAD | NCI-H2085 | SNX-2112 |
| MVP_top60_DNA_seed | MTOR | 2.846558 | KIRC | A498 | WYE-125132 | MVP_top60_DNA+RNA_seed | EGFR | 0.133994 | LUAD | NCI-H2030 | CUDC-101 |
| MVP_top60_DNA_seed | HDAC1 | 2.950869 | KIRC | HA7-RCC | Belinostat | MVP_top60_DNA+RNA_seed | AKT1 | 0.134447 | LUAD | NCI-H522 | AKT inhibitor VIII |
| MVP_top60_DNA_seed | MTOR | 3.018643 | KIRC | A498 | JW-7-52-1 | MVP_top60_DNA+RNA_seed | PIK3CA | 0.134479 | LUAD | EMC-BAC-1 | AZD8835 |
| MVP_top60_DNA_seed | MTOR | 3.086888 | KIRC | A498 | Voxtalisib | MVP_top60_DNA+RNA_seed | HSP90AA1 | 0.135687 | LUAD | HCC-78 | CCT-018159 |
| MVP_top60_DNA_seed | HDAC1 | 3.100667 | KIRC | A498 | Dacinostat | MVP_top60_DNA+RNA_seed | FGFR1 | 0.135954 | LUAD | NCI-H1355 | FGFR_3831 |
| MVP_top60_DNA_seed | HSP90AA1 | 3.151748 | KIRC | RCC10RGB | Elesclomol | MVP_top60_DNA+RNA_seed | EGFR | 0.13668 | LUAD | NCI-H1793 | AST-1306 |
| MVP_top60_DNA_seed | HSP90AB1 | 3.151748 | KIRC | RCC10RGB | Elesclomol | MVP_top60_DNA+RNA_seed | HSP90AA1 | 0.137582 | LUAD | HCC-827 | Luminespib |
| MVP_top60_DNA_seed | HSP90AA1 | 3.177217 | KIRC | A498 | Luminespib | MVP_top60_DNA+RNA_seed | HSP90AA1 | 0.137664 | LUAD | NCI-H1648 | SNX-2112 |
| MVP_top60_DNA_seed | HSP90AB1 | 3.177217 | KIRC | A498 | Luminespib | MVP_top60_DNA+RNA_seed | ATM | 0.13836 | LUAD | NCI-H2009 | KU-55933 |
| MVP_top60_DNA_seed | MTOR | 3.421029 | KIRC | A498 | Torin 2 | MVP_top60_DNA+RNA_seed | AKT1 | 0.13885 | LUAD | NCI-H1563 | GSK690693 |
| MVP_top60_DNA_seed | MTOR | 3.659331 | KIRC | A498 | Apitolisib | MVP_top60_DNA+RNA_seed | MET | 0.139991 | LUAD | Calu-3 | JNJ38877605 |
| MVP_top60_DNA_seed | PIK3CA | -3.218483 | KIRC | RCC-FG2 | AZD8186 | MVP_top60_DNA+RNA_seed | PTK2 | 0.141621 | LUAD | SW1573 | PF-562271 |
| MVP_top60_DNA_seed | PIK3CA | -2.68479 | KIRC | CAL-54 | AZD8186 | MVP_top60_DNA+RNA_seed | BRAF | 0.142884 | LUAD | NCI-H1573 | PLX-4720 |
| MVP_top60_DNA_seed | PIK3CA | -2.420553 | KIRC | A498 | AZD8186 | MVP_top60_DNA+RNA_seed | EGFR | 0.143458 | LUAD | A549 | Gefitinib |
| MVP_top60_DNA_seed | PIK3CA | -2.259398 | KIRC | RCC-JF | AZD8186 | MVP_top60_DNA+RNA_seed | BRAF | 0.143617 | LUAD | NCI-H3122 | HG6-64-1 |
| MVP_top60_DNA_seed | PIK3CA | -2.125832 | KIRC | KMRC-1 | AZD8186 | MVP_top60_DNA+RNA_seed | IGF1R | 0.145148 | LUAD | NCI-H1793 | BMS-536924 |
| MVP_top60_DNA_seed | ATM | -2.07997 | KIRC | RCC-ER | KU-55933 | MVP_top60_DNA+RNA_seed | SRC | 0.145293 | LUAD | NCI-H2347 | NVP-BHG712 |
| MVP_top60_DNA_seed | PIK3CA | -1.724921 | KIRC | RXF393 | AZD8186 | MVP_top60_DNA+RNA_seed | BRAF | 0.145363 | LUAD | NCI-H650 | SB590885 |
| MVP_top60_DNA_seed | PARP1 | -1.580541 | KIRC | RCC-JF | Veliparib | MVP_top60_DNA+RNA_seed | BRAF | 0.14654 | LUAD | NCI-H1568 | HG6-64-1 |
| MVP_top60_DNA_seed | PIK3CA | -1.572785 | KIRC | U031 | AZD8186 | MVP_top60_DNA+RNA_seed | BRAF | 0.147271 | LUAD | HOP-62 | RAF_9304 |
| MVP_top60_DNA_seed | PIK3CA | -1.560281 | KIRC | BFTC-909 | Buparlisib | MVP_top60_DNA+RNA_seed | FGFR1 | 0.147722 | LUAD | RERF-LC-KJ | PD173074 |
| MVP_top60_DNA_seed | PIK3CD | -1.560281 | KIRC | BFTC-909 | Buparlisib | MVP_top60_DNA+RNA_seed | SMARCA4 | 0.148357 | LUAD | H3255 | PFI-3 |
| MVP_top60_DNA_seed | MTOR | -1.510518 | KIRC | RCC-FG2 | Temsirolimus | MVP_top60_DNA+RNA_seed | IGF1R | 0.148908 | LUAD | NCI-H2085 | Linsitinib |
| MVP_top60_DNA_seed | HSP90AA1 | -1.502983 | KIRC | RCC-JF | Luminespib | MVP_top60_DNA+RNA_seed | AURKA | 0.149007 | LUAD | NCI-H1651 | HG-5-113-01 |
| MVP_top60_DNA_seed | HSP90AB1 | -1.502983 | KIRC | RCC-JF | Luminespib | MVP_top60_DNA+RNA_seed | AURKA | 0.149342 | LUAD | EMC-BAC-1 | Alisertib |
| MVP_top60_DNA_seed | HSP90AA1 | -1.458065 | KIRC | RCC-JF | Elesclomol | MVP_top60_DNA+RNA_seed | MAPK14 | 0.149386 | LUAD | NCI-H1792 | TAK-715 |
| MVP_top60_DNA_seed | HSP90AB1 | -1.458065 | KIRC | RCC-JF | Elesclomol | MVP_top60_DNA+RNA_seed | AKT1 | 0.150566 | LUAD | NCI-H3122 | GSK690693 |
| MVP_top60_DNA_seed | HSP90AA1 | -1.427487 | KIRC | RCC-JF | Tanespimycin | MVP_top60_DNA+RNA_seed | IGF1R | 0.151854 | LUAD | NCI-H23 | BMS-536924 |
| MVP_top60_DNA_seed | HSP90AB1 | -1.427487 | KIRC | RCC-JF | Tanespimycin | MVP_top60_DNA+RNA_seed | BRAF | 0.151966 | LUAD | VMRC-LCD | Dabrafenib |
| MVP_top60_DNA_seed | PARP1 | -1.423019 | KIRC | RCC-JF | Rucaparib | MVP_top60_DNA+RNA_seed | BRAF | 0.15212 | LUAD | NCI-H1563 | PLX-4720 |
| MVP_top60_DNA_seed | TP53 | -1.367967 | KIRC | RCC-ER | MIRA-1 | MVP_top60_DNA+RNA_seed | HSP90AA1 | 0.152314 | LUAD | ABC-1 | SNX-2112 |
| MVP_top60_DNA_seed | ESR1 | -1.356483 | KIRC | A498 | Tamoxifen | MVP_top60_DNA+RNA_seed | IGF1R | 0.155229 | LUAD | H3255 | BMS-754807 |
| MVP_top60_DNA_seed | PIK3CA | -1.355301 | KIRC | OS-RC-2 | AZD8186 | MVP_top60_DNA+RNA_seed | MET | 0.156087 | LUAD | EMC-BAC-1 | AZD6094 |

| | | | | | | | | | | | |
|---|---|---|---|---|---|---|---|---|---|---|---|
| MVP_top60_DNA_seed | PIK3CA | -1.348011 | KIRC | NCC021 | AZD8186 | MVP_top60_DNA+RNA_seed | AKT1 | 0.157663 | LUAD | NCI-H3122 | AKT inhibitor VIII |
| MVP_top60_DNA_seed | PIK3CA | -1.337051 | KIRC | RCC-ER | AZD8186 | MVP_top60_DNA+RNA_seed | AURKA | 0.158273 | LUAD | NCI-H1573 | ZM447439 |
| MVP_top60_DNA_seed | PIK3CA | -1.332343 | KIRC | CAL-54 | GNE-317 | MVP_top60_DNA+RNA_seed | AURKA | 0.158969 | LUAD | NCI-H1944 | Alisertib |
| MVP_top60_DNA_seed | PIK3CA | -1.291194 | KIRC | BFTC-909 | GNE-317 | MVP_top60_DNA+RNA_seed | AURKA | 0.159447 | LUAD | NCI-H1734 | Alisertib |
| MVP_top60_DNA_seed | IGF1R | -1.272041 | KIRC | RCC-MF | GSK1904529A | MVP_top60_DNA+RNA_seed | MDM2 | 0.159851 | LUAD | LC-2-ad | Serdemetan |
| MVP_top60_DNA_seed | HSP90AA1 | -1.266588 | KIRC | BFTC-909 | Tanespimycin | MVP_top60_DNA+RNA_seed | FGFR1 | 0.160811 | LUAD | NCI-H2085 | FGFR_3831 |
| MVP_top60_DNA_seed | HSP90AB1 | -1.266588 | KIRC | BFTC-909 | Tanespimycin | MVP_top60_DNA+RNA_seed | BRAF | 0.161044 | LUAD | A549 | PLX-4720 |
| MVP_top60_DNA_seed | ESR1 | -1.241312 | KIRC | BFTC-909 | Tamoxifen | MVP_top60_DNA+RNA_seed | AKT1 | 0.161336 | LUAD | NCI-H1568 | BAY AKT1 |
| MVP_top60_DNA_seed | ESR1 | -1.224183 | KIRC | CAL-54 | Tamoxifen | MVP_top60_DNA+RNA_seed | FGFR1 | 0.161769 | LUAD | NCI-H2122 | PD173074 |
| MVP_top60_DNA_seed | PIK3CA | -1.203648 | KIRC | CAL-54 | Buparlisib | MVP_top60_DNA+RNA_seed | AKT1 | 0.161981 | LUAD | NCI-H2405 | BAY AKT1 |
| MVP_top60_DNA_seed | PIK3CD | -1.203648 | KIRC | CAL-54 | Buparlisib | MVP_top60_DNA+RNA_seed | HDAC1 | 0.162004 | LUAD | HCC-827 | AR-42 |
| MVP_top60_DNA_seed | ATM | -1.184273 | KIRC | A704 | KU-55933 | MVP_top60_DNA+RNA_seed | AURKA | 0.162844 | LUAD | NCI-H3122 | Genentech Cpd 10 |
| MVP_top60_DNA_seed | PIK3CA | -1.184112 | KIRC | A704 | AZD8186 | MVP_top60_DNA+RNA_seed | BRAF | 0.163332 | LUAD | NCI-H1623 | RAF_9304 |
| MVP_top60_DNA_seed | PIK3CA | -1.176332 | KIRC | BFTC-909 | Alpelisib | MVP_top60_DNA+RNA_seed | SRC | 0.163423 | LUAD | LXF-289 | NVP-BHG712 |
| MVP_top60_DNA_seed | MTOR | -1.159736 | KIRC | RCC-JF | Temsirolimus | MVP_top60_DNA+RNA_seed | ATM | 0.163518 | LUAD | NCI-H522 | KU-55933 |
| MVP_top60_DNA_seed | IGF1R | -1.152004 | KIRC | CAL-54 | BMS-536924 | MVP_top60_DNA+RNA_seed | IGF1R | 0.163828 | LUAD | ABC-1 | GSK1904529A |
| MVP_top60_DNA_seed | HDAC1 | -1.140414 | KIRC | RCC-FG2 | PCI-34051 | MVP_top60_DNA+RNA_seed | ATM | 0.165017 | LUAD | NCI-H1792 | KU-60019 |
| MVP_top60_DNA_seed | ATM | -1.134385 | KIRC | NCC010 | KU-55933 | MVP_top60_DNA+RNA_seed | HSP90AA1 | 0.166028 | LUAD | NCI-H1793 | CCT-018159 |
| MVP_top60_DNA_seed | PARP1 | -1.132904 | KIRC | LB996-RCC | Talazoparib | MVP_top60_DNA+RNA_seed | FGFR1 | 0.166604 | LUAD | NCI-H1755 | PD173074 |
| MVP_top60_DNA_seed | PIK3CA | -1.12485 | KIRC | RCC-JF | GNE-317 | MVP_top60_DNA+RNA_seed | EGFR | 0.166765 | LUAD | NCI-H1781 | Gefitinib |
| MVP_top60_DNA_seed | IGF1R | -1.097138 | KIRC | LB996-RCC | BMS-536924 | MVP_top60_DNA+RNA_seed | EGFR | 0.167501 | LUAD | NCI-H1993 | CUDC-101 |
| MVP_top60_DNA_seed | PIK3CA | -1.065702 | KIRC | 786-0 | AZD8186 | MVP_top60_DNA+RNA_seed | EGFR | 0.167645 | LUAD | NCI-H1563 | AST-1306 |
| MVP_top60_DNA_seed | PIK3CA | -1.056549 | KIRC | LB996-RCC | AZD8186 | MVP_top60_DNA+RNA_seed | MET | 0.167709 | LUAD | NCI-H1734 | SU11274 |
| MVP_top60_DNA_seed | PIK3CA | -1.053786 | KIRC | CAKI-1 | AZD8186 | MVP_top60_DNA+RNA_seed | MET | 0.167717 | LUAD | NCI-H1651 | AZD6094 |
| MVP_top60_DNA_seed | HSP90AA1 | -1.016732 | KIRC | BB65-RCC | Tanespimycin | MVP_top60_DNA+RNA_seed | AKT1 | 0.167932 | LUAD | NCI-H1792 | BAY AKT1 |
| MVP_top60_DNA_seed | HSP90AB1 | -1.016732 | KIRC | BB65-RCC | Tanespimycin | MVP_top60_DNA+RNA_seed | AURKA | 0.168099 | LUAD | RERF-LC-MS | CD532 |
| MVP_top60_DNA_seed | PIK3CA | -1.016439 | KIRC | LB996-RCC | GNE-317 | MVP_top60_DNA+RNA_seed | MET | 0.169186 | LUAD | NCI-H1437 | AZD6094 |
| MVP_top60_DNA_seed | HSP90AA1 | -1.008454 | KIRC | RCC-FG2 | Luminespib | MVP_top60_DNA+RNA_seed | HDAC1 | 0.171363 | LUAD | PC-14 | Dacinostat |
| MVP_top60_DNA_seed | HSP90AB1 | -1.008454 | KIRC | RCC-FG2 | Luminespib | MVP_top60_DNA+RNA_seed | AURKA | 0.171861 | LUAD | SW1573 | ZM447439 |
| MVP_top60_DNA_seed | EP300 | -1.003073 | KIRC | 769-P | CPI-637 | MVP_top60_DNA+RNA_seed | EGFR | 0.171946 | LUAD | NCI-H2087 | CI-1033 |
| MVP_top60_DNA_seed | HSP90AA1 | -0.997573 | KIRC | RCC-JF | CCT-018159 | MVP_top60_DNA+RNA_seed | SRC | 0.172039 | LUAD | NCI-H2228 | Bosutinib |
| MVP_top60_DNA_seed | HSP90AB1 | -0.997573 | KIRC | RCC-JF | CCT-018159 | MVP_top60_DNA+RNA_seed | IGF1R | 0.172208 | LUAD | LXF-289 | BMS-536924 |
| MVP_top60_DNA_seed | PIK3CA | -0.993458 | KIRC | RCC-FG2 | GNE-317 | MVP_top60_DNA+RNA_seed | SRC | 0.172723 | LUAD | NCI-H650 | NVP-BHG712 |
| MVP_top60_DNA_seed | PIK3CA | -0.985424 | KIRC | LB996-RCC | Buparlisib | MVP_top60_DNA+RNA_seed | HDAC1 | 0.172726 | LUAD | NCI-H522 | Tubastatin A |
| MVP_top60_DNA_seed | PIK3CD | -0.985424 | KIRC | LB996-RCC | Buparlisib | MVP_top60_DNA+RNA_seed | AURKA | 0.173385 | LUAD | ABC-1 | CD532 |
| MVP_top60_DNA_seed | HSP90AA1 | -0.962962 | KIRC | LB996-RCC | Luminespib | MVP_top60_DNA+RNA_seed | BRAF | 0.173794 | LUAD | SW1573 | PLX-4720 |
| MVP_top60_DNA_seed | HSP90AB1 | -0.962962 | KIRC | LB996-RCC | Luminespib | MVP_top60_DNA+RNA_seed | BRAF | 0.174172 | LUAD | LXF-289 | Dabrafenib |
| MVP_top60_DNA_seed | PARP1 | -0.955883 | KIRC | RCC-JF | Niraparib | MVP_top60_DNA+RNA_seed | AKT1 | 0.174546 | LUAD | SK-LU-1 | AKT inhibitor VIII |
| MVP_top60_DNA_seed | PIK3CA | -0.946664 | KIRC | RCC-ER | Alpelisib | MVP_top60_DNA+RNA_seed | EGFR | 0.175242 | LUAD | ABC-1 | AST-1306 |
| MVP_top60_DNA_seed | IGF1R | -0.941836 | KIRC | RCC-ER | BMS-536924 | MVP_top60_DNA+RNA_seed | SMARCA4 | 0.178408 | LUAD | NCI-H358 | PFI-3 |
| MVP_top60_DNA_seed | HSP90AA1 | -0.940046 | KIRC | LB996-RCC | Tanespimycin | MVP_top60_DNA+RNA_seed | IGF1R | 0.17879 | LUAD | VMRC-LCD | BMS-536924 |

| | | | | | | | | | | | |
|---|---|---|---|---|---|---|---|---|---|---|---|
| MVP_top60_DNA_seed | HSP90AB1 | -0.940046 | KIRC | LB996-RCC | Tanespimycin | MVP_top60_DNA+RNA_seed | IGF1R | 0.179206 | LUAD | NCI-H2342 | BMS-536924 |
| MVP_top60_DNA_seed | PARP1 | -0.91147 | KIRC | RCC-JF | Talazoparib | MVP_top60_DNA+RNA_seed | HDAC1 | 0.179322 | LUAD | NCI-H838 | Tubastatin A |
| MVP_top60_DNA_seed | PIK3CA | -0.911076 | KIRC | LB1047-RCC | Alpelisib | MVP_top60_DNA+RNA_seed | IGF1R | 0.179385 | LUAD | NCI-H1755 | Linsitinib |
| MVP_top60_DNA_seed | IGF1R | -0.890138 | KIRC | RCC-ER | BMS-754807 | MVP_top60_DNA+RNA_seed | SMARCA4 | 0.179965 | LUAD | LC-2-ad | PFI-3 |
| MVP_top60_DNA_seed | ATM | -0.888756 | KIRC | RCC-JW | KU-55933 | MVP_top60_DNA+RNA_seed | IGF1R | 0.181267 | LUAD | NCI-H2087 | GSK1904529A |
| MVP_top60_DNA_seed | PIK3CA | -0.870576 | KIRC | HA7-RCC | AZD8186 | MVP_top60_DNA+RNA_seed | MET | 0.181418 | LUAD | NCI-H1355 | AZD6094 |
| MVP_top60_DNA_seed | IGF1R | -0.863271 | KIRC | U031 | GSK1904529A | MVP_top60_DNA+RNA_seed | PIK3CA | 0.18172 | LUAD | NCI-H23 | AZD8835 |
| MVP_top60_DNA_seed | IGF1R | -0.8624 | KIRC | A704 | BMS-536924 | MVP_top60_DNA+RNA_seed | PIK3CA | 0.182728 | LUAD | NCI-H3122 | PI3Ka_4409 |
| MVP_top60_DNA_seed | PIK3CA | -0.854873 | KIRC | 769-P | GNE-317 | MVP_top60_DNA+RNA_seed | MET | 0.184076 | LUAD | NCI-H1755 | AZD6094 |
| MVP_top60_DNA_seed | PIK3CA | -0.852015 | KIRC | LB2241-RCC | GNE-317 | MVP_top60_DNA+RNA_seed | AURKA | 0.185803 | LUAD | NCI-H522 | ZM447439 |
| MVP_top60_DNA_seed | HSP90AA1 | -0.849202 | KIRC | U031 | CCT-018159 | MVP_top60_DNA+RNA_seed | IGF1R | 0.185863 | LUAD | NCI-H2405 | BMS-536924 |
| MVP_top60_DNA_seed | HSP90AB1 | -0.849202 | KIRC | U031 | CCT-018159 | MVP_top60_DNA+RNA_seed | EGFR | 0.186015 | LUAD | NCI-H1993 | Afatinib |
| MVP_top60_DNA_seed | HSP90AA1 | -0.847173 | KIRC | RCC-FG2 | CCT-018159 | MVP_top60_DNA+RNA_seed | FGFR1 | 0.186421 | LUAD | ABC-1 | FGFR_3831 |
| MVP_top60_DNA_seed | HSP90AB1 | -0.847173 | KIRC | RCC-FG2 | CCT-018159 | MVP_top60_DNA+RNA_seed | AURKA | 0.187163 | LUAD | NCI-H1792 | Alisertib |
| MVP_top60_DNA_seed | IGF1R | -0.816724 | KIRC | NCC021 | BMS-536924 | MVP_top60_DNA+RNA_seed | PIK3CA | 0.187222 | LUAD | NCI-H1650 | PI3Ka_4409 |
| MVP_top60_DNA_seed | HSP90AA1 | -0.805852 | KIRC | RXF393 | Luminespib | MVP_top60_DNA+RNA_seed | IGF1R | 0.187285 | LUAD | NCI-H2087 | Linsitinib |
| MVP_top60_DNA_seed | HSP90AB1 | -0.805852 | KIRC | RXF393 | Luminespib | MVP_top60_DNA+RNA_seed | EGFR | 0.187638 | LUAD | EMC-BAC-2 | AST-1306 |
| MVP_top60_DNA_seed | PIK3CA | -0.803391 | KIRC | RCC10RGB | AZD8186 | MVP_top60_DNA+RNA_seed | HDAC1 | 0.188087 | LUAD | NCI-H2030 | AR-42 |
| MVP_top60_DNA_seed | PIK3CA | -0.803262 | KIRC | ACHN | GNE-317 | MVP_top60_DNA+RNA_seed | HDAC1 | 0.18837 | LUAD | NCI-H322M | Belinostat |
| MVP_top60_DNA_seed | PIK3CA | -0.797416 | KIRC | ACHN | AZD8186 | MVP_top60_DNA+RNA_seed | EGFR | 0.188767 | LUAD | NCI-H441 | AZD8931 |
| MVP_top60_DNA_seed | HDAC1 | -0.795837 | KIRC | RCC-JF | PCI-34051 | MVP_top60_DNA+RNA_seed | AURKA | 0.189375 | LUAD | NCI-H1563 | Genentech Cpd 10 |
| MVP_top60_DNA_seed | HSP90AA1 | -0.789065 | KIRC | RXF393 | CCT-018159 | MVP_top60_DNA+RNA_seed | AKT1 | 0.189506 | LUAD | NCI-H23 | MK-2206 |
| MVP_top60_DNA_seed | HSP90AB1 | -0.789065 | KIRC | RXF393 | CCT-018159 | MVP_top60_DNA+RNA_seed | AKT1 | 0.190701 | LUAD | NCI-H1355 | AKT inhibitor VIII |
| MVP_top60_DNA_seed | AR | -0.785124 | KIRC | U031 | Bicalutamide | MVP_top60_DNA+RNA_seed | EGFR | 0.192345 | LUAD | LC-2-ad | Afatinib |
| MVP_top60_DNA_seed | ATM | -0.783711 | KIRC | RCC10RGB | KU-55933 | MVP_top60_DNA+RNA_seed | FGFR1 | 0.192672 | LUAD | VMRC-LCD | PD173074 |
| MVP_top60_DNA_seed | IGF1R | -0.774009 | KIRC | LB996-RCC | GSK1904529A | MVP_top60_DNA+RNA_seed | EGFR | 0.192772 | LUAD | NCI-H650 | PF-00299804 |
| MVP_top60_DNA_seed | AR | -0.765839 | KIRC | RCC-JF | Bicalutamide | MVP_top60_DNA+RNA_seed | AURKA | 0.193879 | LUAD | NCI-H1568 | ZM447439 |
| MVP_top60_DNA_seed | IGF1R | -0.763919 | KIRC | U031 | BMS-536924 | MVP_top60_DNA+RNA_seed | MET | 0.194184 | LUAD | NCI-H1355 | JNJ38877605 |
| MVP_top60_DNA_seed | MTOR | -0.755409 | KIRC | A498 | Temsirolimus | MVP_top60_DNA+RNA_seed | ATM | 0.194418 | LUAD | NCI-H1568 | CP466722 |
| MVP_top60_DNA_seed | PARP1 | -0.754471 | KIRC | U031 | Niraparib | MVP_top60_DNA+RNA_seed | FGFR1 | 0.194424 | LUAD | NCI-H1437 | FGFR_3831 |
| MVP_top60_DNA_seed | PIK3CA | -0.736097 | KIRC | BFTC-909 | AZD8186 | MVP_top60_DNA+RNA_seed | AURKA | 0.195735 | LUAD | NCI-H2347 | GSK1070916 |
| MVP_top60_DNA_seed | ATM | -0.729976 | KIRC | RCC-FG2 | KU-55933 | MVP_top60_DNA+RNA_seed | FGFR1 | 0.195938 | LUAD | EMC-BAC-1 | PD173074 |
| MVP_top60_DNA_seed | PIK3CA | -0.721635 | KIRC | U031 | GNE-317 | MVP_top60_DNA+RNA_seed | AKT1 | 0.19635 | LUAD | NCI-H1437 | GSK690693 |
| MVP_top60_DNA_seed | HDAC1 | -0.715943 | KIRC | NCC010 | PCI-34051 | MVP_top60_DNA+RNA_seed | EGFR | 0.196811 | LUAD | NCI-H1755 | Afatinib |
| MVP_top60_DNA_seed | IGF1R | -0.708279 | KIRC | NCC010 | BMS-754807 | MVP_top60_DNA+RNA_seed | MET | 0.199832 | LUAD | Calu-6 | JNJ38877605 |
| MVP_top60_DNA_seed | ATM | -0.702862 | KIRC | LB996-RCC | KU-55933 | MVP_top60_DNA+RNA_seed | AURKA | 0.199975 | LUAD | NCI-H1781 | CD532 |
| MVP_top60_DNA_seed | EP300 | -0.699633 | KIRC | CAL-54 | CPI-637 | MVP_top60_DNA+RNA_seed | BRAF | 0.200837 | LUAD | NCI-H2342 | PLX-4720 |
| MVP_top60_DNA_seed | PIK3CA | -0.699149 | KIRC | RCC-ER | Buparlisib | MVP_top60_DNA+RNA_seed | AKT1 | 0.200924 | LUAD | EMC-BAC-1 | BAY AKT1 |
| MVP_top60_DNA_seed | PIK3CD | -0.699149 | KIRC | RCC-ER | Buparlisib | MVP_top60_DNA+RNA_seed | HSP90AA1 | 0.200979 | LUAD | NCI-H1792 | SNX-2112 |
| MVP_top60_DNA_seed | HSP90AA1 | -0.697923 | KIRC | BB65-RCC | Luminespib | MVP_top60_DNA+RNA_seed | EGFR | 0.201012 | LUAD | SK-LU-1 | AST-1306 |
| MVP_top60_DNA_seed | HSP90AB1 | -0.697923 | KIRC | BB65-RCC | Luminespib | MVP_top60_DNA+RNA_seed | HDAC1 | 0.201388 | LUAD | NCI-H2009 | JQ12 |

| | | | | | | | | | | | |
|---|---|---|---|---|---|---|---|---|---|---|---|
| MVP_top60_DNA_seed | IGF1R | -0.695017 | KIRC | A498 | NVP-ADW742 | MVP_top60_DNA+RNA_seed | BRAF | 0.202482 | LUAD | NCI-H1792 | SB590885 |
| MVP_top60_DNA_seed | ESR1 | -0.678958 | KIRC | 786-0 | GDC0810 | MVP_top60_DNA+RNA_seed | SMARCA4 | 0.202512 | LUAD | NCI-H23 | PFI-3 |
| MVP_top60_DNA_seed | HDAC1 | -0.670549 | KIRC | CAKI-1 | PCI-34051 | MVP_top60_DNA+RNA_seed | IGF1R | 0.202535 | LUAD | Calu-3 | Linsitinib |
| MVP_top60_DNA_seed | HSP90AA1 | -0.668507 | KIRC | RCC-JW | Tanespimycin | MVP_top60_DNA+RNA_seed | SRC | 0.203815 | LUAD | NCI-H1648 | Bosutinib |
| MVP_top60_DNA_seed | HSP90AB1 | -0.668507 | KIRC | RCC-JW | Tanespimycin | MVP_top60_DNA+RNA_seed | HDAC1 | 0.203837 | LUAD | NCI-H1573 | Belinostat |
| MVP_top60_DNA_seed | ATM | -0.659454 | KIRC | CAL-54 | KU-55933 | MVP_top60_DNA+RNA_seed | IGF1R | 0.205394 | LUAD | NCI-H2023 | Linsitinib |
| MVP_top60_DNA_seed | MTOR | -0.655868 | KIRC | RCC-ER | Temsirolimus | MVP_top60_DNA+RNA_seed | EGFR | 0.205404 | LUAD | Calu-6 | CI-1033 |
| MVP_top60_DNA_seed | IGF1R | -0.647047 | KIRC | LB996-RCC | NVP-ADW742 | MVP_top60_DNA+RNA_seed | MDM2 | 0.207247 | LUAD | H3255 | Serdemetan |
| MVP_top60_DNA_seed | IGF1R | -0.646483 | KIRC | A704 | NVP-ADW742 | MVP_top60_DNA+RNA_seed | IGF1R | 0.207685 | LUAD | NCI-H1568 | GSK1904529A |
| MVP_top60_DNA_seed | HSP90AA1 | -0.644857 | KIRC | BFTC-909 | Luminespib | MVP_top60_DNA+RNA_seed | AURKA | 0.209002 | LUAD | NCI-H358 | CD532 |
| MVP_top60_DNA_seed | HSP90AB1 | -0.644857 | KIRC | BFTC-909 | Luminespib | MVP_top60_DNA+RNA_seed | AURKA | 0.209612 | LUAD | NCI-H2009 | CD532 |
| MVP_top60_DNA_seed | PIK3CA | -0.641962 | KIRC | RCC-JW | AZD8186 | MVP_top60_DNA+RNA_seed | EGFR | 0.209993 | LUAD | NCI-H2009 | AZD8931 |
| MVP_top60_DNA_seed | TP53 | -0.640654 | KIRC | ACHN | MIRA-1 | MVP_top60_DNA+RNA_seed | MET | 0.211055 | LUAD | NCI-H1693 | AZD6094 |
| MVP_top60_DNA_seed | EP300 | -0.628687 | KIRC | A704 | SGC-CBP30 | MVP_top60_DNA+RNA_seed | EGFR | 0.211771 | LUAD | NCI-H1793 | CI-1033 |
| MVP_top60_DNA_seed | IGF1R | -0.621436 | KIRC | BB65-RCC | NVP-ADW742 | MVP_top60_DNA+RNA_seed | EGFR | 0.211885 | LUAD | HOP-62 | Erlotinib |
| MVP_top60_DNA_seed | PARP1 | -0.617226 | KIRC | RCC-JF | Olaparib | MVP_top60_DNA+RNA_seed | ATM | 0.212281 | LUAD | NCI-H1563 | KU-55933 |
| MVP_top60_DNA_seed | PIK3CA | -0.606246 | KIRC | LB1047-RCC | GNE-317 | MVP_top60_DNA+RNA_seed | MET | 0.212541 | LUAD | ABC-1 | JNJ38877605 |
| MVP_top60_DNA_seed | IGF1R | -0.604646 | KIRC | A498 | BMS-536924 | MVP_top60_DNA+RNA_seed | SMARCA4 | 0.214719 | LUAD | NCI-H2085 | PFI-3 |
| MVP_top60_DNA_seed | IGF1R | -0.604169 | KIRC | RCC-ER | GSK1904529A | MVP_top60_DNA+RNA_seed | HSP90AA1 | 0.21503 | LUAD | NCI-H23 | SNX-2112 |
| MVP_top60_DNA_seed | PIK3CA | -0.592024 | KIRC | RCC-AB | GNE-317 | MVP_top60_DNA+RNA_seed | BRAF | 0.215476 | LUAD | NCI-H2023 | Dabrafenib |
| MVP_top60_DNA_seed | HSP90AA1 | -0.586516 | KIRC | NCC010 | CCT-018159 | MVP_top60_DNA+RNA_seed | FGFR1 | 0.21555 | LUAD | NCI-H23 | FGFR_3831 |
| MVP_top60_DNA_seed | HSP90AB1 | -0.586516 | KIRC | NCC010 | CCT-018159 | MVP_top60_DNA+RNA_seed | AURKA | 0.215798 | LUAD | NCI-H1355 | Tozasertib |
| MVP_top60_DNA_seed | PIK3CA | -0.585482 | KIRC | LB2241-RCC | AZD8186 | MVP_top60_DNA+RNA_seed | AURKA | 0.216129 | LUAD | HOP-62 | Alisertib |
| MVP_top60_DNA_seed | PIK3CA | -0.579237 | KIRC | CAKI-1 | GNE-317 | MVP_top60_DNA+RNA_seed | AKT1 | 0.216239 | LUAD | LC-2-ad | MK-2206 |
| MVP_top60_DNA_seed | HSP90AA1 | -0.578011 | KIRC | RCC-FG2 | Tanespimycin | MVP_top60_DNA+RNA_seed | MET | 0.216392 | LUAD | NCI-H650 | JNJ38877605 |
| MVP_top60_DNA_seed | HSP90AB1 | -0.578011 | KIRC | RCC-FG2 | Tanespimycin | MVP_top60_DNA+RNA_seed | ATM | 0.217734 | LUAD | SK-LU-1 | KU-60019 |
| MVP_top60_DNA_seed | HDAC1 | -0.554545 | KIRC | BFTC-909 | Romidepsin | MVP_top60_DNA+RNA_seed | BRAF | 0.21935 | LUAD | NCI-H1650 | AZ628 |
| MVP_top60_DNA_seed | EP300 | -0.553506 | KIRC | NCC010 | SGC-CBP30 | MVP_top60_DNA+RNA_seed | SMARCA4 | 0.220239 | LUAD | NCI-H2228 | PFI-3 |
| MVP_top60_DNA_seed | TP53 | -0.5454 | KIRC | CAKI-1 | MIRA-1 | MVP_top60_DNA+RNA_seed | EGFR | 0.220298 | LUAD | NCI-H2291 | CI-1033 |
| MVP_top60_DNA_seed | IGF1R | -0.541972 | KIRC | RCC-JF | NVP-ADW742 | MVP_top60_DNA+RNA_seed | PIK3CA | 0.221675 | LUAD | EKVX | PI3Ka_4409 |
| MVP_top60_DNA_seed | ESR1 | -0.541189 | KIRC | U031 | GDC0810 | MVP_top60_DNA+RNA_seed | IGF1R | 0.222188 | LUAD | NCI-H2030 | GSK1904529A |
| MVP_top60_DNA_seed | HSP90AA1 | -0.534673 | KIRC | 769-P | Tanespimycin | MVP_top60_DNA+RNA_seed | BRAF | 0.223477 | LUAD | NCI-H23 | PLX-4720 |
| MVP_top60_DNA_seed | HSP90AB1 | -0.534673 | KIRC | 769-P | Tanespimycin | MVP_top60_DNA+RNA_seed | AURKA | 0.224676 | LUAD | NCI-H2085 | GSK1070916 |
| MVP_top60_DNA_seed | IGF1R | -0.529765 | KIRC | RXF393 | BMS-536924 | MVP_top60_DNA+RNA_seed | EGFR | 0.224853 | LUAD | NCI-H2085 | Cetuximab |
| MVP_top60_DNA_seed | MTOR | -0.527579 | KIRC | RCC-MF | Temsirolimus | MVP_top60_DNA+RNA_seed | IGF1R | 0.225755 | LUAD | NCI-H2342 | Linsitinib |
| MVP_top60_DNA_seed | PIK3CA | -0.527564 | KIRC | 786-0 | GNE-317 | MVP_top60_DNA+RNA_seed | HSP90AA1 | 0.226962 | LUAD | RERF-LC-MS | Luminespib |
| MVP_top60_DNA_seed | IGF1R | -0.526387 | KIRC | NCC010 | GSK1904529A | MVP_top60_DNA+RNA_seed | FGFR1 | 0.227411 | LUAD | EMC-BAC-2 | PD173074 |
| MVP_top60_DNA_seed | HSP90AA1 | -0.520353 | KIRC | 786-0 | Tanespimycin | MVP_top60_DNA+RNA_seed | FGFR1 | 0.227719 | LUAD | HCC-78 | AZD4547 |
| MVP_top60_DNA_seed | HSP90AB1 | -0.520353 | KIRC | 786-0 | Tanespimycin | MVP_top60_DNA+RNA_seed | SMARCA4 | 0.227934 | LUAD | ABC-1 | PFI-3 |
| MVP_top60_DNA_seed | PARP1 | -0.516314 | KIRC | U031 | Rucaparib | MVP_top60_DNA+RNA_seed | HDAC1 | 0.229517 | LUAD | NCI-H1734 | Dacinostat |
| MVP_top60_DNA_seed | IGF1R | -0.515093 | KIRC | RCC-JF | BMS-536924 | MVP_top60_DNA+RNA_seed | MET | 0.229623 | LUAD | LC-2-ad | AZD6094 |

| | | | | | | | | | | | |
|---|---|---|---|---|---|---|---|---|---|---|---|
| MVP_top60_DNA_seed | IGF1R | -0.509704 | KIRC | RCC-FG2 | BMS-536924 | MVP_top60_DNA+RNA_seed | MDM2 | 0.230051 | LUAD | HCC-44 | Serdemetan |
| MVP_top60_DNA_seed | PARP1 | -0.508192 | KIRC | NCC021 | Rucaparib | MVP_top60_DNA+RNA_seed | EGFR | 0.230563 | LUAD | NCI-H2023 | Afatinib |
| MVP_top60_DNA_seed | ESR1 | -0.501132 | KIRC | 786-0 | Tamoxifen | MVP_top60_DNA+RNA_seed | SMARCA4 | 0.230838 | LUAD | NCI-H441 | PFI-3 |
| MVP_top60_DNA_seed | IGF1R | -0.500808 | KIRC | BFTC-909 | NVP-ADW742 | MVP_top60_DNA+RNA_seed | HDAC1 | 0.231683 | LUAD | NCI-H2347 | CAY10603 |
| MVP_top60_DNA_seed | IGF1R | -0.489531 | KIRC | 786-0 | NVP-ADW742 | MVP_top60_DNA+RNA_seed | BRAF | 0.232436 | LUAD | SW1573 | SB590885 |
| MVP_top60_DNA_seed | HDAC1 | -0.47977 | KIRC | RCC-ER | PCI-34051 | MVP_top60_DNA+RNA_seed | ATM | 0.232465 | LUAD | NCI-H1355 | KU-60019 |
| MVP_top60_DNA_seed | PARP1 | -0.478193 | KIRC | RCC-ER | Talazoparib | MVP_top60_DNA+RNA_seed | SMARCA4 | 0.234839 | LUAD | HOP-62 | PFI-3 |
| MVP_top60_DNA_seed | HSP90AA1 | -0.472305 | KIRC | RXF393 | Tanespimycin | MVP_top60_DNA+RNA_seed | IGF1R | 0.235091 | LUAD | HCC-78 | Linsitinib |
| MVP_top60_DNA_seed | HSP90AB1 | -0.472305 | KIRC | RXF393 | Tanespimycin | MVP_top60_DNA+RNA_seed | IGF1R | 0.23597 | LUAD | NCI-H3122 | Linsitinib |
| MVP_top60_DNA_seed | SMARCA4 | -0.472138 | KIRC | A704 | PFI3 | MVP_top60_DNA+RNA_seed | EGFR | 0.237024 | LUAD | NCI-H1734 | AST-1306 |
| MVP_top60_DNA_seed | ESR1 | -0.468891 | KIRC | LB996-RCC | Tamoxifen | MVP_top60_DNA+RNA_seed | ATM | 0.237643 | LUAD | NCI-H1792 | CP466722 |
| MVP_top60_DNA_seed | PARP1 | -0.468231 | KIRC | BFTC-909 | Talazoparib | MVP_top60_DNA+RNA_seed | BRAF | 0.238286 | LUAD | H3255 | Dabrafenib |
| MVP_top60_DNA_seed | PARP1 | -0.465091 | KIRC | RXF393 | Rucaparib | MVP_top60_DNA+RNA_seed | MET | 0.238623 | LUAD | H3255 | SU11274 |
| MVP_top60_DNA_seed | PARP1 | -0.461541 | KIRC | RCC-ER | Veliparib | MVP_top60_DNA+RNA_seed | AURKA | 0.238835 | LUAD | PC-14 | CD532 |
| MVP_top60_DNA_seed | MTOR | -0.460266 | KIRC | RXF393 | Temsirolimus | MVP_top60_DNA+RNA_seed | EGFR | 0.239045 | LUAD | NCI-H838 | Afatinib |
| MVP_top60_DNA_seed | PARP1 | -0.456882 | KIRC | BFTC-909 | Olaparib | MVP_top60_DNA+RNA_seed | HSP90AA1 | 0.241282 | LUAD | Calu-6 | Luminespib |
| MVP_top60_DNA_seed | PIK3CA | -0.456333 | KIRC | RCC-MF | GNE-317 | MVP_top60_DNA+RNA_seed | AURKA | 0.241647 | LUAD | HOP-62 | CD532 |
| MVP_top60_DNA_seed | EP300 | -0.456099 | KIRC | LB996-RCC | CPI-637 | MVP_top60_DNA+RNA_seed | AKT1 | 0.241808 | LUAD | PC-14 | BAY AKT1 |
| MVP_top60_DNA_seed | ESR1 | -0.448495 | KIRC | RCC-ER | Tamoxifen | MVP_top60_DNA+RNA_seed | SRC | 0.242349 | LUAD | HOP-62 | Bosutinib |
| MVP_top60_DNA_seed | TP53 | -0.444953 | KIRC | BFTC-909 | MIRA-1 | MVP_top60_DNA+RNA_seed | FGFR1 | 0.243107 | LUAD | NCI-H1650 | PD173074 |
| MVP_top60_DNA_seed | PIK3CA | -0.435472 | KIRC | RCC-JF | Alpelisib | MVP_top60_DNA+RNA_seed | BRAF | 0.243357 | LUAD | NCI-H2405 | PLX-4720 |
| MVP_top60_DNA_seed | HDAC1 | -0.432309 | KIRC | RCC-MF | PCI-34051 | MVP_top60_DNA+RNA_seed | HDAC1 | 0.243911 | LUAD | NCI-H1792 | JQ12 |
| MVP_top60_DNA_seed | PARP1 | -0.427156 | KIRC | RCC-ER | Rucaparib | MVP_top60_DNA+RNA_seed | MDM2 | 0.243967 | LUAD | EMC-BAC-2 | Serdemetan |
| MVP_top60_DNA_seed | SMARCA4 | -0.427139 | KIRC | NCC010 | PFI3 | MVP_top60_DNA+RNA_seed | AKT1 | 0.243992 | LUAD | NCI-H2122 | BAY AKT1 |
| MVP_top60_DNA_seed | IGF1R | -0.416927 | KIRC | RCC-ER | NVP-ADW742 | MVP_top60_DNA+RNA_seed | HDAC1 | 0.244549 | LUAD | PC-3_[JPC-3] | Belinostat |
| MVP_top60_DNA_seed | IGF1R | -0.410611 | KIRC | LB1047-RCC | BMS-536924 | MVP_top60_DNA+RNA_seed | HDAC1 | 0.245028 | LUAD | NCI-H1755 | Tubastatin A |
| MVP_top60_DNA_seed | HDAC1 | -0.409844 | KIRC | 786-0 | PCI-34051 | MVP_top60_DNA+RNA_seed | IGF1R | 0.245168 | LUAD | NCI-H2347 | GSK1904529A |
| MVP_top60_DNA_seed | PIK3CA | -0.40875 | KIRC | LB1047-RCC | AZD8186 | MVP_top60_DNA+RNA_seed | SMARCA4 | 0.245747 | LUAD | EMC-BAC-2 | PFI-3 |
| MVP_top60_DNA_seed | PARP1 | -0.407898 | KIRC | U031 | Talazoparib | MVP_top60_DNA+RNA_seed | MET | 0.245869 | LUAD | NCI-H1650 | JNJ38877605 |
| MVP_top60_DNA_seed | IGF1R | -0.406177 | KIRC | RCC-ER | Linsitinib | MVP_top60_DNA+RNA_seed | EGFR | 0.246169 | LUAD | NCI-H1648 | Cetuximab |
| MVP_top60_DNA_seed | EP300 | -0.404447 | KIRC | RCC-JW | CPI-637 | MVP_top60_DNA+RNA_seed | HDAC1 | 0.246469 | LUAD | NCI-H441 | JQ12 |
| MVP_top60_DNA_seed | IGF1R | -0.397706 | KIRC | U031 | BMS-754807 | MVP_top60_DNA+RNA_seed | MET | 0.247108 | LUAD | NCI-H358 | AZD6094 |
| MVP_top60_DNA_seed | MTOR | -0.394806 | KIRC | CAL-54 | Temsirolimus | MVP_top60_DNA+RNA_seed | EGFR | 0.247176 | LUAD | NCI-H1734 | Gefitinib |
| MVP_top60_DNA_seed | IGF1R | -0.388324 | KIRC | KMRC-20 | NVP-ADW742 | MVP_top60_DNA+RNA_seed | EGFR | 0.248345 | LUAD | NCI-H2085 | Afatinib |
| MVP_top60_DNA_seed | ESR1 | -0.378197 | KIRC | RCC-ER | GDC0810 | MVP_top60_DNA+RNA_seed | HDAC1 | 0.248443 | LUAD | EMC-BAC-1 | AR-42 |
| MVP_top60_DNA_seed | MTOR | -0.372529 | KIRC | SW156 | Temsirolimus | MVP_top60_DNA+RNA_seed | AKT1 | 0.248464 | LUAD | NCI-H2122 | MK-2206 |
| MVP_top60_DNA_seed | TP53 | -0.370749 | KIRC | RCC-MF | MIRA-1 | MVP_top60_DNA+RNA_seed | BRAF | 0.249717 | LUAD | NCI-H2291 | SB590885 |
| MVP_top60_DNA_seed | PARP1 | -0.370292 | KIRC | 769-P | Niraparib | MVP_top60_DNA+RNA_seed | AURKA | 0.250348 | LUAD | NCI-H1651 | GSK1070916 |
| MVP_top60_DNA_seed | PARP1 | -0.367434 | KIRC | 769-P | Olaparib | MVP_top60_DNA+RNA_seed | HDAC1 | 0.251314 | LUAD | NCI-H1693 | Dacinostat |
| MVP_top60_DNA_seed | IGF1R | -0.364509 | KIRC | LB996-RCC | BMS-754807 | MVP_top60_DNA+RNA_seed | AKT1 | 0.253053 | LUAD | NCI-H1734 | AKT inhibitor VIII |
| MVP_top60_DNA_seed | HDAC1 | -0.364396 | KIRC | RCC-ER | Entinostat | MVP_top60_DNA+RNA_seed | AURKA | 0.253698 | LUAD | EMC-BAC-2 | Genentech Cpd 10 |

| | | | | | | | | | | | |
|---|---|---|---|---|---|---|---|---|---|---|---|
| MVP_top60_DNA_seed | IGF1R | -0.361563 | KIRC | NCC010 | BMS-536924 | MVP_top60_DNA+RNA_seed | AURKA | 0.254645 | LUAD | NCI-H2030 | ZM447439 |
| MVP_top60_DNA_seed | PARP1 | -0.358039 | KIRC | LB1047-RCC | Talazoparib | MVP_top60_DNA+RNA_seed | EGFR | 0.256585 | LUAD | NCI-H1693 | Pelitinib |
| MVP_top60_DNA_seed | MTOR | -0.357142 | KIRC | OS-RC-2 | Temsirolimus | MVP_top60_DNA+RNA_seed | FGFR1 | 0.256856 | LUAD | NCI-H1975 | FGFR_3831 |
| MVP_top60_DNA_seed | PIK3CA | -0.341345 | KIRC | RCC-FG2 | Alpelisib | MVP_top60_DNA+RNA_seed | HDAC1 | 0.257158 | LUAD | NCI-H2405 | AR-42 |
| MVP_top60_DNA_seed | PARP1 | -0.339621 | KIRC | LB2241-RCC | Rucaparib | MVP_top60_DNA+RNA_seed | FGFR1 | 0.257612 | LUAD | RERF-LC-MS | AZD4547 |
| MVP_top60_DNA_seed | HSP90AA1 | -0.331886 | KIRC | KMRC-20 | Tanespimycin | MVP_top60_DNA+RNA_seed | BRAF | 0.258235 | LUAD | NCI-H1355 | AZ628 |
| MVP_top60_DNA_seed | HSP90AB1 | -0.331886 | KIRC | KMRC-20 | Tanespimycin | MVP_top60_DNA+RNA_seed | BRAF | 0.259102 | LUAD | NCI-H1568 | Dabrafenib |
| MVP_top60_DNA_seed | IGF1R | -0.329485 | KIRC | RCC-MF | BMS-536924 | MVP_top60_DNA+RNA_seed | MAPK14 | 0.259821 | LUAD | NCI-H441 | TAK-715 |
| MVP_top60_DNA_seed | HDAC1 | -0.32946 | KIRC | RXF393 | Romidepsin | MVP_top60_DNA+RNA_seed | AURKA | 0.260305 | LUAD | Calu-6 | Alisertib |
| MVP_top60_DNA_seed | MTOR | -0.316733 | KIRC | BFTC-909 | Temsirolimus | MVP_top60_DNA+RNA_seed | AKT1 | 0.260708 | LUAD | ABC-1 | BAY AKT1 |
| MVP_top60_DNA_seed | MTOR | -0.316635 | KIRC | RCC-JW | Temsirolimus | MVP_top60_DNA+RNA_seed | EGFR | 0.261547 | LUAD | NCI-H2228 | AST-1306 |
| MVP_top60_DNA_seed | HSP90AA1 | -0.314135 | KIRC | RCC-ER | CCT-018159 | MVP_top60_DNA+RNA_seed | HDAC1 | 0.2616 | LUAD | HCC-827 | CAY10603 |
| MVP_top60_DNA_seed | HSP90AB1 | -0.314135 | KIRC | RCC-ER | CCT-018159 | MVP_top60_DNA+RNA_seed | IGF1R | 0.261679 | LUAD | NCI-H1563 | BMS-754807 |
| MVP_top60_DNA_seed | HSP90AA1 | -0.311872 | KIRC | CAKI-1 | Elesclomol | MVP_top60_DNA+RNA_seed | HDAC1 | 0.261942 | LUAD | EMC-BAC-1 | Belinostat |
| MVP_top60_DNA_seed | HSP90AB1 | -0.311872 | KIRC | CAKI-1 | Elesclomol | MVP_top60_DNA+RNA_seed | MET | 0.264575 | LUAD | NCI-H2030 | AZD6094 |
| MVP_top60_DNA_seed | PARP1 | -0.304933 | KIRC | KMRC-20 | Veliparib | MVP_top60_DNA+RNA_seed | PIK3CA | 0.267375 | LUAD | NCI-H2405 | PI3Ka_4409 |
| MVP_top60_DNA_seed | ESR1 | -0.303033 | KIRC | ACHN | Tamoxifen | MVP_top60_DNA+RNA_seed | EGFR | 0.268679 | LUAD | EMC-BAC-2 | CI-1033 |
| MVP_top60_DNA_seed | HSP90AA1 | -0.302433 | KIRC | RCC-FG2 | Elesclomol | MVP_top60_DNA+RNA_seed | HSP90AA1 | 0.268714 | LUAD | SK-LU-1 | SNX-2112 |
| MVP_top60_DNA_seed | HSP90AB1 | -0.302433 | KIRC | RCC-FG2 | Elesclomol | MVP_top60_DNA+RNA_seed | MET | 0.26953 | LUAD | NCI-H2085 | AZD6094 |
| MVP_top60_DNA_seed | HDAC1 | -0.301455 | KIRC | RCC-JF | Entinostat | MVP_top60_DNA+RNA_seed | EGFR | 0.270282 | LUAD | NCI-H3122 | AZD8931 |
| MVP_top60_DNA_seed | ATM | -0.300438 | KIRC | KMRC-20 | KU-55933 | MVP_top60_DNA+RNA_seed | EGFR | 0.271199 | LUAD | NCI-H1650 | AZD8931 |
| MVP_top60_DNA_seed | PARP1 | -0.292204 | KIRC | CAKI-1 | Talazoparib | MVP_top60_DNA+RNA_seed | MDM2 | 0.271879 | LUAD | NCI-H2291 | Serdemetan |
| MVP_top60_DNA_seed | MTOR | -0.29049 | KIRC | U031 | Temsirolimus | MVP_top60_DNA+RNA_seed | EGFR | 0.272312 | LUAD | NCI-H1793 | Afatinib |
| MVP_top60_DNA_seed | IGF1R | -0.289694 | KIRC | 769-P | NVP-ADW742 | MVP_top60_DNA+RNA_seed | AKT1 | 0.273024 | LUAD | HOP-62 | MK-2206 |
| MVP_top60_DNA_seed | HDAC1 | -0.287285 | KIRC | BB65-RCC | PCI-34051 | MVP_top60_DNA+RNA_seed | AURKA | 0.273142 | LUAD | NCI-H1693 | Genentech Cpd 10 |
| MVP_top60_DNA_seed | IGF1R | -0.286215 | KIRC | KMRC-20 | BMS-536924 | MVP_top60_DNA+RNA_seed | MDM2 | 0.274105 | LUAD | LC-2-ad | Nutlin-3a (-) |
| MVP_top60_DNA_seed | PARP1 | -0.281727 | KIRC | BFTC-909 | Veliparib | MVP_top60_DNA+RNA_seed | IGF1R | 0.275113 | LUAD | H3255 | Linsitinib |
| MVP_top60_DNA_seed | HDAC1 | -0.275823 | KIRC | U031 | PCI-34051 | MVP_top60_DNA+RNA_seed | HDAC1 | 0.275115 | LUAD | NCI-H2347 | Dacinostat |
| MVP_top60_DNA_seed | TP53 | -0.275164 | KIRC | 769-P | MIRA-1 | MVP_top60_DNA+RNA_seed | EGFR | 0.275273 | LUAD | NCI-H1944 | CI-1033 |
| MVP_top60_DNA_seed | HSP90AA1 | -0.273761 | KIRC | NCC010 | Luminespib | MVP_top60_DNA+RNA_seed | SRC | 0.276463 | LUAD | NCI-H1648 | HG-6-71-01 |
| MVP_top60_DNA_seed | HSP90AB1 | -0.273761 | KIRC | NCC010 | Luminespib | MVP_top60_DNA+RNA_seed | EGFR | 0.276783 | LUAD | NCI-H2087 | AZD8931 |
| MVP_top60_DNA_seed | IGF1R | -0.271285 | KIRC | NCC021 | BMS-754807 | MVP_top60_DNA+RNA_seed | IGF1R | 0.276907 | LUAD | NCI-H1781 | GSK1904529A |
| MVP_top60_DNA_seed | IGF1R | -0.266696 | KIRC | A704 | BMS-754807 | MVP_top60_DNA+RNA_seed | BRAF | 0.277578 | LUAD | NCI-H1755 | SB590885 |
| MVP_top60_DNA_seed | PARP1 | -0.264066 | KIRC | LB2241-RCC | Talazoparib | MVP_top60_DNA+RNA_seed | BRAF | 0.278328 | LUAD | NCI-H1793 | PLX-4720 |
| MVP_top60_DNA_seed | EP300 | -0.258428 | KIRC | 769-P | SGC-CBP30 | MVP_top60_DNA+RNA_seed | MET | 0.278698 | LUAD | HCC-44 | SU11274 |
| MVP_top60_DNA_seed | PARP1 | -0.255388 | KIRC | A704 | Niraparib | MVP_top60_DNA+RNA_seed | EGFR | 0.278978 | LUAD | NCI-H650 | Afatinib |
| MVP_top60_DNA_seed | PIK3CA | -0.253051 | KIRC | U031 | Alpelisib | MVP_top60_DNA+RNA_seed | SRC | 0.279928 | LUAD | NCI-H1944 | Bosutinib |
| MVP_top60_DNA_seed | HDAC1 | -0.25068 | KIRC | RCC-JW | Romidepsin | MVP_top60_DNA+RNA_seed | EGFR | 0.280279 | LUAD | NCI-H2087 | Afatinib |
| MVP_top60_DNA_seed | HDAC1 | -0.25019 | KIRC | RXF393 | PCI-34051 | MVP_top60_DNA+RNA_seed | AURKA | 0.28048 | LUAD | NCI-H2291 | CD532 |
| MVP_top60_DNA_seed | PIK3CA | -0.250163 | KIRC | CAKI-1 | Alpelisib | MVP_top60_DNA+RNA_seed | AKT1 | 0.280534 | LUAD | NCI-H1781 | MK-2206 |
| MVP_top60_DNA_seed | HDAC1 | -0.249587 | KIRC | CAKI-1 | Entinostat | MVP_top60_DNA+RNA_seed | IGF1R | 0.280936 | LUAD | NCI-H1781 | BMS-754807 |

| | | | | | | | | | | | |
|---|---|---|---|---|---|---|---|---|---|---|---|
| MVP_top60_DNA_seed | HSP90AA1 | -0.240579 | KIRC | RXF393 | Elesclomol | MVP_top60_DNA+RNA_seed | EGFR | 0.281767 | LUAD | NCI-H1437 | Pelitinib |
| MVP_top60_DNA_seed | HSP90AB1 | -0.240579 | KIRC | RXF393 | Elesclomol | MVP_top60_DNA+RNA_seed | EGFR | 0.28187 | LUAD | NCI-H1838 | Afatinib |
| MVP_top60_DNA_seed | PIK3CA | -0.237404 | KIRC | A704 | Alpelisib | MVP_top60_DNA+RNA_seed | BRAF | 0.282588 | LUAD | NCI-H1944 | PLX-4720 |
| MVP_top60_DNA_seed | PARP1 | -0.235357 | KIRC | RCC-JW | Niraparib | MVP_top60_DNA+RNA_seed | MDM2 | 0.283097 | LUAD | NCI-H1781 | Serdemetan |
| MVP_top60_DNA_seed | IGF1R | -0.233819 | KIRC | BFTC-909 | Linsitinib | MVP_top60_DNA+RNA_seed | BRAF | 0.283542 | LUAD | NCI-H2085 | SB590885 |
| MVP_top60_DNA_seed | PARP1 | -0.233312 | KIRC | BB65-RCC | Veliparib | MVP_top60_DNA+RNA_seed | FGFR1 | 0.283554 | LUAD | NCI-H441 | FGFR_3831 |
| MVP_top60_DNA_seed | PARP1 | -0.231836 | KIRC | ACHN | Talazoparib | MVP_top60_DNA+RNA_seed | MET | 0.284916 | LUAD | NCI-H2122 | AZD6094 |
| MVP_top60_DNA_seed | SMARCA4 | -0.230155 | KIRC | SN12C | PFI3 | MVP_top60_DNA+RNA_seed | IGF1R | 0.285382 | LUAD | NCI-H2122 | GSK1904529A |
| MVP_top60_DNA_seed | IGF1R | -0.230135 | KIRC | CAKI-1 | BMS-536924 | MVP_top60_DNA+RNA_seed | PIK3CA | 0.285903 | LUAD | NCI-H1781 | PI3Ka_4409 |
| MVP_top60_DNA_seed | IGF1R | -0.223204 | KIRC | CAL-54 | NVP-ADW742 | MVP_top60_DNA+RNA_seed | EGFR | 0.286435 | LUAD | Calu-3 | Cetuximab |
| MVP_top60_DNA_seed | PARP1 | -0.222986 | KIRC | RCC-AB | Talazoparib | MVP_top60_DNA+RNA_seed | MET | 0.287545 | LUAD | LXF-289 | SU11274 |
| MVP_top60_DNA_seed | ESR1 | -0.22078 | KIRC | KMRC-1 | Tamoxifen | MVP_top60_DNA+RNA_seed | SMARCA4 | 0.287923 | LUAD | NCI-H1355 | PFI-3 |
| MVP_top60_DNA_seed | IGF1R | -0.214301 | KIRC | LB2241-RCC | BMS-536924 | MVP_top60_DNA+RNA_seed | PTK2 | 0.288246 | LUAD | NCI-H2405 | PF-562271 |
| MVP_top60_DNA_seed | PIK3CA | -0.210648 | KIRC | NCC021 | Alpelisib | MVP_top60_DNA+RNA_seed | AURKA | 0.288768 | LUAD | NCI-H1975 | ZM447439 |
| MVP_top60_DNA_seed | PIK3CA | -0.210197 | KIRC | RCC-MF | Buparlisib | MVP_top60_DNA+RNA_seed | BRAF | 0.289233 | LUAD | NCI-H1792 | PLX-4720 |
| MVP_top60_DNA_seed | PIK3CD | -0.210197 | KIRC | RCC-MF | Buparlisib | MVP_top60_DNA+RNA_seed | MDM2 | 0.289725 | LUAD | EMC-BAC-1 | Nutlin-3a (-) |
| MVP_top60_DNA_seed | PARP1 | -0.209646 | KIRC | 769-P | Talazoparib | MVP_top60_DNA+RNA_seed | SMARCA4 | 0.289915 | LUAD | RERF-LC-MS | PFI-3 |
| MVP_top60_DNA_seed | HDAC1 | -0.20468 | KIRC | CAL-54 | Romidepsin | MVP_top60_DNA+RNA_seed | ATM | 0.291964 | LUAD | EKVX | KU-55933 |
| MVP_top60_DNA_seed | EP300 | -0.201961 | KIRC | SN12C | SGC-CBP30 | MVP_top60_DNA+RNA_seed | BRAF | 0.292115 | LUAD | NCI-H1975 | RAF_9304 |
| MVP_top60_DNA_seed | IGF1R | -0.200347 | KIRC | BB65-RCC | GSK1904529A | MVP_top60_DNA+RNA_seed | AURKA | 0.293457 | LUAD | NCI-H2347 | CD532 |
| MVP_top60_DNA_seed | ESR1 | -0.198563 | KIRC | KMRC-20 | Tamoxifen | MVP_top60_DNA+RNA_seed | ATM | 0.29535 | LUAD | NCI-H1781 | CP466722 |
| MVP_top60_DNA_seed | PARP1 | -0.197557 | KIRC | LB996-RCC | Olaparib | MVP_top60_DNA+RNA_seed | FGFR1 | 0.297403 | LUAD | NCI-H1650 | FGFR_3831 |
| MVP_top60_DNA_seed | PARP1 | -0.193473 | KIRC | RCC-MF | Rucaparib | MVP_top60_DNA+RNA_seed | AKT1 | 0.298962 | LUAD | NCI-H1648 | BAY AKT1 |
| MVP_top60_DNA_seed | PARP1 | -0.191649 | KIRC | 786-0 | Olaparib | MVP_top60_DNA+RNA_seed | BRAF | 0.299387 | LUAD | NCI-H522 | PLX-4720 |
| MVP_top60_DNA_seed | ATM | -0.189255 | KIRC | RCC-JF | KU-55933 | MVP_top60_DNA+RNA_seed | EGFR | 0.300269 | LUAD | NCI-H1437 | CUDC-101 |
| MVP_top60_DNA_seed | PIK3CA | -0.188336 | KIRC | 786-0 | Buparlisib | MVP_top60_DNA+RNA_seed | FGFR1 | 0.30037 | LUAD | NCI-H2023 | FGFR_3831 |
| MVP_top60_DNA_seed | PIK3CD | -0.188336 | KIRC | 786-0 | Buparlisib | MVP_top60_DNA+RNA_seed | FGFR1 | 0.302223 | LUAD | HCC-827 | PD173074 |
| MVP_top60_DNA_seed | PARP1 | -0.185479 | KIRC | CAL-54 | Talazoparib | MVP_top60_DNA+RNA_seed | HSP90AA1 | 0.303333 | LUAD | NCI-H1781 | Luminespib |
| MVP_top60_DNA_seed | MTOR | -0.185475 | KIRC | TK10 | Temsirolimus | MVP_top60_DNA+RNA_seed | SRC | 0.303414 | LUAD | Calu-6 | Saracatinib |
| MVP_top60_DNA_seed | PIK3CA | -0.173889 | KIRC | RCC-JF | Buparlisib | MVP_top60_DNA+RNA_seed | HDAC1 | 0.304611 | LUAD | NCI-H2085 | CAY10603 |
| MVP_top60_DNA_seed | PIK3CD | -0.173889 | KIRC | RCC-JF | Buparlisib | MVP_top60_DNA+RNA_seed | EGFR | 0.305652 | LUAD | NCI-H1734 | CI-1033 |
| MVP_top60_DNA_seed | IGF1R | -0.168394 | KIRC | CAKI-1 | BMS-754807 | MVP_top60_DNA+RNA_seed | HSP90AA1 | 0.305751 | LUAD | NCI-H1623 | Tanespimycin |
| MVP_top60_DNA_seed | PIK3CA | -0.159832 | KIRC | NCC010 | AZD8186 | MVP_top60_DNA+RNA_seed | PTK2 | 0.305979 | LUAD | NCI-H522 | PF-562271 |
| MVP_top60_DNA_seed | PIK3CA | -0.158216 | KIRC | HA7-RCC | GNE-317 | MVP_top60_DNA+RNA_seed | HSP90AA1 | 0.306737 | LUAD | NCI-H1623 | Elesclomol |
| MVP_top60_DNA_seed | TP53 | -0.157811 | KIRC | RCC-JW | MIRA-1 | MVP_top60_DNA+RNA_seed | AKT1 | 0.306897 | LUAD | NCI-H23 | AKT inhibitor VIII |
| MVP_top60_DNA_seed | ESR1 | -0.156406 | KIRC | A498 | GDC0810 | MVP_top60_DNA+RNA_seed | EGFR | 0.307653 | LUAD | LC-2-ad | Afatinib |
| MVP_top60_DNA_seed | EP300 | -0.156034 | KIRC | HA7-RCC | SGC-CBP30 | MVP_top60_DNA+RNA_seed | EGFR | 0.307721 | LUAD | NCI-H1793 | Afatinib |
| MVP_top60_DNA_seed | IGF1R | -0.153255 | KIRC | ACHN | Linsitinib | MVP_top60_DNA+RNA_seed | MET | 0.307784 | LUAD | SW1573 | SU11274 |
| MVP_top60_DNA_seed | HDAC1 | -0.153153 | KIRC | U031 | Entinostat | MVP_top60_DNA+RNA_seed | PIK3CA | 0.30943 | LUAD | NCI-H358 | PI3Ka_4409 |
| MVP_top60_DNA_seed | HDAC1 | -0.152531 | KIRC | LB1047-RCC | Entinostat | MVP_top60_DNA+RNA_seed | MDM2 | 0.310162 | LUAD | NCI-H1355 | Nutlin-3a (-) |
| MVP_top60_DNA_seed | PARP1 | -0.146702 | KIRC | 769-P | Veliparib | MVP_top60_DNA+RNA_seed | AURKA | 0.310511 | LUAD | EMC-BAC-2 | CD532 |

| | | | | | | | | | | | |
|---|---|---|---|---|---|---|---|---|---|---|---|
| MVP_top60_DNA_seed | PIK3CA | -0.146273 | KIRC | RCC-MF | Alpelisib | MVP_top60_DNA+RNA_seed | HSP90AA1 | 0.310534 | LUAD | Calu-6 | SNX-2112 |
| MVP_top60_DNA_seed | IGF1R | -0.141101 | KIRC | LB2241-RCC | BMS-754807 | MVP_top60_DNA+RNA_seed | BRAF | 0.31078 | LUAD | RERF-LC-KJ | PLX-4720 |
| MVP_top60_DNA_seed | PIK3CA | -0.139308 | KIRC | KMRC-1 | GNE-317 | MVP_top60_DNA+RNA_seed | ATM | 0.310922 | LUAD | NCI-H2228 | CP466722 |
| MVP_top60_DNA_seed | ATM | -0.13407 | KIRC | HA7-RCC | KU-55933 | MVP_top60_DNA+RNA_seed | HSP90AA1 | 0.311178 | LUAD | NCI-H358 | Elesclomol |
| MVP_top60_DNA_seed | HSP90AA1 | -0.131923 | KIRC | SW156 | Luminespib | MVP_top60_DNA+RNA_seed | BRAF | 0.311802 | LUAD | NCI-H1838 | RAF_9304 |
| MVP_top60_DNA_seed | HSP90AB1 | -0.131923 | KIRC | SW156 | Luminespib | MVP_top60_DNA+RNA_seed | EGFR | 0.313807 | LUAD | NCI-H1568 | Gefitinib |
| MVP_top60_DNA_seed | PARP1 | -0.131265 | KIRC | A704 | Talazoparib | MVP_top60_DNA+RNA_seed | EGFR | 0.314094 | LUAD | NCI-H1975 | Lapatinib |
| MVP_top60_DNA_seed | PIK3CA | -0.129105 | KIRC | HA7-RCC | Alpelisib | MVP_top60_DNA+RNA_seed | ATM | 0.315005 | LUAD | NCI-H1975 | KU-55933 |
| MVP_top60_DNA_seed | PARP1 | -0.121325 | KIRC | RCC-MF | Talazoparib | MVP_top60_DNA+RNA_seed | HDAC1 | 0.315659 | LUAD | NCI-H1993 | Dacinostat |
| MVP_top60_DNA_seed | TP53 | -0.118043 | KIRC | RCC-FG2 | MIRA-1 | MVP_top60_DNA+RNA_seed | IGF1R | 0.315751 | LUAD | Calu-6 | BMS-754807 |
| MVP_top60_DNA_seed | IGF1R | -0.108973 | KIRC | SW156 | GSK1904529A | MVP_top60_DNA+RNA_seed | BRAF | 0.315762 | LUAD | NCI-H3122 | Dabrafenib |
| MVP_top60_DNA_seed | ATM | -0.108044 | KIRC | SW156 | KU-55933 | MVP_top60_DNA+RNA_seed | EGFR | 0.316047 | LUAD | NCI-H2030 | PF-00299804 |
| MVP_top60_DNA_seed | PIK3CA | -0.105575 | KIRC | LB2241-RCC | Alpelisib | MVP_top60_DNA+RNA_seed | EGFR | 0.316706 | LUAD | EKVX | AZD8931 |
| MVP_top60_DNA_seed | PARP1 | -0.10226 | KIRC | ACHN | Veliparib | MVP_top60_DNA+RNA_seed | HDAC1 | 0.31697 | LUAD | NCI-H1693 | Tubastatin A |
| MVP_top60_DNA_seed | PIK3CA | -0.10051 | KIRC | RCC-FG2 | Buparlisib | MVP_top60_DNA+RNA_seed | ATM | 0.317537 | LUAD | NCI-H2122 | KU-60019 |
| MVP_top60_DNA_seed | PIK3CD | -0.10051 | KIRC | RCC-FG2 | Buparlisib | MVP_top60_DNA+RNA_seed | EGFR | 0.31847 | LUAD | NCI-H1651 | Afatinib |
| MVP_top60_DNA_seed | AR | -0.092604 | KIRC | RXF393 | Bicalutamide | MVP_top60_DNA+RNA_seed | AURKA | 0.320597 | LUAD | HCC-827 | CD532 |
| MVP_top60_DNA_seed | PARP1 | -0.091432 | KIRC | RCC-MF | Niraparib | MVP_top60_DNA+RNA_seed | EGFR | 0.322121 | LUAD | NCI-H2030 | Afatinib |
| MVP_top60_DNA_seed | IGF1R | -0.09092 | KIRC | NCC010 | NVP-ADW742 | MVP_top60_DNA+RNA_seed | FGFR1 | 0.323147 | LUAD | NCI-H1792 | PD173074 |
| MVP_top60_DNA_seed | HDAC1 | -0.089297 | KIRC | 769-P | PCI-34051 | MVP_top60_DNA+RNA_seed | HSP90AA1 | 0.323887 | LUAD | NCI-H441 | Luminespib |
| MVP_top60_DNA_seed | IGF1R | -0.086369 | KIRC | CAL-54 | Linsitinib | MVP_top60_DNA+RNA_seed | EGFR | 0.323895 | LUAD | NCI-H1734 | Pelitinib |
| MVP_top60_DNA_seed | PIK3CA | -0.085348 | KIRC | A498 | Buparlisib | MVP_top60_DNA+RNA_seed | EGFR | 0.324231 | LUAD | NCI-H1734 | PF-00299804 |
| MVP_top60_DNA_seed | PIK3CD | -0.085348 | KIRC | A498 | Buparlisib | MVP_top60_DNA+RNA_seed | MET | 0.324599 | LUAD | HOP-62 | Crizotinib |
| MVP_top60_DNA_seed | HSP90AA1 | -0.083914 | KIRC | ACHN | Tanespimycin | MVP_top60_DNA+RNA_seed | SMARCA4 | 0.325691 | LUAD | NCI-H838 | PFI-3 |
| MVP_top60_DNA_seed | HSP90AB1 | -0.083914 | KIRC | ACHN | Tanespimycin | MVP_top60_DNA+RNA_seed | MDM2 | 0.325906 | LUAD | COR-L105 | Serdemetan |
| MVP_top60_DNA_seed | MTOR | -0.081958 | KIRC | 769-P | Temsirolimus | MVP_top60_DNA+RNA_seed | AKT1 | 0.326137 | LUAD | NCI-H1568 | AKT inhibitor VIII |
| MVP_top60_DNA_seed | PIK3CA | -0.081313 | KIRC | ACHN | Buparlisib | MVP_top60_DNA+RNA_seed | HSP90AA1 | 0.328214 | LUAD | NCI-H1666 | SNX-2112 |
| MVP_top60_DNA_seed | PIK3CD | -0.081313 | KIRC | ACHN | Buparlisib | MVP_top60_DNA+RNA_seed | AURKA | 0.328294 | LUAD | NCI-H441 | Genentech Cpd 10 |
| MVP_top60_DNA_seed | PIK3CA | -0.079578 | KIRC | U031 | Buparlisib | MVP_top60_DNA+RNA_seed | FGFR1 | 0.328558 | LUAD | NCI-H358 | FGFR_3831 |
| MVP_top60_DNA_seed | PIK3CD | -0.079578 | KIRC | U031 | Buparlisib | MVP_top60_DNA+RNA_seed | HSP90AA1 | 0.328767 | LUAD | NCI-H1651 | Elesclomol |
| MVP_top60_DNA_seed | IGF1R | -0.079044 | KIRC | HA7-RCC | BMS-536924 | MVP_top60_DNA+RNA_seed | EGFR | 0.328855 | LUAD | VMRC-LCD | Gefitinib |
| MVP_top60_DNA_seed | HSP90AA1 | -0.074655 | KIRC | NCC021 | Luminespib | MVP_top60_DNA+RNA_seed | BRAF | 0.328895 | LUAD | H3255 | PLX-4720 |
| MVP_top60_DNA_seed | HSP90AB1 | -0.074655 | KIRC | NCC021 | Luminespib | MVP_top60_DNA+RNA_seed | AKT1 | 0.329214 | LUAD | PC-3_[JPC-3] | AKT inhibitor VIII |
| MVP_top60_DNA_seed | PIK3CA | -0.072205 | KIRC | A498 | GNE-317 | MVP_top60_DNA+RNA_seed | BRAF | 0.330062 | LUAD | LC-2-ad | Dabrafenib |
| MVP_top60_DNA_seed | HSP90AA1 | -0.070257 | KIRC | SW156 | Elesclomol | MVP_top60_DNA+RNA_seed | ATM | 0.330089 | LUAD | NCI-H23 | KU-55933 |
| MVP_top60_DNA_seed | HSP90AB1 | -0.070257 | KIRC | SW156 | Elesclomol | MVP_top60_DNA+RNA_seed | BRAF | 0.330708 | LUAD | NCI-H522 | Dabrafenib |
| MVP_top60_DNA_seed | PIK3CA | -0.069469 | KIRC | CAL-54 | Alpelisib | MVP_top60_DNA+RNA_seed | AURKA | 0.331028 | LUAD | NCI-H3122 | ZM447439 |
| MVP_top60_DNA_seed | SMARCA4 | -0.068164 | KIRC | RCC-ER | PFI3 | MVP_top60_DNA+RNA_seed | BRAF | 0.331504 | LUAD | EMC-BAC-1 | Dabrafenib |
| MVP_top60_DNA_seed | HSP90AA1 | -0.066749 | KIRC | RCC-ER | Luminespib | MVP_top60_DNA+RNA_seed | MET | 0.331839 | LUAD | NCI-H1781 | SU11274 |
| MVP_top60_DNA_seed | HSP90AB1 | -0.066749 | KIRC | RCC-ER | Luminespib | MVP_top60_DNA+RNA_seed | EGFR | 0.332372 | LUAD | NCI-H23 | Afatinib |
| MVP_top60_DNA_seed | HSP90AA1 | -0.065315 | KIRC | A498 | Tanespimycin | MVP_top60_DNA+RNA_seed | AURKA | 0.332614 | LUAD | NCI-H1734 | Genentech Cpd 10 |

| | | | | | | | | | | | |
|---|---|---|---|---|---|---|---|---|---|---|---|
| MVP_top60_DNA_seed | HSP90AB1 | -0.065315 | KIRC | A498 | Tanespimycin | MVP_top60_DNA+RNA_seed | PTK2 | 0.332628 | LUAD | NCI-H1651 | PF-562271 |
| MVP_top60_DNA_seed | EP300 | -0.065191 | KIRC | RCC-JW | SGC-CBP30 | MVP_top60_DNA+RNA_seed | BRAF | 0.332669 | LUAD | NCI-H1651 | PLX-4720 |
| MVP_top60_DNA_seed | IGF1R | -0.062528 | KIRC | A704 | GSK1904529A | MVP_top60_DNA+RNA_seed | IGF1R | 0.334939 | LUAD | PC-3_[JPC-3] | BMS-536924 |
| MVP_top60_DNA_seed | IGF1R | -0.060254 | KIRC | RCC-JF | BMS-754807 | MVP_top60_DNA+RNA_seed | AKT1 | 0.341224 | LUAD | A549 | BAY AKT1 |
| MVP_top60_DNA_seed | PARP1 | -0.060135 | KIRC | NCC010 | Niraparib | MVP_top60_DNA+RNA_seed | BRAF | 0.341313 | LUAD | Calu-3 | RAF_9304 |
| MVP_top60_DNA_seed | PIK3CA | -0.055843 | KIRC | RCC-ER | GNE-317 | MVP_top60_DNA+RNA_seed | HDAC1 | 0.343146 | LUAD | NCI-H1693 | JQ12 |
| MVP_top60_DNA_seed | IGF1R | -0.049247 | KIRC | BFTC-909 | BMS-536924 | MVP_top60_DNA+RNA_seed | AURKA | 0.343929 | LUAD | NCI-H2087 | CD532 |
| MVP_top60_DNA_seed | ESR1 | -0.045743 | KIRC | BFTC-909 | GDC0810 | MVP_top60_DNA+RNA_seed | AURKA | 0.344799 | LUAD | NCI-H1623 | CD532 |
| MVP_top60_DNA_seed | IGF1R | -0.045352 | KIRC | SN12C | BMS-536924 | MVP_top60_DNA+RNA_seed | SMARCA4 | 0.345693 | LUAD | NCI-H2122 | PFI-3 |
| MVP_top60_DNA_seed | HSP90AA1 | -0.040101 | KIRC | RCC-ER | Elesclomol | MVP_top60_DNA+RNA_seed | IGF1R | 0.346202 | LUAD | RERF-LC-MS | BMS-754807 |
| MVP_top60_DNA_seed | HSP90AB1 | -0.040101 | KIRC | RCC-ER | Elesclomol | MVP_top60_DNA+RNA_seed | MET | 0.346429 | LUAD | SW1573 | AZD6094 |
| MVP_top60_DNA_seed | IGF1R | -0.031113 | KIRC | ACHN | BMS-536924 | MVP_top60_DNA+RNA_seed | FGFR1 | 0.346801 | LUAD | NCI-H1568 | FGFR_3831 |
| MVP_top60_DNA_seed | PARP1 | -0.028937 | KIRC | A704 | Olaparib | MVP_top60_DNA+RNA_seed | HDAC1 | 0.347914 | LUAD | HCC-827 | Dacinostat |
| MVP_top60_DNA_seed | ATM | -0.023913 | KIRC | NCC021 | KU-55933 | MVP_top60_DNA+RNA_seed | AURKA | 0.349053 | LUAD | NCI-H1693 | ZM447439 |
| MVP_top60_DNA_seed | PARP1 | -0.019801 | KIRC | 786-0 | Veliparib | MVP_top60_DNA+RNA_seed | HDAC1 | 0.349364 | LUAD | NCI-H1573 | CAY10603 |
| MVP_top60_DNA_seed | ESR1 | -0.019002 | KIRC | RCC10RGB | Tamoxifen | MVP_top60_DNA+RNA_seed | EGFR | 0.349548 | LUAD | NCI-H1355 | AST-1306 |
| MVP_top60_DNA_seed | HSP90AA1 | -0.015529 | KIRC | ACHN | Luminespib | MVP_top60_DNA+RNA_seed | BRAF | 0.351224 | LUAD | HCC-827 | SB590885 |
| MVP_top60_DNA_seed | HSP90AB1 | -0.015529 | KIRC | ACHN | Luminespib | MVP_top60_DNA+RNA_seed | EGFR | 0.351436 | LUAD | SW1573 | Afatinib |
| MVP_top60_DNA_seed | PARP1 | -0.011796 | KIRC | LB996-RCC | Rucaparib | MVP_top60_DNA+RNA_seed | EGFR | 0.353214 | LUAD | HCC-44 | Afatinib |
| MVP_top60_DNA_seed | HSP90AA1 | -0.010222 | KIRC | HA7-RCC | Tanespimycin | MVP_top60_DNA+RNA_seed | HDAC1 | 0.354166 | LUAD | NCI-H1666 | CAY10603 |
| MVP_top60_DNA_seed | HSP90AB1 | -0.010222 | KIRC | HA7-RCC | Tanespimycin | MVP_top60_DNA+RNA_seed | AKT1 | 0.354842 | LUAD | NCI-H1651 | BAY AKT1 |
| MVP_top60_DNA_seed | IGF1R | -0.006827 | KIRC | A704 | Linsitinib | MVP_top60_DNA+RNA_seed | HSP90AA1 | 0.358854 | LUAD | LXF-289 | CCT-018159 |
| MVP_top60_DNA_seed | SMARCA4 | 0.002601 | KIRC | BFTC-909 | PFI3 | MVP_top60_DNA+RNA_seed | EGFR | 0.359013 | LUAD | NCI-H1975 | CUDC-101 |
| MVP_top60_DNA_seed | PARP1 | 0.004185 | KIRC | NCC010 | Veliparib | MVP_top60_DNA+RNA_seed | EGFR | 0.3598 | LUAD | NCI-H838 | Cetuximab |
| MVP_top60_DNA_seed | PARP1 | 0.00523 | KIRC | RXF393 | Niraparib | MVP_top60_DNA+RNA_seed | BRAF | 0.360046 | LUAD | NCI-H1573 | SB590885 |
| MVP_top60_DNA_seed | PIK3CA | 0.007027 | KIRC | RCC10RGB | GNE-317 | MVP_top60_DNA+RNA_seed | AKT1 | 0.36344 | LUAD | NCI-H1568 | AKT inhibitor VIII |
| MVP_top60_DNA_seed | PIK3CA | 0.009887 | KIRC | A704 | Buparlisib | MVP_top60_DNA+RNA_seed | AKT1 | 0.363481 | LUAD | NCI-H2030 | GSK690693 |
| MVP_top60_DNA_seed | PIK3CD | 0.009887 | KIRC | A704 | Buparlisib | MVP_top60_DNA+RNA_seed | HDAC1 | 0.364798 | LUAD | H3255 | JQ12 |
| MVP_top60_DNA_seed | EP300 | 0.013002 | KIRC | RCC10RGB | SGC-CBP30 | MVP_top60_DNA+RNA_seed | AURKA | 0.365042 | LUAD | HCC-827 | ZM447439 |
| MVP_top60_DNA_seed | HSP90AA1 | 0.013567 | KIRC | CAL-54 | Luminespib | MVP_top60_DNA+RNA_seed | IGF1R | 0.365568 | LUAD | PC-14 | Linsitinib |
| MVP_top60_DNA_seed | HSP90AB1 | 0.013567 | KIRC | CAL-54 | Luminespib | MVP_top60_DNA+RNA_seed | HSP90AA1 | 0.36589 | LUAD | NCI-H322M | Tanespimycin |
| MVP_top60_DNA_seed | TP53 | 0.014332 | KIRC | RCC-JF | MIRA-1 | MVP_top60_DNA+RNA_seed | EGFR | 0.366685 | LUAD | NCI-H2023 | AZD8931 |
| MVP_top60_DNA_seed | PARP1 | 0.014728 | KIRC | 769-P | Rucaparib | MVP_top60_DNA+RNA_seed | IGF1R | 0.368296 | LUAD | NCI-H2347 | BMS-754807 |
| MVP_top60_DNA_seed | HSP90AA1 | 0.023327 | KIRC | NCC010 | Tanespimycin | MVP_top60_DNA+RNA_seed | AKT1 | 0.36845 | LUAD | NCI-H1792 | GSK690693 |
| MVP_top60_DNA_seed | HSP90AB1 | 0.023327 | KIRC | NCC010 | Tanespimycin | MVP_top60_DNA+RNA_seed | EGFR | 0.36857 | LUAD | EMC-BAC-2 | PF-00299804 |
| MVP_top60_DNA_seed | PIK3CA | 0.026851 | KIRC | NCC010 | Alpelisib | MVP_top60_DNA+RNA_seed | BRAF | 0.369277 | LUAD | NCI-H3122 | RAF_9304 |
| MVP_top60_DNA_seed | HSP90AA1 | 0.030298 | KIRC | CAKI-1 | Luminespib | MVP_top60_DNA+RNA_seed | HDAC1 | 0.370117 | LUAD | NCI-H650 | JQ12 |
| MVP_top60_DNA_seed | HSP90AB1 | 0.030298 | KIRC | CAKI-1 | Luminespib | MVP_top60_DNA+RNA_seed | EGFR | 0.371778 | LUAD | RERF-LC-MS | AST-1306 |
| MVP_top60_DNA_seed | EP300 | 0.030997 | KIRC | ACHN | CPI-637 | MVP_top60_DNA+RNA_seed | EGFR | 0.372744 | LUAD | NCI-H322M | Afatinib |
| MVP_top60_DNA_seed | TP53 | 0.036065 | KIRC | NCC021 | MIRA-1 | MVP_top60_DNA+RNA_seed | AKT1 | 0.372811 | LUAD | NCI-H1623 | BAY AKT1 |
| MVP_top60_DNA_seed | MTOR | 0.036414 | KIRC | LB996-RCC | Temsirolimus | MVP_top60_DNA+RNA_seed | MET | 0.373257 | LUAD | NCI-H2405 | SU11274 |

| | | | | | | | | | | | |
|---|---|---|---|---|---|---|---|---|---|---|---|
| MVP_top60_DNA_seed | PARP1 | 0.036451 | KIRC | RCC-ER | Olaparib | MVP_top60_DNA+RNA_seed | EGFR | 0.373422 | LUAD | VMRC-LCD | Afatinib |
| MVP_top60_DNA_seed | HDAC1 | 0.037521 | KIRC | A498 | PCI-34051 | MVP_top60_DNA+RNA_seed | AURKA | 0.37366 | LUAD | NCI-H23 | CD532 |
| MVP_top60_DNA_seed | PARP1 | 0.040967 | KIRC | KMRC-20 | Rucaparib | MVP_top60_DNA+RNA_seed | MET | 0.374253 | LUAD | Calu-6 | AZD6094 |
| MVP_top60_DNA_seed | HDAC1 | 0.041232 | KIRC | SW156 | Entinostat | MVP_top60_DNA+RNA_seed | PIK3CA | 0.374923 | LUAD | EKVX | AZD8835 |
| MVP_top60_DNA_seed | TP53 | 0.041486 | KIRC | LB2241-RCC | MIRA-1 | MVP_top60_DNA+RNA_seed | HDAC1 | 0.375007 | LUAD | PC-3_[JPC-3] | CAY10603 |
| MVP_top60_DNA_seed | SMARCA4 | 0.046874 | KIRC | RCC-JF | PFI3 | MVP_top60_DNA+RNA_seed | AKT1 | 0.375139 | LUAD | EMC-BAC-2 | BAY AKT1 |
| MVP_top60_DNA_seed | PARP1 | 0.047616 | KIRC | CAL-54 | Rucaparib | MVP_top60_DNA+RNA_seed | HSP90AA1 | 0.375943 | LUAD | VMRC-LCD | Tanespimycin |
| MVP_top60_DNA_seed | EP300 | 0.051717 | KIRC | 786-0 | CPI-637 | MVP_top60_DNA+RNA_seed | HSP90AA1 | 0.376168 | LUAD | LC-2-ad | CCT-018159 |
| MVP_top60_DNA_seed | IGF1R | 0.053463 | KIRC | 786-0 | Linsitinib | MVP_top60_DNA+RNA_seed | AURKA | 0.378418 | LUAD | NCI-H1651 | Alisertib |
| MVP_top60_DNA_seed | ESR1 | 0.054421 | KIRC | A704 | Tamoxifen | MVP_top60_DNA+RNA_seed | AKT1 | 0.378705 | LUAD | Calu-6 | AKT inhibitor VIII |
| MVP_top60_DNA_seed | HDAC1 | 0.0588 | KIRC | RCC-AB | Entinostat | MVP_top60_DNA+RNA_seed | BRAF | 0.379151 | LUAD | HCC-78 | SB590885 |
| MVP_top60_DNA_seed | PARP1 | 0.060822 | KIRC | RCC10RGB | Niraparib | MVP_top60_DNA+RNA_seed | EGFR | 0.379366 | LUAD | NCI-H441 | Afatinib |
| MVP_top60_DNA_seed | IGF1R | 0.064071 | KIRC | A498 | Linsitinib | MVP_top60_DNA+RNA_seed | HDAC1 | 0.380412 | LUAD | NCI-H1435 | CAY10603 |
| MVP_top60_DNA_seed | PIK3CA | 0.06458 | KIRC | SW156 | AZD8186 | MVP_top60_DNA+RNA_seed | FGFR1 | 0.380916 | LUAD | Calu-3 | AZD4547 |
| MVP_top60_DNA_seed | PIK3CA | 0.065939 | KIRC | RCC-AB | AZD8186 | MVP_top60_DNA+RNA_seed | MDM2 | 0.38131 | LUAD | NCI-H3122 | Serdemetan |
| MVP_top60_DNA_seed | PARP1 | 0.066422 | KIRC | A704 | Rucaparib | MVP_top60_DNA+RNA_seed | FGFR1 | 0.384183 | LUAD | NCI-H2009 | AZD4547 |
| MVP_top60_DNA_seed | IGF1R | 0.06825 | KIRC | RXF393 | Linsitinib | MVP_top60_DNA+RNA_seed | BRAF | 0.384877 | LUAD | NCI-H522 | RAF_9304 |
| MVP_top60_DNA_seed | HDAC1 | 0.070282 | KIRC | KMRC-20 | Entinostat | MVP_top60_DNA+RNA_seed | PIK3CA | 0.385118 | LUAD | NCI-H2087 | AZD8835 |
| MVP_top60_DNA_seed | EP300 | 0.070755 | KIRC | BFTC-909 | CPI-637 | MVP_top60_DNA+RNA_seed | EGFR | 0.385627 | LUAD | NCI-H2085 | CUDC-101 |
| MVP_top60_DNA_seed | SMARCA4 | 0.078614 | KIRC | RCC-JW | PFI3 | MVP_top60_DNA+RNA_seed | PIK3CA | 0.385766 | LUAD | NCI-H650 | AZD8835 |
| MVP_top60_DNA_seed | PARP1 | 0.079421 | KIRC | ACHN | Olaparib | MVP_top60_DNA+RNA_seed | MDM2 | 0.387357 | LUAD | ABC-1 | Serdemetan |
| MVP_top60_DNA_seed | PIK3CA | 0.081977 | KIRC | NCC021 | GNE-317 | MVP_top60_DNA+RNA_seed | AURKA | 0.388187 | LUAD | NCI-H650 | GSK1070916 |
| MVP_top60_DNA_seed | TP53 | 0.082333 | KIRC | NCC010 | MIRA-1 | MVP_top60_DNA+RNA_seed | AURKA | 0.388459 | LUAD | HCC-827 | Alisertib |
| MVP_top60_DNA_seed | PARP1 | 0.084793 | KIRC | A498 | Niraparib | MVP_top60_DNA+RNA_seed | AURKA | 0.388598 | LUAD | NCI-H2087 | Alisertib |
| MVP_top60_DNA_seed | SMARCA4 | 0.084828 | KIRC | 786-0 | PFI3 | MVP_top60_DNA+RNA_seed | AKT1 | 0.389521 | LUAD | NCI-H2291 | MK-2206 |
| MVP_top60_DNA_seed | ESR1 | 0.087509 | KIRC | RCC-JF | Tamoxifen | MVP_top60_DNA+RNA_seed | HDAC1 | 0.390463 | LUAD | H3255 | Dacinostat |
| MVP_top60_DNA_seed | SMARCA4 | 0.093692 | KIRC | 769-P | PFI3 | MVP_top60_DNA+RNA_seed | AURKA | 0.390996 | LUAD | NCI-H1623 | ZM447439 |
| MVP_top60_DNA_seed | HDAC1 | 0.094458 | KIRC | RCC-MF | Entinostat | MVP_top60_DNA+RNA_seed | HDAC1 | 0.391486 | LUAD | NCI-H1993 | AR-42 |
| MVP_top60_DNA_seed | PIK3CA | 0.096705 | KIRC | RXF393 | GNE-317 | MVP_top60_DNA+RNA_seed | HSP90AA1 | 0.391753 | LUAD | NCI-H2122 | SNX-2112 |
| MVP_top60_DNA_seed | IGF1R | 0.097983 | KIRC | CAL-54 | BMS-754807 | MVP_top60_DNA+RNA_seed | BRAF | 0.391972 | LUAD | NCI-H2087 | PLX-4720 |
| MVP_top60_DNA_seed | PIK3CA | 0.103392 | KIRC | RCC-MF | AZD8186 | MVP_top60_DNA+RNA_seed | MET | 0.39239 | LUAD | NCI-H1650 | PHA-665752 |
| MVP_top60_DNA_seed | PARP1 | 0.112661 | KIRC | RCC-FG2 | Niraparib | MVP_top60_DNA+RNA_seed | AKT1 | 0.392818 | LUAD | A549 | AKT inhibitor VIII |
| MVP_top60_DNA_seed | TP53 | 0.113903 | KIRC | U031 | MIRA-1 | MVP_top60_DNA+RNA_seed | EGFR | 0.394744 | LUAD | NCI-H1755 | PF-00299804 |
| MVP_top60_DNA_seed | PARP1 | 0.114057 | KIRC | RCC-FG2 | Veliparib | MVP_top60_DNA+RNA_seed | EGFR | 0.395539 | LUAD | NCI-H441 | AST-1306 |
| MVP_top60_DNA_seed | PARP1 | 0.115824 | KIRC | SN12C | Veliparib | MVP_top60_DNA+RNA_seed | BRAF | 0.396816 | LUAD | NCI-H2085 | PLX-4720 |
| MVP_top60_DNA_seed | ATM | 0.116633 | KIRC | BB65-RCC | KU-55933 | MVP_top60_DNA+RNA_seed | AKT1 | 0.396939 | LUAD | NCI-H1993 | AKT inhibitor VIII |
| MVP_top60_DNA_seed | IGF1R | 0.118136 | KIRC | LB1047-RCC | BMS-754807 | MVP_top60_DNA+RNA_seed | AURKA | 0.397004 | LUAD | NCI-H1355 | GSK1070916 |
| MVP_top60_DNA_seed | PARP1 | 0.124714 | KIRC | A704 | Veliparib | MVP_top60_DNA+RNA_seed | MET | 0.398047 | LUAD | HOP-62 | PHA-665752 |
| MVP_top60_DNA_seed | HSP90AA1 | 0.127148 | KIRC | CAL-54 | Tanespimycin | MVP_top60_DNA+RNA_seed | MDM2 | 0.398386 | LUAD | NCI-H1793 | Serdemetan |
| MVP_top60_DNA_seed | HSP90AB1 | 0.127148 | KIRC | CAL-54 | Tanespimycin | MVP_top60_DNA+RNA_seed | HDAC1 | 0.400107 | LUAD | NCI-H1437 | Tubastatin A |
| MVP_top60_DNA_seed | HSP90AA1 | 0.127691 | KIRC | 786-0 | Luminespib | MVP_top60_DNA+RNA_seed | HDAC1 | 0.400508 | LUAD | NCI-H1693 | Belinostat |

| | | | | | | | | | | | |
|---|---|---|---|---|---|---|---|---|---|---|---|
| MVP_top60_DNA_seed | HSP90AB1 | 0.127691 | KIRC | 786-0 | Luminespib | MVP_top60_DNA+RNA_seed | HDAC1 | 0.400804 | LUAD | NCI-H2085 | JQ12 |
| MVP_top60_DNA_seed | EP300 | 0.129753 | KIRC | ACHN | SGC-CBP30 | MVP_top60_DNA+RNA_seed | PIK3CA | 0.401141 | LUAD | NCI-H2347 | PI3Ka_4409 |
| MVP_top60_DNA_seed | HDAC1 | 0.142093 | KIRC | A704 | PCI-34051 | MVP_top60_DNA+RNA_seed | ATM | 0.401427 | LUAD | NCI-H1734 | CP466722 |
| MVP_top60_DNA_seed | SMARCA4 | 0.142904 | KIRC | LB996-RCC | PFI3 | MVP_top60_DNA+RNA_seed | EGFR | 0.401488 | LUAD | LXF-289 | CI-1033 |
| MVP_top60_DNA_seed | PIK3CA | 0.143317 | KIRC | 769-P | AZD8186 | MVP_top60_DNA+RNA_seed | EGFR | 0.402576 | LUAD | RERF-LC-MS | Afatinib |
| MVP_top60_DNA_seed | HSP90AA1 | 0.143976 | KIRC | LB2241-RCC | Tanespimycin | MVP_top60_DNA+RNA_seed | EGFR | 0.405374 | LUAD | NCI-H2291 | PF-00299804 |
| MVP_top60_DNA_seed | HSP90AB1 | 0.143976 | KIRC | LB2241-RCC | Tanespimycin | MVP_top60_DNA+RNA_seed | HDAC1 | 0.406537 | LUAD | NCI-H322M | CAY10603 |
| MVP_top60_DNA_seed | IGF1R | 0.144922 | KIRC | RCC-FG2 | NVP-ADW742 | MVP_top60_DNA+RNA_seed | EGFR | 0.407697 | LUAD | NCI-H1993 | PF-00299804 |
| MVP_top60_DNA_seed | AR | 0.146122 | KIRC | LB2241-RCC | Bicalutamide | MVP_top60_DNA+RNA_seed | ATM | 0.407946 | LUAD | VMRC-LCD | KU-55933 |
| MVP_top60_DNA_seed | IGF1R | 0.151229 | KIRC | BB65-RCC | BMS-536924 | MVP_top60_DNA+RNA_seed | HSP90AA1 | 0.408308 | LUAD | COR-L105 | CCT-018159 |
| MVP_top60_DNA_seed | HDAC1 | 0.151447 | KIRC | RCC-FG2 | Entinostat | MVP_top60_DNA+RNA_seed | EGFR | 0.408631 | LUAD | NCI-H2291 | Cetuximab |
| MVP_top60_DNA_seed | ATM | 0.151675 | KIRC | RCC-MF | KU-55933 | MVP_top60_DNA+RNA_seed | MET | 0.410695 | LUAD | SK-LU-1 | SU11274 |
| MVP_top60_DNA_seed | HDAC1 | 0.156351 | KIRC | A498 | Romidepsin | MVP_top60_DNA+RNA_seed | SRC | 0.410738 | LUAD | RERF-LC-KJ | NVP-BHG712 |
| MVP_top60_DNA_seed | ESR1 | 0.157262 | KIRC | NCC010 | GDC0810 | MVP_top60_DNA+RNA_seed | MDM2 | 0.412496 | LUAD | NCI-H1993 | Serdemetan |
| MVP_top60_DNA_seed | SMARCA4 | 0.15789 | KIRC | BB65-RCC | PFI3 | MVP_top60_DNA+RNA_seed | BRAF | 0.412541 | LUAD | EKVX | RAF_9304 |
| MVP_top60_DNA_seed | ESR1 | 0.157911 | KIRC | SW156 | GDC0810 | MVP_top60_DNA+RNA_seed | IGF1R | 0.412676 | LUAD | RERF-LC-MS | GSK1904529A |
| MVP_top60_DNA_seed | HSP90AA1 | 0.157954 | KIRC | KMRC-1 | Tanespimycin | MVP_top60_DNA+RNA_seed | EGFR | 0.413185 | LUAD | LXF-289 | AST-1306 |
| MVP_top60_DNA_seed | HSP90AB1 | 0.157954 | KIRC | KMRC-1 | Tanespimycin | MVP_top60_DNA+RNA_seed | EGFR | 0.414291 | LUAD | NCI-H23 | Afatinib |
| MVP_top60_DNA_seed | IGF1R | 0.159967 | KIRC | RCC-MF | Linsitinib | MVP_top60_DNA+RNA_seed | BRAF | 0.414453 | LUAD | NCI-H522 | SB590885 |
| MVP_top60_DNA_seed | PIK3CA | 0.161126 | KIRC | KMRC-20 | AZD8186 | MVP_top60_DNA+RNA_seed | MET | 0.414563 | LUAD | NCI-H1650 | AZD6094 |
| MVP_top60_DNA_seed | IGF1R | 0.166927 | KIRC | 769-P | BMS-754807 | MVP_top60_DNA+RNA_seed | AURKA | 0.41556 | LUAD | NCI-H23 | Alisertib |
| MVP_top60_DNA_seed | IGF1R | 0.167139 | KIRC | U031 | NVP-ADW742 | MVP_top60_DNA+RNA_seed | AKT1 | 0.415652 | LUAD | LXF-289 | GSK690693 |
| MVP_top60_DNA_seed | IGF1R | 0.168683 | KIRC | BFTC-909 | BMS-754807 | MVP_top60_DNA+RNA_seed | AURKA | 0.417359 | LUAD | NCI-H1693 | GSK1070916 |
| MVP_top60_DNA_seed | PIK3CA | 0.171731 | KIRC | 769-P | Buparlisib | MVP_top60_DNA+RNA_seed | BRAF | 0.418035 | LUAD | NCI-H2347 | SB590885 |
| MVP_top60_DNA_seed | PIK3CD | 0.171731 | KIRC | 769-P | Buparlisib | MVP_top60_DNA+RNA_seed | MDM2 | 0.418758 | LUAD | NCI-H522 | Nutlin-3a (-) |
| MVP_top60_DNA_seed | ESR1 | 0.1727 | KIRC | 769-P | Tamoxifen | MVP_top60_DNA+RNA_seed | BRAF | 0.418779 | LUAD | NCI-H2342 | PLX-4720 |
| MVP_top60_DNA_seed | TP53 | 0.175628 | KIRC | HA7-RCC | MIRA-1 | MVP_top60_DNA+RNA_seed | EGFR | 0.419154 | LUAD | NCI-H2347 | Pelitinib |
| MVP_top60_DNA_seed | PIK3CA | 0.17665 | KIRC | VMRC-RCW | AZD8186 | MVP_top60_DNA+RNA_seed | HSP90AA1 | 0.420258 | LUAD | NCI-H1437 | Elesclomol |
| MVP_top60_DNA_seed | ESR1 | 0.177494 | KIRC | SN12C | Tamoxifen | MVP_top60_DNA+RNA_seed | PIK3CA | 0.420377 | LUAD | ABC-1 | AZD8835 |
| MVP_top60_DNA_seed | IGF1R | 0.185537 | KIRC | ACHN | GSK1904529A | MVP_top60_DNA+RNA_seed | EGFR | 0.420476 | LUAD | SK-LU-1 | AZD8931 |
| MVP_top60_DNA_seed | IGF1R | 0.185626 | KIRC | NCC021 | NVP-ADW742 | MVP_top60_DNA+RNA_seed | MDM2 | 0.421239 | LUAD | H3255 | Nutlin-3a (-) |
| MVP_top60_DNA_seed | PIK3CA | 0.18568 | KIRC | SN12C | AZD8186 | MVP_top60_DNA+RNA_seed | BRAF | 0.422675 | LUAD | PC-14 | Dabrafenib |
| MVP_top60_DNA_seed | PIK3CA | 0.188458 | KIRC | NCC010 | Buparlisib | MVP_top60_DNA+RNA_seed | EGFR | 0.424323 | LUAD | EKVX | Pelitinib |
| MVP_top60_DNA_seed | PIK3CD | 0.188458 | KIRC | NCC010 | Buparlisib | MVP_top60_DNA+RNA_seed | MAPK14 | 0.424507 | LUAD | NCI-H1563 | TAK-715 |
| MVP_top60_DNA_seed | HDAC1 | 0.189757 | KIRC | SN12C | Romidepsin | MVP_top60_DNA+RNA_seed | SMARCA4 | 0.424573 | LUAD | NCI-H1993 | PFI-3 |
| MVP_top60_DNA_seed | PIK3CA | 0.190648 | KIRC | SN12C | GNE-317 | MVP_top60_DNA+RNA_seed | SRC | 0.425899 | LUAD | NCI-H1734 | NVP-BHG712 |
| MVP_top60_DNA_seed | PIK3CA | 0.192211 | KIRC | NCC021 | Buparlisib | MVP_top60_DNA+RNA_seed | EGFR | 0.4269 | LUAD | NCI-H441 | PF-00299804 |
| MVP_top60_DNA_seed | PIK3CD | 0.192211 | KIRC | NCC021 | Buparlisib | MVP_top60_DNA+RNA_seed | HDAC1 | 0.427384 | LUAD | SK-LU-1 | JQ12 |
| MVP_top60_DNA_seed | PIK3CA | 0.193138 | KIRC | KMRC-20 | Buparlisib | MVP_top60_DNA+RNA_seed | BRAF | 0.427478 | LUAD | HOP-62 | SB590885 |
| MVP_top60_DNA_seed | PIK3CD | 0.193138 | KIRC | KMRC-20 | Buparlisib | MVP_top60_DNA+RNA_seed | EGFR | 0.427544 | LUAD | NCI-H1651 | AST-1306 |
| MVP_top60_DNA_seed | SMARCA4 | 0.197023 | KIRC | CAL-54 | PFI3 | MVP_top60_DNA+RNA_seed | IGF1R | 0.427607 | LUAD | NCI-H23 | GSK1904529A |

| | | | | | | | | | | | |
|---|---|---|---|---|---|---|---|---|---|---|---|
| MVP_top60_DNA_seed | SMARCA4 | 0.197154 | KIRC | RCC-MF | PFI3 | MVP_top60_DNA+RNA_seed | ATM | 0.427991 | LUAD | COR-L105 | KU-60019 |
| MVP_top60_DNA_seed | PARP1 | 0.198265 | KIRC | U031 | Veliparib | MVP_top60_DNA+RNA_seed | MAPK14 | 0.428059 | LUAD | NCI-H650 | TAK-715 |
| MVP_top60_DNA_seed | PARP1 | 0.198273 | KIRC | RCC-AB | Rucaparib | MVP_top60_DNA+RNA_seed | BRAF | 0.428543 | LUAD | NCI-H1437 | PLX-4720 |
| MVP_top60_DNA_seed | EP300 | 0.202254 | KIRC | SN12C | CPI-637 | MVP_top60_DNA+RNA_seed | BRAF | 0.429244 | LUAD | NCI-H1993 | PLX-4720 |
| MVP_top60_DNA_seed | IGF1R | 0.205418 | KIRC | 786-0 | BMS-754807 | MVP_top60_DNA+RNA_seed | AURKA | 0.430402 | LUAD | NCI-H2023 | ZM447439 |
| MVP_top60_DNA_seed | IGF1R | 0.207612 | KIRC | RCC-AB | GSK1904529A | MVP_top60_DNA+RNA_seed | MDM2 | 0.431784 | LUAD | NCI-H1792 | Nutlin-3a (-) |
| MVP_top60_DNA_seed | PARP1 | 0.207653 | KIRC | U031 | Olaparib | MVP_top60_DNA+RNA_seed | AURKA | 0.431859 | LUAD | NCI-H1666 | Alisertib |
| MVP_top60_DNA_seed | IGF1R | 0.209522 | KIRC | KMRC-20 | GSK1904529A | MVP_top60_DNA+RNA_seed | AKT1 | 0.43242 | LUAD | NCI-H1573 | MK-2206 |
| MVP_top60_DNA_seed | IGF1R | 0.210616 | KIRC | RCC10RGB | BMS-536924 | MVP_top60_DNA+RNA_seed | BRAF | 0.43245 | LUAD | NCI-H1666 | SB590885 |
| MVP_top60_DNA_seed | IGF1R | 0.210986 | KIRC | BFTC-909 | GSK1904529A | MVP_top60_DNA+RNA_seed | BRAF | 0.433497 | LUAD | NCI-H1838 | PLX-4720 |
| MVP_top60_DNA_seed | AR | 0.211074 | KIRC | NCC010 | Bicalutamide | MVP_top60_DNA+RNA_seed | FGFR1 | 0.434691 | LUAD | HOP-62 | AZD4547 |
| MVP_top60_DNA_seed | ESR1 | 0.216767 | KIRC | RCC-JW | GDC0810 | MVP_top60_DNA+RNA_seed | AKT1 | 0.435299 | LUAD | NCI-H2009 | BAY AKT1 |
| MVP_top60_DNA_seed | HDAC1 | 0.217962 | KIRC | OS-RC-2 | Entinostat | MVP_top60_DNA+RNA_seed | EGFR | 0.43555 | LUAD | NCI-H1792 | Afatinib |
| MVP_top60_DNA_seed | IGF1R | 0.222724 | KIRC | U031 | Linsitinib | MVP_top60_DNA+RNA_seed | EGFR | 0.436257 | LUAD | NCI-H1651 | PF-00299804 |
| MVP_top60_DNA_seed | ESR1 | 0.225203 | KIRC | RCC-JF | GDC0810 | MVP_top60_DNA+RNA_seed | EGFR | 0.436492 | LUAD | NCI-H838 | AZD8931 |
| MVP_top60_DNA_seed | PARP1 | 0.228845 | KIRC | A498 | Talazoparib | MVP_top60_DNA+RNA_seed | HSP90AA1 | 0.438402 | LUAD | H3255 | CCT-018159 |
| MVP_top60_DNA_seed | TP53 | 0.22885 | KIRC | TK10 | MIRA-1 | MVP_top60_DNA+RNA_seed | IGF1R | 0.438469 | LUAD | NCI-H1623 | BMS-536924 |
| MVP_top60_DNA_seed | ATM | 0.229116 | KIRC | RXF393 | KU-55933 | MVP_top60_DNA+RNA_seed | AURKA | 0.439025 | LUAD | NCI-H441 | Alisertib |
| MVP_top60_DNA_seed | IGF1R | 0.230284 | KIRC | LB996-RCC | Linsitinib | MVP_top60_DNA+RNA_seed | EGFR | 0.439149 | LUAD | NCI-H2009 | Cetuximab |
| MVP_top60_DNA_seed | AR | 0.236802 | KIRC | LB1047-RCC | Bicalutamide | MVP_top60_DNA+RNA_seed | AURKA | 0.439413 | LUAD | NCI-H1648 | GSK1070916 |
| MVP_top60_DNA_seed | PARP1 | 0.238432 | KIRC | RCC-FG2 | Rucaparib | MVP_top60_DNA+RNA_seed | SRC | 0.439597 | LUAD | NCI-H322M | WH-4-023 |
| MVP_top60_DNA_seed | MTOR | 0.238514 | KIRC | ACHN | Temsirolimus | MVP_top60_DNA+RNA_seed | EGFR | 0.440322 | LUAD | Calu-6 | Erlotinib |
| MVP_top60_DNA_seed | HSP90AA1 | 0.240452 | KIRC | NCC021 | CCT-018159 | MVP_top60_DNA+RNA_seed | IGF1R | 0.442406 | LUAD | NCI-H1650 | GSK1904529A |
| MVP_top60_DNA_seed | HSP90AB1 | 0.240452 | KIRC | NCC021 | CCT-018159 | MVP_top60_DNA+RNA_seed | MET | 0.44253 | LUAD | NCI-H1573 | AZD6094 |
| MVP_top60_DNA_seed | HSP90AA1 | 0.242824 | KIRC | OS-RC-2 | Tanespimycin | MVP_top60_DNA+RNA_seed | AKT1 | 0.442988 | LUAD | SK-LU-1 | BAY AKT1 |
| MVP_top60_DNA_seed | HSP90AB1 | 0.242824 | KIRC | OS-RC-2 | Tanespimycin | MVP_top60_DNA+RNA_seed | HSP90AA1 | 0.44533 | LUAD | H3255 | Luminespib |
| MVP_top60_DNA_seed | PARP1 | 0.243337 | KIRC | SN12C | Talazoparib | MVP_top60_DNA+RNA_seed | AURKA | 0.446395 | LUAD | RERF-LC-MS | HG-5-113-01 |
| MVP_top60_DNA_seed | EP300 | 0.24472 | KIRC | A498 | SGC-CBP30 | MVP_top60_DNA+RNA_seed | EGFR | 0.448467 | LUAD | NCI-H2291 | Afatinib |
| MVP_top60_DNA_seed | HSP90AA1 | 0.245388 | KIRC | U031 | Luminespib | MVP_top60_DNA+RNA_seed | BRAF | 0.448745 | LUAD | HCC-44 | Dabrafenib |
| MVP_top60_DNA_seed | HSP90AB1 | 0.245388 | KIRC | U031 | Luminespib | MVP_top60_DNA+RNA_seed | EGFR | 0.449721 | LUAD | NCI-H1838 | Gefitinib |
| MVP_top60_DNA_seed | IGF1R | 0.258351 | KIRC | RCC-JF | GSK1904529A | MVP_top60_DNA+RNA_seed | EGFR | 0.449859 | LUAD | NCI-H1993 | AZD8931 |
| MVP_top60_DNA_seed | TP53 | 0.258536 | KIRC | BB65-RCC | MIRA-1 | MVP_top60_DNA+RNA_seed | BRAF | 0.451667 | LUAD | Calu-3 | HG6-64-1 |
| MVP_top60_DNA_seed | PIK3CA | 0.259897 | KIRC | RXF393 | Alpelisib | MVP_top60_DNA+RNA_seed | BRAF | 0.454531 | LUAD | EMC-BAC-1 | PLX-4720 |
| MVP_top60_DNA_seed | AR | 0.262855 | KIRC | SW156 | Bicalutamide | MVP_top60_DNA+RNA_seed | HDAC1 | 0.45541 | LUAD | SK-LU-1 | CAY10603 |
| MVP_top60_DNA_seed | HDAC1 | 0.263668 | KIRC | LB1047-RCC | PCI-34051 | MVP_top60_DNA+RNA_seed | IGF1R | 0.455845 | LUAD | NCI-H1563 | Linsitinib |
| MVP_top60_DNA_seed | IGF1R | 0.271211 | KIRC | 786-0 | BMS-536924 | MVP_top60_DNA+RNA_seed | MDM2 | 0.457921 | LUAD | RERF-LC-KJ | Nutlin-3a (-) |
| MVP_top60_DNA_seed | IGF1R | 0.271785 | KIRC | A498 | BMS-754807 | MVP_top60_DNA+RNA_seed | EGFR | 0.457931 | LUAD | RERF-LC-KJ | Pelitinib |
| MVP_top60_DNA_seed | MTOR | 0.276441 | KIRC | RCC-AB | Temsirolimus | MVP_top60_DNA+RNA_seed | HSP90AA1 | 0.458672 | LUAD | NCI-H2228 | CCT-018159 |
| MVP_top60_DNA_seed | PIK3CA | 0.285639 | KIRC | KMRC-20 | GNE-317 | MVP_top60_DNA+RNA_seed | EGFR | 0.460111 | LUAD | NCI-H1563 | Afatinib |
| MVP_top60_DNA_seed | IGF1R | 0.296533 | KIRC | RCC-MF | BMS-754807 | MVP_top60_DNA+RNA_seed | AURKA | 0.460255 | LUAD | NCI-H1651 | CD532 |
| MVP_top60_DNA_seed | IGF1R | 0.298273 | KIRC | LB1047-RCC | GSK1904529A | MVP_top60_DNA+RNA_seed | MET | 0.460526 | LUAD | NCI-H1993 | SU11274 |

| | | | | | | | | | | | |
|---|---|---|---|---|---|---|---|---|---|---|---|
| MVP_top60_DNA_seed | HDAC1 | 0.299265 | KIRC | NCC010 | Entinostat | MVP_top60_DNA+RNA_seed | ATM | 0.461122 | LUAD | NCI-H2087 | KU-60019 |
| MVP_top60_DNA_seed | PARP1 | 0.299742 | KIRC | RCC-MF | Veliparib | MVP_top60_DNA+RNA_seed | HSP90AA1 | 0.461282 | LUAD | Calu-3 | CCT-018159 |
| MVP_top60_DNA_seed | PARP1 | 0.30005 | KIRC | 786-0 | Talazoparib | MVP_top60_DNA+RNA_seed | AURKA | 0.461569 | LUAD | NCI-H2405 | GSK1070916 |
| MVP_top60_DNA_seed | EP300 | 0.3007 | KIRC | KMRC-20 | CPI-637 | MVP_top60_DNA+RNA_seed | EGFR | 0.462667 | LUAD | NCI-H650 | AZD8931 |
| MVP_top60_DNA_seed | HDAC1 | 0.301587 | KIRC | LB996-RCC | Entinostat | MVP_top60_DNA+RNA_seed | ATM | 0.462769 | LUAD | NCI-H1623 | KU-55933 |
| MVP_top60_DNA_seed | TP53 | 0.304421 | KIRC | CAL-54 | MIRA-1 | MVP_top60_DNA+RNA_seed | PTK2 | 0.462964 | LUAD | PC-14 | PF-562271 |
| MVP_top60_DNA_seed | PARP1 | 0.306323 | KIRC | LB996-RCC | Niraparib | MVP_top60_DNA+RNA_seed | FGFR1 | 0.463634 | LUAD | NCI-H522 | AZD4547 |
| MVP_top60_DNA_seed | TP53 | 0.307187 | KIRC | 786-0 | MIRA-1 | MVP_top60_DNA+RNA_seed | HDAC1 | 0.464217 | LUAD | NCI-H1944 | Tubastatin A |
| MVP_top60_DNA_seed | PIK3CA | 0.30787 | KIRC | 769-P | Alpelisib | MVP_top60_DNA+RNA_seed | BRAF | 0.46446 | LUAD | NCI-H2405 | SB590885 |
| MVP_top60_DNA_seed | PARP1 | 0.3108 | KIRC | LB2241-RCC | Niraparib | MVP_top60_DNA+RNA_seed | EGFR | 0.465163 | LUAD | NCI-H2228 | Pelitinib |
| MVP_top60_DNA_seed | IGF1R | 0.311208 | KIRC | KMRC-20 | BMS-754807 | MVP_top60_DNA+RNA_seed | MET | 0.46537 | LUAD | RERF-LC-KJ | SU11274 |
| MVP_top60_DNA_seed | PARP1 | 0.311286 | KIRC | RCC-FG2 | Talazoparib | MVP_top60_DNA+RNA_seed | HDAC1 | 0.465662 | LUAD | EKVX | Entinostat |
| MVP_top60_DNA_seed | HDAC1 | 0.31588 | KIRC | BFTC-909 | Entinostat | MVP_top60_DNA+RNA_seed | AKT1 | 0.465795 | LUAD | PC-3_[JPC-3] | AKT inhibitor VIII |
| MVP_top60_DNA_seed | MTOR | 0.316734 | KIRC | KMRC-1 | Temsirolimus | MVP_top60_DNA+RNA_seed | FGFR1 | 0.466013 | LUAD | SW1573 | AZD4547 |
| MVP_top60_DNA_seed | PARP1 | 0.318218 | KIRC | SW156 | Niraparib | MVP_top60_DNA+RNA_seed | ATM | 0.467663 | LUAD | NCI-H2122 | KU-55933 |
| MVP_top60_DNA_seed | PARP1 | 0.318325 | KIRC | HA7-RCC | Talazoparib | MVP_top60_DNA+RNA_seed | EGFR | 0.468012 | LUAD | NCI-H441 | CI-1033 |
| MVP_top60_DNA_seed | PIK3CA | 0.322928 | KIRC | NCC010 | GNE-317 | MVP_top60_DNA+RNA_seed | EGFR | 0.469634 | LUAD | NCI-H1838 | AZD8931 |
| MVP_top60_DNA_seed | PARP1 | 0.32379 | KIRC | LB1047-RCC | Rucaparib | MVP_top60_DNA+RNA_seed | AKT1 | 0.470904 | LUAD | PC-3_[JPC-3] | GSK690693 |
| MVP_top60_DNA_seed | ATM | 0.325635 | KIRC | ACHN | KU-55933 | MVP_top60_DNA+RNA_seed | BRAF | 0.472367 | LUAD | SW1573 | RAF_9304 |
| MVP_top60_DNA_seed | IGF1R | 0.32936 | KIRC | RCC-MF | NVP-ADW742 | MVP_top60_DNA+RNA_seed | EGFR | 0.474412 | LUAD | RERF-LC-MS | CI-1033 |
| MVP_top60_DNA_seed | PARP1 | 0.330168 | KIRC | SN12C | Rucaparib | MVP_top60_DNA+RNA_seed | AURKA | 0.474566 | LUAD | HOP-62 | ZM447439 |
| MVP_top60_DNA_seed | IGF1R | 0.333165 | KIRC | RCC10RGB | NVP-ADW742 | MVP_top60_DNA+RNA_seed | HDAC1 | 0.475716 | LUAD | Calu-3 | Tubastatin A |
| MVP_top60_DNA_seed | IGF1R | 0.339529 | KIRC | RCC-JW | BMS-536924 | MVP_top60_DNA+RNA_seed | HSP90AA1 | 0.476592 | LUAD | NCI-H1838 | Tanespimycin |
| MVP_top60_DNA_seed | PARP1 | 0.341753 | KIRC | NCC021 | Talazoparib | MVP_top60_DNA+RNA_seed | EGFR | 0.476937 | LUAD | NCI-H2009 | Afatinib |
| MVP_top60_DNA_seed | MTOR | 0.342166 | KIRC | NCC010 | Temsirolimus | MVP_top60_DNA+RNA_seed | FGFR1 | 0.477078 | LUAD | NCI-H2122 | AZD4547 |
| MVP_top60_DNA_seed | HSP90AA1 | 0.345824 | KIRC | 769-P | Luminespib | MVP_top60_DNA+RNA_seed | AURKA | 0.478744 | LUAD | PC-3_[JPC-3] | GSK1070916 |
| MVP_top60_DNA_seed | HSP90AB1 | 0.345824 | KIRC | 769-P | Luminespib | MVP_top60_DNA+RNA_seed | SRC | 0.479197 | LUAD | EKVX | Bosutinib |
| MVP_top60_DNA_seed | HSP90AA1 | 0.348258 | KIRC | LB2241-RCC | Luminespib | MVP_top60_DNA+RNA_seed | MAPK14 | 0.479274 | LUAD | NCI-H1573 | TAK-715 |
| MVP_top60_DNA_seed | HSP90AB1 | 0.348258 | KIRC | LB2241-RCC | Luminespib | MVP_top60_DNA+RNA_seed | EGFR | 0.479737 | LUAD | NCI-H1651 | Cetuximab |
| MVP_top60_DNA_seed | HSP90AA1 | 0.349726 | KIRC | RCC-AB | Luminespib | MVP_top60_DNA+RNA_seed | SRC | 0.481045 | LUAD | Calu-6 | WH-4-023 |
| MVP_top60_DNA_seed | HSP90AB1 | 0.349726 | KIRC | RCC-AB | Luminespib | MVP_top60_DNA+RNA_seed | BRAF | 0.485802 | LUAD | VMRC-LCD | PLX-4720 |
| MVP_top60_DNA_seed | HSP90AA1 | 0.351779 | KIRC | SN12C | Luminespib | MVP_top60_DNA+RNA_seed | AKT1 | 0.485839 | LUAD | NCI-H2087 | AKT inhibitor VIII |
| MVP_top60_DNA_seed | HSP90AB1 | 0.351779 | KIRC | SN12C | Luminespib | MVP_top60_DNA+RNA_seed | EGFR | 0.486252 | LUAD | HCC-78 | AZD8931 |
| MVP_top60_DNA_seed | ATM | 0.352015 | KIRC | BFTC-909 | KU-55933 | MVP_top60_DNA+RNA_seed | AURKA | 0.487057 | LUAD | Calu-6 | Tozasertib |
| MVP_top60_DNA_seed | HDAC1 | 0.353183 | KIRC | RXF393 | Entinostat | MVP_top60_DNA+RNA_seed | AURKA | 0.487295 | LUAD | Calu-3 | GSK1070916 |
| MVP_top60_DNA_seed | HSP90AA1 | 0.355254 | KIRC | RCC-ER | Tanespimycin | MVP_top60_DNA+RNA_seed | BRAF | 0.4876 | LUAD | EMC-BAC-2 | RAF_9304 |
| MVP_top60_DNA_seed | HSP90AB1 | 0.355254 | KIRC | RCC-ER | Tanespimycin | MVP_top60_DNA+RNA_seed | IGF1R | 0.487761 | LUAD | Calu-6 | BMS-536924 |
| MVP_top60_DNA_seed | HDAC1 | 0.355568 | KIRC | RCC10RGB | PCI-34051 | MVP_top60_DNA+RNA_seed | ATM | 0.488647 | LUAD | NCI-H650 | KU-55933 |
| MVP_top60_DNA_seed | PARP1 | 0.355593 | KIRC | KMRC-1 | Olaparib | MVP_top60_DNA+RNA_seed | HDAC1 | 0.48976 | LUAD | NCI-H1648 | Entinostat |
| MVP_top60_DNA_seed | PARP1 | 0.358894 | KIRC | SW156 | Olaparib | MVP_top60_DNA+RNA_seed | IGF1R | 0.490371 | LUAD | NCI-H322M | BMS-536924 |
| MVP_top60_DNA_seed | PARP1 | 0.359136 | KIRC | SW156 | Talazoparib | MVP_top60_DNA+RNA_seed | MET | 0.491171 | LUAD | NCI-H2405 | AZD6094 |

| | | | | | | | | | | | |
|---|---|---|---|---|---|---|---|---|---|---|---|
| MVP_top60_DNA_seed | PIK3CA | 0.359376 | KIRC | OS-RC-2 | GNE-317 | MVP_top60_DNA+RNA_seed | BRAF | 0.491692 | LUAD | PC-3_[JPC-3] | PLX-4720 |
| MVP_top60_DNA_seed | MTOR | 0.363112 | KIRC | CAKI-1 | Temsirolimus | MVP_top60_DNA+RNA_seed | MET | 0.492111 | LUAD | NCI-H1623 | SU11274 |
| MVP_top60_DNA_seed | HSP90AA1 | 0.364493 | KIRC | RCC-MF | Elesclomol | MVP_top60_DNA+RNA_seed | EGFR | 0.492278 | LUAD | NCI-H838 | CUDC-101 |
| MVP_top60_DNA_seed | HSP90AB1 | 0.364493 | KIRC | RCC-MF | Elesclomol | MVP_top60_DNA+RNA_seed | MDM2 | 0.493782 | LUAD | VMRC-LCD | Nutlin-3a (-) |
| MVP_top60_DNA_seed | IGF1R | 0.364926 | KIRC | RXF393 | GSK1904529A | MVP_top60_DNA+RNA_seed | EGFR | 0.494143 | LUAD | NCI-H1437 | Gefitinib |
| MVP_top60_DNA_seed | HDAC1 | 0.366339 | KIRC | ACHN | PCI-34051 | MVP_top60_DNA+RNA_seed | ATM | 0.494755 | LUAD | ABC-1 | KU-60019 |
| MVP_top60_DNA_seed | HDAC1 | 0.366415 | KIRC | NCC021 | Entinostat | MVP_top60_DNA+RNA_seed | EGFR | 0.496306 | LUAD | NCI-H2342 | Gefitinib |
| MVP_top60_DNA_seed | ESR1 | 0.367631 | KIRC | CAL-54 | GDC0810 | MVP_top60_DNA+RNA_seed | FGFR1 | 0.497266 | LUAD | NCI-H2030 | AZD4547 |
| MVP_top60_DNA_seed | PARP1 | 0.368131 | KIRC | 786-0 | Niraparib | MVP_top60_DNA+RNA_seed | ATM | 0.498275 | LUAD | H3255 | KU-55933 |
| MVP_top60_DNA_seed | HSP90AA1 | 0.377435 | KIRC | NCC021 | Elesclomol | MVP_top60_DNA+RNA_seed | BRAF | 0.498886 | LUAD | NCI-H1355 | RAF_9304 |
| MVP_top60_DNA_seed | HSP90AB1 | 0.377435 | KIRC | NCC021 | Elesclomol | MVP_top60_DNA+RNA_seed | HDAC1 | 0.499545 | LUAD | NCI-H322M | AR-42 |
| MVP_top60_DNA_seed | ESR1 | 0.38195 | KIRC | RXF393 | GDC0810 | MVP_top60_DNA+RNA_seed | SMARCA4 | 0.499865 | LUAD | NCI-H1568 | PFI-3 |
| MVP_top60_DNA_seed | SMARCA4 | 0.387165 | KIRC | KMRC-20 | PFI3 | MVP_top60_DNA+RNA_seed | ATM | 0.499968 | LUAD | RERF-LC-MS | KU-55933 |
| MVP_top60_DNA_seed | EP300 | 0.388821 | KIRC | BFTC-909 | SGC-CBP30 | MVP_top60_DNA+RNA_seed | MDM2 | 0.50009 | LUAD | NCI-H2405 | Nutlin-3a (-) |
| MVP_top60_DNA_seed | IGF1R | 0.39041 | KIRC | KMRC-1 | BMS-536924 | MVP_top60_DNA+RNA_seed | AURKA | 0.500558 | LUAD | NCI-H1993 | Genentech Cpd 10 |
| MVP_top60_DNA_seed | HDAC1 | 0.40033 | KIRC | BB65-RCC | Entinostat | MVP_top60_DNA+RNA_seed | MET | 0.500725 | LUAD | NCI-H2342 | SU11274 |
| MVP_top60_DNA_seed | ESR1 | 0.401056 | KIRC | OS-RC-2 | GDC0810 | MVP_top60_DNA+RNA_seed | HDAC1 | 0.50132 | LUAD | SK-LU-1 | Belinostat |
| MVP_top60_DNA_seed | PARP1 | 0.401789 | KIRC | 786-0 | Rucaparib | MVP_top60_DNA+RNA_seed | AURKA | 0.503501 | LUAD | NCI-H650 | Alisertib |
| MVP_top60_DNA_seed | TP53 | 0.403485 | KIRC | SW156 | MIRA-1 | MVP_top60_DNA+RNA_seed | PIK3CA | 0.503865 | LUAD | ABC-1 | PI3Ka_4409 |
| MVP_top60_DNA_seed | ESR1 | 0.403867 | KIRC | ACHN | GDC0810 | MVP_top60_DNA+RNA_seed | HDAC1 | 0.504481 | LUAD | NCI-H2085 | Belinostat |
| MVP_top60_DNA_seed | HSP90AA1 | 0.403962 | KIRC | LB1047-RCC | Tanespimycin | MVP_top60_DNA+RNA_seed | EGFR | 0.504611 | LUAD | NCI-H1650 | CUDC-101 |
| MVP_top60_DNA_seed | HSP90AB1 | 0.403962 | KIRC | LB1047-RCC | Tanespimycin | MVP_top60_DNA+RNA_seed | EGFR | 0.505283 | LUAD | NCI-H1993 | AST-1306 |
| MVP_top60_DNA_seed | SMARCA4 | 0.40752 | KIRC | RCC-FG2 | PFI3 | MVP_top60_DNA+RNA_seed | AURKA | 0.505528 | LUAD | NCI-H1666 | GSK1070916 |
| MVP_top60_DNA_seed | MTOR | 0.415946 | KIRC | RCC10RGB | Temsirolimus | MVP_top60_DNA+RNA_seed | IGF1R | 0.505704 | LUAD | NCI-H2085 | BMS-754807 |
| MVP_top60_DNA_seed | IGF1R | 0.419324 | KIRC | HA7-RCC | BMS-754807 | MVP_top60_DNA+RNA_seed | SRC | 0.505711 | LUAD | EKVX | Saracatinib |
| MVP_top60_DNA_seed | ATM | 0.427488 | KIRC | U031 | KU-55933 | MVP_top60_DNA+RNA_seed | FGFR1 | 0.505803 | LUAD | NCI-H1693 | PD173074 |
| MVP_top60_DNA_seed | PIK3CA | 0.427528 | KIRC | SN12C | Buparlisib | MVP_top60_DNA+RNA_seed | HSP90AA1 | 0.506541 | LUAD | NCI-H1838 | Elesclomol |
| MVP_top60_DNA_seed | PIK3CD | 0.427528 | KIRC | SN12C | Buparlisib | MVP_top60_DNA+RNA_seed | MET | 0.506621 | LUAD | HCC-78 | SU11274 |
| MVP_top60_DNA_seed | PARP1 | 0.432348 | KIRC | KMRC-20 | Talazoparib | MVP_top60_DNA+RNA_seed | ATM | 0.507233 | LUAD | LC-2-ad | KU-60019 |
| MVP_top60_DNA_seed | IGF1R | 0.433283 | KIRC | HA7-RCC | NVP-ADW742 | MVP_top60_DNA+RNA_seed | HSP90AA1 | 0.508474 | LUAD | NCI-H1734 | SNX-2112 |
| MVP_top60_DNA_seed | PARP1 | 0.43587 | KIRC | NCC010 | Talazoparib | MVP_top60_DNA+RNA_seed | BRAF | 0.508785 | LUAD | NCI-H2122 | SB590885 |
| MVP_top60_DNA_seed | HSP90AA1 | 0.438647 | KIRC | KMRC-1 | Luminespib | MVP_top60_DNA+RNA_seed | HDAC1 | 0.508897 | LUAD | SW1573 | Belinostat |
| MVP_top60_DNA_seed | HSP90AB1 | 0.438647 | KIRC | KMRC-1 | Luminespib | MVP_top60_DNA+RNA_seed | SRC | 0.508924 | LUAD | ABC-1 | HG-6-71-01 |
| MVP_top60_DNA_seed | PIK3CA | 0.439711 | KIRC | BB65-RCC | AZD8186 | MVP_top60_DNA+RNA_seed | AKT1 | 0.509073 | LUAD | NCI-H358 | BAY AKT1 |
| MVP_top60_DNA_seed | PARP1 | 0.439735 | KIRC | OS-RC-2 | Niraparib | MVP_top60_DNA+RNA_seed | HDAC1 | 0.509694 | LUAD | NCI-H1435 | Dacinostat |
| MVP_top60_DNA_seed | IGF1R | 0.441179 | KIRC | SW156 | BMS-754807 | MVP_top60_DNA+RNA_seed | PIK3CA | 0.509795 | LUAD | NCI-H522 | AZD8835 |
| MVP_top60_DNA_seed | HSP90AA1 | 0.446857 | KIRC | RCC-JW | Luminespib | MVP_top60_DNA+RNA_seed | EGFR | 0.510279 | LUAD | SK-LU-1 | CUDC-101 |
| MVP_top60_DNA_seed | HSP90AB1 | 0.446857 | KIRC | RCC-JW | Luminespib | MVP_top60_DNA+RNA_seed | EGFR | 0.51045 | LUAD | NCI-H1573 | PF-00299804 |
| MVP_top60_DNA_seed | ESR1 | 0.447783 | KIRC | RCC-JW | Tamoxifen | MVP_top60_DNA+RNA_seed | EGFR | 0.510487 | LUAD | Calu-6 | Lapatinib |
| MVP_top60_DNA_seed | IGF1R | 0.448653 | KIRC | RCC10RGB | Linsitinib | MVP_top60_DNA+RNA_seed | FGFR1 | 0.51076 | LUAD | NCI-H1666 | FGFR_3831 |
| MVP_top60_DNA_seed | PARP1 | 0.449187 | KIRC | BFTC-909 | Rucaparib | MVP_top60_DNA+RNA_seed | EGFR | 0.511534 | LUAD | NCI-H1755 | Gefitinib |

| | | | | | | | | | | | |
|---|---|---|---|---|---|---|---|---|---|---|---|
| MVP_top60_DNA_seed | PARP1 | 0.454882 | KIRC | TK10 | Rucaparib | MVP_top60_DNA+RNA_seed | BRAF | 0.511687 | LUAD | H3255 | PLX-4720 |
| MVP_top60_DNA_seed | PARP1 | 0.461917 | KIRC | BB65-RCC | Olaparib | MVP_top60_DNA+RNA_seed | EGFR | 0.511973 | LUAD | RERF-LC-MS | Cetuximab |
| MVP_top60_DNA_seed | HDAC1 | 0.465005 | KIRC | KMRC-20 | PCI-34051 | MVP_top60_DNA+RNA_seed | BRAF | 0.512696 | LUAD | NCI-H1563 | HG6-64-1 |
| MVP_top60_DNA_seed | HDAC1 | 0.467398 | KIRC | A704 | Entinostat | MVP_top60_DNA+RNA_seed | AKT1 | 0.512834 | LUAD | EMC-BAC-2 | AKT inhibitor VIII |
| MVP_top60_DNA_seed | PIK3CA | 0.469685 | KIRC | RCC-JW | Alpelisib | MVP_top60_DNA+RNA_seed | MET | 0.513053 | LUAD | NCI-H2087 | AZD6094 |
| MVP_top60_DNA_seed | ESR1 | 0.4738 | KIRC | RCC-MF | GDC0810 | MVP_top60_DNA+RNA_seed | MAPK14 | 0.513143 | LUAD | NCI-H1734 | TAK-715 |
| MVP_top60_DNA_seed | PARP1 | 0.480225 | KIRC | A498 | Rucaparib | MVP_top60_DNA+RNA_seed | HDAC1 | 0.514612 | LUAD | NCI-H2085 | Tubastatin A |
| MVP_top60_DNA_seed | EP300 | 0.480629 | KIRC | LB996-RCC | SGC-CBP30 | MVP_top60_DNA+RNA_seed | AURKA | 0.520113 | LUAD | NCI-H3122 | Alisertib |
| MVP_top60_DNA_seed | PIK3CA | 0.481959 | KIRC | LB2241-RCC | Buparlisib | MVP_top60_DNA+RNA_seed | SMARCA4 | 0.520279 | LUAD | NCI-H1781 | PFI-3 |
| MVP_top60_DNA_seed | PIK3CD | 0.481959 | KIRC | LB2241-RCC | Buparlisib | MVP_top60_DNA+RNA_seed | ATM | 0.520475 | LUAD | HOP-62 | KU-55933 |
| MVP_top60_DNA_seed | ESR1 | 0.48518 | KIRC | KMRC-20 | GDC0810 | MVP_top60_DNA+RNA_seed | PIK3CA | 0.520712 | LUAD | NCI-H2291 | AZD8835 |
| MVP_top60_DNA_seed | PARP1 | 0.486263 | KIRC | CAKI-1 | Niraparib | MVP_top60_DNA+RNA_seed | MAPK14 | 0.521036 | LUAD | NCI-H1568 | TAK-715 |
| MVP_top60_DNA_seed | ESR1 | 0.490899 | KIRC | BB65-RCC | GDC0810 | MVP_top60_DNA+RNA_seed | ATM | 0.521038 | LUAD | NCI-H322M | CP466722 |
| MVP_top60_DNA_seed | PARP1 | 0.494226 | KIRC | CAL-54 | Veliparib | MVP_top60_DNA+RNA_seed | AKT1 | 0.521771 | LUAD | SK-LU-1 | AKT inhibitor VIII |
| MVP_top60_DNA_seed | ESR1 | 0.49772 | KIRC | KMRC-1 | GDC0810 | MVP_top60_DNA+RNA_seed | FGFR1 | 0.52226 | LUAD | RERF-LC-KJ | FGFR_3831 |
| MVP_top60_DNA_seed | PIK3CA | 0.498048 | KIRC | SW156 | Alpelisib | MVP_top60_DNA+RNA_seed | BRAF | 0.522804 | LUAD | NCI-H1435 | PLX-4720 |
| MVP_top60_DNA_seed | EP300 | 0.498388 | KIRC | KMRC-1 | CPI-637 | MVP_top60_DNA+RNA_seed | MAPK14 | 0.523521 | LUAD | SK-LU-1 | TAK-715 |
| MVP_top60_DNA_seed | PARP1 | 0.500249 | KIRC | CAL-54 | Niraparib | MVP_top60_DNA+RNA_seed | EGFR | 0.52518 | LUAD | NCI-H1666 | CUDC-101 |
| MVP_top60_DNA_seed | TP53 | 0.50176 | KIRC | OS-RC-2 | MIRA-1 | MVP_top60_DNA+RNA_seed | BRAF | 0.52548 | LUAD | NCI-H2085 | Dabrafenib |
| MVP_top60_DNA_seed | PARP1 | 0.507127 | KIRC | CAL-54 | Olaparib | MVP_top60_DNA+RNA_seed | HDAC1 | 0.526119 | LUAD | NCI-H1792 | Belinostat |
| MVP_top60_DNA_seed | EP300 | 0.511925 | KIRC | 786-0 | SGC-CBP30 | MVP_top60_DNA+RNA_seed | EGFR | 0.52662 | LUAD | NCI-H1563 | PF-00299804 |
| MVP_top60_DNA_seed | EP300 | 0.514848 | KIRC | RXF393 | SGC-CBP30 | MVP_top60_DNA+RNA_seed | IGF1R | 0.527232 | LUAD | PC-3_[JPC-3] | Linsitinib |
| MVP_top60_DNA_seed | HSP90AA1 | 0.51719 | KIRC | A704 | Luminespib | MVP_top60_DNA+RNA_seed | AURKA | 0.527278 | LUAD | NCI-H2228 | ZM447439 |
| MVP_top60_DNA_seed | HSP90AB1 | 0.51719 | KIRC | A704 | Luminespib | MVP_top60_DNA+RNA_seed | HDAC1 | 0.527803 | LUAD | NCI-H1650 | Dacinostat |
| MVP_top60_DNA_seed | HDAC1 | 0.520097 | KIRC | 786-0 | Romidepsin | MVP_top60_DNA+RNA_seed | AKT1 | 0.528128 | LUAD | NCI-H1650 | GSK690693 |
| MVP_top60_DNA_seed | PARP1 | 0.52067 | KIRC | RXF393 | Veliparib | MVP_top60_DNA+RNA_seed | IGF1R | 0.528451 | LUAD | Calu-6 | Linsitinib |
| MVP_top60_DNA_seed | HSP90AA1 | 0.520841 | KIRC | RCC-MF | CCT-018159 | MVP_top60_DNA+RNA_seed | AURKA | 0.528739 | LUAD | RERF-LC-KJ | Alisertib |
| MVP_top60_DNA_seed | HSP90AB1 | 0.520841 | KIRC | RCC-MF | CCT-018159 | MVP_top60_DNA+RNA_seed | BRAF | 0.529692 | LUAD | NCI-H1993 | RAF_9304 |
| MVP_top60_DNA_seed | PIK3CA | 0.521165 | KIRC | LB1047-RCC | Buparlisib | MVP_top60_DNA+RNA_seed | AKT1 | 0.530348 | LUAD | NCI-H2023 | GSK690693 |
| MVP_top60_DNA_seed | PIK3CD | 0.521165 | KIRC | LB1047-RCC | Buparlisib | MVP_top60_DNA+RNA_seed | AKT1 | 0.530471 | LUAD | NCI-H1781 | GSK690693 |
| MVP_top60_DNA_seed | IGF1R | 0.521406 | KIRC | VMRC-RCW | NVP-ADW742 | MVP_top60_DNA+RNA_seed | BRAF | 0.531199 | LUAD | NCI-H1435 | PLX-4720 |
| MVP_top60_DNA_seed | PIK3CA | 0.526806 | KIRC | A704 | GNE-317 | MVP_top60_DNA+RNA_seed | HDAC1 | 0.531311 | LUAD | NCI-H650 | Tubastatin A |
| MVP_top60_DNA_seed | IGF1R | 0.528916 | KIRC | VMRC-RCW | BMS-536924 | MVP_top60_DNA+RNA_seed | IGF1R | 0.531481 | LUAD | NCI-H1755 | GSK1904529A |
| MVP_top60_DNA_seed | IGF1R | 0.536162 | KIRC | OS-RC-2 | GSK1904529A | MVP_top60_DNA+RNA_seed | EGFR | 0.53237 | LUAD | NCI-H838 | Gefitinib |
| MVP_top60_DNA_seed | IGF1R | 0.537402 | KIRC | 769-P | Linsitinib | MVP_top60_DNA+RNA_seed | MDM2 | 0.532374 | LUAD | NCI-H441 | Nutlin-3a (-) |
| MVP_top60_DNA_seed | PARP1 | 0.542838 | KIRC | NCC021 | Olaparib | MVP_top60_DNA+RNA_seed | HDAC1 | 0.533241 | LUAD | Calu-6 | Tubastatin A |
| MVP_top60_DNA_seed | PARP1 | 0.543174 | KIRC | RCC-ER | Niraparib | MVP_top60_DNA+RNA_seed | HSP90AA1 | 0.533587 | LUAD | NCI-H1437 | SNX-2112 |
| MVP_top60_DNA_seed | PIK3CA | 0.548043 | KIRC | LB996-RCC | Alpelisib | MVP_top60_DNA+RNA_seed | AURKA | 0.533891 | LUAD | Calu-6 | Genentech Cpd 10 |
| MVP_top60_DNA_seed | IGF1R | 0.550424 | KIRC | RXF393 | BMS-754807 | MVP_top60_DNA+RNA_seed | PIK3CA | 0.534366 | LUAD | Calu-6 | PI3Ka_4409 |
| MVP_top60_DNA_seed | PARP1 | 0.551669 | KIRC | KMRC-1 | Veliparib | MVP_top60_DNA+RNA_seed | EGFR | 0.534406 | LUAD | NCI-H2405 | CUDC-101 |
| MVP_top60_DNA_seed | HSP90AA1 | 0.552355 | KIRC | BB65-RCC | Elesclomol | MVP_top60_DNA+RNA_seed | MDM2 | 0.534813 | LUAD | NCI-H2009 | Nutlin-3a (-) |

| | | | | | | | | | | | |
|---|---|---|---|---|---|---|---|---|---|---|---|
| MVP_top60_DNA_seed | HSP90AB1 | 0.552355 | KIRC | BB65-RCC | Elesclomol | MVP_top60_DNA+RNA_seed | HSP90AA1 | 0.535888 | LUAD | NCI-H3122 | Elesclomol |
| MVP_top60_DNA_seed | PARP1 | 0.552512 | KIRC | CAKI-1 | Olaparib | MVP_top60_DNA+RNA_seed | FGFR1 | 0.536374 | LUAD | NCI-H358 | AZD4547 |
| MVP_top60_DNA_seed | TP53 | 0.553529 | KIRC | RXF393 | MIRA-1 | MVP_top60_DNA+RNA_seed | BRAF | 0.536937 | LUAD | NCI-H322M | SB590885 |
| MVP_top60_DNA_seed | PARP1 | 0.554823 | KIRC | SN12C | Niraparib | MVP_top60_DNA+RNA_seed | BRAF | 0.537397 | LUAD | NCI-H1755 | RAF_9304 |
| MVP_top60_DNA_seed | IGF1R | 0.556082 | KIRC | NCC021 | GSK1904529A | MVP_top60_DNA+RNA_seed | AKT1 | 0.538857 | LUAD | PC-14 | MK-2206 |
| MVP_top60_DNA_seed | IGF1R | 0.557377 | KIRC | A498 | GSK1904529A | MVP_top60_DNA+RNA_seed | AKT1 | 0.539023 | LUAD | NCI-H2122 | GSK690693 |
| MVP_top60_DNA_seed | MTOR | 0.560461 | KIRC | 786-0 | Temsirolimus | MVP_top60_DNA+RNA_seed | ATM | 0.539599 | LUAD | NCI-H1563 | torin2 |
| MVP_top60_DNA_seed | IGF1R | 0.565225 | KIRC | HA7-RCC | Linsitinib | MVP_top60_DNA+RNA_seed | EGFR | 0.540355 | LUAD | LXF-289 | Gefitinib |
| MVP_top60_DNA_seed | IGF1R | 0.565473 | KIRC | OS-RC-2 | Linsitinib | MVP_top60_DNA+RNA_seed | HDAC1 | 0.541238 | LUAD | NCI-H838 | JQ12 |
| MVP_top60_DNA_seed | PIK3CA | 0.565548 | KIRC | RCC-AB | Buparlisib | MVP_top60_DNA+RNA_seed | BRAF | 0.542069 | LUAD | NCI-H2009 | Dabrafenib |
| MVP_top60_DNA_seed | PIK3CD | 0.565548 | KIRC | RCC-AB | Buparlisib | MVP_top60_DNA+RNA_seed | EGFR | 0.542277 | LUAD | SW1573 | CUDC-101 |
| MVP_top60_DNA_seed | PARP1 | 0.570328 | KIRC | NCC010 | Rucaparib | MVP_top60_DNA+RNA_seed | PIK3CA | 0.543329 | LUAD | HOP-62 | AZD8835 |
| MVP_top60_DNA_seed | PARP1 | 0.570462 | KIRC | ACHN | Rucaparib | MVP_top60_DNA+RNA_seed | IGF1R | 0.544397 | LUAD | NCI-H1568 | BMS-536924 |
| MVP_top60_DNA_seed | PIK3CA | 0.574285 | KIRC | TK10 | GNE-317 | MVP_top60_DNA+RNA_seed | MET | 0.544934 | LUAD | PC-3_[JPC-3] | JNJ38877605 |
| MVP_top60_DNA_seed | AR | 0.575036 | KIRC | RCC-FG2 | Bicalutamide | MVP_top60_DNA+RNA_seed | HDAC1 | 0.545374 | LUAD | NCI-H1734 | Tubastatin A |
| MVP_top60_DNA_seed | HDAC1 | 0.578521 | KIRC | HA7-RCC | Entinostat | MVP_top60_DNA+RNA_seed | SRC | 0.545766 | LUAD | NCI-H1838 | Bosutinib |
| MVP_top60_DNA_seed | MTOR | 0.579023 | KIRC | LB1047-RCC | Temsirolimus | MVP_top60_DNA+RNA_seed | FGFR1 | 0.546828 | LUAD | NCI-H1563 | FGFR_3831 |
| MVP_top60_DNA_seed | PIK3CA | 0.580145 | KIRC | RCC-JW | Buparlisib | MVP_top60_DNA+RNA_seed | MET | 0.549649 | LUAD | NCI-H358 | SU11274 |
| MVP_top60_DNA_seed | PIK3CD | 0.580145 | KIRC | RCC-JW | Buparlisib | MVP_top60_DNA+RNA_seed | EGFR | 0.549665 | LUAD | NCI-H2009 | CI-1033 |
| MVP_top60_DNA_seed | PARP1 | 0.581698 | KIRC | RCC-JW | Veliparib | MVP_top60_DNA+RNA_seed | MET | 0.549937 | LUAD | NCI-H1975 | SU11274 |
| MVP_top60_DNA_seed | IGF1R | 0.584229 | KIRC | RCC-FG2 | BMS-754807 | MVP_top60_DNA+RNA_seed | BRAF | 0.549939 | LUAD | NCI-H1651 | PLX-4720 |
| MVP_top60_DNA_seed | PIK3CA | 0.587686 | KIRC | OS-RC-2 | Alpelisib | MVP_top60_DNA+RNA_seed | BRAF | 0.550298 | LUAD | NCI-H1437 | PLX-4720 |
| MVP_top60_DNA_seed | PIK3CA | 0.590619 | KIRC | RCC-JW | GNE-317 | MVP_top60_DNA+RNA_seed | ATM | 0.55175 | LUAD | NCI-H441 | KU-60019 |
| MVP_top60_DNA_seed | MTOR | 0.59116 | KIRC | KMRC-20 | Temsirolimus | MVP_top60_DNA+RNA_seed | BRAF | 0.552013 | LUAD | ABC-1 | PLX-4720 |
| MVP_top60_DNA_seed | ATM | 0.59152 | KIRC | 786-0 | KU-55933 | MVP_top60_DNA+RNA_seed | MET | 0.553099 | LUAD | NCI-H3122 | AZD6094 |
| MVP_top60_DNA_seed | PIK3CA | 0.592076 | KIRC | CAKI-1 | Buparlisib | MVP_top60_DNA+RNA_seed | HSP90AA1 | 0.553419 | LUAD | PC-3_[JPC-3] | SNX-2112 |
| MVP_top60_DNA_seed | PIK3CD | 0.592076 | KIRC | CAKI-1 | Buparlisib | MVP_top60_DNA+RNA_seed | HDAC1 | 0.555278 | LUAD | A549 | Belinostat |
| MVP_top60_DNA_seed | HDAC1 | 0.595867 | KIRC | LB996-RCC | Romidepsin | MVP_top60_DNA+RNA_seed | AURKA | 0.555386 | LUAD | NCI-H2228 | Alisertib |
| MVP_top60_DNA_seed | HDAC1 | 0.596364 | KIRC | LB2241-RCC | Entinostat | MVP_top60_DNA+RNA_seed | EGFR | 0.555677 | LUAD | NCI-H2009 | PF-00299804 |
| MVP_top60_DNA_seed | PIK3CA | 0.59711 | KIRC | KMRC-1 | Buparlisib | MVP_top60_DNA+RNA_seed | IGF1R | 0.556509 | LUAD | NCI-H358 | GSK1904529A |
| MVP_top60_DNA_seed | PIK3CD | 0.59711 | KIRC | KMRC-1 | Buparlisib | MVP_top60_DNA+RNA_seed | EGFR | 0.556613 | LUAD | NCI-H1666 | Pelitinib |
| MVP_top60_DNA_seed | MTOR | 0.599933 | KIRC | A704 | Temsirolimus | MVP_top60_DNA+RNA_seed | HDAC1 | 0.556762 | LUAD | NCI-H1792 | Tubastatin A |
| MVP_top60_DNA_seed | PARP1 | 0.600654 | KIRC | RCC-AB | Niraparib | MVP_top60_DNA+RNA_seed | AKT1 | 0.556884 | LUAD | NCI-H650 | AKT inhibitor VIII |
| MVP_top60_DNA_seed | MTOR | 0.600924 | KIRC | NCC021 | Temsirolimus | MVP_top60_DNA+RNA_seed | SRC | 0.557319 | LUAD | VMRC-LCD | Bosutinib |
| MVP_top60_DNA_seed | HDAC1 | 0.603041 | KIRC | LB996-RCC | PCI-34051 | MVP_top60_DNA+RNA_seed | PTK2 | 0.558003 | LUAD | NCI-H1944 | PF-562271 |
| MVP_top60_DNA_seed | PARP1 | 0.6043 | KIRC | RXF393 | Talazoparib | MVP_top60_DNA+RNA_seed | BRAF | 0.55814 | LUAD | NCI-H322M | AZ628 |
| MVP_top60_DNA_seed | IGF1R | 0.609547 | KIRC | BB65-RCC | Linsitinib | MVP_top60_DNA+RNA_seed | IGF1R | 0.558765 | LUAD | Calu-3 | BMS-536924 |
| MVP_top60_DNA_seed | IGF1R | 0.609641 | KIRC | ACHN | BMS-754807 | MVP_top60_DNA+RNA_seed | AKT1 | 0.558951 | LUAD | H3255 | GSK690693 |
| MVP_top60_DNA_seed | IGF1R | 0.609994 | KIRC | CAL-54 | GSK1904529A | MVP_top60_DNA+RNA_seed | AURKA | 0.560652 | LUAD | NCI-H1792 | ZM447439 |
| MVP_top60_DNA_seed | SMARCA4 | 0.612045 | KIRC | U031 | PFI3 | MVP_top60_DNA+RNA_seed | HDAC1 | 0.562111 | LUAD | ABC-1 | Tubastatin A |
| MVP_top60_DNA_seed | PARP1 | 0.617799 | KIRC | KMRC-20 | Olaparib | MVP_top60_DNA+RNA_seed | BRAF | 0.56221 | LUAD | NCI-H1650 | RAF_9304 |

| | | | | | | | | | | | |
|---|---|---|---|---|---|---|---|---|---|---|---|
| MVP_top60_DNA_seed | PARP1 | 0.61926 | KIRC | HA7-RCC | Niraparib | MVP_top60_DNA+RNA_seed | HSP90AA1 | 0.562326 | LUAD | NCI-H1568 | Elesclomol |
| MVP_top60_DNA_seed | HSP90AA1 | 0.621203 | KIRC | NCC010 | Elesclomol | MVP_top60_DNA+RNA_seed | EGFR | 0.562436 | LUAD | NCI-H23 | Erlotinib |
| MVP_top60_DNA_seed | HSP90AB1 | 0.621203 | KIRC | NCC010 | Elesclomol | MVP_top60_DNA+RNA_seed | AKT1 | 0.562992 | LUAD | SK-LU-1 | MK-2206 |
| MVP_top60_DNA_seed | HDAC1 | 0.63027 | KIRC | LB2241-RCC | PCI-34051 | MVP_top60_DNA+RNA_seed | PIK3CA | 0.564716 | LUAD | NCI-H1650 | AZD8835 |
| MVP_top60_DNA_seed | IGF1R | 0.630389 | KIRC | 769-P | BMS-536924 | MVP_top60_DNA+RNA_seed | HSP90AA1 | 0.564978 | LUAD | NCI-H1793 | Elesclomol |
| MVP_top60_DNA_seed | HDAC1 | 0.63155 | KIRC | CAL-54 | PCI-34051 | MVP_top60_DNA+RNA_seed | HDAC1 | 0.565586 | LUAD | SW1573 | CAY10603 |
| MVP_top60_DNA_seed | PIK3CA | 0.631708 | KIRC | HA7-RCC | Buparlisib | MVP_top60_DNA+RNA_seed | AURKA | 0.568205 | LUAD | NCI-H2085 | Alisertib |
| MVP_top60_DNA_seed | PIK3CD | 0.631708 | KIRC | HA7-RCC | Buparlisib | MVP_top60_DNA+RNA_seed | MAPK14 | 0.569875 | LUAD | SW1573 | TAK-715 |
| MVP_top60_DNA_seed | TP53 | 0.632368 | KIRC | A704 | MIRA-1 | MVP_top60_DNA+RNA_seed | HSP90AA1 | 0.570675 | LUAD | HCC-827 | Tanespimycin |
| MVP_top60_DNA_seed | HDAC1 | 0.632386 | KIRC | HA7-RCC | Romidepsin | MVP_top60_DNA+RNA_seed | FGFR1 | 0.57072 | LUAD | NCI-H23 | PD173074 |
| MVP_top60_DNA_seed | ESR1 | 0.634383 | KIRC | BB65-RCC | Tamoxifen | MVP_top60_DNA+RNA_seed | MAPK14 | 0.570842 | LUAD | NCI-H2023 | TAK-715 |
| MVP_top60_DNA_seed | EP300 | 0.636231 | KIRC | A704 | CPI-637 | MVP_top60_DNA+RNA_seed | IGF1R | 0.572636 | LUAD | NCI-H1781 | BMS-536924 |
| MVP_top60_DNA_seed | HSP90AA1 | 0.636838 | KIRC | SN12C | Tanespimycin | MVP_top60_DNA+RNA_seed | AKT1 | 0.573545 | LUAD | NCI-H1666 | GSK690693 |
| MVP_top60_DNA_seed | HSP90AB1 | 0.636838 | KIRC | SN12C | Tanespimycin | MVP_top60_DNA+RNA_seed | FGFR1 | 0.573648 | LUAD | NCI-H1437 | PD173074 |
| MVP_top60_DNA_seed | IGF1R | 0.637966 | KIRC | CAKI-1 | GSK1904529A | MVP_top60_DNA+RNA_seed | SMARCA4 | 0.57405 | LUAD | SK-LU-1 | PFI-3 |
| MVP_top60_DNA_seed | AR | 0.638532 | KIRC | HA7-RCC | Bicalutamide | MVP_top60_DNA+RNA_seed | ATM | 0.574413 | LUAD | NCI-H1573 | CP466722 |
| MVP_top60_DNA_seed | HDAC1 | 0.646978 | KIRC | KMRC-20 | Romidepsin | MVP_top60_DNA+RNA_seed | SMARCA4 | 0.574504 | LUAD | NCI-H2087 | PFI-3 |
| MVP_top60_DNA_seed | TP53 | 0.653787 | KIRC | LB1047-RCC | MIRA-1 | MVP_top60_DNA+RNA_seed | EGFR | 0.574742 | LUAD | VMRC-LCD | Afatinib |
| MVP_top60_DNA_seed | ESR1 | 0.656359 | KIRC | 769-P | GDC0810 | MVP_top60_DNA+RNA_seed | HSP90AA1 | 0.576273 | LUAD | NCI-H23 | Luminespib |
| MVP_top60_DNA_seed | ATM | 0.663352 | KIRC | LB1047-RCC | KU-55933 | MVP_top60_DNA+RNA_seed | MDM2 | 0.58081 | LUAD | NCI-H3122 | Nutlin-3a (-) |
| MVP_top60_DNA_seed | AR | 0.664752 | KIRC | RCC-ER | Bicalutamide | MVP_top60_DNA+RNA_seed | PIK3CA | 0.580849 | LUAD | SK-LU-1 | PI3Ka_4409 |
| MVP_top60_DNA_seed | HDAC1 | 0.665242 | KIRC | A498 | Entinostat | MVP_top60_DNA+RNA_seed | AKT1 | 0.583114 | LUAD | A549 | AKT inhibitor VIII |
| MVP_top60_DNA_seed | ATM | 0.675747 | KIRC | A498 | KU-55933 | MVP_top60_DNA+RNA_seed | BRAF | 0.58334 | LUAD | NCI-H1944 | PLX-4720 |
| MVP_top60_DNA_seed | PARP1 | 0.678642 | KIRC | LB2241-RCC | Olaparib | MVP_top60_DNA+RNA_seed | MET | 0.583893 | LUAD | NCI-H23 | Crizotinib |
| MVP_top60_DNA_seed | TP53 | 0.679959 | KIRC | KMRC-1 | MIRA-1 | MVP_top60_DNA+RNA_seed | AURKA | 0.585151 | LUAD | NCI-H1975 | Alisertib |
| MVP_top60_DNA_seed | ESR1 | 0.682241 | KIRC | RCC-FG2 | GDC0810 | MVP_top60_DNA+RNA_seed | HSP90AA1 | 0.585585 | LUAD | Calu-3 | SNX-2112 |
| MVP_top60_DNA_seed | PARP1 | 0.68273 | KIRC | HA7-RCC | Olaparib | MVP_top60_DNA+RNA_seed | AURKA | 0.585869 | LUAD | NCI-H1563 | CD532 |
| MVP_top60_DNA_seed | PARP1 | 0.683074 | KIRC | A498 | Olaparib | MVP_top60_DNA+RNA_seed | BRAF | 0.58669 | LUAD | NCI-H1693 | SB590885 |
| MVP_top60_DNA_seed | PARP1 | 0.684281 | KIRC | HA7-RCC | Veliparib | MVP_top60_DNA+RNA_seed | FGFR1 | 0.586896 | LUAD | SK-LU-1 | FGFR_3831 |
| MVP_top60_DNA_seed | IGF1R | 0.693713 | KIRC | ACHN | NVP-ADW742 | MVP_top60_DNA+RNA_seed | EGFR | 0.588599 | LUAD | NCI-H2405 | Afatinib |
| MVP_top60_DNA_seed | ATM | 0.694802 | KIRC | 769-P | KU-55933 | MVP_top60_DNA+RNA_seed | HDAC1 | 0.588605 | LUAD | NCI-H2405 | Belinostat |
| MVP_top60_DNA_seed | SMARCA4 | 0.695265 | KIRC | LB2241-RCC | PFI3 | MVP_top60_DNA+RNA_seed | AURKA | 0.589176 | LUAD | HCC-44 | Alisertib |
| MVP_top60_DNA_seed | TP53 | 0.706908 | KIRC | SN12C | MIRA-1 | MVP_top60_DNA+RNA_seed | IGF1R | 0.589179 | LUAD | PC-3_[JPC-3] | GSK1904529A |
| MVP_top60_DNA_seed | ESR1 | 0.709393 | KIRC | A704 | GDC0810 | MVP_top60_DNA+RNA_seed | BRAF | 0.589493 | LUAD | NCI-H2087 | SB590885 |
| MVP_top60_DNA_seed | IGF1R | 0.711614 | KIRC | RCC-JW | NVP-ADW742 | MVP_top60_DNA+RNA_seed | SMARCA4 | 0.589571 | LUAD | NCI-H1755 | PFI-3 |
| MVP_top60_DNA_seed | HDAC1 | 0.717405 | KIRC | NCC021 | PCI-34051 | MVP_top60_DNA+RNA_seed | ATM | 0.592273 | LUAD | NCI-H1623 | KU-60019 |
| MVP_top60_DNA_seed | PARP1 | 0.720111 | KIRC | SW156 | Rucaparib | MVP_top60_DNA+RNA_seed | HDAC1 | 0.592667 | LUAD | A549 | Dacinostat |
| MVP_top60_DNA_seed | PIK3CA | 0.720841 | KIRC | RCC-AB | Alpelisib | MVP_top60_DNA+RNA_seed | FGFR1 | 0.593065 | LUAD | COR-L105 | FGFR_3831 |
| MVP_top60_DNA_seed | IGF1R | 0.721112 | KIRC | LB2241-RCC | NVP-ADW742 | MVP_top60_DNA+RNA_seed | HSP90AA1 | 0.593092 | LUAD | NCI-H1573 | SNX-2112 |
| MVP_top60_DNA_seed | PARP1 | 0.724143 | KIRC | LB1047-RCC | Niraparib | MVP_top60_DNA+RNA_seed | AURKA | 0.593119 | LUAD | NCI-H522 | Alisertib |
| MVP_top60_DNA_seed | HDAC1 | 0.726804 | KIRC | RCC-JW | Entinostat | MVP_top60_DNA+RNA_seed | EGFR | 0.593312 | LUAD | NCI-H441 | Gefitinib |

| | | | | | | | | | | | |
|---|---|---|---|---|---|---|---|---|---|---|---|
| MVP_top60_DNA_seed | HDAC1 | 0.728364 | KIRC | ACHN | Entinostat | MVP_top60_DNA+RNA_seed | MET | 0.596322 | LUAD | NCI-H2009 | JNJ38877605 |
| MVP_top60_DNA_seed | PIK3CA | 0.730589 | KIRC | OS-RC-2 | Buparlisib | MVP_top60_DNA+RNA_seed | MDM2 | 0.5969 | LUAD | NCI-H2030 | Serdemetan |
| MVP_top60_DNA_seed | PIK3CD | 0.730589 | KIRC | OS-RC-2 | Buparlisib | MVP_top60_DNA+RNA_seed | EGFR | 0.596901 | LUAD | NCI-H3122 | CI-1033 |
| MVP_top60_DNA_seed | HDAC1 | 0.737516 | KIRC | 769-P | Entinostat | MVP_top60_DNA+RNA_seed | FGFR1 | 0.596924 | LUAD | NCI-H1781 | AZD4547 |
| MVP_top60_DNA_seed | SMARCA4 | 0.738045 | KIRC | OS-RC-2 | PFI3 | MVP_top60_DNA+RNA_seed | MET | 0.597248 | LUAD | A549 | JNJ38877605 |
| MVP_top60_DNA_seed | PARP1 | 0.739728 | KIRC | RCC-JW | Talazoparib | MVP_top60_DNA+RNA_seed | HSP90AA1 | 0.598354 | LUAD | NCI-H1781 | CCT-018159 |
| MVP_top60_DNA_seed | ESR1 | 0.743112 | KIRC | LB996-RCC | GDC0810 | MVP_top60_DNA+RNA_seed | BRAF | 0.598898 | LUAD | NCI-H1838 | AZ628 |
| MVP_top60_DNA_seed | IGF1R | 0.743566 | KIRC | SW156 | BMS-536924 | MVP_top60_DNA+RNA_seed | AURKA | 0.59919 | LUAD | NCI-H2342 | Alisertib |
| MVP_top60_DNA_seed | PARP1 | 0.744849 | KIRC | LB2241-RCC | Veliparib | MVP_top60_DNA+RNA_seed | FGFR1 | 0.599343 | LUAD | RERF-LC-KJ | AZD4547 |
| MVP_top60_DNA_seed | EP300 | 0.745621 | KIRC | A498 | CPI-637 | MVP_top60_DNA+RNA_seed | EGFR | 0.599369 | LUAD | NCI-H1838 | PF-00299804 |
| MVP_top60_DNA_seed | ATM | 0.753981 | KIRC | KMRC-1 | KU-55933 | MVP_top60_DNA+RNA_seed | PIK3CA | 0.599552 | LUAD | NCI-H2405 | AZD8835 |
| MVP_top60_DNA_seed | HSP90AA1 | 0.754848 | KIRC | RCC-AB | Tanespimycin | MVP_top60_DNA+RNA_seed | MET | 0.599864 | LUAD | NCI-H1666 | JNJ38877605 |
| MVP_top60_DNA_seed | HSP90AB1 | 0.754848 | KIRC | RCC-AB | Tanespimycin | MVP_top60_DNA+RNA_seed | EGFR | 0.601051 | LUAD | NCI-H1838 | Cetuximab |
| MVP_top60_DNA_seed | HSP90AA1 | 0.756761 | KIRC | RCC-AB | Elesclomol | MVP_top60_DNA+RNA_seed | FGFR1 | 0.601507 | LUAD | NCI-H1435 | AZD4547 |
| MVP_top60_DNA_seed | HSP90AB1 | 0.756761 | KIRC | RCC-AB | Elesclomol | MVP_top60_DNA+RNA_seed | EGFR | 0.602371 | LUAD | NCI-H1435 | Afatinib |
| MVP_top60_DNA_seed | ESR1 | 0.758063 | KIRC | SN12C | GDC0810 | MVP_top60_DNA+RNA_seed | EGFR | 0.602643 | LUAD | NCI-H2228 | AZD8931 |
| MVP_top60_DNA_seed | SMARCA4 | 0.758393 | KIRC | ACHN | PFI3 | MVP_top60_DNA+RNA_seed | BRAF | 0.603421 | LUAD | NCI-H1944 | SB590885 |
| MVP_top60_DNA_seed | HSP90AA1 | 0.762159 | KIRC | U031 | Tanespimycin | MVP_top60_DNA+RNA_seed | BRAF | 0.603428 | LUAD | NCI-H2122 | HG6-64-1 |
| MVP_top60_DNA_seed | HSP90AB1 | 0.762159 | KIRC | U031 | Tanespimycin | MVP_top60_DNA+RNA_seed | BRAF | 0.603539 | LUAD | RERF-LC-KJ | SB590885 |
| MVP_top60_DNA_seed | IGF1R | 0.771331 | KIRC | RCC-AB | Linsitinib | MVP_top60_DNA+RNA_seed | AURKA | 0.604079 | LUAD | NCI-H358 | Alisertib |
| MVP_top60_DNA_seed | HDAC1 | 0.773843 | KIRC | BFTC-909 | PCI-34051 | MVP_top60_DNA+RNA_seed | IGF1R | 0.604555 | LUAD | NCI-H1355 | GSK1904529A |
| MVP_top60_DNA_seed | IGF1R | 0.776197 | KIRC | HA7-RCC | GSK1904529A | MVP_top60_DNA+RNA_seed | MDM2 | 0.605243 | LUAD | NCI-H322M | Nutlin-3a (-) |
| MVP_top60_DNA_seed | PARP1 | 0.781184 | KIRC | SW156 | Veliparib | MVP_top60_DNA+RNA_seed | AKT1 | 0.605685 | LUAD | H3255 | MK-2206 |
| MVP_top60_DNA_seed | PARP1 | 0.785062 | KIRC | KMRC-1 | Talazoparib | MVP_top60_DNA+RNA_seed | HSP90AA1 | 0.606102 | LUAD | NCI-H1755 | Luminespib |
| MVP_top60_DNA_seed | PARP1 | 0.795836 | KIRC | CAKI-1 | Veliparib | MVP_top60_DNA+RNA_seed | AURKA | 0.60615 | LUAD | NCI-H1781 | GSK1070916 |
| MVP_top60_DNA_seed | HSP90AA1 | 0.796496 | KIRC | TK10 | Tanespimycin | MVP_top60_DNA+RNA_seed | AKT1 | 0.606799 | LUAD | SW1573 | MK-2206 |
| MVP_top60_DNA_seed | HSP90AB1 | 0.796496 | KIRC | TK10 | Tanespimycin | MVP_top60_DNA+RNA_seed | IGF1R | 0.606968 | LUAD | NCI-H1650 | BMS-754807 |
| MVP_top60_DNA_seed | PARP1 | 0.797076 | KIRC | BB65-RCC | Rucaparib | MVP_top60_DNA+RNA_seed | AKT1 | 0.607677 | LUAD | NCI-H1755 | MK-2206 |
| MVP_top60_DNA_seed | TP53 | 0.80106 | KIRC | KMRC-20 | MIRA-1 | MVP_top60_DNA+RNA_seed | BRAF | 0.60836 | LUAD | NCI-H2023 | SB590885 |
| MVP_top60_DNA_seed | PARP1 | 0.803214 | KIRC | LB1047-RCC | Veliparib | MVP_top60_DNA+RNA_seed | HDAC1 | 0.60901 | LUAD | NCI-H1623 | Tubastatin A |
| MVP_top60_DNA_seed | SMARCA4 | 0.809915 | KIRC | RXF393 | PFI3 | MVP_top60_DNA+RNA_seed | HDAC1 | 0.609357 | LUAD | A549 | CAY10603 |
| MVP_top60_DNA_seed | ATM | 0.811201 | KIRC | CAKI-1 | KU-55933 | MVP_top60_DNA+RNA_seed | HDAC1 | 0.610381 | LUAD | NCI-H23 | Entinostat |
| MVP_top60_DNA_seed | ATM | 0.818165 | KIRC | RCC-AB | KU-55933 | MVP_top60_DNA+RNA_seed | HSP90AA1 | 0.611133 | LUAD | NCI-H1792 | Elesclomol |
| MVP_top60_DNA_seed | TP53 | 0.82094 | KIRC | LB996-RCC | MIRA-1 | MVP_top60_DNA+RNA_seed | SMARCA4 | 0.611424 | LUAD | NCI-H2342 | PFI-3 |
| MVP_top60_DNA_seed | PARP1 | 0.822516 | KIRC | BFTC-909 | Niraparib | MVP_top60_DNA+RNA_seed | AURKA | 0.61213 | LUAD | NCI-H1355 | Alisertib |
| MVP_top60_DNA_seed | SMARCA4 | 0.829859 | KIRC | A498 | PFI3 | MVP_top60_DNA+RNA_seed | MET | 0.61219 | LUAD | NCI-H2342 | AZD6094 |
| MVP_top60_DNA_seed | ATM | 0.830618 | KIRC | LB2241-RCC | KU-55933 | MVP_top60_DNA+RNA_seed | MET | 0.612939 | LUAD | NCI-H2347 | SU11274 |
| MVP_top60_DNA_seed | HSP90AA1 | 0.831608 | KIRC | SW156 | Tanespimycin | MVP_top60_DNA+RNA_seed | EGFR | 0.613348 | LUAD | NCI-H2122 | Cetuximab |
| MVP_top60_DNA_seed | HSP90AB1 | 0.831608 | KIRC | SW156 | Tanespimycin | MVP_top60_DNA+RNA_seed | HDAC1 | 0.613763 | LUAD | A549 | AR-42 |
| MVP_top60_DNA_seed | SMARCA4 | 0.833466 | KIRC | SW156 | PFI3 | MVP_top60_DNA+RNA_seed | EGFR | 0.614013 | LUAD | NCI-H2405 | PF-00299804 |
| MVP_top60_DNA_seed | IGF1R | 0.834671 | KIRC | RCC-JF | Linsitinib | MVP_top60_DNA+RNA_seed | IGF1R | 0.614311 | LUAD | Calu-6 | GSK1904529A |

| | | | | | | | | | | | |
|---|---|---|---|---|---|---|---|---|---|---|---|
| MVP_top60_DNA_seed | PARP1 | 0.837168 | KIRC | KMRC-1 | Rucaparib | MVP_top60_DNA+RNA_seed | FGFR1 | 0.618726 | LUAD | NCI-H2087 | AZD4547 |
| MVP_top60_DNA_seed | HDAC1 | 0.839066 | KIRC | RCC-AB | PCI-34051 | MVP_top60_DNA+RNA_seed | BRAF | 0.619248 | LUAD | NCI-H838 | PLX-4720 |
| MVP_top60_DNA_seed | IGF1R | 0.843463 | KIRC | RXF393 | NVP-ADW742 | MVP_top60_DNA+RNA_seed | SRC | 0.6218 | LUAD | NCI-H2122 | NVP-BHG712 |
| MVP_top60_DNA_seed | PARP1 | 0.847539 | KIRC | RCC-AB | Olaparib | MVP_top60_DNA+RNA_seed | ATM | 0.622181 | LUAD | NCI-H838 | CP466722 |
| MVP_top60_DNA_seed | IGF1R | 0.84891 | KIRC | NCC021 | Linsitinib | MVP_top60_DNA+RNA_seed | EGFR | 0.622511 | LUAD | NCI-H1755 | AST-1306 |
| MVP_top60_DNA_seed | EP300 | 0.850651 | KIRC | KMRC-20 | SGC-CBP30 | MVP_top60_DNA+RNA_seed | HDAC1 | 0.622564 | LUAD | NCI-H1563 | CAY10603 |
| MVP_top60_DNA_seed | ESR1 | 0.866638 | KIRC | OS-RC-2 | Tamoxifen | MVP_top60_DNA+RNA_seed | EGFR | 0.623784 | LUAD | NCI-H2009 | Gefitinib |
| MVP_top60_DNA_seed | PARP1 | 0.880042 | KIRC | NCC021 | Niraparib | MVP_top60_DNA+RNA_seed | FGFR1 | 0.624866 | LUAD | NCI-H441 | AZD4547 |
| MVP_top60_DNA_seed | HDAC1 | 0.883983 | KIRC | 786-0 | Entinostat | MVP_top60_DNA+RNA_seed | AURKA | 0.626357 | LUAD | NCI-H1944 | ZM447439 |
| MVP_top60_DNA_seed | IGF1R | 0.89416 | KIRC | KMRC-20 | Linsitinib | MVP_top60_DNA+RNA_seed | SRC | 0.626357 | LUAD | NCI-H1568 | NVP-BHG712 |
| MVP_top60_DNA_seed | AR | 0.895 | KIRC | BB65-RCC | Bicalutamide | MVP_top60_DNA+RNA_seed | EGFR | 0.627227 | LUAD | NCI-H2122 | Afatinib |
| MVP_top60_DNA_seed | IGF1R | 0.895887 | KIRC | SN12C | Linsitinib | MVP_top60_DNA+RNA_seed | BRAF | 0.628586 | LUAD | HOP-62 | PLX-4720 |
| MVP_top60_DNA_seed | PARP1 | 0.899811 | KIRC | NCC010 | Olaparib | MVP_top60_DNA+RNA_seed | EGFR | 0.628603 | LUAD | NCI-H1573 | Afatinib |
| MVP_top60_DNA_seed | PIK3CA | 0.9 | KIRC | BB65-RCC | Buparlisib | MVP_top60_DNA+RNA_seed | AURKA | 0.628976 | LUAD | NCI-H1435 | Alisertib |
| MVP_top60_DNA_seed | PIK3CD | 0.9 | KIRC | BB65-RCC | Buparlisib | MVP_top60_DNA+RNA_seed | MET | 0.629785 | LUAD | NCI-H1975 | Crizotinib |
| MVP_top60_DNA_seed | ESR1 | 0.901768 | KIRC | LB2241-RCC | GDC0810 | MVP_top60_DNA+RNA_seed | EGFR | 0.630638 | LUAD | RERF-LC-KJ | Afatinib |
| MVP_top60_DNA_seed | IGF1R | 0.902805 | KIRC | LB2241-RCC | GSK1904529A | MVP_top60_DNA+RNA_seed | HDAC1 | 0.631434 | LUAD | NCI-H2023 | Dacinostat |
| MVP_top60_DNA_seed | PARP1 | 0.904111 | KIRC | LB996-RCC | Veliparib | MVP_top60_DNA+RNA_seed | HDAC1 | 0.631453 | LUAD | NCI-H2023 | AR-42 |
| MVP_top60_DNA_seed | IGF1R | 0.90767 | KIRC | OS-RC-2 | NVP-ADW742 | MVP_top60_DNA+RNA_seed | HDAC1 | 0.631516 | LUAD | LXF-289 | Tubastatin A |
| MVP_top60_DNA_seed | PARP1 | 0.913282 | KIRC | RCC-JW | Rucaparib | MVP_top60_DNA+RNA_seed | AKT1 | 0.632082 | LUAD | NCI-H1650 | A-443654 |
| MVP_top60_DNA_seed | SMARCA4 | 0.914798 | KIRC | NCC021 | PFI3 | MVP_top60_DNA+RNA_seed | HDAC1 | 0.632484 | LUAD | NCI-H1563 | Belinostat |
| MVP_top60_DNA_seed | HDAC1 | 0.916293 | KIRC | SW156 | PCI-34051 | MVP_top60_DNA+RNA_seed | SRC | 0.632823 | LUAD | NCI-H1975 | NVP-BHG712 |
| MVP_top60_DNA_seed | ESR1 | 0.922205 | KIRC | RCC10RGB | GDC0810 | MVP_top60_DNA+RNA_seed | SMARCA4 | 0.633025 | LUAD | HCC-827 | PFI-3 |
| MVP_top60_DNA_seed | HSP90AA1 | 0.924473 | KIRC | SW156 | CCT-018159 | MVP_top60_DNA+RNA_seed | ATM | 0.634208 | LUAD | NCI-H1651 | CP466722 |
| MVP_top60_DNA_seed | HSP90AB1 | 0.924473 | KIRC | SW156 | CCT-018159 | MVP_top60_DNA+RNA_seed | HDAC1 | 0.634493 | LUAD | NCI-H1781 | Belinostat |
| MVP_top60_DNA_seed | TP53 | 0.925183 | KIRC | RCC-AB | MIRA-1 | MVP_top60_DNA+RNA_seed | PIK3CA | 0.635882 | LUAD | PC-3_[JPC-3] | AZD8835 |
| MVP_top60_DNA_seed | HSP90AA1 | 0.932291 | KIRC | CAKI-1 | Tanespimycin | MVP_top60_DNA+RNA_seed | HSP90AA1 | 0.637913 | LUAD | EMC-BAC-2 | CCT-018159 |
| MVP_top60_DNA_seed | HSP90AB1 | 0.932291 | KIRC | CAKI-1 | Tanespimycin | MVP_top60_DNA+RNA_seed | PIK3CA | 0.638906 | LUAD | NCI-H522 | PI3Ka_4409 |
| MVP_top60_DNA_seed | ATM | 0.933906 | KIRC | OS-RC-2 | KU-55933 | MVP_top60_DNA+RNA_seed | SRC | 0.641287 | LUAD | NCI-H522 | NVP-BHG712 |
| MVP_top60_DNA_seed | HDAC1 | 0.937418 | KIRC | ACHN | Romidepsin | MVP_top60_DNA+RNA_seed | HSP90AA1 | 0.642153 | LUAD | EKVX | Tanespimycin |
| MVP_top60_DNA_seed | TP53 | 0.939831 | KIRC | VMRC-RCW | MIRA-1 | MVP_top60_DNA+RNA_seed | BRAF | 0.642597 | LUAD | NCI-H1781 | SB590885 |
| MVP_top60_DNA_seed | PARP1 | 0.941116 | KIRC | BB65-RCC | Niraparib | MVP_top60_DNA+RNA_seed | BRAF | 0.642958 | LUAD | HCC-78 | RAF_9304 |
| MVP_top60_DNA_seed | HSP90AA1 | 0.942826 | KIRC | RCC-AB | CCT-018159 | MVP_top60_DNA+RNA_seed | HSP90AA1 | 0.643066 | LUAD | NCI-H2122 | Elesclomol |
| MVP_top60_DNA_seed | HSP90AB1 | 0.942826 | KIRC | RCC-AB | CCT-018159 | MVP_top60_DNA+RNA_seed | MET | 0.643817 | LUAD | NCI-H1648 | PHA-665752 |
| MVP_top60_DNA_seed | ESR1 | 0.945693 | KIRC | NCC021 | GDC0810 | MVP_top60_DNA+RNA_seed | BRAF | 0.646688 | LUAD | VMRC-LCD | RAF_9304 |
| MVP_top60_DNA_seed | PARP1 | 0.949428 | KIRC | NCC021 | Veliparib | MVP_top60_DNA+RNA_seed | MDM2 | 0.646972 | LUAD | HCC-78 | Nutlin-3a (-) |
| MVP_top60_DNA_seed | TP53 | 0.954891 | KIRC | A498 | MIRA-1 | MVP_top60_DNA+RNA_seed | ATM | 0.64835 | LUAD | PC-3_[JPC-3] | KU-55933 |
| MVP_top60_DNA_seed | HDAC1 | 0.955875 | KIRC | VMRC-RCZ | Romidepsin | MVP_top60_DNA+RNA_seed | MET | 0.648408 | LUAD | NCI-H2009 | SU11274 |
| MVP_top60_DNA_seed | SMARCA4 | 0.961015 | KIRC | LB1047-RCC | PFI3 | MVP_top60_DNA+RNA_seed | BRAF | 0.64944 | LUAD | NCI-H3122 | PLX-4720 |
| MVP_top60_DNA_seed | HDAC1 | 0.961034 | KIRC | CAL-54 | Entinostat | MVP_top60_DNA+RNA_seed | BRAF | 0.649854 | LUAD | HOP-62 | Dabrafenib |
| MVP_top60_DNA_seed | AR | 0.966853 | KIRC | RCC-MF | Bicalutamide | MVP_top60_DNA+RNA_seed | MDM2 | 0.651283 | LUAD | NCI-H1693 | Nutlin-3a (-) |

| | | | | | | | | | | | |
|---|---|---|---|---|---|---|---|---|---|---|---|
| MVP_top60_DNA_seed | IGF1R | 0.968792 | KIRC | 786-0 | GSK1904529A | MVP_top60_DNA+RNA_seed | MDM2 | 0.651827 | LUAD | NCI-H2228 | Nutlin-3a (-) |
| MVP_top60_DNA_seed | EP300 | 0.969571 | KIRC | VMRC-RCZ | SGC-CBP30 | MVP_top60_DNA+RNA_seed | FGFR1 | 0.652426 | LUAD | NCI-H3122 | AZD4547 |
| MVP_top60_DNA_seed | ESR1 | 0.972005 | KIRC | HA7-RCC | GDC0810 | MVP_top60_DNA+RNA_seed | FGFR1 | 0.652925 | LUAD | H3255 | PD173074 |
| MVP_top60_DNA_seed | ATM | 0.972123 | KIRC | SN12C | KU-55933 | MVP_top60_DNA+RNA_seed | AURKA | 0.653122 | LUAD | NCI-H1650 | GSK1070916 |
| MVP_top60_DNA_seed | PARP1 | 0.978547 | KIRC | ACHN | Niraparib | MVP_top60_DNA+RNA_seed | AURKA | 0.654931 | LUAD | Calu-3 | ZM447439 |
| MVP_top60_DNA_seed | IGF1R | 0.979498 | KIRC | LB1047-RCC | Linsitinib | MVP_top60_DNA+RNA_seed | FGFR1 | 0.656013 | LUAD | HOP-62 | PD173074 |
| MVP_top60_DNA_seed | PARP1 | 0.980198 | KIRC | KMRC-20 | Niraparib | MVP_top60_DNA+RNA_seed | EGFR | 0.65633 | LUAD | NCI-H2087 | Afatinib |
| MVP_top60_DNA_seed | PARP1 | 0.981686 | KIRC | OS-RC-2 | Talazoparib | MVP_top60_DNA+RNA_seed | MDM2 | 0.656779 | LUAD | NCI-H838 | Serdemetan |
| MVP_top60_DNA_seed | PARP1 | 0.986762 | KIRC | A498 | Veliparib | MVP_top60_DNA+RNA_seed | FGFR1 | 0.656869 | LUAD | HCC-827 | FGFR_3831 |
| MVP_top60_DNA_seed | PARP1 | 0.987497 | KIRC | CAKI-1 | Rucaparib | MVP_top60_DNA+RNA_seed | EGFR | 0.657157 | LUAD | NCI-H1563 | AZD8931 |
| MVP_top60_DNA_seed | HSP90AA1 | 0.997302 | KIRC | TK10 | Luminespib | MVP_top60_DNA+RNA_seed | HDAC1 | 0.659073 | LUAD | SW1573 | AR-42 |
| MVP_top60_DNA_seed | HSP90AB1 | 0.997302 | KIRC | TK10 | Luminespib | MVP_top60_DNA+RNA_seed | EGFR | 0.659271 | LUAD | NCI-H1651 | Afatinib |
| MVP_top60_DNA_seed | HSP90AA1 | 0.998496 | KIRC | LB1047-RCC | Elesclomol | MVP_top60_DNA+RNA_seed | IGF1R | 0.659805 | LUAD | NCI-H1993 | BMS-754807 |
| MVP_top60_DNA_seed | HSP90AB1 | 0.998496 | KIRC | LB1047-RCC | Elesclomol | MVP_top60_DNA+RNA_seed | HDAC1 | 0.660538 | LUAD | NCI-H1650 | AR-42 |
| MVP_top60_DNA_seed | EP300 | 0.999916 | KIRC | VMRC-RCZ | CPI-637 | MVP_top60_DNA+RNA_seed | EGFR | 0.660831 | LUAD | SW1573 | AZD8931 |
| MVP_top60_DNA_seed | IGF1R | 1.000174 | KIRC | RCC-JW | GSK1904529A | MVP_top60_DNA+RNA_seed | ATM | 0.661222 | LUAD | SW1573 | CP466722 |
| MVP_top60_DNA_seed | IGF1R | 1.00236 | KIRC | OS-RC-2 | BMS-536924 | MVP_top60_DNA+RNA_seed | SRC | 0.661905 | LUAD | RERF-LC-MS | Bosutinib |
| MVP_top60_DNA_seed | IGF1R | 1.005405 | KIRC | CAKI-1 | NVP-ADW742 | MVP_top60_DNA+RNA_seed | AKT1 | 0.661918 | LUAD | HOP-62 | GSK690693 |
| MVP_top60_DNA_seed | IGF1R | 1.008275 | KIRC | VMRC-RCW | GSK1904529A | MVP_top60_DNA+RNA_seed | AKT1 | 0.663865 | LUAD | SW1573 | GSK690693 |
| MVP_top60_DNA_seed | PARP1 | 1.008975 | KIRC | OS-RC-2 | Veliparib | MVP_top60_DNA+RNA_seed | SMARCA4 | 0.664452 | LUAD | NCI-H522 | PFI-3 |
| MVP_top60_DNA_seed | PARP1 | 1.011598 | KIRC | LB1047-RCC | Olaparib | MVP_top60_DNA+RNA_seed | HDAC1 | 0.66605 | LUAD | NCI-H1650 | CAY10603 |
| MVP_top60_DNA_seed | ESR1 | 1.011666 | KIRC | RCC-MF | Tamoxifen | MVP_top60_DNA+RNA_seed | BRAF | 0.666655 | LUAD | NCI-H2342 | RAF_9304 |
| MVP_top60_DNA_seed | IGF1R | 1.020508 | KIRC | NCC010 | Linsitinib | MVP_top60_DNA+RNA_seed | MET | 0.666677 | LUAD | NCI-H1650 | Crizotinib |
| MVP_top60_DNA_seed | PIK3CA | 1.021881 | KIRC | BB65-RCC | Alpelisib | MVP_top60_DNA+RNA_seed | MDM2 | 0.666825 | LUAD | NCI-H1755 | Nutlin-3a (-) |
| MVP_top60_DNA_seed | PARP1 | 1.027239 | KIRC | BB65-RCC | Talazoparib | MVP_top60_DNA+RNA_seed | MDM2 | 0.666945 | LUAD | HOP-62 | Nutlin-3a (-) |
| MVP_top60_DNA_seed | PARP1 | 1.028791 | KIRC | RCC-MF | Olaparib | MVP_top60_DNA+RNA_seed | BRAF | 0.667395 | LUAD | NCI-H1651 | SB590885 |
| MVP_top60_DNA_seed | PARP1 | 1.039437 | KIRC | RCC10RGB | Talazoparib | MVP_top60_DNA+RNA_seed | MET | 0.667545 | LUAD | NCI-H2228 | AZD6094 |
| MVP_top60_DNA_seed | ESR1 | 1.059703 | KIRC | HA7-RCC | Tamoxifen | MVP_top60_DNA+RNA_seed | PIK3CA | 0.667812 | LUAD | Calu-6 | AZD8835 |
| MVP_top60_DNA_seed | MTOR | 1.063152 | KIRC | BB65-RCC | Temsirolimus | MVP_top60_DNA+RNA_seed | FGFR1 | 0.667924 | LUAD | Calu-6 | AZD4547 |
| MVP_top60_DNA_seed | HSP90AA1 | 1.069167 | KIRC | A704 | Tanespimycin | MVP_top60_DNA+RNA_seed | EGFR | 0.668149 | LUAD | NCI-H2342 | Afatinib |
| MVP_top60_DNA_seed | HSP90AB1 | 1.069167 | KIRC | A704 | Tanespimycin | MVP_top60_DNA+RNA_seed | IGF1R | 0.668197 | LUAD | NCI-H1435 | GSK1904529A |
| MVP_top60_DNA_seed | PIK3CA | 1.072953 | KIRC | KMRC-20 | Alpelisib | MVP_top60_DNA+RNA_seed | BRAF | 0.669367 | LUAD | NCI-H1355 | SB590885 |
| MVP_top60_DNA_seed | IGF1R | 1.076541 | KIRC | 769-P | GSK1904529A | MVP_top60_DNA+RNA_seed | MET | 0.669617 | LUAD | NCI-H2347 | AZD6094 |
| MVP_top60_DNA_seed | MTOR | 1.097443 | KIRC | HA7-RCC | Temsirolimus | MVP_top60_DNA+RNA_seed | BRAF | 0.670816 | LUAD | NCI-H1781 | Dabrafenib |
| MVP_top60_DNA_seed | ESR1 | 1.101727 | KIRC | NCC010 | Tamoxifen | MVP_top60_DNA+RNA_seed | SRC | 0.670843 | LUAD | HCC-827 | NVP-BHG712 |
| MVP_top60_DNA_seed | ESR1 | 1.10184 | KIRC | VMRC-RCZ | Tamoxifen | MVP_top60_DNA+RNA_seed | SMARCA4 | 0.670976 | LUAD | NCI-H2009 | PFI-3 |
| MVP_top60_DNA_seed | PIK3CA | 1.104382 | KIRC | VMRC-RCW | Alpelisib | MVP_top60_DNA+RNA_seed | EGFR | 0.670999 | LUAD | HCC-44 | AST-1306 |
| MVP_top60_DNA_seed | HSP90AA1 | 1.108952 | KIRC | HA7-RCC | Elesclomol | MVP_top60_DNA+RNA_seed | MAPK14 | 0.671119 | LUAD | NCI-H1993 | TAK-715 |
| MVP_top60_DNA_seed | HSP90AB1 | 1.108952 | KIRC | HA7-RCC | Elesclomol | MVP_top60_DNA+RNA_seed | MET | 0.671808 | LUAD | HCC-827 | SU11274 |
| MVP_top60_DNA_seed | ESR1 | 1.11942 | KIRC | LB1047-RCC | GDC0810 | MVP_top60_DNA+RNA_seed | BRAF | 0.671911 | LUAD | A549 | SB590885 |
| MVP_top60_DNA_seed | PIK3CA | 1.121707 | KIRC | BB65-RCC | GNE-317 | MVP_top60_DNA+RNA_seed | AKT1 | 0.672203 | LUAD | NCI-H522 | GSK690693 |

| | | | | | | | | | | | |
|---|---|---|---|---|---|---|---|---|---|---|---|
| MVP_top60_DNA_seed | AR | 1.127689 | KIRC | RCC-AB | Bicalutamide | MVP_top60_DNA+RNA_seed | HSP90AA1 | 0.672529 | LUAD | NCI-H2228 | SNX-2112 |
| MVP_top60_DNA_seed | ESR1 | 1.129094 | KIRC | LB2241-RCC | Tamoxifen | MVP_top60_DNA+RNA_seed | BRAF | 0.672727 | LUAD | NCI-H1666 | PLX-4720 |
| MVP_top60_DNA_seed | PIK3CA | 1.131475 | KIRC | RCC10RGB | Buparlisib | MVP_top60_DNA+RNA_seed | ATM | 0.673168 | LUAD | NCI-H2087 | torin2 |
| MVP_top60_DNA_seed | PIK3CD | 1.131475 | KIRC | RCC10RGB | Buparlisib | MVP_top60_DNA+RNA_seed | AURKA | 0.673563 | LUAD | NCI-H1651 | Genentech Cpd 10 |
| MVP_top60_DNA_seed | IGF1R | 1.133849 | KIRC | RCC-FG2 | GSK1904529A | MVP_top60_DNA+RNA_seed | AURKA | 0.67436 | LUAD | H3255 | Alisertib |
| MVP_top60_DNA_seed | TP53 | 1.134189 | KIRC | RCC10RGB | MIRA-1 | MVP_top60_DNA+RNA_seed | SRC | 0.674369 | LUAD | NCI-H1437 | NVP-BHG712 |
| MVP_top60_DNA_seed | PARP1 | 1.143842 | KIRC | OS-RC-2 | Rucaparib | MVP_top60_DNA+RNA_seed | BRAF | 0.674476 | LUAD | NCI-H1563 | Dabrafenib |
| MVP_top60_DNA_seed | PARP1 | 1.161023 | KIRC | RCC-AB | Veliparib | MVP_top60_DNA+RNA_seed | ATM | 0.675133 | LUAD | NCI-H1651 | KU-55933 |
| MVP_top60_DNA_seed | SMARCA4 | 1.168113 | KIRC | HA7-RCC | PFI3 | MVP_top60_DNA+RNA_seed | FGFR1 | 0.67572 | LUAD | NCI-H2087 | PD173074 |
| MVP_top60_DNA_seed | PIK3CA | 1.168539 | KIRC | ACHN | Alpelisib | MVP_top60_DNA+RNA_seed | AURKA | 0.676929 | LUAD | NCI-H1734 | CD532 |
| MVP_top60_DNA_seed | HDAC1 | 1.169682 | KIRC | HA7-RCC | PCI-34051 | MVP_top60_DNA+RNA_seed | BRAF | 0.678327 | LUAD | NCI-H1944 | Dabrafenib |
| MVP_top60_DNA_seed | SMARCA4 | 1.170472 | KIRC | RCC-AB | PFI3 | MVP_top60_DNA+RNA_seed | HDAC1 | 0.678498 | LUAD | SK-LU-1 | Dacinostat |
| MVP_top60_DNA_seed | HDAC1 | 1.175065 | KIRC | RCC-JW | PCI-34051 | MVP_top60_DNA+RNA_seed | EGFR | 0.678503 | LUAD | Calu-6 | Afatinib |
| MVP_top60_DNA_seed | HDAC1 | 1.175395 | KIRC | A704 | Romidepsin | MVP_top60_DNA+RNA_seed | EGFR | 0.678698 | LUAD | HCC-827 | Cetuximab |
| MVP_top60_DNA_seed | IGF1R | 1.175795 | KIRC | OS-RC-2 | BMS-754807 | MVP_top60_DNA+RNA_seed | FGFR1 | 0.679606 | LUAD | EMC-BAC-1 | AZD4547 |
| MVP_top60_DNA_seed | PARP1 | 1.176243 | KIRC | SN12C | Olaparib | MVP_top60_DNA+RNA_seed | AKT1 | 0.680197 | LUAD | NCI-H2405 | AKT inhibitor VIII |
| MVP_top60_DNA_seed | IGF1R | 1.183257 | KIRC | SN12C | NVP-ADW742 | MVP_top60_DNA+RNA_seed | HDAC1 | 0.6806 | LUAD | NCI-H2009 | Tubastatin A |
| MVP_top60_DNA_seed | PIK3CA | 1.195073 | KIRC | A498 | Alpelisib | MVP_top60_DNA+RNA_seed | BRAF | 0.680907 | LUAD | NCI-H2347 | HG6-64-1 |
| MVP_top60_DNA_seed | IGF1R | 1.196698 | KIRC | LB2241-RCC | Linsitinib | MVP_top60_DNA+RNA_seed | MAPK14 | 0.68117 | LUAD | RERF-LC-MS | TAK-715 |
| MVP_top60_DNA_seed | PIK3CA | 1.218478 | KIRC | SW156 | GNE-317 | MVP_top60_DNA+RNA_seed | AKT1 | 0.683103 | LUAD | NCI-H1793 | BAY AKT1 |
| MVP_top60_DNA_seed | IGF1R | 1.225311 | KIRC | CAKI-1 | Linsitinib | MVP_top60_DNA+RNA_seed | FGFR1 | 0.683585 | LUAD | NCI-H2085 | AZD4547 |
| MVP_top60_DNA_seed | IGF1R | 1.226438 | KIRC | SN12C | BMS-754807 | MVP_top60_DNA+RNA_seed | EGFR | 0.685148 | LUAD | NCI-H1623 | Pelitinib |
| MVP_top60_DNA_seed | IGF1R | 1.22687 | KIRC | RCC-AB | BMS-754807 | MVP_top60_DNA+RNA_seed | AKT1 | 0.685477 | LUAD | NCI-H838 | GSK690693 |
| MVP_top60_DNA_seed | PARP1 | 1.22898 | KIRC | HA7-RCC | Rucaparib | MVP_top60_DNA+RNA_seed | MDM2 | 0.685932 | LUAD | NCI-H441 | Serdemetan |
| MVP_top60_DNA_seed | HSP90AA1 | 1.234003 | KIRC | LB2241-RCC | CCT-018159 | MVP_top60_DNA+RNA_seed | MDM2 | 0.686345 | LUAD | NCI-H1623 | Serdemetan |
| MVP_top60_DNA_seed | HSP90AB1 | 1.234003 | KIRC | LB2241-RCC | CCT-018159 | MVP_top60_DNA+RNA_seed | AURKA | 0.686697 | LUAD | NCI-H441 | HG-5-113-01 |
| MVP_top60_DNA_seed | ESR1 | 1.23426 | KIRC | U031 | Tamoxifen | MVP_top60_DNA+RNA_seed | AURKA | 0.686936 | LUAD | HCC-78 | CD532 |
| MVP_top60_DNA_seed | HSP90AA1 | 1.234402 | KIRC | RCC-MF | Tanespimycin | MVP_top60_DNA+RNA_seed | IGF1R | 0.687022 | LUAD | RERF-LC-KJ | BMS-536924 |
| MVP_top60_DNA_seed | HSP90AB1 | 1.234402 | KIRC | RCC-MF | Tanespimycin | MVP_top60_DNA+RNA_seed | HSP90AA1 | 0.687939 | LUAD | NCI-H2291 | CCT-018159 |
| MVP_top60_DNA_seed | HSP90AA1 | 1.236386 | KIRC | VMRC-RCW | CCT-018159 | MVP_top60_DNA+RNA_seed | BRAF | 0.688028 | LUAD | NCI-H1648 | HG6-64-1 |
| MVP_top60_DNA_seed | HSP90AB1 | 1.236386 | KIRC | VMRC-RCW | CCT-018159 | MVP_top60_DNA+RNA_seed | AURKA | 0.688924 | LUAD | NCI-H2009 | Alisertib |
| MVP_top60_DNA_seed | IGF1R | 1.237731 | KIRC | RCC-AB | BMS-536924 | MVP_top60_DNA+RNA_seed | EGFR | 0.688986 | LUAD | SK-LU-1 | CI-1033 |
| MVP_top60_DNA_seed | PARP1 | 1.238046 | KIRC | VMRC-RCW | Rucaparib | MVP_top60_DNA+RNA_seed | MAPK14 | 0.689158 | LUAD | A549 | TAK-715 |
| MVP_top60_DNA_seed | PIK3CA | 1.239671 | KIRC | RCC10RGB | Alpelisib | MVP_top60_DNA+RNA_seed | BRAF | 0.690136 | LUAD | NCI-H1648 | PLX-4720 |
| MVP_top60_DNA_seed | ATM | 1.247355 | KIRC | VMRC-RCZ | KU-55933 | MVP_top60_DNA+RNA_seed | BRAF | 0.690742 | LUAD | NCI-H3122 | PLX-4720 |
| MVP_top60_DNA_seed | PIK3CA | 1.24889 | KIRC | 786-0 | Alpelisib | MVP_top60_DNA+RNA_seed | EGFR | 0.691048 | LUAD | HOP-62 | Afatinib |
| MVP_top60_DNA_seed | HDAC1 | 1.251491 | KIRC | VMRC-RCW | PCI-34051 | MVP_top60_DNA+RNA_seed | BRAF | 0.692125 | LUAD | NCI-H1437 | SB590885 |
| MVP_top60_DNA_seed | AR | 1.252801 | KIRC | VMRC-RCW | Bicalutamide | MVP_top60_DNA+RNA_seed | PIK3CA | 0.692409 | LUAD | COR-L105 | PI3Ka_4409 |
| MVP_top60_DNA_seed | IGF1R | 1.260442 | KIRC | RCC-JW | Linsitinib | MVP_top60_DNA+RNA_seed | HDAC1 | 0.693036 | LUAD | NCI-H322M | Tubastatin A |
| MVP_top60_DNA_seed | EP300 | 1.269614 | KIRC | CAL-54 | SGC-CBP30 | MVP_top60_DNA+RNA_seed | HSP90AA1 | 0.693214 | LUAD | NCI-H838 | Luminespib |
| MVP_top60_DNA_seed | MTOR | 1.274526 | KIRC | LB2241-RCC | Temsirolimus | MVP_top60_DNA+RNA_seed | FGFR1 | 0.693248 | LUAD | NCI-H522 | PD173074 |

| | | | | | | | | | | | |
|---|---|---|---|---|---|---|---|---|---|---|---|
| MVP_top60_DNA_seed | HSP90AA1 | 1.277113 | KIRC | A498 | Luminespib | MVP_top60_DNA+RNA_seed | HSP90AA1 | 0.693431 | LUAD | NCI-H322M | SNX-2112 |
| MVP_top60_DNA_seed | HSP90AB1 | 1.277113 | KIRC | A498 | Luminespib | MVP_top60_DNA+RNA_seed | ATM | 0.693698 | LUAD | NCI-H2405 | KU-60019 |
| MVP_top60_DNA_seed | HSP90AA1 | 1.277347 | KIRC | U031 | Elesclomol | MVP_top60_DNA+RNA_seed | AURKA | 0.693752 | LUAD | NCI-H3122 | CD532 |
| MVP_top60_DNA_seed | HSP90AB1 | 1.277347 | KIRC | U031 | Elesclomol | MVP_top60_DNA+RNA_seed | SRC | 0.695593 | LUAD | LC-2-ad | Bosutinib |
| MVP_top60_DNA_seed | PARP1 | 1.280049 | KIRC | RCC-JW | Olaparib | MVP_top60_DNA+RNA_seed | AKT1 | 0.696202 | LUAD | NCI-H1993 | GSK690693 |
| MVP_top60_DNA_seed | HDAC1 | 1.281265 | KIRC | SN12C | PCI-34051 | MVP_top60_DNA+RNA_seed | MDM2 | 0.69665 | LUAD | NCI-H2342 | Nutlin-3a (-) |
| MVP_top60_DNA_seed | EP300 | 1.282249 | KIRC | OS-RC-2 | CPI-637 | MVP_top60_DNA+RNA_seed | IGF1R | 0.698289 | LUAD | HOP-62 | GSK1904529A |
| MVP_top60_DNA_seed | IGF1R | 1.283073 | KIRC | VMRC-RCW | BMS-754807 | MVP_top60_DNA+RNA_seed | MDM2 | 0.699467 | LUAD | Calu-3 | Nutlin-3a (-) |
| MVP_top60_DNA_seed | PARP1 | 1.289987 | KIRC | OS-RC-2 | Olaparib | MVP_top60_DNA+RNA_seed | HSP90AA1 | 0.699663 | LUAD | EMC-BAC-1 | CCT-018159 |
| MVP_top60_DNA_seed | MTOR | 1.297446 | KIRC | SN12C | Temsirolimus | MVP_top60_DNA+RNA_seed | ATM | 0.702816 | LUAD | NCI-H1650 | CP466722 |
| MVP_top60_DNA_seed | PARP1 | 1.301085 | KIRC | RXF393 | Olaparib | MVP_top60_DNA+RNA_seed | BRAF | 0.702819 | LUAD | NCI-H1435 | Dabrafenib |
| MVP_top60_DNA_seed | PARP1 | 1.303197 | KIRC | VMRC-RCW | Niraparib | MVP_top60_DNA+RNA_seed | FGFR1 | 0.703768 | LUAD | NCI-H2342 | AZD4547 |
| MVP_top60_DNA_seed | IGF1R | 1.310829 | KIRC | LB1047-RCC | NVP-ADW742 | MVP_top60_DNA+RNA_seed | MET | 0.705187 | LUAD | NCI-H838 | AZD6094 |
| MVP_top60_DNA_seed | IGF1R | 1.318535 | KIRC | KMRC-1 | NVP-ADW742 | MVP_top60_DNA+RNA_seed | AKT1 | 0.705609 | LUAD | NCI-H2405 | MK-2206 |
| MVP_top60_DNA_seed | HDAC1 | 1.318589 | KIRC | VMRC-RCW | Entinostat | MVP_top60_DNA+RNA_seed | EGFR | 0.706322 | LUAD | NCI-H2405 | AZD8931 |
| MVP_top60_DNA_seed | PIK3CA | 1.320015 | KIRC | KMRC-1 | Alpelisib | MVP_top60_DNA+RNA_seed | ATM | 0.70647 | LUAD | NCI-H2085 | KU-60019 |
| MVP_top60_DNA_seed | IGF1R | 1.327066 | KIRC | SW156 | Linsitinib | MVP_top60_DNA+RNA_seed | PIK3CA | 0.706685 | LUAD | H3255 | PI3Ka_4409 |
| MVP_top60_DNA_seed | ESR1 | 1.333047 | KIRC | RCC-AB | GDC0810 | MVP_top60_DNA+RNA_seed | HDAC1 | 0.707176 | LUAD | NCI-H1650 | Tubastatin A |
| MVP_top60_DNA_seed | IGF1R | 1.336044 | KIRC | TK10 | BMS-536924 | MVP_top60_DNA+RNA_seed | EGFR | 0.709277 | LUAD | NCI-H2228 | PF-00299804 |
| MVP_top60_DNA_seed | IGF1R | 1.338909 | KIRC | RCC-FG2 | Linsitinib | MVP_top60_DNA+RNA_seed | BRAF | 0.709432 | LUAD | SK-LU-1 | RAF_9304 |
| MVP_top60_DNA_seed | AR | 1.34485 | KIRC | NCC021 | Bicalutamide | MVP_top60_DNA+RNA_seed | BRAF | 0.709674 | LUAD | NCI-H1437 | HG6-64-1 |
| MVP_top60_DNA_seed | EP300 | 1.346693 | KIRC | OS-RC-2 | SGC-CBP30 | MVP_top60_DNA+RNA_seed | BRAF | 0.710078 | LUAD | HCC-78 | PLX-4720 |
| MVP_top60_DNA_seed | HSP90AA1 | 1.347491 | KIRC | BB65-RCC | CCT-018159 | MVP_top60_DNA+RNA_seed | BRAF | 0.710302 | LUAD | NCI-H2009 | SB590885 |
| MVP_top60_DNA_seed | HSP90AB1 | 1.347491 | KIRC | BB65-RCC | CCT-018159 | MVP_top60_DNA+RNA_seed | MDM2 | 0.711107 | LUAD | NCI-H2228 | Serdemetan |
| MVP_top60_DNA_seed | PIK3CA | 1.351246 | KIRC | VMRC-RCZ | AZD8186 | MVP_top60_DNA+RNA_seed | EGFR | 0.711684 | LUAD | NCI-H2085 | AZD8931 |
| MVP_top60_DNA_seed | HSP90AA1 | 1.385036 | KIRC | OS-RC-2 | Luminespib | MVP_top60_DNA+RNA_seed | ATM | 0.712103 | LUAD | NCI-H522 | KU-60019 |
| MVP_top60_DNA_seed | HSP90AB1 | 1.385036 | KIRC | OS-RC-2 | Luminespib | MVP_top60_DNA+RNA_seed | HDAC1 | 0.713312 | LUAD | NCI-H2122 | Tubastatin A |
| MVP_top60_DNA_seed | ESR1 | 1.389136 | KIRC | RCC-FG2 | Tamoxifen | MVP_top60_DNA+RNA_seed | BRAF | 0.713551 | LUAD | NCI-H1355 | Dabrafenib |
| MVP_top60_DNA_seed | PARP1 | 1.390798 | KIRC | RCC-FG2 | Olaparib | MVP_top60_DNA+RNA_seed | MDM2 | 0.717416 | LUAD | Calu-6 | Serdemetan |
| MVP_top60_DNA_seed | HDAC1 | 1.397825 | KIRC | TK10 | Entinostat | MVP_top60_DNA+RNA_seed | AURKA | 0.720391 | LUAD | SW1573 | Alisertib |
| MVP_top60_DNA_seed | PARP1 | 1.408717 | KIRC | TK10 | Veliparib | MVP_top60_DNA+RNA_seed | BRAF | 0.722368 | LUAD | RERF-LC-MS | SB590885 |
| MVP_top60_DNA_seed | HSP90AA1 | 1.419594 | KIRC | CAKI-1 | CCT-018159 | MVP_top60_DNA+RNA_seed | BRAF | 0.722407 | LUAD | NCI-H1693 | Dabrafenib |
| MVP_top60_DNA_seed | HSP90AB1 | 1.419594 | KIRC | CAKI-1 | CCT-018159 | MVP_top60_DNA+RNA_seed | HDAC1 | 0.72244 | LUAD | NCI-H358 | JQ12 |
| MVP_top60_DNA_seed | ATM | 1.423753 | KIRC | VMRC-RCW | KU-55933 | MVP_top60_DNA+RNA_seed | BRAF | 0.722863 | LUAD | NCI-H2228 | PLX-4720 |
| MVP_top60_DNA_seed | IGF1R | 1.427648 | KIRC | KMRC-1 | BMS-754807 | MVP_top60_DNA+RNA_seed | BRAF | 0.723463 | LUAD | HCC-827 | Dabrafenib |
| MVP_top60_DNA_seed | SMARCA4 | 1.431023 | KIRC | CAKI-1 | PFI3 | MVP_top60_DNA+RNA_seed | AKT1 | 0.72542 | LUAD | NCI-H1651 | GSK690693 |
| MVP_top60_DNA_seed | HDAC1 | 1.44453 | KIRC | TK10 | Romidepsin | MVP_top60_DNA+RNA_seed | EGFR | 0.725663 | LUAD | NCI-H1435 | Gefitinib |
| MVP_top60_DNA_seed | PIK3CA | 1.454868 | KIRC | TK10 | AZD8186 | MVP_top60_DNA+RNA_seed | MET | 0.726015 | LUAD | PC-3_[JPC-3] | SU11274 |
| MVP_top60_DNA_seed | PIK3CA | 1.45568 | KIRC | RXF393 | Buparlisib | MVP_top60_DNA+RNA_seed | BRAF | 0.726995 | LUAD | NCI-H838 | SB590885 |
| MVP_top60_DNA_seed | PIK3CD | 1.45568 | KIRC | RXF393 | Buparlisib | MVP_top60_DNA+RNA_seed | AURKA | 0.727765 | LUAD | NCI-H441 | GSK1070916 |
| MVP_top60_DNA_seed | HSP90AA1 | 1.481733 | KIRC | HA7-RCC | Luminespib | MVP_top60_DNA+RNA_seed | HSP90AA1 | 0.728879 | LUAD | NCI-H1648 | Elesclomol |

| | | | | | | | | | | | |
|---|---|---|---|---|---|---|---|---|---|---|---|
| MVP_top60_DNA_seed | HSP90AB1 | 1.481733 | KIRC | HA7-RCC | Luminespib | MVP_top60_DNA+RNA_seed | EGFR | 0.729725 | LUAD | NCI-H2009 | AST-1306 |
| MVP_top60_DNA_seed | PIK3CA | 1.495973 | KIRC | SN12C | Alpelisib | MVP_top60_DNA+RNA_seed | EGFR | 0.729738 | LUAD | NCI-H2122 | Pelitinib |
| MVP_top60_DNA_seed | HSP90AA1 | 1.506671 | KIRC | KMRC-20 | Luminespib | MVP_top60_DNA+RNA_seed | EGFR | 0.729951 | LUAD | NCI-H23 | Gefitinib |
| MVP_top60_DNA_seed | HSP90AB1 | 1.506671 | KIRC | KMRC-20 | Luminespib | MVP_top60_DNA+RNA_seed | FGFR1 | 0.73089 | LUAD | NCI-H1993 | FGFR_3831 |
| MVP_top60_DNA_seed | HSP90AA1 | 1.507212 | KIRC | LB2241-RCC | Elesclomol | MVP_top60_DNA+RNA_seed | EGFR | 0.731258 | LUAD | NCI-H3122 | AST-1306 |
| MVP_top60_DNA_seed | HSP90AB1 | 1.507212 | KIRC | LB2241-RCC | Elesclomol | MVP_top60_DNA+RNA_seed | EGFR | 0.731424 | LUAD | LXF-289 | PF-00299804 |
| MVP_top60_DNA_seed | ESR1 | 1.509048 | KIRC | NCC021 | Tamoxifen | MVP_top60_DNA+RNA_seed | HDAC1 | 0.731567 | LUAD | NCI-H23 | Dacinostat |
| MVP_top60_DNA_seed | PIK3CA | 1.520458 | KIRC | TK10 | Buparlisib | MVP_top60_DNA+RNA_seed | SRC | 0.731726 | LUAD | NCI-H1563 | Bosutinib |
| MVP_top60_DNA_seed | PIK3CD | 1.520458 | KIRC | TK10 | Buparlisib | MVP_top60_DNA+RNA_seed | HSP90AA1 | 0.732073 | LUAD | HCC-827 | SNX-2112 |
| MVP_top60_DNA_seed | HDAC1 | 1.520889 | KIRC | OS-RC-2 | PCI-34051 | MVP_top60_DNA+RNA_seed | FGFR1 | 0.734283 | LUAD | NCI-H522 | FGFR_3831 |
| MVP_top60_DNA_seed | HDAC1 | 1.521877 | KIRC | 769-P | Romidepsin | MVP_top60_DNA+RNA_seed | EGFR | 0.734982 | LUAD | NCI-H1563 | CUDC-101 |
| MVP_top60_DNA_seed | PARP1 | 1.52898 | KIRC | RCC10RGB | Rucaparib | MVP_top60_DNA+RNA_seed | EGFR | 0.735108 | LUAD | NCI-H1793 | Cetuximab |
| MVP_top60_DNA_seed | PARP1 | 1.536838 | KIRC | VMRC-RCZ | Rucaparib | MVP_top60_DNA+RNA_seed | HDAC1 | 0.735311 | LUAD | Calu-3 | AR-42 |
| MVP_top60_DNA_seed | PARP1 | 1.541467 | KIRC | TK10 | Niraparib | MVP_top60_DNA+RNA_seed | SMARCA4 | 0.736232 | LUAD | Calu-6 | PFI-3 |
| MVP_top60_DNA_seed | PARP1 | 1.543559 | KIRC | RCC10RGB | Veliparib | MVP_top60_DNA+RNA_seed | FGFR1 | 0.736903 | LUAD | NCI-H1975 | AZD4547 |
| MVP_top60_DNA_seed | IGF1R | 1.549533 | KIRC | BB65-RCC | BMS-754807 | MVP_top60_DNA+RNA_seed | AKT1 | 0.73696 | LUAD | HCC-827 | GSK690693 |
| MVP_top60_DNA_seed | IGF1R | 1.553473 | KIRC | SN12C | GSK1904529A | MVP_top60_DNA+RNA_seed | HDAC1 | 0.737508 | LUAD | NCI-H2023 | Belinostat |
| MVP_top60_DNA_seed | HSP90AA1 | 1.559635 | KIRC | LB1047-RCC | Luminespib | MVP_top60_DNA+RNA_seed | AURKA | 0.738294 | LUAD | A549 | Alisertib |
| MVP_top60_DNA_seed | HSP90AB1 | 1.559635 | KIRC | LB1047-RCC | Luminespib | MVP_top60_DNA+RNA_seed | HDAC1 | 0.740176 | LUAD | NCI-H2085 | AR-42 |
| MVP_top60_DNA_seed | AR | 1.572884 | KIRC | CAKI-1 | Bicalutamide | MVP_top60_DNA+RNA_seed | FGFR1 | 0.740315 | LUAD | NCI-H1651 | PD173074 |
| MVP_top60_DNA_seed | MTOR | 1.576607 | KIRC | VMRC-RCZ | Temsirolimus | MVP_top60_DNA+RNA_seed | PTK2 | 0.740883 | LUAD | NCI-H2023 | PF-562271 |
| MVP_top60_DNA_seed | IGF1R | 1.586322 | KIRC | TK10 | Linsitinib | MVP_top60_DNA+RNA_seed | MDM2 | 0.741701 | LUAD | PC-14 | Nutlin-3a (-) |
| MVP_top60_DNA_seed | PARP1 | 1.589189 | KIRC | VMRC-RCZ | Talazoparib | MVP_top60_DNA+RNA_seed | FGFR1 | 0.744307 | LUAD | NCI-H1355 | AZD4547 |
| MVP_top60_DNA_seed | HDAC1 | 1.596102 | KIRC | NCC010 | Romidepsin | MVP_top60_DNA+RNA_seed | FGFR1 | 0.74447 | LUAD | NCI-H1944 | PD173074 |
| MVP_top60_DNA_seed | ESR1 | 1.600789 | KIRC | CAKI-1 | GDC0810 | MVP_top60_DNA+RNA_seed | ATM | 0.744767 | LUAD | NCI-H1666 | KU-60019 |
| MVP_top60_DNA_seed | ESR1 | 1.601588 | KIRC | LB1047-RCC | Tamoxifen | MVP_top60_DNA+RNA_seed | BRAF | 0.745826 | LUAD | NCI-H1650 | SB590885 |
| MVP_top60_DNA_seed | ESR1 | 1.602068 | KIRC | RXF393 | Tamoxifen | MVP_top60_DNA+RNA_seed | FGFR1 | 0.747653 | LUAD | NCI-H1650 | AZD4547 |
| MVP_top60_DNA_seed | PARP1 | 1.605056 | KIRC | VMRC-RCW | Talazoparib | MVP_top60_DNA+RNA_seed | ATM | 0.747876 | LUAD | NCI-H3122 | KU-55933 |
| MVP_top60_DNA_seed | ATM | 1.624105 | KIRC | TK10 | KU-55933 | MVP_top60_DNA+RNA_seed | FGFR1 | 0.748016 | LUAD | NCI-H2291 | FGFR_3831 |
| MVP_top60_DNA_seed | HSP90AA1 | 1.637244 | KIRC | LB1047-RCC | CCT-018159 | MVP_top60_DNA+RNA_seed | AKT1 | 0.74805 | LUAD | ABC-1 | AKT inhibitor VIII |
| MVP_top60_DNA_seed | HSP90AB1 | 1.637244 | KIRC | LB1047-RCC | CCT-018159 | MVP_top60_DNA+RNA_seed | EGFR | 0.748075 | LUAD | NCI-H2009 | Afatinib |
| MVP_top60_DNA_seed | PIK3CA | 1.648698 | KIRC | VMRC-RCW | GNE-317 | MVP_top60_DNA+RNA_seed | MDM2 | 0.748582 | LUAD | NCI-H522 | Serdemetan |
| MVP_top60_DNA_seed | IGF1R | 1.653349 | KIRC | KMRC-1 | Linsitinib | MVP_top60_DNA+RNA_seed | EGFR | 0.748998 | LUAD | NCI-H1650 | PF-00299804 |
| MVP_top60_DNA_seed | HSP90AA1 | 1.679947 | KIRC | NCC021 | Tanespimycin | MVP_top60_DNA+RNA_seed | BRAF | 0.749463 | LUAD | NCI-H2347 | PLX-4720 |
| MVP_top60_DNA_seed | HSP90AB1 | 1.679947 | KIRC | NCC021 | Tanespimycin | MVP_top60_DNA+RNA_seed | ATM | 0.749597 | LUAD | NCI-H1568 | KU-55933 |
| MVP_top60_DNA_seed | TP53 | 1.681904 | KIRC | VMRC-RCZ | MIRA-1 | MVP_top60_DNA+RNA_seed | AKT1 | 0.750086 | LUAD | COR-L105 | BAY AKT1 |
| MVP_top60_DNA_seed | HDAC1 | 1.684918 | KIRC | OS-RC-2 | Romidepsin | MVP_top60_DNA+RNA_seed | HDAC1 | 0.750282 | LUAD | PC-14 | JQ12 |
| MVP_top60_DNA_seed | EP300 | 1.693076 | KIRC | TK10 | CPI-637 | MVP_top60_DNA+RNA_seed | EGFR | 0.750463 | LUAD | NCI-H23 | CUDC-101 |
| MVP_top60_DNA_seed | ESR1 | 1.699914 | KIRC | SW156 | Tamoxifen | MVP_top60_DNA+RNA_seed | FGFR1 | 0.75363 | LUAD | NCI-H1793 | FGFR_3831 |
| MVP_top60_DNA_seed | IGF1R | 1.711974 | KIRC | KMRC-1 | GSK1904529A | MVP_top60_DNA+RNA_seed | HDAC1 | 0.75439 | LUAD | NCI-H1993 | CAY10603 |
| MVP_top60_DNA_seed | EP300 | 1.715111 | KIRC | RCC10RGB | CPI-637 | MVP_top60_DNA+RNA_seed | HSP90AA1 | 0.754425 | LUAD | NCI-H1573 | Luminespib |

| | | | | | | | | | | | |
|---|---|---|---|---|---|---|---|---|---|---|---|
| MVP_top60_DNA_seed | PARP1 | 1.725456 | KIRC | RCC10RGB | Olaparib | MVP_top60_DNA+RNA_seed | EGFR | 0.754637 | LUAD | SW1573 | Pelitinib |
| MVP_top60_DNA_seed | PARP1 | 1.735103 | KIRC | VMRC-RCW | Olaparib | MVP_top60_DNA+RNA_seed | HDAC1 | 0.754933 | LUAD | NCI-H1563 | JQ12 |
| MVP_top60_DNA_seed | PIK3CA | 1.741052 | KIRC | VMRC-RCZ | GNE-317 | MVP_top60_DNA+RNA_seed | AURKA | 0.755228 | LUAD | NCI-H1793 | Alisertib |
| MVP_top60_DNA_seed | HSP90AA1 | 1.757652 | KIRC | HA7-RCC | CCT-018159 | MVP_top60_DNA+RNA_seed | EGFR | 0.75606 | LUAD | NCI-H1573 | Cetuximab |
| MVP_top60_DNA_seed | HSP90AB1 | 1.757652 | KIRC | HA7-RCC | CCT-018159 | MVP_top60_DNA+RNA_seed | HDAC1 | 0.756284 | LUAD | Calu-3 | Belinostat |
| MVP_top60_DNA_seed | PIK3CA | 1.764756 | KIRC | SW156 | Buparlisib | MVP_top60_DNA+RNA_seed | BRAF | 0.756986 | LUAD | NCI-H1573 | PLX-4720 |
| MVP_top60_DNA_seed | PIK3CD | 1.764756 | KIRC | SW156 | Buparlisib | MVP_top60_DNA+RNA_seed | AURKA | 0.75799 | LUAD | RERF-LC-MS | Alisertib |
| MVP_top60_DNA_seed | HSP90AA1 | 1.774957 | KIRC | VMRC-RCZ | Tanespimycin | MVP_top60_DNA+RNA_seed | BRAF | 0.759542 | LUAD | NCI-H1435 | RAF_9304 |
| MVP_top60_DNA_seed | HSP90AB1 | 1.774957 | KIRC | VMRC-RCZ | Tanespimycin | MVP_top60_DNA+RNA_seed | FGFR1 | 0.759761 | LUAD | NCI-H1993 | PD173074 |
| MVP_top60_DNA_seed | PARP1 | 1.779344 | KIRC | VMRC-RCZ | Veliparib | MVP_top60_DNA+RNA_seed | SMARCA4 | 0.759761 | LUAD | RERF-LC-KJ | PFI-3 |
| MVP_top60_DNA_seed | IGF1R | 1.782101 | KIRC | SW156 | NVP-ADW742 | MVP_top60_DNA+RNA_seed | BRAF | 0.760082 | LUAD | COR-L105 | PLX-4720 |
| MVP_top60_DNA_seed | HDAC1 | 1.832891 | KIRC | RCC10RGB | Entinostat | MVP_top60_DNA+RNA_seed | EGFR | 0.760124 | LUAD | ABC-1 | Gefitinib |
| MVP_top60_DNA_seed | IGF1R | 1.835023 | KIRC | RCC-JW | BMS-754807 | MVP_top60_DNA+RNA_seed | BRAF | 0.760176 | LUAD | NCI-H1651 | RAF_9304 |
| MVP_top60_DNA_seed | HSP90AA1 | 1.844277 | KIRC | VMRC-RCW | Luminespib | MVP_top60_DNA+RNA_seed | FGFR1 | 0.760941 | LUAD | NCI-H2023 | AZD4547 |
| MVP_top60_DNA_seed | HSP90AB1 | 1.844277 | KIRC | VMRC-RCW | Luminespib | MVP_top60_DNA+RNA_seed | EGFR | 0.761033 | LUAD | NCI-H2122 | CUDC-101 |
| MVP_top60_DNA_seed | ESR1 | 1.869394 | KIRC | RCC-AB | Tamoxifen | MVP_top60_DNA+RNA_seed | EGFR | 0.7612 | LUAD | RERF-LC-MS | Pelitinib |
| MVP_top60_DNA_seed | IGF1R | 1.884492 | KIRC | RCC10RGB | BMS-754807 | MVP_top60_DNA+RNA_seed | HSP90AA1 | 0.761274 | LUAD | NCI-H2085 | CCT-018159 |
| MVP_top60_DNA_seed | IGF1R | 1.917243 | KIRC | RCC-AB | NVP-ADW742 | MVP_top60_DNA+RNA_seed | HSP90AA1 | 0.762372 | LUAD | NCI-H1573 | Elesclomol |
| MVP_top60_DNA_seed | IGF1R | 1.923712 | KIRC | TK10 | BMS-754807 | MVP_top60_DNA+RNA_seed | EGFR | 0.76361 | LUAD | NCI-H1781 | Cetuximab |
| MVP_top60_DNA_seed | ESR1 | 1.931838 | KIRC | VMRC-RCW | GDC0810 | MVP_top60_DNA+RNA_seed | MDM2 | 0.763852 | LUAD | SK-LU-1 | Serdemetan |
| MVP_top60_DNA_seed | SMARCA4 | 1.986562 | KIRC | TK10 | PFI3 | MVP_top60_DNA+RNA_seed | BRAF | 0.765517 | LUAD | NCI-H2342 | SB590885 |
| MVP_top60_DNA_seed | SMARCA4 | 1.989225 | KIRC | RCC10RGB | PFI3 | MVP_top60_DNA+RNA_seed | PIK3CA | 0.766511 | LUAD | RERF-LC-KJ | AZD8835 |
| MVP_top60_DNA_seed | HDAC1 | 1.995917 | KIRC | RCC10RGB | Romidepsin | MVP_top60_DNA+RNA_seed | IGF1R | 0.766837 | LUAD | HCC-78 | GSK1904529A |
| MVP_top60_DNA_seed | ESR1 | 2.093508 | KIRC | VMRC-RCW | Tamoxifen | MVP_top60_DNA+RNA_seed | FGFR1 | 0.767411 | LUAD | NCI-H1975 | PD173074 |
| MVP_top60_DNA_seed | PARP1 | 2.096802 | KIRC | VMRC-RCW | Veliparib | MVP_top60_DNA+RNA_seed | MET | 0.768618 | LUAD | SK-LU-1 | AZD6094 |
| MVP_top60_DNA_seed | IGF1R | 2.101741 | KIRC | RCC10RGB | GSK1904529A | MVP_top60_DNA+RNA_seed | FGFR1 | 0.769071 | LUAD | SW1573 | PD173074 |
| MVP_top60_DNA_seed | PARP1 | 2.138391 | KIRC | VMRC-RCZ | Olaparib | MVP_top60_DNA+RNA_seed | AURKA | 0.76933 | LUAD | NCI-H441 | ZM447439 |
| MVP_top60_DNA_seed | ESR1 | 2.143903 | KIRC | CAKI-1 | Tamoxifen | MVP_top60_DNA+RNA_seed | AKT1 | 0.769729 | LUAD | EMC-BAC-2 | GSK690693 |
| MVP_top60_DNA_seed | HSP90AA1 | 2.148358 | KIRC | VMRC-RCW | Elesclomol | MVP_top60_DNA+RNA_seed | EGFR | 0.772207 | LUAD | NCI-H2228 | Gefitinib |
| MVP_top60_DNA_seed | HSP90AB1 | 2.148358 | KIRC | VMRC-RCW | Elesclomol | MVP_top60_DNA+RNA_seed | EGFR | 0.772473 | LUAD | NCI-H2405 | Afatinib |
| MVP_top60_DNA_seed | PIK3CA | 2.185823 | KIRC | TK10 | Alpelisib | MVP_top60_DNA+RNA_seed | HDAC1 | 0.77299 | LUAD | Calu-3 | CAY10603 |
| MVP_top60_DNA_seed | ESR1 | 2.191646 | KIRC | VMRC-RCZ | GDC0810 | MVP_top60_DNA+RNA_seed | MDM2 | 0.773315 | LUAD | NCI-H1651 | Nutlin-3a (-) |
| MVP_top60_DNA_seed | SMARCA4 | 2.203211 | KIRC | VMRC-RCZ | PFI3 | MVP_top60_DNA+RNA_seed | MAPK14 | 0.77392 | LUAD | Calu-6 | TAK-715 |
| MVP_top60_DNA_seed | PARP1 | 2.209517 | KIRC | TK10 | Talazoparib | MVP_top60_DNA+RNA_seed | BRAF | 0.773968 | LUAD | RERF-LC-KJ | PLX-4720 |
| MVP_top60_DNA_seed | IGF1R | 2.212351 | KIRC | VMRC-RCZ | GSK1904529A | MVP_top60_DNA+RNA_seed | BRAF | 0.77526 | LUAD | NCI-H650 | PLX-4720 |
| MVP_top60_DNA_seed | PIK3CA | 2.237278 | KIRC | VMRC-RCZ | Buparlisib | MVP_top60_DNA+RNA_seed | IGF1R | 0.775306 | LUAD | NCI-H1975 | GSK1904529A |
| MVP_top60_DNA_seed | PIK3CD | 2.237278 | KIRC | VMRC-RCZ | Buparlisib | MVP_top60_DNA+RNA_seed | EGFR | 0.775334 | LUAD | NCI-H1651 | CI-1033 |
| MVP_top60_DNA_seed | HDAC1 | 2.238131 | KIRC | VMRC-RCZ | PCI-34051 | MVP_top60_DNA+RNA_seed | IGF1R | 0.775697 | LUAD | NCI-H1838 | BMS-536924 |
| MVP_top60_DNA_seed | PARP1 | 2.264141 | KIRC | VMRC-RCZ | Niraparib | MVP_top60_DNA+RNA_seed | SRC | 0.776324 | LUAD | NCI-H23 | Saracatinib |
| MVP_top60_DNA_seed | HSP90AA1 | 2.299331 | KIRC | RCC-MF | Luminespib | MVP_top60_DNA+RNA_seed | MET | 0.777276 | LUAD | H3255 | AZD6094 |
| MVP_top60_DNA_seed | HSP90AB1 | 2.299331 | KIRC | RCC-MF | Luminespib | MVP_top60_DNA+RNA_seed | EGFR | 0.778687 | LUAD | NCI-H322M | AZD8931 |

| | | | | | | | | | | | |
|---|---|---|---|---|---|---|---|---|---|---|---|
| MVP_top60_DNA_seed | HDAC1 | 2.301832 | KIRC | VMRC-RCZ | Entinostat | MVP_top60_DNA+RNA_seed | AURKA | 0.778765 | LUAD | NCI-H2030 | CD532 |
| MVP_top60_DNA_seed | MTOR | 2.324809 | KIRC | VMRC-RCW | Temsirolimus | MVP_top60_DNA+RNA_seed | EGFR | 0.77956 | LUAD | SW1573 | Afatinib |
| MVP_top60_DNA_seed | HDAC1 | 2.335545 | KIRC | TK10 | PCI-34051 | MVP_top60_DNA+RNA_seed | AKT1 | 0.779613 | LUAD | RERF-LC-KJ | GSK690693 |
| MVP_top60_DNA_seed | IGF1R | 2.425755 | KIRC | VMRC-RCZ | BMS-536924 | MVP_top60_DNA+RNA_seed | AURKA | 0.779776 | LUAD | NCI-H2023 | GSK1070916 |
| MVP_top60_DNA_seed | SMARCA4 | 2.435711 | KIRC | VMRC-RCW | PFI3 | MVP_top60_DNA+RNA_seed | MDM2 | 0.780645 | LUAD | NCI-H2347 | Nutlin-3a (-) |
| MVP_top60_DNA_seed | ESR1 | 2.437983 | KIRC | TK10 | GDC0810 | MVP_top60_DNA+RNA_seed | BRAF | 0.782218 | LUAD | ABC-1 | SB590885 |
| MVP_top60_DNA_seed | IGF1R | 2.442234 | KIRC | VMRC-RCZ | Linsitinib | MVP_top60_DNA+RNA_seed | IGF1R | 0.782822 | LUAD | NCI-H650 | BMS-536924 |
| MVP_top60_DNA_seed | IGF1R | 2.446206 | KIRC | VMRC-RCW | Linsitinib | MVP_top60_DNA+RNA_seed | BRAF | 0.782958 | LUAD | NCI-H1568 | PLX-4720 |
| MVP_top60_DNA_seed | HDAC1 | 2.471239 | KIRC | SN12C | Entinostat | MVP_top60_DNA+RNA_seed | HDAC1 | 0.785938 | LUAD | NCI-H1993 | Tubastatin A |
| MVP_top60_DNA_seed | IGF1R | 2.629637 | KIRC | TK10 | NVP-ADW742 | MVP_top60_DNA+RNA_seed | MDM2 | 0.78596 | LUAD | SW1573 | Serdemetan |
| MVP_top60_DNA_seed | EP300 | 2.747749 | KIRC | TK10 | SGC-CBP30 | MVP_top60_DNA+RNA_seed | FGFR1 | 0.786032 | LUAD | NCI-H1781 | PD173074 |
| MVP_top60_DNA_seed | IGF1R | 2.754441 | KIRC | VMRC-RCZ | BMS-754807 | MVP_top60_DNA+RNA_seed | AKT1 | 0.787304 | LUAD | NCI-H522 | MK-2206 |
| MVP_top60_DNA_seed | HSP90AA1 | 2.801677 | KIRC | VMRC-RCW | Tanespimycin | MVP_top60_DNA+RNA_seed | HSP90AA1 | 0.787892 | LUAD | NCI-H1568 | Tanespimycin |
| MVP_top60_DNA_seed | HSP90AB1 | 2.801677 | KIRC | VMRC-RCW | Tanespimycin | MVP_top60_DNA+RNA_seed | AURKA | 0.790867 | LUAD | LXF-289 | CD532 |
| MVP_top60_DNA_seed | HSP90AA1 | 2.810492 | KIRC | RCC10RGB | Tanespimycin | MVP_top60_DNA+RNA_seed | FGFR1 | 0.791474 | LUAD | NCI-H1435 | PD173074 |
| MVP_top60_DNA_seed | HSP90AB1 | 2.810492 | KIRC | RCC10RGB | Tanespimycin | MVP_top60_DNA+RNA_seed | SRC | 0.791899 | LUAD | NCI-H1437 | Bosutinib |
| MVP_top60_DNA_seed | IGF1R | 2.866762 | KIRC | TK10 | GSK1904529A | MVP_top60_DNA+RNA_seed | BRAF | 0.79421 | LUAD | Calu-6 | SB590885 |
| MVP_top60_DNA_seed | PIK3CA | 2.888068 | KIRC | VMRC-RCZ | Alpelisib | MVP_top60_DNA+RNA_seed | HDAC1 | 0.794609 | LUAD | NCI-H1563 | AR-42 |
| MVP_top60_DNA_seed | ESR1 | 2.932502 | KIRC | TK10 | Tamoxifen | MVP_top60_DNA+RNA_seed | SMARCA4 | 0.794778 | LUAD | NCI-H2405 | PFI-3 |
| MVP_top60_DNA_seed | IGF1R | 2.964638 | KIRC | VMRC-RCZ | NVP-ADW742 | MVP_top60_DNA+RNA_seed | BRAF | 0.795398 | LUAD | NCI-H23 | PLX-4720 |
| MVP_top60_DNA_seed | HSP90AA1 | 3.168001 | KIRC | VMRC-RCZ | Luminespib | MVP_top60_DNA+RNA_seed | HDAC1 | 0.797109 | LUAD | H3255 | Belinostat |
| MVP_top60_DNA_seed | HSP90AB1 | 3.168001 | KIRC | VMRC-RCZ | Luminespib | MVP_top60_DNA+RNA_seed | AURKA | 0.797567 | LUAD | NCI-H2030 | Alisertib |
| MVP_top60_DNA_seed | PIK3CA | 3.212868 | KIRC | VMRC-RCW | Buparlisib | MVP_top60_DNA+RNA_seed | PIK3CA | 0.797794 | LUAD | NCI-H1435 | PI3Ka_4409 |
| MVP_top60_DNA_seed | PIK3CD | 3.212868 | KIRC | VMRC-RCW | Buparlisib | MVP_top60_DNA+RNA_seed | FGFR1 | 0.798079 | LUAD | NCI-H2347 | AZD4547 |
| MVP_top60_DNA_seed | HSP90AA1 | 3.345674 | KIRC | RCC10RGB | Luminespib | MVP_top60_DNA+RNA_seed | EGFR | 0.79841 | LUAD | NCI-H838 | Afatinib |
| MVP_top60_DNA_seed | HSP90AB1 | 3.345674 | KIRC | RCC10RGB | Luminespib | MVP_top60_DNA+RNA_seed | BRAF | 0.79909 | LUAD | RERF-LC-KJ | Dabrafenib |
| MVP_top60_DNA_seed | PARP1 | 3.393206 | KIRC | TK10 | Olaparib | MVP_top60_DNA+RNA_seed | HSP90AA1 | 0.79915 | LUAD | NCI-H2009 | CCT-018159 |
| MVP_top60_DNA_seed | MET | -4.837639 | LGG | KINGS-1 | PHA-665752 | MVP_top60_DNA+RNA_seed | BRAF | 0.799203 | LUAD | NCI-H1975 | PLX-4720 |
| MVP_top60_DNA_seed | MET | -3.623891 | LGG | SF539 | Crizotinib | MVP_top60_DNA+RNA_seed | AKT1 | 0.799662 | LUAD | RERF-LC-MS | GSK690693 |
| MVP_top60_DNA_seed | MET | -3.390599 | LGG | MOG-G-UVW | AZD6094 | MVP_top60_DNA+RNA_seed | EGFR | 0.799711 | LUAD | HOP-62 | Afatinib |
| MVP_top60_DNA_seed | MET | -3.106228 | LGG | KINGS-1 | Crizotinib | MVP_top60_DNA+RNA_seed | AKT1 | 0.801137 | LUAD | NCI-H1838 | AKT inhibitor VIII |
| MVP_top60_DNA_seed | MDM2 | -2.743786 | LGG | H4 | Serdemetan | MVP_top60_DNA+RNA_seed | HDAC1 | 0.801623 | LUAD | NCI-H1435 | AR-42 |
| MVP_top60_DNA_seed | MDM2 | -2.548949 | LGG | H4 | Nutlin-3a (-) | MVP_top60_DNA+RNA_seed | HDAC1 | 0.802154 | LUAD | NCI-H1781 | Tubastatin A |
| MVP_top60_DNA_seed | PDGFRB | -2.352912 | LGG | GI-1 | Pazopanib | MVP_top60_DNA+RNA_seed | SMARCA4 | 0.802652 | LUAD | EMC-BAC-1 | PFI-3 |
| MVP_top60_DNA_seed | HSF1 | -2.260876 | LGG | H4 | CCT245232 | MVP_top60_DNA+RNA_seed | HSP90AA1 | 0.802904 | LUAD | NCI-H2342 | CCT-018159 |
| MVP_top60_DNA_seed | HDAC1 | -2.207784 | LGG | no-10 | Parthenolide | MVP_top60_DNA+RNA_seed | AURKA | 0.804246 | LUAD | NCI-H1793 | GSK1070916 |
| MVP_top60_DNA_seed | HDAC1 | -1.8461 | LGG | SF539 | Parthenolide | MVP_top60_DNA+RNA_seed | BRAF | 0.804856 | LUAD | NCI-H522 | PLX-4720 |
| MVP_top60_DNA_seed | CREBBP | -1.821701 | LGG | H4 | I-CBP112 | MVP_top60_DNA+RNA_seed | AURKA | 0.805817 | LUAD | NCI-H1838 | Alisertib |
| MVP_top60_DNA_seed | HDAC1 | -1.81337 | LGG | SF539 | Entinostat | MVP_top60_DNA+RNA_seed | HDAC1 | 0.805953 | LUAD | NCI-H1993 | JQ12 |
| MVP_top60_DNA_seed | EGFR | -1.781339 | LGG | D-502MG | Lapatinib | MVP_top60_DNA+RNA_seed | EGFR | 0.806045 | LUAD | NCI-H1573 | CUDC-101 |
| MVP_top60_DNA_seed | MDM2 | -1.693604 | LGG | D-502MG | Nutlin-3a (-) | MVP_top60_DNA+RNA_seed | HDAC1 | 0.806122 | LUAD | Calu-6 | Entinostat |

| | | | | | | | | | | | |
|---|---|---|---|---|---|---|---|---|---|---|---|
| MVP_top60_DNA_seed | HSP90AA1 | -1.672458 | LGG | MOG-G-UVW | Elesclomol | MVP_top60_DNA+RNA_seed | HDAC1 | 0.806944 | LUAD | H3255 | Tubastatin A |
| MVP_top60_DNA_seed | EGFR | -1.665939 | LGG | D-502MG | Erlotinib | MVP_top60_DNA+RNA_seed | BRAF | 0.806972 | LUAD | NCI-H838 | Dabrafenib |
| MVP_top60_DNA_seed | HSP90AA1 | -1.660513 | LGG | GAMG | Elesclomol | MVP_top60_DNA+RNA_seed | FGFR1 | 0.807423 | LUAD | NCI-H1563 | PD173074 |
| MVP_top60_DNA_seed | HSF1 | -1.643141 | LGG | MOG-G-UVW | CCT245232 | MVP_top60_DNA+RNA_seed | EGFR | 0.808434 | LUAD | NCI-H2342 | Pelitinib |
| MVP_top60_DNA_seed | CDK2 | -1.633484 | LGG | GI-1 | Seliciclib | MVP_top60_DNA+RNA_seed | MET | 0.810252 | LUAD | Calu-3 | AZD6094 |
| MVP_top60_DNA_seed | JAK2 | -1.623002 | LGG | MOG-G-UVW | Ruxolitinib | MVP_top60_DNA+RNA_seed | MDM2 | 0.810913 | LUAD | NCI-H1573 | Serdemetan |
| MVP_top60_DNA_seed | JAK1 | -1.623002 | LGG | MOG-G-UVW | Ruxolitinib | MVP_top60_DNA+RNA_seed | EGFR | 0.812571 | LUAD | RERF-LC-KJ | CUDC-101 |
| MVP_top60_DNA_seed | PIK3CD | -1.61272 | LGG | GAMG | AZD8186 | MVP_top60_DNA+RNA_seed | AURKA | 0.813367 | LUAD | RERF-LC-KJ | CD532 |
| MVP_top60_DNA_seed | MET | -1.600068 | LGG | GI-1 | PHA-665752 | MVP_top60_DNA+RNA_seed | HDAC1 | 0.813411 | LUAD | NCI-H1993 | Belinostat |
| MVP_top60_DNA_seed | HSP90AA1 | -1.572407 | LGG | H4 | Tanespimycin | MVP_top60_DNA+RNA_seed | FGFR1 | 0.814173 | LUAD | PC-14 | AZD4547 |
| MVP_top60_DNA_seed | PDGFRB | -1.563076 | LGG | H4 | Pazopanib | MVP_top60_DNA+RNA_seed | HSP90AA1 | 0.814229 | LUAD | NCI-H1435 | Luminespib |
| MVP_top60_DNA_seed | MET | -1.537092 | LGG | GI-1 | Crizotinib | MVP_top60_DNA+RNA_seed | HDAC1 | 0.815079 | LUAD | NCI-H441 | AR-42 |
| MVP_top60_DNA_seed | IGF1R | -1.47482 | LGG | no-10 | GSK1904529A | MVP_top60_DNA+RNA_seed | ATM | 0.815365 | LUAD | ABC-1 | torin2 |
| MVP_top60_DNA_seed | HSF1 | -1.451036 | LGG | MOG-G-CCM | CCT245467 | MVP_top60_DNA+RNA_seed | ATM | 0.815385 | LUAD | NCI-H1573 | KU-55933 |
| MVP_top60_DNA_seed | JAK3 | -1.448512 | LGG | GI-1 | JAK3_7406 | MVP_top60_DNA+RNA_seed | MET | 0.815674 | LUAD | A549 | AZD6094 |
| MVP_top60_DNA_seed | JAK1 | -1.440348 | LGG | GI-1 | JAK1_3715 | MVP_top60_DNA+RNA_seed | IGF1R | 0.816217 | LUAD | NCI-H1734 | GSK1904529A |
| MVP_top60_DNA_seed | HSP90AA1 | -1.436646 | LGG | MOG-G-CCM | Elesclomol | MVP_top60_DNA+RNA_seed | FGFR1 | 0.817848 | LUAD | H3255 | AZD4547 |
| MVP_top60_DNA_seed | HSP90AA1 | -1.410337 | LGG | D-336MG | Tanespimycin | MVP_top60_DNA+RNA_seed | AURKA | 0.817927 | LUAD | LXF-289 | Alisertib |
| MVP_top60_DNA_seed | PIK3CD | -1.401041 | LGG | KALS-1 | AZD8186 | MVP_top60_DNA+RNA_seed | SRC | 0.818266 | LUAD | ABC-1 | NVP-BHG712 |
| MVP_top60_DNA_seed | PDGFRB | -1.387456 | LGG | GAMG | Pazopanib | MVP_top60_DNA+RNA_seed | HDAC1 | 0.819202 | LUAD | NCI-H1623 | Dacinostat |
| MVP_top60_DNA_seed | MET | -1.378701 | LGG | H4 | JNJ38877605 | MVP_top60_DNA+RNA_seed | BRAF | 0.819332 | LUAD | NCI-H1975 | SB590885 |
| MVP_top60_DNA_seed | HSP90AA1 | -1.36963 | LGG | SF539 | Tanespimycin | MVP_top60_DNA+RNA_seed | EGFR | 0.819354 | LUAD | NCI-H1651 | AZD8931 |
| MVP_top60_DNA_seed | HSP90AA1 | -1.369305 | LGG | GAMG | Tanespimycin | MVP_top60_DNA+RNA_seed | BRAF | 0.820048 | LUAD | NCI-H2085 | RAF_9304 |
| MVP_top60_DNA_seed | PIK3CD | -1.365481 | LGG | D-336MG | AZD8186 | MVP_top60_DNA+RNA_seed | MET | 0.82014 | LUAD | Calu-6 | Crizotinib |
| MVP_top60_DNA_seed | HSP90AA1 | -1.36264 | LGG | SW1783 | Luminespib | MVP_top60_DNA+RNA_seed | IGF1R | 0.821256 | LUAD | NCI-H1793 | BMS-754807 |
| MVP_top60_DNA_seed | EGFR | -1.349885 | LGG | NMC-G1 | Erlotinib | MVP_top60_DNA+RNA_seed | EGFR | 0.821754 | LUAD | NCI-H522 | Pelitinib |
| MVP_top60_DNA_seed | HSP90AA1 | -1.349239 | LGG | Hs-683 | Tanespimycin | MVP_top60_DNA+RNA_seed | EGFR | 0.823436 | LUAD | NCI-H3122 | Cetuximab |
| MVP_top60_DNA_seed | EGFR | -1.348828 | LGG | MOG-G-CCM | Cetuximab | MVP_top60_DNA+RNA_seed | MDM2 | 0.823566 | LUAD | RERF-LC-KJ | Serdemetan |
| MVP_top60_DNA_seed | MET | -1.342467 | LGG | KINGS-1 | JNJ38877605 | MVP_top60_DNA+RNA_seed | AKT1 | 0.825336 | LUAD | NCI-H1568 | GSK690693 |
| MVP_top60_DNA_seed | PIK3CD | -1.338399 | LGG | MOG-G-CCM | AZD8186 | MVP_top60_DNA+RNA_seed | BRAF | 0.826556 | LUAD | NCI-H1437 | Dabrafenib |
| MVP_top60_DNA_seed | IGF1R | -1.33162 | LGG | D-336MG | BMS-536924 | MVP_top60_DNA+RNA_seed | EGFR | 0.826624 | LUAD | LC-2-ad | Gefitinib |
| MVP_top60_DNA_seed | MET | -1.321145 | LGG | MOG-G-CCM | JNJ38877605 | MVP_top60_DNA+RNA_seed | HDAC1 | 0.826857 | LUAD | NCI-H2228 | CAY10603 |
| MVP_top60_DNA_seed | JAK3 | -1.318028 | LGG | H4 | JAK3_7406 | MVP_top60_DNA+RNA_seed | BRAF | 0.827151 | LUAD | NCI-H1666 | AZ628 |
| MVP_top60_DNA_seed | NTRK1 | -1.306101 | LGG | GAMG | AZD1332 | MVP_top60_DNA+RNA_seed | SRC | 0.827339 | LUAD | NCI-H838 | NVP-BHG712 |
| MVP_top60_DNA_seed | PDGFRB | -1.30427 | LGG | NMC-G1 | Pazopanib | MVP_top60_DNA+RNA_seed | FGFR1 | 0.828822 | LUAD | NCI-H1573 | PD173074 |
| MVP_top60_DNA_seed | IGF1R | -1.303804 | LGG | NMC-G1 | BMS-754807 | MVP_top60_DNA+RNA_seed | BRAF | 0.829641 | LUAD | NCI-H1838 | Dabrafenib |
| MVP_top60_DNA_seed | HSP90AA1 | -1.298947 | LGG | MOG-G-UVW | CCT-018159 | MVP_top60_DNA+RNA_seed | SRC | 0.830013 | LUAD | NCI-H23 | WH-4-023 |
| MVP_top60_DNA_seed | HSP90AA1 | -1.273819 | LGG | no-11 | CCT-018159 | MVP_top60_DNA+RNA_seed | HSP90AA1 | 0.830213 | LUAD | PC-3_[JPC-3] | Elesclomol |
| MVP_top60_DNA_seed | MET | -1.259185 | LGG | GI-1 | JNJ38877605 | MVP_top60_DNA+RNA_seed | AURKA | 0.830398 | LUAD | Calu-3 | HG-5-113-01 |
| MVP_top60_DNA_seed | MET | -1.254169 | LGG | MOG-G-UVW | JNJ38877605 | MVP_top60_DNA+RNA_seed | BRAF | 0.831091 | LUAD | RERF-LC-MS | RAF_9304 |
| MVP_top60_DNA_seed | JAK3 | -1.253832 | LGG | SF539 | JAK3_7406 | MVP_top60_DNA+RNA_seed | EGFR | 0.831549 | LUAD | NCI-H1651 | Gefitinib |

| | | | | | | | | | | | |
|---|---|---|---|---|---|---|---|---|---|---|---|
| MVP_top60_DNA_seed | PIK3CD | -1.238452 | LGG | H4 | AZD8186 | MVP_top60_DNA+RNA_seed | AURKA | 0.832222 | LUAD | NCI-H2347 | Genentech Cpd 10 |
| MVP_top60_DNA_seed | SMARCA4 | -1.224278 | LGG | MOG-G-UVW | PFI-3 | MVP_top60_DNA+RNA_seed | HDAC1 | 0.832546 | LUAD | NCI-H1781 | AR-42 |
| MVP_top60_DNA_seed | HSP90AA1 | -1.216265 | LGG | GAMG | SNX-2112 | MVP_top60_DNA+RNA_seed | AURKA | 0.832946 | LUAD | NCI-H23 | Tozasertib |
| MVP_top60_DNA_seed | NTRK1 | -1.174234 | LGG | KINGS-1 | GW441756 | MVP_top60_DNA+RNA_seed | EGFR | 0.83486 | LUAD | NCI-H2405 | AST-1306 |
| MVP_top60_DNA_seed | EGFR | -1.161462 | LGG | GI-1 | Lapatinib | MVP_top60_DNA+RNA_seed | MET | 0.83503 | LUAD | Calu-6 | SU11274 |
| MVP_top60_DNA_seed | JAK3 | -1.1547 | LGG | NMC-G1 | JAK3_7406 | MVP_top60_DNA+RNA_seed | HSP90AA1 | 0.835229 | LUAD | NCI-H2405 | SNX-2112 |
| MVP_top60_DNA_seed | AR | -1.147339 | LGG | H4 | AZD3514 | MVP_top60_DNA+RNA_seed | AURKA | 0.835387 | LUAD | Calu-3 | Genentech Cpd 10 |
| MVP_top60_DNA_seed | AR | -1.137113 | LGG | H4 | Bicalutamide | MVP_top60_DNA+RNA_seed | HDAC1 | 0.835478 | LUAD | NCI-H1792 | Dacinostat |
| MVP_top60_DNA_seed | EGFR | -1.130372 | LGG | NMC-G1 | Lapatinib | MVP_top60_DNA+RNA_seed | MET | 0.836551 | LUAD | HCC-827 | AZD6094 |
| MVP_top60_DNA_seed | HDAC1 | -1.119233 | LGG | NMC-G1 | Entinostat | MVP_top60_DNA+RNA_seed | EGFR | 0.837101 | LUAD | NCI-H1755 | Afatinib |
| MVP_top60_DNA_seed | HSP90AA1 | -1.114903 | LGG | GI-1 | SNX-2112 | MVP_top60_DNA+RNA_seed | IGF1R | 0.837488 | LUAD | PC-14 | BMS-754807 |
| MVP_top60_DNA_seed | HSP90AA1 | -1.093589 | LGG | H4 | Elesclomol | MVP_top60_DNA+RNA_seed | AURKA | 0.837547 | LUAD | NCI-H838 | Genentech Cpd 10 |
| MVP_top60_DNA_seed | MET | -1.093437 | LGG | SF539 | PHA-665752 | MVP_top60_DNA+RNA_seed | HDAC1 | 0.838515 | LUAD | RERF-LC-KJ | JQ12 |
| MVP_top60_DNA_seed | MET | -1.08105 | LGG | GAMG | JNJ38877605 | MVP_top60_DNA+RNA_seed | AURKA | 0.839222 | LUAD | NCI-H1573 | CD532 |
| MVP_top60_DNA_seed | HSP90AA1 | -1.080907 | LGG | GAMG | Luminespib | MVP_top60_DNA+RNA_seed | AKT1 | 0.839409 | LUAD | HCC-78 | AKT inhibitor VIII |
| MVP_top60_DNA_seed | CDK2 | -1.075066 | LGG | MOG-G-UVW | AZD5438 | MVP_top60_DNA+RNA_seed | EGFR | 0.839916 | LUAD | NCI-H3122 | Afatinib |
| MVP_top60_DNA_seed | MET | -1.063293 | LGG | NMC-G1 | JNJ38877605 | MVP_top60_DNA+RNA_seed | SRC | 0.839976 | LUAD | NCI-H2023 | NVP-BHG712 |
| MVP_top60_DNA_seed | IGF1R | -1.06313 | LGG | MOG-G-CCM | GSK1904529A | MVP_top60_DNA+RNA_seed | IGF1R | 0.840642 | LUAD | NCI-H2228 | GSK1904529A |
| MVP_top60_DNA_seed | PDGFRB | -1.060213 | LGG | MOG-G-UVW | Pazopanib | MVP_top60_DNA+RNA_seed | SRC | 0.841284 | LUAD | RERF-LC-MS | NVP-BHG712 |
| MVP_top60_DNA_seed | PIK3CD | -1.054253 | LGG | GI-1 | IC-87114 | MVP_top60_DNA+RNA_seed | HSP90AA1 | 0.842361 | LUAD | NCI-H2009 | SNX-2112 |
| MVP_top60_DNA_seed | JAK2 | -1.03245 | LGG | H4 | AZD1480 | MVP_top60_DNA+RNA_seed | AKT1 | 0.842469 | LUAD | NCI-H1435 | BAY AKT1 |
| MVP_top60_DNA_seed | JAK1 | -1.03245 | LGG | H4 | AZD1480 | MVP_top60_DNA+RNA_seed | AURKA | 0.842575 | LUAD | EKVX | HG-5-113-01 |
| MVP_top60_DNA_seed | IGF1R | -1.023677 | LGG | NMC-G1 | Linsitinib | MVP_top60_DNA+RNA_seed | HDAC1 | 0.842725 | LUAD | NCI-H1651 | Tubastatin A |
| MVP_top60_DNA_seed | MET | -1.021228 | LGG | H4 | AZD6094 | MVP_top60_DNA+RNA_seed | ATM | 0.842743 | LUAD | NCI-H2342 | CP466722 |
| MVP_top60_DNA_seed | IGF1R | -1.016984 | LGG | GI-1 | BMS-536924 | MVP_top60_DNA+RNA_seed | PIK3CA | 0.843485 | LUAD | NCI-H2347 | AZD8835 |
| MVP_top60_DNA_seed | PIK3CD | -1.016392 | LGG | MOG-G-CCM | Idelalisib | MVP_top60_DNA+RNA_seed | EGFR | 0.843616 | LUAD | HCC-44 | PF-00299804 |
| MVP_top60_DNA_seed | AR | -1.014078 | LGG | H4 | Bicalutamide | MVP_top60_DNA+RNA_seed | EGFR | 0.8443 | LUAD | Calu-6 | Cetuximab |
| MVP_top60_DNA_seed | MET | -1.010752 | LGG | GI-1 | AZD6094 | MVP_top60_DNA+RNA_seed | HDAC1 | 0.845933 | LUAD | NCI-H2122 | AR-42 |
| MVP_top60_DNA_seed | HDAC1 | -0.973345 | LGG | GI-1 | Tubastatin A | MVP_top60_DNA+RNA_seed | BRAF | 0.847373 | LUAD | SK-LU-1 | SB590885 |
| MVP_top60_DNA_seed | CDK2 | -0.970511 | LGG | GI-1 | AZD5438 | MVP_top60_DNA+RNA_seed | SMARCA4 | 0.848365 | LUAD | NCI-H1573 | PFI-3 |
| MVP_top60_DNA_seed | HSP90AA1 | -0.950865 | LGG | GI-1 | Elesclomol | MVP_top60_DNA+RNA_seed | HDAC1 | 0.849077 | LUAD | NCI-H2122 | Belinostat |
| MVP_top60_DNA_seed | MET | -0.950305 | LGG | SW1783 | SU11274 | MVP_top60_DNA+RNA_seed | MET | 0.849903 | LUAD | NCI-H1944 | SU11274 |
| MVP_top60_DNA_seed | NTRK1 | -0.942738 | LGG | GI-1 | GW441756 | MVP_top60_DNA+RNA_seed | IGF1R | 0.850172 | LUAD | RERF-LC-MS | Linsitinib |
| MVP_top60_DNA_seed | JAK3 | -0.942255 | LGG | MOG-G-UVW | JAK3_7406 | MVP_top60_DNA+RNA_seed | SMARCA4 | 0.850386 | LUAD | NCI-H2023 | PFI-3 |
| MVP_top60_DNA_seed | CDK2 | -0.927606 | LGG | SF539 | AZD5438 | MVP_top60_DNA+RNA_seed | AKT1 | 0.850466 | LUAD | NCI-H2087 | GSK690693 |
| MVP_top60_DNA_seed | MET | -0.926933 | LGG | D-336MG | JNJ38877605 | MVP_top60_DNA+RNA_seed | AURKA | 0.85192 | LUAD | NCI-H522 | GSK1070916 |
| MVP_top60_DNA_seed | IGF1R | -0.918217 | LGG | D-502MG | GSK1904529A | MVP_top60_DNA+RNA_seed | MET | 0.85194 | LUAD | PC-3_[JPC-3] | AZD6094 |
| MVP_top60_DNA_seed | JAK2 | -0.914868 | LGG | GI-1 | Ruxolitinib | MVP_top60_DNA+RNA_seed | MET | 0.85233 | LUAD | NCI-H1568 | SU11274 |
| MVP_top60_DNA_seed | JAK1 | -0.914868 | LGG | GI-1 | Ruxolitinib | MVP_top60_DNA+RNA_seed | MDM2 | 0.85307 | LUAD | A549 | Serdemetan |
| MVP_top60_DNA_seed | MDM2 | -0.905177 | LGG | SF539 | Serdemetan | MVP_top60_DNA+RNA_seed | EGFR | 0.854444 | LUAD | NCI-H522 | Cetuximab |
| MVP_top60_DNA_seed | HSP90AA1 | -0.902214 | LGG | MOG-G-CCM | SNX-2112 | MVP_top60_DNA+RNA_seed | EGFR | 0.856371 | LUAD | NCI-H1435 | Pelitinib |

| | | | | | | | | | | | |
|---|---|---|---|---|---|---|---|---|---|---|---|
| MVP_top60_DNA_seed | JAK3 | -0.893329 | LGG | KALS-1 | JAK3_7406 | MVP_top60_DNA+RNA_seed | EGFR | 0.858042 | LUAD | RERF-LC-MS | CUDC-101 |
| MVP_top60_DNA_seed | JAK1 | -0.886491 | LGG | KNS-42 | JAK1_3715 | MVP_top60_DNA+RNA_seed | HSP90AA1 | 0.858644 | LUAD | NCI-H1650 | SNX-2112 |
| MVP_top60_DNA_seed | CDK2 | -0.881538 | LGG | NMC-G1 | Seliciclib | MVP_top60_DNA+RNA_seed | BRAF | 0.858686 | LUAD | NCI-H322M | PLX-4720 |
| MVP_top60_DNA_seed | AR | -0.875852 | LGG | Hs-683 | Bicalutamide | MVP_top60_DNA+RNA_seed | AKT1 | 0.859001 | LUAD | PC-14 | AKT inhibitor VIII |
| MVP_top60_DNA_seed | CDK2 | -0.875407 | LGG | GAMG | PHA-793887 | MVP_top60_DNA+RNA_seed | HDAC1 | 0.859444 | LUAD | SK-LU-1 | AR-42 |
| MVP_top60_DNA_seed | HDAC1 | -0.867292 | LGG | GAMG | Belinostat | MVP_top60_DNA+RNA_seed | BRAF | 0.860084 | LUAD | PC-3_[JPC-3] | Dabrafenib |
| MVP_top60_DNA_seed | AR | -0.857091 | LGG | SF539 | AZD3514 | MVP_top60_DNA+RNA_seed | PIK3CA | 0.860304 | LUAD | HOP-62 | PI3Ka_4409 |
| MVP_top60_DNA_seed | HSP90AA1 | -0.855757 | LGG | KINGS-1 | SNX-2112 | MVP_top60_DNA+RNA_seed | HDAC1 | 0.8607 | LUAD | NCI-H838 | Dacinostat |
| MVP_top60_DNA_seed | HDAC1 | -0.853513 | LGG | SF539 | Tubastatin A | MVP_top60_DNA+RNA_seed | SRC | 0.861088 | LUAD | NCI-H1734 | Bosutinib |
| MVP_top60_DNA_seed | MDM2 | -0.845948 | LGG | KNS-81-FD | Nutlin-3a (-) | MVP_top60_DNA+RNA_seed | MET | 0.863324 | LUAD | A549 | SU11274 |
| MVP_top60_DNA_seed | AR | -0.843731 | LGG | GAMG | Bicalutamide | MVP_top60_DNA+RNA_seed | FGFR1 | 0.864414 | LUAD | NCI-H1666 | AZD4547 |
| MVP_top60_DNA_seed | HDAC1 | -0.84248 | LGG | GI-1 | CAY10603 | MVP_top60_DNA+RNA_seed | AKT1 | 0.864507 | LUAD | VMRC-LCD | BAY AKT1 |
| MVP_top60_DNA_seed | HDAC1 | -0.838031 | LGG | MOG-G-UVW | CAY10603 | MVP_top60_DNA+RNA_seed | AURKA | 0.865915 | LUAD | RERF-LC-MS | ZM447439 |
| MVP_top60_DNA_seed | HSP90AA1 | -0.818092 | LGG | GI-1 | CCT-018159 | MVP_top60_DNA+RNA_seed | EGFR | 0.866724 | LUAD | NCI-H1573 | CI-1033 |
| MVP_top60_DNA_seed | HDAC1 | -0.816934 | LGG | GI-1 | Belinostat | MVP_top60_DNA+RNA_seed | MET | 0.867834 | LUAD | Calu-6 | PHA-665752 |
| MVP_top60_DNA_seed | HSP90AA1 | -0.815214 | LGG | H4 | Luminespib | MVP_top60_DNA+RNA_seed | HDAC1 | 0.86841 | LUAD | NCI-H1755 | Dacinostat |
| MVP_top60_DNA_seed | PDGFRB | -0.808254 | LGG | D-502MG | Pazopanib | MVP_top60_DNA+RNA_seed | AKT1 | 0.868454 | LUAD | NCI-H1755 | GSK690693 |
| MVP_top60_DNA_seed | PDGFRA | -0.798891 | LGG | GAMG | Amuvatinib | MVP_top60_DNA+RNA_seed | SRC | 0.869707 | LUAD | H3255 | NVP-BHG712 |
| MVP_top60_DNA_seed | HSP90AA1 | -0.798756 | LGG | GAMG | CCT-018159 | MVP_top60_DNA+RNA_seed | FGFR1 | 0.870663 | LUAD | RERF-LC-MS | PD173074 |
| MVP_top60_DNA_seed | HSP90AA1 | -0.792778 | LGG | SW1783 | CCT-018159 | MVP_top60_DNA+RNA_seed | AURKA | 0.872693 | LUAD | NCI-H322M | GSK1070916 |
| MVP_top60_DNA_seed | HSF1 | -0.792226 | LGG | GAMG | CCT245232 | MVP_top60_DNA+RNA_seed | EGFR | 0.872763 | LUAD | SK-LU-1 | Cetuximab |
| MVP_top60_DNA_seed | HDAC1 | -0.784465 | LGG | GAMG | CAY10603 | MVP_top60_DNA+RNA_seed | AKT1 | 0.87412 | LUAD | NCI-H441 | AKT inhibitor VIII |
| MVP_top60_DNA_seed | NTRK1 | -0.781804 | LGG | GI-1 | AZD1332 | MVP_top60_DNA+RNA_seed | FGFR1 | 0.874736 | LUAD | HCC-827 | AZD4547 |
| MVP_top60_DNA_seed | AR | -0.779093 | LGG | SF539 | Bicalutamide | MVP_top60_DNA+RNA_seed | EGFR | 0.87522 | LUAD | EMC-BAC-2 | Gefitinib |
| MVP_top60_DNA_seed | PDGFRB | -0.778535 | LGG | no-11 | Pazopanib | MVP_top60_DNA+RNA_seed | PTK2 | 0.875995 | LUAD | NCI-H2342 | PF-562271 |
| MVP_top60_DNA_seed | MDM2 | -0.775908 | LGG | NMC-G1 | Serdemetan | MVP_top60_DNA+RNA_seed | EGFR | 0.876815 | LUAD | NCI-H23 | Pelitinib |
| MVP_top60_DNA_seed | CDK2 | -0.768992 | LGG | no-10 | PHA-793887 | MVP_top60_DNA+RNA_seed | ATM | 0.877733 | LUAD | NCI-H2087 | KU-55933 |
| MVP_top60_DNA_seed | MDM2 | -0.764898 | LGG | Hs-683 | Serdemetan | MVP_top60_DNA+RNA_seed | SRC | 0.878006 | LUAD | NCI-H1573 | Bosutinib |
| MVP_top60_DNA_seed | HSP90AA1 | -0.763827 | LGG | KNS-81-FD | Elesclomol | MVP_top60_DNA+RNA_seed | EGFR | 0.878229 | LUAD | NCI-H1437 | Afatinib |
| MVP_top60_DNA_seed | HSP90AA1 | -0.757755 | LGG | KALS-1 | Luminespib | MVP_top60_DNA+RNA_seed | EGFR | 0.879079 | LUAD | HOP-62 | Gefitinib |
| MVP_top60_DNA_seed | NTRK1 | -0.74856 | LGG | MOG-G-CCM | AZD1332 | MVP_top60_DNA+RNA_seed | HSP90AA1 | 0.879398 | LUAD | NCI-H441 | Tanespimycin |
| MVP_top60_DNA_seed | HSP90AA1 | -0.744342 | LGG | SF539 | CCT-018159 | MVP_top60_DNA+RNA_seed | EGFR | 0.879466 | LUAD | NCI-H1651 | CUDC-101 |
| MVP_top60_DNA_seed | MET | -0.743452 | LGG | GI-1 | SU11274 | MVP_top60_DNA+RNA_seed | MET | 0.880069 | LUAD | NCI-H1666 | AZD6094 |
| MVP_top60_DNA_seed | EGFR | -0.738873 | LGG | D-336MG | Lapatinib | MVP_top60_DNA+RNA_seed | MET | 0.880363 | LUAD | NCI-H1975 | PHA-665752 |
| MVP_top60_DNA_seed | MET | -0.720238 | LGG | SW1783 | AZD6094 | MVP_top60_DNA+RNA_seed | EGFR | 0.881841 | LUAD | NCI-H1975 | Gefitinib |
| MVP_top60_DNA_seed | PIK3CD | -0.711306 | LGG | D-336MG | Idelalisib | MVP_top60_DNA+RNA_seed | SRC | 0.882261 | LUAD | HCC-44 | Bosutinib |
| MVP_top60_DNA_seed | HDAC1 | -0.711275 | LGG | GAMG | JQ12 | MVP_top60_DNA+RNA_seed | AKT1 | 0.883131 | LUAD | NCI-H1623 | GSK690693 |
| MVP_top60_DNA_seed | EGFR | -0.710063 | LGG | Hs-683 | Cetuximab | MVP_top60_DNA+RNA_seed | IGF1R | 0.883846 | LUAD | NCI-H1435 | Linsitinib |
| MVP_top60_DNA_seed | MET | -0.705145 | LGG | SF539 | JNJ38877605 | MVP_top60_DNA+RNA_seed | FGFR1 | 0.884137 | LUAD | NCI-H2009 | PD173074 |
| MVP_top60_DNA_seed | PIK3CA | -0.698952 | LGG | H4 | AZD8835 | MVP_top60_DNA+RNA_seed | BRAF | 0.884575 | LUAD | HCC-78 | Dabrafenib |
| MVP_top60_DNA_seed | PIK3CD | -0.698952 | LGG | H4 | AZD8835 | MVP_top60_DNA+RNA_seed | AURKA | 0.885289 | LUAD | Calu-6 | GSK1070916 |

| | | | | | | | | | | | |
|---|---|---|---|---|---|---|---|---|---|---|---|
| MVP_top60_DNA_seed | IGF1R | -0.696871 | LGG | KNS-81-FD | GSK1904529A | MVP_top60_DNA+RNA_seed | SMARCA4 | 0.889656 | LUAD | NCI-H2347 | PFI-3 |
| MVP_top60_DNA_seed | CREBBP | -0.695825 | LGG | GI-1 | I-CBP112 | MVP_top60_DNA+RNA_seed | ATM | 0.890839 | LUAD | SW1573 | KU-60019 |
| MVP_top60_DNA_seed | MDM2 | -0.695641 | LGG | D-336MG | Serdemetan | MVP_top60_DNA+RNA_seed | BRAF | 0.891201 | LUAD | NCI-H1838 | SB590885 |
| MVP_top60_DNA_seed | HSP90AA1 | -0.690561 | LGG | GI-1 | Tanespimycin | MVP_top60_DNA+RNA_seed | EGFR | 0.891446 | LUAD | NCI-H1435 | AZD8931 |
| MVP_top60_DNA_seed | IGF1R | -0.685312 | LGG | NMC-G1 | BMS-536924 | MVP_top60_DNA+RNA_seed | EGFR | 0.892561 | LUAD | NCI-H1993 | CI-1033 |
| MVP_top60_DNA_seed | PIK3CA | -0.68251 | LGG | GAMG | AZD8835 | MVP_top60_DNA+RNA_seed | BRAF | 0.892994 | LUAD | NCI-H2342 | Dabrafenib |
| MVP_top60_DNA_seed | PIK3CD | -0.68251 | LGG | GAMG | AZD8835 | MVP_top60_DNA+RNA_seed | EGFR | 0.893364 | LUAD | NCI-H1650 | Pelitinib |
| MVP_top60_DNA_seed | HDAC1 | -0.680673 | LGG | GI-1 | AR-42 | MVP_top60_DNA+RNA_seed | AURKA | 0.894025 | LUAD | NCI-H1650 | Alisertib |
| MVP_top60_DNA_seed | PIK3CD | -0.674496 | LGG | MOG-G-UVW | AZD8186 | MVP_top60_DNA+RNA_seed | AKT1 | 0.894946 | LUAD | NCI-H1792 | MK-2206 |
| MVP_top60_DNA_seed | EGFR | -0.674168 | LGG | GAMG | Gefitinib | MVP_top60_DNA+RNA_seed | BRAF | 0.89712 | LUAD | NCI-H2405 | PLX-4720 |
| MVP_top60_DNA_seed | JAK3 | -0.670725 | LGG | no-10 | JAK3_7406 | MVP_top60_DNA+RNA_seed | AKT1 | 0.89815 | LUAD | NCI-H2009 | GSK690693 |
| MVP_top60_DNA_seed | JAK2 | -0.663429 | LGG | SF539 | AZD1480 | MVP_top60_DNA+RNA_seed | MET | 0.89904 | LUAD | NCI-H1755 | SU11274 |
| MVP_top60_DNA_seed | JAK1 | -0.663429 | LGG | SF539 | AZD1480 | MVP_top60_DNA+RNA_seed | FGFR1 | 0.89994 | LUAD | A549 | AZD4547 |
| MVP_top60_DNA_seed | HSP90AA1 | -0.662646 | LGG | KALS-1 | CCT-018159 | MVP_top60_DNA+RNA_seed | HDAC1 | 0.900801 | LUAD | SK-LU-1 | Tubastatin A |
| MVP_top60_DNA_seed | PDGFRB | -0.662473 | LGG | SF539 | Masitinib | MVP_top60_DNA+RNA_seed | FGFR1 | 0.901556 | LUAD | NCI-H1648 | AZD4547 |
| MVP_top60_DNA_seed | PDGFRA | -0.662473 | LGG | SF539 | Masitinib | MVP_top60_DNA+RNA_seed | IGF1R | 0.9033 | LUAD | NCI-H23 | Linsitinib |
| MVP_top60_DNA_seed | AR | -0.660166 | LGG | SF539 | Bicalutamide | MVP_top60_DNA+RNA_seed | HDAC1 | 0.903477 | LUAD | NCI-H1781 | JQ12 |
| MVP_top60_DNA_seed | HDAC1 | -0.659762 | LGG | GI-1 | Parthenolide | MVP_top60_DNA+RNA_seed | BRAF | 0.903761 | LUAD | NCI-H1435 | HG6-64-1 |
| MVP_top60_DNA_seed | HDAC1 | -0.653611 | LGG | MOG-G-UVW | Belinostat | MVP_top60_DNA+RNA_seed | HDAC1 | 0.904536 | LUAD | NCI-H1666 | Parthenolide |
| MVP_top60_DNA_seed | PIK3CD | -0.653305 | LGG | GAMG | Idelalisib | MVP_top60_DNA+RNA_seed | AKT1 | 0.904923 | LUAD | NCI-H1568 | MK-2206 |
| MVP_top60_DNA_seed | HDAC1 | -0.653237 | LGG | GAMG | AR-42 | MVP_top60_DNA+RNA_seed | AKT1 | 0.905941 | LUAD | NCI-H2342 | GSK690693 |
| MVP_top60_DNA_seed | NTRK1 | -0.652068 | LGG | H4 | AZD1332 | MVP_top60_DNA+RNA_seed | HSP90AA1 | 0.907162 | LUAD | H3255 | Elesclomol |
| MVP_top60_DNA_seed | AR | -0.648984 | LGG | GAMG | Bicalutamide | MVP_top60_DNA+RNA_seed | BRAF | 0.907875 | LUAD | COR-L105 | Dabrafenib |
| MVP_top60_DNA_seed | HSP90AA1 | -0.648449 | LGG | NMC-G1 | Luminespib | MVP_top60_DNA+RNA_seed | ATM | 0.908892 | LUAD | NCI-H1838 | KU-60019 |
| MVP_top60_DNA_seed | EGFR | -0.636895 | LGG | GI-1 | CUDC-101 | MVP_top60_DNA+RNA_seed | BRAF | 0.90941 | LUAD | NCI-H1355 | PLX-4720 |
| MVP_top60_DNA_seed | JAK2 | -0.635507 | LGG | D-336MG | Fedratinib | MVP_top60_DNA+RNA_seed | ATM | 0.911352 | LUAD | NCI-H441 | CP466722 |
| MVP_top60_DNA_seed | EGFR | -0.631103 | LGG | GAMG | Afatinib | MVP_top60_DNA+RNA_seed | BRAF | 0.911676 | LUAD | NCI-H1435 | SB590885 |
| MVP_top60_DNA_seed | EGFR | -0.629785 | LGG | SF539 | Lapatinib | MVP_top60_DNA+RNA_seed | FGFR1 | 0.912527 | LUAD | NCI-H2405 | PD173074 |
| MVP_top60_DNA_seed | EGFR | -0.627747 | LGG | NMC-G1 | Cetuximab | MVP_top60_DNA+RNA_seed | EGFR | 0.913877 | LUAD | NCI-H2122 | Gefitinib |
| MVP_top60_DNA_seed | MDM2 | -0.624707 | LGG | no-10 | Serdemetan | MVP_top60_DNA+RNA_seed | HDAC1 | 0.914686 | LUAD | H3255 | CAY10603 |
| MVP_top60_DNA_seed | HDAC1 | -0.618569 | LGG | Hs-683 | JQ12 | MVP_top60_DNA+RNA_seed | IGF1R | 0.914868 | LUAD | NCI-H1693 | Linsitinib |
| MVP_top60_DNA_seed | AR | -0.614628 | LGG | GI-1 | Bicalutamide | MVP_top60_DNA+RNA_seed | AURKA | 0.916021 | LUAD | NCI-H2405 | CD532 |
| MVP_top60_DNA_seed | AR | -0.612393 | LGG | GI-1 | AZD3514 | MVP_top60_DNA+RNA_seed | EGFR | 0.916182 | LUAD | NCI-H1993 | Gefitinib |
| MVP_top60_DNA_seed | MDM2 | -0.603874 | LGG | NMC-G1 | Nutlin-3a (-) | MVP_top60_DNA+RNA_seed | FGFR1 | 0.916441 | LUAD | NCI-H2342 | PD173074 |
| MVP_top60_DNA_seed | HDAC1 | -0.600371 | LGG | MOG-G-UVW | AR-42 | MVP_top60_DNA+RNA_seed | MET | 0.916958 | LUAD | NCI-H1568 | JNJ38877605 |
| MVP_top60_DNA_seed | EGFR | -0.597681 | LGG | no-11 | Erlotinib | MVP_top60_DNA+RNA_seed | AURKA | 0.918815 | LUAD | EKVX | CD532 |
| MVP_top60_DNA_seed | PDGFRB | -0.594749 | LGG | GI-1 | Masitinib | MVP_top60_DNA+RNA_seed | FGFR1 | 0.919179 | LUAD | NCI-H2087 | FGFR_3831 |
| MVP_top60_DNA_seed | PDGFRA | -0.594749 | LGG | GI-1 | Masitinib | MVP_top60_DNA+RNA_seed | BRAF | 0.920028 | LUAD | Calu-3 | Dabrafenib |
| MVP_top60_DNA_seed | HSF1 | -0.594091 | LGG | Hs-683 | CCT245232 | MVP_top60_DNA+RNA_seed | EGFR | 0.920788 | LUAD | NCI-H1793 | PF-00299804 |
| MVP_top60_DNA_seed | JAK2 | -0.592728 | LGG | Hs-683 | AZD1480 | MVP_top60_DNA+RNA_seed | SRC | 0.921008 | LUAD | NCI-H1781 | NVP-BHG712 |
| MVP_top60_DNA_seed | JAK1 | -0.592728 | LGG | Hs-683 | AZD1480 | MVP_top60_DNA+RNA_seed | MET | 0.921273 | LUAD | NCI-H1437 | SU11274 |

| | | | | | | | | | | | |
|---|---|---|---|---|---|---|---|---|---|---|---|
| MVP_top60_DNA_seed | MDM2 | -0.588922 | LGG | SW1783 | Serdemetan | MVP_top60_DNA+RNA_seed | EGFR | 0.921467 | LUAD | SK-LU-1 | Afatinib |
| MVP_top60_DNA_seed | HSP90AA1 | -0.588698 | LGG | D-336MG | SNX-2112 | MVP_top60_DNA+RNA_seed | IGF1R | 0.921641 | LUAD | Calu-6 | BMS-536924 |
| MVP_top60_DNA_seed | IGF1R | -0.586355 | LGG | GI-1 | BMS-754807 | MVP_top60_DNA+RNA_seed | BRAF | 0.921815 | LUAD | NCI-H1568 | SB590885 |
| MVP_top60_DNA_seed | MET | -0.584473 | LGG | Hs-683 | JNJ38877605 | MVP_top60_DNA+RNA_seed | MAPK14 | 0.924628 | LUAD | NCI-H1650 | TAK-715 |
| MVP_top60_DNA_seed | HSF1 | -0.58396 | LGG | SF539 | CCT245232 | MVP_top60_DNA+RNA_seed | PIK3CA | 0.926399 | LUAD | NCI-H1573 | AZD8835 |
| MVP_top60_DNA_seed | JAK2 | -0.582256 | LGG | GI-1 | AZD1480 | MVP_top60_DNA+RNA_seed | HSP90AA1 | 0.926415 | LUAD | EKVX | Luminespib |
| MVP_top60_DNA_seed | JAK1 | -0.582256 | LGG | GI-1 | AZD1480 | MVP_top60_DNA+RNA_seed | BRAF | 0.926525 | LUAD | SW1573 | PLX-4720 |
| MVP_top60_DNA_seed | JAK2 | -0.582039 | LGG | NMC-G1 | AZD1480 | MVP_top60_DNA+RNA_seed | EGFR | 0.92702 | LUAD | NCI-H1437 | Afatinib |
| MVP_top60_DNA_seed | JAK1 | -0.582039 | LGG | NMC-G1 | AZD1480 | MVP_top60_DNA+RNA_seed | AKT1 | 0.927586 | LUAD | Calu-6 | GSK690693 |
| MVP_top60_DNA_seed | HDAC1 | -0.581607 | LGG | MOG-G-CCM | Belinostat | MVP_top60_DNA+RNA_seed | AURKA | 0.927754 | LUAD | RERF-LC-KJ | HG-5-113-01 |
| MVP_top60_DNA_seed | IGF1R | -0.580633 | LGG | H4 | BMS-536924 | MVP_top60_DNA+RNA_seed | AURKA | 0.92945 | LUAD | EKVX | ZM447439 |
| MVP_top60_DNA_seed | PIK3CD | -0.579846 | LGG | no-11 | AZD8186 | MVP_top60_DNA+RNA_seed | IGF1R | 0.930459 | LUAD | NCI-H2023 | GSK1904529A |
| MVP_top60_DNA_seed | NTRK1 | -0.574285 | LGG | KALS-1 | GW441756 | MVP_top60_DNA+RNA_seed | MET | 0.930646 | LUAD | EMC-BAC-2 | SU11274 |
| MVP_top60_DNA_seed | JAK2 | -0.572488 | LGG | D-336MG | AZD1480 | MVP_top60_DNA+RNA_seed | FGFR1 | 0.932094 | LUAD | PC-3_[JPC-3] | AZD4547 |
| MVP_top60_DNA_seed | JAK1 | -0.572488 | LGG | D-336MG | AZD1480 | MVP_top60_DNA+RNA_seed | HDAC1 | 0.932469 | LUAD | NCI-H2228 | Belinostat |
| MVP_top60_DNA_seed | HSF1 | -0.567149 | LGG | GI-1 | CCT245232 | MVP_top60_DNA+RNA_seed | SRC | 0.93282 | LUAD | NCI-H1573 | NVP-BHG712 |
| MVP_top60_DNA_seed | EGFR | -0.566105 | LGG | GAMG | CUDC-101 | MVP_top60_DNA+RNA_seed | AKT1 | 0.933064 | LUAD | NCI-H1573 | GSK690693 |
| MVP_top60_DNA_seed | CDK2 | -0.558458 | LGG | NMC-G1 | PHA-793887 | MVP_top60_DNA+RNA_seed | MET | 0.934686 | LUAD | COR-L105 | AZD6094 |
| MVP_top60_DNA_seed | JAK2 | -0.554215 | LGG | D-502MG | Fedratinib | MVP_top60_DNA+RNA_seed | EGFR | 0.935226 | LUAD | NCI-H322M | Cetuximab |
| MVP_top60_DNA_seed | IGF1R | -0.55156 | LGG | SW1783 | GSK1904529A | MVP_top60_DNA+RNA_seed | EGFR | 0.935275 | LUAD | NCI-H1650 | Lapatinib |
| MVP_top60_DNA_seed | HSP90AA1 | -0.548402 | LGG | Hs-683 | Luminespib | MVP_top60_DNA+RNA_seed | FGFR1 | 0.936659 | LUAD | ABC-1 | PD173074 |
| MVP_top60_DNA_seed | JAK2 | -0.547421 | LGG | H4 | Fedratinib | MVP_top60_DNA+RNA_seed | FGFR1 | 0.936928 | LUAD | NCI-H2347 | FGFR_3831 |
| MVP_top60_DNA_seed | EGFR | -0.541582 | LGG | KALS-1 | AZD8931 | MVP_top60_DNA+RNA_seed | EGFR | 0.93745 | LUAD | NCI-H1650 | CI-1033 |
| MVP_top60_DNA_seed | EGFR | -0.538609 | LGG | GI-1 | Cetuximab | MVP_top60_DNA+RNA_seed | HDAC1 | 0.937985 | LUAD | NCI-H1650 | Belinostat |
| MVP_top60_DNA_seed | JAK2 | -0.534685 | LGG | GI-1 | Fedratinib | MVP_top60_DNA+RNA_seed | AURKA | 0.938574 | LUAD | SW1573 | GSK1070916 |
| MVP_top60_DNA_seed | PIK3CA | -0.534211 | LGG | GI-1 | PI3Ka_4409 | MVP_top60_DNA+RNA_seed | IGF1R | 0.940568 | LUAD | NCI-H1650 | BMS-536924 |
| MVP_top60_DNA_seed | PIK3CD | -0.534211 | LGG | GI-1 | PI3Ka_4409 | MVP_top60_DNA+RNA_seed | AURKA | 0.940595 | LUAD | COR-L105 | Alisertib |
| MVP_top60_DNA_seed | AR | -0.533418 | LGG | SW1783 | Bicalutamide | MVP_top60_DNA+RNA_seed | AURKA | 0.940699 | LUAD | NCI-H1355 | CD532 |
| MVP_top60_DNA_seed | HDAC1 | -0.530973 | LGG | MOG-G-CCM | CAY10603 | MVP_top60_DNA+RNA_seed | EGFR | 0.941252 | LUAD | NCI-H2347 | Afatinib |
| MVP_top60_DNA_seed | PIK3CD | -0.529203 | LGG | MOG-G-UVW | Idelalisib | MVP_top60_DNA+RNA_seed | HDAC1 | 0.941503 | LUAD | NCI-H2342 | Tubastatin A |
| MVP_top60_DNA_seed | JAK3 | -0.527706 | LGG | KNS-42 | JAK3_7406 | MVP_top60_DNA+RNA_seed | BRAF | 0.942901 | LUAD | PC-3_[JPC-3] | PLX-4720 |
| MVP_top60_DNA_seed | IGF1R | -0.527631 | LGG | D-336MG | BMS-536924 | MVP_top60_DNA+RNA_seed | BRAF | 0.94359 | LUAD | NCI-H322M | Dabrafenib |
| MVP_top60_DNA_seed | HDAC1 | -0.527554 | LGG | KNS-42 | Tubastatin A | MVP_top60_DNA+RNA_seed | HDAC1 | 0.944524 | LUAD | NCI-H2405 | JQ12 |
| MVP_top60_DNA_seed | HDAC1 | -0.524953 | LGG | GI-1 | JQ12 | MVP_top60_DNA+RNA_seed | HDAC1 | 0.94491 | LUAD | NCI-H1573 | Tubastatin A |
| MVP_top60_DNA_seed | NTRK1 | -0.523561 | LGG | MOG-G-UVW | AZD1332 | MVP_top60_DNA+RNA_seed | HDAC1 | 0.945613 | LUAD | NCI-H1435 | Tubastatin A |
| MVP_top60_DNA_seed | JAK2 | -0.52326 | LGG | GAMG | Ruxolitinib | MVP_top60_DNA+RNA_seed | AKT1 | 0.945842 | LUAD | NCI-H2405 | GSK690693 |
| MVP_top60_DNA_seed | JAK1 | -0.52326 | LGG | GAMG | Ruxolitinib | MVP_top60_DNA+RNA_seed | FGFR1 | 0.946881 | LUAD | Calu-6 | PD173074 |
| MVP_top60_DNA_seed | IGF1R | -0.51883 | LGG | Hs-683 | GSK1904529A | MVP_top60_DNA+RNA_seed | HSP90AA1 | 0.948085 | LUAD | NCI-H322M | CCT-018159 |
| MVP_top60_DNA_seed | PIK3CD | -0.517926 | LGG | SW1783 | AZD8186 | MVP_top60_DNA+RNA_seed | IGF1R | 0.94814 | LUAD | HOP-62 | BMS-536924 |
| MVP_top60_DNA_seed | MAPK1 | -0.517859 | LGG | SF539 | FR-180204 | MVP_top60_DNA+RNA_seed | MET | 0.95018 | LUAD | NCI-H1355 | PHA-665752 |
| MVP_top60_DNA_seed | MAPK3 | -0.517859 | LGG | SF539 | FR-180204 | MVP_top60_DNA+RNA_seed | AURKA | 0.951142 | LUAD | NCI-H2405 | Alisertib |

| | | | | | | | | | | | |
|---|---|---|---|---|---|---|---|---|---|---|---|
| MVP_top60_DNA_seed | MET | -0.517158 | LGG | SW1783 | JNJ38877605 | MVP_top60_DNA+RNA_seed | EGFR | 0.952054 | LUAD | HCC-78 | Gefitinib |
| MVP_top60_DNA_seed | HSP90AA1 | -0.516897 | LGG | MOG-G-CCM | Tanespimycin | MVP_top60_DNA+RNA_seed | HDAC1 | 0.952524 | LUAD | NCI-H23 | Parthenolide |
| MVP_top60_DNA_seed | HSP90AA1 | -0.5153 | LGG | no-11 | Elesclomol | MVP_top60_DNA+RNA_seed | BRAF | 0.953817 | LUAD | NCI-H1563 | SB590885 |
| MVP_top60_DNA_seed | HDAC1 | -0.515299 | LGG | H4 | JQ12 | MVP_top60_DNA+RNA_seed | HDAC1 | 0.954009 | LUAD | NCI-H1792 | AR-42 |
| MVP_top60_DNA_seed | HDAC1 | -0.514579 | LGG | KINGS-1 | CAY10603 | MVP_top60_DNA+RNA_seed | SRC | 0.954527 | LUAD | NCI-H1623 | NVP-BHG712 |
| MVP_top60_DNA_seed | MDM2 | -0.511959 | LGG | MOG-G-UVW | Nutlin-3a (-) | MVP_top60_DNA+RNA_seed | MET | 0.954593 | LUAD | NCI-H2122 | SU11274 |
| MVP_top60_DNA_seed | HSP90AA1 | -0.511529 | LGG | MOG-G-CCM | Luminespib | MVP_top60_DNA+RNA_seed | ATM | 0.956946 | LUAD | NCI-H1693 | KU-55933 |
| MVP_top60_DNA_seed | IGF1R | -0.51114 | LGG | H4 | BMS-754807 | MVP_top60_DNA+RNA_seed | PTK2 | 0.956977 | LUAD | NCI-H1755 | PF-562271 |
| MVP_top60_DNA_seed | PDGFRA | -0.510975 | LGG | Hs-683 | Amuvatinib | MVP_top60_DNA+RNA_seed | BRAF | 0.957459 | LUAD | NCI-H838 | PLX-4720 |
| MVP_top60_DNA_seed | HDAC1 | -0.503689 | LGG | KINGS-1 | AR-42 | MVP_top60_DNA+RNA_seed | AKT1 | 0.957918 | LUAD | NCI-H2347 | GSK690693 |
| MVP_top60_DNA_seed | CDK2 | -0.5014 | LGG | SF539 | PHA-793887 | MVP_top60_DNA+RNA_seed | PIK3CA | 0.959083 | LUAD | NCI-H2087 | PI3Ka_4409 |
| MVP_top60_DNA_seed | EGFR | -0.496939 | LGG | D-502MG | Afatinib | MVP_top60_DNA+RNA_seed | AURKA | 0.959831 | LUAD | NCI-H1568 | Alisertib |
| MVP_top60_DNA_seed | PDGFRB | -0.496163 | LGG | H4 | Masitinib | MVP_top60_DNA+RNA_seed | ATM | 0.959983 | LUAD | SW1573 | KU-55933 |
| MVP_top60_DNA_seed | PDGFRA | -0.496163 | LGG | H4 | Masitinib | MVP_top60_DNA+RNA_seed | FGFR1 | 0.96094 | LUAD | NCI-H2023 | PD173074 |
| MVP_top60_DNA_seed | MDM2 | -0.494603 | LGG | MOG-G-UVW | Serdemetan | MVP_top60_DNA+RNA_seed | HDAC1 | 0.961118 | LUAD | SW1573 | Dacinostat |
| MVP_top60_DNA_seed | EGFR | -0.494186 | LGG | MOG-G-CCM | CUDC-101 | MVP_top60_DNA+RNA_seed | AURKA | 0.961411 | LUAD | NCI-H322M | ZM447439 |
| MVP_top60_DNA_seed | HDAC1 | -0.491805 | LGG | KALS-1 | Entinostat | MVP_top60_DNA+RNA_seed | EGFR | 0.962096 | LUAD | NCI-H1650 | AST-1306 |
| MVP_top60_DNA_seed | JAK2 | -0.490197 | LGG | MOG-G-CCM | AZD1480 | MVP_top60_DNA+RNA_seed | ATM | 0.964407 | LUAD | NCI-H1573 | KU-60019 |
| MVP_top60_DNA_seed | JAK1 | -0.490197 | LGG | MOG-G-CCM | AZD1480 | MVP_top60_DNA+RNA_seed | HDAC1 | 0.964964 | LUAD | LC-2-ad | Parthenolide |
| MVP_top60_DNA_seed | HDAC1 | -0.488706 | LGG | GAMG | Dacinostat | MVP_top60_DNA+RNA_seed | BRAF | 0.964994 | LUAD | NCI-H650 | RAF_9304 |
| MVP_top60_DNA_seed | IGF1R | -0.487432 | LGG | KINGS-1 | BMS-754807 | MVP_top60_DNA+RNA_seed | IGF1R | 0.966032 | LUAD | NCI-H1781 | Linsitinib |
| MVP_top60_DNA_seed | HSP90AA1 | -0.479586 | LGG | KNS-81-FD | Tanespimycin | MVP_top60_DNA+RNA_seed | EGFR | 0.966123 | LUAD | SK-LU-1 | Afatinib |
| MVP_top60_DNA_seed | EGFR | -0.47902 | LGG | MOG-G-CCM | Pelitinib | MVP_top60_DNA+RNA_seed | HDAC1 | 0.967447 | LUAD | NCI-H1792 | CAY10603 |
| MVP_top60_DNA_seed | HSP90AA1 | -0.47662 | LGG | MOG-G-UVW | Luminespib | MVP_top60_DNA+RNA_seed | BRAF | 0.967831 | LUAD | NCI-H838 | RAF_9304 |
| MVP_top60_DNA_seed | EGFR | -0.47516 | LGG | KINGS-1 | CUDC-101 | MVP_top60_DNA+RNA_seed | FGFR1 | 0.970911 | LUAD | NCI-H3122 | PD173074 |
| MVP_top60_DNA_seed | HSP90AA1 | -0.470347 | LGG | no-11 | SNX-2112 | MVP_top60_DNA+RNA_seed | AURKA | 0.973944 | LUAD | NCI-H838 | CD532 |
| MVP_top60_DNA_seed | PDGFRB | -0.469152 | LGG | SW1783 | Masitinib | MVP_top60_DNA+RNA_seed | HDAC1 | 0.97401 | LUAD | RERF-LC-KJ | Tubastatin A |
| MVP_top60_DNA_seed | PDGFRA | -0.469152 | LGG | SW1783 | Masitinib | MVP_top60_DNA+RNA_seed | HDAC1 | 0.974447 | LUAD | NCI-H2023 | CAY10603 |
| MVP_top60_DNA_seed | EGFR | -0.468754 | LGG | KNS-42 | Lapatinib | MVP_top60_DNA+RNA_seed | MET | 0.974448 | LUAD | NCI-H650 | AZD6094 |
| MVP_top60_DNA_seed | HSF1 | -0.467839 | LGG | KALS-1 | CCT245467 | MVP_top60_DNA+RNA_seed | AURKA | 0.974513 | LUAD | NCI-H2342 | CD532 |
| MVP_top60_DNA_seed | NTRK1 | -0.46451 | LGG | NMC-G1 | AZD1332 | MVP_top60_DNA+RNA_seed | FGFR1 | 0.975982 | LUAD | NCI-H2342 | FGFR_3831 |
| MVP_top60_DNA_seed | HSP90AA1 | -0.463572 | LGG | KALS-1 | Tanespimycin | MVP_top60_DNA+RNA_seed | BRAF | 0.97743 | LUAD | NCI-H2030 | PLX-4720 |
| MVP_top60_DNA_seed | NTRK1 | -0.461598 | LGG | D-336MG | AZD1332 | MVP_top60_DNA+RNA_seed | AURKA | 0.978179 | LUAD | NCI-H1781 | Alisertib |
| MVP_top60_DNA_seed | EGFR | -0.460047 | LGG | MOG-G-UVW | Afatinib | MVP_top60_DNA+RNA_seed | FGFR1 | 0.979736 | LUAD | NCI-H650 | AZD4547 |
| MVP_top60_DNA_seed | PIK3CD | -0.458074 | LGG | KALS-1 | Idelalisib | MVP_top60_DNA+RNA_seed | EGFR | 0.980707 | LUAD | RERF-LC-MS | PF-00299804 |
| MVP_top60_DNA_seed | JAK1 | -0.457744 | LGG | SF539 | JAK1_3715 | MVP_top60_DNA+RNA_seed | EGFR | 0.980802 | LUAD | Calu-6 | Gefitinib |
| MVP_top60_DNA_seed | EGFR | -0.456558 | LGG | MOG-G-UVW | CUDC-101 | MVP_top60_DNA+RNA_seed | EGFR | 0.981781 | LUAD | NCI-H2291 | AZD8931 |
| MVP_top60_DNA_seed | HSP90AA1 | -0.454939 | LGG | NMC-G1 | Elesclomol | MVP_top60_DNA+RNA_seed | AKT1 | 0.981845 | LUAD | A549 | GSK690693 |
| MVP_top60_DNA_seed | JAK2 | -0.450427 | LGG | MOG-G-UVW | AZD1480 | MVP_top60_DNA+RNA_seed | PTK2 | 0.983315 | LUAD | NCI-H441 | PF-562271 |
| MVP_top60_DNA_seed | JAK1 | -0.450427 | LGG | MOG-G-UVW | AZD1480 | MVP_top60_DNA+RNA_seed | HSP90AA1 | 0.98345 | LUAD | HCC-78 | SNX-2112 |
| MVP_top60_DNA_seed | JAK2 | -0.448755 | LGG | MOG-G-CCM | Ruxolitinib | MVP_top60_DNA+RNA_seed | PTK2 | 0.983587 | LUAD | PC-3_[JPC-3] | PF-562271 |

| | | | | | | | | | | | |
|---|---|---|---|---|---|---|---|---|---|---|---|
| MVP_top60_DNA_seed | JAK1 | -0.448755 | LGG | MOG-G-CCM | Ruxolitinib | MVP_top60_DNA+RNA_seed | PIK3CA | 0.984511 | LUAD | NCI-H3122 | AZD8835 |
| MVP_top60_DNA_seed | HSP90AA1 | -0.446186 | LGG | D-336MG | Elesclomol | MVP_top60_DNA+RNA_seed | AURKA | 0.984927 | LUAD | NCI-H1755 | CD532 |
| MVP_top60_DNA_seed | HSP90AA1 | -0.442126 | LGG | SW1783 | SNX-2112 | MVP_top60_DNA+RNA_seed | BRAF | 0.986934 | LUAD | Calu-3 | PLX-4720 |
| MVP_top60_DNA_seed | CDK2 | -0.439783 | LGG | no-10 | Seliciclib | MVP_top60_DNA+RNA_seed | HSP90AA1 | 0.987651 | LUAD | RERF-LC-KJ | SNX-2112 |
| MVP_top60_DNA_seed | HSP90AA1 | -0.436448 | LGG | H4 | SNX-2112 | MVP_top60_DNA+RNA_seed | MET | 0.98808 | LUAD | COR-L105 | JNJ38877605 |
| MVP_top60_DNA_seed | HSP90AA1 | -0.433679 | LGG | no-11 | Luminespib | MVP_top60_DNA+RNA_seed | EGFR | 0.988663 | LUAD | NCI-H1355 | CI-1033 |
| MVP_top60_DNA_seed | HSF1 | -0.431512 | LGG | D-336MG | CCT245232 | MVP_top60_DNA+RNA_seed | FGFR1 | 0.990549 | LUAD | NCI-H1651 | FGFR_3831 |
| MVP_top60_DNA_seed | PDGFRB | -0.425799 | LGG | NMC-G1 | Masitinib | MVP_top60_DNA+RNA_seed | AKT1 | 0.99159 | LUAD | NCI-H1693 | AKT inhibitor VIII |
| MVP_top60_DNA_seed | PDGFRA | -0.425799 | LGG | NMC-G1 | Masitinib | MVP_top60_DNA+RNA_seed | PIK3CA | 0.991887 | LUAD | NCI-H1781 | AZD8835 |
| MVP_top60_DNA_seed | MDM2 | -0.425364 | LGG | GAMG | Serdemetan | MVP_top60_DNA+RNA_seed | BRAF | 0.992247 | LUAD | NCI-H1781 | PLX-4720 |
| MVP_top60_DNA_seed | EGFR | -0.424855 | LGG | D-502MG | Pelitinib | MVP_top60_DNA+RNA_seed | EGFR | 0.992257 | LUAD | A549 | AST-1306 |
| MVP_top60_DNA_seed | HDAC1 | -0.42467 | LGG | MOG-G-UVW | Tubastatin A | MVP_top60_DNA+RNA_seed | MDM2 | 0.99241 | LUAD | NCI-H1781 | Nutlin-3a (-) |
| MVP_top60_DNA_seed | HSP90AA1 | -0.423306 | LGG | KNS-81-FD | CCT-018159 | MVP_top60_DNA+RNA_seed | FGFR1 | 0.992511 | LUAD | NCI-H2405 | FGFR_3831 |
| MVP_top60_DNA_seed | PIK3CD | -0.42126 | LGG | SW1783 | Idelalisib | MVP_top60_DNA+RNA_seed | AURKA | 0.992517 | LUAD | RERF-LC-MS | GSK1070916 |
| MVP_top60_DNA_seed | AR | -0.420813 | LGG | MOG-G-UVW | AZD3514 | MVP_top60_DNA+RNA_seed | AKT1 | 0.99275 | LUAD | NCI-H650 | GSK690693 |
| MVP_top60_DNA_seed | EGFR | -0.416291 | LGG | H4 | AZD8931 | MVP_top60_DNA+RNA_seed | BRAF | 0.993986 | LUAD | NCI-H1793 | RAF_9304 |
| MVP_top60_DNA_seed | IGF1R | -0.416285 | LGG | GI-1 | Linsitinib | MVP_top60_DNA+RNA_seed | IGF1R | 0.994898 | LUAD | NCI-H1650 | Linsitinib |
| MVP_top60_DNA_seed | HSF1 | -0.414379 | LGG | NMC-G1 | CCT245232 | MVP_top60_DNA+RNA_seed | EGFR | 0.99823 | LUAD | NCI-H2009 | Pelitinib |
| MVP_top60_DNA_seed | HDAC1 | -0.413486 | LGG | KINGS-1 | Belinostat | MVP_top60_DNA+RNA_seed | PIK3CA | 0.998522 | LUAD | NCI-H1838 | PI3Ka_4409 |
| MVP_top60_DNA_seed | PIK3CA | -0.411424 | LGG | D-336MG | PI3Ka_4409 | MVP_top60_DNA+RNA_seed | SRC | 1.000038 | LUAD | NCI-H1650 | WH-4-023 |
| MVP_top60_DNA_seed | PIK3CD | -0.411424 | LGG | D-336MG | PI3Ka_4409 | MVP_top60_DNA+RNA_seed | MDM2 | 1.000742 | LUAD | NCI-H1563 | Serdemetan |
| MVP_top60_DNA_seed | HSP90AA1 | -0.407872 | LGG | SF539 | Elesclomol | MVP_top60_DNA+RNA_seed | AURKA | 1.000827 | LUAD | NCI-H1568 | HG-5-113-01 |
| MVP_top60_DNA_seed | NTRK1 | -0.407174 | LGG | Hs-683 | AZD1332 | MVP_top60_DNA+RNA_seed | FGFR1 | 1.001329 | LUAD | NCI-H1568 | PD173074 |
| MVP_top60_DNA_seed | HSP90AA1 | -0.403609 | LGG | SF539 | Luminespib | MVP_top60_DNA+RNA_seed | AURKA | 1.002058 | LUAD | NCI-H2342 | ZM447439 |
| MVP_top60_DNA_seed | IGF1R | -0.394485 | LGG | SW1783 | Linsitinib | MVP_top60_DNA+RNA_seed | HSP90AA1 | 1.002477 | LUAD | NCI-H1650 | CCT-018159 |
| MVP_top60_DNA_seed | PIK3CD | -0.393757 | LGG | SW1783 | IC-87114 | MVP_top60_DNA+RNA_seed | FGFR1 | 1.003867 | LUAD | NCI-H1435 | FGFR_3831 |
| MVP_top60_DNA_seed | HSP90AA1 | -0.387926 | LGG | KINGS-1 | Tanespimycin | MVP_top60_DNA+RNA_seed | ATM | 1.004184 | LUAD | NCI-H2405 | KU-55933 |
| MVP_top60_DNA_seed | HDAC1 | -0.384041 | LGG | no-10 | Entinostat | MVP_top60_DNA+RNA_seed | EGFR | 1.004463 | LUAD | NCI-H1838 | Lapatinib |
| MVP_top60_DNA_seed | HSP90AA1 | -0.383438 | LGG | D-502MG | Elesclomol | MVP_top60_DNA+RNA_seed | EGFR | 1.004601 | LUAD | NCI-H2347 | Gefitinib |
| MVP_top60_DNA_seed | JAK2 | -0.37246 | LGG | SW1783 | AZD1480 | MVP_top60_DNA+RNA_seed | FGFR1 | 1.005685 | LUAD | NCI-H2347 | PD173074 |
| MVP_top60_DNA_seed | JAK1 | -0.37246 | LGG | SW1783 | AZD1480 | MVP_top60_DNA+RNA_seed | FGFR1 | 1.005693 | LUAD | A549 | PD173074 |
| MVP_top60_DNA_seed | MET | -0.371562 | LGG | D-336MG | AZD6094 | MVP_top60_DNA+RNA_seed | IGF1R | 1.006802 | LUAD | Calu-3 | BMS-754807 |
| MVP_top60_DNA_seed | SMARCA4 | -0.369949 | LGG | SF539 | PFI-3 | MVP_top60_DNA+RNA_seed | BRAF | 1.008448 | LUAD | RERF-LC-MS | HG6-64-1 |
| MVP_top60_DNA_seed | HDAC1 | -0.368404 | LGG | MOG-G-CCM | Tubastatin A | MVP_top60_DNA+RNA_seed | MET | 1.008848 | LUAD | COR-L105 | SU11274 |
| MVP_top60_DNA_seed | PIK3CA | -0.367911 | LGG | D-336MG | AZD8835 | MVP_top60_DNA+RNA_seed | MET | 1.009157 | LUAD | NCI-H1838 | SU11274 |
| MVP_top60_DNA_seed | PIK3CD | -0.367911 | LGG | D-336MG | AZD8835 | MVP_top60_DNA+RNA_seed | AURKA | 1.010572 | LUAD | NCI-H1650 | Genentech Cpd 10 |
| MVP_top60_DNA_seed | PIK3CA | -0.365229 | LGG | GI-1 | AZD8835 | MVP_top60_DNA+RNA_seed | BRAF | 1.010652 | LUAD | NCI-H1563 | PLX-4720 |
| MVP_top60_DNA_seed | PIK3CD | -0.365229 | LGG | GI-1 | AZD8835 | MVP_top60_DNA+RNA_seed | AURKA | 1.011255 | LUAD | NCI-H1755 | Alisertib |
| MVP_top60_DNA_seed | IGF1R | -0.361601 | LGG | KNS-42 | GSK1904529A | MVP_top60_DNA+RNA_seed | MDM2 | 1.01134 | LUAD | PC-3_[JPC-3] | Serdemetan |
| MVP_top60_DNA_seed | PDGFRB | -0.359974 | LGG | MOG-G-UVW | Masitinib | MVP_top60_DNA+RNA_seed | PTK2 | 1.014873 | LUAD | NCI-H23 | PF-562271 |
| MVP_top60_DNA_seed | PDGFRA | -0.359974 | LGG | MOG-G-UVW | Masitinib | MVP_top60_DNA+RNA_seed | EGFR | 1.015058 | LUAD | ABC-1 | Afatinib |

| | | | | | | | | | | | |
|---|---|---|---|---|---|---|---|---|---|---|---|
| MVP_top60_DNA_seed | HSP90AA1 | -0.356961 | LGG | KINGS-1 | Luminespib | MVP_top60_DNA+RNA_seed | EGFR | 1.015762 | LUAD | NCI-H650 | Gefitinib |
| MVP_top60_DNA_seed | PDGFRB | -0.354652 | LGG | KINGS-1 | Masitinib | MVP_top60_DNA+RNA_seed | EGFR | 1.015941 | LUAD | NCI-H1563 | Gefitinib |
| MVP_top60_DNA_seed | PDGFRA | -0.354652 | LGG | KINGS-1 | Masitinib | MVP_top60_DNA+RNA_seed | BRAF | 1.016349 | LUAD | NCI-H441 | HG6-64-1 |
| MVP_top60_DNA_seed | HSP90AA1 | -0.351225 | LGG | SF539 | SNX-2112 | MVP_top60_DNA+RNA_seed | IGF1R | 1.017024 | LUAD | NCI-H1573 | BMS-754807 |
| MVP_top60_DNA_seed | EGFR | -0.350702 | LGG | SF539 | AZD8931 | MVP_top60_DNA+RNA_seed | AKT1 | 1.017046 | LUAD | VMRC-LCD | MK-2206 |
| MVP_top60_DNA_seed | HSF1 | -0.340787 | LGG | Hs-683 | CCT245467 | MVP_top60_DNA+RNA_seed | MET | 1.017406 | LUAD | NCI-H1435 | SU11274 |
| MVP_top60_DNA_seed | EGFR | -0.340037 | LGG | D-336MG | CI-1033 | MVP_top60_DNA+RNA_seed | MAPK14 | 1.019215 | LUAD | NCI-H838 | TAK-715 |
| MVP_top60_DNA_seed | HSF1 | -0.33906 | LGG | SW1783 | CCT245467 | MVP_top60_DNA+RNA_seed | AKT1 | 1.020157 | LUAD | NCI-H1435 | GSK690693 |
| MVP_top60_DNA_seed | AR | -0.336886 | LGG | MOG-G-UVW | Bicalutamide | MVP_top60_DNA+RNA_seed | AKT1 | 1.020942 | LUAD | RERF-LC-KJ | BAY AKT1 |
| MVP_top60_DNA_seed | HDAC1 | -0.336253 | LGG | MOG-G-CCM | AR-42 | MVP_top60_DNA+RNA_seed | HSP90AA1 | 1.024053 | LUAD | NCI-H2023 | SNX-2112 |
| MVP_top60_DNA_seed | EGFR | -0.335629 | LGG | D-336MG | AST-1306 | MVP_top60_DNA+RNA_seed | MDM2 | 1.025251 | LUAD | NCI-H1623 | Nutlin-3a (-) |
| MVP_top60_DNA_seed | NTRK1 | -0.333787 | LGG | SW1783 | AZD1332 | MVP_top60_DNA+RNA_seed | EGFR | 1.025671 | LUAD | NCI-H1650 | Erlotinib |
| MVP_top60_DNA_seed | CREBBP | -0.327768 | LGG | KNS-42 | I-CBP112 | MVP_top60_DNA+RNA_seed | HDAC1 | 1.026522 | LUAD | H3255 | AR-42 |
| MVP_top60_DNA_seed | CDK2 | -0.3277 | LGG | H4 | AZD5438 | MVP_top60_DNA+RNA_seed | MAPK14 | 1.028325 | LUAD | NCI-H1435 | TAK-715 |
| MVP_top60_DNA_seed | EGFR | -0.327108 | LGG | NMC-G1 | Pelitinib | MVP_top60_DNA+RNA_seed | HDAC1 | 1.029931 | LUAD | RERF-LC-MS | Tubastatin A |
| MVP_top60_DNA_seed | MDM2 | -0.323644 | LGG | KALS-1 | Nutlin-3a (-) | MVP_top60_DNA+RNA_seed | ATM | 1.030683 | LUAD | EKVX | KU-60019 |
| MVP_top60_DNA_seed | EGFR | -0.323543 | LGG | D-336MG | PF-00299804 | MVP_top60_DNA+RNA_seed | HDAC1 | 1.032082 | LUAD | NCI-H2122 | CAY10603 |
| MVP_top60_DNA_seed | EGFR | -0.317082 | LGG | MOG-G-UVW | Cetuximab | MVP_top60_DNA+RNA_seed | EGFR | 1.032886 | LUAD | A549 | PF-00299804 |
| MVP_top60_DNA_seed | HSP90AA1 | -0.314117 | LGG | NMC-G1 | Tanespimycin | MVP_top60_DNA+RNA_seed | BRAF | 1.033739 | LUAD | NCI-H1623 | SB590885 |
| MVP_top60_DNA_seed | NTRK1 | -0.310877 | LGG | H4 | GW441756 | MVP_top60_DNA+RNA_seed | MET | 1.034567 | LUAD | NCI-H1563 | SU11274 |
| MVP_top60_DNA_seed | PIK3CD | -0.308468 | LGG | NMC-G1 | IC-87114 | MVP_top60_DNA+RNA_seed | AKT1 | 1.035559 | LUAD | SK-LU-1 | GSK690693 |
| MVP_top60_DNA_seed | JAK2 | -0.30663 | LGG | SF539 | Fedratinib | MVP_top60_DNA+RNA_seed | BRAF | 1.035902 | LUAD | Calu-3 | SB590885 |
| MVP_top60_DNA_seed | EGFR | -0.306079 | LGG | SW1783 | AST-1306 | MVP_top60_DNA+RNA_seed | ATM | 1.03711 | LUAD | NCI-H1666 | CP466722 |
| MVP_top60_DNA_seed | SMARCA4 | -0.303311 | LGG | Hs-683 | PFI-3 | MVP_top60_DNA+RNA_seed | AURKA | 1.039014 | LUAD | LC-2-ad | CD532 |
| MVP_top60_DNA_seed | HDAC1 | -0.301433 | LGG | NMC-G1 | Tubastatin A | MVP_top60_DNA+RNA_seed | EGFR | 1.04033 | LUAD | NCI-H2347 | AZD8931 |
| MVP_top60_DNA_seed | IGF1R | -0.299929 | LGG | D-502MG | BMS-536924 | MVP_top60_DNA+RNA_seed | AKT1 | 1.041023 | LUAD | Calu-6 | MK-2206 |
| MVP_top60_DNA_seed | HDAC1 | -0.29955 | LGG | MOG-G-CCM | Dacinostat | MVP_top60_DNA+RNA_seed | ATM | 1.042822 | LUAD | NCI-H2023 | CP466722 |
| MVP_top60_DNA_seed | JAK2 | -0.299524 | LGG | D-502MG | AZD1480 | MVP_top60_DNA+RNA_seed | AURKA | 1.04299 | LUAD | NCI-H1623 | HG-5-113-01 |
| MVP_top60_DNA_seed | JAK1 | -0.299524 | LGG | D-502MG | AZD1480 | MVP_top60_DNA+RNA_seed | FGFR1 | 1.043012 | LUAD | COR-L105 | AZD4547 |
| MVP_top60_DNA_seed | HSP90AA1 | -0.299156 | LGG | D-336MG | CCT-018159 | MVP_top60_DNA+RNA_seed | AKT1 | 1.043669 | LUAD | NCI-H1666 | AKT inhibitor VIII |
| MVP_top60_DNA_seed | CDK2 | -0.292523 | LGG | GAMG | AZD5438 | MVP_top60_DNA+RNA_seed | PIK3CA | 1.04412 | LUAD | NCI-H322M | PI3Ka_4409 |
| MVP_top60_DNA_seed | EGFR | -0.289355 | LGG | MOG-G-CCM | CI-1033 | MVP_top60_DNA+RNA_seed | AKT1 | 1.044288 | LUAD | NCI-H2087 | BAY AKT1 |
| MVP_top60_DNA_seed | MDM2 | -0.285322 | LGG | GI-1 | Serdemetan | MVP_top60_DNA+RNA_seed | HDAC1 | 1.045246 | LUAD | NCI-H1563 | Tubastatin A |
| MVP_top60_DNA_seed | MET | -0.285309 | LGG | KNS-81-FD | PHA-665752 | MVP_top60_DNA+RNA_seed | HDAC1 | 1.045299 | LUAD | NCI-H1781 | CAY10603 |
| MVP_top60_DNA_seed | PIK3CD | -0.285147 | LGG | GI-1 | AZD8186 | MVP_top60_DNA+RNA_seed | SRC | 1.045378 | LUAD | NCI-H2087 | HG-6-71-01 |
| MVP_top60_DNA_seed | MET | -0.284271 | LGG | KNS-42 | Crizotinib | MVP_top60_DNA+RNA_seed | AURKA | 1.048357 | LUAD | NCI-H2122 | Alisertib |
| MVP_top60_DNA_seed | HDAC1 | -0.283614 | LGG | SF539 | CAY10603 | MVP_top60_DNA+RNA_seed | SRC | 1.048389 | LUAD | NCI-H2228 | NVP-BHG712 |
| MVP_top60_DNA_seed | HSP90AA1 | -0.28227 | LGG | no-10 | Luminespib | MVP_top60_DNA+RNA_seed | EGFR | 1.050728 | LUAD | NCI-H441 | CUDC-101 |
| MVP_top60_DNA_seed | HSP90AA1 | -0.281765 | LGG | MOG-G-UVW | Tanespimycin | MVP_top60_DNA+RNA_seed | AKT1 | 1.051112 | LUAD | NCI-H1693 | AKT inhibitor VIII |
| MVP_top60_DNA_seed | PIK3CD | -0.28136 | LGG | Hs-683 | Idelalisib | MVP_top60_DNA+RNA_seed | BRAF | 1.052421 | LUAD | NCI-H1693 | PLX-4720 |
| MVP_top60_DNA_seed | EGFR | -0.278218 | LGG | KALS-1 | Erlotinib | MVP_top60_DNA+RNA_seed | AURKA | 1.052634 | LUAD | NCI-H322M | CD532 |

| | | | | | | | | | | | |
|---|---|---|---|---|---|---|---|---|---|---|---|
| MVP_top60_DNA_seed | JAK2 | -0.276988 | LGG | NMC-G1 | Fedratinib | MVP_top60_DNA+RNA_seed | IGF1R | 1.053226 | LUAD | NCI-H322M | Linsitinib |
| MVP_top60_DNA_seed | MAPK1 | -0.276664 | LGG | SF539 | VX-11e | MVP_top60_DNA+RNA_seed | BRAF | 1.053772 | LUAD | NCI-H1650 | PLX-4720 |
| MVP_top60_DNA_seed | MET | -0.269943 | LGG | MOG-G-UVW | SU11274 | MVP_top60_DNA+RNA_seed | MET | 1.053938 | LUAD | NCI-H838 | SU11274 |
| MVP_top60_DNA_seed | MET | -0.269185 | LGG | SF539 | AZD6094 | MVP_top60_DNA+RNA_seed | EGFR | 1.054576 | LUAD | NCI-H2087 | Gefitinib |
| MVP_top60_DNA_seed | HDAC1 | -0.26791 | LGG | SW1783 | JQ12 | MVP_top60_DNA+RNA_seed | MDM2 | 1.055422 | LUAD | NCI-H1435 | Nutlin-3a (-) |
| MVP_top60_DNA_seed | PIK3CD | -0.263489 | LGG | Hs-683 | AZD8186 | MVP_top60_DNA+RNA_seed | EGFR | 1.056975 | LUAD | NCI-H1355 | PF-00299804 |
| MVP_top60_DNA_seed | EGFR | -0.258231 | LGG | D-502MG | Cetuximab | MVP_top60_DNA+RNA_seed | MET | 1.058264 | LUAD | NCI-H2228 | JNJ38877605 |
| MVP_top60_DNA_seed | PDGFRB | -0.25693 | LGG | MOG-G-CCM | Pazopanib | MVP_top60_DNA+RNA_seed | MAPK14 | 1.058299 | LUAD | NCI-H1793 | TAK-715 |
| MVP_top60_DNA_seed | EGFR | -0.256302 | LGG | KNS-42 | Erlotinib | MVP_top60_DNA+RNA_seed | MAPK14 | 1.059016 | LUAD | NCI-H2228 | TAK-715 |
| MVP_top60_DNA_seed | EGFR | -0.254777 | LGG | SF539 | CI-1033 | MVP_top60_DNA+RNA_seed | AURKA | 1.059185 | LUAD | NCI-H1993 | GSK1070916 |
| MVP_top60_DNA_seed | IGF1R | -0.254435 | LGG | GAMG | BMS-754807 | MVP_top60_DNA+RNA_seed | HDAC1 | 1.062302 | LUAD | NCI-H1651 | JQ12 |
| MVP_top60_DNA_seed | NTRK1 | -0.247572 | LGG | MOG-G-UVW | GW441756 | MVP_top60_DNA+RNA_seed | IGF1R | 1.063105 | LUAD | NCI-H322M | BMS-754807 |
| MVP_top60_DNA_seed | AR | -0.241446 | LGG | KALS-1 | Bicalutamide | MVP_top60_DNA+RNA_seed | HDAC1 | 1.063332 | LUAD | NCI-H1648 | Parthenolide |
| MVP_top60_DNA_seed | HSP90AA1 | -0.241149 | LGG | SW1783 | Tanespimycin | MVP_top60_DNA+RNA_seed | MDM2 | 1.063434 | LUAD | RERF-LC-MS | Nutlin-3a (-) |
| MVP_top60_DNA_seed | JAK2 | -0.240489 | LGG | GAMG | Fedratinib | MVP_top60_DNA+RNA_seed | IGF1R | 1.064807 | LUAD | NCI-H1693 | BMS-536924 |
| MVP_top60_DNA_seed | PIK3CA | -0.239207 | LGG | GAMG | PI3Ka_4409 | MVP_top60_DNA+RNA_seed | SRC | 1.065797 | LUAD | NCI-H1666 | NVP-BHG712 |
| MVP_top60_DNA_seed | PIK3CD | -0.239207 | LGG | GAMG | PI3Ka_4409 | MVP_top60_DNA+RNA_seed | IGF1R | 1.066045 | LUAD | Calu-3 | GSK1904529A |
| MVP_top60_DNA_seed | PDGFRB | -0.238102 | LGG | MOG-G-CCM | Masitinib | MVP_top60_DNA+RNA_seed | MET | 1.066741 | LUAD | NCI-H1755 | JNJ38877605 |
| MVP_top60_DNA_seed | PDGFRA | -0.238102 | LGG | MOG-G-CCM | Masitinib | MVP_top60_DNA+RNA_seed | AURKA | 1.067083 | LUAD | NCI-H1437 | CD532 |
| MVP_top60_DNA_seed | HDAC1 | -0.23792 | LGG | GAMG | Tubastatin A | MVP_top60_DNA+RNA_seed | BRAF | 1.067395 | LUAD | RERF-LC-MS | PLX-4720 |
| MVP_top60_DNA_seed | IGF1R | -0.230528 | LGG | GI-1 | BMS-536924 | MVP_top60_DNA+RNA_seed | BRAF | 1.067736 | LUAD | PC-3_[JPC-3] | SB590885 |
| MVP_top60_DNA_seed | EGFR | -0.229225 | LGG | GI-1 | AST-1306 | MVP_top60_DNA+RNA_seed | AURKA | 1.06919 | LUAD | NCI-H650 | ZM447439 |
| MVP_top60_DNA_seed | HDAC1 | -0.226899 | LGG | no-11 | Parthenolide | MVP_top60_DNA+RNA_seed | EGFR | 1.069313 | LUAD | NCI-H1355 | Afatinib |
| MVP_top60_DNA_seed | EGFR | -0.226739 | LGG | SF539 | Afatinib | MVP_top60_DNA+RNA_seed | MDM2 | 1.069513 | LUAD | NCI-H1975 | Nutlin-3a (-) |
| MVP_top60_DNA_seed | HSP90AA1 | -0.224207 | LGG | KALS-1 | SNX-2112 | MVP_top60_DNA+RNA_seed | MDM2 | 1.06964 | LUAD | NCI-H838 | Nutlin-3a (-) |
| MVP_top60_DNA_seed | HDAC1 | -0.222508 | LGG | H4 | AR-42 | MVP_top60_DNA+RNA_seed | AKT1 | 1.070439 | LUAD | Calu-6 | BAY AKT1 |
| MVP_top60_DNA_seed | EGFR | -0.220696 | LGG | SF539 | Pelitinib | MVP_top60_DNA+RNA_seed | AURKA | 1.072425 | LUAD | NCI-H1993 | CD532 |
| MVP_top60_DNA_seed | PIK3CD | -0.219371 | LGG | KNS-81-FD | AZD8186 | MVP_top60_DNA+RNA_seed | BRAF | 1.072544 | LUAD | NCI-H2023 | PLX-4720 |
| MVP_top60_DNA_seed | HSP90AA1 | -0.217983 | LGG | GI-1 | Luminespib | MVP_top60_DNA+RNA_seed | HDAC1 | 1.07317 | LUAD | NCI-H2405 | CAY10603 |
| MVP_top60_DNA_seed | JAK3 | -0.213573 | LGG | SF539 | WHI-P97 | MVP_top60_DNA+RNA_seed | AKT1 | 1.0732 | LUAD | NCI-H1693 | BAY AKT1 |
| MVP_top60_DNA_seed | PDGFRB | -0.212731 | LGG | GAMG | Masitinib | MVP_top60_DNA+RNA_seed | AURKA | 1.075098 | LUAD | EKVX | Alisertib |
| MVP_top60_DNA_seed | PDGFRA | -0.212731 | LGG | GAMG | Masitinib | MVP_top60_DNA+RNA_seed | AURKA | 1.075172 | LUAD | NCI-H2342 | GSK1070916 |
| MVP_top60_DNA_seed | EGFR | -0.210579 | LGG | GAMG | Pelitinib | MVP_top60_DNA+RNA_seed | HSP90AA1 | 1.075897 | LUAD | NCI-H1435 | SNX-2112 |
| MVP_top60_DNA_seed | JAK2 | -0.209278 | LGG | SF539 | Ruxolitinib | MVP_top60_DNA+RNA_seed | HSP90AA1 | 1.077203 | LUAD | NCI-H1993 | Tanespimycin |
| MVP_top60_DNA_seed | JAK1 | -0.209278 | LGG | SF539 | Ruxolitinib | MVP_top60_DNA+RNA_seed | PTK2 | 1.078115 | LUAD | NCI-H1355 | PF-562271 |
| MVP_top60_DNA_seed | EGFR | -0.208433 | LGG | MOG-G-CCM | AST-1306 | MVP_top60_DNA+RNA_seed | SRC | 1.080809 | LUAD | NCI-H1435 | NVP-BHG712 |
| MVP_top60_DNA_seed | IGF1R | -0.207752 | LGG | no-10 | BMS-536924 | MVP_top60_DNA+RNA_seed | MDM2 | 1.081121 | LUAD | NCI-H1568 | Nutlin-3a (-) |
| MVP_top60_DNA_seed | IGF1R | -0.207232 | LGG | no-11 | GSK1904529A | MVP_top60_DNA+RNA_seed | FGFR1 | 1.081756 | LUAD | PC-3_[JPC-3] | FGFR_3831 |
| MVP_top60_DNA_seed | HDAC1 | -0.201079 | LGG | H4 | CAY10603 | MVP_top60_DNA+RNA_seed | AURKA | 1.082671 | LUAD | NCI-H1435 | GSK1070916 |
| MVP_top60_DNA_seed | EGFR | -0.20048 | LGG | D-502MG | CUDC-101 | MVP_top60_DNA+RNA_seed | IGF1R | 1.08468 | LUAD | NCI-H522 | BMS-536924 |
| MVP_top60_DNA_seed | IGF1R | -0.198757 | LGG | KALS-1 | BMS-754807 | MVP_top60_DNA+RNA_seed | PTK2 | 1.087044 | LUAD | RERF-LC-KJ | PF-562271 |

| | | | | | | | | | | | |
|---|---|---|---|---|---|---|---|---|---|---|---|
| MVP_top60_DNA_seed | MET | -0.197549 | LGG | KNS-81-FD | Crizotinib | MVP_top60_DNA+RNA_seed | HSP90AA1 | 1.087114 | LUAD | H3255 | SNX-2112 |
| MVP_top60_DNA_seed | PIK3CA | -0.196107 | LGG | H4 | PI3Ka_4409 | MVP_top60_DNA+RNA_seed | FGFR1 | 1.087179 | LUAD | EKVX | FGFR_3831 |
| MVP_top60_DNA_seed | PIK3CD | -0.196107 | LGG | H4 | PI3Ka_4409 | MVP_top60_DNA+RNA_seed | AKT1 | 1.090283 | LUAD | NCI-H1651 | AKT inhibitor VIII |
| MVP_top60_DNA_seed | PIK3CD | -0.195629 | LGG | GI-1 | Idelalisib | MVP_top60_DNA+RNA_seed | EGFR | 1.09056 | LUAD | EKVX | CUDC-101 |
| MVP_top60_DNA_seed | CREBBP | -0.194042 | LGG | SF539 | I-CBP112 | MVP_top60_DNA+RNA_seed | AKT1 | 1.093007 | LUAD | NCI-H23 | AKT inhibitor VIII |
| MVP_top60_DNA_seed | JAK3 | -0.193025 | LGG | D-336MG | JAK3_7406 | MVP_top60_DNA+RNA_seed | MET | 1.093329 | LUAD | NCI-H2087 | SU11274 |
| MVP_top60_DNA_seed | EGFR | -0.192387 | LGG | GAMG | AST-1306 | MVP_top60_DNA+RNA_seed | BRAF | 1.093706 | LUAD | NCI-H1355 | PLX-4720 |
| MVP_top60_DNA_seed | SMARCA4 | -0.187743 | LGG | GI-1 | PFI-3 | MVP_top60_DNA+RNA_seed | BRAF | 1.096096 | LUAD | HCC-78 | PLX-4720 |
| MVP_top60_DNA_seed | PIK3CD | -0.185602 | LGG | MOG-G-CCM | IC-87114 | MVP_top60_DNA+RNA_seed | SRC | 1.099937 | LUAD | EMC-BAC-2 | NVP-BHG712 |
| MVP_top60_DNA_seed | JAK2 | -0.18462 | LGG | GAMG | AZD1480 | MVP_top60_DNA+RNA_seed | AURKA | 1.100322 | LUAD | NCI-H1568 | CD532 |
| MVP_top60_DNA_seed | JAK1 | -0.18462 | LGG | GAMG | AZD1480 | MVP_top60_DNA+RNA_seed | SRC | 1.100758 | LUAD | NCI-H1838 | Saracatinib |
| MVP_top60_DNA_seed | MET | -0.182894 | LGG | GAMG | SU11274 | MVP_top60_DNA+RNA_seed | SRC | 1.100992 | LUAD | NCI-H1781 | Bosutinib |
| MVP_top60_DNA_seed | PDGFRB | -0.18155 | LGG | Hs-683 | Pazopanib | MVP_top60_DNA+RNA_seed | EGFR | 1.101385 | LUAD | RERF-LC-MS | Gefitinib |
| MVP_top60_DNA_seed | EGFR | -0.180613 | LGG | MOG-G-UVW | AZD8931 | MVP_top60_DNA+RNA_seed | MET | 1.101426 | LUAD | NCI-H1838 | Crizotinib |
| MVP_top60_DNA_seed | JAK3 | -0.178157 | LGG | GI-1 | WHI-P97 | MVP_top60_DNA+RNA_seed | AURKA | 1.101509 | LUAD | HCC-827 | GSK1070916 |
| MVP_top60_DNA_seed | EGFR | -0.170201 | LGG | GI-1 | AZD8931 | MVP_top60_DNA+RNA_seed | HDAC1 | 1.103154 | LUAD | RERF-LC-KJ | CAY10603 |
| MVP_top60_DNA_seed | EGFR | -0.1693 | LGG | D-336MG | Pelitinib | MVP_top60_DNA+RNA_seed | HDAC1 | 1.103951 | LUAD | A549 | Tubastatin A |
| MVP_top60_DNA_seed | EGFR | -0.163842 | LGG | H4 | Pelitinib | MVP_top60_DNA+RNA_seed | SRC | 1.106024 | LUAD | NCI-H1693 | NVP-BHG712 |
| MVP_top60_DNA_seed | JAK2 | -0.160286 | LGG | no-11 | Ruxolitinib | MVP_top60_DNA+RNA_seed | EGFR | 1.106074 | LUAD | NCI-H1666 | Lapatinib |
| MVP_top60_DNA_seed | JAK1 | -0.160286 | LGG | no-11 | Ruxolitinib | MVP_top60_DNA+RNA_seed | AKT1 | 1.108358 | LUAD | NCI-H322M | BAY AKT1 |
| MVP_top60_DNA_seed | CDK2 | -0.159892 | LGG | D-336MG | PHA-793887 | MVP_top60_DNA+RNA_seed | EGFR | 1.108485 | LUAD | NCI-H1573 | Pelitinib |
| MVP_top60_DNA_seed | MET | -0.157504 | LGG | KNS-81-FD | JNJ38877605 | MVP_top60_DNA+RNA_seed | PIK3CA | 1.108915 | LUAD | NCI-H1793 | AZD8835 |
| MVP_top60_DNA_seed | AR | -0.155994 | LGG | GAMG | AZD3514 | MVP_top60_DNA+RNA_seed | AURKA | 1.110766 | LUAD | NCI-H1437 | Alisertib |
| MVP_top60_DNA_seed | JAK1 | -0.152736 | LGG | D-336MG | JAK1_3715 | MVP_top60_DNA+RNA_seed | MAPK14 | 1.110885 | LUAD | NCI-H2009 | TAK-715 |
| MVP_top60_DNA_seed | MAPK1 | -0.149374 | LGG | KALS-1 | VX-11e | MVP_top60_DNA+RNA_seed | MDM2 | 1.115183 | LUAD | NCI-H2405 | Serdemetan |
| MVP_top60_DNA_seed | HSP90AA1 | -0.147447 | LGG | Hs-683 | SNX-2112 | MVP_top60_DNA+RNA_seed | EGFR | 1.116088 | LUAD | NCI-H1666 | Erlotinib |
| MVP_top60_DNA_seed | AR | -0.147358 | LGG | MOG-G-CCM | Bicalutamide | MVP_top60_DNA+RNA_seed | BRAF | 1.116787 | LUAD | HCC-827 | PLX-4720 |
| MVP_top60_DNA_seed | HDAC1 | -0.146023 | LGG | KNS-81-FD | CAY10603 | MVP_top60_DNA+RNA_seed | ATM | 1.116932 | LUAD | NCI-H1435 | CP466722 |
| MVP_top60_DNA_seed | HSP90AA1 | -0.145664 | LGG | no-11 | Tanespimycin | MVP_top60_DNA+RNA_seed | EGFR | 1.117154 | LUAD | NCI-H1435 | Cetuximab |
| MVP_top60_DNA_seed | MET | -0.140649 | LGG | D-336MG | SU11274 | MVP_top60_DNA+RNA_seed | HSP90AA1 | 1.117761 | LUAD | NCI-H1693 | CCT-018159 |
| MVP_top60_DNA_seed | IGF1R | -0.13714 | LGG | GAMG | GSK1904529A | MVP_top60_DNA+RNA_seed | HSP90AA1 | 1.117782 | LUAD | NCI-H1435 | Tanespimycin |
| MVP_top60_DNA_seed | HDAC1 | -0.135378 | LGG | KNS-81-FD | Belinostat | MVP_top60_DNA+RNA_seed | PIK3CA | 1.118268 | LUAD | NCI-H322M | AZD8835 |
| MVP_top60_DNA_seed | JAK3 | -0.134418 | LGG | MOG-G-CCM | JAK3_7406 | MVP_top60_DNA+RNA_seed | PIK3CA | 1.119758 | LUAD | NCI-H1838 | AZD8835 |
| MVP_top60_DNA_seed | HDAC1 | -0.13393 | LGG | MOG-G-CCM | JQ12 | MVP_top60_DNA+RNA_seed | BRAF | 1.120456 | LUAD | COR-L105 | SB590885 |
| MVP_top60_DNA_seed | MDM2 | -0.133356 | LGG | KINGS-1 | Serdemetan | MVP_top60_DNA+RNA_seed | HDAC1 | 1.121056 | LUAD | NCI-H2405 | Tubastatin A |
| MVP_top60_DNA_seed | HDAC1 | -0.127835 | LGG | SF539 | AR-42 | MVP_top60_DNA+RNA_seed | BRAF | 1.121603 | LUAD | Calu-6 | PLX-4720 |
| MVP_top60_DNA_seed | IGF1R | -0.125457 | LGG | MOG-G-UVW | BMS-754807 | MVP_top60_DNA+RNA_seed | HSP90AA1 | 1.121867 | LUAD | Calu-3 | Tanespimycin |
| MVP_top60_DNA_seed | HDAC1 | -0.125259 | LGG | D-336MG | Parthenolide | MVP_top60_DNA+RNA_seed | HDAC1 | 1.122332 | LUAD | NCI-H2023 | Tubastatin A |
| MVP_top60_DNA_seed | SMARCA4 | -0.118443 | LGG | H4 | PFI-3 | MVP_top60_DNA+RNA_seed | EGFR | 1.123169 | LUAD | NCI-H2023 | CUDC-101 |
| MVP_top60_DNA_seed | EGFR | -0.116983 | LGG | KNS-81-FD | Lapatinib | MVP_top60_DNA+RNA_seed | HDAC1 | 1.124975 | LUAD | PC-3_[JPC-3] | Tubastatin A |
| MVP_top60_DNA_seed | PDGFRB | -0.11616 | LGG | Hs-683 | Masitinib | MVP_top60_DNA+RNA_seed | HDAC1 | 1.128467 | LUAD | NCI-H1623 | JQ12 |

| | | | | | | | | | | | |
|---|---|---|---|---|---|---|---|---|---|---|---|
| MVP_top60_DNA_seed | PDGFRA | -0.11616 | LGG | Hs-683 | Masitinib | MVP_top60_DNA+RNA_seed | HSP90AA1 | 1.132215 | LUAD | NCI-H441 | CCT-018159 |
| MVP_top60_DNA_seed | PDGFRB | -0.115943 | LGG | KNS-42 | Pazopanib | MVP_top60_DNA+RNA_seed | MET | 1.133793 | LUAD | NCI-H322M | SU11274 |
| MVP_top60_DNA_seed | SMARCA4 | -0.11527 | LGG | NMC-G1 | PFI-3 | MVP_top60_DNA+RNA_seed | BRAF | 1.133866 | LUAD | NCI-H2347 | PLX-4720 |
| MVP_top60_DNA_seed | CDK2 | -0.112906 | LGG | KNS-81-FD | Seliciclib | MVP_top60_DNA+RNA_seed | SRC | 1.13655 | LUAD | NCI-H2122 | Bosutinib |
| MVP_top60_DNA_seed | HDAC1 | -0.11242 | LGG | D-336MG | CAY10603 | MVP_top60_DNA+RNA_seed | EGFR | 1.136555 | LUAD | RERF-LC-KJ | AST-1306 |
| MVP_top60_DNA_seed | HSF1 | -0.110421 | LGG | KALS-1 | CCT245232 | MVP_top60_DNA+RNA_seed | FGFR1 | 1.13853 | LUAD | NCI-H1838 | FGFR_3831 |
| MVP_top60_DNA_seed | HSF1 | -0.106538 | LGG | SF539 | CCT245467 | MVP_top60_DNA+RNA_seed | BRAF | 1.138736 | LUAD | NCI-H2085 | PLX-4720 |
| MVP_top60_DNA_seed | HDAC1 | -0.105876 | LGG | D-336MG | AR-42 | MVP_top60_DNA+RNA_seed | HDAC1 | 1.140757 | LUAD | NCI-H441 | Belinostat |
| MVP_top60_DNA_seed | EGFR | -0.102049 | LGG | MOG-G-CCM | PF-00299804 | MVP_top60_DNA+RNA_seed | MDM2 | 1.141426 | LUAD | NCI-H1666 | Serdemetan |
| MVP_top60_DNA_seed | PIK3CA | -0.101891 | LGG | MOG-G-UVW | AZD8835 | MVP_top60_DNA+RNA_seed | HSP90AA1 | 1.141735 | LUAD | A549 | Luminespib |
| MVP_top60_DNA_seed | PIK3CD | -0.101891 | LGG | MOG-G-UVW | AZD8835 | MVP_top60_DNA+RNA_seed | MET | 1.142335 | LUAD | NCI-H1623 | AZD6094 |
| MVP_top60_DNA_seed | JAK2 | -0.100842 | LGG | SW1783 | Ruxolitinib | MVP_top60_DNA+RNA_seed | BRAF | 1.142852 | LUAD | NCI-H441 | PLX-4720 |
| MVP_top60_DNA_seed | JAK1 | -0.100842 | LGG | SW1783 | Ruxolitinib | MVP_top60_DNA+RNA_seed | HSP90AA1 | 1.145287 | LUAD | NCI-H2342 | Tanespimycin |
| MVP_top60_DNA_seed | MET | -0.099575 | LGG | Hs-683 | SU11274 | MVP_top60_DNA+RNA_seed | MET | 1.146865 | LUAD | ABC-1 | AZD6094 |
| MVP_top60_DNA_seed | EGFR | -0.099033 | LGG | no-11 | Pelitinib | MVP_top60_DNA+RNA_seed | AKT1 | 1.151738 | LUAD | NCI-H838 | AKT inhibitor VIII |
| MVP_top60_DNA_seed | JAK2 | -0.098593 | LGG | H4 | Ruxolitinib | MVP_top60_DNA+RNA_seed | FGFR1 | 1.154371 | LUAD | NCI-H1573 | FGFR_3831 |
| MVP_top60_DNA_seed | JAK1 | -0.098593 | LGG | H4 | Ruxolitinib | MVP_top60_DNA+RNA_seed | HDAC1 | 1.155033 | LUAD | RERF-LC-MS | Dacinostat |
| MVP_top60_DNA_seed | MDM2 | -0.097984 | LGG | GI-1 | Nutlin-3a (-) | MVP_top60_DNA+RNA_seed | PIK3CA | 1.156105 | LUAD | NCI-H1993 | AZD8835 |
| MVP_top60_DNA_seed | EGFR | -0.096522 | LGG | GAMG | Cetuximab | MVP_top60_DNA+RNA_seed | FGFR1 | 1.156186 | LUAD | Calu-3 | PD173074 |
| MVP_top60_DNA_seed | MET | -0.093858 | LGG | KINGS-1 | AZD6094 | MVP_top60_DNA+RNA_seed | SRC | 1.157476 | LUAD | NCI-H1666 | WH-4-023 |
| MVP_top60_DNA_seed | EGFR | -0.093728 | LGG | GI-1 | CI-1033 | MVP_top60_DNA+RNA_seed | FGFR1 | 1.157557 | LUAD | ABC-1 | AZD4547 |
| MVP_top60_DNA_seed | MDM2 | -0.091549 | LGG | MOG-G-CCM | Serdemetan | MVP_top60_DNA+RNA_seed | MDM2 | 1.158417 | LUAD | PC-3_[JPC-3] | Nutlin-3a (-) |
| MVP_top60_DNA_seed | PIK3CA | -0.088347 | LGG | NMC-G1 | AZD8835 | MVP_top60_DNA+RNA_seed | IGF1R | 1.158795 | LUAD | NCI-H322M | BMS-536924 |
| MVP_top60_DNA_seed | PIK3CD | -0.088347 | LGG | NMC-G1 | AZD8835 | MVP_top60_DNA+RNA_seed | EGFR | 1.158978 | LUAD | NCI-H2228 | Afatinib |
| MVP_top60_DNA_seed | EGFR | -0.086896 | LGG | no-11 | Afatinib | MVP_top60_DNA+RNA_seed | EGFR | 1.161213 | LUAD | NCI-H2342 | CUDC-101 |
| MVP_top60_DNA_seed | IGF1R | -0.083283 | LGG | MOG-G-UVW | GSK1904529A | MVP_top60_DNA+RNA_seed | AURKA | 1.163882 | LUAD | NCI-H1993 | ZM447439 |
| MVP_top60_DNA_seed | IGF1R | -0.082825 | LGG | SF539 | GSK1904529A | MVP_top60_DNA+RNA_seed | EGFR | 1.164977 | LUAD | A549 | CUDC-101 |
| MVP_top60_DNA_seed | EGFR | -0.081845 | LGG | SW1783 | CI-1033 | MVP_top60_DNA+RNA_seed | SRC | 1.165021 | LUAD | HCC-78 | Bosutinib |
| MVP_top60_DNA_seed | JAK1 | -0.081417 | LGG | GAMG | JAK1_3715 | MVP_top60_DNA+RNA_seed | SRC | 1.165229 | LUAD | NCI-H1650 | NVP-BHG712 |
| MVP_top60_DNA_seed | HDAC1 | -0.080449 | LGG | GI-1 | Dacinostat | MVP_top60_DNA+RNA_seed | AKT1 | 1.170507 | LUAD | NCI-H2087 | MK-2206 |
| MVP_top60_DNA_seed | HDAC1 | -0.079693 | LGG | KINGS-1 | Entinostat | MVP_top60_DNA+RNA_seed | AURKA | 1.172044 | LUAD | NCI-H2291 | Alisertib |
| MVP_top60_DNA_seed | NTRK1 | -0.078878 | LGG | SF539 | AZD1332 | MVP_top60_DNA+RNA_seed | BRAF | 1.172307 | LUAD | NCI-H322M | HG6-64-1 |
| MVP_top60_DNA_seed | PDGFRB | -0.077689 | LGG | KALS-1 | Masitinib | MVP_top60_DNA+RNA_seed | EGFR | 1.172583 | LUAD | NCI-H3122 | Gefitinib |
| MVP_top60_DNA_seed | PDGFRA | -0.077689 | LGG | KALS-1 | Masitinib | MVP_top60_DNA+RNA_seed | MET | 1.172594 | LUAD | NCI-H322M | AZD6094 |
| MVP_top60_DNA_seed | CDK2 | -0.072469 | LGG | KALS-1 | PHA-793887 | MVP_top60_DNA+RNA_seed | SRC | 1.176255 | LUAD | NCI-H1435 | Bosutinib |
| MVP_top60_DNA_seed | MET | -0.071437 | LGG | H4 | SU11274 | MVP_top60_DNA+RNA_seed | FGFR1 | 1.176337 | LUAD | NCI-H1793 | PD173074 |
| MVP_top60_DNA_seed | HDAC1 | -0.069356 | LGG | GI-1 | Entinostat | MVP_top60_DNA+RNA_seed | HDAC1 | 1.176401 | LUAD | NCI-H2009 | CAY10603 |
| MVP_top60_DNA_seed | NTRK1 | -0.067733 | LGG | KALS-1 | AZD1332 | MVP_top60_DNA+RNA_seed | AURKA | 1.176713 | LUAD | SW1573 | CD532 |
| MVP_top60_DNA_seed | CDK2 | -0.064496 | LGG | GI-1 | PHA-793887 | MVP_top60_DNA+RNA_seed | HDAC1 | 1.178119 | LUAD | NCI-H2342 | AR-42 |
| MVP_top60_DNA_seed | EGFR | -0.062129 | LGG | GAMG | PF-00299804 | MVP_top60_DNA+RNA_seed | AURKA | 1.179088 | LUAD | NCI-H2291 | ZM447439 |
| MVP_top60_DNA_seed | HDAC1 | -0.06015 | LGG | H4 | Dacinostat | MVP_top60_DNA+RNA_seed | IGF1R | 1.18038 | LUAD | NCI-H2085 | BMS-536924 |

| | | | | | | | | | | | |
|---|---|---|---|---|---|---|---|---|---|---|---|
| MVP_top60_DNA_seed | JAK2 | -0.054349 | LGG | no-10 | AZD1480 | MVP_top60_DNA+RNA_seed | AKT1 | 1.181275 | LUAD | NCI-H1355 | GSK690693 |
| MVP_top60_DNA_seed | JAK1 | -0.054349 | LGG | no-10 | AZD1480 | MVP_top60_DNA+RNA_seed | IGF1R | 1.181537 | LUAD | NCI-H1993 | GSK1904529A |
| MVP_top60_DNA_seed | PIK3CD | -0.052865 | LGG | H4 | Idelalisib | MVP_top60_DNA+RNA_seed | MET | 1.181582 | LUAD | NCI-H1648 | SU11274 |
| MVP_top60_DNA_seed | MAPK1 | -0.05173 | LGG | GAMG | FR-180204 | MVP_top60_DNA+RNA_seed | AURKA | 1.18205 | LUAD | NCI-H2009 | GSK1070916 |
| MVP_top60_DNA_seed | MAPK3 | -0.05173 | LGG | GAMG | FR-180204 | MVP_top60_DNA+RNA_seed | AURKA | 1.182672 | LUAD | NCI-H1838 | CD532 |
| MVP_top60_DNA_seed | EGFR | -0.05156 | LGG | GAMG | CI-1033 | MVP_top60_DNA+RNA_seed | SMARCA4 | 1.183219 | LUAD | PC-3_[JPC-3] | PFI-3 |
| MVP_top60_DNA_seed | CDK2 | -0.051088 | LGG | KINGS-1 | AZD5438 | MVP_top60_DNA+RNA_seed | MET | 1.184237 | LUAD | HCC-827 | JNJ38877605 |
| MVP_top60_DNA_seed | PIK3CD | -0.049303 | LGG | SF539 | IC-87114 | MVP_top60_DNA+RNA_seed | FGFR1 | 1.185836 | LUAD | NCI-H1623 | PD173074 |
| MVP_top60_DNA_seed | CDK2 | -0.04566 | LGG | MOG-G-CCM | AZD5438 | MVP_top60_DNA+RNA_seed | AURKA | 1.186649 | LUAD | NCI-H1573 | Alisertib |
| MVP_top60_DNA_seed | EGFR | -0.044553 | LGG | GI-1 | PF-00299804 | MVP_top60_DNA+RNA_seed | ATM | 1.187008 | LUAD | NCI-H1781 | KU-55933 |
| MVP_top60_DNA_seed | PDGFRB | -0.043774 | LGG | KNS-81-FD | Masitinib | MVP_top60_DNA+RNA_seed | BRAF | 1.192486 | LUAD | NCI-H2023 | HG6-64-1 |
| MVP_top60_DNA_seed | PDGFRA | -0.043774 | LGG | KNS-81-FD | Masitinib | MVP_top60_DNA+RNA_seed | AURKA | 1.192982 | LUAD | NCI-H838 | Alisertib |
| MVP_top60_DNA_seed | EGFR | -0.043042 | LGG | SF539 | Erlotinib | MVP_top60_DNA+RNA_seed | AURKA | 1.193221 | LUAD | NCI-H1623 | Alisertib |
| MVP_top60_DNA_seed | IGF1R | -0.040539 | LGG | GAMG | Linsitinib | MVP_top60_DNA+RNA_seed | MET | 1.193755 | LUAD | NCI-H1793 | AZD6094 |
| MVP_top60_DNA_seed | PIK3CD | -0.039605 | LGG | GAMG | IC-87114 | MVP_top60_DNA+RNA_seed | HSP90AA1 | 1.196928 | LUAD | SW1573 | CCT-018159 |
| MVP_top60_DNA_seed | AR | -0.039421 | LGG | GI-1 | Bicalutamide | MVP_top60_DNA+RNA_seed | AURKA | 1.197466 | LUAD | Calu-3 | CD532 |
| MVP_top60_DNA_seed | EGFR | -0.038619 | LGG | H4 | Afatinib | MVP_top60_DNA+RNA_seed | HSP90AA1 | 1.197576 | LUAD | NCI-H1623 | SNX-2112 |
| MVP_top60_DNA_seed | IGF1R | -0.036527 | LGG | NMC-G1 | GSK1904529A | MVP_top60_DNA+RNA_seed | HDAC1 | 1.198121 | LUAD | NCI-H1666 | Tubastatin A |
| MVP_top60_DNA_seed | EGFR | -0.033075 | LGG | NMC-G1 | PF-00299804 | MVP_top60_DNA+RNA_seed | FGFR1 | 1.198712 | LUAD | NCI-H1993 | AZD4547 |
| MVP_top60_DNA_seed | HDAC1 | -0.029488 | LGG | KNS-42 | CAY10603 | MVP_top60_DNA+RNA_seed | AURKA | 1.201313 | LUAD | NCI-H1993 | Alisertib |
| MVP_top60_DNA_seed | NTRK1 | -0.026218 | LGG | KNS-42 | AZD1332 | MVP_top60_DNA+RNA_seed | EGFR | 1.201904 | LUAD | NCI-H650 | Afatinib |
| MVP_top60_DNA_seed | NTRK1 | -0.022715 | LGG | D-502MG | AZD1332 | MVP_top60_DNA+RNA_seed | AURKA | 1.202304 | LUAD | NCI-H1563 | GSK1070916 |
| MVP_top60_DNA_seed | PIK3CD | -0.019236 | LGG | NMC-G1 | Idelalisib | MVP_top60_DNA+RNA_seed | FGFR1 | 1.202596 | LUAD | NCI-H650 | PD173074 |
| MVP_top60_DNA_seed | EGFR | -0.016892 | LGG | KNS-81-FD | CUDC-101 | MVP_top60_DNA+RNA_seed | IGF1R | 1.204481 | LUAD | NCI-H1573 | GSK1904529A |
| MVP_top60_DNA_seed | HSF1 | -0.013066 | LGG | no-10 | CCT245232 | MVP_top60_DNA+RNA_seed | SRC | 1.208988 | LUAD | NCI-H1651 | Bosutinib |
| MVP_top60_DNA_seed | EGFR | -0.012796 | LGG | no-10 | Lapatinib | MVP_top60_DNA+RNA_seed | IGF1R | 1.209387 | LUAD | NCI-H1651 | BMS-536924 |
| MVP_top60_DNA_seed | HDAC1 | -0.00968 | LGG | NMC-G1 | CAY10603 | MVP_top60_DNA+RNA_seed | EGFR | 1.209552 | LUAD | NCI-H1792 | CUDC-101 |
| MVP_top60_DNA_seed | JAK2 | -0.008265 | LGG | MOG-G-CCM | Fedratinib | MVP_top60_DNA+RNA_seed | SRC | 1.210312 | LUAD | NCI-H2405 | Bosutinib |
| MVP_top60_DNA_seed | AR | -0.00655 | LGG | D-336MG | AZD3514 | MVP_top60_DNA+RNA_seed | HDAC1 | 1.211252 | LUAD | RERF-LC-KJ | Belinostat |
| MVP_top60_DNA_seed | HDAC1 | -0.005671 | LGG | KNS-81-FD | AR-42 | MVP_top60_DNA+RNA_seed | PTK2 | 1.212037 | LUAD | NCI-H2228 | PF-562271 |
| MVP_top60_DNA_seed | PIK3CA | -0.001245 | LGG | SW1783 | PI3Ka_4409 | MVP_top60_DNA+RNA_seed | IGF1R | 1.212867 | LUAD | A549 | GSK1904529A |
| MVP_top60_DNA_seed | PIK3CD | -0.001245 | LGG | SW1783 | PI3Ka_4409 | MVP_top60_DNA+RNA_seed | ATM | 1.215446 | LUAD | HCC-827 | KU-55933 |
| MVP_top60_DNA_seed | EGFR | 0.001096 | LGG | D-336MG | AZD8931 | MVP_top60_DNA+RNA_seed | EGFR | 1.21657 | LUAD | NCI-H1793 | CUDC-101 |
| MVP_top60_DNA_seed | HSF1 | 0.001849 | LGG | KINGS-1 | CCT245232 | MVP_top60_DNA+RNA_seed | HSP90AA1 | 1.216593 | LUAD | EKVX | CCT-018159 |
| MVP_top60_DNA_seed | CDK2 | 0.007232 | LGG | MOG-G-CCM | PHA-793887 | MVP_top60_DNA+RNA_seed | AKT1 | 1.218054 | LUAD | NCI-H1623 | AKT inhibitor VIII |
| MVP_top60_DNA_seed | IGF1R | 0.01021 | LGG | no-11 | Linsitinib | MVP_top60_DNA+RNA_seed | SMARCA4 | 1.219472 | LUAD | COR-L105 | PFI-3 |
| MVP_top60_DNA_seed | IGF1R | 0.010447 | LGG | SW1783 | BMS-754807 | MVP_top60_DNA+RNA_seed | HSP90AA1 | 1.219853 | LUAD | EKVX | SNX-2112 |
| MVP_top60_DNA_seed | EGFR | 0.011479 | LGG | NMC-G1 | AZD8931 | MVP_top60_DNA+RNA_seed | PIK3CA | 1.219951 | LUAD | NCI-H441 | AZD8835 |
| MVP_top60_DNA_seed | HSP90AA1 | 0.011712 | LGG | Hs-683 | Elesclomol | MVP_top60_DNA+RNA_seed | MET | 1.220127 | LUAD | NCI-H2122 | JNJ38877605 |
| MVP_top60_DNA_seed | JAK3 | 0.014215 | LGG | H4 | WHI-P97 | MVP_top60_DNA+RNA_seed | MET | 1.220739 | LUAD | NCI-H1573 | SU11274 |
| MVP_top60_DNA_seed | CDK2 | 0.015496 | LGG | KALS-1 | Seliciclib | MVP_top60_DNA+RNA_seed | BRAF | 1.226464 | LUAD | NCI-H1623 | Dabrafenib |

| | | | | | | | | | | | |
|---|---|---|---|---|---|---|---|---|---|---|---|
| MVP_top60_DNA_seed | PIK3CD | 0.015834 | LGG | D-502MG | AZD8186 | MVP_top60_DNA+RNA_seed | MET | 1.227179 | LUAD | NCI-H2023 | SU11274 |
| MVP_top60_DNA_seed | EGFR | 0.016618 | LGG | SW1783 | PF-00299804 | MVP_top60_DNA+RNA_seed | HDAC1 | 1.227941 | LUAD | NCI-H2228 | AR-42 |
| MVP_top60_DNA_seed | HSF1 | 0.016783 | LGG | D-502MG | CCT245232 | MVP_top60_DNA+RNA_seed | EGFR | 1.230458 | LUAD | SW1573 | PF-00299804 |
| MVP_top60_DNA_seed | EGFR | 0.017897 | LGG | GAMG | AZD8931 | MVP_top60_DNA+RNA_seed | BRAF | 1.230903 | LUAD | ABC-1 | PLX-4720 |
| MVP_top60_DNA_seed | MDM2 | 0.024191 | LGG | Hs-683 | Nutlin-3a (-) | MVP_top60_DNA+RNA_seed | IGF1R | 1.231277 | LUAD | NCI-H1435 | BMS-536924 |
| MVP_top60_DNA_seed | EGFR | 0.026012 | LGG | SF539 | PF-00299804 | MVP_top60_DNA+RNA_seed | PIK3CA | 1.231945 | LUAD | NCI-H1435 | AZD8835 |
| MVP_top60_DNA_seed | AR | 0.028307 | LGG | KNS-42 | AZD3514 | MVP_top60_DNA+RNA_seed | AURKA | 1.233245 | LUAD | SK-LU-1 | Alisertib |
| MVP_top60_DNA_seed | EGFR | 0.028518 | LGG | SF539 | Afatinib | MVP_top60_DNA+RNA_seed | AURKA | 1.233628 | LUAD | NCI-H1650 | Tozasertib |
| MVP_top60_DNA_seed | HDAC1 | 0.030907 | LGG | KALS-1 | CAY10603 | MVP_top60_DNA+RNA_seed | ATM | 1.234763 | LUAD | EMC-BAC-2 | KU-55933 |
| MVP_top60_DNA_seed | EGFR | 0.037276 | LGG | KNS-42 | AZD8931 | MVP_top60_DNA+RNA_seed | AURKA | 1.235035 | LUAD | HCC-44 | CD532 |
| MVP_top60_DNA_seed | EGFR | 0.039318 | LGG | KALS-1 | CUDC-101 | MVP_top60_DNA+RNA_seed | EGFR | 1.236579 | LUAD | SW1573 | CI-1033 |
| MVP_top60_DNA_seed | EGFR | 0.039762 | LGG | no-10 | AZD8931 | MVP_top60_DNA+RNA_seed | FGFR1 | 1.237071 | LUAD | PC-3_[JPC-3] | PD173074 |
| MVP_top60_DNA_seed | HDAC1 | 0.040431 | LGG | no-10 | JQ12 | MVP_top60_DNA+RNA_seed | SRC | 1.237126 | LUAD | NCI-H2087 | Bosutinib |
| MVP_top60_DNA_seed | HDAC1 | 0.042295 | LGG | no-10 | CAY10603 | MVP_top60_DNA+RNA_seed | HSP90AA1 | 1.23936 | LUAD | NCI-H1651 | SNX-2112 |
| MVP_top60_DNA_seed | HDAC1 | 0.042459 | LGG | SW1783 | Belinostat | MVP_top60_DNA+RNA_seed | EGFR | 1.239484 | LUAD | SW1573 | Gefitinib |
| MVP_top60_DNA_seed | EGFR | 0.044501 | LGG | SF539 | AST-1306 | MVP_top60_DNA+RNA_seed | HDAC1 | 1.24026 | LUAD | NCI-H1568 | Tubastatin A |
| MVP_top60_DNA_seed | EGFR | 0.045337 | LGG | KINGS-1 | Afatinib | MVP_top60_DNA+RNA_seed | SRC | 1.240696 | LUAD | NCI-H1623 | Bosutinib |
| MVP_top60_DNA_seed | MAPK1 | 0.048431 | LGG | H4 | FR-180204 | MVP_top60_DNA+RNA_seed | EGFR | 1.241002 | LUAD | RERF-LC-KJ | PF-00299804 |
| MVP_top60_DNA_seed | MAPK3 | 0.048431 | LGG | H4 | FR-180204 | MVP_top60_DNA+RNA_seed | MDM2 | 1.241219 | LUAD | EKVX | Nutlin-3a (-) |
| MVP_top60_DNA_seed | IGF1R | 0.050224 | LGG | GI-1 | GSK1904529A | MVP_top60_DNA+RNA_seed | AURKA | 1.242321 | LUAD | NCI-H2291 | GSK1070916 |
| MVP_top60_DNA_seed | HSF1 | 0.051369 | LGG | GI-1 | CCT245467 | MVP_top60_DNA+RNA_seed | BRAF | 1.242368 | LUAD | EKVX | AZ628 |
| MVP_top60_DNA_seed | PIK3CD | 0.052826 | LGG | SF539 | AZD8186 | MVP_top60_DNA+RNA_seed | SRC | 1.242713 | LUAD | NCI-H1651 | NVP-BHG712 |
| MVP_top60_DNA_seed | EGFR | 0.054718 | LGG | MOG-G-CCM | AZD8931 | MVP_top60_DNA+RNA_seed | AKT1 | 1.24378 | LUAD | NCI-H1651 | MK-2206 |
| MVP_top60_DNA_seed | HDAC1 | 0.055509 | LGG | SF539 | Dacinostat | MVP_top60_DNA+RNA_seed | HDAC1 | 1.244618 | LUAD | NCI-H23 | JQ12 |
| MVP_top60_DNA_seed | EGFR | 0.056061 | LGG | Hs-683 | AZD8931 | MVP_top60_DNA+RNA_seed | FGFR1 | 1.247657 | LUAD | NCI-H441 | PD173074 |
| MVP_top60_DNA_seed | CDK2 | 0.057927 | LGG | SW1783 | AZD5438 | MVP_top60_DNA+RNA_seed | EGFR | 1.250011 | LUAD | SK-LU-1 | PF-00299804 |
| MVP_top60_DNA_seed | HDAC1 | 0.06036 | LGG | SW1783 | CAY10603 | MVP_top60_DNA+RNA_seed | HDAC1 | 1.250412 | LUAD | NCI-H1793 | Belinostat |
| MVP_top60_DNA_seed | PIK3CA | 0.060692 | LGG | SF539 | AZD8835 | MVP_top60_DNA+RNA_seed | FGFR1 | 1.251386 | LUAD | NCI-H1623 | AZD4547 |
| MVP_top60_DNA_seed | PIK3CD | 0.060692 | LGG | SF539 | AZD8835 | MVP_top60_DNA+RNA_seed | BRAF | 1.251558 | LUAD | Calu-3 | PLX-4720 |
| MVP_top60_DNA_seed | JAK2 | 0.061143 | LGG | no-11 | Fedratinib | MVP_top60_DNA+RNA_seed | BRAF | 1.252589 | LUAD | NCI-H1623 | PLX-4720 |
| MVP_top60_DNA_seed | AR | 0.061184 | LGG | MOG-G-UVW | Bicalutamide | MVP_top60_DNA+RNA_seed | MET | 1.254948 | LUAD | NCI-H650 | SU11274 |
| MVP_top60_DNA_seed | PIK3CD | 0.061624 | LGG | H4 | IC-87114 | MVP_top60_DNA+RNA_seed | MET | 1.258353 | LUAD | NCI-H1650 | SU11274 |
| MVP_top60_DNA_seed | EGFR | 0.061681 | LGG | D-336MG | Afatinib | MVP_top60_DNA+RNA_seed | AKT1 | 1.258596 | LUAD | PC-3_[JPC-3] | MK-2206 |
| MVP_top60_DNA_seed | PIK3CA | 0.064094 | LGG | SF539 | PI3Ka_4409 | MVP_top60_DNA+RNA_seed | MDM2 | 1.259967 | LUAD | NCI-H1838 | Nutlin-3a (-) |
| MVP_top60_DNA_seed | PIK3CD | 0.064094 | LGG | SF539 | PI3Ka_4409 | MVP_top60_DNA+RNA_seed | SMARCA4 | 1.260657 | LUAD | NCI-H650 | PFI-3 |
| MVP_top60_DNA_seed | EGFR | 0.070173 | LGG | GAMG | Afatinib | MVP_top60_DNA+RNA_seed | AURKA | 1.260715 | LUAD | NCI-H2023 | Genentech Cpd 10 |
| MVP_top60_DNA_seed | HSP90AA1 | 0.073035 | LGG | no-10 | Tanespimycin | MVP_top60_DNA+RNA_seed | HDAC1 | 1.262397 | LUAD | NCI-H1573 | Dacinostat |
| MVP_top60_DNA_seed | PDGFRB | 0.073579 | LGG | D-336MG | Masitinib | MVP_top60_DNA+RNA_seed | MET | 1.264311 | LUAD | NCI-H2228 | SU11274 |
| MVP_top60_DNA_seed | PDGFRA | 0.073579 | LGG | D-336MG | Masitinib | MVP_top60_DNA+RNA_seed | HDAC1 | 1.265123 | LUAD | HCC-827 | Tubastatin A |
| MVP_top60_DNA_seed | NTRK1 | 0.074976 | LGG | GAMG | GW441756 | MVP_top60_DNA+RNA_seed | BRAF | 1.265144 | LUAD | NCI-H1793 | Dabrafenib |
| MVP_top60_DNA_seed | CREBBP | 0.08268 | LGG | MOG-G-UVW | I-CBP112 | MVP_top60_DNA+RNA_seed | EGFR | 1.266119 | LUAD | NCI-H2228 | CUDC-101 |

| | | | | | | | | | | | |
|---|---|---|---|---|---|---|---|---|---|---|---|
| MVP_top60_DNA_seed | EGFR | 0.083334 | LGG | D-502MG | CI-1033 | MVP_top60_DNA+RNA_seed | AURKA | 1.266812 | LUAD | NCI-H2347 | ZM447439 |
| MVP_top60_DNA_seed | MET | 0.08846 | LGG | GAMG | AZD6094 | MVP_top60_DNA+RNA_seed | SMARCA4 | 1.268707 | LUAD | NCI-H1648 | PFI-3 |
| MVP_top60_DNA_seed | EGFR | 0.103801 | LGG | no-10 | CUDC-101 | MVP_top60_DNA+RNA_seed | HSP90AA1 | 1.268845 | LUAD | PC-3_[JPC-3] | Tanespimycin |
| MVP_top60_DNA_seed | EGFR | 0.109636 | LGG | SW1783 | Afatinib | MVP_top60_DNA+RNA_seed | BRAF | 1.2732 | LUAD | NCI-H1993 | PLX-4720 |
| MVP_top60_DNA_seed | EGFR | 0.111063 | LGG | H4 | PF-00299804 | MVP_top60_DNA+RNA_seed | FGFR1 | 1.274368 | LUAD | NCI-H1838 | PD173074 |
| MVP_top60_DNA_seed | MET | 0.111074 | LGG | no-11 | JNJ38877605 | MVP_top60_DNA+RNA_seed | IGF1R | 1.274881 | LUAD | H3255 | GSK1904529A |
| MVP_top60_DNA_seed | EGFR | 0.113206 | LGG | SW1783 | Gefitinib | MVP_top60_DNA+RNA_seed | SRC | 1.275711 | LUAD | NCI-H1838 | WH-4-023 |
| MVP_top60_DNA_seed | EGFR | 0.115724 | LGG | SW1783 | AZD8931 | MVP_top60_DNA+RNA_seed | IGF1R | 1.277183 | LUAD | NCI-H1573 | Linsitinib |
| MVP_top60_DNA_seed | PIK3CD | 0.118404 | LGG | MOG-G-UVW | IC-87114 | MVP_top60_DNA+RNA_seed | HDAC1 | 1.279632 | LUAD | NCI-H1793 | AR-42 |
| MVP_top60_DNA_seed | CDK2 | 0.119671 | LGG | KNS-42 | Seliciclib | MVP_top60_DNA+RNA_seed | HDAC1 | 1.280853 | LUAD | NCI-H1734 | JQ12 |
| MVP_top60_DNA_seed | HDAC1 | 0.122223 | LGG | SW1783 | AR-42 | MVP_top60_DNA+RNA_seed | AKT1 | 1.282071 | LUAD | NCI-H2342 | AKT inhibitor VIII |
| MVP_top60_DNA_seed | HSP90AA1 | 0.122785 | LGG | NMC-G1 | CCT-018159 | MVP_top60_DNA+RNA_seed | EGFR | 1.28528 | LUAD | NCI-H1993 | Cetuximab |
| MVP_top60_DNA_seed | HDAC1 | 0.123144 | LGG | D-502MG | Belinostat | MVP_top60_DNA+RNA_seed | BRAF | 1.286394 | LUAD | NCI-H1993 | SB590885 |
| MVP_top60_DNA_seed | JAK1 | 0.124847 | LGG | Hs-683 | JAK1_3715 | MVP_top60_DNA+RNA_seed | EGFR | 1.287822 | LUAD | NCI-H1651 | Pelitinib |
| MVP_top60_DNA_seed | PIK3CA | 0.124961 | LGG | MOG-G-CCM | PI3Ka_4409 | MVP_top60_DNA+RNA_seed | MET | 1.289227 | LUAD | NCI-H322M | PHA-665752 |
| MVP_top60_DNA_seed | PIK3CD | 0.124961 | LGG | MOG-G-CCM | PI3Ka_4409 | MVP_top60_DNA+RNA_seed | IGF1R | 1.289297 | LUAD | NCI-H23 | BMS-754807 |
| MVP_top60_DNA_seed | CDK2 | 0.126887 | LGG | no-10 | AZD5438 | MVP_top60_DNA+RNA_seed | PIK3CA | 1.291479 | LUAD | PC-3_[JPC-3] | PI3Ka_4409 |
| MVP_top60_DNA_seed | HSP90AA1 | 0.127529 | LGG | KNS-42 | SNX-2112 | MVP_top60_DNA+RNA_seed | EGFR | 1.291675 | LUAD | NCI-H1355 | Pelitinib |
| MVP_top60_DNA_seed | MAPK1 | 0.127578 | LGG | Hs-683 | FR-180204 | MVP_top60_DNA+RNA_seed | MDM2 | 1.292298 | LUAD | NCI-H2085 | Serdemetan |
| MVP_top60_DNA_seed | MAPK3 | 0.127578 | LGG | Hs-683 | FR-180204 | MVP_top60_DNA+RNA_seed | SRC | 1.293466 | LUAD | NCI-H2291 | Bosutinib |
| MVP_top60_DNA_seed | IGF1R | 0.129962 | LGG | Hs-683 | BMS-754807 | MVP_top60_DNA+RNA_seed | AURKA | 1.293899 | LUAD | NCI-H1573 | Genentech Cpd 10 |
| MVP_top60_DNA_seed | EGFR | 0.130562 | LGG | GI-1 | Pelitinib | MVP_top60_DNA+RNA_seed | AKT1 | 1.294709 | LUAD | NCI-H23 | BAY AKT1 |
| MVP_top60_DNA_seed | EGFR | 0.130695 | LGG | H4 | Cetuximab | MVP_top60_DNA+RNA_seed | MDM2 | 1.295926 | LUAD | NCI-H650 | Nutlin-3a (-) |
| MVP_top60_DNA_seed | JAK2 | 0.131018 | LGG | KNS-81-FD | Ruxolitinib | MVP_top60_DNA+RNA_seed | HSP90AA1 | 1.298267 | LUAD | NCI-H1568 | SNX-2112 |
| MVP_top60_DNA_seed | JAK1 | 0.131018 | LGG | KNS-81-FD | Ruxolitinib | MVP_top60_DNA+RNA_seed | AKT1 | 1.299109 | LUAD | NCI-H522 | BAY AKT1 |
| MVP_top60_DNA_seed | IGF1R | 0.132498 | LGG | GAMG | BMS-536924 | MVP_top60_DNA+RNA_seed | MAPK14 | 1.301362 | LUAD | NCI-H322M | TAK-715 |
| MVP_top60_DNA_seed | HDAC1 | 0.136442 | LGG | NMC-G1 | Dacinostat | MVP_top60_DNA+RNA_seed | AKT1 | 1.303054 | LUAD | NCI-H2023 | AKT inhibitor VIII |
| MVP_top60_DNA_seed | JAK3 | 0.139066 | LGG | D-502MG | WHI-P97 | MVP_top60_DNA+RNA_seed | EGFR | 1.303454 | LUAD | NCI-H441 | Pelitinib |
| MVP_top60_DNA_seed | EGFR | 0.144527 | LGG | KNS-42 | CUDC-101 | MVP_top60_DNA+RNA_seed | HDAC1 | 1.304718 | LUAD | EKVX | AR-42 |
| MVP_top60_DNA_seed | MET | 0.149293 | LGG | KNS-81-FD | AZD6094 | MVP_top60_DNA+RNA_seed | IGF1R | 1.304805 | LUAD | NCI-H1993 | BMS-536924 |
| MVP_top60_DNA_seed | EGFR | 0.155229 | LGG | SF539 | Cetuximab | MVP_top60_DNA+RNA_seed | AKT1 | 1.305485 | LUAD | NCI-H1435 | AKT inhibitor VIII |
| MVP_top60_DNA_seed | PIK3CA | 0.157451 | LGG | KINGS-1 | PI3Ka_4409 | MVP_top60_DNA+RNA_seed | MET | 1.306627 | LUAD | NCI-H1355 | Crizotinib |
| MVP_top60_DNA_seed | PIK3CD | 0.157451 | LGG | KINGS-1 | PI3Ka_4409 | MVP_top60_DNA+RNA_seed | SRC | 1.307892 | LUAD | NCI-H2342 | NVP-BHG712 |
| MVP_top60_DNA_seed | SMARCA4 | 0.16103 | LGG | MOG-G-CCM | PFI-3 | MVP_top60_DNA+RNA_seed | BRAF | 1.309826 | LUAD | RERF-LC-KJ | HG6-64-1 |
| MVP_top60_DNA_seed | PIK3CD | 0.161073 | LGG | no-10 | Idelalisib | MVP_top60_DNA+RNA_seed | FGFR1 | 1.311185 | LUAD | COR-L105 | PD173074 |
| MVP_top60_DNA_seed | CDK2 | 0.166095 | LGG | no-11 | PHA-793887 | MVP_top60_DNA+RNA_seed | SRC | 1.311781 | LUAD | NCI-H3122 | Bosutinib |
| MVP_top60_DNA_seed | JAK2 | 0.166328 | LGG | Hs-683 | Ruxolitinib | MVP_top60_DNA+RNA_seed | EGFR | 1.313376 | LUAD | SW1573 | AST-1306 |
| MVP_top60_DNA_seed | JAK1 | 0.166328 | LGG | Hs-683 | Ruxolitinib | MVP_top60_DNA+RNA_seed | MAPK14 | 1.314011 | LUAD | NCI-H1781 | TAK-715 |
| MVP_top60_DNA_seed | SMARCA4 | 0.167401 | LGG | D-336MG | PFI-3 | MVP_top60_DNA+RNA_seed | HSP90AA1 | 1.317366 | LUAD | NCI-H2085 | Tanespimycin |
| MVP_top60_DNA_seed | EGFR | 0.171176 | LGG | no-10 | Cetuximab | MVP_top60_DNA+RNA_seed | HSP90AA1 | 1.318242 | LUAD | NCI-H1781 | SNX-2112 |
| MVP_top60_DNA_seed | EGFR | 0.172969 | LGG | GI-1 | Afatinib | MVP_top60_DNA+RNA_seed | SRC | 1.320661 | LUAD | NCI-H322M | NVP-BHG712 |

| | | | | | | | | | | | |
|---|---|---|---|---|---|---|---|---|---|---|---|
| MVP_top60_DNA_seed | HSP90AA1 | 0.173306 | LGG | H4 | CCT-018159 | MVP_top60_DNA+RNA_seed | MAPK14 | 1.321587 | LUAD | NCI-H1623 | TAK-715 |
| MVP_top60_DNA_seed | HDAC1 | 0.1739 | LGG | NMC-G1 | Belinostat | MVP_top60_DNA+RNA_seed | AURKA | 1.324399 | LUAD | NCI-H2023 | Alisertib |
| MVP_top60_DNA_seed | PDGFRA | 0.17499 | LGG | MOG-G-CCM | Amuvatinib | MVP_top60_DNA+RNA_seed | BRAF | 1.324822 | LUAD | NCI-H1437 | RAF_9304 |
| MVP_top60_DNA_seed | MET | 0.176994 | LGG | NMC-G1 | AZD6094 | MVP_top60_DNA+RNA_seed | HDAC1 | 1.324855 | LUAD | EKVX | CAY10603 |
| MVP_top60_DNA_seed | MDM2 | 0.177694 | LGG | no-10 | Nutlin-3a (-) | MVP_top60_DNA+RNA_seed | AKT1 | 1.327274 | LUAD | NCI-H2291 | AKT inhibitor VIII |
| MVP_top60_DNA_seed | EGFR | 0.183783 | LGG | GI-1 | Gefitinib | MVP_top60_DNA+RNA_seed | EGFR | 1.328245 | LUAD | NCI-H2405 | CI-1033 |
| MVP_top60_DNA_seed | AR | 0.184791 | LGG | MOG-G-CCM | AZD3514 | MVP_top60_DNA+RNA_seed | IGF1R | 1.331255 | LUAD | PC-3_[JPC-3] | BMS-754807 |
| MVP_top60_DNA_seed | JAK3 | 0.185852 | LGG | SW1783 | JAK3_7406 | MVP_top60_DNA+RNA_seed | MET | 1.332366 | LUAD | NCI-H1648 | AZD6094 |
| MVP_top60_DNA_seed | IGF1R | 0.186751 | LGG | H4 | GSK1904529A | MVP_top60_DNA+RNA_seed | MET | 1.332938 | LUAD | NCI-H1666 | SU11274 |
| MVP_top60_DNA_seed | PDGFRB | 0.187949 | LGG | SF539 | Pazopanib | MVP_top60_DNA+RNA_seed | BRAF | 1.333857 | LUAD | NCI-H1648 | Dabrafenib |
| MVP_top60_DNA_seed | CDK2 | 0.196819 | LGG | H4 | PHA-793887 | MVP_top60_DNA+RNA_seed | HDAC1 | 1.333963 | LUAD | SW1573 | Tubastatin A |
| MVP_top60_DNA_seed | HDAC1 | 0.198714 | LGG | D-502MG | AR-42 | MVP_top60_DNA+RNA_seed | AKT1 | 1.336517 | LUAD | HCC-827 | AKT inhibitor VIII |
| MVP_top60_DNA_seed | HSP90AA1 | 0.20034 | LGG | MOG-G-UVW | SNX-2112 | MVP_top60_DNA+RNA_seed | HSP90AA1 | 1.33746 | LUAD | NCI-H1734 | CCT-018159 |
| MVP_top60_DNA_seed | NTRK1 | 0.200744 | LGG | SF539 | GW441756 | MVP_top60_DNA+RNA_seed | AURKA | 1.339085 | LUAD | NCI-H1435 | CD532 |
| MVP_top60_DNA_seed | JAK2 | 0.206873 | LGG | Hs-683 | Fedratinib | MVP_top60_DNA+RNA_seed | AKT1 | 1.343335 | LUAD | NCI-H1838 | GSK690693 |
| MVP_top60_DNA_seed | NTRK1 | 0.208999 | LGG | no-10 | AZD1332 | MVP_top60_DNA+RNA_seed | SRC | 1.343986 | LUAD | NCI-H1793 | NVP-BHG712 |
| MVP_top60_DNA_seed | HDAC1 | 0.210861 | LGG | KNS-42 | AR-42 | MVP_top60_DNA+RNA_seed | AURKA | 1.34747 | LUAD | NCI-H1573 | GSK1070916 |
| MVP_top60_DNA_seed | PDGFRA | 0.211408 | LGG | GI-1 | Amuvatinib | MVP_top60_DNA+RNA_seed | SRC | 1.34779 | LUAD | Calu-6 | Bosutinib |
| MVP_top60_DNA_seed | EGFR | 0.213495 | LGG | NMC-G1 | CUDC-101 | MVP_top60_DNA+RNA_seed | MET | 1.347876 | LUAD | NCI-H838 | JNJ38877605 |
| MVP_top60_DNA_seed | HDAC1 | 0.214671 | LGG | H4 | Tubastatin A | MVP_top60_DNA+RNA_seed | AKT1 | 1.348315 | LUAD | NCI-H1838 | BAY AKT1 |
| MVP_top60_DNA_seed | IGF1R | 0.215877 | LGG | KINGS-1 | BMS-536924 | MVP_top60_DNA+RNA_seed | EGFR | 1.349207 | LUAD | NCI-H1838 | CUDC-101 |
| MVP_top60_DNA_seed | EGFR | 0.222707 | LGG | no-10 | Afatinib | MVP_top60_DNA+RNA_seed | MET | 1.351211 | LUAD | NCI-H1793 | SU11274 |
| MVP_top60_DNA_seed | EGFR | 0.222872 | LGG | MOG-G-UVW | Gefitinib | MVP_top60_DNA+RNA_seed | SMARCA4 | 1.353313 | LUAD | SW1573 | PFI-3 |
| MVP_top60_DNA_seed | HDAC1 | 0.224656 | LGG | KALS-1 | AR-42 | MVP_top60_DNA+RNA_seed | BRAF | 1.354031 | LUAD | NCI-H1623 | PLX-4720 |
| MVP_top60_DNA_seed | SMARCA4 | 0.225444 | LGG | GAMG | PFI-3 | MVP_top60_DNA+RNA_seed | AURKA | 1.354211 | LUAD | PC-3_[JPC-3] | CD532 |
| MVP_top60_DNA_seed | JAK1 | 0.226431 | LGG | H4 | JAK1_3715 | MVP_top60_DNA+RNA_seed | EGFR | 1.35492 | LUAD | NCI-H3122 | PF-00299804 |
| MVP_top60_DNA_seed | HSP90AA1 | 0.227848 | LGG | Hs-683 | CCT-018159 | MVP_top60_DNA+RNA_seed | AURKA | 1.355437 | LUAD | NCI-H1650 | CD532 |
| MVP_top60_DNA_seed | HDAC1 | 0.230349 | LGG | KNS-42 | Belinostat | MVP_top60_DNA+RNA_seed | HDAC1 | 1.360278 | LUAD | NCI-H1355 | Tubastatin A |
| MVP_top60_DNA_seed | EGFR | 0.230755 | LGG | no-11 | Cetuximab | MVP_top60_DNA+RNA_seed | ATM | 1.362218 | LUAD | Calu-3 | KU-55933 |
| MVP_top60_DNA_seed | PIK3CD | 0.231384 | LGG | D-502MG | IC-87114 | MVP_top60_DNA+RNA_seed | HSP90AA1 | 1.364664 | LUAD | NCI-H1993 | SNX-2112 |
| MVP_top60_DNA_seed | HDAC1 | 0.232024 | LGG | D-336MG | Tubastatin A | MVP_top60_DNA+RNA_seed | EGFR | 1.365092 | LUAD | NCI-H838 | Pelitinib |
| MVP_top60_DNA_seed | PIK3CA | 0.234733 | LGG | MOG-G-UVW | PI3Ka_4409 | MVP_top60_DNA+RNA_seed | AURKA | 1.365345 | LUAD | NCI-H1623 | GSK1070916 |
| MVP_top60_DNA_seed | PIK3CD | 0.234733 | LGG | MOG-G-UVW | PI3Ka_4409 | MVP_top60_DNA+RNA_seed | FGFR1 | 1.368423 | LUAD | LXF-289 | FGFR_3831 |
| MVP_top60_DNA_seed | IGF1R | 0.235643 | LGG | SF539 | BMS-536924 | MVP_top60_DNA+RNA_seed | AURKA | 1.375209 | LUAD | NCI-H1563 | Alisertib |
| MVP_top60_DNA_seed | IGF1R | 0.237749 | LGG | KINGS-1 | Linsitinib | MVP_top60_DNA+RNA_seed | PIK3CA | 1.375567 | LUAD | NCI-H1793 | PI3Ka_4409 |
| MVP_top60_DNA_seed | AR | 0.237875 | LGG | KNS-81-FD | AZD3514 | MVP_top60_DNA+RNA_seed | EGFR | 1.375775 | LUAD | A549 | Pelitinib |
| MVP_top60_DNA_seed | HDAC1 | 0.238353 | LGG | MOG-G-UVW | JQ12 | MVP_top60_DNA+RNA_seed | MET | 1.377565 | LUAD | Calu-3 | SU11274 |
| MVP_top60_DNA_seed | EGFR | 0.240156 | LGG | Hs-683 | AST-1306 | MVP_top60_DNA+RNA_seed | HDAC1 | 1.377679 | LUAD | NCI-H1793 | CAY10603 |
| MVP_top60_DNA_seed | EGFR | 0.246354 | LGG | H4 | CI-1033 | MVP_top60_DNA+RNA_seed | AURKA | 1.380443 | LUAD | NCI-H1838 | GSK1070916 |
| MVP_top60_DNA_seed | AR | 0.249456 | LGG | no-10 | Bicalutamide | MVP_top60_DNA+RNA_seed | MET | 1.380816 | LUAD | NCI-H1437 | JNJ38877605 |
| MVP_top60_DNA_seed | AR | 0.250179 | LGG | KNS-42 | Bicalutamide | MVP_top60_DNA+RNA_seed | AKT1 | 1.381604 | LUAD | PC-3_[JPC-3] | BAY AKT1 |

| | | | | | | | | | | | |
|---|---|---|---|---|---|---|---|---|---|---|---|
| MVP_top60_DNA_seed | HDAC1 | 0.256026 | LGG | KALS-1 | Belinostat | MVP_top60_DNA+RNA_seed | ATM | 1.383735 | LUAD | NCI-H2342 | KU-55933 |
| MVP_top60_DNA_seed | MAPK1 | 0.256074 | LGG | GI-1 | VX-11e | MVP_top60_DNA+RNA_seed | PIK3CA | 1.383771 | LUAD | VMRC-LCD | AZD8835 |
| MVP_top60_DNA_seed | MDM2 | 0.256375 | LGG | SF539 | Nutlin-3a (-) | MVP_top60_DNA+RNA_seed | MDM2 | 1.384006 | LUAD | NCI-H1355 | Serdemetan |
| MVP_top60_DNA_seed | EGFR | 0.258762 | LGG | KINGS-1 | PF-00299804 | MVP_top60_DNA+RNA_seed | AKT1 | 1.384064 | LUAD | Calu-6 | AKT inhibitor VIII |
| MVP_top60_DNA_seed | HDAC1 | 0.259013 | LGG | no-10 | Belinostat | MVP_top60_DNA+RNA_seed | ATM | 1.386201 | LUAD | NCI-H1651 | KU-60019 |
| MVP_top60_DNA_seed | JAK2 | 0.26196 | LGG | KALS-1 | Ruxolitinib | MVP_top60_DNA+RNA_seed | AURKA | 1.386334 | LUAD | NCI-H1666 | Tozasertib |
| MVP_top60_DNA_seed | JAK1 | 0.26196 | LGG | KALS-1 | Ruxolitinib | MVP_top60_DNA+RNA_seed | AURKA | 1.389077 | LUAD | NCI-H1666 | Genentech Cpd 10 |
| MVP_top60_DNA_seed | JAK3 | 0.265819 | LGG | Hs-683 | WHI-P97 | MVP_top60_DNA+RNA_seed | AKT1 | 1.389344 | LUAD | HOP-62 | BAY AKT1 |
| MVP_top60_DNA_seed | EGFR | 0.266573 | LGG | GI-1 | Erlotinib | MVP_top60_DNA+RNA_seed | AURKA | 1.39051 | LUAD | NCI-H1792 | CD532 |
| MVP_top60_DNA_seed | HSF1 | 0.268307 | LGG | GAMG | CCT245467 | MVP_top60_DNA+RNA_seed | HDAC1 | 1.392424 | LUAD | NCI-H1650 | Parthenolide |
| MVP_top60_DNA_seed | HDAC1 | 0.268941 | LGG | SW1783 | Dacinostat | MVP_top60_DNA+RNA_seed | PIK3CA | 1.392587 | LUAD | NCI-H1651 | PI3Ka_4409 |
| MVP_top60_DNA_seed | JAK1 | 0.269524 | LGG | MOG-G-UVW | JAK1_3715 | MVP_top60_DNA+RNA_seed | AURKA | 1.400276 | LUAD | NCI-H2405 | Genentech Cpd 10 |
| MVP_top60_DNA_seed | HDAC1 | 0.269579 | LGG | D-502MG | CAY10603 | MVP_top60_DNA+RNA_seed | MAPK14 | 1.400817 | LUAD | NCI-H1693 | TAK-715 |
| MVP_top60_DNA_seed | JAK1 | 0.278144 | LGG | SW1783 | JAK1_3715 | MVP_top60_DNA+RNA_seed | AKT1 | 1.401164 | LUAD | NCI-H1693 | GSK690693 |
| MVP_top60_DNA_seed | EGFR | 0.278956 | LGG | NMC-G1 | Afatinib | MVP_top60_DNA+RNA_seed | AURKA | 1.402188 | LUAD | NCI-H2085 | CD532 |
| MVP_top60_DNA_seed | CREBBP | 0.279176 | LGG | GAMG | I-CBP112 | MVP_top60_DNA+RNA_seed | FGFR1 | 1.402898 | LUAD | Calu-3 | FGFR_3831 |
| MVP_top60_DNA_seed | AR | 0.279987 | LGG | Hs-683 | Bicalutamide | MVP_top60_DNA+RNA_seed | MET | 1.403986 | LUAD | NCI-H1666 | Crizotinib |
| MVP_top60_DNA_seed | EGFR | 0.28175 | LGG | H4 | AST-1306 | MVP_top60_DNA+RNA_seed | EGFR | 1.404964 | LUAD | NCI-H2009 | CUDC-101 |
| MVP_top60_DNA_seed | PIK3CA | 0.282967 | LGG | D-502MG | PI3Ka_4409 | MVP_top60_DNA+RNA_seed | AURKA | 1.406419 | LUAD | NCI-H1793 | ZM447439 |
| MVP_top60_DNA_seed | PIK3CD | 0.282967 | LGG | D-502MG | PI3Ka_4409 | MVP_top60_DNA+RNA_seed | AKT1 | 1.408539 | LUAD | NCI-H2291 | GSK690693 |
| MVP_top60_DNA_seed | JAK2 | 0.283986 | LGG | D-336MG | Ruxolitinib | MVP_top60_DNA+RNA_seed | HSP90AA1 | 1.408709 | LUAD | NCI-H1435 | CCT-018159 |
| MVP_top60_DNA_seed | JAK1 | 0.283986 | LGG | D-336MG | Ruxolitinib | MVP_top60_DNA+RNA_seed | HDAC1 | 1.41049 | LUAD | NCI-H441 | Tubastatin A |
| MVP_top60_DNA_seed | EGFR | 0.284077 | LGG | NMC-G1 | CI-1033 | MVP_top60_DNA+RNA_seed | HDAC1 | 1.410958 | LUAD | NCI-H2228 | Tubastatin A |
| MVP_top60_DNA_seed | JAK3 | 0.284939 | LGG | KINGS-1 | JAK3_7406 | MVP_top60_DNA+RNA_seed | BRAF | 1.41198 | LUAD | NCI-H2291 | PLX-4720 |
| MVP_top60_DNA_seed | EGFR | 0.289358 | LGG | KINGS-1 | AZD8931 | MVP_top60_DNA+RNA_seed | SRC | 1.413761 | LUAD | NCI-H1838 | NVP-BHG712 |
| MVP_top60_DNA_seed | PIK3CD | 0.2896 | LGG | no-11 | Idelalisib | MVP_top60_DNA+RNA_seed | AKT1 | 1.41423 | LUAD | NCI-H441 | GSK690693 |
| MVP_top60_DNA_seed | MET | 0.292204 | LGG | SF539 | SU11274 | MVP_top60_DNA+RNA_seed | AURKA | 1.415626 | LUAD | NCI-H2342 | Genentech Cpd 10 |
| MVP_top60_DNA_seed | JAK3 | 0.297451 | LGG | KNS-81-FD | JAK3_7406 | MVP_top60_DNA+RNA_seed | PIK3CA | 1.416646 | LUAD | NCI-H650 | PI3Ka_4409 |
| MVP_top60_DNA_seed | HDAC1 | 0.300324 | LGG | NMC-G1 | AR-42 | MVP_top60_DNA+RNA_seed | MAPK14 | 1.42007 | LUAD | NCI-H1651 | TAK-715 |
| MVP_top60_DNA_seed | HSF1 | 0.306013 | LGG | KNS-81-FD | CCT245232 | MVP_top60_DNA+RNA_seed | EGFR | 1.424649 | LUAD | NCI-H2023 | Pelitinib |
| MVP_top60_DNA_seed | MAPK1 | 0.316303 | LGG | GI-1 | FR-180204 | MVP_top60_DNA+RNA_seed | SRC | 1.426138 | LUAD | A549 | NVP-BHG712 |
| MVP_top60_DNA_seed | MAPK3 | 0.316303 | LGG | GI-1 | FR-180204 | MVP_top60_DNA+RNA_seed | EGFR | 1.426197 | LUAD | NCI-H441 | Cetuximab |
| MVP_top60_DNA_seed | IGF1R | 0.31992 | LGG | KALS-1 | Linsitinib | MVP_top60_DNA+RNA_seed | PTK2 | 1.427283 | LUAD | NCI-H1435 | PF-562271 |
| MVP_top60_DNA_seed | HSP90AA1 | 0.326035 | LGG | KNS-81-FD | Luminespib | MVP_top60_DNA+RNA_seed | AKT1 | 1.428344 | LUAD | NCI-H1573 | AKT inhibitor VIII |
| MVP_top60_DNA_seed | MET | 0.327958 | LGG | Hs-683 | AZD6094 | MVP_top60_DNA+RNA_seed | IGF1R | 1.428385 | LUAD | NCI-H1666 | Linsitinib |
| MVP_top60_DNA_seed | NTRK1 | 0.328565 | LGG | KINGS-1 | AZD1332 | MVP_top60_DNA+RNA_seed | EGFR | 1.428748 | LUAD | NCI-H2023 | PF-00299804 |
| MVP_top60_DNA_seed | HSP90AA1 | 0.330346 | LGG | KINGS-1 | CCT-018159 | MVP_top60_DNA+RNA_seed | SMARCA4 | 1.429027 | LUAD | NCI-H1563 | PFI-3 |
| MVP_top60_DNA_seed | CDK2 | 0.332891 | LGG | Hs-683 | PHA-793887 | MVP_top60_DNA+RNA_seed | PIK3CA | 1.429311 | LUAD | NCI-H1651 | AZD8835 |
| MVP_top60_DNA_seed | PIK3CA | 0.332913 | LGG | KALS-1 | PI3Ka_4409 | MVP_top60_DNA+RNA_seed | IGF1R | 1.429544 | LUAD | NCI-H1623 | GSK1904529A |
| MVP_top60_DNA_seed | PIK3CD | 0.332913 | LGG | KALS-1 | PI3Ka_4409 | MVP_top60_DNA+RNA_seed | SRC | 1.430432 | LUAD | NCI-H2347 | Bosutinib |
| MVP_top60_DNA_seed | MAPK1 | 0.333777 | LGG | MOG-G-CCM | VX-11e | MVP_top60_DNA+RNA_seed | ATM | 1.430599 | LUAD | Calu-6 | KU-55933 |

| | | | | | | | | | | | |
|---|---|---|---|---|---|---|---|---|---|---|---|
| MVP_top60_DNA_seed | HDAC1 | 0.334264 | LGG | no-10 | Dacinostat | MVP_top60_DNA+RNA_seed | FGFR1 | 1.43426 | LUAD | NCI-H1838 | AZD4547 |
| MVP_top60_DNA_seed | PDGFRB | 0.337249 | LGG | KNS-81-FD | Pazopanib | MVP_top60_DNA+RNA_seed | AKT1 | 1.43484 | LUAD | SW1573 | AKT inhibitor VIII |
| MVP_top60_DNA_seed | AR | 0.339859 | LGG | NMC-G1 | AZD3514 | MVP_top60_DNA+RNA_seed | BRAF | 1.443016 | LUAD | COR-L105 | PLX-4720 |
| MVP_top60_DNA_seed | JAK2 | 0.341533 | LGG | KINGS-1 | Fedratinib | MVP_top60_DNA+RNA_seed | AURKA | 1.444528 | LUAD | NCI-H322M | Genentech Cpd 10 |
| MVP_top60_DNA_seed | CREBBP | 0.344598 | LGG | Hs-683 | I-CBP112 | MVP_top60_DNA+RNA_seed | AKT1 | 1.445145 | LUAD | HCC-827 | MK-2206 |
| MVP_top60_DNA_seed | AR | 0.348218 | LGG | D-502MG | Bicalutamide | MVP_top60_DNA+RNA_seed | HDAC1 | 1.445772 | LUAD | EKVX | Belinostat |
| MVP_top60_DNA_seed | EGFR | 0.349894 | LGG | SW1783 | Pelitinib | MVP_top60_DNA+RNA_seed | ATM | 1.446998 | LUAD | HCC-827 | CP466722 |
| MVP_top60_DNA_seed | PDGFRA | 0.355171 | LGG | NMC-G1 | Amuvatinib | MVP_top60_DNA+RNA_seed | HDAC1 | 1.448042 | LUAD | NCI-H2009 | AR-42 |
| MVP_top60_DNA_seed | EGFR | 0.356587 | LGG | MOG-G-CCM | Gefitinib | MVP_top60_DNA+RNA_seed | PIK3CA | 1.448167 | LUAD | NCI-H1573 | PI3Ka_4409 |
| MVP_top60_DNA_seed | CDK2 | 0.360743 | LGG | MOG-G-UVW | PHA-793887 | MVP_top60_DNA+RNA_seed | HSP90AA1 | 1.4584 | LUAD | Calu-6 | Tanespimycin |
| MVP_top60_DNA_seed | MET | 0.364471 | LGG | KNS-42 | JNJ38877605 | MVP_top60_DNA+RNA_seed | BRAF | 1.458625 | LUAD | NCI-H1781 | RAF_9304 |
| MVP_top60_DNA_seed | MAPK1 | 0.367936 | LGG | GAMG | VX-11e | MVP_top60_DNA+RNA_seed | FGFR1 | 1.458722 | LUAD | NCI-H2291 | AZD4547 |
| MVP_top60_DNA_seed | JAK2 | 0.369145 | LGG | KALS-1 | AZD1480 | MVP_top60_DNA+RNA_seed | MET | 1.463962 | LUAD | NCI-H1573 | JNJ38877605 |
| MVP_top60_DNA_seed | JAK1 | 0.369145 | LGG | KALS-1 | AZD1480 | MVP_top60_DNA+RNA_seed | AKT1 | 1.464704 | LUAD | NCI-H441 | AKT inhibitor VIII |
| MVP_top60_DNA_seed | NTRK1 | 0.371925 | LGG | NMC-G1 | GW441756 | MVP_top60_DNA+RNA_seed | BRAF | 1.466 | LUAD | NCI-H1993 | HG6-64-1 |
| MVP_top60_DNA_seed | CDK2 | 0.375578 | LGG | SF539 | Seliciclib | MVP_top60_DNA+RNA_seed | HDAC1 | 1.467329 | LUAD | NCI-H2342 | Belinostat |
| MVP_top60_DNA_seed | NTRK1 | 0.378123 | LGG | Hs-683 | GW441756 | MVP_top60_DNA+RNA_seed | AKT1 | 1.468462 | LUAD | NCI-H1793 | GSK690693 |
| MVP_top60_DNA_seed | CREBBP | 0.378143 | LGG | NMC-G1 | I-CBP112 | MVP_top60_DNA+RNA_seed | AKT1 | 1.471681 | LUAD | NCI-H322M | GSK690693 |
| MVP_top60_DNA_seed | HSP90AA1 | 0.386994 | LGG | KNS-42 | Tanespimycin | MVP_top60_DNA+RNA_seed | EGFR | 1.473847 | LUAD | NCI-H1793 | Pelitinib |
| MVP_top60_DNA_seed | JAK3 | 0.38749 | LGG | KNS-42 | WHI-P97 | MVP_top60_DNA+RNA_seed | FGFR1 | 1.476677 | LUAD | NCI-H322M | AZD4547 |
| MVP_top60_DNA_seed | PDGFRA | 0.387975 | LGG | H4 | Amuvatinib | MVP_top60_DNA+RNA_seed | SRC | 1.479589 | LUAD | NCI-H1355 | Bosutinib |
| MVP_top60_DNA_seed | HDAC1 | 0.388608 | LGG | KALS-1 | Tubastatin A | MVP_top60_DNA+RNA_seed | MDM2 | 1.480705 | LUAD | NCI-H1993 | Nutlin-3a (-) |
| MVP_top60_DNA_seed | IGF1R | 0.388686 | LGG | no-11 | BMS-536924 | MVP_top60_DNA+RNA_seed | SMARCA4 | 1.481901 | LUAD | NCI-H1623 | PFI-3 |
| MVP_top60_DNA_seed | JAK2 | 0.390597 | LGG | KNS-42 | Ruxolitinib | MVP_top60_DNA+RNA_seed | ATM | 1.482178 | LUAD | NCI-H1993 | KU-55933 |
| MVP_top60_DNA_seed | JAK1 | 0.390597 | LGG | KNS-42 | Ruxolitinib | MVP_top60_DNA+RNA_seed | PIK3CA | 1.48309 | LUAD | HCC-827 | PI3Ka_4409 |
| MVP_top60_DNA_seed | HSP90AA1 | 0.392055 | LGG | MOG-G-CCM | CCT-018159 | MVP_top60_DNA+RNA_seed | HSP90AA1 | 1.486762 | LUAD | NCI-H2087 | Elesclomol |
| MVP_top60_DNA_seed | JAK2 | 0.393624 | LGG | no-10 | Ruxolitinib | MVP_top60_DNA+RNA_seed | MAPK14 | 1.486937 | LUAD | Calu-3 | TAK-715 |
| MVP_top60_DNA_seed | JAK1 | 0.393624 | LGG | no-10 | Ruxolitinib | MVP_top60_DNA+RNA_seed | MDM2 | 1.487941 | LUAD | EKVX | Serdemetan |
| MVP_top60_DNA_seed | EGFR | 0.393673 | LGG | no-10 | Afatinib | MVP_top60_DNA+RNA_seed | ATM | 1.490753 | LUAD | NCI-H1838 | CP466722 |
| MVP_top60_DNA_seed | HDAC1 | 0.394717 | LGG | Hs-683 | Dacinostat | MVP_top60_DNA+RNA_seed | MET | 1.49142 | LUAD | NCI-H1355 | SU11274 |
| MVP_top60_DNA_seed | EGFR | 0.395668 | LGG | D-502MG | AST-1306 | MVP_top60_DNA+RNA_seed | SMARCA4 | 1.495186 | LUAD | PC-14 | PFI-3 |
| MVP_top60_DNA_seed | MET | 0.398442 | LGG | NMC-G1 | Crizotinib | MVP_top60_DNA+RNA_seed | HSP90AA1 | 1.495901 | LUAD | Calu-6 | Elesclomol |
| MVP_top60_DNA_seed | HDAC1 | 0.398469 | LGG | KINGS-1 | Dacinostat | MVP_top60_DNA+RNA_seed | SRC | 1.496579 | LUAD | Calu-3 | NVP-BHG712 |
| MVP_top60_DNA_seed | PIK3CD | 0.401885 | LGG | Hs-683 | IC-87114 | MVP_top60_DNA+RNA_seed | AKT1 | 1.498574 | LUAD | HCC-827 | BAY AKT1 |
| MVP_top60_DNA_seed | MAPK1 | 0.40371 | LGG | D-336MG | FR-180204 | MVP_top60_DNA+RNA_seed | MET | 1.502235 | LUAD | EKVX | SU11274 |
| MVP_top60_DNA_seed | MAPK3 | 0.40371 | LGG | D-336MG | FR-180204 | MVP_top60_DNA+RNA_seed | BRAF | 1.503912 | LUAD | NCI-H441 | SB590885 |
| MVP_top60_DNA_seed | EGFR | 0.409953 | LGG | KALS-1 | Lapatinib | MVP_top60_DNA+RNA_seed | AURKA | 1.504108 | LUAD | NCI-H1793 | CD532 |
| MVP_top60_DNA_seed | MAPK1 | 0.412061 | LGG | SW1783 | FR-180204 | MVP_top60_DNA+RNA_seed | PIK3CA | 1.50436 | LUAD | NCI-H2085 | PI3Ka_4409 |
| MVP_top60_DNA_seed | MAPK3 | 0.412061 | LGG | SW1783 | FR-180204 | MVP_top60_DNA+RNA_seed | ATM | 1.512469 | LUAD | NCI-H2009 | CP466722 |
| MVP_top60_DNA_seed | HDAC1 | 0.413799 | LGG | KNS-81-FD | Parthenolide | MVP_top60_DNA+RNA_seed | AKT1 | 1.513596 | LUAD | NCI-H322M | A-443654 |
| MVP_top60_DNA_seed | AR | 0.416495 | LGG | no-10 | AZD3514 | MVP_top60_DNA+RNA_seed | HDAC1 | 1.514685 | LUAD | RERF-LC-KJ | AR-42 |

| | | | | | | | | | | | |
|---|---|---|---|---|---|---|---|---|---|---|---|
| MVP_top60_DNA_seed | PIK3CD | 0.419686 | LGG | SF539 | Idelalisib | MVP_top60_DNA+RNA_seed | HSP90AA1 | 1.516398 | LUAD | NCI-H1838 | CCT-018159 |
| MVP_top60_DNA_seed | MAPK1 | 0.41983 | LGG | NMC-G1 | VX-11e | MVP_top60_DNA+RNA_seed | AKT1 | 1.518539 | LUAD | SW1573 | AKT inhibitor VIII |
| MVP_top60_DNA_seed | HSF1 | 0.422368 | LGG | NMC-G1 | CCT245467 | MVP_top60_DNA+RNA_seed | PIK3CA | 1.520595 | LUAD | VMRC-LCD | PI3Ka_4409 |
| MVP_top60_DNA_seed | EGFR | 0.424023 | LGG | Hs-683 | Afatinib | MVP_top60_DNA+RNA_seed | HDAC1 | 1.521377 | LUAD | NCI-H1793 | JQ12 |
| MVP_top60_DNA_seed | EGFR | 0.424677 | LGG | SF539 | Gefitinib | MVP_top60_DNA+RNA_seed | BRAF | 1.521446 | LUAD | NCI-H2291 | PLX-4720 |
| MVP_top60_DNA_seed | EGFR | 0.434767 | LGG | GI-1 | Afatinib | MVP_top60_DNA+RNA_seed | MET | 1.521557 | LUAD | NCI-H1838 | AZD6094 |
| MVP_top60_DNA_seed | EGFR | 0.443061 | LGG | D-336MG | Gefitinib | MVP_top60_DNA+RNA_seed | IGF1R | 1.524332 | LUAD | NCI-H1838 | Linsitinib |
| MVP_top60_DNA_seed | CDK2 | 0.451093 | LGG | Hs-683 | AZD5438 | MVP_top60_DNA+RNA_seed | AURKA | 1.526099 | LUAD | NCI-H1793 | Genentech Cpd 10 |
| MVP_top60_DNA_seed | JAK3 | 0.45255 | LGG | MOG-G-CCM | WHI-P97 | MVP_top60_DNA+RNA_seed | AURKA | 1.527808 | LUAD | PC-3_[JPC-3] | ZM447439 |
| MVP_top60_DNA_seed | JAK2 | 0.453644 | LGG | MOG-G-UVW | Fedratinib | MVP_top60_DNA+RNA_seed | AURKA | 1.528824 | LUAD | NCI-H1975 | CD532 |
| MVP_top60_DNA_seed | IGF1R | 0.455648 | LGG | KINGS-1 | GSK1904529A | MVP_top60_DNA+RNA_seed | BRAF | 1.529359 | LUAD | NCI-H1793 | PLX-4720 |
| MVP_top60_DNA_seed | NTRK1 | 0.456979 | LGG | SW1783 | GW441756 | MVP_top60_DNA+RNA_seed | AURKA | 1.530439 | LUAD | NCI-H2228 | CD532 |
| MVP_top60_DNA_seed | EGFR | 0.456989 | LGG | D-502MG | Afatinib | MVP_top60_DNA+RNA_seed | IGF1R | 1.530975 | LUAD | SW1573 | GSK1904529A |
| MVP_top60_DNA_seed | CDK2 | 0.467317 | LGG | KALS-1 | AZD5438 | MVP_top60_DNA+RNA_seed | BRAF | 1.543202 | LUAD | NCI-H1666 | HG6-64-1 |
| MVP_top60_DNA_seed | HSF1 | 0.471667 | LGG | SW1783 | CCT245232 | MVP_top60_DNA+RNA_seed | MAPK14 | 1.543756 | LUAD | NCI-H2347 | TAK-715 |
| MVP_top60_DNA_seed | PDGFRB | 0.473183 | LGG | KNS-42 | Masitinib | MVP_top60_DNA+RNA_seed | HSP90AA1 | 1.544039 | LUAD | NCI-H1666 | Elesclomol |
| MVP_top60_DNA_seed | PDGFRA | 0.473183 | LGG | KNS-42 | Masitinib | MVP_top60_DNA+RNA_seed | MET | 1.549355 | LUAD | NCI-H2291 | SU11274 |
| MVP_top60_DNA_seed | JAK2 | 0.474307 | LGG | KNS-42 | AZD1480 | MVP_top60_DNA+RNA_seed | SRC | 1.551491 | LUAD | SW1573 | NVP-BHG712 |
| MVP_top60_DNA_seed | JAK1 | 0.474307 | LGG | KNS-42 | AZD1480 | MVP_top60_DNA+RNA_seed | EGFR | 1.556087 | LUAD | NCI-H1993 | Afatinib |
| MVP_top60_DNA_seed | CDK2 | 0.477724 | LGG | D-336MG | AZD5438 | MVP_top60_DNA+RNA_seed | BRAF | 1.556854 | LUAD | EKVX | Dabrafenib |
| MVP_top60_DNA_seed | EGFR | 0.478771 | LGG | KNS-42 | Afatinib | MVP_top60_DNA+RNA_seed | SRC | 1.558977 | LUAD | NCI-H1666 | Saracatinib |
| MVP_top60_DNA_seed | PDGFRB | 0.480065 | LGG | SW1783 | Pazopanib | MVP_top60_DNA+RNA_seed | SRC | 1.56004 | LUAD | Calu-3 | Bosutinib |
| MVP_top60_DNA_seed | IGF1R | 0.483574 | LGG | MOG-G-CCM | BMS-754807 | MVP_top60_DNA+RNA_seed | FGFR1 | 1.561144 | LUAD | NCI-H650 | FGFR_3831 |
| MVP_top60_DNA_seed | EGFR | 0.486965 | LGG | D-502MG | PF-00299804 | MVP_top60_DNA+RNA_seed | BRAF | 1.567386 | LUAD | NCI-H1793 | SB590885 |
| MVP_top60_DNA_seed | HDAC1 | 0.489042 | LGG | no-11 | Dacinostat | MVP_top60_DNA+RNA_seed | IGF1R | 1.568954 | LUAD | NCI-H1666 | BMS-536924 |
| MVP_top60_DNA_seed | CDK2 | 0.490586 | LGG | NMC-G1 | AZD5438 | MVP_top60_DNA+RNA_seed | SRC | 1.569018 | LUAD | NCI-H650 | Bosutinib |
| MVP_top60_DNA_seed | EGFR | 0.490698 | LGG | D-336MG | Erlotinib | MVP_top60_DNA+RNA_seed | AURKA | 1.569425 | LUAD | NCI-H2023 | CD532 |
| MVP_top60_DNA_seed | AR | 0.496831 | LGG | no-10 | Bicalutamide | MVP_top60_DNA+RNA_seed | ATM | 1.569857 | LUAD | ABC-1 | KU-55933 |
| MVP_top60_DNA_seed | PIK3CD | 0.502116 | LGG | KINGS-1 | Idelalisib | MVP_top60_DNA+RNA_seed | PTK2 | 1.57355 | LUAD | HCC-827 | PF-562271 |
| MVP_top60_DNA_seed | HDAC1 | 0.504782 | LGG | Hs-683 | AR-42 | MVP_top60_DNA+RNA_seed | FGFR1 | 1.579167 | LUAD | NCI-H1623 | FGFR_3831 |
| MVP_top60_DNA_seed | HSF1 | 0.505731 | LGG | KINGS-1 | CCT245467 | MVP_top60_DNA+RNA_seed | HDAC1 | 1.583765 | LUAD | EKVX | Parthenolide |
| MVP_top60_DNA_seed | HDAC1 | 0.506336 | LGG | Hs-683 | CAY10603 | MVP_top60_DNA+RNA_seed | AURKA | 1.58794 | LUAD | NCI-H1666 | CD532 |
| MVP_top60_DNA_seed | MET | 0.506356 | LGG | MOG-G-CCM | SU11274 | MVP_top60_DNA+RNA_seed | EGFR | 1.592102 | LUAD | NCI-H1838 | Pelitinib |
| MVP_top60_DNA_seed | EGFR | 0.507585 | LGG | MOG-G-UVW | Afatinib | MVP_top60_DNA+RNA_seed | EGFR | 1.594305 | LUAD | NCI-H2291 | Gefitinib |
| MVP_top60_DNA_seed | EGFR | 0.508347 | LGG | KNS-81-FD | Erlotinib | MVP_top60_DNA+RNA_seed | SRC | 1.597803 | LUAD | NCI-H1355 | NVP-BHG712 |
| MVP_top60_DNA_seed | MET | 0.510986 | LGG | KALS-1 | AZD6094 | MVP_top60_DNA+RNA_seed | AURKA | 1.600124 | LUAD | A549 | CD532 |
| MVP_top60_DNA_seed | MDM2 | 0.512178 | LGG | KNS-81-FD | Serdemetan | MVP_top60_DNA+RNA_seed | EGFR | 1.604902 | LUAD | Calu-6 | AZD8931 |
| MVP_top60_DNA_seed | MAPK1 | 0.512428 | LGG | D-502MG | FR-180204 | MVP_top60_DNA+RNA_seed | HDAC1 | 1.607153 | LUAD | NCI-H1573 | JQ12 |
| MVP_top60_DNA_seed | MAPK3 | 0.512428 | LGG | D-502MG | FR-180204 | MVP_top60_DNA+RNA_seed | HDAC1 | 1.610669 | LUAD | NCI-H1755 | JQ12 |
| MVP_top60_DNA_seed | IGF1R | 0.513125 | LGG | SF539 | BMS-536924 | MVP_top60_DNA+RNA_seed | BRAF | 1.611779 | LUAD | NCI-H1838 | PLX-4720 |
| MVP_top60_DNA_seed | MAPK1 | 0.5143 | LGG | MOG-G-UVW | FR-180204 | MVP_top60_DNA+RNA_seed | MET | 1.613851 | LUAD | EKVX | PHA-665752 |

| | | | | | | | | | | | |
|---|---|---|---|---|---|---|---|---|---|---|---|
| MVP_top60_DNA_seed | MAPK3 | 0.5143 | LGG | MOG-G-UVW | FR-180204 | MVP_top60_DNA+RNA_seed | MAPK14 | 1.616326 | LUAD | NCI-H1666 | TAK-715 |
| MVP_top60_DNA_seed | MET | 0.514803 | LGG | KALS-1 | Crizotinib | MVP_top60_DNA+RNA_seed | AKT1 | 1.618632 | LUAD | RERF-LC-MS | AKT inhibitor VIII |
| MVP_top60_DNA_seed | PIK3CA | 0.515561 | LGG | D-502MG | AZD8835 | MVP_top60_DNA+RNA_seed | AURKA | 1.619338 | LUAD | EMC-BAC-1 | CD532 |
| MVP_top60_DNA_seed | PIK3CD | 0.515561 | LGG | D-502MG | AZD8835 | MVP_top60_DNA+RNA_seed | BRAF | 1.620187 | LUAD | NCI-H1693 | PLX-4720 |
| MVP_top60_DNA_seed | EGFR | 0.516424 | LGG | H4 | Afatinib | MVP_top60_DNA+RNA_seed | AURKA | 1.622022 | LUAD | NCI-H441 | CD532 |
| MVP_top60_DNA_seed | EGFR | 0.519947 | LGG | no-10 | Pelitinib | MVP_top60_DNA+RNA_seed | AKT1 | 1.622874 | LUAD | NCI-H441 | MK-2206 |
| MVP_top60_DNA_seed | JAK1 | 0.524972 | LGG | no-11 | JAK1_3715 | MVP_top60_DNA+RNA_seed | MET | 1.632165 | LUAD | NCI-H1838 | PHA-665752 |
| MVP_top60_DNA_seed | EGFR | 0.525044 | LGG | H4 | Gefitinib | MVP_top60_DNA+RNA_seed | AURKA | 1.637619 | LUAD | NCI-H322M | Alisertib |
| MVP_top60_DNA_seed | PIK3CD | 0.525186 | LGG | KINGS-1 | IC-87114 | MVP_top60_DNA+RNA_seed | AURKA | 1.642485 | LUAD | NCI-H1781 | Genentech Cpd 10 |
| MVP_top60_DNA_seed | HDAC1 | 0.525383 | LGG | KALS-1 | JQ12 | MVP_top60_DNA+RNA_seed | AURKA | 1.647337 | LUAD | NCI-H2122 | CD532 |
| MVP_top60_DNA_seed | EGFR | 0.526874 | LGG | no-10 | AST-1306 | MVP_top60_DNA+RNA_seed | EGFR | 1.650087 | LUAD | NCI-H1693 | Cetuximab |
| MVP_top60_DNA_seed | IGF1R | 0.530973 | LGG | KNS-42 | BMS-536924 | MVP_top60_DNA+RNA_seed | MDM2 | 1.650201 | LUAD | NCI-H2291 | Nutlin-3a (-) |
| MVP_top60_DNA_seed | EGFR | 0.532922 | LGG | MOG-G-UVW | Pelitinib | MVP_top60_DNA+RNA_seed | AURKA | 1.657798 | LUAD | NCI-H1435 | Genentech Cpd 10 |
| MVP_top60_DNA_seed | EGFR | 0.533552 | LGG | SW1783 | Afatinib | MVP_top60_DNA+RNA_seed | AKT1 | 1.660948 | LUAD | NCI-H2347 | MK-2206 |
| MVP_top60_DNA_seed | HDAC1 | 0.53407 | LGG | no-10 | AR-42 | MVP_top60_DNA+RNA_seed | HDAC1 | 1.664309 | LUAD | NCI-H1838 | CAY10603 |
| MVP_top60_DNA_seed | CDK2 | 0.53577 | LGG | no-11 | AZD5438 | MVP_top60_DNA+RNA_seed | FGFR1 | 1.670682 | LUAD | EKVX | PD173074 |
| MVP_top60_DNA_seed | EGFR | 0.539288 | LGG | Hs-683 | Pelitinib | MVP_top60_DNA+RNA_seed | HDAC1 | 1.67207 | LUAD | NCI-H2342 | CAY10603 |
| MVP_top60_DNA_seed | EGFR | 0.540151 | LGG | D-336MG | Afatinib | MVP_top60_DNA+RNA_seed | IGF1R | 1.673572 | LUAD | NCI-H1435 | BMS-754807 |
| MVP_top60_DNA_seed | IGF1R | 0.540187 | LGG | MOG-G-UVW | Linsitinib | MVP_top60_DNA+RNA_seed | PTK2 | 1.674263 | LUAD | NCI-H1993 | PF-562271 |
| MVP_top60_DNA_seed | NTRK1 | 0.551035 | LGG | no-11 | GW441756 | MVP_top60_DNA+RNA_seed | AKT1 | 1.676727 | LUAD | EKVX | A-443654 |
| MVP_top60_DNA_seed | IGF1R | 0.553915 | LGG | SF539 | Linsitinib | MVP_top60_DNA+RNA_seed | AKT1 | 1.684248 | LUAD | NCI-H1573 | BAY AKT1 |
| MVP_top60_DNA_seed | PDGFRB | 0.556625 | LGG | no-11 | Masitinib | MVP_top60_DNA+RNA_seed | EGFR | 1.684419 | LUAD | NCI-H2291 | Pelitinib |
| MVP_top60_DNA_seed | PDGFRA | 0.556625 | LGG | no-11 | Masitinib | MVP_top60_DNA+RNA_seed | AURKA | 1.685601 | LUAD | NCI-H1563 | ZM447439 |
| MVP_top60_DNA_seed | CDK2 | 0.55984 | LGG | KNS-81-FD | AZD5438 | MVP_top60_DNA+RNA_seed | AURKA | 1.686314 | LUAD | NCI-H1838 | Tozasertib |
| MVP_top60_DNA_seed | MDM2 | 0.561538 | LGG | D-502MG | Serdemetan | MVP_top60_DNA+RNA_seed | SRC | 1.686565 | LUAD | EKVX | WH-4-023 |
| MVP_top60_DNA_seed | MET | 0.564271 | LGG | KNS-42 | AZD6094 | MVP_top60_DNA+RNA_seed | AURKA | 1.687343 | LUAD | HCC-827 | Genentech Cpd 10 |
| MVP_top60_DNA_seed | EGFR | 0.564708 | LGG | D-336MG | Cetuximab | MVP_top60_DNA+RNA_seed | AKT1 | 1.690916 | LUAD | NCI-H1793 | AKT inhibitor VIII |
| MVP_top60_DNA_seed | HDAC1 | 0.56926 | LGG | D-502MG | Entinostat | MVP_top60_DNA+RNA_seed | MET | 1.695647 | LUAD | EKVX | JNJ38877605 |
| MVP_top60_DNA_seed | AR | 0.571086 | LGG | NMC-G1 | Bicalutamide | MVP_top60_DNA+RNA_seed | HDAC1 | 1.697951 | LUAD | NCI-H1838 | Tubastatin A |
| MVP_top60_DNA_seed | IGF1R | 0.57539 | LGG | SF539 | BMS-754807 | MVP_top60_DNA+RNA_seed | AURKA | 1.698048 | LUAD | RERF-LC-MS | Genentech Cpd 10 |
| MVP_top60_DNA_seed | MET | 0.577944 | LGG | D-336MG | PHA-665752 | MVP_top60_DNA+RNA_seed | ATM | 1.699962 | LUAD | NCI-H650 | KU-60019 |
| MVP_top60_DNA_seed | HSP90AA1 | 0.58378 | LGG | NMC-G1 | SNX-2112 | MVP_top60_DNA+RNA_seed | HSP90AA1 | 1.700745 | LUAD | NCI-H441 | SNX-2112 |
| MVP_top60_DNA_seed | JAK3 | 0.58727 | LGG | GAMG | JAK3_7406 | MVP_top60_DNA+RNA_seed | AURKA | 1.703742 | LUAD | NCI-H2405 | ZM447439 |
| MVP_top60_DNA_seed | EGFR | 0.589554 | LGG | no-11 | CUDC-101 | MVP_top60_DNA+RNA_seed | EGFR | 1.706033 | LUAD | NCI-H1755 | CI-1033 |
| MVP_top60_DNA_seed | MAPK1 | 0.590481 | LGG | KINGS-1 | VX-11e | MVP_top60_DNA+RNA_seed | PTK2 | 1.712532 | LUAD | NCI-H1623 | PF-562271 |
| MVP_top60_DNA_seed | PDGFRA | 0.594562 | LGG | SW1783 | Amuvatinib | MVP_top60_DNA+RNA_seed | HDAC1 | 1.712739 | LUAD | NCI-H2291 | Tubastatin A |
| MVP_top60_DNA_seed | JAK3 | 0.596082 | LGG | SW1783 | WHI-P97 | MVP_top60_DNA+RNA_seed | AURKA | 1.716566 | LUAD | NCI-H1838 | ZM447439 |
| MVP_top60_DNA_seed | PIK3CD | 0.596434 | LGG | no-10 | AZD8186 | MVP_top60_DNA+RNA_seed | AURKA | 1.717358 | LUAD | NCI-H2009 | Genentech Cpd 10 |
| MVP_top60_DNA_seed | EGFR | 0.596731 | LGG | no-10 | CI-1033 | MVP_top60_DNA+RNA_seed | HDAC1 | 1.718503 | LUAD | NCI-H1793 | Tubastatin A |
| MVP_top60_DNA_seed | MET | 0.597085 | LGG | MOG-G-CCM | AZD6094 | MVP_top60_DNA+RNA_seed | AKT1 | 1.719399 | LUAD | EKVX | GSK690693 |
| MVP_top60_DNA_seed | EGFR | 0.597685 | LGG | D-502MG | AZD8931 | MVP_top60_DNA+RNA_seed | BRAF | 1.726835 | LUAD | EKVX | PLX-4720 |

| | | | | | | | | | | | |
|---|---|---|---|---|---|---|---|---|---|---|---|
| MVP_top60_DNA_seed | MDM2 | 0.598355 | LGG | KALS-1 | Serdemetan | MVP_top60_DNA+RNA_seed | HDAC1 | 1.727161 | LUAD | NCI-H441 | CAY10603 |
| MVP_top60_DNA_seed | PDGFRB | 0.598688 | LGG | KINGS-1 | Pazopanib | MVP_top60_DNA+RNA_seed | PTK2 | 1.727717 | LUAD | NCI-H1693 | PF-562271 |
| MVP_top60_DNA_seed | CDK2 | 0.599819 | LGG | D-336MG | Seliciclib | MVP_top60_DNA+RNA_seed | IGF1R | 1.731836 | LUAD | NCI-H1838 | BMS-536924 |
| MVP_top60_DNA_seed | MAPK1 | 0.602188 | LGG | H4 | VX-11e | MVP_top60_DNA+RNA_seed | HSP90AA1 | 1.737482 | LUAD | A549 | SNX-2112 |
| MVP_top60_DNA_seed | JAK3 | 0.603295 | LGG | D-336MG | WHI-P97 | MVP_top60_DNA+RNA_seed | AURKA | 1.739377 | LUAD | NCI-H322M | Tozasertib |
| MVP_top60_DNA_seed | JAK2 | 0.607839 | LGG | KINGS-1 | AZD1480 | MVP_top60_DNA+RNA_seed | AKT1 | 1.74846 | LUAD | RERF-LC-MS | MK-2206 |
| MVP_top60_DNA_seed | JAK1 | 0.607839 | LGG | KINGS-1 | AZD1480 | MVP_top60_DNA+RNA_seed | HSP90AA1 | 1.750076 | LUAD | NCI-H2291 | Elesclomol |
| MVP_top60_DNA_seed | HDAC1 | 0.610888 | LGG | Hs-683 | Tubastatin A | MVP_top60_DNA+RNA_seed | BRAF | 1.75429 | LUAD | NCI-H1693 | HG6-64-1 |
| MVP_top60_DNA_seed | PDGFRA | 0.618578 | LGG | KNS-81-FD | Amuvatinib | MVP_top60_DNA+RNA_seed | BRAF | 1.757667 | LUAD | ABC-1 | RAF_9304 |
| MVP_top60_DNA_seed | JAK3 | 0.623164 | LGG | D-502MG | JAK3_7406 | MVP_top60_DNA+RNA_seed | SRC | 1.764509 | LUAD | NCI-H441 | NVP-BHG712 |
| MVP_top60_DNA_seed | PIK3CD | 0.623556 | LGG | D-502MG | Idelalisib | MVP_top60_DNA+RNA_seed | SMARCA4 | 1.765601 | LUAD | NCI-H1793 | PFI-3 |
| MVP_top60_DNA_seed | EGFR | 0.624476 | LGG | SW1783 | CUDC-101 | MVP_top60_DNA+RNA_seed | MET | 1.766978 | LUAD | NCI-H441 | SU11274 |
| MVP_top60_DNA_seed | EGFR | 0.628077 | LGG | MOG-G-UVW | AST-1306 | MVP_top60_DNA+RNA_seed | PTK2 | 1.768172 | LUAD | EKVX | PF-562271 |
| MVP_top60_DNA_seed | JAK3 | 0.628821 | LGG | MOG-G-UVW | WHI-P97 | MVP_top60_DNA+RNA_seed | HSP90AA1 | 1.771014 | LUAD | NCI-H1838 | SNX-2112 |
| MVP_top60_DNA_seed | HSF1 | 0.632688 | LGG | no-11 | CCT245232 | MVP_top60_DNA+RNA_seed | IGF1R | 1.772586 | LUAD | NCI-H2291 | Linsitinib |
| MVP_top60_DNA_seed | MDM2 | 0.633955 | LGG | D-336MG | Nutlin-3a (-) | MVP_top60_DNA+RNA_seed | IGF1R | 1.77299 | LUAD | NCI-H1666 | BMS-754807 |
| MVP_top60_DNA_seed | MET | 0.634595 | LGG | no-11 | SU11274 | MVP_top60_DNA+RNA_seed | PIK3CA | 1.779161 | LUAD | COR-L105 | AZD8835 |
| MVP_top60_DNA_seed | CDK2 | 0.635201 | LGG | no-11 | Seliciclib | MVP_top60_DNA+RNA_seed | IGF1R | 1.779629 | LUAD | NCI-H1666 | GSK1904529A |
| MVP_top60_DNA_seed | PIK3CA | 0.63559 | LGG | MOG-G-CCM | AZD8835 | MVP_top60_DNA+RNA_seed | AKT1 | 1.783152 | LUAD | NCI-H650 | BAY AKT1 |
| MVP_top60_DNA_seed | PIK3CD | 0.63559 | LGG | MOG-G-CCM | AZD8835 | MVP_top60_DNA+RNA_seed | AURKA | 1.785861 | LUAD | NCI-H1993 | HG-5-113-01 |
| MVP_top60_DNA_seed | JAK2 | 0.636343 | LGG | KALS-1 | Fedratinib | MVP_top60_DNA+RNA_seed | IGF1R | 1.786217 | LUAD | HCC-827 | BMS-754807 |
| MVP_top60_DNA_seed | HSP90AA1 | 0.638796 | LGG | KINGS-1 | Elesclomol | MVP_top60_DNA+RNA_seed | SMARCA4 | 1.787465 | LUAD | NCI-H322M | PFI-3 |
| MVP_top60_DNA_seed | AR | 0.639445 | LGG | SW1783 | AZD3514 | MVP_top60_DNA+RNA_seed | HDAC1 | 1.790939 | LUAD | NCI-H1838 | Belinostat |
| MVP_top60_DNA_seed | HDAC1 | 0.640212 | LGG | SF539 | Belinostat | MVP_top60_DNA+RNA_seed | AURKA | 1.796747 | LUAD | NCI-H1435 | ZM447439 |
| MVP_top60_DNA_seed | HDAC1 | 0.642354 | LGG | SW1783 | Tubastatin A | MVP_top60_DNA+RNA_seed | MET | 1.798455 | LUAD | NCI-H441 | JNJ38877605 |
| MVP_top60_DNA_seed | HSP90AA1 | 0.642462 | LGG | KALS-1 | Elesclomol | MVP_top60_DNA+RNA_seed | IGF1R | 1.799054 | LUAD | NCI-H1793 | Linsitinib |
| MVP_top60_DNA_seed | EGFR | 0.646058 | LGG | MOG-G-UVW | CI-1033 | MVP_top60_DNA+RNA_seed | AKT1 | 1.799676 | LUAD | NCI-H322M | MK-2206 |
| MVP_top60_DNA_seed | PDGFRA | 0.647222 | LGG | SF539 | Amuvatinib | MVP_top60_DNA+RNA_seed | PTK2 | 1.800382 | LUAD | NCI-H1573 | PF-562271 |
| MVP_top60_DNA_seed | NTRK1 | 0.652532 | LGG | KNS-81-FD | AZD1332 | MVP_top60_DNA+RNA_seed | SRC | 1.802308 | LUAD | EKVX | NVP-BHG712 |
| MVP_top60_DNA_seed | MET | 0.662043 | LGG | D-502MG | PHA-665752 | MVP_top60_DNA+RNA_seed | SRC | 1.803821 | LUAD | NCI-H1993 | Bosutinib |
| MVP_top60_DNA_seed | CDK2 | 0.662239 | LGG | KINGS-1 | PHA-793887 | MVP_top60_DNA+RNA_seed | HSP90AA1 | 1.803824 | LUAD | NCI-H1993 | Luminespib |
| MVP_top60_DNA_seed | MDM2 | 0.663998 | LGG | SW1783 | Nutlin-3a (-) | MVP_top60_DNA+RNA_seed | IGF1R | 1.807654 | LUAD | NCI-H1793 | GSK1904529A |
| MVP_top60_DNA_seed | AR | 0.665049 | LGG | MOG-G-CCM | Bicalutamide | MVP_top60_DNA+RNA_seed | BRAF | 1.828291 | LUAD | NCI-H1838 | HG6-64-1 |
| MVP_top60_DNA_seed | HSP90AA1 | 0.677025 | LGG | SW1783 | Elesclomol | MVP_top60_DNA+RNA_seed | SMARCA4 | 1.828297 | LUAD | NCI-H2291 | PFI-3 |
| MVP_top60_DNA_seed | HDAC1 | 0.679542 | LGG | KALS-1 | Dacinostat | MVP_top60_DNA+RNA_seed | AKT1 | 1.829362 | LUAD | NCI-H2009 | AKT inhibitor VIII |
| MVP_top60_DNA_seed | EGFR | 0.679651 | LGG | no-11 | CI-1033 | MVP_top60_DNA+RNA_seed | BRAF | 1.833713 | LUAD | NCI-H23 | HG6-64-1 |
| MVP_top60_DNA_seed | NTRK1 | 0.680731 | LGG | D-336MG | GW441756 | MVP_top60_DNA+RNA_seed | SRC | 1.834018 | LUAD | NCI-H2291 | NVP-BHG712 |
| MVP_top60_DNA_seed | MET | 0.687242 | LGG | D-502MG | AZD6094 | MVP_top60_DNA+RNA_seed | FGFR1 | 1.848175 | LUAD | NCI-H322M | FGFR_3831 |
| MVP_top60_DNA_seed | EGFR | 0.688353 | LGG | MOG-G-UVW | PF-00299804 | MVP_top60_DNA+RNA_seed | MET | 1.85379 | LUAD | NCI-H322M | Crizotinib |
| MVP_top60_DNA_seed | JAK3 | 0.68877 | LGG | KALS-1 | WHI-P97 | MVP_top60_DNA+RNA_seed | EGFR | 1.858459 | LUAD | NCI-H1355 | CUDC-101 |
| MVP_top60_DNA_seed | CDK2 | 0.691874 | LGG | D-502MG | AZD5438 | MVP_top60_DNA+RNA_seed | BRAF | 1.859683 | LUAD | HCC-78 | HG6-64-1 |

| | | | | | | | | | | | |
|---|---|---|---|---|---|---|---|---|---|---|---|
| MVP_top60_DNA_seed | MAPK1 | 0.692687 | LGG | Hs-683 | VX-11e | MVP_top60_DNA+RNA_seed | MET | 1.8631 | LUAD | EKVX | Crizotinib |
| MVP_top60_DNA_seed | MAPK1 | 0.695593 | LGG | KNS-42 | FR-180204 | MVP_top60_DNA+RNA_seed | AURKA | 1.863414 | LUAD | SW1573 | Genentech Cpd 10 |
| MVP_top60_DNA_seed | MAPK3 | 0.695593 | LGG | KNS-42 | FR-180204 | MVP_top60_DNA+RNA_seed | HDAC1 | 1.864656 | LUAD | NCI-H1355 | AR-42 |
| MVP_top60_DNA_seed | JAK2 | 0.701314 | LGG | KNS-81-FD | AZD1480 | MVP_top60_DNA+RNA_seed | FGFR1 | 1.868144 | LUAD | EKVX | AZD4547 |
| MVP_top60_DNA_seed | JAK1 | 0.701314 | LGG | KNS-81-FD | AZD1480 | MVP_top60_DNA+RNA_seed | HSP90AA1 | 1.873396 | LUAD | NCI-H2342 | SNX-2112 |
| MVP_top60_DNA_seed | AR | 0.701841 | LGG | D-336MG | Bicalutamide | MVP_top60_DNA+RNA_seed | IGF1R | 1.878324 | LUAD | NCI-H1693 | GSK1904529A |
| MVP_top60_DNA_seed | IGF1R | 0.703201 | LGG | KNS-42 | Linsitinib | MVP_top60_DNA+RNA_seed | BRAF | 1.878644 | LUAD | NCI-H1781 | HG6-64-1 |
| MVP_top60_DNA_seed | PIK3CA | 0.703455 | LGG | KINGS-1 | AZD8835 | MVP_top60_DNA+RNA_seed | HDAC1 | 1.879534 | LUAD | NCI-H322M | Parthenolide |
| MVP_top60_DNA_seed | PIK3CD | 0.703455 | LGG | KINGS-1 | AZD8835 | MVP_top60_DNA+RNA_seed | HDAC1 | 1.881398 | LUAD | NCI-H2009 | Belinostat |
| MVP_top60_DNA_seed | MDM2 | 0.705026 | LGG | GAMG | Nutlin-3a (-) | MVP_top60_DNA+RNA_seed | HSP90AA1 | 1.88205 | LUAD | NCI-H1355 | SNX-2112 |
| MVP_top60_DNA_seed | CREBBP | 0.7067 | LGG | D-336MG | I-CBP112 | MVP_top60_DNA+RNA_seed | IGF1R | 1.88319 | LUAD | NCI-H1838 | GSK1904529A |
| MVP_top60_DNA_seed | AR | 0.708113 | LGG | KINGS-1 | Bicalutamide | MVP_top60_DNA+RNA_seed | SRC | 1.88546 | LUAD | NCI-H1437 | HG-6-71-01 |
| MVP_top60_DNA_seed | EGFR | 0.71433 | LGG | no-10 | Erlotinib | MVP_top60_DNA+RNA_seed | AURKA | 1.886977 | LUAD | EKVX | GSK1070916 |
| MVP_top60_DNA_seed | EGFR | 0.716901 | LGG | KALS-1 | Pelitinib | MVP_top60_DNA+RNA_seed | ATM | 1.889199 | LUAD | COR-L105 | KU-55933 |
| MVP_top60_DNA_seed | JAK3 | 0.719945 | LGG | no-10 | WHI-P97 | MVP_top60_DNA+RNA_seed | HDAC1 | 1.890277 | LUAD | NCI-H1838 | Parthenolide |
| MVP_top60_DNA_seed | PIK3CD | 0.721407 | LGG | D-336MG | IC-87114 | MVP_top60_DNA+RNA_seed | SMARCA4 | 1.891292 | LUAD | NCI-H1693 | PFI-3 |
| MVP_top60_DNA_seed | PIK3CD | 0.72634 | LGG | KALS-1 | IC-87114 | MVP_top60_DNA+RNA_seed | EGFR | 1.891895 | LUAD | NCI-H1437 | AST-1306 |
| MVP_top60_DNA_seed | PDGFRA | 0.727253 | LGG | no-10 | Amuvatinib | MVP_top60_DNA+RNA_seed | EGFR | 1.895013 | LUAD | HCC-44 | CI-1033 |
| MVP_top60_DNA_seed | NTRK1 | 0.730293 | LGG | KNS-81-FD | GW441756 | MVP_top60_DNA+RNA_seed | AKT1 | 1.897591 | LUAD | NCI-H1435 | MK-2206 |
| MVP_top60_DNA_seed | HDAC1 | 0.730453 | LGG | SF539 | JQ12 | MVP_top60_DNA+RNA_seed | HDAC1 | 1.898807 | LUAD | NCI-H1838 | AR-42 |
| MVP_top60_DNA_seed | SMARCA4 | 0.733498 | LGG | KALS-1 | PFI-3 | MVP_top60_DNA+RNA_seed | HDAC1 | 1.912496 | LUAD | NCI-H1650 | Entinostat |
| MVP_top60_DNA_seed | IGF1R | 0.735936 | LGG | no-11 | BMS-754807 | MVP_top60_DNA+RNA_seed | BRAF | 1.922278 | LUAD | NCI-H1573 | HG6-64-1 |
| MVP_top60_DNA_seed | PDGFRA | 0.74076 | LGG | D-336MG | Amuvatinib | MVP_top60_DNA+RNA_seed | BRAF | 1.924113 | LUAD | EKVX | SB590885 |
| MVP_top60_DNA_seed | EGFR | 0.741187 | LGG | Hs-683 | Afatinib | MVP_top60_DNA+RNA_seed | AURKA | 1.92505 | LUAD | NCI-H1623 | Genentech Cpd 10 |
| MVP_top60_DNA_seed | HDAC1 | 0.743556 | LGG | no-11 | CAY10603 | MVP_top60_DNA+RNA_seed | HSP90AA1 | 1.937296 | LUAD | NCI-H1648 | CCT-018159 |
| MVP_top60_DNA_seed | HDAC1 | 0.743724 | LGG | no-10 | Tubastatin A | MVP_top60_DNA+RNA_seed | HSP90AA1 | 1.937838 | LUAD | Calu-3 | Elesclomol |
| MVP_top60_DNA_seed | HDAC1 | 0.744461 | LGG | MOG-G-UVW | Dacinostat | MVP_top60_DNA+RNA_seed | ATM | 1.943072 | LUAD | NCI-H2228 | KU-60019 |
| MVP_top60_DNA_seed | NTRK1 | 0.756615 | LGG | no-11 | AZD1332 | MVP_top60_DNA+RNA_seed | PTK2 | 1.944056 | LUAD | NCI-H322M | PF-562271 |
| MVP_top60_DNA_seed | JAK2 | 0.757693 | LGG | no-11 | AZD1480 | MVP_top60_DNA+RNA_seed | HSP90AA1 | 1.946492 | LUAD | NCI-H322M | Elesclomol |
| MVP_top60_DNA_seed | JAK1 | 0.757693 | LGG | no-11 | AZD1480 | MVP_top60_DNA+RNA_seed | IGF1R | 1.951075 | LUAD | NCI-H441 | GSK1904529A |
| MVP_top60_DNA_seed | SMARCA4 | 0.763605 | LGG | KNS-42 | PFI-3 | MVP_top60_DNA+RNA_seed | HDAC1 | 1.955466 | LUAD | NCI-H1355 | CAY10603 |
| MVP_top60_DNA_seed | EGFR | 0.767565 | LGG | NMC-G1 | AST-1306 | MVP_top60_DNA+RNA_seed | AURKA | 1.958492 | LUAD | EKVX | Tozasertib |
| MVP_top60_DNA_seed | PIK3CA | 0.768108 | LGG | SW1783 | AZD8835 | MVP_top60_DNA+RNA_seed | HDAC1 | 1.960504 | LUAD | NCI-H1781 | Dacinostat |
| MVP_top60_DNA_seed | PIK3CD | 0.768108 | LGG | SW1783 | AZD8835 | MVP_top60_DNA+RNA_seed | HDAC1 | 1.962589 | LUAD | RERF-LC-KJ | Dacinostat |
| MVP_top60_DNA_seed | HSF1 | 0.769501 | LGG | KNS-42 | CCT245232 | MVP_top60_DNA+RNA_seed | PTK2 | 1.96495 | LUAD | NCI-H1563 | PF-562271 |
| MVP_top60_DNA_seed | HDAC1 | 0.771197 | LGG | D-502MG | Parthenolide | MVP_top60_DNA+RNA_seed | HSP90AA1 | 1.968778 | LUAD | ABC-1 | CCT-018159 |
| MVP_top60_DNA_seed | MAPK1 | 0.772651 | LGG | SW1783 | VX-11e | MVP_top60_DNA+RNA_seed | HSP90AA1 | 1.9746 | LUAD | NCI-H1693 | Tanespimycin |
| MVP_top60_DNA_seed | CREBBP | 0.772909 | LGG | MOG-G-CCM | I-CBP112 | MVP_top60_DNA+RNA_seed | AURKA | 1.976648 | LUAD | SK-LU-1 | CD532 |
| MVP_top60_DNA_seed | EGFR | 0.774391 | LGG | KNS-42 | Cetuximab | MVP_top60_DNA+RNA_seed | IGF1R | 1.981868 | LUAD | EKVX | GSK1904529A |
| MVP_top60_DNA_seed | MAPK1 | 0.779978 | LGG | MOG-G-UVW | VX-11e | MVP_top60_DNA+RNA_seed | HDAC1 | 1.984077 | LUAD | EKVX | Dacinostat |
| MVP_top60_DNA_seed | MET | 0.782051 | LGG | KALS-1 | PHA-665752 | MVP_top60_DNA+RNA_seed | MDM2 | 1.999421 | LUAD | NCI-H1435 | Serdemetan |

| | | | | | | | | | | | |
|---|---|---|---|---|---|---|---|---|---|---|---|
| MVP_top60_DNA_seed | EGFR | 0.782337 | LGG | no-11 | Gefitinib | MVP_top60_DNA+RNA_seed | EGFR | 2.010551 | LUAD | NCI-H2122 | PF-00299804 |
| MVP_top60_DNA_seed | JAK3 | 0.782638 | LGG | NMC-G1 | WHI-P97 | MVP_top60_DNA+RNA_seed | AKT1 | 2.012874 | LUAD | NCI-H1666 | AKT inhibitor VIII |
| MVP_top60_DNA_seed | PIK3CA | 0.783126 | LGG | no-10 | PI3Ka_4409 | MVP_top60_DNA+RNA_seed | HSP90AA1 | 2.013269 | LUAD | NCI-H1793 | Luminespib |
| MVP_top60_DNA_seed | PIK3CD | 0.783126 | LGG | no-10 | PI3Ka_4409 | MVP_top60_DNA+RNA_seed | BRAF | 2.013966 | LUAD | EKVX | HG6-64-1 |
| MVP_top60_DNA_seed | MET | 0.789522 | LGG | KNS-42 | SU11274 | MVP_top60_DNA+RNA_seed | SRC | 2.014274 | LUAD | NCI-H441 | Bosutinib |
| MVP_top60_DNA_seed | EGFR | 0.789883 | LGG | KNS-42 | Gefitinib | MVP_top60_DNA+RNA_seed | MAPK14 | 2.03145 | LUAD | NCI-H1838 | TAK-715 |
| MVP_top60_DNA_seed | EGFR | 0.790636 | LGG | no-11 | PF-00299804 | MVP_top60_DNA+RNA_seed | MET | 2.033026 | LUAD | EKVX | AZD6094 |
| MVP_top60_DNA_seed | AR | 0.7939 | LGG | KNS-42 | Bicalutamide | MVP_top60_DNA+RNA_seed | HSP90AA1 | 2.036228 | LUAD | NCI-H2347 | SNX-2112 |
| MVP_top60_DNA_seed | IGF1R | 0.795896 | LGG | KNS-81-FD | BMS-754807 | MVP_top60_DNA+RNA_seed | AKT1 | 2.046368 | LUAD | NCI-H1793 | MK-2206 |
| MVP_top60_DNA_seed | EGFR | 0.796652 | LGG | MOG-G-CCM | Afatinib | MVP_top60_DNA+RNA_seed | AKT1 | 2.046969 | LUAD | NCI-H650 | MK-2206 |
| MVP_top60_DNA_seed | MET | 0.798027 | LGG | KINGS-1 | SU11274 | MVP_top60_DNA+RNA_seed | AKT1 | 2.047699 | LUAD | NCI-H322M | AKT inhibitor VIII |
| MVP_top60_DNA_seed | IGF1R | 0.803047 | LGG | no-10 | Linsitinib | MVP_top60_DNA+RNA_seed | AKT1 | 2.048552 | LUAD | NCI-H1838 | AKT inhibitor VIII |
| MVP_top60_DNA_seed | MET | 0.806785 | LGG | no-10 | PHA-665752 | MVP_top60_DNA+RNA_seed | SMARCA4 | 2.056116 | LUAD | NCI-H1838 | PFI-3 |
| MVP_top60_DNA_seed | MET | 0.80932 | LGG | KNS-42 | PHA-665752 | MVP_top60_DNA+RNA_seed | HSP90AA1 | 2.057254 | LUAD | NCI-H1693 | SNX-2112 |
| MVP_top60_DNA_seed | IGF1R | 0.809988 | LGG | H4 | Linsitinib | MVP_top60_DNA+RNA_seed | HDAC1 | 2.058314 | LUAD | EKVX | Tubastatin A |
| MVP_top60_DNA_seed | CDK2 | 0.810102 | LGG | KNS-42 | AZD5438 | MVP_top60_DNA+RNA_seed | HSP90AA1 | 2.062085 | LUAD | ABC-1 | Tanespimycin |
| MVP_top60_DNA_seed | MET | 0.810385 | LGG | no-10 | JNJ38877605 | MVP_top60_DNA+RNA_seed | HDAC1 | 2.094092 | LUAD | SW1573 | JQ12 |
| MVP_top60_DNA_seed | HDAC1 | 0.818311 | LGG | D-336MG | JQ12 | MVP_top60_DNA+RNA_seed | HDAC1 | 2.096667 | LUAD | NCI-H1666 | Entinostat |
| MVP_top60_DNA_seed | EGFR | 0.818872 | LGG | KNS-81-FD | Afatinib | MVP_top60_DNA+RNA_seed | AURKA | 2.112616 | LUAD | NCI-H1838 | Genentech Cpd 10 |
| MVP_top60_DNA_seed | EGFR | 0.821511 | LGG | NMC-G1 | Afatinib | MVP_top60_DNA+RNA_seed | HSP90AA1 | 2.114563 | LUAD | NCI-H1563 | CCT-018159 |
| MVP_top60_DNA_seed | JAK2 | 0.824842 | LGG | no-10 | Fedratinib | MVP_top60_DNA+RNA_seed | HSP90AA1 | 2.122647 | LUAD | NCI-H1563 | SNX-2112 |
| MVP_top60_DNA_seed | MET | 0.829181 | LGG | D-336MG | Crizotinib | MVP_top60_DNA+RNA_seed | HSP90AA1 | 2.12873 | LUAD | SW1573 | SNX-2112 |
| MVP_top60_DNA_seed | EGFR | 0.832433 | LGG | KINGS-1 | Erlotinib | MVP_top60_DNA+RNA_seed | AKT1 | 2.135125 | LUAD | NCI-H2291 | AKT inhibitor VIII |
| MVP_top60_DNA_seed | NTRK1 | 0.833408 | LGG | D-502MG | GW441756 | MVP_top60_DNA+RNA_seed | HSP90AA1 | 2.135751 | LUAD | NCI-H1435 | Elesclomol |
| MVP_top60_DNA_seed | IGF1R | 0.837681 | LGG | Hs-683 | BMS-536924 | MVP_top60_DNA+RNA_seed | EGFR | 2.142759 | LUAD | NCI-H2228 | CI-1033 |
| MVP_top60_DNA_seed | MAPK1 | 0.841288 | LGG | KNS-42 | VX-11e | MVP_top60_DNA+RNA_seed | HDAC1 | 2.164991 | LUAD | NCI-H1666 | JQ12 |
| MVP_top60_DNA_seed | MET | 0.843304 | LGG | KALS-1 | JNJ38877605 | MVP_top60_DNA+RNA_seed | IGF1R | 2.168824 | LUAD | NCI-H2291 | GSK1904529A |
| MVP_top60_DNA_seed | NTRK1 | 0.843486 | LGG | no-10 | GW441756 | MVP_top60_DNA+RNA_seed | AKT1 | 2.17993 | LUAD | NCI-H1993 | MK-2206 |
| MVP_top60_DNA_seed | EGFR | 0.847427 | LGG | KALS-1 | Afatinib | MVP_top60_DNA+RNA_seed | SRC | 2.183864 | LUAD | NCI-H322M | Saracatinib |
| MVP_top60_DNA_seed | PDGFRA | 0.847465 | LGG | no-11 | Amuvatinib | MVP_top60_DNA+RNA_seed | BRAF | 2.184566 | LUAD | NCI-H1623 | HG6-64-1 |
| MVP_top60_DNA_seed | MET | 0.848259 | LGG | NMC-G1 | SU11274 | MVP_top60_DNA+RNA_seed | IGF1R | 2.189054 | LUAD | NCI-H2291 | BMS-754807 |
| MVP_top60_DNA_seed | EGFR | 0.851345 | LGG | Hs-683 | Gefitinib | MVP_top60_DNA+RNA_seed | EGFR | 2.196033 | LUAD | A549 | CI-1033 |
| MVP_top60_DNA_seed | PIK3CA | 0.851969 | LGG | NMC-G1 | PI3Ka_4409 | MVP_top60_DNA+RNA_seed | AURKA | 2.199297 | LUAD | H3255 | CD532 |
| MVP_top60_DNA_seed | PIK3CD | 0.851969 | LGG | NMC-G1 | PI3Ka_4409 | MVP_top60_DNA+RNA_seed | AKT1 | 2.224587 | LUAD | EKVX | AKT inhibitor VIII |
| MVP_top60_DNA_seed | HDAC1 | 0.852089 | LGG | KINGS-1 | Tubastatin A | MVP_top60_DNA+RNA_seed | PTK2 | 2.226134 | LUAD | NCI-H1666 | PF-562271 |
| MVP_top60_DNA_seed | EGFR | 0.853546 | LGG | Hs-683 | CI-1033 | MVP_top60_DNA+RNA_seed | HDAC1 | 2.226534 | LUAD | NCI-H1355 | Belinostat |
| MVP_top60_DNA_seed | AR | 0.85577 | LGG | NMC-G1 | Bicalutamide | MVP_top60_DNA+RNA_seed | HSP90AA1 | 2.228154 | LUAD | COR-L105 | Tanespimycin |
| MVP_top60_DNA_seed | HDAC1 | 0.858198 | LGG | KALS-1 | Parthenolide | MVP_top60_DNA+RNA_seed | MET | 2.230712 | LUAD | NCI-H1793 | JNJ38877605 |
| MVP_top60_DNA_seed | MAPK1 | 0.860392 | LGG | no-10 | VX-11e | MVP_top60_DNA+RNA_seed | IGF1R | 2.254386 | LUAD | NCI-H1838 | BMS-754807 |
| MVP_top60_DNA_seed | EGFR | 0.86105 | LGG | KALS-1 | PF-00299804 | MVP_top60_DNA+RNA_seed | MAPK14 | 2.258166 | LUAD | EKVX | TAK-715 |
| MVP_top60_DNA_seed | AR | 0.86239 | LGG | KINGS-1 | Bicalutamide | MVP_top60_DNA+RNA_seed | ATM | 2.270582 | LUAD | NCI-H1623 | CP466722 |

| | | | | | | | | | | | |
|---|---|---|---|---|---|---|---|---|---|---|---|
| MVP_top60_DNA_seed | MAPK1 | 0.863067 | LGG | NMC-G1 | FR-180204 | MVP_top60_DNA+RNA_seed | ATM | 2.276432 | LUAD | NCI-H2291 | CP466722 |
| MVP_top60_DNA_seed | MAPK3 | 0.863067 | LGG | NMC-G1 | FR-180204 | MVP_top60_DNA+RNA_seed | MDM2 | 2.286987 | LUAD | NCI-H650 | Serdemetan |
| MVP_top60_DNA_seed | IGF1R | 0.868089 | LGG | KNS-42 | BMS-536924 | MVP_top60_DNA+RNA_seed | AKT1 | 2.289173 | LUAD | NCI-H1838 | A-443654 |
| MVP_top60_DNA_seed | EGFR | 0.868104 | LGG | KALS-1 | Gefitinib | MVP_top60_DNA+RNA_seed | HSP90AA1 | 2.293336 | LUAD | HCC-78 | Elesclomol |
| MVP_top60_DNA_seed | EGFR | 0.870005 | LGG | KINGS-1 | Cetuximab | MVP_top60_DNA+RNA_seed | PTK2 | 2.303493 | LUAD | NCI-H1838 | PF-562271 |
| MVP_top60_DNA_seed | EGFR | 0.874775 | LGG | no-10 | PF-00299804 | MVP_top60_DNA+RNA_seed | EGFR | 2.316374 | LUAD | NCI-H2122 | CI-1033 |
| MVP_top60_DNA_seed | JAK1 | 0.879138 | LGG | MOG-G-CCM | JAK1_3715 | MVP_top60_DNA+RNA_seed | EGFR | 2.322947 | LUAD | NCI-H2122 | AST-1306 |
| MVP_top60_DNA_seed | EGFR | 0.881212 | LGG | Hs-683 | PF-00299804 | MVP_top60_DNA+RNA_seed | EGFR | 2.32736 | LUAD | NCI-H2023 | AST-1306 |
| MVP_top60_DNA_seed | HSP90AA1 | 0.881457 | LGG | KNS-81-FD | SNX-2112 | MVP_top60_DNA+RNA_seed | MAPK14 | 2.341349 | LUAD | NCI-H2291 | TAK-715 |
| MVP_top60_DNA_seed | MAPK1 | 0.882235 | LGG | D-336MG | VX-11e | MVP_top60_DNA+RNA_seed | EGFR | 2.343242 | LUAD | NCI-H1435 | CUDC-101 |
| MVP_top60_DNA_seed | JAK1 | 0.895632 | LGG | no-10 | JAK1_3715 | MVP_top60_DNA+RNA_seed | IGF1R | 2.34476 | LUAD | NCI-H1693 | BMS-754807 |
| MVP_top60_DNA_seed | PIK3CA | 0.897285 | LGG | Hs-683 | AZD8835 | MVP_top60_DNA+RNA_seed | EGFR | 2.358305 | LUAD | RERF-LC-KJ | CI-1033 |
| MVP_top60_DNA_seed | PIK3CD | 0.897285 | LGG | Hs-683 | AZD8835 | MVP_top60_DNA+RNA_seed | HDAC1 | 2.36656 | LUAD | NCI-H2291 | JQ12 |
| MVP_top60_DNA_seed | MDM2 | 0.90167 | LGG | KNS-42 | Serdemetan | MVP_top60_DNA+RNA_seed | IGF1R | 2.367299 | LUAD | NCI-H322M | GSK1904529A |
| MVP_top60_DNA_seed | AR | 0.906736 | LGG | KNS-81-FD | Bicalutamide | MVP_top60_DNA+RNA_seed | EGFR | 2.381762 | LUAD | NCI-H2023 | CI-1033 |
| MVP_top60_DNA_seed | PIK3CA | 0.908864 | LGG | KNS-42 | PI3Ka_4409 | MVP_top60_DNA+RNA_seed | ATM | 2.382001 | LUAD | NCI-H441 | KU-55933 |
| MVP_top60_DNA_seed | PIK3CD | 0.908864 | LGG | KNS-42 | PI3Ka_4409 | MVP_top60_DNA+RNA_seed | BRAF | 2.390548 | LUAD | NCI-H1563 | RAF_9304 |
| MVP_top60_DNA_seed | IGF1R | 0.913818 | LGG | KALS-1 | BMS-536924 | MVP_top60_DNA+RNA_seed | EGFR | 2.396535 | LUAD | NCI-H1437 | CI-1033 |
| MVP_top60_DNA_seed | EGFR | 0.916778 | LGG | MOG-G-CCM | Afatinib | MVP_top60_DNA+RNA_seed | HSP90AA1 | 2.400884 | LUAD | NCI-H1793 | SNX-2112 |
| MVP_top60_DNA_seed | CDK2 | 0.918672 | LGG | KNS-81-FD | PHA-793887 | MVP_top60_DNA+RNA_seed | ATM | 2.412826 | LUAD | Calu-3 | KU-60019 |
| MVP_top60_DNA_seed | JAK3 | 0.919756 | LGG | GAMG | WHI-P97 | MVP_top60_DNA+RNA_seed | HDAC1 | 2.416005 | LUAD | NCI-H1666 | Dacinostat |
| MVP_top60_DNA_seed | HSF1 | 0.923046 | LGG | MOG-G-CCM | CCT245232 | MVP_top60_DNA+RNA_seed | AURKA | 2.419294 | LUAD | EKVX | Genentech Cpd 10 |
| MVP_top60_DNA_seed | PIK3CD | 0.923452 | LGG | NMC-G1 | AZD8186 | MVP_top60_DNA+RNA_seed | HSP90AA1 | 2.419708 | LUAD | NCI-H2291 | SNX-2112 |
| MVP_top60_DNA_seed | PIK3CD | 0.927708 | LGG | no-10 | IC-87114 | MVP_top60_DNA+RNA_seed | HDAC1 | 2.433605 | LUAD | NCI-H1563 | Dacinostat |
| MVP_top60_DNA_seed | HSF1 | 0.930243 | LGG | D-502MG | CCT245467 | MVP_top60_DNA+RNA_seed | EGFR | 2.457773 | LUAD | NCI-H838 | CI-1033 |
| MVP_top60_DNA_seed | PDGFRA | 0.930666 | LGG | MOG-G-UVW | Amuvatinib | MVP_top60_DNA+RNA_seed | HDAC1 | 2.45924 | LUAD | NCI-H1838 | JQ12 |
| MVP_top60_DNA_seed | PDGFRB | 0.932948 | LGG | no-10 | Masitinib | MVP_top60_DNA+RNA_seed | AKT1 | 2.476136 | LUAD | NCI-H1793 | AKT inhibitor VIII |
| MVP_top60_DNA_seed | PDGFRA | 0.932948 | LGG | no-10 | Masitinib | MVP_top60_DNA+RNA_seed | HDAC1 | 2.487357 | LUAD | NCI-H322M | JQ12 |
| MVP_top60_DNA_seed | EGFR | 0.936563 | LGG | KNS-81-FD | AST-1306 | MVP_top60_DNA+RNA_seed | AURKA | 2.496792 | LUAD | NCI-H2291 | Genentech Cpd 10 |
| MVP_top60_DNA_seed | PIK3CD | 0.939401 | LGG | KNS-81-FD | Idelalisib | MVP_top60_DNA+RNA_seed | ATM | 2.547208 | LUAD | EKVX | CP466722 |
| MVP_top60_DNA_seed | MET | 0.945665 | LGG | no-11 | Crizotinib | MVP_top60_DNA+RNA_seed | HSP90AA1 | 2.549254 | LUAD | NCI-H1623 | Luminespib |
| MVP_top60_DNA_seed | MET | 0.945859 | LGG | NMC-G1 | PHA-665752 | MVP_top60_DNA+RNA_seed | PTK2 | 2.577374 | LUAD | NCI-H2291 | PF-562271 |
| MVP_top60_DNA_seed | IGF1R | 0.946545 | LGG | Hs-683 | Linsitinib | MVP_top60_DNA+RNA_seed | BRAF | 2.593319 | LUAD | NCI-H2291 | HG6-64-1 |
| MVP_top60_DNA_seed | EGFR | 0.949388 | LGG | KNS-81-FD | Gefitinib | MVP_top60_DNA+RNA_seed | HSP90AA1 | 2.608085 | LUAD | NCI-H1666 | Luminespib |
| MVP_top60_DNA_seed | PIK3CA | 0.952488 | LGG | KALS-1 | AZD8835 | MVP_top60_DNA+RNA_seed | HDAC1 | 2.625685 | LUAD | NCI-H1793 | Dacinostat |
| MVP_top60_DNA_seed | PIK3CD | 0.952488 | LGG | KALS-1 | AZD8835 | MVP_top60_DNA+RNA_seed | AKT1 | 2.650929 | LUAD | NCI-H1666 | A-443654 |
| MVP_top60_DNA_seed | JAK2 | 0.953385 | LGG | KINGS-1 | Ruxolitinib | MVP_top60_DNA+RNA_seed | BRAF | 2.718412 | LUAD | NCI-H322M | RAF_9304 |
| MVP_top60_DNA_seed | JAK1 | 0.953385 | LGG | KINGS-1 | Ruxolitinib | MVP_top60_DNA+RNA_seed | HSP90AA1 | 2.73961 | LUAD | NCI-H1563 | Luminespib |
| MVP_top60_DNA_seed | PIK3CD | 0.955857 | LGG | KNS-42 | IC-87114 | MVP_top60_DNA+RNA_seed | EGFR | 2.792776 | LUAD | NCI-H1568 | CUDC-101 |
| MVP_top60_DNA_seed | EGFR | 0.966507 | LGG | NMC-G1 | Gefitinib | MVP_top60_DNA+RNA_seed | ATM | 2.802996 | LUAD | NCI-H1993 | CP466722 |
| MVP_top60_DNA_seed | PIK3CD | 0.967082 | LGG | no-11 | IC-87114 | MVP_top60_DNA+RNA_seed | ATM | 2.807024 | LUAD | NCI-H322M | KU-60019 |

| | | | | | | | | | | | |
|---|---|---|---|---|---|---|---|---|---|---|---|
| MVP_top60_DNA_seed | NTRK1 | 0.968956 | LGG | MOG-G-CCM | GW441756 | MVP_top60_DNA+RNA_seed | HDAC1 | 2.824876 | LUAD | NCI-H1568 | AR-42 |
| MVP_top60_DNA_seed | HDAC1 | 0.972646 | LGG | KINGS-1 | JQ12 | MVP_top60_DNA+RNA_seed | EGFR | 2.844676 | LUAD | NCI-H838 | AST-1306 |
| MVP_top60_DNA_seed | CREBBP | 0.982842 | LGG | KALS-1 | I-CBP112 | MVP_top60_DNA+RNA_seed | HSP90AA1 | 2.857503 | LUAD | NCI-H1993 | Elesclomol |
| MVP_top60_DNA_seed | PIK3CA | 0.990872 | LGG | KNS-42 | AZD8835 | MVP_top60_DNA+RNA_seed | HSP90AA1 | 2.869074 | LUAD | NCI-H322M | Luminespib |
| MVP_top60_DNA_seed | PIK3CD | 0.990872 | LGG | KNS-42 | AZD8835 | MVP_top60_DNA+RNA_seed | EGFR | 2.886417 | LUAD | NCI-H1437 | PF-00299804 |
| MVP_top60_DNA_seed | JAK1 | 0.992965 | LGG | NMC-G1 | JAK1_3715 | MVP_top60_DNA+RNA_seed | HDAC1 | 2.921016 | LUAD | NCI-H1838 | Entinostat |
| MVP_top60_DNA_seed | EGFR | 0.998428 | LGG | no-11 | Afatinib | MVP_top60_DNA+RNA_seed | HDAC1 | 2.927149 | LUAD | NCI-H1435 | Belinostat |
| MVP_top60_DNA_seed | EGFR | 1.003117 | LGG | KNS-42 | Afatinib | MVP_top60_DNA+RNA_seed | HSP90AA1 | 2.944004 | LUAD | NCI-H1838 | Luminespib |
| MVP_top60_DNA_seed | JAK2 | 1.009866 | LGG | KNS-42 | Fedratinib | MVP_top60_DNA+RNA_seed | HDAC1 | 2.977468 | LUAD | NCI-H322M | Entinostat |
| MVP_top60_DNA_seed | HSF1 | 1.016141 | LGG | KNS-42 | CCT245467 | MVP_top60_DNA+RNA_seed | HDAC1 | 2.982262 | LUAD | NCI-H1568 | CAY10603 |
| MVP_top60_DNA_seed | HSF1 | 1.016795 | LGG | no-10 | CCT245467 | MVP_top60_DNA+RNA_seed | HSP90AA1 | 3.02026 | LUAD | NCI-H441 | Elesclomol |
| MVP_top60_DNA_seed | EGFR | 1.020504 | LGG | KINGS-1 | Afatinib | MVP_top60_DNA+RNA_seed | HDAC1 | 3.066321 | LUAD | NCI-H2291 | CAY10603 |
| MVP_top60_DNA_seed | EGFR | 1.026118 | LGG | KINGS-1 | CI-1033 | MVP_top60_DNA+RNA_seed | HDAC1 | 3.086976 | LUAD | NCI-H1568 | Belinostat |
| MVP_top60_DNA_seed | IGF1R | 1.026583 | LGG | SW1783 | BMS-536924 | MVP_top60_DNA+RNA_seed | HDAC1 | 3.111411 | LUAD | NCI-H2291 | Belinostat |
| MVP_top60_DNA_seed | HDAC1 | 1.036901 | LGG | no-11 | Belinostat | MVP_top60_DNA+RNA_seed | EGFR | 3.111495 | LUAD | NCI-H2291 | CUDC-101 |
| MVP_top60_DNA_seed | MAPK1 | 1.04068 | LGG | MOG-G-CCM | FR-180204 | MVP_top60_DNA+RNA_seed | EGFR | 3.197471 | LUAD | NCI-H838 | PF-00299804 |
| MVP_top60_DNA_seed | MAPK3 | 1.04068 | LGG | MOG-G-CCM | FR-180204 | MVP_top60_DNA+RNA_seed | HDAC1 | 3.251255 | LUAD | NCI-H322M | Dacinostat |
| MVP_top60_DNA_seed | PDGFRB | 1.049618 | LGG | KALS-1 | Pazopanib | MVP_top60_DNA+RNA_seed | HSP90AA1 | 3.292018 | LUAD | NCI-H2291 | Luminespib |
| MVP_top60_DNA_seed | PIK3CA | 1.0522 | LGG | no-10 | AZD8835 | MVP_top60_DNA+RNA_seed | HDAC1 | 3.313231 | LUAD | NCI-H2291 | AR-42 |
| MVP_top60_DNA_seed | PIK3CD | 1.0522 | LGG | no-10 | AZD8835 | MVP_top60_DNA+RNA_seed | MDM2 | 3.439305 | LUAD | NCI-H322M | Serdemetan |
| MVP_top60_DNA_seed | MET | 1.052537 | LGG | no-11 | AZD6094 | MVP_top60_DNA+RNA_seed | HDAC1 | 3.901874 | LUAD | NCI-H2291 | Dacinostat |
| MVP_top60_DNA_seed | IGF1R | 1.057229 | LGG | D-502MG | BMS-536924 | MVP_top60_DNA+RNA_seed | HDAC1 | 4.147478 | LUAD | NCI-H1838 | Dacinostat |
| MVP_top60_DNA_seed | AR | 1.058228 | LGG | D-502MG | AZD3514 | MVP_top60_DNA+RNA_seed | EGFR | -4.854198 | LUAD | HCC-827 | Gefitinib |
| MVP_top60_DNA_seed | MAPK1 | 1.060534 | LGG | D-502MG | VX-11e | MVP_top60_DNA+RNA_seed | MET | -4.82268 | LUAD | NCI-H1993 | Savolitinib |
| MVP_top60_DNA_seed | EGFR | 1.061562 | LGG | KALS-1 | Afatinib | MVP_top60_DNA+RNA_seed | EGFR | -4.716321 | LUAD | HCC-827 | AZD3759 |
| MVP_top60_DNA_seed | EGFR | 1.061812 | LGG | Hs-683 | CUDC-101 | MVP_top60_DNA+RNA_seed | EGFR | -4.534566 | LUAD | PC-14 | AZD3759 |
| MVP_top60_DNA_seed | MDM2 | 1.061853 | LGG | KNS-42 | Nutlin-3a (-) | MVP_top60_DNA+RNA_seed | EGFR | -4.200302 | LUAD | PC-14 | Sapitinib |
| MVP_top60_DNA_seed | HDAC1 | 1.067665 | LGG | KNS-81-FD | Tubastatin A | MVP_top60_DNA+RNA_seed | EGFR | -4.099897 | LUAD | HCC-827 | Osimertinib |
| MVP_top60_DNA_seed | HSP90AA1 | 1.071556 | LGG | KNS-42 | CCT-018159 | MVP_top60_DNA+RNA_seed | EGFR | -4.05738 | LUAD | PC-3_[JPC-3] | Gefitinib |
| MVP_top60_DNA_seed | CDK2 | 1.076836 | LGG | SW1783 | PHA-793887 | MVP_top60_DNA+RNA_seed | EGFR | -3.925376 | LUAD | HCC-827 | Erlotinib |
| MVP_top60_DNA_seed | PIK3CD | 1.08599 | LGG | KNS-81-FD | IC-87114 | MVP_top60_DNA+RNA_seed | EGFR | -3.890432 | LUAD | H3255 | AZD3759 |
| MVP_top60_DNA_seed | MET | 1.088293 | LGG | D-502MG | SU11274 | MVP_top60_DNA+RNA_seed | EGFR | -3.873859 | LUAD | PC-14 | Osimertinib |
| MVP_top60_DNA_seed | IGF1R | 1.093549 | LGG | MOG-G-CCM | BMS-536924 | MVP_top60_DNA+RNA_seed | EGFR | -3.853431 | LUAD | NCI-H1975 | Osimertinib |
| MVP_top60_DNA_seed | IGF1R | 1.095313 | LGG | MOG-G-CCM | Linsitinib | MVP_top60_DNA+RNA_seed | EGFR | -3.747843 | LUAD | PC-14 | Gefitinib |
| MVP_top60_DNA_seed | JAK2 | 1.095511 | LGG | NMC-G1 | Ruxolitinib | MVP_top60_DNA+RNA_seed | EGFR | -3.734321 | LUAD | HCC-827 | Afatinib |
| MVP_top60_DNA_seed | JAK1 | 1.095511 | LGG | NMC-G1 | Ruxolitinib | MVP_top60_DNA+RNA_seed | EGFR | -3.686848 | LUAD | PC-3_[JPC-3] | AZD3759 |
| MVP_top60_DNA_seed | MET | 1.097952 | LGG | D-502MG | Crizotinib | MVP_top60_DNA+RNA_seed | EGFR | -3.551576 | LUAD | PC-14 | Erlotinib |
| MVP_top60_DNA_seed | MAPK1 | 1.100783 | LGG | no-11 | VX-11e | MVP_top60_DNA+RNA_seed | EGFR | -3.473928 | LUAD | NCI-H322M | Sapitinib |
| MVP_top60_DNA_seed | MET | 1.104039 | LGG | no-10 | SU11274 | MVP_top60_DNA+RNA_seed | EGFR | -3.460764 | LUAD | PC-14 | Afatinib |
| MVP_top60_DNA_seed | HSP90AA1 | 1.105953 | LGG | KNS-42 | Elesclomol | MVP_top60_DNA+RNA_seed | EGFR | -3.359337 | LUAD | COR-L105 | Sapitinib |
| MVP_top60_DNA_seed | PIK3CA | 1.106367 | LGG | no-11 | AZD8835 | MVP_top60_DNA+RNA_seed | EGFR | -3.329265 | LUAD | PC-3_[JPC-3] | Osimertinib |

| | | | | | | | | | | | |
|---|---|---|---|---|---|---|---|---|---|---|---|
| MVP_top60_DNA_seed | PIK3CD | 1.106367 | LGG | no-11 | AZD8835 | MVP_top60_DNA+RNA_seed | EGFR | -3.301066 | LUAD | NCI-H322M | Gefitinib |
| MVP_top60_DNA_seed | MET | 1.109998 | LGG | no-10 | Crizotinib | MVP_top60_DNA+RNA_seed | EGFR | -3.27183 | LUAD | COR-L105 | Gefitinib |
| MVP_top60_DNA_seed | MET | 1.111716 | LGG | KNS-81-FD | SU11274 | MVP_top60_DNA+RNA_seed | EGFR | -3.267587 | LUAD | H3255 | Osimertinib |
| MVP_top60_DNA_seed | IGF1R | 1.113075 | LGG | KNS-42 | BMS-754807 | MVP_top60_DNA+RNA_seed | EGFR | -3.259498 | LUAD | H3255 | Afatinib |
| MVP_top60_DNA_seed | HSP90AA1 | 1.11581 | LGG | KNS-42 | Luminespib | MVP_top60_DNA+RNA_seed | EGFR | -3.190589 | LUAD | COR-L105 | Afatinib |
| MVP_top60_DNA_seed | EGFR | 1.117272 | LGG | H4 | CUDC-101 | MVP_top60_DNA+RNA_seed | EGFR | -3.183559 | LUAD | H3255 | Sapitinib |
| MVP_top60_DNA_seed | MET | 1.125976 | LGG | no-10 | AZD6094 | MVP_top60_DNA+RNA_seed | EGFR | -3.108259 | LUAD | PC-3_[JPC-3] | Erlotinib |
| MVP_top60_DNA_seed | EGFR | 1.126234 | LGG | no-10 | Gefitinib | MVP_top60_DNA+RNA_seed | EGFR | -3.061933 | LUAD | PC-3_[JPC-3] | Sapitinib |
| MVP_top60_DNA_seed | EGFR | 1.128556 | LGG | KINGS-1 | Pelitinib | MVP_top60_DNA+RNA_seed | EGFR | -3.043923 | LUAD | NCI-H322M | Afatinib |
| MVP_top60_DNA_seed | HDAC1 | 1.136237 | LGG | KNS-81-FD | Entinostat | MVP_top60_DNA+RNA_seed | EGFR | -2.988821 | LUAD | NCI-H322M | AZD3759 |
| MVP_top60_DNA_seed | JAK3 | 1.138067 | LGG | no-11 | JAK3_7406 | MVP_top60_DNA+RNA_seed | EGFR | -2.95988 | LUAD | H3255 | Gefitinib |
| MVP_top60_DNA_seed | HSF1 | 1.143047 | LGG | D-336MG | CCT245467 | MVP_top60_DNA+RNA_seed | EGFR | -2.869384 | LUAD | NCI-H322M | Erlotinib |
| MVP_top60_DNA_seed | NTRK1 | 1.148471 | LGG | KNS-42 | GW441756 | MVP_top60_DNA+RNA_seed | EGFR | -2.85431 | LUAD | NCI-H1623 | Sapitinib |
| MVP_top60_DNA_seed | PIK3CD | 1.152775 | LGG | KNS-42 | Idelalisib | MVP_top60_DNA+RNA_seed | EGFR | -2.756579 | LUAD | PC-3_[JPC-3] | Afatinib |
| MVP_top60_DNA_seed | EGFR | 1.157726 | LGG | KNS-42 | PF-00299804 | MVP_top60_DNA+RNA_seed | EGFR | -2.60257 | LUAD | COR-L105 | AZD3759 |
| MVP_top60_DNA_seed | JAK1 | 1.163022 | LGG | KINGS-1 | JAK1_3715 | MVP_top60_DNA+RNA_seed | IGF1R | -2.583223 | LUAD | NCI-H2122 | BMS-536924 |
| MVP_top60_DNA_seed | HDAC1 | 1.163441 | LGG | no-11 | AR-42 | MVP_top60_DNA+RNA_seed | EGFR | -2.520692 | LUAD | NCI-H1648 | Sapitinib |
| MVP_top60_DNA_seed | HSF1 | 1.181149 | LGG | MOG-G-UVW | CCT245467 | MVP_top60_DNA+RNA_seed | MET | -2.42713 | LUAD | NCI-H1993 | Crizotinib |
| MVP_top60_DNA_seed | EGFR | 1.190454 | LGG | KINGS-1 | AST-1306 | MVP_top60_DNA+RNA_seed | EGFR | -2.374387 | LUAD | COR-L105 | Osimertinib |
| MVP_top60_DNA_seed | JAK2 | 1.1919 | LGG | D-502MG | Ruxolitinib | MVP_top60_DNA+RNA_seed | EGFR | -2.373031 | LUAD | NCI-H322M | Osimertinib |
| MVP_top60_DNA_seed | JAK1 | 1.1919 | LGG | D-502MG | Ruxolitinib | MVP_top60_DNA+RNA_seed | EGFR | -2.364013 | LUAD | NCI-H1648 | Gefitinib |
| MVP_top60_DNA_seed | CDK2 | 1.194108 | LGG | D-502MG | Seliciclib | MVP_top60_DNA+RNA_seed | EGFR | -2.353975 | LUAD | COR-L105 | Erlotinib |
| MVP_top60_DNA_seed | PDGFRA | 1.201971 | LGG | KNS-42 | Amuvatinib | MVP_top60_DNA+RNA_seed | MET | -2.3503 | LUAD | RERF-LC-MS | Savolitinib |
| MVP_top60_DNA_seed | CREBBP | 1.203669 | LGG | no-10 | I-CBP112 | MVP_top60_DNA+RNA_seed | EGFR | -2.307295 | LUAD | NCI-H1781 | Afatinib |
| MVP_top60_DNA_seed | EGFR | 1.208594 | LGG | KALS-1 | Cetuximab | MVP_top60_DNA+RNA_seed | EGFR | -2.297143 | LUAD | NCI-H1666 | Sapitinib |
| MVP_top60_DNA_seed | MDM2 | 1.210894 | LGG | no-11 | Nutlin-3a (-) | MVP_top60_DNA+RNA_seed | PIK3CA | -2.275126 | LUAD | NCI-H1648 | Alpelisib |
| MVP_top60_DNA_seed | HDAC1 | 1.212105 | LGG | D-502MG | Tubastatin A | MVP_top60_DNA+RNA_seed | EGFR | -2.269518 | LUAD | Calu-3 | Sapitinib |
| MVP_top60_DNA_seed | EGFR | 1.214041 | LGG | no-11 | AZD8931 | MVP_top60_DNA+RNA_seed | EGFR | -2.267607 | LUAD | Calu-3 | Afatinib |
| MVP_top60_DNA_seed | PIK3CD | 1.221336 | LGG | KINGS-1 | AZD8186 | MVP_top60_DNA+RNA_seed | EGFR | -2.255715 | LUAD | NCI-H1623 | Afatinib |
| MVP_top60_DNA_seed | CDK2 | 1.223616 | LGG | D-502MG | PHA-793887 | MVP_top60_DNA+RNA_seed | SRC | -2.169273 | LUAD | PC-14 | Bosutinib |
| MVP_top60_DNA_seed | JAK3 | 1.224175 | LGG | Hs-683 | JAK3_7406 | MVP_top60_DNA+RNA_seed | EGFR | -2.125228 | LUAD | NCI-H1666 | Gefitinib |
| MVP_top60_DNA_seed | EGFR | 1.230171 | LGG | KNS-81-FD | Pelitinib | MVP_top60_DNA+RNA_seed | EGFR | -2.117487 | LUAD | NCI-H1648 | Afatinib |
| MVP_top60_DNA_seed | SMARCA4 | 1.230453 | LGG | no-10 | PFI-3 | MVP_top60_DNA+RNA_seed | EGFR | -2.10784 | LUAD | NCI-H1648 | AZD3759 |
| MVP_top60_DNA_seed | IGF1R | 1.232138 | LGG | no-10 | BMS-754807 | MVP_top60_DNA+RNA_seed | EGFR | -2.096518 | LUAD | EMC-BAC-1 | Afatinib |
| MVP_top60_DNA_seed | PDGFRA | 1.23378 | LGG | KINGS-1 | Amuvatinib | MVP_top60_DNA+RNA_seed | EGFR | -2.087908 | LUAD | NCI-H1648 | Erlotinib |
| MVP_top60_DNA_seed | MAPK1 | 1.241305 | LGG | no-10 | FR-180204 | MVP_top60_DNA+RNA_seed | IGF1R | -2.079693 | LUAD | NCI-H2122 | NVP-ADW742 |
| MVP_top60_DNA_seed | MAPK3 | 1.241305 | LGG | no-10 | FR-180204 | MVP_top60_DNA+RNA_seed | EGFR | -2.060094 | LUAD | NCI-H1648 | Lapatinib |
| MVP_top60_DNA_seed | PDGFRB | 1.252898 | LGG | no-10 | Pazopanib | MVP_top60_DNA+RNA_seed | IGF1R | -2.040206 | LUAD | NCI-H2122 | BMS-754807 |
| MVP_top60_DNA_seed | IGF1R | 1.255866 | LGG | MOG-G-UVW | BMS-536924 | MVP_top60_DNA+RNA_seed | MET | -2.02756 | LUAD | NCI-H3122 | Crizotinib |
| MVP_top60_DNA_seed | PIK3CA | 1.259339 | LGG | KNS-81-FD | AZD8835 | MVP_top60_DNA+RNA_seed | EGFR | -1.995851 | LUAD | NCI-H1568 | Erlotinib |
| MVP_top60_DNA_seed | PIK3CD | 1.259339 | LGG | KNS-81-FD | AZD8835 | MVP_top60_DNA+RNA_seed | HSP90AA1 | -1.975503 | LUAD | A549 | Tanespimycin |

| | | | | | | | | | | | |
|---|---|---|---|---|---|---|---|---|---|---|---|
| MVP_top60_DNA_seed | EGFR | 1.262328 | LGG | KALS-1 | AST-1306 | MVP_top60_DNA+RNA_seed | EGFR | -1.966407 | LUAD | NCI-H1666 | AZD3759 |
| MVP_top60_DNA_seed | MDM2 | 1.266545 | LGG | MOG-G-CCM | Nutlin-3a (-) | MVP_top60_DNA+RNA_seed | EGFR | -1.96437 | LUAD | EMC-BAC-1 | Sapitinib |
| MVP_top60_DNA_seed | CREBBP | 1.271723 | LGG | KINGS-1 | I-CBP112 | MVP_top60_DNA+RNA_seed | HDAC1 | -1.964111 | LUAD | NCI-H2122 | Entinostat |
| MVP_top60_DNA_seed | CDK2 | 1.277353 | LGG | KNS-42 | PHA-793887 | MVP_top60_DNA+RNA_seed | EGFR | -1.928717 | LUAD | EMC-BAC-1 | AZD3759 |
| MVP_top60_DNA_seed | EGFR | 1.282193 | LGG | KNS-81-FD | PF-00299804 | MVP_top60_DNA+RNA_seed | EGFR | -1.928576 | LUAD | NCI-H2030 | AZD3759 |
| MVP_top60_DNA_seed | IGF1R | 1.285342 | LGG | KINGS-1 | BMS-536924 | MVP_top60_DNA+RNA_seed | EGFR | -1.90466 | LUAD | NCI-H1568 | Gefitinib |
| MVP_top60_DNA_seed | HDAC1 | 1.286096 | LGG | NMC-G1 | JQ12 | MVP_top60_DNA+RNA_seed | EZH2 | -1.903149 | LUAD | HCC-78 | GSK343 |
| MVP_top60_DNA_seed | HSF1 | 1.287093 | LGG | H4 | CCT245467 | MVP_top60_DNA+RNA_seed | EGFR | -1.8844 | LUAD | NCI-H1623 | Lapatinib |
| MVP_top60_DNA_seed | PDGFRA | 1.287831 | LGG | KALS-1 | Amuvatinib | MVP_top60_DNA+RNA_seed | EZH2 | -1.85163 | LUAD | Calu-3 | GSK343 |
| MVP_top60_DNA_seed | EGFR | 1.289915 | LGG | no-11 | Lapatinib | MVP_top60_DNA+RNA_seed | EGFR | -1.841329 | LUAD | NCI-H1975 | Afatinib |
| MVP_top60_DNA_seed | EGFR | 1.291035 | LGG | KNS-81-FD | Afatinib | MVP_top60_DNA+RNA_seed | EGFR | -1.841322 | LUAD | NCI-H1648 | Osimertinib |
| MVP_top60_DNA_seed | EGFR | 1.295543 | LGG | KINGS-1 | Gefitinib | MVP_top60_DNA+RNA_seed | EGFR | -1.835846 | LUAD | H3255 | Erlotinib |
| MVP_top60_DNA_seed | IGF1R | 1.304791 | LGG | NMC-G1 | BMS-536924 | MVP_top60_DNA+RNA_seed | EGFR | -1.827677 | LUAD | NCI-H1666 | Erlotinib |
| MVP_top60_DNA_seed | CREBBP | 1.305524 | LGG | no-11 | I-CBP112 | MVP_top60_DNA+RNA_seed | HSP90AA1 | -1.817473 | LUAD | HCC-44 | Tanespimycin |
| MVP_top60_DNA_seed | CREBBP | 1.309189 | LGG | D-502MG | I-CBP112 | MVP_top60_DNA+RNA_seed | AURKA | -1.814104 | LUAD | NCI-H2122 | Alisertib |
| MVP_top60_DNA_seed | HSP90AA1 | 1.311215 | LGG | no-10 | SNX-2112 | MVP_top60_DNA+RNA_seed | SRC | -1.806877 | LUAD | HCC-827 | Bosutinib |
| MVP_top60_DNA_seed | AR | 1.317407 | LGG | no-11 | Bicalutamide | MVP_top60_DNA+RNA_seed | EGFR | -1.788226 | LUAD | NCI-H1568 | Sapitinib |
| MVP_top60_DNA_seed | HDAC1 | 1.322739 | LGG | KNS-42 | Dacinostat | MVP_top60_DNA+RNA_seed | PIK3CA | -1.752711 | LUAD | NCI-H1648 | GNE-317 |
| MVP_top60_DNA_seed | MET | 1.323554 | LGG | no-11 | PHA-665752 | MVP_top60_DNA+RNA_seed | EGFR | -1.742609 | LUAD | H3255 | Lapatinib |
| MVP_top60_DNA_seed | AR | 1.323787 | LGG | KALS-1 | AZD3514 | MVP_top60_DNA+RNA_seed | EGFR | -1.738422 | LUAD | Calu-3 | Lapatinib |
| MVP_top60_DNA_seed | HDAC1 | 1.325815 | LGG | Hs-683 | Belinostat | MVP_top60_DNA+RNA_seed | HSP90AA1 | -1.725962 | LUAD | ABC-1 | Elesclomol |
| MVP_top60_DNA_seed | JAK3 | 1.332564 | LGG | no-11 | WHI-P97 | MVP_top60_DNA+RNA_seed | EGFR | -1.718223 | LUAD | PC-14 | Lapatinib |
| MVP_top60_DNA_seed | EGFR | 1.336917 | LGG | D-502MG | Gefitinib | MVP_top60_DNA+RNA_seed | IGF1R | -1.693025 | LUAD | NCI-H1648 | BMS-536924 |
| MVP_top60_DNA_seed | HDAC1 | 1.340658 | LGG | no-11 | Tubastatin A | MVP_top60_DNA+RNA_seed | EGFR | -1.685023 | LUAD | EMC-BAC-1 | Lapatinib |
| MVP_top60_DNA_seed | JAK1 | 1.343994 | LGG | KALS-1 | JAK1_3715 | MVP_top60_DNA+RNA_seed | EGFR | -1.679859 | LUAD | NCI-H1623 | Erlotinib |
| MVP_top60_DNA_seed | SMARCA4 | 1.347445 | LGG | KINGS-1 | PFI-3 | MVP_top60_DNA+RNA_seed | IGF1R | -1.666667 | LUAD | NCI-H2122 | Linsitinib |
| MVP_top60_DNA_seed | HSF1 | 1.353296 | LGG | KNS-81-FD | CCT245467 | MVP_top60_DNA+RNA_seed | EGFR | -1.662853 | LUAD | NCI-H2122 | Erlotinib |
| MVP_top60_DNA_seed | HSP90AA1 | 1.35569 | LGG | D-502MG | SNX-2112 | MVP_top60_DNA+RNA_seed | KRAS | -1.609231 | LUAD | EMC-BAC-1 | [illegible]RAS (G12C) Inhibitor-1 |
| MVP_top60_DNA_seed | MAPK1 | 1.357089 | LGG | KALS-1 | FR-180204 | MVP_top60_DNA+RNA_seed | EGFR | -1.608489 | LUAD | PC-3_[JPC-3] | Lapatinib |
| MVP_top60_DNA_seed | MAPK3 | 1.357089 | LGG | KALS-1 | FR-180204 | MVP_top60_DNA+RNA_seed | PIK3CA | -1.606638 | LUAD | NCI-H2122 | Alpelisib |
| MVP_top60_DNA_seed | PIK3CD | 1.362629 | LGG | KNS-42 | AZD8186 | MVP_top60_DNA+RNA_seed | EGFR | -1.60601 | LUAD | NCI-H1568 | AZD3759 |
| MVP_top60_DNA_seed | MAPK1 | 1.363923 | LGG | no-11 | FR-180204 | MVP_top60_DNA+RNA_seed | AKT1 | -1.599536 | LUAD | NCI-H1648 | Afuresertib |
| MVP_top60_DNA_seed | MAPK3 | 1.363923 | LGG | no-11 | FR-180204 | MVP_top60_DNA+RNA_seed | HSP90AA1 | -1.577668 | LUAD | PC-14 | Tanespimycin |
| MVP_top60_DNA_seed | SMARCA4 | 1.368213 | LGG | no-11 | PFI-3 | MVP_top60_DNA+RNA_seed | EGFR | -1.565886 | LUAD | NCI-H1781 | Osimertinib |
| MVP_top60_DNA_seed | PIK3CA | 1.372272 | LGG | Hs-683 | PI3Ka_4409 | MVP_top60_DNA+RNA_seed | AKT1 | -1.561795 | LUAD | NCI-H1975 | AZD5363 |
| MVP_top60_DNA_seed | PIK3CD | 1.372272 | LGG | Hs-683 | PI3Ka_4409 | MVP_top60_DNA+RNA_seed | EGFR | -1.561755 | LUAD | NCI-H358 | Sapitinib |
| MVP_top60_DNA_seed | EGFR | 1.379924 | LGG | KINGS-1 | Lapatinib | MVP_top60_DNA+RNA_seed | AKT1 | -1.553685 | LUAD | NCI-H2122 | Ipatasertib |
| MVP_top60_DNA_seed | HDAC1 | 1.392392 | LGG | KNS-42 | Parthenolide | MVP_top60_DNA+RNA_seed | EZH2 | -1.532232 | LUAD | LXF-289 | GSK343 |
| MVP_top60_DNA_seed | MDM2 | 1.418147 | LGG | KINGS-1 | Nutlin-3a (-) | MVP_top60_DNA+RNA_seed | IGF1R | -1.531786 | LUAD | NCI-H1755 | BMS-536924 |
| MVP_top60_DNA_seed | HDAC1 | 1.421033 | LGG | NMC-G1 | Parthenolide | MVP_top60_DNA+RNA_seed | EGFR | -1.516259 | LUAD | NCI-H1781 | Sapitinib |
| MVP_top60_DNA_seed | IGF1R | 1.425803 | LGG | KNS-81-FD | BMS-536924 | MVP_top60_DNA+RNA_seed | EGFR | -1.511946 | LUAD | NCI-H1650 | AZD3759 |

| | | | | | | | | | | | |
|---|---|---|---|---|---|---|---|---|---|---|---|
| MVP_top60_DNA_seed | AR | 1.430927 | LGG | Hs-683 | AZD3514 | MVP_top60_DNA+RNA_seed | IGF1R | -1.502644 | LUAD | NCI-H1975 | BMS-536924 |
| MVP_top60_DNA_seed | AR | 1.4314 | LGG | KNS-81-FD | Bicalutamide | MVP_top60_DNA+RNA_seed | AKT1 | -1.499631 | LUAD | NCI-H1781 | Uprosertib |
| MVP_top60_DNA_seed | EGFR | 1.431837 | LGG | KNS-42 | Pelitinib | MVP_top60_DNA+RNA_seed | PIK3CA | -1.498659 | LUAD | HCC-78 | AZD8186 |
| MVP_top60_DNA_seed | IGF1R | 1.439024 | LGG | D-502MG | Linsitinib | MVP_top60_DNA+RNA_seed | IGF1R | -1.495944 | LUAD | NCI-H1734 | BMS-754807 |
| MVP_top60_DNA_seed | MDM2 | 1.446021 | LGG | no-11 | Serdemetan | MVP_top60_DNA+RNA_seed | PIK3CA | -1.48657 | LUAD | NCI-H1648 | Buparlisib |
| MVP_top60_DNA_seed | JAK2 | 1.463143 | LGG | SW1783 | Fedratinib | MVP_top60_DNA+RNA_seed | IGF1R | -1.48337 | LUAD | NCI-H1651 | GSK1904529A |
| MVP_top60_DNA_seed | MAPK1 | 1.503451 | LGG | KINGS-1 | FR-180204 | MVP_top60_DNA+RNA_seed | MET | -1.478509 | LUAD | RERF-LC-MS | Crizotinib |
| MVP_top60_DNA_seed | MAPK3 | 1.503451 | LGG | KINGS-1 | FR-180204 | MVP_top60_DNA+RNA_seed | HSP90AA1 | -1.476283 | LUAD | NCI-H1734 | Tanespimycin |
| MVP_top60_DNA_seed | MAPK1 | 1.518944 | LGG | KNS-81-FD | VX-11e | MVP_top60_DNA+RNA_seed | HDAC1 | -1.473599 | LUAD | A549 | Romidepsin |
| MVP_top60_DNA_seed | EGFR | 1.52666 | LGG | no-11 | AST-1306 | MVP_top60_DNA+RNA_seed | HSP90AA1 | -1.468469 | LUAD | NCI-H3122 | Tanespimycin |
| MVP_top60_DNA_seed | JAK1 | 1.548438 | LGG | KNS-81-FD | JAK1_3715 | MVP_top60_DNA+RNA_seed | TERT | -1.466218 | LUAD | NCI-H838 | BIBR-1532 |
| MVP_top60_DNA_seed | IGF1R | 1.548943 | LGG | KALS-1 | GSK1904529A | MVP_top60_DNA+RNA_seed | IGF1R | -1.463177 | LUAD | HCC-78 | BMS-536924 |
| MVP_top60_DNA_seed | IGF1R | 1.560831 | LGG | KNS-81-FD | Linsitinib | MVP_top60_DNA+RNA_seed | HSP90AA1 | -1.456579 | LUAD | NCI-H1792 | Tanespimycin |
| MVP_top60_DNA_seed | AR | 1.56942 | LGG | no-11 | Bicalutamide | MVP_top60_DNA+RNA_seed | EGFR | -1.447698 | LUAD | NCI-H358 | Afatinib |
| MVP_top60_DNA_seed | JAK3 | 1.571943 | LGG | KINGS-1 | WHI-P97 | MVP_top60_DNA+RNA_seed | IGF1R | -1.435728 | LUAD | NCI-H2228 | BMS-536924 |
| MVP_top60_DNA_seed | HSP90AA1 | 1.572631 | LGG | D-502MG | Tanespimycin | MVP_top60_DNA+RNA_seed | IGF1R | -1.429392 | LUAD | NCI-H1648 | NVP-ADW742 |
| MVP_top60_DNA_seed | MET | 1.580421 | LGG | D-502MG | JNJ38877605 | MVP_top60_DNA+RNA_seed | SRC | -1.426192 | LUAD | NCI-H322M | Bosutinib |
| MVP_top60_DNA_seed | MAPK1 | 1.604177 | LGG | KNS-81-FD | FR-180204 | MVP_top60_DNA+RNA_seed | HDAC1 | -1.423335 | LUAD | NCI-H1651 | Romidepsin |
| MVP_top60_DNA_seed | MAPK3 | 1.604177 | LGG | KNS-81-FD | FR-180204 | MVP_top60_DNA+RNA_seed | PIK3CA | -1.418419 | LUAD | NCI-H1793 | GNE-317 |
| MVP_top60_DNA_seed | MET | 1.60513 | LGG | KALS-1 | SU11274 | MVP_top60_DNA+RNA_seed | EGFR | -1.417392 | LUAD | NCI-H1666 | Afatinib |
| MVP_top60_DNA_seed | SMARCA4 | 1.616831 | LGG | D-502MG | PFI-3 | MVP_top60_DNA+RNA_seed | IGF1R | -1.41588 | LUAD | NCI-H1623 | Linsitinib |
| MVP_top60_DNA_seed | HSP90AA1 | 1.634333 | LGG | no-10 | CCT-018159 | MVP_top60_DNA+RNA_seed | MDM2 | -1.403221 | LUAD | NCI-H1734 | Serdemetan |
| MVP_top60_DNA_seed | HSP90AA1 | 1.635057 | LGG | D-502MG | Luminespib | MVP_top60_DNA+RNA_seed | PIK3CA | -1.398581 | LUAD | NCI-H1734 | Alpelisib |
| MVP_top60_DNA_seed | PDGFRA | 1.640046 | LGG | D-502MG | Amuvatinib | MVP_top60_DNA+RNA_seed | EGFR | -1.39263 | LUAD | NCI-H1975 | Sapitinib |
| MVP_top60_DNA_seed | JAK2 | 1.640434 | LGG | KNS-81-FD | Fedratinib | MVP_top60_DNA+RNA_seed | MDM2 | -1.391782 | LUAD | A549 | Nutlin-3a (-) |
| MVP_top60_DNA_seed | SMARCA4 | 1.676758 | LGG | KNS-81-FD | PFI-3 | MVP_top60_DNA+RNA_seed | EGFR | -1.391569 | LUAD | RERF-LC-KJ | Erlotinib |
| MVP_top60_DNA_seed | AR | 1.678134 | LGG | KALS-1 | Bicalutamide | MVP_top60_DNA+RNA_seed | EGFR | -1.384082 | LUAD | NCI-H1623 | AZD3759 |
| MVP_top60_DNA_seed | HDAC1 | 1.678203 | LGG | KNS-42 | Entinostat | MVP_top60_DNA+RNA_seed | EZH2 | -1.377759 | LUAD | NCI-H2122 | GSK343 |
| MVP_top60_DNA_seed | EGFR | 1.686139 | LGG | KALS-1 | CI-1033 | MVP_top60_DNA+RNA_seed | IGF1R | -1.375224 | LUAD | NCI-H1437 | BMS-536924 |
| MVP_top60_DNA_seed | HDAC1 | 1.690291 | LGG | KINGS-1 | Parthenolide | MVP_top60_DNA+RNA_seed | PIK3CA | -1.372823 | LUAD | A549 | GNE-317 |
| MVP_top60_DNA_seed | PIK3CA | 1.693859 | LGG | KNS-81-FD | PI3Ka_4409 | MVP_top60_DNA+RNA_seed | AKT1 | -1.37202 | LUAD | NCI-H1975 | AT13148 |
| MVP_top60_DNA_seed | PIK3CD | 1.693859 | LGG | KNS-81-FD | PI3Ka_4409 | MVP_top60_DNA+RNA_seed | EGFR | -1.364414 | LUAD | NCI-H1623 | Gefitinib |
| MVP_top60_DNA_seed | IGF1R | 1.694902 | LGG | KNS-81-FD | BMS-536924 | MVP_top60_DNA+RNA_seed | HSP90AA1 | -1.358527 | LUAD | RERF-LC-MS | Luminespib |
| MVP_top60_DNA_seed | JAK3 | 1.712052 | LGG | KNS-81-FD | WHI-P97 | MVP_top60_DNA+RNA_seed | EGFR | -1.347203 | LUAD | NCI-H2122 | Gefitinib |
| MVP_top60_DNA_seed | EGFR | 1.742203 | LGG | KNS-81-FD | Cetuximab | MVP_top60_DNA+RNA_seed | HDAC1 | -1.341741 | LUAD | NCI-H2122 | Romidepsin |
| MVP_top60_DNA_seed | IGF1R | 1.764359 | LGG | no-10 | BMS-536924 | MVP_top60_DNA+RNA_seed | EGFR | -1.328444 | LUAD | COR-L105 | Lapatinib |
| MVP_top60_DNA_seed | HDAC1 | 1.794333 | LGG | KNS-81-FD | Dacinostat | MVP_top60_DNA+RNA_seed | HDAC1 | -1.326538 | LUAD | NCI-H1437 | Entinostat |
| MVP_top60_DNA_seed | IGF1R | 1.795102 | LGG | no-11 | BMS-536924 | MVP_top60_DNA+RNA_seed | EGFR | -1.326312 | LUAD | EMC-BAC-1 | Osimertinib |
| MVP_top60_DNA_seed | HSP90AA1 | 1.815561 | LGG | no-10 | Elesclomol | MVP_top60_DNA+RNA_seed | PIK3CA | -1.318525 | LUAD | HCC-78 | GNE-317 |
| MVP_top60_DNA_seed | HDAC1 | 1.871194 | LGG | D-336MG | Entinostat | MVP_top60_DNA+RNA_seed | HSP90AA1 | -1.314768 | LUAD | PC-14 | CCT-018159 |
| MVP_top60_DNA_seed | CDK2 | 1.8802 | LGG | KINGS-1 | Seliciclib | MVP_top60_DNA+RNA_seed | EGFR | -1.313911 | LUAD | HCC-827 | Lapatinib |

| | | | | | | | | | | | |
|---|---|---|---|---|---|---|---|---|---|---|---|
| MVP_top60_DNA_seed | HDAC1 | 1.887574 | LGG | D-336MG | Belinostat | MVP_top60_DNA+RNA_seed | SMARCA4 | -1.30627 | LUAD | NCI-H1563 | PFI3 |
| MVP_top60_DNA_seed | PDGFRB | 1.897144 | LGG | D-502MG | Masitinib | MVP_top60_DNA+RNA_seed | EGFR | -1.304226 | LUAD | NCI-H322M | Lapatinib |
| MVP_top60_DNA_seed | PDGFRA | 1.897144 | LGG | D-502MG | Masitinib | MVP_top60_DNA+RNA_seed | IGF1R | -1.302014 | LUAD | HCC-44 | BMS-536924 |
| MVP_top60_DNA_seed | HDAC1 | 1.920137 | LGG | KNS-81-FD | JQ12 | MVP_top60_DNA+RNA_seed | HSP90AA1 | -1.297642 | LUAD | NCI-H1755 | Tanespimycin |
| MVP_top60_DNA_seed | IGF1R | 1.933116 | LGG | KALS-1 | BMS-536924 | MVP_top60_DNA+RNA_seed | EGFR | -1.297525 | LUAD | LC-2-ad | Sapitinib |
| MVP_top60_DNA_seed | AR | 1.937666 | LGG | no-11 | AZD3514 | MVP_top60_DNA+RNA_seed | EGFR | -1.290848 | LUAD | EMC-BAC-1 | Gefitinib |
| MVP_top60_DNA_seed | HDAC1 | 1.943295 | LGG | D-502MG | Dacinostat | MVP_top60_DNA+RNA_seed | EGFR | -1.287555 | LUAD | NCI-H1666 | Lapatinib |
| MVP_top60_DNA_seed | CREBBP | 1.976181 | LGG | KNS-81-FD | I-CBP112 | MVP_top60_DNA+RNA_seed | AKT1 | -1.285676 | LUAD | NCI-H1993 | Uprosertib |
| MVP_top60_DNA_seed | HSF1 | 1.999872 | LGG | no-11 | CCT245467 | MVP_top60_DNA+RNA_seed | HDAC1 | -1.2777 | LUAD | HCC-827 | Entinostat |
| MVP_top60_DNA_seed | AR | 2.033273 | LGG | D-502MG | Bicalutamide | MVP_top60_DNA+RNA_seed | AURKA | -1.265714 | LUAD | LC-2-ad | Tozasertib |
| MVP_top60_DNA_seed | HDAC1 | 2.068112 | LGG | KNS-42 | JQ12 | MVP_top60_DNA+RNA_seed | IGF1R | -1.26433 | LUAD | NCI-H1573 | GSK1904529A |
| MVP_top60_DNA_seed | JAK1 | 2.09747 | LGG | D-502MG | JAK1_3715 | MVP_top60_DNA+RNA_seed | PIK3CA | -1.259063 | LUAD | NCI-H1755 | GNE-317 |
| MVP_top60_DNA_seed | AR | 2.10594 | LGG | KINGS-1 | AZD3514 | MVP_top60_DNA+RNA_seed | PIK3CA | -1.253168 | LUAD | NCI-H1648 | AZD8186 |
| MVP_top60_DNA_seed | HDAC1 | 2.119881 | LGG | no-11 | JQ12 | MVP_top60_DNA+RNA_seed | HDAC1 | -1.249884 | LUAD | LXF-289 | Romidepsin |
| MVP_top60_DNA_seed | HSP90AA1 | 2.197068 | LGG | D-502MG | CCT-018159 | MVP_top60_DNA+RNA_seed | PIK3CA | -1.24466 | LUAD | NCI-H1792 | GNE-317 |
| MVP_top60_DNA_seed | EGFR | 2.352454 | LGG | KNS-42 | AST-1306 | MVP_top60_DNA+RNA_seed | HSP90AA1 | -1.229955 | LUAD | RERF-LC-MS | CCT-018159 |
| MVP_top60_DNA_seed | IGF1R | 2.368743 | LGG | D-502MG | BMS-754807 | MVP_top60_DNA+RNA_seed | PIK3CA | -1.228284 | LUAD | NCI-H2122 | GNE-317 |
| MVP_top60_DNA_seed | HDAC1 | 2.453761 | LGG | D-502MG | JQ12 | MVP_top60_DNA+RNA_seed | PIK3CA | -1.226963 | LUAD | NCI-H1993 | GNE-317 |
| MVP_top60_DNA_seed | EGFR | 2.464475 | LGG | KNS-81-FD | CI-1033 | MVP_top60_DNA+RNA_seed | PIK3CA | -1.225523 | LUAD | COR-L105 | GNE-317 |
| MVP_top60_DNA_seed | EGFR | 2.524426 | LGG | KNS-81-FD | AZD8931 | MVP_top60_DNA+RNA_seed | AURKA | -1.223471 | LUAD | VMRC-LCD | Alisertib |
| MVP_top60_DNA_seed | EGFR | 2.592692 | LGG | KNS-42 | CI-1033 | MVP_top60_DNA+RNA_seed | AKT1 | -1.223328 | LUAD | NCI-H1648 | AZD5363 |
| MVP_top60_DNA_seed | MDM2 | -2.233683 | LGG | H4 | Nutlin-3a (-) | MVP_top60_DNA+RNA_seed | KRAS | -1.223262 | LUAD | NCI-H2122 | RAS (G12C) Inhibitor-1 |
| MVP_top60_DNA_seed | HSP90AA1 | -2.013891 | LGG | H4 | Tanespimycin | MVP_top60_DNA+RNA_seed | HSP90AA1 | -1.216659 | LUAD | NCI-H358 | Tanespimycin |
| MVP_top60_DNA_seed | HSP90AA1 | -2.006503 | LGG | D-336MG | Tanespimycin | MVP_top60_DNA+RNA_seed | EGFR | -1.210974 | LUAD | Calu-3 | Osimertinib |
| MVP_top60_DNA_seed | MET | -1.941204 | LGG | KINGS-1 | Savolitinib | MVP_top60_DNA+RNA_seed | PIK3CA | -1.20902 | LUAD | NCI-H1755 | AZD8186 |
| MVP_top60_DNA_seed | PIK3CA | -1.687761 | LGG | D-336MG | AZD8186 | MVP_top60_DNA+RNA_seed | PIK3CA | -1.20867 | LUAD | RERF-LC-MS | GNE-317 |
| MVP_top60_DNA_seed | HSP90AA1 | -1.628446 | LGG | D-336MG | CCT-018159 | MVP_top60_DNA+RNA_seed | MET | -1.201637 | LUAD | NCI-H23 | Savolitinib |
| MVP_top60_DNA_seed | CREBBP | -1.611296 | LGG | H4 | SGC-CBP30 | MVP_top60_DNA+RNA_seed | PIK3CA | -1.197889 | LUAD | RERF-LC-KJ | Buparlisib |
| MVP_top60_DNA_seed | MET | -1.603405 | LGG | MOG-G-UVW | Savolitinib | MVP_top60_DNA+RNA_seed | AURKA | -1.197243 | LUAD | LC-2-ad | Alisertib |
| MVP_top60_DNA_seed | HSP90AA1 | -1.56758 | LGG | SW1783 | Luminespib | MVP_top60_DNA+RNA_seed | EZH2 | -1.197179 | LUAD | LXF-289 | GSK343 |
| MVP_top60_DNA_seed | CDK2 | -1.539139 | LGG | SF539 | MK-8776 | MVP_top60_DNA+RNA_seed | HDAC1 | -1.195065 | LUAD | HCC-78 | Entinostat |
| MVP_top60_DNA_seed | NTRK1 | -1.537972 | LGG | SF539 | AZD1332 | MVP_top60_DNA+RNA_seed | EGFR | -1.19148 | LUAD | NCI-H1650 | Erlotinib |
| MVP_top60_DNA_seed | PIK3CA | -1.512682 | LGG | MOG-G-CCM | AZD8186 | MVP_top60_DNA+RNA_seed | AKT1 | -1.191153 | LUAD | LXF-289 | AT13148 |
| MVP_top60_DNA_seed | CREBBP | -1.443126 | LGG | SF539 | SGC-CBP30 | MVP_top60_DNA+RNA_seed | SRC | -1.183187 | LUAD | PC-3_[JPC-3] | Bosutinib |
| MVP_top60_DNA_seed | SMARCA4 | -1.418843 | LGG | SF539 | PFI3 | MVP_top60_DNA+RNA_seed | PIK3CA | -1.179015 | LUAD | NCI-H1975 | Buparlisib |
| MVP_top60_DNA_seed | MET | -1.341917 | LGG | H4 | Crizotinib | MVP_top60_DNA+RNA_seed | PIK3CA | -1.174585 | LUAD | RERF-LC-MS | AZD8186 |
| MVP_top60_DNA_seed | IGF1R | -1.337672 | LGG | Hs-683 | GSK1904529A | MVP_top60_DNA+RNA_seed | PIK3CA | -1.168308 | LUAD | RERF-LC-MS | Alpelisib |
| MVP_top60_DNA_seed | HSP90AA1 | -1.333573 | LGG | GAMG | Tanespimycin | MVP_top60_DNA+RNA_seed | PIK3CA | -1.158947 | LUAD | NCI-H1623 | Alpelisib |
| MVP_top60_DNA_seed | MAPK1 | -1.308843 | LGG | H4 | VX-11e | MVP_top60_DNA+RNA_seed | EGFR | -1.144379 | LUAD | NCI-H1573 | Erlotinib |
| MVP_top60_DNA_seed | MDM2 | -1.294905 | LGG | H4 | Serdemetan | MVP_top60_DNA+RNA_seed | IGF1R | -1.143419 | LUAD | NCI-H838 | BMS-536924 |
| MVP_top60_DNA_seed | JAK2 | -1.287951 | LGG | H4 | JAK_8517 | MVP_top60_DNA+RNA_seed | AKT1 | -1.143267 | LUAD | NCI-H2122 | AZD5363 |

| | | | | | | | | | | | |
|---|---|---|---|---|---|---|---|---|---|---|---|
| MVP_top60_DNA_seed | JAK1 | -1.287951 | LGG | H4 | JAK_8517 | MVP_top60_DNA+RNA_seed | EGFR | -1.139855 | LUAD | NCI-H1623 | Osimertinib |
| MVP_top60_DNA_seed | JAK2 | -1.275251 | LGG | SF539 | GSK2276186C | MVP_top60_DNA+RNA_seed | AKT1 | -1.138588 | LUAD | NCI-H2122 | Uprosertib |
| MVP_top60_DNA_seed | JAK1 | -1.275251 | LGG | SF539 | GSK2276186C | MVP_top60_DNA+RNA_seed | HSP90AA1 | -1.131044 | LUAD | NCI-H650 | Luminespib |
| MVP_top60_DNA_seed | JAK3 | -1.275251 | LGG | SF539 | GSK2276186C | MVP_top60_DNA+RNA_seed | KRAS | -1.128535 | LUAD | NCI-H1975 | KRAS (G12C) Inhibitor-1 |
| MVP_top60_DNA_seed | HSP90AA1 | -1.26269 | LGG | MOG-G-CCM | Tanespimycin | MVP_top60_DNA+RNA_seed | AKT1 | -1.125471 | LUAD | NCI-H1975 | Ipatasertib |
| MVP_top60_DNA_seed | PIK3CA | -1.232974 | LGG | SW1783 | GNE-317 | MVP_top60_DNA+RNA_seed | MET | -1.123598 | LUAD | NCI-H1975 | Savolitinib |
| MVP_top60_DNA_seed | JAK2 | -1.230845 | LGG | H4 | AZ960 | MVP_top60_DNA+RNA_seed | AKT1 | -1.120744 | LUAD | NCI-H1781 | Uprosertib |
| MVP_top60_DNA_seed | JAK3 | -1.230845 | LGG | H4 | AZ960 | MVP_top60_DNA+RNA_seed | SMARCA4 | -1.117883 | LUAD | NCI-H23 | PFI3 |
| MVP_top60_DNA_seed | HSP90AA1 | -1.196592 | LGG | D-336MG | Luminespib | MVP_top60_DNA+RNA_seed | IGF1R | -1.112834 | LUAD | COR-L105 | BMS-536924 |
| MVP_top60_DNA_seed | HDAC1 | -1.163551 | LGG | H4 | Romidepsin | MVP_top60_DNA+RNA_seed | ATM | -1.11248 | LUAD | A549 | KU-55933 |
| MVP_top60_DNA_seed | HSP90AA1 | -1.14589 | LGG | Hs-683 | Tanespimycin | MVP_top60_DNA+RNA_seed | EZH2 | -1.110143 | LUAD | NCI-H1651 | GSK343 |
| MVP_top60_DNA_seed | EGFR | -1.141818 | LGG | Hs-683 | Gefitinib | MVP_top60_DNA+RNA_seed | EGFR | -1.109219 | LUAD | RERF-LC-KJ | Gefitinib |
| MVP_top60_DNA_seed | HDAC1 | -1.135802 | LGG | SF539 | PCI-34051 | MVP_top60_DNA+RNA_seed | EGFR | -1.107588 | LUAD | NCI-H1781 | Gefitinib |
| MVP_top60_DNA_seed | SMARCA4 | -1.13386 | LGG | GAMG | PFI3 | MVP_top60_DNA+RNA_seed | AKT1 | -1.098752 | LUAD | NCI-H1993 | AZD5363 |
| MVP_top60_DNA_seed | MAPK1 | -1.127334 | LGG | GAMG | Ulixertinib | MVP_top60_DNA+RNA_seed | MET | -1.088747 | LUAD | NCI-H2122 | Savolitinib |
| MVP_top60_DNA_seed | MAPK3 | -1.127334 | LGG | GAMG | Ulixertinib | MVP_top60_DNA+RNA_seed | AKT1 | -1.087845 | LUAD | NCI-H1993 | MK-2206 |
| MVP_top60_DNA_seed | PIK3CA | -1.121296 | LGG | SW1783 | AZD8186 | MVP_top60_DNA+RNA_seed | EGFR | -1.079003 | LUAD | NCI-H358 | Lapatinib |
| MVP_top60_DNA_seed | JAK1 | -1.117957 | LGG | GI-1 | JAK1_8709 | MVP_top60_DNA+RNA_seed | FGFR1 | -1.076539 | LUAD | NCI-H838 | PD173074 |
| MVP_top60_DNA_seed | MDM2 | -1.117914 | LGG | Hs-683 | Serdemetan | MVP_top60_DNA+RNA_seed | PIK3CA | -1.074684 | LUAD | EMC-BAC-1 | AZD8186 |
| MVP_top60_DNA_seed | HSP90AA1 | -1.111887 | LGG | GAMG | Luminespib | MVP_top60_DNA+RNA_seed | HDAC1 | -1.072302 | LUAD | RERF-LC-MS | PCI-34051 |
| MVP_top60_DNA_seed | JAK2 | -1.110566 | LGG | Hs-683 | AZ960 | MVP_top60_DNA+RNA_seed | ATM | -1.070681 | LUAD | RERF-LC-KJ | KU-55933 |
| MVP_top60_DNA_seed | JAK3 | -1.110566 | LGG | Hs-683 | AZ960 | MVP_top60_DNA+RNA_seed | IGF1R | -1.070253 | LUAD | NCI-H1437 | NVP-ADW742 |
| MVP_top60_DNA_seed | CDK2 | -1.104684 | LGG | D-336MG | Dinaciclib | MVP_top60_DNA+RNA_seed | HSP90AA1 | -1.064813 | LUAD | A549 | Luminespib |
| MVP_top60_DNA_seed | HSP90AA1 | -1.095814 | LGG | H4 | Luminespib | MVP_top60_DNA+RNA_seed | MET | -1.063039 | LUAD | NCI-H1623 | Savolitinib |
| MVP_top60_DNA_seed | EGFR | -1.089063 | LGG | GI-1 | Gefitinib | MVP_top60_DNA+RNA_seed | IGF1R | -1.058671 | LUAD | RERF-LC-MS | NVP-ADW742 |
| MVP_top60_DNA_seed | HSP90AA1 | -1.089046 | LGG | SW1783 | Tanespimycin | MVP_top60_DNA+RNA_seed | BRAF | -1.054628 | LUAD | NCI-H2009 | PLX-4720 |
| MVP_top60_DNA_seed | PIK3CA | -1.060958 | LGG | MOG-G-UVW | GNE-317 | MVP_top60_DNA+RNA_seed | PIK3CA | -1.052842 | LUAD | NCI-H1975 | AZD8186 |
| MVP_top60_DNA_seed | JAK2 | -1.049541 | LGG | D-336MG | AZ960 | MVP_top60_DNA+RNA_seed | IGF1R | -1.051141 | LUAD | NCI-H1666 | BMS-536924 |
| MVP_top60_DNA_seed | JAK3 | -1.049541 | LGG | D-336MG | AZ960 | MVP_top60_DNA+RNA_seed | TP53 | -1.049009 | LUAD | NCI-H1734 | MIRA-1 |
| MVP_top60_DNA_seed | JAK2 | -1.042081 | LGG | GI-1 | GSK2276186C | MVP_top60_DNA+RNA_seed | ATM | -1.048621 | LUAD | NCI-H322M | KU-55933 |
| MVP_top60_DNA_seed | JAK1 | -1.042081 | LGG | GI-1 | GSK2276186C | MVP_top60_DNA+RNA_seed | IGF1R | -1.041825 | LUAD | NCI-H2228 | Linsitinib |
| MVP_top60_DNA_seed | JAK3 | -1.042081 | LGG | GI-1 | GSK2276186C | MVP_top60_DNA+RNA_seed | IGF1R | -1.040294 | LUAD | RERF-LC-MS | BMS-536924 |
| MVP_top60_DNA_seed | HSP90AA1 | -1.034201 | LGG | SF539 | Tanespimycin | MVP_top60_DNA+RNA_seed | FGFR1 | -1.035122 | LUAD | NCI-H2122 | AZD4547 |
| MVP_top60_DNA_seed | JAK2 | -1.027519 | LGG | GI-1 | JAK_8517 | MVP_top60_DNA+RNA_seed | IGF1R | -1.034191 | LUAD | LXF-289 | BMS-754807 |
| MVP_top60_DNA_seed | JAK1 | -1.027519 | LGG | GI-1 | JAK_8517 | MVP_top60_DNA+RNA_seed | MET | -1.028331 | LUAD | NCI-H1944 | Savolitinib |
| MVP_top60_DNA_seed | MDM2 | -1.020447 | LGG | D-502MG | Nutlin-3a (-) | MVP_top60_DNA+RNA_seed | ATM | -1.028141 | LUAD | NCI-H1435 | KU-55933 |
| MVP_top60_DNA_seed | PIK3CA | -1.020332 | LGG | H4 | AZD8186 | MVP_top60_DNA+RNA_seed | EGFR | -1.027958 | LUAD | NCI-H838 | Sapitinib |
| MVP_top60_DNA_seed | MET | -1.003296 | LGG | MOG-G-UVW | Crizotinib | MVP_top60_DNA+RNA_seed | IGF1R | -1.023396 | LUAD | NCI-H1437 | Linsitinib |
| MVP_top60_DNA_seed | CDK2 | -0.995835 | LGG | SW1783 | MK-8776 | MVP_top60_DNA+RNA_seed | HSP90AA1 | -1.021189 | LUAD | NCI-H1792 | Luminespib |
| MVP_top60_DNA_seed | CDK2 | -0.987577 | LGG | KALS-1 | MK-8776 | MVP_top60_DNA+RNA_seed | AKT1 | -1.020747 | LUAD | NCI-H1993 | Ipatasertib |
| MVP_top60_DNA_seed | PIK3CA | -0.981862 | LGG | GAMG | AZD8186 | MVP_top60_DNA+RNA_seed | AKT1 | -1.018807 | LUAD | NCI-H1993 | Afuresertib |

| | | | | | | | | | | | |
|---|---|---|---|---|---|---|---|---|---|---|---|
| MVP_top60_DNA_seed | EGFR | -0.979745 | LGG | Hs-683 | AZD3759 | MVP_top60_DNA+RNA_seed | HSP90AA1 | -1.016678 | LUAD | LXF-289 | Tanespimycin |
| MVP_top60_DNA_seed | PIK3CA | -0.967746 | LGG | D-336MG | Alpelisib | MVP_top60_DNA+RNA_seed | AKT1 | -1.012382 | LUAD | NCI-H838 | AT13148 |
| MVP_top60_DNA_seed | TP53 | -0.959389 | LGG | MOG-G-UVW | MIRA-1 | MVP_top60_DNA+RNA_seed | MET | -1.01205 | LUAD | NCI-H1573 | Savolitinib |
| MVP_top60_DNA_seed | MET | -0.959096 | LGG | H4 | Savolitinib | MVP_top60_DNA+RNA_seed | HSP90AA1 | -1.010772 | LUAD | NCI-H2122 | Tanespimycin |
| MVP_top60_DNA_seed | IGF1R | -0.953325 | LGG | D-336MG | BMS-536924 | MVP_top60_DNA+RNA_seed | AURKA | -1.007577 | LUAD | ABC-1 | ZM447439 |
| MVP_top60_DNA_seed | PIK3CA | -0.940227 | LGG | Hs-683 | AZD8186 | MVP_top60_DNA+RNA_seed | ATM | -1.007379 | LUAD | NCI-H522 | KU-55933 |
| MVP_top60_DNA_seed | HSP90AA1 | -0.921787 | LGG | Hs-683 | Luminespib | MVP_top60_DNA+RNA_seed | AKT1 | -1.004861 | LUAD | NCI-H2122 | GSK2110183B |
| MVP_top60_DNA_seed | HSP90AA1 | -0.902473 | LGG | MOG-G-UVW | Luminespib | MVP_top60_DNA+RNA_seed | AKT1 | -1.002467 | LUAD | NCI-H1781 | AT13148 |
| MVP_top60_DNA_seed | HSP90AA1 | -0.883965 | LGG | Hs-683 | CCT-018159 | MVP_top60_DNA+RNA_seed | AKT1 | -1.002441 | LUAD | HCC-44 | AZD5363 |
| MVP_top60_DNA_seed | MET | -0.868592 | LGG | GI-1 | Crizotinib | MVP_top60_DNA+RNA_seed | AKT1 | -1.00235 | LUAD | NCI-H2122 | Afuresertib |
| MVP_top60_DNA_seed | MET | -0.866465 | LGG | GI-1 | Savolitinib | MVP_top60_DNA+RNA_seed | AURKA | -0.997002 | LUAD | NCI-H1781 | ZM447439 |
| MVP_top60_DNA_seed | MET | -0.852176 | LGG | SF539 | Savolitinib | MVP_top60_DNA+RNA_seed | EGFR | -0.995602 | LUAD | EKVX | AZD3759 |
| MVP_top60_DNA_seed | HDAC1 | -0.850271 | LGG | GI-1 | Romidepsin | MVP_top60_DNA+RNA_seed | HSP90AA1 | -0.99441 | LUAD | NCI-H1755 | Luminespib |
| MVP_top60_DNA_seed | IGF1R | -0.828065 | LGG | H4 | BMS-754807 | MVP_top60_DNA+RNA_seed | HDAC1 | -0.992393 | LUAD | EMC-BAC-1 | Entinostat |
| MVP_top60_DNA_seed | AR | -0.811767 | LGG | D-336MG | Bicalutamide | MVP_top60_DNA+RNA_seed | IGF1R | -0.990169 | LUAD | NCI-H1437 | GSK1904529A |
| MVP_top60_DNA_seed | TP53 | -0.810133 | LGG | SW1783 | MIRA-1 | MVP_top60_DNA+RNA_seed | HSP90AA1 | -0.987039 | LUAD | NCI-H2009 | Luminespib |
| MVP_top60_DNA_seed | MET | -0.797063 | LGG | D-336MG | Savolitinib | MVP_top60_DNA+RNA_seed | AKT1 | -0.986429 | LUAD | A549 | AT13148 |
| MVP_top60_DNA_seed | HSP90AA1 | -0.79415 | LGG | SF539 | Luminespib | MVP_top60_DNA+RNA_seed | BRAF | -0.981614 | LUAD | NCI-H1792 | SB590885 |
| MVP_top60_DNA_seed | IGF1R | -0.783344 | LGG | D-336MG | BMS-754807 | MVP_top60_DNA+RNA_seed | HDAC1 | -0.981466 | LUAD | NCI-H1755 | Romidepsin |
| MVP_top60_DNA_seed | PIK3CA | -0.773063 | LGG | D-336MG | Buparlisib | MVP_top60_DNA+RNA_seed | EGFR | -0.978121 | LUAD | NCI-H1650 | Osimertinib |
| MVP_top60_DNA_seed | PIK3CD | -0.773063 | LGG | D-336MG | Buparlisib | MVP_top60_DNA+RNA_seed | EGFR | -0.973508 | LUAD | RERF-LC-MS | Lapatinib |
| MVP_top60_DNA_seed | EGFR | -0.768215 | LGG | D-336MG | AZD3759 | MVP_top60_DNA+RNA_seed | AURKA | -0.971594 | LUAD | LC-2-ad | ZM447439 |
| MVP_top60_DNA_seed | JAK2 | -0.757137 | LGG | SF539 | JAK_8517 | MVP_top60_DNA+RNA_seed | EZH2 | -0.970408 | LUAD | NCI-H2023 | GSK343 |
| MVP_top60_DNA_seed | JAK1 | -0.757137 | LGG | SF539 | JAK_8517 | MVP_top60_DNA+RNA_seed | AKT1 | -0.969321 | LUAD | NCI-H1623 | Uprosertib |
| MVP_top60_DNA_seed | JAK1 | -0.728464 | LGG | SF539 | JAK1_8709 | MVP_top60_DNA+RNA_seed | EGFR | -0.966727 | LUAD | EMC-BAC-2 | Erlotinib |
| MVP_top60_DNA_seed | CDK2 | -0.722398 | LGG | SF539 | Dinaciclib | MVP_top60_DNA+RNA_seed | IGF1R | -0.966525 | LUAD | LC-2-ad | BMS-754807 |
| MVP_top60_DNA_seed | PIK3CA | -0.688327 | LGG | MOG-G-UVW | Buparlisib | MVP_top60_DNA+RNA_seed | IGF1R | -0.962916 | LUAD | NCI-H1792 | BMS-536924 |
| MVP_top60_DNA_seed | PIK3CD | -0.688327 | LGG | MOG-G-UVW | Buparlisib | MVP_top60_DNA+RNA_seed | IGF1R | -0.96055 | LUAD | LC-2-ad | BMS-536924 |
| MVP_top60_DNA_seed | MDM2 | -0.677117 | LGG | Hs-683 | Nutlin-3a (-) | MVP_top60_DNA+RNA_seed | BRAF | -0.959907 | LUAD | NCI-H2122 | SB590885 |
| MVP_top60_DNA_seed | PIK3CA | -0.676352 | LGG | GI-1 | Buparlisib | MVP_top60_DNA+RNA_seed | EGFR | -0.959136 | LUAD | NCI-H2030 | Sapitinib |
| MVP_top60_DNA_seed | PIK3CD | -0.676352 | LGG | GI-1 | Buparlisib | MVP_top60_DNA+RNA_seed | AKT1 | -0.954396 | LUAD | NCI-H1734 | Ipatasertib |
| MVP_top60_DNA_seed | JAK2 | -0.663281 | LGG | D-502MG | AZ960 | MVP_top60_DNA+RNA_seed | HSP90AA1 | -0.950333 | LUAD | NCI-H1944 | Tanespimycin |
| MVP_top60_DNA_seed | JAK3 | -0.663281 | LGG | D-502MG | AZ960 | MVP_top60_DNA+RNA_seed | HDAC1 | -0.946015 | LUAD | HCC-78 | Romidepsin |
| MVP_top60_DNA_seed | PIK3CA | -0.662721 | LGG | MOG-G-UVW | AZD8186 | MVP_top60_DNA+RNA_seed | SRC | -0.945878 | LUAD | H3255 | Bosutinib |
| MVP_top60_DNA_seed | MAPK1 | -0.654083 | LGG | KALS-1 | SCH772984 | MVP_top60_DNA+RNA_seed | EGFR | -0.944308 | LUAD | NCI-H1666 | Osimertinib |
| MVP_top60_DNA_seed | MAPK3 | -0.654083 | LGG | KALS-1 | SCH772984 | MVP_top60_DNA+RNA_seed | PIK3CA | -0.943231 | LUAD | NCI-H3122 | Buparlisib |
| MVP_top60_DNA_seed | PIK3CA | -0.648271 | LGG | H4 | Buparlisib | MVP_top60_DNA+RNA_seed | AKT1 | -0.940545 | LUAD | HCC-44 | AT13148 |
| MVP_top60_DNA_seed | PIK3CD | -0.648271 | LGG | H4 | Buparlisib | MVP_top60_DNA+RNA_seed | BRAF | -0.939638 | LUAD | RERF-LC-MS | PLX-4720 |
| MVP_top60_DNA_seed | CDK2 | -0.647563 | LGG | SF539 | AZD5438 | MVP_top60_DNA+RNA_seed | EGFR | -0.938908 | LUAD | RERF-LC-KJ | AZD3759 |
| MVP_top60_DNA_seed | PIK3CA | -0.646293 | LGG | MOG-G-CCM | GNE-317 | MVP_top60_DNA+RNA_seed | AKT1 | -0.938658 | LUAD | NCI-H1781 | GSK2110183B |
| MVP_top60_DNA_seed | MAPK1 | -0.644965 | LGG | SF539 | SCH772984 | MVP_top60_DNA+RNA_seed | HSP90AA1 | -0.938482 | LUAD | NCI-H1437 | CCT-018159 |

| | | | | | | | | | | | |
|---|---|---|---|---|---|---|---|---|---|---|---|
| MVP_top60_DNA_seed | MAPK3 | -0.644965 | LGG | SF539 | SCH772984 | MVP_top60_DNA+RNA_seed | HDAC1 | -0.934872 | LUAD | LC-2-ad | Entinostat |
| MVP_top60_DNA_seed | MDM2 | -0.642682 | LGG | KNS-81-FD | Nutlin-3a (-) | MVP_top60_DNA+RNA_seed | ATM | -0.9317 | LUAD | NCI-H2122 | KU-55933 |
| MVP_top60_DNA_seed | IGF1R | -0.640361 | LGG | D-336MG | Linsitinib | MVP_top60_DNA+RNA_seed | PIK3CA | -0.930163 | LUAD | NCI-H2122 | Buparlisib |
| MVP_top60_DNA_seed | IGF1R | -0.626797 | LGG | SF539 | GSK1904529A | MVP_top60_DNA+RNA_seed | EGFR | -0.925628 | LUAD | NCI-H1568 | Osimertinib |
| MVP_top60_DNA_seed | NTRK1 | -0.621357 | LGG | H4 | AZD1332 | MVP_top60_DNA+RNA_seed | HSP90AA1 | -0.92456 | LUAD | EMC-BAC-1 | Tanespimycin |
| MVP_top60_DNA_seed | IGF1R | -0.617368 | LGG | KNS-42 | GSK1904529A | MVP_top60_DNA+RNA_seed | AKT1 | -0.922764 | LUAD | HCC-78 | Ipatasertib |
| MVP_top60_DNA_seed | EGFR | -0.613204 | LGG | Hs-683 | Lapatinib | MVP_top60_DNA+RNA_seed | IGF1R | -0.921882 | LUAD | NCI-H322M | Linsitinib |
| MVP_top60_DNA_seed | JAK2 | -0.612739 | LGG | SF539 | Ruxolitinib | MVP_top60_DNA+RNA_seed | MET | -0.919515 | LUAD | NCI-H2228 | Crizotinib |
| MVP_top60_DNA_seed | JAK1 | -0.612739 | LGG | SF539 | Ruxolitinib | MVP_top60_DNA+RNA_seed | AKT1 | -0.918724 | LUAD | NCI-H1975 | Afuresertib |
| MVP_top60_DNA_seed | JAK2 | -0.61123 | LGG | SF539 | AZ960 | MVP_top60_DNA+RNA_seed | AKT1 | -0.916902 | LUAD | NCI-H1975 | Uprosertib |
| MVP_top60_DNA_seed | JAK3 | -0.61123 | LGG | SF539 | AZ960 | MVP_top60_DNA+RNA_seed | EGFR | -0.91684 | LUAD | EKVX | Sapitinib |
| MVP_top60_DNA_seed | PIK3CA | -0.594503 | LGG | MOG-G-CCM | Buparlisib | MVP_top60_DNA+RNA_seed | EGFR | -0.914333 | LUAD | EMC-BAC-2 | Sapitinib |
| MVP_top60_DNA_seed | PIK3CD | -0.594503 | LGG | MOG-G-CCM | Buparlisib | MVP_top60_DNA+RNA_seed | MET | -0.913338 | LUAD | NCI-H1734 | Savolitinib |
| MVP_top60_DNA_seed | HSP90AA1 | -0.591943 | LGG | GI-1 | Luminespib | MVP_top60_DNA+RNA_seed | TERT | -0.909582 | LUAD | NCI-H1792 | BIBR-1532 |
| MVP_top60_DNA_seed | PIK3CA | -0.577715 | LGG | SF539 | Buparlisib | MVP_top60_DNA+RNA_seed | KRAS | -0.907714 | LUAD | HCC-78 | KRAS (G12C) Inhibitor-1 |
| MVP_top60_DNA_seed | PIK3CD | -0.577715 | LGG | SF539 | Buparlisib | MVP_top60_DNA+RNA_seed | HSP90AA1 | -0.907365 | LUAD | RERF-LC-MS | Tanespimycin |
| MVP_top60_DNA_seed | JAK2 | -0.576559 | LGG | GI-1 | AZ960 | MVP_top60_DNA+RNA_seed | IGF1R | -0.905475 | LUAD | HCC-44 | BMS-754807 |
| MVP_top60_DNA_seed | JAK3 | -0.576559 | LGG | GI-1 | AZ960 | MVP_top60_DNA+RNA_seed | PIK3CA | -0.905297 | LUAD | HCC-44 | AZD8186 |
| MVP_top60_DNA_seed | HSP90AA1 | -0.571869 | LGG | GI-1 | Tanespimycin | MVP_top60_DNA+RNA_seed | IGF1R | -0.905097 | LUAD | COR-L105 | NVP-ADW742 |
| MVP_top60_DNA_seed | MET | -0.568229 | LGG | SF539 | Crizotinib | MVP_top60_DNA+RNA_seed | IGF1R | -0.904804 | LUAD | PC-3_[JPC-3] | BMS-536924 |
| MVP_top60_DNA_seed | EGFR | -0.56726 | LGG | SF539 | Erlotinib | MVP_top60_DNA+RNA_seed | FGFR1 | -0.903348 | LUAD | NCI-H1755 | PD173074 |
| MVP_top60_DNA_seed | EGFR | -0.56455 | LGG | D-502MG | AZD3759 | MVP_top60_DNA+RNA_seed | MDM2 | -0.903095 | LUAD | NCI-H2228 | Serdemetan |
| MVP_top60_DNA_seed | PIK3CA | -0.563741 | LGG | Hs-683 | Buparlisib | MVP_top60_DNA+RNA_seed | EGFR | -0.896158 | LUAD | RERF-LC-MS | AZD3759 |
| MVP_top60_DNA_seed | PIK3CD | -0.563741 | LGG | Hs-683 | Buparlisib | MVP_top60_DNA+RNA_seed | AKT1 | -0.89612 | LUAD | HCC-44 | GSK2110183B |
| MVP_top60_DNA_seed | EGFR | -0.556105 | LGG | D-336MG | Lapatinib | MVP_top60_DNA+RNA_seed | MET | -0.896037 | LUAD | NCI-H2122 | Crizotinib |
| MVP_top60_DNA_seed | NTRK1 | -0.55008 | LGG | GI-1 | AZD1332 | MVP_top60_DNA+RNA_seed | EGFR | -0.895905 | LUAD | EMC-BAC-1 | Erlotinib |
| MVP_top60_DNA_seed | EGFR | -0.540131 | LGG | D-336MG | Osimertinib | MVP_top60_DNA+RNA_seed | AURKA | -0.895412 | LUAD | RERF-LC-MS | Tozasertib |
| MVP_top60_DNA_seed | NTRK1 | -0.524168 | LGG | SF539 | GW441756 | MVP_top60_DNA+RNA_seed | HDAC1 | -0.892282 | LUAD | NCI-H1993 | Romidepsin |
| MVP_top60_DNA_seed | EGFR | -0.517011 | LGG | H4 | Erlotinib | MVP_top60_DNA+RNA_seed | PIK3CA | -0.887305 | LUAD | NCI-H1437 | Alpelisib |
| MVP_top60_DNA_seed | IGF1R | -0.501866 | LGG | MOG-G-UVW | BMS-754807 | MVP_top60_DNA+RNA_seed | TERT | -0.886948 | LUAD | EMC-BAC-1 | BIBR-1532 |
| MVP_top60_DNA_seed | EGFR | -0.479491 | LGG | SF539 | Lapatinib | MVP_top60_DNA+RNA_seed | HSP90AA1 | -0.886648 | LUAD | RERF-LC-MS | Elesclomol |
| MVP_top60_DNA_seed | PIK3CA | -0.477739 | LGG | Hs-683 | Alpelisib | MVP_top60_DNA+RNA_seed | PIK3CA | -0.885205 | LUAD | NCI-H322M | AZD8186 |
| MVP_top60_DNA_seed | IGF1R | -0.476034 | LGG | H4 | GSK1904529A | MVP_top60_DNA+RNA_seed | FGFR1 | -0.885173 | LUAD | SK-LU-1 | PD173074 |
| MVP_top60_DNA_seed | TP53 | -0.460786 | LGG | no-11 | MIRA-1 | MVP_top60_DNA+RNA_seed | HSP90AA1 | -0.884033 | LUAD | NCI-H2009 | Tanespimycin |
| MVP_top60_DNA_seed | MAPK1 | -0.459191 | LGG | Hs-683 | VX-11e | MVP_top60_DNA+RNA_seed | AKT1 | -0.876014 | LUAD | NCI-H1781 | Ipatasertib |
| MVP_top60_DNA_seed | HDAC1 | -0.456593 | LGG | GAMG | Romidepsin | MVP_top60_DNA+RNA_seed | AKT1 | -0.873638 | LUAD | NCI-H2030 | AT13148 |
| MVP_top60_DNA_seed | SMARCA4 | -0.449454 | LGG | H4 | PFI3 | MVP_top60_DNA+RNA_seed | IGF1R | -0.873289 | LUAD | RERF-LC-MS | BMS-754807 |
| MVP_top60_DNA_seed | CDK2 | -0.443816 | LGG | H4 | AZD5438 | MVP_top60_DNA+RNA_seed | PIK3CA | -0.87246 | LUAD | NCI-H1437 | GNE-317 |
| MVP_top60_DNA_seed | TP53 | -0.442532 | LGG | D-502MG | MIRA-1 | MVP_top60_DNA+RNA_seed | EZH2 | -0.870602 | LUAD | RERF-LC-KJ | GSK343 |
| MVP_top60_DNA_seed | MDM2 | -0.434407 | LGG | D-336MG | Serdemetan | MVP_top60_DNA+RNA_seed | EGFR | -0.869284 | LUAD | EKVX | Erlotinib |
| MVP_top60_DNA_seed | PIK3CA | -0.417037 | LGG | SW1783 | Buparlisib | MVP_top60_DNA+RNA_seed | KRAS | -0.868207 | LUAD | LC-2-ad | KRAS (G12C) Inhibitor-1 |

| | | | | | | | | | | | |
|---|---|---|---|---|---|---|---|---|---|---|---|
| MVP_top60_DNA_seed | PIK3CD | -0.417037 | LGG | SW1783 | Buparlisib | MVP_top60_DNA+RNA_seed | IGF1R | -0.863634 | LUAD | A549 | BMS-536924 |
| MVP_top60_DNA_seed | EGFR | -0.410901 | LGG | MOG-G-UVW | AZD3759 | MVP_top60_DNA+RNA_seed | AKT1 | -0.861856 | LUAD | NCI-H1944 | Uprosertib |
| MVP_top60_DNA_seed | IGF1R | -0.409045 | LGG | MOG-G-CCM | BMS-754807 | MVP_top60_DNA+RNA_seed | FGFR1 | -0.85912 | LUAD | RERF-LC-KJ | AZD4547 |
| MVP_top60_DNA_seed | CDK2 | -0.399092 | LGG | H4 | Dinaciclib | MVP_top60_DNA+RNA_seed | HSP90AA1 | -0.859064 | LUAD | HCC-44 | Luminespib |
| MVP_top60_DNA_seed | CDK2 | -0.395722 | LGG | GI-1 | AZD5438 | MVP_top60_DNA+RNA_seed | EGFR | -0.858883 | LUAD | RERF-LC-MS | Sapitinib |
| MVP_top60_DNA_seed | MET | -0.380306 | LGG | KINGS-1 | Crizotinib | MVP_top60_DNA+RNA_seed | ATM | -0.85856 | LUAD | NCI-H1648 | KU-55933 |
| MVP_top60_DNA_seed | HSP90AA1 | -0.376963 | LGG | MOG-G-UVW | Tanespimycin | MVP_top60_DNA+RNA_seed | AKT1 | -0.85829 | LUAD | NCI-H1975 | Uprosertib |
| MVP_top60_DNA_seed | CREBBP | -0.374831 | LGG | GI-1 | SGC-CBP30 | MVP_top60_DNA+RNA_seed | AKT1 | -0.857073 | LUAD | NCI-H3122 | GSK2110183B |
| MVP_top60_DNA_seed | MAPK1 | -0.372414 | LGG | D-336MG | VX-11e | MVP_top60_DNA+RNA_seed | KRAS | -0.854987 | LUAD | NCI-H1792 | KRAS (G12C) Inhibitor-1 |
| MVP_top60_DNA_seed | HDAC1 | -0.371056 | LGG | H4 | PCI-34051 | MVP_top60_DNA+RNA_seed | PIK3CA | -0.854827 | LUAD | NCI-H2030 | AZD8186 |
| MVP_top60_DNA_seed | IGF1R | -0.368394 | LGG | GI-1 | BMS-754807 | MVP_top60_DNA+RNA_seed | EGFR | -0.848727 | LUAD | A549 | Lapatinib |
| MVP_top60_DNA_seed | IGF1R | -0.367611 | LGG | Hs-683 | BMS-754807 | MVP_top60_DNA+RNA_seed | EGFR | -0.846237 | LUAD | NCI-H1650 | Sapitinib |
| MVP_top60_DNA_seed | IDH1 | -0.356654 | LGG | Hs-683 | AGI-5198 | MVP_top60_DNA+RNA_seed | BRAF | -0.845527 | LUAD | NCI-H3122 | SB590885 |
| MVP_top60_DNA_seed | HSP90AA1 | -0.354795 | LGG | MOG-G-CCM | Luminespib | MVP_top60_DNA+RNA_seed | AKT1 | -0.842603 | LUAD | EKVX | AT13148 |
| MVP_top60_DNA_seed | PIK3CA | -0.352235 | LGG | D-502MG | Alpelisib | MVP_top60_DNA+RNA_seed | AKT1 | -0.84054 | LUAD | NCI-H1648 | Ipatasertib |
| MVP_top60_DNA_seed | HSP90AA1 | -0.349427 | LGG | no-10 | Tanespimycin | MVP_top60_DNA+RNA_seed | AKT1 | -0.840355 | LUAD | NCI-H1734 | MK-2206 |
| MVP_top60_DNA_seed | CDK2 | -0.342925 | LGG | Hs-683 | Dinaciclib | MVP_top60_DNA+RNA_seed | IGF1R | -0.839758 | LUAD | HCC-78 | NVP-ADW742 |
| MVP_top60_DNA_seed | JAK2 | -0.33727 | LGG | GAMG | AZ960 | MVP_top60_DNA+RNA_seed | HDAC1 | -0.838397 | LUAD | NCI-H1944 | Entinostat |
| MVP_top60_DNA_seed | JAK3 | -0.33727 | LGG | GAMG | AZ960 | MVP_top60_DNA+RNA_seed | PIK3CA | -0.834628 | LUAD | EMC-BAC-2 | AZD8186 |
| MVP_top60_DNA_seed | PIK3CA | -0.333807 | LGG | KALS-1 | AZD8186 | MVP_top60_DNA+RNA_seed | PIK3CA | -0.832469 | LUAD | NCI-H1944 | Buparlisib |
| MVP_top60_DNA_seed | JAK2 | -0.331334 | LGG | KNS-42 | Ruxolitinib | MVP_top60_DNA+RNA_seed | PIK3CA | -0.829341 | LUAD | NCI-H1993 | AZD8186 |
| MVP_top60_DNA_seed | JAK1 | -0.331334 | LGG | KNS-42 | Ruxolitinib | MVP_top60_DNA+RNA_seed | PIK3CA | -0.826319 | LUAD | EMC-BAC-2 | Alpelisib |
| MVP_top60_DNA_seed | IGF1R | -0.316631 | LGG | GAMG | BMS-536924 | MVP_top60_DNA+RNA_seed | AKT1 | -0.826315 | LUAD | HCC-44 | Uprosertib |
| MVP_top60_DNA_seed | MAPK1 | -0.315064 | LGG | SF539 | Ulixertinib | MVP_top60_DNA+RNA_seed | EGFR | -0.826297 | LUAD | EKVX | Afatinib |
| MVP_top60_DNA_seed | MAPK3 | -0.315064 | LGG | SF539 | Ulixertinib | MVP_top60_DNA+RNA_seed | HDAC1 | -0.826045 | LUAD | NCI-H838 | Entinostat |
| MVP_top60_DNA_seed | EGFR | -0.311257 | LGG | D-502MG | Lapatinib | MVP_top60_DNA+RNA_seed | KRAS | -0.825506 | LUAD | NCI-H1437 | KRAS (G12C) Inhibitor-1 |
| MVP_top60_DNA_seed | EGFR | -0.307399 | LGG | no-10 | AZD3759 | MVP_top60_DNA+RNA_seed | IGF1R | -0.823601 | LUAD | NCI-H1792 | Linsitinib |
| MVP_top60_DNA_seed | HSP90AA1 | -0.301042 | LGG | KINGS-1 | Luminespib | MVP_top60_DNA+RNA_seed | IGF1R | -0.821244 | LUAD | NCI-H3122 | BMS-536924 |
| MVP_top60_DNA_seed | HDAC1 | -0.300157 | LGG | no-10 | PCI-34051 | MVP_top60_DNA+RNA_seed | HDAC1 | -0.819623 | LUAD | NCI-H1993 | PCI-34051 |
| MVP_top60_DNA_seed | PIK3CA | -0.285899 | LGG | SF539 | AZD8186 | MVP_top60_DNA+RNA_seed | PIK3CA | -0.81888 | LUAD | NCI-H2085 | AZD8186 |
| MVP_top60_DNA_seed | SMARCA4 | -0.28166 | LGG | GI-1 | PFI3 | MVP_top60_DNA+RNA_seed | IGF1R | -0.818014 | LUAD | RERF-LC-MS | GSK1904529A |
| MVP_top60_DNA_seed | JAK2 | -0.260695 | LGG | H4 | GSK2276186C | MVP_top60_DNA+RNA_seed | IGF1R | -0.815831 | LUAD | NCI-H838 | Linsitinib |
| MVP_top60_DNA_seed | JAK1 | -0.260695 | LGG | H4 | GSK2276186C | MVP_top60_DNA+RNA_seed | HDAC1 | -0.811205 | LUAD | NCI-H1734 | Entinostat |
| MVP_top60_DNA_seed | JAK3 | -0.260695 | LGG | H4 | GSK2276186C | MVP_top60_DNA+RNA_seed | HSP90AA1 | -0.808601 | LUAD | NCI-H1993 | Tanespimycin |
| MVP_top60_DNA_seed | IGF1R | -0.254046 | LGG | SF539 | BMS-536924 | MVP_top60_DNA+RNA_seed | IGF1R | -0.807448 | LUAD | NCI-H1734 | NVP-ADW742 |
| MVP_top60_DNA_seed | MDM2 | -0.241538 | LGG | no-10 | Serdemetan | MVP_top60_DNA+RNA_seed | EGFR | -0.806559 | LUAD | NCI-H1568 | Afatinib |
| MVP_top60_DNA_seed | CDK2 | -0.239759 | LGG | no-10 | Dinaciclib | MVP_top60_DNA+RNA_seed | SRC | -0.805906 | LUAD | NCI-H1792 | Bosutinib |
| MVP_top60_DNA_seed | HDAC1 | -0.227543 | LGG | GI-1 | PCI-34051 | MVP_top60_DNA+RNA_seed | MDM2 | -0.804969 | LUAD | EMC-BAC-2 | Nutlin-3a (-) |
| MVP_top60_DNA_seed | CDK2 | -0.224702 | LGG | GI-1 | Dinaciclib | MVP_top60_DNA+RNA_seed | HSP90AA1 | -0.804088 | LUAD | PC-14 | Luminespib |
| MVP_top60_DNA_seed | IGF1R | -0.222085 | LGG | MOG-G-CCM | BMS-536924 | MVP_top60_DNA+RNA_seed | EGFR | -0.798745 | LUAD | NCI-H2030 | Lapatinib |
| MVP_top60_DNA_seed | JAK2 | -0.215775 | LGG | H4 | Ruxolitinib | MVP_top60_DNA+RNA_seed | MDM2 | -0.794131 | LUAD | PC-14 | Serdemetan |

| | | | | | | | | | | | |
|---|---|---|---|---|---|---|---|---|---|---|---|
| MVP_top60_DNA_seed | JAK1 | -0.215775 | LGG | H4 | Ruxolitinib | MVP_top60_DNA+RNA_seed | AKT1 | -0.794013 | LUAD | NCI-H2228 | Uprosertib |
| MVP_top60_DNA_seed | PIK3CA | -0.209957 | LGG | H4 | Alpelisib | MVP_top60_DNA+RNA_seed | HSP90AA1 | -0.793277 | LUAD | EMC-BAC-2 | Luminespib |
| MVP_top60_DNA_seed | PIK3CA | -0.196285 | LGG | GI-1 | AZD8186 | MVP_top60_DNA+RNA_seed | PIK3CA | -0.792181 | LUAD | NCI-H838 | AZD8186 |
| MVP_top60_DNA_seed | HDAC1 | -0.190583 | LGG | SF539 | Romidepsin | MVP_top60_DNA+RNA_seed | HDAC1 | -0.789287 | LUAD | NCI-H1975 | Romidepsin |
| MVP_top60_DNA_seed | PIK3CA | -0.186453 | LGG | GAMG | GNE-317 | MVP_top60_DNA+RNA_seed | AKT1 | -0.788663 | LUAD | RERF-LC-KJ | Afuresertib |
| MVP_top60_DNA_seed | HDAC1 | -0.18518 | LGG | D-502MG | PCI-34051 | MVP_top60_DNA+RNA_seed | AKT1 | -0.788326 | LUAD | NCI-H1993 | GSK2110183B |
| MVP_top60_DNA_seed | IGF1R | -0.184771 | LGG | D-502MG | BMS-754807 | MVP_top60_DNA+RNA_seed | EGFR | -0.787682 | LUAD | NCI-H358 | Osimertinib |
| MVP_top60_DNA_seed | NTRK1 | -0.18389 | LGG | GAMG | AZD1332 | MVP_top60_DNA+RNA_seed | EGFR | -0.787111 | LUAD | NCI-H1944 | Erlotinib |
| MVP_top60_DNA_seed | MAPK1 | -0.1824 | LGG | MOG-G-UVW | VX-11e | MVP_top60_DNA+RNA_seed | EGFR | -0.784739 | LUAD | NCI-H1734 | AZD3759 |
| MVP_top60_DNA_seed | MAPK1 | -0.181188 | LGG | MOG-G-UVW | SCH772984 | MVP_top60_DNA+RNA_seed | PIK3CA | -0.784471 | LUAD | NCI-H2291 | AZD8186 |
| MVP_top60_DNA_seed | MAPK3 | -0.181188 | LGG | MOG-G-UVW | SCH772984 | MVP_top60_DNA+RNA_seed | KRAS | -0.781584 | LUAD | NCI-H1993 | KRAS (G12C) Inhibitor-1 |
| MVP_top60_DNA_seed | EGFR | -0.179159 | LGG | H4 | Afatinib | MVP_top60_DNA+RNA_seed | AKT1 | -0.781304 | LUAD | EMC-BAC-2 | AZD5363 |
| MVP_top60_DNA_seed | TP53 | -0.172212 | LGG | GI-1 | MIRA-1 | MVP_top60_DNA+RNA_seed | BRAF | -0.780712 | LUAD | Calu-6 | PLX-4720 |
| MVP_top60_DNA_seed | IDH1 | -0.15915 | LGG | GI-1 | AGI-5198 | MVP_top60_DNA+RNA_seed | ATM | -0.779707 | LUAD | HCC-44 | KU-55933 |
| MVP_top60_DNA_seed | PIK3CA | -0.155749 | LGG | KINGS-1 | AZD8186 | MVP_top60_DNA+RNA_seed | EGFR | -0.776529 | LUAD | LXF-289 | Lapatinib |
| MVP_top60_DNA_seed | IDH1 | -0.155203 | LGG | SF539 | AGI-5198 | MVP_top60_DNA+RNA_seed | PIK3CA | -0.775777 | LUAD | NCI-H1792 | Buparlisib |
| MVP_top60_DNA_seed | HDAC1 | -0.149413 | LGG | no-10 | Entinostat | MVP_top60_DNA+RNA_seed | SRC | -0.775305 | LUAD | NCI-H838 | Bosutinib |
| MVP_top60_DNA_seed | EGFR | -0.149404 | LGG | no-10 | Lapatinib | MVP_top60_DNA+RNA_seed | HDAC1 | -0.775067 | LUAD | HCC-78 | PCI-34051 |
| MVP_top60_DNA_seed | JAK1 | -0.146503 | LGG | H4 | JAK1_8709 | MVP_top60_DNA+RNA_seed | HSP90AA1 | -0.774418 | LUAD | EMC-BAC-1 | Luminespib |
| MVP_top60_DNA_seed | MET | -0.146293 | LGG | D-336MG | Crizotinib | MVP_top60_DNA+RNA_seed | EGFR | -0.774197 | LUAD | EMC-BAC-2 | Osimertinib |
| MVP_top60_DNA_seed | HSP90AA1 | -0.142902 | LGG | no-11 | Luminespib | MVP_top60_DNA+RNA_seed | IGF1R | -0.773649 | LUAD | NCI-H1734 | Linsitinib |
| MVP_top60_DNA_seed | MDM2 | -0.140292 | LGG | no-10 | Nutlin-3a (-) | MVP_top60_DNA+RNA_seed | HDAC1 | -0.772582 | LUAD | NCI-H838 | PCI-34051 |
| MVP_top60_DNA_seed | JAK2 | -0.139351 | LGG | KNS-42 | AZ960 | MVP_top60_DNA+RNA_seed | PIK3CA | -0.769115 | LUAD | HCC-78 | Buparlisib |
| MVP_top60_DNA_seed | JAK3 | -0.139351 | LGG | KNS-42 | AZ960 | MVP_top60_DNA+RNA_seed | TERT | -0.768401 | LUAD | NCI-H2122 | BIBR-1532 |
| MVP_top60_DNA_seed | MAPK1 | -0.13809 | LGG | SF539 | VX-11e | MVP_top60_DNA+RNA_seed | PIK3CA | -0.768246 | LUAD | NCI-H838 | GNE-317 |
| MVP_top60_DNA_seed | MAPK1 | -0.137859 | LGG | H4 | Ulixertinib | MVP_top60_DNA+RNA_seed | EGFR | -0.768056 | LUAD | NCI-H2122 | Afatinib |
| MVP_top60_DNA_seed | MAPK3 | -0.137859 | LGG | H4 | Ulixertinib | MVP_top60_DNA+RNA_seed | SRC | -0.766975 | LUAD | NCI-H1755 | Bosutinib |
| MVP_top60_DNA_seed | MET | -0.136657 | LGG | GAMG | Crizotinib | MVP_top60_DNA+RNA_seed | EGFR | -0.76574 | LUAD | NCI-H2030 | Osimertinib |
| MVP_top60_DNA_seed | EGFR | -0.1312 | LGG | D-502MG | Sapitinib | MVP_top60_DNA+RNA_seed | EGFR | -0.763156 | LUAD | EKVX | Lapatinib |
| MVP_top60_DNA_seed | PIK3CA | -0.130225 | LGG | KINGS-1 | Alpelisib | MVP_top60_DNA+RNA_seed | HSP90AA1 | -0.76203 | LUAD | LC-2-ad | CCT-018159 |
| MVP_top60_DNA_seed | JAK2 | -0.127077 | LGG | no-10 | Ruxolitinib | MVP_top60_DNA+RNA_seed | HDAC1 | -0.761948 | LUAD | NCI-H1623 | Entinostat |
| MVP_top60_DNA_seed | JAK1 | -0.127077 | LGG | no-10 | Ruxolitinib | MVP_top60_DNA+RNA_seed | HSP90AA1 | -0.761902 | LUAD | NCI-H1975 | Tanespimycin |
| MVP_top60_DNA_seed | MDM2 | -0.126379 | LGG | SW1783 | Serdemetan | MVP_top60_DNA+RNA_seed | MDM2 | -0.76162 | LUAD | NCI-H1944 | Nutlin-3a (-) |
| MVP_top60_DNA_seed | EGFR | -0.121743 | LGG | H4 | Sapitinib | MVP_top60_DNA+RNA_seed | IGF1R | -0.760901 | LUAD | NCI-H1355 | Linsitinib |
| MVP_top60_DNA_seed | HSP90AA1 | -0.120805 | LGG | no-11 | CCT-018159 | MVP_top60_DNA+RNA_seed | EGFR | -0.760463 | LUAD | NCI-H2122 | Lapatinib |
| MVP_top60_DNA_seed | IGF1R | -0.113861 | LGG | MOG-G-UVW | NVP-ADW742 | MVP_top60_DNA+RNA_seed | EGFR | -0.758627 | LUAD | NCI-H2122 | AZD3759 |
| MVP_top60_DNA_seed | PIK3CA | -0.112874 | LGG | KNS-81-FD | Alpelisib | MVP_top60_DNA+RNA_seed | KRAS | -0.75525 | LUAD | NCI-H23 | KRAS (G12C) Inhibitor-1 |
| MVP_top60_DNA_seed | AR | -0.112572 | LGG | no-11 | Bicalutamide | MVP_top60_DNA+RNA_seed | PIK3CA | -0.755019 | LUAD | EMC-BAC-1 | Alpelisib |
| MVP_top60_DNA_seed | PIK3CA | -0.112242 | LGG | GAMG | Buparlisib | MVP_top60_DNA+RNA_seed | SRC | -0.753149 | LUAD | NCI-H2030 | Bosutinib |
| MVP_top60_DNA_seed | PIK3CD | -0.112242 | LGG | GAMG | Buparlisib | MVP_top60_DNA+RNA_seed | IGF1R | -0.752205 | LUAD | NCI-H1792 | NVP-ADW742 |
| MVP_top60_DNA_seed | IDH1 | -0.104133 | LGG | KNS-81-FD | AGI-5198 | MVP_top60_DNA+RNA_seed | PIK3CA | -0.752126 | LUAD | NCI-H838 | Alpelisib |

| | | | | | | | | | | | |
|---|---|---|---|---|---|---|---|---|---|---|---|
| MVP_top60_DNA_seed | IGF1R | -0.097656 | LGG | GAMG | GSK1904529A | MVP_top60_DNA+RNA_seed | AURKA | -0.749448 | LUAD | NCI-H1437 | ZM447439 |
| MVP_top60_DNA_seed | IGF1R | -0.09573 | LGG | D-502MG | BMS-536924 | MVP_top60_DNA+RNA_seed | MET | -0.748162 | LUAD | NCI-H2023 | Savolitinib |
| MVP_top60_DNA_seed | EGFR | -0.093755 | LGG | SF539 | Osimertinib | MVP_top60_DNA+RNA_seed | IGF1R | -0.747969 | LUAD | HCC-44 | Linsitinib |
| MVP_top60_DNA_seed | NTRK1 | -0.092539 | LGG | GI-1 | GW441756 | MVP_top60_DNA+RNA_seed | IGF1R | -0.747464 | LUAD | RERF-LC-KJ | GSK1904529A |
| MVP_top60_DNA_seed | EGFR | -0.086449 | LGG | GAMG | Afatinib | MVP_top60_DNA+RNA_seed | MET | -0.744948 | LUAD | HCC-44 | Crizotinib |
| MVP_top60_DNA_seed | NTRK1 | -0.078528 | LGG | D-336MG | GW441756 | MVP_top60_DNA+RNA_seed | PIK3CA | -0.74273 | LUAD | NCI-H2023 | GNE-317 |
| MVP_top60_DNA_seed | NTRK1 | -0.076712 | LGG | KALS-1 | AZD1332 | MVP_top60_DNA+RNA_seed | HDAC1 | -0.741586 | LUAD | NCI-H522 | PCI-34051 |
| MVP_top60_DNA_seed | JAK2 | -0.071193 | LGG | KALS-1 | AZ960 | MVP_top60_DNA+RNA_seed | AKT1 | -0.737062 | LUAD | NCI-H1648 | MK-2206 |
| MVP_top60_DNA_seed | JAK3 | -0.071193 | LGG | KALS-1 | AZ960 | MVP_top60_DNA+RNA_seed | IGF1R | -0.73514 | LUAD | ABC-1 | BMS-754807 |
| MVP_top60_DNA_seed | PIK3CA | -0.066356 | LGG | D-502MG | AZD8186 | MVP_top60_DNA+RNA_seed | HSP90AA1 | -0.734229 | LUAD | NCI-H2030 | Tanespimycin |
| MVP_top60_DNA_seed | IGF1R | -0.061335 | LGG | SW1783 | NVP-ADW742 | MVP_top60_DNA+RNA_seed | PIK3CA | -0.732447 | LUAD | NCI-H3122 | Alpelisib |
| MVP_top60_DNA_seed | MDM2 | -0.060211 | LGG | GAMG | Serdemetan | MVP_top60_DNA+RNA_seed | AKT1 | -0.732278 | LUAD | NCI-H1648 | Uprosertib |
| MVP_top60_DNA_seed | SMARCA4 | -0.05327 | LGG | Hs-683 | PFI3 | MVP_top60_DNA+RNA_seed | TP53 | -0.730298 | LUAD | NCI-H1435 | MIRA-1 |
| MVP_top60_DNA_seed | HSP90AA1 | -0.052969 | LGG | no-10 | Luminespib | MVP_top60_DNA+RNA_seed | PIK3CA | -0.72943 | LUAD | HCC-44 | Buparlisib |
| MVP_top60_DNA_seed | EGFR | -0.048049 | LGG | no-10 | Sapitinib | MVP_top60_DNA+RNA_seed | EGFR | -0.729181 | LUAD | NCI-H1975 | Erlotinib |
| MVP_top60_DNA_seed | EGFR | -0.045606 | LGG | GI-1 | Erlotinib | MVP_top60_DNA+RNA_seed | AKT1 | -0.728317 | LUAD | NCI-H2122 | MK-2206 |
| MVP_top60_DNA_seed | JAK2 | -0.045332 | LGG | KALS-1 | GSK2276186C | MVP_top60_DNA+RNA_seed | PIK3CA | -0.726793 | LUAD | NCI-H1975 | Alpelisib |
| MVP_top60_DNA_seed | JAK1 | -0.045332 | LGG | KALS-1 | GSK2276186C | MVP_top60_DNA+RNA_seed | MET | -0.725844 | LUAD | NCI-H1755 | Crizotinib |
| MVP_top60_DNA_seed | JAK3 | -0.045332 | LGG | KALS-1 | GSK2276186C | MVP_top60_DNA+RNA_seed | EGFR | -0.725677 | LUAD | NCI-H1693 | Afatinib |
| MVP_top60_DNA_seed | MDM2 | -0.037737 | LGG | SF539 | Serdemetan | MVP_top60_DNA+RNA_seed | FGFR1 | -0.7251 | LUAD | NCI-H322M | PD173074 |
| MVP_top60_DNA_seed | EGFR | -0.03322 | LGG | SF539 | Afatinib | MVP_top60_DNA+RNA_seed | IGF1R | -0.724846 | LUAD | A549 | GSK1904529A |
| MVP_top60_DNA_seed | TP53 | -0.030854 | LGG | D-336MG | MIRA-1 | MVP_top60_DNA+RNA_seed | EGFR | -0.723065 | LUAD | NCI-H1437 | Gefitinib |
| MVP_top60_DNA_seed | HSP90AA1 | -0.023553 | LGG | KINGS-1 | Tanespimycin | MVP_top60_DNA+RNA_seed | AKT1 | -0.722518 | LUAD | RERF-LC-MS | AT13148 |
| MVP_top60_DNA_seed | JAK2 | -0.022205 | LGG | GAMG | Ruxolitinib | MVP_top60_DNA+RNA_seed | HSP90AA1 | -0.722228 | LUAD | NCI-H838 | Tanespimycin |
| MVP_top60_DNA_seed | JAK1 | -0.022205 | LGG | GAMG | Ruxolitinib | MVP_top60_DNA+RNA_seed | EGFR | -0.721807 | LUAD | NCI-H1693 | Erlotinib |
| MVP_top60_DNA_seed | PIK3CA | -0.017429 | LGG | SF539 | GNE-317 | MVP_top60_DNA+RNA_seed | AKT1 | -0.721749 | LUAD | NCI-H1993 | Uprosertib |
| MVP_top60_DNA_seed | IGF1R | -0.013415 | LGG | D-502MG | NVP-ADW742 | MVP_top60_DNA+RNA_seed | EZH2 | -0.721317 | LUAD | NCI-H522 | GSK343 |
| MVP_top60_DNA_seed | EGFR | -0.006992 | LGG | KNS-81-FD | Gefitinib | MVP_top60_DNA+RNA_seed | MET | -0.721098 | LUAD | EMC-BAC-1 | Crizotinib |
| MVP_top60_DNA_seed | IGF1R | 0.006927 | LGG | SF539 | BMS-754807 | MVP_top60_DNA+RNA_seed | HSP90AA1 | -0.720347 | LUAD | NCI-H1666 | Tanespimycin |
| MVP_top60_DNA_seed | EGFR | 0.008738 | LGG | GI-1 | Afatinib | MVP_top60_DNA+RNA_seed | EGFR | -0.72027 | LUAD | NCI-H2030 | Erlotinib |
| MVP_top60_DNA_seed | JAK2 | 0.014434 | LGG | KINGS-1 | Ruxolitinib | MVP_top60_DNA+RNA_seed | MET | -0.719839 | LUAD | LXF-289 | Savolitinib |
| MVP_top60_DNA_seed | JAK1 | 0.014434 | LGG | KINGS-1 | Ruxolitinib | MVP_top60_DNA+RNA_seed | PIK3CA | -0.71925 | LUAD | NCI-H1563 | AZD8186 |
| MVP_top60_DNA_seed | EGFR | 0.019251 | LGG | SF539 | Gefitinib | MVP_top60_DNA+RNA_seed | EZH2 | -0.71858 | LUAD | PC-14 | GSK343 |
| MVP_top60_DNA_seed | MAPK1 | 0.020181 | LGG | MOG-G-UVW | Ulixertinib | MVP_top60_DNA+RNA_seed | PIK3CA | -0.71695 | LUAD | A549 | Buparlisib |
| MVP_top60_DNA_seed | MAPK3 | 0.020181 | LGG | MOG-G-UVW | Ulixertinib | MVP_top60_DNA+RNA_seed | IGF1R | -0.716016 | LUAD | NCI-H358 | Linsitinib |
| MVP_top60_DNA_seed | HSP90AA1 | 0.021981 | LGG | D-336MG | Elesclomol | MVP_top60_DNA+RNA_seed | IGF1R | -0.715039 | LUAD | PC-14 | BMS-536924 |
| MVP_top60_DNA_seed | HSP90AA1 | 0.023891 | LGG | KALS-1 | Tanespimycin | MVP_top60_DNA+RNA_seed | HDAC1 | -0.714076 | LUAD | NCI-H2023 | Romidepsin |
| MVP_top60_DNA_seed | IGF1R | 0.025058 | LGG | H4 | BMS-536924 | MVP_top60_DNA+RNA_seed | IGF1R | -0.713322 | LUAD | NCI-H1755 | NVP-ADW742 |
| MVP_top60_DNA_seed | IGF1R | 0.025948 | LGG | GAMG | BMS-754807 | MVP_top60_DNA+RNA_seed | EZH2 | -0.709605 | LUAD | RERF-LC-MS | GSK343 |
| MVP_top60_DNA_seed | MAPK1 | 0.027527 | LGG | D-502MG | SCH772984 | MVP_top60_DNA+RNA_seed | IGF1R | -0.709093 | LUAD | RERF-LC-KJ | BMS-536924 |
| MVP_top60_DNA_seed | MAPK3 | 0.027527 | LGG | D-502MG | SCH772984 | MVP_top60_DNA+RNA_seed | EZH2 | -0.709046 | LUAD | NCI-H1355 | GSK343 |

| | | | | | | | | | | | |
|---|---|---|---|---|---|---|---|---|---|---|---|
| MVP_top60_DNA_seed | PIK3CA | 0.031366 | LGG | GI-1 | Alpelisib | MVP_top60_DNA+RNA_seed | EGFR | -0.707173 | LUAD | NCI-H1781 | Lapatinib |
| MVP_top60_DNA_seed | HSP90AA1 | 0.035518 | LGG | KNS-81-FD | Elesclomol | MVP_top60_DNA+RNA_seed | EGFR | -0.704192 | LUAD | NCI-H2122 | Osimertinib |
| MVP_top60_DNA_seed | TP53 | 0.040713 | LGG | KNS-42 | MIRA-1 | MVP_top60_DNA+RNA_seed | AKT1 | -0.704059 | LUAD | NCI-H3122 | MK-2206 |
| MVP_top60_DNA_seed | PIK3CA | 0.042034 | LGG | KNS-81-FD | GNE-317 | MVP_top60_DNA+RNA_seed | AKT1 | -0.701972 | LUAD | A549 | Uprosertib |
| MVP_top60_DNA_seed | PIK3CA | 0.043355 | LGG | SF539 | Alpelisib | MVP_top60_DNA+RNA_seed | EGFR | -0.701649 | LUAD | NCI-H2122 | Sapitinib |
| MVP_top60_DNA_seed | MAPK1 | 0.044473 | LGG | KALS-1 | VX-11e | MVP_top60_DNA+RNA_seed | MET | -0.699143 | LUAD | NCI-H2023 | Crizotinib |
| MVP_top60_DNA_seed | JAK2 | 0.049194 | LGG | D-336MG | Ruxolitinib | MVP_top60_DNA+RNA_seed | BRAF | -0.697119 | LUAD | EMC-BAC-1 | PLX-4720 |
| MVP_top60_DNA_seed | JAK1 | 0.049194 | LGG | D-336MG | Ruxolitinib | MVP_top60_DNA+RNA_seed | PIK3CA | -0.696165 | LUAD | LC-2-ad | Alpelisib |
| MVP_top60_DNA_seed | HSP90AA1 | 0.061125 | LGG | KALS-1 | Luminespib | MVP_top60_DNA+RNA_seed | AKT1 | -0.695581 | LUAD | HCC-44 | Ipatasertib |
| MVP_top60_DNA_seed | NTRK1 | 0.066354 | LGG | H4 | GW441756 | MVP_top60_DNA+RNA_seed | MDM2 | -0.694639 | LUAD | HCC-44 | Serdemetan |
| MVP_top60_DNA_seed | MDM2 | 0.067884 | LGG | KNS-42 | Nutlin-3a (-) | MVP_top60_DNA+RNA_seed | IGF1R | -0.694518 | LUAD | NCI-H2228 | BMS-754807 |
| MVP_top60_DNA_seed | TP53 | 0.070502 | LGG | SF539 | MIRA-1 | MVP_top60_DNA+RNA_seed | IGF1R | -0.69344 | LUAD | RERF-LC-KJ | NVP-ADW742 |
| MVP_top60_DNA_seed | HSP90AA1 | 0.072799 | LGG | Hs-683 | Elesclomol | MVP_top60_DNA+RNA_seed | AURKA | -0.692987 | LUAD | NCI-H1755 | ZM447439 |
| MVP_top60_DNA_seed | HDAC1 | 0.073463 | LGG | Hs-683 | Entinostat | MVP_top60_DNA+RNA_seed | PIK3CA | -0.688654 | LUAD | NCI-H1755 | Alpelisib |
| MVP_top60_DNA_seed | IGF1R | 0.074096 | LGG | SF539 | NVP-ADW742 | MVP_top60_DNA+RNA_seed | AKT1 | -0.687777 | LUAD | NCI-H2122 | Uprosertib |
| MVP_top60_DNA_seed | IDH1 | 0.078202 | LGG | H4 | AGI-5198 | MVP_top60_DNA+RNA_seed | BRAF | -0.687325 | LUAD | NCI-H1648 | PLX-4720 |
| MVP_top60_DNA_seed | EGFR | 0.080258 | LGG | SW1783 | Osimertinib | MVP_top60_DNA+RNA_seed | BRAF | -0.683802 | LUAD | RERF-LC-MS | Dabrafenib |
| MVP_top60_DNA_seed | IGF1R | 0.08739 | LGG | MOG-G-UVW | BMS-536924 | MVP_top60_DNA+RNA_seed | BRAF | -0.68222 | LUAD | NCI-H2030 | PLX-4720 |
| MVP_top60_DNA_seed | MET | 0.091508 | LGG | Hs-683 | Crizotinib | MVP_top60_DNA+RNA_seed | AKT1 | -0.680964 | LUAD | NCI-H2228 | MK-2206 |
| MVP_top60_DNA_seed | JAK2 | 0.093285 | LGG | KINGS-1 | AZ960 | MVP_top60_DNA+RNA_seed | EGFR | -0.678241 | LUAD | NCI-H1781 | Erlotinib |
| MVP_top60_DNA_seed | JAK3 | 0.093285 | LGG | KINGS-1 | AZ960 | MVP_top60_DNA+RNA_seed | MDM2 | -0.676845 | LUAD | COR-L105 | Nutlin-3a (-) |
| MVP_top60_DNA_seed | JAK2 | 0.095141 | LGG | no-10 | AZ960 | MVP_top60_DNA+RNA_seed | IGF1R | -0.675233 | LUAD | EMC-BAC-2 | BMS-536924 |
| MVP_top60_DNA_seed | JAK3 | 0.095141 | LGG | no-10 | AZ960 | MVP_top60_DNA+RNA_seed | AURKA | -0.673606 | LUAD | EMC-BAC-2 | Alisertib |
| MVP_top60_DNA_seed | JAK2 | 0.097349 | LGG | NMC-G1 | AZ960 | MVP_top60_DNA+RNA_seed | HSP90AA1 | -0.670457 | LUAD | NCI-H838 | Elesclomol |
| MVP_top60_DNA_seed | JAK3 | 0.097349 | LGG | NMC-G1 | AZ960 | MVP_top60_DNA+RNA_seed | AKT1 | -0.664889 | LUAD | NCI-H2030 | MK-2206 |
| MVP_top60_DNA_seed | TP53 | 0.100083 | LGG | Hs-683 | MIRA-1 | MVP_top60_DNA+RNA_seed | TP53 | -0.664239 | LUAD | NCI-H1792 | MIRA-1 |
| MVP_top60_DNA_seed | IGF1R | 0.100854 | LGG | D-336MG | NVP-ADW742 | MVP_top60_DNA+RNA_seed | AURKA | -0.663796 | LUAD | NCI-H838 | ZM447439 |
| MVP_top60_DNA_seed | HSP90AA1 | 0.10473 | LGG | no-10 | CCT-018159 | MVP_top60_DNA+RNA_seed | HSP90AA1 | -0.66342 | LUAD | NCI-H2023 | Tanespimycin |
| MVP_top60_DNA_seed | MDM2 | 0.107415 | LGG | GI-1 | Nutlin-3a (-) | MVP_top60_DNA+RNA_seed | HSP90AA1 | -0.662944 | LUAD | NCI-H2030 | CCT-018159 |
| MVP_top60_DNA_seed | HSP90AA1 | 0.110296 | LGG | KNS-81-FD | Luminespib | MVP_top60_DNA+RNA_seed | IGF1R | -0.658029 | LUAD | LXF-289 | NVP-ADW742 |
| MVP_top60_DNA_seed | HDAC1 | 0.111758 | LGG | GI-1 | Entinostat | MVP_top60_DNA+RNA_seed | SMARCA4 | -0.656587 | LUAD | NCI-H1792 | PFI3 |
| MVP_top60_DNA_seed | NTRK1 | 0.113726 | LGG | SW1783 | GW441756 | MVP_top60_DNA+RNA_seed | IGF1R | -0.655738 | LUAD | NCI-H838 | BMS-754807 |
| MVP_top60_DNA_seed | TP53 | 0.117569 | LGG | no-10 | MIRA-1 | MVP_top60_DNA+RNA_seed | BRAF | -0.654402 | LUAD | NCI-H322M | Dabrafenib |
| MVP_top60_DNA_seed | IGF1R | 0.135355 | LGG | SW1783 | BMS-754807 | MVP_top60_DNA+RNA_seed | IGF1R | -0.652007 | LUAD | NCI-H1437 | BMS-754807 |
| MVP_top60_DNA_seed | CDK2 | 0.136699 | LGG | GAMG | Dinaciclib | MVP_top60_DNA+RNA_seed | AKT1 | -0.650566 | LUAD | NCI-H1944 | AT13148 |
| MVP_top60_DNA_seed | IGF1R | 0.139116 | LGG | SW1783 | BMS-536924 | MVP_top60_DNA+RNA_seed | MET | -0.648689 | LUAD | NCI-H1792 | Crizotinib |
| MVP_top60_DNA_seed | HDAC1 | 0.139322 | LGG | KNS-81-FD | PCI-34051 | MVP_top60_DNA+RNA_seed | EGFR | -0.645534 | LUAD | A549 | AZD3759 |
| MVP_top60_DNA_seed | IDH1 | 0.139609 | LGG | MOG-G-UVW | AGI-5198 | MVP_top60_DNA+RNA_seed | IGF1R | -0.645183 | LUAD | NCI-H2009 | BMS-754807 |
| MVP_top60_DNA_seed | HDAC1 | 0.140868 | LGG | SF539 | Entinostat | MVP_top60_DNA+RNA_seed | SMARCA4 | -0.643857 | LUAD | NCI-H1734 | PFI3 |
| MVP_top60_DNA_seed | EGFR | 0.140876 | LGG | SW1783 | Afatinib | MVP_top60_DNA+RNA_seed | HSP90AA1 | -0.643257 | LUAD | EMC-BAC-2 | Tanespimycin |
| MVP_top60_DNA_seed | PIK3CA | 0.142299 | LGG | MOG-G-CCM | Alpelisib | MVP_top60_DNA+RNA_seed | MET | -0.641633 | LUAD | HCC-78 | Crizotinib |

| | | | | | | | | | | | |
|---|---|---|---|---|---|---|---|---|---|---|---|
| MVP_top60_DNA_seed | HDAC1 | 0.147441 | LGG | D-336MG | PCI-34051 | MVP_top60_DNA+RNA_seed | ATM | -0.637013 | LUAD | NCI-H2030 | KU-55933 |
| MVP_top60_DNA_seed | TP53 | 0.147656 | LGG | H4 | MIRA-1 | MVP_top60_DNA+RNA_seed | SMARCA4 | -0.636456 | LUAD | HCC-78 | PFI3 |
| MVP_top60_DNA_seed | EGFR | 0.151751 | LGG | MOG-G-UVW | Gefitinib | MVP_top60_DNA+RNA_seed | AURKA | -0.636384 | LUAD | NCI-H2085 | ZM447439 |
| MVP_top60_DNA_seed | EGFR | 0.153475 | LGG | MOG-G-UVW | Lapatinib | MVP_top60_DNA+RNA_seed | AURKA | -0.636228 | LUAD | LXF-289 | Alisertib |
| MVP_top60_DNA_seed | IGF1R | 0.157515 | LGG | Hs-683 | BMS-536924 | MVP_top60_DNA+RNA_seed | AKT1 | -0.635213 | LUAD | NCI-H2023 | Uprosertib |
| MVP_top60_DNA_seed | MAPK1 | 0.157746 | LGG | SF539 | Ulixertinib | MVP_top60_DNA+RNA_seed | HDAC1 | -0.634734 | LUAD | Calu-6 | PCI-34051 |
| MVP_top60_DNA_seed | MAPK3 | 0.157746 | LGG | SF539 | Ulixertinib | MVP_top60_DNA+RNA_seed | MDM2 | -0.633027 | LUAD | RERF-LC-KJ | Serdemetan |
| MVP_top60_DNA_seed | EGFR | 0.160787 | LGG | GAMG | AZD3759 | MVP_top60_DNA+RNA_seed | MET | -0.632749 | LUAD | LXF-289 | Crizotinib |
| MVP_top60_DNA_seed | EGFR | 0.161313 | LGG | MOG-G-UVW | Afatinib | MVP_top60_DNA+RNA_seed | BRAF | -0.632672 | LUAD | NCI-H322M | PLX-4720 |
| MVP_top60_DNA_seed | EGFR | 0.164179 | LGG | H4 | Gefitinib | MVP_top60_DNA+RNA_seed | MDM2 | -0.630145 | LUAD | NCI-H1792 | Serdemetan |
| MVP_top60_DNA_seed | EGFR | 0.165006 | LGG | SF539 | Sapitinib | MVP_top60_DNA+RNA_seed | PIK3CA | -0.628623 | LUAD | PC-3_[JPC-3] | GNE-317 |
| MVP_top60_DNA_seed | MET | 0.167905 | LGG | Hs-683 | Savolitinib | MVP_top60_DNA+RNA_seed | HDAC1 | -0.628231 | LUAD | NCI-H322M | Romidepsin |
| MVP_top60_DNA_seed | MAPK1 | 0.168527 | LGG | H4 | SCH772984 | MVP_top60_DNA+RNA_seed | BRAF | -0.626933 | LUAD | COR-L105 | PLX-4720 |
| MVP_top60_DNA_seed | MAPK3 | 0.168527 | LGG | H4 | SCH772984 | MVP_top60_DNA+RNA_seed | PIK3CA | -0.626438 | LUAD | NCI-H1666 | AZD8186 |
| MVP_top60_DNA_seed | AR | 0.173 | LGG | KNS-42 | Bicalutamide | MVP_top60_DNA+RNA_seed | FGFR1 | -0.625612 | LUAD | NCI-H2122 | PD173074 |
| MVP_top60_DNA_seed | CDK2 | 0.174592 | LGG | MOG-G-UVW | MK-8776 | MVP_top60_DNA+RNA_seed | IGF1R | -0.624301 | LUAD | NCI-H358 | BMS-754807 |
| MVP_top60_DNA_seed | EGFR | 0.175121 | LGG | MOG-G-CCM | Lapatinib | MVP_top60_DNA+RNA_seed | IGF1R | -0.622196 | LUAD | NCI-H1792 | BMS-754807 |
| MVP_top60_DNA_seed | IGF1R | 0.175394 | LGG | GI-1 | NVP-ADW742 | MVP_top60_DNA+RNA_seed | PIK3CA | -0.620927 | LUAD | NCI-H1650 | AZD8186 |
| MVP_top60_DNA_seed | TP53 | 0.179887 | LGG | GAMG | MIRA-1 | MVP_top60_DNA+RNA_seed | PIK3CA | -0.618162 | LUAD | NCI-H2122 | AZD8186 |
| MVP_top60_DNA_seed | PIK3CA | 0.183884 | LGG | KINGS-1 | GNE-317 | MVP_top60_DNA+RNA_seed | BRAF | -0.616559 | LUAD | HCC-827 | PLX-4720 |
| MVP_top60_DNA_seed | EGFR | 0.186305 | LGG | D-336MG | Sapitinib | MVP_top60_DNA+RNA_seed | BRAF | -0.615308 | LUAD | NCI-H2122 | PLX-4720 |
| MVP_top60_DNA_seed | MAPK1 | 0.188418 | LGG | GI-1 | VX-11e | MVP_top60_DNA+RNA_seed | KRAS | -0.608473 | LUAD | PC-14 | :RAS (G12C) Inhibitor-1 |
| MVP_top60_DNA_seed | MDM2 | 0.191046 | LGG | SF539 | Nutlin-3a (-) | MVP_top60_DNA+RNA_seed | TERT | -0.607296 | LUAD | NCI-H1573 | BIBR-1532 |
| MVP_top60_DNA_seed | PIK3CA | 0.191285 | LGG | no-10 | AZD8186 | MVP_top60_DNA+RNA_seed | AURKA | -0.606022 | LUAD | NCI-H1781 | Alisertib |
| MVP_top60_DNA_seed | PIK3CA | 0.192235 | LGG | Hs-683 | GNE-317 | MVP_top60_DNA+RNA_seed | AKT1 | -0.60524 | LUAD | RERF-LC-MS | MK-2206 |
| MVP_top60_DNA_seed | IGF1R | 0.194809 | LGG | no-10 | GSK1904529A | MVP_top60_DNA+RNA_seed | EGFR | -0.603694 | LUAD | NCI-H1563 | Osimertinib |
| MVP_top60_DNA_seed | HDAC1 | 0.203171 | LGG | KALS-1 | PCI-34051 | MVP_top60_DNA+RNA_seed | PIK3CA | -0.602351 | LUAD | NCI-H1734 | AZD8186 |
| MVP_top60_DNA_seed | CREBBP | 0.210176 | LGG | KALS-1 | SGC-CBP30 | MVP_top60_DNA+RNA_seed | MET | -0.60022 | LUAD | HCC-44 | Savolitinib |
| MVP_top60_DNA_seed | IGF1R | 0.210586 | LGG | D-336MG | GSK1904529A | MVP_top60_DNA+RNA_seed | AKT1 | -0.600028 | LUAD | NCI-H1734 | Afuresertib |
| MVP_top60_DNA_seed | HDAC1 | 0.216383 | LGG | KALS-1 | Romidepsin | MVP_top60_DNA+RNA_seed | EGFR | -0.599925 | LUAD | NCI-H358 | Erlotinib |
| MVP_top60_DNA_seed | EGFR | 0.216492 | LGG | SW1783 | Gefitinib | MVP_top60_DNA+RNA_seed | EGFR | -0.599418 | LUAD | LC-2-ad | AZD3759 |
| MVP_top60_DNA_seed | HSP90AA1 | 0.219944 | LGG | D-502MG | Luminespib | MVP_top60_DNA+RNA_seed | PIK3CA | -0.598218 | LUAD | NCI-H1623 | Buparlisib |
| MVP_top60_DNA_seed | HDAC1 | 0.232171 | LGG | D-336MG | Entinostat | MVP_top60_DNA+RNA_seed | ATM | -0.597208 | LUAD | NCI-H1792 | KU-55933 |
| MVP_top60_DNA_seed | CDK2 | 0.23851 | LGG | H4 | MK-8776 | MVP_top60_DNA+RNA_seed | AKT1 | -0.595196 | LUAD | EMC-BAC-1 | Ipatasertib |
| MVP_top60_DNA_seed | HSP90AA1 | 0.240221 | LGG | KNS-42 | Tanespimycin | MVP_top60_DNA+RNA_seed | HDAC1 | -0.595179 | LUAD | NCI-H2291 | Entinostat |
| MVP_top60_DNA_seed | EGFR | 0.246484 | LGG | H4 | AZD3759 | MVP_top60_DNA+RNA_seed | AKT1 | -0.593194 | LUAD | NCI-H838 | MK-2206 |
| MVP_top60_DNA_seed | CDK2 | 0.26084 | LGG | no-11 | MK-8776 | MVP_top60_DNA+RNA_seed | AKT1 | -0.592801 | LUAD | NCI-H1781 | MK-2206 |
| MVP_top60_DNA_seed | JAK2 | 0.265798 | LGG | GAMG | GSK2276186C | MVP_top60_DNA+RNA_seed | PIK3CA | -0.590916 | LUAD | NCI-H2030 | GNE-317 |
| MVP_top60_DNA_seed | JAK1 | 0.265798 | LGG | GAMG | GSK2276186C | MVP_top60_DNA+RNA_seed | HSP90AA1 | -0.59049 | LUAD | NCI-H3122 | Luminespib |
| MVP_top60_DNA_seed | JAK3 | 0.265798 | LGG | GAMG | GSK2276186C | MVP_top60_DNA+RNA_seed | EGFR | -0.58924 | LUAD | A549 | Erlotinib |
| MVP_top60_DNA_seed | MET | 0.266456 | LGG | GAMG | Savolitinib | MVP_top60_DNA+RNA_seed | EGFR | -0.588163 | LUAD | LC-2-ad | Erlotinib |

| | | | | | | | | | | | |
|---|---|---|---|---|---|---|---|---|---|---|---|
| MVP_top60_DNA_seed | JAK2 | 0.267275 | LGG | GI-1 | Ruxolitinib | MVP_top60_DNA+RNA_seed | PIK3CA | -0.587139 | LUAD | NCI-H2023 | Alpelisib |
| MVP_top60_DNA_seed | JAK1 | 0.267275 | LGG | GI-1 | Ruxolitinib | MVP_top60_DNA+RNA_seed | AKT1 | -0.585015 | LUAD | NCI-H1781 | Afuresertib |
| MVP_top60_DNA_seed | PIK3CA | 0.267498 | LGG | no-11 | AZD8186 | MVP_top60_DNA+RNA_seed | EZH2 | -0.583697 | LUAD | HCC-44 | GSK343 |
| MVP_top60_DNA_seed | JAK2 | 0.268713 | LGG | GAMG | JAK_8517 | MVP_top60_DNA+RNA_seed | HDAC1 | -0.583352 | LUAD | HCC-44 | Romidepsin |
| MVP_top60_DNA_seed | JAK1 | 0.268713 | LGG | GAMG | JAK_8517 | MVP_top60_DNA+RNA_seed | FGFR1 | -0.58331 | LUAD | LXF-289 | AZD4547 |
| MVP_top60_DNA_seed | EGFR | 0.270554 | LGG | SF539 | AZD3759 | MVP_top60_DNA+RNA_seed | PIK3CA | -0.580946 | LUAD | RERF-LC-MS | Buparlisib |
| MVP_top60_DNA_seed | NTRK1 | 0.271927 | LGG | Hs-683 | GW441756 | MVP_top60_DNA+RNA_seed | PIK3CA | -0.580117 | LUAD | NCI-H1623 | AZD8186 |
| MVP_top60_DNA_seed | PIK3CA | 0.274647 | LGG | GAMG | Alpelisib | MVP_top60_DNA+RNA_seed | MDM2 | -0.578934 | LUAD | NCI-H1437 | Serdemetan |
| MVP_top60_DNA_seed | EGFR | 0.279487 | LGG | H4 | Lapatinib | MVP_top60_DNA+RNA_seed | FGFR1 | -0.578524 | LUAD | EMC-BAC-2 | AZD4547 |
| MVP_top60_DNA_seed | MET | 0.279627 | LGG | D-502MG | Crizotinib | MVP_top60_DNA+RNA_seed | SRC | -0.578392 | LUAD | Calu-3 | Bosutinib |
| MVP_top60_DNA_seed | HSP90AA1 | 0.282723 | LGG | KNS-42 | Elesclomol | MVP_top60_DNA+RNA_seed | MET | -0.578326 | LUAD | A549 | Crizotinib |
| MVP_top60_DNA_seed | HSP90AA1 | 0.289521 | LGG | D-502MG | Elesclomol | MVP_top60_DNA+RNA_seed | EZH2 | -0.578045 | LUAD | NCI-H1792 | GSK343 |
| MVP_top60_DNA_seed | NTRK1 | 0.290459 | LGG | MOG-G-UVW | GW441756 | MVP_top60_DNA+RNA_seed | IGF1R | -0.57765 | LUAD | A549 | Linsitinib |
| MVP_top60_DNA_seed | HSP90AA1 | 0.292117 | LGG | no-10 | Elesclomol | MVP_top60_DNA+RNA_seed | HDAC1 | -0.577474 | LUAD | NCI-H1792 | PCI-34051 |
| MVP_top60_DNA_seed | MAPK1 | 0.298222 | LGG | no-11 | SCH772984 | MVP_top60_DNA+RNA_seed | KRAS | -0.576165 | LUAD | A549 | [illegible]RAS (G12C) Inhibitor-1 |
| MVP_top60_DNA_seed | MAPK3 | 0.298222 | LGG | no-11 | SCH772984 | MVP_top60_DNA+RNA_seed | EZH2 | -0.574844 | LUAD | HCC-44 | GSK343 |
| MVP_top60_DNA_seed | PIK3CA | 0.301169 | LGG | KALS-1 | GNE-317 | MVP_top60_DNA+RNA_seed | EGFR | -0.574442 | LUAD | EMC-BAC-2 | Gefitinib |
| MVP_top60_DNA_seed | EGFR | 0.309872 | LGG | KNS-42 | AZD3759 | MVP_top60_DNA+RNA_seed | EZH2 | -0.573823 | LUAD | HCC-827 | GSK343 |
| MVP_top60_DNA_seed | IGF1R | 0.318602 | LGG | H4 | NVP-ADW742 | MVP_top60_DNA+RNA_seed | AURKA | -0.573581 | LUAD | VMRC-LCD | ZM447439 |
| MVP_top60_DNA_seed | IGF1R | 0.322325 | LGG | GI-1 | BMS-536924 | MVP_top60_DNA+RNA_seed | AURKA | -0.570868 | LUAD | RERF-LC-MS | ZM447439 |
| MVP_top60_DNA_seed | IGF1R | 0.323098 | LGG | GI-1 | Linsitinib | MVP_top60_DNA+RNA_seed | AKT1 | -0.570588 | LUAD | EMC-BAC-2 | Ipatasertib |
| MVP_top60_DNA_seed | HDAC1 | 0.325213 | LGG | GAMG | PCI-34051 | MVP_top60_DNA+RNA_seed | MET | -0.568303 | LUAD | LC-2-ad | Crizotinib |
| MVP_top60_DNA_seed | MAPK1 | 0.329035 | LGG | no-10 | SCH772984 | MVP_top60_DNA+RNA_seed | HDAC1 | -0.567604 | LUAD | COR-L105 | PCI-34051 |
| MVP_top60_DNA_seed | MAPK3 | 0.329035 | LGG | no-10 | SCH772984 | MVP_top60_DNA+RNA_seed | HDAC1 | -0.565808 | LUAD | HCC-827 | Romidepsin |
| MVP_top60_DNA_seed | SMARCA4 | 0.333252 | LGG | D-336MG | PFI3 | MVP_top60_DNA+RNA_seed | AKT1 | -0.564375 | LUAD | NCI-H1975 | MK-2206 |
| MVP_top60_DNA_seed | EGFR | 0.334534 | LGG | D-336MG | Gefitinib | MVP_top60_DNA+RNA_seed | PIK3CA | -0.563451 | LUAD | EMC-BAC-2 | GNE-317 |
| MVP_top60_DNA_seed | EGFR | 0.345412 | LGG | no-10 | Afatinib | MVP_top60_DNA+RNA_seed | IGF1R | -0.561592 | LUAD | NCI-H1755 | Linsitinib |
| MVP_top60_DNA_seed | EGFR | 0.34689 | LGG | GI-1 | Sapitinib | MVP_top60_DNA+RNA_seed | EGFR | -0.561571 | LUAD | A549 | Sapitinib |
| MVP_top60_DNA_seed | MET | 0.349966 | LGG | SW1783 | Crizotinib | MVP_top60_DNA+RNA_seed | BRAF | -0.560148 | LUAD | NCI-H1435 | Dabrafenib |
| MVP_top60_DNA_seed | JAK2 | 0.353721 | LGG | KALS-1 | JAK_8517 | MVP_top60_DNA+RNA_seed | HSP90AA1 | -0.559989 | LUAD | NCI-H1781 | Tanespimycin |
| MVP_top60_DNA_seed | JAK1 | 0.353721 | LGG | KALS-1 | JAK_8517 | MVP_top60_DNA+RNA_seed | PIK3CA | -0.559641 | LUAD | A549 | AZD8186 |
| MVP_top60_DNA_seed | EGFR | 0.354282 | LGG | D-502MG | Erlotinib | MVP_top60_DNA+RNA_seed | MDM2 | -0.559617 | LUAD | RERF-LC-MS | Nutlin-3a (-) |
| MVP_top60_DNA_seed | HDAC1 | 0.357345 | LGG | Hs-683 | PCI-34051 | MVP_top60_DNA+RNA_seed | PIK3CA | -0.55925 | LUAD | NCI-H1734 | Buparlisib |
| MVP_top60_DNA_seed | IGF1R | 0.359556 | LGG | GI-1 | GSK1904529A | MVP_top60_DNA+RNA_seed | IGF1R | -0.558478 | LUAD | NCI-H1666 | Linsitinib |
| MVP_top60_DNA_seed | EGFR | 0.360726 | LGG | GAMG | Gefitinib | MVP_top60_DNA+RNA_seed | IGF1R | -0.555108 | LUAD | A549 | BMS-754807 |
| MVP_top60_DNA_seed | IGF1R | 0.364147 | LGG | D-502MG | Linsitinib | MVP_top60_DNA+RNA_seed | PIK3CA | -0.554255 | LUAD | HCC-78 | Alpelisib |
| MVP_top60_DNA_seed | MDM2 | 0.370709 | LGG | D-502MG | Serdemetan | MVP_top60_DNA+RNA_seed | AKT1 | -0.553987 | LUAD | NCI-H1838 | AT13148 |
| MVP_top60_DNA_seed | MAPK1 | 0.371166 | LGG | D-502MG | VX-11e | MVP_top60_DNA+RNA_seed | PIK3CA | -0.553292 | LUAD | NCI-H2228 | Alpelisib |
| MVP_top60_DNA_seed | EGFR | 0.372986 | LGG | KNS-81-FD | AZD3759 | MVP_top60_DNA+RNA_seed | PIK3CA | -0.552522 | LUAD | NCI-H2009 | Alpelisib |
| MVP_top60_DNA_seed | HSP90AA1 | 0.38062 | LGG | KNS-42 | CCT-018159 | MVP_top60_DNA+RNA_seed | HDAC1 | -0.551923 | LUAD | EMC-BAC-2 | Romidepsin |
| MVP_top60_DNA_seed | CDK2 | 0.380911 | LGG | D-336MG | MK-8776 | MVP_top60_DNA+RNA_seed | SRC | -0.551381 | LUAD | EMC-BAC-1 | Bosutinib |

| | | | | | | | | | | | |
|---|---|---|---|---|---|---|---|---|---|---|---|
| MVP_top60_DNA_seed | EGFR | 0.386058 | LGG | Hs-683 | Sapitinib | MVP_top60_DNA+RNA_seed | EZH2 | -0.551115 | LUAD | NCI-H1435 | GSK343 |
| MVP_top60_DNA_seed | MAPK1 | 0.393264 | LGG | Hs-683 | SCH772984 | MVP_top60_DNA+RNA_seed | PIK3CA | -0.549144 | LUAD | HCC-44 | GNE-317 |
| MVP_top60_DNA_seed | MAPK3 | 0.393264 | LGG | Hs-683 | SCH772984 | MVP_top60_DNA+RNA_seed | SMARCA4 | -0.5489 | LUAD | NCI-H1435 | PFI3 |
| MVP_top60_DNA_seed | EGFR | 0.397765 | LGG | D-502MG | Osimertinib | MVP_top60_DNA+RNA_seed | AKT1 | -0.544665 | LUAD | NCI-H1734 | AZD5363 |
| MVP_top60_DNA_seed | EGFR | 0.40411 | LGG | SW1783 | Sapitinib | MVP_top60_DNA+RNA_seed | EGFR | -0.541517 | LUAD | NCI-H1693 | Sapitinib |
| MVP_top60_DNA_seed | MDM2 | 0.411104 | LGG | GAMG | Nutlin-3a (-) | MVP_top60_DNA+RNA_seed | TERT | -0.540957 | LUAD | NCI-H1623 | BIBR-1532 |
| MVP_top60_DNA_seed | EGFR | 0.411407 | LGG | GAMG | Lapatinib | MVP_top60_DNA+RNA_seed | EGFR | -0.539813 | LUAD | HCC-78 | Osimertinib |
| MVP_top60_DNA_seed | MDM2 | 0.4192 | LGG | no-11 | Nutlin-3a (-) | MVP_top60_DNA+RNA_seed | PIK3CA | -0.537165 | LUAD | NCI-H1666 | Alpelisib |
| MVP_top60_DNA_seed | EGFR | 0.419301 | LGG | SW1783 | AZD3759 | MVP_top60_DNA+RNA_seed | IGF1R | -0.537128 | LUAD | LC-2-ad | NVP-ADW742 |
| MVP_top60_DNA_seed | IDH1 | 0.419509 | LGG | no-10 | AGI-5198 | MVP_top60_DNA+RNA_seed | EGFR | -0.536795 | LUAD | NCI-H650 | Gefitinib |
| MVP_top60_DNA_seed | MET | 0.426146 | LGG | KNS-42 | Crizotinib | MVP_top60_DNA+RNA_seed | EZH2 | -0.536677 | LUAD | NCI-H1944 | GSK343 |
| MVP_top60_DNA_seed | PIK3CA | 0.432726 | LGG | MOG-G-UVW | Alpelisib | MVP_top60_DNA+RNA_seed | PIK3CA | -0.536241 | LUAD | NCI-H1993 | Buparlisib |
| MVP_top60_DNA_seed | MAPK1 | 0.439077 | LGG | KALS-1 | Ulixertinib | MVP_top60_DNA+RNA_seed | PIK3CA | -0.535728 | LUAD | NCI-H2023 | Buparlisib |
| MVP_top60_DNA_seed | MAPK3 | 0.439077 | LGG | KALS-1 | Ulixertinib | MVP_top60_DNA+RNA_seed | MDM2 | -0.535122 | LUAD | NCI-H1651 | Serdemetan |
| MVP_top60_DNA_seed | EGFR | 0.440839 | LGG | H4 | Osimertinib | MVP_top60_DNA+RNA_seed | PIK3CA | -0.53352 | LUAD | COR-L105 | Buparlisib |
| MVP_top60_DNA_seed | EGFR | 0.44375 | LGG | GI-1 | Osimertinib | MVP_top60_DNA+RNA_seed | MET | -0.533176 | LUAD | NCI-H1734 | Crizotinib |
| MVP_top60_DNA_seed | MET | 0.455034 | LGG | SW1783 | Savolitinib | MVP_top60_DNA+RNA_seed | HSP90AA1 | -0.532301 | LUAD | NCI-H322M | Tanespimycin |
| MVP_top60_DNA_seed | HDAC1 | 0.456843 | LGG | H4 | Entinostat | MVP_top60_DNA+RNA_seed | AURKA | -0.532005 | LUAD | NCI-H2122 | ZM447439 |
| MVP_top60_DNA_seed | EGFR | 0.465577 | LGG | KALS-1 | Afatinib | MVP_top60_DNA+RNA_seed | BRAF | -0.530993 | LUAD | A549 | PLX-4720 |
| MVP_top60_DNA_seed | MAPK1 | 0.466278 | LGG | no-11 | VX-11e | MVP_top60_DNA+RNA_seed | EGFR | -0.530947 | LUAD | HCC-78 | Sapitinib |
| MVP_top60_DNA_seed | MET | 0.470143 | LGG | MOG-G-CCM | Crizotinib | MVP_top60_DNA+RNA_seed | IGF1R | -0.53047 | LUAD | NCI-H2009 | Linsitinib |
| MVP_top60_DNA_seed | PIK3CA | 0.477954 | LGG | SW1783 | Alpelisib | MVP_top60_DNA+RNA_seed | AURKA | -0.529581 | LUAD | LXF-289 | ZM447439 |
| MVP_top60_DNA_seed | IGF1R | 0.47917 | LGG | H4 | Linsitinib | MVP_top60_DNA+RNA_seed | EGFR | -0.529259 | LUAD | EMC-BAC-2 | AZD3759 |
| MVP_top60_DNA_seed | EGFR | 0.480667 | LGG | Hs-683 | Erlotinib | MVP_top60_DNA+RNA_seed | AKT1 | -0.528741 | LUAD | LXF-289 | MK-2206 |
| MVP_top60_DNA_seed | CDK2 | 0.487445 | LGG | Hs-683 | MK-8776 | MVP_top60_DNA+RNA_seed | PIK3CA | -0.526357 | LUAD | NCI-H1793 | Buparlisib |
| MVP_top60_DNA_seed | MDM2 | 0.489727 | LGG | MOG-G-CCM | Nutlin-3a (-) | MVP_top60_DNA+RNA_seed | HSP90AA1 | -0.524857 | LUAD | NCI-H1355 | Tanespimycin |
| MVP_top60_DNA_seed | EGFR | 0.490183 | LGG | D-502MG | Afatinib | MVP_top60_DNA+RNA_seed | KRAS | -0.523116 | LUAD | NCI-H650 | KRAS (G12C) Inhibitor-1 |
| MVP_top60_DNA_seed | PIK3CA | 0.490219 | LGG | KINGS-1 | Buparlisib | MVP_top60_DNA+RNA_seed | BRAF | -0.522925 | LUAD | NCI-H1755 | PLX-4720 |
| MVP_top60_DNA_seed | PIK3CD | 0.490219 | LGG | KINGS-1 | Buparlisib | MVP_top60_DNA+RNA_seed | AKT1 | -0.519444 | LUAD | NCI-H2009 | AT13148 |
| MVP_top60_DNA_seed | EGFR | 0.498904 | LGG | Hs-683 | Osimertinib | MVP_top60_DNA+RNA_seed | MET | -0.51892 | LUAD | EMC-BAC-2 | Savolitinib |
| MVP_top60_DNA_seed | JAK2 | 0.501832 | LGG | Hs-683 | Ruxolitinib | MVP_top60_DNA+RNA_seed | EGFR | -0.518054 | LUAD | NCI-H522 | Gefitinib |
| MVP_top60_DNA_seed | JAK1 | 0.501832 | LGG | Hs-683 | Ruxolitinib | MVP_top60_DNA+RNA_seed | HSP90AA1 | -0.516152 | LUAD | NCI-H1734 | Luminespib |
| MVP_top60_DNA_seed | EGFR | 0.505316 | LGG | no-11 | AZD3759 | MVP_top60_DNA+RNA_seed | AKT1 | -0.515056 | LUAD | LXF-289 | GSK2110183B |
| MVP_top60_DNA_seed | MAPK1 | 0.510035 | LGG | KALS-1 | Ulixertinib | MVP_top60_DNA+RNA_seed | PIK3CA | -0.513282 | LUAD | NCI-H1781 | AZD8186 |
| MVP_top60_DNA_seed | MAPK3 | 0.510035 | LGG | KALS-1 | Ulixertinib | MVP_top60_DNA+RNA_seed | HSP90AA1 | -0.511554 | LUAD | NCI-H1437 | Tanespimycin |
| MVP_top60_DNA_seed | IGF1R | 0.514729 | LGG | KINGS-1 | BMS-754807 | MVP_top60_DNA+RNA_seed | HSP90AA1 | -0.511552 | LUAD | NCI-H2228 | Tanespimycin |
| MVP_top60_DNA_seed | IGF1R | 0.516385 | LGG | KINGS-1 | NVP-ADW742 | MVP_top60_DNA+RNA_seed | FGFR1 | -0.509894 | LUAD | LC-2-ad | PD173074 |
| MVP_top60_DNA_seed | CDK2 | 0.519712 | LGG | GAMG | AZD5438 | MVP_top60_DNA+RNA_seed | BRAF | -0.507448 | LUAD | NCI-H1792 | PLX-4720 |
| MVP_top60_DNA_seed | CDK2 | 0.519784 | LGG | D-502MG | Dinaciclib | MVP_top60_DNA+RNA_seed | FGFR1 | -0.503983 | LUAD | RERF-LC-MS | AZD4547 |
| MVP_top60_DNA_seed | MAPK1 | 0.520366 | LGG | MOG-G-CCM | VX-11e | MVP_top60_DNA+RNA_seed | AURKA | -0.503673 | LUAD | NCI-H2085 | Alisertib |
| MVP_top60_DNA_seed | CREBBP | 0.520526 | LGG | KINGS-1 | SGC-CBP30 | MVP_top60_DNA+RNA_seed | FGFR1 | -0.502658 | LUAD | NCI-H1563 | AZD4547 |

| | | | | | | | | | | | |
|---|---|---|---|---|---|---|---|---|---|---|---|
| MVP_top60_DNA_seed | IDH1 | 0.520709 | LGG | MOG-G-CCM | AGI-5198 | MVP_top60_DNA+RNA_seed | ATM | -0.501144 | LUAD | NCI-H1975 | KU-55933 |
| MVP_top60_DNA_seed | HSP90AA1 | 0.522417 | LGG | KNS-42 | Luminespib | MVP_top60_DNA+RNA_seed | AKT1 | -0.501022 | LUAD | NCI-H1755 | AT13148 |
| MVP_top60_DNA_seed | EGFR | 0.531287 | LGG | MOG-G-CCM | Afatinib | MVP_top60_DNA+RNA_seed | AKT1 | -0.49972 | LUAD | NCI-H3122 | Uprosertib |
| MVP_top60_DNA_seed | PIK3CA | 0.534214 | LGG | KNS-81-FD | AZD8186 | MVP_top60_DNA+RNA_seed | IGF1R | -0.49421 | LUAD | EMC-BAC-1 | BMS-754807 |
| MVP_top60_DNA_seed | PIK3CA | 0.535369 | LGG | D-336MG | GNE-317 | MVP_top60_DNA+RNA_seed | HSP90AA1 | -0.493468 | LUAD | LC-2-ad | Elesclomol |
| MVP_top60_DNA_seed | EGFR | 0.53921 | LGG | KALS-1 | Lapatinib | MVP_top60_DNA+RNA_seed | HSP90AA1 | -0.489903 | LUAD | NCI-H838 | Luminespib |
| MVP_top60_DNA_seed | EGFR | 0.539901 | LGG | D-336MG | Afatinib | MVP_top60_DNA+RNA_seed | PIK3CA | -0.488432 | LUAD | NCI-H322M | Buparlisib |
| MVP_top60_DNA_seed | MAPK1 | 0.552011 | LGG | KINGS-1 | Ulixertinib | MVP_top60_DNA+RNA_seed | EGFR | -0.487197 | LUAD | NCI-H1437 | Lapatinib |
| MVP_top60_DNA_seed | MAPK3 | 0.552011 | LGG | KINGS-1 | Ulixertinib | MVP_top60_DNA+RNA_seed | FGFR1 | -0.487101 | LUAD | NCI-H1792 | PD173074 |
| MVP_top60_DNA_seed | MDM2 | 0.558184 | LGG | D-336MG | Nutlin-3a (-) | MVP_top60_DNA+RNA_seed | EZH2 | -0.487004 | LUAD | NCI-H1792 | GSK343 |
| MVP_top60_DNA_seed | EGFR | 0.561184 | LGG | GAMG | Sapitinib | MVP_top60_DNA+RNA_seed | HSP90AA1 | -0.48684 | LUAD | NCI-H1650 | Tanespimycin |
| MVP_top60_DNA_seed | EGFR | 0.571943 | LGG | KNS-42 | Gefitinib | MVP_top60_DNA+RNA_seed | EZH2 | -0.486352 | LUAD | NCI-H1781 | GSK343 |
| MVP_top60_DNA_seed | JAK2 | 0.575721 | LGG | no-11 | Ruxolitinib | MVP_top60_DNA+RNA_seed | AKT1 | -0.48587 | LUAD | NCI-H1437 | MK-2206 |
| MVP_top60_DNA_seed | JAK1 | 0.575721 | LGG | no-11 | Ruxolitinib | MVP_top60_DNA+RNA_seed | MET | -0.484597 | LUAD | VMRC-LCD | Crizotinib |
| MVP_top60_DNA_seed | EGFR | 0.576865 | LGG | GI-1 | AZD3759 | MVP_top60_DNA+RNA_seed | EGFR | -0.484067 | LUAD | NCI-H1563 | Erlotinib |
| MVP_top60_DNA_seed | SMARCA4 | 0.578547 | LGG | KALS-1 | PFI3 | MVP_top60_DNA+RNA_seed | EZH2 | -0.482935 | LUAD | NCI-H1648 | GSK343 |
| MVP_top60_DNA_seed | IGF1R | 0.579641 | LGG | KNS-42 | BMS-536924 | MVP_top60_DNA+RNA_seed | HSP90AA1 | -0.479662 | LUAD | NCI-H2030 | Luminespib |
| MVP_top60_DNA_seed | IGF1R | 0.584779 | LGG | MOG-G-UVW | GSK1904529A | MVP_top60_DNA+RNA_seed | AURKA | -0.479115 | LUAD | RERF-LC-KJ | ZM447439 |
| MVP_top60_DNA_seed | PIK3CA | 0.586555 | LGG | KALS-1 | Buparlisib | MVP_top60_DNA+RNA_seed | HDAC1 | -0.47649 | LUAD | NCI-H2122 | PCI-34051 |
| MVP_top60_DNA_seed | PIK3CD | 0.586555 | LGG | KALS-1 | Buparlisib | MVP_top60_DNA+RNA_seed | IGF1R | -0.475001 | LUAD | NCI-H1781 | GSK1904529A |
| MVP_top60_DNA_seed | EGFR | 0.586737 | LGG | D-336MG | Erlotinib | MVP_top60_DNA+RNA_seed | PIK3CA | -0.471532 | LUAD | A549 | Alpelisib |
| MVP_top60_DNA_seed | MAPK1 | 0.587034 | LGG | KINGS-1 | SCH772984 | MVP_top60_DNA+RNA_seed | AKT1 | -0.471188 | LUAD | NCI-H1734 | AT13148 |
| MVP_top60_DNA_seed | MAPK3 | 0.587034 | LGG | KINGS-1 | SCH772984 | MVP_top60_DNA+RNA_seed | HDAC1 | -0.470349 | LUAD | Calu-6 | Romidepsin |
| MVP_top60_DNA_seed | PIK3CA | 0.590988 | LGG | no-11 | Alpelisib | MVP_top60_DNA+RNA_seed | ATM | -0.470102 | LUAD | NCI-H1568 | KU-55933 |
| MVP_top60_DNA_seed | JAK2 | 0.593351 | LGG | no-11 | AZ960 | MVP_top60_DNA+RNA_seed | AKT1 | -0.46818 | LUAD | NCI-H1623 | Uprosertib |
| MVP_top60_DNA_seed | JAK3 | 0.593351 | LGG | no-11 | AZ960 | MVP_top60_DNA+RNA_seed | SRC | -0.467974 | LUAD | LC-2-ad | Bosutinib |
| MVP_top60_DNA_seed | IGF1R | 0.598804 | LGG | MOG-G-CCM | NVP-ADW742 | MVP_top60_DNA+RNA_seed | PIK3CA | -0.465927 | LUAD | NCI-H2228 | GNE-317 |
| MVP_top60_DNA_seed | MET | 0.599032 | LGG | no-10 | Crizotinib | MVP_top60_DNA+RNA_seed | PIK3CA | -0.465035 | LUAD | EMC-BAC-2 | Buparlisib |
| MVP_top60_DNA_seed | MDM2 | 0.605195 | LGG | NMC-G1 | Nutlin-3a (-) | MVP_top60_DNA+RNA_seed | BRAF | -0.46219 | LUAD | LC-2-ad | PLX-4720 |
| MVP_top60_DNA_seed | MDM2 | 0.606072 | LGG | MOG-G-UVW | Serdemetan | MVP_top60_DNA+RNA_seed | PIK3CA | -0.46151 | LUAD | NCI-H1755 | Buparlisib |
| MVP_top60_DNA_seed | PIK3CA | 0.611713 | LGG | no-10 | Alpelisib | MVP_top60_DNA+RNA_seed | TERT | -0.458878 | LUAD | NCI-H1993 | BIBR-1532 |
| MVP_top60_DNA_seed | EGFR | 0.614726 | LGG | SW1783 | Lapatinib | MVP_top60_DNA+RNA_seed | AKT1 | -0.456887 | LUAD | HCC-78 | AZD5363 |
| MVP_top60_DNA_seed | IDH1 | 0.617818 | LGG | KNS-42 | AGI-5198 | MVP_top60_DNA+RNA_seed | KRAS | -0.455088 | LUAD | EMC-BAC-2 | [illegible]RAS (G12C) Inhibitor-1 |
| MVP_top60_DNA_seed | MAPK1 | 0.618068 | LGG | GI-1 | SCH772984 | MVP_top60_DNA+RNA_seed | AKT1 | -0.453192 | LUAD | PC-14 | Uprosertib |
| MVP_top60_DNA_seed | MAPK3 | 0.618068 | LGG | GI-1 | SCH772984 | MVP_top60_DNA+RNA_seed | IGF1R | -0.451145 | LUAD | LC-2-ad | GSK1904529A |
| MVP_top60_DNA_seed | NTRK1 | 0.624499 | LGG | GAMG | GW441756 | MVP_top60_DNA+RNA_seed | BRAF | -0.450952 | LUAD | Calu-6 | Dabrafenib |
| MVP_top60_DNA_seed | SMARCA4 | 0.637087 | LGG | no-10 | PFI3 | MVP_top60_DNA+RNA_seed | AKT1 | -0.447143 | LUAD | A549 | GSK2110183B |
| MVP_top60_DNA_seed | MDM2 | 0.637138 | LGG | KNS-42 | Serdemetan | MVP_top60_DNA+RNA_seed | EZH2 | -0.443034 | LUAD | NCI-H1734 | GSK343 |
| MVP_top60_DNA_seed | IGF1R | 0.638792 | LGG | KNS-81-FD | BMS-754807 | MVP_top60_DNA+RNA_seed | EGFR | -0.442988 | LUAD | NCI-H1944 | Osimertinib |
| MVP_top60_DNA_seed | MAPK1 | 0.642825 | LGG | KNS-81-FD | SCH772984 | MVP_top60_DNA+RNA_seed | EGFR | -0.442342 | LUAD | A549 | Gefitinib |
| MVP_top60_DNA_seed | MAPK3 | 0.642825 | LGG | KNS-81-FD | SCH772984 | MVP_top60_DNA+RNA_seed | AKT1 | -0.441499 | LUAD | NCI-H322M | GSK2110183B |

| | | | | | | | | | | | |
|---|---|---|---|---|---|---|---|---|---|---|---|
| MVP_top60_DNA_seed | EGFR | 0.643642 | LGG | GI-1 | Lapatinib | MVP_top60_DNA+RNA_seed | HDAC1 | -0.440429 | LUAD | NCI-H1435 | Romidepsin |
| MVP_top60_DNA_seed | CDK2 | 0.65117 | LGG | KNS-42 | Dinaciclib | MVP_top60_DNA+RNA_seed | EGFR | -0.439857 | LUAD | NCI-H1975 | Gefitinib |
| MVP_top60_DNA_seed | MAPK1 | 0.660245 | LGG | KNS-81-FD | VX-11e | MVP_top60_DNA+RNA_seed | EZH2 | -0.439588 | LUAD | NCI-H2087 | GSK343 |
| MVP_top60_DNA_seed | CDK2 | 0.662249 | LGG | no-11 | Dinaciclib | MVP_top60_DNA+RNA_seed | PIK3CA | -0.439283 | LUAD | NCI-H1623 | GNE-317 |
| MVP_top60_DNA_seed | MAPK1 | 0.670797 | LGG | KINGS-1 | VX-11e | MVP_top60_DNA+RNA_seed | AKT1 | -0.438822 | LUAD | COR-L105 | Uprosertib |
| MVP_top60_DNA_seed | MDM2 | 0.672758 | LGG | KALS-1 | Nutlin-3a (-) | MVP_top60_DNA+RNA_seed | PIK3CA | -0.437569 | LUAD | NCI-H1792 | Alpelisib |
| MVP_top60_DNA_seed | EGFR | 0.680258 | LGG | KNS-81-FD | Lapatinib | MVP_top60_DNA+RNA_seed | IGF1R | -0.437178 | LUAD | NCI-H1944 | GSK1904529A |
| MVP_top60_DNA_seed | JAK2 | 0.682173 | LGG | KNS-81-FD | AZ960 | MVP_top60_DNA+RNA_seed | AKT1 | -0.436715 | LUAD | NCI-H2009 | Uprosertib |
| MVP_top60_DNA_seed | JAK3 | 0.682173 | LGG | KNS-81-FD | AZ960 | MVP_top60_DNA+RNA_seed | AKT1 | -0.43662 | LUAD | NCI-H1944 | GSK2110183B |
| MVP_top60_DNA_seed | MAPK1 | 0.684057 | LGG | D-336MG | SCH772984 | MVP_top60_DNA+RNA_seed | AURKA | -0.432636 | LUAD | PC-14 | ZM447439 |
| MVP_top60_DNA_seed | MAPK3 | 0.684057 | LGG | D-336MG | SCH772984 | MVP_top60_DNA+RNA_seed | AURKA | -0.432052 | LUAD | RERF-LC-MS | Alisertib |
| MVP_top60_DNA_seed | HDAC1 | 0.687674 | LGG | KINGS-1 | PCI-34051 | MVP_top60_DNA+RNA_seed | BRAF | -0.427137 | LUAD | NCI-H1435 | SB590885 |
| MVP_top60_DNA_seed | MAPK1 | 0.689237 | LGG | H4 | Ulixertinib | MVP_top60_DNA+RNA_seed | EGFR | -0.426522 | LUAD | NCI-H1693 | Osimertinib |
| MVP_top60_DNA_seed | MAPK3 | 0.689237 | LGG | H4 | Ulixertinib | MVP_top60_DNA+RNA_seed | AKT1 | -0.422941 | LUAD | HCC-44 | MK-2206 |
| MVP_top60_DNA_seed | EGFR | 0.692155 | LGG | no-11 | Erlotinib | MVP_top60_DNA+RNA_seed | PIK3CA | -0.42221 | LUAD | NCI-H1437 | Buparlisib |
| MVP_top60_DNA_seed | IGF1R | 0.698879 | LGG | no-11 | BMS-754807 | MVP_top60_DNA+RNA_seed | IGF1R | -0.419036 | LUAD | NCI-H358 | BMS-536924 |
| MVP_top60_DNA_seed | HSP90AA1 | 0.70047 | LGG | KNS-81-FD | Tanespimycin | MVP_top60_DNA+RNA_seed | ATM | -0.418749 | LUAD | NCI-H3122 | KU-55933 |
| MVP_top60_DNA_seed | MAPK1 | 0.70655 | LGG | KINGS-1 | Ulixertinib | MVP_top60_DNA+RNA_seed | AKT1 | -0.417894 | LUAD | NCI-H1755 | AZD5363 |
| MVP_top60_DNA_seed | MAPK3 | 0.70655 | LGG | KINGS-1 | Ulixertinib | MVP_top60_DNA+RNA_seed | AKT1 | -0.417502 | LUAD | NCI-H1755 | MK-2206 |
| MVP_top60_DNA_seed | CREBBP | 0.708665 | LGG | GAMG | SGC-CBP30 | MVP_top60_DNA+RNA_seed | AKT1 | -0.417351 | LUAD | NCI-H1944 | AZD5363 |
| MVP_top60_DNA_seed | CDK2 | 0.710947 | LGG | KINGS-1 | MK-8776 | MVP_top60_DNA+RNA_seed | PIK3CA | -0.415379 | LUAD | PC-14 | GNE-317 |
| MVP_top60_DNA_seed | EGFR | 0.722897 | LGG | MOG-G-UVW | Erlotinib | MVP_top60_DNA+RNA_seed | KRAS | -0.414707 | LUAD | NCI-H1734 | KRAS (G12C) Inhibitor-1 |
| MVP_top60_DNA_seed | TP53 | 0.723695 | LGG | KALS-1 | MIRA-1 | MVP_top60_DNA+RNA_seed | EGFR | -0.414656 | LUAD | A549 | Afatinib |
| MVP_top60_DNA_seed | IGF1R | 0.723941 | LGG | SF539 | Linsitinib | MVP_top60_DNA+RNA_seed | TERT | -0.414478 | LUAD | NCI-H650 | BIBR-1532 |
| MVP_top60_DNA_seed | PIK3CA | 0.727532 | LGG | KALS-1 | Alpelisib | MVP_top60_DNA+RNA_seed | HDAC1 | -0.412845 | LUAD | NCI-H1651 | PCI-34051 |
| MVP_top60_DNA_seed | AR | 0.735399 | LGG | no-10 | Bicalutamide | MVP_top60_DNA+RNA_seed | HDAC1 | -0.412333 | LUAD | NCI-H1563 | PCI-34051 |
| MVP_top60_DNA_seed | HDAC1 | 0.736119 | LGG | D-502MG | Entinostat | MVP_top60_DNA+RNA_seed | EGFR | -0.411993 | LUAD | NCI-H1693 | AZD3759 |
| MVP_top60_DNA_seed | MET | 0.740069 | LGG | no-10 | Savolitinib | MVP_top60_DNA+RNA_seed | PIK3CA | -0.411127 | LUAD | RERF-LC-KJ | AZD8186 |
| MVP_top60_DNA_seed | EGFR | 0.742559 | LGG | MOG-G-UVW | Osimertinib | MVP_top60_DNA+RNA_seed | AURKA | -0.408736 | LUAD | A549 | Alisertib |
| MVP_top60_DNA_seed | IGF1R | 0.747229 | LGG | SW1783 | Linsitinib | MVP_top60_DNA+RNA_seed | IGF1R | -0.408345 | LUAD | HCC-44 | NVP-ADW742 |
| MVP_top60_DNA_seed | HSP90AA1 | 0.748228 | LGG | D-502MG | CCT-018159 | MVP_top60_DNA+RNA_seed | AKT1 | -0.407921 | LUAD | RERF-LC-KJ | Uprosertib |
| MVP_top60_DNA_seed | SMARCA4 | 0.751453 | LGG | KNS-81-FD | PFI3 | MVP_top60_DNA+RNA_seed | AKT1 | -0.407521 | LUAD | NCI-H1437 | AT13148 |
| MVP_top60_DNA_seed | JAK1 | 0.754211 | LGG | GAMG | JAK1_8709 | MVP_top60_DNA+RNA_seed | IGF1R | -0.407028 | LUAD | NCI-H2122 | GSK1904529A |
| MVP_top60_DNA_seed | MAPK1 | 0.757761 | LGG | no-10 | VX-11e | MVP_top60_DNA+RNA_seed | EGFR | -0.406348 | LUAD | NCI-H1944 | Sapitinib |
| MVP_top60_DNA_seed | IGF1R | 0.762925 | LGG | GAMG | NVP-ADW742 | MVP_top60_DNA+RNA_seed | HSP90AA1 | -0.406218 | LUAD | NCI-H2023 | Luminespib |
| MVP_top60_DNA_seed | EGFR | 0.771658 | LGG | D-502MG | Gefitinib | MVP_top60_DNA+RNA_seed | SMARCA4 | -0.406091 | LUAD | NCI-H2122 | PFI3 |
| MVP_top60_DNA_seed | JAK2 | 0.773665 | LGG | KINGS-1 | JAK_8517 | MVP_top60_DNA+RNA_seed | IGF1R | -0.405163 | LUAD | NCI-H1623 | GSK1904529A |
| MVP_top60_DNA_seed | JAK1 | 0.773665 | LGG | KINGS-1 | JAK_8517 | MVP_top60_DNA+RNA_seed | IGF1R | -0.40173 | LUAD | LXF-289 | BMS-536924 |
| MVP_top60_DNA_seed | TP53 | 0.786286 | LGG | KNS-81-FD | MIRA-1 | MVP_top60_DNA+RNA_seed | IGF1R | -0.401028 | LUAD | NCI-H1781 | BMS-754807 |
| MVP_top60_DNA_seed | IGF1R | 0.787741 | LGG | GAMG | Linsitinib | MVP_top60_DNA+RNA_seed | MET | -0.40006 | LUAD | RERF-LC-KJ | Savolitinib |
| MVP_top60_DNA_seed | NTRK1 | 0.794393 | LGG | no-10 | GW441756 | MVP_top60_DNA+RNA_seed | AURKA | -0.399919 | LUAD | NCI-H1792 | ZM447439 |

| | | | | | | | | | | | |
|---|---|---|---|---|---|---|---|---|---|---|---|
| MVP_top60_DNA_seed | HDAC1 | 0.796849 | LGG | KNS-81-FD | Entinostat | MVP_top60_DNA+RNA_seed | PIK3CA | -0.398005 | LUAD | HCC-44 | Alpelisib |
| MVP_top60_DNA_seed | EGFR | 0.797162 | LGG | no-10 | Gefitinib | MVP_top60_DNA+RNA_seed | EGFR | -0.397211 | LUAD | NCI-H1944 | AZD3759 |
| MVP_top60_DNA_seed | NTRK1 | 0.798759 | LGG | KALS-1 | GW441756 | MVP_top60_DNA+RNA_seed | HDAC1 | -0.396679 | LUAD | PC-14 | Romidepsin |
| MVP_top60_DNA_seed | CDK2 | 0.802007 | LGG | GAMG | MK-8776 | MVP_top60_DNA+RNA_seed | HDAC1 | -0.395466 | LUAD | PC-14 | Entinostat |
| MVP_top60_DNA_seed | PIK3CA | 0.802433 | LGG | D-502MG | GNE-317 | MVP_top60_DNA+RNA_seed | MDM2 | -0.394727 | LUAD | LC-2-ad | Nutlin-3a (-) |
| MVP_top60_DNA_seed | EGFR | 0.803018 | LGG | no-11 | Osimertinib | MVP_top60_DNA+RNA_seed | MET | -0.394085 | LUAD | PC-14 | Crizotinib |
| MVP_top60_DNA_seed | AR | 0.806399 | LGG | Hs-683 | Bicalutamide | MVP_top60_DNA+RNA_seed | TERT | -0.394075 | LUAD | RERF-LC-KJ | BIBR-1532 |
| MVP_top60_DNA_seed | IDH1 | 0.807931 | LGG | D-502MG | AGI-5198 | MVP_top60_DNA+RNA_seed | EGFR | -0.389979 | LUAD | NCI-H2030 | Afatinib |
| MVP_top60_DNA_seed | MET | 0.811632 | LGG | KALS-1 | Crizotinib | MVP_top60_DNA+RNA_seed | HDAC1 | -0.385959 | LUAD | NCI-H1792 | Entinostat |
| MVP_top60_DNA_seed | IDH1 | 0.814015 | LGG | D-336MG | AGI-5198 | MVP_top60_DNA+RNA_seed | HDAC1 | -0.384063 | LUAD | NCI-H1666 | Romidepsin |
| MVP_top60_DNA_seed | EGFR | 0.814218 | LGG | KALS-1 | Sapitinib | MVP_top60_DNA+RNA_seed | EGFR | -0.383381 | LUAD | EKVX | Osimertinib |
| MVP_top60_DNA_seed | MDM2 | 0.817374 | LGG | GI-1 | Serdemetan | MVP_top60_DNA+RNA_seed | HSP90AA1 | -0.382024 | LUAD | NCI-H23 | Tanespimycin |
| MVP_top60_DNA_seed | CDK2 | 0.818499 | LGG | KALS-1 | Dinaciclib | MVP_top60_DNA+RNA_seed | HSP90AA1 | -0.380703 | LUAD | NCI-H838 | CCT-018159 |
| MVP_top60_DNA_seed | JAK2 | 0.858471 | LGG | D-502MG | Ruxolitinib | MVP_top60_DNA+RNA_seed | PIK3CA | -0.380447 | LUAD | COR-L105 | Alpelisib |
| MVP_top60_DNA_seed | JAK1 | 0.858471 | LGG | D-502MG | Ruxolitinib | MVP_top60_DNA+RNA_seed | AKT1 | -0.380179 | LUAD | A549 | Uprosertib |
| MVP_top60_DNA_seed | CDK2 | 0.865512 | LGG | no-10 | MK-8776 | MVP_top60_DNA+RNA_seed | IGF1R | -0.37995 | LUAD | EMC-BAC-1 | BMS-536924 |
| MVP_top60_DNA_seed | EGFR | 0.875275 | LGG | MOG-G-UVW | Sapitinib | MVP_top60_DNA+RNA_seed | AKT1 | -0.377776 | LUAD | PC-14 | Uprosertib |
| MVP_top60_DNA_seed | EGFR | 0.883305 | LGG | SW1783 | Erlotinib | MVP_top60_DNA+RNA_seed | PIK3CA | -0.377532 | LUAD | NCI-H1944 | AZD8186 |
| MVP_top60_DNA_seed | MAPK1 | 0.896822 | LGG | SW1783 | SCH772984 | MVP_top60_DNA+RNA_seed | FGFR1 | -0.37604 | LUAD | NCI-H1437 | PD173074 |
| MVP_top60_DNA_seed | MAPK3 | 0.896822 | LGG | SW1783 | SCH772984 | MVP_top60_DNA+RNA_seed | AURKA | -0.370894 | LUAD | HCC-78 | Alisertib |
| MVP_top60_DNA_seed | PIK3CA | 0.898276 | LGG | D-502MG | Buparlisib | MVP_top60_DNA+RNA_seed | BRAF | -0.368234 | LUAD | A549 | Dabrafenib |
| MVP_top60_DNA_seed | PIK3CD | 0.898276 | LGG | D-502MG | Buparlisib | MVP_top60_DNA+RNA_seed | AKT1 | -0.368233 | LUAD | NCI-H3122 | Uprosertib |
| MVP_top60_DNA_seed | HDAC1 | 0.899412 | LGG | KINGS-1 | Romidepsin | MVP_top60_DNA+RNA_seed | AKT1 | -0.366594 | LUAD | NCI-H1781 | AZD5363 |
| MVP_top60_DNA_seed | EGFR | 0.901063 | LGG | KNS-81-FD | Afatinib | MVP_top60_DNA+RNA_seed | AKT1 | -0.364285 | LUAD | NCI-H3122 | AZD5363 |
| MVP_top60_DNA_seed | MDM2 | 0.907294 | LGG | no-11 | Serdemetan | MVP_top60_DNA+RNA_seed | MET | -0.36361 | LUAD | NCI-H650 | Savolitinib |
| MVP_top60_DNA_seed | EGFR | 0.908559 | LGG | KNS-81-FD | Sapitinib | MVP_top60_DNA+RNA_seed | KRAS | -0.360798 | LUAD | NCI-H1781 | RAS (G12C) Inhibitor-1 |
| MVP_top60_DNA_seed | EGFR | 0.912961 | LGG | KNS-42 | Erlotinib | MVP_top60_DNA+RNA_seed | AKT1 | -0.35713 | LUAD | COR-L105 | Ipatasertib |
| MVP_top60_DNA_seed | MDM2 | 0.923081 | LGG | MOG-G-UVW | Nutlin-3a (-) | MVP_top60_DNA+RNA_seed | SMARCA4 | -0.355079 | LUAD | RERF-LC-KJ | PFI3 |
| MVP_top60_DNA_seed | EGFR | 0.925221 | LGG | MOG-G-CCM | Sapitinib | MVP_top60_DNA+RNA_seed | BRAF | -0.353155 | LUAD | NCI-H1651 | PLX-4720 |
| MVP_top60_DNA_seed | MAPK1 | 0.929413 | LGG | KNS-42 | VX-11e | MVP_top60_DNA+RNA_seed | AKT1 | -0.352144 | LUAD | NCI-H1755 | Ipatasertib |
| MVP_top60_DNA_seed | PIK3CA | 0.93336 | LGG | KNS-42 | Alpelisib | MVP_top60_DNA+RNA_seed | FGFR1 | -0.350758 | LUAD | NCI-H838 | AZD4547 |
| MVP_top60_DNA_seed | EGFR | 0.939677 | LGG | KINGS-1 | Afatinib | MVP_top60_DNA+RNA_seed | PIK3CA | -0.3506 | LUAD | NCI-H2009 | GNE-317 |
| MVP_top60_DNA_seed | IGF1R | 0.94425 | LGG | SW1783 | GSK1904529A | MVP_top60_DNA+RNA_seed | AKT1 | -0.350493 | LUAD | NCI-H1755 | Uprosertib |
| MVP_top60_DNA_seed | EGFR | 0.951583 | LGG | KNS-81-FD | Osimertinib | MVP_top60_DNA+RNA_seed | AKT1 | -0.350168 | LUAD | NCI-H838 | Uprosertib |
| MVP_top60_DNA_seed | MAPK1 | 0.954404 | LGG | no-10 | Ulixertinib | MVP_top60_DNA+RNA_seed | SMARCA4 | -0.348592 | LUAD | NCI-H522 | PFI3 |
| MVP_top60_DNA_seed | MAPK3 | 0.954404 | LGG | no-10 | Ulixertinib | MVP_top60_DNA+RNA_seed | SRC | -0.34845 | LUAD | NCI-H1568 | Bosutinib |
| MVP_top60_DNA_seed | IDH1 | 0.955434 | LGG | SW1783 | AGI-5198 | MVP_top60_DNA+RNA_seed | KRAS | -0.347633 | LUAD | RERF-LC-MS | RAS (G12C) Inhibitor-1 |
| MVP_top60_DNA_seed | EGFR | 0.959616 | LGG | KALS-1 | Gefitinib | MVP_top60_DNA+RNA_seed | AKT1 | -0.346737 | LUAD | NCI-H1944 | Uprosertib |
| MVP_top60_DNA_seed | IGF1R | 0.961793 | LGG | no-11 | GSK1904529A | MVP_top60_DNA+RNA_seed | AKT1 | -0.346317 | LUAD | EMC-BAC-2 | AT13148 |
| MVP_top60_DNA_seed | MAPK1 | 0.965957 | LGG | KNS-81-FD | Ulixertinib | MVP_top60_DNA+RNA_seed | IGF1R | -0.345865 | LUAD | EMC-BAC-2 | Linsitinib |
| MVP_top60_DNA_seed | MAPK3 | 0.965957 | LGG | KNS-81-FD | Ulixertinib | MVP_top60_DNA+RNA_seed | AKT1 | -0.345203 | LUAD | NCI-H1437 | Uprosertib |

| | | | | | | | | | | | |
|---|---|---|---|---|---|---|---|---|---|---|---|
| MVP_top60_DNA_seed | IGF1R | 0.970717 | LGG | MOG-G-CCM | GSK1904529A | MVP_top60_DNA+RNA_seed | EZH2 | -0.344107 | LUAD | COR-L105 | GSK343 |
| MVP_top60_DNA_seed | EGFR | 0.980044 | LGG | KNS-42 | Lapatinib | MVP_top60_DNA+RNA_seed | EGFR | -0.343669 | LUAD | A549 | Osimertinib |
| MVP_top60_DNA_seed | HSP90AA1 | 0.984547 | LGG | no-11 | Tanespimycin | MVP_top60_DNA+RNA_seed | IGF1R | -0.341017 | LUAD | NCI-H322M | NVP-ADW742 |
| MVP_top60_DNA_seed | NTRK1 | 0.987918 | LGG | KNS-42 | GW441756 | MVP_top60_DNA+RNA_seed | AURKA | -0.337657 | LUAD | Calu-6 | ZM447439 |
| MVP_top60_DNA_seed | HSP90AA1 | 0.989697 | LGG | D-502MG | Tanespimycin | MVP_top60_DNA+RNA_seed | MDM2 | -0.337588 | LUAD | NCI-H358 | Serdemetan |
| MVP_top60_DNA_seed | MET | 0.998399 | LGG | KALS-1 | Savolitinib | MVP_top60_DNA+RNA_seed | AKT1 | -0.335884 | LUAD | RERF-LC-MS | Afuresertib |
| MVP_top60_DNA_seed | EGFR | 0.999901 | LGG | KALS-1 | Osimertinib | MVP_top60_DNA+RNA_seed | HSP90AA1 | -0.334286 | LUAD | NCI-H1975 | Luminespib |
| MVP_top60_DNA_seed | IGF1R | 1.003634 | LGG | KALS-1 | Linsitinib | MVP_top60_DNA+RNA_seed | AKT1 | -0.334263 | LUAD | LXF-289 | Uprosertib |
| MVP_top60_DNA_seed | MDM2 | 1.008461 | LGG | SW1783 | Nutlin-3a (-) | MVP_top60_DNA+RNA_seed | EGFR | -0.334096 | LUAD | EKVX | Gefitinib |
| MVP_top60_DNA_seed | NTRK1 | 1.012222 | LGG | MOG-G-CCM | GW441756 | MVP_top60_DNA+RNA_seed | AURKA | -0.334005 | LUAD | A549 | ZM447439 |
| MVP_top60_DNA_seed | HDAC1 | 1.013067 | LGG | no-11 | Entinostat | MVP_top60_DNA+RNA_seed | AKT1 | -0.329797 | LUAD | HCC-44 | Afuresertib |
| MVP_top60_DNA_seed | MDM2 | 1.014852 | LGG | KALS-1 | Serdemetan | MVP_top60_DNA+RNA_seed | IGF1R | -0.329524 | LUAD | NCI-H838 | NVP-ADW742 |
| MVP_top60_DNA_seed | MDM2 | 1.017178 | LGG | KINGS-1 | Serdemetan | MVP_top60_DNA+RNA_seed | HDAC1 | -0.328513 | LUAD | NCI-H650 | Romidepsin |
| MVP_top60_DNA_seed | EGFR | 1.019898 | LGG | no-10 | Erlotinib | MVP_top60_DNA+RNA_seed | MET | -0.328416 | LUAD | NCI-H1437 | Savolitinib |
| MVP_top60_DNA_seed | EGFR | 1.02098 | LGG | KALS-1 | AZD3759 | MVP_top60_DNA+RNA_seed | EZH2 | -0.327654 | LUAD | RERF-LC-KJ | GSK343 |
| MVP_top60_DNA_seed | CDK2 | 1.044159 | LGG | KNS-42 | MK-8776 | MVP_top60_DNA+RNA_seed | TERT | -0.327329 | LUAD | NCI-H23 | BIBR-1532 |
| MVP_top60_DNA_seed | MAPK1 | 1.04757 | LGG | MOG-G-CCM | Ulixertinib | MVP_top60_DNA+RNA_seed | FGFR1 | -0.324083 | LUAD | NCI-H1975 | AZD4547 |
| MVP_top60_DNA_seed | MAPK3 | 1.04757 | LGG | MOG-G-CCM | Ulixertinib | MVP_top60_DNA+RNA_seed | SRC | -0.323756 | LUAD | A549 | Bosutinib |
| MVP_top60_DNA_seed | IGF1R | 1.053237 | LGG | KALS-1 | NVP-ADW742 | MVP_top60_DNA+RNA_seed | IGF1R | -0.323066 | LUAD | NCI-H2030 | BMS-536924 |
| MVP_top60_DNA_seed | IGF1R | 1.065673 | LGG | D-502MG | GSK1904529A | MVP_top60_DNA+RNA_seed | AKT1 | -0.322248 | LUAD | Calu-3 | Uprosertib |
| MVP_top60_DNA_seed | PIK3CA | 1.072966 | LGG | NMC-G1 | AZD8186 | MVP_top60_DNA+RNA_seed | AKT1 | -0.32088 | LUAD | LXF-289 | Uprosertib |
| MVP_top60_DNA_seed | EGFR | 1.076377 | LGG | MOG-G-CCM | Osimertinib | MVP_top60_DNA+RNA_seed | IGF1R | -0.318349 | LUAD | NCI-H1975 | BMS-754807 |
| MVP_top60_DNA_seed | MAPK1 | 1.083612 | LGG | MOG-G-UVW | Ulixertinib | MVP_top60_DNA+RNA_seed | IGF1R | -0.317039 | LUAD | NCI-H1975 | Linsitinib |
| MVP_top60_DNA_seed | MAPK3 | 1.083612 | LGG | MOG-G-UVW | Ulixertinib | MVP_top60_DNA+RNA_seed | ATM | -0.312928 | LUAD | NCI-H1755 | KU-55933 |
| MVP_top60_DNA_seed | MAPK1 | 1.089591 | LGG | D-502MG | Ulixertinib | MVP_top60_DNA+RNA_seed | EGFR | -0.312584 | LUAD | HCC-78 | Afatinib |
| MVP_top60_DNA_seed | MAPK3 | 1.089591 | LGG | D-502MG | Ulixertinib | MVP_top60_DNA+RNA_seed | MET | -0.31253 | LUAD | NCI-H838 | Crizotinib |
| MVP_top60_DNA_seed | IGF1R | 1.091617 | LGG | Hs-683 | Linsitinib | MVP_top60_DNA+RNA_seed | AKT1 | -0.312058 | LUAD | NCI-H1993 | AT13148 |
| MVP_top60_DNA_seed | MDM2 | 1.108658 | LGG | KNS-81-FD | Serdemetan | MVP_top60_DNA+RNA_seed | HDAC1 | -0.31122 | LUAD | NCI-H1781 | PCI-34051 |
| MVP_top60_DNA_seed | EGFR | 1.115263 | LGG | KALS-1 | Erlotinib | MVP_top60_DNA+RNA_seed | IGF1R | -0.310378 | LUAD | Calu-6 | BMS-754807 |
| MVP_top60_DNA_seed | EGFR | 1.121947 | LGG | no-10 | Osimertinib | MVP_top60_DNA+RNA_seed | EGFR | -0.309857 | LUAD | NCI-H1792 | Lapatinib |
| MVP_top60_DNA_seed | NTRK1 | 1.123494 | LGG | D-502MG | GW441756 | MVP_top60_DNA+RNA_seed | SMARCA4 | -0.309242 | LUAD | NCI-H1651 | PFI3 |
| MVP_top60_DNA_seed | MET | 1.133366 | LGG | D-502MG | Savolitinib | MVP_top60_DNA+RNA_seed | AKT1 | -0.30902 | LUAD | NCI-H1435 | AT13148 |
| MVP_top60_DNA_seed | MET | 1.134356 | LGG | KNS-81-FD | Savolitinib | MVP_top60_DNA+RNA_seed | FGFR1 | -0.307087 | LUAD | NCI-H1573 | AZD4547 |
| MVP_top60_DNA_seed | MAPK1 | 1.135408 | LGG | SW1783 | Ulixertinib | MVP_top60_DNA+RNA_seed | AKT1 | -0.305593 | LUAD | EMC-BAC-1 | AT13148 |
| MVP_top60_DNA_seed | MAPK3 | 1.135408 | LGG | SW1783 | Ulixertinib | MVP_top60_DNA+RNA_seed | MDM2 | -0.304836 | LUAD | LC-2-ad | Serdemetan |
| MVP_top60_DNA_seed | MET | 1.136955 | LGG | KNS-42 | Savolitinib | MVP_top60_DNA+RNA_seed | ATM | -0.304213 | LUAD | NCI-H2085 | KU-55933 |
| MVP_top60_DNA_seed | IDH1 | 1.137279 | LGG | GAMG | AGI-5198 | MVP_top60_DNA+RNA_seed | FGFR1 | -0.301253 | LUAD | NCI-H1755 | AZD4547 |
| MVP_top60_DNA_seed | EGFR | 1.148711 | LGG | KNS-42 | Osimertinib | MVP_top60_DNA+RNA_seed | IGF1R | -0.300638 | LUAD | NCI-H2009 | NVP-ADW742 |
| MVP_top60_DNA_seed | EGFR | 1.151089 | LGG | KNS-42 | Afatinib | MVP_top60_DNA+RNA_seed | EGFR | -0.299689 | LUAD | NCI-H1693 | Gefitinib |
| MVP_top60_DNA_seed | EGFR | 1.154416 | LGG | Hs-683 | Afatinib | MVP_top60_DNA+RNA_seed | AURKA | -0.298721 | LUAD | NCI-H1792 | Alisertib |
| MVP_top60_DNA_seed | HDAC1 | 1.161267 | LGG | GAMG | Entinostat | MVP_top60_DNA+RNA_seed | IGF1R | -0.297882 | LUAD | NCI-H1666 | NVP-ADW742 |

| | | | | | | | | | | | |
|---|---|---|---|---|---|---|---|---|---|---|---|
| MVP_top60_DNA_seed | EGFR | 1.163253 | LGG | MOG-G-CCM | Erlotinib | MVP_top60_DNA+RNA_seed | FGFR1 | -0.29747 | LUAD | LXF-289 | PD173074 |
| MVP_top60_DNA_seed | IGF1R | 1.164765 | LGG | KALS-1 | BMS-536924 | MVP_top60_DNA+RNA_seed | EGFR | -0.297008 | LUAD | NCI-H2342 | AZD3759 |
| MVP_top60_DNA_seed | CDK2 | 1.173709 | LGG | MOG-G-CCM | MK-8776 | MVP_top60_DNA+RNA_seed | AKT1 | -0.296447 | LUAD | NCI-H1437 | AZD5363 |
| MVP_top60_DNA_seed | HSP90AA1 | 1.176895 | LGG | no-11 | Elesclomol | MVP_top60_DNA+RNA_seed | AURKA | -0.294825 | LUAD | PC-14 | Alisertib |
| MVP_top60_DNA_seed | PIK3CA | 1.184239 | LGG | no-11 | GNE-317 | MVP_top60_DNA+RNA_seed | BRAF | -0.294686 | LUAD | NCI-H2342 | PLX-4720 |
| MVP_top60_DNA_seed | SMARCA4 | 1.184969 | LGG | KNS-42 | PFI3 | MVP_top60_DNA+RNA_seed | AKT1 | -0.294462 | LUAD | RERF-LC-KJ | AT13148 |
| MVP_top60_DNA_seed | IGF1R | 1.197478 | LGG | NMC-G1 | BMS-754807 | MVP_top60_DNA+RNA_seed | HDAC1 | -0.293966 | LUAD | NCI-H1651 | Entinostat |
| MVP_top60_DNA_seed | IGF1R | 1.198372 | LGG | KINGS-1 | BMS-536924 | MVP_top60_DNA+RNA_seed | KRAS | -0.29333 | LUAD | HCC-44 | :RAS (G12C) Inhibitor-1 |
| MVP_top60_DNA_seed | IGF1R | 1.199393 | LGG | MOG-G-UVW | Linsitinib | MVP_top60_DNA+RNA_seed | AKT1 | -0.291047 | LUAD | RERF-LC-KJ | Ipatasertib |
| MVP_top60_DNA_seed | IGF1R | 1.203018 | LGG | KNS-42 | BMS-754807 | MVP_top60_DNA+RNA_seed | AKT1 | -0.290722 | LUAD | NCI-H1755 | Uprosertib |
| MVP_top60_DNA_seed | SMARCA4 | 1.203649 | LGG | D-502MG | PFI3 | MVP_top60_DNA+RNA_seed | EGFR | -0.287944 | LUAD | NCI-H1650 | Gefitinib |
| MVP_top60_DNA_seed | IGF1R | 1.204453 | LGG | no-10 | Linsitinib | MVP_top60_DNA+RNA_seed | MET | -0.287668 | LUAD | NCI-H2009 | Crizotinib |
| MVP_top60_DNA_seed | EGFR | 1.209865 | LGG | KNS-42 | Sapitinib | MVP_top60_DNA+RNA_seed | AKT1 | -0.2866 | LUAD | PC-14 | Ipatasertib |
| MVP_top60_DNA_seed | IGF1R | 1.219467 | LGG | no-11 | Linsitinib | MVP_top60_DNA+RNA_seed | HSP90AA1 | -0.284152 | LUAD | LXF-289 | Luminespib |
| MVP_top60_DNA_seed | JAK2 | 1.225611 | LGG | KALS-1 | Ruxolitinib | MVP_top60_DNA+RNA_seed | HSP90AA1 | -0.282483 | LUAD | HOP-62 | Tanespimycin |
| MVP_top60_DNA_seed | JAK1 | 1.225611 | LGG | KALS-1 | Ruxolitinib | MVP_top60_DNA+RNA_seed | AKT1 | -0.280823 | LUAD | EMC-BAC-1 | Uprosertib |
| MVP_top60_DNA_seed | IDH1 | 1.226974 | LGG | KINGS-1 | AGI-5198 | MVP_top60_DNA+RNA_seed | MDM2 | -0.279216 | LUAD | NCI-H1666 | Nutlin-3a (-) |
| MVP_top60_DNA_seed | CDK2 | 1.229162 | LGG | KNS-81-FD | Dinaciclib | MVP_top60_DNA+RNA_seed | IGF1R | -0.27861 | LUAD | NCI-H1993 | BMS-754807 |
| MVP_top60_DNA_seed | EGFR | 1.232984 | LGG | GAMG | Erlotinib | MVP_top60_DNA+RNA_seed | BRAF | -0.278561 | LUAD | NCI-H2087 | Dabrafenib |
| MVP_top60_DNA_seed | MAPK1 | 1.262532 | LGG | GAMG | VX-11e | MVP_top60_DNA+RNA_seed | AKT1 | -0.277781 | LUAD | HCC-78 | MK-2206 |
| MVP_top60_DNA_seed | MET | 1.267758 | LGG | KNS-81-FD | Crizotinib | MVP_top60_DNA+RNA_seed | FGFR1 | -0.277235 | LUAD | NCI-H1623 | AZD4547 |
| MVP_top60_DNA_seed | HDAC1 | 1.281261 | LGG | KALS-1 | Entinostat | MVP_top60_DNA+RNA_seed | AKT1 | -0.277097 | LUAD | A549 | Ipatasertib |
| MVP_top60_DNA_seed | PIK3CA | 1.28262 | LGG | KNS-42 | AZD8186 | MVP_top60_DNA+RNA_seed | EZH2 | -0.276916 | LUAD | NCI-H3122 | GSK343 |
| MVP_top60_DNA_seed | MET | 1.283873 | LGG | no-11 | Savolitinib | MVP_top60_DNA+RNA_seed | MDM2 | -0.275424 | LUAD | RERF-LC-MS | Serdemetan |
| MVP_top60_DNA_seed | IGF1R | 1.28555 | LGG | KNS-81-FD | GSK1904529A | MVP_top60_DNA+RNA_seed | MET | -0.274471 | LUAD | NCI-H1435 | Savolitinib |
| MVP_top60_DNA_seed | IGF1R | 1.287068 | LGG | KINGS-1 | Linsitinib | MVP_top60_DNA+RNA_seed | PIK3CA | -0.274163 | LUAD | RERF-LC-KJ | Alpelisib |
| MVP_top60_DNA_seed | MDM2 | 1.291253 | LGG | MOG-G-CCM | Serdemetan | MVP_top60_DNA+RNA_seed | AKT1 | -0.273089 | LUAD | NCI-H322M | Afuresertib |
| MVP_top60_DNA_seed | IGF1R | 1.299138 | LGG | no-10 | BMS-536924 | MVP_top60_DNA+RNA_seed | PIK3CA | -0.27088 | LUAD | PC-14 | Buparlisib |
| MVP_top60_DNA_seed | EGFR | 1.300919 | LGG | KINGS-1 | Sapitinib | MVP_top60_DNA+RNA_seed | SMARCA4 | -0.27023 | LUAD | NCI-H322M | PFI3 |
| MVP_top60_DNA_seed | JAK2 | 1.301292 | LGG | KNS-81-FD | Ruxolitinib | MVP_top60_DNA+RNA_seed | HDAC1 | -0.269725 | LUAD | VMRC-LCD | Entinostat |
| MVP_top60_DNA_seed | JAK1 | 1.301292 | LGG | KNS-81-FD | Ruxolitinib | MVP_top60_DNA+RNA_seed | AKT1 | -0.269722 | LUAD | NCI-H838 | AZD5363 |
| MVP_top60_DNA_seed | EGFR | 1.307215 | LGG | no-11 | Gefitinib | MVP_top60_DNA+RNA_seed | PIK3CA | -0.268148 | LUAD | NCI-H1781 | Buparlisib |
| MVP_top60_DNA_seed | NTRK1 | 1.311351 | LGG | KNS-81-FD | GW441756 | MVP_top60_DNA+RNA_seed | HSP90AA1 | -0.267831 | LUAD | NCI-H522 | Tanespimycin |
| MVP_top60_DNA_seed | IGF1R | 1.318913 | LGG | KALS-1 | BMS-754807 | MVP_top60_DNA+RNA_seed | MDM2 | -0.267588 | LUAD | NCI-H1563 | Nutlin-3a (-) |
| MVP_top60_DNA_seed | IGF1R | 1.331049 | LGG | KNS-42 | Linsitinib | MVP_top60_DNA+RNA_seed | EGFR | -0.266246 | LUAD | RERF-LC-KJ | Sapitinib |
| MVP_top60_DNA_seed | TP53 | 1.333023 | LGG | MOG-G-CCM | MIRA-1 | MVP_top60_DNA+RNA_seed | HSP90AA1 | -0.266191 | LUAD | NCI-H1437 | Elesclomol |
| MVP_top60_DNA_seed | TP53 | 1.334479 | LGG | KINGS-1 | MIRA-1 | MVP_top60_DNA+RNA_seed | TP53 | -0.266147 | LUAD | NCI-H2030 | MIRA-1 |
| MVP_top60_DNA_seed | SMARCA4 | 1.335051 | LGG | KINGS-1 | PFI3 | MVP_top60_DNA+RNA_seed | EGFR | -0.264137 | LUAD | NCI-H1792 | Erlotinib |
| MVP_top60_DNA_seed | EGFR | 1.335166 | LGG | KINGS-1 | AZD3759 | MVP_top60_DNA+RNA_seed | KRAS | -0.263595 | LUAD | LXF-289 | :RAS (G12C) Inhibitor-1 |
| MVP_top60_DNA_seed | MAPK1 | 1.349757 | LGG | GAMG | SCH772984 | MVP_top60_DNA+RNA_seed | EGFR | -0.261681 | LUAD | LC-2-ad | Lapatinib |
| MVP_top60_DNA_seed | MAPK3 | 1.349757 | LGG | GAMG | SCH772984 | MVP_top60_DNA+RNA_seed | HSP90AA1 | -0.259186 | LUAD | NCI-H1650 | Luminespib |

| | | | | | | | | | | | |
|---|---|---|---|---|---|---|---|---|---|---|---|
| MVP_top60_DNA_seed | SMARCA4 | 1.362555 | LGG | no-11 | PFI3 | MVP_top60_DNA+RNA_seed | AKT1 | -0.258985 | LUAD | NCI-H1355 | AT13148 |
| MVP_top60_DNA_seed | PIK3CA | 1.365099 | LGG | no-10 | Buparlisib | MVP_top60_DNA+RNA_seed | EGFR | -0.25795 | LUAD | NCI-H2342 | Sapitinib |
| MVP_top60_DNA_seed | PIK3CD | 1.365099 | LGG | no-10 | Buparlisib | MVP_top60_DNA+RNA_seed | AKT1 | -0.256224 | LUAD | NCI-H2122 | AT13148 |
| MVP_top60_DNA_seed | MET | 1.373876 | LGG | no-11 | Crizotinib | MVP_top60_DNA+RNA_seed | AKT1 | -0.256016 | LUAD | NCI-H1563 | Ipatasertib |
| MVP_top60_DNA_seed | MAPK1 | 1.376468 | LGG | GAMG | Ulixertinib | MVP_top60_DNA+RNA_seed | FGFR1 | -0.255894 | LUAD | LC-2-ad | AZD4547 |
| MVP_top60_DNA_seed | MAPK3 | 1.376468 | LGG | GAMG | Ulixertinib | MVP_top60_DNA+RNA_seed | HDAC1 | -0.255644 | LUAD | NCI-H3122 | Romidepsin |
| MVP_top60_DNA_seed | EGFR | 1.386507 | LGG | GAMG | Osimertinib | MVP_top60_DNA+RNA_seed | AKT1 | -0.254504 | LUAD | NCI-H2023 | Ipatasertib |
| MVP_top60_DNA_seed | IGF1R | 1.405451 | LGG | MOG-G-CCM | Linsitinib | MVP_top60_DNA+RNA_seed | SRC | -0.254038 | LUAD | NCI-H1993 | Bosutinib |
| MVP_top60_DNA_seed | EGFR | 1.414827 | LGG | MOG-G-CCM | Gefitinib | MVP_top60_DNA+RNA_seed | AURKA | -0.252427 | LUAD | NCI-H1734 | ZM447439 |
| MVP_top60_DNA_seed | EGFR | 1.443705 | LGG | MOG-G-CCM | AZD3759 | MVP_top60_DNA+RNA_seed | PIK3CA | -0.251496 | LUAD | COR-L105 | AZD8186 |
| MVP_top60_DNA_seed | PIK3CA | 1.445666 | LGG | no-10 | GNE-317 | MVP_top60_DNA+RNA_seed | AKT1 | -0.250065 | LUAD | NCI-H1944 | MK-2206 |
| MVP_top60_DNA_seed | MAPK1 | 1.448688 | LGG | Hs-683 | Ulixertinib | MVP_top60_DNA+RNA_seed | TP53 | -0.25003 | LUAD | NCI-H2122 | MIRA-1 |
| MVP_top60_DNA_seed | MAPK3 | 1.448688 | LGG | Hs-683 | Ulixertinib | MVP_top60_DNA+RNA_seed | AKT1 | -0.245737 | LUAD | RERF-LC-KJ | AZD5363 |
| MVP_top60_DNA_seed | IGF1R | 1.457879 | LGG | NMC-G1 | BMS-536924 | MVP_top60_DNA+RNA_seed | IGF1R | -0.245726 | LUAD | NCI-H2030 | NVP-ADW742 |
| MVP_top60_DNA_seed | JAK1 | 1.459511 | LGG | KALS-1 | JAK1_8709 | MVP_top60_DNA+RNA_seed | FGFR1 | -0.245156 | LUAD | NCI-H1734 | PD173074 |
| MVP_top60_DNA_seed | MET | 1.470395 | LGG | MOG-G-CCM | Savolitinib | MVP_top60_DNA+RNA_seed | PIK3CA | -0.244183 | LUAD | NCI-H2030 | Alpelisib |
| MVP_top60_DNA_seed | CDK2 | 1.476776 | LGG | D-502MG | MK-8776 | MVP_top60_DNA+RNA_seed | PIK3CA | -0.242331 | LUAD | LXF-289 | Buparlisib |
| MVP_top60_DNA_seed | MDM2 | 1.484019 | LGG | KINGS-1 | Nutlin-3a (-) | MVP_top60_DNA+RNA_seed | EGFR | -0.24091 | LUAD | NCI-H1993 | Afatinib |
| MVP_top60_DNA_seed | EGFR | 1.51225 | LGG | no-11 | Sapitinib | MVP_top60_DNA+RNA_seed | KRAS | -0.239604 | LUAD | NCI-H322M | KRAS (G12C) Inhibitor-1 |
| MVP_top60_DNA_seed | HDAC1 | 1.51684 | LGG | KNS-42 | PCI-34051 | MVP_top60_DNA+RNA_seed | AKT1 | -0.23857 | LUAD | NCI-H3122 | Afuresertib |
| MVP_top60_DNA_seed | CDK2 | 1.528135 | LGG | KNS-81-FD | MK-8776 | MVP_top60_DNA+RNA_seed | MET | -0.23555 | LUAD | NCI-H1437 | Crizotinib |
| MVP_top60_DNA_seed | JAK1 | 1.533643 | LGG | KINGS-1 | JAK1_8709 | MVP_top60_DNA+RNA_seed | HDAC1 | -0.235479 | LUAD | NCI-H1975 | Entinostat |
| MVP_top60_DNA_seed | EGFR | 1.539234 | LGG | NMC-G1 | Erlotinib | MVP_top60_DNA+RNA_seed | IGF1R | -0.235025 | LUAD | NCI-H2009 | BMS-536924 |
| MVP_top60_DNA_seed | IGF1R | 1.548496 | LGG | no-11 | NVP-ADW742 | MVP_top60_DNA+RNA_seed | SRC | -0.234168 | LUAD | NCI-H1648 | Bosutinib |
| MVP_top60_DNA_seed | IGF1R | 1.562746 | LGG | KNS-42 | NVP-ADW742 | MVP_top60_DNA+RNA_seed | IGF1R | -0.234121 | LUAD | NCI-H1755 | BMS-754807 |
| MVP_top60_DNA_seed | PIK3CA | 1.565584 | LGG | NMC-G1 | GNE-317 | MVP_top60_DNA+RNA_seed | EGFR | -0.233537 | LUAD | NCI-H2023 | AZD3759 |
| MVP_top60_DNA_seed | EGFR | 1.566406 | LGG | KINGS-1 | Osimertinib | MVP_top60_DNA+RNA_seed | PIK3CA | -0.228488 | LUAD | NCI-H1437 | AZD8186 |
| MVP_top60_DNA_seed | HDAC1 | 1.579788 | LGG | no-11 | PCI-34051 | MVP_top60_DNA+RNA_seed | TP53 | -0.228242 | LUAD | LC-2-ad | MIRA-1 |
| MVP_top60_DNA_seed | IGF1R | 1.624898 | LGG | KINGS-1 | GSK1904529A | MVP_top60_DNA+RNA_seed | IGF1R | -0.227914 | LUAD | NCI-H322M | BMS-536924 |
| MVP_top60_DNA_seed | MAPK1 | 1.636522 | LGG | GI-1 | Ulixertinib | MVP_top60_DNA+RNA_seed | MDM2 | -0.227815 | LUAD | NCI-H2122 | Nutlin-3a (-) |
| MVP_top60_DNA_seed | MAPK3 | 1.636522 | LGG | GI-1 | Ulixertinib | MVP_top60_DNA+RNA_seed | HSP90AA1 | -0.227219 | LUAD | NCI-H2122 | Luminespib |
| MVP_top60_DNA_seed | MAPK1 | 1.668258 | LGG | MOG-G-CCM | SCH772984 | MVP_top60_DNA+RNA_seed | AKT1 | -0.225638 | LUAD | HCC-44 | Uprosertib |
| MVP_top60_DNA_seed | MAPK3 | 1.668258 | LGG | MOG-G-CCM | SCH772984 | MVP_top60_DNA+RNA_seed | MDM2 | -0.225508 | LUAD | NCI-H1944 | Serdemetan |
| MVP_top60_DNA_seed | IDH1 | 1.678283 | LGG | no-11 | AGI-5198 | MVP_top60_DNA+RNA_seed | IGF1R | -0.224761 | LUAD | NCI-H1563 | BMS-536924 |
| MVP_top60_DNA_seed | MAPK1 | 1.684996 | LGG | MOG-G-CCM | Ulixertinib | MVP_top60_DNA+RNA_seed | ATM | -0.224506 | LUAD | NCI-H1563 | KU-55933 |
| MVP_top60_DNA_seed | MAPK3 | 1.684996 | LGG | MOG-G-CCM | Ulixertinib | MVP_top60_DNA+RNA_seed | HDAC1 | -0.223407 | LUAD | NCI-H1755 | PCI-34051 |
| MVP_top60_DNA_seed | HSP90AA1 | 1.688457 | LGG | NMC-G1 | Elesclomol | MVP_top60_DNA+RNA_seed | HSP90AA1 | -0.222448 | LUAD | NCI-H1993 | Luminespib |
| MVP_top60_DNA_seed | EGFR | 1.69048 | LGG | no-11 | Lapatinib | MVP_top60_DNA+RNA_seed | PIK3CA | -0.222285 | LUAD | Calu-3 | GNE-317 |
| MVP_top60_DNA_seed | NTRK1 | 1.69492 | LGG | KINGS-1 | AZD1332 | MVP_top60_DNA+RNA_seed | ATM | -0.220484 | LUAD | NCI-H358 | KU-55933 |
| MVP_top60_DNA_seed | IGF1R | 1.695092 | LGG | no-11 | BMS-536924 | MVP_top60_DNA+RNA_seed | BRAF | -0.219352 | LUAD | PC-3_[JPC-3] | PLX-4720 |
| MVP_top60_DNA_seed | IDH1 | 1.695498 | LGG | KALS-1 | AGI-5198 | MVP_top60_DNA+RNA_seed | EGFR | -0.218425 | LUAD | EMC-BAC-2 | Afatinib |

| | | | | | | | | | | | |
|---|---|---|---|---|---|---|---|---|---|---|---|
| MVP_top60_DNA_seed | IGF1R | 1.703119 | LGG | Hs-683 | NVP-ADW742 | MVP_top60_DNA+RNA_seed | BRAF | -0.216234 | LUAD | LC-2-ad | Dabrafenib |
| MVP_top60_DNA_seed | CDK2 | 1.703607 | LGG | KALS-1 | AZD5438 | MVP_top60_DNA+RNA_seed | AKT1 | -0.214868 | LUAD | NCI-H1838 | Uprosertib |
| MVP_top60_DNA_seed | PIK3CA | 1.705865 | LGG | KNS-42 | GNE-317 | MVP_top60_DNA+RNA_seed | HSP90AA1 | -0.214178 | LUAD | PC-3_[JPC-3] | Luminespib |
| MVP_top60_DNA_seed | AR | 1.713071 | LGG | D-502MG | Bicalutamide | MVP_top60_DNA+RNA_seed | MET | -0.213508 | LUAD | NCI-H2030 | Crizotinib |
| MVP_top60_DNA_seed | HSP90AA1 | 1.718113 | LGG | NMC-G1 | Luminespib | MVP_top60_DNA+RNA_seed | AURKA | -0.212933 | LUAD | NCI-H1437 | Tozasertib |
| MVP_top60_DNA_seed | MAPK1 | 1.718969 | LGG | SW1783 | VX-11e | MVP_top60_DNA+RNA_seed | TERT | -0.210942 | LUAD | NCI-H1734 | BIBR-1532 |
| MVP_top60_DNA_seed | IGF1R | 1.746644 | LGG | KALS-1 | GSK1904529A | MVP_top60_DNA+RNA_seed | AKT1 | -0.208587 | LUAD | PC-14 | AZD5363 |
| MVP_top60_DNA_seed | IGF1R | 1.783607 | LGG | no-10 | BMS-754807 | MVP_top60_DNA+RNA_seed | EGFR | -0.207757 | LUAD | NCI-H1563 | Gefitinib |
| MVP_top60_DNA_seed | PIK3CA | 1.7939 | LGG | KNS-81-FD | Buparlisib | MVP_top60_DNA+RNA_seed | AKT1 | -0.20669 | LUAD | COR-L105 | MK-2206 |
| MVP_top60_DNA_seed | PIK3CD | 1.7939 | LGG | KNS-81-FD | Buparlisib | MVP_top60_DNA+RNA_seed | HSP90AA1 | -0.205187 | LUAD | Calu-6 | Tanespimycin |
| MVP_top60_DNA_seed | EGFR | 1.79458 | LGG | KINGS-1 | Lapatinib | MVP_top60_DNA+RNA_seed | HDAC1 | -0.205187 | LUAD | A549 | PCI-34051 |
| MVP_top60_DNA_seed | HSP90AA1 | 1.795337 | LGG | KNS-81-FD | CCT-018159 | MVP_top60_DNA+RNA_seed | HDAC1 | -0.204616 | LUAD | NCI-H3122 | Entinostat |
| MVP_top60_DNA_seed | MAPK1 | 1.797113 | LGG | KNS-42 | SCH772984 | MVP_top60_DNA+RNA_seed | PIK3CA | -0.204119 | LUAD | NCI-H838 | Buparlisib |
| MVP_top60_DNA_seed | MAPK3 | 1.797113 | LGG | KNS-42 | SCH772984 | MVP_top60_DNA+RNA_seed | HSP90AA1 | -0.20312 | LUAD | NCI-H2085 | Luminespib |
| MVP_top60_DNA_seed | IGF1R | 1.816401 | LGG | KNS-81-FD | Linsitinib | MVP_top60_DNA+RNA_seed | IGF1R | -0.202586 | LUAD | NCI-H1975 | NVP-ADW742 |
| MVP_top60_DNA_seed | MDM2 | 1.817474 | LGG | NMC-G1 | Serdemetan | MVP_top60_DNA+RNA_seed | MET | -0.201648 | LUAD | NCI-H1650 | Savolitinib |
| MVP_top60_DNA_seed | JAK2 | 1.819837 | LGG | KINGS-1 | GSK2276186C | MVP_top60_DNA+RNA_seed | IGF1R | -0.201112 | LUAD | NCI-H322M | BMS-754807 |
| MVP_top60_DNA_seed | JAK1 | 1.819837 | LGG | KINGS-1 | GSK2276186C | MVP_top60_DNA+RNA_seed | FGFR1 | -0.199558 | LUAD | NCI-H1781 | AZD4547 |
| MVP_top60_DNA_seed | JAK3 | 1.819837 | LGG | KINGS-1 | GSK2276186C | MVP_top60_DNA+RNA_seed | EZH2 | -0.199301 | LUAD | NCI-H1623 | GSK343 |
| MVP_top60_DNA_seed | HDAC1 | 1.833266 | LGG | KNS-42 | Entinostat | MVP_top60_DNA+RNA_seed | PIK3CA | -0.19928 | LUAD | H3255 | AZD8186 |
| MVP_top60_DNA_seed | EGFR | 1.857789 | LGG | KINGS-1 | Gefitinib | MVP_top60_DNA+RNA_seed | BRAF | -0.198889 | LUAD | NCI-H1755 | Dabrafenib |
| MVP_top60_DNA_seed | PIK3CA | 1.861603 | LGG | KNS-42 | Buparlisib | MVP_top60_DNA+RNA_seed | EZH2 | -0.1986 | LUAD | NCI-H1435 | GSK343 |
| MVP_top60_DNA_seed | PIK3CD | 1.861603 | LGG | KNS-42 | Buparlisib | MVP_top60_DNA+RNA_seed | IGF1R | -0.198542 | LUAD | NCI-H1993 | Linsitinib |
| MVP_top60_DNA_seed | EGFR | 1.87069 | LGG | NMC-G1 | Osimertinib | MVP_top60_DNA+RNA_seed | MDM2 | -0.19708 | LUAD | NCI-H322M | Serdemetan |
| MVP_top60_DNA_seed | MAPK1 | 1.893818 | LGG | D-336MG | Ulixertinib | MVP_top60_DNA+RNA_seed | HSP90AA1 | -0.197013 | LUAD | NCI-H1437 | Luminespib |
| MVP_top60_DNA_seed | MAPK3 | 1.893818 | LGG | D-336MG | Ulixertinib | MVP_top60_DNA+RNA_seed | AKT1 | -0.19218 | LUAD | NCI-H2023 | Afuresertib |
| MVP_top60_DNA_seed | NTRK1 | 1.894545 | LGG | KINGS-1 | GW441756 | MVP_top60_DNA+RNA_seed | HSP90AA1 | -0.191238 | LUAD | EMC-BAC-1 | Elesclomol |
| MVP_top60_DNA_seed | AR | 1.905974 | LGG | KNS-81-FD | Bicalutamide | MVP_top60_DNA+RNA_seed | IGF1R | -0.18992 | LUAD | NCI-H1648 | BMS-754807 |
| MVP_top60_DNA_seed | HSP90AA1 | 1.936852 | LGG | NMC-G1 | Tanespimycin | MVP_top60_DNA+RNA_seed | IGF1R | -0.188128 | LUAD | VMRC-LCD | GSK1904529A |
| MVP_top60_DNA_seed | MAPK1 | 1.938406 | LGG | NMC-G1 | VX-11e | MVP_top60_DNA+RNA_seed | PIK3CA | -0.187985 | LUAD | LC-2-ad | Buparlisib |
| MVP_top60_DNA_seed | CDK2 | 1.939496 | LGG | NMC-G1 | MK-8776 | MVP_top60_DNA+RNA_seed | AKT1 | -0.185918 | LUAD | COR-L105 | Afuresertib |
| MVP_top60_DNA_seed | EGFR | 1.941697 | LGG | KINGS-1 | Erlotinib | MVP_top60_DNA+RNA_seed | EGFR | -0.185376 | LUAD | RERF-LC-KJ | Osimertinib |
| MVP_top60_DNA_seed | MAPK1 | 1.950063 | LGG | no-11 | Ulixertinib | MVP_top60_DNA+RNA_seed | MDM2 | -0.184543 | LUAD | NCI-H2347 | Serdemetan |
| MVP_top60_DNA_seed | MAPK3 | 1.950063 | LGG | no-11 | Ulixertinib | MVP_top60_DNA+RNA_seed | MDM2 | -0.18452 | LUAD | NCI-H1437 | Nutlin-3a (-) |
| MVP_top60_DNA_seed | IGF1R | 2.045132 | LGG | no-10 | NVP-ADW742 | MVP_top60_DNA+RNA_seed | MET | -0.182454 | LUAD | NCI-H1648 | Crizotinib |
| MVP_top60_DNA_seed | IDH1 | 2.050594 | LGG | NMC-G1 | AGI-5198 | MVP_top60_DNA+RNA_seed | AKT1 | -0.182337 | LUAD | NCI-H1563 | Uprosertib |
| MVP_top60_DNA_seed | EGFR | 2.072975 | LGG | NMC-G1 | Sapitinib | MVP_top60_DNA+RNA_seed | AKT1 | -0.181232 | LUAD | Calu-3 | MK-2206 |
| MVP_top60_DNA_seed | NTRK1 | 2.081746 | LGG | no-11 | GW441756 | MVP_top60_DNA+RNA_seed | AKT1 | -0.179368 | LUAD | NCI-H1755 | Afuresertib |
| MVP_top60_DNA_seed | MAPK1 | 2.082563 | LGG | SW1783 | Ulixertinib | MVP_top60_DNA+RNA_seed | MET | -0.179254 | LUAD | EMC-BAC-2 | Crizotinib |
| MVP_top60_DNA_seed | MAPK3 | 2.082563 | LGG | SW1783 | Ulixertinib | MVP_top60_DNA+RNA_seed | PIK3CA | -0.178812 | LUAD | NCI-H1693 | GNE-317 |
| MVP_top60_DNA_seed | EGFR | 2.10689 | LGG | KNS-81-FD | Erlotinib | MVP_top60_DNA+RNA_seed | AURKA | -0.178106 | LUAD | NCI-H1648 | ZM447439 |

| | | | | | | | | | | | |
|---|---|---|---|---|---|---|---|---|---|---|---|
| MVP_top60_DNA_seed | MET | 2.113834 | LGG | NMC-G1 | Crizotinib | MVP_top60_DNA+RNA_seed | PIK3CA | -0.177116 | LUAD | LC-2-ad | AZD8186 |
| MVP_top60_DNA_seed | MAPK1 | 2.180075 | LGG | KNS-42 | Ulixertinib | MVP_top60_DNA+RNA_seed | EGFR | -0.175883 | LUAD | NCI-H1563 | Sapitinib |
| MVP_top60_DNA_seed | MAPK3 | 2.180075 | LGG | KNS-42 | Ulixertinib | MVP_top60_DNA+RNA_seed | FGFR1 | -0.174078 | LUAD | NCI-H1568 | AZD4547 |
| MVP_top60_DNA_seed | PIK3CA | 2.257283 | LGG | no-11 | Buparlisib | MVP_top60_DNA+RNA_seed | AKT1 | -0.172351 | LUAD | HCC-78 | Afuresertib |
| MVP_top60_DNA_seed | PIK3CD | 2.257283 | LGG | no-11 | Buparlisib | MVP_top60_DNA+RNA_seed | EGFR | -0.171631 | LUAD | NCI-H1755 | Lapatinib |
| MVP_top60_DNA_seed | EGFR | 2.263084 | LGG | NMC-G1 | Gefitinib | MVP_top60_DNA+RNA_seed | PIK3CA | -0.170387 | LUAD | NCI-H23 | Buparlisib |
| MVP_top60_DNA_seed | TP53 | 2.315779 | LGG | NMC-G1 | MIRA-1 | MVP_top60_DNA+RNA_seed | AKT1 | -0.168826 | LUAD | NCI-H2023 | Uprosertib |
| MVP_top60_DNA_seed | EGFR | 2.325145 | LGG | NMC-G1 | AZD3759 | MVP_top60_DNA+RNA_seed | AKT1 | -0.168825 | LUAD | RERF-LC-MS | AZD5363 |
| MVP_top60_DNA_seed | JAK2 | 2.354717 | LGG | NMC-G1 | Ruxolitinib | MVP_top60_DNA+RNA_seed | FGFR1 | -0.167334 | LUAD | NCI-H3122 | AZD4547 |
| MVP_top60_DNA_seed | JAK1 | 2.354717 | LGG | NMC-G1 | Ruxolitinib | MVP_top60_DNA+RNA_seed | AURKA | -0.165912 | LUAD | HCC-78 | ZM447439 |
| MVP_top60_DNA_seed | IGF1R | 2.36881 | LGG | KNS-81-FD | NVP-ADW742 | MVP_top60_DNA+RNA_seed | EGFR | -0.165431 | LUAD | NCI-H358 | Gefitinib |
| MVP_top60_DNA_seed | HDAC1 | 2.385033 | LGG | NMC-G1 | PCI-34051 | MVP_top60_DNA+RNA_seed | AKT1 | -0.163877 | LUAD | COR-L105 | Uprosertib |
| MVP_top60_DNA_seed | HDAC1 | 2.406989 | LGG | KINGS-1 | Entinostat | MVP_top60_DNA+RNA_seed | EGFR | -0.163454 | LUAD | Calu-3 | Gefitinib |
| MVP_top60_DNA_seed | PIK3CA | 2.538988 | LGG | NMC-G1 | Alpelisib | MVP_top60_DNA+RNA_seed | EGFR | -0.163199 | LUAD | RERF-LC-KJ | Lapatinib |
| MVP_top60_DNA_seed | MAPK1 | 2.539321 | LGG | NMC-G1 | SCH772984 | MVP_top60_DNA+RNA_seed | IGF1R | -0.163181 | LUAD | H3255 | BMS-536924 |
| MVP_top60_DNA_seed | MAPK3 | 2.539321 | LGG | NMC-G1 | SCH772984 | MVP_top60_DNA+RNA_seed | SMARCA4 | -0.162595 | LUAD | LC-2-ad | PFI3 |
| MVP_top60_DNA_seed | CDK2 | 2.557024 | LGG | KINGS-1 | Dinaciclib | MVP_top60_DNA+RNA_seed | AURKA | -0.161895 | LUAD | NCI-H3122 | Alisertib |
| MVP_top60_DNA_seed | HDAC1 | 2.658986 | LGG | NMC-G1 | Entinostat | MVP_top60_DNA+RNA_seed | EGFR | -0.160772 | LUAD | EMC-BAC-2 | Lapatinib |
| MVP_top60_DNA_seed | IGF1R | 2.681031 | LGG | NMC-G1 | NVP-ADW742 | MVP_top60_DNA+RNA_seed | IGF1R | -0.159768 | LUAD | ABC-1 | Linsitinib |
| MVP_top60_DNA_seed | HSP90AA1 | 2.682382 | LGG | NMC-G1 | CCT-018159 | MVP_top60_DNA+RNA_seed | AKT1 | -0.159232 | LUAD | RERF-LC-MS | Ipatasertib |
| MVP_top60_DNA_seed | EGFR | 2.718584 | LGG | NMC-G1 | Lapatinib | MVP_top60_DNA+RNA_seed | SRC | -0.158699 | LUAD | RERF-LC-KJ | Bosutinib |
| MVP_top60_DNA_seed | MAPK1 | 2.796514 | LGG | NMC-G1 | Ulixertinib | MVP_top60_DNA+RNA_seed | HDAC1 | -0.158295 | LUAD | NCI-H1435 | PCI-34051 |
| MVP_top60_DNA_seed | MAPK3 | 2.796514 | LGG | NMC-G1 | Ulixertinib | MVP_top60_DNA+RNA_seed | HDAC1 | -0.15629 | LUAD | RERF-LC-MS | Entinostat |
| MVP_top60_DNA_seed | IGF1R | 2.846875 | LGG | KNS-81-FD | BMS-536924 | MVP_top60_DNA+RNA_seed | AKT1 | -0.156139 | LUAD | A549 | AZD5363 |
| MVP_top60_DNA_seed | EGFR | 2.856642 | LGG | no-11 | Afatinib | MVP_top60_DNA+RNA_seed | HDAC1 | -0.154978 | LUAD | COR-L105 | Romidepsin |
| MVP_top60_DNA_seed | EGFR | 2.954802 | LGG | NMC-G1 | Afatinib | MVP_top60_DNA+RNA_seed | IGF1R | -0.154199 | LUAD | A549 | NVP-ADW742 |
| MVP_top60_DNA_seed | IGF1R | 2.975074 | LGG | NMC-G1 | Linsitinib | MVP_top60_DNA+RNA_seed | EGFR | -0.153439 | LUAD | NCI-H1573 | Gefitinib |
| MVP_top60_DNA_seed | MET | 3.071189 | LGG | NMC-G1 | Savolitinib | MVP_top60_DNA+RNA_seed | AKT1 | -0.153102 | LUAD | NCI-H322M | MK-2206 |
| MVP_top60_DNA_seed | IGF1R | 3.092709 | LGG | NMC-G1 | GSK1904529A | MVP_top60_DNA+RNA_seed | EGFR | -0.152389 | LUAD | NCI-H1651 | Erlotinib |
| MVP_top60_DNA_seed | CDK2 | 3.28446 | LGG | KINGS-1 | AZD5438 | MVP_top60_DNA+RNA_seed | AKT1 | -0.151421 | LUAD | A549 | MK-2206 |
| MVP_top60_DNA_seed | NTRK1 | 3.520693 | LGG | NMC-G1 | GW441756 | MVP_top60_DNA+RNA_seed | AKT1 | -0.150646 | LUAD | NCI-H838 | Afuresertib |
| MVP_top60_DNA_seed | AR | 3.831669 | LGG | NMC-G1 | Bicalutamide | MVP_top60_DNA+RNA_seed | AKT1 | -0.14816 | LUAD | NCI-H1734 | Uprosertib |
| MVP_top60_DNA_seed | SMARCA4 | 3.940776 | LGG | NMC-G1 | PFI3 | MVP_top60_DNA+RNA_seed | HSP90AA1 | -0.148062 | LUAD | LC-2-ad | Luminespib |
| MVP_top60_DNA_seed | CDK2 | 4.680277 | LGG | NMC-G1 | Dinaciclib | MVP_top60_DNA+RNA_seed | FGFR1 | -0.146087 | LUAD | HCC-44 | PD173074 |
| MVP_top60_DNA_seed | PIK3CA | 5.209719 | LGG | NMC-G1 | Buparlisib | MVP_top60_DNA+RNA_seed | HDAC1 | -0.145126 | LUAD | EMC-BAC-2 | PCI-34051 |
| MVP_top60_DNA_seed | PIK3CD | 5.209719 | LGG | NMC-G1 | Buparlisib | MVP_top60_DNA+RNA_seed | IGF1R | -0.144879 | LUAD | NCI-H838 | GSK1904529A |
| MVP_top60_DNA_seed | EGFR | -5.4375 | LUAD | NCI-H1648 | Lapatinib | MVP_top60_DNA+RNA_seed | PIK3CA | -0.14364 | LUAD | NCI-H1781 | GNE-317 |
| MVP_top60_DNA_seed | ERBB2 | -5.4375 | LUAD | NCI-H1648 | Lapatinib | MVP_top60_DNA+RNA_seed | AURKA | -0.143562 | LUAD | NCI-H2342 | Alisertib |
| MVP_top60_DNA_seed | EGFR | -5.341957 | LUAD | HCC-827 | Afatinib | MVP_top60_DNA+RNA_seed | PIK3CA | -0.143288 | LUAD | RERF-LC-KJ | GNE-317 |
| MVP_top60_DNA_seed | ERBB2 | -5.341957 | LUAD | HCC-827 | Afatinib | MVP_top60_DNA+RNA_seed | EGFR | -0.142154 | LUAD | NCI-H1568 | Lapatinib |
| MVP_top60_DNA_seed | MET | -5.097923 | LUAD | NCI-H1993 | AZD6094 | MVP_top60_DNA+RNA_seed | FGFR1 | -0.141793 | LUAD | NCI-H1944 | AZD4547 |

| | | | | | | | | | | | |
|---|---|---|---|---|---|---|---|---|---|---|---|
| MVP_top60_DNA_seed | EGFR | -5.064987 | LUAD | NCI-H1648 | Erlotinib | MVP_top60_DNA+RNA_seed | HSP90AA1 | -0.140696 | LUAD | NCI-H2087 | Tanespimycin |
| MVP_top60_DNA_seed | EGFR | -4.873695 | LUAD | H3255 | Gefitinib | MVP_top60_DNA+RNA_seed | AURKA | -0.139466 | LUAD | NCI-H2023 | ZM447439 |
| MVP_top60_DNA_seed | EGFR | -4.396938 | LUAD | HCC-827 | Gefitinib | MVP_top60_DNA+RNA_seed | HSP90AA1 | -0.139244 | LUAD | NCI-H1648 | Tanespimycin |
| MVP_top60_DNA_seed | EGFR | -4.210085 | LUAD | H3255 | Afatinib | MVP_top60_DNA+RNA_seed | AURKA | -0.138923 | LUAD | NCI-H838 | Tozasertib |
| MVP_top60_DNA_seed | ERBB2 | -4.210085 | LUAD | H3255 | Afatinib | MVP_top60_DNA+RNA_seed | AURKA | -0.138845 | LUAD | HCC-44 | Alisertib |
| MVP_top60_DNA_seed | EGFR | -4.088424 | LUAD | HCC-827 | Afatinib | MVP_top60_DNA+RNA_seed | IGF1R | -0.138716 | LUAD | NCI-H23 | BMS-754807 |
| MVP_top60_DNA_seed | ERBB2 | -4.088424 | LUAD | HCC-827 | Afatinib | MVP_top60_DNA+RNA_seed | PIK3CA | -0.138663 | LUAD | H3255 | GNE-317 |
| MVP_top60_DNA_seed | EGFR | -4.081692 | LUAD | H3255 | Afatinib | MVP_top60_DNA+RNA_seed | PIK3CA | -0.137713 | LUAD | EMC-BAC-1 | Buparlisib |
| MVP_top60_DNA_seed | ERBB2 | -4.081692 | LUAD | H3255 | Afatinib | MVP_top60_DNA+RNA_seed | TP53 | -0.13766 | LUAD | RERF-LC-MS | MIRA-1 |
| MVP_top60_DNA_seed | EGFR | -3.986983 | LUAD | H3255 | Cetuximab | MVP_top60_DNA+RNA_seed | EZH2 | -0.137388 | LUAD | NCI-H1755 | GSK343 |
| MVP_top60_DNA_seed | SMARCA4 | -3.886817 | LUAD | NCI-H1650 | PFI-3 | MVP_top60_DNA+RNA_seed | IGF1R | -0.137199 | LUAD | EMC-BAC-1 | NVP-ADW742 |
| MVP_top60_DNA_seed | EGFR | -3.635979 | LUAD | PC-14 | Afatinib | MVP_top60_DNA+RNA_seed | SMARCA4 | -0.13665 | LUAD | RERF-LC-MS | PFI3 |
| MVP_top60_DNA_seed | ERBB2 | -3.635979 | LUAD | PC-14 | Afatinib | MVP_top60_DNA+RNA_seed | AURKA | -0.13626 | LUAD | NCI-H1944 | Alisertib |
| MVP_top60_DNA_seed | MET | -3.344635 | LUAD | RERF-LC-MS | AZD6094 | MVP_top60_DNA+RNA_seed | IGF1R | -0.136232 | LUAD | EKVX | BMS-536924 |
| MVP_top60_DNA_seed | SRC | -3.312942 | LUAD | HCC-827 | Bosutinib | MVP_top60_DNA+RNA_seed | PIK3CA | -0.135987 | LUAD | NCI-H3122 | GNE-317 |
| MVP_top60_DNA_seed | EGFR | -3.30175 | LUAD | EKVX | Lapatinib | MVP_top60_DNA+RNA_seed | AKT1 | -0.13316 | LUAD | NCI-H2085 | GSK2110183B |
| MVP_top60_DNA_seed | ERBB2 | -3.30175 | LUAD | EKVX | Lapatinib | MVP_top60_DNA+RNA_seed | IGF1R | -0.132611 | LUAD | LXF-289 | Linsitinib |
| MVP_top60_DNA_seed | EGFR | -3.29092 | LUAD | PC-3_[JPC-3] | Afatinib | MVP_top60_DNA+RNA_seed | IGF1R | -0.1326 | LUAD | COR-L105 | Linsitinib |
| MVP_top60_DNA_seed | ERBB2 | -3.29092 | LUAD | PC-3_[JPC-3] | Afatinib | MVP_top60_DNA+RNA_seed | TERT | -0.132516 | LUAD | LXF-289 | BIBR-1532 |
| MVP_top60_DNA_seed | EGFR | -3.289851 | LUAD | NCI-H1568 | Cetuximab | MVP_top60_DNA+RNA_seed | BRAF | -0.130656 | LUAD | EMC-BAC-2 | PLX-4720 |
| MVP_top60_DNA_seed | EGFR | -3.250339 | LUAD | EMC-BAC-1 | Cetuximab | MVP_top60_DNA+RNA_seed | MET | -0.130513 | LUAD | NCI-H1563 | Savolitinib |
| MVP_top60_DNA_seed | EGFR | -3.247407 | LUAD | Calu-3 | PF-00299804 | MVP_top60_DNA+RNA_seed | SRC | -0.130276 | LUAD | NCI-H1666 | Bosutinib |
| MVP_top60_DNA_seed | ERBB2 | -3.247407 | LUAD | Calu-3 | PF-00299804 | MVP_top60_DNA+RNA_seed | HSP90AA1 | -0.130125 | LUAD | RERF-LC-KJ | Tanespimycin |
| MVP_top60_DNA_seed | EGFR | -3.241959 | LUAD | EKVX | Erlotinib | MVP_top60_DNA+RNA_seed | SRC | -0.130046 | LUAD | NCI-H358 | Bosutinib |
| MVP_top60_DNA_seed | GSK3B | -3.232391 | LUAD | HOP-62 | KIN001-042 | MVP_top60_DNA+RNA_seed | AKT1 | -0.129241 | LUAD | RERF-LC-MS | Uprosertib |
| MVP_top60_DNA_seed | EGFR | -3.137964 | LUAD | H3255 | AZD8931 | MVP_top60_DNA+RNA_seed | HDAC1 | -0.127992 | LUAD | NCI-H2030 | PCI-34051 |
| MVP_top60_DNA_seed | ERBB2 | -3.137964 | LUAD | H3255 | AZD8931 | MVP_top60_DNA+RNA_seed | PIK3CA | -0.127475 | LUAD | NCI-H1563 | Buparlisib |
| MVP_top60_DNA_seed | ERBB3 | -3.137964 | LUAD | H3255 | AZD8931 | MVP_top60_DNA+RNA_seed | FGFR1 | -0.12724 | LUAD | NCI-H1648 | AZD4547 |
| MVP_top60_DNA_seed | EGFR | -3.128579 | LUAD | COR-L105 | Pelitinib | MVP_top60_DNA+RNA_seed | PIK3CA | -0.127109 | LUAD | EKVX | AZD8186 |
| MVP_top60_DNA_seed | EGFR | -3.121584 | LUAD | Calu-3 | AST-1306 | MVP_top60_DNA+RNA_seed | PIK3CA | -0.125953 | LUAD | LXF-289 | GNE-317 |
| MVP_top60_DNA_seed | EGFR | -3.094491 | LUAD | HCC-827 | PF-00299804 | MVP_top60_DNA+RNA_seed | TERT | -0.125794 | LUAD | NCI-H322M | BIBR-1532 |
| MVP_top60_DNA_seed | ERBB2 | -3.094491 | LUAD | HCC-827 | PF-00299804 | MVP_top60_DNA+RNA_seed | EZH2 | -0.125613 | LUAD | HCC-78 | GSK343 |
| MVP_top60_DNA_seed | EGFR | -3.061442 | LUAD | PC-3_[JPC-3] | Afatinib | MVP_top60_DNA+RNA_seed | BRAF | -0.124405 | LUAD | NCI-H2405 | SB590885 |
| MVP_top60_DNA_seed | ERBB2 | -3.061442 | LUAD | PC-3_[JPC-3] | Afatinib | MVP_top60_DNA+RNA_seed | AKT1 | -0.124153 | LUAD | Calu-3 | Afuresertib |
| MVP_top60_DNA_seed | EGFR | -3.048994 | LUAD | PC-14 | Afatinib | MVP_top60_DNA+RNA_seed | AKT1 | -0.123179 | LUAD | EMC-BAC-1 | AZD5363 |
| MVP_top60_DNA_seed | ERBB2 | -3.048994 | LUAD | PC-14 | Afatinib | MVP_top60_DNA+RNA_seed | ATM | -0.122111 | LUAD | NCI-H1651 | KU-55933 |
| MVP_top60_DNA_seed | EGFR | -3.006622 | LUAD | HCC-827 | AZD8931 | MVP_top60_DNA+RNA_seed | IGF1R | -0.121883 | LUAD | NCI-H1734 | GSK1904529A |
| MVP_top60_DNA_seed | ERBB2 | -3.006622 | LUAD | HCC-827 | AZD8931 | MVP_top60_DNA+RNA_seed | TERT | -0.119012 | LUAD | LC-2-ad | BIBR-1532 |
| MVP_top60_DNA_seed | ERBB3 | -3.006622 | LUAD | HCC-827 | AZD8931 | MVP_top60_DNA+RNA_seed | SMARCA4 | -0.118711 | LUAD | NCI-H650 | PFI3 |
| MVP_top60_DNA_seed | HSP90AA1 | -2.986095 | LUAD | A549 | Tanespimycin | MVP_top60_DNA+RNA_seed | HSP90AA1 | -0.118463 | LUAD | PC-14 | Elesclomol |
| MVP_top60_DNA_seed | EGFR | -2.962385 | LUAD | PC-14 | AZD8931 | MVP_top60_DNA+RNA_seed | PIK3CA | -0.118434 | LUAD | HCC-827 | AZD8186 |

| | | | | | | | | | | | |
|---|---|---|---|---|---|---|---|---|---|---|---|
| MVP_top60_DNA_seed | ERBB2 | -2.962385 | LUAD | PC-14 | AZD8931 | MVP_top60_DNA+RNA_seed | SMARCA4 | -0.117848 | LUAD | NCI-H1993 | PFI3 |
| MVP_top60_DNA_seed | ERBB3 | -2.962385 | LUAD | PC-14 | AZD8931 | MVP_top60_DNA+RNA_seed | KRAS | -0.117287 | LUAD | NCI-H838 | KRAS (G12C) Inhibitor-1 |
| MVP_top60_DNA_seed | EGFR | -2.961245 | LUAD | NCI-H1623 | PF-00299804 | MVP_top60_DNA+RNA_seed | FGFR1 | -0.114211 | LUAD | NCI-H1975 | PD173074 |
| MVP_top60_DNA_seed | ERBB2 | -2.961245 | LUAD | NCI-H1623 | PF-00299804 | MVP_top60_DNA+RNA_seed | IGF1R | -0.112757 | LUAD | NCI-H1648 | Linsitinib |
| MVP_top60_DNA_seed | EGFR | -2.959705 | LUAD | NCI-H1666 | Cetuximab | MVP_top60_DNA+RNA_seed | AKT1 | -0.112091 | LUAD | PC-14 | Afuresertib |
| MVP_top60_DNA_seed | EGFR | -2.947302 | LUAD | EMC-BAC-1 | Afatinib | MVP_top60_DNA+RNA_seed | AKT1 | -0.111786 | LUAD | PC-14 | MK-2206 |
| MVP_top60_DNA_seed | ERBB2 | -2.947302 | LUAD | EMC-BAC-1 | Afatinib | MVP_top60_DNA+RNA_seed | EGFR | -0.111655 | LUAD | HCC-44 | Afatinib |
| MVP_top60_DNA_seed | EGFR | -2.942942 | LUAD | COR-L105 | Afatinib | MVP_top60_DNA+RNA_seed | HDAC1 | -0.109178 | LUAD | LC-2-ad | PCI-34051 |
| MVP_top60_DNA_seed | ERBB2 | -2.942942 | LUAD | COR-L105 | Afatinib | MVP_top60_DNA+RNA_seed | AKT1 | -0.108312 | LUAD | H3255 | MK-2206 |
| MVP_top60_DNA_seed | MAPK1 | -2.937547 | LUAD | NCI-H1755 | VX-11e | MVP_top60_DNA+RNA_seed | PIK3CA | -0.103037 | LUAD | NCI-H2342 | Alpelisib |
| MVP_top60_DNA_seed | MDM2 | -2.854602 | LUAD | HCC-78 | Serdemetan | MVP_top60_DNA+RNA_seed | PIK3CA | -0.101591 | LUAD | NCI-H1666 | Buparlisib |
| MVP_top60_DNA_seed | ERBB2 | -2.849479 | LUAD | Calu-3 | CP724714 | MVP_top60_DNA+RNA_seed | AKT1 | -0.100328 | LUAD | RERF-LC-KJ | Uprosertib |
| MVP_top60_DNA_seed | IGF1R | -2.847594 | LUAD | NCI-H2342 | GSK1904529A | MVP_top60_DNA+RNA_seed | IGF1R | -0.09898 | LUAD | NCI-H23 | GSK1904529A |
| MVP_top60_DNA_seed | EGFR | -2.79094 | LUAD | Calu-3 | CI-1033 | MVP_top60_DNA+RNA_seed | FGFR1 | -0.098532 | LUAD | NCI-H522 | AZD4547 |
| MVP_top60_DNA_seed | ERBB2 | -2.79094 | LUAD | Calu-3 | CI-1033 | MVP_top60_DNA+RNA_seed | HSP90AA1 | -0.097613 | LUAD | NCI-H650 | Tanespimycin |
| MVP_top60_DNA_seed | SRC | -2.787059 | LUAD | NCI-H1648 | Saracatinib | MVP_top60_DNA+RNA_seed | IGF1R | -0.09395 | LUAD | NCI-H1650 | BMS-536924 |
| MVP_top60_DNA_seed | AKT1 | -2.773876 | LUAD | NCI-H3122 | AKT inhibitor VIII | MVP_top60_DNA+RNA_seed | BRAF | -0.093522 | LUAD | NCI-H2405 | Dabrafenib |
| MVP_top60_DNA_seed | IGF1R | -2.773783 | LUAD | NCI-H2122 | BMS-754807 | MVP_top60_DNA+RNA_seed | IGF1R | -0.092333 | LUAD | PC-14 | NVP-ADW742 |
| MVP_top60_DNA_seed | EGFR | -2.730621 | LUAD | EMC-BAC-1 | Afatinib | MVP_top60_DNA+RNA_seed | HDAC1 | -0.09195 | LUAD | NCI-H2085 | PCI-34051 |
| MVP_top60_DNA_seed | ERBB2 | -2.730621 | LUAD | EMC-BAC-1 | Afatinib | MVP_top60_DNA+RNA_seed | IGF1R | -0.091412 | LUAD | NCI-H1563 | BMS-754807 |
| MVP_top60_DNA_seed | EGFR | -2.717338 | LUAD | EMC-BAC-1 | AZD8931 | MVP_top60_DNA+RNA_seed | ATM | -0.091316 | LUAD | NCI-H1734 | KU-55933 |
| MVP_top60_DNA_seed | ERBB2 | -2.717338 | LUAD | EMC-BAC-1 | AZD8931 | MVP_top60_DNA+RNA_seed | MDM2 | -0.090673 | LUAD | NCI-H838 | Serdemetan |
| MVP_top60_DNA_seed | ERBB3 | -2.717338 | LUAD | EMC-BAC-1 | AZD8931 | MVP_top60_DNA+RNA_seed | ATM | -0.090335 | LUAD | PC-3_[JPC-3] | KU-55933 |
| MVP_top60_DNA_seed | EGFR | -2.707261 | LUAD | NCI-H1781 | PF-00299804 | MVP_top60_DNA+RNA_seed | PIK3CA | -0.089201 | LUAD | NCI-H2291 | Alpelisib |
| MVP_top60_DNA_seed | ERBB2 | -2.707261 | LUAD | NCI-H1781 | PF-00299804 | MVP_top60_DNA+RNA_seed | MDM2 | -0.089055 | LUAD | ABC-1 | Serdemetan |
| MVP_top60_DNA_seed | EGFR | -2.697602 | LUAD | H3255 | PF-00299804 | MVP_top60_DNA+RNA_seed | BRAF | -0.088436 | LUAD | PC-14 | PLX-4720 |
| MVP_top60_DNA_seed | ERBB2 | -2.697602 | LUAD | H3255 | PF-00299804 | MVP_top60_DNA+RNA_seed | IGF1R | -0.085515 | LUAD | NCI-H1693 | NVP-ADW742 |
| MVP_top60_DNA_seed | EGFR | -2.690145 | LUAD | NCI-H1623 | CI-1033 | MVP_top60_DNA+RNA_seed | EZH2 | -0.081768 | LUAD | NCI-H650 | GSK343 |
| MVP_top60_DNA_seed | ERBB2 | -2.690145 | LUAD | NCI-H1623 | CI-1033 | MVP_top60_DNA+RNA_seed | AKT1 | -0.081389 | LUAD | PC-14 | GSK2110183B |
| MVP_top60_DNA_seed | EGFR | -2.651437 | LUAD | NCI-H1568 | AZD8931 | MVP_top60_DNA+RNA_seed | SMARCA4 | -0.08108 | LUAD | LXF-289 | PFI3 |
| MVP_top60_DNA_seed | ERBB2 | -2.651437 | LUAD | NCI-H1568 | AZD8931 | MVP_top60_DNA+RNA_seed | MET | -0.080649 | LUAD | NCI-H358 | Crizotinib |
| MVP_top60_DNA_seed | ERBB3 | -2.651437 | LUAD | NCI-H1568 | AZD8931 | MVP_top60_DNA+RNA_seed | AKT1 | -0.080258 | LUAD | NCI-H1563 | Afuresertib |
| MVP_top60_DNA_seed | EGFR | -2.64536 | LUAD | NCI-H1623 | AST-1306 | MVP_top60_DNA+RNA_seed | EGFR | -0.080005 | LUAD | NCI-H1838 | Gefitinib |
| MVP_top60_DNA_seed | EGFR | -2.630997 | LUAD | HCC-827 | CI-1033 | MVP_top60_DNA+RNA_seed | EGFR | -0.080003 | LUAD | NCI-H1651 | Gefitinib |
| MVP_top60_DNA_seed | ERBB2 | -2.630997 | LUAD | HCC-827 | CI-1033 | MVP_top60_DNA+RNA_seed | PIK3CA | -0.078508 | LUAD | NCI-H1944 | Alpelisib |
| MVP_top60_DNA_seed | EGFR | -2.624419 | LUAD | PC-3_[JPC-3] | Gefitinib | MVP_top60_DNA+RNA_seed | PIK3CA | -0.078505 | LUAD | Calu-6 | GNE-317 |
| MVP_top60_DNA_seed | SRC | -2.604488 | LUAD | LC-2-ad | WH-4-023 | MVP_top60_DNA+RNA_seed | MET | -0.076999 | LUAD | NCI-H23 | Crizotinib |
| MVP_top60_DNA_seed | BRAF | -2.600217 | LUAD | Calu-6 | AZ628 | MVP_top60_DNA+RNA_seed | EGFR | -0.076941 | LUAD | NCI-H838 | Afatinib |
| MVP_top60_DNA_seed | EGFR | -2.576548 | LUAD | NCI-H1623 | Afatinib | MVP_top60_DNA+RNA_seed | TERT | -0.076725 | LUAD | NCI-H2030 | BIBR-1532 |
| MVP_top60_DNA_seed | ERBB2 | -2.576548 | LUAD | NCI-H1623 | Afatinib | MVP_top60_DNA+RNA_seed | BRAF | -0.074879 | LUAD | NCI-H1734 | Dabrafenib |
| MVP_top60_DNA_seed | MAPK1 | -2.544509 | LUAD | Calu-6 | VX-11e | MVP_top60_DNA+RNA_seed | HDAC1 | -0.073644 | LUAD | NCI-H1792 | Romidepsin |

| | | | | | | | | | | | |
|---|---|---|---|---|---|---|---|---|---|---|---|
| MVP_top60_DNA_seed | AKT1 | -2.525469 | LUAD | NCI-H2342 | AKT inhibitor VIII | MVP_top60_DNA+RNA_seed | BRAF | -0.071772 | LUAD | NCI-H1734 | SB590885 |
| MVP_top60_DNA_seed | IGF1R | -2.520279 | LUAD | NCI-H2122 | Linsitinib | MVP_top60_DNA+RNA_seed | AKT1 | -0.071468 | LUAD | PC-14 | AT13148 |
| MVP_top60_DNA_seed | MET | -2.48919 | LUAD | NCI-H2023 | AZD6094 | MVP_top60_DNA+RNA_seed | HDAC1 | -0.071026 | LUAD | NCI-H522 | Entinostat |
| MVP_top60_DNA_seed | IGF1R | -2.475094 | LUAD | NCI-H1355 | BMS-754807 | MVP_top60_DNA+RNA_seed | TP53 | -0.070238 | LUAD | LXF-289 | MIRA-1 |
| MVP_top60_DNA_seed | IGF1R | -2.469308 | LUAD | NCI-H1355 | Linsitinib | MVP_top60_DNA+RNA_seed | AKT1 | -0.069219 | LUAD | NCI-H2228 | Afuresertib |
| MVP_top60_DNA_seed | SRC | -2.458888 | LUAD | NCI-H1755 | NVP-BHG712 | MVP_top60_DNA+RNA_seed | EGFR | -0.06844 | LUAD | NCI-H522 | Sapitinib |
| MVP_top60_DNA_seed | EGFR | -2.449449 | LUAD | NCI-H1666 | AZD8931 | MVP_top60_DNA+RNA_seed | AKT1 | -0.06737 | LUAD | NCI-H1563 | GSK2110183B |
| MVP_top60_DNA_seed | ERBB2 | -2.449449 | LUAD | NCI-H1666 | AZD8931 | MVP_top60_DNA+RNA_seed | HSP90AA1 | -0.066491 | LUAD | NCI-H1944 | Luminespib |
| MVP_top60_DNA_seed | ERBB3 | -2.449449 | LUAD | NCI-H1666 | AZD8931 | MVP_top60_DNA+RNA_seed | BRAF | -0.065928 | LUAD | NCI-H358 | PLX-4720 |
| MVP_top60_DNA_seed | EGFR | -2.431793 | LUAD | H3255 | Pelitinib | MVP_top60_DNA+RNA_seed | AKT1 | -0.063853 | LUAD | NCI-H1792 | Afuresertib |
| MVP_top60_DNA_seed | EGFR | -2.403001 | LUAD | NCI-H1623 | Afatinib | MVP_top60_DNA+RNA_seed | TP53 | -0.063293 | LUAD | NCI-H1437 | MIRA-1 |
| MVP_top60_DNA_seed | ERBB2 | -2.403001 | LUAD | NCI-H1623 | Afatinib | MVP_top60_DNA+RNA_seed | EGFR | -0.062643 | LUAD | NCI-H1435 | AZD3759 |
| MVP_top60_DNA_seed | EGFR | -2.400405 | LUAD | NCI-H1568 | Afatinib | MVP_top60_DNA+RNA_seed | EGFR | -0.062512 | LUAD | NCI-H1563 | Afatinib |
| MVP_top60_DNA_seed | ERBB2 | -2.400405 | LUAD | NCI-H1568 | Afatinib | MVP_top60_DNA+RNA_seed | EGFR | -0.060513 | LUAD | NCI-H1651 | AZD3759 |
| MVP_top60_DNA_seed | EGFR | -2.390909 | LUAD | NCI-H1666 | Afatinib | MVP_top60_DNA+RNA_seed | FGFR1 | -0.058974 | LUAD | NCI-H2009 | PD173074 |
| MVP_top60_DNA_seed | ERBB2 | -2.390909 | LUAD | NCI-H1666 | Afatinib | MVP_top60_DNA+RNA_seed | HSP90AA1 | -0.058974 | LUAD | NCI-H23 | Luminespib |
| MVP_top60_DNA_seed | EGFR | -2.390045 | LUAD | PC-14 | Gefitinib | MVP_top60_DNA+RNA_seed | AURKA | -0.056053 | LUAD | NCI-H2228 | Alisertib |
| MVP_top60_DNA_seed | EGFR | -2.375309 | LUAD | NCI-H1975 | Pelitinib | MVP_top60_DNA+RNA_seed | AKT1 | -0.055584 | LUAD | NCI-H838 | Ipatasertib |
| MVP_top60_DNA_seed | EGFR | -2.373737 | LUAD | HCC-827 | Pelitinib | MVP_top60_DNA+RNA_seed | EGFR | -0.054995 | LUAD | NCI-H1573 | Sapitinib |
| MVP_top60_DNA_seed | HSP90AA1 | -2.371967 | LUAD | ABC-1 | Elesclomol | MVP_top60_DNA+RNA_seed | AKT1 | -0.054084 | LUAD | NCI-H1437 | Ipatasertib |
| MVP_top60_DNA_seed | EGFR | -2.36446 | LUAD | NCI-H1781 | Afatinib | MVP_top60_DNA+RNA_seed | BRAF | -0.053744 | LUAD | NCI-H522 | Dabrafenib |
| MVP_top60_DNA_seed | ERBB2 | -2.36446 | LUAD | NCI-H1781 | Afatinib | MVP_top60_DNA+RNA_seed | EGFR | -0.052991 | LUAD | NCI-H358 | AZD3759 |
| MVP_top60_DNA_seed | AKT1 | -2.358798 | LUAD | NCI-H1755 | AKT inhibitor VIII | MVP_top60_DNA+RNA_seed | EGFR | -0.050957 | LUAD | NCI-H1792 | Gefitinib |
| MVP_top60_DNA_seed | AKT1 | -2.351603 | LUAD | NCI-H1563 | AKT inhibitor VIII | MVP_top60_DNA+RNA_seed | PIK3CA | -0.047645 | LUAD | NCI-H2228 | AZD8186 |
| MVP_top60_DNA_seed | EGFR | -2.333781 | LUAD | H3255 | CI-1033 | MVP_top60_DNA+RNA_seed | BRAF | -0.046643 | LUAD | EMC-BAC-1 | Dabrafenib |
| MVP_top60_DNA_seed | ERBB2 | -2.333781 | LUAD | H3255 | CI-1033 | MVP_top60_DNA+RNA_seed | IGF1R | -0.046488 | LUAD | RERF-LC-KJ | BMS-754807 |
| MVP_top60_DNA_seed | BRAF | -2.314637 | LUAD | LC-2-ad | HG6-64-1 | MVP_top60_DNA+RNA_seed | MDM2 | -0.044598 | LUAD | NCI-H1734 | Nutlin-3a (-) |
| MVP_top60_DNA_seed | EGFR | -2.301404 | LUAD | H3255 | AST-1306 | MVP_top60_DNA+RNA_seed | EGFR | -0.042947 | LUAD | NCI-H1435 | Afatinib |
| MVP_top60_DNA_seed | EGFR | -2.300195 | LUAD | Calu-3 | Afatinib | MVP_top60_DNA+RNA_seed | HSP90AA1 | -0.041669 | LUAD | COR-L105 | Luminespib |
| MVP_top60_DNA_seed | ERBB2 | -2.300195 | LUAD | Calu-3 | Afatinib | MVP_top60_DNA+RNA_seed | SRC | -0.041344 | LUAD | NCI-H2342 | Bosutinib |
| MVP_top60_DNA_seed | MAPK1 | -2.278796 | LUAD | H3255 | FR-180204 | MVP_top60_DNA+RNA_seed | HDAC1 | -0.039471 | LUAD | NCI-H1944 | Romidepsin |
| MVP_top60_DNA_seed | IGF1R | -2.275211 | LUAD | LC-2-ad | BMS-536924 | MVP_top60_DNA+RNA_seed | ATM | -0.039 | LUAD | NCI-H650 | KU-55933 |
| MVP_top60_DNA_seed | HSP90AA1 | -2.268609 | LUAD | NCI-H3122 | CCT-018159 | MVP_top60_DNA+RNA_seed | HSP90AA1 | -0.038665 | LUAD | LC-2-ad | Tanespimycin |
| MVP_top60_DNA_seed | EGFR | -2.236769 | LUAD | NCI-H1666 | Afatinib | MVP_top60_DNA+RNA_seed | MDM2 | -0.038314 | LUAD | Calu-3 | Serdemetan |
| MVP_top60_DNA_seed | ERBB2 | -2.236769 | LUAD | NCI-H1666 | Afatinib | MVP_top60_DNA+RNA_seed | PIK3CA | -0.038234 | LUAD | HOP-62 | GNE-317 |
| MVP_top60_DNA_seed | GSK3B | -2.208405 | LUAD | NCI-H2122 | CHIR-99021 | MVP_top60_DNA+RNA_seed | SRC | -0.03792 | LUAD | NCI-H1781 | Bosutinib |
| MVP_top60_DNA_seed | GSK3B | -2.165265 | LUAD | HOP-62 | CHIR-99021 | MVP_top60_DNA+RNA_seed | SRC | -0.037417 | LUAD | NCI-H2122 | Bosutinib |
| MVP_top60_DNA_seed | BRAF | -2.163684 | LUAD | EMC-BAC-2 | Dabrafenib | MVP_top60_DNA+RNA_seed | MET | -0.03735 | LUAD | NCI-H2009 | Savolitinib |
| MVP_top60_DNA_seed | MAPK1 | -2.145892 | LUAD | PC-3_[JPC-3] | FR-180204 | MVP_top60_DNA+RNA_seed | KRAS | -0.036458 | LUAD | NCI-H2023 | (RAS (G12C) Inhibitor-1 |
| MVP_top60_DNA_seed | AKT1 | -2.125675 | LUAD | NCI-H1437 | AKT inhibitor VIII | MVP_top60_DNA+RNA_seed | FGFR1 | -0.036218 | LUAD | NCI-H650 | PD173074 |
| MVP_top60_DNA_seed | EGFR | -2.11529 | LUAD | Calu-3 | AZD8931 | MVP_top60_DNA+RNA_seed | SRC | -0.035266 | LUAD | NCI-H1435 | Bosutinib |

| | | | | | | | | | | | |
|---|---|---|---|---|---|---|---|---|---|---|---|
| MVP_top60_DNA_seed | ERBB2 | -2.11529 | LUAD | Calu-3 | AZD8931 | MVP_top60_DNA+RNA_seed | BRAF | -0.035156 | LUAD | NCI-H522 | PLX-4720 |
| MVP_top60_DNA_seed | ERBB3 | -2.11529 | LUAD | Calu-3 | AZD8931 | MVP_top60_DNA+RNA_seed | MDM2 | -0.034144 | LUAD | NCI-H2009 | Serdemetan |
| MVP_top60_DNA_seed | AKT1 | -2.112531 | LUAD | NCI-H1975 | BAY AKT1 | MVP_top60_DNA+RNA_seed | HDAC1 | -0.033624 | LUAD | NCI-H23 | PCI-34051 |
| MVP_top60_DNA_seed | EGFR | -2.102305 | LUAD | PC-3_[JPC-3] | Pelitinib | MVP_top60_DNA+RNA_seed | HDAC1 | -0.033596 | LUAD | VMRC-LCD | Romidepsin |
| MVP_top60_DNA_seed | EGFR | -2.096039 | LUAD | PC-3_[JPC-3] | AZD8931 | MVP_top60_DNA+RNA_seed | IGF1R | -0.032573 | LUAD | NCI-H1563 | Linsitinib |
| MVP_top60_DNA_seed | ERBB2 | -2.096039 | LUAD | PC-3_[JPC-3] | AZD8931 | MVP_top60_DNA+RNA_seed | EGFR | -0.032533 | LUAD | NCI-H1944 | Afatinib |
| MVP_top60_DNA_seed | ERBB3 | -2.096039 | LUAD | PC-3_[JPC-3] | AZD8931 | MVP_top60_DNA+RNA_seed | AKT1 | -0.031145 | LUAD | NCI-H1563 | MK-2206 |
| MVP_top60_DNA_seed | AKT1 | -2.080775 | LUAD | LC-2-ad | A-443654 | MVP_top60_DNA+RNA_seed | IGF1R | -0.030377 | LUAD | HCC-827 | BMS-536924 |
| MVP_top60_DNA_seed | GSK3B | -2.074914 | LUAD | NCI-H1993 | SB216763 | MVP_top60_DNA+RNA_seed | HDAC1 | -0.028701 | LUAD | A549 | Entinostat |
| MVP_top60_DNA_seed | GSK3B | -2.06672 | LUAD | NCI-H1792 | CHIR-99021 | MVP_top60_DNA+RNA_seed | AURKA | -0.028556 | LUAD | HOP-62 | Alisertib |
| MVP_top60_DNA_seed | GSK3B | -2.061639 | LUAD | NCI-H2405 | SB216763 | MVP_top60_DNA+RNA_seed | EGFR | -0.026665 | LUAD | NCI-H1437 | AZD3759 |
| MVP_top60_DNA_seed | HSP90AA1 | -2.045994 | LUAD | HCC-44 | Tanespimycin | MVP_top60_DNA+RNA_seed | IGF1R | -0.026585 | LUAD | RERF-LC-MS | Linsitinib |
| MVP_top60_DNA_seed | EGFR | -2.022879 | LUAD | NCI-H1666 | Gefitinib | MVP_top60_DNA+RNA_seed | TERT | -0.02603 | LUAD | NCI-H1435 | BIBR-1532 |
| MVP_top60_DNA_seed | EGFR | -2.022512 | LUAD | NCI-H1781 | Pelitinib | MVP_top60_DNA+RNA_seed | AURKA | -0.024727 | LUAD | NCI-H23 | Alisertib |
| MVP_top60_DNA_seed | AURKA | -2.021489 | LUAD | LC-2-ad | Tozasertib | MVP_top60_DNA+RNA_seed | SMARCA4 | -0.023758 | LUAD | NCI-H2023 | PFI3 |
| MVP_top60_DNA_seed | EGFR | -1.990933 | LUAD | EMC-BAC-1 | Gefitinib | MVP_top60_DNA+RNA_seed | AKT1 | -0.022049 | LUAD | NCI-H2030 | Afuresertib |
| MVP_top60_DNA_seed | EGFR | -1.990647 | LUAD | NCI-H1568 | PF-00299804 | MVP_top60_DNA+RNA_seed | TP53 | -0.021429 | LUAD | NCI-H522 | MIRA-1 |
| MVP_top60_DNA_seed | ERBB2 | -1.990647 | LUAD | NCI-H1568 | PF-00299804 | MVP_top60_DNA+RNA_seed | TP53 | -0.021345 | LUAD | RERF-LC-KJ | MIRA-1 |
| MVP_top60_DNA_seed | ERBB2 | -1.975031 | LUAD | NCI-H1755 | CP724714 | MVP_top60_DNA+RNA_seed | IGF1R | -0.021162 | LUAD | NCI-H1573 | Linsitinib |
| MVP_top60_DNA_seed | EGFR | -1.961863 | LUAD | PC-14 | Pelitinib | MVP_top60_DNA+RNA_seed | TP53 | -0.020545 | LUAD | NCI-H1651 | MIRA-1 |
| MVP_top60_DNA_seed | IGF1R | -1.958465 | LUAD | NCI-H1648 | BMS-536924 | MVP_top60_DNA+RNA_seed | FGFR1 | -0.02 | LUAD | NCI-H522 | PD173074 |
| MVP_top60_DNA_seed | EGFR | -1.944651 | LUAD | NCI-H1781 | AST-1306 | MVP_top60_DNA+RNA_seed | AURKA | -0.018535 | LUAD | ABC-1 | Tozasertib |
| MVP_top60_DNA_seed | EGFR | -1.928812 | LUAD | COR-L105 | Afatinib | MVP_top60_DNA+RNA_seed | AKT1 | -0.018466 | LUAD | H3255 | Afuresertib |
| MVP_top60_DNA_seed | ERBB2 | -1.928812 | LUAD | COR-L105 | Afatinib | MVP_top60_DNA+RNA_seed | HSP90AA1 | -0.018442 | LUAD | NCI-H1781 | Luminespib |
| MVP_top60_DNA_seed | EGFR | -1.913401 | LUAD | NCI-H1623 | AZD8931 | MVP_top60_DNA+RNA_seed | BRAF | -0.018387 | LUAD | EMC-BAC-2 | Dabrafenib |
| MVP_top60_DNA_seed | ERBB2 | -1.913401 | LUAD | NCI-H1623 | AZD8931 | MVP_top60_DNA+RNA_seed | AKT1 | -0.018005 | LUAD | H3255 | Ipatasertib |
| MVP_top60_DNA_seed | ERBB3 | -1.913401 | LUAD | NCI-H1623 | AZD8931 | MVP_top60_DNA+RNA_seed | PIK3CA | -0.015769 | LUAD | NCI-H1563 | Alpelisib |
| MVP_top60_DNA_seed | ATM | -1.905401 | LUAD | HCC-44 | KU-60019 | MVP_top60_DNA+RNA_seed | EGFR | -0.012968 | LUAD | NCI-H1755 | Afatinib |
| MVP_top60_DNA_seed | AKT1 | -1.900726 | LUAD | NCI-H1563 | MK-2206 | MVP_top60_DNA+RNA_seed | EGFR | -0.01269 | LUAD | NCI-H838 | Osimertinib |
| MVP_top60_DNA_seed | IGF1R | -1.89741 | LUAD | NCI-H358 | BMS-754807 | MVP_top60_DNA+RNA_seed | PIK3CA | -0.012626 | LUAD | NCI-H2009 | Buparlisib |
| MVP_top60_DNA_seed | MDM2 | -1.894813 | LUAD | EMC-BAC-2 | Nutlin-3a (-) | MVP_top60_DNA+RNA_seed | EGFR | -0.01189 | LUAD | LC-2-ad | Osimertinib |
| MVP_top60_DNA_seed | BRAF | -1.88687 | LUAD | NCI-H2122 | PLX-4720 | MVP_top60_DNA+RNA_seed | EGFR | -0.011396 | LUAD | NCI-H2085 | Sapitinib |
| MVP_top60_DNA_seed | MET | -1.882533 | LUAD | NCI-H2291 | JNJ38877605 | MVP_top60_DNA+RNA_seed | AKT1 | -0.011382 | LUAD | NCI-H522 | AT13148 |
| MVP_top60_DNA_seed | EGFR | -1.877617 | LUAD | COR-L105 | AZD8931 | MVP_top60_DNA+RNA_seed | PIK3CA | -0.011315 | LUAD | LXF-289 | AZD8186 |
| MVP_top60_DNA_seed | ERBB2 | -1.877617 | LUAD | COR-L105 | AZD8931 | MVP_top60_DNA+RNA_seed | IGF1R | -0.009997 | LUAD | NCI-H1792 | GSK1904529A |
| MVP_top60_DNA_seed | ERBB3 | -1.877617 | LUAD | COR-L105 | AZD8931 | MVP_top60_DNA+RNA_seed | AKT1 | -0.009335 | LUAD | HCC-78 | Uprosertib |
| MVP_top60_DNA_seed | IGF1R | -1.874303 | LUAD | HCC-78 | BMS-536924 | MVP_top60_DNA+RNA_seed | PIK3CA | -0.009014 | LUAD | HCC-827 | GNE-317 |
| MVP_top60_DNA_seed | SRC | -1.8739 | LUAD | H3255 | Bosutinib | MVP_top60_DNA+RNA_seed | IGF1R | -0.008776 | LUAD | NCI-H1975 | GSK1904529A |
| MVP_top60_DNA_seed | GSK3B | -1.869688 | LUAD | NCI-H2342 | KIN001-042 | MVP_top60_DNA+RNA_seed | BRAF | -0.006907 | LUAD | NCI-H1792 | Dabrafenib |
| MVP_top60_DNA_seed | AURKA | -1.865741 | LUAD | PC-14 | Genentech Cpd 10 | MVP_top60_DNA+RNA_seed | MET | -0.003864 | LUAD | NCI-H1792 | Savolitinib |
| MVP_top60_DNA_seed | EGFR | -1.861887 | LUAD | NCI-H1781 | CI-1033 | MVP_top60_DNA+RNA_seed | EGFR | -0.002579 | LUAD | NCI-H838 | Gefitinib |

| | | | | | | | | | | | |
|---|---|---|---|---|---|---|---|---|---|---|---|
| MVP_top60_DNA_seed | ERBB2 | -1.861887 | LUAD | NCI-H1781 | CI-1033 | MVP_top60_DNA+RNA_seed | IGF1R | -0.00174 | LUAD | NCI-H2342 | Linsitinib |
| MVP_top60_DNA_seed | AKT1 | -1.851751 | LUAD | NCI-H1755 | AKT inhibitor VIII | MVP_top60_DNA+RNA_seed | AKT1 | -0.001342 | LUAD | A549 | Afuresertib |
| MVP_top60_DNA_seed | EGFR | -1.846051 | LUAD | NCI-H358 | Pelitinib | MVP_top60_DNA+RNA_seed | MET | -0.000908 | LUAD | NCI-H1648 | Savolitinib |
| MVP_top60_DNA_seed | HDAC1 | -1.831845 | LUAD | Calu-3 | JQ12 | MVP_top60_DNA+RNA_seed | IGF1R | -0.000873 | LUAD | NCI-H650 | NVP-ADW742 |
| MVP_top60_DNA_seed | CREBBP | -1.829394 | LUAD | NCI-H2291 | I-CBP112 | MVP_top60_DNA+RNA_seed | AKT1 | 0.001387 | LUAD | NCI-H1563 | AZD5363 |
| MVP_top60_DNA_seed | IGF1R | -1.82253 | LUAD | NCI-H358 | Linsitinib | MVP_top60_DNA+RNA_seed | HSP90AA1 | 0.003477 | LUAD | NCI-H322M | Luminespib |
| MVP_top60_DNA_seed | HDAC1 | -1.806475 | LUAD | NCI-H1975 | Entinostat | MVP_top60_DNA+RNA_seed | PIK3CA | 0.003942 | LUAD | NCI-H1734 | GNE-317 |
| MVP_top60_DNA_seed | HSP90AA1 | -1.799429 | LUAD | LXF-289 | Tanespimycin | MVP_top60_DNA+RNA_seed | TERT | 0.005902 | LUAD | NCI-H1650 | BIBR-1532 |
| MVP_top60_DNA_seed | EGFR | -1.797091 | LUAD | COR-L105 | PF-00299804 | MVP_top60_DNA+RNA_seed | AKT1 | 0.007821 | LUAD | NCI-H1755 | GSK2110183B |
| MVP_top60_DNA_seed | ERBB2 | -1.797091 | LUAD | COR-L105 | PF-00299804 | MVP_top60_DNA+RNA_seed | IGF1R | 0.008672 | LUAD | NCI-H2009 | GSK1904529A |
| MVP_top60_DNA_seed | GSK3B | -1.789352 | LUAD | NCI-H1755 | CHIR-99021 | MVP_top60_DNA+RNA_seed | IGF1R | 0.00934 | LUAD | NCI-H1944 | Linsitinib |
| MVP_top60_DNA_seed | EGFR | -1.777737 | LUAD | NCI-H1944 | Cetuximab | MVP_top60_DNA+RNA_seed | SRC | 0.009809 | LUAD | HCC-44 | Bosutinib |
| MVP_top60_DNA_seed | MET | -1.763636 | LUAD | NCI-H2347 | JNJ38877605 | MVP_top60_DNA+RNA_seed | AURKA | 0.010821 | LUAD | EMC-BAC-1 | ZM447439 |
| MVP_top60_DNA_seed | HDAC1 | -1.759001 | LUAD | LC-2-ad | JQ12 | MVP_top60_DNA+RNA_seed | SRC | 0.012303 | LUAD | RERF-LC-MS | Bosutinib |
| MVP_top60_DNA_seed | EGFR | -1.758171 | LUAD | NCI-H1568 | CI-1033 | MVP_top60_DNA+RNA_seed | PIK3CA | 0.012533 | LUAD | NCI-H1651 | Buparlisib |
| MVP_top60_DNA_seed | ERBB2 | -1.758171 | LUAD | NCI-H1568 | CI-1033 | MVP_top60_DNA+RNA_seed | EGFR | 0.013265 | LUAD | NCI-H1693 | Lapatinib |
| MVP_top60_DNA_seed | SRC | -1.758019 | LUAD | PC-14 | Bosutinib | MVP_top60_DNA+RNA_seed | KRAS | 0.013546 | LUAD | NCI-H3122 | [illegible]RAS (G12C) Inhibitor-1 |
| MVP_top60_DNA_seed | MET | -1.752393 | LUAD | RERF-LC-MS | JNJ38877605 | MVP_top60_DNA+RNA_seed | EZH2 | 0.014421 | LUAD | NCI-H23 | GSK343 |
| MVP_top60_DNA_seed | AKT1 | -1.750753 | LUAD | NCI-H2228 | AKT inhibitor VIII | MVP_top60_DNA+RNA_seed | PIK3CA | 0.014755 | LUAD | NCI-H1355 | Buparlisib |
| MVP_top60_DNA_seed | GSK3B | -1.735209 | LUAD | NCI-H1792 | AZD7969 | MVP_top60_DNA+RNA_seed | MET | 0.01497 | LUAD | NCI-H2405 | Savolitinib |
| MVP_top60_DNA_seed | IGF1R | -1.730043 | LUAD | NCI-H1944 | GSK1904529A | MVP_top60_DNA+RNA_seed | IGF1R | 0.021004 | LUAD | NCI-H650 | BMS-536924 |
| MVP_top60_DNA_seed | GSK3B | -1.72434 | LUAD | NCI-H1755 | CHIR-99021 | MVP_top60_DNA+RNA_seed | AKT1 | 0.021479 | LUAD | NCI-H2342 | Ipatasertib |
| MVP_top60_DNA_seed | EGFR | -1.724238 | LUAD | PC-3_[JPC-3] | Cetuximab | MVP_top60_DNA+RNA_seed | PIK3CA | 0.022357 | LUAD | NCI-H3122 | AZD8186 |
| MVP_top60_DNA_seed | HSP90AA1 | -1.717851 | LUAD | NCI-H3122 | Tanespimycin | MVP_top60_DNA+RNA_seed | MDM2 | 0.022863 | LUAD | NCI-H1435 | Serdemetan |
| MVP_top60_DNA_seed | IGF1R | -1.715622 | LUAD | HCC-78 | BMS-754807 | MVP_top60_DNA+RNA_seed | MDM2 | 0.024305 | LUAD | NCI-H1993 | Serdemetan |
| MVP_top60_DNA_seed | AKT1 | -1.71489 | LUAD | HCC-44 | MK-2206 | MVP_top60_DNA+RNA_seed | TERT | 0.025029 | LUAD | NCI-H1651 | BIBR-1532 |
| MVP_top60_DNA_seed | GSK3B | -1.707878 | LUAD | NCI-H1648 | CHIR-99021 | MVP_top60_DNA+RNA_seed | EGFR | 0.025665 | LUAD | NCI-H522 | Afatinib |
| MVP_top60_DNA_seed | EGFR | -1.706973 | LUAD | NCI-H1975 | Cetuximab | MVP_top60_DNA+RNA_seed | AKT1 | 0.027672 | LUAD | NCI-H1435 | GSK2110183B |
| MVP_top60_DNA_seed | AKT1 | -1.706946 | LUAD | LXF-289 | AKT inhibitor VIII | MVP_top60_DNA+RNA_seed | HDAC1 | 0.02972 | LUAD | NCI-H1568 | Romidepsin |
| MVP_top60_DNA_seed | SRC | -1.705187 | LUAD | Calu-6 | NVP-BHG712 | MVP_top60_DNA+RNA_seed | HDAC1 | 0.030073 | LUAD | HCC-44 | PCI-34051 |
| MVP_top60_DNA_seed | ATM | -1.694277 | LUAD | NCI-H1838 | KU-55933 | MVP_top60_DNA+RNA_seed | AKT1 | 0.030188 | LUAD | LC-2-ad | AZD5363 |
| MVP_top60_DNA_seed | PIK3CA | -1.691383 | LUAD | NCI-H1944 | AZD8835 | MVP_top60_DNA+RNA_seed | AURKA | 0.031621 | LUAD | NCI-H2030 | ZM447439 |
| MVP_top60_DNA_seed | BRAF | -1.690759 | LUAD | EMC-BAC-2 | PLX-4720 | MVP_top60_DNA+RNA_seed | AKT1 | 0.031621 | LUAD | LXF-289 | Afuresertib |
| MVP_top60_DNA_seed | EGFR | -1.690751 | LUAD | NCI-H1648 | Afatinib | MVP_top60_DNA+RNA_seed | EGFR | 0.031739 | LUAD | LC-2-ad | Afatinib |
| MVP_top60_DNA_seed | ERBB2 | -1.690751 | LUAD | NCI-H1648 | Afatinib | MVP_top60_DNA+RNA_seed | IGF1R | 0.032084 | LUAD | NCI-H2023 | NVP-ADW742 |
| MVP_top60_DNA_seed | ERBB2 | -1.688537 | LUAD | H3255 | CP724714 | MVP_top60_DNA+RNA_seed | EGFR | 0.032678 | LUAD | NCI-H2030 | Gefitinib |
| MVP_top60_DNA_seed | AURKA | -1.679723 | LUAD | NCI-H358 | Genentech Cpd 10 | MVP_top60_DNA+RNA_seed | AURKA | 0.032695 | LUAD | NCI-H1355 | Alisertib |
| MVP_top60_DNA_seed | IGF1R | -1.670658 | LUAD | LC-2-ad | Linsitinib | MVP_top60_DNA+RNA_seed | AKT1 | 0.03296 | LUAD | NCI-H358 | Uprosertib |
| MVP_top60_DNA_seed | EGFR | -1.666768 | LUAD | COR-L105 | AST-1306 | MVP_top60_DNA+RNA_seed | IGF1R | 0.033209 | LUAD | LXF-289 | GSK1904529A |
| MVP_top60_DNA_seed | IGF1R | -1.655968 | LUAD | NCI-H2009 | GSK1904529A | MVP_top60_DNA+RNA_seed | AKT1 | 0.034521 | LUAD | HOP-62 | AT13148 |
| MVP_top60_DNA_seed | IGF1R | -1.651727 | LUAD | NCI-H1437 | BMS-754807 | MVP_top60_DNA+RNA_seed | PIK3CA | 0.034876 | LUAD | Calu-3 | Alpelisib |

| | | | | | | | | | | | |
|---|---|---|---|---|---|---|---|---|---|---|---|
| MVP_top60_DNA_seed | EGFR | -1.649373 | LUAD | NCI-H1755 | Pelitinib | MVP_top60_DNA+RNA_seed | PIK3CA | 0.035645 | LUAD | EKVX | GNE-317 |
| MVP_top60_DNA_seed | EGFR | -1.648424 | LUAD | NCI-H1623 | Gefitinib | MVP_top60_DNA+RNA_seed | MDM2 | 0.037062 | LUAD | NCI-H1693 | Serdemetan |
| MVP_top60_DNA_seed | IGF1R | -1.646996 | LUAD | NCI-H1975 | BMS-536924 | MVP_top60_DNA+RNA_seed | EGFR | 0.03744 | LUAD | NCI-H2009 | AZD3759 |
| MVP_top60_DNA_seed | HSP90AA1 | -1.640404 | LUAD | NCI-H1975 | CCT-018159 | MVP_top60_DNA+RNA_seed | IGF1R | 0.038498 | LUAD | HOP-62 | BMS-536924 |
| MVP_top60_DNA_seed | MET | -1.638251 | LUAD | LC-2-ad | Crizotinib | MVP_top60_DNA+RNA_seed | TERT | 0.040347 | LUAD | NCI-H1781 | BIBR-1532 |
| MVP_top60_DNA_seed | MDM2 | -1.638181 | LUAD | NCI-H1650 | Nutlin-3a (-) | MVP_top60_DNA+RNA_seed | AKT1 | 0.041005 | LUAD | NCI-H2023 | MK-2206 |
| MVP_top60_DNA_seed | EGFR | -1.63695 | LUAD | EMC-BAC-1 | Pelitinib | MVP_top60_DNA+RNA_seed | AURKA | 0.041636 | LUAD | NCI-H358 | ZM447439 |
| MVP_top60_DNA_seed | HSP90AA1 | -1.632857 | LUAD | A549 | CCT-018159 | MVP_top60_DNA+RNA_seed | AURKA | 0.042263 | LUAD | NCI-H1755 | Alisertib |
| MVP_top60_DNA_seed | ERBB2 | -1.628149 | LUAD | NCI-H358 | CP724714 | MVP_top60_DNA+RNA_seed | EZH2 | 0.04233 | LUAD | LC-2-ad | GSK343 |
| MVP_top60_DNA_seed | IGF1R | -1.62212 | LUAD | ABC-1 | Linsitinib | MVP_top60_DNA+RNA_seed | BRAF | 0.043027 | LUAD | NCI-H2122 | Dabrafenib |
| MVP_top60_DNA_seed | EGFR | -1.619433 | LUAD | NCI-H322M | Gefitinib | MVP_top60_DNA+RNA_seed | HSP90AA1 | 0.043841 | LUAD | NCI-H1573 | Tanespimycin |
| MVP_top60_DNA_seed | IGF1R | -1.618078 | LUAD | A549 | BMS-536924 | MVP_top60_DNA+RNA_seed | IGF1R | 0.044606 | LUAD | NCI-H1734 | BMS-536924 |
| MVP_top60_DNA_seed | EGFR | -1.61604 | LUAD | HCC-827 | CUDC-101 | MVP_top60_DNA+RNA_seed | SRC | 0.04476 | LUAD | LXF-289 | Bosutinib |
| MVP_top60_DNA_seed | ERBB2 | -1.61604 | LUAD | HCC-827 | CUDC-101 | MVP_top60_DNA+RNA_seed | AKT1 | 0.045183 | LUAD | EMC-BAC-2 | Afuresertib |
| MVP_top60_DNA_seed | PIK3CA | -1.614816 | LUAD | NCI-H1648 | PI3Ka_4409 | MVP_top60_DNA+RNA_seed | AKT1 | 0.046648 | LUAD | NCI-H2085 | Uprosertib |
| MVP_top60_DNA_seed | IGF1R | -1.614691 | LUAD | NCI-H2228 | BMS-754807 | MVP_top60_DNA+RNA_seed | PIK3CA | 0.046934 | LUAD | PC-3_[JPC-3] | AZD8186 |
| MVP_top60_DNA_seed | MAPK1 | -1.61147 | LUAD | EMC-BAC-1 | FR-180204 | MVP_top60_DNA+RNA_seed | IGF1R | 0.047415 | LUAD | NCI-H1355 | BMS-536924 |
| MVP_top60_DNA_seed | GSK3B | -1.607309 | LUAD | NCI-H1975 | CHIR-99021 | MVP_top60_DNA+RNA_seed | PIK3CA | 0.048077 | LUAD | NCI-H1355 | Alpelisib |
| MVP_top60_DNA_seed | IGF1R | -1.605288 | LUAD | NCI-H1355 | BMS-536924 | MVP_top60_DNA+RNA_seed | IGF1R | 0.049028 | LUAD | NCI-H2030 | Linsitinib |
| MVP_top60_DNA_seed | HSP90AA1 | -1.595321 | LUAD | NCI-H2030 | CCT-018159 | MVP_top60_DNA+RNA_seed | IGF1R | 0.049718 | LUAD | NCI-H2085 | BMS-754807 |
| MVP_top60_DNA_seed | MAPK1 | -1.592688 | LUAD | PC-14 | FR-180204 | MVP_top60_DNA+RNA_seed | HDAC1 | 0.049943 | LUAD | RERF-LC-KJ | PCI-34051 |
| MVP_top60_DNA_seed | EGFR | -1.591446 | LUAD | PC-3_[JPC-3] | AST-1306 | MVP_top60_DNA+RNA_seed | AURKA | 0.050331 | LUAD | NCI-H3122 | ZM447439 |
| MVP_top60_DNA_seed | IGF1R | -1.582989 | LUAD | ABC-1 | BMS-754807 | MVP_top60_DNA+RNA_seed | EGFR | 0.051634 | LUAD | NCI-H1435 | Gefitinib |
| MVP_top60_DNA_seed | HSP90AA1 | -1.579497 | LUAD | NCI-H1755 | CCT-018159 | MVP_top60_DNA+RNA_seed | EZH2 | 0.0518 | LUAD | NCI-H1563 | GSK343 |
| MVP_top60_DNA_seed | EGFR | -1.578569 | LUAD | NCI-H1975 | Afatinib | MVP_top60_DNA+RNA_seed | TERT | 0.052516 | LUAD | HCC-44 | BIBR-1532 |
| MVP_top60_DNA_seed | ERBB2 | -1.578569 | LUAD | NCI-H1975 | Afatinib | MVP_top60_DNA+RNA_seed | AKT1 | 0.052836 | LUAD | NCI-H1975 | GSK2110183B |
| MVP_top60_DNA_seed | GSK3B | -1.57339 | LUAD | NCI-H2122 | CHIR-99021 | MVP_top60_DNA+RNA_seed | PIK3CA | 0.05373 | LUAD | NCI-H1563 | GNE-317 |
| MVP_top60_DNA_seed | ATM | -1.569982 | LUAD | LXF-289 | KU-60019 | MVP_top60_DNA+RNA_seed | EGFR | 0.056497 | LUAD | HOP-62 | Erlotinib |
| MVP_top60_DNA_seed | MAPK1 | -1.564003 | LUAD | NCI-H358 | FR-180204 | MVP_top60_DNA+RNA_seed | FGFR1 | 0.057428 | LUAD | NCI-H441 | PD173074 |
| MVP_top60_DNA_seed | EGFR | -1.56282 | LUAD | NCI-H1648 | PF-00299804 | MVP_top60_DNA+RNA_seed | PIK3CA | 0.061225 | LUAD | NCI-H1792 | AZD8186 |
| MVP_top60_DNA_seed | ERBB2 | -1.56282 | LUAD | NCI-H1648 | PF-00299804 | MVP_top60_DNA+RNA_seed | AURKA | 0.061335 | LUAD | NCI-H2009 | Alisertib |
| MVP_top60_DNA_seed | IGF1R | -1.55855 | LUAD | LC-2-ad | BMS-536924 | MVP_top60_DNA+RNA_seed | IGF1R | 0.062087 | LUAD | NCI-H1793 | BMS-536924 |
| MVP_top60_DNA_seed | AKT1 | -1.552066 | LUAD | NCI-H1437 | AKT inhibitor VIII | MVP_top60_DNA+RNA_seed | IGF1R | 0.062362 | LUAD | NCI-H1838 | BMS-536924 |
| MVP_top60_DNA_seed | EGFR | -1.549928 | LUAD | COR-L105 | CI-1033 | MVP_top60_DNA+RNA_seed | AKT1 | 0.062621 | LUAD | HCC-78 | GSK2110183B |
| MVP_top60_DNA_seed | ERBB2 | -1.549928 | LUAD | COR-L105 | CI-1033 | MVP_top60_DNA+RNA_seed | AURKA | 0.062881 | LUAD | NCI-H2228 | ZM447439 |
| MVP_top60_DNA_seed | ATM | -1.546048 | LUAD | A549 | KU-55933 | MVP_top60_DNA+RNA_seed | IGF1R | 0.064146 | LUAD | PC-3_[JPC-3] | GSK1904529A |
| MVP_top60_DNA_seed | AKT1 | -1.544812 | LUAD | NCI-H2030 | AKT inhibitor VIII | MVP_top60_DNA+RNA_seed | TERT | 0.067007 | LUAD | EMC-BAC-2 | BIBR-1532 |
| MVP_top60_DNA_seed | IGF1R | -1.536601 | LUAD | NCI-H1437 | GSK1904529A | MVP_top60_DNA+RNA_seed | AKT1 | 0.067023 | LUAD | RERF-LC-KJ | MK-2206 |
| MVP_top60_DNA_seed | MDM2 | -1.536493 | LUAD | A549 | Nutlin-3a (-) | MVP_top60_DNA+RNA_seed | AURKA | 0.067836 | LUAD | Calu-6 | Alisertib |
| MVP_top60_DNA_seed | HSP90AA1 | -1.528909 | LUAD | NCI-H3122 | Luminespib | MVP_top60_DNA+RNA_seed | PIK3CA | 0.069132 | LUAD | HOP-62 | Buparlisib |
| MVP_top60_DNA_seed | HDAC1 | -1.525115 | LUAD | NCI-H1944 | Dacinostat | MVP_top60_DNA+RNA_seed | ATM | 0.069168 | LUAD | NCI-H1623 | KU-55933 |

| | | | | | | | | | | | |
|---|---|---|---|---|---|---|---|---|---|---|---|
| MVP_top60_DNA_seed | AKT1 | -1.514808 | LUAD | NCI-H1975 | MK-2206 | MVP_top60_DNA+RNA_seed | HSP90AA1 | 0.069487 | LUAD | NCI-H2228 | Luminespib |
| MVP_top60_DNA_seed | ERBB2 | -1.508208 | LUAD | EMC-BAC-1 | CP724714 | MVP_top60_DNA+RNA_seed | IGF1R | 0.069612 | LUAD | NCI-H1651 | Linsitinib |
| MVP_top60_DNA_seed | EGFR | -1.505636 | LUAD | NCI-H1648 | CI-1033 | MVP_top60_DNA+RNA_seed | FGFR1 | 0.070071 | LUAD | PC-14 | PD173074 |
| MVP_top60_DNA_seed | ERBB2 | -1.505636 | LUAD | NCI-H1648 | CI-1033 | MVP_top60_DNA+RNA_seed | AURKA | 0.070921 | LUAD | NCI-H1693 | ZM447439 |
| MVP_top60_DNA_seed | SRC | -1.503924 | LUAD | COR-L105 | Bosutinib | MVP_top60_DNA+RNA_seed | PIK3CA | 0.071934 | LUAD | NCI-H358 | Alpelisib |
| MVP_top60_DNA_seed | HSP90AA1 | -1.500664 | LUAD | NCI-H23 | Tanespimycin | MVP_top60_DNA+RNA_seed | BRAF | 0.073536 | LUAD | NCI-H838 | PLX-4720 |
| MVP_top60_DNA_seed | IGF1R | -1.495104 | LUAD | NCI-H1437 | BMS-536924 | MVP_top60_DNA+RNA_seed | AKT1 | 0.073686 | LUAD | NCI-H1623 | MK-2206 |
| MVP_top60_DNA_seed | ATM | -1.491805 | LUAD | NCI-H1755 | CP466722 | MVP_top60_DNA+RNA_seed | PIK3CA | 0.073692 | LUAD | NCI-H1993 | Alpelisib |
| MVP_top60_DNA_seed | GSK3B | -1.480274 | LUAD | NCI-H2023 | SB216763 | MVP_top60_DNA+RNA_seed | AKT1 | 0.074104 | LUAD | LC-2-ad | Ipatasertib |
| MVP_top60_DNA_seed | SMARCA4 | -1.475455 | LUAD | NCI-H1666 | PFI-3 | MVP_top60_DNA+RNA_seed | HDAC1 | 0.074211 | LUAD | NCI-H2228 | Romidepsin |
| MVP_top60_DNA_seed | MET | -1.469508 | LUAD | NCI-H1781 | JNJ38877605 | MVP_top60_DNA+RNA_seed | MET | 0.074394 | LUAD | NCI-H1781 | Savolitinib |
| MVP_top60_DNA_seed | IGF1R | -1.46746 | LUAD | NCI-H650 | Linsitinib | MVP_top60_DNA+RNA_seed | SRC | 0.075854 | LUAD | NCI-H1838 | Bosutinib |
| MVP_top60_DNA_seed | ATM | -1.465962 | LUAD | NCI-H2030 | KU-55933 | MVP_top60_DNA+RNA_seed | SRC | 0.076116 | LUAD | COR-L105 | Bosutinib |
| MVP_top60_DNA_seed | IGF1R | -1.46417 | LUAD | NCI-H1666 | BMS-536924 | MVP_top60_DNA+RNA_seed | AURKA | 0.077132 | LUAD | NCI-H322M | ZM447439 |
| MVP_top60_DNA_seed | AURKA | -1.460825 | LUAD | LXF-289 | HG-5-113-01 | MVP_top60_DNA+RNA_seed | EGFR | 0.077419 | LUAD | NCI-H522 | Erlotinib |
| MVP_top60_DNA_seed | HDAC1 | -1.455237 | LUAD | HOP-62 | JQ12 | MVP_top60_DNA+RNA_seed | AKT1 | 0.079151 | LUAD | HCC-827 | AT13148 |
| MVP_top60_DNA_seed | AKT1 | -1.454531 | LUAD | LXF-289 | MK-2206 | MVP_top60_DNA+RNA_seed | IGF1R | 0.079817 | LUAD | NCI-H2030 | BMS-754807 |
| MVP_top60_DNA_seed | MAPK1 | -1.452658 | LUAD | HCC-827 | FR-180204 | MVP_top60_DNA+RNA_seed | FGFR1 | 0.08132 | LUAD | NCI-H650 | AZD4547 |
| MVP_top60_DNA_seed | ATM | -1.448614 | LUAD | LXF-289 | KU-55933 | MVP_top60_DNA+RNA_seed | SMARCA4 | 0.081745 | LUAD | NCI-H838 | PFI3 |
| MVP_top60_DNA_seed | IGF1R | -1.448517 | LUAD | NCI-H838 | BMS-536924 | MVP_top60_DNA+RNA_seed | EZH2 | 0.082741 | LUAD | NCI-H1755 | GSK343 |
| MVP_top60_DNA_seed | EGFR | -1.447333 | LUAD | EKVX | Afatinib | MVP_top60_DNA+RNA_seed | EGFR | 0.08346 | LUAD | NCI-H1944 | Gefitinib |
| MVP_top60_DNA_seed | ERBB2 | -1.447333 | LUAD | EKVX | Afatinib | MVP_top60_DNA+RNA_seed | AURKA | 0.085031 | LUAD | NCI-H838 | Alisertib |
| MVP_top60_DNA_seed | HSP90AA1 | -1.446474 | LUAD | PC-14 | CCT-018159 | MVP_top60_DNA+RNA_seed | AKT1 | 0.085453 | LUAD | NCI-H2228 | Ipatasertib |
| MVP_top60_DNA_seed | HDAC1 | -1.445597 | LUAD | LC-2-ad | Dacinostat | MVP_top60_DNA+RNA_seed | MDM2 | 0.085538 | LUAD | NCI-H3122 | Serdemetan |
| MVP_top60_DNA_seed | HSP90AA1 | -1.428277 | LUAD | NCI-H1563 | Elesclomol | MVP_top60_DNA+RNA_seed | HDAC1 | 0.085555 | LUAD | NCI-H1734 | PCI-34051 |
| MVP_top60_DNA_seed | FGFR1 | -1.426335 | LUAD | LC-2-ad | FGFR_3831 | MVP_top60_DNA+RNA_seed | EGFR | 0.088506 | LUAD | RERF-LC-MS | Erlotinib |
| MVP_top60_DNA_seed | BRAF | -1.415592 | LUAD | NCI-H1781 | PLX-4720 | MVP_top60_DNA+RNA_seed | AURKA | 0.088896 | LUAD | RERF-LC-KJ | Alisertib |
| MVP_top60_DNA_seed | HSP90AA1 | -1.413799 | LUAD | NCI-H1975 | Tanespimycin | MVP_top60_DNA+RNA_seed | HSP90AA1 | 0.089848 | LUAD | Calu-3 | Tanespimycin |
| MVP_top60_DNA_seed | AKT1 | -1.408717 | LUAD | NCI-H1944 | AKT inhibitor VIII | MVP_top60_DNA+RNA_seed | EGFR | 0.091029 | LUAD | NCI-H1792 | Afatinib |
| MVP_top60_DNA_seed | HDAC1 | -1.408523 | LUAD | HCC-44 | Tubastatin A | MVP_top60_DNA+RNA_seed | EGFR | 0.091426 | LUAD | NCI-H1435 | Sapitinib |
| MVP_top60_DNA_seed | IGF1R | -1.402368 | LUAD | LC-2-ad | BMS-754807 | MVP_top60_DNA+RNA_seed | SMARCA4 | 0.092584 | LUAD | NCI-H1437 | PFI3 |
| MVP_top60_DNA_seed | EGFR | -1.401813 | LUAD | NCI-H1355 | Erlotinib | MVP_top60_DNA+RNA_seed | EZH2 | 0.092719 | LUAD | NCI-H1437 | GSK343 |
| MVP_top60_DNA_seed | IGF1R | -1.398164 | LUAD | ABC-1 | BMS-536924 | MVP_top60_DNA+RNA_seed | BRAF | 0.092735 | LUAD | ABC-1 | Dabrafenib |
| MVP_top60_DNA_seed | IGF1R | -1.395073 | LUAD | NCI-H2228 | BMS-536924 | MVP_top60_DNA+RNA_seed | EGFR | 0.094703 | LUAD | NCI-H2085 | Osimertinib |
| MVP_top60_DNA_seed | GSK3B | -1.393308 | LUAD | PC-14 | CHIR-99021 | MVP_top60_DNA+RNA_seed | EGFR | 0.094742 | LUAD | NCI-H1755 | Erlotinib |
| MVP_top60_DNA_seed | HDAC1 | -1.392221 | LUAD | NCI-H522 | Dacinostat | MVP_top60_DNA+RNA_seed | AKT1 | 0.095073 | LUAD | NCI-H322M | AZD5363 |
| MVP_top60_DNA_seed | ATM | -1.392128 | LUAD | NCI-H2030 | KU-60019 | MVP_top60_DNA+RNA_seed | ATM | 0.095384 | LUAD | NCI-H2009 | KU-55933 |
| MVP_top60_DNA_seed | AURKA | -1.387894 | LUAD | LXF-289 | Tozasertib | MVP_top60_DNA+RNA_seed | IGF1R | 0.095522 | LUAD | NCI-H3122 | BMS-754807 |
| MVP_top60_DNA_seed | EGFR | -1.387586 | LUAD | RERF-LC-KJ | Cetuximab | MVP_top60_DNA+RNA_seed | PIK3CA | 0.097448 | LUAD | NCI-H1568 | AZD8186 |
| MVP_top60_DNA_seed | AKT1 | -1.384174 | LUAD | COR-L105 | MK-2206 | MVP_top60_DNA+RNA_seed | IGF1R | 0.097899 | LUAD | NCI-H1355 | GSK1904529A |
| MVP_top60_DNA_seed | EGFR | -1.381905 | LUAD | NCI-H1648 | AZD8931 | MVP_top60_DNA+RNA_seed | EZH2 | 0.098676 | LUAD | NCI-H1651 | GSK343 |

| | | | | | | | | | | | |
|---|---|---|---|---|---|---|---|---|---|---|---|
| MVP_top60_DNA_seed | ERBB2 | -1.381905 | LUAD | NCI-H1648 | AZD8931 | MVP_top60_DNA+RNA_seed | PIK3CA | 0.098733 | LUAD | NCI-H2009 | AZD8186 |
| MVP_top60_DNA_seed | ERBB3 | -1.381905 | LUAD | NCI-H1648 | AZD8931 | MVP_top60_DNA+RNA_seed | EGFR | 0.099261 | LUAD | ABC-1 | Gefitinib |
| MVP_top60_DNA_seed | IGF1R | -1.381452 | LUAD | SK-LU-1 | BMS-754807 | MVP_top60_DNA+RNA_seed | FGFR1 | 0.09973 | LUAD | RERF-LC-MS | PD173074 |
| MVP_top60_DNA_seed | MDM2 | -1.378541 | LUAD | NCI-H2122 | Nutlin-3a (-) | MVP_top60_DNA+RNA_seed | KRAS | 0.100027 | LUAD | NCI-H1651 | (RAS (G12C) Inhibitor-1 |
| MVP_top60_DNA_seed | AURKA | -1.375024 | LUAD | LXF-289 | ZM447439 | MVP_top60_DNA+RNA_seed | EGFR | 0.100266 | LUAD | NCI-H1651 | Osimertinib |
| MVP_top60_DNA_seed | MDM2 | -1.374781 | LUAD | NCI-H1651 | Serdemetan | MVP_top60_DNA+RNA_seed | AKT1 | 0.101856 | LUAD | H3255 | AZD5363 |
| MVP_top60_DNA_seed | HDAC1 | -1.374094 | LUAD | HCC-44 | JQ12 | MVP_top60_DNA+RNA_seed | EGFR | 0.102203 | LUAD | NCI-H2291 | Afatinib |
| MVP_top60_DNA_seed | HSP90AA1 | -1.374089 | LUAD | PC-14 | Tanespimycin | MVP_top60_DNA+RNA_seed | AKT1 | 0.103369 | LUAD | VMRC-LCD | AT13148 |
| MVP_top60_DNA_seed | EGFR | -1.372142 | LUAD | NCI-H1648 | AST-1306 | MVP_top60_DNA+RNA_seed | PIK3CA | 0.104288 | LUAD | NCI-H2228 | Buparlisib |
| MVP_top60_DNA_seed | SRC | -1.36834 | LUAD | NCI-H1648 | WH-4-023 | MVP_top60_DNA+RNA_seed | SMARCA4 | 0.104449 | LUAD | NCI-H1623 | PFI3 |
| MVP_top60_DNA_seed | GSK3B | -1.368122 | LUAD | NCI-H650 | SB216763 | MVP_top60_DNA+RNA_seed | AKT1 | 0.10463 | LUAD | NCI-H2291 | MK-2206 |
| MVP_top60_DNA_seed | HSP90AA1 | -1.366877 | LUAD | SK-LU-1 | Tanespimycin | MVP_top60_DNA+RNA_seed | BRAF | 0.104718 | LUAD | NCI-H3122 | Dabrafenib |
| MVP_top60_DNA_seed | GSK3B | -1.365589 | LUAD | NCI-H2085 | SB216763 | MVP_top60_DNA+RNA_seed | KRAS | 0.105467 | LUAD | H3255 | (RAS (G12C) Inhibitor-1 |
| MVP_top60_DNA_seed | MET | -1.36463 | LUAD | HOP-62 | JNJ38877605 | MVP_top60_DNA+RNA_seed | FGFR1 | 0.10613 | LUAD | NCI-H1781 | PD173074 |
| MVP_top60_DNA_seed | MET | -1.36347 | LUAD | NCI-H2342 | JNJ38877605 | MVP_top60_DNA+RNA_seed | AURKA | 0.107067 | LUAD | NCI-H1437 | Alisertib |
| MVP_top60_DNA_seed | GSK3B | -1.354358 | LUAD | NCI-H1792 | SB216763 | MVP_top60_DNA+RNA_seed | HDAC1 | 0.108466 | LUAD | NCI-H358 | Romidepsin |
| MVP_top60_DNA_seed | HSP90AA1 | -1.35179 | LUAD | NCI-H838 | Tanespimycin | MVP_top60_DNA+RNA_seed | EGFR | 0.10854 | LUAD | HCC-78 | Lapatinib |
| MVP_top60_DNA_seed | MAPK1 | -1.350128 | LUAD | LC-2-ad | VX-11e | MVP_top60_DNA+RNA_seed | TP53 | 0.109622 | LUAD | NCI-H2009 | MIRA-1 |
| MVP_top60_DNA_seed | AKT1 | -1.346078 | LUAD | Calu-3 | AKT inhibitor VIII | MVP_top60_DNA+RNA_seed | EGFR | 0.110253 | LUAD | LXF-289 | AZD3759 |
| MVP_top60_DNA_seed | IGF1R | -1.343221 | LUAD | NCI-H2085 | GSK1904529A | MVP_top60_DNA+RNA_seed | EZH2 | 0.11176 | LUAD | NCI-H1781 | GSK343 |
| MVP_top60_DNA_seed | CREBBP | -1.341935 | LUAD | NCI-H1944 | I-CBP112 | MVP_top60_DNA+RNA_seed | HSP90AA1 | 0.113227 | LUAD | Calu-6 | Luminespib |
| MVP_top60_DNA_seed | HDAC1 | -1.332298 | LUAD | NCI-H1975 | Dacinostat | MVP_top60_DNA+RNA_seed | FGFR1 | 0.113932 | LUAD | HCC-78 | PD173074 |
| MVP_top60_DNA_seed | EGFR | -1.331915 | LUAD | NCI-H1568 | AST-1306 | MVP_top60_DNA+RNA_seed | IGF1R | 0.114046 | LUAD | VMRC-LCD | BMS-754807 |
| MVP_top60_DNA_seed | GSK3B | -1.328385 | LUAD | H3255 | KIN001-042 | MVP_top60_DNA+RNA_seed | FGFR1 | 0.114183 | LUAD | PC-14 | AZD4547 |
| MVP_top60_DNA_seed | AKT1 | -1.318789 | LUAD | NCI-H838 | MK-2206 | MVP_top60_DNA+RNA_seed | AKT1 | 0.114972 | LUAD | NCI-H358 | Ipatasertib |
| MVP_top60_DNA_seed | IGF1R | -1.318446 | LUAD | HCC-44 | GSK1904529A | MVP_top60_DNA+RNA_seed | AKT1 | 0.116562 | LUAD | NCI-H1944 | Ipatasertib |
| MVP_top60_DNA_seed | HSP90AA1 | -1.317377 | LUAD | COR-L105 | Elesclomol | MVP_top60_DNA+RNA_seed | IGF1R | 0.117034 | LUAD | RERF-LC-KJ | Linsitinib |
| MVP_top60_DNA_seed | AURKA | -1.315361 | LUAD | NCI-H1944 | Genentech Cpd 10 | MVP_top60_DNA+RNA_seed | EGFR | 0.118086 | LUAD | NCI-H1781 | AZD3759 |
| MVP_top60_DNA_seed | HDAC1 | -1.311584 | LUAD | NCI-H2122 | JQ12 | MVP_top60_DNA+RNA_seed | AKT1 | 0.118591 | LUAD | EMC-BAC-2 | Uprosertib |
| MVP_top60_DNA_seed | PIK3CA | -1.311541 | LUAD | NCI-H1648 | AZD8835 | MVP_top60_DNA+RNA_seed | AKT1 | 0.11881 | LUAD | NCI-H3122 | Ipatasertib |
| MVP_top60_DNA_seed | IGF1R | -1.309301 | LUAD | NCI-H1975 | Linsitinib | MVP_top60_DNA+RNA_seed | EGFR | 0.119348 | LUAD | NCI-H1573 | Lapatinib |
| MVP_top60_DNA_seed | GSK3B | -1.308885 | LUAD | HCC-78 | KIN001-042 | MVP_top60_DNA+RNA_seed | EGFR | 0.119846 | LUAD | NCI-H2291 | Osimertinib |
| MVP_top60_DNA_seed | IGF1R | -1.305055 | LUAD | NCI-H1355 | BMS-536924 | MVP_top60_DNA+RNA_seed | PIK3CA | 0.120852 | LUAD | NCI-H1666 | GNE-317 |
| MVP_top60_DNA_seed | EGFR | -1.300028 | LUAD | EKVX | Afatinib | MVP_top60_DNA+RNA_seed | IGF1R | 0.121211 | LUAD | NCI-H2023 | BMS-754807 |
| MVP_top60_DNA_seed | ERBB2 | -1.300028 | LUAD | EKVX | Afatinib | MVP_top60_DNA+RNA_seed | TP53 | 0.122093 | LUAD | VMRC-LCD | MIRA-1 |
| MVP_top60_DNA_seed | HSP90AA1 | -1.299512 | LUAD | NCI-H2030 | Tanespimycin | MVP_top60_DNA+RNA_seed | SRC | 0.124534 | LUAD | NCI-H1563 | Bosutinib |
| MVP_top60_DNA_seed | SRC | -1.297763 | LUAD | HOP-62 | WH-4-023 | MVP_top60_DNA+RNA_seed | EGFR | 0.126314 | LUAD | NCI-H838 | Lapatinib |
| MVP_top60_DNA_seed | IGF1R | -1.296557 | LUAD | NCI-H1944 | BMS-536924 | MVP_top60_DNA+RNA_seed | AKT1 | 0.126329 | LUAD | NCI-H322M | Uprosertib |
| MVP_top60_DNA_seed | EGFR | -1.296158 | LUAD | PC-3_[JPC-3] | PF-00299804 | MVP_top60_DNA+RNA_seed | EGFR | 0.126863 | LUAD | NCI-H2085 | Gefitinib |
| MVP_top60_DNA_seed | ERBB2 | -1.296158 | LUAD | PC-3_[JPC-3] | PF-00299804 | MVP_top60_DNA+RNA_seed | EZH2 | 0.127126 | LUAD | NCI-H2085 | GSK343 |
| MVP_top60_DNA_seed | ERBB2 | -1.291878 | LUAD | ABC-1 | CP724714 | MVP_top60_DNA+RNA_seed | TERT | 0.12808 | LUAD | RERF-LC-MS | BIBR-1532 |

| | | | | | | | | | | | |
|---|---|---|---|---|---|---|---|---|---|---|---|
| MVP_top60_DNA_seed | SRC | -1.291483 | LUAD | HOP-62 | Saracatinib | MVP_top60_DNA+RNA_seed | AURKA | 0.128692 | LUAD | NCI-H1623 | ZM447439 |
| MVP_top60_DNA_seed | EGFR | -1.285613 | LUAD | NCI-H2030 | AZD8931 | MVP_top60_DNA+RNA_seed | IGF1R | 0.129808 | LUAD | NCI-H522 | Linsitinib |
| MVP_top60_DNA_seed | ERBB2 | -1.285613 | LUAD | NCI-H2030 | AZD8931 | MVP_top60_DNA+RNA_seed | EGFR | 0.130069 | LUAD | NCI-H1734 | Gefitinib |
| MVP_top60_DNA_seed | ERBB3 | -1.285613 | LUAD | NCI-H2030 | AZD8931 | MVP_top60_DNA+RNA_seed | AKT1 | 0.130914 | LUAD | NCI-H2342 | MK-2206 |
| MVP_top60_DNA_seed | CREBBP | -1.281727 | LUAD | RERF-LC-KJ | I-CBP112 | MVP_top60_DNA+RNA_seed | IGF1R | 0.132113 | LUAD | NCI-H1781 | Linsitinib |
| MVP_top60_DNA_seed | SMARCA4 | -1.280394 | LUAD | VMRC-LCD | PFI-3 | MVP_top60_DNA+RNA_seed | BRAF | 0.133153 | LUAD | SK-LU-1 | PLX-4720 |
| MVP_top60_DNA_seed | EGFR | -1.275361 | LUAD | NCI-H322M | CI-1033 | MVP_top60_DNA+RNA_seed | SMARCA4 | 0.133777 | LUAD | NCI-H1568 | PFI3 |
| MVP_top60_DNA_seed | ERBB2 | -1.275361 | LUAD | NCI-H322M | CI-1033 | MVP_top60_DNA+RNA_seed | ATM | 0.135492 | LUAD | NCI-H1437 | KU-55933 |
| MVP_top60_DNA_seed | EGFR | -1.272352 | LUAD | NCI-H322M | PF-00299804 | MVP_top60_DNA+RNA_seed | IGF1R | 0.136193 | LUAD | NCI-H1944 | BMS-536924 |
| MVP_top60_DNA_seed | ERBB2 | -1.272352 | LUAD | NCI-H322M | PF-00299804 | MVP_top60_DNA+RNA_seed | AKT1 | 0.136211 | LUAD | NCI-H2030 | AZD5363 |
| MVP_top60_DNA_seed | SRC | -1.272267 | LUAD | PC-3_[JPC-3] | Bosutinib | MVP_top60_DNA+RNA_seed | BRAF | 0.137098 | LUAD | NCI-H2030 | Dabrafenib |
| MVP_top60_DNA_seed | AURKA | -1.270062 | LUAD | ABC-1 | ZM447439 | MVP_top60_DNA+RNA_seed | EGFR | 0.137336 | LUAD | NCI-H1650 | Afatinib |
| MVP_top60_DNA_seed | BRAF | -1.26912 | LUAD | Calu-6 | Dabrafenib | MVP_top60_DNA+RNA_seed | AKT1 | 0.137756 | LUAD | NCI-H2342 | AT13148 |
| MVP_top60_DNA_seed | ATM | -1.266769 | LUAD | NCI-H1437 | KU-55933 | MVP_top60_DNA+RNA_seed | IGF1R | 0.138291 | LUAD | NCI-H650 | Linsitinib |
| MVP_top60_DNA_seed | GSK3B | -1.260336 | LUAD | Calu-6 | SB216763 | MVP_top60_DNA+RNA_seed | BRAF | 0.138661 | LUAD | NCI-H23 | SB590885 |
| MVP_top60_DNA_seed | IGF1R | -1.252872 | LUAD | NCI-H1792 | BMS-536924 | MVP_top60_DNA+RNA_seed | PIK3CA | 0.141414 | LUAD | PC-14 | AZD8186 |
| MVP_top60_DNA_seed | SRC | -1.250732 | LUAD | NCI-H1693 | Bosutinib | MVP_top60_DNA+RNA_seed | PIK3CA | 0.143469 | LUAD | PC-14 | Alpelisib |
| MVP_top60_DNA_seed | AKT1 | -1.246958 | LUAD | RERF-LC-KJ | AKT inhibitor VIII | MVP_top60_DNA+RNA_seed | AKT1 | 0.143472 | LUAD | NCI-H2342 | Uprosertib |
| MVP_top60_DNA_seed | EGFR | -1.246743 | LUAD | PC-3_[JPC-3] | CI-1033 | MVP_top60_DNA+RNA_seed | EGFR | 0.144492 | LUAD | HCC-44 | Sapitinib |
| MVP_top60_DNA_seed | ERBB2 | -1.246743 | LUAD | PC-3_[JPC-3] | CI-1033 | MVP_top60_DNA+RNA_seed | MET | 0.14543 | LUAD | NCI-H1568 | Savolitinib |
| MVP_top60_DNA_seed | EGFR | -1.244831 | LUAD | NCI-H358 | Afatinib | MVP_top60_DNA+RNA_seed | AKT1 | 0.147646 | LUAD | NCI-H1666 | Afuresertib |
| MVP_top60_DNA_seed | ERBB2 | -1.244831 | LUAD | NCI-H358 | Afatinib | MVP_top60_DNA+RNA_seed | EGFR | 0.148304 | LUAD | NCI-H2023 | Afatinib |
| MVP_top60_DNA_seed | FGFR1 | -1.244259 | LUAD | HCC-44 | FGFR_3831 | MVP_top60_DNA+RNA_seed | FGFR1 | 0.15023 | LUAD | HCC-44 | AZD4547 |
| MVP_top60_DNA_seed | SRC | -1.243449 | LUAD | NCI-H1975 | WH-4-023 | MVP_top60_DNA+RNA_seed | AURKA | 0.150242 | LUAD | ABC-1 | Alisertib |
| MVP_top60_DNA_seed | EGFR | -1.241932 | LUAD | NCI-H1693 | Afatinib | MVP_top60_DNA+RNA_seed | IGF1R | 0.150999 | LUAD | NCI-H23 | Linsitinib |
| MVP_top60_DNA_seed | ERBB2 | -1.241932 | LUAD | NCI-H1693 | Afatinib | MVP_top60_DNA+RNA_seed | BRAF | 0.151019 | LUAD | NCI-H1781 | PLX-4720 |
| MVP_top60_DNA_seed | IGF1R | -1.24086 | LUAD | NCI-H3122 | GSK1904529A | MVP_top60_DNA+RNA_seed | HDAC1 | 0.151036 | LUAD | NCI-H358 | PCI-34051 |
| MVP_top60_DNA_seed | EGFR | -1.240674 | LUAD | NCI-H322M | Erlotinib | MVP_top60_DNA+RNA_seed | AURKA | 0.154787 | LUAD | H3255 | ZM447439 |
| MVP_top60_DNA_seed | IGF1R | -1.237844 | LUAD | NCI-H650 | BMS-754807 | MVP_top60_DNA+RNA_seed | MDM2 | 0.15482 | LUAD | HCC-827 | Serdemetan |
| MVP_top60_DNA_seed | EGFR | -1.237654 | LUAD | NCI-H1568 | Afatinib | MVP_top60_DNA+RNA_seed | EGFR | 0.155015 | LUAD | NCI-H522 | Osimertinib |
| MVP_top60_DNA_seed | ERBB2 | -1.237654 | LUAD | NCI-H1568 | Afatinib | MVP_top60_DNA+RNA_seed | MDM2 | 0.15502 | LUAD | NCI-H1623 | Serdemetan |
| MVP_top60_DNA_seed | PIK3CA | -1.237086 | LUAD | HCC-78 | AZD8835 | MVP_top60_DNA+RNA_seed | MET | 0.156031 | LUAD | NCI-H1693 | Savolitinib |
| MVP_top60_DNA_seed | FGFR1 | -1.236905 | LUAD | NCI-H2122 | FGFR_3831 | MVP_top60_DNA+RNA_seed | IGF1R | 0.158773 | LUAD | NCI-H23 | NVP-ADW742 |
| MVP_top60_DNA_seed | MDM2 | -1.236763 | LUAD | NCI-H358 | Nutlin-3a (-) | MVP_top60_DNA+RNA_seed | FGFR1 | 0.160446 | LUAD | NCI-H2030 | AZD4547 |
| MVP_top60_DNA_seed | IGF1R | -1.229092 | LUAD | COR-L105 | BMS-754807 | MVP_top60_DNA+RNA_seed | HDAC1 | 0.160706 | LUAD | HCC-44 | Entinostat |
| MVP_top60_DNA_seed | IGF1R | -1.227584 | LUAD | RERF-LC-KJ | Linsitinib | MVP_top60_DNA+RNA_seed | AKT1 | 0.161848 | LUAD | ABC-1 | AT13148 |
| MVP_top60_DNA_seed | MAPK1 | -1.227382 | LUAD | NCI-H1792 | VX-11e | MVP_top60_DNA+RNA_seed | EGFR | 0.163014 | LUAD | NCI-H2009 | Erlotinib |
| MVP_top60_DNA_seed | MET | -1.222975 | LUAD | HOP-62 | SU11274 | MVP_top60_DNA+RNA_seed | PIK3CA | 0.163809 | LUAD | NCI-H1573 | Buparlisib |
| MVP_top60_DNA_seed | HSP90AA1 | -1.222171 | LUAD | NCI-H3122 | SNX-2112 | MVP_top60_DNA+RNA_seed | FGFR1 | 0.164048 | LUAD | NCI-H1435 | AZD4547 |
| MVP_top60_DNA_seed | MAPK1 | -1.221877 | LUAD | NCI-H358 | VX-11e | MVP_top60_DNA+RNA_seed | TERT | 0.164093 | LUAD | NCI-H1437 | BIBR-1532 |
| MVP_top60_DNA_seed | MDM2 | -1.221706 | LUAD | NCI-H1693 | Serdemetan | MVP_top60_DNA+RNA_seed | SMARCA4 | 0.164566 | LUAD | NCI-H2085 | PFI3 |

| | | | | | | | | | | | |
|---|---|---|---|---|---|---|---|---|---|---|---|
| MVP_top60_DNA_seed | MET | -1.217885 | LUAD | EMC-BAC-2 | AZD6094 | MVP_top60_DNA+RNA_seed | BRAF | 0.16549 | LUAD | LC-2-ad | SB590885 |
| MVP_top60_DNA_seed | IGF1R | -1.217236 | LUAD | NCI-H441 | BMS-754807 | MVP_top60_DNA+RNA_seed | SRC | 0.165604 | LUAD | EMC-BAC-2 | Bosutinib |
| MVP_top60_DNA_seed | IGF1R | -1.215199 | LUAD | NCI-H1648 | BMS-754807 | MVP_top60_DNA+RNA_seed | ATM | 0.166237 | LUAD | RERF-LC-MS | KU-55933 |
| MVP_top60_DNA_seed | HDAC1 | -1.215023 | LUAD | NCI-H2122 | Dacinostat | MVP_top60_DNA+RNA_seed | BRAF | 0.167775 | LUAD | RERF-LC-MS | SB590885 |
| MVP_top60_DNA_seed | IGF1R | -1.214805 | LUAD | NCI-H441 | Linsitinib | MVP_top60_DNA+RNA_seed | EGFR | 0.168259 | LUAD | NCI-H2342 | Afatinib |
| MVP_top60_DNA_seed | EGFR | -1.210845 | LUAD | LC-2-ad | Erlotinib | MVP_top60_DNA+RNA_seed | HDAC1 | 0.169077 | LUAD | NCI-H1623 | Romidepsin |
| MVP_top60_DNA_seed | BRAF | -1.206378 | LUAD | HCC-827 | PLX-4720 | MVP_top60_DNA+RNA_seed | IGF1R | 0.169792 | LUAD | LC-2-ad | Linsitinib |
| MVP_top60_DNA_seed | AKT1 | -1.206372 | LUAD | NCI-H1993 | AKT inhibitor VIII | MVP_top60_DNA+RNA_seed | AKT1 | 0.169824 | LUAD | NCI-H1563 | Uprosertib |
| MVP_top60_DNA_seed | HDAC1 | -1.206119 | LUAD | NCI-H1944 | JQ12 | MVP_top60_DNA+RNA_seed | EGFR | 0.172427 | LUAD | NCI-H1734 | Osimertinib |
| MVP_top60_DNA_seed | EGFR | -1.205867 | LUAD | COR-L105 | Gefitinib | MVP_top60_DNA+RNA_seed | AKT1 | 0.172556 | LUAD | NCI-H2009 | Ipatasertib |
| MVP_top60_DNA_seed | HSP90AA1 | -1.203443 | LUAD | HCC-44 | Luminespib | MVP_top60_DNA+RNA_seed | IGF1R | 0.17296 | LUAD | NCI-H1838 | Linsitinib |
| MVP_top60_DNA_seed | AURKA | -1.198895 | LUAD | LC-2-ad | ZM447439 | MVP_top60_DNA+RNA_seed | IGF1R | 0.173024 | LUAD | NCI-H2342 | GSK1904529A |
| MVP_top60_DNA_seed | AKT1 | -1.197261 | LUAD | NCI-H1355 | A-443654 | MVP_top60_DNA+RNA_seed | EGFR | 0.173347 | LUAD | NCI-H1563 | AZD3759 |
| MVP_top60_DNA_seed | HDAC1 | -1.196574 | LUAD | LC-2-ad | Tubastatin A | MVP_top60_DNA+RNA_seed | MDM2 | 0.174225 | LUAD | NCI-H23 | Nutlin-3a (-) |
| MVP_top60_DNA_seed | GSK3B | -1.194133 | LUAD | LXF-289 | CHIR-99021 | MVP_top60_DNA+RNA_seed | AKT1 | 0.17425 | LUAD | NCI-H2291 | Ipatasertib |
| MVP_top60_DNA_seed | SRC | -1.191638 | LUAD | LXF-289 | Saracatinib | MVP_top60_DNA+RNA_seed | MET | 0.174437 | LUAD | LC-2-ad | Savolitinib |
| MVP_top60_DNA_seed | AURKA | -1.187684 | LUAD | EMC-BAC-2 | HG-5-113-01 | MVP_top60_DNA+RNA_seed | AURKA | 0.175629 | LUAD | PC-14 | Tozasertib |
| MVP_top60_DNA_seed | EGFR | -1.186951 | LUAD | NCI-H1975 | Afatinib | MVP_top60_DNA+RNA_seed | FGFR1 | 0.175637 | LUAD | NCI-H1792 | AZD4547 |
| MVP_top60_DNA_seed | ERBB2 | -1.186951 | LUAD | NCI-H1975 | Afatinib | MVP_top60_DNA+RNA_seed | HSP90AA1 | 0.175698 | LUAD | NCI-H1734 | CCT-018159 |
| MVP_top60_DNA_seed | MET | -1.185481 | LUAD | H3255 | JNJ38877605 | MVP_top60_DNA+RNA_seed | IGF1R | 0.176353 | LUAD | NCI-H358 | NVP-ADW742 |
| MVP_top60_DNA_seed | HSP90AA1 | -1.184703 | LUAD | LC-2-ad | Luminespib | MVP_top60_DNA+RNA_seed | KRAS | 0.179106 | LUAD | HOP-62 | KRAS (G12C) Inhibitor-1 |
| MVP_top60_DNA_seed | BRAF | -1.18469 | LUAD | NCI-H1648 | PLX-4720 | MVP_top60_DNA+RNA_seed | AKT1 | 0.181764 | LUAD | NCI-H1666 | Uprosertib |
| MVP_top60_DNA_seed | BRAF | -1.184439 | LUAD | NCI-H1648 | AZ628 | MVP_top60_DNA+RNA_seed | FGFR1 | 0.182429 | LUAD | NCI-H1993 | PD173074 |
| MVP_top60_DNA_seed | HSP90AA1 | -1.183128 | LUAD | NCI-H1975 | Elesclomol | MVP_top60_DNA+RNA_seed | FGFR1 | 0.182714 | LUAD | HCC-827 | AZD4547 |
| MVP_top60_DNA_seed | IGF1R | -1.181323 | LUAD | SK-LU-1 | Linsitinib | MVP_top60_DNA+RNA_seed | FGFR1 | 0.183337 | LUAD | NCI-H358 | PD173074 |
| MVP_top60_DNA_seed | HSP90AA1 | -1.181132 | LUAD | HCC-827 | CCT-018159 | MVP_top60_DNA+RNA_seed | AKT1 | 0.18356 | LUAD | NCI-H1437 | Afuresertib |
| MVP_top60_DNA_seed | AKT1 | -1.179511 | LUAD | ABC-1 | MK-2206 | MVP_top60_DNA+RNA_seed | MDM2 | 0.184045 | LUAD | NCI-H838 | Nutlin-3a (-) |
| MVP_top60_DNA_seed | HDAC1 | -1.178192 | LUAD | NCI-H1755 | CAY10603 | MVP_top60_DNA+RNA_seed | TP53 | 0.184186 | LUAD | NCI-H838 | MIRA-1 |
| MVP_top60_DNA_seed | BRAF | -1.177737 | LUAD | NCI-H1975 | HG6-64-1 | MVP_top60_DNA+RNA_seed | BRAF | 0.186343 | LUAD | NCI-H838 | Dabrafenib |
| MVP_top60_DNA_seed | EGFR | -1.175376 | LUAD | NCI-H1355 | Cetuximab | MVP_top60_DNA+RNA_seed | EGFR | 0.187921 | LUAD | NCI-H1755 | Osimertinib |
| MVP_top60_DNA_seed | EGFR | -1.174421 | LUAD | NCI-H2122 | AZD8931 | MVP_top60_DNA+RNA_seed | IGF1R | 0.18893 | LUAD | NCI-H1355 | BMS-754807 |
| MVP_top60_DNA_seed | ERBB2 | -1.174421 | LUAD | NCI-H2122 | AZD8931 | MVP_top60_DNA+RNA_seed | EGFR | 0.18905 | LUAD | NCI-H23 | Gefitinib |
| MVP_top60_DNA_seed | ERBB3 | -1.174421 | LUAD | NCI-H2122 | AZD8931 | MVP_top60_DNA+RNA_seed | ATM | 0.191098 | LUAD | LC-2-ad | KU-55933 |
| MVP_top60_DNA_seed | ATM | -1.170695 | LUAD | HCC-78 | KU-60019 | MVP_top60_DNA+RNA_seed | EZH2 | 0.191453 | LUAD | EMC-BAC-1 | GSK343 |
| MVP_top60_DNA_seed | EGFR | -1.164955 | LUAD | NCI-H1648 | Afatinib | MVP_top60_DNA+RNA_seed | BRAF | 0.192791 | LUAD | NCI-H1993 | SB590885 |
| MVP_top60_DNA_seed | ERBB2 | -1.164955 | LUAD | NCI-H1648 | Afatinib | MVP_top60_DNA+RNA_seed | TP53 | 0.194821 | LUAD | NCI-H1648 | MIRA-1 |
| MVP_top60_DNA_seed | IGF1R | -1.164914 | LUAD | EKVX | BMS-754807 | MVP_top60_DNA+RNA_seed | HDAC1 | 0.195672 | LUAD | LXF-289 | Entinostat |
| MVP_top60_DNA_seed | HSP90AA1 | -1.164778 | LUAD | NCI-H358 | SNX-2112 | MVP_top60_DNA+RNA_seed | FGFR1 | 0.198604 | LUAD | NCI-H2030 | PD173074 |
| MVP_top60_DNA_seed | AKT1 | -1.16414 | LUAD | NCI-H1975 | AKT inhibitor VIII | MVP_top60_DNA+RNA_seed | EGFR | 0.198991 | LUAD | NCI-H1734 | Lapatinib |
| MVP_top60_DNA_seed | HSP90AA1 | -1.16255 | LUAD | NCI-H1944 | Tanespimycin | MVP_top60_DNA+RNA_seed | AKT1 | 0.199181 | LUAD | NCI-H1792 | Uprosertib |
| MVP_top60_DNA_seed | AURKA | -1.160347 | LUAD | NCI-H2122 | HG-5-113-01 | MVP_top60_DNA+RNA_seed | HSP90AA1 | 0.199386 | LUAD | NCI-H1648 | Luminespib |

| | | | | | | | | | | | |
|---|---|---|---|---|---|---|---|---|---|---|---|
| MVP_top60_DNA_seed | ATM | -1.151655 | LUAD | NCI-H358 | CP466722 | MVP_top60_DNA+RNA_seed | AKT1 | 0.199422 | LUAD | NCI-H322M | Ipatasertib |
| MVP_top60_DNA_seed | HSP90AA1 | -1.151099 | LUAD | NCI-H1975 | Luminespib | MVP_top60_DNA+RNA_seed | TERT | 0.19946 | LUAD | HCC-827 | BIBR-1532 |
| MVP_top60_DNA_seed | AURKA | -1.150993 | LUAD | ABC-1 | HG-5-113-01 | MVP_top60_DNA+RNA_seed | AKT1 | 0.199876 | LUAD | NCI-H1435 | Ipatasertib |
| MVP_top60_DNA_seed | SMARCA4 | -1.147041 | LUAD | HCC-78 | PFI-3 | MVP_top60_DNA+RNA_seed | EGFR | 0.199961 | LUAD | NCI-H1651 | Sapitinib |
| MVP_top60_DNA_seed | HDAC1 | -1.146965 | LUAD | NCI-H1755 | Belinostat | MVP_top60_DNA+RNA_seed | HDAC1 | 0.200021 | LUAD | NCI-H2087 | Romidepsin |
| MVP_top60_DNA_seed | HDAC1 | -1.144915 | LUAD | NCI-H1975 | JQ12 | MVP_top60_DNA+RNA_seed | AKT1 | 0.200071 | LUAD | NCI-H1792 | MK-2206 |
| MVP_top60_DNA_seed | ATM | -1.144747 | LUAD | NCI-H1648 | KU-55933 | MVP_top60_DNA+RNA_seed | AKT1 | 0.200271 | LUAD | NCI-H1792 | GSK2110183B |
| MVP_top60_DNA_seed | IGF1R | -1.142682 | LUAD | HCC-44 | Linsitinib | MVP_top60_DNA+RNA_seed | AKT1 | 0.202565 | LUAD | NCI-H2228 | Uprosertib |
| MVP_top60_DNA_seed | EGFR | -1.142091 | LUAD | Calu-3 | Pelitinib | MVP_top60_DNA+RNA_seed | BRAF | 0.202867 | LUAD | NCI-H2023 | Dabrafenib |
| MVP_top60_DNA_seed | IGF1R | -1.141099 | LUAD | EKVX | Linsitinib | MVP_top60_DNA+RNA_seed | PIK3CA | 0.203638 | LUAD | Calu-6 | Buparlisib |
| MVP_top60_DNA_seed | AURKA | -1.136609 | LUAD | NCI-H1437 | Genentech Cpd 10 | MVP_top60_DNA+RNA_seed | IGF1R | 0.203684 | LUAD | ABC-1 | NVP-ADW742 |
| MVP_top60_DNA_seed | MAPK1 | -1.13507 | LUAD | NCI-H2405 | FR-180204 | MVP_top60_DNA+RNA_seed | SRC | 0.203735 | LUAD | NCI-H1944 | Bosutinib |
| MVP_top60_DNA_seed | BRAF | -1.134449 | LUAD | NCI-H2405 | HG6-64-1 | MVP_top60_DNA+RNA_seed | KRAS | 0.20407 | LUAD | ABC-1 | KRAS (G12C) Inhibitor-1 |
| MVP_top60_DNA_seed | IGF1R | -1.134186 | LUAD | HCC-827 | GSK1904529A | MVP_top60_DNA+RNA_seed | MDM2 | 0.20532 | LUAD | NCI-H2291 | Serdemetan |
| MVP_top60_DNA_seed | AURKA | -1.131602 | LUAD | HCC-78 | Alisertib | MVP_top60_DNA+RNA_seed | HSP90AA1 | 0.205409 | LUAD | NCI-H1651 | Luminespib |
| MVP_top60_DNA_seed | EGFR | -1.131497 | LUAD | NCI-H1975 | PF-00299804 | MVP_top60_DNA+RNA_seed | EZH2 | 0.205414 | LUAD | NCI-H2023 | GSK343 |
| MVP_top60_DNA_seed | ERBB2 | -1.131497 | LUAD | NCI-H1975 | PF-00299804 | MVP_top60_DNA+RNA_seed | BRAF | 0.206037 | LUAD | NCI-H1573 | PLX-4720 |
| MVP_top60_DNA_seed | BRAF | -1.131076 | LUAD | NCI-H522 | HG6-64-1 | MVP_top60_DNA+RNA_seed | AKT1 | 0.206274 | LUAD | NCI-H1651 | Uprosertib |
| MVP_top60_DNA_seed | PIK3CA | -1.130296 | LUAD | NCI-H2122 | PI3Ka_4409 | MVP_top60_DNA+RNA_seed | PIK3CA | 0.206623 | LUAD | NCI-H2342 | AZD8186 |
| MVP_top60_DNA_seed | HDAC1 | -1.1295 | LUAD | LC-2-ad | Belinostat | MVP_top60_DNA+RNA_seed | FGFR1 | 0.206944 | LUAD | Calu-6 | AZD4547 |
| MVP_top60_DNA_seed | IGF1R | -1.128832 | LUAD | HOP-62 | BMS-536924 | MVP_top60_DNA+RNA_seed | BRAF | 0.206968 | LUAD | NCI-H1437 | PLX-4720 |
| MVP_top60_DNA_seed | ATM | -1.127841 | LUAD | RERF-LC-KJ | KU-60019 | MVP_top60_DNA+RNA_seed | EGFR | 0.208743 | LUAD | LXF-289 | Afatinib |
| MVP_top60_DNA_seed | ATM | -1.124553 | LUAD | EMC-BAC-2 | KU-60019 | MVP_top60_DNA+RNA_seed | IGF1R | 0.208878 | LUAD | NCI-H2030 | GSK1904529A |
| MVP_top60_DNA_seed | EGFR | -1.123297 | LUAD | NCI-H1573 | Afatinib | MVP_top60_DNA+RNA_seed | IGF1R | 0.208888 | LUAD | NCI-H2023 | BMS-536924 |
| MVP_top60_DNA_seed | ERBB2 | -1.123297 | LUAD | NCI-H1573 | Afatinib | MVP_top60_DNA+RNA_seed | TERT | 0.208927 | LUAD | NCI-H1944 | BIBR-1532 |
| MVP_top60_DNA_seed | HSP90AA1 | -1.122215 | LUAD | NCI-H1993 | CCT-018159 | MVP_top60_DNA+RNA_seed | AKT1 | 0.209856 | LUAD | NCI-H2023 | GSK2110183B |
| MVP_top60_DNA_seed | HDAC1 | -1.114996 | LUAD | NCI-H1437 | JQ12 | MVP_top60_DNA+RNA_seed | EGFR | 0.210185 | LUAD | NCI-H2023 | Sapitinib |
| MVP_top60_DNA_seed | SRC | -1.113928 | LUAD | SK-LU-1 | Bosutinib | MVP_top60_DNA+RNA_seed | SRC | 0.211163 | LUAD | NCI-H650 | Bosutinib |
| MVP_top60_DNA_seed | PIK3CA | -1.113559 | LUAD | NCI-H1975 | AZD8835 | MVP_top60_DNA+RNA_seed | BRAF | 0.211777 | LUAD | HCC-44 | Dabrafenib |
| MVP_top60_DNA_seed | MDM2 | -1.111722 | LUAD | NCI-H2087 | Nutlin-3a (-) | MVP_top60_DNA+RNA_seed | EGFR | 0.213746 | LUAD | NCI-H3122 | Afatinib |
| MVP_top60_DNA_seed | ATM | -1.111413 | LUAD | A549 | torin2 | MVP_top60_DNA+RNA_seed | EGFR | 0.215366 | LUAD | NCI-H1355 | Sapitinib |
| MVP_top60_DNA_seed | BRAF | -1.109823 | LUAD | HOP-62 | HG6-64-1 | MVP_top60_DNA+RNA_seed | HDAC1 | 0.216866 | LUAD | NCI-H1993 | Entinostat |
| MVP_top60_DNA_seed | AURKA | -1.104659 | LUAD | VMRC-LCD | HG-5-113-01 | MVP_top60_DNA+RNA_seed | FGFR1 | 0.218358 | LUAD | EMC-BAC-1 | PD173074 |
| MVP_top60_DNA_seed | EGFR | -1.102787 | LUAD | NCI-H1975 | AST-1306 | MVP_top60_DNA+RNA_seed | MDM2 | 0.219731 | LUAD | NCI-H1573 | Nutlin-3a (-) |
| MVP_top60_DNA_seed | MET | -1.101569 | LUAD | NCI-H522 | JNJ38877605 | MVP_top60_DNA+RNA_seed | SRC | 0.220595 | LUAD | NCI-H2085 | Bosutinib |
| MVP_top60_DNA_seed | PTK2 | -1.099467 | LUAD | SK-LU-1 | PF-562271 | MVP_top60_DNA+RNA_seed | IGF1R | 0.222408 | LUAD | PC-14 | GSK1904529A |
| MVP_top60_DNA_seed | SRC | -1.09296 | LUAD | PC-14 | HG-6-71-01 | MVP_top60_DNA+RNA_seed | EGFR | 0.224605 | LUAD | NCI-H1792 | Sapitinib |
| MVP_top60_DNA_seed | GSK3B | -1.088733 | LUAD | NCI-H1975 | AZD7969 | MVP_top60_DNA+RNA_seed | MDM2 | 0.227678 | LUAD | HCC-78 | Nutlin-3a (-) |
| MVP_top60_DNA_seed | PIK3CA | -1.087568 | LUAD | NCI-H1944 | PI3Ka_4409 | MVP_top60_DNA+RNA_seed | AKT1 | 0.227925 | LUAD | EMC-BAC-1 | MK-2206 |
| MVP_top60_DNA_seed | PIK3CA | -1.086683 | LUAD | NCI-H441 | PI3Ka_4409 | MVP_top60_DNA+RNA_seed | MDM2 | 0.230052 | LUAD | NCI-H2122 | Serdemetan |
| MVP_top60_DNA_seed | SRC | -1.085245 | LUAD | NCI-H1975 | Saracatinib | MVP_top60_DNA+RNA_seed | ATM | 0.230823 | LUAD | NCI-H2347 | KU-55933 |

| | | | | | | | | | | | |
|---|---|---|---|---|---|---|---|---|---|---|---|
| MVP_top60_DNA_seed | AKT1 | -1.084907 | LUAD | EMC-BAC-2 | MK-2206 | MVP_top60_DNA+RNA_seed | BRAF | 0.231303 | LUAD | NCI-H522 | SB590885 |
| MVP_top60_DNA_seed | EGFR | -1.084571 | LUAD | EKVX | Cetuximab | MVP_top60_DNA+RNA_seed | AKT1 | 0.231715 | LUAD | NCI-H2009 | AZD5363 |
| MVP_top60_DNA_seed | AKT1 | -1.082286 | LUAD | HCC-44 | AKT inhibitor VIII | MVP_top60_DNA+RNA_seed | BRAF | 0.232627 | LUAD | NCI-H2023 | PLX-4720 |
| MVP_top60_DNA_seed | MET | -1.081373 | LUAD | NCI-H23 | AZD6094 | MVP_top60_DNA+RNA_seed | MET | 0.23309 | LUAD | NCI-H2085 | Savolitinib |
| MVP_top60_DNA_seed | PTK2 | -1.079412 | LUAD | LC-2-ad | PF-562271 | MVP_top60_DNA+RNA_seed | EGFR | 0.233724 | LUAD | NCI-H522 | AZD3759 |
| MVP_top60_DNA_seed | HSP90AA1 | -1.079375 | LUAD | NCI-H2085 | Elesclomol | MVP_top60_DNA+RNA_seed | AKT1 | 0.236141 | LUAD | NCI-H2023 | AZD5363 |
| MVP_top60_DNA_seed | HSP90AA1 | -1.078961 | LUAD | NCI-H1792 | CCT-018159 | MVP_top60_DNA+RNA_seed | MET | 0.238613 | LUAD | NCI-H322M | Savolitinib |
| MVP_top60_DNA_seed | ATM | -1.077634 | LUAD | NCI-H2122 | torin2 | MVP_top60_DNA+RNA_seed | EGFR | 0.239435 | LUAD | LXF-289 | Erlotinib |
| MVP_top60_DNA_seed | SRC | -1.075479 | LUAD | LXF-289 | WH-4-023 | MVP_top60_DNA+RNA_seed | AURKA | 0.241089 | LUAD | NCI-H1975 | Alisertib |
| MVP_top60_DNA_seed | IGF1R | -1.07359 | LUAD | NCI-H1755 | BMS-536924 | MVP_top60_DNA+RNA_seed | BRAF | 0.241581 | LUAD | NCI-H650 | SB590885 |
| MVP_top60_DNA_seed | MET | -1.072892 | LUAD | HCC-44 | AZD6094 | MVP_top60_DNA+RNA_seed | BRAF | 0.243461 | LUAD | NCI-H1648 | Dabrafenib |
| MVP_top60_DNA_seed | HSP90AA1 | -1.070715 | LUAD | NCI-H2030 | Luminespib | MVP_top60_DNA+RNA_seed | MDM2 | 0.246866 | LUAD | RERF-LC-KJ | Nutlin-3a (-) |
| MVP_top60_DNA_seed | AURKA | -1.069444 | LUAD | HCC-44 | Genentech Cpd 10 | MVP_top60_DNA+RNA_seed | FGFR1 | 0.247494 | LUAD | A549 | AZD4547 |
| MVP_top60_DNA_seed | HSP90AA1 | -1.069428 | LUAD | NCI-H1755 | Tanespimycin | MVP_top60_DNA+RNA_seed | AURKA | 0.248705 | LUAD | NCI-H1568 | ZM447439 |
| MVP_top60_DNA_seed | ATM | -1.06612 | LUAD | NCI-H1648 | KU-60019 | MVP_top60_DNA+RNA_seed | IGF1R | 0.249786 | LUAD | NCI-H1435 | GSK1904529A |
| MVP_top60_DNA_seed | MAPK1 | -1.063317 | LUAD | LC-2-ad | FR-180204 | MVP_top60_DNA+RNA_seed | ATM | 0.251747 | LUAD | NCI-H23 | KU-55933 |
| MVP_top60_DNA_seed | AKT1 | -1.060228 | LUAD | HCC-78 | AKT inhibitor VIII | MVP_top60_DNA+RNA_seed | MDM2 | 0.251982 | LUAD | NCI-H1648 | Nutlin-3a (-) |
| MVP_top60_DNA_seed | HDAC1 | -1.058888 | LUAD | NCI-H2342 | JQ12 | MVP_top60_DNA+RNA_seed | BRAF | 0.25522 | LUAD | NCI-H1734 | PLX-4720 |
| MVP_top60_DNA_seed | AKT1 | -1.055067 | LUAD | SW1573 | BAY AKT1 | MVP_top60_DNA+RNA_seed | AURKA | 0.256369 | LUAD | NCI-H1650 | Alisertib |
| MVP_top60_DNA_seed | FGFR1 | -1.054026 | LUAD | NCI-H838 | FGFR_3831 | MVP_top60_DNA+RNA_seed | KRAS | 0.257163 | LUAD | RERF-LC-KJ | [illegible]RAS (G12C) Inhibitor-1 |
| MVP_top60_DNA_seed | SRC | -1.051514 | LUAD | ABC-1 | Bosutinib | MVP_top60_DNA+RNA_seed | IGF1R | 0.257593 | LUAD | NCI-H1623 | BMS-754807 |
| MVP_top60_DNA_seed | HSP90AA1 | -1.05091 | LUAD | NCI-H838 | Elesclomol | MVP_top60_DNA+RNA_seed | EGFR | 0.258655 | LUAD | RERF-LC-MS | Osimertinib |
| MVP_top60_DNA_seed | HSP90AA1 | -1.050058 | LUAD | RERF-LC-MS | CCT-018159 | MVP_top60_DNA+RNA_seed | AURKA | 0.258787 | LUAD | NCI-H1944 | ZM447439 |
| MVP_top60_DNA_seed | HDAC1 | -1.043651 | LUAD | NCI-H358 | Tubastatin A | MVP_top60_DNA+RNA_seed | EGFR | 0.258963 | LUAD | NCI-H441 | Erlotinib |
| MVP_top60_DNA_seed | BRAF | -1.043425 | LUAD | NCI-H2030 | HG6-64-1 | MVP_top60_DNA+RNA_seed | EZH2 | 0.2595 | LUAD | NCI-H1993 | GSK343 |
| MVP_top60_DNA_seed | GSK3B | -1.04311 | LUAD | NCI-H1650 | CHIR-99021 | MVP_top60_DNA+RNA_seed | BRAF | 0.259661 | LUAD | RERF-LC-KJ | SB590885 |
| MVP_top60_DNA_seed | HDAC1 | -1.043009 | LUAD | NCI-H1975 | Tubastatin A | MVP_top60_DNA+RNA_seed | AKT1 | 0.260002 | LUAD | EMC-BAC-2 | Uprosertib |
| MVP_top60_DNA_seed | IGF1R | -1.04002 | LUAD | EMC-BAC-1 | BMS-536924 | MVP_top60_DNA+RNA_seed | BRAF | 0.260024 | LUAD | H3255 | Dabrafenib |
| MVP_top60_DNA_seed | AKT1 | -1.039806 | LUAD | NCI-H2228 | MK-2206 | MVP_top60_DNA+RNA_seed | AKT1 | 0.260824 | LUAD | NCI-H23 | AZD5363 |
| MVP_top60_DNA_seed | IGF1R | -1.037951 | LUAD | LXF-289 | BMS-536924 | MVP_top60_DNA+RNA_seed | AKT1 | 0.261546 | LUAD | LC-2-ad | Uprosertib |
| MVP_top60_DNA_seed | HDAC1 | -1.036798 | LUAD | NCI-H3122 | JQ12 | MVP_top60_DNA+RNA_seed | AKT1 | 0.262832 | LUAD | NCI-H2009 | Afuresertib |
| MVP_top60_DNA_seed | HSP90AA1 | -1.036185 | LUAD | NCI-H2122 | Luminespib | MVP_top60_DNA+RNA_seed | EZH2 | 0.264622 | LUAD | NCI-H1993 | GSK343 |
| MVP_top60_DNA_seed | AKT1 | -1.035957 | LUAD | NCI-H1944 | MK-2206 | MVP_top60_DNA+RNA_seed | KRAS | 0.265622 | LUAD | NCI-H1666 | [illegible]RAS (G12C) Inhibitor-1 |
| MVP_top60_DNA_seed | HSP90AA1 | -1.03557 | LUAD | NCI-H1792 | Tanespimycin | MVP_top60_DNA+RNA_seed | AKT1 | 0.265734 | LUAD | NCI-H2291 | Afuresertib |
| MVP_top60_DNA_seed | EGFR | -1.035253 | LUAD | EMC-BAC-2 | Pelitinib | MVP_top60_DNA+RNA_seed | BRAF | 0.26751 | LUAD | NCI-H1993 | Dabrafenib |
| MVP_top60_DNA_seed | ATM | -1.033447 | LUAD | NCI-H1437 | CP466722 | MVP_top60_DNA+RNA_seed | AKT1 | 0.269061 | LUAD | RERF-LC-KJ | GSK2110183B |
| MVP_top60_DNA_seed | MAPK1 | -1.032085 | LUAD | COR-L105 | VX-11e | MVP_top60_DNA+RNA_seed | MET | 0.26999 | LUAD | NCI-H2085 | Crizotinib |
| MVP_top60_DNA_seed | GSK3B | -1.030715 | LUAD | EMC-BAC-2 | AZD7969 | MVP_top60_DNA+RNA_seed | FGFR1 | 0.270559 | LUAD | NCI-H1650 | AZD4547 |
| MVP_top60_DNA_seed | MDM2 | -1.028881 | LUAD | NCI-H2342 | Serdemetan | MVP_top60_DNA+RNA_seed | TP53 | 0.271202 | LUAD | NCI-H2085 | MIRA-1 |
| MVP_top60_DNA_seed | HSP90AA1 | -1.028481 | LUAD | HOP-62 | Luminespib | MVP_top60_DNA+RNA_seed | HDAC1 | 0.27138 | LUAD | RERF-LC-KJ | Entinostat |
| MVP_top60_DNA_seed | AKT1 | -1.028287 | LUAD | NCI-H2009 | AKT inhibitor VIII | MVP_top60_DNA+RNA_seed | AKT1 | 0.271951 | LUAD | NCI-H1623 | AT13148 |

| | | | | | | | | | | | |
|---|---|---|---|---|---|---|---|---|---|---|---|
| MVP_top60_DNA_seed | AURKA | -1.027453 | LUAD | NCI-H2030 | Genentech Cpd 10 | MVP_top60_DNA+RNA_seed | TP53 | 0.273078 | LUAD | NCI-H23 | MIRA-1 |
| MVP_top60_DNA_seed | HSP90AA1 | -1.026593 | LUAD | NCI-H1944 | Elesclomol | MVP_top60_DNA+RNA_seed | FGFR1 | 0.27381 | LUAD | COR-L105 | PD173074 |
| MVP_top60_DNA_seed | SRC | -1.024607 | LUAD | EMC-BAC-1 | Bosutinib | MVP_top60_DNA+RNA_seed | MET | 0.27453 | LUAD | Calu-3 | Crizotinib |
| MVP_top60_DNA_seed | HSP90AA1 | -1.021354 | LUAD | PC-14 | SNX-2112 | MVP_top60_DNA+RNA_seed | TERT | 0.274731 | LUAD | Calu-6 | BIBR-1532 |
| MVP_top60_DNA_seed | PIK3CA | -1.020125 | LUAD | NCI-H1355 | PI3Ka_4409 | MVP_top60_DNA+RNA_seed | PIK3CA | 0.274954 | LUAD | SW1573 | AZD8186 |
| MVP_top60_DNA_seed | CREBBP | -1.014061 | LUAD | HCC-78 | I-CBP112 | MVP_top60_DNA+RNA_seed | AKT1 | 0.27528 | LUAD | NCI-H358 | Afuresertib |
| MVP_top60_DNA_seed | HSP90AA1 | -1.010517 | LUAD | HOP-62 | SNX-2112 | MVP_top60_DNA+RNA_seed | HDAC1 | 0.275315 | LUAD | NCI-H3122 | PCI-34051 |
| MVP_top60_DNA_seed | ATM | -1.008731 | LUAD | NCI-H2085 | KU-55933 | MVP_top60_DNA+RNA_seed | EGFR | 0.275541 | LUAD | NCI-H2009 | Afatinib |
| MVP_top60_DNA_seed | MAPK1 | -1.008585 | LUAD | NCI-H2087 | VX-11e | MVP_top60_DNA+RNA_seed | BRAF | 0.276048 | LUAD | LXF-289 | SB590885 |
| MVP_top60_DNA_seed | HSP90AA1 | -1.008368 | LUAD | HCC-44 | SNX-2112 | MVP_top60_DNA+RNA_seed | MDM2 | 0.277218 | LUAD | EMC-BAC-1 | Serdemetan |
| MVP_top60_DNA_seed | AURKA | -1.006379 | LUAD | NCI-H2087 | Genentech Cpd 10 | MVP_top60_DNA+RNA_seed | BRAF | 0.278245 | LUAD | NCI-H1651 | Dabrafenib |
| MVP_top60_DNA_seed | HSP90AA1 | -1.004941 | LUAD | NCI-H1944 | CCT-018159 | MVP_top60_DNA+RNA_seed | EGFR | 0.278311 | LUAD | HCC-44 | Lapatinib |
| MVP_top60_DNA_seed | AKT1 | -1.004335 | LUAD | NCI-H2342 | BAY AKT1 | MVP_top60_DNA+RNA_seed | IGF1R | 0.278947 | LUAD | NCI-H2087 | NVP-ADW742 |
| MVP_top60_DNA_seed | BRAF | -1.001934 | LUAD | NCI-H1666 | PLX-4720 | MVP_top60_DNA+RNA_seed | EGFR | 0.279115 | LUAD | Calu-3 | Erlotinib |
| MVP_top60_DNA_seed | BRAF | -1.001378 | LUAD | NCI-H2342 | HG6-64-1 | MVP_top60_DNA+RNA_seed | ATM | 0.280303 | LUAD | NCI-H1693 | KU-55933 |
| MVP_top60_DNA_seed | EGFR | -1.001148 | LUAD | NCI-H322M | AST-1306 | MVP_top60_DNA+RNA_seed | BRAF | 0.280881 | LUAD | NCI-H1437 | Dabrafenib |
| MVP_top60_DNA_seed | IGF1R | -1.000676 | LUAD | NCI-H838 | Linsitinib | MVP_top60_DNA+RNA_seed | MDM2 | 0.283916 | LUAD | NCI-H1573 | Serdemetan |
| MVP_top60_DNA_seed | HDAC1 | -0.999116 | LUAD | HCC-44 | Dacinostat | MVP_top60_DNA+RNA_seed | IGF1R | 0.284279 | LUAD | HOP-62 | NVP-ADW742 |
| MVP_top60_DNA_seed | SRC | -0.998977 | LUAD | A549 | HG-6-71-01 | MVP_top60_DNA+RNA_seed | SRC | 0.284795 | LUAD | NCI-H2347 | Bosutinib |
| MVP_top60_DNA_seed | MET | -0.998882 | LUAD | NCI-H1648 | JNJ38877605 | MVP_top60_DNA+RNA_seed | ATM | 0.286418 | LUAD | NCI-H1650 | KU-55933 |
| MVP_top60_DNA_seed | HSP90AA1 | -0.998357 | LUAD | NCI-H1437 | CCT-018159 | MVP_top60_DNA+RNA_seed | EZH2 | 0.286823 | LUAD | EMC-BAC-2 | GSK343 |
| MVP_top60_DNA_seed | SRC | -0.996903 | LUAD | NCI-H2030 | Bosutinib | MVP_top60_DNA+RNA_seed | PIK3CA | 0.287379 | LUAD | NCI-H650 | Buparlisib |
| MVP_top60_DNA_seed | EGFR | -0.992871 | LUAD | NCI-H1648 | Gefitinib | MVP_top60_DNA+RNA_seed | EGFR | 0.288499 | LUAD | NCI-H2228 | Sapitinib |
| MVP_top60_DNA_seed | AURKA | -0.992257 | LUAD | HCC-44 | ZM447439 | MVP_top60_DNA+RNA_seed | AKT1 | 0.289237 | LUAD | NCI-H1435 | Uprosertib |
| MVP_top60_DNA_seed | ATM | -0.992236 | LUAD | NCI-H1648 | torin2 | MVP_top60_DNA+RNA_seed | ATM | 0.290966 | LUAD | NCI-H838 | KU-55933 |
| MVP_top60_DNA_seed | FGFR1 | -0.991899 | LUAD | NCI-H838 | AZD4547 | MVP_top60_DNA+RNA_seed | IGF1R | 0.291363 | LUAD | NCI-H1568 | GSK1904529A |
| MVP_top60_DNA_seed | PIK3CA | -0.989945 | LUAD | SW1573 | AZD8835 | MVP_top60_DNA+RNA_seed | BRAF | 0.291914 | LUAD | NCI-H1437 | SB590885 |
| MVP_top60_DNA_seed | MET | -0.987813 | LUAD | LC-2-ad | JNJ38877605 | MVP_top60_DNA+RNA_seed | PIK3CA | 0.291914 | LUAD | NCI-H1573 | GNE-317 |
| MVP_top60_DNA_seed | PIK3CA | -0.983477 | LUAD | NCI-H1623 | PI3Ka_4409 | MVP_top60_DNA+RNA_seed | IGF1R | 0.296374 | LUAD | Calu-6 | NVP-ADW742 |
| MVP_top60_DNA_seed | PIK3CA | -0.978632 | LUAD | HCC-44 | PI3Ka_4409 | MVP_top60_DNA+RNA_seed | AKT1 | 0.299342 | LUAD | NCI-H1650 | Afuresertib |
| MVP_top60_DNA_seed | AURKA | -0.978187 | LUAD | NCI-H1648 | Genentech Cpd 10 | MVP_top60_DNA+RNA_seed | AKT1 | 0.30122 | LUAD | LC-2-ad | Afuresertib |
| MVP_top60_DNA_seed | EGFR | -0.974108 | LUAD | NCI-H1975 | CI-1033 | MVP_top60_DNA+RNA_seed | AKT1 | 0.303228 | LUAD | NCI-H1666 | MK-2206 |
| MVP_top60_DNA_seed | ERBB2 | -0.974108 | LUAD | NCI-H1975 | CI-1033 | MVP_top60_DNA+RNA_seed | EGFR | 0.303406 | LUAD | RERF-LC-KJ | Afatinib |
| MVP_top60_DNA_seed | IGF1R | -0.973175 | LUAD | NCI-H2030 | BMS-536924 | MVP_top60_DNA+RNA_seed | MDM2 | 0.303465 | LUAD | NCI-H1435 | Nutlin-3a (-) |
| MVP_top60_DNA_seed | AKT1 | -0.971271 | LUAD | HCC-44 | BAY AKT1 | MVP_top60_DNA+RNA_seed | PIK3CA | 0.30589 | LUAD | PC-3_[JPC-3] | Buparlisib |
| MVP_top60_DNA_seed | ATM | -0.970509 | LUAD | NCI-H1734 | KU-60019 | MVP_top60_DNA+RNA_seed | EGFR | 0.306368 | LUAD | NCI-H1435 | Lapatinib |
| MVP_top60_DNA_seed | EGFR | -0.968627 | LUAD | NCI-H1944 | AZD8931 | MVP_top60_DNA+RNA_seed | PIK3CA | 0.306425 | LUAD | EMC-BAC-1 | GNE-317 |
| MVP_top60_DNA_seed | ERBB2 | -0.968627 | LUAD | NCI-H1944 | AZD8931 | MVP_top60_DNA+RNA_seed | BRAF | 0.306714 | LUAD | NCI-H1975 | Dabrafenib |
| MVP_top60_DNA_seed | ERBB3 | -0.968627 | LUAD | NCI-H1944 | AZD8931 | MVP_top60_DNA+RNA_seed | AURKA | 0.306999 | LUAD | HCC-827 | ZM447439 |
| MVP_top60_DNA_seed | HDAC1 | -0.96772 | LUAD | HOP-62 | Tubastatin A | MVP_top60_DNA+RNA_seed | AKT1 | 0.307214 | LUAD | NCI-H2030 | Ipatasertib |
| MVP_top60_DNA_seed | ATM | -0.967298 | LUAD | LXF-289 | torin2 | MVP_top60_DNA+RNA_seed | EGFR | 0.307573 | LUAD | NCI-H1355 | Erlotinib |

| | | | | | | | | | | | |
|---|---|---|---|---|---|---|---|---|---|---|---|
| MVP_top60_DNA_seed | EGFR | -0.966415 | LUAD | EKVX | Gefitinib | MVP_top60_DNA+RNA_seed | BRAF | 0.309539 | LUAD | LXF-289 | PLX-4720 |
| MVP_top60_DNA_seed | ERBB2 | -0.963765 | LUAD | PC-3_[JPC-3] | CP724714 | MVP_top60_DNA+RNA_seed | AKT1 | 0.3096 | LUAD | NCI-H2342 | AZD5363 |
| MVP_top60_DNA_seed | HSP90AA1 | -0.962868 | LUAD | NCI-H2122 | Tanespimycin | MVP_top60_DNA+RNA_seed | HDAC1 | 0.310015 | LUAD | RERF-LC-KJ | Romidepsin |
| MVP_top60_DNA_seed | IGF1R | -0.961389 | LUAD | NCI-H2122 | BMS-536924 | MVP_top60_DNA+RNA_seed | IGF1R | 0.310117 | LUAD | ABC-1 | GSK1904529A |
| MVP_top60_DNA_seed | AURKA | -0.959286 | LUAD | NCI-H1755 | Genentech Cpd 10 | MVP_top60_DNA+RNA_seed | SRC | 0.310274 | LUAD | NCI-H1650 | Bosutinib |
| MVP_top60_DNA_seed | FGFR1 | -0.959115 | LUAD | EMC-BAC-2 | FGFR_3831 | MVP_top60_DNA+RNA_seed | ATM | 0.310577 | LUAD | NCI-H1993 | KU-55933 |
| MVP_top60_DNA_seed | HSP90AA1 | -0.958201 | LUAD | NCI-H2342 | Luminespib | MVP_top60_DNA+RNA_seed | EZH2 | 0.312098 | LUAD | NCI-H838 | GSK343 |
| MVP_top60_DNA_seed | SRC | -0.956777 | LUAD | LC-2-ad | Saracatinib | MVP_top60_DNA+RNA_seed | EGFR | 0.312241 | LUAD | NCI-H441 | Afatinib |
| MVP_top60_DNA_seed | HSP90AA1 | -0.955631 | LUAD | EMC-BAC-2 | Elesclomol | MVP_top60_DNA+RNA_seed | HDAC1 | 0.312803 | LUAD | NCI-H2030 | Entinostat |
| MVP_top60_DNA_seed | BRAF | -0.95384 | LUAD | NCI-H358 | HG6-64-1 | MVP_top60_DNA+RNA_seed | BRAF | 0.312876 | LUAD | NCI-H322M | SB590885 |
| MVP_top60_DNA_seed | MET | -0.953581 | LUAD | NCI-H1975 | AZD6094 | MVP_top60_DNA+RNA_seed | SMARCA4 | 0.312982 | LUAD | HCC-44 | PFI3 |
| MVP_top60_DNA_seed | EGFR | -0.951567 | LUAD | Calu-3 | Afatinib | MVP_top60_DNA+RNA_seed | EZH2 | 0.314116 | LUAD | A549 | GSK343 |
| MVP_top60_DNA_seed | ERBB2 | -0.951567 | LUAD | Calu-3 | Afatinib | MVP_top60_DNA+RNA_seed | EGFR | 0.314177 | LUAD | NCI-H1573 | Afatinib |
| MVP_top60_DNA_seed | MAPK1 | -0.949179 | LUAD | NCI-H1693 | FR-180204 | MVP_top60_DNA+RNA_seed | MDM2 | 0.31581 | LUAD | NCI-H1563 | Serdemetan |
| MVP_top60_DNA_seed | HSP90AA1 | -0.947397 | LUAD | RERF-LC-MS | Elesclomol | MVP_top60_DNA+RNA_seed | MDM2 | 0.315845 | LUAD | NCI-H1755 | Serdemetan |
| MVP_top60_DNA_seed | GSK3B | -0.946058 | LUAD | EMC-BAC-2 | CHIR-99021 | MVP_top60_DNA+RNA_seed | SRC | 0.31614 | LUAD | NCI-H1651 | Bosutinib |
| MVP_top60_DNA_seed | IGF1R | -0.945771 | LUAD | EKVX | BMS-536924 | MVP_top60_DNA+RNA_seed | HDAC1 | 0.316454 | LUAD | NCI-H650 | Entinostat |
| MVP_top60_DNA_seed | PIK3CA | -0.94476 | LUAD | NCI-H2228 | PI3Ka_4409 | MVP_top60_DNA+RNA_seed | AKT1 | 0.317258 | LUAD | LC-2-ad | AT13148 |
| MVP_top60_DNA_seed | GSK3B | -0.938687 | LUAD | HCC-78 | CHIR-99021 | MVP_top60_DNA+RNA_seed | EGFR | 0.317298 | LUAD | NCI-H522 | Lapatinib |
| MVP_top60_DNA_seed | PTK2 | -0.936408 | LUAD | NCI-H2122 | PF-562271 | MVP_top60_DNA+RNA_seed | BRAF | 0.317308 | LUAD | NCI-H2228 | SB590885 |
| MVP_top60_DNA_seed | HSP90AA1 | -0.933845 | LUAD | EMC-BAC-2 | Tanespimycin | MVP_top60_DNA+RNA_seed | BRAF | 0.317629 | LUAD | NCI-H838 | SB590885 |
| MVP_top60_DNA_seed | MDM2 | -0.933725 | LUAD | NCI-H1568 | Serdemetan | MVP_top60_DNA+RNA_seed | AKT1 | 0.318905 | LUAD | EMC-BAC-2 | MK-2206 |
| MVP_top60_DNA_seed | HSP90AA1 | -0.932636 | LUAD | SK-LU-1 | Luminespib | MVP_top60_DNA+RNA_seed | FGFR1 | 0.319649 | LUAD | NCI-H23 | AZD4547 |
| MVP_top60_DNA_seed | EGFR | -0.931851 | LUAD | NCI-H1650 | Gefitinib | MVP_top60_DNA+RNA_seed | PIK3CA | 0.319768 | LUAD | ABC-1 | Alpelisib |
| MVP_top60_DNA_seed | HDAC1 | -0.931749 | LUAD | LC-2-ad | Entinostat | MVP_top60_DNA+RNA_seed | IGF1R | 0.320436 | LUAD | NCI-H23 | BMS-536924 |
| MVP_top60_DNA_seed | AKT1 | -0.931429 | LUAD | NCI-H358 | AKT inhibitor VIII | MVP_top60_DNA+RNA_seed | EGFR | 0.320644 | LUAD | NCI-H2291 | Gefitinib |
| MVP_top60_DNA_seed | PTK2 | -0.930421 | LUAD | NCI-H650 | PF-562271 | MVP_top60_DNA+RNA_seed | MDM2 | 0.32137 | LUAD | A549 | Serdemetan |
| MVP_top60_DNA_seed | MDM2 | -0.929405 | LUAD | RERF-LC-MS | Serdemetan | MVP_top60_DNA+RNA_seed | HSP90AA1 | 0.321549 | LUAD | NCI-H1651 | Tanespimycin |
| MVP_top60_DNA_seed | HSP90AA1 | -0.929309 | LUAD | NCI-H650 | SNX-2112 | MVP_top60_DNA+RNA_seed | SRC | 0.324323 | LUAD | NCI-H522 | Bosutinib |
| MVP_top60_DNA_seed | EGFR | -0.926824 | LUAD | A549 | Afatinib | MVP_top60_DNA+RNA_seed | BRAF | 0.325327 | LUAD | NCI-H1435 | PLX-4720 |
| MVP_top60_DNA_seed | ERBB2 | -0.926824 | LUAD | A549 | Afatinib | MVP_top60_DNA+RNA_seed | EGFR | 0.325384 | LUAD | NCI-H2291 | Sapitinib |
| MVP_top60_DNA_seed | SRC | -0.926307 | LUAD | COR-L105 | NVP-BHG712 | MVP_top60_DNA+RNA_seed | IGF1R | 0.325394 | LUAD | NCI-H1755 | GSK1904529A |
| MVP_top60_DNA_seed | PTK2 | -0.92623 | LUAD | HCC-44 | PF-562271 | MVP_top60_DNA+RNA_seed | BRAF | 0.32556 | LUAD | COR-L105 | Dabrafenib |
| MVP_top60_DNA_seed | ERBB2 | -0.925916 | LUAD | PC-14 | CP724714 | MVP_top60_DNA+RNA_seed | AKT1 | 0.32576 | LUAD | NCI-H1792 | AZD5363 |
| MVP_top60_DNA_seed | MDM2 | -0.925226 | LUAD | NCI-H2087 | Serdemetan | MVP_top60_DNA+RNA_seed | AKT1 | 0.326246 | LUAD | LC-2-ad | MK-2206 |
| MVP_top60_DNA_seed | PTK2 | -0.925092 | LUAD | NCI-H358 | PF-562271 | MVP_top60_DNA+RNA_seed | EGFR | 0.326505 | LUAD | NCI-H1755 | Sapitinib |
| MVP_top60_DNA_seed | HSP90AA1 | -0.923668 | LUAD | NCI-H1437 | Tanespimycin | MVP_top60_DNA+RNA_seed | EZH2 | 0.327683 | LUAD | NCI-H1693 | GSK343 |
| MVP_top60_DNA_seed | MET | -0.923329 | LUAD | NCI-H1993 | JNJ38877605 | MVP_top60_DNA+RNA_seed | IGF1R | 0.32847 | LUAD | NCI-H522 | GSK1904529A |
| MVP_top60_DNA_seed | MET | -0.923318 | LUAD | LXF-289 | PHA-665752 | MVP_top60_DNA+RNA_seed | BRAF | 0.330674 | LUAD | EMC-BAC-2 | SB590885 |
| MVP_top60_DNA_seed | GSK3B | -0.92123 | LUAD | NCI-H2030 | AZD7969 | MVP_top60_DNA+RNA_seed | BRAF | 0.330884 | LUAD | VMRC-LCD | SB590885 |
| MVP_top60_DNA_seed | EGFR | -0.920786 | LUAD | NCI-H650 | Cetuximab | MVP_top60_DNA+RNA_seed | EGFR | 0.330982 | LUAD | NCI-H2405 | Erlotinib |

| | | | | | | | | | | | |
|---|---|---|---|---|---|---|---|---|---|---|---|
| MVP_top60_DNA_seed | IGF1R | -0.919482 | LUAD | RERF-LC-KJ | GSK1904529A | MVP_top60_DNA+RNA_seed | AKT1 | 0.331473 | LUAD | NCI-H2009 | MK-2206 |
| MVP_top60_DNA_seed | HDAC1 | -0.917954 | LUAD | NCI-H1437 | Dacinostat | MVP_top60_DNA+RNA_seed | MDM2 | 0.332898 | LUAD | NCI-H1781 | Serdemetan |
| MVP_top60_DNA_seed | AKT1 | -0.916742 | LUAD | NCI-H2085 | MK-2206 | MVP_top60_DNA+RNA_seed | IGF1R | 0.333397 | LUAD | NCI-H2291 | BMS-536924 |
| MVP_top60_DNA_seed | SMARCA4 | -0.916037 | LUAD | EKVX | PFI-3 | MVP_top60_DNA+RNA_seed | HDAC1 | 0.338056 | LUAD | EMC-BAC-1 | PCI-34051 |
| MVP_top60_DNA_seed | AURKA | -0.914346 | LUAD | A549 | HG-5-113-01 | MVP_top60_DNA+RNA_seed | EGFR | 0.338336 | LUAD | NCI-H2009 | Gefitinib |
| MVP_top60_DNA_seed | IGF1R | -0.913895 | LUAD | HCC-44 | BMS-754807 | MVP_top60_DNA+RNA_seed | MET | 0.338617 | LUAD | NCI-H1651 | Savolitinib |
| MVP_top60_DNA_seed | BRAF | -0.913088 | LUAD | COR-L105 | HG6-64-1 | MVP_top60_DNA+RNA_seed | AKT1 | 0.339169 | LUAD | LXF-289 | AZD5363 |
| MVP_top60_DNA_seed | MAPK1 | -0.910088 | LUAD | NCI-H1944 | FR-180204 | MVP_top60_DNA+RNA_seed | PIK3CA | 0.339773 | LUAD | NCI-H1355 | AZD8186 |
| MVP_top60_DNA_seed | GSK3B | -0.909679 | LUAD | RERF-LC-MS | SB216763 | MVP_top60_DNA+RNA_seed | IGF1R | 0.341253 | LUAD | NCI-H1693 | GSK1904529A |
| MVP_top60_DNA_seed | ATM | -0.909217 | LUAD | LC-2-ad | CP466722 | MVP_top60_DNA+RNA_seed | AURKA | 0.342677 | LUAD | NCI-H1573 | Alisertib |
| MVP_top60_DNA_seed | AURKA | -0.907529 | LUAD | EMC-BAC-1 | HG-5-113-01 | MVP_top60_DNA+RNA_seed | MDM2 | 0.343754 | LUAD | EMC-BAC-1 | Nutlin-3a (-) |
| MVP_top60_DNA_seed | HDAC1 | -0.906758 | LUAD | HOP-62 | Dacinostat | MVP_top60_DNA+RNA_seed | EGFR | 0.344215 | LUAD | NCI-H1993 | Osimertinib |
| MVP_top60_DNA_seed | FGFR1 | -0.906192 | LUAD | NCI-H1355 | PD173074 | MVP_top60_DNA+RNA_seed | MDM2 | 0.344464 | LUAD | NCI-H1792 | Nutlin-3a (-) |
| MVP_top60_DNA_seed | EGFR | -0.904502 | LUAD | NCI-H1781 | AZD8931 | MVP_top60_DNA+RNA_seed | AKT1 | 0.344635 | LUAD | NCI-H2405 | AT13148 |
| MVP_top60_DNA_seed | ERBB2 | -0.904502 | LUAD | NCI-H1781 | AZD8931 | MVP_top60_DNA+RNA_seed | AKT1 | 0.344947 | LUAD | Calu-3 | Uprosertib |
| MVP_top60_DNA_seed | ERBB3 | -0.904502 | LUAD | NCI-H1781 | AZD8931 | MVP_top60_DNA+RNA_seed | PIK3CA | 0.345114 | LUAD | NCI-H2030 | Buparlisib |
| MVP_top60_DNA_seed | HSP90AA1 | -0.901973 | LUAD | NCI-H1693 | Elesclomol | MVP_top60_DNA+RNA_seed | IGF1R | 0.345394 | LUAD | Calu-6 | Linsitinib |
| MVP_top60_DNA_seed | EGFR | -0.901723 | LUAD | NCI-H1755 | CUDC-101 | MVP_top60_DNA+RNA_seed | EGFR | 0.34541 | LUAD | NCI-H1993 | AZD3759 |
| MVP_top60_DNA_seed | ERBB2 | -0.901723 | LUAD | NCI-H1755 | CUDC-101 | MVP_top60_DNA+RNA_seed | FGFR1 | 0.346435 | LUAD | NCI-H1623 | PD173074 |
| MVP_top60_DNA_seed | ATM | -0.899538 | LUAD | NCI-H838 | KU-55933 | MVP_top60_DNA+RNA_seed | AKT1 | 0.346469 | LUAD | NCI-H2023 | AT13148 |
| MVP_top60_DNA_seed | HSP90AA1 | -0.895562 | LUAD | NCI-H1355 | Elesclomol | MVP_top60_DNA+RNA_seed | ATM | 0.346673 | LUAD | NCI-H1666 | KU-55933 |
| MVP_top60_DNA_seed | BRAF | -0.893033 | LUAD | NCI-H1792 | HG6-64-1 | MVP_top60_DNA+RNA_seed | MDM2 | 0.347219 | LUAD | NCI-H522 | Nutlin-3a (-) |
| MVP_top60_DNA_seed | PIK3CA | -0.892877 | LUAD | SW1573 | PI3Ka_4409 | MVP_top60_DNA+RNA_seed | EGFR | 0.347559 | LUAD | NCI-H23 | Erlotinib |
| MVP_top60_DNA_seed | HSP90AA1 | -0.892459 | LUAD | EMC-BAC-1 | Tanespimycin | MVP_top60_DNA+RNA_seed | PIK3CA | 0.347699 | LUAD | HOP-62 | AZD8186 |
| MVP_top60_DNA_seed | AKT1 | -0.891721 | LUAD | NCI-H1648 | AKT inhibitor VIII | MVP_top60_DNA+RNA_seed | BRAF | 0.348081 | LUAD | NCI-H2087 | SB590885 |
| MVP_top60_DNA_seed | EGFR | -0.889269 | LUAD | COR-L105 | CUDC-101 | MVP_top60_DNA+RNA_seed | AKT1 | 0.348915 | LUAD | EMC-BAC-1 | Afuresertib |
| MVP_top60_DNA_seed | ERBB2 | -0.889269 | LUAD | COR-L105 | CUDC-101 | MVP_top60_DNA+RNA_seed | MDM2 | 0.351381 | LUAD | NCI-H2030 | Nutlin-3a (-) |
| MVP_top60_DNA_seed | ERBB2 | -0.888153 | LUAD | LXF-289 | CP724714 | MVP_top60_DNA+RNA_seed | AKT1 | 0.351684 | LUAD | NCI-H1666 | Uprosertib |
| MVP_top60_DNA_seed | HSP90AA1 | -0.88717 | LUAD | NCI-H2023 | CCT-018159 | MVP_top60_DNA+RNA_seed | AKT1 | 0.351788 | LUAD | HCC-78 | Uprosertib |
| MVP_top60_DNA_seed | BRAF | -0.884074 | LUAD | Calu-6 | PLX-4720 | MVP_top60_DNA+RNA_seed | IGF1R | 0.353517 | LUAD | HCC-78 | Linsitinib |
| MVP_top60_DNA_seed | HDAC1 | -0.882527 | LUAD | LC-2-ad | CAY10603 | MVP_top60_DNA+RNA_seed | EGFR | 0.354711 | LUAD | NCI-H2009 | Sapitinib |
| MVP_top60_DNA_seed | AURKA | -0.88191 | LUAD | NCI-H1648 | HG-5-113-01 | MVP_top60_DNA+RNA_seed | AKT1 | 0.354932 | LUAD | HOP-62 | GSK2110183B |
| MVP_top60_DNA_seed | AKT1 | -0.88095 | LUAD | NCI-H2009 | MK-2206 | MVP_top60_DNA+RNA_seed | EGFR | 0.354936 | LUAD | NCI-H1838 | Sapitinib |
| MVP_top60_DNA_seed | HSP90AA1 | -0.880453 | LUAD | NCI-H650 | Luminespib | MVP_top60_DNA+RNA_seed | MET | 0.355221 | LUAD | NCI-H838 | Savolitinib |
| MVP_top60_DNA_seed | MDM2 | -0.877497 | LUAD | LXF-289 | Serdemetan | MVP_top60_DNA+RNA_seed | EGFR | 0.355795 | LUAD | RERF-LC-MS | Gefitinib |
| MVP_top60_DNA_seed | HSP90AA1 | -0.87501 | LUAD | NCI-H2009 | Luminespib | MVP_top60_DNA+RNA_seed | BRAF | 0.357213 | LUAD | NCI-H1781 | SB590885 |
| MVP_top60_DNA_seed | HSP90AA1 | -0.874584 | LUAD | NCI-H2228 | Luminespib | MVP_top60_DNA+RNA_seed | HDAC1 | 0.357475 | LUAD | NCI-H2228 | Entinostat |
| MVP_top60_DNA_seed | PIK3CA | -0.874333 | LUAD | NCI-H1975 | PI3Ka_4409 | MVP_top60_DNA+RNA_seed | AKT1 | 0.35782 | LUAD | NCI-H1650 | AZD5363 |
| MVP_top60_DNA_seed | HSP90AA1 | -0.874079 | LUAD | RERF-LC-KJ | Elesclomol | MVP_top60_DNA+RNA_seed | IGF1R | 0.35878 | LUAD | NCI-H2085 | BMS-536924 |
| MVP_top60_DNA_seed | PIK3CA | -0.872118 | LUAD | RERF-LC-MS | AZD8835 | MVP_top60_DNA+RNA_seed | EGFR | 0.360307 | LUAD | NCI-H1792 | AZD3759 |
| MVP_top60_DNA_seed | AURKA | -0.871966 | LUAD | NCI-H1651 | ZM447439 | MVP_top60_DNA+RNA_seed | BRAF | 0.363396 | LUAD | NCI-H1573 | Dabrafenib |

| | | | | | | | | | | | |
|---|---|---|---|---|---|---|---|---|---|---|---|
| MVP_top60_DNA_seed | AKT1 | -0.870995 | LUAD | NCI-H1993 | BAY AKT1 | MVP_top60_DNA+RNA_seed | BRAF | 0.363838 | LUAD | NCI-H2023 | SB590885 |
| MVP_top60_DNA_seed | AKT1 | -0.869636 | LUAD | HOP-62 | A-443654 | MVP_top60_DNA+RNA_seed | EGFR | 0.364517 | LUAD | NCI-H1734 | Erlotinib |
| MVP_top60_DNA_seed | EGFR | -0.868239 | LUAD | HCC-44 | Afatinib | MVP_top60_DNA+RNA_seed | BRAF | 0.364615 | LUAD | NCI-H2405 | PLX-4720 |
| MVP_top60_DNA_seed | ERBB2 | -0.868239 | LUAD | HCC-44 | Afatinib | MVP_top60_DNA+RNA_seed | PIK3CA | 0.364818 | LUAD | NCI-H2023 | AZD8186 |
| MVP_top60_DNA_seed | MET | -0.86671 | LUAD | LXF-289 | JNJ38877605 | MVP_top60_DNA+RNA_seed | BRAF | 0.365273 | LUAD | NCI-H1975 | PLX-4720 |
| MVP_top60_DNA_seed | HDAC1 | -0.865955 | LUAD | NCI-H1755 | AR-42 | MVP_top60_DNA+RNA_seed | EGFR | 0.365483 | LUAD | NCI-H23 | Afatinib |
| MVP_top60_DNA_seed | AKT1 | -0.86437 | LUAD | ABC-1 | GSK690693 | MVP_top60_DNA+RNA_seed | TP53 | 0.366569 | LUAD | EMC-BAC-1 | MIRA-1 |
| MVP_top60_DNA_seed | IGF1R | -0.864116 | LUAD | NCI-H1648 | BMS-536924 | MVP_top60_DNA+RNA_seed | HSP90AA1 | 0.366938 | LUAD | HOP-62 | Luminespib |
| MVP_top60_DNA_seed | EGFR | -0.861462 | LUAD | H3255 | CUDC-101 | MVP_top60_DNA+RNA_seed | MET | 0.367641 | LUAD | NCI-H1666 | Savolitinib |
| MVP_top60_DNA_seed | ERBB2 | -0.861462 | LUAD | H3255 | CUDC-101 | MVP_top60_DNA+RNA_seed | EGFR | 0.368728 | LUAD | NCI-H2291 | AZD3759 |
| MVP_top60_DNA_seed | PIK3CA | -0.860426 | LUAD | Calu-3 | PI3Ka_4409 | MVP_top60_DNA+RNA_seed | HDAC1 | 0.368921 | LUAD | NCI-H441 | Romidepsin |
| MVP_top60_DNA_seed | EGFR | -0.858535 | LUAD | PC-14 | CUDC-101 | MVP_top60_DNA+RNA_seed | MET | 0.369542 | LUAD | NCI-H650 | Crizotinib |
| MVP_top60_DNA_seed | ERBB2 | -0.858535 | LUAD | PC-14 | CUDC-101 | MVP_top60_DNA+RNA_seed | AKT1 | 0.370265 | LUAD | NCI-H1944 | Afuresertib |
| MVP_top60_DNA_seed | CREBBP | -0.856625 | LUAD | NCI-H1792 | I-CBP112 | MVP_top60_DNA+RNA_seed | EZH2 | 0.37247 | LUAD | NCI-H650 | GSK343 |
| MVP_top60_DNA_seed | ERBB2 | -0.856336 | LUAD | COR-L105 | CP724714 | MVP_top60_DNA+RNA_seed | MET | 0.373413 | LUAD | NCI-H2228 | Savolitinib |
| MVP_top60_DNA_seed | AURKA | -0.855929 | LUAD | HOP-62 | Genentech Cpd 10 | MVP_top60_DNA+RNA_seed | EGFR | 0.374487 | LUAD | NCI-H1734 | Sapitinib |
| MVP_top60_DNA_seed | IGF1R | -0.855027 | LUAD | NCI-H1944 | Linsitinib | MVP_top60_DNA+RNA_seed | HSP90AA1 | 0.379038 | LUAD | NCI-H2342 | Tanespimycin |
| MVP_top60_DNA_seed | SRC | -0.853721 | LUAD | NCI-H1975 | Bosutinib | MVP_top60_DNA+RNA_seed | BRAF | 0.379198 | LUAD | NCI-H1651 | SB590885 |
| MVP_top60_DNA_seed | GSK3B | -0.853435 | LUAD | NCI-H1563 | CHIR-99021 | MVP_top60_DNA+RNA_seed | MDM2 | 0.383577 | LUAD | EMC-BAC-2 | Serdemetan |
| MVP_top60_DNA_seed | BRAF | -0.850744 | LUAD | PC-14 | HG6-64-1 | MVP_top60_DNA+RNA_seed | AKT1 | 0.383772 | LUAD | NCI-H2228 | GSK2110183B |
| MVP_top60_DNA_seed | IGF1R | -0.850236 | LUAD | NCI-H1975 | BMS-754807 | MVP_top60_DNA+RNA_seed | AKT1 | 0.384279 | LUAD | NCI-H322M | Uprosertib |
| MVP_top60_DNA_seed | EGFR | -0.849954 | LUAD | LC-2-ad | Lapatinib | MVP_top60_DNA+RNA_seed | HDAC1 | 0.385286 | LUAD | NCI-H1944 | PCI-34051 |
| MVP_top60_DNA_seed | ERBB2 | -0.849954 | LUAD | LC-2-ad | Lapatinib | MVP_top60_DNA+RNA_seed | IGF1R | 0.386432 | LUAD | HCC-44 | GSK1904529A |
| MVP_top60_DNA_seed | EGFR | -0.849942 | LUAD | NCI-H2122 | Afatinib | MVP_top60_DNA+RNA_seed | PIK3CA | 0.387343 | LUAD | NCI-H522 | Alpelisib |
| MVP_top60_DNA_seed | ERBB2 | -0.849942 | LUAD | NCI-H2122 | Afatinib | MVP_top60_DNA+RNA_seed | IGF1R | 0.388063 | LUAD | NCI-H2291 | BMS-754807 |
| MVP_top60_DNA_seed | EGFR | -0.84962 | LUAD | LC-2-ad | AZD8931 | MVP_top60_DNA+RNA_seed | IGF1R | 0.38899 | LUAD | H3255 | BMS-754807 |
| MVP_top60_DNA_seed | ERBB2 | -0.84962 | LUAD | LC-2-ad | AZD8931 | MVP_top60_DNA+RNA_seed | EZH2 | 0.389964 | LUAD | NCI-H2009 | GSK343 |
| MVP_top60_DNA_seed | ERBB3 | -0.84962 | LUAD | LC-2-ad | AZD8931 | MVP_top60_DNA+RNA_seed | EGFR | 0.39044 | LUAD | NCI-H3122 | Osimertinib |
| MVP_top60_DNA_seed | BRAF | -0.846585 | LUAD | NCI-H358 | PLX-4720 | MVP_top60_DNA+RNA_seed | PIK3CA | 0.390472 | LUAD | NCI-H1793 | AZD8186 |
| MVP_top60_DNA_seed | AKT1 | -0.845301 | LUAD | NCI-H1563 | AKT inhibitor VIII | MVP_top60_DNA+RNA_seed | MET | 0.39217 | LUAD | NCI-H2342 | Savolitinib |
| MVP_top60_DNA_seed | HSP90AA1 | -0.842669 | LUAD | HOP-62 | Tanespimycin | MVP_top60_DNA+RNA_seed | AKT1 | 0.393403 | LUAD | NCI-H1623 | AZD5363 |
| MVP_top60_DNA_seed | AKT1 | -0.839841 | LUAD | LXF-289 | A-443654 | MVP_top60_DNA+RNA_seed | ATM | 0.394137 | LUAD | LXF-289 | KU-55933 |
| MVP_top60_DNA_seed | ATM | -0.839102 | LUAD | SK-LU-1 | CP466722 | MVP_top60_DNA+RNA_seed | AKT1 | 0.394308 | LUAD | NCI-H1838 | MK-2206 |
| MVP_top60_DNA_seed | PIK3CA | -0.839079 | LUAD | HCC-827 | AZD8835 | MVP_top60_DNA+RNA_seed | AURKA | 0.394314 | LUAD | NCI-H1666 | Alisertib |
| MVP_top60_DNA_seed | EGFR | -0.837233 | LUAD | NCI-H2030 | Pelitinib | MVP_top60_DNA+RNA_seed | AURKA | 0.395038 | LUAD | NCI-H358 | Alisertib |
| MVP_top60_DNA_seed | MAPK1 | -0.837223 | LUAD | HCC-44 | VX-11e | MVP_top60_DNA+RNA_seed | EZH2 | 0.396612 | LUAD | NCI-H2122 | GSK343 |
| MVP_top60_DNA_seed | FGFR1 | -0.836622 | LUAD | HCC-44 | PD173074 | MVP_top60_DNA+RNA_seed | IGF1R | 0.397802 | LUAD | EMC-BAC-2 | NVP-ADW742 |
| MVP_top60_DNA_seed | HSP90AA1 | -0.836021 | LUAD | NCI-H23 | Elesclomol | MVP_top60_DNA+RNA_seed | AURKA | 0.39858 | LUAD | H3255 | Alisertib |
| MVP_top60_DNA_seed | AKT1 | -0.833823 | LUAD | RERF-LC-KJ | AKT inhibitor VIII | MVP_top60_DNA+RNA_seed | PIK3CA | 0.39937 | LUAD | NCI-H1435 | AZD8186 |
| MVP_top60_DNA_seed | PTK2 | -0.833327 | LUAD | HOP-62 | PF-562271 | MVP_top60_DNA+RNA_seed | IGF1R | 0.399902 | LUAD | HCC-827 | GSK1904529A |
| MVP_top60_DNA_seed | BRAF | -0.833131 | LUAD | NCI-H3122 | SB590885 | MVP_top60_DNA+RNA_seed | PIK3CA | 0.400589 | LUAD | NCI-H322M | Alpelisib |

| | | | | | | | | | | | |
|---|---|---|---|---|---|---|---|---|---|---|---|
| MVP_top60_DNA_seed | BRAF | -0.832747 | LUAD | NCI-H358 | PLX-4720 | MVP_top60_DNA+RNA_seed | EZH2 | 0.401117 | LUAD | NCI-H23 | GSK343 |
| MVP_top60_DNA_seed | AKT1 | -0.832216 | LUAD | NCI-H2030 | AKT inhibitor VIII | MVP_top60_DNA+RNA_seed | FGFR1 | 0.402116 | LUAD | NCI-H1435 | PD173074 |
| MVP_top60_DNA_seed | MET | -0.828569 | LUAD | NCI-H1734 | JNJ38877605 | MVP_top60_DNA+RNA_seed | EGFR | 0.402792 | LUAD | LC-2-ad | Gefitinib |
| MVP_top60_DNA_seed | IGF1R | -0.82853 | LUAD | NCI-H650 | GSK1904529A | MVP_top60_DNA+RNA_seed | IGF1R | 0.4036 | LUAD | NCI-H2342 | NVP-ADW742 |
| MVP_top60_DNA_seed | HSP90AA1 | -0.82841 | LUAD | SK-LU-1 | Elesclomol | MVP_top60_DNA+RNA_seed | TERT | 0.405246 | LUAD | NCI-H1648 | BIBR-1532 |
| MVP_top60_DNA_seed | AKT1 | -0.828284 | LUAD | LXF-289 | BAY AKT1 | MVP_top60_DNA+RNA_seed | EGFR | 0.405539 | LUAD | NCI-H1437 | Erlotinib |
| MVP_top60_DNA_seed | HSP90AA1 | -0.827909 | LUAD | NCI-H2405 | Luminespib | MVP_top60_DNA+RNA_seed | SMARCA4 | 0.40642 | LUAD | NCI-H1755 | PFI3 |
| MVP_top60_DNA_seed | MAPK1 | -0.826587 | LUAD | RERF-LC-MS | FR-180204 | MVP_top60_DNA+RNA_seed | SRC | 0.412245 | LUAD | NCI-H23 | Bosutinib |
| MVP_top60_DNA_seed | MET | -0.826485 | LUAD | NCI-H1623 | JNJ38877605 | MVP_top60_DNA+RNA_seed | AKT1 | 0.412527 | LUAD | NCI-H1693 | Uprosertib |
| MVP_top60_DNA_seed | AKT1 | -0.826474 | LUAD | NCI-H1975 | A-443654 | MVP_top60_DNA+RNA_seed | MET | 0.413698 | LUAD | RERF-LC-KJ | Crizotinib |
| MVP_top60_DNA_seed | AURKA | -0.825324 | LUAD | HOP-62 | Tozasertib | MVP_top60_DNA+RNA_seed | TERT | 0.415574 | LUAD | NCI-H1975 | BIBR-1532 |
| MVP_top60_DNA_seed | ERBB2 | -0.823224 | LUAD | NCI-H2030 | CP724714 | MVP_top60_DNA+RNA_seed | EZH2 | 0.416819 | LUAD | HOP-62 | GSK343 |
| MVP_top60_DNA_seed | HSP90AA1 | -0.821418 | LUAD | A549 | Elesclomol | MVP_top60_DNA+RNA_seed | EGFR | 0.417187 | LUAD | NCI-H1944 | Lapatinib |
| MVP_top60_DNA_seed | BRAF | -0.821024 | LUAD | SK-LU-1 | HG6-64-1 | MVP_top60_DNA+RNA_seed | IGF1R | 0.417545 | LUAD | NCI-H2228 | NVP-ADW742 |
| MVP_top60_DNA_seed | MET | -0.820218 | LUAD | EMC-BAC-1 | JNJ38877605 | MVP_top60_DNA+RNA_seed | MDM2 | 0.418522 | LUAD | HCC-44 | Nutlin-3a (-) |
| MVP_top60_DNA_seed | EGFR | -0.819897 | LUAD | NCI-H358 | CI-1033 | MVP_top60_DNA+RNA_seed | EZH2 | 0.420148 | LUAD | NCI-H2085 | GSK343 |
| MVP_top60_DNA_seed | ERBB2 | -0.819897 | LUAD | NCI-H358 | CI-1033 | MVP_top60_DNA+RNA_seed | AURKA | 0.420766 | LUAD | NCI-H1648 | Alisertib |
| MVP_top60_DNA_seed | BRAF | -0.81973 | LUAD | NCI-H1792 | RAF_9304 | MVP_top60_DNA+RNA_seed | FGFR1 | 0.421163 | LUAD | RERF-LC-KJ | PD173074 |
| MVP_top60_DNA_seed | PTK2 | -0.817794 | LUAD | NCI-H3122 | PF-562271 | MVP_top60_DNA+RNA_seed | BRAF | 0.422025 | LUAD | NCI-H1838 | Dabrafenib |
| MVP_top60_DNA_seed | AURKA | -0.81424 | LUAD | HCC-78 | GSK1070916 | MVP_top60_DNA+RNA_seed | PIK3CA | 0.422842 | LUAD | H3255 | Alpelisib |
| MVP_top60_DNA_seed | ATM | -0.813508 | LUAD | NCI-H2023 | KU-60019 | MVP_top60_DNA+RNA_seed | MDM2 | 0.424316 | LUAD | SW1573 | Nutlin-3a (-) |
| MVP_top60_DNA_seed | AURKA | -0.812365 | LUAD | HCC-78 | Genentech Cpd 10 | MVP_top60_DNA+RNA_seed | MET | 0.425691 | LUAD | SK-LU-1 | Crizotinib |
| MVP_top60_DNA_seed | BRAF | -0.811809 | LUAD | NCI-H2405 | RAF_9304 | MVP_top60_DNA+RNA_seed | EGFR | 0.426458 | LUAD | NCI-H1792 | Osimertinib |
| MVP_top60_DNA_seed | AKT1 | -0.810689 | LUAD | NCI-H1648 | A-443654 | MVP_top60_DNA+RNA_seed | IGF1R | 0.427617 | LUAD | EMC-BAC-2 | BMS-754807 |
| MVP_top60_DNA_seed | PIK3CA | -0.809024 | LUAD | NCI-H1623 | AZD8835 | MVP_top60_DNA+RNA_seed | IGF1R | 0.428077 | LUAD | ABC-1 | BMS-536924 |
| MVP_top60_DNA_seed | AKT1 | -0.809019 | LUAD | NCI-H522 | AKT inhibitor VIII | MVP_top60_DNA+RNA_seed | AKT1 | 0.428535 | LUAD | NCI-H1568 | AZD5363 |
| MVP_top60_DNA_seed | HSP90AA1 | -0.808824 | LUAD | NCI-H1648 | Luminespib | MVP_top60_DNA+RNA_seed | BRAF | 0.429616 | LUAD | HCC-44 | PLX-4720 |
| MVP_top60_DNA_seed | ATM | -0.807586 | LUAD | NCI-H322M | KU-55933 | MVP_top60_DNA+RNA_seed | AURKA | 0.42967 | LUAD | NCI-H1993 | Alisertib |
| MVP_top60_DNA_seed | HSP90AA1 | -0.807239 | LUAD | SW1573 | Elesclomol | MVP_top60_DNA+RNA_seed | SRC | 0.429782 | LUAD | NCI-H1975 | Bosutinib |
| MVP_top60_DNA_seed | HSP90AA1 | -0.806797 | LUAD | NCI-H1755 | Elesclomol | MVP_top60_DNA+RNA_seed | EGFR | 0.432642 | LUAD | NCI-H838 | AZD3759 |
| MVP_top60_DNA_seed | ATM | -0.806019 | LUAD | COR-L105 | CP466722 | MVP_top60_DNA+RNA_seed | MDM2 | 0.432704 | LUAD | LXF-289 | Serdemetan |
| MVP_top60_DNA_seed | GSK3B | -0.805246 | LUAD | HOP-62 | AZD7969 | MVP_top60_DNA+RNA_seed | SMARCA4 | 0.433167 | LUAD | EMC-BAC-1 | PFI3 |
| MVP_top60_DNA_seed | HDAC1 | -0.804132 | LUAD | NCI-H522 | JQ12 | MVP_top60_DNA+RNA_seed | AURKA | 0.43388 | LUAD | HCC-827 | Alisertib |
| MVP_top60_DNA_seed | AKT1 | -0.803284 | LUAD | Calu-6 | A-443654 | MVP_top60_DNA+RNA_seed | FGFR1 | 0.434337 | LUAD | A549 | PD173074 |
| MVP_top60_DNA_seed | AKT1 | -0.803149 | LUAD | LXF-289 | AKT inhibitor VIII | MVP_top60_DNA+RNA_seed | IGF1R | 0.434682 | LUAD | NCI-H1435 | Linsitinib |
| MVP_top60_DNA_seed | HSP90AA1 | -0.801552 | LUAD | NCI-H1755 | SNX-2112 | MVP_top60_DNA+RNA_seed | PIK3CA | 0.435478 | LUAD | LC-2-ad | GNE-317 |
| MVP_top60_DNA_seed | ATM | -0.801269 | LUAD | RERF-LC-KJ | KU-55933 | MVP_top60_DNA+RNA_seed | MDM2 | 0.43644 | LUAD | NCI-H2023 | Nutlin-3a (-) |
| MVP_top60_DNA_seed | PTK2 | -0.80083 | LUAD | NCI-H2087 | PF-562271 | MVP_top60_DNA+RNA_seed | PIK3CA | 0.436512 | LUAD | NCI-H1435 | Alpelisib |
| MVP_top60_DNA_seed | MET | -0.800421 | LUAD | NCI-H1651 | JNJ38877605 | MVP_top60_DNA+RNA_seed | AKT1 | 0.437616 | LUAD | COR-L105 | GSK2110183B |
| MVP_top60_DNA_seed | HSP90AA1 | -0.800327 | LUAD | COR-L105 | Luminespib | MVP_top60_DNA+RNA_seed | IGF1R | 0.439011 | LUAD | NCI-H2228 | GSK1904529A |
| MVP_top60_DNA_seed | HDAC1 | -0.798706 | LUAD | ABC-1 | JQ12 | MVP_top60_DNA+RNA_seed | TP53 | 0.439079 | LUAD | NCI-H650 | MIRA-1 |

| | | | | | | | | | | | |
|---|---|---|---|---|---|---|---|---|---|---|---|
| MVP_top60_DNA_seed | HDAC1 | -0.798075 | LUAD | LC-2-ad | AR-42 | MVP_top60_DNA+RNA_seed | EGFR | 0.44076 | LUAD | NCI-H1650 | Lapatinib |
| MVP_top60_DNA_seed | MDM2 | -0.797776 | LUAD | NCI-H1755 | Serdemetan | MVP_top60_DNA+RNA_seed | EGFR | 0.440988 | LUAD | NCI-H1435 | Osimertinib |
| MVP_top60_DNA_seed | MDM2 | -0.797588 | LUAD | EMC-BAC-1 | Serdemetan | MVP_top60_DNA+RNA_seed | IGF1R | 0.444181 | LUAD | SW1573 | BMS-754807 |
| MVP_top60_DNA_seed | EGFR | -0.797095 | LUAD | NCI-H322M | Afatinib | MVP_top60_DNA+RNA_seed | TP53 | 0.444214 | LUAD | NCI-H3122 | MIRA-1 |
| MVP_top60_DNA_seed | ERBB2 | -0.797095 | LUAD | NCI-H322M | Afatinib | MVP_top60_DNA+RNA_seed | IGF1R | 0.445258 | LUAD | NCI-H1573 | BMS-536924 |
| MVP_top60_DNA_seed | BRAF | -0.795748 | LUAD | NCI-H1975 | Dabrafenib | MVP_top60_DNA+RNA_seed | HDAC1 | 0.446839 | LUAD | ABC-1 | PCI-34051 |
| MVP_top60_DNA_seed | HSP90AA1 | -0.794547 | LUAD | NCI-H1651 | CCT-018159 | MVP_top60_DNA+RNA_seed | AURKA | 0.448034 | LUAD | H3255 | Tozasertib |
| MVP_top60_DNA_seed | BRAF | -0.793222 | LUAD | LC-2-ad | PLX-4720 | MVP_top60_DNA+RNA_seed | EGFR | 0.450241 | LUAD | NCI-H2023 | Lapatinib |
| MVP_top60_DNA_seed | HDAC1 | -0.792252 | LUAD | HCC-44 | CAY10603 | MVP_top60_DNA+RNA_seed | EGFR | 0.450663 | LUAD | HCC-78 | AZD3759 |
| MVP_top60_DNA_seed | HDAC1 | -0.790653 | LUAD | NCI-H2228 | Dacinostat | MVP_top60_DNA+RNA_seed | PIK3CA | 0.451146 | LUAD | NCI-H1693 | AZD8186 |
| MVP_top60_DNA_seed | IGF1R | -0.789349 | LUAD | NCI-H358 | BMS-536924 | MVP_top60_DNA+RNA_seed | EGFR | 0.453084 | LUAD | Calu-6 | Afatinib |
| MVP_top60_DNA_seed | EGFR | -0.788934 | LUAD | COR-L105 | Cetuximab | MVP_top60_DNA+RNA_seed | BRAF | 0.455351 | LUAD | NCI-H650 | Dabrafenib |
| MVP_top60_DNA_seed | GSK3B | -0.788899 | LUAD | RERF-LC-MS | CHIR-99021 | MVP_top60_DNA+RNA_seed | BRAF | 0.455893 | LUAD | NCI-H23 | PLX-4720 |
| MVP_top60_DNA_seed | BRAF | -0.788367 | LUAD | A549 | HG6-64-1 | MVP_top60_DNA+RNA_seed | EGFR | 0.4563 | LUAD | ABC-1 | AZD3759 |
| MVP_top60_DNA_seed | EGFR | -0.788279 | LUAD | HCC-78 | Afatinib | MVP_top60_DNA+RNA_seed | PIK3CA | 0.456573 | LUAD | NCI-H2085 | GNE-317 |
| MVP_top60_DNA_seed | ERBB2 | -0.788279 | LUAD | HCC-78 | Afatinib | MVP_top60_DNA+RNA_seed | AKT1 | 0.459573 | LUAD | NCI-H1650 | Ipatasertib |
| MVP_top60_DNA_seed | MET | -0.788167 | LUAD | SK-LU-1 | JNJ38877605 | MVP_top60_DNA+RNA_seed | PIK3CA | 0.459856 | LUAD | NCI-H1573 | AZD8186 |
| MVP_top60_DNA_seed | GSK3B | -0.785107 | LUAD | NCI-H1792 | KIN001-042 | MVP_top60_DNA+RNA_seed | ATM | 0.460032 | LUAD | NCI-H1944 | KU-55933 |
| MVP_top60_DNA_seed | CREBBP | -0.78458 | LUAD | NCI-H2122 | I-CBP112 | MVP_top60_DNA+RNA_seed | SRC | 0.460245 | LUAD | HOP-62 | Bosutinib |
| MVP_top60_DNA_seed | SMARCA4 | -0.784166 | LUAD | NCI-H1792 | PFI-3 | MVP_top60_DNA+RNA_seed | EGFR | 0.460661 | LUAD | LXF-289 | Sapitinib |
| MVP_top60_DNA_seed | PIK3CA | -0.784165 | LUAD | Calu-3 | AZD8835 | MVP_top60_DNA+RNA_seed | BRAF | 0.46089 | LUAD | NCI-H1944 | Dabrafenib |
| MVP_top60_DNA_seed | EGFR | -0.782968 | LUAD | NCI-H1693 | CUDC-101 | MVP_top60_DNA+RNA_seed | HSP90AA1 | 0.462044 | LUAD | NCI-H358 | Luminespib |
| MVP_top60_DNA_seed | ERBB2 | -0.782968 | LUAD | NCI-H1693 | CUDC-101 | MVP_top60_DNA+RNA_seed | SRC | 0.462244 | LUAD | NCI-H1734 | Bosutinib |
| MVP_top60_DNA_seed | AKT1 | -0.782192 | LUAD | NCI-H3122 | BAY AKT1 | MVP_top60_DNA+RNA_seed | MDM2 | 0.462428 | LUAD | NCI-H522 | Serdemetan |
| MVP_top60_DNA_seed | ATM | -0.78189 | LUAD | LXF-289 | CP466722 | MVP_top60_DNA+RNA_seed | EGFR | 0.463042 | LUAD | NCI-H2085 | Lapatinib |
| MVP_top60_DNA_seed | IGF1R | -0.780734 | LUAD | NCI-H1563 | GSK1904529A | MVP_top60_DNA+RNA_seed | IGF1R | 0.46317 | LUAD | NCI-H3122 | GSK1904529A |
| MVP_top60_DNA_seed | MAPK1 | -0.780548 | LUAD | NCI-H2030 | FR-180204 | MVP_top60_DNA+RNA_seed | PIK3CA | 0.463421 | LUAD | LXF-289 | Alpelisib |
| MVP_top60_DNA_seed | ATM | -0.779539 | LUAD | NCI-H23 | CP466722 | MVP_top60_DNA+RNA_seed | IGF1R | 0.46507 | LUAD | H3255 | Linsitinib |
| MVP_top60_DNA_seed | EGFR | -0.778734 | LUAD | HCC-78 | CUDC-101 | MVP_top60_DNA+RNA_seed | TERT | 0.46521 | LUAD | NCI-H522 | BIBR-1532 |
| MVP_top60_DNA_seed | ERBB2 | -0.778734 | LUAD | HCC-78 | CUDC-101 | MVP_top60_DNA+RNA_seed | TERT | 0.466496 | LUAD | NCI-H2085 | BIBR-1532 |
| MVP_top60_DNA_seed | BRAF | -0.77557 | LUAD | LXF-289 | HG6-64-1 | MVP_top60_DNA+RNA_seed | PIK3CA | 0.467217 | LUAD | NCI-H1693 | Buparlisib |
| MVP_top60_DNA_seed | HSP90AA1 | -0.772694 | LUAD | NCI-H23 | CCT-018159 | MVP_top60_DNA+RNA_seed | MDM2 | 0.467775 | LUAD | NCI-H2342 | Serdemetan |
| MVP_top60_DNA_seed | AURKA | -0.770918 | LUAD | VMRC-LCD | ZM447439 | MVP_top60_DNA+RNA_seed | AURKA | 0.469075 | LUAD | NCI-H1838 | ZM447439 |
| MVP_top60_DNA_seed | EGFR | -0.770762 | LUAD | PC-14 | PF-00299804 | MVP_top60_DNA+RNA_seed | IGF1R | 0.470001 | LUAD | NCI-H1838 | BMS-754807 |
| MVP_top60_DNA_seed | ERBB2 | -0.770762 | LUAD | PC-14 | PF-00299804 | MVP_top60_DNA+RNA_seed | IGF1R | 0.470075 | LUAD | NCI-H2087 | Linsitinib |
| MVP_top60_DNA_seed | HSP90AA1 | -0.770752 | LUAD | NCI-H1944 | SNX-2112 | MVP_top60_DNA+RNA_seed | PIK3CA | 0.471136 | LUAD | NCI-H1355 | GNE-317 |
| MVP_top60_DNA_seed | HDAC1 | -0.766864 | LUAD | PC-3_[JPC-3] | JQ12 | MVP_top60_DNA+RNA_seed | AURKA | 0.472199 | LUAD | NCI-H2009 | ZM447439 |
| MVP_top60_DNA_seed | MET | -0.76305 | LUAD | LXF-289 | AZD6094 | MVP_top60_DNA+RNA_seed | BRAF | 0.472243 | LUAD | NCI-H1781 | Dabrafenib |
| MVP_top60_DNA_seed | SRC | -0.762393 | LUAD | NCI-H2087 | NVP-BHG712 | MVP_top60_DNA+RNA_seed | IGF1R | 0.473652 | LUAD | PC-14 | Linsitinib |
| MVP_top60_DNA_seed | AKT1 | -0.762254 | LUAD | NCI-H2228 | AKT inhibitor VIII | MVP_top60_DNA+RNA_seed | EZH2 | 0.47381 | LUAD | Calu-6 | GSK343 |
| MVP_top60_DNA_seed | AKT1 | -0.762151 | LUAD | NCI-H1650 | AKT inhibitor VIII | MVP_top60_DNA+RNA_seed | SRC | 0.474074 | LUAD | NCI-H2228 | Bosutinib |

| | | | | | | | | | | | |
|---|---|---|---|---|---|---|---|---|---|---|---|
| MVP_top60_DNA_seed | HSP90AA1 | -0.762084 | LUAD | NCI-H1650 | Luminespib | MVP_top60_DNA+RNA_seed | MET | 0.474964 | LUAD | NCI-H1563 | Crizotinib |
| MVP_top60_DNA_seed | HSP90AA1 | -0.762009 | LUAD | NCI-H2087 | Tanespimycin | MVP_top60_DNA+RNA_seed | AKT1 | 0.476004 | LUAD | NCI-H1792 | Ipatasertib |
| MVP_top60_DNA_seed | BRAF | -0.7615 | LUAD | NCI-H2087 | Dabrafenib | MVP_top60_DNA+RNA_seed | MET | 0.477036 | LUAD | SW1573 | Savolitinib |
| MVP_top60_DNA_seed | PIK3CA | -0.760682 | LUAD | HCC-78 | PI3Ka_4409 | MVP_top60_DNA+RNA_seed | MET | 0.477441 | LUAD | NCI-H1944 | Crizotinib |
| MVP_top60_DNA_seed | CREBBP | -0.757304 | LUAD | NCI-H2030 | I-CBP112 | MVP_top60_DNA+RNA_seed | IGF1R | 0.478872 | LUAD | NCI-H1944 | BMS-754807 |
| MVP_top60_DNA_seed | BRAF | -0.755734 | LUAD | EMC-BAC-2 | PLX-4720 | MVP_top60_DNA+RNA_seed | KRAS | 0.480611 | LUAD | VMRC-LCD | {RAS (G12C) Inhibitor-1 |
| MVP_top60_DNA_seed | FGFR1 | -0.755444 | LUAD | LXF-289 | PD173074 | MVP_top60_DNA+RNA_seed | AURKA | 0.481866 | LUAD | NCI-H650 | Alisertib |
| MVP_top60_DNA_seed | EGFR | -0.755404 | LUAD | NCI-H2405 | Cetuximab | MVP_top60_DNA+RNA_seed | IGF1R | 0.483089 | LUAD | H3255 | GSK1904529A |
| MVP_top60_DNA_seed | ATM | -0.754715 | LUAD | A549 | KU-60019 | MVP_top60_DNA+RNA_seed | EGFR | 0.483998 | LUAD | NCI-H1437 | Sapitinib |
| MVP_top60_DNA_seed | ATM | -0.754364 | LUAD | NCI-H1755 | KU-60019 | MVP_top60_DNA+RNA_seed | MDM2 | 0.487142 | LUAD | NCI-H1648 | Serdemetan |
| MVP_top60_DNA_seed | PTK2 | -0.753482 | LUAD | LXF-289 | PF-562271 | MVP_top60_DNA+RNA_seed | IGF1R | 0.487202 | LUAD | NCI-H1568 | BMS-536924 |
| MVP_top60_DNA_seed | EGFR | -0.753402 | LUAD | LC-2-ad | CUDC-101 | MVP_top60_DNA+RNA_seed | AKT1 | 0.487599 | LUAD | NCI-H23 | Afuresertib |
| MVP_top60_DNA_seed | ERBB2 | -0.753402 | LUAD | LC-2-ad | CUDC-101 | MVP_top60_DNA+RNA_seed | BRAF | 0.489139 | LUAD | NCI-H1573 | SB590885 |
| MVP_top60_DNA_seed | HSP90AA1 | -0.749954 | LUAD | NCI-H1568 | Luminespib | MVP_top60_DNA+RNA_seed | MDM2 | 0.489564 | LUAD | NCI-H1651 | Nutlin-3a (-) |
| MVP_top60_DNA_seed | MET | -0.749332 | LUAD | HCC-78 | JNJ38877605 | MVP_top60_DNA+RNA_seed | AKT1 | 0.490684 | LUAD | NCI-H1650 | Uprosertib |
| MVP_top60_DNA_seed | GSK3B | -0.748926 | LUAD | LC-2-ad | AZD7969 | MVP_top60_DNA+RNA_seed | IGF1R | 0.491428 | LUAD | Calu-6 | GSK1904529A |
| MVP_top60_DNA_seed | FGFR1 | -0.746649 | LUAD | NCI-H1781 | FGFR_3831 | MVP_top60_DNA+RNA_seed | AURKA | 0.491579 | LUAD | EMC-BAC-1 | Alisertib |
| MVP_top60_DNA_seed | HDAC1 | -0.74659 | LUAD | NCI-H1568 | JQ12 | MVP_top60_DNA+RNA_seed | MET | 0.493349 | LUAD | NCI-H1355 | Crizotinib |
| MVP_top60_DNA_seed | HDAC1 | -0.743949 | LUAD | HCC-44 | AR-42 | MVP_top60_DNA+RNA_seed | HDAC1 | 0.494469 | LUAD | NCI-H1666 | Entinostat |
| MVP_top60_DNA_seed | BRAF | -0.742221 | LUAD | H3255 | RAF_9304 | MVP_top60_DNA+RNA_seed | FGFR1 | 0.494507 | LUAD | NCI-H2023 | PD173074 |
| MVP_top60_DNA_seed | MET | -0.74079 | LUAD | NCI-H2087 | JNJ38877605 | MVP_top60_DNA+RNA_seed | EGFR | 0.496589 | LUAD | NCI-H1993 | Erlotinib |
| MVP_top60_DNA_seed | HDAC1 | -0.740112 | LUAD | NCI-H1568 | Dacinostat | MVP_top60_DNA+RNA_seed | AKT1 | 0.496863 | LUAD | NCI-H2228 | AZD5363 |
| MVP_top60_DNA_seed | ATM | -0.73971 | LUAD | NCI-H1648 | CP466722 | MVP_top60_DNA+RNA_seed | KRAS | 0.497351 | LUAD | Calu-6 | {RAS (G12C) Inhibitor-1 |
| MVP_top60_DNA_seed | MDM2 | -0.739485 | LUAD | NCI-H1666 | Nutlin-3a (-) | MVP_top60_DNA+RNA_seed | EGFR | 0.498189 | LUAD | NCI-H1435 | Erlotinib |
| MVP_top60_DNA_seed | BRAF | -0.739242 | LUAD | A549 | Dabrafenib | MVP_top60_DNA+RNA_seed | EGFR | 0.498214 | LUAD | NCI-H650 | Sapitinib |
| MVP_top60_DNA_seed | FGFR1 | -0.737829 | LUAD | SK-LU-1 | PD173074 | MVP_top60_DNA+RNA_seed | EGFR | 0.498421 | LUAD | NCI-H2085 | Afatinib |
| MVP_top60_DNA_seed | ATM | -0.737179 | LUAD | HCC-44 | KU-55933 | MVP_top60_DNA+RNA_seed | BRAF | 0.499074 | LUAD | NCI-H358 | Dabrafenib |
| MVP_top60_DNA_seed | MDM2 | -0.736887 | LUAD | NCI-H1563 | Nutlin-3a (-) | MVP_top60_DNA+RNA_seed | PIK3CA | 0.499505 | LUAD | SW1573 | Alpelisib |
| MVP_top60_DNA_seed | EGFR | -0.735907 | LUAD | NCI-H1693 | Afatinib | MVP_top60_DNA+RNA_seed | TP53 | 0.50233 | LUAD | HCC-44 | MIRA-1 |
| MVP_top60_DNA_seed | ERBB2 | -0.735907 | LUAD | NCI-H1693 | Afatinib | MVP_top60_DNA+RNA_seed | AKT1 | 0.503727 | LUAD | NCI-H1435 | Afuresertib |
| MVP_top60_DNA_seed | MDM2 | -0.735497 | LUAD | ABC-1 | Nutlin-3a (-) | MVP_top60_DNA+RNA_seed | EGFR | 0.504405 | LUAD | NCI-H1975 | Lapatinib |
| MVP_top60_DNA_seed | HSP90AA1 | -0.732356 | LUAD | NCI-H1666 | Tanespimycin | MVP_top60_DNA+RNA_seed | EZH2 | 0.504984 | LUAD | NCI-H1623 | GSK343 |
| MVP_top60_DNA_seed | EGFR | -0.732134 | LUAD | NCI-H1838 | Afatinib | MVP_top60_DNA+RNA_seed | EZH2 | 0.507082 | LUAD | EMC-BAC-2 | GSK343 |
| MVP_top60_DNA_seed | ERBB2 | -0.732134 | LUAD | NCI-H1838 | Afatinib | MVP_top60_DNA+RNA_seed | MDM2 | 0.508387 | LUAD | NCI-H1781 | Nutlin-3a (-) |
| MVP_top60_DNA_seed | AURKA | -0.727087 | LUAD | COR-L105 | Genentech Cpd 10 | MVP_top60_DNA+RNA_seed | HDAC1 | 0.508399 | LUAD | NCI-H1437 | PCI-34051 |
| MVP_top60_DNA_seed | AURKA | -0.726525 | LUAD | NCI-H522 | CD532 | MVP_top60_DNA+RNA_seed | AURKA | 0.510438 | LUAD | NCI-H1563 | Alisertib |
| MVP_top60_DNA_seed | GSK3B | -0.726328 | LUAD | LXF-289 | SB216763 | MVP_top60_DNA+RNA_seed | TP53 | 0.510946 | LUAD | NCI-H1993 | MIRA-1 |
| MVP_top60_DNA_seed | ATM | -0.725217 | LUAD | NCI-H2030 | CP466722 | MVP_top60_DNA+RNA_seed | AKT1 | 0.511628 | LUAD | NCI-H1693 | AT13148 |
| MVP_top60_DNA_seed | EGFR | -0.724683 | LUAD | NCI-H1781 | Afatinib | MVP_top60_DNA+RNA_seed | BRAF | 0.513362 | LUAD | ABC-1 | PLX-4720 |
| MVP_top60_DNA_seed | ERBB2 | -0.724683 | LUAD | NCI-H1781 | Afatinib | MVP_top60_DNA+RNA_seed | AURKA | 0.516417 | LUAD | NCI-H2023 | Alisertib |
| MVP_top60_DNA_seed | HSP90AA1 | -0.724433 | LUAD | NCI-H1568 | CCT-018159 | MVP_top60_DNA+RNA_seed | EGFR | 0.517694 | LUAD | NCI-H2228 | Afatinib |

| | | | | | | | | | | | |
|---|---|---|---|---|---|---|---|---|---|---|---|
| MVP_top60_DNA_seed | GSK3B | -0.723897 | LUAD | A549 | SB216763 | MVP_top60_DNA+RNA_seed | BRAF | 0.518845 | LUAD | LXF-289 | Dabrafenib |
| MVP_top60_DNA_seed | BRAF | -0.723066 | LUAD | NCI-H2228 | SB590885 | MVP_top60_DNA+RNA_seed | HSP90AA1 | 0.519982 | LUAD | HCC-827 | Luminespib |
| MVP_top60_DNA_seed | EGFR | -0.722475 | LUAD | EKVX | AST-1306 | MVP_top60_DNA+RNA_seed | IGF1R | 0.521956 | LUAD | PC-14 | BMS-754807 |
| MVP_top60_DNA_seed | AKT1 | -0.722092 | LUAD | NCI-H2085 | AKT inhibitor VIII | MVP_top60_DNA+RNA_seed | AKT1 | 0.522761 | LUAD | NCI-H23 | GSK2110183B |
| MVP_top60_DNA_seed | PIK3CA | -0.722031 | LUAD | RERF-LC-KJ | PI3Ka_4409 | MVP_top60_DNA+RNA_seed | EGFR | 0.523746 | LUAD | NCI-H1993 | Lapatinib |
| MVP_top60_DNA_seed | HSP90AA1 | -0.721398 | LUAD | NCI-H2023 | Tanespimycin | MVP_top60_DNA+RNA_seed | AKT1 | 0.524217 | LUAD | NCI-H522 | Uprosertib |
| MVP_top60_DNA_seed | HSP90AA1 | -0.718576 | LUAD | EMC-BAC-1 | Elesclomol | MVP_top60_DNA+RNA_seed | IGF1R | 0.5252 | LUAD | Calu-3 | GSK1904529A |
| MVP_top60_DNA_seed | ERBB2 | -0.717202 | LUAD | NCI-H838 | CP724714 | MVP_top60_DNA+RNA_seed | IGF1R | 0.525205 | LUAD | NCI-H1623 | BMS-536924 |
| MVP_top60_DNA_seed | AURKA | -0.716165 | LUAD | Calu-6 | ZM447439 | MVP_top60_DNA+RNA_seed | PIK3CA | 0.525401 | LUAD | NCI-H322M | GNE-317 |
| MVP_top60_DNA_seed | ATM | -0.714051 | LUAD | NCI-H1563 | CP466722 | MVP_top60_DNA+RNA_seed | FGFR1 | 0.52612 | LUAD | NCI-H1563 | PD173074 |
| MVP_top60_DNA_seed | HSP90AA1 | -0.713767 | LUAD | NCI-H522 | Luminespib | MVP_top60_DNA+RNA_seed | MET | 0.526252 | LUAD | NCI-H3122 | Savolitinib |
| MVP_top60_DNA_seed | BRAF | -0.7137 | LUAD | LXF-289 | PLX-4720 | MVP_top60_DNA+RNA_seed | TERT | 0.52758 | LUAD | ABC-1 | BIBR-1532 |
| MVP_top60_DNA_seed | PTK2 | -0.712438 | LUAD | NCI-H1975 | PF-562271 | MVP_top60_DNA+RNA_seed | HDAC1 | 0.527728 | LUAD | Calu-3 | Entinostat |
| MVP_top60_DNA_seed | EGFR | -0.710761 | LUAD | LXF-289 | Cetuximab | MVP_top60_DNA+RNA_seed | AKT1 | 0.527963 | LUAD | NCI-H1666 | AZD5363 |
| MVP_top60_DNA_seed | MAPK1 | -0.710184 | LUAD | EMC-BAC-1 | VX-11e | MVP_top60_DNA+RNA_seed | PIK3CA | 0.529885 | LUAD | PC-3_[JPC-3] | Alpelisib |
| MVP_top60_DNA_seed | HSP90AA1 | -0.708337 | LUAD | NCI-H1355 | Tanespimycin | MVP_top60_DNA+RNA_seed | TP53 | 0.529901 | LUAD | NCI-H1573 | MIRA-1 |
| MVP_top60_DNA_seed | AKT1 | -0.707251 | LUAD | NCI-H2023 | MK-2206 | MVP_top60_DNA+RNA_seed | AKT1 | 0.53014 | LUAD | EKVX | AZD5363 |
| MVP_top60_DNA_seed | SRC | -0.706545 | LUAD | NCI-H23 | Bosutinib | MVP_top60_DNA+RNA_seed | EGFR | 0.530153 | LUAD | NCI-H2023 | Erlotinib |
| MVP_top60_DNA_seed | BRAF | -0.701023 | LUAD | NCI-H2291 | RAF_9304 | MVP_top60_DNA+RNA_seed | MET | 0.530517 | LUAD | NCI-H522 | Savolitinib |
| MVP_top60_DNA_seed | AKT1 | -0.700709 | LUAD | HCC-78 | BAY AKT1 | MVP_top60_DNA+RNA_seed | AKT1 | 0.531035 | LUAD | ABC-1 | Afuresertib |
| MVP_top60_DNA_seed | ATM | -0.700102 | LUAD | PC-14 | CP466722 | MVP_top60_DNA+RNA_seed | AKT1 | 0.531308 | LUAD | NCI-H1623 | Afuresertib |
| MVP_top60_DNA_seed | AURKA | -0.698983 | LUAD | NCI-H1781 | ZM447439 | MVP_top60_DNA+RNA_seed | AKT1 | 0.531847 | LUAD | PC-3_[JPC-3] | AZD5363 |
| MVP_top60_DNA_seed | AURKA | -0.697135 | LUAD | NCI-H1355 | Genentech Cpd 10 | MVP_top60_DNA+RNA_seed | MET | 0.532802 | LUAD | Calu-6 | Crizotinib |
| MVP_top60_DNA_seed | EGFR | -0.696327 | LUAD | ABC-1 | CI-1033 | MVP_top60_DNA+RNA_seed | PIK3CA | 0.534481 | LUAD | NCI-H2291 | GNE-317 |
| MVP_top60_DNA_seed | ERBB2 | -0.696327 | LUAD | ABC-1 | CI-1033 | MVP_top60_DNA+RNA_seed | AKT1 | 0.53583 | LUAD | HOP-62 | AZD5363 |
| MVP_top60_DNA_seed | IGF1R | -0.695728 | LUAD | NCI-H1975 | BMS-536924 | MVP_top60_DNA+RNA_seed | HDAC1 | 0.536 | LUAD | SW1573 | Entinostat |
| MVP_top60_DNA_seed | BRAF | -0.694328 | LUAD | SK-LU-1 | PLX-4720 | MVP_top60_DNA+RNA_seed | BRAF | 0.536581 | LUAD | NCI-H2228 | PLX-4720 |
| MVP_top60_DNA_seed | HDAC1 | -0.691868 | LUAD | HCC-44 | Belinostat | MVP_top60_DNA+RNA_seed | BRAF | 0.537113 | LUAD | EMC-BAC-1 | SB590885 |
| MVP_top60_DNA_seed | BRAF | -0.690777 | LUAD | NCI-H2228 | PLX-4720 | MVP_top60_DNA+RNA_seed | FGFR1 | 0.539245 | LUAD | NCI-H1944 | PD173074 |
| MVP_top60_DNA_seed | IGF1R | -0.689977 | LUAD | SW1573 | BMS-536924 | MVP_top60_DNA+RNA_seed | EGFR | 0.539911 | LUAD | NCI-H650 | Afatinib |
| MVP_top60_DNA_seed | BRAF | -0.689639 | LUAD | VMRC-LCD | PLX-4720 | MVP_top60_DNA+RNA_seed | BRAF | 0.54013 | LUAD | NCI-H1993 | PLX-4720 |
| MVP_top60_DNA_seed | HSP90AA1 | -0.688588 | LUAD | NCI-H2030 | SNX-2112 | MVP_top60_DNA+RNA_seed | SMARCA4 | 0.540359 | LUAD | NCI-H3122 | PFI3 |
| MVP_top60_DNA_seed | ATM | -0.688418 | LUAD | NCI-H2009 | KU-60019 | MVP_top60_DNA+RNA_seed | EGFR | 0.541228 | LUAD | LXF-289 | Osimertinib |
| MVP_top60_DNA_seed | AURKA | -0.686987 | LUAD | HCC-78 | HG-5-113-01 | MVP_top60_DNA+RNA_seed | HSP90AA1 | 0.545184 | LUAD | HCC-78 | Luminespib |
| MVP_top60_DNA_seed | BRAF | -0.686418 | LUAD | NCI-H358 | RAF_9304 | MVP_top60_DNA+RNA_seed | FGFR1 | 0.546248 | LUAD | HOP-62 | AZD4547 |
| MVP_top60_DNA_seed | GSK3B | -0.682946 | LUAD | NCI-H1666 | CHIR-99021 | MVP_top60_DNA+RNA_seed | HDAC1 | 0.546284 | LUAD | NCI-H650 | PCI-34051 |
| MVP_top60_DNA_seed | AURKA | -0.68265 | LUAD | H3255 | Genentech Cpd 10 | MVP_top60_DNA+RNA_seed | TP53 | 0.546422 | LUAD | NCI-H1781 | MIRA-1 |
| MVP_top60_DNA_seed | PIK3CA | -0.681276 | LUAD | NCI-H1568 | PI3Ka_4409 | MVP_top60_DNA+RNA_seed | EGFR | 0.546422 | LUAD | LXF-289 | Gefitinib |
| MVP_top60_DNA_seed | HSP90AA1 | -0.681092 | LUAD | NCI-H2347 | CCT-018159 | MVP_top60_DNA+RNA_seed | PIK3CA | 0.546928 | LUAD | NCI-H522 | Buparlisib |
| MVP_top60_DNA_seed | EGFR | -0.678975 | LUAD | NCI-H2342 | CI-1033 | MVP_top60_DNA+RNA_seed | FGFR1 | 0.548106 | LUAD | NCI-H322M | AZD4547 |
| MVP_top60_DNA_seed | ERBB2 | -0.678975 | LUAD | NCI-H2342 | CI-1033 | MVP_top60_DNA+RNA_seed | HDAC1 | 0.548624 | LUAD | NCI-H1755 | Entinostat |

| | | | | | | | | | | | |
|---|---|---|---|---|---|---|---|---|---|---|---|
| MVP_top60_DNA_seed | HSP90AA1 | -0.678869 | LUAD | LXF-289 | Luminespib | MVP_top60_DNA+RNA_seed | MDM2 | 0.548627 | LUAD | NCI-H1755 | Nutlin-3a (-) |
| MVP_top60_DNA_seed | GSK3B | -0.677573 | LUAD | NCI-H2122 | AZD7969 | MVP_top60_DNA+RNA_seed | MDM2 | 0.549352 | LUAD | NCI-H1975 | Nutlin-3a (-) |
| MVP_top60_DNA_seed | AURKA | -0.677445 | LUAD | ABC-1 | Alisertib | MVP_top60_DNA+RNA_seed | AKT1 | 0.549956 | LUAD | NCI-H322M | AT13148 |
| MVP_top60_DNA_seed | HSP90AA1 | -0.676301 | LUAD | EMC-BAC-2 | SNX-2112 | MVP_top60_DNA+RNA_seed | IGF1R | 0.55195 | LUAD | NCI-H3122 | Linsitinib |
| MVP_top60_DNA_seed | AURKA | -0.676034 | LUAD | SK-LU-1 | HG-5-113-01 | MVP_top60_DNA+RNA_seed | FGFR1 | 0.552816 | LUAD | NCI-H1651 | AZD4547 |
| MVP_top60_DNA_seed | ATM | -0.67364 | LUAD | EMC-BAC-2 | CP466722 | MVP_top60_DNA+RNA_seed | AKT1 | 0.553661 | LUAD | NCI-H23 | Ipatasertib |
| MVP_top60_DNA_seed | MET | -0.673055 | LUAD | NCI-H2405 | JNJ38877605 | MVP_top60_DNA+RNA_seed | BRAF | 0.554123 | LUAD | NCI-H23 | Dabrafenib |
| MVP_top60_DNA_seed | HSP90AA1 | -0.672728 | LUAD | RERF-LC-KJ | CCT-018159 | MVP_top60_DNA+RNA_seed | BRAF | 0.554848 | LUAD | NCI-H1563 | SB590885 |
| MVP_top60_DNA_seed | SRC | -0.672019 | LUAD | EMC-BAC-2 | Bosutinib | MVP_top60_DNA+RNA_seed | AURKA | 0.555526 | LUAD | EMC-BAC-1 | Tozasertib |
| MVP_top60_DNA_seed | IGF1R | -0.668918 | LUAD | NCI-H1648 | Linsitinib | MVP_top60_DNA+RNA_seed | TP53 | 0.556919 | LUAD | ABC-1 | MIRA-1 |
| MVP_top60_DNA_seed | HSP90AA1 | -0.668779 | LUAD | COR-L105 | SNX-2112 | MVP_top60_DNA+RNA_seed | KRAS | 0.557187 | LUAD | NCI-H1563 | KRAS (G12C) Inhibitor-1 |
| MVP_top60_DNA_seed | GSK3B | -0.668775 | LUAD | NCI-H1755 | AZD7969 | MVP_top60_DNA+RNA_seed | IGF1R | 0.557316 | LUAD | NCI-H1693 | Linsitinib |
| MVP_top60_DNA_seed | MET | -0.668265 | LUAD | NCI-H441 | AZD6094 | MVP_top60_DNA+RNA_seed | PIK3CA | 0.559043 | LUAD | NCI-H650 | GNE-317 |
| MVP_top60_DNA_seed | HDAC1 | -0.667757 | LUAD | HOP-62 | CAY10603 | MVP_top60_DNA+RNA_seed | HSP90AA1 | 0.561848 | LUAD | ABC-1 | Luminespib |
| MVP_top60_DNA_seed | ATM | -0.667739 | LUAD | NCI-H358 | KU-55933 | MVP_top60_DNA+RNA_seed | BRAF | 0.562412 | LUAD | PC-14 | Dabrafenib |
| MVP_top60_DNA_seed | AKT1 | -0.666628 | LUAD | EMC-BAC-1 | AKT inhibitor VIII | MVP_top60_DNA+RNA_seed | EGFR | 0.562518 | LUAD | NCI-H2009 | Osimertinib |
| MVP_top60_DNA_seed | EGFR | -0.666492 | LUAD | NCI-H1792 | AZD8931 | MVP_top60_DNA+RNA_seed | BRAF | 0.562526 | LUAD | NCI-H1648 | SB590885 |
| MVP_top60_DNA_seed | ERBB2 | -0.666492 | LUAD | NCI-H1792 | AZD8931 | MVP_top60_DNA+RNA_seed | EGFR | 0.563491 | LUAD | NCI-H838 | Erlotinib |
| MVP_top60_DNA_seed | ERBB3 | -0.666492 | LUAD | NCI-H1792 | AZD8931 | MVP_top60_DNA+RNA_seed | MET | 0.563756 | LUAD | NCI-H1975 | Crizotinib |
| MVP_top60_DNA_seed | MDM2 | -0.666324 | LUAD | NCI-H1944 | Nutlin-3a (-) | MVP_top60_DNA+RNA_seed | EZH2 | 0.564153 | LUAD | NCI-H322M | GSK343 |
| MVP_top60_DNA_seed | CREBBP | -0.665743 | LUAD | NCI-H2347 | I-CBP112 | MVP_top60_DNA+RNA_seed | EZH2 | 0.564188 | LUAD | PC-3_[JPC-3] | GSK343 |
| MVP_top60_DNA_seed | EGFR | -0.664554 | LUAD | NCI-H2030 | Afatinib | MVP_top60_DNA+RNA_seed | PIK3CA | 0.567814 | LUAD | HCC-827 | Alpelisib |
| MVP_top60_DNA_seed | ERBB2 | -0.664554 | LUAD | NCI-H2030 | Afatinib | MVP_top60_DNA+RNA_seed | HSP90AA1 | 0.568702 | LUAD | NCI-H1435 | Tanespimycin |
| MVP_top60_DNA_seed | IGF1R | -0.664243 | LUAD | NCI-H1734 | BMS-754807 | MVP_top60_DNA+RNA_seed | AKT1 | 0.568705 | LUAD | HCC-827 | MK-2206 |
| MVP_top60_DNA_seed | HDAC1 | -0.66281 | LUAD | Calu-6 | Parthenolide | MVP_top60_DNA+RNA_seed | AURKA | 0.56972 | LUAD | NCI-H2291 | ZM447439 |
| MVP_top60_DNA_seed | HDAC1 | -0.662196 | LUAD | LXF-289 | Entinostat | MVP_top60_DNA+RNA_seed | MDM2 | 0.571058 | LUAD | LXF-289 | Nutlin-3a (-) |
| MVP_top60_DNA_seed | PTK2 | -0.662071 | LUAD | Calu-6 | PF-562271 | MVP_top60_DNA+RNA_seed | PIK3CA | 0.57141 | LUAD | ABC-1 | AZD8186 |
| MVP_top60_DNA_seed | HSP90AA1 | -0.660037 | LUAD | LXF-289 | Elesclomol | MVP_top60_DNA+RNA_seed | AKT1 | 0.574826 | LUAD | NCI-H1355 | Uprosertib |
| MVP_top60_DNA_seed | EGFR | -0.659382 | LUAD | PC-14 | AST-1306 | MVP_top60_DNA+RNA_seed | EGFR | 0.576504 | LUAD | NCI-H1838 | Afatinib |
| MVP_top60_DNA_seed | EGFR | -0.657954 | LUAD | NCI-H2085 | Gefitinib | MVP_top60_DNA+RNA_seed | SMARCA4 | 0.577417 | LUAD | EMC-BAC-2 | PFI3 |
| MVP_top60_DNA_seed | BRAF | -0.657135 | LUAD | SK-LU-1 | PLX-4720 | MVP_top60_DNA+RNA_seed | AURKA | 0.578513 | LUAD | NCI-H1623 | Alisertib |
| MVP_top60_DNA_seed | EGFR | -0.656322 | LUAD | NCI-H522 | Afatinib | MVP_top60_DNA+RNA_seed | EGFR | 0.578926 | LUAD | NCI-H2405 | Sapitinib |
| MVP_top60_DNA_seed | ERBB2 | -0.656322 | LUAD | NCI-H522 | Afatinib | MVP_top60_DNA+RNA_seed | AKT1 | 0.579046 | LUAD | NCI-H2291 | AZD5363 |
| MVP_top60_DNA_seed | PIK3CA | -0.6562 | LUAD | EMC-BAC-2 | PI3Ka_4409 | MVP_top60_DNA+RNA_seed | AURKA | 0.579097 | LUAD | Calu-3 | Alisertib |
| MVP_top60_DNA_seed | HSP90AA1 | -0.655859 | LUAD | NCI-H2347 | Tanespimycin | MVP_top60_DNA+RNA_seed | IGF1R | 0.583002 | LUAD | HCC-78 | GSK1904529A |
| MVP_top60_DNA_seed | BRAF | -0.655503 | LUAD | PC-14 | RAF_9304 | MVP_top60_DNA+RNA_seed | EGFR | 0.583439 | LUAD | NCI-H1975 | AZD3759 |
| MVP_top60_DNA_seed | MDM2 | -0.654886 | LUAD | NCI-H23 | Serdemetan | MVP_top60_DNA+RNA_seed | TERT | 0.583715 | LUAD | NCI-H2228 | BIBR-1532 |
| MVP_top60_DNA_seed | HDAC1 | -0.653899 | LUAD | NCI-H358 | Belinostat | MVP_top60_DNA+RNA_seed | AKT1 | 0.58426 | LUAD | Calu-3 | GSK2110183B |
| MVP_top60_DNA_seed | IGF1R | -0.653337 | LUAD | NCI-H1651 | Linsitinib | MVP_top60_DNA+RNA_seed | AKT1 | 0.584392 | LUAD | NCI-H1573 | Ipatasertib |
| MVP_top60_DNA_seed | IGF1R | -0.65133 | LUAD | NCI-H522 | Linsitinib | MVP_top60_DNA+RNA_seed | PIK3CA | 0.584576 | LUAD | NCI-H2087 | Alpelisib |
| MVP_top60_DNA_seed | ATM | -0.651048 | LUAD | LC-2-ad | KU-55933 | MVP_top60_DNA+RNA_seed | PIK3CA | 0.589701 | LUAD | NCI-H23 | AZD8186 |

| | | | | | | | | | | | |
|---|---|---|---|---|---|---|---|---|---|---|---|
| MVP_top60_DNA_seed | GSK3B | -0.649833 | LUAD | HCC-44 | CHIR-99021 | MVP_top60_DNA+RNA_seed | BRAF | 0.590174 | LUAD | A549 | SB590885 |
| MVP_top60_DNA_seed | AURKA | -0.649323 | LUAD | NCI-H23 | Genentech Cpd 10 | MVP_top60_DNA+RNA_seed | MDM2 | 0.59063 | LUAD | ABC-1 | Nutlin-3a (-) |
| MVP_top60_DNA_seed | MET | -0.648208 | LUAD | NCI-H23 | JNJ38877605 | MVP_top60_DNA+RNA_seed | MET | 0.590685 | LUAD | NCI-H2087 | Crizotinib |
| MVP_top60_DNA_seed | AKT1 | -0.6474 | LUAD | NCI-H1648 | AKT inhibitor VIII | MVP_top60_DNA+RNA_seed | IGF1R | 0.590867 | LUAD | NCI-H2087 | BMS-536924 |
| MVP_top60_DNA_seed | PIK3CA | -0.646999 | LUAD | NCI-H1755 | AZD8835 | MVP_top60_DNA+RNA_seed | FGFR1 | 0.591081 | LUAD | NCI-H3122 | PD173074 |
| MVP_top60_DNA_seed | EGFR | -0.646658 | LUAD | LXF-289 | Erlotinib | MVP_top60_DNA+RNA_seed | AKT1 | 0.591214 | LUAD | NCI-H358 | Uprosertib |
| MVP_top60_DNA_seed | HDAC1 | -0.646524 | LUAD | NCI-H1944 | CAY10603 | MVP_top60_DNA+RNA_seed | PIK3CA | 0.592756 | LUAD | NCI-H358 | AZD8186 |
| MVP_top60_DNA_seed | BRAF | -0.645122 | LUAD | NCI-H1792 | PLX-4720 | MVP_top60_DNA+RNA_seed | EGFR | 0.592821 | LUAD | NCI-H1563 | Lapatinib |
| MVP_top60_DNA_seed | BRAF | -0.644732 | LUAD | LXF-289 | SB590885 | MVP_top60_DNA+RNA_seed | HDAC1 | 0.593964 | LUAD | PC-14 | PCI-34051 |
| MVP_top60_DNA_seed | SRC | -0.643993 | LUAD | NCI-H1792 | Bosutinib | MVP_top60_DNA+RNA_seed | BRAF | 0.595342 | LUAD | NCI-H2085 | PLX-4720 |
| MVP_top60_DNA_seed | SRC | -0.642236 | LUAD | NCI-H1355 | WH-4-023 | MVP_top60_DNA+RNA_seed | AURKA | 0.595626 | LUAD | PC-3_[JPC-3] | Alisertib |
| MVP_top60_DNA_seed | MET | -0.641834 | LUAD | PC-14 | SU11274 | MVP_top60_DNA+RNA_seed | BRAF | 0.596116 | LUAD | Calu-6 | SB590885 |
| MVP_top60_DNA_seed | BRAF | -0.641561 | LUAD | NCI-H1650 | PLX-4720 | MVP_top60_DNA+RNA_seed | EGFR | 0.596338 | LUAD | NCI-H2291 | Lapatinib |
| MVP_top60_DNA_seed | SRC | -0.641479 | LUAD | HCC-44 | NVP-BHG712 | MVP_top60_DNA+RNA_seed | EZH2 | 0.596818 | LUAD | NCI-H1975 | GSK343 |
| MVP_top60_DNA_seed | ATM | -0.639384 | LUAD | NCI-H1793 | KU-60019 | MVP_top60_DNA+RNA_seed | BRAF | 0.597619 | LUAD | NCI-H2085 | Dabrafenib |
| MVP_top60_DNA_seed | HDAC1 | -0.638633 | LUAD | NCI-H358 | CAY10603 | MVP_top60_DNA+RNA_seed | MET | 0.598235 | LUAD | EMC-BAC-1 | Savolitinib |
| MVP_top60_DNA_seed | AURKA | -0.638101 | LUAD | NCI-H2085 | ZM447439 | MVP_top60_DNA+RNA_seed | IGF1R | 0.599332 | LUAD | HCC-78 | BMS-754807 |
| MVP_top60_DNA_seed | MAPK1 | -0.638068 | LUAD | NCI-H1437 | VX-11e | MVP_top60_DNA+RNA_seed | EGFR | 0.599903 | LUAD | VMRC-LCD | Afatinib |
| MVP_top60_DNA_seed | MET | -0.634907 | LUAD | NCI-H1792 | JNJ38877605 | MVP_top60_DNA+RNA_seed | EGFR | 0.600094 | LUAD | NCI-H1573 | AZD3759 |
| MVP_top60_DNA_seed | CREBBP | -0.634382 | LUAD | EMC-BAC-2 | I-CBP112 | MVP_top60_DNA+RNA_seed | MDM2 | 0.600156 | LUAD | NCI-H2085 | Nutlin-3a (-) |
| MVP_top60_DNA_seed | PIK3CA | -0.633208 | LUAD | NCI-H2342 | PI3Ka_4409 | MVP_top60_DNA+RNA_seed | TP53 | 0.60034 | LUAD | NCI-H1755 | MIRA-1 |
| MVP_top60_DNA_seed | AURKA | -0.632865 | LUAD | NCI-H358 | ZM447439 | MVP_top60_DNA+RNA_seed | BRAF | 0.601306 | LUAD | H3255 | PLX-4720 |
| MVP_top60_DNA_seed | HSP90AA1 | -0.632264 | LUAD | NCI-H1355 | Luminespib | MVP_top60_DNA+RNA_seed | BRAF | 0.602121 | LUAD | NCI-H1355 | PLX-4720 |
| MVP_top60_DNA_seed | EGFR | -0.632175 | LUAD | NCI-H358 | PF-00299804 | MVP_top60_DNA+RNA_seed | MDM2 | 0.602212 | LUAD | Calu-6 | Serdemetan |
| MVP_top60_DNA_seed | ERBB2 | -0.632175 | LUAD | NCI-H358 | PF-00299804 | MVP_top60_DNA+RNA_seed | AURKA | 0.602494 | LUAD | NCI-H2030 | Alisertib |
| MVP_top60_DNA_seed | MAPK1 | -0.632124 | LUAD | NCI-H3122 | VX-11e | MVP_top60_DNA+RNA_seed | EGFR | 0.602594 | LUAD | HCC-44 | Gefitinib |
| MVP_top60_DNA_seed | BRAF | -0.631858 | LUAD | NCI-H1755 | HG6-64-1 | MVP_top60_DNA+RNA_seed | FGFR1 | 0.605977 | LUAD | NCI-H2291 | PD173074 |
| MVP_top60_DNA_seed | AURKA | -0.630396 | LUAD | LC-2-ad | Genentech Cpd 10 | MVP_top60_DNA+RNA_seed | PIK3CA | 0.606174 | LUAD | HCC-827 | Buparlisib |
| MVP_top60_DNA_seed | ERBB2 | -0.629214 | LUAD | NCI-H1944 | CP724714 | MVP_top60_DNA+RNA_seed | HDAC1 | 0.609312 | LUAD | H3255 | Romidepsin |
| MVP_top60_DNA_seed | HDAC1 | -0.625641 | LUAD | HCC-78 | Belinostat | MVP_top60_DNA+RNA_seed | MDM2 | 0.61031 | LUAD | PC-14 | Nutlin-3a (-) |
| MVP_top60_DNA_seed | HDAC1 | -0.62523 | LUAD | NCI-H1648 | Belinostat | MVP_top60_DNA+RNA_seed | AKT1 | 0.610492 | LUAD | NCI-H522 | Uprosertib |
| MVP_top60_DNA_seed | HSP90AA1 | -0.623574 | LUAD | H3255 | Tanespimycin | MVP_top60_DNA+RNA_seed | PIK3CA | 0.611025 | LUAD | NCI-H1781 | Alpelisib |
| MVP_top60_DNA_seed | IGF1R | -0.623259 | LUAD | NCI-H1651 | BMS-754807 | MVP_top60_DNA+RNA_seed | AKT1 | 0.611646 | LUAD | NCI-H2030 | Uprosertib |
| MVP_top60_DNA_seed | EGFR | -0.622534 | LUAD | NCI-H1563 | Pelitinib | MVP_top60_DNA+RNA_seed | AKT1 | 0.615052 | LUAD | NCI-H1355 | MK-2206 |
| MVP_top60_DNA_seed | MDM2 | -0.620165 | LUAD | NCI-H1650 | Serdemetan | MVP_top60_DNA+RNA_seed | IGF1R | 0.618164 | LUAD | NCI-H1573 | BMS-754807 |
| MVP_top60_DNA_seed | SRC | -0.618727 | LUAD | NCI-H358 | NVP-BHG712 | MVP_top60_DNA+RNA_seed | BRAF | 0.620136 | LUAD | NCI-H2009 | SB590885 |
| MVP_top60_DNA_seed | EGFR | -0.618378 | LUAD | NCI-H358 | Afatinib | MVP_top60_DNA+RNA_seed | AKT1 | 0.620396 | LUAD | NCI-H1563 | AT13148 |
| MVP_top60_DNA_seed | ERBB2 | -0.618378 | LUAD | NCI-H358 | Afatinib | MVP_top60_DNA+RNA_seed | SMARCA4 | 0.623069 | LUAD | VMRC-LCD | PFI3 |
| MVP_top60_DNA_seed | HDAC1 | -0.618053 | LUAD | Calu-6 | Dacinostat | MVP_top60_DNA+RNA_seed | BRAF | 0.624075 | LUAD | VMRC-LCD | PLX-4720 |
| MVP_top60_DNA_seed | SMARCA4 | -0.617358 | LUAD | NCI-H1944 | PFI-3 | MVP_top60_DNA+RNA_seed | HDAC1 | 0.62518 | LUAD | NCI-H1355 | Romidepsin |
| MVP_top60_DNA_seed | EGFR | -0.617067 | LUAD | NCI-H1944 | Pelitinib | MVP_top60_DNA+RNA_seed | TERT | 0.626208 | LUAD | NCI-H1563 | BIBR-1532 |

| | | | | | | | | | | | |
|---|---|---|---|---|---|---|---|---|---|---|---|
| MVP_top60_DNA_seed | BRAF | -0.61523 | LUAD | SW1573 | HG6-64-1 | MVP_top60_DNA+RNA_seed | HSP90AA1 | 0.626784 | LUAD | H3255 | Tanespimycin |
| MVP_top60_DNA_seed | HDAC1 | -0.615097 | LUAD | NCI-H1693 | AR-42 | MVP_top60_DNA+RNA_seed | IGF1R | 0.628832 | LUAD | NCI-H1650 | Linsitinib |
| MVP_top60_DNA_seed | BRAF | -0.61272 | LUAD | NCI-H2030 | PLX-4720 | MVP_top60_DNA+RNA_seed | AKT1 | 0.629469 | LUAD | H3255 | Uprosertib |
| MVP_top60_DNA_seed | PIK3CA | -0.612568 | LUAD | PC-14 | AZD8835 | MVP_top60_DNA+RNA_seed | HDAC1 | 0.63174 | LUAD | H3255 | Entinostat |
| MVP_top60_DNA_seed | MAPK1 | -0.612041 | LUAD | NCI-H1993 | FR-180204 | MVP_top60_DNA+RNA_seed | PIK3CA | 0.631896 | LUAD | NCI-H358 | GNE-317 |
| MVP_top60_DNA_seed | PIK3CA | -0.611118 | LUAD | NCI-H1792 | AZD8835 | MVP_top60_DNA+RNA_seed | MET | 0.632568 | LUAD | NCI-H1651 | Crizotinib |
| MVP_top60_DNA_seed | MET | -0.610997 | LUAD | RERF-LC-MS | SU11274 | MVP_top60_DNA+RNA_seed | AKT1 | 0.634998 | LUAD | NCI-H1792 | AT13148 |
| MVP_top60_DNA_seed | HSP90AA1 | -0.609813 | LUAD | NCI-H1648 | Tanespimycin | MVP_top60_DNA+RNA_seed | AURKA | 0.635389 | LUAD | NCI-H1651 | ZM447439 |
| MVP_top60_DNA_seed | HSP90AA1 | -0.608125 | LUAD | NCI-H650 | Tanespimycin | MVP_top60_DNA+RNA_seed | MDM2 | 0.635966 | LUAD | NCI-H2023 | Serdemetan |
| MVP_top60_DNA_seed | BRAF | -0.60789 | LUAD | NCI-H1975 | PLX-4720 | MVP_top60_DNA+RNA_seed | FGFR1 | 0.636977 | LUAD | NCI-H1437 | AZD4547 |
| MVP_top60_DNA_seed | EGFR | -0.607518 | LUAD | LC-2-ad | Pelitinib | MVP_top60_DNA+RNA_seed | EGFR | 0.637535 | LUAD | NCI-H1437 | Osimertinib |
| MVP_top60_DNA_seed | CREBBP | -0.606151 | LUAD | NCI-H1666 | I-CBP112 | MVP_top60_DNA+RNA_seed | MET | 0.637689 | LUAD | NCI-H1435 | Crizotinib |
| MVP_top60_DNA_seed | IGF1R | -0.605174 | LUAD | NCI-H522 | GSK1904529A | MVP_top60_DNA+RNA_seed | AKT1 | 0.638026 | LUAD | ABC-1 | AZD5363 |
| MVP_top60_DNA_seed | HSP90AA1 | -0.60477 | LUAD | LC-2-ad | Tanespimycin | MVP_top60_DNA+RNA_seed | KRAS | 0.638042 | LUAD | NCI-H522 | KRAS (G12C) Inhibitor-1 |
| MVP_top60_DNA_seed | HDAC1 | -0.604037 | LUAD | PC-14 | Tubastatin A | MVP_top60_DNA+RNA_seed | KRAS | 0.63857 | LUAD | NCI-H2087 | KRAS (G12C) Inhibitor-1 |
| MVP_top60_DNA_seed | IGF1R | -0.603982 | LUAD | NCI-H2228 | Linsitinib | MVP_top60_DNA+RNA_seed | MDM2 | 0.640825 | LUAD | NCI-H650 | Nutlin-3a (-) |
| MVP_top60_DNA_seed | HDAC1 | -0.603575 | LUAD | HCC-78 | Dacinostat | MVP_top60_DNA+RNA_seed | AKT1 | 0.640863 | LUAD | NCI-H2291 | Uprosertib |
| MVP_top60_DNA_seed | HDAC1 | -0.602551 | LUAD | Calu-6 | Belinostat | MVP_top60_DNA+RNA_seed | ATM | 0.641153 | LUAD | EMC-BAC-1 | KU-55933 |
| MVP_top60_DNA_seed | IGF1R | -0.599947 | LUAD | NCI-H1623 | Linsitinib | MVP_top60_DNA+RNA_seed | EGFR | 0.644414 | LUAD | HCC-44 | Osimertinib |
| MVP_top60_DNA_seed | BRAF | -0.599793 | LUAD | NCI-H1666 | RAF_9304 | MVP_top60_DNA+RNA_seed | PIK3CA | 0.644716 | LUAD | NCI-H1651 | Alpelisib |
| MVP_top60_DNA_seed | EGFR | -0.598844 | LUAD | ABC-1 | CUDC-101 | MVP_top60_DNA+RNA_seed | BRAF | 0.647963 | LUAD | NCI-H1623 | Dabrafenib |
| MVP_top60_DNA_seed | ERBB2 | -0.598844 | LUAD | ABC-1 | CUDC-101 | MVP_top60_DNA+RNA_seed | AKT1 | 0.648916 | LUAD | NCI-H1838 | Ipatasertib |
| MVP_top60_DNA_seed | MDM2 | -0.598743 | LUAD | NCI-H1734 | Serdemetan | MVP_top60_DNA+RNA_seed | AKT1 | 0.649513 | LUAD | NCI-H1355 | Uprosertib |
| MVP_top60_DNA_seed | EGFR | -0.597975 | LUAD | NCI-H522 | CUDC-101 | MVP_top60_DNA+RNA_seed | IGF1R | 0.650311 | LUAD | NCI-H2405 | BMS-536924 |
| MVP_top60_DNA_seed | ERBB2 | -0.597975 | LUAD | NCI-H522 | CUDC-101 | MVP_top60_DNA+RNA_seed | EGFR | 0.650505 | LUAD | NCI-H2228 | Erlotinib |
| MVP_top60_DNA_seed | HDAC1 | -0.597717 | LUAD | NCI-H1944 | AR-42 | MVP_top60_DNA+RNA_seed | AKT1 | 0.65206 | LUAD | COR-L105 | AZD5363 |
| MVP_top60_DNA_seed | HSP90AA1 | -0.597047 | LUAD | NCI-H2405 | CCT-018159 | MVP_top60_DNA+RNA_seed | EZH2 | 0.652435 | LUAD | A549 | GSK343 |
| MVP_top60_DNA_seed | ATM | -0.595568 | LUAD | NCI-H2122 | CP466722 | MVP_top60_DNA+RNA_seed | ATM | 0.653319 | LUAD | HOP-62 | KU-55933 |
| MVP_top60_DNA_seed | AURKA | -0.595024 | LUAD | EMC-BAC-1 | GSK1070916 | MVP_top60_DNA+RNA_seed | AKT1 | 0.653655 | LUAD | NCI-H1793 | AT13148 |
| MVP_top60_DNA_seed | SRC | -0.594104 | LUAD | EMC-BAC-1 | NVP-BHG712 | MVP_top60_DNA+RNA_seed | PIK3CA | 0.654089 | LUAD | NCI-H650 | Alpelisib |
| MVP_top60_DNA_seed | HSP90AA1 | -0.592571 | LUAD | NCI-H1944 | Luminespib | MVP_top60_DNA+RNA_seed | EGFR | 0.65424 | LUAD | NCI-H1993 | Gefitinib |
| MVP_top60_DNA_seed | HDAC1 | -0.590629 | LUAD | NCI-H1355 | JQ12 | MVP_top60_DNA+RNA_seed | AKT1 | 0.654981 | LUAD | NCI-H1623 | GSK2110183B |
| MVP_top60_DNA_seed | AKT1 | -0.590362 | LUAD | NCI-H2291 | BAY AKT1 | MVP_top60_DNA+RNA_seed | PIK3CA | 0.65543 | LUAD | NCI-H1650 | Buparlisib |
| MVP_top60_DNA_seed | GSK3B | -0.589654 | LUAD | LC-2-ad | CHIR-99021 | MVP_top60_DNA+RNA_seed | AURKA | 0.655655 | LUAD | NCI-H2030 | Tozasertib |
| MVP_top60_DNA_seed | HDAC1 | -0.589611 | LUAD | PC-14 | Belinostat | MVP_top60_DNA+RNA_seed | BRAF | 0.6565 | LUAD | RERF-LC-KJ | Dabrafenib |
| MVP_top60_DNA_seed | AKT1 | -0.589545 | LUAD | NCI-H2023 | BAY AKT1 | MVP_top60_DNA+RNA_seed | MET | 0.657986 | LUAD | HCC-827 | Crizotinib |
| MVP_top60_DNA_seed | EGFR | -0.588143 | LUAD | NCI-H1648 | CUDC-101 | MVP_top60_DNA+RNA_seed | FGFR1 | 0.658951 | LUAD | NCI-H2405 | AZD4547 |
| MVP_top60_DNA_seed | ERBB2 | -0.588143 | LUAD | NCI-H1648 | CUDC-101 | MVP_top60_DNA+RNA_seed | AKT1 | 0.659749 | LUAD | NCI-H650 | Uprosertib |
| MVP_top60_DNA_seed | IGF1R | -0.587678 | LUAD | NCI-H522 | BMS-754807 | MVP_top60_DNA+RNA_seed | HDAC1 | 0.660331 | LUAD | NCI-H358 | Entinostat |
| MVP_top60_DNA_seed | MDM2 | -0.587523 | LUAD | NCI-H2347 | Serdemetan | MVP_top60_DNA+RNA_seed | HSP90AA1 | 0.661104 | LUAD | NCI-H1568 | Tanespimycin |
| MVP_top60_DNA_seed | BRAF | -0.586351 | LUAD | A549 | RAF_9304 | MVP_top60_DNA+RNA_seed | MDM2 | 0.66126 | LUAD | NCI-H322M | Nutlin-3a (-) |

| | | | | | | | | | | | |
|---|---|---|---|---|---|---|---|---|---|---|---|
| MVP_top60_DNA_seed | SRC | -0.585827 | LUAD | RERF-LC-KJ | Bosutinib | MVP_top60_DNA+RNA_seed | IGF1R | 0.661587 | LUAD | PC-3_[JPC-3] | NVP-ADW742 |
| MVP_top60_DNA_seed | AURKA | -0.584803 | LUAD | PC-3_[JPC-3] | Genentech Cpd 10 | MVP_top60_DNA+RNA_seed | HSP90AA1 | 0.662792 | LUAD | NCI-H1666 | Luminespib |
| MVP_top60_DNA_seed | AURKA | -0.58403 | LUAD | NCI-H358 | GSK1070916 | MVP_top60_DNA+RNA_seed | MET | 0.662799 | LUAD | NCI-H322M | Crizotinib |
| MVP_top60_DNA_seed | HDAC1 | -0.583787 | LUAD | NCI-H1648 | AR-42 | MVP_top60_DNA+RNA_seed | MET | 0.662852 | LUAD | COR-L105 | Crizotinib |
| MVP_top60_DNA_seed | PTK2 | -0.583637 | LUAD | NCI-H838 | PF-562271 | MVP_top60_DNA+RNA_seed | KRAS | 0.664545 | LUAD | NCI-H1944 | (RAS (G12C) Inhibitor-1 |
| MVP_top60_DNA_seed | HSP90AA1 | -0.582992 | LUAD | NCI-H2009 | Tanespimycin | MVP_top60_DNA+RNA_seed | PIK3CA | 0.66601 | LUAD | NCI-H1651 | GNE-317 |
| MVP_top60_DNA_seed | GSK3B | -0.582785 | LUAD | EMC-BAC-1 | KIN001-042 | MVP_top60_DNA+RNA_seed | AKT1 | 0.666867 | LUAD | NCI-H1651 | GSK2110183B |
| MVP_top60_DNA_seed | MDM2 | -0.582278 | LUAD | NCI-H2085 | Nutlin-3a (-) | MVP_top60_DNA+RNA_seed | SRC | 0.667232 | LUAD | NCI-H1693 | Bosutinib |
| MVP_top60_DNA_seed | HSP90AA1 | -0.582159 | LUAD | NCI-H2085 | Luminespib | MVP_top60_DNA+RNA_seed | FGFR1 | 0.667531 | LUAD | NCI-H1666 | AZD4547 |
| MVP_top60_DNA_seed | SRC | -0.581158 | LUAD | NCI-H1650 | Bosutinib | MVP_top60_DNA+RNA_seed | HDAC1 | 0.668035 | LUAD | HOP-62 | Entinostat |
| MVP_top60_DNA_seed | AKT1 | -0.580852 | LUAD | NCI-H2085 | AKT inhibitor VIII | MVP_top60_DNA+RNA_seed | ATM | 0.670889 | LUAD | COR-L105 | KU-55933 |
| MVP_top60_DNA_seed | ERBB2 | -0.580704 | LUAD | NCI-H1648 | CP724714 | MVP_top60_DNA+RNA_seed | MDM2 | 0.671376 | LUAD | NCI-H3122 | Nutlin-3a (-) |
| MVP_top60_DNA_seed | HDAC1 | -0.579974 | LUAD | PC-14 | AR-42 | MVP_top60_DNA+RNA_seed | AKT1 | 0.67156 | LUAD | Calu-6 | MK-2206 |
| MVP_top60_DNA_seed | HSP90AA1 | -0.578678 | LUAD | NCI-H650 | CCT-018159 | MVP_top60_DNA+RNA_seed | BRAF | 0.671589 | LUAD | NCI-H1623 | SB590885 |
| MVP_top60_DNA_seed | EGFR | -0.578358 | LUAD | NCI-H1355 | AZD8931 | MVP_top60_DNA+RNA_seed | AURKA | 0.672937 | LUAD | NCI-H23 | ZM447439 |
| MVP_top60_DNA_seed | ERBB2 | -0.578358 | LUAD | NCI-H1355 | AZD8931 | MVP_top60_DNA+RNA_seed | EGFR | 0.673907 | LUAD | VMRC-LCD | Gefitinib |
| MVP_top60_DNA_seed | ERBB3 | -0.578358 | LUAD | NCI-H1355 | AZD8931 | MVP_top60_DNA+RNA_seed | EZH2 | 0.674217 | LUAD | PC-14 | GSK343 |
| MVP_top60_DNA_seed | ATM | -0.578029 | LUAD | NCI-H1435 | KU-60019 | MVP_top60_DNA+RNA_seed | EGFR | 0.674894 | LUAD | NCI-H23 | AZD3759 |
| MVP_top60_DNA_seed | EGFR | -0.577925 | LUAD | NCI-H1666 | AST-1306 | MVP_top60_DNA+RNA_seed | EZH2 | 0.677033 | LUAD | VMRC-LCD | GSK343 |
| MVP_top60_DNA_seed | BRAF | -0.577473 | LUAD | NCI-H1648 | RAF_9304 | MVP_top60_DNA+RNA_seed | SRC | 0.677186 | LUAD | NCI-H1623 | Bosutinib |
| MVP_top60_DNA_seed | EGFR | -0.575972 | LUAD | NCI-H2342 | AST-1306 | MVP_top60_DNA+RNA_seed | AKT1 | 0.678233 | LUAD | NCI-H650 | GSK2110183B |
| MVP_top60_DNA_seed | SRC | -0.575802 | LUAD | NCI-H2009 | Bosutinib | MVP_top60_DNA+RNA_seed | FGFR1 | 0.678941 | LUAD | NCI-H1666 | PD173074 |
| MVP_top60_DNA_seed | AURKA | -0.574982 | LUAD | NCI-H2009 | ZM447439 | MVP_top60_DNA+RNA_seed | AKT1 | 0.68001 | LUAD | NCI-H358 | GSK2110183B |
| MVP_top60_DNA_seed | MDM2 | -0.574738 | LUAD | PC-14 | Serdemetan | MVP_top60_DNA+RNA_seed | EGFR | 0.6815 | LUAD | Calu-6 | Sapitinib |
| MVP_top60_DNA_seed | CREBBP | -0.572032 | LUAD | NCI-H1650 | I-CBP112 | MVP_top60_DNA+RNA_seed | AURKA | 0.683112 | LUAD | NCI-H1838 | Alisertib |
| MVP_top60_DNA_seed | BRAF | -0.570587 | LUAD | NCI-H1755 | PLX-4720 | MVP_top60_DNA+RNA_seed | EGFR | 0.684061 | LUAD | ABC-1 | Sapitinib |
| MVP_top60_DNA_seed | MET | -0.569336 | LUAD | LC-2-ad | PHA-665752 | MVP_top60_DNA+RNA_seed | EGFR | 0.684482 | LUAD | NCI-H650 | Erlotinib |
| MVP_top60_DNA_seed | PTK2 | -0.568282 | LUAD | NCI-H1650 | PF-562271 | MVP_top60_DNA+RNA_seed | EZH2 | 0.684909 | LUAD | NCI-H1568 | GSK343 |
| MVP_top60_DNA_seed | EGFR | -0.566939 | LUAD | NCI-H358 | AST-1306 | MVP_top60_DNA+RNA_seed | IGF1R | 0.685186 | LUAD | NCI-H650 | BMS-754807 |
| MVP_top60_DNA_seed | GSK3B | -0.566684 | LUAD | PC-14 | AZD7969 | MVP_top60_DNA+RNA_seed | BRAF | 0.686657 | LUAD | NCI-H1666 | Dabrafenib |
| MVP_top60_DNA_seed | MAPK1 | -0.566514 | LUAD | NCI-H1944 | VX-11e | MVP_top60_DNA+RNA_seed | EGFR | 0.687891 | LUAD | NCI-H1755 | AZD3759 |
| MVP_top60_DNA_seed | HSP90AA1 | -0.565504 | LUAD | PC-14 | Luminespib | MVP_top60_DNA+RNA_seed | AKT1 | 0.688797 | LUAD | NCI-H1435 | Uprosertib |
| MVP_top60_DNA_seed | ATM | -0.565224 | LUAD | NCI-H1563 | KU-60019 | MVP_top60_DNA+RNA_seed | FGFR1 | 0.689126 | LUAD | NCI-H1793 | AZD4547 |
| MVP_top60_DNA_seed | AURKA | -0.564473 | LUAD | NCI-H650 | Genentech Cpd 10 | MVP_top60_DNA+RNA_seed | TP53 | 0.690013 | LUAD | NCI-H1944 | MIRA-1 |
| MVP_top60_DNA_seed | SMARCA4 | -0.564417 | LUAD | A549 | PFI-3 | MVP_top60_DNA+RNA_seed | BRAF | 0.690759 | LUAD | SK-LU-1 | Dabrafenib |
| MVP_top60_DNA_seed | MDM2 | -0.564314 | LUAD | HOP-62 | Serdemetan | MVP_top60_DNA+RNA_seed | IGF1R | 0.691104 | LUAD | HCC-827 | NVP-ADW742 |
| MVP_top60_DNA_seed | GSK3B | -0.564291 | LUAD | A549 | AZD7969 | MVP_top60_DNA+RNA_seed | BRAF | 0.691841 | LUAD | NCI-H2228 | Dabrafenib |
| MVP_top60_DNA_seed | ATM | -0.563834 | LUAD | NCI-H1435 | KU-55933 | MVP_top60_DNA+RNA_seed | FGFR1 | 0.693674 | LUAD | NCI-H2085 | PD173074 |
| MVP_top60_DNA_seed | GSK3B | -0.563208 | LUAD | SK-LU-1 | AZD7969 | MVP_top60_DNA+RNA_seed | AKT1 | 0.694874 | LUAD | NCI-H2342 | Afuresertib |
| MVP_top60_DNA_seed | CREBBP | -0.561524 | LUAD | LC-2-ad | I-CBP112 | MVP_top60_DNA+RNA_seed | AKT1 | 0.695611 | LUAD | ABC-1 | Uprosertib |
| MVP_top60_DNA_seed | AURKA | -0.56132 | LUAD | NCI-H1755 | HG-5-113-01 | MVP_top60_DNA+RNA_seed | SMARCA4 | 0.695795 | LUAD | NCI-H358 | PFI3 |

| | | | | | | | | | | | |
|---|---|---|---|---|---|---|---|---|---|---|---|
| MVP_top60_DNA_seed | MAPK1 | -0.561203 | LUAD | COR-L105 | FR-180204 | MVP_top60_DNA+RNA_seed | AKT1 | 0.696011 | LUAD | NCI-H650 | AT13148 |
| MVP_top60_DNA_seed | EGFR | -0.56006 | LUAD | A549 | AZD8931 | MVP_top60_DNA+RNA_seed | HDAC1 | 0.697345 | LUAD | ABC-1 | Entinostat |
| MVP_top60_DNA_seed | ERBB2 | -0.56006 | LUAD | A549 | AZD8931 | MVP_top60_DNA+RNA_seed | HDAC1 | 0.698933 | LUAD | NCI-H2085 | Entinostat |
| MVP_top60_DNA_seed | ERBB3 | -0.56006 | LUAD | A549 | AZD8931 | MVP_top60_DNA+RNA_seed | AKT1 | 0.69903 | LUAD | EKVX | Afuresertib |
| MVP_top60_DNA_seed | IGF1R | -0.559957 | LUAD | NCI-H1734 | Linsitinib | MVP_top60_DNA+RNA_seed | FGFR1 | 0.702511 | LUAD | NCI-H1734 | AZD4547 |
| MVP_top60_DNA_seed | BRAF | -0.559891 | LUAD | HCC-44 | HG6-64-1 | MVP_top60_DNA+RNA_seed | MET | 0.702604 | LUAD | HOP-62 | Savolitinib |
| MVP_top60_DNA_seed | IGF1R | -0.559808 | LUAD | EMC-BAC-2 | BMS-536924 | MVP_top60_DNA+RNA_seed | HSP90AA1 | 0.704094 | LUAD | EKVX | Elesclomol |
| MVP_top60_DNA_seed | PIK3CA | -0.557845 | LUAD | NCI-H838 | AZD8835 | MVP_top60_DNA+RNA_seed | AKT1 | 0.704306 | LUAD | HCC-827 | Ipatasertib |
| MVP_top60_DNA_seed | CREBBP | -0.557741 | LUAD | NCI-H23 | I-CBP112 | MVP_top60_DNA+RNA_seed | AKT1 | 0.705578 | LUAD | NCI-H23 | Uprosertib |
| MVP_top60_DNA_seed | EGFR | -0.557688 | LUAD | LXF-289 | Lapatinib | MVP_top60_DNA+RNA_seed | IGF1R | 0.706995 | LUAD | EMC-BAC-1 | GSK1904529A |
| MVP_top60_DNA_seed | ERBB2 | -0.557688 | LUAD | LXF-289 | Lapatinib | MVP_top60_DNA+RNA_seed | AKT1 | 0.707735 | LUAD | NCI-H2085 | AZD5363 |
| MVP_top60_DNA_seed | ATM | -0.557659 | LUAD | NCI-H1568 | KU-60019 | MVP_top60_DNA+RNA_seed | SRC | 0.707831 | LUAD | NCI-H2087 | Bosutinib |
| MVP_top60_DNA_seed | AKT1 | -0.556928 | LUAD | Calu-3 | BAY AKT1 | MVP_top60_DNA+RNA_seed | AKT1 | 0.709474 | LUAD | NCI-H1666 | Ipatasertib |
| MVP_top60_DNA_seed | AURKA | -0.556336 | LUAD | NCI-H2087 | GSK1070916 | MVP_top60_DNA+RNA_seed | ATM | 0.710379 | LUAD | NCI-H2228 | KU-55933 |
| MVP_top60_DNA_seed | HSP90AA1 | -0.555759 | LUAD | NCI-H2009 | Elesclomol | MVP_top60_DNA+RNA_seed | MDM2 | 0.711229 | LUAD | VMRC-LCD | Serdemetan |
| MVP_top60_DNA_seed | PIK3CA | -0.55444 | LUAD | NCI-H2122 | AZD8835 | MVP_top60_DNA+RNA_seed | EGFR | 0.713687 | LUAD | NCI-H1993 | Sapitinib |
| MVP_top60_DNA_seed | IGF1R | -0.551359 | LUAD | NCI-H2342 | BMS-754807 | MVP_top60_DNA+RNA_seed | IGF1R | 0.715396 | LUAD | Calu-3 | Linsitinib |
| MVP_top60_DNA_seed | HSP90AA1 | -0.550425 | LUAD | NCI-H1650 | Tanespimycin | MVP_top60_DNA+RNA_seed | AKT1 | 0.715435 | LUAD | NCI-H1666 | GSK2110183B |
| MVP_top60_DNA_seed | BRAF | -0.549916 | LUAD | NCI-H441 | PLX-4720 | MVP_top60_DNA+RNA_seed | MET | 0.717704 | LUAD | HOP-62 | Crizotinib |
| MVP_top60_DNA_seed | MAPK1 | -0.54976 | LUAD | NCI-H2405 | VX-11e | MVP_top60_DNA+RNA_seed | MDM2 | 0.717841 | LUAD | NCI-H2030 | Serdemetan |
| MVP_top60_DNA_seed | EGFR | -0.548656 | LUAD | RERF-LC-KJ | AZD8931 | MVP_top60_DNA+RNA_seed | IGF1R | 0.717875 | LUAD | NCI-H1838 | NVP-ADW742 |
| MVP_top60_DNA_seed | ERBB2 | -0.548656 | LUAD | RERF-LC-KJ | AZD8931 | MVP_top60_DNA+RNA_seed | TP53 | 0.718681 | LUAD | EMC-BAC-2 | MIRA-1 |
| MVP_top60_DNA_seed | ERBB3 | -0.548656 | LUAD | RERF-LC-KJ | AZD8931 | MVP_top60_DNA+RNA_seed | AKT1 | 0.718776 | LUAD | NCI-H2085 | MK-2206 |
| MVP_top60_DNA_seed | EGFR | -0.548372 | LUAD | NCI-H2085 | Afatinib | MVP_top60_DNA+RNA_seed | EGFR | 0.71983 | LUAD | NCI-H650 | AZD3759 |
| MVP_top60_DNA_seed | ERBB2 | -0.548372 | LUAD | NCI-H2085 | Afatinib | MVP_top60_DNA+RNA_seed | ATM | 0.720309 | LUAD | NCI-H1573 | KU-55933 |
| MVP_top60_DNA_seed | SRC | -0.547096 | LUAD | NCI-H322M | Bosutinib | MVP_top60_DNA+RNA_seed | SMARCA4 | 0.721451 | LUAD | NCI-H1781 | PFI3 |
| MVP_top60_DNA_seed | FGFR1 | -0.546775 | LUAD | A549 | FGFR_3831 | MVP_top60_DNA+RNA_seed | BRAF | 0.721907 | LUAD | NCI-H1975 | SB590885 |
| MVP_top60_DNA_seed | ATM | -0.544287 | LUAD | NCI-H1355 | CP466722 | MVP_top60_DNA+RNA_seed | PIK3CA | 0.722037 | LUAD | NCI-H358 | Buparlisib |
| MVP_top60_DNA_seed | MDM2 | -0.543896 | LUAD | NCI-H2023 | Serdemetan | MVP_top60_DNA+RNA_seed | HDAC1 | 0.722441 | LUAD | NCI-H522 | Romidepsin |
| MVP_top60_DNA_seed | GSK3B | -0.543638 | LUAD | HCC-44 | AZD7969 | MVP_top60_DNA+RNA_seed | EGFR | 0.723624 | LUAD | NCI-H1793 | Sapitinib |
| MVP_top60_DNA_seed | ATM | -0.543291 | LUAD | PC-3_[JPC-3] | KU-60019 | MVP_top60_DNA+RNA_seed | EGFR | 0.723973 | LUAD | NCI-H2228 | Gefitinib |
| MVP_top60_DNA_seed | AURKA | -0.541447 | LUAD | NCI-H1648 | Tozasertib | MVP_top60_DNA+RNA_seed | IGF1R | 0.72798 | LUAD | NCI-H2342 | BMS-536924 |
| MVP_top60_DNA_seed | EGFR | -0.540564 | LUAD | HCC-78 | AST-1306 | MVP_top60_DNA+RNA_seed | AKT1 | 0.728406 | LUAD | SW1573 | Ipatasertib |
| MVP_top60_DNA_seed | HSP90AA1 | -0.540459 | LUAD | NCI-H1651 | Luminespib | MVP_top60_DNA+RNA_seed | EGFR | 0.728602 | LUAD | HCC-78 | Gefitinib |
| MVP_top60_DNA_seed | ATM | -0.539294 | LUAD | HCC-827 | KU-60019 | MVP_top60_DNA+RNA_seed | HDAC1 | 0.729454 | LUAD | NCI-H23 | Romidepsin |
| MVP_top60_DNA_seed | AURKA | -0.538243 | LUAD | NCI-H2228 | Genentech Cpd 10 | MVP_top60_DNA+RNA_seed | SMARCA4 | 0.730502 | LUAD | A549 | PFI3 |
| MVP_top60_DNA_seed | FGFR1 | -0.538199 | LUAD | H3255 | FGFR_3831 | MVP_top60_DNA+RNA_seed | EZH2 | 0.730699 | LUAD | NCI-H1573 | GSK343 |
| MVP_top60_DNA_seed | EGFR | -0.536147 | LUAD | NCI-H1435 | AST-1306 | MVP_top60_DNA+RNA_seed | BRAF | 0.730791 | LUAD | NCI-H1755 | SB590885 |
| MVP_top60_DNA_seed | FGFR1 | -0.536039 | LUAD | NCI-H1573 | AZD4547 | MVP_top60_DNA+RNA_seed | IGF1R | 0.734011 | LUAD | NCI-H2342 | BMS-754807 |
| MVP_top60_DNA_seed | HSP90AA1 | -0.53594 | LUAD | HOP-62 | Elesclomol | MVP_top60_DNA+RNA_seed | TP53 | 0.734693 | LUAD | NCI-H1693 | MIRA-1 |
| MVP_top60_DNA_seed | AKT1 | -0.535642 | LUAD | NCI-H1435 | AKT inhibitor VIII | MVP_top60_DNA+RNA_seed | AKT1 | 0.736141 | LUAD | PC-3_[JPC-3] | Ipatasertib |

| | | | | | | | | | | | |
|---|---|---|---|---|---|---|---|---|---|---|---|
| MVP_top60_DNA_seed | EGFR | -0.533329 | LUAD | NCI-H1975 | AZD8931 | MVP_top60_DNA+RNA_seed | EGFR | 0.738179 | LUAD | Calu-6 | Lapatinib |
| MVP_top60_DNA_seed | ERBB2 | -0.533329 | LUAD | NCI-H1975 | AZD8931 | MVP_top60_DNA+RNA_seed | TP53 | 0.738409 | LUAD | NCI-H2023 | MIRA-1 |
| MVP_top60_DNA_seed | ERBB3 | -0.533329 | LUAD | NCI-H1975 | AZD8931 | MVP_top60_DNA+RNA_seed | IGF1R | 0.738756 | LUAD | EKVX | Linsitinib |
| MVP_top60_DNA_seed | BRAF | -0.533062 | LUAD | LC-2-ad | RAF_9304 | MVP_top60_DNA+RNA_seed | ATM | 0.741229 | LUAD | NCI-H2342 | KU-55933 |
| MVP_top60_DNA_seed | PTK2 | -0.532298 | LUAD | A549 | PF-562271 | MVP_top60_DNA+RNA_seed | ATM | 0.741313 | LUAD | NCI-H2023 | KU-55933 |
| MVP_top60_DNA_seed | GSK3B | -0.528671 | LUAD | NCI-H1781 | KIN001-042 | MVP_top60_DNA+RNA_seed | SRC | 0.741868 | LUAD | NCI-H1437 | Bosutinib |
| MVP_top60_DNA_seed | ATM | -0.528237 | LUAD | PC-3_[JPC-3] | CP466722 | MVP_top60_DNA+RNA_seed | BRAF | 0.742919 | LUAD | EKVX | PLX-4720 |
| MVP_top60_DNA_seed | HDAC1 | -0.520482 | LUAD | NCI-H1944 | Belinostat | MVP_top60_DNA+RNA_seed | AURKA | 0.744282 | LUAD | COR-L105 | Alisertib |
| MVP_top60_DNA_seed | BRAF | -0.51982 | LUAD | PC-3_[JPC-3] | HG6-64-1 | MVP_top60_DNA+RNA_seed | AKT1 | 0.748213 | LUAD | EMC-BAC-2 | GSK2110183B |
| MVP_top60_DNA_seed | EGFR | -0.519382 | LUAD | NCI-H358 | CUDC-101 | MVP_top60_DNA+RNA_seed | MET | 0.748297 | LUAD | NCI-H2342 | Crizotinib |
| MVP_top60_DNA_seed | ERBB2 | -0.519382 | LUAD | NCI-H358 | CUDC-101 | MVP_top60_DNA+RNA_seed | EGFR | 0.749082 | LUAD | NCI-H2342 | Gefitinib |
| MVP_top60_DNA_seed | MAPK1 | -0.517129 | LUAD | ABC-1 | VX-11e | MVP_top60_DNA+RNA_seed | SMARCA4 | 0.749439 | LUAD | NCI-H1975 | PFI3 |
| MVP_top60_DNA_seed | MET | -0.51645 | LUAD | NCI-H1563 | JNJ38877605 | MVP_top60_DNA+RNA_seed | AKT1 | 0.751414 | LUAD | NCI-H358 | AZD5363 |
| MVP_top60_DNA_seed | PIK3CA | -0.516128 | LUAD | NCI-H1568 | AZD8835 | MVP_top60_DNA+RNA_seed | HDAC1 | 0.753912 | LUAD | EMC-BAC-2 | Entinostat |
| MVP_top60_DNA_seed | IGF1R | -0.515823 | LUAD | NCI-H2087 | BMS-536924 | MVP_top60_DNA+RNA_seed | AKT1 | 0.754303 | LUAD | HCC-827 | Uprosertib |
| MVP_top60_DNA_seed | HSP90AA1 | -0.514611 | LUAD | PC-14 | Elesclomol | MVP_top60_DNA+RNA_seed | IGF1R | 0.755677 | LUAD | NCI-H3122 | NVP-ADW742 |
| MVP_top60_DNA_seed | HDAC1 | -0.51439 | LUAD | COR-L105 | Tubastatin A | MVP_top60_DNA+RNA_seed | HSP90AA1 | 0.756826 | LUAD | NCI-H2087 | Luminespib |
| MVP_top60_DNA_seed | GSK3B | -0.513748 | LUAD | NCI-H1944 | AZD7969 | MVP_top60_DNA+RNA_seed | EGFR | 0.758612 | LUAD | NCI-H2009 | Lapatinib |
| MVP_top60_DNA_seed | HSP90AA1 | -0.512257 | LUAD | Calu-6 | CCT-018159 | MVP_top60_DNA+RNA_seed | TP53 | 0.758629 | LUAD | NCI-H1563 | MIRA-1 |
| MVP_top60_DNA_seed | HSP90AA1 | -0.512241 | LUAD | NCI-H358 | Luminespib | MVP_top60_DNA+RNA_seed | MET | 0.760961 | LUAD | NCI-H1755 | Savolitinib |
| MVP_top60_DNA_seed | BRAF | -0.512051 | LUAD | RERF-LC-MS | PLX-4720 | MVP_top60_DNA+RNA_seed | HDAC1 | 0.761671 | LUAD | NCI-H23 | Entinostat |
| MVP_top60_DNA_seed | ATM | -0.511908 | LUAD | NCI-H1975 | KU-60019 | MVP_top60_DNA+RNA_seed | AKT1 | 0.763262 | LUAD | NCI-H522 | GSK2110183B |
| MVP_top60_DNA_seed | ATM | -0.511369 | LUAD | NCI-H2023 | KU-55933 | MVP_top60_DNA+RNA_seed | AKT1 | 0.763661 | LUAD | SW1573 | AZD5363 |
| MVP_top60_DNA_seed | ATM | -0.510955 | LUAD | NCI-H1437 | KU-60019 | MVP_top60_DNA+RNA_seed | IGF1R | 0.766287 | LUAD | EMC-BAC-2 | GSK1904529A |
| MVP_top60_DNA_seed | GSK3B | -0.509101 | LUAD | NCI-H650 | CHIR-99021 | MVP_top60_DNA+RNA_seed | HDAC1 | 0.76688 | LUAD | HOP-62 | Romidepsin |
| MVP_top60_DNA_seed | FGFR1 | -0.508405 | LUAD | HOP-62 | FGFR_3831 | MVP_top60_DNA+RNA_seed | PIK3CA | 0.768546 | LUAD | NCI-H2405 | GNE-317 |
| MVP_top60_DNA_seed | AURKA | -0.508019 | LUAD | LXF-289 | Genentech Cpd 10 | MVP_top60_DNA+RNA_seed | HDAC1 | 0.768968 | LUAD | NCI-H2023 | Entinostat |
| MVP_top60_DNA_seed | HDAC1 | -0.507902 | LUAD | COR-L105 | CAY10603 | MVP_top60_DNA+RNA_seed | AKT1 | 0.76957 | LUAD | NCI-H2405 | Ipatasertib |
| MVP_top60_DNA_seed | HDAC1 | -0.50664 | LUAD | NCI-H650 | Dacinostat | MVP_top60_DNA+RNA_seed | HSP90AA1 | 0.770846 | LUAD | ABC-1 | CCT-018159 |
| MVP_top60_DNA_seed | BRAF | -0.50553 | LUAD | HCC-44 | RAF_9304 | MVP_top60_DNA+RNA_seed | AURKA | 0.771097 | LUAD | NCI-H522 | Alisertib |
| MVP_top60_DNA_seed | FGFR1 | -0.505281 | LUAD | NCI-H1755 | FGFR_3831 | MVP_top60_DNA+RNA_seed | IGF1R | 0.771693 | LUAD | SW1573 | BMS-536924 |
| MVP_top60_DNA_seed | GSK3B | -0.504131 | LUAD | NCI-H522 | KIN001-042 | MVP_top60_DNA+RNA_seed | EGFR | 0.774301 | LUAD | ABC-1 | Lapatinib |
| MVP_top60_DNA_seed | MDM2 | -0.503327 | LUAD | NCI-H1437 | Serdemetan | MVP_top60_DNA+RNA_seed | TP53 | 0.774438 | LUAD | HCC-78 | MIRA-1 |
| MVP_top60_DNA_seed | ATM | -0.503248 | LUAD | NCI-H838 | KU-60019 | MVP_top60_DNA+RNA_seed | BRAF | 0.77614 | LUAD | NCI-H1563 | Dabrafenib |
| MVP_top60_DNA_seed | HDAC1 | -0.501606 | LUAD | Calu-6 | AR-42 | MVP_top60_DNA+RNA_seed | HSP90AA1 | 0.777057 | LUAD | NCI-H2291 | Luminespib |
| MVP_top60_DNA_seed | ATM | -0.50156 | LUAD | SK-LU-1 | KU-55933 | MVP_top60_DNA+RNA_seed | AKT1 | 0.779293 | LUAD | HOP-62 | Uprosertib |
| MVP_top60_DNA_seed | FGFR1 | -0.499779 | LUAD | NCI-H1944 | FGFR_3831 | MVP_top60_DNA+RNA_seed | EGFR | 0.779507 | LUAD | NCI-H2342 | Osimertinib |
| MVP_top60_DNA_seed | AKT1 | -0.498901 | LUAD | NCI-H1734 | AKT inhibitor VIII | MVP_top60_DNA+RNA_seed | EGFR | 0.779879 | LUAD | NCI-H3122 | Lapatinib |
| MVP_top60_DNA_seed | AKT1 | -0.496511 | LUAD | A549 | MK-2206 | MVP_top60_DNA+RNA_seed | HDAC1 | 0.779987 | LUAD | HOP-62 | PCI-34051 |
| MVP_top60_DNA_seed | PIK3CA | -0.496282 | LUAD | NCI-H1563 | AZD8835 | MVP_top60_DNA+RNA_seed | IGF1R | 0.784109 | LUAD | EKVX | BMS-754807 |
| MVP_top60_DNA_seed | BRAF | -0.496184 | LUAD | HCC-827 | RAF_9304 | MVP_top60_DNA+RNA_seed | TERT | 0.784312 | LUAD | PC-3_[JPC-3] | BIBR-1532 |

| | | | | | | | | | | | |
|---|---|---|---|---|---|---|---|---|---|---|---|
| MVP_top60_DNA_seed | MDM2 | -0.496133 | LUAD | NCI-H1944 | Serdemetan | MVP_top60_DNA+RNA_seed | AKT1 | 0.784618 | LUAD | ABC-1 | GSK2110183B |
| MVP_top60_DNA_seed | IGF1R | -0.496074 | LUAD | COR-L105 | GSK1904529A | MVP_top60_DNA+RNA_seed | MDM2 | 0.784907 | LUAD | NCI-H1993 | Nutlin-3a (-) |
| MVP_top60_DNA_seed | AKT1 | -0.495576 | LUAD | PC-14 | AKT inhibitor VIII | MVP_top60_DNA+RNA_seed | PIK3CA | 0.785271 | LUAD | NCI-H1944 | GNE-317 |
| MVP_top60_DNA_seed | EGFR | -0.493927 | LUAD | EMC-BAC-1 | CUDC-101 | MVP_top60_DNA+RNA_seed | IGF1R | 0.78633 | LUAD | NCI-H1793 | Linsitinib |
| MVP_top60_DNA_seed | ERBB2 | -0.493927 | LUAD | EMC-BAC-1 | CUDC-101 | MVP_top60_DNA+RNA_seed | BRAF | 0.787764 | LUAD | ABC-1 | SB590885 |
| MVP_top60_DNA_seed | SRC | -0.493685 | LUAD | NCI-H2405 | NVP-BHG712 | MVP_top60_DNA+RNA_seed | ATM | 0.789025 | LUAD | SW1573 | KU-55933 |
| MVP_top60_DNA_seed | AURKA | -0.492988 | LUAD | NCI-H1355 | HG-5-113-01 | MVP_top60_DNA+RNA_seed | IGF1R | 0.789942 | LUAD | NCI-H1563 | GSK1904529A |
| MVP_top60_DNA_seed | HDAC1 | -0.492319 | LUAD | ABC-1 | Dacinostat | MVP_top60_DNA+RNA_seed | HDAC1 | 0.791386 | LUAD | SK-LU-1 | Romidepsin |
| MVP_top60_DNA_seed | GSK3B | -0.491257 | LUAD | SK-LU-1 | SB216763 | MVP_top60_DNA+RNA_seed | EZH2 | 0.791926 | LUAD | NCI-H1838 | GSK343 |
| MVP_top60_DNA_seed | EGFR | -0.490492 | LUAD | NCI-H2085 | Pelitinib | MVP_top60_DNA+RNA_seed | IGF1R | 0.793 | LUAD | SK-LU-1 | BMS-536924 |
| MVP_top60_DNA_seed | FGFR1 | -0.489928 | LUAD | VMRC-LCD | FGFR_3831 | MVP_top60_DNA+RNA_seed | AKT1 | 0.794586 | LUAD | NCI-H2085 | Ipatasertib |
| MVP_top60_DNA_seed | HDAC1 | -0.489708 | LUAD | LXF-289 | JQ12 | MVP_top60_DNA+RNA_seed | EGFR | 0.795251 | LUAD | NCI-H1651 | Afatinib |
| MVP_top60_DNA_seed | BRAF | -0.488453 | LUAD | Calu-6 | RAF_9304 | MVP_top60_DNA+RNA_seed | TERT | 0.795957 | LUAD | A549 | BIBR-1532 |
| MVP_top60_DNA_seed | MAPK1 | -0.487417 | LUAD | NCI-H2009 | VX-11e | MVP_top60_DNA+RNA_seed | PIK3CA | 0.79772 | LUAD | VMRC-LCD | Alpelisib |
| MVP_top60_DNA_seed | ATM | -0.48727 | LUAD | RERF-LC-MS | KU-60019 | MVP_top60_DNA+RNA_seed | FGFR1 | 0.798837 | LUAD | NCI-H1993 | AZD4547 |
| MVP_top60_DNA_seed | BRAF | -0.485888 | LUAD | NCI-H2030 | RAF_9304 | MVP_top60_DNA+RNA_seed | MDM2 | 0.801458 | LUAD | HOP-62 | Serdemetan |
| MVP_top60_DNA_seed | AKT1 | -0.485555 | LUAD | EKVX | BAY AKT1 | MVP_top60_DNA+RNA_seed | HDAC1 | 0.80369 | LUAD | NCI-H322M | PCI-34051 |
| MVP_top60_DNA_seed | ATM | -0.485232 | LUAD | RERF-LC-KJ | CP466722 | MVP_top60_DNA+RNA_seed | AKT1 | 0.803698 | LUAD | LXF-289 | Ipatasertib |
| MVP_top60_DNA_seed | AKT1 | -0.485027 | LUAD | HCC-78 | GSK690693 | MVP_top60_DNA+RNA_seed | MDM2 | 0.803918 | LUAD | NCI-H2087 | Nutlin-3a (-) |
| MVP_top60_DNA_seed | HDAC1 | -0.484117 | LUAD | NCI-H3122 | CAY10603 | MVP_top60_DNA+RNA_seed | BRAF | 0.805501 | LUAD | NCI-H2342 | SB590885 |
| MVP_top60_DNA_seed | BRAF | -0.482571 | LUAD | NCI-H2087 | RAF_9304 | MVP_top60_DNA+RNA_seed | EGFR | 0.806373 | LUAD | HOP-62 | Lapatinib |
| MVP_top60_DNA_seed | EGFR | -0.481605 | LUAD | NCI-H322M | Lapatinib | MVP_top60_DNA+RNA_seed | PIK3CA | 0.806887 | LUAD | NCI-H23 | GNE-317 |
| MVP_top60_DNA_seed | ERBB2 | -0.481605 | LUAD | NCI-H322M | Lapatinib | MVP_top60_DNA+RNA_seed | HDAC1 | 0.807011 | LUAD | NCI-H2085 | Romidepsin |
| MVP_top60_DNA_seed | AKT1 | -0.480728 | LUAD | ABC-1 | AKT inhibitor VIII | MVP_top60_DNA+RNA_seed | MDM2 | 0.808288 | LUAD | NCI-H1623 | Nutlin-3a (-) |
| MVP_top60_DNA_seed | BRAF | -0.480584 | LUAD | NCI-H2087 | HG6-64-1 | MVP_top60_DNA+RNA_seed | PIK3CA | 0.808837 | LUAD | NCI-H1651 | AZD8186 |
| MVP_top60_DNA_seed | EGFR | -0.480015 | LUAD | NCI-H1648 | Pelitinib | MVP_top60_DNA+RNA_seed | SMARCA4 | 0.81031 | LUAD | NCI-H1355 | PFI3 |
| MVP_top60_DNA_seed | GSK3B | -0.479987 | LUAD | LXF-289 | AZD7969 | MVP_top60_DNA+RNA_seed | AURKA | 0.810363 | LUAD | SW1573 | Alisertib |
| MVP_top60_DNA_seed | MET | -0.479848 | LUAD | NCI-H2030 | JNJ38877605 | MVP_top60_DNA+RNA_seed | IGF1R | 0.812046 | LUAD | Calu-3 | BMS-754807 |
| MVP_top60_DNA_seed | EGFR | -0.479488 | LUAD | EMC-BAC-1 | PF-00299804 | MVP_top60_DNA+RNA_seed | TP53 | 0.813338 | LUAD | PC-3_[JPC-3] | MIRA-1 |
| MVP_top60_DNA_seed | ERBB2 | -0.479488 | LUAD | EMC-BAC-1 | PF-00299804 | MVP_top60_DNA+RNA_seed | EGFR | 0.813657 | LUAD | NCI-H2291 | Erlotinib |
| MVP_top60_DNA_seed | EGFR | -0.479127 | LUAD | HCC-44 | Gefitinib | MVP_top60_DNA+RNA_seed | EGFR | 0.814658 | LUAD | NCI-H2342 | Erlotinib |
| MVP_top60_DNA_seed | MET | -0.478375 | LUAD | PC-14 | AZD6094 | MVP_top60_DNA+RNA_seed | HDAC1 | 0.81716 | LUAD | COR-L105 | Entinostat |
| MVP_top60_DNA_seed | HDAC1 | -0.474231 | LUAD | NCI-H3122 | Belinostat | MVP_top60_DNA+RNA_seed | MDM2 | 0.817767 | LUAD | H3255 | Nutlin-3a (-) |
| MVP_top60_DNA_seed | HDAC1 | -0.47385 | LUAD | NCI-H1648 | CAY10603 | MVP_top60_DNA+RNA_seed | PIK3CA | 0.819556 | LUAD | NCI-H1650 | GNE-317 |
| MVP_top60_DNA_seed | ATM | -0.473727 | LUAD | NCI-H2087 | CP466722 | MVP_top60_DNA+RNA_seed | FGFR1 | 0.824759 | LUAD | Calu-3 | PD173074 |
| MVP_top60_DNA_seed | MET | -0.47316 | LUAD | LC-2-ad | SU11274 | MVP_top60_DNA+RNA_seed | AURKA | 0.825346 | LUAD | NCI-H1975 | ZM447439 |
| MVP_top60_DNA_seed | EGFR | -0.472183 | LUAD | HCC-44 | Cetuximab | MVP_top60_DNA+RNA_seed | AKT1 | 0.826753 | LUAD | NCI-H1651 | MK-2206 |
| MVP_top60_DNA_seed | HDAC1 | -0.472101 | LUAD | NCI-H2087 | Tubastatin A | MVP_top60_DNA+RNA_seed | IGF1R | 0.828818 | LUAD | NCI-H1793 | GSK1904529A |
| MVP_top60_DNA_seed | EGFR | -0.471048 | LUAD | EMC-BAC-2 | Afatinib | MVP_top60_DNA+RNA_seed | AKT1 | 0.82986 | LUAD | NCI-H2085 | Uprosertib |
| MVP_top60_DNA_seed | ERBB2 | -0.471048 | LUAD | EMC-BAC-2 | Afatinib | MVP_top60_DNA+RNA_seed | IGF1R | 0.82994 | LUAD | NCI-H2405 | BMS-754807 |
| MVP_top60_DNA_seed | EGFR | -0.470369 | LUAD | PC-14 | Cetuximab | MVP_top60_DNA+RNA_seed | EGFR | 0.830972 | LUAD | NCI-H1355 | Afatinib |

| | | | | | | | | | | | |
|---|---|---|---|---|---|---|---|---|---|---|---|
| MVP_top60_DNA_seed | ATM | -0.469854 | LUAD | Calu-6 | KU-60019 | MVP_top60_DNA+RNA_seed | EZH2 | 0.831454 | LUAD | NCI-H1793 | GSK343 |
| MVP_top60_DNA_seed | IGF1R | -0.469304 | LUAD | NCI-H1437 | Linsitinib | MVP_top60_DNA+RNA_seed | AURKA | 0.831531 | LUAD | NCI-H1993 | ZM447439 |
| MVP_top60_DNA_seed | HDAC1 | -0.469182 | LUAD | NCI-H3122 | Tubastatin A | MVP_top60_DNA+RNA_seed | BRAF | 0.833003 | LUAD | NCI-H2085 | SB590885 |
| MVP_top60_DNA_seed | AURKA | -0.468556 | LUAD | H3255 | GSK1070916 | MVP_top60_DNA+RNA_seed | FGFR1 | 0.833099 | LUAD | NCI-H1648 | PD173074 |
| MVP_top60_DNA_seed | MAPK1 | -0.468334 | LUAD | NCI-H1781 | VX-11e | MVP_top60_DNA+RNA_seed | BRAF | 0.836186 | LUAD | NCI-H2009 | Dabrafenib |
| MVP_top60_DNA_seed | EGFR | -0.466621 | LUAD | NCI-H1563 | Cetuximab | MVP_top60_DNA+RNA_seed | HDAC1 | 0.837202 | LUAD | LXF-289 | PCI-34051 |
| MVP_top60_DNA_seed | SRC | -0.465303 | LUAD | LXF-289 | HG-6-71-01 | MVP_top60_DNA+RNA_seed | AKT1 | 0.838818 | LUAD | NCI-H650 | Ipatasertib |
| MVP_top60_DNA_seed | EGFR | -0.465094 | LUAD | ABC-1 | Pelitinib | MVP_top60_DNA+RNA_seed | ATM | 0.839089 | LUAD | NCI-H1781 | KU-55933 |
| MVP_top60_DNA_seed | MET | -0.46465 | LUAD | NCI-H23 | SU11274 | MVP_top60_DNA+RNA_seed | MET | 0.839365 | LUAD | ABC-1 | Savolitinib |
| MVP_top60_DNA_seed | PIK3CA | -0.46407 | LUAD | HCC-44 | AZD8835 | MVP_top60_DNA+RNA_seed | MDM2 | 0.839642 | LUAD | NCI-H650 | Serdemetan |
| MVP_top60_DNA_seed | FGFR1 | -0.463435 | LUAD | NCI-H838 | PD173074 | MVP_top60_DNA+RNA_seed | AURKA | 0.839991 | LUAD | NCI-H2291 | Tozasertib |
| MVP_top60_DNA_seed | HSP90AA1 | -0.461996 | LUAD | HCC-44 | CCT-018159 | MVP_top60_DNA+RNA_seed | TERT | 0.841422 | LUAD | COR-L105 | BIBR-1532 |
| MVP_top60_DNA_seed | AKT1 | -0.461306 | LUAD | LC-2-ad | AKT inhibitor VIII | MVP_top60_DNA+RNA_seed | HDAC1 | 0.843498 | LUAD | NCI-H1355 | Entinostat |
| MVP_top60_DNA_seed | EGFR | -0.459764 | LUAD | NCI-H1755 | Cetuximab | MVP_top60_DNA+RNA_seed | BRAF | 0.84769 | LUAD | RERF-LC-KJ | PLX-4720 |
| MVP_top60_DNA_seed | AURKA | -0.45933 | LUAD | PC-14 | HG-5-113-01 | MVP_top60_DNA+RNA_seed | TP53 | 0.848045 | LUAD | NCI-H1623 | MIRA-1 |
| MVP_top60_DNA_seed | ATM | -0.459248 | LUAD | NCI-H1944 | CP466722 | MVP_top60_DNA+RNA_seed | EGFR | 0.84873 | LUAD | NCI-H2023 | Gefitinib |
| MVP_top60_DNA_seed | AKT1 | -0.458409 | LUAD | H3255 | AKT inhibitor VIII | MVP_top60_DNA+RNA_seed | BRAF | 0.849148 | LUAD | NCI-H2030 | SB590885 |
| MVP_top60_DNA_seed | AKT1 | -0.457943 | LUAD | PC-14 | GSK690693 | MVP_top60_DNA+RNA_seed | KRAS | 0.849262 | LUAD | NCI-H1435 | KRAS (G12C) Inhibitor-1 |
| MVP_top60_DNA_seed | HSP90AA1 | -0.457181 | LUAD | VMRC-LCD | CCT-018159 | MVP_top60_DNA+RNA_seed | EZH2 | 0.85089 | LUAD | NCI-H3122 | GSK343 |
| MVP_top60_DNA_seed | AKT1 | -0.457181 | LUAD | EMC-BAC-1 | GSK690693 | MVP_top60_DNA+RNA_seed | EGFR | 0.852169 | LUAD | HOP-62 | Osimertinib |
| MVP_top60_DNA_seed | GSK3B | -0.456989 | LUAD | EMC-BAC-1 | CHIR-99021 | MVP_top60_DNA+RNA_seed | TERT | 0.852616 | LUAD | NCI-H3122 | BIBR-1532 |
| MVP_top60_DNA_seed | GSK3B | -0.456219 | LUAD | NCI-H322M | SB216763 | MVP_top60_DNA+RNA_seed | AKT1 | 0.855017 | LUAD | NCI-H2085 | AT13148 |
| MVP_top60_DNA_seed | IGF1R | -0.45607 | LUAD | NCI-H1944 | BMS-754807 | MVP_top60_DNA+RNA_seed | AKT1 | 0.855344 | LUAD | SW1573 | MK-2206 |
| MVP_top60_DNA_seed | AKT1 | -0.454312 | LUAD | RERF-LC-KJ | MK-2206 | MVP_top60_DNA+RNA_seed | FGFR1 | 0.855423 | LUAD | NCI-H2009 | AZD4547 |
| MVP_top60_DNA_seed | EGFR | -0.452975 | LUAD | NCI-H1944 | Afatinib | MVP_top60_DNA+RNA_seed | EZH2 | 0.8565 | LUAD | NCI-H522 | GSK343 |
| MVP_top60_DNA_seed | ERBB2 | -0.452975 | LUAD | NCI-H1944 | Afatinib | MVP_top60_DNA+RNA_seed | BRAF | 0.856505 | LUAD | HCC-78 | SB590885 |
| MVP_top60_DNA_seed | HSP90AA1 | -0.449618 | LUAD | HCC-78 | Tanespimycin | MVP_top60_DNA+RNA_seed | HDAC1 | 0.856729 | LUAD | EKVX | Romidepsin |
| MVP_top60_DNA_seed | BRAF | -0.449512 | LUAD | HCC-44 | PLX-4720 | MVP_top60_DNA+RNA_seed | PIK3CA | 0.857441 | LUAD | NCI-H1568 | Buparlisib |
| MVP_top60_DNA_seed | AKT1 | -0.449272 | LUAD | HCC-44 | AKT inhibitor VIII | MVP_top60_DNA+RNA_seed | BRAF | 0.858193 | LUAD | HCC-78 | PLX-4720 |
| MVP_top60_DNA_seed | HDAC1 | -0.448433 | LUAD | NCI-H2228 | JQ12 | MVP_top60_DNA+RNA_seed | AURKA | 0.860309 | LUAD | SK-LU-1 | Alisertib |
| MVP_top60_DNA_seed | ATM | -0.448269 | LUAD | NCI-H2291 | KU-60019 | MVP_top60_DNA+RNA_seed | IGF1R | 0.860822 | LUAD | NCI-H1568 | NVP-ADW742 |
| MVP_top60_DNA_seed | MDM2 | -0.448117 | LUAD | SK-LU-1 | Nutlin-3a (-) | MVP_top60_DNA+RNA_seed | AKT1 | 0.861235 | LUAD | NCI-H1651 | Ipatasertib |
| MVP_top60_DNA_seed | GSK3B | -0.447705 | LUAD | NCI-H2342 | CHIR-99021 | MVP_top60_DNA+RNA_seed | MDM2 | 0.863093 | LUAD | H3255 | Serdemetan |
| MVP_top60_DNA_seed | PIK3CA | -0.44744 | LUAD | LC-2-ad | AZD8835 | MVP_top60_DNA+RNA_seed | AKT1 | 0.863103 | LUAD | SW1573 | Uprosertib |
| MVP_top60_DNA_seed | EGFR | -0.447421 | LUAD | HCC-78 | Cetuximab | MVP_top60_DNA+RNA_seed | EZH2 | 0.863466 | LUAD | NCI-H2030 | GSK343 |
| MVP_top60_DNA_seed | PIK3CA | -0.447255 | LUAD | EMC-BAC-2 | AZD8835 | MVP_top60_DNA+RNA_seed | KRAS | 0.86381 | LUAD | NCI-H358 | KRAS (G12C) Inhibitor-1 |
| MVP_top60_DNA_seed | EGFR | -0.447188 | LUAD | NCI-H1944 | CUDC-101 | MVP_top60_DNA+RNA_seed | IGF1R | 0.863864 | LUAD | HOP-62 | BMS-754807 |
| MVP_top60_DNA_seed | ERBB2 | -0.447188 | LUAD | NCI-H1944 | CUDC-101 | MVP_top60_DNA+RNA_seed | BRAF | 0.864187 | LUAD | HOP-62 | PLX-4720 |
| MVP_top60_DNA_seed | MET | -0.447078 | LUAD | ABC-1 | SU11274 | MVP_top60_DNA+RNA_seed | TERT | 0.865159 | LUAD | NCI-H2023 | BIBR-1532 |
| MVP_top60_DNA_seed | EGFR | -0.446944 | LUAD | NCI-H1568 | Pelitinib | MVP_top60_DNA+RNA_seed | FGFR1 | 0.865231 | LUAD | HCC-827 | PD173074 |
| MVP_top60_DNA_seed | IGF1R | -0.446699 | LUAD | LXF-289 | BMS-754807 | MVP_top60_DNA+RNA_seed | ATM | 0.866654 | LUAD | HCC-78 | KU-55933 |

| | | | | | | | | | | | |
|---|---|---|---|---|---|---|---|---|---|---|---|
| MVP_top60_DNA_seed | AURKA | -0.445561 | LUAD | NCI-H1666 | ZM447439 | MVP_top60_DNA+RNA_seed | TERT | 0.867275 | LUAD | EKVX | BIBR-1532 |
| MVP_top60_DNA_seed | HDAC1 | -0.443942 | LUAD | HOP-62 | Entinostat | MVP_top60_DNA+RNA_seed | BRAF | 0.868314 | LUAD | NCI-H1568 | Dabrafenib |
| MVP_top60_DNA_seed | HDAC1 | -0.443765 | LUAD | HCC-78 | CAY10603 | MVP_top60_DNA+RNA_seed | EGFR | 0.868936 | LUAD | NCI-H23 | Lapatinib |
| MVP_top60_DNA_seed | HSP90AA1 | -0.44359 | LUAD | NCI-H2087 | Luminespib | MVP_top60_DNA+RNA_seed | BRAF | 0.869914 | LUAD | EKVX | Dabrafenib |
| MVP_top60_DNA_seed | IGF1R | -0.443565 | LUAD | RERF-LC-MS | BMS-536924 | MVP_top60_DNA+RNA_seed | EGFR | 0.870559 | LUAD | Calu-3 | AZD3759 |
| MVP_top60_DNA_seed | AURKA | -0.44272 | LUAD | EMC-BAC-1 | ZM447439 | MVP_top60_DNA+RNA_seed | EZH2 | 0.872629 | LUAD | NCI-H1563 | GSK343 |
| MVP_top60_DNA_seed | GSK3B | -0.441885 | LUAD | NCI-H23 | AZD7969 | MVP_top60_DNA+RNA_seed | FGFR1 | 0.874897 | LUAD | Calu-6 | PD173074 |
| MVP_top60_DNA_seed | FGFR1 | -0.440694 | LUAD | NCI-H2228 | AZD4547 | MVP_top60_DNA+RNA_seed | AURKA | 0.877401 | LUAD | NCI-H1651 | Alisertib |
| MVP_top60_DNA_seed | EGFR | -0.440563 | LUAD | NCI-H358 | AZD8931 | MVP_top60_DNA+RNA_seed | TP53 | 0.877456 | LUAD | NCI-H2342 | MIRA-1 |
| MVP_top60_DNA_seed | ERBB2 | -0.440563 | LUAD | NCI-H358 | AZD8931 | MVP_top60_DNA+RNA_seed | PIK3CA | 0.878077 | LUAD | Calu-3 | Buparlisib |
| MVP_top60_DNA_seed | ERBB3 | -0.440563 | LUAD | NCI-H358 | AZD8931 | MVP_top60_DNA+RNA_seed | EZH2 | 0.87872 | LUAD | NCI-H1975 | GSK343 |
| MVP_top60_DNA_seed | IGF1R | -0.440173 | LUAD | EKVX | BMS-536924 | MVP_top60_DNA+RNA_seed | FGFR1 | 0.878764 | LUAD | NCI-H2342 | AZD4547 |
| MVP_top60_DNA_seed | PTK2 | -0.439991 | LUAD | Calu-3 | PF-562271 | MVP_top60_DNA+RNA_seed | FGFR1 | 0.880621 | LUAD | NCI-H2023 | AZD4547 |
| MVP_top60_DNA_seed | MAPK1 | -0.439326 | LUAD | NCI-H1563 | FR-180204 | MVP_top60_DNA+RNA_seed | EGFR | 0.881476 | LUAD | NCI-H1755 | Gefitinib |
| MVP_top60_DNA_seed | AKT1 | -0.43862 | LUAD | COR-L105 | AKT inhibitor VIII | MVP_top60_DNA+RNA_seed | MET | 0.882206 | LUAD | NCI-H1793 | Savolitinib |
| MVP_top60_DNA_seed | MET | -0.438141 | LUAD | HCC-44 | JNJ38877605 | MVP_top60_DNA+RNA_seed | AKT1 | 0.884439 | LUAD | NCI-H1573 | MK-2206 |
| MVP_top60_DNA_seed | HDAC1 | -0.436736 | LUAD | NCI-H358 | AR-42 | MVP_top60_DNA+RNA_seed | IGF1R | 0.885216 | LUAD | VMRC-LCD | NVP-ADW742 |
| MVP_top60_DNA_seed | EGFR | -0.436441 | LUAD | NCI-H2087 | AST-1306 | MVP_top60_DNA+RNA_seed | IGF1R | 0.88563 | LUAD | NCI-H1648 | GSK1904529A |
| MVP_top60_DNA_seed | HDAC1 | -0.435535 | LUAD | HOP-62 | Belinostat | MVP_top60_DNA+RNA_seed | KRAS | 0.886349 | LUAD | NCI-H1623 | :RAS (G12C) Inhibitor-1 |
| MVP_top60_DNA_seed | FGFR1 | -0.434147 | LUAD | HCC-78 | FGFR_3831 | MVP_top60_DNA+RNA_seed | FGFR1 | 0.88757 | LUAD | NCI-H1693 | PD173074 |
| MVP_top60_DNA_seed | AURKA | -0.43412 | LUAD | ABC-1 | Genentech Cpd 10 | MVP_top60_DNA+RNA_seed | AURKA | 0.889618 | LUAD | NCI-H1734 | Tozasertib |
| MVP_top60_DNA_seed | HDAC1 | -0.432696 | LUAD | Calu-3 | Dacinostat | MVP_top60_DNA+RNA_seed | EGFR | 0.889827 | LUAD | VMRC-LCD | Osimertinib |
| MVP_top60_DNA_seed | BRAF | -0.432684 | LUAD | NCI-H1666 | Dabrafenib | MVP_top60_DNA+RNA_seed | MET | 0.890774 | LUAD | NCI-H2347 | Savolitinib |
| MVP_top60_DNA_seed | PIK3CA | -0.43188 | LUAD | NCI-H1666 | AZD8835 | MVP_top60_DNA+RNA_seed | EGFR | 0.891159 | LUAD | NCI-H1355 | AZD3759 |
| MVP_top60_DNA_seed | AKT1 | -0.431308 | LUAD | NCI-H1563 | BAY AKT1 | MVP_top60_DNA+RNA_seed | AURKA | 0.892451 | LUAD | COR-L105 | ZM447439 |
| MVP_top60_DNA_seed | PIK3CA | -0.430608 | LUAD | LXF-289 | PI3Ka_4409 | MVP_top60_DNA+RNA_seed | EGFR | 0.89335 | LUAD | NCI-H1838 | Erlotinib |
| MVP_top60_DNA_seed | HSP90AA1 | -0.42824 | LUAD | LC-2-ad | SNX-2112 | MVP_top60_DNA+RNA_seed | HSP90AA1 | 0.893496 | LUAD | H3255 | CCT-018159 |
| MVP_top60_DNA_seed | HSP90AA1 | -0.427461 | LUAD | SW1573 | Tanespimycin | MVP_top60_DNA+RNA_seed | AKT1 | 0.894983 | LUAD | NCI-H1693 | Uprosertib |
| MVP_top60_DNA_seed | HSP90AA1 | -0.426312 | LUAD | NCI-H1437 | Luminespib | MVP_top60_DNA+RNA_seed | EGFR | 0.895504 | LUAD | NCI-H441 | Sapitinib |
| MVP_top60_DNA_seed | MAPK1 | -0.425349 | LUAD | NCI-H1648 | VX-11e | MVP_top60_DNA+RNA_seed | HDAC1 | 0.896296 | LUAD | NCI-H1563 | Romidepsin |
| MVP_top60_DNA_seed | HDAC1 | -0.421876 | LUAD | Calu-6 | JQ12 | MVP_top60_DNA+RNA_seed | EGFR | 0.898124 | LUAD | HCC-44 | Erlotinib |
| MVP_top60_DNA_seed | MAPK1 | -0.420701 | LUAD | NCI-H522 | FR-180204 | MVP_top60_DNA+RNA_seed | BRAF | 0.899513 | LUAD | NCI-H1793 | PLX-4720 |
| MVP_top60_DNA_seed | BRAF | -0.420679 | LUAD | NCI-H2291 | Dabrafenib | MVP_top60_DNA+RNA_seed | FGFR1 | 0.900424 | LUAD | Calu-3 | AZD4547 |
| MVP_top60_DNA_seed | EGFR | -0.419999 | LUAD | NCI-H2087 | Cetuximab | MVP_top60_DNA+RNA_seed | ATM | 0.901097 | LUAD | EMC-BAC-2 | KU-55933 |
| MVP_top60_DNA_seed | IGF1R | -0.4196 | LUAD | NCI-H838 | BMS-754807 | MVP_top60_DNA+RNA_seed | IGF1R | 0.904785 | LUAD | NCI-H2291 | NVP-ADW742 |
| MVP_top60_DNA_seed | ATM | -0.417586 | LUAD | HCC-44 | CP466722 | MVP_top60_DNA+RNA_seed | IGF1R | 0.905171 | LUAD | NCI-H1435 | NVP-ADW742 |
| MVP_top60_DNA_seed | EGFR | -0.416219 | LUAD | RERF-LC-KJ | Afatinib | MVP_top60_DNA+RNA_seed | HDAC1 | 0.909071 | LUAD | NCI-H2291 | PCI-34051 |
| MVP_top60_DNA_seed | ERBB2 | -0.416219 | LUAD | RERF-LC-KJ | Afatinib | MVP_top60_DNA+RNA_seed | BRAF | 0.909494 | LUAD | HOP-62 | SB590885 |
| MVP_top60_DNA_seed | EGFR | -0.416023 | LUAD | NCI-H1355 | Afatinib | MVP_top60_DNA+RNA_seed | MDM2 | 0.910378 | LUAD | NCI-H2228 | Nutlin-3a (-) |
| MVP_top60_DNA_seed | ERBB2 | -0.416023 | LUAD | NCI-H1355 | Afatinib | MVP_top60_DNA+RNA_seed | SRC | 0.912033 | LUAD | NCI-H1793 | Bosutinib |
| MVP_top60_DNA_seed | HSP90AA1 | -0.415991 | LUAD | HCC-44 | Elesclomol | MVP_top60_DNA+RNA_seed | AKT1 | 0.912764 | LUAD | NCI-H1693 | MK-2206 |

| | | | | | | | | | | | |
|---|---|---|---|---|---|---|---|---|---|---|---|
| MVP_top60_DNA_seed | AKT1 | -0.414767 | LUAD | NCI-H1781 | AKT inhibitor VIII | MVP_top60_DNA+RNA_seed | SMARCA4 | 0.912899 | LUAD | HCC-827 | PFI3 |
| MVP_top60_DNA_seed | HSP90AA1 | -0.414557 | LUAD | PC-3_[JPC-3] | Luminespib | MVP_top60_DNA+RNA_seed | BRAF | 0.912949 | LUAD | NCI-H358 | SB590885 |
| MVP_top60_DNA_seed | BRAF | -0.414052 | LUAD | PC-14 | PLX-4720 | MVP_top60_DNA+RNA_seed | TP53 | 0.914092 | LUAD | Calu-6 | MIRA-1 |
| MVP_top60_DNA_seed | AURKA | -0.411687 | LUAD | RERF-LC-KJ | Genentech Cpd 10 | MVP_top60_DNA+RNA_seed | EGFR | 0.914973 | LUAD | NCI-H23 | Sapitinib |
| MVP_top60_DNA_seed | AURKA | -0.408729 | LUAD | LC-2-ad | Alisertib | MVP_top60_DNA+RNA_seed | MDM2 | 0.917131 | LUAD | NCI-H2009 | Nutlin-3a (-) |
| MVP_top60_DNA_seed | PIK3CA | -0.408583 | LUAD | A549 | AZD8835 | MVP_top60_DNA+RNA_seed | MET | 0.917173 | LUAD | A549 | Savolitinib |
| MVP_top60_DNA_seed | HSP90AA1 | -0.407265 | LUAD | NCI-H2030 | Elesclomol | MVP_top60_DNA+RNA_seed | MET | 0.917527 | LUAD | HCC-78 | Savolitinib |
| MVP_top60_DNA_seed | FGFR1 | -0.40521 | LUAD | NCI-H2228 | PD173074 | MVP_top60_DNA+RNA_seed | HDAC1 | 0.918698 | LUAD | NCI-H1975 | PCI-34051 |
| MVP_top60_DNA_seed | BRAF | -0.405073 | LUAD | HCC-827 | HG6-64-1 | MVP_top60_DNA+RNA_seed | PIK3CA | 0.920335 | LUAD | NCI-H1568 | GNE-317 |
| MVP_top60_DNA_seed | GSK3B | -0.404106 | LUAD | NCI-H23 | KIN001-042 | MVP_top60_DNA+RNA_seed | PIK3CA | 0.921054 | LUAD | NCI-H2342 | GNE-317 |
| MVP_top60_DNA_seed | ATM | -0.403417 | LUAD | SW1573 | torin2 | MVP_top60_DNA+RNA_seed | AKT1 | 0.92106 | LUAD | NCI-H1651 | AZD5363 |
| MVP_top60_DNA_seed | AURKA | -0.402924 | LUAD | NCI-H1437 | HG-5-113-01 | MVP_top60_DNA+RNA_seed | SMARCA4 | 0.921551 | LUAD | Calu-6 | PFI3 |
| MVP_top60_DNA_seed | BRAF | -0.40254 | LUAD | NCI-H1734 | HG6-64-1 | MVP_top60_DNA+RNA_seed | TP53 | 0.92166 | LUAD | NCI-H1975 | MIRA-1 |
| MVP_top60_DNA_seed | AKT1 | -0.402022 | LUAD | NCI-H2122 | AKT inhibitor VIII | MVP_top60_DNA+RNA_seed | HDAC1 | 0.921864 | LUAD | Calu-3 | Romidepsin |
| MVP_top60_DNA_seed | IGF1R | -0.401103 | LUAD | HCC-44 | BMS-536924 | MVP_top60_DNA+RNA_seed | MET | 0.921934 | LUAD | NCI-H2030 | Savolitinib |
| MVP_top60_DNA_seed | ATM | -0.401095 | LUAD | NCI-H2291 | KU-55933 | MVP_top60_DNA+RNA_seed | EZH2 | 0.923147 | LUAD | HCC-827 | GSK343 |
| MVP_top60_DNA_seed | HSP90AA1 | -0.40005 | LUAD | NCI-H2087 | SNX-2112 | MVP_top60_DNA+RNA_seed | EGFR | 0.923693 | LUAD | NCI-H2087 | Afatinib |
| MVP_top60_DNA_seed | AKT1 | -0.399703 | LUAD | NCI-H2122 | AKT inhibitor VIII | MVP_top60_DNA+RNA_seed | MDM2 | 0.924447 | LUAD | Calu-3 | Nutlin-3a (-) |
| MVP_top60_DNA_seed | EGFR | -0.399537 | LUAD | EKVX | CI-1033 | MVP_top60_DNA+RNA_seed | TERT | 0.926156 | LUAD | NCI-H1755 | BIBR-1532 |
| MVP_top60_DNA_seed | ERBB2 | -0.399537 | LUAD | EKVX | CI-1033 | MVP_top60_DNA+RNA_seed | BRAF | 0.926668 | LUAD | NCI-H1944 | SB590885 |
| MVP_top60_DNA_seed | ATM | -0.398691 | LUAD | NCI-H1650 | KU-60019 | MVP_top60_DNA+RNA_seed | BRAF | 0.927375 | LUAD | HOP-62 | Dabrafenib |
| MVP_top60_DNA_seed | PIK3CA | -0.398438 | LUAD | NCI-H23 | PI3Ka_4409 | MVP_top60_DNA+RNA_seed | EGFR | 0.927613 | LUAD | Calu-6 | Erlotinib |
| MVP_top60_DNA_seed | AURKA | -0.39743 | LUAD | NCI-H1568 | Genentech Cpd 10 | MVP_top60_DNA+RNA_seed | IGF1R | 0.927764 | LUAD | Calu-3 | NVP-ADW742 |
| MVP_top60_DNA_seed | PIK3CA | -0.397218 | LUAD | NCI-H1355 | AZD8835 | MVP_top60_DNA+RNA_seed | AURKA | 0.927966 | LUAD | HOP-62 | ZM447439 |
| MVP_top60_DNA_seed | HDAC1 | -0.396289 | LUAD | HOP-62 | AR-42 | MVP_top60_DNA+RNA_seed | BRAF | 0.929629 | LUAD | VMRC-LCD | Dabrafenib |
| MVP_top60_DNA_seed | GSK3B | -0.393912 | LUAD | COR-L105 | CHIR-99021 | MVP_top60_DNA+RNA_seed | PIK3CA | 0.930774 | LUAD | NCI-H2087 | Buparlisib |
| MVP_top60_DNA_seed | AURKA | -0.392518 | LUAD | NCI-H2087 | ZM447439 | MVP_top60_DNA+RNA_seed | PIK3CA | 0.93106 | LUAD | NCI-H2347 | AZD8186 |
| MVP_top60_DNA_seed | HDAC1 | -0.391694 | LUAD | NCI-H1975 | Belinostat | MVP_top60_DNA+RNA_seed | HSP90AA1 | 0.931818 | LUAD | NCI-H522 | Luminespib |
| MVP_top60_DNA_seed | ATM | -0.391139 | LUAD | NCI-H1666 | KU-55933 | MVP_top60_DNA+RNA_seed | AKT1 | 0.932135 | LUAD | NCI-H1650 | MK-2206 |
| MVP_top60_DNA_seed | MET | -0.390652 | LUAD | NCI-H1435 | JNJ38877605 | MVP_top60_DNA+RNA_seed | IGF1R | 0.932998 | LUAD | EMC-BAC-1 | Linsitinib |
| MVP_top60_DNA_seed | BRAF | -0.388667 | LUAD | NCI-H1793 | HG6-64-1 | MVP_top60_DNA+RNA_seed | EGFR | 0.933491 | LUAD | ABC-1 | Osimertinib |
| MVP_top60_DNA_seed | AKT1 | -0.386641 | LUAD | NCI-H1651 | AKT inhibitor VIII | MVP_top60_DNA+RNA_seed | AKT1 | 0.933999 | LUAD | EKVX | MK-2206 |
| MVP_top60_DNA_seed | HDAC1 | -0.386465 | LUAD | HCC-78 | AR-42 | MVP_top60_DNA+RNA_seed | KRAS | 0.936083 | LUAD | NCI-H2030 | RAS (G12C) Inhibitor-1 |
| MVP_top60_DNA_seed | BRAF | -0.385887 | LUAD | NCI-H23 | RAF_9304 | MVP_top60_DNA+RNA_seed | ATM | 0.936511 | LUAD | ABC-1 | KU-55933 |
| MVP_top60_DNA_seed | AKT1 | -0.384716 | LUAD | NCI-H1944 | AKT inhibitor VIII | MVP_top60_DNA+RNA_seed | FGFR1 | 0.936713 | LUAD | NCI-H2228 | PD173074 |
| MVP_top60_DNA_seed | IGF1R | -0.384488 | LUAD | RERF-LC-KJ | BMS-754807 | MVP_top60_DNA+RNA_seed | IGF1R | 0.936912 | LUAD | NCI-H1693 | BMS-536924 |
| MVP_top60_DNA_seed | AURKA | -0.383222 | LUAD | ABC-1 | GSK1070916 | MVP_top60_DNA+RNA_seed | AKT1 | 0.937973 | LUAD | Calu-6 | Uprosertib |
| MVP_top60_DNA_seed | MET | -0.382173 | LUAD | NCI-H1563 | AZD6094 | MVP_top60_DNA+RNA_seed | SMARCA4 | 0.93819 | LUAD | NCI-H2030 | PFI3 |
| MVP_top60_DNA_seed | BRAF | -0.381501 | LUAD | NCI-H2030 | SB590885 | MVP_top60_DNA+RNA_seed | AURKA | 0.940159 | LUAD | EKVX | ZM447439 |
| MVP_top60_DNA_seed | IGF1R | -0.380848 | LUAD | SK-LU-1 | BMS-536924 | MVP_top60_DNA+RNA_seed | EGFR | 0.940165 | LUAD | SK-LU-1 | Sapitinib |
| MVP_top60_DNA_seed | ATM | -0.380128 | LUAD | NCI-H2228 | KU-55933 | MVP_top60_DNA+RNA_seed | IGF1R | 0.940203 | LUAD | HCC-827 | Linsitinib |

| | | | | | | | | | | | |
|---|---|---|---|---|---|---|---|---|---|---|---|
| MVP_top60_DNA_seed | ATM | -0.38011 | LUAD | NCI-H1993 | KU-60019 | MVP_top60_DNA+RNA_seed | PIK3CA | 0.940892 | LUAD | NCI-H2405 | AZD8186 |
| MVP_top60_DNA_seed | PIK3CA | -0.379714 | LUAD | NCI-H2030 | PI3Ka_4409 | MVP_top60_DNA+RNA_seed | AKT1 | 0.942031 | LUAD | ABC-1 | Ipatasertib |
| MVP_top60_DNA_seed | ATM | -0.378262 | LUAD | NCI-H2405 | CP466722 | MVP_top60_DNA+RNA_seed | IGF1R | 0.944484 | LUAD | SW1573 | Linsitinib |
| MVP_top60_DNA_seed | IGF1R | -0.376376 | LUAD | NCI-H2030 | Linsitinib | MVP_top60_DNA+RNA_seed | SMARCA4 | 0.945359 | LUAD | NCI-H1944 | PFI3 |
| MVP_top60_DNA_seed | HDAC1 | -0.376176 | LUAD | NCI-H522 | AR-42 | MVP_top60_DNA+RNA_seed | KRAS | 0.946933 | LUAD | NCI-H2085 | KRAS (G12C) Inhibitor-1 |
| MVP_top60_DNA_seed | IGF1R | -0.376119 | LUAD | NCI-H1573 | BMS-536924 | MVP_top60_DNA+RNA_seed | HSP90AA1 | 0.947345 | LUAD | PC-3_[JPC-3] | Tanespimycin |
| MVP_top60_DNA_seed | GSK3B | -0.372917 | LUAD | NCI-H1568 | KIN001-042 | MVP_top60_DNA+RNA_seed | BRAF | 0.947908 | LUAD | NCI-H1568 | SB590885 |
| MVP_top60_DNA_seed | CREBBP | -0.371881 | LUAD | Calu-3 | I-CBP112 | MVP_top60_DNA+RNA_seed | IGF1R | 0.947925 | LUAD | NCI-H1666 | BMS-754807 |
| MVP_top60_DNA_seed | BRAF | -0.371361 | LUAD | PC-14 | PLX-4720 | MVP_top60_DNA+RNA_seed | TERT | 0.947953 | LUAD | NCI-H2342 | BIBR-1532 |
| MVP_top60_DNA_seed | BRAF | -0.369763 | LUAD | LXF-289 | PLX-4720 | MVP_top60_DNA+RNA_seed | PIK3CA | 0.94951 | LUAD | SK-LU-1 | AZD8186 |
| MVP_top60_DNA_seed | GSK3B | -0.369125 | LUAD | NCI-H23 | SB216763 | MVP_top60_DNA+RNA_seed | HDAC1 | 0.950171 | LUAD | NCI-H322M | Entinostat |
| MVP_top60_DNA_seed | IGF1R | -0.368771 | LUAD | NCI-H838 | GSK1904529A | MVP_top60_DNA+RNA_seed | BRAF | 0.950346 | LUAD | NCI-H1693 | SB590885 |
| MVP_top60_DNA_seed | BRAF | -0.367398 | LUAD | NCI-H650 | Dabrafenib | MVP_top60_DNA+RNA_seed | IGF1R | 0.951134 | LUAD | NCI-H2085 | GSK1904529A |
| MVP_top60_DNA_seed | HSP90AA1 | -0.364479 | LUAD | NCI-H2122 | CCT-018159 | MVP_top60_DNA+RNA_seed | AKT1 | 0.951216 | LUAD | HCC-827 | AZD5363 |
| MVP_top60_DNA_seed | EGFR | -0.362356 | LUAD | NCI-H2342 | Afatinib | MVP_top60_DNA+RNA_seed | EZH2 | 0.951232 | LUAD | ABC-1 | GSK343 |
| MVP_top60_DNA_seed | ERBB2 | -0.362356 | LUAD | NCI-H2342 | Afatinib | MVP_top60_DNA+RNA_seed | EGFR | 0.951415 | LUAD | NCI-H650 | Lapatinib |
| MVP_top60_DNA_seed | IGF1R | -0.362298 | LUAD | COR-L105 | Linsitinib | MVP_top60_DNA+RNA_seed | BRAF | 0.951749 | LUAD | HCC-827 | SB590885 |
| MVP_top60_DNA_seed | GSK3B | -0.362093 | LUAD | LXF-289 | CHIR-99021 | MVP_top60_DNA+RNA_seed | IGF1R | 0.955728 | LUAD | NCI-H2347 | NVP-ADW742 |
| MVP_top60_DNA_seed | IGF1R | -0.361303 | LUAD | H3255 | BMS-536924 | MVP_top60_DNA+RNA_seed | AKT1 | 0.955732 | LUAD | NCI-H1623 | Ipatasertib |
| MVP_top60_DNA_seed | HSP90AA1 | -0.361204 | LUAD | NCI-H1650 | Elesclomol | MVP_top60_DNA+RNA_seed | MET | 0.957671 | LUAD | NCI-H1781 | Crizotinib |
| MVP_top60_DNA_seed | EGFR | -0.361079 | LUAD | HOP-62 | CUDC-101 | MVP_top60_DNA+RNA_seed | EZH2 | 0.957823 | LUAD | NCI-H358 | GSK343 |
| MVP_top60_DNA_seed | ERBB2 | -0.361079 | LUAD | HOP-62 | CUDC-101 | MVP_top60_DNA+RNA_seed | EZH2 | 0.95783 | LUAD | Calu-3 | GSK343 |
| MVP_top60_DNA_seed | GSK3B | -0.360282 | LUAD | NCI-H1838 | SB216763 | MVP_top60_DNA+RNA_seed | PIK3CA | 0.957989 | LUAD | SK-LU-1 | GNE-317 |
| MVP_top60_DNA_seed | PIK3CA | -0.360276 | LUAD | NCI-H2023 | AZD8835 | MVP_top60_DNA+RNA_seed | IGF1R | 0.958939 | LUAD | PC-3_[JPC-3] | Linsitinib |
| MVP_top60_DNA_seed | EGFR | -0.359683 | LUAD | LXF-289 | AZD8931 | MVP_top60_DNA+RNA_seed | PIK3CA | 0.959462 | LUAD | VMRC-LCD | GNE-317 |
| MVP_top60_DNA_seed | ERBB2 | -0.359683 | LUAD | LXF-289 | AZD8931 | MVP_top60_DNA+RNA_seed | EZH2 | 0.95963 | LUAD | NCI-H2087 | GSK343 |
| MVP_top60_DNA_seed | ERBB3 | -0.359683 | LUAD | LXF-289 | AZD8931 | MVP_top60_DNA+RNA_seed | EGFR | 0.960605 | LUAD | NCI-H650 | Osimertinib |
| MVP_top60_DNA_seed | BRAF | -0.358977 | LUAD | LC-2-ad | AZ628 | MVP_top60_DNA+RNA_seed | IGF1R | 0.960727 | LUAD | NCI-H1993 | GSK1904529A |
| MVP_top60_DNA_seed | ATM | -0.358288 | LUAD | NCI-H2347 | KU-60019 | MVP_top60_DNA+RNA_seed | EGFR | 0.960995 | LUAD | NCI-H1437 | Afatinib |
| MVP_top60_DNA_seed | AURKA | -0.357924 | LUAD | NCI-H1437 | GSK1070916 | MVP_top60_DNA+RNA_seed | BRAF | 0.961481 | LUAD | PC-3_[JPC-3] | SB590885 |
| MVP_top60_DNA_seed | MDM2 | -0.357765 | LUAD | NCI-H2009 | Serdemetan | MVP_top60_DNA+RNA_seed | FGFR1 | 0.964134 | LUAD | NCI-H23 | PD173074 |
| MVP_top60_DNA_seed | MDM2 | -0.357401 | LUAD | HCC-827 | Nutlin-3a (-) | MVP_top60_DNA+RNA_seed | HDAC1 | 0.965539 | LUAD | Calu-6 | Entinostat |
| MVP_top60_DNA_seed | HDAC1 | -0.357358 | LUAD | NCI-H3122 | Dacinostat | MVP_top60_DNA+RNA_seed | EGFR | 0.965804 | LUAD | NCI-H1793 | Erlotinib |
| MVP_top60_DNA_seed | HSP90AA1 | -0.356973 | LUAD | LC-2-ad | Elesclomol | MVP_top60_DNA+RNA_seed | FGFR1 | 0.966942 | LUAD | NCI-H2228 | AZD4547 |
| MVP_top60_DNA_seed | FGFR1 | -0.356116 | LUAD | LC-2-ad | PD173074 | MVP_top60_DNA+RNA_seed | AURKA | 0.967196 | LUAD | NCI-H1793 | Alisertib |
| MVP_top60_DNA_seed | PTK2 | -0.355751 | LUAD | NCI-H2009 | PF-562271 | MVP_top60_DNA+RNA_seed | AKT1 | 0.968373 | LUAD | NCI-H1693 | Ipatasertib |
| MVP_top60_DNA_seed | BRAF | -0.35569 | LUAD | NCI-H1650 | HG6-64-1 | MVP_top60_DNA+RNA_seed | MET | 0.968425 | LUAD | NCI-H2291 | Crizotinib |
| MVP_top60_DNA_seed | AKT1 | -0.354109 | LUAD | NCI-H2030 | BAY AKT1 | MVP_top60_DNA+RNA_seed | MDM2 | 0.968829 | LUAD | NCI-H2085 | Serdemetan |
| MVP_top60_DNA_seed | IGF1R | -0.354083 | LUAD | NCI-H2405 | GSK1904529A | MVP_top60_DNA+RNA_seed | KRAS | 0.969335 | LUAD | HCC-827 | KRAS (G12C) Inhibitor-1 |
| MVP_top60_DNA_seed | EGFR | -0.354053 | LUAD | EMC-BAC-2 | AZD8931 | MVP_top60_DNA+RNA_seed | MET | 0.973415 | LUAD | NCI-H2087 | Savolitinib |
| MVP_top60_DNA_seed | ERBB2 | -0.354053 | LUAD | EMC-BAC-2 | AZD8931 | MVP_top60_DNA+RNA_seed | AKT1 | 0.974683 | LUAD | NCI-H358 | MK-2206 |

| | | | | | | | | | | | |
|---|---|---|---|---|---|---|---|---|---|---|---|
| MVP_top60_DNA_seed | ERBB3 | -0.354053 | LUAD | EMC-BAC-2 | AZD8931 | MVP_top60_DNA+RNA_seed | TP53 | 0.977542 | LUAD | NCI-H322M | MIRA-1 |
| MVP_top60_DNA_seed | EGFR | -0.353626 | LUAD | EKVX | PF-00299804 | MVP_top60_DNA+RNA_seed | IGF1R | 0.978539 | LUAD | NCI-H1793 | NVP-ADW742 |
| MVP_top60_DNA_seed | ERBB2 | -0.353626 | LUAD | EKVX | PF-00299804 | MVP_top60_DNA+RNA_seed | IGF1R | 0.97976 | LUAD | HCC-827 | BMS-754807 |
| MVP_top60_DNA_seed | EGFR | -0.353107 | LUAD | NCI-H1792 | Gefitinib | MVP_top60_DNA+RNA_seed | BRAF | 0.979943 | LUAD | Calu-3 | SB590885 |
| MVP_top60_DNA_seed | HDAC1 | -0.351502 | LUAD | EMC-BAC-2 | CAY10603 | MVP_top60_DNA+RNA_seed | AKT1 | 0.980124 | LUAD | NCI-H1650 | GSK2110183B |
| MVP_top60_DNA_seed | SRC | -0.349839 | LUAD | PC-14 | NVP-BHG712 | MVP_top60_DNA+RNA_seed | BRAF | 0.981309 | LUAD | NCI-H1563 | PLX-4720 |
| MVP_top60_DNA_seed | HSP90AA1 | -0.349315 | LUAD | ABC-1 | Luminespib | MVP_top60_DNA+RNA_seed | EGFR | 0.981402 | LUAD | NCI-H3122 | Gefitinib |
| MVP_top60_DNA_seed | ERBB2 | -0.348922 | LUAD | NCI-H1563 | CP724714 | MVP_top60_DNA+RNA_seed | AKT1 | 0.983721 | LUAD | NCI-H1573 | AT13148 |
| MVP_top60_DNA_seed | IGF1R | -0.348873 | LUAD | PC-14 | BMS-536924 | MVP_top60_DNA+RNA_seed | MET | 0.983982 | LUAD | NCI-H522 | Crizotinib |
| MVP_top60_DNA_seed | HDAC1 | -0.348675 | LUAD | EMC-BAC-2 | Belinostat | MVP_top60_DNA+RNA_seed | IGF1R | 0.985523 | LUAD | NCI-H1693 | BMS-754807 |
| MVP_top60_DNA_seed | EGFR | -0.34688 | LUAD | NCI-H23 | Lapatinib | MVP_top60_DNA+RNA_seed | HSP90AA1 | 0.985952 | LUAD | HCC-827 | Tanespimycin |
| MVP_top60_DNA_seed | ERBB2 | -0.34688 | LUAD | NCI-H23 | Lapatinib | MVP_top60_DNA+RNA_seed | TP53 | 0.986453 | LUAD | PC-14 | MIRA-1 |
| MVP_top60_DNA_seed | HSP90AA1 | -0.346649 | LUAD | SK-LU-1 | CCT-018159 | MVP_top60_DNA+RNA_seed | EGFR | 0.989603 | LUAD | NCI-H2085 | AZD3759 |
| MVP_top60_DNA_seed | PIK3CA | -0.346425 | LUAD | NCI-H1666 | PI3Ka_4409 | MVP_top60_DNA+RNA_seed | KRAS | 0.98968 | LUAD | Calu-3 | KRAS (G12C) Inhibitor-1 |
| MVP_top60_DNA_seed | AURKA | -0.344724 | LUAD | EMC-BAC-1 | Genentech Cpd 10 | MVP_top60_DNA+RNA_seed | IGF1R | 0.990332 | LUAD | NCI-H1993 | NVP-ADW742 |
| MVP_top60_DNA_seed | GSK3B | -0.34375 | LUAD | NCI-H358 | AZD7969 | MVP_top60_DNA+RNA_seed | MET | 0.990781 | LUAD | NCI-H1666 | Crizotinib |
| MVP_top60_DNA_seed | ATM | -0.343404 | LUAD | PC-14 | KU-60019 | MVP_top60_DNA+RNA_seed | MDM2 | 0.991437 | LUAD | HCC-827 | Nutlin-3a (-) |
| MVP_top60_DNA_seed | EGFR | -0.342656 | LUAD | NCI-H1792 | Pelitinib | MVP_top60_DNA+RNA_seed | BRAF | 0.994664 | LUAD | NCI-H650 | PLX-4720 |
| MVP_top60_DNA_seed | MAPK1 | -0.342176 | LUAD | NCI-H1792 | FR-180204 | MVP_top60_DNA+RNA_seed | EGFR | 0.996548 | LUAD | NCI-H1793 | Gefitinib |
| MVP_top60_DNA_seed | MET | -0.342002 | LUAD | NCI-H1975 | JNJ38877605 | MVP_top60_DNA+RNA_seed | EGFR | 0.996635 | LUAD | NCI-H1651 | Lapatinib |
| MVP_top60_DNA_seed | HDAC1 | -0.341973 | LUAD | NCI-H1355 | Parthenolide | MVP_top60_DNA+RNA_seed | SRC | 0.99665 | LUAD | VMRC-LCD | Bosutinib |
| MVP_top60_DNA_seed | ATM | -0.341272 | LUAD | NCI-H1793 | KU-55933 | MVP_top60_DNA+RNA_seed | AKT1 | 0.999248 | LUAD | SW1573 | Afuresertib |
| MVP_top60_DNA_seed | MAPK1 | -0.339342 | LUAD | NCI-H23 | VX-11e | MVP_top60_DNA+RNA_seed | AKT1 | 1.002867 | LUAD | NCI-H650 | Afuresertib |
| MVP_top60_DNA_seed | EGFR | -0.338522 | LUAD | NCI-H1734 | Afatinib | MVP_top60_DNA+RNA_seed | EZH2 | 1.003655 | LUAD | COR-L105 | GSK343 |
| MVP_top60_DNA_seed | ERBB2 | -0.338522 | LUAD | NCI-H1734 | Afatinib | MVP_top60_DNA+RNA_seed | HSP90AA1 | 1.003695 | LUAD | HCC-78 | Tanespimycin |
| MVP_top60_DNA_seed | GSK3B | -0.338176 | LUAD | HCC-78 | AZD7969 | MVP_top60_DNA+RNA_seed | IGF1R | 1.004013 | LUAD | HOP-62 | GSK1904529A |
| MVP_top60_DNA_seed | MET | -0.337777 | LUAD | LXF-289 | Crizotinib | MVP_top60_DNA+RNA_seed | MET | 1.004068 | LUAD | PC-14 | Savolitinib |
| MVP_top60_DNA_seed | BRAF | -0.337194 | LUAD | NCI-H1568 | RAF_9304 | MVP_top60_DNA+RNA_seed | FGFR1 | 1.004624 | LUAD | EMC-BAC-2 | PD173074 |
| MVP_top60_DNA_seed | PTK2 | -0.335939 | LUAD | COR-L105 | PF-562271 | MVP_top60_DNA+RNA_seed | MET | 1.004822 | LUAD | VMRC-LCD | Savolitinib |
| MVP_top60_DNA_seed | MET | -0.335149 | LUAD | NCI-H1944 | AZD6094 | MVP_top60_DNA+RNA_seed | EZH2 | 1.007129 | LUAD | VMRC-LCD | GSK343 |
| MVP_top60_DNA_seed | SMARCA4 | -0.334159 | LUAD | NCI-H2030 | PFI-3 | MVP_top60_DNA+RNA_seed | SRC | 1.009268 | LUAD | NCI-H3122 | Bosutinib |
| MVP_top60_DNA_seed | ERBB2 | -0.333151 | LUAD | NCI-H1437 | CP724714 | MVP_top60_DNA+RNA_seed | HSP90AA1 | 1.009437 | LUAD | SK-LU-1 | Luminespib |
| MVP_top60_DNA_seed | EGFR | -0.332907 | LUAD | PC-14 | CI-1033 | MVP_top60_DNA+RNA_seed | IGF1R | 1.009589 | LUAD | NCI-H2023 | Linsitinib |
| MVP_top60_DNA_seed | ERBB2 | -0.332907 | LUAD | PC-14 | CI-1033 | MVP_top60_DNA+RNA_seed | AURKA | 1.0104 | LUAD | NCI-H322M | Alisertib |
| MVP_top60_DNA_seed | IGF1R | -0.332222 | LUAD | NCI-H2405 | Linsitinib | MVP_top60_DNA+RNA_seed | AURKA | 1.010505 | LUAD | EMC-BAC-2 | ZM447439 |
| MVP_top60_DNA_seed | BRAF | -0.332218 | LUAD | LC-2-ad | PLX-4720 | MVP_top60_DNA+RNA_seed | TP53 | 1.010753 | LUAD | HOP-62 | MIRA-1 |
| MVP_top60_DNA_seed | HSP90AA1 | -0.331667 | LUAD | HCC-78 | Luminespib | MVP_top60_DNA+RNA_seed | MDM2 | 1.012476 | LUAD | NCI-H358 | Nutlin-3a (-) |
| MVP_top60_DNA_seed | HDAC1 | -0.330251 | LUAD | NCI-H1975 | Parthenolide | MVP_top60_DNA+RNA_seed | EGFR | 1.013636 | LUAD | NCI-H23 | Osimertinib |
| MVP_top60_DNA_seed | BRAF | -0.330048 | LUAD | NCI-H2009 | HG6-64-1 | MVP_top60_DNA+RNA_seed | TP53 | 1.015556 | LUAD | NCI-H358 | MIRA-1 |
| MVP_top60_DNA_seed | HDAC1 | -0.329632 | LUAD | EMC-BAC-2 | AR-42 | MVP_top60_DNA+RNA_seed | PIK3CA | 1.015869 | LUAD | NCI-H1838 | AZD8186 |
| MVP_top60_DNA_seed | EGFR | -0.329395 | LUAD | NCI-H2342 | Cetuximab | MVP_top60_DNA+RNA_seed | HDAC1 | 1.01639 | LUAD | VMRC-LCD | PCI-34051 |

| | | | | | | | | | | | |
|---|---|---|---|---|---|---|---|---|---|---|---|
| MVP_top60_DNA_seed | MDM2 | -0.329016 | LUAD | NCI-H1648 | Nutlin-3a (-) | MVP_top60_DNA+RNA_seed | HSP90AA1 | 1.017946 | LUAD | RERF-LC-KJ | Luminespib |
| MVP_top60_DNA_seed | ATM | -0.32872 | LUAD | H3255 | KU-60019 | MVP_top60_DNA+RNA_seed | PIK3CA | 1.019241 | LUAD | NCI-H1568 | Alpelisib |
| MVP_top60_DNA_seed | AURKA | -0.328572 | LUAD | COR-L105 | ZM447439 | MVP_top60_DNA+RNA_seed | AURKA | 1.020653 | LUAD | NCI-H2087 | Alisertib |
| MVP_top60_DNA_seed | MET | -0.327084 | LUAD | NCI-H322M | JNJ38877605 | MVP_top60_DNA+RNA_seed | BRAF | 1.020963 | LUAD | NCI-H2342 | Dabrafenib |
| MVP_top60_DNA_seed | HSP90AA1 | -0.326522 | LUAD | NCI-H838 | CCT-018159 | MVP_top60_DNA+RNA_seed | BRAF | 1.021227 | LUAD | NCI-H1623 | PLX-4720 |
| MVP_top60_DNA_seed | AURKA | -0.324213 | LUAD | PC-14 | GSK1070916 | MVP_top60_DNA+RNA_seed | IGF1R | 1.022293 | LUAD | NCI-H1650 | GSK1904529A |
| MVP_top60_DNA_seed | SRC | -0.323515 | LUAD | NCI-H1792 | NVP-BHG712 | MVP_top60_DNA+RNA_seed | FGFR1 | 1.024148 | LUAD | HCC-78 | AZD4547 |
| MVP_top60_DNA_seed | HDAC1 | -0.323348 | LUAD | NCI-H1355 | Dacinostat | MVP_top60_DNA+RNA_seed | IGF1R | 1.024298 | LUAD | NCI-H441 | Linsitinib |
| MVP_top60_DNA_seed | AURKA | -0.322665 | LUAD | NCI-H2122 | ZM447439 | MVP_top60_DNA+RNA_seed | EGFR | 1.024683 | LUAD | RERF-LC-MS | Afatinib |
| MVP_top60_DNA_seed | EGFR | -0.321502 | LUAD | VMRC-LCD | AZD8931 | MVP_top60_DNA+RNA_seed | AKT1 | 1.025872 | LUAD | NCI-H1435 | AZD5363 |
| MVP_top60_DNA_seed | ERBB2 | -0.321502 | LUAD | VMRC-LCD | AZD8931 | MVP_top60_DNA+RNA_seed | PIK3CA | 1.02762 | LUAD | VMRC-LCD | Buparlisib |
| MVP_top60_DNA_seed | ERBB3 | -0.321502 | LUAD | VMRC-LCD | AZD8931 | MVP_top60_DNA+RNA_seed | SMARCA4 | 1.027794 | LUAD | NCI-H1666 | PFI3 |
| MVP_top60_DNA_seed | IGF1R | -0.320594 | LUAD | NCI-H1623 | BMS-754807 | MVP_top60_DNA+RNA_seed | PIK3CA | 1.028748 | LUAD | NCI-H1838 | GNE-317 |
| MVP_top60_DNA_seed | AKT1 | -0.320444 | LUAD | NCI-H2347 | AKT inhibitor VIII | MVP_top60_DNA+RNA_seed | HDAC1 | 1.031691 | LUAD | NCI-H1563 | Entinostat |
| MVP_top60_DNA_seed | IGF1R | -0.319484 | LUAD | LC-2-ad | GSK1904529A | MVP_top60_DNA+RNA_seed | AKT1 | 1.03171 | LUAD | COR-L105 | AT13148 |
| MVP_top60_DNA_seed | MDM2 | -0.319427 | LUAD | SW1573 | Nutlin-3a (-) | MVP_top60_DNA+RNA_seed | KRAS | 1.033303 | LUAD | EKVX | KRAS (G12C) Inhibitor-1 |
| MVP_top60_DNA_seed | IGF1R | -0.318502 | LUAD | NCI-H1792 | BMS-754807 | MVP_top60_DNA+RNA_seed | MDM2 | 1.033423 | LUAD | EKVX | Nutlin-3a (-) |
| MVP_top60_DNA_seed | IGF1R | -0.318294 | LUAD | NCI-H1993 | Linsitinib | MVP_top60_DNA+RNA_seed | MET | 1.034528 | LUAD | NCI-H441 | Crizotinib |
| MVP_top60_DNA_seed | AURKA | -0.31814 | LUAD | NCI-H2228 | GSK1070916 | MVP_top60_DNA+RNA_seed | AKT1 | 1.035306 | LUAD | NCI-H1355 | AZD5363 |
| MVP_top60_DNA_seed | AKT1 | -0.317699 | LUAD | NCI-H1623 | MK-2206 | MVP_top60_DNA+RNA_seed | AKT1 | 1.035421 | LUAD | NCI-H2087 | GSK2110183B |
| MVP_top60_DNA_seed | PIK3CA | -0.317112 | LUAD | NCI-H358 | AZD8835 | MVP_top60_DNA+RNA_seed | EGFR | 1.036243 | LUAD | HCC-44 | AZD3759 |
| MVP_top60_DNA_seed | IGF1R | -0.316427 | LUAD | NCI-H1568 | Linsitinib | MVP_top60_DNA+RNA_seed | MDM2 | 1.038127 | LUAD | PC-3_[JPC-3] | Nutlin-3a (-) |
| MVP_top60_DNA_seed | EGFR | -0.315379 | LUAD | NCI-H441 | Afatinib | MVP_top60_DNA+RNA_seed | AKT1 | 1.041239 | LUAD | NCI-H1355 | Afuresertib |
| MVP_top60_DNA_seed | ERBB2 | -0.315379 | LUAD | NCI-H441 | Afatinib | MVP_top60_DNA+RNA_seed | IGF1R | 1.043226 | LUAD | NCI-H2085 | NVP-ADW742 |
| MVP_top60_DNA_seed | EGFR | -0.315364 | LUAD | NCI-H23 | Cetuximab | MVP_top60_DNA+RNA_seed | TP53 | 1.043648 | LUAD | SK-LU-1 | MIRA-1 |
| MVP_top60_DNA_seed | AURKA | -0.314849 | LUAD | SK-LU-1 | Genentech Cpd 10 | MVP_top60_DNA+RNA_seed | AKT1 | 1.04502 | LUAD | Calu-6 | GSK2110183B |
| MVP_top60_DNA_seed | MET | -0.314713 | LUAD | NCI-H1781 | AZD6094 | MVP_top60_DNA+RNA_seed | BRAF | 1.046126 | LUAD | HCC-78 | Dabrafenib |
| MVP_top60_DNA_seed | SRC | -0.313683 | LUAD | NCI-H2030 | NVP-BHG712 | MVP_top60_DNA+RNA_seed | MET | 1.046588 | LUAD | NCI-H1573 | Crizotinib |
| MVP_top60_DNA_seed | MET | -0.312782 | LUAD | NCI-H1648 | Crizotinib | MVP_top60_DNA+RNA_seed | FGFR1 | 1.046749 | LUAD | NCI-H2087 | PD173074 |
| MVP_top60_DNA_seed | ATM | -0.312659 | LUAD | NCI-H1781 | KU-60019 | MVP_top60_DNA+RNA_seed | MET | 1.047204 | LUAD | PC-3_[JPC-3] | Crizotinib |
| MVP_top60_DNA_seed | MET | -0.31155 | LUAD | NCI-H1944 | JNJ38877605 | MVP_top60_DNA+RNA_seed | AURKA | 1.048085 | LUAD | NCI-H1435 | Alisertib |
| MVP_top60_DNA_seed | SRC | -0.311253 | LUAD | NCI-H1755 | Bosutinib | MVP_top60_DNA+RNA_seed | PIK3CA | 1.051712 | LUAD | NCI-H2347 | GNE-317 |
| MVP_top60_DNA_seed | GSK3B | -0.309146 | LUAD | RERF-LC-KJ | KIN001-042 | MVP_top60_DNA+RNA_seed | PIK3CA | 1.052664 | LUAD | NCI-H1793 | Alpelisib |
| MVP_top60_DNA_seed | EGFR | -0.309056 | LUAD | EMC-BAC-2 | CUDC-101 | MVP_top60_DNA+RNA_seed | FGFR1 | 1.053416 | LUAD | EKVX | PD173074 |
| MVP_top60_DNA_seed | ERBB2 | -0.309056 | LUAD | EMC-BAC-2 | CUDC-101 | MVP_top60_DNA+RNA_seed | TP53 | 1.057105 | LUAD | A549 | MIRA-1 |
| MVP_top60_DNA_seed | FGFR1 | -0.305574 | LUAD | NCI-H2291 | PD173074 | MVP_top60_DNA+RNA_seed | KRAS | 1.058704 | LUAD | NCI-H1355 | KRAS (G12C) Inhibitor-1 |
| MVP_top60_DNA_seed | IGF1R | -0.305561 | LUAD | HOP-62 | BMS-754807 | MVP_top60_DNA+RNA_seed | AKT1 | 1.060136 | LUAD | NCI-H1693 | Afuresertib |
| MVP_top60_DNA_seed | AKT1 | -0.305088 | LUAD | LC-2-ad | GSK690693 | MVP_top60_DNA+RNA_seed | TP53 | 1.060177 | LUAD | NCI-H2087 | MIRA-1 |
| MVP_top60_DNA_seed | ATM | -0.304998 | LUAD | NCI-H1355 | KU-55933 | MVP_top60_DNA+RNA_seed | ATM | 1.060528 | LUAD | VMRC-LCD | KU-55933 |
| MVP_top60_DNA_seed | EGFR | -0.30368 | LUAD | HCC-78 | Pelitinib | MVP_top60_DNA+RNA_seed | AKT1 | 1.06382 | LUAD | VMRC-LCD | GSK2110183B |
| MVP_top60_DNA_seed | HDAC1 | -0.303521 | LUAD | ABC-1 | AR-42 | MVP_top60_DNA+RNA_seed | PIK3CA | 1.064196 | LUAD | Calu-6 | Alpelisib |

| | | | | | | | | | | | |
|---|---|---|---|---|---|---|---|---|---|---|---|
| MVP_top60_DNA_seed | HDAC1 | -0.302572 | LUAD | NCI-H1651 | Dacinostat | MVP_top60_DNA+RNA_seed | IGF1R | 1.064207 | LUAD | NCI-H358 | GSK1904529A |
| MVP_top60_DNA_seed | HDAC1 | -0.302102 | LUAD | NCI-H3122 | AR-42 | MVP_top60_DNA+RNA_seed | IGF1R | 1.064413 | LUAD | NCI-H650 | GSK1904529A |
| MVP_top60_DNA_seed | MDM2 | -0.301973 | LUAD | NCI-H1648 | Serdemetan | MVP_top60_DNA+RNA_seed | MDM2 | 1.065983 | LUAD | Calu-6 | Nutlin-3a (-) |
| MVP_top60_DNA_seed | IGF1R | -0.301722 | LUAD | NCI-H3122 | BMS-754807 | MVP_top60_DNA+RNA_seed | BRAF | 1.066159 | LUAD | SW1573 | SB590885 |
| MVP_top60_DNA_seed | AKT1 | -0.301337 | LUAD | NCI-H1355 | BAY AKT1 | MVP_top60_DNA+RNA_seed | EGFR | 1.066534 | LUAD | HOP-62 | Sapitinib |
| MVP_top60_DNA_seed | PTK2 | -0.300498 | LUAD | NCI-H1792 | PF-562271 | MVP_top60_DNA+RNA_seed | EZH2 | 1.067508 | LUAD | NCI-H1666 | GSK343 |
| MVP_top60_DNA_seed | MAPK1 | -0.300103 | LUAD | NCI-H2085 | FR-180204 | MVP_top60_DNA+RNA_seed | PIK3CA | 1.06766 | LUAD | NCI-H1650 | Alpelisib |
| MVP_top60_DNA_seed | MET | -0.299869 | LUAD | EMC-BAC-1 | SU11274 | MVP_top60_DNA+RNA_seed | EGFR | 1.068391 | LUAD | ABC-1 | Afatinib |
| MVP_top60_DNA_seed | EGFR | -0.299839 | LUAD | NCI-H1734 | CUDC-101 | MVP_top60_DNA+RNA_seed | IGF1R | 1.068636 | LUAD | NCI-H1651 | BMS-536924 |
| MVP_top60_DNA_seed | ERBB2 | -0.299839 | LUAD | NCI-H1734 | CUDC-101 | MVP_top60_DNA+RNA_seed | BRAF | 1.071026 | LUAD | NCI-H1355 | SB590885 |
| MVP_top60_DNA_seed | HSP90AA1 | -0.299531 | LUAD | HOP-62 | CCT-018159 | MVP_top60_DNA+RNA_seed | TP53 | 1.071087 | LUAD | NCI-H2291 | MIRA-1 |
| MVP_top60_DNA_seed | AURKA | -0.299099 | LUAD | SW1573 | HG-5-113-01 | MVP_top60_DNA+RNA_seed | MET | 1.071433 | LUAD | NCI-H1568 | Crizotinib |
| MVP_top60_DNA_seed | AKT1 | -0.297716 | LUAD | H3255 | AKT inhibitor VIII | MVP_top60_DNA+RNA_seed | TP53 | 1.075039 | LUAD | Calu-3 | MIRA-1 |
| MVP_top60_DNA_seed | HDAC1 | -0.296217 | LUAD | ABC-1 | CAY10603 | MVP_top60_DNA+RNA_seed | EGFR | 1.075119 | LUAD | Calu-6 | Gefitinib |
| MVP_top60_DNA_seed | MET | -0.295283 | LUAD | NCI-H2030 | SU11274 | MVP_top60_DNA+RNA_seed | AKT1 | 1.075767 | LUAD | PC-3_[JPC-3] | Afuresertib |
| MVP_top60_DNA_seed | HDAC1 | -0.293775 | LUAD | NCI-H522 | Belinostat | MVP_top60_DNA+RNA_seed | AKT1 | 1.078048 | LUAD | NCI-H1435 | MK-2206 |
| MVP_top60_DNA_seed | HSP90AA1 | -0.293298 | LUAD | RERF-LC-KJ | Tanespimycin | MVP_top60_DNA+RNA_seed | IGF1R | 1.078604 | LUAD | NCI-H2087 | GSK1904529A |
| MVP_top60_DNA_seed | CREBBP | -0.292205 | LUAD | VMRC-LCD | I-CBP112 | MVP_top60_DNA+RNA_seed | HSP90AA1 | 1.078929 | LUAD | NCI-H2291 | Tanespimycin |
| MVP_top60_DNA_seed | EGFR | -0.292033 | LUAD | NCI-H2342 | AZD8931 | MVP_top60_DNA+RNA_seed | AKT1 | 1.082744 | LUAD | NCI-H1651 | AT13148 |
| MVP_top60_DNA_seed | ERBB2 | -0.292033 | LUAD | NCI-H2342 | AZD8931 | MVP_top60_DNA+RNA_seed | SRC | 1.083151 | LUAD | NCI-H1573 | Bosutinib |
| MVP_top60_DNA_seed | ERBB3 | -0.292033 | LUAD | NCI-H2342 | AZD8931 | MVP_top60_DNA+RNA_seed | HSP90AA1 | 1.085614 | LUAD | NCI-H1355 | Luminespib |
| MVP_top60_DNA_seed | AKT1 | -0.291412 | LUAD | NCI-H1693 | MK-2206 | MVP_top60_DNA+RNA_seed | EGFR | 1.08636 | LUAD | VMRC-LCD | Lapatinib |
| MVP_top60_DNA_seed | HSP90AA1 | -0.29078 | LUAD | NCI-H1781 | Elesclomol | MVP_top60_DNA+RNA_seed | IGF1R | 1.090314 | LUAD | NCI-H1568 | BMS-754807 |
| MVP_top60_DNA_seed | BRAF | -0.290296 | LUAD | NCI-H650 | PLX-4720 | MVP_top60_DNA+RNA_seed | BRAF | 1.091162 | LUAD | NCI-H1355 | Dabrafenib |
| MVP_top60_DNA_seed | GSK3B | -0.290275 | LUAD | Calu-6 | AZD7969 | MVP_top60_DNA+RNA_seed | MET | 1.09189 | LUAD | Calu-6 | Savolitinib |
| MVP_top60_DNA_seed | MET | -0.289833 | LUAD | NCI-H1792 | SU11274 | MVP_top60_DNA+RNA_seed | HSP90AA1 | 1.092057 | LUAD | H3255 | Luminespib |
| MVP_top60_DNA_seed | EGFR | -0.289521 | LUAD | NCI-H2291 | Afatinib | MVP_top60_DNA+RNA_seed | FGFR1 | 1.092196 | LUAD | NCI-H2085 | AZD4547 |
| MVP_top60_DNA_seed | ERBB2 | -0.289521 | LUAD | NCI-H2291 | Afatinib | MVP_top60_DNA+RNA_seed | FGFR1 | 1.092953 | LUAD | NCI-H1650 | PD173074 |
| MVP_top60_DNA_seed | BRAF | -0.289088 | LUAD | NCI-H1573 | RAF_9304 | MVP_top60_DNA+RNA_seed | AKT1 | 1.093513 | LUAD | NCI-H1573 | AZD5363 |
| MVP_top60_DNA_seed | HSP90AA1 | -0.286845 | LUAD | NCI-H1792 | Luminespib | MVP_top60_DNA+RNA_seed | AKT1 | 1.096605 | LUAD | NCI-H441 | Uprosertib |
| MVP_top60_DNA_seed | HDAC1 | -0.286826 | LUAD | NCI-H2347 | AR-42 | MVP_top60_DNA+RNA_seed | HDAC1 | 1.098466 | LUAD | NCI-H2087 | Entinostat |
| MVP_top60_DNA_seed | HSP90AA1 | -0.286614 | LUAD | NCI-H2087 | CCT-018159 | MVP_top60_DNA+RNA_seed | IGF1R | 1.100977 | LUAD | VMRC-LCD | Linsitinib |
| MVP_top60_DNA_seed | GSK3B | -0.284986 | LUAD | RERF-LC-MS | AZD7969 | MVP_top60_DNA+RNA_seed | IGF1R | 1.104534 | LUAD | COR-L105 | BMS-754807 |
| MVP_top60_DNA_seed | PTK2 | -0.284743 | LUAD | NCI-H1648 | PF-562271 | MVP_top60_DNA+RNA_seed | EGFR | 1.106349 | LUAD | NCI-H2023 | Osimertinib |
| MVP_top60_DNA_seed | PIK3CA | -0.28472 | LUAD | NCI-H2023 | PI3Ka_4409 | MVP_top60_DNA+RNA_seed | AKT1 | 1.106381 | LUAD | NCI-H23 | AT13148 |
| MVP_top60_DNA_seed | AKT1 | -0.283923 | LUAD | LC-2-ad | AKT inhibitor VIII | MVP_top60_DNA+RNA_seed | PIK3CA | 1.107846 | LUAD | NCI-H2087 | AZD8186 |
| MVP_top60_DNA_seed | IGF1R | -0.281819 | LUAD | NCI-H1650 | BMS-536924 | MVP_top60_DNA+RNA_seed | EGFR | 1.110325 | LUAD | NCI-H2228 | Osimertinib |
| MVP_top60_DNA_seed | AURKA | -0.281556 | LUAD | NCI-H1355 | ZM447439 | MVP_top60_DNA+RNA_seed | AKT1 | 1.111078 | LUAD | NCI-H1792 | Uprosertib |
| MVP_top60_DNA_seed | ATM | -0.281519 | LUAD | NCI-H2347 | CP466722 | MVP_top60_DNA+RNA_seed | FGFR1 | 1.111741 | LUAD | VMRC-LCD | AZD4547 |
| MVP_top60_DNA_seed | SRC | -0.279777 | LUAD | NCI-H1648 | NVP-BHG712 | MVP_top60_DNA+RNA_seed | PIK3CA | 1.116172 | LUAD | ABC-1 | Buparlisib |
| MVP_top60_DNA_seed | FGFR1 | -0.279715 | LUAD | NCI-H23 | AZD4547 | MVP_top60_DNA+RNA_seed | IGF1R | 1.117593 | LUAD | NCI-H1568 | Linsitinib |

| | | | | | | | | | | | |
|---|---|---|---|---|---|---|---|---|---|---|---|
| MVP_top60_DNA_seed | EGFR | -0.279504 | LUAD | NCI-H322M | CUDC-101 | MVP_top60_DNA+RNA_seed | AKT1 | 1.117742 | LUAD | HOP-62 | Afuresertib |
| MVP_top60_DNA_seed | ERBB2 | -0.279504 | LUAD | NCI-H322M | CUDC-101 | MVP_top60_DNA+RNA_seed | IGF1R | 1.118713 | LUAD | NCI-H1993 | BMS-536924 |
| MVP_top60_DNA_seed | HDAC1 | -0.279173 | LUAD | RERF-LC-MS | JQ12 | MVP_top60_DNA+RNA_seed | PIK3CA | 1.121993 | LUAD | NCI-H2087 | GNE-317 |
| MVP_top60_DNA_seed | AURKA | -0.279101 | LUAD | NCI-H1975 | Genentech Cpd 10 | MVP_top60_DNA+RNA_seed | AURKA | 1.124456 | LUAD | NCI-H1573 | ZM447439 |
| MVP_top60_DNA_seed | EGFR | -0.278131 | LUAD | NCI-H1993 | Pelitinib | MVP_top60_DNA+RNA_seed | EZH2 | 1.12697 | LUAD | NCI-H322M | GSK343 |
| MVP_top60_DNA_seed | HDAC1 | -0.276324 | LUAD | NCI-H1975 | CAY10603 | MVP_top60_DNA+RNA_seed | IGF1R | 1.128076 | LUAD | NCI-H1651 | BMS-754807 |
| MVP_top60_DNA_seed | AURKA | -0.276235 | LUAD | NCI-H1944 | GSK1070916 | MVP_top60_DNA+RNA_seed | IGF1R | 1.129236 | LUAD | NCI-H2023 | GSK1904529A |
| MVP_top60_DNA_seed | IGF1R | -0.273746 | LUAD | NCI-H1568 | BMS-754807 | MVP_top60_DNA+RNA_seed | AURKA | 1.129468 | LUAD | NCI-H1563 | ZM447439 |
| MVP_top60_DNA_seed | AURKA | -0.273146 | LUAD | NCI-H650 | CD532 | MVP_top60_DNA+RNA_seed | MDM2 | 1.130567 | LUAD | NCI-H1568 | Serdemetan |
| MVP_top60_DNA_seed | HSP90AA1 | -0.270646 | LUAD | NCI-H1793 | Tanespimycin | MVP_top60_DNA+RNA_seed | AURKA | 1.131258 | LUAD | NCI-H522 | ZM447439 |
| MVP_top60_DNA_seed | ATM | -0.269359 | LUAD | NCI-H1437 | torin2 | MVP_top60_DNA+RNA_seed | AKT1 | 1.131753 | LUAD | NCI-H1568 | Uprosertib |
| MVP_top60_DNA_seed | EGFR | -0.267463 | LUAD | RERF-LC-KJ | Gefitinib | MVP_top60_DNA+RNA_seed | EZH2 | 1.131901 | LUAD | NCI-H1944 | GSK343 |
| MVP_top60_DNA_seed | AURKA | -0.266977 | LUAD | A549 | ZM447439 | MVP_top60_DNA+RNA_seed | MDM2 | 1.132287 | LUAD | NCI-H23 | Serdemetan |
| MVP_top60_DNA_seed | AURKA | -0.266601 | LUAD | NCI-H1755 | GSK1070916 | MVP_top60_DNA+RNA_seed | FGFR1 | 1.134138 | LUAD | SW1573 | AZD4547 |
| MVP_top60_DNA_seed | FGFR1 | -0.266485 | LUAD | NCI-H322M | PD173074 | MVP_top60_DNA+RNA_seed | HSP90AA1 | 1.137607 | LUAD | NCI-H1734 | Elesclomol |
| MVP_top60_DNA_seed | IGF1R | -0.26601 | LUAD | NCI-H2291 | BMS-536924 | MVP_top60_DNA+RNA_seed | TERT | 1.141257 | LUAD | NCI-H2009 | BIBR-1532 |
| MVP_top60_DNA_seed | HDAC1 | -0.265555 | LUAD | NCI-H1623 | Belinostat | MVP_top60_DNA+RNA_seed | HSP90AA1 | 1.141719 | LUAD | NCI-H2405 | Luminespib |
| MVP_top60_DNA_seed | BRAF | -0.264939 | LUAD | NCI-H2009 | PLX-4720 | MVP_top60_DNA+RNA_seed | EZH2 | 1.142225 | LUAD | Calu-6 | GSK343 |
| MVP_top60_DNA_seed | HDAC1 | -0.263539 | LUAD | COR-L105 | AR-42 | MVP_top60_DNA+RNA_seed | TP53 | 1.144447 | LUAD | NCI-H1838 | MIRA-1 |
| MVP_top60_DNA_seed | IGF1R | -0.262586 | LUAD | NCI-H441 | BMS-536924 | MVP_top60_DNA+RNA_seed | SRC | 1.147221 | LUAD | NCI-H2023 | Bosutinib |
| MVP_top60_DNA_seed | EGFR | -0.259758 | LUAD | NCI-H358 | Gefitinib | MVP_top60_DNA+RNA_seed | AKT1 | 1.147313 | LUAD | NCI-H522 | AZD5363 |
| MVP_top60_DNA_seed | ATM | -0.257904 | LUAD | NCI-H1650 | KU-55933 | MVP_top60_DNA+RNA_seed | AURKA | 1.149035 | LUAD | NCI-H1693 | Alisertib |
| MVP_top60_DNA_seed | HSP90AA1 | -0.257085 | LUAD | NCI-H650 | Elesclomol | MVP_top60_DNA+RNA_seed | IGF1R | 1.150477 | LUAD | NCI-H1650 | BMS-754807 |
| MVP_top60_DNA_seed | HDAC1 | -0.256181 | LUAD | NCI-H1975 | AR-42 | MVP_top60_DNA+RNA_seed | BRAF | 1.150579 | LUAD | NCI-H1944 | PLX-4720 |
| MVP_top60_DNA_seed | ATM | -0.25578 | LUAD | NCI-H522 | CP466722 | MVP_top60_DNA+RNA_seed | BRAF | 1.151225 | LUAD | NCI-H3122 | PLX-4720 |
| MVP_top60_DNA_seed | EGFR | -0.255374 | LUAD | HCC-44 | AZD8931 | MVP_top60_DNA+RNA_seed | TERT | 1.155999 | LUAD | H3255 | BIBR-1532 |
| MVP_top60_DNA_seed | ERBB2 | -0.255374 | LUAD | HCC-44 | AZD8931 | MVP_top60_DNA+RNA_seed | SMARCA4 | 1.156127 | LUAD | NCI-H2087 | PFI3 |
| MVP_top60_DNA_seed | ERBB3 | -0.255374 | LUAD | HCC-44 | AZD8931 | MVP_top60_DNA+RNA_seed | AKT1 | 1.15807 | LUAD | HCC-827 | Uprosertib |
| MVP_top60_DNA_seed | AKT1 | -0.251692 | LUAD | NCI-H358 | AKT inhibitor VIII | MVP_top60_DNA+RNA_seed | MDM2 | 1.160978 | LUAD | SK-LU-1 | Nutlin-3a (-) |
| MVP_top60_DNA_seed | BRAF | -0.251225 | LUAD | NCI-H2087 | PLX-4720 | MVP_top60_DNA+RNA_seed | FGFR1 | 1.162131 | LUAD | NCI-H1693 | AZD4547 |
| MVP_top60_DNA_seed | PIK3CA | -0.250988 | LUAD | NCI-H2009 | AZD8835 | MVP_top60_DNA+RNA_seed | EGFR | 1.162746 | LUAD | HCC-78 | Erlotinib |
| MVP_top60_DNA_seed | MDM2 | -0.250532 | LUAD | NCI-H1573 | Nutlin-3a (-) | MVP_top60_DNA+RNA_seed | TP53 | 1.162837 | LUAD | H3255 | MIRA-1 |
| MVP_top60_DNA_seed | FGFR1 | -0.250275 | LUAD | NCI-H2009 | FGFR_3831 | MVP_top60_DNA+RNA_seed | PIK3CA | 1.165026 | LUAD | SW1573 | Buparlisib |
| MVP_top60_DNA_seed | AKT1 | -0.250192 | LUAD | NCI-H1437 | MK-2206 | MVP_top60_DNA+RNA_seed | AKT1 | 1.169195 | LUAD | H3255 | AT13148 |
| MVP_top60_DNA_seed | SRC | -0.248791 | LUAD | NCI-H358 | Bosutinib | MVP_top60_DNA+RNA_seed | MDM2 | 1.171561 | LUAD | NCI-H1666 | Serdemetan |
| MVP_top60_DNA_seed | PIK3CA | -0.248382 | LUAD | NCI-H2030 | AZD8835 | MVP_top60_DNA+RNA_seed | TP53 | 1.172228 | LUAD | NCI-H1355 | MIRA-1 |
| MVP_top60_DNA_seed | HDAC1 | -0.247494 | LUAD | NCI-H358 | Dacinostat | MVP_top60_DNA+RNA_seed | EZH2 | 1.172619 | LUAD | SK-LU-1 | GSK343 |
| MVP_top60_DNA_seed | FGFR1 | -0.247401 | LUAD | LC-2-ad | AZD4547 | MVP_top60_DNA+RNA_seed | HDAC1 | 1.173704 | LUAD | NCI-H1623 | PCI-34051 |
| MVP_top60_DNA_seed | HDAC1 | -0.245823 | LUAD | LXF-289 | Dacinostat | MVP_top60_DNA+RNA_seed | MDM2 | 1.176968 | LUAD | NCI-H1975 | Serdemetan |
| MVP_top60_DNA_seed | PIK3CA | -0.244965 | LUAD | NCI-H1437 | PI3Ka_4409 | MVP_top60_DNA+RNA_seed | IGF1R | 1.177924 | LUAD | SK-LU-1 | Linsitinib |
| MVP_top60_DNA_seed | BRAF | -0.244779 | LUAD | NCI-H1993 | Dabrafenib | MVP_top60_DNA+RNA_seed | MDM2 | 1.178315 | LUAD | VMRC-LCD | Nutlin-3a (-) |

| | | | | | | | | | | | |
|---|---|---|---|---|---|---|---|---|---|---|---|
| MVP_top60_DNA_seed | AURKA | -0.244557 | LUAD | NCI-H2291 | HG-5-113-01 | MVP_top60_DNA+RNA_seed | AKT1 | 1.178429 | LUAD | HCC-827 | Afuresertib |
| MVP_top60_DNA_seed | AURKA | -0.244257 | LUAD | NCI-H2085 | Genentech Cpd 10 | MVP_top60_DNA+RNA_seed | SMARCA4 | 1.180361 | LUAD | ABC-1 | PFI3 |
| MVP_top60_DNA_seed | HSP90AA1 | -0.24408 | LUAD | NCI-H838 | SNX-2112 | MVP_top60_DNA+RNA_seed | EGFR | 1.181664 | LUAD | VMRC-LCD | Sapitinib |
| MVP_top60_DNA_seed | MET | -0.24308 | LUAD | NCI-H2291 | AZD6094 | MVP_top60_DNA+RNA_seed | IGF1R | 1.185303 | LUAD | NCI-H1651 | NVP-ADW742 |
| MVP_top60_DNA_seed | PIK3CA | -0.241774 | LUAD | NCI-H2228 | AZD8835 | MVP_top60_DNA+RNA_seed | BRAF | 1.18532 | LUAD | NCI-H1838 | PLX-4720 |
| MVP_top60_DNA_seed | HDAC1 | -0.241482 | LUAD | PC-14 | CAY10603 | MVP_top60_DNA+RNA_seed | EGFR | 1.185843 | LUAD | NCI-H1573 | Osimertinib |
| MVP_top60_DNA_seed | AKT1 | -0.241002 | LUAD | NCI-H1792 | AKT inhibitor VIII | MVP_top60_DNA+RNA_seed | TERT | 1.186611 | LUAD | HCC-78 | BIBR-1532 |
| MVP_top60_DNA_seed | AKT1 | -0.240609 | LUAD | NCI-H2347 | AKT inhibitor VIII | MVP_top60_DNA+RNA_seed | PIK3CA | 1.186848 | LUAD | NCI-H23 | Alpelisib |
| MVP_top60_DNA_seed | HSP90AA1 | -0.239561 | LUAD | RERF-LC-MS | SNX-2112 | MVP_top60_DNA+RNA_seed | HDAC1 | 1.187097 | LUAD | NCI-H1781 | Entinostat |
| MVP_top60_DNA_seed | HDAC1 | -0.239207 | LUAD | NCI-H1623 | AR-42 | MVP_top60_DNA+RNA_seed | ATM | 1.187124 | LUAD | NCI-H2087 | KU-55933 |
| MVP_top60_DNA_seed | MDM2 | -0.235361 | LUAD | LXF-289 | Nutlin-3a (-) | MVP_top60_DNA+RNA_seed | AKT1 | 1.194185 | LUAD | H3255 | GSK2110183B |
| MVP_top60_DNA_seed | AKT1 | -0.235291 | LUAD | NCI-H838 | AKT inhibitor VIII | MVP_top60_DNA+RNA_seed | AURKA | 1.200749 | LUAD | EKVX | Alisertib |
| MVP_top60_DNA_seed | EGFR | -0.235027 | LUAD | NCI-H23 | CI-1033 | MVP_top60_DNA+RNA_seed | FGFR1 | 1.201416 | LUAD | NCI-H2347 | PD173074 |
| MVP_top60_DNA_seed | ERBB2 | -0.235027 | LUAD | NCI-H23 | CI-1033 | MVP_top60_DNA+RNA_seed | IGF1R | 1.206376 | LUAD | NCI-H1623 | NVP-ADW742 |
| MVP_top60_DNA_seed | EGFR | -0.234702 | LUAD | NCI-H522 | PF-00299804 | MVP_top60_DNA+RNA_seed | BRAF | 1.207166 | LUAD | HCC-827 | Dabrafenib |
| MVP_top60_DNA_seed | ERBB2 | -0.234702 | LUAD | NCI-H522 | PF-00299804 | MVP_top60_DNA+RNA_seed | ATM | 1.208558 | LUAD | HCC-827 | KU-55933 |
| MVP_top60_DNA_seed | ATM | -0.233806 | LUAD | Calu-6 | CP466722 | MVP_top60_DNA+RNA_seed | KRAS | 1.209619 | LUAD | NCI-H2291 | KRAS (G12C) Inhibitor-1 |
| MVP_top60_DNA_seed | HDAC1 | -0.233588 | LUAD | NCI-H2030 | JQ12 | MVP_top60_DNA+RNA_seed | SRC | 1.21069 | LUAD | ABC-1 | Bosutinib |
| MVP_top60_DNA_seed | AKT1 | -0.233523 | LUAD | NCI-H1755 | BAY AKT1 | MVP_top60_DNA+RNA_seed | MDM2 | 1.210848 | LUAD | NCI-H1838 | Nutlin-3a (-) |
| MVP_top60_DNA_seed | ERBB2 | -0.232997 | LUAD | NCI-H1993 | CP724714 | MVP_top60_DNA+RNA_seed | AKT1 | 1.212579 | LUAD | NCI-H1793 | Uprosertib |
| MVP_top60_DNA_seed | HDAC1 | -0.232895 | LUAD | NCI-H1734 | Belinostat | MVP_top60_DNA+RNA_seed | MET | 1.214015 | LUAD | Calu-3 | Savolitinib |
| MVP_top60_DNA_seed | MDM2 | -0.229979 | LUAD | NCI-H2023 | Nutlin-3a (-) | MVP_top60_DNA+RNA_seed | ATM | 1.214093 | LUAD | Calu-6 | KU-55933 |
| MVP_top60_DNA_seed | MDM2 | -0.229418 | LUAD | COR-L105 | Nutlin-3a (-) | MVP_top60_DNA+RNA_seed | BRAF | 1.214278 | LUAD | NCI-H1666 | SB590885 |
| MVP_top60_DNA_seed | HDAC1 | -0.22941 | LUAD | Calu-6 | CAY10603 | MVP_top60_DNA+RNA_seed | BRAF | 1.214825 | LUAD | COR-L105 | SB590885 |
| MVP_top60_DNA_seed | AURKA | -0.227752 | LUAD | NCI-H1792 | Genentech Cpd 10 | MVP_top60_DNA+RNA_seed | IGF1R | 1.219495 | LUAD | NCI-H1573 | NVP-ADW742 |
| MVP_top60_DNA_seed | ATM | -0.226244 | LUAD | HCC-78 | KU-55933 | MVP_top60_DNA+RNA_seed | EGFR | 1.22041 | LUAD | NCI-H3122 | AZD3759 |
| MVP_top60_DNA_seed | BRAF | -0.225985 | LUAD | NCI-H2023 | RAF_9304 | MVP_top60_DNA+RNA_seed | TP53 | 1.221488 | LUAD | COR-L105 | MIRA-1 |
| MVP_top60_DNA_seed | BRAF | -0.225945 | LUAD | NCI-H2405 | Dabrafenib | MVP_top60_DNA+RNA_seed | MDM2 | 1.223567 | LUAD | NCI-H1650 | Nutlin-3a (-) |
| MVP_top60_DNA_seed | AKT1 | -0.22507 | LUAD | NCI-H2087 | AKT inhibitor VIII | MVP_top60_DNA+RNA_seed | IGF1R | 1.223576 | LUAD | NCI-H1944 | NVP-ADW742 |
| MVP_top60_DNA_seed | ATM | -0.224889 | LUAD | NCI-H2085 | CP466722 | MVP_top60_DNA+RNA_seed | HDAC1 | 1.226764 | LUAD | NCI-H1355 | PCI-34051 |
| MVP_top60_DNA_seed | IGF1R | -0.224233 | LUAD | HOP-62 | Linsitinib | MVP_top60_DNA+RNA_seed | AKT1 | 1.227764 | LUAD | NCI-H3122 | AT13148 |
| MVP_top60_DNA_seed | AURKA | -0.222816 | LUAD | NCI-H1437 | ZM447439 | MVP_top60_DNA+RNA_seed | AKT1 | 1.230865 | LUAD | NCI-H2405 | AZD5363 |
| MVP_top60_DNA_seed | GSK3B | -0.222657 | LUAD | NCI-H1568 | CHIR-99021 | MVP_top60_DNA+RNA_seed | IGF1R | 1.23252 | LUAD | NCI-H2085 | Linsitinib |
| MVP_top60_DNA_seed | SRC | -0.222191 | LUAD | NCI-H1793 | Bosutinib | MVP_top60_DNA+RNA_seed | EGFR | 1.235609 | LUAD | NCI-H1355 | Osimertinib |
| MVP_top60_DNA_seed | EGFR | -0.221968 | LUAD | NCI-H358 | Cetuximab | MVP_top60_DNA+RNA_seed | IGF1R | 1.235982 | LUAD | NCI-H2347 | BMS-536924 |
| MVP_top60_DNA_seed | PIK3CA | -0.220916 | LUAD | NCI-H838 | PI3Ka_4409 | MVP_top60_DNA+RNA_seed | MDM2 | 1.238033 | LUAD | NCI-H1793 | Nutlin-3a (-) |
| MVP_top60_DNA_seed | IGF1R | -0.219407 | LUAD | LXF-289 | Linsitinib | MVP_top60_DNA+RNA_seed | FGFR1 | 1.239301 | LUAD | ABC-1 | AZD4547 |
| MVP_top60_DNA_seed | HDAC1 | -0.219281 | LUAD | NCI-H2342 | Dacinostat | MVP_top60_DNA+RNA_seed | MDM2 | 1.240131 | LUAD | NCI-H1568 | Nutlin-3a (-) |
| MVP_top60_DNA_seed | BRAF | -0.219197 | LUAD | SK-LU-1 | Dabrafenib | MVP_top60_DNA+RNA_seed | AKT1 | 1.241163 | LUAD | NCI-H522 | MK-2206 |
| MVP_top60_DNA_seed | HSP90AA1 | -0.218363 | LUAD | PC-3_[JPC-3] | CCT-018159 | MVP_top60_DNA+RNA_seed | HSP90AA1 | 1.242115 | LUAD | EKVX | Luminespib |
| MVP_top60_DNA_seed | AKT1 | -0.217588 | LUAD | NCI-H2228 | GSK690693 | MVP_top60_DNA+RNA_seed | AURKA | 1.242257 | LUAD | NCI-H2291 | Alisertib |

| | | | | | | | | | | | |
|---|---|---|---|---|---|---|---|---|---|---|---|
| MVP_top60_DNA_seed | GSK3B | -0.217121 | LUAD | HCC-78 | SB216763 | MVP_top60_DNA+RNA_seed | IGF1R | 1.245149 | LUAD | NCI-H1781 | NVP-ADW742 |
| MVP_top60_DNA_seed | AKT1 | -0.216813 | LUAD | NCI-H1975 | GSK690693 | MVP_top60_DNA+RNA_seed | MDM2 | 1.247334 | LUAD | NCI-H1355 | Nutlin-3a (-) |
| MVP_top60_DNA_seed | EGFR | -0.215806 | LUAD | Calu-6 | AST-1306 | MVP_top60_DNA+RNA_seed | BRAF | 1.247654 | LUAD | Calu-3 | Dabrafenib |
| MVP_top60_DNA_seed | BRAF | -0.215525 | LUAD | HCC-44 | SB590885 | MVP_top60_DNA+RNA_seed | IGF1R | 1.248273 | LUAD | Calu-6 | BMS-536924 |
| MVP_top60_DNA_seed | EGFR | -0.215374 | LUAD | NCI-H2023 | Cetuximab | MVP_top60_DNA+RNA_seed | FGFR1 | 1.248835 | LUAD | NCI-H358 | AZD4547 |
| MVP_top60_DNA_seed | HSP90AA1 | -0.215281 | LUAD | VMRC-LCD | Elesclomol | MVP_top60_DNA+RNA_seed | AKT1 | 1.248857 | LUAD | EKVX | Ipatasertib |
| MVP_top60_DNA_seed | IGF1R | -0.215138 | LUAD | A549 | BMS-754807 | MVP_top60_DNA+RNA_seed | KRAS | 1.250427 | LUAD | NCI-H1793 | :RAS (G12C) Inhibitor-1 |
| MVP_top60_DNA_seed | BRAF | -0.214903 | LUAD | HOP-62 | AZ628 | MVP_top60_DNA+RNA_seed | BRAF | 1.252297 | LUAD | SW1573 | Dabrafenib |
| MVP_top60_DNA_seed | BRAF | -0.214636 | LUAD | RERF-LC-KJ | RAF_9304 | MVP_top60_DNA+RNA_seed | BRAF | 1.254303 | LUAD | H3255 | SB590885 |
| MVP_top60_DNA_seed | EGFR | -0.213125 | LUAD | ABC-1 | Cetuximab | MVP_top60_DNA+RNA_seed | IGF1R | 1.257779 | LUAD | NCI-H522 | BMS-754807 |
| MVP_top60_DNA_seed | ATM | -0.213007 | LUAD | ABC-1 | CP466722 | MVP_top60_DNA+RNA_seed | AKT1 | 1.258281 | LUAD | NCI-H522 | Ipatasertib |
| MVP_top60_DNA_seed | IGF1R | -0.212976 | LUAD | NCI-H1563 | BMS-536924 | MVP_top60_DNA+RNA_seed | EGFR | 1.25831 | LUAD | NCI-H1793 | Afatinib |
| MVP_top60_DNA_seed | IGF1R | -0.212563 | LUAD | NCI-H1792 | Linsitinib | MVP_top60_DNA+RNA_seed | EGFR | 1.25964 | LUAD | NCI-H2347 | Sapitinib |
| MVP_top60_DNA_seed | GSK3B | -0.212075 | LUAD | PC-14 | CHIR-99021 | MVP_top60_DNA+RNA_seed | EGFR | 1.260114 | LUAD | NCI-H3122 | Erlotinib |
| MVP_top60_DNA_seed | AURKA | -0.211863 | LUAD | NCI-H1755 | ZM447439 | MVP_top60_DNA+RNA_seed | AKT1 | 1.264809 | LUAD | NCI-H1651 | Afuresertib |
| MVP_top60_DNA_seed | EGFR | -0.211454 | LUAD | NCI-H2342 | PF-00299804 | MVP_top60_DNA+RNA_seed | AKT1 | 1.26639 | LUAD | NCI-H23 | MK-2206 |
| MVP_top60_DNA_seed | ERBB2 | -0.211454 | LUAD | NCI-H2342 | PF-00299804 | MVP_top60_DNA+RNA_seed | TERT | 1.27078 | LUAD | PC-14 | BIBR-1532 |
| MVP_top60_DNA_seed | EGFR | -0.211322 | LUAD | NCI-H1666 | PF-00299804 | MVP_top60_DNA+RNA_seed | ATM | 1.27105 | LUAD | NCI-H1793 | KU-55933 |
| MVP_top60_DNA_seed | ERBB2 | -0.211322 | LUAD | NCI-H1666 | PF-00299804 | MVP_top60_DNA+RNA_seed | AKT1 | 1.271304 | LUAD | NCI-H1568 | Ipatasertib |
| MVP_top60_DNA_seed | HDAC1 | -0.211311 | LUAD | PC-3_[JPC-3] | Dacinostat | MVP_top60_DNA+RNA_seed | AKT1 | 1.273734 | LUAD | VMRC-LCD | Ipatasertib |
| MVP_top60_DNA_seed | BRAF | -0.210228 | LUAD | A549 | PLX-4720 | MVP_top60_DNA+RNA_seed | TERT | 1.274898 | LUAD | HOP-62 | BIBR-1532 |
| MVP_top60_DNA_seed | BRAF | -0.208467 | LUAD | NCI-H1755 | Dabrafenib | MVP_top60_DNA+RNA_seed | AKT1 | 1.276877 | LUAD | NCI-H1838 | AZD5363 |
| MVP_top60_DNA_seed | HDAC1 | -0.208076 | LUAD | NCI-H1651 | AR-42 | MVP_top60_DNA+RNA_seed | BRAF | 1.276919 | LUAD | NCI-H2087 | PLX-4720 |
| MVP_top60_DNA_seed | AURKA | -0.207546 | LUAD | NCI-H2347 | Alisertib | MVP_top60_DNA+RNA_seed | EZH2 | 1.279536 | LUAD | NCI-H2342 | GSK343 |
| MVP_top60_DNA_seed | BRAF | -0.205165 | LUAD | NCI-H1755 | PLX-4720 | MVP_top60_DNA+RNA_seed | HDAC1 | 1.281822 | LUAD | NCI-H1781 | Romidepsin |
| MVP_top60_DNA_seed | HDAC1 | -0.205146 | LUAD | NCI-H1734 | CAY10603 | MVP_top60_DNA+RNA_seed | PIK3CA | 1.283535 | LUAD | NCI-H2405 | Buparlisib |
| MVP_top60_DNA_seed | AKT1 | -0.205144 | LUAD | Calu-3 | AKT inhibitor VIII | MVP_top60_DNA+RNA_seed | BRAF | 1.283743 | LUAD | Calu-3 | PLX-4720 |
| MVP_top60_DNA_seed | FGFR1 | -0.204864 | LUAD | SW1573 | FGFR_3831 | MVP_top60_DNA+RNA_seed | HDAC1 | 1.284447 | LUAD | NCI-H1568 | Entinostat |
| MVP_top60_DNA_seed | FGFR1 | -0.203856 | LUAD | EMC-BAC-1 | FGFR_3831 | MVP_top60_DNA+RNA_seed | HDAC1 | 1.284706 | LUAD | Calu-3 | PCI-34051 |
| MVP_top60_DNA_seed | EGFR | -0.20341 | LUAD | ABC-1 | PF-00299804 | MVP_top60_DNA+RNA_seed | SRC | 1.284766 | LUAD | HCC-78 | Bosutinib |
| MVP_top60_DNA_seed | ERBB2 | -0.20341 | LUAD | ABC-1 | PF-00299804 | MVP_top60_DNA+RNA_seed | EGFR | 1.285776 | LUAD | NCI-H2085 | Erlotinib |
| MVP_top60_DNA_seed | BRAF | -0.202006 | LUAD | EMC-BAC-1 | RAF_9304 | MVP_top60_DNA+RNA_seed | HSP90AA1 | 1.289669 | LUAD | NCI-H1838 | Tanespimycin |
| MVP_top60_DNA_seed | SRC | -0.201811 | LUAD | NCI-H1355 | Saracatinib | MVP_top60_DNA+RNA_seed | SRC | 1.291424 | LUAD | NCI-H2009 | Bosutinib |
| MVP_top60_DNA_seed | HSP90AA1 | -0.20147 | LUAD | NCI-H358 | Tanespimycin | MVP_top60_DNA+RNA_seed | HDAC1 | 1.29979 | LUAD | NCI-H441 | Entinostat |
| MVP_top60_DNA_seed | MDM2 | -0.20101 | LUAD | NCI-H1792 | Serdemetan | MVP_top60_DNA+RNA_seed | HDAC1 | 1.300362 | LUAD | NCI-H1650 | PCI-34051 |
| MVP_top60_DNA_seed | AKT1 | -0.198877 | LUAD | NCI-H1792 | AKT inhibitor VIII | MVP_top60_DNA+RNA_seed | IGF1R | 1.301141 | LUAD | NCI-H2087 | BMS-754807 |
| MVP_top60_DNA_seed | HSP90AA1 | -0.197683 | LUAD | NCI-H2228 | Tanespimycin | MVP_top60_DNA+RNA_seed | BRAF | 1.301352 | LUAD | NCI-H1693 | Dabrafenib |
| MVP_top60_DNA_seed | GSK3B | -0.197641 | LUAD | HCC-44 | KIN001-042 | MVP_top60_DNA+RNA_seed | AKT1 | 1.301713 | LUAD | NCI-H2087 | Uprosertib |
| MVP_top60_DNA_seed | EGFR | -0.197594 | LUAD | HCC-78 | CI-1033 | MVP_top60_DNA+RNA_seed | KRAS | 1.30271 | LUAD | NCI-H1755 | :RAS (G12C) Inhibitor-1 |
| MVP_top60_DNA_seed | ERBB2 | -0.197594 | LUAD | HCC-78 | CI-1033 | MVP_top60_DNA+RNA_seed | HSP90AA1 | 1.304098 | LUAD | NCI-H1693 | Luminespib |
| MVP_top60_DNA_seed | HDAC1 | -0.197306 | LUAD | COR-L105 | Belinostat | MVP_top60_DNA+RNA_seed | AKT1 | 1.305034 | LUAD | NCI-H1793 | Ipatasertib |

| | | | | | | | | | | | |
|---|---|---|---|---|---|---|---|---|---|---|---|
| MVP_top60_DNA_seed | AURKA | -0.197242 | LUAD | A549 | Genentech Cpd 10 | MVP_top60_DNA+RNA_seed | AURKA | 1.306098 | LUAD | NCI-H2342 | ZM447439 |
| MVP_top60_DNA_seed | HDAC1 | -0.197104 | LUAD | NCI-H1650 | JQ12 | MVP_top60_DNA+RNA_seed | HDAC1 | 1.30625 | LUAD | NCI-H1666 | PCI-34051 |
| MVP_top60_DNA_seed | EGFR | -0.196598 | LUAD | NCI-H1693 | Gefitinib | MVP_top60_DNA+RNA_seed | SRC | 1.307978 | LUAD | Calu-6 | Bosutinib |
| MVP_top60_DNA_seed | AKT1 | -0.196325 | LUAD | NCI-H1781 | BAY AKT1 | MVP_top60_DNA+RNA_seed | AKT1 | 1.30838 | LUAD | NCI-H650 | MK-2206 |
| MVP_top60_DNA_seed | SRC | -0.196224 | LUAD | A549 | Bosutinib | MVP_top60_DNA+RNA_seed | EGFR | 1.30886 | LUAD | NCI-H2087 | Sapitinib |
| MVP_top60_DNA_seed | BRAF | -0.196063 | LUAD | NCI-H2347 | Dabrafenib | MVP_top60_DNA+RNA_seed | PIK3CA | 1.309286 | LUAD | NCI-H1838 | Alpelisib |
| MVP_top60_DNA_seed | ATM | -0.195846 | LUAD | NCI-H2342 | KU-60019 | MVP_top60_DNA+RNA_seed | IGF1R | 1.310141 | LUAD | NCI-H322M | GSK1904529A |
| MVP_top60_DNA_seed | SMARCA4 | -0.194096 | LUAD | NCI-H1651 | PFI-3 | MVP_top60_DNA+RNA_seed | HSP90AA1 | 1.310422 | LUAD | EKVX | Tanespimycin |
| MVP_top60_DNA_seed | HDAC1 | -0.193826 | LUAD | NCI-H2347 | JQ12 | MVP_top60_DNA+RNA_seed | EGFR | 1.312046 | LUAD | NCI-H2405 | Osimertinib |
| MVP_top60_DNA_seed | EGFR | -0.193496 | LUAD | NCI-H2087 | Pelitinib | MVP_top60_DNA+RNA_seed | FGFR1 | 1.312052 | LUAD | NCI-H2087 | AZD4547 |
| MVP_top60_DNA_seed | HSP90AA1 | -0.192795 | LUAD | NCI-H2347 | Elesclomol | MVP_top60_DNA+RNA_seed | IGF1R | 1.317796 | LUAD | Calu-3 | BMS-536924 |
| MVP_top60_DNA_seed | MAPK1 | -0.19151 | LUAD | NCI-H1993 | VX-11e | MVP_top60_DNA+RNA_seed | IGF1R | 1.317847 | LUAD | NCI-H1435 | BMS-536924 |
| MVP_top60_DNA_seed | FGFR1 | -0.191277 | LUAD | NCI-H2030 | PD173074 | MVP_top60_DNA+RNA_seed | MDM2 | 1.319403 | LUAD | NCI-H2342 | Nutlin-3a (-) |
| MVP_top60_DNA_seed | EGFR | -0.191133 | LUAD | NCI-H1975 | Erlotinib | MVP_top60_DNA+RNA_seed | EGFR | 1.31956 | LUAD | NCI-H2228 | Lapatinib |
| MVP_top60_DNA_seed | EGFR | -0.190391 | LUAD | Calu-6 | CUDC-101 | MVP_top60_DNA+RNA_seed | TP53 | 1.320283 | LUAD | EKVX | MIRA-1 |
| MVP_top60_DNA_seed | ERBB2 | -0.190391 | LUAD | Calu-6 | CUDC-101 | MVP_top60_DNA+RNA_seed | MET | 1.320379 | LUAD | NCI-H358 | Savolitinib |
| MVP_top60_DNA_seed | BRAF | -0.189353 | LUAD | NCI-H23 | AZ628 | MVP_top60_DNA+RNA_seed | FGFR1 | 1.321886 | LUAD | NCI-H1568 | PD173074 |
| MVP_top60_DNA_seed | PIK3CA | -0.188454 | LUAD | H3255 | AZD8835 | MVP_top60_DNA+RNA_seed | AKT1 | 1.324814 | LUAD | VMRC-LCD | MK-2206 |
| MVP_top60_DNA_seed | EGFR | -0.187417 | LUAD | NCI-H1435 | CI-1033 | MVP_top60_DNA+RNA_seed | EGFR | 1.328104 | LUAD | VMRC-LCD | AZD3759 |
| MVP_top60_DNA_seed | ERBB2 | -0.187417 | LUAD | NCI-H1435 | CI-1033 | MVP_top60_DNA+RNA_seed | HSP90AA1 | 1.328407 | LUAD | NCI-H2085 | Tanespimycin |
| MVP_top60_DNA_seed | SRC | -0.186276 | LUAD | NCI-H2122 | HG-6-71-01 | MVP_top60_DNA+RNA_seed | HDAC1 | 1.328981 | LUAD | SW1573 | PCI-34051 |
| MVP_top60_DNA_seed | GSK3B | -0.1862 | LUAD | NCI-H1648 | CHIR-99021 | MVP_top60_DNA+RNA_seed | EGFR | 1.329645 | LUAD | NCI-H2087 | Gefitinib |
| MVP_top60_DNA_seed | ATM | -0.185191 | LUAD | NCI-H1693 | KU-60019 | MVP_top60_DNA+RNA_seed | EGFR | 1.331778 | LUAD | NCI-H1838 | Lapatinib |
| MVP_top60_DNA_seed | HSP90AA1 | -0.185056 | LUAD | NCI-H358 | CCT-018159 | MVP_top60_DNA+RNA_seed | SRC | 1.335851 | LUAD | NCI-H1355 | Bosutinib |
| MVP_top60_DNA_seed | IGF1R | -0.184774 | LUAD | COR-L105 | BMS-536924 | MVP_top60_DNA+RNA_seed | SMARCA4 | 1.338804 | LUAD | HOP-62 | PFI3 |
| MVP_top60_DNA_seed | MET | -0.183608 | LUAD | NCI-H1838 | JNJ38877605 | MVP_top60_DNA+RNA_seed | FGFR1 | 1.341295 | LUAD | PC-3_[JPC-3] | AZD4547 |
| MVP_top60_DNA_seed | HDAC1 | -0.183601 | LUAD | NCI-H650 | CAY10603 | MVP_top60_DNA+RNA_seed | IGF1R | 1.341854 | LUAD | NCI-H1650 | NVP-ADW742 |
| MVP_top60_DNA_seed | AKT1 | -0.183386 | LUAD | Calu-3 | GSK690693 | MVP_top60_DNA+RNA_seed | TP53 | 1.342672 | LUAD | SW1573 | MIRA-1 |
| MVP_top60_DNA_seed | PTK2 | -0.182473 | LUAD | NCI-H2347 | PF-562271 | MVP_top60_DNA+RNA_seed | MET | 1.346259 | LUAD | NCI-H1693 | Crizotinib |
| MVP_top60_DNA_seed | FGFR1 | -0.182266 | LUAD | NCI-H1755 | AZD4547 | MVP_top60_DNA+RNA_seed | EGFR | 1.346517 | LUAD | HOP-62 | Gefitinib |
| MVP_top60_DNA_seed | GSK3B | -0.181826 | LUAD | LXF-289 | KIN001-042 | MVP_top60_DNA+RNA_seed | AKT1 | 1.348592 | LUAD | NCI-H1355 | Ipatasertib |
| MVP_top60_DNA_seed | HSP90AA1 | -0.181576 | LUAD | NCI-H1734 | Luminespib | MVP_top60_DNA+RNA_seed | PIK3CA | 1.348781 | LUAD | NCI-H650 | AZD8186 |
| MVP_top60_DNA_seed | HDAC1 | -0.180821 | LUAD | NCI-H522 | CAY10603 | MVP_top60_DNA+RNA_seed | IGF1R | 1.353082 | LUAD | SW1573 | GSK1904529A |
| MVP_top60_DNA_seed | PIK3CA | -0.179248 | LUAD | LXF-289 | AZD8835 | MVP_top60_DNA+RNA_seed | EGFR | 1.353336 | LUAD | NCI-H2405 | Gefitinib |
| MVP_top60_DNA_seed | EGFR | -0.178781 | LUAD | NCI-H1666 | CI-1033 | MVP_top60_DNA+RNA_seed | HSP90AA1 | 1.354141 | LUAD | EMC-BAC-1 | CCT-018159 |
| MVP_top60_DNA_seed | ERBB2 | -0.178781 | LUAD | NCI-H1666 | CI-1033 | MVP_top60_DNA+RNA_seed | EGFR | 1.354716 | LUAD | NCI-H2087 | Erlotinib |
| MVP_top60_DNA_seed | EGFR | -0.178584 | LUAD | ABC-1 | Afatinib | MVP_top60_DNA+RNA_seed | IGF1R | 1.354913 | LUAD | SK-LU-1 | GSK1904529A |
| MVP_top60_DNA_seed | ERBB2 | -0.178584 | LUAD | ABC-1 | Afatinib | MVP_top60_DNA+RNA_seed | BRAF | 1.357669 | LUAD | NCI-H2347 | Dabrafenib |
| MVP_top60_DNA_seed | SRC | -0.177753 | LUAD | NCI-H1568 | Bosutinib | MVP_top60_DNA+RNA_seed | AKT1 | 1.358761 | LUAD | SW1573 | Uprosertib |
| MVP_top60_DNA_seed | SRC | -0.176779 | LUAD | PC-3_[JPC-3] | NVP-BHG712 | MVP_top60_DNA+RNA_seed | MDM2 | 1.360209 | LUAD | HOP-62 | Nutlin-3a (-) |
| MVP_top60_DNA_seed | MAPK1 | -0.176406 | LUAD | NCI-H1975 | VX-11e | MVP_top60_DNA+RNA_seed | AKT1 | 1.360431 | LUAD | NCI-H1568 | Afuresertib |

| | | | | | | | | | | | |
|---|---|---|---|---|---|---|---|---|---|---|---|
| MVP_top60_DNA_seed | HDAC1 | -0.176288 | LUAD | PC-3_[JPC-3] | AR-42 | MVP_top60_DNA+RNA_seed | MDM2 | 1.361091 | LUAD | NCI-H2291 | Nutlin-3a (-) |
| MVP_top60_DNA_seed | HDAC1 | -0.175075 | LUAD | EMC-BAC-2 | Tubastatin A | MVP_top60_DNA+RNA_seed | HSP90AA1 | 1.365235 | LUAD | NCI-H1563 | Tanespimycin |
| MVP_top60_DNA_seed | EGFR | -0.17504 | LUAD | NCI-H1838 | Erlotinib | MVP_top60_DNA+RNA_seed | AKT1 | 1.365323 | LUAD | SW1573 | GSK2110183B |
| MVP_top60_DNA_seed | HDAC1 | -0.174994 | LUAD | NCI-H2030 | Dacinostat | MVP_top60_DNA+RNA_seed | AKT1 | 1.365437 | LUAD | NCI-H1651 | Uprosertib |
| MVP_top60_DNA_seed | EGFR | -0.17427 | LUAD | SK-LU-1 | Gefitinib | MVP_top60_DNA+RNA_seed | MET | 1.365631 | LUAD | NCI-H1355 | Savolitinib |
| MVP_top60_DNA_seed | SRC | -0.174075 | LUAD | NCI-H1666 | Bosutinib | MVP_top60_DNA+RNA_seed | AKT1 | 1.366518 | LUAD | HOP-62 | Ipatasertib |
| MVP_top60_DNA_seed | EGFR | -0.173758 | LUAD | LXF-289 | Pelitinib | MVP_top60_DNA+RNA_seed | BRAF | 1.370298 | LUAD | NCI-H2347 | PLX-4720 |
| MVP_top60_DNA_seed | HDAC1 | -0.173691 | LUAD | LXF-289 | CAY10603 | MVP_top60_DNA+RNA_seed | AKT1 | 1.37275 | LUAD | HCC-827 | GSK2110183B |
| MVP_top60_DNA_seed | EGFR | -0.173503 | LUAD | HOP-62 | AZD8931 | MVP_top60_DNA+RNA_seed | EGFR | 1.373116 | LUAD | NCI-H1793 | Lapatinib |
| MVP_top60_DNA_seed | ERBB2 | -0.173503 | LUAD | HOP-62 | AZD8931 | MVP_top60_DNA+RNA_seed | AKT1 | 1.376038 | LUAD | NCI-H441 | Uprosertib |
| MVP_top60_DNA_seed | ERBB3 | -0.173503 | LUAD | HOP-62 | AZD8931 | MVP_top60_DNA+RNA_seed | EGFR | 1.376797 | LUAD | NCI-H2087 | Osimertinib |
| MVP_top60_DNA_seed | BRAF | -0.171539 | LUAD | NCI-H2228 | HG6-64-1 | MVP_top60_DNA+RNA_seed | AKT1 | 1.37782 | LUAD | NCI-H23 | Uprosertib |
| MVP_top60_DNA_seed | EGFR | -0.171272 | LUAD | NCI-H1355 | Lapatinib | MVP_top60_DNA+RNA_seed | EGFR | 1.379839 | LUAD | SW1573 | Sapitinib |
| MVP_top60_DNA_seed | ERBB2 | -0.171272 | LUAD | NCI-H1355 | Lapatinib | MVP_top60_DNA+RNA_seed | IGF1R | 1.380565 | LUAD | NCI-H1355 | NVP-ADW742 |
| MVP_top60_DNA_seed | PIK3CA | -0.171097 | LUAD | NCI-H2291 | PI3Ka_4409 | MVP_top60_DNA+RNA_seed | IGF1R | 1.380916 | LUAD | NCI-H522 | NVP-ADW742 |
| MVP_top60_DNA_seed | EGFR | -0.170644 | LUAD | NCI-H3122 | Pelitinib | MVP_top60_DNA+RNA_seed | PIK3CA | 1.382071 | LUAD | EKVX | Alpelisib |
| MVP_top60_DNA_seed | FGFR1 | -0.169764 | LUAD | LXF-289 | AZD4547 | MVP_top60_DNA+RNA_seed | TERT | 1.384092 | LUAD | NCI-H358 | BIBR-1532 |
| MVP_top60_DNA_seed | HDAC1 | -0.168968 | LUAD | RERF-LC-MS | Belinostat | MVP_top60_DNA+RNA_seed | HDAC1 | 1.384361 | LUAD | ABC-1 | Romidepsin |
| MVP_top60_DNA_seed | EGFR | -0.16888 | LUAD | HCC-44 | CUDC-101 | MVP_top60_DNA+RNA_seed | PIK3CA | 1.388825 | LUAD | NCI-H2291 | Buparlisib |
| MVP_top60_DNA_seed | ERBB2 | -0.16888 | LUAD | HCC-44 | CUDC-101 | MVP_top60_DNA+RNA_seed | HDAC1 | 1.389315 | LUAD | NCI-H2023 | PCI-34051 |
| MVP_top60_DNA_seed | MDM2 | -0.16704 | LUAD | Calu-6 | Nutlin-3a (-) | MVP_top60_DNA+RNA_seed | ATM | 1.390152 | LUAD | NCI-H1355 | KU-55933 |
| MVP_top60_DNA_seed | FGFR1 | -0.166974 | LUAD | RERF-LC-MS | FGFR_3831 | MVP_top60_DNA+RNA_seed | IGF1R | 1.391867 | LUAD | NCI-H441 | BMS-754807 |
| MVP_top60_DNA_seed | BRAF | -0.166512 | LUAD | NCI-H322M | PLX-4720 | MVP_top60_DNA+RNA_seed | IGF1R | 1.394338 | LUAD | NCI-H2405 | GSK1904529A |
| MVP_top60_DNA_seed | EGFR | -0.166289 | LUAD | EMC-BAC-1 | CI-1033 | MVP_top60_DNA+RNA_seed | EGFR | 1.394676 | LUAD | HOP-62 | Afatinib |
| MVP_top60_DNA_seed | ERBB2 | -0.166289 | LUAD | EMC-BAC-1 | CI-1033 | MVP_top60_DNA+RNA_seed | AKT1 | 1.39902 | LUAD | NCI-H1648 | AT13148 |
| MVP_top60_DNA_seed | AURKA | -0.165527 | LUAD | NCI-H1944 | CD532 | MVP_top60_DNA+RNA_seed | FGFR1 | 1.399199 | LUAD | EMC-BAC-1 | AZD4547 |
| MVP_top60_DNA_seed | IGF1R | -0.165383 | LUAD | NCI-H2009 | BMS-536924 | MVP_top60_DNA+RNA_seed | MET | 1.399303 | LUAD | NCI-H1838 | Crizotinib |
| MVP_top60_DNA_seed | AKT1 | -0.165204 | LUAD | COR-L105 | AKT inhibitor VIII | MVP_top60_DNA+RNA_seed | AKT1 | 1.399437 | LUAD | NCI-H1693 | GSK2110183B |
| MVP_top60_DNA_seed | PIK3CA | -0.16513 | LUAD | NCI-H1755 | PI3Ka_4409 | MVP_top60_DNA+RNA_seed | EGFR | 1.403831 | LUAD | NCI-H3122 | Sapitinib |
| MVP_top60_DNA_seed | PTK2 | -0.162876 | LUAD | NCI-H1568 | PF-562271 | MVP_top60_DNA+RNA_seed | HSP90AA1 | 1.404259 | LUAD | NCI-H2405 | Tanespimycin |
| MVP_top60_DNA_seed | GSK3B | -0.161619 | LUAD | NCI-H1568 | AZD7969 | MVP_top60_DNA+RNA_seed | HDAC1 | 1.410288 | LUAD | NCI-H1568 | PCI-34051 |
| MVP_top60_DNA_seed | HSP90AA1 | -0.160389 | LUAD | NCI-H2405 | Tanespimycin | MVP_top60_DNA+RNA_seed | PIK3CA | 1.410333 | LUAD | NCI-H1573 | Alpelisib |
| MVP_top60_DNA_seed | HSP90AA1 | -0.160351 | LUAD | NCI-H2291 | Tanespimycin | MVP_top60_DNA+RNA_seed | IGF1R | 1.41099 | LUAD | EKVX | GSK1904529A |
| MVP_top60_DNA_seed | BRAF | -0.160091 | LUAD | H3255 | HG6-64-1 | MVP_top60_DNA+RNA_seed | AURKA | 1.411484 | LUAD | NCI-H2405 | Alisertib |
| MVP_top60_DNA_seed | EGFR | -0.159555 | LUAD | NCI-H2030 | Gefitinib | MVP_top60_DNA+RNA_seed | AKT1 | 1.412291 | LUAD | NCI-H2087 | Uprosertib |
| MVP_top60_DNA_seed | HDAC1 | -0.158975 | LUAD | NCI-H2023 | JQ12 | MVP_top60_DNA+RNA_seed | MDM2 | 1.41243 | LUAD | NCI-H1355 | Serdemetan |
| MVP_top60_DNA_seed | EGFR | -0.158541 | LUAD | HOP-62 | Pelitinib | MVP_top60_DNA+RNA_seed | MET | 1.412749 | LUAD | PC-3_[JPC-3] | Savolitinib |
| MVP_top60_DNA_seed | AKT1 | -0.15815 | LUAD | HOP-62 | AKT inhibitor VIII | MVP_top60_DNA+RNA_seed | FGFR1 | 1.413455 | LUAD | NCI-H1355 | AZD4547 |
| MVP_top60_DNA_seed | IGF1R | -0.157391 | LUAD | NCI-H2087 | BMS-754807 | MVP_top60_DNA+RNA_seed | PIK3CA | 1.413671 | LUAD | NCI-H2085 | Buparlisib |
| MVP_top60_DNA_seed | SRC | -0.156846 | LUAD | HOP-62 | NVP-BHG712 | MVP_top60_DNA+RNA_seed | MET | 1.414504 | LUAD | SW1573 | Crizotinib |
| MVP_top60_DNA_seed | AKT1 | -0.156247 | LUAD | NCI-H1355 | MK-2206 | MVP_top60_DNA+RNA_seed | MET | 1.41557 | LUAD | HCC-827 | Savolitinib |

| | | | | | | | | | | | |
|---|---|---|---|---|---|---|---|---|---|---|---|
| MVP_top60_DNA_seed | EGFR | -0.156022 | LUAD | NCI-H1435 | PF-00299804 | MVP_top60_DNA+RNA_seed | EGFR | 1.416134 | LUAD | NCI-H1838 | Osimertinib |
| MVP_top60_DNA_seed | ERBB2 | -0.156022 | LUAD | NCI-H1435 | PF-00299804 | MVP_top60_DNA+RNA_seed | AKT1 | 1.418869 | LUAD | NCI-H1568 | MK-2206 |
| MVP_top60_DNA_seed | MET | -0.155097 | LUAD | VMRC-LCD | AZD6094 | MVP_top60_DNA+RNA_seed | HDAC1 | 1.422284 | LUAD | EKVX | PCI-34051 |
| MVP_top60_DNA_seed | EGFR | -0.15483 | LUAD | HOP-62 | Lapatinib | MVP_top60_DNA+RNA_seed | FGFR1 | 1.422524 | LUAD | COR-L105 | AZD4547 |
| MVP_top60_DNA_seed | ERBB2 | -0.15483 | LUAD | HOP-62 | Lapatinib | MVP_top60_DNA+RNA_seed | TERT | 1.423055 | LUAD | VMRC-LCD | BIBR-1532 |
| MVP_top60_DNA_seed | IGF1R | -0.152819 | LUAD | SW1573 | BMS-754807 | MVP_top60_DNA+RNA_seed | BRAF | 1.423635 | LUAD | NCI-H1650 | SB590885 |
| MVP_top60_DNA_seed | EGFR | -0.151868 | LUAD | NCI-H1563 | Afatinib | MVP_top60_DNA+RNA_seed | MDM2 | 1.426622 | LUAD | NCI-H441 | Nutlin-3a (-) |
| MVP_top60_DNA_seed | ERBB2 | -0.151868 | LUAD | NCI-H1563 | Afatinib | MVP_top60_DNA+RNA_seed | HSP90AA1 | 1.428228 | LUAD | NCI-H2347 | Tanespimycin |
| MVP_top60_DNA_seed | GSK3B | -0.151784 | LUAD | NCI-H1435 | CHIR-99021 | MVP_top60_DNA+RNA_seed | AKT1 | 1.428663 | LUAD | ABC-1 | MK-2206 |
| MVP_top60_DNA_seed | GSK3B | -0.151257 | LUAD | NCI-H1648 | SB216763 | MVP_top60_DNA+RNA_seed | SRC | 1.431085 | LUAD | SW1573 | Bosutinib |
| MVP_top60_DNA_seed | GSK3B | -0.149761 | LUAD | NCI-H2347 | KIN001-042 | MVP_top60_DNA+RNA_seed | SMARCA4 | 1.4318 | LUAD | COR-L105 | PFI3 |
| MVP_top60_DNA_seed | FGFR1 | -0.149514 | LUAD | PC-14 | PD173074 | MVP_top60_DNA+RNA_seed | AURKA | 1.43194 | LUAD | NCI-H650 | ZM447439 |
| MVP_top60_DNA_seed | EGFR | -0.149422 | LUAD | NCI-H1944 | Gefitinib | MVP_top60_DNA+RNA_seed | EGFR | 1.433437 | LUAD | Calu-6 | Osimertinib |
| MVP_top60_DNA_seed | IGF1R | -0.14871 | LUAD | NCI-H1648 | GSK1904529A | MVP_top60_DNA+RNA_seed | TERT | 1.433595 | LUAD | NCI-H2405 | BIBR-1532 |
| MVP_top60_DNA_seed | ATM | -0.148524 | LUAD | NCI-H1944 | KU-60019 | MVP_top60_DNA+RNA_seed | TERT | 1.43452 | LUAD | NCI-H1568 | BIBR-1532 |
| MVP_top60_DNA_seed | HDAC1 | -0.148386 | LUAD | LXF-289 | Belinostat | MVP_top60_DNA+RNA_seed | AURKA | 1.435071 | LUAD | NCI-H2347 | ZM447439 |
| MVP_top60_DNA_seed | ATM | -0.148223 | LUAD | EMC-BAC-1 | CP466722 | MVP_top60_DNA+RNA_seed | EGFR | 1.435245 | LUAD | HOP-62 | AZD3759 |
| MVP_top60_DNA_seed | BRAF | -0.14743 | LUAD | NCI-H23 | SB590885 | MVP_top60_DNA+RNA_seed | ATM | 1.43625 | LUAD | NCI-H2291 | KU-55933 |
| MVP_top60_DNA_seed | HDAC1 | -0.14683 | LUAD | A549 | JQ12 | MVP_top60_DNA+RNA_seed | AKT1 | 1.43866 | LUAD | Calu-6 | Uprosertib |
| MVP_top60_DNA_seed | FGFR1 | -0.146012 | LUAD | NCI-H2228 | FGFR_3831 | MVP_top60_DNA+RNA_seed | AKT1 | 1.440483 | LUAD | PC-3_[JPC-3] | AT13148 |
| MVP_top60_DNA_seed | EGFR | -0.145112 | LUAD | NCI-H1623 | CUDC-101 | MVP_top60_DNA+RNA_seed | IGF1R | 1.442829 | LUAD | NCI-H1435 | BMS-754807 |
| MVP_top60_DNA_seed | ERBB2 | -0.145112 | LUAD | NCI-H1623 | CUDC-101 | MVP_top60_DNA+RNA_seed | MET | 1.443768 | LUAD | COR-L105 | Savolitinib |
| MVP_top60_DNA_seed | BRAF | -0.144388 | LUAD | NCI-H2347 | RAF_9304 | MVP_top60_DNA+RNA_seed | AKT1 | 1.44533 | LUAD | NCI-H2087 | Ipatasertib |
| MVP_top60_DNA_seed | HDAC1 | -0.143669 | LUAD | EKVX | JQ12 | MVP_top60_DNA+RNA_seed | FGFR1 | 1.445331 | LUAD | NCI-H1573 | PD173074 |
| MVP_top60_DNA_seed | GSK3B | -0.143473 | LUAD | NCI-H358 | CHIR-99021 | MVP_top60_DNA+RNA_seed | EGFR | 1.446492 | LUAD | NCI-H1355 | Gefitinib |
| MVP_top60_DNA_seed | BRAF | -0.143143 | LUAD | LXF-289 | RAF_9304 | MVP_top60_DNA+RNA_seed | AKT1 | 1.448144 | LUAD | NCI-H1568 | GSK2110183B |
| MVP_top60_DNA_seed | BRAF | -0.142131 | LUAD | NCI-H1568 | PLX-4720 | MVP_top60_DNA+RNA_seed | SRC | 1.448639 | LUAD | EKVX | Bosutinib |
| MVP_top60_DNA_seed | AKT1 | -0.141885 | LUAD | NCI-H1944 | GSK690693 | MVP_top60_DNA+RNA_seed | AKT1 | 1.449555 | LUAD | NCI-H2085 | Afuresertib |
| MVP_top60_DNA_seed | EGFR | -0.141755 | LUAD | HCC-78 | Afatinib | MVP_top60_DNA+RNA_seed | HDAC1 | 1.4505 | LUAD | NCI-H2087 | PCI-34051 |
| MVP_top60_DNA_seed | ERBB2 | -0.141755 | LUAD | HCC-78 | Afatinib | MVP_top60_DNA+RNA_seed | HDAC1 | 1.452461 | LUAD | NCI-H1693 | Entinostat |
| MVP_top60_DNA_seed | FGFR1 | -0.141636 | LUAD | PC-14 | FGFR_3831 | MVP_top60_DNA+RNA_seed | MET | 1.453229 | LUAD | NCI-H1838 | Savolitinib |
| MVP_top60_DNA_seed | MDM2 | -0.141423 | LUAD | NCI-H23 | Nutlin-3a (-) | MVP_top60_DNA+RNA_seed | KRAS | 1.453596 | LUAD | NCI-H1693 | KRAS (G12C) Inhibitor-1 |
| MVP_top60_DNA_seed | ATM | -0.141313 | LUAD | PC-14 | torin2 | MVP_top60_DNA+RNA_seed | BRAF | 1.454314 | LUAD | NCI-H1650 | Dabrafenib |
| MVP_top60_DNA_seed | HDAC1 | -0.140213 | LUAD | NCI-H23 | Belinostat | MVP_top60_DNA+RNA_seed | EGFR | 1.457899 | LUAD | NCI-H2087 | Lapatinib |
| MVP_top60_DNA_seed | EGFR | -0.139195 | LUAD | NCI-H1944 | PF-00299804 | MVP_top60_DNA+RNA_seed | AKT1 | 1.459958 | LUAD | NCI-H650 | Uprosertib |
| MVP_top60_DNA_seed | ERBB2 | -0.139195 | LUAD | NCI-H1944 | PF-00299804 | MVP_top60_DNA+RNA_seed | TERT | 1.460395 | LUAD | NCI-H2347 | BIBR-1532 |
| MVP_top60_DNA_seed | EGFR | -0.138089 | LUAD | NCI-H1573 | AZD8931 | MVP_top60_DNA+RNA_seed | AKT1 | 1.46087 | LUAD | Calu-6 | Ipatasertib |
| MVP_top60_DNA_seed | ERBB2 | -0.138089 | LUAD | NCI-H1573 | AZD8931 | MVP_top60_DNA+RNA_seed | AURKA | 1.461505 | LUAD | NCI-H1435 | ZM447439 |
| MVP_top60_DNA_seed | ERBB3 | -0.138089 | LUAD | NCI-H1573 | AZD8931 | MVP_top60_DNA+RNA_seed | HSP90AA1 | 1.46189 | LUAD | H3255 | Elesclomol |
| MVP_top60_DNA_seed | EGFR | -0.138058 | LUAD | Calu-6 | PF-00299804 | MVP_top60_DNA+RNA_seed | EGFR | 1.464976 | LUAD | NCI-H1838 | AZD3759 |
| MVP_top60_DNA_seed | ERBB2 | -0.138058 | LUAD | Calu-6 | PF-00299804 | MVP_top60_DNA+RNA_seed | EGFR | 1.468053 | LUAD | NCI-H2347 | AZD3759 |

| | | | | | | | | | | | |
|---|---|---|---|---|---|---|---|---|---|---|---|
| MVP_top60_DNA_seed | SRC | -0.138032 | LUAD | NCI-H2009 | NVP-BHG712 | MVP_top60_DNA+RNA_seed | EGFR | 1.469333 | LUAD | VMRC-LCD | Erlotinib |
| MVP_top60_DNA_seed | EGFR | -0.137569 | LUAD | NCI-H650 | CUDC-101 | MVP_top60_DNA+RNA_seed | EGFR | 1.469558 | LUAD | NCI-H2228 | AZD3759 |
| MVP_top60_DNA_seed | ERBB2 | -0.137569 | LUAD | NCI-H650 | CUDC-101 | MVP_top60_DNA+RNA_seed | BRAF | 1.470045 | LUAD | NCI-H2347 | SB590885 |
| MVP_top60_DNA_seed | SRC | -0.136877 | LUAD | NCI-H2023 | Bosutinib | MVP_top60_DNA+RNA_seed | EGFR | 1.471242 | LUAD | NCI-H441 | Gefitinib |
| MVP_top60_DNA_seed | AKT1 | -0.13635 | LUAD | NCI-H2405 | AKT inhibitor VIII | MVP_top60_DNA+RNA_seed | FGFR1 | 1.473188 | LUAD | NCI-H1651 | PD173074 |
| MVP_top60_DNA_seed | EGFR | -0.135806 | LUAD | NCI-H1944 | Afatinib | MVP_top60_DNA+RNA_seed | HDAC1 | 1.474664 | LUAD | H3255 | PCI-34051 |
| MVP_top60_DNA_seed | ERBB2 | -0.135806 | LUAD | NCI-H1944 | Afatinib | MVP_top60_DNA+RNA_seed | AKT1 | 1.475109 | LUAD | NCI-H2347 | AZD5363 |
| MVP_top60_DNA_seed | ATM | -0.135725 | LUAD | EMC-BAC-1 | KU-60019 | MVP_top60_DNA+RNA_seed | IGF1R | 1.475151 | LUAD | NCI-H2291 | Linsitinib |
| MVP_top60_DNA_seed | AKT1 | -0.1354 | LUAD | EKVX | MK-2206 | MVP_top60_DNA+RNA_seed | IGF1R | 1.479779 | LUAD | NCI-H441 | BMS-536924 |
| MVP_top60_DNA_seed | AKT1 | -0.134984 | LUAD | RERF-LC-MS | AKT inhibitor VIII | MVP_top60_DNA+RNA_seed | TP53 | 1.479916 | LUAD | NCI-H2228 | MIRA-1 |
| MVP_top60_DNA_seed | GSK3B | -0.134914 | LUAD | NCI-H2087 | SB216763 | MVP_top60_DNA+RNA_seed | AKT1 | 1.483298 | LUAD | Calu-3 | AZD5363 |
| MVP_top60_DNA_seed | ERBB2 | -0.134284 | LUAD | NCI-H2085 | CP724714 | MVP_top60_DNA+RNA_seed | PIK3CA | 1.485284 | LUAD | NCI-H1435 | GNE-317 |
| MVP_top60_DNA_seed | HDAC1 | -0.131321 | LUAD | NCI-H838 | CAY10603 | MVP_top60_DNA+RNA_seed | BRAF | 1.486903 | LUAD | PC-3_[JPC-3] | Dabrafenib |
| MVP_top60_DNA_seed | HDAC1 | -0.129792 | LUAD | NCI-H2405 | Dacinostat | MVP_top60_DNA+RNA_seed | HSP90AA1 | 1.486934 | LUAD | Calu-3 | Luminespib |
| MVP_top60_DNA_seed | HSP90AA1 | -0.12927 | LUAD | NCI-H1693 | Luminespib | MVP_top60_DNA+RNA_seed | EGFR | 1.488139 | LUAD | NCI-H2347 | Osimertinib |
| MVP_top60_DNA_seed | EGFR | -0.128712 | LUAD | NCI-H2087 | PF-00299804 | MVP_top60_DNA+RNA_seed | HDAC1 | 1.491398 | LUAD | NCI-H2405 | Entinostat |
| MVP_top60_DNA_seed | ERBB2 | -0.128712 | LUAD | NCI-H2087 | PF-00299804 | MVP_top60_DNA+RNA_seed | BRAF | 1.495557 | LUAD | NCI-H1838 | SB590885 |
| MVP_top60_DNA_seed | EGFR | -0.128246 | LUAD | LXF-289 | Afatinib | MVP_top60_DNA+RNA_seed | TP53 | 1.497721 | LUAD | NCI-H1666 | MIRA-1 |
| MVP_top60_DNA_seed | ERBB2 | -0.128246 | LUAD | LXF-289 | Afatinib | MVP_top60_DNA+RNA_seed | TP53 | 1.500118 | LUAD | NCI-H1650 | MIRA-1 |
| MVP_top60_DNA_seed | AURKA | -0.126652 | LUAD | EMC-BAC-2 | Alisertib | MVP_top60_DNA+RNA_seed | HSP90AA1 | 1.50074 | LUAD | SW1573 | Luminespib |
| MVP_top60_DNA_seed | MET | -0.125826 | LUAD | NCI-H2023 | JNJ38877605 | MVP_top60_DNA+RNA_seed | MET | 1.505104 | LUAD | H3255 | Crizotinib |
| MVP_top60_DNA_seed | AKT1 | -0.125757 | LUAD | NCI-H2085 | GSK690693 | MVP_top60_DNA+RNA_seed | AKT1 | 1.506064 | LUAD | NCI-H1568 | Uprosertib |
| MVP_top60_DNA_seed | SMARCA4 | -0.124364 | LUAD | HCC-44 | PFI-3 | MVP_top60_DNA+RNA_seed | IGF1R | 1.50765 | LUAD | NCI-H522 | BMS-536924 |
| MVP_top60_DNA_seed | BRAF | -0.122172 | LUAD | NCI-H1355 | HG6-64-1 | MVP_top60_DNA+RNA_seed | AKT1 | 1.510772 | LUAD | NCI-H1355 | GSK2110183B |
| MVP_top60_DNA_seed | BRAF | -0.122092 | LUAD | H3255 | SB590885 | MVP_top60_DNA+RNA_seed | AKT1 | 1.512061 | LUAD | PC-3_[JPC-3] | MK-2206 |
| MVP_top60_DNA_seed | HSP90AA1 | -0.12174 | LUAD | LXF-289 | SNX-2112 | MVP_top60_DNA+RNA_seed | AURKA | 1.513168 | LUAD | NCI-H1734 | Alisertib |
| MVP_top60_DNA_seed | AURKA | -0.121274 | LUAD | HCC-78 | ZM447439 | MVP_top60_DNA+RNA_seed | SMARCA4 | 1.513676 | LUAD | NCI-H1650 | PFI3 |
| MVP_top60_DNA_seed | ERBB2 | -0.120793 | LUAD | LC-2-ad | CP724714 | MVP_top60_DNA+RNA_seed | SRC | 1.514558 | LUAD | NCI-H2291 | Bosutinib |
| MVP_top60_DNA_seed | GSK3B | -0.120726 | LUAD | NCI-H1975 | CHIR-99021 | MVP_top60_DNA+RNA_seed | EZH2 | 1.521444 | LUAD | HOP-62 | GSK343 |
| MVP_top60_DNA_seed | EGFR | -0.12069 | LUAD | NCI-H1355 | Gefitinib | MVP_top60_DNA+RNA_seed | MET | 1.523511 | LUAD | ABC-1 | Crizotinib |
| MVP_top60_DNA_seed | AURKA | -0.119206 | LUAD | HCC-44 | GSK1070916 | MVP_top60_DNA+RNA_seed | FGFR1 | 1.52354 | LUAD | ABC-1 | PD173074 |
| MVP_top60_DNA_seed | EGFR | -0.118356 | LUAD | NCI-H1944 | AST-1306 | MVP_top60_DNA+RNA_seed | HSP90AA1 | 1.524546 | LUAD | NCI-H2291 | CCT-018159 |
| MVP_top60_DNA_seed | HSP90AA1 | -0.11782 | LUAD | Calu-3 | Luminespib | MVP_top60_DNA+RNA_seed | ATM | 1.524624 | LUAD | Calu-3 | KU-55933 |
| MVP_top60_DNA_seed | ATM | -0.11774 | LUAD | EMC-BAC-1 | KU-55933 | MVP_top60_DNA+RNA_seed | IGF1R | 1.527068 | LUAD | NCI-H2347 | Linsitinib |
| MVP_top60_DNA_seed | AKT1 | -0.11683 | LUAD | Calu-3 | MK-2206 | MVP_top60_DNA+RNA_seed | AURKA | 1.53075 | LUAD | NCI-H441 | Alisertib |
| MVP_top60_DNA_seed | HDAC1 | -0.116341 | LUAD | NCI-H1437 | CAY10603 | MVP_top60_DNA+RNA_seed | PIK3CA | 1.532989 | LUAD | NCI-H522 | AZD8186 |
| MVP_top60_DNA_seed | SRC | -0.115851 | LUAD | NCI-H1563 | HG-6-71-01 | MVP_top60_DNA+RNA_seed | FGFR1 | 1.533809 | LUAD | PC-3_[JPC-3] | PD173074 |
| MVP_top60_DNA_seed | ATM | -0.115737 | LUAD | HOP-62 | KU-60019 | MVP_top60_DNA+RNA_seed | AKT1 | 1.534645 | LUAD | NCI-H358 | AT13148 |
| MVP_top60_DNA_seed | EGFR | -0.114885 | LUAD | NCI-H1623 | Cetuximab | MVP_top60_DNA+RNA_seed | FGFR1 | 1.534854 | LUAD | NCI-H2342 | PD173074 |
| MVP_top60_DNA_seed | HDAC1 | -0.113477 | LUAD | NCI-H1573 | AR-42 | MVP_top60_DNA+RNA_seed | EZH2 | 1.536372 | LUAD | NCI-H2291 | GSK343 |
| MVP_top60_DNA_seed | AKT1 | -0.113025 | LUAD | NCI-H1648 | GSK690693 | MVP_top60_DNA+RNA_seed | HSP90AA1 | 1.537239 | LUAD | SW1573 | Tanespimycin |

| | | | | | | | | | | | |
|---|---|---|---|---|---|---|---|---|---|---|---|
| MVP_top60_DNA_seed | IGF1R | -0.112736 | LUAD | A549 | Linsitinib | MVP_top60_DNA+RNA_seed | HSP90AA1 | 1.537521 | LUAD | EKVX | CCT-018159 |
| MVP_top60_DNA_seed | EGFR | -0.11196 | LUAD | NCI-H522 | AZD8931 | MVP_top60_DNA+RNA_seed | TERT | 1.53817 | LUAD | NCI-H2087 | BIBR-1532 |
| MVP_top60_DNA_seed | ERBB2 | -0.11196 | LUAD | NCI-H522 | AZD8931 | MVP_top60_DNA+RNA_seed | ATM | 1.543521 | LUAD | PC-14 | KU-55933 |
| MVP_top60_DNA_seed | ERBB3 | -0.11196 | LUAD | NCI-H522 | AZD8931 | MVP_top60_DNA+RNA_seed | BRAF | 1.546099 | LUAD | NCI-H1793 | Dabrafenib |
| MVP_top60_DNA_seed | EGFR | -0.111926 | LUAD | NCI-H1792 | Cetuximab | MVP_top60_DNA+RNA_seed | IGF1R | 1.548481 | LUAD | SK-LU-1 | NVP-ADW742 |
| MVP_top60_DNA_seed | SRC | -0.111829 | LUAD | LXF-289 | Bosutinib | MVP_top60_DNA+RNA_seed | AKT1 | 1.548559 | LUAD | HCC-78 | AT13148 |
| MVP_top60_DNA_seed | AKT1 | -0.111196 | LUAD | HCC-44 | GSK690693 | MVP_top60_DNA+RNA_seed | IGF1R | 1.551985 | LUAD | PC-3_[JPC-3] | BMS-754807 |
| MVP_top60_DNA_seed | HSP90AA1 | -0.11116 | LUAD | NCI-H1975 | SNX-2112 | MVP_top60_DNA+RNA_seed | AKT1 | 1.554786 | LUAD | NCI-H1838 | Afuresertib |
| MVP_top60_DNA_seed | HDAC1 | -0.110692 | LUAD | COR-L105 | JQ12 | MVP_top60_DNA+RNA_seed | MET | 1.556386 | LUAD | NCI-H1650 | Crizotinib |
| MVP_top60_DNA_seed | EGFR | -0.110083 | LUAD | NCI-H1792 | AST-1306 | MVP_top60_DNA+RNA_seed | IGF1R | 1.556657 | LUAD | NCI-H1793 | BMS-754807 |
| MVP_top60_DNA_seed | PTK2 | -0.109519 | LUAD | RERF-LC-MS | PF-562271 | MVP_top60_DNA+RNA_seed | HDAC1 | 1.556831 | LUAD | SW1573 | Romidepsin |
| MVP_top60_DNA_seed | BRAF | -0.109253 | LUAD | Calu-6 | HG6-64-1 | MVP_top60_DNA+RNA_seed | BRAF | 1.560715 | LUAD | NCI-H2291 | Dabrafenib |
| MVP_top60_DNA_seed | EGFR | -0.108825 | LUAD | LC-2-ad | AST-1306 | MVP_top60_DNA+RNA_seed | AKT1 | 1.561469 | LUAD | NCI-H2087 | Afuresertib |
| MVP_top60_DNA_seed | AKT1 | -0.108126 | LUAD | NCI-H2023 | AKT inhibitor VIII | MVP_top60_DNA+RNA_seed | AKT1 | 1.566781 | LUAD | SK-LU-1 | Uprosertib |
| MVP_top60_DNA_seed | IGF1R | -0.107424 | LUAD | NCI-H2023 | BMS-536924 | MVP_top60_DNA+RNA_seed | TERT | 1.570792 | LUAD | SW1573 | BIBR-1532 |
| MVP_top60_DNA_seed | PTK2 | -0.1074 | LUAD | HCC-78 | PF-562271 | MVP_top60_DNA+RNA_seed | BRAF | 1.575207 | LUAD | NCI-H1666 | PLX-4720 |
| MVP_top60_DNA_seed | MET | -0.107012 | LUAD | NCI-H1568 | AZD6094 | MVP_top60_DNA+RNA_seed | HSP90AA1 | 1.578207 | LUAD | NCI-H1563 | Luminespib |
| MVP_top60_DNA_seed | BRAF | -0.106883 | LUAD | NCI-H1944 | RAF_9304 | MVP_top60_DNA+RNA_seed | PIK3CA | 1.578941 | LUAD | ABC-1 | GNE-317 |
| MVP_top60_DNA_seed | PIK3CA | -0.106684 | LUAD | A549 | PI3Ka_4409 | MVP_top60_DNA+RNA_seed | HSP90AA1 | 1.581275 | LUAD | NCI-H1623 | Tanespimycin |
| MVP_top60_DNA_seed | HDAC1 | -0.106235 | LUAD | HOP-62 | Parthenolide | MVP_top60_DNA+RNA_seed | MET | 1.583766 | LUAD | H3255 | Savolitinib |
| MVP_top60_DNA_seed | MET | -0.105608 | LUAD | NCI-H23 | PHA-665752 | MVP_top60_DNA+RNA_seed | TERT | 1.584771 | LUAD | NCI-H1355 | BIBR-1532 |
| MVP_top60_DNA_seed | BRAF | -0.105565 | LUAD | NCI-H2085 | HG6-64-1 | MVP_top60_DNA+RNA_seed | AKT1 | 1.589153 | LUAD | HOP-62 | Uprosertib |
| MVP_top60_DNA_seed | HDAC1 | -0.105466 | LUAD | NCI-H1734 | AR-42 | MVP_top60_DNA+RNA_seed | AURKA | 1.590916 | LUAD | NCI-H1568 | Alisertib |
| MVP_top60_DNA_seed | AURKA | -0.105292 | LUAD | NCI-H1650 | ZM447439 | MVP_top60_DNA+RNA_seed | EGFR | 1.592716 | LUAD | NCI-H2342 | Lapatinib |
| MVP_top60_DNA_seed | SRC | -0.10519 | LUAD | NCI-H1944 | NVP-BHG712 | MVP_top60_DNA+RNA_seed | PIK3CA | 1.600673 | LUAD | NCI-H2085 | Alpelisib |
| MVP_top60_DNA_seed | BRAF | -0.104639 | LUAD | NCI-H1651 | HG6-64-1 | MVP_top60_DNA+RNA_seed | EGFR | 1.606154 | LUAD | NCI-H1793 | Osimertinib |
| MVP_top60_DNA_seed | ATM | -0.104426 | LUAD | HCC-78 | CP466722 | MVP_top60_DNA+RNA_seed | SMARCA4 | 1.607298 | LUAD | Calu-3 | PFI3 |
| MVP_top60_DNA_seed | EGFR | -0.104168 | LUAD | HCC-78 | PF-00299804 | MVP_top60_DNA+RNA_seed | AURKA | 1.611555 | LUAD | PC-3_[JPC-3] | ZM447439 |
| MVP_top60_DNA_seed | ERBB2 | -0.104168 | LUAD | HCC-78 | PF-00299804 | MVP_top60_DNA+RNA_seed | AKT1 | 1.614298 | LUAD | NCI-H1793 | MK-2206 |
| MVP_top60_DNA_seed | SRC | -0.103125 | LUAD | NCI-H23 | NVP-BHG712 | MVP_top60_DNA+RNA_seed | AURKA | 1.615307 | LUAD | NCI-H2347 | Alisertib |
| MVP_top60_DNA_seed | SMARCA4 | -0.101686 | LUAD | Calu-3 | PFI-3 | MVP_top60_DNA+RNA_seed | SMARCA4 | 1.615845 | LUAD | NCI-H1793 | PFI3 |
| MVP_top60_DNA_seed | IGF1R | -0.101395 | LUAD | NCI-H1792 | GSK1904529A | MVP_top60_DNA+RNA_seed | FGFR1 | 1.62002 | LUAD | SW1573 | PD173074 |
| MVP_top60_DNA_seed | ATM | -0.100832 | LUAD | HOP-62 | CP466722 | MVP_top60_DNA+RNA_seed | AKT1 | 1.621086 | LUAD | EKVX | GSK2110183B |
| MVP_top60_DNA_seed | HDAC1 | -0.100681 | LUAD | NCI-H838 | Belinostat | MVP_top60_DNA+RNA_seed | MDM2 | 1.623235 | LUAD | HCC-78 | Serdemetan |
| MVP_top60_DNA_seed | EGFR | -0.09912 | LUAD | LC-2-ad | Cetuximab | MVP_top60_DNA+RNA_seed | EZH2 | 1.624931 | LUAD | NCI-H358 | GSK343 |
| MVP_top60_DNA_seed | PTK2 | -0.097558 | LUAD | NCI-H2085 | PF-562271 | MVP_top60_DNA+RNA_seed | MDM2 | 1.627616 | LUAD | NCI-H2347 | Nutlin-3a (-) |
| MVP_top60_DNA_seed | EGFR | -0.097104 | LUAD | NCI-H2347 | PF-00299804 | MVP_top60_DNA+RNA_seed | BRAF | 1.628801 | LUAD | NCI-H1793 | SB590885 |
| MVP_top60_DNA_seed | ERBB2 | -0.097104 | LUAD | NCI-H2347 | PF-00299804 | MVP_top60_DNA+RNA_seed | FGFR1 | 1.630316 | LUAD | H3255 | PD173074 |
| MVP_top60_DNA_seed | EGFR | -0.095243 | LUAD | HOP-62 | CI-1033 | MVP_top60_DNA+RNA_seed | KRAS | 1.630854 | LUAD | COR-L105 | (RAS (G12C) Inhibitor-1 |
| MVP_top60_DNA_seed | ERBB2 | -0.095243 | LUAD | HOP-62 | CI-1033 | MVP_top60_DNA+RNA_seed | EGFR | 1.631178 | LUAD | SK-LU-1 | Gefitinib |
| MVP_top60_DNA_seed | FGFR1 | -0.094244 | LUAD | HCC-44 | AZD4547 | MVP_top60_DNA+RNA_seed | IGF1R | 1.633577 | LUAD | NCI-H441 | NVP-ADW742 |

| | | | | | | | | | | | |
|---|---|---|---|---|---|---|---|---|---|---|---|
| MVP_top60_DNA_seed | SRC | -0.093479 | LUAD | LC-2-ad | NVP-BHG712 | MVP_top60_DNA+RNA_seed | FGFR1 | 1.637179 | LUAD | NCI-H2291 | AZD4547 |
| MVP_top60_DNA_seed | MDM2 | -0.093083 | LUAD | NCI-H2030 | Nutlin-3a (-) | MVP_top60_DNA+RNA_seed | AKT1 | 1.637792 | LUAD | Calu-6 | AT13148 |
| MVP_top60_DNA_seed | MET | -0.092929 | LUAD | HCC-78 | AZD6094 | MVP_top60_DNA+RNA_seed | FGFR1 | 1.640935 | LUAD | NCI-H1838 | AZD4547 |
| MVP_top60_DNA_seed | GSK3B | -0.092424 | LUAD | NCI-H1650 | KIN001-042 | MVP_top60_DNA+RNA_seed | MDM2 | 1.641381 | LUAD | NCI-H1693 | Nutlin-3a (-) |
| MVP_top60_DNA_seed | AURKA | -0.090412 | LUAD | NCI-H1792 | GSK1070916 | MVP_top60_DNA+RNA_seed | FGFR1 | 1.64168 | LUAD | NCI-H1793 | PD173074 |
| MVP_top60_DNA_seed | EGFR | -0.08911 | LUAD | SK-LU-1 | Pelitinib | MVP_top60_DNA+RNA_seed | EGFR | 1.641832 | LUAD | SW1573 | Afatinib |
| MVP_top60_DNA_seed | FGFR1 | -0.088425 | LUAD | NCI-H1944 | AZD4547 | MVP_top60_DNA+RNA_seed | BRAF | 1.646322 | LUAD | NCI-H441 | Dabrafenib |
| MVP_top60_DNA_seed | AURKA | -0.088163 | LUAD | NCI-H1944 | HG-5-113-01 | MVP_top60_DNA+RNA_seed | AKT1 | 1.652057 | LUAD | NCI-H1650 | Uprosertib |
| MVP_top60_DNA_seed | MDM2 | -0.088098 | LUAD | VMRC-LCD | Serdemetan | MVP_top60_DNA+RNA_seed | EZH2 | 1.653178 | LUAD | NCI-H1793 | GSK343 |
| MVP_top60_DNA_seed | AKT1 | -0.086511 | LUAD | RERF-LC-MS | BAY AKT1 | MVP_top60_DNA+RNA_seed | HDAC1 | 1.657406 | LUAD | HCC-827 | PCI-34051 |
| MVP_top60_DNA_seed | AKT1 | -0.085574 | LUAD | NCI-H650 | AKT inhibitor VIII | MVP_top60_DNA+RNA_seed | SMARCA4 | 1.658578 | LUAD | NCI-H2228 | PFI3 |
| MVP_top60_DNA_seed | SRC | -0.084794 | LUAD | SW1573 | HG-6-71-01 | MVP_top60_DNA+RNA_seed | IGF1R | 1.662383 | LUAD | HOP-62 | Linsitinib |
| MVP_top60_DNA_seed | HDAC1 | -0.084097 | LUAD | NCI-H1437 | AR-42 | MVP_top60_DNA+RNA_seed | AKT1 | 1.663842 | LUAD | NCI-H1573 | Uprosertib |
| MVP_top60_DNA_seed | GSK3B | -0.084026 | LUAD | NCI-H1651 | KIN001-042 | MVP_top60_DNA+RNA_seed | AKT1 | 1.665035 | LUAD | VMRC-LCD | Uprosertib |
| MVP_top60_DNA_seed | FGFR1 | -0.082982 | LUAD | Calu-6 | FGFR_3831 | MVP_top60_DNA+RNA_seed | AKT1 | 1.667253 | LUAD | NCI-H1693 | AZD5363 |
| MVP_top60_DNA_seed | HDAC1 | -0.082448 | LUAD | NCI-H2087 | Belinostat | MVP_top60_DNA+RNA_seed | EZH2 | 1.669497 | LUAD | NCI-H2228 | GSK343 |
| MVP_top60_DNA_seed | ERBB2 | -0.080802 | LUAD | HOP-62 | CP724714 | MVP_top60_DNA+RNA_seed | EGFR | 1.676934 | LUAD | NCI-H2347 | Afatinib |
| MVP_top60_DNA_seed | HDAC1 | -0.080712 | LUAD | NCI-H1648 | Dacinostat | MVP_top60_DNA+RNA_seed | MDM2 | 1.678429 | LUAD | SW1573 | Serdemetan |
| MVP_top60_DNA_seed | SRC | -0.080599 | LUAD | NCI-H2085 | Bosutinib | MVP_top60_DNA+RNA_seed | ATM | 1.680514 | LUAD | SK-LU-1 | KU-55933 |
| MVP_top60_DNA_seed | HDAC1 | -0.080162 | LUAD | LXF-289 | AR-42 | MVP_top60_DNA+RNA_seed | PIK3CA | 1.681244 | LUAD | NCI-H2347 | Buparlisib |
| MVP_top60_DNA_seed | GSK3B | -0.079858 | LUAD | HOP-62 | CHIR-99021 | MVP_top60_DNA+RNA_seed | ATM | 1.685011 | LUAD | H3255 | KU-55933 |
| MVP_top60_DNA_seed | AURKA | -0.079302 | LUAD | NCI-H838 | GSK1070916 | MVP_top60_DNA+RNA_seed | IGF1R | 1.686758 | LUAD | NCI-H1563 | NVP-ADW742 |
| MVP_top60_DNA_seed | MET | -0.079123 | LUAD | EMC-BAC-2 | JNJ38877605 | MVP_top60_DNA+RNA_seed | EZH2 | 1.689203 | LUAD | NCI-H1568 | GSK343 |
| MVP_top60_DNA_seed | FGFR1 | -0.078476 | LUAD | NCI-H1666 | PD173074 | MVP_top60_DNA+RNA_seed | EGFR | 1.690854 | LUAD | SK-LU-1 | AZD3759 |
| MVP_top60_DNA_seed | EGFR | -0.078303 | LUAD | NCI-H1792 | CI-1033 | MVP_top60_DNA+RNA_seed | FGFR1 | 1.691705 | LUAD | NCI-H441 | AZD4547 |
| MVP_top60_DNA_seed | ERBB2 | -0.078303 | LUAD | NCI-H1792 | CI-1033 | MVP_top60_DNA+RNA_seed | TP53 | 1.691954 | LUAD | NCI-H1568 | MIRA-1 |
| MVP_top60_DNA_seed | MET | -0.077802 | LUAD | PC-14 | JNJ38877605 | MVP_top60_DNA+RNA_seed | EZH2 | 1.694465 | LUAD | NCI-H1650 | GSK343 |
| MVP_top60_DNA_seed | ATM | -0.075855 | LUAD | A549 | CP466722 | MVP_top60_DNA+RNA_seed | AURKA | 1.698161 | LUAD | NCI-H2087 | ZM447439 |
| MVP_top60_DNA_seed | AURKA | -0.075733 | LUAD | SK-LU-1 | GSK1070916 | MVP_top60_DNA+RNA_seed | IGF1R | 1.703473 | LUAD | NCI-H1781 | BMS-536924 |
| MVP_top60_DNA_seed | FGFR1 | -0.075472 | LUAD | NCI-H1693 | AZD4547 | MVP_top60_DNA+RNA_seed | KRAS | 1.703993 | LUAD | NCI-H1650 | KRAS (G12C) Inhibitor-1 |
| MVP_top60_DNA_seed | PTK2 | -0.075042 | LUAD | NCI-H1781 | PF-562271 | MVP_top60_DNA+RNA_seed | HSP90AA1 | 1.705251 | LUAD | NCI-H1568 | Luminespib |
| MVP_top60_DNA_seed | AURKA | -0.073935 | LUAD | NCI-H1648 | ZM447439 | MVP_top60_DNA+RNA_seed | PIK3CA | 1.707488 | LUAD | SK-LU-1 | Alpelisib |
| MVP_top60_DNA_seed | HSP90AA1 | -0.072311 | LUAD | NCI-H2405 | Elesclomol | MVP_top60_DNA+RNA_seed | EZH2 | 1.709431 | LUAD | NCI-H1355 | GSK343 |
| MVP_top60_DNA_seed | AURKA | -0.072279 | LUAD | NCI-H23 | ZM447439 | MVP_top60_DNA+RNA_seed | PIK3CA | 1.713723 | LUAD | NCI-H2342 | Buparlisib |
| MVP_top60_DNA_seed | IGF1R | -0.071287 | LUAD | NCI-H23 | BMS-536924 | MVP_top60_DNA+RNA_seed | SMARCA4 | 1.714755 | LUAD | SK-LU-1 | PFI3 |
| MVP_top60_DNA_seed | AKT1 | -0.069534 | LUAD | NCI-H358 | GSK690693 | MVP_top60_DNA+RNA_seed | BRAF | 1.715931 | LUAD | SW1573 | PLX-4720 |
| MVP_top60_DNA_seed | HDAC1 | -0.068977 | LUAD | NCI-H2087 | JQ12 | MVP_top60_DNA+RNA_seed | AKT1 | 1.716041 | LUAD | NCI-H1666 | AT13148 |
| MVP_top60_DNA_seed | ATM | -0.068203 | LUAD | NCI-H1755 | KU-55933 | MVP_top60_DNA+RNA_seed | TERT | 1.717819 | LUAD | NCI-H1693 | BIBR-1532 |
| MVP_top60_DNA_seed | HSP90AA1 | -0.068129 | LUAD | HCC-827 | Elesclomol | MVP_top60_DNA+RNA_seed | AKT1 | 1.726625 | LUAD | VMRC-LCD | Uprosertib |
| MVP_top60_DNA_seed | HSP90AA1 | -0.068085 | LUAD | EKVX | Elesclomol | MVP_top60_DNA+RNA_seed | TERT | 1.732268 | LUAD | Calu-3 | BIBR-1532 |
| MVP_top60_DNA_seed | HDAC1 | -0.067442 | LUAD | NCI-H1666 | Belinostat | MVP_top60_DNA+RNA_seed | AKT1 | 1.735515 | LUAD | NCI-H1793 | Uprosertib |

| | | | | | | | | | | | |
|---|---|---|---|---|---|---|---|---|---|---|---|
| MVP_top60_DNA_seed | HDAC1 | -0.066834 | LUAD | RERF-LC-MS | AR-42 | MVP_top60_DNA+RNA_seed | HDAC1 | 1.74073 | LUAD | SK-LU-1 | PCI-34051 |
| MVP_top60_DNA_seed | ERBB2 | -0.066446 | LUAD | HCC-827 | CP724714 | MVP_top60_DNA+RNA_seed | MDM2 | 1.742271 | LUAD | EKVX | Serdemetan |
| MVP_top60_DNA_seed | EGFR | -0.065994 | LUAD | NCI-H2291 | AST-1306 | MVP_top60_DNA+RNA_seed | HDAC1 | 1.752217 | LUAD | EKVX | Entinostat |
| MVP_top60_DNA_seed | GSK3B | -0.06553 | LUAD | NCI-H1734 | CHIR-99021 | MVP_top60_DNA+RNA_seed | ATM | 1.754883 | LUAD | EKVX | KU-55933 |
| MVP_top60_DNA_seed | BRAF | -0.064811 | LUAD | VMRC-LCD | SB590885 | MVP_top60_DNA+RNA_seed | EZH2 | 1.758764 | LUAD | SW1573 | GSK343 |
| MVP_top60_DNA_seed | FGFR1 | -0.06452 | LUAD | NCI-H1793 | AZD4547 | MVP_top60_DNA+RNA_seed | FGFR1 | 1.75979 | LUAD | SK-LU-1 | AZD4547 |
| MVP_top60_DNA_seed | SMARCA4 | -0.064269 | LUAD | LXF-289 | PFI-3 | MVP_top60_DNA+RNA_seed | FGFR1 | 1.760949 | LUAD | VMRC-LCD | PD173074 |
| MVP_top60_DNA_seed | EGFR | -0.063764 | LUAD | LXF-289 | CUDC-101 | MVP_top60_DNA+RNA_seed | AKT1 | 1.761526 | LUAD | SK-LU-1 | GSK2110183B |
| MVP_top60_DNA_seed | ERBB2 | -0.063764 | LUAD | LXF-289 | CUDC-101 | MVP_top60_DNA+RNA_seed | TP53 | 1.763346 | LUAD | NCI-H1793 | MIRA-1 |
| MVP_top60_DNA_seed | AKT1 | -0.063253 | LUAD | NCI-H23 | GSK690693 | MVP_top60_DNA+RNA_seed | EGFR | 1.764459 | LUAD | SK-LU-1 | Erlotinib |
| MVP_top60_DNA_seed | EGFR | -0.062741 | LUAD | SW1573 | Cetuximab | MVP_top60_DNA+RNA_seed | FGFR1 | 1.767306 | LUAD | HOP-62 | PD173074 |
| MVP_top60_DNA_seed | AKT1 | -0.062579 | LUAD | HCC-78 | MK-2206 | MVP_top60_DNA+RNA_seed | HDAC1 | 1.768122 | LUAD | NCI-H1793 | PCI-34051 |
| MVP_top60_DNA_seed | MET | -0.06207 | LUAD | NCI-H522 | AZD6094 | MVP_top60_DNA+RNA_seed | EZH2 | 1.768724 | LUAD | NCI-H1666 | GSK343 |
| MVP_top60_DNA_seed | PIK3CA | -0.061745 | LUAD | SK-LU-1 | AZD8835 | MVP_top60_DNA+RNA_seed | EGFR | 1.770075 | LUAD | NCI-H1734 | Afatinib |
| MVP_top60_DNA_seed | AURKA | -0.060406 | LUAD | NCI-H1648 | CD532 | MVP_top60_DNA+RNA_seed | SRC | 1.771157 | LUAD | NCI-H2405 | Bosutinib |
| MVP_top60_DNA_seed | GSK3B | -0.059236 | LUAD | NCI-H650 | AZD7969 | MVP_top60_DNA+RNA_seed | EZH2 | 1.776635 | LUAD | NCI-H2228 | GSK343 |
| MVP_top60_DNA_seed | BRAF | -0.058737 | LUAD | NCI-H1944 | HG6-64-1 | MVP_top60_DNA+RNA_seed | SMARCA4 | 1.777921 | LUAD | SW1573 | PFI3 |
| MVP_top60_DNA_seed | FGFR1 | -0.058704 | LUAD | NCI-H1563 | AZD4547 | MVP_top60_DNA+RNA_seed | EGFR | 1.784245 | LUAD | NCI-H2347 | Erlotinib |
| MVP_top60_DNA_seed | AURKA | -0.058351 | LUAD | PC-14 | Alisertib | MVP_top60_DNA+RNA_seed | MET | 1.785391 | LUAD | EKVX | Savolitinib |
| MVP_top60_DNA_seed | MET | -0.058205 | LUAD | NCI-H1435 | AZD6094 | MVP_top60_DNA+RNA_seed | AKT1 | 1.785968 | LUAD | NCI-H2087 | MK-2206 |
| MVP_top60_DNA_seed | AKT1 | -0.058031 | LUAD | NCI-H1623 | AKT inhibitor VIII | MVP_top60_DNA+RNA_seed | AKT1 | 1.786239 | LUAD | Calu-3 | AT13148 |
| MVP_top60_DNA_seed | BRAF | -0.05781 | LUAD | NCI-H2023 | PLX-4720 | MVP_top60_DNA+RNA_seed | IGF1R | 1.789488 | LUAD | H3255 | NVP-ADW742 |
| MVP_top60_DNA_seed | EGFR | -0.05778 | LUAD | HCC-44 | Pelitinib | MVP_top60_DNA+RNA_seed | AKT1 | 1.790003 | LUAD | NCI-H2347 | Ipatasertib |
| MVP_top60_DNA_seed | HSP90AA1 | -0.056342 | LUAD | NCI-H2228 | Elesclomol | MVP_top60_DNA+RNA_seed | ATM | 1.794129 | LUAD | NCI-H1838 | KU-55933 |
| MVP_top60_DNA_seed | BRAF | -0.055686 | LUAD | NCI-H1573 | Dabrafenib | MVP_top60_DNA+RNA_seed | HSP90AA1 | 1.79805 | LUAD | NCI-H2030 | Elesclomol |
| MVP_top60_DNA_seed | HSP90AA1 | -0.055172 | LUAD | NCI-H1666 | CCT-018159 | MVP_top60_DNA+RNA_seed | AKT1 | 1.79815 | LUAD | NCI-H2405 | Uprosertib |
| MVP_top60_DNA_seed | ERBB2 | -0.054428 | LUAD | NCI-H1975 | CP724714 | MVP_top60_DNA+RNA_seed | SMARCA4 | 1.799288 | LUAD | PC-14 | PFI3 |
| MVP_top60_DNA_seed | PIK3CA | -0.053497 | LUAD | LC-2-ad | PI3Ka_4409 | MVP_top60_DNA+RNA_seed | MDM2 | 1.799551 | LUAD | PC-3_[JPC-3] | Serdemetan |
| MVP_top60_DNA_seed | AKT1 | -0.051924 | LUAD | H3255 | BAY AKT1 | MVP_top60_DNA+RNA_seed | BRAF | 1.809583 | LUAD | PC-14 | SB590885 |
| MVP_top60_DNA_seed | HDAC1 | -0.05048 | LUAD | HCC-78 | Tubastatin A | MVP_top60_DNA+RNA_seed | AKT1 | 1.810585 | LUAD | NCI-H522 | Afuresertib |
| MVP_top60_DNA_seed | BRAF | -0.050473 | LUAD | PC-3_[JPC-3] | RAF_9304 | MVP_top60_DNA+RNA_seed | MET | 1.817294 | LUAD | NCI-H1623 | Crizotinib |
| MVP_top60_DNA_seed | AURKA | -0.050425 | LUAD | NCI-H1563 | HG-5-113-01 | MVP_top60_DNA+RNA_seed | HDAC1 | 1.81901 | LUAD | NCI-H1435 | Entinostat |
| MVP_top60_DNA_seed | HDAC1 | -0.050074 | LUAD | NCI-H23 | AR-42 | MVP_top60_DNA+RNA_seed | AKT1 | 1.819272 | LUAD | NCI-H1793 | AZD5363 |
| MVP_top60_DNA_seed | IGF1R | -0.04824 | LUAD | NCI-H2405 | BMS-754807 | MVP_top60_DNA+RNA_seed | AKT1 | 1.821332 | LUAD | NCI-H2347 | Uprosertib |
| MVP_top60_DNA_seed | PIK3CA | -0.048167 | LUAD | EMC-BAC-1 | PI3Ka_4409 | MVP_top60_DNA+RNA_seed | IGF1R | 1.821663 | LUAD | NCI-H1838 | GSK1904529A |
| MVP_top60_DNA_seed | AURKA | -0.047776 | LUAD | RERF-LC-KJ | ZM447439 | MVP_top60_DNA+RNA_seed | EGFR | 1.821768 | LUAD | NCI-H2087 | AZD3759 |
| MVP_top60_DNA_seed | HDAC1 | -0.047074 | LUAD | NCI-H1437 | Belinostat | MVP_top60_DNA+RNA_seed | EGFR | 1.824114 | LUAD | NCI-H1793 | AZD3759 |
| MVP_top60_DNA_seed | AURKA | -0.046747 | LUAD | NCI-H2030 | GSK1070916 | MVP_top60_DNA+RNA_seed | TP53 | 1.824734 | LUAD | HCC-827 | MIRA-1 |
| MVP_top60_DNA_seed | AURKA | -0.04667 | LUAD | COR-L105 | CD532 | MVP_top60_DNA+RNA_seed | AKT1 | 1.824909 | LUAD | NCI-H1793 | GSK2110183B |
| MVP_top60_DNA_seed | ATM | -0.045788 | LUAD | NCI-H650 | CP466722 | MVP_top60_DNA+RNA_seed | KRAS | 1.826049 | LUAD | NCI-H2405 | KRAS (G12C) Inhibitor-1 |
| MVP_top60_DNA_seed | GSK3B | -0.04506 | LUAD | NCI-H2291 | CHIR-99021 | MVP_top60_DNA+RNA_seed | HDAC1 | 1.83269 | LUAD | NCI-H2228 | PCI-34051 |

| | | | | | | | | | | | |
|---|---|---|---|---|---|---|---|---|---|---|---|
| MVP_top60_DNA_seed | AURKA | -0.044808 | LUAD | COR-L105 | GSK1070916 | MVP_top60_DNA+RNA_seed | MDM2 | 1.83863 | LUAD | NCI-H441 | Serdemetan |
| MVP_top60_DNA_seed | MET | -0.0448 | LUAD | NCI-H358 | JNJ38877605 | MVP_top60_DNA+RNA_seed | EGFR | 1.840042 | LUAD | NCI-H1355 | Lapatinib |
| MVP_top60_DNA_seed | BRAF | -0.043217 | LUAD | NCI-H2122 | PLX-4720 | MVP_top60_DNA+RNA_seed | AKT1 | 1.842697 | LUAD | NCI-H441 | GSK2110183B |
| MVP_top60_DNA_seed | GSK3B | -0.042345 | LUAD | VMRC-LCD | SB216763 | MVP_top60_DNA+RNA_seed | EZH2 | 1.8429 | LUAD | NCI-H2347 | GSK343 |
| MVP_top60_DNA_seed | GSK3B | -0.042053 | LUAD | NCI-H838 | AZD7969 | MVP_top60_DNA+RNA_seed | BRAF | 1.843136 | LUAD | NCI-H1650 | PLX-4720 |
| MVP_top60_DNA_seed | AURKA | -0.041592 | LUAD | NCI-H2087 | HG-5-113-01 | MVP_top60_DNA+RNA_seed | AKT1 | 1.848252 | LUAD | PC-3_[JPC-3] | Uprosertib |
| MVP_top60_DNA_seed | AKT1 | -0.040953 | LUAD | NCI-H2342 | MK-2206 | MVP_top60_DNA+RNA_seed | PIK3CA | 1.852534 | LUAD | HOP-62 | Alpelisib |
| MVP_top60_DNA_seed | BRAF | -0.04087 | LUAD | NCI-H650 | HG6-64-1 | MVP_top60_DNA+RNA_seed | AKT1 | 1.852705 | LUAD | NCI-H2087 | AT13148 |
| MVP_top60_DNA_seed | AURKA | -0.038891 | LUAD | NCI-H1734 | GSK1070916 | MVP_top60_DNA+RNA_seed | MET | 1.853154 | LUAD | NCI-H2291 | Savolitinib |
| MVP_top60_DNA_seed | MET | -0.038381 | LUAD | NCI-H1792 | AZD6094 | MVP_top60_DNA+RNA_seed | EGFR | 1.854342 | LUAD | SW1573 | AZD3759 |
| MVP_top60_DNA_seed | AKT1 | -0.038326 | LUAD | NCI-H1666 | BAY AKT1 | MVP_top60_DNA+RNA_seed | EZH2 | 1.859325 | LUAD | NCI-H1650 | GSK343 |
| MVP_top60_DNA_seed | PIK3CA | -0.037592 | LUAD | NCI-H2342 | AZD8835 | MVP_top60_DNA+RNA_seed | MET | 1.864962 | LUAD | SK-LU-1 | Savolitinib |
| MVP_top60_DNA_seed | CREBBP | -0.035339 | LUAD | HCC-44 | I-CBP112 | MVP_top60_DNA+RNA_seed | AURKA | 1.865375 | LUAD | EKVX | Tozasertib |
| MVP_top60_DNA_seed | MDM2 | -0.034711 | LUAD | NCI-H1793 | Nutlin-3a (-) | MVP_top60_DNA+RNA_seed | AKT1 | 1.866666 | LUAD | NCI-H650 | AZD5363 |
| MVP_top60_DNA_seed | FGFR1 | -0.034512 | LUAD | NCI-H1648 | FGFR_3831 | MVP_top60_DNA+RNA_seed | AKT1 | 1.872643 | LUAD | VMRC-LCD | AZD5363 |
| MVP_top60_DNA_seed | EGFR | -0.03403 | LUAD | NCI-H1838 | AST-1306 | MVP_top60_DNA+RNA_seed | AURKA | 1.873375 | LUAD | NCI-H1650 | ZM447439 |
| MVP_top60_DNA_seed | EGFR | -0.033332 | LUAD | A549 | Cetuximab | MVP_top60_DNA+RNA_seed | AURKA | 1.874314 | LUAD | SW1573 | ZM447439 |
| MVP_top60_DNA_seed | GSK3B | -0.03332 | LUAD | NCI-H1650 | AZD7969 | MVP_top60_DNA+RNA_seed | EGFR | 1.8816 | LUAD | ABC-1 | Erlotinib |
| MVP_top60_DNA_seed | BRAF | -0.032685 | LUAD | NCI-H1975 | AZ628 | MVP_top60_DNA+RNA_seed | EZH2 | 1.881841 | LUAD | NCI-H2405 | GSK343 |
| MVP_top60_DNA_seed | HSP90AA1 | -0.032393 | LUAD | NCI-H1355 | CCT-018159 | MVP_top60_DNA+RNA_seed | EGFR | 1.885326 | LUAD | Calu-6 | AZD3759 |
| MVP_top60_DNA_seed | EGFR | -0.03211 | LUAD | NCI-H23 | PF-00299804 | MVP_top60_DNA+RNA_seed | AKT1 | 1.891077 | LUAD | Calu-6 | Afuresertib |
| MVP_top60_DNA_seed | ERBB2 | -0.03211 | LUAD | NCI-H23 | PF-00299804 | MVP_top60_DNA+RNA_seed | AKT1 | 1.892494 | LUAD | NCI-H441 | Afuresertib |
| MVP_top60_DNA_seed | HDAC1 | -0.031983 | LUAD | NCI-H1435 | JQ12 | MVP_top60_DNA+RNA_seed | FGFR1 | 1.894603 | LUAD | H3255 | AZD4547 |
| MVP_top60_DNA_seed | GSK3B | -0.030962 | LUAD | HCC-827 | SB216763 | MVP_top60_DNA+RNA_seed | PIK3CA | 1.895236 | LUAD | NCI-H441 | AZD8186 |
| MVP_top60_DNA_seed | EGFR | -0.030434 | LUAD | NCI-H2030 | CI-1033 | MVP_top60_DNA+RNA_seed | TERT | 1.901414 | LUAD | NCI-H1793 | BIBR-1532 |
| MVP_top60_DNA_seed | ERBB2 | -0.030434 | LUAD | NCI-H2030 | CI-1033 | MVP_top60_DNA+RNA_seed | TP53 | 1.91054 | LUAD | NCI-H2347 | MIRA-1 |
| MVP_top60_DNA_seed | GSK3B | -0.030264 | LUAD | HCC-44 | SB216763 | MVP_top60_DNA+RNA_seed | IGF1R | 1.911289 | LUAD | SW1573 | NVP-ADW742 |
| MVP_top60_DNA_seed | IGF1R | -0.029919 | LUAD | NCI-H2009 | BMS-754807 | MVP_top60_DNA+RNA_seed | EZH2 | 1.913127 | LUAD | NCI-H441 | GSK343 |
| MVP_top60_DNA_seed | IGF1R | -0.029791 | LUAD | NCI-H2030 | BMS-754807 | MVP_top60_DNA+RNA_seed | AKT1 | 1.917594 | LUAD | HOP-62 | MK-2206 |
| MVP_top60_DNA_seed | SRC | -0.029533 | LUAD | NCI-H2085 | NVP-BHG712 | MVP_top60_DNA+RNA_seed | AKT1 | 1.924887 | LUAD | NCI-H2228 | AT13148 |
| MVP_top60_DNA_seed | AURKA | -0.029091 | LUAD | EMC-BAC-2 | ZM447439 | MVP_top60_DNA+RNA_seed | IGF1R | 1.926888 | LUAD | NCI-H2347 | GSK1904529A |
| MVP_top60_DNA_seed | HDAC1 | -0.029058 | LUAD | NCI-H2087 | AR-42 | MVP_top60_DNA+RNA_seed | EGFR | 1.927509 | LUAD | SW1573 | Gefitinib |
| MVP_top60_DNA_seed | BRAF | -0.027466 | LUAD | EMC-BAC-1 | SB590885 | MVP_top60_DNA+RNA_seed | BRAF | 1.928852 | LUAD | EKVX | SB590885 |
| MVP_top60_DNA_seed | HDAC1 | -0.027446 | LUAD | NCI-H2085 | Dacinostat | MVP_top60_DNA+RNA_seed | KRAS | 1.93085 | LUAD | NCI-H1568 | KRAS (G12C) Inhibitor-1 |
| MVP_top60_DNA_seed | EGFR | -0.026287 | LUAD | NCI-H522 | CI-1033 | MVP_top60_DNA+RNA_seed | TP53 | 1.932986 | LUAD | NCI-H441 | MIRA-1 |
| MVP_top60_DNA_seed | ERBB2 | -0.026287 | LUAD | NCI-H522 | CI-1033 | MVP_top60_DNA+RNA_seed | PIK3CA | 1.94076 | LUAD | VMRC-LCD | AZD8186 |
| MVP_top60_DNA_seed | EGFR | -0.02625 | LUAD | EMC-BAC-2 | Cetuximab | MVP_top60_DNA+RNA_seed | EGFR | 1.942151 | LUAD | SK-LU-1 | Lapatinib |
| MVP_top60_DNA_seed | AURKA | -0.025173 | LUAD | H3255 | ZM447439 | MVP_top60_DNA+RNA_seed | BRAF | 1.946723 | LUAD | NCI-H2291 | PLX-4720 |
| MVP_top60_DNA_seed | AURKA | -0.025067 | LUAD | NCI-H1648 | Alisertib | MVP_top60_DNA+RNA_seed | EZH2 | 1.951714 | LUAD | NCI-H441 | GSK343 |
| MVP_top60_DNA_seed | AURKA | -0.024617 | LUAD | EMC-BAC-2 | GSK1070916 | MVP_top60_DNA+RNA_seed | AKT1 | 1.961121 | LUAD | NCI-H1568 | AT13148 |
| MVP_top60_DNA_seed | EGFR | -0.024421 | LUAD | NCI-H2087 | CUDC-101 | MVP_top60_DNA+RNA_seed | AKT1 | 1.96387 | LUAD | NCI-H441 | MK-2206 |

| | | | | | | | | | | | |
|---|---|---|---|---|---|---|---|---|---|---|---|
| MVP_top60_DNA_seed | ERBB2 | -0.024421 | LUAD | NCI-H2087 | CUDC-101 | MVP_top60_DNA+RNA_seed | PIK3CA | 1.966736 | LUAD | SK-LU-1 | Buparlisib |
| MVP_top60_DNA_seed | EGFR | -0.024176 | LUAD | NCI-H650 | CI-1033 | MVP_top60_DNA+RNA_seed | AURKA | 1.978455 | LUAD | NCI-H1355 | ZM447439 |
| MVP_top60_DNA_seed | ERBB2 | -0.024176 | LUAD | NCI-H650 | CI-1033 | MVP_top60_DNA+RNA_seed | MET | 1.980896 | LUAD | NCI-H441 | Savolitinib |
| MVP_top60_DNA_seed | PIK3CA | -0.023811 | LUAD | NCI-H2009 | PI3Ka_4409 | MVP_top60_DNA+RNA_seed | EGFR | 1.98256 | LUAD | SK-LU-1 | Afatinib |
| MVP_top60_DNA_seed | HSP90AA1 | -0.023167 | LUAD | SW1573 | Luminespib | MVP_top60_DNA+RNA_seed | FGFR1 | 1.986536 | LUAD | NCI-H2347 | AZD4547 |
| MVP_top60_DNA_seed | HDAC1 | -0.022385 | LUAD | NCI-H650 | Belinostat | MVP_top60_DNA+RNA_seed | AKT1 | 1.989425 | LUAD | Calu-3 | Ipatasertib |
| MVP_top60_DNA_seed | EGFR | -0.021439 | LUAD | NCI-H1838 | CI-1033 | MVP_top60_DNA+RNA_seed | TERT | 1.998051 | LUAD | NCI-H1666 | BIBR-1532 |
| MVP_top60_DNA_seed | ERBB2 | -0.021439 | LUAD | NCI-H1838 | CI-1033 | MVP_top60_DNA+RNA_seed | MDM2 | 1.999294 | LUAD | COR-L105 | Serdemetan |
| MVP_top60_DNA_seed | HDAC1 | -0.020088 | LUAD | NCI-H23 | CAY10603 | MVP_top60_DNA+RNA_seed | IGF1R | 2.006281 | LUAD | EKVX | NVP-ADW742 |
| MVP_top60_DNA_seed | MAPK1 | -0.019939 | LUAD | ABC-1 | FR-180204 | MVP_top60_DNA+RNA_seed | HDAC1 | 2.009593 | LUAD | NCI-H441 | PCI-34051 |
| MVP_top60_DNA_seed | HDAC1 | -0.019902 | LUAD | EMC-BAC-1 | CAY10603 | MVP_top60_DNA+RNA_seed | AKT1 | 2.01253 | LUAD | NCI-H2405 | GSK2110183B |
| MVP_top60_DNA_seed | AURKA | -0.018517 | LUAD | RERF-LC-KJ | GSK1070916 | MVP_top60_DNA+RNA_seed | TERT | 2.022634 | LUAD | NCI-H1838 | BIBR-1532 |
| MVP_top60_DNA_seed | HSP90AA1 | -0.018315 | LUAD | NCI-H1781 | Tanespimycin | MVP_top60_DNA+RNA_seed | EZH2 | 2.022846 | LUAD | SW1573 | GSK343 |
| MVP_top60_DNA_seed | HDAC1 | -0.018079 | LUAD | NCI-H23 | Tubastatin A | MVP_top60_DNA+RNA_seed | MDM2 | 2.026136 | LUAD | NCI-H2405 | Nutlin-3a (-) |
| MVP_top60_DNA_seed | EGFR | -0.01726 | LUAD | NCI-H522 | AST-1306 | MVP_top60_DNA+RNA_seed | IGF1R | 2.028363 | LUAD | NCI-H2405 | Linsitinib |
| MVP_top60_DNA_seed | ATM | -0.016552 | LUAD | RERF-LC-MS | CP466722 | MVP_top60_DNA+RNA_seed | FGFR1 | 2.040052 | LUAD | NCI-H2405 | PD173074 |
| MVP_top60_DNA_seed | EGFR | -0.015355 | LUAD | NCI-H3122 | CUDC-101 | MVP_top60_DNA+RNA_seed | KRAS | 2.042551 | LUAD | NCI-H2228 | RAS (G12C) Inhibitor-1 |
| MVP_top60_DNA_seed | ERBB2 | -0.015355 | LUAD | NCI-H3122 | CUDC-101 | MVP_top60_DNA+RNA_seed | IGF1R | 2.043567 | LUAD | NCI-H1666 | GSK1904529A |
| MVP_top60_DNA_seed | HSP90AA1 | -0.014435 | LUAD | NCI-H2023 | Luminespib | MVP_top60_DNA+RNA_seed | TERT | 2.052173 | LUAD | SK-LU-1 | BIBR-1532 |
| MVP_top60_DNA_seed | EGFR | -0.013815 | LUAD | NCI-H2023 | Gefitinib | MVP_top60_DNA+RNA_seed | EZH2 | 2.055256 | LUAD | SK-LU-1 | GSK343 |
| MVP_top60_DNA_seed | AKT1 | -0.012952 | LUAD | NCI-H1734 | GSK690693 | MVP_top60_DNA+RNA_seed | PIK3CA | 2.065836 | LUAD | NCI-H1693 | Alpelisib |
| MVP_top60_DNA_seed | SRC | -0.012772 | LUAD | SK-LU-1 | NVP-BHG712 | MVP_top60_DNA+RNA_seed | EGFR | 2.071103 | LUAD | NCI-H441 | AZD3759 |
| MVP_top60_DNA_seed | AURKA | -0.012542 | LUAD | NCI-H838 | ZM447439 | MVP_top60_DNA+RNA_seed | AKT1 | 2.072122 | LUAD | NCI-H441 | Ipatasertib |
| MVP_top60_DNA_seed | IGF1R | -0.012454 | LUAD | HCC-827 | Linsitinib | MVP_top60_DNA+RNA_seed | FGFR1 | 2.078265 | LUAD | NCI-H1355 | PD173074 |
| MVP_top60_DNA_seed | HDAC1 | -0.011317 | LUAD | NCI-H1623 | CAY10603 | MVP_top60_DNA+RNA_seed | HSP90AA1 | 2.078636 | LUAD | NCI-H1793 | Tanespimycin |
| MVP_top60_DNA_seed | EGFR | -0.011308 | LUAD | PC-3_[JPC-3] | CUDC-101 | MVP_top60_DNA+RNA_seed | EGFR | 2.081775 | LUAD | NCI-H441 | Lapatinib |
| MVP_top60_DNA_seed | ERBB2 | -0.011308 | LUAD | PC-3_[JPC-3] | CUDC-101 | MVP_top60_DNA+RNA_seed | MDM2 | 2.084173 | LUAD | NCI-H1838 | Serdemetan |
| MVP_top60_DNA_seed | EGFR | -0.010331 | LUAD | NCI-H2030 | AST-1306 | MVP_top60_DNA+RNA_seed | AKT1 | 2.084896 | LUAD | EKVX | Uprosertib |
| MVP_top60_DNA_seed | EGFR | -0.009533 | LUAD | HOP-62 | AST-1306 | MVP_top60_DNA+RNA_seed | AKT1 | 2.086222 | LUAD | NCI-H1573 | Afuresertib |
| MVP_top60_DNA_seed | SRC | -0.008841 | LUAD | NCI-H1993 | NVP-BHG712 | MVP_top60_DNA+RNA_seed | HSP90AA1 | 2.093513 | LUAD | NCI-H1693 | Tanespimycin |
| MVP_top60_DNA_seed | BRAF | -0.008536 | LUAD | NCI-H2009 | PLX-4720 | MVP_top60_DNA+RNA_seed | KRAS | 2.098972 | LUAD | SW1573 | RAS (G12C) Inhibitor-1 |
| MVP_top60_DNA_seed | EGFR | -0.008081 | LUAD | LC-2-ad | PF-00299804 | MVP_top60_DNA+RNA_seed | PIK3CA | 2.100023 | LUAD | NCI-H441 | Alpelisib |
| MVP_top60_DNA_seed | ERBB2 | -0.008081 | LUAD | LC-2-ad | PF-00299804 | MVP_top60_DNA+RNA_seed | MDM2 | 2.100282 | LUAD | NCI-H2087 | Serdemetan |
| MVP_top60_DNA_seed | AURKA | -0.006705 | LUAD | NCI-H650 | HG-5-113-01 | MVP_top60_DNA+RNA_seed | AURKA | 2.107135 | LUAD | NCI-H1666 | ZM447439 |
| MVP_top60_DNA_seed | HDAC1 | -0.006091 | LUAD | NCI-H2087 | CAY10603 | MVP_top60_DNA+RNA_seed | SMARCA4 | 2.112213 | LUAD | EKVX | PFI3 |
| MVP_top60_DNA_seed | HDAC1 | -0.005837 | LUAD | NCI-H1693 | CAY10603 | MVP_top60_DNA+RNA_seed | HSP90AA1 | 2.117502 | LUAD | VMRC-LCD | Luminespib |
| MVP_top60_DNA_seed | PIK3CA | -0.005836 | LUAD | NCI-H1563 | PI3Ka_4409 | MVP_top60_DNA+RNA_seed | TERT | 2.124139 | LUAD | NCI-H2291 | BIBR-1532 |
| MVP_top60_DNA_seed | HDAC1 | -0.005655 | LUAD | EMC-BAC-1 | Tubastatin A | MVP_top60_DNA+RNA_seed | PIK3CA | 2.130441 | LUAD | H3255 | Buparlisib |
| MVP_top60_DNA_seed | EGFR | -0.005511 | LUAD | NCI-H23 | AST-1306 | MVP_top60_DNA+RNA_seed | AKT1 | 2.136466 | LUAD | NCI-H1650 | AT13148 |
| MVP_top60_DNA_seed | AKT1 | -0.005169 | LUAD | NCI-H1975 | AKT inhibitor VIII | MVP_top60_DNA+RNA_seed | EGFR | 2.140246 | LUAD | SW1573 | Erlotinib |
| MVP_top60_DNA_seed | BRAF | -0.004932 | LUAD | ABC-1 | HG6-64-1 | MVP_top60_DNA+RNA_seed | PIK3CA | 2.141275 | LUAD | EKVX | Buparlisib |

| | | | | | | | | | | | |
|---|---|---|---|---|---|---|---|---|---|---|---|
| MVP_top60_DNA_seed | IGF1R | -0.003819 | LUAD | NCI-H2347 | Linsitinib | MVP_top60_DNA+RNA_seed | AKT1 | 2.147146 | LUAD | NCI-H2291 | AT13148 |
| MVP_top60_DNA_seed | EGFR | -0.003693 | LUAD | NCI-H322M | Pelitinib | MVP_top60_DNA+RNA_seed | SMARCA4 | 2.151269 | LUAD | H3255 | PFI3 |
| MVP_top60_DNA_seed | AURKA | -0.00367 | LUAD | Calu-3 | Alisertib | MVP_top60_DNA+RNA_seed | HDAC1 | 2.15786 | LUAD | NCI-H1693 | PCI-34051 |
| MVP_top60_DNA_seed | MET | -0.003056 | LUAD | NCI-H522 | SU11274 | MVP_top60_DNA+RNA_seed | PIK3CA | 2.158579 | LUAD | SW1573 | GNE-317 |
| MVP_top60_DNA_seed | AURKA | -0.002764 | LUAD | HOP-62 | GSK1070916 | MVP_top60_DNA+RNA_seed | BRAF | 2.159053 | LUAD | NCI-H441 | PLX-4720 |
| MVP_top60_DNA_seed | EGFR | -0.002066 | LUAD | NCI-H1792 | Afatinib | MVP_top60_DNA+RNA_seed | AKT1 | 2.162631 | LUAD | NCI-H2405 | Uprosertib |
| MVP_top60_DNA_seed | ERBB2 | -0.002066 | LUAD | NCI-H1792 | Afatinib | MVP_top60_DNA+RNA_seed | AKT1 | 2.162741 | LUAD | SK-LU-1 | AT13148 |
| MVP_top60_DNA_seed | HDAC1 | -0.001696 | LUAD | NCI-H2347 | Belinostat | MVP_top60_DNA+RNA_seed | AKT1 | 2.17089 | LUAD | SK-LU-1 | Ipatasertib |
| MVP_top60_DNA_seed | ATM | -0.001618 | LUAD | Calu-3 | CP466722 | MVP_top60_DNA+RNA_seed | AKT1 | 2.174733 | LUAD | SW1573 | AT13148 |
| MVP_top60_DNA_seed | SRC | -0.001381 | LUAD | SW1573 | Bosutinib | MVP_top60_DNA+RNA_seed | MDM2 | 2.179423 | LUAD | NCI-H2405 | Serdemetan |
| MVP_top60_DNA_seed | EGFR | -0.000333 | LUAD | EMC-BAC-1 | AST-1306 | MVP_top60_DNA+RNA_seed | BRAF | 2.182664 | LUAD | NCI-H441 | SB590885 |
| MVP_top60_DNA_seed | ATM | 0.000401 | LUAD | PC-14 | KU-55933 | MVP_top60_DNA+RNA_seed | SMARCA4 | 2.183084 | LUAD | NCI-H2291 | PFI3 |
| MVP_top60_DNA_seed | AKT1 | 0.000643 | LUAD | NCI-H1944 | BAY AKT1 | MVP_top60_DNA+RNA_seed | AURKA | 2.183404 | LUAD | NCI-H1793 | ZM447439 |
| MVP_top60_DNA_seed | IGF1R | 0.001576 | LUAD | SK-LU-1 | GSK1904529A | MVP_top60_DNA+RNA_seed | AKT1 | 2.191459 | LUAD | SK-LU-1 | Uprosertib |
| MVP_top60_DNA_seed | ERBB2 | 0.001765 | LUAD | NCI-H3122 | CP724714 | MVP_top60_DNA+RNA_seed | IGF1R | 2.19161 | LUAD | COR-L105 | GSK1904529A |
| MVP_top60_DNA_seed | FGFR1 | 0.002597 | LUAD | NCI-H1792 | FGFR_3831 | MVP_top60_DNA+RNA_seed | SMARCA4 | 2.192307 | LUAD | NCI-H1693 | PFI3 |
| MVP_top60_DNA_seed | HDAC1 | 0.002935 | LUAD | NCI-H2009 | Dacinostat | MVP_top60_DNA+RNA_seed | HSP90AA1 | 2.19465 | LUAD | NCI-H1573 | Luminespib |
| MVP_top60_DNA_seed | SRC | 0.003275 | LUAD | NCI-H3122 | NVP-BHG712 | MVP_top60_DNA+RNA_seed | BRAF | 2.195264 | LUAD | NCI-H1568 | PLX-4720 |
| MVP_top60_DNA_seed | HDAC1 | 0.004402 | LUAD | NCI-H1648 | Tubastatin A | MVP_top60_DNA+RNA_seed | EZH2 | 2.199728 | LUAD | H3255 | GSK343 |
| MVP_top60_DNA_seed | ATM | 0.004544 | LUAD | NCI-H1792 | KU-55933 | MVP_top60_DNA+RNA_seed | KRAS | 2.199916 | LUAD | SK-LU-1 | KRAS (G12C) Inhibitor-1 |
| MVP_top60_DNA_seed | MAPK1 | 0.004899 | LUAD | NCI-H1755 | FR-180204 | MVP_top60_DNA+RNA_seed | BRAF | 2.201851 | LUAD | NCI-H2291 | SB590885 |
| MVP_top60_DNA_seed | FGFR1 | 0.004972 | LUAD | NCI-H3122 | FGFR_3831 | MVP_top60_DNA+RNA_seed | AKT1 | 2.202921 | LUAD | NCI-H2347 | Afuresertib |
| MVP_top60_DNA_seed | MET | 0.0058 | LUAD | RERF-LC-KJ | AZD6094 | MVP_top60_DNA+RNA_seed | EGFR | 2.204569 | LUAD | NCI-H2347 | Gefitinib |
| MVP_top60_DNA_seed | GSK3B | 0.006121 | LUAD | ABC-1 | KIN001-042 | MVP_top60_DNA+RNA_seed | AURKA | 2.214435 | LUAD | Calu-3 | ZM447439 |
| MVP_top60_DNA_seed | IGF1R | 0.006282 | LUAD | SW1573 | Linsitinib | MVP_top60_DNA+RNA_seed | AKT1 | 2.231771 | LUAD | NCI-H2347 | AT13148 |
| MVP_top60_DNA_seed | AKT1 | 0.006813 | LUAD | HOP-62 | AKT inhibitor VIII | MVP_top60_DNA+RNA_seed | FGFR1 | 2.241299 | LUAD | NCI-H1838 | PD173074 |
| MVP_top60_DNA_seed | HDAC1 | 0.007332 | LUAD | NCI-H650 | AR-42 | MVP_top60_DNA+RNA_seed | HSP90AA1 | 2.241358 | LUAD | VMRC-LCD | Tanespimycin |
| MVP_top60_DNA_seed | EGFR | 0.007981 | LUAD | NCI-H1573 | Gefitinib | MVP_top60_DNA+RNA_seed | EZH2 | 2.244574 | LUAD | NCI-H1693 | GSK343 |
| MVP_top60_DNA_seed | HDAC1 | 0.008029 | LUAD | NCI-H1651 | Belinostat | MVP_top60_DNA+RNA_seed | HDAC1 | 2.250008 | LUAD | NCI-H1693 | Romidepsin |
| MVP_top60_DNA_seed | ATM | 0.008962 | LUAD | VMRC-LCD | KU-60019 | MVP_top60_DNA+RNA_seed | EGFR | 2.283491 | LUAD | SK-LU-1 | Osimertinib |
| MVP_top60_DNA_seed | AURKA | 0.010628 | LUAD | NCI-H1568 | GSK1070916 | MVP_top60_DNA+RNA_seed | SRC | 2.288796 | LUAD | NCI-H441 | Bosutinib |
| MVP_top60_DNA_seed | HDAC1 | 0.011024 | LUAD | COR-L105 | Dacinostat | MVP_top60_DNA+RNA_seed | EZH2 | 2.292879 | LUAD | EKVX | GSK343 |
| MVP_top60_DNA_seed | AKT1 | 0.011316 | LUAD | NCI-H2347 | BAY AKT1 | MVP_top60_DNA+RNA_seed | HSP90AA1 | 2.305945 | LUAD | ABC-1 | Tanespimycin |
| MVP_top60_DNA_seed | HDAC1 | 0.0115 | LUAD | ABC-1 | Belinostat | MVP_top60_DNA+RNA_seed | PIK3CA | 2.306774 | LUAD | NCI-H1435 | Buparlisib |
| MVP_top60_DNA_seed | AKT1 | 0.012084 | LUAD | NCI-H1573 | AKT inhibitor VIII | MVP_top60_DNA+RNA_seed | PIK3CA | 2.318618 | LUAD | Calu-3 | AZD8186 |
| MVP_top60_DNA_seed | EGFR | 0.012487 | LUAD | Calu-6 | Afatinib | MVP_top60_DNA+RNA_seed | HSP90AA1 | 2.318943 | LUAD | COR-L105 | Tanespimycin |
| MVP_top60_DNA_seed | ERBB2 | 0.012487 | LUAD | Calu-6 | Afatinib | MVP_top60_DNA+RNA_seed | EGFR | 2.319069 | LUAD | SW1573 | Osimertinib |
| MVP_top60_DNA_seed | EGFR | 0.01318 | LUAD | NCI-H1792 | PF-00299804 | MVP_top60_DNA+RNA_seed | BRAF | 2.324521 | LUAD | NCI-H1693 | PLX-4720 |
| MVP_top60_DNA_seed | ERBB2 | 0.01318 | LUAD | NCI-H1792 | PF-00299804 | MVP_top60_DNA+RNA_seed | AKT1 | 2.330594 | LUAD | VMRC-LCD | Afuresertib |
| MVP_top60_DNA_seed | FGFR1 | 0.014596 | LUAD | NCI-H1568 | AZD4547 | MVP_top60_DNA+RNA_seed | HDAC1 | 2.335598 | LUAD | NCI-H2405 | PCI-34051 |
| MVP_top60_DNA_seed | HSP90AA1 | 0.015087 | LUAD | NCI-H2342 | Elesclomol | MVP_top60_DNA+RNA_seed | PIK3CA | 2.355811 | LUAD | Calu-6 | AZD8186 |

| | | | | | | | | | | | |
|---|---|---|---|---|---|---|---|---|---|---|---|
| MVP_top60_DNA_seed | ATM | 0.015879 | LUAD | NCI-H358 | KU-60019 | MVP_top60_DNA+RNA_seed | AKT1 | 2.398702 | LUAD | NCI-H2087 | AZD5363 |
| MVP_top60_DNA_seed | GSK3B | 0.016706 | LUAD | NCI-H1755 | SB216763 | MVP_top60_DNA+RNA_seed | HSP90AA1 | 2.414374 | LUAD | NCI-H1793 | Luminespib |
| MVP_top60_DNA_seed | MDM2 | 0.020238 | LUAD | NCI-H1437 | Nutlin-3a (-) | MVP_top60_DNA+RNA_seed | PIK3CA | 2.419897 | LUAD | NCI-H441 | Buparlisib |
| MVP_top60_DNA_seed | PIK3CA | 0.021332 | LUAD | NCI-H2085 | AZD8835 | MVP_top60_DNA+RNA_seed | MDM2 | 2.421773 | LUAD | NCI-H1650 | Serdemetan |
| MVP_top60_DNA_seed | PIK3CA | 0.022361 | LUAD | RERF-LC-MS | PI3Ka_4409 | MVP_top60_DNA+RNA_seed | AKT1 | 2.442261 | LUAD | NCI-H1793 | Afuresertib |
| MVP_top60_DNA_seed | EGFR | 0.022476 | LUAD | NCI-H2347 | Cetuximab | MVP_top60_DNA+RNA_seed | AKT1 | 2.443442 | LUAD | Calu-6 | AZD5363 |
| MVP_top60_DNA_seed | HSP90AA1 | 0.022896 | LUAD | EMC-BAC-1 | SNX-2112 | MVP_top60_DNA+RNA_seed | HSP90AA1 | 2.446973 | LUAD | NCI-H2291 | Elesclomol |
| MVP_top60_DNA_seed | EGFR | 0.023214 | LUAD | A549 | Afatinib | MVP_top60_DNA+RNA_seed | AKT1 | 2.451452 | LUAD | NCI-H2405 | MK-2206 |
| MVP_top60_DNA_seed | ERBB2 | 0.023214 | LUAD | A549 | Afatinib | MVP_top60_DNA+RNA_seed | AKT1 | 2.458361 | LUAD | SK-LU-1 | MK-2206 |
| MVP_top60_DNA_seed | AURKA | 0.023434 | LUAD | SK-LU-1 | ZM447439 | MVP_top60_DNA+RNA_seed | IGF1R | 2.458857 | LUAD | NCI-H2291 | GSK1904529A |
| MVP_top60_DNA_seed | HSP90AA1 | 0.02429 | LUAD | NCI-H1623 | CCT-018159 | MVP_top60_DNA+RNA_seed | TP53 | 2.464529 | LUAD | NCI-H2405 | MIRA-1 |
| MVP_top60_DNA_seed | EGFR | 0.024668 | LUAD | NCI-H650 | Pelitinib | MVP_top60_DNA+RNA_seed | MET | 2.474652 | LUAD | NCI-H2347 | Crizotinib |
| MVP_top60_DNA_seed | HSP90AA1 | 0.024737 | LUAD | NCI-H1734 | Elesclomol | MVP_top60_DNA+RNA_seed | FGFR1 | 2.491065 | LUAD | EKVX | AZD4547 |
| MVP_top60_DNA_seed | EGFR | 0.026267 | LUAD | LC-2-ad | CI-1033 | MVP_top60_DNA+RNA_seed | HDAC1 | 2.497733 | LUAD | NCI-H2405 | Romidepsin |
| MVP_top60_DNA_seed | ERBB2 | 0.026267 | LUAD | LC-2-ad | CI-1033 | MVP_top60_DNA+RNA_seed | PIK3CA | 2.502359 | LUAD | NCI-H522 | GNE-317 |
| MVP_top60_DNA_seed | MET | 0.028147 | LUAD | NCI-H3122 | JNJ38877605 | MVP_top60_DNA+RNA_seed | HSP90AA1 | 2.510571 | LUAD | NCI-H1623 | Luminespib |
| MVP_top60_DNA_seed | GSK3B | 0.028776 | LUAD | NCI-H2023 | AZD7969 | MVP_top60_DNA+RNA_seed | AKT1 | 2.519148 | LUAD | NCI-H2347 | MK-2206 |
| MVP_top60_DNA_seed | ERBB2 | 0.029045 | LUAD | HCC-44 | CP724714 | MVP_top60_DNA+RNA_seed | EGFR | 2.519453 | LUAD | NCI-H2405 | AZD3759 |
| MVP_top60_DNA_seed | AURKA | 0.030073 | LUAD | NCI-H1975 | HG-5-113-01 | MVP_top60_DNA+RNA_seed | MET | 2.565729 | LUAD | NCI-H2405 | Crizotinib |
| MVP_top60_DNA_seed | BRAF | 0.030821 | LUAD | SW1573 | Dabrafenib | MVP_top60_DNA+RNA_seed | EZH2 | 2.595206 | LUAD | NCI-H2405 | GSK343 |
| MVP_top60_DNA_seed | HSP90AA1 | 0.031034 | LUAD | NCI-H2023 | Elesclomol | MVP_top60_DNA+RNA_seed | KRAS | 2.603151 | LUAD | NCI-H441 | RAS (G12C) Inhibitor-1 |
| MVP_top60_DNA_seed | BRAF | 0.031066 | LUAD | EKVX | PLX-4720 | MVP_top60_DNA+RNA_seed | ATM | 2.609157 | LUAD | NCI-H2405 | KU-55933 |
| MVP_top60_DNA_seed | IGF1R | 0.031382 | LUAD | PC-14 | GSK1904529A | MVP_top60_DNA+RNA_seed | AKT1 | 2.616957 | LUAD | NCI-H441 | AZD5363 |
| MVP_top60_DNA_seed | AKT1 | 0.031511 | LUAD | NCI-H1781 | AKT inhibitor VIII | MVP_top60_DNA+RNA_seed | HSP90AA1 | 2.62834 | LUAD | NCI-H1435 | Luminespib |
| MVP_top60_DNA_seed | HDAC1 | 0.032313 | LUAD | NCI-H2030 | CAY10603 | MVP_top60_DNA+RNA_seed | ATM | 2.628708 | LUAD | NCI-H441 | KU-55933 |
| MVP_top60_DNA_seed | ATM | 0.03253 | LUAD | NCI-H1975 | CP466722 | MVP_top60_DNA+RNA_seed | HSP90AA1 | 2.632342 | LUAD | NCI-H2342 | Luminespib |
| MVP_top60_DNA_seed | AKT1 | 0.032543 | LUAD | NCI-H838 | BAY AKT1 | MVP_top60_DNA+RNA_seed | EGFR | 2.654864 | LUAD | NCI-H2347 | Lapatinib |
| MVP_top60_DNA_seed | EGFR | 0.033544 | LUAD | NCI-H2405 | Pelitinib | MVP_top60_DNA+RNA_seed | PIK3CA | 2.656799 | LUAD | NCI-H2347 | Alpelisib |
| MVP_top60_DNA_seed | BRAF | 0.033563 | LUAD | LC-2-ad | SB590885 | MVP_top60_DNA+RNA_seed | MDM2 | 2.718022 | LUAD | NCI-H1793 | Serdemetan |
| MVP_top60_DNA_seed | BRAF | 0.034241 | LUAD | HOP-62 | PLX-4720 | MVP_top60_DNA+RNA_seed | SMARCA4 | 2.718226 | LUAD | NCI-H441 | PFI3 |
| MVP_top60_DNA_seed | EGFR | 0.034841 | LUAD | NCI-H2347 | AST-1306 | MVP_top60_DNA+RNA_seed | IGF1R | 2.722961 | LUAD | NCI-H2347 | BMS-754807 |
| MVP_top60_DNA_seed | BRAF | 0.035379 | LUAD | NCI-H838 | HG6-64-1 | MVP_top60_DNA+RNA_seed | AKT1 | 2.742815 | LUAD | NCI-H2405 | Afuresertib |
| MVP_top60_DNA_seed | GSK3B | 0.03618 | LUAD | NCI-H2085 | CHIR-99021 | MVP_top60_DNA+RNA_seed | AKT1 | 2.749456 | LUAD | SK-LU-1 | AZD5363 |
| MVP_top60_DNA_seed | HDAC1 | 0.037451 | LUAD | HCC-827 | JQ12 | MVP_top60_DNA+RNA_seed | HDAC1 | 2.764612 | LUAD | NCI-H1650 | Entinostat |
| MVP_top60_DNA_seed | HSP90AA1 | 0.037625 | LUAD | NCI-H1573 | CCT-018159 | MVP_top60_DNA+RNA_seed | IGF1R | 2.784329 | LUAD | VMRC-LCD | BMS-536924 |
| MVP_top60_DNA_seed | BRAF | 0.038252 | LUAD | NCI-H441 | RAF_9304 | MVP_top60_DNA+RNA_seed | EGFR | 2.817442 | LUAD | SW1573 | Lapatinib |
| MVP_top60_DNA_seed | HSP90AA1 | 0.039488 | LUAD | NCI-H1573 | Tanespimycin | MVP_top60_DNA+RNA_seed | MET | 2.819847 | LUAD | NCI-H1793 | Crizotinib |
| MVP_top60_DNA_seed | ATM | 0.039839 | LUAD | NCI-H2347 | KU-55933 | MVP_top60_DNA+RNA_seed | AKT1 | 2.831554 | LUAD | NCI-H441 | AT13148 |
| MVP_top60_DNA_seed | HDAC1 | 0.039956 | LUAD | NCI-H2030 | Tubastatin A | MVP_top60_DNA+RNA_seed | AURKA | 2.839326 | LUAD | NCI-H2405 | ZM447439 |
| MVP_top60_DNA_seed | EGFR | 0.040419 | LUAD | NCI-H23 | AZD8931 | MVP_top60_DNA+RNA_seed | MET | 2.878976 | LUAD | EKVX | Crizotinib |
| MVP_top60_DNA_seed | ERBB2 | 0.040419 | LUAD | NCI-H23 | AZD8931 | MVP_top60_DNA+RNA_seed | PIK3CA | 2.886868 | LUAD | NCI-H2405 | Alpelisib |

| | | | | | | | | | | | |
|---|---|---|---|---|---|---|---|---|---|---|---|
| MVP_top60_DNA_seed | ERBB3 | 0.040419 | LUAD | NCI-H23 | AZD8931 | MVP_top60_DNA+RNA_seed | HDAC1 | 2.9272 | LUAD | SK-LU-1 | Entinostat |
| MVP_top60_DNA_seed | GSK3B | 0.041681 | LUAD | PC-3_[JPC-3] | KIN001-042 | MVP_top60_DNA+RNA_seed | EGFR | 2.935743 | LUAD | NCI-H2405 | Lapatinib |
| MVP_top60_DNA_seed | GSK3B | 0.04264 | LUAD | NCI-H3122 | CHIR-99021 | MVP_top60_DNA+RNA_seed | HSP90AA1 | 2.980072 | LUAD | NCI-H441 | Tanespimycin |
| MVP_top60_DNA_seed | AKT1 | 0.042646 | LUAD | NCI-H2085 | BAY AKT1 | MVP_top60_DNA+RNA_seed | EGFR | 2.992514 | LUAD | NCI-H441 | Osimertinib |
| MVP_top60_DNA_seed | IGF1R | 0.042749 | LUAD | LXF-289 | GSK1904529A | MVP_top60_DNA+RNA_seed | TERT | 3.04853 | LUAD | NCI-H441 | BIBR-1532 |
| MVP_top60_DNA_seed | SRC | 0.043018 | LUAD | NCI-H1563 | NVP-BHG712 | MVP_top60_DNA+RNA_seed | AKT1 | 3.107433 | LUAD | SK-LU-1 | Afuresertib |
| MVP_top60_DNA_seed | FGFR1 | 0.043029 | LUAD | NCI-H1792 | AZD4547 | MVP_top60_DNA+RNA_seed | SMARCA4 | 3.153384 | LUAD | NCI-H2405 | PFI3 |
| MVP_top60_DNA_seed | BRAF | 0.043922 | LUAD | COR-L105 | RAF_9304 | MVP_top60_DNA+RNA_seed | PIK3CA | 3.16743 | LUAD | NCI-H441 | GNE-317 |
| MVP_top60_DNA_seed | HDAC1 | 0.044605 | LUAD | NCI-H1355 | Entinostat | MVP_top60_DNA+RNA_seed | EGFR | 3.197082 | LUAD | NCI-H2405 | Afatinib |
| MVP_top60_DNA_seed | ATM | 0.045103 | LUAD | NCI-H3122 | KU-60019 | MVP_top60_DNA+RNA_seed | AURKA | 3.250564 | LUAD | NCI-H441 | ZM447439 |
| MVP_top60_DNA_seed | EGFR | 0.04546 | LUAD | NCI-H2347 | CUDC-101 | MVP_top60_DNA+RNA_seed | IGF1R | 3.281784 | LUAD | NCI-H441 | GSK1904529A |
| MVP_top60_DNA_seed | ERBB2 | 0.04546 | LUAD | NCI-H2347 | CUDC-101 | MVP_top60_DNA+RNA_seed | IGF1R | 3.283733 | LUAD | NCI-H2405 | NVP-ADW742 |
| MVP_top60_DNA_seed | EGFR | 0.046171 | LUAD | Calu-3 | CUDC-101 | MVP_top60_DNA+RNA_seed | HDAC1 | 3.292349 | LUAD | NCI-H1793 | Entinostat |
| MVP_top60_DNA_seed | ERBB2 | 0.046171 | LUAD | Calu-3 | CUDC-101 | MVP_top60_DNA+RNA_seed | HSP90AA1 | 3.313202 | LUAD | NCI-H441 | Luminespib |
| MVP_top60_DNA_seed | HSP90AA1 | 0.04695 | LUAD | NCI-H522 | Tanespimycin | MVP_top60_DNA+RNA_seed | PIK3CA | 3.323685 | LUAD | NCI-H1838 | Buparlisib |
| MVP_top60_DNA_seed | BRAF | 0.047228 | LUAD | NCI-H2030 | Dabrafenib | MVP_top60_DNA+RNA_seed | HSP90AA1 | 3.325474 | LUAD | NCI-H1838 | Luminespib |
| MVP_top60_DNA_seed | MAPK1 | 0.048234 | LUAD | NCI-H2030 | VX-11e | MVP_top60_DNA+RNA_seed | HDAC1 | 3.570933 | LUAD | NCI-H1650 | Romidepsin |
| MVP_top60_DNA_seed | ERBB2 | 0.048955 | LUAD | NCI-H2342 | CP724714 | MVP_top60_DNA+RNA_seed | HDAC1 | 3.572014 | LUAD | NCI-H1793 | Romidepsin |
| MVP_top60_DNA_seed | BRAF | 0.050009 | LUAD | PC-14 | SB590885 | MVP_top60_DNA+RNA_seed | HSP90AA1 | 4.016577 | LUAD | NCI-H2347 | Luminespib |
| MVP_top60_DNA_seed | ATM | 0.050298 | LUAD | H3255 | CP466722 | MVP_top60_DNA+RNA_seed | SMARCA4 | -2.471329 | LUSC | EBC-1 | PFI-3 |
| MVP_top60_DNA_seed | BRAF | 0.052559 | LUAD | EMC-BAC-1 | PLX-4720 | MVP_top60_DNA+RNA_seed | SMARCA2 | -2.471329 | LUSC | EBC-1 | PFI-3 |
| MVP_top60_DNA_seed | HDAC1 | 0.052975 | LUAD | NCI-H441 | Dacinostat | MVP_top60_DNA+RNA_seed | AR | -2.463165 | LUSC | EBC-1 | Bicalutamide |
| MVP_top60_DNA_seed | IGF1R | 0.053062 | LUAD | HCC-827 | BMS-536924 | MVP_top60_DNA+RNA_seed | CREBBP | -2.289563 | LUSC | HARA | I-CBP112 |
| MVP_top60_DNA_seed | MET | 0.054235 | LUAD | NCI-H1666 | PHA-665752 | MVP_top60_DNA+RNA_seed | AR | -2.182747 | LUSC | KNS-62 | AZD3514 |
| MVP_top60_DNA_seed | SRC | 0.055716 | LUAD | NCI-H838 | Bosutinib | MVP_top60_DNA+RNA_seed | AR | -2.102598 | LUSC | EBC-1 | AZD3514 |
| MVP_top60_DNA_seed | FGFR1 | 0.056761 | LUAD | NCI-H1651 | AZD4547 | MVP_top60_DNA+RNA_seed | HSP90AB1 | -2.05752 | LUSC | EBC-1 | Tanespimycin |
| MVP_top60_DNA_seed | GSK3B | 0.056881 | LUAD | NCI-H23 | CHIR-99021 | MVP_top60_DNA+RNA_seed | HDAC1 | -1.700261 | LUSC | EBC-1 | JQ12 |
| MVP_top60_DNA_seed | AKT1 | 0.057221 | LUAD | NCI-H1650 | AKT inhibitor VIII | MVP_top60_DNA+RNA_seed | PIK3CA | -1.665707 | LUSC | SK-MES-1 | PI3Ka_4409 |
| MVP_top60_DNA_seed | MET | 0.057489 | LUAD | RERF-LC-KJ | JNJ38877605 | MVP_top60_DNA+RNA_seed | PIK3CA | -1.469562 | LUSC | RERF-LC-Sq1 | PI3Ka_4409 |
| MVP_top60_DNA_seed | BRAF | 0.058446 | LUAD | NCI-H23 | Dabrafenib | MVP_top60_DNA+RNA_seed | HSP90AB1 | -1.446245 | LUSC | SK-MES-1 | Tanespimycin |
| MVP_top60_DNA_seed | PIK3CA | 0.058583 | LUAD | NCI-H1993 | PI3Ka_4409 | MVP_top60_DNA+RNA_seed | PIK3CA | -1.37393 | LUSC | SK-MES-1 | AZD8835 |
| MVP_top60_DNA_seed | AKT1 | 0.059093 | LUAD | NCI-H322M | AKT inhibitor VIII | MVP_top60_DNA+RNA_seed | HSP90AB1 | -1.349088 | LUSC | LOU-NH91 | Tanespimycin |
| MVP_top60_DNA_seed | HSP90AA1 | 0.059208 | LUAD | NCI-H1563 | Tanespimycin | MVP_top60_DNA+RNA_seed | HDAC1 | -1.276467 | LUSC | LK-2 | JQ12 |
| MVP_top60_DNA_seed | ATM | 0.060047 | LUAD | NCI-H3122 | CP466722 | MVP_top60_DNA+RNA_seed | HSP90AB1 | -1.153069 | LUSC | EPLC-272H | Tanespimycin |
| MVP_top60_DNA_seed | FGFR1 | 0.060867 | LUAD | NCI-H2085 | PD173074 | MVP_top60_DNA+RNA_seed | AR | -1.13906 | LUSC | LK-2 | AZD3514 |
| MVP_top60_DNA_seed | FGFR1 | 0.061006 | LUAD | VMRC-LCD | AZD4547 | MVP_top60_DNA+RNA_seed | HSP90AB1 | -1.117973 | LUSC | SK-MES-1 | Luminespib |
| MVP_top60_DNA_seed | IGF1R | 0.061168 | LUAD | NCI-H1755 | BMS-754807 | MVP_top60_DNA+RNA_seed | HDAC1 | -1.110672 | LUSC | LK-2 | Dacinostat |
| MVP_top60_DNA_seed | MET | 0.062196 | LUAD | NCI-H1651 | SU11274 | MVP_top60_DNA+RNA_seed | HDAC1 | -1.101418 | LUSC | EPLC-272H | Tubastatin A |
| MVP_top60_DNA_seed | MDM2 | 0.062939 | LUAD | HCC-827 | Serdemetan | MVP_top60_DNA+RNA_seed | PIK3CA | -1.09166 | LUSC | RERF-LC-Sq1 | AZD8835 |
| MVP_top60_DNA_seed | EGFR | 0.064074 | LUAD | Calu-6 | Pelitinib | MVP_top60_DNA+RNA_seed | PIK3CA | -1.001354 | LUSC | EPLC-272H | PI3Ka_4409 |
| MVP_top60_DNA_seed | AURKA | 0.065749 | LUAD | Calu-6 | CD532 | MVP_top60_DNA+RNA_seed | HSP90AB1 | -0.979413 | LUSC | KNS-62 | Elesclomol |

| | | | | | | | | | | | |
|---|---|---|---|---|---|---|---|---|---|---|---|
| MVP_top60_DNA_seed | MET | 0.0661 | LUAD | NCI-H2009 | AZD6094 | MVP_top60_DNA+RNA_seed | HSP90AB1 | -0.953538 | LUSC | EPLC-272H | SNX-2112 |
| MVP_top60_DNA_seed | AURKA | 0.066318 | LUAD | NCI-H1975 | Tozasertib | MVP_top60_DNA+RNA_seed | HSP90AB1 | -0.952862 | LUSC | NCI-H2170 | Tanespimycin |
| MVP_top60_DNA_seed | MDM2 | 0.067187 | LUAD | NCI-H1975 | Serdemetan | MVP_top60_DNA+RNA_seed | HDAC1 | -0.952447 | LUSC | EPLC-272H | CAY10603 |
| MVP_top60_DNA_seed | GSK3B | 0.067437 | LUAD | NCI-H1563 | KIN001-042 | MVP_top60_DNA+RNA_seed | AR | -0.941026 | LUSC | SK-MES-1 | Bicalutamide |
| MVP_top60_DNA_seed | AKT1 | 0.068159 | LUAD | LC-2-ad | BAY AKT1 | MVP_top60_DNA+RNA_seed | HSP90AB1 | -0.928431 | LUSC | HCC-15 | Tanespimycin |
| MVP_top60_DNA_seed | SRC | 0.068545 | LUAD | NCI-H2342 | Bosutinib | MVP_top60_DNA+RNA_seed | HDAC1 | -0.920538 | LUSC | EBC-1 | Dacinostat |
| MVP_top60_DNA_seed | MDM2 | 0.068805 | LUAD | HCC-44 | Nutlin-3a (-) | MVP_top60_DNA+RNA_seed | SMARCA4 | -0.913857 | LUSC | KNS-62 | PFI-3 |
| MVP_top60_DNA_seed | EGFR | 0.068918 | LUAD | Calu-3 | Gefitinib | MVP_top60_DNA+RNA_seed | SMARCA2 | -0.913857 | LUSC | KNS-62 | PFI-3 |
| MVP_top60_DNA_seed | GSK3B | 0.068972 | LUAD | NCI-H23 | CHIR-99021 | MVP_top60_DNA+RNA_seed | HSP90AB1 | -0.908413 | LUSC | EPLC-272H | CCT-018159 |
| MVP_top60_DNA_seed | BRAF | 0.068992 | LUAD | NCI-H2009 | RAF_9304 | MVP_top60_DNA+RNA_seed | HSP90AB1 | -0.843278 | LUSC | NCI-H2170 | CCT-018159 |
| MVP_top60_DNA_seed | HDAC1 | 0.069424 | LUAD | RERF-LC-MS | CAY10603 | MVP_top60_DNA+RNA_seed | PIK3CA | -0.834848 | LUSC | EPLC-272H | AZD8835 |
| MVP_top60_DNA_seed | AKT1 | 0.070384 | LUAD | EMC-BAC-1 | MK-2206 | MVP_top60_DNA+RNA_seed | HSP90AB1 | -0.816899 | LUSC | HARA | Tanespimycin |
| MVP_top60_DNA_seed | FGFR1 | 0.070647 | LUAD | NCI-H2405 | AZD4547 | MVP_top60_DNA+RNA_seed | AR | -0.815814 | LUSC | EPLC-272H | AZD3514 |
| MVP_top60_DNA_seed | MET | 0.070683 | LUAD | SW1573 | JNJ38877605 | MVP_top60_DNA+RNA_seed | HDAC1 | -0.814045 | LUSC | EPLC-272H | Belinostat |
| MVP_top60_DNA_seed | ATM | 0.07131 | LUAD | NCI-H1693 | CP466722 | MVP_top60_DNA+RNA_seed | PIK3CA | -0.809777 | LUSC | LOU-NH91 | AZD8835 |
| MVP_top60_DNA_seed | EGFR | 0.071393 | LUAD | EMC-BAC-2 | Afatinib | MVP_top60_DNA+RNA_seed | HSP90AB1 | -0.798116 | LUSC | NCI-H226 | SNX-2112 |
| MVP_top60_DNA_seed | ERBB2 | 0.071393 | LUAD | EMC-BAC-2 | Afatinib | MVP_top60_DNA+RNA_seed | HSP90AB1 | -0.789156 | LUSC | EBC-1 | CCT-018159 |
| MVP_top60_DNA_seed | AURKA | 0.071721 | LUAD | NCI-H1975 | GSK1070916 | MVP_top60_DNA+RNA_seed | HSP90AB1 | -0.760525 | LUSC | HARA | Elesclomol |
| MVP_top60_DNA_seed | SMARCA4 | 0.071758 | LUAD | NCI-H1435 | PFI-3 | MVP_top60_DNA+RNA_seed | CREBBP | -0.75393 | LUSC | LC-1-sq | I-CBP112 |
| MVP_top60_DNA_seed | AKT1 | 0.07226 | LUAD | EKVX | AKT inhibitor VIII | MVP_top60_DNA+RNA_seed | CREBBP | -0.716771 | LUSC | NCI-H226 | I-CBP112 |
| MVP_top60_DNA_seed | ATM | 0.073309 | LUAD | NCI-H1944 | KU-55933 | MVP_top60_DNA+RNA_seed | HSP90AB1 | -0.694522 | LUSC | NCI-H1869 | Tanespimycin |
| MVP_top60_DNA_seed | AURKA | 0.07333 | LUAD | NCI-H23 | GSK1070916 | MVP_top60_DNA+RNA_seed | HDAC1 | -0.673209 | LUSC | EPLC-272H | AR-42 |
| MVP_top60_DNA_seed | MET | 0.073767 | LUAD | HOP-62 | AZD6094 | MVP_top60_DNA+RNA_seed | HDAC1 | -0.64373 | LUSC | NCI-H520 | CAY10603 |
| MVP_top60_DNA_seed | EGFR | 0.074843 | LUAD | NCI-H1650 | Afatinib | MVP_top60_DNA+RNA_seed | HDAC1 | -0.641588 | LUSC | HCC-15 | JQ12 |
| MVP_top60_DNA_seed | ERBB2 | 0.074843 | LUAD | NCI-H1650 | Afatinib | MVP_top60_DNA+RNA_seed | HSP90AB1 | -0.636862 | LUSC | EBC-1 | Luminespib |
| MVP_top60_DNA_seed | ATM | 0.075005 | LUAD | NCI-H1793 | CP466722 | MVP_top60_DNA+RNA_seed | PIK3CA | -0.633699 | LUSC | EBC-1 | PI3Ka_4409 |
| MVP_top60_DNA_seed | HDAC1 | 0.075354 | LUAD | NCI-H2030 | Belinostat | MVP_top60_DNA+RNA_seed | HSP90AB1 | -0.603016 | LUSC | HARA | SNX-2112 |
| MVP_top60_DNA_seed | IGF1R | 0.07543 | LUAD | NCI-H1651 | GSK1904529A | MVP_top60_DNA+RNA_seed | HSP90AB1 | -0.599502 | LUSC | NCI-H520 | SNX-2112 |
| MVP_top60_DNA_seed | HSP90AA1 | 0.076717 | LUAD | RERF-LC-KJ | Luminespib | MVP_top60_DNA+RNA_seed | HDAC1 | -0.596337 | LUSC | NCI-H520 | Tubastatin A |
| MVP_top60_DNA_seed | FGFR1 | 0.077484 | LUAD | NCI-H358 | PD173074 | MVP_top60_DNA+RNA_seed | HSP90AB1 | -0.56938 | LUSC | HARA | CCT-018159 |
| MVP_top60_DNA_seed | HDAC1 | 0.078824 | LUAD | NCI-H838 | AR-42 | MVP_top60_DNA+RNA_seed | SMARCA4 | -0.5672 | LUSC | HARA | PFI-3 |
| MVP_top60_DNA_seed | ATM | 0.079111 | LUAD | NCI-H23 | KU-60019 | MVP_top60_DNA+RNA_seed | SMARCA2 | -0.5672 | LUSC | HARA | PFI-3 |
| MVP_top60_DNA_seed | EGFR | 0.079141 | LUAD | NCI-H3122 | Afatinib | MVP_top60_DNA+RNA_seed | HDAC1 | -0.543127 | LUSC | NCI-H2170 | JQ12 |
| MVP_top60_DNA_seed | ERBB2 | 0.079141 | LUAD | NCI-H3122 | Afatinib | MVP_top60_DNA+RNA_seed | HDAC1 | -0.529467 | LUSC | LK-2 | Belinostat |
| MVP_top60_DNA_seed | AURKA | 0.079601 | LUAD | NCI-H2347 | HG-5-113-01 | MVP_top60_DNA+RNA_seed | HSP90AB1 | -0.523936 | LUSC | NCI-H2170 | Elesclomol |
| MVP_top60_DNA_seed | HDAC1 | 0.080527 | LUAD | NCI-H1651 | CAY10603 | MVP_top60_DNA+RNA_seed | AR | -0.522758 | LUSC | LOU-NH91 | AZD3514 |
| MVP_top60_DNA_seed | PTK2 | 0.080757 | LUAD | NCI-H1437 | PF-562271 | MVP_top60_DNA+RNA_seed | HSP90AB1 | -0.520761 | LUSC | RERF-LC-Sq1 | Tanespimycin |
| MVP_top60_DNA_seed | AKT1 | 0.081894 | LUAD | NCI-H1355 | AKT inhibitor VIII | MVP_top60_DNA+RNA_seed | HSP90AB1 | -0.507348 | LUSC | LK-2 | Tanespimycin |
| MVP_top60_DNA_seed | AKT1 | 0.083347 | LUAD | HCC-827 | AKT inhibitor VIII | MVP_top60_DNA+RNA_seed | SMARCA4 | -0.487779 | LUSC | EPLC-272H | PFI-3 |
| MVP_top60_DNA_seed | BRAF | 0.084937 | LUAD | HCC-44 | PLX-4720 | MVP_top60_DNA+RNA_seed | SMARCA2 | -0.487779 | LUSC | EPLC-272H | PFI-3 |
| MVP_top60_DNA_seed | EGFR | 0.084963 | LUAD | NCI-H1435 | Afatinib | MVP_top60_DNA+RNA_seed | AR | -0.479459 | LUSC | EPLC-272H | Bicalutamide |

| | | | | | | | | | | | |
|---|---|---|---|---|---|---|---|---|---|---|---|
| MVP_top60_DNA_seed | ERBB2 | 0.084963 | LUAD | NCI-H1435 | Afatinib | MVP_top60_DNA+RNA_seed | PIK3CA | -0.455436 | LUSC | KNS-62 | PI3Ka_4409 |
| MVP_top60_DNA_seed | EGFR | 0.085954 | LUAD | NCI-H1573 | AST-1306 | MVP_top60_DNA+RNA_seed | HDAC1 | -0.436315 | LUSC | SK-MES-1 | JQ12 |
| MVP_top60_DNA_seed | EGFR | 0.086348 | LUAD | NCI-H1793 | Gefitinib | MVP_top60_DNA+RNA_seed | HDAC1 | -0.425508 | LUSC | LC-1-sq | Dacinostat |
| MVP_top60_DNA_seed | HSP90AA1 | 0.086513 | LUAD | NCI-H2347 | Luminespib | MVP_top60_DNA+RNA_seed | HSP90AB1 | -0.423257 | LUSC | LK-2 | Elesclomol |
| MVP_top60_DNA_seed | IGF1R | 0.086624 | LUAD | NCI-H2009 | Linsitinib | MVP_top60_DNA+RNA_seed | HDAC1 | -0.416553 | LUSC | NCI-H226 | Belinostat |
| MVP_top60_DNA_seed | AURKA | 0.08672 | LUAD | NCI-H522 | Genentech Cpd 10 | MVP_top60_DNA+RNA_seed | HDAC1 | -0.406793 | LUSC | NCI-H226 | CAY10603 |
| MVP_top60_DNA_seed | HDAC1 | 0.088673 | LUAD | NCI-H2347 | Tubastatin A | MVP_top60_DNA+RNA_seed | HSP90AB1 | -0.402003 | LUSC | SK-MES-1 | CCT-018159 |
| MVP_top60_DNA_seed | ERBB2 | 0.089035 | LUAD | EMC-BAC-2 | CP724714 | MVP_top60_DNA+RNA_seed | HSP90AB1 | -0.388567 | LUSC | NCI-H520 | Luminespib |
| MVP_top60_DNA_seed | GSK3B | 0.089702 | LUAD | NCI-H2347 | AZD7969 | MVP_top60_DNA+RNA_seed | AR | -0.359332 | LUSC | LC-1-sq | AZD3514 |
| MVP_top60_DNA_seed | IGF1R | 0.090597 | LUAD | NCI-H2023 | BMS-754807 | MVP_top60_DNA+RNA_seed | HDAC1 | -0.353983 | LUSC | NCI-H520 | Dacinostat |
| MVP_top60_DNA_seed | GSK3B | 0.092778 | LUAD | NCI-H1623 | AZD7969 | MVP_top60_DNA+RNA_seed | HDAC1 | -0.344443 | LUSC | NCI-H226 | AR-42 |
| MVP_top60_DNA_seed | AURKA | 0.092787 | LUAD | NCI-H1792 | HG-5-113-01 | MVP_top60_DNA+RNA_seed | HDAC1 | -0.341282 | LUSC | NCI-H1869 | Parthenolide |
| MVP_top60_DNA_seed | FGFR1 | 0.094044 | LUAD | NCI-H2030 | FGFR_3831 | MVP_top60_DNA+RNA_seed | HDAC1 | -0.329223 | LUSC | LK-2 | CAY10603 |
| MVP_top60_DNA_seed | BRAF | 0.094367 | LUAD | NCI-H1651 | Dabrafenib | MVP_top60_DNA+RNA_seed | HDAC1 | -0.326793 | LUSC | NCI-H2170 | Dacinostat |
| MVP_top60_DNA_seed | EGFR | 0.094966 | LUAD | RERF-LC-MS | AZD8931 | MVP_top60_DNA+RNA_seed | AR | -0.325215 | LUSC | KNS-62 | Bicalutamide |
| MVP_top60_DNA_seed | ERBB2 | 0.094966 | LUAD | RERF-LC-MS | AZD8931 | MVP_top60_DNA+RNA_seed | HSP90AB1 | -0.324056 | LUSC | NCI-H520 | Tanespimycin |
| MVP_top60_DNA_seed | ERBB3 | 0.094966 | LUAD | RERF-LC-MS | AZD8931 | MVP_top60_DNA+RNA_seed | HSP90AB1 | -0.323072 | LUSC | LOU-NH91 | CCT-018159 |
| MVP_top60_DNA_seed | GSK3B | 0.095427 | LUAD | SK-LU-1 | KIN001-042 | MVP_top60_DNA+RNA_seed | HSP90AB1 | -0.314112 | LUSC | NCI-H226 | Luminespib |
| MVP_top60_DNA_seed | HDAC1 | 0.096712 | LUAD | NCI-H1666 | AR-42 | MVP_top60_DNA+RNA_seed | HDAC1 | -0.301117 | LUSC | LK-2 | AR-42 |
| MVP_top60_DNA_seed | AKT1 | 0.097889 | LUAD | NCI-H3122 | MK-2206 | MVP_top60_DNA+RNA_seed | SMARCA4 | -0.278295 | LUSC | HCC-15 | PFI-3 |
| MVP_top60_DNA_seed | MDM2 | 0.099277 | LUAD | NCI-H2122 | Serdemetan | MVP_top60_DNA+RNA_seed | SMARCA2 | -0.278295 | LUSC | HCC-15 | PFI-3 |
| MVP_top60_DNA_seed | MDM2 | 0.099306 | LUAD | NCI-H358 | Serdemetan | MVP_top60_DNA+RNA_seed | HDAC1 | -0.276912 | LUSC | SK-MES-1 | Dacinostat |
| MVP_top60_DNA_seed | SMARCA4 | 0.099896 | LUAD | NCI-H1975 | PFI-3 | MVP_top60_DNA+RNA_seed | HDAC1 | -0.268619 | LUSC | NCI-H520 | Belinostat |
| MVP_top60_DNA_seed | PIK3CA | 0.100394 | LUAD | NCI-H1792 | PI3Ka_4409 | MVP_top60_DNA+RNA_seed | CREBBP | -0.252649 | LUSC | EBC-1 | I-CBP112 |
| MVP_top60_DNA_seed | BRAF | 0.101724 | LUAD | LXF-289 | AZ628 | MVP_top60_DNA+RNA_seed | PIK3CA | -0.250907 | LUSC | LK-2 | AZD8835 |
| MVP_top60_DNA_seed | MET | 0.102188 | LUAD | NCI-H3122 | SU11274 | MVP_top60_DNA+RNA_seed | PIK3CA | -0.247775 | LUSC | HCC-15 | PI3Ka_4409 |
| MVP_top60_DNA_seed | HDAC1 | 0.10294 | LUAD | LXF-289 | Parthenolide | MVP_top60_DNA+RNA_seed | HSP90AB1 | -0.236974 | LUSC | EBC-1 | Elesclomol |
| MVP_top60_DNA_seed | AURKA | 0.103542 | LUAD | NCI-H1781 | HG-5-113-01 | MVP_top60_DNA+RNA_seed | PIK3CA | -0.233333 | LUSC | NCI-H226 | AZD8835 |
| MVP_top60_DNA_seed | FGFR1 | 0.10405 | LUAD | HCC-78 | PD173074 | MVP_top60_DNA+RNA_seed | PIK3CA | -0.221388 | LUSC | NCI-H1869 | PI3Ka_4409 |
| MVP_top60_DNA_seed | AURKA | 0.105207 | LUAD | PC-3_[JPC-3] | Alisertib | MVP_top60_DNA+RNA_seed | HSP90AB1 | -0.214407 | LUSC | EPLC-272H | Luminespib |
| MVP_top60_DNA_seed | EGFR | 0.105223 | LUAD | NCI-H2030 | Cetuximab | MVP_top60_DNA+RNA_seed | HSP90AB1 | -0.212671 | LUSC | EPLC-272H | Elesclomol |
| MVP_top60_DNA_seed | HDAC1 | 0.10561 | LUAD | NCI-H2087 | Dacinostat | MVP_top60_DNA+RNA_seed | PIK3CA | -0.202814 | LUSC | HARA | AZD8835 |
| MVP_top60_DNA_seed | BRAF | 0.106132 | LUAD | NCI-H2122 | RAF_9304 | MVP_top60_DNA+RNA_seed | HDAC1 | -0.190535 | LUSC | LC-1-sq | Belinostat |
| MVP_top60_DNA_seed | MAPK1 | 0.106166 | LUAD | NCI-H1568 | FR-180204 | MVP_top60_DNA+RNA_seed | HDAC1 | -0.180764 | LUSC | HARA | Tubastatin A |
| MVP_top60_DNA_seed | EGFR | 0.106379 | LUAD | NCI-H2023 | Afatinib | MVP_top60_DNA+RNA_seed | HDAC1 | -0.174574 | LUSC | LC-1-sq | AR-42 |
| MVP_top60_DNA_seed | ERBB2 | 0.106379 | LUAD | NCI-H2023 | Afatinib | MVP_top60_DNA+RNA_seed | HSP90AB1 | -0.156697 | LUSC | LOU-NH91 | Elesclomol |
| MVP_top60_DNA_seed | GSK3B | 0.106942 | LUAD | NCI-H1838 | CHIR-99021 | MVP_top60_DNA+RNA_seed | PIK3CA | -0.155866 | LUSC | HARA | PI3Ka_4409 |
| MVP_top60_DNA_seed | FGFR1 | 0.107135 | LUAD | NCI-H1648 | PD173074 | MVP_top60_DNA+RNA_seed | HSP90AB1 | -0.154184 | LUSC | KNS-62 | Tanespimycin |
| MVP_top60_DNA_seed | MDM2 | 0.107372 | LUAD | Calu-3 | Serdemetan | MVP_top60_DNA+RNA_seed | AR | -0.139412 | LUSC | HCC-15 | AZD3514 |
| MVP_top60_DNA_seed | BRAF | 0.107619 | LUAD | ABC-1 | Dabrafenib | MVP_top60_DNA+RNA_seed | HDAC1 | -0.117491 | LUSC | SK-MES-1 | Belinostat |
| MVP_top60_DNA_seed | EGFR | 0.108768 | LUAD | LXF-289 | Afatinib | MVP_top60_DNA+RNA_seed | HSP90AB1 | -0.116106 | LUSC | LK-2 | SNX-2112 |

| | | | | | | | | | | | |
|---|---|---|---|---|---|---|---|---|---|---|---|
| MVP_top60_DNA_seed | ERBB2 | 0.108768 | LUAD | LXF-289 | Afatinib | MVP_top60_DNA+RNA_seed | AR | -0.105629 | LUSC | LK-2 | Bicalutamide |
| MVP_top60_DNA_seed | AKT1 | 0.109341 | LUAD | NCI-H1650 | MK-2206 | MVP_top60_DNA+RNA_seed | HDAC1 | -0.091381 | LUSC | LC-1-sq | CAY10603 |
| MVP_top60_DNA_seed | FGFR1 | 0.110362 | LUAD | SK-LU-1 | AZD4547 | MVP_top60_DNA+RNA_seed | HDAC1 | -0.087645 | LUSC | NCI-H520 | AR-42 |
| MVP_top60_DNA_seed | MDM2 | 0.111032 | LUAD | NCI-H1838 | Serdemetan | MVP_top60_DNA+RNA_seed | HSP90AB1 | -0.075318 | LUSC | HCC-15 | Elesclomol |
| MVP_top60_DNA_seed | HDAC1 | 0.11186 | LUAD | HCC-827 | Belinostat | MVP_top60_DNA+RNA_seed | HSP90AB1 | -0.074954 | LUSC | HCC-15 | Luminespib |
| MVP_top60_DNA_seed | AKT1 | 0.112574 | LUAD | COR-L105 | GSK690693 | MVP_top60_DNA+RNA_seed | HDAC1 | -0.070004 | LUSC | NCI-H226 | Dacinostat |
| MVP_top60_DNA_seed | EGFR | 0.112963 | LUAD | NCI-H1793 | AZD8931 | MVP_top60_DNA+RNA_seed | HDAC1 | -0.066763 | LUSC | SK-MES-1 | CAY10603 |
| MVP_top60_DNA_seed | ERBB2 | 0.112963 | LUAD | NCI-H1793 | AZD8931 | MVP_top60_DNA+RNA_seed | HSP90AB1 | -0.061017 | LUSC | HARA | Luminespib |
| MVP_top60_DNA_seed | ERBB3 | 0.112963 | LUAD | NCI-H1793 | AZD8931 | MVP_top60_DNA+RNA_seed | AR | -0.053071 | LUSC | RERF-LC-Sq1 | AZD3514 |
| MVP_top60_DNA_seed | EGFR | 0.113173 | LUAD | NCI-H650 | AST-1306 | MVP_top60_DNA+RNA_seed | CREBBP | -0.045072 | LUSC | SK-MES-1 | I-CBP112 |
| MVP_top60_DNA_seed | CREBBP | 0.113327 | LUAD | NCI-H2087 | I-CBP112 | MVP_top60_DNA+RNA_seed | AR | -0.038137 | LUSC | HCC-15 | Bicalutamide |
| MVP_top60_DNA_seed | PTK2 | 0.113588 | LUAD | H3255 | PF-562271 | MVP_top60_DNA+RNA_seed | CREBBP | -0.033738 | LUSC | LK-2 | I-CBP112 |
| MVP_top60_DNA_seed | EGFR | 0.114856 | LUAD | NCI-H1563 | CI-1033 | MVP_top60_DNA+RNA_seed | PIK3CA | -0.032854 | LUSC | NCI-H2170 | PI3Ka_4409 |
| MVP_top60_DNA_seed | ERBB2 | 0.114856 | LUAD | NCI-H1563 | CI-1033 | MVP_top60_DNA+RNA_seed | HDAC1 | -0.014245 | LUSC | LOU-NH91 | Belinostat |
| MVP_top60_DNA_seed | FGFR1 | 0.115337 | LUAD | EMC-BAC-2 | AZD4547 | MVP_top60_DNA+RNA_seed | HDAC1 | -0.008273 | LUSC | NCI-H520 | JQ12 |
| MVP_top60_DNA_seed | BRAF | 0.11597 | LUAD | NCI-H2228 | RAF_9304 | MVP_top60_DNA+RNA_seed | HDAC1 | -0.005246 | LUSC | HARA | Belinostat |
| MVP_top60_DNA_seed | EGFR | 0.118719 | LUAD | VMRC-LCD | Cetuximab | MVP_top60_DNA+RNA_seed | HSP90AB1 | 0.014495 | LUSC | SK-MES-1 | SNX-2112 |
| MVP_top60_DNA_seed | EGFR | 0.119269 | LUAD | HOP-62 | Cetuximab | MVP_top60_DNA+RNA_seed | AR | 0.014828 | LUSC | LK-2 | Bicalutamide |
| MVP_top60_DNA_seed | HSP90AA1 | 0.119605 | LUAD | NCI-H522 | SNX-2112 | MVP_top60_DNA+RNA_seed | CREBBP | 0.016885 | LUSC | EPLC-272H | I-CBP112 |
| MVP_top60_DNA_seed | AURKA | 0.119938 | LUAD | NCI-H3122 | GSK1070916 | MVP_top60_DNA+RNA_seed | HDAC1 | 0.019635 | LUSC | EBC-1 | Belinostat |
| MVP_top60_DNA_seed | EGFR | 0.120587 | LUAD | NCI-H2405 | Gefitinib | MVP_top60_DNA+RNA_seed | HDAC1 | 0.021002 | LUSC | SK-MES-1 | AR-42 |
| MVP_top60_DNA_seed | EGFR | 0.120819 | LUAD | RERF-LC-MS | Afatinib | MVP_top60_DNA+RNA_seed | HSP90AB1 | 0.025448 | LUSC | NCI-H1869 | Luminespib |
| MVP_top60_DNA_seed | ERBB2 | 0.120819 | LUAD | RERF-LC-MS | Afatinib | MVP_top60_DNA+RNA_seed | HDAC1 | 0.029762 | LUSC | LOU-NH91 | CAY10603 |
| MVP_top60_DNA_seed | EGFR | 0.120835 | LUAD | NCI-H2347 | CI-1033 | MVP_top60_DNA+RNA_seed | AR | 0.034641 | LUSC | NCI-H226 | Bicalutamide |
| MVP_top60_DNA_seed | ERBB2 | 0.120835 | LUAD | NCI-H2347 | CI-1033 | MVP_top60_DNA+RNA_seed | PIK3CA | 0.041565 | LUSC | LOU-NH91 | PI3Ka_4409 |
| MVP_top60_DNA_seed | AURKA | 0.122198 | LUAD | PC-14 | ZM447439 | MVP_top60_DNA+RNA_seed | PIK3CA | 0.050047 | LUSC | HCC-15 | AZD8835 |
| MVP_top60_DNA_seed | EGFR | 0.122922 | LUAD | NCI-H1755 | AZD8931 | MVP_top60_DNA+RNA_seed | HDAC1 | 0.051833 | LUSC | LOU-NH91 | JQ12 |
| MVP_top60_DNA_seed | ERBB2 | 0.122922 | LUAD | NCI-H1755 | AZD8931 | MVP_top60_DNA+RNA_seed | HDAC1 | 0.07122 | LUSC | HARA | AR-42 |
| MVP_top60_DNA_seed | ERBB3 | 0.122922 | LUAD | NCI-H1755 | AZD8931 | MVP_top60_DNA+RNA_seed | CREBBP | 0.081399 | LUSC | HCC-15 | I-CBP112 |
| MVP_top60_DNA_seed | HDAC1 | 0.1232 | LUAD | HCC-78 | JQ12 | MVP_top60_DNA+RNA_seed | AR | 0.095524 | LUSC | NCI-H520 | Bicalutamide |
| MVP_top60_DNA_seed | SRC | 0.123697 | LUAD | HCC-78 | NVP-BHG712 | MVP_top60_DNA+RNA_seed | AR | 0.098282 | LUSC | SK-MES-1 | AZD3514 |
| MVP_top60_DNA_seed | GSK3B | 0.124883 | LUAD | NCI-H1792 | CHIR-99021 | MVP_top60_DNA+RNA_seed | PIK3CA | 0.109597 | LUSC | LC-1-sq | AZD8835 |
| MVP_top60_DNA_seed | HDAC1 | 0.125726 | LUAD | NCI-H1648 | JQ12 | MVP_top60_DNA+RNA_seed | HDAC1 | 0.117185 | LUSC | NCI-H2170 | CAY10603 |
| MVP_top60_DNA_seed | EGFR | 0.126278 | LUAD | NCI-H1781 | CUDC-101 | MVP_top60_DNA+RNA_seed | HDAC1 | 0.126375 | LUSC | HARA | CAY10603 |
| MVP_top60_DNA_seed | ERBB2 | 0.126278 | LUAD | NCI-H1781 | CUDC-101 | MVP_top60_DNA+RNA_seed | HDAC1 | 0.139997 | LUSC | EBC-1 | AR-42 |
| MVP_top60_DNA_seed | EGFR | 0.127341 | LUAD | HOP-62 | PF-00299804 | MVP_top60_DNA+RNA_seed | AR | 0.148153 | LUSC | HARA | Bicalutamide |
| MVP_top60_DNA_seed | ERBB2 | 0.127341 | LUAD | HOP-62 | PF-00299804 | MVP_top60_DNA+RNA_seed | HDAC1 | 0.160137 | LUSC | LOU-NH91 | AR-42 |
| MVP_top60_DNA_seed | AKT1 | 0.127859 | LUAD | NCI-H23 | A-443654 | MVP_top60_DNA+RNA_seed | AR | 0.186477 | LUSC | EPLC-272H | Bicalutamide |
| MVP_top60_DNA_seed | HSP90AA1 | 0.129047 | LUAD | NCI-H522 | CCT-018159 | MVP_top60_DNA+RNA_seed | HDAC1 | 0.189457 | LUSC | NCI-H1869 | Entinostat |
| MVP_top60_DNA_seed | HSP90AA1 | 0.129654 | LUAD | RERF-LC-MS | Tanespimycin | MVP_top60_DNA+RNA_seed | HDAC3 | 0.189457 | LUSC | NCI-H1869 | Entinostat |
| MVP_top60_DNA_seed | PIK3CA | 0.13005 | LUAD | PC-14 | PI3Ka_4409 | MVP_top60_DNA+RNA_seed | HSP90AB1 | 0.190544 | LUSC | EBC-1 | SNX-2112 |

| | | | | | | | | | | | |
|---|---|---|---|---|---|---|---|---|---|---|---|
| MVP_top60_DNA_seed | IGF1R | 0.13059 | LUAD | NCI-H2347 | BMS-536924 | MVP_top60_DNA+RNA_seed | SMARCA4 | 0.201605 | LUSC | LK-2 | PFI-3 |
| MVP_top60_DNA_seed | EGFR | 0.132407 | LUAD | NCI-H2347 | Afatinib | MVP_top60_DNA+RNA_seed | SMARCA2 | 0.201605 | LUSC | LK-2 | PFI-3 |
| MVP_top60_DNA_seed | ERBB2 | 0.132407 | LUAD | NCI-H2347 | Afatinib | MVP_top60_DNA+RNA_seed | HDAC1 | 0.204347 | LUSC | LC-1-sq | Tubastatin A |
| MVP_top60_DNA_seed | HSP90AA1 | 0.132426 | LUAD | NCI-H1651 | Tanespimycin | MVP_top60_DNA+RNA_seed | HDAC1 | 0.208178 | LUSC | LK-2 | Tubastatin A |
| MVP_top60_DNA_seed | HSP90AA1 | 0.133602 | LUAD | NCI-H2085 | SNX-2112 | MVP_top60_DNA+RNA_seed | HSP90AB1 | 0.214056 | LUSC | NCI-H2170 | Luminespib |
| MVP_top60_DNA_seed | EGFR | 0.133994 | LUAD | NCI-H2030 | CUDC-101 | MVP_top60_DNA+RNA_seed | SMARCA4 | 0.222003 | LUSC | SK-MES-1 | PFI-3 |
| MVP_top60_DNA_seed | ERBB2 | 0.133994 | LUAD | NCI-H2030 | CUDC-101 | MVP_top60_DNA+RNA_seed | SMARCA2 | 0.222003 | LUSC | SK-MES-1 | PFI-3 |
| MVP_top60_DNA_seed | AKT1 | 0.134447 | LUAD | NCI-H522 | AKT inhibitor VIII | MVP_top60_DNA+RNA_seed | HDAC1 | 0.239963 | LUSC | EBC-1 | CAY10603 |
| MVP_top60_DNA_seed | PIK3CA | 0.134479 | LUAD | EMC-BAC-1 | AZD8835 | MVP_top60_DNA+RNA_seed | HSP90AB1 | 0.258289 | LUSC | NCI-H520 | CCT-018159 |
| MVP_top60_DNA_seed | HSP90AA1 | 0.135687 | LUAD | HCC-78 | CCT-018159 | MVP_top60_DNA+RNA_seed | AR | 0.263244 | LUSC | SK-MES-1 | Bicalutamide |
| MVP_top60_DNA_seed | FGFR1 | 0.135954 | LUAD | NCI-H1355 | FGFR_3831 | MVP_top60_DNA+RNA_seed | HDAC1 | 0.273072 | LUSC | RERF-LC-Sq1 | Belinostat |
| MVP_top60_DNA_seed | EGFR | 0.13668 | LUAD | NCI-H1793 | AST-1306 | MVP_top60_DNA+RNA_seed | HDAC1 | 0.285802 | LUSC | NCI-H1869 | AR-42 |
| MVP_top60_DNA_seed | HSP90AA1 | 0.137582 | LUAD | HCC-827 | Luminespib | MVP_top60_DNA+RNA_seed | HDAC1 | 0.28665 | LUSC | NCI-H2170 | Belinostat |
| MVP_top60_DNA_seed | HSP90AA1 | 0.137664 | LUAD | NCI-H1648 | SNX-2112 | MVP_top60_DNA+RNA_seed | HDAC1 | 0.288484 | LUSC | SK-MES-1 | Tubastatin A |
| MVP_top60_DNA_seed | GSK3B | 0.138151 | LUAD | NCI-H1435 | SB216763 | MVP_top60_DNA+RNA_seed | PIK3CA | 0.293316 | LUSC | NCI-H2170 | AZD8835 |
| MVP_top60_DNA_seed | ATM | 0.13836 | LUAD | NCI-H2009 | KU-55933 | MVP_top60_DNA+RNA_seed | AR | 0.305061 | LUSC | NCI-H2170 | AZD3514 |
| MVP_top60_DNA_seed | GSK3B | 0.138364 | LUAD | SW1573 | CHIR-99021 | MVP_top60_DNA+RNA_seed | AR | 0.305101 | LUSC | LOU-NH91 | Bicalutamide |
| MVP_top60_DNA_seed | AKT1 | 0.13885 | LUAD | NCI-H1563 | GSK690693 | MVP_top60_DNA+RNA_seed | AR | 0.310686 | LUSC | NCI-H1869 | AZD3514 |
| MVP_top60_DNA_seed | MET | 0.139991 | LUAD | Calu-3 | JNJ38877605 | MVP_top60_DNA+RNA_seed | PIK3CA | 0.321961 | LUSC | LC-1-sq | PI3Ka_4409 |
| MVP_top60_DNA_seed | PTK2 | 0.141621 | LUAD | SW1573 | PF-562271 | MVP_top60_DNA+RNA_seed | AR | 0.33193 | LUSC | NCI-H1869 | Bicalutamide |
| MVP_top60_DNA_seed | BRAF | 0.142884 | LUAD | NCI-H1573 | PLX-4720 | MVP_top60_DNA+RNA_seed | HSP90AB1 | 0.333029 | LUSC | RERF-LC-Sq1 | Elesclomol |
| MVP_top60_DNA_seed | EGFR | 0.143458 | LUAD | A549 | Gefitinib | MVP_top60_DNA+RNA_seed | AR | 0.33334 | LUSC | HARA | AZD3514 |
| MVP_top60_DNA_seed | BRAF | 0.143617 | LUAD | NCI-H3122 | HG6-64-1 | MVP_top60_DNA+RNA_seed | HDAC1 | 0.337752 | LUSC | NCI-H226 | JQ12 |
| MVP_top60_DNA_seed | IGF1R | 0.145148 | LUAD | NCI-H1793 | BMS-536924 | MVP_top60_DNA+RNA_seed | PIK3CA | 0.33785 | LUSC | LK-2 | PI3Ka_4409 |
| MVP_top60_DNA_seed | SRC | 0.145293 | LUAD | NCI-H2347 | NVP-BHG712 | MVP_top60_DNA+RNA_seed | HDAC1 | 0.340754 | LUSC | EPLC-272H | Dacinostat |
| MVP_top60_DNA_seed | BRAF | 0.145363 | LUAD | NCI-H650 | SB590885 | MVP_top60_DNA+RNA_seed | HSP90AB1 | 0.341343 | LUSC | HCC-15 | CCT-018159 |
| MVP_top60_DNA_seed | BRAF | 0.14654 | LUAD | NCI-H1568 | HG6-64-1 | MVP_top60_DNA+RNA_seed | HDAC1 | 0.376536 | LUSC | NCI-H226 | Tubastatin A |
| MVP_top60_DNA_seed | BRAF | 0.147271 | LUAD | HOP-62 | RAF_9304 | MVP_top60_DNA+RNA_seed | HSP90AB1 | 0.396326 | LUSC | NCI-H1869 | Elesclomol |
| MVP_top60_DNA_seed | FGFR1 | 0.147722 | LUAD | RERF-LC-KJ | PD173074 | MVP_top60_DNA+RNA_seed | SMARCA4 | 0.40521 | LUSC | NCI-H226 | PFI-3 |
| MVP_top60_DNA_seed | SMARCA4 | 0.148357 | LUAD | H3255 | PFI-3 | MVP_top60_DNA+RNA_seed | SMARCA2 | 0.40521 | LUSC | NCI-H226 | PFI-3 |
| MVP_top60_DNA_seed | IGF1R | 0.148908 | LUAD | NCI-H2085 | Linsitinib | MVP_top60_DNA+RNA_seed | SMARCA4 | 0.405648 | LUSC | NCI-H520 | PFI-3 |
| MVP_top60_DNA_seed | AURKA | 0.149007 | LUAD | NCI-H1651 | HG-5-113-01 | MVP_top60_DNA+RNA_seed | SMARCA2 | 0.405648 | LUSC | NCI-H520 | PFI-3 |
| MVP_top60_DNA_seed | AURKA | 0.149342 | LUAD | EMC-BAC-1 | Alisertib | MVP_top60_DNA+RNA_seed | AR | 0.417828 | LUSC | LC-1-sq | Bicalutamide |
| MVP_top60_DNA_seed | AKT1 | 0.150566 | LUAD | NCI-H3122 | GSK690693 | MVP_top60_DNA+RNA_seed | HDAC1 | 0.429519 | LUSC | RERF-LC-Sq1 | CAY10603 |
| MVP_top60_DNA_seed | IGF1R | 0.151854 | LUAD | NCI-H23 | BMS-536924 | MVP_top60_DNA+RNA_seed | HSP90AB1 | 0.433023 | LUSC | LK-2 | Luminespib |
| MVP_top60_DNA_seed | BRAF | 0.151966 | LUAD | VMRC-LCD | Dabrafenib | MVP_top60_DNA+RNA_seed | HDAC1 | 0.433755 | LUSC | HCC-15 | Dacinostat |
| MVP_top60_DNA_seed | BRAF | 0.15212 | LUAD | NCI-H1563 | PLX-4720 | MVP_top60_DNA+RNA_seed | HDAC1 | 0.437748 | LUSC | RERF-LC-Sq1 | AR-42 |
| MVP_top60_DNA_seed | HSP90AA1 | 0.152314 | LUAD | ABC-1 | SNX-2112 | MVP_top60_DNA+RNA_seed | AR | 0.441245 | LUSC | NCI-H1869 | Bicalutamide |
| MVP_top60_DNA_seed | GSK3B | 0.153123 | LUAD | VMRC-LCD | AZD7969 | MVP_top60_DNA+RNA_seed | AR | 0.44236 | LUSC | NCI-H2170 | Bicalutamide |
| MVP_top60_DNA_seed | GSK3B | 0.153351 | LUAD | NCI-H2405 | CHIR-99021 | MVP_top60_DNA+RNA_seed | HSP90AB1 | 0.445192 | LUSC | NCI-H1869 | CCT-018159 |
| MVP_top60_DNA_seed | GSK3B | 0.155161 | LUAD | NCI-H2122 | SB216763 | MVP_top60_DNA+RNA_seed | CREBBP | 0.449253 | LUSC | NCI-H520 | I-CBP112 |

| | | | | | | | | | | | |
|---|---|---|---|---|---|---|---|---|---|---|---|
| MVP_top60_DNA_seed | IGF1R | 0.155229 | LUAD | H3255 | BMS-754807 | MVP_top60_DNA+RNA_seed | HSP90AB1 | 0.458248 | LUSC | NCI-H520 | Elesclomol |
| MVP_top60_DNA_seed | MET | 0.156087 | LUAD | EMC-BAC-1 | AZD6094 | MVP_top60_DNA+RNA_seed | HDAC1 | 0.463717 | LUSC | NCI-H2170 | AR-42 |
| MVP_top60_DNA_seed | AKT1 | 0.157663 | LUAD | NCI-H3122 | AKT inhibitor VIII | MVP_top60_DNA+RNA_seed | HDAC1 | 0.477411 | LUSC | NCI-H1869 | Belinostat |
| MVP_top60_DNA_seed | AURKA | 0.158273 | LUAD | NCI-H1573 | ZM447439 | MVP_top60_DNA+RNA_seed | HSP90AB1 | 0.499879 | LUSC | NCI-H226 | Tanespimycin |
| MVP_top60_DNA_seed | AURKA | 0.158969 | LUAD | NCI-H1944 | Alisertib | MVP_top60_DNA+RNA_seed | HSP90AB1 | 0.526718 | LUSC | NCI-H226 | Elesclomol |
| MVP_top60_DNA_seed | AURKA | 0.159447 | LUAD | NCI-H1734 | Alisertib | MVP_top60_DNA+RNA_seed | AR | 0.531502 | LUSC | NCI-H226 | AZD3514 |
| MVP_top60_DNA_seed | MDM2 | 0.159851 | LUAD | LC-2-ad | Serdemetan | MVP_top60_DNA+RNA_seed | CREBBP | 0.561683 | LUSC | NCI-H1869 | I-CBP112 |
| MVP_top60_DNA_seed | FGFR1 | 0.160811 | LUAD | NCI-H2085 | FGFR_3831 | MVP_top60_DNA+RNA_seed | AR | 0.568437 | LUSC | LOU-NH91 | Bicalutamide |
| MVP_top60_DNA_seed | BRAF | 0.161044 | LUAD | A549 | PLX-4720 | MVP_top60_DNA+RNA_seed | HSP90AB1 | 0.584051 | LUSC | LC-1-sq | Tanespimycin |
| MVP_top60_DNA_seed | AKT1 | 0.161336 | LUAD | NCI-H1568 | BAY AKT1 | MVP_top60_DNA+RNA_seed | HSP90AB1 | 0.591657 | LUSC | KNS-62 | CCT-018159 |
| MVP_top60_DNA_seed | MAPK1 | 0.161602 | LUAD | NCI-H3122 | FR-180204 | MVP_top60_DNA+RNA_seed | AR | 0.619727 | LUSC | RERF-LC-Sq1 | Bicalutamide |
| MVP_top60_DNA_seed | FGFR1 | 0.161769 | LUAD | NCI-H2122 | PD173074 | MVP_top60_DNA+RNA_seed | AR | 0.642859 | LUSC | NCI-H226 | Bicalutamide |
| MVP_top60_DNA_seed | AKT1 | 0.161981 | LUAD | NCI-H2405 | BAY AKT1 | MVP_top60_DNA+RNA_seed | CREBBP | 0.66458 | LUSC | LOU-NH91 | I-CBP112 |
| MVP_top60_DNA_seed | HDAC1 | 0.162004 | LUAD | HCC-827 | AR-42 | MVP_top60_DNA+RNA_seed | HSP90AB1 | 0.696927 | LUSC | NCI-H226 | CCT-018159 |
| MVP_top60_DNA_seed | AURKA | 0.162844 | LUAD | NCI-H3122 | Genentech Cpd 10 | MVP_top60_DNA+RNA_seed | AR | 0.705625 | LUSC | NCI-H2170 | Bicalutamide |
| MVP_top60_DNA_seed | BRAF | 0.163332 | LUAD | NCI-H1623 | RAF_9304 | MVP_top60_DNA+RNA_seed | HDAC1 | 0.737617 | LUSC | NCI-H1869 | CAY10603 |
| MVP_top60_DNA_seed | SRC | 0.163423 | LUAD | LXF-289 | NVP-BHG712 | MVP_top60_DNA+RNA_seed | HDAC1 | 0.757813 | LUSC | LOU-NH91 | Dacinostat |
| MVP_top60_DNA_seed | ATM | 0.163518 | LUAD | NCI-H522 | KU-55933 | MVP_top60_DNA+RNA_seed | HDAC1 | 0.768894 | LUSC | LOU-NH91 | Tubastatin A |
| MVP_top60_DNA_seed | IGF1R | 0.163828 | LUAD | ABC-1 | GSK1904529A | MVP_top60_DNA+RNA_seed | HDAC1 | 0.798122 | LUSC | HCC-15 | Tubastatin A |
| MVP_top60_DNA_seed | ATM | 0.165017 | LUAD | NCI-H1792 | KU-60019 | MVP_top60_DNA+RNA_seed | HDAC1 | 0.800592 | LUSC | NCI-H1869 | Dacinostat |
| MVP_top60_DNA_seed | HSP90AA1 | 0.166028 | LUAD | NCI-H1793 | CCT-018159 | MVP_top60_DNA+RNA_seed | SMARCA4 | 0.81455 | LUSC | NCI-H2170 | PFI-3 |
| MVP_top60_DNA_seed | FGFR1 | 0.166604 | LUAD | NCI-H1755 | PD173074 | MVP_top60_DNA+RNA_seed | SMARCA2 | 0.81455 | LUSC | NCI-H2170 | PFI-3 |
| MVP_top60_DNA_seed | EGFR | 0.166765 | LUAD | NCI-H1781 | Gefitinib | MVP_top60_DNA+RNA_seed | HDAC1 | 0.815747 | LUSC | HARA | Dacinostat |
| MVP_top60_DNA_seed | EGFR | 0.167501 | LUAD | NCI-H1993 | CUDC-101 | MVP_top60_DNA+RNA_seed | HDAC1 | 0.826961 | LUSC | NCI-H226 | Parthenolide |
| MVP_top60_DNA_seed | ERBB2 | 0.167501 | LUAD | NCI-H1993 | CUDC-101 | MVP_top60_DNA+RNA_seed | HSP90AB1 | 0.845214 | LUSC | NCI-H1869 | SNX-2112 |
| MVP_top60_DNA_seed | EGFR | 0.167645 | LUAD | NCI-H1563 | AST-1306 | MVP_top60_DNA+RNA_seed | PIK3CA | 0.846248 | LUSC | EBC-1 | AZD8835 |
| MVP_top60_DNA_seed | MET | 0.167709 | LUAD | NCI-H1734 | SU11274 | MVP_top60_DNA+RNA_seed | SMARCA4 | 0.849095 | LUSC | RERF-LC-Sq1 | PFI-3 |
| MVP_top60_DNA_seed | MET | 0.167717 | LUAD | NCI-H1651 | AZD6094 | MVP_top60_DNA+RNA_seed | SMARCA2 | 0.849095 | LUSC | RERF-LC-Sq1 | PFI-3 |
| MVP_top60_DNA_seed | AKT1 | 0.167932 | LUAD | NCI-H1792 | BAY AKT1 | MVP_top60_DNA+RNA_seed | HDAC1 | 0.851319 | LUSC | LC-1-sq | JQ12 |
| MVP_top60_DNA_seed | AURKA | 0.168099 | LUAD | RERF-LC-MS | CD532 | MVP_top60_DNA+RNA_seed | HSP90AB1 | 0.852544 | LUSC | LK-2 | CCT-018159 |
| MVP_top60_DNA_seed | MET | 0.169186 | LUAD | NCI-H1437 | AZD6094 | MVP_top60_DNA+RNA_seed | CREBBP | 0.860194 | LUSC | NCI-H2170 | I-CBP112 |
| MVP_top60_DNA_seed | ERBB2 | 0.170403 | LUAD | HCC-78 | CP724714 | MVP_top60_DNA+RNA_seed | HDAC1 | 0.870761 | LUSC | RERF-LC-Sq1 | Tubastatin A |
| MVP_top60_DNA_seed | HDAC1 | 0.171363 | LUAD | PC-14 | Dacinostat | MVP_top60_DNA+RNA_seed | SMARCA4 | 0.886656 | LUSC | NCI-H1869 | PFI-3 |
| MVP_top60_DNA_seed | AURKA | 0.171861 | LUAD | SW1573 | ZM447439 | MVP_top60_DNA+RNA_seed | SMARCA2 | 0.886656 | LUSC | NCI-H1869 | PFI-3 |
| MVP_top60_DNA_seed | EGFR | 0.171946 | LUAD | NCI-H2087 | CI-1033 | MVP_top60_DNA+RNA_seed | AR | 0.909576 | LUSC | NCI-H520 | Bicalutamide |
| MVP_top60_DNA_seed | ERBB2 | 0.171946 | LUAD | NCI-H2087 | CI-1033 | MVP_top60_DNA+RNA_seed | AR | 0.939063 | LUSC | HCC-15 | Bicalutamide |
| MVP_top60_DNA_seed | SRC | 0.172039 | LUAD | NCI-H2228 | Bosutinib | MVP_top60_DNA+RNA_seed | AR | 0.940078 | LUSC | HARA | Bicalutamide |
| MVP_top60_DNA_seed | IGF1R | 0.172208 | LUAD | LXF-289 | BMS-536924 | MVP_top60_DNA+RNA_seed | HDAC1 | 0.944391 | LUSC | NCI-H1869 | Tubastatin A |
| MVP_top60_DNA_seed | SRC | 0.172723 | LUAD | NCI-H650 | NVP-BHG712 | MVP_top60_DNA+RNA_seed | HSP90AB1 | 0.960517 | LUSC | RERF-LC-Sq1 | SNX-2112 |
| MVP_top60_DNA_seed | HDAC1 | 0.172726 | LUAD | NCI-H522 | Tubastatin A | MVP_top60_DNA+RNA_seed | HSP90AB1 | 0.969472 | LUSC | HCC-15 | SNX-2112 |
| MVP_top60_DNA_seed | AURKA | 0.173385 | LUAD | ABC-1 | CD532 | MVP_top60_DNA+RNA_seed | SMARCA4 | 0.989746 | LUSC | LC-1-sq | PFI-3 |

| | | | | | | | | | | | |
|---|---|---|---|---|---|---|---|---|---|---|---|
| MVP_top60_DNA_seed | BRAF | 0.173794 | LUAD | SW1573 | PLX-4720 | MVP_top60_DNA+RNA_seed | SMARCA2 | 0.989746 | LUSC | LC-1-sq | PFI-3 |
| MVP_top60_DNA_seed | BRAF | 0.174172 | LUAD | LXF-289 | Dabrafenib | MVP_top60_DNA+RNA_seed | AR | 1.015018 | LUSC | NCI-H520 | AZD3514 |
| MVP_top60_DNA_seed | AKT1 | 0.174546 | LUAD | SK-LU-1 | AKT inhibitor VIII | MVP_top60_DNA+RNA_seed | HDAC1 | 1.054866 | LUSC | NCI-H2170 | Tubastatin A |
| MVP_top60_DNA_seed | EGFR | 0.175242 | LUAD | ABC-1 | AST-1306 | MVP_top60_DNA+RNA_seed | HDAC1 | 1.064857 | LUSC | EPLC-272H | JQ12 |
| MVP_top60_DNA_seed | SMARCA4 | 0.178408 | LUAD | NCI-H358 | PFI-3 | MVP_top60_DNA+RNA_seed | AR | 1.08513 | LUSC | EBC-1 | Bicalutamide |
| MVP_top60_DNA_seed | IGF1R | 0.17879 | LUAD | VMRC-LCD | BMS-536924 | MVP_top60_DNA+RNA_seed | HDAC1 | 1.102847 | LUSC | HCC-15 | CAY10603 |
| MVP_top60_DNA_seed | IGF1R | 0.179206 | LUAD | NCI-H2342 | BMS-536924 | MVP_top60_DNA+RNA_seed | HSP90AB1 | 1.140622 | LUSC | SK-MES-1 | Elesclomol |
| MVP_top60_DNA_seed | HDAC1 | 0.179322 | LUAD | NCI-H838 | Tubastatin A | MVP_top60_DNA+RNA_seed | PIK3CA | 1.193603 | LUSC | NCI-H226 | PI3Ka_4409 |
| MVP_top60_DNA_seed | IGF1R | 0.179385 | LUAD | NCI-H1755 | Linsitinib | MVP_top60_DNA+RNA_seed | AR | 1.194853 | LUSC | LC-1-sq | Bicalutamide |
| MVP_top60_DNA_seed | SMARCA4 | 0.179965 | LUAD | LC-2-ad | PFI-3 | MVP_top60_DNA+RNA_seed | HDAC1 | 1.205742 | LUSC | HCC-15 | Belinostat |
| MVP_top60_DNA_seed | IGF1R | 0.181267 | LUAD | NCI-H2087 | GSK1904529A | MVP_top60_DNA+RNA_seed | AR | 1.214662 | LUSC | RERF-LC-Sq1 | Bicalutamide |
| MVP_top60_DNA_seed | MET | 0.181418 | LUAD | NCI-H1355 | AZD6094 | MVP_top60_DNA+RNA_seed | HSP90AB1 | 1.2203 | LUSC | NCI-H2170 | SNX-2112 |
| MVP_top60_DNA_seed | PIK3CA | 0.18172 | LUAD | NCI-H23 | AZD8835 | MVP_top60_DNA+RNA_seed | CREBBP | 1.295727 | LUSC | KNS-62 | I-CBP112 |
| MVP_top60_DNA_seed | PIK3CA | 0.182728 | LUAD | NCI-H3122 | PI3Ka_4409 | MVP_top60_DNA+RNA_seed | HSP90AB1 | 1.309972 | LUSC | LOU-NH91 | Luminespib |
| MVP_top60_DNA_seed | MET | 0.184076 | LUAD | NCI-H1755 | AZD6094 | MVP_top60_DNA+RNA_seed | HSP90AB1 | 1.337464 | LUSC | LC-1-sq | Elesclomol |
| MVP_top60_DNA_seed | AURKA | 0.185803 | LUAD | NCI-H522 | ZM447439 | MVP_top60_DNA+RNA_seed | PIK3CA | 1.349994 | LUSC | NCI-H520 | AZD8835 |
| MVP_top60_DNA_seed | IGF1R | 0.185863 | LUAD | NCI-H2405 | BMS-536924 | MVP_top60_DNA+RNA_seed | HDAC1 | 1.382508 | LUSC | EBC-1 | Tubastatin A |
| MVP_top60_DNA_seed | GSK3B | 0.185933 | LUAD | PC-14 | KIN001-042 | MVP_top60_DNA+RNA_seed | HDAC1 | 1.405971 | LUSC | RERF-LC-Sq1 | JQ12 |
| MVP_top60_DNA_seed | EGFR | 0.186015 | LUAD | NCI-H1993 | Afatinib | MVP_top60_DNA+RNA_seed | HSP90AB1 | 1.482617 | LUSC | LC-1-sq | SNX-2112 |
| MVP_top60_DNA_seed | ERBB2 | 0.186015 | LUAD | NCI-H1993 | Afatinib | MVP_top60_DNA+RNA_seed | CREBBP | 1.562696 | LUSC | RERF-LC-Sq1 | I-CBP112 |
| MVP_top60_DNA_seed | FGFR1 | 0.186421 | LUAD | ABC-1 | FGFR_3831 | MVP_top60_DNA+RNA_seed | PIK3CA | 1.576912 | LUSC | NCI-H1869 | AZD8835 |
| MVP_top60_DNA_seed | AURKA | 0.187163 | LUAD | NCI-H1792 | Alisertib | MVP_top60_DNA+RNA_seed | PIK3CA | 1.590637 | LUSC | NCI-H520 | PI3Ka_4409 |
| MVP_top60_DNA_seed | PIK3CA | 0.187222 | LUAD | NCI-H1650 | PI3Ka_4409 | MVP_top60_DNA+RNA_seed | HDAC1 | 1.647033 | LUSC | NCI-H1869 | JQ12 |
| MVP_top60_DNA_seed | IGF1R | 0.187285 | LUAD | NCI-H2087 | Linsitinib | MVP_top60_DNA+RNA_seed | HDAC1 | 1.66459 | LUSC | HCC-15 | AR-42 |
| MVP_top60_DNA_seed | EGFR | 0.187638 | LUAD | EMC-BAC-2 | AST-1306 | MVP_top60_DNA+RNA_seed | HSP90AB1 | 1.668057 | LUSC | LOU-NH91 | SNX-2112 |
| MVP_top60_DNA_seed | HDAC1 | 0.188087 | LUAD | NCI-H2030 | AR-42 | MVP_top60_DNA+RNA_seed | HDAC1 | 1.672272 | LUSC | HARA | JQ12 |
| MVP_top60_DNA_seed | HDAC1 | 0.18837 | LUAD | NCI-H322M | Belinostat | MVP_top60_DNA+RNA_seed | HSP90AB1 | 1.843326 | LUSC | RERF-LC-Sq1 | CCT-018159 |
| MVP_top60_DNA_seed | EGFR | 0.188767 | LUAD | NCI-H441 | AZD8931 | MVP_top60_DNA+RNA_seed | HSP90AB1 | 1.852427 | LUSC | LC-1-sq | Luminespib |
| MVP_top60_DNA_seed | ERBB2 | 0.188767 | LUAD | NCI-H441 | AZD8931 | MVP_top60_DNA+RNA_seed | HSP90AB1 | 2.065733 | LUSC | LC-1-sq | CCT-018159 |
| MVP_top60_DNA_seed | ERBB3 | 0.188767 | LUAD | NCI-H441 | AZD8931 | MVP_top60_DNA+RNA_seed | HDAC1 | 2.100282 | LUSC | NCI-H226 | Entinostat |
| MVP_top60_DNA_seed | GSK3B | 0.189356 | LUAD | NCI-H2009 | AZD7969 | MVP_top60_DNA+RNA_seed | HDAC3 | 2.100282 | LUSC | NCI-H226 | Entinostat |
| MVP_top60_DNA_seed | AURKA | 0.189375 | LUAD | NCI-H1563 | Genentech Cpd 10 | MVP_top60_DNA+RNA_seed | PIK3CA | 2.274945 | LUSC | KNS-62 | AZD8835 |
| MVP_top60_DNA_seed | AKT1 | 0.189506 | LUAD | NCI-H23 | MK-2206 | MVP_top60_DNA+RNA_seed | HDAC1 | 2.494629 | LUSC | RERF-LC-Sq1 | Dacinostat |
| MVP_top60_DNA_seed | ERBB2 | 0.190376 | LUAD | NCI-H1792 | CP724714 | MVP_top60_DNA+RNA_seed | HSP90AB1 | 3.155613 | LUSC | RERF-LC-Sq1 | Luminespib |
| MVP_top60_DNA_seed | AKT1 | 0.190701 | LUAD | NCI-H1355 | AKT inhibitor VIII | MVP_top60_DNA+RNA_seed | HSP90AB1 | -1.807408 | LUSC | NCI-H2170 | Tanespimycin |
| MVP_top60_DNA_seed | EGFR | 0.192345 | LUAD | LC-2-ad | Afatinib | MVP_top60_DNA+RNA_seed | HSP90AB1 | -1.546487 | LUSC | EPLC-272H | Tanespimycin |
| MVP_top60_DNA_seed | ERBB2 | 0.192345 | LUAD | LC-2-ad | Afatinib | MVP_top60_DNA+RNA_seed | PIK3CA | -1.338512 | LUSC | RERF-LC-Sq1 | Alpelisib |
| MVP_top60_DNA_seed | FGFR1 | 0.192672 | LUAD | VMRC-LCD | PD173074 | MVP_top60_DNA+RNA_seed | PIK3CA | -1.220521 | LUSC | EPLC-272H | Alpelisib |
| MVP_top60_DNA_seed | EGFR | 0.192772 | LUAD | NCI-H650 | PF-00299804 | MVP_top60_DNA+RNA_seed | SMARCA4 | -1.163656 | LUSC | EBC-1 | PFI3 |
| MVP_top60_DNA_seed | ERBB2 | 0.192772 | LUAD | NCI-H650 | PF-00299804 | MVP_top60_DNA+RNA_seed | SMARCA2 | -1.163656 | LUSC | EBC-1 | PFI3 |
| MVP_top60_DNA_seed | AURKA | 0.193879 | LUAD | NCI-H1568 | ZM447439 | MVP_top60_DNA+RNA_seed | EZH2 | -1.160375 | LUSC | LC-1-sq | GSK343 |

| | | | | | | | | | | | |
|---|---|---|---|---|---|---|---|---|---|---|---|
| MVP_top60_DNA_seed | MET | 0.194184 | LUAD | NCI-H1355 | JNJ38877605 | MVP_top60_DNA+RNA_seed | PIK3CA | -1.145638 | LUSC | EPLC-272H | Buparlisib |
| MVP_top60_DNA_seed | ATM | 0.194418 | LUAD | NCI-H1568 | CP466722 | MVP_top60_DNA+RNA_seed | PIK3CA | -1.121422 | LUSC | NCI-H2170 | Buparlisib |
| MVP_top60_DNA_seed | FGFR1 | 0.194424 | LUAD | NCI-H1437 | FGFR_3831 | MVP_top60_DNA+RNA_seed | PIK3CA | -1.08911 | LUSC | EPLC-272H | AZD8186 |
| MVP_top60_DNA_seed | AURKA | 0.195735 | LUAD | NCI-H2347 | GSK1070916 | MVP_top60_DNA+RNA_seed | HSP90AB1 | -1.087258 | LUSC | RERF-LC-Sq1 | Tanespimycin |
| MVP_top60_DNA_seed | FGFR1 | 0.195938 | LUAD | EMC-BAC-1 | PD173074 | MVP_top60_DNA+RNA_seed | PIK3CA | -1.036936 | LUSC | EPLC-272H | GNE-317 |
| MVP_top60_DNA_seed | AKT1 | 0.19635 | LUAD | NCI-H1437 | GSK690693 | MVP_top60_DNA+RNA_seed | EZH2 | -1.034054 | LUSC | EBC-1 | GSK343 |
| MVP_top60_DNA_seed | EGFR | 0.196811 | LUAD | NCI-H1755 | Afatinib | MVP_top60_DNA+RNA_seed | CREBBP | -0.985121 | LUSC | LC-1-sq | SGC-CBP30 |
| MVP_top60_DNA_seed | ERBB2 | 0.196811 | LUAD | NCI-H1755 | Afatinib | MVP_top60_DNA+RNA_seed | EZH2 | -0.973241 | LUSC | HARA | GSK343 |
| MVP_top60_DNA_seed | MET | 0.199832 | LUAD | Calu-6 | JNJ38877605 | MVP_top60_DNA+RNA_seed | PIK3CA | -0.955533 | LUSC | HARA | Buparlisib |
| MVP_top60_DNA_seed | AURKA | 0.199975 | LUAD | NCI-H1781 | CD532 | MVP_top60_DNA+RNA_seed | HSP90AB1 | -0.942685 | LUSC | HARA | Luminespib |
| MVP_top60_DNA_seed | BRAF | 0.200837 | LUAD | NCI-H2342 | PLX-4720 | MVP_top60_DNA+RNA_seed | PIK3CA | -0.913577 | LUSC | HARA | Alpelisib |
| MVP_top60_DNA_seed | AKT1 | 0.200924 | LUAD | EMC-BAC-1 | BAY AKT1 | MVP_top60_DNA+RNA_seed | KRAS | -0.911606 | LUSC | EPLC-272H | KRAS (G12C) Inhibitor-1 |
| MVP_top60_DNA_seed | HSP90AA1 | 0.200979 | LUAD | NCI-H1792 | SNX-2112 | MVP_top60_DNA+RNA_seed | HDAC1 | -0.879947 | LUSC | LOU-NH91 | Romidepsin |
| MVP_top60_DNA_seed | EGFR | 0.201012 | LUAD | SK-LU-1 | AST-1306 | MVP_top60_DNA+RNA_seed | HDAC3 | -0.879947 | LUSC | LOU-NH91 | Romidepsin |
| MVP_top60_DNA_seed | HDAC1 | 0.201388 | LUAD | NCI-H2009 | JQ12 | MVP_top60_DNA+RNA_seed | PIK3CA | -0.830417 | LUSC | HARA | GNE-317 |
| MVP_top60_DNA_seed | BRAF | 0.202482 | LUAD | NCI-H1792 | SB590885 | MVP_top60_DNA+RNA_seed | HSP90AB1 | -0.830363 | LUSC | EPLC-272H | Luminespib |
| MVP_top60_DNA_seed | SMARCA4 | 0.202512 | LUAD | NCI-H23 | PFI-3 | MVP_top60_DNA+RNA_seed | PIK3CA | -0.822014 | LUSC | EBC-1 | AZD8186 |
| MVP_top60_DNA_seed | IGF1R | 0.202535 | LUAD | Calu-3 | Linsitinib | MVP_top60_DNA+RNA_seed | EZH2 | -0.817165 | LUSC | EPLC-272H | GSK343 |
| MVP_top60_DNA_seed | SRC | 0.203815 | LUAD | NCI-H1648 | Bosutinib | MVP_top60_DNA+RNA_seed | EZH2 | -0.81452 | LUSC | SK-MES-1 | GSK343 |
| MVP_top60_DNA_seed | HDAC1 | 0.203837 | LUAD | NCI-H1573 | Belinostat | MVP_top60_DNA+RNA_seed | HDAC1 | -0.794417 | LUSC | EBC-1 | Romidepsin |
| MVP_top60_DNA_seed | GSK3B | 0.203905 | LUAD | NCI-H1781 | CHIR-99021 | MVP_top60_DNA+RNA_seed | HDAC3 | -0.794417 | LUSC | EBC-1 | Romidepsin |
| MVP_top60_DNA_seed | IGF1R | 0.205394 | LUAD | NCI-H2023 | Linsitinib | MVP_top60_DNA+RNA_seed | PIK3CA | -0.786418 | LUSC | RERF-LC-Sq1 | GNE-317 |
| MVP_top60_DNA_seed | EGFR | 0.205404 | LUAD | Calu-6 | CI-1033 | MVP_top60_DNA+RNA_seed | PIK3CA | -0.781629 | LUSC | NCI-H2170 | Alpelisib |
| MVP_top60_DNA_seed | ERBB2 | 0.205404 | LUAD | Calu-6 | CI-1033 | MVP_top60_DNA+RNA_seed | HDAC1 | -0.774222 | LUSC | EBC-1 | Entinostat |
| MVP_top60_DNA_seed | MDM2 | 0.207247 | LUAD | H3255 | Serdemetan | MVP_top60_DNA+RNA_seed | HDAC3 | -0.774222 | LUSC | EBC-1 | Entinostat |
| MVP_top60_DNA_seed | IGF1R | 0.207685 | LUAD | NCI-H1568 | GSK1904529A | MVP_top60_DNA+RNA_seed | PIK3CA | -0.729955 | LUSC | EBC-1 | GNE-317 |
| MVP_top60_DNA_seed | AURKA | 0.209002 | LUAD | NCI-H358 | CD532 | MVP_top60_DNA+RNA_seed | PIK3CA | -0.713796 | LUSC | NCI-H520 | AZD8186 |
| MVP_top60_DNA_seed | AURKA | 0.209612 | LUAD | NCI-H2009 | CD532 | MVP_top60_DNA+RNA_seed | HDAC1 | -0.706831 | LUSC | NCI-H520 | Romidepsin |
| MVP_top60_DNA_seed | EGFR | 0.209993 | LUAD | NCI-H2009 | AZD8931 | MVP_top60_DNA+RNA_seed | HDAC3 | -0.706831 | LUSC | NCI-H520 | Romidepsin |
| MVP_top60_DNA_seed | ERBB2 | 0.209993 | LUAD | NCI-H2009 | AZD8931 | MVP_top60_DNA+RNA_seed | HSP90AB1 | -0.690569 | LUSC | HARA | Tanespimycin |
| MVP_top60_DNA_seed | ERBB3 | 0.209993 | LUAD | NCI-H2009 | AZD8931 | MVP_top60_DNA+RNA_seed | EZH2 | -0.632355 | LUSC | EBC-1 | GSK343 |
| MVP_top60_DNA_seed | MET | 0.211055 | LUAD | NCI-H1693 | AZD6094 | MVP_top60_DNA+RNA_seed | HSP90AB1 | -0.592743 | LUSC | HCC-15 | Tanespimycin |
| MVP_top60_DNA_seed | EGFR | 0.211771 | LUAD | NCI-H1793 | CI-1033 | MVP_top60_DNA+RNA_seed | SMARCA4 | -0.568088 | LUSC | SK-MES-1 | PFI3 |
| MVP_top60_DNA_seed | ERBB2 | 0.211771 | LUAD | NCI-H1793 | CI-1033 | MVP_top60_DNA+RNA_seed | SMARCA2 | -0.568088 | LUSC | SK-MES-1 | PFI3 |
| MVP_top60_DNA_seed | EGFR | 0.211885 | LUAD | HOP-62 | Erlotinib | MVP_top60_DNA+RNA_seed | KRAS | -0.540589 | LUSC | NCI-H2170 | KRAS (G12C) Inhibitor-1 |
| MVP_top60_DNA_seed | ATM | 0.212281 | LUAD | NCI-H1563 | KU-55933 | MVP_top60_DNA+RNA_seed | PIK3CA | -0.531937 | LUSC | HCC-15 | Buparlisib |
| MVP_top60_DNA_seed | MET | 0.212541 | LUAD | ABC-1 | JNJ38877605 | MVP_top60_DNA+RNA_seed | CREBBP | -0.518356 | LUSC | KNS-62 | SGC-CBP30 |
| MVP_top60_DNA_seed | GSK3B | 0.213401 | LUAD | NCI-H1563 | AZD7969 | MVP_top60_DNA+RNA_seed | HDAC1 | -0.485702 | LUSC | EPLC-272H | Romidepsin |
| MVP_top60_DNA_seed | MAPK1 | 0.214709 | LUAD | NCI-H1975 | FR-180204 | MVP_top60_DNA+RNA_seed | HDAC3 | -0.485702 | LUSC | EPLC-272H | Romidepsin |
| MVP_top60_DNA_seed | SMARCA4 | 0.214719 | LUAD | NCI-H2085 | PFI-3 | MVP_top60_DNA+RNA_seed | HSP90AB1 | -0.479545 | LUSC | KNS-62 | Luminespib |
| MVP_top60_DNA_seed | HSP90AA1 | 0.21503 | LUAD | NCI-H23 | SNX-2112 | MVP_top60_DNA+RNA_seed | HSP90AB1 | -0.464324 | LUSC | SK-MES-1 | Tanespimycin |

| | | | | | | | | | | | |
|---|---|---|---|---|---|---|---|---|---|---|---|
| MVP_top60_DNA_seed | BRAF | 0.215476 | LUAD | NCI-H2023 | Dabrafenib | MVP_top60_DNA+RNA_seed | PIK3CA | -0.461466 | LUSC | NCI-H2170 | AZD8186 |
| MVP_top60_DNA_seed | FGFR1 | 0.21555 | LUAD | NCI-H23 | FGFR_3831 | MVP_top60_DNA+RNA_seed | CREBBP | -0.459354 | LUSC | EBC-1 | SGC-CBP30 |
| MVP_top60_DNA_seed | AURKA | 0.215798 | LUAD | NCI-H1355 | Tozasertib | MVP_top60_DNA+RNA_seed | CREBBP | -0.453786 | LUSC | EPLC-272H | SGC-CBP30 |
| MVP_top60_DNA_seed | AURKA | 0.216129 | LUAD | HOP-62 | Alisertib | MVP_top60_DNA+RNA_seed | HDAC1 | -0.450813 | LUSC | LK-2 | PCI-34051 |
| MVP_top60_DNA_seed | AKT1 | 0.216239 | LUAD | LC-2-ad | MK-2206 | MVP_top60_DNA+RNA_seed | SMARCA4 | -0.405381 | LUSC | EPLC-272H | PFI3 |
| MVP_top60_DNA_seed | MET | 0.216392 | LUAD | NCI-H650 | JNJ38877605 | MVP_top60_DNA+RNA_seed | SMARCA2 | -0.405381 | LUSC | EPLC-272H | PFI3 |
| MVP_top60_DNA_seed | ATM | 0.217734 | LUAD | SK-LU-1 | KU-60019 | MVP_top60_DNA+RNA_seed | HSP90AB1 | -0.397576 | LUSC | SK-MES-1 | Luminespib |
| MVP_top60_DNA_seed | BRAF | 0.21935 | LUAD | NCI-H1650 | AZ628 | MVP_top60_DNA+RNA_seed | SMARCA4 | -0.358104 | LUSC | RERF-LC-Sq1 | PFI3 |
| MVP_top60_DNA_seed | SMARCA4 | 0.220239 | LUAD | NCI-H2228 | PFI-3 | MVP_top60_DNA+RNA_seed | SMARCA2 | -0.358104 | LUSC | RERF-LC-Sq1 | PFI3 |
| MVP_top60_DNA_seed | EGFR | 0.220298 | LUAD | NCI-H2291 | CI-1033 | MVP_top60_DNA+RNA_seed | SMARCA4 | -0.354313 | LUSC | NCI-H2170 | PFI3 |
| MVP_top60_DNA_seed | ERBB2 | 0.220298 | LUAD | NCI-H2291 | CI-1033 | MVP_top60_DNA+RNA_seed | SMARCA2 | -0.354313 | LUSC | NCI-H2170 | PFI3 |
| MVP_top60_DNA_seed | PIK3CA | 0.221675 | LUAD | EKVX | PI3Ka_4409 | MVP_top60_DNA+RNA_seed | HDAC1 | -0.348232 | LUSC | HARA | Romidepsin |
| MVP_top60_DNA_seed | IGF1R | 0.222188 | LUAD | NCI-H2030 | GSK1904529A | MVP_top60_DNA+RNA_seed | HDAC3 | -0.348232 | LUSC | HARA | Romidepsin |
| MVP_top60_DNA_seed | GSK3B | 0.222978 | LUAD | NCI-H358 | KIN001-042 | MVP_top60_DNA+RNA_seed | PIK3CA | -0.33911 | LUSC | NCI-H520 | GNE-317 |
| MVP_top60_DNA_seed | BRAF | 0.223477 | LUAD | NCI-H23 | PLX-4720 | MVP_top60_DNA+RNA_seed | EZH2 | -0.339083 | LUSC | EPLC-272H | GSK343 |
| MVP_top60_DNA_seed | AURKA | 0.224676 | LUAD | NCI-H2085 | GSK1070916 | MVP_top60_DNA+RNA_seed | PIK3CA | -0.313509 | LUSC | LOU-NH91 | Alpelisib |
| MVP_top60_DNA_seed | EGFR | 0.224853 | LUAD | NCI-H2085 | Cetuximab | MVP_top60_DNA+RNA_seed | PIK3CA | -0.290278 | LUSC | RERF-LC-Sq1 | Buparlisib |
| MVP_top60_DNA_seed | MAPK1 | 0.225491 | LUAD | NCI-H322M | VX-11e | MVP_top60_DNA+RNA_seed | PIK3CA | -0.287494 | LUSC | LOU-NH91 | GNE-317 |
| MVP_top60_DNA_seed | IGF1R | 0.225755 | LUAD | NCI-H2342 | Linsitinib | MVP_top60_DNA+RNA_seed | HSP90AB1 | -0.287353 | LUSC | LOU-NH91 | Tanespimycin |
| MVP_top60_DNA_seed | HSP90AA1 | 0.226962 | LUAD | RERF-LC-MS | Luminespib | MVP_top60_DNA+RNA_seed | PIK3CA | -0.283547 | LUSC | LK-2 | GNE-317 |
| MVP_top60_DNA_seed | FGFR1 | 0.227411 | LUAD | EMC-BAC-2 | PD173074 | MVP_top60_DNA+RNA_seed | PIK3CA | -0.280465 | LUSC | LOU-NH91 | AZD8186 |
| MVP_top60_DNA_seed | FGFR1 | 0.227719 | LUAD | HCC-78 | AZD4547 | MVP_top60_DNA+RNA_seed | PIK3CA | -0.27608 | LUSC | NCI-H1869 | AZD8186 |
| MVP_top60_DNA_seed | SMARCA4 | 0.227934 | LUAD | ABC-1 | PFI-3 | MVP_top60_DNA+RNA_seed | PIK3CA | -0.270405 | LUSC | LK-2 | Buparlisib |
| MVP_top60_DNA_seed | HDAC1 | 0.229517 | LUAD | NCI-H1734 | Dacinostat | MVP_top60_DNA+RNA_seed | HSP90AB1 | -0.259429 | LUSC | EBC-1 | Tanespimycin |
| MVP_top60_DNA_seed | MET | 0.229623 | LUAD | LC-2-ad | AZD6094 | MVP_top60_DNA+RNA_seed | HDAC1 | -0.251449 | LUSC | NCI-H1869 | Romidepsin |
| MVP_top60_DNA_seed | MDM2 | 0.230051 | LUAD | HCC-44 | Serdemetan | MVP_top60_DNA+RNA_seed | HDAC3 | -0.251449 | LUSC | NCI-H1869 | Romidepsin |
| MVP_top60_DNA_seed | GSK3B | 0.230559 | LUAD | NCI-H2405 | CHIR-99021 | MVP_top60_DNA+RNA_seed | EZH2 | -0.248381 | LUSC | KNS-62 | GSK343 |
| MVP_top60_DNA_seed | EGFR | 0.230563 | LUAD | NCI-H2023 | Afatinib | MVP_top60_DNA+RNA_seed | HDAC1 | -0.244018 | LUSC | SK-MES-1 | Romidepsin |
| MVP_top60_DNA_seed | ERBB2 | 0.230563 | LUAD | NCI-H2023 | Afatinib | MVP_top60_DNA+RNA_seed | HDAC3 | -0.244018 | LUSC | SK-MES-1 | Romidepsin |
| MVP_top60_DNA_seed | SMARCA4 | 0.230838 | LUAD | NCI-H441 | PFI-3 | MVP_top60_DNA+RNA_seed | HSP90AB1 | -0.228448 | LUSC | EBC-1 | Luminespib |
| MVP_top60_DNA_seed | HDAC1 | 0.231683 | LUAD | NCI-H2347 | CAY10603 | MVP_top60_DNA+RNA_seed | PIK3CA | -0.210267 | LUSC | EBC-1 | Buparlisib |
| MVP_top60_DNA_seed | BRAF | 0.232436 | LUAD | SW1573 | SB590885 | MVP_top60_DNA+RNA_seed | HSP90AB1 | -0.198681 | LUSC | KNS-62 | Tanespimycin |
| MVP_top60_DNA_seed | ATM | 0.232465 | LUAD | NCI-H1355 | KU-60019 | MVP_top60_DNA+RNA_seed | EZH2 | -0.197828 | LUSC | NCI-H2170 | GSK343 |
| MVP_top60_DNA_seed | SMARCA4 | 0.234839 | LUAD | HOP-62 | PFI-3 | MVP_top60_DNA+RNA_seed | HDAC1 | -0.194467 | LUSC | NCI-H520 | Entinostat |
| MVP_top60_DNA_seed | IGF1R | 0.235091 | LUAD | HCC-78 | Linsitinib | MVP_top60_DNA+RNA_seed | HDAC3 | -0.194467 | LUSC | NCI-H520 | Entinostat |
| MVP_top60_DNA_seed | IGF1R | 0.23597 | LUAD | NCI-H3122 | Linsitinib | MVP_top60_DNA+RNA_seed | HDAC1 | -0.193978 | LUSC | LK-2 | Romidepsin |
| MVP_top60_DNA_seed | EGFR | 0.237024 | LUAD | NCI-H1734 | AST-1306 | MVP_top60_DNA+RNA_seed | HDAC3 | -0.193978 | LUSC | LK-2 | Romidepsin |
| MVP_top60_DNA_seed | ATM | 0.237643 | LUAD | NCI-H1792 | CP466722 | MVP_top60_DNA+RNA_seed | HDAC1 | -0.185329 | LUSC | NCI-H2170 | Entinostat |
| MVP_top60_DNA_seed | BRAF | 0.238286 | LUAD | H3255 | Dabrafenib | MVP_top60_DNA+RNA_seed | HDAC3 | -0.185329 | LUSC | NCI-H2170 | Entinostat |
| MVP_top60_DNA_seed | MET | 0.238623 | LUAD | H3255 | SU11274 | MVP_top60_DNA+RNA_seed | HDAC1 | -0.176826 | LUSC | SK-MES-1 | PCI-34051 |
| MVP_top60_DNA_seed | AURKA | 0.238835 | LUAD | PC-14 | CD532 | MVP_top60_DNA+RNA_seed | HSP90AB1 | -0.170613 | LUSC | LK-2 | Tanespimycin |

| | | | | | | | | | | | |
|---|---|---|---|---|---|---|---|---|---|---|---|
| MVP_top60_DNA_seed | EGFR | 0.239045 | LUAD | NCI-H838 | Afatinib | MVP_top60_DNA+RNA_seed | EZH2 | -0.159379 | LUSC | HARA | GSK343 |
| MVP_top60_DNA_seed | ERBB2 | 0.239045 | LUAD | NCI-H838 | Afatinib | MVP_top60_DNA+RNA_seed | CREBBP | -0.158355 | LUSC | HARA | SGC-CBP30 |
| MVP_top60_DNA_seed | HSP90AA1 | 0.241282 | LUAD | Calu-6 | Luminespib | MVP_top60_DNA+RNA_seed | HSP90AB1 | -0.144757 | LUSC | NCI-H2170 | Luminespib |
| MVP_top60_DNA_seed | AURKA | 0.241647 | LUAD | HOP-62 | CD532 | MVP_top60_DNA+RNA_seed | HSP90AB1 | -0.134436 | LUSC | LC-1-sq | Tanespimycin |
| MVP_top60_DNA_seed | AKT1 | 0.241808 | LUAD | PC-14 | BAY AKT1 | MVP_top60_DNA+RNA_seed | HDAC1 | -0.11281 | LUSC | EPLC-272H | Entinostat |
| MVP_top60_DNA_seed | CREBBP | 0.242344 | LUAD | LXF-289 | I-CBP112 | MVP_top60_DNA+RNA_seed | HDAC3 | -0.11281 | LUSC | EPLC-272H | Entinostat |
| MVP_top60_DNA_seed | SRC | 0.242349 | LUAD | HOP-62 | Bosutinib | MVP_top60_DNA+RNA_seed | PIK3CA | -0.086508 | LUSC | LC-1-sq | Alpelisib |
| MVP_top60_DNA_seed | FGFR1 | 0.243107 | LUAD | NCI-H1650 | PD173074 | MVP_top60_DNA+RNA_seed | HDAC1 | -0.081973 | LUSC | EPLC-272H | PCI-34051 |
| MVP_top60_DNA_seed | BRAF | 0.243357 | LUAD | NCI-H2405 | PLX-4720 | MVP_top60_DNA+RNA_seed | SMARCA4 | -0.070697 | LUSC | HARA | PFI3 |
| MVP_top60_DNA_seed | HDAC1 | 0.243911 | LUAD | NCI-H1792 | JQ12 | MVP_top60_DNA+RNA_seed | SMARCA2 | -0.070697 | LUSC | HARA | PFI3 |
| MVP_top60_DNA_seed | MDM2 | 0.243967 | LUAD | EMC-BAC-2 | Serdemetan | MVP_top60_DNA+RNA_seed | HDAC1 | -0.057777 | LUSC | NCI-H2170 | Romidepsin |
| MVP_top60_DNA_seed | AKT1 | 0.243992 | LUAD | NCI-H2122 | BAY AKT1 | MVP_top60_DNA+RNA_seed | HDAC3 | -0.057777 | LUSC | NCI-H2170 | Romidepsin |
| MVP_top60_DNA_seed | HDAC1 | 0.244549 | LUAD | PC-3_[JPC-3] | Belinostat | MVP_top60_DNA+RNA_seed | PIK3CA | -0.039436 | LUSC | LOU-NH91 | Buparlisib |
| MVP_top60_DNA_seed | HDAC1 | 0.245028 | LUAD | NCI-H1755 | Tubastatin A | MVP_top60_DNA+RNA_seed | CREBBP | -0.037979 | LUSC | LOU-NH91 | SGC-CBP30 |
| MVP_top60_DNA_seed | IGF1R | 0.245168 | LUAD | NCI-H2347 | GSK1904529A | MVP_top60_DNA+RNA_seed | KRAS | -0.016457 | LUSC | KNS-62 | KRAS (G12C) Inhibitor-1 |
| MVP_top60_DNA_seed | SMARCA4 | 0.245747 | LUAD | EMC-BAC-2 | PFI-3 | MVP_top60_DNA+RNA_seed | TP53 | -0.002669 | LUSC | NCI-H1869 | MIRA-1 |
| MVP_top60_DNA_seed | MET | 0.245869 | LUAD | NCI-H1650 | JNJ38877605 | MVP_top60_DNA+RNA_seed | TP53 | 0.006589 | LUSC | NCI-H2170 | MIRA-1 |
| MVP_top60_DNA_seed | EGFR | 0.246169 | LUAD | NCI-H1648 | Cetuximab | MVP_top60_DNA+RNA_seed | KRAS | 0.031088 | LUSC | NCI-H1869 | KRAS (G12C) Inhibitor-1 |
| MVP_top60_DNA_seed | MAPK1 | 0.246313 | LUAD | NCI-H2085 | VX-11e | MVP_top60_DNA+RNA_seed | PIK3CA | 0.035824 | LUSC | SK-MES-1 | AZD8186 |
| MVP_top60_DNA_seed | HDAC1 | 0.246469 | LUAD | NCI-H441 | JQ12 | MVP_top60_DNA+RNA_seed | HDAC1 | 0.057695 | LUSC | LC-1-sq | Romidepsin |
| MVP_top60_DNA_seed | MET | 0.247108 | LUAD | NCI-H358 | AZD6094 | MVP_top60_DNA+RNA_seed | HDAC3 | 0.057695 | LUSC | LC-1-sq | Romidepsin |
| MVP_top60_DNA_seed | EGFR | 0.247176 | LUAD | NCI-H1734 | Gefitinib | MVP_top60_DNA+RNA_seed | TP53 | 0.085711 | LUSC | HCC-15 | MIRA-1 |
| MVP_top60_DNA_seed | EGFR | 0.248345 | LUAD | NCI-H2085 | Afatinib | MVP_top60_DNA+RNA_seed | HDAC1 | 0.086914 | LUSC | NCI-H2170 | PCI-34051 |
| MVP_top60_DNA_seed | ERBB2 | 0.248345 | LUAD | NCI-H2085 | Afatinib | MVP_top60_DNA+RNA_seed | EZH2 | 0.088756 | LUSC | NCI-H2170 | GSK343 |
| MVP_top60_DNA_seed | HDAC1 | 0.248443 | LUAD | EMC-BAC-1 | AR-42 | MVP_top60_DNA+RNA_seed | HDAC1 | 0.112647 | LUSC | HCC-15 | Romidepsin |
| MVP_top60_DNA_seed | AKT1 | 0.248464 | LUAD | NCI-H2122 | MK-2206 | MVP_top60_DNA+RNA_seed | HDAC3 | 0.112647 | LUSC | HCC-15 | Romidepsin |
| MVP_top60_DNA_seed | BRAF | 0.249717 | LUAD | NCI-H2291 | SB590885 | MVP_top60_DNA+RNA_seed | EZH2 | 0.116397 | LUSC | RERF-LC-Sq1 | GSK343 |
| MVP_top60_DNA_seed | AURKA | 0.250348 | LUAD | NCI-H1651 | GSK1070916 | MVP_top60_DNA+RNA_seed | TP53 | 0.117814 | LUSC | NCI-H520 | MIRA-1 |
| MVP_top60_DNA_seed | CREBBP | 0.251016 | LUAD | HOP-62 | I-CBP112 | MVP_top60_DNA+RNA_seed | KRAS | 0.119605 | LUSC | RERF-LC-Sq1 | KRAS (G12C) Inhibitor-1 |
| MVP_top60_DNA_seed | HDAC1 | 0.251314 | LUAD | NCI-H1693 | Dacinostat | MVP_top60_DNA+RNA_seed | TP53 | 0.127705 | LUSC | HARA | MIRA-1 |
| MVP_top60_DNA_seed | AKT1 | 0.253053 | LUAD | NCI-H1734 | AKT inhibitor VIII | MVP_top60_DNA+RNA_seed | KRAS | 0.128121 | LUSC | LK-2 | KRAS (G12C) Inhibitor-1 |
| MVP_top60_DNA_seed | MAPK1 | 0.253324 | LUAD | EMC-BAC-2 | FR-180204 | MVP_top60_DNA+RNA_seed | SMARCA4 | 0.131684 | LUSC | KNS-62 | PFI3 |
| MVP_top60_DNA_seed | AURKA | 0.253698 | LUAD | EMC-BAC-2 | Genentech Cpd 10 | MVP_top60_DNA+RNA_seed | SMARCA2 | 0.131684 | LUSC | KNS-62 | PFI3 |
| MVP_top60_DNA_seed | AURKA | 0.254645 | LUAD | NCI-H2030 | ZM447439 | MVP_top60_DNA+RNA_seed | EZH2 | 0.150594 | LUSC | RERF-LC-Sq1 | GSK343 |
| MVP_top60_DNA_seed | GSK3B | 0.256008 | LUAD | HCC-44 | CHIR-99021 | MVP_top60_DNA+RNA_seed | TP53 | 0.175425 | LUSC | LC-1-sq | MIRA-1 |
| MVP_top60_DNA_seed | EGFR | 0.256585 | LUAD | NCI-H1693 | Pelitinib | MVP_top60_DNA+RNA_seed | PIK3CA | 0.179874 | LUSC | NCI-H520 | Buparlisib |
| MVP_top60_DNA_seed | FGFR1 | 0.256856 | LUAD | NCI-H1975 | FGFR_3831 | MVP_top60_DNA+RNA_seed | HDAC1 | 0.205321 | LUSC | HARA | PCI-34051 |
| MVP_top60_DNA_seed | HDAC1 | 0.257158 | LUAD | NCI-H2405 | AR-42 | MVP_top60_DNA+RNA_seed | CREBBP | 0.208739 | LUSC | NCI-H2170 | SGC-CBP30 |
| MVP_top60_DNA_seed | GSK3B | 0.257273 | LUAD | NCI-H2087 | KIN001-042 | MVP_top60_DNA+RNA_seed | HDAC1 | 0.211623 | LUSC | LOU-NH91 | Entinostat |
| MVP_top60_DNA_seed | FGFR1 | 0.257612 | LUAD | RERF-LC-MS | AZD4547 | MVP_top60_DNA+RNA_seed | HDAC3 | 0.211623 | LUSC | LOU-NH91 | Entinostat |
| MVP_top60_DNA_seed | BRAF | 0.258235 | LUAD | NCI-H1355 | AZ628 | MVP_top60_DNA+RNA_seed | TP53 | 0.215474 | LUSC | LK-2 | MIRA-1 |

| | | | | | | | | | | | |
|---|---|---|---|---|---|---|---|---|---|---|---|
| MVP_top60_DNA_seed | BRAF | 0.259102 | LUAD | NCI-H1568 | Dabrafenib | MVP_top60_DNA+RNA_seed | KRAS | 0.220921 | LUSC | HARA | KRAS (G12C) Inhibitor-1 |
| MVP_top60_DNA_seed | GSK3B | 0.260289 | LUAD | NCI-H1734 | SB216763 | MVP_top60_DNA+RNA_seed | KRAS | 0.232939 | LUSC | LOU-NH91 | KRAS (G12C) Inhibitor-1 |
| MVP_top60_DNA_seed | AURKA | 0.260305 | LUAD | Calu-6 | Alisertib | MVP_top60_DNA+RNA_seed | SMARCA4 | 0.236079 | LUSC | LC-1-sq | PFI3 |
| MVP_top60_DNA_seed | AKT1 | 0.260708 | LUAD | ABC-1 | BAY AKT1 | MVP_top60_DNA+RNA_seed | SMARCA2 | 0.236079 | LUSC | LC-1-sq | PFI3 |
| MVP_top60_DNA_seed | EGFR | 0.261547 | LUAD | NCI-H2228 | AST-1306 | MVP_top60_DNA+RNA_seed | HSP90AB1 | 0.240073 | LUSC | NCI-H520 | Tanespimycin |
| MVP_top60_DNA_seed | HDAC1 | 0.2616 | LUAD | HCC-827 | CAY10603 | MVP_top60_DNA+RNA_seed | SMARCA4 | 0.307263 | LUSC | HCC-15 | PFI3 |
| MVP_top60_DNA_seed | IGF1R | 0.261679 | LUAD | NCI-H1563 | BMS-754807 | MVP_top60_DNA+RNA_seed | SMARCA2 | 0.307263 | LUSC | HCC-15 | PFI3 |
| MVP_top60_DNA_seed | HDAC1 | 0.261942 | LUAD | EMC-BAC-1 | Belinostat | MVP_top60_DNA+RNA_seed | HDAC1 | 0.317533 | LUSC | HCC-15 | PCI-34051 |
| MVP_top60_DNA_seed | CREBBP | 0.263071 | LUAD | NCI-H1568 | I-CBP112 | MVP_top60_DNA+RNA_seed | PIK3CA | 0.328737 | LUSC | SK-MES-1 | GNE-317 |
| MVP_top60_DNA_seed | ERBB2 | 0.264143 | LUAD | RERF-LC-MS | CP724714 | MVP_top60_DNA+RNA_seed | PIK3CA | 0.332925 | LUSC | HARA | AZD8186 |
| MVP_top60_DNA_seed | MET | 0.264575 | LUAD | NCI-H2030 | AZD6094 | MVP_top60_DNA+RNA_seed | HDAC1 | 0.337079 | LUSC | KNS-62 | PCI-34051 |
| MVP_top60_DNA_seed | GSK3B | 0.265986 | LUAD | Calu-6 | CHIR-99021 | MVP_top60_DNA+RNA_seed | HDAC1 | 0.346509 | LUSC | LC-1-sq | Entinostat |
| MVP_top60_DNA_seed | PIK3CA | 0.267375 | LUAD | NCI-H2405 | PI3Ka_4409 | MVP_top60_DNA+RNA_seed | HDAC3 | 0.346509 | LUSC | LC-1-sq | Entinostat |
| MVP_top60_DNA_seed | EGFR | 0.268679 | LUAD | EMC-BAC-2 | CI-1033 | MVP_top60_DNA+RNA_seed | PIK3CA | 0.357542 | LUSC | NCI-H1869 | GNE-317 |
| MVP_top60_DNA_seed | ERBB2 | 0.268679 | LUAD | EMC-BAC-2 | CI-1033 | MVP_top60_DNA+RNA_seed | EZH2 | 0.370852 | LUSC | NCI-H520 | GSK343 |
| MVP_top60_DNA_seed | HSP90AA1 | 0.268714 | LUAD | SK-LU-1 | SNX-2112 | MVP_top60_DNA+RNA_seed | TP53 | 0.371749 | LUSC | EPLC-272H | MIRA-1 |
| MVP_top60_DNA_seed | MET | 0.26953 | LUAD | NCI-H2085 | AZD6094 | MVP_top60_DNA+RNA_seed | HDAC1 | 0.378496 | LUSC | RERF-LC-Sq1 | Entinostat |
| MVP_top60_DNA_seed | EGFR | 0.270282 | LUAD | NCI-H3122 | AZD8931 | MVP_top60_DNA+RNA_seed | HDAC3 | 0.378496 | LUSC | RERF-LC-Sq1 | Entinostat |
| MVP_top60_DNA_seed | ERBB2 | 0.270282 | LUAD | NCI-H3122 | AZD8931 | MVP_top60_DNA+RNA_seed | PIK3CA | 0.384334 | LUSC | RERF-LC-Sq1 | AZD8186 |
| MVP_top60_DNA_seed | ERBB3 | 0.270282 | LUAD | NCI-H3122 | AZD8931 | MVP_top60_DNA+RNA_seed | HDAC1 | 0.415626 | LUSC | LOU-NH91 | PCI-34051 |
| MVP_top60_DNA_seed | EGFR | 0.271199 | LUAD | NCI-H1650 | AZD8931 | MVP_top60_DNA+RNA_seed | EZH2 | 0.444696 | LUSC | LK-2 | GSK343 |
| MVP_top60_DNA_seed | ERBB2 | 0.271199 | LUAD | NCI-H1650 | AZD8931 | MVP_top60_DNA+RNA_seed | SMARCA4 | 0.458455 | LUSC | NCI-H520 | PFI3 |
| MVP_top60_DNA_seed | ERBB3 | 0.271199 | LUAD | NCI-H1650 | AZD8931 | MVP_top60_DNA+RNA_seed | SMARCA2 | 0.458455 | LUSC | NCI-H520 | PFI3 |
| MVP_top60_DNA_seed | MDM2 | 0.271879 | LUAD | NCI-H2291 | Serdemetan | MVP_top60_DNA+RNA_seed | HSP90AB1 | 0.477175 | LUSC | LOU-NH91 | Luminespib |
| MVP_top60_DNA_seed | EGFR | 0.272312 | LUAD | NCI-H1793 | Afatinib | MVP_top60_DNA+RNA_seed | PIK3CA | 0.493276 | LUSC | LC-1-sq | Buparlisib |
| MVP_top60_DNA_seed | ERBB2 | 0.272312 | LUAD | NCI-H1793 | Afatinib | MVP_top60_DNA+RNA_seed | SMARCA4 | 0.498532 | LUSC | LK-2 | PFI3 |
| MVP_top60_DNA_seed | AKT1 | 0.273024 | LUAD | HOP-62 | MK-2206 | MVP_top60_DNA+RNA_seed | SMARCA2 | 0.498532 | LUSC | LK-2 | PFI3 |
| MVP_top60_DNA_seed | AURKA | 0.273142 | LUAD | NCI-H1693 | Genentech Cpd 10 | MVP_top60_DNA+RNA_seed | EZH2 | 0.510513 | LUSC | HCC-15 | GSK343 |
| MVP_top60_DNA_seed | MDM2 | 0.274105 | LUAD | LC-2-ad | Nutlin-3a (-) | MVP_top60_DNA+RNA_seed | HDAC1 | 0.521161 | LUSC | HARA | Entinostat |
| MVP_top60_DNA_seed | MAPK1 | 0.27459 | LUAD | NCI-H2122 | VX-11e | MVP_top60_DNA+RNA_seed | HDAC3 | 0.521161 | LUSC | HARA | Entinostat |
| MVP_top60_DNA_seed | IGF1R | 0.275113 | LUAD | H3255 | Linsitinib | MVP_top60_DNA+RNA_seed | PIK3CA | 0.528927 | LUSC | HCC-15 | GNE-317 |
| MVP_top60_DNA_seed | HDAC1 | 0.275115 | LUAD | NCI-H2347 | Dacinostat | MVP_top60_DNA+RNA_seed | CREBBP | 0.555644 | LUSC | SK-MES-1 | SGC-CBP30 |
| MVP_top60_DNA_seed | EGFR | 0.275273 | LUAD | NCI-H1944 | CI-1033 | MVP_top60_DNA+RNA_seed | HSP90AB1 | 0.557264 | LUSC | LK-2 | Luminespib |
| MVP_top60_DNA_seed | ERBB2 | 0.275273 | LUAD | NCI-H1944 | CI-1033 | MVP_top60_DNA+RNA_seed | PIK3CA | 0.568534 | LUSC | KNS-62 | Alpelisib |
| MVP_top60_DNA_seed | GSK3B | 0.275804 | LUAD | HCC-827 | CHIR-99021 | MVP_top60_DNA+RNA_seed | EZH2 | 0.573817 | LUSC | NCI-H520 | GSK343 |
| MVP_top60_DNA_seed | GSK3B | 0.276437 | LUAD | SW1573 | AZD7969 | MVP_top60_DNA+RNA_seed | TP53 | 0.593917 | LUSC | RERF-LC-Sq1 | MIRA-1 |
| MVP_top60_DNA_seed | SRC | 0.276463 | LUAD | NCI-H1648 | HG-6-71-01 | MVP_top60_DNA+RNA_seed | HDAC1 | 0.60373 | LUSC | KNS-62 | Romidepsin |
| MVP_top60_DNA_seed | EGFR | 0.276783 | LUAD | NCI-H2087 | AZD8931 | MVP_top60_DNA+RNA_seed | HDAC3 | 0.60373 | LUSC | KNS-62 | Romidepsin |
| MVP_top60_DNA_seed | ERBB2 | 0.276783 | LUAD | NCI-H2087 | AZD8931 | MVP_top60_DNA+RNA_seed | EZH2 | 0.609998 | LUSC | KNS-62 | GSK343 |
| MVP_top60_DNA_seed | ERBB3 | 0.276783 | LUAD | NCI-H2087 | AZD8931 | MVP_top60_DNA+RNA_seed | PIK3CA | 0.621815 | LUSC | KNS-62 | GNE-317 |
| MVP_top60_DNA_seed | IGF1R | 0.276907 | LUAD | NCI-H1781 | GSK1904529A | MVP_top60_DNA+RNA_seed | HDAC1 | 0.622254 | LUSC | KNS-62 | Entinostat |

| | | | | | | | | | | | |
|---|---|---|---|---|---|---|---|---|---|---|---|
| MVP_top60_DNA_seed | BRAF | 0.277578 | LUAD | NCI-H1755 | SB590885 | MVP_top60_DNA+RNA_seed | HDAC3 | 0.622254 | LUSC | KNS-62 | Entinostat |
| MVP_top60_DNA_seed | BRAF | 0.278328 | LUAD | NCI-H1793 | PLX-4720 | MVP_top60_DNA+RNA_seed | HDAC1 | 0.623486 | LUSC | NCI-H1869 | Entinostat |
| MVP_top60_DNA_seed | MET | 0.278698 | LUAD | HCC-44 | SU11274 | MVP_top60_DNA+RNA_seed | HDAC3 | 0.623486 | LUSC | NCI-H1869 | Entinostat |
| MVP_top60_DNA_seed | EGFR | 0.278978 | LUAD | NCI-H650 | Afatinib | MVP_top60_DNA+RNA_seed | PIK3CA | 0.637805 | LUSC | NCI-H520 | Alpelisib |
| MVP_top60_DNA_seed | ERBB2 | 0.278978 | LUAD | NCI-H650 | Afatinib | MVP_top60_DNA+RNA_seed | HDAC1 | 0.637975 | LUSC | EBC-1 | PCI-34051 |
| MVP_top60_DNA_seed | SRC | 0.279928 | LUAD | NCI-H1944 | Bosutinib | MVP_top60_DNA+RNA_seed | PIK3CA | 0.638856 | LUSC | SK-MES-1 | Buparlisib |
| MVP_top60_DNA_seed | EGFR | 0.280279 | LUAD | NCI-H2087 | Afatinib | MVP_top60_DNA+RNA_seed | CREBBP | 0.647097 | LUSC | LK-2 | SGC-CBP30 |
| MVP_top60_DNA_seed | ERBB2 | 0.280279 | LUAD | NCI-H2087 | Afatinib | MVP_top60_DNA+RNA_seed | PIK3CA | 0.654019 | LUSC | KNS-62 | Buparlisib |
| MVP_top60_DNA_seed | AURKA | 0.28048 | LUAD | NCI-H2291 | CD532 | MVP_top60_DNA+RNA_seed | SMARCA4 | 0.667792 | LUSC | LOU-NH91 | PFI3 |
| MVP_top60_DNA_seed | AKT1 | 0.280534 | LUAD | NCI-H1781 | MK-2206 | MVP_top60_DNA+RNA_seed | SMARCA2 | 0.667792 | LUSC | LOU-NH91 | PFI3 |
| MVP_top60_DNA_seed | IGF1R | 0.280936 | LUAD | NCI-H1781 | BMS-754807 | MVP_top60_DNA+RNA_seed | EZH2 | 0.667914 | LUSC | HCC-15 | GSK343 |
| MVP_top60_DNA_seed | EGFR | 0.281767 | LUAD | NCI-H1437 | Pelitinib | MVP_top60_DNA+RNA_seed | KRAS | 0.668767 | LUSC | HCC-15 | KRAS (G12C) Inhibitor-1 |
| MVP_top60_DNA_seed | EGFR | 0.28187 | LUAD | NCI-H1838 | Afatinib | MVP_top60_DNA+RNA_seed | EZH2 | 0.681502 | LUSC | LC-1-sq | GSK343 |
| MVP_top60_DNA_seed | ERBB2 | 0.28187 | LUAD | NCI-H1838 | Afatinib | MVP_top60_DNA+RNA_seed | TP53 | 0.691211 | LUSC | KNS-62 | MIRA-1 |
| MVP_top60_DNA_seed | GSK3B | 0.282059 | LUAD | Calu-3 | AZD7969 | MVP_top60_DNA+RNA_seed | HDAC1 | 0.712882 | LUSC | RERF-LC-Sq1 | PCI-34051 |
| MVP_top60_DNA_seed | BRAF | 0.282588 | LUAD | NCI-H1944 | PLX-4720 | MVP_top60_DNA+RNA_seed | TP53 | 0.717299 | LUSC | LOU-NH91 | MIRA-1 |
| MVP_top60_DNA_seed | MDM2 | 0.283097 | LUAD | NCI-H1781 | Serdemetan | MVP_top60_DNA+RNA_seed | CREBBP | 0.723469 | LUSC | NCI-H520 | SGC-CBP30 |
| MVP_top60_DNA_seed | BRAF | 0.283542 | LUAD | NCI-H2085 | SB590885 | MVP_top60_DNA+RNA_seed | PIK3CA | 0.724967 | LUSC | LC-1-sq | AZD8186 |
| MVP_top60_DNA_seed | FGFR1 | 0.283554 | LUAD | NCI-H441 | FGFR_3831 | MVP_top60_DNA+RNA_seed | KRAS | 0.772334 | LUSC | EBC-1 | KRAS (G12C) Inhibitor-1 |
| MVP_top60_DNA_seed | MET | 0.284916 | LUAD | NCI-H2122 | AZD6094 | MVP_top60_DNA+RNA_seed | PIK3CA | 0.786867 | LUSC | LC-1-sq | GNE-317 |
| MVP_top60_DNA_seed | IGF1R | 0.285382 | LUAD | NCI-H2122 | GSK1904529A | MVP_top60_DNA+RNA_seed | EZH2 | 0.807874 | LUSC | NCI-H1869 | GSK343 |
| MVP_top60_DNA_seed | PIK3CA | 0.285903 | LUAD | NCI-H1781 | PI3Ka_4409 | MVP_top60_DNA+RNA_seed | PIK3CA | 0.810293 | LUSC | NCI-H1869 | Alpelisib |
| MVP_top60_DNA_seed | EGFR | 0.286435 | LUAD | Calu-3 | Cetuximab | MVP_top60_DNA+RNA_seed | CREBBP | 0.825808 | LUSC | HCC-15 | SGC-CBP30 |
| MVP_top60_DNA_seed | MET | 0.287545 | LUAD | LXF-289 | SU11274 | MVP_top60_DNA+RNA_seed | EZH2 | 0.832958 | LUSC | LOU-NH91 | GSK343 |
| MVP_top60_DNA_seed | GSK3B | 0.287914 | LUAD | NCI-H1651 | AZD7969 | MVP_top60_DNA+RNA_seed | PIK3CA | 0.833017 | LUSC | HCC-15 | AZD8186 |
| MVP_top60_DNA_seed | SMARCA4 | 0.287923 | LUAD | NCI-H1355 | PFI-3 | MVP_top60_DNA+RNA_seed | TP53 | 0.841218 | LUSC | EBC-1 | MIRA-1 |
| MVP_top60_DNA_seed | PTK2 | 0.288246 | LUAD | NCI-H2405 | PF-562271 | MVP_top60_DNA+RNA_seed | TP53 | 0.849479 | LUSC | SK-MES-1 | MIRA-1 |
| MVP_top60_DNA_seed | AURKA | 0.288768 | LUAD | NCI-H1975 | ZM447439 | MVP_top60_DNA+RNA_seed | EZH2 | 0.896466 | LUSC | SK-MES-1 | GSK343 |
| MVP_top60_DNA_seed | BRAF | 0.289233 | LUAD | NCI-H1792 | PLX-4720 | MVP_top60_DNA+RNA_seed | PIK3CA | 0.906604 | LUSC | LK-2 | AZD8186 |
| MVP_top60_DNA_seed | MDM2 | 0.289725 | LUAD | EMC-BAC-1 | Nutlin-3a (-) | MVP_top60_DNA+RNA_seed | PIK3CA | 0.914289 | LUSC | EBC-1 | Alpelisib |
| MVP_top60_DNA_seed | SMARCA4 | 0.289915 | LUAD | RERF-LC-MS | PFI-3 | MVP_top60_DNA+RNA_seed | SMARCA4 | 0.924267 | LUSC | NCI-H226 | PFI3 |
| MVP_top60_DNA_seed | ATM | 0.291964 | LUAD | EKVX | KU-55933 | MVP_top60_DNA+RNA_seed | SMARCA2 | 0.924267 | LUSC | NCI-H226 | PFI3 |
| MVP_top60_DNA_seed | BRAF | 0.292115 | LUAD | NCI-H1975 | RAF_9304 | MVP_top60_DNA+RNA_seed | HSP90AB1 | 0.953676 | LUSC | NCI-H520 | Luminespib |
| MVP_top60_DNA_seed | GSK3B | 0.292164 | LUAD | NCI-H1573 | SB216763 | MVP_top60_DNA+RNA_seed | EZH2 | 0.967621 | LUSC | LOU-NH91 | GSK343 |
| MVP_top60_DNA_seed | AURKA | 0.293457 | LUAD | NCI-H2347 | CD532 | MVP_top60_DNA+RNA_seed | CREBBP | 0.984065 | LUSC | NCI-H226 | SGC-CBP30 |
| MVP_top60_DNA_seed | ATM | 0.29535 | LUAD | NCI-H1781 | CP466722 | MVP_top60_DNA+RNA_seed | EZH2 | 1.010255 | LUSC | NCI-H1869 | GSK343 |
| MVP_top60_DNA_seed | GSK3B | 0.295488 | LUAD | A549 | CHIR-99021 | MVP_top60_DNA+RNA_seed | HDAC1 | 1.023897 | LUSC | SK-MES-1 | Entinostat |
| MVP_top60_DNA_seed | FGFR1 | 0.297403 | LUAD | NCI-H1650 | FGFR_3831 | MVP_top60_DNA+RNA_seed | HDAC3 | 1.023897 | LUSC | SK-MES-1 | Entinostat |
| MVP_top60_DNA_seed | MAPK1 | 0.297732 | LUAD | NCI-H2087 | FR-180204 | MVP_top60_DNA+RNA_seed | PIK3CA | 1.049427 | LUSC | NCI-H226 | Alpelisib |
| MVP_top60_DNA_seed | GSK3B | 0.297862 | LUAD | NCI-H2030 | SB216763 | MVP_top60_DNA+RNA_seed | CREBBP | 1.051696 | LUSC | NCI-H1869 | SGC-CBP30 |
| MVP_top60_DNA_seed | AKT1 | 0.298962 | LUAD | NCI-H1648 | BAY AKT1 | MVP_top60_DNA+RNA_seed | EZH2 | 1.05326 | LUSC | LK-2 | GSK343 |

| | | | | | | | | | | | |
|---|---|---|---|---|---|---|---|---|---|---|---|
| MVP_top60_DNA_seed | BRAF | 0.299387 | LUAD | NCI-H522 | PLX-4720 | MVP_top60_DNA+RNA_seed | PIK3CA | 1.066896 | LUSC | KNS-62 | AZD8186 |
| MVP_top60_DNA_seed | EGFR | 0.300269 | LUAD | NCI-H1437 | CUDC-101 | MVP_top60_DNA+RNA_seed | SMARCA4 | 1.068902 | LUSC | NCI-H1869 | PFI3 |
| MVP_top60_DNA_seed | ERBB2 | 0.300269 | LUAD | NCI-H1437 | CUDC-101 | MVP_top60_DNA+RNA_seed | SMARCA2 | 1.068902 | LUSC | NCI-H1869 | PFI3 |
| MVP_top60_DNA_seed | FGFR1 | 0.30037 | LUAD | NCI-H2023 | FGFR_3831 | MVP_top60_DNA+RNA_seed | PIK3CA | 1.11521 | LUSC | HCC-15 | Alpelisib |
| MVP_top60_DNA_seed | FGFR1 | 0.302223 | LUAD | HCC-827 | PD173074 | MVP_top60_DNA+RNA_seed | CREBBP | 1.132646 | LUSC | RERF-LC-Sq1 | SGC-CBP30 |
| MVP_top60_DNA_seed | HSP90AA1 | 0.303333 | LUAD | NCI-H1781 | Luminespib | MVP_top60_DNA+RNA_seed | PIK3CA | 1.146915 | LUSC | SK-MES-1 | Alpelisib |
| MVP_top60_DNA_seed | SRC | 0.303414 | LUAD | Calu-6 | Saracatinib | MVP_top60_DNA+RNA_seed | HSP90AB1 | 1.162174 | LUSC | NCI-H226 | Tanespimycin |
| MVP_top60_DNA_seed | HDAC1 | 0.304611 | LUAD | NCI-H2085 | CAY10603 | MVP_top60_DNA+RNA_seed | HDAC1 | 1.16924 | LUSC | NCI-H520 | PCI-34051 |
| MVP_top60_DNA_seed | EGFR | 0.305652 | LUAD | NCI-H1734 | CI-1033 | MVP_top60_DNA+RNA_seed | HSP90AB1 | 1.175011 | LUSC | NCI-H1869 | Luminespib |
| MVP_top60_DNA_seed | ERBB2 | 0.305652 | LUAD | NCI-H1734 | CI-1033 | MVP_top60_DNA+RNA_seed | PIK3CA | 1.175026 | LUSC | LK-2 | Alpelisib |
| MVP_top60_DNA_seed | HSP90AA1 | 0.305751 | LUAD | NCI-H1623 | Tanespimycin | MVP_top60_DNA+RNA_seed | KRAS | 1.183041 | LUSC | NCI-H520 | KRAS (G12C) Inhibitor-1 |
| MVP_top60_DNA_seed | PTK2 | 0.305979 | LUAD | NCI-H522 | PF-562271 | MVP_top60_DNA+RNA_seed | HDAC1 | 1.190399 | LUSC | LK-2 | Entinostat |
| MVP_top60_DNA_seed | HSP90AA1 | 0.306737 | LUAD | NCI-H1623 | Elesclomol | MVP_top60_DNA+RNA_seed | HDAC3 | 1.190399 | LUSC | LK-2 | Entinostat |
| MVP_top60_DNA_seed | AKT1 | 0.306897 | LUAD | NCI-H23 | AKT inhibitor VIII | MVP_top60_DNA+RNA_seed | HDAC1 | 1.250924 | LUSC | NCI-H1869 | PCI-34051 |
| MVP_top60_DNA_seed | EGFR | 0.307653 | LUAD | LC-2-ad | Afatinib | MVP_top60_DNA+RNA_seed | HDAC1 | 1.272295 | LUSC | LC-1-sq | PCI-34051 |
| MVP_top60_DNA_seed | ERBB2 | 0.307653 | LUAD | LC-2-ad | Afatinib | MVP_top60_DNA+RNA_seed | PIK3CA | 1.288456 | LUSC | NCI-H1869 | Buparlisib |
| MVP_top60_DNA_seed | EGFR | 0.307721 | LUAD | NCI-H1793 | Afatinib | MVP_top60_DNA+RNA_seed | HDAC1 | 1.297987 | LUSC | RERF-LC-Sq1 | Romidepsin |
| MVP_top60_DNA_seed | ERBB2 | 0.307721 | LUAD | NCI-H1793 | Afatinib | MVP_top60_DNA+RNA_seed | HDAC3 | 1.297987 | LUSC | RERF-LC-Sq1 | Romidepsin |
| MVP_top60_DNA_seed | MET | 0.307784 | LUAD | SW1573 | SU11274 | MVP_top60_DNA+RNA_seed | KRAS | 1.316273 | LUSC | SK-MES-1 | KRAS (G12C) Inhibitor-1 |
| MVP_top60_DNA_seed | MAPK1 | 0.308461 | LUAD | NCI-H1693 | VX-11e | MVP_top60_DNA+RNA_seed | KRAS | 1.378154 | LUSC | LC-1-sq | KRAS (G12C) Inhibitor-1 |
| MVP_top60_DNA_seed | PIK3CA | 0.30943 | LUAD | NCI-H358 | PI3Ka_4409 | MVP_top60_DNA+RNA_seed | HSP90AB1 | 1.421886 | LUSC | HCC-15 | Luminespib |
| MVP_top60_DNA_seed | MDM2 | 0.310162 | LUAD | NCI-H1355 | Nutlin-3a (-) | MVP_top60_DNA+RNA_seed | EZH2 | 1.536236 | LUSC | NCI-H226 | GSK343 |
| MVP_top60_DNA_seed | AURKA | 0.310511 | LUAD | EMC-BAC-2 | CD532 | MVP_top60_DNA+RNA_seed | HDAC1 | 1.561487 | LUSC | NCI-H226 | Entinostat |
| MVP_top60_DNA_seed | HSP90AA1 | 0.310534 | LUAD | Calu-6 | SNX-2112 | MVP_top60_DNA+RNA_seed | HDAC3 | 1.561487 | LUSC | NCI-H226 | Entinostat |
| MVP_top60_DNA_seed | BRAF | 0.31078 | LUAD | RERF-LC-KJ | PLX-4720 | MVP_top60_DNA+RNA_seed | HSP90AB1 | 1.572441 | LUSC | RERF-LC-Sq1 | Luminespib |
| MVP_top60_DNA_seed | ATM | 0.310922 | LUAD | NCI-H2228 | CP466722 | MVP_top60_DNA+RNA_seed | HSP90AB1 | 1.65739 | LUSC | NCI-H226 | Luminespib |
| MVP_top60_DNA_seed | HSP90AA1 | 0.311178 | LUAD | NCI-H358 | Elesclomol | MVP_top60_DNA+RNA_seed | TP53 | 1.773584 | LUSC | NCI-H226 | MIRA-1 |
| MVP_top60_DNA_seed | BRAF | 0.311802 | LUAD | NCI-H1838 | RAF_9304 | MVP_top60_DNA+RNA_seed | HDAC1 | 1.938385 | LUSC | NCI-H226 | PCI-34051 |
| MVP_top60_DNA_seed | EGFR | 0.313807 | LUAD | NCI-H1568 | Gefitinib | MVP_top60_DNA+RNA_seed | HDAC1 | 1.980422 | LUSC | HCC-15 | Entinostat |
| MVP_top60_DNA_seed | EGFR | 0.314094 | LUAD | NCI-H1975 | Lapatinib | MVP_top60_DNA+RNA_seed | HDAC3 | 1.980422 | LUSC | HCC-15 | Entinostat |
| MVP_top60_DNA_seed | ERBB2 | 0.314094 | LUAD | NCI-H1975 | Lapatinib | MVP_top60_DNA+RNA_seed | HSP90AB1 | 1.996839 | LUSC | LC-1-sq | Luminespib |
| MVP_top60_DNA_seed | ATM | 0.315005 | LUAD | NCI-H1975 | KU-55933 | MVP_top60_DNA+RNA_seed | PIK3CA | 2.103956 | LUSC | NCI-H226 | AZD8186 |
| MVP_top60_DNA_seed | GSK3B | 0.315159 | LUAD | LC-2-ad | KIN001-042 | MVP_top60_DNA+RNA_seed | PIK3CA | 2.336999 | LUSC | NCI-H226 | GNE-317 |
| MVP_top60_DNA_seed | HDAC1 | 0.315659 | LUAD | NCI-H1993 | Dacinostat | MVP_top60_DNA+RNA_seed | PIK3CA | 2.454696 | LUSC | NCI-H226 | Buparlisib |
| MVP_top60_DNA_seed | IGF1R | 0.315751 | LUAD | Calu-6 | BMS-754807 | MVP_top60_DNA+RNA_seed | KRAS | 2.638502 | LUSC | NCI-H226 | KRAS (G12C) Inhibitor-1 |
| MVP_top60_DNA_seed | BRAF | 0.315762 | LUAD | NCI-H3122 | Dabrafenib | MVP_top60_DNA+RNA_seed | EZH2 | 2.732589 | LUSC | NCI-H226 | GSK343 |
| MVP_top60_DNA_seed | EGFR | 0.316047 | LUAD | NCI-H2030 | PF-00299804 | MVP_top60_DNA+RNA_seed | HDAC1 | 3.155465 | LUSC | NCI-H226 | Romidepsin |
| MVP_top60_DNA_seed | ERBB2 | 0.316047 | LUAD | NCI-H2030 | PF-00299804 | MVP_top60_DNA+RNA_seed | HDAC3 | 3.155465 | LUSC | NCI-H226 | Romidepsin |
| MVP_top60_DNA_seed | MAPK1 | 0.316467 | LUAD | NCI-H1437 | FR-180204 | MVP_top60_DNA+RNA_seed | PARP1 | -3.262147 | PRAD | DU-145 | Veliparib |
| MVP_top60_DNA_seed | EGFR | 0.316706 | LUAD | EKVX | AZD8931 | MVP_top60_DNA+RNA_seed | AR | -2.765186 | PRAD | LNCaP-Clone-FGC | Bicalutamide |
| MVP_top60_DNA_seed | ERBB2 | 0.316706 | LUAD | EKVX | AZD8931 | MVP_top60_DNA+RNA_seed | ATM | -2.736602 | PRAD | DU-145 | KU-55933 |

| | | | | | | | | | | | |
|---|---|---|---|---|---|---|---|---|---|---|---|
| MVP_top60_DNA_seed | ERBB3 | 0.316706 | LUAD | EKVX | AZD8931 | MVP_top60_DNA+RNA_seed | GSK3B | -2.692938 | PRAD | DU-145 | SB216763 |
| MVP_top60_DNA_seed | HDAC1 | 0.31697 | LUAD | NCI-H1693 | Tubastatin A | MVP_top60_DNA+RNA_seed | AR | -2.306773 | PRAD | PC-3 | Bicalutamide |
| MVP_top60_DNA_seed | ATM | 0.317537 | LUAD | NCI-H2122 | KU-60019 | MVP_top60_DNA+RNA_seed | ESR1 | -2.152619 | PRAD | LNCaP-Clone-FGC | Fulvestrant |
| MVP_top60_DNA_seed | EGFR | 0.31847 | LUAD | NCI-H1651 | Afatinib | MVP_top60_DNA+RNA_seed | GSK3B | -1.770045 | PRAD | VCaP | SB216763 |
| MVP_top60_DNA_seed | ERBB2 | 0.31847 | LUAD | NCI-H1651 | Afatinib | MVP_top60_DNA+RNA_seed | PARP1 | -1.523428 | PRAD | DU-145 | PARP_9495 |
| MVP_top60_DNA_seed | AURKA | 0.320597 | LUAD | HCC-827 | CD532 | MVP_top60_DNA+RNA_seed | AR | -1.470437 | PRAD | LNCaP-Clone-FGC | AZD3514 |
| MVP_top60_DNA_seed | EGFR | 0.322121 | LUAD | NCI-H2030 | Afatinib | MVP_top60_DNA+RNA_seed | GSK3B | -1.352221 | PRAD | PWR-1E | CHIR-99021 |
| MVP_top60_DNA_seed | ERBB2 | 0.322121 | LUAD | NCI-H2030 | Afatinib | MVP_top60_DNA+RNA_seed | CDK4 | -1.34343 | PRAD | PC-3 | Palbociclib |
| MVP_top60_DNA_seed | FGFR1 | 0.323147 | LUAD | NCI-H1792 | PD173074 | MVP_top60_DNA+RNA_seed | GSK3B | -1.308116 | PRAD | DU-145 | KIN001-042 |
| MVP_top60_DNA_seed | HSP90AA1 | 0.323887 | LUAD | NCI-H441 | Luminespib | MVP_top60_DNA+RNA_seed | HDAC1 | -1.280877 | PRAD | VCaP | AR-42 |
| MVP_top60_DNA_seed | EGFR | 0.323895 | LUAD | NCI-H1734 | Pelitinib | MVP_top60_DNA+RNA_seed | PARP1 | -1.279816 | PRAD | PWR-1E | Olaparib |
| MVP_top60_DNA_seed | EGFR | 0.324231 | LUAD | NCI-H1734 | PF-00299804 | MVP_top60_DNA+RNA_seed | HDAC1 | -1.265822 | PRAD | VCaP | Belinostat |
| MVP_top60_DNA_seed | ERBB2 | 0.324231 | LUAD | NCI-H1734 | PF-00299804 | MVP_top60_DNA+RNA_seed | HDAC1 | -1.179113 | PRAD | LNCaP-Clone-FGC | Belinostat |
| MVP_top60_DNA_seed | MET | 0.324599 | LUAD | HOP-62 | Crizotinib | MVP_top60_DNA+RNA_seed | ESR1 | -1.1777 | PRAD | DU-145 | Tamoxifen |
| MVP_top60_DNA_seed | SMARCA4 | 0.325691 | LUAD | NCI-H838 | PFI-3 | MVP_top60_DNA+RNA_seed | PIK3CA | -1.174944 | PRAD | DU-145 | PI3Ka_4409 |
| MVP_top60_DNA_seed | MDM2 | 0.325906 | LUAD | COR-L105 | Serdemetan | MVP_top60_DNA+RNA_seed | CDK4 | -1.153306 | PRAD | 22RV1 | Palbociclib |
| MVP_top60_DNA_seed | AKT1 | 0.326137 | LUAD | NCI-H1568 | AKT inhibitor VIII | MVP_top60_DNA+RNA_seed | HDAC1 | -1.152706 | PRAD | LNCaP-Clone-FGC | AR-42 |
| MVP_top60_DNA_seed | HSP90AA1 | 0.328214 | LUAD | NCI-H1666 | SNX-2112 | MVP_top60_DNA+RNA_seed | SMARCA4 | -1.145304 | PRAD | DU-145 | PFI-3 |
| MVP_top60_DNA_seed | AURKA | 0.328294 | LUAD | NCI-H441 | Genentech Cpd 10 | MVP_top60_DNA+RNA_seed | HDAC1 | -1.08874 | PRAD | VCaP | Parthenolide |
| MVP_top60_DNA_seed | FGFR1 | 0.328558 | LUAD | NCI-H358 | FGFR_3831 | MVP_top60_DNA+RNA_seed | ESR1 | -1.08327 | PRAD | VCaP | Fulvestrant |
| MVP_top60_DNA_seed | HSP90AA1 | 0.328767 | LUAD | NCI-H1651 | Elesclomol | MVP_top60_DNA+RNA_seed | ESR1 | -1.071545 | PRAD | PWR-1E | Fulvestrant |
| MVP_top60_DNA_seed | EGFR | 0.328855 | LUAD | VMRC-LCD | Gefitinib | MVP_top60_DNA+RNA_seed | PARP1 | -1.05242 | PRAD | PWR-1E | Olaparib |
| MVP_top60_DNA_seed | BRAF | 0.328895 | LUAD | H3255 | PLX-4720 | MVP_top60_DNA+RNA_seed | PARP1 | -1.040411 | PRAD | PC-3 | Veliparib |
| MVP_top60_DNA_seed | AKT1 | 0.329214 | LUAD | PC-3_[JPC-3] | AKT inhibitor VIII | MVP_top60_DNA+RNA_seed | PARP1 | -1.026253 | PRAD | DU-145 | PARP_9482 |
| MVP_top60_DNA_seed | BRAF | 0.330062 | LUAD | LC-2-ad | Dabrafenib | MVP_top60_DNA+RNA_seed | EP300 | -0.997176 | PRAD | LNCaP-Clone-FGC | I-CBP112 |
| MVP_top60_DNA_seed | ATM | 0.330089 | LUAD | NCI-H23 | KU-55933 | MVP_top60_DNA+RNA_seed | HDAC1 | -0.972175 | PRAD | 22RV1 | Dacinostat |
| MVP_top60_DNA_seed | BRAF | 0.330708 | LUAD | NCI-H522 | Dabrafenib | MVP_top60_DNA+RNA_seed | AR | -0.939936 | PRAD | DU-145 | AZD3514 |
| MVP_top60_DNA_seed | AURKA | 0.331028 | LUAD | NCI-H3122 | ZM447439 | MVP_top60_DNA+RNA_seed | AR | -0.938338 | PRAD | PC-3 | Bicalutamide |
| MVP_top60_DNA_seed | BRAF | 0.331504 | LUAD | EMC-BAC-1 | Dabrafenib | MVP_top60_DNA+RNA_seed | HDAC1 | -0.922482 | PRAD | 22RV1 | Belinostat |
| MVP_top60_DNA_seed | MET | 0.331839 | LUAD | NCI-H1781 | SU11274 | MVP_top60_DNA+RNA_seed | ATR | -0.906261 | PRAD | PWR-1E | AZD6738 |
| MVP_top60_DNA_seed | EGFR | 0.332372 | LUAD | NCI-H23 | Afatinib | MVP_top60_DNA+RNA_seed | ATM | -0.891731 | PRAD | DU-145 | KU-60019 |
| MVP_top60_DNA_seed | ERBB2 | 0.332372 | LUAD | NCI-H23 | Afatinib | MVP_top60_DNA+RNA_seed | HDAC1 | -0.881162 | PRAD | 22RV1 | AR-42 |
| MVP_top60_DNA_seed | AURKA | 0.332614 | LUAD | NCI-H1734 | Genentech Cpd 10 | MVP_top60_DNA+RNA_seed | ATR | -0.865979 | PRAD | PWR-1E | AZ20 |
| MVP_top60_DNA_seed | PTK2 | 0.332628 | LUAD | NCI-H1651 | PF-562271 | MVP_top60_DNA+RNA_seed | ATR | -0.865444 | PRAD | 22RV1 | QL-VIII-58 |
| MVP_top60_DNA_seed | BRAF | 0.332669 | LUAD | NCI-H1651 | PLX-4720 | MVP_top60_DNA+RNA_seed | ATM | -0.853892 | PRAD | PWR-1E | KU-60019 |
| MVP_top60_DNA_seed | IGF1R | 0.334939 | LUAD | PC-3_[JPC-3] | BMS-536924 | MVP_top60_DNA+RNA_seed | HDAC1 | -0.84662 | PRAD | VCaP | Tubastatin A |
| MVP_top60_DNA_seed | MAPK1 | 0.336698 | LUAD | RERF-LC-MS | VX-11e | MVP_top60_DNA+RNA_seed | PARP1 | -0.831317 | PRAD | DU-145 | PARP_0108 |
| MVP_top60_DNA_seed | AKT1 | 0.341224 | LUAD | A549 | BAY AKT1 | MVP_top60_DNA+RNA_seed | HDAC1 | -0.77692 | PRAD | LNCaP-Clone-FGC | CAY10603 |
| MVP_top60_DNA_seed | BRAF | 0.341313 | LUAD | Calu-3 | RAF_9304 | MVP_top60_DNA+RNA_seed | PARP1 | -0.759628 | PRAD | 22RV1 | Rucaparib |
| MVP_top60_DNA_seed | HDAC1 | 0.343146 | LUAD | NCI-H1693 | JQ12 | MVP_top60_DNA+RNA_seed | PARP1 | -0.757878 | PRAD | LNCaP-Clone-FGC | Rucaparib |
| MVP_top60_DNA_seed | AURKA | 0.343929 | LUAD | NCI-H2087 | CD532 | MVP_top60_DNA+RNA_seed | HDAC1 | -0.746407 | PRAD | DU-145 | JQ12 |

| | | | | | | | | | | | |
|---|---|---|---|---|---|---|---|---|---|---|---|
| MVP_top60_DNA_seed | AURKA | 0.344799 | LUAD | NCI-H1623 | CD532 | MVP_top60_DNA+RNA_seed | PARP1 | -0.739807 | PRAD | DU-145 | Rucaparib |
| MVP_top60_DNA_seed | GSK3B | 0.345414 | LUAD | NCI-H1437 | CHIR-99021 | MVP_top60_DNA+RNA_seed | HDAC1 | -0.733581 | PRAD | VCaP | CAY10603 |
| MVP_top60_DNA_seed | ERBB2 | 0.345422 | LUAD | NCI-H2087 | CP724714 | MVP_top60_DNA+RNA_seed | PARP1 | -0.730022 | PRAD | DU-145 | Olaparib |
| MVP_top60_DNA_seed | SMARCA4 | 0.345693 | LUAD | NCI-H2122 | PFI-3 | MVP_top60_DNA+RNA_seed | PARP1 | -0.728672 | PRAD | LNCaP-Clone-FGC | PARP_0108 |
| MVP_top60_DNA_seed | IGF1R | 0.346202 | LUAD | RERF-LC-MS | BMS-754807 | MVP_top60_DNA+RNA_seed | HDAC1 | -0.718561 | PRAD | LNCaP-Clone-FGC | Tubastatin A |
| MVP_top60_DNA_seed | MET | 0.346429 | LUAD | SW1573 | AZD6094 | MVP_top60_DNA+RNA_seed | PARP1 | -0.708736 | PRAD | LNCaP-Clone-FGC | Talazoparib |
| MVP_top60_DNA_seed | FGFR1 | 0.346801 | LUAD | NCI-H1568 | FGFR_3831 | MVP_top60_DNA+RNA_seed | ATM | -0.708365 | PRAD | PWR-1E | KU-55933 |
| MVP_top60_DNA_seed | HDAC1 | 0.347914 | LUAD | HCC-827 | Dacinostat | MVP_top60_DNA+RNA_seed | PARP1 | -0.681103 | PRAD | PWR-1E | Talazoparib |
| MVP_top60_DNA_seed | AURKA | 0.349053 | LUAD | NCI-H1693 | ZM447439 | MVP_top60_DNA+RNA_seed | ATM | -0.675106 | PRAD | PC-3 | KU-55933 |
| MVP_top60_DNA_seed | HDAC1 | 0.349364 | LUAD | NCI-H1573 | CAY10603 | MVP_top60_DNA+RNA_seed | HDAC1 | -0.661935 | PRAD | 22RV1 | CAY10603 |
| MVP_top60_DNA_seed | EGFR | 0.349548 | LUAD | NCI-H1355 | AST-1306 | MVP_top60_DNA+RNA_seed | HDAC1 | -0.648199 | PRAD | VCaP | Dacinostat |
| MVP_top60_DNA_seed | BRAF | 0.351224 | LUAD | HCC-827 | SB590885 | MVP_top60_DNA+RNA_seed | PARP1 | -0.622393 | PRAD | LNCaP-Clone-FGC | PARP_9482 |
| MVP_top60_DNA_seed | EGFR | 0.351436 | LUAD | SW1573 | Afatinib | MVP_top60_DNA+RNA_seed | GSK3B | -0.606071 | PRAD | VCaP | CHIR-99021 |
| MVP_top60_DNA_seed | ERBB2 | 0.351436 | LUAD | SW1573 | Afatinib | MVP_top60_DNA+RNA_seed | ATR | -0.567602 | PRAD | DU-145 | AZ20 |
| MVP_top60_DNA_seed | EGFR | 0.353214 | LUAD | HCC-44 | Afatinib | MVP_top60_DNA+RNA_seed | PIK3CA | -0.556787 | PRAD | 22RV1 | PI3Ka_4409 |
| MVP_top60_DNA_seed | ERBB2 | 0.353214 | LUAD | HCC-44 | Afatinib | MVP_top60_DNA+RNA_seed | ATR | -0.545546 | PRAD | LNCaP-Clone-FGC | AZD6738 |
| MVP_top60_DNA_seed | HDAC1 | 0.354166 | LUAD | NCI-H1666 | CAY10603 | MVP_top60_DNA+RNA_seed | HDAC1 | -0.544519 | PRAD | DU-145 | Belinostat |
| MVP_top60_DNA_seed | AKT1 | 0.354842 | LUAD | NCI-H1651 | BAY AKT1 | MVP_top60_DNA+RNA_seed | PARP1 | -0.526103 | PRAD | PWR-1E | PARP_9495 |
| MVP_top60_DNA_seed | HSP90AA1 | 0.358854 | LUAD | LXF-289 | CCT-018159 | MVP_top60_DNA+RNA_seed | AR | -0.50603 | PRAD | 22RV1 | Bicalutamide |
| MVP_top60_DNA_seed | EGFR | 0.359013 | LUAD | NCI-H1975 | CUDC-101 | MVP_top60_DNA+RNA_seed | GSK3B | -0.472573 | PRAD | VCaP | AZD7969 |
| MVP_top60_DNA_seed | ERBB2 | 0.359013 | LUAD | NCI-H1975 | CUDC-101 | MVP_top60_DNA+RNA_seed | PARP1 | -0.470946 | PRAD | 22RV1 | PARP_9482 |
| MVP_top60_DNA_seed | EGFR | 0.3598 | LUAD | NCI-H838 | Cetuximab | MVP_top60_DNA+RNA_seed | HDAC1 | -0.463803 | PRAD | DU-145 | AR-42 |
| MVP_top60_DNA_seed | BRAF | 0.360046 | LUAD | NCI-H1573 | SB590885 | MVP_top60_DNA+RNA_seed | ATM | -0.463026 | PRAD | VCaP | CP466722 |
| MVP_top60_DNA_seed | GSK3B | 0.360416 | LUAD | EMC-BAC-1 | AZD7969 | MVP_top60_DNA+RNA_seed | CDK4 | -0.45584 | PRAD | DU-145 | Palbociclib |
| MVP_top60_DNA_seed | MAPK1 | 0.360572 | LUAD | HCC-44 | FR-180204 | MVP_top60_DNA+RNA_seed | AR | -0.437962 | PRAD | LNCaP-Clone-FGC | Bicalutamide |
| MVP_top60_DNA_seed | GSK3B | 0.362432 | LUAD | NCI-H3122 | AZD7969 | MVP_top60_DNA+RNA_seed | PARP1 | -0.435297 | PRAD | 22RV1 | PARP_0108 |
| MVP_top60_DNA_seed | AKT1 | 0.36344 | LUAD | NCI-H1568 | AKT inhibitor VIII | MVP_top60_DNA+RNA_seed | HDAC1 | -0.435137 | PRAD | DU-145 | Dacinostat |
| MVP_top60_DNA_seed | AKT1 | 0.363481 | LUAD | NCI-H2030 | GSK690693 | MVP_top60_DNA+RNA_seed | GSK3B | -0.425484 | PRAD | 22RV1 | CHIR-99021 |
| MVP_top60_DNA_seed | HDAC1 | 0.364798 | LUAD | H3255 | JQ12 | MVP_top60_DNA+RNA_seed | AR | -0.416302 | PRAD | VCaP | AZD3514 |
| MVP_top60_DNA_seed | AURKA | 0.365042 | LUAD | HCC-827 | ZM447439 | MVP_top60_DNA+RNA_seed | ATM | -0.412618 | PRAD | LNCaP-Clone-FGC | CP466722 |
| MVP_top60_DNA_seed | IGF1R | 0.365568 | LUAD | PC-14 | Linsitinib | MVP_top60_DNA+RNA_seed | PIK3CA | -0.412275 | PRAD | 22RV1 | AZD8835 |
| MVP_top60_DNA_seed | GSK3B | 0.365823 | LUAD | HCC-827 | AZD7969 | MVP_top60_DNA+RNA_seed | HDAC1 | -0.409553 | PRAD | VCaP | Entinostat |
| MVP_top60_DNA_seed | HSP90AA1 | 0.36589 | LUAD | NCI-H322M | Tanespimycin | MVP_top60_DNA+RNA_seed | PARP1 | -0.409432 | PRAD | PC-3 | PARP_0108 |
| MVP_top60_DNA_seed | GSK3B | 0.366481 | LUAD | SW1573 | SB216763 | MVP_top60_DNA+RNA_seed | ATR | -0.407304 | PRAD | LNCaP-Clone-FGC | AZ20 |
| MVP_top60_DNA_seed | EGFR | 0.366685 | LUAD | NCI-H2023 | AZD8931 | MVP_top60_DNA+RNA_seed | HDAC1 | -0.390047 | PRAD | 22RV1 | JQ12 |
| MVP_top60_DNA_seed | ERBB2 | 0.366685 | LUAD | NCI-H2023 | AZD8931 | MVP_top60_DNA+RNA_seed | HDAC1 | -0.379268 | PRAD | DU-145 | CAY10603 |
| MVP_top60_DNA_seed | ERBB3 | 0.366685 | LUAD | NCI-H2023 | AZD8931 | MVP_top60_DNA+RNA_seed | HDAC1 | -0.374469 | PRAD | 22RV1 | Tubastatin A |
| MVP_top60_DNA_seed | IGF1R | 0.368296 | LUAD | NCI-H2347 | BMS-754807 | MVP_top60_DNA+RNA_seed | PIK3CA | -0.367324 | PRAD | DU-145 | AZD8835 |
| MVP_top60_DNA_seed | AKT1 | 0.36845 | LUAD | NCI-H1792 | GSK690693 | MVP_top60_DNA+RNA_seed | PARP1 | -0.364595 | PRAD | PC-3 | PARP_9482 |
| MVP_top60_DNA_seed | EGFR | 0.36857 | LUAD | EMC-BAC-2 | PF-00299804 | MVP_top60_DNA+RNA_seed | GSK3B | -0.353891 | PRAD | LNCaP-Clone-FGC | SB216763 |
| MVP_top60_DNA_seed | ERBB2 | 0.36857 | LUAD | EMC-BAC-2 | PF-00299804 | MVP_top60_DNA+RNA_seed | SMARCA4 | -0.339341 | PRAD | VCaP | PFI-3 |

| | | | | | | | | | | | |
|---|---|---|---|---|---|---|---|---|---|---|---|
| MVP_top60_DNA_seed | BRAF | 0.369277 | LUAD | NCI-H3122 | RAF_9304 | MVP_top60_DNA+RNA_seed | PARP1 | -0.333174 | PRAD | DU-145 | Talazoparib |
| MVP_top60_DNA_seed | HDAC1 | 0.370117 | LUAD | NCI-H650 | JQ12 | MVP_top60_DNA+RNA_seed | PIK3CA | -0.331526 | PRAD | VCaP | AZD8835 |
| MVP_top60_DNA_seed | EGFR | 0.371778 | LUAD | RERF-LC-MS | AST-1306 | MVP_top60_DNA+RNA_seed | EP300 | -0.324634 | PRAD | 22RV1 | I-CBP112 |
| MVP_top60_DNA_seed | EGFR | 0.372744 | LUAD | NCI-H322M | Afatinib | MVP_top60_DNA+RNA_seed | HDAC1 | -0.321626 | PRAD | VCaP | JQ12 |
| MVP_top60_DNA_seed | ERBB2 | 0.372744 | LUAD | NCI-H322M | Afatinib | MVP_top60_DNA+RNA_seed | PARP1 | -0.320227 | PRAD | PWR-1E | Veliparib |
| MVP_top60_DNA_seed | AKT1 | 0.372811 | LUAD | NCI-H1623 | BAY AKT1 | MVP_top60_DNA+RNA_seed | PARP1 | -0.308482 | PRAD | 22RV1 | Talazoparib |
| MVP_top60_DNA_seed | MET | 0.373257 | LUAD | NCI-H2405 | SU11274 | MVP_top60_DNA+RNA_seed | ATR | -0.298652 | PRAD | LNCaP-Clone-FGC | QL-VIII-58 |
| MVP_top60_DNA_seed | EGFR | 0.373422 | LUAD | VMRC-LCD | Afatinib | MVP_top60_DNA+RNA_seed | AR | -0.279002 | PRAD | VCaP | Bicalutamide |
| MVP_top60_DNA_seed | ERBB2 | 0.373422 | LUAD | VMRC-LCD | Afatinib | MVP_top60_DNA+RNA_seed | ESR1 | -0.276456 | PRAD | 22RV1 | Tamoxifen |
| MVP_top60_DNA_seed | AURKA | 0.37366 | LUAD | NCI-H23 | CD532 | MVP_top60_DNA+RNA_seed | PARP1 | -0.243649 | PRAD | LNCaP-Clone-FGC | Olaparib |
| MVP_top60_DNA_seed | MET | 0.374253 | LUAD | Calu-6 | AZD6094 | MVP_top60_DNA+RNA_seed | ATM | -0.237597 | PRAD | LNCaP-Clone-FGC | KU-60019 |
| MVP_top60_DNA_seed | PIK3CA | 0.374923 | LUAD | EKVX | AZD8835 | MVP_top60_DNA+RNA_seed | AR | -0.233028 | PRAD | 22RV1 | AZD3514 |
| MVP_top60_DNA_seed | HDAC1 | 0.375007 | LUAD | PC-3_[JPC-3] | CAY10603 | MVP_top60_DNA+RNA_seed | ATM | -0.228938 | PRAD | LNCaP-Clone-FGC | torin2 |
| MVP_top60_DNA_seed | AKT1 | 0.375139 | LUAD | EMC-BAC-2 | BAY AKT1 | MVP_top60_DNA+RNA_seed | ATR | -0.228938 | PRAD | LNCaP-Clone-FGC | torin2 |
| MVP_top60_DNA_seed | HSP90AA1 | 0.375943 | LUAD | VMRC-LCD | Tanespimycin | MVP_top60_DNA+RNA_seed | PARP1 | -0.207727 | PRAD | PC-3 | Olaparib |
| MVP_top60_DNA_seed | HSP90AA1 | 0.376168 | LUAD | LC-2-ad | CCT-018159 | MVP_top60_DNA+RNA_seed | PARP1 | -0.205763 | PRAD | PC-3 | Talazoparib |
| MVP_top60_DNA_seed | GSK3B | 0.378029 | LUAD | NCI-H838 | CHIR-99021 | MVP_top60_DNA+RNA_seed | HDAC1 | -0.196213 | PRAD | PC-3 | CAY10603 |
| MVP_top60_DNA_seed | AURKA | 0.378418 | LUAD | NCI-H1651 | Alisertib | MVP_top60_DNA+RNA_seed | ATM | -0.18667 | PRAD | 22RV1 | CP466722 |
| MVP_top60_DNA_seed | AKT1 | 0.378705 | LUAD | Calu-6 | AKT inhibitor VIII | MVP_top60_DNA+RNA_seed | AR | -0.177565 | PRAD | DU-145 | Bicalutamide |
| MVP_top60_DNA_seed | BRAF | 0.379151 | LUAD | HCC-78 | SB590885 | MVP_top60_DNA+RNA_seed | ATM | -0.175981 | PRAD | VCaP | KU-55933 |
| MVP_top60_DNA_seed | EGFR | 0.379366 | LUAD | NCI-H441 | Afatinib | MVP_top60_DNA+RNA_seed | ATR | -0.169139 | PRAD | DU-145 | AZD6738 |
| MVP_top60_DNA_seed | ERBB2 | 0.379366 | LUAD | NCI-H441 | Afatinib | MVP_top60_DNA+RNA_seed | EP300 | -0.155857 | PRAD | VCaP | I-CBP112 |
| MVP_top60_DNA_seed | HDAC1 | 0.380412 | LUAD | NCI-H1435 | CAY10603 | MVP_top60_DNA+RNA_seed | HDAC1 | -0.135055 | PRAD | LNCaP-Clone-FGC | Dacinostat |
| MVP_top60_DNA_seed | FGFR1 | 0.380916 | LUAD | Calu-3 | AZD4547 | MVP_top60_DNA+RNA_seed | AR | -0.110202 | PRAD | PC-3 | AZD3514 |
| MVP_top60_DNA_seed | MDM2 | 0.38131 | LUAD | NCI-H3122 | Serdemetan | MVP_top60_DNA+RNA_seed | GSK3B | -0.092599 | PRAD | LNCaP-Clone-FGC | AZD7969 |
| MVP_top60_DNA_seed | FGFR1 | 0.384183 | LUAD | NCI-H2009 | AZD4547 | MVP_top60_DNA+RNA_seed | PARP1 | -0.067815 | PRAD | LNCaP-Clone-FGC | PARP_9495 |
| MVP_top60_DNA_seed | GSK3B | 0.384694 | LUAD | NCI-H2087 | AZD7969 | MVP_top60_DNA+RNA_seed | HDAC1 | -0.061377 | PRAD | PC-3 | Belinostat |
| MVP_top60_DNA_seed | BRAF | 0.384877 | LUAD | NCI-H522 | RAF_9304 | MVP_top60_DNA+RNA_seed | GSK3B | -0.048666 | PRAD | 22RV1 | AZD7969 |
| MVP_top60_DNA_seed | PIK3CA | 0.385118 | LUAD | NCI-H2087 | AZD8835 | MVP_top60_DNA+RNA_seed | SMARCA4 | -0.026262 | PRAD | PWR-1E | PFI-3 |
| MVP_top60_DNA_seed | EGFR | 0.385627 | LUAD | NCI-H2085 | CUDC-101 | MVP_top60_DNA+RNA_seed | PARP1 | -0.01336 | PRAD | PWR-1E | Rucaparib |
| MVP_top60_DNA_seed | ERBB2 | 0.385627 | LUAD | NCI-H2085 | CUDC-101 | MVP_top60_DNA+RNA_seed | PIK3CA | -0.01 | PRAD | VCaP | PI3Ka_4409 |
| MVP_top60_DNA_seed | PIK3CA | 0.385766 | LUAD | NCI-H650 | AZD8835 | MVP_top60_DNA+RNA_seed | SMARCA4 | -0.007397 | PRAD | 22RV1 | PFI-3 |
| MVP_top60_DNA_seed | MDM2 | 0.387357 | LUAD | ABC-1 | Serdemetan | MVP_top60_DNA+RNA_seed | HDAC1 | 0.008543 | PRAD | PC-3 | AR-42 |
| MVP_top60_DNA_seed | AURKA | 0.388187 | LUAD | NCI-H650 | GSK1070916 | MVP_top60_DNA+RNA_seed | ATM | 0.016044 | PRAD | PC-3 | CP466722 |
| MVP_top60_DNA_seed | AURKA | 0.388459 | LUAD | HCC-827 | Alisertib | MVP_top60_DNA+RNA_seed | PIK3CA | 0.022501 | PRAD | PWR-1E | AZD8835 |
| MVP_top60_DNA_seed | AURKA | 0.388598 | LUAD | NCI-H2087 | Alisertib | MVP_top60_DNA+RNA_seed | GSK3B | 0.046287 | PRAD | PC-3 | SB216763 |
| MVP_top60_DNA_seed | AKT1 | 0.389521 | LUAD | NCI-H2291 | MK-2206 | MVP_top60_DNA+RNA_seed | ESR1 | 0.049164 | PRAD | LNCaP-Clone-FGC | Tamoxifen |
| MVP_top60_DNA_seed | HDAC1 | 0.390463 | LUAD | H3255 | Dacinostat | MVP_top60_DNA+RNA_seed | AR | 0.051099 | PRAD | PWR-1E | Bicalutamide |
| MVP_top60_DNA_seed | AURKA | 0.390996 | LUAD | NCI-H1623 | ZM447439 | MVP_top60_DNA+RNA_seed | CDK4 | 0.074618 | PRAD | LNCaP-Clone-FGC | Palbociclib |
| MVP_top60_DNA_seed | HDAC1 | 0.391486 | LUAD | NCI-H1993 | AR-42 | MVP_top60_DNA+RNA_seed | HDAC1 | 0.085457 | PRAD | LNCaP-Clone-FGC | JQ12 |
| MVP_top60_DNA_seed | HSP90AA1 | 0.391753 | LUAD | NCI-H2122 | SNX-2112 | MVP_top60_DNA+RNA_seed | AR | 0.097737 | PRAD | DU-145 | Bicalutamide |

| | | | | | | | | | | | |
|---|---|---|---|---|---|---|---|---|---|---|---|
| MVP_top60_DNA_seed | BRAF | 0.391972 | LUAD | NCI-H2087 | PLX-4720 | MVP_top60_DNA+RNA_seed | HDAC1 | 0.10802 | PRAD | PC-3 | Tubastatin A |
| MVP_top60_DNA_seed | MET | 0.39239 | LUAD | NCI-H1650 | PHA-665752 | MVP_top60_DNA+RNA_seed | GSK3B | 0.109058 | PRAD | DU-145 | CHIR-99021 |
| MVP_top60_DNA_seed | GSK3B | 0.392529 | LUAD | A549 | KIN001-042 | MVP_top60_DNA+RNA_seed | GSK3B | 0.110164 | PRAD | DU-145 | AZD7969 |
| MVP_top60_DNA_seed | AKT1 | 0.392818 | LUAD | A549 | AKT inhibitor VIII | MVP_top60_DNA+RNA_seed | GSK3B | 0.116501 | PRAD | PC-3 | CHIR-99021 |
| MVP_top60_DNA_seed | EGFR | 0.394744 | LUAD | NCI-H1755 | PF-00299804 | MVP_top60_DNA+RNA_seed | CDK4 | 0.137383 | PRAD | VCaP | CGP-082996 |
| MVP_top60_DNA_seed | ERBB2 | 0.394744 | LUAD | NCI-H1755 | PF-00299804 | MVP_top60_DNA+RNA_seed | PIK3CA | 0.144896 | PRAD | LNCaP-Clone-FGC | AZD8835 |
| MVP_top60_DNA_seed | EGFR | 0.395539 | LUAD | NCI-H441 | AST-1306 | MVP_top60_DNA+RNA_seed | PARP1 | 0.16679 | PRAD | PWR-1E | PARP_0108 |
| MVP_top60_DNA_seed | BRAF | 0.396816 | LUAD | NCI-H2085 | PLX-4720 | MVP_top60_DNA+RNA_seed | PARP1 | 0.170597 | PRAD | PWR-1E | PARP_9482 |
| MVP_top60_DNA_seed | AKT1 | 0.396939 | LUAD | NCI-H1993 | AKT inhibitor VIII | MVP_top60_DNA+RNA_seed | EP300 | 0.178589 | PRAD | PWR-1E | I-CBP112 |
| MVP_top60_DNA_seed | AURKA | 0.397004 | LUAD | NCI-H1355 | GSK1070916 | MVP_top60_DNA+RNA_seed | GSK3B | 0.18049 | PRAD | PC-3 | KIN001-042 |
| MVP_top60_DNA_seed | MET | 0.398047 | LUAD | HOP-62 | PHA-665752 | MVP_top60_DNA+RNA_seed | CDK4 | 0.181924 | PRAD | VCaP | Palbociclib |
| MVP_top60_DNA_seed | MDM2 | 0.398386 | LUAD | NCI-H1793 | Serdemetan | MVP_top60_DNA+RNA_seed | HDAC1 | 0.198546 | PRAD | DU-145 | Tubastatin A |
| MVP_top60_DNA_seed | HDAC1 | 0.400107 | LUAD | NCI-H1437 | Tubastatin A | MVP_top60_DNA+RNA_seed | HDAC1 | 0.205203 | PRAD | PC-3 | Dacinostat |
| MVP_top60_DNA_seed | HDAC1 | 0.400508 | LUAD | NCI-H1693 | Belinostat | MVP_top60_DNA+RNA_seed | GSK3B | 0.217229 | PRAD | PWR-1E | KIN001-042 |
| MVP_top60_DNA_seed | HDAC1 | 0.400804 | LUAD | NCI-H2085 | JQ12 | MVP_top60_DNA+RNA_seed | ATR | 0.221189 | PRAD | PC-3 | AZD6738 |
| MVP_top60_DNA_seed | PIK3CA | 0.401141 | LUAD | NCI-H2347 | PI3Ka_4409 | MVP_top60_DNA+RNA_seed | PARP1 | 0.252626 | PRAD | PC-3 | Rucaparib |
| MVP_top60_DNA_seed | ATM | 0.401427 | LUAD | NCI-H1734 | CP466722 | MVP_top60_DNA+RNA_seed | ESR1 | 0.256978 | PRAD | 22RV1 | Fulvestrant |
| MVP_top60_DNA_seed | EGFR | 0.401488 | LUAD | LXF-289 | CI-1033 | MVP_top60_DNA+RNA_seed | GSK3B | 0.290372 | PRAD | LNCaP-Clone-FGC | CHIR-99021 |
| MVP_top60_DNA_seed | ERBB2 | 0.401488 | LUAD | LXF-289 | CI-1033 | MVP_top60_DNA+RNA_seed | ESR1 | 0.32332 | PRAD | DU-145 | Fulvestrant |
| MVP_top60_DNA_seed | EGFR | 0.402576 | LUAD | RERF-LC-MS | Afatinib | MVP_top60_DNA+RNA_seed | ESR1 | 0.339144 | PRAD | PWR-1E | Tamoxifen |
| MVP_top60_DNA_seed | ERBB2 | 0.402576 | LUAD | RERF-LC-MS | Afatinib | MVP_top60_DNA+RNA_seed | GSK3B | 0.349246 | PRAD | PC-3 | AZD7969 |
| MVP_top60_DNA_seed | EGFR | 0.405374 | LUAD | NCI-H2291 | PF-00299804 | MVP_top60_DNA+RNA_seed | PARP1 | 0.35281 | PRAD | VCaP | Talazoparib |
| MVP_top60_DNA_seed | ERBB2 | 0.405374 | LUAD | NCI-H2291 | PF-00299804 | MVP_top60_DNA+RNA_seed | PARP1 | 0.36504 | PRAD | 22RV1 | Olaparib |
| MVP_top60_DNA_seed | GSK3B | 0.406385 | LUAD | RERF-LC-KJ | CHIR-99021 | MVP_top60_DNA+RNA_seed | GSK3B | 0.380022 | PRAD | LNCaP-Clone-FGC | KIN001-042 |
| MVP_top60_DNA_seed | GSK3B | 0.40641 | LUAD | HCC-827 | KIN001-042 | MVP_top60_DNA+RNA_seed | ESR1 | 0.383568 | PRAD | PC-3 | Tamoxifen |
| MVP_top60_DNA_seed | HDAC1 | 0.406537 | LUAD | NCI-H322M | CAY10603 | MVP_top60_DNA+RNA_seed | PARP1 | 0.411417 | PRAD | LNCaP-Clone-FGC | Olaparib |
| MVP_top60_DNA_seed | GSK3B | 0.407106 | LUAD | H3255 | CHIR-99021 | MVP_top60_DNA+RNA_seed | GSK3B | 0.463171 | PRAD | LNCaP-Clone-FGC | CHIR-99021 |
| MVP_top60_DNA_seed | EGFR | 0.407697 | LUAD | NCI-H1993 | PF-00299804 | MVP_top60_DNA+RNA_seed | AR | 0.470589 | PRAD | VCaP | Bicalutamide |
| MVP_top60_DNA_seed | ERBB2 | 0.407697 | LUAD | NCI-H1993 | PF-00299804 | MVP_top60_DNA+RNA_seed | EP300 | 0.485707 | PRAD | DU-145 | I-CBP112 |
| MVP_top60_DNA_seed | ATM | 0.407946 | LUAD | VMRC-LCD | KU-55933 | MVP_top60_DNA+RNA_seed | PIK3CA | 0.489116 | PRAD | PC-3 | AZD8835 |
| MVP_top60_DNA_seed | HSP90AA1 | 0.408308 | LUAD | COR-L105 | CCT-018159 | MVP_top60_DNA+RNA_seed | GSK3B | 0.517366 | PRAD | VCaP | KIN001-042 |
| MVP_top60_DNA_seed | EGFR | 0.408631 | LUAD | NCI-H2291 | Cetuximab | MVP_top60_DNA+RNA_seed | PARP1 | 0.529798 | PRAD | VCaP | Veliparib |
| MVP_top60_DNA_seed | MET | 0.410695 | LUAD | SK-LU-1 | SU11274 | MVP_top60_DNA+RNA_seed | ATR | 0.539113 | PRAD | PC-3 | AZ20 |
| MVP_top60_DNA_seed | SRC | 0.410738 | LUAD | RERF-LC-KJ | NVP-BHG712 | MVP_top60_DNA+RNA_seed | PARP1 | 0.544669 | PRAD | PC-3 | Olaparib |
| MVP_top60_DNA_seed | GSK3B | 0.412376 | LUAD | NCI-H1975 | KIN001-042 | MVP_top60_DNA+RNA_seed | ATR | 0.551016 | PRAD | VCaP | AZ20 |
| MVP_top60_DNA_seed | MDM2 | 0.412496 | LUAD | NCI-H1993 | Serdemetan | MVP_top60_DNA+RNA_seed | PARP1 | 0.556331 | PRAD | PC-3 | PARP_9495 |
| MVP_top60_DNA_seed | BRAF | 0.412541 | LUAD | EKVX | RAF_9304 | MVP_top60_DNA+RNA_seed | ESR1 | 0.563063 | PRAD | VCaP | Tamoxifen |
| MVP_top60_DNA_seed | IGF1R | 0.412676 | LUAD | RERF-LC-MS | GSK1904529A | MVP_top60_DNA+RNA_seed | GSK3B | 0.618646 | PRAD | VCaP | CHIR-99021 |
| MVP_top60_DNA_seed | EGFR | 0.413185 | LUAD | LXF-289 | AST-1306 | MVP_top60_DNA+RNA_seed | ATR | 0.624847 | PRAD | 22RV1 | AZD6738 |
| MVP_top60_DNA_seed | EGFR | 0.414291 | LUAD | NCI-H23 | Afatinib | MVP_top60_DNA+RNA_seed | PARP1 | 0.646756 | PRAD | DU-145 | Olaparib |
| MVP_top60_DNA_seed | ERBB2 | 0.414291 | LUAD | NCI-H23 | Afatinib | MVP_top60_DNA+RNA_seed | PIK3CA | 0.6633 | PRAD | PC-3 | PI3Ka_4409 |

| | | | | | | | | | | | |
|---|---|---|---|---|---|---|---|---|---|---|---|
| MVP_top60_DNA_seed | BRAF | 0.414453 | LUAD | NCI-H522 | SB590885 | MVP_top60_DNA+RNA_seed | GSK3B | 0.694477 | PRAD | PC-3 | CHIR-99021 |
| MVP_top60_DNA_seed | MET | 0.414563 | LUAD | NCI-H1650 | AZD6094 | MVP_top60_DNA+RNA_seed | GSK3B | 0.700894 | PRAD | 22RV1 | CHIR-99021 |
| MVP_top60_DNA_seed | AURKA | 0.41556 | LUAD | NCI-H23 | Alisertib | MVP_top60_DNA+RNA_seed | PIK3CA | 0.701693 | PRAD | LNCaP-Clone-FGC | PI3Ka_4409 |
| MVP_top60_DNA_seed | AKT1 | 0.415652 | LUAD | LXF-289 | GSK690693 | MVP_top60_DNA+RNA_seed | ATR | 0.72534 | PRAD | 22RV1 | AZ20 |
| MVP_top60_DNA_seed | AURKA | 0.417359 | LUAD | NCI-H1693 | GSK1070916 | MVP_top60_DNA+RNA_seed | PARP1 | 0.728695 | PRAD | VCaP | Olaparib |
| MVP_top60_DNA_seed | BRAF | 0.418035 | LUAD | NCI-H2347 | SB590885 | MVP_top60_DNA+RNA_seed | ATM | 0.73058 | PRAD | PC-3 | KU-60019 |
| MVP_top60_DNA_seed | MDM2 | 0.418758 | LUAD | NCI-H522 | Nutlin-3a (-) | MVP_top60_DNA+RNA_seed | PARP1 | 0.731291 | PRAD | VCaP | Olaparib |
| MVP_top60_DNA_seed | BRAF | 0.418779 | LUAD | NCI-H2342 | PLX-4720 | MVP_top60_DNA+RNA_seed | SMARCA4 | 0.753822 | PRAD | LNCaP-Clone-FGC | PFI-3 |
| MVP_top60_DNA_seed | EGFR | 0.419154 | LUAD | NCI-H2347 | Pelitinib | MVP_top60_DNA+RNA_seed | PARP1 | 0.759287 | PRAD | VCaP | Rucaparib |
| MVP_top60_DNA_seed | HSP90AA1 | 0.420258 | LUAD | NCI-H1437 | Elesclomol | MVP_top60_DNA+RNA_seed | SMARCA4 | 0.759298 | PRAD | PC-3 | PFI-3 |
| MVP_top60_DNA_seed | PIK3CA | 0.420377 | LUAD | ABC-1 | AZD8835 | MVP_top60_DNA+RNA_seed | PARP1 | 0.770766 | PRAD | VCaP | PARP_0108 |
| MVP_top60_DNA_seed | EGFR | 0.420476 | LUAD | SK-LU-1 | AZD8931 | MVP_top60_DNA+RNA_seed | ATR | 0.772489 | PRAD | VCaP | AZD6738 |
| MVP_top60_DNA_seed | ERBB2 | 0.420476 | LUAD | SK-LU-1 | AZD8931 | MVP_top60_DNA+RNA_seed | ESR1 | 0.792365 | PRAD | PC-3 | Fulvestrant |
| MVP_top60_DNA_seed | ERBB3 | 0.420476 | LUAD | SK-LU-1 | AZD8931 | MVP_top60_DNA+RNA_seed | HDAC1 | 0.793165 | PRAD | PC-3 | JQ12 |
| MVP_top60_DNA_seed | MDM2 | 0.421239 | LUAD | H3255 | Nutlin-3a (-) | MVP_top60_DNA+RNA_seed | GSK3B | 0.802677 | PRAD | DU-145 | CHIR-99021 |
| MVP_top60_DNA_seed | BRAF | 0.422675 | LUAD | PC-14 | Dabrafenib | MVP_top60_DNA+RNA_seed | ATR | 0.803869 | PRAD | PWR-1E | QL-VIII-58 |
| MVP_top60_DNA_seed | GSK3B | 0.423069 | LUAD | NCI-H2405 | AZD7969 | MVP_top60_DNA+RNA_seed | PARP1 | 0.805579 | PRAD | 22RV1 | Olaparib |
| MVP_top60_DNA_seed | MAPK1 | 0.42414 | LUAD | NCI-H1781 | FR-180204 | MVP_top60_DNA+RNA_seed | CDK4 | 0.807318 | PRAD | LNCaP-Clone-FGC | CGP-082996 |
| MVP_top60_DNA_seed | EGFR | 0.424323 | LUAD | EKVX | Pelitinib | MVP_top60_DNA+RNA_seed | PIK3CA | 0.819393 | PRAD | PWR-1E | PI3Ka_4409 |
| MVP_top60_DNA_seed | SMARCA4 | 0.424573 | LUAD | NCI-H1993 | PFI-3 | MVP_top60_DNA+RNA_seed | GSK3B | 0.884632 | PRAD | 22RV1 | KIN001-042 |
| MVP_top60_DNA_seed | SRC | 0.425899 | LUAD | NCI-H1734 | NVP-BHG712 | MVP_top60_DNA+RNA_seed | GSK3B | 0.886893 | PRAD | 22RV1 | SB216763 |
| MVP_top60_DNA_seed | EGFR | 0.4269 | LUAD | NCI-H441 | PF-00299804 | MVP_top60_DNA+RNA_seed | PARP1 | 0.899247 | PRAD | VCaP | PARP_9495 |
| MVP_top60_DNA_seed | ERBB2 | 0.4269 | LUAD | NCI-H441 | PF-00299804 | MVP_top60_DNA+RNA_seed | EP300 | 0.920034 | PRAD | PC-3 | I-CBP112 |
| MVP_top60_DNA_seed | HDAC1 | 0.427384 | LUAD | SK-LU-1 | JQ12 | MVP_top60_DNA+RNA_seed | PARP1 | 0.932031 | PRAD | 22RV1 | Veliparib |
| MVP_top60_DNA_seed | BRAF | 0.427478 | LUAD | HOP-62 | SB590885 | MVP_top60_DNA+RNA_seed | PARP1 | 0.973581 | PRAD | VCaP | PARP_9482 |
| MVP_top60_DNA_seed | EGFR | 0.427544 | LUAD | NCI-H1651 | AST-1306 | MVP_top60_DNA+RNA_seed | AR | 1.030195 | PRAD | 22RV1 | Bicalutamide |
| MVP_top60_DNA_seed | IGF1R | 0.427607 | LUAD | NCI-H23 | GSK1904529A | MVP_top60_DNA+RNA_seed | HDAC1 | 1.038547 | PRAD | LNCaP-Clone-FGC | Entinostat |
| MVP_top60_DNA_seed | ATM | 0.427991 | LUAD | COR-L105 | KU-60019 | MVP_top60_DNA+RNA_seed | AR | 1.043883 | PRAD | PWR-1E | AZD3514 |
| MVP_top60_DNA_seed | BRAF | 0.428543 | LUAD | NCI-H1437 | PLX-4720 | MVP_top60_DNA+RNA_seed | ATR | 1.163746 | PRAD | VCaP | QL-VIII-58 |
| MVP_top60_DNA_seed | GSK3B | 0.428647 | LUAD | Calu-6 | CHIR-99021 | MVP_top60_DNA+RNA_seed | ATM | 1.204196 | PRAD | 22RV1 | KU-55933 |
| MVP_top60_DNA_seed | BRAF | 0.429244 | LUAD | NCI-H1993 | PLX-4720 | MVP_top60_DNA+RNA_seed | ATM | 1.228193 | PRAD | VCaP | KU-60019 |
| MVP_top60_DNA_seed | AURKA | 0.430402 | LUAD | NCI-H2023 | ZM447439 | MVP_top60_DNA+RNA_seed | HDAC1 | 1.286136 | PRAD | LNCaP-Clone-FGC | Parthenolide |
| MVP_top60_DNA_seed | MDM2 | 0.431784 | LUAD | NCI-H1792 | Nutlin-3a (-) | MVP_top60_DNA+RNA_seed | CDK4 | 1.286301 | PRAD | PWR-1E | Palbociclib |
| MVP_top60_DNA_seed | AURKA | 0.431859 | LUAD | NCI-H1666 | Alisertib | MVP_top60_DNA+RNA_seed | ATM | 1.321879 | PRAD | 22RV1 | KU-60019 |
| MVP_top60_DNA_seed | MAPK1 | 0.4322 | LUAD | NCI-H1648 | FR-180204 | MVP_top60_DNA+RNA_seed | ATM | 1.422892 | PRAD | LNCaP-Clone-FGC | KU-55933 |
| MVP_top60_DNA_seed | AKT1 | 0.43242 | LUAD | NCI-H1573 | MK-2206 | MVP_top60_DNA+RNA_seed | ATM | 1.452518 | PRAD | DU-145 | CP466722 |
| MVP_top60_DNA_seed | BRAF | 0.43245 | LUAD | NCI-H1666 | SB590885 | MVP_top60_DNA+RNA_seed | PARP1 | 1.490444 | PRAD | 22RV1 | PARP_9495 |
| MVP_top60_DNA_seed | BRAF | 0.433497 | LUAD | NCI-H1838 | PLX-4720 | MVP_top60_DNA+RNA_seed | PIK3CA | -3.918726 | PRAD | LNCaP-Clone-FGC | AZD8186 |
| MVP_top60_DNA_seed | FGFR1 | 0.434691 | LUAD | HOP-62 | AZD4547 | MVP_top60_DNA+RNA_seed | ESR1 | -3.122142 | PRAD | LNCaP-Clone-FGC | GDC0810 |
| MVP_top60_DNA_seed | AKT1 | 0.435299 | LUAD | NCI-H2009 | BAY AKT1 | MVP_top60_DNA+RNA_seed | ESR2 | -3.122142 | PRAD | LNCaP-Clone-FGC | GDC0810 |
| MVP_top60_DNA_seed | EGFR | 0.43555 | LUAD | NCI-H1792 | Afatinib | MVP_top60_DNA+RNA_seed | EP300 | -1.937132 | PRAD | LNCaP-Clone-FGC | CPI-637 |

| | | | | | | | | | | | |
|---|---|---|---|---|---|---|---|---|---|---|---|
| MVP_top60_DNA_seed | ERBB2 | 0.43555 | LUAD | NCI-H1792 | Afatinib | MVP_top60_DNA+RNA_seed | CDK4 | -1.667891 | PRAD | LNCaP-Clone-FGC | Ribociclib |
| MVP_top60_DNA_seed | EGFR | 0.436257 | LUAD | NCI-H1651 | PF-00299804 | MVP_top60_DNA+RNA_seed | PIK3CA | -1.521253 | PRAD | LNCaP-Clone-FGC | GNE-317 |
| MVP_top60_DNA_seed | ERBB2 | 0.436257 | LUAD | NCI-H1651 | PF-00299804 | MVP_top60_DNA+RNA_seed | GSK3B | -1.515353 | PRAD | PWR-1E | SB216763 |
| MVP_top60_DNA_seed | EGFR | 0.436492 | LUAD | NCI-H838 | AZD8931 | MVP_top60_DNA+RNA_seed | EP300 | -1.429427 | PRAD | VCaP | CPI-637 |
| MVP_top60_DNA_seed | ERBB2 | 0.436492 | LUAD | NCI-H838 | AZD8931 | MVP_top60_DNA+RNA_seed | EZH2 | -1.380232 | PRAD | PWR-1E | GSK343 |
| MVP_top60_DNA_seed | ERBB3 | 0.436492 | LUAD | NCI-H838 | AZD8931 | MVP_top60_DNA+RNA_seed | EZH2 | -1.353513 | PRAD | LNCaP-Clone-FGC | GSK343 |
| MVP_top60_DNA_seed | GSK3B | 0.437289 | LUAD | NCI-H2023 | CHIR-99021 | MVP_top60_DNA+RNA_seed | CDK4 | -1.25799 | PRAD | LNCaP-Clone-FGC | Palbociclib |
| MVP_top60_DNA_seed | GSK3B | 0.437342 | LUAD | NCI-H1437 | CHIR-99021 | MVP_top60_DNA+RNA_seed | ESR1 | -1.254283 | PRAD | LNCaP-Clone-FGC | Tamoxifen |
| MVP_top60_DNA_seed | MAPK1 | 0.437776 | LUAD | NCI-H1623 | FR-180204 | MVP_top60_DNA+RNA_seed | CDK4 | -1.11126 | PRAD | 22RV1 | Ribociclib |
| MVP_top60_DNA_seed | HSP90AA1 | 0.438402 | LUAD | H3255 | CCT-018159 | MVP_top60_DNA+RNA_seed | PARP1 | -1.070302 | PRAD | LNCaP-Clone-FGC | Olaparib |
| MVP_top60_DNA_seed | GSK3B | 0.438452 | LUAD | NCI-H1437 | SB216763 | MVP_top60_DNA+RNA_seed | HDAC1 | -1.058405 | PRAD | LNCaP-Clone-FGC | Entinostat |
| MVP_top60_DNA_seed | IGF1R | 0.438469 | LUAD | NCI-H1623 | BMS-536924 | MVP_top60_DNA+RNA_seed | PIK3CA | -1.008293 | PRAD | LNCaP-Clone-FGC | Buparlisib |
| MVP_top60_DNA_seed | AURKA | 0.439025 | LUAD | NCI-H441 | Alisertib | MVP_top60_DNA+RNA_seed | HDAC1 | -1.00576 | PRAD | PWR-1E | Romidepsin |
| MVP_top60_DNA_seed | EGFR | 0.439149 | LUAD | NCI-H2009 | Cetuximab | MVP_top60_DNA+RNA_seed | PIK3CA | -0.883386 | PRAD | 22RV1 | Alpelisib |
| MVP_top60_DNA_seed | AURKA | 0.439413 | LUAD | NCI-H1648 | GSK1070916 | MVP_top60_DNA+RNA_seed | ATR | -0.875291 | PRAD | LNCaP-Clone-FGC | VE821 |
| MVP_top60_DNA_seed | SRC | 0.439597 | LUAD | NCI-H322M | WH-4-023 | MVP_top60_DNA+RNA_seed | PARP1 | -0.873991 | PRAD | LNCaP-Clone-FGC | Veliparib |
| MVP_top60_DNA_seed | EGFR | 0.440322 | LUAD | Calu-6 | Erlotinib | MVP_top60_DNA+RNA_seed | ATR | -0.870173 | PRAD | LNCaP-Clone-FGC | AZD6738 |
| MVP_top60_DNA_seed | GSK3B | 0.441399 | LUAD | NCI-H1651 | CHIR-99021 | MVP_top60_DNA+RNA_seed | ATR | -0.830781 | PRAD | PWR-1E | AZD6738 |
| MVP_top60_DNA_seed | IGF1R | 0.442406 | LUAD | NCI-H1650 | GSK1904529A | MVP_top60_DNA+RNA_seed | GSK3B | -0.830021 | PRAD | PC-3 | SB216763 |
| MVP_top60_DNA_seed | MET | 0.44253 | LUAD | NCI-H1573 | AZD6094 | MVP_top60_DNA+RNA_seed | HDAC1 | -0.822506 | PRAD | LNCaP-Clone-FGC | Romidepsin |
| MVP_top60_DNA_seed | AKT1 | 0.442988 | LUAD | SK-LU-1 | BAY AKT1 | MVP_top60_DNA+RNA_seed | EZH2 | -0.805512 | PRAD | LNCaP-Clone-FGC | GSK343 |
| MVP_top60_DNA_seed | HSP90AA1 | 0.44533 | LUAD | H3255 | Luminespib | MVP_top60_DNA+RNA_seed | PARP1 | -0.777444 | PRAD | LNCaP-Clone-FGC | Talazoparib |
| MVP_top60_DNA_seed | AURKA | 0.446395 | LUAD | RERF-LC-MS | HG-5-113-01 | MVP_top60_DNA+RNA_seed | CDK4 | -0.76717 | PRAD | 22RV1 | Palbociclib |
| MVP_top60_DNA_seed | GSK3B | 0.44658 | LUAD | NCI-H1693 | CHIR-99021 | MVP_top60_DNA+RNA_seed | ESR1 | -0.700482 | PRAD | PWR-1E | Tamoxifen |
| MVP_top60_DNA_seed | EGFR | 0.448467 | LUAD | NCI-H2291 | Afatinib | MVP_top60_DNA+RNA_seed | GSK3B | -0.683917 | PRAD | LNCaP-Clone-FGC | SB216763 |
| MVP_top60_DNA_seed | ERBB2 | 0.448467 | LUAD | NCI-H2291 | Afatinib | MVP_top60_DNA+RNA_seed | PIK3CA | -0.654226 | PRAD | 22RV1 | GNE-317 |
| MVP_top60_DNA_seed | BRAF | 0.448745 | LUAD | HCC-44 | Dabrafenib | MVP_top60_DNA+RNA_seed | ATR | -0.619541 | PRAD | LNCaP-Clone-FGC | VE-822 |
| MVP_top60_DNA_seed | ERBB2 | 0.449654 | LUAD | NCI-H1568 | CP724714 | MVP_top60_DNA+RNA_seed | ATM | -0.581479 | PRAD | PWR-1E | KU-55933 |
| MVP_top60_DNA_seed | EGFR | 0.449721 | LUAD | NCI-H1838 | Gefitinib | MVP_top60_DNA+RNA_seed | PIK3CA | -0.541057 | PRAD | PC-3 | AZD8186 |
| MVP_top60_DNA_seed | EGFR | 0.449859 | LUAD | NCI-H1993 | AZD8931 | MVP_top60_DNA+RNA_seed | PARP1 | -0.52367 | PRAD | PWR-1E | Olaparib |
| MVP_top60_DNA_seed | ERBB2 | 0.449859 | LUAD | NCI-H1993 | AZD8931 | MVP_top60_DNA+RNA_seed | PIK3CA | -0.513328 | PRAD | 22RV1 | Buparlisib |
| MVP_top60_DNA_seed | ERBB3 | 0.449859 | LUAD | NCI-H1993 | AZD8931 | MVP_top60_DNA+RNA_seed | EP300 | -0.467554 | PRAD | VCaP | SGC-CBP30 |
| MVP_top60_DNA_seed | BRAF | 0.451667 | LUAD | Calu-3 | HG6-64-1 | MVP_top60_DNA+RNA_seed | CDK4 | -0.456032 | PRAD | PWR-1E | Ribociclib |
| MVP_top60_DNA_seed | BRAF | 0.454531 | LUAD | EMC-BAC-1 | PLX-4720 | MVP_top60_DNA+RNA_seed | SMARCA4 | -0.439419 | PRAD | PWR-1E | PFI3 |
| MVP_top60_DNA_seed | GSK3B | 0.454543 | LUAD | NCI-H2030 | KIN001-042 | MVP_top60_DNA+RNA_seed | PIK3CA | -0.436117 | PRAD | LNCaP-Clone-FGC | Alpelisib |
| MVP_top60_DNA_seed | HDAC1 | 0.45541 | LUAD | SK-LU-1 | CAY10603 | MVP_top60_DNA+RNA_seed | PIK3CA | -0.417913 | PRAD | PWR-1E | Buparlisib |
| MVP_top60_DNA_seed | GSK3B | 0.455613 | LUAD | LC-2-ad | CHIR-99021 | MVP_top60_DNA+RNA_seed | PARP1 | -0.390075 | PRAD | LNCaP-Clone-FGC | Rucaparib |
| MVP_top60_DNA_seed | IGF1R | 0.455845 | LUAD | NCI-H1563 | Linsitinib | MVP_top60_DNA+RNA_seed | PARP1 | -0.384143 | PRAD | LNCaP-Clone-FGC | Niraparib |
| MVP_top60_DNA_seed | MDM2 | 0.457921 | LUAD | RERF-LC-KJ | Nutlin-3a (-) | MVP_top60_DNA+RNA_seed | PARP1 | -0.376943 | PRAD | PWR-1E | Veliparib |
| MVP_top60_DNA_seed | EGFR | 0.457931 | LUAD | RERF-LC-KJ | Pelitinib | MVP_top60_DNA+RNA_seed | PARP1 | -0.36595 | PRAD | 22RV1 | Niraparib |
| MVP_top60_DNA_seed | HSP90AA1 | 0.458672 | LUAD | NCI-H2228 | CCT-018159 | MVP_top60_DNA+RNA_seed | ESR1 | -0.361359 | PRAD | VCaP | Tamoxifen |

| | | | | | | | | | | | |
|---|---|---|---|---|---|---|---|---|---|---|---|
| MVP_top60_DNA_seed | EGFR | 0.460111 | LUAD | NCI-H1563 | Afatinib | MVP_top60_DNA+RNA_seed | HDAC1 | -0.323846 | PRAD | VCaP | Romidepsin |
| MVP_top60_DNA_seed | ERBB2 | 0.460111 | LUAD | NCI-H1563 | Afatinib | MVP_top60_DNA+RNA_seed | PARP1 | -0.238697 | PRAD | VCaP | Rucaparib |
| MVP_top60_DNA_seed | AURKA | 0.460255 | LUAD | NCI-H1651 | CD532 | MVP_top60_DNA+RNA_seed | PIK3CA | -0.229378 | PRAD | PC-3 | GNE-317 |
| MVP_top60_DNA_seed | MET | 0.460526 | LUAD | NCI-H1993 | SU11274 | MVP_top60_DNA+RNA_seed | ESR1 | -0.198261 | PRAD | 22RV1 | Tamoxifen |
| MVP_top60_DNA_seed | ATM | 0.461122 | LUAD | NCI-H2087 | KU-60019 | MVP_top60_DNA+RNA_seed | PARP1 | -0.174937 | PRAD | PWR-1E | Niraparib |
| MVP_top60_DNA_seed | HSP90AA1 | 0.461282 | LUAD | Calu-3 | CCT-018159 | MVP_top60_DNA+RNA_seed | HDAC1 | -0.168062 | PRAD | VCaP | PCI-34051 |
| MVP_top60_DNA_seed | AURKA | 0.461569 | LUAD | NCI-H2405 | GSK1070916 | MVP_top60_DNA+RNA_seed | ATR | -0.120204 | PRAD | PWR-1E | VE821 |
| MVP_top60_DNA_seed | EGFR | 0.462667 | LUAD | NCI-H650 | AZD8931 | MVP_top60_DNA+RNA_seed | PARP1 | -0.111848 | PRAD | 22RV1 | Talazoparib |
| MVP_top60_DNA_seed | ERBB2 | 0.462667 | LUAD | NCI-H650 | AZD8931 | MVP_top60_DNA+RNA_seed | CDK4 | -0.104997 | PRAD | VCaP | Palbociclib |
| MVP_top60_DNA_seed | ERBB3 | 0.462667 | LUAD | NCI-H650 | AZD8931 | MVP_top60_DNA+RNA_seed | HDAC1 | -0.062479 | PRAD | PWR-1E | PCI-34051 |
| MVP_top60_DNA_seed | ATM | 0.462769 | LUAD | NCI-H1623 | KU-55933 | MVP_top60_DNA+RNA_seed | SMARCA4 | -0.04848 | PRAD | LNCaP-Clone-FGC | PFI3 |
| MVP_top60_DNA_seed | PTK2 | 0.462964 | LUAD | PC-14 | PF-562271 | MVP_top60_DNA+RNA_seed | EP300 | -0.029644 | PRAD | PWR-1E | SGC-CBP30 |
| MVP_top60_DNA_seed | FGFR1 | 0.463634 | LUAD | NCI-H522 | AZD4547 | MVP_top60_DNA+RNA_seed | HDAC1 | -0.026224 | PRAD | 22RV1 | Entinostat |
| MVP_top60_DNA_seed | HDAC1 | 0.464217 | LUAD | NCI-H1944 | Tubastatin A | MVP_top60_DNA+RNA_seed | TP53 | -0.014816 | PRAD | PWR-1E | MIRA-1 |
| MVP_top60_DNA_seed | BRAF | 0.46446 | LUAD | NCI-H2405 | SB590885 | MVP_top60_DNA+RNA_seed | EP300 | -0.012331 | PRAD | DU-145 | CPI-637 |
| MVP_top60_DNA_seed | GSK3B | 0.464874 | LUAD | RERF-LC-MS | KIN001-042 | MVP_top60_DNA+RNA_seed | PARP1 | 0.017556 | PRAD | PWR-1E | Talazoparib |
| MVP_top60_DNA_seed | EGFR | 0.465163 | LUAD | NCI-H2228 | Pelitinib | MVP_top60_DNA+RNA_seed | PIK3CA | 0.033091 | PRAD | VCaP | Alpelisib |
| MVP_top60_DNA_seed | MET | 0.46537 | LUAD | RERF-LC-KJ | SU11274 | MVP_top60_DNA+RNA_seed | ATM | 0.046193 | PRAD | PC-3 | KU-55933 |
| MVP_top60_DNA_seed | HDAC1 | 0.465662 | LUAD | EKVX | Entinostat | MVP_top60_DNA+RNA_seed | GSK3B | 0.052881 | PRAD | VCaP | SB216763 |
| MVP_top60_DNA_seed | AKT1 | 0.465795 | LUAD | PC-3_[JPC-3] | AKT inhibitor VIII | MVP_top60_DNA+RNA_seed | PARP1 | 0.057574 | PRAD | VCaP | Olaparib |
| MVP_top60_DNA_seed | GSK3B | 0.465823 | LUAD | NCI-H1437 | KIN001-042 | MVP_top60_DNA+RNA_seed | EZH2 | 0.079193 | PRAD | VCaP | GSK343 |
| MVP_top60_DNA_seed | FGFR1 | 0.466013 | LUAD | SW1573 | AZD4547 | MVP_top60_DNA+RNA_seed | PARP1 | 0.101815 | PRAD | 22RV1 | Rucaparib |
| MVP_top60_DNA_seed | ATM | 0.467663 | LUAD | NCI-H2122 | KU-55933 | MVP_top60_DNA+RNA_seed | PARP1 | 0.131177 | PRAD | PWR-1E | Rucaparib |
| MVP_top60_DNA_seed | MAPK1 | 0.467806 | LUAD | HOP-62 | VX-11e | MVP_top60_DNA+RNA_seed | EP300 | 0.138199 | PRAD | PWR-1E | CPI-637 |
| MVP_top60_DNA_seed | EGFR | 0.468012 | LUAD | NCI-H441 | CI-1033 | MVP_top60_DNA+RNA_seed | PIK3CA | 0.149453 | PRAD | PC-3 | Buparlisib |
| MVP_top60_DNA_seed | ERBB2 | 0.468012 | LUAD | NCI-H441 | CI-1033 | MVP_top60_DNA+RNA_seed | HDAC1 | 0.185731 | PRAD | VCaP | Entinostat |
| MVP_top60_DNA_seed | EGFR | 0.469634 | LUAD | NCI-H1838 | AZD8931 | MVP_top60_DNA+RNA_seed | PARP1 | 0.191115 | PRAD | DU-145 | Veliparib |
| MVP_top60_DNA_seed | ERBB2 | 0.469634 | LUAD | NCI-H1838 | AZD8931 | MVP_top60_DNA+RNA_seed | PARP1 | 0.204799 | PRAD | DU-145 | Talazoparib |
| MVP_top60_DNA_seed | ERBB3 | 0.469634 | LUAD | NCI-H1838 | AZD8931 | MVP_top60_DNA+RNA_seed | ESR1 | 0.213009 | PRAD | VCaP | GDC0810 |
| MVP_top60_DNA_seed | AKT1 | 0.470904 | LUAD | PC-3_[JPC-3] | GSK690693 | MVP_top60_DNA+RNA_seed | ESR2 | 0.213009 | PRAD | VCaP | GDC0810 |
| MVP_top60_DNA_seed | BRAF | 0.472367 | LUAD | SW1573 | RAF_9304 | MVP_top60_DNA+RNA_seed | CDK4 | 0.230549 | PRAD | VCaP | Ribociclib |
| MVP_top60_DNA_seed | EGFR | 0.474412 | LUAD | RERF-LC-MS | CI-1033 | MVP_top60_DNA+RNA_seed | TP53 | 0.243205 | PRAD | VCaP | MIRA-1 |
| MVP_top60_DNA_seed | ERBB2 | 0.474412 | LUAD | RERF-LC-MS | CI-1033 | MVP_top60_DNA+RNA_seed | ESR1 | 0.275997 | PRAD | PWR-1E | GDC0810 |
| MVP_top60_DNA_seed | AURKA | 0.474566 | LUAD | HOP-62 | ZM447439 | MVP_top60_DNA+RNA_seed | ESR2 | 0.275997 | PRAD | PWR-1E | GDC0810 |
| MVP_top60_DNA_seed | HDAC1 | 0.475716 | LUAD | Calu-3 | Tubastatin A | MVP_top60_DNA+RNA_seed | HDAC1 | 0.282483 | PRAD | PWR-1E | Entinostat |
| MVP_top60_DNA_seed | HSP90AA1 | 0.476592 | LUAD | NCI-H1838 | Tanespimycin | MVP_top60_DNA+RNA_seed | PIK3CA | 0.288414 | PRAD | PWR-1E | GNE-317 |
| MVP_top60_DNA_seed | EGFR | 0.476937 | LUAD | NCI-H2009 | Afatinib | MVP_top60_DNA+RNA_seed | ATR | 0.304135 | PRAD | PWR-1E | VE-822 |
| MVP_top60_DNA_seed | ERBB2 | 0.476937 | LUAD | NCI-H2009 | Afatinib | MVP_top60_DNA+RNA_seed | HDAC1 | 0.321559 | PRAD | 22RV1 | Romidepsin |
| MVP_top60_DNA_seed | FGFR1 | 0.477078 | LUAD | NCI-H2122 | AZD4547 | MVP_top60_DNA+RNA_seed | PIK3CA | 0.349073 | PRAD | VCaP | AZD8186 |
| MVP_top60_DNA_seed | AURKA | 0.478744 | LUAD | PC-3_[JPC-3] | GSK1070916 | MVP_top60_DNA+RNA_seed | HDAC1 | 0.381751 | PRAD | LNCaP-Clone-FGC | PCI-34051 |
| MVP_top60_DNA_seed | SRC | 0.479197 | LUAD | EKVX | Bosutinib | MVP_top60_DNA+RNA_seed | EP300 | 0.422506 | PRAD | LNCaP-Clone-FGC | SGC-CBP30 |

| | | | | | | | | | | | |
|---|---|---|---|---|---|---|---|---|---|---|---|
| MVP_top60_DNA_seed | EGFR | 0.479737 | LUAD | NCI-H1651 | Cetuximab | MVP_top60_DNA+RNA_seed | ATM | 0.42586 | PRAD | LNCaP-Clone-FGC | KU-55933 |
| MVP_top60_DNA_seed | SRC | 0.481045 | LUAD | Calu-6 | WH-4-023 | MVP_top60_DNA+RNA_seed | PIK3CA | 0.437684 | PRAD | 22RV1 | AZD8186 |
| MVP_top60_DNA_seed | ERBB2 | 0.482172 | LUAD | NCI-H23 | CP724714 | MVP_top60_DNA+RNA_seed | PARP1 | 0.446962 | PRAD | 22RV1 | Olaparib |
| MVP_top60_DNA_seed | BRAF | 0.485802 | LUAD | VMRC-LCD | PLX-4720 | MVP_top60_DNA+RNA_seed | EZH2 | 0.461793 | PRAD | 22RV1 | GSK343 |
| MVP_top60_DNA_seed | AKT1 | 0.485839 | LUAD | NCI-H2087 | AKT inhibitor VIII | MVP_top60_DNA+RNA_seed | PARP1 | 0.501736 | PRAD | PC-3 | Talazoparib |
| MVP_top60_DNA_seed | EGFR | 0.486252 | LUAD | HCC-78 | AZD8931 | MVP_top60_DNA+RNA_seed | EP300 | 0.522999 | PRAD | 22RV1 | CPI-637 |
| MVP_top60_DNA_seed | ERBB2 | 0.486252 | LUAD | HCC-78 | AZD8931 | MVP_top60_DNA+RNA_seed | HDAC1 | 0.531994 | PRAD | DU-145 | Romidepsin |
| MVP_top60_DNA_seed | ERBB3 | 0.486252 | LUAD | HCC-78 | AZD8931 | MVP_top60_DNA+RNA_seed | ATR | 0.553194 | PRAD | VCaP | VE821 |
| MVP_top60_DNA_seed | AURKA | 0.487057 | LUAD | Calu-6 | Tozasertib | MVP_top60_DNA+RNA_seed | ATR | 0.591904 | PRAD | DU-145 | VE-822 |
| MVP_top60_DNA_seed | MAPK1 | 0.487257 | LUAD | NCI-H1563 | VX-11e | MVP_top60_DNA+RNA_seed | EP300 | 0.602643 | PRAD | DU-145 | SGC-CBP30 |
| MVP_top60_DNA_seed | AURKA | 0.487295 | LUAD | Calu-3 | GSK1070916 | MVP_top60_DNA+RNA_seed | ATR | 0.603229 | PRAD | DU-145 | AZD6738 |
| MVP_top60_DNA_seed | BRAF | 0.4876 | LUAD | EMC-BAC-2 | RAF_9304 | MVP_top60_DNA+RNA_seed | GSK3B | 0.626472 | PRAD | 22RV1 | SB216763 |
| MVP_top60_DNA_seed | IGF1R | 0.487761 | LUAD | Calu-6 | BMS-536924 | MVP_top60_DNA+RNA_seed | ATM | 0.639648 | PRAD | DU-145 | KU-55933 |
| MVP_top60_DNA_seed | ATM | 0.488647 | LUAD | NCI-H650 | KU-55933 | MVP_top60_DNA+RNA_seed | PARP1 | 0.641877 | PRAD | VCaP | Talazoparib |
| MVP_top60_DNA_seed | HDAC1 | 0.48976 | LUAD | NCI-H1648 | Entinostat | MVP_top60_DNA+RNA_seed | ESR1 | 0.655833 | PRAD | DU-145 | GDC0810 |
| MVP_top60_DNA_seed | IGF1R | 0.490371 | LUAD | NCI-H322M | BMS-536924 | MVP_top60_DNA+RNA_seed | ESR2 | 0.655833 | PRAD | DU-145 | GDC0810 |
| MVP_top60_DNA_seed | MET | 0.491171 | LUAD | NCI-H2405 | AZD6094 | MVP_top60_DNA+RNA_seed | EZH2 | 0.674191 | PRAD | PWR-1E | GSK343 |
| MVP_top60_DNA_seed | BRAF | 0.491692 | LUAD | PC-3_[JPC-3] | PLX-4720 | MVP_top60_DNA+RNA_seed | PARP1 | 0.678392 | PRAD | DU-145 | Olaparib |
| MVP_top60_DNA_seed | MET | 0.492111 | LUAD | NCI-H1623 | SU11274 | MVP_top60_DNA+RNA_seed | PIK3CA | 0.678429 | PRAD | PWR-1E | Alpelisib |
| MVP_top60_DNA_seed | EGFR | 0.492278 | LUAD | NCI-H838 | CUDC-101 | MVP_top60_DNA+RNA_seed | EP300 | 0.683503 | PRAD | PC-3 | CPI-637 |
| MVP_top60_DNA_seed | ERBB2 | 0.492278 | LUAD | NCI-H838 | CUDC-101 | MVP_top60_DNA+RNA_seed | TP53 | 0.687703 | PRAD | LNCaP-Clone-FGC | MIRA-1 |
| MVP_top60_DNA_seed | MDM2 | 0.493782 | LUAD | VMRC-LCD | Nutlin-3a (-) | MVP_top60_DNA+RNA_seed | PIK3CA | 0.692404 | PRAD | PWR-1E | AZD8186 |
| MVP_top60_DNA_seed | EGFR | 0.494143 | LUAD | NCI-H1437 | Gefitinib | MVP_top60_DNA+RNA_seed | TP53 | 0.706283 | PRAD | 22RV1 | MIRA-1 |
| MVP_top60_DNA_seed | GSK3B | 0.494419 | LUAD | NCI-H1648 | AZD7969 | MVP_top60_DNA+RNA_seed | PARP1 | 0.72663 | PRAD | VCaP | Niraparib |
| MVP_top60_DNA_seed | ATM | 0.494755 | LUAD | ABC-1 | KU-60019 | MVP_top60_DNA+RNA_seed | ATR | 0.72813 | PRAD | 22RV1 | VE-822 |
| MVP_top60_DNA_seed | GSK3B | 0.495188 | LUAD | NCI-H2009 | KIN001-042 | MVP_top60_DNA+RNA_seed | ATR | 0.752867 | PRAD | PC-3 | AZD6738 |
| MVP_top60_DNA_seed | ERBB2 | 0.495254 | LUAD | NCI-H2405 | CP724714 | MVP_top60_DNA+RNA_seed | SMARCA4 | 0.756082 | PRAD | DU-145 | PFI3 |
| MVP_top60_DNA_seed | MAPK1 | 0.495882 | LUAD | NCI-H1573 | FR-180204 | MVP_top60_DNA+RNA_seed | PIK3CA | 0.757757 | PRAD | PC-3 | Alpelisib |
| MVP_top60_DNA_seed | EGFR | 0.496306 | LUAD | NCI-H2342 | Gefitinib | MVP_top60_DNA+RNA_seed | ESR1 | 0.769845 | PRAD | PC-3 | GDC0810 |
| MVP_top60_DNA_seed | FGFR1 | 0.497266 | LUAD | NCI-H2030 | AZD4547 | MVP_top60_DNA+RNA_seed | ESR2 | 0.769845 | PRAD | PC-3 | GDC0810 |
| MVP_top60_DNA_seed | ATM | 0.498275 | LUAD | H3255 | KU-55933 | MVP_top60_DNA+RNA_seed | SMARCA4 | 0.807282 | PRAD | VCaP | PFI3 |
| MVP_top60_DNA_seed | BRAF | 0.498886 | LUAD | NCI-H1355 | RAF_9304 | MVP_top60_DNA+RNA_seed | SMARCA4 | 0.836821 | PRAD | 22RV1 | PFI3 |
| MVP_top60_DNA_seed | HDAC1 | 0.499545 | LUAD | NCI-H322M | AR-42 | MVP_top60_DNA+RNA_seed | EP300 | 0.844741 | PRAD | 22RV1 | SGC-CBP30 |
| MVP_top60_DNA_seed | SMARCA4 | 0.499865 | LUAD | NCI-H1568 | PFI-3 | MVP_top60_DNA+RNA_seed | PIK3CA | 0.861976 | PRAD | VCaP | Buparlisib |
| MVP_top60_DNA_seed | ATM | 0.499968 | LUAD | RERF-LC-MS | KU-55933 | MVP_top60_DNA+RNA_seed | EP300 | 0.866293 | PRAD | PC-3 | SGC-CBP30 |
| MVP_top60_DNA_seed | MDM2 | 0.50009 | LUAD | NCI-H2405 | Nutlin-3a (-) | MVP_top60_DNA+RNA_seed | PARP1 | 0.872677 | PRAD | 22RV1 | Veliparib |
| MVP_top60_DNA_seed | ERBB2 | 0.500192 | LUAD | NCI-H2009 | CP724714 | MVP_top60_DNA+RNA_seed | PARP1 | 0.874127 | PRAD | PC-3 | Olaparib |
| MVP_top60_DNA_seed | AURKA | 0.500558 | LUAD | NCI-H1993 | Genentech Cpd 10 | MVP_top60_DNA+RNA_seed | EZH2 | 0.889699 | PRAD | VCaP | GSK343 |
| MVP_top60_DNA_seed | MET | 0.500725 | LUAD | NCI-H2342 | SU11274 | MVP_top60_DNA+RNA_seed | ATR | 0.907775 | PRAD | 22RV1 | AZD6738 |
| MVP_top60_DNA_seed | HDAC1 | 0.50132 | LUAD | SK-LU-1 | Belinostat | MVP_top60_DNA+RNA_seed | HDAC1 | 0.90968 | PRAD | PC-3 | Entinostat |
| MVP_top60_DNA_seed | AURKA | 0.503501 | LUAD | NCI-H650 | Alisertib | MVP_top60_DNA+RNA_seed | GSK3B | 0.946724 | PRAD | DU-145 | SB216763 |

| | | | | | | | | | | | |
|---|---|---|---|---|---|---|---|---|---|---|---|
| MVP_top60_DNA_seed | GSK3B | 0.50382 | LUAD | NCI-H2030 | CHIR-99021 | MVP_top60_DNA+RNA_seed | ESR1 | 0.9541 | PRAD | DU-145 | Tamoxifen |
| MVP_top60_DNA_seed | PIK3CA | 0.503865 | LUAD | ABC-1 | PI3Ka_4409 | MVP_top60_DNA+RNA_seed | TP53 | 0.955249 | PRAD | DU-145 | MIRA-1 |
| MVP_top60_DNA_seed | GSK3B | 0.50433 | LUAD | RERF-LC-KJ | CHIR-99021 | MVP_top60_DNA+RNA_seed | CDK4 | 0.956746 | PRAD | PWR-1E | Palbociclib |
| MVP_top60_DNA_seed | HDAC1 | 0.504481 | LUAD | NCI-H2085 | Belinostat | MVP_top60_DNA+RNA_seed | CDK4 | 0.972709 | PRAD | PC-3 | Ribociclib |
| MVP_top60_DNA_seed | EGFR | 0.504611 | LUAD | NCI-H1650 | CUDC-101 | MVP_top60_DNA+RNA_seed | PIK3CA | 1.005586 | PRAD | DU-145 | AZD8186 |
| MVP_top60_DNA_seed | ERBB2 | 0.504611 | LUAD | NCI-H1650 | CUDC-101 | MVP_top60_DNA+RNA_seed | CDK4 | 1.030612 | PRAD | DU-145 | Ribociclib |
| MVP_top60_DNA_seed | EGFR | 0.505283 | LUAD | NCI-H1993 | AST-1306 | MVP_top60_DNA+RNA_seed | EZH2 | 1.037094 | PRAD | 22RV1 | GSK343 |
| MVP_top60_DNA_seed | AURKA | 0.505528 | LUAD | NCI-H1666 | GSK1070916 | MVP_top60_DNA+RNA_seed | HDAC1 | 1.070096 | PRAD | 22RV1 | PCI-34051 |
| MVP_top60_DNA_seed | IGF1R | 0.505704 | LUAD | NCI-H2085 | BMS-754807 | MVP_top60_DNA+RNA_seed | PIK3CA | 1.075058 | PRAD | VCaP | GNE-317 |
| MVP_top60_DNA_seed | SRC | 0.505711 | LUAD | EKVX | Saracatinib | MVP_top60_DNA+RNA_seed | ATM | 1.07653 | PRAD | VCaP | KU-55933 |
| MVP_top60_DNA_seed | FGFR1 | 0.505803 | LUAD | NCI-H1693 | PD173074 | MVP_top60_DNA+RNA_seed | PARP1 | 1.094931 | PRAD | PC-3 | Niraparib |
| MVP_top60_DNA_seed | HSP90AA1 | 0.506541 | LUAD | NCI-H1838 | Elesclomol | MVP_top60_DNA+RNA_seed | PIK3CA | 1.111516 | PRAD | DU-145 | Alpelisib |
| MVP_top60_DNA_seed | MET | 0.506621 | LUAD | HCC-78 | SU11274 | MVP_top60_DNA+RNA_seed | PARP1 | 1.117531 | PRAD | VCaP | Veliparib |
| MVP_top60_DNA_seed | ATM | 0.507233 | LUAD | LC-2-ad | KU-60019 | MVP_top60_DNA+RNA_seed | ATR | 1.131643 | PRAD | PC-3 | VE-822 |
| MVP_top60_DNA_seed | GSK3B | 0.507495 | LUAD | NCI-H358 | SB216763 | MVP_top60_DNA+RNA_seed | PARP1 | 1.145966 | PRAD | PC-3 | Veliparib |
| MVP_top60_DNA_seed | HSP90AA1 | 0.508474 | LUAD | NCI-H1734 | SNX-2112 | MVP_top60_DNA+RNA_seed | EZH2 | 1.198497 | PRAD | DU-145 | GSK343 |
| MVP_top60_DNA_seed | BRAF | 0.508785 | LUAD | NCI-H2122 | SB590885 | MVP_top60_DNA+RNA_seed | HDAC1 | 1.22727 | PRAD | PC-3 | Romidepsin |
| MVP_top60_DNA_seed | HDAC1 | 0.508897 | LUAD | SW1573 | Belinostat | MVP_top60_DNA+RNA_seed | PARP1 | 1.236011 | PRAD | DU-145 | Niraparib |
| MVP_top60_DNA_seed | SRC | 0.508924 | LUAD | ABC-1 | HG-6-71-01 | MVP_top60_DNA+RNA_seed | PARP1 | 1.241793 | PRAD | DU-145 | Rucaparib |
| MVP_top60_DNA_seed | AKT1 | 0.509073 | LUAD | NCI-H358 | BAY AKT1 | MVP_top60_DNA+RNA_seed | PARP1 | 1.262555 | PRAD | PC-3 | Rucaparib |
| MVP_top60_DNA_seed | HDAC1 | 0.509694 | LUAD | NCI-H1435 | Dacinostat | MVP_top60_DNA+RNA_seed | ATM | 1.273546 | PRAD | 22RV1 | KU-55933 |
| MVP_top60_DNA_seed | PIK3CA | 0.509795 | LUAD | NCI-H522 | AZD8835 | MVP_top60_DNA+RNA_seed | ATR | 1.28593 | PRAD | PC-3 | VE821 |
| MVP_top60_DNA_seed | EGFR | 0.510279 | LUAD | SK-LU-1 | CUDC-101 | MVP_top60_DNA+RNA_seed | ATR | 1.303361 | PRAD | VCaP | VE-822 |
| MVP_top60_DNA_seed | ERBB2 | 0.510279 | LUAD | SK-LU-1 | CUDC-101 | MVP_top60_DNA+RNA_seed | ATR | 1.309826 | PRAD | 22RV1 | VE821 |
| MVP_top60_DNA_seed | EGFR | 0.51045 | LUAD | NCI-H1573 | PF-00299804 | MVP_top60_DNA+RNA_seed | SMARCA4 | 1.318882 | PRAD | PC-3 | PFI3 |
| MVP_top60_DNA_seed | ERBB2 | 0.51045 | LUAD | NCI-H1573 | PF-00299804 | MVP_top60_DNA+RNA_seed | EZH2 | 1.340623 | PRAD | DU-145 | GSK343 |
| MVP_top60_DNA_seed | EGFR | 0.510487 | LUAD | Calu-6 | Lapatinib | MVP_top60_DNA+RNA_seed | EZH2 | 1.371107 | PRAD | PC-3 | GSK343 |
| MVP_top60_DNA_seed | ERBB2 | 0.510487 | LUAD | Calu-6 | Lapatinib | MVP_top60_DNA+RNA_seed | PIK3CA | 1.404643 | PRAD | DU-145 | Buparlisib |
| MVP_top60_DNA_seed | FGFR1 | 0.51076 | LUAD | NCI-H1666 | FGFR_3831 | MVP_top60_DNA+RNA_seed | HDAC1 | 1.412924 | PRAD | PC-3 | PCI-34051 |
| MVP_top60_DNA_seed | EGFR | 0.511534 | LUAD | NCI-H1755 | Gefitinib | MVP_top60_DNA+RNA_seed | ESR1 | 1.435482 | PRAD | 22RV1 | GDC0810 |
| MVP_top60_DNA_seed | BRAF | 0.511687 | LUAD | H3255 | PLX-4720 | MVP_top60_DNA+RNA_seed | ESR2 | 1.435482 | PRAD | 22RV1 | GDC0810 |
| MVP_top60_DNA_seed | EGFR | 0.511973 | LUAD | RERF-LC-MS | Cetuximab | MVP_top60_DNA+RNA_seed | ATR | 1.444891 | PRAD | DU-145 | VE821 |
| MVP_top60_DNA_seed | BRAF | 0.512696 | LUAD | NCI-H1563 | HG6-64-1 | MVP_top60_DNA+RNA_seed | HDAC1 | 1.453979 | PRAD | DU-145 | PCI-34051 |
| MVP_top60_DNA_seed | GSK3B | 0.512781 | LUAD | VMRC-LCD | CHIR-99021 | MVP_top60_DNA+RNA_seed | TP53 | 1.463099 | PRAD | PC-3 | MIRA-1 |
| MVP_top60_DNA_seed | AKT1 | 0.512834 | LUAD | EMC-BAC-2 | AKT inhibitor VIII | MVP_top60_DNA+RNA_seed | ESR1 | 1.479545 | PRAD | PC-3 | Tamoxifen |
| MVP_top60_DNA_seed | MET | 0.513053 | LUAD | NCI-H2087 | AZD6094 | MVP_top60_DNA+RNA_seed | CDK4 | 1.540449 | PRAD | PC-3 | Palbociclib |
| MVP_top60_DNA_seed | HDAC1 | 0.514612 | LUAD | NCI-H2085 | Tubastatin A | MVP_top60_DNA+RNA_seed | HDAC1 | 1.544817 | PRAD | DU-145 | Entinostat |
| MVP_top60_DNA_seed | GSK3B | 0.51525 | LUAD | H3255 | CHIR-99021 | MVP_top60_DNA+RNA_seed | CDK4 | 1.564859 | PRAD | DU-145 | Palbociclib |
| MVP_top60_DNA_seed | GSK3B | 0.516563 | LUAD | LC-2-ad | SB216763 | MVP_top60_DNA+RNA_seed | EZH2 | 1.698386 | PRAD | PC-3 | GSK343 |
| MVP_top60_DNA_seed | MAPK1 | 0.51916 | LUAD | NCI-H1651 | FR-180204 | MVP_top60_DNA+RNA_seed | ATR | 1.774057 | PRAD | VCaP | AZD6738 |
| MVP_top60_DNA_seed | AURKA | 0.520113 | LUAD | NCI-H3122 | Alisertib | MVP_top60_DNA+RNA_seed | PIK3CA | 1.796477 | PRAD | DU-145 | GNE-317 |

| | | | | | | | | | | | |
|---|---|---|---|---|---|---|---|---|---|---|---|
| MVP_top60_DNA_seed | SMARCA4 | 0.520279 | LUAD | NCI-H1781 | PFI-3 | MVP_top60_DNA+RNA_seed | MET | -8.579362 | STAD | MKN45 | AZD6094 |
| MVP_top60_DNA_seed | ATM | 0.520475 | LUAD | HOP-62 | KU-55933 | MVP_top60_DNA+RNA_seed | MET | -8.429196 | STAD | SNU-5 | AZD6094 |
| MVP_top60_DNA_seed | PIK3CA | 0.520712 | LUAD | NCI-H2291 | AZD8835 | MVP_top60_DNA+RNA_seed | MET | -6.178313 | STAD | Hs-746T | AZD6094 |
| MVP_top60_DNA_seed | ATM | 0.521038 | LUAD | NCI-H322M | CP466722 | MVP_top60_DNA+RNA_seed | FGFR3 | -5.738215 | STAD | HSC-39 | AZD4547 |
| MVP_top60_DNA_seed | AKT1 | 0.521771 | LUAD | SK-LU-1 | AKT inhibitor VIII | MVP_top60_DNA+RNA_seed | ERBB2 | -5.506139 | STAD | NCI-N87 | CP724714 |
| MVP_top60_DNA_seed | FGFR1 | 0.52226 | LUAD | RERF-LC-KJ | FGFR_3831 | MVP_top60_DNA+RNA_seed | MET | -5.462021 | STAD | SNU-5 | PHA-665752 |
| MVP_top60_DNA_seed | BRAF | 0.522804 | LUAD | NCI-H1435 | PLX-4720 | MVP_top60_DNA+RNA_seed | FGFR3 | -5.404401 | STAD | HSC-39 | PD173074 |
| MVP_top60_DNA_seed | EGFR | 0.52518 | LUAD | NCI-H1666 | CUDC-101 | MVP_top60_DNA+RNA_seed | MET | -5.126293 | STAD | SNU-5 | JNJ38877605 |
| MVP_top60_DNA_seed | ERBB2 | 0.52518 | LUAD | NCI-H1666 | CUDC-101 | MVP_top60_DNA+RNA_seed | FGFR3 | -5.111861 | STAD | KATOIII | PD173074 |
| MVP_top60_DNA_seed | BRAF | 0.52548 | LUAD | NCI-H2085 | Dabrafenib | MVP_top60_DNA+RNA_seed | FGFR3 | -5.015799 | STAD | HSC-39 | AZD4547 |
| MVP_top60_DNA_seed | HDAC1 | 0.526119 | LUAD | NCI-H1792 | Belinostat | MVP_top60_DNA+RNA_seed | MET | -4.798288 | STAD | SNU-5 | Crizotinib |
| MVP_top60_DNA_seed | EGFR | 0.52662 | LUAD | NCI-H1563 | PF-00299804 | MVP_top60_DNA+RNA_seed | MET | -4.659088 | STAD | Hs-746T | JNJ38877605 |
| MVP_top60_DNA_seed | ERBB2 | 0.52662 | LUAD | NCI-H1563 | PF-00299804 | MVP_top60_DNA+RNA_seed | MET | -4.234767 | STAD | MKN45 | JNJ38877605 |
| MVP_top60_DNA_seed | IGF1R | 0.527232 | LUAD | PC-3_[JPC-3] | Linsitinib | MVP_top60_DNA+RNA_seed | FGFR3 | -4.202287 | STAD | KATOIII | AZD4547 |
| MVP_top60_DNA_seed | AURKA | 0.527278 | LUAD | NCI-H2228 | ZM447439 | MVP_top60_DNA+RNA_seed | ERBB2 | -4.174126 | STAD | Hs-746T | CP724714 |
| MVP_top60_DNA_seed | HDAC1 | 0.527803 | LUAD | NCI-H1650 | Dacinostat | MVP_top60_DNA+RNA_seed | ERBB2 | -3.997646 | STAD | SK-GT-2 | Afatinib |
| MVP_top60_DNA_seed | AKT1 | 0.528128 | LUAD | NCI-H1650 | GSK690693 | MVP_top60_DNA+RNA_seed | ERBB2 | -3.708978 | STAD | SK-GT-2 | Afatinib |
| MVP_top60_DNA_seed | IGF1R | 0.528451 | LUAD | Calu-6 | Linsitinib | MVP_top60_DNA+RNA_seed | AR | -3.602727 | STAD | MKN45 | Bicalutamide |
| MVP_top60_DNA_seed | AURKA | 0.528739 | LUAD | RERF-LC-KJ | Alisertib | MVP_top60_DNA+RNA_seed | ERBB2 | -3.579769 | STAD | NUGC-4 | Afatinib |
| MVP_top60_DNA_seed | BRAF | 0.529692 | LUAD | NCI-H1993 | RAF_9304 | MVP_top60_DNA+RNA_seed | FGFR3 | -3.547109 | STAD | SNU-16 | PD173074 |
| MVP_top60_DNA_seed | AKT1 | 0.530348 | LUAD | NCI-H2023 | GSK690693 | MVP_top60_DNA+RNA_seed | ERBB2 | -3.19964 | STAD | NCI-N87 | Afatinib |
| MVP_top60_DNA_seed | AKT1 | 0.530471 | LUAD | NCI-H1781 | GSK690693 | MVP_top60_DNA+RNA_seed | MET | -3.079747 | STAD | MKN45 | SU11274 |
| MVP_top60_DNA_seed | BRAF | 0.531199 | LUAD | NCI-H1435 | PLX-4720 | MVP_top60_DNA+RNA_seed | FGFR3 | -3.047242 | STAD | HSC-39 | FGFR_3831 |
| MVP_top60_DNA_seed | HDAC1 | 0.531311 | LUAD | NCI-H650 | Tubastatin A | MVP_top60_DNA+RNA_seed | ERBB2 | -3.012876 | STAD | NUGC-4 | AZD8931 |
| MVP_top60_DNA_seed | MAPK1 | 0.531357 | LUAD | LXF-289 | FR-180204 | MVP_top60_DNA+RNA_seed | ERBB3 | -3.012876 | STAD | NUGC-4 | AZD8931 |
| MVP_top60_DNA_seed | IGF1R | 0.531481 | LUAD | NCI-H1755 | GSK1904529A | MVP_top60_DNA+RNA_seed | PARP1 | -2.919194 | STAD | HGC-27 | Talazoparib |
| MVP_top60_DNA_seed | EGFR | 0.53237 | LUAD | NCI-H838 | Gefitinib | MVP_top60_DNA+RNA_seed | ERBB2 | -2.840983 | STAD | NCI-N87 | Afatinib |
| MVP_top60_DNA_seed | MDM2 | 0.532374 | LUAD | NCI-H441 | Nutlin-3a (-) | MVP_top60_DNA+RNA_seed | FGFR3 | -2.737636 | STAD | KATOIII | FGFR_3831 |
| MVP_top60_DNA_seed | HDAC1 | 0.533241 | LUAD | Calu-6 | Tubastatin A | MVP_top60_DNA+RNA_seed | PARP1 | -2.725446 | STAD | HGC-27 | Rucaparib |
| MVP_top60_DNA_seed | HSP90AA1 | 0.533587 | LUAD | NCI-H1437 | SNX-2112 | MVP_top60_DNA+RNA_seed | MET | -2.712269 | STAD | IM-95 | JNJ38877605 |
| MVP_top60_DNA_seed | AURKA | 0.533891 | LUAD | Calu-6 | Genentech Cpd 10 | MVP_top60_DNA+RNA_seed | MAPK1 | -2.574266 | STAD | NUGC-4 | FR-180204 |
| MVP_top60_DNA_seed | PIK3CA | 0.534366 | LUAD | Calu-6 | PI3Ka_4409 | MVP_top60_DNA+RNA_seed | MAPK3 | -2.574266 | STAD | NUGC-4 | FR-180204 |
| MVP_top60_DNA_seed | EGFR | 0.534406 | LUAD | NCI-H2405 | CUDC-101 | MVP_top60_DNA+RNA_seed | PIK3CA | -2.507956 | STAD | IM-95 | AZD8835 |
| MVP_top60_DNA_seed | ERBB2 | 0.534406 | LUAD | NCI-H2405 | CUDC-101 | MVP_top60_DNA+RNA_seed | ERBB2 | -2.458722 | STAD | NUGC-4 | PF-00299804 |
| MVP_top60_DNA_seed | GSK3B | 0.534752 | LUAD | NCI-H1651 | SB216763 | MVP_top60_DNA+RNA_seed | ERBB4 | -2.458722 | STAD | NUGC-4 | PF-00299804 |
| MVP_top60_DNA_seed | MDM2 | 0.534813 | LUAD | NCI-H2009 | Nutlin-3a (-) | MVP_top60_DNA+RNA_seed | ESR1 | -2.400741 | STAD | MKN28 | Fulvestrant |
| MVP_top60_DNA_seed | HSP90AA1 | 0.535888 | LUAD | NCI-H3122 | Elesclomol | MVP_top60_DNA+RNA_seed | FGFR3 | -2.29 | STAD | SNU-16 | FGFR_3831 |
| MVP_top60_DNA_seed | FGFR1 | 0.536374 | LUAD | NCI-H358 | AZD4547 | MVP_top60_DNA+RNA_seed | MET | -2.285163 | STAD | Hs-746T | SU11274 |
| MVP_top60_DNA_seed | BRAF | 0.536937 | LUAD | NCI-H322M | SB590885 | MVP_top60_DNA+RNA_seed | ESR1 | -2.256546 | STAD | HGC-27 | Tamoxifen |
| MVP_top60_DNA_seed | BRAF | 0.537397 | LUAD | NCI-H1755 | RAF_9304 | MVP_top60_DNA+RNA_seed | HSP90AA1 | -2.213796 | STAD | AGS | Elesclomol |
| MVP_top60_DNA_seed | GSK3B | 0.53827 | LUAD | NCI-H2087 | CHIR-99021 | MVP_top60_DNA+RNA_seed | HSP90AB1 | -2.213796 | STAD | AGS | Elesclomol |

| | | | | | | | | | | | |
|---|---|---|---|---|---|---|---|---|---|---|---|
| MVP_top60_DNA_seed | AKT1 | 0.538857 | LUAD | PC-14 | MK-2206 | MVP_top60_DNA+RNA_seed | ERBB2 | -2.202897 | STAD | SK-GT-2 | AZD8931 |
| MVP_top60_DNA_seed | AKT1 | 0.539023 | LUAD | NCI-H2122 | GSK690693 | MVP_top60_DNA+RNA_seed | ERBB3 | -2.202897 | STAD | SK-GT-2 | AZD8931 |
| MVP_top60_DNA_seed | ATM | 0.539599 | LUAD | NCI-H1563 | torin2 | MVP_top60_DNA+RNA_seed | HSP90AA1 | -2.199261 | STAD | FU97 | Tanespimycin |
| MVP_top60_DNA_seed | MAPK1 | 0.540144 | LUAD | NCI-H2342 | FR-180204 | MVP_top60_DNA+RNA_seed | HSP90AB1 | -2.199261 | STAD | FU97 | Tanespimycin |
| MVP_top60_DNA_seed | EGFR | 0.540355 | LUAD | LXF-289 | Gefitinib | MVP_top60_DNA+RNA_seed | ERBB2 | -2.197683 | STAD | TGBC11TKB | CP724714 |
| MVP_top60_DNA_seed | CREBBP | 0.540783 | LUAD | NCI-H1781 | I-CBP112 | MVP_top60_DNA+RNA_seed | FGFR3 | -2.175788 | STAD | FU97 | AZD4547 |
| MVP_top60_DNA_seed | HDAC1 | 0.541238 | LUAD | NCI-H838 | JQ12 | MVP_top60_DNA+RNA_seed | PARP1 | -2.154446 | STAD | HGC-27 | Olaparib |
| MVP_top60_DNA_seed | BRAF | 0.542069 | LUAD | NCI-H2009 | Dabrafenib | MVP_top60_DNA+RNA_seed | PARP1 | -2.139724 | STAD | HGC-27 | PARP_9482 |
| MVP_top60_DNA_seed | EGFR | 0.542277 | LUAD | SW1573 | CUDC-101 | MVP_top60_DNA+RNA_seed | ERBB4 | -2.09995 | STAD | AGS | AST-1306 |
| MVP_top60_DNA_seed | ERBB2 | 0.542277 | LUAD | SW1573 | CUDC-101 | MVP_top60_DNA+RNA_seed | ERBB2 | -2.084813 | STAD | AGS | PF-00299804 |
| MVP_top60_DNA_seed | PIK3CA | 0.543329 | LUAD | HOP-62 | AZD8835 | MVP_top60_DNA+RNA_seed | ERBB4 | -2.084813 | STAD | AGS | PF-00299804 |
| MVP_top60_DNA_seed | IGF1R | 0.544397 | LUAD | NCI-H1568 | BMS-536924 | MVP_top60_DNA+RNA_seed | MET | -2.077816 | STAD | HSC-39 | SU11274 |
| MVP_top60_DNA_seed | MET | 0.544934 | LUAD | PC-3_[JPC-3] | JNJ38877605 | MVP_top60_DNA+RNA_seed | EP300 | -2.053495 | STAD | NUGC-4 | I-CBP112 |
| MVP_top60_DNA_seed | HDAC1 | 0.545374 | LUAD | NCI-H1734 | Tubastatin A | MVP_top60_DNA+RNA_seed | CREBBP | -2.053495 | STAD | NUGC-4 | I-CBP112 |
| MVP_top60_DNA_seed | SRC | 0.545766 | LUAD | NCI-H1838 | Bosutinib | MVP_top60_DNA+RNA_seed | MET | -2.033998 | STAD | SNU-5 | SU11274 |
| MVP_top60_DNA_seed | GSK3B | 0.546235 | LUAD | PC-3_[JPC-3] | AZD7969 | MVP_top60_DNA+RNA_seed | PIK3CA | -2.011361 | STAD | AGS | AZD8835 |
| MVP_top60_DNA_seed | ERBB2 | 0.546398 | LUAD | NCI-H1651 | CP724714 | MVP_top60_DNA+RNA_seed | ERBB2 | -1.999928 | STAD | NUGC-3 | Afatinib |
| MVP_top60_DNA_seed | MAPK1 | 0.546673 | LUAD | SK-LU-1 | VX-11e | MVP_top60_DNA+RNA_seed | PARP1 | -1.998801 | STAD | HGC-27 | Olaparib |
| MVP_top60_DNA_seed | FGFR1 | 0.546828 | LUAD | NCI-H1563 | FGFR_3831 | MVP_top60_DNA+RNA_seed | MET | -1.973811 | STAD | KATOIII | SU11274 |
| MVP_top60_DNA_seed | MAPK1 | 0.549055 | LUAD | NCI-H650 | VX-11e | MVP_top60_DNA+RNA_seed | PARP1 | -1.932475 | STAD | HGC-27 | PARP_0108 |
| MVP_top60_DNA_seed | MET | 0.549649 | LUAD | NCI-H358 | SU11274 | MVP_top60_DNA+RNA_seed | HSP90AA1 | -1.922422 | STAD | HGC-27 | Tanespimycin |
| MVP_top60_DNA_seed | EGFR | 0.549665 | LUAD | NCI-H2009 | CI-1033 | MVP_top60_DNA+RNA_seed | HSP90AB1 | -1.922422 | STAD | HGC-27 | Tanespimycin |
| MVP_top60_DNA_seed | ERBB2 | 0.549665 | LUAD | NCI-H2009 | CI-1033 | MVP_top60_DNA+RNA_seed | ERBB2 | -1.907664 | STAD | GCIY | CI-1033 |
| MVP_top60_DNA_seed | MET | 0.549937 | LUAD | NCI-H1975 | SU11274 | MVP_top60_DNA+RNA_seed | ERBB4 | -1.907664 | STAD | GCIY | CI-1033 |
| MVP_top60_DNA_seed | BRAF | 0.549939 | LUAD | NCI-H1651 | PLX-4720 | MVP_top60_DNA+RNA_seed | ERBB2 | -1.847543 | STAD | NCI-N87 | AZD8931 |
| MVP_top60_DNA_seed | BRAF | 0.550298 | LUAD | NCI-H1437 | PLX-4720 | MVP_top60_DNA+RNA_seed | ERBB3 | -1.847543 | STAD | NCI-N87 | AZD8931 |
| MVP_top60_DNA_seed | ATM | 0.55175 | LUAD | NCI-H441 | KU-60019 | MVP_top60_DNA+RNA_seed | EP300 | -1.838052 | STAD | HGC-27 | I-CBP112 |
| MVP_top60_DNA_seed | BRAF | 0.552013 | LUAD | ABC-1 | PLX-4720 | MVP_top60_DNA+RNA_seed | CREBBP | -1.838052 | STAD | HGC-27 | I-CBP112 |
| MVP_top60_DNA_seed | MET | 0.553099 | LUAD | NCI-H3122 | AZD6094 | MVP_top60_DNA+RNA_seed | HSP90AA1 | -1.803018 | STAD | AGS | Tanespimycin |
| MVP_top60_DNA_seed | HSP90AA1 | 0.553419 | LUAD | PC-3_[JPC-3] | SNX-2112 | MVP_top60_DNA+RNA_seed | HSP90AB1 | -1.803018 | STAD | AGS | Tanespimycin |
| MVP_top60_DNA_seed | CREBBP | 0.554584 | LUAD | NCI-H2023 | I-CBP112 | MVP_top60_DNA+RNA_seed | MET | -1.764713 | STAD | HSC-39 | JNJ38877605 |
| MVP_top60_DNA_seed | HDAC1 | 0.555278 | LUAD | A549 | Belinostat | MVP_top60_DNA+RNA_seed | HSP90AA1 | -1.764312 | STAD | OCUM-1 | CCT-018159 |
| MVP_top60_DNA_seed | AURKA | 0.555386 | LUAD | NCI-H2228 | Alisertib | MVP_top60_DNA+RNA_seed | HSP90AB1 | -1.764312 | STAD | OCUM-1 | CCT-018159 |
| MVP_top60_DNA_seed | EGFR | 0.555677 | LUAD | NCI-H2009 | PF-00299804 | MVP_top60_DNA+RNA_seed | ESR1 | -1.75327 | STAD | AGS | Fulvestrant |
| MVP_top60_DNA_seed | ERBB2 | 0.555677 | LUAD | NCI-H2009 | PF-00299804 | MVP_top60_DNA+RNA_seed | ATM | -1.747041 | STAD | HSC-39 | KU-55933 |
| MVP_top60_DNA_seed | IGF1R | 0.556509 | LUAD | NCI-H358 | GSK1904529A | MVP_top60_DNA+RNA_seed | ERBB2 | -1.746646 | STAD | NUGC-3 | Afatinib |
| MVP_top60_DNA_seed | EGFR | 0.556613 | LUAD | NCI-H1666 | Pelitinib | MVP_top60_DNA+RNA_seed | ERBB2 | -1.727366 | STAD | GCIY | PF-00299804 |
| MVP_top60_DNA_seed | HDAC1 | 0.556762 | LUAD | NCI-H1792 | Tubastatin A | MVP_top60_DNA+RNA_seed | ERBB4 | -1.727366 | STAD | GCIY | PF-00299804 |
| MVP_top60_DNA_seed | AKT1 | 0.556884 | LUAD | NCI-H650 | AKT inhibitor VIII | MVP_top60_DNA+RNA_seed | HSP90AA1 | -1.726927 | STAD | RERF-GC-1B | Elesclomol |
| MVP_top60_DNA_seed | SRC | 0.557319 | LUAD | VMRC-LCD | Bosutinib | MVP_top60_DNA+RNA_seed | HSP90AB1 | -1.726927 | STAD | RERF-GC-1B | Elesclomol |
| MVP_top60_DNA_seed | PTK2 | 0.558003 | LUAD | NCI-H1944 | PF-562271 | MVP_top60_DNA+RNA_seed | HDAC1 | -1.712181 | STAD | FU97 | JQ12 |

| | | | | | | | | | | | |
|---|---|---|---|---|---|---|---|---|---|---|---|
| MVP_top60_DNA_seed | BRAF | 0.55814 | LUAD | NCI-H322M | AZ628 | MVP_top60_DNA+RNA_seed | MDM2 | -1.6687 | STAD | NUGC-4 | Serdemetan |
| MVP_top60_DNA_seed | IGF1R | 0.558765 | LUAD | Calu-3 | BMS-536924 | MVP_top60_DNA+RNA_seed | MET | -1.66427 | STAD | GCIY | JNJ38877605 |
| MVP_top60_DNA_seed | AKT1 | 0.558951 | LUAD | H3255 | GSK690693 | MVP_top60_DNA+RNA_seed | MDM2 | -1.636256 | STAD | AGS | Nutlin-3a (-) |
| MVP_top60_DNA_seed | AURKA | 0.560652 | LUAD | NCI-H1792 | ZM447439 | MVP_top60_DNA+RNA_seed | ERBB2 | -1.633241 | STAD | IM-95 | AZD8931 |
| MVP_top60_DNA_seed | HDAC1 | 0.562111 | LUAD | ABC-1 | Tubastatin A | MVP_top60_DNA+RNA_seed | ERBB3 | -1.633241 | STAD | IM-95 | AZD8931 |
| MVP_top60_DNA_seed | BRAF | 0.56221 | LUAD | NCI-H1650 | RAF_9304 | MVP_top60_DNA+RNA_seed | HDAC1 | -1.630624 | STAD | GCIY | Parthenolide |
| MVP_top60_DNA_seed | HSP90AA1 | 0.562326 | LUAD | NCI-H1568 | Elesclomol | MVP_top60_DNA+RNA_seed | FGFR3 | -1.629842 | STAD | RERF-GC-1B | PD173074 |
| MVP_top60_DNA_seed | EGFR | 0.562436 | LUAD | NCI-H23 | Erlotinib | MVP_top60_DNA+RNA_seed | PARP1 | -1.623571 | STAD | NUGC-3 | PARP_0108 |
| MVP_top60_DNA_seed | AKT1 | 0.562992 | LUAD | SK-LU-1 | MK-2206 | MVP_top60_DNA+RNA_seed | MDM2 | -1.621978 | STAD | OCUM-1 | Nutlin-3a (-) |
| MVP_top60_DNA_seed | CREBBP | 0.564273 | LUAD | HCC-827 | I-CBP112 | MVP_top60_DNA+RNA_seed | ESR1 | -1.611139 | STAD | HSC-39 | Fulvestrant |
| MVP_top60_DNA_seed | PIK3CA | 0.564716 | LUAD | NCI-H1650 | AZD8835 | MVP_top60_DNA+RNA_seed | PIK3CA | -1.601479 | STAD | AGS | PI3Ka_4409 |
| MVP_top60_DNA_seed | HSP90AA1 | 0.564978 | LUAD | NCI-H1793 | Elesclomol | MVP_top60_DNA+RNA_seed | SRC | -1.600442 | STAD | NUGC-3 | Bosutinib |
| MVP_top60_DNA_seed | HDAC1 | 0.565586 | LUAD | SW1573 | CAY10603 | MVP_top60_DNA+RNA_seed | PARP1 | -1.583356 | STAD | NUGC-3 | Talazoparib |
| MVP_top60_DNA_seed | AURKA | 0.568205 | LUAD | NCI-H2085 | Alisertib | MVP_top60_DNA+RNA_seed | PARP1 | -1.578801 | STAD | NUGC-3 | PARP_9482 |
| MVP_top60_DNA_seed | GSK3B | 0.570087 | LUAD | NCI-H2405 | KIN001-042 | MVP_top60_DNA+RNA_seed | FGFR3 | -1.561759 | STAD | SNU-16 | AZD4547 |
| MVP_top60_DNA_seed | HSP90AA1 | 0.570675 | LUAD | HCC-827 | Tanespimycin | MVP_top60_DNA+RNA_seed | MAPK1 | -1.544964 | STAD | RERF-GC-1B | FR-180204 |
| MVP_top60_DNA_seed | FGFR1 | 0.57072 | LUAD | NCI-H23 | PD173074 | MVP_top60_DNA+RNA_seed | MAPK3 | -1.544964 | STAD | RERF-GC-1B | FR-180204 |
| MVP_top60_DNA_seed | MAPK1 | 0.571521 | LUAD | NCI-H1734 | FR-180204 | MVP_top60_DNA+RNA_seed | HDAC1 | -1.544682 | STAD | RF-48 | CAY10603 |
| MVP_top60_DNA_seed | IGF1R | 0.572636 | LUAD | NCI-H1781 | BMS-536924 | MVP_top60_DNA+RNA_seed | ESR1 | -1.540627 | STAD | SNU-16 | Tamoxifen |
| MVP_top60_DNA_seed | AKT1 | 0.573545 | LUAD | NCI-H1666 | GSK690693 | MVP_top60_DNA+RNA_seed | ERBB2 | -1.527312 | STAD | NUGC-4 | CI-1033 |
| MVP_top60_DNA_seed | FGFR1 | 0.573648 | LUAD | NCI-H1437 | PD173074 | MVP_top60_DNA+RNA_seed | ERBB4 | -1.527312 | STAD | NUGC-4 | CI-1033 |
| MVP_top60_DNA_seed | GSK3B | 0.573828 | LUAD | NCI-H1944 | KIN001-042 | MVP_top60_DNA+RNA_seed | ERBB4 | -1.522995 | STAD | GCIY | AST-1306 |
| MVP_top60_DNA_seed | SMARCA4 | 0.57405 | LUAD | SK-LU-1 | PFI-3 | MVP_top60_DNA+RNA_seed | PTK2 | -1.522818 | STAD | AGS | PF-562271 |
| MVP_top60_DNA_seed | ATM | 0.574413 | LUAD | NCI-H1573 | CP466722 | MVP_top60_DNA+RNA_seed | PARP1 | -1.522569 | STAD | NUGC-3 | Veliparib |
| MVP_top60_DNA_seed | SMARCA4 | 0.574504 | LUAD | NCI-H2087 | PFI-3 | MVP_top60_DNA+RNA_seed | MET | -1.5209 | STAD | SNU-1 | SU11274 |
| MVP_top60_DNA_seed | EGFR | 0.574742 | LUAD | VMRC-LCD | Afatinib | MVP_top60_DNA+RNA_seed | HSP90AA1 | -1.514956 | STAD | NCI-N87 | Tanespimycin |
| MVP_top60_DNA_seed | ERBB2 | 0.574742 | LUAD | VMRC-LCD | Afatinib | MVP_top60_DNA+RNA_seed | HSP90AB1 | -1.514956 | STAD | NCI-N87 | Tanespimycin |
| MVP_top60_DNA_seed | HSP90AA1 | 0.576273 | LUAD | NCI-H23 | Luminespib | MVP_top60_DNA+RNA_seed | MET | -1.514015 | STAD | TGBC11TKB | JNJ38877605 |
| MVP_top60_DNA_seed | MAPK1 | 0.578478 | LUAD | HOP-62 | FR-180204 | MVP_top60_DNA+RNA_seed | GSK3B | -1.509191 | STAD | TMK-1 | CHIR-99021 |
| MVP_top60_DNA_seed | MAPK1 | 0.57889 | LUAD | NCI-H1650 | FR-180204 | MVP_top60_DNA+RNA_seed | FGFR3 | -1.501861 | STAD | FU97 | FGFR_3831 |
| MVP_top60_DNA_seed | MDM2 | 0.58081 | LUAD | NCI-H3122 | Nutlin-3a (-) | MVP_top60_DNA+RNA_seed | PARP1 | -1.499144 | STAD | NUGC-4 | PARP_9495 |
| MVP_top60_DNA_seed | PIK3CA | 0.580849 | LUAD | SK-LU-1 | PI3Ka_4409 | MVP_top60_DNA+RNA_seed | PIK3CA | -1.49039 | STAD | HSC-39 | PI3Ka_4409 |
| MVP_top60_DNA_seed | ERBB2 | 0.581608 | LUAD | Calu-6 | CP724714 | MVP_top60_DNA+RNA_seed | HDAC1 | -1.469975 | STAD | RF-48 | AR-42 |
| MVP_top60_DNA_seed | AKT1 | 0.583114 | LUAD | A549 | AKT inhibitor VIII | MVP_top60_DNA+RNA_seed | PTK2 | -1.468287 | STAD | HGC-27 | PF-562271 |
| MVP_top60_DNA_seed | BRAF | 0.58334 | LUAD | NCI-H1944 | PLX-4720 | MVP_top60_DNA+RNA_seed | GSK3B | -1.467752 | STAD | HSC-39 | SB216763 |
| MVP_top60_DNA_seed | MET | 0.583893 | LUAD | NCI-H23 | Crizotinib | MVP_top60_DNA+RNA_seed | GSK3B | -1.466871 | STAD | SNU-16 | SB216763 |
| MVP_top60_DNA_seed | AURKA | 0.585151 | LUAD | NCI-H1975 | Alisertib | MVP_top60_DNA+RNA_seed | ERBB2 | -1.456809 | STAD | HSC-39 | CI-1033 |
| MVP_top60_DNA_seed | MAPK1 | 0.585561 | LUAD | NCI-H1666 | FR-180204 | MVP_top60_DNA+RNA_seed | ERBB4 | -1.456809 | STAD | HSC-39 | CI-1033 |
| MVP_top60_DNA_seed | HSP90AA1 | 0.585585 | LUAD | Calu-3 | SNX-2112 | MVP_top60_DNA+RNA_seed | MDM2 | -1.45385 | STAD | RF-48 | Nutlin-3a (-) |
| MVP_top60_DNA_seed | CREBBP | 0.58572 | LUAD | NCI-H1755 | I-CBP112 | MVP_top60_DNA+RNA_seed | SRC | -1.449113 | STAD | SNU-16 | WH-4-023 |
| MVP_top60_DNA_seed | AURKA | 0.585869 | LUAD | NCI-H1563 | CD532 | MVP_top60_DNA+RNA_seed | SRC | -1.445176 | STAD | SNU-1 | NVP-BHG712 |

| | | | | | | | | | | | |
|---|---|---|---|---|---|---|---|---|---|---|---|
| MVP_top60_DNA_seed | BRAF | 0.58669 | LUAD | NCI-H1693 | SB590885 | MVP_top60_DNA+RNA_seed | HDAC1 | -1.441638 | STAD | AGS | JQ12 |
| MVP_top60_DNA_seed | FGFR1 | 0.586896 | LUAD | SK-LU-1 | FGFR_3831 | MVP_top60_DNA+RNA_seed | MAPK1 | -1.420657 | STAD | AGS | VX-11e |
| MVP_top60_DNA_seed | GSK3B | 0.587478 | LUAD | NCI-H2087 | CHIR-99021 | MVP_top60_DNA+RNA_seed | PARP1 | -1.413553 | STAD | HGC-27 | Veliparib |
| MVP_top60_DNA_seed | EGFR | 0.588599 | LUAD | NCI-H2405 | Afatinib | MVP_top60_DNA+RNA_seed | PARP1 | -1.386568 | STAD | SNU-5 | Veliparib |
| MVP_top60_DNA_seed | ERBB2 | 0.588599 | LUAD | NCI-H2405 | Afatinib | MVP_top60_DNA+RNA_seed | MET | -1.386512 | STAD | TMK-1 | AZD6094 |
| MVP_top60_DNA_seed | HDAC1 | 0.588605 | LUAD | NCI-H2405 | Belinostat | MVP_top60_DNA+RNA_seed | ESR1 | -1.383591 | STAD | 23132-87 | Fulvestrant |
| MVP_top60_DNA_seed | AURKA | 0.589176 | LUAD | HCC-44 | Alisertib | MVP_top60_DNA+RNA_seed | GSK3B | -1.380909 | STAD | MKN28 | CHIR-99021 |
| MVP_top60_DNA_seed | IGF1R | 0.589179 | LUAD | PC-3_[JPC-3] | GSK1904529A | MVP_top60_DNA+RNA_seed | HSP90AA1 | -1.346759 | STAD | AGS | CCT-018159 |
| MVP_top60_DNA_seed | BRAF | 0.589493 | LUAD | NCI-H2087 | SB590885 | MVP_top60_DNA+RNA_seed | HSP90AB1 | -1.346759 | STAD | AGS | CCT-018159 |
| MVP_top60_DNA_seed | SMARCA4 | 0.589571 | LUAD | NCI-H1755 | PFI-3 | MVP_top60_DNA+RNA_seed | HDAC1 | -1.346597 | STAD | RF-48 | Parthenolide |
| MVP_top60_DNA_seed | GSK3B | 0.589954 | LUAD | PC-14 | SB216763 | MVP_top60_DNA+RNA_seed | HDAC1 | -1.34113 | STAD | GCIY | JQ12 |
| MVP_top60_DNA_seed | ATM | 0.592273 | LUAD | NCI-H1623 | KU-60019 | MVP_top60_DNA+RNA_seed | HDAC1 | -1.336682 | STAD | HSC-39 | Tubastatin A |
| MVP_top60_DNA_seed | HDAC1 | 0.592667 | LUAD | A549 | Dacinostat | MVP_top60_DNA+RNA_seed | SRC | -1.317775 | STAD | HSC-39 | Bosutinib |
| MVP_top60_DNA_seed | FGFR1 | 0.593065 | LUAD | COR-L105 | FGFR_3831 | MVP_top60_DNA+RNA_seed | HDAC1 | -1.313699 | STAD | RF-48 | Belinostat |
| MVP_top60_DNA_seed | HSP90AA1 | 0.593092 | LUAD | NCI-H1573 | SNX-2112 | MVP_top60_DNA+RNA_seed | HSP90AA1 | -1.312553 | STAD | TMK-1 | Tanespimycin |
| MVP_top60_DNA_seed | AURKA | 0.593119 | LUAD | NCI-H522 | Alisertib | MVP_top60_DNA+RNA_seed | HSP90AB1 | -1.312553 | STAD | TMK-1 | Tanespimycin |
| MVP_top60_DNA_seed | EGFR | 0.593312 | LUAD | NCI-H441 | Gefitinib | MVP_top60_DNA+RNA_seed | ERBB2 | -1.309811 | STAD | RF-48 | CUDC-101 |
| MVP_top60_DNA_seed | GSK3B | 0.595117 | LUAD | NCI-H522 | CHIR-99021 | MVP_top60_DNA+RNA_seed | HSP90AA1 | -1.308562 | STAD | HGC-27 | CCT-018159 |
| MVP_top60_DNA_seed | MET | 0.596322 | LUAD | NCI-H2009 | JNJ38877605 | MVP_top60_DNA+RNA_seed | HSP90AB1 | -1.308562 | STAD | HGC-27 | CCT-018159 |
| MVP_top60_DNA_seed | MDM2 | 0.5969 | LUAD | NCI-H2030 | Serdemetan | MVP_top60_DNA+RNA_seed | GSK3B | -1.296339 | STAD | TMK-1 | CHIR-99021 |
| MVP_top60_DNA_seed | EGFR | 0.596901 | LUAD | NCI-H3122 | CI-1033 | MVP_top60_DNA+RNA_seed | ERBB2 | -1.29334 | STAD | AGS | CI-1033 |
| MVP_top60_DNA_seed | ERBB2 | 0.596901 | LUAD | NCI-H3122 | CI-1033 | MVP_top60_DNA+RNA_seed | ERBB4 | -1.29334 | STAD | AGS | CI-1033 |
| MVP_top60_DNA_seed | FGFR1 | 0.596924 | LUAD | NCI-H1781 | AZD4547 | MVP_top60_DNA+RNA_seed | HDAC1 | -1.292905 | STAD | RF-48 | JQ12 |
| MVP_top60_DNA_seed | MET | 0.597248 | LUAD | A549 | JNJ38877605 | MVP_top60_DNA+RNA_seed | MET | -1.292054 | STAD | TGBC11TKB | SU11274 |
| MVP_top60_DNA_seed | MAPK1 | 0.598201 | LUAD | NCI-H23 | FR-180204 | MVP_top60_DNA+RNA_seed | MAPK1 | -1.290466 | STAD | SNU-1 | VX-11e |
| MVP_top60_DNA_seed | HSP90AA1 | 0.598354 | LUAD | NCI-H1781 | CCT-018159 | MVP_top60_DNA+RNA_seed | MDM2 | -1.284636 | STAD | AGS | Serdemetan |
| MVP_top60_DNA_seed | BRAF | 0.598898 | LUAD | NCI-H1838 | AZ628 | MVP_top60_DNA+RNA_seed | ERBB4 | -1.278249 | STAD | SNU-1 | AST-1306 |
| MVP_top60_DNA_seed | AURKA | 0.59919 | LUAD | NCI-H2342 | Alisertib | MVP_top60_DNA+RNA_seed | HSP90AA1 | -1.274585 | STAD | HSC-39 | Tanespimycin |
| MVP_top60_DNA_seed | FGFR1 | 0.599343 | LUAD | RERF-LC-KJ | AZD4547 | MVP_top60_DNA+RNA_seed | HSP90AB1 | -1.274585 | STAD | HSC-39 | Tanespimycin |
| MVP_top60_DNA_seed | EGFR | 0.599369 | LUAD | NCI-H1838 | PF-00299804 | MVP_top60_DNA+RNA_seed | AR | -1.264694 | STAD | HGC-27 | Bicalutamide |
| MVP_top60_DNA_seed | ERBB2 | 0.599369 | LUAD | NCI-H1838 | PF-00299804 | MVP_top60_DNA+RNA_seed | PARP1 | -1.25944 | STAD | HGC-27 | PARP_9495 |
| MVP_top60_DNA_seed | PIK3CA | 0.599552 | LUAD | NCI-H2405 | AZD8835 | MVP_top60_DNA+RNA_seed | ERBB2 | -1.248274 | STAD | GCIY | Afatinib |
| MVP_top60_DNA_seed | CREBBP | 0.59975 | LUAD | NCI-H1651 | I-CBP112 | MVP_top60_DNA+RNA_seed | PARP1 | -1.242135 | STAD | AGS | PARP_9495 |
| MVP_top60_DNA_seed | MET | 0.599864 | LUAD | NCI-H1666 | JNJ38877605 | MVP_top60_DNA+RNA_seed | PTK2 | -1.240871 | STAD | IM-95 | PF-562271 |
| MVP_top60_DNA_seed | GSK3B | 0.600729 | LUAD | NCI-H3122 | KIN001-042 | MVP_top60_DNA+RNA_seed | SRC | -1.236981 | STAD | SK-GT-2 | Bosutinib |
| MVP_top60_DNA_seed | EGFR | 0.601051 | LUAD | NCI-H1838 | Cetuximab | MVP_top60_DNA+RNA_seed | HDAC1 | -1.229597 | STAD | IM-95 | Dacinostat |
| MVP_top60_DNA_seed | FGFR1 | 0.601507 | LUAD | NCI-H1435 | AZD4547 | MVP_top60_DNA+RNA_seed | MDM2 | -1.216493 | STAD | HSC-39 | Serdemetan |
| MVP_top60_DNA_seed | EGFR | 0.602371 | LUAD | NCI-H1435 | Afatinib | MVP_top60_DNA+RNA_seed | AR | -1.216424 | STAD | FU97 | AZD3514 |
| MVP_top60_DNA_seed | ERBB2 | 0.602371 | LUAD | NCI-H1435 | Afatinib | MVP_top60_DNA+RNA_seed | PARP1 | -1.193517 | STAD | SNU-16 | Olaparib |
| MVP_top60_DNA_seed | GSK3B | 0.602562 | LUAD | NCI-H1666 | KIN001-042 | MVP_top60_DNA+RNA_seed | SRC | -1.192309 | STAD | GCIY | Saracatinib |
| MVP_top60_DNA_seed | EGFR | 0.602643 | LUAD | NCI-H2228 | AZD8931 | MVP_top60_DNA+RNA_seed | PARP1 | -1.191289 | STAD | NUGC-3 | Rucaparib |

| | | | | | | | | | | | |
|---|---|---|---|---|---|---|---|---|---|---|---|
| MVP_top60_DNA_seed | ERBB2 | 0.602643 | LUAD | NCI-H2228 | AZD8931 | MVP_top60_DNA+RNA_seed | HDAC1 | -1.190515 | STAD | RF-48 | Dacinostat |
| MVP_top60_DNA_seed | ERBB3 | 0.602643 | LUAD | NCI-H2228 | AZD8931 | MVP_top60_DNA+RNA_seed | ERBB2 | -1.180036 | STAD | NUGC-4 | CUDC-101 |
| MVP_top60_DNA_seed | BRAF | 0.603421 | LUAD | NCI-H1944 | SB590885 | MVP_top60_DNA+RNA_seed | SRC | -1.179583 | STAD | NUGC-4 | NVP-BHG712 |
| MVP_top60_DNA_seed | BRAF | 0.603428 | LUAD | NCI-H2122 | HG6-64-1 | MVP_top60_DNA+RNA_seed | PARP1 | -1.162049 | STAD | NCI-N87 | PARP_9495 |
| MVP_top60_DNA_seed | BRAF | 0.603539 | LUAD | RERF-LC-KJ | SB590885 | MVP_top60_DNA+RNA_seed | FGFR3 | -1.161364 | STAD | FU97 | AZD4547 |
| MVP_top60_DNA_seed | CREBBP | 0.603861 | LUAD | SW1573 | I-CBP112 | MVP_top60_DNA+RNA_seed | HSP90AA1 | -1.16034 | STAD | MKN45 | Tanespimycin |
| MVP_top60_DNA_seed | AURKA | 0.604079 | LUAD | NCI-H358 | Alisertib | MVP_top60_DNA+RNA_seed | HSP90AB1 | -1.16034 | STAD | MKN45 | Tanespimycin |
| MVP_top60_DNA_seed | IGF1R | 0.604555 | LUAD | NCI-H1355 | GSK1904529A | MVP_top60_DNA+RNA_seed | HSP90AA1 | -1.158613 | STAD | SNU-1 | SNX-2112 |
| MVP_top60_DNA_seed | MDM2 | 0.605243 | LUAD | NCI-H322M | Nutlin-3a (-) | MVP_top60_DNA+RNA_seed | HSP90AB1 | -1.158613 | STAD | SNU-1 | SNX-2112 |
| MVP_top60_DNA_seed | AKT1 | 0.605685 | LUAD | H3255 | MK-2206 | MVP_top60_DNA+RNA_seed | HDAC1 | -1.151827 | STAD | HGC-27 | Dacinostat |
| MVP_top60_DNA_seed | HSP90AA1 | 0.606102 | LUAD | NCI-H1755 | Luminespib | MVP_top60_DNA+RNA_seed | PARP1 | -1.151578 | STAD | RF-48 | Olaparib |
| MVP_top60_DNA_seed | AURKA | 0.60615 | LUAD | NCI-H1781 | GSK1070916 | MVP_top60_DNA+RNA_seed | PARP1 | -1.150262 | STAD | NUGC-3 | Olaparib |
| MVP_top60_DNA_seed | GSK3B | 0.606723 | LUAD | H3255 | AZD7969 | MVP_top60_DNA+RNA_seed | ATM | -1.147537 | STAD | TMK-1 | KU-60019 |
| MVP_top60_DNA_seed | AKT1 | 0.606799 | LUAD | SW1573 | MK-2206 | MVP_top60_DNA+RNA_seed | HSP90AA1 | -1.146672 | STAD | RF-48 | SNX-2112 |
| MVP_top60_DNA_seed | IGF1R | 0.606968 | LUAD | NCI-H1650 | BMS-754807 | MVP_top60_DNA+RNA_seed | HSP90AB1 | -1.146672 | STAD | RF-48 | SNX-2112 |
| MVP_top60_DNA_seed | AKT1 | 0.607677 | LUAD | NCI-H1755 | MK-2206 | MVP_top60_DNA+RNA_seed | MDM2 | -1.143196 | STAD | NUGC-3 | Serdemetan |
| MVP_top60_DNA_seed | BRAF | 0.60836 | LUAD | NCI-H2023 | SB590885 | MVP_top60_DNA+RNA_seed | PARP1 | -1.13811 | STAD | TGBC11TKB | Veliparib |
| MVP_top60_DNA_seed | HDAC1 | 0.60901 | LUAD | NCI-H1623 | Tubastatin A | MVP_top60_DNA+RNA_seed | PARP1 | -1.134671 | STAD | SNU-1 | Olaparib |
| MVP_top60_DNA_seed | HDAC1 | 0.609357 | LUAD | A549 | CAY10603 | MVP_top60_DNA+RNA_seed | ATM | -1.121906 | STAD | GCIY | KU-55933 |
| MVP_top60_DNA_seed | HDAC1 | 0.610381 | LUAD | NCI-H23 | Entinostat | MVP_top60_DNA+RNA_seed | HSP90AA1 | -1.116909 | STAD | NUGC-3 | Elesclomol |
| MVP_top60_DNA_seed | HSP90AA1 | 0.611133 | LUAD | NCI-H1792 | Elesclomol | MVP_top60_DNA+RNA_seed | HSP90AB1 | -1.116909 | STAD | NUGC-3 | Elesclomol |
| MVP_top60_DNA_seed | SMARCA4 | 0.611424 | LUAD | NCI-H2342 | PFI-3 | MVP_top60_DNA+RNA_seed | SRC | -1.111899 | STAD | KATOIII | HG-6-71-01 |
| MVP_top60_DNA_seed | GSK3B | 0.61177 | LUAD | NCI-H1573 | CHIR-99021 | MVP_top60_DNA+RNA_seed | PARP1 | -1.110769 | STAD | KATOIII | Rucaparib |
| MVP_top60_DNA_seed | AURKA | 0.61213 | LUAD | NCI-H1355 | Alisertib | MVP_top60_DNA+RNA_seed | MDM2 | -1.105876 | STAD | IM-95 | Nutlin-3a (-) |
| MVP_top60_DNA_seed | MET | 0.61219 | LUAD | NCI-H2342 | AZD6094 | MVP_top60_DNA+RNA_seed | ERBB2 | -1.09391 | STAD | NCI-N87 | CUDC-101 |
| MVP_top60_DNA_seed | MET | 0.612939 | LUAD | NCI-H2347 | SU11274 | MVP_top60_DNA+RNA_seed | ATM | -1.092186 | STAD | NUGC-3 | KU-55933 |
| MVP_top60_DNA_seed | EGFR | 0.613348 | LUAD | NCI-H2122 | Cetuximab | MVP_top60_DNA+RNA_seed | ERBB2 | -1.08654 | STAD | SNU-1 | PF-00299804 |
| MVP_top60_DNA_seed | ERBB2 | 0.61354 | LUAD | NCI-H1734 | CP724714 | MVP_top60_DNA+RNA_seed | ERBB4 | -1.08654 | STAD | SNU-1 | PF-00299804 |
| MVP_top60_DNA_seed | HDAC1 | 0.613763 | LUAD | A549 | AR-42 | MVP_top60_DNA+RNA_seed | SRC | -1.086301 | STAD | GCIY | WH-4-023 |
| MVP_top60_DNA_seed | EGFR | 0.614013 | LUAD | NCI-H2405 | PF-00299804 | MVP_top60_DNA+RNA_seed | HDAC1 | -1.081741 | STAD | Hs-746T | AR-42 |
| MVP_top60_DNA_seed | ERBB2 | 0.614013 | LUAD | NCI-H2405 | PF-00299804 | MVP_top60_DNA+RNA_seed | HSP90AA1 | -1.080852 | STAD | HGC-27 | Elesclomol |
| MVP_top60_DNA_seed | IGF1R | 0.614311 | LUAD | Calu-6 | GSK1904529A | MVP_top60_DNA+RNA_seed | HSP90AB1 | -1.080852 | STAD | HGC-27 | Elesclomol |
| MVP_top60_DNA_seed | GSK3B | 0.614894 | LUAD | NCI-H2122 | KIN001-042 | MVP_top60_DNA+RNA_seed | PTK2 | -1.08012 | STAD | GCIY | PF-562271 |
| MVP_top60_DNA_seed | ERBB2 | 0.618263 | LUAD | SK-LU-1 | CP724714 | MVP_top60_DNA+RNA_seed | ERBB4 | -1.076841 | STAD | HSC-39 | AST-1306 |
| MVP_top60_DNA_seed | FGFR1 | 0.618726 | LUAD | NCI-H2087 | AZD4547 | MVP_top60_DNA+RNA_seed | SRC | -1.076297 | STAD | HSC-39 | WH-4-023 |
| MVP_top60_DNA_seed | BRAF | 0.619248 | LUAD | NCI-H838 | PLX-4720 | MVP_top60_DNA+RNA_seed | ERBB4 | -1.076218 | STAD | NUGC-4 | AST-1306 |
| MVP_top60_DNA_seed | SRC | 0.6218 | LUAD | NCI-H2122 | NVP-BHG712 | MVP_top60_DNA+RNA_seed | HDAC1 | -1.072515 | STAD | Hs-746T | Tubastatin A |
| MVP_top60_DNA_seed | ATM | 0.622181 | LUAD | NCI-H838 | CP466722 | MVP_top60_DNA+RNA_seed | SRC | -1.069815 | STAD | RF-48 | NVP-BHG712 |
| MVP_top60_DNA_seed | GSK3B | 0.622347 | LUAD | ABC-1 | AZD7969 | MVP_top60_DNA+RNA_seed | MET | -1.066652 | STAD | NUGC-4 | SU11274 |
| MVP_top60_DNA_seed | EGFR | 0.622511 | LUAD | NCI-H1755 | AST-1306 | MVP_top60_DNA+RNA_seed | ESR1 | -1.064554 | STAD | HSC-39 | Tamoxifen |
| MVP_top60_DNA_seed | HDAC1 | 0.622564 | LUAD | NCI-H1563 | CAY10603 | MVP_top60_DNA+RNA_seed | AR | -1.062613 | STAD | TGBC11TKB | Bicalutamide |

| | | | | | | | | | | | |
|---|---|---|---|---|---|---|---|---|---|---|---|
| MVP_top60_DNA_seed | EGFR | 0.623784 | LUAD | NCI-H2009 | Gefitinib | MVP_top60_DNA+RNA_seed | GSK3B | -1.061191 | STAD | NUGC-4 | KIN001-042 |
| MVP_top60_DNA_seed | FGFR1 | 0.624866 | LUAD | NCI-H441 | AZD4547 | MVP_top60_DNA+RNA_seed | HSP90AA1 | -1.057396 | STAD | SNU-5 | Tanespimycin |
| MVP_top60_DNA_seed | AURKA | 0.626357 | LUAD | NCI-H1944 | ZM447439 | MVP_top60_DNA+RNA_seed | HSP90AB1 | -1.057396 | STAD | SNU-5 | Tanespimycin |
| MVP_top60_DNA_seed | SRC | 0.626357 | LUAD | NCI-H1568 | NVP-BHG712 | MVP_top60_DNA+RNA_seed | MET | -1.048543 | STAD | IM-95 | SU11274 |
| MVP_top60_DNA_seed | EGFR | 0.627227 | LUAD | NCI-H2122 | Afatinib | MVP_top60_DNA+RNA_seed | HDAC1 | -1.047527 | STAD | IM-95 | JQ12 |
| MVP_top60_DNA_seed | ERBB2 | 0.627227 | LUAD | NCI-H2122 | Afatinib | MVP_top60_DNA+RNA_seed | MDM2 | -1.042684 | STAD | MKN45 | Nutlin-3a (-) |
| MVP_top60_DNA_seed | BRAF | 0.628586 | LUAD | HOP-62 | PLX-4720 | MVP_top60_DNA+RNA_seed | AR | -1.040687 | STAD | HSC-39 | Bicalutamide |
| MVP_top60_DNA_seed | EGFR | 0.628603 | LUAD | NCI-H1573 | Afatinib | MVP_top60_DNA+RNA_seed | ERBB2 | -1.040137 | STAD | NUGC-4 | Afatinib |
| MVP_top60_DNA_seed | ERBB2 | 0.628603 | LUAD | NCI-H1573 | Afatinib | MVP_top60_DNA+RNA_seed | HSP90AA1 | -1.03792 | STAD | SNU-1 | Luminespib |
| MVP_top60_DNA_seed | AURKA | 0.628976 | LUAD | NCI-H1435 | Alisertib | MVP_top60_DNA+RNA_seed | HSP90AB1 | -1.03792 | STAD | SNU-1 | Luminespib |
| MVP_top60_DNA_seed | MET | 0.629785 | LUAD | NCI-H1975 | Crizotinib | MVP_top60_DNA+RNA_seed | HSP90AA1 | -1.023576 | STAD | HGC-27 | Luminespib |
| MVP_top60_DNA_seed | EGFR | 0.630638 | LUAD | RERF-LC-KJ | Afatinib | MVP_top60_DNA+RNA_seed | HSP90AB1 | -1.023576 | STAD | HGC-27 | Luminespib |
| MVP_top60_DNA_seed | ERBB2 | 0.630638 | LUAD | RERF-LC-KJ | Afatinib | MVP_top60_DNA+RNA_seed | ESR1 | -1.019678 | STAD | FU97 | Tamoxifen |
| MVP_top60_DNA_seed | HDAC1 | 0.631434 | LUAD | NCI-H2023 | Dacinostat | MVP_top60_DNA+RNA_seed | ERBB2 | -1.019476 | STAD | NUGC-3 | AZD8931 |
| MVP_top60_DNA_seed | HDAC1 | 0.631453 | LUAD | NCI-H2023 | AR-42 | MVP_top60_DNA+RNA_seed | ERBB3 | -1.019476 | STAD | NUGC-3 | AZD8931 |
| MVP_top60_DNA_seed | HDAC1 | 0.631516 | LUAD | LXF-289 | Tubastatin A | MVP_top60_DNA+RNA_seed | HSP90AA1 | -1.010232 | STAD | MKN28 | Luminespib |
| MVP_top60_DNA_seed | AKT1 | 0.632082 | LUAD | NCI-H1650 | A-443654 | MVP_top60_DNA+RNA_seed | HSP90AB1 | -1.010232 | STAD | MKN28 | Luminespib |
| MVP_top60_DNA_seed | HDAC1 | 0.632484 | LUAD | NCI-H1563 | Belinostat | MVP_top60_DNA+RNA_seed | HSP90AA1 | -1.00609 | STAD | OCUM-1 | Tanespimycin |
| MVP_top60_DNA_seed | SRC | 0.632823 | LUAD | NCI-H1975 | NVP-BHG712 | MVP_top60_DNA+RNA_seed | HSP90AB1 | -1.00609 | STAD | OCUM-1 | Tanespimycin |
| MVP_top60_DNA_seed | SMARCA4 | 0.633025 | LUAD | HCC-827 | PFI-3 | MVP_top60_DNA+RNA_seed | ATM | -0.996951 | STAD | AGS | KU-60019 |
| MVP_top60_DNA_seed | GSK3B | 0.634184 | LUAD | NCI-H1944 | CHIR-99021 | MVP_top60_DNA+RNA_seed | ERBB2 | -0.995298 | STAD | AGS | AZD8931 |
| MVP_top60_DNA_seed | ATM | 0.634208 | LUAD | NCI-H1651 | CP466722 | MVP_top60_DNA+RNA_seed | ERBB3 | -0.995298 | STAD | AGS | AZD8931 |
| MVP_top60_DNA_seed | HDAC1 | 0.634493 | LUAD | NCI-H1781 | Belinostat | MVP_top60_DNA+RNA_seed | MET | -0.987911 | STAD | GCIY | SU11274 |
| MVP_top60_DNA_seed | PIK3CA | 0.635882 | LUAD | PC-3_[JPC-3] | AZD8835 | MVP_top60_DNA+RNA_seed | MET | -0.984822 | STAD | HSC-39 | AZD6094 |
| MVP_top60_DNA_seed | HSP90AA1 | 0.637913 | LUAD | EMC-BAC-2 | CCT-018159 | MVP_top60_DNA+RNA_seed | ESR1 | -0.980581 | STAD | NUGC-4 | Fulvestrant |
| MVP_top60_DNA_seed | PIK3CA | 0.638906 | LUAD | NCI-H522 | PI3Ka_4409 | MVP_top60_DNA+RNA_seed | HSP90AA1 | -0.972028 | STAD | HSC-39 | CCT-018159 |
| MVP_top60_DNA_seed | GSK3B | 0.638966 | LUAD | NCI-H1573 | AZD7969 | MVP_top60_DNA+RNA_seed | HSP90AB1 | -0.972028 | STAD | HSC-39 | CCT-018159 |
| MVP_top60_DNA_seed | MAPK1 | 0.639694 | LUAD | PC-3_[JPC-3] | VX-11e | MVP_top60_DNA+RNA_seed | ERBB2 | -0.968535 | STAD | HSC-39 | PF-00299804 |
| MVP_top60_DNA_seed | SRC | 0.641287 | LUAD | NCI-H522 | NVP-BHG712 | MVP_top60_DNA+RNA_seed | ERBB4 | -0.968535 | STAD | HSC-39 | PF-00299804 |
| MVP_top60_DNA_seed | HSP90AA1 | 0.642153 | LUAD | EKVX | Tanespimycin | MVP_top60_DNA+RNA_seed | HSP90AA1 | -0.963105 | STAD | NUGC-3 | CCT-018159 |
| MVP_top60_DNA_seed | GSK3B | 0.642455 | LUAD | NCI-H1435 | AZD7969 | MVP_top60_DNA+RNA_seed | HSP90AB1 | -0.963105 | STAD | NUGC-3 | CCT-018159 |
| MVP_top60_DNA_seed | BRAF | 0.642597 | LUAD | NCI-H1781 | SB590885 | MVP_top60_DNA+RNA_seed | AR | -0.960509 | STAD | HGC-27 | Bicalutamide |
| MVP_top60_DNA_seed | BRAF | 0.642958 | LUAD | HCC-78 | RAF_9304 | MVP_top60_DNA+RNA_seed | HSP90AA1 | -0.958702 | STAD | AGS | Luminespib |
| MVP_top60_DNA_seed | HSP90AA1 | 0.643066 | LUAD | NCI-H2122 | Elesclomol | MVP_top60_DNA+RNA_seed | HSP90AB1 | -0.958702 | STAD | AGS | Luminespib |
| MVP_top60_DNA_seed | MET | 0.643817 | LUAD | NCI-H1648 | PHA-665752 | MVP_top60_DNA+RNA_seed | PIK3CA | -0.955002 | STAD | HSC-39 | AZD8835 |
| MVP_top60_DNA_seed | GSK3B | 0.646582 | LUAD | PC-3_[JPC-3] | CHIR-99021 | MVP_top60_DNA+RNA_seed | PARP1 | -0.952909 | STAD | HSC-39 | Olaparib |
| MVP_top60_DNA_seed | BRAF | 0.646688 | LUAD | VMRC-LCD | RAF_9304 | MVP_top60_DNA+RNA_seed | AR | -0.949409 | STAD | HGC-27 | AZD3514 |
| MVP_top60_DNA_seed | MDM2 | 0.646972 | LUAD | HCC-78 | Nutlin-3a (-) | MVP_top60_DNA+RNA_seed | ERBB4 | -0.947594 | STAD | NUGC-3 | AST-1306 |
| MVP_top60_DNA_seed | ATM | 0.64835 | LUAD | PC-3_[JPC-3] | KU-55933 | MVP_top60_DNA+RNA_seed | ERBB2 | -0.940647 | STAD | SNU-1 | CI-1033 |
| MVP_top60_DNA_seed | MET | 0.648408 | LUAD | NCI-H2009 | SU11274 | MVP_top60_DNA+RNA_seed | ERBB4 | -0.940647 | STAD | SNU-1 | CI-1033 |
| MVP_top60_DNA_seed | BRAF | 0.64944 | LUAD | NCI-H3122 | PLX-4720 | MVP_top60_DNA+RNA_seed | HSP90AA1 | -0.935116 | STAD | NUGC-3 | Tanespimycin |

| | | | | | | | | | | | |
|---|---|---|---|---|---|---|---|---|---|---|---|
| MVP_top60_DNA_seed | BRAF | 0.649854 | LUAD | HOP-62 | Dabrafenib | MVP_top60_DNA+RNA_seed | HSP90AB1 | -0.935116 | STAD | NUGC-3 | Tanespimycin |
| MVP_top60_DNA_seed | MDM2 | 0.651283 | LUAD | NCI-H1693 | Nutlin-3a (-) | MVP_top60_DNA+RNA_seed | HSP90AA1 | -0.934868 | STAD | 23132-87 | Tanespimycin |
| MVP_top60_DNA_seed | MDM2 | 0.651827 | LUAD | NCI-H2228 | Nutlin-3a (-) | MVP_top60_DNA+RNA_seed | HSP90AB1 | -0.934868 | STAD | 23132-87 | Tanespimycin |
| MVP_top60_DNA_seed | FGFR1 | 0.652426 | LUAD | NCI-H3122 | AZD4547 | MVP_top60_DNA+RNA_seed | AR | -0.934838 | STAD | FU97 | Bicalutamide |
| MVP_top60_DNA_seed | FGFR1 | 0.652925 | LUAD | H3255 | PD173074 | MVP_top60_DNA+RNA_seed | ERBB2 | -0.930647 | STAD | RERF-GC-1B | CP724714 |
| MVP_top60_DNA_seed | GSK3B | 0.652926 | LUAD | SW1573 | CHIR-99021 | MVP_top60_DNA+RNA_seed | GSK3B | -0.929689 | STAD | HGC-27 | SB216763 |
| MVP_top60_DNA_seed | AURKA | 0.653122 | LUAD | NCI-H1650 | GSK1070916 | MVP_top60_DNA+RNA_seed | MDM2 | -0.926926 | STAD | SNU-1 | Nutlin-3a (-) |
| MVP_top60_DNA_seed | AURKA | 0.654931 | LUAD | Calu-3 | ZM447439 | MVP_top60_DNA+RNA_seed | ATM | -0.916841 | STAD | IM-95 | KU-60019 |
| MVP_top60_DNA_seed | MAPK1 | 0.655617 | LUAD | RERF-LC-KJ | VX-11e | MVP_top60_DNA+RNA_seed | HSP90AA1 | -0.915427 | STAD | MKN28 | Elesclomol |
| MVP_top60_DNA_seed | GSK3B | 0.65578 | LUAD | NCI-H2009 | CHIR-99021 | MVP_top60_DNA+RNA_seed | HSP90AB1 | -0.915427 | STAD | MKN28 | Elesclomol |
| MVP_top60_DNA_seed | ERBB2 | 0.655886 | LUAD | NCI-H2122 | CP724714 | MVP_top60_DNA+RNA_seed | HSP90AA1 | -0.910834 | STAD | TMK-1 | Elesclomol |
| MVP_top60_DNA_seed | FGFR1 | 0.656013 | LUAD | HOP-62 | PD173074 | MVP_top60_DNA+RNA_seed | HSP90AB1 | -0.910834 | STAD | TMK-1 | Elesclomol |
| MVP_top60_DNA_seed | EGFR | 0.65633 | LUAD | NCI-H2087 | Afatinib | MVP_top60_DNA+RNA_seed | ERBB2 | -0.910739 | STAD | GCIY | Lapatinib |
| MVP_top60_DNA_seed | ERBB2 | 0.65633 | LUAD | NCI-H2087 | Afatinib | MVP_top60_DNA+RNA_seed | HDAC1 | -0.910279 | STAD | RF-48 | Tubastatin A |
| MVP_top60_DNA_seed | MDM2 | 0.656779 | LUAD | NCI-H838 | Serdemetan | MVP_top60_DNA+RNA_seed | GSK3B | -0.908126 | STAD | KATOIII | CHIR-99021 |
| MVP_top60_DNA_seed | FGFR1 | 0.656869 | LUAD | HCC-827 | FGFR_3831 | MVP_top60_DNA+RNA_seed | ATM | -0.906994 | STAD | SNU-1 | KU-55933 |
| MVP_top60_DNA_seed | GSK3B | 0.656976 | LUAD | NCI-H1650 | CHIR-99021 | MVP_top60_DNA+RNA_seed | AR | -0.903054 | STAD | MKN28 | AZD3514 |
| MVP_top60_DNA_seed | EGFR | 0.657157 | LUAD | NCI-H1563 | AZD8931 | MVP_top60_DNA+RNA_seed | PARP1 | -0.90247 | STAD | SK-GT-2 | PARP_9495 |
| MVP_top60_DNA_seed | ERBB2 | 0.657157 | LUAD | NCI-H1563 | AZD8931 | MVP_top60_DNA+RNA_seed | HDAC1 | -0.895767 | STAD | AGS | Dacinostat |
| MVP_top60_DNA_seed | ERBB3 | 0.657157 | LUAD | NCI-H1563 | AZD8931 | MVP_top60_DNA+RNA_seed | FGFR3 | -0.888254 | STAD | AGS | FGFR_3831 |
| MVP_top60_DNA_seed | CREBBP | 0.658071 | LUAD | NCI-H1793 | I-CBP112 | MVP_top60_DNA+RNA_seed | FGFR3 | -0.885002 | STAD | SNU-1 | PD173074 |
| MVP_top60_DNA_seed | GSK3B | 0.658195 | LUAD | NCI-H2342 | CHIR-99021 | MVP_top60_DNA+RNA_seed | HDAC1 | -0.884229 | STAD | TMK-1 | JQ12 |
| MVP_top60_DNA_seed | MAPK1 | 0.658389 | LUAD | PC-14 | VX-11e | MVP_top60_DNA+RNA_seed | HDAC1 | -0.880577 | STAD | HGC-27 | JQ12 |
| MVP_top60_DNA_seed | HDAC1 | 0.659073 | LUAD | SW1573 | AR-42 | MVP_top60_DNA+RNA_seed | ERBB2 | -0.876224 | STAD | SNU-1 | CP724714 |
| MVP_top60_DNA_seed | EGFR | 0.659271 | LUAD | NCI-H1651 | Afatinib | MVP_top60_DNA+RNA_seed | MET | -0.875671 | STAD | KATOIII | JNJ38877605 |
| MVP_top60_DNA_seed | ERBB2 | 0.659271 | LUAD | NCI-H1651 | Afatinib | MVP_top60_DNA+RNA_seed | MDM2 | -0.865043 | STAD | TMK-1 | Serdemetan |
| MVP_top60_DNA_seed | IGF1R | 0.659805 | LUAD | NCI-H1993 | BMS-754807 | MVP_top60_DNA+RNA_seed | MET | -0.85186 | STAD | HGC-27 | AZD6094 |
| MVP_top60_DNA_seed | HDAC1 | 0.660538 | LUAD | NCI-H1650 | AR-42 | MVP_top60_DNA+RNA_seed | ERBB2 | -0.845644 | STAD | HGC-27 | CP724714 |
| MVP_top60_DNA_seed | EGFR | 0.660831 | LUAD | SW1573 | AZD8931 | MVP_top60_DNA+RNA_seed | ESR1 | -0.843524 | STAD | FU97 | Fulvestrant |
| MVP_top60_DNA_seed | ERBB2 | 0.660831 | LUAD | SW1573 | AZD8931 | MVP_top60_DNA+RNA_seed | GSK3B | -0.842097 | STAD | FU97 | KIN001-042 |
| MVP_top60_DNA_seed | ERBB3 | 0.660831 | LUAD | SW1573 | AZD8931 | MVP_top60_DNA+RNA_seed | ERBB2 | -0.839761 | STAD | RERF-GC-1B | Afatinib |
| MVP_top60_DNA_seed | ATM | 0.661222 | LUAD | SW1573 | CP466722 | MVP_top60_DNA+RNA_seed | FGFR3 | -0.830987 | STAD | HGC-27 | AZD4547 |
| MVP_top60_DNA_seed | SRC | 0.661905 | LUAD | RERF-LC-MS | Bosutinib | MVP_top60_DNA+RNA_seed | MAPK1 | -0.82765 | STAD | RERF-GC-1B | VX-11e |
| MVP_top60_DNA_seed | AKT1 | 0.661918 | LUAD | HOP-62 | GSK690693 | MVP_top60_DNA+RNA_seed | HSP90AA1 | -0.825112 | STAD | KATOIII | Tanespimycin |
| MVP_top60_DNA_seed | CREBBP | 0.66266 | LUAD | NCI-H522 | I-CBP112 | MVP_top60_DNA+RNA_seed | HSP90AB1 | -0.825112 | STAD | KATOIII | Tanespimycin |
| MVP_top60_DNA_seed | AKT1 | 0.663865 | LUAD | SW1573 | GSK690693 | MVP_top60_DNA+RNA_seed | ATM | -0.82301 | STAD | HSC-39 | KU-60019 |
| MVP_top60_DNA_seed | SMARCA4 | 0.664452 | LUAD | NCI-H522 | PFI-3 | MVP_top60_DNA+RNA_seed | SRC | -0.820092 | STAD | AGS | NVP-BHG712 |
| MVP_top60_DNA_seed | GSK3B | 0.665238 | LUAD | NCI-H650 | CHIR-99021 | MVP_top60_DNA+RNA_seed | HDAC1 | -0.816639 | STAD | Hs-746T | CAY10603 |
| MVP_top60_DNA_seed | HDAC1 | 0.66605 | LUAD | NCI-H1650 | CAY10603 | MVP_top60_DNA+RNA_seed | PARP1 | -0.805281 | STAD | SNU-16 | Veliparib |
| MVP_top60_DNA_seed | BRAF | 0.666655 | LUAD | NCI-H2342 | RAF_9304 | MVP_top60_DNA+RNA_seed | HSP90AA1 | -0.800069 | STAD | SK-GT-2 | CCT-018159 |
| MVP_top60_DNA_seed | MET | 0.666677 | LUAD | NCI-H1650 | Crizotinib | MVP_top60_DNA+RNA_seed | HSP90AB1 | -0.800069 | STAD | SK-GT-2 | CCT-018159 |

| | | | | | | | | | | | |
|---|---|---|---|---|---|---|---|---|---|---|---|
| MVP_top60_DNA_seed | MDM2 | 0.666825 | LUAD | NCI-H1755 | Nutlin-3a (-) | MVP_top60_DNA+RNA_seed | ATM | -0.798055 | STAD | FU97 | KU-55933 |
| MVP_top60_DNA_seed | MDM2 | 0.666945 | LUAD | HOP-62 | Nutlin-3a (-) | MVP_top60_DNA+RNA_seed | HSP90AA1 | -0.796301 | STAD | 23132-87 | CCT-018159 |
| MVP_top60_DNA_seed | BRAF | 0.667395 | LUAD | NCI-H1651 | SB590885 | MVP_top60_DNA+RNA_seed | HSP90AB1 | -0.796301 | STAD | 23132-87 | CCT-018159 |
| MVP_top60_DNA_seed | MET | 0.667545 | LUAD | NCI-H2228 | AZD6094 | MVP_top60_DNA+RNA_seed | GSK3B | -0.793201 | STAD | HGC-27 | KIN001-042 |
| MVP_top60_DNA_seed | PIK3CA | 0.667812 | LUAD | Calu-6 | AZD8835 | MVP_top60_DNA+RNA_seed | HSP90AA1 | -0.786842 | STAD | AGS | SNX-2112 |
| MVP_top60_DNA_seed | FGFR1 | 0.667924 | LUAD | Calu-6 | AZD4547 | MVP_top60_DNA+RNA_seed | HSP90AB1 | -0.786842 | STAD | AGS | SNX-2112 |
| MVP_top60_DNA_seed | EGFR | 0.668149 | LUAD | NCI-H2342 | Afatinib | MVP_top60_DNA+RNA_seed | GSK3B | -0.781636 | STAD | NUGC-4 | CHIR-99021 |
| MVP_top60_DNA_seed | ERBB2 | 0.668149 | LUAD | NCI-H2342 | Afatinib | MVP_top60_DNA+RNA_seed | GSK3B | -0.781365 | STAD | MKN28 | SB216763 |
| MVP_top60_DNA_seed | IGF1R | 0.668197 | LUAD | NCI-H1435 | GSK1904529A | MVP_top60_DNA+RNA_seed | PTK2 | -0.77824 | STAD | RERF-GC-1B | PF-562271 |
| MVP_top60_DNA_seed | BRAF | 0.669367 | LUAD | NCI-H1355 | SB590885 | MVP_top60_DNA+RNA_seed | EP300 | -0.773719 | STAD | TGBC11TKB | I-CBP112 |
| MVP_top60_DNA_seed | MET | 0.669617 | LUAD | NCI-H2347 | AZD6094 | MVP_top60_DNA+RNA_seed | CREBBP | -0.773719 | STAD | TGBC11TKB | I-CBP112 |
| MVP_top60_DNA_seed | BRAF | 0.670816 | LUAD | NCI-H1781 | Dabrafenib | MVP_top60_DNA+RNA_seed | PIK3CA | -0.771988 | STAD | NCI-N87 | PI3Ka_4409 |
| MVP_top60_DNA_seed | SRC | 0.670843 | LUAD | HCC-827 | NVP-BHG712 | MVP_top60_DNA+RNA_seed | HSP90AA1 | -0.765266 | STAD | RF-48 | Luminespib |
| MVP_top60_DNA_seed | SMARCA4 | 0.670976 | LUAD | NCI-H2009 | PFI-3 | MVP_top60_DNA+RNA_seed | HSP90AB1 | -0.765266 | STAD | RF-48 | Luminespib |
| MVP_top60_DNA_seed | EGFR | 0.670999 | LUAD | HCC-44 | AST-1306 | MVP_top60_DNA+RNA_seed | ERBB2 | -0.75817 | STAD | MKN7 | Afatinib |
| MVP_top60_DNA_seed | MET | 0.671808 | LUAD | HCC-827 | SU11274 | MVP_top60_DNA+RNA_seed | PARP1 | -0.757483 | STAD | SK-GT-2 | Veliparib |
| MVP_top60_DNA_seed | BRAF | 0.671911 | LUAD | A549 | SB590885 | MVP_top60_DNA+RNA_seed | MDM2 | -0.751632 | STAD | Hs-746T | Serdemetan |
| MVP_top60_DNA_seed | AKT1 | 0.672203 | LUAD | NCI-H522 | GSK690693 | MVP_top60_DNA+RNA_seed | HDAC1 | -0.750773 | STAD | OCUM-1 | Dacinostat |
| MVP_top60_DNA_seed | HSP90AA1 | 0.672529 | LUAD | NCI-H2228 | SNX-2112 | MVP_top60_DNA+RNA_seed | HSP90AA1 | -0.748233 | STAD | SNU-5 | CCT-018159 |
| MVP_top60_DNA_seed | BRAF | 0.672727 | LUAD | NCI-H1666 | PLX-4720 | MVP_top60_DNA+RNA_seed | HSP90AB1 | -0.748233 | STAD | SNU-5 | CCT-018159 |
| MVP_top60_DNA_seed | ATM | 0.673168 | LUAD | NCI-H2087 | torin2 | MVP_top60_DNA+RNA_seed | ERBB2 | -0.747106 | STAD | RF-48 | Afatinib |
| MVP_top60_DNA_seed | AURKA | 0.673563 | LUAD | NCI-H1651 | Genentech Cpd 10 | MVP_top60_DNA+RNA_seed | GSK3B | -0.74462 | STAD | HSC-39 | AZD7969 |
| MVP_top60_DNA_seed | AURKA | 0.67436 | LUAD | H3255 | Alisertib | MVP_top60_DNA+RNA_seed | FGFR3 | -0.743141 | STAD | MKN45 | PD173074 |
| MVP_top60_DNA_seed | SRC | 0.674369 | LUAD | NCI-H1437 | NVP-BHG712 | MVP_top60_DNA+RNA_seed | MAPK1 | -0.739715 | STAD | IM-95 | FR-180204 |
| MVP_top60_DNA_seed | BRAF | 0.674476 | LUAD | NCI-H1563 | Dabrafenib | MVP_top60_DNA+RNA_seed | MAPK3 | -0.739715 | STAD | IM-95 | FR-180204 |
| MVP_top60_DNA_seed | ATM | 0.675133 | LUAD | NCI-H1651 | KU-55933 | MVP_top60_DNA+RNA_seed | PIK3CA | -0.739382 | STAD | SNU-5 | PI3Ka_4409 |
| MVP_top60_DNA_seed | FGFR1 | 0.67572 | LUAD | NCI-H2087 | PD173074 | MVP_top60_DNA+RNA_seed | MAPK1 | -0.73702 | STAD | IM-95 | VX-11e |
| MVP_top60_DNA_seed | AURKA | 0.676929 | LUAD | NCI-H1734 | CD532 | MVP_top60_DNA+RNA_seed | PARP1 | -0.732341 | STAD | FU97 | Olaparib |
| MVP_top60_DNA_seed | BRAF | 0.678327 | LUAD | NCI-H1944 | Dabrafenib | MVP_top60_DNA+RNA_seed | HSP90AA1 | -0.731187 | STAD | TGBC11TKB | Tanespimycin |
| MVP_top60_DNA_seed | HDAC1 | 0.678498 | LUAD | SK-LU-1 | Dacinostat | MVP_top60_DNA+RNA_seed | HSP90AB1 | -0.731187 | STAD | TGBC11TKB | Tanespimycin |
| MVP_top60_DNA_seed | EGFR | 0.678503 | LUAD | Calu-6 | Afatinib | MVP_top60_DNA+RNA_seed | FYN | -0.730489 | STAD | GCIY | A-770041 |
| MVP_top60_DNA_seed | ERBB2 | 0.678503 | LUAD | Calu-6 | Afatinib | MVP_top60_DNA+RNA_seed | GSK3B | -0.718633 | STAD | SNU-1 | SB216763 |
| MVP_top60_DNA_seed | EGFR | 0.678698 | LUAD | HCC-827 | Cetuximab | MVP_top60_DNA+RNA_seed | PARP1 | -0.718313 | STAD | AGS | Olaparib |
| MVP_top60_DNA_seed | ERBB2 | 0.678854 | LUAD | RERF-LC-KJ | CP724714 | MVP_top60_DNA+RNA_seed | SRC | -0.707845 | STAD | HGC-27 | Bosutinib |
| MVP_top60_DNA_seed | FGFR1 | 0.679606 | LUAD | EMC-BAC-1 | AZD4547 | MVP_top60_DNA+RNA_seed | ATM | -0.707712 | STAD | IM-95 | CP466722 |
| MVP_top60_DNA_seed | AKT1 | 0.680197 | LUAD | NCI-H2405 | AKT inhibitor VIII | MVP_top60_DNA+RNA_seed | FGFR3 | -0.704993 | STAD | 23132-87 | AZD4547 |
| MVP_top60_DNA_seed | HDAC1 | 0.6806 | LUAD | NCI-H2009 | Tubastatin A | MVP_top60_DNA+RNA_seed | HDAC1 | -0.700793 | STAD | RERF-GC-1B | Dacinostat |
| MVP_top60_DNA_seed | BRAF | 0.680907 | LUAD | NCI-H2347 | HG6-64-1 | MVP_top60_DNA+RNA_seed | PARP1 | -0.698158 | STAD | 23132-87 | Olaparib |
| MVP_top60_DNA_seed | ERBB2 | 0.681577 | LUAD | NCI-H522 | CP724714 | MVP_top60_DNA+RNA_seed | ATM | -0.6981 | STAD | HGC-27 | KU-60019 |
| MVP_top60_DNA_seed | AKT1 | 0.683103 | LUAD | NCI-H1793 | BAY AKT1 | MVP_top60_DNA+RNA_seed | SRC | -0.697758 | STAD | IM-95 | NVP-BHG712 |
| MVP_top60_DNA_seed | FGFR1 | 0.683585 | LUAD | NCI-H2085 | AZD4547 | MVP_top60_DNA+RNA_seed | MDM2 | -0.697361 | STAD | FU97 | Serdemetan |

| | | | | | | | | | | | |
|---|---|---|---|---|---|---|---|---|---|---|---|
| MVP_top60_DNA_seed | EGFR | 0.685148 | LUAD | NCI-H1623 | Pelitinib | MVP_top60_DNA+RNA_seed | GSK3B | -0.689565 | STAD | GCIY | KIN001-042 |
| MVP_top60_DNA_seed | AKT1 | 0.685477 | LUAD | NCI-H838 | GSK690693 | MVP_top60_DNA+RNA_seed | HSP90AA1 | -0.68734 | STAD | SK-GT-2 | Elesclomol |
| MVP_top60_DNA_seed | MDM2 | 0.685932 | LUAD | NCI-H441 | Serdemetan | MVP_top60_DNA+RNA_seed | HSP90AB1 | -0.68734 | STAD | SK-GT-2 | Elesclomol |
| MVP_top60_DNA_seed | MDM2 | 0.686345 | LUAD | NCI-H1623 | Serdemetan | MVP_top60_DNA+RNA_seed | ERBB2 | -0.687259 | STAD | TMK-1 | Afatinib |
| MVP_top60_DNA_seed | AURKA | 0.686697 | LUAD | NCI-H441 | HG-5-113-01 | MVP_top60_DNA+RNA_seed | SRC | -0.68444 | STAD | KATOIII | Bosutinib |
| MVP_top60_DNA_seed | AURKA | 0.686936 | LUAD | HCC-78 | CD532 | MVP_top60_DNA+RNA_seed | HDAC1 | -0.68208 | STAD | NCI-N87 | AR-42 |
| MVP_top60_DNA_seed | IGF1R | 0.687022 | LUAD | RERF-LC-KJ | BMS-536924 | MVP_top60_DNA+RNA_seed | HSP90AA1 | -0.680041 | STAD | IM-95 | SNX-2112 |
| MVP_top60_DNA_seed | HSP90AA1 | 0.687939 | LUAD | NCI-H2291 | CCT-018159 | MVP_top60_DNA+RNA_seed | HSP90AB1 | -0.680041 | STAD | IM-95 | SNX-2112 |
| MVP_top60_DNA_seed | BRAF | 0.688028 | LUAD | NCI-H1648 | HG6-64-1 | MVP_top60_DNA+RNA_seed | HSP90AA1 | -0.678794 | STAD | TMK-1 | CCT-018159 |
| MVP_top60_DNA_seed | GSK3B | 0.688685 | LUAD | NCI-H358 | CHIR-99021 | MVP_top60_DNA+RNA_seed | HSP90AB1 | -0.678794 | STAD | TMK-1 | CCT-018159 |
| MVP_top60_DNA_seed | AURKA | 0.688924 | LUAD | NCI-H2009 | Alisertib | MVP_top60_DNA+RNA_seed | SRC | -0.676536 | STAD | RERF-GC-1B | NVP-BHG712 |
| MVP_top60_DNA_seed | EGFR | 0.688986 | LUAD | SK-LU-1 | CI-1033 | MVP_top60_DNA+RNA_seed | AR | -0.673015 | STAD | GCIY | Bicalutamide |
| MVP_top60_DNA_seed | ERBB2 | 0.688986 | LUAD | SK-LU-1 | CI-1033 | MVP_top60_DNA+RNA_seed | HDAC1 | -0.672049 | STAD | MKN28 | JQ12 |
| MVP_top60_DNA_seed | GSK3B | 0.689867 | LUAD | NCI-H2030 | CHIR-99021 | MVP_top60_DNA+RNA_seed | ERBB2 | -0.66765 | STAD | 23132-87 | CUDC-101 |
| MVP_top60_DNA_seed | BRAF | 0.690136 | LUAD | NCI-H1648 | PLX-4720 | MVP_top60_DNA+RNA_seed | MAPK1 | -0.664754 | STAD | MKN28 | FR-180204 |
| MVP_top60_DNA_seed | BRAF | 0.690742 | LUAD | NCI-H3122 | PLX-4720 | MVP_top60_DNA+RNA_seed | MAPK3 | -0.664754 | STAD | MKN28 | FR-180204 |
| MVP_top60_DNA_seed | CREBBP | 0.69077 | LUAD | NCI-H1435 | I-CBP112 | MVP_top60_DNA+RNA_seed | FGFR3 | -0.658334 | STAD | RERF-GC-1B | AZD4547 |
| MVP_top60_DNA_seed | EGFR | 0.691048 | LUAD | HOP-62 | Afatinib | MVP_top60_DNA+RNA_seed | FGFR3 | -0.655139 | STAD | HGC-27 | PD173074 |
| MVP_top60_DNA_seed | ERBB2 | 0.691048 | LUAD | HOP-62 | Afatinib | MVP_top60_DNA+RNA_seed | GSK3B | -0.652984 | STAD | 23132-87 | AZD7969 |
| MVP_top60_DNA_seed | ERBB2 | 0.69181 | LUAD | NCI-H1623 | CP724714 | MVP_top60_DNA+RNA_seed | ERBB2 | -0.650746 | STAD | IM-95 | CP724714 |
| MVP_top60_DNA_seed | BRAF | 0.692125 | LUAD | NCI-H1437 | SB590885 | MVP_top60_DNA+RNA_seed | ERBB2 | -0.648542 | STAD | GCIY | CP724714 |
| MVP_top60_DNA_seed | PIK3CA | 0.692409 | LUAD | COR-L105 | PI3Ka_4409 | MVP_top60_DNA+RNA_seed | HDAC1 | -0.645869 | STAD | NCI-N87 | Belinostat |
| MVP_top60_DNA_seed | HDAC1 | 0.693036 | LUAD | NCI-H322M | Tubastatin A | MVP_top60_DNA+RNA_seed | MET | -0.645854 | STAD | SNU-1 | Crizotinib |
| MVP_top60_DNA_seed | HSP90AA1 | 0.693214 | LUAD | NCI-H838 | Luminespib | MVP_top60_DNA+RNA_seed | FGFR3 | -0.640145 | STAD | FU97 | PD173074 |
| MVP_top60_DNA_seed | FGFR1 | 0.693248 | LUAD | NCI-H522 | PD173074 | MVP_top60_DNA+RNA_seed | MDM2 | -0.632946 | STAD | NUGC-4 | Nutlin-3a (-) |
| MVP_top60_DNA_seed | HSP90AA1 | 0.693431 | LUAD | NCI-H322M | SNX-2112 | MVP_top60_DNA+RNA_seed | ATM | -0.628394 | STAD | RF-48 | CP466722 |
| MVP_top60_DNA_seed | ATM | 0.693698 | LUAD | NCI-H2405 | KU-60019 | MVP_top60_DNA+RNA_seed | AR | -0.62623 | STAD | 23132-87 | Bicalutamide |
| MVP_top60_DNA_seed | AURKA | 0.693752 | LUAD | NCI-H3122 | CD532 | MVP_top60_DNA+RNA_seed | MDM2 | -0.623944 | STAD | HGC-27 | Serdemetan |
| MVP_top60_DNA_seed | SRC | 0.695593 | LUAD | LC-2-ad | Bosutinib | MVP_top60_DNA+RNA_seed | HDAC1 | -0.623534 | STAD | NUGC-3 | JQ12 |
| MVP_top60_DNA_seed | AKT1 | 0.696202 | LUAD | NCI-H1993 | GSK690693 | MVP_top60_DNA+RNA_seed | PARP1 | -0.623179 | STAD | HSC-39 | Veliparib |
| MVP_top60_DNA_seed | MDM2 | 0.69665 | LUAD | NCI-H2342 | Nutlin-3a (-) | MVP_top60_DNA+RNA_seed | GSK3B | -0.616659 | STAD | NCI-N87 | CHIR-99021 |
| MVP_top60_DNA_seed | IGF1R | 0.698289 | LUAD | HOP-62 | GSK1904529A | MVP_top60_DNA+RNA_seed | SRC | -0.615531 | STAD | NUGC-4 | Bosutinib |
| MVP_top60_DNA_seed | MDM2 | 0.699467 | LUAD | Calu-3 | Nutlin-3a (-) | MVP_top60_DNA+RNA_seed | ATM | -0.611452 | STAD | TMK-1 | torin2 |
| MVP_top60_DNA_seed | MAPK1 | 0.699519 | LUAD | NCI-H2023 | FR-180204 | MVP_top60_DNA+RNA_seed | HDAC1 | -0.611059 | STAD | Hs-746T | Belinostat |
| MVP_top60_DNA_seed | HSP90AA1 | 0.699663 | LUAD | EMC-BAC-1 | CCT-018159 | MVP_top60_DNA+RNA_seed | PARP1 | -0.607796 | STAD | TMK-1 | Veliparib |
| MVP_top60_DNA_seed | CREBBP | 0.702038 | LUAD | NCI-H358 | I-CBP112 | MVP_top60_DNA+RNA_seed | ATM | -0.606572 | STAD | TMK-1 | KU-55933 |
| MVP_top60_DNA_seed | CREBBP | 0.702564 | LUAD | NCI-H2009 | I-CBP112 | MVP_top60_DNA+RNA_seed | ERBB2 | -0.605117 | STAD | IM-95 | CI-1033 |
| MVP_top60_DNA_seed | ATM | 0.702816 | LUAD | NCI-H1650 | CP466722 | MVP_top60_DNA+RNA_seed | ERBB4 | -0.605117 | STAD | IM-95 | CI-1033 |
| MVP_top60_DNA_seed | BRAF | 0.702819 | LUAD | NCI-H1435 | Dabrafenib | MVP_top60_DNA+RNA_seed | ERBB4 | -0.604596 | STAD | RERF-GC-1B | AST-1306 |
| MVP_top60_DNA_seed | MAPK1 | 0.703045 | LUAD | NCI-H838 | FR-180204 | MVP_top60_DNA+RNA_seed | GSK3B | -0.603758 | STAD | IM-95 | CHIR-99021 |
| MVP_top60_DNA_seed | FGFR1 | 0.703768 | LUAD | NCI-H2342 | AZD4547 | MVP_top60_DNA+RNA_seed | ERBB2 | -0.602136 | STAD | MKN28 | AZD8931 |

| | | | | | | | | | | | |
|---|---|---|---|---|---|---|---|---|---|---|---|
| MVP_top60_DNA_seed | CREBBP | 0.705049 | LUAD | NCI-H2085 | I-CBP112 | MVP_top60_DNA+RNA_seed | ERBB3 | -0.602136 | STAD | MKN28 | AZD8931 |
| MVP_top60_DNA_seed | MET | 0.705187 | LUAD | NCI-H838 | AZD6094 | MVP_top60_DNA+RNA_seed | HDAC1 | -0.597844 | STAD | FU97 | Dacinostat |
| MVP_top60_DNA_seed | AKT1 | 0.705609 | LUAD | NCI-H2405 | MK-2206 | MVP_top60_DNA+RNA_seed | PTK2 | -0.592193 | STAD | MKN28 | PF-562271 |
| MVP_top60_DNA_seed | EGFR | 0.706322 | LUAD | NCI-H2405 | AZD8931 | MVP_top60_DNA+RNA_seed | ESR1 | -0.585544 | STAD | SNU-16 | Fulvestrant |
| MVP_top60_DNA_seed | ERBB2 | 0.706322 | LUAD | NCI-H2405 | AZD8931 | MVP_top60_DNA+RNA_seed | PARP1 | -0.585243 | STAD | MKN7 | Rucaparib |
| MVP_top60_DNA_seed | ERBB3 | 0.706322 | LUAD | NCI-H2405 | AZD8931 | MVP_top60_DNA+RNA_seed | ESR1 | -0.585061 | STAD | KATOIII | Tamoxifen |
| MVP_top60_DNA_seed | ATM | 0.70647 | LUAD | NCI-H2085 | KU-60019 | MVP_top60_DNA+RNA_seed | PIK3CA | -0.58183 | STAD | 23132-87 | AZD8835 |
| MVP_top60_DNA_seed | GSK3B | 0.70652 | LUAD | NCI-H1781 | CHIR-99021 | MVP_top60_DNA+RNA_seed | ERBB2 | -0.576329 | STAD | Hs-746T | CUDC-101 |
| MVP_top60_DNA_seed | PIK3CA | 0.706685 | LUAD | H3255 | PI3Ka_4409 | MVP_top60_DNA+RNA_seed | ERBB2 | -0.576186 | STAD | KATOIII | PF-00299804 |
| MVP_top60_DNA_seed | HDAC1 | 0.707176 | LUAD | NCI-H1650 | Tubastatin A | MVP_top60_DNA+RNA_seed | ERBB4 | -0.576186 | STAD | KATOIII | PF-00299804 |
| MVP_top60_DNA_seed | EGFR | 0.709277 | LUAD | NCI-H2228 | PF-00299804 | MVP_top60_DNA+RNA_seed | HSP90AA1 | -0.573396 | STAD | HGC-27 | SNX-2112 |
| MVP_top60_DNA_seed | ERBB2 | 0.709277 | LUAD | NCI-H2228 | PF-00299804 | MVP_top60_DNA+RNA_seed | HSP90AB1 | -0.573396 | STAD | HGC-27 | SNX-2112 |
| MVP_top60_DNA_seed | BRAF | 0.709432 | LUAD | SK-LU-1 | RAF_9304 | MVP_top60_DNA+RNA_seed | PARP1 | -0.56865 | STAD | AGS | PARP_0108 |
| MVP_top60_DNA_seed | BRAF | 0.709674 | LUAD | NCI-H1437 | HG6-64-1 | MVP_top60_DNA+RNA_seed | ESR1 | -0.56609 | STAD | TGBC11TKB | Tamoxifen |
| MVP_top60_DNA_seed | BRAF | 0.710078 | LUAD | HCC-78 | PLX-4720 | MVP_top60_DNA+RNA_seed | AR | -0.565261 | STAD | TGBC11TKB | Bicalutamide |
| MVP_top60_DNA_seed | BRAF | 0.710302 | LUAD | NCI-H2009 | SB590885 | MVP_top60_DNA+RNA_seed | MAPK1 | -0.564627 | STAD | MKN45 | VX-11e |
| MVP_top60_DNA_seed | MDM2 | 0.711107 | LUAD | NCI-H2228 | Serdemetan | MVP_top60_DNA+RNA_seed | FGFR3 | -0.558379 | STAD | NUGC-3 | FGFR_3831 |
| MVP_top60_DNA_seed | EGFR | 0.711684 | LUAD | NCI-H2085 | AZD8931 | MVP_top60_DNA+RNA_seed | PARP1 | -0.557984 | STAD | NUGC-3 | PARP_9495 |
| MVP_top60_DNA_seed | ERBB2 | 0.711684 | LUAD | NCI-H2085 | AZD8931 | MVP_top60_DNA+RNA_seed | ATM | -0.556842 | STAD | TGBC11TKB | KU-55933 |
| MVP_top60_DNA_seed | ERBB3 | 0.711684 | LUAD | NCI-H2085 | AZD8931 | MVP_top60_DNA+RNA_seed | FGFR3 | -0.556164 | STAD | RERF-GC-1B | AZD4547 |
| MVP_top60_DNA_seed | ATM | 0.712103 | LUAD | NCI-H522 | KU-60019 | MVP_top60_DNA+RNA_seed | ATM | -0.553314 | STAD | 23132-87 | KU-55933 |
| MVP_top60_DNA_seed | HDAC1 | 0.713312 | LUAD | NCI-H2122 | Tubastatin A | MVP_top60_DNA+RNA_seed | PARP1 | -0.551265 | STAD | MKN28 | PARP_9495 |
| MVP_top60_DNA_seed | BRAF | 0.713551 | LUAD | NCI-H1355 | Dabrafenib | MVP_top60_DNA+RNA_seed | ATM | -0.549809 | STAD | GCIY | CP466722 |
| MVP_top60_DNA_seed | MDM2 | 0.717416 | LUAD | Calu-6 | Serdemetan | MVP_top60_DNA+RNA_seed | ATM | -0.547039 | STAD | MKN7 | KU-55933 |
| MVP_top60_DNA_seed | MAPK1 | 0.719467 | LUAD | Calu-6 | FR-180204 | MVP_top60_DNA+RNA_seed | PIK3CA | -0.544564 | STAD | NCI-N87 | AZD8835 |
| MVP_top60_DNA_seed | AURKA | 0.720391 | LUAD | SW1573 | Alisertib | MVP_top60_DNA+RNA_seed | PARP1 | -0.539058 | STAD | 23132-87 | PARP_9482 |
| MVP_top60_DNA_seed | MAPK1 | 0.721747 | LUAD | NCI-H2342 | VX-11e | MVP_top60_DNA+RNA_seed | AR | -0.537592 | STAD | IM-95 | Bicalutamide |
| MVP_top60_DNA_seed | BRAF | 0.722368 | LUAD | RERF-LC-MS | SB590885 | MVP_top60_DNA+RNA_seed | ERBB2 | -0.53759 | STAD | RERF-GC-1B | CI-1033 |
| MVP_top60_DNA_seed | BRAF | 0.722407 | LUAD | NCI-H1693 | Dabrafenib | MVP_top60_DNA+RNA_seed | ERBB4 | -0.53759 | STAD | RERF-GC-1B | CI-1033 |
| MVP_top60_DNA_seed | HDAC1 | 0.72244 | LUAD | NCI-H358 | JQ12 | MVP_top60_DNA+RNA_seed | MET | -0.537532 | STAD | SNU-16 | AZD6094 |
| MVP_top60_DNA_seed | GSK3B | 0.722529 | LUAD | Calu-6 | KIN001-042 | MVP_top60_DNA+RNA_seed | ATM | -0.533504 | STAD | SNU-16 | KU-55933 |
| MVP_top60_DNA_seed | BRAF | 0.722863 | LUAD | NCI-H2228 | PLX-4720 | MVP_top60_DNA+RNA_seed | GSK3B | -0.528123 | STAD | KATOIII | SB216763 |
| MVP_top60_DNA_seed | BRAF | 0.723463 | LUAD | HCC-827 | Dabrafenib | MVP_top60_DNA+RNA_seed | FGFR3 | -0.524533 | STAD | RF-48 | PD173074 |
| MVP_top60_DNA_seed | AKT1 | 0.72542 | LUAD | NCI-H1651 | GSK690693 | MVP_top60_DNA+RNA_seed | MDM2 | -0.523194 | STAD | SK-GT-2 | Serdemetan |
| MVP_top60_DNA_seed | EGFR | 0.725663 | LUAD | NCI-H1435 | Gefitinib | MVP_top60_DNA+RNA_seed | PARP1 | -0.523022 | STAD | SNU-5 | Rucaparib |
| MVP_top60_DNA_seed | MET | 0.726015 | LUAD | PC-3_[JPC-3] | SU11274 | MVP_top60_DNA+RNA_seed | HSP90AA1 | -0.518581 | STAD | NCI-N87 | SNX-2112 |
| MVP_top60_DNA_seed | MAPK1 | 0.726361 | LUAD | LXF-289 | VX-11e | MVP_top60_DNA+RNA_seed | HSP90AB1 | -0.518581 | STAD | NCI-N87 | SNX-2112 |
| MVP_top60_DNA_seed | BRAF | 0.726995 | LUAD | NCI-H838 | SB590885 | MVP_top60_DNA+RNA_seed | HSP90AA1 | -0.517949 | STAD | TGBC11TKB | SNX-2112 |
| MVP_top60_DNA_seed | AURKA | 0.727765 | LUAD | NCI-H441 | GSK1070916 | MVP_top60_DNA+RNA_seed | HSP90AB1 | -0.517949 | STAD | TGBC11TKB | SNX-2112 |
| MVP_top60_DNA_seed | GSK3B | 0.728513 | LUAD | NCI-H2085 | CHIR-99021 | MVP_top60_DNA+RNA_seed | GSK3B | -0.515859 | STAD | HSC-39 | KIN001-042 |
| MVP_top60_DNA_seed | CREBBP | 0.728861 | LUAD | NCI-H1573 | I-CBP112 | MVP_top60_DNA+RNA_seed | MET | -0.515105 | STAD | MKN7 | SU11274 |

| | | | | | | | | | | | |
|---|---|---|---|---|---|---|---|---|---|---|---|
| MVP_top60_DNA_seed | HSP90AA1 | 0.728879 | LUAD | NCI-H1648 | Elesclomol | MVP_top60_DNA+RNA_seed | SRC | -0.51228 | STAD | HGC-27 | NVP-BHG712 |
| MVP_top60_DNA_seed | EGFR | 0.729725 | LUAD | NCI-H2009 | AST-1306 | MVP_top60_DNA+RNA_seed | PARP1 | -0.512178 | STAD | AGS | PARP_9482 |
| MVP_top60_DNA_seed | EGFR | 0.729738 | LUAD | NCI-H2122 | Pelitinib | MVP_top60_DNA+RNA_seed | ERBB2 | -0.507793 | STAD | NUGC-4 | CP724714 |
| MVP_top60_DNA_seed | EGFR | 0.729951 | LUAD | NCI-H23 | Gefitinib | MVP_top60_DNA+RNA_seed | MET | -0.502076 | STAD | MKN7 | JNJ38877605 |
| MVP_top60_DNA_seed | CREBBP | 0.73078 | LUAD | NCI-H2342 | I-CBP112 | MVP_top60_DNA+RNA_seed | GSK3B | -0.501703 | STAD | HGC-27 | CHIR-99021 |
| MVP_top60_DNA_seed | FGFR1 | 0.73089 | LUAD | NCI-H1993 | FGFR_3831 | MVP_top60_DNA+RNA_seed | FGFR3 | -0.495512 | STAD | NUGC-3 | AZD4547 |
| MVP_top60_DNA_seed | EGFR | 0.731258 | LUAD | NCI-H3122 | AST-1306 | MVP_top60_DNA+RNA_seed | PARP1 | -0.495282 | STAD | 23132-87 | PARP_0108 |
| MVP_top60_DNA_seed | EGFR | 0.731424 | LUAD | LXF-289 | PF-00299804 | MVP_top60_DNA+RNA_seed | GSK3B | -0.495159 | STAD | MKN7 | SB216763 |
| MVP_top60_DNA_seed | ERBB2 | 0.731424 | LUAD | LXF-289 | PF-00299804 | MVP_top60_DNA+RNA_seed | HDAC1 | -0.494925 | STAD | GCIY | Dacinostat |
| MVP_top60_DNA_seed | HDAC1 | 0.731567 | LUAD | NCI-H23 | Dacinostat | MVP_top60_DNA+RNA_seed | ERBB2 | -0.490537 | STAD | SNU-1 | CUDC-101 |
| MVP_top60_DNA_seed | SRC | 0.731726 | LUAD | NCI-H1563 | Bosutinib | MVP_top60_DNA+RNA_seed | ATM | -0.486763 | STAD | SNU-1 | CP466722 |
| MVP_top60_DNA_seed | HSP90AA1 | 0.732073 | LUAD | HCC-827 | SNX-2112 | MVP_top60_DNA+RNA_seed | MET | -0.486593 | STAD | KATOIII | AZD6094 |
| MVP_top60_DNA_seed | FGFR1 | 0.734283 | LUAD | NCI-H522 | FGFR_3831 | MVP_top60_DNA+RNA_seed | GSK3B | -0.480581 | STAD | NUGC-4 | AZD7969 |
| MVP_top60_DNA_seed | MAPK1 | 0.734616 | LUAD | NCI-H2122 | FR-180204 | MVP_top60_DNA+RNA_seed | MDM2 | -0.479252 | STAD | 23132-87 | Nutlin-3a (-) |
| MVP_top60_DNA_seed | EGFR | 0.734982 | LUAD | NCI-H1563 | CUDC-101 | MVP_top60_DNA+RNA_seed | HSP90AA1 | -0.479096 | STAD | GCIY | Luminespib |
| MVP_top60_DNA_seed | ERBB2 | 0.734982 | LUAD | NCI-H1563 | CUDC-101 | MVP_top60_DNA+RNA_seed | HSP90AB1 | -0.479096 | STAD | GCIY | Luminespib |
| MVP_top60_DNA_seed | EGFR | 0.735108 | LUAD | NCI-H1793 | Cetuximab | MVP_top60_DNA+RNA_seed | HSP90AA1 | -0.472617 | STAD | FU97 | CCT-018159 |
| MVP_top60_DNA_seed | HDAC1 | 0.735311 | LUAD | Calu-3 | AR-42 | MVP_top60_DNA+RNA_seed | HSP90AB1 | -0.472617 | STAD | FU97 | CCT-018159 |
| MVP_top60_DNA_seed | SMARCA4 | 0.736232 | LUAD | Calu-6 | PFI-3 | MVP_top60_DNA+RNA_seed | GSK3B | -0.471032 | STAD | TGBC11TKB | KIN001-042 |
| MVP_top60_DNA_seed | FGFR1 | 0.736903 | LUAD | NCI-H1975 | AZD4547 | MVP_top60_DNA+RNA_seed | PARP1 | -0.470017 | STAD | OCUM-1 | Talazoparib |
| MVP_top60_DNA_seed | AKT1 | 0.73696 | LUAD | HCC-827 | GSK690693 | MVP_top60_DNA+RNA_seed | ATM | -0.467569 | STAD | GCIY | KU-60019 |
| MVP_top60_DNA_seed | HDAC1 | 0.737508 | LUAD | NCI-H2023 | Belinostat | MVP_top60_DNA+RNA_seed | AR | -0.466296 | STAD | AGS | Bicalutamide |
| MVP_top60_DNA_seed | AURKA | 0.738294 | LUAD | A549 | Alisertib | MVP_top60_DNA+RNA_seed | AR | -0.462654 | STAD | SK-GT-2 | Bicalutamide |
| MVP_top60_DNA_seed | HDAC1 | 0.740176 | LUAD | NCI-H2085 | AR-42 | MVP_top60_DNA+RNA_seed | HDAC1 | -0.461909 | STAD | NCI-N87 | CAY10603 |
| MVP_top60_DNA_seed | FGFR1 | 0.740315 | LUAD | NCI-H1651 | PD173074 | MVP_top60_DNA+RNA_seed | PTK2 | -0.461073 | STAD | 23132-87 | PF-562271 |
| MVP_top60_DNA_seed | PTK2 | 0.740883 | LUAD | NCI-H2023 | PF-562271 | MVP_top60_DNA+RNA_seed | ERBB2 | -0.458125 | STAD | AGS | CUDC-101 |
| MVP_top60_DNA_seed | MDM2 | 0.741701 | LUAD | PC-14 | Nutlin-3a (-) | MVP_top60_DNA+RNA_seed | ERBB2 | -0.457756 | STAD | MKN28 | CI-1033 |
| MVP_top60_DNA_seed | FGFR1 | 0.744307 | LUAD | NCI-H1355 | AZD4547 | MVP_top60_DNA+RNA_seed | ERBB4 | -0.457756 | STAD | MKN28 | CI-1033 |
| MVP_top60_DNA_seed | FGFR1 | 0.74447 | LUAD | NCI-H1944 | PD173074 | MVP_top60_DNA+RNA_seed | FGFR3 | -0.452904 | STAD | 23132-87 | FGFR_3831 |
| MVP_top60_DNA_seed | ATM | 0.744767 | LUAD | NCI-H1666 | KU-60019 | MVP_top60_DNA+RNA_seed | MET | -0.442766 | STAD | TGBC11TKB | AZD6094 |
| MVP_top60_DNA_seed | BRAF | 0.745826 | LUAD | NCI-H1650 | SB590885 | MVP_top60_DNA+RNA_seed | GSK3B | -0.440756 | STAD | TMK-1 | AZD7969 |
| MVP_top60_DNA_seed | FGFR1 | 0.747653 | LUAD | NCI-H1650 | AZD4547 | MVP_top60_DNA+RNA_seed | MDM2 | -0.437594 | STAD | MKN28 | Serdemetan |
| MVP_top60_DNA_seed | ATM | 0.747876 | LUAD | NCI-H3122 | KU-55933 | MVP_top60_DNA+RNA_seed | GSK3B | -0.437124 | STAD | KATOIII | AZD7969 |
| MVP_top60_DNA_seed | FGFR1 | 0.748016 | LUAD | NCI-H2291 | FGFR_3831 | MVP_top60_DNA+RNA_seed | GSK3B | -0.433852 | STAD | SNU-16 | AZD7969 |
| MVP_top60_DNA_seed | AKT1 | 0.74805 | LUAD | ABC-1 | AKT inhibitor VIII | MVP_top60_DNA+RNA_seed | GSK3B | -0.430945 | STAD | TMK-1 | SB216763 |
| MVP_top60_DNA_seed | EGFR | 0.748075 | LUAD | NCI-H2009 | Afatinib | MVP_top60_DNA+RNA_seed | ERBB2 | -0.428087 | STAD | KATOIII | CI-1033 |
| MVP_top60_DNA_seed | ERBB2 | 0.748075 | LUAD | NCI-H2009 | Afatinib | MVP_top60_DNA+RNA_seed | ERBB4 | -0.428087 | STAD | KATOIII | CI-1033 |
| MVP_top60_DNA_seed | MDM2 | 0.748582 | LUAD | NCI-H522 | Serdemetan | MVP_top60_DNA+RNA_seed | HSP90AA1 | -0.426285 | STAD | SK-GT-2 | Tanespimycin |
| MVP_top60_DNA_seed | EGFR | 0.748998 | LUAD | NCI-H1650 | PF-00299804 | MVP_top60_DNA+RNA_seed | HSP90AB1 | -0.426285 | STAD | SK-GT-2 | Tanespimycin |
| MVP_top60_DNA_seed | ERBB2 | 0.748998 | LUAD | NCI-H1650 | PF-00299804 | MVP_top60_DNA+RNA_seed | FGFR3 | -0.426056 | STAD | MKN7 | PD173074 |
| MVP_top60_DNA_seed | BRAF | 0.749463 | LUAD | NCI-H2347 | PLX-4720 | MVP_top60_DNA+RNA_seed | MAPK1 | -0.424829 | STAD | RF-48 | FR-180204 |

| | | | | | | | | | | | |
|---|---|---|---|---|---|---|---|---|---|---|---|
| MVP_top60_DNA_seed | ATM | 0.749597 | LUAD | NCI-H1568 | KU-55933 | MVP_top60_DNA+RNA_seed | MAPK3 | -0.424829 | STAD | RF-48 | FR-180204 |
| MVP_top60_DNA_seed | AKT1 | 0.750086 | LUAD | COR-L105 | BAY AKT1 | MVP_top60_DNA+RNA_seed | ERBB2 | -0.423921 | STAD | MKN7 | Afatinib |
| MVP_top60_DNA_seed | HDAC1 | 0.750282 | LUAD | PC-14 | JQ12 | MVP_top60_DNA+RNA_seed | HDAC1 | -0.423557 | STAD | MKN28 | Dacinostat |
| MVP_top60_DNA_seed | EGFR | 0.750463 | LUAD | NCI-H23 | CUDC-101 | MVP_top60_DNA+RNA_seed | HSP90AA1 | -0.423536 | STAD | FU97 | Luminespib |
| MVP_top60_DNA_seed | ERBB2 | 0.750463 | LUAD | NCI-H23 | CUDC-101 | MVP_top60_DNA+RNA_seed | HSP90AB1 | -0.423536 | STAD | FU97 | Luminespib |
| MVP_top60_DNA_seed | ERBB2 | 0.75159 | LUAD | NCI-H1355 | CP724714 | MVP_top60_DNA+RNA_seed | HDAC1 | -0.42268 | STAD | TMK-1 | Dacinostat |
| MVP_top60_DNA_seed | MAPK1 | 0.752278 | LUAD | RERF-LC-KJ | FR-180204 | MVP_top60_DNA+RNA_seed | MDM2 | -0.421014 | STAD | 23132-87 | Serdemetan |
| MVP_top60_DNA_seed | GSK3B | 0.753163 | LUAD | EMC-BAC-2 | KIN001-042 | MVP_top60_DNA+RNA_seed | PARP1 | -0.420275 | STAD | TMK-1 | Talazoparib |
| MVP_top60_DNA_seed | FGFR1 | 0.75363 | LUAD | NCI-H1793 | FGFR_3831 | MVP_top60_DNA+RNA_seed | PARP1 | -0.418993 | STAD | FU97 | Veliparib |
| MVP_top60_DNA_seed | CREBBP | 0.753817 | LUAD | Calu-6 | I-CBP112 | MVP_top60_DNA+RNA_seed | HDAC1 | -0.416055 | STAD | NCI-N87 | Dacinostat |
| MVP_top60_DNA_seed | HDAC1 | 0.75439 | LUAD | NCI-H1993 | CAY10603 | MVP_top60_DNA+RNA_seed | MAPK1 | -0.413716 | STAD | MKN28 | VX-11e |
| MVP_top60_DNA_seed | HSP90AA1 | 0.754425 | LUAD | NCI-H1573 | Luminespib | MVP_top60_DNA+RNA_seed | FGFR3 | -0.412827 | STAD | SK-GT-2 | FGFR_3831 |
| MVP_top60_DNA_seed | EGFR | 0.754637 | LUAD | SW1573 | Pelitinib | MVP_top60_DNA+RNA_seed | PARP1 | -0.412779 | STAD | RERF-GC-1B | PARP_9495 |
| MVP_top60_DNA_seed | HDAC1 | 0.754933 | LUAD | NCI-H1563 | JQ12 | MVP_top60_DNA+RNA_seed | HDAC1 | -0.412713 | STAD | SNU-1 | CAY10603 |
| MVP_top60_DNA_seed | AURKA | 0.755228 | LUAD | NCI-H1793 | Alisertib | MVP_top60_DNA+RNA_seed | HDAC1 | -0.409489 | STAD | HGC-27 | CAY10603 |
| MVP_top60_DNA_seed | EGFR | 0.75606 | LUAD | NCI-H1573 | Cetuximab | MVP_top60_DNA+RNA_seed | PIK3CA | -0.406898 | STAD | TMK-1 | PI3Ka_4409 |
| MVP_top60_DNA_seed | HDAC1 | 0.756284 | LUAD | Calu-3 | Belinostat | MVP_top60_DNA+RNA_seed | HDAC1 | -0.406798 | STAD | GCIY | AR-42 |
| MVP_top60_DNA_seed | BRAF | 0.756986 | LUAD | NCI-H1573 | PLX-4720 | MVP_top60_DNA+RNA_seed | ERBB2 | -0.406648 | STAD | FU97 | CI-1033 |
| MVP_top60_DNA_seed | AURKA | 0.75799 | LUAD | RERF-LC-MS | Alisertib | MVP_top60_DNA+RNA_seed | ERBB4 | -0.406648 | STAD | FU97 | CI-1033 |
| MVP_top60_DNA_seed | BRAF | 0.759542 | LUAD | NCI-H1435 | RAF_9304 | MVP_top60_DNA+RNA_seed | AR | -0.403684 | STAD | HSC-39 | AZD3514 |
| MVP_top60_DNA_seed | FGFR1 | 0.759761 | LUAD | NCI-H1993 | PD173074 | MVP_top60_DNA+RNA_seed | AR | -0.398416 | STAD | Hs-746T | AZD3514 |
| MVP_top60_DNA_seed | SMARCA4 | 0.759761 | LUAD | RERF-LC-KJ | PFI-3 | MVP_top60_DNA+RNA_seed | ATM | -0.396905 | STAD | RERF-GC-1B | KU-60019 |
| MVP_top60_DNA_seed | BRAF | 0.760082 | LUAD | COR-L105 | PLX-4720 | MVP_top60_DNA+RNA_seed | AR | -0.395423 | STAD | OCUM-1 | Bicalutamide |
| MVP_top60_DNA_seed | EGFR | 0.760124 | LUAD | ABC-1 | Gefitinib | MVP_top60_DNA+RNA_seed | HDAC1 | -0.39373 | STAD | AGS | AR-42 |
| MVP_top60_DNA_seed | BRAF | 0.760176 | LUAD | NCI-H1651 | RAF_9304 | MVP_top60_DNA+RNA_seed | HSP90AA1 | -0.393635 | STAD | RERF-GC-1B | Luminespib |
| MVP_top60_DNA_seed | FGFR1 | 0.760941 | LUAD | NCI-H2023 | AZD4547 | MVP_top60_DNA+RNA_seed | HSP90AB1 | -0.393635 | STAD | RERF-GC-1B | Luminespib |
| MVP_top60_DNA_seed | EGFR | 0.761033 | LUAD | NCI-H2122 | CUDC-101 | MVP_top60_DNA+RNA_seed | PARP1 | -0.388306 | STAD | Hs-746T | PARP_9482 |
| MVP_top60_DNA_seed | ERBB2 | 0.761033 | LUAD | NCI-H2122 | CUDC-101 | MVP_top60_DNA+RNA_seed | HSP90AA1 | -0.382033 | STAD | GCIY | CCT-018159 |
| MVP_top60_DNA_seed | EGFR | 0.7612 | LUAD | RERF-LC-MS | Pelitinib | MVP_top60_DNA+RNA_seed | HSP90AB1 | -0.382033 | STAD | GCIY | CCT-018159 |
| MVP_top60_DNA_seed | HSP90AA1 | 0.761274 | LUAD | NCI-H2085 | CCT-018159 | MVP_top60_DNA+RNA_seed | SRC | -0.379704 | STAD | MKN28 | Bosutinib |
| MVP_top60_DNA_seed | HSP90AA1 | 0.762372 | LUAD | NCI-H1573 | Elesclomol | MVP_top60_DNA+RNA_seed | PARP1 | -0.377375 | STAD | RERF-GC-1B | Olaparib |
| MVP_top60_DNA_seed | EGFR | 0.76361 | LUAD | NCI-H1781 | Cetuximab | MVP_top60_DNA+RNA_seed | PARP1 | -0.373349 | STAD | RF-48 | Veliparib |
| MVP_top60_DNA_seed | MDM2 | 0.763852 | LUAD | SK-LU-1 | Serdemetan | MVP_top60_DNA+RNA_seed | HSP90AA1 | -0.368181 | STAD | SNU-16 | Tanespimycin |
| MVP_top60_DNA_seed | MAPK1 | 0.76445 | LUAD | NCI-H1573 | VX-11e | MVP_top60_DNA+RNA_seed | HSP90AB1 | -0.368181 | STAD | SNU-16 | Tanespimycin |
| MVP_top60_DNA_seed | BRAF | 0.765517 | LUAD | NCI-H2342 | SB590885 | MVP_top60_DNA+RNA_seed | SRC | -0.367975 | STAD | RF-48 | Bosutinib |
| MVP_top60_DNA_seed | PIK3CA | 0.766511 | LUAD | RERF-LC-KJ | AZD8835 | MVP_top60_DNA+RNA_seed | PIK3CA | -0.366374 | STAD | FU97 | PI3Ka_4409 |
| MVP_top60_DNA_seed | IGF1R | 0.766837 | LUAD | HCC-78 | GSK1904529A | MVP_top60_DNA+RNA_seed | MAPK1 | -0.365888 | STAD | RF-48 | VX-11e |
| MVP_top60_DNA_seed | FGFR1 | 0.767411 | LUAD | NCI-H1975 | PD173074 | MVP_top60_DNA+RNA_seed | ATM | -0.364419 | STAD | SK-GT-2 | KU-60019 |
| MVP_top60_DNA_seed | GSK3B | 0.767994 | LUAD | NCI-H2009 | CHIR-99021 | MVP_top60_DNA+RNA_seed | SRC | -0.364069 | STAD | SNU-16 | Bosutinib |
| MVP_top60_DNA_seed | MET | 0.768618 | LUAD | SK-LU-1 | AZD6094 | MVP_top60_DNA+RNA_seed | GSK3B | -0.362652 | STAD | MKN7 | KIN001-042 |
| MVP_top60_DNA_seed | FGFR1 | 0.769071 | LUAD | SW1573 | PD173074 | MVP_top60_DNA+RNA_seed | HSP90AA1 | -0.362506 | STAD | NUGC-4 | Tanespimycin |

| | | | | | | | | | | | |
|---|---|---|---|---|---|---|---|---|---|---|---|
| MVP_top60_DNA_seed | AURKA | 0.76933 | LUAD | NCI-H441 | ZM447439 | MVP_top60_DNA+RNA_seed | HSP90AB1 | -0.362506 | STAD | NUGC-4 | Tanespimycin |
| MVP_top60_DNA_seed | AKT1 | 0.769729 | LUAD | EMC-BAC-2 | GSK690693 | MVP_top60_DNA+RNA_seed | GSK3B | -0.362202 | STAD | NUGC-4 | CHIR-99021 |
| MVP_top60_DNA_seed | EGFR | 0.772207 | LUAD | NCI-H2228 | Gefitinib | MVP_top60_DNA+RNA_seed | FGFR3 | -0.361398 | STAD | 23132-87 | AZD4547 |
| MVP_top60_DNA_seed | EGFR | 0.772473 | LUAD | NCI-H2405 | Afatinib | MVP_top60_DNA+RNA_seed | HDAC1 | -0.360311 | STAD | 23132-87 | Belinostat |
| MVP_top60_DNA_seed | ERBB2 | 0.772473 | LUAD | NCI-H2405 | Afatinib | MVP_top60_DNA+RNA_seed | AR | -0.359151 | STAD | OCUM-1 | AZD3514 |
| MVP_top60_DNA_seed | HDAC1 | 0.77299 | LUAD | Calu-3 | CAY10603 | MVP_top60_DNA+RNA_seed | FGFR3 | -0.358376 | STAD | TMK-1 | AZD4547 |
| MVP_top60_DNA_seed | MDM2 | 0.773315 | LUAD | NCI-H1651 | Nutlin-3a (-) | MVP_top60_DNA+RNA_seed | HDAC1 | -0.35831 | STAD | AGS | CAY10603 |
| MVP_top60_DNA_seed | BRAF | 0.773968 | LUAD | RERF-LC-KJ | PLX-4720 | MVP_top60_DNA+RNA_seed | MDM2 | -0.357478 | STAD | GCIY | Serdemetan |
| MVP_top60_DNA_seed | CREBBP | 0.775046 | LUAD | NCI-H1975 | I-CBP112 | MVP_top60_DNA+RNA_seed | HDAC1 | -0.357175 | STAD | GCIY | CAY10603 |
| MVP_top60_DNA_seed | BRAF | 0.77526 | LUAD | NCI-H650 | PLX-4720 | MVP_top60_DNA+RNA_seed | GSK3B | -0.355932 | STAD | AGS | AZD7969 |
| MVP_top60_DNA_seed | IGF1R | 0.775306 | LUAD | NCI-H1975 | GSK1904529A | MVP_top60_DNA+RNA_seed | HDAC1 | -0.354964 | STAD | AGS | Belinostat |
| MVP_top60_DNA_seed | EGFR | 0.775334 | LUAD | NCI-H1651 | CI-1033 | MVP_top60_DNA+RNA_seed | GSK3B | -0.353429 | STAD | MKN28 | AZD7969 |
| MVP_top60_DNA_seed | ERBB2 | 0.775334 | LUAD | NCI-H1651 | CI-1033 | MVP_top60_DNA+RNA_seed | MAPK1 | -0.351032 | STAD | NCI-N87 | VX-11e |
| MVP_top60_DNA_seed | IGF1R | 0.775697 | LUAD | NCI-H1838 | BMS-536924 | MVP_top60_DNA+RNA_seed | HDAC1 | -0.348672 | STAD | 23132-87 | AR-42 |
| MVP_top60_DNA_seed | SRC | 0.776324 | LUAD | NCI-H23 | Saracatinib | MVP_top60_DNA+RNA_seed | ATM | -0.34516 | STAD | IM-95 | KU-55933 |
| MVP_top60_DNA_seed | MET | 0.777276 | LUAD | H3255 | AZD6094 | MVP_top60_DNA+RNA_seed | FGFR3 | -0.344998 | STAD | NUGC-3 | AZD4547 |
| MVP_top60_DNA_seed | GSK3B | 0.778259 | LUAD | NCI-H1435 | KIN001-042 | MVP_top60_DNA+RNA_seed | PARP1 | -0.344394 | STAD | NUGC-4 | Talazoparib |
| MVP_top60_DNA_seed | EGFR | 0.778687 | LUAD | NCI-H322M | AZD8931 | MVP_top60_DNA+RNA_seed | ATM | -0.344206 | STAD | FU97 | CP466722 |
| MVP_top60_DNA_seed | ERBB2 | 0.778687 | LUAD | NCI-H322M | AZD8931 | MVP_top60_DNA+RNA_seed | HDAC1 | -0.342954 | STAD | 23132-87 | CAY10603 |
| MVP_top60_DNA_seed | ERBB3 | 0.778687 | LUAD | NCI-H322M | AZD8931 | MVP_top60_DNA+RNA_seed | GSK3B | -0.33502 | STAD | 23132-87 | CHIR-99021 |
| MVP_top60_DNA_seed | AURKA | 0.778765 | LUAD | NCI-H2030 | CD532 | MVP_top60_DNA+RNA_seed | HDAC1 | -0.334021 | STAD | SNU-1 | AR-42 |
| MVP_top60_DNA_seed | EGFR | 0.77956 | LUAD | SW1573 | Afatinib | MVP_top60_DNA+RNA_seed | MAPK1 | -0.332078 | STAD | 23132-87 | VX-11e |
| MVP_top60_DNA_seed | ERBB2 | 0.77956 | LUAD | SW1573 | Afatinib | MVP_top60_DNA+RNA_seed | PARP1 | -0.331209 | STAD | AGS | Rucaparib |
| MVP_top60_DNA_seed | AKT1 | 0.779613 | LUAD | RERF-LC-KJ | GSK690693 | MVP_top60_DNA+RNA_seed | HSP90AA1 | -0.32898 | STAD | IM-95 | Tanespimycin |
| MVP_top60_DNA_seed | GSK3B | 0.779691 | LUAD | RERF-LC-MS | CHIR-99021 | MVP_top60_DNA+RNA_seed | HSP90AB1 | -0.32898 | STAD | IM-95 | Tanespimycin |
| MVP_top60_DNA_seed | AURKA | 0.779776 | LUAD | NCI-H2023 | GSK1070916 | MVP_top60_DNA+RNA_seed | HDAC1 | -0.327405 | STAD | SNU-1 | Belinostat |
| MVP_top60_DNA_seed | MDM2 | 0.780645 | LUAD | NCI-H2347 | Nutlin-3a (-) | MVP_top60_DNA+RNA_seed | FGFR3 | -0.320579 | STAD | OCUM-1 | FGFR_3831 |
| MVP_top60_DNA_seed | MAPK1 | 0.781374 | LUAD | NCI-H2347 | VX-11e | MVP_top60_DNA+RNA_seed | PARP1 | -0.319511 | STAD | 23132-87 | Talazoparib |
| MVP_top60_DNA_seed | BRAF | 0.782218 | LUAD | ABC-1 | SB590885 | MVP_top60_DNA+RNA_seed | PARP1 | -0.319272 | STAD | SNU-5 | PARP_0108 |
| MVP_top60_DNA_seed | IGF1R | 0.782822 | LUAD | NCI-H650 | BMS-536924 | MVP_top60_DNA+RNA_seed | ERBB2 | -0.314995 | STAD | MKN7 | PF-00299804 |
| MVP_top60_DNA_seed | BRAF | 0.782958 | LUAD | NCI-H1568 | PLX-4720 | MVP_top60_DNA+RNA_seed | ERBB4 | -0.314995 | STAD | MKN7 | PF-00299804 |
| MVP_top60_DNA_seed | CREBBP | 0.784039 | LUAD | NCI-H1563 | I-CBP112 | MVP_top60_DNA+RNA_seed | ERBB2 | -0.314747 | STAD | IM-95 | CUDC-101 |
| MVP_top60_DNA_seed | HDAC1 | 0.785938 | LUAD | NCI-H1993 | Tubastatin A | MVP_top60_DNA+RNA_seed | ESR1 | -0.313742 | STAD | SK-GT-2 | Fulvestrant |
| MVP_top60_DNA_seed | MDM2 | 0.78596 | LUAD | SW1573 | Serdemetan | MVP_top60_DNA+RNA_seed | HDAC1 | -0.313686 | STAD | IM-95 | AR-42 |
| MVP_top60_DNA_seed | FGFR1 | 0.786032 | LUAD | NCI-H1781 | PD173074 | MVP_top60_DNA+RNA_seed | PARP1 | -0.308318 | STAD | FU97 | Rucaparib |
| MVP_top60_DNA_seed | AKT1 | 0.787304 | LUAD | NCI-H522 | MK-2206 | MVP_top60_DNA+RNA_seed | ESR1 | -0.307974 | STAD | NUGC-3 | Fulvestrant |
| MVP_top60_DNA_seed | HSP90AA1 | 0.787892 | LUAD | NCI-H1568 | Tanespimycin | MVP_top60_DNA+RNA_seed | ATM | -0.307508 | STAD | 23132-87 | KU-60019 |
| MVP_top60_DNA_seed | AURKA | 0.790867 | LUAD | LXF-289 | CD532 | MVP_top60_DNA+RNA_seed | HSP90AA1 | -0.30662 | STAD | NUGC-4 | Luminespib |
| MVP_top60_DNA_seed | FGFR1 | 0.791474 | LUAD | NCI-H1435 | PD173074 | MVP_top60_DNA+RNA_seed | HSP90AB1 | -0.30662 | STAD | NUGC-4 | Luminespib |
| MVP_top60_DNA_seed | SRC | 0.791899 | LUAD | NCI-H1437 | Bosutinib | MVP_top60_DNA+RNA_seed | PARP1 | -0.305616 | STAD | SNU-5 | Olaparib |
| MVP_top60_DNA_seed | ERBB2 | 0.792386 | LUAD | NCI-H1781 | CP724714 | MVP_top60_DNA+RNA_seed | MDM2 | -0.30519 | STAD | IM-95 | Serdemetan |

| | | | | | | | | | | | |
|---|---|---|---|---|---|---|---|---|---|---|---|
| MVP_top60_DNA_seed | GSK3B | 0.792937 | LUAD | NCI-H1666 | AZD7969 | MVP_top60_DNA+RNA_seed | HSP90AA1 | -0.302075 | STAD | MKN7 | CCT-018159 |
| MVP_top60_DNA_seed | BRAF | 0.79421 | LUAD | Calu-6 | SB590885 | MVP_top60_DNA+RNA_seed | HSP90AB1 | -0.302075 | STAD | MKN7 | CCT-018159 |
| MVP_top60_DNA_seed | HDAC1 | 0.794609 | LUAD | NCI-H1563 | AR-42 | MVP_top60_DNA+RNA_seed | FYN | -0.295457 | STAD | RF-48 | A-770041 |
| MVP_top60_DNA_seed | SMARCA4 | 0.794778 | LUAD | NCI-H2405 | PFI-3 | MVP_top60_DNA+RNA_seed | GSK3B | -0.287943 | STAD | NCI-N87 | SB216763 |
| MVP_top60_DNA_seed | BRAF | 0.795398 | LUAD | NCI-H23 | PLX-4720 | MVP_top60_DNA+RNA_seed | MAPK1 | -0.286058 | STAD | NUGC-4 | VX-11e |
| MVP_top60_DNA_seed | GSK3B | 0.795773 | LUAD | NCI-H838 | KIN001-042 | MVP_top60_DNA+RNA_seed | ERBB4 | -0.286039 | STAD | KATOIII | AST-1306 |
| MVP_top60_DNA_seed | HDAC1 | 0.797109 | LUAD | H3255 | Belinostat | MVP_top60_DNA+RNA_seed | ESR1 | -0.285092 | STAD | RERF-GC-1B | Tamoxifen |
| MVP_top60_DNA_seed | AURKA | 0.797567 | LUAD | NCI-H2030 | Alisertib | MVP_top60_DNA+RNA_seed | HDAC1 | -0.284617 | STAD | GCIY | Belinostat |
| MVP_top60_DNA_seed | PIK3CA | 0.797794 | LUAD | NCI-H1435 | PI3Ka_4409 | MVP_top60_DNA+RNA_seed | ERBB2 | -0.284447 | STAD | 23132-87 | AZD8931 |
| MVP_top60_DNA_seed | FGFR1 | 0.798079 | LUAD | NCI-H2347 | AZD4547 | MVP_top60_DNA+RNA_seed | ERBB3 | -0.284447 | STAD | 23132-87 | AZD8931 |
| MVP_top60_DNA_seed | EGFR | 0.79841 | LUAD | NCI-H838 | Afatinib | MVP_top60_DNA+RNA_seed | SRC | -0.276615 | STAD | TMK-1 | Bosutinib |
| MVP_top60_DNA_seed | ERBB2 | 0.79841 | LUAD | NCI-H838 | Afatinib | MVP_top60_DNA+RNA_seed | ATM | -0.265428 | STAD | FU97 | KU-60019 |
| MVP_top60_DNA_seed | BRAF | 0.79909 | LUAD | RERF-LC-KJ | Dabrafenib | MVP_top60_DNA+RNA_seed | MET | -0.265114 | STAD | SNU-16 | PHA-665752 |
| MVP_top60_DNA_seed | HSP90AA1 | 0.79915 | LUAD | NCI-H2009 | CCT-018159 | MVP_top60_DNA+RNA_seed | AR | -0.263762 | STAD | HSC-39 | Bicalutamide |
| MVP_top60_DNA_seed | BRAF | 0.799203 | LUAD | NCI-H1975 | PLX-4720 | MVP_top60_DNA+RNA_seed | FGFR3 | -0.262097 | STAD | SNU-5 | PD173074 |
| MVP_top60_DNA_seed | GSK3B | 0.799651 | LUAD | NCI-H1993 | CHIR-99021 | MVP_top60_DNA+RNA_seed | PARP1 | -0.260189 | STAD | SNU-5 | Talazoparib |
| MVP_top60_DNA_seed | AKT1 | 0.799662 | LUAD | RERF-LC-MS | GSK690693 | MVP_top60_DNA+RNA_seed | PTK2 | -0.258991 | STAD | FU97 | PF-562271 |
| MVP_top60_DNA_seed | EGFR | 0.799711 | LUAD | HOP-62 | Afatinib | MVP_top60_DNA+RNA_seed | SRC | -0.25301 | STAD | RF-48 | Saracatinib |
| MVP_top60_DNA_seed | ERBB2 | 0.799711 | LUAD | HOP-62 | Afatinib | MVP_top60_DNA+RNA_seed | FGFR3 | -0.251745 | STAD | SNU-5 | AZD4547 |
| MVP_top60_DNA_seed | GSK3B | 0.800158 | LUAD | NCI-H1755 | KIN001-042 | MVP_top60_DNA+RNA_seed | HDAC1 | -0.249618 | STAD | HGC-27 | AR-42 |
| MVP_top60_DNA_seed | AKT1 | 0.801137 | LUAD | NCI-H1838 | AKT inhibitor VIII | MVP_top60_DNA+RNA_seed | ATM | -0.246075 | STAD | NUGC-4 | KU-60019 |
| MVP_top60_DNA_seed | HDAC1 | 0.801623 | LUAD | NCI-H1435 | AR-42 | MVP_top60_DNA+RNA_seed | MDM2 | -0.245297 | STAD | SK-GT-2 | Nutlin-3a (-) |
| MVP_top60_DNA_seed | HDAC1 | 0.802154 | LUAD | NCI-H1781 | Tubastatin A | MVP_top60_DNA+RNA_seed | HDAC1 | -0.244948 | STAD | NUGC-3 | Dacinostat |
| MVP_top60_DNA_seed | SMARCA4 | 0.802652 | LUAD | EMC-BAC-1 | PFI-3 | MVP_top60_DNA+RNA_seed | ERBB2 | -0.24229 | STAD | RERF-GC-1B | AZD8931 |
| MVP_top60_DNA_seed | HSP90AA1 | 0.802904 | LUAD | NCI-H2342 | CCT-018159 | MVP_top60_DNA+RNA_seed | ERBB3 | -0.24229 | STAD | RERF-GC-1B | AZD8931 |
| MVP_top60_DNA_seed | AURKA | 0.804246 | LUAD | NCI-H1793 | GSK1070916 | MVP_top60_DNA+RNA_seed | PARP1 | -0.237646 | STAD | OCUM-1 | Olaparib |
| MVP_top60_DNA_seed | BRAF | 0.804856 | LUAD | NCI-H522 | PLX-4720 | MVP_top60_DNA+RNA_seed | HDAC1 | -0.236947 | STAD | NUGC-4 | Tubastatin A |
| MVP_top60_DNA_seed | MAPK1 | 0.805654 | LUAD | NCI-H1666 | VX-11e | MVP_top60_DNA+RNA_seed | MET | -0.236798 | STAD | FU97 | JNJ38877605 |
| MVP_top60_DNA_seed | AURKA | 0.805817 | LUAD | NCI-H1838 | Alisertib | MVP_top60_DNA+RNA_seed | ERBB2 | -0.23349 | STAD | SNU-16 | AZD8931 |
| MVP_top60_DNA_seed | HDAC1 | 0.805953 | LUAD | NCI-H1993 | JQ12 | MVP_top60_DNA+RNA_seed | ERBB3 | -0.23349 | STAD | SNU-16 | AZD8931 |
| MVP_top60_DNA_seed | EGFR | 0.806045 | LUAD | NCI-H1573 | CUDC-101 | MVP_top60_DNA+RNA_seed | MDM2 | -0.2327 | STAD | HGC-27 | Nutlin-3a (-) |
| MVP_top60_DNA_seed | ERBB2 | 0.806045 | LUAD | NCI-H1573 | CUDC-101 | MVP_top60_DNA+RNA_seed | HDAC1 | -0.226962 | STAD | 23132-87 | Tubastatin A |
| MVP_top60_DNA_seed | HDAC1 | 0.806122 | LUAD | Calu-6 | Entinostat | MVP_top60_DNA+RNA_seed | HDAC1 | -0.223547 | STAD | 23132-87 | JQ12 |
| MVP_top60_DNA_seed | HDAC1 | 0.806944 | LUAD | H3255 | Tubastatin A | MVP_top60_DNA+RNA_seed | HDAC1 | -0.22156 | STAD | HGC-27 | Belinostat |
| MVP_top60_DNA_seed | BRAF | 0.806972 | LUAD | NCI-H838 | Dabrafenib | MVP_top60_DNA+RNA_seed | HDAC1 | -0.22062 | STAD | NUGC-4 | Belinostat |
| MVP_top60_DNA_seed | FGFR1 | 0.807423 | LUAD | NCI-H1563 | PD173074 | MVP_top60_DNA+RNA_seed | ERBB4 | -0.220035 | STAD | TGBC11TKB | AST-1306 |
| MVP_top60_DNA_seed | EGFR | 0.808434 | LUAD | NCI-H2342 | Pelitinib | MVP_top60_DNA+RNA_seed | HSP90AA1 | -0.217351 | STAD | IM-95 | Luminespib |
| MVP_top60_DNA_seed | MET | 0.810252 | LUAD | Calu-3 | AZD6094 | MVP_top60_DNA+RNA_seed | HSP90AB1 | -0.217351 | STAD | IM-95 | Luminespib |
| MVP_top60_DNA_seed | MDM2 | 0.810913 | LUAD | NCI-H1573 | Serdemetan | MVP_top60_DNA+RNA_seed | ERBB2 | -0.21364 | STAD | TGBC11TKB | CI-1033 |
| MVP_top60_DNA_seed | EGFR | 0.812571 | LUAD | RERF-LC-KJ | CUDC-101 | MVP_top60_DNA+RNA_seed | ERBB4 | -0.21364 | STAD | TGBC11TKB | CI-1033 |
| MVP_top60_DNA_seed | ERBB2 | 0.812571 | LUAD | RERF-LC-KJ | CUDC-101 | MVP_top60_DNA+RNA_seed | PARP1 | -0.207898 | STAD | MKN28 | Olaparib |

| | | | | | | | | | | | |
|---|---|---|---|---|---|---|---|---|---|---|---|
| MVP_top60_DNA_seed | AURKA | 0.813367 | LUAD | RERF-LC-KJ | CD532 | MVP_top60_DNA+RNA_seed | MAPK1 | -0.207298 | STAD | TGBC11TKB | VX-11e |
| MVP_top60_DNA_seed | HDAC1 | 0.813411 | LUAD | NCI-H1993 | Belinostat | MVP_top60_DNA+RNA_seed | ERBB4 | -0.206675 | STAD | MKN7 | AST-1306 |
| MVP_top60_DNA_seed | FGFR1 | 0.814173 | LUAD | PC-14 | AZD4547 | MVP_top60_DNA+RNA_seed | PARP1 | -0.206452 | STAD | OCUM-1 | Olaparib |
| MVP_top60_DNA_seed | HSP90AA1 | 0.814229 | LUAD | NCI-H1435 | Luminespib | MVP_top60_DNA+RNA_seed | ESR1 | -0.205362 | STAD | TMK-1 | Fulvestrant |
| MVP_top60_DNA_seed | HDAC1 | 0.815079 | LUAD | NCI-H441 | AR-42 | MVP_top60_DNA+RNA_seed | GSK3B | -0.204024 | STAD | HGC-27 | CHIR-99021 |
| MVP_top60_DNA_seed | ATM | 0.815365 | LUAD | ABC-1 | torin2 | MVP_top60_DNA+RNA_seed | PARP1 | -0.201939 | STAD | TMK-1 | Olaparib |
| MVP_top60_DNA_seed | ATM | 0.815385 | LUAD | NCI-H1573 | KU-55933 | MVP_top60_DNA+RNA_seed | ERBB2 | -0.200245 | STAD | NUGC-3 | CUDC-101 |
| MVP_top60_DNA_seed | MET | 0.815674 | LUAD | A549 | AZD6094 | MVP_top60_DNA+RNA_seed | PARP1 | -0.200034 | STAD | NUGC-4 | Veliparib |
| MVP_top60_DNA_seed | IGF1R | 0.816217 | LUAD | NCI-H1734 | GSK1904529A | MVP_top60_DNA+RNA_seed | HDAC1 | -0.19679 | STAD | 23132-87 | Dacinostat |
| MVP_top60_DNA_seed | FGFR1 | 0.817848 | LUAD | H3255 | AZD4547 | MVP_top60_DNA+RNA_seed | ERBB2 | -0.193304 | STAD | RERF-GC-1B | PF-00299804 |
| MVP_top60_DNA_seed | AURKA | 0.817927 | LUAD | LXF-289 | Alisertib | MVP_top60_DNA+RNA_seed | ERBB4 | -0.193304 | STAD | RERF-GC-1B | PF-00299804 |
| MVP_top60_DNA_seed | SRC | 0.818266 | LUAD | ABC-1 | NVP-BHG712 | MVP_top60_DNA+RNA_seed | SRC | -0.191266 | STAD | SNU-1 | Saracatinib |
| MVP_top60_DNA_seed | HDAC1 | 0.819202 | LUAD | NCI-H1623 | Dacinostat | MVP_top60_DNA+RNA_seed | PARP1 | -0.190851 | STAD | SK-GT-2 | Olaparib |
| MVP_top60_DNA_seed | BRAF | 0.819332 | LUAD | NCI-H1975 | SB590885 | MVP_top60_DNA+RNA_seed | GSK3B | -0.184608 | STAD | Hs-746T | KIN001-042 |
| MVP_top60_DNA_seed | EGFR | 0.819354 | LUAD | NCI-H1651 | AZD8931 | MVP_top60_DNA+RNA_seed | MET | -0.183091 | STAD | IM-95 | AZD6094 |
| MVP_top60_DNA_seed | ERBB2 | 0.819354 | LUAD | NCI-H1651 | AZD8931 | MVP_top60_DNA+RNA_seed | PIK3CA | -0.182956 | STAD | RERF-GC-1B | PI3Ka_4409 |
| MVP_top60_DNA_seed | ERBB3 | 0.819354 | LUAD | NCI-H1651 | AZD8931 | MVP_top60_DNA+RNA_seed | PARP1 | -0.179272 | STAD | MKN28 | Rucaparib |
| MVP_top60_DNA_seed | BRAF | 0.820048 | LUAD | NCI-H2085 | RAF_9304 | MVP_top60_DNA+RNA_seed | FGFR3 | -0.177653 | STAD | RERF-GC-1B | FGFR_3831 |
| MVP_top60_DNA_seed | MET | 0.82014 | LUAD | Calu-6 | Crizotinib | MVP_top60_DNA+RNA_seed | MAPK1 | -0.176446 | STAD | HGC-27 | VX-11e |
| MVP_top60_DNA_seed | IGF1R | 0.821256 | LUAD | NCI-H1793 | BMS-754807 | MVP_top60_DNA+RNA_seed | ERBB2 | -0.175941 | STAD | MKN7 | CI-1033 |
| MVP_top60_DNA_seed | EGFR | 0.821754 | LUAD | NCI-H522 | Pelitinib | MVP_top60_DNA+RNA_seed | ERBB4 | -0.175941 | STAD | MKN7 | CI-1033 |
| MVP_top60_DNA_seed | EGFR | 0.823436 | LUAD | NCI-H3122 | Cetuximab | MVP_top60_DNA+RNA_seed | AR | -0.174018 | STAD | FU97 | Bicalutamide |
| MVP_top60_DNA_seed | MDM2 | 0.823566 | LUAD | RERF-LC-KJ | Serdemetan | MVP_top60_DNA+RNA_seed | GSK3B | -0.169839 | STAD | IM-95 | KIN001-042 |
| MVP_top60_DNA_seed | CREBBP | 0.82431 | LUAD | EMC-BAC-1 | I-CBP112 | MVP_top60_DNA+RNA_seed | HDAC1 | -0.168944 | STAD | TGBC11TKB | AR-42 |
| MVP_top60_DNA_seed | AKT1 | 0.825336 | LUAD | NCI-H1568 | GSK690693 | MVP_top60_DNA+RNA_seed | PIK3CA | -0.166059 | STAD | FU97 | AZD8835 |
| MVP_top60_DNA_seed | BRAF | 0.826556 | LUAD | NCI-H1437 | Dabrafenib | MVP_top60_DNA+RNA_seed | ATM | -0.165111 | STAD | HGC-27 | CP466722 |
| MVP_top60_DNA_seed | EGFR | 0.826624 | LUAD | LC-2-ad | Gefitinib | MVP_top60_DNA+RNA_seed | ERBB4 | -0.163419 | STAD | Hs-746T | AST-1306 |
| MVP_top60_DNA_seed | HDAC1 | 0.826857 | LUAD | NCI-H2228 | CAY10603 | MVP_top60_DNA+RNA_seed | MAPK1 | -0.16321 | STAD | HGC-27 | FR-180204 |
| MVP_top60_DNA_seed | BRAF | 0.827151 | LUAD | NCI-H1666 | AZ628 | MVP_top60_DNA+RNA_seed | MAPK3 | -0.16321 | STAD | HGC-27 | FR-180204 |
| MVP_top60_DNA_seed | SRC | 0.827339 | LUAD | NCI-H838 | NVP-BHG712 | MVP_top60_DNA+RNA_seed | PARP1 | -0.161384 | STAD | TMK-1 | PARP_0108 |
| MVP_top60_DNA_seed | FGFR1 | 0.828822 | LUAD | NCI-H1573 | PD173074 | MVP_top60_DNA+RNA_seed | ERBB2 | -0.160746 | STAD | HGC-27 | CUDC-101 |
| MVP_top60_DNA_seed | BRAF | 0.829641 | LUAD | NCI-H1838 | Dabrafenib | MVP_top60_DNA+RNA_seed | MAPK1 | -0.155294 | STAD | SNU-1 | FR-180204 |
| MVP_top60_DNA_seed | SRC | 0.830013 | LUAD | NCI-H23 | WH-4-023 | MVP_top60_DNA+RNA_seed | MAPK3 | -0.155294 | STAD | SNU-1 | FR-180204 |
| MVP_top60_DNA_seed | HSP90AA1 | 0.830213 | LUAD | PC-3_[JPC-3] | Elesclomol | MVP_top60_DNA+RNA_seed | MET | -0.145601 | STAD | OCUM-1 | AZD6094 |
| MVP_top60_DNA_seed | AURKA | 0.830398 | LUAD | Calu-3 | HG-5-113-01 | MVP_top60_DNA+RNA_seed | SRC | -0.14443 | STAD | TGBC11TKB | NVP-BHG712 |
| MVP_top60_DNA_seed | BRAF | 0.831091 | LUAD | RERF-LC-MS | RAF_9304 | MVP_top60_DNA+RNA_seed | ERBB2 | -0.144342 | STAD | IM-95 | Afatinib |
| MVP_top60_DNA_seed | EGFR | 0.831549 | LUAD | NCI-H1651 | Gefitinib | MVP_top60_DNA+RNA_seed | ERBB2 | -0.138867 | STAD | AGS | CP724714 |
| MVP_top60_DNA_seed | AURKA | 0.832222 | LUAD | NCI-H2347 | Genentech Cpd 10 | MVP_top60_DNA+RNA_seed | SRC | -0.136525 | STAD | NCI-N87 | Bosutinib |
| MVP_top60_DNA_seed | HDAC1 | 0.832546 | LUAD | NCI-H1781 | AR-42 | MVP_top60_DNA+RNA_seed | ATM | -0.134592 | STAD | KATOIII | KU-60019 |
| MVP_top60_DNA_seed | AURKA | 0.832946 | LUAD | NCI-H23 | Tozasertib | MVP_top60_DNA+RNA_seed | ERBB2 | -0.134016 | STAD | FU97 | CP724714 |
| MVP_top60_DNA_seed | EGFR | 0.83486 | LUAD | NCI-H2405 | AST-1306 | MVP_top60_DNA+RNA_seed | FGFR3 | -0.132752 | STAD | OCUM-1 | AZD4547 |

| | | | | | | | | | | | |
|---|---|---|---|---|---|---|---|---|---|---|---|
| MVP_top60_DNA_seed | MET | 0.83503 | LUAD | Calu-6 | SU11274 | MVP_top60_DNA+RNA_seed | PIK3CA | -0.130699 | STAD | 23132-87 | PI3Ka_4409 |
| MVP_top60_DNA_seed | HSP90AA1 | 0.835229 | LUAD | NCI-H2405 | SNX-2112 | MVP_top60_DNA+RNA_seed | ATM | -0.129087 | STAD | HSC-39 | CP466722 |
| MVP_top60_DNA_seed | AURKA | 0.835387 | LUAD | Calu-3 | Genentech Cpd 10 | MVP_top60_DNA+RNA_seed | MAPK1 | -0.128698 | STAD | HSC-39 | FR-180204 |
| MVP_top60_DNA_seed | HDAC1 | 0.835478 | LUAD | NCI-H1792 | Dacinostat | MVP_top60_DNA+RNA_seed | MAPK3 | -0.128698 | STAD | HSC-39 | FR-180204 |
| MVP_top60_DNA_seed | CREBBP | 0.835819 | LUAD | NCI-H2405 | I-CBP112 | MVP_top60_DNA+RNA_seed | PIK3CA | -0.128391 | STAD | RERF-GC-1B | AZD8835 |
| MVP_top60_DNA_seed | MET | 0.836551 | LUAD | HCC-827 | AZD6094 | MVP_top60_DNA+RNA_seed | ERBB2 | -0.128348 | STAD | TGBC11TKB | PF-00299804 |
| MVP_top60_DNA_seed | EGFR | 0.837101 | LUAD | NCI-H1755 | Afatinib | MVP_top60_DNA+RNA_seed | ERBB4 | -0.128348 | STAD | TGBC11TKB | PF-00299804 |
| MVP_top60_DNA_seed | ERBB2 | 0.837101 | LUAD | NCI-H1755 | Afatinib | MVP_top60_DNA+RNA_seed | MAPK1 | -0.127507 | STAD | FU97 | FR-180204 |
| MVP_top60_DNA_seed | IGF1R | 0.837488 | LUAD | PC-14 | BMS-754807 | MVP_top60_DNA+RNA_seed | MAPK3 | -0.127507 | STAD | FU97 | FR-180204 |
| MVP_top60_DNA_seed | AURKA | 0.837547 | LUAD | NCI-H838 | Genentech Cpd 10 | MVP_top60_DNA+RNA_seed | ERBB2 | -0.125099 | STAD | HGC-27 | CI-1033 |
| MVP_top60_DNA_seed | HDAC1 | 0.838515 | LUAD | RERF-LC-KJ | JQ12 | MVP_top60_DNA+RNA_seed | ERBB4 | -0.125099 | STAD | HGC-27 | CI-1033 |
| MVP_top60_DNA_seed | AURKA | 0.839222 | LUAD | NCI-H1573 | CD532 | MVP_top60_DNA+RNA_seed | PARP1 | -0.123764 | STAD | FU97 | Olaparib |
| MVP_top60_DNA_seed | AKT1 | 0.839409 | LUAD | HCC-78 | AKT inhibitor VIII | MVP_top60_DNA+RNA_seed | PARP1 | -0.121965 | STAD | IM-95 | Rucaparib |
| MVP_top60_DNA_seed | EGFR | 0.839916 | LUAD | NCI-H3122 | Afatinib | MVP_top60_DNA+RNA_seed | ERBB2 | -0.120265 | STAD | GCIY | CUDC-101 |
| MVP_top60_DNA_seed | ERBB2 | 0.839916 | LUAD | NCI-H3122 | Afatinib | MVP_top60_DNA+RNA_seed | ERBB2 | -0.120054 | STAD | HSC-39 | Lapatinib |
| MVP_top60_DNA_seed | SRC | 0.839976 | LUAD | NCI-H2023 | NVP-BHG712 | MVP_top60_DNA+RNA_seed | PARP1 | -0.113512 | STAD | SNU-5 | PARP_9482 |
| MVP_top60_DNA_seed | GSK3B | 0.840332 | LUAD | NCI-H1734 | KIN001-042 | MVP_top60_DNA+RNA_seed | HSP90AA1 | -0.110473 | STAD | MKN45 | Elesclomol |
| MVP_top60_DNA_seed | IGF1R | 0.840642 | LUAD | NCI-H2228 | GSK1904529A | MVP_top60_DNA+RNA_seed | HSP90AB1 | -0.110473 | STAD | MKN45 | Elesclomol |
| MVP_top60_DNA_seed | SRC | 0.841284 | LUAD | RERF-LC-MS | NVP-BHG712 | MVP_top60_DNA+RNA_seed | PTK2 | -0.109239 | STAD | NUGC-3 | PF-562271 |
| MVP_top60_DNA_seed | HSP90AA1 | 0.842361 | LUAD | NCI-H2009 | SNX-2112 | MVP_top60_DNA+RNA_seed | FGFR3 | -0.106648 | STAD | HGC-27 | AZD4547 |
| MVP_top60_DNA_seed | AKT1 | 0.842469 | LUAD | NCI-H1435 | BAY AKT1 | MVP_top60_DNA+RNA_seed | HDAC1 | -0.10497 | STAD | TGBC11TKB | Belinostat |
| MVP_top60_DNA_seed | AURKA | 0.842575 | LUAD | EKVX | HG-5-113-01 | MVP_top60_DNA+RNA_seed | ATM | -0.10335 | STAD | MKN45 | KU-60019 |
| MVP_top60_DNA_seed | HDAC1 | 0.842725 | LUAD | NCI-H1651 | Tubastatin A | MVP_top60_DNA+RNA_seed | HSP90AA1 | -0.099072 | STAD | Hs-746T | CCT-018159 |
| MVP_top60_DNA_seed | ATM | 0.842743 | LUAD | NCI-H2342 | CP466722 | MVP_top60_DNA+RNA_seed | HSP90AB1 | -0.099072 | STAD | Hs-746T | CCT-018159 |
| MVP_top60_DNA_seed | PIK3CA | 0.843485 | LUAD | NCI-H2347 | AZD8835 | MVP_top60_DNA+RNA_seed | ERBB2 | -0.093925 | STAD | NUGC-3 | CI-1033 |
| MVP_top60_DNA_seed | EGFR | 0.843616 | LUAD | HCC-44 | PF-00299804 | MVP_top60_DNA+RNA_seed | ERBB4 | -0.093925 | STAD | NUGC-3 | CI-1033 |
| MVP_top60_DNA_seed | ERBB2 | 0.843616 | LUAD | HCC-44 | PF-00299804 | MVP_top60_DNA+RNA_seed | MET | -0.092801 | STAD | SNU-1 | PHA-665752 |
| MVP_top60_DNA_seed | ERBB2 | 0.844253 | LUAD | NCI-H1435 | CP724714 | MVP_top60_DNA+RNA_seed | HSP90AA1 | -0.08514 | STAD | NUGC-3 | Luminespib |
| MVP_top60_DNA_seed | EGFR | 0.8443 | LUAD | Calu-6 | Cetuximab | MVP_top60_DNA+RNA_seed | HSP90AB1 | -0.08514 | STAD | NUGC-3 | Luminespib |
| MVP_top60_DNA_seed | HDAC1 | 0.845933 | LUAD | NCI-H2122 | AR-42 | MVP_top60_DNA+RNA_seed | SRC | -0.083556 | STAD | FU97 | NVP-BHG712 |
| MVP_top60_DNA_seed | GSK3B | 0.846139 | LUAD | HCC-78 | CHIR-99021 | MVP_top60_DNA+RNA_seed | SRC | -0.081569 | STAD | AGS | Bosutinib |
| MVP_top60_DNA_seed | BRAF | 0.847373 | LUAD | SK-LU-1 | SB590885 | MVP_top60_DNA+RNA_seed | PARP1 | -0.080206 | STAD | OCUM-1 | PARP_0108 |
| MVP_top60_DNA_seed | SMARCA4 | 0.848365 | LUAD | NCI-H1573 | PFI-3 | MVP_top60_DNA+RNA_seed | AR | -0.077787 | STAD | NUGC-3 | AZD3514 |
| MVP_top60_DNA_seed | HDAC1 | 0.849077 | LUAD | NCI-H2122 | Belinostat | MVP_top60_DNA+RNA_seed | MET | -0.07549 | STAD | HSC-39 | PHA-665752 |
| MVP_top60_DNA_seed | MET | 0.849903 | LUAD | NCI-H1944 | SU11274 | MVP_top60_DNA+RNA_seed | FGFR3 | -0.074017 | STAD | MKN28 | AZD4547 |
| MVP_top60_DNA_seed | IGF1R | 0.850172 | LUAD | RERF-LC-MS | Linsitinib | MVP_top60_DNA+RNA_seed | HDAC1 | -0.067291 | STAD | IM-95 | Belinostat |
| MVP_top60_DNA_seed | SMARCA4 | 0.850386 | LUAD | NCI-H2023 | PFI-3 | MVP_top60_DNA+RNA_seed | MET | -0.065729 | STAD | AGS | SU11274 |
| MVP_top60_DNA_seed | AKT1 | 0.850466 | LUAD | NCI-H2087 | GSK690693 | MVP_top60_DNA+RNA_seed | ERBB2 | -0.065151 | STAD | AGS | Afatinib |
| MVP_top60_DNA_seed | AURKA | 0.85192 | LUAD | NCI-H522 | GSK1070916 | MVP_top60_DNA+RNA_seed | MET | -0.060469 | STAD | SK-GT-2 | AZD6094 |
| MVP_top60_DNA_seed | MET | 0.85194 | LUAD | PC-3_[JPC-3] | AZD6094 | MVP_top60_DNA+RNA_seed | EP300 | -0.059215 | STAD | RF-48 | I-CBP112 |
| MVP_top60_DNA_seed | GSK3B | 0.852008 | LUAD | NCI-H1355 | AZD7969 | MVP_top60_DNA+RNA_seed | CREBBP | -0.059215 | STAD | RF-48 | I-CBP112 |

| | | | | | | | | | | | |
|---|---|---|---|---|---|---|---|---|---|---|---|
| MVP_top60_DNA_seed | MET | 0.85233 | LUAD | NCI-H1568 | SU11274 | MVP_top60_DNA+RNA_seed | MAPK1 | -0.059028 | STAD | 23132-87 | FR-180204 |
| MVP_top60_DNA_seed | MDM2 | 0.85307 | LUAD | A549 | Serdemetan | MVP_top60_DNA+RNA_seed | MAPK3 | -0.059028 | STAD | 23132-87 | FR-180204 |
| MVP_top60_DNA_seed | EGFR | 0.854444 | LUAD | NCI-H522 | Cetuximab | MVP_top60_DNA+RNA_seed | PTK2 | -0.058158 | STAD | MKN45 | PF-562271 |
| MVP_top60_DNA_seed | MAPK1 | 0.855012 | LUAD | HCC-78 | FR-180204 | MVP_top60_DNA+RNA_seed | AR | -0.057164 | STAD | SNU-16 | AZD3514 |
| MVP_top60_DNA_seed | EGFR | 0.856371 | LUAD | NCI-H1435 | Pelitinib | MVP_top60_DNA+RNA_seed | ATM | -0.053922 | STAD | KATOIII | KU-55933 |
| MVP_top60_DNA_seed | EGFR | 0.858042 | LUAD | RERF-LC-MS | CUDC-101 | MVP_top60_DNA+RNA_seed | HSP90AA1 | -0.051229 | STAD | Hs-746T | Elesclomol |
| MVP_top60_DNA_seed | ERBB2 | 0.858042 | LUAD | RERF-LC-MS | CUDC-101 | MVP_top60_DNA+RNA_seed | HSP90AB1 | -0.051229 | STAD | Hs-746T | Elesclomol |
| MVP_top60_DNA_seed | HSP90AA1 | 0.858644 | LUAD | NCI-H1650 | SNX-2112 | MVP_top60_DNA+RNA_seed | ERBB2 | -0.05119 | STAD | 23132-87 | CI-1033 |
| MVP_top60_DNA_seed | BRAF | 0.858686 | LUAD | NCI-H322M | PLX-4720 | MVP_top60_DNA+RNA_seed | ERBB4 | -0.05119 | STAD | 23132-87 | CI-1033 |
| MVP_top60_DNA_seed | AKT1 | 0.859001 | LUAD | PC-14 | AKT inhibitor VIII | MVP_top60_DNA+RNA_seed | HDAC1 | -0.050676 | STAD | HGC-27 | Tubastatin A |
| MVP_top60_DNA_seed | HDAC1 | 0.859444 | LUAD | SK-LU-1 | AR-42 | MVP_top60_DNA+RNA_seed | PIK3CA | -0.050562 | STAD | Hs-746T | PI3Ka_4409 |
| MVP_top60_DNA_seed | BRAF | 0.860084 | LUAD | PC-3_[JPC-3] | Dabrafenib | MVP_top60_DNA+RNA_seed | ERBB2 | -0.047554 | STAD | HSC-39 | AZD8931 |
| MVP_top60_DNA_seed | PIK3CA | 0.860304 | LUAD | HOP-62 | PI3Ka_4409 | MVP_top60_DNA+RNA_seed | ERBB3 | -0.047554 | STAD | HSC-39 | AZD8931 |
| MVP_top60_DNA_seed | HDAC1 | 0.8607 | LUAD | NCI-H838 | Dacinostat | MVP_top60_DNA+RNA_seed | HDAC1 | -0.034401 | STAD | NUGC-4 | CAY10603 |
| MVP_top60_DNA_seed | SRC | 0.861088 | LUAD | NCI-H1734 | Bosutinib | MVP_top60_DNA+RNA_seed | ERBB2 | -0.032571 | STAD | HSC-39 | CP724714 |
| MVP_top60_DNA_seed | MET | 0.863324 | LUAD | A549 | SU11274 | MVP_top60_DNA+RNA_seed | GSK3B | -0.030606 | STAD | SK-GT-2 | AZD7969 |
| MVP_top60_DNA_seed | FGFR1 | 0.864414 | LUAD | NCI-H1666 | AZD4547 | MVP_top60_DNA+RNA_seed | GSK3B | -0.029111 | STAD | HSC-39 | CHIR-99021 |
| MVP_top60_DNA_seed | AKT1 | 0.864507 | LUAD | VMRC-LCD | BAY AKT1 | MVP_top60_DNA+RNA_seed | EP300 | -0.026335 | STAD | HSC-39 | I-CBP112 |
| MVP_top60_DNA_seed | AURKA | 0.865915 | LUAD | RERF-LC-MS | ZM447439 | MVP_top60_DNA+RNA_seed | CREBBP | -0.026335 | STAD | HSC-39 | I-CBP112 |
| MVP_top60_DNA_seed | EGFR | 0.866724 | LUAD | NCI-H1573 | CI-1033 | MVP_top60_DNA+RNA_seed | AR | -0.023364 | STAD | MKN7 | Bicalutamide |
| MVP_top60_DNA_seed | ERBB2 | 0.866724 | LUAD | NCI-H1573 | CI-1033 | MVP_top60_DNA+RNA_seed | AR | -0.022776 | STAD | RERF-GC-1B | Bicalutamide |
| MVP_top60_DNA_seed | MET | 0.867834 | LUAD | Calu-6 | PHA-665752 | MVP_top60_DNA+RNA_seed | MET | -0.021731 | STAD | AGS | JNJ38877605 |
| MVP_top60_DNA_seed | MAPK1 | 0.86836 | LUAD | H3255 | VX-11e | MVP_top60_DNA+RNA_seed | ERBB2 | -0.020592 | STAD | NUGC-3 | CP724714 |
| MVP_top60_DNA_seed | HDAC1 | 0.86841 | LUAD | NCI-H1755 | Dacinostat | MVP_top60_DNA+RNA_seed | MDM2 | -0.016975 | STAD | MKN45 | Serdemetan |
| MVP_top60_DNA_seed | AKT1 | 0.868454 | LUAD | NCI-H1755 | GSK690693 | MVP_top60_DNA+RNA_seed | HDAC1 | -0.012993 | STAD | NUGC-4 | AR-42 |
| MVP_top60_DNA_seed | SRC | 0.869707 | LUAD | H3255 | NVP-BHG712 | MVP_top60_DNA+RNA_seed | MDM2 | -0.012838 | STAD | MKN7 | Serdemetan |
| MVP_top60_DNA_seed | MAPK1 | 0.869987 | LUAD | NCI-H522 | VX-11e | MVP_top60_DNA+RNA_seed | ATM | -0.00832 | STAD | AGS | KU-55933 |
| MVP_top60_DNA_seed | FGFR1 | 0.870663 | LUAD | RERF-LC-MS | PD173074 | MVP_top60_DNA+RNA_seed | HDAC1 | -0.006399 | STAD | FU97 | Tubastatin A |
| MVP_top60_DNA_seed | AURKA | 0.872693 | LUAD | NCI-H322M | GSK1070916 | MVP_top60_DNA+RNA_seed | GSK3B | -0.004942 | STAD | NUGC-3 | AZD7969 |
| MVP_top60_DNA_seed | EGFR | 0.872763 | LUAD | SK-LU-1 | Cetuximab | MVP_top60_DNA+RNA_seed | ESR1 | -0.004848 | STAD | 23132-87 | Tamoxifen |
| MVP_top60_DNA_seed | AKT1 | 0.87412 | LUAD | NCI-H441 | AKT inhibitor VIII | MVP_top60_DNA+RNA_seed | SRC | -0.002756 | STAD | MKN45 | Bosutinib |
| MVP_top60_DNA_seed | FGFR1 | 0.874736 | LUAD | HCC-827 | AZD4547 | MVP_top60_DNA+RNA_seed | ERBB2 | -0.000572 | STAD | TMK-1 | Afatinib |
| MVP_top60_DNA_seed | EGFR | 0.87522 | LUAD | EMC-BAC-2 | Gefitinib | MVP_top60_DNA+RNA_seed | PARP1 | 0.003599 | STAD | 23132-87 | Rucaparib |
| MVP_top60_DNA_seed | PTK2 | 0.875995 | LUAD | NCI-H2342 | PF-562271 | MVP_top60_DNA+RNA_seed | ERBB2 | 0.004358 | STAD | Hs-746T | CI-1033 |
| MVP_top60_DNA_seed | EGFR | 0.876815 | LUAD | NCI-H23 | Pelitinib | MVP_top60_DNA+RNA_seed | ERBB4 | 0.004358 | STAD | Hs-746T | CI-1033 |
| MVP_top60_DNA_seed | ATM | 0.877733 | LUAD | NCI-H2087 | KU-55933 | MVP_top60_DNA+RNA_seed | ERBB2 | 0.007664 | STAD | HSC-39 | Afatinib |
| MVP_top60_DNA_seed | SRC | 0.878006 | LUAD | NCI-H1573 | Bosutinib | MVP_top60_DNA+RNA_seed | PARP1 | 0.00911 | STAD | Hs-746T | PARP_9495 |
| MVP_top60_DNA_seed | EGFR | 0.878229 | LUAD | NCI-H1437 | Afatinib | MVP_top60_DNA+RNA_seed | HSP90AA1 | 0.009251 | STAD | NCI-N87 | Luminespib |
| MVP_top60_DNA_seed | ERBB2 | 0.878229 | LUAD | NCI-H1437 | Afatinib | MVP_top60_DNA+RNA_seed | HSP90AB1 | 0.009251 | STAD | NCI-N87 | Luminespib |
| MVP_top60_DNA_seed | EGFR | 0.879079 | LUAD | HOP-62 | Gefitinib | MVP_top60_DNA+RNA_seed | AR | 0.00946 | STAD | Hs-746T | Bicalutamide |
| MVP_top60_DNA_seed | HSP90AA1 | 0.879398 | LUAD | NCI-H441 | Tanespimycin | MVP_top60_DNA+RNA_seed | GSK3B | 0.014177 | STAD | HSC-39 | CHIR-99021 |

| | | | | | | | | | | | |
|---|---|---|---|---|---|---|---|---|---|---|---|
| MVP_top60_DNA_seed | EGFR | 0.879466 | LUAD | NCI-H1651 | CUDC-101 | MVP_top60_DNA+RNA_seed | PIK3CA | 0.020711 | STAD | IM-95 | PI3Ka_4409 |
| MVP_top60_DNA_seed | ERBB2 | 0.879466 | LUAD | NCI-H1651 | CUDC-101 | MVP_top60_DNA+RNA_seed | HDAC1 | 0.02156 | STAD | HSC-39 | AR-42 |
| MVP_top60_DNA_seed | MET | 0.880069 | LUAD | NCI-H1666 | AZD6094 | MVP_top60_DNA+RNA_seed | AR | 0.023023 | STAD | AGS | AZD3514 |
| MVP_top60_DNA_seed | MET | 0.880363 | LUAD | NCI-H1975 | PHA-665752 | MVP_top60_DNA+RNA_seed | GSK3B | 0.023117 | STAD | 23132-87 | CHIR-99021 |
| MVP_top60_DNA_seed | EGFR | 0.881841 | LUAD | NCI-H1975 | Gefitinib | MVP_top60_DNA+RNA_seed | MDM2 | 0.025901 | STAD | HSC-39 | Nutlin-3a (-) |
| MVP_top60_DNA_seed | SRC | 0.882261 | LUAD | HCC-44 | Bosutinib | MVP_top60_DNA+RNA_seed | HSP90AA1 | 0.027097 | STAD | OCUM-1 | Elesclomol |
| MVP_top60_DNA_seed | GSK3B | 0.882882 | LUAD | NCI-H838 | CHIR-99021 | MVP_top60_DNA+RNA_seed | HSP90AB1 | 0.027097 | STAD | OCUM-1 | Elesclomol |
| MVP_top60_DNA_seed | AKT1 | 0.883131 | LUAD | NCI-H1623 | GSK690693 | MVP_top60_DNA+RNA_seed | PARP1 | 0.031497 | STAD | OCUM-1 | PARP_9495 |
| MVP_top60_DNA_seed | IGF1R | 0.883846 | LUAD | NCI-H1435 | Linsitinib | MVP_top60_DNA+RNA_seed | ESR1 | 0.035763 | STAD | RERF-GC-1B | Fulvestrant |
| MVP_top60_DNA_seed | FGFR1 | 0.884137 | LUAD | NCI-H2009 | PD173074 | MVP_top60_DNA+RNA_seed | HSP90AA1 | 0.036882 | STAD | NUGC-4 | SNX-2112 |
| MVP_top60_DNA_seed | BRAF | 0.884575 | LUAD | HCC-78 | Dabrafenib | MVP_top60_DNA+RNA_seed | HSP90AB1 | 0.036882 | STAD | NUGC-4 | SNX-2112 |
| MVP_top60_DNA_seed | AURKA | 0.885289 | LUAD | Calu-6 | GSK1070916 | MVP_top60_DNA+RNA_seed | ERBB2 | 0.037018 | STAD | HSC-39 | Afatinib |
| MVP_top60_DNA_seed | SMARCA4 | 0.889656 | LUAD | NCI-H2347 | PFI-3 | MVP_top60_DNA+RNA_seed | SRC | 0.038493 | STAD | NCI-N87 | HG-6-71-01 |
| MVP_top60_DNA_seed | GSK3B | 0.890109 | LUAD | NCI-H1944 | CHIR-99021 | MVP_top60_DNA+RNA_seed | HSP90AA1 | 0.045196 | STAD | TMK-1 | Luminespib |
| MVP_top60_DNA_seed | ATM | 0.890839 | LUAD | SW1573 | KU-60019 | MVP_top60_DNA+RNA_seed | HSP90AB1 | 0.045196 | STAD | TMK-1 | Luminespib |
| MVP_top60_DNA_seed | BRAF | 0.891201 | LUAD | NCI-H1838 | SB590885 | MVP_top60_DNA+RNA_seed | ATM | 0.047239 | STAD | SK-GT-2 | KU-55933 |
| MVP_top60_DNA_seed | EGFR | 0.891446 | LUAD | NCI-H1435 | AZD8931 | MVP_top60_DNA+RNA_seed | FGFR3 | 0.050307 | STAD | MKN28 | FGFR_3831 |
| MVP_top60_DNA_seed | ERBB2 | 0.891446 | LUAD | NCI-H1435 | AZD8931 | MVP_top60_DNA+RNA_seed | HDAC1 | 0.053631 | STAD | FU97 | Belinostat |
| MVP_top60_DNA_seed | ERBB3 | 0.891446 | LUAD | NCI-H1435 | AZD8931 | MVP_top60_DNA+RNA_seed | AR | 0.054807 | STAD | SNU-16 | Bicalutamide |
| MVP_top60_DNA_seed | EGFR | 0.892561 | LUAD | NCI-H1993 | CI-1033 | MVP_top60_DNA+RNA_seed | PARP1 | 0.054836 | STAD | SK-GT-2 | Talazoparib |
| MVP_top60_DNA_seed | ERBB2 | 0.892561 | LUAD | NCI-H1993 | CI-1033 | MVP_top60_DNA+RNA_seed | EP300 | 0.055496 | STAD | FU97 | I-CBP112 |
| MVP_top60_DNA_seed | BRAF | 0.892994 | LUAD | NCI-H2342 | Dabrafenib | MVP_top60_DNA+RNA_seed | CREBBP | 0.055496 | STAD | FU97 | I-CBP112 |
| MVP_top60_DNA_seed | EGFR | 0.893364 | LUAD | NCI-H1650 | Pelitinib | MVP_top60_DNA+RNA_seed | GSK3B | 0.061108 | STAD | SNU-5 | AZD7969 |
| MVP_top60_DNA_seed | AURKA | 0.894025 | LUAD | NCI-H1650 | Alisertib | MVP_top60_DNA+RNA_seed | SRC | 0.064892 | STAD | GCIY | NVP-BHG712 |
| MVP_top60_DNA_seed | AKT1 | 0.894946 | LUAD | NCI-H1792 | MK-2206 | MVP_top60_DNA+RNA_seed | ERBB4 | 0.067172 | STAD | IM-95 | AST-1306 |
| MVP_top60_DNA_seed | GSK3B | 0.895605 | LUAD | ABC-1 | CHIR-99021 | MVP_top60_DNA+RNA_seed | PIK3CA | 0.073779 | STAD | SNU-16 | AZD8835 |
| MVP_top60_DNA_seed | MAPK1 | 0.895678 | LUAD | NCI-H1838 | FR-180204 | MVP_top60_DNA+RNA_seed | PARP1 | 0.075622 | STAD | HSC-39 | Rucaparib |
| MVP_top60_DNA_seed | GSK3B | 0.895826 | LUAD | NCI-H1573 | KIN001-042 | MVP_top60_DNA+RNA_seed | MET | 0.07659 | STAD | HGC-27 | SU11274 |
| MVP_top60_DNA_seed | BRAF | 0.89712 | LUAD | NCI-H2405 | PLX-4720 | MVP_top60_DNA+RNA_seed | AR | 0.089111 | STAD | RERF-GC-1B | AZD3514 |
| MVP_top60_DNA_seed | GSK3B | 0.897135 | LUAD | NCI-H838 | SB216763 | MVP_top60_DNA+RNA_seed | ATM | 0.089537 | STAD | Hs-746T | KU-60019 |
| MVP_top60_DNA_seed | AKT1 | 0.89815 | LUAD | NCI-H2009 | GSK690693 | MVP_top60_DNA+RNA_seed | PARP1 | 0.093078 | STAD | TGBC11TKB | PARP_9495 |
| MVP_top60_DNA_seed | MET | 0.89904 | LUAD | NCI-H1755 | SU11274 | MVP_top60_DNA+RNA_seed | FGFR3 | 0.093478 | STAD | Hs-746T | PD173074 |
| MVP_top60_DNA_seed | FGFR1 | 0.89994 | LUAD | A549 | AZD4547 | MVP_top60_DNA+RNA_seed | ERBB2 | 0.094896 | STAD | FU97 | CUDC-101 |
| MVP_top60_DNA_seed | HDAC1 | 0.900801 | LUAD | SK-LU-1 | Tubastatin A | MVP_top60_DNA+RNA_seed | AR | 0.097163 | STAD | 23132-87 | AZD3514 |
| MVP_top60_DNA_seed | FGFR1 | 0.901556 | LUAD | NCI-H1648 | AZD4547 | MVP_top60_DNA+RNA_seed | PTK2 | 0.09997 | STAD | RF-48 | PF-562271 |
| MVP_top60_DNA_seed | ERBB2 | 0.901932 | LUAD | NCI-H1573 | CP724714 | MVP_top60_DNA+RNA_seed | ESR1 | 0.104338 | STAD | MKN7 | Tamoxifen |
| MVP_top60_DNA_seed | IGF1R | 0.9033 | LUAD | NCI-H23 | Linsitinib | MVP_top60_DNA+RNA_seed | ESR1 | 0.106425 | STAD | OCUM-1 | Fulvestrant |
| MVP_top60_DNA_seed | ERBB2 | 0.903304 | LUAD | NCI-H2023 | CP724714 | MVP_top60_DNA+RNA_seed | PARP1 | 0.107613 | STAD | TMK-1 | PARP_9482 |
| MVP_top60_DNA_seed | HDAC1 | 0.903477 | LUAD | NCI-H1781 | JQ12 | MVP_top60_DNA+RNA_seed | GSK3B | 0.111998 | STAD | FU97 | CHIR-99021 |
| MVP_top60_DNA_seed | GSK3B | 0.903496 | LUAD | NCI-H2347 | CHIR-99021 | MVP_top60_DNA+RNA_seed | MDM2 | 0.113954 | STAD | NUGC-3 | Nutlin-3a (-) |
| MVP_top60_DNA_seed | BRAF | 0.903761 | LUAD | NCI-H1435 | HG6-64-1 | MVP_top60_DNA+RNA_seed | HDAC1 | 0.115686 | STAD | RERF-GC-1B | JQ12 |

| | | | | | | | | | | | |
|---|---|---|---|---|---|---|---|---|---|---|---|
| MVP_top60_DNA_seed | HDAC1 | 0.904536 | LUAD | NCI-H1666 | Parthenolide | MVP_top60_DNA+RNA_seed | ATM | 0.117284 | STAD | OCUM-1 | KU-60019 |
| MVP_top60_DNA_seed | AKT1 | 0.904923 | LUAD | NCI-H1568 | MK-2206 | MVP_top60_DNA+RNA_seed | ATM | 0.117618 | STAD | HGC-27 | KU-55933 |
| MVP_top60_DNA_seed | AKT1 | 0.905941 | LUAD | NCI-H2342 | GSK690693 | MVP_top60_DNA+RNA_seed | PIK3CA | 0.121784 | STAD | TGBC11TKB | AZD8835 |
| MVP_top60_DNA_seed | HSP90AA1 | 0.907162 | LUAD | H3255 | Elesclomol | MVP_top60_DNA+RNA_seed | HSP90AA1 | 0.121889 | STAD | MKN28 | Tanespimycin |
| MVP_top60_DNA_seed | BRAF | 0.907875 | LUAD | COR-L105 | Dabrafenib | MVP_top60_DNA+RNA_seed | HSP90AB1 | 0.121889 | STAD | MKN28 | Tanespimycin |
| MVP_top60_DNA_seed | ATM | 0.908892 | LUAD | NCI-H1838 | KU-60019 | MVP_top60_DNA+RNA_seed | HDAC1 | 0.124559 | STAD | SNU-1 | Tubastatin A |
| MVP_top60_DNA_seed | BRAF | 0.90941 | LUAD | NCI-H1355 | PLX-4720 | MVP_top60_DNA+RNA_seed | AR | 0.125006 | STAD | SK-GT-2 | AZD3514 |
| MVP_top60_DNA_seed | GSK3B | 0.910953 | LUAD | A549 | CHIR-99021 | MVP_top60_DNA+RNA_seed | PARP1 | 0.125134 | STAD | Hs-746T | Olaparib |
| MVP_top60_DNA_seed | ATM | 0.911352 | LUAD | NCI-H441 | CP466722 | MVP_top60_DNA+RNA_seed | PARP1 | 0.125972 | STAD | MKN45 | Rucaparib |
| MVP_top60_DNA_seed | BRAF | 0.911676 | LUAD | NCI-H1435 | SB590885 | MVP_top60_DNA+RNA_seed | PIK3CA | 0.128448 | STAD | NUGC-4 | AZD8835 |
| MVP_top60_DNA_seed | FGFR1 | 0.912527 | LUAD | NCI-H2405 | PD173074 | MVP_top60_DNA+RNA_seed | HSP90AA1 | 0.129645 | STAD | NCI-N87 | CCT-018159 |
| MVP_top60_DNA_seed | MAPK1 | 0.913321 | LUAD | NCI-H2023 | VX-11e | MVP_top60_DNA+RNA_seed | HSP90AB1 | 0.129645 | STAD | NCI-N87 | CCT-018159 |
| MVP_top60_DNA_seed | EGFR | 0.913877 | LUAD | NCI-H2122 | Gefitinib | MVP_top60_DNA+RNA_seed | HDAC1 | 0.137286 | STAD | RF-48 | Entinostat |
| MVP_top60_DNA_seed | HDAC1 | 0.914686 | LUAD | H3255 | CAY10603 | MVP_top60_DNA+RNA_seed | HDAC3 | 0.137286 | STAD | RF-48 | Entinostat |
| MVP_top60_DNA_seed | IGF1R | 0.914868 | LUAD | NCI-H1693 | Linsitinib | MVP_top60_DNA+RNA_seed | PIK3CA | 0.137472 | STAD | MKN28 | AZD8835 |
| MVP_top60_DNA_seed | AURKA | 0.916021 | LUAD | NCI-H2405 | CD532 | MVP_top60_DNA+RNA_seed | ERBB2 | 0.138652 | STAD | SNU-16 | CP724714 |
| MVP_top60_DNA_seed | EGFR | 0.916182 | LUAD | NCI-H1993 | Gefitinib | MVP_top60_DNA+RNA_seed | MET | 0.140019 | STAD | HSC-39 | Crizotinib |
| MVP_top60_DNA_seed | FGFR1 | 0.916441 | LUAD | NCI-H2342 | PD173074 | MVP_top60_DNA+RNA_seed | HSP90AA1 | 0.142344 | STAD | MKN7 | SNX-2112 |
| MVP_top60_DNA_seed | GSK3B | 0.916915 | LUAD | NCI-H1563 | CHIR-99021 | MVP_top60_DNA+RNA_seed | HSP90AB1 | 0.142344 | STAD | MKN7 | SNX-2112 |
| MVP_top60_DNA_seed | MET | 0.916958 | LUAD | NCI-H1568 | JNJ38877605 | MVP_top60_DNA+RNA_seed | HDAC1 | 0.143716 | STAD | NUGC-4 | Dacinostat |
| MVP_top60_DNA_seed | GSK3B | 0.91718 | LUAD | NCI-H1573 | CHIR-99021 | MVP_top60_DNA+RNA_seed | HDAC1 | 0.143809 | STAD | TGBC11TKB | CAY10603 |
| MVP_top60_DNA_seed | AURKA | 0.918815 | LUAD | EKVX | CD532 | MVP_top60_DNA+RNA_seed | GSK3B | 0.145196 | STAD | TGBC11TKB | CHIR-99021 |
| MVP_top60_DNA_seed | FGFR1 | 0.919179 | LUAD | NCI-H2087 | FGFR_3831 | MVP_top60_DNA+RNA_seed | HDAC1 | 0.145259 | STAD | HSC-39 | Entinostat |
| MVP_top60_DNA_seed | BRAF | 0.920028 | LUAD | Calu-3 | Dabrafenib | MVP_top60_DNA+RNA_seed | HDAC3 | 0.145259 | STAD | HSC-39 | Entinostat |
| MVP_top60_DNA_seed | EGFR | 0.920788 | LUAD | NCI-H1793 | PF-00299804 | MVP_top60_DNA+RNA_seed | ERBB2 | 0.146955 | STAD | MKN7 | CP724714 |
| MVP_top60_DNA_seed | ERBB2 | 0.920788 | LUAD | NCI-H1793 | PF-00299804 | MVP_top60_DNA+RNA_seed | PARP1 | 0.147059 | STAD | NCI-N87 | PARP_9482 |
| MVP_top60_DNA_seed | SRC | 0.921008 | LUAD | NCI-H1781 | NVP-BHG712 | MVP_top60_DNA+RNA_seed | HSP90AA1 | 0.154773 | STAD | 23132-87 | Elesclomol |
| MVP_top60_DNA_seed | MET | 0.921273 | LUAD | NCI-H1437 | SU11274 | MVP_top60_DNA+RNA_seed | HSP90AB1 | 0.154773 | STAD | 23132-87 | Elesclomol |
| MVP_top60_DNA_seed | EGFR | 0.921467 | LUAD | SK-LU-1 | Afatinib | MVP_top60_DNA+RNA_seed | MDM2 | 0.155284 | STAD | RERF-GC-1B | Serdemetan |
| MVP_top60_DNA_seed | ERBB2 | 0.921467 | LUAD | SK-LU-1 | Afatinib | MVP_top60_DNA+RNA_seed | ERBB2 | 0.160058 | STAD | MKN7 | AZD8931 |
| MVP_top60_DNA_seed | IGF1R | 0.921641 | LUAD | Calu-6 | BMS-536924 | MVP_top60_DNA+RNA_seed | ERBB3 | 0.160058 | STAD | MKN7 | AZD8931 |
| MVP_top60_DNA_seed | BRAF | 0.921815 | LUAD | NCI-H1568 | SB590885 | MVP_top60_DNA+RNA_seed | MET | 0.1642 | STAD | GCIY | Crizotinib |
| MVP_top60_DNA_seed | CREBBP | 0.924625 | LUAD | SK-LU-1 | I-CBP112 | MVP_top60_DNA+RNA_seed | ERBB2 | 0.167947 | STAD | HGC-27 | Afatinib |
| MVP_top60_DNA_seed | PIK3CA | 0.926399 | LUAD | NCI-H1573 | AZD8835 | MVP_top60_DNA+RNA_seed | MAPK1 | 0.168097 | STAD | FU97 | VX-11e |
| MVP_top60_DNA_seed | HSP90AA1 | 0.926415 | LUAD | EKVX | Luminespib | MVP_top60_DNA+RNA_seed | FYN | 0.171648 | STAD | SNU-1 | A-770041 |
| MVP_top60_DNA_seed | BRAF | 0.926525 | LUAD | SW1573 | PLX-4720 | MVP_top60_DNA+RNA_seed | GSK3B | 0.177124 | STAD | OCUM-1 | AZD7969 |
| MVP_top60_DNA_seed | EGFR | 0.92702 | LUAD | NCI-H1437 | Afatinib | MVP_top60_DNA+RNA_seed | PIK3CA | 0.178054 | STAD | SK-GT-2 | PI3Ka_4409 |
| MVP_top60_DNA_seed | ERBB2 | 0.92702 | LUAD | NCI-H1437 | Afatinib | MVP_top60_DNA+RNA_seed | MAPK1 | 0.181645 | STAD | MKN7 | FR-180204 |
| MVP_top60_DNA_seed | AKT1 | 0.927586 | LUAD | Calu-6 | GSK690693 | MVP_top60_DNA+RNA_seed | MAPK3 | 0.181645 | STAD | MKN7 | FR-180204 |
| MVP_top60_DNA_seed | AURKA | 0.927754 | LUAD | RERF-LC-KJ | HG-5-113-01 | MVP_top60_DNA+RNA_seed | HSP90AA1 | 0.182516 | STAD | IM-95 | CCT-018159 |
| MVP_top60_DNA_seed | AURKA | 0.92945 | LUAD | EKVX | ZM447439 | MVP_top60_DNA+RNA_seed | HSP90AB1 | 0.182516 | STAD | IM-95 | CCT-018159 |

| | | | | | | | | | | | |
|---|---|---|---|---|---|---|---|---|---|---|---|
| MVP_top60_DNA_seed | MAPK1 | 0.929455 | LUAD | NCI-H1435 | FR-180204 | MVP_top60_DNA+RNA_seed | ERBB2 | 0.191637 | STAD | IM-95 | PF-00299804 |
| MVP_top60_DNA_seed | IGF1R | 0.930459 | LUAD | NCI-H2023 | GSK1904529A | MVP_top60_DNA+RNA_seed | ERBB4 | 0.191637 | STAD | IM-95 | PF-00299804 |
| MVP_top60_DNA_seed | MET | 0.930646 | LUAD | EMC-BAC-2 | SU11274 | MVP_top60_DNA+RNA_seed | HSP90AA1 | 0.194683 | STAD | Hs-746T | Tanespimycin |
| MVP_top60_DNA_seed | FGFR1 | 0.932094 | LUAD | PC-3_[JPC-3] | AZD4547 | MVP_top60_DNA+RNA_seed | HSP90AB1 | 0.194683 | STAD | Hs-746T | Tanespimycin |
| MVP_top60_DNA_seed | HDAC1 | 0.932469 | LUAD | NCI-H2228 | Belinostat | MVP_top60_DNA+RNA_seed | FGFR3 | 0.197536 | STAD | AGS | AZD4547 |
| MVP_top60_DNA_seed | SRC | 0.93282 | LUAD | NCI-H1573 | NVP-BHG712 | MVP_top60_DNA+RNA_seed | SRC | 0.197645 | STAD | 23132-87 | Bosutinib |
| MVP_top60_DNA_seed | AKT1 | 0.933064 | LUAD | NCI-H1573 | GSK690693 | MVP_top60_DNA+RNA_seed | MAPK1 | 0.1977 | STAD | TGBC11TKB | FR-180204 |
| MVP_top60_DNA_seed | MET | 0.934686 | LUAD | COR-L105 | AZD6094 | MVP_top60_DNA+RNA_seed | MAPK3 | 0.1977 | STAD | TGBC11TKB | FR-180204 |
| MVP_top60_DNA_seed | EGFR | 0.935226 | LUAD | NCI-H322M | Cetuximab | MVP_top60_DNA+RNA_seed | HDAC1 | 0.198525 | STAD | MKN28 | CAY10603 |
| MVP_top60_DNA_seed | EGFR | 0.935275 | LUAD | NCI-H1650 | Lapatinib | MVP_top60_DNA+RNA_seed | PIK3CA | 0.203951 | STAD | SNU-5 | AZD8835 |
| MVP_top60_DNA_seed | ERBB2 | 0.935275 | LUAD | NCI-H1650 | Lapatinib | MVP_top60_DNA+RNA_seed | SRC | 0.206312 | STAD | TMK-1 | HG-6-71-01 |
| MVP_top60_DNA_seed | FGFR1 | 0.936659 | LUAD | ABC-1 | PD173074 | MVP_top60_DNA+RNA_seed | FGFR3 | 0.207473 | STAD | TGBC11TKB | PD173074 |
| MVP_top60_DNA_seed | FGFR1 | 0.936928 | LUAD | NCI-H2347 | FGFR_3831 | MVP_top60_DNA+RNA_seed | HSP90AA1 | 0.207665 | STAD | MKN45 | SNX-2112 |
| MVP_top60_DNA_seed | MAPK1 | 0.937377 | LUAD | EMC-BAC-2 | VX-11e | MVP_top60_DNA+RNA_seed | HSP90AB1 | 0.207665 | STAD | MKN45 | SNX-2112 |
| MVP_top60_DNA_seed | EGFR | 0.93745 | LUAD | NCI-H1650 | CI-1033 | MVP_top60_DNA+RNA_seed | PARP1 | 0.210012 | STAD | RERF-GC-1B | Olaparib |
| MVP_top60_DNA_seed | ERBB2 | 0.93745 | LUAD | NCI-H1650 | CI-1033 | MVP_top60_DNA+RNA_seed | PTK2 | 0.210308 | STAD | SNU-1 | PF-562271 |
| MVP_top60_DNA_seed | GSK3B | 0.937904 | LUAD | COR-L105 | CHIR-99021 | MVP_top60_DNA+RNA_seed | PARP1 | 0.210667 | STAD | TMK-1 | PARP_9495 |
| MVP_top60_DNA_seed | HDAC1 | 0.937985 | LUAD | NCI-H1650 | Belinostat | MVP_top60_DNA+RNA_seed | GSK3B | 0.21094 | STAD | IM-95 | AZD7969 |
| MVP_top60_DNA_seed | MAPK1 | 0.938403 | LUAD | NCI-H1623 | VX-11e | MVP_top60_DNA+RNA_seed | MET | 0.212317 | STAD | RF-48 | Crizotinib |
| MVP_top60_DNA_seed | AURKA | 0.938574 | LUAD | SW1573 | GSK1070916 | MVP_top60_DNA+RNA_seed | HDAC1 | 0.212807 | STAD | MKN28 | Tubastatin A |
| MVP_top60_DNA_seed | IGF1R | 0.940568 | LUAD | NCI-H1650 | BMS-536924 | MVP_top60_DNA+RNA_seed | GSK3B | 0.213548 | STAD | MKN45 | SB216763 |
| MVP_top60_DNA_seed | AURKA | 0.940595 | LUAD | COR-L105 | Alisertib | MVP_top60_DNA+RNA_seed | GSK3B | 0.216824 | STAD | NUGC-3 | CHIR-99021 |
| MVP_top60_DNA_seed | AURKA | 0.940699 | LUAD | NCI-H1355 | CD532 | MVP_top60_DNA+RNA_seed | PTK2 | 0.217675 | STAD | NUGC-4 | PF-562271 |
| MVP_top60_DNA_seed | CREBBP | 0.941158 | LUAD | H3255 | I-CBP112 | MVP_top60_DNA+RNA_seed | FGFR3 | 0.218777 | STAD | SNU-5 | AZD4547 |
| MVP_top60_DNA_seed | EGFR | 0.941252 | LUAD | NCI-H2347 | Afatinib | MVP_top60_DNA+RNA_seed | HDAC1 | 0.223065 | STAD | NUGC-3 | CAY10603 |
| MVP_top60_DNA_seed | ERBB2 | 0.941252 | LUAD | NCI-H2347 | Afatinib | MVP_top60_DNA+RNA_seed | SRC | 0.224697 | STAD | OCUM-1 | Bosutinib |
| MVP_top60_DNA_seed | HDAC1 | 0.941503 | LUAD | NCI-H2342 | Tubastatin A | MVP_top60_DNA+RNA_seed | EP300 | 0.226234 | STAD | 23132-87 | I-CBP112 |
| MVP_top60_DNA_seed | BRAF | 0.942901 | LUAD | PC-3_[JPC-3] | PLX-4720 | MVP_top60_DNA+RNA_seed | CREBBP | 0.226234 | STAD | 23132-87 | I-CBP112 |
| MVP_top60_DNA_seed | BRAF | 0.94359 | LUAD | NCI-H322M | Dabrafenib | MVP_top60_DNA+RNA_seed | PARP1 | 0.226646 | STAD | TMK-1 | Olaparib |
| MVP_top60_DNA_seed | HDAC1 | 0.944524 | LUAD | NCI-H2405 | JQ12 | MVP_top60_DNA+RNA_seed | PARP1 | 0.235694 | STAD | OCUM-1 | PARP_9482 |
| MVP_top60_DNA_seed | HDAC1 | 0.94491 | LUAD | NCI-H1573 | Tubastatin A | MVP_top60_DNA+RNA_seed | FGFR3 | 0.238331 | STAD | AGS | PD173074 |
| MVP_top60_DNA_seed | HDAC1 | 0.945613 | LUAD | NCI-H1435 | Tubastatin A | MVP_top60_DNA+RNA_seed | PARP1 | 0.238679 | STAD | TGBC11TKB | Rucaparib |
| MVP_top60_DNA_seed | AKT1 | 0.945842 | LUAD | NCI-H2405 | GSK690693 | MVP_top60_DNA+RNA_seed | MET | 0.239117 | STAD | FU97 | AZD6094 |
| MVP_top60_DNA_seed | MAPK1 | 0.946874 | LUAD | NCI-H838 | VX-11e | MVP_top60_DNA+RNA_seed | ATM | 0.23976 | STAD | NCI-N87 | KU-60019 |
| MVP_top60_DNA_seed | FGFR1 | 0.946881 | LUAD | Calu-6 | PD173074 | MVP_top60_DNA+RNA_seed | PARP1 | 0.240103 | STAD | TMK-1 | Rucaparib |
| MVP_top60_DNA_seed | HSP90AA1 | 0.948085 | LUAD | NCI-H322M | CCT-018159 | MVP_top60_DNA+RNA_seed | HSP90AA1 | 0.243622 | STAD | Hs-746T | SNX-2112 |
| MVP_top60_DNA_seed | IGF1R | 0.94814 | LUAD | HOP-62 | BMS-536924 | MVP_top60_DNA+RNA_seed | HSP90AB1 | 0.243622 | STAD | Hs-746T | SNX-2112 |
| MVP_top60_DNA_seed | MET | 0.95018 | LUAD | NCI-H1355 | PHA-665752 | MVP_top60_DNA+RNA_seed | HDAC1 | 0.247638 | STAD | RERF-GC-1B | Tubastatin A |
| MVP_top60_DNA_seed | GSK3B | 0.950982 | LUAD | NCI-H2342 | SB216763 | MVP_top60_DNA+RNA_seed | HDAC1 | 0.247883 | STAD | IM-95 | CAY10603 |
| MVP_top60_DNA_seed | AURKA | 0.951142 | LUAD | NCI-H2405 | Alisertib | MVP_top60_DNA+RNA_seed | PIK3CA | 0.249282 | STAD | KATOIII | AZD8835 |
| MVP_top60_DNA_seed | EGFR | 0.952054 | LUAD | HCC-78 | Gefitinib | MVP_top60_DNA+RNA_seed | GSK3B | 0.250622 | STAD | MKN7 | CHIR-99021 |

| | | | | | | | | | | | |
|---|---|---|---|---|---|---|---|---|---|---|---|
| MVP_top60_DNA_seed | HDAC1 | 0.952524 | LUAD | NCI-H23 | Parthenolide | MVP_top60_DNA+RNA_seed | MET | 0.251625 | STAD | NUGC-4 | JNJ38877605 |
| MVP_top60_DNA_seed | BRAF | 0.953817 | LUAD | NCI-H1563 | SB590885 | MVP_top60_DNA+RNA_seed | PARP1 | 0.255516 | STAD | 23132-87 | PARP_9495 |
| MVP_top60_DNA_seed | HDAC1 | 0.954009 | LUAD | NCI-H1792 | AR-42 | MVP_top60_DNA+RNA_seed | ERBB2 | 0.256146 | STAD | SNU-5 | Afatinib |
| MVP_top60_DNA_seed | SRC | 0.954527 | LUAD | NCI-H1623 | NVP-BHG712 | MVP_top60_DNA+RNA_seed | HDAC1 | 0.257129 | STAD | MKN7 | CAY10603 |
| MVP_top60_DNA_seed | MET | 0.954593 | LUAD | NCI-H2122 | SU11274 | MVP_top60_DNA+RNA_seed | HSP90AA1 | 0.261704 | STAD | OCUM-1 | Luminespib |
| MVP_top60_DNA_seed | ATM | 0.956946 | LUAD | NCI-H1693 | KU-55933 | MVP_top60_DNA+RNA_seed | HSP90AB1 | 0.261704 | STAD | OCUM-1 | Luminespib |
| MVP_top60_DNA_seed | PTK2 | 0.956977 | LUAD | NCI-H1755 | PF-562271 | MVP_top60_DNA+RNA_seed | ERBB4 | 0.262005 | STAD | 23132-87 | AST-1306 |
| MVP_top60_DNA_seed | BRAF | 0.957459 | LUAD | NCI-H838 | PLX-4720 | MVP_top60_DNA+RNA_seed | AR | 0.268343 | STAD | RF-48 | Bicalutamide |
| MVP_top60_DNA_seed | MAPK1 | 0.957655 | LUAD | NCI-H2347 | FR-180204 | MVP_top60_DNA+RNA_seed | ERBB2 | 0.268978 | STAD | SNU-1 | Afatinib |
| MVP_top60_DNA_seed | AKT1 | 0.957918 | LUAD | NCI-H2347 | GSK690693 | MVP_top60_DNA+RNA_seed | PARP1 | 0.269491 | STAD | SNU-16 | PARP_9495 |
| MVP_top60_DNA_seed | PIK3CA | 0.959083 | LUAD | NCI-H2087 | PI3Ka_4409 | MVP_top60_DNA+RNA_seed | MDM2 | 0.270125 | STAD | NCI-N87 | Serdemetan |
| MVP_top60_DNA_seed | AURKA | 0.959831 | LUAD | NCI-H1568 | Alisertib | MVP_top60_DNA+RNA_seed | SRC | 0.274762 | STAD | MKN7 | Bosutinib |
| MVP_top60_DNA_seed | ATM | 0.959983 | LUAD | SW1573 | KU-55933 | MVP_top60_DNA+RNA_seed | MET | 0.276673 | STAD | NUGC-4 | AZD6094 |
| MVP_top60_DNA_seed | FGFR1 | 0.96094 | LUAD | NCI-H2023 | PD173074 | MVP_top60_DNA+RNA_seed | ERBB2 | 0.279114 | STAD | TGBC11TKB | Afatinib |
| MVP_top60_DNA_seed | HDAC1 | 0.961118 | LUAD | SW1573 | Dacinostat | MVP_top60_DNA+RNA_seed | ERBB4 | 0.2807 | STAD | FU97 | AST-1306 |
| MVP_top60_DNA_seed | AURKA | 0.961411 | LUAD | NCI-H322M | ZM447439 | MVP_top60_DNA+RNA_seed | GSK3B | 0.281846 | STAD | AGS | KIN001-042 |
| MVP_top60_DNA_seed | EGFR | 0.962096 | LUAD | NCI-H1650 | AST-1306 | MVP_top60_DNA+RNA_seed | ERBB2 | 0.28185 | STAD | SNU-16 | Lapatinib |
| MVP_top60_DNA_seed | ATM | 0.964407 | LUAD | NCI-H1573 | KU-60019 | MVP_top60_DNA+RNA_seed | AR | 0.282371 | STAD | TGBC11TKB | AZD3514 |
| MVP_top60_DNA_seed | HDAC1 | 0.964964 | LUAD | LC-2-ad | Parthenolide | MVP_top60_DNA+RNA_seed | PARP1 | 0.283499 | STAD | Hs-746T | Talazoparib |
| MVP_top60_DNA_seed | BRAF | 0.964994 | LUAD | NCI-H650 | RAF_9304 | MVP_top60_DNA+RNA_seed | GSK3B | 0.283993 | STAD | AGS | SB216763 |
| MVP_top60_DNA_seed | GSK3B | 0.965476 | LUAD | NCI-H1568 | SB216763 | MVP_top60_DNA+RNA_seed | GSK3B | 0.286311 | STAD | GCIY | CHIR-99021 |
| MVP_top60_DNA_seed | IGF1R | 0.966032 | LUAD | NCI-H1781 | Linsitinib | MVP_top60_DNA+RNA_seed | PARP1 | 0.287025 | STAD | SNU-1 | Veliparib |
| MVP_top60_DNA_seed | EGFR | 0.966123 | LUAD | SK-LU-1 | Afatinib | MVP_top60_DNA+RNA_seed | HDAC1 | 0.288894 | STAD | FU97 | CAY10603 |
| MVP_top60_DNA_seed | ERBB2 | 0.966123 | LUAD | SK-LU-1 | Afatinib | MVP_top60_DNA+RNA_seed | GSK3B | 0.290459 | STAD | OCUM-1 | KIN001-042 |
| MVP_top60_DNA_seed | HDAC1 | 0.967447 | LUAD | NCI-H1792 | CAY10603 | MVP_top60_DNA+RNA_seed | MAPK1 | 0.290598 | STAD | AGS | FR-180204 |
| MVP_top60_DNA_seed | BRAF | 0.967831 | LUAD | NCI-H838 | RAF_9304 | MVP_top60_DNA+RNA_seed | MAPK3 | 0.290598 | STAD | AGS | FR-180204 |
| MVP_top60_DNA_seed | FGFR1 | 0.970911 | LUAD | NCI-H3122 | PD173074 | MVP_top60_DNA+RNA_seed | MET | 0.292771 | STAD | 23132-87 | AZD6094 |
| MVP_top60_DNA_seed | MAPK1 | 0.970923 | LUAD | SW1573 | FR-180204 | MVP_top60_DNA+RNA_seed | HSP90AA1 | 0.294511 | STAD | MKN45 | CCT-018159 |
| MVP_top60_DNA_seed | AURKA | 0.973944 | LUAD | NCI-H838 | CD532 | MVP_top60_DNA+RNA_seed | HSP90AB1 | 0.294511 | STAD | MKN45 | CCT-018159 |
| MVP_top60_DNA_seed | HDAC1 | 0.97401 | LUAD | RERF-LC-KJ | Tubastatin A | MVP_top60_DNA+RNA_seed | GSK3B | 0.297009 | STAD | AGS | CHIR-99021 |
| MVP_top60_DNA_seed | HDAC1 | 0.974447 | LUAD | NCI-H2023 | CAY10603 | MVP_top60_DNA+RNA_seed | PARP1 | 0.300166 | STAD | NCI-N87 | Veliparib |
| MVP_top60_DNA_seed | MET | 0.974448 | LUAD | NCI-H650 | AZD6094 | MVP_top60_DNA+RNA_seed | GSK3B | 0.303219 | STAD | TGBC11TKB | CHIR-99021 |
| MVP_top60_DNA_seed | AURKA | 0.974513 | LUAD | NCI-H2342 | CD532 | MVP_top60_DNA+RNA_seed | ERBB2 | 0.305071 | STAD | SNU-5 | AZD8931 |
| MVP_top60_DNA_seed | FGFR1 | 0.975982 | LUAD | NCI-H2342 | FGFR_3831 | MVP_top60_DNA+RNA_seed | ERBB3 | 0.305071 | STAD | SNU-5 | AZD8931 |
| MVP_top60_DNA_seed | GSK3B | 0.977349 | LUAD | PC-3_[JPC-3] | CHIR-99021 | MVP_top60_DNA+RNA_seed | AR | 0.306706 | STAD | MKN28 | Bicalutamide |
| MVP_top60_DNA_seed | BRAF | 0.97743 | LUAD | NCI-H2030 | PLX-4720 | MVP_top60_DNA+RNA_seed | HDAC1 | 0.313312 | STAD | IM-95 | Tubastatin A |
| MVP_top60_DNA_seed | AURKA | 0.978179 | LUAD | NCI-H1781 | Alisertib | MVP_top60_DNA+RNA_seed | ERBB2 | 0.314203 | STAD | MKN28 | CP724714 |
| MVP_top60_DNA_seed | FGFR1 | 0.979736 | LUAD | NCI-H650 | AZD4547 | MVP_top60_DNA+RNA_seed | HDAC1 | 0.319247 | STAD | TMK-1 | AR-42 |
| MVP_top60_DNA_seed | GSK3B | 0.980034 | LUAD | NCI-H2347 | SB216763 | MVP_top60_DNA+RNA_seed | SRC | 0.319439 | STAD | TGBC11TKB | Bosutinib |
| MVP_top60_DNA_seed | EGFR | 0.980707 | LUAD | RERF-LC-MS | PF-00299804 | MVP_top60_DNA+RNA_seed | HDAC1 | 0.319815 | STAD | SNU-1 | Dacinostat |
| MVP_top60_DNA_seed | ERBB2 | 0.980707 | LUAD | RERF-LC-MS | PF-00299804 | MVP_top60_DNA+RNA_seed | HDAC1 | 0.324896 | STAD | NUGC-3 | Belinostat |

| | | | | | | | | | | | |
|---|---|---|---|---|---|---|---|---|---|---|---|
| MVP_top60_DNA_seed | EGFR | 0.980802 | LUAD | Calu-6 | Gefitinib | MVP_top60_DNA+RNA_seed | PARP1 | 0.324908 | STAD | RERF-GC-1B | Veliparib |
| MVP_top60_DNA_seed | GSK3B | 0.981223 | LUAD | NCI-H3122 | SB216763 | MVP_top60_DNA+RNA_seed | GSK3B | 0.325384 | STAD | MKN45 | CHIR-99021 |
| MVP_top60_DNA_seed | GSK3B | 0.981545 | LUAD | NCI-H2347 | CHIR-99021 | MVP_top60_DNA+RNA_seed | PARP1 | 0.326542 | STAD | 23132-87 | Veliparib |
| MVP_top60_DNA_seed | EGFR | 0.981781 | LUAD | NCI-H2291 | AZD8931 | MVP_top60_DNA+RNA_seed | FGFR3 | 0.327347 | STAD | NUGC-4 | AZD4547 |
| MVP_top60_DNA_seed | ERBB2 | 0.981781 | LUAD | NCI-H2291 | AZD8931 | MVP_top60_DNA+RNA_seed | FGFR3 | 0.327583 | STAD | TGBC11TKB | AZD4547 |
| MVP_top60_DNA_seed | ERBB3 | 0.981781 | LUAD | NCI-H2291 | AZD8931 | MVP_top60_DNA+RNA_seed | PARP1 | 0.330793 | STAD | NCI-N87 | Talazoparib |
| MVP_top60_DNA_seed | AKT1 | 0.981845 | LUAD | A549 | GSK690693 | MVP_top60_DNA+RNA_seed | PARP1 | 0.33521 | STAD | RERF-GC-1B | Talazoparib |
| MVP_top60_DNA_seed | GSK3B | 0.982296 | LUAD | NCI-H2023 | KIN001-042 | MVP_top60_DNA+RNA_seed | ATM | 0.33635 | STAD | Hs-746T | KU-55933 |
| MVP_top60_DNA_seed | GSK3B | 0.982727 | LUAD | NCI-H2228 | KIN001-042 | MVP_top60_DNA+RNA_seed | ERBB2 | 0.340594 | STAD | HGC-27 | Afatinib |
| MVP_top60_DNA_seed | PTK2 | 0.983315 | LUAD | NCI-H441 | PF-562271 | MVP_top60_DNA+RNA_seed | HSP90AA1 | 0.341481 | STAD | TGBC11TKB | CCT-018159 |
| MVP_top60_DNA_seed | HSP90AA1 | 0.98345 | LUAD | HCC-78 | SNX-2112 | MVP_top60_DNA+RNA_seed | HSP90AB1 | 0.341481 | STAD | TGBC11TKB | CCT-018159 |
| MVP_top60_DNA_seed | PTK2 | 0.983587 | LUAD | PC-3_[JPC-3] | PF-562271 | MVP_top60_DNA+RNA_seed | ERBB2 | 0.347888 | STAD | NUGC-3 | PF-00299804 |
| MVP_top60_DNA_seed | CREBBP | 0.98376 | LUAD | COR-L105 | I-CBP112 | MVP_top60_DNA+RNA_seed | ERBB4 | 0.347888 | STAD | NUGC-3 | PF-00299804 |
| MVP_top60_DNA_seed | PIK3CA | 0.984511 | LUAD | NCI-H3122 | AZD8835 | MVP_top60_DNA+RNA_seed | MET | 0.349223 | STAD | HGC-27 | JNJ38877605 |
| MVP_top60_DNA_seed | AURKA | 0.984927 | LUAD | NCI-H1755 | CD532 | MVP_top60_DNA+RNA_seed | SRC | 0.350355 | STAD | HSC-39 | Saracatinib |
| MVP_top60_DNA_seed | BRAF | 0.986934 | LUAD | Calu-3 | PLX-4720 | MVP_top60_DNA+RNA_seed | ATM | 0.352218 | STAD | TGBC11TKB | KU-60019 |
| MVP_top60_DNA_seed | HSP90AA1 | 0.987651 | LUAD | RERF-LC-KJ | SNX-2112 | MVP_top60_DNA+RNA_seed | HSP90AA1 | 0.356116 | STAD | KATOIII | Elesclomol |
| MVP_top60_DNA_seed | MET | 0.98808 | LUAD | COR-L105 | JNJ38877605 | MVP_top60_DNA+RNA_seed | HSP90AB1 | 0.356116 | STAD | KATOIII | Elesclomol |
| MVP_top60_DNA_seed | EGFR | 0.988663 | LUAD | NCI-H1355 | CI-1033 | MVP_top60_DNA+RNA_seed | FGFR3 | 0.356645 | STAD | MKN45 | AZD4547 |
| MVP_top60_DNA_seed | ERBB2 | 0.988663 | LUAD | NCI-H1355 | CI-1033 | MVP_top60_DNA+RNA_seed | HDAC1 | 0.358198 | STAD | NUGC-3 | AR-42 |
| MVP_top60_DNA_seed | GSK3B | 0.989865 | LUAD | NCI-H1355 | CHIR-99021 | MVP_top60_DNA+RNA_seed | ATM | 0.359733 | STAD | RERF-GC-1B | CP466722 |
| MVP_top60_DNA_seed | FGFR1 | 0.990549 | LUAD | NCI-H1651 | FGFR_3831 | MVP_top60_DNA+RNA_seed | PARP1 | 0.360822 | STAD | KATOIII | PARP_9495 |
| MVP_top60_DNA_seed | MAPK1 | 0.990582 | LUAD | NCI-H2009 | FR-180204 | MVP_top60_DNA+RNA_seed | FGFR3 | 0.361168 | STAD | OCUM-1 | AZD4547 |
| MVP_top60_DNA_seed | AKT1 | 0.99159 | LUAD | NCI-H1693 | AKT inhibitor VIII | MVP_top60_DNA+RNA_seed | FGFR3 | 0.361257 | STAD | TGBC11TKB | AZD4547 |
| MVP_top60_DNA_seed | PIK3CA | 0.991887 | LUAD | NCI-H1781 | AZD8835 | MVP_top60_DNA+RNA_seed | HSP90AA1 | 0.36265 | STAD | MKN28 | SNX-2112 |
| MVP_top60_DNA_seed | BRAF | 0.992247 | LUAD | NCI-H1781 | PLX-4720 | MVP_top60_DNA+RNA_seed | HSP90AB1 | 0.36265 | STAD | MKN28 | SNX-2112 |
| MVP_top60_DNA_seed | EGFR | 0.992257 | LUAD | A549 | AST-1306 | MVP_top60_DNA+RNA_seed | AR | 0.364975 | STAD | NUGC-3 | Bicalutamide |
| MVP_top60_DNA_seed | GSK3B | 0.992405 | LUAD | SW1573 | KIN001-042 | MVP_top60_DNA+RNA_seed | PARP1 | 0.368801 | STAD | FU97 | Talazoparib |
| MVP_top60_DNA_seed | MDM2 | 0.99241 | LUAD | NCI-H1781 | Nutlin-3a (-) | MVP_top60_DNA+RNA_seed | HSP90AA1 | 0.375133 | STAD | 23132-87 | Luminespib |
| MVP_top60_DNA_seed | FGFR1 | 0.992511 | LUAD | NCI-H2405 | FGFR_3831 | MVP_top60_DNA+RNA_seed | HSP90AB1 | 0.375133 | STAD | 23132-87 | Luminespib |
| MVP_top60_DNA_seed | AURKA | 0.992517 | LUAD | RERF-LC-MS | GSK1070916 | MVP_top60_DNA+RNA_seed | PARP1 | 0.376367 | STAD | AGS | Veliparib |
| MVP_top60_DNA_seed | AKT1 | 0.99275 | LUAD | NCI-H650 | GSK690693 | MVP_top60_DNA+RNA_seed | PARP1 | 0.377109 | STAD | NUGC-3 | Olaparib |
| MVP_top60_DNA_seed | BRAF | 0.993986 | LUAD | NCI-H1793 | RAF_9304 | MVP_top60_DNA+RNA_seed | MDM2 | 0.380652 | STAD | SNU-16 | Nutlin-3a (-) |
| MVP_top60_DNA_seed | IGF1R | 0.994898 | LUAD | NCI-H1650 | Linsitinib | MVP_top60_DNA+RNA_seed | ESR1 | 0.380847 | STAD | NUGC-4 | Tamoxifen |
| MVP_top60_DNA_seed | GSK3B | 0.995482 | LUAD | NCI-H441 | SB216763 | MVP_top60_DNA+RNA_seed | ERBB2 | 0.381472 | STAD | FU97 | AZD8931 |
| MVP_top60_DNA_seed | MAPK1 | 0.997088 | LUAD | NCI-H1734 | VX-11e | MVP_top60_DNA+RNA_seed | ERBB3 | 0.381472 | STAD | FU97 | AZD8931 |
| MVP_top60_DNA_seed | EGFR | 0.99823 | LUAD | NCI-H2009 | Pelitinib | MVP_top60_DNA+RNA_seed | HDAC1 | 0.383309 | STAD | SNU-1 | Parthenolide |
| MVP_top60_DNA_seed | PIK3CA | 0.998522 | LUAD | NCI-H1838 | PI3Ka_4409 | MVP_top60_DNA+RNA_seed | HDAC1 | 0.383686 | STAD | MKN45 | JQ12 |
| MVP_top60_DNA_seed | SRC | 1.000038 | LUAD | NCI-H1650 | WH-4-023 | MVP_top60_DNA+RNA_seed | EP300 | 0.38651 | STAD | SK-GT-2 | I-CBP112 |
| MVP_top60_DNA_seed | MDM2 | 1.000742 | LUAD | NCI-H1563 | Serdemetan | MVP_top60_DNA+RNA_seed | CREBBP | 0.38651 | STAD | SK-GT-2 | I-CBP112 |
| MVP_top60_DNA_seed | AURKA | 1.000827 | LUAD | NCI-H1568 | HG-5-113-01 | MVP_top60_DNA+RNA_seed | HSP90AA1 | 0.391123 | STAD | GCIY | Tanespimycin |

| | | | | | | | | | | | |
|---|---|---|---|---|---|---|---|---|---|---|---|
| MVP_top60_DNA_seed | FGFR1 | 1.001329 | LUAD | NCI-H1568 | PD173074 | MVP_top60_DNA+RNA_seed | HSP90AB1 | 0.391123 | STAD | GCIY | Tanespimycin |
| MVP_top60_DNA_seed | AURKA | 1.002058 | LUAD | NCI-H2342 | ZM447439 | MVP_top60_DNA+RNA_seed | ERBB2 | 0.394715 | STAD | Hs-746T | PF-00299804 |
| MVP_top60_DNA_seed | HSP90AA1 | 1.002477 | LUAD | NCI-H1650 | CCT-018159 | MVP_top60_DNA+RNA_seed | ERBB4 | 0.394715 | STAD | Hs-746T | PF-00299804 |
| MVP_top60_DNA_seed | GSK3B | 1.002625 | LUAD | NCI-H2023 | CHIR-99021 | MVP_top60_DNA+RNA_seed | HSP90AA1 | 0.398494 | STAD | MKN7 | Elesclomol |
| MVP_top60_DNA_seed | FGFR1 | 1.003867 | LUAD | NCI-H1435 | FGFR_3831 | MVP_top60_DNA+RNA_seed | HSP90AB1 | 0.398494 | STAD | MKN7 | Elesclomol |
| MVP_top60_DNA_seed | ATM | 1.004184 | LUAD | NCI-H2405 | KU-55933 | MVP_top60_DNA+RNA_seed | MET | 0.399172 | STAD | FU97 | SU11274 |
| MVP_top60_DNA_seed | EGFR | 1.004463 | LUAD | NCI-H1838 | Lapatinib | MVP_top60_DNA+RNA_seed | PTK2 | 0.400635 | STAD | OCUM-1 | PF-562271 |
| MVP_top60_DNA_seed | ERBB2 | 1.004463 | LUAD | NCI-H1838 | Lapatinib | MVP_top60_DNA+RNA_seed | SRC | 0.401858 | STAD | NCI-N87 | NVP-BHG712 |
| MVP_top60_DNA_seed | EGFR | 1.004601 | LUAD | NCI-H2347 | Gefitinib | MVP_top60_DNA+RNA_seed | PARP1 | 0.402449 | STAD | SK-GT-2 | PARP_0108 |
| MVP_top60_DNA_seed | ERBB2 | 1.005361 | LUAD | NCI-H650 | CP724714 | MVP_top60_DNA+RNA_seed | PARP1 | 0.406338 | STAD | HSC-39 | Olaparib |
| MVP_top60_DNA_seed | GSK3B | 1.005593 | LUAD | RERF-LC-KJ | SB216763 | MVP_top60_DNA+RNA_seed | SRC | 0.409036 | STAD | GCIY | Bosutinib |
| MVP_top60_DNA_seed | FGFR1 | 1.005685 | LUAD | NCI-H2347 | PD173074 | MVP_top60_DNA+RNA_seed | ATM | 0.410336 | STAD | NUGC-3 | KU-60019 |
| MVP_top60_DNA_seed | FGFR1 | 1.005693 | LUAD | A549 | PD173074 | MVP_top60_DNA+RNA_seed | HDAC1 | 0.410866 | STAD | TGBC11TKB | Tubastatin A |
| MVP_top60_DNA_seed | IGF1R | 1.006802 | LUAD | Calu-3 | BMS-754807 | MVP_top60_DNA+RNA_seed | HDAC1 | 0.41522 | STAD | MKN7 | Belinostat |
| MVP_top60_DNA_seed | GSK3B | 1.007467 | LUAD | EKVX | CHIR-99021 | MVP_top60_DNA+RNA_seed | PARP1 | 0.416395 | STAD | RERF-GC-1B | Rucaparib |
| MVP_top60_DNA_seed | BRAF | 1.008448 | LUAD | RERF-LC-MS | HG6-64-1 | MVP_top60_DNA+RNA_seed | PARP1 | 0.416808 | STAD | FU97 | PARP_0108 |
| MVP_top60_DNA_seed | MET | 1.008848 | LUAD | COR-L105 | SU11274 | MVP_top60_DNA+RNA_seed | MDM2 | 0.420834 | STAD | KATOIII | Serdemetan |
| MVP_top60_DNA_seed | MET | 1.009157 | LUAD | NCI-H1838 | SU11274 | MVP_top60_DNA+RNA_seed | ATM | 0.42355 | STAD | MKN28 | KU-55933 |
| MVP_top60_DNA_seed | AURKA | 1.010572 | LUAD | NCI-H1650 | Genentech Cpd 10 | MVP_top60_DNA+RNA_seed | ATM | 0.423844 | STAD | NCI-N87 | torin2 |
| MVP_top60_DNA_seed | BRAF | 1.010652 | LUAD | NCI-H1563 | PLX-4720 | MVP_top60_DNA+RNA_seed | ERBB2 | 0.424051 | STAD | SNU-5 | Afatinib |
| MVP_top60_DNA_seed | AURKA | 1.011255 | LUAD | NCI-H1755 | Alisertib | MVP_top60_DNA+RNA_seed | GSK3B | 0.425202 | STAD | SK-GT-2 | CHIR-99021 |
| MVP_top60_DNA_seed | MDM2 | 1.01134 | LUAD | PC-3_[JPC-3] | Serdemetan | MVP_top60_DNA+RNA_seed | AR | 0.42524 | STAD | SNU-1 | AZD3514 |
| MVP_top60_DNA_seed | PTK2 | 1.014873 | LUAD | NCI-H23 | PF-562271 | MVP_top60_DNA+RNA_seed | HDAC1 | 0.425624 | STAD | GCIY | Entinostat |
| MVP_top60_DNA_seed | EGFR | 1.015058 | LUAD | ABC-1 | Afatinib | MVP_top60_DNA+RNA_seed | HDAC3 | 0.425624 | STAD | GCIY | Entinostat |
| MVP_top60_DNA_seed | ERBB2 | 1.015058 | LUAD | ABC-1 | Afatinib | MVP_top60_DNA+RNA_seed | ERBB2 | 0.42897 | STAD | MKN45 | CI-1033 |
| MVP_top60_DNA_seed | EGFR | 1.015762 | LUAD | NCI-H650 | Gefitinib | MVP_top60_DNA+RNA_seed | ERBB4 | 0.42897 | STAD | MKN45 | CI-1033 |
| MVP_top60_DNA_seed | EGFR | 1.015941 | LUAD | NCI-H1563 | Gefitinib | MVP_top60_DNA+RNA_seed | MET | 0.432479 | STAD | NUGC-3 | AZD6094 |
| MVP_top60_DNA_seed | BRAF | 1.016349 | LUAD | NCI-H441 | HG6-64-1 | MVP_top60_DNA+RNA_seed | AR | 0.435623 | STAD | NUGC-3 | Bicalutamide |
| MVP_top60_DNA_seed | IGF1R | 1.017024 | LUAD | NCI-H1573 | BMS-754807 | MVP_top60_DNA+RNA_seed | PARP1 | 0.437268 | STAD | MKN7 | PARP_9495 |
| MVP_top60_DNA_seed | AKT1 | 1.017046 | LUAD | VMRC-LCD | MK-2206 | MVP_top60_DNA+RNA_seed | FGFR3 | 0.437522 | STAD | IM-95 | AZD4547 |
| MVP_top60_DNA_seed | MET | 1.017406 | LUAD | NCI-H1435 | SU11274 | MVP_top60_DNA+RNA_seed | ATM | 0.438933 | STAD | 23132-87 | CP466722 |
| MVP_top60_DNA_seed | AKT1 | 1.020157 | LUAD | NCI-H1435 | GSK690693 | MVP_top60_DNA+RNA_seed | MET | 0.444196 | STAD | SNU-16 | Crizotinib |
| MVP_top60_DNA_seed | AKT1 | 1.020942 | LUAD | RERF-LC-KJ | BAY AKT1 | MVP_top60_DNA+RNA_seed | PARP1 | 0.44638 | STAD | IM-95 | Veliparib |
| MVP_top60_DNA_seed | GSK3B | 1.021159 | LUAD | ABC-1 | CHIR-99021 | MVP_top60_DNA+RNA_seed | PARP1 | 0.446594 | STAD | RERF-GC-1B | PARP_0108 |
| MVP_top60_DNA_seed | HSP90AA1 | 1.024053 | LUAD | NCI-H2023 | SNX-2112 | MVP_top60_DNA+RNA_seed | MAPK1 | 0.448173 | STAD | OCUM-1 | VX-11e |
| MVP_top60_DNA_seed | MDM2 | 1.025251 | LUAD | NCI-H1623 | Nutlin-3a (-) | MVP_top60_DNA+RNA_seed | HDAC1 | 0.454467 | STAD | TGBC11TKB | Dacinostat |
| MVP_top60_DNA_seed | EGFR | 1.025671 | LUAD | NCI-H1650 | Erlotinib | MVP_top60_DNA+RNA_seed | ERBB2 | 0.45463 | STAD | RERF-GC-1B | Afatinib |
| MVP_top60_DNA_seed | HDAC1 | 1.026522 | LUAD | H3255 | AR-42 | MVP_top60_DNA+RNA_seed | GSK3B | 0.456557 | STAD | FU97 | CHIR-99021 |
| MVP_top60_DNA_seed | MAPK1 | 1.028918 | LUAD | NCI-H2228 | VX-11e | MVP_top60_DNA+RNA_seed | AR | 0.458698 | STAD | RERF-GC-1B | Bicalutamide |
| MVP_top60_DNA_seed | HDAC1 | 1.029931 | LUAD | RERF-LC-MS | Tubastatin A | MVP_top60_DNA+RNA_seed | FGFR3 | 0.460447 | STAD | NUGC-3 | PD173074 |
| MVP_top60_DNA_seed | ATM | 1.030683 | LUAD | EKVX | KU-60019 | MVP_top60_DNA+RNA_seed | ERBB2 | 0.462993 | STAD | Hs-746T | AZD8931 |

| | | | | | | | | | | | |
|---|---|---|---|---|---|---|---|---|---|---|---|
| MVP_top60_DNA_seed | HDAC1 | 1.032082 | LUAD | NCI-H2122 | CAY10603 | MVP_top60_DNA+RNA_seed | ERBB3 | 0.462993 | STAD | Hs-746T | AZD8931 |
| MVP_top60_DNA_seed | GSK3B | 1.032354 | LUAD | SK-LU-1 | CHIR-99021 | MVP_top60_DNA+RNA_seed | PARP1 | 0.464959 | STAD | HSC-39 | PARP_9495 |
| MVP_top60_DNA_seed | EGFR | 1.032886 | LUAD | A549 | PF-00299804 | MVP_top60_DNA+RNA_seed | HDAC1 | 0.467097 | STAD | FU97 | AR-42 |
| MVP_top60_DNA_seed | ERBB2 | 1.032886 | LUAD | A549 | PF-00299804 | MVP_top60_DNA+RNA_seed | HSP90AA1 | 0.467951 | STAD | SNU-5 | Luminespib |
| MVP_top60_DNA_seed | BRAF | 1.033739 | LUAD | NCI-H1623 | SB590885 | MVP_top60_DNA+RNA_seed | HSP90AB1 | 0.467951 | STAD | SNU-5 | Luminespib |
| MVP_top60_DNA_seed | MET | 1.034567 | LUAD | NCI-H1563 | SU11274 | MVP_top60_DNA+RNA_seed | FGFR3 | 0.468195 | STAD | SK-GT-2 | AZD4547 |
| MVP_top60_DNA_seed | AKT1 | 1.035559 | LUAD | SK-LU-1 | GSK690693 | MVP_top60_DNA+RNA_seed | HSP90AA1 | 0.469142 | STAD | SNU-5 | Elesclomol |
| MVP_top60_DNA_seed | BRAF | 1.035902 | LUAD | Calu-3 | SB590885 | MVP_top60_DNA+RNA_seed | HSP90AB1 | 0.469142 | STAD | SNU-5 | Elesclomol |
| MVP_top60_DNA_seed | ATM | 1.03711 | LUAD | NCI-H1666 | CP466722 | MVP_top60_DNA+RNA_seed | PARP1 | 0.477503 | STAD | Hs-746T | Olaparib |
| MVP_top60_DNA_seed | AURKA | 1.039014 | LUAD | LC-2-ad | CD532 | MVP_top60_DNA+RNA_seed | AR | 0.478677 | STAD | SNU-5 | Bicalutamide |
| MVP_top60_DNA_seed | EGFR | 1.04033 | LUAD | NCI-H2347 | AZD8931 | MVP_top60_DNA+RNA_seed | HDAC1 | 0.479923 | STAD | MKN7 | AR-42 |
| MVP_top60_DNA_seed | ERBB2 | 1.04033 | LUAD | NCI-H2347 | AZD8931 | MVP_top60_DNA+RNA_seed | ERBB2 | 0.48105 | STAD | MKN28 | CUDC-101 |
| MVP_top60_DNA_seed | ERBB3 | 1.04033 | LUAD | NCI-H2347 | AZD8931 | MVP_top60_DNA+RNA_seed | HSP90AA1 | 0.481642 | STAD | NUGC-3 | SNX-2112 |
| MVP_top60_DNA_seed | AKT1 | 1.041023 | LUAD | Calu-6 | MK-2206 | MVP_top60_DNA+RNA_seed | HSP90AB1 | 0.481642 | STAD | NUGC-3 | SNX-2112 |
| MVP_top60_DNA_seed | MAPK1 | 1.041062 | LUAD | NCI-H1435 | VX-11e | MVP_top60_DNA+RNA_seed | PARP1 | 0.483113 | STAD | NCI-N87 | PARP_0108 |
| MVP_top60_DNA_seed | ATM | 1.042822 | LUAD | NCI-H2023 | CP466722 | MVP_top60_DNA+RNA_seed | MAPK1 | 0.484288 | STAD | TMK-1 | VX-11e |
| MVP_top60_DNA_seed | AURKA | 1.04299 | LUAD | NCI-H1623 | HG-5-113-01 | MVP_top60_DNA+RNA_seed | ESR1 | 0.486328 | STAD | IM-95 | Fulvestrant |
| MVP_top60_DNA_seed | FGFR1 | 1.043012 | LUAD | COR-L105 | AZD4547 | MVP_top60_DNA+RNA_seed | AR | 0.487764 | STAD | 23132-87 | Bicalutamide |
| MVP_top60_DNA_seed | CREBBP | 1.04327 | LUAD | RERF-LC-MS | I-CBP112 | MVP_top60_DNA+RNA_seed | ERBB2 | 0.489956 | STAD | 23132-87 | Afatinib |
| MVP_top60_DNA_seed | AKT1 | 1.043669 | LUAD | NCI-H1666 | AKT inhibitor VIII | MVP_top60_DNA+RNA_seed | HSP90AA1 | 0.490265 | STAD | RF-48 | Tanespimycin |
| MVP_top60_DNA_seed | PIK3CA | 1.04412 | LUAD | NCI-H322M | PI3Ka_4409 | MVP_top60_DNA+RNA_seed | HSP90AB1 | 0.490265 | STAD | RF-48 | Tanespimycin |
| MVP_top60_DNA_seed | AKT1 | 1.044288 | LUAD | NCI-H2087 | BAY AKT1 | MVP_top60_DNA+RNA_seed | FGFR3 | 0.495112 | STAD | MKN28 | PD173074 |
| MVP_top60_DNA_seed | HDAC1 | 1.045246 | LUAD | NCI-H1563 | Tubastatin A | MVP_top60_DNA+RNA_seed | FGFR3 | 0.495888 | STAD | IM-95 | FGFR_3831 |
| MVP_top60_DNA_seed | HDAC1 | 1.045299 | LUAD | NCI-H1781 | CAY10603 | MVP_top60_DNA+RNA_seed | ERBB2 | 0.496494 | STAD | KATOIII | AZD8931 |
| MVP_top60_DNA_seed | SRC | 1.045378 | LUAD | NCI-H2087 | HG-6-71-01 | MVP_top60_DNA+RNA_seed | ERBB3 | 0.496494 | STAD | KATOIII | AZD8931 |
| MVP_top60_DNA_seed | AURKA | 1.048357 | LUAD | NCI-H2122 | Alisertib | MVP_top60_DNA+RNA_seed | HSP90AA1 | 0.499024 | STAD | MKN28 | CCT-018159 |
| MVP_top60_DNA_seed | SRC | 1.048389 | LUAD | NCI-H2228 | NVP-BHG712 | MVP_top60_DNA+RNA_seed | HSP90AB1 | 0.499024 | STAD | MKN28 | CCT-018159 |
| MVP_top60_DNA_seed | MAPK1 | 1.048728 | LUAD | NCI-H1838 | VX-11e | MVP_top60_DNA+RNA_seed | PIK3CA | 0.500928 | STAD | MKN45 | PI3Ka_4409 |
| MVP_top60_DNA_seed | EGFR | 1.050728 | LUAD | NCI-H441 | CUDC-101 | MVP_top60_DNA+RNA_seed | HDAC1 | 0.502886 | STAD | NUGC-4 | JQ12 |
| MVP_top60_DNA_seed | ERBB2 | 1.050728 | LUAD | NCI-H441 | CUDC-101 | MVP_top60_DNA+RNA_seed | PARP1 | 0.504494 | STAD | KATOIII | Veliparib |
| MVP_top60_DNA_seed | AKT1 | 1.051112 | LUAD | NCI-H1693 | AKT inhibitor VIII | MVP_top60_DNA+RNA_seed | MDM2 | 0.505255 | STAD | SNU-5 | Nutlin-3a (-) |
| MVP_top60_DNA_seed | BRAF | 1.052421 | LUAD | NCI-H1693 | PLX-4720 | MVP_top60_DNA+RNA_seed | HDAC1 | 0.506164 | STAD | MKN28 | Belinostat |
| MVP_top60_DNA_seed | AURKA | 1.052634 | LUAD | NCI-H322M | CD532 | MVP_top60_DNA+RNA_seed | ERBB2 | 0.507041 | STAD | 23132-87 | Afatinib |
| MVP_top60_DNA_seed | IGF1R | 1.053226 | LUAD | NCI-H322M | Linsitinib | MVP_top60_DNA+RNA_seed | PIK3CA | 0.511737 | STAD | MKN28 | PI3Ka_4409 |
| MVP_top60_DNA_seed | BRAF | 1.053772 | LUAD | NCI-H1650 | PLX-4720 | MVP_top60_DNA+RNA_seed | PARP1 | 0.512887 | STAD | TGBC11TKB | Olaparib |
| MVP_top60_DNA_seed | MET | 1.053938 | LUAD | NCI-H838 | SU11274 | MVP_top60_DNA+RNA_seed | SRC | 0.514907 | STAD | IM-95 | Bosutinib |
| MVP_top60_DNA_seed | EGFR | 1.054576 | LUAD | NCI-H2087 | Gefitinib | MVP_top60_DNA+RNA_seed | ERBB2 | 0.514996 | STAD | RF-48 | CI-1033 |
| MVP_top60_DNA_seed | MDM2 | 1.055422 | LUAD | NCI-H1435 | Nutlin-3a (-) | MVP_top60_DNA+RNA_seed | ERBB4 | 0.514996 | STAD | RF-48 | CI-1033 |
| MVP_top60_DNA_seed | EGFR | 1.056975 | LUAD | NCI-H1355 | PF-00299804 | MVP_top60_DNA+RNA_seed | GSK3B | 0.51567 | STAD | FU97 | SB216763 |
| MVP_top60_DNA_seed | ERBB2 | 1.056975 | LUAD | NCI-H1355 | PF-00299804 | MVP_top60_DNA+RNA_seed | MAPK1 | 0.517562 | STAD | NUGC-3 | VX-11e |
| MVP_top60_DNA_seed | ERBB2 | 1.05826 | LUAD | A549 | CP724714 | MVP_top60_DNA+RNA_seed | HDAC1 | 0.518522 | STAD | MKN45 | Dacinostat |

| | | | | | | | | | | | |
|---|---|---|---|---|---|---|---|---|---|---|---|
| MVP_top60_DNA_seed | MET | 1.058264 | LUAD | NCI-H2228 | JNJ38877605 | MVP_top60_DNA+RNA_seed | PIK3CA | 0.519231 | STAD | NUGC-4 | PI3Ka_4409 |
| MVP_top60_DNA_seed | AURKA | 1.059185 | LUAD | NCI-H1993 | GSK1070916 | MVP_top60_DNA+RNA_seed | ERBB2 | 0.522888 | STAD | MKN7 | CUDC-101 |
| MVP_top60_DNA_seed | GSK3B | 1.06077 | LUAD | SK-LU-1 | CHIR-99021 | MVP_top60_DNA+RNA_seed | PIK3CA | 0.523535 | STAD | SNU-16 | PI3Ka_4409 |
| MVP_top60_DNA_seed | MAPK1 | 1.061257 | LUAD | EKVX | VX-11e | MVP_top60_DNA+RNA_seed | ESR1 | 0.524239 | STAD | MKN7 | Fulvestrant |
| MVP_top60_DNA_seed | HDAC1 | 1.062302 | LUAD | NCI-H1651 | JQ12 | MVP_top60_DNA+RNA_seed | MAPK1 | 0.525733 | STAD | OCUM-1 | FR-180204 |
| MVP_top60_DNA_seed | IGF1R | 1.063105 | LUAD | NCI-H322M | BMS-754807 | MVP_top60_DNA+RNA_seed | MAPK3 | 0.525733 | STAD | OCUM-1 | FR-180204 |
| MVP_top60_DNA_seed | MAPK1 | 1.063178 | LUAD | NCI-H2228 | FR-180204 | MVP_top60_DNA+RNA_seed | MET | 0.525976 | STAD | RERF-GC-1B | SU11274 |
| MVP_top60_DNA_seed | HDAC1 | 1.063332 | LUAD | NCI-H1648 | Parthenolide | MVP_top60_DNA+RNA_seed | MET | 0.529698 | STAD | NCI-N87 | Crizotinib |
| MVP_top60_DNA_seed | MDM2 | 1.063434 | LUAD | RERF-LC-MS | Nutlin-3a (-) | MVP_top60_DNA+RNA_seed | PARP1 | 0.530458 | STAD | SNU-1 | Olaparib |
| MVP_top60_DNA_seed | GSK3B | 1.064528 | LUAD | NCI-H1648 | KIN001-042 | MVP_top60_DNA+RNA_seed | HDAC1 | 0.530899 | STAD | HSC-39 | Parthenolide |
| MVP_top60_DNA_seed | IGF1R | 1.064807 | LUAD | NCI-H1693 | BMS-536924 | MVP_top60_DNA+RNA_seed | AR | 0.531485 | STAD | NUGC-4 | Bicalutamide |
| MVP_top60_DNA_seed | SRC | 1.065797 | LUAD | NCI-H1666 | NVP-BHG712 | MVP_top60_DNA+RNA_seed | GSK3B | 0.53271 | STAD | RERF-GC-1B | CHIR-99021 |
| MVP_top60_DNA_seed | IGF1R | 1.066045 | LUAD | Calu-3 | GSK1904529A | MVP_top60_DNA+RNA_seed | SRC | 0.534293 | STAD | MKN28 | NVP-BHG712 |
| MVP_top60_DNA_seed | MET | 1.066741 | LUAD | NCI-H1755 | JNJ38877605 | MVP_top60_DNA+RNA_seed | GSK3B | 0.534869 | STAD | 23132-87 | KIN001-042 |
| MVP_top60_DNA_seed | AURKA | 1.067083 | LUAD | NCI-H1437 | CD532 | MVP_top60_DNA+RNA_seed | ERBB2 | 0.536215 | STAD | 23132-87 | CP724714 |
| MVP_top60_DNA_seed | BRAF | 1.067395 | LUAD | RERF-LC-MS | PLX-4720 | MVP_top60_DNA+RNA_seed | HDAC1 | 0.544415 | STAD | SNU-16 | Entinostat |
| MVP_top60_DNA_seed | BRAF | 1.067736 | LUAD | PC-3_[JPC-3] | SB590885 | MVP_top60_DNA+RNA_seed | HDAC3 | 0.544415 | STAD | SNU-16 | Entinostat |
| MVP_top60_DNA_seed | AURKA | 1.06919 | LUAD | NCI-H650 | ZM447439 | MVP_top60_DNA+RNA_seed | FGFR3 | 0.544572 | STAD | IM-95 | PD173074 |
| MVP_top60_DNA_seed | EGFR | 1.069313 | LUAD | NCI-H1355 | Afatinib | MVP_top60_DNA+RNA_seed | MAPK1 | 0.547019 | STAD | HSC-39 | VX-11e |
| MVP_top60_DNA_seed | ERBB2 | 1.069313 | LUAD | NCI-H1355 | Afatinib | MVP_top60_DNA+RNA_seed | PARP1 | 0.548019 | STAD | SK-GT-2 | Olaparib |
| MVP_top60_DNA_seed | MDM2 | 1.069513 | LUAD | NCI-H1975 | Nutlin-3a (-) | MVP_top60_DNA+RNA_seed | GSK3B | 0.548838 | STAD | GCIY | SB216763 |
| MVP_top60_DNA_seed | MDM2 | 1.06964 | LUAD | NCI-H838 | Nutlin-3a (-) | MVP_top60_DNA+RNA_seed | GSK3B | 0.55106 | STAD | Hs-746T | CHIR-99021 |
| MVP_top60_DNA_seed | CREBBP | 1.070199 | LUAD | ABC-1 | I-CBP112 | MVP_top60_DNA+RNA_seed | PARP1 | 0.551645 | STAD | 23132-87 | Olaparib |
| MVP_top60_DNA_seed | GSK3B | 1.070289 | LUAD | NCI-H1568 | CHIR-99021 | MVP_top60_DNA+RNA_seed | GSK3B | 0.552627 | STAD | IM-95 | SB216763 |
| MVP_top60_DNA_seed | AKT1 | 1.070439 | LUAD | Calu-6 | BAY AKT1 | MVP_top60_DNA+RNA_seed | AR | 0.553287 | STAD | MKN45 | AZD3514 |
| MVP_top60_DNA_seed | AURKA | 1.072425 | LUAD | NCI-H1993 | CD532 | MVP_top60_DNA+RNA_seed | PIK3CA | 0.554986 | STAD | KATOIII | PI3Ka_4409 |
| MVP_top60_DNA_seed | BRAF | 1.072544 | LUAD | NCI-H2023 | PLX-4720 | MVP_top60_DNA+RNA_seed | MET | 0.55719 | STAD | RF-48 | AZD6094 |
| MVP_top60_DNA_seed | HDAC1 | 1.07317 | LUAD | NCI-H2405 | CAY10603 | MVP_top60_DNA+RNA_seed | FGFR3 | 0.560398 | STAD | RF-48 | AZD4547 |
| MVP_top60_DNA_seed | AKT1 | 1.0732 | LUAD | NCI-H1693 | BAY AKT1 | MVP_top60_DNA+RNA_seed | AR | 0.561141 | STAD | NCI-N87 | AZD3514 |
| MVP_top60_DNA_seed | MAPK1 | 1.073212 | LUAD | NCI-H1568 | VX-11e | MVP_top60_DNA+RNA_seed | ERBB2 | 0.563994 | STAD | OCUM-1 | AZD8931 |
| MVP_top60_DNA_seed | GSK3B | 1.074946 | LUAD | H3255 | SB216763 | MVP_top60_DNA+RNA_seed | ERBB3 | 0.563994 | STAD | OCUM-1 | AZD8931 |
| MVP_top60_DNA_seed | AURKA | 1.075098 | LUAD | EKVX | Alisertib | MVP_top60_DNA+RNA_seed | ERBB2 | 0.564898 | STAD | 23132-87 | PF-00299804 |
| MVP_top60_DNA_seed | AURKA | 1.075172 | LUAD | NCI-H2342 | GSK1070916 | MVP_top60_DNA+RNA_seed | ERBB4 | 0.564898 | STAD | 23132-87 | PF-00299804 |
| MVP_top60_DNA_seed | HSP90AA1 | 1.075897 | LUAD | NCI-H1435 | SNX-2112 | MVP_top60_DNA+RNA_seed | AR | 0.566906 | STAD | NUGC-4 | Bicalutamide |
| MVP_top60_DNA_seed | MAPK1 | 1.076564 | LUAD | NCI-H1651 | VX-11e | MVP_top60_DNA+RNA_seed | GSK3B | 0.567159 | STAD | KATOIII | KIN001-042 |
| MVP_top60_DNA_seed | HSP90AA1 | 1.077203 | LUAD | NCI-H1993 | Tanespimycin | MVP_top60_DNA+RNA_seed | SRC | 0.568032 | STAD | NUGC-3 | NVP-BHG712 |
| MVP_top60_DNA_seed | PTK2 | 1.078115 | LUAD | NCI-H1355 | PF-562271 | MVP_top60_DNA+RNA_seed | ESR1 | 0.568881 | STAD | SNU-1 | Tamoxifen |
| MVP_top60_DNA_seed | CREBBP | 1.079394 | LUAD | NCI-H1993 | I-CBP112 | MVP_top60_DNA+RNA_seed | ERBB2 | 0.569323 | STAD | FU97 | Afatinib |
| MVP_top60_DNA_seed | SRC | 1.080809 | LUAD | NCI-H1435 | NVP-BHG712 | MVP_top60_DNA+RNA_seed | FGFR3 | 0.570671 | STAD | IM-95 | AZD4547 |
| MVP_top60_DNA_seed | GSK3B | 1.080851 | LUAD | NCI-H1435 | CHIR-99021 | MVP_top60_DNA+RNA_seed | PIK3CA | 0.572245 | STAD | OCUM-1 | PI3Ka_4409 |
| MVP_top60_DNA_seed | MDM2 | 1.081121 | LUAD | NCI-H1568 | Nutlin-3a (-) | MVP_top60_DNA+RNA_seed | SRC | 0.572897 | STAD | FU97 | Bosutinib |

| | | | | | | | | | | | |
|---|---|---|---|---|---|---|---|---|---|---|---|
| MVP_top60_DNA_seed | FGFR1 | 1.081756 | LUAD | PC-3_[JPC-3] | FGFR_3831 | MVP_top60_DNA+RNA_seed | MET | 0.573119 | STAD | SNU-1 | AZD6094 |
| MVP_top60_DNA_seed | GSK3B | 1.081991 | LUAD | NCI-H1650 | SB216763 | MVP_top60_DNA+RNA_seed | MET | 0.573554 | STAD | MKN28 | JNJ38877605 |
| MVP_top60_DNA_seed | AURKA | 1.082671 | LUAD | NCI-H1435 | GSK1070916 | MVP_top60_DNA+RNA_seed | FYN | 0.575957 | STAD | HSC-39 | A-770041 |
| MVP_top60_DNA_seed | GSK3B | 1.08442 | LUAD | NCI-H1651 | CHIR-99021 | MVP_top60_DNA+RNA_seed | ERBB2 | 0.577607 | STAD | KATOIII | Afatinib |
| MVP_top60_DNA_seed | IGF1R | 1.08468 | LUAD | NCI-H522 | BMS-536924 | MVP_top60_DNA+RNA_seed | GSK3B | 0.578095 | STAD | MKN45 | CHIR-99021 |
| MVP_top60_DNA_seed | PTK2 | 1.087044 | LUAD | RERF-LC-KJ | PF-562271 | MVP_top60_DNA+RNA_seed | PARP1 | 0.579509 | STAD | FU97 | PARP_9495 |
| MVP_top60_DNA_seed | HSP90AA1 | 1.087114 | LUAD | H3255 | SNX-2112 | MVP_top60_DNA+RNA_seed | ATM | 0.579775 | STAD | MKN7 | CP466722 |
| MVP_top60_DNA_seed | FGFR1 | 1.087179 | LUAD | EKVX | FGFR_3831 | MVP_top60_DNA+RNA_seed | PARP1 | 0.585312 | STAD | FU97 | PARP_9482 |
| MVP_top60_DNA_seed | AKT1 | 1.090283 | LUAD | NCI-H1651 | AKT inhibitor VIII | MVP_top60_DNA+RNA_seed | AR | 0.58645 | STAD | TMK-1 | Bicalutamide |
| MVP_top60_DNA_seed | EGFR | 1.09056 | LUAD | EKVX | CUDC-101 | MVP_top60_DNA+RNA_seed | HSP90AA1 | 0.586509 | STAD | RF-48 | CCT-018159 |
| MVP_top60_DNA_seed | ERBB2 | 1.09056 | LUAD | EKVX | CUDC-101 | MVP_top60_DNA+RNA_seed | HSP90AB1 | 0.586509 | STAD | RF-48 | CCT-018159 |
| MVP_top60_DNA_seed | AKT1 | 1.093007 | LUAD | NCI-H23 | AKT inhibitor VIII | MVP_top60_DNA+RNA_seed | PIK3CA | 0.589523 | STAD | MKN7 | AZD8835 |
| MVP_top60_DNA_seed | ERBB2 | 1.093298 | LUAD | NCI-H2347 | CP724714 | MVP_top60_DNA+RNA_seed | MET | 0.597246 | STAD | AGS | AZD6094 |
| MVP_top60_DNA_seed | MET | 1.093329 | LUAD | NCI-H2087 | SU11274 | MVP_top60_DNA+RNA_seed | PARP1 | 0.598218 | STAD | HSC-39 | PARP_0108 |
| MVP_top60_DNA_seed | BRAF | 1.093706 | LUAD | NCI-H1355 | PLX-4720 | MVP_top60_DNA+RNA_seed | HSP90AA1 | 0.601442 | STAD | KATOIII | CCT-018159 |
| MVP_top60_DNA_seed | BRAF | 1.096096 | LUAD | HCC-78 | PLX-4720 | MVP_top60_DNA+RNA_seed | HSP90AB1 | 0.601442 | STAD | KATOIII | CCT-018159 |
| MVP_top60_DNA_seed | SRC | 1.099937 | LUAD | EMC-BAC-2 | NVP-BHG712 | MVP_top60_DNA+RNA_seed | PARP1 | 0.602563 | STAD | RERF-GC-1B | PARP_9482 |
| MVP_top60_DNA_seed | AURKA | 1.100322 | LUAD | NCI-H1568 | CD532 | MVP_top60_DNA+RNA_seed | AR | 0.605168 | STAD | SNU-1 | Bicalutamide |
| MVP_top60_DNA_seed | SRC | 1.100758 | LUAD | NCI-H1838 | Saracatinib | MVP_top60_DNA+RNA_seed | HSP90AA1 | 0.60624 | STAD | RERF-GC-1B | SNX-2112 |
| MVP_top60_DNA_seed | SRC | 1.100992 | LUAD | NCI-H1781 | Bosutinib | MVP_top60_DNA+RNA_seed | HSP90AB1 | 0.60624 | STAD | RERF-GC-1B | SNX-2112 |
| MVP_top60_DNA_seed | EGFR | 1.101385 | LUAD | RERF-LC-MS | Gefitinib | MVP_top60_DNA+RNA_seed | ESR1 | 0.606326 | STAD | SNU-5 | Tamoxifen |
| MVP_top60_DNA_seed | MET | 1.101426 | LUAD | NCI-H1838 | Crizotinib | MVP_top60_DNA+RNA_seed | MDM2 | 0.612469 | STAD | SNU-16 | Serdemetan |
| MVP_top60_DNA_seed | AURKA | 1.101509 | LUAD | HCC-827 | GSK1070916 | MVP_top60_DNA+RNA_seed | ERBB4 | 0.613193 | STAD | HGC-27 | AST-1306 |
| MVP_top60_DNA_seed | HDAC1 | 1.103154 | LUAD | RERF-LC-KJ | CAY10603 | MVP_top60_DNA+RNA_seed | SRC | 0.618218 | STAD | SNU-1 | Bosutinib |
| MVP_top60_DNA_seed | HDAC1 | 1.103951 | LUAD | A549 | Tubastatin A | MVP_top60_DNA+RNA_seed | GSK3B | 0.624287 | STAD | SNU-1 | CHIR-99021 |
| MVP_top60_DNA_seed | MAPK1 | 1.105672 | LUAD | NCI-H1355 | VX-11e | MVP_top60_DNA+RNA_seed | PARP1 | 0.6243 | STAD | SNU-16 | PARP_0108 |
| MVP_top60_DNA_seed | SRC | 1.106024 | LUAD | NCI-H1693 | NVP-BHG712 | MVP_top60_DNA+RNA_seed | ERBB2 | 0.624443 | STAD | MKN45 | Afatinib |
| MVP_top60_DNA_seed | EGFR | 1.106074 | LUAD | NCI-H1666 | Lapatinib | MVP_top60_DNA+RNA_seed | SRC | 0.624621 | STAD | Hs-746T | NVP-BHG712 |
| MVP_top60_DNA_seed | ERBB2 | 1.106074 | LUAD | NCI-H1666 | Lapatinib | MVP_top60_DNA+RNA_seed | SRC | 0.62529 | STAD | RF-48 | WH-4-023 |
| MVP_top60_DNA_seed | AKT1 | 1.108358 | LUAD | NCI-H322M | BAY AKT1 | MVP_top60_DNA+RNA_seed | ERBB2 | 0.628557 | STAD | TGBC11TKB | CUDC-101 |
| MVP_top60_DNA_seed | EGFR | 1.108485 | LUAD | NCI-H1573 | Pelitinib | MVP_top60_DNA+RNA_seed | FGFR3 | 0.630148 | STAD | 23132-87 | PD173074 |
| MVP_top60_DNA_seed | PIK3CA | 1.108915 | LUAD | NCI-H1793 | AZD8835 | MVP_top60_DNA+RNA_seed | ESR1 | 0.634474 | STAD | SK-GT-2 | Tamoxifen |
| MVP_top60_DNA_seed | MAPK1 | 1.110297 | LUAD | SK-LU-1 | FR-180204 | MVP_top60_DNA+RNA_seed | GSK3B | 0.634734 | STAD | TGBC11TKB | SB216763 |
| MVP_top60_DNA_seed | AURKA | 1.110766 | LUAD | NCI-H1437 | Alisertib | MVP_top60_DNA+RNA_seed | PIK3CA | 0.634969 | STAD | NUGC-3 | AZD8835 |
| MVP_top60_DNA_seed | MAPK1 | 1.111148 | LUAD | NCI-H650 | FR-180204 | MVP_top60_DNA+RNA_seed | HDAC1 | 0.635157 | STAD | SNU-1 | JQ12 |
| MVP_top60_DNA_seed | MDM2 | 1.115183 | LUAD | NCI-H2405 | Serdemetan | MVP_top60_DNA+RNA_seed | HDAC1 | 0.639133 | STAD | AGS | Tubastatin A |
| MVP_top60_DNA_seed | EGFR | 1.116088 | LUAD | NCI-H1666 | Erlotinib | MVP_top60_DNA+RNA_seed | ATM | 0.640117 | STAD | TGBC11TKB | CP466722 |
| MVP_top60_DNA_seed | BRAF | 1.116787 | LUAD | HCC-827 | PLX-4720 | MVP_top60_DNA+RNA_seed | PARP1 | 0.646549 | STAD | MKN28 | Veliparib |
| MVP_top60_DNA_seed | ATM | 1.116932 | LUAD | NCI-H1435 | CP466722 | MVP_top60_DNA+RNA_seed | ATM | 0.647483 | STAD | OCUM-1 | CP466722 |
| MVP_top60_DNA_seed | EGFR | 1.117154 | LUAD | NCI-H1435 | Cetuximab | MVP_top60_DNA+RNA_seed | PARP1 | 0.647902 | STAD | HSC-39 | Talazoparib |
| MVP_top60_DNA_seed | HSP90AA1 | 1.117761 | LUAD | NCI-H1693 | CCT-018159 | MVP_top60_DNA+RNA_seed | PARP1 | 0.651402 | STAD | MKN28 | PARP_0108 |

| | | | | | | | | | | | |
|---|---|---|---|---|---|---|---|---|---|---|---|
| MVP_top60_DNA_seed | HSP90AA1 | 1.117782 | LUAD | NCI-H1435 | Tanespimycin | MVP_top60_DNA+RNA_seed | GSK3B | 0.65237 | STAD | NUGC-4 | SB216763 |
| MVP_top60_DNA_seed | PIK3CA | 1.118268 | LUAD | NCI-H322M | AZD8835 | MVP_top60_DNA+RNA_seed | ERBB2 | 0.65275 | STAD | TMK-1 | CUDC-101 |
| MVP_top60_DNA_seed | PIK3CA | 1.119758 | LUAD | NCI-H1838 | AZD8835 | MVP_top60_DNA+RNA_seed | FYN | 0.659901 | STAD | SNU-16 | A-770041 |
| MVP_top60_DNA_seed | BRAF | 1.120456 | LUAD | COR-L105 | SB590885 | MVP_top60_DNA+RNA_seed | MDM2 | 0.661444 | STAD | RERF-GC-1B | Nutlin-3a (-) |
| MVP_top60_DNA_seed | HDAC1 | 1.121056 | LUAD | NCI-H2405 | Tubastatin A | MVP_top60_DNA+RNA_seed | SRC | 0.665729 | STAD | RERF-GC-1B | Bosutinib |
| MVP_top60_DNA_seed | BRAF | 1.121603 | LUAD | Calu-6 | PLX-4720 | MVP_top60_DNA+RNA_seed | PARP1 | 0.667806 | STAD | MKN45 | PARP_0108 |
| MVP_top60_DNA_seed | HSP90AA1 | 1.121867 | LUAD | Calu-3 | Tanespimycin | MVP_top60_DNA+RNA_seed | PARP1 | 0.671094 | STAD | MKN45 | Veliparib |
| MVP_top60_DNA_seed | HDAC1 | 1.122332 | LUAD | NCI-H2023 | Tubastatin A | MVP_top60_DNA+RNA_seed | GSK3B | 0.671458 | STAD | MKN45 | AZD7969 |
| MVP_top60_DNA_seed | MAPK1 | 1.122885 | LUAD | HCC-78 | VX-11e | MVP_top60_DNA+RNA_seed | ERBB2 | 0.673728 | STAD | SNU-16 | Afatinib |
| MVP_top60_DNA_seed | EGFR | 1.123169 | LUAD | NCI-H2023 | CUDC-101 | MVP_top60_DNA+RNA_seed | ERBB2 | 0.674507 | STAD | OCUM-1 | CP724714 |
| MVP_top60_DNA_seed | ERBB2 | 1.123169 | LUAD | NCI-H2023 | CUDC-101 | MVP_top60_DNA+RNA_seed | HDAC1 | 0.675068 | STAD | MKN28 | AR-42 |
| MVP_top60_DNA_seed | ERBB2 | 1.124652 | LUAD | EKVX | CP724714 | MVP_top60_DNA+RNA_seed | PARP1 | 0.676089 | STAD | Hs-746T | PARP_0108 |
| MVP_top60_DNA_seed | HDAC1 | 1.124975 | LUAD | PC-3_[JPC-3] | Tubastatin A | MVP_top60_DNA+RNA_seed | MET | 0.677291 | STAD | RERF-GC-1B | AZD6094 |
| MVP_top60_DNA_seed | HDAC1 | 1.128467 | LUAD | NCI-H1623 | JQ12 | MVP_top60_DNA+RNA_seed | ESR1 | 0.679352 | STAD | IM-95 | Tamoxifen |
| MVP_top60_DNA_seed | GSK3B | 1.129406 | LUAD | NCI-H522 | AZD7969 | MVP_top60_DNA+RNA_seed | SRC | 0.684412 | STAD | SNU-5 | WH-4-023 |
| MVP_top60_DNA_seed | GSK3B | 1.129717 | LUAD | NCI-H441 | CHIR-99021 | MVP_top60_DNA+RNA_seed | ERBB2 | 0.68628 | STAD | MKN28 | Afatinib |
| MVP_top60_DNA_seed | HSP90AA1 | 1.132215 | LUAD | NCI-H441 | CCT-018159 | MVP_top60_DNA+RNA_seed | GSK3B | 0.686417 | STAD | NUGC-3 | KIN001-042 |
| MVP_top60_DNA_seed | GSK3B | 1.132347 | LUAD | HCC-827 | CHIR-99021 | MVP_top60_DNA+RNA_seed | PARP1 | 0.686795 | STAD | RF-48 | PARP_0108 |
| MVP_top60_DNA_seed | GSK3B | 1.13328 | LUAD | ABC-1 | SB216763 | MVP_top60_DNA+RNA_seed | GSK3B | 0.687524 | STAD | MKN28 | KIN001-042 |
| MVP_top60_DNA_seed | MET | 1.133793 | LUAD | NCI-H322M | SU11274 | MVP_top60_DNA+RNA_seed | PARP1 | 0.688501 | STAD | MKN45 | Talazoparib |
| MVP_top60_DNA_seed | BRAF | 1.133866 | LUAD | NCI-H2347 | PLX-4720 | MVP_top60_DNA+RNA_seed | EP300 | 0.689219 | STAD | NUGC-3 | I-CBP112 |
| MVP_top60_DNA_seed | SRC | 1.13655 | LUAD | NCI-H2122 | Bosutinib | MVP_top60_DNA+RNA_seed | CREBBP | 0.689219 | STAD | NUGC-3 | I-CBP112 |
| MVP_top60_DNA_seed | EGFR | 1.136555 | LUAD | RERF-LC-KJ | AST-1306 | MVP_top60_DNA+RNA_seed | FGFR3 | 0.692894 | STAD | NUGC-4 | PD173074 |
| MVP_top60_DNA_seed | GSK3B | 1.137146 | LUAD | NCI-H2291 | AZD7969 | MVP_top60_DNA+RNA_seed | FGFR3 | 0.696224 | STAD | NCI-N87 | PD173074 |
| MVP_top60_DNA_seed | CREBBP | 1.138044 | LUAD | NCI-H838 | I-CBP112 | MVP_top60_DNA+RNA_seed | EP300 | 0.697561 | STAD | RERF-GC-1B | I-CBP112 |
| MVP_top60_DNA_seed | FGFR1 | 1.13853 | LUAD | NCI-H1838 | FGFR_3831 | MVP_top60_DNA+RNA_seed | CREBBP | 0.697561 | STAD | RERF-GC-1B | I-CBP112 |
| MVP_top60_DNA_seed | BRAF | 1.138736 | LUAD | NCI-H2085 | PLX-4720 | MVP_top60_DNA+RNA_seed | PARP1 | 0.698462 | STAD | NCI-N87 | Rucaparib |
| MVP_top60_DNA_seed | CREBBP | 1.139096 | LUAD | NCI-H1355 | I-CBP112 | MVP_top60_DNA+RNA_seed | ATM | 0.699749 | STAD | NUGC-4 | KU-55933 |
| MVP_top60_DNA_seed | ERBB2 | 1.139187 | LUAD | NCI-H1650 | CP724714 | MVP_top60_DNA+RNA_seed | HDAC1 | 0.700892 | STAD | NCI-N87 | Tubastatin A |
| MVP_top60_DNA_seed | HDAC1 | 1.140757 | LUAD | NCI-H441 | Belinostat | MVP_top60_DNA+RNA_seed | MET | 0.703018 | STAD | MKN7 | AZD6094 |
| MVP_top60_DNA_seed | MDM2 | 1.141426 | LUAD | NCI-H1666 | Serdemetan | MVP_top60_DNA+RNA_seed | HDAC1 | 0.707824 | STAD | TGBC11TKB | JQ12 |
| MVP_top60_DNA_seed | HSP90AA1 | 1.141735 | LUAD | A549 | Luminespib | MVP_top60_DNA+RNA_seed | GSK3B | 0.709362 | STAD | MKN7 | CHIR-99021 |
| MVP_top60_DNA_seed | MET | 1.142335 | LUAD | NCI-H1623 | AZD6094 | MVP_top60_DNA+RNA_seed | PARP1 | 0.710387 | STAD | KATOIII | Olaparib |
| MVP_top60_DNA_seed | BRAF | 1.142852 | LUAD | NCI-H441 | PLX-4720 | MVP_top60_DNA+RNA_seed | MAPK1 | 0.711501 | STAD | Hs-746T | VX-11e |
| MVP_top60_DNA_seed | HSP90AA1 | 1.145287 | LUAD | NCI-H2342 | Tanespimycin | MVP_top60_DNA+RNA_seed | PARP1 | 0.71177 | STAD | RF-48 | Rucaparib |
| MVP_top60_DNA_seed | MET | 1.146865 | LUAD | ABC-1 | AZD6094 | MVP_top60_DNA+RNA_seed | PIK3CA | 0.712438 | STAD | NUGC-3 | PI3Ka_4409 |
| MVP_top60_DNA_seed | CREBBP | 1.149013 | LUAD | NCI-H1648 | I-CBP112 | MVP_top60_DNA+RNA_seed | ERBB2 | 0.713247 | STAD | RF-48 | AZD8931 |
| MVP_top60_DNA_seed | AKT1 | 1.151738 | LUAD | NCI-H838 | AKT inhibitor VIII | MVP_top60_DNA+RNA_seed | ERBB3 | 0.713247 | STAD | RF-48 | AZD8931 |
| MVP_top60_DNA_seed | MAPK1 | 1.152178 | LUAD | A549 | VX-11e | MVP_top60_DNA+RNA_seed | HSP90AA1 | 0.714843 | STAD | GCIY | SNX-2112 |
| MVP_top60_DNA_seed | GSK3B | 1.152276 | LUAD | NCI-H522 | CHIR-99021 | MVP_top60_DNA+RNA_seed | HSP90AB1 | 0.714843 | STAD | GCIY | SNX-2112 |
| MVP_top60_DNA_seed | FGFR1 | 1.154371 | LUAD | NCI-H1573 | FGFR_3831 | MVP_top60_DNA+RNA_seed | PARP1 | 0.716804 | STAD | MKN28 | PARP_9482 |

| | | | | | | | | | | | |
|---|---|---|---|---|---|---|---|---|---|---|---|
| MVP_top60_DNA_seed | HDAC1 | 1.155033 | LUAD | RERF-LC-MS | Dacinostat | MVP_top60_DNA+RNA_seed | GSK3B | 0.723637 | STAD | RF-48 | KIN001-042 |
| MVP_top60_DNA_seed | PIK3CA | 1.156105 | LUAD | NCI-H1993 | AZD8835 | MVP_top60_DNA+RNA_seed | PARP1 | 0.724474 | STAD | SNU-16 | PARP_9482 |
| MVP_top60_DNA_seed | FGFR1 | 1.156186 | LUAD | Calu-3 | PD173074 | MVP_top60_DNA+RNA_seed | HSP90AA1 | 0.72489 | STAD | FU97 | SNX-2112 |
| MVP_top60_DNA_seed | SRC | 1.157476 | LUAD | NCI-H1666 | WH-4-023 | MVP_top60_DNA+RNA_seed | HSP90AB1 | 0.72489 | STAD | FU97 | SNX-2112 |
| MVP_top60_DNA_seed | FGFR1 | 1.157557 | LUAD | ABC-1 | AZD4547 | MVP_top60_DNA+RNA_seed | ERBB2 | 0.725846 | STAD | SNU-5 | Lapatinib |
| MVP_top60_DNA_seed | MDM2 | 1.158417 | LUAD | PC-3_[JPC-3] | Nutlin-3a (-) | MVP_top60_DNA+RNA_seed | PARP1 | 0.728628 | STAD | MKN7 | Talazoparib |
| MVP_top60_DNA_seed | IGF1R | 1.158795 | LUAD | NCI-H322M | BMS-536924 | MVP_top60_DNA+RNA_seed | GSK3B | 0.729397 | STAD | SNU-5 | KIN001-042 |
| MVP_top60_DNA_seed | EGFR | 1.158978 | LUAD | NCI-H2228 | Afatinib | MVP_top60_DNA+RNA_seed | PIK3CA | 0.732466 | STAD | TGBC11TKB | PI3Ka_4409 |
| MVP_top60_DNA_seed | ERBB2 | 1.158978 | LUAD | NCI-H2228 | Afatinib | MVP_top60_DNA+RNA_seed | PARP1 | 0.73517 | STAD | SNU-16 | Rucaparib |
| MVP_top60_DNA_seed | EGFR | 1.161213 | LUAD | NCI-H2342 | CUDC-101 | MVP_top60_DNA+RNA_seed | PARP1 | 0.737408 | STAD | MKN45 | PARP_9482 |
| MVP_top60_DNA_seed | ERBB2 | 1.161213 | LUAD | NCI-H2342 | CUDC-101 | MVP_top60_DNA+RNA_seed | MDM2 | 0.738214 | STAD | SNU-5 | Serdemetan |
| MVP_top60_DNA_seed | AURKA | 1.163882 | LUAD | NCI-H1993 | ZM447439 | MVP_top60_DNA+RNA_seed | ERBB2 | 0.741339 | STAD | KATOIII | Afatinib |
| MVP_top60_DNA_seed | EGFR | 1.164977 | LUAD | A549 | CUDC-101 | MVP_top60_DNA+RNA_seed | SRC | 0.741903 | STAD | SNU-5 | Bosutinib |
| MVP_top60_DNA_seed | ERBB2 | 1.164977 | LUAD | A549 | CUDC-101 | MVP_top60_DNA+RNA_seed | HDAC1 | 0.743373 | STAD | SNU-16 | Parthenolide |
| MVP_top60_DNA_seed | SRC | 1.165021 | LUAD | HCC-78 | Bosutinib | MVP_top60_DNA+RNA_seed | MAPK1 | 0.744045 | STAD | GCIY | FR-180204 |
| MVP_top60_DNA_seed | SRC | 1.165229 | LUAD | NCI-H1650 | NVP-BHG712 | MVP_top60_DNA+RNA_seed | MAPK3 | 0.744045 | STAD | GCIY | FR-180204 |
| MVP_top60_DNA_seed | CREBBP | 1.166272 | LUAD | PC-3_[JPC-3] | I-CBP112 | MVP_top60_DNA+RNA_seed | SRC | 0.744151 | STAD | 23132-87 | NVP-BHG712 |
| MVP_top60_DNA_seed | GSK3B | 1.168901 | LUAD | NCI-H441 | AZD7969 | MVP_top60_DNA+RNA_seed | HDAC1 | 0.745101 | STAD | OCUM-1 | JQ12 |
| MVP_top60_DNA_seed | CREBBP | 1.16938 | LUAD | PC-14 | I-CBP112 | MVP_top60_DNA+RNA_seed | GSK3B | 0.747297 | STAD | SNU-16 | KIN001-042 |
| MVP_top60_DNA_seed | AKT1 | 1.170507 | LUAD | NCI-H2087 | MK-2206 | MVP_top60_DNA+RNA_seed | PARP1 | 0.751056 | STAD | GCIY | Veliparib |
| MVP_top60_DNA_seed | AURKA | 1.172044 | LUAD | NCI-H2291 | Alisertib | MVP_top60_DNA+RNA_seed | ERBB2 | 0.753648 | STAD | FU97 | Afatinib |
| MVP_top60_DNA_seed | BRAF | 1.172307 | LUAD | NCI-H322M | HG6-64-1 | MVP_top60_DNA+RNA_seed | ERBB2 | 0.754714 | STAD | TGBC11TKB | Afatinib |
| MVP_top60_DNA_seed | EGFR | 1.172583 | LUAD | NCI-H3122 | Gefitinib | MVP_top60_DNA+RNA_seed | HDAC1 | 0.754968 | STAD | NCI-N87 | Parthenolide |
| MVP_top60_DNA_seed | MET | 1.172594 | LUAD | NCI-H322M | AZD6094 | MVP_top60_DNA+RNA_seed | MAPK1 | 0.755788 | STAD | GCIY | VX-11e |
| MVP_top60_DNA_seed | SRC | 1.176255 | LUAD | NCI-H1435 | Bosutinib | MVP_top60_DNA+RNA_seed | PIK3CA | 0.758768 | STAD | SK-GT-2 | AZD8835 |
| MVP_top60_DNA_seed | FGFR1 | 1.176337 | LUAD | NCI-H1793 | PD173074 | MVP_top60_DNA+RNA_seed | MDM2 | 0.759175 | STAD | FU97 | Nutlin-3a (-) |
| MVP_top60_DNA_seed | HDAC1 | 1.176401 | LUAD | NCI-H2009 | CAY10603 | MVP_top60_DNA+RNA_seed | ERBB2 | 0.760857 | STAD | HGC-27 | PF-00299804 |
| MVP_top60_DNA_seed | AURKA | 1.176713 | LUAD | SW1573 | CD532 | MVP_top60_DNA+RNA_seed | ERBB4 | 0.760857 | STAD | HGC-27 | PF-00299804 |
| MVP_top60_DNA_seed | HDAC1 | 1.178119 | LUAD | NCI-H2342 | AR-42 | MVP_top60_DNA+RNA_seed | ERBB4 | 0.762208 | STAD | MKN28 | AST-1306 |
| MVP_top60_DNA_seed | AURKA | 1.179088 | LUAD | NCI-H2291 | ZM447439 | MVP_top60_DNA+RNA_seed | HDAC1 | 0.763516 | STAD | SNU-16 | AR-42 |
| MVP_top60_DNA_seed | IGF1R | 1.18038 | LUAD | NCI-H2085 | BMS-536924 | MVP_top60_DNA+RNA_seed | ATM | 0.764145 | STAD | KATOIII | torin2 |
| MVP_top60_DNA_seed | AKT1 | 1.181275 | LUAD | NCI-H1355 | GSK690693 | MVP_top60_DNA+RNA_seed | GSK3B | 0.764976 | STAD | IM-95 | CHIR-99021 |
| MVP_top60_DNA_seed | IGF1R | 1.181537 | LUAD | NCI-H1993 | GSK1904529A | MVP_top60_DNA+RNA_seed | PARP1 | 0.765694 | STAD | HSC-39 | PARP_9482 |
| MVP_top60_DNA_seed | MET | 1.181582 | LUAD | NCI-H1648 | SU11274 | MVP_top60_DNA+RNA_seed | FGFR3 | 0.766972 | STAD | SNU-1 | AZD4547 |
| MVP_top60_DNA_seed | AURKA | 1.18205 | LUAD | NCI-H2009 | GSK1070916 | MVP_top60_DNA+RNA_seed | FGFR3 | 0.767028 | STAD | SK-GT-2 | PD173074 |
| MVP_top60_DNA_seed | AURKA | 1.182672 | LUAD | NCI-H1838 | CD532 | MVP_top60_DNA+RNA_seed | HDAC1 | 0.767679 | STAD | TMK-1 | Tubastatin A |
| MVP_top60_DNA_seed | SMARCA4 | 1.183219 | LUAD | PC-3_[JPC-3] | PFI-3 | MVP_top60_DNA+RNA_seed | MET | 0.769863 | STAD | GCIY | PHA-665752 |
| MVP_top60_DNA_seed | MET | 1.184237 | LUAD | HCC-827 | JNJ38877605 | MVP_top60_DNA+RNA_seed | GSK3B | 0.77531 | STAD | NCI-N87 | AZD7969 |
| MVP_top60_DNA_seed | GSK3B | 1.184672 | LUAD | NCI-H1623 | CHIR-99021 | MVP_top60_DNA+RNA_seed | FGFR3 | 0.777347 | STAD | MKN7 | AZD4547 |
| MVP_top60_DNA_seed | GSK3B | 1.185011 | LUAD | NCI-H1623 | CHIR-99021 | MVP_top60_DNA+RNA_seed | PARP1 | 0.778425 | STAD | NUGC-4 | Rucaparib |
| MVP_top60_DNA_seed | FGFR1 | 1.185836 | LUAD | NCI-H1623 | PD173074 | MVP_top60_DNA+RNA_seed | PARP1 | 0.779419 | STAD | TGBC11TKB | Olaparib |

| | | | | | | | | | | | |
|---|---|---|---|---|---|---|---|---|---|---|---|
| MVP_top60_DNA_seed | AURKA | 1.186649 | LUAD | NCI-H1573 | Alisertib | MVP_top60_DNA+RNA_seed | ESR1 | 0.782169 | STAD | NUGC-3 | Tamoxifen |
| MVP_top60_DNA_seed | ATM | 1.187008 | LUAD | NCI-H1781 | KU-55933 | MVP_top60_DNA+RNA_seed | HDAC1 | 0.783896 | STAD | GCIY | Tubastatin A |
| MVP_top60_DNA_seed | BRAF | 1.192486 | LUAD | NCI-H2023 | HG6-64-1 | MVP_top60_DNA+RNA_seed | AR | 0.786438 | STAD | NUGC-4 | AZD3514 |
| MVP_top60_DNA_seed | AURKA | 1.192982 | LUAD | NCI-H838 | Alisertib | MVP_top60_DNA+RNA_seed | PARP1 | 0.793308 | STAD | OCUM-1 | Rucaparib |
| MVP_top60_DNA_seed | AURKA | 1.193221 | LUAD | NCI-H1623 | Alisertib | MVP_top60_DNA+RNA_seed | MAPK1 | 0.794199 | STAD | NUGC-3 | FR-180204 |
| MVP_top60_DNA_seed | MET | 1.193755 | LUAD | NCI-H1793 | AZD6094 | MVP_top60_DNA+RNA_seed | MAPK3 | 0.794199 | STAD | NUGC-3 | FR-180204 |
| MVP_top60_DNA_seed | ERBB2 | 1.195426 | LUAD | NCI-H1666 | CP724714 | MVP_top60_DNA+RNA_seed | ESR1 | 0.795778 | STAD | NCI-N87 | Fulvestrant |
| MVP_top60_DNA_seed | MAPK1 | 1.19634 | LUAD | NCI-H441 | VX-11e | MVP_top60_DNA+RNA_seed | PARP1 | 0.795868 | STAD | IM-95 | Olaparib |
| MVP_top60_DNA_seed | CREBBP | 1.196853 | LUAD | NCI-H650 | I-CBP112 | MVP_top60_DNA+RNA_seed | GSK3B | 0.799252 | STAD | RERF-GC-1B | SB216763 |
| MVP_top60_DNA_seed | HSP90AA1 | 1.196928 | LUAD | SW1573 | CCT-018159 | MVP_top60_DNA+RNA_seed | ATM | 0.800756 | STAD | MKN45 | CP466722 |
| MVP_top60_DNA_seed | MAPK1 | 1.197331 | LUAD | A549 | FR-180204 | MVP_top60_DNA+RNA_seed | PIK3CA | 0.800803 | STAD | MKN7 | PI3Ka_4409 |
| MVP_top60_DNA_seed | AURKA | 1.197466 | LUAD | Calu-3 | CD532 | MVP_top60_DNA+RNA_seed | HDAC1 | 0.801108 | STAD | MKN7 | Dacinostat |
| MVP_top60_DNA_seed | HSP90AA1 | 1.197576 | LUAD | NCI-H1623 | SNX-2112 | MVP_top60_DNA+RNA_seed | SRC | 0.802908 | STAD | HSC-39 | NVP-BHG712 |
| MVP_top60_DNA_seed | GSK3B | 1.197846 | LUAD | Calu-3 | CHIR-99021 | MVP_top60_DNA+RNA_seed | MDM2 | 0.806473 | STAD | MKN28 | Nutlin-3a (-) |
| MVP_top60_DNA_seed | HDAC1 | 1.198121 | LUAD | NCI-H1666 | Tubastatin A | MVP_top60_DNA+RNA_seed | MAPK1 | 0.806632 | STAD | TMK-1 | FR-180204 |
| MVP_top60_DNA_seed | FGFR1 | 1.198712 | LUAD | NCI-H1993 | AZD4547 | MVP_top60_DNA+RNA_seed | MAPK3 | 0.806632 | STAD | TMK-1 | FR-180204 |
| MVP_top60_DNA_seed | AURKA | 1.201313 | LUAD | NCI-H1993 | Alisertib | MVP_top60_DNA+RNA_seed | ESR1 | 0.807527 | STAD | SNU-1 | Fulvestrant |
| MVP_top60_DNA_seed | EGFR | 1.201904 | LUAD | NCI-H650 | Afatinib | MVP_top60_DNA+RNA_seed | GSK3B | 0.811732 | STAD | SNU-16 | CHIR-99021 |
| MVP_top60_DNA_seed | ERBB2 | 1.201904 | LUAD | NCI-H650 | Afatinib | MVP_top60_DNA+RNA_seed | ERBB2 | 0.81431 | STAD | MKN28 | PF-00299804 |
| MVP_top60_DNA_seed | AURKA | 1.202304 | LUAD | NCI-H1563 | GSK1070916 | MVP_top60_DNA+RNA_seed | ERBB4 | 0.81431 | STAD | MKN28 | PF-00299804 |
| MVP_top60_DNA_seed | FGFR1 | 1.202596 | LUAD | NCI-H650 | PD173074 | MVP_top60_DNA+RNA_seed | PARP1 | 0.817332 | STAD | MKN45 | Olaparib |
| MVP_top60_DNA_seed | IGF1R | 1.204481 | LUAD | NCI-H1573 | GSK1904529A | MVP_top60_DNA+RNA_seed | PARP1 | 0.82579 | STAD | TGBC11TKB | Talazoparib |
| MVP_top60_DNA_seed | GSK3B | 1.206399 | LUAD | NCI-H1666 | SB216763 | MVP_top60_DNA+RNA_seed | HSP90AA1 | 0.82647 | STAD | HSC-39 | Elesclomol |
| MVP_top60_DNA_seed | SRC | 1.208988 | LUAD | NCI-H1651 | Bosutinib | MVP_top60_DNA+RNA_seed | HSP90AB1 | 0.82647 | STAD | HSC-39 | Elesclomol |
| MVP_top60_DNA_seed | IGF1R | 1.209387 | LUAD | NCI-H1651 | BMS-536924 | MVP_top60_DNA+RNA_seed | EP300 | 0.835812 | STAD | MKN7 | I-CBP112 |
| MVP_top60_DNA_seed | EGFR | 1.209552 | LUAD | NCI-H1792 | CUDC-101 | MVP_top60_DNA+RNA_seed | CREBBP | 0.835812 | STAD | MKN7 | I-CBP112 |
| MVP_top60_DNA_seed | ERBB2 | 1.209552 | LUAD | NCI-H1792 | CUDC-101 | MVP_top60_DNA+RNA_seed | HDAC1 | 0.8405 | STAD | SNU-5 | AR-42 |
| MVP_top60_DNA_seed | SRC | 1.210312 | LUAD | NCI-H2405 | Bosutinib | MVP_top60_DNA+RNA_seed | ERBB2 | 0.846034 | STAD | FU97 | PF-00299804 |
| MVP_top60_DNA_seed | HDAC1 | 1.211252 | LUAD | RERF-LC-KJ | Belinostat | MVP_top60_DNA+RNA_seed | ERBB4 | 0.846034 | STAD | FU97 | PF-00299804 |
| MVP_top60_DNA_seed | PTK2 | 1.212037 | LUAD | NCI-H2228 | PF-562271 | MVP_top60_DNA+RNA_seed | PTK2 | 0.848826 | STAD | TGBC11TKB | PF-562271 |
| MVP_top60_DNA_seed | IGF1R | 1.212867 | LUAD | A549 | GSK1904529A | MVP_top60_DNA+RNA_seed | MAPK1 | 0.851745 | STAD | MKN7 | VX-11e |
| MVP_top60_DNA_seed | GSK3B | 1.213736 | LUAD | NCI-H1355 | SB216763 | MVP_top60_DNA+RNA_seed | ESR1 | 0.853524 | STAD | RF-48 | Tamoxifen |
| MVP_top60_DNA_seed | ATM | 1.215446 | LUAD | HCC-827 | KU-55933 | MVP_top60_DNA+RNA_seed | PARP1 | 0.854357 | STAD | RF-48 | PARP_9482 |
| MVP_top60_DNA_seed | GSK3B | 1.215616 | LUAD | NCI-H1563 | SB216763 | MVP_top60_DNA+RNA_seed | MDM2 | 0.861437 | STAD | GCIY | Nutlin-3a (-) |
| MVP_top60_DNA_seed | EGFR | 1.21657 | LUAD | NCI-H1793 | CUDC-101 | MVP_top60_DNA+RNA_seed | HSP90AA1 | 0.865977 | STAD | NCI-N87 | Elesclomol |
| MVP_top60_DNA_seed | ERBB2 | 1.21657 | LUAD | NCI-H1793 | CUDC-101 | MVP_top60_DNA+RNA_seed | HSP90AB1 | 0.865977 | STAD | NCI-N87 | Elesclomol |
| MVP_top60_DNA_seed | HSP90AA1 | 1.216593 | LUAD | EKVX | CCT-018159 | MVP_top60_DNA+RNA_seed | FGFR3 | 0.867024 | STAD | GCIY | PD173074 |
| MVP_top60_DNA_seed | AKT1 | 1.218054 | LUAD | NCI-H1623 | AKT inhibitor VIII | MVP_top60_DNA+RNA_seed | FGFR3 | 0.867664 | STAD | RF-48 | AZD4547 |
| MVP_top60_DNA_seed | ERBB2 | 1.218277 | LUAD | NCI-H322M | CP724714 | MVP_top60_DNA+RNA_seed | AR | 0.867892 | STAD | RF-48 | Bicalutamide |
| MVP_top60_DNA_seed | SMARCA4 | 1.219472 | LUAD | COR-L105 | PFI-3 | MVP_top60_DNA+RNA_seed | PARP1 | 0.868431 | STAD | SK-GT-2 | Rucaparib |
| MVP_top60_DNA_seed | HSP90AA1 | 1.219853 | LUAD | EKVX | SNX-2112 | MVP_top60_DNA+RNA_seed | PARP1 | 0.874247 | STAD | TGBC11TKB | PARP_0108 |

| | | | | | | | | | | | |
|---|---|---|---|---|---|---|---|---|---|---|---|
| MVP_top60_DNA_seed | PIK3CA | 1.219951 | LUAD | NCI-H441 | AZD8835 | MVP_top60_DNA+RNA_seed | HDAC1 | 0.875 | STAD | SNU-16 | Belinostat |
| MVP_top60_DNA_seed | MET | 1.220127 | LUAD | NCI-H2122 | JNJ38877605 | MVP_top60_DNA+RNA_seed | SRC | 0.876367 | STAD | MKN45 | NVP-BHG712 |
| MVP_top60_DNA_seed | MET | 1.220739 | LUAD | NCI-H1573 | SU11274 | MVP_top60_DNA+RNA_seed | PARP1 | 0.87709 | STAD | SK-GT-2 | PARP_9482 |
| MVP_top60_DNA_seed | GSK3B | 1.222497 | LUAD | Calu-3 | CHIR-99021 | MVP_top60_DNA+RNA_seed | HSP90AA1 | 0.877421 | STAD | TMK-1 | SNX-2112 |
| MVP_top60_DNA_seed | BRAF | 1.226464 | LUAD | NCI-H1623 | Dabrafenib | MVP_top60_DNA+RNA_seed | HSP90AB1 | 0.877421 | STAD | TMK-1 | SNX-2112 |
| MVP_top60_DNA_seed | MET | 1.227179 | LUAD | NCI-H2023 | SU11274 | MVP_top60_DNA+RNA_seed | PARP1 | 0.884749 | STAD | TGBC11TKB | PARP_9482 |
| MVP_top60_DNA_seed | HDAC1 | 1.227941 | LUAD | NCI-H2228 | AR-42 | MVP_top60_DNA+RNA_seed | PTK2 | 0.886563 | STAD | HSC-39 | PF-562271 |
| MVP_top60_DNA_seed | EGFR | 1.230458 | LUAD | SW1573 | PF-00299804 | MVP_top60_DNA+RNA_seed | HDAC1 | 0.889929 | STAD | SNU-5 | Entinostat |
| MVP_top60_DNA_seed | ERBB2 | 1.230458 | LUAD | SW1573 | PF-00299804 | MVP_top60_DNA+RNA_seed | HDAC3 | 0.889929 | STAD | SNU-5 | Entinostat |
| MVP_top60_DNA_seed | BRAF | 1.230903 | LUAD | ABC-1 | PLX-4720 | MVP_top60_DNA+RNA_seed | ESR1 | 0.891296 | STAD | TMK-1 | Tamoxifen |
| MVP_top60_DNA_seed | IGF1R | 1.231277 | LUAD | NCI-H1435 | BMS-536924 | MVP_top60_DNA+RNA_seed | PARP1 | 0.89584 | STAD | GCIY | Rucaparib |
| MVP_top60_DNA_seed | PIK3CA | 1.231945 | LUAD | NCI-H1435 | AZD8835 | MVP_top60_DNA+RNA_seed | PARP1 | 0.895889 | STAD | NUGC-4 | PARP_9482 |
| MVP_top60_DNA_seed | AURKA | 1.233245 | LUAD | SK-LU-1 | Alisertib | MVP_top60_DNA+RNA_seed | HDAC1 | 0.897117 | STAD | SNU-5 | CAY10603 |
| MVP_top60_DNA_seed | AURKA | 1.233628 | LUAD | NCI-H1650 | Tozasertib | MVP_top60_DNA+RNA_seed | FGFR3 | 0.900374 | STAD | TMK-1 | FGFR_3831 |
| MVP_top60_DNA_seed | ATM | 1.234763 | LUAD | EMC-BAC-2 | KU-55933 | MVP_top60_DNA+RNA_seed | HSP90AA1 | 0.902853 | STAD | MKN7 | Luminespib |
| MVP_top60_DNA_seed | AURKA | 1.235035 | LUAD | HCC-44 | CD532 | MVP_top60_DNA+RNA_seed | HSP90AB1 | 0.902853 | STAD | MKN7 | Luminespib |
| MVP_top60_DNA_seed | EGFR | 1.236579 | LUAD | SW1573 | CI-1033 | MVP_top60_DNA+RNA_seed | HDAC1 | 0.90912 | STAD | SNU-1 | Entinostat |
| MVP_top60_DNA_seed | ERBB2 | 1.236579 | LUAD | SW1573 | CI-1033 | MVP_top60_DNA+RNA_seed | HDAC3 | 0.90912 | STAD | SNU-1 | Entinostat |
| MVP_top60_DNA_seed | FGFR1 | 1.237071 | LUAD | PC-3_[JPC-3] | PD173074 | MVP_top60_DNA+RNA_seed | MET | 0.910516 | STAD | SNU-1 | JNJ38877605 |
| MVP_top60_DNA_seed | SRC | 1.237126 | LUAD | NCI-H2087 | Bosutinib | MVP_top60_DNA+RNA_seed | ERBB2 | 0.911389 | STAD | TMK-1 | CP724714 |
| MVP_top60_DNA_seed | HSP90AA1 | 1.23936 | LUAD | NCI-H1651 | SNX-2112 | MVP_top60_DNA+RNA_seed | HSP90AA1 | 0.911413 | STAD | FU97 | Elesclomol |
| MVP_top60_DNA_seed | EGFR | 1.239484 | LUAD | SW1573 | Gefitinib | MVP_top60_DNA+RNA_seed | HSP90AB1 | 0.911413 | STAD | FU97 | Elesclomol |
| MVP_top60_DNA_seed | HDAC1 | 1.24026 | LUAD | NCI-H1568 | Tubastatin A | MVP_top60_DNA+RNA_seed | PARP1 | 0.913168 | STAD | GCIY | Olaparib |
| MVP_top60_DNA_seed | SRC | 1.240696 | LUAD | NCI-H1623 | Bosutinib | MVP_top60_DNA+RNA_seed | PARP1 | 0.91451 | STAD | SNU-5 | Olaparib |
| MVP_top60_DNA_seed | EGFR | 1.241002 | LUAD | RERF-LC-KJ | PF-00299804 | MVP_top60_DNA+RNA_seed | MDM2 | 0.916011 | STAD | KATOIII | Nutlin-3a (-) |
| MVP_top60_DNA_seed | ERBB2 | 1.241002 | LUAD | RERF-LC-KJ | PF-00299804 | MVP_top60_DNA+RNA_seed | HSP90AA1 | 0.916055 | STAD | SNU-16 | Elesclomol |
| MVP_top60_DNA_seed | MDM2 | 1.241219 | LUAD | EKVX | Nutlin-3a (-) | MVP_top60_DNA+RNA_seed | HSP90AB1 | 0.916055 | STAD | SNU-16 | Elesclomol |
| MVP_top60_DNA_seed | AURKA | 1.242321 | LUAD | NCI-H2291 | GSK1070916 | MVP_top60_DNA+RNA_seed | ERBB2 | 0.918981 | STAD | RF-48 | Lapatinib |
| MVP_top60_DNA_seed | BRAF | 1.242368 | LUAD | EKVX | AZ628 | MVP_top60_DNA+RNA_seed | MAPK1 | 0.92229 | STAD | NCI-N87 | FR-180204 |
| MVP_top60_DNA_seed | GSK3B | 1.242556 | LUAD | NCI-H1623 | KIN001-042 | MVP_top60_DNA+RNA_seed | MAPK3 | 0.92229 | STAD | NCI-N87 | FR-180204 |
| MVP_top60_DNA_seed | SRC | 1.242713 | LUAD | NCI-H1651 | NVP-BHG712 | MVP_top60_DNA+RNA_seed | HDAC1 | 0.924897 | STAD | SNU-5 | Belinostat |
| MVP_top60_DNA_seed | AKT1 | 1.24378 | LUAD | NCI-H1651 | MK-2206 | MVP_top60_DNA+RNA_seed | GSK3B | 0.927586 | STAD | OCUM-1 | CHIR-99021 |
| MVP_top60_DNA_seed | HDAC1 | 1.244618 | LUAD | NCI-H23 | JQ12 | MVP_top60_DNA+RNA_seed | HDAC1 | 0.930647 | STAD | TMK-1 | CAY10603 |
| MVP_top60_DNA_seed | FGFR1 | 1.247657 | LUAD | NCI-H441 | PD173074 | MVP_top60_DNA+RNA_seed | HDAC1 | 0.930858 | STAD | HSC-39 | JQ12 |
| MVP_top60_DNA_seed | EGFR | 1.250011 | LUAD | SK-LU-1 | PF-00299804 | MVP_top60_DNA+RNA_seed | MDM2 | 0.935312 | STAD | TGBC11TKB | Serdemetan |
| MVP_top60_DNA_seed | ERBB2 | 1.250011 | LUAD | SK-LU-1 | PF-00299804 | MVP_top60_DNA+RNA_seed | GSK3B | 0.937086 | STAD | NUGC-3 | SB216763 |
| MVP_top60_DNA_seed | HDAC1 | 1.250412 | LUAD | NCI-H1793 | Belinostat | MVP_top60_DNA+RNA_seed | PARP1 | 0.937249 | STAD | KATOIII | Olaparib |
| MVP_top60_DNA_seed | FGFR1 | 1.251386 | LUAD | NCI-H1623 | AZD4547 | MVP_top60_DNA+RNA_seed | FGFR3 | 0.941307 | STAD | NCI-N87 | AZD4547 |
| MVP_top60_DNA_seed | BRAF | 1.251558 | LUAD | Calu-3 | PLX-4720 | MVP_top60_DNA+RNA_seed | ATM | 0.94372 | STAD | NUGC-4 | CP466722 |
| MVP_top60_DNA_seed | BRAF | 1.252589 | LUAD | NCI-H1623 | PLX-4720 | MVP_top60_DNA+RNA_seed | PIK3CA | 0.952346 | STAD | RF-48 | AZD8835 |
| MVP_top60_DNA_seed | MET | 1.254948 | LUAD | NCI-H650 | SU11274 | MVP_top60_DNA+RNA_seed | ERBB2 | 0.952632 | STAD | TGBC11TKB | AZD8931 |

| | | | | | | | | | | | |
|---|---|---|---|---|---|---|---|---|---|---|---|
| MVP_top60_DNA_seed | MET | 1.258353 | LUAD | NCI-H1650 | SU11274 | MVP_top60_DNA+RNA_seed | ERBB3 | 0.952632 | STAD | TGBC11TKB | AZD8931 |
| MVP_top60_DNA_seed | AKT1 | 1.258596 | LUAD | PC-3_[JPC-3] | MK-2206 | MVP_top60_DNA+RNA_seed | ERBB2 | 0.953741 | STAD | Hs-746T | Afatinib |
| MVP_top60_DNA_seed | GSK3B | 1.258626 | LUAD | COR-L105 | KIN001-042 | MVP_top60_DNA+RNA_seed | HDAC1 | 0.956902 | STAD | RERF-GC-1B | CAY10603 |
| MVP_top60_DNA_seed | MDM2 | 1.259967 | LUAD | NCI-H1838 | Nutlin-3a (-) | MVP_top60_DNA+RNA_seed | ERBB2 | 0.95716 | STAD | MKN45 | PF-00299804 |
| MVP_top60_DNA_seed | SMARCA4 | 1.260657 | LUAD | NCI-H650 | PFI-3 | MVP_top60_DNA+RNA_seed | ERBB4 | 0.95716 | STAD | MKN45 | PF-00299804 |
| MVP_top60_DNA_seed | AURKA | 1.260715 | LUAD | NCI-H2023 | Genentech Cpd 10 | MVP_top60_DNA+RNA_seed | ERBB2 | 0.957312 | STAD | MKN45 | AZD8931 |
| MVP_top60_DNA_seed | HDAC1 | 1.262397 | LUAD | NCI-H1573 | Dacinostat | MVP_top60_DNA+RNA_seed | ERBB3 | 0.957312 | STAD | MKN45 | AZD8931 |
| MVP_top60_DNA_seed | MAPK1 | 1.262894 | LUAD | NCI-H322M | FR-180204 | MVP_top60_DNA+RNA_seed | ESR1 | 0.957666 | STAD | TGBC11TKB | Fulvestrant |
| MVP_top60_DNA_seed | MET | 1.264311 | LUAD | NCI-H2228 | SU11274 | MVP_top60_DNA+RNA_seed | ERBB2 | 0.961597 | STAD | Hs-746T | Afatinib |
| MVP_top60_DNA_seed | HDAC1 | 1.265123 | LUAD | HCC-827 | Tubastatin A | MVP_top60_DNA+RNA_seed | HDAC1 | 0.962033 | STAD | NUGC-3 | Tubastatin A |
| MVP_top60_DNA_seed | BRAF | 1.265144 | LUAD | NCI-H1793 | Dabrafenib | MVP_top60_DNA+RNA_seed | HSP90AA1 | 0.963145 | STAD | MKN45 | Luminespib |
| MVP_top60_DNA_seed | EGFR | 1.266119 | LUAD | NCI-H2228 | CUDC-101 | MVP_top60_DNA+RNA_seed | HSP90AB1 | 0.963145 | STAD | MKN45 | Luminespib |
| MVP_top60_DNA_seed | ERBB2 | 1.266119 | LUAD | NCI-H2228 | CUDC-101 | MVP_top60_DNA+RNA_seed | ERBB2 | 0.963799 | STAD | OCUM-1 | Afatinib |
| MVP_top60_DNA_seed | AURKA | 1.266812 | LUAD | NCI-H2347 | ZM447439 | MVP_top60_DNA+RNA_seed | MET | 0.965121 | STAD | 23132-87 | SU11274 |
| MVP_top60_DNA_seed | SMARCA4 | 1.268707 | LUAD | NCI-H1648 | PFI-3 | MVP_top60_DNA+RNA_seed | ERBB2 | 0.9741 | STAD | SNU-16 | Afatinib |
| MVP_top60_DNA_seed | HSP90AA1 | 1.268845 | LUAD | PC-3_[JPC-3] | Tanespimycin | MVP_top60_DNA+RNA_seed | MDM2 | 0.97764 | STAD | TMK-1 | Nutlin-3a (-) |
| MVP_top60_DNA_seed | CREBBP | 1.269282 | LUAD | NCI-H1623 | I-CBP112 | MVP_top60_DNA+RNA_seed | MET | 0.978342 | STAD | MKN28 | AZD6094 |
| MVP_top60_DNA_seed | ERBB2 | 1.272465 | LUAD | NCI-H441 | CP724714 | MVP_top60_DNA+RNA_seed | ERBB2 | 0.981788 | STAD | RF-48 | CP724714 |
| MVP_top60_DNA_seed | BRAF | 1.2732 | LUAD | NCI-H1993 | PLX-4720 | MVP_top60_DNA+RNA_seed | FGFR3 | 0.982875 | STAD | MKN45 | AZD4547 |
| MVP_top60_DNA_seed | FGFR1 | 1.274368 | LUAD | NCI-H1838 | PD173074 | MVP_top60_DNA+RNA_seed | HDAC1 | 0.983018 | STAD | HSC-39 | Dacinostat |
| MVP_top60_DNA_seed | IGF1R | 1.274881 | LUAD | H3255 | GSK1904529A | MVP_top60_DNA+RNA_seed | ATM | 0.987829 | STAD | MKN45 | KU-55933 |
| MVP_top60_DNA_seed | GSK3B | 1.27507 | LUAD | PC-3_[JPC-3] | SB216763 | MVP_top60_DNA+RNA_seed | HDAC1 | 0.991616 | STAD | RERF-GC-1B | Belinostat |
| MVP_top60_DNA_seed | SRC | 1.275711 | LUAD | NCI-H1838 | WH-4-023 | MVP_top60_DNA+RNA_seed | HDAC1 | 0.993198 | STAD | SNU-5 | Parthenolide |
| MVP_top60_DNA_seed | IGF1R | 1.277183 | LUAD | NCI-H1573 | Linsitinib | MVP_top60_DNA+RNA_seed | FGFR3 | 0.993217 | STAD | Hs-746T | AZD4547 |
| MVP_top60_DNA_seed | HDAC1 | 1.279632 | LUAD | NCI-H1793 | AR-42 | MVP_top60_DNA+RNA_seed | GSK3B | 0.993845 | STAD | FU97 | AZD7969 |
| MVP_top60_DNA_seed | HDAC1 | 1.280853 | LUAD | NCI-H1734 | JQ12 | MVP_top60_DNA+RNA_seed | PARP1 | 1.000347 | STAD | RF-48 | PARP_9495 |
| MVP_top60_DNA_seed | GSK3B | 1.281849 | LUAD | NCI-H1781 | AZD7969 | MVP_top60_DNA+RNA_seed | MAPK1 | 1.007774 | STAD | SNU-16 | FR-180204 |
| MVP_top60_DNA_seed | AKT1 | 1.282071 | LUAD | NCI-H2342 | AKT inhibitor VIII | MVP_top60_DNA+RNA_seed | MAPK3 | 1.007774 | STAD | SNU-16 | FR-180204 |
| MVP_top60_DNA_seed | GSK3B | 1.282378 | LUAD | NCI-H2228 | SB216763 | MVP_top60_DNA+RNA_seed | ATM | 1.008654 | STAD | SNU-5 | KU-60019 |
| MVP_top60_DNA_seed | GSK3B | 1.283135 | LUAD | Calu-3 | SB216763 | MVP_top60_DNA+RNA_seed | EP300 | 1.013633 | STAD | Hs-746T | I-CBP112 |
| MVP_top60_DNA_seed | EGFR | 1.28528 | LUAD | NCI-H1993 | Cetuximab | MVP_top60_DNA+RNA_seed | CREBBP | 1.013633 | STAD | Hs-746T | I-CBP112 |
| MVP_top60_DNA_seed | BRAF | 1.286394 | LUAD | NCI-H1993 | SB590885 | MVP_top60_DNA+RNA_seed | MDM2 | 1.017186 | STAD | TGBC11TKB | Nutlin-3a (-) |
| MVP_top60_DNA_seed | EGFR | 1.287822 | LUAD | NCI-H1651 | Pelitinib | MVP_top60_DNA+RNA_seed | PARP1 | 1.019075 | STAD | OCUM-1 | Veliparib |
| MVP_top60_DNA_seed | MET | 1.289227 | LUAD | NCI-H322M | PHA-665752 | MVP_top60_DNA+RNA_seed | GSK3B | 1.020996 | STAD | RF-48 | CHIR-99021 |
| MVP_top60_DNA_seed | IGF1R | 1.289297 | LUAD | NCI-H23 | BMS-754807 | MVP_top60_DNA+RNA_seed | AR | 1.022764 | STAD | OCUM-1 | Bicalutamide |
| MVP_top60_DNA_seed | PIK3CA | 1.291479 | LUAD | PC-3_[JPC-3] | PI3Ka_4409 | MVP_top60_DNA+RNA_seed | HSP90AA1 | 1.023274 | STAD | TGBC11TKB | Luminespib |
| MVP_top60_DNA_seed | EGFR | 1.291675 | LUAD | NCI-H1355 | Pelitinib | MVP_top60_DNA+RNA_seed | HSP90AB1 | 1.023274 | STAD | TGBC11TKB | Luminespib |
| MVP_top60_DNA_seed | MDM2 | 1.292298 | LUAD | NCI-H2085 | Serdemetan | MVP_top60_DNA+RNA_seed | FGFR3 | 1.026605 | STAD | OCUM-1 | PD173074 |
| MVP_top60_DNA_seed | SRC | 1.293466 | LUAD | NCI-H2291 | Bosutinib | MVP_top60_DNA+RNA_seed | GSK3B | 1.028067 | STAD | NUGC-3 | CHIR-99021 |
| MVP_top60_DNA_seed | AURKA | 1.293899 | LUAD | NCI-H1573 | Genentech Cpd 10 | MVP_top60_DNA+RNA_seed | PIK3CA | 1.028821 | STAD | OCUM-1 | AZD8835 |
| MVP_top60_DNA_seed | AKT1 | 1.294709 | LUAD | NCI-H23 | BAY AKT1 | MVP_top60_DNA+RNA_seed | ERBB2 | 1.030123 | STAD | MKN45 | Afatinib |

| | | | | | | | | | | | |
|---|---|---|---|---|---|---|---|---|---|---|---|
| MVP_top60_DNA_seed | MDM2 | 1.295926 | LUAD | NCI-H650 | Nutlin-3a (-) | MVP_top60_DNA+RNA_seed | ESR1 | 1.030497 | STAD | KATOIII | Fulvestrant |
| MVP_top60_DNA_seed | HSP90AA1 | 1.298267 | LUAD | NCI-H1568 | SNX-2112 | MVP_top60_DNA+RNA_seed | HDAC1 | 1.034564 | STAD | SNU-16 | Tubastatin A |
| MVP_top60_DNA_seed | AKT1 | 1.299109 | LUAD | NCI-H522 | BAY AKT1 | MVP_top60_DNA+RNA_seed | GSK3B | 1.034863 | STAD | RF-48 | SB216763 |
| MVP_top60_DNA_seed | AKT1 | 1.303054 | LUAD | NCI-H2023 | AKT inhibitor VIII | MVP_top60_DNA+RNA_seed | MET | 1.039295 | STAD | NUGC-3 | SU11274 |
| MVP_top60_DNA_seed | EGFR | 1.303454 | LUAD | NCI-H441 | Pelitinib | MVP_top60_DNA+RNA_seed | ESR1 | 1.039878 | STAD | Hs-746T | Tamoxifen |
| MVP_top60_DNA_seed | HDAC1 | 1.304718 | LUAD | EKVX | AR-42 | MVP_top60_DNA+RNA_seed | ERBB2 | 1.039935 | STAD | SNU-1 | Lapatinib |
| MVP_top60_DNA_seed | IGF1R | 1.304805 | LUAD | NCI-H1993 | BMS-536924 | MVP_top60_DNA+RNA_seed | AR | 1.042086 | STAD | RF-48 | AZD3514 |
| MVP_top60_DNA_seed | AKT1 | 1.305485 | LUAD | NCI-H1435 | AKT inhibitor VIII | MVP_top60_DNA+RNA_seed | MDM2 | 1.046161 | STAD | MKN7 | Nutlin-3a (-) |
| MVP_top60_DNA_seed | MET | 1.306627 | LUAD | NCI-H1355 | Crizotinib | MVP_top60_DNA+RNA_seed | AR | 1.047292 | STAD | SNU-5 | Bicalutamide |
| MVP_top60_DNA_seed | SRC | 1.307892 | LUAD | NCI-H2342 | NVP-BHG712 | MVP_top60_DNA+RNA_seed | EP300 | 1.050984 | STAD | SNU-1 | I-CBP112 |
| MVP_top60_DNA_seed | GSK3B | 1.309614 | LUAD | NCI-H1623 | SB216763 | MVP_top60_DNA+RNA_seed | CREBBP | 1.050984 | STAD | SNU-1 | I-CBP112 |
| MVP_top60_DNA_seed | BRAF | 1.309826 | LUAD | RERF-LC-KJ | HG6-64-1 | MVP_top60_DNA+RNA_seed | PARP1 | 1.066438 | STAD | MKN7 | PARP_0108 |
| MVP_top60_DNA_seed | FGFR1 | 1.311185 | LUAD | COR-L105 | PD173074 | MVP_top60_DNA+RNA_seed | MDM2 | 1.066648 | STAD | RF-48 | Serdemetan |
| MVP_top60_DNA_seed | SRC | 1.311781 | LUAD | NCI-H3122 | Bosutinib | MVP_top60_DNA+RNA_seed | AR | 1.067644 | STAD | MKN7 | Bicalutamide |
| MVP_top60_DNA_seed | EGFR | 1.313376 | LUAD | SW1573 | AST-1306 | MVP_top60_DNA+RNA_seed | GSK3B | 1.072533 | STAD | SNU-5 | CHIR-99021 |
| MVP_top60_DNA_seed | GSK3B | 1.31418 | LUAD | COR-L105 | SB216763 | MVP_top60_DNA+RNA_seed | SRC | 1.073073 | STAD | SNU-16 | NVP-BHG712 |
| MVP_top60_DNA_seed | HSP90AA1 | 1.317366 | LUAD | NCI-H2085 | Tanespimycin | MVP_top60_DNA+RNA_seed | FGFR3 | 1.073296 | STAD | NUGC-4 | FGFR_3831 |
| MVP_top60_DNA_seed | HSP90AA1 | 1.318242 | LUAD | NCI-H1781 | SNX-2112 | MVP_top60_DNA+RNA_seed | PARP1 | 1.074377 | STAD | KATOIII | PARP_9482 |
| MVP_top60_DNA_seed | SRC | 1.320661 | LUAD | NCI-H322M | NVP-BHG712 | MVP_top60_DNA+RNA_seed | HDAC1 | 1.074826 | STAD | OCUM-1 | Tubastatin A |
| MVP_top60_DNA_seed | AURKA | 1.324399 | LUAD | NCI-H2023 | Alisertib | MVP_top60_DNA+RNA_seed | MET | 1.074927 | STAD | RF-48 | PHA-665752 |
| MVP_top60_DNA_seed | BRAF | 1.324822 | LUAD | NCI-H1437 | RAF_9304 | MVP_top60_DNA+RNA_seed | ERBB2 | 1.077495 | STAD | OCUM-1 | Afatinib |
| MVP_top60_DNA_seed | HDAC1 | 1.324855 | LUAD | EKVX | CAY10603 | MVP_top60_DNA+RNA_seed | FGFR3 | 1.081305 | STAD | TGBC11TKB | FGFR_3831 |
| MVP_top60_DNA_seed | GSK3B | 1.326412 | LUAD | NCI-H2228 | AZD7969 | MVP_top60_DNA+RNA_seed | PARP1 | 1.086886 | STAD | SNU-1 | PARP_9482 |
| MVP_top60_DNA_seed | AKT1 | 1.327274 | LUAD | NCI-H2291 | AKT inhibitor VIII | MVP_top60_DNA+RNA_seed | PARP1 | 1.093641 | STAD | NUGC-4 | Olaparib |
| MVP_top60_DNA_seed | EGFR | 1.328245 | LUAD | NCI-H2405 | CI-1033 | MVP_top60_DNA+RNA_seed | ERBB2 | 1.095849 | STAD | RF-48 | PF-00299804 |
| MVP_top60_DNA_seed | ERBB2 | 1.328245 | LUAD | NCI-H2405 | CI-1033 | MVP_top60_DNA+RNA_seed | ERBB4 | 1.095849 | STAD | RF-48 | PF-00299804 |
| MVP_top60_DNA_seed | IGF1R | 1.331255 | LUAD | PC-3_[JPC-3] | BMS-754807 | MVP_top60_DNA+RNA_seed | PARP1 | 1.096766 | STAD | SNU-1 | PARP_0108 |
| MVP_top60_DNA_seed | MET | 1.332366 | LUAD | NCI-H1648 | AZD6094 | MVP_top60_DNA+RNA_seed | HDAC1 | 1.097573 | STAD | MKN7 | Tubastatin A |
| MVP_top60_DNA_seed | MET | 1.332938 | LUAD | NCI-H1666 | SU11274 | MVP_top60_DNA+RNA_seed | ATM | 1.102717 | STAD | MKN7 | KU-60019 |
| MVP_top60_DNA_seed | BRAF | 1.333857 | LUAD | NCI-H1648 | Dabrafenib | MVP_top60_DNA+RNA_seed | PARP1 | 1.10451 | STAD | IM-95 | PARP_9495 |
| MVP_top60_DNA_seed | HDAC1 | 1.333963 | LUAD | SW1573 | Tubastatin A | MVP_top60_DNA+RNA_seed | ERBB2 | 1.106125 | STAD | RF-48 | Afatinib |
| MVP_top60_DNA_seed | MAPK1 | 1.335592 | LUAD | NCI-H1650 | VX-11e | MVP_top60_DNA+RNA_seed | HDAC1 | 1.113724 | STAD | SNU-16 | CAY10603 |
| MVP_top60_DNA_seed | AKT1 | 1.336517 | LUAD | HCC-827 | AKT inhibitor VIII | MVP_top60_DNA+RNA_seed | PARP1 | 1.115564 | STAD | SNU-16 | Talazoparib |
| MVP_top60_DNA_seed | HSP90AA1 | 1.33746 | LUAD | NCI-H1734 | CCT-018159 | MVP_top60_DNA+RNA_seed | MDM2 | 1.11666 | STAD | Hs-746T | Nutlin-3a (-) |
| MVP_top60_DNA_seed | AURKA | 1.339085 | LUAD | NCI-H1435 | CD532 | MVP_top60_DNA+RNA_seed | ERBB2 | 1.116749 | STAD | SNU-5 | CUDC-101 |
| MVP_top60_DNA_seed | GSK3B | 1.340604 | LUAD | NCI-H650 | KIN001-042 | MVP_top60_DNA+RNA_seed | HSP90AA1 | 1.119638 | STAD | TGBC11TKB | Elesclomol |
| MVP_top60_DNA_seed | AKT1 | 1.343335 | LUAD | NCI-H1838 | GSK690693 | MVP_top60_DNA+RNA_seed | HSP90AB1 | 1.119638 | STAD | TGBC11TKB | Elesclomol |
| MVP_top60_DNA_seed | SRC | 1.343986 | LUAD | NCI-H1793 | NVP-BHG712 | MVP_top60_DNA+RNA_seed | EP300 | 1.123118 | STAD | SNU-5 | I-CBP112 |
| MVP_top60_DNA_seed | AURKA | 1.34747 | LUAD | NCI-H1573 | GSK1070916 | MVP_top60_DNA+RNA_seed | CREBBP | 1.123118 | STAD | SNU-5 | I-CBP112 |
| MVP_top60_DNA_seed | SRC | 1.34779 | LUAD | Calu-6 | Bosutinib | MVP_top60_DNA+RNA_seed | MET | 1.123226 | STAD | MKN28 | SU11274 |
| MVP_top60_DNA_seed | MET | 1.347876 | LUAD | NCI-H838 | JNJ38877605 | MVP_top60_DNA+RNA_seed | ESR1 | 1.125403 | STAD | MKN45 | Tamoxifen |

| | | | | | | | | | | | |
|---|---|---|---|---|---|---|---|---|---|---|---|
| MVP_top60_DNA_seed | AKT1 | 1.348315 | LUAD | NCI-H1838 | BAY AKT1 | MVP_top60_DNA+RNA_seed | MAPK1 | 1.12635 | STAD | SNU-16 | VX-11e |
| MVP_top60_DNA_seed | EGFR | 1.349207 | LUAD | NCI-H1838 | CUDC-101 | MVP_top60_DNA+RNA_seed | AR | 1.126367 | STAD | SNU-1 | Bicalutamide |
| MVP_top60_DNA_seed | ERBB2 | 1.349207 | LUAD | NCI-H1838 | CUDC-101 | MVP_top60_DNA+RNA_seed | ATM | 1.128924 | STAD | Hs-746T | CP466722 |
| MVP_top60_DNA_seed | MET | 1.351211 | LUAD | NCI-H1793 | SU11274 | MVP_top60_DNA+RNA_seed | SRC | 1.132299 | STAD | Hs-746T | Bosutinib |
| MVP_top60_DNA_seed | SMARCA4 | 1.353313 | LUAD | SW1573 | PFI-3 | MVP_top60_DNA+RNA_seed | PIK3CA | 1.134076 | STAD | MKN45 | AZD8835 |
| MVP_top60_DNA_seed | BRAF | 1.354031 | LUAD | NCI-H1623 | PLX-4720 | MVP_top60_DNA+RNA_seed | FGFR3 | 1.134314 | STAD | MKN7 | AZD4547 |
| MVP_top60_DNA_seed | AURKA | 1.354211 | LUAD | PC-3_[JPC-3] | CD532 | MVP_top60_DNA+RNA_seed | FGFR3 | 1.137501 | STAD | TMK-1 | PD173074 |
| MVP_top60_DNA_seed | EGFR | 1.35492 | LUAD | NCI-H3122 | PF-00299804 | MVP_top60_DNA+RNA_seed | HDAC1 | 1.139672 | STAD | TMK-1 | Belinostat |
| MVP_top60_DNA_seed | ERBB2 | 1.35492 | LUAD | NCI-H3122 | PF-00299804 | MVP_top60_DNA+RNA_seed | GSK3B | 1.144515 | STAD | RERF-GC-1B | CHIR-99021 |
| MVP_top60_DNA_seed | AURKA | 1.355437 | LUAD | NCI-H1650 | CD532 | MVP_top60_DNA+RNA_seed | PARP1 | 1.148511 | STAD | MKN7 | Olaparib |
| MVP_top60_DNA_seed | GSK3B | 1.358779 | LUAD | NCI-H1666 | CHIR-99021 | MVP_top60_DNA+RNA_seed | PARP1 | 1.152664 | STAD | KATOIII | PARP_0108 |
| MVP_top60_DNA_seed | HDAC1 | 1.360278 | LUAD | NCI-H1355 | Tubastatin A | MVP_top60_DNA+RNA_seed | GSK3B | 1.152683 | STAD | SNU-1 | CHIR-99021 |
| MVP_top60_DNA_seed | ATM | 1.362218 | LUAD | Calu-3 | KU-55933 | MVP_top60_DNA+RNA_seed | PIK3CA | 1.154174 | STAD | RF-48 | PI3Ka_4409 |
| MVP_top60_DNA_seed | GSK3B | 1.36299 | LUAD | NCI-H1838 | KIN001-042 | MVP_top60_DNA+RNA_seed | PARP1 | 1.15656 | STAD | MKN7 | PARP_9482 |
| MVP_top60_DNA_seed | HSP90AA1 | 1.364664 | LUAD | NCI-H1993 | SNX-2112 | MVP_top60_DNA+RNA_seed | ERBB2 | 1.157487 | STAD | SNU-1 | Afatinib |
| MVP_top60_DNA_seed | EGFR | 1.365092 | LUAD | NCI-H838 | Pelitinib | MVP_top60_DNA+RNA_seed | PARP1 | 1.159757 | STAD | MKN7 | Veliparib |
| MVP_top60_DNA_seed | AURKA | 1.365345 | LUAD | NCI-H1623 | GSK1070916 | MVP_top60_DNA+RNA_seed | GSK3B | 1.168373 | STAD | SNU-5 | CHIR-99021 |
| MVP_top60_DNA_seed | GSK3B | 1.368195 | LUAD | NCI-H1693 | CHIR-99021 | MVP_top60_DNA+RNA_seed | PARP1 | 1.181019 | STAD | Hs-746T | Veliparib |
| MVP_top60_DNA_seed | FGFR1 | 1.368423 | LUAD | LXF-289 | FGFR_3831 | MVP_top60_DNA+RNA_seed | HSP90AA1 | 1.186903 | STAD | GCIY | Elesclomol |
| MVP_top60_DNA_seed | GSK3B | 1.373828 | LUAD | NCI-H1437 | AZD7969 | MVP_top60_DNA+RNA_seed | HSP90AB1 | 1.186903 | STAD | GCIY | Elesclomol |
| MVP_top60_DNA_seed | AURKA | 1.375209 | LUAD | NCI-H1563 | Alisertib | MVP_top60_DNA+RNA_seed | HDAC1 | 1.19123 | STAD | MKN7 | JQ12 |
| MVP_top60_DNA_seed | PIK3CA | 1.375567 | LUAD | NCI-H1793 | PI3Ka_4409 | MVP_top60_DNA+RNA_seed | ESR1 | 1.193091 | STAD | NCI-N87 | Tamoxifen |
| MVP_top60_DNA_seed | EGFR | 1.375775 | LUAD | A549 | Pelitinib | MVP_top60_DNA+RNA_seed | FGFR3 | 1.195394 | STAD | Hs-746T | FGFR_3831 |
| MVP_top60_DNA_seed | MET | 1.377565 | LUAD | Calu-3 | SU11274 | MVP_top60_DNA+RNA_seed | HSP90AA1 | 1.200792 | STAD | MKN7 | Tanespimycin |
| MVP_top60_DNA_seed | HDAC1 | 1.377679 | LUAD | NCI-H1793 | CAY10603 | MVP_top60_DNA+RNA_seed | HSP90AB1 | 1.200792 | STAD | MKN7 | Tanespimycin |
| MVP_top60_DNA_seed | AURKA | 1.380443 | LUAD | NCI-H1838 | GSK1070916 | MVP_top60_DNA+RNA_seed | PARP1 | 1.206627 | STAD | NCI-N87 | Olaparib |
| MVP_top60_DNA_seed | MET | 1.380816 | LUAD | NCI-H1437 | JNJ38877605 | MVP_top60_DNA+RNA_seed | PARP1 | 1.209278 | STAD | MKN7 | Olaparib |
| MVP_top60_DNA_seed | AKT1 | 1.381604 | LUAD | PC-3_[JPC-3] | BAY AKT1 | MVP_top60_DNA+RNA_seed | PIK3CA | 1.214622 | STAD | Hs-746T | AZD8835 |
| MVP_top60_DNA_seed | ATM | 1.383735 | LUAD | NCI-H2342 | KU-55933 | MVP_top60_DNA+RNA_seed | ERBB4 | 1.216366 | STAD | RF-48 | AST-1306 |
| MVP_top60_DNA_seed | PIK3CA | 1.383771 | LUAD | VMRC-LCD | AZD8835 | MVP_top60_DNA+RNA_seed | MET | 1.233141 | STAD | NCI-N87 | AZD6094 |
| MVP_top60_DNA_seed | MDM2 | 1.384006 | LUAD | NCI-H1355 | Serdemetan | MVP_top60_DNA+RNA_seed | MET | 1.234687 | STAD | NCI-N87 | SU11274 |
| MVP_top60_DNA_seed | AKT1 | 1.384064 | LUAD | Calu-6 | AKT inhibitor VIII | MVP_top60_DNA+RNA_seed | ERBB2 | 1.235015 | STAD | SNU-16 | CUDC-101 |
| MVP_top60_DNA_seed | ERBB2 | 1.384314 | LUAD | SW1573 | CP724714 | MVP_top60_DNA+RNA_seed | PARP1 | 1.235356 | STAD | RF-48 | Olaparib |
| MVP_top60_DNA_seed | ATM | 1.386201 | LUAD | NCI-H1651 | KU-60019 | MVP_top60_DNA+RNA_seed | PARP1 | 1.238885 | STAD | MKN45 | Olaparib |
| MVP_top60_DNA_seed | AURKA | 1.386334 | LUAD | NCI-H1666 | Tozasertib | MVP_top60_DNA+RNA_seed | HDAC1 | 1.241302 | STAD | MKN45 | Tubastatin A |
| MVP_top60_DNA_seed | AURKA | 1.389077 | LUAD | NCI-H1666 | Genentech Cpd 10 | MVP_top60_DNA+RNA_seed | SRC | 1.243658 | STAD | MKN7 | NVP-BHG712 |
| MVP_top60_DNA_seed | AKT1 | 1.389344 | LUAD | HOP-62 | BAY AKT1 | MVP_top60_DNA+RNA_seed | GSK3B | 1.244341 | STAD | RERF-GC-1B | KIN001-042 |
| MVP_top60_DNA_seed | AURKA | 1.39051 | LUAD | NCI-H1792 | CD532 | MVP_top60_DNA+RNA_seed | AR | 1.24747 | STAD | MKN7 | AZD3514 |
| MVP_top60_DNA_seed | HDAC1 | 1.392424 | LUAD | NCI-H1650 | Parthenolide | MVP_top60_DNA+RNA_seed | MET | 1.251315 | STAD | OCUM-1 | SU11274 |
| MVP_top60_DNA_seed | PIK3CA | 1.392587 | LUAD | NCI-H1651 | PI3Ka_4409 | MVP_top60_DNA+RNA_seed | ERBB2 | 1.254838 | STAD | SNU-5 | CP724714 |
| MVP_top60_DNA_seed | GSK3B | 1.39362 | LUAD | NCI-H1355 | CHIR-99021 | MVP_top60_DNA+RNA_seed | MET | 1.255613 | STAD | 23132-87 | JNJ38877605 |

| | | | | | | | | | | | |
|---|---|---|---|---|---|---|---|---|---|---|---|
| MVP_top60_DNA_seed | MAPK1 | 1.394368 | LUAD | Calu-3 | VX-11e | MVP_top60_DNA+RNA_seed | SRC | 1.258796 | STAD | SNU-1 | WH-4-023 |
| MVP_top60_DNA_seed | AURKA | 1.400276 | LUAD | NCI-H2405 | Genentech Cpd 10 | MVP_top60_DNA+RNA_seed | HDAC1 | 1.264024 | STAD | RERF-GC-1B | AR-42 |
| MVP_top60_DNA_seed | AKT1 | 1.401164 | LUAD | NCI-H1693 | GSK690693 | MVP_top60_DNA+RNA_seed | ATM | 1.264547 | STAD | NUGC-3 | CP466722 |
| MVP_top60_DNA_seed | AURKA | 1.402188 | LUAD | NCI-H2085 | CD532 | MVP_top60_DNA+RNA_seed | PARP1 | 1.271995 | STAD | IM-95 | PARP_9482 |
| MVP_top60_DNA_seed | FGFR1 | 1.402898 | LUAD | Calu-3 | FGFR_3831 | MVP_top60_DNA+RNA_seed | PARP1 | 1.27902 | STAD | SNU-1 | Talazoparib |
| MVP_top60_DNA_seed | MET | 1.403986 | LUAD | NCI-H1666 | Crizotinib | MVP_top60_DNA+RNA_seed | EP300 | 1.279924 | STAD | MKN45 | I-CBP112 |
| MVP_top60_DNA_seed | EGFR | 1.404964 | LUAD | NCI-H2009 | CUDC-101 | MVP_top60_DNA+RNA_seed | CREBBP | 1.279924 | STAD | MKN45 | I-CBP112 |
| MVP_top60_DNA_seed | ERBB2 | 1.404964 | LUAD | NCI-H2009 | CUDC-101 | MVP_top60_DNA+RNA_seed | MET | 1.283609 | STAD | SNU-16 | SU11274 |
| MVP_top60_DNA_seed | AURKA | 1.406419 | LUAD | NCI-H1793 | ZM447439 | MVP_top60_DNA+RNA_seed | HDAC1 | 1.286909 | STAD | SNU-5 | Dacinostat |
| MVP_top60_DNA_seed | AKT1 | 1.408539 | LUAD | NCI-H2291 | GSK690693 | MVP_top60_DNA+RNA_seed | HDAC1 | 1.287335 | STAD | SNU-5 | Tubastatin A |
| MVP_top60_DNA_seed | HSP90AA1 | 1.408709 | LUAD | NCI-H1435 | CCT-018159 | MVP_top60_DNA+RNA_seed | HSP90AA1 | 1.287695 | STAD | NUGC-4 | CCT-018159 |
| MVP_top60_DNA_seed | HDAC1 | 1.41049 | LUAD | NCI-H441 | Tubastatin A | MVP_top60_DNA+RNA_seed | HSP90AB1 | 1.287695 | STAD | NUGC-4 | CCT-018159 |
| MVP_top60_DNA_seed | MAPK1 | 1.410954 | LUAD | HCC-827 | VX-11e | MVP_top60_DNA+RNA_seed | HSP90AA1 | 1.295328 | STAD | HSC-39 | Luminespib |
| MVP_top60_DNA_seed | HDAC1 | 1.410958 | LUAD | NCI-H2228 | Tubastatin A | MVP_top60_DNA+RNA_seed | HSP90AB1 | 1.295328 | STAD | HSC-39 | Luminespib |
| MVP_top60_DNA_seed | GSK3B | 1.411919 | LUAD | NCI-H1993 | CHIR-99021 | MVP_top60_DNA+RNA_seed | ATM | 1.297269 | STAD | RERF-GC-1B | KU-55933 |
| MVP_top60_DNA_seed | BRAF | 1.41198 | LUAD | NCI-H2291 | PLX-4720 | MVP_top60_DNA+RNA_seed | ERBB2 | 1.306524 | STAD | MKN45 | CP724714 |
| MVP_top60_DNA_seed | SRC | 1.413761 | LUAD | NCI-H1838 | NVP-BHG712 | MVP_top60_DNA+RNA_seed | GSK3B | 1.307288 | STAD | SNU-1 | AZD7969 |
| MVP_top60_DNA_seed | AKT1 | 1.41423 | LUAD | NCI-H441 | GSK690693 | MVP_top60_DNA+RNA_seed | SRC | 1.3073 | STAD | SNU-16 | Saracatinib |
| MVP_top60_DNA_seed | AURKA | 1.415626 | LUAD | NCI-H2342 | Genentech Cpd 10 | MVP_top60_DNA+RNA_seed | ATM | 1.309781 | STAD | OCUM-1 | KU-55933 |
| MVP_top60_DNA_seed | PIK3CA | 1.416646 | LUAD | NCI-H650 | PI3Ka_4409 | MVP_top60_DNA+RNA_seed | PARP1 | 1.311284 | STAD | NUGC-4 | PARP_0108 |
| MVP_top60_DNA_seed | ERBB2 | 1.418127 | LUAD | NCI-H1693 | CP724714 | MVP_top60_DNA+RNA_seed | PARP1 | 1.312482 | STAD | KATOIII | Talazoparib |
| MVP_top60_DNA_seed | EGFR | 1.424649 | LUAD | NCI-H2023 | Pelitinib | MVP_top60_DNA+RNA_seed | GSK3B | 1.322174 | STAD | RF-48 | CHIR-99021 |
| MVP_top60_DNA_seed | SRC | 1.426138 | LUAD | A549 | NVP-BHG712 | MVP_top60_DNA+RNA_seed | PARP1 | 1.324956 | STAD | SNU-1 | Rucaparib |
| MVP_top60_DNA_seed | EGFR | 1.426197 | LUAD | NCI-H441 | Cetuximab | MVP_top60_DNA+RNA_seed | SRC | 1.340497 | STAD | TMK-1 | NVP-BHG712 |
| MVP_top60_DNA_seed | GSK3B | 1.426374 | LUAD | NCI-H1993 | KIN001-042 | MVP_top60_DNA+RNA_seed | ATM | 1.342585 | STAD | SNU-16 | KU-60019 |
| MVP_top60_DNA_seed | PTK2 | 1.427283 | LUAD | NCI-H1435 | PF-562271 | MVP_top60_DNA+RNA_seed | AR | 1.345234 | STAD | MKN45 | Bicalutamide |
| MVP_top60_DNA_seed | AKT1 | 1.428344 | LUAD | NCI-H1573 | AKT inhibitor VIII | MVP_top60_DNA+RNA_seed | FYN | 1.350291 | STAD | SNU-5 | A-770041 |
| MVP_top60_DNA_seed | IGF1R | 1.428385 | LUAD | NCI-H1666 | Linsitinib | MVP_top60_DNA+RNA_seed | FGFR3 | 1.351224 | STAD | GCIY | AZD4547 |
| MVP_top60_DNA_seed | EGFR | 1.428748 | LUAD | NCI-H2023 | PF-00299804 | MVP_top60_DNA+RNA_seed | ATM | 1.351348 | STAD | MKN28 | CP466722 |
| MVP_top60_DNA_seed | ERBB2 | 1.428748 | LUAD | NCI-H2023 | PF-00299804 | MVP_top60_DNA+RNA_seed | FGFR3 | 1.351743 | STAD | MKN7 | FGFR_3831 |
| MVP_top60_DNA_seed | SMARCA4 | 1.429027 | LUAD | NCI-H1563 | PFI-3 | MVP_top60_DNA+RNA_seed | HSP90AA1 | 1.359293 | STAD | RF-48 | Elesclomol |
| MVP_top60_DNA_seed | PIK3CA | 1.429311 | LUAD | NCI-H1651 | AZD8835 | MVP_top60_DNA+RNA_seed | HSP90AB1 | 1.359293 | STAD | RF-48 | Elesclomol |
| MVP_top60_DNA_seed | IGF1R | 1.429544 | LUAD | NCI-H1623 | GSK1904529A | MVP_top60_DNA+RNA_seed | MAPK1 | 1.359375 | STAD | SNU-5 | VX-11e |
| MVP_top60_DNA_seed | SRC | 1.430432 | LUAD | NCI-H2347 | Bosutinib | MVP_top60_DNA+RNA_seed | GSK3B | 1.361364 | STAD | Hs-746T | CHIR-99021 |
| MVP_top60_DNA_seed | ATM | 1.430599 | LUAD | Calu-6 | KU-55933 | MVP_top60_DNA+RNA_seed | HSP90AA1 | 1.364521 | STAD | IM-95 | Elesclomol |
| MVP_top60_DNA_seed | CREBBP | 1.430924 | LUAD | NCI-H322M | I-CBP112 | MVP_top60_DNA+RNA_seed | HSP90AB1 | 1.364521 | STAD | IM-95 | Elesclomol |
| MVP_top60_DNA_seed | FGFR1 | 1.43426 | LUAD | NCI-H1838 | AZD4547 | MVP_top60_DNA+RNA_seed | HSP90AA1 | 1.366052 | STAD | NUGC-4 | Elesclomol |
| MVP_top60_DNA_seed | AKT1 | 1.43484 | LUAD | SW1573 | AKT inhibitor VIII | MVP_top60_DNA+RNA_seed | HSP90AB1 | 1.366052 | STAD | NUGC-4 | Elesclomol |
| MVP_top60_DNA_seed | MAPK1 | 1.440228 | LUAD | SW1573 | VX-11e | MVP_top60_DNA+RNA_seed | GSK3B | 1.374242 | STAD | SNU-1 | KIN001-042 |
| MVP_top60_DNA_seed | BRAF | 1.443016 | LUAD | COR-L105 | PLX-4720 | MVP_top60_DNA+RNA_seed | SRC | 1.382787 | STAD | OCUM-1 | NVP-BHG712 |
| MVP_top60_DNA_seed | CREBBP | 1.44452 | LUAD | NCI-H2228 | I-CBP112 | MVP_top60_DNA+RNA_seed | PARP1 | 1.383298 | STAD | IM-95 | PARP_0108 |

| | | | | | | | | | | | |
|---|---|---|---|---|---|---|---|---|---|---|---|
| MVP_top60_DNA_seed | AURKA | 1.444528 | LUAD | NCI-H322M | Genentech Cpd 10 | MVP_top60_DNA+RNA_seed | PARP1 | 1.384827 | STAD | SNU-1 | PARP_9495 |
| MVP_top60_DNA_seed | AKT1 | 1.445145 | LUAD | HCC-827 | MK-2206 | MVP_top60_DNA+RNA_seed | FGFR3 | 1.384828 | STAD | NUGC-4 | AZD4547 |
| MVP_top60_DNA_seed | HDAC1 | 1.445772 | LUAD | EKVX | Belinostat | MVP_top60_DNA+RNA_seed | ATM | 1.392843 | STAD | RF-48 | KU-55933 |
| MVP_top60_DNA_seed | ATM | 1.446998 | LUAD | HCC-827 | CP466722 | MVP_top60_DNA+RNA_seed | MAPK1 | 1.393199 | STAD | SNU-5 | FR-180204 |
| MVP_top60_DNA_seed | HDAC1 | 1.448042 | LUAD | NCI-H2009 | AR-42 | MVP_top60_DNA+RNA_seed | MAPK3 | 1.393199 | STAD | SNU-5 | FR-180204 |
| MVP_top60_DNA_seed | PIK3CA | 1.448167 | LUAD | NCI-H1573 | PI3Ka_4409 | MVP_top60_DNA+RNA_seed | AR | 1.395201 | STAD | KATOIII | AZD3514 |
| MVP_top60_DNA_seed | MAPK1 | 1.452126 | LUAD | Calu-3 | FR-180204 | MVP_top60_DNA+RNA_seed | ERBB2 | 1.395564 | STAD | MKN45 | CUDC-101 |
| MVP_top60_DNA_seed | HSP90AA1 | 1.4584 | LUAD | Calu-6 | Tanespimycin | MVP_top60_DNA+RNA_seed | GSK3B | 1.402784 | STAD | TGBC11TKB | AZD7969 |
| MVP_top60_DNA_seed | BRAF | 1.458625 | LUAD | NCI-H1781 | RAF_9304 | MVP_top60_DNA+RNA_seed | ESR1 | 1.40806 | STAD | Hs-746T | Fulvestrant |
| MVP_top60_DNA_seed | FGFR1 | 1.458722 | LUAD | NCI-H2291 | AZD4547 | MVP_top60_DNA+RNA_seed | MDM2 | 1.409034 | STAD | NCI-N87 | Nutlin-3a (-) |
| MVP_top60_DNA_seed | GSK3B | 1.4618 | LUAD | NCI-H2228 | CHIR-99021 | MVP_top60_DNA+RNA_seed | MET | 1.410312 | STAD | RF-48 | SU11274 |
| MVP_top60_DNA_seed | MET | 1.463962 | LUAD | NCI-H1573 | JNJ38877605 | MVP_top60_DNA+RNA_seed | ESR1 | 1.413893 | STAD | RF-48 | Fulvestrant |
| MVP_top60_DNA_seed | AKT1 | 1.464704 | LUAD | NCI-H441 | AKT inhibitor VIII | MVP_top60_DNA+RNA_seed | MDM2 | 1.438108 | STAD | OCUM-1 | Serdemetan |
| MVP_top60_DNA_seed | BRAF | 1.466 | LUAD | NCI-H1993 | HG6-64-1 | MVP_top60_DNA+RNA_seed | FGFR3 | 1.447633 | STAD | SK-GT-2 | AZD4547 |
| MVP_top60_DNA_seed | HDAC1 | 1.467329 | LUAD | NCI-H2342 | Belinostat | MVP_top60_DNA+RNA_seed | AR | 1.458384 | STAD | SNU-16 | Bicalutamide |
| MVP_top60_DNA_seed | AKT1 | 1.468462 | LUAD | NCI-H1793 | GSK690693 | MVP_top60_DNA+RNA_seed | HSP90AA1 | 1.464536 | STAD | OCUM-1 | SNX-2112 |
| MVP_top60_DNA_seed | GSK3B | 1.470031 | LUAD | NCI-H1693 | SB216763 | MVP_top60_DNA+RNA_seed | HSP90AB1 | 1.464536 | STAD | OCUM-1 | SNX-2112 |
| MVP_top60_DNA_seed | GSK3B | 1.470209 | LUAD | NCI-H1355 | KIN001-042 | MVP_top60_DNA+RNA_seed | HSP90AA1 | 1.466064 | STAD | SNU-16 | SNX-2112 |
| MVP_top60_DNA_seed | AKT1 | 1.471681 | LUAD | NCI-H322M | GSK690693 | MVP_top60_DNA+RNA_seed | HSP90AB1 | 1.466064 | STAD | SNU-16 | SNX-2112 |
| MVP_top60_DNA_seed | EGFR | 1.473847 | LUAD | NCI-H1793 | Pelitinib | MVP_top60_DNA+RNA_seed | PARP1 | 1.466821 | STAD | MKN45 | PARP_9495 |
| MVP_top60_DNA_seed | FGFR1 | 1.476677 | LUAD | NCI-H322M | AZD4547 | MVP_top60_DNA+RNA_seed | ESR1 | 1.469051 | STAD | MKN45 | Fulvestrant |
| MVP_top60_DNA_seed | GSK3B | 1.47827 | LUAD | NCI-H1793 | SB216763 | MVP_top60_DNA+RNA_seed | SRC | 1.479698 | STAD | SNU-5 | NVP-BHG712 |
| MVP_top60_DNA_seed | SRC | 1.479589 | LUAD | NCI-H1355 | Bosutinib | MVP_top60_DNA+RNA_seed | ATM | 1.491787 | STAD | TMK-1 | CP466722 |
| MVP_top60_DNA_seed | MDM2 | 1.480705 | LUAD | NCI-H1993 | Nutlin-3a (-) | MVP_top60_DNA+RNA_seed | GSK3B | 1.497845 | STAD | Hs-746T | SB216763 |
| MVP_top60_DNA_seed | SMARCA4 | 1.481901 | LUAD | NCI-H1623 | PFI-3 | MVP_top60_DNA+RNA_seed | ATM | 1.500206 | STAD | NCI-N87 | CP466722 |
| MVP_top60_DNA_seed | ATM | 1.482178 | LUAD | NCI-H1993 | KU-55933 | MVP_top60_DNA+RNA_seed | ATM | 1.503181 | STAD | SNU-5 | KU-55933 |
| MVP_top60_DNA_seed | PIK3CA | 1.48309 | LUAD | HCC-827 | PI3Ka_4409 | MVP_top60_DNA+RNA_seed | PARP1 | 1.506631 | STAD | SNU-16 | Olaparib |
| MVP_top60_DNA_seed | CREBBP | 1.485092 | LUAD | NCI-H441 | I-CBP112 | MVP_top60_DNA+RNA_seed | MET | 1.51024 | STAD | RERF-GC-1B | JNJ38877605 |
| MVP_top60_DNA_seed | HSP90AA1 | 1.486762 | LUAD | NCI-H2087 | Elesclomol | MVP_top60_DNA+RNA_seed | GSK3B | 1.514643 | STAD | Hs-746T | AZD7969 |
| MVP_top60_DNA_seed | MDM2 | 1.487941 | LUAD | EKVX | Serdemetan | MVP_top60_DNA+RNA_seed | HDAC1 | 1.516312 | STAD | SNU-16 | Dacinostat |
| MVP_top60_DNA_seed | GSK3B | 1.48979 | LUAD | Calu-3 | KIN001-042 | MVP_top60_DNA+RNA_seed | HSP90AA1 | 1.516402 | STAD | RERF-GC-1B | CCT-018159 |
| MVP_top60_DNA_seed | ATM | 1.490753 | LUAD | NCI-H1838 | CP466722 | MVP_top60_DNA+RNA_seed | HSP90AB1 | 1.516402 | STAD | RERF-GC-1B | CCT-018159 |
| MVP_top60_DNA_seed | MET | 1.49142 | LUAD | NCI-H1355 | SU11274 | MVP_top60_DNA+RNA_seed | SRC | 1.517876 | STAD | SNU-5 | Saracatinib |
| MVP_top60_DNA_seed | SMARCA4 | 1.495186 | LUAD | PC-14 | PFI-3 | MVP_top60_DNA+RNA_seed | FGFR3 | 1.520786 | STAD | RF-48 | FGFR_3831 |
| MVP_top60_DNA_seed | HSP90AA1 | 1.495901 | LUAD | Calu-6 | Elesclomol | MVP_top60_DNA+RNA_seed | PARP1 | 1.52318 | STAD | Hs-746T | Rucaparib |
| MVP_top60_DNA_seed | SRC | 1.496579 | LUAD | Calu-3 | NVP-BHG712 | MVP_top60_DNA+RNA_seed | HSP90AA1 | 1.525398 | STAD | SNU-16 | CCT-018159 |
| MVP_top60_DNA_seed | AKT1 | 1.498574 | LUAD | HCC-827 | BAY AKT1 | MVP_top60_DNA+RNA_seed | HSP90AB1 | 1.525398 | STAD | SNU-16 | CCT-018159 |
| MVP_top60_DNA_seed | MET | 1.502235 | LUAD | EKVX | SU11274 | MVP_top60_DNA+RNA_seed | ERBB2 | 1.538234 | STAD | SNU-16 | CI-1033 |
| MVP_top60_DNA_seed | BRAF | 1.503912 | LUAD | NCI-H441 | SB590885 | MVP_top60_DNA+RNA_seed | ERBB4 | 1.538234 | STAD | SNU-16 | CI-1033 |
| MVP_top60_DNA_seed | AURKA | 1.504108 | LUAD | NCI-H1793 | CD532 | MVP_top60_DNA+RNA_seed | HSP90AA1 | 1.561323 | STAD | HSC-39 | SNX-2112 |
| MVP_top60_DNA_seed | PIK3CA | 1.50436 | LUAD | NCI-H2085 | PI3Ka_4409 | MVP_top60_DNA+RNA_seed | HSP90AB1 | 1.561323 | STAD | HSC-39 | SNX-2112 |

| | | | | | | | | | | | |
|---|---|---|---|---|---|---|---|---|---|---|---|
| MVP_top60_DNA_seed | MAPK1 | 1.509338 | LUAD | NCI-H1355 | FR-180204 | MVP_top60_DNA+RNA_seed | MAPK1 | 1.562867 | STAD | MKN45 | FR-180204 |
| MVP_top60_DNA_seed | ATM | 1.512469 | LUAD | NCI-H2009 | CP466722 | MVP_top60_DNA+RNA_seed | MAPK3 | 1.562867 | STAD | MKN45 | FR-180204 |
| MVP_top60_DNA_seed | AKT1 | 1.513596 | LUAD | NCI-H322M | A-443654 | MVP_top60_DNA+RNA_seed | MET | 1.568048 | STAD | NUGC-3 | JNJ38877605 |
| MVP_top60_DNA_seed | HDAC1 | 1.514685 | LUAD | RERF-LC-KJ | AR-42 | MVP_top60_DNA+RNA_seed | FGFR3 | 1.575208 | STAD | Hs-746T | AZD4547 |
| MVP_top60_DNA_seed | HSP90AA1 | 1.516398 | LUAD | NCI-H1838 | CCT-018159 | MVP_top60_DNA+RNA_seed | MAPK1 | 1.579484 | STAD | Hs-746T | FR-180204 |
| MVP_top60_DNA_seed | ERBB2 | 1.518326 | LUAD | NCI-H2228 | CP724714 | MVP_top60_DNA+RNA_seed | MAPK3 | 1.579484 | STAD | Hs-746T | FR-180204 |
| MVP_top60_DNA_seed | AKT1 | 1.518539 | LUAD | SW1573 | AKT inhibitor VIII | MVP_top60_DNA+RNA_seed | GSK3B | 1.581028 | STAD | NCI-N87 | CHIR-99021 |
| MVP_top60_DNA_seed | PIK3CA | 1.520595 | LUAD | VMRC-LCD | PI3Ka_4409 | MVP_top60_DNA+RNA_seed | HDAC1 | 1.591227 | STAD | SNU-5 | JQ12 |
| MVP_top60_DNA_seed | HDAC1 | 1.521377 | LUAD | NCI-H1793 | JQ12 | MVP_top60_DNA+RNA_seed | PARP1 | 1.599957 | STAD | RF-48 | Talazoparib |
| MVP_top60_DNA_seed | BRAF | 1.521446 | LUAD | NCI-H2291 | PLX-4720 | MVP_top60_DNA+RNA_seed | PARP1 | 1.602984 | STAD | SNU-5 | PARP_9495 |
| MVP_top60_DNA_seed | MET | 1.521557 | LUAD | NCI-H1838 | AZD6094 | MVP_top60_DNA+RNA_seed | HDAC1 | 1.603425 | STAD | MKN45 | CAY10603 |
| MVP_top60_DNA_seed | IGF1R | 1.524332 | LUAD | NCI-H1838 | Linsitinib | MVP_top60_DNA+RNA_seed | AR | 1.604421 | STAD | IM-95 | AZD3514 |
| MVP_top60_DNA_seed | AURKA | 1.526099 | LUAD | NCI-H1793 | Genentech Cpd 10 | MVP_top60_DNA+RNA_seed | HDAC1 | 1.641715 | STAD | MKN45 | Belinostat |
| MVP_top60_DNA_seed | AURKA | 1.527808 | LUAD | PC-3_[JPC-3] | ZM447439 | MVP_top60_DNA+RNA_seed | AR | 1.671666 | STAD | NCI-N87 | Bicalutamide |
| MVP_top60_DNA_seed | MAPK1 | 1.528404 | LUAD | EKVX | FR-180204 | MVP_top60_DNA+RNA_seed | GSK3B | 1.675039 | STAD | SNU-5 | SB216763 |
| MVP_top60_DNA_seed | AURKA | 1.528824 | LUAD | NCI-H1975 | CD532 | MVP_top60_DNA+RNA_seed | PTK2 | 1.691122 | STAD | MKN7 | PF-562271 |
| MVP_top60_DNA_seed | BRAF | 1.529359 | LUAD | NCI-H1793 | PLX-4720 | MVP_top60_DNA+RNA_seed | PTK2 | 1.706917 | STAD | SNU-16 | PF-562271 |
| MVP_top60_DNA_seed | AURKA | 1.530439 | LUAD | NCI-H2228 | CD532 | MVP_top60_DNA+RNA_seed | MET | 1.717216 | STAD | OCUM-1 | JNJ38877605 |
| MVP_top60_DNA_seed | IGF1R | 1.530975 | LUAD | SW1573 | GSK1904529A | MVP_top60_DNA+RNA_seed | PARP1 | 1.721144 | STAD | NCI-N87 | Olaparib |
| MVP_top60_DNA_seed | BRAF | 1.543202 | LUAD | NCI-H1666 | HG6-64-1 | MVP_top60_DNA+RNA_seed | FGFR3 | 1.732734 | STAD | NCI-N87 | AZD4547 |
| MVP_top60_DNA_seed | HSP90AA1 | 1.544039 | LUAD | NCI-H1666 | Elesclomol | MVP_top60_DNA+RNA_seed | FGFR3 | 1.740636 | STAD | SNU-1 | AZD4547 |
| MVP_top60_DNA_seed | MET | 1.549355 | LUAD | NCI-H2291 | SU11274 | MVP_top60_DNA+RNA_seed | ERBB2 | 1.742186 | STAD | RERF-GC-1B | CUDC-101 |
| MVP_top60_DNA_seed | SRC | 1.551491 | LUAD | SW1573 | NVP-BHG712 | MVP_top60_DNA+RNA_seed | AR | 1.743914 | STAD | SNU-5 | AZD3514 |
| MVP_top60_DNA_seed | EGFR | 1.556087 | LUAD | NCI-H1993 | Afatinib | MVP_top60_DNA+RNA_seed | AR | 1.763446 | STAD | TMK-1 | AZD3514 |
| MVP_top60_DNA_seed | ERBB2 | 1.556087 | LUAD | NCI-H1993 | Afatinib | MVP_top60_DNA+RNA_seed | HSP90AA1 | 1.775021 | STAD | RERF-GC-1B | Tanespimycin |
| MVP_top60_DNA_seed | BRAF | 1.556854 | LUAD | EKVX | Dabrafenib | MVP_top60_DNA+RNA_seed | HSP90AB1 | 1.775021 | STAD | RERF-GC-1B | Tanespimycin |
| MVP_top60_DNA_seed | SRC | 1.558977 | LUAD | NCI-H1666 | Saracatinib | MVP_top60_DNA+RNA_seed | GSK3B | 1.7764 | STAD | RF-48 | AZD7969 |
| MVP_top60_DNA_seed | SRC | 1.56004 | LUAD | Calu-3 | Bosutinib | MVP_top60_DNA+RNA_seed | HSP90AA1 | 1.780029 | STAD | SNU-5 | SNX-2112 |
| MVP_top60_DNA_seed | FGFR1 | 1.561144 | LUAD | NCI-H650 | FGFR_3831 | MVP_top60_DNA+RNA_seed | HSP90AB1 | 1.780029 | STAD | SNU-5 | SNX-2112 |
| MVP_top60_DNA_seed | BRAF | 1.567386 | LUAD | NCI-H1793 | SB590885 | MVP_top60_DNA+RNA_seed | MET | 1.786694 | STAD | RF-48 | JNJ38877605 |
| MVP_top60_DNA_seed | CREBBP | 1.567711 | LUAD | A549 | I-CBP112 | MVP_top60_DNA+RNA_seed | HDAC1 | 1.787583 | STAD | MKN45 | AR-42 |
| MVP_top60_DNA_seed | IGF1R | 1.568954 | LUAD | NCI-H1666 | BMS-536924 | MVP_top60_DNA+RNA_seed | FGFR3 | 1.800818 | STAD | SNU-1 | FGFR_3831 |
| MVP_top60_DNA_seed | SRC | 1.569018 | LUAD | NCI-H650 | Bosutinib | MVP_top60_DNA+RNA_seed | ATM | 1.856631 | STAD | MKN28 | KU-60019 |
| MVP_top60_DNA_seed | AURKA | 1.569425 | LUAD | NCI-H2023 | CD532 | MVP_top60_DNA+RNA_seed | AR | 1.860389 | STAD | NCI-N87 | Bicalutamide |
| MVP_top60_DNA_seed | ATM | 1.569857 | LUAD | ABC-1 | KU-55933 | MVP_top60_DNA+RNA_seed | ERBB2 | 1.870352 | STAD | SNU-16 | PF-00299804 |
| MVP_top60_DNA_seed | PTK2 | 1.57355 | LUAD | HCC-827 | PF-562271 | MVP_top60_DNA+RNA_seed | ERBB4 | 1.870352 | STAD | SNU-16 | PF-00299804 |
| MVP_top60_DNA_seed | GSK3B | 1.575885 | LUAD | NCI-H441 | KIN001-042 | MVP_top60_DNA+RNA_seed | FGFR3 | 1.871968 | STAD | SNU-5 | FGFR_3831 |
| MVP_top60_DNA_seed | FGFR1 | 1.579167 | LUAD | NCI-H1623 | FGFR_3831 | MVP_top60_DNA+RNA_seed | FGFR3 | 1.889612 | STAD | MKN45 | FGFR_3831 |
| MVP_top60_DNA_seed | HDAC1 | 1.583765 | LUAD | EKVX | Parthenolide | MVP_top60_DNA+RNA_seed | MDM2 | 1.900555 | STAD | SNU-1 | Serdemetan |
| MVP_top60_DNA_seed | AURKA | 1.58794 | LUAD | NCI-H1666 | CD532 | MVP_top60_DNA+RNA_seed | HDAC1 | 1.915128 | STAD | SNU-16 | JQ12 |
| MVP_top60_DNA_seed | EGFR | 1.592102 | LUAD | NCI-H1838 | Pelitinib | MVP_top60_DNA+RNA_seed | PIK3CA | 1.925783 | STAD | SNU-1 | PI3Ka_4409 |

| | | | | | | | | | | | |
|---|---|---|---|---|---|---|---|---|---|---|---|
| MVP_top60_DNA_seed | EGFR | 1.594305 | LUAD | NCI-H2291 | Gefitinib | MVP_top60_DNA+RNA_seed | HSP90AA1 | 1.9342 | STAD | SNU-1 | CCT-018159 |
| MVP_top60_DNA_seed | SRC | 1.597803 | LUAD | NCI-H1355 | NVP-BHG712 | MVP_top60_DNA+RNA_seed | HSP90AB1 | 1.9342 | STAD | SNU-1 | CCT-018159 |
| MVP_top60_DNA_seed | AURKA | 1.600124 | LUAD | A549 | CD532 | MVP_top60_DNA+RNA_seed | ESR1 | 1.943349 | STAD | SNU-5 | Fulvestrant |
| MVP_top60_DNA_seed | EGFR | 1.604902 | LUAD | Calu-6 | AZD8931 | MVP_top60_DNA+RNA_seed | GSK3B | 1.945581 | STAD | RERF-GC-1B | AZD7969 |
| MVP_top60_DNA_seed | ERBB2 | 1.604902 | LUAD | Calu-6 | AZD8931 | MVP_top60_DNA+RNA_seed | ATM | 1.951166 | STAD | AGS | CP466722 |
| MVP_top60_DNA_seed | ERBB3 | 1.604902 | LUAD | Calu-6 | AZD8931 | MVP_top60_DNA+RNA_seed | GSK3B | 1.987441 | STAD | MKN45 | KIN001-042 |
| MVP_top60_DNA_seed | HDAC1 | 1.607153 | LUAD | NCI-H1573 | JQ12 | MVP_top60_DNA+RNA_seed | PTK2 | 2.020085 | STAD | SNU-5 | PF-562271 |
| MVP_top60_DNA_seed | GSK3B | 1.608405 | LUAD | RERF-LC-KJ | AZD7969 | MVP_top60_DNA+RNA_seed | MET | 2.02422 | STAD | SNU-16 | JNJ38877605 |
| MVP_top60_DNA_seed | HDAC1 | 1.610669 | LUAD | NCI-H1755 | JQ12 | MVP_top60_DNA+RNA_seed | GSK3B | 2.049871 | STAD | NCI-N87 | KIN001-042 |
| MVP_top60_DNA_seed | BRAF | 1.611779 | LUAD | NCI-H1838 | PLX-4720 | MVP_top60_DNA+RNA_seed | ATM | 2.055851 | STAD | SNU-1 | KU-60019 |
| MVP_top60_DNA_seed | MET | 1.613851 | LUAD | EKVX | PHA-665752 | MVP_top60_DNA+RNA_seed | ATM | 2.09245 | STAD | NCI-N87 | KU-55933 |
| MVP_top60_DNA_seed | GSK3B | 1.61661 | LUAD | NCI-H1838 | AZD7969 | MVP_top60_DNA+RNA_seed | ATM | 2.104122 | STAD | SNU-16 | CP466722 |
| MVP_top60_DNA_seed | AKT1 | 1.618632 | LUAD | RERF-LC-MS | AKT inhibitor VIII | MVP_top60_DNA+RNA_seed | FGFR3 | 2.110271 | STAD | NCI-N87 | FGFR_3831 |
| MVP_top60_DNA_seed | AURKA | 1.619338 | LUAD | EMC-BAC-1 | CD532 | MVP_top60_DNA+RNA_seed | ATM | 2.140308 | STAD | RF-48 | KU-60019 |
| MVP_top60_DNA_seed | BRAF | 1.620187 | LUAD | NCI-H1693 | PLX-4720 | MVP_top60_DNA+RNA_seed | ERBB4 | 2.175457 | STAD | SNU-16 | AST-1306 |
| MVP_top60_DNA_seed | AURKA | 1.622022 | LUAD | NCI-H441 | CD532 | MVP_top60_DNA+RNA_seed | PIK3CA | 2.177344 | STAD | SNU-1 | AZD8835 |
| MVP_top60_DNA_seed | AKT1 | 1.622874 | LUAD | NCI-H441 | MK-2206 | MVP_top60_DNA+RNA_seed | PTK2 | 2.189078 | STAD | Hs-746T | PF-562271 |
| MVP_top60_DNA_seed | MET | 1.632165 | LUAD | NCI-H1838 | PHA-665752 | MVP_top60_DNA+RNA_seed | HDAC1 | 2.192992 | STAD | Hs-746T | JQ12 |
| MVP_top60_DNA_seed | GSK3B | 1.632844 | LUAD | NCI-H2228 | CHIR-99021 | MVP_top60_DNA+RNA_seed | GSK3B | 2.210329 | STAD | MKN7 | AZD7969 |
| MVP_top60_DNA_seed | AURKA | 1.637619 | LUAD | NCI-H322M | Alisertib | MVP_top60_DNA+RNA_seed | ERBB2 | 2.219959 | STAD | OCUM-1 | CUDC-101 |
| MVP_top60_DNA_seed | GSK3B | 1.639891 | LUAD | NCI-H1793 | CHIR-99021 | MVP_top60_DNA+RNA_seed | ERBB2 | 2.327314 | STAD | SNU-1 | AZD8931 |
| MVP_top60_DNA_seed | AURKA | 1.642485 | LUAD | NCI-H1781 | Genentech Cpd 10 | MVP_top60_DNA+RNA_seed | ERBB3 | 2.327314 | STAD | SNU-1 | AZD8931 |
| MVP_top60_DNA_seed | AURKA | 1.647337 | LUAD | NCI-H2122 | CD532 | MVP_top60_DNA+RNA_seed | ERBB4 | 2.333127 | STAD | OCUM-1 | AST-1306 |
| MVP_top60_DNA_seed | EGFR | 1.650087 | LUAD | NCI-H1693 | Cetuximab | MVP_top60_DNA+RNA_seed | ERBB2 | 2.343604 | STAD | OCUM-1 | CI-1033 |
| MVP_top60_DNA_seed | MDM2 | 1.650201 | LUAD | NCI-H2291 | Nutlin-3a (-) | MVP_top60_DNA+RNA_seed | ERBB4 | 2.343604 | STAD | OCUM-1 | CI-1033 |
| MVP_top60_DNA_seed | AURKA | 1.657798 | LUAD | NCI-H1435 | Genentech Cpd 10 | MVP_top60_DNA+RNA_seed | ATM | 2.430569 | STAD | SNU-5 | CP466722 |
| MVP_top60_DNA_seed | AKT1 | 1.660948 | LUAD | NCI-H2347 | MK-2206 | MVP_top60_DNA+RNA_seed | MET | 2.452519 | STAD | NCI-N87 | JNJ38877605 |
| MVP_top60_DNA_seed | HDAC1 | 1.664309 | LUAD | NCI-H1838 | CAY10603 | MVP_top60_DNA+RNA_seed | ERBB4 | 2.506507 | STAD | MKN45 | AST-1306 |
| MVP_top60_DNA_seed | ERBB2 | 1.664829 | LUAD | NCI-H2291 | CP724714 | MVP_top60_DNA+RNA_seed | ERBB2 | 2.62529 | STAD | SNU-5 | CI-1033 |
| MVP_top60_DNA_seed | FGFR1 | 1.670682 | LUAD | EKVX | PD173074 | MVP_top60_DNA+RNA_seed | ERBB4 | 2.62529 | STAD | SNU-5 | CI-1033 |
| MVP_top60_DNA_seed | HDAC1 | 1.67207 | LUAD | NCI-H2342 | CAY10603 | MVP_top60_DNA+RNA_seed | HDAC1 | 2.639529 | STAD | OCUM-1 | AR-42 |
| MVP_top60_DNA_seed | IGF1R | 1.673572 | LUAD | NCI-H1435 | BMS-754807 | MVP_top60_DNA+RNA_seed | HDAC1 | 2.657814 | STAD | OCUM-1 | Belinostat |
| MVP_top60_DNA_seed | PTK2 | 1.674263 | LUAD | NCI-H1993 | PF-562271 | MVP_top60_DNA+RNA_seed | HSP90AA1 | 2.683601 | STAD | SNU-16 | Luminespib |
| MVP_top60_DNA_seed | AKT1 | 1.676727 | LUAD | EKVX | A-443654 | MVP_top60_DNA+RNA_seed | HSP90AB1 | 2.683601 | STAD | SNU-16 | Luminespib |
| MVP_top60_DNA_seed | AKT1 | 1.684248 | LUAD | NCI-H1573 | BAY AKT1 | MVP_top60_DNA+RNA_seed | PTK2 | 2.698504 | STAD | NCI-N87 | PF-562271 |
| MVP_top60_DNA_seed | EGFR | 1.684419 | LUAD | NCI-H2291 | Pelitinib | MVP_top60_DNA+RNA_seed | HDAC1 | 2.724627 | STAD | OCUM-1 | CAY10603 |
| MVP_top60_DNA_seed | AURKA | 1.685601 | LUAD | NCI-H1563 | ZM447439 | MVP_top60_DNA+RNA_seed | ERBB2 | 3.33718 | STAD | SNU-5 | PF-00299804 |
| MVP_top60_DNA_seed | GSK3B | 1.686055 | LUAD | NCI-H1838 | CHIR-99021 | MVP_top60_DNA+RNA_seed | ERBB4 | 3.33718 | STAD | SNU-5 | PF-00299804 |
| MVP_top60_DNA_seed | AURKA | 1.686314 | LUAD | NCI-H1838 | Tozasertib | MVP_top60_DNA+RNA_seed | ERBB2 | 3.35161 | STAD | OCUM-1 | PF-00299804 |
| MVP_top60_DNA_seed | SRC | 1.686565 | LUAD | EKVX | WH-4-023 | MVP_top60_DNA+RNA_seed | ERBB4 | 3.35161 | STAD | OCUM-1 | PF-00299804 |
| MVP_top60_DNA_seed | AURKA | 1.687343 | LUAD | HCC-827 | Genentech Cpd 10 | MVP_top60_DNA+RNA_seed | ERBB4 | 3.424574 | STAD | SNU-5 | AST-1306 |

| | | | | | | | | | | | |
|---|---|---|---|---|---|---|---|---|---|---|---|
| MVP_top60_DNA_seed | AKT1 | 1.690916 | LUAD | NCI-H1793 | AKT inhibitor VIII | MVP_top60_DNA+RNA_seed | MET | -6.275219 | STAD | MKN45 | Savolitinib |
| MVP_top60_DNA_seed | MET | 1.695647 | LUAD | EKVX | JNJ38877605 | MVP_top60_DNA+RNA_seed | MET | -6.100559 | STAD | SNU-5 | Savolitinib |
| MVP_top60_DNA_seed | GSK3B | 1.696885 | LUAD | NCI-H2291 | KIN001-042 | MVP_top60_DNA+RNA_seed | MET | -5.621499 | STAD | Hs-746T | Savolitinib |
| MVP_top60_DNA_seed | HDAC1 | 1.697951 | LUAD | NCI-H1838 | Tubastatin A | MVP_top60_DNA+RNA_seed | FGFR3 | -5.226735 | STAD | SNU-16 | PD173074 |
| MVP_top60_DNA_seed | AURKA | 1.698048 | LUAD | RERF-LC-MS | Genentech Cpd 10 | MVP_top60_DNA+RNA_seed | ERBB2 | -4.964046 | STAD | NCI-N87 | Afatinib |
| MVP_top60_DNA_seed | ATM | 1.699962 | LUAD | NCI-H650 | KU-60019 | MVP_top60_DNA+RNA_seed | FGFR3 | -4.86989 | STAD | HSC-39 | PD173074 |
| MVP_top60_DNA_seed | GSK3B | 1.700165 | LUAD | NCI-H2342 | AZD7969 | MVP_top60_DNA+RNA_seed | FGFR3 | -4.705369 | STAD | HSC-39 | AZD4547 |
| MVP_top60_DNA_seed | HSP90AA1 | 1.700745 | LUAD | NCI-H441 | SNX-2112 | MVP_top60_DNA+RNA_seed | FGFR3 | -4.610762 | STAD | SNU-16 | AZD4547 |
| MVP_top60_DNA_seed | GSK3B | 1.702811 | LUAD | NCI-H322M | AZD7969 | MVP_top60_DNA+RNA_seed | ERBB2 | -4.104673 | STAD | SK-GT-2 | Afatinib |
| MVP_top60_DNA_seed | AURKA | 1.703742 | LUAD | NCI-H2405 | ZM447439 | MVP_top60_DNA+RNA_seed | ERBB2 | -3.70213 | STAD | NCI-N87 | Lapatinib |
| MVP_top60_DNA_seed | EGFR | 1.706033 | LUAD | NCI-H1755 | CI-1033 | MVP_top60_DNA+RNA_seed | ESR1 | -3.532472 | STAD | FU97 | GDC0810 |
| MVP_top60_DNA_seed | ERBB2 | 1.706033 | LUAD | NCI-H1755 | CI-1033 | MVP_top60_DNA+RNA_seed | MET | -3.373267 | STAD | SNU-5 | Crizotinib |
| MVP_top60_DNA_seed | PTK2 | 1.712532 | LUAD | NCI-H1623 | PF-562271 | MVP_top60_DNA+RNA_seed | FGFR3 | -3.328162 | STAD | KATOIII | AZD4547 |
| MVP_top60_DNA_seed | HDAC1 | 1.712739 | LUAD | NCI-H2291 | Tubastatin A | MVP_top60_DNA+RNA_seed | ERBB2 | -3.096553 | STAD | NCI-N87 | Sapitinib |
| MVP_top60_DNA_seed | AURKA | 1.716566 | LUAD | NCI-H1838 | ZM447439 | MVP_top60_DNA+RNA_seed | ERBB3 | -3.096553 | STAD | NCI-N87 | Sapitinib |
| MVP_top60_DNA_seed | MAPK1 | 1.717322 | LUAD | NCI-H2291 | FR-180204 | MVP_top60_DNA+RNA_seed | MET | -3.039269 | STAD | MKN45 | Crizotinib |
| MVP_top60_DNA_seed | AURKA | 1.717358 | LUAD | NCI-H2009 | Genentech Cpd 10 | MVP_top60_DNA+RNA_seed | MAPK1 | -2.97902 | STAD | AGS | VX-11e |
| MVP_top60_DNA_seed | HDAC1 | 1.718503 | LUAD | NCI-H1793 | Tubastatin A | MVP_top60_DNA+RNA_seed | ERBB2 | -2.940765 | STAD | SK-GT-2 | Sapitinib |
| MVP_top60_DNA_seed | AKT1 | 1.719399 | LUAD | EKVX | GSK690693 | MVP_top60_DNA+RNA_seed | ERBB3 | -2.940765 | STAD | SK-GT-2 | Sapitinib |
| MVP_top60_DNA_seed | BRAF | 1.726835 | LUAD | EKVX | PLX-4720 | MVP_top60_DNA+RNA_seed | MAPK1 | -2.804725 | STAD | AGS | SCH772984 |
| MVP_top60_DNA_seed | HDAC1 | 1.727161 | LUAD | NCI-H441 | CAY10603 | MVP_top60_DNA+RNA_seed | MAPK3 | -2.804725 | STAD | AGS | SCH772984 |
| MVP_top60_DNA_seed | PTK2 | 1.727717 | LUAD | NCI-H1693 | PF-562271 | MVP_top60_DNA+RNA_seed | MAPK1 | -2.791952 | STAD | OCUM-1 | SCH772984 |
| MVP_top60_DNA_seed | IGF1R | 1.731836 | LUAD | NCI-H1838 | BMS-536924 | MVP_top60_DNA+RNA_seed | MAPK3 | -2.791952 | STAD | OCUM-1 | SCH772984 |
| MVP_top60_DNA_seed | GSK3B | 1.733226 | LUAD | NCI-H441 | CHIR-99021 | MVP_top60_DNA+RNA_seed | MAPK1 | -2.726927 | STAD | OCUM-1 | VX-11e |
| MVP_top60_DNA_seed | HSP90AA1 | 1.737482 | LUAD | A549 | SNX-2112 | MVP_top60_DNA+RNA_seed | FGFR3 | -2.695488 | STAD | KATOIII | PD173074 |
| MVP_top60_DNA_seed | AURKA | 1.739377 | LUAD | NCI-H322M | Tozasertib | MVP_top60_DNA+RNA_seed | ERBB2 | -2.431962 | STAD | SK-GT-2 | Lapatinib |
| MVP_top60_DNA_seed | ERBB2 | 1.747601 | LUAD | NCI-H1793 | CP724714 | MVP_top60_DNA+RNA_seed | MAPK1 | -2.237307 | STAD | AGS | Ulixertinib |
| MVP_top60_DNA_seed | AKT1 | 1.74846 | LUAD | RERF-LC-MS | MK-2206 | MVP_top60_DNA+RNA_seed | MAPK3 | -2.237307 | STAD | AGS | Ulixertinib |
| MVP_top60_DNA_seed | HSP90AA1 | 1.750076 | LUAD | NCI-H2291 | Elesclomol | MVP_top60_DNA+RNA_seed | HSP90AA1 | -2.236141 | STAD | HSC-39 | Tanespimycin |
| MVP_top60_DNA_seed | BRAF | 1.75429 | LUAD | NCI-H1693 | HG6-64-1 | MVP_top60_DNA+RNA_seed | HSP90AB1 | -2.236141 | STAD | HSC-39 | Tanespimycin |
| MVP_top60_DNA_seed | BRAF | 1.757667 | LUAD | ABC-1 | RAF_9304 | MVP_top60_DNA+RNA_seed | PIK3CA | -2.195662 | STAD | NCI-N87 | Alpelisib |
| MVP_top60_DNA_seed | SRC | 1.764509 | LUAD | NCI-H441 | NVP-BHG712 | MVP_top60_DNA+RNA_seed | PIK3CA | -2.178419 | STAD | HGC-27 | Alpelisib |
| MVP_top60_DNA_seed | MAPK1 | 1.765249 | LUAD | NCI-H2291 | VX-11e | MVP_top60_DNA+RNA_seed | MAPK1 | -2.14477 | STAD | OCUM-1 | Ulixertinib |
| MVP_top60_DNA_seed | SMARCA4 | 1.765601 | LUAD | NCI-H1793 | PFI-3 | MVP_top60_DNA+RNA_seed | MAPK3 | -2.14477 | STAD | OCUM-1 | Ulixertinib |
| MVP_top60_DNA_seed | MET | 1.766978 | LUAD | NCI-H441 | SU11274 | MVP_top60_DNA+RNA_seed | EZH2 | -2.023559 | STAD | AGS | GSK343 |
| MVP_top60_DNA_seed | PTK2 | 1.768172 | LUAD | EKVX | PF-562271 | MVP_top60_DNA+RNA_seed | PIK3CA | -2.002491 | STAD | HSC-39 | Alpelisib |
| MVP_top60_DNA_seed | HSP90AA1 | 1.771014 | LUAD | NCI-H1838 | SNX-2112 | MVP_top60_DNA+RNA_seed | MAPK1 | -1.999458 | STAD | TGBC11TKB | VX-11e |
| MVP_top60_DNA_seed | IGF1R | 1.772586 | LUAD | NCI-H2291 | Linsitinib | MVP_top60_DNA+RNA_seed | PARP1 | -1.992794 | STAD | HSC-39 | Veliparib |
| MVP_top60_DNA_seed | IGF1R | 1.77299 | LUAD | NCI-H1666 | BMS-754807 | MVP_top60_DNA+RNA_seed | MDM2 | -1.989513 | STAD | OCUM-1 | Nutlin-3a (-) |
| MVP_top60_DNA_seed | PIK3CA | 1.779161 | LUAD | COR-L105 | AZD8835 | MVP_top60_DNA+RNA_seed | HDAC1 | -1.981708 | STAD | TMK-1 | Entinostat |
| MVP_top60_DNA_seed | IGF1R | 1.779629 | LUAD | NCI-H1666 | GSK1904529A | MVP_top60_DNA+RNA_seed | HDAC3 | -1.981708 | STAD | TMK-1 | Entinostat |

| | | | | | | | | | | | |
|---|---|---|---|---|---|---|---|---|---|---|---|
| MVP_top60_DNA_seed | AKT1 | 1.783152 | LUAD | NCI-H650 | BAY AKT1 | MVP_top60_DNA+RNA_seed | ATM | -1.953722 | STAD | HSC-39 | KU-55933 |
| MVP_top60_DNA_seed | AURKA | 1.785861 | LUAD | NCI-H1993 | HG-5-113-01 | MVP_top60_DNA+RNA_seed | MET | -1.95312 | STAD | IM-95 | Savolitinib |
| MVP_top60_DNA_seed | IGF1R | 1.786217 | LUAD | HCC-827 | BMS-754807 | MVP_top60_DNA+RNA_seed | PIK3CA | -1.889177 | STAD | HGC-27 | Buparlisib |
| MVP_top60_DNA_seed | SMARCA4 | 1.787465 | LUAD | NCI-H322M | PFI-3 | MVP_top60_DNA+RNA_seed | PIK3CA | -1.879573 | STAD | 23132-87 | AZD8186 |
| MVP_top60_DNA_seed | HDAC1 | 1.790939 | LUAD | NCI-H1838 | Belinostat | MVP_top60_DNA+RNA_seed | MAPK1 | -1.874616 | STAD | OCUM-1 | Ulixertinib |
| MVP_top60_DNA_seed | AURKA | 1.796747 | LUAD | NCI-H1435 | ZM447439 | MVP_top60_DNA+RNA_seed | MAPK3 | -1.874616 | STAD | OCUM-1 | Ulixertinib |
| MVP_top60_DNA_seed | GSK3B | 1.796957 | LUAD | NCI-H2291 | CHIR-99021 | MVP_top60_DNA+RNA_seed | PIK3CA | -1.873533 | STAD | HGC-27 | GNE-317 |
| MVP_top60_DNA_seed | MET | 1.798455 | LUAD | NCI-H441 | JNJ38877605 | MVP_top60_DNA+RNA_seed | PIK3CA | -1.806595 | STAD | AGS | Alpelisib |
| MVP_top60_DNA_seed | IGF1R | 1.799054 | LUAD | NCI-H1793 | Linsitinib | MVP_top60_DNA+RNA_seed | ESR1 | -1.747615 | STAD | HSC-39 | Tamoxifen |
| MVP_top60_DNA_seed | AKT1 | 1.799676 | LUAD | NCI-H322M | MK-2206 | MVP_top60_DNA+RNA_seed | ESR1 | -1.713851 | STAD | SNU-16 | Tamoxifen |
| MVP_top60_DNA_seed | PTK2 | 1.800382 | LUAD | NCI-H1573 | PF-562271 | MVP_top60_DNA+RNA_seed | ERBB2 | -1.699082 | STAD | NUGC-3 | Sapitinib |
| MVP_top60_DNA_seed | SRC | 1.802308 | LUAD | EKVX | NVP-BHG712 | MVP_top60_DNA+RNA_seed | ERBB3 | -1.699082 | STAD | NUGC-3 | Sapitinib |
| MVP_top60_DNA_seed | SRC | 1.803821 | LUAD | NCI-H1993 | Bosutinib | MVP_top60_DNA+RNA_seed | PIK3CA | -1.690563 | STAD | HGC-27 | AZD8186 |
| MVP_top60_DNA_seed | HSP90AA1 | 1.803824 | LUAD | NCI-H1993 | Luminespib | MVP_top60_DNA+RNA_seed | MET | -1.676809 | STAD | Hs-746T | Crizotinib |
| MVP_top60_DNA_seed | IGF1R | 1.807654 | LUAD | NCI-H1793 | GSK1904529A | MVP_top60_DNA+RNA_seed | PIK3CA | -1.661051 | STAD | FU97 | Alpelisib |
| MVP_top60_DNA_seed | ERBB2 | 1.809032 | LUAD | NCI-H1838 | CP724714 | MVP_top60_DNA+RNA_seed | HSP90AA1 | -1.648706 | STAD | SNU-16 | Tanespimycin |
| MVP_top60_DNA_seed | MAPK1 | 1.81432 | LUAD | NCI-H1793 | VX-11e | MVP_top60_DNA+RNA_seed | HSP90AB1 | -1.648706 | STAD | SNU-16 | Tanespimycin |
| MVP_top60_DNA_seed | BRAF | 1.828291 | LUAD | NCI-H1838 | HG6-64-1 | MVP_top60_DNA+RNA_seed | ESR1 | -1.631682 | STAD | FU97 | Tamoxifen |
| MVP_top60_DNA_seed | SMARCA4 | 1.828297 | LUAD | NCI-H2291 | PFI-3 | MVP_top60_DNA+RNA_seed | EP300 | -1.624699 | STAD | RF-48 | SGC-CBP30 |
| MVP_top60_DNA_seed | AKT1 | 1.829362 | LUAD | NCI-H2009 | AKT inhibitor VIII | MVP_top60_DNA+RNA_seed | CREBBP | -1.624699 | STAD | RF-48 | SGC-CBP30 |
| MVP_top60_DNA_seed | BRAF | 1.833713 | LUAD | NCI-H23 | HG6-64-1 | MVP_top60_DNA+RNA_seed | HSP90AA1 | -1.603973 | STAD | AGS | Tanespimycin |
| MVP_top60_DNA_seed | SRC | 1.834018 | LUAD | NCI-H2291 | NVP-BHG712 | MVP_top60_DNA+RNA_seed | HSP90AB1 | -1.603973 | STAD | AGS | Tanespimycin |
| MVP_top60_DNA_seed | FGFR1 | 1.848175 | LUAD | NCI-H322M | FGFR_3831 | MVP_top60_DNA+RNA_seed | ESR1 | -1.599474 | STAD | RERF-GC-1B | GDC0810 |
| MVP_top60_DNA_seed | MET | 1.85379 | LUAD | NCI-H322M | Crizotinib | MVP_top60_DNA+RNA_seed | PARP1 | -1.593178 | STAD | HSC-39 | Rucaparib |
| MVP_top60_DNA_seed | EGFR | 1.858459 | LUAD | NCI-H1355 | CUDC-101 | MVP_top60_DNA+RNA_seed | SRC | -1.581791 | STAD | RERF-GC-1B | Bosutinib |
| MVP_top60_DNA_seed | ERBB2 | 1.858459 | LUAD | NCI-H1355 | CUDC-101 | MVP_top60_DNA+RNA_seed | MDM2 | -1.580495 | STAD | RF-48 | Nutlin-3a (-) |
| MVP_top60_DNA_seed | BRAF | 1.859683 | LUAD | HCC-78 | HG6-64-1 | MVP_top60_DNA+RNA_seed | HSP90AA1 | -1.545979 | STAD | TGBC11TKB | Tanespimycin |
| MVP_top60_DNA_seed | MET | 1.8631 | LUAD | EKVX | Crizotinib | MVP_top60_DNA+RNA_seed | HSP90AB1 | -1.545979 | STAD | TGBC11TKB | Tanespimycin |
| MVP_top60_DNA_seed | AURKA | 1.863414 | LUAD | SW1573 | Genentech Cpd 10 | MVP_top60_DNA+RNA_seed | HSP90AA1 | -1.539666 | STAD | OCUM-1 | Tanespimycin |
| MVP_top60_DNA_seed | HDAC1 | 1.864656 | LUAD | NCI-H1355 | AR-42 | MVP_top60_DNA+RNA_seed | HSP90AB1 | -1.539666 | STAD | OCUM-1 | Tanespimycin |
| MVP_top60_DNA_seed | GSK3B | 1.865706 | LUAD | NCI-H322M | CHIR-99021 | MVP_top60_DNA+RNA_seed | PIK3CA | -1.513459 | STAD | HSC-39 | GNE-317 |
| MVP_top60_DNA_seed | FGFR1 | 1.868144 | LUAD | EKVX | AZD4547 | MVP_top60_DNA+RNA_seed | HSP90AA1 | -1.504071 | STAD | HSC-39 | Luminespib |
| MVP_top60_DNA_seed | CREBBP | 1.870196 | LUAD | NCI-H1693 | I-CBP112 | MVP_top60_DNA+RNA_seed | HSP90AB1 | -1.504071 | STAD | HSC-39 | Luminespib |
| MVP_top60_DNA_seed | GSK3B | 1.870206 | LUAD | NCI-H322M | CHIR-99021 | MVP_top60_DNA+RNA_seed | FGFR3 | -1.498662 | STAD | SNU-1 | AZD4547 |
| MVP_top60_DNA_seed | HSP90AA1 | 1.873396 | LUAD | NCI-H2342 | SNX-2112 | MVP_top60_DNA+RNA_seed | PARP1 | -1.493874 | STAD | SNU-16 | Olaparib |
| MVP_top60_DNA_seed | IGF1R | 1.878324 | LUAD | NCI-H1693 | GSK1904529A | MVP_top60_DNA+RNA_seed | MDM2 | -1.489373 | STAD | AGS | Nutlin-3a (-) |
| MVP_top60_DNA_seed | BRAF | 1.878644 | LUAD | NCI-H1781 | HG6-64-1 | MVP_top60_DNA+RNA_seed | HDAC1 | -1.48244 | STAD | AGS | PCI-34051 |
| MVP_top60_DNA_seed | HDAC1 | 1.879534 | LUAD | NCI-H322M | Parthenolide | MVP_top60_DNA+RNA_seed | MAPK1 | -1.47265 | STAD | MKN28 | Ulixertinib |
| MVP_top60_DNA_seed | HDAC1 | 1.881398 | LUAD | NCI-H2009 | Belinostat | MVP_top60_DNA+RNA_seed | MAPK3 | -1.47265 | STAD | MKN28 | Ulixertinib |
| MVP_top60_DNA_seed | HSP90AA1 | 1.88205 | LUAD | NCI-H1355 | SNX-2112 | MVP_top60_DNA+RNA_seed | MAPK1 | -1.45617 | STAD | AGS | Ulixertinib |
| MVP_top60_DNA_seed | IGF1R | 1.88319 | LUAD | NCI-H1838 | GSK1904529A | MVP_top60_DNA+RNA_seed | MAPK3 | -1.45617 | STAD | AGS | Ulixertinib |

| | | | | | | | | | | | |
|---|---|---|---|---|---|---|---|---|---|---|---|
| MVP_top60_DNA_seed | SRC | 1.88546 | LUAD | NCI-H1437 | HG-6-71-01 | MVP_top60_DNA+RNA_seed | FGFR3 | -1.438175 | STAD | FU97 | PD173074 |
| MVP_top60_DNA_seed | AURKA | 1.886977 | LUAD | EKVX | GSK1070916 | MVP_top60_DNA+RNA_seed | EP300 | -1.407546 | STAD | SNU-16 | CPI-637 |
| MVP_top60_DNA_seed | ATM | 1.889199 | LUAD | COR-L105 | KU-55933 | MVP_top60_DNA+RNA_seed | MET | -1.406037 | STAD | SNU-16 | Crizotinib |
| MVP_top60_DNA_seed | HDAC1 | 1.890277 | LUAD | NCI-H1838 | Parthenolide | MVP_top60_DNA+RNA_seed | MET | -1.398825 | STAD | HSC-39 | Crizotinib |
| MVP_top60_DNA_seed | SMARCA4 | 1.891292 | LUAD | NCI-H1693 | PFI-3 | MVP_top60_DNA+RNA_seed | ESR1 | -1.384373 | STAD | HGC-27 | GDC0810 |
| MVP_top60_DNA_seed | EGFR | 1.891895 | LUAD | NCI-H1437 | AST-1306 | MVP_top60_DNA+RNA_seed | PARP1 | -1.382899 | STAD | SNU-16 | Veliparib |
| MVP_top60_DNA_seed | EGFR | 1.895013 | LUAD | HCC-44 | CI-1033 | MVP_top60_DNA+RNA_seed | HDAC1 | -1.369778 | STAD | NUGC-4 | Romidepsin |
| MVP_top60_DNA_seed | ERBB2 | 1.895013 | LUAD | HCC-44 | CI-1033 | MVP_top60_DNA+RNA_seed | HDAC3 | -1.369778 | STAD | NUGC-4 | Romidepsin |
| MVP_top60_DNA_seed | AKT1 | 1.897591 | LUAD | NCI-H1435 | MK-2206 | MVP_top60_DNA+RNA_seed | FGFR3 | -1.349506 | STAD | RERF-GC-1B | AZD4547 |
| MVP_top60_DNA_seed | HDAC1 | 1.898807 | LUAD | NCI-H1838 | AR-42 | MVP_top60_DNA+RNA_seed | ATM | -1.34211 | STAD | MKN45 | KU-55933 |
| MVP_top60_DNA_seed | MAPK1 | 1.907285 | LUAD | NCI-H441 | FR-180204 | MVP_top60_DNA+RNA_seed | ERBB2 | -1.340055 | STAD | NUGC-3 | Afatinib |
| MVP_top60_DNA_seed | HDAC1 | 1.912496 | LUAD | NCI-H1650 | Entinostat | MVP_top60_DNA+RNA_seed | PARP1 | -1.321045 | STAD | SNU-16 | Rucaparib |
| MVP_top60_DNA_seed | CREBBP | 1.918098 | LUAD | NCI-H1838 | I-CBP112 | MVP_top60_DNA+RNA_seed | MAPK1 | -1.310121 | STAD | MKN45 | SCH772984 |
| MVP_top60_DNA_seed | BRAF | 1.922278 | LUAD | NCI-H1573 | HG6-64-1 | MVP_top60_DNA+RNA_seed | MAPK3 | -1.310121 | STAD | MKN45 | SCH772984 |
| MVP_top60_DNA_seed | BRAF | 1.924113 | LUAD | EKVX | SB590885 | MVP_top60_DNA+RNA_seed | EZH2 | -1.303093 | STAD | SNU-16 | GSK343 |
| MVP_top60_DNA_seed | AURKA | 1.92505 | LUAD | NCI-H1623 | Genentech Cpd 10 | MVP_top60_DNA+RNA_seed | FGFR3 | -1.274816 | STAD | RERF-GC-1B | PD173074 |
| MVP_top60_DNA_seed | GSK3B | 1.928421 | LUAD | NCI-H2291 | SB216763 | MVP_top60_DNA+RNA_seed | ESR1 | -1.269431 | STAD | IM-95 | GDC0810 |
| MVP_top60_DNA_seed | HSP90AA1 | 1.937296 | LUAD | NCI-H1648 | CCT-018159 | MVP_top60_DNA+RNA_seed | PARP1 | -1.261068 | STAD | HGC-27 | Rucaparib |
| MVP_top60_DNA_seed | HSP90AA1 | 1.937838 | LUAD | Calu-3 | Elesclomol | MVP_top60_DNA+RNA_seed | PARP1 | -1.260067 | STAD | HGC-27 | Olaparib |
| MVP_top60_DNA_seed | ATM | 1.943072 | LUAD | NCI-H2228 | KU-60019 | MVP_top60_DNA+RNA_seed | PIK3CA | -1.255553 | STAD | 23132-87 | Buparlisib |
| MVP_top60_DNA_seed | GSK3B | 1.943544 | LUAD | NCI-H1793 | KIN001-042 | MVP_top60_DNA+RNA_seed | MET | -1.248383 | STAD | HGC-27 | Savolitinib |
| MVP_top60_DNA_seed | PTK2 | 1.944056 | LUAD | NCI-H322M | PF-562271 | MVP_top60_DNA+RNA_seed | MET | -1.230999 | STAD | HSC-39 | Savolitinib |
| MVP_top60_DNA_seed | HSP90AA1 | 1.946492 | LUAD | NCI-H322M | Elesclomol | MVP_top60_DNA+RNA_seed | PARP1 | -1.229143 | STAD | OCUM-1 | Rucaparib |
| MVP_top60_DNA_seed | IGF1R | 1.951075 | LUAD | NCI-H441 | GSK1904529A | MVP_top60_DNA+RNA_seed | MAPK1 | -1.20914 | STAD | NCI-N87 | Ulixertinib |
| MVP_top60_DNA_seed | HDAC1 | 1.955466 | LUAD | NCI-H1355 | CAY10603 | MVP_top60_DNA+RNA_seed | MAPK3 | -1.20914 | STAD | NCI-N87 | Ulixertinib |
| MVP_top60_DNA_seed | AURKA | 1.958492 | LUAD | EKVX | Tozasertib | MVP_top60_DNA+RNA_seed | SRC | -1.207192 | STAD | HGC-27 | Bosutinib |
| MVP_top60_DNA_seed | GSK3B | 1.959799 | LUAD | NCI-H2085 | AZD7969 | MVP_top60_DNA+RNA_seed | MDM2 | -1.206864 | STAD | MKN45 | Nutlin-3a (-) |
| MVP_top60_DNA_seed | HDAC1 | 1.960504 | LUAD | NCI-H1781 | Dacinostat | MVP_top60_DNA+RNA_seed | MDM2 | -1.18676 | STAD | 23132-87 | Nutlin-3a (-) |
| MVP_top60_DNA_seed | HDAC1 | 1.962589 | LUAD | RERF-LC-KJ | Dacinostat | MVP_top60_DNA+RNA_seed | ERBB2 | -1.18579 | STAD | NUGC-3 | Lapatinib |
| MVP_top60_DNA_seed | PTK2 | 1.96495 | LUAD | NCI-H1563 | PF-562271 | MVP_top60_DNA+RNA_seed | HSP90AA1 | -1.182084 | STAD | HGC-27 | Tanespimycin |
| MVP_top60_DNA_seed | HSP90AA1 | 1.968778 | LUAD | ABC-1 | CCT-018159 | MVP_top60_DNA+RNA_seed | HSP90AB1 | -1.182084 | STAD | HGC-27 | Tanespimycin |
| MVP_top60_DNA_seed | GSK3B | 1.970088 | LUAD | NCI-H1793 | CHIR-99021 | MVP_top60_DNA+RNA_seed | MAPK1 | -1.174219 | STAD | TGBC11TKB | SCH772984 |
| MVP_top60_DNA_seed | HSP90AA1 | 1.9746 | LUAD | NCI-H1693 | Tanespimycin | MVP_top60_DNA+RNA_seed | MAPK3 | -1.174219 | STAD | TGBC11TKB | SCH772984 |
| MVP_top60_DNA_seed | AURKA | 1.976648 | LUAD | SK-LU-1 | CD532 | MVP_top60_DNA+RNA_seed | KRAS | -1.16714 | STAD | AGS | KRAS (G12C) Inhibitor-1 |
| MVP_top60_DNA_seed | IGF1R | 1.981868 | LUAD | EKVX | GSK1904529A | MVP_top60_DNA+RNA_seed | EP300 | -1.166446 | STAD | RERF-GC-1B | SGC-CBP30 |
| MVP_top60_DNA_seed | HDAC1 | 1.984077 | LUAD | EKVX | Dacinostat | MVP_top60_DNA+RNA_seed | CREBBP | -1.166446 | STAD | RERF-GC-1B | SGC-CBP30 |
| MVP_top60_DNA_seed | MDM2 | 1.999421 | LUAD | NCI-H1435 | Serdemetan | MVP_top60_DNA+RNA_seed | PIK3CA | -1.166136 | STAD | 23132-87 | GNE-317 |
| MVP_top60_DNA_seed | EGFR | 2.010551 | LUAD | NCI-H2122 | PF-00299804 | MVP_top60_DNA+RNA_seed | EZH2 | -1.165812 | STAD | HSC-39 | GSK343 |
| MVP_top60_DNA_seed | ERBB2 | 2.010551 | LUAD | NCI-H2122 | PF-00299804 | MVP_top60_DNA+RNA_seed | MAPK1 | -1.164394 | STAD | GCIY | Ulixertinib |
| MVP_top60_DNA_seed | AKT1 | 2.012874 | LUAD | NCI-H1666 | AKT inhibitor VIII | MVP_top60_DNA+RNA_seed | MAPK3 | -1.164394 | STAD | GCIY | Ulixertinib |
| MVP_top60_DNA_seed | HSP90AA1 | 2.013269 | LUAD | NCI-H1793 | Luminespib | MVP_top60_DNA+RNA_seed | HDAC1 | -1.158468 | STAD | AGS | Romidepsin |

| | | | | | | | | | | | |
|---|---|---|---|---|---|---|---|---|---|---|---|
| MVP_top60_DNA_seed | BRAF | 2.013966 | LUAD | EKVX | HG6-64-1 | MVP_top60_DNA+RNA_seed | HDAC3 | -1.158468 | STAD | AGS | Romidepsin |
| MVP_top60_DNA_seed | SRC | 2.014274 | LUAD | NCI-H441 | Bosutinib | MVP_top60_DNA+RNA_seed | HDAC1 | -1.152104 | STAD | NUGC-3 | Romidepsin |
| MVP_top60_DNA_seed | MET | 2.033026 | LUAD | EKVX | AZD6094 | MVP_top60_DNA+RNA_seed | HDAC3 | -1.152104 | STAD | NUGC-3 | Romidepsin |
| MVP_top60_DNA_seed | HSP90AA1 | 2.036228 | LUAD | NCI-H2347 | SNX-2112 | MVP_top60_DNA+RNA_seed | ERBB2 | -1.149545 | STAD | GCIY | Afatinib |
| MVP_top60_DNA_seed | AKT1 | 2.046368 | LUAD | NCI-H1793 | MK-2206 | MVP_top60_DNA+RNA_seed | ESR1 | -1.141999 | STAD | HSC-39 | GDC0810 |
| MVP_top60_DNA_seed | AKT1 | 2.046969 | LUAD | NCI-H650 | MK-2206 | MVP_top60_DNA+RNA_seed | EZH2 | -1.136189 | STAD | AGS | GSK343 |
| MVP_top60_DNA_seed | AKT1 | 2.047699 | LUAD | NCI-H322M | AKT inhibitor VIII | MVP_top60_DNA+RNA_seed | FGFR3 | -1.133877 | STAD | 23132-87 | AZD4547 |
| MVP_top60_DNA_seed | AKT1 | 2.048552 | LUAD | NCI-H1838 | AKT inhibitor VIII | MVP_top60_DNA+RNA_seed | EP300 | -1.129626 | STAD | SNU-5 | SGC-CBP30 |
| MVP_top60_DNA_seed | SMARCA4 | 2.056116 | LUAD | NCI-H1838 | PFI-3 | MVP_top60_DNA+RNA_seed | CREBBP | -1.129626 | STAD | SNU-5 | SGC-CBP30 |
| MVP_top60_DNA_seed | HSP90AA1 | 2.057254 | LUAD | NCI-H1693 | SNX-2112 | MVP_top60_DNA+RNA_seed | EP300 | -1.128529 | STAD | NUGC-3 | SGC-CBP30 |
| MVP_top60_DNA_seed | HDAC1 | 2.058314 | LUAD | EKVX | Tubastatin A | MVP_top60_DNA+RNA_seed | CREBBP | -1.128529 | STAD | NUGC-3 | SGC-CBP30 |
| MVP_top60_DNA_seed | GSK3B | 2.059493 | LUAD | COR-L105 | AZD7969 | MVP_top60_DNA+RNA_seed | ERBB2 | -1.120754 | STAD | GCIY | Sapitinib |
| MVP_top60_DNA_seed | HSP90AA1 | 2.062085 | LUAD | ABC-1 | Tanespimycin | MVP_top60_DNA+RNA_seed | ERBB3 | -1.120754 | STAD | GCIY | Sapitinib |
| MVP_top60_DNA_seed | GSK3B | 2.093325 | LUAD | NCI-H1993 | AZD7969 | MVP_top60_DNA+RNA_seed | HSP90AA1 | -1.114071 | STAD | AGS | Luminespib |
| MVP_top60_DNA_seed | HDAC1 | 2.094092 | LUAD | SW1573 | JQ12 | MVP_top60_DNA+RNA_seed | HSP90AB1 | -1.114071 | STAD | AGS | Luminespib |
| MVP_top60_DNA_seed | HDAC1 | 2.096667 | LUAD | NCI-H1666 | Entinostat | MVP_top60_DNA+RNA_seed | PIK3CA | -1.107574 | STAD | HSC-39 | Buparlisib |
| MVP_top60_DNA_seed | AURKA | 2.112616 | LUAD | NCI-H1838 | Genentech Cpd 10 | MVP_top60_DNA+RNA_seed | HSP90AA1 | -1.105157 | STAD | NCI-N87 | Luminespib |
| MVP_top60_DNA_seed | HSP90AA1 | 2.114563 | LUAD | NCI-H1563 | CCT-018159 | MVP_top60_DNA+RNA_seed | HSP90AB1 | -1.105157 | STAD | NCI-N87 | Luminespib |
| MVP_top60_DNA_seed | HSP90AA1 | 2.122647 | LUAD | NCI-H1563 | SNX-2112 | MVP_top60_DNA+RNA_seed | PARP1 | -1.097387 | STAD | HSC-39 | Olaparib |
| MVP_top60_DNA_seed | HSP90AA1 | 2.12873 | LUAD | SW1573 | SNX-2112 | MVP_top60_DNA+RNA_seed | PIK3CA | -1.096937 | STAD | IM-95 | Alpelisib |
| MVP_top60_DNA_seed | MAPK1 | 2.134824 | LUAD | NCI-H1793 | FR-180204 | MVP_top60_DNA+RNA_seed | EZH2 | -1.089951 | STAD | SNU-5 | GSK343 |
| MVP_top60_DNA_seed | AKT1 | 2.135125 | LUAD | NCI-H2291 | AKT inhibitor VIII | MVP_top60_DNA+RNA_seed | MAPK1 | -1.085216 | STAD | MKN45 | Ulixertinib |
| MVP_top60_DNA_seed | HSP90AA1 | 2.135751 | LUAD | NCI-H1435 | Elesclomol | MVP_top60_DNA+RNA_seed | MAPK3 | -1.085216 | STAD | MKN45 | Ulixertinib |
| MVP_top60_DNA_seed | EGFR | 2.142759 | LUAD | NCI-H2228 | CI-1033 | MVP_top60_DNA+RNA_seed | HDAC1 | -1.08404 | STAD | MKN45 | PCI-34051 |
| MVP_top60_DNA_seed | ERBB2 | 2.142759 | LUAD | NCI-H2228 | CI-1033 | MVP_top60_DNA+RNA_seed | MAPK1 | -1.078027 | STAD | NUGC-4 | SCH772984 |
| MVP_top60_DNA_seed | HDAC1 | 2.164991 | LUAD | NCI-H1666 | JQ12 | MVP_top60_DNA+RNA_seed | MAPK3 | -1.078027 | STAD | NUGC-4 | SCH772984 |
| MVP_top60_DNA_seed | IGF1R | 2.168824 | LUAD | NCI-H2291 | GSK1904529A | MVP_top60_DNA+RNA_seed | PIK3CA | -1.074511 | STAD | 23132-87 | Alpelisib |
| MVP_top60_DNA_seed | AKT1 | 2.17993 | LUAD | NCI-H1993 | MK-2206 | MVP_top60_DNA+RNA_seed | ERBB2 | -1.0707 | STAD | NUGC-4 | Sapitinib |
| MVP_top60_DNA_seed | SRC | 2.183864 | LUAD | NCI-H322M | Saracatinib | MVP_top60_DNA+RNA_seed | ERBB3 | -1.0707 | STAD | NUGC-4 | Sapitinib |
| MVP_top60_DNA_seed | BRAF | 2.184566 | LUAD | NCI-H1623 | HG6-64-1 | MVP_top60_DNA+RNA_seed | MAPK1 | -1.069708 | STAD | NCI-N87 | SCH772984 |
| MVP_top60_DNA_seed | IGF1R | 2.189054 | LUAD | NCI-H2291 | BMS-754807 | MVP_top60_DNA+RNA_seed | MAPK3 | -1.069708 | STAD | NCI-N87 | SCH772984 |
| MVP_top60_DNA_seed | EGFR | 2.196033 | LUAD | A549 | CI-1033 | MVP_top60_DNA+RNA_seed | HSP90AA1 | -1.06889 | STAD | SNU-1 | Tanespimycin |
| MVP_top60_DNA_seed | ERBB2 | 2.196033 | LUAD | A549 | CI-1033 | MVP_top60_DNA+RNA_seed | HSP90AB1 | -1.06889 | STAD | SNU-1 | Tanespimycin |
| MVP_top60_DNA_seed | AURKA | 2.199297 | LUAD | H3255 | CD532 | MVP_top60_DNA+RNA_seed | PIK3CA | -1.064823 | STAD | FU97 | GNE-317 |
| MVP_top60_DNA_seed | AKT1 | 2.224587 | LUAD | EKVX | AKT inhibitor VIII | MVP_top60_DNA+RNA_seed | ERBB2 | -1.064414 | STAD | HGC-27 | Lapatinib |
| MVP_top60_DNA_seed | PTK2 | 2.226134 | LUAD | NCI-H1666 | PF-562271 | MVP_top60_DNA+RNA_seed | PIK3CA | -1.052359 | STAD | SNU-16 | Buparlisib |
| MVP_top60_DNA_seed | HDAC1 | 2.226534 | LUAD | NCI-H1355 | Belinostat | MVP_top60_DNA+RNA_seed | MET | -1.03287 | STAD | IM-95 | Crizotinib |
| MVP_top60_DNA_seed | HSP90AA1 | 2.228154 | LUAD | COR-L105 | Tanespimycin | MVP_top60_DNA+RNA_seed | EP300 | -1.032284 | STAD | HSC-39 | CPI-637 |
| MVP_top60_DNA_seed | MET | 2.230712 | LUAD | NCI-H1793 | JNJ38877605 | MVP_top60_DNA+RNA_seed | ERBB2 | -0.992582 | STAD | HSC-39 | Lapatinib |
| MVP_top60_DNA_seed | IGF1R | 2.254386 | LUAD | NCI-H1838 | BMS-754807 | MVP_top60_DNA+RNA_seed | MAPK1 | -0.984578 | STAD | HSC-39 | Ulixertinib |
| MVP_top60_DNA_seed | ATM | 2.270582 | LUAD | NCI-H1623 | CP466722 | MVP_top60_DNA+RNA_seed | MAPK3 | -0.984578 | STAD | HSC-39 | Ulixertinib |

| | | | | | | | | | | | |
|---|---|---|---|---|---|---|---|---|---|---|---|
| MVP_top60_DNA_seed | ATM | 2.276432 | LUAD | NCI-H2291 | CP466722 | MVP_top60_DNA+RNA_seed | ESR1 | -0.983947 | STAD | HGC-27 | Tamoxifen |
| MVP_top60_DNA_seed | MDM2 | 2.286987 | LUAD | NCI-H650 | Serdemetan | MVP_top60_DNA+RNA_seed | MAPK1 | -0.975449 | STAD | HSC-39 | VX-11e |
| MVP_top60_DNA_seed | AKT1 | 2.289173 | LUAD | NCI-H1838 | A-443654 | MVP_top60_DNA+RNA_seed | TP53 | -0.964362 | STAD | SNU-16 | MIRA-1 |
| MVP_top60_DNA_seed | HSP90AA1 | 2.293336 | LUAD | HCC-78 | Elesclomol | MVP_top60_DNA+RNA_seed | ERBB2 | -0.961978 | STAD | RERF-GC-1B | Sapitinib |
| MVP_top60_DNA_seed | PTK2 | 2.303493 | LUAD | NCI-H1838 | PF-562271 | MVP_top60_DNA+RNA_seed | ERBB3 | -0.961978 | STAD | RERF-GC-1B | Sapitinib |
| MVP_top60_DNA_seed | EGFR | 2.316374 | LUAD | NCI-H2122 | CI-1033 | MVP_top60_DNA+RNA_seed | MDM2 | -0.960426 | STAD | RF-48 | Serdemetan |
| MVP_top60_DNA_seed | ERBB2 | 2.316374 | LUAD | NCI-H2122 | CI-1033 | MVP_top60_DNA+RNA_seed | KRAS | -0.957921 | STAD | NCI-N87 | KRAS (G12C) Inhibitor-1 |
| MVP_top60_DNA_seed | EGFR | 2.322947 | LUAD | NCI-H2122 | AST-1306 | MVP_top60_DNA+RNA_seed | PIK3CA | -0.94861 | STAD | HSC-39 | AZD8186 |
| MVP_top60_DNA_seed | GSK3B | 2.324973 | LUAD | EKVX | KIN001-042 | MVP_top60_DNA+RNA_seed | HDAC1 | -0.94795 | STAD | NCI-N87 | Romidepsin |
| MVP_top60_DNA_seed | EGFR | 2.32736 | LUAD | NCI-H2023 | AST-1306 | MVP_top60_DNA+RNA_seed | HDAC3 | -0.94795 | STAD | NCI-N87 | Romidepsin |
| MVP_top60_DNA_seed | GSK3B | 2.330046 | LUAD | EKVX | CHIR-99021 | MVP_top60_DNA+RNA_seed | EP300 | -0.940233 | STAD | 23132-87 | SGC-CBP30 |
| MVP_top60_DNA_seed | GSK3B | 2.34146 | LUAD | NCI-H322M | KIN001-042 | MVP_top60_DNA+RNA_seed | CREBBP | -0.940233 | STAD | 23132-87 | SGC-CBP30 |
| MVP_top60_DNA_seed | EGFR | 2.343242 | LUAD | NCI-H1435 | CUDC-101 | MVP_top60_DNA+RNA_seed | EZH2 | -0.936728 | STAD | MKN28 | GSK343 |
| MVP_top60_DNA_seed | ERBB2 | 2.343242 | LUAD | NCI-H1435 | CUDC-101 | MVP_top60_DNA+RNA_seed | HDAC1 | -0.92607 | STAD | RF-48 | Romidepsin |
| MVP_top60_DNA_seed | IGF1R | 2.34476 | LUAD | NCI-H1693 | BMS-754807 | MVP_top60_DNA+RNA_seed | HDAC3 | -0.92607 | STAD | RF-48 | Romidepsin |
| MVP_top60_DNA_seed | EGFR | 2.358305 | LUAD | RERF-LC-KJ | CI-1033 | MVP_top60_DNA+RNA_seed | FGFR3 | -0.923566 | STAD | AGS | PD173074 |
| MVP_top60_DNA_seed | ERBB2 | 2.358305 | LUAD | RERF-LC-KJ | CI-1033 | MVP_top60_DNA+RNA_seed | GSK3B | -0.91946 | STAD | AGS | SB216763 |
| MVP_top60_DNA_seed | HDAC1 | 2.36656 | LUAD | NCI-H2291 | JQ12 | MVP_top60_DNA+RNA_seed | EZH2 | -0.914135 | STAD | Hs-746T | GSK343 |
| MVP_top60_DNA_seed | IGF1R | 2.367299 | LUAD | NCI-H322M | GSK1904529A | MVP_top60_DNA+RNA_seed | KRAS | -0.911847 | STAD | SNU-5 | KRAS (G12C) Inhibitor-1 |
| MVP_top60_DNA_seed | EGFR | 2.381762 | LUAD | NCI-H2023 | CI-1033 | MVP_top60_DNA+RNA_seed | MAPK1 | -0.91179 | STAD | MKN28 | VX-11e |
| MVP_top60_DNA_seed | ERBB2 | 2.381762 | LUAD | NCI-H2023 | CI-1033 | MVP_top60_DNA+RNA_seed | HDAC1 | -0.911584 | STAD | SNU-5 | PCI-34051 |
| MVP_top60_DNA_seed | ATM | 2.382001 | LUAD | NCI-H441 | KU-55933 | MVP_top60_DNA+RNA_seed | MAPK1 | -0.910393 | STAD | KATOIII | Ulixertinib |
| MVP_top60_DNA_seed | BRAF | 2.390548 | LUAD | NCI-H1563 | RAF_9304 | MVP_top60_DNA+RNA_seed | MAPK3 | -0.910393 | STAD | KATOIII | Ulixertinib |
| MVP_top60_DNA_seed | EGFR | 2.396535 | LUAD | NCI-H1437 | CI-1033 | MVP_top60_DNA+RNA_seed | HSP90AA1 | -0.906857 | STAD | SK-GT-2 | Luminespib |
| MVP_top60_DNA_seed | ERBB2 | 2.396535 | LUAD | NCI-H1437 | CI-1033 | MVP_top60_DNA+RNA_seed | HSP90AB1 | -0.906857 | STAD | SK-GT-2 | Luminespib |
| MVP_top60_DNA_seed | HSP90AA1 | 2.400884 | LUAD | NCI-H1793 | SNX-2112 | MVP_top60_DNA+RNA_seed | MET | -0.903609 | STAD | HGC-27 | Crizotinib |
| MVP_top60_DNA_seed | ATM | 2.412826 | LUAD | Calu-3 | KU-60019 | MVP_top60_DNA+RNA_seed | HSP90AA1 | -0.897585 | STAD | TMK-1 | CCT-018159 |
| MVP_top60_DNA_seed | HDAC1 | 2.416005 | LUAD | NCI-H1666 | Dacinostat | MVP_top60_DNA+RNA_seed | HSP90AB1 | -0.897585 | STAD | TMK-1 | CCT-018159 |
| MVP_top60_DNA_seed | AURKA | 2.419294 | LUAD | EKVX | Genentech Cpd 10 | MVP_top60_DNA+RNA_seed | PIK3CA | -0.895344 | STAD | FU97 | Buparlisib |
| MVP_top60_DNA_seed | HSP90AA1 | 2.419708 | LUAD | NCI-H2291 | SNX-2112 | MVP_top60_DNA+RNA_seed | HDAC1 | -0.883727 | STAD | SK-GT-2 | Romidepsin |
| MVP_top60_DNA_seed | HDAC1 | 2.433605 | LUAD | NCI-H1563 | Dacinostat | MVP_top60_DNA+RNA_seed | HDAC3 | -0.883727 | STAD | SK-GT-2 | Romidepsin |
| MVP_top60_DNA_seed | EGFR | 2.457773 | LUAD | NCI-H838 | CI-1033 | MVP_top60_DNA+RNA_seed | PARP1 | -0.877986 | STAD | NUGC-3 | Talazoparib |
| MVP_top60_DNA_seed | ERBB2 | 2.457773 | LUAD | NCI-H838 | CI-1033 | MVP_top60_DNA+RNA_seed | GSK3B | -0.871876 | STAD | HSC-39 | SB216763 |
| MVP_top60_DNA_seed | HDAC1 | 2.45924 | LUAD | NCI-H1838 | JQ12 | MVP_top60_DNA+RNA_seed | MDM2 | -0.870852 | STAD | HSC-39 | Serdemetan |
| MVP_top60_DNA_seed | AKT1 | 2.476136 | LUAD | NCI-H1793 | AKT inhibitor VIII | MVP_top60_DNA+RNA_seed | HDAC1 | -0.868311 | STAD | NUGC-3 | Entinostat |
| MVP_top60_DNA_seed | HDAC1 | 2.487357 | LUAD | NCI-H322M | JQ12 | MVP_top60_DNA+RNA_seed | HDAC3 | -0.868311 | STAD | NUGC-3 | Entinostat |
| MVP_top60_DNA_seed | AURKA | 2.496792 | LUAD | NCI-H2291 | Genentech Cpd 10 | MVP_top60_DNA+RNA_seed | PIK3CA | -0.860398 | STAD | AGS | GNE-317 |
| MVP_top60_DNA_seed | ATM | 2.547208 | LUAD | EKVX | CP466722 | MVP_top60_DNA+RNA_seed | MET | -0.859577 | STAD | SNU-1 | Savolitinib |
| MVP_top60_DNA_seed | HSP90AA1 | 2.549254 | LUAD | NCI-H1623 | Luminespib | MVP_top60_DNA+RNA_seed | EZH2 | -0.846835 | STAD | 23132-87 | GSK343 |
| MVP_top60_DNA_seed | PTK2 | 2.577374 | LUAD | NCI-H2291 | PF-562271 | MVP_top60_DNA+RNA_seed | PIK3CA | -0.842493 | STAD | NCI-N87 | AZD8186 |
| MVP_top60_DNA_seed | BRAF | 2.593319 | LUAD | NCI-H2291 | HG6-64-1 | MVP_top60_DNA+RNA_seed | ATM | -0.834389 | STAD | SNU-1 | KU-55933 |

| | | | | | | | | | | | |
|---|---|---|---|---|---|---|---|---|---|---|---|
| MVP_top60_DNA_seed | CREBBP | 2.595117 | LUAD | EKVX | I-CBP112 | MVP_top60_DNA+RNA_seed | PIK3CA | -0.832559 | STAD | GCIY | GNE-317 |
| MVP_top60_DNA_seed | HSP90AA1 | 2.608085 | LUAD | NCI-H1666 | Luminespib | MVP_top60_DNA+RNA_seed | HDAC1 | -0.831674 | STAD | MKN7 | Entinostat |
| MVP_top60_DNA_seed | HDAC1 | 2.625685 | LUAD | NCI-H1793 | Dacinostat | MVP_top60_DNA+RNA_seed | HDAC3 | -0.831674 | STAD | MKN7 | Entinostat |
| MVP_top60_DNA_seed | AKT1 | 2.650929 | LUAD | NCI-H1666 | A-443654 | MVP_top60_DNA+RNA_seed | PIK3CA | -0.829642 | STAD | SNU-5 | GNE-317 |
| MVP_top60_DNA_seed | BRAF | 2.718412 | LUAD | NCI-H322M | RAF_9304 | MVP_top60_DNA+RNA_seed | MAPK1 | -0.82574 | STAD | SNU-5 | SCH772984 |
| MVP_top60_DNA_seed | HSP90AA1 | 2.73961 | LUAD | NCI-H1563 | Luminespib | MVP_top60_DNA+RNA_seed | MAPK3 | -0.82574 | STAD | SNU-5 | SCH772984 |
| MVP_top60_DNA_seed | EGFR | 2.792776 | LUAD | NCI-H1568 | CUDC-101 | MVP_top60_DNA+RNA_seed | MAPK1 | -0.817621 | STAD | MKN28 | SCH772984 |
| MVP_top60_DNA_seed | ERBB2 | 2.792776 | LUAD | NCI-H1568 | CUDC-101 | MVP_top60_DNA+RNA_seed | MAPK3 | -0.817621 | STAD | MKN28 | SCH772984 |
| MVP_top60_DNA_seed | ATM | 2.802996 | LUAD | NCI-H1993 | CP466722 | MVP_top60_DNA+RNA_seed | ERBB2 | -0.814856 | STAD | SNU-16 | Lapatinib |
| MVP_top60_DNA_seed | ATM | 2.807024 | LUAD | NCI-H322M | KU-60019 | MVP_top60_DNA+RNA_seed | PIK3CA | -0.814266 | STAD | SNU-5 | AZD8186 |
| MVP_top60_DNA_seed | HDAC1 | 2.824876 | LUAD | NCI-H1568 | AR-42 | MVP_top60_DNA+RNA_seed | EP300 | -0.812311 | STAD | HGC-27 | SGC-CBP30 |
| MVP_top60_DNA_seed | EGFR | 2.844676 | LUAD | NCI-H838 | AST-1306 | MVP_top60_DNA+RNA_seed | CREBBP | -0.812311 | STAD | HGC-27 | SGC-CBP30 |
| MVP_top60_DNA_seed | HSP90AA1 | 2.857503 | LUAD | NCI-H1993 | Elesclomol | MVP_top60_DNA+RNA_seed | ESR1 | -0.812064 | STAD | 23132-87 | Tamoxifen |
| MVP_top60_DNA_seed | HSP90AA1 | 2.869074 | LUAD | NCI-H322M | Luminespib | MVP_top60_DNA+RNA_seed | PARP1 | -0.806521 | STAD | OCUM-1 | Veliparib |
| MVP_top60_DNA_seed | GSK3B | 2.8756 | LUAD | NCI-H1793 | AZD7969 | MVP_top60_DNA+RNA_seed | HSP90AA1 | -0.797028 | STAD | OCUM-1 | Luminespib |
| MVP_top60_DNA_seed | EGFR | 2.886417 | LUAD | NCI-H1437 | PF-00299804 | MVP_top60_DNA+RNA_seed | HSP90AB1 | -0.797028 | STAD | OCUM-1 | Luminespib |
| MVP_top60_DNA_seed | ERBB2 | 2.886417 | LUAD | NCI-H1437 | PF-00299804 | MVP_top60_DNA+RNA_seed | HSP90AA1 | -0.796287 | STAD | RF-48 | Luminespib |
| MVP_top60_DNA_seed | HDAC1 | 2.921016 | LUAD | NCI-H1838 | Entinostat | MVP_top60_DNA+RNA_seed | HSP90AB1 | -0.796287 | STAD | RF-48 | Luminespib |
| MVP_top60_DNA_seed | HDAC1 | 2.927149 | LUAD | NCI-H1435 | Belinostat | MVP_top60_DNA+RNA_seed | PIK3CA | -0.791431 | STAD | AGS | Buparlisib |
| MVP_top60_DNA_seed | HSP90AA1 | 2.944004 | LUAD | NCI-H1838 | Luminespib | MVP_top60_DNA+RNA_seed | TP53 | -0.79102 | STAD | HSC-39 | MIRA-1 |
| MVP_top60_DNA_seed | HDAC1 | 2.977468 | LUAD | NCI-H322M | Entinostat | MVP_top60_DNA+RNA_seed | PIK3CA | -0.790039 | STAD | NUGC-3 | GNE-317 |
| MVP_top60_DNA_seed | HDAC1 | 2.982262 | LUAD | NCI-H1568 | CAY10603 | MVP_top60_DNA+RNA_seed | MAPK1 | -0.789896 | STAD | HGC-27 | Ulixertinib |
| MVP_top60_DNA_seed | HSP90AA1 | 3.02026 | LUAD | NCI-H441 | Elesclomol | MVP_top60_DNA+RNA_seed | MAPK3 | -0.789896 | STAD | HGC-27 | Ulixertinib |
| MVP_top60_DNA_seed | HDAC1 | 3.066321 | LUAD | NCI-H2291 | CAY10603 | MVP_top60_DNA+RNA_seed | PARP1 | -0.788503 | STAD | SNU-1 | Olaparib |
| MVP_top60_DNA_seed | HDAC1 | 3.086976 | LUAD | NCI-H1568 | Belinostat | MVP_top60_DNA+RNA_seed | MAPK1 | -0.78644 | STAD | SNU-5 | Ulixertinib |
| MVP_top60_DNA_seed | HDAC1 | 3.111411 | LUAD | NCI-H2291 | Belinostat | MVP_top60_DNA+RNA_seed | MAPK3 | -0.78644 | STAD | SNU-5 | Ulixertinib |
| MVP_top60_DNA_seed | EGFR | 3.111495 | LUAD | NCI-H2291 | CUDC-101 | MVP_top60_DNA+RNA_seed | MAPK1 | -0.785475 | STAD | SNU-5 | Ulixertinib |
| MVP_top60_DNA_seed | ERBB2 | 3.111495 | LUAD | NCI-H2291 | CUDC-101 | MVP_top60_DNA+RNA_seed | MAPK3 | -0.785475 | STAD | SNU-5 | Ulixertinib |
| MVP_top60_DNA_seed | GSK3B | 3.194575 | LUAD | EKVX | AZD7969 | MVP_top60_DNA+RNA_seed | KRAS | -0.783583 | STAD | SNU-1 | :RAS (G12C) Inhibitor-1 |
| MVP_top60_DNA_seed | EGFR | 3.197471 | LUAD | NCI-H838 | PF-00299804 | MVP_top60_DNA+RNA_seed | MAPK1 | -0.782848 | STAD | HSC-39 | Ulixertinib |
| MVP_top60_DNA_seed | ERBB2 | 3.197471 | LUAD | NCI-H838 | PF-00299804 | MVP_top60_DNA+RNA_seed | MAPK3 | -0.782848 | STAD | HSC-39 | Ulixertinib |
| MVP_top60_DNA_seed | HDAC1 | 3.251255 | LUAD | NCI-H322M | Dacinostat | MVP_top60_DNA+RNA_seed | MAPK1 | -0.778445 | STAD | Hs-746T | SCH772984 |
| MVP_top60_DNA_seed | HSP90AA1 | 3.292018 | LUAD | NCI-H2291 | Luminespib | MVP_top60_DNA+RNA_seed | MAPK3 | -0.778445 | STAD | Hs-746T | SCH772984 |
| MVP_top60_DNA_seed | HDAC1 | 3.313231 | LUAD | NCI-H2291 | AR-42 | MVP_top60_DNA+RNA_seed | PARP1 | -0.773812 | STAD | RF-48 | Olaparib |
| MVP_top60_DNA_seed | MDM2 | 3.439305 | LUAD | NCI-H322M | Serdemetan | MVP_top60_DNA+RNA_seed | HDAC1 | -0.773766 | STAD | 23132-87 | Romidepsin |
| MVP_top60_DNA_seed | HDAC1 | 3.901874 | LUAD | NCI-H2291 | Dacinostat | MVP_top60_DNA+RNA_seed | HDAC3 | -0.773766 | STAD | 23132-87 | Romidepsin |
| MVP_top60_DNA_seed | HDAC1 | 4.147478 | LUAD | NCI-H1838 | Dacinostat | MVP_top60_DNA+RNA_seed | GSK3B | -0.771669 | STAD | RERF-GC-1B | SB216763 |
| MVP_top60_DNA_seed | EGFR | -4.854198 | LUAD | HCC-827 | Gefitinib | MVP_top60_DNA+RNA_seed | HSP90AA1 | -0.770346 | STAD | SNU-5 | Tanespimycin |
| MVP_top60_DNA_seed | MET | -4.82268 | LUAD | NCI-H1993 | Savolitinib | MVP_top60_DNA+RNA_seed | HSP90AB1 | -0.770346 | STAD | SNU-5 | Tanespimycin |
| MVP_top60_DNA_seed | EGFR | -4.716321 | LUAD | HCC-827 | AZD3759 | MVP_top60_DNA+RNA_seed | PIK3CA | -0.770271 | STAD | FU97 | AZD8186 |
| MVP_top60_DNA_seed | EGFR | -4.534566 | LUAD | PC-14 | AZD3759 | MVP_top60_DNA+RNA_seed | PIK3CA | -0.768064 | STAD | Hs-746T | GNE-317 |

| | | | | | | | | | | | |
|---|---|---|---|---|---|---|---|---|---|---|---|
| MVP_top60_DNA_seed | EGFR | -4.200302 | LUAD | PC-14 | Sapitinib | MVP_top60_DNA+RNA_seed | EP300 | -0.750783 | STAD | NUGC-4 | SGC-CBP30 |
| MVP_top60_DNA_seed | ERBB2 | -4.200302 | LUAD | PC-14 | Sapitinib | MVP_top60_DNA+RNA_seed | CREBBP | -0.750783 | STAD | NUGC-4 | SGC-CBP30 |
| MVP_top60_DNA_seed | ERBB3 | -4.200302 | LUAD | PC-14 | Sapitinib | MVP_top60_DNA+RNA_seed | MET | -0.7454 | STAD | SK-GT-2 | Savolitinib |
| MVP_top60_DNA_seed | EGFR | -4.099897 | LUAD | HCC-827 | Osimertinib | MVP_top60_DNA+RNA_seed | HSP90AA1 | -0.744516 | STAD | MKN45 | Tanespimycin |
| MVP_top60_DNA_seed | EGFR | -4.05738 | LUAD | PC-3_[JPC-3] | Gefitinib | MVP_top60_DNA+RNA_seed | HSP90AB1 | -0.744516 | STAD | MKN45 | Tanespimycin |
| MVP_top60_DNA_seed | EGFR | -3.925376 | LUAD | HCC-827 | Erlotinib | MVP_top60_DNA+RNA_seed | PARP1 | -0.743238 | STAD | RF-48 | Veliparib |
| MVP_top60_DNA_seed | EGFR | -3.890432 | LUAD | H3255 | AZD3759 | MVP_top60_DNA+RNA_seed | ESR1 | -0.738363 | STAD | SK-GT-2 | Tamoxifen |
| MVP_top60_DNA_seed | EGFR | -3.873859 | LUAD | PC-14 | Osimertinib | MVP_top60_DNA+RNA_seed | HDAC1 | -0.737076 | STAD | IM-95 | Entinostat |
| MVP_top60_DNA_seed | EGFR | -3.853431 | LUAD | NCI-H1975 | Osimertinib | MVP_top60_DNA+RNA_seed | HDAC3 | -0.737076 | STAD | IM-95 | Entinostat |
| MVP_top60_DNA_seed | EGFR | -3.747843 | LUAD | PC-14 | Gefitinib | MVP_top60_DNA+RNA_seed | ERBB2 | -0.729675 | STAD | NUGC-4 | Afatinib |
| MVP_top60_DNA_seed | EGFR | -3.734321 | LUAD | HCC-827 | Afatinib | MVP_top60_DNA+RNA_seed | ERBB2 | -0.724525 | STAD | MKN7 | Lapatinib |
| MVP_top60_DNA_seed | ERBB2 | -3.734321 | LUAD | HCC-827 | Afatinib | MVP_top60_DNA+RNA_seed | ESR1 | -0.724016 | STAD | SNU-16 | GDC0810 |
| MVP_top60_DNA_seed | EGFR | -3.686848 | LUAD | PC-3_[JPC-3] | AZD3759 | MVP_top60_DNA+RNA_seed | TP53 | -0.720545 | STAD | RF-48 | MIRA-1 |
| MVP_top60_DNA_seed | EGFR | -3.551576 | LUAD | PC-14 | Erlotinib | MVP_top60_DNA+RNA_seed | MAPK1 | -0.708043 | STAD | 23132-87 | VX-11e |
| MVP_top60_DNA_seed | EGFR | -3.473928 | LUAD | NCI-H322M | Sapitinib | MVP_top60_DNA+RNA_seed | ATM | -0.707435 | STAD | SNU-16 | KU-55933 |
| MVP_top60_DNA_seed | ERBB2 | -3.473928 | LUAD | NCI-H322M | Sapitinib | MVP_top60_DNA+RNA_seed | GSK3B | -0.703721 | STAD | 23132-87 | SB216763 |
| MVP_top60_DNA_seed | ERBB3 | -3.473928 | LUAD | NCI-H322M | Sapitinib | MVP_top60_DNA+RNA_seed | MDM2 | -0.701821 | STAD | HGC-27 | Serdemetan |
| MVP_top60_DNA_seed | EGFR | -3.460764 | LUAD | PC-14 | Afatinib | MVP_top60_DNA+RNA_seed | ESR1 | -0.687535 | STAD | NCI-N87 | GDC0810 |
| MVP_top60_DNA_seed | ERBB2 | -3.460764 | LUAD | PC-14 | Afatinib | MVP_top60_DNA+RNA_seed | FGFR3 | -0.675541 | STAD | IM-95 | AZD4547 |
| MVP_top60_DNA_seed | EGFR | -3.359337 | LUAD | COR-L105 | Sapitinib | MVP_top60_DNA+RNA_seed | MAPK1 | -0.672667 | STAD | KATOIII | SCH772984 |
| MVP_top60_DNA_seed | ERBB2 | -3.359337 | LUAD | COR-L105 | Sapitinib | MVP_top60_DNA+RNA_seed | MAPK3 | -0.672667 | STAD | KATOIII | SCH772984 |
| MVP_top60_DNA_seed | ERBB3 | -3.359337 | LUAD | COR-L105 | Sapitinib | MVP_top60_DNA+RNA_seed | EZH2 | -0.66567 | STAD | SNU-1 | GSK343 |
| MVP_top60_DNA_seed | EGFR | -3.329265 | LUAD | PC-3_[JPC-3] | Osimertinib | MVP_top60_DNA+RNA_seed | KRAS | -0.665366 | STAD | MKN45 | (RAS (G12C) Inhibitor-1 |
| MVP_top60_DNA_seed | EGFR | -3.301066 | LUAD | NCI-H322M | Gefitinib | MVP_top60_DNA+RNA_seed | PARP1 | -0.665038 | STAD | SNU-5 | Talazoparib |
| MVP_top60_DNA_seed | EGFR | -3.27183 | LUAD | COR-L105 | Gefitinib | MVP_top60_DNA+RNA_seed | SRC | -0.657963 | STAD | HSC-39 | Bosutinib |
| MVP_top60_DNA_seed | EGFR | -3.267587 | LUAD | H3255 | Osimertinib | MVP_top60_DNA+RNA_seed | MAPK1 | -0.65613 | STAD | TGBC11TKB | Ulixertinib |
| MVP_top60_DNA_seed | EGFR | -3.259498 | LUAD | H3255 | Afatinib | MVP_top60_DNA+RNA_seed | MAPK3 | -0.65613 | STAD | TGBC11TKB | Ulixertinib |
| MVP_top60_DNA_seed | ERBB2 | -3.259498 | LUAD | H3255 | Afatinib | MVP_top60_DNA+RNA_seed | PARP1 | -0.654606 | STAD | TMK-1 | Niraparib |
| MVP_top60_DNA_seed | EGFR | -3.190589 | LUAD | COR-L105 | Afatinib | MVP_top60_DNA+RNA_seed | MAPK1 | -0.645592 | STAD | KATOIII | Ulixertinib |
| MVP_top60_DNA_seed | ERBB2 | -3.190589 | LUAD | COR-L105 | Afatinib | MVP_top60_DNA+RNA_seed | MAPK3 | -0.645592 | STAD | KATOIII | Ulixertinib |
| MVP_top60_DNA_seed | EGFR | -3.183559 | LUAD | H3255 | Sapitinib | MVP_top60_DNA+RNA_seed | MET | -0.645201 | STAD | AGS | Savolitinib |
| MVP_top60_DNA_seed | ERBB2 | -3.183559 | LUAD | H3255 | Sapitinib | MVP_top60_DNA+RNA_seed | GSK3B | -0.6435 | STAD | TMK-1 | CHIR-99021 |
| MVP_top60_DNA_seed | ERBB3 | -3.183559 | LUAD | H3255 | Sapitinib | MVP_top60_DNA+RNA_seed | PARP1 | -0.642907 | STAD | RF-48 | Niraparib |
| MVP_top60_DNA_seed | EGFR | -3.108259 | LUAD | PC-3_[JPC-3] | Erlotinib | MVP_top60_DNA+RNA_seed | ESR1 | -0.641626 | STAD | RERF-GC-1B | Tamoxifen |
| MVP_top60_DNA_seed | EGFR | -3.061933 | LUAD | PC-3_[JPC-3] | Sapitinib | MVP_top60_DNA+RNA_seed | PARP1 | -0.641456 | STAD | HSC-39 | Talazoparib |
| MVP_top60_DNA_seed | ERBB2 | -3.061933 | LUAD | PC-3_[JPC-3] | Sapitinib | MVP_top60_DNA+RNA_seed | PIK3CA | -0.640799 | STAD | MKN45 | GNE-317 |
| MVP_top60_DNA_seed | ERBB3 | -3.061933 | LUAD | PC-3_[JPC-3] | Sapitinib | MVP_top60_DNA+RNA_seed | PIK3CA | -0.636908 | STAD | SNU-5 | Buparlisib |
| MVP_top60_DNA_seed | EGFR | -3.043923 | LUAD | NCI-H322M | Afatinib | MVP_top60_DNA+RNA_seed | ESR1 | -0.633839 | STAD | AGS | Tamoxifen |
| MVP_top60_DNA_seed | ERBB2 | -3.043923 | LUAD | NCI-H322M | Afatinib | MVP_top60_DNA+RNA_seed | PIK3CA | -0.633687 | STAD | SK-GT-2 | Buparlisib |
| MVP_top60_DNA_seed | EGFR | -2.988821 | LUAD | NCI-H322M | AZD3759 | MVP_top60_DNA+RNA_seed | PARP1 | -0.633469 | STAD | OCUM-1 | Niraparib |
| MVP_top60_DNA_seed | EGFR | -2.95988 | LUAD | H3255 | Gefitinib | MVP_top60_DNA+RNA_seed | MDM2 | -0.632532 | STAD | SNU-16 | Serdemetan |

| | | | | | | | | | | | |
|---|---|---|---|---|---|---|---|---|---|---|---|
| MVP_top60_DNA_seed | EGFR | -2.869384 | LUAD | NCI-H322M | Erlotinib | MVP_top60_DNA+RNA_seed | ERBB2 | -0.628949 | STAD | HSC-39 | Sapitinib |
| MVP_top60_DNA_seed | EGFR | -2.85431 | LUAD | NCI-H1623 | Sapitinib | MVP_top60_DNA+RNA_seed | ERBB3 | -0.628949 | STAD | HSC-39 | Sapitinib |
| MVP_top60_DNA_seed | ERBB2 | -2.85431 | LUAD | NCI-H1623 | Sapitinib | MVP_top60_DNA+RNA_seed | MAPK1 | -0.628511 | STAD | MKN45 | VX-11e |
| MVP_top60_DNA_seed | ERBB3 | -2.85431 | LUAD | NCI-H1623 | Sapitinib | MVP_top60_DNA+RNA_seed | MAPK1 | -0.623801 | STAD | SNU-5 | VX-11e |
| MVP_top60_DNA_seed | EGFR | -2.756579 | LUAD | PC-3_[JPC-3] | Afatinib | MVP_top60_DNA+RNA_seed | KRAS | -0.614055 | STAD | RF-48 | RAS (G12C) Inhibitor-1 |
| MVP_top60_DNA_seed | ERBB2 | -2.756579 | LUAD | PC-3_[JPC-3] | Afatinib | MVP_top60_DNA+RNA_seed | PIK3CA | -0.603623 | STAD | SK-GT-2 | Alpelisib |
| MVP_top60_DNA_seed | EGFR | -2.60257 | LUAD | COR-L105 | AZD3759 | MVP_top60_DNA+RNA_seed | MAPK1 | -0.603514 | STAD | TMK-1 | SCH772984 |
| MVP_top60_DNA_seed | IGF1R | -2.583223 | LUAD | NCI-H2122 | BMS-536924 | MVP_top60_DNA+RNA_seed | MAPK3 | -0.603514 | STAD | TMK-1 | SCH772984 |
| MVP_top60_DNA_seed | EGFR | -2.520692 | LUAD | NCI-H1648 | Sapitinib | MVP_top60_DNA+RNA_seed | EZH2 | -0.602076 | STAD | OCUM-1 | GSK343 |
| MVP_top60_DNA_seed | ERBB2 | -2.520692 | LUAD | NCI-H1648 | Sapitinib | MVP_top60_DNA+RNA_seed | MDM2 | -0.60193 | STAD | SNU-1 | Nutlin-3a (-) |
| MVP_top60_DNA_seed | ERBB3 | -2.520692 | LUAD | NCI-H1648 | Sapitinib | MVP_top60_DNA+RNA_seed | MET | -0.598456 | STAD | RERF-GC-1B | Savolitinib |
| MVP_top60_DNA_seed | MET | -2.42713 | LUAD | NCI-H1993 | Crizotinib | MVP_top60_DNA+RNA_seed | MDM2 | -0.597664 | STAD | HSC-39 | Nutlin-3a (-) |
| MVP_top60_DNA_seed | EGFR | -2.374387 | LUAD | COR-L105 | Osimertinib | MVP_top60_DNA+RNA_seed | ERBB2 | -0.596463 | STAD | TMK-1 | Sapitinib |
| MVP_top60_DNA_seed | EGFR | -2.373031 | LUAD | NCI-H322M | Osimertinib | MVP_top60_DNA+RNA_seed | ERBB3 | -0.596463 | STAD | TMK-1 | Sapitinib |
| MVP_top60_DNA_seed | EGFR | -2.364013 | LUAD | NCI-H1648 | Gefitinib | MVP_top60_DNA+RNA_seed | MET | -0.595751 | STAD | NUGC-4 | Savolitinib |
| MVP_top60_DNA_seed | EGFR | -2.353975 | LUAD | COR-L105 | Erlotinib | MVP_top60_DNA+RNA_seed | MDM2 | -0.594361 | STAD | FU97 | Nutlin-3a (-) |
| MVP_top60_DNA_seed | MET | -2.3503 | LUAD | RERF-LC-MS | Savolitinib | MVP_top60_DNA+RNA_seed | HSP90AA1 | -0.593213 | STAD | RERF-GC-1B | Luminespib |
| MVP_top60_DNA_seed | EGFR | -2.307295 | LUAD | NCI-H1781 | Afatinib | MVP_top60_DNA+RNA_seed | HSP90AB1 | -0.593213 | STAD | RERF-GC-1B | Luminespib |
| MVP_top60_DNA_seed | ERBB2 | -2.307295 | LUAD | NCI-H1781 | Afatinib | MVP_top60_DNA+RNA_seed | ESR1 | -0.592299 | STAD | 23132-87 | GDC0810 |
| MVP_top60_DNA_seed | EGFR | -2.297143 | LUAD | NCI-H1666 | Sapitinib | MVP_top60_DNA+RNA_seed | MAPK1 | -0.589577 | STAD | HSC-39 | SCH772984 |
| MVP_top60_DNA_seed | ERBB2 | -2.297143 | LUAD | NCI-H1666 | Sapitinib | MVP_top60_DNA+RNA_seed | MAPK3 | -0.589577 | STAD | HSC-39 | SCH772984 |
| MVP_top60_DNA_seed | ERBB3 | -2.297143 | LUAD | NCI-H1666 | Sapitinib | MVP_top60_DNA+RNA_seed | HDAC1 | -0.585809 | STAD | RF-48 | PCI-34051 |
| MVP_top60_DNA_seed | PIK3CA | -2.275126 | LUAD | NCI-H1648 | Alpelisib | MVP_top60_DNA+RNA_seed | PARP1 | -0.580136 | STAD | TMK-1 | Rucaparib |
| MVP_top60_DNA_seed | EGFR | -2.269518 | LUAD | Calu-3 | Sapitinib | MVP_top60_DNA+RNA_seed | FGFR3 | -0.579611 | STAD | HGC-27 | AZD4547 |
| MVP_top60_DNA_seed | ERBB2 | -2.269518 | LUAD | Calu-3 | Sapitinib | MVP_top60_DNA+RNA_seed | TP53 | -0.578312 | STAD | GCIY | MIRA-1 |
| MVP_top60_DNA_seed | ERBB3 | -2.269518 | LUAD | Calu-3 | Sapitinib | MVP_top60_DNA+RNA_seed | MDM2 | -0.567297 | STAD | TMK-1 | Serdemetan |
| MVP_top60_DNA_seed | EGFR | -2.267607 | LUAD | Calu-3 | Afatinib | MVP_top60_DNA+RNA_seed | HSP90AA1 | -0.5603 | STAD | SNU-5 | Luminespib |
| MVP_top60_DNA_seed | ERBB2 | -2.267607 | LUAD | Calu-3 | Afatinib | MVP_top60_DNA+RNA_seed | HSP90AB1 | -0.5603 | STAD | SNU-5 | Luminespib |
| MVP_top60_DNA_seed | EGFR | -2.255715 | LUAD | NCI-H1623 | Afatinib | MVP_top60_DNA+RNA_seed | EZH2 | -0.559352 | STAD | NUGC-3 | GSK343 |
| MVP_top60_DNA_seed | ERBB2 | -2.255715 | LUAD | NCI-H1623 | Afatinib | MVP_top60_DNA+RNA_seed | PIK3CA | -0.553995 | STAD | IM-95 | Buparlisib |
| MVP_top60_DNA_seed | SRC | -2.169273 | LUAD | PC-14 | Bosutinib | MVP_top60_DNA+RNA_seed | HDAC1 | -0.550885 | STAD | AGS | Entinostat |
| MVP_top60_DNA_seed | EGFR | -2.125228 | LUAD | NCI-H1666 | Gefitinib | MVP_top60_DNA+RNA_seed | HDAC3 | -0.550885 | STAD | AGS | Entinostat |
| MVP_top60_DNA_seed | EGFR | -2.117487 | LUAD | NCI-H1648 | Afatinib | MVP_top60_DNA+RNA_seed | PARP1 | -0.550804 | STAD | HGC-27 | Veliparib |
| MVP_top60_DNA_seed | ERBB2 | -2.117487 | LUAD | NCI-H1648 | Afatinib | MVP_top60_DNA+RNA_seed | MAPK1 | -0.547318 | STAD | 23132-87 | SCH772984 |
| MVP_top60_DNA_seed | EGFR | -2.10784 | LUAD | NCI-H1648 | AZD3759 | MVP_top60_DNA+RNA_seed | MAPK3 | -0.547318 | STAD | 23132-87 | SCH772984 |
| MVP_top60_DNA_seed | EGFR | -2.096518 | LUAD | EMC-BAC-1 | Afatinib | MVP_top60_DNA+RNA_seed | ERBB2 | -0.544321 | STAD | SNU-16 | Afatinib |
| MVP_top60_DNA_seed | ERBB2 | -2.096518 | LUAD | EMC-BAC-1 | Afatinib | MVP_top60_DNA+RNA_seed | HSP90AA1 | -0.540432 | STAD | HGC-27 | Luminespib |
| MVP_top60_DNA_seed | EGFR | -2.087908 | LUAD | NCI-H1648 | Erlotinib | MVP_top60_DNA+RNA_seed | HSP90AB1 | -0.540432 | STAD | HGC-27 | Luminespib |
| MVP_top60_DNA_seed | IGF1R | -2.079693 | LUAD | NCI-H2122 | NVP-ADW742 | MVP_top60_DNA+RNA_seed | KRAS | -0.539066 | STAD | IM-95 | RAS (G12C) Inhibitor-1 |
| MVP_top60_DNA_seed | EGFR | -2.060094 | LUAD | NCI-H1648 | Lapatinib | MVP_top60_DNA+RNA_seed | PIK3CA | -0.536074 | STAD | SK-GT-2 | GNE-317 |
| MVP_top60_DNA_seed | ERBB2 | -2.060094 | LUAD | NCI-H1648 | Lapatinib | MVP_top60_DNA+RNA_seed | HDAC1 | -0.531965 | STAD | NUGC-4 | Entinostat |

| | | | | | | | | | | | |
|---|---|---|---|---|---|---|---|---|---|---|---|
| MVP_top60_DNA_seed | IGF1R | -2.040206 | LUAD | NCI-H2122 | BMS-754807 | MVP_top60_DNA+RNA_seed | HDAC3 | -0.531965 | STAD | NUGC-4 | Entinostat |
| MVP_top60_DNA_seed | MET | -2.02756 | LUAD | NCI-H3122 | Crizotinib | MVP_top60_DNA+RNA_seed | PARP1 | -0.531069 | STAD | AGS | Rucaparib |
| MVP_top60_DNA_seed | EGFR | -1.995851 | LUAD | NCI-H1568 | Erlotinib | MVP_top60_DNA+RNA_seed | MET | -0.530379 | STAD | RERF-GC-1B | Crizotinib |
| MVP_top60_DNA_seed | HSP90AA1 | -1.975503 | LUAD | A549 | Tanespimycin | MVP_top60_DNA+RNA_seed | HDAC1 | -0.528204 | STAD | SK-GT-2 | Entinostat |
| MVP_top60_DNA_seed | EGFR | -1.966407 | LUAD | NCI-H1666 | AZD3759 | MVP_top60_DNA+RNA_seed | HDAC3 | -0.528204 | STAD | SK-GT-2 | Entinostat |
| MVP_top60_DNA_seed | EGFR | -1.96437 | LUAD | EMC-BAC-1 | Sapitinib | MVP_top60_DNA+RNA_seed | ESR1 | -0.527647 | STAD | SK-GT-2 | GDC0810 |
| MVP_top60_DNA_seed | ERBB2 | -1.96437 | LUAD | EMC-BAC-1 | Sapitinib | MVP_top60_DNA+RNA_seed | SRC | -0.526709 | STAD | SNU-16 | Bosutinib |
| MVP_top60_DNA_seed | ERBB3 | -1.96437 | LUAD | EMC-BAC-1 | Sapitinib | MVP_top60_DNA+RNA_seed | PARP1 | -0.525963 | STAD | IM-95 | Veliparib |
| MVP_top60_DNA_seed | HDAC1 | -1.964111 | LUAD | NCI-H2122 | Entinostat | MVP_top60_DNA+RNA_seed | MET | -0.520269 | STAD | SNU-1 | Crizotinib |
| MVP_top60_DNA_seed | MAPK1 | -1.963298 | LUAD | Calu-6 | VX-11e | MVP_top60_DNA+RNA_seed | MDM2 | -0.515019 | STAD | NUGC-3 | Serdemetan |
| MVP_top60_DNA_seed | EGFR | -1.928717 | LUAD | EMC-BAC-1 | AZD3759 | MVP_top60_DNA+RNA_seed | HSP90AA1 | -0.514576 | STAD | IM-95 | Tanespimycin |
| MVP_top60_DNA_seed | EGFR | -1.928576 | LUAD | NCI-H2030 | AZD3759 | MVP_top60_DNA+RNA_seed | HSP90AB1 | -0.514576 | STAD | IM-95 | Tanespimycin |
| MVP_top60_DNA_seed | MAPK1 | -1.922387 | LUAD | Calu-6 | SCH772984 | MVP_top60_DNA+RNA_seed | ATM | -0.513893 | STAD | RERF-GC-1B | KU-55933 |
| MVP_top60_DNA_seed | GSK3B | -1.921231 | LUAD | RERF-LC-MS | SB216763 | MVP_top60_DNA+RNA_seed | EP300 | -0.509442 | STAD | AGS | CPI-637 |
| MVP_top60_DNA_seed | EGFR | -1.90466 | LUAD | NCI-H1568 | Gefitinib | MVP_top60_DNA+RNA_seed | ATM | -0.505711 | STAD | AGS | KU-55933 |
| MVP_top60_DNA_seed | EZH2 | -1.903149 | LUAD | HCC-78 | GSK343 | MVP_top60_DNA+RNA_seed | ESR1 | -0.50288 | STAD | GCIY | GDC0810 |
| MVP_top60_DNA_seed | EGFR | -1.8844 | LUAD | NCI-H1623 | Lapatinib | MVP_top60_DNA+RNA_seed | PIK3CA | -0.502607 | STAD | RERF-GC-1B | Alpelisib |
| MVP_top60_DNA_seed | ERBB2 | -1.8844 | LUAD | NCI-H1623 | Lapatinib | MVP_top60_DNA+RNA_seed | GSK3B | -0.4996 | STAD | SNU-16 | SB216763 |
| MVP_top60_DNA_seed | EZH2 | -1.85163 | LUAD | Calu-3 | GSK343 | MVP_top60_DNA+RNA_seed | MAPK1 | -0.496934 | STAD | SNU-16 | Ulixertinib |
| MVP_top60_DNA_seed | EGFR | -1.841329 | LUAD | NCI-H1975 | Afatinib | MVP_top60_DNA+RNA_seed | MAPK3 | -0.496934 | STAD | SNU-16 | Ulixertinib |
| MVP_top60_DNA_seed | ERBB2 | -1.841329 | LUAD | NCI-H1975 | Afatinib | MVP_top60_DNA+RNA_seed | PIK3CA | -0.494304 | STAD | TMK-1 | Alpelisib |
| MVP_top60_DNA_seed | EGFR | -1.841322 | LUAD | NCI-H1648 | Osimertinib | MVP_top60_DNA+RNA_seed | PIK3CA | -0.492705 | STAD | RF-48 | Buparlisib |
| MVP_top60_DNA_seed | EGFR | -1.835846 | LUAD | H3255 | Erlotinib | MVP_top60_DNA+RNA_seed | ATM | -0.484921 | STAD | HGC-27 | KU-55933 |
| MVP_top60_DNA_seed | EGFR | -1.827677 | LUAD | NCI-H1666 | Erlotinib | MVP_top60_DNA+RNA_seed | MAPK1 | -0.458898 | STAD | MKN45 | Ulixertinib |
| MVP_top60_DNA_seed | HSP90AA1 | -1.817473 | LUAD | HCC-44 | Tanespimycin | MVP_top60_DNA+RNA_seed | MAPK3 | -0.458898 | STAD | MKN45 | Ulixertinib |
| MVP_top60_DNA_seed | AURKA | -1.814104 | LUAD | NCI-H2122 | Alisertib | MVP_top60_DNA+RNA_seed | PIK3CA | -0.458624 | STAD | NCI-N87 | GNE-317 |
| MVP_top60_DNA_seed | SRC | -1.806877 | LUAD | HCC-827 | Bosutinib | MVP_top60_DNA+RNA_seed | HDAC1 | -0.445927 | STAD | KATOIII | Romidepsin |
| MVP_top60_DNA_seed | MAPK1 | -1.794215 | LUAD | NCI-H2291 | SCH772984 | MVP_top60_DNA+RNA_seed | HDAC3 | -0.445927 | STAD | KATOIII | Romidepsin |
| MVP_top60_DNA_seed | EGFR | -1.788226 | LUAD | NCI-H1568 | Sapitinib | MVP_top60_DNA+RNA_seed | ATM | -0.443556 | STAD | GCIY | KU-55933 |
| MVP_top60_DNA_seed | ERBB2 | -1.788226 | LUAD | NCI-H1568 | Sapitinib | MVP_top60_DNA+RNA_seed | MET | -0.442098 | STAD | RF-48 | Savolitinib |
| MVP_top60_DNA_seed | ERBB3 | -1.788226 | LUAD | NCI-H1568 | Sapitinib | MVP_top60_DNA+RNA_seed | ATM | -0.441689 | STAD | RF-48 | KU-55933 |
| MVP_top60_DNA_seed | PIK3CA | -1.752711 | LUAD | NCI-H1648 | GNE-317 | MVP_top60_DNA+RNA_seed | FGFR3 | -0.439307 | STAD | TGBC11TKB | PD173074 |
| MVP_top60_DNA_seed | EGFR | -1.742609 | LUAD | H3255 | Lapatinib | MVP_top60_DNA+RNA_seed | SRC | -0.438876 | STAD | RF-48 | Bosutinib |
| MVP_top60_DNA_seed | ERBB2 | -1.742609 | LUAD | H3255 | Lapatinib | MVP_top60_DNA+RNA_seed | PIK3CA | -0.432008 | STAD | IM-95 | AZD8186 |
| MVP_top60_DNA_seed | EGFR | -1.738422 | LUAD | Calu-3 | Lapatinib | MVP_top60_DNA+RNA_seed | EZH2 | -0.430208 | STAD | NCI-N87 | GSK343 |
| MVP_top60_DNA_seed | ERBB2 | -1.738422 | LUAD | Calu-3 | Lapatinib | MVP_top60_DNA+RNA_seed | EZH2 | -0.428763 | STAD | HGC-27 | GSK343 |
| MVP_top60_DNA_seed | HSP90AA1 | -1.725962 | LUAD | ABC-1 | Elesclomol | MVP_top60_DNA+RNA_seed | ESR1 | -0.425629 | STAD | NUGC-3 | Tamoxifen |
| MVP_top60_DNA_seed | EGFR | -1.718223 | LUAD | PC-14 | Lapatinib | MVP_top60_DNA+RNA_seed | EP300 | -0.413258 | STAD | NCI-N87 | CPI-637 |
| MVP_top60_DNA_seed | ERBB2 | -1.718223 | LUAD | PC-14 | Lapatinib | MVP_top60_DNA+RNA_seed | SRC | -0.404606 | STAD | AGS | Bosutinib |
| MVP_top60_DNA_seed | IGF1R | -1.693025 | LUAD | NCI-H1648 | BMS-536924 | MVP_top60_DNA+RNA_seed | PIK3CA | -0.392938 | STAD | Hs-746T | Buparlisib |
| MVP_top60_DNA_seed | EGFR | -1.685023 | LUAD | EMC-BAC-1 | Lapatinib | MVP_top60_DNA+RNA_seed | HSP90AA1 | -0.392414 | STAD | SNU-1 | Luminespib |

| | | | | | | | | | | | |
|---|---|---|---|---|---|---|---|---|---|---|---|
| MVP_top60_DNA_seed | ERBB2 | -1.685023 | LUAD | EMC-BAC-1 | Lapatinib | MVP_top60_DNA+RNA_seed | HSP90AB1 | -0.392414 | STAD | SNU-1 | Luminespib |
| MVP_top60_DNA_seed | EGFR | -1.679859 | LUAD | NCI-H1623 | Erlotinib | MVP_top60_DNA+RNA_seed | MAPK1 | -0.386403 | STAD | IM-95 | Ulixertinib |
| MVP_top60_DNA_seed | MAPK1 | -1.673425 | LUAD | NCI-H2291 | Ulixertinib | MVP_top60_DNA+RNA_seed | MAPK3 | -0.386403 | STAD | IM-95 | Ulixertinib |
| MVP_top60_DNA_seed | MAPK1 | -1.666908 | LUAD | NCI-H1648 | Ulixertinib | MVP_top60_DNA+RNA_seed | PARP1 | -0.381671 | STAD | MKN45 | Veliparib |
| MVP_top60_DNA_seed | IGF1R | -1.666667 | LUAD | NCI-H2122 | Linsitinib | MVP_top60_DNA+RNA_seed | GSK3B | -0.381661 | STAD | SNU-1 | SB216763 |
| MVP_top60_DNA_seed | EGFR | -1.662853 | LUAD | NCI-H2122 | Erlotinib | MVP_top60_DNA+RNA_seed | ESR1 | -0.380051 | STAD | KATOIII | Tamoxifen |
| MVP_top60_DNA_seed | MAPK1 | -1.658903 | LUAD | NCI-H322M | Ulixertinib | MVP_top60_DNA+RNA_seed | SRC | -0.376421 | STAD | IM-95 | Bosutinib |
| MVP_top60_DNA_seed | KRAS | -1.609231 | LUAD | EMC-BAC-1 | KRAS (G12C) Inhibitor-1: | MVP_top60_DNA+RNA_seed | MAPK1 | -0.375666 | STAD | TMK-1 | Ulixertinib |
| MVP_top60_DNA_seed | EGFR | -1.608489 | LUAD | PC-3_[JPC-3] | Lapatinib | MVP_top60_DNA+RNA_seed | MAPK3 | -0.375666 | STAD | TMK-1 | Ulixertinib |
| MVP_top60_DNA_seed | ERBB2 | -1.608489 | LUAD | PC-3_[JPC-3] | Lapatinib | MVP_top60_DNA+RNA_seed | HSP90AA1 | -0.374191 | STAD | NUGC-3 | Tanespimycin |
| MVP_top60_DNA_seed | PIK3CA | -1.606638 | LUAD | NCI-H2122 | Alpelisib | MVP_top60_DNA+RNA_seed | HSP90AB1 | -0.374191 | STAD | NUGC-3 | Tanespimycin |
| MVP_top60_DNA_seed | EGFR | -1.60601 | LUAD | NCI-H1568 | AZD3759 | MVP_top60_DNA+RNA_seed | PARP1 | -0.368696 | STAD | SNU-5 | Rucaparib |
| MVP_top60_DNA_seed | AKT1 | -1.599536 | LUAD | NCI-H1648 | Afuresertib | MVP_top60_DNA+RNA_seed | MAPK1 | -0.363791 | STAD | Hs-746T | Ulixertinib |
| MVP_top60_DNA_seed | HSP90AA1 | -1.577668 | LUAD | PC-14 | Tanespimycin | MVP_top60_DNA+RNA_seed | MAPK3 | -0.363791 | STAD | Hs-746T | Ulixertinib |
| MVP_top60_DNA_seed | EGFR | -1.565886 | LUAD | NCI-H1781 | Osimertinib | MVP_top60_DNA+RNA_seed | PIK3CA | -0.359993 | STAD | Hs-746T | AZD8186 |
| MVP_top60_DNA_seed | AKT1 | -1.561795 | LUAD | NCI-H1975 | AZD5363 | MVP_top60_DNA+RNA_seed | EZH2 | -0.352909 | STAD | KATOIII | GSK343 |
| MVP_top60_DNA_seed | EGFR | -1.561755 | LUAD | NCI-H358 | Sapitinib | MVP_top60_DNA+RNA_seed | EZH2 | -0.350662 | STAD | 23132-87 | GSK343 |
| MVP_top60_DNA_seed | ERBB2 | -1.561755 | LUAD | NCI-H358 | Sapitinib | MVP_top60_DNA+RNA_seed | FGFR3 | -0.346585 | STAD | NCI-N87 | PD173074 |
| MVP_top60_DNA_seed | ERBB3 | -1.561755 | LUAD | NCI-H358 | Sapitinib | MVP_top60_DNA+RNA_seed | MAPK1 | -0.345874 | STAD | SNU-16 | Ulixertinib |
| MVP_top60_DNA_seed | AKT1 | -1.553685 | LUAD | NCI-H2122 | Ipatasertib | MVP_top60_DNA+RNA_seed | MAPK3 | -0.345874 | STAD | SNU-16 | Ulixertinib |
| MVP_top60_DNA_seed | EZH2 | -1.532232 | LUAD | LXF-289 | GSK343 | MVP_top60_DNA+RNA_seed | FGFR3 | -0.343237 | STAD | SNU-1 | PD173074 |
| MVP_top60_DNA_seed | IGF1R | -1.531786 | LUAD | NCI-H1755 | BMS-536924 | MVP_top60_DNA+RNA_seed | PARP1 | -0.339269 | STAD | SNU-16 | Talazoparib |
| MVP_top60_DNA_seed | EGFR | -1.516259 | LUAD | NCI-H1781 | Sapitinib | MVP_top60_DNA+RNA_seed | ERBB2 | -0.336204 | STAD | HSC-39 | Afatinib |
| MVP_top60_DNA_seed | ERBB2 | -1.516259 | LUAD | NCI-H1781 | Sapitinib | MVP_top60_DNA+RNA_seed | MAPK1 | -0.335714 | STAD | FU97 | VX-11e |
| MVP_top60_DNA_seed | ERBB3 | -1.516259 | LUAD | NCI-H1781 | Sapitinib | MVP_top60_DNA+RNA_seed | PIK3CA | -0.33418 | STAD | SNU-16 | Alpelisib |
| MVP_top60_DNA_seed | EGFR | -1.511946 | LUAD | NCI-H1650 | AZD3759 | MVP_top60_DNA+RNA_seed | PARP1 | -0.320801 | STAD | IM-95 | Niraparib |
| MVP_top60_DNA_seed | IGF1R | -1.502644 | LUAD | NCI-H1975 | BMS-536924 | MVP_top60_DNA+RNA_seed | MAPK1 | -0.316827 | STAD | RERF-GC-1B | VX-11e |
| MVP_top60_DNA_seed | AKT1 | -1.499631 | LUAD | NCI-H1781 | Uprosertib | MVP_top60_DNA+RNA_seed | ESR1 | -0.316824 | STAD | MKN28 | Tamoxifen |
| MVP_top60_DNA_seed | PIK3CA | -1.498659 | LUAD | HCC-78 | AZD8186 | MVP_top60_DNA+RNA_seed | MAPK1 | -0.314581 | STAD | RERF-GC-1B | Ulixertinib |
| MVP_top60_DNA_seed | IGF1R | -1.495944 | LUAD | NCI-H1734 | BMS-754807 | MVP_top60_DNA+RNA_seed | MAPK3 | -0.314581 | STAD | RERF-GC-1B | Ulixertinib |
| MVP_top60_DNA_seed | PIK3CA | -1.48657 | LUAD | NCI-H1648 | Buparlisib | MVP_top60_DNA+RNA_seed | KRAS | -0.312751 | STAD | TMK-1 | KRAS (G12C) Inhibitor-1 |
| MVP_top60_DNA_seed | IGF1R | -1.48337 | LUAD | NCI-H1651 | GSK1904529A | MVP_top60_DNA+RNA_seed | MET | -0.307431 | STAD | SNU-16 | Savolitinib |
| MVP_top60_DNA_seed | MAPK1 | -1.480705 | LUAD | NCI-H1734 | SCH772984 | MVP_top60_DNA+RNA_seed | ERBB2 | -0.293342 | STAD | 23132-87 | Afatinib |
| MVP_top60_DNA_seed | MAPK1 | -1.480067 | LUAD | NCI-H2122 | SCH772984 | MVP_top60_DNA+RNA_seed | HSP90AA1 | -0.293298 | STAD | NUGC-3 | Luminespib |
| MVP_top60_DNA_seed | MET | -1.478509 | LUAD | RERF-LC-MS | Crizotinib | MVP_top60_DNA+RNA_seed | HSP90AB1 | -0.293298 | STAD | NUGC-3 | Luminespib |
| MVP_top60_DNA_seed | HSP90AA1 | -1.476283 | LUAD | NCI-H1734 | Tanespimycin | MVP_top60_DNA+RNA_seed | PARP1 | -0.292667 | STAD | TGBC11TKB | Veliparib |
| MVP_top60_DNA_seed | HDAC1 | -1.473599 | LUAD | A549 | Romidepsin | MVP_top60_DNA+RNA_seed | MAPK1 | -0.292425 | STAD | SNU-1 | Ulixertinib |
| MVP_top60_DNA_seed | HSP90AA1 | -1.468469 | LUAD | NCI-H3122 | Tanespimycin | MVP_top60_DNA+RNA_seed | MAPK3 | -0.292425 | STAD | SNU-1 | Ulixertinib |
| MVP_top60_DNA_seed | TERT | -1.466218 | LUAD | NCI-H838 | BIBR-1532 | MVP_top60_DNA+RNA_seed | ERBB2 | -0.289654 | STAD | NUGC-4 | Lapatinib |
| MVP_top60_DNA_seed | MAPK1 | -1.464964 | LUAD | NCI-H2122 | Ulixertinib | MVP_top60_DNA+RNA_seed | AR | -0.286406 | STAD | TMK-1 | Bicalutamide |
| MVP_top60_DNA_seed | IGF1R | -1.463177 | LUAD | HCC-78 | BMS-536924 | MVP_top60_DNA+RNA_seed | ESR1 | -0.286012 | STAD | RF-48 | GDC0810 |

| | | | | | | | | | | | |
|---|---|---|---|---|---|---|---|---|---|---|---|
| MVP_top60_DNA_seed | HSP90AA1 | -1.456579 | LUAD | NCI-H1792 | Tanespimycin | MVP_top60_DNA+RNA_seed | HSP90AA1 | -0.28304 | STAD | Hs-746T | Luminespib |
| MVP_top60_DNA_seed | EGFR | -1.447698 | LUAD | NCI-H358 | Afatinib | MVP_top60_DNA+RNA_seed | HSP90AB1 | -0.28304 | STAD | Hs-746T | Luminespib |
| MVP_top60_DNA_seed | ERBB2 | -1.447698 | LUAD | NCI-H358 | Afatinib | MVP_top60_DNA+RNA_seed | PARP1 | -0.277943 | STAD | AGS | Olaparib |
| MVP_top60_DNA_seed | MAPK1 | -1.439279 | LUAD | NCI-H1993 | Ulixertinib | MVP_top60_DNA+RNA_seed | KRAS | -0.273421 | STAD | SK-GT-2 | KRAS (G12C) Inhibitor-1 |
| MVP_top60_DNA_seed | IGF1R | -1.435728 | LUAD | NCI-H2228 | BMS-536924 | MVP_top60_DNA+RNA_seed | TP53 | -0.272887 | STAD | HGC-27 | MIRA-1 |
| MVP_top60_DNA_seed | MAPK1 | -1.429738 | LUAD | NCI-H1734 | VX-11e | MVP_top60_DNA+RNA_seed | HDAC1 | -0.268777 | STAD | IM-95 | Romidepsin |
| MVP_top60_DNA_seed | IGF1R | -1.429392 | LUAD | NCI-H1648 | NVP-ADW742 | MVP_top60_DNA+RNA_seed | HDAC3 | -0.268777 | STAD | IM-95 | Romidepsin |
| MVP_top60_DNA_seed | SRC | -1.426192 | LUAD | NCI-H322M | Bosutinib | MVP_top60_DNA+RNA_seed | ESR1 | -0.268664 | STAD | SNU-5 | Tamoxifen |
| MVP_top60_DNA_seed | HDAC1 | -1.423335 | LUAD | NCI-H1651 | Romidepsin | MVP_top60_DNA+RNA_seed | MAPK1 | -0.267497 | STAD | SK-GT-2 | Ulixertinib |
| MVP_top60_DNA_seed | PIK3CA | -1.418419 | LUAD | NCI-H1793 | GNE-317 | MVP_top60_DNA+RNA_seed | MAPK3 | -0.267497 | STAD | SK-GT-2 | Ulixertinib |
| MVP_top60_DNA_seed | EGFR | -1.417392 | LUAD | NCI-H1666 | Afatinib | MVP_top60_DNA+RNA_seed | PARP1 | -0.263871 | STAD | NUGC-3 | Veliparib |
| MVP_top60_DNA_seed | ERBB2 | -1.417392 | LUAD | NCI-H1666 | Afatinib | MVP_top60_DNA+RNA_seed | PIK3CA | -0.263263 | STAD | GCIY | Buparlisib |
| MVP_top60_DNA_seed | IGF1R | -1.41588 | LUAD | NCI-H1623 | Linsitinib | MVP_top60_DNA+RNA_seed | PARP1 | -0.258088 | STAD | SNU-5 | Veliparib |
| MVP_top60_DNA_seed | MAPK1 | -1.40482 | LUAD | NCI-H2291 | VX-11e | MVP_top60_DNA+RNA_seed | MAPK1 | -0.256418 | STAD | Hs-746T | Ulixertinib |
| MVP_top60_DNA_seed | MDM2 | -1.403221 | LUAD | NCI-H1734 | Serdemetan | MVP_top60_DNA+RNA_seed | MAPK3 | -0.256418 | STAD | Hs-746T | Ulixertinib |
| MVP_top60_DNA_seed | MAPK1 | -1.402748 | LUAD | NCI-H2122 | VX-11e | MVP_top60_DNA+RNA_seed | PARP1 | -0.25591 | STAD | 23132-87 | Niraparib |
| MVP_top60_DNA_seed | MAPK1 | -1.400216 | LUAD | RERF-LC-MS | VX-11e | MVP_top60_DNA+RNA_seed | HSP90AA1 | -0.248861 | STAD | Hs-746T | Tanespimycin |
| MVP_top60_DNA_seed | PIK3CA | -1.398581 | LUAD | NCI-H1734 | Alpelisib | MVP_top60_DNA+RNA_seed | HSP90AB1 | -0.248861 | STAD | Hs-746T | Tanespimycin |
| MVP_top60_DNA_seed | MAPK1 | -1.397632 | LUAD | HCC-44 | VX-11e | MVP_top60_DNA+RNA_seed | MET | -0.240744 | STAD | OCUM-1 | Savolitinib |
| MVP_top60_DNA_seed | EGFR | -1.39263 | LUAD | NCI-H1975 | Sapitinib | MVP_top60_DNA+RNA_seed | SRC | -0.238341 | STAD | SNU-5 | Bosutinib |
| MVP_top60_DNA_seed | ERBB2 | -1.39263 | LUAD | NCI-H1975 | Sapitinib | MVP_top60_DNA+RNA_seed | MAPK1 | -0.237803 | STAD | MKN28 | Ulixertinib |
| MVP_top60_DNA_seed | ERBB3 | -1.39263 | LUAD | NCI-H1975 | Sapitinib | MVP_top60_DNA+RNA_seed | MAPK3 | -0.237803 | STAD | MKN28 | Ulixertinib |
| MVP_top60_DNA_seed | MDM2 | -1.391782 | LUAD | A549 | Nutlin-3a (-) | MVP_top60_DNA+RNA_seed | ESR1 | -0.236961 | STAD | KATOIII | GDC0810 |
| MVP_top60_DNA_seed | EGFR | -1.391569 | LUAD | RERF-LC-KJ | Erlotinib | MVP_top60_DNA+RNA_seed | KRAS | -0.232713 | STAD | MKN28 | KRAS (G12C) Inhibitor-1 |
| MVP_top60_DNA_seed | MAPK1 | -1.3881 | LUAD | A549 | VX-11e | MVP_top60_DNA+RNA_seed | EZH2 | -0.22815 | STAD | HGC-27 | GSK343 |
| MVP_top60_DNA_seed | EGFR | -1.384082 | LUAD | NCI-H1623 | AZD3759 | MVP_top60_DNA+RNA_seed | MAPK1 | -0.227401 | STAD | Hs-746T | VX-11e |
| MVP_top60_DNA_seed | EZH2 | -1.377759 | LUAD | NCI-H2122 | GSK343 | MVP_top60_DNA+RNA_seed | HDAC1 | -0.222934 | STAD | RF-48 | Entinostat |
| MVP_top60_DNA_seed | IGF1R | -1.375224 | LUAD | NCI-H1437 | BMS-536924 | MVP_top60_DNA+RNA_seed | HDAC3 | -0.222934 | STAD | RF-48 | Entinostat |
| MVP_top60_DNA_seed | PIK3CA | -1.372823 | LUAD | A549 | GNE-317 | MVP_top60_DNA+RNA_seed | MET | -0.215735 | STAD | TGBC11TKB | Savolitinib |
| MVP_top60_DNA_seed | AKT1 | -1.37202 | LUAD | NCI-H1975 | AT13148 | MVP_top60_DNA+RNA_seed | EZH2 | -0.212438 | STAD | IM-95 | GSK343 |
| MVP_top60_DNA_seed | EGFR | -1.364414 | LUAD | NCI-H1623 | Gefitinib | MVP_top60_DNA+RNA_seed | HDAC1 | -0.201763 | STAD | SNU-5 | Entinostat |
| MVP_top60_DNA_seed | GSK3B | -1.362062 | LUAD | NCI-H2122 | SB216763 | MVP_top60_DNA+RNA_seed | HDAC3 | -0.201763 | STAD | SNU-5 | Entinostat |
| MVP_top60_DNA_seed | HSP90AA1 | -1.358527 | LUAD | RERF-LC-MS | Luminespib | MVP_top60_DNA+RNA_seed | HSP90AA1 | -0.201111 | STAD | FU97 | Luminespib |
| MVP_top60_DNA_seed | EGFR | -1.347203 | LUAD | NCI-H2122 | Gefitinib | MVP_top60_DNA+RNA_seed | HSP90AB1 | -0.201111 | STAD | FU97 | Luminespib |
| MVP_top60_DNA_seed | HDAC1 | -1.341741 | LUAD | NCI-H2122 | Romidepsin | MVP_top60_DNA+RNA_seed | PARP1 | -0.200012 | STAD | AGS | Niraparib |
| MVP_top60_DNA_seed | MAPK1 | -1.328711 | LUAD | NCI-H1993 | VX-11e | MVP_top60_DNA+RNA_seed | HDAC1 | -0.195673 | STAD | MKN7 | PCI-34051 |
| MVP_top60_DNA_seed | EGFR | -1.328444 | LUAD | COR-L105 | Lapatinib | MVP_top60_DNA+RNA_seed | PIK3CA | -0.191993 | STAD | TGBC11TKB | Alpelisib |
| MVP_top60_DNA_seed | ERBB2 | -1.328444 | LUAD | COR-L105 | Lapatinib | MVP_top60_DNA+RNA_seed | ERBB2 | -0.189838 | STAD | AGS | Lapatinib |
| MVP_top60_DNA_seed | HDAC1 | -1.326538 | LUAD | NCI-H1437 | Entinostat | MVP_top60_DNA+RNA_seed | PIK3CA | -0.189207 | STAD | SNU-16 | GNE-317 |
| MVP_top60_DNA_seed | EGFR | -1.326312 | LUAD | EMC-BAC-1 | Osimertinib | MVP_top60_DNA+RNA_seed | KRAS | -0.185502 | STAD | 23132-87 | KRAS (G12C) Inhibitor-1 |
| MVP_top60_DNA_seed | PIK3CA | -1.318525 | LUAD | HCC-78 | GNE-317 | MVP_top60_DNA+RNA_seed | HDAC1 | -0.18429 | STAD | Hs-746T | Entinostat |

| | | | | | | | | | | | |
|---|---|---|---|---|---|---|---|---|---|---|---|
| MVP_top60_DNA_seed | MAPK1 | -1.316756 | LUAD | NCI-H3122 | SCH772984 | MVP_top60_DNA+RNA_seed | HDAC3 | -0.18429 | STAD | Hs-746T | Entinostat |
| MVP_top60_DNA_seed | HSP90AA1 | -1.314768 | LUAD | PC-14 | CCT-018159 | MVP_top60_DNA+RNA_seed | MAPK1 | -0.182584 | STAD | RERF-GC-1B | SCH772984 |
| MVP_top60_DNA_seed | EGFR | -1.313911 | LUAD | HCC-827 | Lapatinib | MVP_top60_DNA+RNA_seed | MAPK3 | -0.182584 | STAD | RERF-GC-1B | SCH772984 |
| MVP_top60_DNA_seed | ERBB2 | -1.313911 | LUAD | HCC-827 | Lapatinib | MVP_top60_DNA+RNA_seed | PIK3CA | -0.181396 | STAD | MKN45 | Alpelisib |
| MVP_top60_DNA_seed | MAPK1 | -1.311611 | LUAD | A549 | Ulixertinib | MVP_top60_DNA+RNA_seed | MAPK1 | -0.17745 | STAD | TMK-1 | VX-11e |
| MVP_top60_DNA_seed | SMARCA4 | -1.30627 | LUAD | NCI-H1563 | PFI3 | MVP_top60_DNA+RNA_seed | SRC | -0.177043 | STAD | SNU-1 | Bosutinib |
| MVP_top60_DNA_seed | GSK3B | -1.304298 | LUAD | A549 | SB216763 | MVP_top60_DNA+RNA_seed | MAPK1 | -0.175867 | STAD | GCIY | VX-11e |
| MVP_top60_DNA_seed | EGFR | -1.304226 | LUAD | NCI-H322M | Lapatinib | MVP_top60_DNA+RNA_seed | GSK3B | -0.171302 | STAD | HGC-27 | SB216763 |
| MVP_top60_DNA_seed | ERBB2 | -1.304226 | LUAD | NCI-H322M | Lapatinib | MVP_top60_DNA+RNA_seed | PIK3CA | -0.170171 | STAD | SNU-1 | Buparlisib |
| MVP_top60_DNA_seed | IGF1R | -1.302014 | LUAD | HCC-44 | BMS-536924 | MVP_top60_DNA+RNA_seed | ERBB2 | -0.169119 | STAD | AGS | Afatinib |
| MVP_top60_DNA_seed | GSK3B | -1.298179 | LUAD | RERF-LC-MS | CHIR-99021 | MVP_top60_DNA+RNA_seed | PARP1 | -0.165446 | STAD | 23132-87 | Talazoparib |
| MVP_top60_DNA_seed | HSP90AA1 | -1.297642 | LUAD | NCI-H1755 | Tanespimycin | MVP_top60_DNA+RNA_seed | TP53 | -0.164612 | STAD | AGS | MIRA-1 |
| MVP_top60_DNA_seed | EGFR | -1.297525 | LUAD | LC-2-ad | Sapitinib | MVP_top60_DNA+RNA_seed | MET | -0.164459 | STAD | AGS | Crizotinib |
| MVP_top60_DNA_seed | ERBB2 | -1.297525 | LUAD | LC-2-ad | Sapitinib | MVP_top60_DNA+RNA_seed | GSK3B | -0.157077 | STAD | RF-48 | SB216763 |
| MVP_top60_DNA_seed | ERBB3 | -1.297525 | LUAD | LC-2-ad | Sapitinib | MVP_top60_DNA+RNA_seed | MAPK1 | -0.148859 | STAD | 23132-87 | Ulixertinib |
| MVP_top60_DNA_seed | EGFR | -1.290848 | LUAD | EMC-BAC-1 | Gefitinib | MVP_top60_DNA+RNA_seed | MAPK3 | -0.148859 | STAD | 23132-87 | Ulixertinib |
| MVP_top60_DNA_seed | EGFR | -1.287555 | LUAD | NCI-H1666 | Lapatinib | MVP_top60_DNA+RNA_seed | PARP1 | -0.146023 | STAD | SK-GT-2 | Talazoparib |
| MVP_top60_DNA_seed | ERBB2 | -1.287555 | LUAD | NCI-H1666 | Lapatinib | MVP_top60_DNA+RNA_seed | ERBB2 | -0.14424 | STAD | 23132-87 | Sapitinib |
| MVP_top60_DNA_seed | AKT1 | -1.285676 | LUAD | NCI-H1993 | Uprosertib | MVP_top60_DNA+RNA_seed | ERBB3 | -0.14424 | STAD | 23132-87 | Sapitinib |
| MVP_top60_DNA_seed | MAPK1 | -1.278117 | LUAD | NCI-H1792 | Ulixertinib | MVP_top60_DNA+RNA_seed | ERBB2 | -0.137624 | STAD | IM-95 | Afatinib |
| MVP_top60_DNA_seed | HDAC1 | -1.2777 | LUAD | HCC-827 | Entinostat | MVP_top60_DNA+RNA_seed | PARP1 | -0.137542 | STAD | SK-GT-2 | Rucaparib |
| MVP_top60_DNA_seed | AURKA | -1.265714 | LUAD | LC-2-ad | Tozasertib | MVP_top60_DNA+RNA_seed | ATM | -0.134883 | STAD | SNU-5 | KU-55933 |
| MVP_top60_DNA_seed | IGF1R | -1.26433 | LUAD | NCI-H1573 | GSK1904529A | MVP_top60_DNA+RNA_seed | MAPK1 | -0.1298 | STAD | RERF-GC-1B | Ulixertinib |
| MVP_top60_DNA_seed | PIK3CA | -1.259063 | LUAD | NCI-H1755 | GNE-317 | MVP_top60_DNA+RNA_seed | MAPK3 | -0.1298 | STAD | RERF-GC-1B | Ulixertinib |
| MVP_top60_DNA_seed | MAPK1 | -1.256618 | LUAD | NCI-H1648 | Ulixertinib | MVP_top60_DNA+RNA_seed | EP300 | -0.129786 | STAD | FU97 | CPI-637 |
| MVP_top60_DNA_seed | PIK3CA | -1.253168 | LUAD | NCI-H1648 | AZD8186 | MVP_top60_DNA+RNA_seed | PARP1 | -0.123559 | STAD | SNU-1 | Talazoparib |
| MVP_top60_DNA_seed | MAPK1 | -1.251548 | LUAD | NCI-H1437 | SCH772984 | MVP_top60_DNA+RNA_seed | ERBB2 | -0.113922 | STAD | TGBC11TKB | Afatinib |
| MVP_top60_DNA_seed | HDAC1 | -1.249884 | LUAD | LXF-289 | Romidepsin | MVP_top60_DNA+RNA_seed | PIK3CA | -0.112712 | STAD | AGS | AZD8186 |
| MVP_top60_DNA_seed | PIK3CA | -1.24466 | LUAD | NCI-H1792 | GNE-317 | MVP_top60_DNA+RNA_seed | ERBB2 | -0.112354 | STAD | SNU-16 | Sapitinib |
| MVP_top60_DNA_seed | HSP90AA1 | -1.229955 | LUAD | RERF-LC-MS | CCT-018159 | MVP_top60_DNA+RNA_seed | ERBB3 | -0.112354 | STAD | SNU-16 | Sapitinib |
| MVP_top60_DNA_seed | PIK3CA | -1.228284 | LUAD | NCI-H2122 | GNE-317 | MVP_top60_DNA+RNA_seed | HDAC1 | -0.110771 | STAD | OCUM-1 | Romidepsin |
| MVP_top60_DNA_seed | PIK3CA | -1.226963 | LUAD | NCI-H1993 | GNE-317 | MVP_top60_DNA+RNA_seed | HDAC3 | -0.110771 | STAD | OCUM-1 | Romidepsin |
| MVP_top60_DNA_seed | PIK3CA | -1.225523 | LUAD | COR-L105 | GNE-317 | MVP_top60_DNA+RNA_seed | PIK3CA | -0.110412 | STAD | SK-GT-2 | AZD8186 |
| MVP_top60_DNA_seed | AURKA | -1.223471 | LUAD | VMRC-LCD | Alisertib | MVP_top60_DNA+RNA_seed | PARP1 | -0.108826 | STAD | SK-GT-2 | Olaparib |
| MVP_top60_DNA_seed | AKT1 | -1.223328 | LUAD | NCI-H1648 | AZD5363 | MVP_top60_DNA+RNA_seed | PIK3CA | -0.108094 | STAD | TMK-1 | Buparlisib |
| MVP_top60_DNA_seed | KRAS | -1.223262 | LUAD | NCI-H2122 | KRAS (G12C) Inhibitor-12 | MVP_top60_DNA+RNA_seed | ERBB2 | -0.103653 | STAD | IM-95 | Sapitinib |
| MVP_top60_DNA_seed | GSK3B | -1.216711 | LUAD | NCI-H1975 | SB216763 | MVP_top60_DNA+RNA_seed | ERBB3 | -0.103653 | STAD | IM-95 | Sapitinib |
| MVP_top60_DNA_seed | HSP90AA1 | -1.216659 | LUAD | NCI-H358 | Tanespimycin | MVP_top60_DNA+RNA_seed | MAPK1 | -0.100731 | STAD | SK-GT-2 | SCH772984 |
| MVP_top60_DNA_seed | EGFR | -1.210974 | LUAD | Calu-3 | Osimertinib | MVP_top60_DNA+RNA_seed | MAPK3 | -0.100731 | STAD | SK-GT-2 | SCH772984 |
| MVP_top60_DNA_seed | PIK3CA | -1.20902 | LUAD | NCI-H1755 | AZD8186 | MVP_top60_DNA+RNA_seed | ESR1 | -0.100388 | STAD | TGBC11TKB | Tamoxifen |
| MVP_top60_DNA_seed | PIK3CA | -1.20867 | LUAD | RERF-LC-MS | GNE-317 | MVP_top60_DNA+RNA_seed | PARP1 | -0.100352 | STAD | FU97 | Talazoparib |

| | | | | | | | | | | | |
|---|---|---|---|---|---|---|---|---|---|---|---|
| MVP_top60_DNA_seed | MET | -1.201637 | LUAD | NCI-H23 | Savolitinib | MVP_top60_DNA+RNA_seed | PIK3CA | -0.100081 | STAD | NCI-N87 | Buparlisib |
| MVP_top60_DNA_seed | PIK3CA | -1.197889 | LUAD | RERF-LC-KJ | Buparlisib | MVP_top60_DNA+RNA_seed | EZH2 | -0.099814 | STAD | MKN45 | GSK343 |
| MVP_top60_DNA_seed | AURKA | -1.197243 | LUAD | LC-2-ad | Alisertib | MVP_top60_DNA+RNA_seed | ATM | -0.094917 | STAD | MKN28 | KU-55933 |
| MVP_top60_DNA_seed | EZH2 | -1.197179 | LUAD | LXF-289 | GSK343 | MVP_top60_DNA+RNA_seed | SRC | -0.092626 | STAD | GCIY | Bosutinib |
| MVP_top60_DNA_seed | HDAC1 | -1.195065 | LUAD | HCC-78 | Entinostat | MVP_top60_DNA+RNA_seed | PIK3CA | -0.091567 | STAD | TMK-1 | AZD8186 |
| MVP_top60_DNA_seed | MAPK1 | -1.194182 | LUAD | NCI-H1437 | VX-11e | MVP_top60_DNA+RNA_seed | HSP90AA1 | -0.091452 | STAD | SK-GT-2 | Tanespimycin |
| MVP_top60_DNA_seed | MAPK1 | -1.193216 | LUAD | NCI-H322M | SCH772984 | MVP_top60_DNA+RNA_seed | HSP90AB1 | -0.091452 | STAD | SK-GT-2 | Tanespimycin |
| MVP_top60_DNA_seed | EGFR | -1.19148 | LUAD | NCI-H1650 | Erlotinib | MVP_top60_DNA+RNA_seed | HSP90AA1 | -0.086544 | STAD | TMK-1 | Elesclomol |
| MVP_top60_DNA_seed | AKT1 | -1.191153 | LUAD | LXF-289 | AT13148 | MVP_top60_DNA+RNA_seed | HSP90AB1 | -0.086544 | STAD | TMK-1 | Elesclomol |
| MVP_top60_DNA_seed | SRC | -1.183187 | LUAD | PC-3_[JPC-3] | Bosutinib | MVP_top60_DNA+RNA_seed | ERBB2 | -0.080522 | STAD | FU97 | Lapatinib |
| MVP_top60_DNA_seed | PIK3CA | -1.179015 | LUAD | NCI-H1975 | Buparlisib | MVP_top60_DNA+RNA_seed | EP300 | -0.078808 | STAD | AGS | SGC-CBP30 |
| MVP_top60_DNA_seed | MAPK1 | -1.176903 | LUAD | Calu-6 | Ulixertinib | MVP_top60_DNA+RNA_seed | CREBBP | -0.078808 | STAD | AGS | SGC-CBP30 |
| MVP_top60_DNA_seed | PIK3CA | -1.174585 | LUAD | RERF-LC-MS | AZD8186 | MVP_top60_DNA+RNA_seed | PIK3CA | -0.072562 | STAD | OCUM-1 | Alpelisib |
| MVP_top60_DNA_seed | MAPK1 | -1.171517 | LUAD | NCI-H1435 | Ulixertinib | MVP_top60_DNA+RNA_seed | PARP1 | -0.071115 | STAD | SNU-1 | Rucaparib |
| MVP_top60_DNA_seed | PIK3CA | -1.168308 | LUAD | RERF-LC-MS | Alpelisib | MVP_top60_DNA+RNA_seed | EZH2 | -0.07009 | STAD | TGBC11TKB | GSK343 |
| MVP_top60_DNA_seed | CREBBP | -1.165161 | LUAD | NCI-H522 | SGC-CBP30 | MVP_top60_DNA+RNA_seed | GSK3B | -0.068422 | STAD | MKN28 | SB216763 |
| MVP_top60_DNA_seed | MAPK1 | -1.161778 | LUAD | Calu-6 | Ulixertinib | MVP_top60_DNA+RNA_seed | ATM | -0.068061 | STAD | FU97 | KU-55933 |
| MVP_top60_DNA_seed | PIK3CA | -1.158947 | LUAD | NCI-H1623 | Alpelisib | MVP_top60_DNA+RNA_seed | MAPK1 | -0.066786 | STAD | SNU-16 | SCH772984 |
| MVP_top60_DNA_seed | MAPK1 | -1.148086 | LUAD | NCI-H1437 | Ulixertinib | MVP_top60_DNA+RNA_seed | MAPK3 | -0.066786 | STAD | SNU-16 | SCH772984 |
| MVP_top60_DNA_seed | EGFR | -1.144379 | LUAD | NCI-H1573 | Erlotinib | MVP_top60_DNA+RNA_seed | ERBB2 | -0.066352 | STAD | MKN7 | Sapitinib |
| MVP_top60_DNA_seed | IGF1R | -1.143419 | LUAD | NCI-H838 | BMS-536924 | MVP_top60_DNA+RNA_seed | ERBB3 | -0.066352 | STAD | MKN7 | Sapitinib |
| MVP_top60_DNA_seed | AKT1 | -1.143267 | LUAD | NCI-H2122 | AZD5363 | MVP_top60_DNA+RNA_seed | TP53 | -0.064001 | STAD | 23132-87 | MIRA-1 |
| MVP_top60_DNA_seed | EGFR | -1.139855 | LUAD | NCI-H1623 | Osimertinib | MVP_top60_DNA+RNA_seed | PIK3CA | -0.061805 | STAD | NUGC-3 | Buparlisib |
| MVP_top60_DNA_seed | AKT1 | -1.138588 | LUAD | NCI-H2122 | Uprosertib | MVP_top60_DNA+RNA_seed | ERBB2 | -0.060068 | STAD | SNU-1 | Lapatinib |
| MVP_top60_DNA_seed | HSP90AA1 | -1.131044 | LUAD | NCI-H650 | Luminespib | MVP_top60_DNA+RNA_seed | ERBB2 | -0.059664 | STAD | SNU-1 | Afatinib |
| MVP_top60_DNA_seed | KRAS | -1.128535 | LUAD | NCI-H1975 | KRAS (G12C) Inhibitor-1 | MVP_top60_DNA+RNA_seed | MAPK1 | -0.05529 | STAD | SK-GT-2 | VX-11e |
| MVP_top60_DNA_seed | AKT1 | -1.125471 | LUAD | NCI-H1975 | Ipatasertib | MVP_top60_DNA+RNA_seed | MAPK1 | -0.053042 | STAD | SNU-1 | Ulixertinib |
| MVP_top60_DNA_seed | MAPK1 | -1.12391 | LUAD | HCC-44 | Ulixertinib | MVP_top60_DNA+RNA_seed | MAPK3 | -0.053042 | STAD | SNU-1 | Ulixertinib |
| MVP_top60_DNA_seed | MET | -1.123598 | LUAD | NCI-H1975 | Savolitinib | MVP_top60_DNA+RNA_seed | KRAS | -0.051228 | STAD | GCIY | KRAS (G12C) Inhibitor-1 |
| MVP_top60_DNA_seed | AKT1 | -1.120744 | LUAD | NCI-H1781 | Uprosertib | MVP_top60_DNA+RNA_seed | PARP1 | -0.047027 | STAD | TMK-1 | Talazoparib |
| MVP_top60_DNA_seed | MAPK1 | -1.117933 | LUAD | NCI-H1648 | SCH772984 | MVP_top60_DNA+RNA_seed | PARP1 | -0.044111 | STAD | KATOIII | Veliparib |
| MVP_top60_DNA_seed | SMARCA4 | -1.117883 | LUAD | NCI-H23 | PFI3 | MVP_top60_DNA+RNA_seed | PARP1 | -0.04201 | STAD | NCI-N87 | Niraparib |
| MVP_top60_DNA_seed | MAPK1 | -1.117005 | LUAD | RERF-LC-MS | Ulixertinib | MVP_top60_DNA+RNA_seed | MDM2 | -0.039627 | STAD | FU97 | Serdemetan |
| MVP_top60_DNA_seed | MAPK1 | -1.116743 | LUAD | NCI-H1792 | VX-11e | MVP_top60_DNA+RNA_seed | HSP90AA1 | -0.037038 | STAD | GCIY | Luminespib |
| MVP_top60_DNA_seed | IGF1R | -1.112834 | LUAD | COR-L105 | BMS-536924 | MVP_top60_DNA+RNA_seed | HSP90AB1 | -0.037038 | STAD | GCIY | Luminespib |
| MVP_top60_DNA_seed | ATM | -1.11248 | LUAD | A549 | KU-55933 | MVP_top60_DNA+RNA_seed | MDM2 | -0.034759 | STAD | IM-95 | Nutlin-3a (-) |
| MVP_top60_DNA_seed | EZH2 | -1.110143 | LUAD | NCI-H1651 | GSK343 | MVP_top60_DNA+RNA_seed | HDAC1 | -0.034086 | STAD | RERF-GC-1B | Romidepsin |
| MVP_top60_DNA_seed | EGFR | -1.109219 | LUAD | RERF-LC-KJ | Gefitinib | MVP_top60_DNA+RNA_seed | HDAC3 | -0.034086 | STAD | RERF-GC-1B | Romidepsin |
| MVP_top60_DNA_seed | EGFR | -1.107588 | LUAD | NCI-H1781 | Gefitinib | MVP_top60_DNA+RNA_seed | ERBB2 | -0.033577 | STAD | HGC-27 | Afatinib |
| MVP_top60_DNA_seed | AKT1 | -1.098752 | LUAD | NCI-H1993 | AZD5363 | MVP_top60_DNA+RNA_seed | HSP90AA1 | -0.032787 | STAD | MKN7 | Elesclomol |
| MVP_top60_DNA_seed | MET | -1.088747 | LUAD | NCI-H2122 | Savolitinib | MVP_top60_DNA+RNA_seed | HSP90AB1 | -0.032787 | STAD | MKN7 | Elesclomol |

| | | | | | | | | | | | |
|---|---|---|---|---|---|---|---|---|---|---|---|
| MVP_top60_DNA_seed | AKT1 | -1.087845 | LUAD | NCI-H1993 | MK-2206 | MVP_top60_DNA+RNA_seed | MAPK1 | -0.029176 | STAD | NUGC-3 | Ulixertinib |
| MVP_top60_DNA_seed | EGFR | -1.079003 | LUAD | NCI-H358 | Lapatinib | MVP_top60_DNA+RNA_seed | MAPK3 | -0.029176 | STAD | NUGC-3 | Ulixertinib |
| MVP_top60_DNA_seed | ERBB2 | -1.079003 | LUAD | NCI-H358 | Lapatinib | MVP_top60_DNA+RNA_seed | PARP1 | -0.023938 | STAD | 23132-87 | Veliparib |
| MVP_top60_DNA_seed | MAPK1 | -1.076551 | LUAD | HCC-44 | SCH772984 | MVP_top60_DNA+RNA_seed | PARP1 | -0.023702 | STAD | TGBC11TKB | Rucaparib |
| MVP_top60_DNA_seed | FGFR1 | -1.076539 | LUAD | NCI-H838 | PD173074 | MVP_top60_DNA+RNA_seed | PIK3CA | -0.023201 | STAD | MKN45 | Buparlisib |
| MVP_top60_DNA_seed | PIK3CA | -1.074684 | LUAD | EMC-BAC-1 | AZD8186 | MVP_top60_DNA+RNA_seed | HDAC1 | -0.022067 | STAD | SNU-5 | Romidepsin |
| MVP_top60_DNA_seed | HDAC1 | -1.072302 | LUAD | RERF-LC-MS | PCI-34051 | MVP_top60_DNA+RNA_seed | HDAC3 | -0.022067 | STAD | SNU-5 | Romidepsin |
| MVP_top60_DNA_seed | ATM | -1.070681 | LUAD | RERF-LC-KJ | KU-55933 | MVP_top60_DNA+RNA_seed | FGFR3 | -0.020568 | STAD | HGC-27 | PD173074 |
| MVP_top60_DNA_seed | IGF1R | -1.070253 | LUAD | NCI-H1437 | NVP-ADW742 | MVP_top60_DNA+RNA_seed | ERBB2 | -0.020032 | STAD | AGS | Sapitinib |
| MVP_top60_DNA_seed | MAPK1 | -1.065512 | LUAD | RERF-LC-MS | SCH772984 | MVP_top60_DNA+RNA_seed | ERBB3 | -0.020032 | STAD | AGS | Sapitinib |
| MVP_top60_DNA_seed | HSP90AA1 | -1.064813 | LUAD | A549 | Luminespib | MVP_top60_DNA+RNA_seed | HSP90AA1 | -0.018806 | STAD | 23132-87 | Tanespimycin |
| MVP_top60_DNA_seed | MET | -1.063039 | LUAD | NCI-H1623 | Savolitinib | MVP_top60_DNA+RNA_seed | HSP90AB1 | -0.018806 | STAD | 23132-87 | Tanespimycin |
| MVP_top60_DNA_seed | IGF1R | -1.058671 | LUAD | RERF-LC-MS | NVP-ADW742 | MVP_top60_DNA+RNA_seed | MET | -0.016684 | STAD | TMK-1 | Crizotinib |
| MVP_top60_DNA_seed | MAPK1 | -1.057018 | LUAD | A549 | SCH772984 | MVP_top60_DNA+RNA_seed | PIK3CA | -0.015433 | STAD | NUGC-3 | Alpelisib |
| MVP_top60_DNA_seed | BRAF | -1.054628 | LUAD | NCI-H2009 | PLX-4720 | MVP_top60_DNA+RNA_seed | PARP1 | -0.009481 | STAD | TGBC11TKB | Niraparib |
| MVP_top60_DNA_seed | PIK3CA | -1.052842 | LUAD | NCI-H1975 | AZD8186 | MVP_top60_DNA+RNA_seed | MAPK1 | 0.001898 | STAD | NUGC-3 | SCH772984 |
| MVP_top60_DNA_seed | IGF1R | -1.051141 | LUAD | NCI-H1666 | BMS-536924 | MVP_top60_DNA+RNA_seed | MAPK3 | 0.001898 | STAD | NUGC-3 | SCH772984 |
| MVP_top60_DNA_seed | TP53 | -1.049009 | LUAD | NCI-H1734 | MIRA-1 | MVP_top60_DNA+RNA_seed | ESR1 | 0.002783 | STAD | Hs-746T | GDC0810 |
| MVP_top60_DNA_seed | ATM | -1.048621 | LUAD | NCI-H322M | KU-55933 | MVP_top60_DNA+RNA_seed | PIK3CA | 0.005655 | STAD | TGBC11TKB | Buparlisib |
| MVP_top60_DNA_seed | IGF1R | -1.041825 | LUAD | NCI-H2228 | Linsitinib | MVP_top60_DNA+RNA_seed | SRC | 0.007134 | STAD | TMK-1 | Bosutinib |
| MVP_top60_DNA_seed | IGF1R | -1.040294 | LUAD | RERF-LC-MS | BMS-536924 | MVP_top60_DNA+RNA_seed | TP53 | 0.008774 | STAD | IM-95 | MIRA-1 |
| MVP_top60_DNA_seed | MAPK1 | -1.039045 | LUAD | NCI-H3122 | Ulixertinib | MVP_top60_DNA+RNA_seed | EZH2 | 0.009195 | STAD | RERF-GC-1B | GSK343 |
| MVP_top60_DNA_seed | FGFR1 | -1.035122 | LUAD | NCI-H2122 | AZD4547 | MVP_top60_DNA+RNA_seed | EZH2 | 0.009494 | STAD | SNU-5 | GSK343 |
| MVP_top60_DNA_seed | IGF1R | -1.034191 | LUAD | LXF-289 | BMS-754807 | MVP_top60_DNA+RNA_seed | ERBB2 | 0.012472 | STAD | MKN28 | Lapatinib |
| MVP_top60_DNA_seed | MAPK1 | -1.033773 | LUAD | NCI-H2085 | Ulixertinib | MVP_top60_DNA+RNA_seed | HDAC1 | 0.013655 | STAD | Hs-746T | Romidepsin |
| MVP_top60_DNA_seed | MET | -1.028331 | LUAD | NCI-H1944 | Savolitinib | MVP_top60_DNA+RNA_seed | HDAC3 | 0.013655 | STAD | Hs-746T | Romidepsin |
| MVP_top60_DNA_seed | ATM | -1.028141 | LUAD | NCI-H1435 | KU-55933 | MVP_top60_DNA+RNA_seed | KRAS | 0.015757 | STAD | OCUM-1 | KRAS (G12C) Inhibitor-1 |
| MVP_top60_DNA_seed | EGFR | -1.027958 | LUAD | NCI-H838 | Sapitinib | MVP_top60_DNA+RNA_seed | SRC | 0.018182 | STAD | NUGC-3 | Bosutinib |
| MVP_top60_DNA_seed | ERBB2 | -1.027958 | LUAD | NCI-H838 | Sapitinib | MVP_top60_DNA+RNA_seed | HSP90AA1 | 0.022141 | STAD | MKN28 | Tanespimycin |
| MVP_top60_DNA_seed | ERBB3 | -1.027958 | LUAD | NCI-H838 | Sapitinib | MVP_top60_DNA+RNA_seed | HSP90AB1 | 0.022141 | STAD | MKN28 | Tanespimycin |
| MVP_top60_DNA_seed | IGF1R | -1.023396 | LUAD | NCI-H1437 | Linsitinib | MVP_top60_DNA+RNA_seed | MAPK1 | 0.022994 | STAD | 23132-87 | Ulixertinib |
| MVP_top60_DNA_seed | HSP90AA1 | -1.021189 | LUAD | NCI-H1792 | Luminespib | MVP_top60_DNA+RNA_seed | MAPK3 | 0.022994 | STAD | 23132-87 | Ulixertinib |
| MVP_top60_DNA_seed | AKT1 | -1.020747 | LUAD | NCI-H1993 | Ipatasertib | MVP_top60_DNA+RNA_seed | ESR1 | 0.023539 | STAD | IM-95 | Tamoxifen |
| MVP_top60_DNA_seed | AKT1 | -1.018807 | LUAD | NCI-H1993 | Afuresertib | MVP_top60_DNA+RNA_seed | MAPK1 | 0.024982 | STAD | IM-95 | SCH772984 |
| MVP_top60_DNA_seed | MAPK1 | -1.017031 | LUAD | NCI-H1975 | Ulixertinib | MVP_top60_DNA+RNA_seed | MAPK3 | 0.024982 | STAD | IM-95 | SCH772984 |
| MVP_top60_DNA_seed | HSP90AA1 | -1.016678 | LUAD | LXF-289 | Tanespimycin | MVP_top60_DNA+RNA_seed | FGFR3 | 0.025966 | STAD | NUGC-3 | PD173074 |
| MVP_top60_DNA_seed | AKT1 | -1.012382 | LUAD | NCI-H838 | AT13148 | MVP_top60_DNA+RNA_seed | ERBB2 | 0.027895 | STAD | MKN28 | Sapitinib |
| MVP_top60_DNA_seed | MET | -1.01205 | LUAD | NCI-H1573 | Savolitinib | MVP_top60_DNA+RNA_seed | ERBB3 | 0.027895 | STAD | MKN28 | Sapitinib |
| MVP_top60_DNA_seed | HSP90AA1 | -1.010772 | LUAD | NCI-H2122 | Tanespimycin | MVP_top60_DNA+RNA_seed | ESR1 | 0.028829 | STAD | TMK-1 | GDC0810 |
| MVP_top60_DNA_seed | AURKA | -1.007577 | LUAD | ABC-1 | ZM447439 | MVP_top60_DNA+RNA_seed | PARP1 | 0.030553 | STAD | Hs-746T | Olaparib |
| MVP_top60_DNA_seed | ATM | -1.007379 | LUAD | NCI-H522 | KU-55933 | MVP_top60_DNA+RNA_seed | FGFR3 | 0.031612 | STAD | SK-GT-2 | AZD4547 |

| | | | | | | | | | | | |
|---|---|---|---|---|---|---|---|---|---|---|---|
| MVP_top60_DNA_seed | AKT1 | -1.004861 | LUAD | NCI-H2122 | GSK2110183B | MVP_top60_DNA+RNA_seed | HSP90AA1 | 0.033424 | STAD | RF-48 | Tanespimycin |
| MVP_top60_DNA_seed | AKT1 | -1.002467 | LUAD | NCI-H1781 | AT13148 | MVP_top60_DNA+RNA_seed | HSP90AB1 | 0.033424 | STAD | RF-48 | Tanespimycin |
| MVP_top60_DNA_seed | AKT1 | -1.002441 | LUAD | HCC-44 | AZD5363 | MVP_top60_DNA+RNA_seed | MAPK1 | 0.033804 | STAD | FU97 | Ulixertinib |
| MVP_top60_DNA_seed | AKT1 | -1.00235 | LUAD | NCI-H2122 | Afuresertib | MVP_top60_DNA+RNA_seed | MAPK3 | 0.033804 | STAD | FU97 | Ulixertinib |
| MVP_top60_DNA_seed | MAPK1 | -0.997032 | LUAD | NCI-H2087 | SCH772984 | MVP_top60_DNA+RNA_seed | PIK3CA | 0.036404 | STAD | MKN7 | AZD8186 |
| MVP_top60_DNA_seed | AURKA | -0.997002 | LUAD | NCI-H1781 | ZM447439 | MVP_top60_DNA+RNA_seed | FGFR3 | 0.038038 | STAD | AGS | AZD4547 |
| MVP_top60_DNA_seed | EGFR | -0.995602 | LUAD | EKVX | AZD3759 | MVP_top60_DNA+RNA_seed | ERBB2 | 0.03859 | STAD | Hs-746T | Lapatinib |
| MVP_top60_DNA_seed | MAPK1 | -0.995143 | LUAD | NCI-H1993 | Ulixertinib | MVP_top60_DNA+RNA_seed | PARP1 | 0.042462 | STAD | FU97 | Olaparib |
| MVP_top60_DNA_seed | HSP90AA1 | -0.99441 | LUAD | NCI-H1755 | Luminespib | MVP_top60_DNA+RNA_seed | EZH2 | 0.043467 | STAD | SNU-1 | GSK343 |
| MVP_top60_DNA_seed | HDAC1 | -0.992393 | LUAD | EMC-BAC-1 | Entinostat | MVP_top60_DNA+RNA_seed | PIK3CA | 0.043728 | STAD | MKN7 | GNE-317 |
| MVP_top60_DNA_seed | IGF1R | -0.990169 | LUAD | NCI-H1437 | GSK1904529A | MVP_top60_DNA+RNA_seed | ESR1 | 0.044685 | STAD | NUGC-4 | GDC0810 |
| MVP_top60_DNA_seed | MAPK1 | -0.987395 | LUAD | NCI-H2405 | Ulixertinib | MVP_top60_DNA+RNA_seed | MAPK1 | 0.046051 | STAD | GCIY | Ulixertinib |
| MVP_top60_DNA_seed | HSP90AA1 | -0.987039 | LUAD | NCI-H2009 | Luminespib | MVP_top60_DNA+RNA_seed | MAPK3 | 0.046051 | STAD | GCIY | Ulixertinib |
| MVP_top60_DNA_seed | AKT1 | -0.986429 | LUAD | A549 | AT13148 | MVP_top60_DNA+RNA_seed | ERBB2 | 0.046841 | STAD | RERF-GC-1B | Afatinib |
| MVP_top60_DNA_seed | MAPK1 | -0.985587 | LUAD | NCI-H838 | Ulixertinib | MVP_top60_DNA+RNA_seed | EP300 | 0.047649 | STAD | GCIY | SGC-CBP30 |
| MVP_top60_DNA_seed | BRAF | -0.981614 | LUAD | NCI-H1792 | SB590885 | MVP_top60_DNA+RNA_seed | CREBBP | 0.047649 | STAD | GCIY | SGC-CBP30 |
| MVP_top60_DNA_seed | HDAC1 | -0.981466 | LUAD | NCI-H1755 | Romidepsin | MVP_top60_DNA+RNA_seed | ATM | 0.0506 | STAD | Hs-746T | KU-55933 |
| MVP_top60_DNA_seed | MAPK1 | -0.978603 | LUAD | NCI-H1755 | VX-11e | MVP_top60_DNA+RNA_seed | ERBB2 | 0.050663 | STAD | MKN28 | Afatinib |
| MVP_top60_DNA_seed | EGFR | -0.978121 | LUAD | NCI-H1650 | Osimertinib | MVP_top60_DNA+RNA_seed | EP300 | 0.059794 | STAD | HGC-27 | CPI-637 |
| MVP_top60_DNA_seed | EGFR | -0.973508 | LUAD | RERF-LC-MS | Lapatinib | MVP_top60_DNA+RNA_seed | PIK3CA | 0.060791 | STAD | TGBC11TKB | GNE-317 |
| MVP_top60_DNA_seed | ERBB2 | -0.973508 | LUAD | RERF-LC-MS | Lapatinib | MVP_top60_DNA+RNA_seed | PIK3CA | 0.062867 | STAD | GCIY | AZD8186 |
| MVP_top60_DNA_seed | AURKA | -0.971594 | LUAD | LC-2-ad | ZM447439 | MVP_top60_DNA+RNA_seed | PARP1 | 0.072455 | STAD | SNU-1 | Veliparib |
| MVP_top60_DNA_seed | EZH2 | -0.970408 | LUAD | NCI-H2023 | GSK343 | MVP_top60_DNA+RNA_seed | PIK3CA | 0.0727 | STAD | RF-48 | AZD8186 |
| MVP_top60_DNA_seed | AKT1 | -0.969321 | LUAD | NCI-H1623 | Uprosertib | MVP_top60_DNA+RNA_seed | ERBB2 | 0.073149 | STAD | RERF-GC-1B | Lapatinib |
| MVP_top60_DNA_seed | EGFR | -0.966727 | LUAD | EMC-BAC-2 | Erlotinib | MVP_top60_DNA+RNA_seed | MDM2 | 0.074679 | STAD | TGBC11TKB | Serdemetan |
| MVP_top60_DNA_seed | IGF1R | -0.966525 | LUAD | LC-2-ad | BMS-754807 | MVP_top60_DNA+RNA_seed | PARP1 | 0.078822 | STAD | RF-48 | Rucaparib |
| MVP_top60_DNA_seed | IGF1R | -0.962916 | LUAD | NCI-H1792 | BMS-536924 | MVP_top60_DNA+RNA_seed | MDM2 | 0.07984 | STAD | HGC-27 | Nutlin-3a (-) |
| MVP_top60_DNA_seed | MAPK1 | -0.962386 | LUAD | NCI-H1993 | SCH772984 | MVP_top60_DNA+RNA_seed | PARP1 | 0.086451 | STAD | AGS | Veliparib |
| MVP_top60_DNA_seed | IGF1R | -0.96055 | LUAD | LC-2-ad | BMS-536924 | MVP_top60_DNA+RNA_seed | GSK3B | 0.091457 | STAD | IM-95 | SB216763 |
| MVP_top60_DNA_seed | BRAF | -0.959907 | LUAD | NCI-H2122 | SB590885 | MVP_top60_DNA+RNA_seed | SRC | 0.093718 | STAD | NUGC-4 | Bosutinib |
| MVP_top60_DNA_seed | EGFR | -0.959136 | LUAD | NCI-H2030 | Sapitinib | MVP_top60_DNA+RNA_seed | ESR1 | 0.093965 | STAD | TGBC11TKB | GDC0810 |
| MVP_top60_DNA_seed | ERBB2 | -0.959136 | LUAD | NCI-H2030 | Sapitinib | MVP_top60_DNA+RNA_seed | FGFR3 | 0.094442 | STAD | TMK-1 | PD173074 |
| MVP_top60_DNA_seed | ERBB3 | -0.959136 | LUAD | NCI-H2030 | Sapitinib | MVP_top60_DNA+RNA_seed | MAPK1 | 0.097508 | STAD | NUGC-3 | Ulixertinib |
| MVP_top60_DNA_seed | AKT1 | -0.954396 | LUAD | NCI-H1734 | Ipatasertib | MVP_top60_DNA+RNA_seed | MAPK3 | 0.097508 | STAD | NUGC-3 | Ulixertinib |
| MVP_top60_DNA_seed | HSP90AA1 | -0.950333 | LUAD | NCI-H1944 | Tanespimycin | MVP_top60_DNA+RNA_seed | HDAC1 | 0.107099 | STAD | 23132-87 | Entinostat |
| MVP_top60_DNA_seed | HDAC1 | -0.946015 | LUAD | HCC-78 | Romidepsin | MVP_top60_DNA+RNA_seed | HDAC3 | 0.107099 | STAD | 23132-87 | Entinostat |
| MVP_top60_DNA_seed | SRC | -0.945878 | LUAD | H3255 | Bosutinib | MVP_top60_DNA+RNA_seed | PARP1 | 0.107179 | STAD | KATOIII | Olaparib |
| MVP_top60_DNA_seed | MAPK1 | -0.945769 | LUAD | NCI-H2122 | Ulixertinib | MVP_top60_DNA+RNA_seed | EP300 | 0.107301 | STAD | OCUM-1 | SGC-CBP30 |
| MVP_top60_DNA_seed | MAPK1 | -0.945618 | LUAD | NCI-H322M | VX-11e | MVP_top60_DNA+RNA_seed | CREBBP | 0.107301 | STAD | OCUM-1 | SGC-CBP30 |
| MVP_top60_DNA_seed | EGFR | -0.944308 | LUAD | NCI-H1666 | Osimertinib | MVP_top60_DNA+RNA_seed | PARP1 | 0.111616 | STAD | RERF-GC-1B | Olaparib |
| MVP_top60_DNA_seed | PIK3CA | -0.943231 | LUAD | NCI-H3122 | Buparlisib | MVP_top60_DNA+RNA_seed | PARP1 | 0.111725 | STAD | IM-95 | Olaparib |

| | | | | | | | | | | | |
|---|---|---|---|---|---|---|---|---|---|---|---|
| MVP_top60_DNA_seed | AKT1 | -0.940545 | LUAD | HCC-44 | AT13148 | MVP_top60_DNA+RNA_seed | MAPK1 | 0.113167 | STAD | TGBC11TKB | Ulixertinib |
| MVP_top60_DNA_seed | BRAF | -0.939638 | LUAD | RERF-LC-MS | PLX-4720 | MVP_top60_DNA+RNA_seed | MAPK3 | 0.113167 | STAD | TGBC11TKB | Ulixertinib |
| MVP_top60_DNA_seed | EGFR | -0.938908 | LUAD | RERF-LC-KJ | AZD3759 | MVP_top60_DNA+RNA_seed | FGFR3 | 0.114731 | STAD | SNU-5 | AZD4547 |
| MVP_top60_DNA_seed | AKT1 | -0.938658 | LUAD | NCI-H1781 | GSK2110183B | MVP_top60_DNA+RNA_seed | PIK3CA | 0.116808 | STAD | RF-48 | Alpelisib |
| MVP_top60_DNA_seed | HSP90AA1 | -0.938482 | LUAD | NCI-H1437 | CCT-018159 | MVP_top60_DNA+RNA_seed | HSP90AA1 | 0.118825 | STAD | MKN45 | Luminespib |
| MVP_top60_DNA_seed | HDAC1 | -0.934872 | LUAD | LC-2-ad | Entinostat | MVP_top60_DNA+RNA_seed | HSP90AB1 | 0.118825 | STAD | MKN45 | Luminespib |
| MVP_top60_DNA_seed | MAPK1 | -0.933265 | LUAD | COR-L105 | Ulixertinib | MVP_top60_DNA+RNA_seed | MET | 0.123733 | STAD | OCUM-1 | Crizotinib |
| MVP_top60_DNA_seed | ATM | -0.9317 | LUAD | NCI-H2122 | KU-55933 | MVP_top60_DNA+RNA_seed | HDAC1 | 0.130555 | STAD | MKN28 | PCI-34051 |
| MVP_top60_DNA_seed | PIK3CA | -0.930163 | LUAD | NCI-H2122 | Buparlisib | MVP_top60_DNA+RNA_seed | FGFR3 | 0.13309 | STAD | RF-48 | AZD4547 |
| MVP_top60_DNA_seed | EGFR | -0.925628 | LUAD | NCI-H1568 | Osimertinib | MVP_top60_DNA+RNA_seed | ERBB2 | 0.133165 | STAD | TGBC11TKB | Lapatinib |
| MVP_top60_DNA_seed | HSP90AA1 | -0.92456 | LUAD | EMC-BAC-1 | Tanespimycin | MVP_top60_DNA+RNA_seed | PARP1 | 0.138421 | STAD | TMK-1 | Olaparib |
| MVP_top60_DNA_seed | AKT1 | -0.922764 | LUAD | HCC-78 | Ipatasertib | MVP_top60_DNA+RNA_seed | PARP1 | 0.139336 | STAD | RF-48 | Talazoparib |
| MVP_top60_DNA_seed | IGF1R | -0.921882 | LUAD | NCI-H322M | Linsitinib | MVP_top60_DNA+RNA_seed | MDM2 | 0.140719 | STAD | GCIY | Serdemetan |
| MVP_top60_DNA_seed | MET | -0.919515 | LUAD | NCI-H2228 | Crizotinib | MVP_top60_DNA+RNA_seed | HSP90AA1 | 0.145553 | STAD | 23132-87 | Luminespib |
| MVP_top60_DNA_seed | AKT1 | -0.918724 | LUAD | NCI-H1975 | Afuresertib | MVP_top60_DNA+RNA_seed | HSP90AB1 | 0.145553 | STAD | 23132-87 | Luminespib |
| MVP_top60_DNA_seed | AKT1 | -0.916902 | LUAD | NCI-H1975 | Uprosertib | MVP_top60_DNA+RNA_seed | TP53 | 0.145707 | STAD | FU97 | MIRA-1 |
| MVP_top60_DNA_seed | EGFR | -0.91684 | LUAD | EKVX | Sapitinib | MVP_top60_DNA+RNA_seed | EP300 | 0.147874 | STAD | SNU-5 | CPI-637 |
| MVP_top60_DNA_seed | ERBB2 | -0.91684 | LUAD | EKVX | Sapitinib | MVP_top60_DNA+RNA_seed | GSK3B | 0.150724 | STAD | SK-GT-2 | SB216763 |
| MVP_top60_DNA_seed | ERBB3 | -0.91684 | LUAD | EKVX | Sapitinib | MVP_top60_DNA+RNA_seed | ERBB2 | 0.15512 | STAD | HGC-27 | Sapitinib |
| MVP_top60_DNA_seed | EGFR | -0.914333 | LUAD | EMC-BAC-2 | Sapitinib | MVP_top60_DNA+RNA_seed | ERBB3 | 0.15512 | STAD | HGC-27 | Sapitinib |
| MVP_top60_DNA_seed | ERBB2 | -0.914333 | LUAD | EMC-BAC-2 | Sapitinib | MVP_top60_DNA+RNA_seed | HDAC1 | 0.162536 | STAD | HGC-27 | PCI-34051 |
| MVP_top60_DNA_seed | ERBB3 | -0.914333 | LUAD | EMC-BAC-2 | Sapitinib | MVP_top60_DNA+RNA_seed | TP53 | 0.163433 | STAD | MKN7 | MIRA-1 |
| MVP_top60_DNA_seed | MET | -0.913338 | LUAD | NCI-H1734 | Savolitinib | MVP_top60_DNA+RNA_seed | ATM | 0.165988 | STAD | NCI-N87 | KU-55933 |
| MVP_top60_DNA_seed | TERT | -0.909582 | LUAD | NCI-H1792 | BIBR-1532 | MVP_top60_DNA+RNA_seed | MAPK1 | 0.166255 | STAD | HGC-27 | Ulixertinib |
| MVP_top60_DNA_seed | KRAS | -0.907714 | LUAD | HCC-78 | KRAS (G12C) Inhibitor-1 | MVP_top60_DNA+RNA_seed | MAPK3 | 0.166255 | STAD | HGC-27 | Ulixertinib |
| MVP_top60_DNA_seed | HSP90AA1 | -0.907365 | LUAD | RERF-LC-MS | Tanespimycin | MVP_top60_DNA+RNA_seed | HDAC1 | 0.174872 | STAD | HGC-27 | Romidepsin |
| MVP_top60_DNA_seed | IGF1R | -0.905475 | LUAD | HCC-44 | BMS-754807 | MVP_top60_DNA+RNA_seed | HDAC3 | 0.174872 | STAD | HGC-27 | Romidepsin |
| MVP_top60_DNA_seed | PIK3CA | -0.905297 | LUAD | HCC-44 | AZD8186 | MVP_top60_DNA+RNA_seed | MET | 0.17519 | STAD | NCI-N87 | Savolitinib |
| MVP_top60_DNA_seed | IGF1R | -0.905097 | LUAD | COR-L105 | NVP-ADW742 | MVP_top60_DNA+RNA_seed | HSP90AA1 | 0.177348 | STAD | SNU-16 | Luminespib |
| MVP_top60_DNA_seed | IGF1R | -0.904804 | LUAD | PC-3_[JPC-3] | BMS-536924 | MVP_top60_DNA+RNA_seed | HSP90AB1 | 0.177348 | STAD | SNU-16 | Luminespib |
| MVP_top60_DNA_seed | FGFR1 | -0.903348 | LUAD | NCI-H1755 | PD173074 | MVP_top60_DNA+RNA_seed | HSP90AA1 | 0.179068 | STAD | GCIY | Tanespimycin |
| MVP_top60_DNA_seed | MDM2 | -0.903095 | LUAD | NCI-H2228 | Serdemetan | MVP_top60_DNA+RNA_seed | HSP90AB1 | 0.179068 | STAD | GCIY | Tanespimycin |
| MVP_top60_DNA_seed | MAPK1 | -0.89779 | LUAD | NCI-H522 | Ulixertinib | MVP_top60_DNA+RNA_seed | MAPK1 | 0.179324 | STAD | RF-48 | SCH772984 |
| MVP_top60_DNA_seed | EGFR | -0.896158 | LUAD | RERF-LC-MS | AZD3759 | MVP_top60_DNA+RNA_seed | MAPK3 | 0.179324 | STAD | RF-48 | SCH772984 |
| MVP_top60_DNA_seed | AKT1 | -0.89612 | LUAD | HCC-44 | GSK2110183B | MVP_top60_DNA+RNA_seed | EP300 | 0.182026 | STAD | IM-95 | SGC-CBP30 |
| MVP_top60_DNA_seed | MET | -0.896037 | LUAD | NCI-H2122 | Crizotinib | MVP_top60_DNA+RNA_seed | CREBBP | 0.182026 | STAD | IM-95 | SGC-CBP30 |
| MVP_top60_DNA_seed | EGFR | -0.895905 | LUAD | EMC-BAC-1 | Erlotinib | MVP_top60_DNA+RNA_seed | GSK3B | 0.188284 | STAD | SNU-5 | SB216763 |
| MVP_top60_DNA_seed | AURKA | -0.895412 | LUAD | RERF-LC-MS | Tozasertib | MVP_top60_DNA+RNA_seed | EP300 | 0.188979 | STAD | TGBC11TKB | CPI-637 |
| MVP_top60_DNA_seed | HDAC1 | -0.892282 | LUAD | NCI-H1993 | Romidepsin | MVP_top60_DNA+RNA_seed | HDAC1 | 0.189704 | STAD | 23132-87 | PCI-34051 |
| MVP_top60_DNA_seed | PIK3CA | -0.887305 | LUAD | NCI-H1437 | Alpelisib | MVP_top60_DNA+RNA_seed | MDM2 | 0.193363 | STAD | RERF-GC-1B | Serdemetan |
| MVP_top60_DNA_seed | TERT | -0.886948 | LUAD | EMC-BAC-1 | BIBR-1532 | MVP_top60_DNA+RNA_seed | MET | 0.198369 | STAD | RF-48 | Crizotinib |

| | | | | | | | | | | | |
|---|---|---|---|---|---|---|---|---|---|---|---|
| MVP_top60_DNA_seed | HSP90AA1 | -0.886648 | LUAD | RERF-LC-MS | Elesclomol | MVP_top60_DNA+RNA_seed | HSP90AA1 | 0.208561 | STAD | KATOIII | Tanespimycin |
| MVP_top60_DNA_seed | PIK3CA | -0.885205 | LUAD | NCI-H322M | AZD8186 | MVP_top60_DNA+RNA_seed | HSP90AB1 | 0.208561 | STAD | KATOIII | Tanespimycin |
| MVP_top60_DNA_seed | FGFR1 | -0.885173 | LUAD | SK-LU-1 | PD173074 | MVP_top60_DNA+RNA_seed | EP300 | 0.209651 | STAD | SNU-1 | CPI-637 |
| MVP_top60_DNA_seed | HSP90AA1 | -0.884033 | LUAD | NCI-H2009 | Tanespimycin | MVP_top60_DNA+RNA_seed | ESR1 | 0.21113 | STAD | AGS | GDC0810 |
| MVP_top60_DNA_seed | GSK3B | -0.878616 | LUAD | Calu-6 | SB216763 | MVP_top60_DNA+RNA_seed | MDM2 | 0.22093 | STAD | SNU-16 | Nutlin-3a (-) |
| MVP_top60_DNA_seed | AKT1 | -0.876014 | LUAD | NCI-H1781 | Ipatasertib | MVP_top60_DNA+RNA_seed | MAPK1 | 0.221079 | STAD | NUGC-3 | VX-11e |
| MVP_top60_DNA_seed | AKT1 | -0.873638 | LUAD | NCI-H2030 | AT13148 | MVP_top60_DNA+RNA_seed | MAPK1 | 0.222337 | STAD | NUGC-4 | Ulixertinib |
| MVP_top60_DNA_seed | IGF1R | -0.873289 | LUAD | RERF-LC-MS | BMS-754807 | MVP_top60_DNA+RNA_seed | MAPK3 | 0.222337 | STAD | NUGC-4 | Ulixertinib |
| MVP_top60_DNA_seed | PIK3CA | -0.87246 | LUAD | NCI-H1437 | GNE-317 | MVP_top60_DNA+RNA_seed | PIK3CA | 0.222733 | STAD | SNU-5 | Alpelisib |
| MVP_top60_DNA_seed | EZH2 | -0.870602 | LUAD | RERF-LC-KJ | GSK343 | MVP_top60_DNA+RNA_seed | PARP1 | 0.226749 | STAD | SK-GT-2 | Veliparib |
| MVP_top60_DNA_seed | EGFR | -0.869284 | LUAD | EKVX | Erlotinib | MVP_top60_DNA+RNA_seed | HDAC1 | 0.22766 | STAD | OCUM-1 | PCI-34051 |
| MVP_top60_DNA_seed | KRAS | -0.868207 | LUAD | LC-2-ad | KRAS (G12C) Inhibitor-1 | MVP_top60_DNA+RNA_seed | PARP1 | 0.227916 | STAD | AGS | Talazoparib |
| MVP_top60_DNA_seed | MAPK1 | -0.864135 | LUAD | NCI-H1666 | Ulixertinib | MVP_top60_DNA+RNA_seed | EZH2 | 0.228916 | STAD | FU97 | GSK343 |
| MVP_top60_DNA_seed | IGF1R | -0.863634 | LUAD | A549 | BMS-536924 | MVP_top60_DNA+RNA_seed | MDM2 | 0.233225 | STAD | SNU-5 | Nutlin-3a (-) |
| MVP_top60_DNA_seed | AKT1 | -0.861856 | LUAD | NCI-H1944 | Uprosertib | MVP_top60_DNA+RNA_seed | MDM2 | 0.233994 | STAD | SNU-5 | Serdemetan |
| MVP_top60_DNA_seed | FGFR1 | -0.85912 | LUAD | RERF-LC-KJ | AZD4547 | MVP_top60_DNA+RNA_seed | SRC | 0.23633 | STAD | OCUM-1 | Bosutinib |
| MVP_top60_DNA_seed | HSP90AA1 | -0.859064 | LUAD | HCC-44 | Luminespib | MVP_top60_DNA+RNA_seed | PARP1 | 0.237019 | STAD | NUGC-3 | Niraparib |
| MVP_top60_DNA_seed | EGFR | -0.858883 | LUAD | RERF-LC-MS | Sapitinib | MVP_top60_DNA+RNA_seed | MDM2 | 0.242162 | STAD | AGS | Serdemetan |
| MVP_top60_DNA_seed | ERBB2 | -0.858883 | LUAD | RERF-LC-MS | Sapitinib | MVP_top60_DNA+RNA_seed | PARP1 | 0.245962 | STAD | SNU-5 | Niraparib |
| MVP_top60_DNA_seed | ERBB3 | -0.858883 | LUAD | RERF-LC-MS | Sapitinib | MVP_top60_DNA+RNA_seed | ERBB2 | 0.246427 | STAD | MKN7 | Afatinib |
| MVP_top60_DNA_seed | ATM | -0.85856 | LUAD | NCI-H1648 | KU-55933 | MVP_top60_DNA+RNA_seed | PIK3CA | 0.246863 | STAD | RF-48 | GNE-317 |
| MVP_top60_DNA_seed | AKT1 | -0.85829 | LUAD | NCI-H1975 | Uprosertib | MVP_top60_DNA+RNA_seed | HSP90AA1 | 0.248015 | STAD | FU97 | Tanespimycin |
| MVP_top60_DNA_seed | AKT1 | -0.857073 | LUAD | NCI-H3122 | GSK2110183B | MVP_top60_DNA+RNA_seed | HSP90AB1 | 0.248015 | STAD | FU97 | Tanespimycin |
| MVP_top60_DNA_seed | KRAS | -0.854987 | LUAD | NCI-H1792 | KRAS (G12C) Inhibitor-1 | MVP_top60_DNA+RNA_seed | ESR1 | 0.248251 | STAD | SNU-1 | GDC0810 |
| MVP_top60_DNA_seed | PIK3CA | -0.854827 | LUAD | NCI-H2030 | AZD8186 | MVP_top60_DNA+RNA_seed | SRC | 0.250155 | STAD | SK-GT-2 | Bosutinib |
| MVP_top60_DNA_seed | MAPK1 | -0.849033 | LUAD | NCI-H1666 | SCH772984 | MVP_top60_DNA+RNA_seed | GSK3B | 0.252279 | STAD | MKN45 | SB216763 |
| MVP_top60_DNA_seed | EGFR | -0.848727 | LUAD | A549 | Lapatinib | MVP_top60_DNA+RNA_seed | PARP1 | 0.258452 | STAD | 23132-87 | Olaparib |
| MVP_top60_DNA_seed | ERBB2 | -0.848727 | LUAD | A549 | Lapatinib | MVP_top60_DNA+RNA_seed | MET | 0.258961 | STAD | 23132-87 | Savolitinib |
| MVP_top60_DNA_seed | EGFR | -0.846237 | LUAD | NCI-H1650 | Sapitinib | MVP_top60_DNA+RNA_seed | EZH2 | 0.260477 | STAD | NUGC-3 | GSK343 |
| MVP_top60_DNA_seed | ERBB2 | -0.846237 | LUAD | NCI-H1650 | Sapitinib | MVP_top60_DNA+RNA_seed | HDAC1 | 0.260893 | STAD | NUGC-3 | PCI-34051 |
| MVP_top60_DNA_seed | ERBB3 | -0.846237 | LUAD | NCI-H1650 | Sapitinib | MVP_top60_DNA+RNA_seed | ERBB2 | 0.261371 | STAD | RF-48 | Afatinib |
| MVP_top60_DNA_seed | BRAF | -0.845527 | LUAD | NCI-H3122 | SB590885 | MVP_top60_DNA+RNA_seed | PIK3CA | 0.261699 | STAD | NUGC-3 | AZD8186 |
| MVP_top60_DNA_seed | AKT1 | -0.842603 | LUAD | EKVX | AT13148 | MVP_top60_DNA+RNA_seed | FGFR3 | 0.261821 | STAD | MKN28 | PD173074 |
| MVP_top60_DNA_seed | AKT1 | -0.84054 | LUAD | NCI-H1648 | Ipatasertib | MVP_top60_DNA+RNA_seed | MAPK1 | 0.265918 | STAD | NCI-N87 | VX-11e |
| MVP_top60_DNA_seed | AKT1 | -0.840355 | LUAD | NCI-H1734 | MK-2206 | MVP_top60_DNA+RNA_seed | MET | 0.276314 | STAD | MKN28 | Crizotinib |
| MVP_top60_DNA_seed | IGF1R | -0.839758 | LUAD | HCC-78 | NVP-ADW742 | MVP_top60_DNA+RNA_seed | EZH2 | 0.276845 | STAD | RF-48 | GSK343 |
| MVP_top60_DNA_seed | HDAC1 | -0.838397 | LUAD | NCI-H1944 | Entinostat | MVP_top60_DNA+RNA_seed | TP53 | 0.278515 | STAD | TMK-1 | MIRA-1 |
| MVP_top60_DNA_seed | MAPK1 | -0.835191 | LUAD | NCI-H1755 | Ulixertinib | MVP_top60_DNA+RNA_seed | HSP90AA1 | 0.287888 | STAD | TMK-1 | Luminespib |
| MVP_top60_DNA_seed | PIK3CA | -0.834628 | LUAD | EMC-BAC-2 | AZD8186 | MVP_top60_DNA+RNA_seed | HSP90AB1 | 0.287888 | STAD | TMK-1 | Luminespib |
| MVP_top60_DNA_seed | PIK3CA | -0.832469 | LUAD | NCI-H1944 | Buparlisib | MVP_top60_DNA+RNA_seed | SRC | 0.292812 | STAD | MKN28 | Bosutinib |
| MVP_top60_DNA_seed | PIK3CA | -0.829341 | LUAD | NCI-H1993 | AZD8186 | MVP_top60_DNA+RNA_seed | PARP1 | 0.293604 | STAD | MKN45 | Niraparib |

| | | | | | | | | | | | |
|---|---|---|---|---|---|---|---|---|---|---|---|
| MVP_top60_DNA_seed | PIK3CA | -0.826319 | LUAD | EMC-BAC-2 | Alpelisib | MVP_top60_DNA+RNA_seed | ATM | 0.293658 | STAD | NUGC-3 | KU-55933 |
| MVP_top60_DNA_seed | AKT1 | -0.826315 | LUAD | HCC-44 | Uprosertib | MVP_top60_DNA+RNA_seed | PARP1 | 0.29819 | STAD | SNU-5 | Olaparib |
| MVP_top60_DNA_seed | EGFR | -0.826297 | LUAD | EKVX | Afatinib | MVP_top60_DNA+RNA_seed | ATM | 0.298397 | STAD | IM-95 | KU-55933 |
| MVP_top60_DNA_seed | ERBB2 | -0.826297 | LUAD | EKVX | Afatinib | MVP_top60_DNA+RNA_seed | PIK3CA | 0.299203 | STAD | MKN7 | Alpelisib |
| MVP_top60_DNA_seed | HDAC1 | -0.826045 | LUAD | NCI-H838 | Entinostat | MVP_top60_DNA+RNA_seed | HSP90AA1 | 0.299986 | STAD | MKN28 | Luminespib |
| MVP_top60_DNA_seed | KRAS | -0.825506 | LUAD | NCI-H1437 | KRAS (G12C) Inhibitor-12 | MVP_top60_DNA+RNA_seed | HSP90AB1 | 0.299986 | STAD | MKN28 | Luminespib |
| MVP_top60_DNA_seed | IGF1R | -0.823601 | LUAD | NCI-H1792 | Linsitinib | MVP_top60_DNA+RNA_seed | MET | 0.301869 | STAD | TGBC11TKB | Crizotinib |
| MVP_top60_DNA_seed | IGF1R | -0.821244 | LUAD | NCI-H3122 | BMS-536924 | MVP_top60_DNA+RNA_seed | ERBB2 | 0.305537 | STAD | GCIY | Lapatinib |
| MVP_top60_DNA_seed | HDAC1 | -0.819623 | LUAD | NCI-H1993 | PCI-34051 | MVP_top60_DNA+RNA_seed | ESR1 | 0.310481 | STAD | SNU-5 | GDC0810 |
| MVP_top60_DNA_seed | PIK3CA | -0.81888 | LUAD | NCI-H2085 | AZD8186 | MVP_top60_DNA+RNA_seed | MAPK1 | 0.310979 | STAD | MKN7 | VX-11e |
| MVP_top60_DNA_seed | IGF1R | -0.818014 | LUAD | RERF-LC-MS | GSK1904529A | MVP_top60_DNA+RNA_seed | ERBB2 | 0.312197 | STAD | TMK-1 | Lapatinib |
| MVP_top60_DNA_seed | IGF1R | -0.815831 | LUAD | NCI-H838 | Linsitinib | MVP_top60_DNA+RNA_seed | PARP1 | 0.31232 | STAD | GCIY | Niraparib |
| MVP_top60_DNA_seed | HDAC1 | -0.811205 | LUAD | NCI-H1734 | Entinostat | MVP_top60_DNA+RNA_seed | EP300 | 0.318274 | STAD | RF-48 | CPI-637 |
| MVP_top60_DNA_seed | HSP90AA1 | -0.808601 | LUAD | NCI-H1993 | Tanespimycin | MVP_top60_DNA+RNA_seed | HDAC1 | 0.321258 | STAD | SK-GT-2 | PCI-34051 |
| MVP_top60_DNA_seed | IGF1R | -0.807448 | LUAD | NCI-H1734 | NVP-ADW742 | MVP_top60_DNA+RNA_seed | MDM2 | 0.329854 | STAD | RERF-GC-1B | Nutlin-3a (-) |
| MVP_top60_DNA_seed | EGFR | -0.806559 | LUAD | NCI-H1568 | Afatinib | MVP_top60_DNA+RNA_seed | HSP90AA1 | 0.340552 | STAD | IM-95 | Luminespib |
| MVP_top60_DNA_seed | ERBB2 | -0.806559 | LUAD | NCI-H1568 | Afatinib | MVP_top60_DNA+RNA_seed | HSP90AB1 | 0.340552 | STAD | IM-95 | Luminespib |
| MVP_top60_DNA_seed | SRC | -0.805906 | LUAD | NCI-H1792 | Bosutinib | MVP_top60_DNA+RNA_seed | MET | 0.343877 | STAD | TMK-1 | Savolitinib |
| MVP_top60_DNA_seed | MDM2 | -0.804969 | LUAD | EMC-BAC-2 | Nutlin-3a (-) | MVP_top60_DNA+RNA_seed | ESR1 | 0.34605 | STAD | MKN45 | GDC0810 |
| MVP_top60_DNA_seed | HSP90AA1 | -0.804088 | LUAD | PC-14 | Luminespib | MVP_top60_DNA+RNA_seed | TP53 | 0.34861 | STAD | SNU-1 | MIRA-1 |
| MVP_top60_DNA_seed | EGFR | -0.798745 | LUAD | NCI-H2030 | Lapatinib | MVP_top60_DNA+RNA_seed | MAPK1 | 0.358626 | STAD | KATOIII | VX-11e |
| MVP_top60_DNA_seed | ERBB2 | -0.798745 | LUAD | NCI-H2030 | Lapatinib | MVP_top60_DNA+RNA_seed | PARP1 | 0.359144 | STAD | RERF-GC-1B | Talazoparib |
| MVP_top60_DNA_seed | MDM2 | -0.794131 | LUAD | PC-14 | Serdemetan | MVP_top60_DNA+RNA_seed | MAPK1 | 0.359989 | STAD | HGC-27 | VX-11e |
| MVP_top60_DNA_seed | AKT1 | -0.794013 | LUAD | NCI-H2228 | Uprosertib | MVP_top60_DNA+RNA_seed | PARP1 | 0.36064 | STAD | TMK-1 | Veliparib |
| MVP_top60_DNA_seed | HSP90AA1 | -0.793277 | LUAD | EMC-BAC-2 | Luminespib | MVP_top60_DNA+RNA_seed | ATM | 0.363082 | STAD | SK-GT-2 | KU-55933 |
| MVP_top60_DNA_seed | PIK3CA | -0.792181 | LUAD | NCI-H838 | AZD8186 | MVP_top60_DNA+RNA_seed | MAPK1 | 0.368653 | STAD | NCI-N87 | Ulixertinib |
| MVP_top60_DNA_seed | HDAC1 | -0.789287 | LUAD | NCI-H1975 | Romidepsin | MVP_top60_DNA+RNA_seed | MAPK3 | 0.368653 | STAD | NCI-N87 | Ulixertinib |
| MVP_top60_DNA_seed | AKT1 | -0.788663 | LUAD | RERF-LC-KJ | Afuresertib | MVP_top60_DNA+RNA_seed | PIK3CA | 0.36905 | STAD | MKN28 | Alpelisib |
| MVP_top60_DNA_seed | AKT1 | -0.788326 | LUAD | NCI-H1993 | GSK2110183B | MVP_top60_DNA+RNA_seed | MAPK1 | 0.37124 | STAD | SNU-1 | SCH772984 |
| MVP_top60_DNA_seed | EGFR | -0.787682 | LUAD | NCI-H358 | Osimertinib | MVP_top60_DNA+RNA_seed | MAPK3 | 0.37124 | STAD | SNU-1 | SCH772984 |
| MVP_top60_DNA_seed | EGFR | -0.787111 | LUAD | NCI-H1944 | Erlotinib | MVP_top60_DNA+RNA_seed | MAPK1 | 0.374746 | STAD | MKN7 | SCH772984 |
| MVP_top60_DNA_seed | EGFR | -0.784739 | LUAD | NCI-H1734 | AZD3759 | MVP_top60_DNA+RNA_seed | MAPK3 | 0.374746 | STAD | MKN7 | SCH772984 |
| MVP_top60_DNA_seed | PIK3CA | -0.784471 | LUAD | NCI-H2291 | AZD8186 | MVP_top60_DNA+RNA_seed | HDAC1 | 0.375703 | STAD | OCUM-1 | Entinostat |
| MVP_top60_DNA_seed | KRAS | -0.781584 | LUAD | NCI-H1993 | KRAS (G12C) Inhibitor-12 | MVP_top60_DNA+RNA_seed | HDAC3 | 0.375703 | STAD | OCUM-1 | Entinostat |
| MVP_top60_DNA_seed | AKT1 | -0.781304 | LUAD | EMC-BAC-2 | AZD5363 | MVP_top60_DNA+RNA_seed | FGFR3 | 0.377844 | STAD | Hs-746T | PD173074 |
| MVP_top60_DNA_seed | BRAF | -0.780712 | LUAD | Calu-6 | PLX-4720 | MVP_top60_DNA+RNA_seed | PARP1 | 0.379974 | STAD | HGC-27 | Niraparib |
| MVP_top60_DNA_seed | ATM | -0.779707 | LUAD | HCC-44 | KU-55933 | MVP_top60_DNA+RNA_seed | MET | 0.381507 | STAD | FU97 | Crizotinib |
| MVP_top60_DNA_seed | MAPK1 | -0.779643 | LUAD | NCI-H1666 | VX-11e | MVP_top60_DNA+RNA_seed | TP53 | 0.383155 | STAD | SNU-5 | MIRA-1 |
| MVP_top60_DNA_seed | EGFR | -0.776529 | LUAD | LXF-289 | Lapatinib | MVP_top60_DNA+RNA_seed | PARP1 | 0.39381 | STAD | NUGC-3 | Olaparib |
| MVP_top60_DNA_seed | ERBB2 | -0.776529 | LUAD | LXF-289 | Lapatinib | MVP_top60_DNA+RNA_seed | ERBB2 | 0.396143 | STAD | MKN45 | Lapatinib |
| MVP_top60_DNA_seed | PIK3CA | -0.775777 | LUAD | NCI-H1792 | Buparlisib | MVP_top60_DNA+RNA_seed | PIK3CA | 0.396228 | STAD | MKN28 | Buparlisib |

| | | | | | | | | | | | |
|---|---|---|---|---|---|---|---|---|---|---|---|
| MVP_top60_DNA_seed | SRC | -0.775305 | LUAD | NCI-H838 | Bosutinib | MVP_top60_DNA+RNA_seed | SRC | 0.39658 | STAD | FU97 | Bosutinib |
| MVP_top60_DNA_seed | HDAC1 | -0.775067 | LUAD | HCC-78 | PCI-34051 | MVP_top60_DNA+RNA_seed | GSK3B | 0.402097 | STAD | TMK-1 | SB216763 |
| MVP_top60_DNA_seed | HSP90AA1 | -0.774418 | LUAD | EMC-BAC-1 | Luminespib | MVP_top60_DNA+RNA_seed | MAPK1 | 0.402851 | STAD | SK-GT-2 | Ulixertinib |
| MVP_top60_DNA_seed | EGFR | -0.774197 | LUAD | EMC-BAC-2 | Osimertinib | MVP_top60_DNA+RNA_seed | MAPK3 | 0.402851 | STAD | SK-GT-2 | Ulixertinib |
| MVP_top60_DNA_seed | IGF1R | -0.773649 | LUAD | NCI-H1734 | Linsitinib | MVP_top60_DNA+RNA_seed | EP300 | 0.405531 | STAD | Hs-746T | CPI-637 |
| MVP_top60_DNA_seed | HDAC1 | -0.772582 | LUAD | NCI-H838 | PCI-34051 | MVP_top60_DNA+RNA_seed | TP53 | 0.406069 | STAD | RERF-GC-1B | MIRA-1 |
| MVP_top60_DNA_seed | PIK3CA | -0.769115 | LUAD | HCC-78 | Buparlisib | MVP_top60_DNA+RNA_seed | MAPK1 | 0.413718 | STAD | SNU-1 | VX-11e |
| MVP_top60_DNA_seed | TERT | -0.768401 | LUAD | NCI-H2122 | BIBR-1532 | MVP_top60_DNA+RNA_seed | PIK3CA | 0.415397 | STAD | MKN45 | AZD8186 |
| MVP_top60_DNA_seed | PIK3CA | -0.768246 | LUAD | NCI-H838 | GNE-317 | MVP_top60_DNA+RNA_seed | ERBB2 | 0.41743 | STAD | IM-95 | Lapatinib |
| MVP_top60_DNA_seed | EGFR | -0.768056 | LUAD | NCI-H2122 | Afatinib | MVP_top60_DNA+RNA_seed | PARP1 | 0.420565 | STAD | NCI-N87 | Talazoparib |
| MVP_top60_DNA_seed | ERBB2 | -0.768056 | LUAD | NCI-H2122 | Afatinib | MVP_top60_DNA+RNA_seed | MET | 0.424269 | STAD | NUGC-3 | Crizotinib |
| MVP_top60_DNA_seed | SRC | -0.766975 | LUAD | NCI-H1755 | Bosutinib | MVP_top60_DNA+RNA_seed | MAPK1 | 0.424381 | STAD | FU97 | SCH772984 |
| MVP_top60_DNA_seed | EGFR | -0.76574 | LUAD | NCI-H2030 | Osimertinib | MVP_top60_DNA+RNA_seed | MAPK3 | 0.424381 | STAD | FU97 | SCH772984 |
| MVP_top60_DNA_seed | EGFR | -0.763156 | LUAD | EKVX | Lapatinib | MVP_top60_DNA+RNA_seed | MAPK1 | 0.425057 | STAD | HGC-27 | SCH772984 |
| MVP_top60_DNA_seed | ERBB2 | -0.763156 | LUAD | EKVX | Lapatinib | MVP_top60_DNA+RNA_seed | MAPK3 | 0.425057 | STAD | HGC-27 | SCH772984 |
| MVP_top60_DNA_seed | HSP90AA1 | -0.76203 | LUAD | LC-2-ad | CCT-018159 | MVP_top60_DNA+RNA_seed | HDAC1 | 0.429843 | STAD | KATOIII | Entinostat |
| MVP_top60_DNA_seed | HDAC1 | -0.761948 | LUAD | NCI-H1623 | Entinostat | MVP_top60_DNA+RNA_seed | HDAC3 | 0.429843 | STAD | KATOIII | Entinostat |
| MVP_top60_DNA_seed | HSP90AA1 | -0.761902 | LUAD | NCI-H1975 | Tanespimycin | MVP_top60_DNA+RNA_seed | TP53 | 0.433549 | STAD | SK-GT-2 | MIRA-1 |
| MVP_top60_DNA_seed | MDM2 | -0.76162 | LUAD | NCI-H1944 | Nutlin-3a (-) | MVP_top60_DNA+RNA_seed | FGFR3 | 0.438476 | STAD | FU97 | AZD4547 |
| MVP_top60_DNA_seed | IGF1R | -0.760901 | LUAD | NCI-H1355 | Linsitinib | MVP_top60_DNA+RNA_seed | PARP1 | 0.440392 | STAD | RERF-GC-1B | Veliparib |
| MVP_top60_DNA_seed | EGFR | -0.760463 | LUAD | NCI-H2122 | Lapatinib | MVP_top60_DNA+RNA_seed | PARP1 | 0.441055 | STAD | OCUM-1 | Talazoparib |
| MVP_top60_DNA_seed | ERBB2 | -0.760463 | LUAD | NCI-H2122 | Lapatinib | MVP_top60_DNA+RNA_seed | PIK3CA | 0.442209 | STAD | GCIY | Alpelisib |
| MVP_top60_DNA_seed | EGFR | -0.758627 | LUAD | NCI-H2122 | AZD3759 | MVP_top60_DNA+RNA_seed | EZH2 | 0.447403 | STAD | SK-GT-2 | GSK343 |
| MVP_top60_DNA_seed | MAPK1 | -0.756574 | LUAD | A549 | Ulixertinib | MVP_top60_DNA+RNA_seed | EP300 | 0.454674 | STAD | SNU-1 | SGC-CBP30 |
| MVP_top60_DNA_seed | KRAS | -0.75525 | LUAD | NCI-H23 | KRAS (G12C) Inhibitor-1 | MVP_top60_DNA+RNA_seed | CREBBP | 0.454674 | STAD | SNU-1 | SGC-CBP30 |
| MVP_top60_DNA_seed | PIK3CA | -0.755019 | LUAD | EMC-BAC-1 | Alpelisib | MVP_top60_DNA+RNA_seed | KRAS | 0.457661 | STAD | HGC-27 | RAS (G12C) Inhibitor-1 |
| MVP_top60_DNA_seed | SRC | -0.753149 | LUAD | NCI-H2030 | Bosutinib | MVP_top60_DNA+RNA_seed | EP300 | 0.461379 | STAD | MKN28 | SGC-CBP30 |
| MVP_top60_DNA_seed | IGF1R | -0.752205 | LUAD | NCI-H1792 | NVP-ADW742 | MVP_top60_DNA+RNA_seed | CREBBP | 0.461379 | STAD | MKN28 | SGC-CBP30 |
| MVP_top60_DNA_seed | PIK3CA | -0.752126 | LUAD | NCI-H838 | Alpelisib | MVP_top60_DNA+RNA_seed | EZH2 | 0.466528 | STAD | TMK-1 | GSK343 |
| MVP_top60_DNA_seed | AURKA | -0.749448 | LUAD | NCI-H1437 | ZM447439 | MVP_top60_DNA+RNA_seed | MAPK1 | 0.468294 | STAD | GCIY | SCH772984 |
| MVP_top60_DNA_seed | MET | -0.748162 | LUAD | NCI-H2023 | Savolitinib | MVP_top60_DNA+RNA_seed | MAPK3 | 0.468294 | STAD | GCIY | SCH772984 |
| MVP_top60_DNA_seed | IGF1R | -0.747969 | LUAD | HCC-44 | Linsitinib | MVP_top60_DNA+RNA_seed | PARP1 | 0.468892 | STAD | MKN28 | Rucaparib |
| MVP_top60_DNA_seed | IGF1R | -0.747464 | LUAD | RERF-LC-KJ | GSK1904529A | MVP_top60_DNA+RNA_seed | EP300 | 0.471067 | STAD | NUGC-3 | CPI-637 |
| MVP_top60_DNA_seed | MET | -0.744948 | LUAD | HCC-44 | Crizotinib | MVP_top60_DNA+RNA_seed | FGFR3 | 0.474056 | STAD | 23132-87 | PD173074 |
| MVP_top60_DNA_seed | PIK3CA | -0.74273 | LUAD | NCI-H2023 | GNE-317 | MVP_top60_DNA+RNA_seed | EP300 | 0.476173 | STAD | MKN45 | CPI-637 |
| MVP_top60_DNA_seed | HDAC1 | -0.741586 | LUAD | NCI-H522 | PCI-34051 | MVP_top60_DNA+RNA_seed | ERBB2 | 0.479741 | STAD | TGBC11TKB | Sapitinib |
| MVP_top60_DNA_seed | AKT1 | -0.737062 | LUAD | NCI-H1648 | MK-2206 | MVP_top60_DNA+RNA_seed | ERBB3 | 0.479741 | STAD | TGBC11TKB | Sapitinib |
| MVP_top60_DNA_seed | IGF1R | -0.73514 | LUAD | ABC-1 | BMS-754807 | MVP_top60_DNA+RNA_seed | ATM | 0.479952 | STAD | 23132-87 | KU-55933 |
| MVP_top60_DNA_seed | HSP90AA1 | -0.734229 | LUAD | NCI-H2030 | Tanespimycin | MVP_top60_DNA+RNA_seed | FGFR3 | 0.481047 | STAD | IM-95 | PD173074 |
| MVP_top60_DNA_seed | PIK3CA | -0.732447 | LUAD | NCI-H3122 | Alpelisib | MVP_top60_DNA+RNA_seed | PIK3CA | 0.481446 | STAD | RERF-GC-1B | AZD8186 |
| MVP_top60_DNA_seed | AKT1 | -0.732278 | LUAD | NCI-H1648 | Uprosertib | MVP_top60_DNA+RNA_seed | MDM2 | 0.484299 | STAD | NUGC-4 | Nutlin-3a (-) |

| | | | | | | | | | | | |
|---|---|---|---|---|---|---|---|---|---|---|---|
| MVP_top60_DNA_seed | MAPK1 | -0.73216 | LUAD | NCI-H358 | SCH772984 | MVP_top60_DNA+RNA_seed | MET | 0.491468 | STAD | NUGC-3 | Savolitinib |
| MVP_top60_DNA_seed | TP53 | -0.730298 | LUAD | NCI-H1435 | MIRA-1 | MVP_top60_DNA+RNA_seed | ATM | 0.494666 | STAD | TMK-1 | KU-55933 |
| MVP_top60_DNA_seed | PIK3CA | -0.72943 | LUAD | HCC-44 | Buparlisib | MVP_top60_DNA+RNA_seed | HDAC1 | 0.495248 | STAD | IM-95 | PCI-34051 |
| MVP_top60_DNA_seed | EGFR | -0.729181 | LUAD | NCI-H1975 | Erlotinib | MVP_top60_DNA+RNA_seed | HDAC1 | 0.497644 | STAD | RERF-GC-1B | Entinostat |
| MVP_top60_DNA_seed | AKT1 | -0.728317 | LUAD | NCI-H2122 | MK-2206 | MVP_top60_DNA+RNA_seed | HDAC3 | 0.497644 | STAD | RERF-GC-1B | Entinostat |
| MVP_top60_DNA_seed | GSK3B | -0.726963 | LUAD | NCI-H1755 | SB216763 | MVP_top60_DNA+RNA_seed | GSK3B | 0.499467 | STAD | NUGC-3 | SB216763 |
| MVP_top60_DNA_seed | PIK3CA | -0.726793 | LUAD | NCI-H1975 | Alpelisib | MVP_top60_DNA+RNA_seed | PARP1 | 0.501902 | STAD | MKN45 | Talazoparib |
| MVP_top60_DNA_seed | MET | -0.725844 | LUAD | NCI-H1755 | Crizotinib | MVP_top60_DNA+RNA_seed | PARP1 | 0.510591 | STAD | NUGC-4 | Niraparib |
| MVP_top60_DNA_seed | EGFR | -0.725677 | LUAD | NCI-H1693 | Afatinib | MVP_top60_DNA+RNA_seed | ERBB2 | 0.512615 | STAD | KATOIII | Sapitinib |
| MVP_top60_DNA_seed | ERBB2 | -0.725677 | LUAD | NCI-H1693 | Afatinib | MVP_top60_DNA+RNA_seed | ERBB3 | 0.512615 | STAD | KATOIII | Sapitinib |
| MVP_top60_DNA_seed | FGFR1 | -0.7251 | LUAD | NCI-H322M | PD173074 | MVP_top60_DNA+RNA_seed | ESR1 | 0.516392 | STAD | MKN28 | GDC0810 |
| MVP_top60_DNA_seed | IGF1R | -0.724846 | LUAD | A549 | GSK1904529A | MVP_top60_DNA+RNA_seed | FGFR3 | 0.521063 | STAD | OCUM-1 | PD173074 |
| MVP_top60_DNA_seed | EGFR | -0.723065 | LUAD | NCI-H1437 | Gefitinib | MVP_top60_DNA+RNA_seed | MDM2 | 0.53308 | STAD | 23132-87 | Serdemetan |
| MVP_top60_DNA_seed | AKT1 | -0.722518 | LUAD | RERF-LC-MS | AT13148 | MVP_top60_DNA+RNA_seed | GSK3B | 0.533731 | STAD | MKN7 | CHIR-99021 |
| MVP_top60_DNA_seed | GSK3B | -0.722291 | LUAD | NCI-H650 | SB216763 | MVP_top60_DNA+RNA_seed | EP300 | 0.53914 | STAD | SK-GT-2 | SGC-CBP30 |
| MVP_top60_DNA_seed | HSP90AA1 | -0.722228 | LUAD | NCI-H838 | Tanespimycin | MVP_top60_DNA+RNA_seed | CREBBP | 0.53914 | STAD | SK-GT-2 | SGC-CBP30 |
| MVP_top60_DNA_seed | EGFR | -0.721807 | LUAD | NCI-H1693 | Erlotinib | MVP_top60_DNA+RNA_seed | HDAC1 | 0.541107 | STAD | GCIY | Romidepsin |
| MVP_top60_DNA_seed | AKT1 | -0.721749 | LUAD | NCI-H1993 | Uprosertib | MVP_top60_DNA+RNA_seed | HDAC3 | 0.541107 | STAD | GCIY | Romidepsin |
| MVP_top60_DNA_seed | EZH2 | -0.721317 | LUAD | NCI-H522 | GSK343 | MVP_top60_DNA+RNA_seed | FGFR3 | 0.551204 | STAD | TGBC11TKB | AZD4547 |
| MVP_top60_DNA_seed | MET | -0.721098 | LUAD | EMC-BAC-1 | Crizotinib | MVP_top60_DNA+RNA_seed | PARP1 | 0.5526 | STAD | GCIY | Olaparib |
| MVP_top60_DNA_seed | HSP90AA1 | -0.720347 | LUAD | NCI-H1666 | Tanespimycin | MVP_top60_DNA+RNA_seed | MAPK1 | 0.553461 | STAD | IM-95 | VX-11e |
| MVP_top60_DNA_seed | EGFR | -0.72027 | LUAD | NCI-H2030 | Erlotinib | MVP_top60_DNA+RNA_seed | FGFR3 | 0.555995 | STAD | MKN45 | PD173074 |
| MVP_top60_DNA_seed | MET | -0.719839 | LUAD | LXF-289 | Savolitinib | MVP_top60_DNA+RNA_seed | HDAC1 | 0.564163 | STAD | SNU-1 | Entinostat |
| MVP_top60_DNA_seed | PIK3CA | -0.71925 | LUAD | NCI-H1563 | AZD8186 | MVP_top60_DNA+RNA_seed | HDAC3 | 0.564163 | STAD | SNU-1 | Entinostat |
| MVP_top60_DNA_seed | EZH2 | -0.71858 | LUAD | PC-14 | GSK343 | MVP_top60_DNA+RNA_seed | ESR1 | 0.574937 | STAD | Hs-746T | Tamoxifen |
| MVP_top60_DNA_seed | PIK3CA | -0.71695 | LUAD | A549 | Buparlisib | MVP_top60_DNA+RNA_seed | GSK3B | 0.578226 | STAD | TGBC11TKB | SB216763 |
| MVP_top60_DNA_seed | IGF1R | -0.716016 | LUAD | NCI-H358 | Linsitinib | MVP_top60_DNA+RNA_seed | GSK3B | 0.583817 | STAD | NCI-N87 | SB216763 |
| MVP_top60_DNA_seed | IGF1R | -0.715039 | LUAD | PC-14 | BMS-536924 | MVP_top60_DNA+RNA_seed | MAPK1 | 0.585739 | STAD | NUGC-4 | Ulixertinib |
| MVP_top60_DNA_seed | HDAC1 | -0.714076 | LUAD | NCI-H2023 | Romidepsin | MVP_top60_DNA+RNA_seed | MAPK3 | 0.585739 | STAD | NUGC-4 | Ulixertinib |
| MVP_top60_DNA_seed | IGF1R | -0.713322 | LUAD | NCI-H1755 | NVP-ADW742 | MVP_top60_DNA+RNA_seed | TP53 | 0.588419 | STAD | TGBC11TKB | MIRA-1 |
| MVP_top60_DNA_seed | EZH2 | -0.709605 | LUAD | RERF-LC-MS | GSK343 | MVP_top60_DNA+RNA_seed | EP300 | 0.590533 | STAD | MKN45 | SGC-CBP30 |
| MVP_top60_DNA_seed | IGF1R | -0.709093 | LUAD | RERF-LC-KJ | BMS-536924 | MVP_top60_DNA+RNA_seed | CREBBP | 0.590533 | STAD | MKN45 | SGC-CBP30 |
| MVP_top60_DNA_seed | EZH2 | -0.709046 | LUAD | NCI-H1355 | GSK343 | MVP_top60_DNA+RNA_seed | MDM2 | 0.590792 | STAD | TMK-1 | Nutlin-3a (-) |
| MVP_top60_DNA_seed | EGFR | -0.707173 | LUAD | NCI-H1781 | Lapatinib | MVP_top60_DNA+RNA_seed | EP300 | 0.596401 | STAD | GCIY | CPI-637 |
| MVP_top60_DNA_seed | ERBB2 | -0.707173 | LUAD | NCI-H1781 | Lapatinib | MVP_top60_DNA+RNA_seed | PIK3CA | 0.598927 | STAD | SNU-1 | AZD8186 |
| MVP_top60_DNA_seed | EGFR | -0.704192 | LUAD | NCI-H2122 | Osimertinib | MVP_top60_DNA+RNA_seed | EP300 | 0.599162 | STAD | NCI-N87 | SGC-CBP30 |
| MVP_top60_DNA_seed | AKT1 | -0.704059 | LUAD | NCI-H3122 | MK-2206 | MVP_top60_DNA+RNA_seed | CREBBP | 0.599162 | STAD | NCI-N87 | SGC-CBP30 |
| MVP_top60_DNA_seed | AKT1 | -0.701972 | LUAD | A549 | Uprosertib | MVP_top60_DNA+RNA_seed | ERBB2 | 0.601589 | STAD | Hs-746T | Afatinib |
| MVP_top60_DNA_seed | EGFR | -0.701649 | LUAD | NCI-H2122 | Sapitinib | MVP_top60_DNA+RNA_seed | ERBB2 | 0.603182 | STAD | RF-48 | Sapitinib |
| MVP_top60_DNA_seed | ERBB2 | -0.701649 | LUAD | NCI-H2122 | Sapitinib | MVP_top60_DNA+RNA_seed | ERBB3 | 0.603182 | STAD | RF-48 | Sapitinib |
| MVP_top60_DNA_seed | ERBB3 | -0.701649 | LUAD | NCI-H2122 | Sapitinib | MVP_top60_DNA+RNA_seed | HDAC1 | 0.607904 | STAD | HGC-27 | Entinostat |

| | | | | | | | | | | | |
|---|---|---|---|---|---|---|---|---|---|---|---|
| MVP_top60_DNA_seed | MAPK1 | -0.700037 | LUAD | LC-2-ad | VX-11e | MVP_top60_DNA+RNA_seed | HDAC3 | 0.607904 | STAD | HGC-27 | Entinostat |
| MVP_top60_DNA_seed | MET | -0.699143 | LUAD | NCI-H2023 | Crizotinib | MVP_top60_DNA+RNA_seed | SRC | 0.612194 | STAD | KATOIII | Bosutinib |
| MVP_top60_DNA_seed | BRAF | -0.697119 | LUAD | EMC-BAC-1 | PLX-4720 | MVP_top60_DNA+RNA_seed | ESR1 | 0.613222 | STAD | SNU-1 | Tamoxifen |
| MVP_top60_DNA_seed | PIK3CA | -0.696165 | LUAD | LC-2-ad | Alpelisib | MVP_top60_DNA+RNA_seed | PARP1 | 0.613311 | STAD | Hs-746T | Talazoparib |
| MVP_top60_DNA_seed | AKT1 | -0.695581 | LUAD | HCC-44 | Ipatasertib | MVP_top60_DNA+RNA_seed | PARP1 | 0.616044 | STAD | RERF-GC-1B | Rucaparib |
| MVP_top60_DNA_seed | MDM2 | -0.694639 | LUAD | HCC-44 | Serdemetan | MVP_top60_DNA+RNA_seed | PIK3CA | 0.616749 | STAD | RERF-GC-1B | Buparlisib |
| MVP_top60_DNA_seed | IGF1R | -0.694518 | LUAD | NCI-H2228 | BMS-754807 | MVP_top60_DNA+RNA_seed | MET | 0.617708 | STAD | KATOIII | Crizotinib |
| MVP_top60_DNA_seed | IGF1R | -0.69344 | LUAD | RERF-LC-KJ | NVP-ADW742 | MVP_top60_DNA+RNA_seed | MET | 0.619445 | STAD | SK-GT-2 | Crizotinib |
| MVP_top60_DNA_seed | AURKA | -0.692987 | LUAD | NCI-H1755 | ZM447439 | MVP_top60_DNA+RNA_seed | ESR1 | 0.623097 | STAD | MKN7 | GDC0810 |
| MVP_top60_DNA_seed | MAPK1 | -0.692314 | LUAD | NCI-H2405 | Ulixertinib | MVP_top60_DNA+RNA_seed | ESR1 | 0.633735 | STAD | MKN45 | Tamoxifen |
| MVP_top60_DNA_seed | PIK3CA | -0.688654 | LUAD | NCI-H1755 | Alpelisib | MVP_top60_DNA+RNA_seed | PARP1 | 0.6348 | STAD | NCI-N87 | Olaparib |
| MVP_top60_DNA_seed | GSK3B | -0.688475 | LUAD | LXF-289 | SB216763 | MVP_top60_DNA+RNA_seed | MDM2 | 0.642 | STAD | IM-95 | Serdemetan |
| MVP_top60_DNA_seed | AKT1 | -0.687777 | LUAD | NCI-H2122 | Uprosertib | MVP_top60_DNA+RNA_seed | MET | 0.642108 | STAD | GCIY | Crizotinib |
| MVP_top60_DNA_seed | BRAF | -0.687325 | LUAD | NCI-H1648 | PLX-4720 | MVP_top60_DNA+RNA_seed | PARP1 | 0.655058 | STAD | Hs-746T | Niraparib |
| MVP_top60_DNA_seed | BRAF | -0.683802 | LUAD | RERF-LC-MS | Dabrafenib | MVP_top60_DNA+RNA_seed | PARP1 | 0.658834 | STAD | SK-GT-2 | Niraparib |
| MVP_top60_DNA_seed | BRAF | -0.68222 | LUAD | NCI-H2030 | PLX-4720 | MVP_top60_DNA+RNA_seed | ERBB2 | 0.663193 | STAD | 23132-87 | Lapatinib |
| MVP_top60_DNA_seed | AKT1 | -0.680964 | LUAD | NCI-H2228 | MK-2206 | MVP_top60_DNA+RNA_seed | PARP1 | 0.66387 | STAD | NUGC-3 | Rucaparib |
| MVP_top60_DNA_seed | CREBBP | -0.679072 | LUAD | NCI-H1781 | SGC-CBP30 | MVP_top60_DNA+RNA_seed | ESR1 | 0.663927 | STAD | GCIY | Tamoxifen |
| MVP_top60_DNA_seed | EGFR | -0.678241 | LUAD | NCI-H1781 | Erlotinib | MVP_top60_DNA+RNA_seed | ATM | 0.66448 | STAD | TGBC11TKB | KU-55933 |
| MVP_top60_DNA_seed | MDM2 | -0.676845 | LUAD | COR-L105 | Nutlin-3a (-) | MVP_top60_DNA+RNA_seed | HDAC1 | 0.667302 | STAD | SNU-1 | PCI-34051 |
| MVP_top60_DNA_seed | IGF1R | -0.675233 | LUAD | EMC-BAC-2 | BMS-536924 | MVP_top60_DNA+RNA_seed | TP53 | 0.67081 | STAD | OCUM-1 | MIRA-1 |
| MVP_top60_DNA_seed | AURKA | -0.673606 | LUAD | EMC-BAC-2 | Alisertib | MVP_top60_DNA+RNA_seed | ERBB2 | 0.670885 | STAD | SNU-1 | Sapitinib |
| MVP_top60_DNA_seed | HSP90AA1 | -0.670457 | LUAD | NCI-H838 | Elesclomol | MVP_top60_DNA+RNA_seed | ERBB3 | 0.670885 | STAD | SNU-1 | Sapitinib |
| MVP_top60_DNA_seed | MAPK1 | -0.669467 | LUAD | NCI-H322M | Ulixertinib | MVP_top60_DNA+RNA_seed | MAPK1 | 0.672457 | STAD | NUGC-4 | VX-11e |
| MVP_top60_DNA_seed | GSK3B | -0.665667 | LUAD | NCI-H1792 | SB216763 | MVP_top60_DNA+RNA_seed | EP300 | 0.677818 | STAD | KATOIII | SGC-CBP30 |
| MVP_top60_DNA_seed | AKT1 | -0.664889 | LUAD | NCI-H2030 | MK-2206 | MVP_top60_DNA+RNA_seed | CREBBP | 0.677818 | STAD | KATOIII | SGC-CBP30 |
| MVP_top60_DNA_seed | TP53 | -0.664239 | LUAD | NCI-H1792 | MIRA-1 | MVP_top60_DNA+RNA_seed | SRC | 0.680894 | STAD | 23132-87 | Bosutinib |
| MVP_top60_DNA_seed | AURKA | -0.663796 | LUAD | NCI-H838 | ZM447439 | MVP_top60_DNA+RNA_seed | KRAS | 0.682097 | STAD | NUGC-4 | KRAS (G12C) Inhibitor-1 |
| MVP_top60_DNA_seed | HSP90AA1 | -0.66342 | LUAD | NCI-H2023 | Tanespimycin | MVP_top60_DNA+RNA_seed | PIK3CA | 0.69062 | STAD | RERF-GC-1B | GNE-317 |
| MVP_top60_DNA_seed | HSP90AA1 | -0.662944 | LUAD | NCI-H2030 | CCT-018159 | MVP_top60_DNA+RNA_seed | MDM2 | 0.692716 | STAD | SNU-1 | Serdemetan |
| MVP_top60_DNA_seed | IGF1R | -0.658029 | LUAD | LXF-289 | NVP-ADW742 | MVP_top60_DNA+RNA_seed | MET | 0.699545 | STAD | FU97 | Savolitinib |
| MVP_top60_DNA_seed | CREBBP | -0.657493 | LUAD | RERF-LC-KJ | SGC-CBP30 | MVP_top60_DNA+RNA_seed | FGFR3 | 0.703335 | STAD | RF-48 | PD173074 |
| MVP_top60_DNA_seed | MAPK1 | -0.656946 | LUAD | NCI-H1734 | Ulixertinib | MVP_top60_DNA+RNA_seed | EZH2 | 0.707938 | STAD | RERF-GC-1B | GSK343 |
| MVP_top60_DNA_seed | SMARCA4 | -0.656587 | LUAD | NCI-H1792 | PFI3 | MVP_top60_DNA+RNA_seed | PARP1 | 0.718571 | STAD | GCIY | Veliparib |
| MVP_top60_DNA_seed | IGF1R | -0.655738 | LUAD | NCI-H838 | BMS-754807 | MVP_top60_DNA+RNA_seed | TP53 | 0.719516 | STAD | NUGC-3 | MIRA-1 |
| MVP_top60_DNA_seed | BRAF | -0.654402 | LUAD | NCI-H322M | Dabrafenib | MVP_top60_DNA+RNA_seed | HDAC1 | 0.729204 | STAD | MKN28 | Romidepsin |
| MVP_top60_DNA_seed | IGF1R | -0.652007 | LUAD | NCI-H1437 | BMS-754807 | MVP_top60_DNA+RNA_seed | HDAC3 | 0.729204 | STAD | MKN28 | Romidepsin |
| MVP_top60_DNA_seed | AKT1 | -0.650566 | LUAD | NCI-H1944 | AT13148 | MVP_top60_DNA+RNA_seed | PIK3CA | 0.73458 | STAD | TMK-1 | GNE-317 |
| MVP_top60_DNA_seed | MET | -0.648689 | LUAD | NCI-H1792 | Crizotinib | MVP_top60_DNA+RNA_seed | HDAC1 | 0.736435 | STAD | NCI-N87 | Entinostat |
| MVP_top60_DNA_seed | EGFR | -0.645534 | LUAD | A549 | AZD3759 | MVP_top60_DNA+RNA_seed | HDAC3 | 0.736435 | STAD | NCI-N87 | Entinostat |
| MVP_top60_DNA_seed | IGF1R | -0.645183 | LUAD | NCI-H2009 | BMS-754807 | MVP_top60_DNA+RNA_seed | PARP1 | 0.73943 | STAD | Hs-746T | Veliparib |

| | | | | | | | | | | | |
|---|---|---|---|---|---|---|---|---|---|---|---|
| MVP_top60_DNA_seed | SMARCA4 | -0.643857 | LUAD | NCI-H1734 | PFI3 | MVP_top60_DNA+RNA_seed | HSP90AA1 | 0.743032 | STAD | TGBC11TKB | Luminespib |
| MVP_top60_DNA_seed | HSP90AA1 | -0.643257 | LUAD | EMC-BAC-2 | Tanespimycin | MVP_top60_DNA+RNA_seed | HSP90AB1 | 0.743032 | STAD | TGBC11TKB | Luminespib |
| MVP_top60_DNA_seed | MET | -0.641633 | LUAD | HCC-78 | Crizotinib | MVP_top60_DNA+RNA_seed | PARP1 | 0.749619 | STAD | RERF-GC-1B | Niraparib |
| MVP_top60_DNA_seed | ATM | -0.637013 | LUAD | NCI-H2030 | KU-55933 | MVP_top60_DNA+RNA_seed | TP53 | 0.752842 | STAD | NCI-N87 | MIRA-1 |
| MVP_top60_DNA_seed | SMARCA4 | -0.636456 | LUAD | HCC-78 | PFI3 | MVP_top60_DNA+RNA_seed | EP300 | 0.759056 | STAD | NUGC-4 | CPI-637 |
| MVP_top60_DNA_seed | AURKA | -0.636384 | LUAD | NCI-H2085 | ZM447439 | MVP_top60_DNA+RNA_seed | HDAC1 | 0.763636 | STAD | GCIY | PCI-34051 |
| MVP_top60_DNA_seed | AURKA | -0.636228 | LUAD | LXF-289 | Alisertib | MVP_top60_DNA+RNA_seed | MDM2 | 0.76536 | STAD | NCI-N87 | Nutlin-3a (-) |
| MVP_top60_DNA_seed | AKT1 | -0.635213 | LUAD | NCI-H2023 | Uprosertib | MVP_top60_DNA+RNA_seed | FGFR3 | 0.770224 | STAD | GCIY | PD173074 |
| MVP_top60_DNA_seed | HDAC1 | -0.634734 | LUAD | Calu-6 | PCI-34051 | MVP_top60_DNA+RNA_seed | MAPK1 | 0.770431 | STAD | MKN7 | Ulixertinib |
| MVP_top60_DNA_seed | MDM2 | -0.633027 | LUAD | RERF-LC-KJ | Serdemetan | MVP_top60_DNA+RNA_seed | MAPK3 | 0.770431 | STAD | MKN7 | Ulixertinib |
| MVP_top60_DNA_seed | MET | -0.632749 | LUAD | LXF-289 | Crizotinib | MVP_top60_DNA+RNA_seed | PARP1 | 0.776116 | STAD | MKN28 | Veliparib |
| MVP_top60_DNA_seed | BRAF | -0.632672 | LUAD | NCI-H322M | PLX-4720 | MVP_top60_DNA+RNA_seed | KRAS | 0.778574 | STAD | NUGC-3 | KRAS (G12C) Inhibitor-1 |
| MVP_top60_DNA_seed | MDM2 | -0.630145 | LUAD | NCI-H1792 | Serdemetan | MVP_top60_DNA+RNA_seed | EP300 | 0.779691 | STAD | SK-GT-2 | CPI-637 |
| MVP_top60_DNA_seed | PIK3CA | -0.628623 | LUAD | PC-3_[JPC-3] | GNE-317 | MVP_top60_DNA+RNA_seed | PARP1 | 0.782376 | STAD | SNU-1 | Niraparib |
| MVP_top60_DNA_seed | HDAC1 | -0.628231 | LUAD | NCI-H322M | Romidepsin | MVP_top60_DNA+RNA_seed | HDAC1 | 0.790381 | STAD | MKN45 | Entinostat |
| MVP_top60_DNA_seed | BRAF | -0.626933 | LUAD | COR-L105 | PLX-4720 | MVP_top60_DNA+RNA_seed | HDAC3 | 0.790381 | STAD | MKN45 | Entinostat |
| MVP_top60_DNA_seed | PIK3CA | -0.626438 | LUAD | NCI-H1666 | AZD8186 | MVP_top60_DNA+RNA_seed | ERBB2 | 0.793279 | STAD | SNU-5 | Afatinib |
| MVP_top60_DNA_seed | FGFR1 | -0.625612 | LUAD | NCI-H2122 | PD173074 | MVP_top60_DNA+RNA_seed | PIK3CA | 0.7938 | STAD | SNU-16 | AZD8186 |
| MVP_top60_DNA_seed | IGF1R | -0.624301 | LUAD | NCI-H358 | BMS-754807 | MVP_top60_DNA+RNA_seed | HDAC1 | 0.800946 | STAD | Hs-746T | PCI-34051 |
| MVP_top60_DNA_seed | IGF1R | -0.622196 | LUAD | NCI-H1792 | BMS-754807 | MVP_top60_DNA+RNA_seed | PARP1 | 0.801737 | STAD | IM-95 | Talazoparib |
| MVP_top60_DNA_seed | PIK3CA | -0.620927 | LUAD | NCI-H1650 | AZD8186 | MVP_top60_DNA+RNA_seed | ESR1 | 0.80209 | STAD | NUGC-3 | GDC0810 |
| MVP_top60_DNA_seed | PIK3CA | -0.618162 | LUAD | NCI-H2122 | AZD8186 | MVP_top60_DNA+RNA_seed | HDAC1 | 0.80635 | STAD | NCI-N87 | PCI-34051 |
| MVP_top60_DNA_seed | MAPK1 | -0.617451 | LUAD | EMC-BAC-2 | SCH772984 | MVP_top60_DNA+RNA_seed | AR | 0.808352 | STAD | MKN7 | Bicalutamide |
| MVP_top60_DNA_seed | MAPK1 | -0.617043 | LUAD | NCI-H2030 | VX-11e | MVP_top60_DNA+RNA_seed | ERBB2 | 0.817424 | STAD | FU97 | Sapitinib |
| MVP_top60_DNA_seed | BRAF | -0.616559 | LUAD | HCC-827 | PLX-4720 | MVP_top60_DNA+RNA_seed | ERBB3 | 0.817424 | STAD | FU97 | Sapitinib |
| MVP_top60_DNA_seed | BRAF | -0.615308 | LUAD | NCI-H2122 | PLX-4720 | MVP_top60_DNA+RNA_seed | FGFR3 | 0.819547 | STAD | NUGC-3 | AZD4547 |
| MVP_top60_DNA_seed | MAPK1 | -0.615241 | LUAD | NCI-H1755 | SCH772984 | MVP_top60_DNA+RNA_seed | EP300 | 0.819904 | STAD | IM-95 | CPI-637 |
| MVP_top60_DNA_seed | MAPK1 | -0.611977 | LUAD | NCI-H1944 | VX-11e | MVP_top60_DNA+RNA_seed | PARP1 | 0.825619 | STAD | MKN28 | Niraparib |
| MVP_top60_DNA_seed | MAPK1 | -0.609124 | LUAD | NCI-H1944 | Ulixertinib | MVP_top60_DNA+RNA_seed | MDM2 | 0.827733 | STAD | SK-GT-2 | Serdemetan |
| MVP_top60_DNA_seed | KRAS | -0.608473 | LUAD | PC-14 | KRAS (G12C) Inhibitor-1 | MVP_top60_DNA+RNA_seed | FGFR3 | 0.833458 | STAD | Hs-746T | AZD4547 |
| MVP_top60_DNA_seed | TERT | -0.607296 | LUAD | NCI-H1573 | BIBR-1532 | MVP_top60_DNA+RNA_seed | FGFR3 | 0.834795 | STAD | MKN45 | AZD4547 |
| MVP_top60_DNA_seed | AURKA | -0.606022 | LUAD | NCI-H1781 | Alisertib | MVP_top60_DNA+RNA_seed | TP53 | 0.838111 | STAD | MKN28 | MIRA-1 |
| MVP_top60_DNA_seed | MAPK1 | -0.605748 | LUAD | NCI-H1648 | VX-11e | MVP_top60_DNA+RNA_seed | PARP1 | 0.839379 | STAD | GCIY | Rucaparib |
| MVP_top60_DNA_seed | AKT1 | -0.60524 | LUAD | RERF-LC-MS | MK-2206 | MVP_top60_DNA+RNA_seed | PARP1 | 0.846367 | STAD | 23132-87 | Rucaparib |
| MVP_top60_DNA_seed | CREBBP | -0.604581 | LUAD | NCI-H23 | SGC-CBP30 | MVP_top60_DNA+RNA_seed | EZH2 | 0.851509 | STAD | MKN45 | GSK343 |
| MVP_top60_DNA_seed | EGFR | -0.603694 | LUAD | NCI-H1563 | Osimertinib | MVP_top60_DNA+RNA_seed | MDM2 | 0.852634 | STAD | GCIY | Nutlin-3a (-) |
| MVP_top60_DNA_seed | PIK3CA | -0.602351 | LUAD | NCI-H1734 | AZD8186 | MVP_top60_DNA+RNA_seed | PARP1 | 0.852889 | STAD | IM-95 | Rucaparib |
| MVP_top60_DNA_seed | MAPK1 | -0.600555 | LUAD | NCI-H358 | Ulixertinib | MVP_top60_DNA+RNA_seed | PIK3CA | 0.855342 | STAD | SNU-1 | Alpelisib |
| MVP_top60_DNA_seed | MET | -0.60022 | LUAD | HCC-44 | Savolitinib | MVP_top60_DNA+RNA_seed | PIK3CA | 0.855649 | STAD | NUGC-4 | Alpelisib |
| MVP_top60_DNA_seed | AKT1 | -0.600028 | LUAD | NCI-H1734 | Afuresertib | MVP_top60_DNA+RNA_seed | MET | 0.855978 | STAD | GCIY | Savolitinib |
| MVP_top60_DNA_seed | EGFR | -0.599925 | LUAD | NCI-H358 | Erlotinib | MVP_top60_DNA+RNA_seed | HSP90AA1 | 0.85794 | STAD | TMK-1 | Tanespimycin |

| | | | | | | | | | | | |
|---|---|---|---|---|---|---|---|---|---|---|---|
| MVP_top60_DNA_seed | EGFR | -0.599418 | LUAD | LC-2-ad | AZD3759 | MVP_top60_DNA+RNA_seed | HSP90AB1 | 0.85794 | STAD | TMK-1 | Tanespimycin |
| MVP_top60_DNA_seed | PIK3CA | -0.598218 | LUAD | NCI-H1623 | Buparlisib | MVP_top60_DNA+RNA_seed | EZH2 | 0.862819 | STAD | SK-GT-2 | GSK343 |
| MVP_top60_DNA_seed | ATM | -0.597208 | LUAD | NCI-H1792 | KU-55933 | MVP_top60_DNA+RNA_seed | FGFR3 | 0.869434 | STAD | MKN28 | AZD4547 |
| MVP_top60_DNA_seed | AKT1 | -0.595196 | LUAD | EMC-BAC-1 | Ipatasertib | MVP_top60_DNA+RNA_seed | PARP1 | 0.871256 | STAD | MKN45 | Olaparib |
| MVP_top60_DNA_seed | HDAC1 | -0.595179 | LUAD | NCI-H2291 | Entinostat | MVP_top60_DNA+RNA_seed | MAPK1 | 0.872168 | STAD | FU97 | Ulixertinib |
| MVP_top60_DNA_seed | AKT1 | -0.593194 | LUAD | NCI-H838 | MK-2206 | MVP_top60_DNA+RNA_seed | MAPK3 | 0.872168 | STAD | FU97 | Ulixertinib |
| MVP_top60_DNA_seed | AKT1 | -0.592801 | LUAD | NCI-H1781 | MK-2206 | MVP_top60_DNA+RNA_seed | ESR1 | 0.872895 | STAD | NCI-N87 | Tamoxifen |
| MVP_top60_DNA_seed | PIK3CA | -0.590916 | LUAD | NCI-H2030 | GNE-317 | MVP_top60_DNA+RNA_seed | ERBB2 | 0.876586 | STAD | MKN45 | Afatinib |
| MVP_top60_DNA_seed | HSP90AA1 | -0.59049 | LUAD | NCI-H3122 | Luminespib | MVP_top60_DNA+RNA_seed | PARP1 | 0.879927 | STAD | OCUM-1 | Olaparib |
| MVP_top60_DNA_seed | EGFR | -0.58924 | LUAD | A549 | Erlotinib | MVP_top60_DNA+RNA_seed | EP300 | 0.883488 | STAD | KATOIII | CPI-637 |
| MVP_top60_DNA_seed | EGFR | -0.588163 | LUAD | LC-2-ad | Erlotinib | MVP_top60_DNA+RNA_seed | MAPK1 | 0.883756 | STAD | IM-95 | Ulixertinib |
| MVP_top60_DNA_seed | PIK3CA | -0.587139 | LUAD | NCI-H2023 | Alpelisib | MVP_top60_DNA+RNA_seed | MAPK3 | 0.883756 | STAD | IM-95 | Ulixertinib |
| MVP_top60_DNA_seed | AKT1 | -0.585015 | LUAD | NCI-H1781 | Afuresertib | MVP_top60_DNA+RNA_seed | EZH2 | 0.889557 | STAD | IM-95 | GSK343 |
| MVP_top60_DNA_seed | EZH2 | -0.583697 | LUAD | HCC-44 | GSK343 | MVP_top60_DNA+RNA_seed | FGFR3 | 0.892925 | STAD | SK-GT-2 | PD173074 |
| MVP_top60_DNA_seed | HDAC1 | -0.583352 | LUAD | HCC-44 | Romidepsin | MVP_top60_DNA+RNA_seed | ATM | 0.89813 | STAD | NUGC-4 | KU-55933 |
| MVP_top60_DNA_seed | FGFR1 | -0.58331 | LUAD | LXF-289 | AZD4547 | MVP_top60_DNA+RNA_seed | PARP1 | 0.90933 | STAD | MKN7 | Niraparib |
| MVP_top60_DNA_seed | PIK3CA | -0.580946 | LUAD | RERF-LC-MS | Buparlisib | MVP_top60_DNA+RNA_seed | EP300 | 0.909502 | STAD | 23132-87 | CPI-637 |
| MVP_top60_DNA_seed | PIK3CA | -0.580117 | LUAD | NCI-H1623 | AZD8186 | MVP_top60_DNA+RNA_seed | EZH2 | 0.911102 | STAD | MKN28 | GSK343 |
| MVP_top60_DNA_seed | MDM2 | -0.578934 | LUAD | NCI-H1437 | Serdemetan | MVP_top60_DNA+RNA_seed | KRAS | 0.913606 | STAD | RERF-GC-1B | KRAS (G12C) Inhibitor-1 |
| MVP_top60_DNA_seed | FGFR1 | -0.578524 | LUAD | EMC-BAC-2 | AZD4547 | MVP_top60_DNA+RNA_seed | PIK3CA | 0.924553 | STAD | MKN28 | AZD8186 |
| MVP_top60_DNA_seed | SRC | -0.578392 | LUAD | Calu-3 | Bosutinib | MVP_top60_DNA+RNA_seed | PARP1 | 0.929083 | STAD | FU97 | Rucaparib |
| MVP_top60_DNA_seed | MET | -0.578326 | LUAD | A549 | Crizotinib | MVP_top60_DNA+RNA_seed | GSK3B | 0.92934 | STAD | Hs-746T | SB216763 |
| MVP_top60_DNA_seed | EZH2 | -0.578045 | LUAD | NCI-H1792 | GSK343 | MVP_top60_DNA+RNA_seed | FGFR3 | 0.930064 | STAD | SNU-5 | PD173074 |
| MVP_top60_DNA_seed | IGF1R | -0.57765 | LUAD | A549 | Linsitinib | MVP_top60_DNA+RNA_seed | HDAC1 | 0.932902 | STAD | TMK-1 | PCI-34051 |
| MVP_top60_DNA_seed | HDAC1 | -0.577474 | LUAD | NCI-H1792 | PCI-34051 | MVP_top60_DNA+RNA_seed | FGFR3 | 0.935127 | STAD | OCUM-1 | AZD4547 |
| MVP_top60_DNA_seed | KRAS | -0.576165 | LUAD | A549 | KRAS (G12C) Inhibitor-1 | MVP_top60_DNA+RNA_seed | MET | 0.945278 | STAD | NCI-N87 | Crizotinib |
| MVP_top60_DNA_seed | EZH2 | -0.574844 | LUAD | HCC-44 | GSK343 | MVP_top60_DNA+RNA_seed | ERBB2 | 0.945989 | STAD | FU97 | Afatinib |
| MVP_top60_DNA_seed | EGFR | -0.574442 | LUAD | EMC-BAC-2 | Gefitinib | MVP_top60_DNA+RNA_seed | ESR1 | 0.958655 | STAD | RF-48 | Tamoxifen |
| MVP_top60_DNA_seed | EZH2 | -0.573823 | LUAD | HCC-827 | GSK343 | MVP_top60_DNA+RNA_seed | MDM2 | 0.966061 | STAD | SK-GT-2 | Nutlin-3a (-) |
| MVP_top60_DNA_seed | AURKA | -0.573581 | LUAD | VMRC-LCD | ZM447439 | MVP_top60_DNA+RNA_seed | EP300 | 0.967239 | STAD | OCUM-1 | CPI-637 |
| MVP_top60_DNA_seed | AURKA | -0.570868 | LUAD | RERF-LC-MS | ZM447439 | MVP_top60_DNA+RNA_seed | HDAC1 | 0.972412 | STAD | SNU-1 | Romidepsin |
| MVP_top60_DNA_seed | AKT1 | -0.570588 | LUAD | EMC-BAC-2 | Ipatasertib | MVP_top60_DNA+RNA_seed | HDAC3 | 0.972412 | STAD | SNU-1 | Romidepsin |
| MVP_top60_DNA_seed | MET | -0.568303 | LUAD | LC-2-ad | Crizotinib | MVP_top60_DNA+RNA_seed | HSP90AA1 | 0.973899 | STAD | RERF-GC-1B | Tanespimycin |
| MVP_top60_DNA_seed | HDAC1 | -0.567604 | LUAD | COR-L105 | PCI-34051 | MVP_top60_DNA+RNA_seed | HSP90AB1 | 0.973899 | STAD | RERF-GC-1B | Tanespimycin |
| MVP_top60_DNA_seed | HDAC1 | -0.565808 | LUAD | HCC-827 | Romidepsin | MVP_top60_DNA+RNA_seed | PARP1 | 0.976224 | STAD | MKN7 | Olaparib |
| MVP_top60_DNA_seed | AKT1 | -0.564375 | LUAD | NCI-H1975 | MK-2206 | MVP_top60_DNA+RNA_seed | PIK3CA | 0.976605 | STAD | OCUM-1 | Buparlisib |
| MVP_top60_DNA_seed | PIK3CA | -0.563451 | LUAD | EMC-BAC-2 | GNE-317 | MVP_top60_DNA+RNA_seed | HDAC1 | 0.97777 | STAD | RERF-GC-1B | PCI-34051 |
| MVP_top60_DNA_seed | IGF1R | -0.561592 | LUAD | NCI-H1755 | Linsitinib | MVP_top60_DNA+RNA_seed | ESR1 | 0.978548 | STAD | MKN7 | Tamoxifen |
| MVP_top60_DNA_seed | EGFR | -0.561571 | LUAD | A549 | Sapitinib | MVP_top60_DNA+RNA_seed | FGFR3 | 0.98229 | STAD | TMK-1 | AZD4547 |
| MVP_top60_DNA_seed | ERBB2 | -0.561571 | LUAD | A549 | Sapitinib | MVP_top60_DNA+RNA_seed | MDM2 | 0.986817 | STAD | MKN28 | Nutlin-3a (-) |
| MVP_top60_DNA_seed | ERBB3 | -0.561571 | LUAD | A549 | Sapitinib | MVP_top60_DNA+RNA_seed | PARP1 | 0.987638 | STAD | GCIY | Talazoparib |

| | | | | | | | | | | | |
|---|---|---|---|---|---|---|---|---|---|---|---|
| MVP_top60_DNA_seed | BRAF | -0.560148 | LUAD | NCI-H1435 | Dabrafenib | MVP_top60_DNA+RNA_seed | MDM2 | 0.996569 | STAD | MKN7 | Serdemetan |
| MVP_top60_DNA_seed | HSP90AA1 | -0.559989 | LUAD | NCI-H1781 | Tanespimycin | MVP_top60_DNA+RNA_seed | FGFR3 | 0.996848 | STAD | NCI-N87 | AZD4547 |
| MVP_top60_DNA_seed | PIK3CA | -0.559641 | LUAD | A549 | AZD8186 | MVP_top60_DNA+RNA_seed | EZH2 | 1.004534 | STAD | Hs-746T | GSK343 |
| MVP_top60_DNA_seed | MDM2 | -0.559617 | LUAD | RERF-LC-MS | Nutlin-3a (-) | MVP_top60_DNA+RNA_seed | MET | 1.010332 | STAD | MKN28 | Savolitinib |
| MVP_top60_DNA_seed | PIK3CA | -0.55925 | LUAD | NCI-H1734 | Buparlisib | MVP_top60_DNA+RNA_seed | EZH2 | 1.011769 | STAD | NCI-N87 | GSK343 |
| MVP_top60_DNA_seed | IGF1R | -0.558478 | LUAD | NCI-H1666 | Linsitinib | MVP_top60_DNA+RNA_seed | TP53 | 1.053476 | STAD | MKN45 | MIRA-1 |
| MVP_top60_DNA_seed | IGF1R | -0.555108 | LUAD | A549 | BMS-754807 | MVP_top60_DNA+RNA_seed | PARP1 | 1.056646 | STAD | MKN45 | Rucaparib |
| MVP_top60_DNA_seed | PIK3CA | -0.554255 | LUAD | HCC-78 | Alpelisib | MVP_top60_DNA+RNA_seed | EZH2 | 1.056917 | STAD | KATOIII | GSK343 |
| MVP_top60_DNA_seed | AKT1 | -0.553987 | LUAD | NCI-H1838 | AT13148 | MVP_top60_DNA+RNA_seed | MDM2 | 1.059129 | STAD | MKN28 | Serdemetan |
| MVP_top60_DNA_seed | PIK3CA | -0.553292 | LUAD | NCI-H2228 | Alpelisib | MVP_top60_DNA+RNA_seed | PARP1 | 1.059471 | STAD | TGBC11TKB | Olaparib |
| MVP_top60_DNA_seed | PIK3CA | -0.552522 | LUAD | NCI-H2009 | Alpelisib | MVP_top60_DNA+RNA_seed | ERBB2 | 1.059574 | STAD | SNU-5 | Sapitinib |
| MVP_top60_DNA_seed | HDAC1 | -0.551923 | LUAD | EMC-BAC-2 | Romidepsin | MVP_top60_DNA+RNA_seed | ERBB3 | 1.059574 | STAD | SNU-5 | Sapitinib |
| MVP_top60_DNA_seed | SRC | -0.551381 | LUAD | EMC-BAC-1 | Bosutinib | MVP_top60_DNA+RNA_seed | EP300 | 1.071057 | STAD | RERF-GC-1B | CPI-637 |
| MVP_top60_DNA_seed | EZH2 | -0.551115 | LUAD | NCI-H1435 | GSK343 | MVP_top60_DNA+RNA_seed | MDM2 | 1.073453 | STAD | TGBC11TKB | Nutlin-3a (-) |
| MVP_top60_DNA_seed | MAPK1 | -0.550671 | LUAD | COR-L105 | SCH772984 | MVP_top60_DNA+RNA_seed | GSK3B | 1.075772 | STAD | GCIY | SB216763 |
| MVP_top60_DNA_seed | PIK3CA | -0.549144 | LUAD | HCC-44 | GNE-317 | MVP_top60_DNA+RNA_seed | MET | 1.079609 | STAD | 23132-87 | Crizotinib |
| MVP_top60_DNA_seed | SMARCA4 | -0.5489 | LUAD | NCI-H1435 | PFI3 | MVP_top60_DNA+RNA_seed | PIK3CA | 1.080422 | STAD | OCUM-1 | AZD8186 |
| MVP_top60_DNA_seed | AKT1 | -0.544665 | LUAD | NCI-H1734 | AZD5363 | MVP_top60_DNA+RNA_seed | ERBB2 | 1.085179 | STAD | SNU-5 | Lapatinib |
| MVP_top60_DNA_seed | EGFR | -0.541517 | LUAD | NCI-H1693 | Sapitinib | MVP_top60_DNA+RNA_seed | PARP1 | 1.08855 | STAD | MKN7 | Veliparib |
| MVP_top60_DNA_seed | ERBB2 | -0.541517 | LUAD | NCI-H1693 | Sapitinib | MVP_top60_DNA+RNA_seed | ERBB2 | 1.093234 | STAD | OCUM-1 | Sapitinib |
| MVP_top60_DNA_seed | ERBB3 | -0.541517 | LUAD | NCI-H1693 | Sapitinib | MVP_top60_DNA+RNA_seed | ERBB3 | 1.093234 | STAD | OCUM-1 | Sapitinib |
| MVP_top60_DNA_seed | TERT | -0.540957 | LUAD | NCI-H1623 | BIBR-1532 | MVP_top60_DNA+RNA_seed | KRAS | 1.096434 | STAD | MKN7 | RAS (G12C) Inhibitor-1 |
| MVP_top60_DNA_seed | EGFR | -0.539813 | LUAD | HCC-78 | Osimertinib | MVP_top60_DNA+RNA_seed | MDM2 | 1.096944 | STAD | Hs-746T | Nutlin-3a (-) |
| MVP_top60_DNA_seed | PIK3CA | -0.537165 | LUAD | NCI-H1666 | Alpelisib | MVP_top60_DNA+RNA_seed | PARP1 | 1.104811 | STAD | MKN7 | Talazoparib |
| MVP_top60_DNA_seed | IGF1R | -0.537128 | LUAD | LC-2-ad | NVP-ADW742 | MVP_top60_DNA+RNA_seed | HDAC1 | 1.109871 | STAD | KATOIII | PCI-34051 |
| MVP_top60_DNA_seed | EGFR | -0.536795 | LUAD | NCI-H650 | Gefitinib | MVP_top60_DNA+RNA_seed | EZH2 | 1.129435 | STAD | MKN7 | GSK343 |
| MVP_top60_DNA_seed | EZH2 | -0.536677 | LUAD | NCI-H1944 | GSK343 | MVP_top60_DNA+RNA_seed | PARP1 | 1.135467 | STAD | NUGC-4 | Talazoparib |
| MVP_top60_DNA_seed | PIK3CA | -0.536241 | LUAD | NCI-H1993 | Buparlisib | MVP_top60_DNA+RNA_seed | FGFR3 | 1.150755 | STAD | GCIY | AZD4547 |
| MVP_top60_DNA_seed | PIK3CA | -0.535728 | LUAD | NCI-H2023 | Buparlisib | MVP_top60_DNA+RNA_seed | ESR1 | 1.152807 | STAD | OCUM-1 | Tamoxifen |
| MVP_top60_DNA_seed | MDM2 | -0.535122 | LUAD | NCI-H1651 | Serdemetan | MVP_top60_DNA+RNA_seed | KRAS | 1.159427 | STAD | Hs-746T | RAS (G12C) Inhibitor-1 |
| MVP_top60_DNA_seed | PIK3CA | -0.53352 | LUAD | COR-L105 | Buparlisib | MVP_top60_DNA+RNA_seed | ESR1 | 1.161804 | STAD | OCUM-1 | GDC0810 |
| MVP_top60_DNA_seed | MET | -0.533176 | LUAD | NCI-H1734 | Crizotinib | MVP_top60_DNA+RNA_seed | MDM2 | 1.165518 | STAD | MKN45 | Serdemetan |
| MVP_top60_DNA_seed | MAPK1 | -0.532531 | LUAD | HCC-827 | SCH772984 | MVP_top60_DNA+RNA_seed | SRC | 1.167276 | STAD | MKN7 | Bosutinib |
| MVP_top60_DNA_seed | MAPK1 | -0.532378 | LUAD | NCI-H2087 | Ulixertinib | MVP_top60_DNA+RNA_seed | PIK3CA | 1.187297 | STAD | IM-95 | GNE-317 |
| MVP_top60_DNA_seed | HSP90AA1 | -0.532301 | LUAD | NCI-H322M | Tanespimycin | MVP_top60_DNA+RNA_seed | HDAC1 | 1.188913 | STAD | GCIY | Entinostat |
| MVP_top60_DNA_seed | AURKA | -0.532005 | LUAD | NCI-H2122 | ZM447439 | MVP_top60_DNA+RNA_seed | HDAC3 | 1.188913 | STAD | GCIY | Entinostat |
| MVP_top60_DNA_seed | BRAF | -0.530993 | LUAD | A549 | PLX-4720 | MVP_top60_DNA+RNA_seed | SRC | 1.190962 | STAD | TGBC11TKB | Bosutinib |
| MVP_top60_DNA_seed | EGFR | -0.530947 | LUAD | HCC-78 | Sapitinib | MVP_top60_DNA+RNA_seed | PARP1 | 1.200796 | STAD | KATOIII | Talazoparib |
| MVP_top60_DNA_seed | ERBB2 | -0.530947 | LUAD | HCC-78 | Sapitinib | MVP_top60_DNA+RNA_seed | MDM2 | 1.205417 | STAD | KATOIII | Serdemetan |
| MVP_top60_DNA_seed | ERBB3 | -0.530947 | LUAD | HCC-78 | Sapitinib | MVP_top60_DNA+RNA_seed | ESR1 | 1.216674 | STAD | TMK-1 | Tamoxifen |
| MVP_top60_DNA_seed | IGF1R | -0.53047 | LUAD | NCI-H2009 | Linsitinib | MVP_top60_DNA+RNA_seed | MDM2 | 1.220894 | STAD | NUGC-3 | Nutlin-3a (-) |

| | | | | | | | | | | | |
|---|---|---|---|---|---|---|---|---|---|---|---|
| MVP_top60_DNA_seed | AURKA | -0.529581 | LUAD | LXF-289 | ZM447439 | MVP_top60_DNA+RNA_seed | ERBB2 | 1.223624 | STAD | KATOIII | Lapatinib |
| MVP_top60_DNA_seed | EGFR | -0.529259 | LUAD | EMC-BAC-2 | AZD3759 | MVP_top60_DNA+RNA_seed | PIK3CA | 1.223944 | STAD | MKN7 | Buparlisib |
| MVP_top60_DNA_seed | AKT1 | -0.528741 | LUAD | LXF-289 | MK-2206 | MVP_top60_DNA+RNA_seed | GSK3B | 1.225564 | STAD | OCUM-1 | SB216763 |
| MVP_top60_DNA_seed | MAPK1 | -0.527116 | LUAD | EMC-BAC-1 | SCH772984 | MVP_top60_DNA+RNA_seed | PIK3CA | 1.234249 | STAD | OCUM-1 | GNE-317 |
| MVP_top60_DNA_seed | PIK3CA | -0.526357 | LUAD | NCI-H1793 | Buparlisib | MVP_top60_DNA+RNA_seed | TP53 | 1.240448 | STAD | NUGC-4 | MIRA-1 |
| MVP_top60_DNA_seed | HSP90AA1 | -0.524857 | LUAD | NCI-H1355 | Tanespimycin | MVP_top60_DNA+RNA_seed | EP300 | 1.254091 | STAD | MKN28 | CPI-637 |
| MVP_top60_DNA_seed | KRAS | -0.523116 | LUAD | NCI-H650 | KRAS (G12C) Inhibitor-1 | MVP_top60_DNA+RNA_seed | ATM | 1.264994 | STAD | OCUM-1 | KU-55933 |
| MVP_top60_DNA_seed | BRAF | -0.522925 | LUAD | NCI-H1755 | PLX-4720 | MVP_top60_DNA+RNA_seed | TP53 | 1.27456 | STAD | Hs-746T | MIRA-1 |
| MVP_top60_DNA_seed | AKT1 | -0.519444 | LUAD | NCI-H2009 | AT13148 | MVP_top60_DNA+RNA_seed | PARP1 | 1.282248 | STAD | FU97 | Veliparib |
| MVP_top60_DNA_seed | MET | -0.51892 | LUAD | EMC-BAC-2 | Savolitinib | MVP_top60_DNA+RNA_seed | MDM2 | 1.294502 | STAD | MKN7 | Nutlin-3a (-) |
| MVP_top60_DNA_seed | EGFR | -0.518054 | LUAD | NCI-H522 | Gefitinib | MVP_top60_DNA+RNA_seed | HSP90AA1 | 1.299079 | STAD | NUGC-4 | Tanespimycin |
| MVP_top60_DNA_seed | HSP90AA1 | -0.516152 | LUAD | NCI-H1734 | Luminespib | MVP_top60_DNA+RNA_seed | HSP90AB1 | 1.299079 | STAD | NUGC-4 | Tanespimycin |
| MVP_top60_DNA_seed | AKT1 | -0.515056 | LUAD | LXF-289 | GSK2110183B | MVP_top60_DNA+RNA_seed | EZH2 | 1.33071 | STAD | GCIY | GSK343 |
| MVP_top60_DNA_seed | PIK3CA | -0.513282 | LUAD | NCI-H1781 | AZD8186 | MVP_top60_DNA+RNA_seed | GSK3B | 1.33164 | STAD | MKN7 | SB216763 |
| MVP_top60_DNA_seed | HSP90AA1 | -0.511554 | LUAD | NCI-H1437 | Tanespimycin | MVP_top60_DNA+RNA_seed | PIK3CA | 1.33347 | STAD | KATOIII | Alpelisib |
| MVP_top60_DNA_seed | HSP90AA1 | -0.511552 | LUAD | NCI-H2228 | Tanespimycin | MVP_top60_DNA+RNA_seed | HDAC1 | 1.333627 | STAD | MKN28 | Entinostat |
| MVP_top60_DNA_seed | FGFR1 | -0.509894 | LUAD | LC-2-ad | PD173074 | MVP_top60_DNA+RNA_seed | HDAC3 | 1.333627 | STAD | MKN28 | Entinostat |
| MVP_top60_DNA_seed | MAPK1 | -0.507662 | LUAD | NCI-H1666 | Ulixertinib | MVP_top60_DNA+RNA_seed | PIK3CA | 1.34511 | STAD | TGBC11TKB | AZD8186 |
| MVP_top60_DNA_seed | BRAF | -0.507448 | LUAD | NCI-H1792 | PLX-4720 | MVP_top60_DNA+RNA_seed | SRC | 1.349176 | STAD | MKN45 | Bosutinib |
| MVP_top60_DNA_seed | FGFR1 | -0.503983 | LUAD | RERF-LC-MS | AZD4547 | MVP_top60_DNA+RNA_seed | EZH2 | 1.35059 | STAD | GCIY | GSK343 |
| MVP_top60_DNA_seed | AURKA | -0.503673 | LUAD | NCI-H2085 | Alisertib | MVP_top60_DNA+RNA_seed | MDM2 | 1.357746 | STAD | OCUM-1 | Serdemetan |
| MVP_top60_DNA_seed | MAPK1 | -0.503372 | LUAD | NCI-H1792 | SCH772984 | MVP_top60_DNA+RNA_seed | TP53 | 1.362835 | STAD | KATOIII | MIRA-1 |
| MVP_top60_DNA_seed | FGFR1 | -0.502658 | LUAD | NCI-H1563 | AZD4547 | MVP_top60_DNA+RNA_seed | MET | 1.363142 | STAD | MKN7 | Crizotinib |
| MVP_top60_DNA_seed | ATM | -0.501144 | LUAD | NCI-H1975 | KU-55933 | MVP_top60_DNA+RNA_seed | FGFR3 | 1.363982 | STAD | MKN7 | PD173074 |
| MVP_top60_DNA_seed | AKT1 | -0.501022 | LUAD | NCI-H1755 | AT13148 | MVP_top60_DNA+RNA_seed | PARP1 | 1.364628 | STAD | TGBC11TKB | Talazoparib |
| MVP_top60_DNA_seed | MAPK1 | -0.49983 | LUAD | NCI-H1792 | Ulixertinib | MVP_top60_DNA+RNA_seed | ATM | 1.369068 | STAD | MKN7 | KU-55933 |
| MVP_top60_DNA_seed | AKT1 | -0.49972 | LUAD | NCI-H3122 | Uprosertib | MVP_top60_DNA+RNA_seed | ATM | 1.379442 | STAD | KATOIII | KU-55933 |
| MVP_top60_DNA_seed | IGF1R | -0.49421 | LUAD | EMC-BAC-1 | BMS-754807 | MVP_top60_DNA+RNA_seed | SRC | 1.419095 | STAD | Hs-746T | Bosutinib |
| MVP_top60_DNA_seed | HSP90AA1 | -0.493468 | LUAD | LC-2-ad | Elesclomol | MVP_top60_DNA+RNA_seed | PARP1 | 1.429855 | STAD | MKN28 | Talazoparib |
| MVP_top60_DNA_seed | HSP90AA1 | -0.489903 | LUAD | NCI-H838 | Luminespib | MVP_top60_DNA+RNA_seed | PARP1 | 1.437628 | STAD | KATOIII | Niraparib |
| MVP_top60_DNA_seed | PIK3CA | -0.488432 | LUAD | NCI-H322M | Buparlisib | MVP_top60_DNA+RNA_seed | MET | 1.447934 | STAD | KATOIII | Savolitinib |
| MVP_top60_DNA_seed | EGFR | -0.487197 | LUAD | NCI-H1437 | Lapatinib | MVP_top60_DNA+RNA_seed | KRAS | 1.453123 | STAD | KATOIII | KRAS (G12C) Inhibitor-1 |
| MVP_top60_DNA_seed | ERBB2 | -0.487197 | LUAD | NCI-H1437 | Lapatinib | MVP_top60_DNA+RNA_seed | PIK3CA | 1.466011 | STAD | Hs-746T | Alpelisib |
| MVP_top60_DNA_seed | FGFR1 | -0.487101 | LUAD | NCI-H1792 | PD173074 | MVP_top60_DNA+RNA_seed | EZH2 | 1.466522 | STAD | OCUM-1 | GSK343 |
| MVP_top60_DNA_seed | EZH2 | -0.487004 | LUAD | NCI-H1792 | GSK343 | MVP_top60_DNA+RNA_seed | ERBB2 | 1.474767 | STAD | Hs-746T | Sapitinib |
| MVP_top60_DNA_seed | HSP90AA1 | -0.48684 | LUAD | NCI-H1650 | Tanespimycin | MVP_top60_DNA+RNA_seed | ERBB3 | 1.474767 | STAD | Hs-746T | Sapitinib |
| MVP_top60_DNA_seed | EZH2 | -0.486352 | LUAD | NCI-H1781 | GSK343 | MVP_top60_DNA+RNA_seed | GSK3B | 1.477037 | STAD | FU97 | SB216763 |
| MVP_top60_DNA_seed | AKT1 | -0.48587 | LUAD | NCI-H1437 | MK-2206 | MVP_top60_DNA+RNA_seed | PARP1 | 1.517544 | STAD | Hs-746T | Rucaparib |
| MVP_top60_DNA_seed | MET | -0.484597 | LUAD | VMRC-LCD | Crizotinib | MVP_top60_DNA+RNA_seed | HSP90AA1 | 1.542322 | STAD | MKN7 | Luminespib |
| MVP_top60_DNA_seed | EGFR | -0.484067 | LUAD | NCI-H1563 | Erlotinib | MVP_top60_DNA+RNA_seed | HSP90AB1 | 1.542322 | STAD | MKN7 | Luminespib |
| MVP_top60_DNA_seed | EZH2 | -0.482935 | LUAD | NCI-H1648 | GSK343 | MVP_top60_DNA+RNA_seed | MAPK1 | 1.544501 | STAD | RF-48 | VX-11e |

| | | | | | | | | | | | |
|---|---|---|---|---|---|---|---|---|---|---|---|
| MVP_top60_DNA_seed | HSP90AA1 | -0.479662 | LUAD | NCI-H2030 | Luminespib | MVP_top60_DNA+RNA_seed | HDAC1 | 1.581781 | STAD | MKN45 | Romidepsin |
| MVP_top60_DNA_seed | MAPK1 | -0.479565 | LUAD | COR-L105 | VX-11e | MVP_top60_DNA+RNA_seed | HDAC3 | 1.581781 | STAD | MKN45 | Romidepsin |
| MVP_top60_DNA_seed | AURKA | -0.479115 | LUAD | RERF-LC-KJ | ZM447439 | MVP_top60_DNA+RNA_seed | EP300 | 1.589912 | STAD | Hs-746T | SGC-CBP30 |
| MVP_top60_DNA_seed | HDAC1 | -0.47649 | LUAD | NCI-H2122 | PCI-34051 | MVP_top60_DNA+RNA_seed | CREBBP | 1.589912 | STAD | Hs-746T | SGC-CBP30 |
| MVP_top60_DNA_seed | IGF1R | -0.475001 | LUAD | NCI-H1781 | GSK1904529A | MVP_top60_DNA+RNA_seed | MET | 1.613309 | STAD | MKN7 | Savolitinib |
| MVP_top60_DNA_seed | MAPK1 | -0.474652 | LUAD | NCI-H1623 | SCH772984 | MVP_top60_DNA+RNA_seed | PARP1 | 1.647467 | STAD | KATOIII | Rucaparib |
| MVP_top60_DNA_seed | GSK3B | -0.474045 | LUAD | EMC-BAC-1 | CHIR-99021 | MVP_top60_DNA+RNA_seed | PARP1 | 1.649013 | STAD | NUGC-4 | Olaparib |
| MVP_top60_DNA_seed | PIK3CA | -0.471532 | LUAD | A549 | Alpelisib | MVP_top60_DNA+RNA_seed | MAPK1 | 1.650232 | STAD | RF-48 | Ulixertinib |
| MVP_top60_DNA_seed | MAPK1 | -0.471355 | LUAD | NCI-H3122 | VX-11e | MVP_top60_DNA+RNA_seed | MAPK3 | 1.650232 | STAD | RF-48 | Ulixertinib |
| MVP_top60_DNA_seed | AKT1 | -0.471188 | LUAD | NCI-H1734 | AT13148 | MVP_top60_DNA+RNA_seed | MDM2 | 1.651727 | STAD | KATOIII | Nutlin-3a (-) |
| MVP_top60_DNA_seed | HDAC1 | -0.470349 | LUAD | Calu-6 | Romidepsin | MVP_top60_DNA+RNA_seed | ERBB2 | 1.655498 | STAD | RF-48 | Lapatinib |
| MVP_top60_DNA_seed | ATM | -0.470102 | LUAD | NCI-H1568 | KU-55933 | MVP_top60_DNA+RNA_seed | PIK3CA | 1.678729 | STAD | KATOIII | AZD8186 |
| MVP_top60_DNA_seed | AKT1 | -0.46818 | LUAD | NCI-H1623 | Uprosertib | MVP_top60_DNA+RNA_seed | ERBB2 | 1.687067 | STAD | MKN45 | Sapitinib |
| MVP_top60_DNA_seed | SRC | -0.467974 | LUAD | LC-2-ad | Bosutinib | MVP_top60_DNA+RNA_seed | ERBB3 | 1.687067 | STAD | MKN45 | Sapitinib |
| MVP_top60_DNA_seed | PIK3CA | -0.465927 | LUAD | NCI-H2228 | GNE-317 | MVP_top60_DNA+RNA_seed | PIK3CA | 1.709535 | STAD | NUGC-4 | AZD8186 |
| MVP_top60_DNA_seed | PIK3CA | -0.465035 | LUAD | EMC-BAC-2 | Buparlisib | MVP_top60_DNA+RNA_seed | EZH2 | 1.7156 | STAD | NUGC-4 | GSK343 |
| MVP_top60_DNA_seed | BRAF | -0.46219 | LUAD | LC-2-ad | PLX-4720 | MVP_top60_DNA+RNA_seed | ERBB2 | 1.715738 | STAD | TMK-1 | Afatinib |
| MVP_top60_DNA_seed | PIK3CA | -0.46151 | LUAD | NCI-H1755 | Buparlisib | MVP_top60_DNA+RNA_seed | GSK3B | 1.71904 | STAD | KATOIII | SB216763 |
| MVP_top60_DNA_seed | TERT | -0.458878 | LUAD | NCI-H1993 | BIBR-1532 | MVP_top60_DNA+RNA_seed | FGFR3 | 1.724345 | STAD | NUGC-4 | AZD4547 |
| MVP_top60_DNA_seed | GSK3B | -0.457544 | LUAD | NCI-H1435 | SB216763 | MVP_top60_DNA+RNA_seed | PARP1 | 1.726646 | STAD | MKN28 | Olaparib |
| MVP_top60_DNA_seed | AKT1 | -0.456887 | LUAD | HCC-78 | AZD5363 | MVP_top60_DNA+RNA_seed | HSP90AA1 | 1.738538 | STAD | KATOIII | Luminespib |
| MVP_top60_DNA_seed | KRAS | -0.455088 | LUAD | EMC-BAC-2 | KRAS (G12C) Inhibitor-12 | MVP_top60_DNA+RNA_seed | HSP90AB1 | 1.738538 | STAD | KATOIII | Luminespib |
| MVP_top60_DNA_seed | AKT1 | -0.453192 | LUAD | PC-14 | Uprosertib | MVP_top60_DNA+RNA_seed | ERBB2 | 1.755136 | STAD | OCUM-1 | Afatinib |
| MVP_top60_DNA_seed | IGF1R | -0.451145 | LUAD | LC-2-ad | GSK1904529A | MVP_top60_DNA+RNA_seed | PIK3CA | 1.757653 | STAD | KATOIII | GNE-317 |
| MVP_top60_DNA_seed | BRAF | -0.450952 | LUAD | Calu-6 | Dabrafenib | MVP_top60_DNA+RNA_seed | HDAC1 | 1.762681 | STAD | NUGC-4 | PCI-34051 |
| MVP_top60_DNA_seed | AKT1 | -0.447143 | LUAD | A549 | GSK2110183B | MVP_top60_DNA+RNA_seed | HSP90AA1 | 1.769519 | STAD | MKN7 | CCT-018159 |
| MVP_top60_DNA_seed | EZH2 | -0.443034 | LUAD | NCI-H1734 | GSK343 | MVP_top60_DNA+RNA_seed | HSP90AB1 | 1.769519 | STAD | MKN7 | CCT-018159 |
| MVP_top60_DNA_seed | EGFR | -0.442988 | LUAD | NCI-H1944 | Osimertinib | MVP_top60_DNA+RNA_seed | PARP1 | 1.777769 | STAD | MKN7 | Rucaparib |
| MVP_top60_DNA_seed | EGFR | -0.442342 | LUAD | A549 | Gefitinib | MVP_top60_DNA+RNA_seed | PARP1 | 1.780279 | STAD | NUGC-4 | Rucaparib |
| MVP_top60_DNA_seed | AKT1 | -0.441499 | LUAD | NCI-H322M | GSK2110183B | MVP_top60_DNA+RNA_seed | MDM2 | 1.791178 | STAD | NUGC-4 | Serdemetan |
| MVP_top60_DNA_seed | MAPK1 | -0.441212 | LUAD | NCI-H838 | VX-11e | MVP_top60_DNA+RNA_seed | EZH2 | 1.821195 | STAD | NUGC-4 | GSK343 |
| MVP_top60_DNA_seed | HDAC1 | -0.440429 | LUAD | NCI-H1435 | Romidepsin | MVP_top60_DNA+RNA_seed | GSK3B | 1.855118 | STAD | NUGC-4 | SB216763 |
| MVP_top60_DNA_seed | EGFR | -0.439857 | LUAD | NCI-H1975 | Gefitinib | MVP_top60_DNA+RNA_seed | MDM2 | 1.944473 | STAD | Hs-746T | Serdemetan |
| MVP_top60_DNA_seed | EZH2 | -0.439588 | LUAD | NCI-H2087 | GSK343 | MVP_top60_DNA+RNA_seed | HSP90AA1 | 1.952751 | STAD | MKN7 | Tanespimycin |
| MVP_top60_DNA_seed | PIK3CA | -0.439283 | LUAD | NCI-H1623 | GNE-317 | MVP_top60_DNA+RNA_seed | HSP90AB1 | 1.952751 | STAD | MKN7 | Tanespimycin |
| MVP_top60_DNA_seed | AKT1 | -0.438822 | LUAD | COR-L105 | Uprosertib | MVP_top60_DNA+RNA_seed | PIK3CA | 1.960357 | STAD | NUGC-4 | Buparlisib |
| MVP_top60_DNA_seed | PIK3CA | -0.437569 | LUAD | NCI-H1792 | Alpelisib | MVP_top60_DNA+RNA_seed | ERBB2 | 1.972143 | STAD | OCUM-1 | Lapatinib |
| MVP_top60_DNA_seed | IGF1R | -0.437178 | LUAD | NCI-H1944 | GSK1904529A | MVP_top60_DNA+RNA_seed | PIK3CA | 1.980186 | STAD | NUGC-4 | GNE-317 |
| MVP_top60_DNA_seed | AKT1 | -0.436715 | LUAD | NCI-H2009 | Uprosertib | MVP_top60_DNA+RNA_seed | ESR1 | 2.096074 | STAD | NUGC-4 | Tamoxifen |
| MVP_top60_DNA_seed | AKT1 | -0.43662 | LUAD | NCI-H1944 | GSK2110183B | MVP_top60_DNA+RNA_seed | HSP90AA1 | 2.101438 | STAD | NUGC-4 | Luminespib |
| MVP_top60_DNA_seed | AURKA | -0.432636 | LUAD | PC-14 | ZM447439 | MVP_top60_DNA+RNA_seed | HSP90AB1 | 2.101438 | STAD | NUGC-4 | Luminespib |

| | | | | | | | | | | | |
|---|---|---|---|---|---|---|---|---|---|---|---|
| MVP_top60_DNA_seed | AURKA | -0.432052 | LUAD | RERF-LC-MS | Alisertib | MVP_top60_DNA+RNA_seed | PARP1 | 2.141636 | STAD | NUGC-4 | Veliparib |
| MVP_top60_DNA_seed | CREBBP | -0.427187 | LUAD | NCI-H1792 | SGC-CBP30 | MVP_top60_DNA+RNA_seed | FGFR3 | 2.16447 | STAD | MKN7 | AZD4547 |
| MVP_top60_DNA_seed | BRAF | -0.427137 | LUAD | NCI-H1435 | SB590885 | MVP_top60_DNA+RNA_seed | FGFR3 | 2.280056 | STAD | NUGC-4 | PD173074 |
| MVP_top60_DNA_seed | EGFR | -0.426522 | LUAD | NCI-H1693 | Osimertinib | MVP_top60_DNA+RNA_seed | PIK3CA | 2.305225 | STAD | KATOIII | Buparlisib |
| MVP_top60_DNA_seed | AKT1 | -0.422941 | LUAD | HCC-44 | MK-2206 | MVP_top60_DNA+RNA_seed | ERBB2 | 2.393403 | STAD | KATOIII | Afatinib |
| MVP_top60_DNA_seed | PIK3CA | -0.42221 | LUAD | NCI-H1437 | Buparlisib | MVP_top60_DNA+RNA_seed | PIK3CA | 2.561985 | STAD | MKN28 | GNE-317 |
| MVP_top60_DNA_seed | IGF1R | -0.419036 | LUAD | NCI-H358 | BMS-536924 | MVP_top60_DNA+RNA_seed | MET | 2.845476 | STAD | NUGC-4 | Crizotinib |
| MVP_top60_DNA_seed | ATM | -0.418749 | LUAD | NCI-H3122 | KU-55933 | MVP_top60_DNA+RNA_seed | GSK3B | -4.461638 | THCA | IHH-4 | KIN001-042 |
| MVP_top60_DNA_seed | AKT1 | -0.417894 | LUAD | NCI-H1755 | AZD5363 | MVP_top60_DNA+RNA_seed | RET | -3.946255 | THCA | CGTH-W-1 | Motesanib |
| MVP_top60_DNA_seed | AKT1 | -0.417502 | LUAD | NCI-H1755 | MK-2206 | MVP_top60_DNA+RNA_seed | BRAF | -3.729294 | THCA | K5 | Dabrafenib |
| MVP_top60_DNA_seed | AKT1 | -0.417351 | LUAD | NCI-H1944 | AZD5363 | MVP_top60_DNA+RNA_seed | BRAF | -3.412404 | THCA | BHT-101 | Dabrafenib |
| MVP_top60_DNA_seed | MAPK1 | -0.417176 | LUAD | NCI-H2009 | VX-11e | MVP_top60_DNA+RNA_seed | BRAF | -3.190105 | THCA | BHT-101 | PLX-4720 |
| MVP_top60_DNA_seed | PIK3CA | -0.415379 | LUAD | PC-14 | GNE-317 | MVP_top60_DNA+RNA_seed | BRAF | -3.135102 | THCA | IHH-4 | Dabrafenib |
| MVP_top60_DNA_seed | KRAS | -0.414707 | LUAD | NCI-H1734 | KRAS (G12C) Inhibitor-12 | MVP_top60_DNA+RNA_seed | BRAF | -3.105461 | THCA | HTC-C3 | Dabrafenib |
| MVP_top60_DNA_seed | EGFR | -0.414656 | LUAD | A549 | Afatinib | MVP_top60_DNA+RNA_seed | BRAF | -2.966357 | THCA | CGTH-W-1 | HG6-64-1 |
| MVP_top60_DNA_seed | ERBB2 | -0.414656 | LUAD | A549 | Afatinib | MVP_top60_DNA+RNA_seed | BRAF | -2.961541 | THCA | K5 | PLX-4720 |
| MVP_top60_DNA_seed | TERT | -0.414478 | LUAD | NCI-H650 | BIBR-1532 | MVP_top60_DNA+RNA_seed | BRAF | -2.905795 | THCA | BHT-101 | PLX-4720 |
| MVP_top60_DNA_seed | MAPK1 | -0.414043 | LUAD | HCC-78 | VX-11e | MVP_top60_DNA+RNA_seed | BRAF | -2.830382 | THCA | HTC-C3 | PLX-4720 |
| MVP_top60_DNA_seed | HDAC1 | -0.412845 | LUAD | NCI-H1651 | PCI-34051 | MVP_top60_DNA+RNA_seed | BRAF | -2.827309 | THCA | K5 | PLX-4720 |
| MVP_top60_DNA_seed | HDAC1 | -0.412333 | LUAD | NCI-H1563 | PCI-34051 | MVP_top60_DNA+RNA_seed | CHEK2 | -2.727623 | THCA | CGTH-W-1 | AZD7762 |
| MVP_top60_DNA_seed | EGFR | -0.411993 | LUAD | NCI-H1693 | AZD3759 | MVP_top60_DNA+RNA_seed | BRAF | -2.688852 | THCA | HTC-C3 | RAF_9304 |
| MVP_top60_DNA_seed | PIK3CA | -0.411127 | LUAD | RERF-LC-KJ | AZD8186 | MVP_top60_DNA+RNA_seed | BRAF | -2.604383 | THCA | BHT-101 | SB590885 |
| MVP_top60_DNA_seed | AURKA | -0.408736 | LUAD | A549 | Alisertib | MVP_top60_DNA+RNA_seed | BRAF | -2.492753 | THCA | HTC-C3 | PLX-4720 |
| MVP_top60_DNA_seed | IGF1R | -0.408345 | LUAD | HCC-44 | NVP-ADW742 | MVP_top60_DNA+RNA_seed | BRAF | -2.480329 | THCA | B-CPAP | Dabrafenib |
| MVP_top60_DNA_seed | MAPK1 | -0.408304 | LUAD | NCI-H1944 | SCH772984 | MVP_top60_DNA+RNA_seed | GSK3B | -2.431374 | THCA | B-CPAP | CHIR-99021 |
| MVP_top60_DNA_seed | AKT1 | -0.407921 | LUAD | RERF-LC-KJ | Uprosertib | MVP_top60_DNA+RNA_seed | BRAF | -2.430714 | THCA | K5 | RAF_9304 |
| MVP_top60_DNA_seed | AKT1 | -0.407521 | LUAD | NCI-H1437 | AT13148 | MVP_top60_DNA+RNA_seed | BRAF | -2.371192 | THCA | BHT-101 | RAF_9304 |
| MVP_top60_DNA_seed | IGF1R | -0.407028 | LUAD | NCI-H2122 | GSK1904529A | MVP_top60_DNA+RNA_seed | GSK3B | -2.310407 | THCA | BHT-101 | CHIR-99021 |
| MVP_top60_DNA_seed | EGFR | -0.406348 | LUAD | NCI-H1944 | Sapitinib | MVP_top60_DNA+RNA_seed | GSK3B | -2.250906 | THCA | ASH-3 | CHIR-99021 |
| MVP_top60_DNA_seed | ERBB2 | -0.406348 | LUAD | NCI-H1944 | Sapitinib | MVP_top60_DNA+RNA_seed | GSK3B | -2.250659 | THCA | B-CPAP | CHIR-99021 |
| MVP_top60_DNA_seed | ERBB3 | -0.406348 | LUAD | NCI-H1944 | Sapitinib | MVP_top60_DNA+RNA_seed | MAPK1 | -2.176373 | THCA | BHT-101 | VX-11e |
| MVP_top60_DNA_seed | HSP90AA1 | -0.406218 | LUAD | NCI-H2023 | Luminespib | MVP_top60_DNA+RNA_seed | BRAF | -2.172551 | THCA | 8305C | Dabrafenib |
| MVP_top60_DNA_seed | SMARCA4 | -0.406091 | LUAD | NCI-H2122 | PFI3 | MVP_top60_DNA+RNA_seed | HDAC1 | -2.1696 | THCA | CGTH-W-1 | JQ12 |
| MVP_top60_DNA_seed | IGF1R | -0.405163 | LUAD | NCI-H1623 | GSK1904529A | MVP_top60_DNA+RNA_seed | BRAF | -2.151843 | THCA | K5 | AZ628 |
| MVP_top60_DNA_seed | IGF1R | -0.40173 | LUAD | LXF-289 | BMS-536924 | MVP_top60_DNA+RNA_seed | RET | -2.119921 | THCA | HTC-C3 | Motesanib |
| MVP_top60_DNA_seed | GSK3B | -0.401089 | LUAD | NCI-H1693 | SB216763 | MVP_top60_DNA+RNA_seed | GSK3B | -2.047005 | THCA | BHT-101 | SB216763 |
| MVP_top60_DNA_seed | IGF1R | -0.401028 | LUAD | NCI-H1781 | BMS-754807 | MVP_top60_DNA+RNA_seed | BRAF | -2.037898 | THCA | 8305C | PLX-4720 |
| MVP_top60_DNA_seed | MET | -0.40006 | LUAD | RERF-LC-KJ | Savolitinib | MVP_top60_DNA+RNA_seed | BRAF | -2.027428 | THCA | CGTH-W-1 | AZ628 |
| MVP_top60_DNA_seed | AURKA | -0.399919 | LUAD | NCI-H1792 | ZM447439 | MVP_top60_DNA+RNA_seed | HDAC1 | -1.95383 | THCA | ML-1 | JQ12 |
| MVP_top60_DNA_seed | PIK3CA | -0.398005 | LUAD | HCC-44 | Alpelisib | MVP_top60_DNA+RNA_seed | BRAF | -1.905405 | THCA | IHH-4 | PLX-4720 |
| MVP_top60_DNA_seed | EGFR | -0.397211 | LUAD | NCI-H1944 | AZD3759 | MVP_top60_DNA+RNA_seed | BRAF | -1.828709 | THCA | TT2609-C02 | HG6-64-1 |

| | | | | | | | | | | | |
|---|---|---|---|---|---|---|---|---|---|---|---|
| MVP_top60_DNA_seed | HDAC1 | -0.396679 | LUAD | PC-14 | Romidepsin | MVP_top60_DNA+RNA_seed | RET | -1.811525 | THCA | TT | Motesanib |
| MVP_top60_DNA_seed | HDAC1 | -0.395466 | LUAD | PC-14 | Entinostat | MVP_top60_DNA+RNA_seed | PDPK1 | -1.799658 | THCA | IHH-4 | OSU-03012 |
| MVP_top60_DNA_seed | MDM2 | -0.394727 | LUAD | LC-2-ad | Nutlin-3a (-) | MVP_top60_DNA+RNA_seed | ATM | -1.797046 | THCA | IHH-4 | KU-60019 |
| MVP_top60_DNA_seed | MET | -0.394085 | LUAD | PC-14 | Crizotinib | MVP_top60_DNA+RNA_seed | GSK3B | -1.793115 | THCA | CGTH-W-1 | CHIR-99021 |
| MVP_top60_DNA_seed | TERT | -0.394075 | LUAD | RERF-LC-KJ | BIBR-1532 | MVP_top60_DNA+RNA_seed | BRAF | -1.773559 | THCA | 8305C | PLX-4720 |
| MVP_top60_DNA_seed | EGFR | -0.389979 | LUAD | NCI-H2030 | Afatinib | MVP_top60_DNA+RNA_seed | PTK2 | -1.758295 | THCA | B-CPAP | PF-562271 |
| MVP_top60_DNA_seed | ERBB2 | -0.389979 | LUAD | NCI-H2030 | Afatinib | MVP_top60_DNA+RNA_seed | PTK2 | -1.743982 | THCA | 8305C | PF-562271 |
| MVP_top60_DNA_seed | HDAC1 | -0.385959 | LUAD | NCI-H1792 | Entinostat | MVP_top60_DNA+RNA_seed | PTK2 | -1.742409 | THCA | KMH-2 | PF-562271 |
| MVP_top60_DNA_seed | HDAC1 | -0.384063 | LUAD | NCI-H1666 | Romidepsin | MVP_top60_DNA+RNA_seed | AKT1 | -1.742069 | THCA | BHT-101 | MK-2206 |
| MVP_top60_DNA_seed | EGFR | -0.383381 | LUAD | EKVX | Osimertinib | MVP_top60_DNA+RNA_seed | AKT2 | -1.742069 | THCA | BHT-101 | MK-2206 |
| MVP_top60_DNA_seed | HSP90AA1 | -0.382024 | LUAD | NCI-H23 | Tanespimycin | MVP_top60_DNA+RNA_seed | BRAF | -1.672569 | THCA | IHH-4 | SB590885 |
| MVP_top60_DNA_seed | HSP90AA1 | -0.380703 | LUAD | NCI-H838 | CCT-018159 | MVP_top60_DNA+RNA_seed | PDPK1 | -1.668786 | THCA | TT2609-C02 | OSU-03012 |
| MVP_top60_DNA_seed | PIK3CA | -0.380447 | LUAD | COR-L105 | Alpelisib | MVP_top60_DNA+RNA_seed | GSK3B | -1.638434 | THCA | HTC-C3 | KIN001-042 |
| MVP_top60_DNA_seed | AKT1 | -0.380179 | LUAD | A549 | Uprosertib | MVP_top60_DNA+RNA_seed | HDAC1 | -1.616837 | THCA | TT2609-C02 | JQ12 |
| MVP_top60_DNA_seed | IGF1R | -0.37995 | LUAD | EMC-BAC-1 | BMS-536924 | MVP_top60_DNA+RNA_seed | GSK3B | -1.607433 | THCA | CGTH-W-1 | CHIR-99021 |
| MVP_top60_DNA_seed | AKT1 | -0.377776 | LUAD | PC-14 | Uprosertib | MVP_top60_DNA+RNA_seed | GSK3B | -1.604268 | THCA | ML-1 | CHIR-99021 |
| MVP_top60_DNA_seed | PIK3CA | -0.377532 | LUAD | NCI-H1944 | AZD8186 | MVP_top60_DNA+RNA_seed | BRAF | -1.554192 | THCA | B-CPAP | PLX-4720 |
| MVP_top60_DNA_seed | FGFR1 | -0.37604 | LUAD | NCI-H1437 | PD173074 | MVP_top60_DNA+RNA_seed | BRAF | -1.542311 | THCA | FTC-133 | PLX-4720 |
| MVP_top60_DNA_seed | GSK3B | -0.373946 | LUAD | NCI-H1623 | SB216763 | MVP_top60_DNA+RNA_seed | GSK3B | -1.535477 | THCA | K5 | CHIR-99021 |
| MVP_top60_DNA_seed | MAPK1 | -0.372104 | LUAD | NCI-H2087 | Ulixertinib | MVP_top60_DNA+RNA_seed | BRAF | -1.51961 | THCA | B-CPAP | PLX-4720 |
| MVP_top60_DNA_seed | MAPK1 | -0.371559 | LUAD | PC-14 | VX-11e | MVP_top60_DNA+RNA_seed | PTK2 | -1.455613 | THCA | TT2609-C02 | PF-562271 |
| MVP_top60_DNA_seed | MAPK1 | -0.371154 | LUAD | NCI-H2405 | SCH772984 | MVP_top60_DNA+RNA_seed | HDAC1 | -1.447506 | THCA | IHH-4 | JQ12 |
| MVP_top60_DNA_seed | AURKA | -0.370894 | LUAD | HCC-78 | Alisertib | MVP_top60_DNA+RNA_seed | ATM | -1.435817 | THCA | IHH-4 | KU-55933 |
| MVP_top60_DNA_seed | BRAF | -0.368234 | LUAD | A549 | Dabrafenib | MVP_top60_DNA+RNA_seed | BRAF | -1.423171 | THCA | IHH-4 | PLX-4720 |
| MVP_top60_DNA_seed | AKT1 | -0.368233 | LUAD | NCI-H3122 | Uprosertib | MVP_top60_DNA+RNA_seed | GSK3B | -1.386729 | THCA | K5 | AZD7969 |
| MVP_top60_DNA_seed | AKT1 | -0.366594 | LUAD | NCI-H1781 | AZD5363 | MVP_top60_DNA+RNA_seed | BRAF | -1.362543 | THCA | 8305C | SB590885 |
| MVP_top60_DNA_seed | MAPK1 | -0.366394 | LUAD | HCC-44 | Ulixertinib | MVP_top60_DNA+RNA_seed | PTK2 | -1.348531 | THCA | CAL-62 | PF-562271 |
| MVP_top60_DNA_seed | AKT1 | -0.364285 | LUAD | NCI-H3122 | AZD5363 | MVP_top60_DNA+RNA_seed | HDAC1 | -1.338739 | THCA | TT2609-C02 | Dacinostat |
| MVP_top60_DNA_seed | MAPK1 | -0.36368 | LUAD | LXF-289 | VX-11e | MVP_top60_DNA+RNA_seed | AKT1 | -1.335264 | THCA | TT2609-C02 | AKT inhibitor VIII |
| MVP_top60_DNA_seed | MET | -0.36361 | LUAD | NCI-H650 | Savolitinib | MVP_top60_DNA+RNA_seed | AKT2 | -1.335264 | THCA | TT2609-C02 | AKT inhibitor VIII |
| MVP_top60_DNA_seed | KRAS | -0.360798 | LUAD | NCI-H1781 | KRAS (G12C) Inhibitor-12 | MVP_top60_DNA+RNA_seed | AKT3 | -1.335264 | THCA | TT2609-C02 | AKT inhibitor VIII |
| MVP_top60_DNA_seed | AKT1 | -0.35713 | LUAD | COR-L105 | Ipatasertib | MVP_top60_DNA+RNA_seed | GSK3B | -1.329082 | THCA | 8505C | CHIR-99021 |
| MVP_top60_DNA_seed | SMARCA4 | -0.355079 | LUAD | RERF-LC-KJ | PFI3 | MVP_top60_DNA+RNA_seed | BRAF | -1.308841 | THCA | IHH-4 | RAF_9304 |
| MVP_top60_DNA_seed | BRAF | -0.353155 | LUAD | NCI-H1651 | PLX-4720 | MVP_top60_DNA+RNA_seed | CHEK2 | -1.308715 | THCA | IHH-4 | AZD7762 |
| MVP_top60_DNA_seed | AKT1 | -0.352144 | LUAD | NCI-H1755 | Ipatasertib | MVP_top60_DNA+RNA_seed | ATM | -1.291053 | THCA | BHT-101 | KU-55933 |
| MVP_top60_DNA_seed | FGFR1 | -0.350758 | LUAD | NCI-H838 | AZD4547 | MVP_top60_DNA+RNA_seed | MAPK1 | -1.283128 | THCA | HTC-C3 | VX-11e |
| MVP_top60_DNA_seed | PIK3CA | -0.3506 | LUAD | NCI-H2009 | GNE-317 | MVP_top60_DNA+RNA_seed | BRAF | -1.282376 | THCA | B-CPAP | HG6-64-1 |
| MVP_top60_DNA_seed | AKT1 | -0.350493 | LUAD | NCI-H1755 | Uprosertib | MVP_top60_DNA+RNA_seed | ATM | -1.265003 | THCA | K5 | torin2 |
| MVP_top60_DNA_seed | MAPK1 | -0.350207 | LUAD | EKVX | SCH772984 | MVP_top60_DNA+RNA_seed | AKT1 | -1.262621 | THCA | BHT-101 | AKT inhibitor VIII |
| MVP_top60_DNA_seed | AKT1 | -0.350168 | LUAD | NCI-H838 | Uprosertib | MVP_top60_DNA+RNA_seed | AKT2 | -1.262621 | THCA | BHT-101 | AKT inhibitor VIII |
| MVP_top60_DNA_seed | SMARCA4 | -0.348592 | LUAD | NCI-H522 | PFI3 | MVP_top60_DNA+RNA_seed | AKT3 | -1.262621 | THCA | BHT-101 | AKT inhibitor VIII |

| | | | | | | | | | | | |
|---|---|---|---|---|---|---|---|---|---|---|---|
| MVP_top60_DNA_seed | SRC | -0.34845 | LUAD | NCI-H1568 | Bosutinib | MVP_top60_DNA+RNA_seed | GSK3B | -1.239157 | THCA | B-CPAP | AZD7969 |
| MVP_top60_DNA_seed | KRAS | -0.347633 | LUAD | RERF-LC-MS | KRAS (G12C) Inhibitor-1[illegible] | MVP_top60_DNA+RNA_seed | PTK2 | -1.231651 | THCA | CGTH-W-1 | PF-562271 |
| MVP_top60_DNA_seed | AKT1 | -0.346737 | LUAD | NCI-H1944 | Uprosertib | MVP_top60_DNA+RNA_seed | RET | -1.225834 | THCA | TT2609-C02 | Motesanib |
| MVP_top60_DNA_seed | AKT1 | -0.346317 | LUAD | EMC-BAC-2 | AT13148 | MVP_top60_DNA+RNA_seed | BRAF | -1.224839 | THCA | KMH-2 | SB590885 |
| MVP_top60_DNA_seed | IGF1R | -0.345865 | LUAD | EMC-BAC-2 | Linsitinib | MVP_top60_DNA+RNA_seed | ATM | -1.218818 | THCA | CAL-62 | torin2 |
| MVP_top60_DNA_seed | AKT1 | -0.345203 | LUAD | NCI-H1437 | Uprosertib | MVP_top60_DNA+RNA_seed | ATM | -1.205962 | THCA | 8305C | KU-60019 |
| MVP_top60_DNA_seed | EZH2 | -0.344107 | LUAD | COR-L105 | GSK343 | MVP_top60_DNA+RNA_seed | BRAF | -1.201393 | THCA | KMH-2 | Dabrafenib |
| MVP_top60_DNA_seed | EGFR | -0.343669 | LUAD | A549 | Osimertinib | MVP_top60_DNA+RNA_seed | GSK3B | -1.192884 | THCA | KMH-2 | SB216763 |
| MVP_top60_DNA_seed | IGF1R | -0.341017 | LUAD | NCI-H322M | NVP-ADW742 | MVP_top60_DNA+RNA_seed | ATM | -1.178386 | THCA | B-CPAP | KU-60019 |
| MVP_top60_DNA_seed | AURKA | -0.337657 | LUAD | Calu-6 | ZM447439 | MVP_top60_DNA+RNA_seed | BRAF | -1.173142 | THCA | ML-1 | HG6-64-1 |
| MVP_top60_DNA_seed | MDM2 | -0.337588 | LUAD | NCI-H358 | Serdemetan | MVP_top60_DNA+RNA_seed | BRAF | -1.163381 | THCA | 8305C | HG6-64-1 |
| MVP_top60_DNA_seed | AKT1 | -0.335884 | LUAD | RERF-LC-MS | Afuresertib | MVP_top60_DNA+RNA_seed | PDPK1 | -1.162175 | THCA | 8505C | OSU-03012 |
| MVP_top60_DNA_seed | HSP90AA1 | -0.334286 | LUAD | NCI-H1975 | Luminespib | MVP_top60_DNA+RNA_seed | HDAC1 | -1.160062 | THCA | HTC-C3 | JQ12 |
| MVP_top60_DNA_seed | AKT1 | -0.334263 | LUAD | LXF-289 | Uprosertib | MVP_top60_DNA+RNA_seed | RET | -1.158994 | THCA | CAL-62 | Motesanib |
| MVP_top60_DNA_seed | EGFR | -0.334096 | LUAD | EKVX | Gefitinib | MVP_top60_DNA+RNA_seed | AKT1 | -1.15598 | THCA | ML-1 | AKT inhibitor VIII |
| MVP_top60_DNA_seed | AURKA | -0.334005 | LUAD | A549 | ZM447439 | MVP_top60_DNA+RNA_seed | AKT2 | -1.15598 | THCA | ML-1 | AKT inhibitor VIII |
| MVP_top60_DNA_seed | AKT1 | -0.329797 | LUAD | HCC-44 | Afuresertib | MVP_top60_DNA+RNA_seed | AKT3 | -1.15598 | THCA | ML-1 | AKT inhibitor VIII |
| MVP_top60_DNA_seed | IGF1R | -0.329524 | LUAD | NCI-H838 | NVP-ADW742 | MVP_top60_DNA+RNA_seed | BRAF | -1.154696 | THCA | ML-1 | PLX-4720 |
| MVP_top60_DNA_seed | HDAC1 | -0.328513 | LUAD | NCI-H650 | Romidepsin | MVP_top60_DNA+RNA_seed | HDAC1 | -1.152329 | THCA | CGTH-W-1 | Dacinostat |
| MVP_top60_DNA_seed | MET | -0.328416 | LUAD | NCI-H1437 | Savolitinib | MVP_top60_DNA+RNA_seed | AKT1 | -1.135161 | THCA | BHT-101 | BAY AKT1 |
| MVP_top60_DNA_seed | GSK3B | -0.32784 | LUAD | RERF-LC-KJ | SB216763 | MVP_top60_DNA+RNA_seed | AKT2 | -1.135161 | THCA | BHT-101 | BAY AKT1 |
| MVP_top60_DNA_seed | EZH2 | -0.327654 | LUAD | RERF-LC-KJ | GSK343 | MVP_top60_DNA+RNA_seed | AKT3 | -1.135161 | THCA | BHT-101 | BAY AKT1 |
| MVP_top60_DNA_seed | TERT | -0.327329 | LUAD | NCI-H23 | BIBR-1532 | MVP_top60_DNA+RNA_seed | GSK3B | -1.131743 | THCA | CAL-62 | AZD7969 |
| MVP_top60_DNA_seed | FGFR1 | -0.324083 | LUAD | NCI-H1975 | AZD4547 | MVP_top60_DNA+RNA_seed | GSK3B | -1.12833 | THCA | IHH-4 | AZD7969 |
| MVP_top60_DNA_seed | SRC | -0.323756 | LUAD | A549 | Bosutinib | MVP_top60_DNA+RNA_seed | PTK2 | -1.124678 | THCA | IHH-4 | PF-562271 |
| MVP_top60_DNA_seed | IGF1R | -0.323066 | LUAD | NCI-H2030 | BMS-536924 | MVP_top60_DNA+RNA_seed | PTK2 | -1.110141 | THCA | ASH-3 | PF-562271 |
| MVP_top60_DNA_seed | AKT1 | -0.322248 | LUAD | Calu-3 | Uprosertib | MVP_top60_DNA+RNA_seed | PDPK1 | -1.100357 | THCA | CAL-62 | OSU-03012 |
| MVP_top60_DNA_seed | AKT1 | -0.32088 | LUAD | LXF-289 | Uprosertib | MVP_top60_DNA+RNA_seed | GSK3B | -1.091751 | THCA | IHH-4 | SB216763 |
| MVP_top60_DNA_seed | MAPK1 | -0.320552 | LUAD | NCI-H3122 | Ulixertinib | MVP_top60_DNA+RNA_seed | ATM | -1.08938 | THCA | 8305C | KU-55933 |
| MVP_top60_DNA_seed | CREBBP | -0.318636 | LUAD | NCI-H1435 | SGC-CBP30 | MVP_top60_DNA+RNA_seed | BRAF | -1.088167 | THCA | 8305C | RAF_9304 |
| MVP_top60_DNA_seed | MAPK1 | -0.318421 | LUAD | NCI-H23 | Ulixertinib | MVP_top60_DNA+RNA_seed | GSK3B | -1.079866 | THCA | 8505C | CHIR-99021 |
| MVP_top60_DNA_seed | IGF1R | -0.318349 | LUAD | NCI-H1975 | BMS-754807 | MVP_top60_DNA+RNA_seed | AKT1 | -1.066972 | THCA | TT | AKT inhibitor VIII |
| MVP_top60_DNA_seed | GSK3B | -0.317591 | LUAD | NCI-H1437 | SB216763 | MVP_top60_DNA+RNA_seed | AKT2 | -1.066972 | THCA | TT | AKT inhibitor VIII |
| MVP_top60_DNA_seed | IGF1R | -0.317039 | LUAD | NCI-H1975 | Linsitinib | MVP_top60_DNA+RNA_seed | AKT3 | -1.066972 | THCA | TT | AKT inhibitor VIII |
| MVP_top60_DNA_seed | GSK3B | -0.316424 | LUAD | NCI-H322M | SB216763 | MVP_top60_DNA+RNA_seed | GSK3B | -1.054235 | THCA | KMH-2 | CHIR-99021 |
| MVP_top60_DNA_seed | ATM | -0.312928 | LUAD | NCI-H1755 | KU-55933 | MVP_top60_DNA+RNA_seed | BRAF | -1.031488 | THCA | TT2609-C02 | PLX-4720 |
| MVP_top60_DNA_seed | EGFR | -0.312584 | LUAD | HCC-78 | Afatinib | MVP_top60_DNA+RNA_seed | BRAF | -1.023462 | THCA | IHH-4 | HG6-64-1 |
| MVP_top60_DNA_seed | ERBB2 | -0.312584 | LUAD | HCC-78 | Afatinib | MVP_top60_DNA+RNA_seed | CHEK2 | -1.013809 | THCA | CAL-62 | AZD7762 |
| MVP_top60_DNA_seed | MET | -0.31253 | LUAD | NCI-H838 | Crizotinib | MVP_top60_DNA+RNA_seed | HDAC1 | -1.009404 | THCA | HTC-C3 | Belinostat |
| MVP_top60_DNA_seed | AKT1 | -0.312058 | LUAD | NCI-H1993 | AT13148 | MVP_top60_DNA+RNA_seed | PTK2 | -1.005164 | THCA | ML-1 | PF-562271 |
| MVP_top60_DNA_seed | HDAC1 | -0.31122 | LUAD | NCI-H1781 | PCI-34051 | MVP_top60_DNA+RNA_seed | GSK3B | -1.001871 | THCA | K5 | CHIR-99021 |

| | | | | | | | | | | | |
|---|---|---|---|---|---|---|---|---|---|---|---|
| MVP_top60_DNA_seed | IGF1R | -0.310378 | LUAD | Calu-6 | BMS-754807 | MVP_top60_DNA+RNA_seed | HDAC1 | -0.992644 | THCA | B-CPAP | JQ12 |
| MVP_top60_DNA_seed | EGFR | -0.309857 | LUAD | NCI-H1792 | Lapatinib | MVP_top60_DNA+RNA_seed | PTK2 | -0.983662 | THCA | BHT-101 | PF-562271 |
| MVP_top60_DNA_seed | ERBB2 | -0.309857 | LUAD | NCI-H1792 | Lapatinib | MVP_top60_DNA+RNA_seed | RET | -0.982642 | THCA | BHT-101 | Motesanib |
| MVP_top60_DNA_seed | SMARCA4 | -0.309242 | LUAD | NCI-H1651 | PFI3 | MVP_top60_DNA+RNA_seed | MAPK1 | -0.950621 | THCA | K5 | FR-180204 |
| MVP_top60_DNA_seed | AKT1 | -0.30902 | LUAD | NCI-H1435 | AT13148 | MVP_top60_DNA+RNA_seed | MAPK3 | -0.950621 | THCA | K5 | FR-180204 |
| MVP_top60_DNA_seed | FGFR1 | -0.307087 | LUAD | NCI-H1573 | AZD4547 | MVP_top60_DNA+RNA_seed | BRAF | -0.94498 | THCA | CAL-62 | RAF_9304 |
| MVP_top60_DNA_seed | GSK3B | -0.30684 | LUAD | LC-2-ad | CHIR-99021 | MVP_top60_DNA+RNA_seed | ATM | -0.944882 | THCA | HTC-C3 | CP466722 |
| MVP_top60_DNA_seed | AKT1 | -0.305593 | LUAD | EMC-BAC-1 | AT13148 | MVP_top60_DNA+RNA_seed | ATM | -0.941654 | THCA | 8505C | KU-60019 |
| MVP_top60_DNA_seed | MAPK1 | -0.305471 | LUAD | NCI-H1568 | SCH772984 | MVP_top60_DNA+RNA_seed | GSK3B | -0.93908 | THCA | RO82-W-1 | CHIR-99021 |
| MVP_top60_DNA_seed | MDM2 | -0.304836 | LUAD | LC-2-ad | Serdemetan | MVP_top60_DNA+RNA_seed | MAPK1 | -0.933707 | THCA | B-CPAP | VX-11e |
| MVP_top60_DNA_seed | ATM | -0.304213 | LUAD | NCI-H2085 | KU-55933 | MVP_top60_DNA+RNA_seed | MAPK1 | -0.933083 | THCA | TT2609-C02 | VX-11e |
| MVP_top60_DNA_seed | FGFR1 | -0.301253 | LUAD | NCI-H1755 | AZD4547 | MVP_top60_DNA+RNA_seed | GSK3B | -0.925468 | THCA | TT2609-C02 | CHIR-99021 |
| MVP_top60_DNA_seed | IGF1R | -0.300638 | LUAD | NCI-H2009 | NVP-ADW742 | MVP_top60_DNA+RNA_seed | HDAC1 | -0.92111 | THCA | B-CPAP | CAY10603 |
| MVP_top60_DNA_seed | EGFR | -0.299689 | LUAD | NCI-H1693 | Gefitinib | MVP_top60_DNA+RNA_seed | MAPK1 | -0.915528 | THCA | 8505C | VX-11e |
| MVP_top60_DNA_seed | AURKA | -0.298721 | LUAD | NCI-H1792 | Alisertib | MVP_top60_DNA+RNA_seed | GSK3B | -0.904701 | THCA | B-CPAP | KIN001-042 |
| MVP_top60_DNA_seed | IGF1R | -0.297882 | LUAD | NCI-H1666 | NVP-ADW742 | MVP_top60_DNA+RNA_seed | MAPK1 | -0.899585 | THCA | IHH-4 | VX-11e |
| MVP_top60_DNA_seed | FGFR1 | -0.29747 | LUAD | LXF-289 | PD173074 | MVP_top60_DNA+RNA_seed | ATM | -0.892202 | THCA | CGTH-W-1 | torin2 |
| MVP_top60_DNA_seed | EGFR | -0.297008 | LUAD | NCI-H2342 | AZD3759 | MVP_top60_DNA+RNA_seed | HDAC1 | -0.888667 | THCA | 8505C | Dacinostat |
| MVP_top60_DNA_seed | AKT1 | -0.296447 | LUAD | NCI-H1437 | AZD5363 | MVP_top60_DNA+RNA_seed | HDAC1 | -0.878096 | THCA | 8505C | JQ12 |
| MVP_top60_DNA_seed | AURKA | -0.294825 | LUAD | PC-14 | Alisertib | MVP_top60_DNA+RNA_seed | BRAF | -0.870308 | THCA | TT2609-C02 | PLX-4720 |
| MVP_top60_DNA_seed | BRAF | -0.294686 | LUAD | NCI-H2342 | PLX-4720 | MVP_top60_DNA+RNA_seed | ATM | -0.868791 | THCA | ML-1 | KU-55933 |
| MVP_top60_DNA_seed | AKT1 | -0.294462 | LUAD | RERF-LC-KJ | AT13148 | MVP_top60_DNA+RNA_seed | HDAC1 | -0.85957 | THCA | BHT-101 | AR-42 |
| MVP_top60_DNA_seed | HDAC1 | -0.293966 | LUAD | NCI-H1651 | Entinostat | MVP_top60_DNA+RNA_seed | RET | -0.85325 | THCA | RO82-W-1 | Motesanib |
| MVP_top60_DNA_seed | KRAS | -0.29333 | LUAD | HCC-44 | KRAS (G12C) Inhibitor-1 | MVP_top60_DNA+RNA_seed | BRAF | -0.842455 | THCA | HTC-C3 | HG6-64-1 |
| MVP_top60_DNA_seed | AKT1 | -0.291047 | LUAD | RERF-LC-KJ | Ipatasertib | MVP_top60_DNA+RNA_seed | ATM | -0.825398 | THCA | KMH-2 | CP466722 |
| MVP_top60_DNA_seed | AKT1 | -0.290722 | LUAD | NCI-H1755 | Uprosertib | MVP_top60_DNA+RNA_seed | HDAC1 | -0.817087 | THCA | ML-1 | Dacinostat |
| MVP_top60_DNA_seed | EGFR | -0.287944 | LUAD | NCI-H1650 | Gefitinib | MVP_top60_DNA+RNA_seed | BRAF | -0.813015 | THCA | HTC-C3 | SB590885 |
| MVP_top60_DNA_seed | MET | -0.287668 | LUAD | NCI-H2009 | Crizotinib | MVP_top60_DNA+RNA_seed | PDPK1 | -0.808843 | THCA | CGTH-W-1 | BX-912 |
| MVP_top60_DNA_seed | AKT1 | -0.2866 | LUAD | PC-14 | Ipatasertib | MVP_top60_DNA+RNA_seed | GSK3B | -0.803248 | THCA | ASH-3 | CHIR-99021 |
| MVP_top60_DNA_seed | HSP90AA1 | -0.284152 | LUAD | LXF-289 | Luminespib | MVP_top60_DNA+RNA_seed | CHEK2 | -0.802362 | THCA | FTC-133 | AZD7762 |
| MVP_top60_DNA_seed | HSP90AA1 | -0.282483 | LUAD | HOP-62 | Tanespimycin | MVP_top60_DNA+RNA_seed | PIK3CG | -0.800033 | THCA | B-CPAP | PIK-93 |
| MVP_top60_DNA_seed | AKT1 | -0.280823 | LUAD | EMC-BAC-1 | Uprosertib | MVP_top60_DNA+RNA_seed | GSK3B | -0.799404 | THCA | 8305C | CHIR-99021 |
| MVP_top60_DNA_seed | MDM2 | -0.279216 | LUAD | NCI-H1666 | Nutlin-3a (-) | MVP_top60_DNA+RNA_seed | CHEK2 | -0.791571 | THCA | ML-1 | AZD7762 |
| MVP_top60_DNA_seed | IGF1R | -0.27861 | LUAD | NCI-H1993 | BMS-754807 | MVP_top60_DNA+RNA_seed | RET | -0.791025 | THCA | 8505C | Motesanib |
| MVP_top60_DNA_seed | BRAF | -0.278561 | LUAD | NCI-H2087 | Dabrafenib | MVP_top60_DNA+RNA_seed | ATM | -0.773274 | THCA | TT2609-C02 | KU-60019 |
| MVP_top60_DNA_seed | AKT1 | -0.277781 | LUAD | HCC-78 | MK-2206 | MVP_top60_DNA+RNA_seed | AKT1 | -0.773118 | THCA | FTC-133 | AKT inhibitor VIII |
| MVP_top60_DNA_seed | FGFR1 | -0.277235 | LUAD | NCI-H1623 | AZD4547 | MVP_top60_DNA+RNA_seed | AKT2 | -0.773118 | THCA | FTC-133 | AKT inhibitor VIII |
| MVP_top60_DNA_seed | AKT1 | -0.277097 | LUAD | A549 | Ipatasertib | MVP_top60_DNA+RNA_seed | AKT3 | -0.773118 | THCA | FTC-133 | AKT inhibitor VIII |
| MVP_top60_DNA_seed | EZH2 | -0.276916 | LUAD | NCI-H3122 | GSK343 | MVP_top60_DNA+RNA_seed | GSK3B | -0.771928 | THCA | RO82-W-1 | AZD7969 |
| MVP_top60_DNA_seed | MDM2 | -0.275424 | LUAD | RERF-LC-MS | Serdemetan | MVP_top60_DNA+RNA_seed | HDAC1 | -0.76804 | THCA | B-CPAP | Belinostat |
| MVP_top60_DNA_seed | MET | -0.274471 | LUAD | NCI-H1435 | Savolitinib | MVP_top60_DNA+RNA_seed | PDPK1 | -0.765822 | THCA | 8305C | OSU-03012 |

| | | | | | | | | | | | |
|---|---|---|---|---|---|---|---|---|---|---|---|
| MVP_top60_DNA_seed | PIK3CA | -0.274163 | LUAD | RERF-LC-KJ | Alpelisib | MVP_top60_DNA+RNA_seed | BRAF | -0.763833 | THCA | B-CPAP | RAF_9304 |
| MVP_top60_DNA_seed | GSK3B | -0.273202 | LUAD | NCI-H1734 | SB216763 | MVP_top60_DNA+RNA_seed | HDAC1 | -0.763123 | THCA | BHT-101 | Belinostat |
| MVP_top60_DNA_seed | AKT1 | -0.273089 | LUAD | NCI-H322M | Afuresertib | MVP_top60_DNA+RNA_seed | BRAF | -0.758091 | THCA | RO82-W-1 | PLX-4720 |
| MVP_top60_DNA_seed | PIK3CA | -0.27088 | LUAD | PC-14 | Buparlisib | MVP_top60_DNA+RNA_seed | HDAC1 | -0.747378 | THCA | K5 | Entinostat |
| MVP_top60_DNA_seed | SMARCA4 | -0.27023 | LUAD | NCI-H322M | PFI3 | MVP_top60_DNA+RNA_seed | HDAC1 | -0.728411 | THCA | KMH-2 | Dacinostat |
| MVP_top60_DNA_seed | HDAC1 | -0.269725 | LUAD | VMRC-LCD | Entinostat | MVP_top60_DNA+RNA_seed | BRAF | -0.71913 | THCA | 8505C | Dabrafenib |
| MVP_top60_DNA_seed | AKT1 | -0.269722 | LUAD | NCI-H838 | AZD5363 | MVP_top60_DNA+RNA_seed | GSK3B | -0.718318 | THCA | K5 | SB216763 |
| MVP_top60_DNA_seed | PIK3CA | -0.268148 | LUAD | NCI-H1781 | Buparlisib | MVP_top60_DNA+RNA_seed | MAPK1 | -0.715936 | THCA | B-CPAP | FR-180204 |
| MVP_top60_DNA_seed | HSP90AA1 | -0.267831 | LUAD | NCI-H522 | Tanespimycin | MVP_top60_DNA+RNA_seed | MAPK3 | -0.715936 | THCA | B-CPAP | FR-180204 |
| MVP_top60_DNA_seed | MDM2 | -0.267588 | LUAD | NCI-H1563 | Nutlin-3a (-) | MVP_top60_DNA+RNA_seed | ATM | -0.711223 | THCA | ML-1 | CP466722 |
| MVP_top60_DNA_seed | MAPK1 | -0.266353 | LUAD | NCI-H2009 | SCH772984 | MVP_top60_DNA+RNA_seed | AKT1 | -0.710688 | THCA | IHH-4 | MK-2206 |
| MVP_top60_DNA_seed | EGFR | -0.266246 | LUAD | RERF-LC-KJ | Sapitinib | MVP_top60_DNA+RNA_seed | AKT2 | -0.710688 | THCA | IHH-4 | MK-2206 |
| MVP_top60_DNA_seed | ERBB2 | -0.266246 | LUAD | RERF-LC-KJ | Sapitinib | MVP_top60_DNA+RNA_seed | HDAC1 | -0.70789 | THCA | 8305C | Dacinostat |
| MVP_top60_DNA_seed | ERBB3 | -0.266246 | LUAD | RERF-LC-KJ | Sapitinib | MVP_top60_DNA+RNA_seed | ATM | -0.707837 | THCA | K5 | KU-60019 |
| MVP_top60_DNA_seed | HSP90AA1 | -0.266191 | LUAD | NCI-H1437 | Elesclomol | MVP_top60_DNA+RNA_seed | BRAF | -0.706258 | THCA | CAL-62 | Dabrafenib |
| MVP_top60_DNA_seed | TP53 | -0.266147 | LUAD | NCI-H2030 | MIRA-1 | MVP_top60_DNA+RNA_seed | HDAC1 | -0.69876 | THCA | HTC-C3 | CAY10603 |
| MVP_top60_DNA_seed | EGFR | -0.264137 | LUAD | NCI-H1792 | Erlotinib | MVP_top60_DNA+RNA_seed | PIK3CG | -0.691587 | THCA | BHT-101 | PIK-93 |
| MVP_top60_DNA_seed | KRAS | -0.263595 | LUAD | LXF-289 | KRAS (G12C) Inhibitor-12 | MVP_top60_DNA+RNA_seed | PDPK1 | -0.687044 | THCA | ML-1 | OSU-03012 |
| MVP_top60_DNA_seed | EGFR | -0.261681 | LUAD | LC-2-ad | Lapatinib | MVP_top60_DNA+RNA_seed | BRAF | -0.686113 | THCA | TT2609-C02 | SB590885 |
| MVP_top60_DNA_seed | ERBB2 | -0.261681 | LUAD | LC-2-ad | Lapatinib | MVP_top60_DNA+RNA_seed | HDAC1 | -0.68595 | THCA | ML-1 | Belinostat |
| MVP_top60_DNA_seed | HSP90AA1 | -0.259186 | LUAD | NCI-H1650 | Luminespib | MVP_top60_DNA+RNA_seed | HDAC1 | -0.683656 | THCA | HTC-C3 | AR-42 |
| MVP_top60_DNA_seed | AKT1 | -0.258985 | LUAD | NCI-H1355 | AT13148 | MVP_top60_DNA+RNA_seed | ATM | -0.674507 | THCA | ASH-3 | KU-60019 |
| MVP_top60_DNA_seed | EGFR | -0.25795 | LUAD | NCI-H2342 | Sapitinib | MVP_top60_DNA+RNA_seed | GSK3B | -0.671053 | THCA | CGTH-W-1 | SB216763 |
| MVP_top60_DNA_seed | ERBB2 | -0.25795 | LUAD | NCI-H2342 | Sapitinib | MVP_top60_DNA+RNA_seed | PDPK1 | -0.667673 | THCA | ASH-3 | OSU-03012 |
| MVP_top60_DNA_seed | ERBB3 | -0.25795 | LUAD | NCI-H2342 | Sapitinib | MVP_top60_DNA+RNA_seed | HDAC1 | -0.665707 | THCA | BHT-101 | JQ12 |
| MVP_top60_DNA_seed | AKT1 | -0.256224 | LUAD | NCI-H2122 | AT13148 | MVP_top60_DNA+RNA_seed | MAPK1 | -0.650267 | THCA | BHT-101 | FR-180204 |
| MVP_top60_DNA_seed | AKT1 | -0.256016 | LUAD | NCI-H1563 | Ipatasertib | MVP_top60_DNA+RNA_seed | MAPK3 | -0.650267 | THCA | BHT-101 | FR-180204 |
| MVP_top60_DNA_seed | FGFR1 | -0.255894 | LUAD | LC-2-ad | AZD4547 | MVP_top60_DNA+RNA_seed | PTK2 | -0.641648 | THCA | 8505C | PF-562271 |
| MVP_top60_DNA_seed | HDAC1 | -0.255644 | LUAD | NCI-H3122 | Romidepsin | MVP_top60_DNA+RNA_seed | ATM | -0.632848 | THCA | IHH-4 | CP466722 |
| MVP_top60_DNA_seed | AKT1 | -0.254504 | LUAD | NCI-H2023 | Ipatasertib | MVP_top60_DNA+RNA_seed | HDAC1 | -0.632105 | THCA | BHT-101 | CAY10603 |
| MVP_top60_DNA_seed | SRC | -0.254038 | LUAD | NCI-H1993 | Bosutinib | MVP_top60_DNA+RNA_seed | PDPK1 | -0.627212 | THCA | KMH-2 | BX-912 |
| MVP_top60_DNA_seed | AURKA | -0.252427 | LUAD | NCI-H1734 | ZM447439 | MVP_top60_DNA+RNA_seed | HDAC1 | -0.613768 | THCA | K5 | JQ12 |
| MVP_top60_DNA_seed | PIK3CA | -0.251496 | LUAD | COR-L105 | AZD8186 | MVP_top60_DNA+RNA_seed | HDAC1 | -0.612377 | THCA | B-CPAP | AR-42 |
| MVP_top60_DNA_seed | AKT1 | -0.250065 | LUAD | NCI-H1944 | MK-2206 | MVP_top60_DNA+RNA_seed | BRAF | -0.606979 | THCA | KMH-2 | RAF_9304 |
| MVP_top60_DNA_seed | TP53 | -0.25003 | LUAD | NCI-H2122 | MIRA-1 | MVP_top60_DNA+RNA_seed | MAPK1 | -0.594281 | THCA | HTC-C3 | FR-180204 |
| MVP_top60_DNA_seed | AKT1 | -0.245737 | LUAD | RERF-LC-KJ | AZD5363 | MVP_top60_DNA+RNA_seed | MAPK3 | -0.594281 | THCA | HTC-C3 | FR-180204 |
| MVP_top60_DNA_seed | IGF1R | -0.245726 | LUAD | NCI-H2030 | NVP-ADW742 | MVP_top60_DNA+RNA_seed | CHEK2 | -0.592674 | THCA | FTC-133 | AZD7762 |
| MVP_top60_DNA_seed | FGFR1 | -0.245156 | LUAD | NCI-H1734 | PD173074 | MVP_top60_DNA+RNA_seed | BRAF | -0.5883 | THCA | TT | SB590885 |
| MVP_top60_DNA_seed | PIK3CA | -0.244183 | LUAD | NCI-H2030 | Alpelisib | MVP_top60_DNA+RNA_seed | PTK2 | -0.587439 | THCA | HTC-C3 | PF-562271 |
| MVP_top60_DNA_seed | PIK3CA | -0.242331 | LUAD | LXF-289 | Buparlisib | MVP_top60_DNA+RNA_seed | HDAC1 | -0.587288 | THCA | ML-1 | CAY10603 |
| MVP_top60_DNA_seed | EGFR | -0.24091 | LUAD | NCI-H1993 | Afatinib | MVP_top60_DNA+RNA_seed | HDAC1 | -0.585637 | THCA | K5 | Parthenolide |

| | | | | | | | | | | | |
|---|---|---|---|---|---|---|---|---|---|---|---|
| MVP_top60_DNA_seed | ERBB2 | -0.24091 | LUAD | NCI-H1993 | Afatinib | MVP_top60_DNA+RNA_seed | CHEK2 | -0.575681 | THCA | 8305C | AZD7762 |
| MVP_top60_DNA_seed | MAPK1 | -0.240839 | LUAD | PC-14 | Ulixertinib | MVP_top60_DNA+RNA_seed | GSK3B | -0.564997 | THCA | RO82-W-1 | CHIR-99021 |
| MVP_top60_DNA_seed | KRAS | -0.239604 | LUAD | NCI-H322M | KRAS (G12C) Inhibitor-12 | MVP_top60_DNA+RNA_seed | AKT1 | -0.56319 | THCA | TT | MK-2206 |
| MVP_top60_DNA_seed | AKT1 | -0.23857 | LUAD | NCI-H3122 | Afuresertib | MVP_top60_DNA+RNA_seed | AKT2 | -0.56319 | THCA | TT | MK-2206 |
| MVP_top60_DNA_seed | MET | -0.23555 | LUAD | NCI-H1437 | Crizotinib | MVP_top60_DNA+RNA_seed | CHEK2 | -0.555376 | THCA | KMH-2 | AZD7762 |
| MVP_top60_DNA_seed | HDAC1 | -0.235479 | LUAD | NCI-H1975 | Entinostat | MVP_top60_DNA+RNA_seed | RET | -0.553464 | THCA | FTC-133 | Motesanib |
| MVP_top60_DNA_seed | MAPK1 | -0.235449 | LUAD | NCI-H1755 | Ulixertinib | MVP_top60_DNA+RNA_seed | HDAC1 | -0.552855 | THCA | TT2609-C02 | Tubastatin A |
| MVP_top60_DNA_seed | IGF1R | -0.235025 | LUAD | NCI-H2009 | BMS-536924 | MVP_top60_DNA+RNA_seed | HDAC1 | -0.552481 | THCA | B-CPAP | Dacinostat |
| MVP_top60_DNA_seed | SRC | -0.234168 | LUAD | NCI-H1648 | Bosutinib | MVP_top60_DNA+RNA_seed | RET | -0.550059 | THCA | B-CPAP | Motesanib |
| MVP_top60_DNA_seed | IGF1R | -0.234121 | LUAD | NCI-H1755 | BMS-754807 | MVP_top60_DNA+RNA_seed | AKT1 | -0.546823 | THCA | ASH-3 | AKT inhibitor VIII |
| MVP_top60_DNA_seed | EGFR | -0.233537 | LUAD | NCI-H2023 | AZD3759 | MVP_top60_DNA+RNA_seed | AKT2 | -0.546823 | THCA | ASH-3 | AKT inhibitor VIII |
| MVP_top60_DNA_seed | CREBBP | -0.233493 | LUAD | HCC-78 | SGC-CBP30 | MVP_top60_DNA+RNA_seed | AKT3 | -0.546823 | THCA | ASH-3 | AKT inhibitor VIII |
| MVP_top60_DNA_seed | PIK3CA | -0.228488 | LUAD | NCI-H1437 | AZD8186 | MVP_top60_DNA+RNA_seed | ATM | -0.545485 | THCA | TT2609-C02 | KU-55933 |
| MVP_top60_DNA_seed | TP53 | -0.228242 | LUAD | LC-2-ad | MIRA-1 | MVP_top60_DNA+RNA_seed | CHEK2 | -0.537076 | THCA | KMH-2 | AZD7762 |
| MVP_top60_DNA_seed | IGF1R | -0.227914 | LUAD | NCI-H322M | BMS-536924 | MVP_top60_DNA+RNA_seed | GSK3B | -0.535206 | THCA | 8505C | KIN001-042 |
| MVP_top60_DNA_seed | MDM2 | -0.227815 | LUAD | NCI-H2122 | Nutlin-3a (-) | MVP_top60_DNA+RNA_seed | BRAF | -0.53173 | THCA | ASH-3 | RAF_9304 |
| MVP_top60_DNA_seed | HSP90AA1 | -0.227219 | LUAD | NCI-H2122 | Luminespib | MVP_top60_DNA+RNA_seed | CHEK2 | -0.528478 | THCA | IHH-4 | AZD7762 |
| MVP_top60_DNA_seed | AKT1 | -0.225638 | LUAD | HCC-44 | Uprosertib | MVP_top60_DNA+RNA_seed | GSK3B | -0.525849 | THCA | KMH-2 | CHIR-99021 |
| MVP_top60_DNA_seed | MDM2 | -0.225508 | LUAD | NCI-H1944 | Serdemetan | MVP_top60_DNA+RNA_seed | AKT1 | -0.521863 | THCA | CGTH-W-1 | AKT inhibitor VIII |
| MVP_top60_DNA_seed | IGF1R | -0.224761 | LUAD | NCI-H1563 | BMS-536924 | MVP_top60_DNA+RNA_seed | AKT2 | -0.521863 | THCA | CGTH-W-1 | AKT inhibitor VIII |
| MVP_top60_DNA_seed | ATM | -0.224506 | LUAD | NCI-H1563 | KU-55933 | MVP_top60_DNA+RNA_seed | AKT3 | -0.521863 | THCA | CGTH-W-1 | AKT inhibitor VIII |
| MVP_top60_DNA_seed | HDAC1 | -0.223407 | LUAD | NCI-H1755 | PCI-34051 | MVP_top60_DNA+RNA_seed | ATM | -0.515829 | THCA | TT | KU-60019 |
| MVP_top60_DNA_seed | HSP90AA1 | -0.222448 | LUAD | NCI-H1993 | Luminespib | MVP_top60_DNA+RNA_seed | CHEK2 | -0.515183 | THCA | 8505C | AZD7762 |
| MVP_top60_DNA_seed | PIK3CA | -0.222285 | LUAD | Calu-3 | GNE-317 | MVP_top60_DNA+RNA_seed | ATM | -0.513165 | THCA | HTC-C3 | KU-60019 |
| MVP_top60_DNA_seed | CREBBP | -0.221252 | LUAD | NCI-H1666 | SGC-CBP30 | MVP_top60_DNA+RNA_seed | GSK3B | -0.512483 | THCA | BHT-101 | AZD7969 |
| MVP_top60_DNA_seed | ATM | -0.220484 | LUAD | NCI-H358 | KU-55933 | MVP_top60_DNA+RNA_seed | PDPK1 | -0.509294 | THCA | TT2609-C02 | BX-912 |
| MVP_top60_DNA_seed | BRAF | -0.219352 | LUAD | PC-3_[JPC-3] | PLX-4720 | MVP_top60_DNA+RNA_seed | BRAF | -0.489515 | THCA | 8505C | SB590885 |
| MVP_top60_DNA_seed | EGFR | -0.218425 | LUAD | EMC-BAC-2 | Afatinib | MVP_top60_DNA+RNA_seed | AKT1 | -0.481397 | THCA | CGTH-W-1 | A-443654 |
| MVP_top60_DNA_seed | ERBB2 | -0.218425 | LUAD | EMC-BAC-2 | Afatinib | MVP_top60_DNA+RNA_seed | AKT2 | -0.481397 | THCA | CGTH-W-1 | A-443654 |
| MVP_top60_DNA_seed | BRAF | -0.216234 | LUAD | LC-2-ad | Dabrafenib | MVP_top60_DNA+RNA_seed | AKT3 | -0.481397 | THCA | CGTH-W-1 | A-443654 |
| MVP_top60_DNA_seed | AKT1 | -0.214868 | LUAD | NCI-H1838 | Uprosertib | MVP_top60_DNA+RNA_seed | ATM | -0.479085 | THCA | BHT-101 | KU-60019 |
| MVP_top60_DNA_seed | MAPK1 | -0.214569 | LUAD | HOP-62 | VX-11e | MVP_top60_DNA+RNA_seed | HDAC1 | -0.473664 | THCA | TT2609-C02 | Belinostat |
| MVP_top60_DNA_seed | HSP90AA1 | -0.214178 | LUAD | PC-3_[JPC-3] | Luminespib | MVP_top60_DNA+RNA_seed | HDAC1 | -0.473279 | THCA | IHH-4 | Belinostat |
| MVP_top60_DNA_seed | MET | -0.213508 | LUAD | NCI-H2030 | Crizotinib | MVP_top60_DNA+RNA_seed | PDPK1 | -0.471103 | THCA | IHH-4 | BX-912 |
| MVP_top60_DNA_seed | AURKA | -0.212933 | LUAD | NCI-H1437 | Tozasertib | MVP_top60_DNA+RNA_seed | GSK3B | -0.467068 | THCA | CAL-62 | CHIR-99021 |
| MVP_top60_DNA_seed | TERT | -0.210942 | LUAD | NCI-H1734 | BIBR-1532 | MVP_top60_DNA+RNA_seed | AKT1 | -0.465209 | THCA | IHH-4 | AKT inhibitor VIII |
| MVP_top60_DNA_seed | AKT1 | -0.208587 | LUAD | PC-14 | AZD5363 | MVP_top60_DNA+RNA_seed | AKT2 | -0.465209 | THCA | IHH-4 | AKT inhibitor VIII |
| MVP_top60_DNA_seed | EGFR | -0.207757 | LUAD | NCI-H1563 | Gefitinib | MVP_top60_DNA+RNA_seed | AKT3 | -0.465209 | THCA | IHH-4 | AKT inhibitor VIII |
| MVP_top60_DNA_seed | AKT1 | -0.20669 | LUAD | COR-L105 | MK-2206 | MVP_top60_DNA+RNA_seed | HDAC1 | -0.462103 | THCA | TT2609-C02 | CAY10603 |
| MVP_top60_DNA_seed | HSP90AA1 | -0.205187 | LUAD | Calu-6 | Tanespimycin | MVP_top60_DNA+RNA_seed | PDPK1 | -0.459865 | THCA | KMH-2 | OSU-03012 |
| MVP_top60_DNA_seed | HDAC1 | -0.205187 | LUAD | A549 | PCI-34051 | MVP_top60_DNA+RNA_seed | HDAC1 | -0.458914 | THCA | ML-1 | AR-42 |

| | | | | | | | | | | | |
|---|---|---|---|---|---|---|---|---|---|---|---|
| MVP_top60_DNA_seed | HDAC1 | -0.204616 | LUAD | NCI-H3122 | Entinostat | MVP_top60_DNA+RNA_seed | AKT1 | -0.454229 | THCA | K5 | BAY AKT1 |
| MVP_top60_DNA_seed | PIK3CA | -0.204119 | LUAD | NCI-H838 | Buparlisib | MVP_top60_DNA+RNA_seed | AKT2 | -0.454229 | THCA | K5 | BAY AKT1 |
| MVP_top60_DNA_seed | HSP90AA1 | -0.20312 | LUAD | NCI-H2085 | Luminespib | MVP_top60_DNA+RNA_seed | AKT3 | -0.454229 | THCA | K5 | BAY AKT1 |
| MVP_top60_DNA_seed | IGF1R | -0.202586 | LUAD | NCI-H1975 | NVP-ADW742 | MVP_top60_DNA+RNA_seed | MAPK1 | -0.453868 | THCA | CAL-62 | VX-11e |
| MVP_top60_DNA_seed | MET | -0.201648 | LUAD | NCI-H1650 | Savolitinib | MVP_top60_DNA+RNA_seed | ATM | -0.452208 | THCA | CGTH-W-1 | KU-55933 |
| MVP_top60_DNA_seed | IGF1R | -0.201112 | LUAD | NCI-H322M | BMS-754807 | MVP_top60_DNA+RNA_seed | HDAC1 | -0.443166 | THCA | IHH-4 | AR-42 |
| MVP_top60_DNA_seed | FGFR1 | -0.199558 | LUAD | NCI-H1781 | AZD4547 | MVP_top60_DNA+RNA_seed | HDAC1 | -0.440462 | THCA | BHT-101 | Tubastatin A |
| MVP_top60_DNA_seed | EZH2 | -0.199301 | LUAD | NCI-H1623 | GSK343 | MVP_top60_DNA+RNA_seed | PDPK1 | -0.43414 | THCA | 8505C | BX-912 |
| MVP_top60_DNA_seed | PIK3CA | -0.19928 | LUAD | H3255 | AZD8186 | MVP_top60_DNA+RNA_seed | PIK3CG | -0.425946 | THCA | K5 | AS605240 |
| MVP_top60_DNA_seed | BRAF | -0.198889 | LUAD | NCI-H1755 | Dabrafenib | MVP_top60_DNA+RNA_seed | RET | -0.422726 | THCA | ML-1 | Motesanib |
| MVP_top60_DNA_seed | EZH2 | -0.1986 | LUAD | NCI-H1435 | GSK343 | MVP_top60_DNA+RNA_seed | ATM | -0.421715 | THCA | KMH-2 | KU-60019 |
| MVP_top60_DNA_seed | IGF1R | -0.198542 | LUAD | NCI-H1993 | Linsitinib | MVP_top60_DNA+RNA_seed | MAPK1 | -0.414665 | THCA | IHH-4 | FR-180204 |
| MVP_top60_DNA_seed | MAPK1 | -0.197237 | LUAD | NCI-H838 | SCH772984 | MVP_top60_DNA+RNA_seed | MAPK3 | -0.414665 | THCA | IHH-4 | FR-180204 |
| MVP_top60_DNA_seed | MDM2 | -0.19708 | LUAD | NCI-H322M | Serdemetan | MVP_top60_DNA+RNA_seed | RET | -0.413693 | THCA | IHH-4 | Motesanib |
| MVP_top60_DNA_seed | HSP90AA1 | -0.197013 | LUAD | NCI-H1437 | Luminespib | MVP_top60_DNA+RNA_seed | PDPK1 | -0.413168 | THCA | CGTH-W-1 | OSU-03012 |
| MVP_top60_DNA_seed | MAPK1 | -0.195984 | LUAD | EMC-BAC-1 | VX-11e | MVP_top60_DNA+RNA_seed | GSK3B | -0.39953 | THCA | RO82-W-1 | SB216763 |
| MVP_top60_DNA_seed | AKT1 | -0.19218 | LUAD | NCI-H2023 | Afuresertib | MVP_top60_DNA+RNA_seed | HDAC1 | -0.398658 | THCA | 8505C | CAY10603 |
| MVP_top60_DNA_seed | HSP90AA1 | -0.191238 | LUAD | EMC-BAC-1 | Elesclomol | MVP_top60_DNA+RNA_seed | HDAC1 | -0.396079 | THCA | TT2609-C02 | AR-42 |
| MVP_top60_DNA_seed | IGF1R | -0.18992 | LUAD | NCI-H1648 | BMS-754807 | MVP_top60_DNA+RNA_seed | BRAF | -0.392081 | THCA | ASH-3 | Dabrafenib |
| MVP_top60_DNA_seed | IGF1R | -0.188128 | LUAD | VMRC-LCD | GSK1904529A | MVP_top60_DNA+RNA_seed | ATM | -0.391261 | THCA | ASH-3 | KU-55933 |
| MVP_top60_DNA_seed | PIK3CA | -0.187985 | LUAD | LC-2-ad | Buparlisib | MVP_top60_DNA+RNA_seed | ATM | -0.374302 | THCA | BHT-101 | CP466722 |
| MVP_top60_DNA_seed | AKT1 | -0.185918 | LUAD | COR-L105 | Afuresertib | MVP_top60_DNA+RNA_seed | GSK3B | -0.372682 | THCA | IHH-4 | CHIR-99021 |
| MVP_top60_DNA_seed | EGFR | -0.185376 | LUAD | RERF-LC-KJ | Osimertinib | MVP_top60_DNA+RNA_seed | BRAF | -0.370822 | THCA | K5 | HG6-64-1 |
| MVP_top60_DNA_seed | MDM2 | -0.184543 | LUAD | NCI-H2347 | Serdemetan | MVP_top60_DNA+RNA_seed | BRAF | -0.366295 | THCA | CAL-62 | PLX-4720 |
| MVP_top60_DNA_seed | MDM2 | -0.18452 | LUAD | NCI-H1437 | Nutlin-3a (-) | MVP_top60_DNA+RNA_seed | GSK3B | -0.366067 | THCA | B-CPAP | SB216763 |
| MVP_top60_DNA_seed | MET | -0.182454 | LUAD | NCI-H1648 | Crizotinib | MVP_top60_DNA+RNA_seed | HDAC1 | -0.345252 | THCA | HTC-C3 | Tubastatin A |
| MVP_top60_DNA_seed | AKT1 | -0.182337 | LUAD | NCI-H1563 | Uprosertib | MVP_top60_DNA+RNA_seed | HDAC1 | -0.33994 | THCA | KMH-2 | JQ12 |
| MVP_top60_DNA_seed | AKT1 | -0.181232 | LUAD | Calu-3 | MK-2206 | MVP_top60_DNA+RNA_seed | HDAC1 | -0.332175 | THCA | CAL-62 | Tubastatin A |
| MVP_top60_DNA_seed | MAPK1 | -0.180387 | LUAD | NCI-H2023 | Ulixertinib | MVP_top60_DNA+RNA_seed | HDAC1 | -0.331273 | THCA | IHH-4 | Dacinostat |
| MVP_top60_DNA_seed | AKT1 | -0.179368 | LUAD | NCI-H1755 | Afuresertib | MVP_top60_DNA+RNA_seed | HDAC1 | -0.327175 | THCA | IHH-4 | CAY10603 |
| MVP_top60_DNA_seed | MET | -0.179254 | LUAD | EMC-BAC-2 | Crizotinib | MVP_top60_DNA+RNA_seed | BRAF | -0.324975 | THCA | B-CPAP | SB590885 |
| MVP_top60_DNA_seed | PIK3CA | -0.178812 | LUAD | NCI-H1693 | GNE-317 | MVP_top60_DNA+RNA_seed | CHEK2 | -0.308969 | THCA | HTC-C3 | AZD7762 |
| MVP_top60_DNA_seed | AURKA | -0.178106 | LUAD | NCI-H1648 | ZM447439 | MVP_top60_DNA+RNA_seed | PDPK1 | -0.303598 | THCA | B-CPAP | BX-912 |
| MVP_top60_DNA_seed | PIK3CA | -0.177116 | LUAD | LC-2-ad | AZD8186 | MVP_top60_DNA+RNA_seed | HDAC1 | -0.303126 | THCA | CAL-62 | Belinostat |
| MVP_top60_DNA_seed | EGFR | -0.175883 | LUAD | NCI-H1563 | Sapitinib | MVP_top60_DNA+RNA_seed | HDAC1 | -0.297678 | THCA | B-CPAP | Tubastatin A |
| MVP_top60_DNA_seed | ERBB2 | -0.175883 | LUAD | NCI-H1563 | Sapitinib | MVP_top60_DNA+RNA_seed | PIK3CG | -0.29514 | THCA | HTC-C3 | PIK-93 |
| MVP_top60_DNA_seed | ERBB3 | -0.175883 | LUAD | NCI-H1563 | Sapitinib | MVP_top60_DNA+RNA_seed | AKT1 | -0.288646 | THCA | KMH-2 | GSK690693 |
| MVP_top60_DNA_seed | MAPK1 | -0.174893 | LUAD | LXF-289 | SCH772984 | MVP_top60_DNA+RNA_seed | AKT2 | -0.288646 | THCA | KMH-2 | GSK690693 |
| MVP_top60_DNA_seed | FGFR1 | -0.174078 | LUAD | NCI-H1568 | AZD4547 | MVP_top60_DNA+RNA_seed | AKT3 | -0.288646 | THCA | KMH-2 | GSK690693 |
| MVP_top60_DNA_seed | AKT1 | -0.172351 | LUAD | HCC-78 | Afuresertib | MVP_top60_DNA+RNA_seed | HDAC1 | -0.287986 | THCA | 8305C | JQ12 |
| MVP_top60_DNA_seed | EGFR | -0.171631 | LUAD | NCI-H1755 | Lapatinib | MVP_top60_DNA+RNA_seed | PIK3CG | -0.284458 | THCA | BHT-101 | AS605240 |

| | | | | | | | | | | | |
|---|---|---|---|---|---|---|---|---|---|---|---|
| MVP_top60_DNA_seed | ERBB2 | -0.171631 | LUAD | NCI-H1755 | Lapatinib | MVP_top60_DNA+RNA_seed | HDAC1 | -0.281563 | THCA | 8505C | Belinostat |
| MVP_top60_DNA_seed | PIK3CA | -0.170387 | LUAD | NCI-H23 | Buparlisib | MVP_top60_DNA+RNA_seed | BRAF | -0.272961 | THCA | 8505C | PLX-4720 |
| MVP_top60_DNA_seed | AKT1 | -0.168826 | LUAD | NCI-H2023 | Uprosertib | MVP_top60_DNA+RNA_seed | PIK3CG | -0.268929 | THCA | 8505C | AS605240 |
| MVP_top60_DNA_seed | AKT1 | -0.168825 | LUAD | RERF-LC-MS | AZD5363 | MVP_top60_DNA+RNA_seed | PIK3CG | -0.266371 | THCA | 8505C | PIK-93 |
| MVP_top60_DNA_seed | FGFR1 | -0.167334 | LUAD | NCI-H3122 | AZD4547 | MVP_top60_DNA+RNA_seed | GSK3B | -0.265778 | THCA | HTC-C3 | AZD7969 |
| MVP_top60_DNA_seed | AURKA | -0.165912 | LUAD | HCC-78 | ZM447439 | MVP_top60_DNA+RNA_seed | GSK3B | -0.255147 | THCA | ASH-3 | AZD7969 |
| MVP_top60_DNA_seed | EGFR | -0.165431 | LUAD | NCI-H358 | Gefitinib | MVP_top60_DNA+RNA_seed | HDAC1 | -0.254665 | THCA | K5 | Dacinostat |
| MVP_top60_DNA_seed | AKT1 | -0.163877 | LUAD | COR-L105 | Uprosertib | MVP_top60_DNA+RNA_seed | CHEK2 | -0.245498 | THCA | K5 | AZD7762 |
| MVP_top60_DNA_seed | EGFR | -0.163454 | LUAD | Calu-3 | Gefitinib | MVP_top60_DNA+RNA_seed | PDPK1 | -0.243332 | THCA | ML-1 | BX-912 |
| MVP_top60_DNA_seed | EGFR | -0.163199 | LUAD | RERF-LC-KJ | Lapatinib | MVP_top60_DNA+RNA_seed | BRAF | -0.242036 | THCA | CAL-62 | HG6-64-1 |
| MVP_top60_DNA_seed | ERBB2 | -0.163199 | LUAD | RERF-LC-KJ | Lapatinib | MVP_top60_DNA+RNA_seed | ATM | -0.239267 | THCA | 8505C | KU-55933 |
| MVP_top60_DNA_seed | IGF1R | -0.163181 | LUAD | H3255 | BMS-536924 | MVP_top60_DNA+RNA_seed | PIK3CG | -0.236601 | THCA | K5 | PIK-93 |
| MVP_top60_DNA_seed | SMARCA4 | -0.162595 | LUAD | LC-2-ad | PFI3 | MVP_top60_DNA+RNA_seed | BRAF | -0.23467 | THCA | 8505C | HG6-64-1 |
| MVP_top60_DNA_seed | AURKA | -0.161895 | LUAD | NCI-H3122 | Alisertib | MVP_top60_DNA+RNA_seed | PDPK1 | -0.213444 | THCA | B-CPAP | OSU-03012 |
| MVP_top60_DNA_seed | MAPK1 | -0.161727 | LUAD | PC-14 | SCH772984 | MVP_top60_DNA+RNA_seed | ATM | -0.212464 | THCA | 8505C | torin2 |
| MVP_top60_DNA_seed | MAPK1 | -0.161591 | LUAD | HOP-62 | SCH772984 | MVP_top60_DNA+RNA_seed | AKT1 | -0.208708 | THCA | B-CPAP | MK-2206 |
| MVP_top60_DNA_seed | MAPK1 | -0.161551 | LUAD | NCI-H2087 | VX-11e | MVP_top60_DNA+RNA_seed | AKT2 | -0.208708 | THCA | B-CPAP | MK-2206 |
| MVP_top60_DNA_seed | EGFR | -0.160772 | LUAD | EMC-BAC-2 | Lapatinib | MVP_top60_DNA+RNA_seed | ATM | -0.201777 | THCA | ASH-3 | CP466722 |
| MVP_top60_DNA_seed | ERBB2 | -0.160772 | LUAD | EMC-BAC-2 | Lapatinib | MVP_top60_DNA+RNA_seed | GSK3B | -0.200796 | THCA | IHH-4 | CHIR-99021 |
| MVP_top60_DNA_seed | CREBBP | -0.160366 | LUAD | NCI-H2023 | SGC-CBP30 | MVP_top60_DNA+RNA_seed | HDAC1 | -0.192304 | THCA | BHT-101 | Dacinostat |
| MVP_top60_DNA_seed | IGF1R | -0.159768 | LUAD | ABC-1 | Linsitinib | MVP_top60_DNA+RNA_seed | BRAF | -0.186451 | THCA | CAL-62 | PLX-4720 |
| MVP_top60_DNA_seed | AKT1 | -0.159232 | LUAD | RERF-LC-MS | Ipatasertib | MVP_top60_DNA+RNA_seed | AKT1 | -0.182832 | THCA | KMH-2 | AKT inhibitor VIII |
| MVP_top60_DNA_seed | SRC | -0.158699 | LUAD | RERF-LC-KJ | Bosutinib | MVP_top60_DNA+RNA_seed | AKT2 | -0.182832 | THCA | KMH-2 | AKT inhibitor VIII |
| MVP_top60_DNA_seed | HDAC1 | -0.158295 | LUAD | NCI-H1435 | PCI-34051 | MVP_top60_DNA+RNA_seed | AKT3 | -0.182832 | THCA | KMH-2 | AKT inhibitor VIII |
| MVP_top60_DNA_seed | HDAC1 | -0.15629 | LUAD | RERF-LC-MS | Entinostat | MVP_top60_DNA+RNA_seed | CHEK2 | -0.174044 | THCA | TT | AZD7762 |
| MVP_top60_DNA_seed | AKT1 | -0.156139 | LUAD | A549 | AZD5363 | MVP_top60_DNA+RNA_seed | GSK3B | -0.170817 | THCA | KMH-2 | AZD7969 |
| MVP_top60_DNA_seed | HDAC1 | -0.154978 | LUAD | COR-L105 | Romidepsin | MVP_top60_DNA+RNA_seed | AKT1 | -0.170145 | THCA | IHH-4 | BAY AKT1 |
| MVP_top60_DNA_seed | IGF1R | -0.154199 | LUAD | A549 | NVP-ADW742 | MVP_top60_DNA+RNA_seed | AKT2 | -0.170145 | THCA | IHH-4 | BAY AKT1 |
| MVP_top60_DNA_seed | EGFR | -0.153439 | LUAD | NCI-H1573 | Gefitinib | MVP_top60_DNA+RNA_seed | AKT3 | -0.170145 | THCA | IHH-4 | BAY AKT1 |
| MVP_top60_DNA_seed | AKT1 | -0.153102 | LUAD | NCI-H322M | MK-2206 | MVP_top60_DNA+RNA_seed | BRAF | -0.164558 | THCA | KMH-2 | PLX-4720 |
| MVP_top60_DNA_seed | EGFR | -0.152389 | LUAD | NCI-H1651 | Erlotinib | MVP_top60_DNA+RNA_seed | BRAF | -0.164224 | THCA | CGTH-W-1 | Dabrafenib |
| MVP_top60_DNA_seed | AKT1 | -0.151421 | LUAD | A549 | MK-2206 | MVP_top60_DNA+RNA_seed | AKT1 | -0.158883 | THCA | 8505C | MK-2206 |
| MVP_top60_DNA_seed | AKT1 | -0.150646 | LUAD | NCI-H838 | Afuresertib | MVP_top60_DNA+RNA_seed | AKT2 | -0.158883 | THCA | 8505C | MK-2206 |
| MVP_top60_DNA_seed | MAPK1 | -0.149149 | LUAD | NCI-H1623 | Ulixertinib | MVP_top60_DNA+RNA_seed | GSK3B | -0.15719 | THCA | 8505C | AZD7969 |
| MVP_top60_DNA_seed | AKT1 | -0.14816 | LUAD | NCI-H1734 | Uprosertib | MVP_top60_DNA+RNA_seed | HDAC1 | -0.156541 | THCA | K5 | AR-42 |
| MVP_top60_DNA_seed | HSP90AA1 | -0.148062 | LUAD | LC-2-ad | Luminespib | MVP_top60_DNA+RNA_seed | GSK3B | -0.152963 | THCA | TT2609-C02 | CHIR-99021 |
| MVP_top60_DNA_seed | MAPK1 | -0.146615 | LUAD | NCI-H358 | VX-11e | MVP_top60_DNA+RNA_seed | ATM | -0.145785 | THCA | FTC-133 | KU-60019 |
| MVP_top60_DNA_seed | FGFR1 | -0.146087 | LUAD | HCC-44 | PD173074 | MVP_top60_DNA+RNA_seed | CHEK2 | -0.142249 | THCA | ASH-3 | AZD7762 |
| MVP_top60_DNA_seed | HDAC1 | -0.145126 | LUAD | EMC-BAC-2 | PCI-34051 | MVP_top60_DNA+RNA_seed | HDAC1 | -0.133908 | THCA | CAL-62 | Dacinostat |
| MVP_top60_DNA_seed | IGF1R | -0.144879 | LUAD | NCI-H838 | GSK1904529A | MVP_top60_DNA+RNA_seed | AKT1 | -0.129599 | THCA | HTC-C3 | AKT inhibitor VIII |
| MVP_top60_DNA_seed | MAPK1 | -0.144282 | LUAD | NCI-H1781 | VX-11e | MVP_top60_DNA+RNA_seed | AKT2 | -0.129599 | THCA | HTC-C3 | AKT inhibitor VIII |

| | | | | | | | | | | | |
|---|---|---|---|---|---|---|---|---|---|---|---|
| MVP_top60_DNA_seed | PIK3CA | -0.14364 | LUAD | NCI-H1781 | GNE-317 | MVP_top60_DNA+RNA_seed | AKT3 | -0.129599 | THCA | HTC-C3 | AKT inhibitor VIII |
| MVP_top60_DNA_seed | AURKA | -0.143562 | LUAD | NCI-H2342 | Alisertib | MVP_top60_DNA+RNA_seed | HDAC1 | -0.11734 | THCA | K5 | CAY10603 |
| MVP_top60_DNA_seed | PIK3CA | -0.143288 | LUAD | RERF-LC-KJ | GNE-317 | MVP_top60_DNA+RNA_seed | GSK3B | -0.113085 | THCA | KMH-2 | KIN001-042 |
| MVP_top60_DNA_seed | EGFR | -0.142154 | LUAD | NCI-H1568 | Lapatinib | MVP_top60_DNA+RNA_seed | HDAC1 | -0.106354 | THCA | CAL-62 | AR-42 |
| MVP_top60_DNA_seed | ERBB2 | -0.142154 | LUAD | NCI-H1568 | Lapatinib | MVP_top60_DNA+RNA_seed | PTK2 | -0.091913 | THCA | K5 | PF-562271 |
| MVP_top60_DNA_seed | FGFR1 | -0.141793 | LUAD | NCI-H1944 | AZD4547 | MVP_top60_DNA+RNA_seed | BRAF | -0.085211 | THCA | RO82-W-1 | HG6-64-1 |
| MVP_top60_DNA_seed | MAPK1 | -0.141767 | LUAD | LC-2-ad | SCH772984 | MVP_top60_DNA+RNA_seed | BRAF | -0.081779 | THCA | CGTH-W-1 | PLX-4720 |
| MVP_top60_DNA_seed | HSP90AA1 | -0.140696 | LUAD | NCI-H2087 | Tanespimycin | MVP_top60_DNA+RNA_seed | HDAC1 | -0.078662 | THCA | 8505C | AR-42 |
| MVP_top60_DNA_seed | AURKA | -0.139466 | LUAD | NCI-H2023 | ZM447439 | MVP_top60_DNA+RNA_seed | PIK3CG | -0.077083 | THCA | CAL-62 | AS605240 |
| MVP_top60_DNA_seed | HSP90AA1 | -0.139244 | LUAD | NCI-H1648 | Tanespimycin | MVP_top60_DNA+RNA_seed | GSK3B | -0.073815 | THCA | CAL-62 | CHIR-99021 |
| MVP_top60_DNA_seed | AURKA | -0.138923 | LUAD | NCI-H838 | Tozasertib | MVP_top60_DNA+RNA_seed | CHEK2 | -0.063791 | THCA | B-CPAP | AZD7762 |
| MVP_top60_DNA_seed | AURKA | -0.138845 | LUAD | HCC-44 | Alisertib | MVP_top60_DNA+RNA_seed | ATM | -0.054763 | THCA | K5 | KU-55933 |
| MVP_top60_DNA_seed | IGF1R | -0.138716 | LUAD | NCI-H23 | BMS-754807 | MVP_top60_DNA+RNA_seed | MAPK1 | -0.051215 | THCA | 8305C | FR-180204 |
| MVP_top60_DNA_seed | PIK3CA | -0.138663 | LUAD | H3255 | GNE-317 | MVP_top60_DNA+RNA_seed | MAPK3 | -0.051215 | THCA | 8305C | FR-180204 |
| MVP_top60_DNA_seed | PIK3CA | -0.137713 | LUAD | EMC-BAC-1 | Buparlisib | MVP_top60_DNA+RNA_seed | AKT1 | -0.042584 | THCA | HTC-C3 | AKT inhibitor VIII |
| MVP_top60_DNA_seed | TP53 | -0.13766 | LUAD | RERF-LC-MS | MIRA-1 | MVP_top60_DNA+RNA_seed | AKT2 | -0.042584 | THCA | HTC-C3 | AKT inhibitor VIII |
| MVP_top60_DNA_seed | EZH2 | -0.137388 | LUAD | NCI-H1755 | GSK343 | MVP_top60_DNA+RNA_seed | AKT3 | -0.042584 | THCA | HTC-C3 | AKT inhibitor VIII |
| MVP_top60_DNA_seed | IGF1R | -0.137199 | LUAD | EMC-BAC-1 | NVP-ADW742 | MVP_top60_DNA+RNA_seed | BRAF | -0.04043 | THCA | ML-1 | Dabrafenib |
| MVP_top60_DNA_seed | SMARCA4 | -0.13665 | LUAD | RERF-LC-MS | PFI3 | MVP_top60_DNA+RNA_seed | MAPK1 | -0.036196 | THCA | 8505C | FR-180204 |
| MVP_top60_DNA_seed | AURKA | -0.13626 | LUAD | NCI-H1944 | Alisertib | MVP_top60_DNA+RNA_seed | MAPK3 | -0.036196 | THCA | 8505C | FR-180204 |
| MVP_top60_DNA_seed | IGF1R | -0.136232 | LUAD | EKVX | BMS-536924 | MVP_top60_DNA+RNA_seed | HDAC1 | -0.035633 | THCA | 8505C | Tubastatin A |
| MVP_top60_DNA_seed | PIK3CA | -0.135987 | LUAD | NCI-H3122 | GNE-317 | MVP_top60_DNA+RNA_seed | BRAF | -0.035248 | THCA | KMH-2 | HG6-64-1 |
| MVP_top60_DNA_seed | AKT1 | -0.13316 | LUAD | NCI-H2085 | GSK2110183B | MVP_top60_DNA+RNA_seed | GSK3B | -0.0272 | THCA | TT2609-C02 | SB216763 |
| MVP_top60_DNA_seed | IGF1R | -0.132611 | LUAD | LXF-289 | Linsitinib | MVP_top60_DNA+RNA_seed | AKT1 | -0.025516 | THCA | K5 | A-443654 |
| MVP_top60_DNA_seed | IGF1R | -0.1326 | LUAD | COR-L105 | Linsitinib | MVP_top60_DNA+RNA_seed | AKT2 | -0.025516 | THCA | K5 | A-443654 |
| MVP_top60_DNA_seed | TERT | -0.132516 | LUAD | LXF-289 | BIBR-1532 | MVP_top60_DNA+RNA_seed | AKT3 | -0.025516 | THCA | K5 | A-443654 |
| MVP_top60_DNA_seed | BRAF | -0.130656 | LUAD | EMC-BAC-2 | PLX-4720 | MVP_top60_DNA+RNA_seed | AKT1 | -0.024435 | THCA | CGTH-W-1 | MK-2206 |
| MVP_top60_DNA_seed | MET | -0.130513 | LUAD | NCI-H1563 | Savolitinib | MVP_top60_DNA+RNA_seed | AKT2 | -0.024435 | THCA | CGTH-W-1 | MK-2206 |
| MVP_top60_DNA_seed | SRC | -0.130276 | LUAD | NCI-H1666 | Bosutinib | MVP_top60_DNA+RNA_seed | HDAC1 | -0.023376 | THCA | IHH-4 | Tubastatin A |
| MVP_top60_DNA_seed | HSP90AA1 | -0.130125 | LUAD | RERF-LC-KJ | Tanespimycin | MVP_top60_DNA+RNA_seed | AKT1 | -0.019583 | THCA | FTC-133 | BAY AKT1 |
| MVP_top60_DNA_seed | SRC | -0.130046 | LUAD | NCI-H358 | Bosutinib | MVP_top60_DNA+RNA_seed | AKT2 | -0.019583 | THCA | FTC-133 | BAY AKT1 |
| MVP_top60_DNA_seed | AKT1 | -0.129241 | LUAD | RERF-LC-MS | Uprosertib | MVP_top60_DNA+RNA_seed | AKT3 | -0.019583 | THCA | FTC-133 | BAY AKT1 |
| MVP_top60_DNA_seed | MAPK1 | -0.128113 | LUAD | EMC-BAC-1 | Ulixertinib | MVP_top60_DNA+RNA_seed | HDAC1 | -0.016061 | THCA | HTC-C3 | Dacinostat |
| MVP_top60_DNA_seed | HDAC1 | -0.127992 | LUAD | NCI-H2030 | PCI-34051 | MVP_top60_DNA+RNA_seed | CHEK2 | -0.015845 | THCA | K5 | AZD7762 |
| MVP_top60_DNA_seed | PIK3CA | -0.127475 | LUAD | NCI-H1563 | Buparlisib | MVP_top60_DNA+RNA_seed | CHEK2 | -0.009868 | THCA | CAL-62 | AZD7762 |
| MVP_top60_DNA_seed | FGFR1 | -0.12724 | LUAD | NCI-H1648 | AZD4547 | MVP_top60_DNA+RNA_seed | HDAC1 | -0.008344 | THCA | CAL-62 | JQ12 |
| MVP_top60_DNA_seed | PIK3CA | -0.127109 | LUAD | EKVX | AZD8186 | MVP_top60_DNA+RNA_seed | CHEK2 | -0.005508 | THCA | 8305C | AZD7762 |
| MVP_top60_DNA_seed | PIK3CA | -0.125953 | LUAD | LXF-289 | GNE-317 | MVP_top60_DNA+RNA_seed | HDAC1 | -0.005206 | THCA | RO82-W-1 | CAY10603 |
| MVP_top60_DNA_seed | TERT | -0.125794 | LUAD | NCI-H322M | BIBR-1532 | MVP_top60_DNA+RNA_seed | ATM | -0.001181 | THCA | CAL-62 | KU-60019 |
| MVP_top60_DNA_seed | EZH2 | -0.125613 | LUAD | HCC-78 | GSK343 | MVP_top60_DNA+RNA_seed | BRAF | 0.001482 | THCA | 8505C | RAF_9304 |
| MVP_top60_DNA_seed | BRAF | -0.124405 | LUAD | NCI-H2405 | SB590885 | MVP_top60_DNA+RNA_seed | HDAC1 | 0.003584 | THCA | CGTH-W-1 | Entinostat |

| | | | | | | | | | | | |
|---|---|---|---|---|---|---|---|---|---|---|---|
| MVP_top60_DNA_seed | AKT1 | -0.124153 | LUAD | Calu-3 | Afuresertib | MVP_top60_DNA+RNA_seed | ATM | 0.004693 | THCA | HTC-C3 | KU-55933 |
| MVP_top60_DNA_seed | AKT1 | -0.123179 | LUAD | EMC-BAC-1 | AZD5363 | MVP_top60_DNA+RNA_seed | HDAC1 | 0.009976 | THCA | RO82-W-1 | Belinostat |
| MVP_top60_DNA_seed | GSK3B | -0.122991 | LUAD | NCI-H2085 | SB216763 | MVP_top60_DNA+RNA_seed | AKT1 | 0.012071 | THCA | IHH-4 | AKT inhibitor VIII |
| MVP_top60_DNA_seed | ATM | -0.122111 | LUAD | NCI-H1651 | KU-55933 | MVP_top60_DNA+RNA_seed | AKT2 | 0.012071 | THCA | IHH-4 | AKT inhibitor VIII |
| MVP_top60_DNA_seed | IGF1R | -0.121883 | LUAD | NCI-H1734 | GSK1904529A | MVP_top60_DNA+RNA_seed | AKT3 | 0.012071 | THCA | IHH-4 | AKT inhibitor VIII |
| MVP_top60_DNA_seed | TERT | -0.119012 | LUAD | LC-2-ad | BIBR-1532 | MVP_top60_DNA+RNA_seed | HDAC1 | 0.016729 | THCA | TT | JQ12 |
| MVP_top60_DNA_seed | SMARCA4 | -0.118711 | LUAD | NCI-H650 | PFI3 | MVP_top60_DNA+RNA_seed | HDAC1 | 0.02288 | THCA | K5 | Belinostat |
| MVP_top60_DNA_seed | HSP90AA1 | -0.118463 | LUAD | PC-14 | Elesclomol | MVP_top60_DNA+RNA_seed | ATM | 0.039882 | THCA | K5 | CP466722 |
| MVP_top60_DNA_seed | PIK3CA | -0.118434 | LUAD | HCC-827 | AZD8186 | MVP_top60_DNA+RNA_seed | PDPK1 | 0.04981 | THCA | HTC-C3 | OSU-03012 |
| MVP_top60_DNA_seed | SMARCA4 | -0.117848 | LUAD | NCI-H1993 | PFI3 | MVP_top60_DNA+RNA_seed | PIK3CG | 0.064785 | THCA | KMH-2 | AS605240 |
| MVP_top60_DNA_seed | KRAS | -0.117287 | LUAD | NCI-H838 | KRAS (G12C) Inhibitor-1: | MVP_top60_DNA+RNA_seed | AKT1 | 0.071723 | THCA | KMH-2 | BAY AKT1 |
| MVP_top60_DNA_seed | FGFR1 | -0.114211 | LUAD | NCI-H1975 | PD173074 | MVP_top60_DNA+RNA_seed | AKT2 | 0.071723 | THCA | KMH-2 | BAY AKT1 |
| MVP_top60_DNA_seed | IGF1R | -0.112757 | LUAD | NCI-H1648 | Linsitinib | MVP_top60_DNA+RNA_seed | AKT3 | 0.071723 | THCA | KMH-2 | BAY AKT1 |
| MVP_top60_DNA_seed | AKT1 | -0.112091 | LUAD | PC-14 | Afuresertib | MVP_top60_DNA+RNA_seed | HDAC1 | 0.073299 | THCA | RO82-W-1 | AR-42 |
| MVP_top60_DNA_seed | AKT1 | -0.111786 | LUAD | PC-14 | MK-2206 | MVP_top60_DNA+RNA_seed | AKT1 | 0.074453 | THCA | BHT-101 | AKT inhibitor VIII |
| MVP_top60_DNA_seed | EGFR | -0.111655 | LUAD | HCC-44 | Afatinib | MVP_top60_DNA+RNA_seed | AKT2 | 0.074453 | THCA | BHT-101 | AKT inhibitor VIII |
| MVP_top60_DNA_seed | ERBB2 | -0.111655 | LUAD | HCC-44 | Afatinib | MVP_top60_DNA+RNA_seed | AKT3 | 0.074453 | THCA | BHT-101 | AKT inhibitor VIII |
| MVP_top60_DNA_seed | HDAC1 | -0.109178 | LUAD | LC-2-ad | PCI-34051 | MVP_top60_DNA+RNA_seed | HDAC1 | 0.078315 | THCA | K5 | Tubastatin A |
| MVP_top60_DNA_seed | AKT1 | -0.108312 | LUAD | H3255 | MK-2206 | MVP_top60_DNA+RNA_seed | PIK3CG | 0.081648 | THCA | IHH-4 | PIK-93 |
| MVP_top60_DNA_seed | PIK3CA | -0.103037 | LUAD | NCI-H2342 | Alpelisib | MVP_top60_DNA+RNA_seed | HDAC1 | 0.093076 | THCA | CGTH-W-1 | AR-42 |
| MVP_top60_DNA_seed | PIK3CA | -0.101591 | LUAD | NCI-H1666 | Buparlisib | MVP_top60_DNA+RNA_seed | PDPK1 | 0.096586 | THCA | ASH-3 | BX-912 |
| MVP_top60_DNA_seed | AKT1 | -0.100328 | LUAD | RERF-LC-KJ | Uprosertib | MVP_top60_DNA+RNA_seed | ATM | 0.100739 | THCA | RO82-W-1 | KU-60019 |
| MVP_top60_DNA_seed | IGF1R | -0.09898 | LUAD | NCI-H23 | GSK1904529A | MVP_top60_DNA+RNA_seed | ATM | 0.106576 | THCA | CAL-62 | KU-55933 |
| MVP_top60_DNA_seed | FGFR1 | -0.098532 | LUAD | NCI-H522 | AZD4547 | MVP_top60_DNA+RNA_seed | RET | 0.108057 | THCA | K5 | Motesanib |
| MVP_top60_DNA_seed | HSP90AA1 | -0.097613 | LUAD | NCI-H650 | Tanespimycin | MVP_top60_DNA+RNA_seed | BRAF | 0.111801 | THCA | RO82-W-1 | Dabrafenib |
| MVP_top60_DNA_seed | IGF1R | -0.09395 | LUAD | NCI-H1650 | BMS-536924 | MVP_top60_DNA+RNA_seed | AKT1 | 0.112808 | THCA | KMH-2 | AKT inhibitor VIII |
| MVP_top60_DNA_seed | BRAF | -0.093522 | LUAD | NCI-H2405 | Dabrafenib | MVP_top60_DNA+RNA_seed | AKT2 | 0.112808 | THCA | KMH-2 | AKT inhibitor VIII |
| MVP_top60_DNA_seed | IGF1R | -0.092333 | LUAD | PC-14 | NVP-ADW742 | MVP_top60_DNA+RNA_seed | AKT3 | 0.112808 | THCA | KMH-2 | AKT inhibitor VIII |
| MVP_top60_DNA_seed | HDAC1 | -0.09195 | LUAD | NCI-H2085 | PCI-34051 | MVP_top60_DNA+RNA_seed | PIK3CG | 0.116703 | THCA | ML-1 | PIK-93 |
| MVP_top60_DNA_seed | IGF1R | -0.091412 | LUAD | NCI-H1563 | BMS-754807 | MVP_top60_DNA+RNA_seed | AKT1 | 0.118602 | THCA | K5 | AKT inhibitor VIII |
| MVP_top60_DNA_seed | ATM | -0.091316 | LUAD | NCI-H1734 | KU-55933 | MVP_top60_DNA+RNA_seed | AKT2 | 0.118602 | THCA | K5 | AKT inhibitor VIII |
| MVP_top60_DNA_seed | MDM2 | -0.090673 | LUAD | NCI-H838 | Serdemetan | MVP_top60_DNA+RNA_seed | AKT3 | 0.118602 | THCA | K5 | AKT inhibitor VIII |
| MVP_top60_DNA_seed | ATM | -0.090335 | LUAD | PC-3_[JPC-3] | KU-55933 | MVP_top60_DNA+RNA_seed | AKT1 | 0.123679 | THCA | RO82-W-1 | BAY AKT1 |
| MVP_top60_DNA_seed | PIK3CA | -0.089201 | LUAD | NCI-H2291 | Alpelisib | MVP_top60_DNA+RNA_seed | AKT2 | 0.123679 | THCA | RO82-W-1 | BAY AKT1 |
| MVP_top60_DNA_seed | MDM2 | -0.089055 | LUAD | ABC-1 | Serdemetan | MVP_top60_DNA+RNA_seed | AKT3 | 0.123679 | THCA | RO82-W-1 | BAY AKT1 |
| MVP_top60_DNA_seed | BRAF | -0.088436 | LUAD | PC-14 | PLX-4720 | MVP_top60_DNA+RNA_seed | GSK3B | 0.124593 | THCA | FTC-133 | SB216763 |
| MVP_top60_DNA_seed | IGF1R | -0.085515 | LUAD | NCI-H1693 | NVP-ADW742 | MVP_top60_DNA+RNA_seed | AKT1 | 0.128532 | THCA | K5 | GSK690693 |
| MVP_top60_DNA_seed | CREBBP | -0.084277 | LUAD | NCI-H1563 | SGC-CBP30 | MVP_top60_DNA+RNA_seed | AKT2 | 0.128532 | THCA | K5 | GSK690693 |
| MVP_top60_DNA_seed | EZH2 | -0.081768 | LUAD | NCI-H650 | GSK343 | MVP_top60_DNA+RNA_seed | AKT3 | 0.128532 | THCA | K5 | GSK690693 |
| MVP_top60_DNA_seed | AKT1 | -0.081389 | LUAD | PC-14 | GSK2110183B | MVP_top60_DNA+RNA_seed | BRAF | 0.13551 | THCA | ML-1 | PLX-4720 |
| MVP_top60_DNA_seed | SMARCA4 | -0.08108 | LUAD | LXF-289 | PFI3 | MVP_top60_DNA+RNA_seed | PIK3CG | 0.140116 | THCA | CAL-62 | PIK-93 |

| | | | | | | | | | | | |
|---|---|---|---|---|---|---|---|---|---|---|---|
| MVP_top60_DNA_seed | MET | -0.080649 | LUAD | NCI-H358 | Crizotinib | MVP_top60_DNA+RNA_seed | AKT1 | 0.140238 | THCA | TT2609-C02 | MK-2206 |
| MVP_top60_DNA_seed | AKT1 | -0.080258 | LUAD | NCI-H1563 | Afuresertib | MVP_top60_DNA+RNA_seed | AKT2 | 0.140238 | THCA | TT2609-C02 | MK-2206 |
| MVP_top60_DNA_seed | EGFR | -0.080005 | LUAD | NCI-H1838 | Gefitinib | MVP_top60_DNA+RNA_seed | CHEK2 | 0.154721 | THCA | HTC-C3 | AZD7762 |
| MVP_top60_DNA_seed | EGFR | -0.080003 | LUAD | NCI-H1651 | Gefitinib | MVP_top60_DNA+RNA_seed | HDAC1 | 0.155716 | THCA | CGTH-W-1 | Belinostat |
| MVP_top60_DNA_seed | PIK3CA | -0.078508 | LUAD | NCI-H1944 | Alpelisib | MVP_top60_DNA+RNA_seed | BRAF | 0.161866 | THCA | TT | Dabrafenib |
| MVP_top60_DNA_seed | PIK3CA | -0.078505 | LUAD | Calu-6 | GNE-317 | MVP_top60_DNA+RNA_seed | PTK2 | 0.164523 | THCA | RO82-W-1 | PF-562271 |
| MVP_top60_DNA_seed | MET | -0.076999 | LUAD | NCI-H23 | Crizotinib | MVP_top60_DNA+RNA_seed | ATM | 0.164551 | THCA | CGTH-W-1 | KU-60019 |
| MVP_top60_DNA_seed | EGFR | -0.076941 | LUAD | NCI-H838 | Afatinib | MVP_top60_DNA+RNA_seed | AKT1 | 0.165025 | THCA | 8305C | MK-2206 |
| MVP_top60_DNA_seed | ERBB2 | -0.076941 | LUAD | NCI-H838 | Afatinib | MVP_top60_DNA+RNA_seed | AKT2 | 0.165025 | THCA | 8305C | MK-2206 |
| MVP_top60_DNA_seed | TERT | -0.076725 | LUAD | NCI-H2030 | BIBR-1532 | MVP_top60_DNA+RNA_seed | PDPK1 | 0.165865 | THCA | CAL-62 | BX-912 |
| MVP_top60_DNA_seed | BRAF | -0.074879 | LUAD | NCI-H1734 | Dabrafenib | MVP_top60_DNA+RNA_seed | GSK3B | 0.173615 | THCA | FTC-133 | CHIR-99021 |
| MVP_top60_DNA_seed | HDAC1 | -0.073644 | LUAD | NCI-H1792 | Romidepsin | MVP_top60_DNA+RNA_seed | PIK3CG | 0.17854 | THCA | CGTH-W-1 | PIK-93 |
| MVP_top60_DNA_seed | BRAF | -0.071772 | LUAD | NCI-H1734 | SB590885 | MVP_top60_DNA+RNA_seed | AKT1 | 0.180136 | THCA | RO82-W-1 | AKT inhibitor VIII |
| MVP_top60_DNA_seed | AKT1 | -0.071468 | LUAD | PC-14 | AT13148 | MVP_top60_DNA+RNA_seed | AKT2 | 0.180136 | THCA | RO82-W-1 | AKT inhibitor VIII |
| MVP_top60_DNA_seed | HDAC1 | -0.071026 | LUAD | NCI-H522 | Entinostat | MVP_top60_DNA+RNA_seed | AKT3 | 0.180136 | THCA | RO82-W-1 | AKT inhibitor VIII |
| MVP_top60_DNA_seed | TP53 | -0.070238 | LUAD | LXF-289 | MIRA-1 | MVP_top60_DNA+RNA_seed | MAPK1 | 0.18019 | THCA | TT2609-C02 | FR-180204 |
| MVP_top60_DNA_seed | AKT1 | -0.069219 | LUAD | NCI-H2228 | Afuresertib | MVP_top60_DNA+RNA_seed | MAPK3 | 0.18019 | THCA | TT2609-C02 | FR-180204 |
| MVP_top60_DNA_seed | EGFR | -0.06844 | LUAD | NCI-H522 | Sapitinib | MVP_top60_DNA+RNA_seed | AKT1 | 0.18882 | THCA | 8505C | GSK690693 |
| MVP_top60_DNA_seed | ERBB2 | -0.06844 | LUAD | NCI-H522 | Sapitinib | MVP_top60_DNA+RNA_seed | AKT2 | 0.18882 | THCA | 8505C | GSK690693 |
| MVP_top60_DNA_seed | ERBB3 | -0.06844 | LUAD | NCI-H522 | Sapitinib | MVP_top60_DNA+RNA_seed | AKT3 | 0.18882 | THCA | 8505C | GSK690693 |
| MVP_top60_DNA_seed | AKT1 | -0.06737 | LUAD | NCI-H1563 | GSK2110183B | MVP_top60_DNA+RNA_seed | RET | 0.203243 | THCA | KMH-2 | Motesanib |
| MVP_top60_DNA_seed | HSP90AA1 | -0.066491 | LUAD | NCI-H1944 | Luminespib | MVP_top60_DNA+RNA_seed | AKT1 | 0.215195 | THCA | 8505C | AKT inhibitor VIII |
| MVP_top60_DNA_seed | BRAF | -0.065928 | LUAD | NCI-H358 | PLX-4720 | MVP_top60_DNA+RNA_seed | AKT2 | 0.215195 | THCA | 8505C | AKT inhibitor VIII |
| MVP_top60_DNA_seed | MAPK1 | -0.065092 | LUAD | NCI-H2023 | VX-11e | MVP_top60_DNA+RNA_seed | AKT3 | 0.215195 | THCA | 8505C | AKT inhibitor VIII |
| MVP_top60_DNA_seed | MAPK1 | -0.064385 | LUAD | EMC-BAC-2 | Ulixertinib | MVP_top60_DNA+RNA_seed | CHEK2 | 0.221166 | THCA | TT2609-C02 | AZD7762 |
| MVP_top60_DNA_seed | AKT1 | -0.063853 | LUAD | NCI-H1792 | Afuresertib | MVP_top60_DNA+RNA_seed | MAPK1 | 0.222853 | THCA | KMH-2 | FR-180204 |
| MVP_top60_DNA_seed | TP53 | -0.063293 | LUAD | NCI-H1437 | MIRA-1 | MVP_top60_DNA+RNA_seed | MAPK3 | 0.222853 | THCA | KMH-2 | FR-180204 |
| MVP_top60_DNA_seed | EGFR | -0.062643 | LUAD | NCI-H1435 | AZD3759 | MVP_top60_DNA+RNA_seed | PDPK1 | 0.230659 | THCA | K5 | BX-912 |
| MVP_top60_DNA_seed | EGFR | -0.062512 | LUAD | NCI-H1563 | Afatinib | MVP_top60_DNA+RNA_seed | AKT1 | 0.233858 | THCA | IHH-4 | GSK690693 |
| MVP_top60_DNA_seed | ERBB2 | -0.062512 | LUAD | NCI-H1563 | Afatinib | MVP_top60_DNA+RNA_seed | AKT2 | 0.233858 | THCA | IHH-4 | GSK690693 |
| MVP_top60_DNA_seed | EGFR | -0.060513 | LUAD | NCI-H1651 | AZD3759 | MVP_top60_DNA+RNA_seed | AKT3 | 0.233858 | THCA | IHH-4 | GSK690693 |
| MVP_top60_DNA_seed | FGFR1 | -0.058974 | LUAD | NCI-H2009 | PD173074 | MVP_top60_DNA+RNA_seed | PDPK1 | 0.23749 | THCA | K5 | OSU-03012 |
| MVP_top60_DNA_seed | HSP90AA1 | -0.058974 | LUAD | NCI-H23 | Luminespib | MVP_top60_DNA+RNA_seed | PIK3CG | 0.239262 | THCA | 8305C | PIK-93 |
| MVP_top60_DNA_seed | AURKA | -0.056053 | LUAD | NCI-H2228 | Alisertib | MVP_top60_DNA+RNA_seed | RET | 0.241375 | THCA | ASH-3 | Motesanib |
| MVP_top60_DNA_seed | AKT1 | -0.055584 | LUAD | NCI-H838 | Ipatasertib | MVP_top60_DNA+RNA_seed | BRAF | 0.24194 | THCA | TT2609-C02 | Dabrafenib |
| MVP_top60_DNA_seed | EGFR | -0.054995 | LUAD | NCI-H1573 | Sapitinib | MVP_top60_DNA+RNA_seed | BRAF | 0.241945 | THCA | RO82-W-1 | RAF_9304 |
| MVP_top60_DNA_seed | ERBB2 | -0.054995 | LUAD | NCI-H1573 | Sapitinib | MVP_top60_DNA+RNA_seed | MAPK1 | 0.249067 | THCA | CAL-62 | FR-180204 |
| MVP_top60_DNA_seed | ERBB3 | -0.054995 | LUAD | NCI-H1573 | Sapitinib | MVP_top60_DNA+RNA_seed | MAPK3 | 0.249067 | THCA | CAL-62 | FR-180204 |
| MVP_top60_DNA_seed | MAPK1 | -0.054823 | LUAD | NCI-H358 | Ulixertinib | MVP_top60_DNA+RNA_seed | BRAF | 0.25163 | THCA | 8505C | PLX-4720 |
| MVP_top60_DNA_seed | AKT1 | -0.054084 | LUAD | NCI-H1437 | Ipatasertib | MVP_top60_DNA+RNA_seed | BRAF | 0.252942 | THCA | TT | RAF_9304 |
| MVP_top60_DNA_seed | BRAF | -0.053744 | LUAD | NCI-H522 | Dabrafenib | MVP_top60_DNA+RNA_seed | HDAC1 | 0.258404 | THCA | KMH-2 | CAY10603 |

| | | | | | | | | | | | |
|---|---|---|---|---|---|---|---|---|---|---|---|
| MVP_top60_DNA_seed | MAPK1 | -0.053177 | LUAD | NCI-H2023 | SCH772984 | MVP_top60_DNA+RNA_seed | MAPK1 | 0.264829 | THCA | CGTH-W-1 | FR-180204 |
| MVP_top60_DNA_seed | EGFR | -0.052991 | LUAD | NCI-H358 | AZD3759 | MVP_top60_DNA+RNA_seed | MAPK3 | 0.264829 | THCA | CGTH-W-1 | FR-180204 |
| MVP_top60_DNA_seed | EGFR | -0.050957 | LUAD | NCI-H1792 | Gefitinib | MVP_top60_DNA+RNA_seed | GSK3B | 0.265169 | THCA | CAL-62 | KIN001-042 |
| MVP_top60_DNA_seed | PIK3CA | -0.047645 | LUAD | NCI-H2228 | AZD8186 | MVP_top60_DNA+RNA_seed | HDAC1 | 0.265439 | THCA | CGTH-W-1 | CAY10603 |
| MVP_top60_DNA_seed | MAPK1 | -0.047333 | LUAD | RERF-LC-KJ | SCH772984 | MVP_top60_DNA+RNA_seed | ATM | 0.26645 | THCA | ML-1 | KU-60019 |
| MVP_top60_DNA_seed | BRAF | -0.046643 | LUAD | EMC-BAC-1 | Dabrafenib | MVP_top60_DNA+RNA_seed | AKT1 | 0.268555 | THCA | ASH-3 | MK-2206 |
| MVP_top60_DNA_seed | IGF1R | -0.046488 | LUAD | RERF-LC-KJ | BMS-754807 | MVP_top60_DNA+RNA_seed | AKT2 | 0.268555 | THCA | ASH-3 | MK-2206 |
| MVP_top60_DNA_seed | MDM2 | -0.044598 | LUAD | NCI-H1734 | Nutlin-3a (-) | MVP_top60_DNA+RNA_seed | HDAC1 | 0.278697 | THCA | FTC-133 | Dacinostat |
| MVP_top60_DNA_seed | MAPK1 | -0.043573 | LUAD | NCI-H1975 | SCH772984 | MVP_top60_DNA+RNA_seed | GSK3B | 0.292474 | THCA | BHT-101 | CHIR-99021 |
| MVP_top60_DNA_seed | EGFR | -0.042947 | LUAD | NCI-H1435 | Afatinib | MVP_top60_DNA+RNA_seed | HDAC1 | 0.29621 | THCA | CAL-62 | CAY10603 |
| MVP_top60_DNA_seed | ERBB2 | -0.042947 | LUAD | NCI-H1435 | Afatinib | MVP_top60_DNA+RNA_seed | GSK3B | 0.296564 | THCA | 8305C | AZD7969 |
| MVP_top60_DNA_seed | HSP90AA1 | -0.041669 | LUAD | COR-L105 | Luminespib | MVP_top60_DNA+RNA_seed | MAPK1 | 0.298846 | THCA | 8305C | VX-11e |
| MVP_top60_DNA_seed | SRC | -0.041344 | LUAD | NCI-H2342 | Bosutinib | MVP_top60_DNA+RNA_seed | GSK3B | 0.305956 | THCA | HTC-C3 | CHIR-99021 |
| MVP_top60_DNA_seed | HDAC1 | -0.039471 | LUAD | NCI-H1944 | Romidepsin | MVP_top60_DNA+RNA_seed | BRAF | 0.307688 | THCA | FTC-133 | SB590885 |
| MVP_top60_DNA_seed | ATM | -0.039 | LUAD | NCI-H650 | KU-55933 | MVP_top60_DNA+RNA_seed | GSK3B | 0.308457 | THCA | BHT-101 | KIN001-042 |
| MVP_top60_DNA_seed | HSP90AA1 | -0.038665 | LUAD | LC-2-ad | Tanespimycin | MVP_top60_DNA+RNA_seed | PIK3CG | 0.3097 | THCA | TT2609-C02 | AS605240 |
| MVP_top60_DNA_seed | MDM2 | -0.038314 | LUAD | Calu-3 | Serdemetan | MVP_top60_DNA+RNA_seed | PDPK1 | 0.314436 | THCA | 8305C | BX-912 |
| MVP_top60_DNA_seed | PIK3CA | -0.038234 | LUAD | HOP-62 | GNE-317 | MVP_top60_DNA+RNA_seed | ATM | 0.317324 | THCA | CAL-62 | CP466722 |
| MVP_top60_DNA_seed | SRC | -0.03792 | LUAD | NCI-H1781 | Bosutinib | MVP_top60_DNA+RNA_seed | AKT1 | 0.318722 | THCA | 8305C | BAY AKT1 |
| MVP_top60_DNA_seed | SRC | -0.037417 | LUAD | NCI-H2122 | Bosutinib | MVP_top60_DNA+RNA_seed | AKT2 | 0.318722 | THCA | 8305C | BAY AKT1 |
| MVP_top60_DNA_seed | MET | -0.03735 | LUAD | NCI-H2009 | Savolitinib | MVP_top60_DNA+RNA_seed | AKT3 | 0.318722 | THCA | 8305C | BAY AKT1 |
| MVP_top60_DNA_seed | MAPK1 | -0.036926 | LUAD | EKVX | VX-11e | MVP_top60_DNA+RNA_seed | CHEK2 | 0.318755 | THCA | B-CPAP | AZD7762 |
| MVP_top60_DNA_seed | KRAS | -0.036458 | LUAD | NCI-H2023 | KRAS (G12C) Inhibitor-1[illegible] | MVP_top60_DNA+RNA_seed | GSK3B | 0.32138 | THCA | K5 | KIN001-042 |
| MVP_top60_DNA_seed | FGFR1 | -0.036218 | LUAD | NCI-H650 | PD173074 | MVP_top60_DNA+RNA_seed | BRAF | 0.3238 | THCA | FTC-133 | PLX-4720 |
| MVP_top60_DNA_seed | SRC | -0.035266 | LUAD | NCI-H1435 | Bosutinib | MVP_top60_DNA+RNA_seed | PDPK1 | 0.326596 | THCA | TT | BX-912 |
| MVP_top60_DNA_seed | BRAF | -0.035156 | LUAD | NCI-H522 | PLX-4720 | MVP_top60_DNA+RNA_seed | ATM | 0.338034 | THCA | B-CPAP | CP466722 |
| MVP_top60_DNA_seed | MDM2 | -0.034144 | LUAD | NCI-H2009 | Serdemetan | MVP_top60_DNA+RNA_seed | ATM | 0.340556 | THCA | CGTH-W-1 | CP466722 |
| MVP_top60_DNA_seed | HDAC1 | -0.033624 | LUAD | NCI-H23 | PCI-34051 | MVP_top60_DNA+RNA_seed | GSK3B | 0.34145 | THCA | ASH-3 | KIN001-042 |
| MVP_top60_DNA_seed | HDAC1 | -0.033596 | LUAD | VMRC-LCD | Romidepsin | MVP_top60_DNA+RNA_seed | AKT1 | 0.345478 | THCA | CGTH-W-1 | AKT inhibitor VIII |
| MVP_top60_DNA_seed | IGF1R | -0.032573 | LUAD | NCI-H1563 | Linsitinib | MVP_top60_DNA+RNA_seed | AKT2 | 0.345478 | THCA | CGTH-W-1 | AKT inhibitor VIII |
| MVP_top60_DNA_seed | EGFR | -0.032533 | LUAD | NCI-H1944 | Afatinib | MVP_top60_DNA+RNA_seed | AKT3 | 0.345478 | THCA | CGTH-W-1 | AKT inhibitor VIII |
| MVP_top60_DNA_seed | ERBB2 | -0.032533 | LUAD | NCI-H1944 | Afatinib | MVP_top60_DNA+RNA_seed | ATM | 0.353416 | THCA | TT | KU-55933 |
| MVP_top60_DNA_seed | AKT1 | -0.031145 | LUAD | NCI-H1563 | MK-2206 | MVP_top60_DNA+RNA_seed | PDPK1 | 0.376134 | THCA | BHT-101 | OSU-03012 |
| MVP_top60_DNA_seed | IGF1R | -0.030377 | LUAD | HCC-827 | BMS-536924 | MVP_top60_DNA+RNA_seed | ATM | 0.377689 | THCA | 8505C | CP466722 |
| MVP_top60_DNA_seed | HDAC1 | -0.028701 | LUAD | A549 | Entinostat | MVP_top60_DNA+RNA_seed | BRAF | 0.380782 | THCA | ASH-3 | SB590885 |
| MVP_top60_DNA_seed | AURKA | -0.028556 | LUAD | HOP-62 | Alisertib | MVP_top60_DNA+RNA_seed | CHEK2 | 0.391805 | THCA | BHT-101 | AZD7762 |
| MVP_top60_DNA_seed | EGFR | -0.026665 | LUAD | NCI-H1437 | AZD3759 | MVP_top60_DNA+RNA_seed | AKT1 | 0.398747 | THCA | TT | BAY AKT1 |
| MVP_top60_DNA_seed | IGF1R | -0.026585 | LUAD | RERF-LC-MS | Linsitinib | MVP_top60_DNA+RNA_seed | AKT2 | 0.398747 | THCA | TT | BAY AKT1 |
| MVP_top60_DNA_seed | TERT | -0.02603 | LUAD | NCI-H1435 | BIBR-1532 | MVP_top60_DNA+RNA_seed | AKT3 | 0.398747 | THCA | TT | BAY AKT1 |
| MVP_top60_DNA_seed | AURKA | -0.024727 | LUAD | NCI-H23 | Alisertib | MVP_top60_DNA+RNA_seed | CHEK2 | 0.413405 | THCA | RO82-W-1 | AZD7762 |
| MVP_top60_DNA_seed | SMARCA4 | -0.023758 | LUAD | NCI-H2023 | PFI3 | MVP_top60_DNA+RNA_seed | GSK3B | 0.413921 | THCA | CGTH-W-1 | KIN001-042 |

| | | | | | | | | | | | |
|---|---|---|---|---|---|---|---|---|---|---|---|
| MVP_top60_DNA_seed | GSK3B | -0.023224 | LUAD | NCI-H1734 | CHIR-99021 | MVP_top60_DNA+RNA_seed | HDAC1 | 0.425525 | THCA | RO82-W-1 | Tubastatin A |
| MVP_top60_DNA_seed | AKT1 | -0.022049 | LUAD | NCI-H2030 | Afuresertib | MVP_top60_DNA+RNA_seed | BRAF | 0.427087 | THCA | CGTH-W-1 | PLX-4720 |
| MVP_top60_DNA_seed | TP53 | -0.021429 | LUAD | NCI-H522 | MIRA-1 | MVP_top60_DNA+RNA_seed | HDAC1 | 0.434938 | THCA | ASH-3 | Dacinostat |
| MVP_top60_DNA_seed | TP53 | -0.021345 | LUAD | RERF-LC-KJ | MIRA-1 | MVP_top60_DNA+RNA_seed | CHEK2 | 0.437393 | THCA | 8505C | AZD7762 |
| MVP_top60_DNA_seed | IGF1R | -0.021162 | LUAD | NCI-H1573 | Linsitinib | MVP_top60_DNA+RNA_seed | AKT1 | 0.441879 | THCA | B-CPAP | GSK690693 |
| MVP_top60_DNA_seed | TP53 | -0.020545 | LUAD | NCI-H1651 | MIRA-1 | MVP_top60_DNA+RNA_seed | AKT2 | 0.441879 | THCA | B-CPAP | GSK690693 |
| MVP_top60_DNA_seed | FGFR1 | -0.02 | LUAD | NCI-H522 | PD173074 | MVP_top60_DNA+RNA_seed | AKT3 | 0.441879 | THCA | B-CPAP | GSK690693 |
| MVP_top60_DNA_seed | MAPK1 | -0.019164 | LUAD | NCI-H1573 | Ulixertinib | MVP_top60_DNA+RNA_seed | ATM | 0.448252 | THCA | B-CPAP | KU-55933 |
| MVP_top60_DNA_seed | AURKA | -0.018535 | LUAD | ABC-1 | Tozasertib | MVP_top60_DNA+RNA_seed | GSK3B | 0.449242 | THCA | TT | AZD7969 |
| MVP_top60_DNA_seed | AKT1 | -0.018466 | LUAD | H3255 | Afuresertib | MVP_top60_DNA+RNA_seed | PIK3CG | 0.454936 | THCA | HTC-C3 | AS605240 |
| MVP_top60_DNA_seed | HSP90AA1 | -0.018442 | LUAD | NCI-H1781 | Luminespib | MVP_top60_DNA+RNA_seed | AKT1 | 0.46 | THCA | FTC-133 | MK-2206 |
| MVP_top60_DNA_seed | BRAF | -0.018387 | LUAD | EMC-BAC-2 | Dabrafenib | MVP_top60_DNA+RNA_seed | AKT2 | 0.46 | THCA | FTC-133 | MK-2206 |
| MVP_top60_DNA_seed | AKT1 | -0.018005 | LUAD | H3255 | Ipatasertib | MVP_top60_DNA+RNA_seed | MAPK1 | 0.460452 | THCA | KMH-2 | VX-11e |
| MVP_top60_DNA_seed | MAPK1 | -0.017889 | LUAD | NCI-H650 | SCH772984 | MVP_top60_DNA+RNA_seed | AKT1 | 0.464012 | THCA | CAL-62 | AKT inhibitor VIII |
| MVP_top60_DNA_seed | PIK3CA | -0.015769 | LUAD | NCI-H1563 | Alpelisib | MVP_top60_DNA+RNA_seed | AKT2 | 0.464012 | THCA | CAL-62 | AKT inhibitor VIII |
| MVP_top60_DNA_seed | MAPK1 | -0.015382 | LUAD | SK-LU-1 | SCH772984 | MVP_top60_DNA+RNA_seed | AKT3 | 0.464012 | THCA | CAL-62 | AKT inhibitor VIII |
| MVP_top60_DNA_seed | CREBBP | -0.014929 | LUAD | NCI-H2122 | SGC-CBP30 | MVP_top60_DNA+RNA_seed | BRAF | 0.47435 | THCA | KMH-2 | PLX-4720 |
| MVP_top60_DNA_seed | MAPK1 | -0.013411 | LUAD | NCI-H2030 | Ulixertinib | MVP_top60_DNA+RNA_seed | BRAF | 0.476929 | THCA | ASH-3 | PLX-4720 |
| MVP_top60_DNA_seed | EGFR | -0.012968 | LUAD | NCI-H1755 | Afatinib | MVP_top60_DNA+RNA_seed | BRAF | 0.478727 | THCA | FTC-133 | Dabrafenib |
| MVP_top60_DNA_seed | ERBB2 | -0.012968 | LUAD | NCI-H1755 | Afatinib | MVP_top60_DNA+RNA_seed | AKT1 | 0.482172 | THCA | ASH-3 | GSK690693 |
| MVP_top60_DNA_seed | EGFR | -0.01269 | LUAD | NCI-H838 | Osimertinib | MVP_top60_DNA+RNA_seed | AKT2 | 0.482172 | THCA | ASH-3 | GSK690693 |
| MVP_top60_DNA_seed | PIK3CA | -0.012626 | LUAD | NCI-H2009 | Buparlisib | MVP_top60_DNA+RNA_seed | AKT3 | 0.482172 | THCA | ASH-3 | GSK690693 |
| MVP_top60_DNA_seed | EGFR | -0.01189 | LUAD | LC-2-ad | Osimertinib | MVP_top60_DNA+RNA_seed | AKT1 | 0.496458 | THCA | RO82-W-1 | MK-2206 |
| MVP_top60_DNA_seed | EGFR | -0.011396 | LUAD | NCI-H2085 | Sapitinib | MVP_top60_DNA+RNA_seed | AKT2 | 0.496458 | THCA | RO82-W-1 | MK-2206 |
| MVP_top60_DNA_seed | ERBB2 | -0.011396 | LUAD | NCI-H2085 | Sapitinib | MVP_top60_DNA+RNA_seed | BRAF | 0.503209 | THCA | FTC-133 | RAF_9304 |
| MVP_top60_DNA_seed | ERBB3 | -0.011396 | LUAD | NCI-H2085 | Sapitinib | MVP_top60_DNA+RNA_seed | AKT1 | 0.513617 | THCA | ASH-3 | AKT inhibitor VIII |
| MVP_top60_DNA_seed | AKT1 | -0.011382 | LUAD | NCI-H522 | AT13148 | MVP_top60_DNA+RNA_seed | AKT2 | 0.513617 | THCA | ASH-3 | AKT inhibitor VIII |
| MVP_top60_DNA_seed | PIK3CA | -0.011315 | LUAD | LXF-289 | AZD8186 | MVP_top60_DNA+RNA_seed | AKT3 | 0.513617 | THCA | ASH-3 | AKT inhibitor VIII |
| MVP_top60_DNA_seed | CREBBP | -0.010388 | LUAD | NCI-H1993 | SGC-CBP30 | MVP_top60_DNA+RNA_seed | AKT1 | 0.518521 | THCA | CGTH-W-1 | GSK690693 |
| MVP_top60_DNA_seed | IGF1R | -0.009997 | LUAD | NCI-H1792 | GSK1904529A | MVP_top60_DNA+RNA_seed | AKT2 | 0.518521 | THCA | CGTH-W-1 | GSK690693 |
| MVP_top60_DNA_seed | AKT1 | -0.009335 | LUAD | HCC-78 | Uprosertib | MVP_top60_DNA+RNA_seed | AKT3 | 0.518521 | THCA | CGTH-W-1 | GSK690693 |
| MVP_top60_DNA_seed | PIK3CA | -0.009014 | LUAD | HCC-827 | GNE-317 | MVP_top60_DNA+RNA_seed | PDPK1 | 0.518904 | THCA | BHT-101 | BX-912 |
| MVP_top60_DNA_seed | IGF1R | -0.008776 | LUAD | NCI-H1975 | GSK1904529A | MVP_top60_DNA+RNA_seed | HDAC1 | 0.530875 | THCA | KMH-2 | Tubastatin A |
| MVP_top60_DNA_seed | BRAF | -0.006907 | LUAD | NCI-H1792 | Dabrafenib | MVP_top60_DNA+RNA_seed | AKT1 | 0.537145 | THCA | ASH-3 | BAY AKT1 |
| MVP_top60_DNA_seed | MAPK1 | -0.003918 | LUAD | NCI-H1781 | Ulixertinib | MVP_top60_DNA+RNA_seed | AKT2 | 0.537145 | THCA | ASH-3 | BAY AKT1 |
| MVP_top60_DNA_seed | MET | -0.003864 | LUAD | NCI-H1792 | Savolitinib | MVP_top60_DNA+RNA_seed | AKT3 | 0.537145 | THCA | ASH-3 | BAY AKT1 |
| MVP_top60_DNA_seed | EGFR | -0.002579 | LUAD | NCI-H838 | Gefitinib | MVP_top60_DNA+RNA_seed | GSK3B | 0.542286 | THCA | HTC-C3 | SB216763 |
| MVP_top60_DNA_seed | IGF1R | -0.00174 | LUAD | NCI-H2342 | Linsitinib | MVP_top60_DNA+RNA_seed | BRAF | 0.542429 | THCA | TT2609-C02 | RAF_9304 |
| MVP_top60_DNA_seed | AKT1 | -0.001342 | LUAD | A549 | Afuresertib | MVP_top60_DNA+RNA_seed | AKT1 | 0.546883 | THCA | HTC-C3 | MK-2206 |
| MVP_top60_DNA_seed | MET | -0.000908 | LUAD | NCI-H1648 | Savolitinib | MVP_top60_DNA+RNA_seed | AKT2 | 0.546883 | THCA | HTC-C3 | MK-2206 |
| MVP_top60_DNA_seed | IGF1R | -0.000873 | LUAD | NCI-H650 | NVP-ADW742 | MVP_top60_DNA+RNA_seed | MAPK1 | 0.547965 | THCA | ML-1 | FR-180204 |

| | | | | | | | | | | | |
|---|---|---|---|---|---|---|---|---|---|---|---|
| MVP_top60_DNA_seed | AKT1 | 0.001387 | LUAD | NCI-H1563 | AZD5363 | MVP_top60_DNA+RNA_seed | MAPK3 | 0.547965 | THCA | ML-1 | FR-180204 |
| MVP_top60_DNA_seed | HSP90AA1 | 0.003477 | LUAD | NCI-H322M | Luminespib | MVP_top60_DNA+RNA_seed | CHEK2 | 0.551334 | THCA | ASH-3 | AZD7762 |
| MVP_top60_DNA_seed | PIK3CA | 0.003942 | LUAD | NCI-H1734 | GNE-317 | MVP_top60_DNA+RNA_seed | AKT1 | 0.555869 | THCA | B-CPAP | AKT inhibitor VIII |
| MVP_top60_DNA_seed | TERT | 0.005902 | LUAD | NCI-H1650 | BIBR-1532 | MVP_top60_DNA+RNA_seed | AKT2 | 0.555869 | THCA | B-CPAP | AKT inhibitor VIII |
| MVP_top60_DNA_seed | MAPK1 | 0.006191 | LUAD | NCI-H1573 | Ulixertinib | MVP_top60_DNA+RNA_seed | AKT3 | 0.555869 | THCA | B-CPAP | AKT inhibitor VIII |
| MVP_top60_DNA_seed | AKT1 | 0.007821 | LUAD | NCI-H1755 | GSK2110183B | MVP_top60_DNA+RNA_seed | BRAF | 0.558517 | THCA | ASH-3 | HG6-64-1 |
| MVP_top60_DNA_seed | IGF1R | 0.008672 | LUAD | NCI-H2009 | GSK1904529A | MVP_top60_DNA+RNA_seed | AKT1 | 0.560545 | THCA | B-CPAP | BAY AKT1 |
| MVP_top60_DNA_seed | IGF1R | 0.00934 | LUAD | NCI-H1944 | Linsitinib | MVP_top60_DNA+RNA_seed | AKT2 | 0.560545 | THCA | B-CPAP | BAY AKT1 |
| MVP_top60_DNA_seed | SRC | 0.009809 | LUAD | HCC-44 | Bosutinib | MVP_top60_DNA+RNA_seed | AKT3 | 0.560545 | THCA | B-CPAP | BAY AKT1 |
| MVP_top60_DNA_seed | AURKA | 0.010821 | LUAD | EMC-BAC-1 | ZM447439 | MVP_top60_DNA+RNA_seed | PIK3CG | 0.570249 | THCA | IHH-4 | AS605240 |
| MVP_top60_DNA_seed | SRC | 0.012303 | LUAD | RERF-LC-MS | Bosutinib | MVP_top60_DNA+RNA_seed | HDAC1 | 0.576176 | THCA | FTC-133 | Belinostat |
| MVP_top60_DNA_seed | PIK3CA | 0.012533 | LUAD | NCI-H1651 | Buparlisib | MVP_top60_DNA+RNA_seed | GSK3B | 0.579491 | THCA | HTC-C3 | CHIR-99021 |
| MVP_top60_DNA_seed | EGFR | 0.013265 | LUAD | NCI-H1693 | Lapatinib | MVP_top60_DNA+RNA_seed | PIK3CG | 0.583872 | THCA | ML-1 | AS605240 |
| MVP_top60_DNA_seed | ERBB2 | 0.013265 | LUAD | NCI-H1693 | Lapatinib | MVP_top60_DNA+RNA_seed | AKT1 | 0.598072 | THCA | TT2609-C02 | AKT inhibitor VIII |
| MVP_top60_DNA_seed | KRAS | 0.013546 | LUAD | NCI-H3122 | KRAS (G12C) Inhibitor-1[illegible] | MVP_top60_DNA+RNA_seed | AKT2 | 0.598072 | THCA | TT2609-C02 | AKT inhibitor VIII |
| MVP_top60_DNA_seed | CREBBP | 0.013998 | LUAD | NCI-H650 | SGC-CBP30 | MVP_top60_DNA+RNA_seed | AKT3 | 0.598072 | THCA | TT2609-C02 | AKT inhibitor VIII |
| MVP_top60_DNA_seed | EZH2 | 0.014421 | LUAD | NCI-H23 | GSK343 | MVP_top60_DNA+RNA_seed | BRAF | 0.604345 | THCA | BHT-101 | HG6-64-1 |
| MVP_top60_DNA_seed | GSK3B | 0.014575 | LUAD | LC-2-ad | SB216763 | MVP_top60_DNA+RNA_seed | ATM | 0.604481 | THCA | KMH-2 | KU-55933 |
| MVP_top60_DNA_seed | PIK3CA | 0.014755 | LUAD | NCI-H1355 | Buparlisib | MVP_top60_DNA+RNA_seed | AKT1 | 0.60813 | THCA | CAL-62 | GSK690693 |
| MVP_top60_DNA_seed | MET | 0.01497 | LUAD | NCI-H2405 | Savolitinib | MVP_top60_DNA+RNA_seed | AKT2 | 0.60813 | THCA | CAL-62 | GSK690693 |
| MVP_top60_DNA_seed | IGF1R | 0.021004 | LUAD | NCI-H650 | BMS-536924 | MVP_top60_DNA+RNA_seed | AKT3 | 0.60813 | THCA | CAL-62 | GSK690693 |
| MVP_top60_DNA_seed | AKT1 | 0.021479 | LUAD | NCI-H2342 | Ipatasertib | MVP_top60_DNA+RNA_seed | BRAF | 0.61027 | THCA | CAL-62 | SB590885 |
| MVP_top60_DNA_seed | PIK3CA | 0.022357 | LUAD | NCI-H3122 | AZD8186 | MVP_top60_DNA+RNA_seed | BRAF | 0.623824 | THCA | RO82-W-1 | PLX-4720 |
| MVP_top60_DNA_seed | MDM2 | 0.022863 | LUAD | NCI-H1435 | Serdemetan | MVP_top60_DNA+RNA_seed | AKT1 | 0.630547 | THCA | BHT-101 | GSK690693 |
| MVP_top60_DNA_seed | MDM2 | 0.024305 | LUAD | NCI-H1993 | Serdemetan | MVP_top60_DNA+RNA_seed | AKT2 | 0.630547 | THCA | BHT-101 | GSK690693 |
| MVP_top60_DNA_seed | TERT | 0.025029 | LUAD | NCI-H1651 | BIBR-1532 | MVP_top60_DNA+RNA_seed | AKT3 | 0.630547 | THCA | BHT-101 | GSK690693 |
| MVP_top60_DNA_seed | EGFR | 0.025665 | LUAD | NCI-H522 | Afatinib | MVP_top60_DNA+RNA_seed | AKT1 | 0.632176 | THCA | 8505C | AKT inhibitor VIII |
| MVP_top60_DNA_seed | ERBB2 | 0.025665 | LUAD | NCI-H522 | Afatinib | MVP_top60_DNA+RNA_seed | AKT2 | 0.632176 | THCA | 8505C | AKT inhibitor VIII |
| MVP_top60_DNA_seed | AKT1 | 0.027672 | LUAD | NCI-H1435 | GSK2110183B | MVP_top60_DNA+RNA_seed | AKT3 | 0.632176 | THCA | 8505C | AKT inhibitor VIII |
| MVP_top60_DNA_seed | HDAC1 | 0.02972 | LUAD | NCI-H1568 | Romidepsin | MVP_top60_DNA+RNA_seed | AKT1 | 0.642781 | THCA | KMH-2 | MK-2206 |
| MVP_top60_DNA_seed | HDAC1 | 0.030073 | LUAD | HCC-44 | PCI-34051 | MVP_top60_DNA+RNA_seed | AKT2 | 0.642781 | THCA | KMH-2 | MK-2206 |
| MVP_top60_DNA_seed | AKT1 | 0.030188 | LUAD | LC-2-ad | AZD5363 | MVP_top60_DNA+RNA_seed | GSK3B | 0.647225 | THCA | ML-1 | KIN001-042 |
| MVP_top60_DNA_seed | AURKA | 0.031621 | LUAD | NCI-H2030 | ZM447439 | MVP_top60_DNA+RNA_seed | AKT1 | 0.647524 | THCA | K5 | AKT inhibitor VIII |
| MVP_top60_DNA_seed | AKT1 | 0.031621 | LUAD | LXF-289 | Afuresertib | MVP_top60_DNA+RNA_seed | AKT2 | 0.647524 | THCA | K5 | AKT inhibitor VIII |
| MVP_top60_DNA_seed | EGFR | 0.031739 | LUAD | LC-2-ad | Afatinib | MVP_top60_DNA+RNA_seed | AKT3 | 0.647524 | THCA | K5 | AKT inhibitor VIII |
| MVP_top60_DNA_seed | ERBB2 | 0.031739 | LUAD | LC-2-ad | Afatinib | MVP_top60_DNA+RNA_seed | PDPK1 | 0.650017 | THCA | RO82-W-1 | BX-912 |
| MVP_top60_DNA_seed | IGF1R | 0.032084 | LUAD | NCI-H2023 | NVP-ADW742 | MVP_top60_DNA+RNA_seed | BRAF | 0.653543 | THCA | CGTH-W-1 | SB590885 |
| MVP_top60_DNA_seed | EGFR | 0.032678 | LUAD | NCI-H2030 | Gefitinib | MVP_top60_DNA+RNA_seed | GSK3B | 0.657198 | THCA | CAL-62 | SB216763 |
| MVP_top60_DNA_seed | AURKA | 0.032695 | LUAD | NCI-H1355 | Alisertib | MVP_top60_DNA+RNA_seed | HDAC1 | 0.660448 | THCA | RO82-W-1 | Dacinostat |
| MVP_top60_DNA_seed | AKT1 | 0.03296 | LUAD | NCI-H358 | Uprosertib | MVP_top60_DNA+RNA_seed | CHEK2 | 0.669386 | THCA | RO82-W-1 | AZD7762 |
| MVP_top60_DNA_seed | IGF1R | 0.033209 | LUAD | LXF-289 | GSK1904529A | MVP_top60_DNA+RNA_seed | BRAF | 0.670799 | THCA | TT | HG6-64-1 |

| | | | | | | | | | | | |
|---|---|---|---|---|---|---|---|---|---|---|---|
| MVP_top60_DNA_seed | GSK3B | 0.033569 | LUAD | NCI-H1648 | SB216763 | MVP_top60_DNA+RNA_seed | MAPK1 | 0.673209 | THCA | ASH-3 | FR-180204 |
| MVP_top60_DNA_seed | AKT1 | 0.034521 | LUAD | HOP-62 | AT13148 | MVP_top60_DNA+RNA_seed | MAPK3 | 0.673209 | THCA | ASH-3 | FR-180204 |
| MVP_top60_DNA_seed | PIK3CA | 0.034876 | LUAD | Calu-3 | Alpelisib | MVP_top60_DNA+RNA_seed | AKT1 | 0.67578 | THCA | TT2609-C02 | GSK690693 |
| MVP_top60_DNA_seed | PIK3CA | 0.035645 | LUAD | EKVX | GNE-317 | MVP_top60_DNA+RNA_seed | AKT2 | 0.67578 | THCA | TT2609-C02 | GSK690693 |
| MVP_top60_DNA_seed | MDM2 | 0.037062 | LUAD | NCI-H1693 | Serdemetan | MVP_top60_DNA+RNA_seed | AKT3 | 0.67578 | THCA | TT2609-C02 | GSK690693 |
| MVP_top60_DNA_seed | EGFR | 0.03744 | LUAD | NCI-H2009 | AZD3759 | MVP_top60_DNA+RNA_seed | PIK3CG | 0.676735 | THCA | ASH-3 | AS605240 |
| MVP_top60_DNA_seed | CREBBP | 0.03794 | LUAD | NCI-H322M | SGC-CBP30 | MVP_top60_DNA+RNA_seed | HDAC1 | 0.683032 | THCA | FTC-133 | JQ12 |
| MVP_top60_DNA_seed | IGF1R | 0.038498 | LUAD | HOP-62 | BMS-536924 | MVP_top60_DNA+RNA_seed | HDAC1 | 0.694364 | THCA | CGTH-W-1 | Tubastatin A |
| MVP_top60_DNA_seed | MAPK1 | 0.039706 | LUAD | NCI-H1355 | SCH772984 | MVP_top60_DNA+RNA_seed | HDAC1 | 0.707075 | THCA | ASH-3 | JQ12 |
| MVP_top60_DNA_seed | TERT | 0.040347 | LUAD | NCI-H1781 | BIBR-1532 | MVP_top60_DNA+RNA_seed | AKT1 | 0.710134 | THCA | FTC-133 | GSK690693 |
| MVP_top60_DNA_seed | AKT1 | 0.041005 | LUAD | NCI-H2023 | MK-2206 | MVP_top60_DNA+RNA_seed | AKT2 | 0.710134 | THCA | FTC-133 | GSK690693 |
| MVP_top60_DNA_seed | AURKA | 0.041636 | LUAD | NCI-H358 | ZM447439 | MVP_top60_DNA+RNA_seed | AKT3 | 0.710134 | THCA | FTC-133 | GSK690693 |
| MVP_top60_DNA_seed | AURKA | 0.042263 | LUAD | NCI-H1755 | Alisertib | MVP_top60_DNA+RNA_seed | GSK3B | 0.713104 | THCA | TT2609-C02 | KIN001-042 |
| MVP_top60_DNA_seed | EZH2 | 0.04233 | LUAD | LC-2-ad | GSK343 | MVP_top60_DNA+RNA_seed | MAPK1 | 0.713488 | THCA | FTC-133 | VX-11e |
| MVP_top60_DNA_seed | BRAF | 0.043027 | LUAD | NCI-H2122 | Dabrafenib | MVP_top60_DNA+RNA_seed | HDAC1 | 0.720626 | THCA | ASH-3 | Tubastatin A |
| MVP_top60_DNA_seed | HSP90AA1 | 0.043841 | LUAD | NCI-H1573 | Tanespimycin | MVP_top60_DNA+RNA_seed | AKT1 | 0.721096 | THCA | HTC-C3 | GSK690693 |
| MVP_top60_DNA_seed | IGF1R | 0.044606 | LUAD | NCI-H1734 | BMS-536924 | MVP_top60_DNA+RNA_seed | AKT2 | 0.721096 | THCA | HTC-C3 | GSK690693 |
| MVP_top60_DNA_seed | SRC | 0.04476 | LUAD | LXF-289 | Bosutinib | MVP_top60_DNA+RNA_seed | AKT3 | 0.721096 | THCA | HTC-C3 | GSK690693 |
| MVP_top60_DNA_seed | AKT1 | 0.045183 | LUAD | EMC-BAC-2 | Afuresertib | MVP_top60_DNA+RNA_seed | HDAC1 | 0.740788 | THCA | FTC-133 | CAY10603 |
| MVP_top60_DNA_seed | AKT1 | 0.046648 | LUAD | NCI-H2085 | Uprosertib | MVP_top60_DNA+RNA_seed | PIK3CG | 0.745687 | THCA | KMH-2 | PIK-93 |
| MVP_top60_DNA_seed | PIK3CA | 0.046934 | LUAD | PC-3_[JPC-3] | AZD8186 | MVP_top60_DNA+RNA_seed | MAPK1 | 0.765539 | THCA | K5 | VX-11e |
| MVP_top60_DNA_seed | MAPK1 | 0.047236 | LUAD | LXF-289 | Ulixertinib | MVP_top60_DNA+RNA_seed | MAPK1 | 0.775514 | THCA | FTC-133 | FR-180204 |
| MVP_top60_DNA_seed | IGF1R | 0.047415 | LUAD | NCI-H1355 | BMS-536924 | MVP_top60_DNA+RNA_seed | MAPK3 | 0.775514 | THCA | FTC-133 | FR-180204 |
| MVP_top60_DNA_seed | PIK3CA | 0.048077 | LUAD | NCI-H1355 | Alpelisib | MVP_top60_DNA+RNA_seed | PDPK1 | 0.777246 | THCA | HTC-C3 | BX-912 |
| MVP_top60_DNA_seed | IGF1R | 0.049028 | LUAD | NCI-H2030 | Linsitinib | MVP_top60_DNA+RNA_seed | AKT1 | 0.777805 | THCA | HTC-C3 | BAY AKT1 |
| MVP_top60_DNA_seed | IGF1R | 0.049718 | LUAD | NCI-H2085 | BMS-754807 | MVP_top60_DNA+RNA_seed | AKT2 | 0.777805 | THCA | HTC-C3 | BAY AKT1 |
| MVP_top60_DNA_seed | HDAC1 | 0.049943 | LUAD | RERF-LC-KJ | PCI-34051 | MVP_top60_DNA+RNA_seed | AKT3 | 0.777805 | THCA | HTC-C3 | BAY AKT1 |
| MVP_top60_DNA_seed | AURKA | 0.050331 | LUAD | NCI-H3122 | ZM447439 | MVP_top60_DNA+RNA_seed | HDAC1 | 0.778947 | THCA | FTC-133 | AR-42 |
| MVP_top60_DNA_seed | EGFR | 0.051634 | LUAD | NCI-H1435 | Gefitinib | MVP_top60_DNA+RNA_seed | MAPK1 | 0.779093 | THCA | ML-1 | VX-11e |
| MVP_top60_DNA_seed | EZH2 | 0.0518 | LUAD | NCI-H1563 | GSK343 | MVP_top60_DNA+RNA_seed | AKT1 | 0.798693 | THCA | 8305C | AKT inhibitor VIII |
| MVP_top60_DNA_seed | TERT | 0.052516 | LUAD | HCC-44 | BIBR-1532 | MVP_top60_DNA+RNA_seed | AKT2 | 0.798693 | THCA | 8305C | AKT inhibitor VIII |
| MVP_top60_DNA_seed | AKT1 | 0.052836 | LUAD | NCI-H1975 | GSK2110183B | MVP_top60_DNA+RNA_seed | AKT3 | 0.798693 | THCA | 8305C | AKT inhibitor VIII |
| MVP_top60_DNA_seed | PIK3CA | 0.05373 | LUAD | NCI-H1563 | GNE-317 | MVP_top60_DNA+RNA_seed | AKT1 | 0.805927 | THCA | B-CPAP | AKT inhibitor VIII |
| MVP_top60_DNA_seed | EGFR | 0.056497 | LUAD | HOP-62 | Erlotinib | MVP_top60_DNA+RNA_seed | AKT2 | 0.805927 | THCA | B-CPAP | AKT inhibitor VIII |
| MVP_top60_DNA_seed | FGFR1 | 0.057428 | LUAD | NCI-H441 | PD173074 | MVP_top60_DNA+RNA_seed | AKT3 | 0.805927 | THCA | B-CPAP | AKT inhibitor VIII |
| MVP_top60_DNA_seed | GSK3B | 0.060486 | LUAD | NCI-H23 | SB216763 | MVP_top60_DNA+RNA_seed | GSK3B | 0.807173 | THCA | FTC-133 | KIN001-042 |
| MVP_top60_DNA_seed | PIK3CA | 0.061225 | LUAD | NCI-H1792 | AZD8186 | MVP_top60_DNA+RNA_seed | PIK3CG | 0.809936 | THCA | TT2609-C02 | PIK-93 |
| MVP_top60_DNA_seed | AURKA | 0.061335 | LUAD | NCI-H2009 | Alisertib | MVP_top60_DNA+RNA_seed | ATM | 0.815754 | THCA | RO82-W-1 | KU-55933 |
| MVP_top60_DNA_seed | IGF1R | 0.062087 | LUAD | NCI-H1793 | BMS-536924 | MVP_top60_DNA+RNA_seed | CHEK2 | 0.822093 | THCA | BHT-101 | AZD7762 |
| MVP_top60_DNA_seed | IGF1R | 0.062362 | LUAD | NCI-H1838 | BMS-536924 | MVP_top60_DNA+RNA_seed | BRAF | 0.831109 | THCA | ASH-3 | PLX-4720 |
| MVP_top60_DNA_seed | AKT1 | 0.062621 | LUAD | HCC-78 | GSK2110183B | MVP_top60_DNA+RNA_seed | RET | 0.846469 | THCA | 8305C | Motesanib |

| | | | | | | | | | | | |
|---|---|---|---|---|---|---|---|---|---|---|---|
| MVP_top60_DNA_seed | AURKA | 0.062881 | LUAD | NCI-H2228 | ZM447439 | MVP_top60_DNA+RNA_seed | MAPK1 | 0.865998 | THCA | CGTH-W-1 | VX-11e |
| MVP_top60_DNA_seed | IGF1R | 0.064146 | LUAD | PC-3_[JPC-3] | GSK1904529A | MVP_top60_DNA+RNA_seed | ATM | 0.876813 | THCA | FTC-133 | KU-55933 |
| MVP_top60_DNA_seed | TERT | 0.067007 | LUAD | EMC-BAC-2 | BIBR-1532 | MVP_top60_DNA+RNA_seed | GSK3B | 0.879238 | THCA | TT2609-C02 | AZD7969 |
| MVP_top60_DNA_seed | AKT1 | 0.067023 | LUAD | RERF-LC-KJ | MK-2206 | MVP_top60_DNA+RNA_seed | HDAC1 | 0.883865 | THCA | FTC-133 | Tubastatin A |
| MVP_top60_DNA_seed | AURKA | 0.067836 | LUAD | Calu-6 | Alisertib | MVP_top60_DNA+RNA_seed | PIK3CG | 0.884857 | THCA | CGTH-W-1 | AS605240 |
| MVP_top60_DNA_seed | PIK3CA | 0.069132 | LUAD | HOP-62 | Buparlisib | MVP_top60_DNA+RNA_seed | GSK3B | 0.885482 | THCA | 8305C | SB216763 |
| MVP_top60_DNA_seed | ATM | 0.069168 | LUAD | NCI-H1623 | KU-55933 | MVP_top60_DNA+RNA_seed | PIK3CG | 0.886255 | THCA | B-CPAP | AS605240 |
| MVP_top60_DNA_seed | MAPK1 | 0.069206 | LUAD | NCI-H522 | VX-11e | MVP_top60_DNA+RNA_seed | MAPK1 | 0.899212 | THCA | ASH-3 | VX-11e |
| MVP_top60_DNA_seed | HSP90AA1 | 0.069487 | LUAD | NCI-H2228 | Luminespib | MVP_top60_DNA+RNA_seed | BRAF | 0.916491 | THCA | TT | PLX-4720 |
| MVP_top60_DNA_seed | IGF1R | 0.069612 | LUAD | NCI-H1651 | Linsitinib | MVP_top60_DNA+RNA_seed | PDPK1 | 0.92546 | THCA | RO82-W-1 | OSU-03012 |
| MVP_top60_DNA_seed | FGFR1 | 0.070071 | LUAD | PC-14 | PD173074 | MVP_top60_DNA+RNA_seed | HDAC1 | 0.928272 | THCA | CGTH-W-1 | Parthenolide |
| MVP_top60_DNA_seed | AURKA | 0.070921 | LUAD | NCI-H1693 | ZM447439 | MVP_top60_DNA+RNA_seed | GSK3B | 0.934547 | THCA | 8305C | CHIR-99021 |
| MVP_top60_DNA_seed | GSK3B | 0.071382 | LUAD | SW1573 | SB216763 | MVP_top60_DNA+RNA_seed | HDAC1 | 0.942386 | THCA | TT | Dacinostat |
| MVP_top60_DNA_seed | PIK3CA | 0.071934 | LUAD | NCI-H358 | Alpelisib | MVP_top60_DNA+RNA_seed | AKT1 | 0.953518 | THCA | CAL-62 | MK-2206 |
| MVP_top60_DNA_seed | BRAF | 0.073536 | LUAD | NCI-H838 | PLX-4720 | MVP_top60_DNA+RNA_seed | AKT2 | 0.953518 | THCA | CAL-62 | MK-2206 |
| MVP_top60_DNA_seed | AKT1 | 0.073686 | LUAD | NCI-H1623 | MK-2206 | MVP_top60_DNA+RNA_seed | AKT1 | 0.984208 | THCA | ML-1 | AKT inhibitor VIII |
| MVP_top60_DNA_seed | PIK3CA | 0.073692 | LUAD | NCI-H1993 | Alpelisib | MVP_top60_DNA+RNA_seed | AKT2 | 0.984208 | THCA | ML-1 | AKT inhibitor VIII |
| MVP_top60_DNA_seed | AKT1 | 0.074104 | LUAD | LC-2-ad | Ipatasertib | MVP_top60_DNA+RNA_seed | AKT3 | 0.984208 | THCA | ML-1 | AKT inhibitor VIII |
| MVP_top60_DNA_seed | HDAC1 | 0.074211 | LUAD | NCI-H2228 | Romidepsin | MVP_top60_DNA+RNA_seed | AKT1 | 1.017811 | THCA | ML-1 | GSK690693 |
| MVP_top60_DNA_seed | MET | 0.074394 | LUAD | NCI-H1781 | Savolitinib | MVP_top60_DNA+RNA_seed | AKT2 | 1.017811 | THCA | ML-1 | GSK690693 |
| MVP_top60_DNA_seed | SRC | 0.075854 | LUAD | NCI-H1838 | Bosutinib | MVP_top60_DNA+RNA_seed | AKT3 | 1.017811 | THCA | ML-1 | GSK690693 |
| MVP_top60_DNA_seed | SRC | 0.076116 | LUAD | COR-L105 | Bosutinib | MVP_top60_DNA+RNA_seed | ATM | 1.023871 | THCA | 8305C | CP466722 |
| MVP_top60_DNA_seed | AURKA | 0.077132 | LUAD | NCI-H322M | ZM447439 | MVP_top60_DNA+RNA_seed | BRAF | 1.027277 | THCA | RO82-W-1 | SB590885 |
| MVP_top60_DNA_seed | EGFR | 0.077419 | LUAD | NCI-H522 | Erlotinib | MVP_top60_DNA+RNA_seed | GSK3B | 1.030288 | THCA | ML-1 | SB216763 |
| MVP_top60_DNA_seed | AKT1 | 0.079151 | LUAD | HCC-827 | AT13148 | MVP_top60_DNA+RNA_seed | HDAC1 | 1.032466 | THCA | KMH-2 | Belinostat |
| MVP_top60_DNA_seed | IGF1R | 0.079817 | LUAD | NCI-H2030 | BMS-754807 | MVP_top60_DNA+RNA_seed | PIK3CG | 1.036995 | THCA | RO82-W-1 | PIK-93 |
| MVP_top60_DNA_seed | FGFR1 | 0.08132 | LUAD | NCI-H650 | AZD4547 | MVP_top60_DNA+RNA_seed | HDAC1 | 1.039646 | THCA | ML-1 | Tubastatin A |
| MVP_top60_DNA_seed | SMARCA4 | 0.081745 | LUAD | NCI-H838 | PFI3 | MVP_top60_DNA+RNA_seed | CHEK2 | 1.046318 | THCA | TT2609-C02 | AZD7762 |
| MVP_top60_DNA_seed | EZH2 | 0.082741 | LUAD | NCI-H1755 | GSK343 | MVP_top60_DNA+RNA_seed | AKT1 | 1.050863 | THCA | 8305C | GSK690693 |
| MVP_top60_DNA_seed | EGFR | 0.08346 | LUAD | NCI-H1944 | Gefitinib | MVP_top60_DNA+RNA_seed | AKT2 | 1.050863 | THCA | 8305C | GSK690693 |
| MVP_top60_DNA_seed | GSK3B | 0.084672 | LUAD | NCI-H1781 | SB216763 | MVP_top60_DNA+RNA_seed | AKT3 | 1.050863 | THCA | 8305C | GSK690693 |
| MVP_top60_DNA_seed | AURKA | 0.085031 | LUAD | NCI-H838 | Alisertib | MVP_top60_DNA+RNA_seed | PDPK1 | 1.057225 | THCA | FTC-133 | OSU-03012 |
| MVP_top60_DNA_seed | AKT1 | 0.085453 | LUAD | NCI-H2228 | Ipatasertib | MVP_top60_DNA+RNA_seed | GSK3B | 1.059125 | THCA | ML-1 | CHIR-99021 |
| MVP_top60_DNA_seed | MDM2 | 0.085538 | LUAD | NCI-H3122 | Serdemetan | MVP_top60_DNA+RNA_seed | GSK3B | 1.06425 | THCA | FTC-133 | CHIR-99021 |
| MVP_top60_DNA_seed | HDAC1 | 0.085555 | LUAD | NCI-H1734 | PCI-34051 | MVP_top60_DNA+RNA_seed | AKT1 | 1.085726 | THCA | RO82-W-1 | GSK690693 |
| MVP_top60_DNA_seed | MAPK1 | 0.088278 | LUAD | NCI-H2228 | Ulixertinib | MVP_top60_DNA+RNA_seed | AKT2 | 1.085726 | THCA | RO82-W-1 | GSK690693 |
| MVP_top60_DNA_seed | EGFR | 0.088506 | LUAD | RERF-LC-MS | Erlotinib | MVP_top60_DNA+RNA_seed | AKT3 | 1.085726 | THCA | RO82-W-1 | GSK690693 |
| MVP_top60_DNA_seed | AURKA | 0.088896 | LUAD | RERF-LC-KJ | Alisertib | MVP_top60_DNA+RNA_seed | AKT1 | 1.106976 | THCA | CAL-62 | AKT inhibitor VIII |
| MVP_top60_DNA_seed | HSP90AA1 | 0.089848 | LUAD | Calu-3 | Tanespimycin | MVP_top60_DNA+RNA_seed | AKT2 | 1.106976 | THCA | CAL-62 | AKT inhibitor VIII |
| MVP_top60_DNA_seed | EGFR | 0.091029 | LUAD | NCI-H1792 | Afatinib | MVP_top60_DNA+RNA_seed | AKT3 | 1.106976 | THCA | CAL-62 | AKT inhibitor VIII |
| MVP_top60_DNA_seed | ERBB2 | 0.091029 | LUAD | NCI-H1792 | Afatinib | MVP_top60_DNA+RNA_seed | PDPK1 | 1.131757 | THCA | TT | OSU-03012 |

| | | | | | | | | | | | |
|---|---|---|---|---|---|---|---|---|---|---|---|
| MVP_top60_DNA_seed | EGFR | 0.091426 | LUAD | NCI-H1435 | Sapitinib | MVP_top60_DNA+RNA_seed | PTK2 | 1.222335 | THCA | FTC-133 | PF-562271 |
| MVP_top60_DNA_seed | ERBB2 | 0.091426 | LUAD | NCI-H1435 | Sapitinib | MVP_top60_DNA+RNA_seed | GSK3B | 1.240107 | THCA | FTC-133 | AZD7969 |
| MVP_top60_DNA_seed | ERBB3 | 0.091426 | LUAD | NCI-H1435 | Sapitinib | MVP_top60_DNA+RNA_seed | HDAC1 | 1.262085 | THCA | 8305C | Tubastatin A |
| MVP_top60_DNA_seed | SMARCA4 | 0.092584 | LUAD | NCI-H1437 | PFI3 | MVP_top60_DNA+RNA_seed | AKT1 | 1.295044 | THCA | 8305C | AKT inhibitor VIII |
| MVP_top60_DNA_seed | EZH2 | 0.092719 | LUAD | NCI-H1437 | GSK343 | MVP_top60_DNA+RNA_seed | AKT2 | 1.295044 | THCA | 8305C | AKT inhibitor VIII |
| MVP_top60_DNA_seed | BRAF | 0.092735 | LUAD | ABC-1 | Dabrafenib | MVP_top60_DNA+RNA_seed | AKT3 | 1.295044 | THCA | 8305C | AKT inhibitor VIII |
| MVP_top60_DNA_seed | GSK3B | 0.093595 | LUAD | NCI-H1437 | CHIR-99021 | MVP_top60_DNA+RNA_seed | GSK3B | 1.29885 | THCA | 8305C | KIN001-042 |
| MVP_top60_DNA_seed | MAPK1 | 0.094436 | LUAD | NCI-H1623 | VX-11e | MVP_top60_DNA+RNA_seed | BRAF | 1.301277 | THCA | FTC-133 | HG6-64-1 |
| MVP_top60_DNA_seed | EGFR | 0.094703 | LUAD | NCI-H2085 | Osimertinib | MVP_top60_DNA+RNA_seed | PIK3CG | 1.318563 | THCA | FTC-133 | AS605240 |
| MVP_top60_DNA_seed | EGFR | 0.094742 | LUAD | NCI-H1755 | Erlotinib | MVP_top60_DNA+RNA_seed | MAPK1 | 1.33145 | THCA | RO82-W-1 | VX-11e |
| MVP_top60_DNA_seed | AKT1 | 0.095073 | LUAD | NCI-H322M | AZD5363 | MVP_top60_DNA+RNA_seed | CHEK2 | 1.341407 | THCA | TT | AZD7762 |
| MVP_top60_DNA_seed | ATM | 0.095384 | LUAD | NCI-H2009 | KU-55933 | MVP_top60_DNA+RNA_seed | MAPK1 | 1.344383 | THCA | TT | VX-11e |
| MVP_top60_DNA_seed | IGF1R | 0.095522 | LUAD | NCI-H3122 | BMS-754807 | MVP_top60_DNA+RNA_seed | HDAC1 | 1.344893 | THCA | RO82-W-1 | JQ12 |
| MVP_top60_DNA_seed | GSK3B | 0.096356 | LUAD | NCI-H2023 | SB216763 | MVP_top60_DNA+RNA_seed | BRAF | 1.355515 | THCA | TT | PLX-4720 |
| MVP_top60_DNA_seed | PIK3CA | 0.097448 | LUAD | NCI-H1568 | AZD8186 | MVP_top60_DNA+RNA_seed | PIK3CG | 1.371806 | THCA | 8305C | AS605240 |
| MVP_top60_DNA_seed | IGF1R | 0.097899 | LUAD | NCI-H1355 | GSK1904529A | MVP_top60_DNA+RNA_seed | GSK3B | 1.381463 | THCA | RO82-W-1 | KIN001-042 |
| MVP_top60_DNA_seed | EZH2 | 0.098676 | LUAD | NCI-H1651 | GSK343 | MVP_top60_DNA+RNA_seed | PDPK1 | 1.383265 | THCA | FTC-133 | BX-912 |
| MVP_top60_DNA_seed | PIK3CA | 0.098733 | LUAD | NCI-H2009 | AZD8186 | MVP_top60_DNA+RNA_seed | PIK3CG | 1.400985 | THCA | ASH-3 | PIK-93 |
| MVP_top60_DNA_seed | EGFR | 0.099261 | LUAD | ABC-1 | Gefitinib | MVP_top60_DNA+RNA_seed | AKT1 | 1.413432 | THCA | TT | GSK690693 |
| MVP_top60_DNA_seed | FGFR1 | 0.09973 | LUAD | RERF-LC-MS | PD173074 | MVP_top60_DNA+RNA_seed | AKT2 | 1.413432 | THCA | TT | GSK690693 |
| MVP_top60_DNA_seed | KRAS | 0.100027 | LUAD | NCI-H1651 | KRAS (G12C) Inhibitor-12 | MVP_top60_DNA+RNA_seed | AKT3 | 1.413432 | THCA | TT | GSK690693 |
| MVP_top60_DNA_seed | EGFR | 0.100266 | LUAD | NCI-H1651 | Osimertinib | MVP_top60_DNA+RNA_seed | MAPK1 | 1.414238 | THCA | RO82-W-1 | FR-180204 |
| MVP_top60_DNA_seed | AKT1 | 0.101856 | LUAD | H3255 | AZD5363 | MVP_top60_DNA+RNA_seed | MAPK3 | 1.414238 | THCA | RO82-W-1 | FR-180204 |
| MVP_top60_DNA_seed | EGFR | 0.102203 | LUAD | NCI-H2291 | Afatinib | MVP_top60_DNA+RNA_seed | HDAC1 | 1.509265 | THCA | ASH-3 | CAY10603 |
| MVP_top60_DNA_seed | ERBB2 | 0.102203 | LUAD | NCI-H2291 | Afatinib | MVP_top60_DNA+RNA_seed | AKT1 | 1.564793 | THCA | TT2609-C02 | BAY AKT1 |
| MVP_top60_DNA_seed | MAPK1 | 0.102761 | LUAD | NCI-H1975 | VX-11e | MVP_top60_DNA+RNA_seed | AKT2 | 1.564793 | THCA | TT2609-C02 | BAY AKT1 |
| MVP_top60_DNA_seed | AKT1 | 0.103369 | LUAD | VMRC-LCD | AT13148 | MVP_top60_DNA+RNA_seed | AKT3 | 1.564793 | THCA | TT2609-C02 | BAY AKT1 |
| MVP_top60_DNA_seed | PIK3CA | 0.104288 | LUAD | NCI-H2228 | Buparlisib | MVP_top60_DNA+RNA_seed | GSK3B | 1.569962 | THCA | TT | CHIR-99021 |
| MVP_top60_DNA_seed | SMARCA4 | 0.104449 | LUAD | NCI-H1623 | PFI3 | MVP_top60_DNA+RNA_seed | HDAC1 | 1.577116 | THCA | ASH-3 | AR-42 |
| MVP_top60_DNA_seed | AKT1 | 0.10463 | LUAD | NCI-H2291 | MK-2206 | MVP_top60_DNA+RNA_seed | AKT1 | 1.58178 | THCA | ML-1 | BAY AKT1 |
| MVP_top60_DNA_seed | BRAF | 0.104718 | LUAD | NCI-H3122 | Dabrafenib | MVP_top60_DNA+RNA_seed | AKT2 | 1.58178 | THCA | ML-1 | BAY AKT1 |
| MVP_top60_DNA_seed | MAPK1 | 0.10535 | LUAD | HCC-78 | SCH772984 | MVP_top60_DNA+RNA_seed | AKT3 | 1.58178 | THCA | ML-1 | BAY AKT1 |
| MVP_top60_DNA_seed | KRAS | 0.105467 | LUAD | H3255 | KRAS (G12C) Inhibitor-12 | MVP_top60_DNA+RNA_seed | GSK3B | 1.654439 | THCA | TT | SB216763 |
| MVP_top60_DNA_seed | FGFR1 | 0.10613 | LUAD | NCI-H1781 | PD173074 | MVP_top60_DNA+RNA_seed | HDAC1 | 1.69053 | THCA | KMH-2 | AR-42 |
| MVP_top60_DNA_seed | AURKA | 0.107067 | LUAD | NCI-H1437 | Alisertib | MVP_top60_DNA+RNA_seed | HDAC1 | 1.701735 | THCA | TT | Tubastatin A |
| MVP_top60_DNA_seed | HDAC1 | 0.108466 | LUAD | NCI-H358 | Romidepsin | MVP_top60_DNA+RNA_seed | GSK3B | 1.715476 | THCA | TT | CHIR-99021 |
| MVP_top60_DNA_seed | EGFR | 0.10854 | LUAD | HCC-78 | Lapatinib | MVP_top60_DNA+RNA_seed | PIK3CG | 1.731511 | THCA | FTC-133 | PIK-93 |
| MVP_top60_DNA_seed | ERBB2 | 0.10854 | LUAD | HCC-78 | Lapatinib | MVP_top60_DNA+RNA_seed | AKT1 | 1.752216 | THCA | FTC-133 | AKT inhibitor VIII |
| MVP_top60_DNA_seed | TP53 | 0.109622 | LUAD | NCI-H2009 | MIRA-1 | MVP_top60_DNA+RNA_seed | AKT2 | 1.752216 | THCA | FTC-133 | AKT inhibitor VIII |
| MVP_top60_DNA_seed | EGFR | 0.110253 | LUAD | LXF-289 | AZD3759 | MVP_top60_DNA+RNA_seed | AKT3 | 1.752216 | THCA | FTC-133 | AKT inhibitor VIII |
| MVP_top60_DNA_seed | EZH2 | 0.11176 | LUAD | NCI-H1781 | GSK343 | MVP_top60_DNA+RNA_seed | HDAC1 | 1.910143 | THCA | ASH-3 | Belinostat |

| | | | | | | | | | | | |
|---|---|---|---|---|---|---|---|---|---|---|---|
| MVP_top60_DNA_seed | HSP90AA1 | 0.113227 | LUAD | Calu-6 | Luminespib | MVP_top60_DNA+RNA_seed | PIK3CG | 1.958764 | THCA | RO82-W-1 | AS605240 |
| MVP_top60_DNA_seed | FGFR1 | 0.113932 | LUAD | HCC-78 | PD173074 | MVP_top60_DNA+RNA_seed | PIK3CG | 1.96255 | THCA | TT | PIK-93 |
| MVP_top60_DNA_seed | IGF1R | 0.114046 | LUAD | VMRC-LCD | BMS-754807 | MVP_top60_DNA+RNA_seed | PIK3CG | 1.985059 | THCA | TT | AS605240 |
| MVP_top60_DNA_seed | FGFR1 | 0.114183 | LUAD | PC-14 | AZD4547 | MVP_top60_DNA+RNA_seed | AKT1 | 2.049624 | THCA | RO82-W-1 | AKT inhibitor VIII |
| MVP_top60_DNA_seed | AKT1 | 0.114972 | LUAD | NCI-H358 | Ipatasertib | MVP_top60_DNA+RNA_seed | AKT2 | 2.049624 | THCA | RO82-W-1 | AKT inhibitor VIII |
| MVP_top60_DNA_seed | AKT1 | 0.116562 | LUAD | NCI-H1944 | Ipatasertib | MVP_top60_DNA+RNA_seed | AKT3 | 2.049624 | THCA | RO82-W-1 | AKT inhibitor VIII |
| MVP_top60_DNA_seed | IGF1R | 0.117034 | LUAD | RERF-LC-KJ | Linsitinib | MVP_top60_DNA+RNA_seed | ATM | 2.089043 | THCA | TT2609-C02 | CP466722 |
| MVP_top60_DNA_seed | EGFR | 0.118086 | LUAD | NCI-H1781 | AZD3759 | MVP_top60_DNA+RNA_seed | MAPK1 | 2.094196 | THCA | TT | FR-180204 |
| MVP_top60_DNA_seed | AKT1 | 0.118591 | LUAD | EMC-BAC-2 | Uprosertib | MVP_top60_DNA+RNA_seed | MAPK3 | 2.094196 | THCA | TT | FR-180204 |
| MVP_top60_DNA_seed | AKT1 | 0.11881 | LUAD | NCI-H3122 | Ipatasertib | MVP_top60_DNA+RNA_seed | GSK3B | 2.131041 | THCA | TT | KIN001-042 |
| MVP_top60_DNA_seed | EGFR | 0.119348 | LUAD | NCI-H1573 | Lapatinib | MVP_top60_DNA+RNA_seed | PTK2 | 2.198954 | THCA | TT | PF-562271 |
| MVP_top60_DNA_seed | ERBB2 | 0.119348 | LUAD | NCI-H1573 | Lapatinib | MVP_top60_DNA+RNA_seed | HDAC1 | 2.312474 | THCA | 8305C | AR-42 |
| MVP_top60_DNA_seed | EGFR | 0.119846 | LUAD | NCI-H2291 | Osimertinib | MVP_top60_DNA+RNA_seed | ATM | 2.326309 | THCA | RO82-W-1 | CP466722 |
| MVP_top60_DNA_seed | MAPK1 | 0.119902 | LUAD | NCI-H1568 | Ulixertinib | MVP_top60_DNA+RNA_seed | AKT1 | 2.457782 | THCA | TT | AKT inhibitor VIII |
| MVP_top60_DNA_seed | PIK3CA | 0.120852 | LUAD | NCI-H1666 | GNE-317 | MVP_top60_DNA+RNA_seed | AKT2 | 2.457782 | THCA | TT | AKT inhibitor VIII |
| MVP_top60_DNA_seed | IGF1R | 0.121211 | LUAD | NCI-H2023 | BMS-754807 | MVP_top60_DNA+RNA_seed | AKT3 | 2.457782 | THCA | TT | AKT inhibitor VIII |
| MVP_top60_DNA_seed | TP53 | 0.122093 | LUAD | VMRC-LCD | MIRA-1 | MVP_top60_DNA+RNA_seed | ATM | 2.462246 | THCA | FTC-133 | CP466722 |
| MVP_top60_DNA_seed | GSK3B | 0.123615 | LUAD | NCI-H2342 | SB216763 | MVP_top60_DNA+RNA_seed | ATM | 2.506771 | THCA | TT | CP466722 |
| MVP_top60_DNA_seed | SRC | 0.124534 | LUAD | NCI-H1563 | Bosutinib | MVP_top60_DNA+RNA_seed | HDAC1 | 2.509226 | THCA | TT | AR-42 |
| MVP_top60_DNA_seed | EGFR | 0.126314 | LUAD | NCI-H838 | Lapatinib | MVP_top60_DNA+RNA_seed | HDAC1 | 2.532809 | THCA | TT | CAY10603 |
| MVP_top60_DNA_seed | ERBB2 | 0.126314 | LUAD | NCI-H838 | Lapatinib | MVP_top60_DNA+RNA_seed | HDAC1 | 2.784592 | THCA | TT | Belinostat |
| MVP_top60_DNA_seed | AKT1 | 0.126329 | LUAD | NCI-H322M | Uprosertib | MVP_top60_DNA+RNA_seed | HDAC1 | 2.998205 | THCA | 8305C | CAY10603 |
| MVP_top60_DNA_seed | EGFR | 0.126863 | LUAD | NCI-H2085 | Gefitinib | MVP_top60_DNA+RNA_seed | HDAC1 | 3.113187 | THCA | 8305C | Belinostat |
| MVP_top60_DNA_seed | EZH2 | 0.127126 | LUAD | NCI-H2085 | GSK343 | MVP_top60_DNA+RNA_seed | BRAF | -2.904049 | THCA | K5 | Dabrafenib |
| MVP_top60_DNA_seed | GSK3B | 0.127832 | LUAD | NCI-H522 | SB216763 | MVP_top60_DNA+RNA_seed | GSK3B | -2.306073 | THCA | B-CPAP | SB216763 |
| MVP_top60_DNA_seed | TERT | 0.12808 | LUAD | RERF-LC-MS | BIBR-1532 | MVP_top60_DNA+RNA_seed | BRAF | -2.070305 | THCA | BHT-101 | Dabrafenib |
| MVP_top60_DNA_seed | AURKA | 0.128692 | LUAD | NCI-H1623 | ZM447439 | MVP_top60_DNA+RNA_seed | BRAF | -2.008943 | THCA | K5 | PLX-4720 |
| MVP_top60_DNA_seed | IGF1R | 0.129808 | LUAD | NCI-H522 | Linsitinib | MVP_top60_DNA+RNA_seed | MAPK1 | -1.972739 | THCA | KMH-2 | VX-11e |
| MVP_top60_DNA_seed | EGFR | 0.130069 | LUAD | NCI-H1734 | Gefitinib | MVP_top60_DNA+RNA_seed | BRAF | -1.889949 | THCA | HTC-C3 | PLX-4720 |
| MVP_top60_DNA_seed | AKT1 | 0.130914 | LUAD | NCI-H2342 | MK-2206 | MVP_top60_DNA+RNA_seed | MAPK1 | -1.76979 | THCA | KMH-2 | SCH772984 |
| MVP_top60_DNA_seed | IGF1R | 0.132113 | LUAD | NCI-H1781 | Linsitinib | MVP_top60_DNA+RNA_seed | MAPK3 | -1.76979 | THCA | KMH-2 | SCH772984 |
| MVP_top60_DNA_seed | BRAF | 0.133153 | LUAD | SK-LU-1 | PLX-4720 | MVP_top60_DNA+RNA_seed | MAPK1 | -1.757059 | THCA | BHT-101 | VX-11e |
| MVP_top60_DNA_seed | MAPK1 | 0.133268 | LUAD | NCI-H650 | Ulixertinib | MVP_top60_DNA+RNA_seed | MAPK1 | -1.755432 | THCA | HTC-C3 | Ulixertinib |
| MVP_top60_DNA_seed | SMARCA4 | 0.133777 | LUAD | NCI-H1568 | PFI3 | MVP_top60_DNA+RNA_seed | MAPK3 | -1.755432 | THCA | HTC-C3 | Ulixertinib |
| MVP_top60_DNA_seed | MAPK1 | 0.13379 | LUAD | NCI-H1781 | SCH772984 | MVP_top60_DNA+RNA_seed | MAPK1 | -1.730831 | THCA | KMH-2 | Ulixertinib |
| MVP_top60_DNA_seed | ATM | 0.135492 | LUAD | NCI-H1437 | KU-55933 | MVP_top60_DNA+RNA_seed | MAPK3 | -1.730831 | THCA | KMH-2 | Ulixertinib |
| MVP_top60_DNA_seed | IGF1R | 0.136193 | LUAD | NCI-H1944 | BMS-536924 | MVP_top60_DNA+RNA_seed | BRAF | -1.699237 | THCA | K5 | SB590885 |
| MVP_top60_DNA_seed | AKT1 | 0.136211 | LUAD | NCI-H2030 | AZD5363 | MVP_top60_DNA+RNA_seed | HDAC1 | -1.694317 | THCA | K5 | Romidepsin |
| MVP_top60_DNA_seed | BRAF | 0.137098 | LUAD | NCI-H2030 | Dabrafenib | MVP_top60_DNA+RNA_seed | MAPK1 | -1.683262 | THCA | CAL-62 | Ulixertinib |
| MVP_top60_DNA_seed | EGFR | 0.137336 | LUAD | NCI-H1650 | Afatinib | MVP_top60_DNA+RNA_seed | MAPK3 | -1.683262 | THCA | CAL-62 | Ulixertinib |
| MVP_top60_DNA_seed | ERBB2 | 0.137336 | LUAD | NCI-H1650 | Afatinib | MVP_top60_DNA+RNA_seed | MAPK1 | -1.556041 | THCA | BHT-101 | SCH772984 |

| | | | | | | | | | | | |
|---|---|---|---|---|---|---|---|---|---|---|---|
| MVP_top60_DNA_seed | MAPK1 | 0.137373 | LUAD | NCI-H650 | Ulixertinib | MVP_top60_DNA+RNA_seed | MAPK3 | -1.556041 | THCA | BHT-101 | SCH772984 |
| MVP_top60_DNA_seed | AKT1 | 0.137756 | LUAD | NCI-H2342 | AT13148 | MVP_top60_DNA+RNA_seed | BRAF | -1.548644 | THCA | HTC-C3 | Dabrafenib |
| MVP_top60_DNA_seed | IGF1R | 0.138291 | LUAD | NCI-H650 | Linsitinib | MVP_top60_DNA+RNA_seed | MAPK1 | -1.431794 | THCA | IHH-4 | Ulixertinib |
| MVP_top60_DNA_seed | BRAF | 0.138661 | LUAD | NCI-H23 | SB590885 | MVP_top60_DNA+RNA_seed | MAPK3 | -1.431794 | THCA | IHH-4 | Ulixertinib |
| MVP_top60_DNA_seed | PIK3CA | 0.141414 | LUAD | PC-14 | AZD8186 | MVP_top60_DNA+RNA_seed | MAPK1 | -1.365373 | THCA | HTC-C3 | Ulixertinib |
| MVP_top60_DNA_seed | MAPK1 | 0.142806 | LUAD | SK-LU-1 | VX-11e | MVP_top60_DNA+RNA_seed | MAPK3 | -1.365373 | THCA | HTC-C3 | Ulixertinib |
| MVP_top60_DNA_seed | PIK3CA | 0.143469 | LUAD | PC-14 | Alpelisib | MVP_top60_DNA+RNA_seed | BRAF | -1.363286 | THCA | BHT-101 | PLX-4720 |
| MVP_top60_DNA_seed | AKT1 | 0.143472 | LUAD | NCI-H2342 | Uprosertib | MVP_top60_DNA+RNA_seed | MAPK1 | -1.343518 | THCA | BHT-101 | Ulixertinib |
| MVP_top60_DNA_seed | MAPK1 | 0.143573 | LUAD | HOP-62 | Ulixertinib | MVP_top60_DNA+RNA_seed | MAPK3 | -1.343518 | THCA | BHT-101 | Ulixertinib |
| MVP_top60_DNA_seed | EGFR | 0.144492 | LUAD | HCC-44 | Sapitinib | MVP_top60_DNA+RNA_seed | MAPK1 | -1.338722 | THCA | IHH-4 | VX-11e |
| MVP_top60_DNA_seed | ERBB2 | 0.144492 | LUAD | HCC-44 | Sapitinib | MVP_top60_DNA+RNA_seed | BRAF | -1.326271 | THCA | 8305C | Dabrafenib |
| MVP_top60_DNA_seed | ERBB3 | 0.144492 | LUAD | HCC-44 | Sapitinib | MVP_top60_DNA+RNA_seed | BRAF | -1.308531 | THCA | IHH-4 | Dabrafenib |
| MVP_top60_DNA_seed | MET | 0.14543 | LUAD | NCI-H1568 | Savolitinib | MVP_top60_DNA+RNA_seed | MAPK1 | -1.278527 | THCA | IHH-4 | SCH772984 |
| MVP_top60_DNA_seed | CREBBP | 0.147297 | LUAD | NCI-H2087 | SGC-CBP30 | MVP_top60_DNA+RNA_seed | MAPK3 | -1.278527 | THCA | IHH-4 | SCH772984 |
| MVP_top60_DNA_seed | AKT1 | 0.147646 | LUAD | NCI-H1666 | Afuresertib | MVP_top60_DNA+RNA_seed | HDAC1 | -1.250755 | THCA | IHH-4 | Romidepsin |
| MVP_top60_DNA_seed | EGFR | 0.148304 | LUAD | NCI-H2023 | Afatinib | MVP_top60_DNA+RNA_seed | BRAF | -1.24812 | THCA | IHH-4 | PLX-4720 |
| MVP_top60_DNA_seed | ERBB2 | 0.148304 | LUAD | NCI-H2023 | Afatinib | MVP_top60_DNA+RNA_seed | AKT1 | -1.24793 | THCA | KMH-2 | AT13148 |
| MVP_top60_DNA_seed | FGFR1 | 0.15023 | LUAD | HCC-44 | AZD4547 | MVP_top60_DNA+RNA_seed | MAPK1 | -1.238702 | THCA | BHT-101 | Ulixertinib |
| MVP_top60_DNA_seed | AURKA | 0.150242 | LUAD | ABC-1 | Alisertib | MVP_top60_DNA+RNA_seed | MAPK3 | -1.238702 | THCA | BHT-101 | Ulixertinib |
| MVP_top60_DNA_seed | IGF1R | 0.150999 | LUAD | NCI-H23 | Linsitinib | MVP_top60_DNA+RNA_seed | BRAF | -1.182661 | THCA | B-CPAP | Dabrafenib |
| MVP_top60_DNA_seed | BRAF | 0.151019 | LUAD | NCI-H1781 | PLX-4720 | MVP_top60_DNA+RNA_seed | RET | -1.146579 | THCA | B-CPAP | Motesanib |
| MVP_top60_DNA_seed | HDAC1 | 0.151036 | LUAD | NCI-H358 | PCI-34051 | MVP_top60_DNA+RNA_seed | MAPK1 | -1.131306 | THCA | K5 | Ulixertinib |
| MVP_top60_DNA_seed | GSK3B | 0.153766 | LUAD | HOP-62 | SB216763 | MVP_top60_DNA+RNA_seed | MAPK3 | -1.131306 | THCA | K5 | Ulixertinib |
| MVP_top60_DNA_seed | AURKA | 0.154787 | LUAD | H3255 | ZM447439 | MVP_top60_DNA+RNA_seed | MAPK1 | -1.114038 | THCA | HTC-C3 | VX-11e |
| MVP_top60_DNA_seed | MDM2 | 0.15482 | LUAD | HCC-827 | Serdemetan | MVP_top60_DNA+RNA_seed | AKT1 | -1.105186 | THCA | FTC-133 | AZD5363 |
| MVP_top60_DNA_seed | MAPK1 | 0.154947 | LUAD | NCI-H1355 | Ulixertinib | MVP_top60_DNA+RNA_seed | AKT2 | -1.105186 | THCA | FTC-133 | AZD5363 |
| MVP_top60_DNA_seed | EGFR | 0.155015 | LUAD | NCI-H522 | Osimertinib | MVP_top60_DNA+RNA_seed | AKT3 | -1.105186 | THCA | FTC-133 | AZD5363 |
| MVP_top60_DNA_seed | MDM2 | 0.15502 | LUAD | NCI-H1623 | Serdemetan | MVP_top60_DNA+RNA_seed | CHEK2 | -1.092126 | THCA | CAL-62 | AZD7762 |
| MVP_top60_DNA_seed | MET | 0.156031 | LUAD | NCI-H1693 | Savolitinib | MVP_top60_DNA+RNA_seed | GSK3B | -1.089368 | THCA | KMH-2 | SB216763 |
| MVP_top60_DNA_seed | IGF1R | 0.158773 | LUAD | NCI-H23 | NVP-ADW742 | MVP_top60_DNA+RNA_seed | BRAF | -1.081835 | THCA | ML-1 | PLX-4720 |
| MVP_top60_DNA_seed | FGFR1 | 0.160446 | LUAD | NCI-H2030 | AZD4547 | MVP_top60_DNA+RNA_seed | GSK3B | -1.078602 | THCA | KMH-2 | CHIR-99021 |
| MVP_top60_DNA_seed | HDAC1 | 0.160706 | LUAD | HCC-44 | Entinostat | MVP_top60_DNA+RNA_seed | HDAC1 | -1.070706 | THCA | KMH-2 | PCI-34051 |
| MVP_top60_DNA_seed | AKT1 | 0.161848 | LUAD | ABC-1 | AT13148 | MVP_top60_DNA+RNA_seed | PIK3CG | -1.048896 | THCA | BHT-101 | CZC24832 |
| MVP_top60_DNA_seed | EGFR | 0.163014 | LUAD | NCI-H2009 | Erlotinib | MVP_top60_DNA+RNA_seed | HDAC1 | -1.048756 | THCA | 8505C | PCI-34051 |
| MVP_top60_DNA_seed | PIK3CA | 0.163809 | LUAD | NCI-H1573 | Buparlisib | MVP_top60_DNA+RNA_seed | MAPK1 | -1.038659 | THCA | K5 | SCH772984 |
| MVP_top60_DNA_seed | FGFR1 | 0.164048 | LUAD | NCI-H1435 | AZD4547 | MVP_top60_DNA+RNA_seed | MAPK3 | -1.038659 | THCA | K5 | SCH772984 |
| MVP_top60_DNA_seed | TERT | 0.164093 | LUAD | NCI-H1437 | BIBR-1532 | MVP_top60_DNA+RNA_seed | AKT1 | -1.013336 | THCA | FTC-133 | Afuresertib |
| MVP_top60_DNA_seed | SMARCA4 | 0.164566 | LUAD | NCI-H2085 | PFI3 | MVP_top60_DNA+RNA_seed | AKT2 | -1.013336 | THCA | FTC-133 | Afuresertib |
| MVP_top60_DNA_seed | BRAF | 0.16549 | LUAD | LC-2-ad | SB590885 | MVP_top60_DNA+RNA_seed | AKT3 | -1.013336 | THCA | FTC-133 | Afuresertib |
| MVP_top60_DNA_seed | SRC | 0.165604 | LUAD | EMC-BAC-2 | Bosutinib | MVP_top60_DNA+RNA_seed | AKT1 | -1.004826 | THCA | K5 | AT13148 |
| MVP_top60_DNA_seed | ATM | 0.166237 | LUAD | RERF-LC-MS | KU-55933 | MVP_top60_DNA+RNA_seed | AKT1 | -0.953548 | THCA | K5 | Afuresertib |

| | | | | | | | | | | | |
|---|---|---|---|---|---|---|---|---|---|---|---|
| MVP_top60_DNA_seed | BRAF | 0.167775 | LUAD | RERF-LC-MS | SB590885 | MVP_top60_DNA+RNA_seed | AKT2 | -0.953548 | THCA | K5 | Afuresertib |
| MVP_top60_DNA_seed | EGFR | 0.168259 | LUAD | NCI-H2342 | Afatinib | MVP_top60_DNA+RNA_seed | AKT3 | -0.953548 | THCA | K5 | Afuresertib |
| MVP_top60_DNA_seed | ERBB2 | 0.168259 | LUAD | NCI-H2342 | Afatinib | MVP_top60_DNA+RNA_seed | KRAS | -0.951678 | THCA | IHH-4 | (RAS (G12C) Inhibitor-1 |
| MVP_top60_DNA_seed | HDAC1 | 0.169077 | LUAD | NCI-H1623 | Romidepsin | MVP_top60_DNA+RNA_seed | KRAS | -0.948296 | THCA | ASH-3 | (RAS (G12C) Inhibitor-1 |
| MVP_top60_DNA_seed | IGF1R | 0.169792 | LUAD | LC-2-ad | Linsitinib | MVP_top60_DNA+RNA_seed | GSK3B | -0.862785 | THCA | K5 | SB216763 |
| MVP_top60_DNA_seed | AKT1 | 0.169824 | LUAD | NCI-H1563 | Uprosertib | MVP_top60_DNA+RNA_seed | HDAC1 | -0.8621 | THCA | BHT-101 | PCI-34051 |
| MVP_top60_DNA_seed | EGFR | 0.172427 | LUAD | NCI-H1734 | Osimertinib | MVP_top60_DNA+RNA_seed | BRAF | -0.841682 | THCA | KMH-2 | SB590885 |
| MVP_top60_DNA_seed | AKT1 | 0.172556 | LUAD | NCI-H2009 | Ipatasertib | MVP_top60_DNA+RNA_seed | BRAF | -0.811101 | THCA | BHT-101 | SB590885 |
| MVP_top60_DNA_seed | IGF1R | 0.17296 | LUAD | NCI-H1838 | Linsitinib | MVP_top60_DNA+RNA_seed | KRAS | -0.791263 | THCA | BHT-101 | (RAS (G12C) Inhibitor-1 |
| MVP_top60_DNA_seed | IGF1R | 0.173024 | LUAD | NCI-H2342 | GSK1904529A | MVP_top60_DNA+RNA_seed | HDAC1 | -0.787006 | THCA | KMH-2 | Entinostat |
| MVP_top60_DNA_seed | EGFR | 0.173347 | LUAD | NCI-H1563 | AZD3759 | MVP_top60_DNA+RNA_seed | BRAF | -0.779147 | THCA | 8305C | PLX-4720 |
| MVP_top60_DNA_seed | MDM2 | 0.174225 | LUAD | NCI-H23 | Nutlin-3a (-) | MVP_top60_DNA+RNA_seed | HDAC1 | -0.744797 | THCA | 8305C | Romidepsin |
| MVP_top60_DNA_seed | AKT1 | 0.17425 | LUAD | NCI-H2291 | Ipatasertib | MVP_top60_DNA+RNA_seed | BRAF | -0.729044 | THCA | B-CPAP | PLX-4720 |
| MVP_top60_DNA_seed | MET | 0.174437 | LUAD | LC-2-ad | Savolitinib | MVP_top60_DNA+RNA_seed | BRAF | -0.727578 | THCA | KMH-2 | Dabrafenib |
| MVP_top60_DNA_seed | AURKA | 0.175629 | LUAD | PC-14 | Tozasertib | MVP_top60_DNA+RNA_seed | PIK3CG | -0.727389 | THCA | BHT-101 | Buparlisib |
| MVP_top60_DNA_seed | FGFR1 | 0.175637 | LUAD | NCI-H1792 | AZD4547 | MVP_top60_DNA+RNA_seed | AKT1 | -0.727366 | THCA | K5 | AZD5363 |
| MVP_top60_DNA_seed | HSP90AA1 | 0.175698 | LUAD | NCI-H1734 | CCT-018159 | MVP_top60_DNA+RNA_seed | AKT2 | -0.727366 | THCA | K5 | AZD5363 |
| MVP_top60_DNA_seed | IGF1R | 0.176353 | LUAD | NCI-H358 | NVP-ADW742 | MVP_top60_DNA+RNA_seed | AKT3 | -0.727366 | THCA | K5 | AZD5363 |
| MVP_top60_DNA_seed | KRAS | 0.179106 | LUAD | HOP-62 | (RAS (G12C) Inhibitor-1: | MVP_top60_DNA+RNA_seed | BRAF | -0.724427 | THCA | KMH-2 | PLX-4720 |
| MVP_top60_DNA_seed | AKT1 | 0.181764 | LUAD | NCI-H1666 | Uprosertib | MVP_top60_DNA+RNA_seed | MAPK1 | -0.715407 | THCA | IHH-4 | Ulixertinib |
| MVP_top60_DNA_seed | FGFR1 | 0.182429 | LUAD | NCI-H1993 | PD173074 | MVP_top60_DNA+RNA_seed | MAPK3 | -0.715407 | THCA | IHH-4 | Ulixertinib |
| MVP_top60_DNA_seed | FGFR1 | 0.182714 | LUAD | HCC-827 | AZD4547 | MVP_top60_DNA+RNA_seed | BRAF | -0.706696 | THCA | 8505C | Dabrafenib |
| MVP_top60_DNA_seed | FGFR1 | 0.183337 | LUAD | NCI-H358 | PD173074 | MVP_top60_DNA+RNA_seed | PIK3CG | -0.702152 | THCA | FTC-133 | Buparlisib |
| MVP_top60_DNA_seed | AKT1 | 0.18356 | LUAD | NCI-H1437 | Afuresertib | MVP_top60_DNA+RNA_seed | AKT1 | -0.700062 | THCA | K5 | GSK2110183B |
| MVP_top60_DNA_seed | MDM2 | 0.184045 | LUAD | NCI-H838 | Nutlin-3a (-) | MVP_top60_DNA+RNA_seed | AKT2 | -0.700062 | THCA | K5 | GSK2110183B |
| MVP_top60_DNA_seed | TP53 | 0.184186 | LUAD | NCI-H838 | MIRA-1 | MVP_top60_DNA+RNA_seed | AKT3 | -0.700062 | THCA | K5 | GSK2110183B |
| MVP_top60_DNA_seed | BRAF | 0.186343 | LUAD | NCI-H838 | Dabrafenib | MVP_top60_DNA+RNA_seed | PIK3CG | -0.685384 | THCA | IHH-4 | Buparlisib |
| MVP_top60_DNA_seed | MAPK1 | 0.186742 | LUAD | NCI-H1975 | Ulixertinib | MVP_top60_DNA+RNA_seed | PIK3CG | -0.678075 | THCA | K5 | Buparlisib |
| MVP_top60_DNA_seed | MAPK1 | 0.187084 | LUAD | NCI-H2228 | SCH772984 | MVP_top60_DNA+RNA_seed | ATM | -0.677234 | THCA | ASH-3 | KU-55933 |
| MVP_top60_DNA_seed | EGFR | 0.187921 | LUAD | NCI-H1755 | Osimertinib | MVP_top60_DNA+RNA_seed | MAPK1 | -0.673281 | THCA | HTC-C3 | SCH772984 |
| MVP_top60_DNA_seed | IGF1R | 0.18893 | LUAD | NCI-H1355 | BMS-754807 | MVP_top60_DNA+RNA_seed | MAPK3 | -0.673281 | THCA | HTC-C3 | SCH772984 |
| MVP_top60_DNA_seed | EGFR | 0.18905 | LUAD | NCI-H23 | Gefitinib | MVP_top60_DNA+RNA_seed | MAPK1 | -0.665421 | THCA | K5 | Ulixertinib |
| MVP_top60_DNA_seed | MAPK1 | 0.190508 | LUAD | NCI-H2085 | VX-11e | MVP_top60_DNA+RNA_seed | MAPK3 | -0.665421 | THCA | K5 | Ulixertinib |
| MVP_top60_DNA_seed | ATM | 0.191098 | LUAD | LC-2-ad | KU-55933 | MVP_top60_DNA+RNA_seed | AKT1 | -0.634041 | THCA | ASH-3 | AT13148 |
| MVP_top60_DNA_seed | EZH2 | 0.191453 | LUAD | EMC-BAC-1 | GSK343 | MVP_top60_DNA+RNA_seed | AKT1 | -0.626325 | THCA | BHT-101 | AT13148 |
| MVP_top60_DNA_seed | BRAF | 0.192791 | LUAD | NCI-H1993 | SB590885 | MVP_top60_DNA+RNA_seed | AKT1 | -0.614942 | THCA | FTC-133 | Ipatasertib |
| MVP_top60_DNA_seed | TP53 | 0.194821 | LUAD | NCI-H1648 | MIRA-1 | MVP_top60_DNA+RNA_seed | AKT3 | -0.614942 | THCA | FTC-133 | Ipatasertib |
| MVP_top60_DNA_seed | HDAC1 | 0.195672 | LUAD | LXF-289 | Entinostat | MVP_top60_DNA+RNA_seed | MAPK1 | -0.600139 | THCA | TT | Ulixertinib |
| MVP_top60_DNA_seed | MAPK1 | 0.195718 | LUAD | HCC-827 | VX-11e | MVP_top60_DNA+RNA_seed | MAPK3 | -0.600139 | THCA | TT | Ulixertinib |
| MVP_top60_DNA_seed | FGFR1 | 0.198604 | LUAD | NCI-H2030 | PD173074 | MVP_top60_DNA+RNA_seed | AKT1 | -0.586595 | THCA | K5 | Ipatasertib |
| MVP_top60_DNA_seed | CREBBP | 0.198754 | LUAD | VMRC-LCD | SGC-CBP30 | MVP_top60_DNA+RNA_seed | AKT3 | -0.586595 | THCA | K5 | Ipatasertib |

| | | | | | | | | | | | |
|---|---|---|---|---|---|---|---|---|---|---|---|
| MVP_top60_DNA_seed | EGFR | 0.198991 | LUAD | NCI-H1734 | Lapatinib | MVP_top60_DNA+RNA_seed | AKT1 | -0.584098 | THCA | K5 | Uprosertib |
| MVP_top60_DNA_seed | ERBB2 | 0.198991 | LUAD | NCI-H1734 | Lapatinib | MVP_top60_DNA+RNA_seed | AKT2 | -0.584098 | THCA | K5 | Uprosertib |
| MVP_top60_DNA_seed | AKT1 | 0.199181 | LUAD | NCI-H1792 | Uprosertib | MVP_top60_DNA+RNA_seed | AKT3 | -0.584098 | THCA | K5 | Uprosertib |
| MVP_top60_DNA_seed | HSP90AA1 | 0.199386 | LUAD | NCI-H1648 | Luminespib | MVP_top60_DNA+RNA_seed | HDAC1 | -0.57334 | THCA | ASH-3 | Romidepsin |
| MVP_top60_DNA_seed | AKT1 | 0.199422 | LUAD | NCI-H322M | Ipatasertib | MVP_top60_DNA+RNA_seed | AKT1 | -0.562402 | THCA | IHH-4 | Uprosertib |
| MVP_top60_DNA_seed | TERT | 0.19946 | LUAD | HCC-827 | BIBR-1532 | MVP_top60_DNA+RNA_seed | AKT2 | -0.562402 | THCA | IHH-4 | Uprosertib |
| MVP_top60_DNA_seed | AKT1 | 0.199876 | LUAD | NCI-H1435 | Ipatasertib | MVP_top60_DNA+RNA_seed | AKT3 | -0.562402 | THCA | IHH-4 | Uprosertib |
| MVP_top60_DNA_seed | EGFR | 0.199961 | LUAD | NCI-H1651 | Sapitinib | MVP_top60_DNA+RNA_seed | MAPK1 | -0.549286 | THCA | 8505C | SCH772984 |
| MVP_top60_DNA_seed | ERBB2 | 0.199961 | LUAD | NCI-H1651 | Sapitinib | MVP_top60_DNA+RNA_seed | MAPK3 | -0.549286 | THCA | 8505C | SCH772984 |
| MVP_top60_DNA_seed | ERBB3 | 0.199961 | LUAD | NCI-H1651 | Sapitinib | MVP_top60_DNA+RNA_seed | BRAF | -0.507576 | THCA | ML-1 | Dabrafenib |
| MVP_top60_DNA_seed | HDAC1 | 0.200021 | LUAD | NCI-H2087 | Romidepsin | MVP_top60_DNA+RNA_seed | PIK3CG | -0.49927 | THCA | B-CPAP | Buparlisib |
| MVP_top60_DNA_seed | AKT1 | 0.200071 | LUAD | NCI-H1792 | MK-2206 | MVP_top60_DNA+RNA_seed | MAPK1 | -0.491213 | THCA | TT2609-C02 | SCH772984 |
| MVP_top60_DNA_seed | AKT1 | 0.200271 | LUAD | NCI-H1792 | GSK2110183B | MVP_top60_DNA+RNA_seed | MAPK3 | -0.491213 | THCA | TT2609-C02 | SCH772984 |
| MVP_top60_DNA_seed | AKT1 | 0.202565 | LUAD | NCI-H2228 | Uprosertib | MVP_top60_DNA+RNA_seed | AKT1 | -0.488621 | THCA | ASH-3 | GSK2110183B |
| MVP_top60_DNA_seed | BRAF | 0.202867 | LUAD | NCI-H2023 | Dabrafenib | MVP_top60_DNA+RNA_seed | AKT2 | -0.488621 | THCA | ASH-3 | GSK2110183B |
| MVP_top60_DNA_seed | PIK3CA | 0.203638 | LUAD | Calu-6 | Buparlisib | MVP_top60_DNA+RNA_seed | AKT3 | -0.488621 | THCA | ASH-3 | GSK2110183B |
| MVP_top60_DNA_seed | IGF1R | 0.203684 | LUAD | ABC-1 | NVP-ADW742 | MVP_top60_DNA+RNA_seed | MAPK1 | -0.476449 | THCA | ASH-3 | SCH772984 |
| MVP_top60_DNA_seed | SRC | 0.203735 | LUAD | NCI-H1944 | Bosutinib | MVP_top60_DNA+RNA_seed | MAPK3 | -0.476449 | THCA | ASH-3 | SCH772984 |
| MVP_top60_DNA_seed | MAPK1 | 0.203867 | LUAD | NCI-H650 | VX-11e | MVP_top60_DNA+RNA_seed | HDAC1 | -0.472437 | THCA | FTC-133 | Romidepsin |
| MVP_top60_DNA_seed | KRAS | 0.20407 | LUAD | ABC-1 | KRAS (G12C) Inhibitor-12 | MVP_top60_DNA+RNA_seed | GSK3B | -0.463637 | THCA | BHT-101 | SB216763 |
| MVP_top60_DNA_seed | MAPK1 | 0.205317 | LUAD | LC-2-ad | Ulixertinib | MVP_top60_DNA+RNA_seed | ATM | -0.461576 | THCA | IHH-4 | KU-55933 |
| MVP_top60_DNA_seed | MDM2 | 0.20532 | LUAD | NCI-H2291 | Serdemetan | MVP_top60_DNA+RNA_seed | AKT1 | -0.461444 | THCA | KMH-2 | AZD5363 |
| MVP_top60_DNA_seed | HSP90AA1 | 0.205409 | LUAD | NCI-H1651 | Luminespib | MVP_top60_DNA+RNA_seed | AKT2 | -0.461444 | THCA | KMH-2 | AZD5363 |
| MVP_top60_DNA_seed | EZH2 | 0.205414 | LUAD | NCI-H2023 | GSK343 | MVP_top60_DNA+RNA_seed | AKT3 | -0.461444 | THCA | KMH-2 | AZD5363 |
| MVP_top60_DNA_seed | BRAF | 0.206037 | LUAD | NCI-H1573 | PLX-4720 | MVP_top60_DNA+RNA_seed | PIK3CG | -0.441235 | THCA | ASH-3 | Buparlisib |
| MVP_top60_DNA_seed | AKT1 | 0.206274 | LUAD | NCI-H1651 | Uprosertib | MVP_top60_DNA+RNA_seed | AKT1 | -0.422296 | THCA | FTC-133 | Uprosertib |
| MVP_top60_DNA_seed | PIK3CA | 0.206623 | LUAD | NCI-H2342 | AZD8186 | MVP_top60_DNA+RNA_seed | AKT2 | -0.422296 | THCA | FTC-133 | Uprosertib |
| MVP_top60_DNA_seed | FGFR1 | 0.206944 | LUAD | Calu-6 | AZD4547 | MVP_top60_DNA+RNA_seed | AKT3 | -0.422296 | THCA | FTC-133 | Uprosertib |
| MVP_top60_DNA_seed | BRAF | 0.206968 | LUAD | NCI-H1437 | PLX-4720 | MVP_top60_DNA+RNA_seed | HDAC1 | -0.417698 | THCA | CAL-62 | Romidepsin |
| MVP_top60_DNA_seed | EGFR | 0.208743 | LUAD | LXF-289 | Afatinib | MVP_top60_DNA+RNA_seed | MAPK1 | -0.413029 | THCA | CAL-62 | Ulixertinib |
| MVP_top60_DNA_seed | ERBB2 | 0.208743 | LUAD | LXF-289 | Afatinib | MVP_top60_DNA+RNA_seed | MAPK3 | -0.413029 | THCA | CAL-62 | Ulixertinib |
| MVP_top60_DNA_seed | IGF1R | 0.208878 | LUAD | NCI-H2030 | GSK1904529A | MVP_top60_DNA+RNA_seed | PIK3CG | -0.412696 | THCA | 8305C | Buparlisib |
| MVP_top60_DNA_seed | IGF1R | 0.208888 | LUAD | NCI-H2023 | BMS-536924 | MVP_top60_DNA+RNA_seed | AKT1 | -0.408279 | THCA | FTC-133 | GSK2110183B |
| MVP_top60_DNA_seed | TERT | 0.208927 | LUAD | NCI-H1944 | BIBR-1532 | MVP_top60_DNA+RNA_seed | AKT2 | -0.408279 | THCA | FTC-133 | GSK2110183B |
| MVP_top60_DNA_seed | AKT1 | 0.209856 | LUAD | NCI-H2023 | GSK2110183B | MVP_top60_DNA+RNA_seed | AKT3 | -0.408279 | THCA | FTC-133 | GSK2110183B |
| MVP_top60_DNA_seed | EGFR | 0.210185 | LUAD | NCI-H2023 | Sapitinib | MVP_top60_DNA+RNA_seed | CHEK2 | -0.405347 | THCA | FTC-133 | AZD7762 |
| MVP_top60_DNA_seed | ERBB2 | 0.210185 | LUAD | NCI-H2023 | Sapitinib | MVP_top60_DNA+RNA_seed | AKT1 | -0.385085 | THCA | K5 | MK-2206 |
| MVP_top60_DNA_seed | ERBB3 | 0.210185 | LUAD | NCI-H2023 | Sapitinib | MVP_top60_DNA+RNA_seed | AKT2 | -0.385085 | THCA | K5 | MK-2206 |
| MVP_top60_DNA_seed | SRC | 0.211163 | LUAD | NCI-H650 | Bosutinib | MVP_top60_DNA+RNA_seed | AKT1 | -0.384849 | THCA | IHH-4 | Ipatasertib |
| MVP_top60_DNA_seed | BRAF | 0.211777 | LUAD | HCC-44 | Dabrafenib | MVP_top60_DNA+RNA_seed | AKT3 | -0.384849 | THCA | IHH-4 | Ipatasertib |
| MVP_top60_DNA_seed | EGFR | 0.213746 | LUAD | NCI-H3122 | Afatinib | MVP_top60_DNA+RNA_seed | BRAF | -0.380654 | THCA | B-CPAP | SB590885 |

| | | | | | | | | | | | |
|---|---|---|---|---|---|---|---|---|---|---|---|
| MVP_top60_DNA_seed | ERBB2 | 0.213746 | LUAD | NCI-H3122 | Afatinib | MVP_top60_DNA+RNA_seed | BRAF | -0.377407 | THCA | IHH-4 | SB590885 |
| MVP_top60_DNA_seed | EGFR | 0.215366 | LUAD | NCI-H1355 | Sapitinib | MVP_top60_DNA+RNA_seed | AKT1 | -0.372713 | THCA | BHT-101 | MK-2206 |
| MVP_top60_DNA_seed | ERBB2 | 0.215366 | LUAD | NCI-H1355 | Sapitinib | MVP_top60_DNA+RNA_seed | AKT2 | -0.372713 | THCA | BHT-101 | MK-2206 |
| MVP_top60_DNA_seed | ERBB3 | 0.215366 | LUAD | NCI-H1355 | Sapitinib | MVP_top60_DNA+RNA_seed | KRAS | -0.36196 | THCA | K5 | KRAS (G12C) Inhibitor-1 |
| MVP_top60_DNA_seed | HDAC1 | 0.216866 | LUAD | NCI-H1993 | Entinostat | MVP_top60_DNA+RNA_seed | HDAC1 | -0.358899 | THCA | B-CPAP | Romidepsin |
| MVP_top60_DNA_seed | FGFR1 | 0.218358 | LUAD | EMC-BAC-1 | PD173074 | MVP_top60_DNA+RNA_seed | AKT1 | -0.350262 | THCA | B-CPAP | Afuresertib |
| MVP_top60_DNA_seed | MDM2 | 0.219731 | LUAD | NCI-H1573 | Nutlin-3a (-) | MVP_top60_DNA+RNA_seed | AKT2 | -0.350262 | THCA | B-CPAP | Afuresertib |
| MVP_top60_DNA_seed | SRC | 0.220595 | LUAD | NCI-H2085 | Bosutinib | MVP_top60_DNA+RNA_seed | AKT3 | -0.350262 | THCA | B-CPAP | Afuresertib |
| MVP_top60_DNA_seed | MAPK1 | 0.22067 | LUAD | NCI-H1573 | SCH772984 | MVP_top60_DNA+RNA_seed | PIK3CG | -0.318633 | THCA | 8505C | CZC24832 |
| MVP_top60_DNA_seed | IGF1R | 0.222408 | LUAD | PC-14 | GSK1904529A | MVP_top60_DNA+RNA_seed | PIK3CG | -0.316575 | THCA | CAL-62 | Buparlisib |
| MVP_top60_DNA_seed | EGFR | 0.224605 | LUAD | NCI-H1792 | Sapitinib | MVP_top60_DNA+RNA_seed | HDAC1 | -0.304496 | THCA | K5 | PCI-34051 |
| MVP_top60_DNA_seed | ERBB2 | 0.224605 | LUAD | NCI-H1792 | Sapitinib | MVP_top60_DNA+RNA_seed | CHEK2 | -0.303186 | THCA | HTC-C3 | AZD7762 |
| MVP_top60_DNA_seed | ERBB3 | 0.224605 | LUAD | NCI-H1792 | Sapitinib | MVP_top60_DNA+RNA_seed | MAPK1 | -0.300332 | THCA | ASH-3 | VX-11e |
| MVP_top60_DNA_seed | GSK3B | 0.225405 | LUAD | HCC-44 | SB216763 | MVP_top60_DNA+RNA_seed | CHEK2 | -0.296783 | THCA | KMH-2 | AZD7762 |
| MVP_top60_DNA_seed | MDM2 | 0.227678 | LUAD | HCC-78 | Nutlin-3a (-) | MVP_top60_DNA+RNA_seed | PIK3CG | -0.284821 | THCA | KMH-2 | Buparlisib |
| MVP_top60_DNA_seed | AKT1 | 0.227925 | LUAD | EMC-BAC-1 | MK-2206 | MVP_top60_DNA+RNA_seed | GSK3B | -0.282 | THCA | IHH-4 | SB216763 |
| MVP_top60_DNA_seed | MDM2 | 0.230052 | LUAD | NCI-H2122 | Serdemetan | MVP_top60_DNA+RNA_seed | AKT1 | -0.265183 | THCA | KMH-2 | MK-2206 |
| MVP_top60_DNA_seed | ATM | 0.230823 | LUAD | NCI-H2347 | KU-55933 | MVP_top60_DNA+RNA_seed | AKT2 | -0.265183 | THCA | KMH-2 | MK-2206 |
| MVP_top60_DNA_seed | BRAF | 0.231303 | LUAD | NCI-H522 | SB590885 | MVP_top60_DNA+RNA_seed | AKT1 | -0.259045 | THCA | KMH-2 | Afuresertib |
| MVP_top60_DNA_seed | AKT1 | 0.231715 | LUAD | NCI-H2009 | AZD5363 | MVP_top60_DNA+RNA_seed | AKT2 | -0.259045 | THCA | KMH-2 | Afuresertib |
| MVP_top60_DNA_seed | BRAF | 0.232627 | LUAD | NCI-H2023 | PLX-4720 | MVP_top60_DNA+RNA_seed | AKT3 | -0.259045 | THCA | KMH-2 | Afuresertib |
| MVP_top60_DNA_seed | MET | 0.23309 | LUAD | NCI-H2085 | Savolitinib | MVP_top60_DNA+RNA_seed | AKT1 | -0.254152 | THCA | B-CPAP | AT13148 |
| MVP_top60_DNA_seed | EGFR | 0.233724 | LUAD | NCI-H522 | AZD3759 | MVP_top60_DNA+RNA_seed | MAPK1 | -0.242485 | THCA | CAL-62 | SCH772984 |
| MVP_top60_DNA_seed | AKT1 | 0.236141 | LUAD | NCI-H2023 | AZD5363 | MVP_top60_DNA+RNA_seed | MAPK3 | -0.242485 | THCA | CAL-62 | SCH772984 |
| MVP_top60_DNA_seed | MET | 0.238613 | LUAD | NCI-H322M | Savolitinib | MVP_top60_DNA+RNA_seed | AKT1 | -0.238933 | THCA | B-CPAP | Uprosertib |
| MVP_top60_DNA_seed | EGFR | 0.239435 | LUAD | LXF-289 | Erlotinib | MVP_top60_DNA+RNA_seed | AKT2 | -0.238933 | THCA | B-CPAP | Uprosertib |
| MVP_top60_DNA_seed | AURKA | 0.241089 | LUAD | NCI-H1975 | Alisertib | MVP_top60_DNA+RNA_seed | AKT3 | -0.238933 | THCA | B-CPAP | Uprosertib |
| MVP_top60_DNA_seed | BRAF | 0.241581 | LUAD | NCI-H650 | SB590885 | MVP_top60_DNA+RNA_seed | BRAF | -0.238721 | THCA | CGTH-W-1 | SB590885 |
| MVP_top60_DNA_seed | BRAF | 0.243461 | LUAD | NCI-H1648 | Dabrafenib | MVP_top60_DNA+RNA_seed | HDAC1 | -0.2379 | THCA | FTC-133 | Entinostat |
| MVP_top60_DNA_seed | MDM2 | 0.246866 | LUAD | RERF-LC-KJ | Nutlin-3a (-) | MVP_top60_DNA+RNA_seed | HDAC1 | -0.232194 | THCA | HTC-C3 | PCI-34051 |
| MVP_top60_DNA_seed | FGFR1 | 0.247494 | LUAD | A549 | AZD4547 | MVP_top60_DNA+RNA_seed | CHEK2 | -0.219384 | THCA | FTC-133 | MK-8776 |
| MVP_top60_DNA_seed | AURKA | 0.248705 | LUAD | NCI-H1568 | ZM447439 | MVP_top60_DNA+RNA_seed | HDAC1 | -0.215744 | THCA | IHH-4 | Entinostat |
| MVP_top60_DNA_seed | IGF1R | 0.249786 | LUAD | NCI-H1435 | GSK1904529A | MVP_top60_DNA+RNA_seed | MAPK1 | -0.203577 | THCA | 8505C | Ulixertinib |
| MVP_top60_DNA_seed | ATM | 0.251747 | LUAD | NCI-H23 | KU-55933 | MVP_top60_DNA+RNA_seed | MAPK3 | -0.203577 | THCA | 8505C | Ulixertinib |
| MVP_top60_DNA_seed | MDM2 | 0.251982 | LUAD | NCI-H1648 | Nutlin-3a (-) | MVP_top60_DNA+RNA_seed | MAPK1 | -0.20344 | THCA | ASH-3 | Ulixertinib |
| MVP_top60_DNA_seed | BRAF | 0.25522 | LUAD | NCI-H1734 | PLX-4720 | MVP_top60_DNA+RNA_seed | MAPK3 | -0.20344 | THCA | ASH-3 | Ulixertinib |
| MVP_top60_DNA_seed | AURKA | 0.256369 | LUAD | NCI-H1650 | Alisertib | MVP_top60_DNA+RNA_seed | BRAF | -0.188294 | THCA | CGTH-W-1 | PLX-4720 |
| MVP_top60_DNA_seed | KRAS | 0.257163 | LUAD | RERF-LC-KJ | KRAS (G12C) Inhibitor-1 | MVP_top60_DNA+RNA_seed | PIK3CG | -0.187949 | THCA | KMH-2 | CZC24832 |
| MVP_top60_DNA_seed | IGF1R | 0.257593 | LUAD | NCI-H1623 | BMS-754807 | MVP_top60_DNA+RNA_seed | HDAC1 | -0.181671 | THCA | K5 | Entinostat |
| MVP_top60_DNA_seed | CREBBP | 0.258503 | LUAD | NCI-H1651 | SGC-CBP30 | MVP_top60_DNA+RNA_seed | PIK3CG | -0.175235 | THCA | HTC-C3 | Buparlisib |
| MVP_top60_DNA_seed | EGFR | 0.258655 | LUAD | RERF-LC-MS | Osimertinib | MVP_top60_DNA+RNA_seed | BRAF | -0.168751 | THCA | FTC-133 | Dabrafenib |

| | | | | | | | | | | | |
|---|---|---|---|---|---|---|---|---|---|---|---|
| MVP_top60_DNA_seed | AURKA | 0.258787 | LUAD | NCI-H1944 | ZM447439 | MVP_top60_DNA+RNA_seed | RET | -0.154526 | THCA | KMH-2 | Motesanib |
| MVP_top60_DNA_seed | EGFR | 0.258963 | LUAD | NCI-H441 | Erlotinib | MVP_top60_DNA+RNA_seed | MAPK1 | -0.151983 | THCA | FTC-133 | Ulixertinib |
| MVP_top60_DNA_seed | EZH2 | 0.2595 | LUAD | NCI-H1993 | GSK343 | MVP_top60_DNA+RNA_seed | MAPK3 | -0.151983 | THCA | FTC-133 | Ulixertinib |
| MVP_top60_DNA_seed | BRAF | 0.259661 | LUAD | RERF-LC-KJ | SB590885 | MVP_top60_DNA+RNA_seed | RET | -0.139767 | THCA | IHH-4 | Motesanib |
| MVP_top60_DNA_seed | AKT1 | 0.260002 | LUAD | EMC-BAC-2 | Uprosertib | MVP_top60_DNA+RNA_seed | MAPK1 | -0.132502 | THCA | 8305C | SCH772984 |
| MVP_top60_DNA_seed | BRAF | 0.260024 | LUAD | H3255 | Dabrafenib | MVP_top60_DNA+RNA_seed | MAPK3 | -0.132502 | THCA | 8305C | SCH772984 |
| MVP_top60_DNA_seed | AKT1 | 0.260824 | LUAD | NCI-H23 | AZD5363 | MVP_top60_DNA+RNA_seed | AKT1 | -0.132233 | THCA | CAL-62 | AT13148 |
| MVP_top60_DNA_seed | AKT1 | 0.261546 | LUAD | LC-2-ad | Uprosertib | MVP_top60_DNA+RNA_seed | HDAC1 | -0.130585 | THCA | BHT-101 | Entinostat |
| MVP_top60_DNA_seed | AKT1 | 0.262832 | LUAD | NCI-H2009 | Afuresertib | MVP_top60_DNA+RNA_seed | GSK3B | -0.125859 | THCA | TT2609-C02 | SB216763 |
| MVP_top60_DNA_seed | EZH2 | 0.264622 | LUAD | NCI-H1993 | GSK343 | MVP_top60_DNA+RNA_seed | MAPK1 | -0.123184 | THCA | TT2609-C02 | Ulixertinib |
| MVP_top60_DNA_seed | KRAS | 0.265622 | LUAD | NCI-H1666 | KRAS (G12C) Inhibitor-1[illegible] | MVP_top60_DNA+RNA_seed | MAPK3 | -0.123184 | THCA | TT2609-C02 | Ulixertinib |
| MVP_top60_DNA_seed | AKT1 | 0.265734 | LUAD | NCI-H2291 | Afuresertib | MVP_top60_DNA+RNA_seed | HDAC1 | -0.118586 | THCA | 8505C | Entinostat |
| MVP_top60_DNA_seed | BRAF | 0.26751 | LUAD | NCI-H1993 | Dabrafenib | MVP_top60_DNA+RNA_seed | CHEK2 | -0.095069 | THCA | IHH-4 | AZD7762 |
| MVP_top60_DNA_seed | AKT1 | 0.269061 | LUAD | RERF-LC-KJ | GSK2110183B | MVP_top60_DNA+RNA_seed | MAPK1 | -0.09403 | THCA | ASH-3 | Ulixertinib |
| MVP_top60_DNA_seed | MET | 0.26999 | LUAD | NCI-H2085 | Crizotinib | MVP_top60_DNA+RNA_seed | MAPK3 | -0.09403 | THCA | ASH-3 | Ulixertinib |
| MVP_top60_DNA_seed | FGFR1 | 0.270559 | LUAD | NCI-H1650 | AZD4547 | MVP_top60_DNA+RNA_seed | MAPK1 | -0.089216 | THCA | K5 | VX-11e |
| MVP_top60_DNA_seed | TP53 | 0.271202 | LUAD | NCI-H2085 | MIRA-1 | MVP_top60_DNA+RNA_seed | MAPK1 | -0.073967 | THCA | B-CPAP | Ulixertinib |
| MVP_top60_DNA_seed | HDAC1 | 0.27138 | LUAD | RERF-LC-KJ | Entinostat | MVP_top60_DNA+RNA_seed | MAPK3 | -0.073967 | THCA | B-CPAP | Ulixertinib |
| MVP_top60_DNA_seed | AKT1 | 0.271951 | LUAD | NCI-H1623 | AT13148 | MVP_top60_DNA+RNA_seed | HDAC1 | -0.073074 | THCA | HTC-C3 | Entinostat |
| MVP_top60_DNA_seed | GSK3B | 0.272976 | LUAD | NCI-H838 | SB216763 | MVP_top60_DNA+RNA_seed | AKT1 | -0.070647 | THCA | 8305C | MK-2206 |
| MVP_top60_DNA_seed | TP53 | 0.273078 | LUAD | NCI-H23 | MIRA-1 | MVP_top60_DNA+RNA_seed | AKT2 | -0.070647 | THCA | 8305C | MK-2206 |
| MVP_top60_DNA_seed | FGFR1 | 0.27381 | LUAD | COR-L105 | PD173074 | MVP_top60_DNA+RNA_seed | MAPK1 | -0.065603 | THCA | 8505C | VX-11e |
| MVP_top60_DNA_seed | MET | 0.27453 | LUAD | Calu-3 | Crizotinib | MVP_top60_DNA+RNA_seed | CHEK2 | -0.062307 | THCA | CGTH-W-1 | MK-8776 |
| MVP_top60_DNA_seed | TERT | 0.274731 | LUAD | Calu-6 | BIBR-1532 | MVP_top60_DNA+RNA_seed | AKT1 | -0.059484 | THCA | K5 | Uprosertib |
| MVP_top60_DNA_seed | PIK3CA | 0.274954 | LUAD | SW1573 | AZD8186 | MVP_top60_DNA+RNA_seed | AKT2 | -0.059484 | THCA | K5 | Uprosertib |
| MVP_top60_DNA_seed | AKT1 | 0.27528 | LUAD | NCI-H358 | Afuresertib | MVP_top60_DNA+RNA_seed | AKT3 | -0.059484 | THCA | K5 | Uprosertib |
| MVP_top60_DNA_seed | HDAC1 | 0.275315 | LUAD | NCI-H3122 | PCI-34051 | MVP_top60_DNA+RNA_seed | AKT1 | -0.050801 | THCA | KMH-2 | Ipatasertib |
| MVP_top60_DNA_seed | EGFR | 0.275541 | LUAD | NCI-H2009 | Afatinib | MVP_top60_DNA+RNA_seed | AKT3 | -0.050801 | THCA | KMH-2 | Ipatasertib |
| MVP_top60_DNA_seed | ERBB2 | 0.275541 | LUAD | NCI-H2009 | Afatinib | MVP_top60_DNA+RNA_seed | GSK3B | -0.050169 | THCA | FTC-133 | SB216763 |
| MVP_top60_DNA_seed | BRAF | 0.276048 | LUAD | LXF-289 | SB590885 | MVP_top60_DNA+RNA_seed | AKT1 | -0.035112 | THCA | B-CPAP | AZD5363 |
| MVP_top60_DNA_seed | MDM2 | 0.277218 | LUAD | EMC-BAC-1 | Serdemetan | MVP_top60_DNA+RNA_seed | AKT2 | -0.035112 | THCA | B-CPAP | AZD5363 |
| MVP_top60_DNA_seed | BRAF | 0.278245 | LUAD | NCI-H1651 | Dabrafenib | MVP_top60_DNA+RNA_seed | AKT3 | -0.035112 | THCA | B-CPAP | AZD5363 |
| MVP_top60_DNA_seed | EGFR | 0.278311 | LUAD | HCC-44 | Lapatinib | MVP_top60_DNA+RNA_seed | MAPK1 | -0.020558 | THCA | B-CPAP | SCH772984 |
| MVP_top60_DNA_seed | ERBB2 | 0.278311 | LUAD | HCC-44 | Lapatinib | MVP_top60_DNA+RNA_seed | MAPK3 | -0.020558 | THCA | B-CPAP | SCH772984 |
| MVP_top60_DNA_seed | IGF1R | 0.278947 | LUAD | NCI-H2087 | NVP-ADW742 | MVP_top60_DNA+RNA_seed | AKT1 | -0.012276 | THCA | ASH-3 | Afuresertib |
| MVP_top60_DNA_seed | EGFR | 0.279115 | LUAD | Calu-3 | Erlotinib | MVP_top60_DNA+RNA_seed | AKT2 | -0.012276 | THCA | ASH-3 | Afuresertib |
| MVP_top60_DNA_seed | ATM | 0.280303 | LUAD | NCI-H1693 | KU-55933 | MVP_top60_DNA+RNA_seed | AKT3 | -0.012276 | THCA | ASH-3 | Afuresertib |
| MVP_top60_DNA_seed | BRAF | 0.280881 | LUAD | NCI-H1437 | Dabrafenib | MVP_top60_DNA+RNA_seed | GSK3B | -0.005432 | THCA | HTC-C3 | SB216763 |
| MVP_top60_DNA_seed | MDM2 | 0.283916 | LUAD | NCI-H1573 | Serdemetan | MVP_top60_DNA+RNA_seed | CHEK2 | -0.003886 | THCA | KMH-2 | MK-8776 |
| MVP_top60_DNA_seed | IGF1R | 0.284279 | LUAD | HOP-62 | NVP-ADW742 | MVP_top60_DNA+RNA_seed | HDAC1 | -0.000759 | THCA | 8505C | Romidepsin |
| MVP_top60_DNA_seed | SRC | 0.284795 | LUAD | NCI-H2347 | Bosutinib | MVP_top60_DNA+RNA_seed | AKT1 | 0.006181 | THCA | 8305C | AT13148 |

| | | | | | | | | | | | |
|---|---|---|---|---|---|---|---|---|---|---|---|
| MVP_top60_DNA_seed | ATM | 0.286418 | LUAD | NCI-H1650 | KU-55933 | MVP_top60_DNA+RNA_seed | CHEK2 | 0.012385 | THCA | 8305C | AZD7762 |
| MVP_top60_DNA_seed | EZH2 | 0.286823 | LUAD | EMC-BAC-2 | GSK343 | MVP_top60_DNA+RNA_seed | AKT1 | 0.022588 | THCA | KMH-2 | Uprosertib |
| MVP_top60_DNA_seed | PIK3CA | 0.287379 | LUAD | NCI-H650 | Buparlisib | MVP_top60_DNA+RNA_seed | AKT2 | 0.022588 | THCA | KMH-2 | Uprosertib |
| MVP_top60_DNA_seed | EGFR | 0.288499 | LUAD | NCI-H2228 | Sapitinib | MVP_top60_DNA+RNA_seed | AKT3 | 0.022588 | THCA | KMH-2 | Uprosertib |
| MVP_top60_DNA_seed | ERBB2 | 0.288499 | LUAD | NCI-H2228 | Sapitinib | MVP_top60_DNA+RNA_seed | CHEK2 | 0.022619 | THCA | CGTH-W-1 | AZD7762 |
| MVP_top60_DNA_seed | ERBB3 | 0.288499 | LUAD | NCI-H2228 | Sapitinib | MVP_top60_DNA+RNA_seed | CHEK2 | 0.024054 | THCA | HTC-C3 | MK-8776 |
| MVP_top60_DNA_seed | AKT1 | 0.289237 | LUAD | NCI-H1435 | Uprosertib | MVP_top60_DNA+RNA_seed | MAPK1 | 0.034688 | THCA | B-CPAP | VX-11e |
| MVP_top60_DNA_seed | ATM | 0.290966 | LUAD | NCI-H838 | KU-55933 | MVP_top60_DNA+RNA_seed | CHEK2 | 0.0574 | THCA | IHH-4 | MK-8776 |
| MVP_top60_DNA_seed | IGF1R | 0.291363 | LUAD | NCI-H1568 | GSK1904529A | MVP_top60_DNA+RNA_seed | MAPK1 | 0.058668 | THCA | B-CPAP | Ulixertinib |
| MVP_top60_DNA_seed | BRAF | 0.291914 | LUAD | NCI-H1437 | SB590885 | MVP_top60_DNA+RNA_seed | MAPK3 | 0.058668 | THCA | B-CPAP | Ulixertinib |
| MVP_top60_DNA_seed | PIK3CA | 0.291914 | LUAD | NCI-H1573 | GNE-317 | MVP_top60_DNA+RNA_seed | AKT1 | 0.07147 | THCA | ASH-3 | AZD5363 |
| MVP_top60_DNA_seed | MAPK1 | 0.292022 | LUAD | NCI-H2030 | SCH772984 | MVP_top60_DNA+RNA_seed | AKT2 | 0.07147 | THCA | ASH-3 | AZD5363 |
| MVP_top60_DNA_seed | IGF1R | 0.296374 | LUAD | Calu-6 | NVP-ADW742 | MVP_top60_DNA+RNA_seed | AKT3 | 0.07147 | THCA | ASH-3 | AZD5363 |
| MVP_top60_DNA_seed | AKT1 | 0.299342 | LUAD | NCI-H1650 | Afuresertib | MVP_top60_DNA+RNA_seed | ATM | 0.082146 | THCA | CGTH-W-1 | KU-55933 |
| MVP_top60_DNA_seed | AKT1 | 0.30122 | LUAD | LC-2-ad | Afuresertib | MVP_top60_DNA+RNA_seed | AKT1 | 0.082997 | THCA | IHH-4 | MK-2206 |
| MVP_top60_DNA_seed | AKT1 | 0.303228 | LUAD | NCI-H1666 | MK-2206 | MVP_top60_DNA+RNA_seed | AKT2 | 0.082997 | THCA | IHH-4 | MK-2206 |
| MVP_top60_DNA_seed | EGFR | 0.303406 | LUAD | RERF-LC-KJ | Afatinib | MVP_top60_DNA+RNA_seed | AKT1 | 0.088719 | THCA | IHH-4 | Afuresertib |
| MVP_top60_DNA_seed | ERBB2 | 0.303406 | LUAD | RERF-LC-KJ | Afatinib | MVP_top60_DNA+RNA_seed | AKT2 | 0.088719 | THCA | IHH-4 | Afuresertib |
| MVP_top60_DNA_seed | MDM2 | 0.303465 | LUAD | NCI-H1435 | Nutlin-3a (-) | MVP_top60_DNA+RNA_seed | AKT3 | 0.088719 | THCA | IHH-4 | Afuresertib |
| MVP_top60_DNA_seed | PIK3CA | 0.30589 | LUAD | PC-3_[JPC-3] | Buparlisib | MVP_top60_DNA+RNA_seed | HDAC1 | 0.090957 | THCA | ASH-3 | Entinostat |
| MVP_top60_DNA_seed | EGFR | 0.306368 | LUAD | NCI-H1435 | Lapatinib | MVP_top60_DNA+RNA_seed | AKT1 | 0.096746 | THCA | 8305C | Ipatasertib |
| MVP_top60_DNA_seed | ERBB2 | 0.306368 | LUAD | NCI-H1435 | Lapatinib | MVP_top60_DNA+RNA_seed | AKT3 | 0.096746 | THCA | 8305C | Ipatasertib |
| MVP_top60_DNA_seed | PIK3CA | 0.306425 | LUAD | EMC-BAC-1 | GNE-317 | MVP_top60_DNA+RNA_seed | CHEK2 | 0.099675 | THCA | B-CPAP | AZD7762 |
| MVP_top60_DNA_seed | BRAF | 0.306714 | LUAD | NCI-H1975 | Dabrafenib | MVP_top60_DNA+RNA_seed | ATM | 0.104186 | THCA | HTC-C3 | KU-55933 |
| MVP_top60_DNA_seed | AURKA | 0.306999 | LUAD | HCC-827 | ZM447439 | MVP_top60_DNA+RNA_seed | ATM | 0.11089 | THCA | 8305C | KU-55933 |
| MVP_top60_DNA_seed | AKT1 | 0.307214 | LUAD | NCI-H2030 | Ipatasertib | MVP_top60_DNA+RNA_seed | AKT1 | 0.116142 | THCA | CAL-62 | Ipatasertib |
| MVP_top60_DNA_seed | EGFR | 0.307573 | LUAD | NCI-H1355 | Erlotinib | MVP_top60_DNA+RNA_seed | AKT3 | 0.116142 | THCA | CAL-62 | Ipatasertib |
| MVP_top60_DNA_seed | BRAF | 0.309539 | LUAD | LXF-289 | PLX-4720 | MVP_top60_DNA+RNA_seed | RET | 0.121121 | THCA | CAL-62 | Motesanib |
| MVP_top60_DNA_seed | AKT1 | 0.3096 | LUAD | NCI-H2342 | AZD5363 | MVP_top60_DNA+RNA_seed | HDAC1 | 0.125716 | THCA | B-CPAP | PCI-34051 |
| MVP_top60_DNA_seed | HDAC1 | 0.310015 | LUAD | RERF-LC-KJ | Romidepsin | MVP_top60_DNA+RNA_seed | MAPK1 | 0.134393 | THCA | CAL-62 | VX-11e |
| MVP_top60_DNA_seed | IGF1R | 0.310117 | LUAD | ABC-1 | GSK1904529A | MVP_top60_DNA+RNA_seed | PIK3CG | 0.139202 | THCA | B-CPAP | CZC24832 |
| MVP_top60_DNA_seed | SRC | 0.310274 | LUAD | NCI-H1650 | Bosutinib | MVP_top60_DNA+RNA_seed | MAPK1 | 0.151567 | THCA | 8505C | Ulixertinib |
| MVP_top60_DNA_seed | ATM | 0.310577 | LUAD | NCI-H1993 | KU-55933 | MVP_top60_DNA+RNA_seed | MAPK3 | 0.151567 | THCA | 8505C | Ulixertinib |
| MVP_top60_DNA_seed | EZH2 | 0.312098 | LUAD | NCI-H838 | GSK343 | MVP_top60_DNA+RNA_seed | PIK3CG | 0.154241 | THCA | IHH-4 | CZC24832 |
| MVP_top60_DNA_seed | EGFR | 0.312241 | LUAD | NCI-H441 | Afatinib | MVP_top60_DNA+RNA_seed | GSK3B | 0.157933 | THCA | CGTH-W-1 | SB216763 |
| MVP_top60_DNA_seed | ERBB2 | 0.312241 | LUAD | NCI-H441 | Afatinib | MVP_top60_DNA+RNA_seed | RET | 0.158754 | THCA | ASH-3 | Motesanib |
| MVP_top60_DNA_seed | HDAC1 | 0.312803 | LUAD | NCI-H2030 | Entinostat | MVP_top60_DNA+RNA_seed | KRAS | 0.164964 | THCA | KMH-2 | KRAS (G12C) Inhibitor-1 |
| MVP_top60_DNA_seed | BRAF | 0.312876 | LUAD | NCI-H322M | SB590885 | MVP_top60_DNA+RNA_seed | ATM | 0.16506 | THCA | KMH-2 | KU-55933 |
| MVP_top60_DNA_seed | SMARCA4 | 0.312982 | LUAD | HCC-44 | PFI3 | MVP_top60_DNA+RNA_seed | KRAS | 0.165603 | THCA | CAL-62 | KRAS (G12C) Inhibitor-1 |
| MVP_top60_DNA_seed | EZH2 | 0.314116 | LUAD | A549 | GSK343 | MVP_top60_DNA+RNA_seed | HDAC1 | 0.166609 | THCA | HTC-C3 | Romidepsin |
| MVP_top60_DNA_seed | EGFR | 0.314177 | LUAD | NCI-H1573 | Afatinib | MVP_top60_DNA+RNA_seed | AKT1 | 0.172248 | THCA | ASH-3 | Ipatasertib |

| | | | | | | | | | | | |
|---|---|---|---|---|---|---|---|---|---|---|---|
| MVP_top60_DNA_seed | ERBB2 | 0.314177 | LUAD | NCI-H1573 | Afatinib | MVP_top60_DNA+RNA_seed | AKT3 | 0.172248 | THCA | ASH-3 | Ipatasertib |
| MVP_top60_DNA_seed | GSK3B | 0.314897 | LUAD | VMRC-LCD | SB216763 | MVP_top60_DNA+RNA_seed | AKT1 | 0.182526 | THCA | B-CPAP | MK-2206 |
| MVP_top60_DNA_seed | GSK3B | 0.315584 | LUAD | NCI-H1993 | SB216763 | MVP_top60_DNA+RNA_seed | AKT2 | 0.182526 | THCA | B-CPAP | MK-2206 |
| MVP_top60_DNA_seed | MDM2 | 0.31581 | LUAD | NCI-H1563 | Serdemetan | MVP_top60_DNA+RNA_seed | PIK3CG | 0.190162 | THCA | HTC-C3 | CZC24832 |
| MVP_top60_DNA_seed | MDM2 | 0.315845 | LUAD | NCI-H1755 | Serdemetan | MVP_top60_DNA+RNA_seed | AKT1 | 0.193557 | THCA | BHT-101 | Afuresertib |
| MVP_top60_DNA_seed | SRC | 0.31614 | LUAD | NCI-H1651 | Bosutinib | MVP_top60_DNA+RNA_seed | AKT2 | 0.193557 | THCA | BHT-101 | Afuresertib |
| MVP_top60_DNA_seed | HDAC1 | 0.316454 | LUAD | NCI-H650 | Entinostat | MVP_top60_DNA+RNA_seed | AKT3 | 0.193557 | THCA | BHT-101 | Afuresertib |
| MVP_top60_DNA_seed | AKT1 | 0.317258 | LUAD | LC-2-ad | AT13148 | MVP_top60_DNA+RNA_seed | HDAC1 | 0.197236 | THCA | 8305C | PCI-34051 |
| MVP_top60_DNA_seed | EGFR | 0.317298 | LUAD | NCI-H522 | Lapatinib | MVP_top60_DNA+RNA_seed | AKT1 | 0.197588 | THCA | CAL-62 | Afuresertib |
| MVP_top60_DNA_seed | ERBB2 | 0.317298 | LUAD | NCI-H522 | Lapatinib | MVP_top60_DNA+RNA_seed | AKT2 | 0.197588 | THCA | CAL-62 | Afuresertib |
| MVP_top60_DNA_seed | BRAF | 0.317308 | LUAD | NCI-H2228 | SB590885 | MVP_top60_DNA+RNA_seed | AKT3 | 0.197588 | THCA | CAL-62 | Afuresertib |
| MVP_top60_DNA_seed | BRAF | 0.317629 | LUAD | NCI-H838 | SB590885 | MVP_top60_DNA+RNA_seed | CHEK2 | 0.248206 | THCA | ML-1 | MK-8776 |
| MVP_top60_DNA_seed | AKT1 | 0.318905 | LUAD | EMC-BAC-2 | MK-2206 | MVP_top60_DNA+RNA_seed | RET | 0.250691 | THCA | FTC-133 | Motesanib |
| MVP_top60_DNA_seed | FGFR1 | 0.319649 | LUAD | NCI-H23 | AZD4547 | MVP_top60_DNA+RNA_seed | ATM | 0.251063 | THCA | K5 | KU-55933 |
| MVP_top60_DNA_seed | PIK3CA | 0.319768 | LUAD | ABC-1 | Alpelisib | MVP_top60_DNA+RNA_seed | RET | 0.25247 | THCA | K5 | Motesanib |
| MVP_top60_DNA_seed | IGF1R | 0.320436 | LUAD | NCI-H23 | BMS-536924 | MVP_top60_DNA+RNA_seed | CHEK2 | 0.258476 | THCA | K5 | AZD7762 |
| MVP_top60_DNA_seed | EGFR | 0.320644 | LUAD | NCI-H2291 | Gefitinib | MVP_top60_DNA+RNA_seed | ATM | 0.260663 | THCA | BHT-101 | KU-55933 |
| MVP_top60_DNA_seed | MDM2 | 0.32137 | LUAD | A549 | Serdemetan | MVP_top60_DNA+RNA_seed | BRAF | 0.268676 | THCA | 8505C | PLX-4720 |
| MVP_top60_DNA_seed | HSP90AA1 | 0.321549 | LUAD | NCI-H1651 | Tanespimycin | MVP_top60_DNA+RNA_seed | BRAF | 0.269089 | THCA | CAL-62 | SB590885 |
| MVP_top60_DNA_seed | CREBBP | 0.321754 | LUAD | NCI-H2085 | SGC-CBP30 | MVP_top60_DNA+RNA_seed | GSK3B | 0.271067 | THCA | CAL-62 | SB216763 |
| MVP_top60_DNA_seed | SRC | 0.324323 | LUAD | NCI-H522 | Bosutinib | MVP_top60_DNA+RNA_seed | AKT1 | 0.279299 | THCA | 8305C | Afuresertib |
| MVP_top60_DNA_seed | BRAF | 0.325327 | LUAD | NCI-H1435 | PLX-4720 | MVP_top60_DNA+RNA_seed | AKT2 | 0.279299 | THCA | 8305C | Afuresertib |
| MVP_top60_DNA_seed | EGFR | 0.325384 | LUAD | NCI-H2291 | Sapitinib | MVP_top60_DNA+RNA_seed | AKT3 | 0.279299 | THCA | 8305C | Afuresertib |
| MVP_top60_DNA_seed | ERBB2 | 0.325384 | LUAD | NCI-H2291 | Sapitinib | MVP_top60_DNA+RNA_seed | MAPK1 | 0.296411 | THCA | CGTH-W-1 | SCH772984 |
| MVP_top60_DNA_seed | ERBB3 | 0.325384 | LUAD | NCI-H2291 | Sapitinib | MVP_top60_DNA+RNA_seed | MAPK3 | 0.296411 | THCA | CGTH-W-1 | SCH772984 |
| MVP_top60_DNA_seed | IGF1R | 0.325394 | LUAD | NCI-H1755 | GSK1904529A | MVP_top60_DNA+RNA_seed | MAPK1 | 0.297869 | THCA | 8305C | VX-11e |
| MVP_top60_DNA_seed | BRAF | 0.32556 | LUAD | COR-L105 | Dabrafenib | MVP_top60_DNA+RNA_seed | RET | 0.303606 | THCA | BHT-101 | Motesanib |
| MVP_top60_DNA_seed | AKT1 | 0.32576 | LUAD | NCI-H1792 | AZD5363 | MVP_top60_DNA+RNA_seed | AKT1 | 0.309416 | THCA | BHT-101 | Uprosertib |
| MVP_top60_DNA_seed | AKT1 | 0.326246 | LUAD | LC-2-ad | MK-2206 | MVP_top60_DNA+RNA_seed | AKT2 | 0.309416 | THCA | BHT-101 | Uprosertib |
| MVP_top60_DNA_seed | EGFR | 0.326505 | LUAD | NCI-H1755 | Sapitinib | MVP_top60_DNA+RNA_seed | AKT3 | 0.309416 | THCA | BHT-101 | Uprosertib |
| MVP_top60_DNA_seed | ERBB2 | 0.326505 | LUAD | NCI-H1755 | Sapitinib | MVP_top60_DNA+RNA_seed | PIK3CG | 0.324424 | THCA | ASH-3 | CZC24832 |
| MVP_top60_DNA_seed | ERBB3 | 0.326505 | LUAD | NCI-H1755 | Sapitinib | MVP_top60_DNA+RNA_seed | AKT1 | 0.346773 | THCA | FTC-133 | MK-2206 |
| MVP_top60_DNA_seed | GSK3B | 0.326635 | LUAD | HCC-827 | SB216763 | MVP_top60_DNA+RNA_seed | AKT2 | 0.346773 | THCA | FTC-133 | MK-2206 |
| MVP_top60_DNA_seed | EZH2 | 0.327683 | LUAD | NCI-H1693 | GSK343 | MVP_top60_DNA+RNA_seed | AKT1 | 0.350825 | THCA | BHT-101 | Ipatasertib |
| MVP_top60_DNA_seed | IGF1R | 0.32847 | LUAD | NCI-H522 | GSK1904529A | MVP_top60_DNA+RNA_seed | AKT3 | 0.350825 | THCA | BHT-101 | Ipatasertib |
| MVP_top60_DNA_seed | BRAF | 0.330674 | LUAD | EMC-BAC-2 | SB590885 | MVP_top60_DNA+RNA_seed | MAPK1 | 0.357318 | THCA | CGTH-W-1 | VX-11e |
| MVP_top60_DNA_seed | BRAF | 0.330884 | LUAD | VMRC-LCD | SB590885 | MVP_top60_DNA+RNA_seed | HDAC1 | 0.367898 | THCA | FTC-133 | PCI-34051 |
| MVP_top60_DNA_seed | EGFR | 0.330982 | LUAD | NCI-H2405 | Erlotinib | MVP_top60_DNA+RNA_seed | RET | 0.368546 | THCA | HTC-C3 | Motesanib |
| MVP_top60_DNA_seed | AKT1 | 0.331473 | LUAD | NCI-H2009 | MK-2206 | MVP_top60_DNA+RNA_seed | BRAF | 0.372867 | THCA | FTC-133 | PLX-4720 |
| MVP_top60_DNA_seed | MDM2 | 0.332898 | LUAD | NCI-H1781 | Serdemetan | MVP_top60_DNA+RNA_seed | PIK3CG | 0.373617 | THCA | 8505C | Buparlisib |
| MVP_top60_DNA_seed | IGF1R | 0.333397 | LUAD | NCI-H2291 | BMS-536924 | MVP_top60_DNA+RNA_seed | MAPK1 | 0.378 | THCA | RO82-W-1 | Ulixertinib |

| | | | | | | | | | | | |
|---|---|---|---|---|---|---|---|---|---|---|---|
| MVP_top60_DNA_seed | HDAC1 | 0.338056 | LUAD | EMC-BAC-1 | PCI-34051 | MVP_top60_DNA+RNA_seed | MAPK3 | 0.378 | THCA | RO82-W-1 | Ulixertinib |
| MVP_top60_DNA_seed | EGFR | 0.338336 | LUAD | NCI-H2009 | Gefitinib | MVP_top60_DNA+RNA_seed | BRAF | 0.386036 | THCA | ASH-3 | SB590885 |
| MVP_top60_DNA_seed | MET | 0.338617 | LUAD | NCI-H1651 | Savolitinib | MVP_top60_DNA+RNA_seed | CHEK2 | 0.397495 | THCA | TT | MK-8776 |
| MVP_top60_DNA_seed | AKT1 | 0.339169 | LUAD | LXF-289 | AZD5363 | MVP_top60_DNA+RNA_seed | AKT1 | 0.423141 | THCA | ASH-3 | Uprosertib |
| MVP_top60_DNA_seed | PIK3CA | 0.339773 | LUAD | NCI-H1355 | AZD8186 | MVP_top60_DNA+RNA_seed | AKT2 | 0.423141 | THCA | ASH-3 | Uprosertib |
| MVP_top60_DNA_seed | IGF1R | 0.341253 | LUAD | NCI-H1693 | GSK1904529A | MVP_top60_DNA+RNA_seed | AKT3 | 0.423141 | THCA | ASH-3 | Uprosertib |
| MVP_top60_DNA_seed | AURKA | 0.342677 | LUAD | NCI-H1573 | Alisertib | MVP_top60_DNA+RNA_seed | RET | 0.425459 | THCA | CGTH-W-1 | Motesanib |
| MVP_top60_DNA_seed | MDM2 | 0.343754 | LUAD | EMC-BAC-1 | Nutlin-3a (-) | MVP_top60_DNA+RNA_seed | CHEK2 | 0.430461 | THCA | CAL-62 | MK-8776 |
| MVP_top60_DNA_seed | EGFR | 0.344215 | LUAD | NCI-H1993 | Osimertinib | MVP_top60_DNA+RNA_seed | KRAS | 0.432497 | THCA | 8505C | RAS (G12C) Inhibitor-1 |
| MVP_top60_DNA_seed | MDM2 | 0.344464 | LUAD | NCI-H1792 | Nutlin-3a (-) | MVP_top60_DNA+RNA_seed | AKT1 | 0.434285 | THCA | IHH-4 | GSK2110183B |
| MVP_top60_DNA_seed | AKT1 | 0.344635 | LUAD | NCI-H2405 | AT13148 | MVP_top60_DNA+RNA_seed | AKT2 | 0.434285 | THCA | IHH-4 | GSK2110183B |
| MVP_top60_DNA_seed | AKT1 | 0.344947 | LUAD | Calu-3 | Uprosertib | MVP_top60_DNA+RNA_seed | AKT3 | 0.434285 | THCA | IHH-4 | GSK2110183B |
| MVP_top60_DNA_seed | PIK3CA | 0.345114 | LUAD | NCI-H2030 | Buparlisib | MVP_top60_DNA+RNA_seed | HDAC1 | 0.451747 | THCA | BHT-101 | Romidepsin |
| MVP_top60_DNA_seed | IGF1R | 0.345394 | LUAD | Calu-6 | Linsitinib | MVP_top60_DNA+RNA_seed | CHEK2 | 0.45675 | THCA | 8505C | MK-8776 |
| MVP_top60_DNA_seed | EGFR | 0.34541 | LUAD | NCI-H1993 | AZD3759 | MVP_top60_DNA+RNA_seed | AKT1 | 0.459115 | THCA | BHT-101 | Uprosertib |
| MVP_top60_DNA_seed | FGFR1 | 0.346435 | LUAD | NCI-H1623 | PD173074 | MVP_top60_DNA+RNA_seed | AKT2 | 0.459115 | THCA | BHT-101 | Uprosertib |
| MVP_top60_DNA_seed | AKT1 | 0.346469 | LUAD | NCI-H2023 | AT13148 | MVP_top60_DNA+RNA_seed | AKT3 | 0.459115 | THCA | BHT-101 | Uprosertib |
| MVP_top60_DNA_seed | ATM | 0.346673 | LUAD | NCI-H1666 | KU-55933 | MVP_top60_DNA+RNA_seed | GSK3B | 0.460811 | THCA | ASH-3 | SB216763 |
| MVP_top60_DNA_seed | MDM2 | 0.347219 | LUAD | NCI-H522 | Nutlin-3a (-) | MVP_top60_DNA+RNA_seed | HDAC1 | 0.467791 | THCA | B-CPAP | Entinostat |
| MVP_top60_DNA_seed | EGFR | 0.347559 | LUAD | NCI-H23 | Erlotinib | MVP_top60_DNA+RNA_seed | MAPK1 | 0.47052 | THCA | ML-1 | Ulixertinib |
| MVP_top60_DNA_seed | PIK3CA | 0.347699 | LUAD | HOP-62 | AZD8186 | MVP_top60_DNA+RNA_seed | MAPK3 | 0.47052 | THCA | ML-1 | Ulixertinib |
| MVP_top60_DNA_seed | BRAF | 0.348081 | LUAD | NCI-H2087 | SB590885 | MVP_top60_DNA+RNA_seed | AKT1 | 0.475187 | THCA | BHT-101 | AZD5363 |
| MVP_top60_DNA_seed | AKT1 | 0.348915 | LUAD | EMC-BAC-1 | Afuresertib | MVP_top60_DNA+RNA_seed | AKT2 | 0.475187 | THCA | BHT-101 | AZD5363 |
| MVP_top60_DNA_seed | MAPK1 | 0.350731 | LUAD | RERF-LC-KJ | Ulixertinib | MVP_top60_DNA+RNA_seed | AKT3 | 0.475187 | THCA | BHT-101 | AZD5363 |
| MVP_top60_DNA_seed | GSK3B | 0.350824 | LUAD | NCI-H1651 | SB216763 | MVP_top60_DNA+RNA_seed | MAPK1 | 0.478319 | THCA | TT2609-C02 | Ulixertinib |
| MVP_top60_DNA_seed | MDM2 | 0.351381 | LUAD | NCI-H2030 | Nutlin-3a (-) | MVP_top60_DNA+RNA_seed | MAPK3 | 0.478319 | THCA | TT2609-C02 | Ulixertinib |
| MVP_top60_DNA_seed | MAPK1 | 0.351477 | LUAD | PC-14 | Ulixertinib | MVP_top60_DNA+RNA_seed | ATM | 0.494355 | THCA | CAL-62 | KU-55933 |
| MVP_top60_DNA_seed | AKT1 | 0.351684 | LUAD | NCI-H1666 | Uprosertib | MVP_top60_DNA+RNA_seed | AKT1 | 0.512286 | THCA | ASH-3 | MK-2206 |
| MVP_top60_DNA_seed | AKT1 | 0.351788 | LUAD | HCC-78 | Uprosertib | MVP_top60_DNA+RNA_seed | AKT2 | 0.512286 | THCA | ASH-3 | MK-2206 |
| MVP_top60_DNA_seed | IGF1R | 0.353517 | LUAD | HCC-78 | Linsitinib | MVP_top60_DNA+RNA_seed | AKT1 | 0.517844 | THCA | TT | MK-2206 |
| MVP_top60_DNA_seed | EGFR | 0.354711 | LUAD | NCI-H2009 | Sapitinib | MVP_top60_DNA+RNA_seed | AKT2 | 0.517844 | THCA | TT | MK-2206 |
| MVP_top60_DNA_seed | ERBB2 | 0.354711 | LUAD | NCI-H2009 | Sapitinib | MVP_top60_DNA+RNA_seed | CHEK2 | 0.519508 | THCA | ML-1 | AZD7762 |
| MVP_top60_DNA_seed | ERBB3 | 0.354711 | LUAD | NCI-H2009 | Sapitinib | MVP_top60_DNA+RNA_seed | BRAF | 0.543479 | THCA | ASH-3 | Dabrafenib |
| MVP_top60_DNA_seed | AKT1 | 0.354932 | LUAD | HOP-62 | GSK2110183B | MVP_top60_DNA+RNA_seed | MAPK1 | 0.543589 | THCA | ML-1 | SCH772984 |
| MVP_top60_DNA_seed | EGFR | 0.354936 | LUAD | NCI-H1838 | Sapitinib | MVP_top60_DNA+RNA_seed | MAPK3 | 0.543589 | THCA | ML-1 | SCH772984 |
| MVP_top60_DNA_seed | ERBB2 | 0.354936 | LUAD | NCI-H1838 | Sapitinib | MVP_top60_DNA+RNA_seed | AKT1 | 0.551077 | THCA | HTC-C3 | Uprosertib |
| MVP_top60_DNA_seed | ERBB3 | 0.354936 | LUAD | NCI-H1838 | Sapitinib | MVP_top60_DNA+RNA_seed | AKT2 | 0.551077 | THCA | HTC-C3 | Uprosertib |
| MVP_top60_DNA_seed | MET | 0.355221 | LUAD | NCI-H838 | Savolitinib | MVP_top60_DNA+RNA_seed | AKT3 | 0.551077 | THCA | HTC-C3 | Uprosertib |
| MVP_top60_DNA_seed | EGFR | 0.355795 | LUAD | RERF-LC-MS | Gefitinib | MVP_top60_DNA+RNA_seed | KRAS | 0.55186 | THCA | HTC-C3 | RAS (G12C) Inhibitor-1 |
| MVP_top60_DNA_seed | MAPK1 | 0.356539 | LUAD | NCI-H2405 | VX-11e | MVP_top60_DNA+RNA_seed | HDAC1 | 0.553659 | THCA | RO82-W-1 | Romidepsin |
| MVP_top60_DNA_seed | BRAF | 0.357213 | LUAD | NCI-H1781 | SB590885 | MVP_top60_DNA+RNA_seed | CHEK2 | 0.556117 | THCA | ASH-3 | AZD7762 |

| | | | | | | | | | | | |
|---|---|---|---|---|---|---|---|---|---|---|---|
| MVP_top60_DNA_seed | HDAC1 | 0.357475 | LUAD | NCI-H2228 | Entinostat | MVP_top60_DNA+RNA_seed | HDAC1 | 0.572394 | THCA | ML-1 | PCI-34051 |
| MVP_top60_DNA_seed | AKT1 | 0.35782 | LUAD | NCI-H1650 | AZD5363 | MVP_top60_DNA+RNA_seed | PIK3CG | 0.579816 | THCA | CAL-62 | CZC24832 |
| MVP_top60_DNA_seed | IGF1R | 0.35878 | LUAD | NCI-H2085 | BMS-536924 | MVP_top60_DNA+RNA_seed | BRAF | 0.581894 | THCA | FTC-133 | SB590885 |
| MVP_top60_DNA_seed | EGFR | 0.360307 | LUAD | NCI-H1792 | AZD3759 | MVP_top60_DNA+RNA_seed | AKT1 | 0.587831 | THCA | ML-1 | AT13148 |
| MVP_top60_DNA_seed | BRAF | 0.363396 | LUAD | NCI-H1573 | Dabrafenib | MVP_top60_DNA+RNA_seed | PIK3CG | 0.592532 | THCA | K5 | CZC24832 |
| MVP_top60_DNA_seed | BRAF | 0.363838 | LUAD | NCI-H2023 | SB590885 | MVP_top60_DNA+RNA_seed | BRAF | 0.605339 | THCA | 8305C | SB590885 |
| MVP_top60_DNA_seed | EGFR | 0.364517 | LUAD | NCI-H1734 | Erlotinib | MVP_top60_DNA+RNA_seed | AKT1 | 0.605455 | THCA | HTC-C3 | Afuresertib |
| MVP_top60_DNA_seed | BRAF | 0.364615 | LUAD | NCI-H2405 | PLX-4720 | MVP_top60_DNA+RNA_seed | AKT2 | 0.605455 | THCA | HTC-C3 | Afuresertib |
| MVP_top60_DNA_seed | PIK3CA | 0.364818 | LUAD | NCI-H2023 | AZD8186 | MVP_top60_DNA+RNA_seed | AKT3 | 0.605455 | THCA | HTC-C3 | Afuresertib |
| MVP_top60_DNA_seed | BRAF | 0.365273 | LUAD | NCI-H1975 | PLX-4720 | MVP_top60_DNA+RNA_seed | AKT1 | 0.611823 | THCA | B-CPAP | Ipatasertib |
| MVP_top60_DNA_seed | EGFR | 0.365483 | LUAD | NCI-H23 | Afatinib | MVP_top60_DNA+RNA_seed | AKT3 | 0.611823 | THCA | B-CPAP | Ipatasertib |
| MVP_top60_DNA_seed | ERBB2 | 0.365483 | LUAD | NCI-H23 | Afatinib | MVP_top60_DNA+RNA_seed | BRAF | 0.615266 | THCA | HTC-C3 | SB590885 |
| MVP_top60_DNA_seed | GSK3B | 0.366255 | LUAD | EMC-BAC-2 | SB216763 | MVP_top60_DNA+RNA_seed | CHEK2 | 0.619692 | THCA | 8305C | MK-8776 |
| MVP_top60_DNA_seed | TP53 | 0.366569 | LUAD | EMC-BAC-1 | MIRA-1 | MVP_top60_DNA+RNA_seed | AKT1 | 0.622402 | THCA | IHH-4 | AT13148 |
| MVP_top60_DNA_seed | HSP90AA1 | 0.366938 | LUAD | HOP-62 | Luminespib | MVP_top60_DNA+RNA_seed | MAPK1 | 0.624351 | THCA | CGTH-W-1 | Ulixertinib |
| MVP_top60_DNA_seed | MET | 0.367641 | LUAD | NCI-H1666 | Savolitinib | MVP_top60_DNA+RNA_seed | MAPK3 | 0.624351 | THCA | CGTH-W-1 | Ulixertinib |
| MVP_top60_DNA_seed | EGFR | 0.368728 | LUAD | NCI-H2291 | AZD3759 | MVP_top60_DNA+RNA_seed | CHEK2 | 0.626977 | THCA | ASH-3 | MK-8776 |
| MVP_top60_DNA_seed | HDAC1 | 0.368921 | LUAD | NCI-H441 | Romidepsin | MVP_top60_DNA+RNA_seed | AKT1 | 0.632152 | THCA | IHH-4 | AZD5363 |
| MVP_top60_DNA_seed | MET | 0.369542 | LUAD | NCI-H650 | Crizotinib | MVP_top60_DNA+RNA_seed | AKT2 | 0.632152 | THCA | IHH-4 | AZD5363 |
| MVP_top60_DNA_seed | AKT1 | 0.370265 | LUAD | NCI-H1944 | Afuresertib | MVP_top60_DNA+RNA_seed | AKT3 | 0.632152 | THCA | IHH-4 | AZD5363 |
| MVP_top60_DNA_seed | EZH2 | 0.37247 | LUAD | NCI-H650 | GSK343 | MVP_top60_DNA+RNA_seed | AKT1 | 0.63469 | THCA | ASH-3 | Uprosertib |
| MVP_top60_DNA_seed | MET | 0.373413 | LUAD | NCI-H2228 | Savolitinib | MVP_top60_DNA+RNA_seed | AKT2 | 0.63469 | THCA | ASH-3 | Uprosertib |
| MVP_top60_DNA_seed | EGFR | 0.374487 | LUAD | NCI-H1734 | Sapitinib | MVP_top60_DNA+RNA_seed | AKT3 | 0.63469 | THCA | ASH-3 | Uprosertib |
| MVP_top60_DNA_seed | ERBB2 | 0.374487 | LUAD | NCI-H1734 | Sapitinib | MVP_top60_DNA+RNA_seed | MAPK1 | 0.636907 | THCA | CGTH-W-1 | Ulixertinib |
| MVP_top60_DNA_seed | ERBB3 | 0.374487 | LUAD | NCI-H1734 | Sapitinib | MVP_top60_DNA+RNA_seed | MAPK3 | 0.636907 | THCA | CGTH-W-1 | Ulixertinib |
| MVP_top60_DNA_seed | HSP90AA1 | 0.379038 | LUAD | NCI-H2342 | Tanespimycin | MVP_top60_DNA+RNA_seed | AKT1 | 0.667776 | THCA | 8305C | AZD5363 |
| MVP_top60_DNA_seed | BRAF | 0.379198 | LUAD | NCI-H1651 | SB590885 | MVP_top60_DNA+RNA_seed | AKT2 | 0.667776 | THCA | 8305C | AZD5363 |
| MVP_top60_DNA_seed | CREBBP | 0.379849 | LUAD | HCC-44 | SGC-CBP30 | MVP_top60_DNA+RNA_seed | AKT3 | 0.667776 | THCA | 8305C | AZD5363 |
| MVP_top60_DNA_seed | CREBBP | 0.380504 | LUAD | NCI-H1975 | SGC-CBP30 | MVP_top60_DNA+RNA_seed | GSK3B | 0.680266 | THCA | ML-1 | CHIR-99021 |
| MVP_top60_DNA_seed | MAPK1 | 0.381292 | LUAD | LXF-289 | Ulixertinib | MVP_top60_DNA+RNA_seed | HDAC1 | 0.681284 | THCA | IHH-4 | PCI-34051 |
| MVP_top60_DNA_seed | MDM2 | 0.383577 | LUAD | EMC-BAC-2 | Serdemetan | MVP_top60_DNA+RNA_seed | AKT1 | 0.682358 | THCA | IHH-4 | Uprosertib |
| MVP_top60_DNA_seed | AKT1 | 0.383772 | LUAD | NCI-H2228 | GSK2110183B | MVP_top60_DNA+RNA_seed | AKT2 | 0.682358 | THCA | IHH-4 | Uprosertib |
| MVP_top60_DNA_seed | AKT1 | 0.384279 | LUAD | NCI-H322M | Uprosertib | MVP_top60_DNA+RNA_seed | AKT3 | 0.682358 | THCA | IHH-4 | Uprosertib |
| MVP_top60_DNA_seed | HDAC1 | 0.385286 | LUAD | NCI-H1944 | PCI-34051 | MVP_top60_DNA+RNA_seed | ATM | 0.695164 | THCA | FTC-133 | KU-55933 |
| MVP_top60_DNA_seed | IGF1R | 0.386432 | LUAD | HCC-44 | GSK1904529A | MVP_top60_DNA+RNA_seed | PIK3CG | 0.696872 | THCA | FTC-133 | CZC24832 |
| MVP_top60_DNA_seed | MAPK1 | 0.386751 | LUAD | ABC-1 | VX-11e | MVP_top60_DNA+RNA_seed | KRAS | 0.703899 | THCA | 8305C | KRAS (G12C) Inhibitor-1 |
| MVP_top60_DNA_seed | PIK3CA | 0.387343 | LUAD | NCI-H522 | Alpelisib | MVP_top60_DNA+RNA_seed | CHEK2 | 0.711308 | THCA | 8505C | AZD7762 |
| MVP_top60_DNA_seed | IGF1R | 0.388063 | LUAD | NCI-H2291 | BMS-754807 | MVP_top60_DNA+RNA_seed | MAPK1 | 0.712598 | THCA | 8305C | Ulixertinib |
| MVP_top60_DNA_seed | IGF1R | 0.38899 | LUAD | H3255 | BMS-754807 | MVP_top60_DNA+RNA_seed | MAPK3 | 0.712598 | THCA | 8305C | Ulixertinib |
| MVP_top60_DNA_seed | EZH2 | 0.389964 | LUAD | NCI-H2009 | GSK343 | MVP_top60_DNA+RNA_seed | AKT1 | 0.71606 | THCA | BHT-101 | GSK2110183B |
| MVP_top60_DNA_seed | EGFR | 0.39044 | LUAD | NCI-H3122 | Osimertinib | MVP_top60_DNA+RNA_seed | AKT2 | 0.71606 | THCA | BHT-101 | GSK2110183B |

| | | | | | | | | | | | |
|---|---|---|---|---|---|---|---|---|---|---|---|
| MVP_top60_DNA_seed | PIK3CA | 0.390472 | LUAD | NCI-H1793 | AZD8186 | MVP_top60_DNA+RNA_seed | AKT3 | 0.71606 | THCA | BHT-101 | GSK2110183B |
| MVP_top60_DNA_seed | MET | 0.39217 | LUAD | NCI-H2342 | Savolitinib | MVP_top60_DNA+RNA_seed | MAPK1 | 0.716426 | THCA | TT | Ulixertinib |
| MVP_top60_DNA_seed | AKT1 | 0.393403 | LUAD | NCI-H1623 | AZD5363 | MVP_top60_DNA+RNA_seed | MAPK3 | 0.716426 | THCA | TT | Ulixertinib |
| MVP_top60_DNA_seed | ATM | 0.394137 | LUAD | LXF-289 | KU-55933 | MVP_top60_DNA+RNA_seed | AKT1 | 0.729228 | THCA | CAL-62 | AZD5363 |
| MVP_top60_DNA_seed | AKT1 | 0.394308 | LUAD | NCI-H1838 | MK-2206 | MVP_top60_DNA+RNA_seed | AKT2 | 0.729228 | THCA | CAL-62 | AZD5363 |
| MVP_top60_DNA_seed | AURKA | 0.394314 | LUAD | NCI-H1666 | Alisertib | MVP_top60_DNA+RNA_seed | AKT3 | 0.729228 | THCA | CAL-62 | AZD5363 |
| MVP_top60_DNA_seed | AURKA | 0.395038 | LUAD | NCI-H358 | Alisertib | MVP_top60_DNA+RNA_seed | BRAF | 0.730547 | THCA | CAL-62 | Dabrafenib |
| MVP_top60_DNA_seed | EZH2 | 0.396612 | LUAD | NCI-H2122 | GSK343 | MVP_top60_DNA+RNA_seed | CHEK2 | 0.733719 | THCA | TT2609-C02 | AZD7762 |
| MVP_top60_DNA_seed | IGF1R | 0.397802 | LUAD | EMC-BAC-2 | NVP-ADW742 | MVP_top60_DNA+RNA_seed | RET | 0.739204 | THCA | 8505C | Motesanib |
| MVP_top60_DNA_seed | AURKA | 0.39858 | LUAD | H3255 | Alisertib | MVP_top60_DNA+RNA_seed | ATM | 0.756658 | THCA | B-CPAP | KU-55933 |
| MVP_top60_DNA_seed | PIK3CA | 0.39937 | LUAD | NCI-H1435 | AZD8186 | MVP_top60_DNA+RNA_seed | AKT1 | 0.763665 | THCA | CAL-62 | GSK2110183B |
| MVP_top60_DNA_seed | IGF1R | 0.399902 | LUAD | HCC-827 | GSK1904529A | MVP_top60_DNA+RNA_seed | AKT2 | 0.763665 | THCA | CAL-62 | GSK2110183B |
| MVP_top60_DNA_seed | PIK3CA | 0.400589 | LUAD | NCI-H322M | Alpelisib | MVP_top60_DNA+RNA_seed | AKT3 | 0.763665 | THCA | CAL-62 | GSK2110183B |
| MVP_top60_DNA_seed | EZH2 | 0.401117 | LUAD | NCI-H23 | GSK343 | MVP_top60_DNA+RNA_seed | AKT1 | 0.776184 | THCA | HTC-C3 | AZD5363 |
| MVP_top60_DNA_seed | FGFR1 | 0.402116 | LUAD | NCI-H1435 | PD173074 | MVP_top60_DNA+RNA_seed | AKT2 | 0.776184 | THCA | HTC-C3 | AZD5363 |
| MVP_top60_DNA_seed | EGFR | 0.402792 | LUAD | LC-2-ad | Gefitinib | MVP_top60_DNA+RNA_seed | AKT3 | 0.776184 | THCA | HTC-C3 | AZD5363 |
| MVP_top60_DNA_seed | IGF1R | 0.4036 | LUAD | NCI-H2342 | NVP-ADW742 | MVP_top60_DNA+RNA_seed | HDAC1 | 0.794174 | THCA | ASH-3 | PCI-34051 |
| MVP_top60_DNA_seed | MAPK1 | 0.404561 | LUAD | NCI-H2009 | Ulixertinib | MVP_top60_DNA+RNA_seed | AKT1 | 0.809369 | THCA | 8505C | Afuresertib |
| MVP_top60_DNA_seed | TERT | 0.405246 | LUAD | NCI-H1648 | BIBR-1532 | MVP_top60_DNA+RNA_seed | AKT2 | 0.809369 | THCA | 8505C | Afuresertib |
| MVP_top60_DNA_seed | MAPK1 | 0.40542 | LUAD | NCI-H1693 | SCH772984 | MVP_top60_DNA+RNA_seed | AKT3 | 0.809369 | THCA | 8505C | Afuresertib |
| MVP_top60_DNA_seed | EGFR | 0.405539 | LUAD | NCI-H1437 | Erlotinib | MVP_top60_DNA+RNA_seed | AKT1 | 0.820935 | THCA | HTC-C3 | MK-2206 |
| MVP_top60_DNA_seed | SMARCA4 | 0.40642 | LUAD | NCI-H1755 | PFI3 | MVP_top60_DNA+RNA_seed | AKT2 | 0.820935 | THCA | HTC-C3 | MK-2206 |
| MVP_top60_DNA_seed | CREBBP | 0.407675 | LUAD | NCI-H1793 | SGC-CBP30 | MVP_top60_DNA+RNA_seed | AKT1 | 0.833195 | THCA | TT | AZD5363 |
| MVP_top60_DNA_seed | SRC | 0.412245 | LUAD | NCI-H23 | Bosutinib | MVP_top60_DNA+RNA_seed | AKT2 | 0.833195 | THCA | TT | AZD5363 |
| MVP_top60_DNA_seed | AKT1 | 0.412527 | LUAD | NCI-H1693 | Uprosertib | MVP_top60_DNA+RNA_seed | AKT3 | 0.833195 | THCA | TT | AZD5363 |
| MVP_top60_DNA_seed | MET | 0.413698 | LUAD | RERF-LC-KJ | Crizotinib | MVP_top60_DNA+RNA_seed | BRAF | 0.844063 | THCA | TT2609-C02 | SB590885 |
| MVP_top60_DNA_seed | TERT | 0.415574 | LUAD | NCI-H1975 | BIBR-1532 | MVP_top60_DNA+RNA_seed | BRAF | 0.846021 | THCA | CGTH-W-1 | Dabrafenib |
| MVP_top60_DNA_seed | EZH2 | 0.416819 | LUAD | HOP-62 | GSK343 | MVP_top60_DNA+RNA_seed | GSK3B | 0.848699 | THCA | 8505C | SB216763 |
| MVP_top60_DNA_seed | EGFR | 0.417187 | LUAD | NCI-H1944 | Lapatinib | MVP_top60_DNA+RNA_seed | CHEK2 | 0.855055 | THCA | BHT-101 | AZD7762 |
| MVP_top60_DNA_seed | ERBB2 | 0.417187 | LUAD | NCI-H1944 | Lapatinib | MVP_top60_DNA+RNA_seed | KRAS | 0.863473 | THCA | FTC-133 | :RAS (G12C) Inhibitor-1 |
| MVP_top60_DNA_seed | IGF1R | 0.417545 | LUAD | NCI-H2228 | NVP-ADW742 | MVP_top60_DNA+RNA_seed | AKT1 | 0.865932 | THCA | HTC-C3 | GSK2110183B |
| MVP_top60_DNA_seed | MDM2 | 0.418522 | LUAD | HCC-44 | Nutlin-3a (-) | MVP_top60_DNA+RNA_seed | AKT2 | 0.865932 | THCA | HTC-C3 | GSK2110183B |
| MVP_top60_DNA_seed | EZH2 | 0.420148 | LUAD | NCI-H2085 | GSK343 | MVP_top60_DNA+RNA_seed | AKT3 | 0.865932 | THCA | HTC-C3 | GSK2110183B |
| MVP_top60_DNA_seed | AURKA | 0.420766 | LUAD | NCI-H1648 | Alisertib | MVP_top60_DNA+RNA_seed | MAPK1 | 0.877 | THCA | ML-1 | VX-11e |
| MVP_top60_DNA_seed | FGFR1 | 0.421163 | LUAD | RERF-LC-KJ | PD173074 | MVP_top60_DNA+RNA_seed | HDAC1 | 0.88349 | THCA | TT2609-C02 | Entinostat |
| MVP_top60_DNA_seed | BRAF | 0.422025 | LUAD | NCI-H1838 | Dabrafenib | MVP_top60_DNA+RNA_seed | MAPK1 | 0.886116 | THCA | FTC-133 | Ulixertinib |
| MVP_top60_DNA_seed | PIK3CA | 0.422842 | LUAD | H3255 | Alpelisib | MVP_top60_DNA+RNA_seed | MAPK3 | 0.886116 | THCA | FTC-133 | Ulixertinib |
| MVP_top60_DNA_seed | MDM2 | 0.424316 | LUAD | SW1573 | Nutlin-3a (-) | MVP_top60_DNA+RNA_seed | AKT1 | 0.887738 | THCA | HTC-C3 | Uprosertib |
| MVP_top60_DNA_seed | CREBBP | 0.42467 | LUAD | PC-14 | SGC-CBP30 | MVP_top60_DNA+RNA_seed | AKT2 | 0.887738 | THCA | HTC-C3 | Uprosertib |
| MVP_top60_DNA_seed | MET | 0.425691 | LUAD | SK-LU-1 | Crizotinib | MVP_top60_DNA+RNA_seed | AKT3 | 0.887738 | THCA | HTC-C3 | Uprosertib |
| MVP_top60_DNA_seed | EGFR | 0.426458 | LUAD | NCI-H1792 | Osimertinib | MVP_top60_DNA+RNA_seed | RET | 0.90392 | THCA | 8305C | Motesanib |

| | | | | | | | | | | | |
|---|---|---|---|---|---|---|---|---|---|---|---|
| MVP_top60_DNA_seed | IGF1R | 0.427617 | LUAD | EMC-BAC-2 | BMS-754807 | MVP_top60_DNA+RNA_seed | AKT1 | 0.914714 | THCA | TT2609-C02 | AT13148 |
| MVP_top60_DNA_seed | IGF1R | 0.428077 | LUAD | ABC-1 | BMS-536924 | MVP_top60_DNA+RNA_seed | BRAF | 0.919604 | THCA | 8505C | SB590885 |
| MVP_top60_DNA_seed | GSK3B | 0.428519 | LUAD | NCI-H1573 | SB216763 | MVP_top60_DNA+RNA_seed | AKT1 | 0.927061 | THCA | 8505C | Uprosertib |
| MVP_top60_DNA_seed | AKT1 | 0.428535 | LUAD | NCI-H1568 | AZD5363 | MVP_top60_DNA+RNA_seed | AKT2 | 0.927061 | THCA | 8505C | Uprosertib |
| MVP_top60_DNA_seed | BRAF | 0.429616 | LUAD | HCC-44 | PLX-4720 | MVP_top60_DNA+RNA_seed | AKT3 | 0.927061 | THCA | 8505C | Uprosertib |
| MVP_top60_DNA_seed | AURKA | 0.42967 | LUAD | NCI-H1993 | Alisertib | MVP_top60_DNA+RNA_seed | AKT1 | 0.927763 | THCA | HTC-C3 | AT13148 |
| MVP_top60_DNA_seed | SRC | 0.429782 | LUAD | NCI-H1975 | Bosutinib | MVP_top60_DNA+RNA_seed | MAPK1 | 0.931795 | THCA | TT | SCH772984 |
| MVP_top60_DNA_seed | EGFR | 0.432642 | LUAD | NCI-H838 | AZD3759 | MVP_top60_DNA+RNA_seed | MAPK3 | 0.931795 | THCA | TT | SCH772984 |
| MVP_top60_DNA_seed | MDM2 | 0.432704 | LUAD | LXF-289 | Serdemetan | MVP_top60_DNA+RNA_seed | BRAF | 0.939749 | THCA | TT2609-C02 | Dabrafenib |
| MVP_top60_DNA_seed | SMARCA4 | 0.433167 | LUAD | EMC-BAC-1 | PFI3 | MVP_top60_DNA+RNA_seed | AKT1 | 0.955885 | THCA | CAL-62 | Uprosertib |
| MVP_top60_DNA_seed | AURKA | 0.43388 | LUAD | HCC-827 | Alisertib | MVP_top60_DNA+RNA_seed | AKT2 | 0.955885 | THCA | CAL-62 | Uprosertib |
| MVP_top60_DNA_seed | FGFR1 | 0.434337 | LUAD | A549 | PD173074 | MVP_top60_DNA+RNA_seed | AKT3 | 0.955885 | THCA | CAL-62 | Uprosertib |
| MVP_top60_DNA_seed | IGF1R | 0.434682 | LUAD | NCI-H1435 | Linsitinib | MVP_top60_DNA+RNA_seed | CHEK2 | 0.959975 | THCA | BHT-101 | MK-8776 |
| MVP_top60_DNA_seed | PIK3CA | 0.435478 | LUAD | LC-2-ad | GNE-317 | MVP_top60_DNA+RNA_seed | AKT1 | 0.980526 | THCA | CGTH-W-1 | MK-2206 |
| MVP_top60_DNA_seed | MDM2 | 0.43644 | LUAD | NCI-H2023 | Nutlin-3a (-) | MVP_top60_DNA+RNA_seed | AKT2 | 0.980526 | THCA | CGTH-W-1 | MK-2206 |
| MVP_top60_DNA_seed | PIK3CA | 0.436512 | LUAD | NCI-H1435 | Alpelisib | MVP_top60_DNA+RNA_seed | GSK3B | 0.998085 | THCA | 8305C | SB216763 |
| MVP_top60_DNA_seed | AKT1 | 0.437616 | LUAD | COR-L105 | GSK2110183B | MVP_top60_DNA+RNA_seed | MAPK1 | 1.002336 | THCA | 8305C | Ulixertinib |
| MVP_top60_DNA_seed | IGF1R | 0.439011 | LUAD | NCI-H2228 | GSK1904529A | MVP_top60_DNA+RNA_seed | MAPK3 | 1.002336 | THCA | 8305C | Ulixertinib |
| MVP_top60_DNA_seed | TP53 | 0.439079 | LUAD | NCI-H650 | MIRA-1 | MVP_top60_DNA+RNA_seed | HDAC1 | 1.003123 | THCA | CAL-62 | PCI-34051 |
| MVP_top60_DNA_seed | EGFR | 0.44076 | LUAD | NCI-H1650 | Lapatinib | MVP_top60_DNA+RNA_seed | BRAF | 1.008652 | THCA | TT | Dabrafenib |
| MVP_top60_DNA_seed | ERBB2 | 0.44076 | LUAD | NCI-H1650 | Lapatinib | MVP_top60_DNA+RNA_seed | AKT1 | 1.042761 | THCA | CGTH-W-1 | Uprosertib |
| MVP_top60_DNA_seed | EGFR | 0.440988 | LUAD | NCI-H1435 | Osimertinib | MVP_top60_DNA+RNA_seed | AKT2 | 1.042761 | THCA | CGTH-W-1 | Uprosertib |
| MVP_top60_DNA_seed | IGF1R | 0.444181 | LUAD | SW1573 | BMS-754807 | MVP_top60_DNA+RNA_seed | AKT3 | 1.042761 | THCA | CGTH-W-1 | Uprosertib |
| MVP_top60_DNA_seed | TP53 | 0.444214 | LUAD | NCI-H3122 | MIRA-1 | MVP_top60_DNA+RNA_seed | AKT1 | 1.047035 | THCA | B-CPAP | Uprosertib |
| MVP_top60_DNA_seed | IGF1R | 0.445258 | LUAD | NCI-H1573 | BMS-536924 | MVP_top60_DNA+RNA_seed | AKT2 | 1.047035 | THCA | B-CPAP | Uprosertib |
| MVP_top60_DNA_seed | HDAC1 | 0.446839 | LUAD | ABC-1 | PCI-34051 | MVP_top60_DNA+RNA_seed | AKT3 | 1.047035 | THCA | B-CPAP | Uprosertib |
| MVP_top60_DNA_seed | AURKA | 0.448034 | LUAD | H3255 | Tozasertib | MVP_top60_DNA+RNA_seed | AKT1 | 1.057772 | THCA | TT | Afuresertib |
| MVP_top60_DNA_seed | MAPK1 | 0.448601 | LUAD | SK-LU-1 | Ulixertinib | MVP_top60_DNA+RNA_seed | AKT2 | 1.057772 | THCA | TT | Afuresertib |
| MVP_top60_DNA_seed | MAPK1 | 0.450238 | LUAD | EMC-BAC-2 | Ulixertinib | MVP_top60_DNA+RNA_seed | AKT3 | 1.057772 | THCA | TT | Afuresertib |
| MVP_top60_DNA_seed | EGFR | 0.450241 | LUAD | NCI-H2023 | Lapatinib | MVP_top60_DNA+RNA_seed | CHEK2 | 1.062999 | THCA | K5 | MK-8776 |
| MVP_top60_DNA_seed | ERBB2 | 0.450241 | LUAD | NCI-H2023 | Lapatinib | MVP_top60_DNA+RNA_seed | BRAF | 1.073604 | THCA | ASH-3 | PLX-4720 |
| MVP_top60_DNA_seed | EGFR | 0.450663 | LUAD | HCC-78 | AZD3759 | MVP_top60_DNA+RNA_seed | CHEK2 | 1.080986 | THCA | B-CPAP | MK-8776 |
| MVP_top60_DNA_seed | PIK3CA | 0.451146 | LUAD | NCI-H1693 | AZD8186 | MVP_top60_DNA+RNA_seed | AKT1 | 1.083041 | THCA | 8505C | MK-2206 |
| MVP_top60_DNA_seed | EGFR | 0.453084 | LUAD | Calu-6 | Afatinib | MVP_top60_DNA+RNA_seed | AKT2 | 1.083041 | THCA | 8505C | MK-2206 |
| MVP_top60_DNA_seed | ERBB2 | 0.453084 | LUAD | Calu-6 | Afatinib | MVP_top60_DNA+RNA_seed | AKT1 | 1.100716 | THCA | CGTH-W-1 | Afuresertib |
| MVP_top60_DNA_seed | BRAF | 0.455351 | LUAD | NCI-H650 | Dabrafenib | MVP_top60_DNA+RNA_seed | AKT2 | 1.100716 | THCA | CGTH-W-1 | Afuresertib |
| MVP_top60_DNA_seed | BRAF | 0.455893 | LUAD | NCI-H23 | PLX-4720 | MVP_top60_DNA+RNA_seed | AKT3 | 1.100716 | THCA | CGTH-W-1 | Afuresertib |
| MVP_top60_DNA_seed | EGFR | 0.4563 | LUAD | ABC-1 | AZD3759 | MVP_top60_DNA+RNA_seed | HDAC1 | 1.103791 | THCA | CAL-62 | Entinostat |
| MVP_top60_DNA_seed | PIK3CA | 0.456573 | LUAD | NCI-H2085 | GNE-317 | MVP_top60_DNA+RNA_seed | AKT1 | 1.120207 | THCA | 8505C | Ipatasertib |
| MVP_top60_DNA_seed | AKT1 | 0.459573 | LUAD | NCI-H1650 | Ipatasertib | MVP_top60_DNA+RNA_seed | AKT3 | 1.120207 | THCA | 8505C | Ipatasertib |
| MVP_top60_DNA_seed | PIK3CA | 0.459856 | LUAD | NCI-H1573 | AZD8186 | MVP_top60_DNA+RNA_seed | CHEK2 | 1.120942 | THCA | TT2609-C02 | MK-8776 |

| | | | | | | | | | | | |
|---|---|---|---|---|---|---|---|---|---|---|---|
| MVP_top60_DNA_seed | ATM | 0.460032 | LUAD | NCI-H1944 | KU-55933 | MVP_top60_DNA+RNA_seed | AKT1 | 1.132721 | THCA | CAL-62 | MK-2206 |
| MVP_top60_DNA_seed | SRC | 0.460245 | LUAD | HOP-62 | Bosutinib | MVP_top60_DNA+RNA_seed | AKT2 | 1.132721 | THCA | CAL-62 | MK-2206 |
| MVP_top60_DNA_seed | EGFR | 0.460661 | LUAD | LXF-289 | Sapitinib | MVP_top60_DNA+RNA_seed | ATM | 1.184934 | THCA | 8505C | KU-55933 |
| MVP_top60_DNA_seed | ERBB2 | 0.460661 | LUAD | LXF-289 | Sapitinib | MVP_top60_DNA+RNA_seed | AKT1 | 1.186441 | THCA | 8505C | AZD5363 |
| MVP_top60_DNA_seed | ERBB3 | 0.460661 | LUAD | LXF-289 | Sapitinib | MVP_top60_DNA+RNA_seed | AKT2 | 1.186441 | THCA | 8505C | AZD5363 |
| MVP_top60_DNA_seed | BRAF | 0.46089 | LUAD | NCI-H1944 | Dabrafenib | MVP_top60_DNA+RNA_seed | AKT3 | 1.186441 | THCA | 8505C | AZD5363 |
| MVP_top60_DNA_seed | HSP90AA1 | 0.462044 | LUAD | NCI-H358 | Luminespib | MVP_top60_DNA+RNA_seed | RET | 1.194374 | THCA | ML-1 | Motesanib |
| MVP_top60_DNA_seed | SRC | 0.462244 | LUAD | NCI-H1734 | Bosutinib | MVP_top60_DNA+RNA_seed | AKT1 | 1.201658 | THCA | 8505C | GSK2110183B |
| MVP_top60_DNA_seed | MDM2 | 0.462428 | LUAD | NCI-H522 | Serdemetan | MVP_top60_DNA+RNA_seed | AKT2 | 1.201658 | THCA | 8505C | GSK2110183B |
| MVP_top60_DNA_seed | EGFR | 0.463042 | LUAD | NCI-H2085 | Lapatinib | MVP_top60_DNA+RNA_seed | AKT3 | 1.201658 | THCA | 8505C | GSK2110183B |
| MVP_top60_DNA_seed | ERBB2 | 0.463042 | LUAD | NCI-H2085 | Lapatinib | MVP_top60_DNA+RNA_seed | AKT1 | 1.211165 | THCA | HTC-C3 | Ipatasertib |
| MVP_top60_DNA_seed | IGF1R | 0.46317 | LUAD | NCI-H3122 | GSK1904529A | MVP_top60_DNA+RNA_seed | AKT3 | 1.211165 | THCA | HTC-C3 | Ipatasertib |
| MVP_top60_DNA_seed | PIK3CA | 0.463421 | LUAD | LXF-289 | Alpelisib | MVP_top60_DNA+RNA_seed | AKT1 | 1.212533 | THCA | B-CPAP | GSK2110183B |
| MVP_top60_DNA_seed | MAPK1 | 0.464968 | LUAD | NCI-H23 | Ulixertinib | MVP_top60_DNA+RNA_seed | AKT2 | 1.212533 | THCA | B-CPAP | GSK2110183B |
| MVP_top60_DNA_seed | IGF1R | 0.46507 | LUAD | H3255 | Linsitinib | MVP_top60_DNA+RNA_seed | AKT3 | 1.212533 | THCA | B-CPAP | GSK2110183B |
| MVP_top60_DNA_seed | TERT | 0.46521 | LUAD | NCI-H522 | BIBR-1532 | MVP_top60_DNA+RNA_seed | AKT1 | 1.214275 | THCA | TT2609-C02 | Afuresertib |
| MVP_top60_DNA_seed | TERT | 0.466496 | LUAD | NCI-H2085 | BIBR-1532 | MVP_top60_DNA+RNA_seed | AKT2 | 1.214275 | THCA | TT2609-C02 | Afuresertib |
| MVP_top60_DNA_seed | PIK3CA | 0.467217 | LUAD | NCI-H1693 | Buparlisib | MVP_top60_DNA+RNA_seed | AKT3 | 1.214275 | THCA | TT2609-C02 | Afuresertib |
| MVP_top60_DNA_seed | MDM2 | 0.467775 | LUAD | NCI-H2342 | Serdemetan | MVP_top60_DNA+RNA_seed | BRAF | 1.232725 | THCA | CAL-62 | PLX-4720 |
| MVP_top60_DNA_seed | AURKA | 0.469075 | LUAD | NCI-H1838 | ZM447439 | MVP_top60_DNA+RNA_seed | KRAS | 1.239146 | THCA | B-CPAP | :RAS (G12C) Inhibitor-1 |
| MVP_top60_DNA_seed | IGF1R | 0.470001 | LUAD | NCI-H1838 | BMS-754807 | MVP_top60_DNA+RNA_seed | HDAC1 | 1.267906 | THCA | 8305C | Entinostat |
| MVP_top60_DNA_seed | IGF1R | 0.470075 | LUAD | NCI-H2087 | Linsitinib | MVP_top60_DNA+RNA_seed | PIK3CG | 1.279192 | THCA | CGTH-W-1 | Buparlisib |
| MVP_top60_DNA_seed | MAPK1 | 0.470906 | LUAD | PC-3_[JPC-3] | Ulixertinib | MVP_top60_DNA+RNA_seed | AKT1 | 1.284222 | THCA | CAL-62 | Uprosertib |
| MVP_top60_DNA_seed | PIK3CA | 0.471136 | LUAD | NCI-H1355 | GNE-317 | MVP_top60_DNA+RNA_seed | AKT2 | 1.284222 | THCA | CAL-62 | Uprosertib |
| MVP_top60_DNA_seed | AURKA | 0.472199 | LUAD | NCI-H2009 | ZM447439 | MVP_top60_DNA+RNA_seed | AKT3 | 1.284222 | THCA | CAL-62 | Uprosertib |
| MVP_top60_DNA_seed | BRAF | 0.472243 | LUAD | NCI-H1781 | Dabrafenib | MVP_top60_DNA+RNA_seed | HDAC1 | 1.295316 | THCA | ML-1 | Entinostat |
| MVP_top60_DNA_seed | MAPK1 | 0.472601 | LUAD | NCI-H23 | VX-11e | MVP_top60_DNA+RNA_seed | HDAC1 | 1.295599 | THCA | TT2609-C02 | Romidepsin |
| MVP_top60_DNA_seed | IGF1R | 0.473652 | LUAD | PC-14 | Linsitinib | MVP_top60_DNA+RNA_seed | PIK3CG | 1.2998 | THCA | ML-1 | CZC24832 |
| MVP_top60_DNA_seed | EZH2 | 0.47381 | LUAD | Calu-6 | GSK343 | MVP_top60_DNA+RNA_seed | AKT1 | 1.323573 | THCA | CGTH-W-1 | AZD5363 |
| MVP_top60_DNA_seed | SRC | 0.474074 | LUAD | NCI-H2228 | Bosutinib | MVP_top60_DNA+RNA_seed | AKT2 | 1.323573 | THCA | CGTH-W-1 | AZD5363 |
| MVP_top60_DNA_seed | MET | 0.474964 | LUAD | NCI-H1563 | Crizotinib | MVP_top60_DNA+RNA_seed | AKT3 | 1.323573 | THCA | CGTH-W-1 | AZD5363 |
| MVP_top60_DNA_seed | AKT1 | 0.476004 | LUAD | NCI-H1792 | Ipatasertib | MVP_top60_DNA+RNA_seed | AKT1 | 1.327396 | THCA | 8305C | GSK2110183B |
| MVP_top60_DNA_seed | MET | 0.477036 | LUAD | SW1573 | Savolitinib | MVP_top60_DNA+RNA_seed | AKT2 | 1.327396 | THCA | 8305C | GSK2110183B |
| MVP_top60_DNA_seed | MET | 0.477441 | LUAD | NCI-H1944 | Crizotinib | MVP_top60_DNA+RNA_seed | AKT3 | 1.327396 | THCA | 8305C | GSK2110183B |
| MVP_top60_DNA_seed | IGF1R | 0.478872 | LUAD | NCI-H1944 | BMS-754807 | MVP_top60_DNA+RNA_seed | BRAF | 1.327751 | THCA | ML-1 | SB590885 |
| MVP_top60_DNA_seed | KRAS | 0.480611 | LUAD | VMRC-LCD | KRAS (G12C) Inhibitor-1: | MVP_top60_DNA+RNA_seed | AKT1 | 1.342937 | THCA | RO82-W-1 | Uprosertib |
| MVP_top60_DNA_seed | MAPK1 | 0.48086 | LUAD | Calu-3 | VX-11e | MVP_top60_DNA+RNA_seed | AKT2 | 1.342937 | THCA | RO82-W-1 | Uprosertib |
| MVP_top60_DNA_seed | AURKA | 0.481866 | LUAD | NCI-H650 | Alisertib | MVP_top60_DNA+RNA_seed | AKT3 | 1.342937 | THCA | RO82-W-1 | Uprosertib |
| MVP_top60_DNA_seed | IGF1R | 0.483089 | LUAD | H3255 | GSK1904529A | MVP_top60_DNA+RNA_seed | PIK3CG | 1.353847 | THCA | TT2609-C02 | Buparlisib |
| MVP_top60_DNA_seed | EGFR | 0.483998 | LUAD | NCI-H1437 | Sapitinib | MVP_top60_DNA+RNA_seed | GSK3B | 1.382225 | THCA | ML-1 | SB216763 |
| MVP_top60_DNA_seed | ERBB2 | 0.483998 | LUAD | NCI-H1437 | Sapitinib | MVP_top60_DNA+RNA_seed | PIK3CG | 1.442314 | THCA | 8305C | CZC24832 |

| | | | | | | | | | | | |
|---|---|---|---|---|---|---|---|---|---|---|---|
| MVP_top60_DNA_seed | ERBB3 | 0.483998 | LUAD | NCI-H1437 | Sapitinib | MVP_top60_DNA+RNA_seed | AKT1 | 1.444889 | THCA | CGTH-W-1 | Ipatasertib |
| MVP_top60_DNA_seed | MDM2 | 0.487142 | LUAD | NCI-H1648 | Serdemetan | MVP_top60_DNA+RNA_seed | AKT3 | 1.444889 | THCA | CGTH-W-1 | Ipatasertib |
| MVP_top60_DNA_seed | IGF1R | 0.487202 | LUAD | NCI-H1568 | BMS-536924 | MVP_top60_DNA+RNA_seed | AKT1 | 1.464269 | THCA | RO82-W-1 | AZD5363 |
| MVP_top60_DNA_seed | AKT1 | 0.487599 | LUAD | NCI-H23 | Afuresertib | MVP_top60_DNA+RNA_seed | AKT2 | 1.464269 | THCA | RO82-W-1 | AZD5363 |
| MVP_top60_DNA_seed | BRAF | 0.489139 | LUAD | NCI-H1573 | SB590885 | MVP_top60_DNA+RNA_seed | AKT3 | 1.464269 | THCA | RO82-W-1 | AZD5363 |
| MVP_top60_DNA_seed | MDM2 | 0.489564 | LUAD | NCI-H1651 | Nutlin-3a (-) | MVP_top60_DNA+RNA_seed | CHEK2 | 1.468932 | THCA | RO82-W-1 | MK-8776 |
| MVP_top60_DNA_seed | AKT1 | 0.490684 | LUAD | NCI-H1650 | Uprosertib | MVP_top60_DNA+RNA_seed | HDAC1 | 1.47177 | THCA | TT2609-C02 | PCI-34051 |
| MVP_top60_DNA_seed | IGF1R | 0.491428 | LUAD | Calu-6 | GSK1904529A | MVP_top60_DNA+RNA_seed | CHEK2 | 1.472233 | THCA | TT | AZD7762 |
| MVP_top60_DNA_seed | AURKA | 0.491579 | LUAD | EMC-BAC-1 | Alisertib | MVP_top60_DNA+RNA_seed | AKT1 | 1.483532 | THCA | CGTH-W-1 | AT13148 |
| MVP_top60_DNA_seed | MET | 0.493349 | LUAD | NCI-H1355 | Crizotinib | MVP_top60_DNA+RNA_seed | BRAF | 1.50131 | THCA | TT | SB590885 |
| MVP_top60_DNA_seed | HDAC1 | 0.494469 | LUAD | NCI-H1666 | Entinostat | MVP_top60_DNA+RNA_seed | AKT1 | 1.532642 | THCA | 8305C | Uprosertib |
| MVP_top60_DNA_seed | FGFR1 | 0.494507 | LUAD | NCI-H2023 | PD173074 | MVP_top60_DNA+RNA_seed | AKT2 | 1.532642 | THCA | 8305C | Uprosertib |
| MVP_top60_DNA_seed | EGFR | 0.496589 | LUAD | NCI-H1993 | Erlotinib | MVP_top60_DNA+RNA_seed | AKT3 | 1.532642 | THCA | 8305C | Uprosertib |
| MVP_top60_DNA_seed | AKT1 | 0.496863 | LUAD | NCI-H2228 | AZD5363 | MVP_top60_DNA+RNA_seed | MAPK1 | 1.53516 | THCA | FTC-133 | SCH772984 |
| MVP_top60_DNA_seed | KRAS | 0.497351 | LUAD | Calu-6 | KRAS (G12C) Inhibitor-12 | MVP_top60_DNA+RNA_seed | MAPK3 | 1.53516 | THCA | FTC-133 | SCH772984 |
| MVP_top60_DNA_seed | EGFR | 0.498189 | LUAD | NCI-H1435 | Erlotinib | MVP_top60_DNA+RNA_seed | AKT1 | 1.552267 | THCA | TT | Uprosertib |
| MVP_top60_DNA_seed | EGFR | 0.498214 | LUAD | NCI-H650 | Sapitinib | MVP_top60_DNA+RNA_seed | AKT2 | 1.552267 | THCA | TT | Uprosertib |
| MVP_top60_DNA_seed | ERBB2 | 0.498214 | LUAD | NCI-H650 | Sapitinib | MVP_top60_DNA+RNA_seed | AKT3 | 1.552267 | THCA | TT | Uprosertib |
| MVP_top60_DNA_seed | ERBB3 | 0.498214 | LUAD | NCI-H650 | Sapitinib | MVP_top60_DNA+RNA_seed | AKT1 | 1.556382 | THCA | TT | Ipatasertib |
| MVP_top60_DNA_seed | EGFR | 0.498421 | LUAD | NCI-H2085 | Afatinib | MVP_top60_DNA+RNA_seed | AKT3 | 1.556382 | THCA | TT | Ipatasertib |
| MVP_top60_DNA_seed | ERBB2 | 0.498421 | LUAD | NCI-H2085 | Afatinib | MVP_top60_DNA+RNA_seed | ATM | 1.574088 | THCA | ML-1 | KU-55933 |
| MVP_top60_DNA_seed | BRAF | 0.499074 | LUAD | NCI-H358 | Dabrafenib | MVP_top60_DNA+RNA_seed | RET | 1.5804 | THCA | TT2609-C02 | Motesanib |
| MVP_top60_DNA_seed | PIK3CA | 0.499505 | LUAD | SW1573 | Alpelisib | MVP_top60_DNA+RNA_seed | AKT1 | 1.605319 | THCA | ML-1 | Ipatasertib |
| MVP_top60_DNA_seed | TP53 | 0.50233 | LUAD | HCC-44 | MIRA-1 | MVP_top60_DNA+RNA_seed | AKT3 | 1.605319 | THCA | ML-1 | Ipatasertib |
| MVP_top60_DNA_seed | AKT1 | 0.503727 | LUAD | NCI-H1435 | Afuresertib | MVP_top60_DNA+RNA_seed | AKT1 | 1.606235 | THCA | TT2609-C02 | GSK2110183B |
| MVP_top60_DNA_seed | EGFR | 0.504405 | LUAD | NCI-H1975 | Lapatinib | MVP_top60_DNA+RNA_seed | AKT2 | 1.606235 | THCA | TT2609-C02 | GSK2110183B |
| MVP_top60_DNA_seed | ERBB2 | 0.504405 | LUAD | NCI-H1975 | Lapatinib | MVP_top60_DNA+RNA_seed | AKT3 | 1.606235 | THCA | TT2609-C02 | GSK2110183B |
| MVP_top60_DNA_seed | CREBBP | 0.504676 | LUAD | Calu-3 | SGC-CBP30 | MVP_top60_DNA+RNA_seed | BRAF | 1.632561 | THCA | RO82-W-1 | PLX-4720 |
| MVP_top60_DNA_seed | EZH2 | 0.504984 | LUAD | NCI-H1623 | GSK343 | MVP_top60_DNA+RNA_seed | AKT1 | 1.634191 | THCA | 8505C | Uprosertib |
| MVP_top60_DNA_seed | EZH2 | 0.507082 | LUAD | EMC-BAC-2 | GSK343 | MVP_top60_DNA+RNA_seed | AKT2 | 1.634191 | THCA | 8505C | Uprosertib |
| MVP_top60_DNA_seed | MDM2 | 0.508387 | LUAD | NCI-H1781 | Nutlin-3a (-) | MVP_top60_DNA+RNA_seed | AKT3 | 1.634191 | THCA | 8505C | Uprosertib |
| MVP_top60_DNA_seed | HDAC1 | 0.508399 | LUAD | NCI-H1437 | PCI-34051 | MVP_top60_DNA+RNA_seed | PIK3CG | 1.651604 | THCA | RO82-W-1 | CZC24832 |
| MVP_top60_DNA_seed | CREBBP | 0.508964 | LUAD | COR-L105 | SGC-CBP30 | MVP_top60_DNA+RNA_seed | ATM | 1.660709 | THCA | TT2609-C02 | KU-55933 |
| MVP_top60_DNA_seed | AURKA | 0.510438 | LUAD | NCI-H1563 | Alisertib | MVP_top60_DNA+RNA_seed | AKT1 | 1.674566 | THCA | ML-1 | AZD5363 |
| MVP_top60_DNA_seed | TP53 | 0.510946 | LUAD | NCI-H1993 | MIRA-1 | MVP_top60_DNA+RNA_seed | AKT2 | 1.674566 | THCA | ML-1 | AZD5363 |
| MVP_top60_DNA_seed | AKT1 | 0.511628 | LUAD | NCI-H1693 | AT13148 | MVP_top60_DNA+RNA_seed | AKT3 | 1.674566 | THCA | ML-1 | AZD5363 |
| MVP_top60_DNA_seed | MAPK1 | 0.512368 | LUAD | NCI-H1435 | Ulixertinib | MVP_top60_DNA+RNA_seed | AKT1 | 1.713256 | THCA | TT2609-C02 | Ipatasertib |
| MVP_top60_DNA_seed | BRAF | 0.513362 | LUAD | ABC-1 | PLX-4720 | MVP_top60_DNA+RNA_seed | AKT3 | 1.713256 | THCA | TT2609-C02 | Ipatasertib |
| MVP_top60_DNA_seed | MAPK1 | 0.513378 | LUAD | H3255 | VX-11e | MVP_top60_DNA+RNA_seed | PIK3CG | 1.753511 | THCA | RO82-W-1 | Buparlisib |
| MVP_top60_DNA_seed | AURKA | 0.516417 | LUAD | NCI-H2023 | Alisertib | MVP_top60_DNA+RNA_seed | AKT1 | 1.758311 | THCA | 8505C | AT13148 |
| MVP_top60_DNA_seed | EGFR | 0.517694 | LUAD | NCI-H2228 | Afatinib | MVP_top60_DNA+RNA_seed | AKT1 | 1.766555 | THCA | 8305C | Uprosertib |

| | | | | | | | | | | | |
|---|---|---|---|---|---|---|---|---|---|---|---|
| MVP_top60_DNA_seed | ERBB2 | 0.517694 | LUAD | NCI-H2228 | Afatinib | MVP_top60_DNA+RNA_seed | AKT2 | 1.766555 | THCA | 8305C | Uprosertib |
| MVP_top60_DNA_seed | BRAF | 0.518845 | LUAD | LXF-289 | Dabrafenib | MVP_top60_DNA+RNA_seed | AKT3 | 1.766555 | THCA | 8305C | Uprosertib |
| MVP_top60_DNA_seed | HSP90AA1 | 0.519982 | LUAD | HCC-827 | Luminespib | MVP_top60_DNA+RNA_seed | AKT1 | 1.802675 | THCA | TT2609-C02 | Uprosertib |
| MVP_top60_DNA_seed | IGF1R | 0.521956 | LUAD | PC-14 | BMS-754807 | MVP_top60_DNA+RNA_seed | AKT2 | 1.802675 | THCA | TT2609-C02 | Uprosertib |
| MVP_top60_DNA_seed | AKT1 | 0.522761 | LUAD | NCI-H23 | GSK2110183B | MVP_top60_DNA+RNA_seed | AKT3 | 1.802675 | THCA | TT2609-C02 | Uprosertib |
| MVP_top60_DNA_seed | EGFR | 0.523746 | LUAD | NCI-H1993 | Lapatinib | MVP_top60_DNA+RNA_seed | AKT1 | 1.812247 | THCA | ML-1 | MK-2206 |
| MVP_top60_DNA_seed | ERBB2 | 0.523746 | LUAD | NCI-H1993 | Lapatinib | MVP_top60_DNA+RNA_seed | AKT2 | 1.812247 | THCA | ML-1 | MK-2206 |
| MVP_top60_DNA_seed | AKT1 | 0.524217 | LUAD | NCI-H522 | Uprosertib | MVP_top60_DNA+RNA_seed | RET | 1.819942 | THCA | TT | Motesanib |
| MVP_top60_DNA_seed | IGF1R | 0.5252 | LUAD | Calu-3 | GSK1904529A | MVP_top60_DNA+RNA_seed | HDAC1 | 1.835974 | THCA | RO82-W-1 | PCI-34051 |
| MVP_top60_DNA_seed | IGF1R | 0.525205 | LUAD | NCI-H1623 | BMS-536924 | MVP_top60_DNA+RNA_seed | ATM | 1.880356 | THCA | RO82-W-1 | KU-55933 |
| MVP_top60_DNA_seed | PIK3CA | 0.525401 | LUAD | NCI-H322M | GNE-317 | MVP_top60_DNA+RNA_seed | KRAS | 1.913729 | THCA | ML-1 | KRAS (G12C) Inhibitor-1 |
| MVP_top60_DNA_seed | FGFR1 | 0.52612 | LUAD | NCI-H1563 | PD173074 | MVP_top60_DNA+RNA_seed | BRAF | 1.917072 | THCA | TT2609-C02 | PLX-4720 |
| MVP_top60_DNA_seed | MET | 0.526252 | LUAD | NCI-H3122 | Savolitinib | MVP_top60_DNA+RNA_seed | PIK3CG | 1.922315 | THCA | ML-1 | Buparlisib |
| MVP_top60_DNA_seed | TERT | 0.52758 | LUAD | ABC-1 | BIBR-1532 | MVP_top60_DNA+RNA_seed | AKT1 | 1.933148 | THCA | RO82-W-1 | AT13148 |
| MVP_top60_DNA_seed | HDAC1 | 0.527728 | LUAD | Calu-3 | Entinostat | MVP_top60_DNA+RNA_seed | AKT1 | 1.960196 | THCA | RO82-W-1 | GSK2110183B |
| MVP_top60_DNA_seed | AKT1 | 0.527963 | LUAD | NCI-H1666 | AZD5363 | MVP_top60_DNA+RNA_seed | AKT2 | 1.960196 | THCA | RO82-W-1 | GSK2110183B |
| MVP_top60_DNA_seed | GSK3B | 0.529682 | LUAD | PC-14 | CHIR-99021 | MVP_top60_DNA+RNA_seed | AKT3 | 1.960196 | THCA | RO82-W-1 | GSK2110183B |
| MVP_top60_DNA_seed | PIK3CA | 0.529885 | LUAD | PC-3_[JPC-3] | Alpelisib | MVP_top60_DNA+RNA_seed | AKT1 | 1.96079 | THCA | RO82-W-1 | MK-2206 |
| MVP_top60_DNA_seed | TP53 | 0.529901 | LUAD | NCI-H1573 | MIRA-1 | MVP_top60_DNA+RNA_seed | AKT2 | 1.96079 | THCA | RO82-W-1 | MK-2206 |
| MVP_top60_DNA_seed | AKT1 | 0.53014 | LUAD | EKVX | AZD5363 | MVP_top60_DNA+RNA_seed | AKT1 | 1.993555 | THCA | RO82-W-1 | Afuresertib |
| MVP_top60_DNA_seed | EGFR | 0.530153 | LUAD | NCI-H2023 | Erlotinib | MVP_top60_DNA+RNA_seed | AKT2 | 1.993555 | THCA | RO82-W-1 | Afuresertib |
| MVP_top60_DNA_seed | MET | 0.530517 | LUAD | NCI-H522 | Savolitinib | MVP_top60_DNA+RNA_seed | AKT3 | 1.993555 | THCA | RO82-W-1 | Afuresertib |
| MVP_top60_DNA_seed | AKT1 | 0.531035 | LUAD | ABC-1 | Afuresertib | MVP_top60_DNA+RNA_seed | BRAF | 2.011685 | THCA | TT | PLX-4720 |
| MVP_top60_DNA_seed | AKT1 | 0.531308 | LUAD | NCI-H1623 | Afuresertib | MVP_top60_DNA+RNA_seed | AKT1 | 2.025232 | THCA | TT2609-C02 | AZD5363 |
| MVP_top60_DNA_seed | AKT1 | 0.531847 | LUAD | PC-3_[JPC-3] | AZD5363 | MVP_top60_DNA+RNA_seed | AKT2 | 2.025232 | THCA | TT2609-C02 | AZD5363 |
| MVP_top60_DNA_seed | MET | 0.532802 | LUAD | Calu-6 | Crizotinib | MVP_top60_DNA+RNA_seed | AKT3 | 2.025232 | THCA | TT2609-C02 | AZD5363 |
| MVP_top60_DNA_seed | MAPK1 | 0.532867 | LUAD | NCI-H1651 | VX-11e | MVP_top60_DNA+RNA_seed | BRAF | 2.030313 | THCA | RO82-W-1 | Dabrafenib |
| MVP_top60_DNA_seed | PIK3CA | 0.534481 | LUAD | NCI-H2291 | GNE-317 | MVP_top60_DNA+RNA_seed | GSK3B | 2.07841 | THCA | TT | SB216763 |
| MVP_top60_DNA_seed | AKT1 | 0.53583 | LUAD | HOP-62 | AZD5363 | MVP_top60_DNA+RNA_seed | PIK3CG | 2.135251 | THCA | TT2609-C02 | CZC24832 |
| MVP_top60_DNA_seed | HDAC1 | 0.536 | LUAD | SW1573 | Entinostat | MVP_top60_DNA+RNA_seed | ATM | 2.159676 | THCA | TT | KU-55933 |
| MVP_top60_DNA_seed | BRAF | 0.536581 | LUAD | NCI-H2228 | PLX-4720 | MVP_top60_DNA+RNA_seed | KRAS | 2.161661 | THCA | TT2609-C02 | KRAS (G12C) Inhibitor-1 |
| MVP_top60_DNA_seed | BRAF | 0.537113 | LUAD | EMC-BAC-1 | SB590885 | MVP_top60_DNA+RNA_seed | MAPK1 | 2.217249 | THCA | RO82-W-1 | SCH772984 |
| MVP_top60_DNA_seed | FGFR1 | 0.539245 | LUAD | NCI-H1944 | PD173074 | MVP_top60_DNA+RNA_seed | MAPK3 | 2.217249 | THCA | RO82-W-1 | SCH772984 |
| MVP_top60_DNA_seed | EGFR | 0.539911 | LUAD | NCI-H650 | Afatinib | MVP_top60_DNA+RNA_seed | AKT1 | 2.230667 | THCA | TT2609-C02 | Uprosertib |
| MVP_top60_DNA_seed | ERBB2 | 0.539911 | LUAD | NCI-H650 | Afatinib | MVP_top60_DNA+RNA_seed | AKT2 | 2.230667 | THCA | TT2609-C02 | Uprosertib |
| MVP_top60_DNA_seed | BRAF | 0.54013 | LUAD | NCI-H1993 | PLX-4720 | MVP_top60_DNA+RNA_seed | AKT3 | 2.230667 | THCA | TT2609-C02 | Uprosertib |
| MVP_top60_DNA_seed | SMARCA4 | 0.540359 | LUAD | NCI-H3122 | PFI3 | MVP_top60_DNA+RNA_seed | MAPK1 | 2.240985 | THCA | TT2609-C02 | VX-11e |
| MVP_top60_DNA_seed | CREBBP | 0.540847 | LUAD | NCI-H1944 | SGC-CBP30 | MVP_top60_DNA+RNA_seed | AKT1 | 2.300894 | THCA | ML-1 | Uprosertib |
| MVP_top60_DNA_seed | EGFR | 0.541228 | LUAD | LXF-289 | Osimertinib | MVP_top60_DNA+RNA_seed | AKT2 | 2.300894 | THCA | ML-1 | Uprosertib |
| MVP_top60_DNA_seed | HSP90AA1 | 0.545184 | LUAD | HCC-78 | Luminespib | MVP_top60_DNA+RNA_seed | AKT3 | 2.300894 | THCA | ML-1 | Uprosertib |
| MVP_top60_DNA_seed | FGFR1 | 0.546248 | LUAD | HOP-62 | AZD4547 | MVP_top60_DNA+RNA_seed | BRAF | 2.359152 | THCA | RO82-W-1 | SB590885 |

| | | | | | | | | | | | |
|---|---|---|---|---|---|---|---|---|---|---|---|
| MVP_top60_DNA_seed | HDAC1 | 0.546284 | LUAD | NCI-H650 | PCI-34051 | MVP_top60_DNA+RNA_seed | AKT1 | 2.420424 | THCA | TT2609-C02 | MK-2206 |
| MVP_top60_DNA_seed | TP53 | 0.546422 | LUAD | NCI-H1781 | MIRA-1 | MVP_top60_DNA+RNA_seed | AKT2 | 2.420424 | THCA | TT2609-C02 | MK-2206 |
| MVP_top60_DNA_seed | EGFR | 0.546422 | LUAD | LXF-289 | Gefitinib | MVP_top60_DNA+RNA_seed | AKT1 | 2.431171 | THCA | TT | AT13148 |
| MVP_top60_DNA_seed | PIK3CA | 0.546928 | LUAD | NCI-H522 | Buparlisib | MVP_top60_DNA+RNA_seed | AKT1 | 2.510429 | THCA | RO82-W-1 | Ipatasertib |
| MVP_top60_DNA_seed | FGFR1 | 0.548106 | LUAD | NCI-H322M | AZD4547 | MVP_top60_DNA+RNA_seed | AKT3 | 2.510429 | THCA | RO82-W-1 | Ipatasertib |
| MVP_top60_DNA_seed | HDAC1 | 0.548624 | LUAD | NCI-H1755 | Entinostat | MVP_top60_DNA+RNA_seed | MAPK1 | 2.518446 | THCA | TT | VX-11e |
| MVP_top60_DNA_seed | MDM2 | 0.548627 | LUAD | NCI-H1755 | Nutlin-3a (-) | MVP_top60_DNA+RNA_seed | RET | 2.530202 | THCA | RO82-W-1 | Motesanib |
| MVP_top60_DNA_seed | MDM2 | 0.549352 | LUAD | NCI-H1975 | Nutlin-3a (-) | MVP_top60_DNA+RNA_seed | CHEK2 | 2.539113 | THCA | RO82-W-1 | AZD7762 |
| MVP_top60_DNA_seed | AKT1 | 0.549956 | LUAD | NCI-H322M | AT13148 | MVP_top60_DNA+RNA_seed | MAPK1 | 2.699379 | THCA | RO82-W-1 | VX-11e |
| MVP_top60_DNA_seed | IGF1R | 0.55195 | LUAD | NCI-H3122 | Linsitinib | MVP_top60_DNA+RNA_seed | MAPK1 | 2.723591 | THCA | RO82-W-1 | Ulixertinib |
| MVP_top60_DNA_seed | FGFR1 | 0.552816 | LUAD | NCI-H1651 | AZD4547 | MVP_top60_DNA+RNA_seed | MAPK3 | 2.723591 | THCA | RO82-W-1 | Ulixertinib |
| MVP_top60_DNA_seed | AKT1 | 0.553661 | LUAD | NCI-H23 | Ipatasertib | MVP_top60_DNA+RNA_seed | GSK3B | 2.745417 | THCA | RO82-W-1 | SB216763 |
| MVP_top60_DNA_seed | BRAF | 0.554123 | LUAD | NCI-H23 | Dabrafenib | MVP_top60_DNA+RNA_seed | PIK3CG | 2.751355 | THCA | TT | Buparlisib |
| MVP_top60_DNA_seed | BRAF | 0.554848 | LUAD | NCI-H1563 | SB590885 | MVP_top60_DNA+RNA_seed | AKT1 | 2.83111 | THCA | RO82-W-1 | Uprosertib |
| MVP_top60_DNA_seed | AURKA | 0.555526 | LUAD | EMC-BAC-1 | Tozasertib | MVP_top60_DNA+RNA_seed | AKT2 | 2.83111 | THCA | RO82-W-1 | Uprosertib |
| MVP_top60_DNA_seed | TP53 | 0.556919 | LUAD | ABC-1 | MIRA-1 | MVP_top60_DNA+RNA_seed | AKT3 | 2.83111 | THCA | RO82-W-1 | Uprosertib |
| MVP_top60_DNA_seed | KRAS | 0.557187 | LUAD | NCI-H1563 | KRAS (G12C) Inhibitor-1 | MVP_top60_DNA+RNA_seed | AKT1 | 2.837742 | THCA | ML-1 | Afuresertib |
| MVP_top60_DNA_seed | IGF1R | 0.557316 | LUAD | NCI-H1693 | Linsitinib | MVP_top60_DNA+RNA_seed | AKT2 | 2.837742 | THCA | ML-1 | Afuresertib |
| MVP_top60_DNA_seed | MAPK1 | 0.557451 | LUAD | H3255 | SCH772984 | MVP_top60_DNA+RNA_seed | AKT3 | 2.837742 | THCA | ML-1 | Afuresertib |
| MVP_top60_DNA_seed | PIK3CA | 0.559043 | LUAD | NCI-H650 | GNE-317 | MVP_top60_DNA+RNA_seed | KRAS | 2.906905 | THCA | RO82-W-1 | KRAS (G12C) Inhibitor-1 |
| MVP_top60_DNA_seed | MAPK1 | 0.56085 | LUAD | HCC-78 | Ulixertinib | MVP_top60_DNA+RNA_seed | HDAC1 | 3.081271 | THCA | RO82-W-1 | Entinostat |
| MVP_top60_DNA_seed | GSK3B | 0.560968 | LUAD | EMC-BAC-1 | SB216763 | MVP_top60_DNA+RNA_seed | FGFR2 | -3.839158 | UCEC | MFE-280 | AZD4547 |
| MVP_top60_DNA_seed | MAPK1 | 0.561811 | LUAD | NCI-H1568 | VX-11e | MVP_top60_DNA+RNA_seed | FGFR2 | -3.193966 | UCEC | AN3-CA | PD173074 |
| MVP_top60_DNA_seed | HSP90AA1 | 0.561848 | LUAD | ABC-1 | Luminespib | MVP_top60_DNA+RNA_seed | FGFR2 | -3.138837 | UCEC | AN3-CA | AZD4547 |
| MVP_top60_DNA_seed | GSK3B | 0.562071 | LUAD | NCI-H838 | CHIR-99021 | MVP_top60_DNA+RNA_seed | ESR1 | -2.903364 | UCEC | MFE-280 | Tamoxifen |
| MVP_top60_DNA_seed | BRAF | 0.562412 | LUAD | PC-14 | Dabrafenib | MVP_top60_DNA+RNA_seed | FGFR2 | -2.831998 | UCEC | AN3-CA | AZD4547 |
| MVP_top60_DNA_seed | EGFR | 0.562518 | LUAD | NCI-H2009 | Osimertinib | MVP_top60_DNA+RNA_seed | JAK1 | -2.681385 | UCEC | AN3-CA | AZD1480 |
| MVP_top60_DNA_seed | BRAF | 0.562526 | LUAD | NCI-H1648 | SB590885 | MVP_top60_DNA+RNA_seed | JAK1 | -2.574905 | UCEC | MFE-280 | AZD1480 |
| MVP_top60_DNA_seed | EGFR | 0.563491 | LUAD | NCI-H838 | Erlotinib | MVP_top60_DNA+RNA_seed | AKT3 | -2.540572 | UCEC | COLO-684 | GSK690693 |
| MVP_top60_DNA_seed | MET | 0.563756 | LUAD | NCI-H1975 | Crizotinib | MVP_top60_DNA+RNA_seed | AKT1 | -2.540572 | UCEC | COLO-684 | GSK690693 |
| MVP_top60_DNA_seed | EZH2 | 0.564153 | LUAD | NCI-H322M | GSK343 | MVP_top60_DNA+RNA_seed | FGFR2 | -2.286659 | UCEC | MFE-296 | AZD4547 |
| MVP_top60_DNA_seed | EZH2 | 0.564188 | LUAD | PC-3_[JPC-3] | GSK343 | MVP_top60_DNA+RNA_seed | EGFR | -2.253594 | UCEC | SNG-M | PF-00299804 |
| MVP_top60_DNA_seed | PIK3CA | 0.567814 | LUAD | HCC-827 | Alpelisib | MVP_top60_DNA+RNA_seed | ERBB2 | -2.253594 | UCEC | SNG-M | PF-00299804 |
| MVP_top60_DNA_seed | HSP90AA1 | 0.568702 | LUAD | NCI-H1435 | Tanespimycin | MVP_top60_DNA+RNA_seed | FGFR2 | -2.095604 | UCEC | MFE-280 | FGFR_3831 |
| MVP_top60_DNA_seed | AKT1 | 0.568705 | LUAD | HCC-827 | MK-2206 | MVP_top60_DNA+RNA_seed | AKT3 | -2.045663 | UCEC | AN3-CA | AKT inhibitor VIII |
| MVP_top60_DNA_seed | AURKA | 0.56972 | LUAD | NCI-H2291 | ZM447439 | MVP_top60_DNA+RNA_seed | AKT1 | -2.045663 | UCEC | AN3-CA | AKT inhibitor VIII |
| MVP_top60_DNA_seed | MDM2 | 0.571058 | LUAD | LXF-289 | Nutlin-3a (-) | MVP_top60_DNA+RNA_seed | AKT3 | -2.045604 | UCEC | MFE-296 | AKT inhibitor VIII |
| MVP_top60_DNA_seed | PIK3CA | 0.57141 | LUAD | ABC-1 | AZD8186 | MVP_top60_DNA+RNA_seed | AKT1 | -2.045604 | UCEC | MFE-296 | AKT inhibitor VIII |
| MVP_top60_DNA_seed | AKT1 | 0.574826 | LUAD | NCI-H1355 | Uprosertib | MVP_top60_DNA+RNA_seed | FGFR2 | -1.92556 | UCEC | MFE-296 | AZD4547 |
| MVP_top60_DNA_seed | EGFR | 0.576504 | LUAD | NCI-H1838 | Afatinib | MVP_top60_DNA+RNA_seed | FGFR2 | -1.813677 | UCEC | MFE-296 | PD173074 |
| MVP_top60_DNA_seed | ERBB2 | 0.576504 | LUAD | NCI-H1838 | Afatinib | MVP_top60_DNA+RNA_seed | EGFR | -1.791614 | UCEC | SNG-M | AZD8931 |

| | | | | | | | | | | | |
|---|---|---|---|---|---|---|---|---|---|---|---|
| MVP_top60_DNA_seed | SMARCA4 | 0.577417 | LUAD | EMC-BAC-2 | PFI3 | MVP_top60_DNA+RNA_seed | ERBB2 | -1.791614 | UCEC | SNG-M | AZD8931 |
| MVP_top60_DNA_seed | AURKA | 0.578513 | LUAD | NCI-H1623 | Alisertib | MVP_top60_DNA+RNA_seed | AKT3 | -1.667957 | UCEC | COLO-684 | AKT inhibitor VIII |
| MVP_top60_DNA_seed | EGFR | 0.578926 | LUAD | NCI-H2405 | Sapitinib | MVP_top60_DNA+RNA_seed | AKT1 | -1.667957 | UCEC | COLO-684 | AKT inhibitor VIII |
| MVP_top60_DNA_seed | ERBB2 | 0.578926 | LUAD | NCI-H2405 | Sapitinib | MVP_top60_DNA+RNA_seed | MDM2 | -1.665184 | UCEC | SNG-M | Nutlin-3a (-) |
| MVP_top60_DNA_seed | ERBB3 | 0.578926 | LUAD | NCI-H2405 | Sapitinib | MVP_top60_DNA+RNA_seed | EGFR | -1.6008 | UCEC | SNG-M | CI-1033 |
| MVP_top60_DNA_seed | AKT1 | 0.579046 | LUAD | NCI-H2291 | AZD5363 | MVP_top60_DNA+RNA_seed | ERBB2 | -1.6008 | UCEC | SNG-M | CI-1033 |
| MVP_top60_DNA_seed | AURKA | 0.579097 | LUAD | Calu-3 | Alisertib | MVP_top60_DNA+RNA_seed | EGFR | -1.598851 | UCEC | MFE-319 | Cetuximab |
| MVP_top60_DNA_seed | IGF1R | 0.583002 | LUAD | HCC-78 | GSK1904529A | MVP_top60_DNA+RNA_seed | AKT1 | -1.586867 | UCEC | COLO-684 | MK-2206 |
| MVP_top60_DNA_seed | EGFR | 0.583439 | LUAD | NCI-H1975 | AZD3759 | MVP_top60_DNA+RNA_seed | EGFR | -1.560385 | UCEC | SNG-M | Pelitinib |
| MVP_top60_DNA_seed | TERT | 0.583715 | LUAD | NCI-H2228 | BIBR-1532 | MVP_top60_DNA+RNA_seed | JAK1 | -1.509739 | UCEC | MFE-296 | AZD1480 |
| MVP_top60_DNA_seed | AKT1 | 0.58426 | LUAD | Calu-3 | GSK2110183B | MVP_top60_DNA+RNA_seed | PIK3CA | -1.502812 | UCEC | MFE-280 | PI3Ka_4409 |
| MVP_top60_DNA_seed | AKT1 | 0.584392 | LUAD | NCI-H1573 | Ipatasertib | MVP_top60_DNA+RNA_seed | PIK3CD | -1.502812 | UCEC | MFE-280 | PI3Ka_4409 |
| MVP_top60_DNA_seed | PIK3CA | 0.584576 | LUAD | NCI-H2087 | Alpelisib | MVP_top60_DNA+RNA_seed | HSP90AA1 | -1.450464 | UCEC | SNG-M | CCT-018159 |
| MVP_top60_DNA_seed | MAPK1 | 0.585068 | LUAD | NCI-H1573 | VX-11e | MVP_top60_DNA+RNA_seed | HSP90AB1 | -1.450464 | UCEC | SNG-M | CCT-018159 |
| MVP_top60_DNA_seed | MAPK1 | 0.5868 | LUAD | NCI-H1838 | VX-11e | MVP_top60_DNA+RNA_seed | PIK3CB | -1.407155 | UCEC | MFE-296 | AZD6482 |
| MVP_top60_DNA_seed | PIK3CA | 0.589701 | LUAD | NCI-H23 | AZD8186 | MVP_top60_DNA+RNA_seed | AKT3 | -1.406257 | UCEC | MFE-296 | BAY AKT1 |
| MVP_top60_DNA_seed | BRAF | 0.590174 | LUAD | A549 | SB590885 | MVP_top60_DNA+RNA_seed | AKT1 | -1.406257 | UCEC | MFE-296 | BAY AKT1 |
| MVP_top60_DNA_seed | MDM2 | 0.59063 | LUAD | ABC-1 | Nutlin-3a (-) | MVP_top60_DNA+RNA_seed | AKT3 | -1.390049 | UCEC | SNG-M | GSK690693 |
| MVP_top60_DNA_seed | MET | 0.590685 | LUAD | NCI-H2087 | Crizotinib | MVP_top60_DNA+RNA_seed | AKT1 | -1.390049 | UCEC | SNG-M | GSK690693 |
| MVP_top60_DNA_seed | IGF1R | 0.590867 | LUAD | NCI-H2087 | BMS-536924 | MVP_top60_DNA+RNA_seed | HSP90AA1 | -1.362448 | UCEC | SNG-M | Luminespib |
| MVP_top60_DNA_seed | FGFR1 | 0.591081 | LUAD | NCI-H3122 | PD173074 | MVP_top60_DNA+RNA_seed | HSP90AB1 | -1.362448 | UCEC | SNG-M | Luminespib |
| MVP_top60_DNA_seed | AKT1 | 0.591214 | LUAD | NCI-H358 | Uprosertib | MVP_top60_DNA+RNA_seed | PIK3CA | -1.306172 | UCEC | SNG-M | AZD8835 |
| MVP_top60_DNA_seed | PIK3CA | 0.592756 | LUAD | NCI-H358 | AZD8186 | MVP_top60_DNA+RNA_seed | PIK3CD | -1.306172 | UCEC | SNG-M | AZD8835 |
| MVP_top60_DNA_seed | EGFR | 0.592821 | LUAD | NCI-H1563 | Lapatinib | MVP_top60_DNA+RNA_seed | EGFR | -1.305625 | UCEC | AN3-CA | CI-1033 |
| MVP_top60_DNA_seed | ERBB2 | 0.592821 | LUAD | NCI-H1563 | Lapatinib | MVP_top60_DNA+RNA_seed | ERBB2 | -1.305625 | UCEC | AN3-CA | CI-1033 |
| MVP_top60_DNA_seed | HDAC1 | 0.593964 | LUAD | PC-14 | PCI-34051 | MVP_top60_DNA+RNA_seed | PIK3CB | -1.301199 | UCEC | COLO-684 | AZD6482 |
| MVP_top60_DNA_seed | BRAF | 0.595342 | LUAD | NCI-H2085 | PLX-4720 | MVP_top60_DNA+RNA_seed | ERBB2 | -1.289139 | UCEC | SNG-M | CP724714 |
| MVP_top60_DNA_seed | AURKA | 0.595626 | LUAD | PC-3_[JPC-3] | Alisertib | MVP_top60_DNA+RNA_seed | EGFR | -1.286658 | UCEC | MFE-319 | Afatinib |
| MVP_top60_DNA_seed | BRAF | 0.596116 | LUAD | Calu-6 | SB590885 | MVP_top60_DNA+RNA_seed | ERBB2 | -1.286658 | UCEC | MFE-319 | Afatinib |
| MVP_top60_DNA_seed | EGFR | 0.596338 | LUAD | NCI-H2291 | Lapatinib | MVP_top60_DNA+RNA_seed | HSP90AA1 | -1.280748 | UCEC | AN3-CA | Tanespimycin |
| MVP_top60_DNA_seed | ERBB2 | 0.596338 | LUAD | NCI-H2291 | Lapatinib | MVP_top60_DNA+RNA_seed | HSP90AB1 | -1.280748 | UCEC | AN3-CA | Tanespimycin |
| MVP_top60_DNA_seed | EZH2 | 0.596818 | LUAD | NCI-H1975 | GSK343 | MVP_top60_DNA+RNA_seed | HSP90AA1 | -1.274147 | UCEC | HEC-1 | Tanespimycin |
| MVP_top60_DNA_seed | BRAF | 0.597619 | LUAD | NCI-H2085 | Dabrafenib | MVP_top60_DNA+RNA_seed | HSP90AB1 | -1.274147 | UCEC | HEC-1 | Tanespimycin |
| MVP_top60_DNA_seed | MET | 0.598235 | LUAD | EMC-BAC-1 | Savolitinib | MVP_top60_DNA+RNA_seed | AKT3 | -1.264695 | UCEC | MFE-296 | GSK690693 |
| MVP_top60_DNA_seed | IGF1R | 0.599332 | LUAD | HCC-78 | BMS-754807 | MVP_top60_DNA+RNA_seed | AKT1 | -1.264695 | UCEC | MFE-296 | GSK690693 |
| MVP_top60_DNA_seed | EGFR | 0.599903 | LUAD | VMRC-LCD | Afatinib | MVP_top60_DNA+RNA_seed | AKT3 | -1.262466 | UCEC | AN3-CA | BAY AKT1 |
| MVP_top60_DNA_seed | ERBB2 | 0.599903 | LUAD | VMRC-LCD | Afatinib | MVP_top60_DNA+RNA_seed | AKT1 | -1.262466 | UCEC | AN3-CA | BAY AKT1 |
| MVP_top60_DNA_seed | EGFR | 0.600094 | LUAD | NCI-H1573 | AZD3759 | MVP_top60_DNA+RNA_seed | AKT3 | -1.253918 | UCEC | COLO-684 | A-443654 |
| MVP_top60_DNA_seed | MDM2 | 0.600156 | LUAD | NCI-H2085 | Nutlin-3a (-) | MVP_top60_DNA+RNA_seed | AKT1 | -1.253918 | UCEC | COLO-684 | A-443654 |
| MVP_top60_DNA_seed | TP53 | 0.60034 | LUAD | NCI-H1755 | MIRA-1 | MVP_top60_DNA+RNA_seed | AR | -1.242198 | UCEC | SNG-M | AZD3514 |
| MVP_top60_DNA_seed | BRAF | 0.601306 | LUAD | H3255 | PLX-4720 | MVP_top60_DNA+RNA_seed | EGFR | -1.206584 | UCEC | SNG-M | AST-1306 |

| | | | | | | | | | | | |
|---|---|---|---|---|---|---|---|---|---|---|---|
| MVP_top60_DNA_seed | MAPK1 | 0.601745 | LUAD | HCC-78 | Ulixertinib | MVP_top60_DNA+RNA_seed | PIK3CA | -1.182536 | UCEC | MFE-280 | AZD8835 |
| MVP_top60_DNA_seed | BRAF | 0.602121 | LUAD | NCI-H1355 | PLX-4720 | MVP_top60_DNA+RNA_seed | PIK3CD | -1.182536 | UCEC | MFE-280 | AZD8835 |
| MVP_top60_DNA_seed | MDM2 | 0.602212 | LUAD | Calu-6 | Serdemetan | MVP_top60_DNA+RNA_seed | EGFR | -1.180624 | UCEC | MFE-319 | Gefitinib |
| MVP_top60_DNA_seed | AURKA | 0.602494 | LUAD | NCI-H2030 | Alisertib | MVP_top60_DNA+RNA_seed | HSP90AA1 | -1.165754 | UCEC | AN3-CA | Luminespib |
| MVP_top60_DNA_seed | EGFR | 0.602594 | LUAD | HCC-44 | Gefitinib | MVP_top60_DNA+RNA_seed | HSP90AB1 | -1.165754 | UCEC | AN3-CA | Luminespib |
| MVP_top60_DNA_seed | MAPK1 | 0.603471 | LUAD | NCI-H1435 | VX-11e | MVP_top60_DNA+RNA_seed | AKT1 | -1.155082 | UCEC | MFE-296 | MK-2206 |
| MVP_top60_DNA_seed | FGFR1 | 0.605977 | LUAD | NCI-H2291 | PD173074 | MVP_top60_DNA+RNA_seed | SRC | -1.15495 | UCEC | KLE | WH-4-023 |
| MVP_top60_DNA_seed | PIK3CA | 0.606174 | LUAD | HCC-827 | Buparlisib | MVP_top60_DNA+RNA_seed | FGFR2 | -1.154504 | UCEC | MFE-280 | PD173074 |
| MVP_top60_DNA_seed | HDAC1 | 0.609312 | LUAD | H3255 | Romidepsin | MVP_top60_DNA+RNA_seed | ATM | -1.138256 | UCEC | MFE-319 | KU-55933 |
| MVP_top60_DNA_seed | MDM2 | 0.61031 | LUAD | PC-14 | Nutlin-3a (-) | MVP_top60_DNA+RNA_seed | AKT3 | -1.136416 | UCEC | MFE-280 | GSK690693 |
| MVP_top60_DNA_seed | AKT1 | 0.610492 | LUAD | NCI-H522 | Uprosertib | MVP_top60_DNA+RNA_seed | AKT1 | -1.136416 | UCEC | MFE-280 | GSK690693 |
| MVP_top60_DNA_seed | PIK3CA | 0.611025 | LUAD | NCI-H1781 | Alpelisib | MVP_top60_DNA+RNA_seed | PIK3CD | -1.124603 | UCEC | COLO-684 | Idelalisib |
| MVP_top60_DNA_seed | MAPK1 | 0.611508 | LUAD | RERF-LC-KJ | Ulixertinib | MVP_top60_DNA+RNA_seed | MAPK1 | -1.080554 | UCEC | SNG-M | VX-11e |
| MVP_top60_DNA_seed | AKT1 | 0.611646 | LUAD | NCI-H2030 | Uprosertib | MVP_top60_DNA+RNA_seed | MDM2 | -1.071433 | UCEC | MFE-296 | Nutlin-3a (-) |
| MVP_top60_DNA_seed | AKT1 | 0.615052 | LUAD | NCI-H1355 | MK-2206 | MVP_top60_DNA+RNA_seed | PIK3CA | -1.065015 | UCEC | SNG-M | PI3Ka_4409 |
| MVP_top60_DNA_seed | CREBBP | 0.617467 | LUAD | ABC-1 | SGC-CBP30 | MVP_top60_DNA+RNA_seed | PIK3CD | -1.065015 | UCEC | SNG-M | PI3Ka_4409 |
| MVP_top60_DNA_seed | IGF1R | 0.618164 | LUAD | NCI-H1573 | BMS-754807 | MVP_top60_DNA+RNA_seed | EGFR | -1.062848 | UCEC | KLE | AST-1306 |
| MVP_top60_DNA_seed | MAPK1 | 0.618194 | LUAD | NCI-H1693 | Ulixertinib | MVP_top60_DNA+RNA_seed | AR | -1.050465 | UCEC | MFE-319 | AZD3514 |
| MVP_top60_DNA_seed | BRAF | 0.620136 | LUAD | NCI-H2009 | SB590885 | MVP_top60_DNA+RNA_seed | EGFR | -1.038138 | UCEC | MFE-319 | AZD8931 |
| MVP_top60_DNA_seed | AKT1 | 0.620396 | LUAD | NCI-H1563 | AT13148 | MVP_top60_DNA+RNA_seed | ERBB2 | -1.038138 | UCEC | MFE-319 | AZD8931 |
| MVP_top60_DNA_seed | SMARCA4 | 0.623069 | LUAD | VMRC-LCD | PFI3 | MVP_top60_DNA+RNA_seed | FGFR2 | -1.007139 | UCEC | AN3-CA | FGFR_3831 |
| MVP_top60_DNA_seed | BRAF | 0.624075 | LUAD | VMRC-LCD | PLX-4720 | MVP_top60_DNA+RNA_seed | ATM | -0.98435 | UCEC | SNG-M | CP466722 |
| MVP_top60_DNA_seed | HDAC1 | 0.62518 | LUAD | NCI-H1355 | Romidepsin | MVP_top60_DNA+RNA_seed | ATM | -0.967779 | UCEC | MFE-319 | KU-60019 |
| MVP_top60_DNA_seed | TERT | 0.626208 | LUAD | NCI-H1563 | BIBR-1532 | MVP_top60_DNA+RNA_seed | EGFR | -0.965278 | UCEC | MFE-280 | AST-1306 |
| MVP_top60_DNA_seed | HSP90AA1 | 0.626784 | LUAD | H3255 | Tanespimycin | MVP_top60_DNA+RNA_seed | HSP90AA1 | -0.942022 | UCEC | HEC-1 | Elesclomol |
| MVP_top60_DNA_seed | IGF1R | 0.628832 | LUAD | NCI-H1650 | Linsitinib | MVP_top60_DNA+RNA_seed | HSP90AB1 | -0.942022 | UCEC | HEC-1 | Elesclomol |
| MVP_top60_DNA_seed | GSK3B | 0.629259 | LUAD | NCI-H2009 | SB216763 | MVP_top60_DNA+RNA_seed | ATM | -0.941115 | UCEC | MFE-319 | CP466722 |
| MVP_top60_DNA_seed | AKT1 | 0.629469 | LUAD | H3255 | Uprosertib | MVP_top60_DNA+RNA_seed | AKT1 | -0.938485 | UCEC | MFE-280 | MK-2206 |
| MVP_top60_DNA_seed | HDAC1 | 0.63174 | LUAD | H3255 | Entinostat | MVP_top60_DNA+RNA_seed | SRC | -0.932293 | UCEC | SNG-M | NVP-BHG712 |
| MVP_top60_DNA_seed | PIK3CA | 0.631896 | LUAD | NCI-H358 | GNE-317 | MVP_top60_DNA+RNA_seed | EGFR | -0.922846 | UCEC | MFE-280 | CI-1033 |
| MVP_top60_DNA_seed | MET | 0.632568 | LUAD | NCI-H1651 | Crizotinib | MVP_top60_DNA+RNA_seed | ERBB2 | -0.922846 | UCEC | MFE-280 | CI-1033 |
| MVP_top60_DNA_seed | MAPK1 | 0.634823 | LUAD | HOP-62 | Ulixertinib | MVP_top60_DNA+RNA_seed | ESR1 | -0.92168 | UCEC | AN3-CA | Tamoxifen |
| MVP_top60_DNA_seed | AKT1 | 0.634998 | LUAD | NCI-H1792 | AT13148 | MVP_top60_DNA+RNA_seed | AKT1 | -0.911742 | UCEC | AN3-CA | MK-2206 |
| MVP_top60_DNA_seed | AURKA | 0.635389 | LUAD | NCI-H1651 | ZM447439 | MVP_top60_DNA+RNA_seed | EGFR | -0.904027 | UCEC | MFE-319 | Afatinib |
| MVP_top60_DNA_seed | MDM2 | 0.635966 | LUAD | NCI-H2023 | Serdemetan | MVP_top60_DNA+RNA_seed | ERBB2 | -0.904027 | UCEC | MFE-319 | Afatinib |
| MVP_top60_DNA_seed | MAPK1 | 0.6362 | LUAD | NCI-H1355 | VX-11e | MVP_top60_DNA+RNA_seed | AKT3 | -0.883088 | UCEC | MFE-319 | AKT inhibitor VIII |
| MVP_top60_DNA_seed | FGFR1 | 0.636977 | LUAD | NCI-H1437 | AZD4547 | MVP_top60_DNA+RNA_seed | AKT1 | -0.883088 | UCEC | MFE-319 | AKT inhibitor VIII |
| MVP_top60_DNA_seed | EGFR | 0.637535 | LUAD | NCI-H1437 | Osimertinib | MVP_top60_DNA+RNA_seed | HSP90AA1 | -0.867371 | UCEC | COLO-684 | CCT-018159 |
| MVP_top60_DNA_seed | MET | 0.637689 | LUAD | NCI-H1435 | Crizotinib | MVP_top60_DNA+RNA_seed | HSP90AB1 | -0.867371 | UCEC | COLO-684 | CCT-018159 |
| MVP_top60_DNA_seed | AKT1 | 0.638026 | LUAD | ABC-1 | AZD5363 | MVP_top60_DNA+RNA_seed | ESR1 | -0.866579 | UCEC | SNG-M | Fulvestrant |
| MVP_top60_DNA_seed | KRAS | 0.638042 | LUAD | NCI-H522 | KRAS (G12C) Inhibitor-1: | MVP_top60_DNA+RNA_seed | AKT3 | -0.863762 | UCEC | COLO-684 | BAY AKT1 |

| | | | | | | | | | | | |
|---|---|---|---|---|---|---|---|---|---|---|---|
| MVP_top60_DNA_seed | KRAS | 0.63857 | LUAD | NCI-H2087 | KRAS (G12C) Inhibitor-12 | MVP_top60_DNA+RNA_seed | AKT1 | -0.863762 | UCEC | COLO-684 | BAY AKT1 |
| MVP_top60_DNA_seed | CREBBP | 0.638787 | LUAD | LXF-289 | SGC-CBP30 | MVP_top60_DNA+RNA_seed | PIK3CB | -0.861903 | UCEC | KLE | AZD8186 |
| MVP_top60_DNA_seed | MAPK1 | 0.638902 | LUAD | NCI-H2085 | SCH772984 | MVP_top60_DNA+RNA_seed | PIK3CD | -0.861903 | UCEC | KLE | AZD8186 |
| MVP_top60_DNA_seed | MDM2 | 0.640825 | LUAD | NCI-H650 | Nutlin-3a (-) | MVP_top60_DNA+RNA_seed | EGFR | -0.856102 | UCEC | HEC-1 | Cetuximab |
| MVP_top60_DNA_seed | AKT1 | 0.640863 | LUAD | NCI-H2291 | Uprosertib | MVP_top60_DNA+RNA_seed | PIK3CB | -0.856077 | UCEC | COLO-684 | AZD8186 |
| MVP_top60_DNA_seed | ATM | 0.641153 | LUAD | EMC-BAC-1 | KU-55933 | MVP_top60_DNA+RNA_seed | PIK3CD | -0.856077 | UCEC | COLO-684 | AZD8186 |
| MVP_top60_DNA_seed | EGFR | 0.644414 | LUAD | HCC-44 | Osimertinib | MVP_top60_DNA+RNA_seed | FGFR2 | -0.837374 | UCEC | MFE-296 | FGFR_3831 |
| MVP_top60_DNA_seed | PIK3CA | 0.644716 | LUAD | NCI-H1651 | Alpelisib | MVP_top60_DNA+RNA_seed | PIK3CD | -0.829826 | UCEC | COLO-684 | IC-87114 |
| MVP_top60_DNA_seed | CREBBP | 0.646789 | LUAD | NCI-H1568 | SGC-CBP30 | MVP_top60_DNA+RNA_seed | HSP90AA1 | -0.820435 | UCEC | COLO-684 | Elesclomol |
| MVP_top60_DNA_seed | BRAF | 0.647963 | LUAD | NCI-H1623 | Dabrafenib | MVP_top60_DNA+RNA_seed | HSP90AB1 | -0.820435 | UCEC | COLO-684 | Elesclomol |
| MVP_top60_DNA_seed | MAPK1 | 0.648519 | LUAD | NCI-H1623 | Ulixertinib | MVP_top60_DNA+RNA_seed | JAK1 | -0.801292 | UCEC | SNG-M | AZD1480 |
| MVP_top60_DNA_seed | AKT1 | 0.648916 | LUAD | NCI-H1838 | Ipatasertib | MVP_top60_DNA+RNA_seed | HSP90AA1 | -0.797967 | UCEC | AN3-CA | Elesclomol |
| MVP_top60_DNA_seed | MAPK1 | 0.649072 | LUAD | NCI-H1568 | Ulixertinib | MVP_top60_DNA+RNA_seed | HSP90AB1 | -0.797967 | UCEC | AN3-CA | Elesclomol |
| MVP_top60_DNA_seed | AKT1 | 0.649513 | LUAD | NCI-H1355 | Uprosertib | MVP_top60_DNA+RNA_seed | ERBB2 | -0.796951 | UCEC | MFE-280 | CP724714 |
| MVP_top60_DNA_seed | IGF1R | 0.650311 | LUAD | NCI-H2405 | BMS-536924 | MVP_top60_DNA+RNA_seed | ESR1 | -0.794746 | UCEC | SNG-M | Tamoxifen |
| MVP_top60_DNA_seed | EGFR | 0.650505 | LUAD | NCI-H2228 | Erlotinib | MVP_top60_DNA+RNA_seed | HSP90AA1 | -0.794576 | UCEC | SNG-M | SNX-2112 |
| MVP_top60_DNA_seed | AKT1 | 0.65206 | LUAD | COR-L105 | AZD5363 | MVP_top60_DNA+RNA_seed | HSP90AB1 | -0.794576 | UCEC | SNG-M | SNX-2112 |
| MVP_top60_DNA_seed | EZH2 | 0.652435 | LUAD | A549 | GSK343 | MVP_top60_DNA+RNA_seed | HSP90AA1 | -0.792124 | UCEC | MFE-296 | Tanespimycin |
| MVP_top60_DNA_seed | ATM | 0.653319 | LUAD | HOP-62 | KU-55933 | MVP_top60_DNA+RNA_seed | HSP90AB1 | -0.792124 | UCEC | MFE-296 | Tanespimycin |
| MVP_top60_DNA_seed | AKT1 | 0.653655 | LUAD | NCI-H1793 | AT13148 | MVP_top60_DNA+RNA_seed | AR | -0.786412 | UCEC | SNG-M | Bicalutamide |
| MVP_top60_DNA_seed | PIK3CA | 0.654089 | LUAD | NCI-H650 | Alpelisib | MVP_top60_DNA+RNA_seed | AKT3 | -0.778096 | UCEC | KLE | AKT inhibitor VIII |
| MVP_top60_DNA_seed | EGFR | 0.65424 | LUAD | NCI-H1993 | Gefitinib | MVP_top60_DNA+RNA_seed | AKT1 | -0.778096 | UCEC | KLE | AKT inhibitor VIII |
| MVP_top60_DNA_seed | AKT1 | 0.654981 | LUAD | NCI-H1623 | GSK2110183B | MVP_top60_DNA+RNA_seed | ATM | -0.757481 | UCEC | HEC-1 | KU-60019 |
| MVP_top60_DNA_seed | PIK3CA | 0.65543 | LUAD | NCI-H1650 | Buparlisib | MVP_top60_DNA+RNA_seed | AKT3 | -0.750188 | UCEC | HEC-1 | AKT inhibitor VIII |
| MVP_top60_DNA_seed | AURKA | 0.655655 | LUAD | NCI-H2030 | Tozasertib | MVP_top60_DNA+RNA_seed | AKT1 | -0.750188 | UCEC | HEC-1 | AKT inhibitor VIII |
| MVP_top60_DNA_seed | BRAF | 0.6565 | LUAD | RERF-LC-KJ | Dabrafenib | MVP_top60_DNA+RNA_seed | PIK3CB | -0.746986 | UCEC | AN3-CA | AZD8186 |
| MVP_top60_DNA_seed | CREBBP | 0.657911 | LUAD | Calu-6 | SGC-CBP30 | MVP_top60_DNA+RNA_seed | PIK3CD | -0.746986 | UCEC | AN3-CA | AZD8186 |
| MVP_top60_DNA_seed | MET | 0.657986 | LUAD | HCC-827 | Crizotinib | MVP_top60_DNA+RNA_seed | EGFR | -0.743217 | UCEC | MFE-280 | PF-00299804 |
| MVP_top60_DNA_seed | FGFR1 | 0.658951 | LUAD | NCI-H2405 | AZD4547 | MVP_top60_DNA+RNA_seed | ERBB2 | -0.743217 | UCEC | MFE-280 | PF-00299804 |
| MVP_top60_DNA_seed | AKT1 | 0.659749 | LUAD | NCI-H650 | Uprosertib | MVP_top60_DNA+RNA_seed | AR | -0.729614 | UCEC | HEC-1 | AZD3514 |
| MVP_top60_DNA_seed | HDAC1 | 0.660331 | LUAD | NCI-H358 | Entinostat | MVP_top60_DNA+RNA_seed | AR | -0.721671 | UCEC | MFE-319 | Bicalutamide |
| MVP_top60_DNA_seed | HSP90AA1 | 0.661104 | LUAD | NCI-H1568 | Tanespimycin | MVP_top60_DNA+RNA_seed | PIK3CB | -0.702514 | UCEC | AN3-CA | AZD6482 |
| MVP_top60_DNA_seed | MDM2 | 0.66126 | LUAD | NCI-H322M | Nutlin-3a (-) | MVP_top60_DNA+RNA_seed | SRC | -0.697529 | UCEC | SNG-M | Bosutinib |
| MVP_top60_DNA_seed | IGF1R | 0.661587 | LUAD | PC-3_[JPC-3] | NVP-ADW742 | MVP_top60_DNA+RNA_seed | PIK3CA | -0.695396 | UCEC | AN3-CA | AZD8835 |
| MVP_top60_DNA_seed | HSP90AA1 | 0.662792 | LUAD | NCI-H1666 | Luminespib | MVP_top60_DNA+RNA_seed | PIK3CD | -0.695396 | UCEC | AN3-CA | AZD8835 |
| MVP_top60_DNA_seed | MET | 0.662799 | LUAD | NCI-H322M | Crizotinib | MVP_top60_DNA+RNA_seed | AKT3 | -0.694469 | UCEC | KLE | A-443654 |
| MVP_top60_DNA_seed | MET | 0.662852 | LUAD | COR-L105 | Crizotinib | MVP_top60_DNA+RNA_seed | AKT1 | -0.694469 | UCEC | KLE | A-443654 |
| MVP_top60_DNA_seed | KRAS | 0.664545 | LUAD | NCI-H1944 | KRAS (G12C) Inhibitor-12 | MVP_top60_DNA+RNA_seed | AKT3 | -0.686103 | UCEC | MFE-280 | AKT inhibitor VIII |
| MVP_top60_DNA_seed | PIK3CA | 0.66601 | LUAD | NCI-H1651 | GNE-317 | MVP_top60_DNA+RNA_seed | AKT1 | -0.686103 | UCEC | MFE-280 | AKT inhibitor VIII |
| MVP_top60_DNA_seed | AKT1 | 0.666867 | LUAD | NCI-H1651 | GSK2110183B | MVP_top60_DNA+RNA_seed | EP300 | -0.666443 | UCEC | SNG-M | I-CBP112 |
| MVP_top60_DNA_seed | SRC | 0.667232 | LUAD | NCI-H1693 | Bosutinib | MVP_top60_DNA+RNA_seed | PIK3CB | -0.659503 | UCEC | SNG-M | AZD8186 |

| | | | | | | | | | | | |
|---|---|---|---|---|---|---|---|---|---|---|---|
| MVP_top60_DNA_seed | FGFR1 | 0.667531 | LUAD | NCI-H1666 | AZD4547 | MVP_top60_DNA+RNA_seed | PIK3CD | -0.659503 | UCEC | SNG-M | AZD8186 |
| MVP_top60_DNA_seed | HDAC1 | 0.668035 | LUAD | HOP-62 | Entinostat | MVP_top60_DNA+RNA_seed | EGFR | -0.650466 | UCEC | KLE | Pelitinib |
| MVP_top60_DNA_seed | ATM | 0.670889 | LUAD | COR-L105 | KU-55933 | MVP_top60_DNA+RNA_seed | HSP90AA1 | -0.6457 | UCEC | COLO-684 | SNX-2112 |
| MVP_top60_DNA_seed | MDM2 | 0.671376 | LUAD | NCI-H3122 | Nutlin-3a (-) | MVP_top60_DNA+RNA_seed | HSP90AB1 | -0.6457 | UCEC | COLO-684 | SNX-2112 |
| MVP_top60_DNA_seed | AKT1 | 0.67156 | LUAD | Calu-6 | MK-2206 | MVP_top60_DNA+RNA_seed | HDAC1 | -0.638229 | UCEC | SNG-M | JQ12 |
| MVP_top60_DNA_seed | BRAF | 0.671589 | LUAD | NCI-H1623 | SB590885 | MVP_top60_DNA+RNA_seed | HDAC2 | -0.638229 | UCEC | SNG-M | JQ12 |
| MVP_top60_DNA_seed | AURKA | 0.672937 | LUAD | NCI-H23 | ZM447439 | MVP_top60_DNA+RNA_seed | MAPK1 | -0.625057 | UCEC | KLE | FR-180204 |
| MVP_top60_DNA_seed | EGFR | 0.673907 | LUAD | VMRC-LCD | Gefitinib | MVP_top60_DNA+RNA_seed | MAPK3 | -0.625057 | UCEC | KLE | FR-180204 |
| MVP_top60_DNA_seed | EZH2 | 0.674217 | LUAD | PC-14 | GSK343 | MVP_top60_DNA+RNA_seed | EGFR | -0.622425 | UCEC | MFE-280 | CUDC-101 |
| MVP_top60_DNA_seed | EGFR | 0.674894 | LUAD | NCI-H23 | AZD3759 | MVP_top60_DNA+RNA_seed | ERBB2 | -0.622425 | UCEC | MFE-280 | CUDC-101 |
| MVP_top60_DNA_seed | EZH2 | 0.677033 | LUAD | VMRC-LCD | GSK343 | MVP_top60_DNA+RNA_seed | EGFR | -0.622161 | UCEC | MFE-280 | Pelitinib |
| MVP_top60_DNA_seed | SRC | 0.677186 | LUAD | NCI-H1623 | Bosutinib | MVP_top60_DNA+RNA_seed | ATM | -0.608482 | UCEC | COLO-684 | KU-55933 |
| MVP_top60_DNA_seed | AKT1 | 0.678233 | LUAD | NCI-H650 | GSK2110183B | MVP_top60_DNA+RNA_seed | HSP90AA1 | -0.598157 | UCEC | HEC-1 | CCT-018159 |
| MVP_top60_DNA_seed | FGFR1 | 0.678941 | LUAD | NCI-H1666 | PD173074 | MVP_top60_DNA+RNA_seed | HSP90AB1 | -0.598157 | UCEC | HEC-1 | CCT-018159 |
| MVP_top60_DNA_seed | AKT1 | 0.68001 | LUAD | NCI-H358 | GSK2110183B | MVP_top60_DNA+RNA_seed | HSP90AA1 | -0.59084 | UCEC | MFE-296 | Luminespib |
| MVP_top60_DNA_seed | EGFR | 0.6815 | LUAD | Calu-6 | Sapitinib | MVP_top60_DNA+RNA_seed | HSP90AB1 | -0.59084 | UCEC | MFE-296 | Luminespib |
| MVP_top60_DNA_seed | ERBB2 | 0.6815 | LUAD | Calu-6 | Sapitinib | MVP_top60_DNA+RNA_seed | ATM | -0.589341 | UCEC | MFE-296 | KU-60019 |
| MVP_top60_DNA_seed | ERBB3 | 0.6815 | LUAD | Calu-6 | Sapitinib | MVP_top60_DNA+RNA_seed | HSP90AA1 | -0.582916 | UCEC | AN3-CA | CCT-018159 |
| MVP_top60_DNA_seed | MAPK1 | 0.681793 | LUAD | COR-L105 | Ulixertinib | MVP_top60_DNA+RNA_seed | HSP90AB1 | -0.582916 | UCEC | AN3-CA | CCT-018159 |
| MVP_top60_DNA_seed | AURKA | 0.683112 | LUAD | NCI-H1838 | Alisertib | MVP_top60_DNA+RNA_seed | JAK1 | -0.566585 | UCEC | SNG-M | Ruxolitinib |
| MVP_top60_DNA_seed | EGFR | 0.684061 | LUAD | ABC-1 | Sapitinib | MVP_top60_DNA+RNA_seed | ESR1 | -0.559523 | UCEC | MFE-319 | Tamoxifen |
| MVP_top60_DNA_seed | ERBB2 | 0.684061 | LUAD | ABC-1 | Sapitinib | MVP_top60_DNA+RNA_seed | AR | -0.558596 | UCEC | AN3-CA | Bicalutamide |
| MVP_top60_DNA_seed | ERBB3 | 0.684061 | LUAD | ABC-1 | Sapitinib | MVP_top60_DNA+RNA_seed | PIK3CD | -0.555003 | UCEC | MFE-280 | Idelalisib |
| MVP_top60_DNA_seed | EGFR | 0.684482 | LUAD | NCI-H650 | Erlotinib | MVP_top60_DNA+RNA_seed | JAK1 | -0.550619 | UCEC | SNG-M | JAK1_3715 |
| MVP_top60_DNA_seed | EZH2 | 0.684909 | LUAD | NCI-H1568 | GSK343 | MVP_top60_DNA+RNA_seed | ATM | -0.540231 | UCEC | AN3-CA | KU-55933 |
| MVP_top60_DNA_seed | IGF1R | 0.685186 | LUAD | NCI-H650 | BMS-754807 | MVP_top60_DNA+RNA_seed | HDAC1 | -0.53721 | UCEC | SNG-M | Dacinostat |
| MVP_top60_DNA_seed | MAPK1 | 0.685684 | LUAD | NCI-H1793 | Ulixertinib | MVP_top60_DNA+RNA_seed | ATM | -0.532342 | UCEC | COLO-684 | CP466722 |
| MVP_top60_DNA_seed | MAPK1 | 0.685857 | LUAD | NCI-H1435 | SCH772984 | MVP_top60_DNA+RNA_seed | AR | -0.529467 | UCEC | HEC-1 | Bicalutamide |
| MVP_top60_DNA_seed | BRAF | 0.686657 | LUAD | NCI-H1666 | Dabrafenib | MVP_top60_DNA+RNA_seed | AKT1 | -0.525196 | UCEC | EN | MK-2206 |
| MVP_top60_DNA_seed | GSK3B | 0.687864 | LUAD | NCI-H2228 | SB216763 | MVP_top60_DNA+RNA_seed | HSP90AA1 | -0.523921 | UCEC | EN | Elesclomol |
| MVP_top60_DNA_seed | EGFR | 0.687891 | LUAD | NCI-H1755 | AZD3759 | MVP_top60_DNA+RNA_seed | HSP90AB1 | -0.523921 | UCEC | EN | Elesclomol |
| MVP_top60_DNA_seed | AKT1 | 0.688797 | LUAD | NCI-H1435 | Uprosertib | MVP_top60_DNA+RNA_seed | ESR1 | -0.517665 | UCEC | HEC-1 | Fulvestrant |
| MVP_top60_DNA_seed | FGFR1 | 0.689126 | LUAD | NCI-H1793 | AZD4547 | MVP_top60_DNA+RNA_seed | PIK3CB | -0.517357 | UCEC | MFE-280 | AZD8186 |
| MVP_top60_DNA_seed | TP53 | 0.690013 | LUAD | NCI-H1944 | MIRA-1 | MVP_top60_DNA+RNA_seed | PIK3CD | -0.517357 | UCEC | MFE-280 | AZD8186 |
| MVP_top60_DNA_seed | BRAF | 0.690759 | LUAD | SK-LU-1 | Dabrafenib | MVP_top60_DNA+RNA_seed | HDAC1 | -0.517153 | UCEC | AN3-CA | Dacinostat |
| MVP_top60_DNA_seed | IGF1R | 0.691104 | LUAD | HCC-827 | NVP-ADW742 | MVP_top60_DNA+RNA_seed | HSP90AA1 | -0.510537 | UCEC | MFE-319 | Luminespib |
| MVP_top60_DNA_seed | BRAF | 0.691841 | LUAD | NCI-H2228 | Dabrafenib | MVP_top60_DNA+RNA_seed | HSP90AB1 | -0.510537 | UCEC | MFE-319 | Luminespib |
| MVP_top60_DNA_seed | FGFR1 | 0.693674 | LUAD | NCI-H2085 | PD173074 | MVP_top60_DNA+RNA_seed | AKT3 | -0.505071 | UCEC | AN3-CA | GSK690693 |
| MVP_top60_DNA_seed | AKT1 | 0.694874 | LUAD | NCI-H2342 | Afuresertib | MVP_top60_DNA+RNA_seed | AKT1 | -0.505071 | UCEC | AN3-CA | GSK690693 |
| MVP_top60_DNA_seed | AKT1 | 0.695611 | LUAD | ABC-1 | Uprosertib | MVP_top60_DNA+RNA_seed | AKT3 | -0.483919 | UCEC | MFE-319 | AKT inhibitor VIII |
| MVP_top60_DNA_seed | SMARCA4 | 0.695795 | LUAD | NCI-H358 | PFI3 | MVP_top60_DNA+RNA_seed | AKT1 | -0.483919 | UCEC | MFE-319 | AKT inhibitor VIII |

| | | | | | | | | | | | |
|---|---|---|---|---|---|---|---|---|---|---|---|
| MVP_top60_DNA_seed | AKT1 | 0.696011 | LUAD | NCI-H650 | AT13148 | MVP_top60_DNA+RNA_seed | EGFR | -0.482508 | UCEC | COLO-684 | Gefitinib |
| MVP_top60_DNA_seed | MAPK1 | 0.696321 | LUAD | NCI-H2347 | Ulixertinib | MVP_top60_DNA+RNA_seed | EGFR | -0.473667 | UCEC | SNG-M | Afatinib |
| MVP_top60_DNA_seed | HDAC1 | 0.697345 | LUAD | ABC-1 | Entinostat | MVP_top60_DNA+RNA_seed | ERBB2 | -0.473667 | UCEC | SNG-M | Afatinib |
| MVP_top60_DNA_seed | MAPK1 | 0.69736 | LUAD | NCI-H522 | Ulixertinib | MVP_top60_DNA+RNA_seed | AKT3 | -0.469321 | UCEC | EN | GSK690693 |
| MVP_top60_DNA_seed | GSK3B | 0.698437 | LUAD | NCI-H1563 | SB216763 | MVP_top60_DNA+RNA_seed | AKT1 | -0.469321 | UCEC | EN | GSK690693 |
| MVP_top60_DNA_seed | HDAC1 | 0.698933 | LUAD | NCI-H2085 | Entinostat | MVP_top60_DNA+RNA_seed | HSP90AA1 | -0.466099 | UCEC | MFE-319 | Tanespimycin |
| MVP_top60_DNA_seed | AKT1 | 0.69903 | LUAD | EKVX | Afuresertib | MVP_top60_DNA+RNA_seed | HSP90AB1 | -0.466099 | UCEC | MFE-319 | Tanespimycin |
| MVP_top60_DNA_seed | FGFR1 | 0.702511 | LUAD | NCI-H1734 | AZD4547 | MVP_top60_DNA+RNA_seed | SRC | -0.464 | UCEC | KLE | Saracatinib |
| MVP_top60_DNA_seed | MET | 0.702604 | LUAD | HOP-62 | Savolitinib | MVP_top60_DNA+RNA_seed | ATM | -0.458963 | UCEC | AN3-CA | KU-60019 |
| MVP_top60_DNA_seed | HSP90AA1 | 0.704094 | LUAD | EKVX | Elesclomol | MVP_top60_DNA+RNA_seed | EP300 | -0.453855 | UCEC | KLE | I-CBP112 |
| MVP_top60_DNA_seed | AKT1 | 0.704306 | LUAD | HCC-827 | Ipatasertib | MVP_top60_DNA+RNA_seed | MDM2 | -0.442765 | UCEC | HEC-1 | Nutlin-3a (-) |
| MVP_top60_DNA_seed | AKT1 | 0.705578 | LUAD | NCI-H23 | Uprosertib | MVP_top60_DNA+RNA_seed | HDAC1 | -0.425759 | UCEC | COLO-684 | CAY10603 |
| MVP_top60_DNA_seed | IGF1R | 0.706995 | LUAD | EMC-BAC-1 | GSK1904529A | MVP_top60_DNA+RNA_seed | EGFR | -0.416391 | UCEC | MFE-296 | AZD8931 |
| MVP_top60_DNA_seed | AKT1 | 0.707735 | LUAD | NCI-H2085 | AZD5363 | MVP_top60_DNA+RNA_seed | ERBB2 | -0.416391 | UCEC | MFE-296 | AZD8931 |
| MVP_top60_DNA_seed | SRC | 0.707831 | LUAD | NCI-H2087 | Bosutinib | MVP_top60_DNA+RNA_seed | EGFR | -0.413687 | UCEC | HEC-1 | Afatinib |
| MVP_top60_DNA_seed | MAPK1 | 0.708247 | LUAD | NCI-H1651 | Ulixertinib | MVP_top60_DNA+RNA_seed | ERBB2 | -0.413687 | UCEC | HEC-1 | Afatinib |
| MVP_top60_DNA_seed | MAPK1 | 0.709469 | LUAD | NCI-H2228 | VX-11e | MVP_top60_DNA+RNA_seed | PIK3CB | -0.405042 | UCEC | HEC-1 | AZD6482 |
| MVP_top60_DNA_seed | AKT1 | 0.709474 | LUAD | NCI-H1666 | Ipatasertib | MVP_top60_DNA+RNA_seed | AKT3 | -0.400349 | UCEC | KLE | BAY AKT1 |
| MVP_top60_DNA_seed | ATM | 0.710379 | LUAD | NCI-H2228 | KU-55933 | MVP_top60_DNA+RNA_seed | AKT1 | -0.400349 | UCEC | KLE | BAY AKT1 |
| MVP_top60_DNA_seed | MAPK1 | 0.710781 | LUAD | EKVX | Ulixertinib | MVP_top60_DNA+RNA_seed | HSP90AA1 | -0.397375 | UCEC | MFE-296 | Elesclomol |
| MVP_top60_DNA_seed | MDM2 | 0.711229 | LUAD | VMRC-LCD | Serdemetan | MVP_top60_DNA+RNA_seed | HSP90AB1 | -0.397375 | UCEC | MFE-296 | Elesclomol |
| MVP_top60_DNA_seed | MAPK1 | 0.711257 | LUAD | NCI-H1944 | Ulixertinib | MVP_top60_DNA+RNA_seed | PIK3CA | -0.392543 | UCEC | AN3-CA | PI3Ka_4409 |
| MVP_top60_DNA_seed | EGFR | 0.713687 | LUAD | NCI-H1993 | Sapitinib | MVP_top60_DNA+RNA_seed | PIK3CD | -0.392543 | UCEC | AN3-CA | PI3Ka_4409 |
| MVP_top60_DNA_seed | ERBB2 | 0.713687 | LUAD | NCI-H1993 | Sapitinib | MVP_top60_DNA+RNA_seed | AKT3 | -0.381262 | UCEC | SNG-M | AKT inhibitor VIII |
| MVP_top60_DNA_seed | ERBB3 | 0.713687 | LUAD | NCI-H1993 | Sapitinib | MVP_top60_DNA+RNA_seed | AKT1 | -0.381262 | UCEC | SNG-M | AKT inhibitor VIII |
| MVP_top60_DNA_seed | GSK3B | 0.715034 | LUAD | PC-3_[JPC-3] | SB216763 | MVP_top60_DNA+RNA_seed | MAPK1 | -0.375542 | UCEC | SNG-M | FR-180204 |
| MVP_top60_DNA_seed | IGF1R | 0.715396 | LUAD | Calu-3 | Linsitinib | MVP_top60_DNA+RNA_seed | MAPK3 | -0.375542 | UCEC | SNG-M | FR-180204 |
| MVP_top60_DNA_seed | AKT1 | 0.715435 | LUAD | NCI-H1666 | GSK2110183B | MVP_top60_DNA+RNA_seed | PIK3CD | -0.374953 | UCEC | MFE-280 | IC-87114 |
| MVP_top60_DNA_seed | MET | 0.717704 | LUAD | HOP-62 | Crizotinib | MVP_top60_DNA+RNA_seed | SRC | -0.374751 | UCEC | MFE-280 | NVP-BHG712 |
| MVP_top60_DNA_seed | MDM2 | 0.717841 | LUAD | NCI-H2030 | Serdemetan | MVP_top60_DNA+RNA_seed | AKT3 | -0.366585 | UCEC | AN3-CA | AKT inhibitor VIII |
| MVP_top60_DNA_seed | IGF1R | 0.717875 | LUAD | NCI-H1838 | NVP-ADW742 | MVP_top60_DNA+RNA_seed | AKT1 | -0.366585 | UCEC | AN3-CA | AKT inhibitor VIII |
| MVP_top60_DNA_seed | MAPK1 | 0.718598 | LUAD | NCI-H2009 | Ulixertinib | MVP_top60_DNA+RNA_seed | FGFR2 | -0.355229 | UCEC | SNG-M | FGFR_3831 |
| MVP_top60_DNA_seed | TP53 | 0.718681 | LUAD | EMC-BAC-2 | MIRA-1 | MVP_top60_DNA+RNA_seed | HDAC1 | -0.3448 | UCEC | MFE-280 | CAY10603 |
| MVP_top60_DNA_seed | AKT1 | 0.718776 | LUAD | NCI-H2085 | MK-2206 | MVP_top60_DNA+RNA_seed | FGFR2 | -0.342315 | UCEC | COLO-684 | AZD4547 |
| MVP_top60_DNA_seed | EGFR | 0.71983 | LUAD | NCI-H650 | AZD3759 | MVP_top60_DNA+RNA_seed | HSP90AA1 | -0.333341 | UCEC | MFE-319 | CCT-018159 |
| MVP_top60_DNA_seed | ATM | 0.720309 | LUAD | NCI-H1573 | KU-55933 | MVP_top60_DNA+RNA_seed | HSP90AB1 | -0.333341 | UCEC | MFE-319 | CCT-018159 |
| MVP_top60_DNA_seed | SMARCA4 | 0.721451 | LUAD | NCI-H1781 | PFI3 | MVP_top60_DNA+RNA_seed | AKT1 | -0.327348 | UCEC | KLE | MK-2206 |
| MVP_top60_DNA_seed | BRAF | 0.721907 | LUAD | NCI-H1975 | SB590885 | MVP_top60_DNA+RNA_seed | HSP90AA1 | -0.31414 | UCEC | MFE-319 | Elesclomol |
| MVP_top60_DNA_seed | PIK3CA | 0.722037 | LUAD | NCI-H358 | Buparlisib | MVP_top60_DNA+RNA_seed | HSP90AB1 | -0.31414 | UCEC | MFE-319 | Elesclomol |
| MVP_top60_DNA_seed | HDAC1 | 0.722441 | LUAD | NCI-H522 | Romidepsin | MVP_top60_DNA+RNA_seed | AKT3 | -0.310089 | UCEC | KLE | GSK690693 |
| MVP_top60_DNA_seed | EGFR | 0.723624 | LUAD | NCI-H1793 | Sapitinib | MVP_top60_DNA+RNA_seed | AKT1 | -0.310089 | UCEC | KLE | GSK690693 |

| | | | | | | | | | | | |
|---|---|---|---|---|---|---|---|---|---|---|---|
| MVP_top60_DNA_seed | ERBB2 | 0.723624 | LUAD | NCI-H1793 | Sapitinib | MVP_top60_DNA+RNA_seed | AR | -0.300636 | UCEC | MFE-280 | AZD3514 |
| MVP_top60_DNA_seed | ERBB3 | 0.723624 | LUAD | NCI-H1793 | Sapitinib | MVP_top60_DNA+RNA_seed | ESR1 | -0.299305 | UCEC | MFE-296 | Tamoxifen |
| MVP_top60_DNA_seed | EGFR | 0.723973 | LUAD | NCI-H2228 | Gefitinib | MVP_top60_DNA+RNA_seed | EGFR | -0.294777 | UCEC | MFE-319 | PF-00299804 |
| MVP_top60_DNA_seed | IGF1R | 0.72798 | LUAD | NCI-H2342 | BMS-536924 | MVP_top60_DNA+RNA_seed | ERBB2 | -0.294777 | UCEC | MFE-319 | PF-00299804 |
| MVP_top60_DNA_seed | GSK3B | 0.728388 | LUAD | NCI-H2030 | SB216763 | MVP_top60_DNA+RNA_seed | ESR1 | -0.291264 | UCEC | MFE-319 | Fulvestrant |
| MVP_top60_DNA_seed | AKT1 | 0.728406 | LUAD | SW1573 | Ipatasertib | MVP_top60_DNA+RNA_seed | HDAC1 | -0.275283 | UCEC | MFE-280 | Belinostat |
| MVP_top60_DNA_seed | EGFR | 0.728602 | LUAD | HCC-78 | Gefitinib | MVP_top60_DNA+RNA_seed | EGFR | -0.274487 | UCEC | KLE | Afatinib |
| MVP_top60_DNA_seed | HDAC1 | 0.729454 | LUAD | NCI-H23 | Romidepsin | MVP_top60_DNA+RNA_seed | ERBB2 | -0.274487 | UCEC | KLE | Afatinib |
| MVP_top60_DNA_seed | SMARCA4 | 0.730502 | LUAD | A549 | PFI3 | MVP_top60_DNA+RNA_seed | AKT3 | -0.273754 | UCEC | COLO-684 | AKT inhibitor VIII |
| MVP_top60_DNA_seed | GSK3B | 0.730525 | LUAD | COR-L105 | SB216763 | MVP_top60_DNA+RNA_seed | AKT1 | -0.273754 | UCEC | COLO-684 | AKT inhibitor VIII |
| MVP_top60_DNA_seed | EZH2 | 0.730699 | LUAD | NCI-H1573 | GSK343 | MVP_top60_DNA+RNA_seed | EGFR | -0.263096 | UCEC | COLO-684 | CUDC-101 |
| MVP_top60_DNA_seed | BRAF | 0.730791 | LUAD | NCI-H1755 | SB590885 | MVP_top60_DNA+RNA_seed | ERBB2 | -0.263096 | UCEC | COLO-684 | CUDC-101 |
| MVP_top60_DNA_seed | IGF1R | 0.734011 | LUAD | NCI-H2342 | BMS-754807 | MVP_top60_DNA+RNA_seed | AKT3 | -0.254252 | UCEC | KLE | AKT inhibitor VIII |
| MVP_top60_DNA_seed | TP53 | 0.734693 | LUAD | NCI-H1693 | MIRA-1 | MVP_top60_DNA+RNA_seed | AKT1 | -0.254252 | UCEC | KLE | AKT inhibitor VIII |
| MVP_top60_DNA_seed | MAPK1 | 0.73612 | LUAD | SW1573 | Ulixertinib | MVP_top60_DNA+RNA_seed | AKT3 | -0.248601 | UCEC | EN | BAY AKT1 |
| MVP_top60_DNA_seed | AKT1 | 0.736141 | LUAD | PC-3_[JPC-3] | Ipatasertib | MVP_top60_DNA+RNA_seed | AKT1 | -0.248601 | UCEC | EN | BAY AKT1 |
| MVP_top60_DNA_seed | EGFR | 0.738179 | LUAD | Calu-6 | Lapatinib | MVP_top60_DNA+RNA_seed | ATM | -0.244126 | UCEC | HEC-1 | KU-55933 |
| MVP_top60_DNA_seed | ERBB2 | 0.738179 | LUAD | Calu-6 | Lapatinib | MVP_top60_DNA+RNA_seed | HDAC1 | -0.234194 | UCEC | AN3-CA | JQ12 |
| MVP_top60_DNA_seed | TP53 | 0.738409 | LUAD | NCI-H2023 | MIRA-1 | MVP_top60_DNA+RNA_seed | HDAC2 | -0.234194 | UCEC | AN3-CA | JQ12 |
| MVP_top60_DNA_seed | IGF1R | 0.738756 | LUAD | EKVX | Linsitinib | MVP_top60_DNA+RNA_seed | HDAC1 | -0.226359 | UCEC | HEC-1 | Dacinostat |
| MVP_top60_DNA_seed | ATM | 0.741229 | LUAD | NCI-H2342 | KU-55933 | MVP_top60_DNA+RNA_seed | MDM2 | -0.22556 | UCEC | MFE-319 | Nutlin-3a (-) |
| MVP_top60_DNA_seed | ATM | 0.741313 | LUAD | NCI-H2023 | KU-55933 | MVP_top60_DNA+RNA_seed | HDAC1 | -0.221093 | UCEC | COLO-684 | Tubastatin A |
| MVP_top60_DNA_seed | SRC | 0.741868 | LUAD | NCI-H1437 | Bosutinib | MVP_top60_DNA+RNA_seed | EGFR | -0.213487 | UCEC | KLE | CUDC-101 |
| MVP_top60_DNA_seed | BRAF | 0.742919 | LUAD | EKVX | PLX-4720 | MVP_top60_DNA+RNA_seed | ERBB2 | -0.213487 | UCEC | KLE | CUDC-101 |
| MVP_top60_DNA_seed | AURKA | 0.744282 | LUAD | COR-L105 | Alisertib | MVP_top60_DNA+RNA_seed | MAPK1 | -0.210891 | UCEC | MFE-280 | VX-11e |
| MVP_top60_DNA_seed | MAPK1 | 0.74797 | LUAD | RERF-LC-KJ | VX-11e | MVP_top60_DNA+RNA_seed | HDAC1 | -0.210836 | UCEC | MFE-296 | Dacinostat |
| MVP_top60_DNA_seed | AKT1 | 0.748213 | LUAD | EMC-BAC-2 | GSK2110183B | MVP_top60_DNA+RNA_seed | AKT3 | -0.204435 | UCEC | SNG-M | BAY AKT1 |
| MVP_top60_DNA_seed | MET | 0.748297 | LUAD | NCI-H2342 | Crizotinib | MVP_top60_DNA+RNA_seed | AKT1 | -0.204435 | UCEC | SNG-M | BAY AKT1 |
| MVP_top60_DNA_seed | EGFR | 0.749082 | LUAD | NCI-H2342 | Gefitinib | MVP_top60_DNA+RNA_seed | HDAC1 | -0.195598 | UCEC | EN | Belinostat |
| MVP_top60_DNA_seed | SMARCA4 | 0.749439 | LUAD | NCI-H1975 | PFI3 | MVP_top60_DNA+RNA_seed | EGFR | -0.190407 | UCEC | KLE | PF-00299804 |
| MVP_top60_DNA_seed | AKT1 | 0.751414 | LUAD | NCI-H358 | AZD5363 | MVP_top60_DNA+RNA_seed | ERBB2 | -0.190407 | UCEC | KLE | PF-00299804 |
| MVP_top60_DNA_seed | MAPK1 | 0.75299 | LUAD | NCI-H2347 | SCH772984 | MVP_top60_DNA+RNA_seed | HDAC1 | -0.189258 | UCEC | COLO-684 | Dacinostat |
| MVP_top60_DNA_seed | HDAC1 | 0.753912 | LUAD | EMC-BAC-2 | Entinostat | MVP_top60_DNA+RNA_seed | ERBB2 | -0.187817 | UCEC | KLE | CP724714 |
| MVP_top60_DNA_seed | AKT1 | 0.754303 | LUAD | HCC-827 | Uprosertib | MVP_top60_DNA+RNA_seed | MAPK1 | -0.187804 | UCEC | COLO-684 | VX-11e |
| MVP_top60_DNA_seed | IGF1R | 0.755677 | LUAD | NCI-H3122 | NVP-ADW742 | MVP_top60_DNA+RNA_seed | PIK3CD | -0.168739 | UCEC | SNG-M | Idelalisib |
| MVP_top60_DNA_seed | HSP90AA1 | 0.756826 | LUAD | NCI-H2087 | Luminespib | MVP_top60_DNA+RNA_seed | PIK3CD | -0.163341 | UCEC | KLE | Idelalisib |
| MVP_top60_DNA_seed | EGFR | 0.758612 | LUAD | NCI-H2009 | Lapatinib | MVP_top60_DNA+RNA_seed | JAK1 | -0.159153 | UCEC | MFE-319 | AZD1480 |
| MVP_top60_DNA_seed | ERBB2 | 0.758612 | LUAD | NCI-H2009 | Lapatinib | MVP_top60_DNA+RNA_seed | AKT3 | -0.152611 | UCEC | SNG-M | AKT inhibitor VIII |
| MVP_top60_DNA_seed | TP53 | 0.758629 | LUAD | NCI-H1563 | MIRA-1 | MVP_top60_DNA+RNA_seed | AKT1 | -0.152611 | UCEC | SNG-M | AKT inhibitor VIII |
| MVP_top60_DNA_seed | MET | 0.760961 | LUAD | NCI-H1755 | Savolitinib | MVP_top60_DNA+RNA_seed | PIK3CB | -0.150755 | UCEC | MFE-319 | AZD6482 |
| MVP_top60_DNA_seed | HDAC1 | 0.761671 | LUAD | NCI-H23 | Entinostat | MVP_top60_DNA+RNA_seed | EGFR | -0.148695 | UCEC | MFE-319 | AST-1306 |

| | | | | | | | | | | | |
|---|---|---|---|---|---|---|---|---|---|---|---|
| MVP_top60_DNA_seed | AKT1 | 0.763262 | LUAD | NCI-H522 | GSK2110183B | MVP_top60_DNA+RNA_seed | EGFR | -0.13801 | UCEC | SNG-M | Cetuximab |
| MVP_top60_DNA_seed | AKT1 | 0.763661 | LUAD | SW1573 | AZD5363 | MVP_top60_DNA+RNA_seed | PIK3CA | -0.137169 | UCEC | COLO-684 | AZD8835 |
| MVP_top60_DNA_seed | IGF1R | 0.766287 | LUAD | EMC-BAC-2 | GSK1904529A | MVP_top60_DNA+RNA_seed | PIK3CD | -0.137169 | UCEC | COLO-684 | AZD8835 |
| MVP_top60_DNA_seed | HDAC1 | 0.76688 | LUAD | HOP-62 | Romidepsin | MVP_top60_DNA+RNA_seed | AKT3 | -0.132474 | UCEC | MFE-319 | BAY AKT1 |
| MVP_top60_DNA_seed | PIK3CA | 0.768546 | LUAD | NCI-H2405 | GNE-317 | MVP_top60_DNA+RNA_seed | AKT1 | -0.132474 | UCEC | MFE-319 | BAY AKT1 |
| MVP_top60_DNA_seed | HDAC1 | 0.768968 | LUAD | NCI-H2023 | Entinostat | MVP_top60_DNA+RNA_seed | AR | -0.128718 | UCEC | MFE-296 | AZD3514 |
| MVP_top60_DNA_seed | AKT1 | 0.76957 | LUAD | NCI-H2405 | Ipatasertib | MVP_top60_DNA+RNA_seed | HSP90AA1 | -0.125845 | UCEC | MFE-296 | CCT-018159 |
| MVP_top60_DNA_seed | HSP90AA1 | 0.770846 | LUAD | ABC-1 | CCT-018159 | MVP_top60_DNA+RNA_seed | HSP90AB1 | -0.125845 | UCEC | MFE-296 | CCT-018159 |
| MVP_top60_DNA_seed | AURKA | 0.771097 | LUAD | NCI-H522 | Alisertib | MVP_top60_DNA+RNA_seed | JAK1 | -0.120985 | UCEC | COLO-684 | Ruxolitinib |
| MVP_top60_DNA_seed | IGF1R | 0.771693 | LUAD | SW1573 | BMS-536924 | MVP_top60_DNA+RNA_seed | EGFR | -0.119893 | UCEC | HEC-1 | AZD8931 |
| MVP_top60_DNA_seed | GSK3B | 0.773945 | LUAD | NCI-H2030 | CHIR-99021 | MVP_top60_DNA+RNA_seed | ERBB2 | -0.119893 | UCEC | HEC-1 | AZD8931 |
| MVP_top60_DNA_seed | EGFR | 0.774301 | LUAD | ABC-1 | Lapatinib | MVP_top60_DNA+RNA_seed | HDAC1 | -0.113898 | UCEC | EN | CAY10603 |
| MVP_top60_DNA_seed | ERBB2 | 0.774301 | LUAD | ABC-1 | Lapatinib | MVP_top60_DNA+RNA_seed | EP300 | -0.107449 | UCEC | HEC-1 | I-CBP112 |
| MVP_top60_DNA_seed | TP53 | 0.774438 | LUAD | HCC-78 | MIRA-1 | MVP_top60_DNA+RNA_seed | MDM2 | -0.103957 | UCEC | HEC-1 | Serdemetan |
| MVP_top60_DNA_seed | BRAF | 0.77614 | LUAD | NCI-H1563 | Dabrafenib | MVP_top60_DNA+RNA_seed | JAK1 | -0.103402 | UCEC | HEC-1 | AZD1480 |
| MVP_top60_DNA_seed | HSP90AA1 | 0.777057 | LUAD | NCI-H2291 | Luminespib | MVP_top60_DNA+RNA_seed | HDAC1 | -0.10036 | UCEC | MFE-296 | Tubastatin A |
| MVP_top60_DNA_seed | AKT1 | 0.779293 | LUAD | HOP-62 | Uprosertib | MVP_top60_DNA+RNA_seed | SRC | -0.099624 | UCEC | COLO-684 | NVP-BHG712 |
| MVP_top60_DNA_seed | EGFR | 0.779507 | LUAD | NCI-H2342 | Osimertinib | MVP_top60_DNA+RNA_seed | ERBB2 | -0.091649 | UCEC | MFE-319 | CP724714 |
| MVP_top60_DNA_seed | EGFR | 0.779879 | LUAD | NCI-H3122 | Lapatinib | MVP_top60_DNA+RNA_seed | PIK3CB | -0.090714 | UCEC | MFE-296 | AZD8186 |
| MVP_top60_DNA_seed | ERBB2 | 0.779879 | LUAD | NCI-H3122 | Lapatinib | MVP_top60_DNA+RNA_seed | PIK3CD | -0.090714 | UCEC | MFE-296 | AZD8186 |
| MVP_top60_DNA_seed | HDAC1 | 0.779987 | LUAD | HOP-62 | PCI-34051 | MVP_top60_DNA+RNA_seed | HSP90AA1 | -0.088273 | UCEC | COLO-684 | Luminespib |
| MVP_top60_DNA_seed | GSK3B | 0.781998 | LUAD | NCI-H1650 | SB216763 | MVP_top60_DNA+RNA_seed | HSP90AB1 | -0.088273 | UCEC | COLO-684 | Luminespib |
| MVP_top60_DNA_seed | IGF1R | 0.784109 | LUAD | EKVX | BMS-754807 | MVP_top60_DNA+RNA_seed | HDAC1 | -0.082807 | UCEC | MFE-296 | JQ12 |
| MVP_top60_DNA_seed | TERT | 0.784312 | LUAD | PC-3_[JPC-3] | BIBR-1532 | MVP_top60_DNA+RNA_seed | HDAC2 | -0.082807 | UCEC | MFE-296 | JQ12 |
| MVP_top60_DNA_seed | AKT1 | 0.784618 | LUAD | ABC-1 | GSK2110183B | MVP_top60_DNA+RNA_seed | MDM2 | -0.079099 | UCEC | SNG-M | Serdemetan |
| MVP_top60_DNA_seed | MDM2 | 0.784907 | LUAD | NCI-H1993 | Nutlin-3a (-) | MVP_top60_DNA+RNA_seed | SRC | -0.074364 | UCEC | KLE | NVP-BHG712 |
| MVP_top60_DNA_seed | PIK3CA | 0.785271 | LUAD | NCI-H1944 | GNE-317 | MVP_top60_DNA+RNA_seed | FGFR2 | -0.071005 | UCEC | HEC-1 | AZD4547 |
| MVP_top60_DNA_seed | CREBBP | 0.786261 | LUAD | NCI-H1755 | SGC-CBP30 | MVP_top60_DNA+RNA_seed | EGFR | -0.063469 | UCEC | EN | CI-1033 |
| MVP_top60_DNA_seed | IGF1R | 0.78633 | LUAD | NCI-H1793 | Linsitinib | MVP_top60_DNA+RNA_seed | ERBB2 | -0.063469 | UCEC | EN | CI-1033 |
| MVP_top60_DNA_seed | BRAF | 0.787764 | LUAD | ABC-1 | SB590885 | MVP_top60_DNA+RNA_seed | FGFR2 | -0.061881 | UCEC | HEC-1 | PD173074 |
| MVP_top60_DNA_seed | ATM | 0.789025 | LUAD | SW1573 | KU-55933 | MVP_top60_DNA+RNA_seed | ATM | -0.054838 | UCEC | KLE | CP466722 |
| MVP_top60_DNA_seed | MAPK1 | 0.789912 | LUAD | NCI-H1781 | Ulixertinib | MVP_top60_DNA+RNA_seed | HDAC1 | -0.051517 | UCEC | MFE-319 | Dacinostat |
| MVP_top60_DNA_seed | IGF1R | 0.789942 | LUAD | NCI-H1563 | GSK1904529A | MVP_top60_DNA+RNA_seed | ATM | -0.049526 | UCEC | COLO-684 | KU-60019 |
| MVP_top60_DNA_seed | HDAC1 | 0.791386 | LUAD | SK-LU-1 | Romidepsin | MVP_top60_DNA+RNA_seed | HDAC1 | -0.045742 | UCEC | EN | Tubastatin A |
| MVP_top60_DNA_seed | EZH2 | 0.791926 | LUAD | NCI-H1838 | GSK343 | MVP_top60_DNA+RNA_seed | EGFR | -0.04525 | UCEC | KLE | AZD8931 |
| MVP_top60_DNA_seed | IGF1R | 0.793 | LUAD | SK-LU-1 | BMS-536924 | MVP_top60_DNA+RNA_seed | ERBB2 | -0.04525 | UCEC | KLE | AZD8931 |
| MVP_top60_DNA_seed | AKT1 | 0.794586 | LUAD | NCI-H2085 | Ipatasertib | MVP_top60_DNA+RNA_seed | EGFR | -0.044238 | UCEC | EN | CUDC-101 |
| MVP_top60_DNA_seed | EGFR | 0.795251 | LUAD | NCI-H1651 | Afatinib | MVP_top60_DNA+RNA_seed | ERBB2 | -0.044238 | UCEC | EN | CUDC-101 |
| MVP_top60_DNA_seed | ERBB2 | 0.795251 | LUAD | NCI-H1651 | Afatinib | MVP_top60_DNA+RNA_seed | AKT3 | -0.04317 | UCEC | EN | AKT inhibitor VIII |
| MVP_top60_DNA_seed | TERT | 0.795957 | LUAD | A549 | BIBR-1532 | MVP_top60_DNA+RNA_seed | AKT1 | -0.04317 | UCEC | EN | AKT inhibitor VIII |
| MVP_top60_DNA_seed | PIK3CA | 0.79772 | LUAD | VMRC-LCD | Alpelisib | MVP_top60_DNA+RNA_seed | JAK1 | -0.029314 | UCEC | MFE-296 | JAK1_3715 |

| | | | | | | | | | | | |
|---|---|---|---|---|---|---|---|---|---|---|---|
| MVP_top60_DNA_seed | FGFR1 | 0.798837 | LUAD | NCI-H1993 | AZD4547 | MVP_top60_DNA+RNA_seed | HDAC1 | -0.026238 | UCEC | HEC-1 | Belinostat |
| MVP_top60_DNA_seed | MDM2 | 0.801458 | LUAD | HOP-62 | Serdemetan | MVP_top60_DNA+RNA_seed | AKT1 | -0.025501 | UCEC | HEC-1 | MK-2206 |
| MVP_top60_DNA_seed | HDAC1 | 0.80369 | LUAD | NCI-H322M | PCI-34051 | MVP_top60_DNA+RNA_seed | JAK1 | -0.02547 | UCEC | HEC-1 | JAK1_3715 |
| MVP_top60_DNA_seed | AKT1 | 0.803698 | LUAD | LXF-289 | Ipatasertib | MVP_top60_DNA+RNA_seed | HSP90AA1 | -0.019557 | UCEC | MFE-280 | SNX-2112 |
| MVP_top60_DNA_seed | MDM2 | 0.803918 | LUAD | NCI-H2087 | Nutlin-3a (-) | MVP_top60_DNA+RNA_seed | HSP90AB1 | -0.019557 | UCEC | MFE-280 | SNX-2112 |
| MVP_top60_DNA_seed | BRAF | 0.805501 | LUAD | NCI-H2342 | SB590885 | MVP_top60_DNA+RNA_seed | MAPK1 | -0.018591 | UCEC | MFE-280 | FR-180204 |
| MVP_top60_DNA_seed | EGFR | 0.806373 | LUAD | HOP-62 | Lapatinib | MVP_top60_DNA+RNA_seed | MAPK3 | -0.018591 | UCEC | MFE-280 | FR-180204 |
| MVP_top60_DNA_seed | ERBB2 | 0.806373 | LUAD | HOP-62 | Lapatinib | MVP_top60_DNA+RNA_seed | EP300 | -0.009974 | UCEC | MFE-296 | I-CBP112 |
| MVP_top60_DNA_seed | PIK3CA | 0.806887 | LUAD | NCI-H23 | GNE-317 | MVP_top60_DNA+RNA_seed | PIK3CA | -0.004274 | UCEC | KLE | PI3Ka_4409 |
| MVP_top60_DNA_seed | HDAC1 | 0.807011 | LUAD | NCI-H2085 | Romidepsin | MVP_top60_DNA+RNA_seed | PIK3CD | -0.004274 | UCEC | KLE | PI3Ka_4409 |
| MVP_top60_DNA_seed | MDM2 | 0.808288 | LUAD | NCI-H1623 | Nutlin-3a (-) | MVP_top60_DNA+RNA_seed | PIK3CA | 0.002161 | UCEC | MFE-296 | AZD8835 |
| MVP_top60_DNA_seed | PIK3CA | 0.808837 | LUAD | NCI-H1651 | AZD8186 | MVP_top60_DNA+RNA_seed | PIK3CD | 0.002161 | UCEC | MFE-296 | AZD8835 |
| MVP_top60_DNA_seed | SMARCA4 | 0.81031 | LUAD | NCI-H1355 | PFI3 | MVP_top60_DNA+RNA_seed | EGFR | 0.004627 | UCEC | HEC-1 | CI-1033 |
| MVP_top60_DNA_seed | AURKA | 0.810363 | LUAD | SW1573 | Alisertib | MVP_top60_DNA+RNA_seed | ERBB2 | 0.004627 | UCEC | HEC-1 | CI-1033 |
| MVP_top60_DNA_seed | IGF1R | 0.812046 | LUAD | Calu-3 | BMS-754807 | MVP_top60_DNA+RNA_seed | ESR1 | 0.007057 | UCEC | KLE | Fulvestrant |
| MVP_top60_DNA_seed | TP53 | 0.813338 | LUAD | PC-3_[JPC-3] | MIRA-1 | MVP_top60_DNA+RNA_seed | PIK3CA | 0.007765 | UCEC | HEC-1 | PI3Ka_4409 |
| MVP_top60_DNA_seed | EGFR | 0.813657 | LUAD | NCI-H2291 | Erlotinib | MVP_top60_DNA+RNA_seed | PIK3CD | 0.007765 | UCEC | HEC-1 | PI3Ka_4409 |
| MVP_top60_DNA_seed | EGFR | 0.814658 | LUAD | NCI-H2342 | Erlotinib | MVP_top60_DNA+RNA_seed | HDAC1 | 0.008111 | UCEC | EN | AR-42 |
| MVP_top60_DNA_seed | HDAC1 | 0.81716 | LUAD | COR-L105 | Entinostat | MVP_top60_DNA+RNA_seed | EGFR | 0.010594 | UCEC | COLO-684 | CI-1033 |
| MVP_top60_DNA_seed | MDM2 | 0.817767 | LUAD | H3255 | Nutlin-3a (-) | MVP_top60_DNA+RNA_seed | ERBB2 | 0.010594 | UCEC | COLO-684 | CI-1033 |
| MVP_top60_DNA_seed | CREBBP | 0.818512 | LUAD | EKVX | SGC-CBP30 | MVP_top60_DNA+RNA_seed | EGFR | 0.010679 | UCEC | COLO-684 | Pelitinib |
| MVP_top60_DNA_seed | PIK3CA | 0.819556 | LUAD | NCI-H1650 | GNE-317 | MVP_top60_DNA+RNA_seed | EGFR | 0.01629 | UCEC | HEC-1 | Gefitinib |
| MVP_top60_DNA_seed | CREBBP | 0.820648 | LUAD | NCI-H3122 | SGC-CBP30 | MVP_top60_DNA+RNA_seed | HDAC1 | 0.018214 | UCEC | HEC-1 | AR-42 |
| MVP_top60_DNA_seed | FGFR1 | 0.824759 | LUAD | Calu-3 | PD173074 | MVP_top60_DNA+RNA_seed | ESR1 | 0.023711 | UCEC | MFE-296 | Fulvestrant |
| MVP_top60_DNA_seed | AURKA | 0.825346 | LUAD | NCI-H1975 | ZM447439 | MVP_top60_DNA+RNA_seed | SRC | 0.026441 | UCEC | EN | NVP-BHG712 |
| MVP_top60_DNA_seed | AKT1 | 0.826753 | LUAD | NCI-H1651 | MK-2206 | MVP_top60_DNA+RNA_seed | EGFR | 0.034598 | UCEC | MFE-319 | CI-1033 |
| MVP_top60_DNA_seed | IGF1R | 0.828818 | LUAD | NCI-H1793 | GSK1904529A | MVP_top60_DNA+RNA_seed | ERBB2 | 0.034598 | UCEC | MFE-319 | CI-1033 |
| MVP_top60_DNA_seed | AKT1 | 0.82986 | LUAD | NCI-H2085 | Uprosertib | MVP_top60_DNA+RNA_seed | HDAC1 | 0.035617 | UCEC | MFE-280 | Tubastatin A |
| MVP_top60_DNA_seed | IGF1R | 0.82994 | LUAD | NCI-H2405 | BMS-754807 | MVP_top60_DNA+RNA_seed | ERBB2 | 0.04091 | UCEC | COLO-684 | CP724714 |
| MVP_top60_DNA_seed | EGFR | 0.830972 | LUAD | NCI-H1355 | Afatinib | MVP_top60_DNA+RNA_seed | HDAC1 | 0.048415 | UCEC | MFE-280 | AR-42 |
| MVP_top60_DNA_seed | ERBB2 | 0.830972 | LUAD | NCI-H1355 | Afatinib | MVP_top60_DNA+RNA_seed | EGFR | 0.053618 | UCEC | MFE-296 | AST-1306 |
| MVP_top60_DNA_seed | EZH2 | 0.831454 | LUAD | NCI-H1793 | GSK343 | MVP_top60_DNA+RNA_seed | HDAC1 | 0.058416 | UCEC | MFE-319 | JQ12 |
| MVP_top60_DNA_seed | AURKA | 0.831531 | LUAD | NCI-H1993 | ZM447439 | MVP_top60_DNA+RNA_seed | HDAC2 | 0.058416 | UCEC | MFE-319 | JQ12 |
| MVP_top60_DNA_seed | BRAF | 0.833003 | LUAD | NCI-H2085 | SB590885 | MVP_top60_DNA+RNA_seed | SRC | 0.0597 | UCEC | HEC-1 | Bosutinib |
| MVP_top60_DNA_seed | FGFR1 | 0.833099 | LUAD | NCI-H1648 | PD173074 | MVP_top60_DNA+RNA_seed | EGFR | 0.062381 | UCEC | MFE-296 | CI-1033 |
| MVP_top60_DNA_seed | GSK3B | 0.836066 | LUAD | NCI-H358 | SB216763 | MVP_top60_DNA+RNA_seed | ERBB2 | 0.062381 | UCEC | MFE-296 | CI-1033 |
| MVP_top60_DNA_seed | BRAF | 0.836186 | LUAD | NCI-H2009 | Dabrafenib | MVP_top60_DNA+RNA_seed | JAK1 | 0.063622 | UCEC | MFE-319 | Ruxolitinib |
| MVP_top60_DNA_seed | HDAC1 | 0.837202 | LUAD | LXF-289 | PCI-34051 | MVP_top60_DNA+RNA_seed | HDAC1 | 0.06482 | UCEC | KLE | CAY10603 |
| MVP_top60_DNA_seed | AKT1 | 0.838818 | LUAD | NCI-H650 | Ipatasertib | MVP_top60_DNA+RNA_seed | HSP90AA1 | 0.070505 | UCEC | COLO-684 | Tanespimycin |
| MVP_top60_DNA_seed | ATM | 0.839089 | LUAD | NCI-H1781 | KU-55933 | MVP_top60_DNA+RNA_seed | HSP90AB1 | 0.070505 | UCEC | COLO-684 | Tanespimycin |
| MVP_top60_DNA_seed | MET | 0.839365 | LUAD | ABC-1 | Savolitinib | MVP_top60_DNA+RNA_seed | HDAC1 | 0.079084 | UCEC | KLE | Dacinostat |

| | | | | | | | | | | | |
|---|---|---|---|---|---|---|---|---|---|---|---|
| MVP_top60_DNA_seed | MDM2 | 0.839642 | LUAD | NCI-H650 | Serdemetan | MVP_top60_DNA+RNA_seed | HSP90AA1 | 0.094957 | UCEC | HEC-1 | Luminespib |
| MVP_top60_DNA_seed | AURKA | 0.839991 | LUAD | NCI-H2291 | Tozasertib | MVP_top60_DNA+RNA_seed | HSP90AB1 | 0.094957 | UCEC | HEC-1 | Luminespib |
| MVP_top60_DNA_seed | TERT | 0.841422 | LUAD | COR-L105 | BIBR-1532 | MVP_top60_DNA+RNA_seed | HDAC1 | 0.101031 | UCEC | HEC-1 | JQ12 |
| MVP_top60_DNA_seed | HDAC1 | 0.843498 | LUAD | NCI-H1355 | Entinostat | MVP_top60_DNA+RNA_seed | HDAC2 | 0.101031 | UCEC | HEC-1 | JQ12 |
| MVP_top60_DNA_seed | BRAF | 0.84769 | LUAD | RERF-LC-KJ | PLX-4720 | MVP_top60_DNA+RNA_seed | FGFR2 | 0.102188 | UCEC | COLO-684 | PD173074 |
| MVP_top60_DNA_seed | TP53 | 0.848045 | LUAD | NCI-H1623 | MIRA-1 | MVP_top60_DNA+RNA_seed | FGFR2 | 0.105396 | UCEC | SNG-M | AZD4547 |
| MVP_top60_DNA_seed | EGFR | 0.84873 | LUAD | NCI-H2023 | Gefitinib | MVP_top60_DNA+RNA_seed | ATM | 0.113108 | UCEC | MFE-296 | KU-55933 |
| MVP_top60_DNA_seed | MAPK1 | 0.849074 | LUAD | NCI-H522 | SCH772984 | MVP_top60_DNA+RNA_seed | HDAC1 | 0.126755 | UCEC | KLE | Belinostat |
| MVP_top60_DNA_seed | BRAF | 0.849148 | LUAD | NCI-H2030 | SB590885 | MVP_top60_DNA+RNA_seed | HDAC1 | 0.128478 | UCEC | HEC-1 | CAY10603 |
| MVP_top60_DNA_seed | KRAS | 0.849262 | LUAD | NCI-H1435 | KRAS (G12C) Inhibitor-12 | MVP_top60_DNA+RNA_seed | SRC | 0.136586 | UCEC | MFE-319 | NVP-BHG712 |
| MVP_top60_DNA_seed | EZH2 | 0.85089 | LUAD | NCI-H3122 | GSK343 | MVP_top60_DNA+RNA_seed | EGFR | 0.149855 | UCEC | HEC-1 | CUDC-101 |
| MVP_top60_DNA_seed | EGFR | 0.852169 | LUAD | HOP-62 | Osimertinib | MVP_top60_DNA+RNA_seed | ERBB2 | 0.149855 | UCEC | HEC-1 | CUDC-101 |
| MVP_top60_DNA_seed | TERT | 0.852616 | LUAD | NCI-H3122 | BIBR-1532 | MVP_top60_DNA+RNA_seed | HDAC1 | 0.151208 | UCEC | COLO-684 | Belinostat |
| MVP_top60_DNA_seed | AKT1 | 0.855017 | LUAD | NCI-H2085 | AT13148 | MVP_top60_DNA+RNA_seed | PIK3CA | 0.154596 | UCEC | HEC-1 | AZD8835 |
| MVP_top60_DNA_seed | AKT1 | 0.855344 | LUAD | SW1573 | MK-2206 | MVP_top60_DNA+RNA_seed | PIK3CD | 0.154596 | UCEC | HEC-1 | AZD8835 |
| MVP_top60_DNA_seed | FGFR1 | 0.855423 | LUAD | NCI-H2009 | AZD4547 | MVP_top60_DNA+RNA_seed | PIK3CB | 0.167623 | UCEC | MFE-319 | AZD8186 |
| MVP_top60_DNA_seed | EZH2 | 0.8565 | LUAD | NCI-H522 | GSK343 | MVP_top60_DNA+RNA_seed | PIK3CD | 0.167623 | UCEC | MFE-319 | AZD8186 |
| MVP_top60_DNA_seed | BRAF | 0.856505 | LUAD | HCC-78 | SB590885 | MVP_top60_DNA+RNA_seed | PIK3CD | 0.183443 | UCEC | MFE-319 | Idelalisib |
| MVP_top60_DNA_seed | HDAC1 | 0.856729 | LUAD | EKVX | Romidepsin | MVP_top60_DNA+RNA_seed | AR | 0.186985 | UCEC | COLO-684 | Bicalutamide |
| MVP_top60_DNA_seed | PIK3CA | 0.857441 | LUAD | NCI-H1568 | Buparlisib | MVP_top60_DNA+RNA_seed | ATM | 0.192287 | UCEC | EN | CP466722 |
| MVP_top60_DNA_seed | BRAF | 0.858193 | LUAD | HCC-78 | PLX-4720 | MVP_top60_DNA+RNA_seed | EGFR | 0.207992 | UCEC | MFE-296 | Cetuximab |
| MVP_top60_DNA_seed | AURKA | 0.860309 | LUAD | SK-LU-1 | Alisertib | MVP_top60_DNA+RNA_seed | EP300 | 0.208765 | UCEC | MFE-319 | I-CBP112 |
| MVP_top60_DNA_seed | IGF1R | 0.860822 | LUAD | NCI-H1568 | NVP-ADW742 | MVP_top60_DNA+RNA_seed | EGFR | 0.211331 | UCEC | AN3-CA | AZD8931 |
| MVP_top60_DNA_seed | AKT1 | 0.861235 | LUAD | NCI-H1651 | Ipatasertib | MVP_top60_DNA+RNA_seed | ERBB2 | 0.211331 | UCEC | AN3-CA | AZD8931 |
| MVP_top60_DNA_seed | MDM2 | 0.863093 | LUAD | H3255 | Serdemetan | MVP_top60_DNA+RNA_seed | FGFR2 | 0.220404 | UCEC | EN | FGFR_3831 |
| MVP_top60_DNA_seed | AKT1 | 0.863103 | LUAD | SW1573 | Uprosertib | MVP_top60_DNA+RNA_seed | EGFR | 0.221284 | UCEC | AN3-CA | CUDC-101 |
| MVP_top60_DNA_seed | EZH2 | 0.863466 | LUAD | NCI-H2030 | GSK343 | MVP_top60_DNA+RNA_seed | ERBB2 | 0.221284 | UCEC | AN3-CA | CUDC-101 |
| MVP_top60_DNA_seed | KRAS | 0.86381 | LUAD | NCI-H358 | KRAS (G12C) Inhibitor-12 | MVP_top60_DNA+RNA_seed | PIK3CD | 0.223966 | UCEC | HEC-1 | IC-87114 |
| MVP_top60_DNA_seed | IGF1R | 0.863864 | LUAD | HOP-62 | BMS-754807 | MVP_top60_DNA+RNA_seed | FGFR2 | 0.233763 | UCEC | MFE-319 | PD173074 |
| MVP_top60_DNA_seed | BRAF | 0.864187 | LUAD | HOP-62 | PLX-4720 | MVP_top60_DNA+RNA_seed | JAK1 | 0.239651 | UCEC | HEC-1 | Ruxolitinib |
| MVP_top60_DNA_seed | TERT | 0.865159 | LUAD | NCI-H2023 | BIBR-1532 | MVP_top60_DNA+RNA_seed | HSP90AA1 | 0.239798 | UCEC | MFE-280 | Tanespimycin |
| MVP_top60_DNA_seed | FGFR1 | 0.865231 | LUAD | HCC-827 | PD173074 | MVP_top60_DNA+RNA_seed | HSP90AB1 | 0.239798 | UCEC | MFE-280 | Tanespimycin |
| MVP_top60_DNA_seed | ATM | 0.866654 | LUAD | HCC-78 | KU-55933 | MVP_top60_DNA+RNA_seed | MAPK1 | 0.253668 | UCEC | MFE-319 | VX-11e |
| MVP_top60_DNA_seed | TERT | 0.867275 | LUAD | EKVX | BIBR-1532 | MVP_top60_DNA+RNA_seed | FGFR2 | 0.254212 | UCEC | MFE-319 | FGFR_3831 |
| MVP_top60_DNA_seed | BRAF | 0.868314 | LUAD | NCI-H1568 | Dabrafenib | MVP_top60_DNA+RNA_seed | PIK3CB | 0.263521 | UCEC | MFE-280 | AZD6482 |
| MVP_top60_DNA_seed | EGFR | 0.868936 | LUAD | NCI-H23 | Lapatinib | MVP_top60_DNA+RNA_seed | EGFR | 0.263839 | UCEC | KLE | CI-1033 |
| MVP_top60_DNA_seed | ERBB2 | 0.868936 | LUAD | NCI-H23 | Lapatinib | MVP_top60_DNA+RNA_seed | ERBB2 | 0.263839 | UCEC | KLE | CI-1033 |
| MVP_top60_DNA_seed | BRAF | 0.869914 | LUAD | EKVX | Dabrafenib | MVP_top60_DNA+RNA_seed | AR | 0.274026 | UCEC | MFE-319 | Bicalutamide |
| MVP_top60_DNA_seed | EGFR | 0.870559 | LUAD | Calu-3 | AZD3759 | MVP_top60_DNA+RNA_seed | AKT3 | 0.277979 | UCEC | MFE-296 | AKT inhibitor VIII |
| MVP_top60_DNA_seed | GSK3B | 0.871758 | LUAD | ABC-1 | SB216763 | MVP_top60_DNA+RNA_seed | AKT1 | 0.277979 | UCEC | MFE-296 | AKT inhibitor VIII |
| MVP_top60_DNA_seed | EZH2 | 0.872629 | LUAD | NCI-H1563 | GSK343 | MVP_top60_DNA+RNA_seed | ESR1 | 0.282681 | UCEC | HEC-1 | Tamoxifen |

| | | | | | | | | | | | |
|---|---|---|---|---|---|---|---|---|---|---|---|
| MVP_top60_DNA_seed | FGFR1 | 0.874897 | LUAD | Calu-6 | PD173074 | MVP_top60_DNA+RNA_seed | MDM2 | 0.289948 | UCEC | AN3-CA | Nutlin-3a (-) |
| MVP_top60_DNA_seed | AURKA | 0.877401 | LUAD | NCI-H1651 | Alisertib | MVP_top60_DNA+RNA_seed | ATM | 0.291618 | UCEC | AN3-CA | CP466722 |
| MVP_top60_DNA_seed | TP53 | 0.877456 | LUAD | NCI-H2342 | MIRA-1 | MVP_top60_DNA+RNA_seed | AR | 0.291829 | UCEC | HEC-1 | Bicalutamide |
| MVP_top60_DNA_seed | PIK3CA | 0.878077 | LUAD | Calu-3 | Buparlisib | MVP_top60_DNA+RNA_seed | ESR1 | 0.299525 | UCEC | MFE-280 | Fulvestrant |
| MVP_top60_DNA_seed | EZH2 | 0.87872 | LUAD | NCI-H1975 | GSK343 | MVP_top60_DNA+RNA_seed | JAK1 | 0.299954 | UCEC | MFE-280 | Ruxolitinib |
| MVP_top60_DNA_seed | FGFR1 | 0.878764 | LUAD | NCI-H2342 | AZD4547 | MVP_top60_DNA+RNA_seed | FGFR2 | 0.300365 | UCEC | HEC-1 | AZD4547 |
| MVP_top60_DNA_seed | FGFR1 | 0.880621 | LUAD | NCI-H2023 | AZD4547 | MVP_top60_DNA+RNA_seed | AR | 0.31215 | UCEC | KLE | AZD3514 |
| MVP_top60_DNA_seed | EGFR | 0.881476 | LUAD | NCI-H1755 | Gefitinib | MVP_top60_DNA+RNA_seed | PIK3CB | 0.322234 | UCEC | SNG-M | AZD6482 |
| MVP_top60_DNA_seed | MET | 0.882206 | LUAD | NCI-H1793 | Savolitinib | MVP_top60_DNA+RNA_seed | AKT3 | 0.327086 | UCEC | MFE-280 | AKT inhibitor VIII |
| MVP_top60_DNA_seed | AKT1 | 0.884439 | LUAD | NCI-H1573 | MK-2206 | MVP_top60_DNA+RNA_seed | AKT1 | 0.327086 | UCEC | MFE-280 | AKT inhibitor VIII |
| MVP_top60_DNA_seed | IGF1R | 0.885216 | LUAD | VMRC-LCD | NVP-ADW742 | MVP_top60_DNA+RNA_seed | AKT3 | 0.327418 | UCEC | MFE-319 | GSK690693 |
| MVP_top60_DNA_seed | IGF1R | 0.88563 | LUAD | NCI-H1648 | GSK1904529A | MVP_top60_DNA+RNA_seed | AKT1 | 0.327418 | UCEC | MFE-319 | GSK690693 |
| MVP_top60_DNA_seed | KRAS | 0.886349 | LUAD | NCI-H1623 | KRAS (G12C) Inhibitor-1: | MVP_top60_DNA+RNA_seed | ATM | 0.330904 | UCEC | MFE-280 | CP466722 |
| MVP_top60_DNA_seed | FGFR1 | 0.88757 | LUAD | NCI-H1693 | PD173074 | MVP_top60_DNA+RNA_seed | PIK3CD | 0.332221 | UCEC | AN3-CA | Idelalisib |
| MVP_top60_DNA_seed | GSK3B | 0.888882 | LUAD | NCI-H1568 | SB216763 | MVP_top60_DNA+RNA_seed | AR | 0.334759 | UCEC | AN3-CA | Bicalutamide |
| MVP_top60_DNA_seed | GSK3B | 0.889213 | LUAD | EKVX | SB216763 | MVP_top60_DNA+RNA_seed | PIK3CD | 0.335865 | UCEC | MFE-319 | IC-87114 |
| MVP_top60_DNA_seed | AURKA | 0.889618 | LUAD | NCI-H1734 | Tozasertib | MVP_top60_DNA+RNA_seed | ATM | 0.340159 | UCEC | MFE-296 | CP466722 |
| MVP_top60_DNA_seed | EGFR | 0.889827 | LUAD | VMRC-LCD | Osimertinib | MVP_top60_DNA+RNA_seed | MAPK1 | 0.340857 | UCEC | MFE-319 | FR-180204 |
| MVP_top60_DNA_seed | MET | 0.890774 | LUAD | NCI-H2347 | Savolitinib | MVP_top60_DNA+RNA_seed | MAPK3 | 0.340857 | UCEC | MFE-319 | FR-180204 |
| MVP_top60_DNA_seed | EGFR | 0.891159 | LUAD | NCI-H1355 | AZD3759 | MVP_top60_DNA+RNA_seed | AR | 0.342183 | UCEC | KLE | Bicalutamide |
| MVP_top60_DNA_seed | AURKA | 0.892451 | LUAD | COR-L105 | ZM447439 | MVP_top60_DNA+RNA_seed | JAK1 | 0.342569 | UCEC | AN3-CA | Ruxolitinib |
| MVP_top60_DNA_seed | EGFR | 0.89335 | LUAD | NCI-H1838 | Erlotinib | MVP_top60_DNA+RNA_seed | FGFR2 | 0.344305 | UCEC | COLO-684 | FGFR_3831 |
| MVP_top60_DNA_seed | HSP90AA1 | 0.893496 | LUAD | H3255 | CCT-018159 | MVP_top60_DNA+RNA_seed | PIK3CB | 0.347108 | UCEC | KLE | AZD6482 |
| MVP_top60_DNA_seed | AKT1 | 0.894983 | LUAD | NCI-H1693 | Uprosertib | MVP_top60_DNA+RNA_seed | PIK3CA | 0.352447 | UCEC | COLO-684 | PI3Ka_4409 |
| MVP_top60_DNA_seed | EGFR | 0.895504 | LUAD | NCI-H441 | Sapitinib | MVP_top60_DNA+RNA_seed | PIK3CD | 0.352447 | UCEC | COLO-684 | PI3Ka_4409 |
| MVP_top60_DNA_seed | ERBB2 | 0.895504 | LUAD | NCI-H441 | Sapitinib | MVP_top60_DNA+RNA_seed | EGFR | 0.35841 | UCEC | AN3-CA | Afatinib |
| MVP_top60_DNA_seed | ERBB3 | 0.895504 | LUAD | NCI-H441 | Sapitinib | MVP_top60_DNA+RNA_seed | ERBB2 | 0.35841 | UCEC | AN3-CA | Afatinib |
| MVP_top60_DNA_seed | HDAC1 | 0.896296 | LUAD | NCI-H1563 | Romidepsin | MVP_top60_DNA+RNA_seed | HDAC1 | 0.361733 | UCEC | KLE | AR-42 |
| MVP_top60_DNA_seed | EGFR | 0.898124 | LUAD | HCC-44 | Erlotinib | MVP_top60_DNA+RNA_seed | FGFR2 | 0.369629 | UCEC | MFE-319 | AZD4547 |
| MVP_top60_DNA_seed | BRAF | 0.899513 | LUAD | NCI-H1793 | PLX-4720 | MVP_top60_DNA+RNA_seed | EGFR | 0.370158 | UCEC | EN | Cetuximab |
| MVP_top60_DNA_seed | FGFR1 | 0.900424 | LUAD | Calu-3 | AZD4547 | MVP_top60_DNA+RNA_seed | ATM | 0.375809 | UCEC | SNG-M | KU-60019 |
| MVP_top60_DNA_seed | ATM | 0.901097 | LUAD | EMC-BAC-2 | KU-55933 | MVP_top60_DNA+RNA_seed | HDAC1 | 0.382997 | UCEC | AN3-CA | CAY10603 |
| MVP_top60_DNA_seed | MAPK1 | 0.901227 | LUAD | NCI-H23 | SCH772984 | MVP_top60_DNA+RNA_seed | PIK3CA | 0.384848 | UCEC | MFE-296 | PI3Ka_4409 |
| MVP_top60_DNA_seed | MAPK1 | 0.902933 | LUAD | NCI-H1355 | Ulixertinib | MVP_top60_DNA+RNA_seed | PIK3CD | 0.384848 | UCEC | MFE-296 | PI3Ka_4409 |
| MVP_top60_DNA_seed | IGF1R | 0.904785 | LUAD | NCI-H2291 | NVP-ADW742 | MVP_top60_DNA+RNA_seed | SRC | 0.391633 | UCEC | MFE-319 | Bosutinib |
| MVP_top60_DNA_seed | IGF1R | 0.905171 | LUAD | NCI-H1435 | NVP-ADW742 | MVP_top60_DNA+RNA_seed | HDAC1 | 0.395963 | UCEC | AN3-CA | AR-42 |
| MVP_top60_DNA_seed | GSK3B | 0.905399 | LUAD | Calu-3 | SB216763 | MVP_top60_DNA+RNA_seed | JAK1 | 0.426205 | UCEC | MFE-319 | JAK1_3715 |
| MVP_top60_DNA_seed | MAPK1 | 0.905866 | LUAD | NCI-H1651 | SCH772984 | MVP_top60_DNA+RNA_seed | MDM2 | 0.426412 | UCEC | EN | Serdemetan |
| MVP_top60_DNA_seed | GSK3B | 0.906786 | LUAD | NCI-H3122 | SB216763 | MVP_top60_DNA+RNA_seed | EGFR | 0.428356 | UCEC | EN | AST-1306 |
| MVP_top60_DNA_seed | HDAC1 | 0.909071 | LUAD | NCI-H2291 | PCI-34051 | MVP_top60_DNA+RNA_seed | HDAC1 | 0.431141 | UCEC | KLE | Tubastatin A |
| MVP_top60_DNA_seed | BRAF | 0.909494 | LUAD | HOP-62 | SB590885 | MVP_top60_DNA+RNA_seed | FGFR2 | 0.441923 | UCEC | SNG-M | PD173074 |

| | | | | | | | | | | | |
|---|---|---|---|---|---|---|---|---|---|---|---|
| MVP_top60_DNA_seed | MDM2 | 0.910378 | LUAD | NCI-H2228 | Nutlin-3a (-) | MVP_top60_DNA+RNA_seed | HSP90AA1 | 0.445405 | UCEC | EN | CCT-018159 |
| MVP_top60_DNA_seed | SRC | 0.912033 | LUAD | NCI-H1793 | Bosutinib | MVP_top60_DNA+RNA_seed | HSP90AB1 | 0.445405 | UCEC | EN | CCT-018159 |
| MVP_top60_DNA_seed | AKT1 | 0.912764 | LUAD | NCI-H1693 | MK-2206 | MVP_top60_DNA+RNA_seed | MDM2 | 0.45042 | UCEC | AN3-CA | Serdemetan |
| MVP_top60_DNA_seed | SMARCA4 | 0.912899 | LUAD | HCC-827 | PFI3 | MVP_top60_DNA+RNA_seed | AKT1 | 0.455757 | UCEC | SNG-M | MK-2206 |
| MVP_top60_DNA_seed | BRAF | 0.912949 | LUAD | NCI-H358 | SB590885 | MVP_top60_DNA+RNA_seed | HSP90AA1 | 0.457367 | UCEC | SNG-M | Elesclomol |
| MVP_top60_DNA_seed | TP53 | 0.914092 | LUAD | Calu-6 | MIRA-1 | MVP_top60_DNA+RNA_seed | HSP90AB1 | 0.457367 | UCEC | SNG-M | Elesclomol |
| MVP_top60_DNA_seed | MAPK1 | 0.914629 | LUAD | ABC-1 | Ulixertinib | MVP_top60_DNA+RNA_seed | ATM | 0.458843 | UCEC | SNG-M | KU-55933 |
| MVP_top60_DNA_seed | EGFR | 0.914973 | LUAD | NCI-H23 | Sapitinib | MVP_top60_DNA+RNA_seed | MDM2 | 0.463917 | UCEC | KLE | Serdemetan |
| MVP_top60_DNA_seed | ERBB2 | 0.914973 | LUAD | NCI-H23 | Sapitinib | MVP_top60_DNA+RNA_seed | EGFR | 0.48369 | UCEC | KLE | Afatinib |
| MVP_top60_DNA_seed | ERBB3 | 0.914973 | LUAD | NCI-H23 | Sapitinib | MVP_top60_DNA+RNA_seed | ERBB2 | 0.48369 | UCEC | KLE | Afatinib |
| MVP_top60_DNA_seed | MAPK1 | 0.91604 | LUAD | EMC-BAC-2 | VX-11e | MVP_top60_DNA+RNA_seed | HSP90AA1 | 0.484913 | UCEC | EN | SNX-2112 |
| MVP_top60_DNA_seed | MDM2 | 0.917131 | LUAD | NCI-H2009 | Nutlin-3a (-) | MVP_top60_DNA+RNA_seed | HSP90AB1 | 0.484913 | UCEC | EN | SNX-2112 |
| MVP_top60_DNA_seed | MET | 0.917173 | LUAD | A549 | Savolitinib | MVP_top60_DNA+RNA_seed | EGFR | 0.486628 | UCEC | AN3-CA | AST-1306 |
| MVP_top60_DNA_seed | MET | 0.917527 | LUAD | HCC-78 | Savolitinib | MVP_top60_DNA+RNA_seed | ATM | 0.493472 | UCEC | MFE-280 | KU-55933 |
| MVP_top60_DNA_seed | HDAC1 | 0.918698 | LUAD | NCI-H1975 | PCI-34051 | MVP_top60_DNA+RNA_seed | JAK1 | 0.494036 | UCEC | MFE-296 | Ruxolitinib |
| MVP_top60_DNA_seed | PIK3CA | 0.920335 | LUAD | NCI-H1568 | GNE-317 | MVP_top60_DNA+RNA_seed | MAPK1 | 0.500664 | UCEC | HEC-1 | FR-180204 |
| MVP_top60_DNA_seed | PIK3CA | 0.921054 | LUAD | NCI-H2342 | GNE-317 | MVP_top60_DNA+RNA_seed | MAPK3 | 0.500664 | UCEC | HEC-1 | FR-180204 |
| MVP_top60_DNA_seed | AKT1 | 0.92106 | LUAD | NCI-H1651 | AZD5363 | MVP_top60_DNA+RNA_seed | HDAC1 | 0.504916 | UCEC | MFE-280 | Dacinostat |
| MVP_top60_DNA_seed | SMARCA4 | 0.921551 | LUAD | Calu-6 | PFI3 | MVP_top60_DNA+RNA_seed | AR | 0.507807 | UCEC | COLO-684 | AZD3514 |
| MVP_top60_DNA_seed | TP53 | 0.92166 | LUAD | NCI-H1975 | MIRA-1 | MVP_top60_DNA+RNA_seed | PIK3CD | 0.508062 | UCEC | AN3-CA | IC-87114 |
| MVP_top60_DNA_seed | GSK3B | 0.921822 | LUAD | HCC-78 | SB216763 | MVP_top60_DNA+RNA_seed | SRC | 0.511928 | UCEC | AN3-CA | NVP-BHG712 |
| MVP_top60_DNA_seed | HDAC1 | 0.921864 | LUAD | Calu-3 | Romidepsin | MVP_top60_DNA+RNA_seed | AR | 0.516299 | UCEC | MFE-296 | Bicalutamide |
| MVP_top60_DNA_seed | MET | 0.921934 | LUAD | NCI-H2030 | Savolitinib | MVP_top60_DNA+RNA_seed | JAK1 | 0.52226 | UCEC | MFE-280 | JAK1_3715 |
| MVP_top60_DNA_seed | EZH2 | 0.923147 | LUAD | HCC-827 | GSK343 | MVP_top60_DNA+RNA_seed | ESR1 | 0.536283 | UCEC | KLE | Tamoxifen |
| MVP_top60_DNA_seed | EGFR | 0.923693 | LUAD | NCI-H2087 | Afatinib | MVP_top60_DNA+RNA_seed | HSP90AA1 | 0.537936 | UCEC | KLE | SNX-2112 |
| MVP_top60_DNA_seed | ERBB2 | 0.923693 | LUAD | NCI-H2087 | Afatinib | MVP_top60_DNA+RNA_seed | HSP90AB1 | 0.537936 | UCEC | KLE | SNX-2112 |
| MVP_top60_DNA_seed | MDM2 | 0.924447 | LUAD | Calu-3 | Nutlin-3a (-) | MVP_top60_DNA+RNA_seed | SRC | 0.547493 | UCEC | HEC-1 | NVP-BHG712 |
| MVP_top60_DNA_seed | TERT | 0.926156 | LUAD | NCI-H1755 | BIBR-1532 | MVP_top60_DNA+RNA_seed | PIK3CA | 0.563782 | UCEC | MFE-319 | AZD8835 |
| MVP_top60_DNA_seed | BRAF | 0.926668 | LUAD | NCI-H1944 | SB590885 | MVP_top60_DNA+RNA_seed | PIK3CD | 0.563782 | UCEC | MFE-319 | AZD8835 |
| MVP_top60_DNA_seed | BRAF | 0.927375 | LUAD | HOP-62 | Dabrafenib | MVP_top60_DNA+RNA_seed | MAPK1 | 0.57192 | UCEC | AN3-CA | VX-11e |
| MVP_top60_DNA_seed | EGFR | 0.927613 | LUAD | Calu-6 | Erlotinib | MVP_top60_DNA+RNA_seed | EP300 | 0.573295 | UCEC | AN3-CA | I-CBP112 |
| MVP_top60_DNA_seed | IGF1R | 0.927764 | LUAD | Calu-3 | NVP-ADW742 | MVP_top60_DNA+RNA_seed | EP300 | 0.576443 | UCEC | COLO-684 | I-CBP112 |
| MVP_top60_DNA_seed | AURKA | 0.927966 | LUAD | HOP-62 | ZM447439 | MVP_top60_DNA+RNA_seed | HDAC1 | 0.576983 | UCEC | MFE-296 | CAY10603 |
| MVP_top60_DNA_seed | BRAF | 0.929629 | LUAD | VMRC-LCD | Dabrafenib | MVP_top60_DNA+RNA_seed | EGFR | 0.582641 | UCEC | AN3-CA | Afatinib |
| MVP_top60_DNA_seed | PIK3CA | 0.930774 | LUAD | NCI-H2087 | Buparlisib | MVP_top60_DNA+RNA_seed | ERBB2 | 0.582641 | UCEC | AN3-CA | Afatinib |
| MVP_top60_DNA_seed | PIK3CA | 0.93106 | LUAD | NCI-H2347 | AZD8186 | MVP_top60_DNA+RNA_seed | AR | 0.586072 | UCEC | EN | Bicalutamide |
| MVP_top60_DNA_seed | HSP90AA1 | 0.931818 | LUAD | NCI-H522 | Luminespib | MVP_top60_DNA+RNA_seed | HDAC1 | 0.587212 | UCEC | KLE | JQ12 |
| MVP_top60_DNA_seed | AKT1 | 0.932135 | LUAD | NCI-H1650 | MK-2206 | MVP_top60_DNA+RNA_seed | HDAC2 | 0.587212 | UCEC | KLE | JQ12 |
| MVP_top60_DNA_seed | IGF1R | 0.932998 | LUAD | EMC-BAC-1 | Linsitinib | MVP_top60_DNA+RNA_seed | HSP90AA1 | 0.593566 | UCEC | MFE-280 | Elesclomol |
| MVP_top60_DNA_seed | EGFR | 0.933491 | LUAD | ABC-1 | Osimertinib | MVP_top60_DNA+RNA_seed | HSP90AB1 | 0.593566 | UCEC | MFE-280 | Elesclomol |
| MVP_top60_DNA_seed | AKT1 | 0.933999 | LUAD | EKVX | MK-2206 | MVP_top60_DNA+RNA_seed | EGFR | 0.596501 | UCEC | AN3-CA | PF-00299804 |

| | | | | | | | | | | | |
|---|---|---|---|---|---|---|---|---|---|---|---|
| MVP_top60_DNA_seed | MAPK1 | 0.93517 | LUAD | NCI-H441 | VX-11e | MVP_top60_DNA+RNA_seed | ERBB2 | 0.596501 | UCEC | AN3-CA | PF-00299804 |
| MVP_top60_DNA_seed | KRAS | 0.936083 | LUAD | NCI-H2030 | KRAS (G12C) Inhibitor-1[illegible] | MVP_top60_DNA+RNA_seed | HSP90AA1 | 0.59667 | UCEC | EN | Tanespimycin |
| MVP_top60_DNA_seed | ATM | 0.936511 | LUAD | ABC-1 | KU-55933 | MVP_top60_DNA+RNA_seed | HSP90AB1 | 0.59667 | UCEC | EN | Tanespimycin |
| MVP_top60_DNA_seed | FGFR1 | 0.936713 | LUAD | NCI-H2228 | PD173074 | MVP_top60_DNA+RNA_seed | ATM | 0.599189 | UCEC | KLE | KU-60019 |
| MVP_top60_DNA_seed | IGF1R | 0.936912 | LUAD | NCI-H1693 | BMS-536924 | MVP_top60_DNA+RNA_seed | ERBB2 | 0.6102 | UCEC | HEC-1 | CP724714 |
| MVP_top60_DNA_seed | AKT1 | 0.937973 | LUAD | Calu-6 | Uprosertib | MVP_top60_DNA+RNA_seed | EGFR | 0.610908 | UCEC | MFE-280 | AZD8931 |
| MVP_top60_DNA_seed | SMARCA4 | 0.93819 | LUAD | NCI-H2030 | PFI3 | MVP_top60_DNA+RNA_seed | ERBB2 | 0.610908 | UCEC | MFE-280 | AZD8931 |
| MVP_top60_DNA_seed | MAPK1 | 0.939697 | LUAD | VMRC-LCD | Ulixertinib | MVP_top60_DNA+RNA_seed | FGFR2 | 0.612214 | UCEC | EN | AZD4547 |
| MVP_top60_DNA_seed | AURKA | 0.940159 | LUAD | EKVX | ZM447439 | MVP_top60_DNA+RNA_seed | SRC | 0.616683 | UCEC | MFE-296 | Bosutinib |
| MVP_top60_DNA_seed | EGFR | 0.940165 | LUAD | SK-LU-1 | Sapitinib | MVP_top60_DNA+RNA_seed | HSP90AA1 | 0.624883 | UCEC | MFE-319 | SNX-2112 |
| MVP_top60_DNA_seed | ERBB2 | 0.940165 | LUAD | SK-LU-1 | Sapitinib | MVP_top60_DNA+RNA_seed | HSP90AB1 | 0.624883 | UCEC | MFE-319 | SNX-2112 |
| MVP_top60_DNA_seed | ERBB3 | 0.940165 | LUAD | SK-LU-1 | Sapitinib | MVP_top60_DNA+RNA_seed | EGFR | 0.639637 | UCEC | MFE-296 | Afatinib |
| MVP_top60_DNA_seed | IGF1R | 0.940203 | LUAD | HCC-827 | Linsitinib | MVP_top60_DNA+RNA_seed | ERBB2 | 0.639637 | UCEC | MFE-296 | Afatinib |
| MVP_top60_DNA_seed | PIK3CA | 0.940892 | LUAD | NCI-H2405 | AZD8186 | MVP_top60_DNA+RNA_seed | FGFR2 | 0.647232 | UCEC | SNG-M | AZD4547 |
| MVP_top60_DNA_seed | AKT1 | 0.942031 | LUAD | ABC-1 | Ipatasertib | MVP_top60_DNA+RNA_seed | JAK1 | 0.650458 | UCEC | EN | AZD1480 |
| MVP_top60_DNA_seed | IGF1R | 0.944484 | LUAD | SW1573 | Linsitinib | MVP_top60_DNA+RNA_seed | EGFR | 0.653766 | UCEC | MFE-319 | Pelitinib |
| MVP_top60_DNA_seed | SMARCA4 | 0.945359 | LUAD | NCI-H1944 | PFI3 | MVP_top60_DNA+RNA_seed | SRC | 0.654226 | UCEC | COLO-684 | Bosutinib |
| MVP_top60_DNA_seed | KRAS | 0.946933 | LUAD | NCI-H2085 | KRAS (G12C) Inhibitor-1[illegible] | MVP_top60_DNA+RNA_seed | ERBB2 | 0.654458 | UCEC | MFE-296 | CP724714 |
| MVP_top60_DNA_seed | MAPK1 | 0.946978 | LUAD | NCI-H2342 | VX-11e | MVP_top60_DNA+RNA_seed | ERBB2 | 0.655068 | UCEC | AN3-CA | CP724714 |
| MVP_top60_DNA_seed | HSP90AA1 | 0.947345 | LUAD | PC-3_[JPC-3] | Tanespimycin | MVP_top60_DNA+RNA_seed | EGFR | 0.659161 | UCEC | KLE | Gefitinib |
| MVP_top60_DNA_seed | MAPK1 | 0.947726 | LUAD | ABC-1 | SCH772984 | MVP_top60_DNA+RNA_seed | EGFR | 0.663197 | UCEC | AN3-CA | Cetuximab |
| MVP_top60_DNA_seed | BRAF | 0.947908 | LUAD | NCI-H1568 | SB590885 | MVP_top60_DNA+RNA_seed | MDM2 | 0.677846 | UCEC | COLO-684 | Nutlin-3a (-) |
| MVP_top60_DNA_seed | IGF1R | 0.947925 | LUAD | NCI-H1666 | BMS-754807 | MVP_top60_DNA+RNA_seed | EGFR | 0.678647 | UCEC | MFE-296 | Gefitinib |
| MVP_top60_DNA_seed | TERT | 0.947953 | LUAD | NCI-H2342 | BIBR-1532 | MVP_top60_DNA+RNA_seed | EGFR | 0.685851 | UCEC | HEC-1 | Afatinib |
| MVP_top60_DNA_seed | PIK3CA | 0.94951 | LUAD | SK-LU-1 | AZD8186 | MVP_top60_DNA+RNA_seed | ERBB2 | 0.685851 | UCEC | HEC-1 | Afatinib |
| MVP_top60_DNA_seed | HDAC1 | 0.950171 | LUAD | NCI-H322M | Entinostat | MVP_top60_DNA+RNA_seed | HDAC1 | 0.686067 | UCEC | MFE-319 | Tubastatin A |
| MVP_top60_DNA_seed | BRAF | 0.950346 | LUAD | NCI-H1693 | SB590885 | MVP_top60_DNA+RNA_seed | MAPK1 | 0.687686 | UCEC | EN | VX-11e |
| MVP_top60_DNA_seed | IGF1R | 0.951134 | LUAD | NCI-H2085 | GSK1904529A | MVP_top60_DNA+RNA_seed | EGFR | 0.691755 | UCEC | HEC-1 | Pelitinib |
| MVP_top60_DNA_seed | AKT1 | 0.951216 | LUAD | HCC-827 | AZD5363 | MVP_top60_DNA+RNA_seed | EGFR | 0.699062 | UCEC | COLO-684 | AST-1306 |
| MVP_top60_DNA_seed | EZH2 | 0.951232 | LUAD | ABC-1 | GSK343 | MVP_top60_DNA+RNA_seed | HDAC1 | 0.699207 | UCEC | AN3-CA | Belinostat |
| MVP_top60_DNA_seed | EGFR | 0.951415 | LUAD | NCI-H650 | Lapatinib | MVP_top60_DNA+RNA_seed | HDAC1 | 0.703336 | UCEC | MFE-296 | Belinostat |
| MVP_top60_DNA_seed | ERBB2 | 0.951415 | LUAD | NCI-H650 | Lapatinib | MVP_top60_DNA+RNA_seed | JAK1 | 0.708809 | UCEC | AN3-CA | JAK1_3715 |
| MVP_top60_DNA_seed | BRAF | 0.951749 | LUAD | HCC-827 | SB590885 | MVP_top60_DNA+RNA_seed | PIK3CD | 0.709942 | UCEC | MFE-296 | IC-87114 |
| MVP_top60_DNA_seed | IGF1R | 0.955728 | LUAD | NCI-H2347 | NVP-ADW742 | MVP_top60_DNA+RNA_seed | EGFR | 0.711985 | UCEC | COLO-684 | Lapatinib |
| MVP_top60_DNA_seed | AKT1 | 0.955732 | LUAD | NCI-H1623 | Ipatasertib | MVP_top60_DNA+RNA_seed | ERBB2 | 0.711985 | UCEC | COLO-684 | Lapatinib |
| MVP_top60_DNA_seed | MET | 0.957671 | LUAD | NCI-H1781 | Crizotinib | MVP_top60_DNA+RNA_seed | HSP90AA1 | 0.721388 | UCEC | AN3-CA | SNX-2112 |
| MVP_top60_DNA_seed | EZH2 | 0.957823 | LUAD | NCI-H358 | GSK343 | MVP_top60_DNA+RNA_seed | HSP90AB1 | 0.721388 | UCEC | AN3-CA | SNX-2112 |
| MVP_top60_DNA_seed | EZH2 | 0.95783 | LUAD | Calu-3 | GSK343 | MVP_top60_DNA+RNA_seed | HDAC1 | 0.724498 | UCEC | HEC-1 | Tubastatin A |
| MVP_top60_DNA_seed | PIK3CA | 0.957989 | LUAD | SK-LU-1 | GNE-317 | MVP_top60_DNA+RNA_seed | MAPK1 | 0.726595 | UCEC | AN3-CA | FR-180204 |
| MVP_top60_DNA_seed | IGF1R | 0.958939 | LUAD | PC-3_[JPC-3] | Linsitinib | MVP_top60_DNA+RNA_seed | MAPK3 | 0.726595 | UCEC | AN3-CA | FR-180204 |
| MVP_top60_DNA_seed | CREBBP | 0.958965 | LUAD | EMC-BAC-2 | SGC-CBP30 | MVP_top60_DNA+RNA_seed | EGFR | 0.728492 | UCEC | MFE-280 | Gefitinib |

| | | | | | | | | | | | |
|---|---|---|---|---|---|---|---|---|---|---|---|
| MVP_top60_DNA_seed | PIK3CA | 0.959462 | LUAD | VMRC-LCD | GNE-317 | MVP_top60_DNA+RNA_seed | HDAC1 | 0.733862 | UCEC | COLO-684 | AR-42 |
| MVP_top60_DNA_seed | EZH2 | 0.95963 | LUAD | NCI-H2087 | GSK343 | MVP_top60_DNA+RNA_seed | EGFR | 0.737688 | UCEC | HEC-1 | AST-1306 |
| MVP_top60_DNA_seed | EGFR | 0.960605 | LUAD | NCI-H650 | Osimertinib | MVP_top60_DNA+RNA_seed | MDM2 | 0.749433 | UCEC | COLO-684 | Serdemetan |
| MVP_top60_DNA_seed | IGF1R | 0.960727 | LUAD | NCI-H1993 | GSK1904529A | MVP_top60_DNA+RNA_seed | PIK3CB | 0.750803 | UCEC | COLO-684 | TGX221 |
| MVP_top60_DNA_seed | EGFR | 0.960995 | LUAD | NCI-H1437 | Afatinib | MVP_top60_DNA+RNA_seed | FGFR2 | 0.75276 | UCEC | COLO-684 | AZD4547 |
| MVP_top60_DNA_seed | ERBB2 | 0.960995 | LUAD | NCI-H1437 | Afatinib | MVP_top60_DNA+RNA_seed | ATM | 0.765162 | UCEC | EN | KU-55933 |
| MVP_top60_DNA_seed | BRAF | 0.961481 | LUAD | PC-3_[JPC-3] | SB590885 | MVP_top60_DNA+RNA_seed | AR | 0.774001 | UCEC | AN3-CA | AZD3514 |
| MVP_top60_DNA_seed | MAPK1 | 0.963991 | LUAD | VMRC-LCD | VX-11e | MVP_top60_DNA+RNA_seed | EGFR | 0.77468 | UCEC | SNG-M | Gefitinib |
| MVP_top60_DNA_seed | FGFR1 | 0.964134 | LUAD | NCI-H23 | PD173074 | MVP_top60_DNA+RNA_seed | SRC | 0.776491 | UCEC | COLO-684 | WH-4-023 |
| MVP_top60_DNA_seed | HDAC1 | 0.965539 | LUAD | Calu-6 | Entinostat | MVP_top60_DNA+RNA_seed | HDAC1 | 0.778957 | UCEC | MFE-280 | JQ12 |
| MVP_top60_DNA_seed | EGFR | 0.965804 | LUAD | NCI-H1793 | Erlotinib | MVP_top60_DNA+RNA_seed | HDAC2 | 0.778957 | UCEC | MFE-280 | JQ12 |
| MVP_top60_DNA_seed | FGFR1 | 0.966942 | LUAD | NCI-H2228 | AZD4547 | MVP_top60_DNA+RNA_seed | HDAC1 | 0.780934 | UCEC | SNG-M | Tubastatin A |
| MVP_top60_DNA_seed | AURKA | 0.967196 | LUAD | NCI-H1793 | Alisertib | MVP_top60_DNA+RNA_seed | AKT3 | 0.782152 | UCEC | HEC-1 | GSK690693 |
| MVP_top60_DNA_seed | CREBBP | 0.967589 | LUAD | NCI-H1623 | SGC-CBP30 | MVP_top60_DNA+RNA_seed | AKT1 | 0.782152 | UCEC | HEC-1 | GSK690693 |
| MVP_top60_DNA_seed | MAPK1 | 0.967755 | LUAD | NCI-H1693 | VX-11e | MVP_top60_DNA+RNA_seed | FGFR2 | 0.786376 | UCEC | MFE-319 | AZD4547 |
| MVP_top60_DNA_seed | AKT1 | 0.968373 | LUAD | NCI-H1693 | Ipatasertib | MVP_top60_DNA+RNA_seed | JAK1 | 0.794613 | UCEC | EN | JAK1_3715 |
| MVP_top60_DNA_seed | MET | 0.968425 | LUAD | NCI-H2291 | Crizotinib | MVP_top60_DNA+RNA_seed | AR | 0.804154 | UCEC | MFE-280 | Bicalutamide |
| MVP_top60_DNA_seed | MDM2 | 0.968829 | LUAD | NCI-H2085 | Serdemetan | MVP_top60_DNA+RNA_seed | ATM | 0.811403 | UCEC | EN | KU-60019 |
| MVP_top60_DNA_seed | KRAS | 0.969335 | LUAD | HCC-827 | (RAS (G12C) Inhibitor-1: | MVP_top60_DNA+RNA_seed | HSP90AA1 | 0.812381 | UCEC | KLE | Luminespib |
| MVP_top60_DNA_seed | MET | 0.973415 | LUAD | NCI-H2087 | Savolitinib | MVP_top60_DNA+RNA_seed | HSP90AB1 | 0.812381 | UCEC | KLE | Luminespib |
| MVP_top60_DNA_seed | AKT1 | 0.974683 | LUAD | NCI-H358 | MK-2206 | MVP_top60_DNA+RNA_seed | MAPK1 | 0.814761 | UCEC | MFE-296 | FR-180204 |
| MVP_top60_DNA_seed | TP53 | 0.977542 | LUAD | NCI-H322M | MIRA-1 | MVP_top60_DNA+RNA_seed | MAPK3 | 0.814761 | UCEC | MFE-296 | FR-180204 |
| MVP_top60_DNA_seed | IGF1R | 0.978539 | LUAD | NCI-H1793 | NVP-ADW742 | MVP_top60_DNA+RNA_seed | EGFR | 0.81677 | UCEC | HEC-1 | PF-00299804 |
| MVP_top60_DNA_seed | IGF1R | 0.97976 | LUAD | HCC-827 | BMS-754807 | MVP_top60_DNA+RNA_seed | ERBB2 | 0.81677 | UCEC | HEC-1 | PF-00299804 |
| MVP_top60_DNA_seed | BRAF | 0.979943 | LUAD | Calu-3 | SB590885 | MVP_top60_DNA+RNA_seed | EGFR | 0.823994 | UCEC | MFE-296 | PF-00299804 |
| MVP_top60_DNA_seed | AKT1 | 0.980124 | LUAD | NCI-H1650 | GSK2110183B | MVP_top60_DNA+RNA_seed | ERBB2 | 0.823994 | UCEC | MFE-296 | PF-00299804 |
| MVP_top60_DNA_seed | BRAF | 0.981309 | LUAD | NCI-H1563 | PLX-4720 | MVP_top60_DNA+RNA_seed | MDM2 | 0.831763 | UCEC | MFE-280 | Nutlin-3a (-) |
| MVP_top60_DNA_seed | EGFR | 0.981402 | LUAD | NCI-H3122 | Gefitinib | MVP_top60_DNA+RNA_seed | PIK3CD | 0.832741 | UCEC | HEC-1 | Idelalisib |
| MVP_top60_DNA_seed | AKT1 | 0.983721 | LUAD | NCI-H1573 | AT13148 | MVP_top60_DNA+RNA_seed | AR | 0.842284 | UCEC | SNG-M | Bicalutamide |
| MVP_top60_DNA_seed | MET | 0.983982 | LUAD | NCI-H522 | Crizotinib | MVP_top60_DNA+RNA_seed | MAPK1 | 0.842517 | UCEC | HEC-1 | VX-11e |
| MVP_top60_DNA_seed | IGF1R | 0.985523 | LUAD | NCI-H1693 | BMS-754807 | MVP_top60_DNA+RNA_seed | MDM2 | 0.845624 | UCEC | MFE-319 | Serdemetan |
| MVP_top60_DNA_seed | HSP90AA1 | 0.985952 | LUAD | HCC-827 | Tanespimycin | MVP_top60_DNA+RNA_seed | EGFR | 0.847915 | UCEC | EN | PF-00299804 |
| MVP_top60_DNA_seed | GSK3B | 0.986433 | LUAD | ABC-1 | CHIR-99021 | MVP_top60_DNA+RNA_seed | ERBB2 | 0.847915 | UCEC | EN | PF-00299804 |
| MVP_top60_DNA_seed | TP53 | 0.986453 | LUAD | PC-14 | MIRA-1 | MVP_top60_DNA+RNA_seed | HDAC1 | 0.851064 | UCEC | COLO-684 | Entinostat |
| MVP_top60_DNA_seed | GSK3B | 0.989316 | LUAD | NCI-H1944 | SB216763 | MVP_top60_DNA+RNA_seed | JAK1 | 0.853787 | UCEC | KLE | Ruxolitinib |
| MVP_top60_DNA_seed | EGFR | 0.989603 | LUAD | NCI-H2085 | AZD3759 | MVP_top60_DNA+RNA_seed | ESR1 | 0.854879 | UCEC | COLO-684 | Tamoxifen |
| MVP_top60_DNA_seed | KRAS | 0.98968 | LUAD | Calu-3 | (RAS (G12C) Inhibitor-1: | MVP_top60_DNA+RNA_seed | ERBB2 | 0.876731 | UCEC | EN | CP724714 |
| MVP_top60_DNA_seed | IGF1R | 0.990332 | LUAD | NCI-H1993 | NVP-ADW742 | MVP_top60_DNA+RNA_seed | EGFR | 0.884773 | UCEC | KLE | Cetuximab |
| MVP_top60_DNA_seed | MET | 0.990781 | LUAD | NCI-H1666 | Crizotinib | MVP_top60_DNA+RNA_seed | JAK1 | 0.884858 | UCEC | COLO-684 | AZD1480 |
| MVP_top60_DNA_seed | MDM2 | 0.991437 | LUAD | HCC-827 | Nutlin-3a (-) | MVP_top60_DNA+RNA_seed | EGFR | 0.885137 | UCEC | MFE-280 | Afatinib |
| MVP_top60_DNA_seed | BRAF | 0.994664 | LUAD | NCI-H650 | PLX-4720 | MVP_top60_DNA+RNA_seed | ERBB2 | 0.885137 | UCEC | MFE-280 | Afatinib |

| | | | | | | | | | | | |
|---|---|---|---|---|---|---|---|---|---|---|---|
| MVP_top60_DNA_seed | EGFR | 0.996548 | LUAD | NCI-H1793 | Gefitinib | MVP_top60_DNA+RNA_seed | AKT3 | 0.887171 | UCEC | HEC-1 | AKT inhibitor VIII |
| MVP_top60_DNA_seed | EGFR | 0.996635 | LUAD | NCI-H1651 | Lapatinib | MVP_top60_DNA+RNA_seed | AKT1 | 0.887171 | UCEC | HEC-1 | AKT inhibitor VIII |
| MVP_top60_DNA_seed | ERBB2 | 0.996635 | LUAD | NCI-H1651 | Lapatinib | MVP_top60_DNA+RNA_seed | MAPK1 | 0.894344 | UCEC | MFE-296 | VX-11e |
| MVP_top60_DNA_seed | SRC | 0.99665 | LUAD | VMRC-LCD | Bosutinib | MVP_top60_DNA+RNA_seed | HDAC1 | 0.900099 | UCEC | AN3-CA | Tubastatin A |
| MVP_top60_DNA_seed | AKT1 | 0.999248 | LUAD | SW1573 | Afuresertib | MVP_top60_DNA+RNA_seed | SRC | 0.90174 | UCEC | AN3-CA | Bosutinib |
| MVP_top60_DNA_seed | AKT1 | 1.002867 | LUAD | NCI-H650 | Afuresertib | MVP_top60_DNA+RNA_seed | FGFR2 | 0.917464 | UCEC | HEC-1 | FGFR_3831 |
| MVP_top60_DNA_seed | EZH2 | 1.003655 | LUAD | COR-L105 | GSK343 | MVP_top60_DNA+RNA_seed | AR | 0.917979 | UCEC | MFE-296 | Bicalutamide |
| MVP_top60_DNA_seed | HSP90AA1 | 1.003695 | LUAD | HCC-78 | Tanespimycin | MVP_top60_DNA+RNA_seed | EGFR | 0.919158 | UCEC | COLO-684 | Cetuximab |
| MVP_top60_DNA_seed | IGF1R | 1.004013 | LUAD | HOP-62 | GSK1904529A | MVP_top60_DNA+RNA_seed | EGFR | 0.9262 | UCEC | COLO-684 | Erlotinib |
| MVP_top60_DNA_seed | MET | 1.004068 | LUAD | PC-14 | Savolitinib | MVP_top60_DNA+RNA_seed | PIK3CD | 0.928035 | UCEC | MFE-296 | Idelalisib |
| MVP_top60_DNA_seed | FGFR1 | 1.004624 | LUAD | EMC-BAC-2 | PD173074 | MVP_top60_DNA+RNA_seed | JAK1 | 0.938225 | UCEC | COLO-684 | JAK1_3715 |
| MVP_top60_DNA_seed | MET | 1.004822 | LUAD | VMRC-LCD | Savolitinib | MVP_top60_DNA+RNA_seed | EGFR | 0.960282 | UCEC | COLO-684 | PF-00299804 |
| MVP_top60_DNA_seed | EZH2 | 1.007129 | LUAD | VMRC-LCD | GSK343 | MVP_top60_DNA+RNA_seed | ERBB2 | 0.960282 | UCEC | COLO-684 | PF-00299804 |
| MVP_top60_DNA_seed | CREBBP | 1.009031 | LUAD | SK-LU-1 | SGC-CBP30 | MVP_top60_DNA+RNA_seed | ATM | 0.972207 | UCEC | KLE | KU-55933 |
| MVP_top60_DNA_seed | SRC | 1.009268 | LUAD | NCI-H3122 | Bosutinib | MVP_top60_DNA+RNA_seed | EGFR | 0.984534 | UCEC | MFE-280 | Afatinib |
| MVP_top60_DNA_seed | HSP90AA1 | 1.009437 | LUAD | SK-LU-1 | Luminespib | MVP_top60_DNA+RNA_seed | ERBB2 | 0.984534 | UCEC | MFE-280 | Afatinib |
| MVP_top60_DNA_seed | IGF1R | 1.009589 | LUAD | NCI-H2023 | Linsitinib | MVP_top60_DNA+RNA_seed | HSP90AA1 | 0.985196 | UCEC | SNG-M | Tanespimycin |
| MVP_top60_DNA_seed | AURKA | 1.0104 | LUAD | NCI-H322M | Alisertib | MVP_top60_DNA+RNA_seed | HSP90AB1 | 0.985196 | UCEC | SNG-M | Tanespimycin |
| MVP_top60_DNA_seed | AURKA | 1.010505 | LUAD | EMC-BAC-2 | ZM447439 | MVP_top60_DNA+RNA_seed | SRC | 0.990642 | UCEC | COLO-684 | Saracatinib |
| MVP_top60_DNA_seed | TP53 | 1.010753 | LUAD | HOP-62 | MIRA-1 | MVP_top60_DNA+RNA_seed | EGFR | 0.993025 | UCEC | MFE-296 | Afatinib |
| MVP_top60_DNA_seed | MDM2 | 1.012476 | LUAD | NCI-H358 | Nutlin-3a (-) | MVP_top60_DNA+RNA_seed | ERBB2 | 0.993025 | UCEC | MFE-296 | Afatinib |
| MVP_top60_DNA_seed | EGFR | 1.013636 | LUAD | NCI-H23 | Osimertinib | MVP_top60_DNA+RNA_seed | AR | 0.99526 | UCEC | COLO-684 | Bicalutamide |
| MVP_top60_DNA_seed | TP53 | 1.015556 | LUAD | NCI-H358 | MIRA-1 | MVP_top60_DNA+RNA_seed | EGFR | 1.003911 | UCEC | COLO-684 | Afatinib |
| MVP_top60_DNA_seed | PIK3CA | 1.015869 | LUAD | NCI-H1838 | AZD8186 | MVP_top60_DNA+RNA_seed | ERBB2 | 1.003911 | UCEC | COLO-684 | Afatinib |
| MVP_top60_DNA_seed | HDAC1 | 1.01639 | LUAD | VMRC-LCD | PCI-34051 | MVP_top60_DNA+RNA_seed | EGFR | 1.010634 | UCEC | COLO-684 | Afatinib |
| MVP_top60_DNA_seed | MAPK1 | 1.016473 | LUAD | NCI-H1693 | Ulixertinib | MVP_top60_DNA+RNA_seed | ERBB2 | 1.010634 | UCEC | COLO-684 | Afatinib |
| MVP_top60_DNA_seed | HSP90AA1 | 1.017946 | LUAD | RERF-LC-KJ | Luminespib | MVP_top60_DNA+RNA_seed | FGFR2 | 1.011234 | UCEC | KLE | AZD4547 |
| MVP_top60_DNA_seed | PIK3CA | 1.019241 | LUAD | NCI-H1568 | Alpelisib | MVP_top60_DNA+RNA_seed | HSP90AA1 | 1.020393 | UCEC | MFE-296 | SNX-2112 |
| MVP_top60_DNA_seed | MAPK1 | 1.019381 | LUAD | NCI-H1651 | Ulixertinib | MVP_top60_DNA+RNA_seed | HSP90AB1 | 1.020393 | UCEC | MFE-296 | SNX-2112 |
| MVP_top60_DNA_seed | AURKA | 1.020653 | LUAD | NCI-H2087 | Alisertib | MVP_top60_DNA+RNA_seed | EGFR | 1.02499 | UCEC | AN3-CA | Gefitinib |
| MVP_top60_DNA_seed | BRAF | 1.020963 | LUAD | NCI-H2342 | Dabrafenib | MVP_top60_DNA+RNA_seed | MAPK1 | 1.045057 | UCEC | COLO-684 | FR-180204 |
| MVP_top60_DNA_seed | BRAF | 1.021227 | LUAD | NCI-H1623 | PLX-4720 | MVP_top60_DNA+RNA_seed | MAPK3 | 1.045057 | UCEC | COLO-684 | FR-180204 |
| MVP_top60_DNA_seed | IGF1R | 1.022293 | LUAD | NCI-H1650 | GSK1904529A | MVP_top60_DNA+RNA_seed | MAPK1 | 1.047686 | UCEC | KLE | VX-11e |
| MVP_top60_DNA_seed | FGFR1 | 1.024148 | LUAD | HCC-78 | AZD4547 | MVP_top60_DNA+RNA_seed | FGFR2 | 1.067797 | UCEC | EN | AZD4547 |
| MVP_top60_DNA_seed | IGF1R | 1.024298 | LUAD | NCI-H441 | Linsitinib | MVP_top60_DNA+RNA_seed | MDM2 | 1.068278 | UCEC | MFE-296 | Serdemetan |
| MVP_top60_DNA_seed | EGFR | 1.024683 | LUAD | RERF-LC-MS | Afatinib | MVP_top60_DNA+RNA_seed | PIK3CB | 1.068292 | UCEC | HEC-1 | AZD8186 |
| MVP_top60_DNA_seed | ERBB2 | 1.024683 | LUAD | RERF-LC-MS | Afatinib | MVP_top60_DNA+RNA_seed | PIK3CD | 1.068292 | UCEC | HEC-1 | AZD8186 |
| MVP_top60_DNA_seed | AKT1 | 1.025872 | LUAD | NCI-H1435 | AZD5363 | MVP_top60_DNA+RNA_seed | EP300 | 1.095465 | UCEC | MFE-280 | I-CBP112 |
| MVP_top60_DNA_seed | PIK3CA | 1.02762 | LUAD | VMRC-LCD | Buparlisib | MVP_top60_DNA+RNA_seed | EGFR | 1.103031 | UCEC | AN3-CA | Pelitinib |
| MVP_top60_DNA_seed | SMARCA4 | 1.027794 | LUAD | NCI-H1666 | PFI3 | MVP_top60_DNA+RNA_seed | HSP90AA1 | 1.106335 | UCEC | KLE | Elesclomol |
| MVP_top60_DNA_seed | PIK3CA | 1.028748 | LUAD | NCI-H1838 | GNE-317 | MVP_top60_DNA+RNA_seed | HSP90AB1 | 1.106335 | UCEC | KLE | Elesclomol |

| | | | | | | | | | | | |
|---|---|---|---|---|---|---|---|---|---|---|---|
| MVP_top60_DNA_seed | MAPK1 | 1.029536 | LUAD | NCI-H1650 | Ulixertinib | MVP_top60_DNA+RNA_seed | SRC | 1.121127 | UCEC | MFE-296 | NVP-BHG712 |
| MVP_top60_DNA_seed | HDAC1 | 1.031691 | LUAD | NCI-H1563 | Entinostat | MVP_top60_DNA+RNA_seed | FGFR2 | 1.159924 | UCEC | EN | PD173074 |
| MVP_top60_DNA_seed | AKT1 | 1.03171 | LUAD | COR-L105 | AT13148 | MVP_top60_DNA+RNA_seed | ESR1 | 1.172355 | UCEC | AN3-CA | Fulvestrant |
| MVP_top60_DNA_seed | KRAS | 1.033303 | LUAD | EKVX | KRAS (G12C) Inhibitor-12 | MVP_top60_DNA+RNA_seed | HDAC1 | 1.177407 | UCEC | COLO-684 | JQ12 |
| MVP_top60_DNA_seed | MDM2 | 1.033423 | LUAD | EKVX | Nutlin-3a (-) | MVP_top60_DNA+RNA_seed | HDAC2 | 1.177407 | UCEC | COLO-684 | JQ12 |
| MVP_top60_DNA_seed | MET | 1.034528 | LUAD | NCI-H441 | Crizotinib | MVP_top60_DNA+RNA_seed | PIK3CA | 1.18858 | UCEC | MFE-319 | PI3Ka_4409 |
| MVP_top60_DNA_seed | AKT1 | 1.035306 | LUAD | NCI-H1355 | AZD5363 | MVP_top60_DNA+RNA_seed | PIK3CD | 1.18858 | UCEC | MFE-319 | PI3Ka_4409 |
| MVP_top60_DNA_seed | AKT1 | 1.035421 | LUAD | NCI-H2087 | GSK2110183B | MVP_top60_DNA+RNA_seed | EGFR | 1.188775 | UCEC | EN | Afatinib |
| MVP_top60_DNA_seed | EGFR | 1.036243 | LUAD | HCC-44 | AZD3759 | MVP_top60_DNA+RNA_seed | ERBB2 | 1.188775 | UCEC | EN | Afatinib |
| MVP_top60_DNA_seed | MDM2 | 1.038127 | LUAD | PC-3_[JPC-3] | Nutlin-3a (-) | MVP_top60_DNA+RNA_seed | PIK3CD | 1.191703 | UCEC | EN | IC-87114 |
| MVP_top60_DNA_seed | AKT1 | 1.041239 | LUAD | NCI-H1355 | Afuresertib | MVP_top60_DNA+RNA_seed | HDAC1 | 1.195439 | UCEC | EN | Dacinostat |
| MVP_top60_DNA_seed | IGF1R | 1.043226 | LUAD | NCI-H2085 | NVP-ADW742 | MVP_top60_DNA+RNA_seed | JAK1 | 1.217774 | UCEC | EN | Ruxolitinib |
| MVP_top60_DNA_seed | TP53 | 1.043648 | LUAD | SK-LU-1 | MIRA-1 | MVP_top60_DNA+RNA_seed | EGFR | 1.218919 | UCEC | MFE-296 | Pelitinib |
| MVP_top60_DNA_seed | AKT1 | 1.04502 | LUAD | Calu-6 | GSK2110183B | MVP_top60_DNA+RNA_seed | PIK3CA | 1.254999 | UCEC | KLE | AZD8835 |
| MVP_top60_DNA_seed | BRAF | 1.046126 | LUAD | HCC-78 | Dabrafenib | MVP_top60_DNA+RNA_seed | PIK3CD | 1.254999 | UCEC | KLE | AZD8835 |
| MVP_top60_DNA_seed | MET | 1.046588 | LUAD | NCI-H1573 | Crizotinib | MVP_top60_DNA+RNA_seed | HDAC1 | 1.269922 | UCEC | MFE-296 | AR-42 |
| MVP_top60_DNA_seed | FGFR1 | 1.046749 | LUAD | NCI-H2087 | PD173074 | MVP_top60_DNA+RNA_seed | HSP90AA1 | 1.286752 | UCEC | EN | Luminespib |
| MVP_top60_DNA_seed | MET | 1.047204 | LUAD | PC-3_[JPC-3] | Crizotinib | MVP_top60_DNA+RNA_seed | HSP90AB1 | 1.286752 | UCEC | EN | Luminespib |
| MVP_top60_DNA_seed | AURKA | 1.048085 | LUAD | NCI-H1435 | Alisertib | MVP_top60_DNA+RNA_seed | FGFR2 | 1.300353 | UCEC | KLE | FGFR_3831 |
| MVP_top60_DNA_seed | PIK3CA | 1.051712 | LUAD | NCI-H2347 | GNE-317 | MVP_top60_DNA+RNA_seed | AR | 1.30838 | UCEC | EN | Bicalutamide |
| MVP_top60_DNA_seed | PIK3CA | 1.052664 | LUAD | NCI-H1793 | Alpelisib | MVP_top60_DNA+RNA_seed | HDAC1 | 1.318465 | UCEC | COLO-684 | Parthenolide |
| MVP_top60_DNA_seed | FGFR1 | 1.053416 | LUAD | EKVX | PD173074 | MVP_top60_DNA+RNA_seed | EGFR | 1.320902 | UCEC | COLO-684 | AZD8931 |
| MVP_top60_DNA_seed | TP53 | 1.057105 | LUAD | A549 | MIRA-1 | MVP_top60_DNA+RNA_seed | ERBB2 | 1.320902 | UCEC | COLO-684 | AZD8931 |
| MVP_top60_DNA_seed | KRAS | 1.058704 | LUAD | NCI-H1355 | KRAS (G12C) Inhibitor-12 | MVP_top60_DNA+RNA_seed | EGFR | 1.322524 | UCEC | MFE-296 | CUDC-101 |
| MVP_top60_DNA_seed | AKT1 | 1.060136 | LUAD | NCI-H1693 | Afuresertib | MVP_top60_DNA+RNA_seed | ERBB2 | 1.322524 | UCEC | MFE-296 | CUDC-101 |
| MVP_top60_DNA_seed | TP53 | 1.060177 | LUAD | NCI-H2087 | MIRA-1 | MVP_top60_DNA+RNA_seed | AKT3 | 1.329107 | UCEC | HEC-1 | BAY AKT1 |
| MVP_top60_DNA_seed | ATM | 1.060528 | LUAD | VMRC-LCD | KU-55933 | MVP_top60_DNA+RNA_seed | AKT1 | 1.329107 | UCEC | HEC-1 | BAY AKT1 |
| MVP_top60_DNA_seed | AKT1 | 1.06382 | LUAD | VMRC-LCD | GSK2110183B | MVP_top60_DNA+RNA_seed | PIK3CD | 1.333835 | UCEC | EN | Idelalisib |
| MVP_top60_DNA_seed | PIK3CA | 1.064196 | LUAD | Calu-6 | Alpelisib | MVP_top60_DNA+RNA_seed | HSP90AA1 | 1.344395 | UCEC | MFE-280 | Luminespib |
| MVP_top60_DNA_seed | IGF1R | 1.064207 | LUAD | NCI-H358 | GSK1904529A | MVP_top60_DNA+RNA_seed | HSP90AB1 | 1.344395 | UCEC | MFE-280 | Luminespib |
| MVP_top60_DNA_seed | IGF1R | 1.064413 | LUAD | NCI-H650 | GSK1904529A | MVP_top60_DNA+RNA_seed | ESR1 | 1.360619 | UCEC | EN | Tamoxifen |
| MVP_top60_DNA_seed | MDM2 | 1.065983 | LUAD | Calu-6 | Nutlin-3a (-) | MVP_top60_DNA+RNA_seed | HSP90AA1 | 1.386121 | UCEC | KLE | Tanespimycin |
| MVP_top60_DNA_seed | BRAF | 1.066159 | LUAD | SW1573 | SB590885 | MVP_top60_DNA+RNA_seed | HSP90AB1 | 1.386121 | UCEC | KLE | Tanespimycin |
| MVP_top60_DNA_seed | EGFR | 1.066534 | LUAD | HOP-62 | Sapitinib | MVP_top60_DNA+RNA_seed | ATM | 1.411894 | UCEC | MFE-280 | KU-60019 |
| MVP_top60_DNA_seed | ERBB2 | 1.066534 | LUAD | HOP-62 | Sapitinib | MVP_top60_DNA+RNA_seed | ATM | 1.419228 | UCEC | HEC-1 | CP466722 |
| MVP_top60_DNA_seed | ERBB3 | 1.066534 | LUAD | HOP-62 | Sapitinib | MVP_top60_DNA+RNA_seed | HDAC1 | 1.427128 | UCEC | MFE-319 | AR-42 |
| MVP_top60_DNA_seed | MAPK1 | 1.067057 | LUAD | NCI-H2228 | Ulixertinib | MVP_top60_DNA+RNA_seed | EGFR | 1.427347 | UCEC | EN | Afatinib |
| MVP_top60_DNA_seed | EZH2 | 1.067508 | LUAD | NCI-H1666 | GSK343 | MVP_top60_DNA+RNA_seed | ERBB2 | 1.427347 | UCEC | EN | Afatinib |
| MVP_top60_DNA_seed | PIK3CA | 1.06766 | LUAD | NCI-H1650 | Alpelisib | MVP_top60_DNA+RNA_seed | MDM2 | 1.431443 | UCEC | EN | Nutlin-3a (-) |
| MVP_top60_DNA_seed | EGFR | 1.068391 | LUAD | ABC-1 | Afatinib | MVP_top60_DNA+RNA_seed | FGFR2 | 1.445917 | UCEC | MFE-280 | AZD4547 |
| MVP_top60_DNA_seed | ERBB2 | 1.068391 | LUAD | ABC-1 | Afatinib | MVP_top60_DNA+RNA_seed | EGFR | 1.455239 | UCEC | MFE-319 | CUDC-101 |

| | | | | | | | | | | | |
|---|---|---|---|---|---|---|---|---|---|---|---|
| MVP_top60_DNA_seed | IGF1R | 1.068636 | LUAD | NCI-H1651 | BMS-536924 | MVP_top60_DNA+RNA_seed | ERBB2 | 1.455239 | UCEC | MFE-319 | CUDC-101 |
| MVP_top60_DNA_seed | CREBBP | 1.070575 | LUAD | NCI-H1355 | SGC-CBP30 | MVP_top60_DNA+RNA_seed | MAPK1 | 1.461863 | UCEC | EN | FR-180204 |
| MVP_top60_DNA_seed | BRAF | 1.071026 | LUAD | NCI-H1355 | SB590885 | MVP_top60_DNA+RNA_seed | MAPK3 | 1.461863 | UCEC | EN | FR-180204 |
| MVP_top60_DNA_seed | TP53 | 1.071087 | LUAD | NCI-H2291 | MIRA-1 | MVP_top60_DNA+RNA_seed | PIK3CD | 1.47506 | UCEC | KLE | IC-87114 |
| MVP_top60_DNA_seed | MET | 1.071433 | LUAD | NCI-H1568 | Crizotinib | MVP_top60_DNA+RNA_seed | EGFR | 1.483304 | UCEC | EN | Pelitinib |
| MVP_top60_DNA_seed | TP53 | 1.075039 | LUAD | Calu-3 | MIRA-1 | MVP_top60_DNA+RNA_seed | MDM2 | 1.518861 | UCEC | KLE | Nutlin-3a (-) |
| MVP_top60_DNA_seed | EGFR | 1.075119 | LUAD | Calu-6 | Gefitinib | MVP_top60_DNA+RNA_seed | SRC | 1.528067 | UCEC | KLE | Bosutinib |
| MVP_top60_DNA_seed | MAPK1 | 1.075654 | LUAD | NCI-H441 | SCH772984 | MVP_top60_DNA+RNA_seed | SRC | 1.551759 | UCEC | EN | Bosutinib |
| MVP_top60_DNA_seed | AKT1 | 1.075767 | LUAD | PC-3_[JPC-3] | Afuresertib | MVP_top60_DNA+RNA_seed | HDAC1 | 1.553633 | UCEC | MFE-319 | CAY10603 |
| MVP_top60_DNA_seed | AKT1 | 1.078048 | LUAD | NCI-H1435 | MK-2206 | MVP_top60_DNA+RNA_seed | PIK3CB | 1.571729 | UCEC | EN | AZD8186 |
| MVP_top60_DNA_seed | IGF1R | 1.078604 | LUAD | NCI-H2087 | GSK1904529A | MVP_top60_DNA+RNA_seed | PIK3CD | 1.571729 | UCEC | EN | AZD8186 |
| MVP_top60_DNA_seed | HSP90AA1 | 1.078929 | LUAD | NCI-H2291 | Tanespimycin | MVP_top60_DNA+RNA_seed | HSP90AA1 | 1.606551 | UCEC | HEC-1 | SNX-2112 |
| MVP_top60_DNA_seed | AKT1 | 1.082744 | LUAD | NCI-H1651 | AT13148 | MVP_top60_DNA+RNA_seed | HSP90AB1 | 1.606551 | UCEC | HEC-1 | SNX-2112 |
| MVP_top60_DNA_seed | SRC | 1.083151 | LUAD | NCI-H1573 | Bosutinib | MVP_top60_DNA+RNA_seed | JAK1 | 1.615952 | UCEC | KLE | JAK1_3715 |
| MVP_top60_DNA_seed | HSP90AA1 | 1.085614 | LUAD | NCI-H1355 | Luminespib | MVP_top60_DNA+RNA_seed | FGFR2 | 1.631694 | UCEC | KLE | PD173074 |
| MVP_top60_DNA_seed | EGFR | 1.08636 | LUAD | VMRC-LCD | Lapatinib | MVP_top60_DNA+RNA_seed | ESR1 | 1.67379 | UCEC | EN | Fulvestrant |
| MVP_top60_DNA_seed | ERBB2 | 1.08636 | LUAD | VMRC-LCD | Lapatinib | MVP_top60_DNA+RNA_seed | HDAC1 | 1.679567 | UCEC | SNG-M | CAY10603 |
| MVP_top60_DNA_seed | IGF1R | 1.090314 | LUAD | NCI-H1568 | BMS-754807 | MVP_top60_DNA+RNA_seed | ESR1 | 1.726912 | UCEC | COLO-684 | Fulvestrant |
| MVP_top60_DNA_seed | BRAF | 1.091162 | LUAD | NCI-H1355 | Dabrafenib | MVP_top60_DNA+RNA_seed | PIK3CA | 1.727071 | UCEC | EN | PI3Ka_4409 |
| MVP_top60_DNA_seed | MET | 1.09189 | LUAD | Calu-6 | Savolitinib | MVP_top60_DNA+RNA_seed | PIK3CD | 1.727071 | UCEC | EN | PI3Ka_4409 |
| MVP_top60_DNA_seed | MAPK1 | 1.091984 | LUAD | NCI-H1563 | VX-11e | MVP_top60_DNA+RNA_seed | HDAC1 | 1.744332 | UCEC | MFE-319 | Belinostat |
| MVP_top60_DNA_seed | HSP90AA1 | 1.092057 | LUAD | H3255 | Luminespib | MVP_top60_DNA+RNA_seed | SRC | 1.746463 | UCEC | MFE-280 | Bosutinib |
| MVP_top60_DNA_seed | FGFR1 | 1.092196 | LUAD | NCI-H2085 | AZD4547 | MVP_top60_DNA+RNA_seed | HSP90AA1 | 1.808225 | UCEC | KLE | CCT-018159 |
| MVP_top60_DNA_seed | FGFR1 | 1.092953 | LUAD | NCI-H1650 | PD173074 | MVP_top60_DNA+RNA_seed | HSP90AB1 | 1.808225 | UCEC | KLE | CCT-018159 |
| MVP_top60_DNA_seed | AKT1 | 1.093513 | LUAD | NCI-H1573 | AZD5363 | MVP_top60_DNA+RNA_seed | HDAC1 | 1.832348 | UCEC | EN | JQ12 |
| MVP_top60_DNA_seed | AKT1 | 1.096605 | LUAD | NCI-H441 | Uprosertib | MVP_top60_DNA+RNA_seed | HDAC2 | 1.832348 | UCEC | EN | JQ12 |
| MVP_top60_DNA_seed | HDAC1 | 1.098466 | LUAD | NCI-H2087 | Entinostat | MVP_top60_DNA+RNA_seed | PIK3CB | 1.860746 | UCEC | EN | AZD6482 |
| MVP_top60_DNA_seed | IGF1R | 1.100977 | LUAD | VMRC-LCD | Linsitinib | MVP_top60_DNA+RNA_seed | EGFR | 1.863248 | UCEC | EN | Gefitinib |
| MVP_top60_DNA_seed | IGF1R | 1.104534 | LUAD | COR-L105 | BMS-754807 | MVP_top60_DNA+RNA_seed | AKT3 | 1.884926 | UCEC | EN | AKT inhibitor VIII |
| MVP_top60_DNA_seed | EGFR | 1.106349 | LUAD | NCI-H2023 | Osimertinib | MVP_top60_DNA+RNA_seed | AKT1 | 1.884926 | UCEC | EN | AKT inhibitor VIII |
| MVP_top60_DNA_seed | AKT1 | 1.106381 | LUAD | NCI-H23 | AT13148 | MVP_top60_DNA+RNA_seed | AR | 2.015584 | UCEC | KLE | Bicalutamide |
| MVP_top60_DNA_seed | PIK3CA | 1.107846 | LUAD | NCI-H2087 | AZD8186 | MVP_top60_DNA+RNA_seed | FGFR2 | 2.092606 | UCEC | KLE | AZD4547 |
| MVP_top60_DNA_seed | MAPK1 | 1.108681 | LUAD | Calu-3 | Ulixertinib | MVP_top60_DNA+RNA_seed | JAK1 | 2.101062 | UCEC | KLE | AZD1480 |
| MVP_top60_DNA_seed | EGFR | 1.110325 | LUAD | NCI-H2228 | Osimertinib | MVP_top60_DNA+RNA_seed | HSP90AA1 | 2.13042 | UCEC | MFE-280 | CCT-018159 |
| MVP_top60_DNA_seed | AKT1 | 1.111078 | LUAD | NCI-H1792 | Uprosertib | MVP_top60_DNA+RNA_seed | HSP90AB1 | 2.13042 | UCEC | MFE-280 | CCT-018159 |
| MVP_top60_DNA_seed | FGFR1 | 1.111741 | LUAD | VMRC-LCD | AZD4547 | MVP_top60_DNA+RNA_seed | AR | 2.13383 | UCEC | EN | AZD3514 |
| MVP_top60_DNA_seed | PIK3CA | 1.116172 | LUAD | ABC-1 | Buparlisib | MVP_top60_DNA+RNA_seed | MDM2 | 2.430501 | UCEC | MFE-280 | Serdemetan |
| MVP_top60_DNA_seed | IGF1R | 1.117593 | LUAD | NCI-H1568 | Linsitinib | MVP_top60_DNA+RNA_seed | EP300 | -1.900587 | UCEC | EN | SGC-CBP30 |
| MVP_top60_DNA_seed | AKT1 | 1.117742 | LUAD | HOP-62 | Afuresertib | MVP_top60_DNA+RNA_seed | MAPK1 | -1.850765 | UCEC | EN | Ulixertinib |
| MVP_top60_DNA_seed | IGF1R | 1.118713 | LUAD | NCI-H1993 | BMS-536924 | MVP_top60_DNA+RNA_seed | MAPK3 | -1.850765 | UCEC | EN | Ulixertinib |
| MVP_top60_DNA_seed | PIK3CA | 1.121993 | LUAD | NCI-H2087 | GNE-317 | MVP_top60_DNA+RNA_seed | JAK1 | -1.754477 | UCEC | AN3-CA | GSK2276186C |

| | | | | | | | | | | | |
|---|---|---|---|---|---|---|---|---|---|---|---|
| MVP_top60_DNA_seed | AURKA | 1.124456 | LUAD | NCI-H1573 | ZM447439 | MVP_top60_DNA+RNA_seed | FGFR2 | -1.681912 | UCEC | AN3-CA | PD173074 |
| MVP_top60_DNA_seed | EZH2 | 1.12697 | LUAD | NCI-H322M | GSK343 | MVP_top60_DNA+RNA_seed | AKT3 | -1.538701 | UCEC | AN3-CA | GSK2110183B |
| MVP_top60_DNA_seed | IGF1R | 1.128076 | LUAD | NCI-H1651 | BMS-754807 | MVP_top60_DNA+RNA_seed | AKT1 | -1.538701 | UCEC | AN3-CA | GSK2110183B |
| MVP_top60_DNA_seed | IGF1R | 1.129236 | LUAD | NCI-H2023 | GSK1904529A | MVP_top60_DNA+RNA_seed | AKT3 | -1.461307 | UCEC | COLO-684 | GSK2110183B |
| MVP_top60_DNA_seed | AURKA | 1.129468 | LUAD | NCI-H1563 | ZM447439 | MVP_top60_DNA+RNA_seed | AKT1 | -1.461307 | UCEC | COLO-684 | GSK2110183B |
| MVP_top60_DNA_seed | MAPK1 | 1.129559 | LUAD | NCI-H2023 | Ulixertinib | MVP_top60_DNA+RNA_seed | FGFR2 | -1.455099 | UCEC | AN3-CA | AZD4547 |
| MVP_top60_DNA_seed | GSK3B | 1.129818 | LUAD | PC-14 | SB216763 | MVP_top60_DNA+RNA_seed | AKT3 | -1.441089 | UCEC | AN3-CA | Uprosertib |
| MVP_top60_DNA_seed | MDM2 | 1.130567 | LUAD | NCI-H1568 | Serdemetan | MVP_top60_DNA+RNA_seed | AKT1 | -1.441089 | UCEC | AN3-CA | Uprosertib |
| MVP_top60_DNA_seed | AURKA | 1.131258 | LUAD | NCI-H522 | ZM447439 | MVP_top60_DNA+RNA_seed | PIK3CA | -1.433814 | UCEC | MFE-280 | Alpelisib |
| MVP_top60_DNA_seed | AKT1 | 1.131753 | LUAD | NCI-H1568 | Uprosertib | MVP_top60_DNA+RNA_seed | JAK1 | -1.401798 | UCEC | AN3-CA | JAK_8517 |
| MVP_top60_DNA_seed | EZH2 | 1.131901 | LUAD | NCI-H1944 | GSK343 | MVP_top60_DNA+RNA_seed | AKT3 | -1.393069 | UCEC | EN | AZD5363 |
| MVP_top60_DNA_seed | MDM2 | 1.132287 | LUAD | NCI-H23 | Serdemetan | MVP_top60_DNA+RNA_seed | AKT1 | -1.393069 | UCEC | EN | AZD5363 |
| MVP_top60_DNA_seed | FGFR1 | 1.134138 | LUAD | SW1573 | AZD4547 | MVP_top60_DNA+RNA_seed | AKT3 | -1.363351 | UCEC | AN3-CA | Ipatasertib |
| MVP_top60_DNA_seed | HSP90AA1 | 1.137607 | LUAD | NCI-H1734 | Elesclomol | MVP_top60_DNA+RNA_seed | AKT1 | -1.363351 | UCEC | AN3-CA | Ipatasertib |
| MVP_top60_DNA_seed | TERT | 1.141257 | LUAD | NCI-H2009 | BIBR-1532 | MVP_top60_DNA+RNA_seed | AKT3 | -1.33634 | UCEC | EN | GSK2110183B |
| MVP_top60_DNA_seed | HSP90AA1 | 1.141719 | LUAD | NCI-H2405 | Luminespib | MVP_top60_DNA+RNA_seed | AKT1 | -1.33634 | UCEC | EN | GSK2110183B |
| MVP_top60_DNA_seed | EZH2 | 1.142225 | LUAD | Calu-6 | GSK343 | MVP_top60_DNA+RNA_seed | AKT3 | -1.322297 | UCEC | COLO-684 | Uprosertib |
| MVP_top60_DNA_seed | TP53 | 1.144447 | LUAD | NCI-H1838 | MIRA-1 | MVP_top60_DNA+RNA_seed | AKT1 | -1.322297 | UCEC | COLO-684 | Uprosertib |
| MVP_top60_DNA_seed | SRC | 1.147221 | LUAD | NCI-H2023 | Bosutinib | MVP_top60_DNA+RNA_seed | HSP90AA1 | -1.268945 | UCEC | AN3-CA | Tanespimycin |
| MVP_top60_DNA_seed | AKT1 | 1.147313 | LUAD | NCI-H522 | AZD5363 | MVP_top60_DNA+RNA_seed | HSP90AB1 | -1.268945 | UCEC | AN3-CA | Tanespimycin |
| MVP_top60_DNA_seed | AURKA | 1.149035 | LUAD | NCI-H1693 | Alisertib | MVP_top60_DNA+RNA_seed | AKT3 | -1.246177 | UCEC | COLO-684 | Afuresertib |
| MVP_top60_DNA_seed | IGF1R | 1.150477 | LUAD | NCI-H1650 | BMS-754807 | MVP_top60_DNA+RNA_seed | AKT1 | -1.246177 | UCEC | COLO-684 | Afuresertib |
| MVP_top60_DNA_seed | BRAF | 1.150579 | LUAD | NCI-H1944 | PLX-4720 | MVP_top60_DNA+RNA_seed | AKT3 | -1.225323 | UCEC | EN | Afuresertib |
| MVP_top60_DNA_seed | BRAF | 1.151225 | LUAD | NCI-H3122 | PLX-4720 | MVP_top60_DNA+RNA_seed | AKT1 | -1.225323 | UCEC | EN | Afuresertib |
| MVP_top60_DNA_seed | TERT | 1.155999 | LUAD | H3255 | BIBR-1532 | MVP_top60_DNA+RNA_seed | AKT3 | -1.197812 | UCEC | EN | Uprosertib |
| MVP_top60_DNA_seed | SMARCA4 | 1.156127 | LUAD | NCI-H2087 | PFI3 | MVP_top60_DNA+RNA_seed | AKT1 | -1.197812 | UCEC | EN | Uprosertib |
| MVP_top60_DNA_seed | AKT1 | 1.15807 | LUAD | HCC-827 | Uprosertib | MVP_top60_DNA+RNA_seed | AKT3 | -1.173212 | UCEC | AN3-CA | AZD5363 |
| MVP_top60_DNA_seed | MAPK1 | 1.159179 | LUAD | NCI-H2347 | VX-11e | MVP_top60_DNA+RNA_seed | AKT1 | -1.173212 | UCEC | AN3-CA | AZD5363 |
| MVP_top60_DNA_seed | MDM2 | 1.160978 | LUAD | SK-LU-1 | Nutlin-3a (-) | MVP_top60_DNA+RNA_seed | PIK3CA | -1.129591 | UCEC | AN3-CA | Alpelisib |
| MVP_top60_DNA_seed | FGFR1 | 1.162131 | LUAD | NCI-H1693 | AZD4547 | MVP_top60_DNA+RNA_seed | PIK3CA | -1.128491 | UCEC | KLE | AZD8186 |
| MVP_top60_DNA_seed | EGFR | 1.162746 | LUAD | HCC-78 | Erlotinib | MVP_top60_DNA+RNA_seed | PIK3CB | -1.128491 | UCEC | KLE | AZD8186 |
| MVP_top60_DNA_seed | TP53 | 1.162837 | LUAD | H3255 | MIRA-1 | MVP_top60_DNA+RNA_seed | AKT1 | -1.121104 | UCEC | AN3-CA | MK-2206 |
| MVP_top60_DNA_seed | PIK3CA | 1.165026 | LUAD | SW1573 | Buparlisib | MVP_top60_DNA+RNA_seed | AKT1 | -1.098586 | UCEC | COLO-684 | MK-2206 |
| MVP_top60_DNA_seed | AKT1 | 1.169195 | LUAD | H3255 | AT13148 | MVP_top60_DNA+RNA_seed | KRAS | -1.093533 | UCEC | SNG-M | (RAS (G12C) Inhibitor-1 |
| MVP_top60_DNA_seed | MDM2 | 1.171561 | LUAD | NCI-H1666 | Serdemetan | MVP_top60_DNA+RNA_seed | ATM | -1.082897 | UCEC | MFE-319 | KU-55933 |
| MVP_top60_DNA_seed | TP53 | 1.172228 | LUAD | NCI-H1355 | MIRA-1 | MVP_top60_DNA+RNA_seed | AKT1 | -1.07994 | UCEC | MFE-280 | MK-2206 |
| MVP_top60_DNA_seed | EZH2 | 1.172619 | LUAD | SK-LU-1 | GSK343 | MVP_top60_DNA+RNA_seed | AKT3 | -1.069252 | UCEC | AN3-CA | Uprosertib |
| MVP_top60_DNA_seed | HDAC1 | 1.173704 | LUAD | NCI-H1623 | PCI-34051 | MVP_top60_DNA+RNA_seed | AKT1 | -1.069252 | UCEC | AN3-CA | Uprosertib |
| MVP_top60_DNA_seed | MDM2 | 1.176968 | LUAD | NCI-H1975 | Serdemetan | MVP_top60_DNA+RNA_seed | PIK3CA | -1.05072 | UCEC | AN3-CA | GNE-317 |
| MVP_top60_DNA_seed | IGF1R | 1.177924 | LUAD | SK-LU-1 | Linsitinib | MVP_top60_DNA+RNA_seed | AKT3 | -1.034058 | UCEC | MFE-280 | Uprosertib |
| MVP_top60_DNA_seed | MDM2 | 1.178315 | LUAD | VMRC-LCD | Nutlin-3a (-) | MVP_top60_DNA+RNA_seed | AKT1 | -1.034058 | UCEC | MFE-280 | Uprosertib |

| | | | | | | | | | | | |
|---|---|---|---|---|---|---|---|---|---|---|---|
| MVP_top60_DNA_seed | AKT1 | 1.178429 | LUAD | HCC-827 | Afuresertib | MVP_top60_DNA+RNA_seed | AKT1 | -1.033309 | UCEC | KLE | MK-2206 |
| MVP_top60_DNA_seed | SMARCA4 | 1.180361 | LUAD | ABC-1 | PFI3 | MVP_top60_DNA+RNA_seed | AKT3 | -1.029426 | UCEC | EN | Ipatasertib |
| MVP_top60_DNA_seed | EGFR | 1.181664 | LUAD | VMRC-LCD | Sapitinib | MVP_top60_DNA+RNA_seed | AKT1 | -1.029426 | UCEC | EN | Ipatasertib |
| MVP_top60_DNA_seed | ERBB2 | 1.181664 | LUAD | VMRC-LCD | Sapitinib | MVP_top60_DNA+RNA_seed | PIK3CA | -0.986437 | UCEC | AN3-CA | AZD8186 |
| MVP_top60_DNA_seed | ERBB3 | 1.181664 | LUAD | VMRC-LCD | Sapitinib | MVP_top60_DNA+RNA_seed | PIK3CB | -0.986437 | UCEC | AN3-CA | AZD8186 |
| MVP_top60_DNA_seed | IGF1R | 1.185303 | LUAD | NCI-H1651 | NVP-ADW742 | MVP_top60_DNA+RNA_seed | ESR1 | -0.986386 | UCEC | MFE-319 | Tamoxifen |
| MVP_top60_DNA_seed | BRAF | 1.18532 | LUAD | NCI-H1838 | PLX-4720 | MVP_top60_DNA+RNA_seed | ATM | -0.980089 | UCEC | AN3-CA | KU-55933 |
| MVP_top60_DNA_seed | EGFR | 1.185843 | LUAD | NCI-H1573 | Osimertinib | MVP_top60_DNA+RNA_seed | ESR1 | -0.97847 | UCEC | COLO-684 | Tamoxifen |
| MVP_top60_DNA_seed | TERT | 1.186611 | LUAD | HCC-78 | BIBR-1532 | MVP_top60_DNA+RNA_seed | FGFR2 | -0.977111 | UCEC | MFE-280 | AZD4547 |
| MVP_top60_DNA_seed | PIK3CA | 1.186848 | LUAD | NCI-H23 | Alpelisib | MVP_top60_DNA+RNA_seed | AKT1 | -0.967928 | UCEC | SNG-M | AT13148 |
| MVP_top60_DNA_seed | HDAC1 | 1.187097 | LUAD | NCI-H1781 | Entinostat | MVP_top60_DNA+RNA_seed | JAK1 | -0.957908 | UCEC | COLO-684 | GSK2276186C |
| MVP_top60_DNA_seed | ATM | 1.187124 | LUAD | NCI-H2087 | KU-55933 | MVP_top60_DNA+RNA_seed | ESR1 | -0.955704 | UCEC | SNG-M | GDC0810 |
| MVP_top60_DNA_seed | MAPK1 | 1.1878 | LUAD | SW1573 | Ulixertinib | MVP_top60_DNA+RNA_seed | AKT3 | -0.948204 | UCEC | COLO-684 | Ipatasertib |
| MVP_top60_DNA_seed | CREBBP | 1.190798 | LUAD | A549 | SGC-CBP30 | MVP_top60_DNA+RNA_seed | AKT1 | -0.948204 | UCEC | COLO-684 | Ipatasertib |
| MVP_top60_DNA_seed | AKT1 | 1.194185 | LUAD | H3255 | GSK2110183B | MVP_top60_DNA+RNA_seed | AKT3 | -0.939066 | UCEC | COLO-684 | Uprosertib |
| MVP_top60_DNA_seed | MAPK1 | 1.196934 | LUAD | NCI-H1563 | Ulixertinib | MVP_top60_DNA+RNA_seed | AKT1 | -0.939066 | UCEC | COLO-684 | Uprosertib |
| MVP_top60_DNA_seed | MAPK1 | 1.197012 | LUAD | PC-3_[JPC-3] | SCH772984 | MVP_top60_DNA+RNA_seed | MDM2 | -0.928289 | UCEC | SNG-M | Nutlin-3a (-) |
| MVP_top60_DNA_seed | AURKA | 1.200749 | LUAD | EKVX | Alisertib | MVP_top60_DNA+RNA_seed | AKT3 | -0.918815 | UCEC | MFE-296 | Uprosertib |
| MVP_top60_DNA_seed | FGFR1 | 1.201416 | LUAD | NCI-H2347 | PD173074 | MVP_top60_DNA+RNA_seed | AKT1 | -0.918815 | UCEC | MFE-296 | Uprosertib |
| MVP_top60_DNA_seed | IGF1R | 1.206376 | LUAD | NCI-H1623 | NVP-ADW742 | MVP_top60_DNA+RNA_seed | EP300 | -0.907567 | UCEC | HEC-1 | SGC-CBP30 |
| MVP_top60_DNA_seed | BRAF | 1.207166 | LUAD | HCC-827 | Dabrafenib | MVP_top60_DNA+RNA_seed | AKT3 | -0.877355 | UCEC | COLO-684 | AZD5363 |
| MVP_top60_DNA_seed | ATM | 1.208558 | LUAD | HCC-827 | KU-55933 | MVP_top60_DNA+RNA_seed | AKT1 | -0.877355 | UCEC | COLO-684 | AZD5363 |
| MVP_top60_DNA_seed | KRAS | 1.209619 | LUAD | NCI-H2291 | KRAS (G12C) Inhibitor-1: | MVP_top60_DNA+RNA_seed | EP300 | -0.863046 | UCEC | KLE | SGC-CBP30 |
| MVP_top60_DNA_seed | SRC | 1.21069 | LUAD | ABC-1 | Bosutinib | MVP_top60_DNA+RNA_seed | ESR1 | -0.862256 | UCEC | EN | GDC0810 |
| MVP_top60_DNA_seed | MDM2 | 1.210848 | LUAD | NCI-H1838 | Nutlin-3a (-) | MVP_top60_DNA+RNA_seed | EGFR | -0.803748 | UCEC | KLE | Sapitinib |
| MVP_top60_DNA_seed | AKT1 | 1.212579 | LUAD | NCI-H1793 | Uprosertib | MVP_top60_DNA+RNA_seed | ERBB2 | -0.803748 | UCEC | KLE | Sapitinib |
| MVP_top60_DNA_seed | MAPK1 | 1.213698 | LUAD | NCI-H2342 | Ulixertinib | MVP_top60_DNA+RNA_seed | PIK3CA | -0.799668 | UCEC | AN3-CA | Buparlisib |
| MVP_top60_DNA_seed | MET | 1.214015 | LUAD | Calu-3 | Savolitinib | MVP_top60_DNA+RNA_seed | PIK3CB | -0.799668 | UCEC | AN3-CA | Buparlisib |
| MVP_top60_DNA_seed | ATM | 1.214093 | LUAD | Calu-6 | KU-55933 | MVP_top60_DNA+RNA_seed | PIK3CD | -0.799668 | UCEC | AN3-CA | Buparlisib |
| MVP_top60_DNA_seed | BRAF | 1.214278 | LUAD | NCI-H1666 | SB590885 | MVP_top60_DNA+RNA_seed | FGFR2 | -0.787283 | UCEC | MFE-296 | AZD4547 |
| MVP_top60_DNA_seed | BRAF | 1.214825 | LUAD | COR-L105 | SB590885 | MVP_top60_DNA+RNA_seed | JAK1 | -0.779399 | UCEC | AN3-CA | Ruxolitinib |
| MVP_top60_DNA_seed | IGF1R | 1.219495 | LUAD | NCI-H1573 | NVP-ADW742 | MVP_top60_DNA+RNA_seed | PIK3CA | -0.776155 | UCEC | MFE-319 | GNE-317 |
| MVP_top60_DNA_seed | EGFR | 1.22041 | LUAD | NCI-H3122 | AZD3759 | MVP_top60_DNA+RNA_seed | AKT3 | -0.771595 | UCEC | MFE-280 | Uprosertib |
| MVP_top60_DNA_seed | TP53 | 1.221488 | LUAD | COR-L105 | MIRA-1 | MVP_top60_DNA+RNA_seed | AKT1 | -0.771595 | UCEC | MFE-280 | Uprosertib |
| MVP_top60_DNA_seed | MDM2 | 1.223567 | LUAD | NCI-H1650 | Nutlin-3a (-) | MVP_top60_DNA+RNA_seed | EGFR | -0.765043 | UCEC | SNG-M | Sapitinib |
| MVP_top60_DNA_seed | IGF1R | 1.223576 | LUAD | NCI-H1944 | NVP-ADW742 | MVP_top60_DNA+RNA_seed | ERBB2 | -0.765043 | UCEC | SNG-M | Sapitinib |
| MVP_top60_DNA_seed | HDAC1 | 1.226764 | LUAD | NCI-H1355 | PCI-34051 | MVP_top60_DNA+RNA_seed | AKT3 | -0.752001 | UCEC | AN3-CA | Afuresertib |
| MVP_top60_DNA_seed | AKT1 | 1.227764 | LUAD | NCI-H3122 | AT13148 | MVP_top60_DNA+RNA_seed | AKT1 | -0.752001 | UCEC | AN3-CA | Afuresertib |
| MVP_top60_DNA_seed | AKT1 | 1.230865 | LUAD | NCI-H2405 | AZD5363 | MVP_top60_DNA+RNA_seed | EGFR | -0.740338 | UCEC | MFE-319 | Erlotinib |
| MVP_top60_DNA_seed | IGF1R | 1.23252 | LUAD | NCI-H2085 | Linsitinib | MVP_top60_DNA+RNA_seed | AKT3 | -0.693632 | UCEC | MFE-319 | Uprosertib |
| MVP_top60_DNA_seed | EGFR | 1.235609 | LUAD | NCI-H1355 | Osimertinib | MVP_top60_DNA+RNA_seed | AKT1 | -0.693632 | UCEC | MFE-319 | Uprosertib |

| | | | | | | | | | | | |
|---|---|---|---|---|---|---|---|---|---|---|---|
| MVP_top60_DNA_seed | IGF1R | 1.235982 | LUAD | NCI-H2347 | BMS-536924 | MVP_top60_DNA+RNA_seed | PIK3CA | -0.682345 | UCEC | COLO-684 | GNE-317 |
| MVP_top60_DNA_seed | MDM2 | 1.238033 | LUAD | NCI-H1793 | Nutlin-3a (-) | MVP_top60_DNA+RNA_seed | ATM | -0.677619 | UCEC | MFE-280 | KU-55933 |
| MVP_top60_DNA_seed | FGFR1 | 1.239301 | LUAD | ABC-1 | AZD4547 | MVP_top60_DNA+RNA_seed | AKT1 | -0.650013 | UCEC | AN3-CA | AT13148 |
| MVP_top60_DNA_seed | MDM2 | 1.240131 | LUAD | NCI-H1568 | Nutlin-3a (-) | MVP_top60_DNA+RNA_seed | AKT3 | -0.647068 | UCEC | MFE-319 | Ipatasertib |
| MVP_top60_DNA_seed | AKT1 | 1.241163 | LUAD | NCI-H522 | MK-2206 | MVP_top60_DNA+RNA_seed | AKT1 | -0.647068 | UCEC | MFE-319 | Ipatasertib |
| MVP_top60_DNA_seed | HSP90AA1 | 1.242115 | LUAD | EKVX | Luminespib | MVP_top60_DNA+RNA_seed | AKT1 | -0.631172 | UCEC | MFE-319 | AT13148 |
| MVP_top60_DNA_seed | AURKA | 1.242257 | LUAD | NCI-H2291 | Alisertib | MVP_top60_DNA+RNA_seed | AKT3 | -0.623315 | UCEC | MFE-296 | Afuresertib |
| MVP_top60_DNA_seed | IGF1R | 1.245149 | LUAD | NCI-H1781 | NVP-ADW742 | MVP_top60_DNA+RNA_seed | AKT1 | -0.623315 | UCEC | MFE-296 | Afuresertib |
| MVP_top60_DNA_seed | MDM2 | 1.247334 | LUAD | NCI-H1355 | Nutlin-3a (-) | MVP_top60_DNA+RNA_seed | JAK1 | -0.620585 | UCEC | MFE-280 | JAK_8517 |
| MVP_top60_DNA_seed | BRAF | 1.247654 | LUAD | Calu-3 | Dabrafenib | MVP_top60_DNA+RNA_seed | EGFR | -0.614033 | UCEC | AN3-CA | Sapitinib |
| MVP_top60_DNA_seed | IGF1R | 1.248273 | LUAD | Calu-6 | BMS-536924 | MVP_top60_DNA+RNA_seed | ERBB2 | -0.614033 | UCEC | AN3-CA | Sapitinib |
| MVP_top60_DNA_seed | FGFR1 | 1.248835 | LUAD | NCI-H358 | AZD4547 | MVP_top60_DNA+RNA_seed | AKT3 | -0.609343 | UCEC | MFE-296 | Uprosertib |
| MVP_top60_DNA_seed | AKT1 | 1.248857 | LUAD | EKVX | Ipatasertib | MVP_top60_DNA+RNA_seed | AKT1 | -0.609343 | UCEC | MFE-296 | Uprosertib |
| MVP_top60_DNA_seed | KRAS | 1.250427 | LUAD | NCI-H1793 | KRAS (G12C) Inhibitor-1[illegible] | MVP_top60_DNA+RNA_seed | TP53 | -0.609038 | UCEC | AN3-CA | MIRA-1 |
| MVP_top60_DNA_seed | BRAF | 1.252297 | LUAD | SW1573 | Dabrafenib | MVP_top60_DNA+RNA_seed | EP300 | -0.60117 | UCEC | MFE-296 | SGC-CBP30 |
| MVP_top60_DNA_seed | BRAF | 1.254303 | LUAD | H3255 | SB590885 | MVP_top60_DNA+RNA_seed | AKT3 | -0.595333 | UCEC | EN | Uprosertib |
| MVP_top60_DNA_seed | GSK3B | 1.255876 | LUAD | NCI-H2087 | SB216763 | MVP_top60_DNA+RNA_seed | AKT1 | -0.595333 | UCEC | EN | Uprosertib |
| MVP_top60_DNA_seed | IGF1R | 1.257779 | LUAD | NCI-H522 | BMS-754807 | MVP_top60_DNA+RNA_seed | MAPK1 | -0.590423 | UCEC | COLO-684 | Ulixertinib |
| MVP_top60_DNA_seed | AKT1 | 1.258281 | LUAD | NCI-H522 | Ipatasertib | MVP_top60_DNA+RNA_seed | MAPK3 | -0.590423 | UCEC | COLO-684 | Ulixertinib |
| MVP_top60_DNA_seed | EGFR | 1.25831 | LUAD | NCI-H1793 | Afatinib | MVP_top60_DNA+RNA_seed | AKT3 | -0.587729 | UCEC | MFE-280 | GSK2110183B |
| MVP_top60_DNA_seed | ERBB2 | 1.25831 | LUAD | NCI-H1793 | Afatinib | MVP_top60_DNA+RNA_seed | AKT1 | -0.587729 | UCEC | MFE-280 | GSK2110183B |
| MVP_top60_DNA_seed | EGFR | 1.25964 | LUAD | NCI-H2347 | Sapitinib | MVP_top60_DNA+RNA_seed | ESR1 | -0.580206 | UCEC | MFE-296 | Tamoxifen |
| MVP_top60_DNA_seed | ERBB2 | 1.25964 | LUAD | NCI-H2347 | Sapitinib | MVP_top60_DNA+RNA_seed | JAK1 | -0.578613 | UCEC | AN3-CA | JAK1_8709 |
| MVP_top60_DNA_seed | ERBB3 | 1.25964 | LUAD | NCI-H2347 | Sapitinib | MVP_top60_DNA+RNA_seed | HDAC1 | -0.568278 | UCEC | AN3-CA | PCI-34051 |
| MVP_top60_DNA_seed | EGFR | 1.260114 | LUAD | NCI-H3122 | Erlotinib | MVP_top60_DNA+RNA_seed | AKT3 | -0.556221 | UCEC | MFE-280 | Ipatasertib |
| MVP_top60_DNA_seed | AKT1 | 1.264809 | LUAD | NCI-H1651 | Afuresertib | MVP_top60_DNA+RNA_seed | AKT1 | -0.556221 | UCEC | MFE-280 | Ipatasertib |
| MVP_top60_DNA_seed | AKT1 | 1.26639 | LUAD | NCI-H23 | MK-2206 | MVP_top60_DNA+RNA_seed | PIK3CA | -0.553514 | UCEC | COLO-684 | Alpelisib |
| MVP_top60_DNA_seed | TERT | 1.27078 | LUAD | PC-14 | BIBR-1532 | MVP_top60_DNA+RNA_seed | AKT1 | -0.526036 | UCEC | EN | MK-2206 |
| MVP_top60_DNA_seed | ATM | 1.27105 | LUAD | NCI-H1793 | KU-55933 | MVP_top60_DNA+RNA_seed | AKT3 | -0.515101 | UCEC | MFE-319 | Uprosertib |
| MVP_top60_DNA_seed | AKT1 | 1.271304 | LUAD | NCI-H1568 | Ipatasertib | MVP_top60_DNA+RNA_seed | AKT1 | -0.515101 | UCEC | MFE-319 | Uprosertib |
| MVP_top60_DNA_seed | AKT1 | 1.273734 | LUAD | VMRC-LCD | Ipatasertib | MVP_top60_DNA+RNA_seed | JAK1 | -0.49145 | UCEC | COLO-684 | JAK1_8709 |
| MVP_top60_DNA_seed | TERT | 1.274898 | LUAD | HOP-62 | BIBR-1532 | MVP_top60_DNA+RNA_seed | HSP90AA1 | -0.485312 | UCEC | MFE-319 | Tanespimycin |
| MVP_top60_DNA_seed | AKT1 | 1.276877 | LUAD | NCI-H1838 | AZD5363 | MVP_top60_DNA+RNA_seed | HSP90AB1 | -0.485312 | UCEC | MFE-319 | Tanespimycin |
| MVP_top60_DNA_seed | BRAF | 1.276919 | LUAD | NCI-H2087 | PLX-4720 | MVP_top60_DNA+RNA_seed | JAK1 | -0.483644 | UCEC | MFE-296 | Ruxolitinib |
| MVP_top60_DNA_seed | EZH2 | 1.279536 | LUAD | NCI-H2342 | GSK343 | MVP_top60_DNA+RNA_seed | ESR1 | -0.474905 | UCEC | MFE-319 | GDC0810 |
| MVP_top60_DNA_seed | HDAC1 | 1.281822 | LUAD | NCI-H1781 | Romidepsin | MVP_top60_DNA+RNA_seed | HDAC1 | -0.468427 | UCEC | COLO-684 | Romidepsin |
| MVP_top60_DNA_seed | PIK3CA | 1.283535 | LUAD | NCI-H2405 | Buparlisib | MVP_top60_DNA+RNA_seed | HDAC2 | -0.468427 | UCEC | COLO-684 | Romidepsin |
| MVP_top60_DNA_seed | BRAF | 1.283743 | LUAD | Calu-3 | PLX-4720 | MVP_top60_DNA+RNA_seed | FGFR2 | -0.441738 | UCEC | MFE-319 | AZD4547 |
| MVP_top60_DNA_seed | HDAC1 | 1.284447 | LUAD | NCI-H1568 | Entinostat | MVP_top60_DNA+RNA_seed | EGFR | -0.427167 | UCEC | KLE | Afatinib |
| MVP_top60_DNA_seed | HDAC1 | 1.284706 | LUAD | Calu-3 | PCI-34051 | MVP_top60_DNA+RNA_seed | ERBB2 | -0.427167 | UCEC | KLE | Afatinib |
| MVP_top60_DNA_seed | SRC | 1.284766 | LUAD | HCC-78 | Bosutinib | MVP_top60_DNA+RNA_seed | PIK3CA | -0.402583 | UCEC | COLO-684 | AZD8186 |

| | | | | | | | | | | | |
|---|---|---|---|---|---|---|---|---|---|---|---|
| MVP_top60_DNA_seed | EGFR | 1.285776 | LUAD | NCI-H2085 | Erlotinib | MVP_top60_DNA+RNA_seed | PIK3CB | -0.402583 | UCEC | COLO-684 | AZD8186 |
| MVP_top60_DNA_seed | HSP90AA1 | 1.289669 | LUAD | NCI-H1838 | Tanespimycin | MVP_top60_DNA+RNA_seed | AKT3 | -0.385786 | UCEC | MFE-296 | Ipatasertib |
| MVP_top60_DNA_seed | SRC | 1.291424 | LUAD | NCI-H2009 | Bosutinib | MVP_top60_DNA+RNA_seed | AKT1 | -0.385786 | UCEC | MFE-296 | Ipatasertib |
| MVP_top60_DNA_seed | HDAC1 | 1.29979 | LUAD | NCI-H441 | Entinostat | MVP_top60_DNA+RNA_seed | HDAC1 | -0.381355 | UCEC | MFE-319 | Entinostat |
| MVP_top60_DNA_seed | HDAC1 | 1.300362 | LUAD | NCI-H1650 | PCI-34051 | MVP_top60_DNA+RNA_seed | AKT3 | -0.375179 | UCEC | MFE-319 | AZD5363 |
| MVP_top60_DNA_seed | IGF1R | 1.301141 | LUAD | NCI-H2087 | BMS-754807 | MVP_top60_DNA+RNA_seed | AKT1 | -0.375179 | UCEC | MFE-319 | AZD5363 |
| MVP_top60_DNA_seed | BRAF | 1.301352 | LUAD | NCI-H1693 | Dabrafenib | MVP_top60_DNA+RNA_seed | KRAS | -0.368617 | UCEC | HEC-1 | (RAS (G12C) Inhibitor-1 |
| MVP_top60_DNA_seed | AKT1 | 1.301713 | LUAD | NCI-H2087 | Uprosertib | MVP_top60_DNA+RNA_seed | EP300 | -0.361843 | UCEC | MFE-319 | CPI-637 |
| MVP_top60_DNA_seed | KRAS | 1.30271 | LUAD | NCI-H1755 | KRAS (G12C) Inhibitor-1: | MVP_top60_DNA+RNA_seed | EGFR | -0.344944 | UCEC | MFE-280 | Sapitinib |
| MVP_top60_DNA_seed | HSP90AA1 | 1.304098 | LUAD | NCI-H1693 | Luminespib | MVP_top60_DNA+RNA_seed | ERBB2 | -0.344944 | UCEC | MFE-280 | Sapitinib |
| MVP_top60_DNA_seed | AKT1 | 1.305034 | LUAD | NCI-H1793 | Ipatasertib | MVP_top60_DNA+RNA_seed | AKT3 | -0.338625 | UCEC | MFE-319 | Afuresertib |
| MVP_top60_DNA_seed | AURKA | 1.306098 | LUAD | NCI-H2342 | ZM447439 | MVP_top60_DNA+RNA_seed | AKT1 | -0.338625 | UCEC | MFE-319 | Afuresertib |
| MVP_top60_DNA_seed | HDAC1 | 1.30625 | LUAD | NCI-H1666 | PCI-34051 | MVP_top60_DNA+RNA_seed | FGFR2 | -0.337154 | UCEC | MFE-280 | PD173074 |
| MVP_top60_DNA_seed | SRC | 1.307978 | LUAD | Calu-6 | Bosutinib | MVP_top60_DNA+RNA_seed | ESR1 | -0.33711 | UCEC | KLE | GDC0810 |
| MVP_top60_DNA_seed | AKT1 | 1.30838 | LUAD | NCI-H650 | MK-2206 | MVP_top60_DNA+RNA_seed | JAK1 | -0.335557 | UCEC | SNG-M | JAK_8517 |
| MVP_top60_DNA_seed | EGFR | 1.30886 | LUAD | NCI-H2087 | Sapitinib | MVP_top60_DNA+RNA_seed | EP300 | -0.326335 | UCEC | COLO-684 | SGC-CBP30 |
| MVP_top60_DNA_seed | ERBB2 | 1.30886 | LUAD | NCI-H2087 | Sapitinib | MVP_top60_DNA+RNA_seed | EP300 | -0.309197 | UCEC | SNG-M | SGC-CBP30 |
| MVP_top60_DNA_seed | ERBB3 | 1.30886 | LUAD | NCI-H2087 | Sapitinib | MVP_top60_DNA+RNA_seed | HSP90AA1 | -0.30519 | UCEC | AN3-CA | Luminespib |
| MVP_top60_DNA_seed | PIK3CA | 1.309286 | LUAD | NCI-H1838 | Alpelisib | MVP_top60_DNA+RNA_seed | HSP90AB1 | -0.30519 | UCEC | AN3-CA | Luminespib |
| MVP_top60_DNA_seed | IGF1R | 1.310141 | LUAD | NCI-H322M | GSK1904529A | MVP_top60_DNA+RNA_seed | PIK3CA | -0.300771 | UCEC | KLE | GNE-317 |
| MVP_top60_DNA_seed | HSP90AA1 | 1.310422 | LUAD | EKVX | Tanespimycin | MVP_top60_DNA+RNA_seed | PIK3CA | -0.277586 | UCEC | EN | Alpelisib |
| MVP_top60_DNA_seed | EGFR | 1.312046 | LUAD | NCI-H2405 | Osimertinib | MVP_top60_DNA+RNA_seed | HDAC1 | -0.272126 | UCEC | MFE-280 | PCI-34051 |
| MVP_top60_DNA_seed | FGFR1 | 1.312052 | LUAD | NCI-H2087 | AZD4547 | MVP_top60_DNA+RNA_seed | KRAS | -0.264657 | UCEC | EN | (RAS (G12C) Inhibitor-1 |
| MVP_top60_DNA_seed | IGF1R | 1.317796 | LUAD | Calu-3 | BMS-536924 | MVP_top60_DNA+RNA_seed | MAPK1 | -0.261615 | UCEC | SNG-M | VX-11e |
| MVP_top60_DNA_seed | IGF1R | 1.317847 | LUAD | NCI-H1435 | BMS-536924 | MVP_top60_DNA+RNA_seed | EGFR | -0.257983 | UCEC | MFE-319 | AZD3759 |
| MVP_top60_DNA_seed | MDM2 | 1.319403 | LUAD | NCI-H2342 | Nutlin-3a (-) | MVP_top60_DNA+RNA_seed | ESR1 | -0.253878 | UCEC | COLO-684 | GDC0810 |
| MVP_top60_DNA_seed | EGFR | 1.31956 | LUAD | NCI-H2228 | Lapatinib | MVP_top60_DNA+RNA_seed | AKT3 | -0.24562 | UCEC | SNG-M | AZD5363 |
| MVP_top60_DNA_seed | ERBB2 | 1.31956 | LUAD | NCI-H2228 | Lapatinib | MVP_top60_DNA+RNA_seed | AKT1 | -0.24562 | UCEC | SNG-M | AZD5363 |
| MVP_top60_DNA_seed | GSK3B | 1.31991 | LUAD | H3255 | CHIR-99021 | MVP_top60_DNA+RNA_seed | AKT3 | -0.239528 | UCEC | MFE-280 | AZD5363 |
| MVP_top60_DNA_seed | TP53 | 1.320283 | LUAD | EKVX | MIRA-1 | MVP_top60_DNA+RNA_seed | AKT1 | -0.239528 | UCEC | MFE-280 | AZD5363 |
| MVP_top60_DNA_seed | MET | 1.320379 | LUAD | NCI-H358 | Savolitinib | MVP_top60_DNA+RNA_seed | HDAC1 | -0.235431 | UCEC | HEC-1 | Romidepsin |
| MVP_top60_DNA_seed | FGFR1 | 1.321886 | LUAD | NCI-H1568 | PD173074 | MVP_top60_DNA+RNA_seed | HDAC2 | -0.235431 | UCEC | HEC-1 | Romidepsin |
| MVP_top60_DNA_seed | AKT1 | 1.324814 | LUAD | VMRC-LCD | MK-2206 | MVP_top60_DNA+RNA_seed | PIK3CA | -0.22301 | UCEC | MFE-280 | GNE-317 |
| MVP_top60_DNA_seed | EGFR | 1.328104 | LUAD | VMRC-LCD | AZD3759 | MVP_top60_DNA+RNA_seed | KRAS | -0.204616 | UCEC | COLO-684 | (RAS (G12C) Inhibitor-1 |
| MVP_top60_DNA_seed | HSP90AA1 | 1.328407 | LUAD | NCI-H2085 | Tanespimycin | MVP_top60_DNA+RNA_seed | EGFR | -0.201322 | UCEC | KLE | Osimertinib |
| MVP_top60_DNA_seed | HDAC1 | 1.328981 | LUAD | SW1573 | PCI-34051 | MVP_top60_DNA+RNA_seed | HDAC1 | -0.19928 | UCEC | KLE | PCI-34051 |
| MVP_top60_DNA_seed | EGFR | 1.329645 | LUAD | NCI-H2087 | Gefitinib | MVP_top60_DNA+RNA_seed | AKT3 | -0.187275 | UCEC | MFE-280 | Afuresertib |
| MVP_top60_DNA_seed | EGFR | 1.331778 | LUAD | NCI-H1838 | Lapatinib | MVP_top60_DNA+RNA_seed | AKT1 | -0.187275 | UCEC | MFE-280 | Afuresertib |
| MVP_top60_DNA_seed | ERBB2 | 1.331778 | LUAD | NCI-H1838 | Lapatinib | MVP_top60_DNA+RNA_seed | JAK1 | -0.184399 | UCEC | SNG-M | Ruxolitinib |
| MVP_top60_DNA_seed | SRC | 1.335851 | LUAD | NCI-H1355 | Bosutinib | MVP_top60_DNA+RNA_seed | HDAC1 | -0.165932 | UCEC | COLO-684 | PCI-34051 |
| MVP_top60_DNA_seed | SMARCA4 | 1.338804 | LUAD | HOP-62 | PFI3 | MVP_top60_DNA+RNA_seed | AKT3 | -0.156282 | UCEC | KLE | Uprosertib |

| | | | | | | | | | | | |
|---|---|---|---|---|---|---|---|---|---|---|---|
| MVP_top60_DNA_seed | FGFR1 | 1.341295 | LUAD | PC-3_[JPC-3] | AZD4547 | MVP_top60_DNA+RNA_seed | AKT1 | -0.156282 | UCEC | KLE | Uprosertib |
| MVP_top60_DNA_seed | IGF1R | 1.341854 | LUAD | NCI-H1650 | NVP-ADW742 | MVP_top60_DNA+RNA_seed | ATM | -0.154584 | UCEC | MFE-296 | KU-55933 |
| MVP_top60_DNA_seed | TP53 | 1.342672 | LUAD | SW1573 | MIRA-1 | MVP_top60_DNA+RNA_seed | EP300 | -0.152548 | UCEC | SNG-M | CPI-637 |
| MVP_top60_DNA_seed | MET | 1.346259 | LUAD | NCI-H1693 | Crizotinib | MVP_top60_DNA+RNA_seed | FGFR2 | -0.151758 | UCEC | SNG-M | PD173074 |
| MVP_top60_DNA_seed | EGFR | 1.346517 | LUAD | HOP-62 | Gefitinib | MVP_top60_DNA+RNA_seed | MDM2 | -0.141892 | UCEC | AN3-CA | Nutlin-3a (-) |
| MVP_top60_DNA_seed | AKT1 | 1.348592 | LUAD | NCI-H1355 | Ipatasertib | MVP_top60_DNA+RNA_seed | EGFR | -0.139967 | UCEC | MFE-296 | Erlotinib |
| MVP_top60_DNA_seed | PIK3CA | 1.348781 | LUAD | NCI-H650 | AZD8186 | MVP_top60_DNA+RNA_seed | JAK1 | -0.129582 | UCEC | SNG-M | JAK1_8709 |
| MVP_top60_DNA_seed | IGF1R | 1.353082 | LUAD | SW1573 | GSK1904529A | MVP_top60_DNA+RNA_seed | ATM | -0.127018 | UCEC | SNG-M | KU-55933 |
| MVP_top60_DNA_seed | EGFR | 1.353336 | LUAD | NCI-H2405 | Gefitinib | MVP_top60_DNA+RNA_seed | PIK3CA | -0.117276 | UCEC | MFE-319 | Buparlisib |
| MVP_top60_DNA_seed | HSP90AA1 | 1.354141 | LUAD | EMC-BAC-1 | CCT-018159 | MVP_top60_DNA+RNA_seed | PIK3CB | -0.117276 | UCEC | MFE-319 | Buparlisib |
| MVP_top60_DNA_seed | MAPK1 | 1.354426 | LUAD | NCI-H1563 | Ulixertinib | MVP_top60_DNA+RNA_seed | PIK3CD | -0.117276 | UCEC | MFE-319 | Buparlisib |
| MVP_top60_DNA_seed | EGFR | 1.354716 | LUAD | NCI-H2087 | Erlotinib | MVP_top60_DNA+RNA_seed | JAK1 | -0.11364 | UCEC | MFE-280 | Ruxolitinib |
| MVP_top60_DNA_seed | IGF1R | 1.354913 | LUAD | SK-LU-1 | GSK1904529A | MVP_top60_DNA+RNA_seed | JAK1 | -0.095985 | UCEC | MFE-319 | JAK1_8709 |
| MVP_top60_DNA_seed | BRAF | 1.357669 | LUAD | NCI-H2347 | Dabrafenib | MVP_top60_DNA+RNA_seed | AKT3 | -0.095928 | UCEC | MFE-296 | AZD5363 |
| MVP_top60_DNA_seed | AKT1 | 1.358761 | LUAD | SW1573 | Uprosertib | MVP_top60_DNA+RNA_seed | AKT1 | -0.095928 | UCEC | MFE-296 | AZD5363 |
| MVP_top60_DNA_seed | MDM2 | 1.360209 | LUAD | HOP-62 | Nutlin-3a (-) | MVP_top60_DNA+RNA_seed | EGFR | -0.094472 | UCEC | KLE | AZD3759 |
| MVP_top60_DNA_seed | AKT1 | 1.360431 | LUAD | NCI-H1568 | Afuresertib | MVP_top60_DNA+RNA_seed | PIK3CA | -0.092167 | UCEC | MFE-280 | AZD8186 |
| MVP_top60_DNA_seed | MDM2 | 1.361091 | LUAD | NCI-H2291 | Nutlin-3a (-) | MVP_top60_DNA+RNA_seed | PIK3CB | -0.092167 | UCEC | MFE-280 | AZD8186 |
| MVP_top60_DNA_seed | MAPK1 | 1.362223 | LUAD | SK-LU-1 | Ulixertinib | MVP_top60_DNA+RNA_seed | MDM2 | -0.09068 | UCEC | MFE-296 | Serdemetan |
| MVP_top60_DNA_seed | HSP90AA1 | 1.365235 | LUAD | NCI-H1563 | Tanespimycin | MVP_top60_DNA+RNA_seed | EGFR | -0.082744 | UCEC | MFE-319 | Gefitinib |
| MVP_top60_DNA_seed | AKT1 | 1.365323 | LUAD | SW1573 | GSK2110183B | MVP_top60_DNA+RNA_seed | EGFR | -0.082371 | UCEC | AN3-CA | Erlotinib |
| MVP_top60_DNA_seed | AKT1 | 1.365437 | LUAD | NCI-H1651 | Uprosertib | MVP_top60_DNA+RNA_seed | MDM2 | -0.068658 | UCEC | MFE-319 | Serdemetan |
| MVP_top60_DNA_seed | MET | 1.365631 | LUAD | NCI-H1355 | Savolitinib | MVP_top60_DNA+RNA_seed | EGFR | -0.058552 | UCEC | MFE-319 | Osimertinib |
| MVP_top60_DNA_seed | AKT1 | 1.366518 | LUAD | HOP-62 | Ipatasertib | MVP_top60_DNA+RNA_seed | MDM2 | -0.056338 | UCEC | COLO-684 | Nutlin-3a (-) |
| MVP_top60_DNA_seed | BRAF | 1.370298 | LUAD | NCI-H2347 | PLX-4720 | MVP_top60_DNA+RNA_seed | HSP90AA1 | -0.05537 | UCEC | MFE-319 | Luminespib |
| MVP_top60_DNA_seed | AKT1 | 1.37275 | LUAD | HCC-827 | GSK2110183B | MVP_top60_DNA+RNA_seed | HSP90AB1 | -0.05537 | UCEC | MFE-319 | Luminespib |
| MVP_top60_DNA_seed | EGFR | 1.373116 | LUAD | NCI-H1793 | Lapatinib | MVP_top60_DNA+RNA_seed | EP300 | -0.049097 | UCEC | MFE-280 | SGC-CBP30 |
| MVP_top60_DNA_seed | ERBB2 | 1.373116 | LUAD | NCI-H1793 | Lapatinib | MVP_top60_DNA+RNA_seed | MDM2 | -0.047125 | UCEC | MFE-319 | Nutlin-3a (-) |
| MVP_top60_DNA_seed | MAPK1 | 1.373967 | LUAD | VMRC-LCD | SCH772984 | MVP_top60_DNA+RNA_seed | ESR1 | -0.046219 | UCEC | MFE-280 | Tamoxifen |
| MVP_top60_DNA_seed | AKT1 | 1.376038 | LUAD | NCI-H441 | Uprosertib | MVP_top60_DNA+RNA_seed | EGFR | -0.044788 | UCEC | SNG-M | AZD3759 |
| MVP_top60_DNA_seed | EGFR | 1.376797 | LUAD | NCI-H2087 | Osimertinib | MVP_top60_DNA+RNA_seed | PIK3CA | -0.035895 | UCEC | COLO-684 | Buparlisib |
| MVP_top60_DNA_seed | AKT1 | 1.37782 | LUAD | NCI-H23 | Uprosertib | MVP_top60_DNA+RNA_seed | PIK3CB | -0.035895 | UCEC | COLO-684 | Buparlisib |
| MVP_top60_DNA_seed | EGFR | 1.379839 | LUAD | SW1573 | Sapitinib | MVP_top60_DNA+RNA_seed | PIK3CD | -0.035895 | UCEC | COLO-684 | Buparlisib |
| MVP_top60_DNA_seed | ERBB2 | 1.379839 | LUAD | SW1573 | Sapitinib | MVP_top60_DNA+RNA_seed | KRAS | -0.031796 | UCEC | AN3-CA | (RAS (G12C) Inhibitor-1 |
| MVP_top60_DNA_seed | ERBB3 | 1.379839 | LUAD | SW1573 | Sapitinib | MVP_top60_DNA+RNA_seed | MAPK1 | -0.031319 | UCEC | AN3-CA | VX-11e |
| MVP_top60_DNA_seed | IGF1R | 1.380565 | LUAD | NCI-H1355 | NVP-ADW742 | MVP_top60_DNA+RNA_seed | EGFR | -0.010186 | UCEC | MFE-280 | Afatinib |
| MVP_top60_DNA_seed | IGF1R | 1.380916 | LUAD | NCI-H522 | NVP-ADW742 | MVP_top60_DNA+RNA_seed | ERBB2 | -0.010186 | UCEC | MFE-280 | Afatinib |
| MVP_top60_DNA_seed | PIK3CA | 1.382071 | LUAD | EKVX | Alpelisib | MVP_top60_DNA+RNA_seed | PIK3CA | 0.004545 | UCEC | EN | Buparlisib |
| MVP_top60_DNA_seed | TERT | 1.384092 | LUAD | NCI-H358 | BIBR-1532 | MVP_top60_DNA+RNA_seed | PIK3CB | 0.004545 | UCEC | EN | Buparlisib |
| MVP_top60_DNA_seed | HDAC1 | 1.384361 | LUAD | ABC-1 | Romidepsin | MVP_top60_DNA+RNA_seed | PIK3CD | 0.004545 | UCEC | EN | Buparlisib |
| MVP_top60_DNA_seed | PIK3CA | 1.388825 | LUAD | NCI-H2291 | Buparlisib | MVP_top60_DNA+RNA_seed | JAK1 | 0.009319 | UCEC | EN | Ruxolitinib |

| | | | | | | | | | | | |
|---|---|---|---|---|---|---|---|---|---|---|---|
| MVP_top60_DNA_seed | HDAC1 | 1.389315 | LUAD | NCI-H2023 | PCI-34051 | MVP_top60_DNA+RNA_seed | FGFR2 | 0.011596 | UCEC | MFE-296 | PD173074 |
| MVP_top60_DNA_seed | ATM | 1.390152 | LUAD | NCI-H1355 | KU-55933 | MVP_top60_DNA+RNA_seed | HSP90AA1 | 0.016781 | UCEC | SNG-M | Luminespib |
| MVP_top60_DNA_seed | IGF1R | 1.391867 | LUAD | NCI-H441 | BMS-754807 | MVP_top60_DNA+RNA_seed | HSP90AB1 | 0.016781 | UCEC | SNG-M | Luminespib |
| MVP_top60_DNA_seed | MAPK1 | 1.394269 | LUAD | NCI-H2342 | Ulixertinib | MVP_top60_DNA+RNA_seed | AKT3 | 0.028776 | UCEC | SNG-M | Uprosertib |
| MVP_top60_DNA_seed | IGF1R | 1.394338 | LUAD | NCI-H2405 | GSK1904529A | MVP_top60_DNA+RNA_seed | AKT1 | 0.028776 | UCEC | SNG-M | Uprosertib |
| MVP_top60_DNA_seed | EGFR | 1.394676 | LUAD | HOP-62 | Afatinib | MVP_top60_DNA+RNA_seed | HSP90AA1 | 0.034726 | UCEC | COLO-684 | Tanespimycin |
| MVP_top60_DNA_seed | ERBB2 | 1.394676 | LUAD | HOP-62 | Afatinib | MVP_top60_DNA+RNA_seed | HSP90AB1 | 0.034726 | UCEC | COLO-684 | Tanespimycin |
| MVP_top60_DNA_seed | AKT1 | 1.39902 | LUAD | NCI-H1648 | AT13148 | MVP_top60_DNA+RNA_seed | ESR1 | 0.048635 | UCEC | SNG-M | Tamoxifen |
| MVP_top60_DNA_seed | FGFR1 | 1.399199 | LUAD | EMC-BAC-1 | AZD4547 | MVP_top60_DNA+RNA_seed | EGFR | 0.048945 | UCEC | COLO-684 | Afatinib |
| MVP_top60_DNA_seed | MET | 1.399303 | LUAD | NCI-H1838 | Crizotinib | MVP_top60_DNA+RNA_seed | ERBB2 | 0.048945 | UCEC | COLO-684 | Afatinib |
| MVP_top60_DNA_seed | AKT1 | 1.399437 | LUAD | NCI-H1693 | GSK2110183B | MVP_top60_DNA+RNA_seed | HSP90AA1 | 0.066108 | UCEC | MFE-280 | Tanespimycin |
| MVP_top60_DNA_seed | EGFR | 1.403831 | LUAD | NCI-H3122 | Sapitinib | MVP_top60_DNA+RNA_seed | HSP90AB1 | 0.066108 | UCEC | MFE-280 | Tanespimycin |
| MVP_top60_DNA_seed | ERBB2 | 1.403831 | LUAD | NCI-H3122 | Sapitinib | MVP_top60_DNA+RNA_seed | EGFR | 0.072826 | UCEC | MFE-280 | Lapatinib |
| MVP_top60_DNA_seed | ERBB3 | 1.403831 | LUAD | NCI-H3122 | Sapitinib | MVP_top60_DNA+RNA_seed | ERBB2 | 0.072826 | UCEC | MFE-280 | Lapatinib |
| MVP_top60_DNA_seed | HSP90AA1 | 1.404259 | LUAD | NCI-H2405 | Tanespimycin | MVP_top60_DNA+RNA_seed | AKT3 | 0.073645 | UCEC | SNG-M | Ipatasertib |
| MVP_top60_DNA_seed | HDAC1 | 1.410288 | LUAD | NCI-H1568 | PCI-34051 | MVP_top60_DNA+RNA_seed | AKT1 | 0.073645 | UCEC | SNG-M | Ipatasertib |
| MVP_top60_DNA_seed | PIK3CA | 1.410333 | LUAD | NCI-H1573 | Alpelisib | MVP_top60_DNA+RNA_seed | EGFR | 0.079518 | UCEC | COLO-684 | AZD3759 |
| MVP_top60_DNA_seed | IGF1R | 1.41099 | LUAD | EKVX | GSK1904529A | MVP_top60_DNA+RNA_seed | MDM2 | 0.079842 | UCEC | COLO-684 | Serdemetan |
| MVP_top60_DNA_seed | AURKA | 1.411484 | LUAD | NCI-H2405 | Alisertib | MVP_top60_DNA+RNA_seed | AKT1 | 0.085373 | UCEC | COLO-684 | AT13148 |
| MVP_top60_DNA_seed | AKT1 | 1.412291 | LUAD | NCI-H2087 | Uprosertib | MVP_top60_DNA+RNA_seed | EGFR | 0.098884 | UCEC | SNG-M | Afatinib |
| MVP_top60_DNA_seed | MDM2 | 1.41243 | LUAD | NCI-H1355 | Serdemetan | MVP_top60_DNA+RNA_seed | ERBB2 | 0.098884 | UCEC | SNG-M | Afatinib |
| MVP_top60_DNA_seed | MET | 1.412749 | LUAD | PC-3_[JPC-3] | Savolitinib | MVP_top60_DNA+RNA_seed | ESR1 | 0.099888 | UCEC | KLE | Tamoxifen |
| MVP_top60_DNA_seed | FGFR1 | 1.413455 | LUAD | NCI-H1355 | AZD4547 | MVP_top60_DNA+RNA_seed | MAPK1 | 0.105911 | UCEC | COLO-684 | VX-11e |
| MVP_top60_DNA_seed | PIK3CA | 1.413671 | LUAD | NCI-H2085 | Buparlisib | MVP_top60_DNA+RNA_seed | EGFR | 0.112023 | UCEC | SNG-M | Osimertinib |
| MVP_top60_DNA_seed | MET | 1.414504 | LUAD | SW1573 | Crizotinib | MVP_top60_DNA+RNA_seed | HDAC1 | 0.117694 | UCEC | SNG-M | PCI-34051 |
| MVP_top60_DNA_seed | MET | 1.41557 | LUAD | HCC-827 | Savolitinib | MVP_top60_DNA+RNA_seed | PIK3CA | 0.121978 | UCEC | KLE | Buparlisib |
| MVP_top60_DNA_seed | EGFR | 1.416134 | LUAD | NCI-H1838 | Osimertinib | MVP_top60_DNA+RNA_seed | PIK3CB | 0.121978 | UCEC | KLE | Buparlisib |
| MVP_top60_DNA_seed | MAPK1 | 1.417401 | LUAD | NCI-H1793 | VX-11e | MVP_top60_DNA+RNA_seed | PIK3CD | 0.121978 | UCEC | KLE | Buparlisib |
| MVP_top60_DNA_seed | AKT1 | 1.418869 | LUAD | NCI-H1568 | MK-2206 | MVP_top60_DNA+RNA_seed | ESR1 | 0.122817 | UCEC | AN3-CA | GDC0810 |
| MVP_top60_DNA_seed | HDAC1 | 1.422284 | LUAD | EKVX | PCI-34051 | MVP_top60_DNA+RNA_seed | TP53 | 0.125506 | UCEC | SNG-M | MIRA-1 |
| MVP_top60_DNA_seed | FGFR1 | 1.422524 | LUAD | COR-L105 | AZD4547 | MVP_top60_DNA+RNA_seed | MDM2 | 0.127979 | UCEC | MFE-296 | Nutlin-3a (-) |
| MVP_top60_DNA_seed | TERT | 1.423055 | LUAD | VMRC-LCD | BIBR-1532 | MVP_top60_DNA+RNA_seed | PIK3CA | 0.139196 | UCEC | EN | GNE-317 |
| MVP_top60_DNA_seed | BRAF | 1.423635 | LUAD | NCI-H1650 | SB590885 | MVP_top60_DNA+RNA_seed | MAPK1 | 0.142764 | UCEC | SNG-M | Ulixertinib |
| MVP_top60_DNA_seed | MDM2 | 1.426622 | LUAD | NCI-H441 | Nutlin-3a (-) | MVP_top60_DNA+RNA_seed | MAPK3 | 0.142764 | UCEC | SNG-M | Ulixertinib |
| MVP_top60_DNA_seed | HSP90AA1 | 1.428228 | LUAD | NCI-H2347 | Tanespimycin | MVP_top60_DNA+RNA_seed | AKT1 | 0.143375 | UCEC | EN | AT13148 |
| MVP_top60_DNA_seed | GSK3B | 1.428387 | LUAD | NCI-H1355 | SB216763 | MVP_top60_DNA+RNA_seed | EGFR | 0.147381 | UCEC | SNG-M | Erlotinib |
| MVP_top60_DNA_seed | AKT1 | 1.428663 | LUAD | ABC-1 | MK-2206 | MVP_top60_DNA+RNA_seed | TP53 | 0.157663 | UCEC | MFE-296 | MIRA-1 |
| MVP_top60_DNA_seed | SRC | 1.431085 | LUAD | SW1573 | Bosutinib | MVP_top60_DNA+RNA_seed | MAPK1 | 0.15838 | UCEC | MFE-319 | Ulixertinib |
| MVP_top60_DNA_seed | SMARCA4 | 1.4318 | LUAD | COR-L105 | PFI3 | MVP_top60_DNA+RNA_seed | MAPK3 | 0.15838 | UCEC | MFE-319 | Ulixertinib |
| MVP_top60_DNA_seed | AURKA | 1.43194 | LUAD | NCI-H650 | ZM447439 | MVP_top60_DNA+RNA_seed | EGFR | 0.164156 | UCEC | MFE-319 | Lapatinib |
| MVP_top60_DNA_seed | EGFR | 1.433437 | LUAD | Calu-6 | Osimertinib | MVP_top60_DNA+RNA_seed | ERBB2 | 0.164156 | UCEC | MFE-319 | Lapatinib |

| | | | | | | | | | | | |
|---|---|---|---|---|---|---|---|---|---|---|---|
| MVP_top60_DNA_seed | TERT | 1.433595 | LUAD | NCI-H2405 | BIBR-1532 | MVP_top60_DNA+RNA_seed | ESR1 | 0.167485 | UCEC | EN | Tamoxifen |
| MVP_top60_DNA_seed | TERT | 1.43452 | LUAD | NCI-H1568 | BIBR-1532 | MVP_top60_DNA+RNA_seed | EGFR | 0.168569 | UCEC | KLE | Erlotinib |
| MVP_top60_DNA_seed | AURKA | 1.435071 | LUAD | NCI-H2347 | ZM447439 | MVP_top60_DNA+RNA_seed | TP53 | 0.169217 | UCEC | COLO-684 | MIRA-1 |
| MVP_top60_DNA_seed | EGFR | 1.435245 | LUAD | HOP-62 | AZD3759 | MVP_top60_DNA+RNA_seed | EP300 | 0.171773 | UCEC | EN | CPI-637 |
| MVP_top60_DNA_seed | ATM | 1.43625 | LUAD | NCI-H2291 | KU-55933 | MVP_top60_DNA+RNA_seed | JAK1 | 0.17188 | UCEC | SNG-M | GSK2276186C |
| MVP_top60_DNA_seed | AKT1 | 1.43866 | LUAD | Calu-6 | Uprosertib | MVP_top60_DNA+RNA_seed | ESR1 | 0.176844 | UCEC | HEC-1 | Tamoxifen |
| MVP_top60_DNA_seed | AKT1 | 1.440483 | LUAD | PC-3_[JPC-3] | AT13148 | MVP_top60_DNA+RNA_seed | EGFR | 0.176851 | UCEC | COLO-684 | Lapatinib |
| MVP_top60_DNA_seed | IGF1R | 1.442829 | LUAD | NCI-H1435 | BMS-754807 | MVP_top60_DNA+RNA_seed | ERBB2 | 0.176851 | UCEC | COLO-684 | Lapatinib |
| MVP_top60_DNA_seed | MET | 1.443768 | LUAD | COR-L105 | Savolitinib | MVP_top60_DNA+RNA_seed | EGFR | 0.18344 | UCEC | MFE-319 | Sapitinib |
| MVP_top60_DNA_seed | AKT1 | 1.44533 | LUAD | NCI-H2087 | Ipatasertib | MVP_top60_DNA+RNA_seed | ERBB2 | 0.18344 | UCEC | MFE-319 | Sapitinib |
| MVP_top60_DNA_seed | FGFR1 | 1.445331 | LUAD | NCI-H1573 | PD173074 | MVP_top60_DNA+RNA_seed | EP300 | 0.189186 | UCEC | KLE | CPI-637 |
| MVP_top60_DNA_seed | EGFR | 1.446492 | LUAD | NCI-H1355 | Gefitinib | MVP_top60_DNA+RNA_seed | HDAC1 | 0.191042 | UCEC | AN3-CA | Romidepsin |
| MVP_top60_DNA_seed | AKT1 | 1.448144 | LUAD | NCI-H1568 | GSK2110183B | MVP_top60_DNA+RNA_seed | HDAC2 | 0.191042 | UCEC | AN3-CA | Romidepsin |
| MVP_top60_DNA_seed | SRC | 1.448639 | LUAD | EKVX | Bosutinib | MVP_top60_DNA+RNA_seed | EP300 | 0.191117 | UCEC | COLO-684 | CPI-637 |
| MVP_top60_DNA_seed | AKT1 | 1.449555 | LUAD | NCI-H2085 | Afuresertib | MVP_top60_DNA+RNA_seed | HDAC1 | 0.194166 | UCEC | MFE-319 | PCI-34051 |
| MVP_top60_DNA_seed | HDAC1 | 1.4505 | LUAD | NCI-H2087 | PCI-34051 | MVP_top60_DNA+RNA_seed | JAK1 | 0.19547 | UCEC | MFE-280 | GSK2276186C |
| MVP_top60_DNA_seed | HDAC1 | 1.452461 | LUAD | NCI-H1693 | Entinostat | MVP_top60_DNA+RNA_seed | HDAC1 | 0.198837 | UCEC | SNG-M | Romidepsin |
| MVP_top60_DNA_seed | MET | 1.453229 | LUAD | NCI-H1838 | Savolitinib | MVP_top60_DNA+RNA_seed | HDAC2 | 0.198837 | UCEC | SNG-M | Romidepsin |
| MVP_top60_DNA_seed | KRAS | 1.453596 | LUAD | NCI-H1693 | KRAS (G12C) Inhibitor-1 | MVP_top60_DNA+RNA_seed | FGFR2 | 0.204251 | UCEC | KLE | PD173074 |
| MVP_top60_DNA_seed | BRAF | 1.454314 | LUAD | NCI-H1650 | Dabrafenib | MVP_top60_DNA+RNA_seed | EGFR | 0.212163 | UCEC | MFE-296 | AZD3759 |
| MVP_top60_DNA_seed | EGFR | 1.457899 | LUAD | NCI-H2087 | Lapatinib | MVP_top60_DNA+RNA_seed | PIK3CA | 0.230603 | UCEC | SNG-M | Alpelisib |
| MVP_top60_DNA_seed | ERBB2 | 1.457899 | LUAD | NCI-H2087 | Lapatinib | MVP_top60_DNA+RNA_seed | EP300 | 0.231413 | UCEC | MFE-280 | CPI-637 |
| MVP_top60_DNA_seed | MAPK1 | 1.459351 | LUAD | PC-3_[JPC-3] | VX-11e | MVP_top60_DNA+RNA_seed | KRAS | 0.232402 | UCEC | MFE-319 | RAS (G12C) Inhibitor-1 |
| MVP_top60_DNA_seed | AKT1 | 1.459958 | LUAD | NCI-H650 | Uprosertib | MVP_top60_DNA+RNA_seed | JAK1 | 0.239616 | UCEC | COLO-684 | Ruxolitinib |
| MVP_top60_DNA_seed | TERT | 1.460395 | LUAD | NCI-H2347 | BIBR-1532 | MVP_top60_DNA+RNA_seed | EP300 | 0.240869 | UCEC | AN3-CA | SGC-CBP30 |
| MVP_top60_DNA_seed | AKT1 | 1.46087 | LUAD | Calu-6 | Ipatasertib | MVP_top60_DNA+RNA_seed | MAPK1 | 0.258765 | UCEC | MFE-319 | SCH772984 |
| MVP_top60_DNA_seed | AURKA | 1.461505 | LUAD | NCI-H1435 | ZM447439 | MVP_top60_DNA+RNA_seed | MAPK3 | 0.258765 | UCEC | MFE-319 | SCH772984 |
| MVP_top60_DNA_seed | HSP90AA1 | 1.46189 | LUAD | H3255 | Elesclomol | MVP_top60_DNA+RNA_seed | ESR1 | 0.260571 | UCEC | MFE-280 | GDC0810 |
| MVP_top60_DNA_seed | EGFR | 1.464976 | LUAD | NCI-H1838 | AZD3759 | MVP_top60_DNA+RNA_seed | JAK1 | 0.282599 | UCEC | EN | JAK1_8709 |
| MVP_top60_DNA_seed | EGFR | 1.468053 | LUAD | NCI-H2347 | AZD3759 | MVP_top60_DNA+RNA_seed | EGFR | 0.293901 | UCEC | MFE-296 | Osimertinib |
| MVP_top60_DNA_seed | EGFR | 1.469333 | LUAD | VMRC-LCD | Erlotinib | MVP_top60_DNA+RNA_seed | HDAC1 | 0.295173 | UCEC | MFE-296 | PCI-34051 |
| MVP_top60_DNA_seed | EGFR | 1.469558 | LUAD | NCI-H2228 | AZD3759 | MVP_top60_DNA+RNA_seed | EGFR | 0.297354 | UCEC | COLO-684 | Erlotinib |
| MVP_top60_DNA_seed | BRAF | 1.470045 | LUAD | NCI-H2347 | SB590885 | MVP_top60_DNA+RNA_seed | SRC | 0.300929 | UCEC | SNG-M | Bosutinib |
| MVP_top60_DNA_seed | EGFR | 1.471242 | LUAD | NCI-H441 | Gefitinib | MVP_top60_DNA+RNA_seed | HDAC1 | 0.304622 | UCEC | MFE-319 | Romidepsin |
| MVP_top60_DNA_seed | FGFR1 | 1.473188 | LUAD | NCI-H1651 | PD173074 | MVP_top60_DNA+RNA_seed | HDAC2 | 0.304622 | UCEC | MFE-319 | Romidepsin |
| MVP_top60_DNA_seed | HDAC1 | 1.474664 | LUAD | H3255 | PCI-34051 | MVP_top60_DNA+RNA_seed | EGFR | 0.311072 | UCEC | HEC-1 | Sapitinib |
| MVP_top60_DNA_seed | AKT1 | 1.475109 | LUAD | NCI-H2347 | AZD5363 | MVP_top60_DNA+RNA_seed | ERBB2 | 0.311072 | UCEC | HEC-1 | Sapitinib |
| MVP_top60_DNA_seed | IGF1R | 1.475151 | LUAD | NCI-H2291 | Linsitinib | MVP_top60_DNA+RNA_seed | MAPK1 | 0.315083 | UCEC | MFE-296 | SCH772984 |
| MVP_top60_DNA_seed | IGF1R | 1.479779 | LUAD | NCI-H441 | BMS-536924 | MVP_top60_DNA+RNA_seed | MAPK3 | 0.315083 | UCEC | MFE-296 | SCH772984 |
| MVP_top60_DNA_seed | TP53 | 1.479916 | LUAD | NCI-H2228 | MIRA-1 | MVP_top60_DNA+RNA_seed | PIK3CA | 0.320685 | UCEC | MFE-319 | AZD8186 |
| MVP_top60_DNA_seed | AKT1 | 1.483298 | LUAD | Calu-3 | AZD5363 | MVP_top60_DNA+RNA_seed | PIK3CB | 0.320685 | UCEC | MFE-319 | AZD8186 |

| | | | | | | | | | | | |
|---|---|---|---|---|---|---|---|---|---|---|---|
| MVP_top60_DNA_seed | PIK3CA | 1.485284 | LUAD | NCI-H1435 | GNE-317 | MVP_top60_DNA+RNA_seed | MAPK1 | 0.3282 | UCEC | MFE-296 | Ulixertinib |
| MVP_top60_DNA_seed | BRAF | 1.486903 | LUAD | PC-3_[JPC-3] | Dabrafenib | MVP_top60_DNA+RNA_seed | MAPK3 | 0.3282 | UCEC | MFE-296 | Ulixertinib |
| MVP_top60_DNA_seed | HSP90AA1 | 1.486934 | LUAD | Calu-3 | Luminespib | MVP_top60_DNA+RNA_seed | AKT3 | 0.336334 | UCEC | SNG-M | GSK2110183B |
| MVP_top60_DNA_seed | EGFR | 1.488139 | LUAD | NCI-H2347 | Osimertinib | MVP_top60_DNA+RNA_seed | AKT1 | 0.336334 | UCEC | SNG-M | GSK2110183B |
| MVP_top60_DNA_seed | MAPK1 | 1.489314 | LUAD | NCI-H2085 | Ulixertinib | MVP_top60_DNA+RNA_seed | MAPK1 | 0.341619 | UCEC | SNG-M | SCH772984 |
| MVP_top60_DNA_seed | HDAC1 | 1.491398 | LUAD | NCI-H2405 | Entinostat | MVP_top60_DNA+RNA_seed | MAPK3 | 0.341619 | UCEC | SNG-M | SCH772984 |
| MVP_top60_DNA_seed | MAPK1 | 1.494903 | LUAD | SW1573 | VX-11e | MVP_top60_DNA+RNA_seed | HSP90AA1 | 0.34164 | UCEC | COLO-684 | Luminespib |
| MVP_top60_DNA_seed | BRAF | 1.495557 | LUAD | NCI-H1838 | SB590885 | MVP_top60_DNA+RNA_seed | HSP90AB1 | 0.34164 | UCEC | COLO-684 | Luminespib |
| MVP_top60_DNA_seed | TP53 | 1.497721 | LUAD | NCI-H1666 | MIRA-1 | MVP_top60_DNA+RNA_seed | EGFR | 0.348403 | UCEC | AN3-CA | Gefitinib |
| MVP_top60_DNA_seed | TP53 | 1.500118 | LUAD | NCI-H1650 | MIRA-1 | MVP_top60_DNA+RNA_seed | MAPK1 | 0.348671 | UCEC | AN3-CA | SCH772984 |
| MVP_top60_DNA_seed | HSP90AA1 | 1.50074 | LUAD | SW1573 | Luminespib | MVP_top60_DNA+RNA_seed | MAPK3 | 0.348671 | UCEC | AN3-CA | SCH772984 |
| MVP_top60_DNA_seed | MAPK1 | 1.500884 | LUAD | Calu-3 | SCH772984 | MVP_top60_DNA+RNA_seed | ATM | 0.35817 | UCEC | COLO-684 | KU-55933 |
| MVP_top60_DNA_seed | CREBBP | 1.503523 | LUAD | H3255 | SGC-CBP30 | MVP_top60_DNA+RNA_seed | EGFR | 0.371226 | UCEC | AN3-CA | Afatinib |
| MVP_top60_DNA_seed | MET | 1.505104 | LUAD | H3255 | Crizotinib | MVP_top60_DNA+RNA_seed | ERBB2 | 0.371226 | UCEC | AN3-CA | Afatinib |
| MVP_top60_DNA_seed | AKT1 | 1.506064 | LUAD | NCI-H1568 | Uprosertib | MVP_top60_DNA+RNA_seed | PIK3CA | 0.376584 | UCEC | SNG-M | GNE-317 |
| MVP_top60_DNA_seed | IGF1R | 1.50765 | LUAD | NCI-H522 | BMS-536924 | MVP_top60_DNA+RNA_seed | EGFR | 0.380384 | UCEC | KLE | Lapatinib |
| MVP_top60_DNA_seed | AKT1 | 1.510772 | LUAD | NCI-H1355 | GSK2110183B | MVP_top60_DNA+RNA_seed | ERBB2 | 0.380384 | UCEC | KLE | Lapatinib |
| MVP_top60_DNA_seed | AKT1 | 1.512061 | LUAD | PC-3_[JPC-3] | MK-2206 | MVP_top60_DNA+RNA_seed | EGFR | 0.384107 | UCEC | SNG-M | Gefitinib |
| MVP_top60_DNA_seed | CREBBP | 1.51309 | LUAD | NCI-H1650 | SGC-CBP30 | MVP_top60_DNA+RNA_seed | ESR1 | 0.389647 | UCEC | HEC-1 | GDC0810 |
| MVP_top60_DNA_seed | AURKA | 1.513168 | LUAD | NCI-H1734 | Alisertib | MVP_top60_DNA+RNA_seed | AKT1 | 0.393859 | UCEC | MFE-280 | AT13148 |
| MVP_top60_DNA_seed | MAPK1 | 1.513301 | LUAD | NCI-H1563 | SCH772984 | MVP_top60_DNA+RNA_seed | SRC | 0.394768 | UCEC | COLO-684 | Bosutinib |
| MVP_top60_DNA_seed | SMARCA4 | 1.513676 | LUAD | NCI-H1650 | PFI3 | MVP_top60_DNA+RNA_seed | EGFR | 0.406535 | UCEC | SNG-M | Lapatinib |
| MVP_top60_DNA_seed | SRC | 1.514558 | LUAD | NCI-H2291 | Bosutinib | MVP_top60_DNA+RNA_seed | ERBB2 | 0.406535 | UCEC | SNG-M | Lapatinib |
| MVP_top60_DNA_seed | EZH2 | 1.521444 | LUAD | HOP-62 | GSK343 | MVP_top60_DNA+RNA_seed | PIK3CA | 0.414572 | UCEC | SNG-M | Buparlisib |
| MVP_top60_DNA_seed | MET | 1.523511 | LUAD | ABC-1 | Crizotinib | MVP_top60_DNA+RNA_seed | PIK3CB | 0.414572 | UCEC | SNG-M | Buparlisib |
| MVP_top60_DNA_seed | FGFR1 | 1.52354 | LUAD | ABC-1 | PD173074 | MVP_top60_DNA+RNA_seed | PIK3CD | 0.414572 | UCEC | SNG-M | Buparlisib |
| MVP_top60_DNA_seed | HSP90AA1 | 1.524546 | LUAD | NCI-H2291 | CCT-018159 | MVP_top60_DNA+RNA_seed | JAK1 | 0.425695 | UCEC | MFE-319 | Ruxolitinib |
| MVP_top60_DNA_seed | ATM | 1.524624 | LUAD | Calu-3 | KU-55933 | MVP_top60_DNA+RNA_seed | KRAS | 0.426664 | UCEC | MFE-280 | KRAS (G12C) Inhibitor-1 |
| MVP_top60_DNA_seed | GSK3B | 1.526006 | LUAD | H3255 | SB216763 | MVP_top60_DNA+RNA_seed | ESR1 | 0.42959 | UCEC | MFE-296 | GDC0810 |
| MVP_top60_DNA_seed | IGF1R | 1.527068 | LUAD | NCI-H2347 | Linsitinib | MVP_top60_DNA+RNA_seed | EGFR | 0.430831 | UCEC | MFE-319 | Afatinib |
| MVP_top60_DNA_seed | AURKA | 1.53075 | LUAD | NCI-H441 | Alisertib | MVP_top60_DNA+RNA_seed | ERBB2 | 0.430831 | UCEC | MFE-319 | Afatinib |
| MVP_top60_DNA_seed | PIK3CA | 1.532989 | LUAD | NCI-H522 | AZD8186 | MVP_top60_DNA+RNA_seed | EP300 | 0.432925 | UCEC | MFE-296 | CPI-637 |
| MVP_top60_DNA_seed | FGFR1 | 1.533809 | LUAD | PC-3_[JPC-3] | PD173074 | MVP_top60_DNA+RNA_seed | EGFR | 0.433326 | UCEC | KLE | Gefitinib |
| MVP_top60_DNA_seed | AKT1 | 1.534645 | LUAD | NCI-H358 | AT13148 | MVP_top60_DNA+RNA_seed | AKT3 | 0.437204 | UCEC | MFE-319 | GSK2110183B |
| MVP_top60_DNA_seed | FGFR1 | 1.534854 | LUAD | NCI-H2342 | PD173074 | MVP_top60_DNA+RNA_seed | AKT1 | 0.437204 | UCEC | MFE-319 | GSK2110183B |
| MVP_top60_DNA_seed | EZH2 | 1.536372 | LUAD | NCI-H2291 | GSK343 | MVP_top60_DNA+RNA_seed | FGFR2 | 0.454732 | UCEC | SNG-M | AZD4547 |
| MVP_top60_DNA_seed | HSP90AA1 | 1.537239 | LUAD | SW1573 | Tanespimycin | MVP_top60_DNA+RNA_seed | JAK1 | 0.471991 | UCEC | MFE-296 | GSK2276186C |
| MVP_top60_DNA_seed | HSP90AA1 | 1.537521 | LUAD | EKVX | CCT-018159 | MVP_top60_DNA+RNA_seed | EGFR | 0.484221 | UCEC | MFE-296 | Sapitinib |
| MVP_top60_DNA_seed | TERT | 1.53817 | LUAD | NCI-H2087 | BIBR-1532 | MVP_top60_DNA+RNA_seed | ERBB2 | 0.484221 | UCEC | MFE-296 | Sapitinib |
| MVP_top60_DNA_seed | ATM | 1.543521 | LUAD | PC-14 | KU-55933 | MVP_top60_DNA+RNA_seed | PIK3CA | 0.485906 | UCEC | HEC-1 | GNE-317 |
| MVP_top60_DNA_seed | BRAF | 1.546099 | LUAD | NCI-H1793 | Dabrafenib | MVP_top60_DNA+RNA_seed | JAK1 | 0.487375 | UCEC | MFE-319 | GSK2276186C |

| | | | | | | | | | | | |
|---|---|---|---|---|---|---|---|---|---|---|---|
| MVP_top60_DNA_seed | IGF1R | 1.548481 | LUAD | SK-LU-1 | NVP-ADW742 | MVP_top60_DNA+RNA_seed | HSP90AA1 | 0.489232 | UCEC | MFE-296 | Tanespimycin |
| MVP_top60_DNA_seed | AKT1 | 1.548559 | LUAD | HCC-78 | AT13148 | MVP_top60_DNA+RNA_seed | HSP90AB1 | 0.489232 | UCEC | MFE-296 | Tanespimycin |
| MVP_top60_DNA_seed | IGF1R | 1.551985 | LUAD | PC-3_[JPC-3] | BMS-754807 | MVP_top60_DNA+RNA_seed | HSP90AA1 | 0.493502 | UCEC | MFE-280 | Luminespib |
| MVP_top60_DNA_seed | AKT1 | 1.554786 | LUAD | NCI-H1838 | Afuresertib | MVP_top60_DNA+RNA_seed | HSP90AB1 | 0.493502 | UCEC | MFE-280 | Luminespib |
| MVP_top60_DNA_seed | MET | 1.556386 | LUAD | NCI-H1650 | Crizotinib | MVP_top60_DNA+RNA_seed | EGFR | 0.499133 | UCEC | AN3-CA | AZD3759 |
| MVP_top60_DNA_seed | IGF1R | 1.556657 | LUAD | NCI-H1793 | BMS-754807 | MVP_top60_DNA+RNA_seed | JAK1 | 0.506222 | UCEC | EN | JAK_8517 |
| MVP_top60_DNA_seed | HDAC1 | 1.556831 | LUAD | SW1573 | Romidepsin | MVP_top60_DNA+RNA_seed | AKT3 | 0.538987 | UCEC | KLE | AZD5363 |
| MVP_top60_DNA_seed | BRAF | 1.560715 | LUAD | NCI-H2291 | Dabrafenib | MVP_top60_DNA+RNA_seed | AKT1 | 0.538987 | UCEC | KLE | AZD5363 |
| MVP_top60_DNA_seed | AKT1 | 1.561469 | LUAD | NCI-H2087 | Afuresertib | MVP_top60_DNA+RNA_seed | AKT1 | 0.542338 | UCEC | MFE-296 | AT13148 |
| MVP_top60_DNA_seed | MAPK1 | 1.562262 | LUAD | HCC-827 | Ulixertinib | MVP_top60_DNA+RNA_seed | PIK3CA | 0.543186 | UCEC | MFE-280 | Buparlisib |
| MVP_top60_DNA_seed | CREBBP | 1.566448 | LUAD | HOP-62 | SGC-CBP30 | MVP_top60_DNA+RNA_seed | PIK3CB | 0.543186 | UCEC | MFE-280 | Buparlisib |
| MVP_top60_DNA_seed | AKT1 | 1.566781 | LUAD | SK-LU-1 | Uprosertib | MVP_top60_DNA+RNA_seed | PIK3CD | 0.543186 | UCEC | MFE-280 | Buparlisib |
| MVP_top60_DNA_seed | MAPK1 | 1.568388 | LUAD | NCI-H1650 | SCH772984 | MVP_top60_DNA+RNA_seed | TP53 | 0.56048 | UCEC | MFE-319 | MIRA-1 |
| MVP_top60_DNA_seed | TERT | 1.570792 | LUAD | SW1573 | BIBR-1532 | MVP_top60_DNA+RNA_seed | AKT3 | 0.56375 | UCEC | SNG-M | Afuresertib |
| MVP_top60_DNA_seed | BRAF | 1.575207 | LUAD | NCI-H1666 | PLX-4720 | MVP_top60_DNA+RNA_seed | AKT1 | 0.56375 | UCEC | SNG-M | Afuresertib |
| MVP_top60_DNA_seed | GSK3B | 1.575581 | LUAD | NCI-H1666 | SB216763 | MVP_top60_DNA+RNA_seed | EGFR | 0.564521 | UCEC | MFE-280 | AZD3759 |
| MVP_top60_DNA_seed | HSP90AA1 | 1.578207 | LUAD | NCI-H1563 | Luminespib | MVP_top60_DNA+RNA_seed | JAK1 | 0.578082 | UCEC | MFE-319 | JAK_8517 |
| MVP_top60_DNA_seed | PIK3CA | 1.578941 | LUAD | ABC-1 | GNE-317 | MVP_top60_DNA+RNA_seed | HDAC1 | 0.600748 | UCEC | MFE-296 | Romidepsin |
| MVP_top60_DNA_seed | HSP90AA1 | 1.581275 | LUAD | NCI-H1623 | Tanespimycin | MVP_top60_DNA+RNA_seed | HDAC2 | 0.600748 | UCEC | MFE-296 | Romidepsin |
| MVP_top60_DNA_seed | MET | 1.583766 | LUAD | H3255 | Savolitinib | MVP_top60_DNA+RNA_seed | AKT1 | 0.605984 | UCEC | SNG-M | MK-2206 |
| MVP_top60_DNA_seed | TERT | 1.584771 | LUAD | NCI-H1355 | BIBR-1532 | MVP_top60_DNA+RNA_seed | MAPK1 | 0.606594 | UCEC | EN | SCH772984 |
| MVP_top60_DNA_seed | CREBBP | 1.587429 | LUAD | HCC-827 | SGC-CBP30 | MVP_top60_DNA+RNA_seed | MAPK3 | 0.606594 | UCEC | EN | SCH772984 |
| MVP_top60_DNA_seed | AKT1 | 1.589153 | LUAD | HOP-62 | Uprosertib | MVP_top60_DNA+RNA_seed | HDAC1 | 0.608743 | UCEC | MFE-280 | Romidepsin |
| MVP_top60_DNA_seed | AURKA | 1.590916 | LUAD | NCI-H1568 | Alisertib | MVP_top60_DNA+RNA_seed | HDAC2 | 0.608743 | UCEC | MFE-280 | Romidepsin |
| MVP_top60_DNA_seed | EGFR | 1.592716 | LUAD | NCI-H2342 | Lapatinib | MVP_top60_DNA+RNA_seed | FGFR2 | 0.620923 | UCEC | HEC-1 | AZD4547 |
| MVP_top60_DNA_seed | ERBB2 | 1.592716 | LUAD | NCI-H2342 | Lapatinib | MVP_top60_DNA+RNA_seed | EGFR | 0.62759 | UCEC | AN3-CA | Lapatinib |
| MVP_top60_DNA_seed | MAPK1 | 1.597288 | LUAD | Calu-3 | Ulixertinib | MVP_top60_DNA+RNA_seed | ERBB2 | 0.62759 | UCEC | AN3-CA | Lapatinib |
| MVP_top60_DNA_seed | MAPK1 | 1.598327 | LUAD | H3255 | Ulixertinib | MVP_top60_DNA+RNA_seed | HDAC1 | 0.633137 | UCEC | HEC-1 | Entinostat |
| MVP_top60_DNA_seed | PIK3CA | 1.600673 | LUAD | NCI-H2085 | Alpelisib | MVP_top60_DNA+RNA_seed | SRC | 0.650923 | UCEC | AN3-CA | Bosutinib |
| MVP_top60_DNA_seed | EGFR | 1.606154 | LUAD | NCI-H1793 | Osimertinib | MVP_top60_DNA+RNA_seed | SRC | 0.653856 | UCEC | HEC-1 | Bosutinib |
| MVP_top60_DNA_seed | SMARCA4 | 1.607298 | LUAD | Calu-3 | PFI3 | MVP_top60_DNA+RNA_seed | ATM | 0.65422 | UCEC | EN | KU-55933 |
| MVP_top60_DNA_seed | AURKA | 1.611555 | LUAD | PC-3_[JPC-3] | ZM447439 | MVP_top60_DNA+RNA_seed | HDAC1 | 0.656557 | UCEC | AN3-CA | Entinostat |
| MVP_top60_DNA_seed | AKT1 | 1.614298 | LUAD | NCI-H1793 | MK-2206 | MVP_top60_DNA+RNA_seed | MDM2 | 0.659737 | UCEC | HEC-1 | Nutlin-3a (-) |
| MVP_top60_DNA_seed | AURKA | 1.615307 | LUAD | NCI-H2347 | Alisertib | MVP_top60_DNA+RNA_seed | EGFR | 0.667135 | UCEC | EN | Afatinib |
| MVP_top60_DNA_seed | SMARCA4 | 1.615845 | LUAD | NCI-H1793 | PFI3 | MVP_top60_DNA+RNA_seed | ERBB2 | 0.667135 | UCEC | EN | Afatinib |
| MVP_top60_DNA_seed | FGFR1 | 1.62002 | LUAD | SW1573 | PD173074 | MVP_top60_DNA+RNA_seed | EGFR | 0.668489 | UCEC | HEC-1 | AZD3759 |
| MVP_top60_DNA_seed | AKT1 | 1.621086 | LUAD | EKVX | GSK2110183B | MVP_top60_DNA+RNA_seed | MAPK1 | 0.668563 | UCEC | AN3-CA | Ulixertinib |
| MVP_top60_DNA_seed | CREBBP | 1.622068 | LUAD | NCI-H358 | SGC-CBP30 | MVP_top60_DNA+RNA_seed | MAPK3 | 0.668563 | UCEC | AN3-CA | Ulixertinib |
| MVP_top60_DNA_seed | MDM2 | 1.623235 | LUAD | HCC-78 | Serdemetan | MVP_top60_DNA+RNA_seed | MDM2 | 0.671411 | UCEC | SNG-M | Serdemetan |
| MVP_top60_DNA_seed | EZH2 | 1.624931 | LUAD | NCI-H358 | GSK343 | MVP_top60_DNA+RNA_seed | EGFR | 0.672078 | UCEC | MFE-296 | Gefitinib |
| MVP_top60_DNA_seed | MDM2 | 1.627616 | LUAD | NCI-H2347 | Nutlin-3a (-) | MVP_top60_DNA+RNA_seed | AKT3 | 0.675819 | UCEC | MFE-296 | GSK2110183B |

| | | | | | | | | | | | |
|---|---|---|---|---|---|---|---|---|---|---|---|
| MVP_top60_DNA_seed | GSK3B | 1.627729 | LUAD | NCI-H1838 | SB216763 | MVP_top60_DNA+RNA_seed | AKT1 | 0.675819 | UCEC | MFE-296 | GSK2110183B |
| MVP_top60_DNA_seed | BRAF | 1.628801 | LUAD | NCI-H1793 | SB590885 | MVP_top60_DNA+RNA_seed | MAPK1 | 0.69056 | UCEC | HEC-1 | Ulixertinib |
| MVP_top60_DNA_seed | FGFR1 | 1.630316 | LUAD | H3255 | PD173074 | MVP_top60_DNA+RNA_seed | MAPK3 | 0.69056 | UCEC | HEC-1 | Ulixertinib |
| MVP_top60_DNA_seed | KRAS | 1.630854 | LUAD | COR-L105 | KRAS (G12C) Inhibitor-1: | MVP_top60_DNA+RNA_seed | AKT1 | 0.692956 | UCEC | HEC-1 | AT13148 |
| MVP_top60_DNA_seed | EGFR | 1.631178 | LUAD | SK-LU-1 | Gefitinib | MVP_top60_DNA+RNA_seed | SRC | 0.694887 | UCEC | MFE-296 | Bosutinib |
| MVP_top60_DNA_seed | IGF1R | 1.633577 | LUAD | NCI-H441 | NVP-ADW742 | MVP_top60_DNA+RNA_seed | EGFR | 0.699991 | UCEC | MFE-280 | Osimertinib |
| MVP_top60_DNA_seed | GSK3B | 1.635198 | LUAD | NCI-H1793 | SB216763 | MVP_top60_DNA+RNA_seed | AKT3 | 0.702212 | UCEC | HEC-1 | AZD5363 |
| MVP_top60_DNA_seed | FGFR1 | 1.637179 | LUAD | NCI-H2291 | AZD4547 | MVP_top60_DNA+RNA_seed | AKT1 | 0.702212 | UCEC | HEC-1 | AZD5363 |
| MVP_top60_DNA_seed | AKT1 | 1.637792 | LUAD | Calu-6 | AT13148 | MVP_top60_DNA+RNA_seed | EP300 | 0.711783 | UCEC | AN3-CA | CPI-637 |
| MVP_top60_DNA_seed | FGFR1 | 1.640935 | LUAD | NCI-H1838 | AZD4547 | MVP_top60_DNA+RNA_seed | MAPK1 | 0.712296 | UCEC | MFE-280 | SCH772984 |
| MVP_top60_DNA_seed | MDM2 | 1.641381 | LUAD | NCI-H1693 | Nutlin-3a (-) | MVP_top60_DNA+RNA_seed | MAPK3 | 0.712296 | UCEC | MFE-280 | SCH772984 |
| MVP_top60_DNA_seed | FGFR1 | 1.64168 | LUAD | NCI-H1793 | PD173074 | MVP_top60_DNA+RNA_seed | MDM2 | 0.712865 | UCEC | EN | Nutlin-3a (-) |
| MVP_top60_DNA_seed | EGFR | 1.641832 | LUAD | SW1573 | Afatinib | MVP_top60_DNA+RNA_seed | AKT3 | 0.713039 | UCEC | SNG-M | Uprosertib |
| MVP_top60_DNA_seed | ERBB2 | 1.641832 | LUAD | SW1573 | Afatinib | MVP_top60_DNA+RNA_seed | AKT1 | 0.713039 | UCEC | SNG-M | Uprosertib |
| MVP_top60_DNA_seed | BRAF | 1.646322 | LUAD | NCI-H441 | Dabrafenib | MVP_top60_DNA+RNA_seed | EGFR | 0.716971 | UCEC | AN3-CA | Osimertinib |
| MVP_top60_DNA_seed | MAPK1 | 1.646986 | LUAD | NCI-H441 | Ulixertinib | MVP_top60_DNA+RNA_seed | EP300 | 0.722915 | UCEC | HEC-1 | CPI-637 |
| MVP_top60_DNA_seed | AKT1 | 1.652057 | LUAD | NCI-H1650 | Uprosertib | MVP_top60_DNA+RNA_seed | TP53 | 0.730928 | UCEC | EN | MIRA-1 |
| MVP_top60_DNA_seed | EZH2 | 1.653178 | LUAD | NCI-H1793 | GSK343 | MVP_top60_DNA+RNA_seed | SRC | 0.742165 | UCEC | MFE-319 | Bosutinib |
| MVP_top60_DNA_seed | MAPK1 | 1.653726 | LUAD | NCI-H1793 | SCH772984 | MVP_top60_DNA+RNA_seed | FGFR2 | 0.750459 | UCEC | COLO-684 | AZD4547 |
| MVP_top60_DNA_seed | HDAC1 | 1.657406 | LUAD | HCC-827 | PCI-34051 | MVP_top60_DNA+RNA_seed | HDAC1 | 0.767931 | UCEC | EN | Entinostat |
| MVP_top60_DNA_seed | SMARCA4 | 1.658578 | LUAD | NCI-H2228 | PFI3 | MVP_top60_DNA+RNA_seed | HDAC1 | 0.790518 | UCEC | HEC-1 | PCI-34051 |
| MVP_top60_DNA_seed | IGF1R | 1.662383 | LUAD | HOP-62 | Linsitinib | MVP_top60_DNA+RNA_seed | HSP90AA1 | 0.791403 | UCEC | EN | Tanespimycin |
| MVP_top60_DNA_seed | AKT1 | 1.663842 | LUAD | NCI-H1573 | Uprosertib | MVP_top60_DNA+RNA_seed | HSP90AB1 | 0.791403 | UCEC | EN | Tanespimycin |
| MVP_top60_DNA_seed | AKT1 | 1.665035 | LUAD | VMRC-LCD | Uprosertib | MVP_top60_DNA+RNA_seed | ESR1 | 0.793876 | UCEC | AN3-CA | Tamoxifen |
| MVP_top60_DNA_seed | AKT1 | 1.667253 | LUAD | NCI-H1693 | AZD5363 | MVP_top60_DNA+RNA_seed | HSP90AA1 | 0.807676 | UCEC | SNG-M | Tanespimycin |
| MVP_top60_DNA_seed | EZH2 | 1.669497 | LUAD | NCI-H2228 | GSK343 | MVP_top60_DNA+RNA_seed | HSP90AB1 | 0.807676 | UCEC | SNG-M | Tanespimycin |
| MVP_top60_DNA_seed | EGFR | 1.676934 | LUAD | NCI-H2347 | Afatinib | MVP_top60_DNA+RNA_seed | AKT3 | 0.812305 | UCEC | HEC-1 | GSK2110183B |
| MVP_top60_DNA_seed | ERBB2 | 1.676934 | LUAD | NCI-H2347 | Afatinib | MVP_top60_DNA+RNA_seed | AKT1 | 0.812305 | UCEC | HEC-1 | GSK2110183B |
| MVP_top60_DNA_seed | MDM2 | 1.678429 | LUAD | SW1573 | Serdemetan | MVP_top60_DNA+RNA_seed | PIK3CA | 0.816956 | UCEC | MFE-296 | Buparlisib |
| MVP_top60_DNA_seed | ATM | 1.680514 | LUAD | SK-LU-1 | KU-55933 | MVP_top60_DNA+RNA_seed | PIK3CB | 0.816956 | UCEC | MFE-296 | Buparlisib |
| MVP_top60_DNA_seed | PIK3CA | 1.681244 | LUAD | NCI-H2347 | Buparlisib | MVP_top60_DNA+RNA_seed | PIK3CD | 0.816956 | UCEC | MFE-296 | Buparlisib |
| MVP_top60_DNA_seed | ATM | 1.685011 | LUAD | H3255 | KU-55933 | MVP_top60_DNA+RNA_seed | HDAC1 | 0.817593 | UCEC | EN | PCI-34051 |
| MVP_top60_DNA_seed | IGF1R | 1.686758 | LUAD | NCI-H1563 | NVP-ADW742 | MVP_top60_DNA+RNA_seed | MAPK1 | 0.820356 | UCEC | EN | Ulixertinib |
| MVP_top60_DNA_seed | EZH2 | 1.689203 | LUAD | NCI-H1568 | GSK343 | MVP_top60_DNA+RNA_seed | MAPK3 | 0.820356 | UCEC | EN | Ulixertinib |
| MVP_top60_DNA_seed | EGFR | 1.690854 | LUAD | SK-LU-1 | AZD3759 | MVP_top60_DNA+RNA_seed | KRAS | 0.824168 | UCEC | KLE | KRAS (G12C) Inhibitor-1 |
| MVP_top60_DNA_seed | FGFR1 | 1.691705 | LUAD | NCI-H441 | AZD4547 | MVP_top60_DNA+RNA_seed | HDAC1 | 0.834873 | UCEC | EN | Romidepsin |
| MVP_top60_DNA_seed | TP53 | 1.691954 | LUAD | NCI-H1568 | MIRA-1 | MVP_top60_DNA+RNA_seed | HDAC2 | 0.834873 | UCEC | EN | Romidepsin |
| MVP_top60_DNA_seed | EZH2 | 1.694465 | LUAD | NCI-H1650 | GSK343 | MVP_top60_DNA+RNA_seed | JAK1 | 0.841807 | UCEC | MFE-280 | JAK1_8709 |
| MVP_top60_DNA_seed | AURKA | 1.698161 | LUAD | NCI-H2087 | ZM447439 | MVP_top60_DNA+RNA_seed | MAPK1 | 0.851823 | UCEC | HEC-1 | VX-11e |
| MVP_top60_DNA_seed | IGF1R | 1.703473 | LUAD | NCI-H1781 | BMS-536924 | MVP_top60_DNA+RNA_seed | EP300 | 0.853598 | UCEC | MFE-319 | SGC-CBP30 |
| MVP_top60_DNA_seed | KRAS | 1.703993 | LUAD | NCI-H1650 | KRAS (G12C) Inhibitor-1: | MVP_top60_DNA+RNA_seed | TP53 | 0.860405 | UCEC | MFE-280 | MIRA-1 |

| | | | | | | | | | | | |
|---|---|---|---|---|---|---|---|---|---|---|---|
| MVP_top60_DNA_seed | HSP90AA1 | 1.705251 | LUAD | NCI-H1568 | Luminespib | MVP_top60_DNA+RNA_seed | EGFR | 0.863622 | UCEC | EN | Osimertinib |
| MVP_top60_DNA_seed | MAPK1 | 1.706446 | LUAD | NCI-H2347 | Ulixertinib | MVP_top60_DNA+RNA_seed | HDAC1 | 0.864204 | UCEC | MFE-296 | Entinostat |
| MVP_top60_DNA_seed | PIK3CA | 1.707488 | LUAD | SK-LU-1 | Alpelisib | MVP_top60_DNA+RNA_seed | MAPK1 | 0.87711 | UCEC | KLE | VX-11e |
| MVP_top60_DNA_seed | EZH2 | 1.709431 | LUAD | NCI-H1355 | GSK343 | MVP_top60_DNA+RNA_seed | JAK1 | 0.888952 | UCEC | HEC-1 | GSK2276186C |
| MVP_top60_DNA_seed | MAPK1 | 1.712134 | LUAD | VMRC-LCD | Ulixertinib | MVP_top60_DNA+RNA_seed | ATM | 0.891619 | UCEC | HEC-1 | KU-55933 |
| MVP_top60_DNA_seed | GSK3B | 1.713486 | LUAD | EKVX | CHIR-99021 | MVP_top60_DNA+RNA_seed | KRAS | 0.892833 | UCEC | MFE-296 | KRAS (G12C) Inhibitor-1 |
| MVP_top60_DNA_seed | PIK3CA | 1.713723 | LUAD | NCI-H2342 | Buparlisib | MVP_top60_DNA+RNA_seed | JAK1 | 0.896522 | UCEC | EN | GSK2276186C |
| MVP_top60_DNA_seed | SMARCA4 | 1.714755 | LUAD | SK-LU-1 | PFI3 | MVP_top60_DNA+RNA_seed | MAPK1 | 0.899826 | UCEC | SNG-M | Ulixertinib |
| MVP_top60_DNA_seed | BRAF | 1.715931 | LUAD | SW1573 | PLX-4720 | MVP_top60_DNA+RNA_seed | MAPK3 | 0.899826 | UCEC | SNG-M | Ulixertinib |
| MVP_top60_DNA_seed | AKT1 | 1.716041 | LUAD | NCI-H1666 | AT13148 | MVP_top60_DNA+RNA_seed | EGFR | 0.902998 | UCEC | HEC-1 | Erlotinib |
| MVP_top60_DNA_seed | TERT | 1.717819 | LUAD | NCI-H1693 | BIBR-1532 | MVP_top60_DNA+RNA_seed | MAPK1 | 0.913964 | UCEC | MFE-319 | VX-11e |
| MVP_top60_DNA_seed | MAPK1 | 1.725318 | LUAD | NCI-H2342 | SCH772984 | MVP_top60_DNA+RNA_seed | MAPK1 | 0.915343 | UCEC | MFE-280 | VX-11e |
| MVP_top60_DNA_seed | MAPK1 | 1.725905 | LUAD | HCC-827 | Ulixertinib | MVP_top60_DNA+RNA_seed | PIK3CA | 0.916666 | UCEC | HEC-1 | Buparlisib |
| MVP_top60_DNA_seed | AKT1 | 1.726625 | LUAD | VMRC-LCD | Uprosertib | MVP_top60_DNA+RNA_seed | PIK3CB | 0.916666 | UCEC | HEC-1 | Buparlisib |
| MVP_top60_DNA_seed | TERT | 1.732268 | LUAD | Calu-3 | BIBR-1532 | MVP_top60_DNA+RNA_seed | PIK3CD | 0.916666 | UCEC | HEC-1 | Buparlisib |
| MVP_top60_DNA_seed | AKT1 | 1.735515 | LUAD | NCI-H1793 | Uprosertib | MVP_top60_DNA+RNA_seed | MAPK1 | 0.935447 | UCEC | AN3-CA | Ulixertinib |
| MVP_top60_DNA_seed | HDAC1 | 1.74073 | LUAD | SK-LU-1 | PCI-34051 | MVP_top60_DNA+RNA_seed | MAPK3 | 0.935447 | UCEC | AN3-CA | Ulixertinib |
| MVP_top60_DNA_seed | MDM2 | 1.742271 | LUAD | EKVX | Serdemetan | MVP_top60_DNA+RNA_seed | AKT1 | 0.937382 | UCEC | MFE-296 | MK-2206 |
| MVP_top60_DNA_seed | HDAC1 | 1.752217 | LUAD | EKVX | Entinostat | MVP_top60_DNA+RNA_seed | EGFR | 0.938034 | UCEC | EN | AZD3759 |
| MVP_top60_DNA_seed | ATM | 1.754883 | LUAD | EKVX | KU-55933 | MVP_top60_DNA+RNA_seed | PIK3CA | 0.94508 | UCEC | EN | AZD8186 |
| MVP_top60_DNA_seed | EZH2 | 1.758764 | LUAD | SW1573 | GSK343 | MVP_top60_DNA+RNA_seed | PIK3CB | 0.94508 | UCEC | EN | AZD8186 |
| MVP_top60_DNA_seed | FGFR1 | 1.75979 | LUAD | SK-LU-1 | AZD4547 | MVP_top60_DNA+RNA_seed | FGFR2 | 0.946602 | UCEC | EN | AZD4547 |
| MVP_top60_DNA_seed | FGFR1 | 1.760949 | LUAD | VMRC-LCD | PD173074 | MVP_top60_DNA+RNA_seed | PIK3CA | 0.953357 | UCEC | MFE-319 | Alpelisib |
| MVP_top60_DNA_seed | AKT1 | 1.761526 | LUAD | SK-LU-1 | GSK2110183B | MVP_top60_DNA+RNA_seed | HSP90AA1 | 0.954938 | UCEC | MFE-296 | Luminespib |
| MVP_top60_DNA_seed | TP53 | 1.763346 | LUAD | NCI-H1793 | MIRA-1 | MVP_top60_DNA+RNA_seed | HSP90AB1 | 0.954938 | UCEC | MFE-296 | Luminespib |
| MVP_top60_DNA_seed | EGFR | 1.764459 | LUAD | SK-LU-1 | Erlotinib | MVP_top60_DNA+RNA_seed | SRC | 0.974677 | UCEC | KLE | Bosutinib |
| MVP_top60_DNA_seed | FGFR1 | 1.767306 | LUAD | HOP-62 | PD173074 | MVP_top60_DNA+RNA_seed | AKT3 | 0.976671 | UCEC | KLE | Ipatasertib |
| MVP_top60_DNA_seed | GSK3B | 1.768041 | LUAD | NCI-H2291 | CHIR-99021 | MVP_top60_DNA+RNA_seed | AKT1 | 0.976671 | UCEC | KLE | Ipatasertib |
| MVP_top60_DNA_seed | HDAC1 | 1.768122 | LUAD | NCI-H1793 | PCI-34051 | MVP_top60_DNA+RNA_seed | EGFR | 1.004285 | UCEC | EN | Gefitinib |
| MVP_top60_DNA_seed | EZH2 | 1.768724 | LUAD | NCI-H1666 | GSK343 | MVP_top60_DNA+RNA_seed | EGFR | 1.017563 | UCEC | EN | Sapitinib |
| MVP_top60_DNA_seed | EGFR | 1.770075 | LUAD | NCI-H1734 | Afatinib | MVP_top60_DNA+RNA_seed | ERBB2 | 1.017563 | UCEC | EN | Sapitinib |
| MVP_top60_DNA_seed | ERBB2 | 1.770075 | LUAD | NCI-H1734 | Afatinib | MVP_top60_DNA+RNA_seed | EGFR | 1.021104 | UCEC | EN | Lapatinib |
| MVP_top60_DNA_seed | SRC | 1.771157 | LUAD | NCI-H2405 | Bosutinib | MVP_top60_DNA+RNA_seed | ERBB2 | 1.021104 | UCEC | EN | Lapatinib |
| MVP_top60_DNA_seed | EZH2 | 1.776635 | LUAD | NCI-H2228 | GSK343 | MVP_top60_DNA+RNA_seed | PIK3CA | 1.02174 | UCEC | HEC-1 | AZD8186 |
| MVP_top60_DNA_seed | SMARCA4 | 1.777921 | LUAD | SW1573 | PFI3 | MVP_top60_DNA+RNA_seed | PIK3CB | 1.02174 | UCEC | HEC-1 | AZD8186 |
| MVP_top60_DNA_seed | EGFR | 1.784245 | LUAD | NCI-H2347 | Erlotinib | MVP_top60_DNA+RNA_seed | AKT3 | 1.043738 | UCEC | HEC-1 | Afuresertib |
| MVP_top60_DNA_seed | MET | 1.785391 | LUAD | EKVX | Savolitinib | MVP_top60_DNA+RNA_seed | AKT1 | 1.043738 | UCEC | HEC-1 | Afuresertib |
| MVP_top60_DNA_seed | AKT1 | 1.785968 | LUAD | NCI-H2087 | MK-2206 | MVP_top60_DNA+RNA_seed | PIK3CA | 1.043925 | UCEC | SNG-M | AZD8186 |
| MVP_top60_DNA_seed | AKT1 | 1.786239 | LUAD | Calu-3 | AT13148 | MVP_top60_DNA+RNA_seed | PIK3CB | 1.043925 | UCEC | SNG-M | AZD8186 |
| MVP_top60_DNA_seed | IGF1R | 1.789488 | LUAD | H3255 | NVP-ADW742 | MVP_top60_DNA+RNA_seed | EGFR | 1.044043 | UCEC | COLO-684 | Gefitinib |
| MVP_top60_DNA_seed | AKT1 | 1.790003 | LUAD | NCI-H2347 | Ipatasertib | MVP_top60_DNA+RNA_seed | JAK1 | 1.059244 | UCEC | HEC-1 | Ruxolitinib |

| | | | | | | | | | | | |
|---|---|---|---|---|---|---|---|---|---|---|---|
| MVP_top60_DNA_seed | GSK3B | 1.792691 | LUAD | SK-LU-1 | SB216763 | MVP_top60_DNA+RNA_seed | AKT1 | 1.065309 | UCEC | HEC-1 | MK-2206 |
| MVP_top60_DNA_seed | ATM | 1.794129 | LUAD | NCI-H1838 | KU-55933 | MVP_top60_DNA+RNA_seed | TP53 | 1.069404 | UCEC | HEC-1 | MIRA-1 |
| MVP_top60_DNA_seed | HSP90AA1 | 1.79805 | LUAD | NCI-H2030 | Elesclomol | MVP_top60_DNA+RNA_seed | PIK3CA | 1.070358 | UCEC | MFE-296 | Alpelisib |
| MVP_top60_DNA_seed | AKT1 | 1.79815 | LUAD | NCI-H2405 | Uprosertib | MVP_top60_DNA+RNA_seed | HDAC1 | 1.08261 | UCEC | MFE-280 | Entinostat |
| MVP_top60_DNA_seed | SMARCA4 | 1.799288 | LUAD | PC-14 | PFI3 | MVP_top60_DNA+RNA_seed | MDM2 | 1.084862 | UCEC | KLE | Nutlin-3a (-) |
| MVP_top60_DNA_seed | MDM2 | 1.799551 | LUAD | PC-3_[JPC-3] | Serdemetan | MVP_top60_DNA+RNA_seed | MAPK1 | 1.089382 | UCEC | KLE | SCH772984 |
| MVP_top60_DNA_seed | BRAF | 1.809583 | LUAD | PC-14 | SB590885 | MVP_top60_DNA+RNA_seed | MAPK3 | 1.089382 | UCEC | KLE | SCH772984 |
| MVP_top60_DNA_seed | AKT1 | 1.810585 | LUAD | NCI-H522 | Afuresertib | MVP_top60_DNA+RNA_seed | PIK3CA | 1.09639 | UCEC | MFE-296 | AZD8186 |
| MVP_top60_DNA_seed | MET | 1.817294 | LUAD | NCI-H1623 | Crizotinib | MVP_top60_DNA+RNA_seed | PIK3CB | 1.09639 | UCEC | MFE-296 | AZD8186 |
| MVP_top60_DNA_seed | HDAC1 | 1.81901 | LUAD | NCI-H1435 | Entinostat | MVP_top60_DNA+RNA_seed | MAPK1 | 1.10766 | UCEC | MFE-319 | Ulixertinib |
| MVP_top60_DNA_seed | AKT1 | 1.819272 | LUAD | NCI-H1793 | AZD5363 | MVP_top60_DNA+RNA_seed | MAPK3 | 1.10766 | UCEC | MFE-319 | Ulixertinib |
| MVP_top60_DNA_seed | AKT1 | 1.821332 | LUAD | NCI-H2347 | Uprosertib | MVP_top60_DNA+RNA_seed | JAK1 | 1.109738 | UCEC | MFE-296 | JAK1_8709 |
| MVP_top60_DNA_seed | IGF1R | 1.821663 | LUAD | NCI-H1838 | GSK1904529A | MVP_top60_DNA+RNA_seed | MDM2 | 1.122706 | UCEC | EN | Serdemetan |
| MVP_top60_DNA_seed | EGFR | 1.821768 | LUAD | NCI-H2087 | AZD3759 | MVP_top60_DNA+RNA_seed | SRC | 1.124856 | UCEC | MFE-280 | Bosutinib |
| MVP_top60_DNA_seed | EGFR | 1.824114 | LUAD | NCI-H1793 | AZD3759 | MVP_top60_DNA+RNA_seed | EGFR | 1.128682 | UCEC | HEC-1 | Osimertinib |
| MVP_top60_DNA_seed | TP53 | 1.824734 | LUAD | HCC-827 | MIRA-1 | MVP_top60_DNA+RNA_seed | JAK1 | 1.130173 | UCEC | MFE-296 | JAK_8517 |
| MVP_top60_DNA_seed | AKT1 | 1.824909 | LUAD | NCI-H1793 | GSK2110183B | MVP_top60_DNA+RNA_seed | EGFR | 1.136334 | UCEC | HEC-1 | Gefitinib |
| MVP_top60_DNA_seed | KRAS | 1.826049 | LUAD | NCI-H2405 | KRAS (G12C) Inhibitor-12 | MVP_top60_DNA+RNA_seed | MAPK1 | 1.155624 | UCEC | COLO-684 | SCH772984 |
| MVP_top60_DNA_seed | HDAC1 | 1.83269 | LUAD | NCI-H2228 | PCI-34051 | MVP_top60_DNA+RNA_seed | MAPK3 | 1.155624 | UCEC | COLO-684 | SCH772984 |
| MVP_top60_DNA_seed | MDM2 | 1.83863 | LUAD | NCI-H441 | Serdemetan | MVP_top60_DNA+RNA_seed | MAPK1 | 1.160351 | UCEC | MFE-296 | Ulixertinib |
| MVP_top60_DNA_seed | EGFR | 1.840042 | LUAD | NCI-H1355 | Lapatinib | MVP_top60_DNA+RNA_seed | MAPK3 | 1.160351 | UCEC | MFE-296 | Ulixertinib |
| MVP_top60_DNA_seed | ERBB2 | 1.840042 | LUAD | NCI-H1355 | Lapatinib | MVP_top60_DNA+RNA_seed | MDM2 | 1.180671 | UCEC | MFE-280 | Nutlin-3a (-) |
| MVP_top60_DNA_seed | AKT1 | 1.842697 | LUAD | NCI-H441 | GSK2110183B | MVP_top60_DNA+RNA_seed | AKT3 | 1.182879 | UCEC | KLE | Uprosertib |
| MVP_top60_DNA_seed | EZH2 | 1.8429 | LUAD | NCI-H2347 | GSK343 | MVP_top60_DNA+RNA_seed | AKT1 | 1.182879 | UCEC | KLE | Uprosertib |
| MVP_top60_DNA_seed | BRAF | 1.843136 | LUAD | NCI-H1650 | PLX-4720 | MVP_top60_DNA+RNA_seed | HDAC1 | 1.183379 | UCEC | SNG-M | Entinostat |
| MVP_top60_DNA_seed | AKT1 | 1.848252 | LUAD | PC-3_[JPC-3] | Uprosertib | MVP_top60_DNA+RNA_seed | PIK3CA | 1.200719 | UCEC | HEC-1 | Alpelisib |
| MVP_top60_DNA_seed | PIK3CA | 1.852534 | LUAD | HOP-62 | Alpelisib | MVP_top60_DNA+RNA_seed | AKT1 | 1.206756 | UCEC | MFE-319 | MK-2206 |
| MVP_top60_DNA_seed | AKT1 | 1.852705 | LUAD | NCI-H2087 | AT13148 | MVP_top60_DNA+RNA_seed | EGFR | 1.228972 | UCEC | EN | Erlotinib |
| MVP_top60_DNA_seed | MET | 1.853154 | LUAD | NCI-H2291 | Savolitinib | MVP_top60_DNA+RNA_seed | AKT3 | 1.229678 | UCEC | HEC-1 | Uprosertib |
| MVP_top60_DNA_seed | MAPK1 | 1.853288 | LUAD | PC-3_[JPC-3] | Ulixertinib | MVP_top60_DNA+RNA_seed | AKT1 | 1.229678 | UCEC | HEC-1 | Uprosertib |
| MVP_top60_DNA_seed | EGFR | 1.854342 | LUAD | SW1573 | AZD3759 | MVP_top60_DNA+RNA_seed | FGFR2 | 1.238673 | UCEC | EN | PD173074 |
| MVP_top60_DNA_seed | EZH2 | 1.859325 | LUAD | NCI-H1650 | GSK343 | MVP_top60_DNA+RNA_seed | MAPK1 | 1.239816 | UCEC | KLE | Ulixertinib |
| MVP_top60_DNA_seed | MET | 1.864962 | LUAD | SK-LU-1 | Savolitinib | MVP_top60_DNA+RNA_seed | MAPK3 | 1.239816 | UCEC | KLE | Ulixertinib |
| MVP_top60_DNA_seed | AURKA | 1.865375 | LUAD | EKVX | Tozasertib | MVP_top60_DNA+RNA_seed | MDM2 | 1.241378 | UCEC | AN3-CA | Serdemetan |
| MVP_top60_DNA_seed | AKT1 | 1.866666 | LUAD | NCI-H650 | AZD5363 | MVP_top60_DNA+RNA_seed | SRC | 1.241598 | UCEC | EN | Bosutinib |
| MVP_top60_DNA_seed | AKT1 | 1.872643 | LUAD | VMRC-LCD | AZD5363 | MVP_top60_DNA+RNA_seed | FGFR2 | 1.252341 | UCEC | KLE | AZD4547 |
| MVP_top60_DNA_seed | AURKA | 1.873375 | LUAD | NCI-H1650 | ZM447439 | MVP_top60_DNA+RNA_seed | AKT3 | 1.264935 | UCEC | HEC-1 | Ipatasertib |
| MVP_top60_DNA_seed | AURKA | 1.874314 | LUAD | SW1573 | ZM447439 | MVP_top60_DNA+RNA_seed | AKT1 | 1.264935 | UCEC | HEC-1 | Ipatasertib |
| MVP_top60_DNA_seed | MAPK1 | 1.877676 | LUAD | NCI-H1838 | SCH772984 | MVP_top60_DNA+RNA_seed | EGFR | 1.265159 | UCEC | HEC-1 | Afatinib |
| MVP_top60_DNA_seed | EGFR | 1.8816 | LUAD | ABC-1 | Erlotinib | MVP_top60_DNA+RNA_seed | ERBB2 | 1.265159 | UCEC | HEC-1 | Afatinib |
| MVP_top60_DNA_seed | EZH2 | 1.881841 | LUAD | NCI-H2405 | GSK343 | MVP_top60_DNA+RNA_seed | MAPK1 | 1.276293 | UCEC | HEC-1 | Ulixertinib |

| | | | | | | | | | | | |
|---|---|---|---|---|---|---|---|---|---|---|---|
| MVP_top60_DNA_seed | EGFR | 1.885326 | LUAD | Calu-6 | AZD3759 | MVP_top60_DNA+RNA_seed | MAPK3 | 1.276293 | UCEC | HEC-1 | Ulixertinib |
| MVP_top60_DNA_seed | AKT1 | 1.891077 | LUAD | Calu-6 | Afuresertib | MVP_top60_DNA+RNA_seed | EGFR | 1.286195 | UCEC | COLO-684 | Sapitinib |
| MVP_top60_DNA_seed | AKT1 | 1.892494 | LUAD | NCI-H441 | Afuresertib | MVP_top60_DNA+RNA_seed | ERBB2 | 1.286195 | UCEC | COLO-684 | Sapitinib |
| MVP_top60_DNA_seed | FGFR1 | 1.894603 | LUAD | H3255 | AZD4547 | MVP_top60_DNA+RNA_seed | MAPK1 | 1.287893 | UCEC | KLE | Ulixertinib |
| MVP_top60_DNA_seed | PIK3CA | 1.895236 | LUAD | NCI-H441 | AZD8186 | MVP_top60_DNA+RNA_seed | MAPK3 | 1.287893 | UCEC | KLE | Ulixertinib |
| MVP_top60_DNA_seed | TERT | 1.901414 | LUAD | NCI-H1793 | BIBR-1532 | MVP_top60_DNA+RNA_seed | EGFR | 1.290504 | UCEC | MFE-280 | Gefitinib |
| MVP_top60_DNA_seed | TP53 | 1.91054 | LUAD | NCI-H2347 | MIRA-1 | MVP_top60_DNA+RNA_seed | MAPK1 | 1.316997 | UCEC | COLO-684 | Ulixertinib |
| MVP_top60_DNA_seed | IGF1R | 1.911289 | LUAD | SW1573 | NVP-ADW742 | MVP_top60_DNA+RNA_seed | MAPK3 | 1.316997 | UCEC | COLO-684 | Ulixertinib |
| MVP_top60_DNA_seed | EZH2 | 1.913127 | LUAD | NCI-H441 | GSK343 | MVP_top60_DNA+RNA_seed | HSP90AA1 | 1.317908 | UCEC | KLE | Luminespib |
| MVP_top60_DNA_seed | AKT1 | 1.917594 | LUAD | HOP-62 | MK-2206 | MVP_top60_DNA+RNA_seed | HSP90AB1 | 1.317908 | UCEC | KLE | Luminespib |
| MVP_top60_DNA_seed | AKT1 | 1.924887 | LUAD | NCI-H2228 | AT13148 | MVP_top60_DNA+RNA_seed | AKT3 | 1.368983 | UCEC | KLE | Afuresertib |
| MVP_top60_DNA_seed | IGF1R | 1.926888 | LUAD | NCI-H2347 | GSK1904529A | MVP_top60_DNA+RNA_seed | AKT1 | 1.368983 | UCEC | KLE | Afuresertib |
| MVP_top60_DNA_seed | EGFR | 1.927509 | LUAD | SW1573 | Gefitinib | MVP_top60_DNA+RNA_seed | HSP90AA1 | 1.369409 | UCEC | EN | Luminespib |
| MVP_top60_DNA_seed | BRAF | 1.928852 | LUAD | EKVX | SB590885 | MVP_top60_DNA+RNA_seed | HSP90AB1 | 1.369409 | UCEC | EN | Luminespib |
| MVP_top60_DNA_seed | KRAS | 1.93085 | LUAD | NCI-H1568 | KRAS (G12C) Inhibitor-1 | MVP_top60_DNA+RNA_seed | EGFR | 1.378763 | UCEC | MFE-296 | Afatinib |
| MVP_top60_DNA_seed | TP53 | 1.932986 | LUAD | NCI-H441 | MIRA-1 | MVP_top60_DNA+RNA_seed | ERBB2 | 1.378763 | UCEC | MFE-296 | Afatinib |
| MVP_top60_DNA_seed | GSK3B | 1.935306 | LUAD | NCI-H2347 | SB216763 | MVP_top60_DNA+RNA_seed | MDM2 | 1.381257 | UCEC | MFE-280 | Serdemetan |
| MVP_top60_DNA_seed | PIK3CA | 1.94076 | LUAD | VMRC-LCD | AZD8186 | MVP_top60_DNA+RNA_seed | JAK1 | 1.386109 | UCEC | HEC-1 | JAK1_8709 |
| MVP_top60_DNA_seed | MAPK1 | 1.941893 | LUAD | SW1573 | SCH772984 | MVP_top60_DNA+RNA_seed | HDAC1 | 1.394134 | UCEC | COLO-684 | Entinostat |
| MVP_top60_DNA_seed | EGFR | 1.942151 | LUAD | SK-LU-1 | Lapatinib | MVP_top60_DNA+RNA_seed | HSP90AA1 | 1.40746 | UCEC | KLE | Tanespimycin |
| MVP_top60_DNA_seed | ERBB2 | 1.942151 | LUAD | SK-LU-1 | Lapatinib | MVP_top60_DNA+RNA_seed | HSP90AB1 | 1.40746 | UCEC | KLE | Tanespimycin |
| MVP_top60_DNA_seed | CREBBP | 1.943634 | LUAD | NCI-H441 | SGC-CBP30 | MVP_top60_DNA+RNA_seed | MAPK1 | 1.440149 | UCEC | MFE-280 | Ulixertinib |
| MVP_top60_DNA_seed | BRAF | 1.946723 | LUAD | NCI-H2291 | PLX-4720 | MVP_top60_DNA+RNA_seed | MAPK3 | 1.440149 | UCEC | MFE-280 | Ulixertinib |
| MVP_top60_DNA_seed | EZH2 | 1.951714 | LUAD | NCI-H441 | GSK343 | MVP_top60_DNA+RNA_seed | FGFR2 | 1.456139 | UCEC | MFE-319 | PD173074 |
| MVP_top60_DNA_seed | CREBBP | 1.959093 | LUAD | SW1573 | SGC-CBP30 | MVP_top60_DNA+RNA_seed | ATM | 1.460351 | UCEC | KLE | KU-55933 |
| MVP_top60_DNA_seed | AKT1 | 1.961121 | LUAD | NCI-H1568 | AT13148 | MVP_top60_DNA+RNA_seed | AKT1 | 1.480473 | UCEC | KLE | AT13148 |
| MVP_top60_DNA_seed | MAPK1 | 1.961487 | LUAD | NCI-H1650 | Ulixertinib | MVP_top60_DNA+RNA_seed | MAPK1 | 1.485205 | UCEC | HEC-1 | SCH772984 |
| MVP_top60_DNA_seed | AKT1 | 1.96387 | LUAD | NCI-H441 | MK-2206 | MVP_top60_DNA+RNA_seed | MAPK3 | 1.485205 | UCEC | HEC-1 | SCH772984 |
| MVP_top60_DNA_seed | PIK3CA | 1.966736 | LUAD | SK-LU-1 | Buparlisib | MVP_top60_DNA+RNA_seed | HDAC1 | 1.500964 | UCEC | KLE | Romidepsin |
| MVP_top60_DNA_seed | MAPK1 | 1.967123 | LUAD | NCI-H441 | Ulixertinib | MVP_top60_DNA+RNA_seed | HDAC2 | 1.500964 | UCEC | KLE | Romidepsin |
| MVP_top60_DNA_seed | AURKA | 1.978455 | LUAD | NCI-H1355 | ZM447439 | MVP_top60_DNA+RNA_seed | EGFR | 1.524505 | UCEC | MFE-280 | Erlotinib |
| MVP_top60_DNA_seed | MET | 1.980896 | LUAD | NCI-H441 | Savolitinib | MVP_top60_DNA+RNA_seed | MAPK1 | 1.545871 | UCEC | MFE-296 | VX-11e |
| MVP_top60_DNA_seed | EGFR | 1.98256 | LUAD | SK-LU-1 | Afatinib | MVP_top60_DNA+RNA_seed | MAPK1 | 1.555213 | UCEC | EN | VX-11e |
| MVP_top60_DNA_seed | ERBB2 | 1.98256 | LUAD | SK-LU-1 | Afatinib | MVP_top60_DNA+RNA_seed | AKT3 | 1.572434 | UCEC | KLE | GSK2110183B |
| MVP_top60_DNA_seed | FGFR1 | 1.986536 | LUAD | NCI-H2347 | AZD4547 | MVP_top60_DNA+RNA_seed | AKT1 | 1.572434 | UCEC | KLE | GSK2110183B |
| MVP_top60_DNA_seed | AKT1 | 1.989425 | LUAD | Calu-3 | Ipatasertib | MVP_top60_DNA+RNA_seed | TP53 | 1.5776 | UCEC | KLE | MIRA-1 |
| MVP_top60_DNA_seed | TERT | 1.998051 | LUAD | NCI-H1666 | BIBR-1532 | MVP_top60_DNA+RNA_seed | JAK1 | 1.620162 | UCEC | HEC-1 | JAK_8517 |
| MVP_top60_DNA_seed | MDM2 | 1.999294 | LUAD | COR-L105 | Serdemetan | MVP_top60_DNA+RNA_seed | EGFR | 1.622469 | UCEC | COLO-684 | Osimertinib |
| MVP_top60_DNA_seed | IGF1R | 2.006281 | LUAD | EKVX | NVP-ADW742 | MVP_top60_DNA+RNA_seed | JAK1 | 1.641106 | UCEC | KLE | Ruxolitinib |
| MVP_top60_DNA_seed | HDAC1 | 2.009593 | LUAD | NCI-H441 | PCI-34051 | MVP_top60_DNA+RNA_seed | PIK3CA | 1.64178 | UCEC | KLE | Alpelisib |
| MVP_top60_DNA_seed | AKT1 | 2.01253 | LUAD | NCI-H2405 | GSK2110183B | MVP_top60_DNA+RNA_seed | PIK3CA | 1.651226 | UCEC | MFE-296 | GNE-317 |

| | | | | | | | | | | | |
|---|---|---|---|---|---|---|---|---|---|---|---|
| MVP_top60_DNA_seed | TERT | 2.022634 | LUAD | NCI-H1838 | BIBR-1532 | MVP_top60_DNA+RNA_seed | JAK1 | 1.666591 | UCEC | KLE | GSK2276186C |
| MVP_top60_DNA_seed | EZH2 | 2.022846 | LUAD | SW1573 | GSK343 | MVP_top60_DNA+RNA_seed | HDAC1 | 1.698881 | UCEC | KLE | Entinostat |
| MVP_top60_DNA_seed | MDM2 | 2.026136 | LUAD | NCI-H2405 | Nutlin-3a (-) | MVP_top60_DNA+RNA_seed | MDM2 | 1.726795 | UCEC | KLE | Serdemetan |
| MVP_top60_DNA_seed | IGF1R | 2.028363 | LUAD | NCI-H2405 | Linsitinib | MVP_top60_DNA+RNA_seed | EGFR | 1.72751 | UCEC | MFE-296 | Lapatinib |
| MVP_top60_DNA_seed | FGFR1 | 2.040052 | LUAD | NCI-H2405 | PD173074 | MVP_top60_DNA+RNA_seed | ERBB2 | 1.72751 | UCEC | MFE-296 | Lapatinib |
| MVP_top60_DNA_seed | KRAS | 2.042551 | LUAD | NCI-H2228 | KRAS (G12C) Inhibitor-1 | MVP_top60_DNA+RNA_seed | AKT3 | 1.729271 | UCEC | HEC-1 | Uprosertib |
| MVP_top60_DNA_seed | IGF1R | 2.043567 | LUAD | NCI-H1666 | GSK1904529A | MVP_top60_DNA+RNA_seed | AKT1 | 1.729271 | UCEC | HEC-1 | Uprosertib |
| MVP_top60_DNA_seed | CREBBP | 2.049884 | LUAD | NCI-H2228 | SGC-CBP30 | MVP_top60_DNA+RNA_seed | JAK1 | 1.782162 | UCEC | COLO-684 | JAK_8517 |
| MVP_top60_DNA_seed | TERT | 2.052173 | LUAD | SK-LU-1 | BIBR-1532 | MVP_top60_DNA+RNA_seed | FGFR2 | 1.811944 | UCEC | COLO-684 | PD173074 |
| MVP_top60_DNA_seed | EZH2 | 2.055256 | LUAD | SK-LU-1 | GSK343 | MVP_top60_DNA+RNA_seed | MAPK1 | 1.858141 | UCEC | MFE-280 | Ulixertinib |
| MVP_top60_DNA_seed | PIK3CA | 2.065836 | LUAD | NCI-H1693 | Alpelisib | MVP_top60_DNA+RNA_seed | MAPK3 | 1.858141 | UCEC | MFE-280 | Ulixertinib |
| MVP_top60_DNA_seed | EGFR | 2.071103 | LUAD | NCI-H441 | AZD3759 | MVP_top60_DNA+RNA_seed | HSP90AA1 | 1.863317 | UCEC | HEC-1 | Tanespimycin |
| MVP_top60_DNA_seed | AKT1 | 2.072122 | LUAD | NCI-H441 | Ipatasertib | MVP_top60_DNA+RNA_seed | HSP90AB1 | 1.863317 | UCEC | HEC-1 | Tanespimycin |
| MVP_top60_DNA_seed | FGFR1 | 2.078265 | LUAD | NCI-H1355 | PD173074 | MVP_top60_DNA+RNA_seed | FGFR2 | 1.915425 | UCEC | HEC-1 | PD173074 |
| MVP_top60_DNA_seed | HSP90AA1 | 2.078636 | LUAD | NCI-H1793 | Tanespimycin | MVP_top60_DNA+RNA_seed | MDM2 | 1.915843 | UCEC | HEC-1 | Serdemetan |
| MVP_top60_DNA_seed | MAPK1 | 2.081009 | LUAD | NCI-H1793 | Ulixertinib | MVP_top60_DNA+RNA_seed | EGFR | 1.971861 | UCEC | HEC-1 | Lapatinib |
| MVP_top60_DNA_seed | EGFR | 2.081775 | LUAD | NCI-H441 | Lapatinib | MVP_top60_DNA+RNA_seed | ERBB2 | 1.971861 | UCEC | HEC-1 | Lapatinib |
| MVP_top60_DNA_seed | ERBB2 | 2.081775 | LUAD | NCI-H441 | Lapatinib | MVP_top60_DNA+RNA_seed | HSP90AA1 | 1.980393 | UCEC | HEC-1 | Luminespib |
| MVP_top60_DNA_seed | MDM2 | 2.084173 | LUAD | NCI-H1838 | Serdemetan | MVP_top60_DNA+RNA_seed | HSP90AB1 | 1.980393 | UCEC | HEC-1 | Luminespib |
| MVP_top60_DNA_seed | AKT1 | 2.084896 | LUAD | EKVX | Uprosertib | MVP_top60_DNA+RNA_seed | JAK1 | 2.248089 | UCEC | KLE | JAK1_8709 |
| MVP_top60_DNA_seed | AKT1 | 2.086222 | LUAD | NCI-H1573 | Afuresertib | MVP_top60_DNA+RNA_seed | JAK1 | 2.426792 | UCEC | KLE | JAK_8517 |
| MVP_top60_DNA_seed | HSP90AA1 | 2.093513 | LUAD | NCI-H1693 | Tanespimycin | top60_altered_DNA+RNA_seed | PIK3CA | -2.363975 | BLCA | CAL-29 | AZD8835 |
| MVP_top60_DNA_seed | MAPK1 | 2.098189 | LUAD | NCI-H1838 | Ulixertinib | top60_altered_DNA+RNA_seed | PIK3CA | -2.256468 | BLCA | CAL-29 | PI3Ka_4409 |
| MVP_top60_DNA_seed | KRAS | 2.098972 | LUAD | SW1573 | KRAS (G12C) Inhibitor-1 | top60_altered_DNA+RNA_seed | PIK3CA | -1.581022 | BLCA | BFTC-905 | AZD8835 |
| MVP_top60_DNA_seed | PIK3CA | 2.100023 | LUAD | NCI-H441 | Alpelisib | top60_altered_DNA+RNA_seed | PIK3CA | -1.503629 | BLCA | BFTC-905 | PI3Ka_4409 |
| MVP_top60_DNA_seed | MDM2 | 2.100282 | LUAD | NCI-H2087 | Serdemetan | top60_altered_DNA+RNA_seed | PIK3CA | -1.067843 | BLCA | HT-1197 | AZD8835 |
| MVP_top60_DNA_seed | AURKA | 2.107135 | LUAD | NCI-H1666 | ZM447439 | top60_altered_DNA+RNA_seed | PIK3CA | -0.997563 | BLCA | RT-112 | PI3Ka_4409 |
| MVP_top60_DNA_seed | SMARCA4 | 2.112213 | LUAD | EKVX | PFI3 | top60_altered_DNA+RNA_seed | PIK3CA | -0.625709 | BLCA | RT-112 | AZD8835 |
| MVP_top60_DNA_seed | HSP90AA1 | 2.117502 | LUAD | VMRC-LCD | Luminespib | top60_altered_DNA+RNA_seed | PIK3CA | -0.57933 | BLCA | TCCSUP | AZD8835 |
| MVP_top60_DNA_seed | TERT | 2.124139 | LUAD | NCI-H2291 | BIBR-1532 | top60_altered_DNA+RNA_seed | PIK3CA | -0.521185 | BLCA | SW1710 | AZD8835 |
| MVP_top60_DNA_seed | PIK3CA | 2.130441 | LUAD | H3255 | Buparlisib | top60_altered_DNA+RNA_seed | PIK3CA | -0.475212 | BLCA | 5637 | PI3Ka_4409 |
| MVP_top60_DNA_seed | AKT1 | 2.136466 | LUAD | NCI-H1650 | AT13148 | top60_altered_DNA+RNA_seed | PIK3CA | -0.367211 | BLCA | 639-V | AZD8835 |
| MVP_top60_DNA_seed | EGFR | 2.140246 | LUAD | SW1573 | Erlotinib | top60_altered_DNA+RNA_seed | PIK3CA | -0.302125 | BLCA | SW780 | PI3Ka_4409 |
| MVP_top60_DNA_seed | PIK3CA | 2.141275 | LUAD | EKVX | Buparlisib | top60_altered_DNA+RNA_seed | PIK3CA | -0.295805 | BLCA | LB831-BLC | AZD8835 |
| MVP_top60_DNA_seed | AKT1 | 2.147146 | LUAD | NCI-H2291 | AT13148 | top60_altered_DNA+RNA_seed | PIK3CA | -0.227711 | BLCA | HT-1197 | PI3Ka_4409 |
| MVP_top60_DNA_seed | SMARCA4 | 2.151269 | LUAD | H3255 | PFI3 | top60_altered_DNA+RNA_seed | PIK3CA | -0.226616 | BLCA | 647-V | PI3Ka_4409 |
| MVP_top60_DNA_seed | HDAC1 | 2.15786 | LUAD | NCI-H1693 | PCI-34051 | top60_altered_DNA+RNA_seed | PIK3CA | 0.049539 | BLCA | SW1710 | PI3Ka_4409 |
| MVP_top60_DNA_seed | PIK3CA | 2.158579 | LUAD | SW1573 | GNE-317 | top60_altered_DNA+RNA_seed | PIK3CA | 0.050942 | BLCA | J82 | PI3Ka_4409 |
| MVP_top60_DNA_seed | BRAF | 2.159053 | LUAD | NCI-H441 | PLX-4720 | top60_altered_DNA+RNA_seed | PIK3CA | 0.120483 | BLCA | DSH1 | AZD8835 |
| MVP_top60_DNA_seed | AKT1 | 2.162631 | LUAD | NCI-H2405 | Uprosertib | top60_altered_DNA+RNA_seed | PIK3CA | 0.123523 | BLCA | 5637 | AZD8835 |
| MVP_top60_DNA_seed | AKT1 | 2.162741 | LUAD | SK-LU-1 | AT13148 | top60_altered_DNA+RNA_seed | PIK3CA | 0.212412 | BLCA | DSH1 | PI3Ka_4409 |

| | | | | | | | | | | | |
|---|---|---|---|---|---|---|---|---|---|---|---|
| MVP_top60_DNA_seed | AKT1 | 2.17089 | LUAD | SK-LU-1 | Ipatasertib | top60_altered_DNA+RNA_seed | PIK3CA | 0.217414 | BLCA | T24 | AZD8835 |
| MVP_top60_DNA_seed | AKT1 | 2.174733 | LUAD | SW1573 | AT13148 | top60_altered_DNA+RNA_seed | PIK3CA | 0.230402 | BLCA | 639-V | PI3Ka_4409 |
| MVP_top60_DNA_seed | MDM2 | 2.179423 | LUAD | NCI-H2405 | Serdemetan | top60_altered_DNA+RNA_seed | PIK3CA | 0.306539 | BLCA | UM-UC-3 | AZD8835 |
| MVP_top60_DNA_seed | BRAF | 2.182664 | LUAD | NCI-H441 | SB590885 | top60_altered_DNA+RNA_seed | PIK3CA | 0.378449 | BLCA | 647-V | AZD8835 |
| MVP_top60_DNA_seed | SMARCA4 | 2.183084 | LUAD | NCI-H2291 | PFI3 | top60_altered_DNA+RNA_seed | PIK3CA | 0.379359 | BLCA | LB831-BLC | PI3Ka_4409 |
| MVP_top60_DNA_seed | AURKA | 2.183404 | LUAD | NCI-H1793 | ZM447439 | top60_altered_DNA+RNA_seed | PIK3CA | 0.485958 | BLCA | J82 | AZD8835 |
| MVP_top60_DNA_seed | AKT1 | 2.191459 | LUAD | SK-LU-1 | Uprosertib | top60_altered_DNA+RNA_seed | PIK3CA | 0.670531 | BLCA | HT-1376 | AZD8835 |
| MVP_top60_DNA_seed | IGF1R | 2.19161 | LUAD | COR-L105 | GSK1904529A | top60_altered_DNA+RNA_seed | PIK3CA | 0.758114 | BLCA | KU-19-19 | AZD8835 |
| MVP_top60_DNA_seed | SMARCA4 | 2.192307 | LUAD | NCI-H1693 | PFI3 | top60_altered_DNA+RNA_seed | PIK3CA | 0.811732 | BLCA | UM-UC-3 | PI3Ka_4409 |
| MVP_top60_DNA_seed | HSP90AA1 | 2.19465 | LUAD | NCI-H1573 | Luminespib | top60_altered_DNA+RNA_seed | PIK3CA | 0.831832 | BLCA | HT-1376 | PI3Ka_4409 |
| MVP_top60_DNA_seed | BRAF | 2.195264 | LUAD | NCI-H1568 | PLX-4720 | top60_altered_DNA+RNA_seed | PIK3CA | 0.840047 | BLCA | RT4 | AZD8835 |
| MVP_top60_DNA_seed | EZH2 | 2.199728 | LUAD | H3255 | GSK343 | top60_altered_DNA+RNA_seed | PIK3CA | 0.936587 | BLCA | SW780 | AZD8835 |
| MVP_top60_DNA_seed | KRAS | 2.199916 | LUAD | SK-LU-1 | KRAS (G12C) Inhibitor-1 | top60_altered_DNA+RNA_seed | PIK3CA | 0.983085 | BLCA | T24 | PI3Ka_4409 |
| MVP_top60_DNA_seed | BRAF | 2.201851 | LUAD | NCI-H2291 | SB590885 | top60_altered_DNA+RNA_seed | PIK3CA | 1.110391 | BLCA | KU-19-19 | PI3Ka_4409 |
| MVP_top60_DNA_seed | AKT1 | 2.202921 | LUAD | NCI-H2347 | Afuresertib | top60_altered_DNA+RNA_seed | PIK3CA | 1.846686 | BLCA | RT4 | PI3Ka_4409 |
| MVP_top60_DNA_seed | EGFR | 2.204569 | LUAD | NCI-H2347 | Gefitinib | top60_altered_DNA+RNA_seed | PIK3CA | -1.984079 | BLCA | CAL-29 | Alpelisib |
| MVP_top60_DNA_seed | AURKA | 2.214435 | LUAD | Calu-3 | ZM447439 | top60_altered_DNA+RNA_seed | TP53 | -1.730724 | BLCA | BFTC-905 | MIRA-1 |
| MVP_top60_DNA_seed | AKT1 | 2.231771 | LUAD | NCI-H2347 | AT13148 | top60_altered_DNA+RNA_seed | PIK3CA | -1.147597 | BLCA | CAL-29 | GNE-317 |
| MVP_top60_DNA_seed | FGFR1 | 2.241299 | LUAD | NCI-H1838 | PD173074 | top60_altered_DNA+RNA_seed | PIK3CA | -1.094133 | BLCA | CAL-29 | Buparlisib |
| MVP_top60_DNA_seed | HSP90AA1 | 2.241358 | LUAD | VMRC-LCD | Tanespimycin | top60_altered_DNA+RNA_seed | TERT | -1.023961 | BLCA | 639-V | BIBR-1532 |
| MVP_top60_DNA_seed | EZH2 | 2.244574 | LUAD | NCI-H1693 | GSK343 | top60_altered_DNA+RNA_seed | TERT | -1.011537 | BLCA | DSH1 | BIBR-1532 |
| MVP_top60_DNA_seed | HDAC1 | 2.250008 | LUAD | NCI-H1693 | Romidepsin | top60_altered_DNA+RNA_seed | TERT | -0.902284 | BLCA | HT-1376 | BIBR-1532 |
| MVP_top60_DNA_seed | GSK3B | 2.281729 | LUAD | NCI-H2405 | SB216763 | top60_altered_DNA+RNA_seed | PIK3CA | -0.884619 | BLCA | CAL-29 | AZD8186 |
| MVP_top60_DNA_seed | EGFR | 2.283491 | LUAD | SK-LU-1 | Osimertinib | top60_altered_DNA+RNA_seed | PIK3CA | -0.837054 | BLCA | BFTC-905 | Buparlisib |
| MVP_top60_DNA_seed | SRC | 2.288796 | LUAD | NCI-H441 | Bosutinib | top60_altered_DNA+RNA_seed | PIK3CA | -0.799839 | BLCA | BFTC-905 | Alpelisib |
| MVP_top60_DNA_seed | EZH2 | 2.292879 | LUAD | EKVX | GSK343 | top60_altered_DNA+RNA_seed | PIK3CA | -0.777686 | BLCA | 5637 | Buparlisib |
| MVP_top60_DNA_seed | HSP90AA1 | 2.305945 | LUAD | ABC-1 | Tanespimycin | top60_altered_DNA+RNA_seed | TERT | -0.77393 | BLCA | HT-1197 | BIBR-1532 |
| MVP_top60_DNA_seed | PIK3CA | 2.306774 | LUAD | NCI-H1435 | Buparlisib | top60_altered_DNA+RNA_seed | PIK3CA | -0.69325 | BLCA | HT-1197 | Buparlisib |
| MVP_top60_DNA_seed | MAPK1 | 2.309493 | LUAD | NCI-H1650 | VX-11e | top60_altered_DNA+RNA_seed | PIK3CA | -0.670616 | BLCA | RT-112 | Alpelisib |
| MVP_top60_DNA_seed | PIK3CA | 2.318618 | LUAD | Calu-3 | AZD8186 | top60_altered_DNA+RNA_seed | PIK3CA | -0.562267 | BLCA | DSH1 | GNE-317 |
| MVP_top60_DNA_seed | HSP90AA1 | 2.318943 | LUAD | COR-L105 | Tanespimycin | top60_altered_DNA+RNA_seed | PIK3CA | -0.533608 | BLCA | HT-1197 | GNE-317 |
| MVP_top60_DNA_seed | EGFR | 2.319069 | LUAD | SW1573 | Osimertinib | top60_altered_DNA+RNA_seed | TERT | -0.520061 | BLCA | 647-V | BIBR-1532 |
| MVP_top60_DNA_seed | BRAF | 2.324521 | LUAD | NCI-H1693 | PLX-4720 | top60_altered_DNA+RNA_seed | PIK3CA | -0.481732 | BLCA | KU-19-19 | Buparlisib |
| MVP_top60_DNA_seed | AKT1 | 2.330594 | LUAD | VMRC-LCD | Afuresertib | top60_altered_DNA+RNA_seed | PIK3CA | -0.476776 | BLCA | KU-19-19 | GNE-317 |
| MVP_top60_DNA_seed | HDAC1 | 2.335598 | LUAD | NCI-H2405 | PCI-34051 | top60_altered_DNA+RNA_seed | TERT | -0.410889 | BLCA | SW780 | BIBR-1532 |
| MVP_top60_DNA_seed | PIK3CA | 2.355811 | LUAD | Calu-6 | AZD8186 | top60_altered_DNA+RNA_seed | PIK3CA | -0.335329 | BLCA | 647-V | AZD8186 |
| MVP_top60_DNA_seed | CREBBP | 2.370258 | LUAD | NCI-H1693 | SGC-CBP30 | top60_altered_DNA+RNA_seed | TERT | -0.327465 | BLCA | RT-112 | BIBR-1532 |
| MVP_top60_DNA_seed | AKT1 | 2.398702 | LUAD | NCI-H2087 | AZD5363 | top60_altered_DNA+RNA_seed | PIK3CA | -0.311178 | BLCA | 639-V | AZD8186 |
| MVP_top60_DNA_seed | HSP90AA1 | 2.414374 | LUAD | NCI-H1793 | Luminespib | top60_altered_DNA+RNA_seed | PIK3CA | -0.297558 | BLCA | 639-V | Buparlisib |
| MVP_top60_DNA_seed | PIK3CA | 2.419897 | LUAD | NCI-H441 | Buparlisib | top60_altered_DNA+RNA_seed | PIK3CA | -0.28342 | BLCA | RT-112 | GNE-317 |
| MVP_top60_DNA_seed | MDM2 | 2.421773 | LUAD | NCI-H1650 | Serdemetan | top60_altered_DNA+RNA_seed | PIK3CA | -0.273018 | BLCA | BFTC-905 | GNE-317 |

| | | | | | | | | | | | |
|---|---|---|---|---|---|---|---|---|---|---|---|
| MVP_top60_DNA_seed | AKT1 | 2.442261 | LUAD | NCI-H1793 | Afuresertib | top60_altered_DNA+RNA_seed | PIK3CA | -0.270785 | BLCA | TCCSUP | GNE-317 |
| MVP_top60_DNA_seed | AKT1 | 2.443442 | LUAD | Calu-6 | AZD5363 | top60_altered_DNA+RNA_seed | PIK3CA | -0.22432 | BLCA | UM-UC-3 | GNE-317 |
| MVP_top60_DNA_seed | HSP90AA1 | 2.446973 | LUAD | NCI-H2291 | Elesclomol | top60_altered_DNA+RNA_seed | TP53 | -0.214167 | BLCA | 639-V | MIRA-1 |
| MVP_top60_DNA_seed | AKT1 | 2.451452 | LUAD | NCI-H2405 | MK-2206 | top60_altered_DNA+RNA_seed | PIK3CA | -0.202548 | BLCA | HT-1197 | AZD8186 |
| MVP_top60_DNA_seed | AKT1 | 2.458361 | LUAD | SK-LU-1 | MK-2206 | top60_altered_DNA+RNA_seed | PIK3CA | -0.167433 | BLCA | RT-112 | Buparlisib |
| MVP_top60_DNA_seed | IGF1R | 2.458857 | LUAD | NCI-H2291 | GSK1904529A | top60_altered_DNA+RNA_seed | PIK3CA | -0.141735 | BLCA | UM-UC-3 | AZD8186 |
| MVP_top60_DNA_seed | TP53 | 2.464529 | LUAD | NCI-H2405 | MIRA-1 | top60_altered_DNA+RNA_seed | TP53 | -0.130501 | BLCA | RT-112 | MIRA-1 |
| MVP_top60_DNA_seed | MET | 2.474652 | LUAD | NCI-H2347 | Crizotinib | top60_altered_DNA+RNA_seed | PIK3CA | -0.11969 | BLCA | 647-V | GNE-317 |
| MVP_top60_DNA_seed | FGFR1 | 2.491065 | LUAD | EKVX | AZD4547 | top60_altered_DNA+RNA_seed | PIK3CA | -0.110066 | BLCA | TCCSUP | Buparlisib |
| MVP_top60_DNA_seed | HDAC1 | 2.497733 | LUAD | NCI-H2405 | Romidepsin | top60_altered_DNA+RNA_seed | PIK3CA | -0.101739 | BLCA | BFTC-905 | AZD8186 |
| MVP_top60_DNA_seed | PIK3CA | 2.502359 | LUAD | NCI-H522 | GNE-317 | top60_altered_DNA+RNA_seed | PIK3CA | -0.087925 | BLCA | LB831-BLC | Alpelisib |
| MVP_top60_DNA_seed | HSP90AA1 | 2.510571 | LUAD | NCI-H1623 | Luminespib | top60_altered_DNA+RNA_seed | PIK3CA | -0.070269 | BLCA | DSH1 | Buparlisib |
| MVP_top60_DNA_seed | AKT1 | 2.519148 | LUAD | NCI-H2347 | MK-2206 | top60_altered_DNA+RNA_seed | TERT | -0.031656 | BLCA | 5637 | BIBR-1532 |
| MVP_top60_DNA_seed | EGFR | 2.519453 | LUAD | NCI-H2405 | AZD3759 | top60_altered_DNA+RNA_seed | PIK3CA | -0.013456 | BLCA | 647-V | Buparlisib |
| MVP_top60_DNA_seed | CREBBP | 2.527151 | LUAD | NCI-H2405 | SGC-CBP30 | top60_altered_DNA+RNA_seed | PIK3CA | -0.010391 | BLCA | 5637 | GNE-317 |
| MVP_top60_DNA_seed | MET | 2.565729 | LUAD | NCI-H2405 | Crizotinib | top60_altered_DNA+RNA_seed | PIK3CA | -0.005178 | BLCA | HT-1197 | Alpelisib |
| MVP_top60_DNA_seed | EZH2 | 2.595206 | LUAD | NCI-H2405 | GSK343 | top60_altered_DNA+RNA_seed | PIK3CA | 0.010687 | BLCA | 5637 | Alpelisib |
| MVP_top60_DNA_seed | KRAS | 2.603151 | LUAD | NCI-H441 | KRAS (G12C) Inhibitor-1 | top60_altered_DNA+RNA_seed | PIK3CA | 0.019742 | BLCA | UM-UC-3 | Buparlisib |
| MVP_top60_DNA_seed | ATM | 2.609157 | LUAD | NCI-H2405 | KU-55933 | top60_altered_DNA+RNA_seed | TP53 | 0.039981 | BLCA | KU-19-19 | MIRA-1 |
| MVP_top60_DNA_seed | AKT1 | 2.616957 | LUAD | NCI-H441 | AZD5363 | top60_altered_DNA+RNA_seed | PIK3CA | 0.070222 | BLCA | DSH1 | Alpelisib |
| MVP_top60_DNA_seed | HSP90AA1 | 2.62834 | LUAD | NCI-H1435 | Luminespib | top60_altered_DNA+RNA_seed | PIK3CA | 0.077521 | BLCA | 639-V | GNE-317 |
| MVP_top60_DNA_seed | ATM | 2.628708 | LUAD | NCI-H441 | KU-55933 | top60_altered_DNA+RNA_seed | TP53 | 0.082077 | BLCA | CAL-29 | MIRA-1 |
| MVP_top60_DNA_seed | HSP90AA1 | 2.632342 | LUAD | NCI-H2342 | Luminespib | top60_altered_DNA+RNA_seed | PIK3CA | 0.095867 | BLCA | SW1710 | Buparlisib |
| MVP_top60_DNA_seed | EGFR | 2.654864 | LUAD | NCI-H2347 | Lapatinib | top60_altered_DNA+RNA_seed | TERT | 0.113943 | BLCA | KU-19-19 | BIBR-1532 |
| MVP_top60_DNA_seed | ERBB2 | 2.654864 | LUAD | NCI-H2347 | Lapatinib | top60_altered_DNA+RNA_seed | TERT | 0.117448 | BLCA | SW1710 | BIBR-1532 |
| MVP_top60_DNA_seed | PIK3CA | 2.656799 | LUAD | NCI-H2347 | Alpelisib | top60_altered_DNA+RNA_seed | TERT | 0.129428 | BLCA | BFTC-905 | BIBR-1532 |
| MVP_top60_DNA_seed | GSK3B | 2.658274 | LUAD | NCI-H441 | SB216763 | top60_altered_DNA+RNA_seed | TERT | 0.165043 | BLCA | CAL-29 | BIBR-1532 |
| MVP_top60_DNA_seed | MDM2 | 2.718022 | LUAD | NCI-H1793 | Serdemetan | top60_altered_DNA+RNA_seed | PIK3CA | 0.16847 | BLCA | SW780 | GNE-317 |
| MVP_top60_DNA_seed | SMARCA4 | 2.718226 | LUAD | NCI-H441 | PFI3 | top60_altered_DNA+RNA_seed | TP53 | 0.170108 | BLCA | UM-UC-3 | MIRA-1 |
| MVP_top60_DNA_seed | IGF1R | 2.722961 | LUAD | NCI-H2347 | BMS-754807 | top60_altered_DNA+RNA_seed | TP53 | 0.209842 | BLCA | DSH1 | MIRA-1 |
| MVP_top60_DNA_seed | GSK3B | 2.73389 | LUAD | NCI-H2291 | SB216763 | top60_altered_DNA+RNA_seed | PIK3CA | 0.220832 | BLCA | SW1710 | GNE-317 |
| MVP_top60_DNA_seed | AKT1 | 2.742815 | LUAD | NCI-H2405 | Afuresertib | top60_altered_DNA+RNA_seed | PIK3CA | 0.255993 | BLCA | LB831-BLC | GNE-317 |
| MVP_top60_DNA_seed | AKT1 | 2.749456 | LUAD | SK-LU-1 | AZD5363 | top60_altered_DNA+RNA_seed | TERT | 0.318581 | BLCA | T24 | BIBR-1532 |
| MVP_top60_DNA_seed | HDAC1 | 2.764612 | LUAD | NCI-H1650 | Entinostat | top60_altered_DNA+RNA_seed | PIK3CA | 0.336452 | BLCA | LB831-BLC | AZD8186 |
| MVP_top60_DNA_seed | IGF1R | 2.784329 | LUAD | VMRC-LCD | BMS-536924 | top60_altered_DNA+RNA_seed | PIK3CA | 0.349232 | BLCA | SW1710 | AZD8186 |
| MVP_top60_DNA_seed | EGFR | 2.817442 | LUAD | SW1573 | Lapatinib | top60_altered_DNA+RNA_seed | PIK3CA | 0.380056 | BLCA | DSH1 | AZD8186 |
| MVP_top60_DNA_seed | ERBB2 | 2.817442 | LUAD | SW1573 | Lapatinib | top60_altered_DNA+RNA_seed | TP53 | 0.382708 | BLCA | 647-V | MIRA-1 |
| MVP_top60_DNA_seed | MET | 2.819847 | LUAD | NCI-H1793 | Crizotinib | top60_altered_DNA+RNA_seed | PIK3CA | 0.404888 | BLCA | 5637 | AZD8186 |
| MVP_top60_DNA_seed | AKT1 | 2.831554 | LUAD | NCI-H441 | AT13148 | top60_altered_DNA+RNA_seed | TP53 | 0.417231 | BLCA | SW1710 | MIRA-1 |
| MVP_top60_DNA_seed | AURKA | 2.839326 | LUAD | NCI-H2405 | ZM447439 | top60_altered_DNA+RNA_seed | PIK3CA | 0.435335 | BLCA | HT-1376 | AZD8186 |
| MVP_top60_DNA_seed | MET | 2.878976 | LUAD | EKVX | Crizotinib | top60_altered_DNA+RNA_seed | PIK3CA | 0.44544 | BLCA | SW780 | Buparlisib |

| | | | | | | | | | | | |
|---|---|---|---|---|---|---|---|---|---|---|---|
| MVP_top60_DNA_seed | PIK3CA | 2.886868 | LUAD | NCI-H2405 | Alpelisib | top60_altered_DNA+RNA_seed | TP53 | 0.480846 | BLCA | T24 | MIRA-1 |
| MVP_top60_DNA_seed | HDAC1 | 2.9272 | LUAD | SK-LU-1 | Entinostat | top60_altered_DNA+RNA_seed | TERT | 0.486949 | BLCA | RT4 | BIBR-1532 |
| MVP_top60_DNA_seed | EGFR | 2.935743 | LUAD | NCI-H2405 | Lapatinib | top60_altered_DNA+RNA_seed | PIK3CA | 0.491344 | BLCA | RT4 | Buparlisib |
| MVP_top60_DNA_seed | ERBB2 | 2.935743 | LUAD | NCI-H2405 | Lapatinib | top60_altered_DNA+RNA_seed | PIK3CA | 0.505059 | BLCA | TCCSUP | Alpelisib |
| MVP_top60_DNA_seed | MAPK1 | 2.962967 | LUAD | NCI-H1838 | Ulixertinib | top60_altered_DNA+RNA_seed | TP53 | 0.520183 | BLCA | SW780 | MIRA-1 |
| MVP_top60_DNA_seed | HSP90AA1 | 2.980072 | LUAD | NCI-H441 | Tanespimycin | top60_altered_DNA+RNA_seed | TP53 | 0.520968 | BLCA | RT4 | MIRA-1 |
| MVP_top60_DNA_seed | EGFR | 2.992514 | LUAD | NCI-H441 | Osimertinib | top60_altered_DNA+RNA_seed | PIK3CA | 0.557464 | BLCA | RT-112 | AZD8186 |
| MVP_top60_DNA_seed | TERT | 3.04853 | LUAD | NCI-H441 | BIBR-1532 | top60_altered_DNA+RNA_seed | PIK3CA | 0.599658 | BLCA | 639-V | Alpelisib |
| MVP_top60_DNA_seed | AKT1 | 3.107433 | LUAD | SK-LU-1 | Afuresertib | top60_altered_DNA+RNA_seed | PIK3CA | 0.61902 | BLCA | LB831-BLC | Buparlisib |
| MVP_top60_DNA_seed | SMARCA4 | 3.153384 | LUAD | NCI-H2405 | PFI3 | top60_altered_DNA+RNA_seed | TP53 | 0.670544 | BLCA | 5637 | MIRA-1 |
| MVP_top60_DNA_seed | PIK3CA | 3.16743 | LUAD | NCI-H441 | GNE-317 | top60_altered_DNA+RNA_seed | PIK3CA | 0.672557 | BLCA | TCCSUP | AZD8186 |
| MVP_top60_DNA_seed | EGFR | 3.197082 | LUAD | NCI-H2405 | Afatinib | top60_altered_DNA+RNA_seed | TP53 | 0.706531 | BLCA | HT-1197 | MIRA-1 |
| MVP_top60_DNA_seed | ERBB2 | 3.197082 | LUAD | NCI-H2405 | Afatinib | top60_altered_DNA+RNA_seed | TERT | 0.718228 | BLCA | UM-UC-3 | BIBR-1532 |
| MVP_top60_DNA_seed | AURKA | 3.250564 | LUAD | NCI-H441 | ZM447439 | top60_altered_DNA+RNA_seed | TP53 | 0.720144 | BLCA | TCCSUP | MIRA-1 |
| MVP_top60_DNA_seed | IGF1R | 3.281784 | LUAD | NCI-H441 | GSK1904529A | top60_altered_DNA+RNA_seed | TP53 | 0.736355 | BLCA | LB831-BLC | MIRA-1 |
| MVP_top60_DNA_seed | IGF1R | 3.283733 | LUAD | NCI-H2405 | NVP-ADW742 | top60_altered_DNA+RNA_seed | PIK3CA | 0.770447 | BLCA | RT4 | Alpelisib |
| MVP_top60_DNA_seed | HDAC1 | 3.292349 | LUAD | NCI-H1793 | Entinostat | top60_altered_DNA+RNA_seed | TP53 | 0.778608 | BLCA | J82 | MIRA-1 |
| MVP_top60_DNA_seed | HSP90AA1 | 3.313202 | LUAD | NCI-H441 | Luminespib | top60_altered_DNA+RNA_seed | PIK3CA | 0.807117 | BLCA | T24 | AZD8186 |
| MVP_top60_DNA_seed | PIK3CA | 3.323685 | LUAD | NCI-H1838 | Buparlisib | top60_altered_DNA+RNA_seed | TERT | 0.853057 | BLCA | LB831-BLC | BIBR-1532 |
| MVP_top60_DNA_seed | HSP90AA1 | 3.325474 | LUAD | NCI-H1838 | Luminespib | top60_altered_DNA+RNA_seed | TERT | 0.853201 | BLCA | TCCSUP | BIBR-1532 |
| MVP_top60_DNA_seed | HDAC1 | 3.570933 | LUAD | NCI-H1650 | Romidepsin | top60_altered_DNA+RNA_seed | PIK3CA | 0.864754 | BLCA | KU-19-19 | AZD8186 |
| MVP_top60_DNA_seed | HDAC1 | 3.572014 | LUAD | NCI-H1793 | Romidepsin | top60_altered_DNA+RNA_seed | PIK3CA | 0.933356 | BLCA | T24 | GNE-317 |
| MVP_top60_DNA_seed | HSP90AA1 | 4.016577 | LUAD | NCI-H2347 | Luminespib | top60_altered_DNA+RNA_seed | TP53 | 0.988377 | BLCA | HT-1376 | MIRA-1 |
| MVP_top60_DNA_seed | EGFR | -2.71814 | LUSC | NCI-H2170 | Afatinib | top60_altered_DNA+RNA_seed | PIK3CA | 1.024332 | BLCA | J82 | AZD8186 |
| MVP_top60_DNA_seed | EGFR | -2.517836 | LUSC | HARA | Gefitinib | top60_altered_DNA+RNA_seed | TERT | 1.043692 | BLCA | J82 | BIBR-1532 |
| MVP_top60_DNA_seed | SMARCA4 | -2.471329 | LUSC | EBC-1 | PFI-3 | top60_altered_DNA+RNA_seed | PIK3CA | 1.077254 | BLCA | KU-19-19 | Alpelisib |
| MVP_top60_DNA_seed | SMARCA2 | -2.471329 | LUSC | EBC-1 | PFI-3 | top60_altered_DNA+RNA_seed | PIK3CA | 1.103866 | BLCA | T24 | Alpelisib |
| MVP_top60_DNA_seed | AR | -2.463165 | LUSC | EBC-1 | Bicalutamide | top60_altered_DNA+RNA_seed | PIK3CA | 1.110764 | BLCA | RT4 | GNE-317 |
| MVP_top60_DNA_seed | EGFR | -2.436966 | LUSC | SK-MES-1 | Cetuximab | top60_altered_DNA+RNA_seed | PIK3CA | 1.13296 | BLCA | SW1710 | Alpelisib |
| MVP_top60_DNA_seed | EGFR | -2.389588 | LUSC | NCI-H2170 | Afatinib | top60_altered_DNA+RNA_seed | PIK3CA | 1.261823 | BLCA | 647-V | Alpelisib |
| MVP_top60_DNA_seed | CREBBP | -2.289563 | LUSC | HARA | I-CBP112 | top60_altered_DNA+RNA_seed | PIK3CA | 1.29491 | BLCA | UM-UC-3 | Alpelisib |
| MVP_top60_DNA_seed | AR | -2.182747 | LUSC | KNS-62 | AZD3514 | top60_altered_DNA+RNA_seed | PIK3CA | 1.309147 | BLCA | J82 | Alpelisib |
| MVP_top60_DNA_seed | EGFR | -2.146963 | LUSC | RERF-LC-Sq1 | AZD8931 | top60_altered_DNA+RNA_seed | PIK3CA | 1.405059 | BLCA | SW780 | Alpelisib |
| MVP_top60_DNA_seed | EGFR | -2.137225 | LUSC | SK-MES-1 | Pelitinib | top60_altered_DNA+RNA_seed | PIK3CA | 1.680588 | BLCA | T24 | Buparlisib |
| MVP_top60_DNA_seed | AR | -2.102598 | LUSC | EBC-1 | AZD3514 | top60_altered_DNA+RNA_seed | PIK3CA | 1.706332 | BLCA | RT4 | AZD8186 |
| MVP_top60_DNA_seed | EGFR | -1.847581 | LUSC | RERF-LC-Sq1 | Afatinib | top60_altered_DNA+RNA_seed | PIK3CA | 1.715415 | BLCA | HT-1376 | GNE-317 |
| MVP_top60_DNA_seed | EGFR | -1.841615 | LUSC | RERF-LC-Sq1 | Afatinib | top60_altered_DNA+RNA_seed | PIK3CA | 1.716928 | BLCA | SW780 | AZD8186 |
| MVP_top60_DNA_seed | EGFR | -1.792281 | LUSC | NCI-H1869 | Erlotinib | top60_altered_DNA+RNA_seed | PIK3CA | 1.756588 | BLCA | HT-1376 | Alpelisib |
| MVP_top60_DNA_seed | ESR1 | -1.701233 | LUSC | EBC-1 | Tamoxifen | top60_altered_DNA+RNA_seed | PIK3CA | 1.951983 | BLCA | J82 | Buparlisib |
| MVP_top60_DNA_seed | HDAC1 | -1.700261 | LUSC | EBC-1 | JQ12 | top60_altered_DNA+RNA_seed | PIK3CA | 2.716316 | BLCA | HT-1376 | Buparlisib |
| MVP_top60_DNA_seed | HDAC2 | -1.700261 | LUSC | EBC-1 | JQ12 | top60_altered_DNA+RNA_seed | PIK3CA | 2.971117 | BLCA | J82 | GNE-317 |

| | | | | | | | | | | | |
|---|---|---|---|---|---|---|---|---|---|---|---|
| MVP_top60_DNA_seed | PIK3CA | -1.665707 | LUSC | SK-MES-1 | PI3Ka_4409 | top60_altered_DNA+RNA_seed | PIK3CA | -2.548715 | BRCA | OCUB-M | PI3Ka_4409 |
| MVP_top60_DNA_seed | EGFR | -1.593722 | LUSC | LOU-NH91 | Afatinib | top60_altered_DNA+RNA_seed | PIK3CA | -2.357662 | BRCA | MDA-MB-453 | PI3Ka_4409 |
| MVP_top60_DNA_seed | GSK3B | -1.579193 | LUSC | EPLC-272H | KIN001-042 | top60_altered_DNA+RNA_seed | PIK3CA | -2.077815 | BRCA | MDA-MB-453 | AZD8835 |
| MVP_top60_DNA_seed | EGFR | -1.561145 | LUSC | LOU-NH91 | AZD8931 | top60_altered_DNA+RNA_seed | PIK3CA | -2.033158 | BRCA | HCC202 | PI3Ka_4409 |
| MVP_top60_DNA_seed | EGFR | -1.491791 | LUSC | RERF-LC-Sq1 | Gefitinib | top60_altered_DNA+RNA_seed | PIK3CA | -2.01571 | BRCA | MDA-MB-361 | PI3Ka_4409 |
| MVP_top60_DNA_seed | EGFR | -1.490837 | LUSC | NCI-H2170 | AZD8931 | top60_altered_DNA+RNA_seed | PIK3CA | -1.993881 | BRCA | MDA-MB-361 | AZD8835 |
| MVP_top60_DNA_seed | PIK3CA | -1.469562 | LUSC | RERF-LC-Sq1 | PI3Ka_4409 | top60_altered_DNA+RNA_seed | PIK3CA | -1.923421 | BRCA | HCC1954 | PI3Ka_4409 |
| MVP_top60_DNA_seed | ESR1 | -1.407604 | LUSC | HARA | Fulvestrant | top60_altered_DNA+RNA_seed | PIK3CA | -1.888956 | BRCA | MDA-MB-330 | AZD8835 |
| MVP_top60_DNA_seed | ESR1 | -1.405606 | LUSC | NCI-H1869 | Fulvestrant | top60_altered_DNA+RNA_seed | PIK3CA | -1.819452 | BRCA | OCUB-M | AZD8835 |
| MVP_top60_DNA_seed | PIK3CA | -1.37393 | LUSC | SK-MES-1 | AZD8835 | top60_altered_DNA+RNA_seed | PIK3CA | -1.72137 | BRCA | HCC1599 | PI3Ka_4409 |
| MVP_top60_DNA_seed | EGFR | -1.355352 | LUSC | RERF-LC-Sq1 | Cetuximab | top60_altered_DNA+RNA_seed | PIK3CA | -1.437506 | BRCA | JIMT-1 | AZD8835 |
| MVP_top60_DNA_seed | EGFR | -1.343181 | LUSC | EPLC-272H | Afatinib | top60_altered_DNA+RNA_seed | PIK3CA | -1.311004 | BRCA | EFM-19 | PI3Ka_4409 |
| MVP_top60_DNA_seed | EGFR | -1.338893 | LUSC | SK-MES-1 | Afatinib | top60_altered_DNA+RNA_seed | PIK3CA | -1.288787 | BRCA | EFM-192A | PI3Ka_4409 |
| MVP_top60_DNA_seed | ESR1 | -1.337827 | LUSC | RERF-LC-Sq1 | Fulvestrant | top60_altered_DNA+RNA_seed | PLK1 | -1.265636 | BRCA | COLO-824 | PLK_6522 |
| MVP_top60_DNA_seed | EGFR | -1.329189 | LUSC | LC-1-sq | Gefitinib | top60_altered_DNA+RNA_seed | PIK3CA | -1.199455 | BRCA | BT-20 | AZD8835 |
| MVP_top60_DNA_seed | ESR1 | -1.303025 | LUSC | KNS-62 | Tamoxifen | top60_altered_DNA+RNA_seed | PIK3CA | -1.194497 | BRCA | EFM-19 | AZD8835 |
| MVP_top60_DNA_seed | HDAC1 | -1.276467 | LUSC | LK-2 | JQ12 | top60_altered_DNA+RNA_seed | PLK1 | -1.179885 | BRCA | OCUB-M | BI-2536 |
| MVP_top60_DNA_seed | HDAC2 | -1.276467 | LUSC | LK-2 | JQ12 | top60_altered_DNA+RNA_seed | PLK1 | -1.173377 | BRCA | AU565 | PLK_6522 |
| MVP_top60_DNA_seed | EGFR | -1.266214 | LUSC | LOU-NH91 | Gefitinib | top60_altered_DNA+RNA_seed | PIK3CA | -1.156943 | BRCA | HCC1599 | AZD8835 |
| MVP_top60_DNA_seed | EGFR | -1.195642 | LUSC | NCI-H1869 | Lapatinib | top60_altered_DNA+RNA_seed | PIK3CA | -1.1376 | BRCA | UACC-893 | PI3Ka_4409 |
| MVP_top60_DNA_seed | GSK3B | -1.151212 | LUSC | LOU-NH91 | AZD7969 | top60_altered_DNA+RNA_seed | PIK3CA | -1.130069 | BRCA | T47D | PI3Ka_4409 |
| MVP_top60_DNA_seed | EGFR | -1.143583 | LUSC | HARA | Afatinib | top60_altered_DNA+RNA_seed | PIK3CA | -1.12156 | BRCA | MCF7 | AZD8835 |
| MVP_top60_DNA_seed | AR | -1.13906 | LUSC | LK-2 | AZD3514 | top60_altered_DNA+RNA_seed | PLK1 | -1.116155 | BRCA | OCUB-M | GW843682X |
| MVP_top60_DNA_seed | EGFR | -1.124315 | LUSC | NCI-H1869 | Cetuximab | top60_altered_DNA+RNA_seed | PIK3CA | -1.072273 | BRCA | JIMT-1 | PI3Ka_4409 |
| MVP_top60_DNA_seed | HDAC1 | -1.110672 | LUSC | LK-2 | Dacinostat | top60_altered_DNA+RNA_seed | PLK1 | -0.969456 | BRCA | BT-549 | PLK_6522 |
| MVP_top60_DNA_seed | HDAC1 | -1.101418 | LUSC | EPLC-272H | Tubastatin A | top60_altered_DNA+RNA_seed | PIK3CA | -0.962684 | BRCA | EVSA-T | PI3Ka_4409 |
| MVP_top60_DNA_seed | PIK3CA | -1.09166 | LUSC | RERF-LC-Sq1 | AZD8835 | top60_altered_DNA+RNA_seed | PLK1 | -0.948112 | BRCA | MFM-223 | PLK_6522 |
| MVP_top60_DNA_seed | GSK3B | -1.058428 | LUSC | EPLC-272H | CHIR-99021 | top60_altered_DNA+RNA_seed | PIK3CA | -0.937617 | BRCA | HCC1954 | AZD8835 |
| MVP_top60_DNA_seed | GSK3B | -1.038421 | LUSC | LC-1-sq | CHIR-99021 | top60_altered_DNA+RNA_seed | PIK3CA | -0.895769 | BRCA | BT-20 | PI3Ka_4409 |
| MVP_top60_DNA_seed | GSK3B | -1.031 | LUSC | NCI-H226 | CHIR-99021 | top60_altered_DNA+RNA_seed | PIK3CA | -0.8937 | BRCA | BT-474 | PI3Ka_4409 |
| MVP_top60_DNA_seed | EGFR | -1.023936 | LUSC | EPLC-272H | CI-1033 | top60_altered_DNA+RNA_seed | PLK1 | -0.869562 | BRCA | MDA-MB-361 | PLK_6522 |
| MVP_top60_DNA_seed | EGFR | -1.022259 | LUSC | EPLC-272H | Pelitinib | top60_altered_DNA+RNA_seed | PLK1 | -0.857366 | BRCA | OCUB-M | PLK_6522 |
| MVP_top60_DNA_seed | RAC1 | -1.016352 | LUSC | LC-1-sq | EHT-1864 | top60_altered_DNA+RNA_seed | PLK1 | -0.853843 | BRCA | MDA-MB-468 | PLK_6522 |
| MVP_top60_DNA_seed | PIK3CA | -1.001354 | LUSC | EPLC-272H | PI3Ka_4409 | top60_altered_DNA+RNA_seed | PIK3CA | -0.852086 | BRCA | UACC-893 | AZD8835 |
| MVP_top60_DNA_seed | EGFR | -0.996358 | LUSC | EPLC-272H | Afatinib | top60_altered_DNA+RNA_seed | PLK1 | -0.833838 | BRCA | HCC1599 | GW843682X |
| MVP_top60_DNA_seed | EGFR | -0.978311 | LUSC | NCI-H1869 | CI-1033 | top60_altered_DNA+RNA_seed | PIK3CA | -0.832895 | BRCA | MFM-223 | AZD8835 |
| MVP_top60_DNA_seed | HDAC1 | -0.952447 | LUSC | EPLC-272H | CAY10603 | top60_altered_DNA+RNA_seed | PLK1 | -0.821775 | BRCA | HCC1187 | PLK_6522 |
| MVP_top60_DNA_seed | AR | -0.941026 | LUSC | SK-MES-1 | Bicalutamide | top60_altered_DNA+RNA_seed | PLK1 | -0.796427 | BRCA | CAL-120 | PLK_6522 |
| MVP_top60_DNA_seed | EGFR | -0.937233 | LUSC | EPLC-272H | AZD8931 | top60_altered_DNA+RNA_seed | PIK3CA | -0.787269 | BRCA | MCF7 | PI3Ka_4409 |
| MVP_top60_DNA_seed | EGFR | -0.928276 | LUSC | RERF-LC-Sq1 | CI-1033 | top60_altered_DNA+RNA_seed | PIK3CA | -0.782566 | BRCA | EFM-192A | AZD8835 |
| MVP_top60_DNA_seed | EGFR | -0.925958 | LUSC | SK-MES-1 | Afatinib | top60_altered_DNA+RNA_seed | PIK3CA | -0.778535 | BRCA | BT-474 | AZD8835 |

| | | | | | | | | | | | |
|---|---|---|---|---|---|---|---|---|---|---|---|
| MVP_top60_DNA_seed | HDAC1 | -0.920538 | LUSC | EBC-1 | Dacinostat | top60_altered_DNA+RNA_seed | PIK3CA | -0.767928 | BRCA | HCC202 | AZD8835 |
| MVP_top60_DNA_seed | SMARCA4 | -0.913857 | LUSC | KNS-62 | PFI-3 | top60_altered_DNA+RNA_seed | PLK1 | -0.733683 | BRCA | HCC1599 | BI-2536 |
| MVP_top60_DNA_seed | SMARCA2 | -0.913857 | LUSC | KNS-62 | PFI-3 | top60_altered_DNA+RNA_seed | PLK1 | -0.698038 | BRCA | EFM-19 | PLK_6522 |
| MVP_top60_DNA_seed | GSK3B | -0.904919 | LUSC | SK-MES-1 | SB216763 | top60_altered_DNA+RNA_seed | PLK1 | -0.689839 | BRCA | HCC38 | PLK_6522 |
| MVP_top60_DNA_seed | EGFR | -0.87797 | LUSC | EPLC-272H | CUDC-101 | top60_altered_DNA+RNA_seed | PLK1 | -0.653011 | BRCA | HCC1143 | PLK_6522 |
| MVP_top60_DNA_seed | EGFR | -0.874985 | LUSC | NCI-H1869 | PF-00299804 | top60_altered_DNA+RNA_seed | PLK1 | -0.637338 | BRCA | MRK-nu-1 | PLK_6522 |
| MVP_top60_DNA_seed | EGFR | -0.861098 | LUSC | EPLC-272H | AST-1306 | top60_altered_DNA+RNA_seed | PLK1 | -0.578569 | BRCA | JIMT-1 | PLK_6522 |
| MVP_top60_DNA_seed | EGFR | -0.835148 | LUSC | HARA | AZD8931 | top60_altered_DNA+RNA_seed | PLK1 | -0.576057 | BRCA | HCC1395 | PLK_6522 |
| MVP_top60_DNA_seed | PIK3CA | -0.834848 | LUSC | EPLC-272H | AZD8835 | top60_altered_DNA+RNA_seed | PIK3CA | -0.574712 | BRCA | HCC1187 | AZD8835 |
| MVP_top60_DNA_seed | ESR1 | -0.830783 | LUSC | LK-2 | Fulvestrant | top60_altered_DNA+RNA_seed | PIK3CA | -0.573288 | BRCA | T47D | AZD8835 |
| MVP_top60_DNA_seed | EGFR | -0.826362 | LUSC | LC-1-sq | AZD8931 | top60_altered_DNA+RNA_seed | PIK3CA | -0.568687 | BRCA | MFM-223 | PI3Ka_4409 |
| MVP_top60_DNA_seed | AR | -0.815814 | LUSC | EPLC-272H | AZD3514 | top60_altered_DNA+RNA_seed | PIK3CA | -0.565849 | BRCA | HCC1187 | PI3Ka_4409 |
| MVP_top60_DNA_seed | HDAC1 | -0.814045 | LUSC | EPLC-272H | Belinostat | top60_altered_DNA+RNA_seed | PLK1 | -0.516691 | BRCA | CAL-51 | PLK_6522 |
| MVP_top60_DNA_seed | PIK3CA | -0.809777 | LUSC | LOU-NH91 | AZD8835 | top60_altered_DNA+RNA_seed | PIK3CA | -0.453369 | BRCA | MDA-MB-330 | PI3Ka_4409 |
| MVP_top60_DNA_seed | EGFR | -0.777122 | LUSC | NCI-H2170 | Gefitinib | top60_altered_DNA+RNA_seed | PIK3CA | -0.43744 | BRCA | EVSA-T | AZD8835 |
| MVP_top60_DNA_seed | EGFR | -0.774567 | LUSC | EPLC-272H | Gefitinib | top60_altered_DNA+RNA_seed | PLK1 | -0.425524 | BRCA | EVSA-T | PLK_6522 |
| MVP_top60_DNA_seed | EGFR | -0.771381 | LUSC | LC-1-sq | Afatinib | top60_altered_DNA+RNA_seed | PLK1 | -0.420428 | BRCA | HCC1954 | PLK_6522 |
| MVP_top60_DNA_seed | CREBBP | -0.75393 | LUSC | LC-1-sq | I-CBP112 | top60_altered_DNA+RNA_seed | PLK1 | -0.408664 | BRCA | DU-4475 | GW843682X |
| MVP_top60_DNA_seed | EGFR | -0.749354 | LUSC | HARA | Pelitinib | top60_altered_DNA+RNA_seed | PLK1 | -0.334662 | BRCA | HCC1187 | GW843682X |
| MVP_top60_DNA_seed | RAC1 | -0.734201 | LUSC | NCI-H1869 | EHT-1864 | top60_altered_DNA+RNA_seed | PIK3CA | -0.297693 | BRCA | MDA-MB-175-VII | AZD8835 |
| MVP_top60_DNA_seed | ESR1 | -0.724432 | LUSC | LOU-NH91 | Fulvestrant | top60_altered_DNA+RNA_seed | PLK1 | -0.292947 | BRCA | HCC1187 | BI-2536 |
| MVP_top60_DNA_seed | GSK3B | -0.719088 | LUSC | SK-MES-1 | AZD7969 | top60_altered_DNA+RNA_seed | PLK1 | -0.271905 | BRCA | MCF7 | PLK_6522 |
| MVP_top60_DNA_seed | CREBBP | -0.716771 | LUSC | NCI-H226 | I-CBP112 | top60_altered_DNA+RNA_seed | PLK1 | -0.253283 | BRCA | MDA-MB-231 | PLK_6522 |
| MVP_top60_DNA_seed | EGFR | -0.686041 | LUSC | SK-MES-1 | AZD8931 | top60_altered_DNA+RNA_seed | PIK3CA | -0.24184 | BRCA | AU565 | AZD8835 |
| MVP_top60_DNA_seed | HDAC1 | -0.673209 | LUSC | EPLC-272H | AR-42 | top60_altered_DNA+RNA_seed | PIK3CA | -0.234688 | BRCA | MDA-MB-175-VII | PI3Ka_4409 |
| MVP_top60_DNA_seed | EGFR | -0.671443 | LUSC | NCI-H1869 | AZD8931 | top60_altered_DNA+RNA_seed | PLK1 | -0.197318 | BRCA | CAL-148 | BI-2536 |
| MVP_top60_DNA_seed | RAC1 | -0.657535 | LUSC | HARA | EHT-1864 | top60_altered_DNA+RNA_seed | PLK1 | -0.18996 | BRCA | CAL-148 | GW843682X |
| MVP_top60_DNA_seed | HDAC1 | -0.64373 | LUSC | NCI-H520 | CAY10603 | top60_altered_DNA+RNA_seed | PIK3CA | -0.185494 | BRCA | HCC1569 | PI3Ka_4409 |
| MVP_top60_DNA_seed | HDAC1 | -0.641588 | LUSC | HCC-15 | JQ12 | top60_altered_DNA+RNA_seed | PIK3CA | -0.182922 | BRCA | CAL-51 | PI3Ka_4409 |
| MVP_top60_DNA_seed | HDAC2 | -0.641588 | LUSC | HCC-15 | JQ12 | top60_altered_DNA+RNA_seed | PIK3CA | -0.171098 | BRCA | CAL-120 | AZD8835 |
| MVP_top60_DNA_seed | GSK3B | -0.63458 | LUSC | HARA | KIN001-042 | top60_altered_DNA+RNA_seed | PIK3CA | -0.14942 | BRCA | CAL-85-1 | PI3Ka_4409 |
| MVP_top60_DNA_seed | PIK3CA | -0.633699 | LUSC | EBC-1 | PI3Ka_4409 | top60_altered_DNA+RNA_seed | PIK3CA | -0.149209 | BRCA | AU565 | PI3Ka_4409 |
| MVP_top60_DNA_seed | ESR1 | -0.629856 | LUSC | EPLC-272H | Fulvestrant | top60_altered_DNA+RNA_seed | PLK1 | -0.137538 | BRCA | Hs-578-T | PLK_6522 |
| MVP_top60_DNA_seed | EGFR | -0.629384 | LUSC | KNS-62 | Afatinib | top60_altered_DNA+RNA_seed | PIK3CA | -0.111097 | BRCA | HCC2218 | PI3Ka_4409 |
| MVP_top60_DNA_seed | ESR1 | -0.609709 | LUSC | EPLC-272H | Tamoxifen | top60_altered_DNA+RNA_seed | PLK1 | -0.096615 | BRCA | MDA-MB-436 | PLK_6522 |
| MVP_top60_DNA_seed | EGFR | -0.59923 | LUSC | NCI-H1869 | Pelitinib | top60_altered_DNA+RNA_seed | PLK1 | -0.095186 | BRCA | DU-4475 | BI-2536 |
| MVP_top60_DNA_seed | HDAC1 | -0.596337 | LUSC | NCI-H520 | Tubastatin A | top60_altered_DNA+RNA_seed | PIK3CA | -0.057415 | BRCA | HCC70 | PI3Ka_4409 |
| MVP_top60_DNA_seed | EGFR | -0.590916 | LUSC | NCI-H1869 | AST-1306 | top60_altered_DNA+RNA_seed | PIK3CA | -0.021242 | BRCA | HCC1143 | AZD8835 |
| MVP_top60_DNA_seed | EGFR | -0.588844 | LUSC | RERF-LC-Sq1 | PF-00299804 | top60_altered_DNA+RNA_seed | PIK3CA | 0.017213 | BRCA | HCC2157 | AZD8835 |
| MVP_top60_DNA_seed | SMARCA4 | -0.5672 | LUSC | HARA | PFI-3 | top60_altered_DNA+RNA_seed | PLK1 | 0.018303 | BRCA | MDA-MB-453 | PLK_6522 |
| MVP_top60_DNA_seed | SMARCA2 | -0.5672 | LUSC | HARA | PFI-3 | top60_altered_DNA+RNA_seed | PIK3CA | 0.04396 | BRCA | HCC1143 | PI3Ka_4409 |

| | | | | | | | | | | | |
|---|---|---|---|---|---|---|---|---|---|---|---|
| MVP_top60_DNA_seed | EGFR | -0.557407 | LUSC | LOU-NH91 | Cetuximab | top60_altered_DNA+RNA_seed | PLK1 | 0.04863 | BRCA | EVSA-T | BI-2536 |
| MVP_top60_DNA_seed | HDAC1 | -0.543127 | LUSC | NCI-H2170 | JQ12 | top60_altered_DNA+RNA_seed | PIK3CA | 0.062903 | BRCA | MDA-MB-415 | PI3Ka_4409 |
| MVP_top60_DNA_seed | HDAC2 | -0.543127 | LUSC | NCI-H2170 | JQ12 | top60_altered_DNA+RNA_seed | PIK3CA | 0.070522 | BRCA | HCC2157 | PI3Ka_4409 |
| MVP_top60_DNA_seed | EGFR | -0.543019 | LUSC | HARA | Afatinib | top60_altered_DNA+RNA_seed | PIK3CA | 0.077335 | BRCA | COLO-824 | AZD8835 |
| MVP_top60_DNA_seed | HDAC1 | -0.529467 | LUSC | LK-2 | Belinostat | top60_altered_DNA+RNA_seed | PIK3CA | 0.089588 | BRCA | BT-549 | PI3Ka_4409 |
| MVP_top60_DNA_seed | GSK3B | -0.52887 | LUSC | LK-2 | KIN001-042 | top60_altered_DNA+RNA_seed | PIK3CA | 0.096349 | BRCA | HCC1569 | AZD8835 |
| MVP_top60_DNA_seed | RAC1 | -0.527973 | LUSC | LK-2 | EHT-1864 | top60_altered_DNA+RNA_seed | PIK3CA | 0.140748 | BRCA | COLO-824 | PI3Ka_4409 |
| MVP_top60_DNA_seed | AR | -0.522758 | LUSC | LOU-NH91 | AZD3514 | top60_altered_DNA+RNA_seed | PIK3CA | 0.195146 | BRCA | CAL-120 | PI3Ka_4409 |
| MVP_top60_DNA_seed | EGFR | -0.508225 | LUSC | NCI-H520 | CUDC-101 | top60_altered_DNA+RNA_seed | PIK3CA | 0.20947 | BRCA | DU-4475 | AZD8835 |
| MVP_top60_DNA_seed | GSK3B | -0.507467 | LUSC | NCI-H2170 | CHIR-99021 | top60_altered_DNA+RNA_seed | PLK1 | 0.215164 | BRCA | MRK-nu-1 | BI-2536 |
| MVP_top60_DNA_seed | EGFR | -0.498153 | LUSC | EPLC-272H | Cetuximab | top60_altered_DNA+RNA_seed | PIK3CA | 0.263913 | BRCA | Hs-578-T | PI3Ka_4409 |
| MVP_top60_DNA_seed | SMARCA4 | -0.487779 | LUSC | EPLC-272H | PFI-3 | top60_altered_DNA+RNA_seed | PIK3CA | 0.273674 | BRCA | MDA-MB-231 | AZD8835 |
| MVP_top60_DNA_seed | SMARCA2 | -0.487779 | LUSC | EPLC-272H | PFI-3 | top60_altered_DNA+RNA_seed | PLK1 | 0.287728 | BRCA | BT-20 | PLK_6522 |
| MVP_top60_DNA_seed | AR | -0.479459 | LUSC | EPLC-272H | Bicalutamide | top60_altered_DNA+RNA_seed | PIK3CA | 0.357184 | BRCA | BT-483 | PI3Ka_4409 |
| MVP_top60_DNA_seed | GSK3B | -0.47076 | LUSC | LC-1-sq | CHIR-99021 | top60_altered_DNA+RNA_seed | PLK1 | 0.374765 | BRCA | EVSA-T | GW843682X |
| MVP_top60_DNA_seed | EGFR | -0.468166 | LUSC | LC-1-sq | Pelitinib | top60_altered_DNA+RNA_seed | PIK3CA | 0.494257 | BRCA | HDQ-P1 | PI3Ka_4409 |
| MVP_top60_DNA_seed | PIK3CA | -0.455436 | LUSC | KNS-62 | PI3Ka_4409 | top60_altered_DNA+RNA_seed | PIK3CA | 0.513118 | BRCA | HCC2218 | AZD8835 |
| MVP_top60_DNA_seed | ESR1 | -0.439128 | LUSC | SK-MES-1 | Fulvestrant | top60_altered_DNA+RNA_seed | PIK3CA | 0.51515 | BRCA | DU-4475 | PI3Ka_4409 |
| MVP_top60_DNA_seed | HDAC1 | -0.436315 | LUSC | SK-MES-1 | JQ12 | top60_altered_DNA+RNA_seed | PIK3CA | 0.551516 | BRCA | HCC1419 | AZD8835 |
| MVP_top60_DNA_seed | HDAC2 | -0.436315 | LUSC | SK-MES-1 | JQ12 | top60_altered_DNA+RNA_seed | PLK1 | 0.560414 | BRCA | MDA-MB-157 | PLK_6522 |
| MVP_top60_DNA_seed | HDAC1 | -0.425508 | LUSC | LC-1-sq | Dacinostat | top60_altered_DNA+RNA_seed | PLK1 | 0.582692 | BRCA | DU-4475 | PLK_6522 |
| MVP_top60_DNA_seed | HDAC1 | -0.416553 | LUSC | NCI-H226 | Belinostat | top60_altered_DNA+RNA_seed | PIK3CA | 0.586171 | BRCA | CAL-85-1 | AZD8835 |
| MVP_top60_DNA_seed | EGFR | -0.408762 | LUSC | NCI-H520 | Pelitinib | top60_altered_DNA+RNA_seed | PIK3CA | 0.605227 | BRCA | HDQ-P1 | AZD8835 |
| MVP_top60_DNA_seed | HDAC1 | -0.406793 | LUSC | NCI-H226 | CAY10603 | top60_altered_DNA+RNA_seed | PIK3CA | 0.650719 | BRCA | HCC70 | AZD8835 |
| MVP_top60_DNA_seed | EGFR | -0.395706 | LUSC | EPLC-272H | PF-00299804 | top60_altered_DNA+RNA_seed | PIK3CA | 0.674489 | BRCA | BT-549 | AZD8835 |
| MVP_top60_DNA_seed | EGFR | -0.380808 | LUSC | NCI-H226 | Cetuximab | top60_altered_DNA+RNA_seed | PIK3CA | 0.676875 | BRCA | HCC1419 | PI3Ka_4409 |
| MVP_top60_DNA_seed | ESR1 | -0.372579 | LUSC | HCC-15 | Tamoxifen | top60_altered_DNA+RNA_seed | PIK3CA | 0.754134 | BRCA | MDA-MB-468 | PI3Ka_4409 |
| MVP_top60_DNA_seed | EGFR | -0.365694 | LUSC | NCI-H1869 | Gefitinib | top60_altered_DNA+RNA_seed | PLK1 | 0.771832 | BRCA | BT-474 | BI-2536 |
| MVP_top60_DNA_seed | AR | -0.359332 | LUSC | LC-1-sq | AZD3514 | top60_altered_DNA+RNA_seed | PLK1 | 0.782784 | BRCA | HCC2218 | GW843682X |
| MVP_top60_DNA_seed | HDAC1 | -0.353983 | LUSC | NCI-H520 | Dacinostat | top60_altered_DNA+RNA_seed | PLK1 | 0.818127 | BRCA | HCC1569 | PLK_6522 |
| MVP_top60_DNA_seed | HDAC1 | -0.344443 | LUSC | NCI-H226 | AR-42 | top60_altered_DNA+RNA_seed | PLK1 | 0.87786 | BRCA | HCC1937 | PLK_6522 |
| MVP_top60_DNA_seed | HDAC1 | -0.341282 | LUSC | NCI-H1869 | Parthenolide | top60_altered_DNA+RNA_seed | PLK1 | 0.917323 | BRCA | HCC2218 | BI-2536 |
| MVP_top60_DNA_seed | EGFR | -0.33099 | LUSC | HARA | Cetuximab | top60_altered_DNA+RNA_seed | PIK3CA | 0.936188 | BRCA | MDA-MB-415 | AZD8835 |
| MVP_top60_DNA_seed | HDAC1 | -0.329223 | LUSC | LK-2 | CAY10603 | top60_altered_DNA+RNA_seed | PLK1 | 0.970128 | BRCA | HDQ-P1 | PLK_6522 |
| MVP_top60_DNA_seed | EGFR | -0.328744 | LUSC | EBC-1 | Afatinib | top60_altered_DNA+RNA_seed | PIK3CA | 0.972751 | BRCA | Hs-578-T | AZD8835 |
| MVP_top60_DNA_seed | HDAC1 | -0.326793 | LUSC | NCI-H2170 | Dacinostat | top60_altered_DNA+RNA_seed | PIK3CA | 0.987493 | BRCA | HCC1395 | AZD8835 |
| MVP_top60_DNA_seed | AR | -0.325215 | LUSC | KNS-62 | Bicalutamide | top60_altered_DNA+RNA_seed | PIK3CA | 0.997241 | BRCA | ZR-75-30 | AZD8835 |
| MVP_top60_DNA_seed | EGFR | -0.319172 | LUSC | SK-MES-1 | Gefitinib | top60_altered_DNA+RNA_seed | PIK3CA | 1.012949 | BRCA | HCC38 | PI3Ka_4409 |
| MVP_top60_DNA_seed | GSK3B | -0.319026 | LUSC | NCI-H226 | AZD7969 | top60_altered_DNA+RNA_seed | PLK1 | 1.026712 | BRCA | HCC2157 | BI-2536 |
| MVP_top60_DNA_seed | HDAC1 | -0.301117 | LUSC | LK-2 | AR-42 | top60_altered_DNA+RNA_seed | PIK3CA | 1.082358 | BRCA | HCC38 | AZD8835 |
| MVP_top60_DNA_seed | GSK3B | -0.300652 | LUSC | HARA | AZD7969 | top60_altered_DNA+RNA_seed | PLK1 | 1.083181 | BRCA | COLO-824 | GW843682X |

| | | | | | | | | | | | |
|---|---|---|---|---|---|---|---|---|---|---|---|
| MVP_top60_DNA_seed | SMARCA4 | -0.278295 | LUSC | HCC-15 | PFI-3 | top60_altered_DNA+RNA_seed | PIK3CA | 1.15106 | BRCA | BT-483 | AZD8835 |
| MVP_top60_DNA_seed | SMARCA2 | -0.278295 | LUSC | HCC-15 | PFI-3 | top60_altered_DNA+RNA_seed | PIK3CA | 1.15663 | BRCA | MDA-MB-468 | AZD8835 |
| MVP_top60_DNA_seed | HDAC1 | -0.276912 | LUSC | SK-MES-1 | Dacinostat | top60_altered_DNA+RNA_seed | PLK1 | 1.158757 | BRCA | MDA-MB-415 | PLK_6522 |
| MVP_top60_DNA_seed | HDAC1 | -0.268619 | LUSC | NCI-H520 | Belinostat | top60_altered_DNA+RNA_seed | PLK1 | 1.172116 | BRCA | COLO-824 | BI-2536 |
| MVP_top60_DNA_seed | RAC1 | -0.265676 | LUSC | HCC-15 | EHT-1864 | top60_altered_DNA+RNA_seed | PIK3CA | 1.175306 | BRCA | CAL-51 | AZD8835 |
| MVP_top60_DNA_seed | CREBBP | -0.252649 | LUSC | EBC-1 | I-CBP112 | top60_altered_DNA+RNA_seed | PIK3CA | 1.217205 | BRCA | HCC1937 | PI3Ka_4409 |
| MVP_top60_DNA_seed | PIK3CA | -0.250907 | LUSC | LK-2 | AZD8835 | top60_altered_DNA+RNA_seed | PLK1 | 1.227898 | BRCA | MFM-223 | GW843682X |
| MVP_top60_DNA_seed | PIK3CA | -0.247775 | LUSC | HCC-15 | PI3Ka_4409 | top60_altered_DNA+RNA_seed | PIK3CA | 1.260121 | BRCA | MDA-MB-231 | PI3Ka_4409 |
| MVP_top60_DNA_seed | EGFR | -0.239382 | LUSC | NCI-H1869 | Afatinib | top60_altered_DNA+RNA_seed | PLK1 | 1.309873 | BRCA | CAL-85-1 | PLK_6522 |
| MVP_top60_DNA_seed | GSK3B | -0.238382 | LUSC | NCI-H1869 | CHIR-99021 | top60_altered_DNA+RNA_seed | PIK3CA | 1.317085 | BRCA | MDA-MB-157 | AZD8835 |
| MVP_top60_DNA_seed | ESR1 | -0.235672 | LUSC | EBC-1 | Fulvestrant | top60_altered_DNA+RNA_seed | PIK3CA | 1.320549 | BRCA | HCC1428 | AZD8835 |
| MVP_top60_DNA_seed | PIK3CA | -0.233333 | LUSC | NCI-H226 | AZD8835 | top60_altered_DNA+RNA_seed | PLK1 | 1.340377 | BRCA | MFM-223 | BI-2536 |
| MVP_top60_DNA_seed | GSK3B | -0.232769 | LUSC | NCI-H2170 | KIN001-042 | top60_altered_DNA+RNA_seed | PIK3CA | 1.344327 | BRCA | MDA-MB-436 | AZD8835 |
| MVP_top60_DNA_seed | EGFR | -0.229857 | LUSC | LK-2 | AZD8931 | top60_altered_DNA+RNA_seed | PLK1 | 1.357253 | BRCA | BT-474 | GW843682X |
| MVP_top60_DNA_seed | EGFR | -0.222436 | LUSC | LK-2 | CUDC-101 | top60_altered_DNA+RNA_seed | PIK3CA | 1.386546 | BRCA | HCC1428 | PI3Ka_4409 |
| MVP_top60_DNA_seed | PIK3CA | -0.221388 | LUSC | NCI-H1869 | PI3Ka_4409 | top60_altered_DNA+RNA_seed | PLK1 | 1.415635 | BRCA | HCC2157 | GW843682X |
| MVP_top60_DNA_seed | GSK3B | -0.217893 | LUSC | EPLC-272H | AZD7969 | top60_altered_DNA+RNA_seed | PIK3CA | 1.450804 | BRCA | ZR-75-30 | PI3Ka_4409 |
| MVP_top60_DNA_seed | PIK3CA | -0.202814 | LUSC | HARA | AZD8835 | top60_altered_DNA+RNA_seed | PLK1 | 1.465972 | BRCA | HCC1599 | PLK_6522 |
| MVP_top60_DNA_seed | HDAC1 | -0.190535 | LUSC | LC-1-sq | Belinostat | top60_altered_DNA+RNA_seed | PIK3CA | 1.479792 | BRCA | HCC1937 | AZD8835 |
| MVP_top60_DNA_seed | HDAC1 | -0.180764 | LUSC | HARA | Tubastatin A | top60_altered_DNA+RNA_seed | PLK1 | 1.482361 | BRCA | CAMA-1 | PLK_6522 |
| MVP_top60_DNA_seed | HDAC1 | -0.174574 | LUSC | LC-1-sq | AR-42 | top60_altered_DNA+RNA_seed | PIK3CA | 1.482499 | BRCA | MDA-MB-157 | PI3Ka_4409 |
| MVP_top60_DNA_seed | EGFR | -0.159153 | LUSC | HARA | CUDC-101 | top60_altered_DNA+RNA_seed | PLK1 | 1.575889 | BRCA | HCC1428 | PLK_6522 |
| MVP_top60_DNA_seed | EGFR | -0.159054 | LUSC | NCI-H1869 | Afatinib | top60_altered_DNA+RNA_seed | PIK3CA | 1.682452 | BRCA | HCC1500 | AZD8835 |
| MVP_top60_DNA_seed | PIK3CA | -0.155866 | LUSC | HARA | PI3Ka_4409 | top60_altered_DNA+RNA_seed | PLK1 | 1.72722 | BRCA | MDA-MB-330 | PLK_6522 |
| MVP_top60_DNA_seed | GSK3B | -0.150141 | LUSC | NCI-H2170 | AZD7969 | top60_altered_DNA+RNA_seed | PLK1 | 1.753902 | BRCA | T47D | PLK_6522 |
| MVP_top60_DNA_seed | GSK3B | -0.142127 | LUSC | SK-MES-1 | CHIR-99021 | top60_altered_DNA+RNA_seed | PLK1 | 1.827528 | BRCA | HCC202 | PLK_6522 |
| MVP_top60_DNA_seed | AR | -0.139412 | LUSC | HCC-15 | AZD3514 | top60_altered_DNA+RNA_seed | PIK3CA | 1.886279 | BRCA | MRK-nu-1 | PI3Ka_4409 |
| MVP_top60_DNA_seed | GSK3B | -0.124244 | LUSC | LOU-NH91 | KIN001-042 | top60_altered_DNA+RNA_seed | PIK3CA | 1.903586 | BRCA | MRK-nu-1 | AZD8835 |
| MVP_top60_DNA_seed | EGFR | -0.123208 | LUSC | SK-MES-1 | CUDC-101 | top60_altered_DNA+RNA_seed | PLK1 | 1.919401 | BRCA | UACC-893 | PLK_6522 |
| MVP_top60_DNA_seed | EGFR | -0.118687 | LUSC | LC-1-sq | Cetuximab | top60_altered_DNA+RNA_seed | PIK3CA | 1.962095 | BRCA | HCC1500 | PI3Ka_4409 |
| MVP_top60_DNA_seed | HDAC1 | -0.117491 | LUSC | SK-MES-1 | Belinostat | top60_altered_DNA+RNA_seed | PLK1 | 2.016825 | BRCA | HCC1500 | PLK_6522 |
| MVP_top60_DNA_seed | GSK3B | -0.11046 | LUSC | LOU-NH91 | CHIR-99021 | top60_altered_DNA+RNA_seed | PIK3CA | 2.047064 | BRCA | HCC1395 | PI3Ka_4409 |
| MVP_top60_DNA_seed | AR | -0.105629 | LUSC | LK-2 | Bicalutamide | top60_altered_DNA+RNA_seed | PIK3CA | 2.180415 | BRCA | CAMA-1 | AZD8835 |
| MVP_top60_DNA_seed | EGFR | -0.102951 | LUSC | LC-1-sq | Afatinib | top60_altered_DNA+RNA_seed | PIK3CA | 2.290532 | BRCA | CAMA-1 | PI3Ka_4409 |
| MVP_top60_DNA_seed | EGFR | -0.095902 | LUSC | LOU-NH91 | CUDC-101 | top60_altered_DNA+RNA_seed | PLK1 | 2.296477 | BRCA | HCC70 | PLK_6522 |
| MVP_top60_DNA_seed | GSK3B | -0.093246 | LUSC | EBC-1 | AZD7969 | top60_altered_DNA+RNA_seed | PIK3CA | 2.332789 | BRCA | MDA-MB-436 | PI3Ka_4409 |
| MVP_top60_DNA_seed | HDAC1 | -0.091381 | LUSC | LC-1-sq | CAY10603 | top60_altered_DNA+RNA_seed | PLK1 | 2.396567 | BRCA | HCC2218 | PLK_6522 |
| MVP_top60_DNA_seed | HDAC1 | -0.087645 | LUSC | NCI-H520 | AR-42 | top60_altered_DNA+RNA_seed | PLK1 | 2.488824 | BRCA | HCC1419 | PLK_6522 |
| MVP_top60_DNA_seed | HDAC1 | -0.070004 | LUSC | NCI-H226 | Dacinostat | top60_altered_DNA+RNA_seed | PLK1 | 2.498751 | BRCA | BT-474 | PLK_6522 |
| MVP_top60_DNA_seed | HDAC1 | -0.066763 | LUSC | SK-MES-1 | CAY10603 | top60_altered_DNA+RNA_seed | PLK1 | 2.618082 | BRCA | MDA-MB-175-VII | PLK_6522 |
| MVP_top60_DNA_seed | EGFR | -0.060777 | LUSC | KNS-62 | Gefitinib | top60_altered_DNA+RNA_seed | PLK1 | 2.642613 | BRCA | ZR-75-30 | PLK_6522 |

| | | | | | | | | | | | |
|---|---|---|---|---|---|---|---|---|---|---|---|
| MVP_top60_DNA_seed | EGFR | -0.05582 | LUSC | NCI-H226 | PF-00299804 | top60_altered_DNA+RNA_seed | PLK1 | 2.659119 | BRCA | BT-483 | PLK_6522 |
| MVP_top60_DNA_seed | AR | -0.053071 | LUSC | RERF-LC-Sq1 | AZD3514 | top60_altered_DNA+RNA_seed | PIK3CA | -4.087508 | BRCA | YMB-1-E | AZD8186 |
| MVP_top60_DNA_seed | CREBBP | -0.045072 | LUSC | SK-MES-1 | I-CBP112 | top60_altered_DNA+RNA_seed | PIK3CA | -3.846124 | BRCA | EVSA-T | AZD8186 |
| MVP_top60_DNA_seed | EGFR | -0.043968 | LUSC | NCI-H226 | CUDC-101 | top60_altered_DNA+RNA_seed | PIK3CA | -3.378294 | BRCA | MDA-MB-415 | AZD8186 |
| MVP_top60_DNA_seed | AR | -0.038137 | LUSC | HCC-15 | Bicalutamide | top60_altered_DNA+RNA_seed | PIK3CA | -2.478395 | BRCA | MDA-MB-361 | Alpelisib |
| MVP_top60_DNA_seed | CREBBP | -0.033738 | LUSC | LK-2 | I-CBP112 | top60_altered_DNA+RNA_seed | PIK3CA | -2.467028 | BRCA | MDA-MB-330 | Alpelisib |
| MVP_top60_DNA_seed | PIK3CA | -0.032854 | LUSC | NCI-H2170 | PI3Ka_4409 | top60_altered_DNA+RNA_seed | PIK3CA | -2.349952 | BRCA | OCUB-M | Alpelisib |
| MVP_top60_DNA_seed | EGFR | -0.025233 | LUSC | HCC-15 | Afatinib | top60_altered_DNA+RNA_seed | TP53 | -2.331311 | BRCA | HCC1599 | MIRA-1 |
| MVP_top60_DNA_seed | EGFR | -0.020839 | LUSC | NCI-H520 | PF-00299804 | top60_altered_DNA+RNA_seed | PIK3CA | -2.200339 | BRCA | UACC-893 | Alpelisib |
| MVP_top60_DNA_seed | HDAC1 | -0.014245 | LUSC | LOU-NH91 | Belinostat | top60_altered_DNA+RNA_seed | PIK3CA | -2.198675 | BRCA | MDA-MB-453 | Alpelisib |
| MVP_top60_DNA_seed | HDAC1 | -0.008273 | LUSC | NCI-H520 | JQ12 | top60_altered_DNA+RNA_seed | PIK3CA | -2.109527 | BRCA | HCC70 | AZD8186 |
| MVP_top60_DNA_seed | HDAC2 | -0.008273 | LUSC | NCI-H520 | JQ12 | top60_altered_DNA+RNA_seed | TP53 | -2.017852 | BRCA | MRK-nu-1 | MIRA-1 |
| MVP_top60_DNA_seed | HDAC1 | -0.005246 | LUSC | HARA | Belinostat | top60_altered_DNA+RNA_seed | PIK3CA | -1.979001 | BRCA | HCC2157 | AZD8186 |
| MVP_top60_DNA_seed | RAC1 | 0.005383 | LUSC | RERF-LC-Sq1 | EHT-1864 | top60_altered_DNA+RNA_seed | PIK3CA | -1.88702 | BRCA | MDA-MB-330 | AZD8186 |
| MVP_top60_DNA_seed | EGFR | 0.012334 | LUSC | NCI-H2170 | Pelitinib | top60_altered_DNA+RNA_seed | PIK3CA | -1.882454 | BRCA | CAMA-1 | AZD8186 |
| MVP_top60_DNA_seed | AR | 0.014828 | LUSC | LK-2 | Bicalutamide | top60_altered_DNA+RNA_seed | PIK3CA | -1.729419 | BRCA | CAL-148 | Alpelisib |
| MVP_top60_DNA_seed | CREBBP | 0.016885 | LUSC | EPLC-272H | I-CBP112 | top60_altered_DNA+RNA_seed | PLK1 | -1.686304 | BRCA | MRK-nu-1 | BI-2536 |
| MVP_top60_DNA_seed | EGFR | 0.018382 | LUSC | NCI-H226 | Pelitinib | top60_altered_DNA+RNA_seed | PIK3CA | -1.675579 | BRCA | BT-20 | Alpelisib |
| MVP_top60_DNA_seed | HDAC1 | 0.019635 | LUSC | EBC-1 | Belinostat | top60_altered_DNA+RNA_seed | PIK3CA | -1.673389 | BRCA | OCUB-M | GNE-317 |
| MVP_top60_DNA_seed | HDAC1 | 0.021002 | LUSC | SK-MES-1 | AR-42 | top60_altered_DNA+RNA_seed | PIK3CA | -1.607008 | BRCA | EFM-19 | Alpelisib |
| MVP_top60_DNA_seed | EGFR | 0.022852 | LUSC | NCI-H226 | Gefitinib | top60_altered_DNA+RNA_seed | PIK3CA | -1.544994 | BRCA | MRK-nu-1 | GNE-317 |
| MVP_top60_DNA_seed | HDAC1 | 0.029762 | LUSC | LOU-NH91 | CAY10603 | top60_altered_DNA+RNA_seed | PIK3CA | -1.452375 | BRCA | MDA-MB-415 | GNE-317 |
| MVP_top60_DNA_seed | AR | 0.034641 | LUSC | NCI-H226 | Bicalutamide | top60_altered_DNA+RNA_seed | PIK3CA | -1.424528 | BRCA | HCC2157 | GNE-317 |
| MVP_top60_DNA_seed | RAC1 | 0.035068 | LUSC | NCI-H520 | EHT-1864 | top60_altered_DNA+RNA_seed | PIK3CA | -1.401092 | BRCA | HCC1954 | Alpelisib |
| MVP_top60_DNA_seed | PIK3CA | 0.041565 | LUSC | LOU-NH91 | PI3Ka_4409 | top60_altered_DNA+RNA_seed | PIK3CA | -1.397942 | BRCA | MDA-MB-175-VII | GNE-317 |
| MVP_top60_DNA_seed | PIK3CA | 0.050047 | LUSC | HCC-15 | AZD8835 | top60_altered_DNA+RNA_seed | PIK3CA | -1.37047 | BRCA | MDA-MB-361 | GNE-317 |
| MVP_top60_DNA_seed | HDAC1 | 0.051833 | LUSC | LOU-NH91 | JQ12 | top60_altered_DNA+RNA_seed | PIK3CA | -1.319474 | BRCA | EVSA-T | GNE-317 |
| MVP_top60_DNA_seed | HDAC2 | 0.051833 | LUSC | LOU-NH91 | JQ12 | top60_altered_DNA+RNA_seed | PLK1 | -1.306051 | BRCA | Hs-578-T | BI-2536 |
| MVP_top60_DNA_seed | GSK3B | 0.060861 | LUSC | NCI-H1869 | CHIR-99021 | top60_altered_DNA+RNA_seed | PIK3CA | -1.305722 | BRCA | T47D | Alpelisib |
| MVP_top60_DNA_seed | EGFR | 0.061958 | LUSC | LK-2 | PF-00299804 | top60_altered_DNA+RNA_seed | PIK3CA | -1.30157 | BRCA | HDQ-P1 | GNE-317 |
| MVP_top60_DNA_seed | EGFR | 0.068269 | LUSC | LOU-NH91 | AST-1306 | top60_altered_DNA+RNA_seed | PLK1 | -1.299101 | BRCA | OCUB-M | BI-2536 |
| MVP_top60_DNA_seed | HDAC1 | 0.07122 | LUSC | HARA | AR-42 | top60_altered_DNA+RNA_seed | PIK3CA | -1.298818 | BRCA | HCC2157 | Buparlisib |
| MVP_top60_DNA_seed | RAC1 | 0.073751 | LUSC | SK-MES-1 | EHT-1864 | top60_altered_DNA+RNA_seed | PIK3CA | -1.282233 | BRCA | MDA-MB-330 | GNE-317 |
| MVP_top60_DNA_seed | EGFR | 0.074059 | LUSC | NCI-H226 | Lapatinib | top60_altered_DNA+RNA_seed | PIK3CA | -1.243567 | BRCA | HDQ-P1 | AZD8186 |
| MVP_top60_DNA_seed | RAC1 | 0.074877 | LUSC | EBC-1 | EHT-1864 | top60_altered_DNA+RNA_seed | PIK3CA | -1.240029 | BRCA | EFM-19 | GNE-317 |
| MVP_top60_DNA_seed | EGFR | 0.079498 | LUSC | LK-2 | AST-1306 | top60_altered_DNA+RNA_seed | PLK1 | -1.238342 | BRCA | HCC1599 | BI-2536 |
| MVP_top60_DNA_seed | CREBBP | 0.081399 | LUSC | HCC-15 | I-CBP112 | top60_altered_DNA+RNA_seed | PIK3CA | -1.236731 | BRCA | CAL-51 | GNE-317 |
| MVP_top60_DNA_seed | AR | 0.095524 | LUSC | NCI-H520 | Bicalutamide | top60_altered_DNA+RNA_seed | PIK3CA | -1.223717 | BRCA | EVSA-T | Buparlisib |
| MVP_top60_DNA_seed | AR | 0.098282 | LUSC | SK-MES-1 | AZD3514 | top60_altered_DNA+RNA_seed | PIK3CA | -1.217177 | BRCA | BT-474 | Alpelisib |
| MVP_top60_DNA_seed | EGFR | 0.109192 | LUSC | RERF-LC-Sq1 | AST-1306 | top60_altered_DNA+RNA_seed | PIK3CA | -1.207284 | BRCA | UACC-893 | GNE-317 |
| MVP_top60_DNA_seed | PIK3CA | 0.109597 | LUSC | LC-1-sq | AZD8835 | top60_altered_DNA+RNA_seed | TP53 | -1.185478 | BRCA | HCC2157 | MIRA-1 |

| | | | | | | | | | | | |
|---|---|---|---|---|---|---|---|---|---|---|---|
| MVP_top60_DNA_seed | HDAC1 | 0.117185 | LUSC | NCI-H2170 | CAY10603 | top60_altered_DNA+RNA_seed | TP53 | -1.155661 | BRCA | HCC1419 | MIRA-1 |
| MVP_top60_DNA_seed | HDAC1 | 0.126375 | LUSC | HARA | CAY10603 | top60_altered_DNA+RNA_seed | PIK3CA | -1.128218 | BRCA | MDA-MB-175-VII | Alpelisib |
| MVP_top60_DNA_seed | ESR1 | 0.128759 | LUSC | NCI-H2170 | Fulvestrant | top60_altered_DNA+RNA_seed | PIK3CA | -1.098943 | BRCA | CAL-148 | GNE-317 |
| MVP_top60_DNA_seed | EGFR | 0.132605 | LUSC | NCI-H520 | Cetuximab | top60_altered_DNA+RNA_seed | PIK3CA | -1.078618 | BRCA | HCC1395 | AZD8186 |
| MVP_top60_DNA_seed | HDAC1 | 0.139997 | LUSC | EBC-1 | AR-42 | top60_altered_DNA+RNA_seed | PIK3CA | -1.076969 | BRCA | CAMA-1 | GNE-317 |
| MVP_top60_DNA_seed | AR | 0.148153 | LUSC | HARA | Bicalutamide | top60_altered_DNA+RNA_seed | PIK3CA | -1.074375 | BRCA | T47D | GNE-317 |
| MVP_top60_DNA_seed | GSK3B | 0.151437 | LUSC | NCI-H520 | KIN001-042 | top60_altered_DNA+RNA_seed | PIK3CA | -1.073624 | BRCA | HCC1599 | Alpelisib |
| MVP_top60_DNA_seed | HDAC1 | 0.160137 | LUSC | LOU-NH91 | AR-42 | top60_altered_DNA+RNA_seed | PLK1 | -1.068707 | BRCA | HCC2157 | BI-2536 |
| MVP_top60_DNA_seed | EGFR | 0.161354 | LUSC | HARA | PF-00299804 | top60_altered_DNA+RNA_seed | PIK3CA | -1.026325 | BRCA | CAL-51 | Buparlisib |
| MVP_top60_DNA_seed | ESR1 | 0.165591 | LUSC | RERF-LC-Sq1 | Tamoxifen | top60_altered_DNA+RNA_seed | TP53 | -1.02495 | BRCA | OCUB-M | MIRA-1 |
| MVP_top60_DNA_seed | ESR1 | 0.166707 | LUSC | NCI-H1869 | Tamoxifen | top60_altered_DNA+RNA_seed | PIK3CA | -1.022631 | BRCA | EFM-19 | AZD8186 |
| MVP_top60_DNA_seed | EGFR | 0.181102 | LUSC | EBC-1 | AZD8931 | top60_altered_DNA+RNA_seed | PIK3CA | -1.019348 | BRCA | MDA-MB-361 | Buparlisib |
| MVP_top60_DNA_seed | AR | 0.186477 | LUSC | EPLC-272H | Bicalutamide | top60_altered_DNA+RNA_seed | PIK3CA | -1.018997 | BRCA | MDA-MB-175-VII | Buparlisib |
| MVP_top60_DNA_seed | HDAC1 | 0.189457 | LUSC | NCI-H1869 | Entinostat | top60_altered_DNA+RNA_seed | PIK3CA | -1.016139 | BRCA | HCC1569 | Alpelisib |
| MVP_top60_DNA_seed | HDAC3 | 0.189457 | LUSC | NCI-H1869 | Entinostat | top60_altered_DNA+RNA_seed | PLK1 | -0.952557 | BRCA | CAL-51 | BI-2536 |
| MVP_top60_DNA_seed | ESR1 | 0.192421 | LUSC | HCC-15 | Fulvestrant | top60_altered_DNA+RNA_seed | PLK1 | -0.935033 | BRCA | COLO-824 | BI-2536 |
| MVP_top60_DNA_seed | GSK3B | 0.197491 | LUSC | NCI-H2170 | CHIR-99021 | top60_altered_DNA+RNA_seed | PIK3CA | -0.932406 | BRCA | HDQ-P1 | Buparlisib |
| MVP_top60_DNA_seed | SMARCA4 | 0.201605 | LUSC | LK-2 | PFI-3 | top60_altered_DNA+RNA_seed | PIK3CA | -0.931641 | BRCA | AU565 | Alpelisib |
| MVP_top60_DNA_seed | SMARCA2 | 0.201605 | LUSC | LK-2 | PFI-3 | top60_altered_DNA+RNA_seed | PIK3CA | -0.927964 | BRCA | T47D | AZD8186 |
| MVP_top60_DNA_seed | HDAC1 | 0.204347 | LUSC | LC-1-sq | Tubastatin A | top60_altered_DNA+RNA_seed | TP53 | -0.920382 | BRCA | MDA-MB-415 | MIRA-1 |
| MVP_top60_DNA_seed | GSK3B | 0.205212 | LUSC | EPLC-272H | CHIR-99021 | top60_altered_DNA+RNA_seed | PIK3CA | -0.917627 | BRCA | CAL-120 | GNE-317 |
| MVP_top60_DNA_seed | HDAC1 | 0.208178 | LUSC | LK-2 | Tubastatin A | top60_altered_DNA+RNA_seed | PIK3CA | -0.892798 | BRCA | T47D | Buparlisib |
| MVP_top60_DNA_seed | SMARCA4 | 0.222003 | LUSC | SK-MES-1 | PFI-3 | top60_altered_DNA+RNA_seed | PIK3CA | -0.882716 | BRCA | AU565 | AZD8186 |
| MVP_top60_DNA_seed | SMARCA2 | 0.222003 | LUSC | SK-MES-1 | PFI-3 | top60_altered_DNA+RNA_seed | PIK3CA | -0.880401 | BRCA | Hs-578-T | AZD8186 |
| MVP_top60_DNA_seed | EGFR | 0.227284 | LUSC | LK-2 | Afatinib | top60_altered_DNA+RNA_seed | PIK3CA | -0.853556 | BRCA | MCF7 | Alpelisib |
| MVP_top60_DNA_seed | HDAC1 | 0.239963 | LUSC | EBC-1 | CAY10603 | top60_altered_DNA+RNA_seed | PIK3CA | -0.850261 | BRCA | EFM-192A | Alpelisib |
| MVP_top60_DNA_seed | GSK3B | 0.244941 | LUSC | EBC-1 | KIN001-042 | top60_altered_DNA+RNA_seed | PIK3CA | -0.840747 | BRCA | MDA-MB-361 | AZD8186 |
| MVP_top60_DNA_seed | EGFR | 0.251709 | LUSC | HCC-15 | Gefitinib | top60_altered_DNA+RNA_seed | PIK3CA | -0.819784 | BRCA | COLO-824 | AZD8186 |
| MVP_top60_DNA_seed | EGFR | 0.261545 | LUSC | NCI-H520 | AZD8931 | top60_altered_DNA+RNA_seed | PIK3CA | -0.809161 | BRCA | MRK-nu-1 | Buparlisib |
| MVP_top60_DNA_seed | AR | 0.263244 | LUSC | SK-MES-1 | Bicalutamide | top60_altered_DNA+RNA_seed | PLK1 | -0.780478 | BRCA | EVSA-T | BI-2536 |
| MVP_top60_DNA_seed | EGFR | 0.270292 | LUSC | EBC-1 | CUDC-101 | top60_altered_DNA+RNA_seed | PIK3CA | -0.772239 | BRCA | OCUB-M | Buparlisib |
| MVP_top60_DNA_seed | GSK3B | 0.27056 | LUSC | NCI-H226 | KIN001-042 | top60_altered_DNA+RNA_seed | PIK3CA | -0.745503 | BRCA | HCC1143 | AZD8186 |
| MVP_top60_DNA_seed | HDAC1 | 0.273072 | LUSC | RERF-LC-Sq1 | Belinostat | top60_altered_DNA+RNA_seed | PIK3CA | -0.743736 | BRCA | Hs-578-T | Buparlisib |
| MVP_top60_DNA_seed | EGFR | 0.280073 | LUSC | LOU-NH91 | CI-1033 | top60_altered_DNA+RNA_seed | PIK3CA | -0.741553 | BRCA | MDA-MB-468 | AZD8186 |
| MVP_top60_DNA_seed | HDAC1 | 0.285802 | LUSC | NCI-H1869 | AR-42 | top60_altered_DNA+RNA_seed | PIK3CA | -0.73116 | BRCA | JIMT-1 | GNE-317 |
| MVP_top60_DNA_seed | HDAC1 | 0.28665 | LUSC | NCI-H2170 | Belinostat | top60_altered_DNA+RNA_seed | PIK3CA | -0.730409 | BRCA | Hs-578-T | Alpelisib |
| MVP_top60_DNA_seed | ESR1 | 0.28685 | LUSC | NCI-H226 | Fulvestrant | top60_altered_DNA+RNA_seed | PLK1 | -0.711158 | BRCA | HCC1599 | MK-1775 |
| MVP_top60_DNA_seed | HDAC1 | 0.288484 | LUSC | SK-MES-1 | Tubastatin A | top60_altered_DNA+RNA_seed | PIK3CA | -0.704781 | BRCA | CAL-148 | AZD8186 |
| MVP_top60_DNA_seed | PIK3CA | 0.293316 | LUSC | NCI-H2170 | AZD8835 | top60_altered_DNA+RNA_seed | PIK3CA | -0.702855 | BRCA | HCC1395 | Buparlisib |
| MVP_top60_DNA_seed | AR | 0.305061 | LUSC | NCI-H2170 | AZD3514 | top60_altered_DNA+RNA_seed | PLK1 | -0.700967 | BRCA | JIMT-1 | BI-2536 |
| MVP_top60_DNA_seed | AR | 0.305101 | LUSC | LOU-NH91 | Bicalutamide | top60_altered_DNA+RNA_seed | PLK1 | -0.699233 | BRCA | DU-4475 | BI-2536 |

| | | | | | | | | | | | |
|---|---|---|---|---|---|---|---|---|---|---|---|
| MVP_top60_DNA_seed | AR | 0.310686 | LUSC | NCI-H1869 | AZD3514 | top60_altered_DNA+RNA_seed | PIK3CA | -0.691065 | BRCA | EFM-192A | GNE-317 |
| MVP_top60_DNA_seed | EGFR | 0.312275 | LUSC | NCI-H520 | CI-1033 | top60_altered_DNA+RNA_seed | PIK3CA | -0.67704 | BRCA | UACC-812 | AZD8186 |
| MVP_top60_DNA_seed | EGFR | 0.31889 | LUSC | HCC-15 | AZD8931 | top60_altered_DNA+RNA_seed | PIK3CA | -0.660445 | BRCA | EFM-192A | Buparlisib |
| MVP_top60_DNA_seed | EGFR | 0.319449 | LUSC | NCI-H226 | AST-1306 | top60_altered_DNA+RNA_seed | PIK3CA | -0.653864 | BRCA | AU565 | GNE-317 |
| MVP_top60_DNA_seed | PIK3CA | 0.321961 | LUSC | LC-1-sq | PI3Ka_4409 | top60_altered_DNA+RNA_seed | PIK3CA | -0.652526 | BRCA | MDA-MB-330 | Buparlisib |
| MVP_top60_DNA_seed | EGFR | 0.325973 | LUSC | KNS-62 | Afatinib | top60_altered_DNA+RNA_seed | PIK3CA | -0.652473 | BRCA | UACC-812 | Alpelisib |
| MVP_top60_DNA_seed | AR | 0.33193 | LUSC | NCI-H1869 | Bicalutamide | top60_altered_DNA+RNA_seed | TP53 | -0.647937 | BRCA | EVSA-T | MIRA-1 |
| MVP_top60_DNA_seed | AR | 0.33334 | LUSC | HARA | AZD3514 | top60_altered_DNA+RNA_seed | PIK3CA | -0.607739 | BRCA | Hs-578-T | GNE-317 |
| MVP_top60_DNA_seed | ESR1 | 0.334162 | LUSC | NCI-H2170 | Tamoxifen | top60_altered_DNA+RNA_seed | PIK3CA | -0.601721 | BRCA | COLO-824 | Alpelisib |
| MVP_top60_DNA_seed | HDAC1 | 0.337752 | LUSC | NCI-H226 | JQ12 | top60_altered_DNA+RNA_seed | PIK3CA | -0.596498 | BRCA | DU-4475 | Buparlisib |
| MVP_top60_DNA_seed | HDAC2 | 0.337752 | LUSC | NCI-H226 | JQ12 | top60_altered_DNA+RNA_seed | PIK3CA | -0.595752 | BRCA | HCC1143 | Buparlisib |
| MVP_top60_DNA_seed | PIK3CA | 0.33785 | LUSC | LK-2 | PI3Ka_4409 | top60_altered_DNA+RNA_seed | PIK3CA | -0.592111 | BRCA | MDA-MB-175-VII | AZD8186 |
| MVP_top60_DNA_seed | HDAC1 | 0.340754 | LUSC | EPLC-272H | Dacinostat | top60_altered_DNA+RNA_seed | PIK3CA | -0.57039 | BRCA | HCC1569 | AZD8186 |
| MVP_top60_DNA_seed | EGFR | 0.341694 | LUSC | RERF-LC-Sq1 | CUDC-101 | top60_altered_DNA+RNA_seed | PIK3CA | -0.568248 | BRCA | CAL-148 | Buparlisib |
| MVP_top60_DNA_seed | GSK3B | 0.342609 | LUSC | RERF-LC-Sq1 | KIN001-042 | top60_altered_DNA+RNA_seed | TP53 | -0.527752 | BRCA | HCC70 | MIRA-1 |
| MVP_top60_DNA_seed | RAC1 | 0.343428 | LUSC | EPLC-272H | EHT-1864 | top60_altered_DNA+RNA_seed | PLK1 | -0.525683 | BRCA | CAL-148 | MK-1775 |
| MVP_top60_DNA_seed | GSK3B | 0.344113 | LUSC | LC-1-sq | AZD7969 | top60_altered_DNA+RNA_seed | PLK1 | -0.524674 | BRCA | Hs-578-T | MK-1775 |
| MVP_top60_DNA_seed | EGFR | 0.370835 | LUSC | NCI-H2170 | CUDC-101 | top60_altered_DNA+RNA_seed | PIK3CA | -0.515105 | BRCA | AU565 | Buparlisib |
| MVP_top60_DNA_seed | HDAC1 | 0.376536 | LUSC | NCI-H226 | Tubastatin A | top60_altered_DNA+RNA_seed | PIK3CA | -0.514386 | BRCA | HCC1143 | GNE-317 |
| MVP_top60_DNA_seed | ESR1 | 0.377092 | LUSC | LC-1-sq | Fulvestrant | top60_altered_DNA+RNA_seed | PIK3CA | -0.503639 | BRCA | HCC70 | GNE-317 |
| MVP_top60_DNA_seed | EGFR | 0.383653 | LUSC | NCI-H520 | Afatinib | top60_altered_DNA+RNA_seed | PLK1 | -0.502995 | BRCA | DU-4475 | MK-1775 |
| MVP_top60_DNA_seed | EGFR | 0.386856 | LUSC | NCI-H226 | AZD8931 | top60_altered_DNA+RNA_seed | TP53 | -0.499832 | BRCA | CAL-148 | MIRA-1 |
| MVP_top60_DNA_seed | EGFR | 0.389426 | LUSC | NCI-H226 | Afatinib | top60_altered_DNA+RNA_seed | PIK3CA | -0.486361 | BRCA | HCC1954 | Buparlisib |
| MVP_top60_DNA_seed | SMARCA4 | 0.40521 | LUSC | NCI-H226 | PFI-3 | top60_altered_DNA+RNA_seed | PIK3CA | -0.464579 | BRCA | MDA-MB-231 | GNE-317 |
| MVP_top60_DNA_seed | SMARCA2 | 0.40521 | LUSC | NCI-H226 | PFI-3 | top60_altered_DNA+RNA_seed | TP53 | -0.454454 | BRCA | DU-4475 | MIRA-1 |
| MVP_top60_DNA_seed | SMARCA4 | 0.405648 | LUSC | NCI-H520 | PFI-3 | top60_altered_DNA+RNA_seed | PIK3CA | -0.441574 | BRCA | COLO-824 | Buparlisib |
| MVP_top60_DNA_seed | SMARCA2 | 0.405648 | LUSC | NCI-H520 | PFI-3 | top60_altered_DNA+RNA_seed | PIK3CA | -0.441497 | BRCA | HCC1143 | Alpelisib |
| MVP_top60_DNA_seed | ESR1 | 0.411571 | LUSC | SK-MES-1 | Tamoxifen | top60_altered_DNA+RNA_seed | PIK3CA | -0.438056 | BRCA | HCC1954 | GNE-317 |
| MVP_top60_DNA_seed | AR | 0.417828 | LUSC | LC-1-sq | Bicalutamide | top60_altered_DNA+RNA_seed | PIK3CA | -0.43752 | BRCA | MCF7 | Buparlisib |
| MVP_top60_DNA_seed | EGFR | 0.417973 | LUSC | LOU-NH91 | PF-00299804 | top60_altered_DNA+RNA_seed | PIK3CA | -0.436057 | BRCA | HDQ-P1 | Alpelisib |
| MVP_top60_DNA_seed | HDAC1 | 0.429519 | LUSC | RERF-LC-Sq1 | CAY10603 | top60_altered_DNA+RNA_seed | PIK3CA | -0.413514 | BRCA | EFM-192A | AZD8186 |
| MVP_top60_DNA_seed | HDAC1 | 0.433755 | LUSC | HCC-15 | Dacinostat | top60_altered_DNA+RNA_seed | PIK3CA | -0.410153 | BRCA | YMB-1-E | GNE-317 |
| MVP_top60_DNA_seed | HDAC1 | 0.437748 | LUSC | RERF-LC-Sq1 | AR-42 | top60_altered_DNA+RNA_seed | PLK1 | -0.406585 | BRCA | CAL-51 | MK-1775 |
| MVP_top60_DNA_seed | EGFR | 0.441048 | LUSC | RERF-LC-Sq1 | Pelitinib | top60_altered_DNA+RNA_seed | PIK3CA | -0.402635 | BRCA | EFM-19 | Buparlisib |
| MVP_top60_DNA_seed | AR | 0.441245 | LUSC | NCI-H1869 | Bicalutamide | top60_altered_DNA+RNA_seed | TP53 | -0.401514 | BRCA | AU565 | MIRA-1 |
| MVP_top60_DNA_seed | AR | 0.44236 | LUSC | NCI-H2170 | Bicalutamide | top60_altered_DNA+RNA_seed | PIK3CA | -0.372092 | BRCA | ZR-75-30 | Alpelisib |
| MVP_top60_DNA_seed | GSK3B | 0.444556 | LUSC | LK-2 | CHIR-99021 | top60_altered_DNA+RNA_seed | PIK3CA | -0.37003 | BRCA | HCC1599 | Buparlisib |
| MVP_top60_DNA_seed | EGFR | 0.446848 | LUSC | NCI-H226 | Erlotinib | top60_altered_DNA+RNA_seed | PLK1 | -0.348205 | BRCA | HCC1395 | MK-1775 |
| MVP_top60_DNA_seed | ESR1 | 0.448263 | LUSC | LK-2 | Tamoxifen | top60_altered_DNA+RNA_seed | PIK3CA | -0.332338 | BRCA | HCC70 | Buparlisib |
| MVP_top60_DNA_seed | CREBBP | 0.449253 | LUSC | NCI-H520 | I-CBP112 | top60_altered_DNA+RNA_seed | PIK3CA | -0.32986 | BRCA | MFM-223 | Alpelisib |
| MVP_top60_DNA_seed | HDAC1 | 0.463717 | LUSC | NCI-H2170 | AR-42 | top60_altered_DNA+RNA_seed | PIK3CA | -0.328263 | BRCA | CAL-85-1 | GNE-317 |

| | | | | | | | | | | | |
|---|---|---|---|---|---|---|---|---|---|---|---|
| MVP_top60_DNA_seed | HDAC1 | 0.477411 | LUSC | NCI-H1869 | Belinostat | top60_altered_DNA+RNA_seed | PIK3CA | -0.321277 | BRCA | MDA-MB-453 | Buparlisib |
| MVP_top60_DNA_seed | EGFR | 0.48258 | LUSC | NCI-H520 | AST-1306 | top60_altered_DNA+RNA_seed | PIK3CA | -0.320262 | BRCA | BT-20 | Buparlisib |
| MVP_top60_DNA_seed | EGFR | 0.513495 | LUSC | NCI-H226 | CI-1033 | top60_altered_DNA+RNA_seed | PIK3CA | -0.312548 | BRCA | BT-20 | GNE-317 |
| MVP_top60_DNA_seed | GSK3B | 0.516564 | LUSC | HARA | CHIR-99021 | top60_altered_DNA+RNA_seed | PIK3CA | -0.261162 | BRCA | JIMT-1 | Alpelisib |
| MVP_top60_DNA_seed | GSK3B | 0.529754 | LUSC | NCI-H2170 | SB216763 | top60_altered_DNA+RNA_seed | PIK3CA | -0.238073 | BRCA | CAL-85-1 | Alpelisib |
| MVP_top60_DNA_seed | AR | 0.531502 | LUSC | NCI-H226 | AZD3514 | top60_altered_DNA+RNA_seed | PIK3CA | -0.212837 | BRCA | HCC1937 | AZD8186 |
| MVP_top60_DNA_seed | CREBBP | 0.561683 | LUSC | NCI-H1869 | I-CBP112 | top60_altered_DNA+RNA_seed | PIK3CA | -0.208771 | BRCA | MFM-223 | GNE-317 |
| MVP_top60_DNA_seed | AR | 0.568437 | LUSC | LOU-NH91 | Bicalutamide | top60_altered_DNA+RNA_seed | PIK3CA | -0.208619 | BRCA | HCC1569 | GNE-317 |
| MVP_top60_DNA_seed | GSK3B | 0.570337 | LUSC | NCI-H226 | SB216763 | top60_altered_DNA+RNA_seed | TP53 | -0.205086 | BRCA | HCC1187 | MIRA-1 |
| MVP_top60_DNA_seed | GSK3B | 0.572063 | LUSC | LK-2 | SB216763 | top60_altered_DNA+RNA_seed | PLK1 | -0.203687 | BRCA | HCC1569 | MK-1775 |
| MVP_top60_DNA_seed | ESR1 | 0.5819 | LUSC | NCI-H226 | Tamoxifen | top60_altered_DNA+RNA_seed | TP53 | -0.201608 | BRCA | HCC1500 | MIRA-1 |
| MVP_top60_DNA_seed | EGFR | 0.585062 | LUSC | HARA | AST-1306 | top60_altered_DNA+RNA_seed | PIK3CA | -0.175846 | BRCA | UACC-812 | GNE-317 |
| MVP_top60_DNA_seed | EGFR | 0.585682 | LUSC | LOU-NH91 | Pelitinib | top60_altered_DNA+RNA_seed | PIK3CA | -0.175299 | BRCA | UACC-893 | AZD8186 |
| MVP_top60_DNA_seed | ESR1 | 0.586323 | LUSC | NCI-H520 | Tamoxifen | top60_altered_DNA+RNA_seed | PIK3CA | -0.168908 | BRCA | MDA-MB-453 | AZD8186 |
| MVP_top60_DNA_seed | EGFR | 0.59893 | LUSC | HCC-15 | Pelitinib | top60_altered_DNA+RNA_seed | TP53 | -0.167296 | BRCA | BT-20 | MIRA-1 |
| MVP_top60_DNA_seed | ESR1 | 0.604023 | LUSC | LOU-NH91 | Tamoxifen | top60_altered_DNA+RNA_seed | TP53 | -0.164861 | BRCA | MDA-MB-453 | MIRA-1 |
| MVP_top60_DNA_seed | EGFR | 0.612179 | LUSC | NCI-H2170 | AST-1306 | top60_altered_DNA+RNA_seed | PIK3CA | -0.147878 | BRCA | MDA-MB-468 | GNE-317 |
| MVP_top60_DNA_seed | GSK3B | 0.618538 | LUSC | NCI-H226 | CHIR-99021 | top60_altered_DNA+RNA_seed | PIK3CA | -0.142764 | BRCA | CAL-51 | Alpelisib |
| MVP_top60_DNA_seed | AR | 0.619727 | LUSC | RERF-LC-Sq1 | Bicalutamide | top60_altered_DNA+RNA_seed | PIK3CA | -0.132617 | BRCA | JIMT-1 | Buparlisib |
| MVP_top60_DNA_seed | GSK3B | 0.620706 | LUSC | EPLC-272H | SB216763 | top60_altered_DNA+RNA_seed | PIK3CA | -0.13028 | BRCA | YMB-1-E | Buparlisib |
| MVP_top60_DNA_seed | GSK3B | 0.624045 | LUSC | SK-MES-1 | KIN001-042 | top60_altered_DNA+RNA_seed | PLK1 | -0.129855 | BRCA | BT-549 | MK-1775 |
| MVP_top60_DNA_seed | GSK3B | 0.62458 | LUSC | NCI-H520 | CHIR-99021 | top60_altered_DNA+RNA_seed | TP53 | -0.128637 | BRCA | HCC2218 | MIRA-1 |
| MVP_top60_DNA_seed | EGFR | 0.627333 | LUSC | LK-2 | Afatinib | top60_altered_DNA+RNA_seed | PIK3CA | -0.122867 | BRCA | HCC1187 | AZD8186 |
| MVP_top60_DNA_seed | AR | 0.642859 | LUSC | NCI-H226 | Bicalutamide | top60_altered_DNA+RNA_seed | PIK3CA | -0.105804 | BRCA | MDA-MB-453 | GNE-317 |
| MVP_top60_DNA_seed | EGFR | 0.64977 | LUSC | HCC-15 | AST-1306 | top60_altered_DNA+RNA_seed | PIK3CA | -0.096342 | BRCA | HCC2218 | Alpelisib |
| MVP_top60_DNA_seed | EGFR | 0.655347 | LUSC | HCC-15 | Afatinib | top60_altered_DNA+RNA_seed | PIK3CA | -0.09236 | BRCA | HCC70 | Alpelisib |
| MVP_top60_DNA_seed | GSK3B | 0.66191 | LUSC | NCI-H520 | AZD7969 | top60_altered_DNA+RNA_seed | PIK3CA | -0.086838 | BRCA | DU-4475 | GNE-317 |
| MVP_top60_DNA_seed | CREBBP | 0.66458 | LUSC | LOU-NH91 | I-CBP112 | top60_altered_DNA+RNA_seed | PLK1 | -0.086005 | BRCA | BT-549 | BI-2536 |
| MVP_top60_DNA_seed | GSK3B | 0.677727 | LUSC | NCI-H520 | SB216763 | top60_altered_DNA+RNA_seed | PLK1 | -0.078954 | BRCA | HCC38 | MK-1775 |
| MVP_top60_DNA_seed | EGFR | 0.681761 | LUSC | LK-2 | CI-1033 | top60_altered_DNA+RNA_seed | PIK3CA | -0.077314 | BRCA | HCC2157 | Alpelisib |
| MVP_top60_DNA_seed | EGFR | 0.686117 | LUSC | NCI-H2170 | PF-00299804 | top60_altered_DNA+RNA_seed | PIK3CA | -0.064745 | BRCA | MDA-MB-415 | Buparlisib |
| MVP_top60_DNA_seed | AR | 0.705625 | LUSC | NCI-H2170 | Bicalutamide | top60_altered_DNA+RNA_seed | PLK1 | -0.06018 | BRCA | JIMT-1 | MK-1775 |
| MVP_top60_DNA_seed | GSK3B | 0.730788 | LUSC | LOU-NH91 | CHIR-99021 | top60_altered_DNA+RNA_seed | PLK1 | -0.037378 | BRCA | MDA-MB-453 | BI-2536 |
| MVP_top60_DNA_seed | HDAC1 | 0.737617 | LUSC | NCI-H1869 | CAY10603 | top60_altered_DNA+RNA_seed | PIK3CA | -0.014331 | BRCA | BT-549 | GNE-317 |
| MVP_top60_DNA_seed | GSK3B | 0.74223 | LUSC | NCI-H1869 | AZD7969 | top60_altered_DNA+RNA_seed | PLK1 | -0.010128 | BRCA | MDA-MB-231 | BI-2536 |
| MVP_top60_DNA_seed | EGFR | 0.742982 | LUSC | NCI-H226 | Afatinib | top60_altered_DNA+RNA_seed | PIK3CA | -0.006998 | BRCA | MCF7 | AZD8186 |
| MVP_top60_DNA_seed | EGFR | 0.75569 | LUSC | EBC-1 | Cetuximab | top60_altered_DNA+RNA_seed | PIK3CA | -0.003606 | BRCA | COLO-824 | GNE-317 |
| MVP_top60_DNA_seed | EGFR | 0.757769 | LUSC | NCI-H520 | Gefitinib | top60_altered_DNA+RNA_seed | PIK3CA | -0.003436 | BRCA | EVSA-T | Alpelisib |
| MVP_top60_DNA_seed | HDAC1 | 0.757813 | LUSC | LOU-NH91 | Dacinostat | top60_altered_DNA+RNA_seed | PLK1 | 0.006015 | BRCA | MCF7 | BI-2536 |
| MVP_top60_DNA_seed | EGFR | 0.760335 | LUSC | NCI-H2170 | Cetuximab | top60_altered_DNA+RNA_seed | TP53 | 0.010507 | BRCA | MDA-MB-330 | MIRA-1 |
| MVP_top60_DNA_seed | HDAC1 | 0.768894 | LUSC | LOU-NH91 | Tubastatin A | top60_altered_DNA+RNA_seed | PIK3CA | 0.031282 | BRCA | HCC1954 | AZD8186 |

| | | | | | | | | | | | |
|---|---|---|---|---|---|---|---|---|---|---|---|
| MVP_top60_DNA_seed | EGFR | 0.780336 | LUSC | EBC-1 | Gefitinib | top60_altered_DNA+RNA_seed | PIK3CA | 0.044886 | BRCA | BT-549 | AZD8186 |
| MVP_top60_DNA_seed | EGFR | 0.788466 | LUSC | EBC-1 | Pelitinib | top60_altered_DNA+RNA_seed | PIK3CA | 0.053041 | BRCA | CAMA-1 | Buparlisib |
| MVP_top60_DNA_seed | GSK3B | 0.793099 | LUSC | HARA | CHIR-99021 | top60_altered_DNA+RNA_seed | PIK3CA | 0.05419 | BRCA | MDA-MB-468 | Buparlisib |
| MVP_top60_DNA_seed | EGFR | 0.796768 | LUSC | KNS-62 | CI-1033 | top60_altered_DNA+RNA_seed | PLK1 | 0.067211 | BRCA | HCC1143 | MK-1775 |
| MVP_top60_DNA_seed | HDAC1 | 0.798122 | LUSC | HCC-15 | Tubastatin A | top60_altered_DNA+RNA_seed | TP53 | 0.067588 | BRCA | CAL-85-1 | MIRA-1 |
| MVP_top60_DNA_seed | HDAC1 | 0.800592 | LUSC | NCI-H1869 | Dacinostat | top60_altered_DNA+RNA_seed | PIK3CA | 0.072001 | BRCA | HCC1395 | Alpelisib |
| MVP_top60_DNA_seed | EGFR | 0.808393 | LUSC | LC-1-sq | CUDC-101 | top60_altered_DNA+RNA_seed | TP53 | 0.079871 | BRCA | CAMA-1 | MIRA-1 |
| MVP_top60_DNA_seed | GSK3B | 0.810188 | LUSC | NCI-H520 | CHIR-99021 | top60_altered_DNA+RNA_seed | PLK1 | 0.087046 | BRCA | MDA-MB-157 | MK-1775 |
| MVP_top60_DNA_seed | EGFR | 0.811609 | LUSC | NCI-H1869 | CUDC-101 | top60_altered_DNA+RNA_seed | TP53 | 0.09638 | BRCA | HDQ-P1 | MIRA-1 |
| MVP_top60_DNA_seed | SMARCA4 | 0.81455 | LUSC | NCI-H2170 | PFI-3 | top60_altered_DNA+RNA_seed | PIK3CA | 0.103107 | BRCA | UACC-893 | Buparlisib |
| MVP_top60_DNA_seed | SMARCA2 | 0.81455 | LUSC | NCI-H2170 | PFI-3 | top60_altered_DNA+RNA_seed | TP53 | 0.112399 | BRCA | HCC1569 | MIRA-1 |
| MVP_top60_DNA_seed | HDAC1 | 0.815747 | LUSC | HARA | Dacinostat | top60_altered_DNA+RNA_seed | TP53 | 0.124896 | BRCA | Hs-578-T | MIRA-1 |
| MVP_top60_DNA_seed | EGFR | 0.823646 | LUSC | LK-2 | Cetuximab | top60_altered_DNA+RNA_seed | TP53 | 0.129468 | BRCA | JIMT-1 | MIRA-1 |
| MVP_top60_DNA_seed | HDAC1 | 0.826961 | LUSC | NCI-H226 | Parthenolide | top60_altered_DNA+RNA_seed | TP53 | 0.15232 | BRCA | MDA-MB-231 | MIRA-1 |
| MVP_top60_DNA_seed | ESR1 | 0.832648 | LUSC | HARA | Tamoxifen | top60_altered_DNA+RNA_seed | PIK3CA | 0.163942 | BRCA | CAL-51 | AZD8186 |
| MVP_top60_DNA_seed | PIK3CA | 0.846248 | LUSC | EBC-1 | AZD8835 | top60_altered_DNA+RNA_seed | PLK1 | 0.176091 | BRCA | HCC1143 | BI-2536 |
| MVP_top60_DNA_seed | SMARCA4 | 0.849095 | LUSC | RERF-LC-Sq1 | PFI-3 | top60_altered_DNA+RNA_seed | PIK3CA | 0.19012 | BRCA | MDA-MB-436 | AZD8186 |
| MVP_top60_DNA_seed | SMARCA2 | 0.849095 | LUSC | RERF-LC-Sq1 | PFI-3 | top60_altered_DNA+RNA_seed | PLK1 | 0.207684 | BRCA | HDQ-P1 | MK-1775 |
| MVP_top60_DNA_seed | HDAC1 | 0.851319 | LUSC | LC-1-sq | JQ12 | top60_altered_DNA+RNA_seed | PLK1 | 0.212815 | BRCA | EVSA-T | MK-1775 |
| MVP_top60_DNA_seed | HDAC2 | 0.851319 | LUSC | LC-1-sq | JQ12 | top60_altered_DNA+RNA_seed | PIK3CA | 0.213926 | BRCA | MDA-MB-157 | Buparlisib |
| MVP_top60_DNA_seed | EGFR | 0.852472 | LUSC | LK-2 | Gefitinib | top60_altered_DNA+RNA_seed | TP53 | 0.221206 | BRCA | UACC-812 | MIRA-1 |
| MVP_top60_DNA_seed | EGFR | 0.85632 | LUSC | SK-MES-1 | AST-1306 | top60_altered_DNA+RNA_seed | PLK1 | 0.224669 | BRCA | COLO-824 | MK-1775 |
| MVP_top60_DNA_seed | CREBBP | 0.860194 | LUSC | NCI-H2170 | I-CBP112 | top60_altered_DNA+RNA_seed | PIK3CA | 0.227552 | BRCA | CAL-85-1 | Buparlisib |
| MVP_top60_DNA_seed | EGFR | 0.87026 | LUSC | HCC-15 | Cetuximab | top60_altered_DNA+RNA_seed | PLK1 | 0.230955 | BRCA | MDA-MB-231 | MK-1775 |
| MVP_top60_DNA_seed | GSK3B | 0.870638 | LUSC | HCC-15 | AZD7969 | top60_altered_DNA+RNA_seed | PLK1 | 0.231566 | BRCA | MFM-223 | MK-1775 |
| MVP_top60_DNA_seed | HDAC1 | 0.870761 | LUSC | RERF-LC-Sq1 | Tubastatin A | top60_altered_DNA+RNA_seed | PLK1 | 0.237452 | BRCA | MDA-MB-468 | MK-1775 |
| MVP_top60_DNA_seed | GSK3B | 0.872481 | LUSC | NCI-H1869 | KIN001-042 | top60_altered_DNA+RNA_seed | TP53 | 0.238802 | BRCA | HCC1954 | MIRA-1 |
| MVP_top60_DNA_seed | SMARCA4 | 0.886656 | LUSC | NCI-H1869 | PFI-3 | top60_altered_DNA+RNA_seed | TP53 | 0.246462 | BRCA | MDA-MB-361 | MIRA-1 |
| MVP_top60_DNA_seed | SMARCA2 | 0.886656 | LUSC | NCI-H1869 | PFI-3 | top60_altered_DNA+RNA_seed | PIK3CA | 0.268441 | BRCA | YMB-1-E | Alpelisib |
| MVP_top60_DNA_seed | GSK3B | 0.892331 | LUSC | LK-2 | CHIR-99021 | top60_altered_DNA+RNA_seed | PIK3CA | 0.273017 | BRCA | BT-483 | Alpelisib |
| MVP_top60_DNA_seed | AR | 0.909576 | LUSC | NCI-H520 | Bicalutamide | top60_altered_DNA+RNA_seed | PIK3CA | 0.274275 | BRCA | CAL-120 | Buparlisib |
| MVP_top60_DNA_seed | EGFR | 0.93335 | LUSC | LK-2 | Pelitinib | top60_altered_DNA+RNA_seed | PIK3CA | 0.280119 | BRCA | DU-4475 | Alpelisib |
| MVP_top60_DNA_seed | AR | 0.939063 | LUSC | HCC-15 | Bicalutamide | top60_altered_DNA+RNA_seed | PIK3CA | 0.286683 | BRCA | BT-474 | AZD8186 |
| MVP_top60_DNA_seed | AR | 0.940078 | LUSC | HARA | Bicalutamide | top60_altered_DNA+RNA_seed | TP53 | 0.294139 | BRCA | MDA-MB-157 | MIRA-1 |
| MVP_top60_DNA_seed | HDAC1 | 0.944391 | LUSC | NCI-H1869 | Tubastatin A | top60_altered_DNA+RNA_seed | TP53 | 0.311105 | BRCA | UACC-893 | MIRA-1 |
| MVP_top60_DNA_seed | ESR1 | 0.946758 | LUSC | NCI-H520 | Fulvestrant | top60_altered_DNA+RNA_seed | PIK3CA | 0.315274 | BRCA | DU-4475 | AZD8186 |
| MVP_top60_DNA_seed | SMARCA4 | 0.989746 | LUSC | LC-1-sq | PFI-3 | top60_altered_DNA+RNA_seed | PIK3CA | 0.319258 | BRCA | HCC1395 | GNE-317 |
| MVP_top60_DNA_seed | SMARCA2 | 0.989746 | LUSC | LC-1-sq | PFI-3 | top60_altered_DNA+RNA_seed | PLK1 | 0.341469 | BRCA | CAL-85-1 | BI-2536 |
| MVP_top60_DNA_seed | EGFR | 0.995471 | LUSC | EBC-1 | Afatinib | top60_altered_DNA+RNA_seed | PIK3CA | 0.342549 | BRCA | HCC1599 | GNE-317 |
| MVP_top60_DNA_seed | AR | 1.015018 | LUSC | NCI-H520 | AZD3514 | top60_altered_DNA+RNA_seed | PIK3CA | 0.349377 | BRCA | HCC38 | GNE-317 |
| MVP_top60_DNA_seed | ESR1 | 1.048906 | LUSC | LC-1-sq | Tamoxifen | top60_altered_DNA+RNA_seed | PIK3CA | 0.358545 | BRCA | HCC2218 | AZD8186 |

| | | | | | | | | | | | |
|---|---|---|---|---|---|---|---|---|---|---|---|
| MVP_top60_DNA_seed | GSK3B | 1.049155 | LUSC | SK-MES-1 | CHIR-99021 | top60_altered_DNA+RNA_seed | PIK3CA | 0.373971 | BRCA | BT-20 | AZD8186 |
| MVP_top60_DNA_seed | EGFR | 1.051407 | LUSC | NCI-H520 | Afatinib | top60_altered_DNA+RNA_seed | TP53 | 0.389817 | BRCA | CAL-51 | MIRA-1 |
| MVP_top60_DNA_seed | HDAC1 | 1.054866 | LUSC | NCI-H2170 | Tubastatin A | top60_altered_DNA+RNA_seed | TP53 | 0.393963 | BRCA | YMB-1-E | MIRA-1 |
| MVP_top60_DNA_seed | EGFR | 1.064489 | LUSC | KNS-62 | AZD8931 | top60_altered_DNA+RNA_seed | PIK3CA | 0.396797 | BRCA | OCUB-M | AZD8186 |
| MVP_top60_DNA_seed | HDAC1 | 1.064857 | LUSC | EPLC-272H | JQ12 | top60_altered_DNA+RNA_seed | PIK3CA | 0.405435 | BRCA | UACC-812 | Buparlisib |
| MVP_top60_DNA_seed | HDAC2 | 1.064857 | LUSC | EPLC-272H | JQ12 | top60_altered_DNA+RNA_seed | PIK3CA | 0.422393 | BRCA | HCC1419 | AZD8186 |
| MVP_top60_DNA_seed | GSK3B | 1.067995 | LUSC | NCI-H1869 | SB216763 | top60_altered_DNA+RNA_seed | PLK1 | 0.448416 | BRCA | MDA-MB-361 | MK-1775 |
| MVP_top60_DNA_seed | RAC1 | 1.083852 | LUSC | KNS-62 | EHT-1864 | top60_altered_DNA+RNA_seed | TP53 | 0.453655 | BRCA | MDA-MB-468 | MIRA-1 |
| MVP_top60_DNA_seed | AR | 1.08513 | LUSC | EBC-1 | Bicalutamide | top60_altered_DNA+RNA_seed | PIK3CA | 0.459333 | BRCA | BT-549 | Buparlisib |
| MVP_top60_DNA_seed | GSK3B | 1.086718 | LUSC | EBC-1 | SB216763 | top60_altered_DNA+RNA_seed | PLK1 | 0.462976 | BRCA | UACC-893 | BI-2536 |
| MVP_top60_DNA_seed | EGFR | 1.101244 | LUSC | HARA | CI-1033 | top60_altered_DNA+RNA_seed | PIK3CA | 0.4689 | BRCA | MDA-MB-415 | Alpelisib |
| MVP_top60_DNA_seed | HDAC1 | 1.102847 | LUSC | HCC-15 | CAY10603 | top60_altered_DNA+RNA_seed | PLK1 | 0.482187 | BRCA | MDA-MB-453 | MK-1775 |
| MVP_top60_DNA_seed | EGFR | 1.122653 | LUSC | KNS-62 | AST-1306 | top60_altered_DNA+RNA_seed | PLK1 | 0.483204 | BRCA | OCUB-M | MK-1775 |
| MVP_top60_DNA_seed | GSK3B | 1.134153 | LUSC | HCC-15 | CHIR-99021 | top60_altered_DNA+RNA_seed | PIK3CA | 0.487822 | BRCA | CAL-85-1 | AZD8186 |
| MVP_top60_DNA_seed | GSK3B | 1.134506 | LUSC | HCC-15 | KIN001-042 | top60_altered_DNA+RNA_seed | TP53 | 0.499196 | BRCA | T47D | MIRA-1 |
| MVP_top60_DNA_seed | GSK3B | 1.154785 | LUSC | HCC-15 | CHIR-99021 | top60_altered_DNA+RNA_seed | PIK3CA | 0.501662 | BRCA | HCC1937 | Alpelisib |
| MVP_top60_DNA_seed | GSK3B | 1.178022 | LUSC | EBC-1 | CHIR-99021 | top60_altered_DNA+RNA_seed | PLK1 | 0.506409 | BRCA | CAL-120 | MK-1775 |
| MVP_top60_DNA_seed | EGFR | 1.184737 | LUSC | EBC-1 | AST-1306 | top60_altered_DNA+RNA_seed | PIK3CA | 0.522996 | BRCA | MDA-MB-157 | GNE-317 |
| MVP_top60_DNA_seed | RAC1 | 1.18549 | LUSC | NCI-H2170 | EHT-1864 | top60_altered_DNA+RNA_seed | PIK3CA | 0.530461 | BRCA | HCC38 | Alpelisib |
| MVP_top60_DNA_seed | PIK3CA | 1.193603 | LUSC | NCI-H226 | PI3Ka_4409 | top60_altered_DNA+RNA_seed | PIK3CA | 0.533957 | BRCA | HCC1419 | Alpelisib |
| MVP_top60_DNA_seed | AR | 1.194853 | LUSC | LC-1-sq | Bicalutamide | top60_altered_DNA+RNA_seed | PIK3CA | 0.535915 | BRCA | HCC38 | AZD8186 |
| MVP_top60_DNA_seed | EGFR | 1.201283 | LUSC | HCC-15 | CI-1033 | top60_altered_DNA+RNA_seed | PLK1 | 0.53622 | BRCA | HCC70 | MK-1775 |
| MVP_top60_DNA_seed | HDAC1 | 1.205742 | LUSC | HCC-15 | Belinostat | top60_altered_DNA+RNA_seed | PIK3CA | 0.540628 | BRCA | JIMT-1 | AZD8186 |
| MVP_top60_DNA_seed | AR | 1.214662 | LUSC | RERF-LC-Sq1 | Bicalutamide | top60_altered_DNA+RNA_seed | PIK3CA | 0.541288 | BRCA | BT-549 | Alpelisib |
| MVP_top60_DNA_seed | GSK3B | 1.254587 | LUSC | EBC-1 | CHIR-99021 | top60_altered_DNA+RNA_seed | PLK1 | 0.545301 | BRCA | CAL-85-1 | MK-1775 |
| MVP_top60_DNA_seed | EGFR | 1.259781 | LUSC | HCC-15 | PF-00299804 | top60_altered_DNA+RNA_seed | TP53 | 0.557608 | BRCA | ZR-75-30 | MIRA-1 |
| MVP_top60_DNA_seed | EGFR | 1.269245 | LUSC | KNS-62 | Cetuximab | top60_altered_DNA+RNA_seed | PIK3CA | 0.561574 | BRCA | MFM-223 | Buparlisib |
| MVP_top60_DNA_seed | CREBBP | 1.295727 | LUSC | KNS-62 | I-CBP112 | top60_altered_DNA+RNA_seed | PIK3CA | 0.56285 | BRCA | MDA-MB-231 | Buparlisib |
| MVP_top60_DNA_seed | EGFR | 1.32694 | LUSC | EBC-1 | PF-00299804 | top60_altered_DNA+RNA_seed | PLK1 | 0.596821 | BRCA | HCC2157 | MK-1775 |
| MVP_top60_DNA_seed | PIK3CA | 1.349994 | LUSC | NCI-H520 | AZD8835 | top60_altered_DNA+RNA_seed | TP53 | 0.597125 | BRCA | COLO-824 | MIRA-1 |
| MVP_top60_DNA_seed | RAC1 | 1.353848 | LUSC | NCI-H226 | EHT-1864 | top60_altered_DNA+RNA_seed | PIK3CA | 0.60026 | BRCA | ZR-75-30 | AZD8186 |
| MVP_top60_DNA_seed | HDAC1 | 1.382508 | LUSC | EBC-1 | Tubastatin A | top60_altered_DNA+RNA_seed | PIK3CA | 0.600468 | BRCA | MCF7 | GNE-317 |
| MVP_top60_DNA_seed | GSK3B | 1.403683 | LUSC | RERF-LC-Sq1 | CHIR-99021 | top60_altered_DNA+RNA_seed | PIK3CA | 0.604254 | BRCA | MDA-MB-157 | Alpelisib |
| MVP_top60_DNA_seed | HDAC1 | 1.405971 | LUSC | RERF-LC-Sq1 | JQ12 | top60_altered_DNA+RNA_seed | PIK3CA | 0.611107 | BRCA | CAMA-1 | Alpelisib |
| MVP_top60_DNA_seed | HDAC2 | 1.405971 | LUSC | RERF-LC-Sq1 | JQ12 | top60_altered_DNA+RNA_seed | TP53 | 0.612158 | BRCA | EFM-19 | MIRA-1 |
| MVP_top60_DNA_seed | ESR1 | 1.492669 | LUSC | KNS-62 | Fulvestrant | top60_altered_DNA+RNA_seed | PLK1 | 0.61745 | BRCA | EFM-19 | BI-2536 |
| MVP_top60_DNA_seed | EGFR | 1.503045 | LUSC | EBC-1 | CI-1033 | top60_altered_DNA+RNA_seed | PIK3CA | 0.618825 | BRCA | HCC1187 | Buparlisib |
| MVP_top60_DNA_seed | GSK3B | 1.51208 | LUSC | KNS-62 | KIN001-042 | top60_altered_DNA+RNA_seed | PLK1 | 0.646539 | BRCA | MRK-nu-1 | MK-1775 |
| MVP_top60_DNA_seed | EGFR | 1.51472 | LUSC | HCC-15 | CUDC-101 | top60_altered_DNA+RNA_seed | PLK1 | 0.654826 | BRCA | CAL-120 | BI-2536 |
| MVP_top60_DNA_seed | GSK3B | 1.51823 | LUSC | KNS-62 | CHIR-99021 | top60_altered_DNA+RNA_seed | TP53 | 0.671396 | BRCA | EFM-192A | MIRA-1 |
| MVP_top60_DNA_seed | CREBBP | 1.562696 | LUSC | RERF-LC-Sq1 | I-CBP112 | top60_altered_DNA+RNA_seed | PLK1 | 0.677287 | BRCA | HCC1395 | BI-2536 |

| | | | | | | | | | | | |
|---|---|---|---|---|---|---|---|---|---|---|---|
| MVP_top60_DNA_seed | PIK3CA | 1.576912 | LUSC | NCI-H1869 | AZD8835 | top60_altered_DNA+RNA_seed | PIK3CA | 0.678447 | BRCA | MDA-MB-231 | Alpelisib |
| MVP_top60_DNA_seed | EGFR | 1.58722 | LUSC | KNS-62 | PF-00299804 | top60_altered_DNA+RNA_seed | PIK3CA | 0.685429 | BRCA | MRK-nu-1 | Alpelisib |
| MVP_top60_DNA_seed | PIK3CA | 1.590637 | LUSC | NCI-H520 | PI3Ka_4409 | top60_altered_DNA+RNA_seed | PLK1 | 0.691406 | BRCA | HCC1954 | MK-1775 |
| MVP_top60_DNA_seed | HDAC1 | 1.647033 | LUSC | NCI-H1869 | JQ12 | top60_altered_DNA+RNA_seed | PLK1 | 0.70649 | BRCA | AU565 | MK-1775 |
| MVP_top60_DNA_seed | HDAC2 | 1.647033 | LUSC | NCI-H1869 | JQ12 | top60_altered_DNA+RNA_seed | PIK3CA | 0.706951 | BRCA | MDA-MB-468 | Alpelisib |
| MVP_top60_DNA_seed | HDAC1 | 1.66459 | LUSC | HCC-15 | AR-42 | top60_altered_DNA+RNA_seed | PLK1 | 0.712753 | BRCA | MDA-MB-157 | BI-2536 |
| MVP_top60_DNA_seed | HDAC1 | 1.672272 | LUSC | HARA | JQ12 | top60_altered_DNA+RNA_seed | PLK1 | 0.733164 | BRCA | HCC1187 | MK-1775 |
| MVP_top60_DNA_seed | HDAC2 | 1.672272 | LUSC | HARA | JQ12 | top60_altered_DNA+RNA_seed | TP53 | 0.756287 | BRCA | HCC38 | MIRA-1 |
| MVP_top60_DNA_seed | GSK3B | 1.688594 | LUSC | RERF-LC-Sq1 | CHIR-99021 | top60_altered_DNA+RNA_seed | TP53 | 0.759119 | BRCA | HCC1395 | MIRA-1 |
| MVP_top60_DNA_seed | RAC1 | 2.024044 | LUSC | LOU-NH91 | EHT-1864 | top60_altered_DNA+RNA_seed | PIK3CA | 0.776656 | BRCA | HCC1937 | GNE-317 |
| MVP_top60_DNA_seed | GSK3B | 2.032697 | LUSC | KNS-62 | AZD7969 | top60_altered_DNA+RNA_seed | PIK3CA | 0.776945 | BRCA | HCC1187 | Alpelisib |
| MVP_top60_DNA_seed | EGFR | 2.064351 | LUSC | NCI-H2170 | CI-1033 | top60_altered_DNA+RNA_seed | PLK1 | 0.801968 | BRCA | YMB-1-E | BI-2536 |
| MVP_top60_DNA_seed | EGFR | 2.096853 | LUSC | SK-MES-1 | CI-1033 | top60_altered_DNA+RNA_seed | PLK1 | 0.852642 | BRCA | BT-20 | MK-1775 |
| MVP_top60_DNA_seed | HDAC1 | 2.100282 | LUSC | NCI-H226 | Entinostat | top60_altered_DNA+RNA_seed | PLK1 | 0.854893 | BRCA | MDA-MB-361 | BI-2536 |
| MVP_top60_DNA_seed | HDAC3 | 2.100282 | LUSC | NCI-H226 | Entinostat | top60_altered_DNA+RNA_seed | PIK3CA | 0.857775 | BRCA | HCC2218 | Buparlisib |
| MVP_top60_DNA_seed | EGFR | 2.122355 | LUSC | SK-MES-1 | PF-00299804 | top60_altered_DNA+RNA_seed | PLK1 | 0.857858 | BRCA | HCC1937 | BI-2536 |
| MVP_top60_DNA_seed | GSK3B | 2.18796 | LUSC | RERF-LC-Sq1 | AZD7969 | top60_altered_DNA+RNA_seed | TP53 | 0.860765 | BRCA | MCF7 | MIRA-1 |
| MVP_top60_DNA_seed | PIK3CA | 2.274945 | LUSC | KNS-62 | AZD8835 | top60_altered_DNA+RNA_seed | PLK1 | 0.863951 | BRCA | HCC1569 | BI-2536 |
| MVP_top60_DNA_seed | HDAC1 | 2.494629 | LUSC | RERF-LC-Sq1 | Dacinostat | top60_altered_DNA+RNA_seed | PLK1 | 0.870907 | BRCA | HCC1428 | BI-2536 |
| MVP_top60_DNA_seed | EGFR | -3.132005 | LUSC | NCI-H2170 | Afatinib | top60_altered_DNA+RNA_seed | PIK3CA | 0.8721 | BRCA | HCC1599 | AZD8186 |
| MVP_top60_DNA_seed | EGFR | -3.069377 | LUSC | NCI-H2170 | Osimertinib | top60_altered_DNA+RNA_seed | PLK1 | 0.891807 | BRCA | MDA-MB-330 | BI-2536 |
| MVP_top60_DNA_seed | EGFR | -2.612939 | LUSC | NCI-H2170 | Lapatinib | top60_altered_DNA+RNA_seed | PLK1 | 0.897235 | BRCA | EFM-19 | MK-1775 |
| MVP_top60_DNA_seed | EGFR | -2.373387 | LUSC | NCI-H2170 | Sapitinib | top60_altered_DNA+RNA_seed | PIK3CA | 0.901775 | BRCA | BT-474 | GNE-317 |
| MVP_top60_DNA_seed | EGFR | -2.303236 | LUSC | RERF-LC-Sq1 | Gefitinib | top60_altered_DNA+RNA_seed | PIK3CA | 0.911682 | BRCA | MRK-nu-1 | AZD8186 |
| MVP_top60_DNA_seed | EGFR | -2.240216 | LUSC | RERF-LC-Sq1 | Sapitinib | top60_altered_DNA+RNA_seed | PLK1 | 0.913904 | BRCA | HCC1500 | BI-2536 |
| MVP_top60_DNA_seed | EGFR | -1.926747 | LUSC | NCI-H2170 | Gefitinib | top60_altered_DNA+RNA_seed | PIK3CA | 0.919742 | BRCA | HCC1569 | Buparlisib |
| MVP_top60_DNA_seed | EGFR | -1.759858 | LUSC | LOU-NH91 | Erlotinib | top60_altered_DNA+RNA_seed | PIK3CA | 0.9207 | BRCA | MFM-223 | AZD8186 |
| MVP_top60_DNA_seed | EGFR | -1.753993 | LUSC | LC-1-sq | Gefitinib | top60_altered_DNA+RNA_seed | PLK1 | 0.941195 | BRCA | HCC70 | BI-2536 |
| MVP_top60_DNA_seed | EGFR | -1.738314 | LUSC | NCI-H1869 | Gefitinib | top60_altered_DNA+RNA_seed | PIK3CA | 0.951276 | BRCA | MDA-MB-436 | Alpelisib |
| MVP_top60_DNA_seed | EGFR | -1.718778 | LUSC | RERF-LC-Sq1 | Afatinib | top60_altered_DNA+RNA_seed | TP53 | 1.036629 | BRCA | HCC1937 | MIRA-1 |
| MVP_top60_DNA_seed | EGFR | -1.710952 | LUSC | NCI-H1869 | Erlotinib | top60_altered_DNA+RNA_seed | PIK3CA | 1.040492 | BRCA | HCC38 | Buparlisib |
| MVP_top60_DNA_seed | EGFR | -1.687019 | LUSC | RERF-LC-Sq1 | AZD3759 | top60_altered_DNA+RNA_seed | PIK3CA | 1.045803 | BRCA | MDA-MB-231 | AZD8186 |
| MVP_top60_DNA_seed | EGFR | -1.672699 | LUSC | RERF-LC-Sq1 | Erlotinib | top60_altered_DNA+RNA_seed | PLK1 | 1.051407 | BRCA | HCC38 | BI-2536 |
| MVP_top60_DNA_seed | EGFR | -1.641911 | LUSC | NCI-H2170 | Erlotinib | top60_altered_DNA+RNA_seed | PLK1 | 1.052519 | BRCA | T47D | BI-2536 |
| MVP_top60_DNA_seed | EGFR | -1.636937 | LUSC | LOU-NH91 | AZD3759 | top60_altered_DNA+RNA_seed | PLK1 | 1.063651 | BRCA | HCC1937 | MK-1775 |
| MVP_top60_DNA_seed | GSK3B | -1.520108 | LUSC | EBC-1 | SB216763 | top60_altered_DNA+RNA_seed | PLK1 | 1.065279 | BRCA | MDA-MB-436 | MK-1775 |
| MVP_top60_DNA_seed | EGFR | -1.4507 | LUSC | RERF-LC-Sq1 | Lapatinib | top60_altered_DNA+RNA_seed | PIK3CA | 1.071977 | BRCA | MDA-MB-436 | GNE-317 |
| MVP_top60_DNA_seed | EGFR | -1.438509 | LUSC | RERF-LC-Sq1 | Osimertinib | top60_altered_DNA+RNA_seed | PLK1 | 1.072475 | BRCA | HDQ-P1 | BI-2536 |
| MVP_top60_DNA_seed | EGFR | -1.428391 | LUSC | LOU-NH91 | Gefitinib | top60_altered_DNA+RNA_seed | PLK1 | 1.078397 | BRCA | HCC1500 | MK-1775 |
| MVP_top60_DNA_seed | EGFR | -1.41677 | LUSC | EPLC-272H | Gefitinib | top60_altered_DNA+RNA_seed | PIK3CA | 1.098639 | BRCA | HCC1419 | Buparlisib |
| MVP_top60_DNA_seed | EGFR | -1.387167 | LUSC | LC-1-sq | Erlotinib | top60_altered_DNA+RNA_seed | TP53 | 1.099032 | BRCA | BT-549 | MIRA-1 |

| | | | | | | | | | | | |
|---|---|---|---|---|---|---|---|---|---|---|---|
| MVP_top60_DNA_seed | EGFR | -1.363386 | LUSC | LC-1-sq | Sapitinib | top60_altered_DNA+RNA_seed | PIK3CA | 1.099167 | BRCA | BT-474 | Buparlisib |
| MVP_top60_DNA_seed | PIK3CA | -1.338512 | LUSC | RERF-LC-Sq1 | Alpelisib | top60_altered_DNA+RNA_seed | PIK3CA | 1.100603 | BRCA | ZR-75-30 | GNE-317 |
| MVP_top60_DNA_seed | ESR1 | -1.258784 | LUSC | HARA | Tamoxifen | top60_altered_DNA+RNA_seed | PIK3CA | 1.140993 | BRCA | HCC1937 | Buparlisib |
| MVP_top60_DNA_seed | ESR1 | -1.258168 | LUSC | EBC-1 | GDC0810 | top60_altered_DNA+RNA_seed | TP53 | 1.142254 | BRCA | HCC1143 | MIRA-1 |
| MVP_top60_DNA_seed | EGFR | -1.225478 | LUSC | NCI-H1869 | Osimertinib | top60_altered_DNA+RNA_seed | PIK3CA | 1.149817 | BRCA | HCC1500 | Alpelisib |
| MVP_top60_DNA_seed | PIK3CA | -1.220521 | LUSC | EPLC-272H | Alpelisib | top60_altered_DNA+RNA_seed | PLK1 | 1.158134 | BRCA | T47D | MK-1775 |
| MVP_top60_DNA_seed | SMARCA4 | -1.163656 | LUSC | EBC-1 | PFI3 | top60_altered_DNA+RNA_seed | PIK3CA | 1.15996 | BRCA | MDA-MB-157 | AZD8186 |
| MVP_top60_DNA_seed | SMARCA2 | -1.163656 | LUSC | EBC-1 | PFI3 | top60_altered_DNA+RNA_seed | PIK3CA | 1.160204 | BRCA | MDA-MB-436 | Buparlisib |
| MVP_top60_DNA_seed | EZH2 | -1.160375 | LUSC | LC-1-sq | GSK343 | top60_altered_DNA+RNA_seed | TP53 | 1.167902 | BRCA | MFM-223 | MIRA-1 |
| MVP_top60_DNA_seed | PIK3CA | -1.145638 | LUSC | EPLC-272H | Buparlisib | top60_altered_DNA+RNA_seed | PLK1 | 1.196486 | BRCA | CAMA-1 | MK-1775 |
| MVP_top60_DNA_seed | PIK3CA | -1.121422 | LUSC | NCI-H2170 | Buparlisib | top60_altered_DNA+RNA_seed | TP53 | 1.204967 | BRCA | MDA-MB-436 | MIRA-1 |
| MVP_top60_DNA_seed | EGFR | -1.110884 | LUSC | EPLC-272H | Sapitinib | top60_altered_DNA+RNA_seed | TP53 | 1.22708 | BRCA | CAL-120 | MIRA-1 |
| MVP_top60_DNA_seed | EGFR | -1.096208 | LUSC | SK-MES-1 | Sapitinib | top60_altered_DNA+RNA_seed | PIK3CA | 1.246289 | BRCA | BT-483 | GNE-317 |
| MVP_top60_DNA_seed | PIK3CA | -1.08911 | LUSC | EPLC-272H | AZD8186 | top60_altered_DNA+RNA_seed | PIK3CA | 1.275595 | BRCA | CAL-120 | AZD8186 |
| MVP_top60_DNA_seed | EGFR | -1.066494 | LUSC | HARA | Sapitinib | top60_altered_DNA+RNA_seed | PLK1 | 1.276223 | BRCA | MDA-MB-175-VII | MK-1775 |
| MVP_top60_DNA_seed | EGFR | -1.0663 | LUSC | SK-MES-1 | Erlotinib | top60_altered_DNA+RNA_seed | PLK1 | 1.287501 | BRCA | HCC1428 | MK-1775 |
| MVP_top60_DNA_seed | EGFR | -1.063689 | LUSC | LOU-NH91 | Osimertinib | top60_altered_DNA+RNA_seed | TP53 | 1.29189 | BRCA | BT-474 | MIRA-1 |
| MVP_top60_DNA_seed | EGFR | -1.05362 | LUSC | NCI-H1869 | AZD3759 | top60_altered_DNA+RNA_seed | PLK1 | 1.308867 | BRCA | MDA-MB-415 | BI-2536 |
| MVP_top60_DNA_seed | PIK3CA | -1.036936 | LUSC | EPLC-272H | GNE-317 | top60_altered_DNA+RNA_seed | PLK1 | 1.315157 | BRCA | BT-474 | BI-2536 |
| MVP_top60_DNA_seed | EZH2 | -1.034054 | LUSC | EBC-1 | GSK343 | top60_altered_DNA+RNA_seed | PLK1 | 1.348728 | BRCA | CAMA-1 | BI-2536 |
| MVP_top60_DNA_seed | EGFR | -0.997257 | LUSC | HARA | Gefitinib | top60_altered_DNA+RNA_seed | PLK1 | 1.354522 | BRCA | MDA-MB-330 | MK-1775 |
| MVP_top60_DNA_seed | CREBBP | -0.985121 | LUSC | LC-1-sq | SGC-CBP30 | top60_altered_DNA+RNA_seed | PLK1 | 1.359822 | BRCA | HCC2218 | BI-2536 |
| MVP_top60_DNA_seed | EZH2 | -0.973241 | LUSC | HARA | GSK343 | top60_altered_DNA+RNA_seed | PLK1 | 1.382404 | BRCA | HCC1954 | BI-2536 |
| MVP_top60_DNA_seed | PIK3CA | -0.955533 | LUSC | HARA | Buparlisib | top60_altered_DNA+RNA_seed | PLK1 | 1.38815 | BRCA | HCC1187 | BI-2536 |
| MVP_top60_DNA_seed | EGFR | -0.943743 | LUSC | HARA | Lapatinib | top60_altered_DNA+RNA_seed | TP53 | 1.444911 | BRCA | HCC1428 | MIRA-1 |
| MVP_top60_DNA_seed | EGFR | -0.940591 | LUSC | NCI-H1869 | Lapatinib | top60_altered_DNA+RNA_seed | PIK3CA | 1.481992 | BRCA | HCC1500 | Buparlisib |
| MVP_top60_DNA_seed | EGFR | -0.940366 | LUSC | LOU-NH91 | Sapitinib | top60_altered_DNA+RNA_seed | PLK1 | 1.546905 | BRCA | BT-483 | BI-2536 |
| MVP_top60_DNA_seed | EGFR | -0.922265 | LUSC | EPLC-272H | Afatinib | top60_altered_DNA+RNA_seed | PLK1 | 1.552509 | BRCA | UACC-812 | MK-1775 |
| MVP_top60_DNA_seed | PIK3CA | -0.913577 | LUSC | HARA | Alpelisib | top60_altered_DNA+RNA_seed | PIK3CA | 1.553813 | BRCA | HCC1187 | GNE-317 |
| MVP_top60_DNA_seed | EGFR | -0.891851 | LUSC | LC-1-sq | AZD3759 | top60_altered_DNA+RNA_seed | PLK1 | 1.569675 | BRCA | BT-474 | MK-1775 |
| MVP_top60_DNA_seed | EGFR | -0.887201 | LUSC | NCI-H1869 | Sapitinib | top60_altered_DNA+RNA_seed | PIK3CA | 1.582609 | BRCA | HCC1419 | GNE-317 |
| MVP_top60_DNA_seed | HDAC1 | -0.879947 | LUSC | LOU-NH91 | Romidepsin | top60_altered_DNA+RNA_seed | PLK1 | 1.585866 | BRCA | UACC-893 | MK-1775 |
| MVP_top60_DNA_seed | HDAC3 | -0.879947 | LUSC | LOU-NH91 | Romidepsin | top60_altered_DNA+RNA_seed | PLK1 | 1.598239 | BRCA | MDA-MB-436 | BI-2536 |
| MVP_top60_DNA_seed | HDAC2 | -0.879947 | LUSC | LOU-NH91 | Romidepsin | top60_altered_DNA+RNA_seed | PLK1 | 1.600018 | BRCA | EFM-192A | BI-2536 |
| MVP_top60_DNA_seed | EGFR | -0.872074 | LUSC | SK-MES-1 | Gefitinib | top60_altered_DNA+RNA_seed | PIK3CA | 1.601177 | BRCA | HCC1500 | AZD8186 |
| MVP_top60_DNA_seed | EGFR | -0.858463 | LUSC | LOU-NH91 | Afatinib | top60_altered_DNA+RNA_seed | PLK1 | 1.61371 | BRCA | MDA-MB-175-VII | BI-2536 |
| MVP_top60_DNA_seed | EGFR | -0.857846 | LUSC | SK-MES-1 | Afatinib | top60_altered_DNA+RNA_seed | PLK1 | 1.659986 | BRCA | HCC2218 | MK-1775 |
| MVP_top60_DNA_seed | PIK3CA | -0.830417 | LUSC | HARA | GNE-317 | top60_altered_DNA+RNA_seed | PLK1 | 1.672698 | BRCA | MCF7 | MK-1775 |
| MVP_top60_DNA_seed | PIK3CA | -0.822014 | LUSC | EBC-1 | AZD8186 | top60_altered_DNA+RNA_seed | PLK1 | 1.682074 | BRCA | MDA-MB-415 | MK-1775 |
| MVP_top60_DNA_seed | EGFR | -0.820673 | LUSC | EPLC-272H | Lapatinib | top60_altered_DNA+RNA_seed | PIK3CA | 1.70168 | BRCA | ZR-75-30 | Buparlisib |
| MVP_top60_DNA_seed | EZH2 | -0.817165 | LUSC | EPLC-272H | GSK343 | top60_altered_DNA+RNA_seed | PLK1 | 1.820042 | BRCA | HCC1419 | MK-1775 |

| | | | | | | | | | | | |
|---|---|---|---|---|---|---|---|---|---|---|---|
| MVP_top60_DNA_seed | EZH2 | -0.81452 | LUSC | SK-MES-1 | GSK343 | top60_altered_DNA+RNA_seed | PLK1 | 1.826433 | BRCA | ZR-75-30 | BI-2536 |
| MVP_top60_DNA_seed | RAC1 | -0.796256 | LUSC | NCI-H520 | EHT-1864 | top60_altered_DNA+RNA_seed | TP53 | 1.829054 | BRCA | MDA-MB-175-VII | MIRA-1 |
| MVP_top60_DNA_seed | HDAC1 | -0.794417 | LUSC | EBC-1 | Romidepsin | top60_altered_DNA+RNA_seed | PLK1 | 1.851474 | BRCA | HCC1419 | BI-2536 |
| MVP_top60_DNA_seed | HDAC3 | -0.794417 | LUSC | EBC-1 | Romidepsin | top60_altered_DNA+RNA_seed | TP53 | 1.908261 | BRCA | BT-483 | MIRA-1 |
| MVP_top60_DNA_seed | HDAC2 | -0.794417 | LUSC | EBC-1 | Romidepsin | top60_altered_DNA+RNA_seed | PIK3CA | 1.92054 | BRCA | HCC1500 | GNE-317 |
| MVP_top60_DNA_seed | PIK3CA | -0.786418 | LUSC | RERF-LC-Sq1 | GNE-317 | top60_altered_DNA+RNA_seed | PLK1 | 2.00218 | BRCA | UACC-812 | BI-2536 |
| MVP_top60_DNA_seed | GSK3B | -0.78493 | LUSC | HARA | SB216763 | top60_altered_DNA+RNA_seed | PIK3CA | 2.052789 | BRCA | CAL-120 | Alpelisib |
| MVP_top60_DNA_seed | PIK3CA | -0.781629 | LUSC | NCI-H2170 | Alpelisib | top60_altered_DNA+RNA_seed | PIK3CA | 2.131534 | BRCA | BT-483 | Buparlisib |
| MVP_top60_DNA_seed | HDAC1 | -0.774222 | LUSC | EBC-1 | Entinostat | top60_altered_DNA+RNA_seed | PIK3CA | 2.322717 | BRCA | HCC1428 | GNE-317 |
| MVP_top60_DNA_seed | HDAC3 | -0.774222 | LUSC | EBC-1 | Entinostat | top60_altered_DNA+RNA_seed | PLK1 | 2.39159 | BRCA | EFM-192A | MK-1775 |
| MVP_top60_DNA_seed | EGFR | -0.755037 | LUSC | HARA | Afatinib | top60_altered_DNA+RNA_seed | PLK1 | 2.463964 | BRCA | YMB-1-E | MK-1775 |
| MVP_top60_DNA_seed | EGFR | -0.754414 | LUSC | HARA | Erlotinib | top60_altered_DNA+RNA_seed | PIK3CA | 2.57169 | BRCA | HCC1428 | Buparlisib |
| MVP_top60_DNA_seed | PIK3CA | -0.729955 | LUSC | EBC-1 | GNE-317 | top60_altered_DNA+RNA_seed | PIK3CA | 2.676332 | BRCA | HCC1428 | AZD8186 |
| MVP_top60_DNA_seed | ESR1 | -0.719565 | LUSC | EPLC-272H | GDC0810 | top60_altered_DNA+RNA_seed | PLK1 | 2.788994 | BRCA | BT-483 | MK-1775 |
| MVP_top60_DNA_seed | PIK3CA | -0.713796 | LUSC | NCI-H520 | AZD8186 | top60_altered_DNA+RNA_seed | PIK3CA | 2.896574 | BRCA | HCC1428 | Alpelisib |
| MVP_top60_DNA_seed | EGFR | -0.706849 | LUSC | EPLC-272H | Osimertinib | top60_altered_DNA+RNA_seed | PIK3CA | 2.896804 | BRCA | BT-483 | AZD8186 |
| MVP_top60_DNA_seed | HDAC1 | -0.706831 | LUSC | NCI-H520 | Romidepsin | top60_altered_DNA+RNA_seed | PLK1 | 3.008957 | BRCA | ZR-75-30 | MK-1775 |
| MVP_top60_DNA_seed | HDAC3 | -0.706831 | LUSC | NCI-H520 | Romidepsin | top60_altered_DNA+RNA_seed | BRAF | -4.363396 | COADREAD | COLO-205 | SB590885 |
| MVP_top60_DNA_seed | HDAC2 | -0.706831 | LUSC | NCI-H520 | Romidepsin | top60_altered_DNA+RNA_seed | BRAF | -4.227096 | COADREAD | COLO-205 | PLX-4720 |
| MVP_top60_DNA_seed | EGFR | -0.681676 | LUSC | HARA | Osimertinib | top60_altered_DNA+RNA_seed | BRAF | -2.966581 | COADREAD | MDST8 | Dabrafenib |
| MVP_top60_DNA_seed | EZH2 | -0.632355 | LUSC | EBC-1 | GSK343 | top60_altered_DNA+RNA_seed | PIK3CA | -2.885395 | COADREAD | SNU-407 | AZD8835 |
| MVP_top60_DNA_seed | EGFR | -0.599268 | LUSC | SK-MES-1 | AZD3759 | top60_altered_DNA+RNA_seed | BRAF | -2.486896 | COADREAD | RKO | AZ628 |
| MVP_top60_DNA_seed | ESR1 | -0.590708 | LUSC | KNS-62 | GDC0810 | top60_altered_DNA+RNA_seed | BRAF | -2.479845 | COADREAD | COLO-205 | PLX-4720 |
| MVP_top60_DNA_seed | EGFR | -0.579935 | LUSC | EPLC-272H | Erlotinib | top60_altered_DNA+RNA_seed | BRAF | -2.286426 | COADREAD | HCT-116 | RAF_9304 |
| MVP_top60_DNA_seed | SMARCA4 | -0.568088 | LUSC | SK-MES-1 | PFI3 | top60_altered_DNA+RNA_seed | ATM | -2.26219 | COADREAD | LoVo | KU-55933 |
| MVP_top60_DNA_seed | SMARCA2 | -0.568088 | LUSC | SK-MES-1 | PFI3 | top60_altered_DNA+RNA_seed | BRAF | -2.16582 | COADREAD | CL-11 | RAF_9304 |
| MVP_top60_DNA_seed | EGFR | -0.563444 | LUSC | SK-MES-1 | Lapatinib | top60_altered_DNA+RNA_seed | BRAF | -2.150984 | COADREAD | LS-411N | AZ628 |
| MVP_top60_DNA_seed | ESR1 | -0.560603 | LUSC | SK-MES-1 | GDC0810 | top60_altered_DNA+RNA_seed | BRAF | -2.064105 | COADREAD | MDST8 | RAF_9304 |
| MVP_top60_DNA_seed | PIK3CA | -0.531937 | LUSC | HCC-15 | Buparlisib | top60_altered_DNA+RNA_seed | BRAF | -1.973443 | COADREAD | C2BBe1 | SB590885 |
| MVP_top60_DNA_seed | CREBBP | -0.518356 | LUSC | KNS-62 | SGC-CBP30 | top60_altered_DNA+RNA_seed | BRAF | -1.791122 | COADREAD | MDST8 | PLX-4720 |
| MVP_top60_DNA_seed | HDAC1 | -0.485702 | LUSC | EPLC-272H | Romidepsin | top60_altered_DNA+RNA_seed | BRAF | -1.761566 | COADREAD | HT-29 | PLX-4720 |
| MVP_top60_DNA_seed | HDAC3 | -0.485702 | LUSC | EPLC-272H | Romidepsin | top60_altered_DNA+RNA_seed | BRAF | -1.707595 | COADREAD | HT-29 | PLX-4720 |
| MVP_top60_DNA_seed | HDAC2 | -0.485702 | LUSC | EPLC-272H | Romidepsin | top60_altered_DNA+RNA_seed | BRAF | -1.697462 | COADREAD | RKO | RAF_9304 |
| MVP_top60_DNA_seed | EGFR | -0.479051 | LUSC | SK-MES-1 | Osimertinib | top60_altered_DNA+RNA_seed | BRAF | -1.692775 | COADREAD | HT-29 | SB590885 |
| MVP_top60_DNA_seed | EGFR | -0.476759 | LUSC | HCC-15 | Erlotinib | top60_altered_DNA+RNA_seed | BRAF | -1.676996 | COADREAD | LS-411N | PLX-4720 |
| MVP_top60_DNA_seed | PIK3CA | -0.461466 | LUSC | NCI-H2170 | AZD8186 | top60_altered_DNA+RNA_seed | BRAF | -1.67516 | COADREAD | HCC2998 | AZ628 |
| MVP_top60_DNA_seed | CREBBP | -0.459354 | LUSC | EBC-1 | SGC-CBP30 | top60_altered_DNA+RNA_seed | PIK3CA | -1.570969 | COADREAD | CaR-1 | PI3Ka_4409 |
| MVP_top60_DNA_seed | CREBBP | -0.453786 | LUSC | EPLC-272H | SGC-CBP30 | top60_altered_DNA+RNA_seed | ATM | -1.540048 | COADREAD | LS-411N | KU-60019 |
| MVP_top60_DNA_seed | HDAC1 | -0.450813 | LUSC | LK-2 | PCI-34051 | top60_altered_DNA+RNA_seed | PIK3CA | -1.527487 | COADREAD | T84 | AZD8835 |
| MVP_top60_DNA_seed | SMARCA4 | -0.405381 | LUSC | EPLC-272H | PFI3 | top60_altered_DNA+RNA_seed | BRAF | -1.517703 | COADREAD | SK-CO-1 | SB590885 |
| MVP_top60_DNA_seed | SMARCA2 | -0.405381 | LUSC | EPLC-272H | PFI3 | top60_altered_DNA+RNA_seed | BRAF | -1.481452 | COADREAD | HT-29 | RAF_9304 |

| | | | | | | | | | | | |
|---|---|---|---|---|---|---|---|---|---|---|---|
| MVP_top60_DNA_seed | GSK3B | -0.383184 | LUSC | KNS-62 | SB216763 | top60_altered_DNA+RNA_seed | PIK3CA | -1.479836 | COADREAD | SW48 | AZD8835 |
| MVP_top60_DNA_seed | EGFR | -0.380056 | LUSC | HARA | AZD3759 | top60_altered_DNA+RNA_seed | BRAF | -1.402812 | COADREAD | NCI-H747 | AZ628 |
| MVP_top60_DNA_seed | EGFR | -0.377992 | LUSC | EBC-1 | Osimertinib | top60_altered_DNA+RNA_seed | BRAF | -1.385014 | COADREAD | RKO | Dabrafenib |
| MVP_top60_DNA_seed | RAC1 | -0.373581 | LUSC | LK-2 | EHT-1864 | top60_altered_DNA+RNA_seed | BRAF | -1.331183 | COADREAD | LS-411N | PLX-4720 |
| MVP_top60_DNA_seed | RAC1 | -0.373311 | LUSC | EPLC-272H | EHT-1864 | top60_altered_DNA+RNA_seed | ATM | -1.289227 | COADREAD | HCT-116 | KU-60019 |
| MVP_top60_DNA_seed | EGFR | -0.365011 | LUSC | LC-1-sq | Afatinib | top60_altered_DNA+RNA_seed | BRAF | -1.25549 | COADREAD | RKO | SB590885 |
| MVP_top60_DNA_seed | ESR1 | -0.364402 | LUSC | KNS-62 | Tamoxifen | top60_altered_DNA+RNA_seed | BRAF | -1.246299 | COADREAD | COLO-205 | RAF_9304 |
| MVP_top60_DNA_seed | SMARCA4 | -0.358104 | LUSC | RERF-LC-Sq1 | PFI3 | top60_altered_DNA+RNA_seed | BRAF | -1.215661 | COADREAD | SNU-C5 | PLX-4720 |
| MVP_top60_DNA_seed | SMARCA2 | -0.358104 | LUSC | RERF-LC-Sq1 | PFI3 | top60_altered_DNA+RNA_seed | BRAF | -1.207046 | COADREAD | CCK-81 | SB590885 |
| MVP_top60_DNA_seed | ESR1 | -0.355397 | LUSC | EBC-1 | Tamoxifen | top60_altered_DNA+RNA_seed | ATM | -1.191718 | COADREAD | MDST8 | torin2 |
| MVP_top60_DNA_seed | SMARCA4 | -0.354313 | LUSC | NCI-H2170 | PFI3 | top60_altered_DNA+RNA_seed | PIK3CA | -1.178649 | COADREAD | SK-CO-1 | AZD8835 |
| MVP_top60_DNA_seed | SMARCA2 | -0.354313 | LUSC | NCI-H2170 | PFI3 | top60_altered_DNA+RNA_seed | ATM | -1.134419 | COADREAD | NCI-H716 | KU-60019 |
| MVP_top60_DNA_seed | HDAC1 | -0.348232 | LUSC | HARA | Romidepsin | top60_altered_DNA+RNA_seed | BRAF | -1.132251 | COADREAD | LS-513 | AZ628 |
| MVP_top60_DNA_seed | HDAC3 | -0.348232 | LUSC | HARA | Romidepsin | top60_altered_DNA+RNA_seed | ATM | -1.124209 | COADREAD | SNU-175 | CP466722 |
| MVP_top60_DNA_seed | HDAC2 | -0.348232 | LUSC | HARA | Romidepsin | top60_altered_DNA+RNA_seed | BRAF | -1.097259 | COADREAD | LoVo | HG6-64-1 |
| MVP_top60_DNA_seed | ESR1 | -0.347209 | LUSC | HCC-15 | Tamoxifen | top60_altered_DNA+RNA_seed | PIK3CA | -1.086516 | COADREAD | GP5d | AZD8835 |
| MVP_top60_DNA_seed | PIK3CA | -0.33911 | LUSC | NCI-H520 | GNE-317 | top60_altered_DNA+RNA_seed | PIK3CA | -1.084276 | COADREAD | DiFi | PI3Ka_4409 |
| MVP_top60_DNA_seed | EZH2 | -0.339083 | LUSC | EPLC-272H | GSK343 | top60_altered_DNA+RNA_seed | PIK3CA | -1.081583 | COADREAD | SNU-407 | PI3Ka_4409 |
| MVP_top60_DNA_seed | EGFR | -0.32884 | LUSC | LOU-NH91 | Lapatinib | top60_altered_DNA+RNA_seed | BRAF | -1.077068 | COADREAD | COLO-205 | HG6-64-1 |
| MVP_top60_DNA_seed | EGFR | -0.326951 | LUSC | LC-1-sq | Osimertinib | top60_altered_DNA+RNA_seed | BRAF | -1.074232 | COADREAD | LS-513 | RAF_9304 |
| MVP_top60_DNA_seed | ESR1 | -0.314204 | LUSC | LK-2 | GDC0810 | top60_altered_DNA+RNA_seed | BRAF | -1.066656 | COADREAD | LS-411N | SB590885 |
| MVP_top60_DNA_seed | PIK3CA | -0.313509 | LUSC | LOU-NH91 | Alpelisib | top60_altered_DNA+RNA_seed | PIK3CA | -1.060381 | COADREAD | NCI-H747 | AZD8835 |
| MVP_top60_DNA_seed | ESR1 | -0.306721 | LUSC | HARA | GDC0810 | top60_altered_DNA+RNA_seed | BRAF | -1.048822 | COADREAD | LoVo | PLX-4720 |
| MVP_top60_DNA_seed | ESR1 | -0.304732 | LUSC | HCC-15 | GDC0810 | top60_altered_DNA+RNA_seed | ATM | -1.042063 | COADREAD | NCI-H716 | KU-55933 |
| MVP_top60_DNA_seed | RAC1 | -0.296971 | LUSC | HCC-15 | EHT-1864 | top60_altered_DNA+RNA_seed | BRAF | -1.031048 | COADREAD | MDST8 | PLX-4720 |
| MVP_top60_DNA_seed | PIK3CA | -0.290278 | LUSC | RERF-LC-Sq1 | Buparlisib | top60_altered_DNA+RNA_seed | PIK3CA | -1.013241 | COADREAD | HCT-116 | PI3Ka_4409 |
| MVP_top60_DNA_seed | PIK3CA | -0.287494 | LUSC | LOU-NH91 | GNE-317 | top60_altered_DNA+RNA_seed | ATM | -1.012839 | COADREAD | SNU-C5 | KU-55933 |
| MVP_top60_DNA_seed | PIK3CA | -0.283547 | LUSC | LK-2 | GNE-317 | top60_altered_DNA+RNA_seed | BRAF | -0.99188 | COADREAD | LS-411N | Dabrafenib |
| MVP_top60_DNA_seed | RAC1 | -0.283352 | LUSC | KNS-62 | EHT-1864 | top60_altered_DNA+RNA_seed | BRAF | -0.990068 | COADREAD | HCT-116 | Dabrafenib |
| MVP_top60_DNA_seed | PIK3CA | -0.280465 | LUSC | LOU-NH91 | AZD8186 | top60_altered_DNA+RNA_seed | PIK3CA | -0.984215 | COADREAD | GP5d | PI3Ka_4409 |
| MVP_top60_DNA_seed | PIK3CA | -0.27608 | LUSC | NCI-H1869 | AZD8186 | top60_altered_DNA+RNA_seed | ATM | -0.98135 | COADREAD | KM12 | KU-60019 |
| MVP_top60_DNA_seed | PIK3CA | -0.270405 | LUSC | LK-2 | Buparlisib | top60_altered_DNA+RNA_seed | BRAF | -0.953793 | COADREAD | LS-411N | RAF_9304 |
| MVP_top60_DNA_seed | HDAC1 | -0.251449 | LUSC | NCI-H1869 | Romidepsin | top60_altered_DNA+RNA_seed | PIK3CA | -0.928663 | COADREAD | HCT-116 | AZD8835 |
| MVP_top60_DNA_seed | HDAC3 | -0.251449 | LUSC | NCI-H1869 | Romidepsin | top60_altered_DNA+RNA_seed | BRAF | -0.905346 | COADREAD | SNU-C5 | Dabrafenib |
| MVP_top60_DNA_seed | HDAC2 | -0.251449 | LUSC | NCI-H1869 | Romidepsin | top60_altered_DNA+RNA_seed | ATM | -0.890736 | COADREAD | LoVo | KU-60019 |
| MVP_top60_DNA_seed | EZH2 | -0.248381 | LUSC | KNS-62 | GSK343 | top60_altered_DNA+RNA_seed | PIK3CA | -0.869761 | COADREAD | RKO | AZD8835 |
| MVP_top60_DNA_seed | EGFR | -0.247437 | LUSC | HCC-15 | Gefitinib | top60_altered_DNA+RNA_seed | BRAF | -0.866334 | COADREAD | HCC2998 | RAF_9304 |
| MVP_top60_DNA_seed | HDAC1 | -0.244018 | LUSC | SK-MES-1 | Romidepsin | top60_altered_DNA+RNA_seed | BRAF | -0.858547 | COADREAD | SW948 | SB590885 |
| MVP_top60_DNA_seed | HDAC3 | -0.244018 | LUSC | SK-MES-1 | Romidepsin | top60_altered_DNA+RNA_seed | PIK3CA | -0.844898 | COADREAD | SNU-C1 | AZD8835 |
| MVP_top60_DNA_seed | HDAC2 | -0.244018 | LUSC | SK-MES-1 | Romidepsin | top60_altered_DNA+RNA_seed | PIK3CA | -0.843751 | COADREAD | SW48 | PI3Ka_4409 |
| MVP_top60_DNA_seed | PIK3CA | -0.210267 | LUSC | EBC-1 | Buparlisib | top60_altered_DNA+RNA_seed | ATM | -0.825948 | COADREAD | CL-11 | KU-60019 |

| | | | | | | | | | | | |
|---|---|---|---|---|---|---|---|---|---|---|---|
| MVP_top60_DNA_seed | EZH2 | -0.197828 | LUSC | NCI-H2170 | GSK343 | top60_altered_DNA+RNA_seed | BRAF | -0.801322 | COADREAD | NCI-H747 | RAF_9304 |
| MVP_top60_DNA_seed | HDAC1 | -0.194467 | LUSC | NCI-H520 | Entinostat | top60_altered_DNA+RNA_seed | BRAF | -0.800854 | COADREAD | HCT-116 | HG6-64-1 |
| MVP_top60_DNA_seed | HDAC3 | -0.194467 | LUSC | NCI-H520 | Entinostat | top60_altered_DNA+RNA_seed | ATM | -0.790348 | COADREAD | HCT-116 | torin2 |
| MVP_top60_DNA_seed | HDAC1 | -0.193978 | LUSC | LK-2 | Romidepsin | top60_altered_DNA+RNA_seed | ATM | -0.790196 | COADREAD | RKO | KU-60019 |
| MVP_top60_DNA_seed | HDAC3 | -0.193978 | LUSC | LK-2 | Romidepsin | top60_altered_DNA+RNA_seed | BRAF | -0.77774 | COADREAD | HCT-15 | HG6-64-1 |
| MVP_top60_DNA_seed | HDAC2 | -0.193978 | LUSC | LK-2 | Romidepsin | top60_altered_DNA+RNA_seed | ATM | -0.776531 | COADREAD | SNU-81 | CP466722 |
| MVP_top60_DNA_seed | HDAC1 | -0.185329 | LUSC | NCI-H2170 | Entinostat | top60_altered_DNA+RNA_seed | BRAF | -0.743223 | COADREAD | LoVo | Dabrafenib |
| MVP_top60_DNA_seed | HDAC3 | -0.185329 | LUSC | NCI-H2170 | Entinostat | top60_altered_DNA+RNA_seed | BRAF | -0.740234 | COADREAD | GP5d | HG6-64-1 |
| MVP_top60_DNA_seed | HDAC1 | -0.176826 | LUSC | SK-MES-1 | PCI-34051 | top60_altered_DNA+RNA_seed | PIK3CA | -0.72489 | COADREAD | COLO-320-HSR | AZD8835 |
| MVP_top60_DNA_seed | EGFR | -0.166191 | LUSC | LK-2 | Osimertinib | top60_altered_DNA+RNA_seed | PIK3CA | -0.715688 | COADREAD | RKO | PI3Ka_4409 |
| MVP_top60_DNA_seed | EZH2 | -0.159379 | LUSC | HARA | GSK343 | top60_altered_DNA+RNA_seed | BRAF | -0.709088 | COADREAD | LoVo | RAF_9304 |
| MVP_top60_DNA_seed | CREBBP | -0.158355 | LUSC | HARA | SGC-CBP30 | top60_altered_DNA+RNA_seed | ATM | -0.709043 | COADREAD | HCT-116 | CP466722 |
| MVP_top60_DNA_seed | EGFR | -0.156562 | LUSC | NCI-H1869 | Afatinib | top60_altered_DNA+RNA_seed | ATM | -0.698908 | COADREAD | LS-180 | CP466722 |
| MVP_top60_DNA_seed | ESR1 | -0.116985 | LUSC | RERF-LC-Sq1 | Tamoxifen | top60_altered_DNA+RNA_seed | PIK3CA | -0.694177 | COADREAD | COLO-205 | PI3Ka_4409 |
| MVP_top60_DNA_seed | GSK3B | -0.115766 | LUSC | SK-MES-1 | SB216763 | top60_altered_DNA+RNA_seed | BRAF | -0.684369 | COADREAD | KM12 | AZ628 |
| MVP_top60_DNA_seed | HDAC1 | -0.11281 | LUSC | EPLC-272H | Entinostat | top60_altered_DNA+RNA_seed | BRAF | -0.675173 | COADREAD | CL-11 | AZ628 |
| MVP_top60_DNA_seed | HDAC3 | -0.11281 | LUSC | EPLC-272H | Entinostat | top60_altered_DNA+RNA_seed | ATM | -0.673183 | COADREAD | SNU-81 | KU-55933 |
| MVP_top60_DNA_seed | ESR1 | -0.096543 | LUSC | LK-2 | Tamoxifen | top60_altered_DNA+RNA_seed | BRAF | -0.662747 | COADREAD | SK-CO-1 | RAF_9304 |
| MVP_top60_DNA_seed | PIK3CA | -0.086508 | LUSC | LC-1-sq | Alpelisib | top60_altered_DNA+RNA_seed | ATM | -0.635834 | COADREAD | CW-2 | KU-60019 |
| MVP_top60_DNA_seed | HDAC1 | -0.081973 | LUSC | EPLC-272H | PCI-34051 | top60_altered_DNA+RNA_seed | PIK3CA | -0.627517 | COADREAD | DiFi | AZD8835 |
| MVP_top60_DNA_seed | SMARCA4 | -0.070697 | LUSC | HARA | PFI3 | top60_altered_DNA+RNA_seed | PIK3CA | -0.624684 | COADREAD | SNU-C5 | AZD8835 |
| MVP_top60_DNA_seed | SMARCA2 | -0.070697 | LUSC | HARA | PFI3 | top60_altered_DNA+RNA_seed | ATM | -0.61809 | COADREAD | NCI-H747 | CP466722 |
| MVP_top60_DNA_seed | HDAC1 | -0.057777 | LUSC | NCI-H2170 | Romidepsin | top60_altered_DNA+RNA_seed | BRAF | -0.614838 | COADREAD | NCI-H508 | PLX-4720 |
| MVP_top60_DNA_seed | HDAC3 | -0.057777 | LUSC | NCI-H2170 | Romidepsin | top60_altered_DNA+RNA_seed | PIK3CA | -0.61323 | COADREAD | SK-CO-1 | PI3Ka_4409 |
| MVP_top60_DNA_seed | HDAC2 | -0.057777 | LUSC | NCI-H2170 | Romidepsin | top60_altered_DNA+RNA_seed | BRAF | -0.603733 | COADREAD | RKO | PLX-4720 |
| MVP_top60_DNA_seed | PIK3CA | -0.039436 | LUSC | LOU-NH91 | Buparlisib | top60_altered_DNA+RNA_seed | BRAF | -0.598427 | COADREAD | SNU-C5 | PLX-4720 |
| MVP_top60_DNA_seed | CREBBP | -0.037979 | LUSC | LOU-NH91 | SGC-CBP30 | top60_altered_DNA+RNA_seed | BRAF | -0.592205 | COADREAD | NCI-H716 | SB590885 |
| MVP_top60_DNA_seed | TP53 | -0.002669 | LUSC | NCI-H1869 | MIRA-1 | top60_altered_DNA+RNA_seed | ATM | -0.587908 | COADREAD | CCK-81 | KU-60019 |
| MVP_top60_DNA_seed | TP53 | 0.006589 | LUSC | NCI-H2170 | MIRA-1 | top60_altered_DNA+RNA_seed | PIK3CA | -0.586716 | COADREAD | SNU-C5 | PI3Ka_4409 |
| MVP_top60_DNA_seed | PIK3CA | 0.035824 | LUSC | SK-MES-1 | AZD8186 | top60_altered_DNA+RNA_seed | ATM | -0.584066 | COADREAD | COLO-320-HSR | KU-60019 |
| MVP_top60_DNA_seed | EGFR | 0.042899 | LUSC | KNS-62 | Gefitinib | top60_altered_DNA+RNA_seed | PIK3CA | -0.572921 | COADREAD | SNU-C1 | PI3Ka_4409 |
| MVP_top60_DNA_seed | EGFR | 0.057401 | LUSC | NCI-H520 | Afatinib | top60_altered_DNA+RNA_seed | BRAF | -0.572903 | COADREAD | LS-411N | HG6-64-1 |
| MVP_top60_DNA_seed | HDAC1 | 0.057695 | LUSC | LC-1-sq | Romidepsin | top60_altered_DNA+RNA_seed | BRAF | -0.571538 | COADREAD | HCT-116 | SB590885 |
| MVP_top60_DNA_seed | HDAC3 | 0.057695 | LUSC | LC-1-sq | Romidepsin | top60_altered_DNA+RNA_seed | BRAF | -0.557702 | COADREAD | LS-180 | SB590885 |
| MVP_top60_DNA_seed | HDAC2 | 0.057695 | LUSC | LC-1-sq | Romidepsin | top60_altered_DNA+RNA_seed | BRAF | -0.544497 | COADREAD | SW948 | RAF_9304 |
| MVP_top60_DNA_seed | TP53 | 0.085711 | LUSC | HCC-15 | MIRA-1 | top60_altered_DNA+RNA_seed | PIK3CA | -0.542571 | COADREAD | LoVo | AZD8835 |
| MVP_top60_DNA_seed | HDAC1 | 0.086914 | LUSC | NCI-H2170 | PCI-34051 | top60_altered_DNA+RNA_seed | PIK3CA | -0.539297 | COADREAD | NCI-H747 | PI3Ka_4409 |
| MVP_top60_DNA_seed | EZH2 | 0.088756 | LUSC | NCI-H2170 | GSK343 | top60_altered_DNA+RNA_seed | PIK3CA | -0.534642 | COADREAD | HT-29 | AZD8835 |
| MVP_top60_DNA_seed | EGFR | 0.108767 | LUSC | NCI-H2170 | AZD3759 | top60_altered_DNA+RNA_seed | PIK3CA | -0.521684 | COADREAD | NCI-H716 | PI3Ka_4409 |
| MVP_top60_DNA_seed | HDAC1 | 0.112647 | LUSC | HCC-15 | Romidepsin | top60_altered_DNA+RNA_seed | ATM | -0.519922 | COADREAD | CL-11 | KU-55933 |
| MVP_top60_DNA_seed | HDAC3 | 0.112647 | LUSC | HCC-15 | Romidepsin | top60_altered_DNA+RNA_seed | ATM | -0.518202 | COADREAD | SK-CO-1 | KU-55933 |

| | | | | | | | | | | | |
|---|---|---|---|---|---|---|---|---|---|---|---|
| MVP_top60_DNA_seed | HDAC2 | 0.112647 | LUSC | HCC-15 | Romidepsin | top60_altered_DNA+RNA_seed | BRAF | -0.513367 | COADREAD | CaR-1 | RAF_9304 |
| MVP_top60_DNA_seed | EZH2 | 0.116397 | LUSC | RERF-LC-Sq1 | GSK343 | top60_altered_DNA+RNA_seed | ATM | -0.511225 | COADREAD | T84 | CP466722 |
| MVP_top60_DNA_seed | TP53 | 0.117814 | LUSC | NCI-H520 | MIRA-1 | top60_altered_DNA+RNA_seed | ATM | -0.510789 | COADREAD | SNU-1040 | CP466722 |
| MVP_top60_DNA_seed | EGFR | 0.118382 | LUSC | EBC-1 | Erlotinib | top60_altered_DNA+RNA_seed | BRAF | -0.495341 | COADREAD | MDST8 | AZ628 |
| MVP_top60_DNA_seed | TP53 | 0.127705 | LUSC | HARA | MIRA-1 | top60_altered_DNA+RNA_seed | BRAF | -0.48826 | COADREAD | SNU-407 | RAF_9304 |
| MVP_top60_DNA_seed | SMARCA4 | 0.131684 | LUSC | KNS-62 | PFI3 | top60_altered_DNA+RNA_seed | BRAF | -0.474294 | COADREAD | NCI-H716 | RAF_9304 |
| MVP_top60_DNA_seed | SMARCA2 | 0.131684 | LUSC | KNS-62 | PFI3 | top60_altered_DNA+RNA_seed | PIK3CA | -0.474138 | COADREAD | SW948 | PI3Ka_4409 |
| MVP_top60_DNA_seed | EZH2 | 0.150594 | LUSC | RERF-LC-Sq1 | GSK343 | top60_altered_DNA+RNA_seed | ATM | -0.463427 | COADREAD | COLO-320-HSR | CP466722 |
| MVP_top60_DNA_seed | GSK3B | 0.164155 | LUSC | EPLC-272H | SB216763 | top60_altered_DNA+RNA_seed | ATM | -0.461765 | COADREAD | CCK-81 | CP466722 |
| MVP_top60_DNA_seed | TP53 | 0.175425 | LUSC | LC-1-sq | MIRA-1 | top60_altered_DNA+RNA_seed | PIK3CA | -0.433547 | COADREAD | NCI-H508 | PI3Ka_4409 |
| MVP_top60_DNA_seed | EGFR | 0.177649 | LUSC | LC-1-sq | Lapatinib | top60_altered_DNA+RNA_seed | ATM | -0.432508 | COADREAD | HCT-15 | CP466722 |
| MVP_top60_DNA_seed | PIK3CA | 0.179874 | LUSC | NCI-H520 | Buparlisib | top60_altered_DNA+RNA_seed | ATM | -0.432033 | COADREAD | SNU-81 | KU-60019 |
| MVP_top60_DNA_seed | ESR1 | 0.180197 | LUSC | NCI-H520 | Tamoxifen | top60_altered_DNA+RNA_seed | BRAF | -0.417936 | COADREAD | MDST8 | SB590885 |
| MVP_top60_DNA_seed | HDAC1 | 0.205321 | LUSC | HARA | PCI-34051 | top60_altered_DNA+RNA_seed | ATM | -0.414445 | COADREAD | KM12 | CP466722 |
| MVP_top60_DNA_seed | CREBBP | 0.208739 | LUSC | NCI-H2170 | SGC-CBP30 | top60_altered_DNA+RNA_seed | BRAF | -0.383245 | COADREAD | HCT-15 | PLX-4720 |
| MVP_top60_DNA_seed | HDAC1 | 0.211623 | LUSC | LOU-NH91 | Entinostat | top60_altered_DNA+RNA_seed | ATM | -0.38166 | COADREAD | COLO-205 | CP466722 |
| MVP_top60_DNA_seed | HDAC3 | 0.211623 | LUSC | LOU-NH91 | Entinostat | top60_altered_DNA+RNA_seed | PIK3CA | -0.36886 | COADREAD | LS-513 | PI3Ka_4409 |
| MVP_top60_DNA_seed | TP53 | 0.215474 | LUSC | LK-2 | MIRA-1 | top60_altered_DNA+RNA_seed | BRAF | -0.367373 | COADREAD | LS-180 | HG6-64-1 |
| MVP_top60_DNA_seed | SMARCA4 | 0.236079 | LUSC | LC-1-sq | PFI3 | top60_altered_DNA+RNA_seed | BRAF | -0.36118 | COADREAD | SW48 | RAF_9304 |
| MVP_top60_DNA_seed | SMARCA2 | 0.236079 | LUSC | LC-1-sq | PFI3 | top60_altered_DNA+RNA_seed | ATM | -0.360689 | COADREAD | HCT-15 | KU-60019 |
| MVP_top60_DNA_seed | EGFR | 0.237191 | LUSC | EBC-1 | Lapatinib | top60_altered_DNA+RNA_seed | ATM | -0.353132 | COADREAD | RKO | CP466722 |
| MVP_top60_DNA_seed | ESR1 | 0.264243 | LUSC | NCI-H1869 | Tamoxifen | top60_altered_DNA+RNA_seed | ATM | -0.353111 | COADREAD | SW620 | CP466722 |
| MVP_top60_DNA_seed | GSK3B | 0.283736 | LUSC | NCI-H2170 | SB216763 | top60_altered_DNA+RNA_seed | BRAF | -0.348715 | COADREAD | NCI-H747 | HG6-64-1 |
| MVP_top60_DNA_seed | SMARCA4 | 0.307263 | LUSC | HCC-15 | PFI3 | top60_altered_DNA+RNA_seed | PIK3CA | -0.347499 | COADREAD | COLO-320-HSR | PI3Ka_4409 |
| MVP_top60_DNA_seed | SMARCA2 | 0.307263 | LUSC | HCC-15 | PFI3 | top60_altered_DNA+RNA_seed | BRAF | -0.345525 | COADREAD | KM12 | HG6-64-1 |
| MVP_top60_DNA_seed | EGFR | 0.309128 | LUSC | KNS-62 | AZD3759 | top60_altered_DNA+RNA_seed | PIK3CA | -0.344309 | COADREAD | NCI-H716 | AZD8835 |
| MVP_top60_DNA_seed | EGFR | 0.31637 | LUSC | LK-2 | Afatinib | top60_altered_DNA+RNA_seed | PIK3CA | -0.336371 | COADREAD | SW1463 | PI3Ka_4409 |
| MVP_top60_DNA_seed | HDAC1 | 0.317533 | LUSC | HCC-15 | PCI-34051 | top60_altered_DNA+RNA_seed | BRAF | -0.335172 | COADREAD | GP5d | RAF_9304 |
| MVP_top60_DNA_seed | ESR1 | 0.317926 | LUSC | NCI-H2170 | Tamoxifen | top60_altered_DNA+RNA_seed | PIK3CA | -0.319794 | COADREAD | COLO-205 | AZD8835 |
| MVP_top60_DNA_seed | GSK3B | 0.322646 | LUSC | LK-2 | SB216763 | top60_altered_DNA+RNA_seed | PIK3CA | -0.318476 | COADREAD | MDST8 | AZD8835 |
| MVP_top60_DNA_seed | PIK3CA | 0.328737 | LUSC | SK-MES-1 | GNE-317 | top60_altered_DNA+RNA_seed | BRAF | -0.30831 | COADREAD | SW1417 | PLX-4720 |
| MVP_top60_DNA_seed | PIK3CA | 0.332925 | LUSC | HARA | AZD8186 | top60_altered_DNA+RNA_seed | BRAF | -0.303669 | COADREAD | SNU-C5 | RAF_9304 |
| MVP_top60_DNA_seed | HDAC1 | 0.337079 | LUSC | KNS-62 | PCI-34051 | top60_altered_DNA+RNA_seed | ATM | -0.283284 | COADREAD | SW48 | KU-55933 |
| MVP_top60_DNA_seed | HDAC1 | 0.346509 | LUSC | LC-1-sq | Entinostat | top60_altered_DNA+RNA_seed | ATM | -0.277802 | COADREAD | MDST8 | KU-60019 |
| MVP_top60_DNA_seed | HDAC3 | 0.346509 | LUSC | LC-1-sq | Entinostat | top60_altered_DNA+RNA_seed | ATM | -0.273401 | COADREAD | SK-CO-1 | KU-60019 |
| MVP_top60_DNA_seed | PIK3CA | 0.357542 | LUSC | NCI-H1869 | GNE-317 | top60_altered_DNA+RNA_seed | PIK3CA | -0.2684 | COADREAD | CCK-81 | AZD8835 |
| MVP_top60_DNA_seed | ESR1 | 0.362385 | LUSC | NCI-H2170 | GDC0810 | top60_altered_DNA+RNA_seed | BRAF | -0.265451 | COADREAD | SNU-407 | Dabrafenib |
| MVP_top60_DNA_seed | EZH2 | 0.370852 | LUSC | NCI-H520 | GSK343 | top60_altered_DNA+RNA_seed | BRAF | -0.265331 | COADREAD | HT-115 | RAF_9304 |
| MVP_top60_DNA_seed | TP53 | 0.371749 | LUSC | EPLC-272H | MIRA-1 | top60_altered_DNA+RNA_seed | ATM | -0.243581 | COADREAD | LS-411N | CP466722 |
| MVP_top60_DNA_seed | HDAC1 | 0.378496 | LUSC | RERF-LC-Sq1 | Entinostat | top60_altered_DNA+RNA_seed | BRAF | -0.242675 | COADREAD | SNU-C2B | RAF_9304 |
| MVP_top60_DNA_seed | HDAC3 | 0.378496 | LUSC | RERF-LC-Sq1 | Entinostat | top60_altered_DNA+RNA_seed | BRAF | -0.238157 | COADREAD | NCI-H716 | PLX-4720 |

| | | | | | | | | | | | |
|---|---|---|---|---|---|---|---|---|---|---|---|
| MVP_top60_DNA_seed | PIK3CA | 0.384334 | LUSC | RERF-LC-Sq1 | AZD8186 | top60_altered_DNA+RNA_seed | BRAF | -0.220269 | COADREAD | SW837 | RAF_9304 |
| MVP_top60_DNA_seed | RAC1 | 0.386393 | LUSC | HARA | EHT-1864 | top60_altered_DNA+RNA_seed | BRAF | -0.191986 | COADREAD | RCM-1 | Dabrafenib |
| MVP_top60_DNA_seed | EGFR | 0.40832 | LUSC | LK-2 | AZD3759 | top60_altered_DNA+RNA_seed | PIK3CA | -0.191277 | COADREAD | LS-180 | AZD8835 |
| MVP_top60_DNA_seed | HDAC1 | 0.415626 | LUSC | LOU-NH91 | PCI-34051 | top60_altered_DNA+RNA_seed | PIK3CA | -0.182657 | COADREAD | LoVo | PI3Ka_4409 |
| MVP_top60_DNA_seed | EGFR | 0.431164 | LUSC | KNS-62 | Osimertinib | top60_altered_DNA+RNA_seed | ATM | -0.182176 | COADREAD | SW837 | CP466722 |
| MVP_top60_DNA_seed | EGFR | 0.442611 | LUSC | KNS-62 | Erlotinib | top60_altered_DNA+RNA_seed | ATM | -0.173003 | COADREAD | LoVo | torin2 |
| MVP_top60_DNA_seed | EZH2 | 0.444696 | LUSC | LK-2 | GSK343 | top60_altered_DNA+RNA_seed | PIK3CA | -0.168674 | COADREAD | HCT-15 | AZD8835 |
| MVP_top60_DNA_seed | RAC1 | 0.448091 | LUSC | NCI-H2170 | EHT-1864 | top60_altered_DNA+RNA_seed | BRAF | -0.154587 | COADREAD | LS-180 | RAF_9304 |
| MVP_top60_DNA_seed | EGFR | 0.449123 | LUSC | NCI-H520 | Lapatinib | top60_altered_DNA+RNA_seed | ATM | -0.151994 | COADREAD | HCT-15 | KU-55933 |
| MVP_top60_DNA_seed | SMARCA4 | 0.458455 | LUSC | NCI-H520 | PFI3 | top60_altered_DNA+RNA_seed | BRAF | -0.147022 | COADREAD | SW1417 | RAF_9304 |
| MVP_top60_DNA_seed | SMARCA2 | 0.458455 | LUSC | NCI-H520 | PFI3 | top60_altered_DNA+RNA_seed | PIK3CA | -0.130352 | COADREAD | T84 | PI3Ka_4409 |
| MVP_top60_DNA_seed | PIK3CA | 0.493276 | LUSC | LC-1-sq | Buparlisib | top60_altered_DNA+RNA_seed | BRAF | -0.125091 | COADREAD | LS-180 | PLX-4720 |
| MVP_top60_DNA_seed | SMARCA4 | 0.498532 | LUSC | LK-2 | PFI3 | top60_altered_DNA+RNA_seed | PIK3CA | -0.118668 | COADREAD | NCI-H508 | AZD8835 |
| MVP_top60_DNA_seed | SMARCA2 | 0.498532 | LUSC | LK-2 | PFI3 | top60_altered_DNA+RNA_seed | PIK3CA | -0.113655 | COADREAD | HT-115 | PI3Ka_4409 |
| MVP_top60_DNA_seed | GSK3B | 0.509331 | LUSC | NCI-H520 | SB216763 | top60_altered_DNA+RNA_seed | BRAF | -0.109933 | COADREAD | SNU-C5 | SB590885 |
| MVP_top60_DNA_seed | EZH2 | 0.510513 | LUSC | HCC-15 | GSK343 | top60_altered_DNA+RNA_seed | BRAF | -0.106926 | COADREAD | C2BBe1 | HG6-64-1 |
| MVP_top60_DNA_seed | EGFR | 0.516684 | LUSC | KNS-62 | Afatinib | top60_altered_DNA+RNA_seed | PIK3CA | -0.099566 | COADREAD | LS-513 | AZD8835 |
| MVP_top60_DNA_seed | HDAC1 | 0.521161 | LUSC | HARA | Entinostat | top60_altered_DNA+RNA_seed | BRAF | -0.092861 | COADREAD | CL-11 | Dabrafenib |
| MVP_top60_DNA_seed | HDAC3 | 0.521161 | LUSC | HARA | Entinostat | top60_altered_DNA+RNA_seed | PIK3CA | -0.092356 | COADREAD | HT-115 | AZD8835 |
| MVP_top60_DNA_seed | EGFR | 0.524351 | LUSC | LK-2 | Lapatinib | top60_altered_DNA+RNA_seed | BRAF | -0.088897 | COADREAD | SK-CO-1 | Dabrafenib |
| MVP_top60_DNA_seed | PIK3CA | 0.528927 | LUSC | HCC-15 | GNE-317 | top60_altered_DNA+RNA_seed | PIK3CA | -0.086547 | COADREAD | LS-180 | PI3Ka_4409 |
| MVP_top60_DNA_seed | ESR1 | 0.52982 | LUSC | SK-MES-1 | Tamoxifen | top60_altered_DNA+RNA_seed | PIK3CA | -0.077205 | COADREAD | CCK-81 | PI3Ka_4409 |
| MVP_top60_DNA_seed | GSK3B | 0.533185 | LUSC | RERF-LC-Sq1 | SB216763 | top60_altered_DNA+RNA_seed | BRAF | -0.076556 | COADREAD | RKO | HG6-64-1 |
| MVP_top60_DNA_seed | GSK3B | 0.537321 | LUSC | LC-1-sq | SB216763 | top60_altered_DNA+RNA_seed | ATM | -0.067091 | COADREAD | SNU-407 | CP466722 |
| MVP_top60_DNA_seed | EGFR | 0.554849 | LUSC | EBC-1 | Sapitinib | top60_altered_DNA+RNA_seed | BRAF | -0.064654 | COADREAD | HT-29 | HG6-64-1 |
| MVP_top60_DNA_seed | CREBBP | 0.555644 | LUSC | SK-MES-1 | SGC-CBP30 | top60_altered_DNA+RNA_seed | BRAF | -0.063906 | COADREAD | HT55 | RAF_9304 |
| MVP_top60_DNA_seed | ESR1 | 0.558151 | LUSC | LC-1-sq | Tamoxifen | top60_altered_DNA+RNA_seed | ATM | -0.057984 | COADREAD | SW48 | torin2 |
| MVP_top60_DNA_seed | PIK3CA | 0.568534 | LUSC | KNS-62 | Alpelisib | top60_altered_DNA+RNA_seed | BRAF | -0.056508 | COADREAD | RCM-1 | SB590885 |
| MVP_top60_DNA_seed | EZH2 | 0.573817 | LUSC | NCI-H520 | GSK343 | top60_altered_DNA+RNA_seed | BRAF | -0.052428 | COADREAD | SW620 | HG6-64-1 |
| MVP_top60_DNA_seed | GSK3B | 0.57578 | LUSC | HCC-15 | SB216763 | top60_altered_DNA+RNA_seed | BRAF | -0.052056 | COADREAD | NCI-H716 | HG6-64-1 |
| MVP_top60_DNA_seed | EGFR | 0.57822 | LUSC | NCI-H520 | Erlotinib | top60_altered_DNA+RNA_seed | BRAF | -0.051633 | COADREAD | LoVo | SB590885 |
| MVP_top60_DNA_seed | EGFR | 0.5917 | LUSC | KNS-62 | Lapatinib | top60_altered_DNA+RNA_seed | BRAF | -0.046015 | COADREAD | SW620 | RAF_9304 |
| MVP_top60_DNA_seed | TP53 | 0.593917 | LUSC | RERF-LC-Sq1 | MIRA-1 | top60_altered_DNA+RNA_seed | BRAF | -0.038668 | COADREAD | RCM-1 | RAF_9304 |
| MVP_top60_DNA_seed | HDAC1 | 0.60373 | LUSC | KNS-62 | Romidepsin | top60_altered_DNA+RNA_seed | ATM | -0.037897 | COADREAD | LS-180 | KU-55933 |
| MVP_top60_DNA_seed | HDAC3 | 0.60373 | LUSC | KNS-62 | Romidepsin | top60_altered_DNA+RNA_seed | ATM | -0.035626 | COADREAD | GP5d | CP466722 |
| MVP_top60_DNA_seed | HDAC2 | 0.60373 | LUSC | KNS-62 | Romidepsin | top60_altered_DNA+RNA_seed | BRAF | -0.024449 | COADREAD | COLO-678 | HG6-64-1 |
| MVP_top60_DNA_seed | EZH2 | 0.609998 | LUSC | KNS-62 | GSK343 | top60_altered_DNA+RNA_seed | ATM | -0.012139 | COADREAD | RKO | KU-55933 |
| MVP_top60_DNA_seed | PIK3CA | 0.621815 | LUSC | KNS-62 | GNE-317 | top60_altered_DNA+RNA_seed | BRAF | 0.005611 | COADREAD | LS-123 | RAF_9304 |
| MVP_top60_DNA_seed | HDAC1 | 0.622254 | LUSC | KNS-62 | Entinostat | top60_altered_DNA+RNA_seed | BRAF | 0.008408 | COADREAD | C2BBe1 | RAF_9304 |
| MVP_top60_DNA_seed | HDAC3 | 0.622254 | LUSC | KNS-62 | Entinostat | top60_altered_DNA+RNA_seed | BRAF | 0.014266 | COADREAD | SW1417 | PLX-4720 |
| MVP_top60_DNA_seed | HDAC1 | 0.623486 | LUSC | NCI-H1869 | Entinostat | top60_altered_DNA+RNA_seed | BRAF | 0.021297 | COADREAD | SW1116 | RAF_9304 |

| | | | | | | | | | | | |
|---|---|---|---|---|---|---|---|---|---|---|---|
| MVP_top60_DNA_seed | HDAC3 | 0.623486 | LUSC | NCI-H1869 | Entinostat | top60_altered_DNA+RNA_seed | PIK3CA | 0.022367 | COADREAD | SNU-81 | AZD8835 |
| MVP_top60_DNA_seed | EGFR | 0.624659 | LUSC | HCC-15 | Lapatinib | top60_altered_DNA+RNA_seed | PIK3CA | 0.028046 | COADREAD | CL-11 | PI3Ka_4409 |
| MVP_top60_DNA_seed | EGFR | 0.625961 | LUSC | KNS-62 | Sapitinib | top60_altered_DNA+RNA_seed | BRAF | 0.036466 | COADREAD | SW1463 | SB590885 |
| MVP_top60_DNA_seed | PIK3CA | 0.637805 | LUSC | NCI-H520 | Alpelisib | top60_altered_DNA+RNA_seed | PIK3CA | 0.037754 | COADREAD | LS-123 | PI3Ka_4409 |
| MVP_top60_DNA_seed | HDAC1 | 0.637975 | LUSC | EBC-1 | PCI-34051 | top60_altered_DNA+RNA_seed | PIK3CA | 0.038105 | COADREAD | SW837 | PI3Ka_4409 |
| MVP_top60_DNA_seed | PIK3CA | 0.638856 | LUSC | SK-MES-1 | Buparlisib | top60_altered_DNA+RNA_seed | PIK3CA | 0.039911 | COADREAD | SW948 | AZD8835 |
| MVP_top60_DNA_seed | CREBBP | 0.647097 | LUSC | LK-2 | SGC-CBP30 | top60_altered_DNA+RNA_seed | BRAF | 0.046935 | COADREAD | CCK-81 | RAF_9304 |
| MVP_top60_DNA_seed | PIK3CA | 0.654019 | LUSC | KNS-62 | Buparlisib | top60_altered_DNA+RNA_seed | BRAF | 0.050947 | COADREAD | GP5d | PLX-4720 |
| MVP_top60_DNA_seed | SMARCA4 | 0.667792 | LUSC | LOU-NH91 | PFI3 | top60_altered_DNA+RNA_seed | BRAF | 0.053849 | COADREAD | COLO-320-HSR | HG6-64-1 |
| MVP_top60_DNA_seed | SMARCA2 | 0.667792 | LUSC | LOU-NH91 | PFI3 | top60_altered_DNA+RNA_seed | PIK3CA | 0.057106 | COADREAD | HT-29 | PI3Ka_4409 |
| MVP_top60_DNA_seed | EZH2 | 0.667914 | LUSC | HCC-15 | GSK343 | top60_altered_DNA+RNA_seed | ATM | 0.062083 | COADREAD | SW620 | KU-60019 |
| MVP_top60_DNA_seed | EZH2 | 0.681502 | LUSC | LC-1-sq | GSK343 | top60_altered_DNA+RNA_seed | BRAF | 0.063099 | COADREAD | SK-CO-1 | PLX-4720 |
| MVP_top60_DNA_seed | EGFR | 0.68342 | LUSC | HCC-15 | Osimertinib | top60_altered_DNA+RNA_seed | PIK3CA | 0.064424 | COADREAD | LS-1034 | PI3Ka_4409 |
| MVP_top60_DNA_seed | TP53 | 0.691211 | LUSC | KNS-62 | MIRA-1 | top60_altered_DNA+RNA_seed | BRAF | 0.06462 | COADREAD | SW1417 | Dabrafenib |
| MVP_top60_DNA_seed | EGFR | 0.691763 | LUSC | NCI-H520 | Gefitinib | top60_altered_DNA+RNA_seed | BRAF | 0.067297 | COADREAD | NCI-H747 | Dabrafenib |
| MVP_top60_DNA_seed | HDAC1 | 0.712882 | LUSC | RERF-LC-Sq1 | PCI-34051 | top60_altered_DNA+RNA_seed | BRAF | 0.074205 | COADREAD | SW48 | PLX-4720 |
| MVP_top60_DNA_seed | RAC1 | 0.714193 | LUSC | EBC-1 | EHT-1864 | top60_altered_DNA+RNA_seed | PIK3CA | 0.078169 | COADREAD | SNU-1040 | PI3Ka_4409 |
| MVP_top60_DNA_seed | EGFR | 0.715945 | LUSC | HCC-15 | Afatinib | top60_altered_DNA+RNA_seed | BRAF | 0.078758 | COADREAD | RKO | PLX-4720 |
| MVP_top60_DNA_seed | TP53 | 0.717299 | LUSC | LOU-NH91 | MIRA-1 | top60_altered_DNA+RNA_seed | ATM | 0.079666 | COADREAD | HT-29 | CP466722 |
| MVP_top60_DNA_seed | CREBBP | 0.723469 | LUSC | NCI-H520 | SGC-CBP30 | top60_altered_DNA+RNA_seed | ATM | 0.083235 | COADREAD | C2BBe1 | KU-60019 |
| MVP_top60_DNA_seed | PIK3CA | 0.724967 | LUSC | LC-1-sq | AZD8186 | top60_altered_DNA+RNA_seed | ATM | 0.090307 | COADREAD | CL-11 | torin2 |
| MVP_top60_DNA_seed | EGFR | 0.744094 | LUSC | EBC-1 | Afatinib | top60_altered_DNA+RNA_seed | BRAF | 0.094188 | COADREAD | HT-115 | Dabrafenib |
| MVP_top60_DNA_seed | GSK3B | 0.769577 | LUSC | LOU-NH91 | SB216763 | top60_altered_DNA+RNA_seed | BRAF | 0.103934 | COADREAD | HCT-15 | Dabrafenib |
| MVP_top60_DNA_seed | PIK3CA | 0.786867 | LUSC | LC-1-sq | GNE-317 | top60_altered_DNA+RNA_seed | BRAF | 0.109116 | COADREAD | C2BBe1 | PLX-4720 |
| MVP_top60_DNA_seed | EGFR | 0.797979 | LUSC | NCI-H520 | Osimertinib | top60_altered_DNA+RNA_seed | ATM | 0.114673 | COADREAD | GP5d | torin2 |
| MVP_top60_DNA_seed | EZH2 | 0.807874 | LUSC | NCI-H1869 | GSK343 | top60_altered_DNA+RNA_seed | BRAF | 0.116283 | COADREAD | KM12 | Dabrafenib |
| MVP_top60_DNA_seed | PIK3CA | 0.810293 | LUSC | NCI-H1869 | Alpelisib | top60_altered_DNA+RNA_seed | ATM | 0.125947 | COADREAD | HT-115 | torin2 |
| MVP_top60_DNA_seed | EGFR | 0.811197 | LUSC | EBC-1 | Gefitinib | top60_altered_DNA+RNA_seed | BRAF | 0.141169 | COADREAD | NCI-H716 | PLX-4720 |
| MVP_top60_DNA_seed | ESR1 | 0.817596 | LUSC | EPLC-272H | Tamoxifen | top60_altered_DNA+RNA_seed | PIK3CA | 0.148652 | COADREAD | LS-1034 | AZD8835 |
| MVP_top60_DNA_seed | CREBBP | 0.825808 | LUSC | HCC-15 | SGC-CBP30 | top60_altered_DNA+RNA_seed | PIK3CA | 0.151439 | COADREAD | HT55 | PI3Ka_4409 |
| MVP_top60_DNA_seed | EZH2 | 0.832958 | LUSC | LOU-NH91 | GSK343 | top60_altered_DNA+RNA_seed | PIK3CA | 0.165612 | COADREAD | CL-11 | AZD8835 |
| MVP_top60_DNA_seed | PIK3CA | 0.833017 | LUSC | HCC-15 | AZD8186 | top60_altered_DNA+RNA_seed | ATM | 0.166745 | COADREAD | HCC2998 | CP466722 |
| MVP_top60_DNA_seed | TP53 | 0.841218 | LUSC | EBC-1 | MIRA-1 | top60_altered_DNA+RNA_seed | BRAF | 0.167257 | COADREAD | GP5d | SB590885 |
| MVP_top60_DNA_seed | TP53 | 0.849479 | LUSC | SK-MES-1 | MIRA-1 | top60_altered_DNA+RNA_seed | BRAF | 0.182679 | COADREAD | SW48 | Dabrafenib |
| MVP_top60_DNA_seed | EGFR | 0.855926 | LUSC | LK-2 | Erlotinib | top60_altered_DNA+RNA_seed | ATM | 0.191918 | COADREAD | HCT-116 | KU-55933 |
| MVP_top60_DNA_seed | EGFR | 0.863873 | LUSC | EBC-1 | AZD3759 | top60_altered_DNA+RNA_seed | BRAF | 0.195223 | COADREAD | SW48 | HG6-64-1 |
| MVP_top60_DNA_seed | ESR1 | 0.871464 | LUSC | NCI-H1869 | GDC0810 | top60_altered_DNA+RNA_seed | PIK3CA | 0.200779 | COADREAD | LS-123 | AZD8835 |
| MVP_top60_DNA_seed | ESR1 | 0.873256 | LUSC | NCI-H226 | Tamoxifen | top60_altered_DNA+RNA_seed | BRAF | 0.218369 | COADREAD | COLO-320-HSR | SB590885 |
| MVP_top60_DNA_seed | ESR1 | 0.894158 | LUSC | RERF-LC-Sq1 | GDC0810 | top60_altered_DNA+RNA_seed | ATM | 0.220984 | COADREAD | CaR-1 | KU-55933 |
| MVP_top60_DNA_seed | EZH2 | 0.896466 | LUSC | SK-MES-1 | GSK343 | top60_altered_DNA+RNA_seed | ATM | 0.232157 | COADREAD | LS-123 | CP466722 |
| MVP_top60_DNA_seed | ESR1 | 0.905891 | LUSC | NCI-H520 | GDC0810 | top60_altered_DNA+RNA_seed | BRAF | 0.242776 | COADREAD | HCT-116 | PLX-4720 |

| | | | | | | | | | | | |
|---|---|---|---|---|---|---|---|---|---|---|---|
| MVP_top60_DNA_seed | PIK3CA | 0.906604 | LUSC | LK-2 | AZD8186 | top60_altered_DNA+RNA_seed | BRAF | 0.251086 | COADREAD | COLO-678 | RAF_9304 |
| MVP_top60_DNA_seed | PIK3CA | 0.914289 | LUSC | EBC-1 | Alpelisib | top60_altered_DNA+RNA_seed | BRAF | 0.25152 | COADREAD | SW837 | PLX-4720 |
| MVP_top60_DNA_seed | SMARCA4 | 0.924267 | LUSC | NCI-H226 | PFI3 | top60_altered_DNA+RNA_seed | BRAF | 0.260894 | COADREAD | SW1417 | SB590885 |
| MVP_top60_DNA_seed | SMARCA2 | 0.924267 | LUSC | NCI-H226 | PFI3 | top60_altered_DNA+RNA_seed | PIK3CA | 0.265577 | COADREAD | HT55 | AZD8835 |
| MVP_top60_DNA_seed | EGFR | 0.930107 | LUSC | LK-2 | Gefitinib | top60_altered_DNA+RNA_seed | ATM | 0.267102 | COADREAD | COLO-320-HSR | KU-55933 |
| MVP_top60_DNA_seed | EGFR | 0.961985 | LUSC | EPLC-272H | AZD3759 | top60_altered_DNA+RNA_seed | BRAF | 0.267861 | COADREAD | HCC2998 | PLX-4720 |
| MVP_top60_DNA_seed | EZH2 | 0.967621 | LUSC | LOU-NH91 | GSK343 | top60_altered_DNA+RNA_seed | BRAF | 0.26976 | COADREAD | SW1417 | HG6-64-1 |
| MVP_top60_DNA_seed | CREBBP | 0.984065 | LUSC | NCI-H226 | SGC-CBP30 | top60_altered_DNA+RNA_seed | PIK3CA | 0.271829 | COADREAD | SW1463 | AZD8835 |
| MVP_top60_DNA_seed | ESR1 | 1.007554 | LUSC | LC-1-sq | GDC0810 | top60_altered_DNA+RNA_seed | BRAF | 0.279003 | COADREAD | HCT-15 | PLX-4720 |
| MVP_top60_DNA_seed | EZH2 | 1.010255 | LUSC | NCI-H1869 | GSK343 | top60_altered_DNA+RNA_seed | ATM | 0.279929 | COADREAD | SNU-61 | CP466722 |
| MVP_top60_DNA_seed | HDAC1 | 1.023897 | LUSC | SK-MES-1 | Entinostat | top60_altered_DNA+RNA_seed | PIK3CA | 0.286264 | COADREAD | SNU-C2B | PI3Ka_4409 |
| MVP_top60_DNA_seed | HDAC3 | 1.023897 | LUSC | SK-MES-1 | Entinostat | top60_altered_DNA+RNA_seed | ATM | 0.287877 | COADREAD | GP5d | KU-60019 |
| MVP_top60_DNA_seed | PIK3CA | 1.049427 | LUSC | NCI-H226 | Alpelisib | top60_altered_DNA+RNA_seed | ATM | 0.289545 | COADREAD | NCI-H508 | KU-55933 |
| MVP_top60_DNA_seed | CREBBP | 1.051696 | LUSC | NCI-H1869 | SGC-CBP30 | top60_altered_DNA+RNA_seed | BRAF | 0.289821 | COADREAD | KM12 | SB590885 |
| MVP_top60_DNA_seed | EZH2 | 1.05326 | LUSC | LK-2 | GSK343 | top60_altered_DNA+RNA_seed | BRAF | 0.294653 | COADREAD | LS-123 | AZ628 |
| MVP_top60_DNA_seed | PIK3CA | 1.066896 | LUSC | KNS-62 | AZD8186 | top60_altered_DNA+RNA_seed | ATM | 0.299476 | COADREAD | LS-1034 | KU-60019 |
| MVP_top60_DNA_seed | SMARCA4 | 1.068902 | LUSC | NCI-H1869 | PFI3 | top60_altered_DNA+RNA_seed | BRAF | 0.30351 | COADREAD | T84 | RAF_9304 |
| MVP_top60_DNA_seed | SMARCA2 | 1.068902 | LUSC | NCI-H1869 | PFI3 | top60_altered_DNA+RNA_seed | BRAF | 0.304312 | COADREAD | NCI-H716 | Dabrafenib |
| MVP_top60_DNA_seed | EGFR | 1.113786 | LUSC | HCC-15 | Sapitinib | top60_altered_DNA+RNA_seed | BRAF | 0.31342 | COADREAD | HCC2998 | SB590885 |
| MVP_top60_DNA_seed | PIK3CA | 1.11521 | LUSC | HCC-15 | Alpelisib | top60_altered_DNA+RNA_seed | PIK3CA | 0.315191 | COADREAD | HCC2998 | PI3Ka_4409 |
| MVP_top60_DNA_seed | ESR1 | 1.122618 | LUSC | LOU-NH91 | Tamoxifen | top60_altered_DNA+RNA_seed | BRAF | 0.318441 | COADREAD | SW620 | Dabrafenib |
| MVP_top60_DNA_seed | CREBBP | 1.132646 | LUSC | RERF-LC-Sq1 | SGC-CBP30 | top60_altered_DNA+RNA_seed | PIK3CA | 0.320958 | COADREAD | HCT-15 | PI3Ka_4409 |
| MVP_top60_DNA_seed | PIK3CA | 1.146915 | LUSC | SK-MES-1 | Alpelisib | top60_altered_DNA+RNA_seed | PIK3CA | 0.331059 | COADREAD | SW1417 | PI3Ka_4409 |
| MVP_top60_DNA_seed | HDAC1 | 1.16924 | LUSC | NCI-H520 | PCI-34051 | top60_altered_DNA+RNA_seed | BRAF | 0.332525 | COADREAD | GP5d | PLX-4720 |
| MVP_top60_DNA_seed | PIK3CA | 1.175026 | LUSC | LK-2 | Alpelisib | top60_altered_DNA+RNA_seed | BRAF | 0.340068 | COADREAD | NCI-H508 | RAF_9304 |
| MVP_top60_DNA_seed | RAC1 | 1.180865 | LUSC | SK-MES-1 | EHT-1864 | top60_altered_DNA+RNA_seed | ATM | 0.340794 | COADREAD | SW48 | CP466722 |
| MVP_top60_DNA_seed | HDAC1 | 1.190399 | LUSC | LK-2 | Entinostat | top60_altered_DNA+RNA_seed | ATM | 0.345977 | COADREAD | NCI-H747 | KU-60019 |
| MVP_top60_DNA_seed | HDAC3 | 1.190399 | LUSC | LK-2 | Entinostat | top60_altered_DNA+RNA_seed | BRAF | 0.347518 | COADREAD | NCI-H508 | Dabrafenib |
| MVP_top60_DNA_seed | EGFR | 1.202981 | LUSC | LK-2 | Sapitinib | top60_altered_DNA+RNA_seed | ATM | 0.356093 | COADREAD | HT-115 | KU-60019 |
| MVP_top60_DNA_seed | ESR1 | 1.248155 | LUSC | LOU-NH91 | GDC0810 | top60_altered_DNA+RNA_seed | ATM | 0.364793 | COADREAD | NCI-H716 | CP466722 |
| MVP_top60_DNA_seed | HDAC1 | 1.250924 | LUSC | NCI-H1869 | PCI-34051 | top60_altered_DNA+RNA_seed | BRAF | 0.36533 | COADREAD | DiFi | Dabrafenib |
| MVP_top60_DNA_seed | EGFR | 1.262909 | LUSC | HCC-15 | AZD3759 | top60_altered_DNA+RNA_seed | ATM | 0.366626 | COADREAD | LS-513 | CP466722 |
| MVP_top60_DNA_seed | HDAC1 | 1.272295 | LUSC | LC-1-sq | PCI-34051 | top60_altered_DNA+RNA_seed | BRAF | 0.368817 | COADREAD | NCI-H747 | SB590885 |
| MVP_top60_DNA_seed | RAC1 | 1.284342 | LUSC | LC-1-sq | EHT-1864 | top60_altered_DNA+RNA_seed | BRAF | 0.371569 | COADREAD | CaR-1 | SB590885 |
| MVP_top60_DNA_seed | PIK3CA | 1.288456 | LUSC | NCI-H1869 | Buparlisib | top60_altered_DNA+RNA_seed | ATM | 0.374839 | COADREAD | LS-1034 | CP466722 |
| MVP_top60_DNA_seed | HDAC1 | 1.297987 | LUSC | RERF-LC-Sq1 | Romidepsin | top60_altered_DNA+RNA_seed | ATM | 0.38181 | COADREAD | C2BBe1 | torin2 |
| MVP_top60_DNA_seed | HDAC3 | 1.297987 | LUSC | RERF-LC-Sq1 | Romidepsin | top60_altered_DNA+RNA_seed | ATM | 0.382052 | COADREAD | CaR-1 | torin2 |
| MVP_top60_DNA_seed | HDAC2 | 1.297987 | LUSC | RERF-LC-Sq1 | Romidepsin | top60_altered_DNA+RNA_seed | PIK3CA | 0.388994 | COADREAD | SNU-81 | PI3Ka_4409 |
| MVP_top60_DNA_seed | EGFR | 1.320047 | LUSC | NCI-H520 | AZD3759 | top60_altered_DNA+RNA_seed | BRAF | 0.392273 | COADREAD | CCK-81 | HG6-64-1 |
| MVP_top60_DNA_seed | EGFR | 1.393748 | LUSC | NCI-H226 | Gefitinib | top60_altered_DNA+RNA_seed | BRAF | 0.392488 | COADREAD | SW620 | PLX-4720 |
| MVP_top60_DNA_seed | EGFR | 1.401717 | LUSC | NCI-H520 | Sapitinib | top60_altered_DNA+RNA_seed | BRAF | 0.392974 | COADREAD | C2BBe1 | AZ628 |

| | | | | | | | | | | | |
|---|---|---|---|---|---|---|---|---|---|---|---|
| MVP_top60_DNA_seed | EZH2 | 1.536236 | LUSC | NCI-H226 | GSK343 | top60_altered_DNA+RNA_seed | ATM | 0.395235 | COADREAD | HCT-15 | torin2 |
| MVP_top60_DNA_seed | HDAC1 | 1.561487 | LUSC | NCI-H226 | Entinostat | top60_altered_DNA+RNA_seed | BRAF | 0.397356 | COADREAD | DiFi | RAF_9304 |
| MVP_top60_DNA_seed | HDAC3 | 1.561487 | LUSC | NCI-H226 | Entinostat | top60_altered_DNA+RNA_seed | ATM | 0.397829 | COADREAD | HT-115 | KU-55933 |
| MVP_top60_DNA_seed | RAC1 | 1.587087 | LUSC | LOU-NH91 | EHT-1864 | top60_altered_DNA+RNA_seed | ATM | 0.403881 | COADREAD | C2BBe1 | KU-55933 |
| MVP_top60_DNA_seed | TP53 | 1.773584 | LUSC | NCI-H226 | MIRA-1 | top60_altered_DNA+RNA_seed | BRAF | 0.403922 | COADREAD | SW1463 | RAF_9304 |
| MVP_top60_DNA_seed | EGFR | 1.922212 | LUSC | NCI-H226 | Sapitinib | top60_altered_DNA+RNA_seed | ATM | 0.404426 | COADREAD | MDST8 | KU-55933 |
| MVP_top60_DNA_seed | HDAC1 | 1.938385 | LUSC | NCI-H226 | PCI-34051 | top60_altered_DNA+RNA_seed | BRAF | 0.410079 | COADREAD | SW1116 | Dabrafenib |
| MVP_top60_DNA_seed | EGFR | 1.965884 | LUSC | NCI-H226 | Erlotinib | top60_altered_DNA+RNA_seed | BRAF | 0.412899 | COADREAD | NCI-H716 | AZ628 |
| MVP_top60_DNA_seed | RAC1 | 1.968116 | LUSC | NCI-H226 | EHT-1864 | top60_altered_DNA+RNA_seed | PIK3CA | 0.416232 | COADREAD | MDST8 | PI3Ka_4409 |
| MVP_top60_DNA_seed | HDAC1 | 1.980422 | LUSC | HCC-15 | Entinostat | top60_altered_DNA+RNA_seed | BRAF | 0.427269 | COADREAD | COLO-320-HSR | RAF_9304 |
| MVP_top60_DNA_seed | HDAC3 | 1.980422 | LUSC | HCC-15 | Entinostat | top60_altered_DNA+RNA_seed | BRAF | 0.441962 | COADREAD | LS-123 | Dabrafenib |
| MVP_top60_DNA_seed | PIK3CA | 2.103956 | LUSC | NCI-H226 | AZD8186 | top60_altered_DNA+RNA_seed | BRAF | 0.444489 | COADREAD | CW-2 | RAF_9304 |
| MVP_top60_DNA_seed | EGFR | 2.119627 | LUSC | NCI-H226 | Afatinib | top60_altered_DNA+RNA_seed | BRAF | 0.448024 | COADREAD | SK-CO-1 | PLX-4720 |
| MVP_top60_DNA_seed | EGFR | 2.248937 | LUSC | NCI-H226 | Osimertinib | top60_altered_DNA+RNA_seed | BRAF | 0.450682 | COADREAD | RCM-1 | PLX-4720 |
| MVP_top60_DNA_seed | RAC1 | 2.274074 | LUSC | RERF-LC-Sq1 | EHT-1864 | top60_altered_DNA+RNA_seed | ATM | 0.450843 | COADREAD | SW620 | torin2 |
| MVP_top60_DNA_seed | PIK3CA | 2.336999 | LUSC | NCI-H226 | GNE-317 | top60_altered_DNA+RNA_seed | ATM | 0.459969 | COADREAD | SW837 | KU-60019 |
| MVP_top60_DNA_seed | PIK3CA | 2.454696 | LUSC | NCI-H226 | Buparlisib | top60_altered_DNA+RNA_seed | PIK3CA | 0.465554 | COADREAD | SW837 | AZD8835 |
| MVP_top60_DNA_seed | ESR1 | 2.456959 | LUSC | NCI-H226 | GDC0810 | top60_altered_DNA+RNA_seed | BRAF | 0.469949 | COADREAD | COLO-678 | Dabrafenib |
| MVP_top60_DNA_seed | GSK3B | 2.535895 | LUSC | NCI-H226 | SB216763 | top60_altered_DNA+RNA_seed | BRAF | 0.472184 | COADREAD | COLO-320-HSR | PLX-4720 |
| MVP_top60_DNA_seed | EGFR | 2.62472 | LUSC | NCI-H226 | Lapatinib | top60_altered_DNA+RNA_seed | BRAF | 0.493704 | COADREAD | HCC2998 | Dabrafenib |
| MVP_top60_DNA_seed | EZH2 | 2.732589 | LUSC | NCI-H226 | GSK343 | top60_altered_DNA+RNA_seed | BRAF | 0.495644 | COADREAD | LS-1034 | AZ628 |
| MVP_top60_DNA_seed | HDAC1 | 3.155465 | LUSC | NCI-H226 | Romidepsin | top60_altered_DNA+RNA_seed | BRAF | 0.514964 | COADREAD | LoVo | PLX-4720 |
| MVP_top60_DNA_seed | HDAC3 | 3.155465 | LUSC | NCI-H226 | Romidepsin | top60_altered_DNA+RNA_seed | ATM | 0.532861 | COADREAD | COLO-205 | KU-55933 |
| MVP_top60_DNA_seed | HDAC2 | 3.155465 | LUSC | NCI-H226 | Romidepsin | top60_altered_DNA+RNA_seed | BRAF | 0.533077 | COADREAD | HCC2998 | PLX-4720 |
| MVP_top60_DNA_seed | EGFR | 3.249286 | LUSC | NCI-H226 | AZD3759 | top60_altered_DNA+RNA_seed | ATM | 0.540154 | COADREAD | CaR-1 | CP466722 |
| MVP_top60_DNA_seed | EGFR | -3.504502 | OV | OVMIU | Afatinib | top60_altered_DNA+RNA_seed | ATM | 0.556719 | COADREAD | SW48 | KU-60019 |
| MVP_top60_DNA_seed | ERBB2 | -3.504502 | OV | OVMIU | Afatinib | top60_altered_DNA+RNA_seed | ATM | 0.564396 | COADREAD | SNU-C2B | torin2 |
| MVP_top60_DNA_seed | EGFR | -3.452544 | OV | OVMIU | Afatinib | top60_altered_DNA+RNA_seed | BRAF | 0.57794 | COADREAD | HT-115 | PLX-4720 |
| MVP_top60_DNA_seed | ERBB2 | -3.452544 | OV | OVMIU | Afatinib | top60_altered_DNA+RNA_seed | BRAF | 0.581441 | COADREAD | LS-513 | Dabrafenib |
| MVP_top60_DNA_seed | EGFR | -3.389491 | OV | OVCA420 | Afatinib | top60_altered_DNA+RNA_seed | PIK3CA | 0.59458 | COADREAD | SNU-C2B | AZD8835 |
| MVP_top60_DNA_seed | ERBB2 | -3.389491 | OV | OVCA420 | Afatinib | top60_altered_DNA+RNA_seed | BRAF | 0.59687 | COADREAD | COLO-320-HSR | PLX-4720 |
| MVP_top60_DNA_seed | EGFR | -3.249644 | OV | OVMIU | AZD8931 | top60_altered_DNA+RNA_seed | BRAF | 0.620732 | COADREAD | SW48 | SB590885 |
| MVP_top60_DNA_seed | ERBB2 | -3.249644 | OV | OVMIU | AZD8931 | top60_altered_DNA+RNA_seed | BRAF | 0.623059 | COADREAD | HCT-116 | PLX-4720 |
| MVP_top60_DNA_seed | EGFR | -3.223649 | OV | OVCA420 | AZD8931 | top60_altered_DNA+RNA_seed | BRAF | 0.62713 | COADREAD | LS-513 | SB590885 |
| MVP_top60_DNA_seed | ERBB2 | -3.223649 | OV | OVCA420 | AZD8931 | top60_altered_DNA+RNA_seed | BRAF | 0.628952 | COADREAD | SNU-1040 | Dabrafenib |
| MVP_top60_DNA_seed | EGFR | -3.20211 | OV | OVMIU | Cetuximab | top60_altered_DNA+RNA_seed | BRAF | 0.631429 | COADREAD | GP5d | Dabrafenib |
| MVP_top60_DNA_seed | EGFR | -2.844187 | OV | OVTOKO | Cetuximab | top60_altered_DNA+RNA_seed | BRAF | 0.64165 | COADREAD | DiFi | PLX-4720 |
| MVP_top60_DNA_seed | ERBB2 | -2.687266 | OV | SW626 | CP724714 | top60_altered_DNA+RNA_seed | BRAF | 0.645082 | COADREAD | HT-115 | SB590885 |
| MVP_top60_DNA_seed | EGFR | -2.676136 | OV | OVMIU | Gefitinib | top60_altered_DNA+RNA_seed | BRAF | 0.650184 | COADREAD | LS-513 | PLX-4720 |
| MVP_top60_DNA_seed | EGFR | -2.511985 | OV | OVCA420 | Afatinib | top60_altered_DNA+RNA_seed | ATM | 0.652341 | COADREAD | SNU-C2B | KU-60019 |
| MVP_top60_DNA_seed | ERBB2 | -2.511985 | OV | OVCA420 | Afatinib | top60_altered_DNA+RNA_seed | ATM | 0.654055 | COADREAD | SNU-407 | KU-55933 |

| | | | | | | | | | | | |
|---|---|---|---|---|---|---|---|---|---|---|---|
| MVP_top60_DNA_seed | EGFR | -2.505204 | OV | OVMIU | PF-00299804 | top60_altered_DNA+RNA_seed | BRAF | 0.663698 | COADREAD | SNU-407 | PLX-4720 |
| MVP_top60_DNA_seed | ERBB2 | -2.505204 | OV | OVMIU | PF-00299804 | top60_altered_DNA+RNA_seed | BRAF | 0.664534 | COADREAD | SNU-C2B | PLX-4720 |
| MVP_top60_DNA_seed | EGFR | -2.501474 | OV | OVCA420 | Cetuximab | top60_altered_DNA+RNA_seed | ATM | 0.677636 | COADREAD | SW1463 | KU-55933 |
| MVP_top60_DNA_seed | EGFR | -2.293661 | OV | OVMIU | Pelitinib | top60_altered_DNA+RNA_seed | PIK3CA | 0.686876 | COADREAD | HCC2998 | AZD8835 |
| MVP_top60_DNA_seed | PIK3CA | -2.212427 | OV | OAW-42 | AZD8835 | top60_altered_DNA+RNA_seed | BRAF | 0.695822 | COADREAD | SNU-C2B | HG6-64-1 |
| MVP_top60_DNA_seed | EGFR | -2.176996 | OV | FU-OV-1 | Cetuximab | top60_altered_DNA+RNA_seed | BRAF | 0.698147 | COADREAD | COLO-320-HSR | Dabrafenib |
| MVP_top60_DNA_seed | PIK3CA | -1.965628 | OV | OAW-42 | PI3Ka_4409 | top60_altered_DNA+RNA_seed | ATM | 0.705629 | COADREAD | SW1417 | CP466722 |
| MVP_top60_DNA_seed | ERBB2 | -1.959643 | OV | OVMIU | CP724714 | top60_altered_DNA+RNA_seed | BRAF | 0.709396 | COADREAD | SNU-81 | SB590885 |
| MVP_top60_DNA_seed | EGFR | -1.955357 | OV | JHOS-2 | Afatinib | top60_altered_DNA+RNA_seed | BRAF | 0.71386 | COADREAD | CCK-81 | PLX-4720 |
| MVP_top60_DNA_seed | ERBB2 | -1.955357 | OV | JHOS-2 | Afatinib | top60_altered_DNA+RNA_seed | ATM | 0.713882 | COADREAD | CaR-1 | KU-60019 |
| MVP_top60_DNA_seed | EGFR | -1.933563 | OV | OVMIU | AST-1306 | top60_altered_DNA+RNA_seed | BRAF | 0.718315 | COADREAD | SNU-C1 | PLX-4720 |
| MVP_top60_DNA_seed | EGFR | -1.925823 | OV | JHOS-2 | AZD8931 | top60_altered_DNA+RNA_seed | BRAF | 0.729073 | COADREAD | DiFi | SB590885 |
| MVP_top60_DNA_seed | ERBB2 | -1.925823 | OV | JHOS-2 | AZD8931 | top60_altered_DNA+RNA_seed | PIK3CA | 0.737672 | COADREAD | CW-2 | AZD8835 |
| MVP_top60_DNA_seed | EGFR | -1.899328 | OV | OVCA433 | Cetuximab | top60_altered_DNA+RNA_seed | BRAF | 0.737854 | COADREAD | KM12 | PLX-4720 |
| MVP_top60_DNA_seed | EGFR | -1.84639 | OV | UWB1.289 | PF-00299804 | top60_altered_DNA+RNA_seed | ATM | 0.742282 | COADREAD | HCC2998 | KU-60019 |
| MVP_top60_DNA_seed | ERBB2 | -1.84639 | OV | UWB1.289 | PF-00299804 | top60_altered_DNA+RNA_seed | PIK3CA | 0.746074 | COADREAD | RCM-1 | AZD8835 |
| MVP_top60_DNA_seed | EGFR | -1.818531 | OV | OVTOKO | Gefitinib | top60_altered_DNA+RNA_seed | BRAF | 0.758533 | COADREAD | T84 | Dabrafenib |
| MVP_top60_DNA_seed | PTK2 | -1.808301 | OV | IGROV-1 | PF-562271 | top60_altered_DNA+RNA_seed | BRAF | 0.762312 | COADREAD | HCC2998 | HG6-64-1 |
| MVP_top60_DNA_seed | PTK2 | -1.766052 | OV | OVTOKO | PF-562271 | top60_altered_DNA+RNA_seed | ATM | 0.772567 | COADREAD | GP5d | KU-55933 |
| MVP_top60_DNA_seed | ATR | -1.746032 | OV | TOV-21G | AZ20 | top60_altered_DNA+RNA_seed | BRAF | 0.774553 | COADREAD | SNU-C1 | Dabrafenib |
| MVP_top60_DNA_seed | MAPK1 | -1.744475 | OV | TOV-112D | FR-180204 | top60_altered_DNA+RNA_seed | BRAF | 0.775816 | COADREAD | SNU-407 | SB590885 |
| MVP_top60_DNA_seed | MAPK3 | -1.744475 | OV | TOV-112D | FR-180204 | top60_altered_DNA+RNA_seed | ATM | 0.776763 | COADREAD | HCC2998 | KU-55933 |
| MVP_top60_DNA_seed | PIK3CA | -1.719603 | OV | JHOS-4 | PI3Ka_4409 | top60_altered_DNA+RNA_seed | ATM | 0.782786 | COADREAD | LoVo | CP466722 |
| MVP_top60_DNA_seed | PIK3CA | -1.706503 | OV | OVCA420 | AZD8835 | top60_altered_DNA+RNA_seed | BRAF | 0.791578 | COADREAD | T84 | PLX-4720 |
| MVP_top60_DNA_seed | CDK2 | -1.656108 | OV | OC-314 | AZD5438 | top60_altered_DNA+RNA_seed | BRAF | 0.793588 | COADREAD | SNU-81 | RAF_9304 |
| MVP_top60_DNA_seed | CDK2 | -1.653905 | OV | OAW-42 | AZD5438 | top60_altered_DNA+RNA_seed | ATM | 0.796034 | COADREAD | SNU-C2B | CP466722 |
| MVP_top60_DNA_seed | CDK1 | -1.582676 | OV | OVK-18 | RO-3306 | top60_altered_DNA+RNA_seed | BRAF | 0.80362 | COADREAD | SW1463 | PLX-4720 |
| MVP_top60_DNA_seed | EGFR | -1.574138 | OV | OVMIU | CI-1033 | top60_altered_DNA+RNA_seed | ATM | 0.806017 | COADREAD | SW1417 | KU-60019 |
| MVP_top60_DNA_seed | ERBB2 | -1.574138 | OV | OVMIU | CI-1033 | top60_altered_DNA+RNA_seed | BRAF | 0.807135 | COADREAD | T84 | SB590885 |
| MVP_top60_DNA_seed | EGFR | -1.547228 | OV | OVCA420 | Gefitinib | top60_altered_DNA+RNA_seed | ATM | 0.810344 | COADREAD | LS-123 | KU-60019 |
| MVP_top60_DNA_seed | EGFR | -1.544256 | OV | UWB1.289 | Cetuximab | top60_altered_DNA+RNA_seed | ATM | 0.811286 | COADREAD | LS-411N | KU-55933 |
| MVP_top60_DNA_seed | CREBBP | -1.512957 | OV | OC-314 | I-CBP112 | top60_altered_DNA+RNA_seed | BRAF | 0.811666 | COADREAD | SW948 | PLX-4720 |
| MVP_top60_DNA_seed | CDK2 | -1.508726 | OV | OVCAR-5 | AZD5438 | top60_altered_DNA+RNA_seed | BRAF | 0.813107 | COADREAD | LS-1034 | PLX-4720 |
| MVP_top60_DNA_seed | PIK3CA | -1.497561 | OV | OVMIU | PI3Ka_4409 | top60_altered_DNA+RNA_seed | BRAF | 0.813415 | COADREAD | SW620 | SB590885 |
| MVP_top60_DNA_seed | EGFR | -1.492782 | OV | OVTOKO | Afatinib | top60_altered_DNA+RNA_seed | BRAF | 0.832385 | COADREAD | HT55 | SB590885 |
| MVP_top60_DNA_seed | ERBB2 | -1.492782 | OV | OVTOKO | Afatinib | top60_altered_DNA+RNA_seed | BRAF | 0.836697 | COADREAD | CL-11 | SB590885 |
| MVP_top60_DNA_seed | EGFR | -1.490607 | OV | UWB1.289 | AST-1306 | top60_altered_DNA+RNA_seed | ATM | 0.840684 | COADREAD | SNU-C5 | KU-60019 |
| MVP_top60_DNA_seed | PTK2 | -1.466293 | OV | OVCAR-5 | PF-562271 | top60_altered_DNA+RNA_seed | ATM | 0.84323 | COADREAD | CCK-81 | KU-55933 |
| MVP_top60_DNA_seed | EGFR | -1.464812 | OV | TOV-112D | AZD8931 | top60_altered_DNA+RNA_seed | BRAF | 0.848974 | COADREAD | SNU-C2B | Dabrafenib |
| MVP_top60_DNA_seed | ERBB2 | -1.464812 | OV | TOV-112D | AZD8931 | top60_altered_DNA+RNA_seed | BRAF | 0.851612 | COADREAD | LS-1034 | SB590885 |
| MVP_top60_DNA_seed | ATR | -1.464491 | OV | Hey | AZ20 | top60_altered_DNA+RNA_seed | BRAF | 0.852871 | COADREAD | LS-123 | HG6-64-1 |

| | | | | | | | | | | | |
|---|---|---|---|---|---|---|---|---|---|---|---|
| MVP_top60_DNA_seed | PIK3CA | -1.458712 | OV | SK-OV-3 | AZD8835 | top60_altered_DNA+RNA_seed | ATM | 0.853584 | COADREAD | C2BBe1 | CP466722 |
| MVP_top60_DNA_seed | CDK1 | -1.447133 | OV | Hey | RO-3306 | top60_altered_DNA+RNA_seed | BRAF | 0.85741 | COADREAD | CCK-81 | Dabrafenib |
| MVP_top60_DNA_seed | EGFR | -1.431757 | OV | OVISE | Gefitinib | top60_altered_DNA+RNA_seed | BRAF | 0.864044 | COADREAD | NCI-H508 | PLX-4720 |
| MVP_top60_DNA_seed | PIK3CA | -1.431707 | OV | OVCA420 | PI3Ka_4409 | top60_altered_DNA+RNA_seed | BRAF | 0.870694 | COADREAD | SNU-1040 | PLX-4720 |
| MVP_top60_DNA_seed | ATR | -1.417569 | OV | OC-314 | QL-VIII-58 | top60_altered_DNA+RNA_seed | ATM | 0.871659 | COADREAD | LS-1034 | KU-55933 |
| MVP_top60_DNA_seed | EGFR | -1.415459 | OV | UWB1.289 | CI-1033 | top60_altered_DNA+RNA_seed | BRAF | 0.882413 | COADREAD | HT-115 | HG6-64-1 |
| MVP_top60_DNA_seed | ERBB2 | -1.415459 | OV | UWB1.289 | CI-1033 | top60_altered_DNA+RNA_seed | BRAF | 0.886147 | COADREAD | SNU-81 | PLX-4720 |
| MVP_top60_DNA_seed | ERBB2 | -1.414995 | OV | TOV-112D | CP724714 | top60_altered_DNA+RNA_seed | BRAF | 0.889052 | COADREAD | HCT-15 | RAF_9304 |
| MVP_top60_DNA_seed | EGFR | -1.397004 | OV | JHOS-2 | Afatinib | top60_altered_DNA+RNA_seed | PIK3CA | 0.898392 | COADREAD | SW620 | AZD8835 |
| MVP_top60_DNA_seed | ERBB2 | -1.397004 | OV | JHOS-2 | Afatinib | top60_altered_DNA+RNA_seed | ATM | 0.90616 | COADREAD | KM12 | KU-55933 |
| MVP_top60_DNA_seed | PIK3CA | -1.396413 | OV | OVMIU | AZD8835 | top60_altered_DNA+RNA_seed | BRAF | 0.906177 | COADREAD | SW48 | PLX-4720 |
| MVP_top60_DNA_seed | EGFR | -1.395021 | OV | TOV-112D | Afatinib | top60_altered_DNA+RNA_seed | BRAF | 0.914824 | COADREAD | C2BBe1 | PLX-4720 |
| MVP_top60_DNA_seed | ERBB2 | -1.395021 | OV | TOV-112D | Afatinib | top60_altered_DNA+RNA_seed | PIK3CA | 0.918071 | COADREAD | SW1116 | PI3Ka_4409 |
| MVP_top60_DNA_seed | ATR | -1.389386 | OV | TOV-21G | AZD6738 | top60_altered_DNA+RNA_seed | BRAF | 0.919444 | COADREAD | SW1463 | Dabrafenib |
| MVP_top60_DNA_seed | EGFR | -1.362403 | OV | OAW-42 | Gefitinib | top60_altered_DNA+RNA_seed | BRAF | 0.946948 | COADREAD | LS-1034 | Dabrafenib |
| MVP_top60_DNA_seed | EGFR | -1.361183 | OV | OVISE | Afatinib | top60_altered_DNA+RNA_seed | BRAF | 0.965513 | COADREAD | CCK-81 | PLX-4720 |
| MVP_top60_DNA_seed | ERBB2 | -1.361183 | OV | OVISE | Afatinib | top60_altered_DNA+RNA_seed | BRAF | 0.966486 | COADREAD | CaR-1 | HG6-64-1 |
| MVP_top60_DNA_seed | MAPK1 | -1.345372 | OV | OVMIU | FR-180204 | top60_altered_DNA+RNA_seed | BRAF | 0.969409 | COADREAD | DiFi | PLX-4720 |
| MVP_top60_DNA_seed | MAPK3 | -1.345372 | OV | OVMIU | FR-180204 | top60_altered_DNA+RNA_seed | ATM | 0.975841 | COADREAD | DiFi | KU-55933 |
| MVP_top60_DNA_seed | CREBBP | -1.334749 | OV | OVCA420 | I-CBP112 | top60_altered_DNA+RNA_seed | ATM | 0.979241 | COADREAD | SNU-C1 | KU-55933 |
| MVP_top60_DNA_seed | EGFR | -1.327917 | OV | OVISE | Pelitinib | top60_altered_DNA+RNA_seed | BRAF | 0.988457 | COADREAD | HCT-15 | SB590885 |
| MVP_top60_DNA_seed | CDK2 | -1.321659 | OV | IGROV-1 | PHA-793887 | top60_altered_DNA+RNA_seed | BRAF | 0.989371 | COADREAD | LS-513 | HG6-64-1 |
| MVP_top60_DNA_seed | CDK2 | -1.321499 | OV | OV-7 | AZD5438 | top60_altered_DNA+RNA_seed | BRAF | 0.989568 | COADREAD | CL-11 | PLX-4720 |
| MVP_top60_DNA_seed | PLK1 | -1.320188 | OV | TOV-112D | PLK_6522 | top60_altered_DNA+RNA_seed | BRAF | 0.999727 | COADREAD | C2BBe1 | Dabrafenib |
| MVP_top60_DNA_seed | PIK3CA | -1.299272 | OV | OV-7 | PI3Ka_4409 | top60_altered_DNA+RNA_seed | ATM | 1.00267 | COADREAD | HT-115 | CP466722 |
| MVP_top60_DNA_seed | EGFR | -1.296688 | OV | JHOS-4 | Afatinib | top60_altered_DNA+RNA_seed | PIK3CA | 1.012971 | COADREAD | COLO-678 | AZD8835 |
| MVP_top60_DNA_seed | ERBB2 | -1.296688 | OV | JHOS-4 | Afatinib | top60_altered_DNA+RNA_seed | BRAF | 1.017936 | COADREAD | SW1463 | HG6-64-1 |
| MVP_top60_DNA_seed | CDK2 | -1.295189 | OV | OC-314 | PHA-793887 | top60_altered_DNA+RNA_seed | ATM | 1.01836 | COADREAD | DiFi | KU-60019 |
| MVP_top60_DNA_seed | EGFR | -1.28691 | OV | OVTOKO | AZD8931 | top60_altered_DNA+RNA_seed | BRAF | 1.02012 | COADREAD | SNU-81 | PLX-4720 |
| MVP_top60_DNA_seed | ERBB2 | -1.28691 | OV | OVTOKO | AZD8931 | top60_altered_DNA+RNA_seed | BRAF | 1.025394 | COADREAD | LS-123 | PLX-4720 |
| MVP_top60_DNA_seed | CDK2 | -1.274937 | OV | OVCAR-8 | AZD5438 | top60_altered_DNA+RNA_seed | PIK3CA | 1.026844 | COADREAD | CaR-1 | AZD8835 |
| MVP_top60_DNA_seed | EGFR | -1.269775 | OV | OC-314 | Pelitinib | top60_altered_DNA+RNA_seed | BRAF | 1.027412 | COADREAD | LS-1034 | RAF_9304 |
| MVP_top60_DNA_seed | CDK2 | -1.261499 | OV | Hey | AZD5438 | top60_altered_DNA+RNA_seed | BRAF | 1.02833 | COADREAD | KM12 | PLX-4720 |
| MVP_top60_DNA_seed | SIRT1 | -1.251509 | OV | Caov-3 | Selisistat | top60_altered_DNA+RNA_seed | BRAF | 1.042429 | COADREAD | RCM-1 | PLX-4720 |
| MVP_top60_DNA_seed | EGFR | -1.244922 | OV | JHOS-2 | Gefitinib | top60_altered_DNA+RNA_seed | BRAF | 1.043 | COADREAD | SNU-C2B | PLX-4720 |
| MVP_top60_DNA_seed | ERBB2 | -1.244635 | OV | OV-17R | CP724714 | top60_altered_DNA+RNA_seed | BRAF | 1.045255 | COADREAD | SNU-C2B | SB590885 |
| MVP_top60_DNA_seed | PIK3CA | -1.239267 | OV | JHOS-2 | AZD8835 | top60_altered_DNA+RNA_seed | BRAF | 1.055755 | COADREAD | LS-180 | PLX-4720 |
| MVP_top60_DNA_seed | MAPK1 | -1.235047 | OV | FU-OV-1 | FR-180204 | top60_altered_DNA+RNA_seed | BRAF | 1.06569 | COADREAD | LS-180 | Dabrafenib |
| MVP_top60_DNA_seed | MAPK3 | -1.235047 | OV | FU-OV-1 | FR-180204 | top60_altered_DNA+RNA_seed | BRAF | 1.08096 | COADREAD | LS-123 | PLX-4720 |
| MVP_top60_DNA_seed | EGFR | -1.223719 | OV | OAW-28 | CUDC-101 | top60_altered_DNA+RNA_seed | BRAF | 1.083645 | COADREAD | T84 | PLX-4720 |
| MVP_top60_DNA_seed | ERBB2 | -1.223719 | OV | OAW-28 | CUDC-101 | top60_altered_DNA+RNA_seed | BRAF | 1.083854 | COADREAD | SNU-81 | Dabrafenib |

| | | | | | | | | | | | |
|---|---|---|---|---|---|---|---|---|---|---|---|
| MVP_top60_DNA_seed | CREBBP | -1.220332 | OV | TOV-112D | I-CBP112 | top60_altered_DNA+RNA_seed | BRAF | 1.09231 | COADREAD | T84 | HG6-64-1 |
| MVP_top60_DNA_seed | EGFR | -1.215193 | OV | OVCAR-4 | Erlotinib | top60_altered_DNA+RNA_seed | BRAF | 1.102298 | COADREAD | HT55 | PLX-4720 |
| MVP_top60_DNA_seed | PIK3CA | -1.208861 | OV | SK-OV-3 | PI3Ka_4409 | top60_altered_DNA+RNA_seed | ATM | 1.112672 | COADREAD | RCM-1 | torin2 |
| MVP_top60_DNA_seed | EGFR | -1.207818 | OV | TOV-112D | Gefitinib | top60_altered_DNA+RNA_seed | BRAF | 1.125855 | COADREAD | SNU-61 | Dabrafenib |
| MVP_top60_DNA_seed | CDK2 | -1.196376 | OV | Hey | PHA-793887 | top60_altered_DNA+RNA_seed | BRAF | 1.145662 | COADREAD | COLO-678 | SB590885 |
| MVP_top60_DNA_seed | HDAC2 | -1.183018 | OV | OVCAR-5 | JQ12 | top60_altered_DNA+RNA_seed | BRAF | 1.148894 | COADREAD | CW-2 | Dabrafenib |
| MVP_top60_DNA_seed | MAPK1 | -1.1774 | OV | SW626 | VX-11e | top60_altered_DNA+RNA_seed | BRAF | 1.149149 | COADREAD | CaR-1 | Dabrafenib |
| MVP_top60_DNA_seed | HDAC2 | -1.171945 | OV | OVTOKO | JQ12 | top60_altered_DNA+RNA_seed | ATM | 1.149447 | COADREAD | HT-29 | KU-55933 |
| MVP_top60_DNA_seed | ATR | -1.16231 | OV | TOV-21G | QL-VIII-58 | top60_altered_DNA+RNA_seed | BRAF | 1.15232 | COADREAD | LS-1034 | PLX-4720 |
| MVP_top60_DNA_seed | MAPK1 | -1.159616 | OV | OVMIU | VX-11e | top60_altered_DNA+RNA_seed | BRAF | 1.155179 | COADREAD | HT-115 | PLX-4720 |
| MVP_top60_DNA_seed | EGFR | -1.157088 | OV | TOV-112D | CI-1033 | top60_altered_DNA+RNA_seed | BRAF | 1.169008 | COADREAD | CW-2 | PLX-4720 |
| MVP_top60_DNA_seed | ERBB2 | -1.157088 | OV | TOV-112D | CI-1033 | top60_altered_DNA+RNA_seed | ATM | 1.170588 | COADREAD | SW1417 | KU-55933 |
| MVP_top60_DNA_seed | EGFR | -1.157041 | OV | JHOS-4 | PF-00299804 | top60_altered_DNA+RNA_seed | BRAF | 1.181593 | COADREAD | CW-2 | PLX-4720 |
| MVP_top60_DNA_seed | ERBB2 | -1.157041 | OV | JHOS-4 | PF-00299804 | top60_altered_DNA+RNA_seed | BRAF | 1.184731 | COADREAD | SW837 | Dabrafenib |
| MVP_top60_DNA_seed | EGFR | -1.155473 | OV | DOV13 | Afatinib | top60_altered_DNA+RNA_seed | BRAF | 1.192472 | COADREAD | MDST8 | HG6-64-1 |
| MVP_top60_DNA_seed | ERBB2 | -1.155473 | OV | DOV13 | Afatinib | top60_altered_DNA+RNA_seed | ATM | 1.192943 | COADREAD | COLO-678 | KU-60019 |
| MVP_top60_DNA_seed | HDAC2 | -1.14853 | OV | OVCAR-4 | JQ12 | top60_altered_DNA+RNA_seed | BRAF | 1.195714 | COADREAD | CW-2 | AZ628 |
| MVP_top60_DNA_seed | MAPK1 | -1.138765 | OV | IGROV-1 | FR-180204 | top60_altered_DNA+RNA_seed | BRAF | 1.198266 | COADREAD | SNU-407 | PLX-4720 |
| MVP_top60_DNA_seed | MAPK3 | -1.138765 | OV | IGROV-1 | FR-180204 | top60_altered_DNA+RNA_seed | ATM | 1.199291 | COADREAD | CL-11 | CP466722 |
| MVP_top60_DNA_seed | PLK1 | -1.131978 | OV | OC-314 | PLK_6522 | top60_altered_DNA+RNA_seed | BRAF | 1.200588 | COADREAD | SNU-C1 | HG6-64-1 |
| MVP_top60_DNA_seed | ATR | -1.127962 | OV | OC-314 | AZ20 | top60_altered_DNA+RNA_seed | BRAF | 1.20171 | COADREAD | SW948 | HG6-64-1 |
| MVP_top60_DNA_seed | CDK2 | -1.124565 | OV | Caov-3 | PHA-793887 | top60_altered_DNA+RNA_seed | PIK3CA | 1.211845 | COADREAD | C2BBe1 | AZD8835 |
| MVP_top60_DNA_seed | EGFR | -1.12043 | OV | TOV-21G | AZD8931 | top60_altered_DNA+RNA_seed | BRAF | 1.222679 | COADREAD | CW-2 | SB590885 |
| MVP_top60_DNA_seed | ERBB2 | -1.12043 | OV | TOV-21G | AZD8931 | top60_altered_DNA+RNA_seed | ATM | 1.224194 | COADREAD | SNU-C5 | CP466722 |
| MVP_top60_DNA_seed | HDAC2 | -1.103019 | OV | OAW-28 | JQ12 | top60_altered_DNA+RNA_seed | PIK3CA | 1.228973 | COADREAD | SW1417 | AZD8835 |
| MVP_top60_DNA_seed | EGFR | -1.102202 | OV | IGROV-1 | Afatinib | top60_altered_DNA+RNA_seed | ATM | 1.232944 | COADREAD | SNU-407 | KU-60019 |
| MVP_top60_DNA_seed | ERBB2 | -1.102202 | OV | IGROV-1 | Afatinib | top60_altered_DNA+RNA_seed | ATM | 1.24555 | COADREAD | T84 | torin2 |
| MVP_top60_DNA_seed | MAPK1 | -1.100074 | OV | Hey | VX-11e | top60_altered_DNA+RNA_seed | BRAF | 1.262356 | COADREAD | SW948 | Dabrafenib |
| MVP_top60_DNA_seed | EGFR | -1.086844 | OV | TOV-112D | Afatinib | top60_altered_DNA+RNA_seed | BRAF | 1.280658 | COADREAD | LS-123 | SB590885 |
| MVP_top60_DNA_seed | ERBB2 | -1.086844 | OV | TOV-112D | Afatinib | top60_altered_DNA+RNA_seed | ATM | 1.280746 | COADREAD | SW948 | CP466722 |
| MVP_top60_DNA_seed | EGFR | -1.081656 | OV | OVISE | Afatinib | top60_altered_DNA+RNA_seed | ATM | 1.283206 | COADREAD | T84 | KU-55933 |
| MVP_top60_DNA_seed | ERBB2 | -1.081656 | OV | OVISE | Afatinib | top60_altered_DNA+RNA_seed | PIK3CA | 1.290498 | COADREAD | RCM-1 | PI3Ka_4409 |
| MVP_top60_DNA_seed | CDK2 | -1.074864 | OV | OVMIU | AZD5438 | top60_altered_DNA+RNA_seed | BRAF | 1.294629 | COADREAD | LS-1034 | HG6-64-1 |
| MVP_top60_DNA_seed | PLK1 | -1.074588 | OV | OVCAR-5 | PLK_6522 | top60_altered_DNA+RNA_seed | ATM | 1.300873 | COADREAD | COLO-205 | KU-60019 |
| MVP_top60_DNA_seed | MAPK1 | -1.058951 | OV | IGROV-1 | VX-11e | top60_altered_DNA+RNA_seed | BRAF | 1.303785 | COADREAD | NCI-H747 | PLX-4720 |
| MVP_top60_DNA_seed | ATR | -1.056402 | OV | Hey | torin2 | top60_altered_DNA+RNA_seed | BRAF | 1.307196 | COADREAD | COLO-320-HSR | AZ628 |
| MVP_top60_DNA_seed | EGFR | -1.037635 | OV | Caov-4 | Cetuximab | top60_altered_DNA+RNA_seed | ATM | 1.330659 | COADREAD | RCM-1 | CP466722 |
| MVP_top60_DNA_seed | MAPK1 | -1.036857 | OV | OAW-28 | FR-180204 | top60_altered_DNA+RNA_seed | PIK3CA | 1.336968 | COADREAD | LS-411N | AZD8835 |
| MVP_top60_DNA_seed | MAPK3 | -1.036857 | OV | OAW-28 | FR-180204 | top60_altered_DNA+RNA_seed | ATM | 1.362202 | COADREAD | SNU-C1 | KU-60019 |
| MVP_top60_DNA_seed | EGFR | -1.022697 | OV | PEO1 | Afatinib | top60_altered_DNA+RNA_seed | PIK3CA | 1.377467 | COADREAD | CW-2 | PI3Ka_4409 |
| MVP_top60_DNA_seed | ERBB2 | -1.022697 | OV | PEO1 | Afatinib | top60_altered_DNA+RNA_seed | BRAF | 1.384328 | COADREAD | LS-513 | PLX-4720 |

| | | | | | | | | | | | |
|---|---|---|---|---|---|---|---|---|---|---|---|
| MVP_top60_DNA_seed | PTK2 | -1.010748 | OV | TOV-112D | PF-562271 | top60_altered_DNA+RNA_seed | ATM | 1.385027 | COADREAD | NCI-H747 | KU-55933 |
| MVP_top60_DNA_seed | PLK1 | -0.998567 | OV | OVCAR-8 | PLK_6522 | top60_altered_DNA+RNA_seed | ATM | 1.389542 | COADREAD | SW1116 | CP466722 |
| MVP_top60_DNA_seed | ATR | -0.994535 | OV | OVK-18 | AZ20 | top60_altered_DNA+RNA_seed | BRAF | 1.394983 | COADREAD | CaR-1 | PLX-4720 |
| MVP_top60_DNA_seed | PIK3CA | -0.993002 | OV | JHOS-4 | AZD8835 | top60_altered_DNA+RNA_seed | ATM | 1.398281 | COADREAD | CW-2 | torin2 |
| MVP_top60_DNA_seed | HDAC2 | -0.99253 | OV | IGROV-1 | JQ12 | top60_altered_DNA+RNA_seed | ATM | 1.398285 | COADREAD | SNU-C1 | CP466722 |
| MVP_top60_DNA_seed | HDAC2 | -0.992307 | OV | OVCAR-8 | JQ12 | top60_altered_DNA+RNA_seed | BRAF | 1.412692 | COADREAD | SNU-61 | SB590885 |
| MVP_top60_DNA_seed | EGFR | -0.987409 | OV | JHOS-4 | CI-1033 | top60_altered_DNA+RNA_seed | ATM | 1.416189 | COADREAD | CW-2 | KU-55933 |
| MVP_top60_DNA_seed | ERBB2 | -0.987409 | OV | JHOS-4 | CI-1033 | top60_altered_DNA+RNA_seed | ATM | 1.417218 | COADREAD | HT55 | KU-55933 |
| MVP_top60_DNA_seed | EGFR | -0.985174 | OV | JHOS-4 | AST-1306 | top60_altered_DNA+RNA_seed | BRAF | 1.423385 | COADREAD | SNU-C1 | AZ628 |
| MVP_top60_DNA_seed | MAPK1 | -0.979244 | OV | OVCAR-8 | VX-11e | top60_altered_DNA+RNA_seed | BRAF | 1.43449 | COADREAD | SNU-1040 | RAF_9304 |
| MVP_top60_DNA_seed | CDK1 | -0.979209 | OV | PEO1 | RO-3306 | top60_altered_DNA+RNA_seed | BRAF | 1.435047 | COADREAD | SW948 | PLX-4720 |
| MVP_top60_DNA_seed | EGFR | -0.976441 | OV | DOV13 | Afatinib | top60_altered_DNA+RNA_seed | ATM | 1.45201 | COADREAD | SW620 | KU-55933 |
| MVP_top60_DNA_seed | ERBB2 | -0.976441 | OV | DOV13 | Afatinib | top60_altered_DNA+RNA_seed | BRAF | 1.466158 | COADREAD | COLO-678 | PLX-4720 |
| MVP_top60_DNA_seed | ATR | -0.975289 | OV | OVCAR-5 | AZ20 | top60_altered_DNA+RNA_seed | ATM | 1.501923 | COADREAD | HT55 | KU-60019 |
| MVP_top60_DNA_seed | ATR | -0.974947 | OV | PEO1 | AZD6738 | top60_altered_DNA+RNA_seed | ATM | 1.50684 | COADREAD | RCM-1 | KU-60019 |
| MVP_top60_DNA_seed | MAPK1 | -0.974303 | OV | OAW-28 | VX-11e | top60_altered_DNA+RNA_seed | BRAF | 1.525145 | COADREAD | SW837 | HG6-64-1 |
| MVP_top60_DNA_seed | ERBB2 | -0.969459 | OV | OC-314 | CP724714 | top60_altered_DNA+RNA_seed | BRAF | 1.531545 | COADREAD | NCI-H747 | PLX-4720 |
| MVP_top60_DNA_seed | EGFR | -0.967934 | OV | OV-56 | Cetuximab | top60_altered_DNA+RNA_seed | BRAF | 1.53931 | COADREAD | CL-11 | HG6-64-1 |
| MVP_top60_DNA_seed | MAPK1 | -0.955965 | OV | OVCAR-8 | FR-180204 | top60_altered_DNA+RNA_seed | PIK3CA | 1.564331 | COADREAD | SNU-61 | AZD8835 |
| MVP_top60_DNA_seed | MAPK3 | -0.955965 | OV | OVCAR-8 | FR-180204 | top60_altered_DNA+RNA_seed | BRAF | 1.56492 | COADREAD | NCI-H508 | SB590885 |
| MVP_top60_DNA_seed | MAPK1 | -0.955212 | OV | JHOS-2 | FR-180204 | top60_altered_DNA+RNA_seed | PIK3CA | 1.574111 | COADREAD | SW1116 | AZD8835 |
| MVP_top60_DNA_seed | MAPK3 | -0.955212 | OV | JHOS-2 | FR-180204 | top60_altered_DNA+RNA_seed | ATM | 1.598014 | COADREAD | RCM-1 | KU-55933 |
| MVP_top60_DNA_seed | EGFR | -0.954487 | OV | OVTOKO | Afatinib | top60_altered_DNA+RNA_seed | BRAF | 1.599263 | COADREAD | CL-11 | PLX-4720 |
| MVP_top60_DNA_seed | ERBB2 | -0.954487 | OV | OVTOKO | Afatinib | top60_altered_DNA+RNA_seed | PIK3CA | 1.61439 | COADREAD | COLO-678 | PI3Ka_4409 |
| MVP_top60_DNA_seed | PIK3CA | -0.933535 | OV | JHOS-2 | PI3Ka_4409 | top60_altered_DNA+RNA_seed | BRAF | 1.627943 | COADREAD | RCM-1 | HG6-64-1 |
| MVP_top60_DNA_seed | PLK1 | -0.928599 | OV | JHOS-2 | PLK_6522 | top60_altered_DNA+RNA_seed | BRAF | 1.633544 | COADREAD | SW1116 | PLX-4720 |
| MVP_top60_DNA_seed | ATR | -0.926995 | OV | DOV13 | AZD6738 | top60_altered_DNA+RNA_seed | ATM | 1.635518 | COADREAD | CW-2 | CP466722 |
| MVP_top60_DNA_seed | EGFR | -0.917313 | OV | UWB1.289 | Afatinib | top60_altered_DNA+RNA_seed | BRAF | 1.648702 | COADREAD | COLO-678 | PLX-4720 |
| MVP_top60_DNA_seed | ERBB2 | -0.917313 | OV | UWB1.289 | Afatinib | top60_altered_DNA+RNA_seed | BRAF | 1.65166 | COADREAD | KM12 | RAF_9304 |
| MVP_top60_DNA_seed | CDK1 | -0.908444 | OV | TOV-21G | RO-3306 | top60_altered_DNA+RNA_seed | BRAF | 1.660652 | COADREAD | SW1116 | SB590885 |
| MVP_top60_DNA_seed | EGFR | -0.901759 | OV | TOV-112D | AST-1306 | top60_altered_DNA+RNA_seed | BRAF | 1.666703 | COADREAD | SW620 | PLX-4720 |
| MVP_top60_DNA_seed | CDK2 | -0.894975 | OV | TOV-21G | AZD5438 | top60_altered_DNA+RNA_seed | PIK3CA | 1.673282 | COADREAD | SW620 | PI3Ka_4409 |
| MVP_top60_DNA_seed | EGFR | -0.890952 | OV | OV-7 | Gefitinib | top60_altered_DNA+RNA_seed | ATM | 1.693951 | COADREAD | SW1463 | CP466722 |
| MVP_top60_DNA_seed | EGFR | -0.887644 | OV | UWB1.289 | Afatinib | top60_altered_DNA+RNA_seed | ATM | 1.695339 | COADREAD | MDST8 | CP466722 |
| MVP_top60_DNA_seed | ERBB2 | -0.887644 | OV | UWB1.289 | Afatinib | top60_altered_DNA+RNA_seed | BRAF | 1.706309 | COADREAD | SNU-C1 | SB590885 |
| MVP_top60_DNA_seed | ATR | -0.884442 | OV | DOV13 | torin2 | top60_altered_DNA+RNA_seed | ATM | 1.724064 | COADREAD | SNU-C2B | KU-55933 |
| MVP_top60_DNA_seed | EGFR | -0.87383 | OV | PEO1 | CUDC-101 | top60_altered_DNA+RNA_seed | BRAF | 1.725127 | COADREAD | SNU-61 | PLX-4720 |
| MVP_top60_DNA_seed | ERBB2 | -0.87383 | OV | PEO1 | CUDC-101 | top60_altered_DNA+RNA_seed | PIK3CA | 1.773409 | COADREAD | LS-411N | PI3Ka_4409 |
| MVP_top60_DNA_seed | CDK2 | -0.870739 | OV | TOV-112D | PHA-793887 | top60_altered_DNA+RNA_seed | ATM | 1.782405 | COADREAD | LS-513 | KU-55933 |
| MVP_top60_DNA_seed | ATR | -0.870066 | OV | OVCAR-3 | QL-VIII-58 | top60_altered_DNA+RNA_seed | ATM | 1.796726 | COADREAD | COLO-678 | KU-55933 |
| MVP_top60_DNA_seed | PLK1 | -0.859263 | OV | JHOS-4 | PLK_6522 | top60_altered_DNA+RNA_seed | BRAF | 1.79791 | COADREAD | SNU-61 | RAF_9304 |

| | | | | | | | | | | | |
|---|---|---|---|---|---|---|---|---|---|---|---|
| MVP_top60_DNA_seed | ATR | -0.854396 | OV | OC-314 | AZD6738 | top60_altered_DNA+RNA_seed | BRAF | 1.798255 | COADREAD | SW837 | PLX-4720 |
| MVP_top60_DNA_seed | ATR | -0.849217 | OV | Hey | QL-VIII-58 | top60_altered_DNA+RNA_seed | PIK3CA | 1.799843 | COADREAD | SNU-61 | PI3Ka_4409 |
| MVP_top60_DNA_seed | ATR | -0.847937 | OV | OVCA433 | QL-VIII-58 | top60_altered_DNA+RNA_seed | BRAF | 1.809291 | COADREAD | SNU-61 | PLX-4720 |
| MVP_top60_DNA_seed | ATR | -0.844993 | OV | OVCAR-5 | AZD6738 | top60_altered_DNA+RNA_seed | PIK3CA | 1.826066 | COADREAD | KM12 | AZD8835 |
| MVP_top60_DNA_seed | PIK3CA | -0.834445 | OV | PEO1 | AZD8835 | top60_altered_DNA+RNA_seed | PIK3CA | 1.869052 | COADREAD | C2BBe1 | PI3Ka_4409 |
| MVP_top60_DNA_seed | ATR | -0.827462 | OV | OV-56 | AZD6738 | top60_altered_DNA+RNA_seed | BRAF | 1.89441 | COADREAD | SNU-C1 | PLX-4720 |
| MVP_top60_DNA_seed | MAPK1 | -0.826009 | OV | OVCAR-5 | FR-180204 | top60_altered_DNA+RNA_seed | BRAF | 1.90181 | COADREAD | HT55 | HG6-64-1 |
| MVP_top60_DNA_seed | MAPK3 | -0.826009 | OV | OVCAR-5 | FR-180204 | top60_altered_DNA+RNA_seed | PIK3CA | 1.928324 | COADREAD | SNU-1040 | AZD8835 |
| MVP_top60_DNA_seed | EGFR | -0.821005 | OV | PEO1 | AST-1306 | top60_altered_DNA+RNA_seed | ATM | 1.9308 | COADREAD | SW837 | KU-55933 |
| MVP_top60_DNA_seed | CDK2 | -0.808262 | OV | OVCA433 | AZD5438 | top60_altered_DNA+RNA_seed | ATM | 2.012473 | COADREAD | COLO-678 | CP466722 |
| MVP_top60_DNA_seed | PIK3CA | -0.808092 | OV | OVCA433 | AZD8835 | top60_altered_DNA+RNA_seed | ATM | 2.01646 | COADREAD | T84 | KU-60019 |
| MVP_top60_DNA_seed | MAPK1 | -0.807823 | OV | DOV13 | FR-180204 | top60_altered_DNA+RNA_seed | PIK3CA | 2.09473 | COADREAD | KM12 | PI3Ka_4409 |
| MVP_top60_DNA_seed | MAPK3 | -0.807823 | OV | DOV13 | FR-180204 | top60_altered_DNA+RNA_seed | ATM | 2.112754 | COADREAD | LS-123 | KU-55933 |
| MVP_top60_DNA_seed | EGFR | -0.804813 | OV | OVCAR-8 | CUDC-101 | top60_altered_DNA+RNA_seed | ATM | 2.116676 | COADREAD | HT55 | torin2 |
| MVP_top60_DNA_seed | ERBB2 | -0.804813 | OV | OVCAR-8 | CUDC-101 | top60_altered_DNA+RNA_seed | ATM | 2.148201 | COADREAD | HT-29 | KU-60019 |
| MVP_top60_DNA_seed | ATR | -0.804809 | OV | TOV-112D | torin2 | top60_altered_DNA+RNA_seed | ATM | 2.230647 | COADREAD | SW948 | KU-55933 |
| MVP_top60_DNA_seed | ATR | -0.804071 | OV | TOV-112D | AZ20 | top60_altered_DNA+RNA_seed | ATM | 2.230884 | COADREAD | LS-513 | KU-60019 |
| MVP_top60_DNA_seed | EGFR | -0.800549 | OV | TOV-112D | Pelitinib | top60_altered_DNA+RNA_seed | ATM | 2.296439 | COADREAD | KM12 | torin2 |
| MVP_top60_DNA_seed | EGFR | -0.795922 | OV | JHOS-4 | Afatinib | top60_altered_DNA+RNA_seed | ATM | 2.299113 | COADREAD | SW948 | KU-60019 |
| MVP_top60_DNA_seed | ERBB2 | -0.795922 | OV | JHOS-4 | Afatinib | top60_altered_DNA+RNA_seed | BRAF | 2.320515 | COADREAD | SW1116 | PLX-4720 |
| MVP_top60_DNA_seed | SIRT1 | -0.793659 | OV | TOV-112D | Selisistat | top60_altered_DNA+RNA_seed | ATM | 2.384769 | COADREAD | LS-180 | KU-60019 |
| MVP_top60_DNA_seed | MAPK1 | -0.790554 | OV | Caov-3 | FR-180204 | top60_altered_DNA+RNA_seed | BRAF | 2.439146 | COADREAD | CW-2 | HG6-64-1 |
| MVP_top60_DNA_seed | MAPK3 | -0.790554 | OV | Caov-3 | FR-180204 | top60_altered_DNA+RNA_seed | ATM | 2.446635 | COADREAD | SNU-61 | KU-55933 |
| MVP_top60_DNA_seed | ATR | -0.785925 | OV | PEO1 | QL-VIII-58 | top60_altered_DNA+RNA_seed | ATM | 2.534752 | COADREAD | LS-180 | torin2 |
| MVP_top60_DNA_seed | EGFR | -0.779181 | OV | OC-314 | AZD8931 | top60_altered_DNA+RNA_seed | BRAF | 2.574624 | COADREAD | SW1463 | PLX-4720 |
| MVP_top60_DNA_seed | ERBB2 | -0.779181 | OV | OC-314 | AZD8931 | top60_altered_DNA+RNA_seed | ATM | 2.624244 | COADREAD | HT-29 | torin2 |
| MVP_top60_DNA_seed | ATR | -0.778611 | OV | DOV13 | QL-VIII-58 | top60_altered_DNA+RNA_seed | ATM | 2.62843 | COADREAD | HT55 | CP466722 |
| MVP_top60_DNA_seed | CDK1 | -0.769702 | OV | OVCAR-5 | RO-3306 | top60_altered_DNA+RNA_seed | ATM | 2.713114 | COADREAD | SNU-61 | KU-60019 |
| MVP_top60_DNA_seed | EGFR | -0.769295 | OV | OV-17R | CUDC-101 | top60_altered_DNA+RNA_seed | ATM | 2.716752 | COADREAD | SW1116 | KU-60019 |
| MVP_top60_DNA_seed | ERBB2 | -0.769295 | OV | OV-17R | CUDC-101 | top60_altered_DNA+RNA_seed | ATM | 2.752819 | COADREAD | SW1116 | KU-55933 |
| MVP_top60_DNA_seed | ATR | -0.762748 | OV | OVISE | AZ20 | top60_altered_DNA+RNA_seed | BRAF | 2.958328 | COADREAD | SNU-C1 | RAF_9304 |
| MVP_top60_DNA_seed | ATR | -0.757883 | OV | OVK-18 | QL-VIII-58 | top60_altered_DNA+RNA_seed | ATM | 3.090585 | COADREAD | SW1463 | KU-60019 |
| MVP_top60_DNA_seed | CDK2 | -0.733933 | OV | JHOS-4 | AZD5438 | top60_altered_DNA+RNA_seed | ATM | 3.099592 | COADREAD | SW948 | torin2 |
| MVP_top60_DNA_seed | EGFR | -0.733762 | OV | DOV13 | Gefitinib | top60_altered_DNA+RNA_seed | ATM | 3.150686 | COADREAD | NCI-H508 | KU-60019 |
| MVP_top60_DNA_seed | HDAC2 | -0.728114 | OV | OC-314 | JQ12 | top60_altered_DNA+RNA_seed | BRAF | -3.715734 | COADREAD | COLO-205 | SB590885 |
| MVP_top60_DNA_seed | MAPK1 | -0.72598 | OV | OVCA420 | FR-180204 | top60_altered_DNA+RNA_seed | BRAF | -2.42515 | COADREAD | CCK-81 | SB590885 |
| MVP_top60_DNA_seed | MAPK3 | -0.72598 | OV | OVCA420 | FR-180204 | top60_altered_DNA+RNA_seed | BRAF | -2.424628 | COADREAD | COLO-205 | PLX-4720 |
| MVP_top60_DNA_seed | EGFR | -0.72132 | OV | TOV-21G | Afatinib | top60_altered_DNA+RNA_seed | BRAF | -2.336353 | COADREAD | COLO-205 | Dabrafenib |
| MVP_top60_DNA_seed | ERBB2 | -0.72132 | OV | TOV-21G | Afatinib | top60_altered_DNA+RNA_seed | PIK3CA | -2.306434 | COADREAD | C2BBe1 | AZD8186 |
| MVP_top60_DNA_seed | ATR | -0.720911 | OV | Hey | AZD6738 | top60_altered_DNA+RNA_seed | BRAF | -2.208516 | COADREAD | HT-29 | Dabrafenib |
| MVP_top60_DNA_seed | ERBB2 | -0.720549 | OV | OV-90 | CP724714 | top60_altered_DNA+RNA_seed | PIK3CA | -2.162493 | COADREAD | NCI-H508 | Alpelisib |

| | | | | | | | | | | | |
|---|---|---|---|---|---|---|---|---|---|---|---|
| MVP_top60_DNA_seed | PLK1 | -0.719285 | OV | PEO1 | PLK_6522 | top60_altered_DNA+RNA_seed | PIK3CA | -2.120441 | COADREAD | LoVo | AZD8186 |
| MVP_top60_DNA_seed | CDK2 | -0.717152 | OV | OVISE | PHA-793887 | top60_altered_DNA+RNA_seed | BRAF | -1.782926 | COADREAD | MDST8 | Dabrafenib |
| MVP_top60_DNA_seed | PLK1 | -0.703159 | OV | TOV-21G | PLK_6522 | top60_altered_DNA+RNA_seed | BRAF | -1.585375 | COADREAD | HT-29 | PLX-4720 |
| MVP_top60_DNA_seed | ATR | -0.695646 | OV | DOV13 | AZ20 | top60_altered_DNA+RNA_seed | ATM | -1.525765 | COADREAD | SK-CO-1 | KU-55933 |
| MVP_top60_DNA_seed | ATR | -0.692719 | OV | TOV-112D | QL-VIII-58 | top60_altered_DNA+RNA_seed | BRAF | -1.518973 | COADREAD | RKO | Dabrafenib |
| MVP_top60_DNA_seed | ATR | -0.673438 | OV | OVK-18 | AZD6738 | top60_altered_DNA+RNA_seed | PIK3CA | -1.501377 | COADREAD | GP5d | Alpelisib |
| MVP_top60_DNA_seed | ATR | -0.672778 | OV | OVCAR-4 | AZD6738 | top60_altered_DNA+RNA_seed | ATM | -1.481905 | COADREAD | RKO | KU-55933 |
| MVP_top60_DNA_seed | ATR | -0.671129 | OV | OVCAR-4 | QL-VIII-58 | top60_altered_DNA+RNA_seed | BRAF | -1.427691 | COADREAD | LS-411N | PLX-4720 |
| MVP_top60_DNA_seed | EGFR | -0.670123 | OV | EFO-21 | Pelitinib | top60_altered_DNA+RNA_seed | KRAS | -1.426956 | COADREAD | KM12 | :RAS (G12C) Inhibitor-1 |
| MVP_top60_DNA_seed | ERBB2 | -0.666775 | OV | OVTOKO | CP724714 | top60_altered_DNA+RNA_seed | ATM | -1.418079 | COADREAD | CCK-81 | KU-55933 |
| MVP_top60_DNA_seed | CDK2 | -0.66469 | OV | JHOS-2 | AZD5438 | top60_altered_DNA+RNA_seed | BRAF | -1.412943 | COADREAD | HT-29 | SB590885 |
| MVP_top60_DNA_seed | EGFR | -0.658399 | OV | OVCA420 | AST-1306 | top60_altered_DNA+RNA_seed | BRAF | -1.3808 | COADREAD | RCM-1 | SB590885 |
| MVP_top60_DNA_seed | EGFR | -0.656851 | OV | OVTOKO | Pelitinib | top60_altered_DNA+RNA_seed | ATM | -1.3741 | COADREAD | LoVo | KU-55933 |
| MVP_top60_DNA_seed | PLK1 | -0.655019 | OV | OVCAR-4 | BI-2536 | top60_altered_DNA+RNA_seed | BRAF | -1.369627 | COADREAD | MDST8 | PLX-4720 |
| MVP_top60_DNA_seed | ATR | -0.652403 | OV | UWB1.289 | AZD6738 | top60_altered_DNA+RNA_seed | BRAF | -1.313137 | COADREAD | MDST8 | SB590885 |
| MVP_top60_DNA_seed | MAPK1 | -0.651249 | OV | OV-90 | VX-11e | top60_altered_DNA+RNA_seed | ATM | -1.312324 | COADREAD | T84 | KU-55933 |
| MVP_top60_DNA_seed | EGFR | -0.646575 | OV | Hey | CUDC-101 | top60_altered_DNA+RNA_seed | KRAS | -1.301712 | COADREAD | HCT-15 | :RAS (G12C) Inhibitor-1 |
| MVP_top60_DNA_seed | ERBB2 | -0.646575 | OV | Hey | CUDC-101 | top60_altered_DNA+RNA_seed | BRAF | -1.232931 | COADREAD | SW1417 | Dabrafenib |
| MVP_top60_DNA_seed | CDK1 | -0.642556 | OV | TOV-112D | RO-3306 | top60_altered_DNA+RNA_seed | PIK3CA | -1.201292 | COADREAD | SW48 | AZD8186 |
| MVP_top60_DNA_seed | PIK3CA | -0.640021 | OV | OV-90 | PI3Ka_4409 | top60_altered_DNA+RNA_seed | PIK3CA | -1.197354 | COADREAD | HCT-15 | GNE-317 |
| MVP_top60_DNA_seed | EGFR | -0.63855 | OV | TOV-21G | Pelitinib | top60_altered_DNA+RNA_seed | PIK3CA | -1.158287 | COADREAD | NCI-H508 | AZD8186 |
| MVP_top60_DNA_seed | PIK3CA | -0.632105 | OV | OVCAR-5 | AZD8835 | top60_altered_DNA+RNA_seed | BRAF | -1.156491 | COADREAD | LS-411N | Dabrafenib |
| MVP_top60_DNA_seed | PLK1 | -0.629037 | OV | Hey | PLK_6522 | top60_altered_DNA+RNA_seed | PIK3CA | -1.125492 | COADREAD | SNU-81 | AZD8186 |
| MVP_top60_DNA_seed | PIK3CA | -0.625477 | OV | TOV-21G | AZD8835 | top60_altered_DNA+RNA_seed | BRAF | -1.125398 | COADREAD | LoVo | Dabrafenib |
| MVP_top60_DNA_seed | CDK1 | -0.624096 | OV | OAW-28 | RO-3306 | top60_altered_DNA+RNA_seed | PIK3CA | -1.117837 | COADREAD | SW48 | GNE-317 |
| MVP_top60_DNA_seed | EGFR | -0.620778 | OV | OC-314 | Cetuximab | top60_altered_DNA+RNA_seed | PIK3CA | -1.104252 | COADREAD | C2BBe1 | GNE-317 |
| MVP_top60_DNA_seed | EGFR | -0.61899 | OV | IGROV-1 | Afatinib | top60_altered_DNA+RNA_seed | TP53 | -1.101466 | COADREAD | COLO-320-HSR | MIRA-1 |
| MVP_top60_DNA_seed | ERBB2 | -0.61899 | OV | IGROV-1 | Afatinib | top60_altered_DNA+RNA_seed | BRAF | -1.062641 | COADREAD | LoVo | SB590885 |
| MVP_top60_DNA_seed | EGFR | -0.613523 | OV | SK-OV-3 | Pelitinib | top60_altered_DNA+RNA_seed | BRAF | -1.05669 | COADREAD | HCT-116 | Dabrafenib |
| MVP_top60_DNA_seed | CDK2 | -0.613249 | OV | OVTOKO | AZD5438 | top60_altered_DNA+RNA_seed | PIK3CA | -1.052704 | COADREAD | SW48 | Alpelisib |
| MVP_top60_DNA_seed | ERBB2 | -0.601852 | OV | OVCAR-8 | CP724714 | top60_altered_DNA+RNA_seed | PIK3CA | -1.051702 | COADREAD | COLO-320-HSR | Buparlisib |
| MVP_top60_DNA_seed | PIK3CA | -0.599678 | OV | PEO1 | PI3Ka_4409 | top60_altered_DNA+RNA_seed | PIK3CA | -1.051678 | COADREAD | T84 | Alpelisib |
| MVP_top60_DNA_seed | MAPK1 | -0.595204 | OV | OVKATE | VX-11e | top60_altered_DNA+RNA_seed | PIK3CA | -1.026602 | COADREAD | HCT-15 | Alpelisib |
| MVP_top60_DNA_seed | ATR | -0.590559 | OV | PEO1 | AZ20 | top60_altered_DNA+RNA_seed | ATM | -1.019802 | COADREAD | HCC2998 | KU-55933 |
| MVP_top60_DNA_seed | CDK2 | -0.589302 | OV | OVCA420 | AZD5438 | top60_altered_DNA+RNA_seed | PIK3CA | -1.008135 | COADREAD | DiFi | Alpelisib |
| MVP_top60_DNA_seed | EGFR | -0.584936 | OV | UWB1.289 | AZD8931 | top60_altered_DNA+RNA_seed | PIK3CA | -1.004211 | COADREAD | SK-CO-1 | AZD8186 |
| MVP_top60_DNA_seed | ERBB2 | -0.584936 | OV | UWB1.289 | AZD8931 | top60_altered_DNA+RNA_seed | BRAF | -0.994267 | COADREAD | RKO | SB590885 |
| MVP_top60_DNA_seed | CDK1 | -0.584431 | OV | OAW-42 | RO-3306 | top60_altered_DNA+RNA_seed | PIK3CA | -0.993072 | COADREAD | MDST8 | GNE-317 |
| MVP_top60_DNA_seed | MAPK1 | -0.576446 | OV | JHOS-3 | FR-180204 | top60_altered_DNA+RNA_seed | PIK3CA | -0.977944 | COADREAD | SK-CO-1 | Buparlisib |
| MVP_top60_DNA_seed | MAPK3 | -0.576446 | OV | JHOS-3 | FR-180204 | top60_altered_DNA+RNA_seed | PIK3CA | -0.958464 | COADREAD | NCI-H716 | AZD8186 |
| MVP_top60_DNA_seed | EGFR | -0.573651 | OV | SK-OV-3 | Afatinib | top60_altered_DNA+RNA_seed | KRAS | -0.948158 | COADREAD | HT55 | :RAS (G12C) Inhibitor-1 |

| | | | | | | | | | | | |
|---|---|---|---|---|---|---|---|---|---|---|---|
| MVP_top60_DNA_seed | ERBB2 | -0.573651 | OV | SK-OV-3 | Afatinib | top60_altered_DNA+RNA_seed | PIK3CA | -0.935631 | COADREAD | NCI-H747 | Alpelisib |
| MVP_top60_DNA_seed | EGFR | -0.573368 | OV | Caov-3 | CI-1033 | top60_altered_DNA+RNA_seed | BRAF | -0.93309 | COADREAD | NCI-H716 | SB590885 |
| MVP_top60_DNA_seed | ERBB2 | -0.573368 | OV | Caov-3 | CI-1033 | top60_altered_DNA+RNA_seed | PIK3CA | -0.924875 | COADREAD | CaR-1 | GNE-317 |
| MVP_top60_DNA_seed | MAPK1 | -0.568936 | OV | TOV-112D | VX-11e | top60_altered_DNA+RNA_seed | PIK3CA | -0.911292 | COADREAD | COLO-205 | GNE-317 |
| MVP_top60_DNA_seed | PIK3CA | -0.562 | OV | Hey | AZD8835 | top60_altered_DNA+RNA_seed | TP53 | -0.910323 | COADREAD | COLO-205 | MIRA-1 |
| MVP_top60_DNA_seed | SIRT1 | -0.559864 | OV | OAW-42 | Selisistat | top60_altered_DNA+RNA_seed | PIK3CA | -0.897857 | COADREAD | HT55 | Alpelisib |
| MVP_top60_DNA_seed | PLK1 | -0.559411 | OV | OVCAR-3 | PLK_6522 | top60_altered_DNA+RNA_seed | PIK3CA | -0.889066 | COADREAD | CCK-81 | Alpelisib |
| MVP_top60_DNA_seed | ATR | -0.551134 | OV | OVTOKO | AZD6738 | top60_altered_DNA+RNA_seed | BRAF | -0.8602 | COADREAD | RKO | PLX-4720 |
| MVP_top60_DNA_seed | MAPK1 | -0.547932 | OV | Caov-4 | VX-11e | top60_altered_DNA+RNA_seed | BRAF | -0.839029 | COADREAD | KM12 | SB590885 |
| MVP_top60_DNA_seed | PIK3CA | -0.538582 | OV | OVTOKO | AZD8835 | top60_altered_DNA+RNA_seed | PIK3CA | -0.83054 | COADREAD | HT-29 | Alpelisib |
| MVP_top60_DNA_seed | EGFR | -0.538335 | OV | Hey | PF-00299804 | top60_altered_DNA+RNA_seed | BRAF | -0.827317 | COADREAD | SNU-C5 | Dabrafenib |
| MVP_top60_DNA_seed | ERBB2 | -0.538335 | OV | Hey | PF-00299804 | top60_altered_DNA+RNA_seed | PIK3CA | -0.827186 | COADREAD | HCT-15 | Buparlisib |
| MVP_top60_DNA_seed | CDK1 | -0.534285 | OV | DOV13 | RO-3306 | top60_altered_DNA+RNA_seed | PIK3CA | -0.826032 | COADREAD | SK-CO-1 | GNE-317 |
| MVP_top60_DNA_seed | PIK3CA | -0.532522 | OV | OV-7 | AZD8835 | top60_altered_DNA+RNA_seed | PIK3CA | -0.80982 | COADREAD | GP5d | Buparlisib |
| MVP_top60_DNA_seed | ATR | -0.523104 | OV | OVMIU | AZ20 | top60_altered_DNA+RNA_seed | BRAF | -0.809263 | COADREAD | LoVo | PLX-4720 |
| MVP_top60_DNA_seed | SIRT1 | -0.520943 | OV | Hey | Selisistat | top60_altered_DNA+RNA_seed | PIK3CA | -0.808737 | COADREAD | COLO-320-HSR | Alpelisib |
| MVP_top60_DNA_seed | EGFR | -0.512663 | OV | SK-OV-3 | AZD8931 | top60_altered_DNA+RNA_seed | PIK3CA | -0.807439 | COADREAD | MDST8 | Buparlisib |
| MVP_top60_DNA_seed | ERBB2 | -0.512663 | OV | SK-OV-3 | AZD8931 | top60_altered_DNA+RNA_seed | PIK3CA | -0.797717 | COADREAD | NCI-H747 | Buparlisib |
| MVP_top60_DNA_seed | EGFR | -0.510918 | OV | TOV-21G | Afatinib | top60_altered_DNA+RNA_seed | PIK3CA | -0.795901 | COADREAD | NCI-H747 | GNE-317 |
| MVP_top60_DNA_seed | ERBB2 | -0.510918 | OV | TOV-21G | Afatinib | top60_altered_DNA+RNA_seed | BRAF | -0.785509 | COADREAD | SNU-C5 | SB590885 |
| MVP_top60_DNA_seed | EGFR | -0.508057 | OV | OVKATE | CUDC-101 | top60_altered_DNA+RNA_seed | PIK3CA | -0.783071 | COADREAD | SW948 | Alpelisib |
| MVP_top60_DNA_seed | ERBB2 | -0.508057 | OV | OVKATE | CUDC-101 | top60_altered_DNA+RNA_seed | PIK3CA | -0.772557 | COADREAD | LoVo | Buparlisib |
| MVP_top60_DNA_seed | ATR | -0.507371 | OV | JHOS-2 | torin2 | top60_altered_DNA+RNA_seed | PIK3CA | -0.765258 | COADREAD | COLO-205 | Buparlisib |
| MVP_top60_DNA_seed | EGFR | -0.505804 | OV | OVCAR-4 | Lapatinib | top60_altered_DNA+RNA_seed | BRAF | -0.763068 | COADREAD | LS-513 | SB590885 |
| MVP_top60_DNA_seed | ERBB2 | -0.505804 | OV | OVCAR-4 | Lapatinib | top60_altered_DNA+RNA_seed | PIK3CA | -0.758564 | COADREAD | SNU-175 | Buparlisib |
| MVP_top60_DNA_seed | MAPK1 | -0.49899 | OV | PEO1 | VX-11e | top60_altered_DNA+RNA_seed | PIK3CA | -0.755599 | COADREAD | SW48 | Buparlisib |
| MVP_top60_DNA_seed | EGFR | -0.494932 | OV | PEO1 | Afatinib | top60_altered_DNA+RNA_seed | PIK3CA | -0.753669 | COADREAD | LoVo | GNE-317 |
| MVP_top60_DNA_seed | ERBB2 | -0.494932 | OV | PEO1 | Afatinib | top60_altered_DNA+RNA_seed | TP53 | -0.749332 | COADREAD | LS-513 | MIRA-1 |
| MVP_top60_DNA_seed | EGFR | -0.494125 | OV | OVCAR-8 | AZD8931 | top60_altered_DNA+RNA_seed | BRAF | -0.747864 | COADREAD | SNU-C5 | PLX-4720 |
| MVP_top60_DNA_seed | ERBB2 | -0.494125 | OV | OVCAR-8 | AZD8931 | top60_altered_DNA+RNA_seed | BRAF | -0.739477 | COADREAD | SK-CO-1 | SB590885 |
| MVP_top60_DNA_seed | EGFR | -0.491329 | OV | PEO1 | AZD8931 | top60_altered_DNA+RNA_seed | PIK3CA | -0.735045 | COADREAD | SW948 | GNE-317 |
| MVP_top60_DNA_seed | ERBB2 | -0.491329 | OV | PEO1 | AZD8931 | top60_altered_DNA+RNA_seed | PIK3CA | -0.732431 | COADREAD | C2BBe1 | Buparlisib |
| MVP_top60_DNA_seed | HDAC2 | -0.481002 | OV | TOV-112D | JQ12 | top60_altered_DNA+RNA_seed | PIK3CA | -0.728024 | COADREAD | KM12 | Buparlisib |
| MVP_top60_DNA_seed | HDAC2 | -0.478038 | OV | SW626 | JQ12 | top60_altered_DNA+RNA_seed | PIK3CA | -0.698622 | COADREAD | COLO-320-HSR | GNE-317 |
| MVP_top60_DNA_seed | PTK2 | -0.473914 | OV | OVMIU | PF-562271 | top60_altered_DNA+RNA_seed | PIK3CA | -0.691121 | COADREAD | DiFi | GNE-317 |
| MVP_top60_DNA_seed | PIK3CA | -0.470036 | OV | OVISE | PI3Ka_4409 | top60_altered_DNA+RNA_seed | ATM | -0.679532 | COADREAD | SNU-C5 | KU-55933 |
| MVP_top60_DNA_seed | EGFR | -0.463047 | OV | OAW-42 | AZD8931 | top60_altered_DNA+RNA_seed | BRAF | -0.66488 | COADREAD | KM12 | Dabrafenib |
| MVP_top60_DNA_seed | ERBB2 | -0.463047 | OV | OAW-42 | AZD8931 | top60_altered_DNA+RNA_seed | PIK3CA | -0.661258 | COADREAD | DiFi | Buparlisib |
| MVP_top60_DNA_seed | EGFR | -0.458684 | OV | JHOS-4 | AZD8931 | top60_altered_DNA+RNA_seed | ATM | -0.660589 | COADREAD | MDST8 | KU-55933 |
| MVP_top60_DNA_seed | ERBB2 | -0.458684 | OV | JHOS-4 | AZD8931 | top60_altered_DNA+RNA_seed | BRAF | -0.658788 | COADREAD | NCI-H508 | Dabrafenib |
| MVP_top60_DNA_seed | PIK3CA | -0.458504 | OV | OC-314 | PI3Ka_4409 | top60_altered_DNA+RNA_seed | KRAS | -0.652878 | COADREAD | HCC2998 | RAS (G12C) Inhibitor-1 |

| | | | | | | | | | | | |
|---|---|---|---|---|---|---|---|---|---|---|---|
| MVP_top60_DNA_seed | EGFR | -0.458263 | OV | Caov-3 | AST-1306 | top60_altered_DNA+RNA_seed | BRAF | -0.647974 | COADREAD | COLO-320-HSR | PLX-4720 |
| MVP_top60_DNA_seed | MAPK1 | -0.454837 | OV | OVCA433 | FR-180204 | top60_altered_DNA+RNA_seed | ATM | -0.620347 | COADREAD | SW837 | KU-55933 |
| MVP_top60_DNA_seed | MAPK3 | -0.454837 | OV | OVCA433 | FR-180204 | top60_altered_DNA+RNA_seed | PIK3CA | -0.60258 | COADREAD | SK-CO-1 | Alpelisib |
| MVP_top60_DNA_seed | ATR | -0.454471 | OV | TOV-112D | AZD6738 | top60_altered_DNA+RNA_seed | PIK3CA | -0.602487 | COADREAD | SNU-C5 | Buparlisib |
| MVP_top60_DNA_seed | MAPK1 | -0.454322 | OV | UWB1.289 | FR-180204 | top60_altered_DNA+RNA_seed | PIK3CA | -0.594705 | COADREAD | HCT-116 | Buparlisib |
| MVP_top60_DNA_seed | MAPK3 | -0.454322 | OV | UWB1.289 | FR-180204 | top60_altered_DNA+RNA_seed | BRAF | -0.585591 | COADREAD | HT-115 | SB590885 |
| MVP_top60_DNA_seed | CDK2 | -0.45206 | OV | DOV13 | AZD5438 | top60_altered_DNA+RNA_seed | BRAF | -0.577844 | COADREAD | HCT-15 | PLX-4720 |
| MVP_top60_DNA_seed | EGFR | -0.45051 | OV | OVCAR-4 | Afatinib | top60_altered_DNA+RNA_seed | PIK3CA | -0.551992 | COADREAD | COLO-320-HSR | AZD8186 |
| MVP_top60_DNA_seed | ERBB2 | -0.45051 | OV | OVCAR-4 | Afatinib | top60_altered_DNA+RNA_seed | BRAF | -0.54219 | COADREAD | GP5d | SB590885 |
| MVP_top60_DNA_seed | ATR | -0.437915 | OV | OV-7 | torin2 | top60_altered_DNA+RNA_seed | PIK3CA | -0.527602 | COADREAD | HT-115 | Buparlisib |
| MVP_top60_DNA_seed | ERBB2 | -0.430945 | OV | Caov-3 | CP724714 | top60_altered_DNA+RNA_seed | PIK3CA | -0.508767 | COADREAD | HCT-116 | Alpelisib |
| MVP_top60_DNA_seed | CDK1 | -0.429268 | OV | OVCA433 | RO-3306 | top60_altered_DNA+RNA_seed | TP53 | -0.502326 | COADREAD | GP5d | MIRA-1 |
| MVP_top60_DNA_seed | EGFR | -0.428237 | OV | DOV13 | AZD8931 | top60_altered_DNA+RNA_seed | PIK3CA | -0.484905 | COADREAD | CCK-81 | Buparlisib |
| MVP_top60_DNA_seed | ERBB2 | -0.428237 | OV | DOV13 | AZD8931 | top60_altered_DNA+RNA_seed | PIK3CA | -0.48204 | COADREAD | CaR-1 | AZD8186 |
| MVP_top60_DNA_seed | PLK1 | -0.426391 | OV | SK-OV-3 | PLK_6522 | top60_altered_DNA+RNA_seed | PIK3CA | -0.478742 | COADREAD | NCI-H747 | AZD8186 |
| MVP_top60_DNA_seed | SIRT1 | -0.424002 | OV | TOV-21G | Selisistat | top60_altered_DNA+RNA_seed | KRAS | -0.460015 | COADREAD | GP5d | KRAS (G12C) Inhibitor-1 |
| MVP_top60_DNA_seed | EGFR | -0.420237 | OV | OVCAR-5 | Pelitinib | top60_altered_DNA+RNA_seed | KRAS | -0.452548 | COADREAD | LoVo | KRAS (G12C) Inhibitor-1 |
| MVP_top60_DNA_seed | PLK1 | -0.417625 | OV | OVCAR-4 | GW843682X | top60_altered_DNA+RNA_seed | ATM | -0.441003 | COADREAD | CaR-1 | KU-55933 |
| MVP_top60_DNA_seed | CREBBP | -0.414213 | OV | OVTOKO | I-CBP112 | top60_altered_DNA+RNA_seed | KRAS | -0.43604 | COADREAD | SW620 | KRAS (G12C) Inhibitor-1 |
| MVP_top60_DNA_seed | EGFR | -0.411905 | OV | TOV-21G | Cetuximab | top60_altered_DNA+RNA_seed | PIK3CA | -0.432524 | COADREAD | SNU-407 | Buparlisib |
| MVP_top60_DNA_seed | EGFR | -0.410339 | OV | TOV-112D | CUDC-101 | top60_altered_DNA+RNA_seed | TP53 | -0.424729 | COADREAD | DiFi | MIRA-1 |
| MVP_top60_DNA_seed | ERBB2 | -0.410339 | OV | TOV-112D | CUDC-101 | top60_altered_DNA+RNA_seed | PIK3CA | -0.42315 | COADREAD | RKO | Alpelisib |
| MVP_top60_DNA_seed | ATR | -0.409953 | OV | OV-56 | AZ20 | top60_altered_DNA+RNA_seed | PIK3CA | -0.408004 | COADREAD | HCT-116 | GNE-317 |
| MVP_top60_DNA_seed | PTK2 | -0.409326 | OV | OC-314 | PF-562271 | top60_altered_DNA+RNA_seed | BRAF | -0.404416 | COADREAD | HT55 | PLX-4720 |
| MVP_top60_DNA_seed | EGFR | -0.40362 | OV | OVCAR-8 | Cetuximab | top60_altered_DNA+RNA_seed | PIK3CA | -0.39723 | COADREAD | SNU-407 | Alpelisib |
| MVP_top60_DNA_seed | EGFR | -0.402394 | OV | OAW-28 | AST-1306 | top60_altered_DNA+RNA_seed | BRAF | -0.394189 | COADREAD | HCC2998 | SB590885 |
| MVP_top60_DNA_seed | HDAC2 | -0.401567 | OV | SK-OV-3 | JQ12 | top60_altered_DNA+RNA_seed | ATM | -0.391766 | COADREAD | C2BBe1 | KU-55933 |
| MVP_top60_DNA_seed | EGFR | -0.399535 | OV | Hey | Afatinib | top60_altered_DNA+RNA_seed | PIK3CA | -0.389519 | COADREAD | LS-180 | GNE-317 |
| MVP_top60_DNA_seed | ERBB2 | -0.399535 | OV | Hey | Afatinib | top60_altered_DNA+RNA_seed | BRAF | -0.380895 | COADREAD | HCT-15 | Dabrafenib |
| MVP_top60_DNA_seed | EGFR | -0.399066 | OV | OVCA433 | Afatinib | top60_altered_DNA+RNA_seed | PIK3CA | -0.380122 | COADREAD | COLO-205 | AZD8186 |
| MVP_top60_DNA_seed | ERBB2 | -0.399066 | OV | OVCA433 | Afatinib | top60_altered_DNA+RNA_seed | KRAS | -0.379246 | COADREAD | CW-2 | KRAS (G12C) Inhibitor-1 |
| MVP_top60_DNA_seed | CDK1 | -0.398886 | OV | OVCAR-4 | RO-3306 | top60_altered_DNA+RNA_seed | ATM | -0.378328 | COADREAD | SNU-C1 | KU-55933 |
| MVP_top60_DNA_seed | EGFR | -0.392937 | OV | OC-314 | Afatinib | top60_altered_DNA+RNA_seed | BRAF | -0.375272 | COADREAD | SNU-81 | PLX-4720 |
| MVP_top60_DNA_seed | ERBB2 | -0.392937 | OV | OC-314 | Afatinib | top60_altered_DNA+RNA_seed | TP53 | -0.365414 | COADREAD | CW-2 | MIRA-1 |
| MVP_top60_DNA_seed | ATR | -0.39252 | OV | IGROV-1 | AZ20 | top60_altered_DNA+RNA_seed | TP53 | -0.364305 | COADREAD | HT55 | MIRA-1 |
| MVP_top60_DNA_seed | PIK3CA | -0.390209 | OV | OC-314 | AZD8835 | top60_altered_DNA+RNA_seed | TP53 | -0.337806 | COADREAD | CCK-81 | MIRA-1 |
| MVP_top60_DNA_seed | ATR | -0.389893 | OV | OV-90 | AZ20 | top60_altered_DNA+RNA_seed | BRAF | -0.337173 | COADREAD | SW1417 | PLX-4720 |
| MVP_top60_DNA_seed | PIK3CA | -0.38871 | OV | OVISE | AZD8835 | top60_altered_DNA+RNA_seed | PIK3CA | -0.337026 | COADREAD | COLO-205 | Alpelisib |
| MVP_top60_DNA_seed | EGFR | -0.384349 | OV | JHOS-2 | CUDC-101 | top60_altered_DNA+RNA_seed | KRAS | -0.334025 | COADREAD | RKO | KRAS (G12C) Inhibitor-1 |
| MVP_top60_DNA_seed | ERBB2 | -0.384349 | OV | JHOS-2 | CUDC-101 | top60_altered_DNA+RNA_seed | PIK3CA | -0.321428 | COADREAD | CaR-1 | Alpelisib |
| MVP_top60_DNA_seed | EGFR | -0.378614 | OV | OAW-28 | PF-00299804 | top60_altered_DNA+RNA_seed | KRAS | -0.297829 | COADREAD | SW48 | KRAS (G12C) Inhibitor-1 |

| | | | | | | | | | | | |
|---|---|---|---|---|---|---|---|---|---|---|---|
| MVP_top60_DNA_seed | ERBB2 | -0.378614 | OV | OAW-28 | PF-00299804 | top60_altered_DNA+RNA_seed | BRAF | -0.294989 | COADREAD | SK-CO-1 | Dabrafenib |
| MVP_top60_DNA_seed | ATR | -0.377995 | OV | OV-90 | torin2 | top60_altered_DNA+RNA_seed | BRAF | -0.292846 | COADREAD | LS-1034 | Dabrafenib |
| MVP_top60_DNA_seed | EGFR | -0.375952 | OV | OVCAR-8 | Pelitinib | top60_altered_DNA+RNA_seed | PIK3CA | -0.290618 | COADREAD | LS-411N | GNE-317 |
| MVP_top60_DNA_seed | ATR | -0.37169 | OV | OVTOKO | AZ20 | top60_altered_DNA+RNA_seed | ATM | -0.278066 | COADREAD | CW-2 | KU-55933 |
| MVP_top60_DNA_seed | ERBB2 | -0.370724 | OV | OVCAR-4 | CP724714 | top60_altered_DNA+RNA_seed | KRAS | -0.259001 | COADREAD | NCI-H716 | KRAS (G12C) Inhibitor-1 |
| MVP_top60_DNA_seed | PTK2 | -0.369307 | OV | OVCAR-3 | PF-562271 | top60_altered_DNA+RNA_seed | KRAS | -0.249931 | COADREAD | SNU-C5 | KRAS (G12C) Inhibitor-1 |
| MVP_top60_DNA_seed | ATR | -0.36796 | OV | OV-90 | QL-VIII-58 | top60_altered_DNA+RNA_seed | PIK3CA | -0.24948 | COADREAD | KM12 | GNE-317 |
| MVP_top60_DNA_seed | PLK1 | -0.36771 | OV | OV-90 | PLK_6522 | top60_altered_DNA+RNA_seed | BRAF | -0.237639 | COADREAD | HT55 | Dabrafenib |
| MVP_top60_DNA_seed | EGFR | -0.367545 | OV | OVCA433 | Afatinib | top60_altered_DNA+RNA_seed | PIK3CA | -0.233439 | COADREAD | CaR-1 | Buparlisib |
| MVP_top60_DNA_seed | ERBB2 | -0.367545 | OV | OVCA433 | Afatinib | top60_altered_DNA+RNA_seed | PIK3CA | -0.227941 | COADREAD | LS-123 | GNE-317 |
| MVP_top60_DNA_seed | EGFR | -0.364101 | OV | OVMIU | CUDC-101 | top60_altered_DNA+RNA_seed | BRAF | -0.213705 | COADREAD | HCT-116 | PLX-4720 |
| MVP_top60_DNA_seed | ERBB2 | -0.364101 | OV | OVMIU | CUDC-101 | top60_altered_DNA+RNA_seed | KRAS | -0.211738 | COADREAD | LS-513 | KRAS (G12C) Inhibitor-1 |
| MVP_top60_DNA_seed | EGFR | -0.359361 | OV | JHOS-2 | AST-1306 | top60_altered_DNA+RNA_seed | BRAF | -0.204938 | COADREAD | DiFi | SB590885 |
| MVP_top60_DNA_seed | PTK2 | -0.353918 | OV | SK-OV-3 | PF-562271 | top60_altered_DNA+RNA_seed | PIK3CA | -0.194916 | COADREAD | HT-29 | GNE-317 |
| MVP_top60_DNA_seed | EGFR | -0.350057 | OV | FU-OV-1 | PF-00299804 | top60_altered_DNA+RNA_seed | PIK3CA | -0.192095 | COADREAD | LS-513 | Alpelisib |
| MVP_top60_DNA_seed | ERBB2 | -0.350057 | OV | FU-OV-1 | PF-00299804 | top60_altered_DNA+RNA_seed | TP53 | -0.191951 | COADREAD | MDST8 | MIRA-1 |
| MVP_top60_DNA_seed | CDK2 | -0.348316 | OV | SW626 | PHA-793887 | top60_altered_DNA+RNA_seed | PIK3CA | -0.190287 | COADREAD | HCC2998 | Buparlisib |
| MVP_top60_DNA_seed | CREBBP | -0.348048 | OV | TOV-21G | I-CBP112 | top60_altered_DNA+RNA_seed | ATM | -0.186846 | COADREAD | NCI-H716 | KU-55933 |
| MVP_top60_DNA_seed | MAPK1 | -0.345991 | OV | SW626 | FR-180204 | top60_altered_DNA+RNA_seed | ATM | -0.184286 | COADREAD | HT55 | KU-55933 |
| MVP_top60_DNA_seed | MAPK3 | -0.345991 | OV | SW626 | FR-180204 | top60_altered_DNA+RNA_seed | BRAF | -0.18041 | COADREAD | HCT-116 | SB590885 |
| MVP_top60_DNA_seed | EGFR | -0.338642 | OV | OVCAR-4 | Pelitinib | top60_altered_DNA+RNA_seed | PIK3CA | -0.172189 | COADREAD | LS-180 | Alpelisib |
| MVP_top60_DNA_seed | EGFR | -0.338095 | OV | IGROV-1 | Cetuximab | top60_altered_DNA+RNA_seed | PIK3CA | -0.170209 | COADREAD | SNU-407 | AZD8186 |
| MVP_top60_DNA_seed | PIK3CA | -0.327897 | OV | OVTOKO | PI3Ka_4409 | top60_altered_DNA+RNA_seed | PIK3CA | -0.164333 | COADREAD | HT-29 | AZD8186 |
| MVP_top60_DNA_seed | EGFR | -0.321757 | OV | OVISE | CI-1033 | top60_altered_DNA+RNA_seed | BRAF | -0.163102 | COADREAD | CaR-1 | Dabrafenib |
| MVP_top60_DNA_seed | ERBB2 | -0.321757 | OV | OVISE | CI-1033 | top60_altered_DNA+RNA_seed | PIK3CA | -0.158809 | COADREAD | HT-115 | Alpelisib |
| MVP_top60_DNA_seed | EGFR | -0.320122 | OV | OC-314 | Afatinib | top60_altered_DNA+RNA_seed | PIK3CA | -0.156407 | COADREAD | MDST8 | AZD8186 |
| MVP_top60_DNA_seed | ERBB2 | -0.320122 | OV | OC-314 | Afatinib | top60_altered_DNA+RNA_seed | PIK3CA | -0.155478 | COADREAD | SW948 | Buparlisib |
| MVP_top60_DNA_seed | PIK3CA | -0.313719 | OV | FU-OV-1 | AZD8835 | top60_altered_DNA+RNA_seed | PIK3CA | -0.13814 | COADREAD | CW-2 | Alpelisib |
| MVP_top60_DNA_seed | PIK3CA | -0.310992 | OV | JHOS-3 | AZD8835 | top60_altered_DNA+RNA_seed | PIK3CA | -0.136585 | COADREAD | LS-123 | AZD8186 |
| MVP_top60_DNA_seed | EGFR | -0.309073 | OV | SK-OV-3 | CUDC-101 | top60_altered_DNA+RNA_seed | ATM | -0.125136 | COADREAD | KM12 | KU-55933 |
| MVP_top60_DNA_seed | ERBB2 | -0.309073 | OV | SK-OV-3 | CUDC-101 | top60_altered_DNA+RNA_seed | PIK3CA | -0.115279 | COADREAD | SNU-C5 | Alpelisib |
| MVP_top60_DNA_seed | PLK1 | -0.304214 | OV | OV-7 | PLK_6522 | top60_altered_DNA+RNA_seed | PIK3CA | -0.115211 | COADREAD | LoVo | Alpelisib |
| MVP_top60_DNA_seed | MAPK1 | -0.300943 | OV | OVK-18 | VX-11e | top60_altered_DNA+RNA_seed | PIK3CA | -0.09687 | COADREAD | HT55 | AZD8186 |
| MVP_top60_DNA_seed | MAPK1 | -0.295116 | OV | JHOS-3 | VX-11e | top60_altered_DNA+RNA_seed | PIK3CA | -0.085083 | COADREAD | T84 | AZD8186 |
| MVP_top60_DNA_seed | EGFR | -0.292905 | OV | OVCAR-3 | PF-00299804 | top60_altered_DNA+RNA_seed | PIK3CA | -0.080345 | COADREAD | HCC2998 | GNE-317 |
| MVP_top60_DNA_seed | ERBB2 | -0.292905 | OV | OVCAR-3 | PF-00299804 | top60_altered_DNA+RNA_seed | BRAF | -0.073861 | COADREAD | COLO-320-HSR | SB590885 |
| MVP_top60_DNA_seed | ERBB2 | -0.292664 | OV | Caov-4 | CP724714 | top60_altered_DNA+RNA_seed | PIK3CA | -0.072293 | COADREAD | RCM-1 | Alpelisib |
| MVP_top60_DNA_seed | CDK2 | -0.292641 | OV | TOV-112D | AZD5438 | top60_altered_DNA+RNA_seed | PIK3CA | -0.072201 | COADREAD | GP5d | GNE-317 |
| MVP_top60_DNA_seed | EGFR | -0.289967 | OV | OAW-28 | Gefitinib | top60_altered_DNA+RNA_seed | KRAS | -0.072151 | COADREAD | NCI-H747 | KRAS (G12C) Inhibitor-1 |
| MVP_top60_DNA_seed | EGFR | -0.285147 | OV | Hey | Pelitinib | top60_altered_DNA+RNA_seed | KRAS | -0.069158 | COADREAD | HT-29 | KRAS (G12C) Inhibitor-1 |
| MVP_top60_DNA_seed | EGFR | -0.281826 | OV | OVCAR-4 | AZD8931 | top60_altered_DNA+RNA_seed | PIK3CA | -0.066991 | COADREAD | CW-2 | Buparlisib |

| | | | | | | | | | | | |
|---|---|---|---|---|---|---|---|---|---|---|---|
| MVP_top60_DNA_seed | ERBB2 | -0.281826 | OV | OVCAR-4 | AZD8931 | top60_altered_DNA+RNA_seed | BRAF | -0.06537 | COADREAD | CW-2 | SB590885 |
| MVP_top60_DNA_seed | EGFR | -0.28104 | OV | FU-OV-1 | Pelitinib | top60_altered_DNA+RNA_seed | PIK3CA | -0.060572 | COADREAD | SW1463 | GNE-317 |
| MVP_top60_DNA_seed | SIRT1 | -0.27679 | OV | OVCAR-8 | Selisistat | top60_altered_DNA+RNA_seed | ATM | -0.059558 | COADREAD | HT-115 | KU-55933 |
| MVP_top60_DNA_seed | EGFR | -0.272535 | OV | OVISE | CUDC-101 | top60_altered_DNA+RNA_seed | PIK3CA | -0.058033 | COADREAD | HCC2998 | AZD8186 |
| MVP_top60_DNA_seed | ERBB2 | -0.272535 | OV | OVISE | CUDC-101 | top60_altered_DNA+RNA_seed | KRAS | -0.055151 | COADREAD | HCT-116 | KRAS (G12C) Inhibitor-1 |
| MVP_top60_DNA_seed | EGFR | -0.260657 | OV | OVCA433 | Gefitinib | top60_altered_DNA+RNA_seed | TP53 | -0.052856 | COADREAD | HCT-116 | MIRA-1 |
| MVP_top60_DNA_seed | EGFR | -0.259992 | OV | OVCA420 | PF-00299804 | top60_altered_DNA+RNA_seed | ATM | -0.0517 | COADREAD | GP5d | KU-55933 |
| MVP_top60_DNA_seed | ERBB2 | -0.259992 | OV | OVCA420 | PF-00299804 | top60_altered_DNA+RNA_seed | BRAF | -0.046993 | COADREAD | NCI-H716 | Dabrafenib |
| MVP_top60_DNA_seed | CDK2 | -0.250428 | OV | OVCAR-8 | PHA-793887 | top60_altered_DNA+RNA_seed | KRAS | -0.034769 | COADREAD | SNU-1040 | KRAS (G12C) Inhibitor-1 |
| MVP_top60_DNA_seed | EGFR | -0.245188 | OV | OAW-28 | Pelitinib | top60_altered_DNA+RNA_seed | BRAF | -0.032983 | COADREAD | SW948 | Dabrafenib |
| MVP_top60_DNA_seed | MAPK1 | -0.244719 | OV | OC-314 | FR-180204 | top60_altered_DNA+RNA_seed | BRAF | -0.032273 | COADREAD | HT55 | SB590885 |
| MVP_top60_DNA_seed | MAPK3 | -0.244719 | OV | OC-314 | FR-180204 | top60_altered_DNA+RNA_seed | PIK3CA | -0.027099 | COADREAD | RKO | Buparlisib |
| MVP_top60_DNA_seed | CDK2 | -0.243671 | OV | OV-90 | PHA-793887 | top60_altered_DNA+RNA_seed | PIK3CA | -0.026078 | COADREAD | SNU-C5 | AZD8186 |
| MVP_top60_DNA_seed | EGFR | -0.23894 | OV | OVCAR-5 | CUDC-101 | top60_altered_DNA+RNA_seed | PIK3CA | -0.019827 | COADREAD | NCI-H508 | Buparlisib |
| MVP_top60_DNA_seed | ERBB2 | -0.23894 | OV | OVCAR-5 | CUDC-101 | top60_altered_DNA+RNA_seed | PIK3CA | -0.000728 | COADREAD | SNU-407 | GNE-317 |
| MVP_top60_DNA_seed | EGFR | -0.234373 | OV | SK-OV-3 | Afatinib | top60_altered_DNA+RNA_seed | PIK3CA | 0.002855 | COADREAD | SNU-175 | AZD8186 |
| MVP_top60_DNA_seed | ERBB2 | -0.234373 | OV | SK-OV-3 | Afatinib | top60_altered_DNA+RNA_seed | PIK3CA | 0.006593 | COADREAD | SNU-175 | GNE-317 |
| MVP_top60_DNA_seed | PIK3CA | -0.234136 | OV | OAW-28 | PI3Ka_4409 | top60_altered_DNA+RNA_seed | PIK3CA | 0.0087 | COADREAD | HT-29 | Buparlisib |
| MVP_top60_DNA_seed | ATR | -0.233404 | OV | SK-OV-3 | AZD6738 | top60_altered_DNA+RNA_seed | PIK3CA | 0.017624 | COADREAD | CCK-81 | AZD8186 |
| MVP_top60_DNA_seed | EGFR | -0.232302 | OV | UWB1.289 | Gefitinib | top60_altered_DNA+RNA_seed | ATM | 0.018433 | COADREAD | COLO-320-HSR | KU-55933 |
| MVP_top60_DNA_seed | ATR | -0.23188 | OV | SW626 | AZD6738 | top60_altered_DNA+RNA_seed | TP53 | 0.031989 | COADREAD | HCC2998 | MIRA-1 |
| MVP_top60_DNA_seed | EGFR | -0.231222 | OV | PEO1 | CI-1033 | top60_altered_DNA+RNA_seed | BRAF | 0.032811 | COADREAD | LS-411N | SB590885 |
| MVP_top60_DNA_seed | ERBB2 | -0.231222 | OV | PEO1 | CI-1033 | top60_altered_DNA+RNA_seed | KRAS | 0.053306 | COADREAD | LS-411N | KRAS (G12C) Inhibitor-1 |
| MVP_top60_DNA_seed | EGFR | -0.224454 | OV | SW626 | Pelitinib | top60_altered_DNA+RNA_seed | BRAF | 0.057755 | COADREAD | SNU-C1 | Dabrafenib |
| MVP_top60_DNA_seed | PLK1 | -0.219924 | OV | DOV13 | PLK_6522 | top60_altered_DNA+RNA_seed | BRAF | 0.058484 | COADREAD | CaR-1 | PLX-4720 |
| MVP_top60_DNA_seed | PIK3CA | -0.217999 | OV | FU-OV-1 | PI3Ka_4409 | top60_altered_DNA+RNA_seed | PIK3CA | 0.06485 | COADREAD | DiFi | AZD8186 |
| MVP_top60_DNA_seed | HDAC2 | -0.215612 | OV | OVMIU | JQ12 | top60_altered_DNA+RNA_seed | TP53 | 0.065601 | COADREAD | SK-CO-1 | MIRA-1 |
| MVP_top60_DNA_seed | CDK2 | -0.214283 | OV | OVCAR-5 | PHA-793887 | top60_altered_DNA+RNA_seed | PIK3CA | 0.067512 | COADREAD | SNU-175 | Alpelisib |
| MVP_top60_DNA_seed | EGFR | -0.209945 | OV | UWB1.289 | CUDC-101 | top60_altered_DNA+RNA_seed | BRAF | 0.079847 | COADREAD | LS-513 | Dabrafenib |
| MVP_top60_DNA_seed | ERBB2 | -0.209945 | OV | UWB1.289 | CUDC-101 | top60_altered_DNA+RNA_seed | PIK3CA | 0.085727 | COADREAD | GP5d | AZD8186 |
| MVP_top60_DNA_seed | ATR | -0.209737 | OV | OVCA433 | AZD6738 | top60_altered_DNA+RNA_seed | PIK3CA | 0.087524 | COADREAD | SW837 | GNE-317 |
| MVP_top60_DNA_seed | EGFR | -0.207891 | OV | OC-314 | CUDC-101 | top60_altered_DNA+RNA_seed | TP53 | 0.089289 | COADREAD | HT-115 | MIRA-1 |
| MVP_top60_DNA_seed | ERBB2 | -0.207891 | OV | OC-314 | CUDC-101 | top60_altered_DNA+RNA_seed | BRAF | 0.095294 | COADREAD | SW948 | PLX-4720 |
| MVP_top60_DNA_seed | EGFR | -0.205825 | OV | OVCAR-4 | AST-1306 | top60_altered_DNA+RNA_seed | BRAF | 0.101765 | COADREAD | DiFi | PLX-4720 |
| MVP_top60_DNA_seed | CDK2 | -0.205808 | OV | OVMIU | PHA-793887 | top60_altered_DNA+RNA_seed | KRAS | 0.104894 | COADREAD | SNU-175 | KRAS (G12C) Inhibitor-1 |
| MVP_top60_DNA_seed | EGFR | -0.204223 | OV | OVCAR-8 | Afatinib | top60_altered_DNA+RNA_seed | TP53 | 0.111455 | COADREAD | SNU-175 | MIRA-1 |
| MVP_top60_DNA_seed | ERBB2 | -0.204223 | OV | OVCAR-8 | Afatinib | top60_altered_DNA+RNA_seed | PIK3CA | 0.117066 | COADREAD | HCC2998 | Alpelisib |
| MVP_top60_DNA_seed | EGFR | -0.203367 | OV | TOV-21G | Gefitinib | top60_altered_DNA+RNA_seed | BRAF | 0.133763 | COADREAD | SK-CO-1 | PLX-4720 |
| MVP_top60_DNA_seed | PIK3CA | -0.203367 | OV | JHOS-3 | PI3Ka_4409 | top60_altered_DNA+RNA_seed | PIK3CA | 0.140505 | COADREAD | T84 | Buparlisib |
| MVP_top60_DNA_seed | ATR | -0.203205 | OV | JHOS-2 | QL-VIII-58 | top60_altered_DNA+RNA_seed | PIK3CA | 0.152698 | COADREAD | HT55 | Buparlisib |
| MVP_top60_DNA_seed | PLK1 | -0.202991 | OV | OVMIU | PLK_6522 | top60_altered_DNA+RNA_seed | PIK3CA | 0.152922 | COADREAD | SNU-C5 | GNE-317 |

| | | | | | | | | | | | |
|---|---|---|---|---|---|---|---|---|---|---|---|
| MVP_top60_DNA_seed | CDK1 | -0.192644 | OV | SK-OV-3 | RO-3306 | top60_altered_DNA+RNA_seed | PIK3CA | 0.153117 | COADREAD | NCI-H508 | GNE-317 |
| MVP_top60_DNA_seed | PIK3CA | -0.181188 | OV | UWB1.289 | AZD8835 | top60_altered_DNA+RNA_seed | KRAS | 0.172869 | COADREAD | SNU-407 | KRAS (G12C) Inhibitor-1 |
| MVP_top60_DNA_seed | EGFR | -0.180348 | OV | IGROV-1 | CUDC-101 | top60_altered_DNA+RNA_seed | PIK3CA | 0.178998 | COADREAD | LS-180 | AZD8186 |
| MVP_top60_DNA_seed | ERBB2 | -0.180348 | OV | IGROV-1 | CUDC-101 | top60_altered_DNA+RNA_seed | BRAF | 0.188235 | COADREAD | T84 | PLX-4720 |
| MVP_top60_DNA_seed | MAPK1 | -0.170161 | OV | FU-OV-1 | VX-11e | top60_altered_DNA+RNA_seed | BRAF | 0.192778 | COADREAD | HCC2998 | Dabrafenib |
| MVP_top60_DNA_seed | PIK3CA | -0.168138 | OV | DOV13 | AZD8835 | top60_altered_DNA+RNA_seed | ATM | 0.195901 | COADREAD | LS-1034 | KU-55933 |
| MVP_top60_DNA_seed | ATR | -0.164886 | OV | OVCAR-8 | AZD6738 | top60_altered_DNA+RNA_seed | TP53 | 0.205136 | COADREAD | T84 | MIRA-1 |
| MVP_top60_DNA_seed | MAPK1 | -0.163935 | OV | Hey | FR-180204 | top60_altered_DNA+RNA_seed | KRAS | 0.207918 | COADREAD | SK-CO-1 | KRAS (G12C) Inhibitor-1 |
| MVP_top60_DNA_seed | MAPK3 | -0.163935 | OV | Hey | FR-180204 | top60_altered_DNA+RNA_seed | PIK3CA | 0.220829 | COADREAD | SNU-C1 | GNE-317 |
| MVP_top60_DNA_seed | EGFR | -0.162833 | OV | Caov-3 | Pelitinib | top60_altered_DNA+RNA_seed | PIK3CA | 0.22992 | COADREAD | HCT-15 | AZD8186 |
| MVP_top60_DNA_seed | CDK1 | -0.161981 | OV | OVTOKO | RO-3306 | top60_altered_DNA+RNA_seed | ATM | 0.233929 | COADREAD | SW1463 | KU-55933 |
| MVP_top60_DNA_seed | ATR | -0.159534 | OV | OVCAR-4 | AZ20 | top60_altered_DNA+RNA_seed | PIK3CA | 0.24029 | COADREAD | SW620 | GNE-317 |
| MVP_top60_DNA_seed | EGFR | -0.158349 | OV | FU-OV-1 | AST-1306 | top60_altered_DNA+RNA_seed | TP53 | 0.240471 | COADREAD | LS-180 | MIRA-1 |
| MVP_top60_DNA_seed | PLK1 | -0.156934 | OV | OAW-28 | PLK_6522 | top60_altered_DNA+RNA_seed | TP53 | 0.240883 | COADREAD | SW48 | MIRA-1 |
| MVP_top60_DNA_seed | EGFR | -0.144374 | OV | OAW-42 | Afatinib | top60_altered_DNA+RNA_seed | PIK3CA | 0.246012 | COADREAD | HT55 | GNE-317 |
| MVP_top60_DNA_seed | ERBB2 | -0.144374 | OV | OAW-42 | Afatinib | top60_altered_DNA+RNA_seed | BRAF | 0.248667 | COADREAD | COLO-320-HSR | Dabrafenib |
| MVP_top60_DNA_seed | EGFR | -0.143192 | OV | IGROV-1 | Gefitinib | top60_altered_DNA+RNA_seed | ATM | 0.257987 | COADREAD | HCT-116 | KU-55933 |
| MVP_top60_DNA_seed | CDK1 | -0.142103 | OV | JHOS-2 | RO-3306 | top60_altered_DNA+RNA_seed | BRAF | 0.26104 | COADREAD | SNU-407 | Dabrafenib |
| MVP_top60_DNA_seed | ATR | -0.139533 | OV | OVISE | AZD6738 | top60_altered_DNA+RNA_seed | BRAF | 0.262301 | COADREAD | LS-1034 | SB590885 |
| MVP_top60_DNA_seed | EGFR | -0.13232 | OV | JHOS-3 | Afatinib | top60_altered_DNA+RNA_seed | TP53 | 0.268349 | COADREAD | SNU-81 | MIRA-1 |
| MVP_top60_DNA_seed | ERBB2 | -0.13232 | OV | JHOS-3 | Afatinib | top60_altered_DNA+RNA_seed | TP53 | 0.268778 | COADREAD | RKO | MIRA-1 |
| MVP_top60_DNA_seed | EGFR | -0.128062 | OV | OAW-28 | CI-1033 | top60_altered_DNA+RNA_seed | ATM | 0.269069 | COADREAD | SNU-81 | KU-55933 |
| MVP_top60_DNA_seed | ERBB2 | -0.128062 | OV | OAW-28 | CI-1033 | top60_altered_DNA+RNA_seed | BRAF | 0.272616 | COADREAD | NCI-H716 | PLX-4720 |
| MVP_top60_DNA_seed | ATR | -0.126766 | OV | OV-17R | AZD6738 | top60_altered_DNA+RNA_seed | TP53 | 0.280323 | COADREAD | HCT-15 | MIRA-1 |
| MVP_top60_DNA_seed | ATR | -0.118046 | OV | OV-7 | QL-VIII-58 | top60_altered_DNA+RNA_seed | BRAF | 0.286585 | COADREAD | SNU-175 | SB590885 |
| MVP_top60_DNA_seed | MAPK1 | -0.112744 | OV | OVISE | FR-180204 | top60_altered_DNA+RNA_seed | BRAF | 0.29153 | COADREAD | GP5d | PLX-4720 |
| MVP_top60_DNA_seed | MAPK3 | -0.112744 | OV | OVISE | FR-180204 | top60_altered_DNA+RNA_seed | BRAF | 0.300419 | COADREAD | RCM-1 | Dabrafenib |
| MVP_top60_DNA_seed | ATR | -0.107979 | OV | OAW-28 | AZD6738 | top60_altered_DNA+RNA_seed | PIK3CA | 0.310936 | COADREAD | T84 | GNE-317 |
| MVP_top60_DNA_seed | ERBB2 | -0.105358 | OV | IGROV-1 | CP724714 | top60_altered_DNA+RNA_seed | BRAF | 0.314626 | COADREAD | HT-115 | PLX-4720 |
| MVP_top60_DNA_seed | EGFR | -0.100379 | OV | OVCAR-5 | Afatinib | top60_altered_DNA+RNA_seed | BRAF | 0.328334 | COADREAD | HCT-15 | SB590885 |
| MVP_top60_DNA_seed | ERBB2 | -0.100379 | OV | OVCAR-5 | Afatinib | top60_altered_DNA+RNA_seed | ATM | 0.339776 | COADREAD | SNU-175 | KU-55933 |
| MVP_top60_DNA_seed | CREBBP | -0.099707 | OV | PEO1 | I-CBP112 | top60_altered_DNA+RNA_seed | BRAF | 0.346097 | COADREAD | SNU-61 | SB590885 |
| MVP_top60_DNA_seed | PIK3CA | -0.097192 | OV | OVCAR-3 | AZD8835 | top60_altered_DNA+RNA_seed | PIK3CA | 0.348843 | COADREAD | SNU-81 | GNE-317 |
| MVP_top60_DNA_seed | EGFR | -0.095798 | OV | OVISE | AZD8931 | top60_altered_DNA+RNA_seed | BRAF | 0.356624 | COADREAD | SW1463 | SB590885 |
| MVP_top60_DNA_seed | ERBB2 | -0.095798 | OV | OVISE | AZD8931 | top60_altered_DNA+RNA_seed | BRAF | 0.358405 | COADREAD | SW837 | SB590885 |
| MVP_top60_DNA_seed | PLK1 | -0.094969 | OV | OVCA433 | PLK_6522 | top60_altered_DNA+RNA_seed | PIK3CA | 0.364046 | COADREAD | LS-123 | Buparlisib |
| MVP_top60_DNA_seed | CDK2 | -0.090218 | OV | OV-56 | AZD5438 | top60_altered_DNA+RNA_seed | BRAF | 0.379977 | COADREAD | CW-2 | Dabrafenib |
| MVP_top60_DNA_seed | EGFR | -0.089183 | OV | OVCAR-3 | AZD8931 | top60_altered_DNA+RNA_seed | ATM | 0.380863 | COADREAD | CL-11 | KU-55933 |
| MVP_top60_DNA_seed | ERBB2 | -0.089183 | OV | OVCAR-3 | AZD8931 | top60_altered_DNA+RNA_seed | PIK3CA | 0.381011 | COADREAD | SW620 | Buparlisib |
| MVP_top60_DNA_seed | MAPK1 | -0.086536 | OV | DOV13 | VX-11e | top60_altered_DNA+RNA_seed | TP53 | 0.382935 | COADREAD | CaR-1 | MIRA-1 |
| MVP_top60_DNA_seed | MAPK1 | -0.081423 | OV | OVCAR-5 | VX-11e | top60_altered_DNA+RNA_seed | PIK3CA | 0.389427 | COADREAD | NCI-H716 | Buparlisib |

| | | | | | | | | | | | |
|---|---|---|---|---|---|---|---|---|---|---|---|
| MVP_top60_DNA_seed | CREBBP | -0.08011 | OV | OVCAR-8 | I-CBP112 | top60_altered_DNA+RNA_seed | KRAS | 0.394184 | COADREAD | DiFi | KRAS (G12C) Inhibitor-1 |
| MVP_top60_DNA_seed | EGFR | -0.074794 | OV | OVCAR-4 | CI-1033 | top60_altered_DNA+RNA_seed | TP53 | 0.396854 | COADREAD | NCI-H508 | MIRA-1 |
| MVP_top60_DNA_seed | ERBB2 | -0.074794 | OV | OVCAR-4 | CI-1033 | top60_altered_DNA+RNA_seed | PIK3CA | 0.409103 | COADREAD | SW837 | AZD8186 |
| MVP_top60_DNA_seed | EGFR | -0.073211 | OV | Caov-3 | PF-00299804 | top60_altered_DNA+RNA_seed | PIK3CA | 0.411115 | COADREAD | HT-115 | GNE-317 |
| MVP_top60_DNA_seed | ERBB2 | -0.073211 | OV | Caov-3 | PF-00299804 | top60_altered_DNA+RNA_seed | PIK3CA | 0.414408 | COADREAD | CCK-81 | GNE-317 |
| MVP_top60_DNA_seed | CDK1 | -0.071137 | OV | OVISE | RO-3306 | top60_altered_DNA+RNA_seed | BRAF | 0.421528 | COADREAD | SW837 | PLX-4720 |
| MVP_top60_DNA_seed | PLK1 | -0.066653 | OV | OVCAR-4 | PLK_6522 | top60_altered_DNA+RNA_seed | PIK3CA | 0.425256 | COADREAD | SW1463 | AZD8186 |
| MVP_top60_DNA_seed | HDAC2 | -0.058636 | OV | OV-90 | JQ12 | top60_altered_DNA+RNA_seed | PIK3CA | 0.426361 | COADREAD | HCT-116 | AZD8186 |
| MVP_top60_DNA_seed | EGFR | -0.055038 | OV | Hey | Cetuximab | top60_altered_DNA+RNA_seed | BRAF | 0.427606 | COADREAD | CCK-81 | PLX-4720 |
| MVP_top60_DNA_seed | EGFR | -0.05389 | OV | PEO1 | PF-00299804 | top60_altered_DNA+RNA_seed | BRAF | 0.432668 | COADREAD | SW948 | SB590885 |
| MVP_top60_DNA_seed | ERBB2 | -0.05389 | OV | PEO1 | PF-00299804 | top60_altered_DNA+RNA_seed | ATM | 0.433564 | COADREAD | HCT-15 | KU-55933 |
| MVP_top60_DNA_seed | EGFR | -0.053756 | OV | OAW-28 | Afatinib | top60_altered_DNA+RNA_seed | BRAF | 0.434516 | COADREAD | CaR-1 | SB590885 |
| MVP_top60_DNA_seed | ERBB2 | -0.053756 | OV | OAW-28 | Afatinib | top60_altered_DNA+RNA_seed | TP53 | 0.436215 | COADREAD | KM12 | MIRA-1 |
| MVP_top60_DNA_seed | MAPK1 | -0.049253 | OV | UWB1.289 | VX-11e | top60_altered_DNA+RNA_seed | BRAF | 0.441368 | COADREAD | SNU-81 | Dabrafenib |
| MVP_top60_DNA_seed | EGFR | -0.045096 | OV | TOV-112D | PF-00299804 | top60_altered_DNA+RNA_seed | PIK3CA | 0.447664 | COADREAD | LS-180 | Buparlisib |
| MVP_top60_DNA_seed | ERBB2 | -0.045096 | OV | TOV-112D | PF-00299804 | top60_altered_DNA+RNA_seed | BRAF | 0.448205 | COADREAD | LS-180 | SB590885 |
| MVP_top60_DNA_seed | CDK2 | -0.040005 | OV | OVISE | AZD5438 | top60_altered_DNA+RNA_seed | BRAF | 0.450449 | COADREAD | HT-115 | Dabrafenib |
| MVP_top60_DNA_seed | EGFR | -0.037062 | OV | OAW-42 | Afatinib | top60_altered_DNA+RNA_seed | PIK3CA | 0.45689 | COADREAD | LS-411N | Buparlisib |
| MVP_top60_DNA_seed | ERBB2 | -0.037062 | OV | OAW-42 | Afatinib | top60_altered_DNA+RNA_seed | PIK3CA | 0.460474 | COADREAD | SNU-C1 | Alpelisib |
| MVP_top60_DNA_seed | EGFR | -0.036486 | OV | OV-90 | Afatinib | top60_altered_DNA+RNA_seed | BRAF | 0.472625 | COADREAD | CL-11 | Dabrafenib |
| MVP_top60_DNA_seed | ERBB2 | -0.036486 | OV | OV-90 | Afatinib | top60_altered_DNA+RNA_seed | BRAF | 0.473968 | COADREAD | SW620 | Dabrafenib |
| MVP_top60_DNA_seed | SIRT1 | -0.033818 | OV | OVCA433 | Selisistat | top60_altered_DNA+RNA_seed | TP53 | 0.484948 | COADREAD | LS-411N | MIRA-1 |
| MVP_top60_DNA_seed | CDK2 | -0.031396 | OV | PEO1 | AZD5438 | top60_altered_DNA+RNA_seed | BRAF | 0.495745 | COADREAD | SNU-1040 | Dabrafenib |
| MVP_top60_DNA_seed | EGFR | -0.030438 | OV | Caov-4 | CI-1033 | top60_altered_DNA+RNA_seed | PIK3CA | 0.504079 | COADREAD | SNU-81 | Buparlisib |
| MVP_top60_DNA_seed | ERBB2 | -0.030438 | OV | Caov-4 | CI-1033 | top60_altered_DNA+RNA_seed | BRAF | 0.50889 | COADREAD | C2BBe1 | SB590885 |
| MVP_top60_DNA_seed | PLK1 | -0.023742 | OV | OVTOKO | PLK_6522 | top60_altered_DNA+RNA_seed | BRAF | 0.515803 | COADREAD | SNU-175 | Dabrafenib |
| MVP_top60_DNA_seed | CDK1 | -0.023376 | OV | OC-314 | RO-3306 | top60_altered_DNA+RNA_seed | TP53 | 0.519311 | COADREAD | LoVo | MIRA-1 |
| MVP_top60_DNA_seed | EGFR | -0.021862 | OV | IGROV-1 | Pelitinib | top60_altered_DNA+RNA_seed | PIK3CA | 0.522049 | COADREAD | C2BBe1 | Alpelisib |
| MVP_top60_DNA_seed | CDK2 | -0.019482 | OV | SK-OV-3 | PHA-793887 | top60_altered_DNA+RNA_seed | ATM | 0.522708 | COADREAD | DiFi | KU-55933 |
| MVP_top60_DNA_seed | PIK3CA | -0.015978 | OV | TOV-112D | AZD8835 | top60_altered_DNA+RNA_seed | PIK3CA | 0.523605 | COADREAD | SW1417 | AZD8186 |
| MVP_top60_DNA_seed | PLK1 | -0.01542 | OV | UWB1.289 | PLK_6522 | top60_altered_DNA+RNA_seed | PIK3CA | 0.523866 | COADREAD | SNU-C2B | GNE-317 |
| MVP_top60_DNA_seed | SIRT1 | -0.010394 | OV | OV-56 | Selisistat | top60_altered_DNA+RNA_seed | BRAF | 0.542197 | COADREAD | SNU-81 | SB590885 |
| MVP_top60_DNA_seed | EGFR | -0.009389 | OV | IGROV-1 | AST-1306 | top60_altered_DNA+RNA_seed | BRAF | 0.544978 | COADREAD | NCI-H508 | SB590885 |
| MVP_top60_DNA_seed | EGFR | -0.008859 | OV | OVCA433 | AST-1306 | top60_altered_DNA+RNA_seed | PIK3CA | 0.546419 | COADREAD | SW1116 | Alpelisib |
| MVP_top60_DNA_seed | PIK3CA | -0.002644 | OV | IGROV-1 | AZD8835 | top60_altered_DNA+RNA_seed | KRAS | 0.556824 | COADREAD | CaR-1 | KRAS (G12C) Inhibitor-1 |
| MVP_top60_DNA_seed | MAPK1 | 0.001262 | OV | OVKATE | FR-180204 | top60_altered_DNA+RNA_seed | BRAF | 0.574168 | COADREAD | LS-513 | PLX-4720 |
| MVP_top60_DNA_seed | MAPK3 | 0.001262 | OV | OVKATE | FR-180204 | top60_altered_DNA+RNA_seed | BRAF | 0.574511 | COADREAD | T84 | SB590885 |
| MVP_top60_DNA_seed | ATR | 0.00292 | OV | OVCA420 | AZD6738 | top60_altered_DNA+RNA_seed | PIK3CA | 0.57669 | COADREAD | LS-123 | Alpelisib |
| MVP_top60_DNA_seed | ATR | 0.004029 | OV | SW626 | torin2 | top60_altered_DNA+RNA_seed | BRAF | 0.582661 | COADREAD | SW1417 | SB590885 |
| MVP_top60_DNA_seed | PIK3CA | 0.005346 | OV | OVCAR-3 | PI3Ka_4409 | top60_altered_DNA+RNA_seed | BRAF | 0.585585 | COADREAD | SW48 | PLX-4720 |
| MVP_top60_DNA_seed | ATR | 0.008689 | OV | OVMIU | AZD6738 | top60_altered_DNA+RNA_seed | ATM | 0.59262 | COADREAD | NCI-H747 | KU-55933 |

| | | | | | | | | | | | |
|---|---|---|---|---|---|---|---|---|---|---|---|
| MVP_top60_DNA_seed | EGFR | 0.01009 | OV | OC-314 | Gefitinib | top60_altered_DNA+RNA_seed | ATM | 0.596974 | COADREAD | SNU-407 | KU-55933 |
| MVP_top60_DNA_seed | EGFR | 0.011684 | OV | DOV13 | AST-1306 | top60_altered_DNA+RNA_seed | PIK3CA | 0.603429 | COADREAD | NCI-H716 | Alpelisib |
| MVP_top60_DNA_seed | EGFR | 0.017362 | OV | FU-OV-1 | Afatinib | top60_altered_DNA+RNA_seed | BRAF | 0.60628 | COADREAD | SW1463 | Dabrafenib |
| MVP_top60_DNA_seed | ERBB2 | 0.017362 | OV | FU-OV-1 | Afatinib | top60_altered_DNA+RNA_seed | BRAF | 0.610297 | COADREAD | SNU-407 | SB590885 |
| MVP_top60_DNA_seed | CDK1 | 0.020775 | OV | Caov-4 | RO-3306 | top60_altered_DNA+RNA_seed | TP53 | 0.611766 | COADREAD | LS-1034 | MIRA-1 |
| MVP_top60_DNA_seed | EGFR | 0.022974 | OV | OV-7 | Afatinib | top60_altered_DNA+RNA_seed | ATM | 0.629547 | COADREAD | NCI-H508 | KU-55933 |
| MVP_top60_DNA_seed | ERBB2 | 0.022974 | OV | OV-7 | Afatinib | top60_altered_DNA+RNA_seed | BRAF | 0.641139 | COADREAD | NCI-H747 | SB590885 |
| MVP_top60_DNA_seed | EGFR | 0.025629 | OV | DOV13 | CI-1033 | top60_altered_DNA+RNA_seed | BRAF | 0.643356 | COADREAD | T84 | Dabrafenib |
| MVP_top60_DNA_seed | ERBB2 | 0.025629 | OV | DOV13 | CI-1033 | top60_altered_DNA+RNA_seed | PIK3CA | 0.645643 | COADREAD | CL-11 | Alpelisib |
| MVP_top60_DNA_seed | CREBBP | 0.028339 | OV | OVCA433 | I-CBP112 | top60_altered_DNA+RNA_seed | TP53 | 0.646022 | COADREAD | SW1463 | MIRA-1 |
| MVP_top60_DNA_seed | CDK2 | 0.034543 | OV | OV-7 | PHA-793887 | top60_altered_DNA+RNA_seed | ATM | 0.651683 | COADREAD | LS-513 | KU-55933 |
| MVP_top60_DNA_seed | PIK3CA | 0.034709 | OV | OV-17R | PI3Ka_4409 | top60_altered_DNA+RNA_seed | PIK3CA | 0.651935 | COADREAD | HT-115 | AZD8186 |
| MVP_top60_DNA_seed | PIK3CA | 0.040835 | OV | TOV-112D | PI3Ka_4409 | top60_altered_DNA+RNA_seed | ATM | 0.668588 | COADREAD | COLO-205 | KU-55933 |
| MVP_top60_DNA_seed | ERBB2 | 0.041592 | OV | EFO-21 | CP724714 | top60_altered_DNA+RNA_seed | KRAS | 0.679288 | COADREAD | SW948 | RAS (G12C) Inhibitor-1 |
| MVP_top60_DNA_seed | PTK2 | 0.042812 | OV | TOV-21G | PF-562271 | top60_altered_DNA+RNA_seed | PIK3CA | 0.683489 | COADREAD | CL-11 | AZD8186 |
| MVP_top60_DNA_seed | EGFR | 0.043274 | OV | OVCAR-8 | Afatinib | top60_altered_DNA+RNA_seed | BRAF | 0.685516 | COADREAD | SW48 | SB590885 |
| MVP_top60_DNA_seed | ERBB2 | 0.043274 | OV | OVCAR-8 | Afatinib | top60_altered_DNA+RNA_seed | KRAS | 0.687276 | COADREAD | HT-115 | RAS (G12C) Inhibitor-1 |
| MVP_top60_DNA_seed | MAPK1 | 0.050413 | OV | Caov-3 | VX-11e | top60_altered_DNA+RNA_seed | BRAF | 0.693735 | COADREAD | CW-2 | PLX-4720 |
| MVP_top60_DNA_seed | EGFR | 0.051464 | OV | OVCA433 | CUDC-101 | top60_altered_DNA+RNA_seed | ATM | 0.694096 | COADREAD | SW948 | KU-55933 |
| MVP_top60_DNA_seed | ERBB2 | 0.051464 | OV | OVCA433 | CUDC-101 | top60_altered_DNA+RNA_seed | ATM | 0.704125 | COADREAD | HT-29 | KU-55933 |
| MVP_top60_DNA_seed | CDK1 | 0.055634 | OV | OVCAR-3 | RO-3306 | top60_altered_DNA+RNA_seed | PIK3CA | 0.716953 | COADREAD | RCM-1 | AZD8186 |
| MVP_top60_DNA_seed | MAPK1 | 0.056373 | OV | SK-OV-3 | FR-180204 | top60_altered_DNA+RNA_seed | BRAF | 0.729667 | COADREAD | C2BBe1 | PLX-4720 |
| MVP_top60_DNA_seed | MAPK3 | 0.056373 | OV | SK-OV-3 | FR-180204 | top60_altered_DNA+RNA_seed | PIK3CA | 0.732518 | COADREAD | CL-11 | Buparlisib |
| MVP_top60_DNA_seed | CDK2 | 0.056393 | OV | FU-OV-1 | PHA-793887 | top60_altered_DNA+RNA_seed | ATM | 0.740156 | COADREAD | SW48 | KU-55933 |
| MVP_top60_DNA_seed | MAPK1 | 0.058108 | OV | OVCAR-4 | VX-11e | top60_altered_DNA+RNA_seed | PIK3CA | 0.743821 | COADREAD | RKO | AZD8186 |
| MVP_top60_DNA_seed | PIK3CA | 0.05963 | OV | DOV13 | PI3Ka_4409 | top60_altered_DNA+RNA_seed | TP53 | 0.752339 | COADREAD | CL-11 | MIRA-1 |
| MVP_top60_DNA_seed | EGFR | 0.065222 | OV | OVCAR-5 | AZD8931 | top60_altered_DNA+RNA_seed | BRAF | 0.76091 | COADREAD | CCK-81 | Dabrafenib |
| MVP_top60_DNA_seed | ERBB2 | 0.065222 | OV | OVCAR-5 | AZD8931 | top60_altered_DNA+RNA_seed | PIK3CA | 0.769725 | COADREAD | SW620 | Alpelisib |
| MVP_top60_DNA_seed | CDK2 | 0.066075 | OV | SK-OV-3 | AZD5438 | top60_altered_DNA+RNA_seed | BRAF | 0.785674 | COADREAD | CL-11 | SB590885 |
| MVP_top60_DNA_seed | PIK3CA | 0.066184 | OV | UWB1.289 | PI3Ka_4409 | top60_altered_DNA+RNA_seed | BRAF | 0.788804 | COADREAD | LS-180 | Dabrafenib |
| MVP_top60_DNA_seed | MAPK1 | 0.067431 | OV | OVTOKO | FR-180204 | top60_altered_DNA+RNA_seed | PIK3CA | 0.794784 | COADREAD | SNU-C2B | Buparlisib |
| MVP_top60_DNA_seed | MAPK3 | 0.067431 | OV | OVTOKO | FR-180204 | top60_altered_DNA+RNA_seed | BRAF | 0.803115 | COADREAD | RCM-1 | PLX-4720 |
| MVP_top60_DNA_seed | EGFR | 0.070956 | OV | OVCAR-4 | PF-00299804 | top60_altered_DNA+RNA_seed | ATM | 0.804114 | COADREAD | RCM-1 | KU-55933 |
| MVP_top60_DNA_seed | ERBB2 | 0.070956 | OV | OVCAR-4 | PF-00299804 | top60_altered_DNA+RNA_seed | ATM | 0.808598 | COADREAD | SW1116 | KU-55933 |
| MVP_top60_DNA_seed | PTK2 | 0.071001 | OV | OVCAR-4 | PF-562271 | top60_altered_DNA+RNA_seed | PIK3CA | 0.811062 | COADREAD | SW1417 | GNE-317 |
| MVP_top60_DNA_seed | EGFR | 0.071245 | OV | UWB1.289 | Pelitinib | top60_altered_DNA+RNA_seed | BRAF | 0.812777 | COADREAD | SW620 | PLX-4720 |
| MVP_top60_DNA_seed | PIK3CA | 0.072523 | OV | OVCA433 | PI3Ka_4409 | top60_altered_DNA+RNA_seed | BRAF | 0.83446 | COADREAD | SNU-C1 | SB590885 |
| MVP_top60_DNA_seed | EGFR | 0.073307 | OV | JHOS-2 | CI-1033 | top60_altered_DNA+RNA_seed | BRAF | 0.838636 | COADREAD | KM12 | PLX-4720 |
| MVP_top60_DNA_seed | ERBB2 | 0.073307 | OV | JHOS-2 | CI-1033 | top60_altered_DNA+RNA_seed | BRAF | 0.839014 | COADREAD | GP5d | Dabrafenib |
| MVP_top60_DNA_seed | EGFR | 0.07349 | OV | OVCAR-4 | Gefitinib | top60_altered_DNA+RNA_seed | PIK3CA | 0.843262 | COADREAD | SNU-61 | Alpelisib |
| MVP_top60_DNA_seed | EGFR | 0.073745 | OV | OV-90 | CUDC-101 | top60_altered_DNA+RNA_seed | PIK3CA | 0.846979 | COADREAD | SNU-C1 | Buparlisib |

| | | | | | | | | | | | |
|---|---|---|---|---|---|---|---|---|---|---|---|
| MVP_top60_DNA_seed | ERBB2 | 0.073745 | OV | OV-90 | CUDC-101 | top60_altered_DNA+RNA_seed | PIK3CA | 0.849273 | COADREAD | SW837 | Buparlisib |
| MVP_top60_DNA_seed | EGFR | 0.077021 | OV | JHOS-2 | PF-00299804 | top60_altered_DNA+RNA_seed | KRAS | 0.850839 | COADREAD | COLO-320-HSR | RAS (G12C) Inhibitor-1 |
| MVP_top60_DNA_seed | ERBB2 | 0.077021 | OV | JHOS-2 | PF-00299804 | top60_altered_DNA+RNA_seed | BRAF | 0.855065 | COADREAD | SNU-175 | PLX-4720 |
| MVP_top60_DNA_seed | CDK2 | 0.077107 | OV | OV-56 | PHA-793887 | top60_altered_DNA+RNA_seed | PIK3CA | 0.87246 | COADREAD | SW948 | AZD8186 |
| MVP_top60_DNA_seed | EGFR | 0.078483 | OV | Hey | AZD8931 | top60_altered_DNA+RNA_seed | TP53 | 0.872614 | COADREAD | HT-29 | MIRA-1 |
| MVP_top60_DNA_seed | ERBB2 | 0.078483 | OV | Hey | AZD8931 | top60_altered_DNA+RNA_seed | PIK3CA | 0.87469 | COADREAD | SW1463 | Alpelisib |
| MVP_top60_DNA_seed | CREBBP | 0.081388 | OV | OV-17R | I-CBP112 | top60_altered_DNA+RNA_seed | TP53 | 0.877092 | COADREAD | SW948 | MIRA-1 |
| MVP_top60_DNA_seed | ATR | 0.084415 | OV | FU-OV-1 | AZ20 | top60_altered_DNA+RNA_seed | TP53 | 0.892991 | COADREAD | SNU-407 | MIRA-1 |
| MVP_top60_DNA_seed | EGFR | 0.085082 | OV | OV-7 | AST-1306 | top60_altered_DNA+RNA_seed | PIK3CA | 0.893689 | COADREAD | LS-1034 | Alpelisib |
| MVP_top60_DNA_seed | MAPK1 | 0.086274 | OV | OVCA420 | VX-11e | top60_altered_DNA+RNA_seed | TP53 | 0.920579 | COADREAD | SNU-61 | MIRA-1 |
| MVP_top60_DNA_seed | EGFR | 0.090987 | OV | OC-314 | PF-00299804 | top60_altered_DNA+RNA_seed | BRAF | 0.922635 | COADREAD | SW1116 | SB590885 |
| MVP_top60_DNA_seed | ERBB2 | 0.090987 | OV | OC-314 | PF-00299804 | top60_altered_DNA+RNA_seed | BRAF | 0.949576 | COADREAD | SW48 | Dabrafenib |
| MVP_top60_DNA_seed | EGFR | 0.096494 | OV | OVCA433 | AZD8931 | top60_altered_DNA+RNA_seed | BRAF | 0.958502 | COADREAD | SW1463 | PLX-4720 |
| MVP_top60_DNA_seed | ERBB2 | 0.096494 | OV | OVCA433 | AZD8931 | top60_altered_DNA+RNA_seed | ATM | 0.966502 | COADREAD | SNU-61 | KU-55933 |
| MVP_top60_DNA_seed | EGFR | 0.097343 | OV | Caov-4 | AST-1306 | top60_altered_DNA+RNA_seed | PIK3CA | 0.97764 | COADREAD | SW1417 | Alpelisib |
| MVP_top60_DNA_seed | EGFR | 0.101357 | OV | OC-314 | AST-1306 | top60_altered_DNA+RNA_seed | PIK3CA | 0.98394 | COADREAD | SNU-61 | Buparlisib |
| MVP_top60_DNA_seed | CDK2 | 0.101982 | OV | OVK-18 | AZD5438 | top60_altered_DNA+RNA_seed | BRAF | 0.988845 | COADREAD | HCC2998 | PLX-4720 |
| MVP_top60_DNA_seed | PTK2 | 0.1039 | OV | OVISE | PF-562271 | top60_altered_DNA+RNA_seed | KRAS | 0.989445 | COADREAD | CL-11 | RAS (G12C) Inhibitor-1 |
| MVP_top60_DNA_seed | ATR | 0.104305 | OV | OVCAR-3 | AZ20 | top60_altered_DNA+RNA_seed | BRAF | 1.013611 | COADREAD | DiFi | Dabrafenib |
| MVP_top60_DNA_seed | EGFR | 0.109526 | OV | OVCAR-8 | AST-1306 | top60_altered_DNA+RNA_seed | ATM | 1.018111 | COADREAD | LS-411N | KU-55933 |
| MVP_top60_DNA_seed | PTK2 | 0.115421 | OV | OVCAR-8 | PF-562271 | top60_altered_DNA+RNA_seed | BRAF | 1.02829 | COADREAD | NCI-H508 | PLX-4720 |
| MVP_top60_DNA_seed | EGFR | 0.117145 | OV | OV-17R | Cetuximab | top60_altered_DNA+RNA_seed | ATM | 1.028655 | COADREAD | LS-180 | KU-55933 |
| MVP_top60_DNA_seed | CDK2 | 0.11818 | OV | OVCAR-3 | AZD5438 | top60_altered_DNA+RNA_seed | PIK3CA | 1.029105 | COADREAD | MDST8 | Alpelisib |
| MVP_top60_DNA_seed | SIRT1 | 0.119977 | OV | OC-314 | Selisistat | top60_altered_DNA+RNA_seed | TP53 | 1.043725 | COADREAD | SNU-C5 | MIRA-1 |
| MVP_top60_DNA_seed | CDK1 | 0.126016 | OV | JHOS-4 | RO-3306 | top60_altered_DNA+RNA_seed | TP53 | 1.048788 | COADREAD | SNU-C1 | MIRA-1 |
| MVP_top60_DNA_seed | EGFR | 0.127154 | OV | OVCAR-3 | Gefitinib | top60_altered_DNA+RNA_seed | KRAS | 1.051334 | COADREAD | CCK-81 | RAS (G12C) Inhibitor-1 |
| MVP_top60_DNA_seed | EGFR | 0.127813 | OV | OV-17R | AST-1306 | top60_altered_DNA+RNA_seed | KRAS | 1.053421 | COADREAD | MDST8 | RAS (G12C) Inhibitor-1 |
| MVP_top60_DNA_seed | PTK2 | 0.130294 | OV | SW626 | PF-562271 | top60_altered_DNA+RNA_seed | PIK3CA | 1.054854 | COADREAD | SNU-C2B | Alpelisib |
| MVP_top60_DNA_seed | EGFR | 0.130754 | OV | Caov-4 | PF-00299804 | top60_altered_DNA+RNA_seed | PIK3CA | 1.068569 | COADREAD | SNU-C2B | AZD8186 |
| MVP_top60_DNA_seed | ERBB2 | 0.130754 | OV | Caov-4 | PF-00299804 | top60_altered_DNA+RNA_seed | TP53 | 1.072246 | COADREAD | SW837 | MIRA-1 |
| MVP_top60_DNA_seed | ATR | 0.131934 | OV | OVCAR-8 | QL-VIII-58 | top60_altered_DNA+RNA_seed | TP53 | 1.083472 | COADREAD | NCI-H716 | MIRA-1 |
| MVP_top60_DNA_seed | EGFR | 0.133356 | OV | FU-OV-1 | Afatinib | top60_altered_DNA+RNA_seed | TP53 | 1.084915 | COADREAD | SW620 | MIRA-1 |
| MVP_top60_DNA_seed | ERBB2 | 0.133356 | OV | FU-OV-1 | Afatinib | top60_altered_DNA+RNA_seed | BRAF | 1.089549 | COADREAD | C2BBe1 | Dabrafenib |
| MVP_top60_DNA_seed | ATR | 0.133853 | OV | OV-7 | AZD6738 | top60_altered_DNA+RNA_seed | BRAF | 1.094671 | COADREAD | SNU-61 | PLX-4720 |
| MVP_top60_DNA_seed | EGFR | 0.134288 | OV | PEO1 | Cetuximab | top60_altered_DNA+RNA_seed | BRAF | 1.104791 | COADREAD | SNU-1040 | PLX-4720 |
| MVP_top60_DNA_seed | ATR | 0.136355 | OV | SW626 | QL-VIII-58 | top60_altered_DNA+RNA_seed | BRAF | 1.108184 | COADREAD | SNU-407 | PLX-4720 |
| MVP_top60_DNA_seed | CREBBP | 0.138543 | OV | OAW-42 | I-CBP112 | top60_altered_DNA+RNA_seed | KRAS | 1.118692 | COADREAD | SW837 | RAS (G12C) Inhibitor-1 |
| MVP_top60_DNA_seed | CDK1 | 0.14163 | OV | OV-17R | RO-3306 | top60_altered_DNA+RNA_seed | KRAS | 1.119802 | COADREAD | LS-1034 | RAS (G12C) Inhibitor-1 |
| MVP_top60_DNA_seed | PLK1 | 0.142191 | OV | OVK-18 | PLK_6522 | top60_altered_DNA+RNA_seed | KRAS | 1.127681 | COADREAD | SNU-61 | RAS (G12C) Inhibitor-1 |
| MVP_top60_DNA_seed | ERBB2 | 0.142349 | OV | SK-OV-3 | CP724714 | top60_altered_DNA+RNA_seed | KRAS | 1.130554 | COADREAD | NCI-H508 | RAS (G12C) Inhibitor-1 |
| MVP_top60_DNA_seed | PLK1 | 0.142359 | OV | SW626 | PLK_6522 | top60_altered_DNA+RNA_seed | PIK3CA | 1.142895 | COADREAD | LS-1034 | GNE-317 |

| | | | | | | | | | | | |
|---|---|---|---|---|---|---|---|---|---|---|---|
| MVP_top60_DNA_seed | ATR | 0.151121 | OV | OVCAR-8 | AZ20 | top60_altered_DNA+RNA_seed | PIK3CA | 1.160921 | COADREAD | SNU-1040 | Alpelisib |
| MVP_top60_DNA_seed | PTK2 | 0.151912 | OV | EFO-21 | PF-562271 | top60_altered_DNA+RNA_seed | BRAF | 1.162674 | COADREAD | LS-1034 | PLX-4720 |
| MVP_top60_DNA_seed | MAPK1 | 0.153562 | OV | OVCAR-4 | FR-180204 | top60_altered_DNA+RNA_seed | KRAS | 1.198722 | COADREAD | C2BBe1 | KRAS (G12C) Inhibitor-1 |
| MVP_top60_DNA_seed | MAPK3 | 0.153562 | OV | OVCAR-4 | FR-180204 | top60_altered_DNA+RNA_seed | KRAS | 1.199411 | COADREAD | COLO-205 | KRAS (G12C) Inhibitor-1 |
| MVP_top60_DNA_seed | MAPK1 | 0.158495 | OV | JHOS-2 | VX-11e | top60_altered_DNA+RNA_seed | KRAS | 1.200548 | COADREAD | T84 | KRAS (G12C) Inhibitor-1 |
| MVP_top60_DNA_seed | ATR | 0.15981 | OV | OV-17R | AZ20 | top60_altered_DNA+RNA_seed | BRAF | 1.202437 | COADREAD | SW837 | Dabrafenib |
| MVP_top60_DNA_seed | EGFR | 0.16041 | OV | JHOS-3 | AST-1306 | top60_altered_DNA+RNA_seed | PIK3CA | 1.217937 | COADREAD | NCI-H716 | GNE-317 |
| MVP_top60_DNA_seed | CDK2 | 0.172198 | OV | OVCAR-4 | PHA-793887 | top60_altered_DNA+RNA_seed | KRAS | 1.230805 | COADREAD | SNU-81 | KRAS (G12C) Inhibitor-1 |
| MVP_top60_DNA_seed | EGFR | 0.178825 | OV | OVCAR-3 | Afatinib | top60_altered_DNA+RNA_seed | TP53 | 1.236705 | COADREAD | LS-123 | MIRA-1 |
| MVP_top60_DNA_seed | ERBB2 | 0.178825 | OV | OVCAR-3 | Afatinib | top60_altered_DNA+RNA_seed | KRAS | 1.240603 | COADREAD | SNU-C1 | KRAS (G12C) Inhibitor-1 |
| MVP_top60_DNA_seed | SIRT1 | 0.184124 | OV | PEO1 | Selisistat | top60_altered_DNA+RNA_seed | BRAF | 1.244345 | COADREAD | LS-123 | Dabrafenib |
| MVP_top60_DNA_seed | EGFR | 0.186542 | OV | OVCA420 | CUDC-101 | top60_altered_DNA+RNA_seed | TP53 | 1.244492 | COADREAD | SW1116 | MIRA-1 |
| MVP_top60_DNA_seed | ERBB2 | 0.186542 | OV | OVCA420 | CUDC-101 | top60_altered_DNA+RNA_seed | PIK3CA | 1.245301 | COADREAD | CW-2 | AZD8186 |
| MVP_top60_DNA_seed | EGFR | 0.187545 | OV | JHOS-3 | Cetuximab | top60_altered_DNA+RNA_seed | PIK3CA | 1.253307 | COADREAD | RCM-1 | GNE-317 |
| MVP_top60_DNA_seed | ATR | 0.191348 | OV | OV-90 | AZD6738 | top60_altered_DNA+RNA_seed | PIK3CA | 1.257325 | COADREAD | SNU-61 | AZD8186 |
| MVP_top60_DNA_seed | EGFR | 0.194659 | OV | Hey | AST-1306 | top60_altered_DNA+RNA_seed | PIK3CA | 1.270247 | COADREAD | SW1463 | Buparlisib |
| MVP_top60_DNA_seed | CREBBP | 0.20518 | OV | OV-90 | I-CBP112 | top60_altered_DNA+RNA_seed | KRAS | 1.278234 | COADREAD | RCM-1 | KRAS (G12C) Inhibitor-1 |
| MVP_top60_DNA_seed | ERBB2 | 0.206327 | OV | OVCA433 | CP724714 | top60_altered_DNA+RNA_seed | BRAF | 1.289872 | COADREAD | SNU-C2B | SB590885 |
| MVP_top60_DNA_seed | EGFR | 0.211481 | OV | OV-90 | Pelitinib | top60_altered_DNA+RNA_seed | KRAS | 1.302596 | COADREAD | LS-180 | KRAS (G12C) Inhibitor-1 |
| MVP_top60_DNA_seed | CDK1 | 0.216762 | OV | OVCAR-8 | RO-3306 | top60_altered_DNA+RNA_seed | BRAF | 1.329114 | COADREAD | SW1116 | Dabrafenib |
| MVP_top60_DNA_seed | EGFR | 0.218704 | OV | EFO-21 | AST-1306 | top60_altered_DNA+RNA_seed | BRAF | 1.347972 | COADREAD | LS-180 | PLX-4720 |
| MVP_top60_DNA_seed | PIK3CA | 0.221642 | OV | OV-90 | AZD8835 | top60_altered_DNA+RNA_seed | ATM | 1.357096 | COADREAD | SW1417 | KU-55933 |
| MVP_top60_DNA_seed | EGFR | 0.22499 | OV | JHOS-2 | Pelitinib | top60_altered_DNA+RNA_seed | PIK3CA | 1.368682 | COADREAD | CL-11 | GNE-317 |
| MVP_top60_DNA_seed | PIK3CA | 0.229529 | OV | SW626 | AZD8835 | top60_altered_DNA+RNA_seed | TP53 | 1.369924 | COADREAD | RCM-1 | MIRA-1 |
| MVP_top60_DNA_seed | EGFR | 0.232655 | OV | SW626 | AZD8931 | top60_altered_DNA+RNA_seed | PIK3CA | 1.369951 | COADREAD | LS-411N | Alpelisib |
| MVP_top60_DNA_seed | ERBB2 | 0.232655 | OV | SW626 | AZD8931 | top60_altered_DNA+RNA_seed | PIK3CA | 1.385943 | COADREAD | SW620 | AZD8186 |
| MVP_top60_DNA_seed | CDK1 | 0.238682 | OV | OV-56 | RO-3306 | top60_altered_DNA+RNA_seed | PIK3CA | 1.393056 | COADREAD | SW1417 | Buparlisib |
| MVP_top60_DNA_seed | PIK3CA | 0.239303 | OV | OVCAR-5 | PI3Ka_4409 | top60_altered_DNA+RNA_seed | PIK3CA | 1.412744 | COADREAD | SNU-81 | Alpelisib |
| MVP_top60_DNA_seed | EGFR | 0.244128 | OV | OVK-18 | AZD8931 | top60_altered_DNA+RNA_seed | KRAS | 1.442541 | COADREAD | SW1463 | KRAS (G12C) Inhibitor-1 |
| MVP_top60_DNA_seed | ERBB2 | 0.244128 | OV | OVK-18 | AZD8931 | top60_altered_DNA+RNA_seed | BRAF | 1.455466 | COADREAD | NCI-H747 | Dabrafenib |
| MVP_top60_DNA_seed | PLK1 | 0.245106 | OV | OVCA420 | PLK_6522 | top60_altered_DNA+RNA_seed | BRAF | 1.457877 | COADREAD | SNU-C1 | PLX-4720 |
| MVP_top60_DNA_seed | PIK3CA | 0.25014 | OV | OVCAR-8 | AZD8835 | top60_altered_DNA+RNA_seed | KRAS | 1.460753 | COADREAD | SNU-C2B | KRAS (G12C) Inhibitor-1 |
| MVP_top60_DNA_seed | EGFR | 0.251424 | OV | OV-56 | Afatinib | top60_altered_DNA+RNA_seed | PIK3CA | 1.474853 | COADREAD | SW837 | Alpelisib |
| MVP_top60_DNA_seed | ERBB2 | 0.251424 | OV | OV-56 | Afatinib | top60_altered_DNA+RNA_seed | PIK3CA | 1.489188 | COADREAD | KM12 | Alpelisib |
| MVP_top60_DNA_seed | EGFR | 0.253739 | OV | OVISE | AST-1306 | top60_altered_DNA+RNA_seed | PIK3CA | 1.493804 | COADREAD | RKO | GNE-317 |
| MVP_top60_DNA_seed | EGFR | 0.2563 | OV | OVCAR-3 | CI-1033 | top60_altered_DNA+RNA_seed | PIK3CA | 1.499855 | COADREAD | LS-1034 | Buparlisib |
| MVP_top60_DNA_seed | ERBB2 | 0.2563 | OV | OVCAR-3 | CI-1033 | top60_altered_DNA+RNA_seed | BRAF | 1.516738 | COADREAD | LS-123 | SB590885 |
| MVP_top60_DNA_seed | ERBB2 | 0.256319 | OV | OAW-42 | CP724714 | top60_altered_DNA+RNA_seed | PIK3CA | 1.548009 | COADREAD | LS-411N | AZD8186 |
| MVP_top60_DNA_seed | EGFR | 0.25663 | OV | DOV13 | Cetuximab | top60_altered_DNA+RNA_seed | BRAF | 1.602098 | COADREAD | CL-11 | PLX-4720 |
| MVP_top60_DNA_seed | EGFR | 0.259778 | OV | OAW-28 | Afatinib | top60_altered_DNA+RNA_seed | PIK3CA | 1.603978 | COADREAD | SNU-C1 | AZD8186 |
| MVP_top60_DNA_seed | ERBB2 | 0.259778 | OV | OAW-28 | Afatinib | top60_altered_DNA+RNA_seed | KRAS | 1.604202 | COADREAD | SW1116 | KRAS (G12C) Inhibitor-1 |

| | | | | | | | | | | | |
|---|---|---|---|---|---|---|---|---|---|---|---|
| MVP_top60_DNA_seed | CREBBP | 0.262398 | OV | OV-56 | I-CBP112 | top60_altered_DNA+RNA_seed | ATM | 1.637887 | COADREAD | LS-123 | KU-55933 |
| MVP_top60_DNA_seed | ERBB2 | 0.263336 | OV | Hey | CP724714 | top60_altered_DNA+RNA_seed | PIK3CA | 1.652514 | COADREAD | LS-1034 | AZD8186 |
| MVP_top60_DNA_seed | EGFR | 0.264362 | OV | JHOS-3 | PF-00299804 | top60_altered_DNA+RNA_seed | BRAF | 1.658252 | COADREAD | SW1116 | PLX-4720 |
| MVP_top60_DNA_seed | ERBB2 | 0.264362 | OV | JHOS-3 | PF-00299804 | top60_altered_DNA+RNA_seed | TP53 | 1.663562 | COADREAD | C2BBe1 | MIRA-1 |
| MVP_top60_DNA_seed | SIRT1 | 0.266534 | OV | OVK-18 | Selisistat | top60_altered_DNA+RNA_seed | PIK3CA | 1.670726 | COADREAD | KM12 | AZD8186 |
| MVP_top60_DNA_seed | ATR | 0.266869 | OV | SK-OV-3 | AZ20 | top60_altered_DNA+RNA_seed | BRAF | 1.692513 | COADREAD | SNU-C2B | PLX-4720 |
| MVP_top60_DNA_seed | PIK3CA | 0.268818 | OV | OAW-28 | AZD8835 | top60_altered_DNA+RNA_seed | TP53 | 1.727207 | COADREAD | NCI-H747 | MIRA-1 |
| MVP_top60_DNA_seed | PIK3CA | 0.272268 | OV | OV-17R | AZD8835 | top60_altered_DNA+RNA_seed | BRAF | 1.729052 | COADREAD | LS-123 | PLX-4720 |
| MVP_top60_DNA_seed | EGFR | 0.273708 | OV | TOV-112D | Cetuximab | top60_altered_DNA+RNA_seed | BRAF | 1.746145 | COADREAD | NCI-H747 | PLX-4720 |
| MVP_top60_DNA_seed | CDK1 | 0.276053 | OV | SW626 | RO-3306 | top60_altered_DNA+RNA_seed | ATM | 1.762167 | COADREAD | SNU-C2B | KU-55933 |
| MVP_top60_DNA_seed | ATR | 0.281213 | OV | OAW-28 | AZ20 | top60_altered_DNA+RNA_seed | PIK3CA | 1.771889 | COADREAD | SW1116 | Buparlisib |
| MVP_top60_DNA_seed | CDK2 | 0.281311 | OV | OVCAR-4 | Seliciclib | top60_altered_DNA+RNA_seed | PIK3CA | 1.863039 | COADREAD | CW-2 | GNE-317 |
| MVP_top60_DNA_seed | EGFR | 0.282769 | OV | JHOS-3 | Pelitinib | top60_altered_DNA+RNA_seed | PIK3CA | 1.876554 | COADREAD | SW1116 | GNE-317 |
| MVP_top60_DNA_seed | EGFR | 0.285291 | OV | OVCAR-3 | Afatinib | top60_altered_DNA+RNA_seed | BRAF | 1.88081 | COADREAD | SNU-C2B | Dabrafenib |
| MVP_top60_DNA_seed | ERBB2 | 0.285291 | OV | OVCAR-3 | Afatinib | top60_altered_DNA+RNA_seed | PIK3CA | 1.907945 | COADREAD | COLO-678 | AZD8186 |
| MVP_top60_DNA_seed | EGFR | 0.286862 | OV | Hey | Gefitinib | top60_altered_DNA+RNA_seed | PIK3CA | 1.98122 | COADREAD | SW1116 | AZD8186 |
| MVP_top60_DNA_seed | EGFR | 0.287507 | OV | EFO-21 | PF-00299804 | top60_altered_DNA+RNA_seed | KRAS | 2.035954 | COADREAD | LS-123 | KRAS (G12C) Inhibitor-1 |
| MVP_top60_DNA_seed | ERBB2 | 0.287507 | OV | EFO-21 | PF-00299804 | top60_altered_DNA+RNA_seed | TP53 | 2.051736 | COADREAD | SNU-C2B | MIRA-1 |
| MVP_top60_DNA_seed | ATR | 0.290767 | OV | OV-7 | AZ20 | top60_altered_DNA+RNA_seed | BRAF | 2.158822 | COADREAD | COLO-678 | Dabrafenib |
| MVP_top60_DNA_seed | CDK1 | 0.290795 | OV | FU-OV-1 | RO-3306 | top60_altered_DNA+RNA_seed | PIK3CA | 2.296675 | COADREAD | RCM-1 | Buparlisib |
| MVP_top60_DNA_seed | ATR | 0.290944 | OV | JHOS-3 | QL-VIII-58 | top60_altered_DNA+RNA_seed | BRAF | 2.373089 | COADREAD | SW620 | SB590885 |
| MVP_top60_DNA_seed | CDK2 | 0.297104 | OV | OAW-42 | PHA-793887 | top60_altered_DNA+RNA_seed | ATM | 2.380035 | COADREAD | SW620 | KU-55933 |
| MVP_top60_DNA_seed | PIK3CA | 0.298793 | OV | OVCAR-8 | PI3Ka_4409 | top60_altered_DNA+RNA_seed | BRAF | 2.411923 | COADREAD | COLO-678 | SB590885 |
| MVP_top60_DNA_seed | EGFR | 0.30384 | OV | DOV13 | PF-00299804 | top60_altered_DNA+RNA_seed | KRAS | 2.416192 | COADREAD | SW1417 | KRAS (G12C) Inhibitor-1 |
| MVP_top60_DNA_seed | ERBB2 | 0.30384 | OV | DOV13 | PF-00299804 | top60_altered_DNA+RNA_seed | TP53 | 2.502555 | COADREAD | SW1417 | MIRA-1 |
| MVP_top60_DNA_seed | ATR | 0.304117 | OV | OVCA433 | AZ20 | top60_altered_DNA+RNA_seed | PIK3CA | 2.581429 | COADREAD | COLO-678 | Alpelisib |
| MVP_top60_DNA_seed | CDK2 | 0.307842 | OV | OVTOKO | PHA-793887 | top60_altered_DNA+RNA_seed | KRAS | 2.680078 | COADREAD | COLO-678 | KRAS (G12C) Inhibitor-1 |
| MVP_top60_DNA_seed | MAPK1 | 0.308762 | OV | OV-7 | FR-180204 | top60_altered_DNA+RNA_seed | PIK3CA | 2.855048 | COADREAD | SNU-61 | GNE-317 |
| MVP_top60_DNA_seed | MAPK3 | 0.308762 | OV | OV-7 | FR-180204 | top60_altered_DNA+RNA_seed | ATM | 2.874474 | COADREAD | COLO-678 | KU-55933 |
| MVP_top60_DNA_seed | EGFR | 0.311567 | OV | OAW-42 | Pelitinib | top60_altered_DNA+RNA_seed | TP53 | 3.209116 | COADREAD | COLO-678 | MIRA-1 |
| MVP_top60_DNA_seed | EGFR | 0.312248 | OV | OV-17R | Afatinib | top60_altered_DNA+RNA_seed | BRAF | 3.295513 | COADREAD | COLO-678 | PLX-4720 |
| MVP_top60_DNA_seed | ERBB2 | 0.312248 | OV | OV-17R | Afatinib | top60_altered_DNA+RNA_seed | PIK3CA | 3.578788 | COADREAD | COLO-678 | Buparlisib |
| MVP_top60_DNA_seed | EGFR | 0.314219 | OV | OVCA433 | PF-00299804 | top60_altered_DNA+RNA_seed | PIK3CA | 3.798474 | COADREAD | COLO-678 | GNE-317 |
| MVP_top60_DNA_seed | ERBB2 | 0.314219 | OV | OVCA433 | PF-00299804 | top60_altered_DNA+RNA_seed | PDGFRA | -3.050319 | GBM | D-247MG | Amuvatinib |
| MVP_top60_DNA_seed | PIK3CA | 0.31749 | OV | OVK-18 | PI3Ka_4409 | top60_altered_DNA+RNA_seed | PDGFRA | -2.850486 | GBM | D-247MG | Masitinib |
| MVP_top60_DNA_seed | MAPK1 | 0.317719 | OV | OC-314 | VX-11e | top60_altered_DNA+RNA_seed | PDGFRA | -2.358508 | GBM | KS-1 | Amuvatinib |
| MVP_top60_DNA_seed | EGFR | 0.317937 | OV | OVCAR-3 | AST-1306 | top60_altered_DNA+RNA_seed | PDGFRA | -1.687812 | GBM | AM-38 | Amuvatinib |
| MVP_top60_DNA_seed | ATR | 0.321825 | OV | OAW-42 | AZD6738 | top60_altered_DNA+RNA_seed | CDK4 | -1.652208 | GBM | D-263MG | Palbociclib |
| MVP_top60_DNA_seed | EGFR | 0.328001 | OV | OV-17R | PF-00299804 | top60_altered_DNA+RNA_seed | EGFR | -1.546145 | GBM | KS-1 | Erlotinib |
| MVP_top60_DNA_seed | ERBB2 | 0.328001 | OV | OV-17R | PF-00299804 | top60_altered_DNA+RNA_seed | CDK4 | -1.453012 | GBM | U251 | Palbociclib |
| MVP_top60_DNA_seed | EGFR | 0.329557 | OV | EFO-21 | AZD8931 | top60_altered_DNA+RNA_seed | EGFR | -1.189085 | GBM | U251 | CUDC-101 |

| | | | | | | | | | | | |
|---|---|---|---|---|---|---|---|---|---|---|---|
| MVP_top60_DNA_seed | ERBB2 | 0.329557 | OV | EFO-21 | AZD8931 | top60_altered_DNA+RNA_seed | EGFR | -1.16436 | GBM | DK-MG | Pelitinib |
| MVP_top60_DNA_seed | PIK3CA | 0.338485 | OV | Caov-4 | PI3Ka_4409 | top60_altered_DNA+RNA_seed | CDK4 | -1.153023 | GBM | YH-13 | Palbociclib |
| MVP_top60_DNA_seed | ATR | 0.34037 | OV | JHOS-2 | AZ20 | top60_altered_DNA+RNA_seed | CDK4 | -1.122478 | GBM | KS-1 | CGP-082996 |
| MVP_top60_DNA_seed | ERBB2 | 0.341413 | OV | OVISE | CP724714 | top60_altered_DNA+RNA_seed | PDGFRA | -1.114523 | GBM | DK-MG | Masitinib |
| MVP_top60_DNA_seed | EGFR | 0.343516 | OV | OVCAR-8 | Gefitinib | top60_altered_DNA+RNA_seed | CDK4 | -1.097164 | GBM | GB-1 | CGP-082996 |
| MVP_top60_DNA_seed | EGFR | 0.34883 | OV | OVISE | PF-00299804 | top60_altered_DNA+RNA_seed | EGFR | -1.028082 | GBM | Becker | Erlotinib |
| MVP_top60_DNA_seed | ERBB2 | 0.34883 | OV | OVISE | PF-00299804 | top60_altered_DNA+RNA_seed | CDK4 | -1.011535 | GBM | SF268 | Palbociclib |
| MVP_top60_DNA_seed | SIRT1 | 0.350829 | OV | DOV13 | Selisistat | top60_altered_DNA+RNA_seed | EGFR | -0.954878 | GBM | U-87-MG | Afatinib |
| MVP_top60_DNA_seed | EGFR | 0.356433 | OV | OV-7 | CUDC-101 | top60_altered_DNA+RNA_seed | PDGFRA | -0.941443 | GBM | D-423MG | Amuvatinib |
| MVP_top60_DNA_seed | ERBB2 | 0.356433 | OV | OV-7 | CUDC-101 | top60_altered_DNA+RNA_seed | EGFR | -0.941235 | GBM | KS-1 | Lapatinib |
| MVP_top60_DNA_seed | EGFR | 0.357495 | OV | FU-OV-1 | Gefitinib | top60_altered_DNA+RNA_seed | CDK4 | -0.93056 | GBM | YKG-1 | Palbociclib |
| MVP_top60_DNA_seed | MAPK1 | 0.362808 | OV | TOV-21G | FR-180204 | top60_altered_DNA+RNA_seed | EGFR | -0.924194 | GBM | SF268 | CUDC-101 |
| MVP_top60_DNA_seed | MAPK3 | 0.362808 | OV | TOV-21G | FR-180204 | top60_altered_DNA+RNA_seed | EGFR | -0.919244 | GBM | A172 | Cetuximab |
| MVP_top60_DNA_seed | EGFR | 0.3707 | OV | OVK-18 | AST-1306 | top60_altered_DNA+RNA_seed | CDK4 | -0.9049 | GBM | Becker | Palbociclib |
| MVP_top60_DNA_seed | EGFR | 0.373749 | OV | OV-56 | CUDC-101 | top60_altered_DNA+RNA_seed | CDK4 | -0.881495 | GBM | T98G | Palbociclib |
| MVP_top60_DNA_seed | ERBB2 | 0.373749 | OV | OV-56 | CUDC-101 | top60_altered_DNA+RNA_seed | CDK4 | -0.874117 | GBM | SW1088 | Palbociclib |
| MVP_top60_DNA_seed | EGFR | 0.376141 | OV | EFO-21 | Afatinib | top60_altered_DNA+RNA_seed | EGFR | -0.859529 | GBM | GB-1 | CUDC-101 |
| MVP_top60_DNA_seed | ERBB2 | 0.376141 | OV | EFO-21 | Afatinib | top60_altered_DNA+RNA_seed | PDGFRA | -0.835034 | GBM | U251 | Amuvatinib |
| MVP_top60_DNA_seed | EGFR | 0.376204 | OV | OAW-28 | AZD8931 | top60_altered_DNA+RNA_seed | CDK4 | -0.764434 | GBM | SF295 | Palbociclib |
| MVP_top60_DNA_seed | ERBB2 | 0.376204 | OV | OAW-28 | AZD8931 | top60_altered_DNA+RNA_seed | EGFR | -0.747323 | GBM | Becker | AZD8931 |
| MVP_top60_DNA_seed | EGFR | 0.382121 | OV | JHOS-3 | AZD8931 | top60_altered_DNA+RNA_seed | CDK4 | -0.730447 | GBM | D-423MG | Palbociclib |
| MVP_top60_DNA_seed | ERBB2 | 0.382121 | OV | JHOS-3 | AZD8931 | top60_altered_DNA+RNA_seed | EGFR | -0.730374 | GBM | D-542MG | Cetuximab |
| MVP_top60_DNA_seed | ERBB2 | 0.383915 | OV | OV-56 | CP724714 | top60_altered_DNA+RNA_seed | EGFR | -0.724267 | GBM | KS-1 | CI-1033 |
| MVP_top60_DNA_seed | PIK3CA | 0.384149 | OV | IGROV-1 | PI3Ka_4409 | top60_altered_DNA+RNA_seed | CDK4 | -0.720499 | GBM | D-247MG | Palbociclib |
| MVP_top60_DNA_seed | CREBBP | 0.385181 | OV | SW626 | I-CBP112 | top60_altered_DNA+RNA_seed | PDGFRA | -0.712378 | GBM | U251 | Masitinib |
| MVP_top60_DNA_seed | EGFR | 0.385408 | OV | OVK-18 | Gefitinib | top60_altered_DNA+RNA_seed | CDK4 | -0.709831 | GBM | DK-MG | Palbociclib |
| MVP_top60_DNA_seed | EGFR | 0.38625 | OV | OVCAR-5 | Cetuximab | top60_altered_DNA+RNA_seed | EGFR | -0.704283 | GBM | YH-13 | AZD8931 |
| MVP_top60_DNA_seed | EGFR | 0.388411 | OV | SW626 | PF-00299804 | top60_altered_DNA+RNA_seed | CDK4 | -0.680791 | GBM | SNB75 | CGP-082996 |
| MVP_top60_DNA_seed | ERBB2 | 0.388411 | OV | SW626 | PF-00299804 | top60_altered_DNA+RNA_seed | EGFR | -0.671917 | GBM | U-87-MG | Cetuximab |
| MVP_top60_DNA_seed | EGFR | 0.399193 | OV | OVCAR-4 | Afatinib | top60_altered_DNA+RNA_seed | EGFR | -0.652341 | GBM | SW1088 | Gefitinib |
| MVP_top60_DNA_seed | ERBB2 | 0.399193 | OV | OVCAR-4 | Afatinib | top60_altered_DNA+RNA_seed | PDGFRA | -0.646932 | GBM | SK-MG-1 | Masitinib |
| MVP_top60_DNA_seed | PIK3CA | 0.401536 | OV | OVCAR-4 | PI3Ka_4409 | top60_altered_DNA+RNA_seed | EGFR | -0.645346 | GBM | 8-MG-BA | Pelitinib |
| MVP_top60_DNA_seed | PTK2 | 0.403877 | OV | FU-OV-1 | PF-562271 | top60_altered_DNA+RNA_seed | EGFR | -0.630226 | GBM | D-566MG | CUDC-101 |
| MVP_top60_DNA_seed | PIK3CA | 0.407055 | OV | SW626 | PI3Ka_4409 | top60_altered_DNA+RNA_seed | PDGFRA | -0.624878 | GBM | 8-MG-BA | Masitinib |
| MVP_top60_DNA_seed | EGFR | 0.410146 | OV | EFO-21 | Afatinib | top60_altered_DNA+RNA_seed | EGFR | -0.622044 | GBM | YKG-1 | Afatinib |
| MVP_top60_DNA_seed | ERBB2 | 0.410146 | OV | EFO-21 | Afatinib | top60_altered_DNA+RNA_seed | EGFR | -0.621896 | GBM | D-423MG | CUDC-101 |
| MVP_top60_DNA_seed | CDK2 | 0.410315 | OV | SW626 | AZD5438 | top60_altered_DNA+RNA_seed | CDK4 | -0.589324 | GBM | GB-1 | Palbociclib |
| MVP_top60_DNA_seed | EGFR | 0.410327 | OV | SW626 | Afatinib | top60_altered_DNA+RNA_seed | EGFR | -0.55904 | GBM | SF295 | Afatinib |
| MVP_top60_DNA_seed | ERBB2 | 0.410327 | OV | SW626 | Afatinib | top60_altered_DNA+RNA_seed | EGFR | -0.556108 | GBM | KS-1 | Pelitinib |
| MVP_top60_DNA_seed | EGFR | 0.412504 | OV | OV-17R | Afatinib | top60_altered_DNA+RNA_seed | PDGFRA | -0.548783 | GBM | D-566MG | Masitinib |
| MVP_top60_DNA_seed | ERBB2 | 0.412504 | OV | OV-17R | Afatinib | top60_altered_DNA+RNA_seed | EGFR | -0.539459 | GBM | D-542MG | Erlotinib |

| | | | | | | | | | | | |
|---|---|---|---|---|---|---|---|---|---|---|---|
| MVP_top60_DNA_seed | PIK3CA | 0.413785 | OV | OVCAR-4 | AZD8835 | top60_altered_DNA+RNA_seed | EGFR | -0.53299 | GBM | CAS-1 | Pelitinib |
| MVP_top60_DNA_seed | EGFR | 0.419071 | OV | IGROV-1 | AZD8931 | top60_altered_DNA+RNA_seed | EGFR | -0.528168 | GBM | YH-13 | AST-1306 |
| MVP_top60_DNA_seed | ERBB2 | 0.419071 | OV | IGROV-1 | AZD8931 | top60_altered_DNA+RNA_seed | CDK4 | -0.522497 | GBM | KS-1 | Palbociclib |
| MVP_top60_DNA_seed | EGFR | 0.419247 | OV | SK-OV-3 | AST-1306 | top60_altered_DNA+RNA_seed | EGFR | -0.521341 | GBM | D-392MG | Lapatinib |
| MVP_top60_DNA_seed | MAPK1 | 0.41925 | OV | OVTOKO | VX-11e | top60_altered_DNA+RNA_seed | PDGFRA | -0.516043 | GBM | YKG-1 | Masitinib |
| MVP_top60_DNA_seed | EGFR | 0.419505 | OV | OVTOKO | CUDC-101 | top60_altered_DNA+RNA_seed | EGFR | -0.501998 | GBM | A172 | Gefitinib |
| MVP_top60_DNA_seed | ERBB2 | 0.419505 | OV | OVTOKO | CUDC-101 | top60_altered_DNA+RNA_seed | EGFR | -0.498351 | GBM | GB-1 | Cetuximab |
| MVP_top60_DNA_seed | EGFR | 0.420455 | OV | Hey | CI-1033 | top60_altered_DNA+RNA_seed | CDK4 | -0.488739 | GBM | SK-MG-1 | Palbociclib |
| MVP_top60_DNA_seed | ERBB2 | 0.420455 | OV | Hey | CI-1033 | top60_altered_DNA+RNA_seed | EGFR | -0.464147 | GBM | DK-MG | CI-1033 |
| MVP_top60_DNA_seed | EGFR | 0.423843 | OV | OVCAR-3 | Cetuximab | top60_altered_DNA+RNA_seed | EGFR | -0.463366 | GBM | SF268 | Cetuximab |
| MVP_top60_DNA_seed | PIK3CA | 0.425958 | OV | OVK-18 | AZD8835 | top60_altered_DNA+RNA_seed | PDGFRA | -0.456348 | GBM | A172 | Masitinib |
| MVP_top60_DNA_seed | SIRT1 | 0.426947 | OV | OVCAR-3 | Selisistat | top60_altered_DNA+RNA_seed | EGFR | -0.42457 | GBM | Becker | Lapatinib |
| MVP_top60_DNA_seed | EGFR | 0.429438 | OV | OVCA420 | Pelitinib | top60_altered_DNA+RNA_seed | EGFR | -0.42128 | GBM | D-247MG | Lapatinib |
| MVP_top60_DNA_seed | HDAC2 | 0.431629 | OV | OAW-42 | JQ12 | top60_altered_DNA+RNA_seed | EGFR | -0.40314 | GBM | YH-13 | Afatinib |
| MVP_top60_DNA_seed | EGFR | 0.434066 | OV | OVCA420 | CI-1033 | top60_altered_DNA+RNA_seed | EGFR | -0.398414 | GBM | SF126 | Cetuximab |
| MVP_top60_DNA_seed | ERBB2 | 0.434066 | OV | OVCA420 | CI-1033 | top60_altered_DNA+RNA_seed | EGFR | -0.397437 | GBM | D-247MG | CUDC-101 |
| MVP_top60_DNA_seed | EGFR | 0.439751 | OV | OV-56 | AST-1306 | top60_altered_DNA+RNA_seed | CDK4 | -0.396022 | GBM | LN-18 | Palbociclib |
| MVP_top60_DNA_seed | CDK1 | 0.440665 | OV | EFO-21 | RO-3306 | top60_altered_DNA+RNA_seed | EGFR | -0.392801 | GBM | U251 | AZD8931 |
| MVP_top60_DNA_seed | CREBBP | 0.448724 | OV | FU-OV-1 | I-CBP112 | top60_altered_DNA+RNA_seed | CDK4 | -0.384323 | GBM | 8-MG-BA | CGP-082996 |
| MVP_top60_DNA_seed | EGFR | 0.448966 | OV | TOV-21G | CI-1033 | top60_altered_DNA+RNA_seed | EGFR | -0.383351 | GBM | SF295 | Gefitinib |
| MVP_top60_DNA_seed | ERBB2 | 0.448966 | OV | TOV-21G | CI-1033 | top60_altered_DNA+RNA_seed | EGFR | -0.378426 | GBM | KS-1 | CUDC-101 |
| MVP_top60_DNA_seed | EGFR | 0.449242 | OV | SW626 | Cetuximab | top60_altered_DNA+RNA_seed | EGFR | -0.373229 | GBM | D-566MG | Gefitinib |
| MVP_top60_DNA_seed | ATR | 0.453137 | OV | SW626 | AZ20 | top60_altered_DNA+RNA_seed | EGFR | -0.366672 | GBM | SK-MG-1 | AZD8931 |
| MVP_top60_DNA_seed | EGFR | 0.455572 | OV | JHOS-3 | CI-1033 | top60_altered_DNA+RNA_seed | EGFR | -0.36401 | GBM | T98G | Cetuximab |
| MVP_top60_DNA_seed | ERBB2 | 0.455572 | OV | JHOS-3 | CI-1033 | top60_altered_DNA+RNA_seed | EGFR | -0.357658 | GBM | SF268 | AZD8931 |
| MVP_top60_DNA_seed | ERBB2 | 0.46008 | OV | FU-OV-1 | CP724714 | top60_altered_DNA+RNA_seed | PDGFRA | -0.345152 | GBM | SF268 | Masitinib |
| MVP_top60_DNA_seed | PLK1 | 0.461587 | OV | EFO-21 | PLK_6522 | top60_altered_DNA+RNA_seed | EGFR | -0.341445 | GBM | D-542MG | AZD8931 |
| MVP_top60_DNA_seed | EGFR | 0.478294 | OV | OVCAR-5 | AST-1306 | top60_altered_DNA+RNA_seed | EGFR | -0.335944 | GBM | D-566MG | Afatinib |
| MVP_top60_DNA_seed | EGFR | 0.48548 | OV | OVCAR-5 | Afatinib | top60_altered_DNA+RNA_seed | EGFR | -0.331832 | GBM | AM-38 | Pelitinib |
| MVP_top60_DNA_seed | ERBB2 | 0.48548 | OV | OVCAR-5 | Afatinib | top60_altered_DNA+RNA_seed | PDGFRA | -0.317828 | GBM | D-423MG | Masitinib |
| MVP_top60_DNA_seed | ATR | 0.48761 | OV | UWB1.289 | QL-VIII-58 | top60_altered_DNA+RNA_seed | PDGFRA | -0.311239 | GBM | AM-38 | Masitinib |
| MVP_top60_DNA_seed | EGFR | 0.490034 | OV | Caov-3 | CUDC-101 | top60_altered_DNA+RNA_seed | EGFR | -0.309137 | GBM | 8-MG-BA | Gefitinib |
| MVP_top60_DNA_seed | ERBB2 | 0.490034 | OV | Caov-3 | CUDC-101 | top60_altered_DNA+RNA_seed | CDK4 | -0.302098 | GBM | D-566MG | Palbociclib |
| MVP_top60_DNA_seed | EGFR | 0.492054 | OV | OV-17R | AZD8931 | top60_altered_DNA+RNA_seed | PDGFRA | -0.299402 | GBM | YH-13 | Masitinib |
| MVP_top60_DNA_seed | ERBB2 | 0.492054 | OV | OV-17R | AZD8931 | top60_altered_DNA+RNA_seed | EGFR | -0.297176 | GBM | D-392MG | Erlotinib |
| MVP_top60_DNA_seed | EGFR | 0.493721 | OV | OAW-42 | CUDC-101 | top60_altered_DNA+RNA_seed | PDGFRA | -0.296604 | GBM | SF295 | Masitinib |
| MVP_top60_DNA_seed | ERBB2 | 0.493721 | OV | OAW-42 | CUDC-101 | top60_altered_DNA+RNA_seed | PDGFRA | -0.294946 | GBM | LNZTA3WT4 | Amuvatinib |
| MVP_top60_DNA_seed | ERBB2 | 0.499022 | OV | PEO1 | CP724714 | top60_altered_DNA+RNA_seed | EGFR | -0.290201 | GBM | SF268 | Lapatinib |
| MVP_top60_DNA_seed | SIRT1 | 0.507668 | OV | SW626 | Selisistat | top60_altered_DNA+RNA_seed | EGFR | -0.283792 | GBM | U-87-MG | Afatinib |
| MVP_top60_DNA_seed | CDK2 | 0.507763 | OV | EFO-21 | PHA-793887 | top60_altered_DNA+RNA_seed | CDK4 | -0.283675 | GBM | SF268 | CGP-082996 |
| MVP_top60_DNA_seed | EGFR | 0.507827 | OV | Hey | Afatinib | top60_altered_DNA+RNA_seed | EGFR | -0.277359 | GBM | D-247MG | Erlotinib |

| | | | | | | | | | | | |
|---|---|---|---|---|---|---|---|---|---|---|---|
| MVP_top60_DNA_seed | ERBB2 | 0.507827 | OV | Hey | Afatinib | top60_altered_DNA+RNA_seed | EGFR | -0.27486 | GBM | KS-1 | AZD8931 |
| MVP_top60_DNA_seed | ATR | 0.511671 | OV | JHOS-4 | AZD6738 | top60_altered_DNA+RNA_seed | PDGFRA | -0.27437 | GBM | SF126 | Masitinib |
| MVP_top60_DNA_seed | CDK2 | 0.512121 | OV | FU-OV-1 | AZD5438 | top60_altered_DNA+RNA_seed | EGFR | -0.270597 | GBM | KS-1 | PF-00299804 |
| MVP_top60_DNA_seed | HDAC2 | 0.512782 | OV | TOV-21G | JQ12 | top60_altered_DNA+RNA_seed | EGFR | -0.269198 | GBM | SF126 | AZD8931 |
| MVP_top60_DNA_seed | MAPK1 | 0.513124 | OV | OV-56 | VX-11e | top60_altered_DNA+RNA_seed | EGFR | -0.259102 | GBM | D-542MG | Lapatinib |
| MVP_top60_DNA_seed | EGFR | 0.517032 | OV | OV-56 | AZD8931 | top60_altered_DNA+RNA_seed | EGFR | -0.252138 | GBM | SNB75 | AZD8931 |
| MVP_top60_DNA_seed | ERBB2 | 0.517032 | OV | OV-56 | AZD8931 | top60_altered_DNA+RNA_seed | PDGFRA | -0.250136 | GBM | GB-1 | Amuvatinib |
| MVP_top60_DNA_seed | ERBB2 | 0.518705 | OV | OV-7 | CP724714 | top60_altered_DNA+RNA_seed | EGFR | -0.249357 | GBM | 8-MG-BA | CUDC-101 |
| MVP_top60_DNA_seed | EGFR | 0.520386 | OV | OV-7 | AZD8931 | top60_altered_DNA+RNA_seed | EGFR | -0.233569 | GBM | DBTRG-05MG | Pelitinib |
| MVP_top60_DNA_seed | ERBB2 | 0.520386 | OV | OV-7 | AZD8931 | top60_altered_DNA+RNA_seed | PDGFRA | -0.226488 | GBM | KS-1 | Masitinib |
| MVP_top60_DNA_seed | CDK1 | 0.52225 | OV | OV-90 | RO-3306 | top60_altered_DNA+RNA_seed | CDK4 | -0.222332 | GBM | U-118-MG | Palbociclib |
| MVP_top60_DNA_seed | EGFR | 0.522787 | OV | OV-7 | Afatinib | top60_altered_DNA+RNA_seed | EGFR | -0.218535 | GBM | D-423MG | Pelitinib |
| MVP_top60_DNA_seed | ERBB2 | 0.522787 | OV | OV-7 | Afatinib | top60_altered_DNA+RNA_seed | EGFR | -0.217538 | GBM | SK-MG-1 | Cetuximab |
| MVP_top60_DNA_seed | ATR | 0.530359 | OV | Caov-4 | AZD6738 | top60_altered_DNA+RNA_seed | EGFR | -0.208843 | GBM | LN-229 | Afatinib |
| MVP_top60_DNA_seed | SIRT1 | 0.535034 | OV | SK-OV-3 | Selisistat | top60_altered_DNA+RNA_seed | EGFR | -0.207617 | GBM | LN-405 | Gefitinib |
| MVP_top60_DNA_seed | ERBB2 | 0.541631 | OV | OVCA420 | CP724714 | top60_altered_DNA+RNA_seed | EGFR | -0.192046 | GBM | GB-1 | AST-1306 |
| MVP_top60_DNA_seed | HDAC2 | 0.553916 | OV | OVISE | JQ12 | top60_altered_DNA+RNA_seed | EGFR | -0.192031 | GBM | U-87-MG | AZD8931 |
| MVP_top60_DNA_seed | EGFR | 0.563602 | OV | Caov-4 | Afatinib | top60_altered_DNA+RNA_seed | EGFR | -0.190368 | GBM | KS-1 | Afatinib |
| MVP_top60_DNA_seed | ERBB2 | 0.563602 | OV | Caov-4 | Afatinib | top60_altered_DNA+RNA_seed | EGFR | -0.186879 | GBM | YKG-1 | CI-1033 |
| MVP_top60_DNA_seed | ATR | 0.566979 | OV | FU-OV-1 | AZD6738 | top60_altered_DNA+RNA_seed | EGFR | -0.180533 | GBM | KS-1 | Afatinib |
| MVP_top60_DNA_seed | EGFR | 0.577851 | OV | TOV-21G | PF-00299804 | top60_altered_DNA+RNA_seed | EGFR | -0.179381 | GBM | DBTRG-05MG | Gefitinib |
| MVP_top60_DNA_seed | ERBB2 | 0.577851 | OV | TOV-21G | PF-00299804 | top60_altered_DNA+RNA_seed | EGFR | -0.171645 | GBM | U251 | Pelitinib |
| MVP_top60_DNA_seed | EGFR | 0.580209 | OV | EFO-21 | Gefitinib | top60_altered_DNA+RNA_seed | EGFR | -0.158119 | GBM | D-542MG | CUDC-101 |
| MVP_top60_DNA_seed | EGFR | 0.581071 | OV | OVK-18 | CI-1033 | top60_altered_DNA+RNA_seed | EGFR | -0.157381 | GBM | 8-MG-BA | Cetuximab |
| MVP_top60_DNA_seed | ERBB2 | 0.581071 | OV | OVK-18 | CI-1033 | top60_altered_DNA+RNA_seed | EGFR | -0.146351 | GBM | A172 | CUDC-101 |
| MVP_top60_DNA_seed | EGFR | 0.583217 | OV | SW626 | AST-1306 | top60_altered_DNA+RNA_seed | EGFR | -0.143505 | GBM | GB-1 | CI-1033 |
| MVP_top60_DNA_seed | CREBBP | 0.5862 | OV | OVCAR-4 | I-CBP112 | top60_altered_DNA+RNA_seed | EGFR | -0.142824 | GBM | YKG-1 | Cetuximab |
| MVP_top60_DNA_seed | EGFR | 0.588764 | OV | OVKATE | AZD8931 | top60_altered_DNA+RNA_seed | EGFR | -0.13723 | GBM | DK-MG | PF-00299804 |
| MVP_top60_DNA_seed | ERBB2 | 0.588764 | OV | OVKATE | AZD8931 | top60_altered_DNA+RNA_seed | CDK4 | -0.130802 | GBM | AM-38 | Palbociclib |
| MVP_top60_DNA_seed | CDK2 | 0.589295 | OV | OVKATE | PHA-793887 | top60_altered_DNA+RNA_seed | PDGFRA | -0.12509 | GBM | U-87-MG | Masitinib |
| MVP_top60_DNA_seed | EGFR | 0.592142 | OV | OVCAR-4 | CUDC-101 | top60_altered_DNA+RNA_seed | EGFR | -0.119082 | GBM | T98G | AZD8931 |
| MVP_top60_DNA_seed | ERBB2 | 0.592142 | OV | OVCAR-4 | CUDC-101 | top60_altered_DNA+RNA_seed | EGFR | -0.116551 | GBM | SF126 | CI-1033 |
| MVP_top60_DNA_seed | EGFR | 0.593542 | OV | OV-17R | Pelitinib | top60_altered_DNA+RNA_seed | PDGFRA | -0.110574 | GBM | CAS-1 | Amuvatinib |
| MVP_top60_DNA_seed | ERBB2 | 0.59566 | OV | DOV13 | CP724714 | top60_altered_DNA+RNA_seed | EGFR | -0.108835 | GBM | M059J | AZD8931 |
| MVP_top60_DNA_seed | EGFR | 0.598017 | OV | OV-56 | Gefitinib | top60_altered_DNA+RNA_seed | PDGFRA | -0.107572 | GBM | SF268 | Amuvatinib |
| MVP_top60_DNA_seed | MAPK1 | 0.602227 | OV | OVK-18 | FR-180204 | top60_altered_DNA+RNA_seed | EGFR | -0.095753 | GBM | CAS-1 | AST-1306 |
| MVP_top60_DNA_seed | MAPK3 | 0.602227 | OV | OVK-18 | FR-180204 | top60_altered_DNA+RNA_seed | EGFR | -0.085668 | GBM | AM-38 | Gefitinib |
| MVP_top60_DNA_seed | EGFR | 0.606708 | OV | OV-7 | Pelitinib | top60_altered_DNA+RNA_seed | EGFR | -0.078794 | GBM | M059J | CUDC-101 |
| MVP_top60_DNA_seed | EGFR | 0.613324 | OV | OVK-18 | Afatinib | top60_altered_DNA+RNA_seed | CDK4 | -0.07251 | GBM | D-392MG | Palbociclib |
| MVP_top60_DNA_seed | ERBB2 | 0.613324 | OV | OVK-18 | Afatinib | top60_altered_DNA+RNA_seed | EGFR | -0.057445 | GBM | AM-38 | AST-1306 |
| MVP_top60_DNA_seed | EGFR | 0.616663 | OV | OVCA433 | CI-1033 | top60_altered_DNA+RNA_seed | EGFR | -0.046977 | GBM | AM-38 | CUDC-101 |

| | | | | | | | | | | | |
|---|---|---|---|---|---|---|---|---|---|---|---|
| MVP_top60_DNA_seed | ERBB2 | 0.616663 | OV | OVCA433 | CI-1033 | top60_altered_DNA+RNA_seed | EGFR | -0.038738 | GBM | D-423MG | Cetuximab |
| MVP_top60_DNA_seed | PLK1 | 0.617904 | OV | OV-56 | PLK_6522 | top60_altered_DNA+RNA_seed | EGFR | -0.033848 | GBM | YH-13 | CI-1033 |
| MVP_top60_DNA_seed | EGFR | 0.621653 | OV | IGROV-1 | PF-00299804 | top60_altered_DNA+RNA_seed | EGFR | -0.029853 | GBM | 42-MG-BA | CUDC-101 |
| MVP_top60_DNA_seed | ERBB2 | 0.621653 | OV | IGROV-1 | PF-00299804 | top60_altered_DNA+RNA_seed | EGFR | -0.024533 | GBM | KS-1 | AST-1306 |
| MVP_top60_DNA_seed | MAPK1 | 0.622199 | OV | OV-90 | FR-180204 | top60_altered_DNA+RNA_seed | EGFR | -0.021607 | GBM | 8-MG-BA | Erlotinib |
| MVP_top60_DNA_seed | MAPK3 | 0.622199 | OV | OV-90 | FR-180204 | top60_altered_DNA+RNA_seed | EGFR | -0.013575 | GBM | SF126 | AST-1306 |
| MVP_top60_DNA_seed | EGFR | 0.624245 | OV | SW626 | CI-1033 | top60_altered_DNA+RNA_seed | EGFR | -0.011909 | GBM | YKG-1 | PF-00299804 |
| MVP_top60_DNA_seed | ERBB2 | 0.624245 | OV | SW626 | CI-1033 | top60_altered_DNA+RNA_seed | EGFR | -0.010827 | GBM | LN-229 | AST-1306 |
| MVP_top60_DNA_seed | EGFR | 0.624729 | OV | SK-OV-3 | PF-00299804 | top60_altered_DNA+RNA_seed | PDGFRA | -0.009646 | GBM | LN-18 | Masitinib |
| MVP_top60_DNA_seed | ERBB2 | 0.624729 | OV | SK-OV-3 | PF-00299804 | top60_altered_DNA+RNA_seed | CDK4 | -0.000924 | GBM | LNZTA3WT4 | Palbociclib |
| MVP_top60_DNA_seed | EGFR | 0.629718 | OV | OAW-42 | Cetuximab | top60_altered_DNA+RNA_seed | PDGFRA | -0.000135 | GBM | 8-MG-BA | Amuvatinib |
| MVP_top60_DNA_seed | ERBB2 | 0.635727 | OV | JHOS-2 | CP724714 | top60_altered_DNA+RNA_seed | PDGFRA | -0.000123 | GBM | D-542MG | Amuvatinib |
| MVP_top60_DNA_seed | MAPK1 | 0.636668 | OV | OV-56 | FR-180204 | top60_altered_DNA+RNA_seed | EGFR | 0.006771 | GBM | SNB75 | Lapatinib |
| MVP_top60_DNA_seed | MAPK3 | 0.636668 | OV | OV-56 | FR-180204 | top60_altered_DNA+RNA_seed | EGFR | 0.007005 | GBM | YKG-1 | Pelitinib |
| MVP_top60_DNA_seed | EGFR | 0.638623 | OV | EFO-21 | Cetuximab | top60_altered_DNA+RNA_seed | PDGFRA | 0.009774 | GBM | LN-405 | Masitinib |
| MVP_top60_DNA_seed | PIK3CA | 0.644558 | OV | Hey | PI3Ka_4409 | top60_altered_DNA+RNA_seed | EGFR | 0.012495 | GBM | U251 | Cetuximab |
| MVP_top60_DNA_seed | EGFR | 0.656271 | OV | OV-56 | Afatinib | top60_altered_DNA+RNA_seed | EGFR | 0.01419 | GBM | U251 | PF-00299804 |
| MVP_top60_DNA_seed | ERBB2 | 0.656271 | OV | OV-56 | Afatinib | top60_altered_DNA+RNA_seed | EGFR | 0.01544 | GBM | 8-MG-BA | CI-1033 |
| MVP_top60_DNA_seed | ATR | 0.657401 | OV | JHOS-2 | AZD6738 | top60_altered_DNA+RNA_seed | EGFR | 0.020324 | GBM | D-423MG | Afatinib |
| MVP_top60_DNA_seed | EGFR | 0.663077 | OV | PEO1 | Gefitinib | top60_altered_DNA+RNA_seed | EGFR | 0.029027 | GBM | 8-MG-BA | Lapatinib |
| MVP_top60_DNA_seed | EGFR | 0.664689 | OV | Caov-4 | AZD8931 | top60_altered_DNA+RNA_seed | EGFR | 0.032605 | GBM | LN-18 | AST-1306 |
| MVP_top60_DNA_seed | ERBB2 | 0.664689 | OV | Caov-4 | AZD8931 | top60_altered_DNA+RNA_seed | EGFR | 0.037946 | GBM | LNZTA3WT4 | AST-1306 |
| MVP_top60_DNA_seed | CDK2 | 0.673438 | OV | Caov-4 | PHA-793887 | top60_altered_DNA+RNA_seed | EGFR | 0.04348 | GBM | D-423MG | PF-00299804 |
| MVP_top60_DNA_seed | CDK2 | 0.678272 | OV | OAW-28 | PHA-793887 | top60_altered_DNA+RNA_seed | EGFR | 0.044373 | GBM | U251 | AST-1306 |
| MVP_top60_DNA_seed | PIK3CA | 0.680012 | OV | TOV-21G | PI3Ka_4409 | top60_altered_DNA+RNA_seed | EGFR | 0.044568 | GBM | D-263MG | AST-1306 |
| MVP_top60_DNA_seed | CDK2 | 0.680335 | OV | OVK-18 | PHA-793887 | top60_altered_DNA+RNA_seed | EGFR | 0.045027 | GBM | LN-229 | CI-1033 |
| MVP_top60_DNA_seed | SIRT1 | 0.684861 | OV | OVTOKO | Selisistat | top60_altered_DNA+RNA_seed | PDGFRA | 0.047682 | GBM | A172 | Amuvatinib |
| MVP_top60_DNA_seed | MAPK1 | 0.690436 | OV | EFO-21 | FR-180204 | top60_altered_DNA+RNA_seed | EGFR | 0.04783 | GBM | SK-MG-1 | CUDC-101 |
| MVP_top60_DNA_seed | MAPK3 | 0.690436 | OV | EFO-21 | FR-180204 | top60_altered_DNA+RNA_seed | EGFR | 0.053342 | GBM | LN-18 | Cetuximab |
| MVP_top60_DNA_seed | CREBBP | 0.692837 | OV | OVISE | I-CBP112 | top60_altered_DNA+RNA_seed | EGFR | 0.055296 | GBM | LN-18 | AZD8931 |
| MVP_top60_DNA_seed | CDK1 | 0.694921 | OV | OVCA420 | RO-3306 | top60_altered_DNA+RNA_seed | EGFR | 0.066144 | GBM | SF295 | AZD8931 |
| MVP_top60_DNA_seed | CREBBP | 0.69629 | OV | OVK-18 | I-CBP112 | top60_altered_DNA+RNA_seed | EGFR | 0.067477 | GBM | LN-18 | PF-00299804 |
| MVP_top60_DNA_seed | CDK1 | 0.699219 | OV | OVMIU | RO-3306 | top60_altered_DNA+RNA_seed | EGFR | 0.078088 | GBM | 8-MG-BA | Afatinib |
| MVP_top60_DNA_seed | CDK2 | 0.69958 | OV | OVCA433 | PHA-793887 | top60_altered_DNA+RNA_seed | EGFR | 0.083535 | GBM | U251 | CI-1033 |
| MVP_top60_DNA_seed | MAPK1 | 0.705145 | OV | OVISE | VX-11e | top60_altered_DNA+RNA_seed | PDGFRA | 0.085676 | GBM | YKG-1 | Amuvatinib |
| MVP_top60_DNA_seed | CDK2 | 0.705505 | OV | UWB1.289 | PHA-793887 | top60_altered_DNA+RNA_seed | EGFR | 0.087284 | GBM | 8-MG-BA | AST-1306 |
| MVP_top60_DNA_seed | ATR | 0.707593 | OV | JHOS-3 | AZD6738 | top60_altered_DNA+RNA_seed | CDK4 | 0.09599 | GBM | DBTRG-05MG | Palbociclib |
| MVP_top60_DNA_seed | PLK1 | 0.707941 | OV | OVISE | PLK_6522 | top60_altered_DNA+RNA_seed | EGFR | 0.100039 | GBM | SK-MG-1 | PF-00299804 |
| MVP_top60_DNA_seed | EGFR | 0.710458 | OV | OAW-28 | Cetuximab | top60_altered_DNA+RNA_seed | EGFR | 0.100493 | GBM | D-566MG | CI-1033 |
| MVP_top60_DNA_seed | PTK2 | 0.711747 | OV | Caov-3 | PF-562271 | top60_altered_DNA+RNA_seed | EGFR | 0.101348 | GBM | YKG-1 | AZD8931 |
| MVP_top60_DNA_seed | ERBB2 | 0.720163 | OV | OAW-28 | CP724714 | top60_altered_DNA+RNA_seed | EGFR | 0.101937 | GBM | LN-229 | AZD8931 |

| | | | | | | | | | | | |
|---|---|---|---|---|---|---|---|---|---|---|---|
| MVP_top60_DNA_seed | EGFR | 0.73249 | OV | OVCA433 | Pelitinib | top60_altered_DNA+RNA_seed | EGFR | 0.106911 | GBM | D-542MG | AST-1306 |
| MVP_top60_DNA_seed | SIRT1 | 0.736952 | OV | EFO-21 | Selisistat | top60_altered_DNA+RNA_seed | EGFR | 0.109626 | GBM | LN-229 | PF-00299804 |
| MVP_top60_DNA_seed | EGFR | 0.738285 | OV | IGROV-1 | CI-1033 | top60_altered_DNA+RNA_seed | EGFR | 0.120889 | GBM | AM-38 | CI-1033 |
| MVP_top60_DNA_seed | ERBB2 | 0.738285 | OV | IGROV-1 | CI-1033 | top60_altered_DNA+RNA_seed | EGFR | 0.122608 | GBM | 42-MG-BA | CI-1033 |
| MVP_top60_DNA_seed | CDK2 | 0.738645 | OV | JHOS-3 | AZD5438 | top60_altered_DNA+RNA_seed | EGFR | 0.122964 | GBM | YH-13 | Pelitinib |
| MVP_top60_DNA_seed | EGFR | 0.740485 | OV | SW626 | Afatinib | top60_altered_DNA+RNA_seed | EGFR | 0.126075 | GBM | D-247MG | AZD8931 |
| MVP_top60_DNA_seed | ERBB2 | 0.740485 | OV | SW626 | Afatinib | top60_altered_DNA+RNA_seed | EGFR | 0.129469 | GBM | LN-229 | Afatinib |
| MVP_top60_DNA_seed | ATR | 0.747277 | OV | OV-56 | QL-VIII-58 | top60_altered_DNA+RNA_seed | PDGFRA | 0.132873 | GBM | Becker | Amuvatinib |
| MVP_top60_DNA_seed | EGFR | 0.752121 | OV | OVK-18 | Pelitinib | top60_altered_DNA+RNA_seed | EGFR | 0.133898 | GBM | 8-MG-BA | PF-00299804 |
| MVP_top60_DNA_seed | ERBB2 | 0.763122 | OV | TOV-21G | CP724714 | top60_altered_DNA+RNA_seed | EGFR | 0.134567 | GBM | D-423MG | AZD8931 |
| MVP_top60_DNA_seed | SIRT1 | 0.777233 | OV | IGROV-1 | Selisistat | top60_altered_DNA+RNA_seed | EGFR | 0.140854 | GBM | D-566MG | Pelitinib |
| MVP_top60_DNA_seed | MAPK1 | 0.781306 | OV | SK-OV-3 | VX-11e | top60_altered_DNA+RNA_seed | EGFR | 0.156294 | GBM | D-566MG | AST-1306 |
| MVP_top60_DNA_seed | MAPK1 | 0.782732 | OV | OV-17R | VX-11e | top60_altered_DNA+RNA_seed | EGFR | 0.160378 | GBM | AM-38 | PF-00299804 |
| MVP_top60_DNA_seed | MAPK1 | 0.788442 | OV | OV-17R | FR-180204 | top60_altered_DNA+RNA_seed | EGFR | 0.160936 | GBM | Becker | Pelitinib |
| MVP_top60_DNA_seed | MAPK3 | 0.788442 | OV | OV-17R | FR-180204 | top60_altered_DNA+RNA_seed | EGFR | 0.165266 | GBM | U-87-MG | AST-1306 |
| MVP_top60_DNA_seed | EGFR | 0.803309 | OV | DOV13 | Pelitinib | top60_altered_DNA+RNA_seed | EGFR | 0.167431 | GBM | SNB75 | AST-1306 |
| MVP_top60_DNA_seed | EGFR | 0.809043 | OV | TOV-21G | CUDC-101 | top60_altered_DNA+RNA_seed | EGFR | 0.167731 | GBM | DBTRG-05MG | CUDC-101 |
| MVP_top60_DNA_seed | ERBB2 | 0.809043 | OV | TOV-21G | CUDC-101 | top60_altered_DNA+RNA_seed | EGFR | 0.171606 | GBM | D-392MG | Cetuximab |
| MVP_top60_DNA_seed | CREBBP | 0.809929 | OV | IGROV-1 | I-CBP112 | top60_altered_DNA+RNA_seed | PDGFRA | 0.176476 | GBM | DBTRG-05MG | Masitinib |
| MVP_top60_DNA_seed | MAPK1 | 0.810676 | OV | OAW-42 | FR-180204 | top60_altered_DNA+RNA_seed | EGFR | 0.176952 | GBM | SF126 | Erlotinib |
| MVP_top60_DNA_seed | MAPK3 | 0.810676 | OV | OAW-42 | FR-180204 | top60_altered_DNA+RNA_seed | EGFR | 0.185456 | GBM | A172 | AZD8931 |
| MVP_top60_DNA_seed | CREBBP | 0.813446 | OV | JHOS-2 | I-CBP112 | top60_altered_DNA+RNA_seed | EGFR | 0.188649 | GBM | SF295 | CUDC-101 |
| MVP_top60_DNA_seed | CDK2 | 0.813636 | OV | JHOS-3 | PHA-793887 | top60_altered_DNA+RNA_seed | EGFR | 0.193134 | GBM | YH-13 | CUDC-101 |
| MVP_top60_DNA_seed | EGFR | 0.818465 | OV | JHOS-3 | Afatinib | top60_altered_DNA+RNA_seed | EGFR | 0.203313 | GBM | LN-229 | CUDC-101 |
| MVP_top60_DNA_seed | ERBB2 | 0.818465 | OV | JHOS-3 | Afatinib | top60_altered_DNA+RNA_seed | EGFR | 0.207045 | GBM | GB-1 | Lapatinib |
| MVP_top60_DNA_seed | MAPK1 | 0.821878 | OV | OVCA433 | VX-11e | top60_altered_DNA+RNA_seed | EGFR | 0.216187 | GBM | SK-MG-1 | AST-1306 |
| MVP_top60_DNA_seed | HDAC2 | 0.8293 | OV | OVCAR-3 | JQ12 | top60_altered_DNA+RNA_seed | EGFR | 0.22013 | GBM | YKG-1 | AST-1306 |
| MVP_top60_DNA_seed | EGFR | 0.831506 | OV | FU-OV-1 | CUDC-101 | top60_altered_DNA+RNA_seed | EGFR | 0.220175 | GBM | LN-229 | Gefitinib |
| MVP_top60_DNA_seed | ERBB2 | 0.831506 | OV | FU-OV-1 | CUDC-101 | top60_altered_DNA+RNA_seed | PDGFRA | 0.225253 | GBM | M059J | Masitinib |
| MVP_top60_DNA_seed | EGFR | 0.834946 | OV | JHOS-4 | Cetuximab | top60_altered_DNA+RNA_seed | EGFR | 0.231418 | GBM | GB-1 | Afatinib |
| MVP_top60_DNA_seed | ATR | 0.836926 | OV | OAW-42 | AZ20 | top60_altered_DNA+RNA_seed | EGFR | 0.239334 | GBM | D-263MG | Erlotinib |
| MVP_top60_DNA_seed | SIRT1 | 0.840872 | OV | JHOS-2 | Selisistat | top60_altered_DNA+RNA_seed | EGFR | 0.241185 | GBM | YH-13 | Gefitinib |
| MVP_top60_DNA_seed | ATR | 0.841585 | OV | EFO-21 | AZD6738 | top60_altered_DNA+RNA_seed | EGFR | 0.248324 | GBM | LN-18 | CI-1033 |
| MVP_top60_DNA_seed | CREBBP | 0.842755 | OV | OVMIU | I-CBP112 | top60_altered_DNA+RNA_seed | CDK4 | 0.248666 | GBM | SF126 | CGP-082996 |
| MVP_top60_DNA_seed | EGFR | 0.844522 | OV | OVISE | Cetuximab | top60_altered_DNA+RNA_seed | EGFR | 0.255388 | GBM | Becker | Cetuximab |
| MVP_top60_DNA_seed | MAPK1 | 0.847293 | OV | PEO1 | FR-180204 | top60_altered_DNA+RNA_seed | EGFR | 0.25594 | GBM | SF295 | Cetuximab |
| MVP_top60_DNA_seed | MAPK3 | 0.847293 | OV | PEO1 | FR-180204 | top60_altered_DNA+RNA_seed | EGFR | 0.261073 | GBM | KS-1 | Gefitinib |
| MVP_top60_DNA_seed | EGFR | 0.855153 | OV | OAW-42 | AST-1306 | top60_altered_DNA+RNA_seed | CDK4 | 0.267169 | GBM | D-247MG | CGP-082996 |
| MVP_top60_DNA_seed | EGFR | 0.855926 | OV | FU-OV-1 | AZD8931 | top60_altered_DNA+RNA_seed | EGFR | 0.269698 | GBM | LN-18 | Afatinib |
| MVP_top60_DNA_seed | ERBB2 | 0.855926 | OV | FU-OV-1 | AZD8931 | top60_altered_DNA+RNA_seed | EGFR | 0.274048 | GBM | SF126 | CUDC-101 |
| MVP_top60_DNA_seed | PIK3CA | 0.856435 | OV | EFO-21 | PI3Ka_4409 | top60_altered_DNA+RNA_seed | EGFR | 0.276723 | GBM | DBTRG-05MG | AZD8931 |

| | | | | | | | | | | | |
|---|---|---|---|---|---|---|---|---|---|---|---|
| MVP_top60_DNA_seed | EGFR | 0.857398 | OV | EFO-21 | CUDC-101 | top60_altered_DNA+RNA_seed | EGFR | 0.276843 | GBM | SW1088 | Afatinib |
| MVP_top60_DNA_seed | ERBB2 | 0.857398 | OV | EFO-21 | CUDC-101 | top60_altered_DNA+RNA_seed | EGFR | 0.279396 | GBM | 8-MG-BA | AZD8931 |
| MVP_top60_DNA_seed | EGFR | 0.859797 | OV | OVCAR-8 | PF-00299804 | top60_altered_DNA+RNA_seed | EGFR | 0.282952 | GBM | CAS-1 | PF-00299804 |
| MVP_top60_DNA_seed | ERBB2 | 0.859797 | OV | OVCAR-8 | PF-00299804 | top60_altered_DNA+RNA_seed | EGFR | 0.286063 | GBM | SNB75 | Afatinib |
| MVP_top60_DNA_seed | ATR | 0.860601 | OV | OVCA420 | AZ20 | top60_altered_DNA+RNA_seed | EGFR | 0.287584 | GBM | D-263MG | AZD8931 |
| MVP_top60_DNA_seed | PLK1 | 0.863045 | OV | FU-OV-1 | PLK_6522 | top60_altered_DNA+RNA_seed | EGFR | 0.290069 | GBM | U251 | Gefitinib |
| MVP_top60_DNA_seed | SIRT1 | 0.863411 | OV | OVMIU | Selisistat | top60_altered_DNA+RNA_seed | EGFR | 0.292605 | GBM | D-247MG | AST-1306 |
| MVP_top60_DNA_seed | EGFR | 0.869206 | OV | OVK-18 | Cetuximab | top60_altered_DNA+RNA_seed | EGFR | 0.294226 | GBM | GB-1 | Pelitinib |
| MVP_top60_DNA_seed | EGFR | 0.873814 | OV | OV-7 | PF-00299804 | top60_altered_DNA+RNA_seed | EGFR | 0.3009 | GBM | Becker | Afatinib |
| MVP_top60_DNA_seed | ERBB2 | 0.873814 | OV | OV-7 | PF-00299804 | top60_altered_DNA+RNA_seed | EGFR | 0.306633 | GBM | LN-18 | Gefitinib |
| MVP_top60_DNA_seed | PIK3CA | 0.874359 | OV | OV-56 | AZD8835 | top60_altered_DNA+RNA_seed | EGFR | 0.309742 | GBM | D-392MG | CUDC-101 |
| MVP_top60_DNA_seed | CDK2 | 0.877049 | OV | TOV-21G | PHA-793887 | top60_altered_DNA+RNA_seed | EGFR | 0.310051 | GBM | LN-18 | Afatinib |
| MVP_top60_DNA_seed | PIK3CA | 0.883747 | OV | OV-56 | PI3Ka_4409 | top60_altered_DNA+RNA_seed | EGFR | 0.311172 | GBM | M059J | CI-1033 |
| MVP_top60_DNA_seed | ATR | 0.884575 | OV | OVISE | QL-VIII-58 | top60_altered_DNA+RNA_seed | EGFR | 0.313652 | GBM | CAS-1 | CI-1033 |
| MVP_top60_DNA_seed | CREBBP | 0.891538 | OV | OVCAR-3 | I-CBP112 | top60_altered_DNA+RNA_seed | EGFR | 0.315478 | GBM | KS-1 | Cetuximab |
| MVP_top60_DNA_seed | EGFR | 0.895468 | OV | OV-90 | Cetuximab | top60_altered_DNA+RNA_seed | EGFR | 0.317128 | GBM | GB-1 | PF-00299804 |
| MVP_top60_DNA_seed | EGFR | 0.895836 | OV | JHOS-4 | Gefitinib | top60_altered_DNA+RNA_seed | EGFR | 0.324796 | GBM | U251 | Afatinib |
| MVP_top60_DNA_seed | EGFR | 0.896317 | OV | TOV-21G | AST-1306 | top60_altered_DNA+RNA_seed | EGFR | 0.32485 | GBM | D-566MG | Afatinib |
| MVP_top60_DNA_seed | ATR | 0.901385 | OV | OVCAR-3 | AZD6738 | top60_altered_DNA+RNA_seed | EGFR | 0.325862 | GBM | D-263MG | Lapatinib |
| MVP_top60_DNA_seed | SIRT1 | 0.90318 | OV | OV-90 | Selisistat | top60_altered_DNA+RNA_seed | EGFR | 0.328818 | GBM | D-423MG | CI-1033 |
| MVP_top60_DNA_seed | CDK1 | 0.903298 | OV | UWB1.289 | RO-3306 | top60_altered_DNA+RNA_seed | PDGFRA | 0.331779 | GBM | T98G | Amuvatinib |
| MVP_top60_DNA_seed | EGFR | 0.903918 | OV | OV-90 | PF-00299804 | top60_altered_DNA+RNA_seed | EGFR | 0.34101 | GBM | DK-MG | AST-1306 |
| MVP_top60_DNA_seed | ERBB2 | 0.903918 | OV | OV-90 | PF-00299804 | top60_altered_DNA+RNA_seed | EGFR | 0.349775 | GBM | T98G | Afatinib |
| MVP_top60_DNA_seed | PIK3CA | 0.904176 | OV | OVKATE | AZD8835 | top60_altered_DNA+RNA_seed | PDGFRA | 0.353126 | GBM | D-392MG | Masitinib |
| MVP_top60_DNA_seed | EGFR | 0.906141 | OV | OVTOKO | PF-00299804 | top60_altered_DNA+RNA_seed | EGFR | 0.359435 | GBM | T98G | Pelitinib |
| MVP_top60_DNA_seed | ERBB2 | 0.906141 | OV | OVTOKO | PF-00299804 | top60_altered_DNA+RNA_seed | PDGFRA | 0.361447 | GBM | T98G | Masitinib |
| MVP_top60_DNA_seed | EGFR | 0.909097 | OV | OV-56 | PF-00299804 | top60_altered_DNA+RNA_seed | EGFR | 0.361453 | GBM | D-392MG | AST-1306 |
| MVP_top60_DNA_seed | ERBB2 | 0.909097 | OV | OV-56 | PF-00299804 | top60_altered_DNA+RNA_seed | EGFR | 0.362003 | GBM | U251 | Afatinib |
| MVP_top60_DNA_seed | ATR | 0.912303 | OV | EFO-21 | AZ20 | top60_altered_DNA+RNA_seed | EGFR | 0.365374 | GBM | M059J | Cetuximab |
| MVP_top60_DNA_seed | EGFR | 0.914679 | OV | SK-OV-3 | Cetuximab | top60_altered_DNA+RNA_seed | EGFR | 0.36579 | GBM | SF126 | PF-00299804 |
| MVP_top60_DNA_seed | ERBB2 | 0.929386 | OV | JHOS-3 | CP724714 | top60_altered_DNA+RNA_seed | EGFR | 0.368807 | GBM | D-566MG | AZD8931 |
| MVP_top60_DNA_seed | EGFR | 0.933966 | OV | FU-OV-1 | CI-1033 | top60_altered_DNA+RNA_seed | EGFR | 0.374295 | GBM | YH-13 | PF-00299804 |
| MVP_top60_DNA_seed | ERBB2 | 0.933966 | OV | FU-OV-1 | CI-1033 | top60_altered_DNA+RNA_seed | EGFR | 0.376762 | GBM | U-118-MG | CUDC-101 |
| MVP_top60_DNA_seed | CREBBP | 0.93423 | OV | OVCAR-5 | I-CBP112 | top60_altered_DNA+RNA_seed | EGFR | 0.376843 | GBM | M059J | Gefitinib |
| MVP_top60_DNA_seed | EGFR | 0.948544 | OV | OC-314 | CI-1033 | top60_altered_DNA+RNA_seed | EGFR | 0.379354 | GBM | 42-MG-BA | Cetuximab |
| MVP_top60_DNA_seed | ERBB2 | 0.948544 | OV | OC-314 | CI-1033 | top60_altered_DNA+RNA_seed | EGFR | 0.39038 | GBM | A172 | Pelitinib |
| MVP_top60_DNA_seed | SIRT1 | 0.950118 | OV | OVCAR-5 | Selisistat | top60_altered_DNA+RNA_seed | EGFR | 0.390957 | GBM | D-392MG | Gefitinib |
| MVP_top60_DNA_seed | EGFR | 0.95063 | OV | OVK-18 | Afatinib | top60_altered_DNA+RNA_seed | CDK4 | 0.398742 | GBM | U-87-MG | Palbociclib |
| MVP_top60_DNA_seed | ERBB2 | 0.95063 | OV | OVK-18 | Afatinib | top60_altered_DNA+RNA_seed | EGFR | 0.3991 | GBM | D-392MG | AZD8931 |
| MVP_top60_DNA_seed | EGFR | 0.967018 | OV | JHOS-2 | Cetuximab | top60_altered_DNA+RNA_seed | EGFR | 0.399554 | GBM | M059J | AST-1306 |
| MVP_top60_DNA_seed | EGFR | 0.969627 | OV | OV-7 | CI-1033 | top60_altered_DNA+RNA_seed | EGFR | 0.403308 | GBM | SF268 | Pelitinib |

| | | | | | | | | | | | |
|---|---|---|---|---|---|---|---|---|---|---|---|
| MVP_top60_DNA_seed | ERBB2 | 0.969627 | OV | OV-7 | CI-1033 | top60_altered_DNA+RNA_seed | PDGFRA | 0.408351 | GBM | D-392MG | Amuvatinib |
| MVP_top60_DNA_seed | EGFR | 0.991125 | OV | JHOS-3 | CUDC-101 | top60_altered_DNA+RNA_seed | EGFR | 0.41024 | GBM | 8-MG-BA | Afatinib |
| MVP_top60_DNA_seed | ERBB2 | 0.991125 | OV | JHOS-3 | CUDC-101 | top60_altered_DNA+RNA_seed | EGFR | 0.427033 | GBM | SK-MG-1 | Afatinib |
| MVP_top60_DNA_seed | MAPK1 | 0.993033 | OV | TOV-21G | VX-11e | top60_altered_DNA+RNA_seed | EGFR | 0.433777 | GBM | SK-MG-1 | Afatinib |
| MVP_top60_DNA_seed | EGFR | 0.995952 | OV | OVCAR-4 | Cetuximab | top60_altered_DNA+RNA_seed | EGFR | 0.436719 | GBM | SF268 | Afatinib |
| MVP_top60_DNA_seed | CDK2 | 1.000002 | OV | JHOS-2 | PHA-793887 | top60_altered_DNA+RNA_seed | PDGFRA | 0.440862 | GBM | SW1088 | Masitinib |
| MVP_top60_DNA_seed | ATR | 1.000023 | OV | OVTOKO | QL-VIII-58 | top60_altered_DNA+RNA_seed | EGFR | 0.45681 | GBM | LNZTA3WT4 | CI-1033 |
| MVP_top60_DNA_seed | EGFR | 1.0058 | OV | OV-7 | Cetuximab | top60_altered_DNA+RNA_seed | EGFR | 0.462306 | GBM | D-423MG | AST-1306 |
| MVP_top60_DNA_seed | EGFR | 1.009324 | OV | JHOS-3 | Gefitinib | top60_altered_DNA+RNA_seed | PDGFRA | 0.466775 | GBM | SF295 | Amuvatinib |
| MVP_top60_DNA_seed | CREBBP | 1.013559 | OV | Caov-4 | I-CBP112 | top60_altered_DNA+RNA_seed | EGFR | 0.468353 | GBM | DK-MG | CUDC-101 |
| MVP_top60_DNA_seed | EGFR | 1.017603 | OV | OV-56 | Pelitinib | top60_altered_DNA+RNA_seed | EGFR | 0.46971 | GBM | DK-MG | Cetuximab |
| MVP_top60_DNA_seed | PTK2 | 1.025922 | OV | OAW-42 | PF-562271 | top60_altered_DNA+RNA_seed | EGFR | 0.474493 | GBM | 42-MG-BA | PF-00299804 |
| MVP_top60_DNA_seed | EGFR | 1.029584 | OV | OV-90 | Afatinib | top60_altered_DNA+RNA_seed | EGFR | 0.476711 | GBM | SNB75 | CUDC-101 |
| MVP_top60_DNA_seed | ERBB2 | 1.029584 | OV | OV-90 | Afatinib | top60_altered_DNA+RNA_seed | EGFR | 0.480983 | GBM | LN-229 | Cetuximab |
| MVP_top60_DNA_seed | EGFR | 1.038749 | OV | OAW-42 | CI-1033 | top60_altered_DNA+RNA_seed | EGFR | 0.481954 | GBM | LN-405 | AST-1306 |
| MVP_top60_DNA_seed | ERBB2 | 1.038749 | OV | OAW-42 | CI-1033 | top60_altered_DNA+RNA_seed | EGFR | 0.485131 | GBM | SW1088 | AST-1306 |
| MVP_top60_DNA_seed | SIRT1 | 1.038798 | OV | OVCAR-4 | Selisistat | top60_altered_DNA+RNA_seed | EGFR | 0.498474 | GBM | D-542MG | PF-00299804 |
| MVP_top60_DNA_seed | SIRT1 | 1.042903 | OV | OV-17R | Selisistat | top60_altered_DNA+RNA_seed | EGFR | 0.500836 | GBM | LN-229 | Pelitinib |
| MVP_top60_DNA_seed | MAPK1 | 1.045191 | OV | OAW-42 | VX-11e | top60_altered_DNA+RNA_seed | EGFR | 0.51882 | GBM | U-118-MG | Gefitinib |
| MVP_top60_DNA_seed | CDK2 | 1.051243 | OV | Caov-4 | AZD5438 | top60_altered_DNA+RNA_seed | EGFR | 0.525061 | GBM | YKG-1 | Gefitinib |
| MVP_top60_DNA_seed | MAPK1 | 1.061977 | OV | OVCAR-3 | FR-180204 | top60_altered_DNA+RNA_seed | PDGFRA | 0.525591 | GBM | D-566MG | Amuvatinib |
| MVP_top60_DNA_seed | MAPK3 | 1.061977 | OV | OVCAR-3 | FR-180204 | top60_altered_DNA+RNA_seed | EGFR | 0.528014 | GBM | D-263MG | Afatinib |
| MVP_top60_DNA_seed | SIRT1 | 1.062914 | OV | OVCA420 | Selisistat | top60_altered_DNA+RNA_seed | EGFR | 0.530853 | GBM | D-423MG | Gefitinib |
| MVP_top60_DNA_seed | ERBB2 | 1.071243 | OV | UWB1.289 | CP724714 | top60_altered_DNA+RNA_seed | EGFR | 0.534881 | GBM | SF295 | Afatinib |
| MVP_top60_DNA_seed | CREBBP | 1.071784 | OV | SK-OV-3 | I-CBP112 | top60_altered_DNA+RNA_seed | EGFR | 0.535548 | GBM | A172 | Afatinib |
| MVP_top60_DNA_seed | EGFR | 1.071988 | OV | OVKATE | CI-1033 | top60_altered_DNA+RNA_seed | CDK4 | 0.537366 | GBM | SF126 | Palbociclib |
| MVP_top60_DNA_seed | ERBB2 | 1.071988 | OV | OVKATE | CI-1033 | top60_altered_DNA+RNA_seed | EGFR | 0.545301 | GBM | LN-18 | CUDC-101 |
| MVP_top60_DNA_seed | CDK1 | 1.07238 | OV | OVKATE | RO-3306 | top60_altered_DNA+RNA_seed | PDGFRA | 0.545993 | GBM | LN-18 | Amuvatinib |
| MVP_top60_DNA_seed | CDK2 | 1.075199 | OV | OV-17R | AZD5438 | top60_altered_DNA+RNA_seed | EGFR | 0.546202 | GBM | Becker | Afatinib |
| MVP_top60_DNA_seed | EGFR | 1.07647 | OV | OVK-18 | PF-00299804 | top60_altered_DNA+RNA_seed | EGFR | 0.548579 | GBM | SW1088 | AZD8931 |
| MVP_top60_DNA_seed | ERBB2 | 1.07647 | OV | OVK-18 | PF-00299804 | top60_altered_DNA+RNA_seed | PDGFRA | 0.551922 | GBM | CAS-1 | Masitinib |
| MVP_top60_DNA_seed | ERBB2 | 1.079518 | OV | OVCAR-3 | CP724714 | top60_altered_DNA+RNA_seed | PDGFRA | 0.556644 | GBM | U-118-MG | Masitinib |
| MVP_top60_DNA_seed | SIRT1 | 1.102974 | OV | JHOS-3 | Selisistat | top60_altered_DNA+RNA_seed | EGFR | 0.557431 | GBM | SF126 | Pelitinib |
| MVP_top60_DNA_seed | CREBBP | 1.109743 | OV | OAW-28 | I-CBP112 | top60_altered_DNA+RNA_seed | EGFR | 0.562046 | GBM | DBTRG-05MG | AST-1306 |
| MVP_top60_DNA_seed | SIRT1 | 1.117972 | OV | FU-OV-1 | Selisistat | top60_altered_DNA+RNA_seed | PDGFRA | 0.562627 | GBM | YH-13 | Amuvatinib |
| MVP_top60_DNA_seed | SIRT1 | 1.130195 | OV | OAW-28 | Selisistat | top60_altered_DNA+RNA_seed | EGFR | 0.57615 | GBM | D-263MG | CI-1033 |
| MVP_top60_DNA_seed | PTK2 | 1.132375 | OV | OV-90 | PF-562271 | top60_altered_DNA+RNA_seed | EGFR | 0.578175 | GBM | 42-MG-BA | Afatinib |
| MVP_top60_DNA_seed | EGFR | 1.134516 | OV | OVCAR-5 | Gefitinib | top60_altered_DNA+RNA_seed | EGFR | 0.584384 | GBM | D-566MG | PF-00299804 |
| MVP_top60_DNA_seed | EGFR | 1.144733 | OV | PEO1 | Pelitinib | top60_altered_DNA+RNA_seed | EGFR | 0.588596 | GBM | T98G | Afatinib |
| MVP_top60_DNA_seed | EGFR | 1.146451 | OV | OVCAR-5 | PF-00299804 | top60_altered_DNA+RNA_seed | EGFR | 0.59153 | GBM | SK-MG-1 | CI-1033 |
| MVP_top60_DNA_seed | ERBB2 | 1.146451 | OV | OVCAR-5 | PF-00299804 | top60_altered_DNA+RNA_seed | EGFR | 0.592515 | GBM | D-263MG | PF-00299804 |

| | | | | | | | | | | | |
|---|---|---|---|---|---|---|---|---|---|---|---|
| MVP_top60_DNA_seed | PLK1 | 1.147771 | OV | Caov-4 | PLK_6522 | top60_altered_DNA+RNA_seed | EGFR | 0.593466 | GBM | GB-1 | Afatinib |
| MVP_top60_DNA_seed | EGFR | 1.157004 | OV | SW626 | CUDC-101 | top60_altered_DNA+RNA_seed | EGFR | 0.596205 | GBM | LN-405 | Cetuximab |
| MVP_top60_DNA_seed | ERBB2 | 1.157004 | OV | SW626 | CUDC-101 | top60_altered_DNA+RNA_seed | EGFR | 0.599575 | GBM | D-263MG | Cetuximab |
| MVP_top60_DNA_seed | EGFR | 1.161196 | OV | OVKATE | Afatinib | top60_altered_DNA+RNA_seed | EGFR | 0.603259 | GBM | Becker | AST-1306 |
| MVP_top60_DNA_seed | ERBB2 | 1.161196 | OV | OVKATE | Afatinib | top60_altered_DNA+RNA_seed | PDGFRA | 0.604991 | GBM | 42-MG-BA | Masitinib |
| MVP_top60_DNA_seed | CDK2 | 1.162349 | OV | OV-90 | AZD5438 | top60_altered_DNA+RNA_seed | PDGFRA | 0.608412 | GBM | U-118-MG | Amuvatinib |
| MVP_top60_DNA_seed | ERBB2 | 1.171862 | OV | OVK-18 | CP724714 | top60_altered_DNA+RNA_seed | EGFR | 0.611763 | GBM | DK-MG | Afatinib |
| MVP_top60_DNA_seed | PIK3CA | 1.180401 | OV | EFO-21 | AZD8835 | top60_altered_DNA+RNA_seed | EGFR | 0.613775 | GBM | T98G | Gefitinib |
| MVP_top60_DNA_seed | MAPK1 | 1.184026 | OV | OVCAR-3 | VX-11e | top60_altered_DNA+RNA_seed | PDGFRA | 0.619007 | GBM | SF126 | Amuvatinib |
| MVP_top60_DNA_seed | EGFR | 1.18536 | OV | OVTOKO | AST-1306 | top60_altered_DNA+RNA_seed | EGFR | 0.620391 | GBM | 42-MG-BA | AST-1306 |
| MVP_top60_DNA_seed | EGFR | 1.189854 | OV | OV-17R | Gefitinib | top60_altered_DNA+RNA_seed | EGFR | 0.62599 | GBM | SF268 | PF-00299804 |
| MVP_top60_DNA_seed | MAPK1 | 1.19005 | OV | OV-7 | VX-11e | top60_altered_DNA+RNA_seed | EGFR | 0.627451 | GBM | A172 | Afatinib |
| MVP_top60_DNA_seed | EGFR | 1.190791 | OV | OVKATE | Pelitinib | top60_altered_DNA+RNA_seed | EGFR | 0.636266 | GBM | LNZTA3WT4 | Cetuximab |
| MVP_top60_DNA_seed | CDK2 | 1.194528 | OV | OVCAR-4 | AZD5438 | top60_altered_DNA+RNA_seed | EGFR | 0.639196 | GBM | D-392MG | PF-00299804 |
| MVP_top60_DNA_seed | CDK2 | 1.198341 | OV | EFO-21 | AZD5438 | top60_altered_DNA+RNA_seed | EGFR | 0.643621 | GBM | Becker | CI-1033 |
| MVP_top60_DNA_seed | PLK1 | 1.199736 | OV | JHOS-3 | PLK_6522 | top60_altered_DNA+RNA_seed | EGFR | 0.645694 | GBM | CAS-1 | CUDC-101 |
| MVP_top60_DNA_seed | PLK1 | 1.201251 | OV | OAW-42 | PLK_6522 | top60_altered_DNA+RNA_seed | EGFR | 0.648066 | GBM | U-118-MG | PF-00299804 |
| MVP_top60_DNA_seed | EGFR | 1.20888 | OV | OV-90 | AZD8931 | top60_altered_DNA+RNA_seed | EGFR | 0.648765 | GBM | D-263MG | Gefitinib |
| MVP_top60_DNA_seed | ERBB2 | 1.20888 | OV | OV-90 | AZD8931 | top60_altered_DNA+RNA_seed | EGFR | 0.651567 | GBM | D-247MG | Cetuximab |
| MVP_top60_DNA_seed | EGFR | 1.215426 | OV | OV-90 | Gefitinib | top60_altered_DNA+RNA_seed | EGFR | 0.654301 | GBM | SF295 | PF-00299804 |
| MVP_top60_DNA_seed | CDK1 | 1.222013 | OV | IGROV-1 | RO-3306 | top60_altered_DNA+RNA_seed | EGFR | 0.660387 | GBM | U-87-MG | PF-00299804 |
| MVP_top60_DNA_seed | CREBBP | 1.223831 | OV | EFO-21 | I-CBP112 | top60_altered_DNA+RNA_seed | EGFR | 0.661416 | GBM | 42-MG-BA | Gefitinib |
| MVP_top60_DNA_seed | EGFR | 1.231738 | OV | OAW-42 | PF-00299804 | top60_altered_DNA+RNA_seed | EGFR | 0.666765 | GBM | CAS-1 | Lapatinib |
| MVP_top60_DNA_seed | ERBB2 | 1.231738 | OV | OAW-42 | PF-00299804 | top60_altered_DNA+RNA_seed | EGFR | 0.674141 | GBM | A172 | AST-1306 |
| MVP_top60_DNA_seed | PTK2 | 1.234536 | OV | OAW-28 | PF-562271 | top60_altered_DNA+RNA_seed | EGFR | 0.675234 | GBM | M059J | PF-00299804 |
| MVP_top60_DNA_seed | CDK1 | 1.237233 | OV | OV-7 | RO-3306 | top60_altered_DNA+RNA_seed | EGFR | 0.677006 | GBM | D-263MG | Afatinib |
| MVP_top60_DNA_seed | ATR | 1.245692 | OV | UWB1.289 | AZ20 | top60_altered_DNA+RNA_seed | EGFR | 0.6825 | GBM | D-392MG | Afatinib |
| MVP_top60_DNA_seed | CDK2 | 1.281315 | OV | PEO1 | PHA-793887 | top60_altered_DNA+RNA_seed | EGFR | 0.686034 | GBM | U-87-MG | CUDC-101 |
| MVP_top60_DNA_seed | CDK2 | 1.289969 | OV | UWB1.289 | AZD5438 | top60_altered_DNA+RNA_seed | EGFR | 0.689103 | GBM | M059J | Afatinib |
| MVP_top60_DNA_seed | EGFR | 1.290009 | OV | OV-90 | AST-1306 | top60_altered_DNA+RNA_seed | EGFR | 0.695419 | GBM | U-118-MG | AZD8931 |
| MVP_top60_DNA_seed | EGFR | 1.300384 | OV | SW626 | Gefitinib | top60_altered_DNA+RNA_seed | EGFR | 0.695518 | GBM | Becker | Gefitinib |
| MVP_top60_DNA_seed | CREBBP | 1.301005 | OV | OV-7 | I-CBP112 | top60_altered_DNA+RNA_seed | EGFR | 0.698169 | GBM | 42-MG-BA | Afatinib |
| MVP_top60_DNA_seed | PLK1 | 1.310764 | OV | OV-17R | PLK_6522 | top60_altered_DNA+RNA_seed | EGFR | 0.698915 | GBM | 42-MG-BA | AZD8931 |
| MVP_top60_DNA_seed | CDK2 | 1.332341 | OV | DOV13 | PHA-793887 | top60_altered_DNA+RNA_seed | EGFR | 0.699406 | GBM | YKG-1 | Afatinib |
| MVP_top60_DNA_seed | EGFR | 1.339641 | OV | Caov-4 | Pelitinib | top60_altered_DNA+RNA_seed | EGFR | 0.699493 | GBM | CAS-1 | Cetuximab |
| MVP_top60_DNA_seed | CREBBP | 1.357867 | OV | JHOS-3 | I-CBP112 | top60_altered_DNA+RNA_seed | EGFR | 0.699883 | GBM | SK-MG-1 | Pelitinib |
| MVP_top60_DNA_seed | EGFR | 1.364733 | OV | Caov-4 | Afatinib | top60_altered_DNA+RNA_seed | PDGFRA | 0.701246 | GBM | GB-1 | Masitinib |
| MVP_top60_DNA_seed | ERBB2 | 1.364733 | OV | Caov-4 | Afatinib | top60_altered_DNA+RNA_seed | PDGFRA | 0.706838 | GBM | D-263MG | Amuvatinib |
| MVP_top60_DNA_seed | ATR | 1.390102 | OV | JHOS-3 | AZ20 | top60_altered_DNA+RNA_seed | EGFR | 0.708314 | GBM | T98G | AST-1306 |
| MVP_top60_DNA_seed | CDK2 | 1.400008 | OV | OVCAR-3 | PHA-793887 | top60_altered_DNA+RNA_seed | EGFR | 0.709652 | GBM | SW1088 | Pelitinib |
| MVP_top60_DNA_seed | SIRT1 | 1.400225 | OV | OV-7 | Selisistat | top60_altered_DNA+RNA_seed | EGFR | 0.715007 | GBM | U-87-MG | CI-1033 |

| | | | | | | | | | | | |
|---|---|---|---|---|---|---|---|---|---|---|---|
| MVP_top60_DNA_seed | SIRT1 | 1.403929 | OV | OVISE | Selisistat | top60_altered_DNA+RNA_seed | EGFR | 0.716797 | GBM | YH-13 | Afatinib |
| MVP_top60_DNA_seed | ATR | 1.404032 | OV | OVKATE | QL-VIII-58 | top60_altered_DNA+RNA_seed | EGFR | 0.730349 | GBM | 42-MG-BA | Pelitinib |
| MVP_top60_DNA_seed | PIK3CA | 1.404989 | OV | Caov-4 | AZD8835 | top60_altered_DNA+RNA_seed | EGFR | 0.732461 | GBM | SF126 | Gefitinib |
| MVP_top60_DNA_seed | ATR | 1.425687 | OV | Caov-4 | AZ20 | top60_altered_DNA+RNA_seed | EGFR | 0.741936 | GBM | D-392MG | CI-1033 |
| MVP_top60_DNA_seed | MAPK1 | 1.431124 | OV | EFO-21 | VX-11e | top60_altered_DNA+RNA_seed | PDGFRA | 0.742057 | GBM | DBTRG-05MG | Amuvatinib |
| MVP_top60_DNA_seed | ERBB2 | 1.438801 | OV | OVCAR-5 | CP724714 | top60_altered_DNA+RNA_seed | EGFR | 0.748635 | GBM | AM-38 | Lapatinib |
| MVP_top60_DNA_seed | EGFR | 1.462138 | OV | EFO-21 | CI-1033 | top60_altered_DNA+RNA_seed | EGFR | 0.753517 | GBM | SF126 | Afatinib |
| MVP_top60_DNA_seed | ERBB2 | 1.462138 | OV | EFO-21 | CI-1033 | top60_altered_DNA+RNA_seed | EGFR | 0.755547 | GBM | U-87-MG | Lapatinib |
| MVP_top60_DNA_seed | EGFR | 1.471723 | OV | OVCAR-3 | Pelitinib | top60_altered_DNA+RNA_seed | EGFR | 0.774173 | GBM | SF295 | Pelitinib |
| MVP_top60_DNA_seed | HDAC2 | 1.473147 | OV | EFO-21 | JQ12 | top60_altered_DNA+RNA_seed | EGFR | 0.775129 | GBM | AM-38 | Afatinib |
| MVP_top60_DNA_seed | CREBBP | 1.477652 | OV | UWB1.289 | I-CBP112 | top60_altered_DNA+RNA_seed | EGFR | 0.778509 | GBM | D-392MG | Pelitinib |
| MVP_top60_DNA_seed | HDAC2 | 1.496014 | OV | Caov-3 | JQ12 | top60_altered_DNA+RNA_seed | EGFR | 0.780786 | GBM | AM-38 | Erlotinib |
| MVP_top60_DNA_seed | CDK2 | 1.500956 | OV | OVCA420 | PHA-793887 | top60_altered_DNA+RNA_seed | EGFR | 0.782544 | GBM | LN-405 | CUDC-101 |
| MVP_top60_DNA_seed | EGFR | 1.515091 | OV | OVKATE | Gefitinib | top60_altered_DNA+RNA_seed | EGFR | 0.782733 | GBM | SNB75 | Cetuximab |
| MVP_top60_DNA_seed | CDK1 | 1.530862 | OV | JHOS-3 | RO-3306 | top60_altered_DNA+RNA_seed | EGFR | 0.785877 | GBM | SF126 | Lapatinib |
| MVP_top60_DNA_seed | EGFR | 1.549411 | OV | OV-17R | CI-1033 | top60_altered_DNA+RNA_seed | EGFR | 0.786171 | GBM | LN-405 | AZD8931 |
| MVP_top60_DNA_seed | ERBB2 | 1.549411 | OV | OV-17R | CI-1033 | top60_altered_DNA+RNA_seed | EGFR | 0.792512 | GBM | LN-405 | Afatinib |
| MVP_top60_DNA_seed | PIK3CA | 1.588414 | OV | OVKATE | PI3Ka_4409 | top60_altered_DNA+RNA_seed | EGFR | 0.798119 | GBM | SF126 | Afatinib |
| MVP_top60_DNA_seed | EGFR | 1.596128 | OV | SK-OV-3 | CI-1033 | top60_altered_DNA+RNA_seed | EGFR | 0.799122 | GBM | U-87-MG | Gefitinib |
| MVP_top60_DNA_seed | ERBB2 | 1.596128 | OV | SK-OV-3 | CI-1033 | top60_altered_DNA+RNA_seed | EGFR | 0.803099 | GBM | GB-1 | AZD8931 |
| MVP_top60_DNA_seed | ATR | 1.601575 | OV | OVCA420 | QL-VIII-58 | top60_altered_DNA+RNA_seed | EGFR | 0.81115 | GBM | SF295 | AST-1306 |
| MVP_top60_DNA_seed | EGFR | 1.646104 | OV | OVKATE | Afatinib | top60_altered_DNA+RNA_seed | EGFR | 0.813821 | GBM | LNZTA3WT4 | PF-00299804 |
| MVP_top60_DNA_seed | ERBB2 | 1.646104 | OV | OVKATE | Afatinib | top60_altered_DNA+RNA_seed | EGFR | 0.824329 | GBM | SF268 | Erlotinib |
| MVP_top60_DNA_seed | ATR | 1.665497 | OV | JHOS-4 | AZ20 | top60_altered_DNA+RNA_seed | EGFR | 0.848985 | GBM | T98G | PF-00299804 |
| MVP_top60_DNA_seed | SIRT1 | 1.666463 | OV | UWB1.289 | Selisistat | top60_altered_DNA+RNA_seed | EGFR | 0.85078 | GBM | D-423MG | Afatinib |
| MVP_top60_DNA_seed | EGFR | 1.680505 | OV | OVCAR-8 | CI-1033 | top60_altered_DNA+RNA_seed | EGFR | 0.852691 | GBM | LNZTA3WT4 | Afatinib |
| MVP_top60_DNA_seed | ERBB2 | 1.680505 | OV | OVCAR-8 | CI-1033 | top60_altered_DNA+RNA_seed | EGFR | 0.854566 | GBM | GB-1 | Gefitinib |
| MVP_top60_DNA_seed | CDK2 | 1.698282 | OV | OV-17R | PHA-793887 | top60_altered_DNA+RNA_seed | PDGFRA | 0.861017 | GBM | SNB75 | Masitinib |
| MVP_top60_DNA_seed | CDK2 | 1.716718 | OV | IGROV-1 | AZD5438 | top60_altered_DNA+RNA_seed | EGFR | 0.868661 | GBM | M059J | Afatinib |
| MVP_top60_DNA_seed | ATR | 1.725267 | OV | OVKATE | AZ20 | top60_altered_DNA+RNA_seed | PDGFRA | 0.869844 | GBM | LNZTA3WT4 | Masitinib |
| MVP_top60_DNA_seed | EGFR | 1.765705 | OV | Caov-4 | Gefitinib | top60_altered_DNA+RNA_seed | EGFR | 0.871061 | GBM | SK-MG-1 | Gefitinib |
| MVP_top60_DNA_seed | EGFR | 1.767556 | OV | OVKATE | AST-1306 | top60_altered_DNA+RNA_seed | EGFR | 0.875322 | GBM | LNZTA3WT4 | Afatinib |
| MVP_top60_DNA_seed | MAPK1 | 1.799396 | OV | Caov-4 | FR-180204 | top60_altered_DNA+RNA_seed | EGFR | 0.893863 | GBM | SF268 | AST-1306 |
| MVP_top60_DNA_seed | MAPK3 | 1.799396 | OV | Caov-4 | FR-180204 | top60_altered_DNA+RNA_seed | PDGFRA | 0.896973 | GBM | 42-MG-BA | Amuvatinib |
| MVP_top60_DNA_seed | EGFR | 1.814328 | OV | OVTOKO | CI-1033 | top60_altered_DNA+RNA_seed | EGFR | 0.905008 | GBM | CAS-1 | AZD8931 |
| MVP_top60_DNA_seed | ERBB2 | 1.814328 | OV | OVTOKO | CI-1033 | top60_altered_DNA+RNA_seed | EGFR | 0.905612 | GBM | LNZTA3WT4 | Pelitinib |
| MVP_top60_DNA_seed | HDAC2 | 1.877844 | OV | Caov-4 | JQ12 | top60_altered_DNA+RNA_seed | EGFR | 0.907562 | GBM | U-118-MG | AST-1306 |
| MVP_top60_DNA_seed | ATR | 1.897527 | OV | OVKATE | torin2 | top60_altered_DNA+RNA_seed | PDGFRA | 0.907839 | GBM | D-542MG | Masitinib |
| MVP_top60_DNA_seed | ERBB2 | 1.948335 | OV | OVKATE | CP724714 | top60_altered_DNA+RNA_seed | EGFR | 0.911172 | GBM | Becker | CUDC-101 |
| MVP_top60_DNA_seed | ATR | 2.017703 | OV | OV-17R | QL-VIII-58 | top60_altered_DNA+RNA_seed | EGFR | 0.918206 | GBM | LN-405 | PF-00299804 |
| MVP_top60_DNA_seed | EGFR | 2.022766 | OV | OV-90 | CI-1033 | top60_altered_DNA+RNA_seed | EGFR | 0.922185 | GBM | A172 | PF-00299804 |

| | | | | | | | | | | | |
|---|---|---|---|---|---|---|---|---|---|---|---|
| MVP_top60_DNA_seed | ERBB2 | 2.022766 | OV | OV-90 | CI-1033 | top60_altered_DNA+RNA_seed | EGFR | 0.922255 | GBM | SNB75 | PF-00299804 |
| MVP_top60_DNA_seed | SIRT1 | 2.038062 | OV | OVKATE | Selisistat | top60_altered_DNA+RNA_seed | EGFR | 0.928215 | GBM | SW1088 | PF-00299804 |
| MVP_top60_DNA_seed | CDK2 | 2.057971 | OV | OAW-28 | AZD5438 | top60_altered_DNA+RNA_seed | EGFR | 0.930239 | GBM | SF268 | Afatinib |
| MVP_top60_DNA_seed | EGFR | 2.109053 | OV | OVCAR-5 | CI-1033 | top60_altered_DNA+RNA_seed | EGFR | 0.930559 | GBM | DBTRG-05MG | Cetuximab |
| MVP_top60_DNA_seed | ERBB2 | 2.109053 | OV | OVCAR-5 | CI-1033 | top60_altered_DNA+RNA_seed | EGFR | 0.934676 | GBM | D-247MG | Gefitinib |
| MVP_top60_DNA_seed | HDAC2 | 2.18599 | OV | FU-OV-1 | JQ12 | top60_altered_DNA+RNA_seed | EGFR | 0.948982 | GBM | DBTRG-05MG | PF-00299804 |
| MVP_top60_DNA_seed | EGFR | 2.231858 | OV | DOV13 | CUDC-101 | top60_altered_DNA+RNA_seed | EGFR | 0.950111 | GBM | LN-18 | Pelitinib |
| MVP_top60_DNA_seed | ERBB2 | 2.231858 | OV | DOV13 | CUDC-101 | top60_altered_DNA+RNA_seed | EGFR | 0.950739 | GBM | D-263MG | CUDC-101 |
| MVP_top60_DNA_seed | EGFR | 2.23706 | OV | OV-56 | CI-1033 | top60_altered_DNA+RNA_seed | EGFR | 0.963154 | GBM | D-247MG | PF-00299804 |
| MVP_top60_DNA_seed | ERBB2 | 2.23706 | OV | OV-56 | CI-1033 | top60_altered_DNA+RNA_seed | EGFR | 0.966872 | GBM | LN-405 | Afatinib |
| MVP_top60_DNA_seed | EGFR | 2.30949 | OV | OVK-18 | CUDC-101 | top60_altered_DNA+RNA_seed | EGFR | 0.973124 | GBM | GB-1 | Erlotinib |
| MVP_top60_DNA_seed | ERBB2 | 2.30949 | OV | OVK-18 | CUDC-101 | top60_altered_DNA+RNA_seed | CDK4 | 0.97739 | GBM | 42-MG-BA | Palbociclib |
| MVP_top60_DNA_seed | ATR | 2.311137 | OV | OVKATE | AZD6738 | top60_altered_DNA+RNA_seed | PDGFRA | 0.9916 | GBM | M059J | Amuvatinib |
| MVP_top60_DNA_seed | PTK2 | 2.324968 | OV | Caov-4 | PF-562271 | top60_altered_DNA+RNA_seed | EGFR | 0.997767 | GBM | U-118-MG | Cetuximab |
| MVP_top60_DNA_seed | HDAC2 | 2.400782 | OV | OVKATE | JQ12 | top60_altered_DNA+RNA_seed | EGFR | 1.007573 | GBM | DK-MG | Gefitinib |
| MVP_top60_DNA_seed | EGFR | 2.49227 | OV | OVCAR-3 | CUDC-101 | top60_altered_DNA+RNA_seed | EGFR | 1.013694 | GBM | U-118-MG | Afatinib |
| MVP_top60_DNA_seed | ERBB2 | 2.49227 | OV | OVCAR-3 | CUDC-101 | top60_altered_DNA+RNA_seed | EGFR | 1.023486 | GBM | DBTRG-05MG | Afatinib |
| MVP_top60_DNA_seed | PLK1 | 2.578193 | OV | IGROV-1 | PLK_6522 | top60_altered_DNA+RNA_seed | EGFR | 1.028719 | GBM | D-392MG | Afatinib |
| MVP_top60_DNA_seed | PTK2 | 2.985457 | OV | OVKATE | PF-562271 | top60_altered_DNA+RNA_seed | EGFR | 1.030523 | GBM | SW1088 | Afatinib |
| MVP_top60_DNA_seed | PLK1 | 3.129011 | OV | OVKATE | PLK_6522 | top60_altered_DNA+RNA_seed | EGFR | 1.034087 | GBM | Becker | PF-00299804 |
| MVP_top60_DNA_seed | EGFR | 3.346423 | OV | OVKATE | PF-00299804 | top60_altered_DNA+RNA_seed | EGFR | 1.041416 | GBM | DBTRG-05MG | Afatinib |
| MVP_top60_DNA_seed | ERBB2 | 3.346423 | OV | OVKATE | PF-00299804 | top60_altered_DNA+RNA_seed | EGFR | 1.041725 | GBM | SF268 | Gefitinib |
| MVP_top60_DNA_seed | CDK2 | 3.607598 | OV | OVKATE | AZD5438 | top60_altered_DNA+RNA_seed | EGFR | 1.047505 | GBM | CAS-1 | Erlotinib |
| MVP_top60_DNA_seed | EGFR | -3.133954 | OV | OVCA420 | Gefitinib | top60_altered_DNA+RNA_seed | CDK4 | 1.050188 | GBM | 8-MG-BA | Palbociclib |
| MVP_top60_DNA_seed | EGFR | -3.118836 | OV | OVMIU | Afatinib | top60_altered_DNA+RNA_seed | EGFR | 1.051861 | GBM | D-542MG | Afatinib |
| MVP_top60_DNA_seed | ERBB2 | -3.118836 | OV | OVMIU | Afatinib | top60_altered_DNA+RNA_seed | PDGFRA | 1.054111 | GBM | LN-405 | Amuvatinib |
| MVP_top60_DNA_seed | EGFR | -2.968273 | OV | OVCA420 | Sapitinib | top60_altered_DNA+RNA_seed | EGFR | 1.060722 | GBM | DK-MG | Afatinib |
| MVP_top60_DNA_seed | ERBB2 | -2.968273 | OV | OVCA420 | Sapitinib | top60_altered_DNA+RNA_seed | EGFR | 1.06613 | GBM | D-542MG | Afatinib |
| MVP_top60_DNA_seed | PIK3CA | -2.937083 | OV | IGROV-1 | Alpelisib | top60_altered_DNA+RNA_seed | EGFR | 1.074244 | GBM | AM-38 | Cetuximab |
| MVP_top60_DNA_seed | EGFR | -2.734318 | OV | OVCA420 | Erlotinib | top60_altered_DNA+RNA_seed | CDK4 | 1.080758 | GBM | CAS-1 | Palbociclib |
| MVP_top60_DNA_seed | EGFR | -2.693684 | OV | OVMIU | Sapitinib | top60_altered_DNA+RNA_seed | EGFR | 1.084594 | GBM | CAS-1 | Afatinib |
| MVP_top60_DNA_seed | ERBB2 | -2.693684 | OV | OVMIU | Sapitinib | top60_altered_DNA+RNA_seed | EGFR | 1.088397 | GBM | SNB75 | Pelitinib |
| MVP_top60_DNA_seed | EGFR | -2.576517 | OV | OVCA420 | Afatinib | top60_altered_DNA+RNA_seed | EGFR | 1.091108 | GBM | M059J | Pelitinib |
| MVP_top60_DNA_seed | ERBB2 | -2.576517 | OV | OVCA420 | Afatinib | top60_altered_DNA+RNA_seed | EGFR | 1.092916 | GBM | T98G | CUDC-101 |
| MVP_top60_DNA_seed | EGFR | -2.513178 | OV | OVMIU | Gefitinib | top60_altered_DNA+RNA_seed | EGFR | 1.093055 | GBM | SW1088 | Cetuximab |
| MVP_top60_DNA_seed | PIK3CA | -2.347388 | OV | IGROV-1 | Buparlisib | top60_altered_DNA+RNA_seed | EGFR | 1.093282 | GBM | D-542MG | CI-1033 |
| MVP_top60_DNA_seed | EGFR | -2.296658 | OV | OVCA420 | AZD3759 | top60_altered_DNA+RNA_seed | EGFR | 1.114354 | GBM | SNB75 | CI-1033 |
| MVP_top60_DNA_seed | PIK3CA | -2.20993 | OV | IGROV-1 | AZD8186 | top60_altered_DNA+RNA_seed | EGFR | 1.11661 | GBM | LNZTA3WT4 | Gefitinib |
| MVP_top60_DNA_seed | EGFR | -2.157637 | OV | IGROV-1 | Sapitinib | top60_altered_DNA+RNA_seed | EGFR | 1.123878 | GBM | AM-38 | Afatinib |
| MVP_top60_DNA_seed | ERBB2 | -2.157637 | OV | IGROV-1 | Sapitinib | top60_altered_DNA+RNA_seed | CDK4 | 1.124655 | GBM | U-87-MG | CGP-082996 |
| MVP_top60_DNA_seed | EGFR | -2.088229 | OV | OVCA420 | Osimertinib | top60_altered_DNA+RNA_seed | EGFR | 1.128897 | GBM | D-247MG | Afatinib |

| | | | | | | | | | | | |
|---|---|---|---|---|---|---|---|---|---|---|---|
| MVP_top60_DNA_seed | EGFR | -2.028156 | OV | IGROV-1 | Erlotinib | top60_altered_DNA+RNA_seed | EGFR | 1.164147 | GBM | LN-405 | Pelitinib |
| MVP_top60_DNA_seed | EGFR | -1.995398 | OV | IGROV-1 | Gefitinib | top60_altered_DNA+RNA_seed | EGFR | 1.169363 | GBM | D-247MG | Afatinib |
| MVP_top60_DNA_seed | PIK3CA | -1.980327 | OV | JHOS-4 | Alpelisib | top60_altered_DNA+RNA_seed | EGFR | 1.183565 | GBM | D-542MG | Gefitinib |
| MVP_top60_DNA_seed | PIK3CA | -1.974779 | OV | IGROV-1 | GNE-317 | top60_altered_DNA+RNA_seed | EGFR | 1.217418 | GBM | U-87-MG | Erlotinib |
| MVP_top60_DNA_seed | EGFR | -1.960451 | OV | OVMIU | Erlotinib | top60_altered_DNA+RNA_seed | CDK4 | 1.232426 | GBM | D-263MG | CGP-082996 |
| MVP_top60_DNA_seed | EGFR | -1.941037 | OV | OVCA420 | Lapatinib | top60_altered_DNA+RNA_seed | EGFR | 1.234474 | GBM | CAS-1 | Afatinib |
| MVP_top60_DNA_seed | ERBB2 | -1.941037 | OV | OVCA420 | Lapatinib | top60_altered_DNA+RNA_seed | EGFR | 1.248653 | GBM | SNB75 | Afatinib |
| MVP_top60_DNA_seed | EGFR | -1.856838 | OV | IGROV-1 | AZD3759 | top60_altered_DNA+RNA_seed | CDK4 | 1.249184 | GBM | AM-38 | CGP-082996 |
| MVP_top60_DNA_seed | EGFR | -1.836583 | OV | IGROV-1 | Afatinib | top60_altered_DNA+RNA_seed | EGFR | 1.261068 | GBM | U-118-MG | Afatinib |
| MVP_top60_DNA_seed | ERBB2 | -1.836583 | OV | IGROV-1 | Afatinib | top60_altered_DNA+RNA_seed | EGFR | 1.280577 | GBM | CAS-1 | Gefitinib |
| MVP_top60_DNA_seed | EGFR | -1.814445 | OV | IGROV-1 | Lapatinib | top60_altered_DNA+RNA_seed | PDGFRA | 1.280743 | GBM | LN-229 | Amuvatinib |
| MVP_top60_DNA_seed | ERBB2 | -1.814445 | OV | IGROV-1 | Lapatinib | top60_altered_DNA+RNA_seed | PDGFRA | 1.317396 | GBM | Becker | Masitinib |
| MVP_top60_DNA_seed | EGFR | -1.74378 | OV | IGROV-1 | Osimertinib | top60_altered_DNA+RNA_seed | EGFR | 1.318872 | GBM | DK-MG | AZD8931 |
| MVP_top60_DNA_seed | MAPK1 | -1.705442 | OV | IGROV-1 | Ulixertinib | top60_altered_DNA+RNA_seed | EGFR | 1.321979 | GBM | SNB75 | Erlotinib |
| MVP_top60_DNA_seed | MAPK3 | -1.705442 | OV | IGROV-1 | Ulixertinib | top60_altered_DNA+RNA_seed | PDGFRA | 1.330843 | GBM | SW1088 | Amuvatinib |
| MVP_top60_DNA_seed | EGFR | -1.646551 | OV | UWB1.289 | AZD3759 | top60_altered_DNA+RNA_seed | PDGFRA | 1.340726 | GBM | SNB75 | Amuvatinib |
| MVP_top60_DNA_seed | EGFR | -1.615976 | OV | OVMIU | Lapatinib | top60_altered_DNA+RNA_seed | PDGFRA | 1.356297 | GBM | DK-MG | Amuvatinib |
| MVP_top60_DNA_seed | ERBB2 | -1.615976 | OV | OVMIU | Lapatinib | top60_altered_DNA+RNA_seed | CDK4 | 1.357365 | GBM | LN-405 | Palbociclib |
| MVP_top60_DNA_seed | PIK3CA | -1.59291 | OV | JHOS-4 | AZD8186 | top60_altered_DNA+RNA_seed | EGFR | 1.364496 | GBM | U-118-MG | Pelitinib |
| MVP_top60_DNA_seed | MAPK1 | -1.552045 | OV | IGROV-1 | VX-11e | top60_altered_DNA+RNA_seed | CDK4 | 1.396558 | GBM | Becker | CGP-082996 |
| MVP_top60_DNA_seed | EGFR | -1.546529 | OV | JHOS-4 | AZD3759 | top60_altered_DNA+RNA_seed | EGFR | 1.39991 | GBM | U-87-MG | Pelitinib |
| MVP_top60_DNA_seed | MAPK1 | -1.538248 | OV | Hey | Ulixertinib | top60_altered_DNA+RNA_seed | EGFR | 1.433478 | GBM | SF268 | CI-1033 |
| MVP_top60_DNA_seed | MAPK3 | -1.538248 | OV | Hey | Ulixertinib | top60_altered_DNA+RNA_seed | EGFR | 1.439725 | GBM | D-263MG | Pelitinib |
| MVP_top60_DNA_seed | EGFR | -1.528977 | OV | OVMIU | AZD3759 | top60_altered_DNA+RNA_seed | PDGFRA | 1.501964 | GBM | U-87-MG | Amuvatinib |
| MVP_top60_DNA_seed | MAPK1 | -1.521224 | OV | IGROV-1 | SCH772984 | top60_altered_DNA+RNA_seed | EGFR | 1.516762 | GBM | D-247MG | Pelitinib |
| MVP_top60_DNA_seed | MAPK3 | -1.521224 | OV | IGROV-1 | SCH772984 | top60_altered_DNA+RNA_seed | EGFR | 1.520523 | GBM | SNB75 | Gefitinib |
| MVP_top60_DNA_seed | EGFR | -1.501336 | OV | JHOS-3 | AZD3759 | top60_altered_DNA+RNA_seed | PDGFRA | 1.533048 | GBM | LN-229 | Masitinib |
| MVP_top60_DNA_seed | ATR | -1.475445 | OV | UWB1.289 | VE821 | top60_altered_DNA+RNA_seed | EGFR | 1.541826 | GBM | SW1088 | CI-1033 |
| MVP_top60_DNA_seed | EGFR | -1.43416 | OV | UWB1.289 | Sapitinib | top60_altered_DNA+RNA_seed | EGFR | 1.544437 | GBM | U-118-MG | CI-1033 |
| MVP_top60_DNA_seed | ERBB2 | -1.43416 | OV | UWB1.289 | Sapitinib | top60_altered_DNA+RNA_seed | EGFR | 1.553352 | GBM | D-542MG | Pelitinib |
| MVP_top60_DNA_seed | MAPK1 | -1.417974 | OV | OV-90 | SCH772984 | top60_altered_DNA+RNA_seed | CDK4 | 1.554434 | GBM | M059J | Palbociclib |
| MVP_top60_DNA_seed | MAPK3 | -1.417974 | OV | OV-90 | SCH772984 | top60_altered_DNA+RNA_seed | EGFR | 1.571145 | GBM | D-247MG | CI-1033 |
| MVP_top60_DNA_seed | MAPK1 | -1.394059 | OV | Caov-4 | SCH772984 | top60_altered_DNA+RNA_seed | EGFR | 1.672108 | GBM | AM-38 | AZD8931 |
| MVP_top60_DNA_seed | MAPK3 | -1.394059 | OV | Caov-4 | SCH772984 | top60_altered_DNA+RNA_seed | CDK4 | 1.678389 | GBM | D-542MG | Palbociclib |
| MVP_top60_DNA_seed | PLK1 | -1.383747 | OV | OC-314 | BI-2536 | top60_altered_DNA+RNA_seed | CDK4 | 1.67935 | GBM | CAS-1 | CGP-082996 |
| MVP_top60_DNA_seed | ATR | -1.380449 | OV | JHOS-4 | VE821 | top60_altered_DNA+RNA_seed | PDGFRA | 1.747289 | GBM | D-263MG | Masitinib |
| MVP_top60_DNA_seed | EGFR | -1.318335 | OV | JHOS-4 | Lapatinib | top60_altered_DNA+RNA_seed | EGFR | 1.79721 | GBM | SF295 | CI-1033 |
| MVP_top60_DNA_seed | ERBB2 | -1.318335 | OV | JHOS-4 | Lapatinib | top60_altered_DNA+RNA_seed | EGFR | 1.824225 | GBM | A172 | CI-1033 |
| MVP_top60_DNA_seed | MAPK1 | -1.295009 | OV | Hey | VX-11e | top60_altered_DNA+RNA_seed | EGFR | 1.873966 | GBM | T98G | CI-1033 |
| MVP_top60_DNA_seed | EGFR | -1.265364 | OV | OVMIU | Osimertinib | top60_altered_DNA+RNA_seed | EGFR | 1.886643 | GBM | LNZTA3WT4 | AZD8931 |
| MVP_top60_DNA_seed | PIK3CA | -1.259542 | OV | OV-7 | AZD8186 | top60_altered_DNA+RNA_seed | EGFR | 1.894947 | GBM | DBTRG-05MG | CI-1033 |

| | | | | | | | | | | | |
|---|---|---|---|---|---|---|---|---|---|---|---|
| MVP_top60_DNA_seed | ATR | -1.255075 | OV | UWB1.289 | VE-822 | top60_altered_DNA+RNA_seed | EGFR | 1.929988 | GBM | LN-405 | CI-1033 |
| MVP_top60_DNA_seed | PIK3CA | -1.24505 | OV | UWB1.289 | Alpelisib | top60_altered_DNA+RNA_seed | EGFR | 1.939787 | GBM | SW1088 | CUDC-101 |
| MVP_top60_DNA_seed | EGFR | -1.223736 | OV | OVTOKO | Sapitinib | top60_altered_DNA+RNA_seed | CDK4 | 1.965836 | GBM | SNB75 | Palbociclib |
| MVP_top60_DNA_seed | ERBB2 | -1.223736 | OV | OVTOKO | Sapitinib | top60_altered_DNA+RNA_seed | EGFR | 2.206699 | GBM | LNZTA3WT4 | CUDC-101 |
| MVP_top60_DNA_seed | EGFR | -1.203697 | OV | UWB1.289 | Gefitinib | top60_altered_DNA+RNA_seed | CDK4 | -1.409729 | GBM | D-245MG | Ribociclib |
| MVP_top60_DNA_seed | MAPK1 | -1.203455 | OV | JHOS-3 | SCH772984 | top60_altered_DNA+RNA_seed | TP53 | -1.090076 | GBM | LNZTA3WT4 | MIRA-1 |
| MVP_top60_DNA_seed | MAPK3 | -1.203455 | OV | JHOS-3 | SCH772984 | top60_altered_DNA+RNA_seed | EGFR | -1.068129 | GBM | D-423MG | AZD3759 |
| MVP_top60_DNA_seed | CDK2 | -1.19679 | OV | DOV13 | AZD5438 | top60_altered_DNA+RNA_seed | CDK4 | -1.023605 | GBM | YH-13 | Ribociclib |
| MVP_top60_DNA_seed | MAPK1 | -1.193496 | OV | JHOS-3 | Ulixertinib | top60_altered_DNA+RNA_seed | CDK4 | -1.003435 | GBM | A172 | Ribociclib |
| MVP_top60_DNA_seed | MAPK3 | -1.193496 | OV | JHOS-3 | Ulixertinib | top60_altered_DNA+RNA_seed | TP53 | -0.962093 | GBM | D-566MG | MIRA-1 |
| MVP_top60_DNA_seed | HDAC2 | -1.193233 | OV | DOV13 | Romidepsin | top60_altered_DNA+RNA_seed | EGFR | -0.844443 | GBM | LN-229 | Erlotinib |
| MVP_top60_DNA_seed | EGFR | -1.186493 | OV | JHOS-4 | Osimertinib | top60_altered_DNA+RNA_seed | EGFR | -0.783918 | GBM | 8-MG-BA | Lapatinib |
| MVP_top60_DNA_seed | MAPK1 | -1.141669 | OV | IGROV-1 | Ulixertinib | top60_altered_DNA+RNA_seed | EGFR | -0.74154 | GBM | D-566MG | Lapatinib |
| MVP_top60_DNA_seed | MAPK3 | -1.141669 | OV | IGROV-1 | Ulixertinib | top60_altered_DNA+RNA_seed | EGFR | -0.69609 | GBM | LN-18 | Lapatinib |
| MVP_top60_DNA_seed | MAPK1 | -1.141542 | OV | Hey | SCH772984 | top60_altered_DNA+RNA_seed | EGFR | -0.690795 | GBM | SF126 | Osimertinib |
| MVP_top60_DNA_seed | MAPK3 | -1.141542 | OV | Hey | SCH772984 | top60_altered_DNA+RNA_seed | CDK4 | -0.676501 | GBM | LNZTA3WT4 | Palbociclib |
| MVP_top60_DNA_seed | PLK1 | -1.13958 | OV | IGROV-1 | MK-1775 | top60_altered_DNA+RNA_seed | CDK4 | -0.663907 | GBM | DK-MG | Ribociclib |
| MVP_top60_DNA_seed | CREBBP | -1.134426 | OV | OVCA420 | SGC-CBP30 | top60_altered_DNA+RNA_seed | EGFR | -0.641273 | GBM | D-423MG | Lapatinib |
| MVP_top60_DNA_seed | PIK3CA | -1.124618 | OV | JHOS-2 | AZD8186 | top60_altered_DNA+RNA_seed | TP53 | -0.55726 | GBM | SF295 | MIRA-1 |
| MVP_top60_DNA_seed | EGFR | -1.113572 | OV | OV-17R | AZD3759 | top60_altered_DNA+RNA_seed | EGFR | -0.514164 | GBM | SF126 | Lapatinib |
| MVP_top60_DNA_seed | EGFR | -1.111674 | OV | JHOS-4 | Gefitinib | top60_altered_DNA+RNA_seed | CDK4 | -0.506462 | GBM | D-263MG | Ribociclib |
| MVP_top60_DNA_seed | EGFR | -1.108936 | OV | UWB1.289 | Osimertinib | top60_altered_DNA+RNA_seed | CDK4 | -0.498988 | GBM | A172 | Palbociclib |
| MVP_top60_DNA_seed | ATR | -1.106741 | OV | IGROV-1 | VE821 | top60_altered_DNA+RNA_seed | TP53 | -0.487817 | GBM | GB-1 | MIRA-1 |
| MVP_top60_DNA_seed | ATR | -1.101243 | OV | IGROV-1 | VE-822 | top60_altered_DNA+RNA_seed | EGFR | -0.478769 | GBM | SF295 | AZD3759 |
| MVP_top60_DNA_seed | EGFR | -1.081521 | OV | OVCA433 | Lapatinib | top60_altered_DNA+RNA_seed | EGFR | -0.470561 | GBM | LN-18 | Erlotinib |
| MVP_top60_DNA_seed | ERBB2 | -1.081521 | OV | OVCA433 | Lapatinib | top60_altered_DNA+RNA_seed | EGFR | -0.439711 | GBM | 8-MG-BA | Gefitinib |
| MVP_top60_DNA_seed | EGFR | -1.056988 | OV | JHOS-4 | Sapitinib | top60_altered_DNA+RNA_seed | EGFR | -0.409178 | GBM | YKG-1 | Sapitinib |
| MVP_top60_DNA_seed | ERBB2 | -1.056988 | OV | JHOS-4 | Sapitinib | top60_altered_DNA+RNA_seed | TP53 | -0.396213 | GBM | D-423MG | MIRA-1 |
| MVP_top60_DNA_seed | ATR | -1.05282 | OV | OV-17R | VE-822 | top60_altered_DNA+RNA_seed | EGFR | -0.393865 | GBM | GB-1 | Lapatinib |
| MVP_top60_DNA_seed | PIK3CA | -1.046228 | OV | OVCAR-5 | Alpelisib | top60_altered_DNA+RNA_seed | EGFR | -0.377618 | GBM | YKG-1 | Lapatinib |
| MVP_top60_DNA_seed | ATR | -1.045771 | OV | OVTOKO | VE-822 | top60_altered_DNA+RNA_seed | CDK4 | -0.373544 | GBM | LNZTA3WT4 | Ribociclib |
| MVP_top60_DNA_seed | MAPK1 | -1.030877 | OV | OVCAR-5 | SCH772984 | top60_altered_DNA+RNA_seed | CDK4 | -0.369037 | GBM | Becker | Palbociclib |
| MVP_top60_DNA_seed | MAPK3 | -1.030877 | OV | OVCAR-5 | SCH772984 | top60_altered_DNA+RNA_seed | EGFR | -0.358775 | GBM | SNB75 | AZD3759 |
| MVP_top60_DNA_seed | CDK1 | -1.030729 | OV | Caov-3 | RO-3306 | top60_altered_DNA+RNA_seed | EGFR | -0.356905 | GBM | LN-229 | Sapitinib |
| MVP_top60_DNA_seed | ATR | -1.023467 | OV | OVISE | VE-822 | top60_altered_DNA+RNA_seed | CDK4 | -0.355037 | GBM | YH-13 | Palbociclib |
| MVP_top60_DNA_seed | EGFR | -1.019114 | OV | UWB1.289 | Afatinib | top60_altered_DNA+RNA_seed | EGFR | -0.339388 | GBM | GB-1 | Erlotinib |
| MVP_top60_DNA_seed | ERBB2 | -1.019114 | OV | UWB1.289 | Afatinib | top60_altered_DNA+RNA_seed | EGFR | -0.334036 | GBM | YKG-1 | AZD3759 |
| MVP_top60_DNA_seed | ATR | -1.005852 | OV | OVK-18 | VE821 | top60_altered_DNA+RNA_seed | TP53 | -0.319202 | GBM | KS-1 | MIRA-1 |
| MVP_top60_DNA_seed | CDK2 | -1.00144 | OV | OVK-18 | Dinaciclib | top60_altered_DNA+RNA_seed | EGFR | -0.317899 | GBM | D-423MG | Osimertinib |
| MVP_top60_DNA_seed | CDK1 | -1.00144 | OV | OVK-18 | Dinaciclib | top60_altered_DNA+RNA_seed | TP53 | -0.309237 | GBM | LN-18 | MIRA-1 |
| MVP_top60_DNA_seed | PIK3CA | -0.999203 | OV | OVTOKO | AZD8186 | top60_altered_DNA+RNA_seed | TP53 | -0.290196 | GBM | D-247MG | MIRA-1 |

| | | | | | | | | | | | |
|---|---|---|---|---|---|---|---|---|---|---|---|
| MVP_top60_DNA_seed | CDK2 | -0.984231 | OV | TOV-112D | Dinaciclib | top60_altered_DNA+RNA_seed | CDK4 | -0.271687 | GBM | 42-MG-BA | Ribociclib |
| MVP_top60_DNA_seed | CDK1 | -0.984231 | OV | TOV-112D | Dinaciclib | top60_altered_DNA+RNA_seed | EGFR | -0.26647 | GBM | DK-MG | Osimertinib |
| MVP_top60_DNA_seed | ATR | -0.982576 | OV | OVCAR-5 | VE-822 | top60_altered_DNA+RNA_seed | CDK4 | -0.262916 | GBM | D-263MG | Palbociclib |
| MVP_top60_DNA_seed | CDK2 | -0.980008 | OV | JHOS-4 | Dinaciclib | top60_altered_DNA+RNA_seed | EGFR | -0.258488 | GBM | SF126 | Gefitinib |
| MVP_top60_DNA_seed | CDK1 | -0.980008 | OV | JHOS-4 | Dinaciclib | top60_altered_DNA+RNA_seed | EGFR | -0.252165 | GBM | U-87-MG | Erlotinib |
| MVP_top60_DNA_seed | EGFR | -0.975808 | OV | UWB1.289 | Erlotinib | top60_altered_DNA+RNA_seed | TP53 | -0.246912 | GBM | A172 | MIRA-1 |
| MVP_top60_DNA_seed | ATR | -0.9729 | OV | OV-17R | AZD6738 | top60_altered_DNA+RNA_seed | TP53 | -0.241901 | GBM | YH-13 | MIRA-1 |
| MVP_top60_DNA_seed | MAPK1 | -0.969957 | OV | SW626 | SCH772984 | top60_altered_DNA+RNA_seed | EGFR | -0.235615 | GBM | LN-229 | Osimertinib |
| MVP_top60_DNA_seed | MAPK3 | -0.969957 | OV | SW626 | SCH772984 | top60_altered_DNA+RNA_seed | CDK4 | -0.231495 | GBM | D-566MG | Palbociclib |
| MVP_top60_DNA_seed | EGFR | -0.969223 | OV | OV-17R | Lapatinib | top60_altered_DNA+RNA_seed | EGFR | -0.23003 | GBM | AM-38 | Erlotinib |
| MVP_top60_DNA_seed | ERBB2 | -0.969223 | OV | OV-17R | Lapatinib | top60_altered_DNA+RNA_seed | EGFR | -0.225836 | GBM | 42-MG-BA | Lapatinib |
| MVP_top60_DNA_seed | MAPK1 | -0.968864 | OV | JHOS-4 | VX-11e | top60_altered_DNA+RNA_seed | CDK4 | -0.208943 | GBM | SF295 | Palbociclib |
| MVP_top60_DNA_seed | ATR | -0.955839 | OV | Caov-3 | VE-822 | top60_altered_DNA+RNA_seed | TP53 | -0.206846 | GBM | 8-MG-BA | MIRA-1 |
| MVP_top60_DNA_seed | CDK2 | -0.95154 | OV | TOV-21G | AZD5438 | top60_altered_DNA+RNA_seed | EGFR | -0.19335 | GBM | D-245MG | Sapitinib |
| MVP_top60_DNA_seed | CDK2 | -0.948231 | OV | UWB1.289 | MK-8776 | top60_altered_DNA+RNA_seed | CDK4 | -0.181636 | GBM | YKG-1 | Palbociclib |
| MVP_top60_DNA_seed | PIK3CA | -0.935568 | OV | OVCA420 | Alpelisib | top60_altered_DNA+RNA_seed | EGFR | -0.17646 | GBM | SW1088 | Sapitinib |
| MVP_top60_DNA_seed | PLK1 | -0.918473 | OV | TOV-112D | BI-2536 | top60_altered_DNA+RNA_seed | TP53 | -0.170488 | GBM | 42-MG-BA | MIRA-1 |
| MVP_top60_DNA_seed | EGFR | -0.907491 | OV | OC-314 | Afatinib | top60_altered_DNA+RNA_seed | TP53 | -0.162285 | GBM | D-263MG | MIRA-1 |
| MVP_top60_DNA_seed | ERBB2 | -0.907491 | OV | OC-314 | Afatinib | top60_altered_DNA+RNA_seed | EGFR | -0.138111 | GBM | 42-MG-BA | Osimertinib |
| MVP_top60_DNA_seed | PLK1 | -0.899786 | OV | UWB1.289 | MK-1775 | top60_altered_DNA+RNA_seed | CDK4 | -0.137955 | GBM | AM-38 | Palbociclib |
| MVP_top60_DNA_seed | PIK3CA | -0.899656 | OV | TOV-112D | Buparlisib | top60_altered_DNA+RNA_seed | EGFR | -0.12271 | GBM | 8-MG-BA | Erlotinib |
| MVP_top60_DNA_seed | EGFR | -0.880526 | OV | TOV-21G | Erlotinib | top60_altered_DNA+RNA_seed | CDK4 | -0.12219 | GBM | D-566MG | Ribociclib |
| MVP_top60_DNA_seed | EGFR | -0.880265 | OV | OV-7 | AZD3759 | top60_altered_DNA+RNA_seed | EGFR | -0.116719 | GBM | A172 | Afatinib |
| MVP_top60_DNA_seed | EGFR | -0.875835 | OV | OVISE | Sapitinib | top60_altered_DNA+RNA_seed | CDK4 | -0.102853 | GBM | D-247MG | Palbociclib |
| MVP_top60_DNA_seed | ERBB2 | -0.875835 | OV | OVISE | Sapitinib | top60_altered_DNA+RNA_seed | EGFR | -0.0997 | GBM | CAS-1 | Lapatinib |
| MVP_top60_DNA_seed | MAPK1 | -0.874792 | OV | JHOS-3 | VX-11e | top60_altered_DNA+RNA_seed | CDK4 | -0.086184 | GBM | KS-1 | Ribociclib |
| MVP_top60_DNA_seed | ATR | -0.873599 | OV | JHOS-4 | AZD6738 | top60_altered_DNA+RNA_seed | EGFR | -0.082638 | GBM | SNB75 | Osimertinib |
| MVP_top60_DNA_seed | TP53 | -0.87162 | OV | OVCA433 | MIRA-1 | top60_altered_DNA+RNA_seed | EGFR | -0.06101 | GBM | LN-229 | Gefitinib |
| MVP_top60_DNA_seed | EGFR | -0.870149 | OV | SK-OV-3 | Osimertinib | top60_altered_DNA+RNA_seed | EGFR | -0.059066 | GBM | 8-MG-BA | Sapitinib |
| MVP_top60_DNA_seed | PLK1 | -0.867592 | OV | JHOS-4 | BI-2536 | top60_altered_DNA+RNA_seed | EGFR | -0.057422 | GBM | A172 | Gefitinib |
| MVP_top60_DNA_seed | EGFR | -0.864306 | OV | UWB1.289 | Lapatinib | top60_altered_DNA+RNA_seed | EGFR | -0.055395 | GBM | D-542MG | Lapatinib |
| MVP_top60_DNA_seed | ERBB2 | -0.864306 | OV | UWB1.289 | Lapatinib | top60_altered_DNA+RNA_seed | CDK4 | -0.053038 | GBM | U251 | Palbociclib |
| MVP_top60_DNA_seed | PIK3CA | -0.862955 | OV | OC-314 | AZD8186 | top60_altered_DNA+RNA_seed | EGFR | -0.052 | GBM | 42-MG-BA | Afatinib |
| MVP_top60_DNA_seed | ATR | -0.861319 | OV | OC-314 | VE821 | top60_altered_DNA+RNA_seed | EGFR | -0.050785 | GBM | A172 | Osimertinib |
| MVP_top60_DNA_seed | EGFR | -0.85374 | OV | OC-314 | Gefitinib | top60_altered_DNA+RNA_seed | EGFR | -0.042264 | GBM | D-566MG | Erlotinib |
| MVP_top60_DNA_seed | MAPK1 | -0.847011 | OV | Hey | Ulixertinib | top60_altered_DNA+RNA_seed | TP53 | -0.03173 | GBM | M059J | MIRA-1 |
| MVP_top60_DNA_seed | MAPK3 | -0.847011 | OV | Hey | Ulixertinib | top60_altered_DNA+RNA_seed | EGFR | -0.0309 | GBM | SF295 | Osimertinib |
| MVP_top60_DNA_seed | PIK3CA | -0.845269 | OV | JHOS-4 | GNE-317 | top60_altered_DNA+RNA_seed | CDK4 | -0.030002 | GBM | U-87-MG | Palbociclib |
| MVP_top60_DNA_seed | EGFR | -0.842177 | OV | SK-OV-3 | Sapitinib | top60_altered_DNA+RNA_seed | CDK4 | -0.021426 | GBM | 8-MG-BA | Palbociclib |
| MVP_top60_DNA_seed | ERBB2 | -0.842177 | OV | SK-OV-3 | Sapitinib | top60_altered_DNA+RNA_seed | EGFR | -0.015156 | GBM | U-87-MG | AZD3759 |
| MVP_top60_DNA_seed | EGFR | -0.837256 | OV | OVISE | Erlotinib | top60_altered_DNA+RNA_seed | EGFR | -0.014364 | GBM | YH-13 | Erlotinib |

| | | | | | | | | | | | |
|---|---|---|---|---|---|---|---|---|---|---|---|
| MVP_top60_DNA_seed | ATR | -0.835629 | OV | Caov-3 | VE821 | top60_altered_DNA+RNA_seed | CDK4 | -0.012447 | GBM | 8-MG-BA | Ribociclib |
| MVP_top60_DNA_seed | EGFR | -0.831773 | OV | JHOS-2 | Erlotinib | top60_altered_DNA+RNA_seed | EGFR | -0.011375 | GBM | YKG-1 | Osimertinib |
| MVP_top60_DNA_seed | PLK1 | -0.830113 | OV | Caov-4 | MK-1775 | top60_altered_DNA+RNA_seed | EGFR | -0.007482 | GBM | YH-13 | Osimertinib |
| MVP_top60_DNA_seed | ATR | -0.829194 | OV | OVISE | AZD6738 | top60_altered_DNA+RNA_seed | EGFR | -0.005381 | GBM | SF126 | Sapitinib |
| MVP_top60_DNA_seed | MAPK1 | -0.829057 | OV | OVMIU | Ulixertinib | top60_altered_DNA+RNA_seed | EGFR | 0.002614 | GBM | M059J | Erlotinib |
| MVP_top60_DNA_seed | MAPK3 | -0.829057 | OV | OVMIU | Ulixertinib | top60_altered_DNA+RNA_seed | EGFR | 0.004394 | GBM | U251 | Erlotinib |
| MVP_top60_DNA_seed | EGFR | -0.81937 | OV | OVCAR-8 | Erlotinib | top60_altered_DNA+RNA_seed | TP53 | 0.007679 | GBM | T98G | MIRA-1 |
| MVP_top60_DNA_seed | CREBBP | -0.811154 | OV | TOV-112D | SGC-CBP30 | top60_altered_DNA+RNA_seed | EGFR | 0.008988 | GBM | KS-1 | Sapitinib |
| MVP_top60_DNA_seed | MAPK1 | -0.811027 | OV | Caov-4 | Ulixertinib | top60_altered_DNA+RNA_seed | EGFR | 0.010215 | GBM | SF268 | Erlotinib |
| MVP_top60_DNA_seed | MAPK3 | -0.811027 | OV | Caov-4 | Ulixertinib | top60_altered_DNA+RNA_seed | EGFR | 0.027993 | GBM | A172 | Lapatinib |
| MVP_top60_DNA_seed | PIK3CA | -0.808473 | OV | OC-314 | GNE-317 | top60_altered_DNA+RNA_seed | EGFR | 0.029904 | GBM | YKG-1 | Gefitinib |
| MVP_top60_DNA_seed | PIK3CA | -0.788485 | OV | JHOS-3 | Alpelisib | top60_altered_DNA+RNA_seed | EGFR | 0.030789 | GBM | 8-MG-BA | Afatinib |
| MVP_top60_DNA_seed | PIK3CA | -0.787397 | OV | JHOS-2 | GNE-317 | top60_altered_DNA+RNA_seed | CDK4 | 0.031313 | GBM | SW1088 | Palbociclib |
| MVP_top60_DNA_seed | MAPK1 | -0.778775 | OV | OV-90 | Ulixertinib | top60_altered_DNA+RNA_seed | EGFR | 0.035055 | GBM | LN-18 | Sapitinib |
| MVP_top60_DNA_seed | MAPK3 | -0.778775 | OV | OV-90 | Ulixertinib | top60_altered_DNA+RNA_seed | EGFR | 0.036472 | GBM | DK-MG | Erlotinib |
| MVP_top60_DNA_seed | PIK3CA | -0.778659 | OV | OVCAR-3 | AZD8186 | top60_altered_DNA+RNA_seed | TP53 | 0.063766 | GBM | SNB75 | MIRA-1 |
| MVP_top60_DNA_seed | PLK1 | -0.777975 | OV | OVCAR-8 | MK-1775 | top60_altered_DNA+RNA_seed | EGFR | 0.065361 | GBM | D-566MG | Gefitinib |
| MVP_top60_DNA_seed | ATR | -0.7643 | OV | JHOS-4 | VE-822 | top60_altered_DNA+RNA_seed | EGFR | 0.067748 | GBM | LNZTA3WT4 | Afatinib |
| MVP_top60_DNA_seed | PIK3CA | -0.761504 | OV | OV-17R | Alpelisib | top60_altered_DNA+RNA_seed | CDK4 | 0.076872 | GBM | D-423MG | Palbociclib |
| MVP_top60_DNA_seed | ATR | -0.761042 | OV | IGROV-1 | AZD6738 | top60_altered_DNA+RNA_seed | CDK4 | 0.080189 | GBM | D-245MG | Palbociclib |
| MVP_top60_DNA_seed | TP53 | -0.752262 | OV | JHOS-4 | MIRA-1 | top60_altered_DNA+RNA_seed | TP53 | 0.090294 | GBM | SF126 | MIRA-1 |
| MVP_top60_DNA_seed | CDK2 | -0.747499 | OV | Caov-3 | MK-8776 | top60_altered_DNA+RNA_seed | EGFR | 0.091768 | GBM | U-118-MG | AZD3759 |
| MVP_top60_DNA_seed | PLK1 | -0.740897 | OV | OV-17R | MK-1775 | top60_altered_DNA+RNA_seed | EGFR | 0.100732 | GBM | SF295 | Lapatinib |
| MVP_top60_DNA_seed | EGFR | -0.730247 | OV | OC-314 | Lapatinib | top60_altered_DNA+RNA_seed | EGFR | 0.110759 | GBM | U-118-MG | Osimertinib |
| MVP_top60_DNA_seed | ERBB2 | -0.730247 | OV | OC-314 | Lapatinib | top60_altered_DNA+RNA_seed | EGFR | 0.114846 | GBM | SK-MG-1 | Afatinib |
| MVP_top60_DNA_seed | PLK1 | -0.72156 | OV | OVCA433 | MK-1775 | top60_altered_DNA+RNA_seed | EGFR | 0.115143 | GBM | DK-MG | Sapitinib |
| MVP_top60_DNA_seed | MAPK1 | -0.717897 | OV | JHOS-4 | SCH772984 | top60_altered_DNA+RNA_seed | EGFR | 0.118834 | GBM | D-542MG | Osimertinib |
| MVP_top60_DNA_seed | MAPK3 | -0.717897 | OV | JHOS-4 | SCH772984 | top60_altered_DNA+RNA_seed | EGFR | 0.124166 | GBM | SF126 | AZD3759 |
| MVP_top60_DNA_seed | EGFR | -0.7148 | OV | JHOS-4 | Afatinib | top60_altered_DNA+RNA_seed | EGFR | 0.12491 | GBM | AM-38 | AZD3759 |
| MVP_top60_DNA_seed | ERBB2 | -0.7148 | OV | JHOS-4 | Afatinib | top60_altered_DNA+RNA_seed | CDK4 | 0.128484 | GBM | SF268 | Ribociclib |
| MVP_top60_DNA_seed | PIK3CA | -0.712748 | OV | JHOS-4 | Buparlisib | top60_altered_DNA+RNA_seed | EGFR | 0.132447 | GBM | D-566MG | AZD3759 |
| MVP_top60_DNA_seed | ATR | -0.712223 | OV | TOV-21G | VE-822 | top60_altered_DNA+RNA_seed | EGFR | 0.134693 | GBM | LNZTA3WT4 | Lapatinib |
| MVP_top60_DNA_seed | PIK3CA | -0.701671 | OV | OV-17R | AZD8186 | top60_altered_DNA+RNA_seed | EGFR | 0.135413 | GBM | T98G | Gefitinib |
| MVP_top60_DNA_seed | PIK3CA | -0.699598 | OV | OVISE | Alpelisib | top60_altered_DNA+RNA_seed | EGFR | 0.148283 | GBM | DK-MG | Afatinib |
| MVP_top60_DNA_seed | PLK1 | -0.696988 | OV | DOV13 | MK-1775 | top60_altered_DNA+RNA_seed | EGFR | 0.149432 | GBM | D-263MG | AZD3759 |
| MVP_top60_DNA_seed | EGFR | -0.693524 | OV | JHOS-4 | Erlotinib | top60_altered_DNA+RNA_seed | CDK4 | 0.152862 | GBM | SF126 | Ribociclib |
| MVP_top60_DNA_seed | EGFR | -0.68574 | OV | DOV13 | Sapitinib | top60_altered_DNA+RNA_seed | CDK4 | 0.169151 | GBM | D-423MG | Ribociclib |
| MVP_top60_DNA_seed | ERBB2 | -0.68574 | OV | DOV13 | Sapitinib | top60_altered_DNA+RNA_seed | CDK4 | 0.169831 | GBM | LN-18 | Ribociclib |
| MVP_top60_DNA_seed | MAPK1 | -0.683593 | OV | EFO-21 | Ulixertinib | top60_altered_DNA+RNA_seed | EGFR | 0.17414 | GBM | U-118-MG | Erlotinib |
| MVP_top60_DNA_seed | MAPK3 | -0.683593 | OV | EFO-21 | Ulixertinib | top60_altered_DNA+RNA_seed | EGFR | 0.175123 | GBM | D-423MG | Gefitinib |
| MVP_top60_DNA_seed | MAPK1 | -0.677495 | OV | JHOS-4 | Ulixertinib | top60_altered_DNA+RNA_seed | EGFR | 0.177191 | GBM | SF268 | Afatinib |

| | | | | | | | | | | | |
|---|---|---|---|---|---|---|---|---|---|---|---|
| MVP_top60_DNA_seed | MAPK3 | -0.677495 | OV | JHOS-4 | Ulixertinib | top60_altered_DNA+RNA_seed | EGFR | 0.180797 | GBM | A172 | AZD3759 |
| MVP_top60_DNA_seed | PIK3CA | -0.677243 | OV | Hey | GNE-317 | top60_altered_DNA+RNA_seed | EGFR | 0.181391 | GBM | A172 | Erlotinib |
| MVP_top60_DNA_seed | CDK2 | -0.674952 | OV | OVCA433 | Dinaciclib | top60_altered_DNA+RNA_seed | EGFR | 0.183624 | GBM | D-245MG | Afatinib |
| MVP_top60_DNA_seed | CDK1 | -0.674952 | OV | OVCA433 | Dinaciclib | top60_altered_DNA+RNA_seed | CDK4 | 0.18412 | GBM | DK-MG | Palbociclib |
| MVP_top60_DNA_seed | ATR | -0.674189 | OV | OV-17R | VE821 | top60_altered_DNA+RNA_seed | EGFR | 0.192616 | GBM | D-245MG | Osimertinib |
| MVP_top60_DNA_seed | ATR | -0.673238 | OV | OC-314 | AZD6738 | top60_altered_DNA+RNA_seed | EGFR | 0.192717 | GBM | SF268 | Gefitinib |
| MVP_top60_DNA_seed | PIK3CA | -0.671725 | OV | OVCA433 | Alpelisib | top60_altered_DNA+RNA_seed | CDK4 | 0.195795 | GBM | U-118-MG | Palbociclib |
| MVP_top60_DNA_seed | ATR | -0.669794 | OV | Caov-3 | AZD6738 | top60_altered_DNA+RNA_seed | EGFR | 0.197386 | GBM | D-247MG | Osimertinib |
| MVP_top60_DNA_seed | EGFR | -0.668982 | OV | TOV-21G | Sapitinib | top60_altered_DNA+RNA_seed | EGFR | 0.200155 | GBM | AM-38 | Lapatinib |
| MVP_top60_DNA_seed | ERBB2 | -0.668982 | OV | TOV-21G | Sapitinib | top60_altered_DNA+RNA_seed | EGFR | 0.209204 | GBM | U251 | AZD3759 |
| MVP_top60_DNA_seed | ATR | -0.665142 | OV | TOV-21G | AZD6738 | top60_altered_DNA+RNA_seed | EGFR | 0.20971 | GBM | YH-13 | Afatinib |
| MVP_top60_DNA_seed | CDK2 | -0.660164 | OV | TOV-21G | Dinaciclib | top60_altered_DNA+RNA_seed | EGFR | 0.225471 | GBM | M059J | Afatinib |
| MVP_top60_DNA_seed | CDK1 | -0.660164 | OV | TOV-21G | Dinaciclib | top60_altered_DNA+RNA_seed | EGFR | 0.232195 | GBM | M059J | Lapatinib |
| MVP_top60_DNA_seed | ATR | -0.655233 | OV | DOV13 | VE821 | top60_altered_DNA+RNA_seed | EGFR | 0.233071 | GBM | DK-MG | AZD3759 |
| MVP_top60_DNA_seed | CDK2 | -0.653564 | OV | Caov-4 | MK-8776 | top60_altered_DNA+RNA_seed | EGFR | 0.237068 | GBM | U-87-MG | Gefitinib |
| MVP_top60_DNA_seed | PIK3CA | -0.651845 | OV | OVCAR-5 | GNE-317 | top60_altered_DNA+RNA_seed | EGFR | 0.241009 | GBM | U-87-MG | Sapitinib |
| MVP_top60_DNA_seed | ATR | -0.6407 | OV | TOV-112D | VE821 | top60_altered_DNA+RNA_seed | EGFR | 0.255549 | GBM | LN-229 | AZD3759 |
| MVP_top60_DNA_seed | ATR | -0.640093 | OV | Caov-4 | AZD6738 | top60_altered_DNA+RNA_seed | CDK4 | 0.261538 | GBM | SF295 | Ribociclib |
| MVP_top60_DNA_seed | PIK3CA | -0.631969 | OV | Hey | Buparlisib | top60_altered_DNA+RNA_seed | TP53 | 0.264647 | GBM | YKG-1 | MIRA-1 |
| MVP_top60_DNA_seed | PIK3CA | -0.629394 | OV | OVK-18 | GNE-317 | top60_altered_DNA+RNA_seed | EGFR | 0.269102 | GBM | 42-MG-BA | Sapitinib |
| MVP_top60_DNA_seed | ATR | -0.628828 | OV | OVK-18 | AZD6738 | top60_altered_DNA+RNA_seed | EGFR | 0.27457 | GBM | SW1088 | Erlotinib |
| MVP_top60_DNA_seed | EGFR | -0.62548 | OV | OVISE | Osimertinib | top60_altered_DNA+RNA_seed | EGFR | 0.275207 | GBM | D-542MG | Gefitinib |
| MVP_top60_DNA_seed | PLK1 | -0.624533 | OV | JHOS-4 | MK-1775 | top60_altered_DNA+RNA_seed | EGFR | 0.280485 | GBM | Becker | AZD3759 |
| MVP_top60_DNA_seed | HDAC2 | -0.621595 | OV | SW626 | Romidepsin | top60_altered_DNA+RNA_seed | CDK4 | 0.290798 | GBM | U-87-MG | Ribociclib |
| MVP_top60_DNA_seed | PLK1 | -0.618885 | OV | OVCAR-8 | BI-2536 | top60_altered_DNA+RNA_seed | EGFR | 0.293897 | GBM | SW1088 | Osimertinib |
| MVP_top60_DNA_seed | PLK1 | -0.617369 | OV | Caov-3 | MK-1775 | top60_altered_DNA+RNA_seed | EGFR | 0.294884 | GBM | T98G | Lapatinib |
| MVP_top60_DNA_seed | CDK2 | -0.602219 | OV | OVCAR-8 | MK-8776 | top60_altered_DNA+RNA_seed | EGFR | 0.296246 | GBM | CAS-1 | Sapitinib |
| MVP_top60_DNA_seed | HDAC2 | -0.601464 | OV | TOV-21G | Romidepsin | top60_altered_DNA+RNA_seed | TP53 | 0.299196 | GBM | AM-38 | MIRA-1 |
| MVP_top60_DNA_seed | PIK3CA | -0.600582 | OV | OV-7 | Alpelisib | top60_altered_DNA+RNA_seed | CDK4 | 0.305782 | GBM | CAS-1 | Palbociclib |
| MVP_top60_DNA_seed | EGFR | -0.595363 | OV | Caov-4 | Lapatinib | top60_altered_DNA+RNA_seed | TP53 | 0.3112 | GBM | D-542MG | MIRA-1 |
| MVP_top60_DNA_seed | ERBB2 | -0.595363 | OV | Caov-4 | Lapatinib | top60_altered_DNA+RNA_seed | TP53 | 0.313675 | GBM | Becker | MIRA-1 |
| MVP_top60_DNA_seed | HDAC2 | -0.595187 | OV | OVCAR-5 | Romidepsin | top60_altered_DNA+RNA_seed | EGFR | 0.327969 | GBM | D-423MG | Erlotinib |
| MVP_top60_DNA_seed | ATR | -0.593255 | OV | DOV13 | AZD6738 | top60_altered_DNA+RNA_seed | EGFR | 0.334174 | GBM | 8-MG-BA | AZD3759 |
| MVP_top60_DNA_seed | MAPK1 | -0.592917 | OV | OV-7 | VX-11e | top60_altered_DNA+RNA_seed | EGFR | 0.334651 | GBM | D-423MG | Afatinib |
| MVP_top60_DNA_seed | CDK2 | -0.590846 | OV | TOV-112D | AZD5438 | top60_altered_DNA+RNA_seed | EGFR | 0.337738 | GBM | D-566MG | Osimertinib |
| MVP_top60_DNA_seed | EGFR | -0.585671 | OV | OC-314 | Osimertinib | top60_altered_DNA+RNA_seed | EGFR | 0.364198 | GBM | SF268 | Sapitinib |
| MVP_top60_DNA_seed | PLK1 | -0.582124 | OV | OC-314 | MK-1775 | top60_altered_DNA+RNA_seed | EGFR | 0.364904 | GBM | D-247MG | Sapitinib |
| MVP_top60_DNA_seed | HDAC2 | -0.576194 | OV | Hey | Romidepsin | top60_altered_DNA+RNA_seed | EGFR | 0.366709 | GBM | AM-38 | Gefitinib |
| MVP_top60_DNA_seed | ATR | -0.571755 | OV | UWB1.289 | AZD6738 | top60_altered_DNA+RNA_seed | CDK4 | 0.375127 | GBM | D-392MG | Ribociclib |
| MVP_top60_DNA_seed | EGFR | -0.571298 | OV | OVTOKO | Erlotinib | top60_altered_DNA+RNA_seed | EGFR | 0.37903 | GBM | SF295 | Gefitinib |
| MVP_top60_DNA_seed | MAPK1 | -0.56809 | OV | Caov-4 | VX-11e | top60_altered_DNA+RNA_seed | EGFR | 0.379162 | GBM | YKG-1 | Erlotinib |

| | | | | | | | | | | | |
|---|---|---|---|---|---|---|---|---|---|---|---|
| MVP_top60_DNA_seed | EGFR | -0.56631 | OV | JHOS-3 | Sapitinib | top60_altered_DNA+RNA_seed | EGFR | 0.382277 | GBM | D-247MG | Lapatinib |
| MVP_top60_DNA_seed | ERBB2 | -0.56631 | OV | JHOS-3 | Sapitinib | top60_altered_DNA+RNA_seed | EGFR | 0.384935 | GBM | 42-MG-BA | AZD3759 |
| MVP_top60_DNA_seed | EGFR | -0.563138 | OV | OVCAR-8 | Gefitinib | top60_altered_DNA+RNA_seed | CDK4 | 0.385871 | GBM | SF126 | Palbociclib |
| MVP_top60_DNA_seed | EGFR | -0.562711 | OV | JHOS-2 | Afatinib | top60_altered_DNA+RNA_seed | EGFR | 0.3876 | GBM | KS-1 | Afatinib |
| MVP_top60_DNA_seed | ERBB2 | -0.562711 | OV | JHOS-2 | Afatinib | top60_altered_DNA+RNA_seed | EGFR | 0.393013 | GBM | YH-13 | Sapitinib |
| MVP_top60_DNA_seed | ATR | -0.558595 | OV | OV-7 | VE821 | top60_altered_DNA+RNA_seed | EGFR | 0.393487 | GBM | SK-MG-1 | Osimertinib |
| MVP_top60_DNA_seed | MAPK1 | -0.558441 | OV | OVCA420 | VX-11e | top60_altered_DNA+RNA_seed | CDK4 | 0.394151 | GBM | D-542MG | Ribociclib |
| MVP_top60_DNA_seed | HDAC2 | -0.554065 | OV | OVCAR-8 | Romidepsin | top60_altered_DNA+RNA_seed | EGFR | 0.400711 | GBM | D-245MG | Gefitinib |
| MVP_top60_DNA_seed | PIK3CA | -0.550588 | OV | UWB1.289 | AZD8186 | top60_altered_DNA+RNA_seed | EGFR | 0.402606 | GBM | SK-MG-1 | Erlotinib |
| MVP_top60_DNA_seed | MAPK1 | -0.549852 | OV | OVCA433 | SCH772984 | top60_altered_DNA+RNA_seed | CDK4 | 0.410973 | GBM | Becker | Ribociclib |
| MVP_top60_DNA_seed | MAPK3 | -0.549852 | OV | OVCA433 | SCH772984 | top60_altered_DNA+RNA_seed | EGFR | 0.412974 | GBM | D-566MG | Sapitinib |
| MVP_top60_DNA_seed | PLK1 | -0.546608 | OV | TOV-21G | MK-1775 | top60_altered_DNA+RNA_seed | EGFR | 0.417341 | GBM | U-118-MG | Gefitinib |
| MVP_top60_DNA_seed | EGFR | -0.544116 | OV | OVCAR-8 | Lapatinib | top60_altered_DNA+RNA_seed | EGFR | 0.431461 | GBM | LNZTA3WT4 | AZD3759 |
| MVP_top60_DNA_seed | ERBB2 | -0.544116 | OV | OVCAR-8 | Lapatinib | top60_altered_DNA+RNA_seed | EGFR | 0.447714 | GBM | D-423MG | Sapitinib |
| MVP_top60_DNA_seed | EGFR | -0.537828 | OV | DOV13 | Afatinib | top60_altered_DNA+RNA_seed | EGFR | 0.455249 | GBM | M059J | AZD3759 |
| MVP_top60_DNA_seed | ERBB2 | -0.537828 | OV | DOV13 | Afatinib | top60_altered_DNA+RNA_seed | TP53 | 0.47018 | GBM | D-245MG | MIRA-1 |
| MVP_top60_DNA_seed | TP53 | -0.53417 | OV | TOV-21G | MIRA-1 | top60_altered_DNA+RNA_seed | CDK4 | 0.472011 | GBM | D-392MG | Palbociclib |
| MVP_top60_DNA_seed | EGFR | -0.532761 | OV | OVCA433 | Gefitinib | top60_altered_DNA+RNA_seed | EGFR | 0.475655 | GBM | A172 | Sapitinib |
| MVP_top60_DNA_seed | PLK1 | -0.532313 | OV | IGROV-1 | BI-2536 | top60_altered_DNA+RNA_seed | CDK4 | 0.480546 | GBM | SK-MG-1 | Palbociclib |
| MVP_top60_DNA_seed | MAPK1 | -0.520294 | OV | OVCAR-8 | Ulixertinib | top60_altered_DNA+RNA_seed | EGFR | 0.4854 | GBM | KS-1 | Erlotinib |
| MVP_top60_DNA_seed | MAPK3 | -0.520294 | OV | OVCAR-8 | Ulixertinib | top60_altered_DNA+RNA_seed | TP53 | 0.49485 | GBM | U251 | MIRA-1 |
| MVP_top60_DNA_seed | PLK1 | -0.519958 | OV | Caov-3 | BI-2536 | top60_altered_DNA+RNA_seed | EGFR | 0.503098 | GBM | D-263MG | Gefitinib |
| MVP_top60_DNA_seed | CDK2 | -0.515419 | OV | DOV13 | Dinaciclib | top60_altered_DNA+RNA_seed | CDK4 | 0.505729 | GBM | KS-1 | Palbociclib |
| MVP_top60_DNA_seed | CDK1 | -0.515419 | OV | DOV13 | Dinaciclib | top60_altered_DNA+RNA_seed | EGFR | 0.506116 | GBM | LN-229 | Afatinib |
| MVP_top60_DNA_seed | EGFR | -0.512875 | OV | OVCAR-3 | Erlotinib | top60_altered_DNA+RNA_seed | EGFR | 0.509287 | GBM | U-87-MG | Osimertinib |
| MVP_top60_DNA_seed | EGFR | -0.51085 | OV | Caov-4 | Erlotinib | top60_altered_DNA+RNA_seed | EGFR | 0.510367 | GBM | SNB75 | Gefitinib |
| MVP_top60_DNA_seed | EGFR | -0.508799 | OV | OC-314 | AZD3759 | top60_altered_DNA+RNA_seed | EGFR | 0.514688 | GBM | SW1088 | Afatinib |
| MVP_top60_DNA_seed | ATR | -0.508452 | OV | OVCA433 | VE821 | top60_altered_DNA+RNA_seed | CDK4 | 0.515567 | GBM | T98G | Palbociclib |
| MVP_top60_DNA_seed | CDK1 | -0.506704 | OV | OV-7 | RO-3306 | top60_altered_DNA+RNA_seed | EGFR | 0.520009 | GBM | LN-405 | Gefitinib |
| MVP_top60_DNA_seed | EGFR | -0.505699 | OV | JHOS-3 | Gefitinib | top60_altered_DNA+RNA_seed | EGFR | 0.526224 | GBM | D-542MG | Afatinib |
| MVP_top60_DNA_seed | PLK1 | -0.504499 | OV | OVCAR-3 | MK-1775 | top60_altered_DNA+RNA_seed | CDK4 | 0.527481 | GBM | AM-38 | Ribociclib |
| MVP_top60_DNA_seed | PLK1 | -0.50092 | OV | DOV13 | BI-2536 | top60_altered_DNA+RNA_seed | EGFR | 0.530148 | GBM | D-542MG | Sapitinib |
| MVP_top60_DNA_seed | EGFR | -0.498893 | OV | OC-314 | Sapitinib | top60_altered_DNA+RNA_seed | CDK4 | 0.533868 | GBM | LN-18 | Palbociclib |
| MVP_top60_DNA_seed | ERBB2 | -0.498893 | OV | OC-314 | Sapitinib | top60_altered_DNA+RNA_seed | EGFR | 0.537031 | GBM | 42-MG-BA | Erlotinib |
| MVP_top60_DNA_seed | ATR | -0.495835 | OV | JHOS-3 | VE821 | top60_altered_DNA+RNA_seed | CDK4 | 0.537833 | GBM | D-247MG | Ribociclib |
| MVP_top60_DNA_seed | EGFR | -0.482246 | OV | JHOS-2 | Osimertinib | top60_altered_DNA+RNA_seed | EGFR | 0.546951 | GBM | DK-MG | Lapatinib |
| MVP_top60_DNA_seed | PIK3CA | -0.479701 | OV | JHOS-2 | Alpelisib | top60_altered_DNA+RNA_seed | EGFR | 0.547239 | GBM | SF268 | Lapatinib |
| MVP_top60_DNA_seed | CDK1 | -0.474284 | OV | OV-17R | RO-3306 | top60_altered_DNA+RNA_seed | CDK4 | 0.557772 | GBM | SK-MG-1 | Ribociclib |
| MVP_top60_DNA_seed | EGFR | -0.46862 | OV | Hey | Erlotinib | top60_altered_DNA+RNA_seed | EGFR | 0.56236 | GBM | 8-MG-BA | Osimertinib |
| MVP_top60_DNA_seed | EGFR | -0.458849 | OV | TOV-112D | Osimertinib | top60_altered_DNA+RNA_seed | EGFR | 0.575883 | GBM | D-542MG | AZD3759 |
| MVP_top60_DNA_seed | PIK3CA | -0.457196 | OV | OAW-42 | Alpelisib | top60_altered_DNA+RNA_seed | EGFR | 0.579175 | GBM | YH-13 | Lapatinib |

| | | | | | | | | | | | |
|---|---|---|---|---|---|---|---|---|---|---|---|
| MVP_top60_DNA_seed | EGFR | -0.455596 | OV | JHOS-2 | Sapitinib | top60_altered_DNA+RNA_seed | EGFR | 0.587528 | GBM | 42-MG-BA | Gefitinib |
| MVP_top60_DNA_seed | ERBB2 | -0.455596 | OV | JHOS-2 | Sapitinib | top60_altered_DNA+RNA_seed | EGFR | 0.590257 | GBM | M059J | Sapitinib |
| MVP_top60_DNA_seed | EGFR | -0.452952 | OV | TOV-112D | AZD3759 | top60_altered_DNA+RNA_seed | EGFR | 0.595524 | GBM | SK-MG-1 | AZD3759 |
| MVP_top60_DNA_seed | PLK1 | -0.449972 | OV | OVISE | MK-1775 | top60_altered_DNA+RNA_seed | EGFR | 0.595763 | GBM | D-542MG | Erlotinib |
| MVP_top60_DNA_seed | MAPK1 | -0.438041 | OV | Caov-4 | Ulixertinib | top60_altered_DNA+RNA_seed | EGFR | 0.595809 | GBM | D-245MG | Lapatinib |
| MVP_top60_DNA_seed | MAPK3 | -0.438041 | OV | Caov-4 | Ulixertinib | top60_altered_DNA+RNA_seed | CDK4 | 0.605292 | GBM | GB-1 | Palbociclib |
| MVP_top60_DNA_seed | CDK1 | -0.436699 | OV | TOV-112D | RO-3306 | top60_altered_DNA+RNA_seed | CDK4 | 0.61874 | GBM | YKG-1 | Ribociclib |
| MVP_top60_DNA_seed | PIK3CA | -0.43021 | OV | OV-7 | GNE-317 | top60_altered_DNA+RNA_seed | EGFR | 0.621027 | GBM | LN-18 | Gefitinib |
| MVP_top60_DNA_seed | MAPK1 | -0.429453 | OV | OVCAR-5 | Ulixertinib | top60_altered_DNA+RNA_seed | EGFR | 0.627379 | GBM | Becker | Osimertinib |
| MVP_top60_DNA_seed | MAPK3 | -0.429453 | OV | OVCAR-5 | Ulixertinib | top60_altered_DNA+RNA_seed | EGFR | 0.628974 | GBM | GB-1 | Sapitinib |
| MVP_top60_DNA_seed | EGFR | -0.423926 | OV | OVTOKO | Afatinib | top60_altered_DNA+RNA_seed | EGFR | 0.629459 | GBM | Becker | Afatinib |
| MVP_top60_DNA_seed | ERBB2 | -0.423926 | OV | OVTOKO | Afatinib | top60_altered_DNA+RNA_seed | EGFR | 0.635474 | GBM | D-247MG | Afatinib |
| MVP_top60_DNA_seed | MAPK1 | -0.423218 | OV | OVCAR-5 | VX-11e | top60_altered_DNA+RNA_seed | EGFR | 0.639898 | GBM | T98G | AZD3759 |
| MVP_top60_DNA_seed | ATR | -0.419272 | OV | OVCAR-4 | VE-822 | top60_altered_DNA+RNA_seed | TP53 | 0.645376 | GBM | LN-229 | MIRA-1 |
| MVP_top60_DNA_seed | PLK1 | -0.417173 | OV | OV-7 | BI-2536 | top60_altered_DNA+RNA_seed | TP53 | 0.648846 | GBM | U-118-MG | MIRA-1 |
| MVP_top60_DNA_seed | PIK3CA | -0.413093 | OV | OVCA420 | AZD8186 | top60_altered_DNA+RNA_seed | EGFR | 0.662326 | GBM | YH-13 | AZD3759 |
| MVP_top60_DNA_seed | ATR | -0.40687 | OV | TOV-112D | AZD6738 | top60_altered_DNA+RNA_seed | CDK4 | 0.663048 | GBM | SW1088 | Ribociclib |
| MVP_top60_DNA_seed | MAPK1 | -0.397765 | OV | OVK-18 | Ulixertinib | top60_altered_DNA+RNA_seed | EGFR | 0.666117 | GBM | DBTRG-05MG | Erlotinib |
| MVP_top60_DNA_seed | MAPK3 | -0.397765 | OV | OVK-18 | Ulixertinib | top60_altered_DNA+RNA_seed | EGFR | 0.667379 | GBM | AM-38 | Osimertinib |
| MVP_top60_DNA_seed | CDK2 | -0.395464 | OV | JHOS-4 | MK-8776 | top60_altered_DNA+RNA_seed | TP53 | 0.673706 | GBM | D-392MG | MIRA-1 |
| MVP_top60_DNA_seed | MAPK1 | -0.395229 | OV | OVCA433 | VX-11e | top60_altered_DNA+RNA_seed | EGFR | 0.686248 | GBM | AM-38 | Afatinib |
| MVP_top60_DNA_seed | EGFR | -0.393171 | OV | OVTOKO | Lapatinib | top60_altered_DNA+RNA_seed | EGFR | 0.688348 | GBM | CAS-1 | Osimertinib |
| MVP_top60_DNA_seed | ERBB2 | -0.393171 | OV | OVTOKO | Lapatinib | top60_altered_DNA+RNA_seed | CDK4 | 0.691742 | GBM | M059J | Ribociclib |
| MVP_top60_DNA_seed | EGFR | -0.390831 | OV | TOV-112D | Gefitinib | top60_altered_DNA+RNA_seed | EGFR | 0.702379 | GBM | D-392MG | Osimertinib |
| MVP_top60_DNA_seed | MAPK1 | -0.386203 | OV | UWB1.289 | SCH772984 | top60_altered_DNA+RNA_seed | CDK4 | 0.703697 | GBM | U251 | Ribociclib |
| MVP_top60_DNA_seed | MAPK3 | -0.386203 | OV | UWB1.289 | SCH772984 | top60_altered_DNA+RNA_seed | CDK4 | 0.704074 | GBM | SNB75 | Ribociclib |
| MVP_top60_DNA_seed | HDAC2 | -0.382116 | OV | OVK-18 | Romidepsin | top60_altered_DNA+RNA_seed | EGFR | 0.706663 | GBM | M059J | Osimertinib |
| MVP_top60_DNA_seed | PIK3CA | -0.38052 | OV | OVCAR-8 | Buparlisib | top60_altered_DNA+RNA_seed | TP53 | 0.714854 | GBM | U-87-MG | MIRA-1 |
| MVP_top60_DNA_seed | PIK3CA | -0.377635 | OV | OC-314 | Buparlisib | top60_altered_DNA+RNA_seed | EGFR | 0.71768 | GBM | T98G | Afatinib |
| MVP_top60_DNA_seed | CDK2 | -0.374381 | OV | OV-7 | Dinaciclib | top60_altered_DNA+RNA_seed | EGFR | 0.722859 | GBM | SW1088 | Lapatinib |
| MVP_top60_DNA_seed | CDK1 | -0.374381 | OV | OV-7 | Dinaciclib | top60_altered_DNA+RNA_seed | EGFR | 0.7261 | GBM | Becker | Gefitinib |
| MVP_top60_DNA_seed | HDAC2 | -0.373973 | OV | TOV-112D | Romidepsin | top60_altered_DNA+RNA_seed | EGFR | 0.738527 | GBM | LN-229 | Lapatinib |
| MVP_top60_DNA_seed | EGFR | -0.371985 | OV | OVTOKO | AZD3759 | top60_altered_DNA+RNA_seed | CDK4 | 0.744679 | GBM | 42-MG-BA | Palbociclib |
| MVP_top60_DNA_seed | CDK1 | -0.367366 | OV | JHOS-4 | RO-3306 | top60_altered_DNA+RNA_seed | EGFR | 0.746715 | GBM | CAS-1 | Gefitinib |
| MVP_top60_DNA_seed | MAPK1 | -0.364788 | OV | OVMIU | SCH772984 | top60_altered_DNA+RNA_seed | EGFR | 0.758851 | GBM | SK-MG-1 | Lapatinib |
| MVP_top60_DNA_seed | MAPK3 | -0.364788 | OV | OVMIU | SCH772984 | top60_altered_DNA+RNA_seed | EGFR | 0.760432 | GBM | D-392MG | AZD3759 |
| MVP_top60_DNA_seed | MAPK1 | -0.364464 | OV | Caov-3 | VX-11e | top60_altered_DNA+RNA_seed | EGFR | 0.780566 | GBM | D-392MG | Afatinib |
| MVP_top60_DNA_seed | MAPK1 | -0.363752 | OV | OV-17R | VX-11e | top60_altered_DNA+RNA_seed | EGFR | 0.787238 | GBM | SW1088 | AZD3759 |
| MVP_top60_DNA_seed | MAPK1 | -0.361106 | OV | PEO1 | Ulixertinib | top60_altered_DNA+RNA_seed | CDK4 | 0.792326 | GBM | SNB75 | Palbociclib |
| MVP_top60_DNA_seed | MAPK3 | -0.361106 | OV | PEO1 | Ulixertinib | top60_altered_DNA+RNA_seed | EGFR | 0.803738 | GBM | CAS-1 | Afatinib |
| MVP_top60_DNA_seed | MAPK1 | -0.35858 | OV | OV-17R | SCH772984 | top60_altered_DNA+RNA_seed | EGFR | 0.809603 | GBM | LN-405 | Erlotinib |

| | | | | | | | | | | | |
|---|---|---|---|---|---|---|---|---|---|---|---|
| MVP_top60_DNA_seed | MAPK3 | -0.35858 | OV | OV-17R | SCH772984 | top60_altered_DNA+RNA_seed | EGFR | 0.810487 | GBM | U251 | Gefitinib |
| MVP_top60_DNA_seed | MAPK1 | -0.355889 | OV | OVCA433 | Ulixertinib | top60_altered_DNA+RNA_seed | EGFR | 0.811865 | GBM | SNB75 | Sapitinib |
| MVP_top60_DNA_seed | MAPK3 | -0.355889 | OV | OVCA433 | Ulixertinib | top60_altered_DNA+RNA_seed | EGFR | 0.820737 | GBM | D-245MG | AZD3759 |
| MVP_top60_DNA_seed | CDK2 | -0.355841 | OV | Caov-3 | Dinaciclib | top60_altered_DNA+RNA_seed | EGFR | 0.822151 | GBM | D-247MG | Gefitinib |
| MVP_top60_DNA_seed | CDK1 | -0.355841 | OV | Caov-3 | Dinaciclib | top60_altered_DNA+RNA_seed | EGFR | 0.823413 | GBM | KS-1 | Osimertinib |
| MVP_top60_DNA_seed | EGFR | -0.352552 | OV | OVISE | AZD3759 | top60_altered_DNA+RNA_seed | CDK4 | 0.825781 | GBM | T98G | Ribociclib |
| MVP_top60_DNA_seed | MAPK1 | -0.351637 | OV | SW626 | Ulixertinib | top60_altered_DNA+RNA_seed | EGFR | 0.835318 | GBM | YH-13 | Gefitinib |
| MVP_top60_DNA_seed | MAPK3 | -0.351637 | OV | SW626 | Ulixertinib | top60_altered_DNA+RNA_seed | EGFR | 0.849635 | GBM | T98G | Sapitinib |
| MVP_top60_DNA_seed | PLK1 | -0.349701 | OV | UWB1.289 | BI-2536 | top60_altered_DNA+RNA_seed | TP53 | 0.851192 | GBM | SK-MG-1 | MIRA-1 |
| MVP_top60_DNA_seed | MAPK1 | -0.349429 | OV | DOV13 | VX-11e | top60_altered_DNA+RNA_seed | EGFR | 0.860964 | GBM | SF295 | Sapitinib |
| MVP_top60_DNA_seed | CDK1 | -0.34808 | OV | JHOS-2 | RO-3306 | top60_altered_DNA+RNA_seed | EGFR | 0.871233 | GBM | LN-405 | AZD3759 |
| MVP_top60_DNA_seed | EGFR | -0.346865 | OV | FU-OV-1 | Sapitinib | top60_altered_DNA+RNA_seed | TP53 | 0.871374 | GBM | SF268 | MIRA-1 |
| MVP_top60_DNA_seed | ERBB2 | -0.346865 | OV | FU-OV-1 | Sapitinib | top60_altered_DNA+RNA_seed | EGFR | 0.87195 | GBM | D-263MG | Erlotinib |
| MVP_top60_DNA_seed | PIK3CA | -0.343246 | OV | SK-OV-3 | Alpelisib | top60_altered_DNA+RNA_seed | EGFR | 0.884814 | GBM | D-392MG | Gefitinib |
| MVP_top60_DNA_seed | PIK3CA | -0.338385 | OV | Caov-4 | GNE-317 | top60_altered_DNA+RNA_seed | EGFR | 0.884914 | GBM | KS-1 | Gefitinib |
| MVP_top60_DNA_seed | EGFR | -0.337572 | OV | JHOS-3 | Lapatinib | top60_altered_DNA+RNA_seed | EGFR | 0.888304 | GBM | SK-MG-1 | Sapitinib |
| MVP_top60_DNA_seed | ERBB2 | -0.337572 | OV | JHOS-3 | Lapatinib | top60_altered_DNA+RNA_seed | EGFR | 0.895519 | GBM | SF126 | Erlotinib |
| MVP_top60_DNA_seed | MAPK1 | -0.336081 | OV | TOV-112D | Ulixertinib | top60_altered_DNA+RNA_seed | CDK4 | 0.899147 | GBM | SF268 | Palbociclib |
| MVP_top60_DNA_seed | MAPK3 | -0.336081 | OV | TOV-112D | Ulixertinib | top60_altered_DNA+RNA_seed | EGFR | 0.908304 | GBM | SF268 | AZD3759 |
| MVP_top60_DNA_seed | EGFR | -0.335796 | OV | TOV-21G | Afatinib | top60_altered_DNA+RNA_seed | EGFR | 0.910745 | GBM | LNZTA3WT4 | Sapitinib |
| MVP_top60_DNA_seed | ERBB2 | -0.335796 | OV | TOV-21G | Afatinib | top60_altered_DNA+RNA_seed | CDK4 | 0.917941 | GBM | LN-229 | Ribociclib |
| MVP_top60_DNA_seed | PIK3CA | -0.328832 | OV | UWB1.289 | GNE-317 | top60_altered_DNA+RNA_seed | EGFR | 0.931245 | GBM | U251 | Osimertinib |
| MVP_top60_DNA_seed | EGFR | -0.328476 | OV | TOV-112D | Erlotinib | top60_altered_DNA+RNA_seed | EGFR | 0.931524 | GBM | SF126 | Afatinib |
| MVP_top60_DNA_seed | MAPK1 | -0.327813 | OV | OVMIU | Ulixertinib | top60_altered_DNA+RNA_seed | EGFR | 0.932418 | GBM | CAS-1 | Erlotinib |
| MVP_top60_DNA_seed | MAPK3 | -0.327813 | OV | OVMIU | Ulixertinib | top60_altered_DNA+RNA_seed | EGFR | 0.938563 | GBM | T98G | Erlotinib |
| MVP_top60_DNA_seed | PIK3CA | -0.323738 | OV | DOV13 | Alpelisib | top60_altered_DNA+RNA_seed | EGFR | 0.941274 | GBM | D-566MG | Afatinib |
| MVP_top60_DNA_seed | MAPK1 | -0.322975 | OV | OV-90 | VX-11e | top60_altered_DNA+RNA_seed | EGFR | 0.945968 | GBM | D-247MG | Erlotinib |
| MVP_top60_DNA_seed | MAPK1 | -0.315509 | OV | UWB1.289 | VX-11e | top60_altered_DNA+RNA_seed | EGFR | 0.949568 | GBM | U-118-MG | Afatinib |
| MVP_top60_DNA_seed | EGFR | -0.31519 | OV | OV-7 | Sapitinib | top60_altered_DNA+RNA_seed | EGFR | 0.955973 | GBM | T98G | Osimertinib |
| MVP_top60_DNA_seed | ERBB2 | -0.31519 | OV | OV-7 | Sapitinib | top60_altered_DNA+RNA_seed | EGFR | 0.956074 | GBM | D-263MG | Sapitinib |
| MVP_top60_DNA_seed | ATR | -0.315187 | OV | Hey | AZD6738 | top60_altered_DNA+RNA_seed | EGFR | 0.965185 | GBM | SNB75 | Lapatinib |
| MVP_top60_DNA_seed | PIK3CA | -0.313284 | OV | OVCAR-8 | GNE-317 | top60_altered_DNA+RNA_seed | EGFR | 0.972104 | GBM | LN-18 | AZD3759 |
| MVP_top60_DNA_seed | PIK3CA | -0.312622 | OV | OV-17R | GNE-317 | top60_altered_DNA+RNA_seed | EGFR | 0.973538 | GBM | SF295 | Afatinib |
| MVP_top60_DNA_seed | ATR | -0.304809 | OV | TOV-21G | VE821 | top60_altered_DNA+RNA_seed | TP53 | 0.975512 | GBM | DK-MG | MIRA-1 |
| MVP_top60_DNA_seed | ATR | -0.300775 | OV | OAW-42 | VE-822 | top60_altered_DNA+RNA_seed | EGFR | 0.979684 | GBM | SK-MG-1 | Gefitinib |
| MVP_top60_DNA_seed | CDK2 | -0.300764 | OV | IGROV-1 | AZD5438 | top60_altered_DNA+RNA_seed | EGFR | 0.980727 | GBM | LN-405 | Sapitinib |
| MVP_top60_DNA_seed | PLK1 | -0.298357 | OV | OVCAR-3 | BI-2536 | top60_altered_DNA+RNA_seed | EGFR | 0.986548 | GBM | D-263MG | Osimertinib |
| MVP_top60_DNA_seed | EGFR | -0.292651 | OV | OV-17R | Osimertinib | top60_altered_DNA+RNA_seed | EGFR | 0.9873 | GBM | D-263MG | Afatinib |
| MVP_top60_DNA_seed | MAPK1 | -0.290673 | OV | JHOS-2 | VX-11e | top60_altered_DNA+RNA_seed | EGFR | 0.991396 | GBM | M059J | Gefitinib |
| MVP_top60_DNA_seed | PIK3CA | -0.290111 | OV | OVCA420 | GNE-317 | top60_altered_DNA+RNA_seed | EGFR | 1.000318 | GBM | YKG-1 | Afatinib |
| MVP_top60_DNA_seed | PIK3CA | -0.290079 | OV | OAW-42 | GNE-317 | top60_altered_DNA+RNA_seed | CDK4 | 1.00091 | GBM | LN-405 | Ribociclib |

| | | | | | | | | | | | |
|---|---|---|---|---|---|---|---|---|---|---|---|
| MVP_top60_DNA_seed | PLK1 | -0.288643 | OV | OVK-18 | BI-2536 | top60_altered_DNA+RNA_seed | EGFR | 1.023988 | GBM | Becker | Sapitinib |
| MVP_top60_DNA_seed | CDK2 | -0.285532 | OV | OVCAR-8 | AZD5438 | top60_altered_DNA+RNA_seed | EGFR | 1.027722 | GBM | U251 | Afatinib |
| MVP_top60_DNA_seed | PIK3CA | -0.284163 | OV | Caov-3 | AZD8186 | top60_altered_DNA+RNA_seed | EGFR | 1.039273 | GBM | LN-18 | Afatinib |
| MVP_top60_DNA_seed | TP53 | -0.279165 | OV | OVCAR-8 | MIRA-1 | top60_altered_DNA+RNA_seed | EGFR | 1.039408 | GBM | KS-1 | AZD3759 |
| MVP_top60_DNA_seed | MAPK1 | -0.279006 | OV | OC-314 | VX-11e | top60_altered_DNA+RNA_seed | EGFR | 1.04507 | GBM | LN-18 | Osimertinib |
| MVP_top60_DNA_seed | MAPK1 | -0.271499 | OV | OVCA420 | Ulixertinib | top60_altered_DNA+RNA_seed | EGFR | 1.049049 | GBM | D-247MG | AZD3759 |
| MVP_top60_DNA_seed | MAPK3 | -0.271499 | OV | OVCA420 | Ulixertinib | top60_altered_DNA+RNA_seed | EGFR | 1.056207 | GBM | SW1088 | Gefitinib |
| MVP_top60_DNA_seed | ATR | -0.270435 | OV | OVCAR-8 | VE-822 | top60_altered_DNA+RNA_seed | EGFR | 1.063211 | GBM | GB-1 | Gefitinib |
| MVP_top60_DNA_seed | CDK2 | -0.270034 | OV | OVTOKO | AZD5438 | top60_altered_DNA+RNA_seed | EGFR | 1.086889 | GBM | GB-1 | Afatinib |
| MVP_top60_DNA_seed | CDK2 | -0.269554 | OV | JHOS-2 | MK-8776 | top60_altered_DNA+RNA_seed | EGFR | 1.088469 | GBM | Becker | Erlotinib |
| MVP_top60_DNA_seed | CDK2 | -0.268568 | OV | OVK-18 | AZD5438 | top60_altered_DNA+RNA_seed | TP53 | 1.104613 | GBM | LN-405 | MIRA-1 |
| MVP_top60_DNA_seed | ATR | -0.266311 | OV | SK-OV-3 | AZD6738 | top60_altered_DNA+RNA_seed | EGFR | 1.128677 | GBM | AM-38 | Sapitinib |
| MVP_top60_DNA_seed | PIK3CA | -0.264715 | OV | OAW-42 | Buparlisib | top60_altered_DNA+RNA_seed | EGFR | 1.130392 | GBM | D-392MG | Erlotinib |
| MVP_top60_DNA_seed | EGFR | -0.261898 | OV | JHOS-3 | Osimertinib | top60_altered_DNA+RNA_seed | EGFR | 1.14068 | GBM | D-392MG | Sapitinib |
| MVP_top60_DNA_seed | EGFR | -0.261719 | OV | DOV13 | AZD3759 | top60_altered_DNA+RNA_seed | CDK4 | 1.157249 | GBM | U-118-MG | Ribociclib |
| MVP_top60_DNA_seed | PIK3CA | -0.261182 | OV | OVTOKO | GNE-317 | top60_altered_DNA+RNA_seed | CDK4 | 1.178543 | GBM | M059J | Palbociclib |
| MVP_top60_DNA_seed | EGFR | -0.259079 | OV | SK-OV-3 | Erlotinib | top60_altered_DNA+RNA_seed | CDK4 | 1.180443 | GBM | CAS-1 | Ribociclib |
| MVP_top60_DNA_seed | CDK1 | -0.257936 | OV | IGROV-1 | RO-3306 | top60_altered_DNA+RNA_seed | EGFR | 1.180972 | GBM | U251 | Lapatinib |
| MVP_top60_DNA_seed | TP53 | -0.252356 | OV | OV-7 | MIRA-1 | top60_altered_DNA+RNA_seed | TP53 | 1.185029 | GBM | SW1088 | MIRA-1 |
| MVP_top60_DNA_seed | EGFR | -0.252306 | OV | FU-OV-1 | Osimertinib | top60_altered_DNA+RNA_seed | EGFR | 1.186611 | GBM | SF268 | Osimertinib |
| MVP_top60_DNA_seed | CDK1 | -0.250423 | OV | OC-314 | RO-3306 | top60_altered_DNA+RNA_seed | EGFR | 1.187698 | GBM | DBTRG-05MG | Osimertinib |
| MVP_top60_DNA_seed | MAPK1 | -0.248751 | OV | FU-OV-1 | Ulixertinib | top60_altered_DNA+RNA_seed | EGFR | 1.201006 | GBM | LN-405 | Osimertinib |
| MVP_top60_DNA_seed | MAPK3 | -0.248751 | OV | FU-OV-1 | Ulixertinib | top60_altered_DNA+RNA_seed | EGFR | 1.222906 | GBM | U-118-MG | Lapatinib |
| MVP_top60_DNA_seed | PIK3CA | -0.24486 | OV | OVCA433 | GNE-317 | top60_altered_DNA+RNA_seed | EGFR | 1.234905 | GBM | DK-MG | Gefitinib |
| MVP_top60_DNA_seed | CDK2 | -0.240953 | OV | OV-17R | MK-8776 | top60_altered_DNA+RNA_seed | EGFR | 1.247878 | GBM | CAS-1 | AZD3759 |
| MVP_top60_DNA_seed | CREBBP | -0.239878 | OV | OV-56 | SGC-CBP30 | top60_altered_DNA+RNA_seed | TP53 | 1.249454 | GBM | CAS-1 | MIRA-1 |
| MVP_top60_DNA_seed | CDK2 | -0.239534 | OV | JHOS-3 | Dinaciclib | top60_altered_DNA+RNA_seed | EGFR | 1.260517 | GBM | SNB75 | Erlotinib |
| MVP_top60_DNA_seed | CDK1 | -0.239534 | OV | JHOS-3 | Dinaciclib | top60_altered_DNA+RNA_seed | CDK4 | 1.263292 | GBM | LN-405 | Palbociclib |
| MVP_top60_DNA_seed | EGFR | -0.237971 | OV | Caov-4 | Gefitinib | top60_altered_DNA+RNA_seed | EGFR | 1.283699 | GBM | D-263MG | Lapatinib |
| MVP_top60_DNA_seed | CDK2 | -0.235295 | OV | Hey | AZD5438 | top60_altered_DNA+RNA_seed | EGFR | 1.288037 | GBM | D-245MG | Erlotinib |
| MVP_top60_DNA_seed | MAPK1 | -0.234218 | OV | OV-90 | Ulixertinib | top60_altered_DNA+RNA_seed | CDK4 | 1.299051 | GBM | D-542MG | Palbociclib |
| MVP_top60_DNA_seed | MAPK3 | -0.234218 | OV | OV-90 | Ulixertinib | top60_altered_DNA+RNA_seed | CDK4 | 1.331047 | GBM | GB-1 | Ribociclib |
| MVP_top60_DNA_seed | ATR | -0.231782 | OV | OV-56 | VE-822 | top60_altered_DNA+RNA_seed | EGFR | 1.375176 | GBM | SNB75 | Afatinib |
| MVP_top60_DNA_seed | EGFR | -0.2232 | OV | OVK-18 | Osimertinib | top60_altered_DNA+RNA_seed | EGFR | 1.378183 | GBM | DBTRG-05MG | AZD3759 |
| MVP_top60_DNA_seed | CDK2 | -0.218232 | OV | TOV-21G | MK-8776 | top60_altered_DNA+RNA_seed | EGFR | 1.39833 | GBM | GB-1 | AZD3759 |
| MVP_top60_DNA_seed | EGFR | -0.215025 | OV | OVCAR-3 | Gefitinib | top60_altered_DNA+RNA_seed | EGFR | 1.401945 | GBM | U251 | Sapitinib |
| MVP_top60_DNA_seed | EGFR | -0.210089 | OV | OC-314 | Erlotinib | top60_altered_DNA+RNA_seed | EGFR | 1.425539 | GBM | U-87-MG | Afatinib |
| MVP_top60_DNA_seed | PIK3CA | -0.208394 | OV | OVISE | Buparlisib | top60_altered_DNA+RNA_seed | EGFR | 1.430242 | GBM | U-87-MG | Lapatinib |
| MVP_top60_DNA_seed | EGFR | -0.206868 | OV | Caov-4 | Osimertinib | top60_altered_DNA+RNA_seed | EGFR | 1.446664 | GBM | DBTRG-05MG | Afatinib |
| MVP_top60_DNA_seed | EGFR | -0.203308 | OV | OVK-18 | AZD3759 | top60_altered_DNA+RNA_seed | EGFR | 1.4674 | GBM | Becker | Lapatinib |
| MVP_top60_DNA_seed | MAPK1 | -0.201958 | OV | PEO1 | Ulixertinib | top60_altered_DNA+RNA_seed | CDK4 | 1.489772 | GBM | LN-229 | Palbociclib |

| | | | | | | | | | | | |
|---|---|---|---|---|---|---|---|---|---|---|---|
| MVP_top60_DNA_seed | MAPK3 | -0.201958 | OV | PEO1 | Ulixertinib | top60_altered_DNA+RNA_seed | EGFR | 1.52125 | GBM | LN-405 | Afatinib |
| MVP_top60_DNA_seed | CDK2 | -0.199985 | OV | JHOS-2 | Dinaciclib | top60_altered_DNA+RNA_seed | EGFR | 1.562033 | GBM | KS-1 | Lapatinib |
| MVP_top60_DNA_seed | CDK1 | -0.199985 | OV | JHOS-2 | Dinaciclib | top60_altered_DNA+RNA_seed | CDK4 | 1.691322 | GBM | DBTRG-05MG | Ribociclib |
| MVP_top60_DNA_seed | PLK1 | -0.193507 | OV | OVMIU | MK-1775 | top60_altered_DNA+RNA_seed | EGFR | 1.716475 | GBM | SF295 | Erlotinib |
| MVP_top60_DNA_seed | ATR | -0.192324 | OV | PEO1 | VE821 | top60_altered_DNA+RNA_seed | EGFR | 1.747985 | GBM | LN-405 | Lapatinib |
| MVP_top60_DNA_seed | PIK3CA | -0.189805 | OV | TOV-21G | GNE-317 | top60_altered_DNA+RNA_seed | EGFR | 1.750427 | GBM | U-118-MG | Sapitinib |
| MVP_top60_DNA_seed | CREBBP | -0.186346 | OV | DOV13 | SGC-CBP30 | top60_altered_DNA+RNA_seed | EGFR | 1.767129 | GBM | DBTRG-05MG | Sapitinib |
| MVP_top60_DNA_seed | PIK3CA | -0.185794 | OV | OVCAR-8 | AZD8186 | top60_altered_DNA+RNA_seed | EGFR | 1.805704 | GBM | GB-1 | Osimertinib |
| MVP_top60_DNA_seed | PLK1 | -0.185345 | OV | PEO1 | BI-2536 | top60_altered_DNA+RNA_seed | CDK4 | 2.065252 | GBM | DBTRG-05MG | Palbociclib |
| MVP_top60_DNA_seed | PIK3CA | -0.181595 | OV | OVISE | AZD8186 | top60_altered_DNA+RNA_seed | EGFR | 2.079235 | GBM | D-392MG | Lapatinib |
| MVP_top60_DNA_seed | PIK3CA | -0.173488 | OV | OVCA433 | AZD8186 | top60_altered_DNA+RNA_seed | EGFR | 2.094157 | GBM | DBTRG-05MG | Gefitinib |
| MVP_top60_DNA_seed | ATR | -0.171541 | OV | Caov-4 | VE821 | top60_altered_DNA+RNA_seed | TP53 | 2.125063 | GBM | DBTRG-05MG | MIRA-1 |
| MVP_top60_DNA_seed | CDK2 | -0.1708 | OV | UWB1.289 | Dinaciclib | top60_altered_DNA+RNA_seed | EGFR | 2.576791 | GBM | DBTRG-05MG | Lapatinib |
| MVP_top60_DNA_seed | CDK1 | -0.1708 | OV | UWB1.289 | Dinaciclib | top60_altered_DNA+RNA_seed | FGFR4 | -2.4591 | KIRC | U031 | FGFR_0939 |
| MVP_top60_DNA_seed | ATR | -0.166759 | OV | OVCA433 | AZD6738 | top60_altered_DNA+RNA_seed | FGFR4 | -1.657996 | KIRC | RCC-JW | FGFR_0939 |
| MVP_top60_DNA_seed | EGFR | -0.16618 | OV | OVCAR-3 | Osimertinib | top60_altered_DNA+RNA_seed | FGFR4 | -1.281831 | KIRC | HA7-RCC | FGFR_0939 |
| MVP_top60_DNA_seed | PIK3CA | -0.166083 | OV | OVMIU | Alpelisib | top60_altered_DNA+RNA_seed | FGFR4 | -0.9546 | KIRC | LB1047-RCC | FGFR_0939 |
| MVP_top60_DNA_seed | MAPK1 | -0.164819 | OV | SW626 | VX-11e | top60_altered_DNA+RNA_seed | FGFR4 | -0.919515 | KIRC | RCC-AB | FGFR_0939 |
| MVP_top60_DNA_seed | EGFR | -0.161102 | OV | OVISE | Gefitinib | top60_altered_DNA+RNA_seed | FGFR4 | -0.892812 | KIRC | 786-0 | FGFR_0939 |
| MVP_top60_DNA_seed | CDK2 | -0.159256 | OV | IGROV-1 | Dinaciclib | top60_altered_DNA+RNA_seed | FGFR4 | -0.875187 | KIRC | A704 | FGFR_0939 |
| MVP_top60_DNA_seed | CDK1 | -0.159256 | OV | IGROV-1 | Dinaciclib | top60_altered_DNA+RNA_seed | FGFR4 | -0.775996 | KIRC | NCC010 | FGFR_3831 |
| MVP_top60_DNA_seed | EGFR | -0.158794 | OV | OV-7 | Lapatinib | top60_altered_DNA+RNA_seed | FGFR4 | -0.749345 | KIRC | BFTC-909 | FGFR_0939 |
| MVP_top60_DNA_seed | ERBB2 | -0.158794 | OV | OV-7 | Lapatinib | top60_altered_DNA+RNA_seed | FGFR4 | -0.73585 | KIRC | RCC10RGB | FGFR_0939 |
| MVP_top60_DNA_seed | PIK3CA | -0.157264 | OV | OVCA420 | Buparlisib | top60_altered_DNA+RNA_seed | FGFR4 | -0.713748 | KIRC | A704 | FGFR_3831 |
| MVP_top60_DNA_seed | PIK3CA | -0.156449 | OV | TOV-112D | GNE-317 | top60_altered_DNA+RNA_seed | FGFR4 | -0.655322 | KIRC | RCC-FG2 | FGFR_0939 |
| MVP_top60_DNA_seed | PIK3CA | -0.156147 | OV | SK-OV-3 | GNE-317 | top60_altered_DNA+RNA_seed | FGFR4 | -0.65324 | KIRC | RXF393 | FGFR_3831 |
| MVP_top60_DNA_seed | ATR | -0.154518 | OV | OVK-18 | VE-822 | top60_altered_DNA+RNA_seed | FGFR4 | -0.652829 | KIRC | LB996-RCC | FGFR_0939 |
| MVP_top60_DNA_seed | HDAC2 | -0.153835 | OV | OVMIU | Romidepsin | top60_altered_DNA+RNA_seed | FGFR4 | -0.609385 | KIRC | RCC-JF | FGFR_0939 |
| MVP_top60_DNA_seed | PLK1 | -0.151794 | OV | OV-7 | MK-1775 | top60_altered_DNA+RNA_seed | FGFR4 | -0.494367 | KIRC | 769-P | FGFR_0939 |
| MVP_top60_DNA_seed | PIK3CA | -0.147607 | OV | OAW-42 | AZD8186 | top60_altered_DNA+RNA_seed | FGFR4 | -0.486649 | KIRC | NCC021 | FGFR_0939 |
| MVP_top60_DNA_seed | EGFR | -0.144205 | OV | SK-OV-3 | Gefitinib | top60_altered_DNA+RNA_seed | FGFR4 | -0.461804 | KIRC | SN12C | FGFR_0939 |
| MVP_top60_DNA_seed | MAPK1 | -0.143153 | OV | OVCAR-5 | Ulixertinib | top60_altered_DNA+RNA_seed | FGFR4 | -0.453204 | KIRC | LB2241-RCC | FGFR_3831 |
| MVP_top60_DNA_seed | MAPK3 | -0.143153 | OV | OVCAR-5 | Ulixertinib | top60_altered_DNA+RNA_seed | FGFR4 | -0.405782 | KIRC | BFTC-909 | FGFR_3831 |
| MVP_top60_DNA_seed | PLK1 | -0.141657 | OV | JHOS-2 | BI-2536 | top60_altered_DNA+RNA_seed | FGFR4 | -0.374645 | KIRC | HA7-RCC | FGFR_3831 |
| MVP_top60_DNA_seed | PIK3CA | -0.134733 | OV | OVK-18 | AZD8186 | top60_altered_DNA+RNA_seed | FGFR4 | -0.3568 | KIRC | LB1047-RCC | FGFR_3831 |
| MVP_top60_DNA_seed | EGFR | -0.134446 | OV | OV-7 | Gefitinib | top60_altered_DNA+RNA_seed | FGFR4 | -0.302501 | KIRC | NCC010 | FGFR_0939 |
| MVP_top60_DNA_seed | EGFR | -0.132675 | OV | OVCAR-5 | Sapitinib | top60_altered_DNA+RNA_seed | FGFR4 | -0.296539 | KIRC | LB2241-RCC | FGFR_0939 |
| MVP_top60_DNA_seed | ERBB2 | -0.132675 | OV | OVCAR-5 | Sapitinib | top60_altered_DNA+RNA_seed | FGFR4 | -0.262488 | KIRC | 769-P | FGFR_3831 |
| MVP_top60_DNA_seed | MAPK1 | -0.132617 | OV | OVCAR-8 | SCH772984 | top60_altered_DNA+RNA_seed | FGFR4 | -0.261901 | KIRC | KMRC-20 | FGFR_0939 |
| MVP_top60_DNA_seed | MAPK3 | -0.132617 | OV | OVCAR-8 | SCH772984 | top60_altered_DNA+RNA_seed | FGFR4 | -0.258943 | KIRC | RCC-JW | FGFR_3831 |
| MVP_top60_DNA_seed | PLK1 | -0.130273 | OV | TOV-21G | BI-2536 | top60_altered_DNA+RNA_seed | FGFR4 | -0.248438 | KIRC | VMRC-RCZ | FGFR_0939 |

| | | | | | | | | | | | |
|---|---|---|---|---|---|---|---|---|---|---|---|
| MVP_top60_DNA_seed | EGFR | -0.12785 | OV | Caov-3 | Gefitinib | top60_altered_DNA+RNA_seed | FGFR4 | -0.241968 | KIRC | RCC-MF | FGFR_0939 |
| MVP_top60_DNA_seed | CDK1 | -0.127128 | OV | Hey | RO-3306 | top60_altered_DNA+RNA_seed | FGFR4 | -0.219742 | KIRC | RXF393 | FGFR_0939 |
| MVP_top60_DNA_seed | MAPK1 | -0.123439 | OV | TOV-112D | SCH772984 | top60_altered_DNA+RNA_seed | FGFR4 | -0.199676 | KIRC | SW156 | FGFR_0939 |
| MVP_top60_DNA_seed | MAPK3 | -0.123439 | OV | TOV-112D | SCH772984 | top60_altered_DNA+RNA_seed | FGFR4 | -0.116518 | KIRC | A498 | FGFR_3831 |
| MVP_top60_DNA_seed | MAPK1 | -0.119392 | OV | OV-7 | SCH772984 | top60_altered_DNA+RNA_seed | FGFR4 | -0.112227 | KIRC | 786-0 | FGFR_3831 |
| MVP_top60_DNA_seed | MAPK3 | -0.119392 | OV | OV-7 | SCH772984 | top60_altered_DNA+RNA_seed | FGFR4 | -0.109773 | KIRC | BB65-RCC | FGFR_0939 |
| MVP_top60_DNA_seed | EGFR | -0.117674 | OV | OVCAR-8 | Sapitinib | top60_altered_DNA+RNA_seed | FGFR4 | -0.080865 | KIRC | SN12C | FGFR_3831 |
| MVP_top60_DNA_seed | ERBB2 | -0.117674 | OV | OVCAR-8 | Sapitinib | top60_altered_DNA+RNA_seed | FGFR4 | -0.052158 | KIRC | KMRC-1 | FGFR_0939 |
| MVP_top60_DNA_seed | MAPK1 | -0.116254 | OV | TOV-112D | Ulixertinib | top60_altered_DNA+RNA_seed | FGFR4 | -0.044434 | KIRC | LB996-RCC | FGFR_3831 |
| MVP_top60_DNA_seed | MAPK3 | -0.116254 | OV | TOV-112D | Ulixertinib | top60_altered_DNA+RNA_seed | FGFR4 | 0.022847 | KIRC | RCC-AB | FGFR_3831 |
| MVP_top60_DNA_seed | MAPK1 | -0.116234 | OV | OVCAR-8 | Ulixertinib | top60_altered_DNA+RNA_seed | FGFR4 | 0.035524 | KIRC | CAKI-1 | FGFR_0939 |
| MVP_top60_DNA_seed | MAPK3 | -0.116234 | OV | OVCAR-8 | Ulixertinib | top60_altered_DNA+RNA_seed | FGFR4 | 0.06514 | KIRC | ACHN | FGFR_3831 |
| MVP_top60_DNA_seed | EGFR | -0.113702 | OV | SK-OV-3 | Lapatinib | top60_altered_DNA+RNA_seed | FGFR4 | 0.088982 | KIRC | RCC-ER | FGFR_0939 |
| MVP_top60_DNA_seed | ERBB2 | -0.113702 | OV | SK-OV-3 | Lapatinib | top60_altered_DNA+RNA_seed | FGFR4 | 0.152314 | KIRC | RCC-JF | FGFR_3831 |
| MVP_top60_DNA_seed | TP53 | -0.111277 | OV | OV-17R | MIRA-1 | top60_altered_DNA+RNA_seed | FGFR4 | 0.175746 | KIRC | NCC021 | FGFR_3831 |
| MVP_top60_DNA_seed | EGFR | -0.109964 | OV | OVCAR-8 | Osimertinib | top60_altered_DNA+RNA_seed | FGFR4 | 0.242805 | KIRC | CAKI-1 | FGFR_3831 |
| MVP_top60_DNA_seed | EGFR | -0.107045 | OV | OVISE | Lapatinib | top60_altered_DNA+RNA_seed | FGFR4 | 0.253419 | KIRC | TK10 | FGFR_0939 |
| MVP_top60_DNA_seed | ERBB2 | -0.107045 | OV | OVISE | Lapatinib | top60_altered_DNA+RNA_seed | FGFR4 | 0.284177 | KIRC | KMRC-20 | FGFR_3831 |
| MVP_top60_DNA_seed | ATR | -0.105929 | OV | OVCAR-5 | AZD6738 | top60_altered_DNA+RNA_seed | FGFR4 | 0.386611 | KIRC | KMRC-1 | FGFR_3831 |
| MVP_top60_DNA_seed | CDK2 | -0.105713 | OV | JHOS-3 | MK-8776 | top60_altered_DNA+RNA_seed | FGFR4 | 0.407777 | KIRC | CAL-54 | FGFR_3831 |
| MVP_top60_DNA_seed | EGFR | -0.097373 | OV | DOV13 | Lapatinib | top60_altered_DNA+RNA_seed | FGFR4 | 0.476178 | KIRC | BB65-RCC | FGFR_3831 |
| MVP_top60_DNA_seed | ERBB2 | -0.097373 | OV | DOV13 | Lapatinib | top60_altered_DNA+RNA_seed | FGFR4 | 0.522441 | KIRC | A498 | FGFR_0939 |
| MVP_top60_DNA_seed | MAPK1 | -0.097169 | OV | TOV-21G | SCH772984 | top60_altered_DNA+RNA_seed | FGFR4 | 0.558079 | KIRC | U031 | FGFR_3831 |
| MVP_top60_DNA_seed | MAPK3 | -0.097169 | OV | TOV-21G | SCH772984 | top60_altered_DNA+RNA_seed | FGFR4 | 0.621739 | KIRC | VMRC-RCZ | FGFR_3831 |
| MVP_top60_DNA_seed | CDK2 | -0.091991 | OV | IGROV-1 | MK-8776 | top60_altered_DNA+RNA_seed | FGFR4 | 0.668249 | KIRC | OS-RC-2 | FGFR_0939 |
| MVP_top60_DNA_seed | PIK3CA | -0.09046 | OV | OC-314 | Alpelisib | top60_altered_DNA+RNA_seed | FGFR4 | 0.736576 | KIRC | ACHN | FGFR_0939 |
| MVP_top60_DNA_seed | EGFR | -0.089418 | OV | OVISE | Afatinib | top60_altered_DNA+RNA_seed | FGFR4 | 0.824612 | KIRC | TK10 | FGFR_3831 |
| MVP_top60_DNA_seed | ERBB2 | -0.089418 | OV | OVISE | Afatinib | top60_altered_DNA+RNA_seed | FGFR4 | 0.932604 | KIRC | CAL-54 | FGFR_0939 |
| MVP_top60_DNA_seed | PIK3CA | -0.086084 | OV | OV-7 | Buparlisib | top60_altered_DNA+RNA_seed | FGFR4 | 1.151913 | KIRC | SW156 | FGFR_3831 |
| MVP_top60_DNA_seed | ATR | -0.082502 | OV | TOV-112D | VE-822 | top60_altered_DNA+RNA_seed | FGFR4 | 1.15443 | KIRC | OS-RC-2 | FGFR_3831 |
| MVP_top60_DNA_seed | PLK1 | -0.071695 | OV | OVCAR-4 | MK-1775 | top60_altered_DNA+RNA_seed | FGFR4 | 1.190802 | KIRC | RCC-FG2 | FGFR_3831 |
| MVP_top60_DNA_seed | ATR | -0.070267 | OV | OVCAR-8 | VE821 | top60_altered_DNA+RNA_seed | FGFR4 | 1.559144 | KIRC | RCC-MF | FGFR_3831 |
| MVP_top60_DNA_seed | EGFR | -0.070219 | OV | OVCAR-3 | AZD3759 | top60_altered_DNA+RNA_seed | FGFR4 | 1.7371 | KIRC | RCC-ER | FGFR_3831 |
| MVP_top60_DNA_seed | PIK3CA | -0.07014 | OV | OVCAR-3 | Buparlisib | top60_altered_DNA+RNA_seed | FGFR4 | 2.045145 | KIRC | RCC10RGB | FGFR_3831 |
| MVP_top60_DNA_seed | MAPK1 | -0.066918 | OV | FU-OV-1 | VX-11e | top60_altered_DNA+RNA_seed | KRAS | -1.609231 | LUAD | EMC-BAC-1 | KRAS (G12C) Inhibitor-1 |
| MVP_top60_DNA_seed | CDK2 | -0.065475 | OV | OV-56 | MK-8776 | top60_altered_DNA+RNA_seed | TERT | -1.466218 | LUAD | NCI-H838 | BIBR-1532 |
| MVP_top60_DNA_seed | TP53 | -0.063357 | OV | DOV13 | MIRA-1 | top60_altered_DNA+RNA_seed | BRD9 | -1.314061 | LUAD | RERF-LC-KJ | Bromosporine |
| MVP_top60_DNA_seed | EGFR | -0.061661 | OV | SK-OV-3 | Afatinib | top60_altered_DNA+RNA_seed | BRD9 | -1.230202 | LUAD | NCI-H1734 | I-BRD9 |
| MVP_top60_DNA_seed | ERBB2 | -0.061661 | OV | SK-OV-3 | Afatinib | top60_altered_DNA+RNA_seed | KRAS | -1.223262 | LUAD | NCI-H2122 | KRAS (G12C) Inhibitor-1 |
| MVP_top60_DNA_seed | CREBBP | -0.060077 | OV | PEO1 | SGC-CBP30 | top60_altered_DNA+RNA_seed | BRD9 | -1.173616 | LUAD | HCC-78 | I-BRD9 |
| MVP_top60_DNA_seed | ATR | -0.058029 | OV | OVCAR-8 | AZD6738 | top60_altered_DNA+RNA_seed | BRD9 | -1.164588 | LUAD | HCC-44 | Bromosporine |

| | | | | | | | | | | | |
|---|---|---|---|---|---|---|---|---|---|---|---|
| MVP_top60_DNA_seed | EGFR | -0.055679 | OV | FU-OV-1 | AZD3759 | top60_altered_DNA+RNA_seed | BRD9 | -1.152894 | LUAD | COR-L105 | Bromosporine |
| MVP_top60_DNA_seed | EGFR | -0.055493 | OV | TOV-112D | Lapatinib | top60_altered_DNA+RNA_seed | KRAS | -1.128535 | LUAD | NCI-H1975 | RAS (G12C) Inhibitor-1 |
| MVP_top60_DNA_seed | ERBB2 | -0.055493 | OV | TOV-112D | Lapatinib | top60_altered_DNA+RNA_seed | TP53 | -1.049009 | LUAD | NCI-H1734 | MIRA-1 |
| MVP_top60_DNA_seed | MAPK1 | -0.053611 | OV | TOV-112D | VX-11e | top60_altered_DNA+RNA_seed | BRD9 | -0.988706 | LUAD | COR-L105 | I-BRD9 |
| MVP_top60_DNA_seed | CDK2 | -0.044341 | OV | OVISE | AZD5438 | top60_altered_DNA+RNA_seed | TERT | -0.909582 | LUAD | NCI-H1792 | BIBR-1532 |
| MVP_top60_DNA_seed | EGFR | -0.042259 | OV | Caov-3 | Erlotinib | top60_altered_DNA+RNA_seed | KRAS | -0.907714 | LUAD | HCC-78 | RAS (G12C) Inhibitor-1 |
| MVP_top60_DNA_seed | CDK2 | -0.041877 | OV | OVCAR-5 | AZD5438 | top60_altered_DNA+RNA_seed | TERT | -0.886948 | LUAD | EMC-BAC-1 | BIBR-1532 |
| MVP_top60_DNA_seed | EGFR | -0.041507 | OV | JHOS-2 | AZD3759 | top60_altered_DNA+RNA_seed | KRAS | -0.868207 | LUAD | LC-2-ad | RAS (G12C) Inhibitor-1 |
| MVP_top60_DNA_seed | MAPK1 | -0.035018 | OV | JHOS-2 | SCH772984 | top60_altered_DNA+RNA_seed | KRAS | -0.854987 | LUAD | NCI-H1792 | RAS (G12C) Inhibitor-1 |
| MVP_top60_DNA_seed | MAPK3 | -0.035018 | OV | JHOS-2 | SCH772984 | top60_altered_DNA+RNA_seed | KRAS | -0.825506 | LUAD | NCI-H1437 | RAS (G12C) Inhibitor-1 |
| MVP_top60_DNA_seed | TP53 | -0.033943 | OV | JHOS-2 | MIRA-1 | top60_altered_DNA+RNA_seed | KRAS | -0.781584 | LUAD | NCI-H1993 | RAS (G12C) Inhibitor-1 |
| MVP_top60_DNA_seed | MAPK1 | -0.033927 | OV | OVCA420 | SCH772984 | top60_altered_DNA+RNA_seed | TERT | -0.768401 | LUAD | NCI-H2122 | BIBR-1532 |
| MVP_top60_DNA_seed | MAPK3 | -0.033927 | OV | OVCA420 | SCH772984 | top60_altered_DNA+RNA_seed | KRAS | -0.75525 | LUAD | NCI-H23 | RAS (G12C) Inhibitor-1 |
| MVP_top60_DNA_seed | EGFR | -0.03246 | OV | JHOS-3 | Afatinib | top60_altered_DNA+RNA_seed | TP53 | -0.730298 | LUAD | NCI-H1435 | MIRA-1 |
| MVP_top60_DNA_seed | ERBB2 | -0.03246 | OV | JHOS-3 | Afatinib | top60_altered_DNA+RNA_seed | BRD9 | -0.689971 | LUAD | LXF-289 | Bromosporine |
| MVP_top60_DNA_seed | CDK2 | -0.030031 | OV | OVCAR-8 | Dinaciclib | top60_altered_DNA+RNA_seed | TP53 | -0.664239 | LUAD | NCI-H1792 | MIRA-1 |
| MVP_top60_DNA_seed | CDK1 | -0.030031 | OV | OVCAR-8 | Dinaciclib | top60_altered_DNA+RNA_seed | BRD9 | -0.659039 | LUAD | RERF-LC-MS | I-BRD9 |
| MVP_top60_DNA_seed | EGFR | -0.027943 | OV | JHOS-2 | Gefitinib | top60_altered_DNA+RNA_seed | BRD9 | -0.628006 | LUAD | HCC-78 | Bromosporine |
| MVP_top60_DNA_seed | CDK2 | -0.026302 | OV | OVCA433 | MK-8776 | top60_altered_DNA+RNA_seed | KRAS | -0.608473 | LUAD | PC-14 | RAS (G12C) Inhibitor-1 |
| MVP_top60_DNA_seed | TP53 | -0.024273 | OV | OC-314 | MIRA-1 | top60_altered_DNA+RNA_seed | TERT | -0.607296 | LUAD | NCI-H1573 | BIBR-1532 |
| MVP_top60_DNA_seed | PIK3CA | -0.024098 | OV | OVCAR-3 | GNE-317 | top60_altered_DNA+RNA_seed | BRD9 | -0.58437 | LUAD | NCI-H522 | Bromosporine |
| MVP_top60_DNA_seed | CREBBP | -0.023308 | OV | OVK-18 | SGC-CBP30 | top60_altered_DNA+RNA_seed | BRD9 | -0.583015 | LUAD | EMC-BAC-2 | Bromosporine |
| MVP_top60_DNA_seed | MAPK1 | -0.021259 | OV | SW626 | Ulixertinib | top60_altered_DNA+RNA_seed | KRAS | -0.576165 | LUAD | A549 | RAS (G12C) Inhibitor-1 |
| MVP_top60_DNA_seed | MAPK3 | -0.021259 | OV | SW626 | Ulixertinib | top60_altered_DNA+RNA_seed | BRD9 | -0.572938 | LUAD | NCI-H1648 | I-BRD9 |
| MVP_top60_DNA_seed | CDK2 | -0.020154 | OV | OVMIU | Dinaciclib | top60_altered_DNA+RNA_seed | TERT | -0.540957 | LUAD | NCI-H1623 | BIBR-1532 |
| MVP_top60_DNA_seed | CDK1 | -0.020154 | OV | OVMIU | Dinaciclib | top60_altered_DNA+RNA_seed | BRD9 | -0.529817 | LUAD | NCI-H838 | I-BRD9 |
| MVP_top60_DNA_seed | PLK1 | -0.014341 | OV | JHOS-3 | MK-1775 | top60_altered_DNA+RNA_seed | KRAS | -0.523116 | LUAD | NCI-H650 | RAS (G12C) Inhibitor-1 |
| MVP_top60_DNA_seed | CDK2 | -0.013888 | OV | OVCAR-5 | Dinaciclib | top60_altered_DNA+RNA_seed | BRD9 | -0.489473 | LUAD | NCI-H2122 | I-BRD9 |
| MVP_top60_DNA_seed | CDK1 | -0.013888 | OV | OVCAR-5 | Dinaciclib | top60_altered_DNA+RNA_seed | BRD9 | -0.460898 | LUAD | NCI-H1944 | Bromosporine |
| MVP_top60_DNA_seed | CDK2 | -0.010936 | OV | OVTOKO | MK-8776 | top60_altered_DNA+RNA_seed | TERT | -0.458878 | LUAD | NCI-H1993 | BIBR-1532 |
| MVP_top60_DNA_seed | TP53 | -0.005617 | OV | OVCAR-4 | MIRA-1 | top60_altered_DNA+RNA_seed | KRAS | -0.455088 | LUAD | EMC-BAC-2 | RAS (G12C) Inhibitor-1 |
| MVP_top60_DNA_seed | PIK3CA | -0.000185 | OV | TOV-21G | AZD8186 | top60_altered_DNA+RNA_seed | KRAS | -0.414707 | LUAD | NCI-H1734 | RAS (G12C) Inhibitor-1 |
| MVP_top60_DNA_seed | CDK2 | 0.004147 | OV | OVK-18 | MK-8776 | top60_altered_DNA+RNA_seed | TERT | -0.414478 | LUAD | NCI-H650 | BIBR-1532 |
| MVP_top60_DNA_seed | EGFR | 0.008779 | OV | OVCAR-8 | Afatinib | top60_altered_DNA+RNA_seed | TERT | -0.394075 | LUAD | RERF-LC-KJ | BIBR-1532 |
| MVP_top60_DNA_seed | ERBB2 | 0.008779 | OV | OVCAR-8 | Afatinib | top60_altered_DNA+RNA_seed | BRD9 | -0.365915 | LUAD | NCI-H1792 | Bromosporine |
| MVP_top60_DNA_seed | EGFR | 0.010403 | OV | OVKATE | Erlotinib | top60_altered_DNA+RNA_seed | KRAS | -0.360798 | LUAD | NCI-H1781 | RAS (G12C) Inhibitor-1 |
| MVP_top60_DNA_seed | EGFR | 0.010667 | OV | Hey | Gefitinib | top60_altered_DNA+RNA_seed | KRAS | -0.347633 | LUAD | RERF-LC-MS | RAS (G12C) Inhibitor-1 |
| MVP_top60_DNA_seed | MAPK1 | 0.011635 | OV | TOV-21G | Ulixertinib | top60_altered_DNA+RNA_seed | BRD9 | -0.336778 | LUAD | NCI-H1666 | Bromosporine |
| MVP_top60_DNA_seed | MAPK3 | 0.011635 | OV | TOV-21G | Ulixertinib | top60_altered_DNA+RNA_seed | TERT | -0.327329 | LUAD | NCI-H23 | BIBR-1532 |
| MVP_top60_DNA_seed | EGFR | 0.017402 | OV | OV-17R | Sapitinib | top60_altered_DNA+RNA_seed | KRAS | -0.29333 | LUAD | HCC-44 | RAS (G12C) Inhibitor-1 |
| MVP_top60_DNA_seed | ERBB2 | 0.017402 | OV | OV-17R | Sapitinib | top60_altered_DNA+RNA_seed | BRD9 | -0.276848 | LUAD | NCI-H1437 | I-BRD9 |

| | | | | | | | | | | | |
|---|---|---|---|---|---|---|---|---|---|---|---|
| MVP_top60_DNA_seed | CDK1 | 0.018918 | OV | OVCA433 | RO-3306 | top60_altered_DNA+RNA_seed | TP53 | -0.266147 | LUAD | NCI-H2030 | MIRA-1 |
| MVP_top60_DNA_seed | MAPK1 | 0.020037 | OV | OV-7 | Ulixertinib | top60_altered_DNA+RNA_seed | KRAS | -0.263595 | LUAD | LXF-289 | KRAS (G12C) Inhibitor-1 |
| MVP_top60_DNA_seed | MAPK3 | 0.020037 | OV | OV-7 | Ulixertinib | top60_altered_DNA+RNA_seed | TP53 | -0.25003 | LUAD | NCI-H2122 | MIRA-1 |
| MVP_top60_DNA_seed | PIK3CA | 0.020126 | OV | Caov-4 | Alpelisib | top60_altered_DNA+RNA_seed | KRAS | -0.239604 | LUAD | NCI-H322M | KRAS (G12C) Inhibitor-1 |
| MVP_top60_DNA_seed | PLK1 | 0.024417 | OV | SK-OV-3 | MK-1775 | top60_altered_DNA+RNA_seed | BRD9 | -0.236018 | LUAD | EMC-BAC-2 | I-BRD9 |
| MVP_top60_DNA_seed | MAPK1 | 0.024607 | OV | OVCAR-3 | Ulixertinib | top60_altered_DNA+RNA_seed | TP53 | -0.228242 | LUAD | LC-2-ad | MIRA-1 |
| MVP_top60_DNA_seed | MAPK3 | 0.024607 | OV | OVCAR-3 | Ulixertinib | top60_altered_DNA+RNA_seed | BRD9 | -0.21285 | LUAD | NCI-H1993 | Bromosporine |
| MVP_top60_DNA_seed | HDAC2 | 0.034343 | OV | SK-OV-3 | Romidepsin | top60_altered_DNA+RNA_seed | TERT | -0.210942 | LUAD | NCI-H1734 | BIBR-1532 |
| MVP_top60_DNA_seed | EGFR | 0.037227 | OV | OVCAR-4 | Sapitinib | top60_altered_DNA+RNA_seed | BRD9 | -0.202968 | LUAD | NCI-H1563 | I-BRD9 |
| MVP_top60_DNA_seed | ERBB2 | 0.037227 | OV | OVCAR-4 | Sapitinib | top60_altered_DNA+RNA_seed | BRD9 | -0.202866 | LUAD | NCI-H2122 | Bromosporine |
| MVP_top60_DNA_seed | PIK3CA | 0.039057 | OV | Caov-3 | Alpelisib | top60_altered_DNA+RNA_seed | BRD9 | -0.187309 | LUAD | NCI-H1666 | I-BRD9 |
| MVP_top60_DNA_seed | MAPK1 | 0.040995 | OV | JHOS-2 | Ulixertinib | top60_altered_DNA+RNA_seed | BRD9 | -0.185913 | LUAD | NCI-H1792 | I-BRD9 |
| MVP_top60_DNA_seed | MAPK3 | 0.040995 | OV | JHOS-2 | Ulixertinib | top60_altered_DNA+RNA_seed | BRD9 | -0.156733 | LUAD | NCI-H1651 | Bromosporine |
| MVP_top60_DNA_seed | CDK1 | 0.043156 | OV | DOV13 | RO-3306 | top60_altered_DNA+RNA_seed | BRD9 | -0.14542 | LUAD | NCI-H2009 | I-BRD9 |
| MVP_top60_DNA_seed | MAPK1 | 0.048844 | OV | OVK-18 | VX-11e | top60_altered_DNA+RNA_seed | TP53 | -0.13766 | LUAD | RERF-LC-MS | MIRA-1 |
| MVP_top60_DNA_seed | ATR | 0.052328 | OV | DOV13 | VE-822 | top60_altered_DNA+RNA_seed | TERT | -0.132516 | LUAD | LXF-289 | BIBR-1532 |
| MVP_top60_DNA_seed | PLK1 | 0.052957 | OV | OVTOKO | MK-1775 | top60_altered_DNA+RNA_seed | TERT | -0.125794 | LUAD | NCI-H322M | BIBR-1532 |
| MVP_top60_DNA_seed | MAPK1 | 0.05528 | OV | OC-314 | SCH772984 | top60_altered_DNA+RNA_seed | TERT | -0.119012 | LUAD | LC-2-ad | BIBR-1532 |
| MVP_top60_DNA_seed | MAPK3 | 0.05528 | OV | OC-314 | SCH772984 | top60_altered_DNA+RNA_seed | KRAS | -0.117287 | LUAD | NCI-H838 | KRAS (G12C) Inhibitor-1 |
| MVP_top60_DNA_seed | TP53 | 0.056099 | OV | OVISE | MIRA-1 | top60_altered_DNA+RNA_seed | BRD9 | -0.114776 | LUAD | A549 | Bromosporine |
| MVP_top60_DNA_seed | ATR | 0.056907 | OV | JHOS-3 | AZD6738 | top60_altered_DNA+RNA_seed | BRD9 | -0.099113 | LUAD | NCI-H1651 | I-BRD9 |
| MVP_top60_DNA_seed | CDK2 | 0.061533 | OV | OVISE | MK-8776 | top60_altered_DNA+RNA_seed | TERT | -0.076725 | LUAD | NCI-H2030 | BIBR-1532 |
| MVP_top60_DNA_seed | EGFR | 0.067263 | OV | OV-17R | Gefitinib | top60_altered_DNA+RNA_seed | TP53 | -0.070238 | LUAD | LXF-289 | MIRA-1 |
| MVP_top60_DNA_seed | MAPK1 | 0.074288 | OV | TOV-21G | VX-11e | top60_altered_DNA+RNA_seed | TP53 | -0.063293 | LUAD | NCI-H1437 | MIRA-1 |
| MVP_top60_DNA_seed | MAPK1 | 0.076398 | OV | OVK-18 | SCH772984 | top60_altered_DNA+RNA_seed | BRD9 | -0.038769 | LUAD | NCI-H1755 | I-BRD9 |
| MVP_top60_DNA_seed | MAPK3 | 0.076398 | OV | OVK-18 | SCH772984 | top60_altered_DNA+RNA_seed | BRD9 | -0.037207 | LUAD | NCI-H1975 | Bromosporine |
| MVP_top60_DNA_seed | MAPK1 | 0.08052 | OV | DOV13 | Ulixertinib | top60_altered_DNA+RNA_seed | KRAS | -0.036458 | LUAD | NCI-H2023 | KRAS (G12C) Inhibitor-1 |
| MVP_top60_DNA_seed | MAPK3 | 0.08052 | OV | DOV13 | Ulixertinib | top60_altered_DNA+RNA_seed | TERT | -0.02603 | LUAD | NCI-H1435 | BIBR-1532 |
| MVP_top60_DNA_seed | PIK3CA | 0.082336 | OV | OVTOKO | Buparlisib | top60_altered_DNA+RNA_seed | TP53 | -0.021429 | LUAD | NCI-H522 | MIRA-1 |
| MVP_top60_DNA_seed | ATR | 0.082344 | OV | OV-7 | AZD6738 | top60_altered_DNA+RNA_seed | TP53 | -0.021345 | LUAD | RERF-LC-KJ | MIRA-1 |
| MVP_top60_DNA_seed | MAPK1 | 0.082954 | OV | UWB1.289 | Ulixertinib | top60_altered_DNA+RNA_seed | TP53 | -0.020545 | LUAD | NCI-H1651 | MIRA-1 |
| MVP_top60_DNA_seed | MAPK3 | 0.082954 | OV | UWB1.289 | Ulixertinib | top60_altered_DNA+RNA_seed | TERT | 0.005902 | LUAD | NCI-H1650 | BIBR-1532 |
| MVP_top60_DNA_seed | CDK2 | 0.083604 | OV | OAW-28 | MK-8776 | top60_altered_DNA+RNA_seed | KRAS | 0.013546 | LUAD | NCI-H3122 | KRAS (G12C) Inhibitor-1 |
| MVP_top60_DNA_seed | PIK3CA | 0.083968 | OV | DOV13 | Buparlisib | top60_altered_DNA+RNA_seed | TERT | 0.025029 | LUAD | NCI-H1651 | BIBR-1532 |
| MVP_top60_DNA_seed | PLK1 | 0.090571 | OV | JHOS-2 | MK-1775 | top60_altered_DNA+RNA_seed | BRD9 | 0.03762 | LUAD | NCI-H1975 | I-BRD9 |
| MVP_top60_DNA_seed | EGFR | 0.09127 | OV | OAW-28 | Gefitinib | top60_altered_DNA+RNA_seed | TERT | 0.040347 | LUAD | NCI-H1781 | BIBR-1532 |
| MVP_top60_DNA_seed | TP53 | 0.091522 | OV | IGROV-1 | MIRA-1 | top60_altered_DNA+RNA_seed | BRD9 | 0.045812 | LUAD | HCC-44 | I-BRD9 |
| MVP_top60_DNA_seed | PIK3CA | 0.091785 | OV | OVCA433 | Buparlisib | top60_altered_DNA+RNA_seed | TERT | 0.052516 | LUAD | HCC-44 | BIBR-1532 |
| MVP_top60_DNA_seed | PIK3CA | 0.09816 | OV | Caov-4 | Buparlisib | top60_altered_DNA+RNA_seed | TERT | 0.067007 | LUAD | EMC-BAC-2 | BIBR-1532 |
| MVP_top60_DNA_seed | PIK3CA | 0.104986 | OV | OAW-28 | Buparlisib | top60_altered_DNA+RNA_seed | BRD9 | 0.073841 | LUAD | NCI-H322M | Bromosporine |
| MVP_top60_DNA_seed | EGFR | 0.11171 | OV | Caov-3 | AZD3759 | top60_altered_DNA+RNA_seed | KRAS | 0.100027 | LUAD | NCI-H1651 | KRAS (G12C) Inhibitor-1 |

| | | | | | | | | | | | |
|---|---|---|---|---|---|---|---|---|---|---|---|
| MVP_top60_DNA_seed | PIK3CA | 0.112797 | OV | FU-OV-1 | AZD8186 | top60_altered_DNA+RNA_seed | KRAS | 0.105467 | LUAD | H3255 | RAS (G12C) Inhibitor-1 |
| MVP_top60_DNA_seed | MAPK1 | 0.116989 | OV | Caov-3 | Ulixertinib | top60_altered_DNA+RNA_seed | TP53 | 0.109622 | LUAD | NCI-H2009 | MIRA-1 |
| MVP_top60_DNA_seed | MAPK3 | 0.116989 | OV | Caov-3 | Ulixertinib | top60_altered_DNA+RNA_seed | BRD9 | 0.115613 | LUAD | NCI-H1755 | Bromosporine |
| MVP_top60_DNA_seed | TP53 | 0.117296 | OV | PEO1 | MIRA-1 | top60_altered_DNA+RNA_seed | TP53 | 0.122093 | LUAD | VMRC-LCD | MIRA-1 |
| MVP_top60_DNA_seed | ATR | 0.11942 | OV | PEO1 | AZD6738 | top60_altered_DNA+RNA_seed | TERT | 0.12808 | LUAD | RERF-LC-MS | BIBR-1532 |
| MVP_top60_DNA_seed | CDK2 | 0.119934 | OV | OVMIU | AZD5438 | top60_altered_DNA+RNA_seed | TERT | 0.164093 | LUAD | NCI-H1437 | BIBR-1532 |
| MVP_top60_DNA_seed | EGFR | 0.124286 | OV | OVCAR-4 | Erlotinib | top60_altered_DNA+RNA_seed | KRAS | 0.179106 | LUAD | HOP-62 | RAS (G12C) Inhibitor-1 |
| MVP_top60_DNA_seed | CREBBP | 0.128693 | OV | OVCAR-8 | SGC-CBP30 | top60_altered_DNA+RNA_seed | TP53 | 0.184186 | LUAD | NCI-H838 | MIRA-1 |
| MVP_top60_DNA_seed | PIK3CA | 0.136027 | OV | OVMIU | GNE-317 | top60_altered_DNA+RNA_seed | TP53 | 0.194821 | LUAD | NCI-H1648 | MIRA-1 |
| MVP_top60_DNA_seed | ATR | 0.137407 | OV | JHOS-2 | AZD6738 | top60_altered_DNA+RNA_seed | BRD9 | 0.198308 | LUAD | NCI-H2087 | Bromosporine |
| MVP_top60_DNA_seed | EGFR | 0.137588 | OV | DOV13 | Erlotinib | top60_altered_DNA+RNA_seed | TERT | 0.19946 | LUAD | HCC-827 | BIBR-1532 |
| MVP_top60_DNA_seed | PLK1 | 0.138561 | OV | TOV-112D | MK-1775 | top60_altered_DNA+RNA_seed | BRD9 | 0.203063 | LUAD | NCI-H23 | Bromosporine |
| MVP_top60_DNA_seed | MAPK1 | 0.138613 | OV | OC-314 | Ulixertinib | top60_altered_DNA+RNA_seed | KRAS | 0.20407 | LUAD | ABC-1 | RAS (G12C) Inhibitor-1 |
| MVP_top60_DNA_seed | MAPK3 | 0.138613 | OV | OC-314 | Ulixertinib | top60_altered_DNA+RNA_seed | TERT | 0.208927 | LUAD | NCI-H1944 | BIBR-1532 |
| MVP_top60_DNA_seed | EGFR | 0.145869 | OV | TOV-21G | Osimertinib | top60_altered_DNA+RNA_seed | BRD9 | 0.218902 | LUAD | A549 | I-BRD9 |
| MVP_top60_DNA_seed | EGFR | 0.147127 | OV | OVCAR-3 | Afatinib | top60_altered_DNA+RNA_seed | KRAS | 0.257163 | LUAD | RERF-LC-KJ | RAS (G12C) Inhibitor-1 |
| MVP_top60_DNA_seed | ERBB2 | 0.147127 | OV | OVCAR-3 | Afatinib | top60_altered_DNA+RNA_seed | BRD9 | 0.261197 | LUAD | NCI-H358 | I-BRD9 |
| MVP_top60_DNA_seed | EGFR | 0.152615 | OV | OVCA433 | Osimertinib | top60_altered_DNA+RNA_seed | KRAS | 0.265622 | LUAD | NCI-H1666 | RAS (G12C) Inhibitor-1 |
| MVP_top60_DNA_seed | PIK3CA | 0.154344 | OV | EFO-21 | GNE-317 | top60_altered_DNA+RNA_seed | TP53 | 0.271202 | LUAD | NCI-H2085 | MIRA-1 |
| MVP_top60_DNA_seed | MAPK1 | 0.157819 | OV | OVMIU | VX-11e | top60_altered_DNA+RNA_seed | TP53 | 0.273078 | LUAD | NCI-H23 | MIRA-1 |
| MVP_top60_DNA_seed | PIK3CA | 0.158628 | OV | JHOS-3 | AZD8186 | top60_altered_DNA+RNA_seed | BRD9 | 0.274441 | LUAD | Calu-6 | Bromosporine |
| MVP_top60_DNA_seed | PIK3CA | 0.158952 | OV | OVISE | GNE-317 | top60_altered_DNA+RNA_seed | TERT | 0.274731 | LUAD | Calu-6 | BIBR-1532 |
| MVP_top60_DNA_seed | PIK3CA | 0.159528 | OV | OVTOKO | Alpelisib | top60_altered_DNA+RNA_seed | BRD9 | 0.306316 | LUAD | NCI-H1944 | I-BRD9 |
| MVP_top60_DNA_seed | EGFR | 0.160546 | OV | OAW-28 | Erlotinib | top60_altered_DNA+RNA_seed | BRD9 | 0.31203 | LUAD | NCI-H1993 | I-BRD9 |
| MVP_top60_DNA_seed | EGFR | 0.171285 | OV | OVCAR-3 | Sapitinib | top60_altered_DNA+RNA_seed | BRD9 | 0.355059 | LUAD | NCI-H2030 | I-BRD9 |
| MVP_top60_DNA_seed | ERBB2 | 0.171285 | OV | OVCAR-3 | Sapitinib | top60_altered_DNA+RNA_seed | BRD9 | 0.35988 | LUAD | ABC-1 | I-BRD9 |
| MVP_top60_DNA_seed | PIK3CA | 0.176384 | OV | OVCAR-5 | Buparlisib | top60_altered_DNA+RNA_seed | TP53 | 0.366569 | LUAD | EMC-BAC-1 | MIRA-1 |
| MVP_top60_DNA_seed | CDK2 | 0.177396 | OV | OAW-42 | AZD5438 | top60_altered_DNA+RNA_seed | BRD9 | 0.382116 | LUAD | PC-3_[JPC-3] | I-BRD9 |
| MVP_top60_DNA_seed | EGFR | 0.180072 | OV | TOV-21G | AZD3759 | top60_altered_DNA+RNA_seed | BRD9 | 0.392292 | LUAD | NCI-H2405 | I-BRD9 |
| MVP_top60_DNA_seed | MAPK1 | 0.182318 | OV | Caov-3 | SCH772984 | top60_altered_DNA+RNA_seed | TERT | 0.405246 | LUAD | NCI-H1648 | BIBR-1532 |
| MVP_top60_DNA_seed | MAPK3 | 0.182318 | OV | Caov-3 | SCH772984 | top60_altered_DNA+RNA_seed | TERT | 0.415574 | LUAD | NCI-H1975 | BIBR-1532 |
| MVP_top60_DNA_seed | TP53 | 0.182884 | OV | Caov-3 | MIRA-1 | top60_altered_DNA+RNA_seed | TP53 | 0.439079 | LUAD | NCI-H650 | MIRA-1 |
| MVP_top60_DNA_seed | EGFR | 0.186212 | OV | PEO1 | Sapitinib | top60_altered_DNA+RNA_seed | BRD9 | 0.442087 | LUAD | LC-2-ad | I-BRD9 |
| MVP_top60_DNA_seed | ERBB2 | 0.186212 | OV | PEO1 | Sapitinib | top60_altered_DNA+RNA_seed | TP53 | 0.444214 | LUAD | NCI-H3122 | MIRA-1 |
| MVP_top60_DNA_seed | CDK2 | 0.188975 | OV | OV-17R | Dinaciclib | top60_altered_DNA+RNA_seed | BRD9 | 0.453932 | LUAD | NCI-H650 | Bromosporine |
| MVP_top60_DNA_seed | CDK1 | 0.188975 | OV | OV-17R | Dinaciclib | top60_altered_DNA+RNA_seed | TERT | 0.46521 | LUAD | NCI-H522 | BIBR-1532 |
| MVP_top60_DNA_seed | PLK1 | 0.189239 | OV | OVTOKO | BI-2536 | top60_altered_DNA+RNA_seed | TERT | 0.466496 | LUAD | NCI-H2085 | BIBR-1532 |
| MVP_top60_DNA_seed | PIK3CA | 0.189843 | OV | SK-OV-3 | AZD8186 | top60_altered_DNA+RNA_seed | BRD9 | 0.467879 | LUAD | NCI-H1435 | I-BRD9 |
| MVP_top60_DNA_seed | MAPK1 | 0.192521 | OV | OVISE | SCH772984 | top60_altered_DNA+RNA_seed | BRD9 | 0.477396 | LUAD | VMRC-LCD | Bromosporine |
| MVP_top60_DNA_seed | MAPK3 | 0.192521 | OV | OVISE | SCH772984 | top60_altered_DNA+RNA_seed | KRAS | 0.480611 | LUAD | VMRC-LCD | RAS (G12C) Inhibitor-1 |
| MVP_top60_DNA_seed | ATR | 0.194866 | OV | FU-OV-1 | VE821 | top60_altered_DNA+RNA_seed | KRAS | 0.497351 | LUAD | Calu-6 | RAS (G12C) Inhibitor-1 |

| | | | | | | | | | | | |
|---|---|---|---|---|---|---|---|---|---|---|---|
| MVP_top60_DNA_seed | TP53 | 0.207213 | OV | OAW-28 | MIRA-1 | top60_altered_DNA+RNA_seed | TP53 | 0.50233 | LUAD | HCC-44 | MIRA-1 |
| MVP_top60_DNA_seed | TP53 | 0.208587 | OV | TOV-112D | MIRA-1 | top60_altered_DNA+RNA_seed | BRD9 | 0.504722 | LUAD | NCI-H23 | I-BRD9 |
| MVP_top60_DNA_seed | CDK2 | 0.213191 | OV | SW626 | AZD5438 | top60_altered_DNA+RNA_seed | TP53 | 0.510946 | LUAD | NCI-H1993 | MIRA-1 |
| MVP_top60_DNA_seed | MAPK1 | 0.214267 | OV | OAW-28 | Ulixertinib | top60_altered_DNA+RNA_seed | BRD9 | 0.514473 | LUAD | VMRC-LCD | I-BRD9 |
| MVP_top60_DNA_seed | MAPK3 | 0.214267 | OV | OAW-28 | Ulixertinib | top60_altered_DNA+RNA_seed | TERT | 0.52758 | LUAD | ABC-1 | BIBR-1532 |
| MVP_top60_DNA_seed | MAPK1 | 0.221031 | OV | FU-OV-1 | SCH772984 | top60_altered_DNA+RNA_seed | TP53 | 0.529901 | LUAD | NCI-H1573 | MIRA-1 |
| MVP_top60_DNA_seed | MAPK3 | 0.221031 | OV | FU-OV-1 | SCH772984 | top60_altered_DNA+RNA_seed | BRD9 | 0.539094 | LUAD | NCI-H1623 | Bromosporine |
| MVP_top60_DNA_seed | EGFR | 0.22453 | OV | DOV13 | Gefitinib | top60_altered_DNA+RNA_seed | BRD9 | 0.54331 | LUAD | H3255 | I-BRD9 |
| MVP_top60_DNA_seed | EGFR | 0.226219 | OV | Caov-4 | Afatinib | top60_altered_DNA+RNA_seed | TP53 | 0.546422 | LUAD | NCI-H1781 | MIRA-1 |
| MVP_top60_DNA_seed | ERBB2 | 0.226219 | OV | Caov-4 | Afatinib | top60_altered_DNA+RNA_seed | BRD9 | 0.549555 | LUAD | NCI-H322M | I-BRD9 |
| MVP_top60_DNA_seed | PLK1 | 0.228285 | OV | OVCA433 | BI-2536 | top60_altered_DNA+RNA_seed | TP53 | 0.556919 | LUAD | ABC-1 | MIRA-1 |
| MVP_top60_DNA_seed | CDK1 | 0.22865 | OV | OVK-18 | RO-3306 | top60_altered_DNA+RNA_seed | KRAS | 0.557187 | LUAD | NCI-H1563 | RAS (G12C) Inhibitor-1 |
| MVP_top60_DNA_seed | EGFR | 0.236076 | OV | OAW-28 | Lapatinib | top60_altered_DNA+RNA_seed | TERT | 0.583715 | LUAD | NCI-H2228 | BIBR-1532 |
| MVP_top60_DNA_seed | ERBB2 | 0.236076 | OV | OAW-28 | Lapatinib | top60_altered_DNA+RNA_seed | BRD9 | 0.597292 | LUAD | NCI-H1793 | I-BRD9 |
| MVP_top60_DNA_seed | EGFR | 0.239452 | OV | OVKATE | Osimertinib | top60_altered_DNA+RNA_seed | TP53 | 0.60034 | LUAD | NCI-H1755 | MIRA-1 |
| MVP_top60_DNA_seed | EGFR | 0.241538 | OV | FU-OV-1 | Lapatinib | top60_altered_DNA+RNA_seed | BRD9 | 0.62427 | LUAD | NCI-H1563 | Bromosporine |
| MVP_top60_DNA_seed | ERBB2 | 0.241538 | OV | FU-OV-1 | Lapatinib | top60_altered_DNA+RNA_seed | TERT | 0.626208 | LUAD | NCI-H1563 | BIBR-1532 |
| MVP_top60_DNA_seed | MAPK1 | 0.24362 | OV | OVCAR-4 | VX-11e | top60_altered_DNA+RNA_seed | KRAS | 0.638042 | LUAD | NCI-H522 | RAS (G12C) Inhibitor-1 |
| MVP_top60_DNA_seed | EGFR | 0.244486 | OV | OVTOKO | Gefitinib | top60_altered_DNA+RNA_seed | KRAS | 0.63857 | LUAD | NCI-H2087 | RAS (G12C) Inhibitor-1 |
| MVP_top60_DNA_seed | EGFR | 0.25082 | OV | OV-17R | Afatinib | top60_altered_DNA+RNA_seed | KRAS | 0.664545 | LUAD | NCI-H1944 | RAS (G12C) Inhibitor-1 |
| MVP_top60_DNA_seed | ERBB2 | 0.25082 | OV | OV-17R | Afatinib | top60_altered_DNA+RNA_seed | BRD9 | 0.670687 | LUAD | PC-14 | I-BRD9 |
| MVP_top60_DNA_seed | EGFR | 0.252805 | OV | OAW-42 | Erlotinib | top60_altered_DNA+RNA_seed | BRD9 | 0.682399 | LUAD | NCI-H1573 | I-BRD9 |
| MVP_top60_DNA_seed | EGFR | 0.259345 | OV | OVTOKO | Osimertinib | top60_altered_DNA+RNA_seed | TP53 | 0.690013 | LUAD | NCI-H1944 | MIRA-1 |
| MVP_top60_DNA_seed | EGFR | 0.266819 | OV | OVCAR-5 | Afatinib | top60_altered_DNA+RNA_seed | BRD9 | 0.714904 | LUAD | NCI-H650 | I-BRD9 |
| MVP_top60_DNA_seed | ERBB2 | 0.266819 | OV | OVCAR-5 | Afatinib | top60_altered_DNA+RNA_seed | TP53 | 0.718681 | LUAD | EMC-BAC-2 | MIRA-1 |
| MVP_top60_DNA_seed | EGFR | 0.267207 | OV | OAW-28 | Osimertinib | top60_altered_DNA+RNA_seed | BRD9 | 0.719418 | LUAD | NCI-H1435 | Bromosporine |
| MVP_top60_DNA_seed | EGFR | 0.267503 | OV | Caov-4 | AZD3759 | top60_altered_DNA+RNA_seed | BRD9 | 0.723754 | LUAD | EMC-BAC-1 | I-BRD9 |
| MVP_top60_DNA_seed | EGFR | 0.26798 | OV | OVCA433 | AZD3759 | top60_altered_DNA+RNA_seed | TP53 | 0.734693 | LUAD | NCI-H1693 | MIRA-1 |
| MVP_top60_DNA_seed | PIK3CA | 0.270717 | OV | PEO1 | Alpelisib | top60_altered_DNA+RNA_seed | TP53 | 0.738409 | LUAD | NCI-H2023 | MIRA-1 |
| MVP_top60_DNA_seed | PIK3CA | 0.273956 | OV | UWB1.289 | Buparlisib | top60_altered_DNA+RNA_seed | TP53 | 0.758629 | LUAD | NCI-H1563 | MIRA-1 |
| MVP_top60_DNA_seed | MAPK1 | 0.274644 | OV | OC-314 | Ulixertinib | top60_altered_DNA+RNA_seed | TP53 | 0.774438 | LUAD | HCC-78 | MIRA-1 |
| MVP_top60_DNA_seed | MAPK3 | 0.274644 | OV | OC-314 | Ulixertinib | top60_altered_DNA+RNA_seed | TERT | 0.784312 | LUAD | PC-3_[JPC-3] | BIBR-1532 |
| MVP_top60_DNA_seed | CREBBP | 0.274942 | OV | OVCAR-3 | SGC-CBP30 | top60_altered_DNA+RNA_seed | BRD9 | 0.789368 | LUAD | NCI-H3122 | Bromosporine |
| MVP_top60_DNA_seed | ATR | 0.279162 | OV | OVCAR-3 | VE821 | top60_altered_DNA+RNA_seed | BRD9 | 0.792468 | LUAD | NCI-H2023 | Bromosporine |
| MVP_top60_DNA_seed | PIK3CA | 0.290924 | OV | JHOS-2 | Buparlisib | top60_altered_DNA+RNA_seed | TERT | 0.795957 | LUAD | A549 | BIBR-1532 |
| MVP_top60_DNA_seed | TP53 | 0.298653 | OV | OVCAR-3 | MIRA-1 | top60_altered_DNA+RNA_seed | BRD9 | 0.799301 | LUAD | NCI-H522 | I-BRD9 |
| MVP_top60_DNA_seed | CDK2 | 0.303444 | OV | OVTOKO | Dinaciclib | top60_altered_DNA+RNA_seed | BRD9 | 0.801363 | LUAD | Calu-3 | Bromosporine |
| MVP_top60_DNA_seed | CDK1 | 0.303444 | OV | OVTOKO | Dinaciclib | top60_altered_DNA+RNA_seed | BRD9 | 0.804124 | LUAD | NCI-H1568 | Bromosporine |
| MVP_top60_DNA_seed | EGFR | 0.304161 | OV | JHOS-3 | Erlotinib | top60_altered_DNA+RNA_seed | TP53 | 0.813338 | LUAD | PC-3_[JPC-3] | MIRA-1 |
| MVP_top60_DNA_seed | EGFR | 0.305324 | OV | OVCA433 | Erlotinib | top60_altered_DNA+RNA_seed | BRD9 | 0.830891 | LUAD | RERF-LC-KJ | I-BRD9 |
| MVP_top60_DNA_seed | PIK3CA | 0.309134 | OV | JHOS-3 | GNE-317 | top60_altered_DNA+RNA_seed | TERT | 0.841422 | LUAD | COR-L105 | BIBR-1532 |

| | | | | | | | | | | | |
|---|---|---|---|---|---|---|---|---|---|---|---|
| MVP_top60_DNA_seed | EGFR | 0.31049 | OV | OVCAR-8 | AZD3759 | top60_altered_DNA+RNA_seed | TP53 | 0.848045 | LUAD | NCI-H1623 | MIRA-1 |
| MVP_top60_DNA_seed | MAPK1 | 0.311697 | OV | OVCAR-3 | Ulixertinib | top60_altered_DNA+RNA_seed | KRAS | 0.849262 | LUAD | NCI-H1435 | :RAS (G12C) Inhibitor-1 |
| MVP_top60_DNA_seed | MAPK3 | 0.311697 | OV | OVCAR-3 | Ulixertinib | top60_altered_DNA+RNA_seed | TERT | 0.852616 | LUAD | NCI-H3122 | BIBR-1532 |
| MVP_top60_DNA_seed | EGFR | 0.311732 | OV | Caov-3 | Lapatinib | top60_altered_DNA+RNA_seed | KRAS | 0.86381 | LUAD | NCI-H358 | :RAS (G12C) Inhibitor-1 |
| MVP_top60_DNA_seed | ERBB2 | 0.311732 | OV | Caov-3 | Lapatinib | top60_altered_DNA+RNA_seed | TERT | 0.865159 | LUAD | NCI-H2023 | BIBR-1532 |
| MVP_top60_DNA_seed | PIK3CA | 0.312313 | OV | OVMIU | AZD8186 | top60_altered_DNA+RNA_seed | TERT | 0.867275 | LUAD | EKVX | BIBR-1532 |
| MVP_top60_DNA_seed | PLK1 | 0.313473 | OV | OV-17R | BI-2536 | top60_altered_DNA+RNA_seed | BRD9 | 0.869967 | LUAD | NCI-H1355 | Bromosporine |
| MVP_top60_DNA_seed | TP53 | 0.315496 | OV | SW626 | MIRA-1 | top60_altered_DNA+RNA_seed | BRD9 | 0.871911 | LUAD | NCI-H2087 | I-BRD9 |
| MVP_top60_DNA_seed | EGFR | 0.318129 | OV | OV-7 | Afatinib | top60_altered_DNA+RNA_seed | TP53 | 0.877456 | LUAD | NCI-H2342 | MIRA-1 |
| MVP_top60_DNA_seed | ERBB2 | 0.318129 | OV | OV-7 | Afatinib | top60_altered_DNA+RNA_seed | KRAS | 0.886349 | LUAD | NCI-H1623 | :RAS (G12C) Inhibitor-1 |
| MVP_top60_DNA_seed | ATR | 0.31815 | OV | OAW-42 | AZD6738 | top60_altered_DNA+RNA_seed | BRD9 | 0.891852 | LUAD | LXF-289 | I-BRD9 |
| MVP_top60_DNA_seed | PIK3CA | 0.318488 | OV | OV-56 | GNE-317 | top60_altered_DNA+RNA_seed | TP53 | 0.914092 | LUAD | Calu-6 | MIRA-1 |
| MVP_top60_DNA_seed | PIK3CA | 0.319101 | OV | OVCAR-8 | Alpelisib | top60_altered_DNA+RNA_seed | TP53 | 0.92166 | LUAD | NCI-H1975 | MIRA-1 |
| MVP_top60_DNA_seed | PLK1 | 0.328451 | OV | PEO1 | MK-1775 | top60_altered_DNA+RNA_seed | TERT | 0.926156 | LUAD | NCI-H1755 | BIBR-1532 |
| MVP_top60_DNA_seed | MAPK1 | 0.336232 | OV | OAW-28 | Ulixertinib | top60_altered_DNA+RNA_seed | KRAS | 0.936083 | LUAD | NCI-H2030 | :RAS (G12C) Inhibitor-1 |
| MVP_top60_DNA_seed | MAPK3 | 0.336232 | OV | OAW-28 | Ulixertinib | top60_altered_DNA+RNA_seed | BRD9 | 0.938286 | LUAD | HCC-827 | Bromosporine |
| MVP_top60_DNA_seed | CDK1 | 0.33662 | OV | UWB1.289 | RO-3306 | top60_altered_DNA+RNA_seed | KRAS | 0.946933 | LUAD | NCI-H2085 | :RAS (G12C) Inhibitor-1 |
| MVP_top60_DNA_seed | ATR | 0.344679 | OV | OVCAR-4 | AZD6738 | top60_altered_DNA+RNA_seed | TERT | 0.947953 | LUAD | NCI-H2342 | BIBR-1532 |
| MVP_top60_DNA_seed | PIK3CA | 0.345481 | OV | Caov-3 | GNE-317 | top60_altered_DNA+RNA_seed | KRAS | 0.969335 | LUAD | HCC-827 | :RAS (G12C) Inhibitor-1 |
| MVP_top60_DNA_seed | CDK1 | 0.346979 | OV | OVCAR-8 | RO-3306 | top60_altered_DNA+RNA_seed | TP53 | 0.977542 | LUAD | NCI-H322M | MIRA-1 |
| MVP_top60_DNA_seed | EGFR | 0.348579 | OV | OAW-42 | Sapitinib | top60_altered_DNA+RNA_seed | BRD9 | 0.98021 | LUAD | NCI-H2342 | I-BRD9 |
| MVP_top60_DNA_seed | ERBB2 | 0.348579 | OV | OAW-42 | Sapitinib | top60_altered_DNA+RNA_seed | TP53 | 0.986453 | LUAD | PC-14 | MIRA-1 |
| MVP_top60_DNA_seed | HDAC2 | 0.349571 | OV | OVTOKO | Romidepsin | top60_altered_DNA+RNA_seed | KRAS | 0.98968 | LUAD | Calu-3 | :RAS (G12C) Inhibitor-1 |
| MVP_top60_DNA_seed | EGFR | 0.354037 | OV | TOV-112D | Sapitinib | top60_altered_DNA+RNA_seed | BRD9 | 1.000166 | LUAD | NCI-H1781 | Bromosporine |
| MVP_top60_DNA_seed | ERBB2 | 0.354037 | OV | TOV-112D | Sapitinib | top60_altered_DNA+RNA_seed | TP53 | 1.010753 | LUAD | HOP-62 | MIRA-1 |
| MVP_top60_DNA_seed | PIK3CA | 0.368077 | OV | EFO-21 | AZD8186 | top60_altered_DNA+RNA_seed | TP53 | 1.015556 | LUAD | NCI-H358 | MIRA-1 |
| MVP_top60_DNA_seed | EGFR | 0.380393 | OV | PEO1 | Afatinib | top60_altered_DNA+RNA_seed | KRAS | 1.033303 | LUAD | EKVX | :RAS (G12C) Inhibitor-1 |
| MVP_top60_DNA_seed | ERBB2 | 0.380393 | OV | PEO1 | Afatinib | top60_altered_DNA+RNA_seed | TP53 | 1.043648 | LUAD | SK-LU-1 | MIRA-1 |
| MVP_top60_DNA_seed | MAPK1 | 0.383381 | OV | OVCAR-4 | SCH772984 | top60_altered_DNA+RNA_seed | TP53 | 1.057105 | LUAD | A549 | MIRA-1 |
| MVP_top60_DNA_seed | MAPK3 | 0.383381 | OV | OVCAR-4 | SCH772984 | top60_altered_DNA+RNA_seed | KRAS | 1.058704 | LUAD | NCI-H1355 | :RAS (G12C) Inhibitor-1 |
| MVP_top60_DNA_seed | ATR | 0.383462 | OV | OVCA420 | VE-822 | top60_altered_DNA+RNA_seed | BRD9 | 1.058707 | LUAD | NCI-H1781 | I-BRD9 |
| MVP_top60_DNA_seed | EGFR | 0.384329 | OV | OVCAR-4 | Afatinib | top60_altered_DNA+RNA_seed | TP53 | 1.060177 | LUAD | NCI-H2087 | MIRA-1 |
| MVP_top60_DNA_seed | ERBB2 | 0.384329 | OV | OVCAR-4 | Afatinib | top60_altered_DNA+RNA_seed | BRD9 | 1.069834 | LUAD | NCI-H3122 | I-BRD9 |
| MVP_top60_DNA_seed | ATR | 0.386103 | OV | SW626 | AZD6738 | top60_altered_DNA+RNA_seed | TP53 | 1.071087 | LUAD | NCI-H2291 | MIRA-1 |
| MVP_top60_DNA_seed | MAPK1 | 0.388164 | OV | OVISE | Ulixertinib | top60_altered_DNA+RNA_seed | TP53 | 1.075039 | LUAD | Calu-3 | MIRA-1 |
| MVP_top60_DNA_seed | MAPK3 | 0.388164 | OV | OVISE | Ulixertinib | top60_altered_DNA+RNA_seed | BRD9 | 1.115935 | LUAD | NCI-H1623 | I-BRD9 |
| MVP_top60_DNA_seed | EGFR | 0.394212 | OV | TOV-112D | Afatinib | top60_altered_DNA+RNA_seed | BRD9 | 1.121407 | LUAD | NCI-H1355 | I-BRD9 |
| MVP_top60_DNA_seed | ERBB2 | 0.394212 | OV | TOV-112D | Afatinib | top60_altered_DNA+RNA_seed | TERT | 1.141257 | LUAD | NCI-H2009 | BIBR-1532 |
| MVP_top60_DNA_seed | TP53 | 0.403741 | OV | OVK-18 | MIRA-1 | top60_altered_DNA+RNA_seed | TP53 | 1.144447 | LUAD | NCI-H1838 | MIRA-1 |
| MVP_top60_DNA_seed | EGFR | 0.407777 | OV | OVCA433 | Sapitinib | top60_altered_DNA+RNA_seed | TERT | 1.155999 | LUAD | H3255 | BIBR-1532 |
| MVP_top60_DNA_seed | ERBB2 | 0.407777 | OV | OVCA433 | Sapitinib | top60_altered_DNA+RNA_seed | BRD9 | 1.156979 | LUAD | ABC-1 | Bromosporine |

| | | | | | | | | | | | |
|---|---|---|---|---|---|---|---|---|---|---|---|
| MVP_top60_DNA_seed | CDK2 | 0.407934 | OV | OAW-42 | MK-8776 | top60_altered_DNA+RNA_seed | TP53 | 1.162837 | LUAD | H3255 | MIRA-1 |
| MVP_top60_DNA_seed | CDK1 | 0.408611 | OV | TOV-21G | RO-3306 | top60_altered_DNA+RNA_seed | BRD9 | 1.169214 | LUAD | NCI-H2228 | I-BRD9 |
| MVP_top60_DNA_seed | PLK1 | 0.411119 | OV | Hey | BI-2536 | top60_altered_DNA+RNA_seed | TP53 | 1.172228 | LUAD | NCI-H1355 | MIRA-1 |
| MVP_top60_DNA_seed | PIK3CA | 0.415692 | OV | OVK-18 | Buparlisib | top60_altered_DNA+RNA_seed | TERT | 1.186611 | LUAD | HCC-78 | BIBR-1532 |
| MVP_top60_DNA_seed | MAPK1 | 0.424012 | OV | TOV-21G | Ulixertinib | top60_altered_DNA+RNA_seed | BRD9 | 1.187279 | LUAD | NCI-H358 | Bromosporine |
| MVP_top60_DNA_seed | MAPK3 | 0.424012 | OV | TOV-21G | Ulixertinib | top60_altered_DNA+RNA_seed | KRAS | 1.209619 | LUAD | NCI-H2291 | KRAS (G12C) Inhibitor-1 |
| MVP_top60_DNA_seed | PLK1 | 0.424561 | OV | Hey | MK-1775 | top60_altered_DNA+RNA_seed | BRD9 | 1.221117 | LUAD | PC-14 | Bromosporine |
| MVP_top60_DNA_seed | EGFR | 0.42545 | OV | OVCA433 | Afatinib | top60_altered_DNA+RNA_seed | TP53 | 1.221488 | LUAD | COR-L105 | MIRA-1 |
| MVP_top60_DNA_seed | ERBB2 | 0.42545 | OV | OVCA433 | Afatinib | top60_altered_DNA+RNA_seed | BRD9 | 1.248195 | LUAD | NCI-H2085 | I-BRD9 |
| MVP_top60_DNA_seed | PIK3CA | 0.426306 | OV | Hey | AZD8186 | top60_altered_DNA+RNA_seed | KRAS | 1.250427 | LUAD | NCI-H1793 | KRAS (G12C) Inhibitor-1 |
| MVP_top60_DNA_seed | CDK2 | 0.431246 | OV | TOV-112D | MK-8776 | top60_altered_DNA+RNA_seed | TERT | 1.27078 | LUAD | PC-14 | BIBR-1532 |
| MVP_top60_DNA_seed | PLK1 | 0.433384 | OV | OVCAR-5 | BI-2536 | top60_altered_DNA+RNA_seed | TERT | 1.274898 | LUAD | HOP-62 | BIBR-1532 |
| MVP_top60_DNA_seed | MAPK1 | 0.434274 | OV | PEO1 | SCH772984 | top60_altered_DNA+RNA_seed | KRAS | 1.30271 | LUAD | NCI-H1755 | KRAS (G12C) Inhibitor-1 |
| MVP_top60_DNA_seed | MAPK3 | 0.434274 | OV | PEO1 | SCH772984 | top60_altered_DNA+RNA_seed | TP53 | 1.320283 | LUAD | EKVX | MIRA-1 |
| MVP_top60_DNA_seed | CREBBP | 0.435889 | OV | SW626 | SGC-CBP30 | top60_altered_DNA+RNA_seed | BRD9 | 1.329192 | LUAD | SW1573 | Bromosporine |
| MVP_top60_DNA_seed | EGFR | 0.437123 | OV | FU-OV-1 | Afatinib | top60_altered_DNA+RNA_seed | TP53 | 1.342672 | LUAD | SW1573 | MIRA-1 |
| MVP_top60_DNA_seed | ERBB2 | 0.437123 | OV | FU-OV-1 | Afatinib | top60_altered_DNA+RNA_seed | BRD9 | 1.344005 | LUAD | Calu-3 | I-BRD9 |
| MVP_top60_DNA_seed | PIK3CA | 0.440805 | OV | OVCAR-5 | AZD8186 | top60_altered_DNA+RNA_seed | BRD9 | 1.347902 | LUAD | NCI-H1693 | I-BRD9 |
| MVP_top60_DNA_seed | PIK3CA | 0.441456 | OV | OV-56 | AZD8186 | top60_altered_DNA+RNA_seed | BRD9 | 1.380537 | LUAD | HOP-62 | Bromosporine |
| MVP_top60_DNA_seed | EGFR | 0.443917 | OV | OVCAR-4 | Osimertinib | top60_altered_DNA+RNA_seed | TERT | 1.384092 | LUAD | NCI-H358 | BIBR-1532 |
| MVP_top60_DNA_seed | EGFR | 0.44598 | OV | Caov-4 | Sapitinib | top60_altered_DNA+RNA_seed | BRD9 | 1.396357 | LUAD | HOP-62 | I-BRD9 |
| MVP_top60_DNA_seed | ERBB2 | 0.44598 | OV | Caov-4 | Sapitinib | top60_altered_DNA+RNA_seed | BRD9 | 1.410531 | LUAD | NCI-H2085 | Bromosporine |
| MVP_top60_DNA_seed | EGFR | 0.454038 | OV | OVCAR-5 | Lapatinib | top60_altered_DNA+RNA_seed | TERT | 1.423055 | LUAD | VMRC-LCD | BIBR-1532 |
| MVP_top60_DNA_seed | ERBB2 | 0.454038 | OV | OVCAR-5 | Lapatinib | top60_altered_DNA+RNA_seed | TERT | 1.433595 | LUAD | NCI-H2405 | BIBR-1532 |
| MVP_top60_DNA_seed | ATR | 0.456795 | OV | OAW-28 | VE-822 | top60_altered_DNA+RNA_seed | TERT | 1.43452 | LUAD | NCI-H1568 | BIBR-1532 |
| MVP_top60_DNA_seed | PIK3CA | 0.465026 | OV | PEO1 | AZD8186 | top60_altered_DNA+RNA_seed | BRD9 | 1.445571 | LUAD | EKVX | I-BRD9 |
| MVP_top60_DNA_seed | EGFR | 0.467883 | OV | OAW-42 | Afatinib | top60_altered_DNA+RNA_seed | KRAS | 1.453596 | LUAD | NCI-H1693 | KRAS (G12C) Inhibitor-1 |
| MVP_top60_DNA_seed | ERBB2 | 0.467883 | OV | OAW-42 | Afatinib | top60_altered_DNA+RNA_seed | TERT | 1.460395 | LUAD | NCI-H2347 | BIBR-1532 |
| MVP_top60_DNA_seed | CDK2 | 0.469452 | OV | OVISE | Dinaciclib | top60_altered_DNA+RNA_seed | TP53 | 1.479916 | LUAD | NCI-H2228 | MIRA-1 |
| MVP_top60_DNA_seed | CDK1 | 0.469452 | OV | OVISE | Dinaciclib | top60_altered_DNA+RNA_seed | TP53 | 1.497721 | LUAD | NCI-H1666 | MIRA-1 |
| MVP_top60_DNA_seed | EGFR | 0.48386 | OV | PEO1 | Erlotinib | top60_altered_DNA+RNA_seed | TP53 | 1.500118 | LUAD | NCI-H1650 | MIRA-1 |
| MVP_top60_DNA_seed | PIK3CA | 0.503464 | OV | DOV13 | AZD8186 | top60_altered_DNA+RNA_seed | BRD9 | 1.527926 | LUAD | NCI-H2023 | I-BRD9 |
| MVP_top60_DNA_seed | PLK1 | 0.503873 | OV | OVCA420 | MK-1775 | top60_altered_DNA+RNA_seed | TERT | 1.53817 | LUAD | NCI-H2087 | BIBR-1532 |
| MVP_top60_DNA_seed | PLK1 | 0.508448 | OV | OVK-18 | MK-1775 | top60_altered_DNA+RNA_seed | BRD9 | 1.544361 | LUAD | SK-LU-1 | I-BRD9 |
| MVP_top60_DNA_seed | TP53 | 0.514513 | OV | UWB1.289 | MIRA-1 | top60_altered_DNA+RNA_seed | TERT | 1.570792 | LUAD | SW1573 | BIBR-1532 |
| MVP_top60_DNA_seed | EGFR | 0.517848 | OV | OV-17R | Erlotinib | top60_altered_DNA+RNA_seed | BRD9 | 1.578414 | LUAD | NCI-H1650 | I-BRD9 |
| MVP_top60_DNA_seed | HDAC2 | 0.524996 | OV | IGROV-1 | Romidepsin | top60_altered_DNA+RNA_seed | TERT | 1.584771 | LUAD | NCI-H1355 | BIBR-1532 |
| MVP_top60_DNA_seed | ATR | 0.526936 | OV | OV-90 | AZD6738 | top60_altered_DNA+RNA_seed | BRD9 | 1.588914 | LUAD | SK-LU-1 | Bromosporine |
| MVP_top60_DNA_seed | CREBBP | 0.527058 | OV | OAW-42 | SGC-CBP30 | top60_altered_DNA+RNA_seed | BRD9 | 1.602406 | LUAD | Calu-6 | I-BRD9 |
| MVP_top60_DNA_seed | MAPK1 | 0.528733 | OV | EFO-21 | SCH772984 | top60_altered_DNA+RNA_seed | BRD9 | 1.61943 | LUAD | NCI-H1838 | I-BRD9 |
| MVP_top60_DNA_seed | MAPK3 | 0.528733 | OV | EFO-21 | SCH772984 | top60_altered_DNA+RNA_seed | KRAS | 1.630854 | LUAD | COR-L105 | KRAS (G12C) Inhibitor-1 |

| | | | | | | | | | | | |
|---|---|---|---|---|---|---|---|---|---|---|---|
| MVP_top60_DNA_seed | PIK3CA | 0.530888 | OV | FU-OV-1 | GNE-317 | top60_altered_DNA+RNA_seed | BRD9 | 1.673066 | LUAD | NCI-H2291 | I-BRD9 |
| MVP_top60_DNA_seed | PLK1 | 0.540098 | OV | SK-OV-3 | BI-2536 | top60_altered_DNA+RNA_seed | TP53 | 1.691954 | LUAD | NCI-H1568 | MIRA-1 |
| MVP_top60_DNA_seed | MAPK1 | 0.551168 | OV | OVCAR-4 | Ulixertinib | top60_altered_DNA+RNA_seed | KRAS | 1.703993 | LUAD | NCI-H1650 | RAS (G12C) Inhibitor-1 |
| MVP_top60_DNA_seed | MAPK3 | 0.551168 | OV | OVCAR-4 | Ulixertinib | top60_altered_DNA+RNA_seed | TERT | 1.717819 | LUAD | NCI-H1693 | BIBR-1532 |
| MVP_top60_DNA_seed | PIK3CA | 0.555958 | OV | Caov-4 | AZD8186 | top60_altered_DNA+RNA_seed | TERT | 1.732268 | LUAD | Calu-3 | BIBR-1532 |
| MVP_top60_DNA_seed | PLK1 | 0.556638 | OV | OAW-42 | MK-1775 | top60_altered_DNA+RNA_seed | BRD9 | 1.73555 | LUAD | NCI-H2405 | Bromosporine |
| MVP_top60_DNA_seed | PLK1 | 0.557664 | OV | OVCAR-4 | BI-2536 | top60_altered_DNA+RNA_seed | BRD9 | 1.742991 | LUAD | NCI-H441 | Bromosporine |
| MVP_top60_DNA_seed | EGFR | 0.559364 | OV | OV-7 | Erlotinib | top60_altered_DNA+RNA_seed | TP53 | 1.763346 | LUAD | NCI-H1793 | MIRA-1 |
| MVP_top60_DNA_seed | MAPK1 | 0.561644 | OV | PEO1 | VX-11e | top60_altered_DNA+RNA_seed | BRD9 | 1.789527 | LUAD | NCI-H2228 | Bromosporine |
| MVP_top60_DNA_seed | EGFR | 0.562147 | OV | EFO-21 | AZD3759 | top60_altered_DNA+RNA_seed | TP53 | 1.824734 | LUAD | HCC-827 | MIRA-1 |
| MVP_top60_DNA_seed | EGFR | 0.567645 | OV | OAW-28 | Afatinib | top60_altered_DNA+RNA_seed | KRAS | 1.826049 | LUAD | NCI-H2405 | RAS (G12C) Inhibitor-1 |
| MVP_top60_DNA_seed | ERBB2 | 0.567645 | OV | OAW-28 | Afatinib | top60_altered_DNA+RNA_seed | TERT | 1.901414 | LUAD | NCI-H1793 | BIBR-1532 |
| MVP_top60_DNA_seed | ATR | 0.570872 | OV | PEO1 | VE-822 | top60_altered_DNA+RNA_seed | TP53 | 1.91054 | LUAD | NCI-H2347 | MIRA-1 |
| MVP_top60_DNA_seed | CREBBP | 0.582908 | OV | OAW-28 | SGC-CBP30 | top60_altered_DNA+RNA_seed | KRAS | 1.93085 | LUAD | NCI-H1568 | RAS (G12C) Inhibitor-1 |
| MVP_top60_DNA_seed | EGFR | 0.585179 | OV | OAW-28 | AZD3759 | top60_altered_DNA+RNA_seed | TP53 | 1.932986 | LUAD | NCI-H441 | MIRA-1 |
| MVP_top60_DNA_seed | CDK2 | 0.585441 | OV | OVCAR-3 | AZD5438 | top60_altered_DNA+RNA_seed | TERT | 1.998051 | LUAD | NCI-H1666 | BIBR-1532 |
| MVP_top60_DNA_seed | ATR | 0.589343 | OV | OVCAR-3 | AZD6738 | top60_altered_DNA+RNA_seed | BRD9 | 2.004657 | LUAD | NCI-H1568 | I-BRD9 |
| MVP_top60_DNA_seed | CDK1 | 0.591599 | OV | OAW-42 | RO-3306 | top60_altered_DNA+RNA_seed | TERT | 2.022634 | LUAD | NCI-H1838 | BIBR-1532 |
| MVP_top60_DNA_seed | EGFR | 0.596143 | OV | Hey | Osimertinib | top60_altered_DNA+RNA_seed | KRAS | 2.042551 | LUAD | NCI-H2228 | RAS (G12C) Inhibitor-1 |
| MVP_top60_DNA_seed | EGFR | 0.599316 | OV | OVCAR-5 | Gefitinib | top60_altered_DNA+RNA_seed | TERT | 2.052173 | LUAD | SK-LU-1 | BIBR-1532 |
| MVP_top60_DNA_seed | PIK3CA | 0.608699 | OV | TOV-112D | Alpelisib | top60_altered_DNA+RNA_seed | BRD9 | 2.066195 | LUAD | NCI-H2347 | I-BRD9 |
| MVP_top60_DNA_seed | MAPK1 | 0.615538 | OV | DOV13 | SCH772984 | top60_altered_DNA+RNA_seed | BRD9 | 2.07942 | LUAD | NCI-H1650 | Bromosporine |
| MVP_top60_DNA_seed | MAPK3 | 0.615538 | OV | DOV13 | SCH772984 | top60_altered_DNA+RNA_seed | BRD9 | 2.093849 | LUAD | HCC-827 | I-BRD9 |
| MVP_top60_DNA_seed | ATR | 0.622789 | OV | OAW-28 | VE821 | top60_altered_DNA+RNA_seed | KRAS | 2.098972 | LUAD | SW1573 | RAS (G12C) Inhibitor-1 |
| MVP_top60_DNA_seed | MAPK1 | 0.624783 | OV | OVISE | Ulixertinib | top60_altered_DNA+RNA_seed | BRD9 | 2.105804 | LUAD | NCI-H1693 | Bromosporine |
| MVP_top60_DNA_seed | MAPK3 | 0.624783 | OV | OVISE | Ulixertinib | top60_altered_DNA+RNA_seed | TERT | 2.124139 | LUAD | NCI-H2291 | BIBR-1532 |
| MVP_top60_DNA_seed | PIK3CA | 0.625535 | OV | OVMIU | Buparlisib | top60_altered_DNA+RNA_seed | BRD9 | 2.139753 | LUAD | SW1573 | I-BRD9 |
| MVP_top60_DNA_seed | EGFR | 0.62562 | OV | EFO-21 | Sapitinib | top60_altered_DNA+RNA_seed | BRD9 | 2.148125 | LUAD | EKVX | Bromosporine |
| MVP_top60_DNA_seed | ERBB2 | 0.62562 | OV | EFO-21 | Sapitinib | top60_altered_DNA+RNA_seed | BRD9 | 2.156061 | LUAD | NCI-H1793 | Bromosporine |
| MVP_top60_DNA_seed | MAPK1 | 0.632702 | OV | EFO-21 | Ulixertinib | top60_altered_DNA+RNA_seed | KRAS | 2.199916 | LUAD | SK-LU-1 | RAS (G12C) Inhibitor-1 |
| MVP_top60_DNA_seed | MAPK3 | 0.632702 | OV | EFO-21 | Ulixertinib | top60_altered_DNA+RNA_seed | BRD9 | 2.222534 | LUAD | H3255 | Bromosporine |
| MVP_top60_DNA_seed | PLK1 | 0.636074 | OV | OVMIU | BI-2536 | top60_altered_DNA+RNA_seed | BRD9 | 2.398814 | LUAD | NCI-H441 | I-BRD9 |
| MVP_top60_DNA_seed | PLK1 | 0.637006 | OV | OAW-28 | BI-2536 | top60_altered_DNA+RNA_seed | TP53 | 2.464529 | LUAD | NCI-H2405 | MIRA-1 |
| MVP_top60_DNA_seed | PIK3CA | 0.637138 | OV | OV-90 | GNE-317 | top60_altered_DNA+RNA_seed | KRAS | 2.603151 | LUAD | NCI-H441 | RAS (G12C) Inhibitor-1 |
| MVP_top60_DNA_seed | CDK1 | 0.639744 | OV | Caov-4 | RO-3306 | top60_altered_DNA+RNA_seed | TERT | 3.04853 | LUAD | NCI-H441 | BIBR-1532 |
| MVP_top60_DNA_seed | EGFR | 0.64377 | OV | Hey | Sapitinib | top60_altered_DNA+RNA_seed | PIK3CA | -1.665707 | LUSC | SK-MES-1 | PI3Ka_4409 |
| MVP_top60_DNA_seed | ERBB2 | 0.64377 | OV | Hey | Sapitinib | top60_altered_DNA+RNA_seed | PIK3CA | -1.469562 | LUSC | RERF-LC-Sq1 | PI3Ka_4409 |
| MVP_top60_DNA_seed | EGFR | 0.644157 | OV | PEO1 | Lapatinib | top60_altered_DNA+RNA_seed | PIK3CA | -1.37393 | LUSC | SK-MES-1 | AZD8835 |
| MVP_top60_DNA_seed | ERBB2 | 0.644157 | OV | PEO1 | Lapatinib | top60_altered_DNA+RNA_seed | PIK3CA | -1.09166 | LUSC | RERF-LC-Sq1 | AZD8835 |
| MVP_top60_DNA_seed | HDAC2 | 0.649048 | OV | OVCAR-3 | Romidepsin | top60_altered_DNA+RNA_seed | PIK3CA | -1.001354 | LUSC | EPLC-272H | PI3Ka_4409 |
| MVP_top60_DNA_seed | EGFR | 0.659365 | OV | OVCAR-4 | Gefitinib | top60_altered_DNA+RNA_seed | PIK3CA | -0.834848 | LUSC | EPLC-272H | AZD8835 |

| | | | | | | | | | | | |
|---|---|---|---|---|---|---|---|---|---|---|---|
| MVP_top60_DNA_seed | ATR | 0.668172 | OV | OVCAR-4 | VE821 | top60_altered_DNA+RNA_seed | PIK3CA | -0.809777 | LUSC | LOU-NH91 | AZD8835 |
| MVP_top60_DNA_seed | PLK1 | 0.668206 | OV | EFO-21 | MK-1775 | top60_altered_DNA+RNA_seed | PIK3CA | -0.633699 | LUSC | EBC-1 | PI3Ka_4409 |
| MVP_top60_DNA_seed | EGFR | 0.669822 | OV | OAW-42 | Osimertinib | top60_altered_DNA+RNA_seed | PIK3CA | -0.455436 | LUSC | KNS-62 | PI3Ka_4409 |
| MVP_top60_DNA_seed | ATR | 0.674262 | OV | OV-7 | VE-822 | top60_altered_DNA+RNA_seed | PIK3CA | -0.250907 | LUSC | LK-2 | AZD8835 |
| MVP_top60_DNA_seed | EGFR | 0.674837 | OV | OVKATE | Sapitinib | top60_altered_DNA+RNA_seed | PIK3CA | -0.247775 | LUSC | HCC-15 | PI3Ka_4409 |
| MVP_top60_DNA_seed | ERBB2 | 0.674837 | OV | OVKATE | Sapitinib | top60_altered_DNA+RNA_seed | PIK3CA | -0.233333 | LUSC | NCI-H226 | AZD8835 |
| MVP_top60_DNA_seed | PLK1 | 0.67553 | OV | FU-OV-1 | MK-1775 | top60_altered_DNA+RNA_seed | PIK3CA | -0.221388 | LUSC | NCI-H1869 | PI3Ka_4409 |
| MVP_top60_DNA_seed | CDK2 | 0.67558 | OV | OC-314 | MK-8776 | top60_altered_DNA+RNA_seed | PIK3CA | -0.202814 | LUSC | HARA | AZD8835 |
| MVP_top60_DNA_seed | TP53 | 0.677733 | OV | Caov-4 | MIRA-1 | top60_altered_DNA+RNA_seed | PIK3CA | -0.155866 | LUSC | HARA | PI3Ka_4409 |
| MVP_top60_DNA_seed | PIK3CA | 0.678552 | OV | OAW-28 | AZD8186 | top60_altered_DNA+RNA_seed | PIK3CA | -0.032854 | LUSC | NCI-H2170 | PI3Ka_4409 |
| MVP_top60_DNA_seed | PIK3CA | 0.685968 | OV | OV-90 | Alpelisib | top60_altered_DNA+RNA_seed | PIK3CA | 0.041565 | LUSC | LOU-NH91 | PI3Ka_4409 |
| MVP_top60_DNA_seed | EGFR | 0.689226 | OV | OVCAR-5 | AZD3759 | top60_altered_DNA+RNA_seed | PIK3CA | 0.050047 | LUSC | HCC-15 | AZD8835 |
| MVP_top60_DNA_seed | PIK3CA | 0.689699 | OV | Caov-3 | Buparlisib | top60_altered_DNA+RNA_seed | PIK3CA | 0.109597 | LUSC | LC-1-sq | AZD8835 |
| MVP_top60_DNA_seed | PIK3CA | 0.69064 | OV | SK-OV-3 | Buparlisib | top60_altered_DNA+RNA_seed | PIK3CA | 0.293316 | LUSC | NCI-H2170 | AZD8835 |
| MVP_top60_DNA_seed | EGFR | 0.69179 | OV | Hey | Afatinib | top60_altered_DNA+RNA_seed | PIK3CA | 0.321961 | LUSC | LC-1-sq | PI3Ka_4409 |
| MVP_top60_DNA_seed | ERBB2 | 0.69179 | OV | Hey | Afatinib | top60_altered_DNA+RNA_seed | PIK3CA | 0.33785 | LUSC | LK-2 | PI3Ka_4409 |
| MVP_top60_DNA_seed | PLK1 | 0.694648 | OV | OVCAR-5 | MK-1775 | top60_altered_DNA+RNA_seed | PIK3CA | 0.846248 | LUSC | EBC-1 | AZD8835 |
| MVP_top60_DNA_seed | EGFR | 0.697177 | OV | OV-7 | Osimertinib | top60_altered_DNA+RNA_seed | PIK3CA | 1.193603 | LUSC | NCI-H226 | PI3Ka_4409 |
| MVP_top60_DNA_seed | CDK2 | 0.697382 | OV | OVCAR-3 | MK-8776 | top60_altered_DNA+RNA_seed | PIK3CA | 1.349994 | LUSC | NCI-H520 | AZD8835 |
| MVP_top60_DNA_seed | ATR | 0.701252 | OV | OAW-28 | AZD6738 | top60_altered_DNA+RNA_seed | PIK3CA | 1.576912 | LUSC | NCI-H1869 | AZD8835 |
| MVP_top60_DNA_seed | EGFR | 0.701983 | OV | JHOS-2 | Lapatinib | top60_altered_DNA+RNA_seed | PIK3CA | 1.590637 | LUSC | NCI-H520 | PI3Ka_4409 |
| MVP_top60_DNA_seed | ERBB2 | 0.701983 | OV | JHOS-2 | Lapatinib | top60_altered_DNA+RNA_seed | PIK3CA | 2.274945 | LUSC | KNS-62 | AZD8835 |
| MVP_top60_DNA_seed | PLK1 | 0.705055 | OV | SW626 | MK-1775 | top60_altered_DNA+RNA_seed | PIK3CA | -1.338512 | LUSC | RERF-LC-Sq1 | Alpelisib |
| MVP_top60_DNA_seed | PLK1 | 0.705911 | OV | OV-90 | MK-1775 | top60_altered_DNA+RNA_seed | PIK3CA | -1.220521 | LUSC | EPLC-272H | Alpelisib |
| MVP_top60_DNA_seed | PLK1 | 0.706774 | OV | OVISE | BI-2536 | top60_altered_DNA+RNA_seed | PIK3CA | -1.145638 | LUSC | EPLC-272H | Buparlisib |
| MVP_top60_DNA_seed | ATR | 0.71314 | OV | OVTOKO | VE821 | top60_altered_DNA+RNA_seed | PIK3CA | -1.121422 | LUSC | NCI-H2170 | Buparlisib |
| MVP_top60_DNA_seed | CREBBP | 0.7183 | OV | IGROV-1 | SGC-CBP30 | top60_altered_DNA+RNA_seed | PIK3CA | -1.08911 | LUSC | EPLC-272H | AZD8186 |
| MVP_top60_DNA_seed | EGFR | 0.724061 | OV | DOV13 | Osimertinib | top60_altered_DNA+RNA_seed | PIK3CA | -1.036936 | LUSC | EPLC-272H | GNE-317 |
| MVP_top60_DNA_seed | EGFR | 0.724746 | OV | OAW-28 | Sapitinib | top60_altered_DNA+RNA_seed | PIK3CA | -0.955533 | LUSC | HARA | Buparlisib |
| MVP_top60_DNA_seed | ERBB2 | 0.724746 | OV | OAW-28 | Sapitinib | top60_altered_DNA+RNA_seed | PIK3CA | -0.913577 | LUSC | HARA | Alpelisib |
| MVP_top60_DNA_seed | CDK1 | 0.727073 | OV | OVCA420 | RO-3306 | top60_altered_DNA+RNA_seed | PIK3CA | -0.830417 | LUSC | HARA | GNE-317 |
| MVP_top60_DNA_seed | CDK2 | 0.732001 | OV | OVCAR-4 | MK-8776 | top60_altered_DNA+RNA_seed | PIK3CA | -0.822014 | LUSC | EBC-1 | AZD8186 |
| MVP_top60_DNA_seed | ATR | 0.733493 | OV | OVCA420 | AZD6738 | top60_altered_DNA+RNA_seed | PIK3CA | -0.786418 | LUSC | RERF-LC-Sq1 | GNE-317 |
| MVP_top60_DNA_seed | EGFR | 0.74451 | OV | EFO-21 | Osimertinib | top60_altered_DNA+RNA_seed | PIK3CA | -0.781629 | LUSC | NCI-H2170 | Alpelisib |
| MVP_top60_DNA_seed | PLK1 | 0.746856 | OV | SW626 | BI-2536 | top60_altered_DNA+RNA_seed | PIK3CA | -0.729955 | LUSC | EBC-1 | GNE-317 |
| MVP_top60_DNA_seed | MAPK1 | 0.747943 | OV | OVCAR-3 | VX-11e | top60_altered_DNA+RNA_seed | PIK3CA | -0.713796 | LUSC | NCI-H520 | AZD8186 |
| MVP_top60_DNA_seed | MAPK1 | 0.749637 | OV | FU-OV-1 | Ulixertinib | top60_altered_DNA+RNA_seed | PIK3CA | -0.531937 | LUSC | HCC-15 | Buparlisib |
| MVP_top60_DNA_seed | MAPK3 | 0.749637 | OV | FU-OV-1 | Ulixertinib | top60_altered_DNA+RNA_seed | PIK3CA | -0.461466 | LUSC | NCI-H2170 | AZD8186 |
| MVP_top60_DNA_seed | CDK2 | 0.75144 | OV | OVMIU | MK-8776 | top60_altered_DNA+RNA_seed | PIK3CA | -0.33911 | LUSC | NCI-H520 | GNE-317 |
| MVP_top60_DNA_seed | CDK2 | 0.757646 | OV | SW626 | Dinaciclib | top60_altered_DNA+RNA_seed | PIK3CA | -0.313509 | LUSC | LOU-NH91 | Alpelisib |
| MVP_top60_DNA_seed | CDK1 | 0.757646 | OV | SW626 | Dinaciclib | top60_altered_DNA+RNA_seed | PIK3CA | -0.290278 | LUSC | RERF-LC-Sq1 | Buparlisib |

| | | | | | | | | | | | |
|---|---|---|---|---|---|---|---|---|---|---|---|
| MVP_top60_DNA_seed | ATR | 0.760458 | OV | OV-90 | VE821 | top60_altered_DNA+RNA_seed | PIK3CA | -0.287494 | LUSC | LOU-NH91 | GNE-317 |
| MVP_top60_DNA_seed | ATR | 0.768533 | OV | OVCAR-3 | VE-822 | top60_altered_DNA+RNA_seed | PIK3CA | -0.283547 | LUSC | LK-2 | GNE-317 |
| MVP_top60_DNA_seed | EGFR | 0.770683 | OV | OAW-42 | Gefitinib | top60_altered_DNA+RNA_seed | PIK3CA | -0.280465 | LUSC | LOU-NH91 | AZD8186 |
| MVP_top60_DNA_seed | TP53 | 0.77188 | OV | OAW-42 | MIRA-1 | top60_altered_DNA+RNA_seed | PIK3CA | -0.27608 | LUSC | NCI-H1869 | AZD8186 |
| MVP_top60_DNA_seed | EGFR | 0.772278 | OV | SK-OV-3 | AZD3759 | top60_altered_DNA+RNA_seed | PIK3CA | -0.270405 | LUSC | LK-2 | Buparlisib |
| MVP_top60_DNA_seed | CDK2 | 0.778411 | OV | OVCAR-3 | Dinaciclib | top60_altered_DNA+RNA_seed | PIK3CA | -0.210267 | LUSC | EBC-1 | Buparlisib |
| MVP_top60_DNA_seed | CDK1 | 0.778411 | OV | OVCAR-3 | Dinaciclib | top60_altered_DNA+RNA_seed | PIK3CA | -0.086508 | LUSC | LC-1-sq | Alpelisib |
| MVP_top60_DNA_seed | EGFR | 0.778437 | OV | TOV-21G | Gefitinib | top60_altered_DNA+RNA_seed | PIK3CA | -0.039436 | LUSC | LOU-NH91 | Buparlisib |
| MVP_top60_DNA_seed | EGFR | 0.781674 | OV | SW626 | Erlotinib | top60_altered_DNA+RNA_seed | TP53 | -0.002669 | LUSC | NCI-H1869 | MIRA-1 |
| MVP_top60_DNA_seed | EGFR | 0.781798 | OV | OV-56 | AZD3759 | top60_altered_DNA+RNA_seed | TP53 | 0.006589 | LUSC | NCI-H2170 | MIRA-1 |
| MVP_top60_DNA_seed | PIK3CA | 0.78181 | OV | FU-OV-1 | Buparlisib | top60_altered_DNA+RNA_seed | PIK3CA | 0.035824 | LUSC | SK-MES-1 | AZD8186 |
| MVP_top60_DNA_seed | EGFR | 0.78353 | OV | OV-56 | Gefitinib | top60_altered_DNA+RNA_seed | TP53 | 0.085711 | LUSC | HCC-15 | MIRA-1 |
| MVP_top60_DNA_seed | EGFR | 0.792114 | OV | Caov-3 | Sapitinib | top60_altered_DNA+RNA_seed | TP53 | 0.117814 | LUSC | NCI-H520 | MIRA-1 |
| MVP_top60_DNA_seed | ERBB2 | 0.792114 | OV | Caov-3 | Sapitinib | top60_altered_DNA+RNA_seed | TP53 | 0.127705 | LUSC | HARA | MIRA-1 |
| MVP_top60_DNA_seed | EGFR | 0.792888 | OV | Caov-3 | Afatinib | top60_altered_DNA+RNA_seed | TP53 | 0.175425 | LUSC | LC-1-sq | MIRA-1 |
| MVP_top60_DNA_seed | ERBB2 | 0.792888 | OV | Caov-3 | Afatinib | top60_altered_DNA+RNA_seed | PIK3CA | 0.179874 | LUSC | NCI-H520 | Buparlisib |
| MVP_top60_DNA_seed | PIK3CA | 0.793259 | OV | OV-17R | Buparlisib | top60_altered_DNA+RNA_seed | TP53 | 0.215474 | LUSC | LK-2 | MIRA-1 |
| MVP_top60_DNA_seed | PIK3CA | 0.793972 | OV | FU-OV-1 | Alpelisib | top60_altered_DNA+RNA_seed | PIK3CA | 0.328737 | LUSC | SK-MES-1 | GNE-317 |
| MVP_top60_DNA_seed | EGFR | 0.801624 | OV | OVCAR-4 | Lapatinib | top60_altered_DNA+RNA_seed | PIK3CA | 0.332925 | LUSC | HARA | AZD8186 |
| MVP_top60_DNA_seed | ERBB2 | 0.801624 | OV | OVCAR-4 | Lapatinib | top60_altered_DNA+RNA_seed | PIK3CA | 0.357542 | LUSC | NCI-H1869 | GNE-317 |
| MVP_top60_DNA_seed | EGFR | 0.814524 | OV | Hey | Lapatinib | top60_altered_DNA+RNA_seed | TP53 | 0.371749 | LUSC | EPLC-272H | MIRA-1 |
| MVP_top60_DNA_seed | ERBB2 | 0.814524 | OV | Hey | Lapatinib | top60_altered_DNA+RNA_seed | PIK3CA | 0.384334 | LUSC | RERF-LC-Sq1 | AZD8186 |
| MVP_top60_DNA_seed | EGFR | 0.816678 | OV | OAW-42 | AZD3759 | top60_altered_DNA+RNA_seed | PIK3CA | 0.493276 | LUSC | LC-1-sq | Buparlisib |
| MVP_top60_DNA_seed | ATR | 0.816996 | OV | OVTOKO | AZD6738 | top60_altered_DNA+RNA_seed | PIK3CA | 0.528927 | LUSC | HCC-15 | GNE-317 |
| MVP_top60_DNA_seed | CDK2 | 0.82782 | OV | SW626 | MK-8776 | top60_altered_DNA+RNA_seed | PIK3CA | 0.568534 | LUSC | KNS-62 | Alpelisib |
| MVP_top60_DNA_seed | EGFR | 0.830073 | OV | Caov-3 | Osimertinib | top60_altered_DNA+RNA_seed | TP53 | 0.593917 | LUSC | RERF-LC-Sq1 | MIRA-1 |
| MVP_top60_DNA_seed | PIK3CA | 0.831355 | OV | TOV-21G | Buparlisib | top60_altered_DNA+RNA_seed | PIK3CA | 0.621815 | LUSC | KNS-62 | GNE-317 |
| MVP_top60_DNA_seed | PIK3CA | 0.836084 | OV | OVCAR-3 | Alpelisib | top60_altered_DNA+RNA_seed | PIK3CA | 0.637805 | LUSC | NCI-H520 | Alpelisib |
| MVP_top60_DNA_seed | MAPK1 | 0.836602 | OV | DOV13 | Ulixertinib | top60_altered_DNA+RNA_seed | PIK3CA | 0.638856 | LUSC | SK-MES-1 | Buparlisib |
| MVP_top60_DNA_seed | MAPK3 | 0.836602 | OV | DOV13 | Ulixertinib | top60_altered_DNA+RNA_seed | PIK3CA | 0.654019 | LUSC | KNS-62 | Buparlisib |
| MVP_top60_DNA_seed | HDAC2 | 0.837352 | OV | OVISE | Romidepsin | top60_altered_DNA+RNA_seed | TP53 | 0.691211 | LUSC | KNS-62 | MIRA-1 |
| MVP_top60_DNA_seed | CDK2 | 0.837568 | OV | PEO1 | Dinaciclib | top60_altered_DNA+RNA_seed | TP53 | 0.717299 | LUSC | LOU-NH91 | MIRA-1 |
| MVP_top60_DNA_seed | CDK1 | 0.837568 | OV | PEO1 | Dinaciclib | top60_altered_DNA+RNA_seed | PIK3CA | 0.724967 | LUSC | LC-1-sq | AZD8186 |
| MVP_top60_DNA_seed | CDK2 | 0.840847 | OV | SK-OV-3 | MK-8776 | top60_altered_DNA+RNA_seed | PIK3CA | 0.786867 | LUSC | LC-1-sq | GNE-317 |
| MVP_top60_DNA_seed | PIK3CA | 0.85254 | OV | SW626 | Alpelisib | top60_altered_DNA+RNA_seed | PIK3CA | 0.810293 | LUSC | NCI-H1869 | Alpelisib |
| MVP_top60_DNA_seed | ATR | 0.853702 | OV | OVISE | VE821 | top60_altered_DNA+RNA_seed | PIK3CA | 0.833017 | LUSC | HCC-15 | AZD8186 |
| MVP_top60_DNA_seed | EGFR | 0.859868 | OV | PEO1 | Osimertinib | top60_altered_DNA+RNA_seed | TP53 | 0.841218 | LUSC | EBC-1 | MIRA-1 |
| MVP_top60_DNA_seed | CREBBP | 0.865919 | OV | OVCAR-5 | SGC-CBP30 | top60_altered_DNA+RNA_seed | TP53 | 0.849479 | LUSC | SK-MES-1 | MIRA-1 |
| MVP_top60_DNA_seed | PIK3CA | 0.872908 | OV | OVK-18 | Alpelisib | top60_altered_DNA+RNA_seed | PIK3CA | 0.906604 | LUSC | LK-2 | AZD8186 |
| MVP_top60_DNA_seed | PIK3CA | 0.877878 | OV | OAW-28 | GNE-317 | top60_altered_DNA+RNA_seed | PIK3CA | 0.914289 | LUSC | EBC-1 | Alpelisib |
| MVP_top60_DNA_seed | CDK2 | 0.8819 | OV | DOV13 | MK-8776 | top60_altered_DNA+RNA_seed | PIK3CA | 1.049427 | LUSC | NCI-H226 | Alpelisib |

| | | | | | | | | | | | |
|---|---|---|---|---|---|---|---|---|---|---|---|
| MVP_top60_DNA_seed | CDK2 | 0.881912 | OV | Hey | MK-8776 | top60_altered_DNA+RNA_seed | PIK3CA | 1.066896 | LUSC | KNS-62 | AZD8186 |
| MVP_top60_DNA_seed | CDK2 | 0.882902 | OV | PEO1 | MK-8776 | top60_altered_DNA+RNA_seed | PIK3CA | 1.11521 | LUSC | HCC-15 | Alpelisib |
| MVP_top60_DNA_seed | EGFR | 0.882938 | OV | OVK-18 | Lapatinib | top60_altered_DNA+RNA_seed | PIK3CA | 1.146915 | LUSC | SK-MES-1 | Alpelisib |
| MVP_top60_DNA_seed | ERBB2 | 0.882938 | OV | OVK-18 | Lapatinib | top60_altered_DNA+RNA_seed | PIK3CA | 1.175026 | LUSC | LK-2 | Alpelisib |
| MVP_top60_DNA_seed | TP53 | 0.888744 | OV | OV-56 | MIRA-1 | top60_altered_DNA+RNA_seed | PIK3CA | 1.288456 | LUSC | NCI-H1869 | Buparlisib |
| MVP_top60_DNA_seed | CDK2 | 0.892834 | OV | OAW-42 | Dinaciclib | top60_altered_DNA+RNA_seed | TP53 | 1.773584 | LUSC | NCI-H226 | MIRA-1 |
| MVP_top60_DNA_seed | CDK1 | 0.892834 | OV | OAW-42 | Dinaciclib | top60_altered_DNA+RNA_seed | PIK3CA | 2.103956 | LUSC | NCI-H226 | AZD8186 |
| MVP_top60_DNA_seed | ATR | 0.893273 | OV | JHOS-2 | VE-822 | top60_altered_DNA+RNA_seed | PIK3CA | 2.336999 | LUSC | NCI-H226 | GNE-317 |
| MVP_top60_DNA_seed | CDK2 | 0.894909 | OV | OVCA420 | MK-8776 | top60_altered_DNA+RNA_seed | PIK3CA | 2.454696 | LUSC | NCI-H226 | Buparlisib |
| MVP_top60_DNA_seed | ATR | 0.913409 | OV | OVMIU | AZD6738 | top60_altered_DNA+RNA_seed | TP53 | -0.014816 | PRAD | PWR-1E | MIRA-1 |
| MVP_top60_DNA_seed | PLK1 | 0.916338 | OV | OAW-28 | MK-1775 | top60_altered_DNA+RNA_seed | TP53 | 0.243205 | PRAD | VCaP | MIRA-1 |
| MVP_top60_DNA_seed | PIK3CA | 0.920256 | OV | OVCAR-4 | GNE-317 | top60_altered_DNA+RNA_seed | TP53 | 0.687703 | PRAD | LNCaP-Clone-FGC | MIRA-1 |
| MVP_top60_DNA_seed | MAPK1 | 0.923247 | OV | OVISE | VX-11e | top60_altered_DNA+RNA_seed | TP53 | 0.706283 | PRAD | 22RV1 | MIRA-1 |
| MVP_top60_DNA_seed | TP53 | 0.92833 | OV | OVCA420 | MIRA-1 | top60_altered_DNA+RNA_seed | TP53 | 0.955249 | PRAD | DU-145 | MIRA-1 |
| MVP_top60_DNA_seed | MAPK1 | 0.935237 | OV | OVTOKO | Ulixertinib | top60_altered_DNA+RNA_seed | TP53 | 1.463099 | PRAD | PC-3 | MIRA-1 |
| MVP_top60_DNA_seed | MAPK3 | 0.935237 | OV | OVTOKO | Ulixertinib | top60_altered_DNA+RNA_seed | ERBB2 | -5.506139 | STAD | NCI-N87 | CP724714 |
| MVP_top60_DNA_seed | MAPK1 | 0.936502 | OV | OVTOKO | Ulixertinib | top60_altered_DNA+RNA_seed | ERBB2 | -4.174126 | STAD | Hs-746T | CP724714 |
| MVP_top60_DNA_seed | MAPK3 | 0.936502 | OV | OVTOKO | Ulixertinib | top60_altered_DNA+RNA_seed | ERBB2 | -3.997646 | STAD | SK-GT-2 | Afatinib |
| MVP_top60_DNA_seed | MAPK1 | 0.9369 | OV | OVCAR-3 | SCH772984 | top60_altered_DNA+RNA_seed | ERBB2 | -3.708978 | STAD | SK-GT-2 | Afatinib |
| MVP_top60_DNA_seed | MAPK3 | 0.9369 | OV | OVCAR-3 | SCH772984 | top60_altered_DNA+RNA_seed | ERBB2 | -3.579769 | STAD | NUGC-4 | Afatinib |
| MVP_top60_DNA_seed | CREBBP | 0.939377 | OV | TOV-21G | SGC-CBP30 | top60_altered_DNA+RNA_seed | ERBB2 | -3.19964 | STAD | NCI-N87 | Afatinib |
| MVP_top60_DNA_seed | ATR | 0.939732 | OV | Hey | VE821 | top60_altered_DNA+RNA_seed | ERBB2 | -3.012876 | STAD | NUGC-4 | AZD8931 |
| MVP_top60_DNA_seed | EGFR | 0.944285 | OV | OVK-18 | Gefitinib | top60_altered_DNA+RNA_seed | ERBB2 | -2.840983 | STAD | NCI-N87 | Afatinib |
| MVP_top60_DNA_seed | CDK2 | 0.950015 | OV | OV-90 | MK-8776 | top60_altered_DNA+RNA_seed | PIK3CA | -2.507956 | STAD | IM-95 | AZD8835 |
| MVP_top60_DNA_seed | EGFR | 0.950293 | OV | OVK-18 | Sapitinib | top60_altered_DNA+RNA_seed | ERBB2 | -2.458722 | STAD | NUGC-4 | PF-00299804 |
| MVP_top60_DNA_seed | ERBB2 | 0.950293 | OV | OVK-18 | Sapitinib | top60_altered_DNA+RNA_seed | ERBB4 | -2.458722 | STAD | NUGC-4 | PF-00299804 |
| MVP_top60_DNA_seed | TP53 | 0.954593 | OV | FU-OV-1 | MIRA-1 | top60_altered_DNA+RNA_seed | ERBB2 | -2.202897 | STAD | SK-GT-2 | AZD8931 |
| MVP_top60_DNA_seed | EGFR | 0.960304 | OV | TOV-21G | Lapatinib | top60_altered_DNA+RNA_seed | ERBB2 | -2.197683 | STAD | TGBC11TKB | CP724714 |
| MVP_top60_DNA_seed | ERBB2 | 0.960304 | OV | TOV-21G | Lapatinib | top60_altered_DNA+RNA_seed | ERBB4 | -2.09995 | STAD | AGS | AST-1306 |
| MVP_top60_DNA_seed | TP53 | 0.960346 | OV | OV-90 | MIRA-1 | top60_altered_DNA+RNA_seed | ERBB2 | -2.084813 | STAD | AGS | PF-00299804 |
| MVP_top60_DNA_seed | PIK3CA | 0.961998 | OV | PEO1 | Buparlisib | top60_altered_DNA+RNA_seed | ERBB4 | -2.084813 | STAD | AGS | PF-00299804 |
| MVP_top60_DNA_seed | TP53 | 0.962155 | OV | JHOS-3 | MIRA-1 | top60_altered_DNA+RNA_seed | PIK3CA | -2.011361 | STAD | AGS | AZD8835 |
| MVP_top60_DNA_seed | TP53 | 0.967064 | OV | OVMIU | MIRA-1 | top60_altered_DNA+RNA_seed | ERBB2 | -1.999928 | STAD | NUGC-3 | Afatinib |
| MVP_top60_DNA_seed | MAPK1 | 0.967149 | OV | SK-OV-3 | SCH772984 | top60_altered_DNA+RNA_seed | ERBB2 | -1.907664 | STAD | GCIY | CI-1033 |
| MVP_top60_DNA_seed | MAPK3 | 0.967149 | OV | SK-OV-3 | SCH772984 | top60_altered_DNA+RNA_seed | ERBB4 | -1.907664 | STAD | GCIY | CI-1033 |
| MVP_top60_DNA_seed | EGFR | 0.970792 | OV | OV-56 | Sapitinib | top60_altered_DNA+RNA_seed | ERBB2 | -1.847543 | STAD | NCI-N87 | AZD8931 |
| MVP_top60_DNA_seed | ERBB2 | 0.970792 | OV | OV-56 | Sapitinib | top60_altered_DNA+RNA_seed | ERBB2 | -1.746646 | STAD | NUGC-3 | Afatinib |
| MVP_top60_DNA_seed | ATR | 0.975446 | OV | FU-OV-1 | AZD6738 | top60_altered_DNA+RNA_seed | ERBB2 | -1.727366 | STAD | GCIY | PF-00299804 |
| MVP_top60_DNA_seed | CDK2 | 0.980615 | OV | OV-7 | MK-8776 | top60_altered_DNA+RNA_seed | ERBB4 | -1.727366 | STAD | GCIY | PF-00299804 |
| MVP_top60_DNA_seed | ATR | 0.981207 | OV | OVCA433 | VE-822 | top60_altered_DNA+RNA_seed | ERBB2 | -1.633241 | STAD | IM-95 | AZD8931 |
| MVP_top60_DNA_seed | PIK3CA | 0.987509 | OV | OAW-28 | Alpelisib | top60_altered_DNA+RNA_seed | PIK3CA | -1.601479 | STAD | AGS | PI3Ka_4409 |

| | | | | | | | | | | | |
|---|---|---|---|---|---|---|---|---|---|---|---|
| MVP_top60_DNA_seed | CDK2 | 0.98896 | OV | OV-56 | Dinaciclib | top60_altered_DNA+RNA_seed | ERBB2 | -1.527312 | STAD | NUGC-4 | CI-1033 |
| MVP_top60_DNA_seed | CDK1 | 0.98896 | OV | OV-56 | Dinaciclib | top60_altered_DNA+RNA_seed | ERBB4 | -1.527312 | STAD | NUGC-4 | CI-1033 |
| MVP_top60_DNA_seed | MAPK1 | 0.992451 | OV | OVCAR-4 | Ulixertinib | top60_altered_DNA+RNA_seed | ERBB4 | -1.522995 | STAD | GCIY | AST-1306 |
| MVP_top60_DNA_seed | MAPK3 | 0.992451 | OV | OVCAR-4 | Ulixertinib | top60_altered_DNA+RNA_seed | PIK3CA | -1.49039 | STAD | HSC-39 | PI3Ka_4409 |
| MVP_top60_DNA_seed | MAPK1 | 1.009598 | OV | OVCA420 | Ulixertinib | top60_altered_DNA+RNA_seed | ERBB2 | -1.456809 | STAD | HSC-39 | CI-1033 |
| MVP_top60_DNA_seed | MAPK3 | 1.009598 | OV | OVCA420 | Ulixertinib | top60_altered_DNA+RNA_seed | ERBB4 | -1.456809 | STAD | HSC-39 | CI-1033 |
| MVP_top60_DNA_seed | ATR | 1.012375 | OV | JHOS-3 | VE-822 | top60_altered_DNA+RNA_seed | ERBB2 | -1.309811 | STAD | RF-48 | CUDC-101 |
| MVP_top60_DNA_seed | CREBBP | 1.021635 | OV | OVTOKO | SGC-CBP30 | top60_altered_DNA+RNA_seed | ERBB2 | -1.29334 | STAD | AGS | CI-1033 |
| MVP_top60_DNA_seed | PLK1 | 1.030871 | OV | FU-OV-1 | BI-2536 | top60_altered_DNA+RNA_seed | ERBB4 | -1.29334 | STAD | AGS | CI-1033 |
| MVP_top60_DNA_seed | CREBBP | 1.033161 | OV | OV-90 | SGC-CBP30 | top60_altered_DNA+RNA_seed | ERBB4 | -1.278249 | STAD | SNU-1 | AST-1306 |
| MVP_top60_DNA_seed | CREBBP | 1.037695 | OV | OVCAR-4 | SGC-CBP30 | top60_altered_DNA+RNA_seed | ERBB2 | -1.248274 | STAD | GCIY | Afatinib |
| MVP_top60_DNA_seed | EGFR | 1.041304 | OV | OVKATE | Gefitinib | top60_altered_DNA+RNA_seed | ERBB2 | -1.180036 | STAD | NUGC-4 | CUDC-101 |
| MVP_top60_DNA_seed | MAPK1 | 1.05795 | OV | OAW-28 | SCH772984 | top60_altered_DNA+RNA_seed | ERBB2 | -1.09391 | STAD | NCI-N87 | CUDC-101 |
| MVP_top60_DNA_seed | MAPK3 | 1.05795 | OV | OAW-28 | SCH772984 | top60_altered_DNA+RNA_seed | ERBB2 | -1.08654 | STAD | SNU-1 | PF-00299804 |
| MVP_top60_DNA_seed | PIK3CA | 1.061637 | OV | Hey | Alpelisib | top60_altered_DNA+RNA_seed | ERBB4 | -1.08654 | STAD | SNU-1 | PF-00299804 |
| MVP_top60_DNA_seed | PLK1 | 1.062788 | OV | OV-90 | BI-2536 | top60_altered_DNA+RNA_seed | ERBB4 | -1.076841 | STAD | HSC-39 | AST-1306 |
| MVP_top60_DNA_seed | EGFR | 1.071326 | OV | EFO-21 | Erlotinib | top60_altered_DNA+RNA_seed | ERBB4 | -1.076218 | STAD | NUGC-4 | AST-1306 |
| MVP_top60_DNA_seed | PLK1 | 1.077414 | OV | JHOS-3 | BI-2536 | top60_altered_DNA+RNA_seed | ERBB2 | -1.040137 | STAD | NUGC-4 | Afatinib |
| MVP_top60_DNA_seed | CREBBP | 1.079566 | OV | Hey | SGC-CBP30 | top60_altered_DNA+RNA_seed | ERBB2 | -1.019476 | STAD | NUGC-3 | AZD8931 |
| MVP_top60_DNA_seed | PLK1 | 1.086393 | OV | OVCA420 | BI-2536 | top60_altered_DNA+RNA_seed | ERBB2 | -0.995298 | STAD | AGS | AZD8931 |
| MVP_top60_DNA_seed | CDK1 | 1.093098 | OV | OVCAR-4 | RO-3306 | top60_altered_DNA+RNA_seed | ERBB2 | -0.968535 | STAD | HSC-39 | PF-00299804 |
| MVP_top60_DNA_seed | ATR | 1.100956 | OV | OV-90 | VE-822 | top60_altered_DNA+RNA_seed | ERBB4 | -0.968535 | STAD | HSC-39 | PF-00299804 |
| MVP_top60_DNA_seed | PIK3CA | 1.103096 | OV | PEO1 | GNE-317 | top60_altered_DNA+RNA_seed | PIK3CA | -0.955002 | STAD | HSC-39 | AZD8835 |
| MVP_top60_DNA_seed | EGFR | 1.103196 | OV | OVKATE | Afatinib | top60_altered_DNA+RNA_seed | ERBB4 | -0.947594 | STAD | NUGC-3 | AST-1306 |
| MVP_top60_DNA_seed | ERBB2 | 1.103196 | OV | OVKATE | Afatinib | top60_altered_DNA+RNA_seed | ERBB2 | -0.940647 | STAD | SNU-1 | CI-1033 |
| MVP_top60_DNA_seed | CDK1 | 1.10465 | OV | OAW-28 | RO-3306 | top60_altered_DNA+RNA_seed | ERBB4 | -0.940647 | STAD | SNU-1 | CI-1033 |
| MVP_top60_DNA_seed | MAPK1 | 1.113879 | OV | OAW-42 | Ulixertinib | top60_altered_DNA+RNA_seed | ERBB2 | -0.930647 | STAD | RERF-GC-1B | CP724714 |
| MVP_top60_DNA_seed | MAPK3 | 1.113879 | OV | OAW-42 | Ulixertinib | top60_altered_DNA+RNA_seed | ERBB2 | -0.910739 | STAD | GCIY | Lapatinib |
| MVP_top60_DNA_seed | CDK2 | 1.119525 | OV | OAW-28 | AZD5438 | top60_altered_DNA+RNA_seed | ERBB2 | -0.876224 | STAD | SNU-1 | CP724714 |
| MVP_top60_DNA_seed | PIK3CA | 1.120648 | OV | OVCAR-4 | Buparlisib | top60_altered_DNA+RNA_seed | ERBB2 | -0.845644 | STAD | HGC-27 | CP724714 |
| MVP_top60_DNA_seed | PIK3CA | 1.125647 | OV | TOV-21G | Alpelisib | top60_altered_DNA+RNA_seed | ERBB2 | -0.839761 | STAD | RERF-GC-1B | Afatinib |
| MVP_top60_DNA_seed | TP53 | 1.132044 | OV | OVTOKO | MIRA-1 | top60_altered_DNA+RNA_seed | PIK3CA | -0.771988 | STAD | NCI-N87 | PI3Ka_4409 |
| MVP_top60_DNA_seed | EGFR | 1.132715 | OV | FU-OV-1 | Erlotinib | top60_altered_DNA+RNA_seed | ERBB2 | -0.75817 | STAD | MKN7 | Afatinib |
| MVP_top60_DNA_seed | HDAC2 | 1.144626 | OV | OV-90 | Romidepsin | top60_altered_DNA+RNA_seed | ERBB2 | -0.747106 | STAD | RF-48 | Afatinib |
| MVP_top60_DNA_seed | CDK2 | 1.153832 | OV | Hey | Dinaciclib | top60_altered_DNA+RNA_seed | PIK3CA | -0.739382 | STAD | SNU-5 | PI3Ka_4409 |
| MVP_top60_DNA_seed | CDK1 | 1.153832 | OV | Hey | Dinaciclib | top60_altered_DNA+RNA_seed | ERBB2 | -0.687259 | STAD | TMK-1 | Afatinib |
| MVP_top60_DNA_seed | EGFR | 1.155033 | OV | OVCAR-4 | AZD3759 | top60_altered_DNA+RNA_seed | ERBB2 | -0.66765 | STAD | 23132-87 | CUDC-101 |
| MVP_top60_DNA_seed | CDK1 | 1.156057 | OV | OVMIU | RO-3306 | top60_altered_DNA+RNA_seed | ERBB2 | -0.650746 | STAD | IM-95 | CP724714 |
| MVP_top60_DNA_seed | CDK1 | 1.160813 | OV | OVCAR-5 | RO-3306 | top60_altered_DNA+RNA_seed | ERBB2 | -0.648542 | STAD | GCIY | CP724714 |
| MVP_top60_DNA_seed | PIK3CA | 1.171585 | OV | TOV-112D | AZD8186 | top60_altered_DNA+RNA_seed | ERBB2 | -0.605117 | STAD | IM-95 | CI-1033 |
| MVP_top60_DNA_seed | EGFR | 1.172017 | OV | OVK-18 | Erlotinib | top60_altered_DNA+RNA_seed | ERBB4 | -0.605117 | STAD | IM-95 | CI-1033 |

| | | | | | | | | | | | |
|---|---|---|---|---|---|---|---|---|---|---|---|
| MVP_top60_DNA_seed | EGFR | 1.184508 | OV | SW626 | Afatinib | top60_altered_DNA+RNA_seed | ERBB4 | -0.604596 | STAD | RERF-GC-1B | AST-1306 |
| MVP_top60_DNA_seed | ERBB2 | 1.184508 | OV | SW626 | Afatinib | top60_altered_DNA+RNA_seed | ERBB2 | -0.602136 | STAD | MKN28 | AZD8931 |
| MVP_top60_DNA_seed | PIK3CA | 1.184693 | OV | OVKATE | AZD8186 | top60_altered_DNA+RNA_seed | PIK3CA | -0.58183 | STAD | 23132-87 | AZD8835 |
| MVP_top60_DNA_seed | CDK2 | 1.188538 | OV | FU-OV-1 | MK-8776 | top60_altered_DNA+RNA_seed | ERBB2 | -0.576329 | STAD | Hs-746T | CUDC-101 |
| MVP_top60_DNA_seed | EGFR | 1.189712 | OV | OVKATE | AZD3759 | top60_altered_DNA+RNA_seed | ERBB2 | -0.576186 | STAD | KATOIII | PF-00299804 |
| MVP_top60_DNA_seed | CDK1 | 1.199338 | OV | OVISE | RO-3306 | top60_altered_DNA+RNA_seed | ERBB4 | -0.576186 | STAD | KATOIII | PF-00299804 |
| MVP_top60_DNA_seed | ATR | 1.201095 | OV | JHOS-2 | VE821 | top60_altered_DNA+RNA_seed | PIK3CA | -0.544564 | STAD | NCI-N87 | AZD8835 |
| MVP_top60_DNA_seed | CREBBP | 1.207917 | OV | OVISE | SGC-CBP30 | top60_altered_DNA+RNA_seed | ERBB2 | -0.53759 | STAD | RERF-GC-1B | CI-1033 |
| MVP_top60_DNA_seed | MAPK1 | 1.21074 | OV | OAW-42 | Ulixertinib | top60_altered_DNA+RNA_seed | ERBB4 | -0.53759 | STAD | RERF-GC-1B | CI-1033 |
| MVP_top60_DNA_seed | MAPK3 | 1.21074 | OV | OAW-42 | Ulixertinib | top60_altered_DNA+RNA_seed | ERBB2 | -0.507793 | STAD | NUGC-4 | CP724714 |
| MVP_top60_DNA_seed | TP53 | 1.211391 | OV | Hey | MIRA-1 | top60_altered_DNA+RNA_seed | ERBB2 | -0.490537 | STAD | SNU-1 | CUDC-101 |
| MVP_top60_DNA_seed | MAPK1 | 1.225528 | OV | OAW-28 | VX-11e | top60_altered_DNA+RNA_seed | ERBB2 | -0.458125 | STAD | AGS | CUDC-101 |
| MVP_top60_DNA_seed | PLK1 | 1.22863 | OV | Caov-4 | BI-2536 | top60_altered_DNA+RNA_seed | ERBB2 | -0.457756 | STAD | MKN28 | CI-1033 |
| MVP_top60_DNA_seed | CDK1 | 1.231961 | OV | JHOS-3 | RO-3306 | top60_altered_DNA+RNA_seed | ERBB4 | -0.457756 | STAD | MKN28 | CI-1033 |
| MVP_top60_DNA_seed | MAPK1 | 1.2332 | OV | OVTOKO | VX-11e | top60_altered_DNA+RNA_seed | ERBB2 | -0.428087 | STAD | KATOIII | CI-1033 |
| MVP_top60_DNA_seed | EGFR | 1.239986 | OV | OV-56 | Erlotinib | top60_altered_DNA+RNA_seed | ERBB4 | -0.428087 | STAD | KATOIII | CI-1033 |
| MVP_top60_DNA_seed | ATR | 1.241047 | OV | SK-OV-3 | VE821 | top60_altered_DNA+RNA_seed | ERBB2 | -0.423921 | STAD | MKN7 | Afatinib |
| MVP_top60_DNA_seed | MAPK1 | 1.242756 | OV | OVTOKO | SCH772984 | top60_altered_DNA+RNA_seed | PIK3CA | -0.406898 | STAD | TMK-1 | PI3Ka_4409 |
| MVP_top60_DNA_seed | MAPK3 | 1.242756 | OV | OVTOKO | SCH772984 | top60_altered_DNA+RNA_seed | ERBB2 | -0.406648 | STAD | FU97 | CI-1033 |
| MVP_top60_DNA_seed | HDAC2 | 1.24551 | OV | OV-56 | Romidepsin | top60_altered_DNA+RNA_seed | ERBB4 | -0.406648 | STAD | FU97 | CI-1033 |
| MVP_top60_DNA_seed | EGFR | 1.248838 | OV | OVCAR-3 | Lapatinib | top60_altered_DNA+RNA_seed | PIK3CA | -0.366374 | STAD | FU97 | PI3Ka_4409 |
| MVP_top60_DNA_seed | ERBB2 | 1.248838 | OV | OVCAR-3 | Lapatinib | top60_altered_DNA+RNA_seed | ERBB2 | -0.314995 | STAD | MKN7 | PF-00299804 |
| MVP_top60_DNA_seed | PIK3CA | 1.250227 | OV | SW626 | GNE-317 | top60_altered_DNA+RNA_seed | ERBB4 | -0.314995 | STAD | MKN7 | PF-00299804 |
| MVP_top60_DNA_seed | EGFR | 1.262003 | OV | EFO-21 | Afatinib | top60_altered_DNA+RNA_seed | ERBB2 | -0.314747 | STAD | IM-95 | CUDC-101 |
| MVP_top60_DNA_seed | ERBB2 | 1.262003 | OV | EFO-21 | Afatinib | top60_altered_DNA+RNA_seed | ERBB4 | -0.286039 | STAD | KATOIII | AST-1306 |
| MVP_top60_DNA_seed | ATR | 1.276742 | OV | OVMIU | VE821 | top60_altered_DNA+RNA_seed | ERBB2 | -0.284447 | STAD | 23132-87 | AZD8931 |
| MVP_top60_DNA_seed | ATR | 1.278417 | OV | OAW-42 | VE821 | top60_altered_DNA+RNA_seed | ERBB2 | -0.24229 | STAD | RERF-GC-1B | AZD8931 |
| MVP_top60_DNA_seed | CDK2 | 1.279014 | OV | OAW-28 | Dinaciclib | top60_altered_DNA+RNA_seed | ERBB2 | -0.23349 | STAD | SNU-16 | AZD8931 |
| MVP_top60_DNA_seed | CDK1 | 1.279014 | OV | OAW-28 | Dinaciclib | top60_altered_DNA+RNA_seed | ERBB4 | -0.220035 | STAD | TGBC11TKB | AST-1306 |
| MVP_top60_DNA_seed | EGFR | 1.284414 | OV | OVKATE | Lapatinib | top60_altered_DNA+RNA_seed | ERBB2 | -0.21364 | STAD | TGBC11TKB | CI-1033 |
| MVP_top60_DNA_seed | ERBB2 | 1.284414 | OV | OVKATE | Lapatinib | top60_altered_DNA+RNA_seed | ERBB4 | -0.21364 | STAD | TGBC11TKB | CI-1033 |
| MVP_top60_DNA_seed | CDK2 | 1.286833 | OV | EFO-21 | MK-8776 | top60_altered_DNA+RNA_seed | ERBB4 | -0.206675 | STAD | MKN7 | AST-1306 |
| MVP_top60_DNA_seed | MAPK1 | 1.288101 | OV | OV-17R | Ulixertinib | top60_altered_DNA+RNA_seed | ERBB2 | -0.200245 | STAD | NUGC-3 | CUDC-101 |
| MVP_top60_DNA_seed | MAPK3 | 1.288101 | OV | OV-17R | Ulixertinib | top60_altered_DNA+RNA_seed | ERBB2 | -0.193304 | STAD | RERF-GC-1B | PF-00299804 |
| MVP_top60_DNA_seed | CDK2 | 1.291227 | OV | OVCAR-5 | MK-8776 | top60_altered_DNA+RNA_seed | ERBB4 | -0.193304 | STAD | RERF-GC-1B | PF-00299804 |
| MVP_top60_DNA_seed | EGFR | 1.300841 | OV | OV-56 | Afatinib | top60_altered_DNA+RNA_seed | PIK3CA | -0.182956 | STAD | RERF-GC-1B | PI3Ka_4409 |
| MVP_top60_DNA_seed | ERBB2 | 1.300841 | OV | OV-56 | Afatinib | top60_altered_DNA+RNA_seed | ERBB2 | -0.175941 | STAD | MKN7 | CI-1033 |
| MVP_top60_DNA_seed | PLK1 | 1.307273 | OV | OV-56 | MK-1775 | top60_altered_DNA+RNA_seed | ERBB4 | -0.175941 | STAD | MKN7 | CI-1033 |
| MVP_top60_DNA_seed | CDK1 | 1.307978 | OV | SK-OV-3 | RO-3306 | top60_altered_DNA+RNA_seed | PIK3CA | -0.166059 | STAD | FU97 | AZD8835 |
| MVP_top60_DNA_seed | EGFR | 1.312166 | OV | PEO1 | Gefitinib | top60_altered_DNA+RNA_seed | ERBB4 | -0.163419 | STAD | Hs-746T | AST-1306 |
| MVP_top60_DNA_seed | MAPK1 | 1.318788 | OV | EFO-21 | VX-11e | top60_altered_DNA+RNA_seed | ERBB2 | -0.160746 | STAD | HGC-27 | CUDC-101 |

| | | | | | | | | | | | |
|---|---|---|---|---|---|---|---|---|---|---|---|
| MVP_top60_DNA_seed | EGFR | 1.324072 | OV | SW626 | Sapitinib | top60_altered_DNA+RNA_seed | ERBB2 | -0.144342 | STAD | IM-95 | Afatinib |
| MVP_top60_DNA_seed | ERBB2 | 1.324072 | OV | SW626 | Sapitinib | top60_altered_DNA+RNA_seed | ERBB2 | -0.138867 | STAD | AGS | CP724714 |
| MVP_top60_DNA_seed | PIK3CA | 1.348812 | OV | OV-90 | Buparlisib | top60_altered_DNA+RNA_seed | ERBB2 | -0.134016 | STAD | FU97 | CP724714 |
| MVP_top60_DNA_seed | CDK1 | 1.360186 | OV | OVCAR-3 | RO-3306 | top60_altered_DNA+RNA_seed | PIK3CA | -0.130699 | STAD | 23132-87 | PI3Ka_4409 |
| MVP_top60_DNA_seed | ATR | 1.364334 | OV | Hey | VE-822 | top60_altered_DNA+RNA_seed | PIK3CA | -0.128391 | STAD | RERF-GC-1B | AZD8835 |
| MVP_top60_DNA_seed | ATR | 1.36502 | OV | OV-56 | VE821 | top60_altered_DNA+RNA_seed | ERBB2 | -0.128348 | STAD | TGBC11TKB | PF-00299804 |
| MVP_top60_DNA_seed | ATR | 1.384674 | OV | OVCA420 | VE821 | top60_altered_DNA+RNA_seed | ERBB4 | -0.128348 | STAD | TGBC11TKB | PF-00299804 |
| MVP_top60_DNA_seed | TP53 | 1.387695 | OV | OVCAR-5 | MIRA-1 | top60_altered_DNA+RNA_seed | ERBB2 | -0.125099 | STAD | HGC-27 | CI-1033 |
| MVP_top60_DNA_seed | EGFR | 1.396589 | OV | OV-90 | Erlotinib | top60_altered_DNA+RNA_seed | ERBB4 | -0.125099 | STAD | HGC-27 | CI-1033 |
| MVP_top60_DNA_seed | ATR | 1.403126 | OV | SW626 | VE821 | top60_altered_DNA+RNA_seed | ERBB2 | -0.120265 | STAD | GCIY | CUDC-101 |
| MVP_top60_DNA_seed | EGFR | 1.411217 | OV | OV-90 | AZD3759 | top60_altered_DNA+RNA_seed | ERBB2 | -0.120054 | STAD | HSC-39 | Lapatinib |
| MVP_top60_DNA_seed | CDK2 | 1.412598 | OV | OVCA420 | Dinaciclib | top60_altered_DNA+RNA_seed | ERBB2 | -0.093925 | STAD | NUGC-3 | CI-1033 |
| MVP_top60_DNA_seed | CDK1 | 1.412598 | OV | OVCA420 | Dinaciclib | top60_altered_DNA+RNA_seed | ERBB4 | -0.093925 | STAD | NUGC-3 | CI-1033 |
| MVP_top60_DNA_seed | TP53 | 1.422896 | OV | SK-OV-3 | MIRA-1 | top60_altered_DNA+RNA_seed | ERBB2 | -0.065151 | STAD | AGS | Afatinib |
| MVP_top60_DNA_seed | PIK3CA | 1.437695 | OV | JHOS-3 | Buparlisib | top60_altered_DNA+RNA_seed | ERBB2 | -0.05119 | STAD | 23132-87 | CI-1033 |
| MVP_top60_DNA_seed | PIK3CA | 1.445814 | OV | EFO-21 | Buparlisib | top60_altered_DNA+RNA_seed | ERBB4 | -0.05119 | STAD | 23132-87 | CI-1033 |
| MVP_top60_DNA_seed | EGFR | 1.449047 | OV | SW626 | Gefitinib | top60_altered_DNA+RNA_seed | PIK3CA | -0.050562 | STAD | Hs-746T | PI3Ka_4409 |
| MVP_top60_DNA_seed | MAPK1 | 1.467614 | OV | OVKATE | Ulixertinib | top60_altered_DNA+RNA_seed | ERBB2 | -0.047554 | STAD | HSC-39 | AZD8931 |
| MVP_top60_DNA_seed | MAPK3 | 1.467614 | OV | OVKATE | Ulixertinib | top60_altered_DNA+RNA_seed | ERBB2 | -0.032571 | STAD | HSC-39 | CP724714 |
| MVP_top60_DNA_seed | MAPK1 | 1.470404 | OV | OAW-42 | SCH772984 | top60_altered_DNA+RNA_seed | ERBB2 | -0.020592 | STAD | NUGC-3 | CP724714 |
| MVP_top60_DNA_seed | MAPK3 | 1.470404 | OV | OAW-42 | SCH772984 | top60_altered_DNA+RNA_seed | ERBB2 | -0.000572 | STAD | TMK-1 | Afatinib |
| MVP_top60_DNA_seed | CDK2 | 1.479749 | OV | EFO-21 | Dinaciclib | top60_altered_DNA+RNA_seed | ERBB2 | 0.004358 | STAD | Hs-746T | CI-1033 |
| MVP_top60_DNA_seed | CDK1 | 1.479749 | OV | EFO-21 | Dinaciclib | top60_altered_DNA+RNA_seed | ERBB4 | 0.004358 | STAD | Hs-746T | CI-1033 |
| MVP_top60_DNA_seed | EGFR | 1.481052 | OV | PEO1 | AZD3759 | top60_altered_DNA+RNA_seed | ERBB2 | 0.007664 | STAD | HSC-39 | Afatinib |
| MVP_top60_DNA_seed | PIK3CA | 1.48927 | OV | SW626 | Buparlisib | top60_altered_DNA+RNA_seed | PIK3CA | 0.020711 | STAD | IM-95 | PI3Ka_4409 |
| MVP_top60_DNA_seed | EGFR | 1.493503 | OV | SW626 | Osimertinib | top60_altered_DNA+RNA_seed | ERBB2 | 0.037018 | STAD | HSC-39 | Afatinib |
| MVP_top60_DNA_seed | EGFR | 1.494488 | OV | OV-90 | Afatinib | top60_altered_DNA+RNA_seed | ERBB4 | 0.067172 | STAD | IM-95 | AST-1306 |
| MVP_top60_DNA_seed | ERBB2 | 1.494488 | OV | OV-90 | Afatinib | top60_altered_DNA+RNA_seed | PIK3CA | 0.073779 | STAD | SNU-16 | AZD8835 |
| MVP_top60_DNA_seed | PLK1 | 1.495418 | OV | EFO-21 | BI-2536 | top60_altered_DNA+RNA_seed | ERBB2 | 0.094896 | STAD | FU97 | CUDC-101 |
| MVP_top60_DNA_seed | MAPK1 | 1.501427 | OV | OV-56 | Ulixertinib | top60_altered_DNA+RNA_seed | PIK3CA | 0.121784 | STAD | TGBC11TKB | AZD8835 |
| MVP_top60_DNA_seed | MAPK3 | 1.501427 | OV | OV-56 | Ulixertinib | top60_altered_DNA+RNA_seed | PIK3CA | 0.128448 | STAD | NUGC-4 | AZD8835 |
| MVP_top60_DNA_seed | EGFR | 1.503579 | OV | FU-OV-1 | Gefitinib | top60_altered_DNA+RNA_seed | PIK3CA | 0.137472 | STAD | MKN28 | AZD8835 |
| MVP_top60_DNA_seed | PIK3CA | 1.51291 | OV | OVKATE | GNE-317 | top60_altered_DNA+RNA_seed | ERBB2 | 0.138652 | STAD | SNU-16 | CP724714 |
| MVP_top60_DNA_seed | EGFR | 1.527364 | OV | EFO-21 | Gefitinib | top60_altered_DNA+RNA_seed | ERBB2 | 0.146955 | STAD | MKN7 | CP724714 |
| MVP_top60_DNA_seed | ATR | 1.529522 | OV | OVMIU | VE-822 | top60_altered_DNA+RNA_seed | ERBB2 | 0.160058 | STAD | MKN7 | AZD8931 |
| MVP_top60_DNA_seed | PIK3CA | 1.536942 | OV | OVCAR-4 | AZD8186 | top60_altered_DNA+RNA_seed | ERBB2 | 0.167947 | STAD | HGC-27 | Afatinib |
| MVP_top60_DNA_seed | MAPK1 | 1.543985 | OV | OVCAR-8 | VX-11e | top60_altered_DNA+RNA_seed | PIK3CA | 0.178054 | STAD | SK-GT-2 | PI3Ka_4409 |
| MVP_top60_DNA_seed | CDK2 | 1.558421 | OV | OVCAR-4 | Dinaciclib | top60_altered_DNA+RNA_seed | ERBB2 | 0.191637 | STAD | IM-95 | PF-00299804 |
| MVP_top60_DNA_seed | CDK1 | 1.558421 | OV | OVCAR-4 | Dinaciclib | top60_altered_DNA+RNA_seed | ERBB4 | 0.191637 | STAD | IM-95 | PF-00299804 |
| MVP_top60_DNA_seed | EGFR | 1.564358 | OV | EFO-21 | Lapatinib | top60_altered_DNA+RNA_seed | PIK3CA | 0.203951 | STAD | SNU-5 | AZD8835 |
| MVP_top60_DNA_seed | ERBB2 | 1.564358 | OV | EFO-21 | Lapatinib | top60_altered_DNA+RNA_seed | PIK3CA | 0.249282 | STAD | KATOIII | AZD8835 |

| | | | | | | | | | | | |
|---|---|---|---|---|---|---|---|---|---|---|---|
| MVP_top60_DNA_seed | EGFR | 1.566351 | OV | Hey | AZD3759 | top60_altered_DNA+RNA_seed | ERBB2 | 0.256146 | STAD | SNU-5 | Afatinib |
| MVP_top60_DNA_seed | CDK2 | 1.56827 | OV | OV-90 | AZD5438 | top60_altered_DNA+RNA_seed | ERBB4 | 0.262005 | STAD | 23132-87 | AST-1306 |
| MVP_top60_DNA_seed | MAPK1 | 1.575187 | OV | OV-56 | VX-11e | top60_altered_DNA+RNA_seed | ERBB2 | 0.268978 | STAD | SNU-1 | Afatinib |
| MVP_top60_DNA_seed | PIK3CA | 1.583205 | OV | OV-56 | Buparlisib | top60_altered_DNA+RNA_seed | ERBB2 | 0.279114 | STAD | TGBC11TKB | Afatinib |
| MVP_top60_DNA_seed | ATR | 1.612715 | OV | EFO-21 | AZD6738 | top60_altered_DNA+RNA_seed | ERBB4 | 0.2807 | STAD | FU97 | AST-1306 |
| MVP_top60_DNA_seed | HDAC2 | 1.615375 | OV | OAW-28 | Romidepsin | top60_altered_DNA+RNA_seed | ERBB2 | 0.28185 | STAD | SNU-16 | Lapatinib |
| MVP_top60_DNA_seed | CDK2 | 1.616995 | OV | SK-OV-3 | Dinaciclib | top60_altered_DNA+RNA_seed | ERBB2 | 0.305071 | STAD | SNU-5 | AZD8931 |
| MVP_top60_DNA_seed | CDK1 | 1.616995 | OV | SK-OV-3 | Dinaciclib | top60_altered_DNA+RNA_seed | ERBB2 | 0.314203 | STAD | MKN28 | CP724714 |
| MVP_top60_DNA_seed | CREBBP | 1.62136 | OV | OVMIU | SGC-CBP30 | top60_altered_DNA+RNA_seed | ERBB2 | 0.340594 | STAD | HGC-27 | Afatinib |
| MVP_top60_DNA_seed | EGFR | 1.624466 | OV | OAW-42 | Lapatinib | top60_altered_DNA+RNA_seed | ERBB2 | 0.347888 | STAD | NUGC-3 | PF-00299804 |
| MVP_top60_DNA_seed | ERBB2 | 1.624466 | OV | OAW-42 | Lapatinib | top60_altered_DNA+RNA_seed | ERBB4 | 0.347888 | STAD | NUGC-3 | PF-00299804 |
| MVP_top60_DNA_seed | EGFR | 1.635413 | OV | OV-90 | Sapitinib | top60_altered_DNA+RNA_seed | ERBB2 | 0.381472 | STAD | FU97 | AZD8931 |
| MVP_top60_DNA_seed | ERBB2 | 1.635413 | OV | OV-90 | Sapitinib | top60_altered_DNA+RNA_seed | ERBB2 | 0.394715 | STAD | Hs-746T | PF-00299804 |
| MVP_top60_DNA_seed | PIK3CA | 1.642875 | OV | OVCAR-4 | Alpelisib | top60_altered_DNA+RNA_seed | ERBB4 | 0.394715 | STAD | Hs-746T | PF-00299804 |
| MVP_top60_DNA_seed | CDK1 | 1.656661 | OV | OV-56 | RO-3306 | top60_altered_DNA+RNA_seed | ERBB2 | 0.424051 | STAD | SNU-5 | Afatinib |
| MVP_top60_DNA_seed | EGFR | 1.656998 | OV | OVCAR-5 | Erlotinib | top60_altered_DNA+RNA_seed | ERBB2 | 0.42897 | STAD | MKN45 | CI-1033 |
| MVP_top60_DNA_seed | ATR | 1.664987 | OV | SW626 | VE-822 | top60_altered_DNA+RNA_seed | ERBB4 | 0.42897 | STAD | MKN45 | CI-1033 |
| MVP_top60_DNA_seed | CDK1 | 1.675011 | OV | OV-90 | RO-3306 | top60_altered_DNA+RNA_seed | ERBB2 | 0.45463 | STAD | RERF-GC-1B | Afatinib |
| MVP_top60_DNA_seed | EGFR | 1.676269 | OV | OV-56 | Osimertinib | top60_altered_DNA+RNA_seed | ERBB2 | 0.462993 | STAD | Hs-746T | AZD8931 |
| MVP_top60_DNA_seed | ATR | 1.676673 | OV | OVCAR-5 | VE821 | top60_altered_DNA+RNA_seed | ERBB2 | 0.48105 | STAD | MKN28 | CUDC-101 |
| MVP_top60_DNA_seed | CREBBP | 1.686987 | OV | SK-OV-3 | SGC-CBP30 | top60_altered_DNA+RNA_seed | ERBB2 | 0.489956 | STAD | 23132-87 | Afatinib |
| MVP_top60_DNA_seed | PIK3CA | 1.690777 | OV | OV-56 | Alpelisib | top60_altered_DNA+RNA_seed | ERBB2 | 0.496494 | STAD | KATOIII | AZD8931 |
| MVP_top60_DNA_seed | HDAC2 | 1.696267 | OV | OVCA420 | Romidepsin | top60_altered_DNA+RNA_seed | PIK3CA | 0.500928 | STAD | MKN45 | PI3Ka_4409 |
| MVP_top60_DNA_seed | MAPK1 | 1.698266 | OV | OVKATE | SCH772984 | top60_altered_DNA+RNA_seed | ERBB2 | 0.507041 | STAD | 23132-87 | Afatinib |
| MVP_top60_DNA_seed | MAPK3 | 1.698266 | OV | OVKATE | SCH772984 | top60_altered_DNA+RNA_seed | PIK3CA | 0.511737 | STAD | MKN28 | PI3Ka_4409 |
| MVP_top60_DNA_seed | CDK1 | 1.704197 | OV | SW626 | RO-3306 | top60_altered_DNA+RNA_seed | ERBB2 | 0.514996 | STAD | RF-48 | CI-1033 |
| MVP_top60_DNA_seed | EGFR | 1.715002 | OV | OV-90 | Lapatinib | top60_altered_DNA+RNA_seed | ERBB4 | 0.514996 | STAD | RF-48 | CI-1033 |
| MVP_top60_DNA_seed | ERBB2 | 1.715002 | OV | OV-90 | Lapatinib | top60_altered_DNA+RNA_seed | PIK3CA | 0.519231 | STAD | NUGC-4 | PI3Ka_4409 |
| MVP_top60_DNA_seed | ATR | 1.717195 | OV | SK-OV-3 | VE-822 | top60_altered_DNA+RNA_seed | ERBB2 | 0.522888 | STAD | MKN7 | CUDC-101 |
| MVP_top60_DNA_seed | CDK2 | 1.742357 | OV | OV-90 | Dinaciclib | top60_altered_DNA+RNA_seed | PIK3CA | 0.523535 | STAD | SNU-16 | PI3Ka_4409 |
| MVP_top60_DNA_seed | CDK1 | 1.742357 | OV | OV-90 | Dinaciclib | top60_altered_DNA+RNA_seed | ERBB2 | 0.536215 | STAD | 23132-87 | CP724714 |
| MVP_top60_DNA_seed | PIK3CA | 1.743283 | OV | SW626 | AZD8186 | top60_altered_DNA+RNA_seed | PIK3CA | 0.554986 | STAD | KATOIII | PI3Ka_4409 |
| MVP_top60_DNA_seed | CDK1 | 1.758703 | OV | OVTOKO | RO-3306 | top60_altered_DNA+RNA_seed | ERBB2 | 0.563994 | STAD | OCUM-1 | AZD8931 |
| MVP_top60_DNA_seed | CDK2 | 1.760814 | OV | PEO1 | AZD5438 | top60_altered_DNA+RNA_seed | ERBB2 | 0.564898 | STAD | 23132-87 | PF-00299804 |
| MVP_top60_DNA_seed | PLK1 | 1.772337 | OV | OV-56 | BI-2536 | top60_altered_DNA+RNA_seed | ERBB4 | 0.564898 | STAD | 23132-87 | PF-00299804 |
| MVP_top60_DNA_seed | MAPK1 | 1.780926 | OV | OAW-42 | VX-11e | top60_altered_DNA+RNA_seed | ERBB2 | 0.569323 | STAD | FU97 | Afatinib |
| MVP_top60_DNA_seed | EGFR | 1.800697 | OV | OV-90 | Gefitinib | top60_altered_DNA+RNA_seed | PIK3CA | 0.572245 | STAD | OCUM-1 | PI3Ka_4409 |
| MVP_top60_DNA_seed | CDK2 | 1.833921 | OV | OVCAR-4 | AZD5438 | top60_altered_DNA+RNA_seed | ERBB2 | 0.577607 | STAD | KATOIII | Afatinib |
| MVP_top60_DNA_seed | PIK3CA | 1.854031 | OV | EFO-21 | Alpelisib | top60_altered_DNA+RNA_seed | PIK3CA | 0.589523 | STAD | MKN7 | AZD8835 |
| MVP_top60_DNA_seed | ATR | 1.870468 | OV | OV-56 | AZD6738 | top60_altered_DNA+RNA_seed | ERBB4 | 0.613193 | STAD | HGC-27 | AST-1306 |
| MVP_top60_DNA_seed | EGFR | 1.876163 | OV | SW626 | Lapatinib | top60_altered_DNA+RNA_seed | ERBB2 | 0.624443 | STAD | MKN45 | Afatinib |

| | | | | | | | | | | | |
|---|---|---|---|---|---|---|---|---|---|---|---|
| MVP_top60_DNA_seed | ERBB2 | 1.876163 | OV | SW626 | Lapatinib | top60_altered_DNA+RNA_seed | ERBB2 | 0.628557 | STAD | TGBC11TKB | CUDC-101 |
| MVP_top60_DNA_seed | CDK1 | 1.889445 | OV | FU-OV-1 | RO-3306 | top60_altered_DNA+RNA_seed | PIK3CA | 0.634969 | STAD | NUGC-3 | AZD8835 |
| MVP_top60_DNA_seed | EGFR | 1.898484 | OV | OVCAR-5 | Osimertinib | top60_altered_DNA+RNA_seed | ERBB2 | 0.65275 | STAD | TMK-1 | CUDC-101 |
| MVP_top60_DNA_seed | MAPK1 | 1.913957 | OV | SK-OV-3 | VX-11e | top60_altered_DNA+RNA_seed | ERBB2 | 0.673728 | STAD | SNU-16 | Afatinib |
| MVP_top60_DNA_seed | PIK3CA | 1.919447 | OV | OV-90 | AZD8186 | top60_altered_DNA+RNA_seed | ERBB2 | 0.674507 | STAD | OCUM-1 | CP724714 |
| MVP_top60_DNA_seed | MAPK1 | 1.939801 | OV | OV-56 | SCH772984 | top60_altered_DNA+RNA_seed | ERBB2 | 0.68628 | STAD | MKN28 | Afatinib |
| MVP_top60_DNA_seed | MAPK3 | 1.939801 | OV | OV-56 | SCH772984 | top60_altered_DNA+RNA_seed | PIK3CA | 0.712438 | STAD | NUGC-3 | PI3Ka_4409 |
| MVP_top60_DNA_seed | MAPK1 | 1.956783 | OV | SK-OV-3 | Ulixertinib | top60_altered_DNA+RNA_seed | ERBB2 | 0.713247 | STAD | RF-48 | AZD8931 |
| MVP_top60_DNA_seed | MAPK3 | 1.956783 | OV | SK-OV-3 | Ulixertinib | top60_altered_DNA+RNA_seed | ERBB2 | 0.725846 | STAD | SNU-5 | Lapatinib |
| MVP_top60_DNA_seed | MAPK1 | 1.960825 | OV | SK-OV-3 | Ulixertinib | top60_altered_DNA+RNA_seed | PIK3CA | 0.732466 | STAD | TGBC11TKB | PI3Ka_4409 |
| MVP_top60_DNA_seed | MAPK3 | 1.960825 | OV | SK-OV-3 | Ulixertinib | top60_altered_DNA+RNA_seed | ERBB2 | 0.741339 | STAD | KATOIII | Afatinib |
| MVP_top60_DNA_seed | PIK3CA | 1.964508 | OV | OVKATE | Alpelisib | top60_altered_DNA+RNA_seed | ERBB2 | 0.753648 | STAD | FU97 | Afatinib |
| MVP_top60_DNA_seed | CDK1 | 1.965873 | OV | PEO1 | RO-3306 | top60_altered_DNA+RNA_seed | ERBB2 | 0.754714 | STAD | TGBC11TKB | Afatinib |
| MVP_top60_DNA_seed | ATR | 1.987183 | OV | EFO-21 | VE821 | top60_altered_DNA+RNA_seed | PIK3CA | 0.758768 | STAD | SK-GT-2 | AZD8835 |
| MVP_top60_DNA_seed | CDK2 | 2.008173 | OV | EFO-21 | AZD5438 | top60_altered_DNA+RNA_seed | ERBB2 | 0.760857 | STAD | HGC-27 | PF-00299804 |
| MVP_top60_DNA_seed | EGFR | 2.015888 | OV | OV-56 | Lapatinib | top60_altered_DNA+RNA_seed | ERBB4 | 0.760857 | STAD | HGC-27 | PF-00299804 |
| MVP_top60_DNA_seed | ERBB2 | 2.015888 | OV | OV-56 | Lapatinib | top60_altered_DNA+RNA_seed | ERBB4 | 0.762208 | STAD | MKN28 | AST-1306 |
| MVP_top60_DNA_seed | CDK2 | 2.049569 | OV | SK-OV-3 | AZD5438 | top60_altered_DNA+RNA_seed | PIK3CA | 0.800803 | STAD | MKN7 | PI3Ka_4409 |
| MVP_top60_DNA_seed | CREBBP | 2.052992 | OV | EFO-21 | SGC-CBP30 | top60_altered_DNA+RNA_seed | ERBB2 | 0.81431 | STAD | MKN28 | PF-00299804 |
| MVP_top60_DNA_seed | ATR | 2.168268 | OV | EFO-21 | VE-822 | top60_altered_DNA+RNA_seed | ERBB4 | 0.81431 | STAD | MKN28 | PF-00299804 |
| MVP_top60_DNA_seed | HDAC2 | 2.241086 | OV | PEO1 | Romidepsin | top60_altered_DNA+RNA_seed | ERBB2 | 0.846034 | STAD | FU97 | PF-00299804 |
| MVP_top60_DNA_seed | PIK3CA | 2.282247 | OV | OVKATE | Buparlisib | top60_altered_DNA+RNA_seed | ERBB4 | 0.846034 | STAD | FU97 | PF-00299804 |
| MVP_top60_DNA_seed | TP53 | 2.285099 | OV | EFO-21 | MIRA-1 | top60_altered_DNA+RNA_seed | ERBB2 | 0.911389 | STAD | TMK-1 | CP724714 |
| MVP_top60_DNA_seed | EGFR | 2.320427 | OV | OV-90 | Osimertinib | top60_altered_DNA+RNA_seed | ERBB2 | 0.918981 | STAD | RF-48 | Lapatinib |
| MVP_top60_DNA_seed | MAPK1 | 2.338687 | OV | OV-56 | Ulixertinib | top60_altered_DNA+RNA_seed | PIK3CA | 0.952346 | STAD | RF-48 | AZD8835 |
| MVP_top60_DNA_seed | MAPK3 | 2.338687 | OV | OV-56 | Ulixertinib | top60_altered_DNA+RNA_seed | ERBB2 | 0.952632 | STAD | TGBC11TKB | AZD8931 |
| MVP_top60_DNA_seed | EGFR | 2.348204 | OV | SW626 | AZD3759 | top60_altered_DNA+RNA_seed | ERBB2 | 0.953741 | STAD | Hs-746T | Afatinib |
| MVP_top60_DNA_seed | PLK1 | 2.391882 | OV | OVKATE | MK-1775 | top60_altered_DNA+RNA_seed | ERBB2 | 0.95716 | STAD | MKN45 | PF-00299804 |
| MVP_top60_DNA_seed | HDAC2 | 2.394554 | OV | OVCAR-4 | Romidepsin | top60_altered_DNA+RNA_seed | ERBB4 | 0.95716 | STAD | MKN45 | PF-00299804 |
| MVP_top60_DNA_seed | ATR | 2.492328 | OV | OVKATE | VE821 | top60_altered_DNA+RNA_seed | ERBB2 | 0.957312 | STAD | MKN45 | AZD8931 |
| MVP_top60_DNA_seed | CDK2 | 2.502915 | OV | OV-56 | AZD5438 | top60_altered_DNA+RNA_seed | ERBB2 | 0.961597 | STAD | Hs-746T | Afatinib |
| MVP_top60_DNA_seed | EGFR | 2.505243 | OV | OVK-18 | Afatinib | top60_altered_DNA+RNA_seed | ERBB2 | 0.963799 | STAD | OCUM-1 | Afatinib |
| MVP_top60_DNA_seed | ERBB2 | 2.505243 | OV | OVK-18 | Afatinib | top60_altered_DNA+RNA_seed | ERBB2 | 0.9741 | STAD | SNU-16 | Afatinib |
| MVP_top60_DNA_seed | CDK2 | 2.627391 | OV | OVKATE | MK-8776 | top60_altered_DNA+RNA_seed | ERBB2 | 0.981788 | STAD | RF-48 | CP724714 |
| MVP_top60_DNA_seed | CDK1 | 2.639398 | OV | EFO-21 | RO-3306 | top60_altered_DNA+RNA_seed | PIK3CA | 1.028821 | STAD | OCUM-1 | AZD8835 |
| MVP_top60_DNA_seed | MAPK1 | 2.669309 | OV | OVKATE | Ulixertinib | top60_altered_DNA+RNA_seed | ERBB2 | 1.030123 | STAD | MKN45 | Afatinib |
| MVP_top60_DNA_seed | MAPK3 | 2.669309 | OV | OVKATE | Ulixertinib | top60_altered_DNA+RNA_seed | ERBB2 | 1.039935 | STAD | SNU-1 | Lapatinib |
| MVP_top60_DNA_seed | PLK1 | 2.713057 | OV | OVKATE | BI-2536 | top60_altered_DNA+RNA_seed | ERBB2 | 1.077495 | STAD | OCUM-1 | Afatinib |
| MVP_top60_DNA_seed | CDK1 | 2.724575 | OV | OVKATE | RO-3306 | top60_altered_DNA+RNA_seed | ERBB2 | 1.095849 | STAD | RF-48 | PF-00299804 |
| MVP_top60_DNA_seed | HDAC2 | 2.735783 | OV | EFO-21 | Romidepsin | top60_altered_DNA+RNA_seed | ERBB4 | 1.095849 | STAD | RF-48 | PF-00299804 |
| MVP_top60_DNA_seed | TP53 | 2.834131 | OV | OVKATE | MIRA-1 | top60_altered_DNA+RNA_seed | ERBB2 | 1.106125 | STAD | RF-48 | Afatinib |

| | | | | | | | | | | | |
|---|---|---|---|---|---|---|---|---|---|---|---|
| MVP_top60_DNA_seed | MAPK1 | 2.867698 | OV | OVKATE | VX-11e | top60_altered_DNA+RNA_seed | ERBB2 | 1.116749 | STAD | SNU-5 | CUDC-101 |
| MVP_top60_DNA_seed | ATR | 2.953747 | OV | OVKATE | AZD6738 | top60_altered_DNA+RNA_seed | PIK3CA | 1.134076 | STAD | MKN45 | AZD8835 |
| MVP_top60_DNA_seed | HDAC2 | 3.667555 | OV | OAW-42 | Romidepsin | top60_altered_DNA+RNA_seed | PIK3CA | 1.154174 | STAD | RF-48 | PI3Ka_4409 |
| MVP_top60_DNA_seed | PARP1 | -3.262147 | PRAD | DU-145 | Veliparib | top60_altered_DNA+RNA_seed | ERBB2 | 1.157487 | STAD | SNU-1 | Afatinib |
| MVP_top60_DNA_seed | AR | -2.765186 | PRAD | LNCaP-Clone-FGC | Bicalutamide | top60_altered_DNA+RNA_seed | PIK3CA | 1.214622 | STAD | Hs-746T | AZD8835 |
| MVP_top60_DNA_seed | ATM | -2.736602 | PRAD | DU-145 | KU-55933 | top60_altered_DNA+RNA_seed | ERBB4 | 1.216366 | STAD | RF-48 | AST-1306 |
| MVP_top60_DNA_seed | GSK3B | -2.692938 | PRAD | DU-145 | SB216763 | top60_altered_DNA+RNA_seed | ERBB2 | 1.235015 | STAD | SNU-16 | CUDC-101 |
| MVP_top60_DNA_seed | AR | -2.306773 | PRAD | PC-3 | Bicalutamide | top60_altered_DNA+RNA_seed | ERBB2 | 1.254838 | STAD | SNU-5 | CP724714 |
| MVP_top60_DNA_seed | ESR1 | -2.152619 | PRAD | LNCaP-Clone-FGC | Fulvestrant | top60_altered_DNA+RNA_seed | ERBB2 | 1.306524 | STAD | MKN45 | CP724714 |
| MVP_top60_DNA_seed | GSK3B | -1.770045 | PRAD | VCaP | SB216763 | top60_altered_DNA+RNA_seed | ERBB2 | 1.395564 | STAD | MKN45 | CUDC-101 |
| MVP_top60_DNA_seed | PARP1 | -1.523428 | PRAD | DU-145 | PARP_9495 | top60_altered_DNA+RNA_seed | ERBB2 | 1.538234 | STAD | SNU-16 | CI-1033 |
| MVP_top60_DNA_seed | AR | -1.470437 | PRAD | LNCaP-Clone-FGC | AZD3514 | top60_altered_DNA+RNA_seed | ERBB4 | 1.538234 | STAD | SNU-16 | CI-1033 |
| MVP_top60_DNA_seed | GSK3B | -1.352221 | PRAD | PWR-1E | CHIR-99021 | top60_altered_DNA+RNA_seed | ERBB2 | 1.742186 | STAD | RERF-GC-1B | CUDC-101 |
| MVP_top60_DNA_seed | CDK4 | -1.34343 | PRAD | PC-3 | Palbociclib | top60_altered_DNA+RNA_seed | ERBB2 | 1.870352 | STAD | SNU-16 | PF-00299804 |
| MVP_top60_DNA_seed | GSK3B | -1.308116 | PRAD | DU-145 | KIN001-042 | top60_altered_DNA+RNA_seed | ERBB4 | 1.870352 | STAD | SNU-16 | PF-00299804 |
| MVP_top60_DNA_seed | HDAC1 | -1.280877 | PRAD | VCaP | AR-42 | top60_altered_DNA+RNA_seed | PIK3CA | 1.925783 | STAD | SNU-1 | PI3Ka_4409 |
| MVP_top60_DNA_seed | PARP1 | -1.279816 | PRAD | PWR-1E | Olaparib | top60_altered_DNA+RNA_seed | ERBB4 | 2.175457 | STAD | SNU-16 | AST-1306 |
| MVP_top60_DNA_seed | HDAC1 | -1.265822 | PRAD | VCaP | Belinostat | top60_altered_DNA+RNA_seed | PIK3CA | 2.177344 | STAD | SNU-1 | AZD8835 |
| MVP_top60_DNA_seed | MAPK3 | -1.197452 | PRAD | 22RV1 | FR-180204 | top60_altered_DNA+RNA_seed | ERBB2 | 2.219959 | STAD | OCUM-1 | CUDC-101 |
| MVP_top60_DNA_seed | PTK2 | -1.179388 | PRAD | DU-145 | PF-562271 | top60_altered_DNA+RNA_seed | ERBB2 | 2.327314 | STAD | SNU-1 | AZD8931 |
| MVP_top60_DNA_seed | HDAC1 | -1.179113 | PRAD | LNCaP-Clone-FGC | Belinostat | top60_altered_DNA+RNA_seed | ERBB4 | 2.333127 | STAD | OCUM-1 | AST-1306 |
| MVP_top60_DNA_seed | ESR1 | -1.1777 | PRAD | DU-145 | Tamoxifen | top60_altered_DNA+RNA_seed | ERBB2 | 2.343604 | STAD | OCUM-1 | CI-1033 |
| MVP_top60_DNA_seed | PIK3CA | -1.174944 | PRAD | DU-145 | PI3Ka_4409 | top60_altered_DNA+RNA_seed | ERBB4 | 2.343604 | STAD | OCUM-1 | CI-1033 |
| MVP_top60_DNA_seed | CDK4 | -1.153306 | PRAD | 22RV1 | Palbociclib | top60_altered_DNA+RNA_seed | ERBB4 | 2.506507 | STAD | MKN45 | AST-1306 |
| MVP_top60_DNA_seed | HDAC1 | -1.152706 | PRAD | LNCaP-Clone-FGC | AR-42 | top60_altered_DNA+RNA_seed | ERBB2 | 2.62529 | STAD | SNU-5 | CI-1033 |
| MVP_top60_DNA_seed | MAPK3 | -1.113497 | PRAD | VCaP | FR-180204 | top60_altered_DNA+RNA_seed | ERBB4 | 2.62529 | STAD | SNU-5 | CI-1033 |
| MVP_top60_DNA_seed | HDAC1 | -1.08874 | PRAD | VCaP | Parthenolide | top60_altered_DNA+RNA_seed | ERBB2 | 3.33718 | STAD | SNU-5 | PF-00299804 |
| MVP_top60_DNA_seed | ESR1 | -1.08327 | PRAD | VCaP | Fulvestrant | top60_altered_DNA+RNA_seed | ERBB4 | 3.33718 | STAD | SNU-5 | PF-00299804 |
| MVP_top60_DNA_seed | ESR1 | -1.071545 | PRAD | PWR-1E | Fulvestrant | top60_altered_DNA+RNA_seed | ERBB2 | 3.35161 | STAD | OCUM-1 | PF-00299804 |
| MVP_top60_DNA_seed | PARP1 | -1.05242 | PRAD | PWR-1E | Olaparib | top60_altered_DNA+RNA_seed | ERBB4 | 3.35161 | STAD | OCUM-1 | PF-00299804 |
| MVP_top60_DNA_seed | PARP1 | -1.040411 | PRAD | PC-3 | Veliparib | top60_altered_DNA+RNA_seed | ERBB4 | 3.424574 | STAD | SNU-5 | AST-1306 |
| MVP_top60_DNA_seed | PARP1 | -1.026253 | PRAD | DU-145 | PARP_9482 | top60_altered_DNA+RNA_seed | ERBB2 | -4.964046 | STAD | NCI-N87 | Afatinib |
| MVP_top60_DNA_seed | EP300 | -0.997176 | PRAD | LNCaP-Clone-FGC | I-CBP112 | top60_altered_DNA+RNA_seed | ERBB2 | -4.104673 | STAD | SK-GT-2 | Afatinib |
| MVP_top60_DNA_seed | HDAC1 | -0.972175 | PRAD | 22RV1 | Dacinostat | top60_altered_DNA+RNA_seed | ERBB2 | -3.70213 | STAD | NCI-N87 | Lapatinib |
| MVP_top60_DNA_seed | AR | -0.939936 | PRAD | DU-145 | AZD3514 | top60_altered_DNA+RNA_seed | ERBB2 | -3.096553 | STAD | NCI-N87 | Sapitinib |
| MVP_top60_DNA_seed | AR | -0.938338 | PRAD | PC-3 | Bicalutamide | top60_altered_DNA+RNA_seed | ERBB2 | -2.940765 | STAD | SK-GT-2 | Sapitinib |
| MVP_top60_DNA_seed | HDAC1 | -0.922482 | PRAD | 22RV1 | Belinostat | top60_altered_DNA+RNA_seed | ERBB2 | -2.431962 | STAD | SK-GT-2 | Lapatinib |
| MVP_top60_DNA_seed | ATR | -0.906261 | PRAD | PWR-1E | AZD6738 | top60_altered_DNA+RNA_seed | PIK3CA | -2.195662 | STAD | NCI-N87 | Alpelisib |
| MVP_top60_DNA_seed | ATM | -0.891731 | PRAD | DU-145 | KU-60019 | top60_altered_DNA+RNA_seed | PIK3CA | -2.178419 | STAD | HGC-27 | Alpelisib |
| MVP_top60_DNA_seed | HDAC1 | -0.881162 | PRAD | 22RV1 | AR-42 | top60_altered_DNA+RNA_seed | PIK3CA | -2.002491 | STAD | HSC-39 | Alpelisib |
| MVP_top60_DNA_seed | ATR | -0.865979 | PRAD | PWR-1E | AZ20 | top60_altered_DNA+RNA_seed | PIK3CA | -1.889177 | STAD | HGC-27 | Buparlisib |

| | | | | | | | | | | | |
|---|---|---|---|---|---|---|---|---|---|---|---|
| MVP_top60_DNA_seed | ATR | -0.865444 | PRAD | 22RV1 | QL-VIII-58 | top60_altered_DNA+RNA_seed | PIK3CA | -1.879573 | STAD | 23132-87 | AZD8186 |
| MVP_top60_DNA_seed | ATM | -0.853892 | PRAD | PWR-1E | KU-60019 | top60_altered_DNA+RNA_seed | PIK3CA | -1.873533 | STAD | HGC-27 | GNE-317 |
| MVP_top60_DNA_seed | HDAC1 | -0.84662 | PRAD | VCaP | Tubastatin A | top60_altered_DNA+RNA_seed | PIK3CA | -1.806595 | STAD | AGS | Alpelisib |
| MVP_top60_DNA_seed | PARP1 | -0.831317 | PRAD | DU-145 | PARP_0108 | top60_altered_DNA+RNA_seed | ERBB2 | -1.699082 | STAD | NUGC-3 | Sapitinib |
| MVP_top60_DNA_seed | BRAF | -0.814215 | PRAD | DU-145 | PLX-4720 | top60_altered_DNA+RNA_seed | PIK3CA | -1.690563 | STAD | HGC-27 | AZD8186 |
| MVP_top60_DNA_seed | HDAC1 | -0.77692 | PRAD | LNCaP-Clone-FGC | CAY10603 | top60_altered_DNA+RNA_seed | PIK3CA | -1.661051 | STAD | FU97 | Alpelisib |
| MVP_top60_DNA_seed | PARP1 | -0.759628 | PRAD | 22RV1 | Rucaparib | top60_altered_DNA+RNA_seed | PIK3CA | -1.513459 | STAD | HSC-39 | GNE-317 |
| MVP_top60_DNA_seed | PARP1 | -0.757878 | PRAD | LNCaP-Clone-FGC | Rucaparib | top60_altered_DNA+RNA_seed | ERBB2 | -1.340055 | STAD | NUGC-3 | Afatinib |
| MVP_top60_DNA_seed | HDAC1 | -0.746407 | PRAD | DU-145 | JQ12 | top60_altered_DNA+RNA_seed | PIK3CA | -1.255553 | STAD | 23132-87 | Buparlisib |
| MVP_top60_DNA_seed | PARP1 | -0.739807 | PRAD | DU-145 | Rucaparib | top60_altered_DNA+RNA_seed | ERBB2 | -1.18579 | STAD | NUGC-3 | Lapatinib |
| MVP_top60_DNA_seed | HDAC1 | -0.733581 | PRAD | VCaP | CAY10603 | top60_altered_DNA+RNA_seed | PIK3CA | -1.166136 | STAD | 23132-87 | GNE-317 |
| MVP_top60_DNA_seed | PARP1 | -0.730022 | PRAD | DU-145 | Olaparib | top60_altered_DNA+RNA_seed | ERBB2 | -1.149545 | STAD | GCIY | Afatinib |
| MVP_top60_DNA_seed | PARP1 | -0.728672 | PRAD | LNCaP-Clone-FGC | PARP_0108 | top60_altered_DNA+RNA_seed | ERBB2 | -1.120754 | STAD | GCIY | Sapitinib |
| MVP_top60_DNA_seed | HDAC1 | -0.718561 | PRAD | LNCaP-Clone-FGC | Tubastatin A | top60_altered_DNA+RNA_seed | PIK3CA | -1.107574 | STAD | HSC-39 | Buparlisib |
| MVP_top60_DNA_seed | PARP1 | -0.708736 | PRAD | LNCaP-Clone-FGC | Talazoparib | top60_altered_DNA+RNA_seed | PIK3CA | -1.096937 | STAD | IM-95 | Alpelisib |
| MVP_top60_DNA_seed | ATM | -0.708365 | PRAD | PWR-1E | KU-55933 | top60_altered_DNA+RNA_seed | PIK3CA | -1.074511 | STAD | 23132-87 | Alpelisib |
| MVP_top60_DNA_seed | PARP1 | -0.681103 | PRAD | PWR-1E | Talazoparib | top60_altered_DNA+RNA_seed | ERBB2 | -1.0707 | STAD | NUGC-4 | Sapitinib |
| MVP_top60_DNA_seed | ATM | -0.675106 | PRAD | PC-3 | KU-55933 | top60_altered_DNA+RNA_seed | PIK3CA | -1.064823 | STAD | FU97 | GNE-317 |
| MVP_top60_DNA_seed | HDAC1 | -0.661935 | PRAD | 22RV1 | CAY10603 | top60_altered_DNA+RNA_seed | ERBB2 | -1.064414 | STAD | HGC-27 | Lapatinib |
| MVP_top60_DNA_seed | HDAC1 | -0.648199 | PRAD | VCaP | Dacinostat | top60_altered_DNA+RNA_seed | PIK3CA | -1.052359 | STAD | SNU-16 | Buparlisib |
| MVP_top60_DNA_seed | BRAF | -0.633081 | PRAD | DU-145 | HG6-64-1 | top60_altered_DNA+RNA_seed | ERBB2 | -0.992582 | STAD | HSC-39 | Lapatinib |
| MVP_top60_DNA_seed | PARP1 | -0.622393 | PRAD | LNCaP-Clone-FGC | PARP_9482 | top60_altered_DNA+RNA_seed | TP53 | -0.964362 | STAD | SNU-16 | MIRA-1 |
| MVP_top60_DNA_seed | GSK3B | -0.606071 | PRAD | VCaP | CHIR-99021 | top60_altered_DNA+RNA_seed | ERBB2 | -0.961978 | STAD | RERF-GC-1B | Sapitinib |
| MVP_top60_DNA_seed | BRAF | -0.602904 | PRAD | DU-145 | SB590885 | top60_altered_DNA+RNA_seed | PIK3CA | -0.94861 | STAD | HSC-39 | AZD8186 |
| MVP_top60_DNA_seed | ATR | -0.567602 | PRAD | DU-145 | AZ20 | top60_altered_DNA+RNA_seed | PIK3CA | -0.895344 | STAD | FU97 | Buparlisib |
| MVP_top60_DNA_seed | PIK3CA | -0.556787 | PRAD | 22RV1 | PI3Ka_4409 | top60_altered_DNA+RNA_seed | PIK3CA | -0.860398 | STAD | AGS | GNE-317 |
| MVP_top60_DNA_seed | BRAF | -0.552257 | PRAD | PC-3 | PLX-4720 | top60_altered_DNA+RNA_seed | PIK3CA | -0.842493 | STAD | NCI-N87 | AZD8186 |
| MVP_top60_DNA_seed | ATR | -0.545546 | PRAD | LNCaP-Clone-FGC | AZD6738 | top60_altered_DNA+RNA_seed | PIK3CA | -0.832559 | STAD | GCIY | GNE-317 |
| MVP_top60_DNA_seed | HDAC1 | -0.544519 | PRAD | DU-145 | Belinostat | top60_altered_DNA+RNA_seed | PIK3CA | -0.829642 | STAD | SNU-5 | GNE-317 |
| MVP_top60_DNA_seed | PARP1 | -0.526103 | PRAD | PWR-1E | PARP_9495 | top60_altered_DNA+RNA_seed | ERBB2 | -0.814856 | STAD | SNU-16 | Lapatinib |
| MVP_top60_DNA_seed | AR | -0.50603 | PRAD | 22RV1 | Bicalutamide | top60_altered_DNA+RNA_seed | PIK3CA | -0.814266 | STAD | SNU-5 | AZD8186 |
| MVP_top60_DNA_seed | GSK3B | -0.472573 | PRAD | VCaP | AZD7969 | top60_altered_DNA+RNA_seed | PIK3CA | -0.791431 | STAD | AGS | Buparlisib |
| MVP_top60_DNA_seed | PARP1 | -0.470946 | PRAD | 22RV1 | PARP_9482 | top60_altered_DNA+RNA_seed | TP53 | -0.79102 | STAD | HSC-39 | MIRA-1 |
| MVP_top60_DNA_seed | HDAC1 | -0.463803 | PRAD | DU-145 | AR-42 | top60_altered_DNA+RNA_seed | PIK3CA | -0.790039 | STAD | NUGC-3 | GNE-317 |
| MVP_top60_DNA_seed | ATM | -0.463026 | PRAD | VCaP | CP466722 | top60_altered_DNA+RNA_seed | PIK3CA | -0.770271 | STAD | FU97 | AZD8186 |
| MVP_top60_DNA_seed | CDK4 | -0.45584 | PRAD | DU-145 | Palbociclib | top60_altered_DNA+RNA_seed | PIK3CA | -0.768064 | STAD | Hs-746T | GNE-317 |
| MVP_top60_DNA_seed | AR | -0.437962 | PRAD | LNCaP-Clone-FGC | Bicalutamide | top60_altered_DNA+RNA_seed | ERBB2 | -0.729675 | STAD | NUGC-4 | Afatinib |
| MVP_top60_DNA_seed | PARP1 | -0.435297 | PRAD | 22RV1 | PARP_0108 | top60_altered_DNA+RNA_seed | ERBB2 | -0.724525 | STAD | MKN7 | Lapatinib |
| MVP_top60_DNA_seed | HDAC1 | -0.435137 | PRAD | DU-145 | Dacinostat | top60_altered_DNA+RNA_seed | TP53 | -0.720545 | STAD | RF-48 | MIRA-1 |
| MVP_top60_DNA_seed | GSK3B | -0.425484 | PRAD | 22RV1 | CHIR-99021 | top60_altered_DNA+RNA_seed | PIK3CA | -0.640799 | STAD | MKN45 | GNE-317 |
| MVP_top60_DNA_seed | AR | -0.416302 | PRAD | VCaP | AZD3514 | top60_altered_DNA+RNA_seed | PIK3CA | -0.636908 | STAD | SNU-5 | Buparlisib |

| | | | | | | | | | | | |
|---|---|---|---|---|---|---|---|---|---|---|---|
| MVP_top60_DNA_seed | ATM | -0.412618 | PRAD | LNCaP-Clone-FGC | CP466722 | top60_altered_DNA+RNA_seed | PIK3CA | -0.633687 | STAD | SK-GT-2 | Buparlisib |
| MVP_top60_DNA_seed | PIK3CA | -0.412275 | PRAD | 22RV1 | AZD8835 | top60_altered_DNA+RNA_seed | ERBB2 | -0.628949 | STAD | HSC-39 | Sapitinib |
| MVP_top60_DNA_seed | HDAC1 | -0.409553 | PRAD | VCaP | Entinostat | top60_altered_DNA+RNA_seed | PIK3CA | -0.603623 | STAD | SK-GT-2 | Alpelisib |
| MVP_top60_DNA_seed | PARP1 | -0.409432 | PRAD | PC-3 | PARP_0108 | top60_altered_DNA+RNA_seed | ERBB2 | -0.596463 | STAD | TMK-1 | Sapitinib |
| MVP_top60_DNA_seed | ATR | -0.407304 | PRAD | LNCaP-Clone-FGC | AZ20 | top60_altered_DNA+RNA_seed | TP53 | -0.578312 | STAD | GCIY | MIRA-1 |
| MVP_top60_DNA_seed | BRAF | -0.392579 | PRAD | VCaP | SB590885 | top60_altered_DNA+RNA_seed | PIK3CA | -0.553995 | STAD | IM-95 | Buparlisib |
| MVP_top60_DNA_seed | HDAC1 | -0.390047 | PRAD | 22RV1 | JQ12 | top60_altered_DNA+RNA_seed | ERBB2 | -0.544321 | STAD | SNU-16 | Afatinib |
| MVP_top60_DNA_seed | HDAC1 | -0.379268 | PRAD | DU-145 | CAY10603 | top60_altered_DNA+RNA_seed | PIK3CA | -0.536074 | STAD | SK-GT-2 | GNE-317 |
| MVP_top60_DNA_seed | HDAC1 | -0.374469 | PRAD | 22RV1 | Tubastatin A | top60_altered_DNA+RNA_seed | PIK3CA | -0.502607 | STAD | RERF-GC-1B | Alpelisib |
| MVP_top60_DNA_seed | PIK3CA | -0.367324 | PRAD | DU-145 | AZD8835 | top60_altered_DNA+RNA_seed | PIK3CA | -0.494304 | STAD | TMK-1 | Alpelisib |
| MVP_top60_DNA_seed | PARP1 | -0.364595 | PRAD | PC-3 | PARP_9482 | top60_altered_DNA+RNA_seed | PIK3CA | -0.492705 | STAD | RF-48 | Buparlisib |
| MVP_top60_DNA_seed | BRAF | -0.357259 | PRAD | LNCaP-Clone-FGC | PLX-4720 | top60_altered_DNA+RNA_seed | PIK3CA | -0.458624 | STAD | NCI-N87 | GNE-317 |
| MVP_top60_DNA_seed | GSK3B | -0.353891 | PRAD | LNCaP-Clone-FGC | SB216763 | top60_altered_DNA+RNA_seed | PIK3CA | -0.432008 | STAD | IM-95 | AZD8186 |
| MVP_top60_DNA_seed | PARP1 | -0.333174 | PRAD | DU-145 | Talazoparib | top60_altered_DNA+RNA_seed | PIK3CA | -0.392938 | STAD | Hs-746T | Buparlisib |
| MVP_top60_DNA_seed | PIK3CA | -0.331526 | PRAD | VCaP | AZD8835 | top60_altered_DNA+RNA_seed | PIK3CA | -0.359993 | STAD | Hs-746T | AZD8186 |
| MVP_top60_DNA_seed | EP300 | -0.324634 | PRAD | 22RV1 | I-CBP112 | top60_altered_DNA+RNA_seed | ERBB2 | -0.336204 | STAD | HSC-39 | Afatinib |
| MVP_top60_DNA_seed | HDAC1 | -0.321626 | PRAD | VCaP | JQ12 | top60_altered_DNA+RNA_seed | PIK3CA | -0.33418 | STAD | SNU-16 | Alpelisib |
| MVP_top60_DNA_seed | PARP1 | -0.320227 | PRAD | PWR-1E | Veliparib | top60_altered_DNA+RNA_seed | ERBB2 | -0.293342 | STAD | 23132-87 | Afatinib |
| MVP_top60_DNA_seed | PARP1 | -0.308482 | PRAD | 22RV1 | Talazoparib | top60_altered_DNA+RNA_seed | ERBB2 | -0.289654 | STAD | NUGC-4 | Lapatinib |
| MVP_top60_DNA_seed | ATR | -0.298652 | PRAD | LNCaP-Clone-FGC | QL-VIII-58 | top60_altered_DNA+RNA_seed | TP53 | -0.272887 | STAD | HGC-27 | MIRA-1 |
| MVP_top60_DNA_seed | BRAF | -0.298345 | PRAD | LNCaP-Clone-FGC | Dabrafenib | top60_altered_DNA+RNA_seed | PIK3CA | -0.263263 | STAD | GCIY | Buparlisib |
| MVP_top60_DNA_seed | AR | -0.279002 | PRAD | VCaP | Bicalutamide | top60_altered_DNA+RNA_seed | PIK3CA | -0.191993 | STAD | TGBC11TKB | Alpelisib |
| MVP_top60_DNA_seed | ESR1 | -0.276456 | PRAD | 22RV1 | Tamoxifen | top60_altered_DNA+RNA_seed | ERBB2 | -0.189838 | STAD | AGS | Lapatinib |
| MVP_top60_DNA_seed | PARP1 | -0.243649 | PRAD | LNCaP-Clone-FGC | Olaparib | top60_altered_DNA+RNA_seed | PIK3CA | -0.189207 | STAD | SNU-16 | GNE-317 |
| MVP_top60_DNA_seed | PTK2 | -0.243334 | PRAD | 22RV1 | PF-562271 | top60_altered_DNA+RNA_seed | PIK3CA | -0.181396 | STAD | MKN45 | Alpelisib |
| MVP_top60_DNA_seed | ATM | -0.237597 | PRAD | LNCaP-Clone-FGC | KU-60019 | top60_altered_DNA+RNA_seed | PIK3CA | -0.170171 | STAD | SNU-1 | Buparlisib |
| MVP_top60_DNA_seed | BRAF | -0.235269 | PRAD | DU-145 | RAF_9304 | top60_altered_DNA+RNA_seed | ERBB2 | -0.169119 | STAD | AGS | Afatinib |
| MVP_top60_DNA_seed | AR | -0.233028 | PRAD | 22RV1 | AZD3514 | top60_altered_DNA+RNA_seed | TP53 | -0.164612 | STAD | AGS | MIRA-1 |
| MVP_top60_DNA_seed | ATM | -0.228938 | PRAD | LNCaP-Clone-FGC | torin2 | top60_altered_DNA+RNA_seed | ERBB2 | -0.14424 | STAD | 23132-87 | Sapitinib |
| MVP_top60_DNA_seed | ATR | -0.228938 | PRAD | LNCaP-Clone-FGC | torin2 | top60_altered_DNA+RNA_seed | ERBB2 | -0.137624 | STAD | IM-95 | Afatinib |
| MVP_top60_DNA_seed | BRAF | -0.215364 | PRAD | DU-145 | PLX-4720 | top60_altered_DNA+RNA_seed | ERBB2 | -0.113922 | STAD | TGBC11TKB | Afatinib |
| MVP_top60_DNA_seed | PARP1 | -0.207727 | PRAD | PC-3 | Olaparib | top60_altered_DNA+RNA_seed | PIK3CA | -0.112712 | STAD | AGS | AZD8186 |
| MVP_top60_DNA_seed | PARP1 | -0.205763 | PRAD | PC-3 | Talazoparib | top60_altered_DNA+RNA_seed | ERBB2 | -0.112354 | STAD | SNU-16 | Sapitinib |
| MVP_top60_DNA_seed | HDAC1 | -0.196213 | PRAD | PC-3 | CAY10603 | top60_altered_DNA+RNA_seed | PIK3CA | -0.110412 | STAD | SK-GT-2 | AZD8186 |
| MVP_top60_DNA_seed | ATM | -0.18667 | PRAD | 22RV1 | CP466722 | top60_altered_DNA+RNA_seed | PIK3CA | -0.108094 | STAD | TMK-1 | Buparlisib |
| MVP_top60_DNA_seed | BRAF | -0.179587 | PRAD | VCaP | Dabrafenib | top60_altered_DNA+RNA_seed | ERBB2 | -0.103653 | STAD | IM-95 | Sapitinib |
| MVP_top60_DNA_seed | AR | -0.177565 | PRAD | DU-145 | Bicalutamide | top60_altered_DNA+RNA_seed | PIK3CA | -0.100081 | STAD | NCI-N87 | Buparlisib |
| MVP_top60_DNA_seed | ATM | -0.175981 | PRAD | VCaP | KU-55933 | top60_altered_DNA+RNA_seed | PIK3CA | -0.091567 | STAD | TMK-1 | AZD8186 |
| MVP_top60_DNA_seed | PTK2 | -0.172219 | PRAD | PC-3 | PF-562271 | top60_altered_DNA+RNA_seed | ERBB2 | -0.080522 | STAD | FU97 | Lapatinib |
| MVP_top60_DNA_seed | ATR | -0.169139 | PRAD | DU-145 | AZD6738 | top60_altered_DNA+RNA_seed | PIK3CA | -0.072562 | STAD | OCUM-1 | Alpelisib |
| MVP_top60_DNA_seed | MAPK3 | -0.164128 | PRAD | LNCaP-Clone-FGC | FR-180204 | top60_altered_DNA+RNA_seed | ERBB2 | -0.066352 | STAD | MKN7 | Sapitinib |

| | | | | | | | | | | | |
|---|---|---|---|---|---|---|---|---|---|---|---|
| MVP_top60_DNA_seed | EP300 | -0.155857 | PRAD | VCaP | I-CBP112 | top60_altered_DNA+RNA_seed | TP53 | -0.064001 | STAD | 23132-87 | MIRA-1 |
| MVP_top60_DNA_seed | BRAF | -0.135525 | PRAD | DU-145 | Dabrafenib | top60_altered_DNA+RNA_seed | PIK3CA | -0.061805 | STAD | NUGC-3 | Buparlisib |
| MVP_top60_DNA_seed | HDAC1 | -0.135055 | PRAD | LNCaP-Clone-FGC | Dacinostat | top60_altered_DNA+RNA_seed | ERBB2 | -0.060068 | STAD | SNU-1 | Lapatinib |
| MVP_top60_DNA_seed | AR | -0.110202 | PRAD | PC-3 | AZD3514 | top60_altered_DNA+RNA_seed | ERBB2 | -0.059664 | STAD | SNU-1 | Afatinib |
| MVP_top60_DNA_seed | GSK3B | -0.092599 | PRAD | LNCaP-Clone-FGC | AZD7969 | top60_altered_DNA+RNA_seed | ERBB2 | -0.033577 | STAD | HGC-27 | Afatinib |
| MVP_top60_DNA_seed | PTK2 | -0.083354 | PRAD | VCaP | PF-562271 | top60_altered_DNA+RNA_seed | PIK3CA | -0.023201 | STAD | MKN45 | Buparlisib |
| MVP_top60_DNA_seed | PARP1 | -0.067815 | PRAD | LNCaP-Clone-FGC | PARP_9495 | top60_altered_DNA+RNA_seed | ERBB2 | -0.020032 | STAD | AGS | Sapitinib |
| MVP_top60_DNA_seed | HDAC1 | -0.061377 | PRAD | PC-3 | Belinostat | top60_altered_DNA+RNA_seed | PIK3CA | -0.015433 | STAD | NUGC-3 | Alpelisib |
| MVP_top60_DNA_seed | GSK3B | -0.048666 | PRAD | 22RV1 | AZD7969 | top60_altered_DNA+RNA_seed | PIK3CA | 0.005655 | STAD | TGBC11TKB | Buparlisib |
| MVP_top60_DNA_seed | BRAF | -0.041804 | PRAD | 22RV1 | SB590885 | top60_altered_DNA+RNA_seed | TP53 | 0.008774 | STAD | IM-95 | MIRA-1 |
| MVP_top60_DNA_seed | PARP1 | -0.01336 | PRAD | PWR-1E | Rucaparib | top60_altered_DNA+RNA_seed | ERBB2 | 0.012472 | STAD | MKN28 | Lapatinib |
| MVP_top60_DNA_seed | PIK3CA | -0.01 | PRAD | VCaP | PI3Ka_4409 | top60_altered_DNA+RNA_seed | ERBB2 | 0.027895 | STAD | MKN28 | Sapitinib |
| MVP_top60_DNA_seed | HDAC1 | 0.008543 | PRAD | PC-3 | AR-42 | top60_altered_DNA+RNA_seed | PIK3CA | 0.036404 | STAD | MKN7 | AZD8186 |
| MVP_top60_DNA_seed | ATM | 0.016044 | PRAD | PC-3 | CP466722 | top60_altered_DNA+RNA_seed | ERBB2 | 0.03859 | STAD | Hs-746T | Lapatinib |
| MVP_top60_DNA_seed | PIK3CA | 0.022501 | PRAD | PWR-1E | AZD8835 | top60_altered_DNA+RNA_seed | PIK3CA | 0.043728 | STAD | MKN7 | GNE-317 |
| MVP_top60_DNA_seed | BRAF | 0.035323 | PRAD | PWR-1E | PLX-4720 | top60_altered_DNA+RNA_seed | ERBB2 | 0.046841 | STAD | RERF-GC-1B | Afatinib |
| MVP_top60_DNA_seed | GSK3B | 0.046287 | PRAD | PC-3 | SB216763 | top60_altered_DNA+RNA_seed | ERBB2 | 0.050663 | STAD | MKN28 | Afatinib |
| MVP_top60_DNA_seed | ESR1 | 0.049164 | PRAD | LNCaP-Clone-FGC | Tamoxifen | top60_altered_DNA+RNA_seed | PIK3CA | 0.060791 | STAD | TGBC11TKB | GNE-317 |
| MVP_top60_DNA_seed | AR | 0.051099 | PRAD | PWR-1E | Bicalutamide | top60_altered_DNA+RNA_seed | PIK3CA | 0.062867 | STAD | GCIY | AZD8186 |
| MVP_top60_DNA_seed | CDK4 | 0.074618 | PRAD | LNCaP-Clone-FGC | Palbociclib | top60_altered_DNA+RNA_seed | PIK3CA | 0.0727 | STAD | RF-48 | AZD8186 |
| MVP_top60_DNA_seed | HDAC1 | 0.085457 | PRAD | LNCaP-Clone-FGC | JQ12 | top60_altered_DNA+RNA_seed | ERBB2 | 0.073149 | STAD | RERF-GC-1B | Lapatinib |
| MVP_top60_DNA_seed | AR | 0.097737 | PRAD | DU-145 | Bicalutamide | top60_altered_DNA+RNA_seed | PIK3CA | 0.116808 | STAD | RF-48 | Alpelisib |
| MVP_top60_DNA_seed | BRAF | 0.098459 | PRAD | PWR-1E | SB590885 | top60_altered_DNA+RNA_seed | ERBB2 | 0.133165 | STAD | TGBC11TKB | Lapatinib |
| MVP_top60_DNA_seed | HDAC1 | 0.10802 | PRAD | PC-3 | Tubastatin A | top60_altered_DNA+RNA_seed | TP53 | 0.145707 | STAD | FU97 | MIRA-1 |
| MVP_top60_DNA_seed | GSK3B | 0.109058 | PRAD | DU-145 | CHIR-99021 | top60_altered_DNA+RNA_seed | ERBB2 | 0.15512 | STAD | HGC-27 | Sapitinib |
| MVP_top60_DNA_seed | GSK3B | 0.110164 | PRAD | DU-145 | AZD7969 | top60_altered_DNA+RNA_seed | TP53 | 0.163433 | STAD | MKN7 | MIRA-1 |
| MVP_top60_DNA_seed | GSK3B | 0.116501 | PRAD | PC-3 | CHIR-99021 | top60_altered_DNA+RNA_seed | PIK3CA | 0.222733 | STAD | SNU-5 | Alpelisib |
| MVP_top60_DNA_seed | BRAF | 0.122962 | PRAD | 22RV1 | Dabrafenib | top60_altered_DNA+RNA_seed | ERBB2 | 0.246427 | STAD | MKN7 | Afatinib |
| MVP_top60_DNA_seed | BRAF | 0.134685 | PRAD | PC-3 | SB590885 | top60_altered_DNA+RNA_seed | PIK3CA | 0.246863 | STAD | RF-48 | GNE-317 |
| MVP_top60_DNA_seed | BRAF | 0.135765 | PRAD | 22RV1 | PLX-4720 | top60_altered_DNA+RNA_seed | ERBB2 | 0.261371 | STAD | RF-48 | Afatinib |
| MVP_top60_DNA_seed | CDK4 | 0.137383 | PRAD | VCaP | CGP-082996 | top60_altered_DNA+RNA_seed | PIK3CA | 0.261699 | STAD | NUGC-3 | AZD8186 |
| MVP_top60_DNA_seed | BRAF | 0.14097 | PRAD | LNCaP-Clone-FGC | RAF_9304 | top60_altered_DNA+RNA_seed | TP53 | 0.278515 | STAD | TMK-1 | MIRA-1 |
| MVP_top60_DNA_seed | PIK3CA | 0.144896 | PRAD | LNCaP-Clone-FGC | AZD8835 | top60_altered_DNA+RNA_seed | PIK3CA | 0.299203 | STAD | MKN7 | Alpelisib |
| MVP_top60_DNA_seed | PARP1 | 0.16679 | PRAD | PWR-1E | PARP_0108 | top60_altered_DNA+RNA_seed | ERBB2 | 0.305537 | STAD | GCIY | Lapatinib |
| MVP_top60_DNA_seed | PARP1 | 0.170597 | PRAD | PWR-1E | PARP_9482 | top60_altered_DNA+RNA_seed | ERBB2 | 0.312197 | STAD | TMK-1 | Lapatinib |
| MVP_top60_DNA_seed | EP300 | 0.178589 | PRAD | PWR-1E | I-CBP112 | top60_altered_DNA+RNA_seed | TP53 | 0.34861 | STAD | SNU-1 | MIRA-1 |
| MVP_top60_DNA_seed | GSK3B | 0.18049 | PRAD | PC-3 | KIN001-042 | top60_altered_DNA+RNA_seed | PIK3CA | 0.36905 | STAD | MKN28 | Alpelisib |
| MVP_top60_DNA_seed | CDK4 | 0.181924 | PRAD | VCaP | Palbociclib | top60_altered_DNA+RNA_seed | TP53 | 0.383155 | STAD | SNU-5 | MIRA-1 |
| MVP_top60_DNA_seed | HDAC1 | 0.198546 | PRAD | DU-145 | Tubastatin A | top60_altered_DNA+RNA_seed | ERBB2 | 0.396143 | STAD | MKN45 | Lapatinib |
| MVP_top60_DNA_seed | HDAC1 | 0.205203 | PRAD | PC-3 | Dacinostat | top60_altered_DNA+RNA_seed | PIK3CA | 0.396228 | STAD | MKN28 | Buparlisib |
| MVP_top60_DNA_seed | BRAF | 0.209381 | PRAD | PC-3 | PLX-4720 | top60_altered_DNA+RNA_seed | TP53 | 0.406069 | STAD | RERF-GC-1B | MIRA-1 |

| | | | | | | | | | | | |
|---|---|---|---|---|---|---|---|---|---|---|---|
| MVP_top60_DNA_seed | GSK3B | 0.217229 | PRAD | PWR-1E | KIN001-042 | top60_altered_DNA+RNA_seed | PIK3CA | 0.415397 | STAD | MKN45 | AZD8186 |
| MVP_top60_DNA_seed | ATR | 0.221189 | PRAD | PC-3 | AZD6738 | top60_altered_DNA+RNA_seed | ERBB2 | 0.41743 | STAD | IM-95 | Lapatinib |
| MVP_top60_DNA_seed | PARP1 | 0.252626 | PRAD | PC-3 | Rucaparib | top60_altered_DNA+RNA_seed | TP53 | 0.433549 | STAD | SK-GT-2 | MIRA-1 |
| MVP_top60_DNA_seed | ESR1 | 0.256978 | PRAD | 22RV1 | Fulvestrant | top60_altered_DNA+RNA_seed | PIK3CA | 0.442209 | STAD | GCIY | Alpelisib |
| MVP_top60_DNA_seed | GSK3B | 0.290372 | PRAD | LNCaP-Clone-FGC | CHIR-99021 | top60_altered_DNA+RNA_seed | ERBB2 | 0.479741 | STAD | TGBC11TKB | Sapitinib |
| MVP_top60_DNA_seed | MAPK3 | 0.31184 | PRAD | PC-3 | FR-180204 | top60_altered_DNA+RNA_seed | PIK3CA | 0.481446 | STAD | RERF-GC-1B | AZD8186 |
| MVP_top60_DNA_seed | ESR1 | 0.32332 | PRAD | DU-145 | Fulvestrant | top60_altered_DNA+RNA_seed | ERBB2 | 0.512615 | STAD | KATOIII | Sapitinib |
| MVP_top60_DNA_seed | ESR1 | 0.339144 | PRAD | PWR-1E | Tamoxifen | top60_altered_DNA+RNA_seed | TP53 | 0.588419 | STAD | TGBC11TKB | MIRA-1 |
| MVP_top60_DNA_seed | BRAF | 0.341279 | PRAD | 22RV1 | RAF_9304 | top60_altered_DNA+RNA_seed | PIK3CA | 0.598927 | STAD | SNU-1 | AZD8186 |
| MVP_top60_DNA_seed | GSK3B | 0.349246 | PRAD | PC-3 | AZD7969 | top60_altered_DNA+RNA_seed | ERBB2 | 0.601589 | STAD | Hs-746T | Afatinib |
| MVP_top60_DNA_seed | PARP1 | 0.35281 | PRAD | VCaP | Talazoparib | top60_altered_DNA+RNA_seed | ERBB2 | 0.603182 | STAD | RF-48 | Sapitinib |
| MVP_top60_DNA_seed | PARP1 | 0.36504 | PRAD | 22RV1 | Olaparib | top60_altered_DNA+RNA_seed | PIK3CA | 0.616749 | STAD | RERF-GC-1B | Buparlisib |
| MVP_top60_DNA_seed | GSK3B | 0.380022 | PRAD | LNCaP-Clone-FGC | KIN001-042 | top60_altered_DNA+RNA_seed | ERBB2 | 0.663193 | STAD | 23132-87 | Lapatinib |
| MVP_top60_DNA_seed | ESR1 | 0.383568 | PRAD | PC-3 | Tamoxifen | top60_altered_DNA+RNA_seed | TP53 | 0.67081 | STAD | OCUM-1 | MIRA-1 |
| MVP_top60_DNA_seed | PARP1 | 0.411417 | PRAD | LNCaP-Clone-FGC | Olaparib | top60_altered_DNA+RNA_seed | ERBB2 | 0.670885 | STAD | SNU-1 | Sapitinib |
| MVP_top60_DNA_seed | BRAF | 0.427131 | PRAD | PWR-1E | PLX-4720 | top60_altered_DNA+RNA_seed | PIK3CA | 0.69062 | STAD | RERF-GC-1B | GNE-317 |
| MVP_top60_DNA_seed | MAPK3 | 0.42764 | PRAD | DU-145 | FR-180204 | top60_altered_DNA+RNA_seed | TP53 | 0.719516 | STAD | NUGC-3 | MIRA-1 |
| MVP_top60_DNA_seed | GSK3B | 0.463171 | PRAD | LNCaP-Clone-FGC | CHIR-99021 | top60_altered_DNA+RNA_seed | PIK3CA | 0.73458 | STAD | TMK-1 | GNE-317 |
| MVP_top60_DNA_seed | AR | 0.470589 | PRAD | VCaP | Bicalutamide | top60_altered_DNA+RNA_seed | TP53 | 0.752842 | STAD | NCI-N87 | MIRA-1 |
| MVP_top60_DNA_seed | BRAF | 0.47927 | PRAD | 22RV1 | HG6-64-1 | top60_altered_DNA+RNA_seed | ERBB2 | 0.793279 | STAD | SNU-5 | Afatinib |
| MVP_top60_DNA_seed | EP300 | 0.485707 | PRAD | DU-145 | I-CBP112 | top60_altered_DNA+RNA_seed | PIK3CA | 0.7938 | STAD | SNU-16 | AZD8186 |
| MVP_top60_DNA_seed | PIK3CA | 0.489116 | PRAD | PC-3 | AZD8835 | top60_altered_DNA+RNA_seed | ERBB2 | 0.817424 | STAD | FU97 | Sapitinib |
| MVP_top60_DNA_seed | GSK3B | 0.517366 | PRAD | VCaP | KIN001-042 | top60_altered_DNA+RNA_seed | TP53 | 0.838111 | STAD | MKN28 | MIRA-1 |
| MVP_top60_DNA_seed | PARP1 | 0.529798 | PRAD | VCaP | Veliparib | top60_altered_DNA+RNA_seed | PIK3CA | 0.855342 | STAD | SNU-1 | Alpelisib |
| MVP_top60_DNA_seed | ATR | 0.539113 | PRAD | PC-3 | AZ20 | top60_altered_DNA+RNA_seed | PIK3CA | 0.855649 | STAD | NUGC-4 | Alpelisib |
| MVP_top60_DNA_seed | PARP1 | 0.544669 | PRAD | PC-3 | Olaparib | top60_altered_DNA+RNA_seed | ERBB2 | 0.876586 | STAD | MKN45 | Afatinib |
| MVP_top60_DNA_seed | ATR | 0.551016 | PRAD | VCaP | AZ20 | top60_altered_DNA+RNA_seed | PIK3CA | 0.924553 | STAD | MKN28 | AZD8186 |
| MVP_top60_DNA_seed | PARP1 | 0.556331 | PRAD | PC-3 | PARP_9495 | top60_altered_DNA+RNA_seed | ERBB2 | 0.945989 | STAD | FU97 | Afatinib |
| MVP_top60_DNA_seed | ESR1 | 0.563063 | PRAD | VCaP | Tamoxifen | top60_altered_DNA+RNA_seed | PIK3CA | 0.976605 | STAD | OCUM-1 | Buparlisib |
| MVP_top60_DNA_seed | BRAF | 0.570707 | PRAD | VCaP | PLX-4720 | top60_altered_DNA+RNA_seed | TP53 | 1.053476 | STAD | MKN45 | MIRA-1 |
| MVP_top60_DNA_seed | BRAF | 0.578012 | PRAD | VCaP | RAF_9304 | top60_altered_DNA+RNA_seed | ERBB2 | 1.059574 | STAD | SNU-5 | Sapitinib |
| MVP_top60_DNA_seed | GSK3B | 0.618646 | PRAD | VCaP | CHIR-99021 | top60_altered_DNA+RNA_seed | PIK3CA | 1.080422 | STAD | OCUM-1 | AZD8186 |
| MVP_top60_DNA_seed | ATR | 0.624847 | PRAD | 22RV1 | AZD6738 | top60_altered_DNA+RNA_seed | ERBB2 | 1.085179 | STAD | SNU-5 | Lapatinib |
| MVP_top60_DNA_seed | BRAF | 0.642236 | PRAD | PWR-1E | Dabrafenib | top60_altered_DNA+RNA_seed | ERBB2 | 1.093234 | STAD | OCUM-1 | Sapitinib |
| MVP_top60_DNA_seed | PARP1 | 0.646756 | PRAD | DU-145 | Olaparib | top60_altered_DNA+RNA_seed | PIK3CA | 1.187297 | STAD | IM-95 | GNE-317 |
| MVP_top60_DNA_seed | PIK3CA | 0.6633 | PRAD | PC-3 | PI3Ka_4409 | top60_altered_DNA+RNA_seed | ERBB2 | 1.223624 | STAD | KATOIII | Lapatinib |
| MVP_top60_DNA_seed | GSK3B | 0.694477 | PRAD | PC-3 | CHIR-99021 | top60_altered_DNA+RNA_seed | PIK3CA | 1.223944 | STAD | MKN7 | Buparlisib |
| MVP_top60_DNA_seed | GSK3B | 0.700894 | PRAD | 22RV1 | CHIR-99021 | top60_altered_DNA+RNA_seed | PIK3CA | 1.234249 | STAD | OCUM-1 | GNE-317 |
| MVP_top60_DNA_seed | PIK3CA | 0.701693 | PRAD | LNCaP-Clone-FGC | PI3Ka_4409 | top60_altered_DNA+RNA_seed | TP53 | 1.240448 | STAD | NUGC-4 | MIRA-1 |
| MVP_top60_DNA_seed | ATR | 0.72534 | PRAD | 22RV1 | AZ20 | top60_altered_DNA+RNA_seed | TP53 | 1.27456 | STAD | Hs-746T | MIRA-1 |
| MVP_top60_DNA_seed | PARP1 | 0.728695 | PRAD | VCaP | Olaparib | top60_altered_DNA+RNA_seed | PIK3CA | 1.33347 | STAD | KATOIII | Alpelisib |

| | | | | | | | | | | | |
|---|---|---|---|---|---|---|---|---|---|---|---|
| MVP_top60_DNA_seed | ATM | 0.73058 | PRAD | PC-3 | KU-60019 | top60_altered_DNA+RNA_seed | PIK3CA | 1.34511 | STAD | TGBC11TKB | AZD8186 |
| MVP_top60_DNA_seed | PARP1 | 0.731291 | PRAD | VCaP | Olaparib | top60_altered_DNA+RNA_seed | TP53 | 1.362835 | STAD | KATOIII | MIRA-1 |
| MVP_top60_DNA_seed | PARP1 | 0.759287 | PRAD | VCaP | Rucaparib | top60_altered_DNA+RNA_seed | PIK3CA | 1.466011 | STAD | Hs-746T | Alpelisib |
| MVP_top60_DNA_seed | PARP1 | 0.770766 | PRAD | VCaP | PARP_0108 | top60_altered_DNA+RNA_seed | ERBB2 | 1.474767 | STAD | Hs-746T | Sapitinib |
| MVP_top60_DNA_seed | ATR | 0.772489 | PRAD | VCaP | AZD6738 | top60_altered_DNA+RNA_seed | ERBB2 | 1.655498 | STAD | RF-48 | Lapatinib |
| MVP_top60_DNA_seed | BRAF | 0.77415 | PRAD | LNCaP-Clone-FGC | SB590885 | top60_altered_DNA+RNA_seed | PIK3CA | 1.678729 | STAD | KATOIII | AZD8186 |
| MVP_top60_DNA_seed | ESR1 | 0.792365 | PRAD | PC-3 | Fulvestrant | top60_altered_DNA+RNA_seed | ERBB2 | 1.687067 | STAD | MKN45 | Sapitinib |
| MVP_top60_DNA_seed | HDAC1 | 0.793165 | PRAD | PC-3 | JQ12 | top60_altered_DNA+RNA_seed | PIK3CA | 1.709535 | STAD | NUGC-4 | AZD8186 |
| MVP_top60_DNA_seed | GSK3B | 0.802677 | PRAD | DU-145 | CHIR-99021 | top60_altered_DNA+RNA_seed | ERBB2 | 1.715738 | STAD | TMK-1 | Afatinib |
| MVP_top60_DNA_seed | ATR | 0.803869 | PRAD | PWR-1E | QL-VIII-58 | top60_altered_DNA+RNA_seed | ERBB2 | 1.755136 | STAD | OCUM-1 | Afatinib |
| MVP_top60_DNA_seed | PARP1 | 0.805579 | PRAD | 22RV1 | Olaparib | top60_altered_DNA+RNA_seed | PIK3CA | 1.757653 | STAD | KATOIII | GNE-317 |
| MVP_top60_DNA_seed | CDK4 | 0.807318 | PRAD | LNCaP-Clone-FGC | CGP-082996 | top60_altered_DNA+RNA_seed | PIK3CA | 1.960357 | STAD | NUGC-4 | Buparlisib |
| MVP_top60_DNA_seed | BRAF | 0.818377 | PRAD | VCaP | AZ628 | top60_altered_DNA+RNA_seed | ERBB2 | 1.972143 | STAD | OCUM-1 | Lapatinib |
| MVP_top60_DNA_seed | PTK2 | 0.819074 | PRAD | LNCaP-Clone-FGC | PF-562271 | top60_altered_DNA+RNA_seed | PIK3CA | 1.980186 | STAD | NUGC-4 | GNE-317 |
| MVP_top60_DNA_seed | PIK3CA | 0.819393 | PRAD | PWR-1E | PI3Ka_4409 | top60_altered_DNA+RNA_seed | PIK3CA | 2.305225 | STAD | KATOIII | Buparlisib |
| MVP_top60_DNA_seed | GSK3B | 0.884632 | PRAD | 22RV1 | KIN001-042 | top60_altered_DNA+RNA_seed | ERBB2 | 2.393403 | STAD | KATOIII | Afatinib |
| MVP_top60_DNA_seed | GSK3B | 0.886893 | PRAD | 22RV1 | SB216763 | top60_altered_DNA+RNA_seed | PIK3CA | 2.561985 | STAD | MKN28 | GNE-317 |
| MVP_top60_DNA_seed | PARP1 | 0.899247 | PRAD | VCaP | PARP_9495 | top60_altered_DNA+RNA_seed | RET | -3.946255 | THCA | CGTH-W-1 | Motesanib |
| MVP_top60_DNA_seed | EP300 | 0.920034 | PRAD | PC-3 | I-CBP112 | top60_altered_DNA+RNA_seed | BRAF | -3.729294 | THCA | K5 | Dabrafenib |
| MVP_top60_DNA_seed | BRAF | 0.923904 | PRAD | VCaP | HG6-64-1 | top60_altered_DNA+RNA_seed | BRAF | -3.412404 | THCA | BHT-101 | Dabrafenib |
| MVP_top60_DNA_seed | BRAF | 0.927436 | PRAD | VCaP | PLX-4720 | top60_altered_DNA+RNA_seed | BRAF | -3.190105 | THCA | BHT-101 | PLX-4720 |
| MVP_top60_DNA_seed | PARP1 | 0.932031 | PRAD | 22RV1 | Veliparib | top60_altered_DNA+RNA_seed | BRAF | -3.135102 | THCA | IHH-4 | Dabrafenib |
| MVP_top60_DNA_seed | BRAF | 0.956961 | PRAD | PC-3 | Dabrafenib | top60_altered_DNA+RNA_seed | BRAF | -3.105461 | THCA | HTC-C3 | Dabrafenib |
| MVP_top60_DNA_seed | PARP1 | 0.973581 | PRAD | VCaP | PARP_9482 | top60_altered_DNA+RNA_seed | BRAF | -2.966357 | THCA | CGTH-W-1 | HG6-64-1 |
| MVP_top60_DNA_seed | BRAF | 1.021972 | PRAD | LNCaP-Clone-FGC | AZ628 | top60_altered_DNA+RNA_seed | BRAF | -2.961541 | THCA | K5 | PLX-4720 |
| MVP_top60_DNA_seed | BRAF | 1.02543 | PRAD | PC-3 | RAF_9304 | top60_altered_DNA+RNA_seed | BRAF | -2.905795 | THCA | BHT-101 | PLX-4720 |
| MVP_top60_DNA_seed | AR | 1.030195 | PRAD | 22RV1 | Bicalutamide | top60_altered_DNA+RNA_seed | BRAF | -2.830382 | THCA | HTC-C3 | PLX-4720 |
| MVP_top60_DNA_seed | HDAC1 | 1.038547 | PRAD | LNCaP-Clone-FGC | Entinostat | top60_altered_DNA+RNA_seed | BRAF | -2.827309 | THCA | K5 | PLX-4720 |
| MVP_top60_DNA_seed | AR | 1.043883 | PRAD | PWR-1E | AZD3514 | top60_altered_DNA+RNA_seed | BRAF | -2.688852 | THCA | HTC-C3 | RAF_9304 |
| MVP_top60_DNA_seed | BRAF | 1.094727 | PRAD | 22RV1 | PLX-4720 | top60_altered_DNA+RNA_seed | BRAF | -2.604383 | THCA | BHT-101 | SB590885 |
| MVP_top60_DNA_seed | BRAF | 1.129819 | PRAD | LNCaP-Clone-FGC | PLX-4720 | top60_altered_DNA+RNA_seed | BRAF | -2.492753 | THCA | HTC-C3 | PLX-4720 |
| MVP_top60_DNA_seed | ATR | 1.163746 | PRAD | VCaP | QL-VIII-58 | top60_altered_DNA+RNA_seed | BRAF | -2.480329 | THCA | B-CPAP | Dabrafenib |
| MVP_top60_DNA_seed | ATM | 1.204196 | PRAD | 22RV1 | KU-55933 | top60_altered_DNA+RNA_seed | BRAF | -2.430714 | THCA | K5 | RAF_9304 |
| MVP_top60_DNA_seed | ATM | 1.228193 | PRAD | VCaP | KU-60019 | top60_altered_DNA+RNA_seed | BRAF | -2.371192 | THCA | BHT-101 | RAF_9304 |
| MVP_top60_DNA_seed | HDAC1 | 1.286136 | PRAD | LNCaP-Clone-FGC | Parthenolide | top60_altered_DNA+RNA_seed | BRAF | -2.172551 | THCA | 8305C | Dabrafenib |
| MVP_top60_DNA_seed | CDK4 | 1.286301 | PRAD | PWR-1E | Palbociclib | top60_altered_DNA+RNA_seed | BRAF | -2.151843 | THCA | K5 | AZ628 |
| MVP_top60_DNA_seed | BRAF | 1.289507 | PRAD | PWR-1E | RAF_9304 | top60_altered_DNA+RNA_seed | RET | -2.119921 | THCA | HTC-C3 | Motesanib |
| MVP_top60_DNA_seed | ATM | 1.321879 | PRAD | 22RV1 | KU-60019 | top60_altered_DNA+RNA_seed | BRAF | -2.037898 | THCA | 8305C | PLX-4720 |
| MVP_top60_DNA_seed | ATM | 1.422892 | PRAD | LNCaP-Clone-FGC | KU-55933 | top60_altered_DNA+RNA_seed | BRAF | -2.027428 | THCA | CGTH-W-1 | AZ628 |
| MVP_top60_DNA_seed | ATM | 1.452518 | PRAD | DU-145 | CP466722 | top60_altered_DNA+RNA_seed | BRAF | -1.905405 | THCA | IHH-4 | PLX-4720 |
| MVP_top60_DNA_seed | PARP1 | 1.490444 | PRAD | 22RV1 | PARP_9495 | top60_altered_DNA+RNA_seed | BRAF | -1.828709 | THCA | TT2609-C02 | HG6-64-1 |

| | | | | | | | | | | | |
|---|---|---|---|---|---|---|---|---|---|---|---|
| MVP_top60_DNA_seed | BRAF | 1.612236 | PRAD | LNCaP-Clone-FGC | HG6-64-1 | top60_altered_DNA+RNA_seed | RET | -1.811525 | THCA | TT | Motesanib |
| MVP_top60_DNA_seed | BRAF | 1.639419 | PRAD | PC-3 | HG6-64-1 | top60_altered_DNA+RNA_seed | ATM | -1.797046 | THCA | IHH-4 | KU-60019 |
| MVP_top60_DNA_seed | PIK3CA | -3.918726 | PRAD | LNCaP-Clone-FGC | AZD8186 | top60_altered_DNA+RNA_seed | BRAF | -1.773559 | THCA | 8305C | PLX-4720 |
| MVP_top60_DNA_seed | ESR1 | -3.122142 | PRAD | LNCaP-Clone-FGC | GDC0810 | top60_altered_DNA+RNA_seed | AKT1 | -1.742069 | THCA | BHT-101 | MK-2206 |
| MVP_top60_DNA_seed | ESR2 | -3.122142 | PRAD | LNCaP-Clone-FGC | GDC0810 | top60_altered_DNA+RNA_seed | BRAF | -1.672569 | THCA | IHH-4 | SB590885 |
| MVP_top60_DNA_seed | EP300 | -1.937132 | PRAD | LNCaP-Clone-FGC | CPI-637 | top60_altered_DNA+RNA_seed | BRAF | -1.554192 | THCA | B-CPAP | PLX-4720 |
| MVP_top60_DNA_seed | USP7 | -1.816793 | PRAD | LNCaP-Clone-FGC | P22077 | top60_altered_DNA+RNA_seed | BRAF | -1.542311 | THCA | FTC-133 | PLX-4720 |
| MVP_top60_DNA_seed | CDK4 | -1.667891 | PRAD | LNCaP-Clone-FGC | Ribociclib | top60_altered_DNA+RNA_seed | BRAF | -1.51961 | THCA | B-CPAP | PLX-4720 |
| MVP_top60_DNA_seed | PIK3CA | -1.521253 | PRAD | LNCaP-Clone-FGC | GNE-317 | top60_altered_DNA+RNA_seed | ATM | -1.435817 | THCA | IHH-4 | KU-55933 |
| MVP_top60_DNA_seed | GSK3B | -1.515353 | PRAD | PWR-1E | SB216763 | top60_altered_DNA+RNA_seed | BRAF | -1.423171 | THCA | IHH-4 | PLX-4720 |
| MVP_top60_DNA_seed | EP300 | -1.429427 | PRAD | VCaP | CPI-637 | top60_altered_DNA+RNA_seed | BRAF | -1.362543 | THCA | 8305C | SB590885 |
| MVP_top60_DNA_seed | EZH2 | -1.380232 | PRAD | PWR-1E | GSK343 | top60_altered_DNA+RNA_seed | AKT1 | -1.335264 | THCA | TT2609-C02 | AKT inhibitor VIII |
| MVP_top60_DNA_seed | EZH2 | -1.353513 | PRAD | LNCaP-Clone-FGC | GSK343 | top60_altered_DNA+RNA_seed | BRAF | -1.308841 | THCA | IHH-4 | RAF_9304 |
| MVP_top60_DNA_seed | CDK4 | -1.25799 | PRAD | LNCaP-Clone-FGC | Palbociclib | top60_altered_DNA+RNA_seed | ATM | -1.291053 | THCA | BHT-101 | KU-55933 |
| MVP_top60_DNA_seed | ESR1 | -1.254283 | PRAD | LNCaP-Clone-FGC | Tamoxifen | top60_altered_DNA+RNA_seed | BRAF | -1.282376 | THCA | B-CPAP | HG6-64-1 |
| MVP_top60_DNA_seed | CDK4 | -1.11126 | PRAD | 22RV1 | Ribociclib | top60_altered_DNA+RNA_seed | ATM | -1.265003 | THCA | K5 | torin2 |
| MVP_top60_DNA_seed | PARP1 | -1.070302 | PRAD | LNCaP-Clone-FGC | Olaparib | top60_altered_DNA+RNA_seed | AKT1 | -1.262621 | THCA | BHT-101 | AKT inhibitor VIII |
| MVP_top60_DNA_seed | HDAC1 | -1.058405 | PRAD | LNCaP-Clone-FGC | Entinostat | top60_altered_DNA+RNA_seed | RET | -1.225834 | THCA | TT2609-C02 | Motesanib |
| MVP_top60_DNA_seed | PIK3CA | -1.008293 | PRAD | LNCaP-Clone-FGC | Buparlisib | top60_altered_DNA+RNA_seed | BRAF | -1.224839 | THCA | KMH-2 | SB590885 |
| MVP_top60_DNA_seed | HDAC1 | -1.00576 | PRAD | PWR-1E | Romidepsin | top60_altered_DNA+RNA_seed | ATM | -1.218818 | THCA | CAL-62 | torin2 |
| MVP_top60_DNA_seed | PIK3CA | -0.883386 | PRAD | 22RV1 | Alpelisib | top60_altered_DNA+RNA_seed | ATM | -1.205962 | THCA | 8305C | KU-60019 |
| MVP_top60_DNA_seed | ATR | -0.875291 | PRAD | LNCaP-Clone-FGC | VE821 | top60_altered_DNA+RNA_seed | BRAF | -1.201393 | THCA | KMH-2 | Dabrafenib |
| MVP_top60_DNA_seed | PARP1 | -0.873991 | PRAD | LNCaP-Clone-FGC | Veliparib | top60_altered_DNA+RNA_seed | ATM | -1.178386 | THCA | B-CPAP | KU-60019 |
| MVP_top60_DNA_seed | ATR | -0.870173 | PRAD | LNCaP-Clone-FGC | AZD6738 | top60_altered_DNA+RNA_seed | BRAF | -1.173142 | THCA | ML-1 | HG6-64-1 |
| MVP_top60_DNA_seed | ATR | -0.830781 | PRAD | PWR-1E | AZD6738 | top60_altered_DNA+RNA_seed | BRAF | -1.163381 | THCA | 8305C | HG6-64-1 |
| MVP_top60_DNA_seed | GSK3B | -0.830021 | PRAD | PC-3 | SB216763 | top60_altered_DNA+RNA_seed | RET | -1.158994 | THCA | CAL-62 | Motesanib |
| MVP_top60_DNA_seed | HDAC1 | -0.822506 | PRAD | LNCaP-Clone-FGC | Romidepsin | top60_altered_DNA+RNA_seed | AKT1 | -1.15598 | THCA | ML-1 | AKT inhibitor VIII |
| MVP_top60_DNA_seed | EZH2 | -0.805512 | PRAD | LNCaP-Clone-FGC | GSK343 | top60_altered_DNA+RNA_seed | BRAF | -1.154696 | THCA | ML-1 | PLX-4720 |
| MVP_top60_DNA_seed | PARP1 | -0.777444 | PRAD | LNCaP-Clone-FGC | Talazoparib | top60_altered_DNA+RNA_seed | AKT1 | -1.135161 | THCA | BHT-101 | BAY AKT1 |
| MVP_top60_DNA_seed | CDK4 | -0.76717 | PRAD | 22RV1 | Palbociclib | top60_altered_DNA+RNA_seed | ATM | -1.08938 | THCA | 8305C | KU-55933 |
| MVP_top60_DNA_seed | ESR1 | -0.700482 | PRAD | PWR-1E | Tamoxifen | top60_altered_DNA+RNA_seed | BRAF | -1.088167 | THCA | 8305C | RAF_9304 |
| MVP_top60_DNA_seed | GSK3B | -0.683917 | PRAD | LNCaP-Clone-FGC | SB216763 | top60_altered_DNA+RNA_seed | AKT1 | -1.066972 | THCA | TT | AKT inhibitor VIII |
| MVP_top60_DNA_seed | PIK3CA | -0.654226 | PRAD | 22RV1 | GNE-317 | top60_altered_DNA+RNA_seed | BRAF | -1.031488 | THCA | TT2609-C02 | PLX-4720 |
| MVP_top60_DNA_seed | ATR | -0.619541 | PRAD | LNCaP-Clone-FGC | VE-822 | top60_altered_DNA+RNA_seed | BRAF | -1.023462 | THCA | IHH-4 | HG6-64-1 |
| MVP_top60_DNA_seed | ATM | -0.581479 | PRAD | PWR-1E | KU-55933 | top60_altered_DNA+RNA_seed | RET | -0.982642 | THCA | BHT-101 | Motesanib |
| MVP_top60_DNA_seed | PIK3CA | -0.541057 | PRAD | PC-3 | AZD8186 | top60_altered_DNA+RNA_seed | BRAF | -0.94498 | THCA | CAL-62 | RAF_9304 |
| MVP_top60_DNA_seed | PARP1 | -0.52367 | PRAD | PWR-1E | Olaparib | top60_altered_DNA+RNA_seed | ATM | -0.944882 | THCA | HTC-C3 | CP466722 |
| MVP_top60_DNA_seed | PIK3CA | -0.513328 | PRAD | 22RV1 | Buparlisib | top60_altered_DNA+RNA_seed | ATM | -0.941654 | THCA | 8505C | KU-60019 |
| MVP_top60_DNA_seed | EP300 | -0.467554 | PRAD | VCaP | SGC-CBP30 | top60_altered_DNA+RNA_seed | ATM | -0.892202 | THCA | CGTH-W-1 | torin2 |
| MVP_top60_DNA_seed | MAPK3 | -0.45826 | PRAD | VCaP | Ulixertinib | top60_altered_DNA+RNA_seed | BRAF | -0.870308 | THCA | TT2609-C02 | PLX-4720 |
| MVP_top60_DNA_seed | CDK4 | -0.456032 | PRAD | PWR-1E | Ribociclib | top60_altered_DNA+RNA_seed | ATM | -0.868791 | THCA | ML-1 | KU-55933 |

| | | | | | | | | | | | |
|---|---|---|---|---|---|---|---|---|---|---|---|
| MVP_top60_DNA_seed | MAPK3 | -0.438599 | PRAD | LNCaP-Clone-FGC | Ulixertinib | top60_altered_DNA+RNA_seed | RET | -0.85325 | THCA | RO82-W-1 | Motesanib |
| MVP_top60_DNA_seed | PIK3CA | -0.436117 | PRAD | LNCaP-Clone-FGC | Alpelisib | top60_altered_DNA+RNA_seed | BRAF | -0.842455 | THCA | HTC-C3 | HG6-64-1 |
| MVP_top60_DNA_seed | PIK3CA | -0.417913 | PRAD | PWR-1E | Buparlisib | top60_altered_DNA+RNA_seed | ATM | -0.825398 | THCA | KMH-2 | CP466722 |
| MVP_top60_DNA_seed | PARP1 | -0.390075 | PRAD | LNCaP-Clone-FGC | Rucaparib | top60_altered_DNA+RNA_seed | BRAF | -0.813015 | THCA | HTC-C3 | SB590885 |
| MVP_top60_DNA_seed | PARP1 | -0.384143 | PRAD | LNCaP-Clone-FGC | Niraparib | top60_altered_DNA+RNA_seed | RET | -0.791025 | THCA | 8505C | Motesanib |
| MVP_top60_DNA_seed | PARP1 | -0.376943 | PRAD | PWR-1E | Veliparib | top60_altered_DNA+RNA_seed | ATM | -0.773274 | THCA | TT2609-C02 | KU-60019 |
| MVP_top60_DNA_seed | PARP1 | -0.36595 | PRAD | 22RV1 | Niraparib | top60_altered_DNA+RNA_seed | AKT1 | -0.773118 | THCA | FTC-133 | AKT inhibitor VIII |
| MVP_top60_DNA_seed | ESR1 | -0.361359 | PRAD | VCaP | Tamoxifen | top60_altered_DNA+RNA_seed | BRAF | -0.763833 | THCA | B-CPAP | RAF_9304 |
| MVP_top60_DNA_seed | HDAC1 | -0.323846 | PRAD | VCaP | Romidepsin | top60_altered_DNA+RNA_seed | BRAF | -0.758091 | THCA | RO82-W-1 | PLX-4720 |
| MVP_top60_DNA_seed | BRAF | -0.31693 | PRAD | LNCaP-Clone-FGC | SB590885 | top60_altered_DNA+RNA_seed | BRAF | -0.71913 | THCA | 8505C | Dabrafenib |
| MVP_top60_DNA_seed | PARP1 | -0.238697 | PRAD | VCaP | Rucaparib | top60_altered_DNA+RNA_seed | ATM | -0.711223 | THCA | ML-1 | CP466722 |
| MVP_top60_DNA_seed | MAPK3 | -0.237906 | PRAD | LNCaP-Clone-FGC | Ulixertinib | top60_altered_DNA+RNA_seed | AKT1 | -0.710688 | THCA | IHH-4 | MK-2206 |
| MVP_top60_DNA_seed | PIK3CA | -0.229378 | PRAD | PC-3 | GNE-317 | top60_altered_DNA+RNA_seed | ATM | -0.707837 | THCA | K5 | KU-60019 |
| MVP_top60_DNA_seed | BRAF | -0.218562 | PRAD | VCaP | SB590885 | top60_altered_DNA+RNA_seed | BRAF | -0.706258 | THCA | CAL-62 | Dabrafenib |
| MVP_top60_DNA_seed | ESR1 | -0.198261 | PRAD | 22RV1 | Tamoxifen | top60_altered_DNA+RNA_seed | BRAF | -0.686113 | THCA | TT2609-C02 | SB590885 |
| MVP_top60_DNA_seed | PARP1 | -0.174937 | PRAD | PWR-1E | Niraparib | top60_altered_DNA+RNA_seed | ATM | -0.674507 | THCA | ASH-3 | KU-60019 |
| MVP_top60_DNA_seed | HDAC1 | -0.168062 | PRAD | VCaP | PCI-34051 | top60_altered_DNA+RNA_seed | ATM | -0.632848 | THCA | IHH-4 | CP466722 |
| MVP_top60_DNA_seed | BRAF | -0.158114 | PRAD | VCaP | Dabrafenib | top60_altered_DNA+RNA_seed | BRAF | -0.606979 | THCA | KMH-2 | RAF_9304 |
| MVP_top60_DNA_seed | USP7 | -0.139823 | PRAD | PWR-1E | P22077 | top60_altered_DNA+RNA_seed | BRAF | -0.5883 | THCA | TT | SB590885 |
| MVP_top60_DNA_seed | ATR | -0.120204 | PRAD | PWR-1E | VE821 | top60_altered_DNA+RNA_seed | AKT1 | -0.56319 | THCA | TT | MK-2206 |
| MVP_top60_DNA_seed | PARP1 | -0.111848 | PRAD | 22RV1 | Talazoparib | top60_altered_DNA+RNA_seed | RET | -0.553464 | THCA | FTC-133 | Motesanib |
| MVP_top60_DNA_seed | CDK4 | -0.104997 | PRAD | VCaP | Palbociclib | top60_altered_DNA+RNA_seed | RET | -0.550059 | THCA | B-CPAP | Motesanib |
| MVP_top60_DNA_seed | HDAC1 | -0.062479 | PRAD | PWR-1E | PCI-34051 | top60_altered_DNA+RNA_seed | AKT1 | -0.546823 | THCA | ASH-3 | AKT inhibitor VIII |
| MVP_top60_DNA_seed | BRAF | -0.046353 | PRAD | LNCaP-Clone-FGC | Dabrafenib | top60_altered_DNA+RNA_seed | ATM | -0.545485 | THCA | TT2609-C02 | KU-55933 |
| MVP_top60_DNA_seed | EP300 | -0.029644 | PRAD | PWR-1E | SGC-CBP30 | top60_altered_DNA+RNA_seed | BRAF | -0.53173 | THCA | ASH-3 | RAF_9304 |
| MVP_top60_DNA_seed | HDAC1 | -0.026224 | PRAD | 22RV1 | Entinostat | top60_altered_DNA+RNA_seed | AKT1 | -0.521863 | THCA | CGTH-W-1 | AKT inhibitor VIII |
| MVP_top60_DNA_seed | TP53 | -0.014816 | PRAD | PWR-1E | MIRA-1 | top60_altered_DNA+RNA_seed | ATM | -0.515829 | THCA | TT | KU-60019 |
| MVP_top60_DNA_seed | EP300 | -0.012331 | PRAD | DU-145 | CPI-637 | top60_altered_DNA+RNA_seed | ATM | -0.513165 | THCA | HTC-C3 | KU-60019 |
| MVP_top60_DNA_seed | PARP1 | 0.017556 | PRAD | PWR-1E | Talazoparib | top60_altered_DNA+RNA_seed | BRAF | -0.489515 | THCA | 8505C | SB590885 |
| MVP_top60_DNA_seed | PIK3CA | 0.033091 | PRAD | VCaP | Alpelisib | top60_altered_DNA+RNA_seed | AKT1 | -0.481397 | THCA | CGTH-W-1 | A-443654 |
| MVP_top60_DNA_seed | ATM | 0.046193 | PRAD | PC-3 | KU-55933 | top60_altered_DNA+RNA_seed | ATM | -0.479085 | THCA | BHT-101 | KU-60019 |
| MVP_top60_DNA_seed | GSK3B | 0.052881 | PRAD | VCaP | SB216763 | top60_altered_DNA+RNA_seed | AKT1 | -0.465209 | THCA | IHH-4 | AKT inhibitor VIII |
| MVP_top60_DNA_seed | PARP1 | 0.057574 | PRAD | VCaP | Olaparib | top60_altered_DNA+RNA_seed | AKT1 | -0.454229 | THCA | K5 | BAY AKT1 |
| MVP_top60_DNA_seed | EZH2 | 0.079193 | PRAD | VCaP | GSK343 | top60_altered_DNA+RNA_seed | ATM | -0.452208 | THCA | CGTH-W-1 | KU-55933 |
| MVP_top60_DNA_seed | PARP1 | 0.101815 | PRAD | 22RV1 | Rucaparib | top60_altered_DNA+RNA_seed | RET | -0.422726 | THCA | ML-1 | Motesanib |
| MVP_top60_DNA_seed | PARP1 | 0.131177 | PRAD | PWR-1E | Rucaparib | top60_altered_DNA+RNA_seed | ATM | -0.421715 | THCA | KMH-2 | KU-60019 |
| MVP_top60_DNA_seed | EP300 | 0.138199 | PRAD | PWR-1E | CPI-637 | top60_altered_DNA+RNA_seed | RET | -0.413693 | THCA | IHH-4 | Motesanib |
| MVP_top60_DNA_seed | PIK3CA | 0.149453 | PRAD | PC-3 | Buparlisib | top60_altered_DNA+RNA_seed | BRAF | -0.392081 | THCA | ASH-3 | Dabrafenib |
| MVP_top60_DNA_seed | HDAC1 | 0.185731 | PRAD | VCaP | Entinostat | top60_altered_DNA+RNA_seed | ATM | -0.391261 | THCA | ASH-3 | KU-55933 |
| MVP_top60_DNA_seed | MAPK3 | 0.186089 | PRAD | PWR-1E | SCH772984 | top60_altered_DNA+RNA_seed | ATM | -0.374302 | THCA | BHT-101 | CP466722 |
| MVP_top60_DNA_seed | PARP1 | 0.191115 | PRAD | DU-145 | Veliparib | top60_altered_DNA+RNA_seed | BRAF | -0.370822 | THCA | K5 | HG6-64-1 |

| | | | | | | | | | | | |
|---|---|---|---|---|---|---|---|---|---|---|---|
| MVP_top60_DNA_seed | PARP1 | 0.204799 | PRAD | DU-145 | Talazoparib | top60_altered_DNA+RNA_seed | BRAF | -0.366295 | THCA | CAL-62 | PLX-4720 |
| MVP_top60_DNA_seed | ESR1 | 0.213009 | PRAD | VCaP | GDC0810 | top60_altered_DNA+RNA_seed | BRAF | -0.324975 | THCA | B-CPAP | SB590885 |
| MVP_top60_DNA_seed | ESR2 | 0.213009 | PRAD | VCaP | GDC0810 | top60_altered_DNA+RNA_seed | AKT1 | -0.288646 | THCA | KMH-2 | GSK690693 |
| MVP_top60_DNA_seed | CDK4 | 0.230549 | PRAD | VCaP | Ribociclib | top60_altered_DNA+RNA_seed | BRAF | -0.272961 | THCA | 8505C | PLX-4720 |
| MVP_top60_DNA_seed | TP53 | 0.243205 | PRAD | VCaP | MIRA-1 | top60_altered_DNA+RNA_seed | BRAF | -0.242036 | THCA | CAL-62 | HG6-64-1 |
| MVP_top60_DNA_seed | MAPK3 | 0.246707 | PRAD | 22RV1 | Ulixertinib | top60_altered_DNA+RNA_seed | ATM | -0.239267 | THCA | 8505C | KU-55933 |
| MVP_top60_DNA_seed | BRAF | 0.248985 | PRAD | PWR-1E | PLX-4720 | top60_altered_DNA+RNA_seed | BRAF | -0.23467 | THCA | 8505C | HG6-64-1 |
| MVP_top60_DNA_seed | BRAF | 0.250083 | PRAD | PWR-1E | SB590885 | top60_altered_DNA+RNA_seed | ATM | -0.212464 | THCA | 8505C | torin2 |
| MVP_top60_DNA_seed | USP7 | 0.275383 | PRAD | PC-3 | P22077 | top60_altered_DNA+RNA_seed | AKT1 | -0.208708 | THCA | B-CPAP | MK-2206 |
| MVP_top60_DNA_seed | ESR1 | 0.275997 | PRAD | PWR-1E | GDC0810 | top60_altered_DNA+RNA_seed | ATM | -0.201777 | THCA | ASH-3 | CP466722 |
| MVP_top60_DNA_seed | ESR2 | 0.275997 | PRAD | PWR-1E | GDC0810 | top60_altered_DNA+RNA_seed | BRAF | -0.186451 | THCA | CAL-62 | PLX-4720 |
| MVP_top60_DNA_seed | HDAC1 | 0.282483 | PRAD | PWR-1E | Entinostat | top60_altered_DNA+RNA_seed | AKT1 | -0.182832 | THCA | KMH-2 | AKT inhibitor VIII |
| MVP_top60_DNA_seed | PIK3CA | 0.288414 | PRAD | PWR-1E | GNE-317 | top60_altered_DNA+RNA_seed | AKT1 | -0.170145 | THCA | IHH-4 | BAY AKT1 |
| MVP_top60_DNA_seed | ATR | 0.304135 | PRAD | PWR-1E | VE-822 | top60_altered_DNA+RNA_seed | BRAF | -0.164558 | THCA | KMH-2 | PLX-4720 |
| MVP_top60_DNA_seed | MAPK3 | 0.304985 | PRAD | 22RV1 | SCH772984 | top60_altered_DNA+RNA_seed | BRAF | -0.164224 | THCA | CGTH-W-1 | Dabrafenib |
| MVP_top60_DNA_seed | HDAC1 | 0.321559 | PRAD | 22RV1 | Romidepsin | top60_altered_DNA+RNA_seed | AKT1 | -0.158883 | THCA | 8505C | MK-2206 |
| MVP_top60_DNA_seed | PIK3CA | 0.349073 | PRAD | VCaP | AZD8186 | top60_altered_DNA+RNA_seed | ATM | -0.145785 | THCA | FTC-133 | KU-60019 |
| MVP_top60_DNA_seed | HDAC1 | 0.381751 | PRAD | LNCaP-Clone-FGC | PCI-34051 | top60_altered_DNA+RNA_seed | AKT1 | -0.129599 | THCA | HTC-C3 | AKT inhibitor VIII |
| MVP_top60_DNA_seed | BRAF | 0.398518 | PRAD | LNCaP-Clone-FGC | PLX-4720 | top60_altered_DNA+RNA_seed | BRAF | -0.085211 | THCA | RO82-W-1 | HG6-64-1 |
| MVP_top60_DNA_seed | MAPK3 | 0.418259 | PRAD | PWR-1E | Ulixertinib | top60_altered_DNA+RNA_seed | BRAF | -0.081779 | THCA | CGTH-W-1 | PLX-4720 |
| MVP_top60_DNA_seed | EP300 | 0.422506 | PRAD | LNCaP-Clone-FGC | SGC-CBP30 | top60_altered_DNA+RNA_seed | ATM | -0.054763 | THCA | K5 | KU-55933 |
| MVP_top60_DNA_seed | ATM | 0.42586 | PRAD | LNCaP-Clone-FGC | KU-55933 | top60_altered_DNA+RNA_seed | AKT1 | -0.042584 | THCA | HTC-C3 | AKT inhibitor VIII |
| MVP_top60_DNA_seed | PIK3CA | 0.437684 | PRAD | 22RV1 | AZD8186 | top60_altered_DNA+RNA_seed | BRAF | -0.04043 | THCA | ML-1 | Dabrafenib |
| MVP_top60_DNA_seed | PARP1 | 0.446962 | PRAD | 22RV1 | Olaparib | top60_altered_DNA+RNA_seed | BRAF | -0.035248 | THCA | KMH-2 | HG6-64-1 |
| MVP_top60_DNA_seed | MAPK3 | 0.448432 | PRAD | 22RV1 | Ulixertinib | top60_altered_DNA+RNA_seed | AKT1 | -0.025516 | THCA | K5 | A-443654 |
| MVP_top60_DNA_seed | EZH2 | 0.461793 | PRAD | 22RV1 | GSK343 | top60_altered_DNA+RNA_seed | AKT1 | -0.024435 | THCA | CGTH-W-1 | MK-2206 |
| MVP_top60_DNA_seed | PARP1 | 0.501736 | PRAD | PC-3 | Talazoparib | top60_altered_DNA+RNA_seed | AKT1 | -0.019583 | THCA | FTC-133 | BAY AKT1 |
| MVP_top60_DNA_seed | EP300 | 0.522999 | PRAD | 22RV1 | CPI-637 | top60_altered_DNA+RNA_seed | ATM | -0.001181 | THCA | CAL-62 | KU-60019 |
| MVP_top60_DNA_seed | HDAC1 | 0.531994 | PRAD | DU-145 | Romidepsin | top60_altered_DNA+RNA_seed | BRAF | 0.001482 | THCA | 8505C | RAF_9304 |
| MVP_top60_DNA_seed | BRAF | 0.537231 | PRAD | PWR-1E | Dabrafenib | top60_altered_DNA+RNA_seed | ATM | 0.004693 | THCA | HTC-C3 | KU-55933 |
| MVP_top60_DNA_seed | ATR | 0.553194 | PRAD | VCaP | VE821 | top60_altered_DNA+RNA_seed | AKT1 | 0.012071 | THCA | IHH-4 | AKT inhibitor VIII |
| MVP_top60_DNA_seed | ATR | 0.591904 | PRAD | DU-145 | VE-822 | top60_altered_DNA+RNA_seed | ATM | 0.039882 | THCA | K5 | CP466722 |
| MVP_top60_DNA_seed | EP300 | 0.602643 | PRAD | DU-145 | SGC-CBP30 | top60_altered_DNA+RNA_seed | AKT1 | 0.071723 | THCA | KMH-2 | BAY AKT1 |
| MVP_top60_DNA_seed | ATR | 0.603229 | PRAD | DU-145 | AZD6738 | top60_altered_DNA+RNA_seed | AKT1 | 0.074453 | THCA | BHT-101 | AKT inhibitor VIII |
| MVP_top60_DNA_seed | GSK3B | 0.626472 | PRAD | 22RV1 | SB216763 | top60_altered_DNA+RNA_seed | ATM | 0.100739 | THCA | RO82-W-1 | KU-60019 |
| MVP_top60_DNA_seed | ATM | 0.639648 | PRAD | DU-145 | KU-55933 | top60_altered_DNA+RNA_seed | ATM | 0.106576 | THCA | CAL-62 | KU-55933 |
| MVP_top60_DNA_seed | PARP1 | 0.641877 | PRAD | VCaP | Talazoparib | top60_altered_DNA+RNA_seed | RET | 0.108057 | THCA | K5 | Motesanib |
| MVP_top60_DNA_seed | ESR1 | 0.655833 | PRAD | DU-145 | GDC0810 | top60_altered_DNA+RNA_seed | BRAF | 0.111801 | THCA | RO82-W-1 | Dabrafenib |
| MVP_top60_DNA_seed | ESR2 | 0.655833 | PRAD | DU-145 | GDC0810 | top60_altered_DNA+RNA_seed | AKT1 | 0.112808 | THCA | KMH-2 | AKT inhibitor VIII |
| MVP_top60_DNA_seed | EZH2 | 0.674191 | PRAD | PWR-1E | GSK343 | top60_altered_DNA+RNA_seed | AKT1 | 0.118602 | THCA | K5 | AKT inhibitor VIII |
| MVP_top60_DNA_seed | PARP1 | 0.678392 | PRAD | DU-145 | Olaparib | top60_altered_DNA+RNA_seed | AKT1 | 0.123679 | THCA | RO82-W-1 | BAY AKT1 |

| | | | | | | | | | | | |
|---|---|---|---|---|---|---|---|---|---|---|---|
| MVP_top60_DNA_seed | PIK3CA | 0.678429 | PRAD | PWR-1E | Alpelisib | top60_altered_DNA+RNA_seed | AKT1 | 0.128532 | THCA | K5 | GSK690693 |
| MVP_top60_DNA_seed | EP300 | 0.683503 | PRAD | PC-3 | CPI-637 | top60_altered_DNA+RNA_seed | BRAF | 0.13551 | THCA | ML-1 | PLX-4720 |
| MVP_top60_DNA_seed | TP53 | 0.687703 | PRAD | LNCaP-Clone-FGC | MIRA-1 | top60_altered_DNA+RNA_seed | AKT1 | 0.140238 | THCA | TT2609-C02 | MK-2206 |
| MVP_top60_DNA_seed | PIK3CA | 0.692404 | PRAD | PWR-1E | AZD8186 | top60_altered_DNA+RNA_seed | BRAF | 0.161866 | THCA | TT | Dabrafenib |
| MVP_top60_DNA_seed | TP53 | 0.706283 | PRAD | 22RV1 | MIRA-1 | top60_altered_DNA+RNA_seed | ATM | 0.164551 | THCA | CGTH-W-1 | KU-60019 |
| MVP_top60_DNA_seed | MAPK3 | 0.712034 | PRAD | VCaP | Ulixertinib | top60_altered_DNA+RNA_seed | AKT1 | 0.165025 | THCA | 8305C | MK-2206 |
| MVP_top60_DNA_seed | PARP1 | 0.72663 | PRAD | VCaP | Niraparib | top60_altered_DNA+RNA_seed | AKT1 | 0.180136 | THCA | RO82-W-1 | AKT inhibitor VIII |
| MVP_top60_DNA_seed | ATR | 0.72813 | PRAD | 22RV1 | VE-822 | top60_altered_DNA+RNA_seed | AKT1 | 0.18882 | THCA | 8505C | GSK690693 |
| MVP_top60_DNA_seed | MAPK3 | 0.752388 | PRAD | LNCaP-Clone-FGC | SCH772984 | top60_altered_DNA+RNA_seed | RET | 0.203243 | THCA | KMH-2 | Motesanib |
| MVP_top60_DNA_seed | ATR | 0.752867 | PRAD | PC-3 | AZD6738 | top60_altered_DNA+RNA_seed | AKT1 | 0.215195 | THCA | 8505C | AKT inhibitor VIII |
| MVP_top60_DNA_seed | PIK3CA | 0.757757 | PRAD | PC-3 | Alpelisib | top60_altered_DNA+RNA_seed | AKT1 | 0.233858 | THCA | IHH-4 | GSK690693 |
| MVP_top60_DNA_seed | ESR1 | 0.769845 | PRAD | PC-3 | GDC0810 | top60_altered_DNA+RNA_seed | RET | 0.241375 | THCA | ASH-3 | Motesanib |
| MVP_top60_DNA_seed | ESR2 | 0.769845 | PRAD | PC-3 | GDC0810 | top60_altered_DNA+RNA_seed | BRAF | 0.24194 | THCA | TT2609-C02 | Dabrafenib |
| MVP_top60_DNA_seed | BRAF | 0.806339 | PRAD | DU-145 | Dabrafenib | top60_altered_DNA+RNA_seed | BRAF | 0.241945 | THCA | RO82-W-1 | RAF_9304 |
| MVP_top60_DNA_seed | EP300 | 0.844741 | PRAD | 22RV1 | SGC-CBP30 | top60_altered_DNA+RNA_seed | BRAF | 0.25163 | THCA | 8505C | PLX-4720 |
| MVP_top60_DNA_seed | PIK3CA | 0.861976 | PRAD | VCaP | Buparlisib | top60_altered_DNA+RNA_seed | BRAF | 0.252942 | THCA | TT | RAF_9304 |
| MVP_top60_DNA_seed | EP300 | 0.866293 | PRAD | PC-3 | SGC-CBP30 | top60_altered_DNA+RNA_seed | ATM | 0.26645 | THCA | ML-1 | KU-60019 |
| MVP_top60_DNA_seed | PARP1 | 0.872677 | PRAD | 22RV1 | Veliparib | top60_altered_DNA+RNA_seed | AKT1 | 0.268555 | THCA | ASH-3 | MK-2206 |
| MVP_top60_DNA_seed | PARP1 | 0.874127 | PRAD | PC-3 | Olaparib | top60_altered_DNA+RNA_seed | BRAF | 0.307688 | THCA | FTC-133 | SB590885 |
| MVP_top60_DNA_seed | BRAF | 0.87528 | PRAD | 22RV1 | PLX-4720 | top60_altered_DNA+RNA_seed | ATM | 0.317324 | THCA | CAL-62 | CP466722 |
| MVP_top60_DNA_seed | EZH2 | 0.889699 | PRAD | VCaP | GSK343 | top60_altered_DNA+RNA_seed | AKT1 | 0.318722 | THCA | 8305C | BAY AKT1 |
| MVP_top60_DNA_seed | BRAF | 0.902922 | PRAD | DU-145 | SB590885 | top60_altered_DNA+RNA_seed | BRAF | 0.3238 | THCA | FTC-133 | PLX-4720 |
| MVP_top60_DNA_seed | ATR | 0.907775 | PRAD | 22RV1 | AZD6738 | top60_altered_DNA+RNA_seed | ATM | 0.338034 | THCA | B-CPAP | CP466722 |
| MVP_top60_DNA_seed | HDAC1 | 0.90968 | PRAD | PC-3 | Entinostat | top60_altered_DNA+RNA_seed | ATM | 0.340556 | THCA | CGTH-W-1 | CP466722 |
| MVP_top60_DNA_seed | MAPK3 | 0.935252 | PRAD | DU-145 | Ulixertinib | top60_altered_DNA+RNA_seed | AKT1 | 0.345478 | THCA | CGTH-W-1 | AKT inhibitor VIII |
| MVP_top60_DNA_seed | GSK3B | 0.946724 | PRAD | DU-145 | SB216763 | top60_altered_DNA+RNA_seed | ATM | 0.353416 | THCA | TT | KU-55933 |
| MVP_top60_DNA_seed | BRAF | 0.952669 | PRAD | PC-3 | Dabrafenib | top60_altered_DNA+RNA_seed | ATM | 0.377689 | THCA | 8505C | CP466722 |
| MVP_top60_DNA_seed | ESR1 | 0.9541 | PRAD | DU-145 | Tamoxifen | top60_altered_DNA+RNA_seed | BRAF | 0.380782 | THCA | ASH-3 | SB590885 |
| MVP_top60_DNA_seed | TP53 | 0.955249 | PRAD | DU-145 | MIRA-1 | top60_altered_DNA+RNA_seed | AKT1 | 0.398747 | THCA | TT | BAY AKT1 |
| MVP_top60_DNA_seed | CDK4 | 0.956746 | PRAD | PWR-1E | Palbociclib | top60_altered_DNA+RNA_seed | BRAF | 0.427087 | THCA | CGTH-W-1 | PLX-4720 |
| MVP_top60_DNA_seed | CDK4 | 0.972709 | PRAD | PC-3 | Ribociclib | top60_altered_DNA+RNA_seed | AKT1 | 0.441879 | THCA | B-CPAP | GSK690693 |
| MVP_top60_DNA_seed | PIK3CA | 1.005586 | PRAD | DU-145 | AZD8186 | top60_altered_DNA+RNA_seed | ATM | 0.448252 | THCA | B-CPAP | KU-55933 |
| MVP_top60_DNA_seed | BRAF | 1.018457 | PRAD | 22RV1 | SB590885 | top60_altered_DNA+RNA_seed | AKT1 | 0.46 | THCA | FTC-133 | MK-2206 |
| MVP_top60_DNA_seed | CDK4 | 1.030612 | PRAD | DU-145 | Ribociclib | top60_altered_DNA+RNA_seed | AKT1 | 0.464012 | THCA | CAL-62 | AKT inhibitor VIII |
| MVP_top60_DNA_seed | EZH2 | 1.037094 | PRAD | 22RV1 | GSK343 | top60_altered_DNA+RNA_seed | BRAF | 0.47435 | THCA | KMH-2 | PLX-4720 |
| MVP_top60_DNA_seed | HDAC1 | 1.070096 | PRAD | 22RV1 | PCI-34051 | top60_altered_DNA+RNA_seed | BRAF | 0.476929 | THCA | ASH-3 | PLX-4720 |
| MVP_top60_DNA_seed | PIK3CA | 1.075058 | PRAD | VCaP | GNE-317 | top60_altered_DNA+RNA_seed | BRAF | 0.478727 | THCA | FTC-133 | Dabrafenib |
| MVP_top60_DNA_seed | ATM | 1.07653 | PRAD | VCaP | KU-55933 | top60_altered_DNA+RNA_seed | AKT1 | 0.482172 | THCA | ASH-3 | GSK690693 |
| MVP_top60_DNA_seed | PARP1 | 1.094931 | PRAD | PC-3 | Niraparib | top60_altered_DNA+RNA_seed | AKT1 | 0.496458 | THCA | RO82-W-1 | MK-2206 |
| MVP_top60_DNA_seed | PIK3CA | 1.111516 | PRAD | DU-145 | Alpelisib | top60_altered_DNA+RNA_seed | BRAF | 0.503209 | THCA | FTC-133 | RAF_9304 |
| MVP_top60_DNA_seed | PARP1 | 1.117531 | PRAD | VCaP | Veliparib | top60_altered_DNA+RNA_seed | AKT1 | 0.513617 | THCA | ASH-3 | AKT inhibitor VIII |

| | | | | | | | | | | | |
|---|---|---|---|---|---|---|---|---|---|---|---|
| MVP_top60_DNA_seed | ATR | 1.131643 | PRAD | PC-3 | VE-822 | top60_altered_DNA+RNA_seed | AKT1 | 0.518521 | THCA | CGTH-W-1 | GSK690693 |
| MVP_top60_DNA_seed | PARP1 | 1.145966 | PRAD | PC-3 | Veliparib | top60_altered_DNA+RNA_seed | AKT1 | 0.537145 | THCA | ASH-3 | BAY AKT1 |
| MVP_top60_DNA_seed | BRAF | 1.177123 | PRAD | 22RV1 | Dabrafenib | top60_altered_DNA+RNA_seed | BRAF | 0.542429 | THCA | TT2609-C02 | RAF_9304 |
| MVP_top60_DNA_seed | MAPK3 | 1.193568 | PRAD | PWR-1E | Ulixertinib | top60_altered_DNA+RNA_seed | AKT1 | 0.546883 | THCA | HTC-C3 | MK-2206 |
| MVP_top60_DNA_seed | EZH2 | 1.198497 | PRAD | DU-145 | GSK343 | top60_altered_DNA+RNA_seed | AKT1 | 0.555869 | THCA | B-CPAP | AKT inhibitor VIII |
| MVP_top60_DNA_seed | BRAF | 1.199828 | PRAD | PC-3 | SB590885 | top60_altered_DNA+RNA_seed | BRAF | 0.558517 | THCA | ASH-3 | HG6-64-1 |
| MVP_top60_DNA_seed | USP7 | 1.221179 | PRAD | VCaP | P22077 | top60_altered_DNA+RNA_seed | AKT1 | 0.560545 | THCA | B-CPAP | BAY AKT1 |
| MVP_top60_DNA_seed | HDAC1 | 1.22727 | PRAD | PC-3 | Romidepsin | top60_altered_DNA+RNA_seed | AKT1 | 0.598072 | THCA | TT2609-C02 | AKT inhibitor VIII |
| MVP_top60_DNA_seed | USP7 | 1.229009 | PRAD | DU-145 | P22077 | top60_altered_DNA+RNA_seed | BRAF | 0.604345 | THCA | BHT-101 | HG6-64-1 |
| MVP_top60_DNA_seed | BRAF | 1.233131 | PRAD | PC-3 | PLX-4720 | top60_altered_DNA+RNA_seed | ATM | 0.604481 | THCA | KMH-2 | KU-55933 |
| MVP_top60_DNA_seed | MAPK3 | 1.233322 | PRAD | PC-3 | Ulixertinib | top60_altered_DNA+RNA_seed | AKT1 | 0.60813 | THCA | CAL-62 | GSK690693 |
| MVP_top60_DNA_seed | PARP1 | 1.236011 | PRAD | DU-145 | Niraparib | top60_altered_DNA+RNA_seed | BRAF | 0.61027 | THCA | CAL-62 | SB590885 |
| MVP_top60_DNA_seed | PARP1 | 1.241793 | PRAD | DU-145 | Rucaparib | top60_altered_DNA+RNA_seed | BRAF | 0.623824 | THCA | RO82-W-1 | PLX-4720 |
| MVP_top60_DNA_seed | PARP1 | 1.262555 | PRAD | PC-3 | Rucaparib | top60_altered_DNA+RNA_seed | AKT1 | 0.630547 | THCA | BHT-101 | GSK690693 |
| MVP_top60_DNA_seed | ATM | 1.273546 | PRAD | 22RV1 | KU-55933 | top60_altered_DNA+RNA_seed | AKT1 | 0.632176 | THCA | 8505C | AKT inhibitor VIII |
| MVP_top60_DNA_seed | ATR | 1.28593 | PRAD | PC-3 | VE821 | top60_altered_DNA+RNA_seed | AKT1 | 0.642781 | THCA | KMH-2 | MK-2206 |
| MVP_top60_DNA_seed | ATR | 1.303361 | PRAD | VCaP | VE-822 | top60_altered_DNA+RNA_seed | AKT1 | 0.647524 | THCA | K5 | AKT inhibitor VIII |
| MVP_top60_DNA_seed | ATR | 1.309826 | PRAD | 22RV1 | VE821 | top60_altered_DNA+RNA_seed | BRAF | 0.653543 | THCA | CGTH-W-1 | SB590885 |
| MVP_top60_DNA_seed | EZH2 | 1.340623 | PRAD | DU-145 | GSK343 | top60_altered_DNA+RNA_seed | BRAF | 0.670799 | THCA | TT | HG6-64-1 |
| MVP_top60_DNA_seed | USP7 | 1.364387 | PRAD | 22RV1 | P22077 | top60_altered_DNA+RNA_seed | AKT1 | 0.67578 | THCA | TT2609-C02 | GSK690693 |
| MVP_top60_DNA_seed | EZH2 | 1.371107 | PRAD | PC-3 | GSK343 | top60_altered_DNA+RNA_seed | AKT1 | 0.710134 | THCA | FTC-133 | GSK690693 |
| MVP_top60_DNA_seed | PIK3CA | 1.404643 | PRAD | DU-145 | Buparlisib | top60_altered_DNA+RNA_seed | AKT1 | 0.721096 | THCA | HTC-C3 | GSK690693 |
| MVP_top60_DNA_seed | HDAC1 | 1.412924 | PRAD | PC-3 | PCI-34051 | top60_altered_DNA+RNA_seed | AKT1 | 0.777805 | THCA | HTC-C3 | BAY AKT1 |
| MVP_top60_DNA_seed | ESR1 | 1.435482 | PRAD | 22RV1 | GDC0810 | top60_altered_DNA+RNA_seed | AKT1 | 0.798693 | THCA | 8305C | AKT inhibitor VIII |
| MVP_top60_DNA_seed | ESR2 | 1.435482 | PRAD | 22RV1 | GDC0810 | top60_altered_DNA+RNA_seed | AKT1 | 0.805927 | THCA | B-CPAP | AKT inhibitor VIII |
| MVP_top60_DNA_seed | ATR | 1.444891 | PRAD | DU-145 | VE821 | top60_altered_DNA+RNA_seed | ATM | 0.815754 | THCA | RO82-W-1 | KU-55933 |
| MVP_top60_DNA_seed | HDAC1 | 1.453979 | PRAD | DU-145 | PCI-34051 | top60_altered_DNA+RNA_seed | BRAF | 0.831109 | THCA | ASH-3 | PLX-4720 |
| MVP_top60_DNA_seed | TP53 | 1.463099 | PRAD | PC-3 | MIRA-1 | top60_altered_DNA+RNA_seed | RET | 0.846469 | THCA | 8305C | Motesanib |
| MVP_top60_DNA_seed | ESR1 | 1.479545 | PRAD | PC-3 | Tamoxifen | top60_altered_DNA+RNA_seed | ATM | 0.876813 | THCA | FTC-133 | KU-55933 |
| MVP_top60_DNA_seed | BRAF | 1.484246 | PRAD | VCaP | PLX-4720 | top60_altered_DNA+RNA_seed | BRAF | 0.916491 | THCA | TT | PLX-4720 |
| MVP_top60_DNA_seed | BRAF | 1.522732 | PRAD | DU-145 | PLX-4720 | top60_altered_DNA+RNA_seed | AKT1 | 0.953518 | THCA | CAL-62 | MK-2206 |
| MVP_top60_DNA_seed | CDK4 | 1.540449 | PRAD | PC-3 | Palbociclib | top60_altered_DNA+RNA_seed | AKT1 | 0.984208 | THCA | ML-1 | AKT inhibitor VIII |
| MVP_top60_DNA_seed | HDAC1 | 1.544817 | PRAD | DU-145 | Entinostat | top60_altered_DNA+RNA_seed | AKT1 | 1.017811 | THCA | ML-1 | GSK690693 |
| MVP_top60_DNA_seed | CDK4 | 1.564859 | PRAD | DU-145 | Palbociclib | top60_altered_DNA+RNA_seed | ATM | 1.023871 | THCA | 8305C | CP466722 |
| MVP_top60_DNA_seed | MAPK3 | 1.638994 | PRAD | VCaP | SCH772984 | top60_altered_DNA+RNA_seed | BRAF | 1.027277 | THCA | RO82-W-1 | SB590885 |
| MVP_top60_DNA_seed | MAPK3 | 1.660256 | PRAD | DU-145 | SCH772984 | top60_altered_DNA+RNA_seed | AKT1 | 1.050863 | THCA | 8305C | GSK690693 |
| MVP_top60_DNA_seed | MAPK3 | 1.682992 | PRAD | PC-3 | Ulixertinib | top60_altered_DNA+RNA_seed | AKT1 | 1.085726 | THCA | RO82-W-1 | GSK690693 |
| MVP_top60_DNA_seed | EZH2 | 1.698386 | PRAD | PC-3 | GSK343 | top60_altered_DNA+RNA_seed | AKT1 | 1.106976 | THCA | CAL-62 | AKT inhibitor VIII |
| MVP_top60_DNA_seed | ATR | 1.774057 | PRAD | VCaP | AZD6738 | top60_altered_DNA+RNA_seed | AKT1 | 1.295044 | THCA | 8305C | AKT inhibitor VIII |
| MVP_top60_DNA_seed | PIK3CA | 1.796477 | PRAD | DU-145 | GNE-317 | top60_altered_DNA+RNA_seed | BRAF | 1.301277 | THCA | FTC-133 | HG6-64-1 |
| MVP_top60_DNA_seed | MAPK3 | 1.798863 | PRAD | DU-145 | Ulixertinib | top60_altered_DNA+RNA_seed | BRAF | 1.355515 | THCA | TT | PLX-4720 |

| | | | | | | | | | | | |
|---|---|---|---|---|---|---|---|---|---|---|---|
| MVP_top60_DNA_seed | MAPK3 | 2.081677 | PRAD | PC-3 | SCH772984 | top60_altered_DNA+RNA_seed | AKT1 | 1.413432 | THCA | TT | GSK690693 |
| MVP_top60_DNA_seed | MET | -8.579362 | STAD | MKN45 | AZD6094 | top60_altered_DNA+RNA_seed | AKT1 | 1.564793 | THCA | TT2609-C02 | BAY AKT1 |
| MVP_top60_DNA_seed | MET | -8.429196 | STAD | SNU-5 | AZD6094 | top60_altered_DNA+RNA_seed | AKT1 | 1.58178 | THCA | ML-1 | BAY AKT1 |
| MVP_top60_DNA_seed | EGFR | -7.349604 | STAD | NUGC-4 | Cetuximab | top60_altered_DNA+RNA_seed | AKT1 | 1.752216 | THCA | FTC-133 | AKT inhibitor VIII |
| MVP_top60_DNA_seed | MET | -6.178313 | STAD | Hs-746T | AZD6094 | top60_altered_DNA+RNA_seed | AKT1 | 2.049624 | THCA | RO82-W-1 | AKT inhibitor VIII |
| MVP_top60_DNA_seed | FGFR3 | -5.738215 | STAD | HSC-39 | AZD4547 | top60_altered_DNA+RNA_seed | ATM | 2.089043 | THCA | TT2609-C02 | CP466722 |
| MVP_top60_DNA_seed | ERBB2 | -5.506139 | STAD | NCI-N87 | CP724714 | top60_altered_DNA+RNA_seed | ATM | 2.326309 | THCA | RO82-W-1 | CP466722 |
| MVP_top60_DNA_seed | MET | -5.462021 | STAD | SNU-5 | PHA-665752 | top60_altered_DNA+RNA_seed | AKT1 | 2.457782 | THCA | TT | AKT inhibitor VIII |
| MVP_top60_DNA_seed | FGFR3 | -5.404401 | STAD | HSC-39 | PD173074 | top60_altered_DNA+RNA_seed | ATM | 2.462246 | THCA | FTC-133 | CP466722 |
| MVP_top60_DNA_seed | MET | -5.126293 | STAD | SNU-5 | JNJ38877605 | top60_altered_DNA+RNA_seed | ATM | 2.506771 | THCA | TT | CP466722 |
| MVP_top60_DNA_seed | FGFR3 | -5.111861 | STAD | KATOIII | PD173074 | top60_altered_DNA+RNA_seed | BRAF | -2.904049 | THCA | K5 | Dabrafenib |
| MVP_top60_DNA_seed | FGFR3 | -5.015799 | STAD | HSC-39 | AZD4547 | top60_altered_DNA+RNA_seed | BRAF | -2.070305 | THCA | BHT-101 | Dabrafenib |
| MVP_top60_DNA_seed | MET | -4.798288 | STAD | SNU-5 | Crizotinib | top60_altered_DNA+RNA_seed | BRAF | -2.008943 | THCA | K5 | PLX-4720 |
| MVP_top60_DNA_seed | MET | -4.659088 | STAD | Hs-746T | JNJ38877605 | top60_altered_DNA+RNA_seed | BRAF | -1.889949 | THCA | HTC-C3 | PLX-4720 |
| MVP_top60_DNA_seed | PDGFRA | -4.265569 | STAD | Hs-746T | Amuvatinib | top60_altered_DNA+RNA_seed | BRAF | -1.699237 | THCA | K5 | SB590885 |
| MVP_top60_DNA_seed | MET | -4.234767 | STAD | MKN45 | JNJ38877605 | top60_altered_DNA+RNA_seed | BRAF | -1.548644 | THCA | HTC-C3 | Dabrafenib |
| MVP_top60_DNA_seed | FGFR3 | -4.202287 | STAD | KATOIII | AZD4547 | top60_altered_DNA+RNA_seed | BRAF | -1.363286 | THCA | BHT-101 | PLX-4720 |
| MVP_top60_DNA_seed | ERBB2 | -4.174126 | STAD | Hs-746T | CP724714 | top60_altered_DNA+RNA_seed | BRAF | -1.326271 | THCA | 8305C | Dabrafenib |
| MVP_top60_DNA_seed | AKT1 | -4.021365 | STAD | HGC-27 | AKT inhibitor VIII | top60_altered_DNA+RNA_seed | BRAF | -1.308531 | THCA | IHH-4 | Dabrafenib |
| MVP_top60_DNA_seed | ERBB2 | -3.997646 | STAD | SK-GT-2 | Afatinib | top60_altered_DNA+RNA_seed | BRAF | -1.24812 | THCA | IHH-4 | PLX-4720 |
| MVP_top60_DNA_seed | EGFR | -3.997646 | STAD | SK-GT-2 | Afatinib | top60_altered_DNA+RNA_seed | AKT1 | -1.24793 | THCA | KMH-2 | AT13148 |
| MVP_top60_DNA_seed | ERBB2 | -3.708978 | STAD | SK-GT-2 | Afatinib | top60_altered_DNA+RNA_seed | BRAF | -1.182661 | THCA | B-CPAP | Dabrafenib |
| MVP_top60_DNA_seed | EGFR | -3.708978 | STAD | SK-GT-2 | Afatinib | top60_altered_DNA+RNA_seed | RET | -1.146579 | THCA | B-CPAP | Motesanib |
| MVP_top60_DNA_seed | IGF1R | -3.673674 | STAD | NUGC-3 | GSK1904529A | top60_altered_DNA+RNA_seed | AKT1 | -1.105186 | THCA | FTC-133 | AZD5363 |
| MVP_top60_DNA_seed | AR | -3.602727 | STAD | MKN45 | Bicalutamide | top60_altered_DNA+RNA_seed | BRAF | -1.081835 | THCA | ML-1 | PLX-4720 |
| MVP_top60_DNA_seed | ERBB2 | -3.579769 | STAD | NUGC-4 | Afatinib | top60_altered_DNA+RNA_seed | AKT1 | -1.013336 | THCA | FTC-133 | Afuresertib |
| MVP_top60_DNA_seed | EGFR | -3.579769 | STAD | NUGC-4 | Afatinib | top60_altered_DNA+RNA_seed | AKT1 | -1.004826 | THCA | K5 | AT13148 |
| MVP_top60_DNA_seed | FGFR3 | -3.547109 | STAD | SNU-16 | PD173074 | top60_altered_DNA+RNA_seed | AKT1 | -0.953548 | THCA | K5 | Afuresertib |
| MVP_top60_DNA_seed | ERBB2 | -3.19964 | STAD | NCI-N87 | Afatinib | top60_altered_DNA+RNA_seed | BRAF | -0.841682 | THCA | KMH-2 | SB590885 |
| MVP_top60_DNA_seed | EGFR | -3.19964 | STAD | NCI-N87 | Afatinib | top60_altered_DNA+RNA_seed | BRAF | -0.811101 | THCA | BHT-101 | SB590885 |
| MVP_top60_DNA_seed | MET | -3.079747 | STAD | MKN45 | SU11274 | top60_altered_DNA+RNA_seed | BRAF | -0.779147 | THCA | 8305C | PLX-4720 |
| MVP_top60_DNA_seed | FGFR3 | -3.047242 | STAD | HSC-39 | FGFR_3831 | top60_altered_DNA+RNA_seed | EHMT1 | -0.759624 | THCA | K5 | A-366 |
| MVP_top60_DNA_seed | ERBB2 | -3.012876 | STAD | NUGC-4 | AZD8931 | top60_altered_DNA+RNA_seed | EHMT1 | -0.743298 | THCA | CAL-62 | A-366 |
| MVP_top60_DNA_seed | ERBB3 | -3.012876 | STAD | NUGC-4 | AZD8931 | top60_altered_DNA+RNA_seed | BRAF | -0.729044 | THCA | B-CPAP | PLX-4720 |
| MVP_top60_DNA_seed | EGFR | -3.012876 | STAD | NUGC-4 | AZD8931 | top60_altered_DNA+RNA_seed | BRAF | -0.727578 | THCA | KMH-2 | Dabrafenib |
| MVP_top60_DNA_seed | AKT1 | -2.981395 | STAD | HGC-27 | BAY AKT1 | top60_altered_DNA+RNA_seed | AKT1 | -0.727366 | THCA | K5 | AZD5363 |
| MVP_top60_DNA_seed | EGFR | -2.858724 | STAD | NCI-N87 | Pelitinib | top60_altered_DNA+RNA_seed | BRAF | -0.724427 | THCA | KMH-2 | PLX-4720 |
| MVP_top60_DNA_seed | ERBB2 | -2.840983 | STAD | NCI-N87 | Afatinib | top60_altered_DNA+RNA_seed | BRAF | -0.706696 | THCA | 8505C | Dabrafenib |
| MVP_top60_DNA_seed | EGFR | -2.840983 | STAD | NCI-N87 | Afatinib | top60_altered_DNA+RNA_seed | AKT1 | -0.700062 | THCA | K5 | GSK2110183B |
| MVP_top60_DNA_seed | IGF1R | -2.822499 | STAD | OCUM-1 | GSK1904529A | top60_altered_DNA+RNA_seed | ATM | -0.677234 | THCA | ASH-3 | KU-55933 |
| MVP_top60_DNA_seed | IGF1R | -2.820374 | STAD | HGC-27 | GSK1904529A | top60_altered_DNA+RNA_seed | EHMT1 | -0.653728 | THCA | CGTH-W-1 | A-366 |

| | | | | | | | | | | | |
|---|---|---|---|---|---|---|---|---|---|---|---|
| MVP_top60_DNA_seed | AKT1 | -2.759774 | STAD | 23132-87 | AKT inhibitor VIII | top60_altered_DNA+RNA_seed | AKT1 | -0.634041 | THCA | ASH-3 | AT13148 |
| MVP_top60_DNA_seed | FGFR3 | -2.737636 | STAD | KATOIII | FGFR_3831 | top60_altered_DNA+RNA_seed | AKT1 | -0.626325 | THCA | BHT-101 | AT13148 |
| MVP_top60_DNA_seed | MET | -2.712269 | STAD | IM-95 | JNJ38877605 | top60_altered_DNA+RNA_seed | AKT1 | -0.614942 | THCA | FTC-133 | Ipatasertib |
| MVP_top60_DNA_seed | MAPK1 | -2.574266 | STAD | NUGC-4 | FR-180204 | top60_altered_DNA+RNA_seed | AKT1 | -0.586595 | THCA | K5 | Ipatasertib |
| MVP_top60_DNA_seed | MAPK3 | -2.574266 | STAD | NUGC-4 | FR-180204 | top60_altered_DNA+RNA_seed | AKT1 | -0.584098 | THCA | K5 | Uprosertib |
| MVP_top60_DNA_seed | PIK3CA | -2.507956 | STAD | IM-95 | AZD8835 | top60_altered_DNA+RNA_seed | AKT1 | -0.562402 | THCA | IHH-4 | Uprosertib |
| MVP_top60_DNA_seed | ERBB2 | -2.458722 | STAD | NUGC-4 | PF-00299804 | top60_altered_DNA+RNA_seed | BRAF | -0.507576 | THCA | ML-1 | Dabrafenib |
| MVP_top60_DNA_seed | ERBB4 | -2.458722 | STAD | NUGC-4 | PF-00299804 | top60_altered_DNA+RNA_seed | AKT1 | -0.488621 | THCA | ASH-3 | GSK2110183B |
| MVP_top60_DNA_seed | EGFR | -2.458722 | STAD | NUGC-4 | PF-00299804 | top60_altered_DNA+RNA_seed | ATM | -0.461576 | THCA | IHH-4 | KU-55933 |
| MVP_top60_DNA_seed | ESR1 | -2.400741 | STAD | MKN28 | Fulvestrant | top60_altered_DNA+RNA_seed | AKT1 | -0.461444 | THCA | KMH-2 | AZD5363 |
| MVP_top60_DNA_seed | EGFR | -2.291194 | STAD | GCIY | Erlotinib | top60_altered_DNA+RNA_seed | AKT1 | -0.422296 | THCA | FTC-133 | Uprosertib |
| MVP_top60_DNA_seed | FGFR3 | -2.29 | STAD | SNU-16 | FGFR_3831 | top60_altered_DNA+RNA_seed | AKT1 | -0.408279 | THCA | FTC-133 | GSK2110183B |
| MVP_top60_DNA_seed | MET | -2.285163 | STAD | Hs-746T | SU11274 | top60_altered_DNA+RNA_seed | EHMT1 | -0.400559 | THCA | IHH-4 | A-366 |
| MVP_top60_DNA_seed | ESR1 | -2.256546 | STAD | HGC-27 | Tamoxifen | top60_altered_DNA+RNA_seed | AKT1 | -0.385085 | THCA | K5 | MK-2206 |
| MVP_top60_DNA_seed | AKT1 | -2.235648 | STAD | IM-95 | AKT inhibitor VIII | top60_altered_DNA+RNA_seed | AKT1 | -0.384849 | THCA | IHH-4 | Ipatasertib |
| MVP_top60_DNA_seed | HSP90AA1 | -2.213796 | STAD | AGS | Elesclomol | top60_altered_DNA+RNA_seed | BRAF | -0.380654 | THCA | B-CPAP | SB590885 |
| MVP_top60_DNA_seed | ERBB2 | -2.202897 | STAD | SK-GT-2 | AZD8931 | top60_altered_DNA+RNA_seed | BRAF | -0.377407 | THCA | IHH-4 | SB590885 |
| MVP_top60_DNA_seed | ERBB3 | -2.202897 | STAD | SK-GT-2 | AZD8931 | top60_altered_DNA+RNA_seed | AKT1 | -0.372713 | THCA | BHT-101 | MK-2206 |
| MVP_top60_DNA_seed | EGFR | -2.202897 | STAD | SK-GT-2 | AZD8931 | top60_altered_DNA+RNA_seed | AKT1 | -0.350262 | THCA | B-CPAP | Afuresertib |
| MVP_top60_DNA_seed | HSP90AA1 | -2.199261 | STAD | FU97 | Tanespimycin | top60_altered_DNA+RNA_seed | AKT1 | -0.265183 | THCA | KMH-2 | MK-2206 |
| MVP_top60_DNA_seed | ERBB2 | -2.197683 | STAD | TGBC11TKB | CP724714 | top60_altered_DNA+RNA_seed | AKT1 | -0.259045 | THCA | KMH-2 | Afuresertib |
| MVP_top60_DNA_seed | FGFR3 | -2.175788 | STAD | FU97 | AZD4547 | top60_altered_DNA+RNA_seed | AKT1 | -0.254152 | THCA | B-CPAP | AT13148 |
| MVP_top60_DNA_seed | ERBB4 | -2.09995 | STAD | AGS | AST-1306 | top60_altered_DNA+RNA_seed | AKT1 | -0.238933 | THCA | B-CPAP | Uprosertib |
| MVP_top60_DNA_seed | EGFR | -2.09995 | STAD | AGS | AST-1306 | top60_altered_DNA+RNA_seed | BRAF | -0.238721 | THCA | CGTH-W-1 | SB590885 |
| MVP_top60_DNA_seed | EGFR | -2.099271 | STAD | NUGC-3 | Cetuximab | top60_altered_DNA+RNA_seed | BRAF | -0.188294 | THCA | CGTH-W-1 | PLX-4720 |
| MVP_top60_DNA_seed | ERBB2 | -2.084813 | STAD | AGS | PF-00299804 | top60_altered_DNA+RNA_seed | BRAF | -0.168751 | THCA | FTC-133 | Dabrafenib |
| MVP_top60_DNA_seed | ERBB4 | -2.084813 | STAD | AGS | PF-00299804 | top60_altered_DNA+RNA_seed | RET | -0.154526 | THCA | KMH-2 | Motesanib |
| MVP_top60_DNA_seed | EGFR | -2.084813 | STAD | AGS | PF-00299804 | top60_altered_DNA+RNA_seed | RET | -0.139767 | THCA | IHH-4 | Motesanib |
| MVP_top60_DNA_seed | MET | -2.077816 | STAD | HSC-39 | SU11274 | top60_altered_DNA+RNA_seed | AKT1 | -0.132233 | THCA | CAL-62 | AT13148 |
| MVP_top60_DNA_seed | EP300 | -2.053495 | STAD | NUGC-4 | I-CBP112 | top60_altered_DNA+RNA_seed | EHMT1 | -0.107764 | THCA | ASH-3 | A-366 |
| MVP_top60_DNA_seed | CREBBP | -2.053495 | STAD | NUGC-4 | I-CBP112 | top60_altered_DNA+RNA_seed | AKT1 | -0.070647 | THCA | 8305C | MK-2206 |
| MVP_top60_DNA_seed | AKT1 | -2.037163 | STAD | 23132-87 | BAY AKT1 | top60_altered_DNA+RNA_seed | AKT1 | -0.059484 | THCA | K5 | Uprosertib |
| MVP_top60_DNA_seed | MET | -2.033998 | STAD | SNU-5 | SU11274 | top60_altered_DNA+RNA_seed | AKT1 | -0.050801 | THCA | KMH-2 | Ipatasertib |
| MVP_top60_DNA_seed | AKT1 | -2.017287 | STAD | FU97 | BAY AKT1 | top60_altered_DNA+RNA_seed | AKT1 | -0.035112 | THCA | B-CPAP | AZD5363 |
| MVP_top60_DNA_seed | PIK3CA | -2.011361 | STAD | AGS | AZD8835 | top60_altered_DNA+RNA_seed | EHMT1 | -0.026972 | THCA | B-CPAP | A-366 |
| MVP_top60_DNA_seed | ERBB2 | -1.999928 | STAD | NUGC-3 | Afatinib | top60_altered_DNA+RNA_seed | AKT1 | -0.012276 | THCA | ASH-3 | Afuresertib |
| MVP_top60_DNA_seed | EGFR | -1.999928 | STAD | NUGC-3 | Afatinib | top60_altered_DNA+RNA_seed | AKT1 | 0.006181 | THCA | 8305C | AT13148 |
| MVP_top60_DNA_seed | MET | -1.973811 | STAD | KATOIII | SU11274 | top60_altered_DNA+RNA_seed | AKT1 | 0.022588 | THCA | KMH-2 | Uprosertib |
| MVP_top60_DNA_seed | PDGFRA | -1.960612 | STAD | MKN7 | Amuvatinib | top60_altered_DNA+RNA_seed | AKT1 | 0.07147 | THCA | ASH-3 | AZD5363 |
| MVP_top60_DNA_seed | EGFR | -1.938283 | STAD | NUGC-3 | Gefitinib | top60_altered_DNA+RNA_seed | ATM | 0.082146 | THCA | CGTH-W-1 | KU-55933 |
| MVP_top60_DNA_seed | HSP90AA1 | -1.922422 | STAD | HGC-27 | Tanespimycin | top60_altered_DNA+RNA_seed | AKT1 | 0.082997 | THCA | IHH-4 | MK-2206 |

| | | | | | | | | | | | |
|---|---|---|---|---|---|---|---|---|---|---|---|
| MVP_top60_DNA_seed | ERBB2 | -1.907664 | STAD | GCIY | CI-1033 | top60_altered_DNA+RNA_seed | AKT1 | 0.088719 | THCA | IHH-4 | Afuresertib |
| MVP_top60_DNA_seed | ERBB4 | -1.907664 | STAD | GCIY | CI-1033 | top60_altered_DNA+RNA_seed | AKT1 | 0.096746 | THCA | 8305C | Ipatasertib |
| MVP_top60_DNA_seed | EGFR | -1.907664 | STAD | GCIY | CI-1033 | top60_altered_DNA+RNA_seed | ATM | 0.104186 | THCA | HTC-C3 | KU-55933 |
| MVP_top60_DNA_seed | AKT1 | -1.849627 | STAD | FU97 | AKT inhibitor VIII | top60_altered_DNA+RNA_seed | ATM | 0.11089 | THCA | 8305C | KU-55933 |
| MVP_top60_DNA_seed | ERBB2 | -1.847543 | STAD | NCI-N87 | AZD8931 | top60_altered_DNA+RNA_seed | AKT1 | 0.116142 | THCA | CAL-62 | Ipatasertib |
| MVP_top60_DNA_seed | ERBB3 | -1.847543 | STAD | NCI-N87 | AZD8931 | top60_altered_DNA+RNA_seed | RET | 0.121121 | THCA | CAL-62 | Motesanib |
| MVP_top60_DNA_seed | EGFR | -1.847543 | STAD | NCI-N87 | AZD8931 | top60_altered_DNA+RNA_seed | RET | 0.158754 | THCA | ASH-3 | Motesanib |
| MVP_top60_DNA_seed | EP300 | -1.838052 | STAD | HGC-27 | I-CBP112 | top60_altered_DNA+RNA_seed | ATM | 0.16506 | THCA | KMH-2 | KU-55933 |
| MVP_top60_DNA_seed | CREBBP | -1.838052 | STAD | HGC-27 | I-CBP112 | top60_altered_DNA+RNA_seed | AKT1 | 0.172248 | THCA | ASH-3 | Ipatasertib |
| MVP_top60_DNA_seed | AKT1 | -1.818354 | STAD | FU97 | MK-2206 | top60_altered_DNA+RNA_seed | AKT1 | 0.182526 | THCA | B-CPAP | MK-2206 |
| MVP_top60_DNA_seed | IGF1R | -1.815888 | STAD | RERF-GC-1B | BMS-754807 | top60_altered_DNA+RNA_seed | AKT1 | 0.193557 | THCA | BHT-101 | Afuresertib |
| MVP_top60_DNA_seed | IGF1R | -1.812214 | STAD | SNU-16 | BMS-536924 | top60_altered_DNA+RNA_seed | AKT1 | 0.197588 | THCA | CAL-62 | Afuresertib |
| MVP_top60_DNA_seed | HSP90AA1 | -1.803018 | STAD | AGS | Tanespimycin | top60_altered_DNA+RNA_seed | EHMT1 | 0.230574 | THCA | 8505C | A-366 |
| MVP_top60_DNA_seed | IGF1R | -1.797944 | STAD | IM-95 | GSK1904529A | top60_altered_DNA+RNA_seed | RET | 0.250691 | THCA | FTC-133 | Motesanib |
| MVP_top60_DNA_seed | MET | -1.764713 | STAD | HSC-39 | JNJ38877605 | top60_altered_DNA+RNA_seed | ATM | 0.251063 | THCA | K5 | KU-55933 |
| MVP_top60_DNA_seed | HSP90AA1 | -1.764312 | STAD | OCUM-1 | CCT-018159 | top60_altered_DNA+RNA_seed | RET | 0.25247 | THCA | K5 | Motesanib |
| MVP_top60_DNA_seed | ESR1 | -1.75327 | STAD | AGS | Fulvestrant | top60_altered_DNA+RNA_seed | ATM | 0.260663 | THCA | BHT-101 | KU-55933 |
| MVP_top60_DNA_seed | ATM | -1.747041 | STAD | HSC-39 | KU-55933 | top60_altered_DNA+RNA_seed | BRAF | 0.268676 | THCA | 8505C | PLX-4720 |
| MVP_top60_DNA_seed | ERBB2 | -1.746646 | STAD | NUGC-3 | Afatinib | top60_altered_DNA+RNA_seed | BRAF | 0.269089 | THCA | CAL-62 | SB590885 |
| MVP_top60_DNA_seed | EGFR | -1.746646 | STAD | NUGC-3 | Afatinib | top60_altered_DNA+RNA_seed | AKT1 | 0.279299 | THCA | 8305C | Afuresertib |
| MVP_top60_DNA_seed | ERBB2 | -1.727366 | STAD | GCIY | PF-00299804 | top60_altered_DNA+RNA_seed | RET | 0.303606 | THCA | BHT-101 | Motesanib |
| MVP_top60_DNA_seed | ERBB4 | -1.727366 | STAD | GCIY | PF-00299804 | top60_altered_DNA+RNA_seed | AKT1 | 0.309416 | THCA | BHT-101 | Uprosertib |
| MVP_top60_DNA_seed | EGFR | -1.727366 | STAD | GCIY | PF-00299804 | top60_altered_DNA+RNA_seed | AKT1 | 0.346773 | THCA | FTC-133 | MK-2206 |
| MVP_top60_DNA_seed | HSP90AA1 | -1.726927 | STAD | RERF-GC-1B | Elesclomol | top60_altered_DNA+RNA_seed | AKT1 | 0.350825 | THCA | BHT-101 | Ipatasertib |
| MVP_top60_DNA_seed | HDAC1 | -1.712181 | STAD | FU97 | JQ12 | top60_altered_DNA+RNA_seed | EHMT1 | 0.351451 | THCA | BHT-101 | A-366 |
| MVP_top60_DNA_seed | AKT1 | -1.688103 | STAD | HGC-27 | AKT inhibitor VIII | top60_altered_DNA+RNA_seed | RET | 0.368546 | THCA | HTC-C3 | Motesanib |
| MVP_top60_DNA_seed | MDM2 | -1.6687 | STAD | NUGC-4 | Serdemetan | top60_altered_DNA+RNA_seed | BRAF | 0.372867 | THCA | FTC-133 | PLX-4720 |
| MVP_top60_DNA_seed | MET | -1.66427 | STAD | GCIY | JNJ38877605 | top60_altered_DNA+RNA_seed | BRAF | 0.386036 | THCA | ASH-3 | SB590885 |
| MVP_top60_DNA_seed | IGF1R | -1.663404 | STAD | RERF-GC-1B | BMS-536924 | top60_altered_DNA+RNA_seed | AKT1 | 0.423141 | THCA | ASH-3 | Uprosertib |
| MVP_top60_DNA_seed | AKT1 | -1.661144 | STAD | 23132-87 | MK-2206 | top60_altered_DNA+RNA_seed | RET | 0.425459 | THCA | CGTH-W-1 | Motesanib |
| MVP_top60_DNA_seed | MDM2 | -1.636256 | STAD | AGS | Nutlin-3a (-) | top60_altered_DNA+RNA_seed | AKT1 | 0.434285 | THCA | IHH-4 | GSK2110183B |
| MVP_top60_DNA_seed | ERBB2 | -1.633241 | STAD | IM-95 | AZD8931 | top60_altered_DNA+RNA_seed | AKT1 | 0.459115 | THCA | BHT-101 | Uprosertib |
| MVP_top60_DNA_seed | ERBB3 | -1.633241 | STAD | IM-95 | AZD8931 | top60_altered_DNA+RNA_seed | AKT1 | 0.475187 | THCA | BHT-101 | AZD5363 |
| MVP_top60_DNA_seed | EGFR | -1.633241 | STAD | IM-95 | AZD8931 | top60_altered_DNA+RNA_seed | ATM | 0.494355 | THCA | CAL-62 | KU-55933 |
| MVP_top60_DNA_seed | HDAC1 | -1.630624 | STAD | GCIY | Parthenolide | top60_altered_DNA+RNA_seed | AKT1 | 0.512286 | THCA | ASH-3 | MK-2206 |
| MVP_top60_DNA_seed | FGFR3 | -1.629842 | STAD | RERF-GC-1B | PD173074 | top60_altered_DNA+RNA_seed | AKT1 | 0.517844 | THCA | TT | MK-2206 |
| MVP_top60_DNA_seed | AKT1 | -1.622226 | STAD | IM-95 | AKT inhibitor VIII | top60_altered_DNA+RNA_seed | BRAF | 0.543479 | THCA | ASH-3 | Dabrafenib |
| MVP_top60_DNA_seed | MDM2 | -1.621978 | STAD | OCUM-1 | Nutlin-3a (-) | top60_altered_DNA+RNA_seed | AKT1 | 0.551077 | THCA | HTC-C3 | Uprosertib |
| MVP_top60_DNA_seed | ESR1 | -1.611139 | STAD | HSC-39 | Fulvestrant | top60_altered_DNA+RNA_seed | BRAF | 0.581894 | THCA | FTC-133 | SB590885 |
| MVP_top60_DNA_seed | EGFR | -1.611112 | STAD | NUGC-4 | Gefitinib | top60_altered_DNA+RNA_seed | AKT1 | 0.587831 | THCA | ML-1 | AT13148 |
| MVP_top60_DNA_seed | PIK3CA | -1.601479 | STAD | AGS | PI3Ka_4409 | top60_altered_DNA+RNA_seed | EHMT1 | 0.601777 | THCA | 8305C | A-366 |

| | | | | | | | | | | | |
|---|---|---|---|---|---|---|---|---|---|---|---|
| MVP_top60_DNA_seed | SRC | -1.600442 | STAD | NUGC-3 | Bosutinib | top60_altered_DNA+RNA_seed | BRAF | 0.605339 | THCA | 8305C | SB590885 |
| MVP_top60_DNA_seed | AKT1 | -1.600319 | STAD | HGC-27 | MK-2206 | top60_altered_DNA+RNA_seed | AKT1 | 0.605455 | THCA | HTC-C3 | Afuresertib |
| MVP_top60_DNA_seed | FGFR3 | -1.561759 | STAD | SNU-16 | AZD4547 | top60_altered_DNA+RNA_seed | AKT1 | 0.611823 | THCA | B-CPAP | Ipatasertib |
| MVP_top60_DNA_seed | MAPK1 | -1.544964 | STAD | RERF-GC-1B | FR-180204 | top60_altered_DNA+RNA_seed | BRAF | 0.615266 | THCA | HTC-C3 | SB590885 |
| MVP_top60_DNA_seed | MAPK3 | -1.544964 | STAD | RERF-GC-1B | FR-180204 | top60_altered_DNA+RNA_seed | AKT1 | 0.622402 | THCA | IHH-4 | AT13148 |
| MVP_top60_DNA_seed | HDAC1 | -1.544682 | STAD | RF-48 | CAY10603 | top60_altered_DNA+RNA_seed | AKT1 | 0.632152 | THCA | IHH-4 | AZD5363 |
| MVP_top60_DNA_seed | ESR1 | -1.540627 | STAD | SNU-16 | Tamoxifen | top60_altered_DNA+RNA_seed | AKT1 | 0.63469 | THCA | ASH-3 | Uprosertib |
| MVP_top60_DNA_seed | IGF1R | -1.53435 | STAD | GCIY | BMS-536924 | top60_altered_DNA+RNA_seed | AKT1 | 0.667776 | THCA | 8305C | AZD5363 |
| MVP_top60_DNA_seed | ERBB2 | -1.527312 | STAD | NUGC-4 | CI-1033 | top60_altered_DNA+RNA_seed | AKT1 | 0.682358 | THCA | IHH-4 | Uprosertib |
| MVP_top60_DNA_seed | ERBB4 | -1.527312 | STAD | NUGC-4 | CI-1033 | top60_altered_DNA+RNA_seed | ATM | 0.695164 | THCA | FTC-133 | KU-55933 |
| MVP_top60_DNA_seed | EGFR | -1.527312 | STAD | NUGC-4 | CI-1033 | top60_altered_DNA+RNA_seed | AKT1 | 0.71606 | THCA | BHT-101 | GSK2110183B |
| MVP_top60_DNA_seed | IGF1R | -1.527013 | STAD | RERF-GC-1B | Linsitinib | top60_altered_DNA+RNA_seed | AKT1 | 0.729228 | THCA | CAL-62 | AZD5363 |
| MVP_top60_DNA_seed | ERBB4 | -1.522995 | STAD | GCIY | AST-1306 | top60_altered_DNA+RNA_seed | BRAF | 0.730547 | THCA | CAL-62 | Dabrafenib |
| MVP_top60_DNA_seed | EGFR | -1.522995 | STAD | GCIY | AST-1306 | top60_altered_DNA+RNA_seed | RET | 0.739204 | THCA | 8505C | Motesanib |
| MVP_top60_DNA_seed | MET | -1.5209 | STAD | SNU-1 | SU11274 | top60_altered_DNA+RNA_seed | ATM | 0.756658 | THCA | B-CPAP | KU-55933 |
| MVP_top60_DNA_seed | HSP90AA1 | -1.514956 | STAD | NCI-N87 | Tanespimycin | top60_altered_DNA+RNA_seed | AKT1 | 0.763665 | THCA | CAL-62 | GSK2110183B |
| MVP_top60_DNA_seed | MET | -1.514015 | STAD | TGBC11TKB | JNJ38877605 | top60_altered_DNA+RNA_seed | AKT1 | 0.776184 | THCA | HTC-C3 | AZD5363 |
| MVP_top60_DNA_seed | GSK3B | -1.509191 | STAD | TMK-1 | CHIR-99021 | top60_altered_DNA+RNA_seed | EHMT1 | 0.801091 | THCA | HTC-C3 | A-366 |
| MVP_top60_DNA_seed | FGFR3 | -1.501861 | STAD | FU97 | FGFR_3831 | top60_altered_DNA+RNA_seed | AKT1 | 0.809369 | THCA | 8505C | Afuresertib |
| MVP_top60_DNA_seed | PIK3CA | -1.49039 | STAD | HSC-39 | PI3Ka_4409 | top60_altered_DNA+RNA_seed | AKT1 | 0.820935 | THCA | HTC-C3 | MK-2206 |
| MVP_top60_DNA_seed | IGF1R | -1.477709 | STAD | FU97 | GSK1904529A | top60_altered_DNA+RNA_seed | AKT1 | 0.833195 | THCA | TT | AZD5363 |
| MVP_top60_DNA_seed | HDAC1 | -1.469975 | STAD | RF-48 | AR-42 | top60_altered_DNA+RNA_seed | BRAF | 0.844063 | THCA | TT2609-C02 | SB590885 |
| MVP_top60_DNA_seed | GSK3B | -1.467752 | STAD | HSC-39 | SB216763 | top60_altered_DNA+RNA_seed | BRAF | 0.846021 | THCA | CGTH-W-1 | Dabrafenib |
| MVP_top60_DNA_seed | GSK3B | -1.466871 | STAD | SNU-16 | SB216763 | top60_altered_DNA+RNA_seed | AKT1 | 0.865932 | THCA | HTC-C3 | GSK2110183B |
| MVP_top60_DNA_seed | AKT1 | -1.461182 | STAD | SK-GT-2 | MK-2206 | top60_altered_DNA+RNA_seed | AKT1 | 0.887738 | THCA | HTC-C3 | Uprosertib |
| MVP_top60_DNA_seed | ERBB2 | -1.456809 | STAD | HSC-39 | CI-1033 | top60_altered_DNA+RNA_seed | EHMT1 | 0.898935 | THCA | FTC-133 | A-366 |
| MVP_top60_DNA_seed | ERBB4 | -1.456809 | STAD | HSC-39 | CI-1033 | top60_altered_DNA+RNA_seed | RET | 0.90392 | THCA | 8305C | Motesanib |
| MVP_top60_DNA_seed | EGFR | -1.456809 | STAD | HSC-39 | CI-1033 | top60_altered_DNA+RNA_seed | AKT1 | 0.914714 | THCA | TT2609-C02 | AT13148 |
| MVP_top60_DNA_seed | MDM2 | -1.45385 | STAD | RF-48 | Nutlin-3a (-) | top60_altered_DNA+RNA_seed | BRAF | 0.919604 | THCA | 8505C | SB590885 |
| MVP_top60_DNA_seed | SRC | -1.449113 | STAD | SNU-16 | WH-4-023 | top60_altered_DNA+RNA_seed | AKT1 | 0.927061 | THCA | 8505C | Uprosertib |
| MVP_top60_DNA_seed | SRC | -1.445176 | STAD | SNU-1 | NVP-BHG712 | top60_altered_DNA+RNA_seed | AKT1 | 0.927763 | THCA | HTC-C3 | AT13148 |
| MVP_top60_DNA_seed | HDAC1 | -1.441638 | STAD | AGS | JQ12 | top60_altered_DNA+RNA_seed | BRAF | 0.939749 | THCA | TT2609-C02 | Dabrafenib |
| MVP_top60_DNA_seed | MAPK1 | -1.420657 | STAD | AGS | VX-11e | top60_altered_DNA+RNA_seed | AKT1 | 0.955885 | THCA | CAL-62 | Uprosertib |
| MVP_top60_DNA_seed | MET | -1.386512 | STAD | TMK-1 | AZD6094 | top60_altered_DNA+RNA_seed | AKT1 | 0.980526 | THCA | CGTH-W-1 | MK-2206 |
| MVP_top60_DNA_seed | ESR1 | -1.383591 | STAD | 23132-87 | Fulvestrant | top60_altered_DNA+RNA_seed | BRAF | 1.008652 | THCA | TT | Dabrafenib |
| MVP_top60_DNA_seed | IGF1R | -1.382387 | STAD | KATOIII | BMS-536924 | top60_altered_DNA+RNA_seed | AKT1 | 1.042761 | THCA | CGTH-W-1 | Uprosertib |
| MVP_top60_DNA_seed | AKT1 | -1.382261 | STAD | NUGC-3 | AKT inhibitor VIII | top60_altered_DNA+RNA_seed | AKT1 | 1.047035 | THCA | B-CPAP | Uprosertib |
| MVP_top60_DNA_seed | GSK3B | -1.380909 | STAD | MKN28 | CHIR-99021 | top60_altered_DNA+RNA_seed | AKT1 | 1.057772 | THCA | TT | Afuresertib |
| MVP_top60_DNA_seed | HSP90AA1 | -1.346759 | STAD | AGS | CCT-018159 | top60_altered_DNA+RNA_seed | BRAF | 1.073604 | THCA | ASH-3 | PLX-4720 |
| MVP_top60_DNA_seed | HDAC1 | -1.346597 | STAD | RF-48 | Parthenolide | top60_altered_DNA+RNA_seed | AKT1 | 1.083041 | THCA | 8505C | MK-2206 |
| MVP_top60_DNA_seed | HDAC1 | -1.34113 | STAD | GCIY | JQ12 | top60_altered_DNA+RNA_seed | AKT1 | 1.100716 | THCA | CGTH-W-1 | Afuresertib |

| | | | | | | | | | | | |
|---|---|---|---|---|---|---|---|---|---|---|---|
| MVP_top60_DNA_seed | AKT1 | -1.336956 | STAD | IM-95 | GSK690693 | top60_altered_DNA+RNA_seed | AKT1 | 1.120207 | THCA | 8505C | Ipatasertib |
| MVP_top60_DNA_seed | HDAC1 | -1.336682 | STAD | HSC-39 | Tubastatin A | top60_altered_DNA+RNA_seed | AKT1 | 1.132721 | THCA | CAL-62 | MK-2206 |
| MVP_top60_DNA_seed | AKT1 | -1.33382 | STAD | SK-GT-2 | BAY AKT1 | top60_altered_DNA+RNA_seed | ATM | 1.184934 | THCA | 8505C | KU-55933 |
| MVP_top60_DNA_seed | AKT1 | -1.333458 | STAD | RF-48 | GSK690693 | top60_altered_DNA+RNA_seed | AKT1 | 1.186441 | THCA | 8505C | AZD5363 |
| MVP_top60_DNA_seed | IGF1R | -1.326116 | STAD | GCIY | BMS-754807 | top60_altered_DNA+RNA_seed | RET | 1.194374 | THCA | ML-1 | Motesanib |
| MVP_top60_DNA_seed | SRC | -1.317775 | STAD | HSC-39 | Bosutinib | top60_altered_DNA+RNA_seed | AKT1 | 1.201658 | THCA | 8505C | GSK2110183B |
| MVP_top60_DNA_seed | HDAC1 | -1.313699 | STAD | RF-48 | Belinostat | top60_altered_DNA+RNA_seed | AKT1 | 1.211165 | THCA | HTC-C3 | Ipatasertib |
| MVP_top60_DNA_seed | HSP90AA1 | -1.312553 | STAD | TMK-1 | Tanespimycin | top60_altered_DNA+RNA_seed | AKT1 | 1.212533 | THCA | B-CPAP | GSK2110183B |
| MVP_top60_DNA_seed | ERBB2 | -1.309811 | STAD | RF-48 | CUDC-101 | top60_altered_DNA+RNA_seed | AKT1 | 1.214275 | THCA | TT2609-C02 | Afuresertib |
| MVP_top60_DNA_seed | EGFR | -1.309811 | STAD | RF-48 | CUDC-101 | top60_altered_DNA+RNA_seed | BRAF | 1.232725 | THCA | CAL-62 | PLX-4720 |
| MVP_top60_DNA_seed | HSP90AA1 | -1.308562 | STAD | HGC-27 | CCT-018159 | top60_altered_DNA+RNA_seed | AKT1 | 1.284222 | THCA | CAL-62 | Uprosertib |
| MVP_top60_DNA_seed | IGF1R | -1.302914 | STAD | HSC-39 | BMS-536924 | top60_altered_DNA+RNA_seed | AKT1 | 1.323573 | THCA | CGTH-W-1 | AZD5363 |
| MVP_top60_DNA_seed | GSK3B | -1.296339 | STAD | TMK-1 | CHIR-99021 | top60_altered_DNA+RNA_seed | AKT1 | 1.327396 | THCA | 8305C | GSK2110183B |
| MVP_top60_DNA_seed | ERBB2 | -1.29334 | STAD | AGS | CI-1033 | top60_altered_DNA+RNA_seed | BRAF | 1.327751 | THCA | ML-1 | SB590885 |
| MVP_top60_DNA_seed | ERBB4 | -1.29334 | STAD | AGS | CI-1033 | top60_altered_DNA+RNA_seed | AKT1 | 1.342937 | THCA | RO82-W-1 | Uprosertib |
| MVP_top60_DNA_seed | EGFR | -1.29334 | STAD | AGS | CI-1033 | top60_altered_DNA+RNA_seed | AKT1 | 1.444889 | THCA | CGTH-W-1 | Ipatasertib |
| MVP_top60_DNA_seed | HDAC1 | -1.292905 | STAD | RF-48 | JQ12 | top60_altered_DNA+RNA_seed | AKT1 | 1.464269 | THCA | RO82-W-1 | AZD5363 |
| MVP_top60_DNA_seed | MET | -1.292054 | STAD | TGBC11TKB | SU11274 | top60_altered_DNA+RNA_seed | AKT1 | 1.483532 | THCA | CGTH-W-1 | AT13148 |
| MVP_top60_DNA_seed | MAPK1 | -1.290466 | STAD | SNU-1 | VX-11e | top60_altered_DNA+RNA_seed | BRAF | 1.50131 | THCA | TT | SB590885 |
| MVP_top60_DNA_seed | MDM2 | -1.284636 | STAD | AGS | Serdemetan | top60_altered_DNA+RNA_seed | EHMT1 | 1.504114 | THCA | TT2609-C02 | A-366 |
| MVP_top60_DNA_seed | ERBB4 | -1.278249 | STAD | SNU-1 | AST-1306 | top60_altered_DNA+RNA_seed | AKT1 | 1.532642 | THCA | 8305C | Uprosertib |
| MVP_top60_DNA_seed | EGFR | -1.278249 | STAD | SNU-1 | AST-1306 | top60_altered_DNA+RNA_seed | AKT1 | 1.552267 | THCA | TT | Uprosertib |
| MVP_top60_DNA_seed | HSP90AA1 | -1.274585 | STAD | HSC-39 | Tanespimycin | top60_altered_DNA+RNA_seed | AKT1 | 1.556382 | THCA | TT | Ipatasertib |
| MVP_top60_DNA_seed | IGF1R | -1.271007 | STAD | TMK-1 | BMS-536924 | top60_altered_DNA+RNA_seed | ATM | 1.574088 | THCA | ML-1 | KU-55933 |
| MVP_top60_DNA_seed | AR | -1.264694 | STAD | HGC-27 | Bicalutamide | top60_altered_DNA+RNA_seed | RET | 1.5804 | THCA | TT2609-C02 | Motesanib |
| MVP_top60_DNA_seed | EGFR | -1.262881 | STAD | SNU-16 | Gefitinib | top60_altered_DNA+RNA_seed | AKT1 | 1.605319 | THCA | ML-1 | Ipatasertib |
| MVP_top60_DNA_seed | AKT1 | -1.262656 | STAD | RF-48 | AKT inhibitor VIII | top60_altered_DNA+RNA_seed | AKT1 | 1.606235 | THCA | TT2609-C02 | GSK2110183B |
| MVP_top60_DNA_seed | ERBB2 | -1.248274 | STAD | GCIY | Afatinib | top60_altered_DNA+RNA_seed | BRAF | 1.632561 | THCA | RO82-W-1 | PLX-4720 |
| MVP_top60_DNA_seed | EGFR | -1.248274 | STAD | GCIY | Afatinib | top60_altered_DNA+RNA_seed | AKT1 | 1.634191 | THCA | 8505C | Uprosertib |
| MVP_top60_DNA_seed | SRC | -1.236981 | STAD | SK-GT-2 | Bosutinib | top60_altered_DNA+RNA_seed | ATM | 1.660709 | THCA | TT2609-C02 | KU-55933 |
| MVP_top60_DNA_seed | HDAC1 | -1.229597 | STAD | IM-95 | Dacinostat | top60_altered_DNA+RNA_seed | AKT1 | 1.674566 | THCA | ML-1 | AZD5363 |
| MVP_top60_DNA_seed | AKT1 | -1.221597 | STAD | OCUM-1 | MK-2206 | top60_altered_DNA+RNA_seed | AKT1 | 1.713256 | THCA | TT2609-C02 | Ipatasertib |
| MVP_top60_DNA_seed | MDM2 | -1.216493 | STAD | HSC-39 | Serdemetan | top60_altered_DNA+RNA_seed | AKT1 | 1.758311 | THCA | 8505C | AT13148 |
| MVP_top60_DNA_seed | AR | -1.216424 | STAD | FU97 | AZD3514 | top60_altered_DNA+RNA_seed | AKT1 | 1.766555 | THCA | 8305C | Uprosertib |
| MVP_top60_DNA_seed | SRC | -1.192309 | STAD | GCIY | Saracatinib | top60_altered_DNA+RNA_seed | AKT1 | 1.802675 | THCA | TT2609-C02 | Uprosertib |
| MVP_top60_DNA_seed | HDAC1 | -1.190515 | STAD | RF-48 | Dacinostat | top60_altered_DNA+RNA_seed | AKT1 | 1.812247 | THCA | ML-1 | MK-2206 |
| MVP_top60_DNA_seed | ERBB2 | -1.180036 | STAD | NUGC-4 | CUDC-101 | top60_altered_DNA+RNA_seed | RET | 1.819942 | THCA | TT | Motesanib |
| MVP_top60_DNA_seed | EGFR | -1.180036 | STAD | NUGC-4 | CUDC-101 | top60_altered_DNA+RNA_seed | ATM | 1.880356 | THCA | RO82-W-1 | KU-55933 |
| MVP_top60_DNA_seed | SRC | -1.179583 | STAD | NUGC-4 | NVP-BHG712 | top60_altered_DNA+RNA_seed | BRAF | 1.917072 | THCA | TT2609-C02 | PLX-4720 |
| MVP_top60_DNA_seed | FGFR3 | -1.161364 | STAD | FU97 | AZD4547 | top60_altered_DNA+RNA_seed | AKT1 | 1.933148 | THCA | RO82-W-1 | AT13148 |
| MVP_top60_DNA_seed | HSP90AA1 | -1.16034 | STAD | MKN45 | Tanespimycin | top60_altered_DNA+RNA_seed | AKT1 | 1.960196 | THCA | RO82-W-1 | GSK2110183B |

| | | | | | | | | | | | |
|---|---|---|---|---|---|---|---|---|---|---|---|
| MVP_top60_DNA_seed | HSP90AA1 | -1.158613 | STAD | SNU-1 | SNX-2112 | top60_altered_DNA+RNA_seed | AKT1 | 1.96079 | THCA | RO82-W-1 | MK-2206 |
| MVP_top60_DNA_seed | HDAC1 | -1.151827 | STAD | HGC-27 | Dacinostat | top60_altered_DNA+RNA_seed | AKT1 | 1.993555 | THCA | RO82-W-1 | Afuresertib |
| MVP_top60_DNA_seed | ATM | -1.147537 | STAD | TMK-1 | KU-60019 | top60_altered_DNA+RNA_seed | BRAF | 2.011685 | THCA | TT | PLX-4720 |
| MVP_top60_DNA_seed | HSP90AA1 | -1.146672 | STAD | RF-48 | SNX-2112 | top60_altered_DNA+RNA_seed | AKT1 | 2.025232 | THCA | TT2609-C02 | AZD5363 |
| MVP_top60_DNA_seed | MDM2 | -1.143196 | STAD | NUGC-3 | Serdemetan | top60_altered_DNA+RNA_seed | BRAF | 2.030313 | THCA | RO82-W-1 | Dabrafenib |
| MVP_top60_DNA_seed | ATM | -1.121906 | STAD | GCIY | KU-55933 | top60_altered_DNA+RNA_seed | EHMT1 | 2.122321 | THCA | TT | A-366 |
| MVP_top60_DNA_seed | HSP90AA1 | -1.116909 | STAD | NUGC-3 | Elesclomol | top60_altered_DNA+RNA_seed | ATM | 2.159676 | THCA | TT | KU-55933 |
| MVP_top60_DNA_seed | SRC | -1.111899 | STAD | KATOIII | HG-6-71-01 | top60_altered_DNA+RNA_seed | AKT1 | 2.230667 | THCA | TT2609-C02 | Uprosertib |
| MVP_top60_DNA_seed | IGF1R | -1.11013 | STAD | NUGC-4 | GSK1904529A | top60_altered_DNA+RNA_seed | AKT1 | 2.300894 | THCA | ML-1 | Uprosertib |
| MVP_top60_DNA_seed | MDM2 | -1.105876 | STAD | IM-95 | Nutlin-3a (-) | top60_altered_DNA+RNA_seed | BRAF | 2.359152 | THCA | RO82-W-1 | SB590885 |
| MVP_top60_DNA_seed | AKT1 | -1.094234 | STAD | TMK-1 | MK-2206 | top60_altered_DNA+RNA_seed | AKT1 | 2.420424 | THCA | TT2609-C02 | MK-2206 |
| MVP_top60_DNA_seed | ERBB2 | -1.09391 | STAD | NCI-N87 | CUDC-101 | top60_altered_DNA+RNA_seed | AKT1 | 2.431171 | THCA | TT | AT13148 |
| MVP_top60_DNA_seed | EGFR | -1.09391 | STAD | NCI-N87 | CUDC-101 | top60_altered_DNA+RNA_seed | AKT1 | 2.510429 | THCA | RO82-W-1 | Ipatasertib |
| MVP_top60_DNA_seed | ATM | -1.092186 | STAD | NUGC-3 | KU-55933 | top60_altered_DNA+RNA_seed | RET | 2.530202 | THCA | RO82-W-1 | Motesanib |
| MVP_top60_DNA_seed | ERBB2 | -1.08654 | STAD | SNU-1 | PF-00299804 | top60_altered_DNA+RNA_seed | AKT1 | 2.83111 | THCA | RO82-W-1 | Uprosertib |
| MVP_top60_DNA_seed | ERBB4 | -1.08654 | STAD | SNU-1 | PF-00299804 | top60_altered_DNA+RNA_seed | AKT1 | 2.837742 | THCA | ML-1 | Afuresertib |
| MVP_top60_DNA_seed | EGFR | -1.08654 | STAD | SNU-1 | PF-00299804 | top60_altered_DNA+RNA_seed | EHMT1 | 3.201287 | THCA | RO82-W-1 | A-366 |
| MVP_top60_DNA_seed | SRC | -1.086301 | STAD | GCIY | WH-4-023 | top60_altered_DNA+RNA_seed | FGFR2 | -3.839158 | UCEC | MFE-280 | AZD4547 |
| MVP_top60_DNA_seed | AKT1 | -1.085057 | STAD | AGS | MK-2206 | top60_altered_DNA+RNA_seed | FGFR2 | -3.193966 | UCEC | AN3-CA | PD173074 |
| MVP_top60_DNA_seed | EGFR | -1.08236 | STAD | IM-95 | Gefitinib | top60_altered_DNA+RNA_seed | FGFR2 | -3.138837 | UCEC | AN3-CA | AZD4547 |
| MVP_top60_DNA_seed | HDAC1 | -1.081741 | STAD | Hs-746T | AR-42 | top60_altered_DNA+RNA_seed | FGFR2 | -2.831998 | UCEC | AN3-CA | AZD4547 |
| MVP_top60_DNA_seed | HSP90AA1 | -1.080852 | STAD | HGC-27 | Elesclomol | top60_altered_DNA+RNA_seed | AKT3 | -2.540572 | UCEC | COLO-684 | GSK690693 |
| MVP_top60_DNA_seed | ERBB4 | -1.076841 | STAD | HSC-39 | AST-1306 | top60_altered_DNA+RNA_seed | FGFR2 | -2.286659 | UCEC | MFE-296 | AZD4547 |
| MVP_top60_DNA_seed | EGFR | -1.076841 | STAD | HSC-39 | AST-1306 | top60_altered_DNA+RNA_seed | FGFR2 | -2.095604 | UCEC | MFE-280 | FGFR_3831 |
| MVP_top60_DNA_seed | SRC | -1.076297 | STAD | HSC-39 | WH-4-023 | top60_altered_DNA+RNA_seed | AKT3 | -2.045663 | UCEC | AN3-CA | AKT inhibitor VIII |
| MVP_top60_DNA_seed | ERBB4 | -1.076218 | STAD | NUGC-4 | AST-1306 | top60_altered_DNA+RNA_seed | AKT3 | -2.045604 | UCEC | MFE-296 | AKT inhibitor VIII |
| MVP_top60_DNA_seed | EGFR | -1.076218 | STAD | NUGC-4 | AST-1306 | top60_altered_DNA+RNA_seed | FGFR2 | -1.92556 | UCEC | MFE-296 | AZD4547 |
| MVP_top60_DNA_seed | HDAC1 | -1.072515 | STAD | Hs-746T | Tubastatin A | top60_altered_DNA+RNA_seed | FGFR2 | -1.813677 | UCEC | MFE-296 | PD173074 |
| MVP_top60_DNA_seed | SRC | -1.069815 | STAD | RF-48 | NVP-BHG712 | top60_altered_DNA+RNA_seed | AKT3 | -1.667957 | UCEC | COLO-684 | AKT inhibitor VIII |
| MVP_top60_DNA_seed | MET | -1.066652 | STAD | NUGC-4 | SU11274 | top60_altered_DNA+RNA_seed | PIK3CA | -1.502812 | UCEC | MFE-280 | PI3Ka_4409 |
| MVP_top60_DNA_seed | ESR1 | -1.064554 | STAD | HSC-39 | Tamoxifen | top60_altered_DNA+RNA_seed | AKT3 | -1.406257 | UCEC | MFE-296 | BAY AKT1 |
| MVP_top60_DNA_seed | AR | -1.062613 | STAD | TGBC11TKB | Bicalutamide | top60_altered_DNA+RNA_seed | AKT3 | -1.390049 | UCEC | SNG-M | GSK690693 |
| MVP_top60_DNA_seed | GSK3B | -1.061191 | STAD | NUGC-4 | KIN001-042 | top60_altered_DNA+RNA_seed | PIK3CA | -1.306172 | UCEC | SNG-M | AZD8835 |
| MVP_top60_DNA_seed | HSP90AA1 | -1.057396 | STAD | SNU-5 | Tanespimycin | top60_altered_DNA+RNA_seed | AKT3 | -1.264695 | UCEC | MFE-296 | GSK690693 |
| MVP_top60_DNA_seed | MET | -1.048543 | STAD | IM-95 | SU11274 | top60_altered_DNA+RNA_seed | AKT3 | -1.262466 | UCEC | AN3-CA | BAY AKT1 |
| MVP_top60_DNA_seed | HDAC1 | -1.047527 | STAD | IM-95 | JQ12 | top60_altered_DNA+RNA_seed | AKT3 | -1.253918 | UCEC | COLO-684 | A-443654 |
| MVP_top60_DNA_seed | MDM2 | -1.042684 | STAD | MKN45 | Nutlin-3a (-) | top60_altered_DNA+RNA_seed | PIK3CA | -1.182536 | UCEC | MFE-280 | AZD8835 |
| MVP_top60_DNA_seed | EGFR | -1.041277 | STAD | IM-95 | Pelitinib | top60_altered_DNA+RNA_seed | FGFR2 | -1.154504 | UCEC | MFE-280 | PD173074 |
| MVP_top60_DNA_seed | AR | -1.040687 | STAD | HSC-39 | Bicalutamide | top60_altered_DNA+RNA_seed | ATM | -1.138256 | UCEC | MFE-319 | KU-55933 |
| MVP_top60_DNA_seed | ERBB2 | -1.040137 | STAD | NUGC-4 | Afatinib | top60_altered_DNA+RNA_seed | AKT3 | -1.136416 | UCEC | MFE-280 | GSK690693 |
| MVP_top60_DNA_seed | EGFR | -1.040137 | STAD | NUGC-4 | Afatinib | top60_altered_DNA+RNA_seed | PIK3CA | -1.065015 | UCEC | SNG-M | PI3Ka_4409 |

| | | | | | | | | | | | |
|---|---|---|---|---|---|---|---|---|---|---|---|
| MVP_top60_DNA_seed | HSP90AA1 | -1.03792 | STAD | SNU-1 | Luminespib | top60_altered_DNA+RNA_seed | FGFR2 | -1.007139 | UCEC | AN3-CA | FGFR_3831 |
| MVP_top60_DNA_seed | HSP90AA1 | -1.023576 | STAD | HGC-27 | Luminespib | top60_altered_DNA+RNA_seed | ATM | -0.98435 | UCEC | SNG-M | CP466722 |
| MVP_top60_DNA_seed | ESR1 | -1.019678 | STAD | FU97 | Tamoxifen | top60_altered_DNA+RNA_seed | ATM | -0.967779 | UCEC | MFE-319 | KU-60019 |
| MVP_top60_DNA_seed | ERBB2 | -1.019476 | STAD | NUGC-3 | AZD8931 | top60_altered_DNA+RNA_seed | ATM | -0.941115 | UCEC | MFE-319 | CP466722 |
| MVP_top60_DNA_seed | ERBB3 | -1.019476 | STAD | NUGC-3 | AZD8931 | top60_altered_DNA+RNA_seed | AKT3 | -0.883088 | UCEC | MFE-319 | AKT inhibitor VIII |
| MVP_top60_DNA_seed | EGFR | -1.019476 | STAD | NUGC-3 | AZD8931 | top60_altered_DNA+RNA_seed | AKT3 | -0.863762 | UCEC | COLO-684 | BAY AKT1 |
| MVP_top60_DNA_seed | AKT1 | -1.012918 | STAD | 23132-87 | AKT inhibitor VIII | top60_altered_DNA+RNA_seed | FGFR2 | -0.837374 | UCEC | MFE-296 | FGFR_3831 |
| MVP_top60_DNA_seed | HSP90AA1 | -1.010232 | STAD | MKN28 | Luminespib | top60_altered_DNA+RNA_seed | AKT3 | -0.778096 | UCEC | KLE | AKT inhibitor VIII |
| MVP_top60_DNA_seed | HSP90AA1 | -1.00609 | STAD | OCUM-1 | Tanespimycin | top60_altered_DNA+RNA_seed | ATM | -0.757481 | UCEC | HEC-1 | KU-60019 |
| MVP_top60_DNA_seed | IGF1R | -1.005483 | STAD | HSC-39 | BMS-536924 | top60_altered_DNA+RNA_seed | AKT3 | -0.750188 | UCEC | HEC-1 | AKT inhibitor VIII |
| MVP_top60_DNA_seed | ATM | -0.996951 | STAD | AGS | KU-60019 | top60_altered_DNA+RNA_seed | PIK3CA | -0.695396 | UCEC | AN3-CA | AZD8835 |
| MVP_top60_DNA_seed | ERBB2 | -0.995298 | STAD | AGS | AZD8931 | top60_altered_DNA+RNA_seed | AKT3 | -0.694469 | UCEC | KLE | A-443654 |
| MVP_top60_DNA_seed | ERBB3 | -0.995298 | STAD | AGS | AZD8931 | top60_altered_DNA+RNA_seed | AKT3 | -0.686103 | UCEC | MFE-280 | AKT inhibitor VIII |
| MVP_top60_DNA_seed | EGFR | -0.995298 | STAD | AGS | AZD8931 | top60_altered_DNA+RNA_seed | ATM | -0.608482 | UCEC | COLO-684 | KU-55933 |
| MVP_top60_DNA_seed | MET | -0.987911 | STAD | GCIY | SU11274 | top60_altered_DNA+RNA_seed | ATM | -0.589341 | UCEC | MFE-296 | KU-60019 |
| MVP_top60_DNA_seed | MET | -0.984822 | STAD | HSC-39 | AZD6094 | top60_altered_DNA+RNA_seed | ATM | -0.540231 | UCEC | AN3-CA | KU-55933 |
| MVP_top60_DNA_seed | ESR1 | -0.980581 | STAD | NUGC-4 | Fulvestrant | top60_altered_DNA+RNA_seed | ATM | -0.532342 | UCEC | COLO-684 | CP466722 |
| MVP_top60_DNA_seed | EGFR | -0.975714 | STAD | SNU-1 | Gefitinib | top60_altered_DNA+RNA_seed | AKT3 | -0.505071 | UCEC | AN3-CA | GSK690693 |
| MVP_top60_DNA_seed | PDGFRA | -0.975156 | STAD | HSC-39 | Masitinib | top60_altered_DNA+RNA_seed | AKT3 | -0.483919 | UCEC | MFE-319 | AKT inhibitor VIII |
| MVP_top60_DNA_seed | EGFR | -0.974989 | STAD | RF-48 | Gefitinib | top60_altered_DNA+RNA_seed | AKT3 | -0.469321 | UCEC | EN | GSK690693 |
| MVP_top60_DNA_seed | HSP90AA1 | -0.972028 | STAD | HSC-39 | CCT-018159 | top60_altered_DNA+RNA_seed | ATM | -0.458963 | UCEC | AN3-CA | KU-60019 |
| MVP_top60_DNA_seed | ERBB2 | -0.968535 | STAD | HSC-39 | PF-00299804 | top60_altered_DNA+RNA_seed | AKT3 | -0.400349 | UCEC | KLE | BAY AKT1 |
| MVP_top60_DNA_seed | ERBB4 | -0.968535 | STAD | HSC-39 | PF-00299804 | top60_altered_DNA+RNA_seed | PIK3CA | -0.392543 | UCEC | AN3-CA | PI3Ka_4409 |
| MVP_top60_DNA_seed | EGFR | -0.968535 | STAD | HSC-39 | PF-00299804 | top60_altered_DNA+RNA_seed | AKT3 | -0.381262 | UCEC | SNG-M | AKT inhibitor VIII |
| MVP_top60_DNA_seed | AKT1 | -0.964502 | STAD | HSC-39 | BAY AKT1 | top60_altered_DNA+RNA_seed | AKT3 | -0.366585 | UCEC | AN3-CA | AKT inhibitor VIII |
| MVP_top60_DNA_seed | HSP90AA1 | -0.963105 | STAD | NUGC-3 | CCT-018159 | top60_altered_DNA+RNA_seed | FGFR2 | -0.355229 | UCEC | SNG-M | FGFR_3831 |
| MVP_top60_DNA_seed | AR | -0.960509 | STAD | HGC-27 | Bicalutamide | top60_altered_DNA+RNA_seed | FGFR2 | -0.342315 | UCEC | COLO-684 | AZD4547 |
| MVP_top60_DNA_seed | HSP90AA1 | -0.958702 | STAD | AGS | Luminespib | top60_altered_DNA+RNA_seed | AKT3 | -0.310089 | UCEC | KLE | GSK690693 |
| MVP_top60_DNA_seed | PIK3CA | -0.955002 | STAD | HSC-39 | AZD8835 | top60_altered_DNA+RNA_seed | AKT3 | -0.273754 | UCEC | COLO-684 | AKT inhibitor VIII |
| MVP_top60_DNA_seed | EGFR | -0.954623 | STAD | GCIY | Gefitinib | top60_altered_DNA+RNA_seed | AKT3 | -0.254252 | UCEC | KLE | AKT inhibitor VIII |
| MVP_top60_DNA_seed | AR | -0.949409 | STAD | HGC-27 | AZD3514 | top60_altered_DNA+RNA_seed | AKT3 | -0.248601 | UCEC | EN | BAY AKT1 |
| MVP_top60_DNA_seed | ERBB4 | -0.947594 | STAD | NUGC-3 | AST-1306 | top60_altered_DNA+RNA_seed | ATM | -0.244126 | UCEC | HEC-1 | KU-55933 |
| MVP_top60_DNA_seed | EGFR | -0.947594 | STAD | NUGC-3 | AST-1306 | top60_altered_DNA+RNA_seed | AKT3 | -0.204435 | UCEC | SNG-M | BAY AKT1 |
| MVP_top60_DNA_seed | ERBB2 | -0.940647 | STAD | SNU-1 | CI-1033 | top60_altered_DNA+RNA_seed | AKT3 | -0.152611 | UCEC | SNG-M | AKT inhibitor VIII |
| MVP_top60_DNA_seed | ERBB4 | -0.940647 | STAD | SNU-1 | CI-1033 | top60_altered_DNA+RNA_seed | PIK3CA | -0.137169 | UCEC | COLO-684 | AZD8835 |
| MVP_top60_DNA_seed | EGFR | -0.940647 | STAD | SNU-1 | CI-1033 | top60_altered_DNA+RNA_seed | AKT3 | -0.132474 | UCEC | MFE-319 | BAY AKT1 |
| MVP_top60_DNA_seed | HSP90AA1 | -0.935116 | STAD | NUGC-3 | Tanespimycin | top60_altered_DNA+RNA_seed | FGFR2 | -0.071005 | UCEC | HEC-1 | AZD4547 |
| MVP_top60_DNA_seed | HSP90AA1 | -0.934868 | STAD | 23132-87 | Tanespimycin | top60_altered_DNA+RNA_seed | FGFR2 | -0.061881 | UCEC | HEC-1 | PD173074 |
| MVP_top60_DNA_seed | AR | -0.934838 | STAD | FU97 | Bicalutamide | top60_altered_DNA+RNA_seed | ATM | -0.054838 | UCEC | KLE | CP466722 |
| MVP_top60_DNA_seed | ERBB2 | -0.930647 | STAD | RERF-GC-1B | CP724714 | top60_altered_DNA+RNA_seed | ATM | -0.049526 | UCEC | COLO-684 | KU-60019 |
| MVP_top60_DNA_seed | GSK3B | -0.929689 | STAD | HGC-27 | SB216763 | top60_altered_DNA+RNA_seed | AKT3 | -0.04317 | UCEC | EN | AKT inhibitor VIII |

| | | | | | | | | | | | |
|---|---|---|---|---|---|---|---|---|---|---|---|
| MVP_top60_DNA_seed | MDM2 | -0.926926 | STAD | SNU-1 | Nutlin-3a (-) | top60_altered_DNA+RNA_seed | PIK3CA | -0.004274 | UCEC | KLE | PI3Ka_4409 |
| MVP_top60_DNA_seed | ATM | -0.916841 | STAD | IM-95 | KU-60019 | top60_altered_DNA+RNA_seed | PIK3CA | 0.002161 | UCEC | MFE-296 | AZD8835 |
| MVP_top60_DNA_seed | HSP90AA1 | -0.915427 | STAD | MKN28 | Elesclomol | top60_altered_DNA+RNA_seed | PIK3CA | 0.007765 | UCEC | HEC-1 | PI3Ka_4409 |
| MVP_top60_DNA_seed | HSP90AA1 | -0.910834 | STAD | TMK-1 | Elesclomol | top60_altered_DNA+RNA_seed | FGFR2 | 0.102188 | UCEC | COLO-684 | PD173074 |
| MVP_top60_DNA_seed | ERBB2 | -0.910739 | STAD | GCIY | Lapatinib | top60_altered_DNA+RNA_seed | FGFR2 | 0.105396 | UCEC | SNG-M | AZD4547 |
| MVP_top60_DNA_seed | EGFR | -0.910739 | STAD | GCIY | Lapatinib | top60_altered_DNA+RNA_seed | ATM | 0.113108 | UCEC | MFE-296 | KU-55933 |
| MVP_top60_DNA_seed | HDAC1 | -0.910279 | STAD | RF-48 | Tubastatin A | top60_altered_DNA+RNA_seed | PIK3CA | 0.154596 | UCEC | HEC-1 | AZD8835 |
| MVP_top60_DNA_seed | GSK3B | -0.908126 | STAD | KATOIII | CHIR-99021 | top60_altered_DNA+RNA_seed | ATM | 0.192287 | UCEC | EN | CP466722 |
| MVP_top60_DNA_seed | AKT1 | -0.907878 | STAD | FU97 | AKT inhibitor VIII | top60_altered_DNA+RNA_seed | FGFR2 | 0.220404 | UCEC | EN | FGFR_3831 |
| MVP_top60_DNA_seed | ATM | -0.906994 | STAD | SNU-1 | KU-55933 | top60_altered_DNA+RNA_seed | FGFR2 | 0.233763 | UCEC | MFE-319 | PD173074 |
| MVP_top60_DNA_seed | AKT1 | -0.903383 | STAD | HSC-39 | MK-2206 | top60_altered_DNA+RNA_seed | FGFR2 | 0.254212 | UCEC | MFE-319 | FGFR_3831 |
| MVP_top60_DNA_seed | AR | -0.903054 | STAD | MKN28 | AZD3514 | top60_altered_DNA+RNA_seed | AKT3 | 0.277979 | UCEC | MFE-296 | AKT inhibitor VIII |
| MVP_top60_DNA_seed | HDAC1 | -0.895767 | STAD | AGS | Dacinostat | top60_altered_DNA+RNA_seed | ATM | 0.291618 | UCEC | AN3-CA | CP466722 |
| MVP_top60_DNA_seed | EGFR | -0.891589 | STAD | SNU-5 | Gefitinib | top60_altered_DNA+RNA_seed | FGFR2 | 0.300365 | UCEC | HEC-1 | AZD4547 |
| MVP_top60_DNA_seed | FGFR3 | -0.888254 | STAD | AGS | FGFR_3831 | top60_altered_DNA+RNA_seed | AKT3 | 0.327086 | UCEC | MFE-280 | AKT inhibitor VIII |
| MVP_top60_DNA_seed | FGFR3 | -0.885002 | STAD | SNU-1 | PD173074 | top60_altered_DNA+RNA_seed | AKT3 | 0.327418 | UCEC | MFE-319 | GSK690693 |
| MVP_top60_DNA_seed | HDAC1 | -0.884229 | STAD | TMK-1 | JQ12 | top60_altered_DNA+RNA_seed | ATM | 0.330904 | UCEC | MFE-280 | CP466722 |
| MVP_top60_DNA_seed | HDAC1 | -0.880577 | STAD | HGC-27 | JQ12 | top60_altered_DNA+RNA_seed | ATM | 0.340159 | UCEC | MFE-296 | CP466722 |
| MVP_top60_DNA_seed | ERBB2 | -0.876224 | STAD | SNU-1 | CP724714 | top60_altered_DNA+RNA_seed | FGFR2 | 0.344305 | UCEC | COLO-684 | FGFR_3831 |
| MVP_top60_DNA_seed | MET | -0.875671 | STAD | KATOIII | JNJ38877605 | top60_altered_DNA+RNA_seed | PIK3CA | 0.352447 | UCEC | COLO-684 | PI3Ka_4409 |
| MVP_top60_DNA_seed | AKT1 | -0.868331 | STAD | AGS | AKT inhibitor VIII | top60_altered_DNA+RNA_seed | FGFR2 | 0.369629 | UCEC | MFE-319 | AZD4547 |
| MVP_top60_DNA_seed | MDM2 | -0.865043 | STAD | TMK-1 | Serdemetan | top60_altered_DNA+RNA_seed | ATM | 0.375809 | UCEC | SNG-M | KU-60019 |
| MVP_top60_DNA_seed | AKT1 | -0.861828 | STAD | Hs-746T | GSK690693 | top60_altered_DNA+RNA_seed | PIK3CA | 0.384848 | UCEC | MFE-296 | PI3Ka_4409 |
| MVP_top60_DNA_seed | IGF1R | -0.860508 | STAD | IM-95 | BMS-754807 | top60_altered_DNA+RNA_seed | FGFR2 | 0.441923 | UCEC | SNG-M | PD173074 |
| MVP_top60_DNA_seed | AKT1 | -0.859178 | STAD | TMK-1 | AKT inhibitor VIII | top60_altered_DNA+RNA_seed | ATM | 0.458843 | UCEC | SNG-M | KU-55933 |
| MVP_top60_DNA_seed | MET | -0.85186 | STAD | HGC-27 | AZD6094 | top60_altered_DNA+RNA_seed | ATM | 0.493472 | UCEC | MFE-280 | KU-55933 |
| MVP_top60_DNA_seed | ERBB2 | -0.845644 | STAD | HGC-27 | CP724714 | top60_altered_DNA+RNA_seed | PIK3CA | 0.563782 | UCEC | MFE-319 | AZD8835 |
| MVP_top60_DNA_seed | ESR1 | -0.843524 | STAD | FU97 | Fulvestrant | top60_altered_DNA+RNA_seed | ATM | 0.599189 | UCEC | KLE | KU-60019 |
| MVP_top60_DNA_seed | GSK3B | -0.842097 | STAD | FU97 | KIN001-042 | top60_altered_DNA+RNA_seed | FGFR2 | 0.612214 | UCEC | EN | AZD4547 |
| MVP_top60_DNA_seed | ERBB2 | -0.839761 | STAD | RERF-GC-1B | Afatinib | top60_altered_DNA+RNA_seed | FGFR2 | 0.647232 | UCEC | SNG-M | AZD4547 |
| MVP_top60_DNA_seed | EGFR | -0.839761 | STAD | RERF-GC-1B | Afatinib | top60_altered_DNA+RNA_seed | FGFR2 | 0.75276 | UCEC | COLO-684 | AZD4547 |
| MVP_top60_DNA_seed | IGF1R | -0.832431 | STAD | SNU-16 | BMS-536924 | top60_altered_DNA+RNA_seed | ATM | 0.765162 | UCEC | EN | KU-55933 |
| MVP_top60_DNA_seed | FGFR3 | -0.830987 | STAD | HGC-27 | AZD4547 | top60_altered_DNA+RNA_seed | AKT3 | 0.782152 | UCEC | HEC-1 | GSK690693 |
| MVP_top60_DNA_seed | MAPK1 | -0.82765 | STAD | RERF-GC-1B | VX-11e | top60_altered_DNA+RNA_seed | FGFR2 | 0.786376 | UCEC | MFE-319 | AZD4547 |
| MVP_top60_DNA_seed | HSP90AA1 | -0.825112 | STAD | KATOIII | Tanespimycin | top60_altered_DNA+RNA_seed | ATM | 0.811403 | UCEC | EN | KU-60019 |
| MVP_top60_DNA_seed | ATM | -0.82301 | STAD | HSC-39 | KU-60019 | top60_altered_DNA+RNA_seed | AKT3 | 0.887171 | UCEC | HEC-1 | AKT inhibitor VIII |
| MVP_top60_DNA_seed | SRC | -0.820092 | STAD | AGS | NVP-BHG712 | top60_altered_DNA+RNA_seed | FGFR2 | 0.917464 | UCEC | HEC-1 | FGFR_3831 |
| MVP_top60_DNA_seed | PDGFRA | -0.8197 | STAD | GCIY | Amuvatinib | top60_altered_DNA+RNA_seed | ATM | 0.972207 | UCEC | KLE | KU-55933 |
| MVP_top60_DNA_seed | HDAC1 | -0.816639 | STAD | Hs-746T | CAY10603 | top60_altered_DNA+RNA_seed | FGFR2 | 1.011234 | UCEC | KLE | AZD4547 |
| MVP_top60_DNA_seed | AKT1 | -0.808116 | STAD | RF-48 | AKT inhibitor VIII | top60_altered_DNA+RNA_seed | FGFR2 | 1.067797 | UCEC | EN | AZD4547 |
| MVP_top60_DNA_seed | HSP90AA1 | -0.800069 | STAD | SK-GT-2 | CCT-018159 | top60_altered_DNA+RNA_seed | FGFR2 | 1.159924 | UCEC | EN | PD173074 |

| | | | | | | | | | | | |
|---|---|---|---|---|---|---|---|---|---|---|---|
| MVP_top60_DNA_seed | ATM | -0.798055 | STAD | FU97 | KU-55933 | top60_altered_DNA+RNA_seed | PIK3CA | 1.18858 | UCEC | MFE-319 | PI3Ka_4409 |
| MVP_top60_DNA_seed | HSP90AA1 | -0.796301 | STAD | 23132-87 | CCT-018159 | top60_altered_DNA+RNA_seed | PIK3CA | 1.254999 | UCEC | KLE | AZD8835 |
| MVP_top60_DNA_seed | GSK3B | -0.793201 | STAD | HGC-27 | KIN001-042 | top60_altered_DNA+RNA_seed | FGFR2 | 1.300353 | UCEC | KLE | FGFR_3831 |
| MVP_top60_DNA_seed | HSP90AA1 | -0.786842 | STAD | AGS | SNX-2112 | top60_altered_DNA+RNA_seed | AKT3 | 1.329107 | UCEC | HEC-1 | BAY AKT1 |
| MVP_top60_DNA_seed | GSK3B | -0.781636 | STAD | NUGC-4 | CHIR-99021 | top60_altered_DNA+RNA_seed | ATM | 1.411894 | UCEC | MFE-280 | KU-60019 |
| MVP_top60_DNA_seed | GSK3B | -0.781365 | STAD | MKN28 | SB216763 | top60_altered_DNA+RNA_seed | ATM | 1.419228 | UCEC | HEC-1 | CP466722 |
| MVP_top60_DNA_seed | EP300 | -0.773719 | STAD | TGBC11TKB | I-CBP112 | top60_altered_DNA+RNA_seed | FGFR2 | 1.445917 | UCEC | MFE-280 | AZD4547 |
| MVP_top60_DNA_seed | CREBBP | -0.773719 | STAD | TGBC11TKB | I-CBP112 | top60_altered_DNA+RNA_seed | FGFR2 | 1.631694 | UCEC | KLE | PD173074 |
| MVP_top60_DNA_seed | PIK3CA | -0.771988 | STAD | NCI-N87 | PI3Ka_4409 | top60_altered_DNA+RNA_seed | PIK3CA | 1.727071 | UCEC | EN | PI3Ka_4409 |
| MVP_top60_DNA_seed | HSP90AA1 | -0.765266 | STAD | RF-48 | Luminespib | top60_altered_DNA+RNA_seed | AKT3 | 1.884926 | UCEC | EN | AKT inhibitor VIII |
| MVP_top60_DNA_seed | IGF1R | -0.764149 | STAD | TMK-1 | Linsitinib | top60_altered_DNA+RNA_seed | FGFR2 | 2.092606 | UCEC | KLE | AZD4547 |
| MVP_top60_DNA_seed | ERBB2 | -0.75817 | STAD | MKN7 | Afatinib | top60_altered_DNA+RNA_seed | FGFR2 | -1.681912 | UCEC | AN3-CA | PD173074 |
| MVP_top60_DNA_seed | EGFR | -0.75817 | STAD | MKN7 | Afatinib | top60_altered_DNA+RNA_seed | AKT3 | -1.538701 | UCEC | AN3-CA | GSK2110183B |
| MVP_top60_DNA_seed | MDM2 | -0.751632 | STAD | Hs-746T | Serdemetan | top60_altered_DNA+RNA_seed | AKT3 | -1.461307 | UCEC | COLO-684 | GSK2110183B |
| MVP_top60_DNA_seed | HDAC1 | -0.750773 | STAD | OCUM-1 | Dacinostat | top60_altered_DNA+RNA_seed | FGFR2 | -1.455099 | UCEC | AN3-CA | AZD4547 |
| MVP_top60_DNA_seed | HSP90AA1 | -0.748233 | STAD | SNU-5 | CCT-018159 | top60_altered_DNA+RNA_seed | AKT3 | -1.441089 | UCEC | AN3-CA | Uprosertib |
| MVP_top60_DNA_seed | ERBB2 | -0.747106 | STAD | RF-48 | Afatinib | top60_altered_DNA+RNA_seed | PIK3CA | -1.433814 | UCEC | MFE-280 | Alpelisib |
| MVP_top60_DNA_seed | EGFR | -0.747106 | STAD | RF-48 | Afatinib | top60_altered_DNA+RNA_seed | AKT3 | -1.393069 | UCEC | EN | AZD5363 |
| MVP_top60_DNA_seed | IGF1R | -0.746269 | STAD | FU97 | BMS-536924 | top60_altered_DNA+RNA_seed | AKT3 | -1.363351 | UCEC | AN3-CA | Ipatasertib |
| MVP_top60_DNA_seed | GSK3B | -0.74462 | STAD | HSC-39 | AZD7969 | top60_altered_DNA+RNA_seed | AKT3 | -1.33634 | UCEC | EN | GSK2110183B |
| MVP_top60_DNA_seed | FGFR3 | -0.743141 | STAD | MKN45 | PD173074 | top60_altered_DNA+RNA_seed | AKT3 | -1.322297 | UCEC | COLO-684 | Uprosertib |
| MVP_top60_DNA_seed | AKT1 | -0.742596 | STAD | 23132-87 | GSK690693 | top60_altered_DNA+RNA_seed | AKT3 | -1.246177 | UCEC | COLO-684 | Afuresertib |
| MVP_top60_DNA_seed | MAPK1 | -0.739715 | STAD | IM-95 | FR-180204 | top60_altered_DNA+RNA_seed | AKT3 | -1.225323 | UCEC | EN | Afuresertib |
| MVP_top60_DNA_seed | MAPK3 | -0.739715 | STAD | IM-95 | FR-180204 | top60_altered_DNA+RNA_seed | AKT3 | -1.197812 | UCEC | EN | Uprosertib |
| MVP_top60_DNA_seed | PIK3CA | -0.739382 | STAD | SNU-5 | PI3Ka_4409 | top60_altered_DNA+RNA_seed | AKT3 | -1.173212 | UCEC | AN3-CA | AZD5363 |
| MVP_top60_DNA_seed | MAPK1 | -0.73702 | STAD | IM-95 | VX-11e | top60_altered_DNA+RNA_seed | PIK3CA | -1.129591 | UCEC | AN3-CA | Alpelisib |
| MVP_top60_DNA_seed | PDGFRA | -0.733506 | STAD | RERF-GC-1B | Masitinib | top60_altered_DNA+RNA_seed | PIK3CA | -1.128491 | UCEC | KLE | AZD8186 |
| MVP_top60_DNA_seed | HSP90AA1 | -0.731187 | STAD | TGBC11TKB | Tanespimycin | top60_altered_DNA+RNA_seed | KRAS | -1.093533 | UCEC | SNG-M | [illegible]RAS (G12C) Inhibitor-1 |
| MVP_top60_DNA_seed | AKT1 | -0.725855 | STAD | HSC-39 | AKT inhibitor VIII | top60_altered_DNA+RNA_seed | ATM | -1.082897 | UCEC | MFE-319 | KU-55933 |
| MVP_top60_DNA_seed | EGFR | -0.721985 | STAD | GCIY | Pelitinib | top60_altered_DNA+RNA_seed | AKT3 | -1.069252 | UCEC | AN3-CA | Uprosertib |
| MVP_top60_DNA_seed | GSK3B | -0.718633 | STAD | SNU-1 | SB216763 | top60_altered_DNA+RNA_seed | PIK3CA | -1.05072 | UCEC | AN3-CA | GNE-317 |
| MVP_top60_DNA_seed | EGFR | -0.715991 | STAD | Hs-746T | Pelitinib | top60_altered_DNA+RNA_seed | AKT3 | -1.034058 | UCEC | MFE-280 | Uprosertib |
| MVP_top60_DNA_seed | AKT1 | -0.710851 | STAD | Hs-746T | AKT inhibitor VIII | top60_altered_DNA+RNA_seed | AKT3 | -1.029426 | UCEC | EN | Ipatasertib |
| MVP_top60_DNA_seed | SRC | -0.707845 | STAD | HGC-27 | Bosutinib | top60_altered_DNA+RNA_seed | PIK3CA | -0.986437 | UCEC | AN3-CA | AZD8186 |
| MVP_top60_DNA_seed | ATM | -0.707712 | STAD | IM-95 | CP466722 | top60_altered_DNA+RNA_seed | ATM | -0.980089 | UCEC | AN3-CA | KU-55933 |
| MVP_top60_DNA_seed | FGFR3 | -0.704993 | STAD | 23132-87 | AZD4547 | top60_altered_DNA+RNA_seed | FGFR2 | -0.977111 | UCEC | MFE-280 | AZD4547 |
| MVP_top60_DNA_seed | HDAC1 | -0.700793 | STAD | RERF-GC-1B | Dacinostat | top60_altered_DNA+RNA_seed | AKT3 | -0.948204 | UCEC | COLO-684 | Ipatasertib |
| MVP_top60_DNA_seed | AKT1 | -0.699987 | STAD | HSC-39 | GSK690693 | top60_altered_DNA+RNA_seed | AKT3 | -0.939066 | UCEC | COLO-684 | Uprosertib |
| MVP_top60_DNA_seed | ATM | -0.6981 | STAD | HGC-27 | KU-60019 | top60_altered_DNA+RNA_seed | AKT3 | -0.918815 | UCEC | MFE-296 | Uprosertib |
| MVP_top60_DNA_seed | SRC | -0.697758 | STAD | IM-95 | NVP-BHG712 | top60_altered_DNA+RNA_seed | AKT3 | -0.877355 | UCEC | COLO-684 | AZD5363 |
| MVP_top60_DNA_seed | MDM2 | -0.697361 | STAD | FU97 | Serdemetan | top60_altered_DNA+RNA_seed | PIK3CA | -0.799668 | UCEC | AN3-CA | Buparlisib |

| | | | | | | | | | | | |
|---|---|---|---|---|---|---|---|---|---|---|---|
| MVP_top60_DNA_seed | PDGFRA | -0.696992 | STAD | HGC-27 | Amuvatinib | top60_altered_DNA+RNA_seed | FGFR2 | -0.787283 | UCEC | MFE-296 | AZD4547 |
| MVP_top60_DNA_seed | GSK3B | -0.689565 | STAD | GCIY | KIN001-042 | top60_altered_DNA+RNA_seed | PIK3CA | -0.776155 | UCEC | MFE-319 | GNE-317 |
| MVP_top60_DNA_seed | HSP90AA1 | -0.68734 | STAD | SK-GT-2 | Elesclomol | top60_altered_DNA+RNA_seed | AKT3 | -0.771595 | UCEC | MFE-280 | Uprosertib |
| MVP_top60_DNA_seed | ERBB2 | -0.687259 | STAD | TMK-1 | Afatinib | top60_altered_DNA+RNA_seed | AKT3 | -0.752001 | UCEC | AN3-CA | Afuresertib |
| MVP_top60_DNA_seed | EGFR | -0.687259 | STAD | TMK-1 | Afatinib | top60_altered_DNA+RNA_seed | AKT3 | -0.693632 | UCEC | MFE-319 | Uprosertib |
| MVP_top60_DNA_seed | AKT1 | -0.686 | STAD | SNU-1 | AKT inhibitor VIII | top60_altered_DNA+RNA_seed | PIK3CA | -0.682345 | UCEC | COLO-684 | GNE-317 |
| MVP_top60_DNA_seed | SRC | -0.68444 | STAD | KATOIII | Bosutinib | top60_altered_DNA+RNA_seed | ATM | -0.677619 | UCEC | MFE-280 | KU-55933 |
| MVP_top60_DNA_seed | HDAC1 | -0.68208 | STAD | NCI-N87 | AR-42 | top60_altered_DNA+RNA_seed | AKT3 | -0.647068 | UCEC | MFE-319 | Ipatasertib |
| MVP_top60_DNA_seed | HSP90AA1 | -0.680041 | STAD | IM-95 | SNX-2112 | top60_altered_DNA+RNA_seed | AKT3 | -0.623315 | UCEC | MFE-296 | Afuresertib |
| MVP_top60_DNA_seed | HSP90AA1 | -0.678794 | STAD | TMK-1 | CCT-018159 | top60_altered_DNA+RNA_seed | AKT3 | -0.609343 | UCEC | MFE-296 | Uprosertib |
| MVP_top60_DNA_seed | PDGFRA | -0.677449 | STAD | HGC-27 | Masitinib | top60_altered_DNA+RNA_seed | TP53 | -0.609038 | UCEC | AN3-CA | MIRA-1 |
| MVP_top60_DNA_seed | SRC | -0.676536 | STAD | RERF-GC-1B | NVP-BHG712 | top60_altered_DNA+RNA_seed | AKT3 | -0.595333 | UCEC | EN | Uprosertib |
| MVP_top60_DNA_seed | AR | -0.673015 | STAD | GCIY | Bicalutamide | top60_altered_DNA+RNA_seed | AKT3 | -0.587729 | UCEC | MFE-280 | GSK2110183B |
| MVP_top60_DNA_seed | HDAC1 | -0.672049 | STAD | MKN28 | JQ12 | top60_altered_DNA+RNA_seed | AKT3 | -0.556221 | UCEC | MFE-280 | Ipatasertib |
| MVP_top60_DNA_seed | IGF1R | -0.671629 | STAD | MKN28 | GSK1904529A | top60_altered_DNA+RNA_seed | PIK3CA | -0.553514 | UCEC | COLO-684 | Alpelisib |
| MVP_top60_DNA_seed | ERBB2 | -0.66765 | STAD | 23132-87 | CUDC-101 | top60_altered_DNA+RNA_seed | AKT3 | -0.515101 | UCEC | MFE-319 | Uprosertib |
| MVP_top60_DNA_seed | EGFR | -0.66765 | STAD | 23132-87 | CUDC-101 | top60_altered_DNA+RNA_seed | FGFR2 | -0.441738 | UCEC | MFE-319 | AZD4547 |
| MVP_top60_DNA_seed | MAPK1 | -0.664754 | STAD | MKN28 | FR-180204 | top60_altered_DNA+RNA_seed | PIK3CA | -0.402583 | UCEC | COLO-684 | AZD8186 |
| MVP_top60_DNA_seed | MAPK3 | -0.664754 | STAD | MKN28 | FR-180204 | top60_altered_DNA+RNA_seed | AKT3 | -0.385786 | UCEC | MFE-296 | Ipatasertib |
| MVP_top60_DNA_seed | IGF1R | -0.660689 | STAD | HGC-27 | BMS-754807 | top60_altered_DNA+RNA_seed | AKT3 | -0.375179 | UCEC | MFE-319 | AZD5363 |
| MVP_top60_DNA_seed | AKT1 | -0.660567 | STAD | SNU-1 | A-443654 | top60_altered_DNA+RNA_seed | KRAS | -0.368617 | UCEC | HEC-1 | :RAS (G12C) Inhibitor-1 |
| MVP_top60_DNA_seed | FGFR3 | -0.658334 | STAD | RERF-GC-1B | AZD4547 | top60_altered_DNA+RNA_seed | AKT3 | -0.338625 | UCEC | MFE-319 | Afuresertib |
| MVP_top60_DNA_seed | FGFR3 | -0.655139 | STAD | HGC-27 | PD173074 | top60_altered_DNA+RNA_seed | FGFR2 | -0.337154 | UCEC | MFE-280 | PD173074 |
| MVP_top60_DNA_seed | GSK3B | -0.652984 | STAD | 23132-87 | AZD7969 | top60_altered_DNA+RNA_seed | PIK3CA | -0.300771 | UCEC | KLE | GNE-317 |
| MVP_top60_DNA_seed | ERBB2 | -0.650746 | STAD | IM-95 | CP724714 | top60_altered_DNA+RNA_seed | PIK3CA | -0.277586 | UCEC | EN | Alpelisib |
| MVP_top60_DNA_seed | IGF1R | -0.650201 | STAD | NUGC-4 | BMS-536924 | top60_altered_DNA+RNA_seed | KRAS | -0.264657 | UCEC | EN | :RAS (G12C) Inhibitor-1 |
| MVP_top60_DNA_seed | ERBB2 | -0.648542 | STAD | GCIY | CP724714 | top60_altered_DNA+RNA_seed | AKT3 | -0.24562 | UCEC | SNG-M | AZD5363 |
| MVP_top60_DNA_seed | HDAC1 | -0.645869 | STAD | NCI-N87 | Belinostat | top60_altered_DNA+RNA_seed | AKT3 | -0.239528 | UCEC | MFE-280 | AZD5363 |
| MVP_top60_DNA_seed | MET | -0.645854 | STAD | SNU-1 | Crizotinib | top60_altered_DNA+RNA_seed | PIK3CA | -0.22301 | UCEC | MFE-280 | GNE-317 |
| MVP_top60_DNA_seed | FGFR3 | -0.640145 | STAD | FU97 | PD173074 | top60_altered_DNA+RNA_seed | KRAS | -0.204616 | UCEC | COLO-684 | :RAS (G12C) Inhibitor-1 |
| MVP_top60_DNA_seed | EGFR | -0.637361 | STAD | TMK-1 | Gefitinib | top60_altered_DNA+RNA_seed | AKT3 | -0.187275 | UCEC | MFE-280 | Afuresertib |
| MVP_top60_DNA_seed | MDM2 | -0.632946 | STAD | NUGC-4 | Nutlin-3a (-) | top60_altered_DNA+RNA_seed | AKT3 | -0.156282 | UCEC | KLE | Uprosertib |
| MVP_top60_DNA_seed | ATM | -0.628394 | STAD | RF-48 | CP466722 | top60_altered_DNA+RNA_seed | ATM | -0.154584 | UCEC | MFE-296 | KU-55933 |
| MVP_top60_DNA_seed | AR | -0.62623 | STAD | 23132-87 | Bicalutamide | top60_altered_DNA+RNA_seed | FGFR2 | -0.151758 | UCEC | SNG-M | PD173074 |
| MVP_top60_DNA_seed | MDM2 | -0.623944 | STAD | HGC-27 | Serdemetan | top60_altered_DNA+RNA_seed | ATM | -0.127018 | UCEC | SNG-M | KU-55933 |
| MVP_top60_DNA_seed | HDAC1 | -0.623534 | STAD | NUGC-3 | JQ12 | top60_altered_DNA+RNA_seed | PIK3CA | -0.117276 | UCEC | MFE-319 | Buparlisib |
| MVP_top60_DNA_seed | GSK3B | -0.616659 | STAD | NCI-N87 | CHIR-99021 | top60_altered_DNA+RNA_seed | AKT3 | -0.095928 | UCEC | MFE-296 | AZD5363 |
| MVP_top60_DNA_seed | SRC | -0.615531 | STAD | NUGC-4 | Bosutinib | top60_altered_DNA+RNA_seed | PIK3CA | -0.092167 | UCEC | MFE-280 | AZD8186 |
| MVP_top60_DNA_seed | ATM | -0.611452 | STAD | TMK-1 | torin2 | top60_altered_DNA+RNA_seed | PIK3CA | -0.035895 | UCEC | COLO-684 | Buparlisib |
| MVP_top60_DNA_seed | HDAC1 | -0.611059 | STAD | Hs-746T | Belinostat | top60_altered_DNA+RNA_seed | KRAS | -0.031796 | UCEC | AN3-CA | :RAS (G12C) Inhibitor-1 |
| MVP_top60_DNA_seed | ATM | -0.606572 | STAD | TMK-1 | KU-55933 | top60_altered_DNA+RNA_seed | PIK3CA | 0.004545 | UCEC | EN | Buparlisib |

| | | | | | | | | | | | |
|---|---|---|---|---|---|---|---|---|---|---|---|
| MVP_top60_DNA_seed | EGFR | -0.605785 | STAD | SK-GT-2 | Gefitinib | top60_altered_DNA+RNA_seed | FGFR2 | 0.011596 | UCEC | MFE-296 | PD173074 |
| MVP_top60_DNA_seed | ERBB2 | -0.605117 | STAD | IM-95 | CI-1033 | top60_altered_DNA+RNA_seed | AKT3 | 0.028776 | UCEC | SNG-M | Uprosertib |
| MVP_top60_DNA_seed | ERBB4 | -0.605117 | STAD | IM-95 | CI-1033 | top60_altered_DNA+RNA_seed | AKT3 | 0.073645 | UCEC | SNG-M | Ipatasertib |
| MVP_top60_DNA_seed | EGFR | -0.605117 | STAD | IM-95 | CI-1033 | top60_altered_DNA+RNA_seed | PIK3CA | 0.121978 | UCEC | KLE | Buparlisib |
| MVP_top60_DNA_seed | ERBB4 | -0.604596 | STAD | RERF-GC-1B | AST-1306 | top60_altered_DNA+RNA_seed | TP53 | 0.125506 | UCEC | SNG-M | MIRA-1 |
| MVP_top60_DNA_seed | EGFR | -0.604596 | STAD | RERF-GC-1B | AST-1306 | top60_altered_DNA+RNA_seed | PIK3CA | 0.139196 | UCEC | EN | GNE-317 |
| MVP_top60_DNA_seed | GSK3B | -0.603758 | STAD | IM-95 | CHIR-99021 | top60_altered_DNA+RNA_seed | TP53 | 0.157663 | UCEC | MFE-296 | MIRA-1 |
| MVP_top60_DNA_seed | ERBB2 | -0.602136 | STAD | MKN28 | AZD8931 | top60_altered_DNA+RNA_seed | TP53 | 0.169217 | UCEC | COLO-684 | MIRA-1 |
| MVP_top60_DNA_seed | ERBB3 | -0.602136 | STAD | MKN28 | AZD8931 | top60_altered_DNA+RNA_seed | FGFR2 | 0.204251 | UCEC | KLE | PD173074 |
| MVP_top60_DNA_seed | EGFR | -0.602136 | STAD | MKN28 | AZD8931 | top60_altered_DNA+RNA_seed | PIK3CA | 0.230603 | UCEC | SNG-M | Alpelisib |
| MVP_top60_DNA_seed | HDAC1 | -0.597844 | STAD | FU97 | Dacinostat | top60_altered_DNA+RNA_seed | KRAS | 0.232402 | UCEC | MFE-319 | KRAS (G12C) Inhibitor-1 |
| MVP_top60_DNA_seed | ESR1 | -0.585544 | STAD | SNU-16 | Fulvestrant | top60_altered_DNA+RNA_seed | PIK3CA | 0.320685 | UCEC | MFE-319 | AZD8186 |
| MVP_top60_DNA_seed | ESR1 | -0.585061 | STAD | KATOIII | Tamoxifen | top60_altered_DNA+RNA_seed | AKT3 | 0.336334 | UCEC | SNG-M | GSK2110183B |
| MVP_top60_DNA_seed | PIK3CA | -0.58183 | STAD | 23132-87 | AZD8835 | top60_altered_DNA+RNA_seed | ATM | 0.35817 | UCEC | COLO-684 | KU-55933 |
| MVP_top60_DNA_seed | ERBB2 | -0.576329 | STAD | Hs-746T | CUDC-101 | top60_altered_DNA+RNA_seed | PIK3CA | 0.376584 | UCEC | SNG-M | GNE-317 |
| MVP_top60_DNA_seed | EGFR | -0.576329 | STAD | Hs-746T | CUDC-101 | top60_altered_DNA+RNA_seed | PIK3CA | 0.414572 | UCEC | SNG-M | Buparlisib |
| MVP_top60_DNA_seed | ERBB2 | -0.576186 | STAD | KATOIII | PF-00299804 | top60_altered_DNA+RNA_seed | KRAS | 0.426664 | UCEC | MFE-280 | KRAS (G12C) Inhibitor-1 |
| MVP_top60_DNA_seed | ERBB4 | -0.576186 | STAD | KATOIII | PF-00299804 | top60_altered_DNA+RNA_seed | AKT3 | 0.437204 | UCEC | MFE-319 | GSK2110183B |
| MVP_top60_DNA_seed | EGFR | -0.576186 | STAD | KATOIII | PF-00299804 | top60_altered_DNA+RNA_seed | FGFR2 | 0.454732 | UCEC | SNG-M | AZD4547 |
| MVP_top60_DNA_seed | HSP90AA1 | -0.573396 | STAD | HGC-27 | SNX-2112 | top60_altered_DNA+RNA_seed | PIK3CA | 0.485906 | UCEC | HEC-1 | GNE-317 |
| MVP_top60_DNA_seed | IGF1R | -0.569122 | STAD | 23132-87 | GSK1904529A | top60_altered_DNA+RNA_seed | AKT3 | 0.538987 | UCEC | KLE | AZD5363 |
| MVP_top60_DNA_seed | ESR1 | -0.56609 | STAD | TGBC11TKB | Tamoxifen | top60_altered_DNA+RNA_seed | PIK3CA | 0.543186 | UCEC | MFE-280 | Buparlisib |
| MVP_top60_DNA_seed | AR | -0.565261 | STAD | TGBC11TKB | Bicalutamide | top60_altered_DNA+RNA_seed | TP53 | 0.56048 | UCEC | MFE-319 | MIRA-1 |
| MVP_top60_DNA_seed | MAPK1 | -0.564627 | STAD | MKN45 | VX-11e | top60_altered_DNA+RNA_seed | AKT3 | 0.56375 | UCEC | SNG-M | Afuresertib |
| MVP_top60_DNA_seed | FGFR3 | -0.558379 | STAD | NUGC-3 | FGFR_3831 | top60_altered_DNA+RNA_seed | FGFR2 | 0.620923 | UCEC | HEC-1 | AZD4547 |
| MVP_top60_DNA_seed | ATM | -0.556842 | STAD | TGBC11TKB | KU-55933 | top60_altered_DNA+RNA_seed | ATM | 0.65422 | UCEC | EN | KU-55933 |
| MVP_top60_DNA_seed | FGFR3 | -0.556164 | STAD | RERF-GC-1B | AZD4547 | top60_altered_DNA+RNA_seed | AKT3 | 0.675819 | UCEC | MFE-296 | GSK2110183B |
| MVP_top60_DNA_seed | ATM | -0.553314 | STAD | 23132-87 | KU-55933 | top60_altered_DNA+RNA_seed | AKT3 | 0.702212 | UCEC | HEC-1 | AZD5363 |
| MVP_top60_DNA_seed | ATM | -0.549809 | STAD | GCIY | CP466722 | top60_altered_DNA+RNA_seed | AKT3 | 0.713039 | UCEC | SNG-M | Uprosertib |
| MVP_top60_DNA_seed | ATM | -0.547039 | STAD | MKN7 | KU-55933 | top60_altered_DNA+RNA_seed | TP53 | 0.730928 | UCEC | EN | MIRA-1 |
| MVP_top60_DNA_seed | PIK3CA | -0.544564 | STAD | NCI-N87 | AZD8835 | top60_altered_DNA+RNA_seed | FGFR2 | 0.750459 | UCEC | COLO-684 | AZD4547 |
| MVP_top60_DNA_seed | PDGFRA | -0.53974 | STAD | FU97 | Amuvatinib | top60_altered_DNA+RNA_seed | AKT3 | 0.812305 | UCEC | HEC-1 | GSK2110183B |
| MVP_top60_DNA_seed | AR | -0.537592 | STAD | IM-95 | Bicalutamide | top60_altered_DNA+RNA_seed | PIK3CA | 0.816956 | UCEC | MFE-296 | Buparlisib |
| MVP_top60_DNA_seed | ERBB2 | -0.53759 | STAD | RERF-GC-1B | CI-1033 | top60_altered_DNA+RNA_seed | KRAS | 0.824168 | UCEC | KLE | KRAS (G12C) Inhibitor-1 |
| MVP_top60_DNA_seed | ERBB4 | -0.53759 | STAD | RERF-GC-1B | CI-1033 | top60_altered_DNA+RNA_seed | TP53 | 0.860405 | UCEC | MFE-280 | MIRA-1 |
| MVP_top60_DNA_seed | EGFR | -0.53759 | STAD | RERF-GC-1B | CI-1033 | top60_altered_DNA+RNA_seed | ATM | 0.891619 | UCEC | HEC-1 | KU-55933 |
| MVP_top60_DNA_seed | MET | -0.537532 | STAD | SNU-16 | AZD6094 | top60_altered_DNA+RNA_seed | KRAS | 0.892833 | UCEC | MFE-296 | KRAS (G12C) Inhibitor-1 |
| MVP_top60_DNA_seed | ATM | -0.533504 | STAD | SNU-16 | KU-55933 | top60_altered_DNA+RNA_seed | PIK3CA | 0.916666 | UCEC | HEC-1 | Buparlisib |
| MVP_top60_DNA_seed | GSK3B | -0.528123 | STAD | KATOIII | SB216763 | top60_altered_DNA+RNA_seed | PIK3CA | 0.94508 | UCEC | EN | AZD8186 |
| MVP_top60_DNA_seed | FGFR3 | -0.524533 | STAD | RF-48 | PD173074 | top60_altered_DNA+RNA_seed | FGFR2 | 0.946602 | UCEC | EN | AZD4547 |
| MVP_top60_DNA_seed | MDM2 | -0.523194 | STAD | SK-GT-2 | Serdemetan | top60_altered_DNA+RNA_seed | PIK3CA | 0.953357 | UCEC | MFE-319 | Alpelisib |

| | | | | | | | | | | | |
|---|---|---|---|---|---|---|---|---|---|---|---|
| MVP_top60_DNA_seed | HSP90AA1 | -0.518581 | STAD | NCI-N87 | SNX-2112 | top60_altered_DNA+RNA_seed | AKT3 | 0.976671 | UCEC | KLE | Ipatasertib |
| MVP_top60_DNA_seed | HSP90AA1 | -0.517949 | STAD | TGBC11TKB | SNX-2112 | top60_altered_DNA+RNA_seed | PIK3CA | 1.02174 | UCEC | HEC-1 | AZD8186 |
| MVP_top60_DNA_seed | GSK3B | -0.515859 | STAD | HSC-39 | KIN001-042 | top60_altered_DNA+RNA_seed | AKT3 | 1.043738 | UCEC | HEC-1 | Afuresertib |
| MVP_top60_DNA_seed | AKT1 | -0.515213 | STAD | RF-48 | A-443654 | top60_altered_DNA+RNA_seed | PIK3CA | 1.043925 | UCEC | SNG-M | AZD8186 |
| MVP_top60_DNA_seed | MET | -0.515105 | STAD | MKN7 | SU11274 | top60_altered_DNA+RNA_seed | TP53 | 1.069404 | UCEC | HEC-1 | MIRA-1 |
| MVP_top60_DNA_seed | SRC | -0.51228 | STAD | HGC-27 | NVP-BHG712 | top60_altered_DNA+RNA_seed | PIK3CA | 1.070358 | UCEC | MFE-296 | Alpelisib |
| MVP_top60_DNA_seed | ERBB2 | -0.507793 | STAD | NUGC-4 | CP724714 | top60_altered_DNA+RNA_seed | PIK3CA | 1.09639 | UCEC | MFE-296 | AZD8186 |
| MVP_top60_DNA_seed | MET | -0.502076 | STAD | MKN7 | JNJ38877605 | top60_altered_DNA+RNA_seed | AKT3 | 1.182879 | UCEC | KLE | Uprosertib |
| MVP_top60_DNA_seed | GSK3B | -0.501703 | STAD | HGC-27 | CHIR-99021 | top60_altered_DNA+RNA_seed | PIK3CA | 1.200719 | UCEC | HEC-1 | Alpelisib |
| MVP_top60_DNA_seed | FGFR3 | -0.495512 | STAD | NUGC-3 | AZD4547 | top60_altered_DNA+RNA_seed | AKT3 | 1.229678 | UCEC | HEC-1 | Uprosertib |
| MVP_top60_DNA_seed | GSK3B | -0.495159 | STAD | MKN7 | SB216763 | top60_altered_DNA+RNA_seed | FGFR2 | 1.238673 | UCEC | EN | PD173074 |
| MVP_top60_DNA_seed | HDAC1 | -0.494925 | STAD | GCIY | Dacinostat | top60_altered_DNA+RNA_seed | FGFR2 | 1.252341 | UCEC | KLE | AZD4547 |
| MVP_top60_DNA_seed | ERBB2 | -0.490537 | STAD | SNU-1 | CUDC-101 | top60_altered_DNA+RNA_seed | AKT3 | 1.264935 | UCEC | HEC-1 | Ipatasertib |
| MVP_top60_DNA_seed | EGFR | -0.490537 | STAD | SNU-1 | CUDC-101 | top60_altered_DNA+RNA_seed | AKT3 | 1.368983 | UCEC | KLE | Afuresertib |
| MVP_top60_DNA_seed | ATM | -0.486763 | STAD | SNU-1 | CP466722 | top60_altered_DNA+RNA_seed | FGFR2 | 1.456139 | UCEC | MFE-319 | PD173074 |
| MVP_top60_DNA_seed | MET | -0.486593 | STAD | KATOIII | AZD6094 | top60_altered_DNA+RNA_seed | ATM | 1.460351 | UCEC | KLE | KU-55933 |
| MVP_top60_DNA_seed | AKT1 | -0.482983 | STAD | NCI-N87 | AKT inhibitor VIII | top60_altered_DNA+RNA_seed | AKT3 | 1.572434 | UCEC | KLE | GSK2110183B |
| MVP_top60_DNA_seed | GSK3B | -0.480581 | STAD | NUGC-4 | AZD7969 | top60_altered_DNA+RNA_seed | TP53 | 1.5776 | UCEC | KLE | MIRA-1 |
| MVP_top60_DNA_seed | MDM2 | -0.479252 | STAD | 23132-87 | Nutlin-3a (-) | top60_altered_DNA+RNA_seed | PIK3CA | 1.64178 | UCEC | KLE | Alpelisib |
| MVP_top60_DNA_seed | HSP90AA1 | -0.479096 | STAD | GCIY | Luminespib | top60_altered_DNA+RNA_seed | PIK3CA | 1.651226 | UCEC | MFE-296 | GNE-317 |
| MVP_top60_DNA_seed | HSP90AA1 | -0.472617 | STAD | FU97 | CCT-018159 | top60_altered_DNA+RNA_seed | AKT3 | 1.729271 | UCEC | HEC-1 | Uprosertib |
| MVP_top60_DNA_seed | GSK3B | -0.471032 | STAD | TGBC11TKB | KIN001-042 | top60_altered_DNA+RNA_seed | FGFR2 | 1.811944 | UCEC | COLO-684 | PD173074 |
| MVP_top60_DNA_seed | AKT1 | -0.469945 | STAD | MKN28 | AKT inhibitor VIII | top60_altered_DNA+RNA_seed | FGFR2 | 1.915425 | UCEC | HEC-1 | PD173074 |
| MVP_top60_DNA_seed | ATM | -0.467569 | STAD | GCIY | KU-60019 | | | | | | |
| MVP_top60_DNA_seed | IGF1R | -0.466854 | STAD | IM-95 | Linsitinib | | | | | | |
| MVP_top60_DNA_seed | AR | -0.466296 | STAD | AGS | Bicalutamide | | | | | | |
| MVP_top60_DNA_seed | AR | -0.462654 | STAD | SK-GT-2 | Bicalutamide | | | | | | |
| MVP_top60_DNA_seed | HDAC1 | -0.461909 | STAD | NCI-N87 | CAY10603 | | | | | | |
| MVP_top60_DNA_seed | ERBB2 | -0.458125 | STAD | AGS | CUDC-101 | | | | | | |
| MVP_top60_DNA_seed | EGFR | -0.458125 | STAD | AGS | CUDC-101 | | | | | | |
| MVP_top60_DNA_seed | ERBB2 | -0.457756 | STAD | MKN28 | CI-1033 | | | | | | |
| MVP_top60_DNA_seed | ERBB4 | -0.457756 | STAD | MKN28 | CI-1033 | | | | | | |
| MVP_top60_DNA_seed | EGFR | -0.457756 | STAD | MKN28 | CI-1033 | | | | | | |
| MVP_top60_DNA_seed | IGF1R | -0.457534 | STAD | GCIY | Linsitinib | | | | | | |
| MVP_top60_DNA_seed | IGF1R | -0.457308 | STAD | MKN45 | BMS-754807 | | | | | | |
| MVP_top60_DNA_seed | FGFR3 | -0.452904 | STAD | 23132-87 | FGFR_3831 | | | | | | |
| MVP_top60_DNA_seed | EGFR | -0.449358 | STAD | RF-48 | Pelitinib | | | | | | |
| MVP_top60_DNA_seed | EGFR | -0.447518 | STAD | SNU-1 | Pelitinib | | | | | | |
| MVP_top60_DNA_seed | AKT1 | -0.443938 | STAD | RF-48 | MK-2206 | | | | | | |
| MVP_top60_DNA_seed | MET | -0.442766 | STAD | TGBC11TKB | AZD6094 | | | | | | |
| MVP_top60_DNA_seed | GSK3B | -0.440756 | STAD | TMK-1 | AZD7969 | | | | | | |

| | | | | | |
|---|---|---|---|---|---|
| MVP_top60_DNA_seed | MDM2 | -0.437594 | STAD | MKN28 | Serdemetan |
| MVP_top60_DNA_seed | GSK3B | -0.437124 | STAD | KATOIII | AZD7969 |
| MVP_top60_DNA_seed | GSK3B | -0.433852 | STAD | SNU-16 | AZD7969 |
| MVP_top60_DNA_seed | IGF1R | -0.431737 | STAD | 23132-87 | BMS-536924 |
| MVP_top60_DNA_seed | GSK3B | -0.430945 | STAD | TMK-1 | SB216763 |
| MVP_top60_DNA_seed | AKT1 | -0.429835 | STAD | MKN45 | BAY AKT1 |
| MVP_top60_DNA_seed | ERBB2 | -0.428087 | STAD | KATOIII | CI-1033 |
| MVP_top60_DNA_seed | ERBB4 | -0.428087 | STAD | KATOIII | CI-1033 |
| MVP_top60_DNA_seed | EGFR | -0.428087 | STAD | KATOIII | CI-1033 |
| MVP_top60_DNA_seed | HSP90AA1 | -0.426285 | STAD | SK-GT-2 | Tanespimycin |
| MVP_top60_DNA_seed | FGFR3 | -0.426056 | STAD | MKN7 | PD173074 |
| MVP_top60_DNA_seed | MAPK1 | -0.424829 | STAD | RF-48 | FR-180204 |
| MVP_top60_DNA_seed | MAPK3 | -0.424829 | STAD | RF-48 | FR-180204 |
| MVP_top60_DNA_seed | ERBB2 | -0.423921 | STAD | MKN7 | Afatinib |
| MVP_top60_DNA_seed | EGFR | -0.423921 | STAD | MKN7 | Afatinib |
| MVP_top60_DNA_seed | HDAC1 | -0.423557 | STAD | MKN28 | Dacinostat |
| MVP_top60_DNA_seed | HSP90AA1 | -0.423536 | STAD | FU97 | Luminespib |
| MVP_top60_DNA_seed | HDAC1 | -0.42268 | STAD | TMK-1 | Dacinostat |
| MVP_top60_DNA_seed | MDM2 | -0.421014 | STAD | 23132-87 | Serdemetan |
| MVP_top60_DNA_seed | HDAC1 | -0.416055 | STAD | NCI-N87 | Dacinostat |
| MVP_top60_DNA_seed | MAPK1 | -0.413716 | STAD | MKN28 | VX-11e |
| MVP_top60_DNA_seed | FGFR3 | -0.412827 | STAD | SK-GT-2 | FGFR_3831 |
| MVP_top60_DNA_seed | HDAC1 | -0.412713 | STAD | SNU-1 | CAY10603 |
| MVP_top60_DNA_seed | HDAC1 | -0.409489 | STAD | HGC-27 | CAY10603 |
| MVP_top60_DNA_seed | IGF1R | -0.409407 | STAD | AGS | BMS-754807 |
| MVP_top60_DNA_seed | PIK3CA | -0.406898 | STAD | TMK-1 | PI3Ka_4409 |
| MVP_top60_DNA_seed | HDAC1 | -0.406798 | STAD | GCIY | AR-42 |
| MVP_top60_DNA_seed | ERBB2 | -0.406648 | STAD | FU97 | CI-1033 |
| MVP_top60_DNA_seed | ERBB4 | -0.406648 | STAD | FU97 | CI-1033 |
| MVP_top60_DNA_seed | EGFR | -0.406648 | STAD | FU97 | CI-1033 |
| MVP_top60_DNA_seed | AR | -0.403684 | STAD | HSC-39 | AZD3514 |
| MVP_top60_DNA_seed | AR | -0.398416 | STAD | Hs-746T | AZD3514 |
| MVP_top60_DNA_seed | EGFR | -0.397649 | STAD | HSC-39 | Gefitinib |
| MVP_top60_DNA_seed | ATM | -0.396905 | STAD | RERF-GC-1B | KU-60019 |
| MVP_top60_DNA_seed | AR | -0.395423 | STAD | OCUM-1 | Bicalutamide |
| MVP_top60_DNA_seed | HDAC1 | -0.39373 | STAD | AGS | AR-42 |
| MVP_top60_DNA_seed | HSP90AA1 | -0.393635 | STAD | RERF-GC-1B | Luminespib |
| MVP_top60_DNA_seed | HSP90AA1 | -0.382033 | STAD | GCIY | CCT-018159 |
| MVP_top60_DNA_seed | EGFR | -0.381789 | STAD | MKN7 | Gefitinib |
| MVP_top60_DNA_seed | SRC | -0.379704 | STAD | MKN28 | Bosutinib |
| MVP_top60_DNA_seed | PDGFRA | -0.37388 | STAD | GCIY | Masitinib |

| | | | | | |
|---|---|---|---|---|---|
| MVP_top60_DNA_seed | HSP90AA1 | -0.368181 | STAD | SNU-16 | Tanespimycin |
| MVP_top60_DNA_seed | SRC | -0.367975 | STAD | RF-48 | Bosutinib |
| MVP_top60_DNA_seed | PIK3CA | -0.366374 | STAD | FU97 | PI3Ka_4409 |
| MVP_top60_DNA_seed | MAPK1 | -0.365888 | STAD | RF-48 | VX-11e |
| MVP_top60_DNA_seed | ATM | -0.364419 | STAD | SK-GT-2 | KU-60019 |
| MVP_top60_DNA_seed | SRC | -0.364069 | STAD | SNU-16 | Bosutinib |
| MVP_top60_DNA_seed | GSK3B | -0.362652 | STAD | MKN7 | KIN001-042 |
| MVP_top60_DNA_seed | HSP90AA1 | -0.362506 | STAD | NUGC-4 | Tanespimycin |
| MVP_top60_DNA_seed | GSK3B | -0.362202 | STAD | NUGC-4 | CHIR-99021 |
| MVP_top60_DNA_seed | FGFR3 | -0.361398 | STAD | 23132-87 | AZD4547 |
| MVP_top60_DNA_seed | HDAC1 | -0.360311 | STAD | 23132-87 | Belinostat |
| MVP_top60_DNA_seed | AR | -0.359151 | STAD | OCUM-1 | AZD3514 |
| MVP_top60_DNA_seed | FGFR3 | -0.358376 | STAD | TMK-1 | AZD4547 |
| MVP_top60_DNA_seed | HDAC1 | -0.35831 | STAD | AGS | CAY10603 |
| MVP_top60_DNA_seed | MDM2 | -0.357478 | STAD | GCIY | Serdemetan |
| MVP_top60_DNA_seed | HDAC1 | -0.357175 | STAD | GCIY | CAY10603 |
| MVP_top60_DNA_seed | EGFR | -0.356797 | STAD | HGC-27 | Gefitinib |
| MVP_top60_DNA_seed | GSK3B | -0.355932 | STAD | AGS | AZD7969 |
| MVP_top60_DNA_seed | HDAC1 | -0.354964 | STAD | AGS | Belinostat |
| MVP_top60_DNA_seed | GSK3B | -0.353429 | STAD | MKN28 | AZD7969 |
| MVP_top60_DNA_seed | AKT1 | -0.351674 | STAD | RERF-GC-1B | AKT inhibitor VIII |
| MVP_top60_DNA_seed | MAPK1 | -0.351032 | STAD | NCI-N87 | VX-11e |
| MVP_top60_DNA_seed | HDAC1 | -0.348672 | STAD | 23132-87 | AR-42 |
| MVP_top60_DNA_seed | ATM | -0.34516 | STAD | IM-95 | KU-55933 |
| MVP_top60_DNA_seed | FGFR3 | -0.344998 | STAD | NUGC-3 | AZD4547 |
| MVP_top60_DNA_seed | ATM | -0.344206 | STAD | FU97 | CP466722 |
| MVP_top60_DNA_seed | HDAC1 | -0.342954 | STAD | 23132-87 | CAY10603 |
| MVP_top60_DNA_seed | PDGFRA | -0.338331 | STAD | OCUM-1 | Amuvatinib |
| MVP_top60_DNA_seed | GSK3B | -0.33502 | STAD | 23132-87 | CHIR-99021 |
| MVP_top60_DNA_seed | HDAC1 | -0.334021 | STAD | SNU-1 | AR-42 |
| MVP_top60_DNA_seed | MAPK1 | -0.332078 | STAD | 23132-87 | VX-11e |
| MVP_top60_DNA_seed | HSP90AA1 | -0.32898 | STAD | IM-95 | Tanespimycin |
| MVP_top60_DNA_seed | AKT1 | -0.328138 | STAD | GCIY | A-443654 |
| MVP_top60_DNA_seed | HDAC1 | -0.327405 | STAD | SNU-1 | Belinostat |
| MVP_top60_DNA_seed | FGFR3 | -0.320579 | STAD | OCUM-1 | FGFR_3831 |
| MVP_top60_DNA_seed | ERBB2 | -0.314995 | STAD | MKN7 | PF-00299804 |
| MVP_top60_DNA_seed | ERBB4 | -0.314995 | STAD | MKN7 | PF-00299804 |
| MVP_top60_DNA_seed | EGFR | -0.314995 | STAD | MKN7 | PF-00299804 |
| MVP_top60_DNA_seed | ERBB2 | -0.314747 | STAD | IM-95 | CUDC-101 |
| MVP_top60_DNA_seed | EGFR | -0.314747 | STAD | IM-95 | CUDC-101 |
| MVP_top60_DNA_seed | ESR1 | -0.313742 | STAD | SK-GT-2 | Fulvestrant |

| | | | | | |
|---|---|---|---|---|---|
| MVP_top60_DNA_seed | HDAC1 | -0.313686 | STAD | IM-95 | AR-42 |
| MVP_top60_DNA_seed | PDGFRA | -0.312262 | STAD | SNU-1 | Masitinib |
| MVP_top60_DNA_seed | IGF1R | -0.309885 | STAD | TMK-1 | BMS-754807 |
| MVP_top60_DNA_seed | ESR1 | -0.307974 | STAD | NUGC-3 | Fulvestrant |
| MVP_top60_DNA_seed | ATM | -0.307508 | STAD | 23132-87 | KU-60019 |
| MVP_top60_DNA_seed | HSP90AA1 | -0.30662 | STAD | NUGC-4 | Luminespib |
| MVP_top60_DNA_seed | MDM2 | -0.30519 | STAD | IM-95 | Serdemetan |
| MVP_top60_DNA_seed | HSP90AA1 | -0.302075 | STAD | MKN7 | CCT-018159 |
| MVP_top60_DNA_seed | AKT1 | -0.293573 | STAD | SNU-16 | BAY AKT1 |
| MVP_top60_DNA_seed | IGF1R | -0.292153 | STAD | OCUM-1 | BMS-754807 |
| MVP_top60_DNA_seed | GSK3B | -0.287943 | STAD | NCI-N87 | SB216763 |
| MVP_top60_DNA_seed | MAPK1 | -0.286058 | STAD | NUGC-4 | VX-11e |
| MVP_top60_DNA_seed | ERBB4 | -0.286039 | STAD | KATOIII | AST-1306 |
| MVP_top60_DNA_seed | EGFR | -0.286039 | STAD | KATOIII | AST-1306 |
| MVP_top60_DNA_seed | ESR1 | -0.285092 | STAD | RERF-GC-1B | Tamoxifen |
| MVP_top60_DNA_seed | HDAC1 | -0.284617 | STAD | GCIY | Belinostat |
| MVP_top60_DNA_seed | ERBB2 | -0.284447 | STAD | 23132-87 | AZD8931 |
| MVP_top60_DNA_seed | ERBB3 | -0.284447 | STAD | 23132-87 | AZD8931 |
| MVP_top60_DNA_seed | EGFR | -0.284447 | STAD | 23132-87 | AZD8931 |
| MVP_top60_DNA_seed | AKT1 | -0.277269 | STAD | MKN28 | MK-2206 |
| MVP_top60_DNA_seed | SRC | -0.276615 | STAD | TMK-1 | Bosutinib |
| MVP_top60_DNA_seed | AKT1 | -0.269116 | STAD | NCI-N87 | MK-2206 |
| MVP_top60_DNA_seed | EGFR | -0.265827 | STAD | HSC-39 | Pelitinib |
| MVP_top60_DNA_seed | ATM | -0.265428 | STAD | FU97 | KU-60019 |
| MVP_top60_DNA_seed | MET | -0.265114 | STAD | SNU-16 | PHA-665752 |
| MVP_top60_DNA_seed | AR | -0.263762 | STAD | HSC-39 | Bicalutamide |
| MVP_top60_DNA_seed | FGFR3 | -0.262097 | STAD | SNU-5 | PD173074 |
| MVP_top60_DNA_seed | EGFR | -0.256264 | STAD | RERF-GC-1B | Gefitinib |
| MVP_top60_DNA_seed | EGFR | -0.255527 | STAD | NUGC-4 | Pelitinib |
| MVP_top60_DNA_seed | IGF1R | -0.253787 | STAD | HSC-39 | GSK1904529A |
| MVP_top60_DNA_seed | SRC | -0.25301 | STAD | RF-48 | Saracatinib |
| MVP_top60_DNA_seed | FGFR3 | -0.251745 | STAD | SNU-5 | AZD4547 |
| MVP_top60_DNA_seed | HDAC1 | -0.249618 | STAD | HGC-27 | AR-42 |
| MVP_top60_DNA_seed | ATM | -0.246075 | STAD | NUGC-4 | KU-60019 |
| MVP_top60_DNA_seed | MDM2 | -0.245297 | STAD | SK-GT-2 | Nutlin-3a (-) |
| MVP_top60_DNA_seed | HDAC1 | -0.244948 | STAD | NUGC-3 | Dacinostat |
| MVP_top60_DNA_seed | ERBB2 | -0.24229 | STAD | RERF-GC-1B | AZD8931 |
| MVP_top60_DNA_seed | ERBB3 | -0.24229 | STAD | RERF-GC-1B | AZD8931 |
| MVP_top60_DNA_seed | EGFR | -0.24229 | STAD | RERF-GC-1B | AZD8931 |
| MVP_top60_DNA_seed | HDAC1 | -0.236947 | STAD | NUGC-4 | Tubastatin A |
| MVP_top60_DNA_seed | MET | -0.236798 | STAD | FU97 | JNJ38877605 |

| | | | | | |
|---|---|---|---|---|---|
| MVP_top60_DNA_seed | ERBB2 | -0.23349 | STAD | SNU-16 | AZD8931 |
| MVP_top60_DNA_seed | ERBB3 | -0.23349 | STAD | SNU-16 | AZD8931 |
| MVP_top60_DNA_seed | EGFR | -0.23349 | STAD | SNU-16 | AZD8931 |
| MVP_top60_DNA_seed | MDM2 | -0.2327 | STAD | HGC-27 | Nutlin-3a (-) |
| MVP_top60_DNA_seed | HDAC1 | -0.226962 | STAD | 23132-87 | Tubastatin A |
| MVP_top60_DNA_seed | HDAC1 | -0.223547 | STAD | 23132-87 | JQ12 |
| MVP_top60_DNA_seed | HDAC1 | -0.22156 | STAD | HGC-27 | Belinostat |
| MVP_top60_DNA_seed | HDAC1 | -0.22062 | STAD | NUGC-4 | Belinostat |
| MVP_top60_DNA_seed | ERBB4 | -0.220035 | STAD | TGBC11TKB | AST-1306 |
| MVP_top60_DNA_seed | EGFR | -0.220035 | STAD | TGBC11TKB | AST-1306 |
| MVP_top60_DNA_seed | PDGFRA | -0.218372 | STAD | MKN7 | Masitinib |
| MVP_top60_DNA_seed | HSP90AA1 | -0.217351 | STAD | IM-95 | Luminespib |
| MVP_top60_DNA_seed | EGFR | -0.215431 | STAD | AGS | Gefitinib |
| MVP_top60_DNA_seed | ERBB2 | -0.21364 | STAD | TGBC11TKB | CI-1033 |
| MVP_top60_DNA_seed | ERBB4 | -0.21364 | STAD | TGBC11TKB | CI-1033 |
| MVP_top60_DNA_seed | EGFR | -0.21364 | STAD | TGBC11TKB | CI-1033 |
| MVP_top60_DNA_seed | MAPK1 | -0.207298 | STAD | TGBC11TKB | VX-11e |
| MVP_top60_DNA_seed | ERBB4 | -0.206675 | STAD | MKN7 | AST-1306 |
| MVP_top60_DNA_seed | EGFR | -0.206675 | STAD | MKN7 | AST-1306 |
| MVP_top60_DNA_seed | AKT1 | -0.205625 | STAD | RERF-GC-1B | GSK690693 |
| MVP_top60_DNA_seed | ESR1 | -0.205362 | STAD | TMK-1 | Fulvestrant |
| MVP_top60_DNA_seed | GSK3B | -0.204024 | STAD | HGC-27 | CHIR-99021 |
| MVP_top60_DNA_seed | IGF1R | -0.201036 | STAD | HGC-27 | BMS-536924 |
| MVP_top60_DNA_seed | ERBB2 | -0.200245 | STAD | NUGC-3 | CUDC-101 |
| MVP_top60_DNA_seed | EGFR | -0.200245 | STAD | NUGC-3 | CUDC-101 |
| MVP_top60_DNA_seed | IGF1R | -0.198799 | STAD | TMK-1 | GSK1904529A |
| MVP_top60_DNA_seed | HDAC1 | -0.19679 | STAD | 23132-87 | Dacinostat |
| MVP_top60_DNA_seed | ERBB2 | -0.193304 | STAD | RERF-GC-1B | PF-00299804 |
| MVP_top60_DNA_seed | ERBB4 | -0.193304 | STAD | RERF-GC-1B | PF-00299804 |
| MVP_top60_DNA_seed | EGFR | -0.193304 | STAD | RERF-GC-1B | PF-00299804 |
| MVP_top60_DNA_seed | SRC | -0.191266 | STAD | SNU-1 | Saracatinib |
| MVP_top60_DNA_seed | GSK3B | -0.184608 | STAD | Hs-746T | KIN001-042 |
| MVP_top60_DNA_seed | MET | -0.183091 | STAD | IM-95 | AZD6094 |
| MVP_top60_DNA_seed | PIK3CA | -0.182956 | STAD | RERF-GC-1B | PI3Ka_4409 |
| MVP_top60_DNA_seed | FGFR3 | -0.177653 | STAD | RERF-GC-1B | FGFR_3831 |
| MVP_top60_DNA_seed | MAPK1 | -0.176446 | STAD | HGC-27 | VX-11e |
| MVP_top60_DNA_seed | ERBB2 | -0.175941 | STAD | MKN7 | CI-1033 |
| MVP_top60_DNA_seed | ERBB4 | -0.175941 | STAD | MKN7 | CI-1033 |
| MVP_top60_DNA_seed | EGFR | -0.175941 | STAD | MKN7 | CI-1033 |
| MVP_top60_DNA_seed | AR | -0.174018 | STAD | FU97 | Bicalutamide |
| MVP_top60_DNA_seed | GSK3B | -0.169839 | STAD | IM-95 | KIN001-042 |

| | | | | | |
|---|---|---|---|---|---|
| MVP_top60_DNA_seed | HDAC1 | -0.168944 | STAD | TGBC11TKB | AR-42 |
| MVP_top60_DNA_seed | PIK3CA | -0.166059 | STAD | FU97 | AZD8835 |
| MVP_top60_DNA_seed | ATM | -0.165111 | STAD | HGC-27 | CP466722 |
| MVP_top60_DNA_seed | ERBB4 | -0.163419 | STAD | Hs-746T | AST-1306 |
| MVP_top60_DNA_seed | EGFR | -0.163419 | STAD | Hs-746T | AST-1306 |
| MVP_top60_DNA_seed | MAPK1 | -0.16321 | STAD | HGC-27 | FR-180204 |
| MVP_top60_DNA_seed | MAPK3 | -0.16321 | STAD | HGC-27 | FR-180204 |
| MVP_top60_DNA_seed | IGF1R | -0.163085 | STAD | GCIY | GSK1904529A |
| MVP_top60_DNA_seed | ERBB2 | -0.160746 | STAD | HGC-27 | CUDC-101 |
| MVP_top60_DNA_seed | EGFR | -0.160746 | STAD | HGC-27 | CUDC-101 |
| MVP_top60_DNA_seed | IGF1R | -0.156819 | STAD | SNU-5 | BMS-754807 |
| MVP_top60_DNA_seed | MAPK1 | -0.155294 | STAD | SNU-1 | FR-180204 |
| MVP_top60_DNA_seed | MAPK3 | -0.155294 | STAD | SNU-1 | FR-180204 |
| MVP_top60_DNA_seed | PDGFRA | -0.152662 | STAD | HSC-39 | Amuvatinib |
| MVP_top60_DNA_seed | AKT1 | -0.152583 | STAD | MKN45 | AKT inhibitor VIII |
| MVP_top60_DNA_seed | AKT1 | -0.151994 | STAD | FU97 | GSK690693 |
| MVP_top60_DNA_seed | MET | -0.145601 | STAD | OCUM-1 | AZD6094 |
| MVP_top60_DNA_seed | SRC | -0.14443 | STAD | TGBC11TKB | NVP-BHG712 |
| MVP_top60_DNA_seed | ERBB2 | -0.144342 | STAD | IM-95 | Afatinib |
| MVP_top60_DNA_seed | EGFR | -0.144342 | STAD | IM-95 | Afatinib |
| MVP_top60_DNA_seed | ERBB2 | -0.138867 | STAD | AGS | CP724714 |
| MVP_top60_DNA_seed | SRC | -0.136525 | STAD | NCI-N87 | Bosutinib |
| MVP_top60_DNA_seed | ATM | -0.134592 | STAD | KATOIII | KU-60019 |
| MVP_top60_DNA_seed | EGFR | -0.134148 | STAD | SK-GT-2 | Cetuximab |
| MVP_top60_DNA_seed | ERBB2 | -0.134016 | STAD | FU97 | CP724714 |
| MVP_top60_DNA_seed | FGFR3 | -0.132752 | STAD | OCUM-1 | AZD4547 |
| MVP_top60_DNA_seed | PIK3CA | -0.130699 | STAD | 23132-87 | PI3Ka_4409 |
| MVP_top60_DNA_seed | ATM | -0.129087 | STAD | HSC-39 | CP466722 |
| MVP_top60_DNA_seed | MAPK1 | -0.128698 | STAD | HSC-39 | FR-180204 |
| MVP_top60_DNA_seed | MAPK3 | -0.128698 | STAD | HSC-39 | FR-180204 |
| MVP_top60_DNA_seed | PIK3CA | -0.128391 | STAD | RERF-GC-1B | AZD8835 |
| MVP_top60_DNA_seed | ERBB2 | -0.128348 | STAD | TGBC11TKB | PF-00299804 |
| MVP_top60_DNA_seed | ERBB4 | -0.128348 | STAD | TGBC11TKB | PF-00299804 |
| MVP_top60_DNA_seed | EGFR | -0.128348 | STAD | TGBC11TKB | PF-00299804 |
| MVP_top60_DNA_seed | MAPK1 | -0.127507 | STAD | FU97 | FR-180204 |
| MVP_top60_DNA_seed | MAPK3 | -0.127507 | STAD | FU97 | FR-180204 |
| MVP_top60_DNA_seed | ERBB2 | -0.125099 | STAD | HGC-27 | CI-1033 |
| MVP_top60_DNA_seed | ERBB4 | -0.125099 | STAD | HGC-27 | CI-1033 |
| MVP_top60_DNA_seed | EGFR | -0.125099 | STAD | HGC-27 | CI-1033 |
| MVP_top60_DNA_seed | ERBB2 | -0.120265 | STAD | GCIY | CUDC-101 |
| MVP_top60_DNA_seed | EGFR | -0.120265 | STAD | GCIY | CUDC-101 |

| | | | | | |
|---|---|---|---|---|---|
| MVP_top60_DNA_seed | ERBB2 | -0.120054 | STAD | HSC-39 | Lapatinib |
| MVP_top60_DNA_seed | EGFR | -0.120054 | STAD | HSC-39 | Lapatinib |
| MVP_top60_DNA_seed | AKT1 | -0.112506 | STAD | OCUM-1 | BAY AKT1 |
| MVP_top60_DNA_seed | HSP90AA1 | -0.110473 | STAD | MKN45 | Elesclomol |
| MVP_top60_DNA_seed | EGFR | -0.109712 | STAD | RERF-GC-1B | Pelitinib |
| MVP_top60_DNA_seed | FGFR3 | -0.106648 | STAD | HGC-27 | AZD4547 |
| MVP_top60_DNA_seed | EGFR | -0.105705 | STAD | NUGC-3 | Pelitinib |
| MVP_top60_DNA_seed | HDAC1 | -0.10497 | STAD | TGBC11TKB | Belinostat |
| MVP_top60_DNA_seed | ATM | -0.10335 | STAD | MKN45 | KU-60019 |
| MVP_top60_DNA_seed | PDGFRA | -0.100945 | STAD | RF-48 | Masitinib |
| MVP_top60_DNA_seed | HSP90AA1 | -0.099072 | STAD | Hs-746T | CCT-018159 |
| MVP_top60_DNA_seed | PDGFRA | -0.096982 | STAD | TGBC11TKB | Amuvatinib |
| MVP_top60_DNA_seed | ERBB2 | -0.093925 | STAD | NUGC-3 | CI-1033 |
| MVP_top60_DNA_seed | ERBB4 | -0.093925 | STAD | NUGC-3 | CI-1033 |
| MVP_top60_DNA_seed | EGFR | -0.093925 | STAD | NUGC-3 | CI-1033 |
| MVP_top60_DNA_seed | MET | -0.092801 | STAD | SNU-1 | PHA-665752 |
| MVP_top60_DNA_seed | HSP90AA1 | -0.08514 | STAD | NUGC-3 | Luminespib |
| MVP_top60_DNA_seed | AKT1 | -0.084674 | STAD | Hs-746T | MK-2206 |
| MVP_top60_DNA_seed | SRC | -0.083556 | STAD | FU97 | NVP-BHG712 |
| MVP_top60_DNA_seed | SRC | -0.081569 | STAD | AGS | Bosutinib |
| MVP_top60_DNA_seed | AR | -0.077787 | STAD | NUGC-3 | AZD3514 |
| MVP_top60_DNA_seed | EGFR | -0.075942 | STAD | SNU-16 | Erlotinib |
| MVP_top60_DNA_seed | MET | -0.07549 | STAD | HSC-39 | PHA-665752 |
| MVP_top60_DNA_seed | FGFR3 | -0.074017 | STAD | MKN28 | AZD4547 |
| MVP_top60_DNA_seed | AKT1 | -0.070877 | STAD | TGBC11TKB | GSK690693 |
| MVP_top60_DNA_seed | HDAC1 | -0.067291 | STAD | IM-95 | Belinostat |
| MVP_top60_DNA_seed | MET | -0.065729 | STAD | AGS | SU11274 |
| MVP_top60_DNA_seed | ERBB2 | -0.065151 | STAD | AGS | Afatinib |
| MVP_top60_DNA_seed | EGFR | -0.065151 | STAD | AGS | Afatinib |
| MVP_top60_DNA_seed | PDGFRA | -0.064315 | STAD | Hs-746T | Masitinib |
| MVP_top60_DNA_seed | MET | -0.060469 | STAD | SK-GT-2 | AZD6094 |
| MVP_top60_DNA_seed | EP300 | -0.059215 | STAD | RF-48 | I-CBP112 |
| MVP_top60_DNA_seed | CREBBP | -0.059215 | STAD | RF-48 | I-CBP112 |
| MVP_top60_DNA_seed | MAPK1 | -0.059028 | STAD | 23132-87 | FR-180204 |
| MVP_top60_DNA_seed | MAPK3 | -0.059028 | STAD | 23132-87 | FR-180204 |
| MVP_top60_DNA_seed | AR | -0.057164 | STAD | SNU-16 | AZD3514 |
| MVP_top60_DNA_seed | ATM | -0.053922 | STAD | KATOIII | KU-55933 |
| MVP_top60_DNA_seed | IGF1R | -0.052014 | STAD | SNU-1 | Linsitinib |
| MVP_top60_DNA_seed | HSP90AA1 | -0.051229 | STAD | Hs-746T | Elesclomol |
| MVP_top60_DNA_seed | ERBB2 | -0.05119 | STAD | 23132-87 | CI-1033 |
| MVP_top60_DNA_seed | ERBB4 | -0.05119 | STAD | 23132-87 | CI-1033 |

| | | | | | |
|---|---|---|---|---|---|
| MVP_top60_DNA_seed | EGFR | -0.05119 | STAD | 23132-87 | CI-1033 |
| MVP_top60_DNA_seed | HDAC1 | -0.050676 | STAD | HGC-27 | Tubastatin A |
| MVP_top60_DNA_seed | PIK3CA | -0.050562 | STAD | Hs-746T | PI3Ka_4409 |
| MVP_top60_DNA_seed | ERBB2 | -0.047554 | STAD | HSC-39 | AZD8931 |
| MVP_top60_DNA_seed | ERBB3 | -0.047554 | STAD | HSC-39 | AZD8931 |
| MVP_top60_DNA_seed | EGFR | -0.047554 | STAD | HSC-39 | AZD8931 |
| MVP_top60_DNA_seed | AKT1 | -0.035029 | STAD | KATOIII | MK-2206 |
| MVP_top60_DNA_seed | HDAC1 | -0.034401 | STAD | NUGC-4 | CAY10603 |
| MVP_top60_DNA_seed | ERBB2 | -0.032571 | STAD | HSC-39 | CP724714 |
| MVP_top60_DNA_seed | GSK3B | -0.030606 | STAD | SK-GT-2 | AZD7969 |
| MVP_top60_DNA_seed | GSK3B | -0.029111 | STAD | HSC-39 | CHIR-99021 |
| MVP_top60_DNA_seed | EP300 | -0.026335 | STAD | HSC-39 | I-CBP112 |
| MVP_top60_DNA_seed | CREBBP | -0.026335 | STAD | HSC-39 | I-CBP112 |
| MVP_top60_DNA_seed | AR | -0.023364 | STAD | MKN7 | Bicalutamide |
| MVP_top60_DNA_seed | AR | -0.022776 | STAD | RERF-GC-1B | Bicalutamide |
| MVP_top60_DNA_seed | MET | -0.021731 | STAD | AGS | JNJ38877605 |
| MVP_top60_DNA_seed | ERBB2 | -0.020592 | STAD | NUGC-3 | CP724714 |
| MVP_top60_DNA_seed | MDM2 | -0.016975 | STAD | MKN45 | Serdemetan |
| MVP_top60_DNA_seed | HDAC1 | -0.012993 | STAD | NUGC-4 | AR-42 |
| MVP_top60_DNA_seed | MDM2 | -0.012838 | STAD | MKN7 | Serdemetan |
| MVP_top60_DNA_seed | ATM | -0.00832 | STAD | AGS | KU-55933 |
| MVP_top60_DNA_seed | HDAC1 | -0.006399 | STAD | FU97 | Tubastatin A |
| MVP_top60_DNA_seed | GSK3B | -0.004942 | STAD | NUGC-3 | AZD7969 |
| MVP_top60_DNA_seed | ESR1 | -0.004848 | STAD | 23132-87 | Tamoxifen |
| MVP_top60_DNA_seed | SRC | -0.002756 | STAD | MKN45 | Bosutinib |
| MVP_top60_DNA_seed | EGFR | -0.00259 | STAD | MKN28 | Pelitinib |
| MVP_top60_DNA_seed | ERBB2 | -0.000572 | STAD | TMK-1 | Afatinib |
| MVP_top60_DNA_seed | EGFR | -0.000572 | STAD | TMK-1 | Afatinib |
| MVP_top60_DNA_seed | ERBB2 | 0.004358 | STAD | Hs-746T | CI-1033 |
| MVP_top60_DNA_seed | ERBB4 | 0.004358 | STAD | Hs-746T | CI-1033 |
| MVP_top60_DNA_seed | EGFR | 0.004358 | STAD | Hs-746T | CI-1033 |
| MVP_top60_DNA_seed | ERBB2 | 0.007664 | STAD | HSC-39 | Afatinib |
| MVP_top60_DNA_seed | EGFR | 0.007664 | STAD | HSC-39 | Afatinib |
| MVP_top60_DNA_seed | HSP90AA1 | 0.009251 | STAD | NCI-N87 | Luminespib |
| MVP_top60_DNA_seed | AR | 0.00946 | STAD | Hs-746T | Bicalutamide |
| MVP_top60_DNA_seed | EGFR | 0.0135 | STAD | TGBC11TKB | Cetuximab |
| MVP_top60_DNA_seed | GSK3B | 0.014177 | STAD | HSC-39 | CHIR-99021 |
| MVP_top60_DNA_seed | PDGFRA | 0.014935 | STAD | SNU-1 | Amuvatinib |
| MVP_top60_DNA_seed | IGF1R | 0.015512 | STAD | MKN28 | Linsitinib |
| MVP_top60_DNA_seed | EGFR | 0.018384 | STAD | HGC-27 | Cetuximab |
| MVP_top60_DNA_seed | PIK3CA | 0.020711 | STAD | IM-95 | PI3Ka_4409 |

| | | | | | |
|---|---|---|---|---|---|
| MVP_top60_DNA_seed | HDAC1 | 0.02156 | STAD | HSC-39 | AR-42 |
| MVP_top60_DNA_seed | AR | 0.023023 | STAD | AGS | AZD3514 |
| MVP_top60_DNA_seed | GSK3B | 0.023117 | STAD | 23132-87 | CHIR-99021 |
| MVP_top60_DNA_seed | PDGFRA | 0.024025 | STAD | 23132-87 | Amuvatinib |
| MVP_top60_DNA_seed | MDM2 | 0.025901 | STAD | HSC-39 | Nutlin-3a (-) |
| MVP_top60_DNA_seed | PDGFRA | 0.026272 | STAD | IM-95 | Masitinib |
| MVP_top60_DNA_seed | HSP90AA1 | 0.027097 | STAD | OCUM-1 | Elesclomol |
| MVP_top60_DNA_seed | IGF1R | 0.033387 | STAD | SNU-1 | BMS-536924 |
| MVP_top60_DNA_seed | ESR1 | 0.035763 | STAD | RERF-GC-1B | Fulvestrant |
| MVP_top60_DNA_seed | HSP90AA1 | 0.036882 | STAD | NUGC-4 | SNX-2112 |
| MVP_top60_DNA_seed | ERBB2 | 0.037018 | STAD | HSC-39 | Afatinib |
| MVP_top60_DNA_seed | EGFR | 0.037018 | STAD | HSC-39 | Afatinib |
| MVP_top60_DNA_seed | SRC | 0.038493 | STAD | NCI-N87 | HG-6-71-01 |
| MVP_top60_DNA_seed | AKT1 | 0.042074 | STAD | RERF-GC-1B | BAY AKT1 |
| MVP_top60_DNA_seed | HSP90AA1 | 0.045196 | STAD | TMK-1 | Luminespib |
| MVP_top60_DNA_seed | ATM | 0.047239 | STAD | SK-GT-2 | KU-55933 |
| MVP_top60_DNA_seed | FGFR3 | 0.050307 | STAD | MKN28 | FGFR_3831 |
| MVP_top60_DNA_seed | HDAC1 | 0.053631 | STAD | FU97 | Belinostat |
| MVP_top60_DNA_seed | AKT1 | 0.054176 | STAD | Hs-746T | BAY AKT1 |
| MVP_top60_DNA_seed | AR | 0.054807 | STAD | SNU-16 | Bicalutamide |
| MVP_top60_DNA_seed | EP300 | 0.055496 | STAD | FU97 | I-CBP112 |
| MVP_top60_DNA_seed | CREBBP | 0.055496 | STAD | FU97 | I-CBP112 |
| MVP_top60_DNA_seed | GSK3B | 0.061108 | STAD | SNU-5 | AZD7969 |
| MVP_top60_DNA_seed | SRC | 0.064892 | STAD | GCIY | NVP-BHG712 |
| MVP_top60_DNA_seed | ERBB4 | 0.067172 | STAD | IM-95 | AST-1306 |
| MVP_top60_DNA_seed | EGFR | 0.067172 | STAD | IM-95 | AST-1306 |
| MVP_top60_DNA_seed | PIK3CA | 0.073779 | STAD | SNU-16 | AZD8835 |
| MVP_top60_DNA_seed | MET | 0.07659 | STAD | HGC-27 | SU11274 |
| MVP_top60_DNA_seed | PDGFRA | 0.083786 | STAD | RF-48 | Amuvatinib |
| MVP_top60_DNA_seed | EGFR | 0.084699 | STAD | SNU-1 | Erlotinib |
| MVP_top60_DNA_seed | AR | 0.089111 | STAD | RERF-GC-1B | AZD3514 |
| MVP_top60_DNA_seed | ATM | 0.089537 | STAD | Hs-746T | KU-60019 |
| MVP_top60_DNA_seed | FGFR3 | 0.093478 | STAD | Hs-746T | PD173074 |
| MVP_top60_DNA_seed | ERBB2 | 0.094896 | STAD | FU97 | CUDC-101 |
| MVP_top60_DNA_seed | EGFR | 0.094896 | STAD | FU97 | CUDC-101 |
| MVP_top60_DNA_seed | IGF1R | 0.096761 | STAD | FU97 | Linsitinib |
| MVP_top60_DNA_seed | AR | 0.097163 | STAD | 23132-87 | AZD3514 |
| MVP_top60_DNA_seed | IGF1R | 0.09757 | STAD | NCI-N87 | GSK1904529A |
| MVP_top60_DNA_seed | ESR1 | 0.104338 | STAD | MKN7 | Tamoxifen |
| MVP_top60_DNA_seed | ESR1 | 0.106425 | STAD | OCUM-1 | Fulvestrant |
| MVP_top60_DNA_seed | AKT1 | 0.111714 | STAD | AGS | GSK690693 |

| | | | | | |
|---|---|---|---|---|---|
| MVP_top60_DNA_seed | GSK3B | 0.111998 | STAD | FU97 | CHIR-99021 |
| MVP_top60_DNA_seed | MDM2 | 0.113954 | STAD | NUGC-3 | Nutlin-3a (-) |
| MVP_top60_DNA_seed | PDGFRA | 0.11419 | STAD | AGS | Amuvatinib |
| MVP_top60_DNA_seed | EGFR | 0.114711 | STAD | HSC-39 | Cetuximab |
| MVP_top60_DNA_seed | HDAC1 | 0.115686 | STAD | RERF-GC-1B | JQ12 |
| MVP_top60_DNA_seed | ATM | 0.117284 | STAD | OCUM-1 | KU-60019 |
| MVP_top60_DNA_seed | ATM | 0.117618 | STAD | HGC-27 | KU-55933 |
| MVP_top60_DNA_seed | PIK3CA | 0.121784 | STAD | TGBC11TKB | AZD8835 |
| MVP_top60_DNA_seed | HSP90AA1 | 0.121889 | STAD | MKN28 | Tanespimycin |
| MVP_top60_DNA_seed | IGF1R | 0.12417 | STAD | MKN28 | BMS-754807 |
| MVP_top60_DNA_seed | HDAC1 | 0.124559 | STAD | SNU-1 | Tubastatin A |
| MVP_top60_DNA_seed | AR | 0.125006 | STAD | SK-GT-2 | AZD3514 |
| MVP_top60_DNA_seed | PIK3CA | 0.128448 | STAD | NUGC-4 | AZD8835 |
| MVP_top60_DNA_seed | HSP90AA1 | 0.129645 | STAD | NCI-N87 | CCT-018159 |
| MVP_top60_DNA_seed | IGF1R | 0.136913 | STAD | HSC-39 | BMS-754807 |
| MVP_top60_DNA_seed | HDAC1 | 0.137286 | STAD | RF-48 | Entinostat |
| MVP_top60_DNA_seed | PIK3CA | 0.137472 | STAD | MKN28 | AZD8835 |
| MVP_top60_DNA_seed | ERBB2 | 0.138652 | STAD | SNU-16 | CP724714 |
| MVP_top60_DNA_seed | MET | 0.140019 | STAD | HSC-39 | Crizotinib |
| MVP_top60_DNA_seed | HSP90AA1 | 0.142344 | STAD | MKN7 | SNX-2112 |
| MVP_top60_DNA_seed | HDAC1 | 0.143716 | STAD | NUGC-4 | Dacinostat |
| MVP_top60_DNA_seed | HDAC1 | 0.143809 | STAD | TGBC11TKB | CAY10603 |
| MVP_top60_DNA_seed | GSK3B | 0.145196 | STAD | TGBC11TKB | CHIR-99021 |
| MVP_top60_DNA_seed | HDAC1 | 0.145259 | STAD | HSC-39 | Entinostat |
| MVP_top60_DNA_seed | EGFR | 0.146634 | STAD | HSC-39 | Erlotinib |
| MVP_top60_DNA_seed | PDGFRA | 0.146856 | STAD | SNU-16 | Amuvatinib |
| MVP_top60_DNA_seed | ERBB2 | 0.146955 | STAD | MKN7 | CP724714 |
| MVP_top60_DNA_seed | IGF1R | 0.153969 | STAD | RF-48 | BMS-536924 |
| MVP_top60_DNA_seed | HSP90AA1 | 0.154773 | STAD | 23132-87 | Elesclomol |
| MVP_top60_DNA_seed | MDM2 | 0.155284 | STAD | RERF-GC-1B | Serdemetan |
| MVP_top60_DNA_seed | ERBB2 | 0.160058 | STAD | MKN7 | AZD8931 |
| MVP_top60_DNA_seed | ERBB3 | 0.160058 | STAD | MKN7 | AZD8931 |
| MVP_top60_DNA_seed | EGFR | 0.160058 | STAD | MKN7 | AZD8931 |
| MVP_top60_DNA_seed | AKT1 | 0.161817 | STAD | NCI-N87 | AKT inhibitor VIII |
| MVP_top60_DNA_seed | MET | 0.1642 | STAD | GCIY | Crizotinib |
| MVP_top60_DNA_seed | ERBB2 | 0.167947 | STAD | HGC-27 | Afatinib |
| MVP_top60_DNA_seed | EGFR | 0.167947 | STAD | HGC-27 | Afatinib |
| MVP_top60_DNA_seed | MAPK1 | 0.168097 | STAD | FU97 | VX-11e |
| MVP_top60_DNA_seed | IGF1R | 0.171694 | STAD | RF-48 | BMS-754807 |
| MVP_top60_DNA_seed | IGF1R | 0.173062 | STAD | HGC-27 | Linsitinib |
| MVP_top60_DNA_seed | GSK3B | 0.177124 | STAD | OCUM-1 | AZD7969 |

| | | | | | |
|---|---|---|---|---|---|
| MVP_top60_DNA_seed | PIK3CA | 0.178054 | STAD | SK-GT-2 | PI3Ka_4409 |
| MVP_top60_DNA_seed | MAPK1 | 0.181645 | STAD | MKN7 | FR-180204 |
| MVP_top60_DNA_seed | MAPK3 | 0.181645 | STAD | MKN7 | FR-180204 |
| MVP_top60_DNA_seed | HSP90AA1 | 0.182516 | STAD | IM-95 | CCT-018159 |
| MVP_top60_DNA_seed | ERBB2 | 0.191637 | STAD | IM-95 | PF-00299804 |
| MVP_top60_DNA_seed | ERBB4 | 0.191637 | STAD | IM-95 | PF-00299804 |
| MVP_top60_DNA_seed | EGFR | 0.191637 | STAD | IM-95 | PF-00299804 |
| MVP_top60_DNA_seed | HSP90AA1 | 0.194683 | STAD | Hs-746T | Tanespimycin |
| MVP_top60_DNA_seed | IGF1R | 0.195008 | STAD | AGS | GSK1904529A |
| MVP_top60_DNA_seed | FGFR3 | 0.197536 | STAD | AGS | AZD4547 |
| MVP_top60_DNA_seed | SRC | 0.197645 | STAD | 23132-87 | Bosutinib |
| MVP_top60_DNA_seed | MAPK1 | 0.1977 | STAD | TGBC11TKB | FR-180204 |
| MVP_top60_DNA_seed | MAPK3 | 0.1977 | STAD | TGBC11TKB | FR-180204 |
| MVP_top60_DNA_seed | HDAC1 | 0.198525 | STAD | MKN28 | CAY10603 |
| MVP_top60_DNA_seed | IGF1R | 0.199956 | STAD | 23132-87 | Linsitinib |
| MVP_top60_DNA_seed | PIK3CA | 0.203951 | STAD | SNU-5 | AZD8835 |
| MVP_top60_DNA_seed | SRC | 0.206312 | STAD | TMK-1 | HG-6-71-01 |
| MVP_top60_DNA_seed | FGFR3 | 0.207473 | STAD | TGBC11TKB | PD173074 |
| MVP_top60_DNA_seed | HSP90AA1 | 0.207665 | STAD | MKN45 | SNX-2112 |
| MVP_top60_DNA_seed | GSK3B | 0.21094 | STAD | IM-95 | AZD7969 |
| MVP_top60_DNA_seed | MET | 0.212317 | STAD | RF-48 | Crizotinib |
| MVP_top60_DNA_seed | HDAC1 | 0.212807 | STAD | MKN28 | Tubastatin A |
| MVP_top60_DNA_seed | GSK3B | 0.213548 | STAD | MKN45 | SB216763 |
| MVP_top60_DNA_seed | GSK3B | 0.216824 | STAD | NUGC-3 | CHIR-99021 |
| MVP_top60_DNA_seed | FGFR3 | 0.218777 | STAD | SNU-5 | AZD4547 |
| MVP_top60_DNA_seed | HDAC1 | 0.223065 | STAD | NUGC-3 | CAY10603 |
| MVP_top60_DNA_seed | SRC | 0.224697 | STAD | OCUM-1 | Bosutinib |
| MVP_top60_DNA_seed | EP300 | 0.226234 | STAD | 23132-87 | I-CBP112 |
| MVP_top60_DNA_seed | CREBBP | 0.226234 | STAD | 23132-87 | I-CBP112 |
| MVP_top60_DNA_seed | AKT1 | 0.235142 | STAD | HSC-39 | A-443654 |
| MVP_top60_DNA_seed | EGFR | 0.23612 | STAD | HGC-27 | Pelitinib |
| MVP_top60_DNA_seed | EGFR | 0.23747 | STAD | MKN7 | Cetuximab |
| MVP_top60_DNA_seed | FGFR3 | 0.238331 | STAD | AGS | PD173074 |
| MVP_top60_DNA_seed | MET | 0.239117 | STAD | FU97 | AZD6094 |
| MVP_top60_DNA_seed | ATM | 0.23976 | STAD | NCI-N87 | KU-60019 |
| MVP_top60_DNA_seed | PDGFRA | 0.240904 | STAD | 23132-87 | Masitinib |
| MVP_top60_DNA_seed | HSP90AA1 | 0.243622 | STAD | Hs-746T | SNX-2112 |
| MVP_top60_DNA_seed | HDAC1 | 0.247638 | STAD | RERF-GC-1B | Tubastatin A |
| MVP_top60_DNA_seed | HDAC1 | 0.247883 | STAD | IM-95 | CAY10603 |
| MVP_top60_DNA_seed | PIK3CA | 0.249282 | STAD | KATOIII | AZD8835 |
| MVP_top60_DNA_seed | GSK3B | 0.250622 | STAD | MKN7 | CHIR-99021 |

| | | | | | |
|---|---|---|---|---|---|
| MVP_top60_DNA_seed | MET | 0.251625 | STAD | NUGC-4 | JNJ38877605 |
| MVP_top60_DNA_seed | EGFR | 0.252753 | STAD | TGBC11TKB | Gefitinib |
| MVP_top60_DNA_seed | ERBB2 | 0.256146 | STAD | SNU-5 | Afatinib |
| MVP_top60_DNA_seed | EGFR | 0.256146 | STAD | SNU-5 | Afatinib |
| MVP_top60_DNA_seed | HDAC1 | 0.257129 | STAD | MKN7 | CAY10603 |
| MVP_top60_DNA_seed | AKT1 | 0.260779 | STAD | SNU-5 | BAY AKT1 |
| MVP_top60_DNA_seed | HSP90AA1 | 0.261704 | STAD | OCUM-1 | Luminespib |
| MVP_top60_DNA_seed | ERBB4 | 0.262005 | STAD | 23132-87 | AST-1306 |
| MVP_top60_DNA_seed | EGFR | 0.262005 | STAD | 23132-87 | AST-1306 |
| MVP_top60_DNA_seed | AKT1 | 0.264889 | STAD | SNU-5 | MK-2206 |
| MVP_top60_DNA_seed | AR | 0.268343 | STAD | RF-48 | Bicalutamide |
| MVP_top60_DNA_seed | ERBB2 | 0.268978 | STAD | SNU-1 | Afatinib |
| MVP_top60_DNA_seed | EGFR | 0.268978 | STAD | SNU-1 | Afatinib |
| MVP_top60_DNA_seed | MDM2 | 0.270125 | STAD | NCI-N87 | Serdemetan |
| MVP_top60_DNA_seed | SRC | 0.274762 | STAD | MKN7 | Bosutinib |
| MVP_top60_DNA_seed | PDGFRA | 0.275417 | STAD | IM-95 | Amuvatinib |
| MVP_top60_DNA_seed | MET | 0.276673 | STAD | NUGC-4 | AZD6094 |
| MVP_top60_DNA_seed | EGFR | 0.2783 | STAD | MKN28 | Gefitinib |
| MVP_top60_DNA_seed | ERBB2 | 0.279114 | STAD | TGBC11TKB | Afatinib |
| MVP_top60_DNA_seed | EGFR | 0.279114 | STAD | TGBC11TKB | Afatinib |
| MVP_top60_DNA_seed | ERBB4 | 0.2807 | STAD | FU97 | AST-1306 |
| MVP_top60_DNA_seed | EGFR | 0.2807 | STAD | FU97 | AST-1306 |
| MVP_top60_DNA_seed | GSK3B | 0.281846 | STAD | AGS | KIN001-042 |
| MVP_top60_DNA_seed | ERBB2 | 0.28185 | STAD | SNU-16 | Lapatinib |
| MVP_top60_DNA_seed | EGFR | 0.28185 | STAD | SNU-16 | Lapatinib |
| MVP_top60_DNA_seed | AR | 0.282371 | STAD | TGBC11TKB | AZD3514 |
| MVP_top60_DNA_seed | GSK3B | 0.283993 | STAD | AGS | SB216763 |
| MVP_top60_DNA_seed | GSK3B | 0.286311 | STAD | GCIY | CHIR-99021 |
| MVP_top60_DNA_seed | PDGFRA | 0.287735 | STAD | NUGC-3 | Amuvatinib |
| MVP_top60_DNA_seed | HDAC1 | 0.288894 | STAD | FU97 | CAY10603 |
| MVP_top60_DNA_seed | GSK3B | 0.290459 | STAD | OCUM-1 | KIN001-042 |
| MVP_top60_DNA_seed | MAPK1 | 0.290598 | STAD | AGS | FR-180204 |
| MVP_top60_DNA_seed | MAPK3 | 0.290598 | STAD | AGS | FR-180204 |
| MVP_top60_DNA_seed | MET | 0.292771 | STAD | 23132-87 | AZD6094 |
| MVP_top60_DNA_seed | HSP90AA1 | 0.294511 | STAD | MKN45 | CCT-018159 |
| MVP_top60_DNA_seed | GSK3B | 0.297009 | STAD | AGS | CHIR-99021 |
| MVP_top60_DNA_seed | IGF1R | 0.298186 | STAD | RF-48 | BMS-536924 |
| MVP_top60_DNA_seed | AKT1 | 0.298993 | STAD | NUGC-3 | GSK690693 |
| MVP_top60_DNA_seed | GSK3B | 0.303219 | STAD | TGBC11TKB | CHIR-99021 |
| MVP_top60_DNA_seed | PDGFRA | 0.30461 | STAD | SNU-5 | Amuvatinib |
| MVP_top60_DNA_seed | ERBB2 | 0.305071 | STAD | SNU-5 | AZD8931 |

| | | | | | |
|---|---|---|---|---|---|
| MVP_top60_DNA_seed | ERBB3 | 0.305071 | STAD | SNU-5 | AZD8931 |
| MVP_top60_DNA_seed | EGFR | 0.305071 | STAD | SNU-5 | AZD8931 |
| MVP_top60_DNA_seed | AR | 0.306706 | STAD | MKN28 | Bicalutamide |
| MVP_top60_DNA_seed | EGFR | 0.308943 | STAD | 23132-87 | Cetuximab |
| MVP_top60_DNA_seed | HDAC1 | 0.313312 | STAD | IM-95 | Tubastatin A |
| MVP_top60_DNA_seed | ERBB2 | 0.314203 | STAD | MKN28 | CP724714 |
| MVP_top60_DNA_seed | IGF1R | 0.317619 | STAD | RF-48 | Linsitinib |
| MVP_top60_DNA_seed | HDAC1 | 0.319247 | STAD | TMK-1 | AR-42 |
| MVP_top60_DNA_seed | SRC | 0.319439 | STAD | TGBC11TKB | Bosutinib |
| MVP_top60_DNA_seed | HDAC1 | 0.319815 | STAD | SNU-1 | Dacinostat |
| MVP_top60_DNA_seed | HDAC1 | 0.324896 | STAD | NUGC-3 | Belinostat |
| MVP_top60_DNA_seed | GSK3B | 0.325384 | STAD | MKN45 | CHIR-99021 |
| MVP_top60_DNA_seed | FGFR3 | 0.327347 | STAD | NUGC-4 | AZD4547 |
| MVP_top60_DNA_seed | FGFR3 | 0.327583 | STAD | TGBC11TKB | AZD4547 |
| MVP_top60_DNA_seed | ATM | 0.33635 | STAD | Hs-746T | KU-55933 |
| MVP_top60_DNA_seed | ERBB2 | 0.340594 | STAD | HGC-27 | Afatinib |
| MVP_top60_DNA_seed | EGFR | 0.340594 | STAD | HGC-27 | Afatinib |
| MVP_top60_DNA_seed | HSP90AA1 | 0.341481 | STAD | TGBC11TKB | CCT-018159 |
| MVP_top60_DNA_seed | AKT1 | 0.341562 | STAD | RERF-GC-1B | AKT inhibitor VIII |
| MVP_top60_DNA_seed | ERBB2 | 0.347888 | STAD | NUGC-3 | PF-00299804 |
| MVP_top60_DNA_seed | ERBB4 | 0.347888 | STAD | NUGC-3 | PF-00299804 |
| MVP_top60_DNA_seed | EGFR | 0.347888 | STAD | NUGC-3 | PF-00299804 |
| MVP_top60_DNA_seed | MET | 0.349223 | STAD | HGC-27 | JNJ38877605 |
| MVP_top60_DNA_seed | IGF1R | 0.349661 | STAD | TGBC11TKB | GSK1904529A |
| MVP_top60_DNA_seed | SRC | 0.350355 | STAD | HSC-39 | Saracatinib |
| MVP_top60_DNA_seed | ATM | 0.352218 | STAD | TGBC11TKB | KU-60019 |
| MVP_top60_DNA_seed | PDGFRA | 0.353112 | STAD | MKN28 | Amuvatinib |
| MVP_top60_DNA_seed | PDGFRA | 0.355783 | STAD | RERF-GC-1B | Amuvatinib |
| MVP_top60_DNA_seed | HSP90AA1 | 0.356116 | STAD | KATOIII | Elesclomol |
| MVP_top60_DNA_seed | FGFR3 | 0.356645 | STAD | MKN45 | AZD4547 |
| MVP_top60_DNA_seed | HDAC1 | 0.358198 | STAD | NUGC-3 | AR-42 |
| MVP_top60_DNA_seed | ATM | 0.359733 | STAD | RERF-GC-1B | CP466722 |
| MVP_top60_DNA_seed | FGFR3 | 0.361168 | STAD | OCUM-1 | AZD4547 |
| MVP_top60_DNA_seed | FGFR3 | 0.361257 | STAD | TGBC11TKB | AZD4547 |
| MVP_top60_DNA_seed | EGFR | 0.362285 | STAD | FU97 | Pelitinib |
| MVP_top60_DNA_seed | HSP90AA1 | 0.36265 | STAD | MKN28 | SNX-2112 |
| MVP_top60_DNA_seed | AR | 0.364975 | STAD | NUGC-3 | Bicalutamide |
| MVP_top60_DNA_seed | EGFR | 0.365901 | STAD | FU97 | Cetuximab |
| MVP_top60_DNA_seed | IGF1R | 0.366091 | STAD | TGBC11TKB | Linsitinib |
| MVP_top60_DNA_seed | EGFR | 0.373634 | STAD | RERF-GC-1B | Cetuximab |
| MVP_top60_DNA_seed | HSP90AA1 | 0.375133 | STAD | 23132-87 | Luminespib |

| | | | | | |
|---|---|---|---|---|---|
| MVP_top60_DNA_seed | AKT1 | 0.375585 | STAD | NUGC-3 | BAY AKT1 |
| MVP_top60_DNA_seed | PDGFRA | 0.378357 | STAD | FU97 | Masitinib |
| MVP_top60_DNA_seed | MDM2 | 0.380652 | STAD | SNU-16 | Nutlin-3a (-) |
| MVP_top60_DNA_seed | ESR1 | 0.380847 | STAD | NUGC-4 | Tamoxifen |
| MVP_top60_DNA_seed | ERBB2 | 0.381472 | STAD | FU97 | AZD8931 |
| MVP_top60_DNA_seed | ERBB3 | 0.381472 | STAD | FU97 | AZD8931 |
| MVP_top60_DNA_seed | EGFR | 0.381472 | STAD | FU97 | AZD8931 |
| MVP_top60_DNA_seed | AKT1 | 0.382474 | STAD | GCIY | AKT inhibitor VIII |
| MVP_top60_DNA_seed | IGF1R | 0.382898 | STAD | NUGC-4 | BMS-754807 |
| MVP_top60_DNA_seed | HDAC1 | 0.383309 | STAD | SNU-1 | Parthenolide |
| MVP_top60_DNA_seed | HDAC1 | 0.383686 | STAD | MKN45 | JQ12 |
| MVP_top60_DNA_seed | EP300 | 0.38651 | STAD | SK-GT-2 | I-CBP112 |
| MVP_top60_DNA_seed | CREBBP | 0.38651 | STAD | SK-GT-2 | I-CBP112 |
| MVP_top60_DNA_seed | AKT1 | 0.388978 | STAD | NUGC-3 | MK-2206 |
| MVP_top60_DNA_seed | PDGFRA | 0.389858 | STAD | NUGC-4 | Masitinib |
| MVP_top60_DNA_seed | HSP90AA1 | 0.391123 | STAD | GCIY | Tanespimycin |
| MVP_top60_DNA_seed | AKT1 | 0.39428 | STAD | AGS | AKT inhibitor VIII |
| MVP_top60_DNA_seed | ERBB2 | 0.394715 | STAD | Hs-746T | PF-00299804 |
| MVP_top60_DNA_seed | ERBB4 | 0.394715 | STAD | Hs-746T | PF-00299804 |
| MVP_top60_DNA_seed | EGFR | 0.394715 | STAD | Hs-746T | PF-00299804 |
| MVP_top60_DNA_seed | HSP90AA1 | 0.398494 | STAD | MKN7 | Elesclomol |
| MVP_top60_DNA_seed | EGFR | 0.398772 | STAD | 23132-87 | Pelitinib |
| MVP_top60_DNA_seed | MET | 0.399172 | STAD | FU97 | SU11274 |
| MVP_top60_DNA_seed | SRC | 0.401858 | STAD | NCI-N87 | NVP-BHG712 |
| MVP_top60_DNA_seed | IGF1R | 0.402289 | STAD | NUGC-4 | Linsitinib |
| MVP_top60_DNA_seed | EGFR | 0.408043 | STAD | KATOIII | Gefitinib |
| MVP_top60_DNA_seed | SRC | 0.409036 | STAD | GCIY | Bosutinib |
| MVP_top60_DNA_seed | ATM | 0.410336 | STAD | NUGC-3 | KU-60019 |
| MVP_top60_DNA_seed | IGF1R | 0.41052 | STAD | IM-95 | BMS-536924 |
| MVP_top60_DNA_seed | HDAC1 | 0.410866 | STAD | TGBC11TKB | Tubastatin A |
| MVP_top60_DNA_seed | HDAC1 | 0.41522 | STAD | MKN7 | Belinostat |
| MVP_top60_DNA_seed | AKT1 | 0.42067 | STAD | NUGC-4 | GSK690693 |
| MVP_top60_DNA_seed | MDM2 | 0.420834 | STAD | KATOIII | Serdemetan |
| MVP_top60_DNA_seed | AKT1 | 0.420867 | STAD | GCIY | MK-2206 |
| MVP_top60_DNA_seed | ATM | 0.42355 | STAD | MKN28 | KU-55933 |
| MVP_top60_DNA_seed | ATM | 0.423844 | STAD | NCI-N87 | torin2 |
| MVP_top60_DNA_seed | ERBB2 | 0.424051 | STAD | SNU-5 | Afatinib |
| MVP_top60_DNA_seed | EGFR | 0.424051 | STAD | SNU-5 | Afatinib |
| MVP_top60_DNA_seed | GSK3B | 0.425202 | STAD | SK-GT-2 | CHIR-99021 |
| MVP_top60_DNA_seed | AR | 0.42524 | STAD | SNU-1 | AZD3514 |
| MVP_top60_DNA_seed | HDAC1 | 0.425624 | STAD | GCIY | Entinostat |

| | | | | | |
|---|---|---|---|---|---|
| MVP_top60_DNA_seed | AKT1 | 0.426446 | STAD | TGBC11TKB | AKT inhibitor VIII |
| MVP_top60_DNA_seed | ERBB2 | 0.42897 | STAD | MKN45 | CI-1033 |
| MVP_top60_DNA_seed | ERBB4 | 0.42897 | STAD | MKN45 | CI-1033 |
| MVP_top60_DNA_seed | EGFR | 0.42897 | STAD | MKN45 | CI-1033 |
| MVP_top60_DNA_seed | MET | 0.432479 | STAD | NUGC-3 | AZD6094 |
| MVP_top60_DNA_seed | AR | 0.435623 | STAD | NUGC-3 | Bicalutamide |
| MVP_top60_DNA_seed | FGFR3 | 0.437522 | STAD | IM-95 | AZD4547 |
| MVP_top60_DNA_seed | ATM | 0.438933 | STAD | 23132-87 | CP466722 |
| MVP_top60_DNA_seed | MET | 0.444196 | STAD | SNU-16 | Crizotinib |
| MVP_top60_DNA_seed | MAPK1 | 0.448173 | STAD | OCUM-1 | VX-11e |
| MVP_top60_DNA_seed | HDAC1 | 0.454467 | STAD | TGBC11TKB | Dacinostat |
| MVP_top60_DNA_seed | ERBB2 | 0.45463 | STAD | RERF-GC-1B | Afatinib |
| MVP_top60_DNA_seed | EGFR | 0.45463 | STAD | RERF-GC-1B | Afatinib |
| MVP_top60_DNA_seed | GSK3B | 0.456557 | STAD | FU97 | CHIR-99021 |
| MVP_top60_DNA_seed | EGFR | 0.457378 | STAD | AGS | Pelitinib |
| MVP_top60_DNA_seed | AR | 0.458698 | STAD | RERF-GC-1B | Bicalutamide |
| MVP_top60_DNA_seed | FGFR3 | 0.460447 | STAD | NUGC-3 | PD173074 |
| MVP_top60_DNA_seed | ERBB2 | 0.462993 | STAD | Hs-746T | AZD8931 |
| MVP_top60_DNA_seed | ERBB3 | 0.462993 | STAD | Hs-746T | AZD8931 |
| MVP_top60_DNA_seed | EGFR | 0.462993 | STAD | Hs-746T | AZD8931 |
| MVP_top60_DNA_seed | AKT1 | 0.463223 | STAD | MKN28 | GSK690693 |
| MVP_top60_DNA_seed | HDAC1 | 0.467097 | STAD | FU97 | AR-42 |
| MVP_top60_DNA_seed | HSP90AA1 | 0.467951 | STAD | SNU-5 | Luminespib |
| MVP_top60_DNA_seed | FGFR3 | 0.468195 | STAD | SK-GT-2 | AZD4547 |
| MVP_top60_DNA_seed | HSP90AA1 | 0.469142 | STAD | SNU-5 | Elesclomol |
| MVP_top60_DNA_seed | AKT1 | 0.471694 | STAD | TMK-1 | GSK690693 |
| MVP_top60_DNA_seed | AR | 0.478677 | STAD | SNU-5 | Bicalutamide |
| MVP_top60_DNA_seed | HDAC1 | 0.479923 | STAD | MKN7 | AR-42 |
| MVP_top60_DNA_seed | ERBB2 | 0.48105 | STAD | MKN28 | CUDC-101 |
| MVP_top60_DNA_seed | EGFR | 0.48105 | STAD | MKN28 | CUDC-101 |
| MVP_top60_DNA_seed | HSP90AA1 | 0.481642 | STAD | NUGC-3 | SNX-2112 |
| MVP_top60_DNA_seed | MAPK1 | 0.484288 | STAD | TMK-1 | VX-11e |
| MVP_top60_DNA_seed | AKT1 | 0.485455 | STAD | MKN45 | GSK690693 |
| MVP_top60_DNA_seed | ESR1 | 0.486328 | STAD | IM-95 | Fulvestrant |
| MVP_top60_DNA_seed | AR | 0.487764 | STAD | 23132-87 | Bicalutamide |
| MVP_top60_DNA_seed | AKT1 | 0.488048 | STAD | GCIY | GSK690693 |
| MVP_top60_DNA_seed | ERBB2 | 0.489956 | STAD | 23132-87 | Afatinib |
| MVP_top60_DNA_seed | EGFR | 0.489956 | STAD | 23132-87 | Afatinib |
| MVP_top60_DNA_seed | HSP90AA1 | 0.490265 | STAD | RF-48 | Tanespimycin |
| MVP_top60_DNA_seed | EGFR | 0.494435 | STAD | NCI-N87 | Cetuximab |
| MVP_top60_DNA_seed | FGFR3 | 0.495112 | STAD | MKN28 | PD173074 |

| | | | | | |
|---|---|---|---|---|---|
| MVP_top60_DNA_seed | FGFR3 | 0.495888 | STAD | IM-95 | FGFR_3831 |
| MVP_top60_DNA_seed | ERBB2 | 0.496494 | STAD | KATOIII | AZD8931 |
| MVP_top60_DNA_seed | ERBB3 | 0.496494 | STAD | KATOIII | AZD8931 |
| MVP_top60_DNA_seed | EGFR | 0.496494 | STAD | KATOIII | AZD8931 |
| MVP_top60_DNA_seed | AKT1 | 0.497842 | STAD | NUGC-4 | AKT inhibitor VIII |
| MVP_top60_DNA_seed | HSP90AA1 | 0.499024 | STAD | MKN28 | CCT-018159 |
| MVP_top60_DNA_seed | PIK3CA | 0.500928 | STAD | MKN45 | PI3Ka_4409 |
| MVP_top60_DNA_seed | HDAC1 | 0.502886 | STAD | NUGC-4 | JQ12 |
| MVP_top60_DNA_seed | MDM2 | 0.505255 | STAD | SNU-5 | Nutlin-3a (-) |
| MVP_top60_DNA_seed | HDAC1 | 0.506164 | STAD | MKN28 | Belinostat |
| MVP_top60_DNA_seed | ERBB2 | 0.507041 | STAD | 23132-87 | Afatinib |
| MVP_top60_DNA_seed | EGFR | 0.507041 | STAD | 23132-87 | Afatinib |
| MVP_top60_DNA_seed | PIK3CA | 0.511737 | STAD | MKN28 | PI3Ka_4409 |
| MVP_top60_DNA_seed | SRC | 0.514907 | STAD | IM-95 | Bosutinib |
| MVP_top60_DNA_seed | ERBB2 | 0.514996 | STAD | RF-48 | CI-1033 |
| MVP_top60_DNA_seed | ERBB4 | 0.514996 | STAD | RF-48 | CI-1033 |
| MVP_top60_DNA_seed | EGFR | 0.514996 | STAD | RF-48 | CI-1033 |
| MVP_top60_DNA_seed | GSK3B | 0.51567 | STAD | FU97 | SB216763 |
| MVP_top60_DNA_seed | MAPK1 | 0.517562 | STAD | NUGC-3 | VX-11e |
| MVP_top60_DNA_seed | PDGFRA | 0.517755 | STAD | MKN28 | Masitinib |
| MVP_top60_DNA_seed | HDAC1 | 0.518522 | STAD | MKN45 | Dacinostat |
| MVP_top60_DNA_seed | PIK3CA | 0.519231 | STAD | NUGC-4 | PI3Ka_4409 |
| MVP_top60_DNA_seed | ERBB2 | 0.522888 | STAD | MKN7 | CUDC-101 |
| MVP_top60_DNA_seed | EGFR | 0.522888 | STAD | MKN7 | CUDC-101 |
| MVP_top60_DNA_seed | PIK3CA | 0.523535 | STAD | SNU-16 | PI3Ka_4409 |
| MVP_top60_DNA_seed | ESR1 | 0.524239 | STAD | MKN7 | Fulvestrant |
| MVP_top60_DNA_seed | MAPK1 | 0.525733 | STAD | OCUM-1 | FR-180204 |
| MVP_top60_DNA_seed | MAPK3 | 0.525733 | STAD | OCUM-1 | FR-180204 |
| MVP_top60_DNA_seed | MET | 0.525976 | STAD | RERF-GC-1B | SU11274 |
| MVP_top60_DNA_seed | MET | 0.529698 | STAD | NCI-N87 | Crizotinib |
| MVP_top60_DNA_seed | HDAC1 | 0.530899 | STAD | HSC-39 | Parthenolide |
| MVP_top60_DNA_seed | AR | 0.531485 | STAD | NUGC-4 | Bicalutamide |
| MVP_top60_DNA_seed | GSK3B | 0.53271 | STAD | RERF-GC-1B | CHIR-99021 |
| MVP_top60_DNA_seed | SRC | 0.534293 | STAD | MKN28 | NVP-BHG712 |
| MVP_top60_DNA_seed | GSK3B | 0.534869 | STAD | 23132-87 | KIN001-042 |
| MVP_top60_DNA_seed | ERBB2 | 0.536215 | STAD | 23132-87 | CP724714 |
| MVP_top60_DNA_seed | IGF1R | 0.538549 | STAD | OCUM-1 | Linsitinib |
| MVP_top60_DNA_seed | HDAC1 | 0.544415 | STAD | SNU-16 | Entinostat |
| MVP_top60_DNA_seed | FGFR3 | 0.544572 | STAD | IM-95 | PD173074 |
| MVP_top60_DNA_seed | MAPK1 | 0.547019 | STAD | HSC-39 | VX-11e |
| MVP_top60_DNA_seed | GSK3B | 0.548838 | STAD | GCIY | SB216763 |

| | | | | | |
|---|---|---|---|---|---|
| MVP_top60_DNA_seed | GSK3B | 0.55106 | STAD | Hs-746T | CHIR-99021 |
| MVP_top60_DNA_seed | GSK3B | 0.552627 | STAD | IM-95 | SB216763 |
| MVP_top60_DNA_seed | AR | 0.553287 | STAD | MKN45 | AZD3514 |
| MVP_top60_DNA_seed | PIK3CA | 0.554986 | STAD | KATOIII | PI3Ka_4409 |
| MVP_top60_DNA_seed | MET | 0.55719 | STAD | RF-48 | AZD6094 |
| MVP_top60_DNA_seed | FGFR3 | 0.560398 | STAD | RF-48 | AZD4547 |
| MVP_top60_DNA_seed | AR | 0.561141 | STAD | NCI-N87 | AZD3514 |
| MVP_top60_DNA_seed | PDGFRA | 0.561969 | STAD | NUGC-4 | Amuvatinib |
| MVP_top60_DNA_seed | EGFR | 0.563138 | STAD | FU97 | Gefitinib |
| MVP_top60_DNA_seed | ERBB2 | 0.563994 | STAD | OCUM-1 | AZD8931 |
| MVP_top60_DNA_seed | ERBB3 | 0.563994 | STAD | OCUM-1 | AZD8931 |
| MVP_top60_DNA_seed | EGFR | 0.563994 | STAD | OCUM-1 | AZD8931 |
| MVP_top60_DNA_seed | ERBB2 | 0.564898 | STAD | 23132-87 | PF-00299804 |
| MVP_top60_DNA_seed | ERBB4 | 0.564898 | STAD | 23132-87 | PF-00299804 |
| MVP_top60_DNA_seed | EGFR | 0.564898 | STAD | 23132-87 | PF-00299804 |
| MVP_top60_DNA_seed | AR | 0.566906 | STAD | NUGC-4 | Bicalutamide |
| MVP_top60_DNA_seed | GSK3B | 0.567159 | STAD | KATOIII | KIN001-042 |
| MVP_top60_DNA_seed | SRC | 0.568032 | STAD | NUGC-3 | NVP-BHG712 |
| MVP_top60_DNA_seed | ESR1 | 0.568881 | STAD | SNU-1 | Tamoxifen |
| MVP_top60_DNA_seed | ERBB2 | 0.569323 | STAD | FU97 | Afatinib |
| MVP_top60_DNA_seed | EGFR | 0.569323 | STAD | FU97 | Afatinib |
| MVP_top60_DNA_seed | FGFR3 | 0.570671 | STAD | IM-95 | AZD4547 |
| MVP_top60_DNA_seed | PIK3CA | 0.572245 | STAD | OCUM-1 | PI3Ka_4409 |
| MVP_top60_DNA_seed | SRC | 0.572897 | STAD | FU97 | Bosutinib |
| MVP_top60_DNA_seed | MET | 0.573119 | STAD | SNU-1 | AZD6094 |
| MVP_top60_DNA_seed | MET | 0.573554 | STAD | MKN28 | JNJ38877605 |
| MVP_top60_DNA_seed | IGF1R | 0.573642 | STAD | HSC-39 | Linsitinib |
| MVP_top60_DNA_seed | ERBB2 | 0.577607 | STAD | KATOIII | Afatinib |
| MVP_top60_DNA_seed | EGFR | 0.577607 | STAD | KATOIII | Afatinib |
| MVP_top60_DNA_seed | GSK3B | 0.578095 | STAD | MKN45 | CHIR-99021 |
| MVP_top60_DNA_seed | ATM | 0.579775 | STAD | MKN7 | CP466722 |
| MVP_top60_DNA_seed | AR | 0.58645 | STAD | TMK-1 | Bicalutamide |
| MVP_top60_DNA_seed | HSP90AA1 | 0.586509 | STAD | RF-48 | CCT-018159 |
| MVP_top60_DNA_seed | PIK3CA | 0.589523 | STAD | MKN7 | AZD8835 |
| MVP_top60_DNA_seed | MET | 0.597246 | STAD | AGS | AZD6094 |
| MVP_top60_DNA_seed | HSP90AA1 | 0.601442 | STAD | KATOIII | CCT-018159 |
| MVP_top60_DNA_seed | AR | 0.605168 | STAD | SNU-1 | Bicalutamide |
| MVP_top60_DNA_seed | HSP90AA1 | 0.60624 | STAD | RERF-GC-1B | SNX-2112 |
| MVP_top60_DNA_seed | ESR1 | 0.606326 | STAD | SNU-5 | Tamoxifen |
| MVP_top60_DNA_seed | MDM2 | 0.612469 | STAD | SNU-16 | Serdemetan |
| MVP_top60_DNA_seed | EGFR | 0.612701 | STAD | Hs-746T | Cetuximab |

| | | | | | |
|---|---|---|---|---|---|
| MVP_top60_DNA_seed | ERBB4 | 0.613193 | STAD | HGC-27 | AST-1306 |
| MVP_top60_DNA_seed | EGFR | 0.613193 | STAD | HGC-27 | AST-1306 |
| MVP_top60_DNA_seed | SRC | 0.618218 | STAD | SNU-1 | Bosutinib |
| MVP_top60_DNA_seed | AKT1 | 0.620135 | STAD | SNU-1 | AKT inhibitor VIII |
| MVP_top60_DNA_seed | GSK3B | 0.624287 | STAD | SNU-1 | CHIR-99021 |
| MVP_top60_DNA_seed | ERBB2 | 0.624443 | STAD | MKN45 | Afatinib |
| MVP_top60_DNA_seed | EGFR | 0.624443 | STAD | MKN45 | Afatinib |
| MVP_top60_DNA_seed | SRC | 0.624621 | STAD | Hs-746T | NVP-BHG712 |
| MVP_top60_DNA_seed | SRC | 0.62529 | STAD | RF-48 | WH-4-023 |
| MVP_top60_DNA_seed | PDGFRA | 0.625708 | STAD | NCI-N87 | Masitinib |
| MVP_top60_DNA_seed | ERBB2 | 0.628557 | STAD | TGBC11TKB | CUDC-101 |
| MVP_top60_DNA_seed | EGFR | 0.628557 | STAD | TGBC11TKB | CUDC-101 |
| MVP_top60_DNA_seed | FGFR3 | 0.630148 | STAD | 23132-87 | PD173074 |
| MVP_top60_DNA_seed | AKT1 | 0.63353 | STAD | NUGC-4 | MK-2206 |
| MVP_top60_DNA_seed | ESR1 | 0.634474 | STAD | SK-GT-2 | Tamoxifen |
| MVP_top60_DNA_seed | GSK3B | 0.634734 | STAD | TGBC11TKB | SB216763 |
| MVP_top60_DNA_seed | PIK3CA | 0.634969 | STAD | NUGC-3 | AZD8835 |
| MVP_top60_DNA_seed | HDAC1 | 0.635157 | STAD | SNU-1 | JQ12 |
| MVP_top60_DNA_seed | AKT1 | 0.637226 | STAD | HSC-39 | AKT inhibitor VIII |
| MVP_top60_DNA_seed | HDAC1 | 0.639133 | STAD | AGS | Tubastatin A |
| MVP_top60_DNA_seed | ATM | 0.640117 | STAD | TGBC11TKB | CP466722 |
| MVP_top60_DNA_seed | IGF1R | 0.643202 | STAD | SNU-1 | BMS-754807 |
| MVP_top60_DNA_seed | ATM | 0.647483 | STAD | OCUM-1 | CP466722 |
| MVP_top60_DNA_seed | AKT1 | 0.649262 | STAD | SNU-1 | GSK690693 |
| MVP_top60_DNA_seed | GSK3B | 0.65237 | STAD | NUGC-4 | SB216763 |
| MVP_top60_DNA_seed | ERBB2 | 0.65275 | STAD | TMK-1 | CUDC-101 |
| MVP_top60_DNA_seed | EGFR | 0.65275 | STAD | TMK-1 | CUDC-101 |
| MVP_top60_DNA_seed | EGFR | 0.65499 | STAD | MKN45 | Pelitinib |
| MVP_top60_DNA_seed | AKT1 | 0.658592 | STAD | SNU-16 | A-443654 |
| MVP_top60_DNA_seed | MDM2 | 0.661444 | STAD | RERF-GC-1B | Nutlin-3a (-) |
| MVP_top60_DNA_seed | SRC | 0.665729 | STAD | RERF-GC-1B | Bosutinib |
| MVP_top60_DNA_seed | GSK3B | 0.671458 | STAD | MKN45 | AZD7969 |
| MVP_top60_DNA_seed | ERBB2 | 0.673728 | STAD | SNU-16 | Afatinib |
| MVP_top60_DNA_seed | EGFR | 0.673728 | STAD | SNU-16 | Afatinib |
| MVP_top60_DNA_seed | ERBB2 | 0.674507 | STAD | OCUM-1 | CP724714 |
| MVP_top60_DNA_seed | PDGFRA | 0.674727 | STAD | TGBC11TKB | Masitinib |
| MVP_top60_DNA_seed | HDAC1 | 0.675068 | STAD | MKN28 | AR-42 |
| MVP_top60_DNA_seed | MET | 0.677291 | STAD | RERF-GC-1B | AZD6094 |
| MVP_top60_DNA_seed | ESR1 | 0.679352 | STAD | IM-95 | Tamoxifen |
| MVP_top60_DNA_seed | SRC | 0.684412 | STAD | SNU-5 | WH-4-023 |
| MVP_top60_DNA_seed | ERBB2 | 0.68628 | STAD | MKN28 | Afatinib |

| | | | | | |
|---|---|---|---|---|---|
| MVP_top60_DNA_seed | EGFR | 0.68628 | STAD | MKN28 | Afatinib |
| MVP_top60_DNA_seed | GSK3B | 0.686417 | STAD | NUGC-3 | KIN001-042 |
| MVP_top60_DNA_seed | GSK3B | 0.687524 | STAD | MKN28 | KIN001-042 |
| MVP_top60_DNA_seed | EP300 | 0.689219 | STAD | NUGC-3 | I-CBP112 |
| MVP_top60_DNA_seed | CREBBP | 0.689219 | STAD | NUGC-3 | I-CBP112 |
| MVP_top60_DNA_seed | FGFR3 | 0.692894 | STAD | NUGC-4 | PD173074 |
| MVP_top60_DNA_seed | FGFR3 | 0.696224 | STAD | NCI-N87 | PD173074 |
| MVP_top60_DNA_seed | EGFR | 0.697077 | STAD | RF-48 | Erlotinib |
| MVP_top60_DNA_seed | EP300 | 0.697561 | STAD | RERF-GC-1B | I-CBP112 |
| MVP_top60_DNA_seed | CREBBP | 0.697561 | STAD | RERF-GC-1B | I-CBP112 |
| MVP_top60_DNA_seed | ATM | 0.699749 | STAD | NUGC-4 | KU-55933 |
| MVP_top60_DNA_seed | HDAC1 | 0.700892 | STAD | NCI-N87 | Tubastatin A |
| MVP_top60_DNA_seed | MET | 0.703018 | STAD | MKN7 | AZD6094 |
| MVP_top60_DNA_seed | PDGFRA | 0.706911 | STAD | AGS | Masitinib |
| MVP_top60_DNA_seed | HDAC1 | 0.707824 | STAD | TGBC11TKB | JQ12 |
| MVP_top60_DNA_seed | GSK3B | 0.709362 | STAD | MKN7 | CHIR-99021 |
| MVP_top60_DNA_seed | MAPK1 | 0.711501 | STAD | Hs-746T | VX-11e |
| MVP_top60_DNA_seed | PIK3CA | 0.712438 | STAD | NUGC-3 | PI3Ka_4409 |
| MVP_top60_DNA_seed | ERBB2 | 0.713247 | STAD | RF-48 | AZD8931 |
| MVP_top60_DNA_seed | ERBB3 | 0.713247 | STAD | RF-48 | AZD8931 |
| MVP_top60_DNA_seed | EGFR | 0.713247 | STAD | RF-48 | AZD8931 |
| MVP_top60_DNA_seed | HSP90AA1 | 0.714843 | STAD | GCIY | SNX-2112 |
| MVP_top60_DNA_seed | IGF1R | 0.721255 | STAD | AGS | Linsitinib |
| MVP_top60_DNA_seed | GSK3B | 0.723637 | STAD | RF-48 | KIN001-042 |
| MVP_top60_DNA_seed | HSP90AA1 | 0.72489 | STAD | FU97 | SNX-2112 |
| MVP_top60_DNA_seed | ERBB2 | 0.725846 | STAD | SNU-5 | Lapatinib |
| MVP_top60_DNA_seed | EGFR | 0.725846 | STAD | SNU-5 | Lapatinib |
| MVP_top60_DNA_seed | GSK3B | 0.729397 | STAD | SNU-5 | KIN001-042 |
| MVP_top60_DNA_seed | PIK3CA | 0.732466 | STAD | TGBC11TKB | PI3Ka_4409 |
| MVP_top60_DNA_seed | MDM2 | 0.738214 | STAD | SNU-5 | Serdemetan |
| MVP_top60_DNA_seed | ERBB2 | 0.741339 | STAD | KATOIII | Afatinib |
| MVP_top60_DNA_seed | EGFR | 0.741339 | STAD | KATOIII | Afatinib |
| MVP_top60_DNA_seed | SRC | 0.741903 | STAD | SNU-5 | Bosutinib |
| MVP_top60_DNA_seed | HDAC1 | 0.743373 | STAD | SNU-16 | Parthenolide |
| MVP_top60_DNA_seed | MAPK1 | 0.744045 | STAD | GCIY | FR-180204 |
| MVP_top60_DNA_seed | MAPK3 | 0.744045 | STAD | GCIY | FR-180204 |
| MVP_top60_DNA_seed | SRC | 0.744151 | STAD | 23132-87 | NVP-BHG712 |
| MVP_top60_DNA_seed | HDAC1 | 0.745101 | STAD | OCUM-1 | JQ12 |
| MVP_top60_DNA_seed | GSK3B | 0.747297 | STAD | SNU-16 | KIN001-042 |
| MVP_top60_DNA_seed | IGF1R | 0.74801 | STAD | MKN7 | Linsitinib |
| MVP_top60_DNA_seed | ERBB2 | 0.753648 | STAD | FU97 | Afatinib |

| | | | | | |
|---|---|---|---|---|---|
| MVP_top60_DNA_seed | EGFR | 0.753648 | STAD | FU97 | Afatinib |
| MVP_top60_DNA_seed | ERBB2 | 0.754714 | STAD | TGBC11TKB | Afatinib |
| MVP_top60_DNA_seed | EGFR | 0.754714 | STAD | TGBC11TKB | Afatinib |
| MVP_top60_DNA_seed | HDAC1 | 0.754968 | STAD | NCI-N87 | Parthenolide |
| MVP_top60_DNA_seed | MAPK1 | 0.755788 | STAD | GCIY | VX-11e |
| MVP_top60_DNA_seed | PIK3CA | 0.758768 | STAD | SK-GT-2 | AZD8835 |
| MVP_top60_DNA_seed | MDM2 | 0.759175 | STAD | FU97 | Nutlin-3a (-) |
| MVP_top60_DNA_seed | ERBB2 | 0.760857 | STAD | HGC-27 | PF-00299804 |
| MVP_top60_DNA_seed | ERBB4 | 0.760857 | STAD | HGC-27 | PF-00299804 |
| MVP_top60_DNA_seed | EGFR | 0.760857 | STAD | HGC-27 | PF-00299804 |
| MVP_top60_DNA_seed | ERBB4 | 0.762208 | STAD | MKN28 | AST-1306 |
| MVP_top60_DNA_seed | EGFR | 0.762208 | STAD | MKN28 | AST-1306 |
| MVP_top60_DNA_seed | HDAC1 | 0.763516 | STAD | SNU-16 | AR-42 |
| MVP_top60_DNA_seed | ATM | 0.764145 | STAD | KATOIII | torin2 |
| MVP_top60_DNA_seed | GSK3B | 0.764976 | STAD | IM-95 | CHIR-99021 |
| MVP_top60_DNA_seed | FGFR3 | 0.766972 | STAD | SNU-1 | AZD4547 |
| MVP_top60_DNA_seed | FGFR3 | 0.767028 | STAD | SK-GT-2 | PD173074 |
| MVP_top60_DNA_seed | HDAC1 | 0.767679 | STAD | TMK-1 | Tubastatin A |
| MVP_top60_DNA_seed | MET | 0.769863 | STAD | GCIY | PHA-665752 |
| MVP_top60_DNA_seed | GSK3B | 0.77531 | STAD | NCI-N87 | AZD7969 |
| MVP_top60_DNA_seed | FGFR3 | 0.777347 | STAD | MKN7 | AZD4547 |
| MVP_top60_DNA_seed | PDGFRA | 0.777823 | STAD | TMK-1 | Amuvatinib |
| MVP_top60_DNA_seed | AKT1 | 0.781248 | STAD | RF-48 | BAY AKT1 |
| MVP_top60_DNA_seed | ESR1 | 0.782169 | STAD | NUGC-3 | Tamoxifen |
| MVP_top60_DNA_seed | HDAC1 | 0.783896 | STAD | GCIY | Tubastatin A |
| MVP_top60_DNA_seed | AKT1 | 0.784931 | STAD | GCIY | AKT inhibitor VIII |
| MVP_top60_DNA_seed | AR | 0.786438 | STAD | NUGC-4 | AZD3514 |
| MVP_top60_DNA_seed | IGF1R | 0.788819 | STAD | SNU-16 | BMS-754807 |
| MVP_top60_DNA_seed | MAPK1 | 0.794199 | STAD | NUGC-3 | FR-180204 |
| MVP_top60_DNA_seed | MAPK3 | 0.794199 | STAD | NUGC-3 | FR-180204 |
| MVP_top60_DNA_seed | ESR1 | 0.795778 | STAD | NCI-N87 | Fulvestrant |
| MVP_top60_DNA_seed | GSK3B | 0.799252 | STAD | RERF-GC-1B | SB216763 |
| MVP_top60_DNA_seed | ATM | 0.800756 | STAD | MKN45 | CP466722 |
| MVP_top60_DNA_seed | PIK3CA | 0.800803 | STAD | MKN7 | PI3Ka_4409 |
| MVP_top60_DNA_seed | HDAC1 | 0.801108 | STAD | MKN7 | Dacinostat |
| MVP_top60_DNA_seed | IGF1R | 0.801831 | STAD | TGBC11TKB | BMS-536924 |
| MVP_top60_DNA_seed | SRC | 0.802908 | STAD | HSC-39 | NVP-BHG712 |
| MVP_top60_DNA_seed | MDM2 | 0.806473 | STAD | MKN28 | Nutlin-3a (-) |
| MVP_top60_DNA_seed | MAPK1 | 0.806632 | STAD | TMK-1 | FR-180204 |
| MVP_top60_DNA_seed | MAPK3 | 0.806632 | STAD | TMK-1 | FR-180204 |
| MVP_top60_DNA_seed | ESR1 | 0.807527 | STAD | SNU-1 | Fulvestrant |

| | | | | | |
|---|---|---|---|---|---|
| MVP_top60_DNA_seed | IGF1R | 0.808872 | STAD | MKN45 | GSK1904529A |
| MVP_top60_DNA_seed | GSK3B | 0.811732 | STAD | SNU-16 | CHIR-99021 |
| MVP_top60_DNA_seed | ERBB2 | 0.81431 | STAD | MKN28 | PF-00299804 |
| MVP_top60_DNA_seed | ERBB4 | 0.81431 | STAD | MKN28 | PF-00299804 |
| MVP_top60_DNA_seed | EGFR | 0.81431 | STAD | MKN28 | PF-00299804 |
| MVP_top60_DNA_seed | PDGFRA | 0.816393 | STAD | MKN45 | Amuvatinib |
| MVP_top60_DNA_seed | AKT1 | 0.818167 | STAD | SNU-5 | A-443654 |
| MVP_top60_DNA_seed | HSP90AA1 | 0.82647 | STAD | HSC-39 | Elesclomol |
| MVP_top60_DNA_seed | IGF1R | 0.829994 | STAD | FU97 | BMS-754807 |
| MVP_top60_DNA_seed | EP300 | 0.835812 | STAD | MKN7 | I-CBP112 |
| MVP_top60_DNA_seed | CREBBP | 0.835812 | STAD | MKN7 | I-CBP112 |
| MVP_top60_DNA_seed | HDAC1 | 0.8405 | STAD | SNU-5 | AR-42 |
| MVP_top60_DNA_seed | EGFR | 0.842127 | STAD | OCUM-1 | Cetuximab |
| MVP_top60_DNA_seed | ERBB2 | 0.846034 | STAD | FU97 | PF-00299804 |
| MVP_top60_DNA_seed | ERBB4 | 0.846034 | STAD | FU97 | PF-00299804 |
| MVP_top60_DNA_seed | EGFR | 0.846034 | STAD | FU97 | PF-00299804 |
| MVP_top60_DNA_seed | AKT1 | 0.848417 | STAD | MKN28 | AKT inhibitor VIII |
| MVP_top60_DNA_seed | MAPK1 | 0.851745 | STAD | MKN7 | VX-11e |
| MVP_top60_DNA_seed | EGFR | 0.852291 | STAD | 23132-87 | Gefitinib |
| MVP_top60_DNA_seed | ESR1 | 0.853524 | STAD | RF-48 | Tamoxifen |
| MVP_top60_DNA_seed | MDM2 | 0.861437 | STAD | GCIY | Nutlin-3a (-) |
| MVP_top60_DNA_seed | IGF1R | 0.863254 | STAD | NUGC-3 | BMS-536924 |
| MVP_top60_DNA_seed | HSP90AA1 | 0.865977 | STAD | NCI-N87 | Elesclomol |
| MVP_top60_DNA_seed | FGFR3 | 0.867024 | STAD | GCIY | PD173074 |
| MVP_top60_DNA_seed | FGFR3 | 0.867664 | STAD | RF-48 | AZD4547 |
| MVP_top60_DNA_seed | AR | 0.867892 | STAD | RF-48 | Bicalutamide |
| MVP_top60_DNA_seed | HDAC1 | 0.875 | STAD | SNU-16 | Belinostat |
| MVP_top60_DNA_seed | SRC | 0.876367 | STAD | MKN45 | NVP-BHG712 |
| MVP_top60_DNA_seed | HSP90AA1 | 0.877421 | STAD | TMK-1 | SNX-2112 |
| MVP_top60_DNA_seed | AKT1 | 0.880606 | STAD | SNU-16 | GSK690693 |
| MVP_top60_DNA_seed | IGF1R | 0.881746 | STAD | 23132-87 | BMS-754807 |
| MVP_top60_DNA_seed | HDAC1 | 0.889929 | STAD | SNU-5 | Entinostat |
| MVP_top60_DNA_seed | ESR1 | 0.891296 | STAD | TMK-1 | Tamoxifen |
| MVP_top60_DNA_seed | IGF1R | 0.891849 | STAD | SNU-16 | GSK1904529A |
| MVP_top60_DNA_seed | AKT1 | 0.892712 | STAD | SNU-16 | AKT inhibitor VIII |
| MVP_top60_DNA_seed | HDAC1 | 0.897117 | STAD | SNU-5 | CAY10603 |
| MVP_top60_DNA_seed | IGF1R | 0.8976 | STAD | NCI-N87 | BMS-536924 |
| MVP_top60_DNA_seed | FGFR3 | 0.900374 | STAD | TMK-1 | FGFR_3831 |
| MVP_top60_DNA_seed | AKT1 | 0.900601 | STAD | SNU-1 | BAY AKT1 |
| MVP_top60_DNA_seed | HSP90AA1 | 0.902853 | STAD | MKN7 | Luminespib |
| MVP_top60_DNA_seed | HDAC1 | 0.90912 | STAD | SNU-1 | Entinostat |

| | | | | | |
|---|---|---|---|---|---|
| MVP_top60_DNA_seed | MET | 0.910516 | STAD | SNU-1 | JNJ38877605 |
| MVP_top60_DNA_seed | ERBB2 | 0.911389 | STAD | TMK-1 | CP724714 |
| MVP_top60_DNA_seed | HSP90AA1 | 0.911413 | STAD | FU97 | Elesclomol |
| MVP_top60_DNA_seed | MDM2 | 0.916011 | STAD | KATOIII | Nutlin-3a (-) |
| MVP_top60_DNA_seed | HSP90AA1 | 0.916055 | STAD | SNU-16 | Elesclomol |
| MVP_top60_DNA_seed | ERBB2 | 0.918981 | STAD | RF-48 | Lapatinib |
| MVP_top60_DNA_seed | EGFR | 0.918981 | STAD | RF-48 | Lapatinib |
| MVP_top60_DNA_seed | MAPK1 | 0.92229 | STAD | NCI-N87 | FR-180204 |
| MVP_top60_DNA_seed | MAPK3 | 0.92229 | STAD | NCI-N87 | FR-180204 |
| MVP_top60_DNA_seed | HDAC1 | 0.924897 | STAD | SNU-5 | Belinostat |
| MVP_top60_DNA_seed | AKT1 | 0.926086 | STAD | MKN7 | AKT inhibitor VIII |
| MVP_top60_DNA_seed | GSK3B | 0.927586 | STAD | OCUM-1 | CHIR-99021 |
| MVP_top60_DNA_seed | HDAC1 | 0.930647 | STAD | TMK-1 | CAY10603 |
| MVP_top60_DNA_seed | HDAC1 | 0.930858 | STAD | HSC-39 | JQ12 |
| MVP_top60_DNA_seed | EGFR | 0.931296 | STAD | SNU-5 | Cetuximab |
| MVP_top60_DNA_seed | AKT1 | 0.934279 | STAD | MKN7 | MK-2206 |
| MVP_top60_DNA_seed | MDM2 | 0.935312 | STAD | TGBC11TKB | Serdemetan |
| MVP_top60_DNA_seed | GSK3B | 0.937086 | STAD | NUGC-3 | SB216763 |
| MVP_top60_DNA_seed | FGFR3 | 0.941307 | STAD | NCI-N87 | AZD4547 |
| MVP_top60_DNA_seed | AKT1 | 0.943295 | STAD | SNU-1 | MK-2206 |
| MVP_top60_DNA_seed | ATM | 0.94372 | STAD | NUGC-4 | CP466722 |
| MVP_top60_DNA_seed | AKT1 | 0.944729 | STAD | MKN7 | GSK690693 |
| MVP_top60_DNA_seed | IGF1R | 0.94694 | STAD | RERF-GC-1B | GSK1904529A |
| MVP_top60_DNA_seed | AKT1 | 0.950219 | STAD | OCUM-1 | AKT inhibitor VIII |
| MVP_top60_DNA_seed | PIK3CA | 0.952346 | STAD | RF-48 | AZD8835 |
| MVP_top60_DNA_seed | ERBB2 | 0.952632 | STAD | TGBC11TKB | AZD8931 |
| MVP_top60_DNA_seed | ERBB3 | 0.952632 | STAD | TGBC11TKB | AZD8931 |
| MVP_top60_DNA_seed | EGFR | 0.952632 | STAD | TGBC11TKB | AZD8931 |
| MVP_top60_DNA_seed | ERBB2 | 0.953741 | STAD | Hs-746T | Afatinib |
| MVP_top60_DNA_seed | EGFR | 0.953741 | STAD | Hs-746T | Afatinib |
| MVP_top60_DNA_seed | HDAC1 | 0.956902 | STAD | RERF-GC-1B | CAY10603 |
| MVP_top60_DNA_seed | ERBB2 | 0.95716 | STAD | MKN45 | PF-00299804 |
| MVP_top60_DNA_seed | ERBB4 | 0.95716 | STAD | MKN45 | PF-00299804 |
| MVP_top60_DNA_seed | EGFR | 0.95716 | STAD | MKN45 | PF-00299804 |
| MVP_top60_DNA_seed | ERBB2 | 0.957312 | STAD | MKN45 | AZD8931 |
| MVP_top60_DNA_seed | ERBB3 | 0.957312 | STAD | MKN45 | AZD8931 |
| MVP_top60_DNA_seed | EGFR | 0.957312 | STAD | MKN45 | AZD8931 |
| MVP_top60_DNA_seed | ESR1 | 0.957666 | STAD | TGBC11TKB | Fulvestrant |
| MVP_top60_DNA_seed | ERBB2 | 0.961597 | STAD | Hs-746T | Afatinib |
| MVP_top60_DNA_seed | EGFR | 0.961597 | STAD | Hs-746T | Afatinib |
| MVP_top60_DNA_seed | HDAC1 | 0.962033 | STAD | NUGC-3 | Tubastatin A |

| | | | | | |
|---|---|---|---|---|---|
| MVP_top60_DNA_seed | HSP90AA1 | 0.963145 | STAD | MKN45 | Luminespib |
| MVP_top60_DNA_seed | ERBB2 | 0.963799 | STAD | OCUM-1 | Afatinib |
| MVP_top60_DNA_seed | EGFR | 0.963799 | STAD | OCUM-1 | Afatinib |
| MVP_top60_DNA_seed | MET | 0.965121 | STAD | 23132-87 | SU11274 |
| MVP_top60_DNA_seed | AKT1 | 0.965362 | STAD | TGBC11TKB | AKT inhibitor VIII |
| MVP_top60_DNA_seed | AKT1 | 0.971556 | STAD | OCUM-1 | GSK690693 |
| MVP_top60_DNA_seed | ERBB2 | 0.9741 | STAD | SNU-16 | Afatinib |
| MVP_top60_DNA_seed | EGFR | 0.9741 | STAD | SNU-16 | Afatinib |
| MVP_top60_DNA_seed | IGF1R | 0.975447 | STAD | MKN45 | Linsitinib |
| MVP_top60_DNA_seed | MDM2 | 0.97764 | STAD | TMK-1 | Nutlin-3a (-) |
| MVP_top60_DNA_seed | MET | 0.978342 | STAD | MKN28 | AZD6094 |
| MVP_top60_DNA_seed | ERBB2 | 0.981788 | STAD | RF-48 | CP724714 |
| MVP_top60_DNA_seed | FGFR3 | 0.982875 | STAD | MKN45 | AZD4547 |
| MVP_top60_DNA_seed | HDAC1 | 0.983018 | STAD | HSC-39 | Dacinostat |
| MVP_top60_DNA_seed | IGF1R | 0.98364 | STAD | NUGC-3 | BMS-754807 |
| MVP_top60_DNA_seed | ATM | 0.987829 | STAD | MKN45 | KU-55933 |
| MVP_top60_DNA_seed | HDAC1 | 0.991616 | STAD | RERF-GC-1B | Belinostat |
| MVP_top60_DNA_seed | HDAC1 | 0.993198 | STAD | SNU-5 | Parthenolide |
| MVP_top60_DNA_seed | FGFR3 | 0.993217 | STAD | Hs-746T | AZD4547 |
| MVP_top60_DNA_seed | GSK3B | 0.993845 | STAD | FU97 | AZD7969 |
| MVP_top60_DNA_seed | MAPK1 | 1.007774 | STAD | SNU-16 | FR-180204 |
| MVP_top60_DNA_seed | MAPK3 | 1.007774 | STAD | SNU-16 | FR-180204 |
| MVP_top60_DNA_seed | ATM | 1.008654 | STAD | SNU-5 | KU-60019 |
| MVP_top60_DNA_seed | EP300 | 1.013633 | STAD | Hs-746T | I-CBP112 |
| MVP_top60_DNA_seed | CREBBP | 1.013633 | STAD | Hs-746T | I-CBP112 |
| MVP_top60_DNA_seed | EGFR | 1.016793 | STAD | SNU-1 | Cetuximab |
| MVP_top60_DNA_seed | MDM2 | 1.017186 | STAD | TGBC11TKB | Nutlin-3a (-) |
| MVP_top60_DNA_seed | GSK3B | 1.020996 | STAD | RF-48 | CHIR-99021 |
| MVP_top60_DNA_seed | AR | 1.022764 | STAD | OCUM-1 | Bicalutamide |
| MVP_top60_DNA_seed | HSP90AA1 | 1.023274 | STAD | TGBC11TKB | Luminespib |
| MVP_top60_DNA_seed | FGFR3 | 1.026605 | STAD | OCUM-1 | PD173074 |
| MVP_top60_DNA_seed | GSK3B | 1.028067 | STAD | NUGC-3 | CHIR-99021 |
| MVP_top60_DNA_seed | PIK3CA | 1.028821 | STAD | OCUM-1 | AZD8835 |
| MVP_top60_DNA_seed | ERBB2 | 1.030123 | STAD | MKN45 | Afatinib |
| MVP_top60_DNA_seed | EGFR | 1.030123 | STAD | MKN45 | Afatinib |
| MVP_top60_DNA_seed | ESR1 | 1.030497 | STAD | KATOIII | Fulvestrant |
| MVP_top60_DNA_seed | HDAC1 | 1.034564 | STAD | SNU-16 | Tubastatin A |
| MVP_top60_DNA_seed | GSK3B | 1.034863 | STAD | RF-48 | SB216763 |
| MVP_top60_DNA_seed | MET | 1.039295 | STAD | NUGC-3 | SU11274 |
| MVP_top60_DNA_seed | ESR1 | 1.039878 | STAD | Hs-746T | Tamoxifen |
| MVP_top60_DNA_seed | ERBB2 | 1.039935 | STAD | SNU-1 | Lapatinib |

| | | | | | |
|---|---|---|---|---|---|
| MVP_top60_DNA_seed | EGFR | 1.039935 | STAD | SNU-1 | Lapatinib |
| MVP_top60_DNA_seed | AR | 1.042086 | STAD | RF-48 | AZD3514 |
| MVP_top60_DNA_seed | MDM2 | 1.046161 | STAD | MKN7 | Nutlin-3a (-) |
| MVP_top60_DNA_seed | AR | 1.047292 | STAD | SNU-5 | Bicalutamide |
| MVP_top60_DNA_seed | EP300 | 1.050984 | STAD | SNU-1 | I-CBP112 |
| MVP_top60_DNA_seed | CREBBP | 1.050984 | STAD | SNU-1 | I-CBP112 |
| MVP_top60_DNA_seed | EGFR | 1.055442 | STAD | RF-48 | Cetuximab |
| MVP_top60_DNA_seed | IGF1R | 1.056589 | STAD | MKN7 | GSK1904529A |
| MVP_top60_DNA_seed | EGFR | 1.059487 | STAD | SNU-5 | Erlotinib |
| MVP_top60_DNA_seed | AKT1 | 1.059655 | STAD | NUGC-4 | AKT inhibitor VIII |
| MVP_top60_DNA_seed | MDM2 | 1.066648 | STAD | RF-48 | Serdemetan |
| MVP_top60_DNA_seed | AR | 1.067644 | STAD | MKN7 | Bicalutamide |
| MVP_top60_DNA_seed | AKT1 | 1.070062 | STAD | SNU-5 | GSK690693 |
| MVP_top60_DNA_seed | GSK3B | 1.072533 | STAD | SNU-5 | CHIR-99021 |
| MVP_top60_DNA_seed | SRC | 1.073073 | STAD | SNU-16 | NVP-BHG712 |
| MVP_top60_DNA_seed | FGFR3 | 1.073296 | STAD | NUGC-4 | FGFR_3831 |
| MVP_top60_DNA_seed | HDAC1 | 1.074826 | STAD | OCUM-1 | Tubastatin A |
| MVP_top60_DNA_seed | MET | 1.074927 | STAD | RF-48 | PHA-665752 |
| MVP_top60_DNA_seed | ERBB2 | 1.077495 | STAD | OCUM-1 | Afatinib |
| MVP_top60_DNA_seed | EGFR | 1.077495 | STAD | OCUM-1 | Afatinib |
| MVP_top60_DNA_seed | FGFR3 | 1.081305 | STAD | TGBC11TKB | FGFR_3831 |
| MVP_top60_DNA_seed | AKT1 | 1.093208 | STAD | MKN7 | AKT inhibitor VIII |
| MVP_top60_DNA_seed | ERBB2 | 1.095849 | STAD | RF-48 | PF-00299804 |
| MVP_top60_DNA_seed | ERBB4 | 1.095849 | STAD | RF-48 | PF-00299804 |
| MVP_top60_DNA_seed | EGFR | 1.095849 | STAD | RF-48 | PF-00299804 |
| MVP_top60_DNA_seed | HDAC1 | 1.097573 | STAD | MKN7 | Tubastatin A |
| MVP_top60_DNA_seed | ATM | 1.102717 | STAD | MKN7 | KU-60019 |
| MVP_top60_DNA_seed | IGF1R | 1.10573 | STAD | SNU-16 | Linsitinib |
| MVP_top60_DNA_seed | PDGFRA | 1.105921 | STAD | TMK-1 | Masitinib |
| MVP_top60_DNA_seed | ERBB2 | 1.106125 | STAD | RF-48 | Afatinib |
| MVP_top60_DNA_seed | EGFR | 1.106125 | STAD | RF-48 | Afatinib |
| MVP_top60_DNA_seed | IGF1R | 1.110929 | STAD | NUGC-3 | Linsitinib |
| MVP_top60_DNA_seed | PDGFRA | 1.111995 | STAD | SNU-16 | Masitinib |
| MVP_top60_DNA_seed | HDAC1 | 1.113724 | STAD | SNU-16 | CAY10603 |
| MVP_top60_DNA_seed | MDM2 | 1.11666 | STAD | Hs-746T | Nutlin-3a (-) |
| MVP_top60_DNA_seed | ERBB2 | 1.116749 | STAD | SNU-5 | CUDC-101 |
| MVP_top60_DNA_seed | EGFR | 1.116749 | STAD | SNU-5 | CUDC-101 |
| MVP_top60_DNA_seed | HSP90AA1 | 1.119638 | STAD | TGBC11TKB | Elesclomol |
| MVP_top60_DNA_seed | EP300 | 1.123118 | STAD | SNU-5 | I-CBP112 |
| MVP_top60_DNA_seed | CREBBP | 1.123118 | STAD | SNU-5 | I-CBP112 |
| MVP_top60_DNA_seed | MET | 1.123226 | STAD | MKN28 | SU11274 |

| | | | | | |
|---|---|---|---|---|---|
| MVP_top60_DNA_seed | ESR1 | 1.125403 | STAD | MKN45 | Tamoxifen |
| MVP_top60_DNA_seed | MAPK1 | 1.12635 | STAD | SNU-16 | VX-11e |
| MVP_top60_DNA_seed | AR | 1.126367 | STAD | SNU-1 | Bicalutamide |
| MVP_top60_DNA_seed | ATM | 1.128924 | STAD | Hs-746T | CP466722 |
| MVP_top60_DNA_seed | SRC | 1.132299 | STAD | Hs-746T | Bosutinib |
| MVP_top60_DNA_seed | EGFR | 1.132996 | STAD | OCUM-1 | Gefitinib |
| MVP_top60_DNA_seed | PIK3CA | 1.134076 | STAD | MKN45 | AZD8835 |
| MVP_top60_DNA_seed | FGFR3 | 1.134314 | STAD | MKN7 | AZD4547 |
| MVP_top60_DNA_seed | FGFR3 | 1.137501 | STAD | TMK-1 | PD173074 |
| MVP_top60_DNA_seed | HDAC1 | 1.139672 | STAD | TMK-1 | Belinostat |
| MVP_top60_DNA_seed | GSK3B | 1.144515 | STAD | RERF-GC-1B | CHIR-99021 |
| MVP_top60_DNA_seed | GSK3B | 1.152683 | STAD | SNU-1 | CHIR-99021 |
| MVP_top60_DNA_seed | PIK3CA | 1.154174 | STAD | RF-48 | PI3Ka_4409 |
| MVP_top60_DNA_seed | ERBB2 | 1.157487 | STAD | SNU-1 | Afatinib |
| MVP_top60_DNA_seed | EGFR | 1.157487 | STAD | SNU-1 | Afatinib |
| MVP_top60_DNA_seed | IGF1R | 1.160714 | STAD | TGBC11TKB | BMS-754807 |
| MVP_top60_DNA_seed | GSK3B | 1.168373 | STAD | SNU-5 | CHIR-99021 |
| MVP_top60_DNA_seed | PDGFRA | 1.179917 | STAD | OCUM-1 | Masitinib |
| MVP_top60_DNA_seed | AKT1 | 1.186697 | STAD | MKN45 | AKT inhibitor VIII |
| MVP_top60_DNA_seed | HSP90AA1 | 1.186903 | STAD | GCIY | Elesclomol |
| MVP_top60_DNA_seed | HDAC1 | 1.19123 | STAD | MKN7 | JQ12 |
| MVP_top60_DNA_seed | ESR1 | 1.193091 | STAD | NCI-N87 | Tamoxifen |
| MVP_top60_DNA_seed | FGFR3 | 1.195394 | STAD | Hs-746T | FGFR_3831 |
| MVP_top60_DNA_seed | HSP90AA1 | 1.200792 | STAD | MKN7 | Tanespimycin |
| MVP_top60_DNA_seed | IGF1R | 1.201183 | STAD | SK-GT-2 | BMS-536924 |
| MVP_top60_DNA_seed | IGF1R | 1.202969 | STAD | MKN7 | BMS-754807 |
| MVP_top60_DNA_seed | PIK3CA | 1.214622 | STAD | Hs-746T | AZD8835 |
| MVP_top60_DNA_seed | ERBB4 | 1.216366 | STAD | RF-48 | AST-1306 |
| MVP_top60_DNA_seed | EGFR | 1.216366 | STAD | RF-48 | AST-1306 |
| MVP_top60_DNA_seed | EGFR | 1.224546 | STAD | TGBC11TKB | Pelitinib |
| MVP_top60_DNA_seed | MET | 1.233141 | STAD | NCI-N87 | AZD6094 |
| MVP_top60_DNA_seed | MET | 1.234687 | STAD | NCI-N87 | SU11274 |
| MVP_top60_DNA_seed | ERBB2 | 1.235015 | STAD | SNU-16 | CUDC-101 |
| MVP_top60_DNA_seed | EGFR | 1.235015 | STAD | SNU-16 | CUDC-101 |
| MVP_top60_DNA_seed | HDAC1 | 1.241302 | STAD | MKN45 | Tubastatin A |
| MVP_top60_DNA_seed | SRC | 1.243658 | STAD | MKN7 | NVP-BHG712 |
| MVP_top60_DNA_seed | GSK3B | 1.244341 | STAD | RERF-GC-1B | KIN001-042 |
| MVP_top60_DNA_seed | AR | 1.24747 | STAD | MKN7 | AZD3514 |
| MVP_top60_DNA_seed | EGFR | 1.250439 | STAD | OCUM-1 | Pelitinib |
| MVP_top60_DNA_seed | MET | 1.251315 | STAD | OCUM-1 | SU11274 |
| MVP_top60_DNA_seed | IGF1R | 1.253537 | STAD | RF-48 | GSK1904529A |

| | | | | | |
|---|---|---|---|---|---|
| MVP_top60_DNA_seed | ERBB2 | 1.254838 | STAD | SNU-5 | CP724714 |
| MVP_top60_DNA_seed | MET | 1.255613 | STAD | 23132-87 | JNJ38877605 |
| MVP_top60_DNA_seed | SRC | 1.258796 | STAD | SNU-1 | WH-4-023 |
| MVP_top60_DNA_seed | EGFR | 1.261047 | STAD | MKN45 | Cetuximab |
| MVP_top60_DNA_seed | HDAC1 | 1.264024 | STAD | RERF-GC-1B | AR-42 |
| MVP_top60_DNA_seed | ATM | 1.264547 | STAD | NUGC-3 | CP466722 |
| MVP_top60_DNA_seed | AKT1 | 1.267448 | STAD | OCUM-1 | AKT inhibitor VIII |
| MVP_top60_DNA_seed | EP300 | 1.279924 | STAD | MKN45 | I-CBP112 |
| MVP_top60_DNA_seed | CREBBP | 1.279924 | STAD | MKN45 | I-CBP112 |
| MVP_top60_DNA_seed | MET | 1.283609 | STAD | SNU-16 | SU11274 |
| MVP_top60_DNA_seed | HDAC1 | 1.286909 | STAD | SNU-5 | Dacinostat |
| MVP_top60_DNA_seed | HDAC1 | 1.287335 | STAD | SNU-5 | Tubastatin A |
| MVP_top60_DNA_seed | HSP90AA1 | 1.287695 | STAD | NUGC-4 | CCT-018159 |
| MVP_top60_DNA_seed | AKT1 | 1.292285 | STAD | TMK-1 | AKT inhibitor VIII |
| MVP_top60_DNA_seed | HSP90AA1 | 1.295328 | STAD | HSC-39 | Luminespib |
| MVP_top60_DNA_seed | ATM | 1.297269 | STAD | RERF-GC-1B | KU-55933 |
| MVP_top60_DNA_seed | EGFR | 1.300367 | STAD | SNU-16 | Cetuximab |
| MVP_top60_DNA_seed | ERBB2 | 1.306524 | STAD | MKN45 | CP724714 |
| MVP_top60_DNA_seed | GSK3B | 1.307288 | STAD | SNU-1 | AZD7969 |
| MVP_top60_DNA_seed | SRC | 1.3073 | STAD | SNU-16 | Saracatinib |
| MVP_top60_DNA_seed | PDGFRA | 1.309532 | STAD | MKN45 | Masitinib |
| MVP_top60_DNA_seed | ATM | 1.309781 | STAD | OCUM-1 | KU-55933 |
| MVP_top60_DNA_seed | EGFR | 1.312675 | STAD | MKN45 | Gefitinib |
| MVP_top60_DNA_seed | GSK3B | 1.322174 | STAD | RF-48 | CHIR-99021 |
| MVP_top60_DNA_seed | PDGFRA | 1.329032 | STAD | NUGC-3 | Masitinib |
| MVP_top60_DNA_seed | SRC | 1.340497 | STAD | TMK-1 | NVP-BHG712 |
| MVP_top60_DNA_seed | AKT1 | 1.3423 | STAD | SNU-16 | AKT inhibitor VIII |
| MVP_top60_DNA_seed | ATM | 1.342585 | STAD | SNU-16 | KU-60019 |
| MVP_top60_DNA_seed | AKT1 | 1.342707 | STAD | NCI-N87 | GSK690693 |
| MVP_top60_DNA_seed | AR | 1.345234 | STAD | MKN45 | Bicalutamide |
| MVP_top60_DNA_seed | FGFR3 | 1.351224 | STAD | GCIY | AZD4547 |
| MVP_top60_DNA_seed | ATM | 1.351348 | STAD | MKN28 | CP466722 |
| MVP_top60_DNA_seed | FGFR3 | 1.351743 | STAD | MKN7 | FGFR_3831 |
| MVP_top60_DNA_seed | IGF1R | 1.355878 | STAD | SNU-5 | Linsitinib |
| MVP_top60_DNA_seed | HSP90AA1 | 1.359293 | STAD | RF-48 | Elesclomol |
| MVP_top60_DNA_seed | MAPK1 | 1.359375 | STAD | SNU-5 | VX-11e |
| MVP_top60_DNA_seed | GSK3B | 1.361364 | STAD | Hs-746T | CHIR-99021 |
| MVP_top60_DNA_seed | AKT1 | 1.364001 | STAD | Hs-746T | AKT inhibitor VIII |
| MVP_top60_DNA_seed | HSP90AA1 | 1.364521 | STAD | IM-95 | Elesclomol |
| MVP_top60_DNA_seed | HSP90AA1 | 1.366052 | STAD | NUGC-4 | Elesclomol |
| MVP_top60_DNA_seed | GSK3B | 1.374242 | STAD | SNU-1 | KIN001-042 |

| | | | | | |
|---|---|---|---|---|---|
| MVP_top60_DNA_seed | IGF1R | 1.376614 | STAD | SNU-1 | GSK1904529A |
| MVP_top60_DNA_seed | IGF1R | 1.377586 | STAD | SNU-5 | GSK1904529A |
| MVP_top60_DNA_seed | AKT1 | 1.38167 | STAD | NUGC-3 | AKT inhibitor VIII |
| MVP_top60_DNA_seed | SRC | 1.382787 | STAD | OCUM-1 | NVP-BHG712 |
| MVP_top60_DNA_seed | FGFR3 | 1.384828 | STAD | NUGC-4 | AZD4547 |
| MVP_top60_DNA_seed | ATM | 1.392843 | STAD | RF-48 | KU-55933 |
| MVP_top60_DNA_seed | MAPK1 | 1.393199 | STAD | SNU-5 | FR-180204 |
| MVP_top60_DNA_seed | MAPK3 | 1.393199 | STAD | SNU-5 | FR-180204 |
| MVP_top60_DNA_seed | AR | 1.395201 | STAD | KATOIII | AZD3514 |
| MVP_top60_DNA_seed | ERBB2 | 1.395564 | STAD | MKN45 | CUDC-101 |
| MVP_top60_DNA_seed | EGFR | 1.395564 | STAD | MKN45 | CUDC-101 |
| MVP_top60_DNA_seed | EGFR | 1.401933 | STAD | TMK-1 | Pelitinib |
| MVP_top60_DNA_seed | GSK3B | 1.402784 | STAD | TGBC11TKB | AZD7969 |
| MVP_top60_DNA_seed | AKT1 | 1.407707 | STAD | TGBC11TKB | BAY AKT1 |
| MVP_top60_DNA_seed | ESR1 | 1.40806 | STAD | Hs-746T | Fulvestrant |
| MVP_top60_DNA_seed | MDM2 | 1.409034 | STAD | NCI-N87 | Nutlin-3a (-) |
| MVP_top60_DNA_seed | MET | 1.410312 | STAD | RF-48 | SU11274 |
| MVP_top60_DNA_seed | ESR1 | 1.413893 | STAD | RF-48 | Fulvestrant |
| MVP_top60_DNA_seed | EGFR | 1.427712 | STAD | SNU-16 | Pelitinib |
| MVP_top60_DNA_seed | MDM2 | 1.438108 | STAD | OCUM-1 | Serdemetan |
| MVP_top60_DNA_seed | IGF1R | 1.442478 | STAD | SNU-1 | BMS-536924 |
| MVP_top60_DNA_seed | AKT1 | 1.44525 | STAD | MKN7 | BAY AKT1 |
| MVP_top60_DNA_seed | FGFR3 | 1.447633 | STAD | SK-GT-2 | AZD4547 |
| MVP_top60_DNA_seed | AR | 1.458384 | STAD | SNU-16 | Bicalutamide |
| MVP_top60_DNA_seed | HSP90AA1 | 1.464536 | STAD | OCUM-1 | SNX-2112 |
| MVP_top60_DNA_seed | HSP90AA1 | 1.466064 | STAD | SNU-16 | SNX-2112 |
| MVP_top60_DNA_seed | ESR1 | 1.469051 | STAD | MKN45 | Fulvestrant |
| MVP_top60_DNA_seed | SRC | 1.479698 | STAD | SNU-5 | NVP-BHG712 |
| MVP_top60_DNA_seed | AKT1 | 1.481516 | STAD | NUGC-4 | BAY AKT1 |
| MVP_top60_DNA_seed | IGF1R | 1.486068 | STAD | MKN45 | BMS-536924 |
| MVP_top60_DNA_seed | ATM | 1.491787 | STAD | TMK-1 | CP466722 |
| MVP_top60_DNA_seed | GSK3B | 1.497845 | STAD | Hs-746T | SB216763 |
| MVP_top60_DNA_seed | ATM | 1.500206 | STAD | NCI-N87 | CP466722 |
| MVP_top60_DNA_seed | ATM | 1.503181 | STAD | SNU-5 | KU-55933 |
| MVP_top60_DNA_seed | MET | 1.51024 | STAD | RERF-GC-1B | JNJ38877605 |
| MVP_top60_DNA_seed | GSK3B | 1.514643 | STAD | Hs-746T | AZD7969 |
| MVP_top60_DNA_seed | HDAC1 | 1.516312 | STAD | SNU-16 | Dacinostat |
| MVP_top60_DNA_seed | HSP90AA1 | 1.516402 | STAD | RERF-GC-1B | CCT-018159 |
| MVP_top60_DNA_seed | SRC | 1.517876 | STAD | SNU-5 | Saracatinib |
| MVP_top60_DNA_seed | FGFR3 | 1.520786 | STAD | RF-48 | FGFR_3831 |
| MVP_top60_DNA_seed | HSP90AA1 | 1.525398 | STAD | SNU-16 | CCT-018159 |

| | | | | | |
|---|---|---|---|---|---|
| MVP_top60_DNA_seed | EGFR | 1.532345 | STAD | MKN7 | Pelitinib |
| MVP_top60_DNA_seed | ERBB2 | 1.538234 | STAD | SNU-16 | CI-1033 |
| MVP_top60_DNA_seed | ERBB4 | 1.538234 | STAD | SNU-16 | CI-1033 |
| MVP_top60_DNA_seed | EGFR | 1.538234 | STAD | SNU-16 | CI-1033 |
| MVP_top60_DNA_seed | EGFR | 1.545938 | STAD | Hs-746T | Gefitinib |
| MVP_top60_DNA_seed | HSP90AA1 | 1.561323 | STAD | HSC-39 | SNX-2112 |
| MVP_top60_DNA_seed | MAPK1 | 1.562867 | STAD | MKN45 | FR-180204 |
| MVP_top60_DNA_seed | MAPK3 | 1.562867 | STAD | MKN45 | FR-180204 |
| MVP_top60_DNA_seed | MET | 1.568048 | STAD | NUGC-3 | JNJ38877605 |
| MVP_top60_DNA_seed | FGFR3 | 1.575208 | STAD | Hs-746T | AZD4547 |
| MVP_top60_DNA_seed | MAPK1 | 1.579484 | STAD | Hs-746T | FR-180204 |
| MVP_top60_DNA_seed | MAPK3 | 1.579484 | STAD | Hs-746T | FR-180204 |
| MVP_top60_DNA_seed | GSK3B | 1.581028 | STAD | NCI-N87 | CHIR-99021 |
| MVP_top60_DNA_seed | PDGFRA | 1.58717 | STAD | SNU-5 | Masitinib |
| MVP_top60_DNA_seed | IGF1R | 1.590636 | STAD | SNU-5 | BMS-536924 |
| MVP_top60_DNA_seed | HDAC1 | 1.591227 | STAD | SNU-5 | JQ12 |
| MVP_top60_DNA_seed | EGFR | 1.599433 | STAD | NCI-N87 | Gefitinib |
| MVP_top60_DNA_seed | HDAC1 | 1.603425 | STAD | MKN45 | CAY10603 |
| MVP_top60_DNA_seed | AR | 1.604421 | STAD | IM-95 | AZD3514 |
| MVP_top60_DNA_seed | HDAC1 | 1.641715 | STAD | MKN45 | Belinostat |
| MVP_top60_DNA_seed | EGFR | 1.646496 | STAD | SNU-5 | Pelitinib |
| MVP_top60_DNA_seed | AR | 1.671666 | STAD | NCI-N87 | Bicalutamide |
| MVP_top60_DNA_seed | GSK3B | 1.675039 | STAD | SNU-5 | SB216763 |
| MVP_top60_DNA_seed | IGF1R | 1.708497 | STAD | MKN7 | BMS-536924 |
| MVP_top60_DNA_seed | MET | 1.717216 | STAD | OCUM-1 | JNJ38877605 |
| MVP_top60_DNA_seed | AKT1 | 1.731044 | STAD | TGBC11TKB | MK-2206 |
| MVP_top60_DNA_seed | FGFR3 | 1.732734 | STAD | NCI-N87 | AZD4547 |
| MVP_top60_DNA_seed | FGFR3 | 1.740636 | STAD | SNU-1 | AZD4547 |
| MVP_top60_DNA_seed | ERBB2 | 1.742186 | STAD | RERF-GC-1B | CUDC-101 |
| MVP_top60_DNA_seed | EGFR | 1.742186 | STAD | RERF-GC-1B | CUDC-101 |
| MVP_top60_DNA_seed | AR | 1.743914 | STAD | SNU-5 | AZD3514 |
| MVP_top60_DNA_seed | AKT1 | 1.746835 | STAD | SNU-5 | AKT inhibitor VIII |
| MVP_top60_DNA_seed | AR | 1.763446 | STAD | TMK-1 | AZD3514 |
| MVP_top60_DNA_seed | HSP90AA1 | 1.775021 | STAD | RERF-GC-1B | Tanespimycin |
| MVP_top60_DNA_seed | GSK3B | 1.7764 | STAD | RF-48 | AZD7969 |
| MVP_top60_DNA_seed | HSP90AA1 | 1.780029 | STAD | SNU-5 | SNX-2112 |
| MVP_top60_DNA_seed | MET | 1.786694 | STAD | RF-48 | JNJ38877605 |
| MVP_top60_DNA_seed | HDAC1 | 1.787583 | STAD | MKN45 | AR-42 |
| MVP_top60_DNA_seed | FGFR3 | 1.800818 | STAD | SNU-1 | FGFR_3831 |
| MVP_top60_DNA_seed | ATM | 1.856631 | STAD | MKN28 | KU-60019 |
| MVP_top60_DNA_seed | AR | 1.860389 | STAD | NCI-N87 | Bicalutamide |

| | | | | | |
|---|---|---|---|---|---|
| MVP_top60_DNA_seed | AKT1 | 1.868179 | STAD | SNU-16 | MK-2206 |
| MVP_top60_DNA_seed | ERBB2 | 1.870352 | STAD | SNU-16 | PF-00299804 |
| MVP_top60_DNA_seed | ERBB4 | 1.870352 | STAD | SNU-16 | PF-00299804 |
| MVP_top60_DNA_seed | EGFR | 1.870352 | STAD | SNU-16 | PF-00299804 |
| MVP_top60_DNA_seed | FGFR3 | 1.871968 | STAD | SNU-5 | FGFR_3831 |
| MVP_top60_DNA_seed | IGF1R | 1.873926 | STAD | SNU-5 | BMS-536924 |
| MVP_top60_DNA_seed | FGFR3 | 1.889612 | STAD | MKN45 | FGFR_3831 |
| MVP_top60_DNA_seed | IGF1R | 1.893876 | STAD | NCI-N87 | Linsitinib |
| MVP_top60_DNA_seed | MDM2 | 1.900555 | STAD | SNU-1 | Serdemetan |
| MVP_top60_DNA_seed | HDAC1 | 1.915128 | STAD | SNU-16 | JQ12 |
| MVP_top60_DNA_seed | PDGFRA | 1.92009 | STAD | NCI-N87 | Amuvatinib |
| MVP_top60_DNA_seed | PIK3CA | 1.925783 | STAD | SNU-1 | PI3Ka_4409 |
| MVP_top60_DNA_seed | HSP90AA1 | 1.9342 | STAD | SNU-1 | CCT-018159 |
| MVP_top60_DNA_seed | ESR1 | 1.943349 | STAD | SNU-5 | Fulvestrant |
| MVP_top60_DNA_seed | GSK3B | 1.945581 | STAD | RERF-GC-1B | AZD7969 |
| MVP_top60_DNA_seed | ATM | 1.951166 | STAD | AGS | CP466722 |
| MVP_top60_DNA_seed | AKT1 | 1.966951 | STAD | SNU-5 | AKT inhibitor VIII |
| MVP_top60_DNA_seed | GSK3B | 1.987441 | STAD | MKN45 | KIN001-042 |
| MVP_top60_DNA_seed | IGF1R | 1.998483 | STAD | Hs-746T | BMS-536924 |
| MVP_top60_DNA_seed | MET | 2.02422 | STAD | SNU-16 | JNJ38877605 |
| MVP_top60_DNA_seed | GSK3B | 2.049871 | STAD | NCI-N87 | KIN001-042 |
| MVP_top60_DNA_seed | ATM | 2.055851 | STAD | SNU-1 | KU-60019 |
| MVP_top60_DNA_seed | ATM | 2.09245 | STAD | NCI-N87 | KU-55933 |
| MVP_top60_DNA_seed | ATM | 2.104122 | STAD | SNU-16 | CP466722 |
| MVP_top60_DNA_seed | FGFR3 | 2.110271 | STAD | NCI-N87 | FGFR_3831 |
| MVP_top60_DNA_seed | ATM | 2.140308 | STAD | RF-48 | KU-60019 |
| MVP_top60_DNA_seed | ERBB4 | 2.175457 | STAD | SNU-16 | AST-1306 |
| MVP_top60_DNA_seed | EGFR | 2.175457 | STAD | SNU-16 | AST-1306 |
| MVP_top60_DNA_seed | PIK3CA | 2.177344 | STAD | SNU-1 | AZD8835 |
| MVP_top60_DNA_seed | HDAC1 | 2.192992 | STAD | Hs-746T | JQ12 |
| MVP_top60_DNA_seed | GSK3B | 2.210329 | STAD | MKN7 | AZD7969 |
| MVP_top60_DNA_seed | ERBB2 | 2.219959 | STAD | OCUM-1 | CUDC-101 |
| MVP_top60_DNA_seed | EGFR | 2.219959 | STAD | OCUM-1 | CUDC-101 |
| MVP_top60_DNA_seed | IGF1R | 2.225572 | STAD | NCI-N87 | BMS-754807 |
| MVP_top60_DNA_seed | ERBB2 | 2.327314 | STAD | SNU-1 | AZD8931 |
| MVP_top60_DNA_seed | ERBB3 | 2.327314 | STAD | SNU-1 | AZD8931 |
| MVP_top60_DNA_seed | EGFR | 2.327314 | STAD | SNU-1 | AZD8931 |
| MVP_top60_DNA_seed | ERBB4 | 2.333127 | STAD | OCUM-1 | AST-1306 |
| MVP_top60_DNA_seed | EGFR | 2.333127 | STAD | OCUM-1 | AST-1306 |
| MVP_top60_DNA_seed | ERBB2 | 2.343604 | STAD | OCUM-1 | CI-1033 |
| MVP_top60_DNA_seed | ERBB4 | 2.343604 | STAD | OCUM-1 | CI-1033 |

| | | | | | |
|---|---|---|---|---|---|
| MVP_top60_DNA_seed | EGFR | 2.343604 | STAD | OCUM-1 | CI-1033 |
| MVP_top60_DNA_seed | ATM | 2.430569 | STAD | SNU-5 | CP466722 |
| MVP_top60_DNA_seed | MET | 2.452519 | STAD | NCI-N87 | JNJ38877605 |
| MVP_top60_DNA_seed | ERBB4 | 2.506507 | STAD | MKN45 | AST-1306 |
| MVP_top60_DNA_seed | EGFR | 2.506507 | STAD | MKN45 | AST-1306 |
| MVP_top60_DNA_seed | ERBB2 | 2.62529 | STAD | SNU-5 | CI-1033 |
| MVP_top60_DNA_seed | ERBB4 | 2.62529 | STAD | SNU-5 | CI-1033 |
| MVP_top60_DNA_seed | EGFR | 2.62529 | STAD | SNU-5 | CI-1033 |
| MVP_top60_DNA_seed | HDAC1 | 2.639529 | STAD | OCUM-1 | AR-42 |
| MVP_top60_DNA_seed | HDAC1 | 2.657814 | STAD | OCUM-1 | Belinostat |
| MVP_top60_DNA_seed | HSP90AA1 | 2.683601 | STAD | SNU-16 | Luminespib |
| MVP_top60_DNA_seed | HDAC1 | 2.724627 | STAD | OCUM-1 | CAY10603 |
| MVP_top60_DNA_seed | ERBB2 | 3.33718 | STAD | SNU-5 | PF-00299804 |
| MVP_top60_DNA_seed | ERBB4 | 3.33718 | STAD | SNU-5 | PF-00299804 |
| MVP_top60_DNA_seed | EGFR | 3.33718 | STAD | SNU-5 | PF-00299804 |
| MVP_top60_DNA_seed | ERBB2 | 3.35161 | STAD | OCUM-1 | PF-00299804 |
| MVP_top60_DNA_seed | ERBB4 | 3.35161 | STAD | OCUM-1 | PF-00299804 |
| MVP_top60_DNA_seed | EGFR | 3.35161 | STAD | OCUM-1 | PF-00299804 |
| MVP_top60_DNA_seed | ERBB4 | 3.424574 | STAD | SNU-5 | AST-1306 |
| MVP_top60_DNA_seed | EGFR | 3.424574 | STAD | SNU-5 | AST-1306 |
| MVP_top60_DNA_seed | MET | -6.275219 | STAD | MKN45 | Savolitinib |
| MVP_top60_DNA_seed | MET | -6.100559 | STAD | SNU-5 | Savolitinib |
| MVP_top60_DNA_seed | MET | -5.621499 | STAD | Hs-746T | Savolitinib |
| MVP_top60_DNA_seed | FGFR3 | -5.226735 | STAD | SNU-16 | PD173074 |
| MVP_top60_DNA_seed | ERBB2 | -4.964046 | STAD | NCI-N87 | Afatinib |
| MVP_top60_DNA_seed | EGFR | -4.964046 | STAD | NCI-N87 | Afatinib |
| MVP_top60_DNA_seed | FGFR3 | -4.86989 | STAD | HSC-39 | PD173074 |
| MVP_top60_DNA_seed | FGFR3 | -4.705369 | STAD | HSC-39 | AZD4547 |
| MVP_top60_DNA_seed | FGFR3 | -4.610762 | STAD | SNU-16 | AZD4547 |
| MVP_top60_DNA_seed | ERBB2 | -4.104673 | STAD | SK-GT-2 | Afatinib |
| MVP_top60_DNA_seed | EGFR | -4.104673 | STAD | SK-GT-2 | Afatinib |
| MVP_top60_DNA_seed | ERBB2 | -3.70213 | STAD | NCI-N87 | Lapatinib |
| MVP_top60_DNA_seed | EGFR | -3.70213 | STAD | NCI-N87 | Lapatinib |
| MVP_top60_DNA_seed | EGFR | -3.6903 | STAD | SK-GT-2 | Osimertinib |
| MVP_top60_DNA_seed | ESR1 | -3.532472 | STAD | FU97 | GDC0810 |
| MVP_top60_DNA_seed | EGFR | -3.476284 | STAD | NCI-N87 | Osimertinib |
| MVP_top60_DNA_seed | MET | -3.373267 | STAD | SNU-5 | Crizotinib |
| MVP_top60_DNA_seed | FGFR3 | -3.328162 | STAD | KATOIII | AZD4547 |
| MVP_top60_DNA_seed | ERBB2 | -3.096553 | STAD | NCI-N87 | Sapitinib |
| MVP_top60_DNA_seed | ERBB3 | -3.096553 | STAD | NCI-N87 | Sapitinib |
| MVP_top60_DNA_seed | EGFR | -3.096553 | STAD | NCI-N87 | Sapitinib |

| | | | | | |
|---|---|---|---|---|---|
| MVP_top60_DNA_seed | MET | -3.039269 | STAD | MKN45 | Crizotinib |
| MVP_top60_DNA_seed | EGFR | -3.029148 | STAD | SK-GT-2 | Gefitinib |
| MVP_top60_DNA_seed | MAPK1 | -2.97902 | STAD | AGS | VX-11e |
| MVP_top60_DNA_seed | ERBB2 | -2.940765 | STAD | SK-GT-2 | Sapitinib |
| MVP_top60_DNA_seed | ERBB3 | -2.940765 | STAD | SK-GT-2 | Sapitinib |
| MVP_top60_DNA_seed | EGFR | -2.940765 | STAD | SK-GT-2 | Sapitinib |
| MVP_top60_DNA_seed | MAPK1 | -2.804725 | STAD | AGS | SCH772984 |
| MVP_top60_DNA_seed | MAPK3 | -2.804725 | STAD | AGS | SCH772984 |
| MVP_top60_DNA_seed | MAPK1 | -2.791952 | STAD | OCUM-1 | SCH772984 |
| MVP_top60_DNA_seed | MAPK3 | -2.791952 | STAD | OCUM-1 | SCH772984 |
| MVP_top60_DNA_seed | AKT1 | -2.738816 | STAD | HGC-27 | Afuresertib |
| MVP_top60_DNA_seed | MAPK1 | -2.726927 | STAD | OCUM-1 | VX-11e |
| MVP_top60_DNA_seed | FGFR3 | -2.695488 | STAD | KATOIII | PD173074 |
| MVP_top60_DNA_seed | AKT1 | -2.611303 | STAD | HGC-27 | Ipatasertib |
| MVP_top60_DNA_seed | EGFR | -2.523602 | STAD | SK-GT-2 | Erlotinib |
| MVP_top60_DNA_seed | ERBB2 | -2.431962 | STAD | SK-GT-2 | Lapatinib |
| MVP_top60_DNA_seed | EGFR | -2.431962 | STAD | SK-GT-2 | Lapatinib |
| MVP_top60_DNA_seed | AKT1 | -2.425695 | STAD | HGC-27 | AZD5363 |
| MVP_top60_DNA_seed | IGF1R | -2.377363 | STAD | HSC-39 | BMS-536924 |
| MVP_top60_DNA_seed | MAPK1 | -2.237307 | STAD | AGS | Ulixertinib |
| MVP_top60_DNA_seed | MAPK3 | -2.237307 | STAD | AGS | Ulixertinib |
| MVP_top60_DNA_seed | HSP90AA1 | -2.236141 | STAD | HSC-39 | Tanespimycin |
| MVP_top60_DNA_seed | AKT1 | -2.234097 | STAD | HGC-27 | GSK2110183B |
| MVP_top60_DNA_seed | AKT1 | -2.232916 | STAD | HGC-27 | MK-2206 |
| MVP_top60_DNA_seed | PIK3CA | -2.195662 | STAD | NCI-N87 | Alpelisib |
| MVP_top60_DNA_seed | AKT1 | -2.185663 | STAD | HGC-27 | Uprosertib |
| MVP_top60_DNA_seed | PIK3CA | -2.178419 | STAD | HGC-27 | Alpelisib |
| MVP_top60_DNA_seed | MAPK1 | -2.14477 | STAD | OCUM-1 | Ulixertinib |
| MVP_top60_DNA_seed | MAPK3 | -2.14477 | STAD | OCUM-1 | Ulixertinib |
| MVP_top60_DNA_seed | IGF1R | -2.115001 | STAD | RERF-GC-1B | BMS-754807 |
| MVP_top60_DNA_seed | EGFR | -2.107292 | STAD | NUGC-3 | Gefitinib |
| MVP_top60_DNA_seed | EGFR | -2.058122 | STAD | NUGC-3 | Erlotinib |
| MVP_top60_DNA_seed | PIK3CA | -2.002491 | STAD | HSC-39 | Alpelisib |
| MVP_top60_DNA_seed | MAPK1 | -1.999458 | STAD | TGBC11TKB | VX-11e |
| MVP_top60_DNA_seed | MDM2 | -1.989513 | STAD | OCUM-1 | Nutlin-3a (-) |
| MVP_top60_DNA_seed | EGFR | -1.986916 | STAD | SK-GT-2 | AZD3759 |
| MVP_top60_DNA_seed | HDAC1 | -1.981708 | STAD | TMK-1 | Entinostat |
| MVP_top60_DNA_seed | ATM | -1.953722 | STAD | HSC-39 | KU-55933 |
| MVP_top60_DNA_seed | MET | -1.95312 | STAD | IM-95 | Savolitinib |
| MVP_top60_DNA_seed | IGF1R | -1.929294 | STAD | RERF-GC-1B | BMS-536924 |
| MVP_top60_DNA_seed | AKT1 | -1.925667 | STAD | IM-95 | GSK2110183B |

| | | | | | |
|---|---|---|---|---|---|
| MVP_top60_DNA_seed | IGF1R | -1.907502 | STAD | RERF-GC-1B | NVP-ADW742 |
| MVP_top60_DNA_seed | AKT1 | -1.903326 | STAD | IM-95 | Uprosertib |
| MVP_top60_DNA_seed | PIK3CA | -1.889177 | STAD | HGC-27 | Buparlisib |
| MVP_top60_DNA_seed | PIK3CA | -1.879573 | STAD | 23132-87 | AZD8186 |
| MVP_top60_DNA_seed | MAPK1 | -1.874616 | STAD | OCUM-1 | Ulixertinib |
| MVP_top60_DNA_seed | MAPK3 | -1.874616 | STAD | OCUM-1 | Ulixertinib |
| MVP_top60_DNA_seed | PIK3CA | -1.873533 | STAD | HGC-27 | GNE-317 |
| MVP_top60_DNA_seed | AKT1 | -1.836172 | STAD | HGC-27 | Uprosertib |
| MVP_top60_DNA_seed | PIK3CA | -1.806595 | STAD | AGS | Alpelisib |
| MVP_top60_DNA_seed | ESR1 | -1.747615 | STAD | HSC-39 | Tamoxifen |
| MVP_top60_DNA_seed | IGF1R | -1.738342 | STAD | SNU-16 | BMS-536924 |
| MVP_top60_DNA_seed | IGF1R | -1.731946 | STAD | RERF-GC-1B | Linsitinib |
| MVP_top60_DNA_seed | EGFR | -1.71851 | STAD | SNU-1 | Erlotinib |
| MVP_top60_DNA_seed | ESR1 | -1.713851 | STAD | SNU-16 | Tamoxifen |
| MVP_top60_DNA_seed | ERBB2 | -1.699082 | STAD | NUGC-3 | Sapitinib |
| MVP_top60_DNA_seed | ERBB3 | -1.699082 | STAD | NUGC-3 | Sapitinib |
| MVP_top60_DNA_seed | EGFR | -1.699082 | STAD | NUGC-3 | Sapitinib |
| MVP_top60_DNA_seed | PIK3CA | -1.690563 | STAD | HGC-27 | AZD8186 |
| MVP_top60_DNA_seed | MET | -1.676809 | STAD | Hs-746T | Crizotinib |
| MVP_top60_DNA_seed | PIK3CA | -1.661051 | STAD | FU97 | Alpelisib |
| MVP_top60_DNA_seed | HSP90AA1 | -1.648706 | STAD | SNU-16 | Tanespimycin |
| MVP_top60_DNA_seed | AKT1 | -1.644049 | STAD | AGS | MK-2206 |
| MVP_top60_DNA_seed | ESR1 | -1.631682 | STAD | FU97 | Tamoxifen |
| MVP_top60_DNA_seed | EP300 | -1.624699 | STAD | RF-48 | SGC-CBP30 |
| MVP_top60_DNA_seed | CREBBP | -1.624699 | STAD | RF-48 | SGC-CBP30 |
| MVP_top60_DNA_seed | HSP90AA1 | -1.603973 | STAD | AGS | Tanespimycin |
| MVP_top60_DNA_seed | ESR1 | -1.599474 | STAD | RERF-GC-1B | GDC0810 |
| MVP_top60_DNA_seed | SRC | -1.581791 | STAD | RERF-GC-1B | Bosutinib |
| MVP_top60_DNA_seed | MDM2 | -1.580495 | STAD | RF-48 | Nutlin-3a (-) |
| MVP_top60_DNA_seed | AKT1 | -1.575102 | STAD | 23132-87 | Uprosertib |
| MVP_top60_DNA_seed | HSP90AA1 | -1.545979 | STAD | TGBC11TKB | Tanespimycin |
| MVP_top60_DNA_seed | HSP90AA1 | -1.539666 | STAD | OCUM-1 | Tanespimycin |
| MVP_top60_DNA_seed | AKT1 | -1.516725 | STAD | 23132-87 | Uprosertib |
| MVP_top60_DNA_seed | PIK3CA | -1.513459 | STAD | HSC-39 | GNE-317 |
| MVP_top60_DNA_seed | HSP90AA1 | -1.504071 | STAD | HSC-39 | Luminespib |
| MVP_top60_DNA_seed | FGFR3 | -1.498662 | STAD | SNU-1 | AZD4547 |
| MVP_top60_DNA_seed | IGF1R | -1.495139 | STAD | SNU-16 | GSK1904529A |
| MVP_top60_DNA_seed | MDM2 | -1.489373 | STAD | AGS | Nutlin-3a (-) |
| MVP_top60_DNA_seed | IGF1R | -1.483885 | STAD | HSC-39 | Linsitinib |
| MVP_top60_DNA_seed | HDAC1 | -1.48244 | STAD | AGS | PCI-34051 |
| MVP_top60_DNA_seed | MAPK1 | -1.47265 | STAD | MKN28 | Ulixertinib |

| | | | | | |
|---|---|---|---|---|---|
| MVP_top60_DNA_seed | MAPK3 | -1.47265 | STAD | MKN28 | Ulixertinib |
| MVP_top60_DNA_seed | MAPK1 | -1.45617 | STAD | AGS | Ulixertinib |
| MVP_top60_DNA_seed | MAPK3 | -1.45617 | STAD | AGS | Ulixertinib |
| MVP_top60_DNA_seed | IGF1R | -1.452679 | STAD | SNU-16 | Linsitinib |
| MVP_top60_DNA_seed | EGFR | -1.439239 | STAD | NUGC-3 | AZD3759 |
| MVP_top60_DNA_seed | FGFR3 | -1.438175 | STAD | FU97 | PD173074 |
| MVP_top60_DNA_seed | AKT1 | -1.431153 | STAD | AGS | Uprosertib |
| MVP_top60_DNA_seed | AKT1 | -1.430333 | STAD | NCI-N87 | AT13148 |
| MVP_top60_DNA_seed | EP300 | -1.407546 | STAD | SNU-16 | CPI-637 |
| MVP_top60_DNA_seed | MET | -1.406037 | STAD | SNU-16 | Crizotinib |
| MVP_top60_DNA_seed | MET | -1.398825 | STAD | HSC-39 | Crizotinib |
| MVP_top60_DNA_seed | IGF1R | -1.384697 | STAD | HSC-39 | NVP-ADW742 |
| MVP_top60_DNA_seed | ESR1 | -1.384373 | STAD | HGC-27 | GDC0810 |
| MVP_top60_DNA_seed | AKT1 | -1.373034 | STAD | NCI-N87 | Uprosertib |
| MVP_top60_DNA_seed | HDAC1 | -1.369778 | STAD | NUGC-4 | Romidepsin |
| MVP_top60_DNA_seed | AKT1 | -1.349839 | STAD | HGC-27 | AT13148 |
| MVP_top60_DNA_seed | FGFR3 | -1.349506 | STAD | RERF-GC-1B | AZD4547 |
| MVP_top60_DNA_seed | AKT1 | -1.347719 | STAD | 23132-87 | AT13148 |
| MVP_top60_DNA_seed | IGF1R | -1.34305 | STAD | HSC-39 | BMS-754807 |
| MVP_top60_DNA_seed | EGFR | -1.343018 | STAD | HSC-39 | AZD3759 |
| MVP_top60_DNA_seed | ATM | -1.34211 | STAD | MKN45 | KU-55933 |
| MVP_top60_DNA_seed | ERBB2 | -1.340055 | STAD | NUGC-3 | Afatinib |
| MVP_top60_DNA_seed | EGFR | -1.340055 | STAD | NUGC-3 | Afatinib |
| MVP_top60_DNA_seed | AKT1 | -1.322319 | STAD | TMK-1 | Ipatasertib |
| MVP_top60_DNA_seed | MAPK1 | -1.310121 | STAD | MKN45 | SCH772984 |
| MVP_top60_DNA_seed | MAPK3 | -1.310121 | STAD | MKN45 | SCH772984 |
| MVP_top60_DNA_seed | EGFR | -1.296907 | STAD | NUGC-4 | AZD3759 |
| MVP_top60_DNA_seed | AKT1 | -1.285219 | STAD | 23132-87 | AZD5363 |
| MVP_top60_DNA_seed | AKT1 | -1.28451 | STAD | SK-GT-2 | AT13148 |
| MVP_top60_DNA_seed | FGFR3 | -1.274816 | STAD | RERF-GC-1B | PD173074 |
| MVP_top60_DNA_seed | AKT1 | -1.273648 | STAD | 23132-87 | GSK2110183B |
| MVP_top60_DNA_seed | ESR1 | -1.269431 | STAD | IM-95 | GDC0810 |
| MVP_top60_DNA_seed | AKT1 | -1.257823 | STAD | AGS | GSK2110183B |
| MVP_top60_DNA_seed | PIK3CA | -1.255553 | STAD | 23132-87 | Buparlisib |
| MVP_top60_DNA_seed | MET | -1.248383 | STAD | HGC-27 | Savolitinib |
| MVP_top60_DNA_seed | IGF1R | -1.247327 | STAD | HGC-27 | GSK1904529A |
| MVP_top60_DNA_seed | AKT1 | -1.239847 | STAD | HSC-39 | AT13148 |
| MVP_top60_DNA_seed | MET | -1.230999 | STAD | HSC-39 | Savolitinib |
| MVP_top60_DNA_seed | AKT1 | -1.209787 | STAD | HSC-39 | Ipatasertib |
| MVP_top60_DNA_seed | MAPK1 | -1.20914 | STAD | NCI-N87 | Ulixertinib |
| MVP_top60_DNA_seed | MAPK3 | -1.20914 | STAD | NCI-N87 | Ulixertinib |

| | | | | | |
|---|---|---|---|---|---|
| MVP_top60_DNA_seed | SRC | -1.207192 | STAD | HGC-27 | Bosutinib |
| MVP_top60_DNA_seed | MDM2 | -1.206864 | STAD | MKN45 | Nutlin-3a (-) |
| MVP_top60_DNA_seed | AKT1 | -1.194714 | STAD | MKN45 | Uprosertib |
| MVP_top60_DNA_seed | MDM2 | -1.18676 | STAD | 23132-87 | Nutlin-3a (-) |
| MVP_top60_DNA_seed | ERBB2 | -1.18579 | STAD | NUGC-3 | Lapatinib |
| MVP_top60_DNA_seed | EGFR | -1.18579 | STAD | NUGC-3 | Lapatinib |
| MVP_top60_DNA_seed | HSP90AA1 | -1.182084 | STAD | HGC-27 | Tanespimycin |
| MVP_top60_DNA_seed | IGF1R | -1.177343 | STAD | SNU-16 | BMS-754807 |
| MVP_top60_DNA_seed | AKT1 | -1.176424 | STAD | HSC-39 | Afuresertib |
| MVP_top60_DNA_seed | MAPK1 | -1.174219 | STAD | TGBC11TKB | SCH772984 |
| MVP_top60_DNA_seed | MAPK3 | -1.174219 | STAD | TGBC11TKB | SCH772984 |
| MVP_top60_DNA_seed | KRAS | -1.16714 | STAD | AGS | KRAS (G12C) Inhibitor-12 |
| MVP_top60_DNA_seed | EP300 | -1.166446 | STAD | RERF-GC-1B | SGC-CBP30 |
| MVP_top60_DNA_seed | CREBBP | -1.166446 | STAD | RERF-GC-1B | SGC-CBP30 |
| MVP_top60_DNA_seed | PIK3CA | -1.166136 | STAD | 23132-87 | GNE-317 |
| MVP_top60_DNA_seed | MAPK1 | -1.164394 | STAD | GCIY | Ulixertinib |
| MVP_top60_DNA_seed | MAPK3 | -1.164394 | STAD | GCIY | Ulixertinib |
| MVP_top60_DNA_seed | AKT1 | -1.158885 | STAD | TMK-1 | AZD5363 |
| MVP_top60_DNA_seed | HDAC1 | -1.158468 | STAD | AGS | Romidepsin |
| MVP_top60_DNA_seed | HDAC1 | -1.152104 | STAD | NUGC-3 | Romidepsin |
| MVP_top60_DNA_seed | ERBB2 | -1.149545 | STAD | GCIY | Afatinib |
| MVP_top60_DNA_seed | EGFR | -1.149545 | STAD | GCIY | Afatinib |
| MVP_top60_DNA_seed | AKT1 | -1.14792 | STAD | 23132-87 | Ipatasertib |
| MVP_top60_DNA_seed | ESR1 | -1.141999 | STAD | HSC-39 | GDC0810 |
| MVP_top60_DNA_seed | FGFR3 | -1.133877 | STAD | 23132-87 | AZD4547 |
| MVP_top60_DNA_seed | EP300 | -1.129626 | STAD | SNU-5 | SGC-CBP30 |
| MVP_top60_DNA_seed | CREBBP | -1.129626 | STAD | SNU-5 | SGC-CBP30 |
| MVP_top60_DNA_seed | EP300 | -1.128529 | STAD | NUGC-3 | SGC-CBP30 |
| MVP_top60_DNA_seed | CREBBP | -1.128529 | STAD | NUGC-3 | SGC-CBP30 |
| MVP_top60_DNA_seed | ERBB2 | -1.120754 | STAD | GCIY | Sapitinib |
| MVP_top60_DNA_seed | ERBB3 | -1.120754 | STAD | GCIY | Sapitinib |
| MVP_top60_DNA_seed | EGFR | -1.120754 | STAD | GCIY | Sapitinib |
| MVP_top60_DNA_seed | AKT1 | -1.115329 | STAD | AGS | Uprosertib |
| MVP_top60_DNA_seed | HSP90AA1 | -1.114071 | STAD | AGS | Luminespib |
| MVP_top60_DNA_seed | PIK3CA | -1.107574 | STAD | HSC-39 | Buparlisib |
| MVP_top60_DNA_seed | HSP90AA1 | -1.105157 | STAD | NCI-N87 | Luminespib |
| MVP_top60_DNA_seed | EGFR | -1.101097 | STAD | 23132-87 | Gefitinib |
| MVP_top60_DNA_seed | PIK3CA | -1.096937 | STAD | IM-95 | Alpelisib |
| MVP_top60_DNA_seed | EGFR | -1.09315 | STAD | HSC-39 | Erlotinib |
| MVP_top60_DNA_seed | MAPK1 | -1.085216 | STAD | MKN45 | Ulixertinib |
| MVP_top60_DNA_seed | MAPK3 | -1.085216 | STAD | MKN45 | Ulixertinib |

| | | | | | |
|---|---|---|---|---|---|
| MVP_top60_DNA_seed | HDAC1 | -1.08404 | STAD | MKN45 | PCI-34051 |
| MVP_top60_DNA_seed | MAPK1 | -1.078027 | STAD | NUGC-4 | SCH772984 |
| MVP_top60_DNA_seed | MAPK3 | -1.078027 | STAD | NUGC-4 | SCH772984 |
| MVP_top60_DNA_seed | PIK3CA | -1.074511 | STAD | 23132-87 | Alpelisib |
| MVP_top60_DNA_seed | ERBB2 | -1.0707 | STAD | NUGC-4 | Sapitinib |
| MVP_top60_DNA_seed | ERBB3 | -1.0707 | STAD | NUGC-4 | Sapitinib |
| MVP_top60_DNA_seed | EGFR | -1.0707 | STAD | NUGC-4 | Sapitinib |
| MVP_top60_DNA_seed | MAPK1 | -1.069708 | STAD | NCI-N87 | SCH772984 |
| MVP_top60_DNA_seed | MAPK3 | -1.069708 | STAD | NCI-N87 | SCH772984 |
| MVP_top60_DNA_seed | EGFR | -1.069024 | STAD | HSC-39 | Osimertinib |
| MVP_top60_DNA_seed | HSP90AA1 | -1.06889 | STAD | SNU-1 | Tanespimycin |
| MVP_top60_DNA_seed | AKT1 | -1.068491 | STAD | 23132-87 | MK-2206 |
| MVP_top60_DNA_seed | AKT1 | -1.065711 | STAD | IM-95 | Afuresertib |
| MVP_top60_DNA_seed | PIK3CA | -1.064823 | STAD | FU97 | GNE-317 |
| MVP_top60_DNA_seed | ERBB2 | -1.064414 | STAD | HGC-27 | Lapatinib |
| MVP_top60_DNA_seed | EGFR | -1.064414 | STAD | HGC-27 | Lapatinib |
| MVP_top60_DNA_seed | IGF1R | -1.05591 | STAD | HGC-27 | BMS-754807 |
| MVP_top60_DNA_seed | IGF1R | -1.054155 | STAD | HSC-39 | GSK1904529A |
| MVP_top60_DNA_seed | PIK3CA | -1.052359 | STAD | SNU-16 | Buparlisib |
| MVP_top60_DNA_seed | MET | -1.03287 | STAD | IM-95 | Crizotinib |
| MVP_top60_DNA_seed | EP300 | -1.032284 | STAD | HSC-39 | CPI-637 |
| MVP_top60_DNA_seed | AKT1 | -1.028039 | STAD | 23132-87 | Afuresertib |
| MVP_top60_DNA_seed | AKT1 | -1.023256 | STAD | AGS | Afuresertib |
| MVP_top60_DNA_seed | AKT1 | -1.012419 | STAD | HSC-39 | AZD5363 |
| MVP_top60_DNA_seed | AKT1 | -1.007262 | STAD | AGS | AT13148 |
| MVP_top60_DNA_seed | AKT1 | -1.004213 | STAD | TMK-1 | Afuresertib |
| MVP_top60_DNA_seed | AKT1 | -0.995071 | STAD | IM-95 | Ipatasertib |
| MVP_top60_DNA_seed | ERBB2 | -0.992582 | STAD | HSC-39 | Lapatinib |
| MVP_top60_DNA_seed | EGFR | -0.992582 | STAD | HSC-39 | Lapatinib |
| MVP_top60_DNA_seed | MAPK1 | -0.984578 | STAD | HSC-39 | Ulixertinib |
| MVP_top60_DNA_seed | MAPK3 | -0.984578 | STAD | HSC-39 | Ulixertinib |
| MVP_top60_DNA_seed | ESR1 | -0.983947 | STAD | HGC-27 | Tamoxifen |
| MVP_top60_DNA_seed | MAPK1 | -0.975449 | STAD | HSC-39 | VX-11e |
| MVP_top60_DNA_seed | EGFR | -0.96723 | STAD | NCI-N87 | Gefitinib |
| MVP_top60_DNA_seed | EGFR | -0.964959 | STAD | HGC-27 | Erlotinib |
| MVP_top60_DNA_seed | AKT1 | -0.964844 | STAD | MKN45 | Uprosertib |
| MVP_top60_DNA_seed | TP53 | -0.964362 | STAD | SNU-16 | MIRA-1 |
| MVP_top60_DNA_seed | ERBB2 | -0.961978 | STAD | RERF-GC-1B | Sapitinib |
| MVP_top60_DNA_seed | ERBB3 | -0.961978 | STAD | RERF-GC-1B | Sapitinib |
| MVP_top60_DNA_seed | EGFR | -0.961978 | STAD | RERF-GC-1B | Sapitinib |
| MVP_top60_DNA_seed | MDM2 | -0.960426 | STAD | RF-48 | Serdemetan |

| | | | | | |
|---|---|---|---|---|---|
| MVP_top60_DNA_seed | KRAS | -0.957921 | STAD | NCI-N87 | KRAS (G12C) Inhibitor-12 |
| MVP_top60_DNA_seed | PIK3CA | -0.94861 | STAD | HSC-39 | AZD8186 |
| MVP_top60_DNA_seed | HDAC1 | -0.94795 | STAD | NCI-N87 | Romidepsin |
| MVP_top60_DNA_seed | EP300 | -0.940233 | STAD | 23132-87 | SGC-CBP30 |
| MVP_top60_DNA_seed | CREBBP | -0.940233 | STAD | 23132-87 | SGC-CBP30 |
| MVP_top60_DNA_seed | AKT1 | -0.939247 | STAD | HSC-39 | MK-2206 |
| MVP_top60_DNA_seed | IGF1R | -0.938207 | STAD | 23132-87 | GSK1904529A |
| MVP_top60_DNA_seed | EGFR | -0.92994 | STAD | NUGC-3 | Osimertinib |
| MVP_top60_DNA_seed | HDAC1 | -0.92607 | STAD | RF-48 | Romidepsin |
| MVP_top60_DNA_seed | IGF1R | -0.924343 | STAD | SNU-5 | GSK1904529A |
| MVP_top60_DNA_seed | AKT1 | -0.923776 | STAD | TMK-1 | Uprosertib |
| MVP_top60_DNA_seed | FGFR3 | -0.923566 | STAD | AGS | PD173074 |
| MVP_top60_DNA_seed | GSK3B | -0.91946 | STAD | AGS | SB216763 |
| MVP_top60_DNA_seed | KRAS | -0.911847 | STAD | SNU-5 | KRAS (G12C) Inhibitor-12 |
| MVP_top60_DNA_seed | MAPK1 | -0.91179 | STAD | MKN28 | VX-11e |
| MVP_top60_DNA_seed | HDAC1 | -0.911584 | STAD | SNU-5 | PCI-34051 |
| MVP_top60_DNA_seed | AKT1 | -0.910502 | STAD | IM-95 | AZD5363 |
| MVP_top60_DNA_seed | MAPK1 | -0.910393 | STAD | KATOIII | Ulixertinib |
| MVP_top60_DNA_seed | MAPK3 | -0.910393 | STAD | KATOIII | Ulixertinib |
| MVP_top60_DNA_seed | HSP90AA1 | -0.906857 | STAD | SK-GT-2 | Luminespib |
| MVP_top60_DNA_seed | MET | -0.903609 | STAD | HGC-27 | Crizotinib |
| MVP_top60_DNA_seed | AKT1 | -0.902486 | STAD | FU97 | Ipatasertib |
| MVP_top60_DNA_seed | HSP90AA1 | -0.897585 | STAD | TMK-1 | CCT-018159 |
| MVP_top60_DNA_seed | PIK3CA | -0.895344 | STAD | FU97 | Buparlisib |
| MVP_top60_DNA_seed | HDAC1 | -0.883727 | STAD | SK-GT-2 | Romidepsin |
| MVP_top60_DNA_seed | IGF1R | -0.878678 | STAD | SNU-16 | NVP-ADW742 |
| MVP_top60_DNA_seed | AKT1 | -0.875407 | STAD | NCI-N87 | Ipatasertib |
| MVP_top60_DNA_seed | IGF1R | -0.871944 | STAD | SNU-1 | NVP-ADW742 |
| MVP_top60_DNA_seed | GSK3B | -0.871876 | STAD | HSC-39 | SB216763 |
| MVP_top60_DNA_seed | MDM2 | -0.870852 | STAD | HSC-39 | Serdemetan |
| MVP_top60_DNA_seed | HDAC1 | -0.868311 | STAD | NUGC-3 | Entinostat |
| MVP_top60_DNA_seed | AKT1 | -0.865999 | STAD | FU97 | Afuresertib |
| MVP_top60_DNA_seed | PIK3CA | -0.860398 | STAD | AGS | GNE-317 |
| MVP_top60_DNA_seed | MET | -0.859577 | STAD | SNU-1 | Savolitinib |
| MVP_top60_DNA_seed | AKT1 | -0.859384 | STAD | FU97 | AZD5363 |
| MVP_top60_DNA_seed | AKT1 | -0.851785 | STAD | AGS | Ipatasertib |
| MVP_top60_DNA_seed | PIK3CA | -0.842493 | STAD | NCI-N87 | AZD8186 |
| MVP_top60_DNA_seed | ATM | -0.834389 | STAD | SNU-1 | KU-55933 |
| MVP_top60_DNA_seed | PIK3CA | -0.832559 | STAD | GCIY | GNE-317 |
| MVP_top60_DNA_seed | HDAC1 | -0.831674 | STAD | MKN7 | Entinostat |
| MVP_top60_DNA_seed | PIK3CA | -0.829642 | STAD | SNU-5 | GNE-317 |

| | | | | | |
|---|---|---|---|---|---|
| MVP_top60_DNA_seed | MAPK1 | -0.82574 | STAD | SNU-5 | SCH772984 |
| MVP_top60_DNA_seed | MAPK3 | -0.82574 | STAD | SNU-5 | SCH772984 |
| MVP_top60_DNA_seed | AKT1 | -0.818769 | STAD | OCUM-1 | Uprosertib |
| MVP_top60_DNA_seed | MAPK1 | -0.817621 | STAD | MKN28 | SCH772984 |
| MVP_top60_DNA_seed | MAPK3 | -0.817621 | STAD | MKN28 | SCH772984 |
| MVP_top60_DNA_seed | ERBB2 | -0.814856 | STAD | SNU-16 | Lapatinib |
| MVP_top60_DNA_seed | EGFR | -0.814856 | STAD | SNU-16 | Lapatinib |
| MVP_top60_DNA_seed | PIK3CA | -0.814266 | STAD | SNU-5 | AZD8186 |
| MVP_top60_DNA_seed | EP300 | -0.812311 | STAD | HGC-27 | SGC-CBP30 |
| MVP_top60_DNA_seed | CREBBP | -0.812311 | STAD | HGC-27 | SGC-CBP30 |
| MVP_top60_DNA_seed | ESR1 | -0.812064 | STAD | 23132-87 | Tamoxifen |
| MVP_top60_DNA_seed | AKT1 | -0.805596 | STAD | RF-48 | Uprosertib |
| MVP_top60_DNA_seed | HSP90AA1 | -0.797028 | STAD | OCUM-1 | Luminespib |
| MVP_top60_DNA_seed | HSP90AA1 | -0.796287 | STAD | RF-48 | Luminespib |
| MVP_top60_DNA_seed | PIK3CA | -0.791431 | STAD | AGS | Buparlisib |
| MVP_top60_DNA_seed | TP53 | -0.79102 | STAD | HSC-39 | MIRA-1 |
| MVP_top60_DNA_seed | PIK3CA | -0.790039 | STAD | NUGC-3 | GNE-317 |
| MVP_top60_DNA_seed | MAPK1 | -0.789896 | STAD | HGC-27 | Ulixertinib |
| MVP_top60_DNA_seed | MAPK3 | -0.789896 | STAD | HGC-27 | Ulixertinib |
| MVP_top60_DNA_seed | EGFR | -0.788562 | STAD | NCI-N87 | AZD3759 |
| MVP_top60_DNA_seed | MAPK1 | -0.78644 | STAD | SNU-5 | Ulixertinib |
| MVP_top60_DNA_seed | MAPK3 | -0.78644 | STAD | SNU-5 | Ulixertinib |
| MVP_top60_DNA_seed | MAPK1 | -0.785475 | STAD | SNU-5 | Ulixertinib |
| MVP_top60_DNA_seed | MAPK3 | -0.785475 | STAD | SNU-5 | Ulixertinib |
| MVP_top60_DNA_seed | KRAS | -0.783583 | STAD | SNU-1 | KRAS (G12C) Inhibitor-12 |
| MVP_top60_DNA_seed | MAPK1 | -0.782848 | STAD | HSC-39 | Ulixertinib |
| MVP_top60_DNA_seed | MAPK3 | -0.782848 | STAD | HSC-39 | Ulixertinib |
| MVP_top60_DNA_seed | EGFR | -0.781258 | STAD | GCIY | AZD3759 |
| MVP_top60_DNA_seed | MAPK1 | -0.778445 | STAD | Hs-746T | SCH772984 |
| MVP_top60_DNA_seed | MAPK3 | -0.778445 | STAD | Hs-746T | SCH772984 |
| MVP_top60_DNA_seed | HDAC1 | -0.773766 | STAD | 23132-87 | Romidepsin |
| MVP_top60_DNA_seed | AKT1 | -0.772267 | STAD | MKN45 | GSK2110183B |
| MVP_top60_DNA_seed | GSK3B | -0.771669 | STAD | RERF-GC-1B | SB216763 |
| MVP_top60_DNA_seed | HSP90AA1 | -0.770346 | STAD | SNU-5 | Tanespimycin |
| MVP_top60_DNA_seed | PIK3CA | -0.770271 | STAD | FU97 | AZD8186 |
| MVP_top60_DNA_seed | PIK3CA | -0.768064 | STAD | Hs-746T | GNE-317 |
| MVP_top60_DNA_seed | AKT1 | -0.767251 | STAD | NCI-N87 | Uprosertib |
| MVP_top60_DNA_seed | IGF1R | -0.761767 | STAD | OCUM-1 | BMS-536924 |
| MVP_top60_DNA_seed | EP300 | -0.750783 | STAD | NUGC-4 | SGC-CBP30 |
| MVP_top60_DNA_seed | CREBBP | -0.750783 | STAD | NUGC-4 | SGC-CBP30 |
| MVP_top60_DNA_seed | MET | -0.7454 | STAD | SK-GT-2 | Savolitinib |

| | | | | | |
|---|---|---|---|---|---|
| MVP_top60_DNA_seed | HSP90AA1 | -0.744516 | STAD | MKN45 | Tanespimycin |
| MVP_top60_DNA_seed | EGFR | -0.744344 | STAD | GCIY | Gefitinib |
| MVP_top60_DNA_seed | ESR1 | -0.738363 | STAD | SK-GT-2 | Tamoxifen |
| MVP_top60_DNA_seed | HDAC1 | -0.737076 | STAD | IM-95 | Entinostat |
| MVP_top60_DNA_seed | ERBB2 | -0.729675 | STAD | NUGC-4 | Afatinib |
| MVP_top60_DNA_seed | EGFR | -0.729675 | STAD | NUGC-4 | Afatinib |
| MVP_top60_DNA_seed | IGF1R | -0.729587 | STAD | FU97 | NVP-ADW742 |
| MVP_top60_DNA_seed | ERBB2 | -0.724525 | STAD | MKN7 | Lapatinib |
| MVP_top60_DNA_seed | EGFR | -0.724525 | STAD | MKN7 | Lapatinib |
| MVP_top60_DNA_seed | ESR1 | -0.724016 | STAD | SNU-16 | GDC0810 |
| MVP_top60_DNA_seed | TP53 | -0.720545 | STAD | RF-48 | MIRA-1 |
| MVP_top60_DNA_seed | EGFR | -0.717928 | STAD | 23132-87 | Erlotinib |
| MVP_top60_DNA_seed | EGFR | -0.712181 | STAD | SNU-16 | Osimertinib |
| MVP_top60_DNA_seed | MAPK1 | -0.708043 | STAD | 23132-87 | VX-11e |
| MVP_top60_DNA_seed | ATM | -0.707435 | STAD | SNU-16 | KU-55933 |
| MVP_top60_DNA_seed | GSK3B | -0.703721 | STAD | 23132-87 | SB216763 |
| MVP_top60_DNA_seed | MDM2 | -0.701821 | STAD | HGC-27 | Serdemetan |
| MVP_top60_DNA_seed | AKT1 | -0.697813 | STAD | AGS | AZD5363 |
| MVP_top60_DNA_seed | ESR1 | -0.687535 | STAD | NCI-N87 | GDC0810 |
| MVP_top60_DNA_seed | AKT1 | -0.686143 | STAD | SNU-5 | AT13148 |
| MVP_top60_DNA_seed | EGFR | -0.67953 | STAD | RERF-GC-1B | Erlotinib |
| MVP_top60_DNA_seed | AKT1 | -0.678676 | STAD | SK-GT-2 | Afuresertib |
| MVP_top60_DNA_seed | FGFR3 | -0.675541 | STAD | IM-95 | AZD4547 |
| MVP_top60_DNA_seed | MAPK1 | -0.672667 | STAD | KATOIII | SCH772984 |
| MVP_top60_DNA_seed | MAPK3 | -0.672667 | STAD | KATOIII | SCH772984 |
| MVP_top60_DNA_seed | KRAS | -0.665366 | STAD | MKN45 | KRAS (G12C) Inhibitor-12 |
| MVP_top60_DNA_seed | AKT1 | -0.664113 | STAD | SNU-16 | Ipatasertib |
| MVP_top60_DNA_seed | AKT1 | -0.664085 | STAD | MKN45 | Afuresertib |
| MVP_top60_DNA_seed | SRC | -0.657963 | STAD | HSC-39 | Bosutinib |
| MVP_top60_DNA_seed | MAPK1 | -0.65613 | STAD | TGBC11TKB | Ulixertinib |
| MVP_top60_DNA_seed | MAPK3 | -0.65613 | STAD | TGBC11TKB | Ulixertinib |
| MVP_top60_DNA_seed | EGFR | -0.646859 | STAD | TMK-1 | Osimertinib |
| MVP_top60_DNA_seed | MAPK1 | -0.645592 | STAD | KATOIII | Ulixertinib |
| MVP_top60_DNA_seed | MAPK3 | -0.645592 | STAD | KATOIII | Ulixertinib |
| MVP_top60_DNA_seed | MET | -0.645201 | STAD | AGS | Savolitinib |
| MVP_top60_DNA_seed | GSK3B | -0.6435 | STAD | TMK-1 | CHIR-99021 |
| MVP_top60_DNA_seed | ESR1 | -0.641626 | STAD | RERF-GC-1B | Tamoxifen |
| MVP_top60_DNA_seed | PIK3CA | -0.640799 | STAD | MKN45 | GNE-317 |
| MVP_top60_DNA_seed | PIK3CA | -0.636908 | STAD | SNU-5 | Buparlisib |
| MVP_top60_DNA_seed | ESR1 | -0.633839 | STAD | AGS | Tamoxifen |
| MVP_top60_DNA_seed | PIK3CA | -0.633687 | STAD | SK-GT-2 | Buparlisib |

| | | | | | |
|---|---|---|---|---|---|
| MVP_top60_DNA_seed | MDM2 | -0.632532 | STAD | SNU-16 | Serdemetan |
| MVP_top60_DNA_seed | AKT1 | -0.63233 | STAD | NCI-N87 | MK-2206 |
| MVP_top60_DNA_seed | ERBB2 | -0.628949 | STAD | HSC-39 | Sapitinib |
| MVP_top60_DNA_seed | ERBB3 | -0.628949 | STAD | HSC-39 | Sapitinib |
| MVP_top60_DNA_seed | EGFR | -0.628949 | STAD | HSC-39 | Sapitinib |
| MVP_top60_DNA_seed | MAPK1 | -0.628511 | STAD | MKN45 | VX-11e |
| MVP_top60_DNA_seed | MAPK1 | -0.623801 | STAD | SNU-5 | VX-11e |
| MVP_top60_DNA_seed | KRAS | -0.614055 | STAD | RF-48 | KRAS (G12C) Inhibitor-12 |
| MVP_top60_DNA_seed | PIK3CA | -0.603623 | STAD | SK-GT-2 | Alpelisib |
| MVP_top60_DNA_seed | MAPK1 | -0.603514 | STAD | TMK-1 | SCH772984 |
| MVP_top60_DNA_seed | MAPK3 | -0.603514 | STAD | TMK-1 | SCH772984 |
| MVP_top60_DNA_seed | MDM2 | -0.60193 | STAD | SNU-1 | Nutlin-3a (-) |
| MVP_top60_DNA_seed | EGFR | -0.598918 | STAD | RERF-GC-1B | Gefitinib |
| MVP_top60_DNA_seed | MET | -0.598456 | STAD | RERF-GC-1B | Savolitinib |
| MVP_top60_DNA_seed | MDM2 | -0.597664 | STAD | HSC-39 | Nutlin-3a (-) |
| MVP_top60_DNA_seed | ERBB2 | -0.596463 | STAD | TMK-1 | Sapitinib |
| MVP_top60_DNA_seed | ERBB3 | -0.596463 | STAD | TMK-1 | Sapitinib |
| MVP_top60_DNA_seed | EGFR | -0.596463 | STAD | TMK-1 | Sapitinib |
| MVP_top60_DNA_seed | MET | -0.595751 | STAD | NUGC-4 | Savolitinib |
| MVP_top60_DNA_seed | MDM2 | -0.594361 | STAD | FU97 | Nutlin-3a (-) |
| MVP_top60_DNA_seed | HSP90AA1 | -0.593213 | STAD | RERF-GC-1B | Luminespib |
| MVP_top60_DNA_seed | ESR1 | -0.592299 | STAD | 23132-87 | GDC0810 |
| MVP_top60_DNA_seed | EGFR | -0.590912 | STAD | HSC-39 | Gefitinib |
| MVP_top60_DNA_seed | AKT1 | -0.590559 | STAD | RERF-GC-1B | GSK2110183B |
| MVP_top60_DNA_seed | MAPK1 | -0.589577 | STAD | HSC-39 | SCH772984 |
| MVP_top60_DNA_seed | MAPK3 | -0.589577 | STAD | HSC-39 | SCH772984 |
| MVP_top60_DNA_seed | AKT1 | -0.585844 | STAD | SK-GT-2 | MK-2206 |
| MVP_top60_DNA_seed | HDAC1 | -0.585809 | STAD | RF-48 | PCI-34051 |
| MVP_top60_DNA_seed | IGF1R | -0.579911 | STAD | KATOIII | GSK1904529A |
| MVP_top60_DNA_seed | FGFR3 | -0.579611 | STAD | HGC-27 | AZD4547 |
| MVP_top60_DNA_seed | TP53 | -0.578312 | STAD | GCIY | MIRA-1 |
| MVP_top60_DNA_seed | MDM2 | -0.567297 | STAD | TMK-1 | Serdemetan |
| MVP_top60_DNA_seed | IGF1R | -0.565347 | STAD | SNU-5 | BMS-754807 |
| MVP_top60_DNA_seed | AKT1 | -0.560976 | STAD | SNU-5 | AZD5363 |
| MVP_top60_DNA_seed | HSP90AA1 | -0.5603 | STAD | SNU-5 | Luminespib |
| MVP_top60_DNA_seed | EGFR | -0.554187 | STAD | AGS | Erlotinib |
| MVP_top60_DNA_seed | PIK3CA | -0.553995 | STAD | IM-95 | Buparlisib |
| MVP_top60_DNA_seed | HDAC1 | -0.550885 | STAD | AGS | Entinostat |
| MVP_top60_DNA_seed | AKT1 | -0.549007 | STAD | GCIY | Uprosertib |
| MVP_top60_DNA_seed | MAPK1 | -0.547318 | STAD | 23132-87 | SCH772984 |
| MVP_top60_DNA_seed | MAPK3 | -0.547318 | STAD | 23132-87 | SCH772984 |

| | | | | | |
|---|---|---|---|---|---|
| MVP_top60_DNA_seed | ERBB2 | -0.544321 | STAD | SNU-16 | Afatinib |
| MVP_top60_DNA_seed | EGFR | -0.544321 | STAD | SNU-16 | Afatinib |
| MVP_top60_DNA_seed | HSP90AA1 | -0.540432 | STAD | HGC-27 | Luminespib |
| MVP_top60_DNA_seed | KRAS | -0.539066 | STAD | IM-95 | KRAS (G12C) Inhibitor-12 |
| MVP_top60_DNA_seed | PIK3CA | -0.536074 | STAD | SK-GT-2 | GNE-317 |
| MVP_top60_DNA_seed | AKT1 | -0.534309 | STAD | NCI-N87 | GSK2110183B |
| MVP_top60_DNA_seed | HDAC1 | -0.531965 | STAD | NUGC-4 | Entinostat |
| MVP_top60_DNA_seed | EGFR | -0.531315 | STAD | MKN7 | Osimertinib |
| MVP_top60_DNA_seed | MET | -0.530379 | STAD | RERF-GC-1B | Crizotinib |
| MVP_top60_DNA_seed | HDAC1 | -0.528204 | STAD | SK-GT-2 | Entinostat |
| MVP_top60_DNA_seed | ESR1 | -0.527647 | STAD | SK-GT-2 | GDC0810 |
| MVP_top60_DNA_seed | EGFR | -0.527474 | STAD | TGBC11TKB | AZD3759 |
| MVP_top60_DNA_seed | AKT1 | -0.527459 | STAD | SNU-1 | Uprosertib |
| MVP_top60_DNA_seed | SRC | -0.526709 | STAD | SNU-16 | Bosutinib |
| MVP_top60_DNA_seed | IGF1R | -0.525031 | STAD | HGC-27 | NVP-ADW742 |
| MVP_top60_DNA_seed | MET | -0.520269 | STAD | SNU-1 | Crizotinib |
| MVP_top60_DNA_seed | AKT1 | -0.518175 | STAD | IM-95 | MK-2206 |
| MVP_top60_DNA_seed | MDM2 | -0.515019 | STAD | NUGC-3 | Serdemetan |
| MVP_top60_DNA_seed | HSP90AA1 | -0.514576 | STAD | IM-95 | Tanespimycin |
| MVP_top60_DNA_seed | ATM | -0.513893 | STAD | RERF-GC-1B | KU-55933 |
| MVP_top60_DNA_seed | IGF1R | -0.51346 | STAD | KATOIII | BMS-536924 |
| MVP_top60_DNA_seed | EP300 | -0.509442 | STAD | AGS | CPI-637 |
| MVP_top60_DNA_seed | AKT1 | -0.506538 | STAD | MKN45 | Ipatasertib |
| MVP_top60_DNA_seed | IGF1R | -0.506035 | STAD | MKN28 | Linsitinib |
| MVP_top60_DNA_seed | ATM | -0.505711 | STAD | AGS | KU-55933 |
| MVP_top60_DNA_seed | ESR1 | -0.50288 | STAD | GCIY | GDC0810 |
| MVP_top60_DNA_seed | PIK3CA | -0.502607 | STAD | RERF-GC-1B | Alpelisib |
| MVP_top60_DNA_seed | GSK3B | -0.4996 | STAD | SNU-16 | SB216763 |
| MVP_top60_DNA_seed | MAPK1 | -0.496934 | STAD | SNU-16 | Ulixertinib |
| MVP_top60_DNA_seed | MAPK3 | -0.496934 | STAD | SNU-16 | Ulixertinib |
| MVP_top60_DNA_seed | PIK3CA | -0.494304 | STAD | TMK-1 | Alpelisib |
| MVP_top60_DNA_seed | PIK3CA | -0.492705 | STAD | RF-48 | Buparlisib |
| MVP_top60_DNA_seed | IGF1R | -0.487912 | STAD | OCUM-1 | Linsitinib |
| MVP_top60_DNA_seed | ATM | -0.484921 | STAD | HGC-27 | KU-55933 |
| MVP_top60_DNA_seed | IGF1R | -0.478903 | STAD | FU97 | GSK1904529A |
| MVP_top60_DNA_seed | IGF1R | -0.473767 | STAD | GCIY | BMS-536924 |
| MVP_top60_DNA_seed | AKT1 | -0.469298 | STAD | SNU-5 | GSK2110183B |
| MVP_top60_DNA_seed | IGF1R | -0.4668 | STAD | MKN45 | BMS-754807 |
| MVP_top60_DNA_seed | MAPK1 | -0.458898 | STAD | MKN45 | Ulixertinib |
| MVP_top60_DNA_seed | MAPK3 | -0.458898 | STAD | MKN45 | Ulixertinib |
| MVP_top60_DNA_seed | PIK3CA | -0.458624 | STAD | NCI-N87 | GNE-317 |

| | | | | | |
|---|---|---|---|---|---|
| MVP_top60_DNA_seed | AKT1 | -0.458105 | STAD | MKN45 | AZD5363 |
| MVP_top60_DNA_seed | IGF1R | -0.456437 | STAD | HGC-27 | BMS-536924 |
| MVP_top60_DNA_seed | IGF1R | -0.446697 | STAD | IM-95 | BMS-754807 |
| MVP_top60_DNA_seed | HDAC1 | -0.445927 | STAD | KATOIII | Romidepsin |
| MVP_top60_DNA_seed | ATM | -0.443556 | STAD | GCIY | KU-55933 |
| MVP_top60_DNA_seed | AKT1 | -0.443199 | STAD | SNU-16 | AZD5363 |
| MVP_top60_DNA_seed | MET | -0.442098 | STAD | RF-48 | Savolitinib |
| MVP_top60_DNA_seed | ATM | -0.441689 | STAD | RF-48 | KU-55933 |
| MVP_top60_DNA_seed | FGFR3 | -0.439307 | STAD | TGBC11TKB | PD173074 |
| MVP_top60_DNA_seed | SRC | -0.438876 | STAD | RF-48 | Bosutinib |
| MVP_top60_DNA_seed | PIK3CA | -0.432008 | STAD | IM-95 | AZD8186 |
| MVP_top60_DNA_seed | AKT1 | -0.428792 | STAD | NUGC-3 | Ipatasertib |
| MVP_top60_DNA_seed | IGF1R | -0.428461 | STAD | TMK-1 | NVP-ADW742 |
| MVP_top60_DNA_seed | ESR1 | -0.425629 | STAD | NUGC-3 | Tamoxifen |
| MVP_top60_DNA_seed | EP300 | -0.413258 | STAD | NCI-N87 | CPI-637 |
| MVP_top60_DNA_seed | SRC | -0.404606 | STAD | AGS | Bosutinib |
| MVP_top60_DNA_seed | IGF1R | -0.402743 | STAD | 23132-87 | NVP-ADW742 |
| MVP_top60_DNA_seed | IGF1R | -0.401925 | STAD | AGS | NVP-ADW742 |
| MVP_top60_DNA_seed | AKT1 | -0.401152 | STAD | OCUM-1 | AZD5363 |
| MVP_top60_DNA_seed | IGF1R | -0.395323 | STAD | TMK-1 | BMS-536924 |
| MVP_top60_DNA_seed | PIK3CA | -0.392938 | STAD | Hs-746T | Buparlisib |
| MVP_top60_DNA_seed | HSP90AA1 | -0.392414 | STAD | SNU-1 | Luminespib |
| MVP_top60_DNA_seed | IGF1R | -0.390436 | STAD | KATOIII | BMS-754807 |
| MVP_top60_DNA_seed | MAPK1 | -0.386403 | STAD | IM-95 | Ulixertinib |
| MVP_top60_DNA_seed | MAPK3 | -0.386403 | STAD | IM-95 | Ulixertinib |
| MVP_top60_DNA_seed | AKT1 | -0.385877 | STAD | SK-GT-2 | Uprosertib |
| MVP_top60_DNA_seed | IGF1R | -0.383355 | STAD | TGBC11TKB | Linsitinib |
| MVP_top60_DNA_seed | GSK3B | -0.381661 | STAD | SNU-1 | SB216763 |
| MVP_top60_DNA_seed | ESR1 | -0.380051 | STAD | KATOIII | Tamoxifen |
| MVP_top60_DNA_seed | SRC | -0.376421 | STAD | IM-95 | Bosutinib |
| MVP_top60_DNA_seed | MAPK1 | -0.375666 | STAD | TMK-1 | Ulixertinib |
| MVP_top60_DNA_seed | MAPK3 | -0.375666 | STAD | TMK-1 | Ulixertinib |
| MVP_top60_DNA_seed | HSP90AA1 | -0.374191 | STAD | NUGC-3 | Tanespimycin |
| MVP_top60_DNA_seed | IGF1R | -0.369864 | STAD | TMK-1 | BMS-754807 |
| MVP_top60_DNA_seed | EGFR | -0.36942 | STAD | HGC-27 | Osimertinib |
| MVP_top60_DNA_seed | AKT1 | -0.368926 | STAD | NCI-N87 | AZD5363 |
| MVP_top60_DNA_seed | AKT1 | -0.368098 | STAD | SK-GT-2 | Ipatasertib |
| MVP_top60_DNA_seed | EGFR | -0.367993 | STAD | AGS | Gefitinib |
| MVP_top60_DNA_seed | AKT1 | -0.363889 | STAD | SNU-16 | Afuresertib |
| MVP_top60_DNA_seed | MAPK1 | -0.363791 | STAD | Hs-746T | Ulixertinib |
| MVP_top60_DNA_seed | MAPK3 | -0.363791 | STAD | Hs-746T | Ulixertinib |

| | | | | | |
|---|---|---|---|---|---|
| MVP_top60_DNA_seed | PIK3CA | -0.359993 | STAD | Hs-746T | AZD8186 |
| MVP_top60_DNA_seed | AKT1 | -0.353568 | STAD | RF-48 | GSK2110183B |
| MVP_top60_DNA_seed | FGFR3 | -0.346585 | STAD | NCI-N87 | PD173074 |
| MVP_top60_DNA_seed | MAPK1 | -0.345874 | STAD | SNU-16 | Ulixertinib |
| MVP_top60_DNA_seed | MAPK3 | -0.345874 | STAD | SNU-16 | Ulixertinib |
| MVP_top60_DNA_seed | AKT1 | -0.343667 | STAD | SNU-16 | MK-2206 |
| MVP_top60_DNA_seed | FGFR3 | -0.343237 | STAD | SNU-1 | PD173074 |
| MVP_top60_DNA_seed | IGF1R | -0.341117 | STAD | Hs-746T | GSK1904529A |
| MVP_top60_DNA_seed | ERBB2 | -0.336204 | STAD | HSC-39 | Afatinib |
| MVP_top60_DNA_seed | EGFR | -0.336204 | STAD | HSC-39 | Afatinib |
| MVP_top60_DNA_seed | MAPK1 | -0.335714 | STAD | FU97 | VX-11e |
| MVP_top60_DNA_seed | PIK3CA | -0.33418 | STAD | SNU-16 | Alpelisib |
| MVP_top60_DNA_seed | MAPK1 | -0.316827 | STAD | RERF-GC-1B | VX-11e |
| MVP_top60_DNA_seed | ESR1 | -0.316824 | STAD | MKN28 | Tamoxifen |
| MVP_top60_DNA_seed | AKT1 | -0.315209 | STAD | TMK-1 | AT13148 |
| MVP_top60_DNA_seed | MAPK1 | -0.314581 | STAD | RERF-GC-1B | Ulixertinib |
| MVP_top60_DNA_seed | MAPK3 | -0.314581 | STAD | RERF-GC-1B | Ulixertinib |
| MVP_top60_DNA_seed | EGFR | -0.313668 | STAD | NUGC-4 | Erlotinib |
| MVP_top60_DNA_seed | KRAS | -0.312751 | STAD | TMK-1 | KRAS (G12C) Inhibitor-12 |
| MVP_top60_DNA_seed | AKT1 | -0.312153 | STAD | FU97 | Uprosertib |
| MVP_top60_DNA_seed | EGFR | -0.311995 | STAD | IM-95 | Osimertinib |
| MVP_top60_DNA_seed | MET | -0.307431 | STAD | SNU-16 | Savolitinib |
| MVP_top60_DNA_seed | IGF1R | -0.303875 | STAD | TMK-1 | GSK1904529A |
| MVP_top60_DNA_seed | AKT1 | -0.303238 | STAD | RERF-GC-1B | Uprosertib |
| MVP_top60_DNA_seed | IGF1R | -0.29984 | STAD | SNU-1 | GSK1904529A |
| MVP_top60_DNA_seed | IGF1R | -0.294561 | STAD | IM-95 | GSK1904529A |
| MVP_top60_DNA_seed | ERBB2 | -0.293342 | STAD | 23132-87 | Afatinib |
| MVP_top60_DNA_seed | EGFR | -0.293342 | STAD | 23132-87 | Afatinib |
| MVP_top60_DNA_seed | HSP90AA1 | -0.293298 | STAD | NUGC-3 | Luminespib |
| MVP_top60_DNA_seed | MAPK1 | -0.292425 | STAD | SNU-1 | Ulixertinib |
| MVP_top60_DNA_seed | MAPK3 | -0.292425 | STAD | SNU-1 | Ulixertinib |
| MVP_top60_DNA_seed | ERBB2 | -0.289654 | STAD | NUGC-4 | Lapatinib |
| MVP_top60_DNA_seed | EGFR | -0.289654 | STAD | NUGC-4 | Lapatinib |
| MVP_top60_DNA_seed | AR | -0.286406 | STAD | TMK-1 | Bicalutamide |
| MVP_top60_DNA_seed | ESR1 | -0.286012 | STAD | RF-48 | GDC0810 |
| MVP_top60_DNA_seed | AKT1 | -0.284079 | STAD | Hs-746T | Uprosertib |
| MVP_top60_DNA_seed | HSP90AA1 | -0.28304 | STAD | Hs-746T | Luminespib |
| MVP_top60_DNA_seed | EGFR | -0.279593 | STAD | GCIY | Erlotinib |
| MVP_top60_DNA_seed | EGFR | -0.277349 | STAD | TMK-1 | Erlotinib |
| MVP_top60_DNA_seed | KRAS | -0.273421 | STAD | SK-GT-2 | KRAS (G12C) Inhibitor-12 |
| MVP_top60_DNA_seed | TP53 | -0.272887 | STAD | HGC-27 | MIRA-1 |

| | | | | | |
|---|---|---|---|---|---|
| MVP_top60_DNA_seed | HDAC1 | -0.268777 | STAD | IM-95 | Romidepsin |
| MVP_top60_DNA_seed | ESR1 | -0.268664 | STAD | SNU-5 | Tamoxifen |
| MVP_top60_DNA_seed | AKT1 | -0.268201 | STAD | IM-95 | Uprosertib |
| MVP_top60_DNA_seed | AKT1 | -0.268092 | STAD | SK-GT-2 | AZD5363 |
| MVP_top60_DNA_seed | MAPK1 | -0.267497 | STAD | SK-GT-2 | Ulixertinib |
| MVP_top60_DNA_seed | MAPK3 | -0.267497 | STAD | SK-GT-2 | Ulixertinib |
| MVP_top60_DNA_seed | EGFR | -0.264183 | STAD | NUGC-4 | Osimertinib |
| MVP_top60_DNA_seed | PIK3CA | -0.263263 | STAD | GCIY | Buparlisib |
| MVP_top60_DNA_seed | AKT1 | -0.260163 | STAD | Hs-746T | MK-2206 |
| MVP_top60_DNA_seed | MAPK1 | -0.256418 | STAD | Hs-746T | Ulixertinib |
| MVP_top60_DNA_seed | MAPK3 | -0.256418 | STAD | Hs-746T | Ulixertinib |
| MVP_top60_DNA_seed | IGF1R | -0.251082 | STAD | MKN28 | BMS-536924 |
| MVP_top60_DNA_seed | IGF1R | -0.250416 | STAD | SK-GT-2 | NVP-ADW742 |
| MVP_top60_DNA_seed | AKT1 | -0.249218 | STAD | Hs-746T | Afuresertib |
| MVP_top60_DNA_seed | HSP90AA1 | -0.248861 | STAD | Hs-746T | Tanespimycin |
| MVP_top60_DNA_seed | MET | -0.240744 | STAD | OCUM-1 | Savolitinib |
| MVP_top60_DNA_seed | SRC | -0.238341 | STAD | SNU-5 | Bosutinib |
| MVP_top60_DNA_seed | MAPK1 | -0.237803 | STAD | MKN28 | Ulixertinib |
| MVP_top60_DNA_seed | MAPK3 | -0.237803 | STAD | MKN28 | Ulixertinib |
| MVP_top60_DNA_seed | ESR1 | -0.236961 | STAD | KATOIII | GDC0810 |
| MVP_top60_DNA_seed | EGFR | -0.235631 | STAD | SNU-16 | Gefitinib |
| MVP_top60_DNA_seed | AKT1 | -0.235456 | STAD | NCI-N87 | Afuresertib |
| MVP_top60_DNA_seed | KRAS | -0.232713 | STAD | MKN28 | KRAS (G12C) Inhibitor-12 |
| MVP_top60_DNA_seed | EGFR | -0.232404 | STAD | SNU-16 | AZD3759 |
| MVP_top60_DNA_seed | MAPK1 | -0.227401 | STAD | Hs-746T | VX-11e |
| MVP_top60_DNA_seed | HDAC1 | -0.222934 | STAD | RF-48 | Entinostat |
| MVP_top60_DNA_seed | MET | -0.215735 | STAD | TGBC11TKB | Savolitinib |
| MVP_top60_DNA_seed | IGF1R | -0.207395 | STAD | SNU-1 | BMS-754807 |
| MVP_top60_DNA_seed | AKT1 | -0.204172 | STAD | Hs-746T | GSK2110183B |
| MVP_top60_DNA_seed | HDAC1 | -0.201763 | STAD | SNU-5 | Entinostat |
| MVP_top60_DNA_seed | HSP90AA1 | -0.201111 | STAD | FU97 | Luminespib |
| MVP_top60_DNA_seed | AKT1 | -0.20004 | STAD | MKN45 | MK-2206 |
| MVP_top60_DNA_seed | AKT1 | -0.196703 | STAD | KATOIII | GSK2110183B |
| MVP_top60_DNA_seed | HDAC1 | -0.195673 | STAD | MKN7 | PCI-34051 |
| MVP_top60_DNA_seed | PIK3CA | -0.191993 | STAD | TGBC11TKB | Alpelisib |
| MVP_top60_DNA_seed | ERBB2 | -0.189838 | STAD | AGS | Lapatinib |
| MVP_top60_DNA_seed | EGFR | -0.189838 | STAD | AGS | Lapatinib |
| MVP_top60_DNA_seed | PIK3CA | -0.189207 | STAD | SNU-16 | GNE-317 |
| MVP_top60_DNA_seed | EGFR | -0.187189 | STAD | TGBC11TKB | Gefitinib |
| MVP_top60_DNA_seed | KRAS | -0.185502 | STAD | 23132-87 | KRAS (G12C) Inhibitor-12 |
| MVP_top60_DNA_seed | HDAC1 | -0.18429 | STAD | Hs-746T | Entinostat |

| | | | | | |
|---|---|---|---|---|---|
| MVP_top60_DNA_seed | MAPK1 | -0.182584 | STAD | RERF-GC-1B | SCH772984 |
| MVP_top60_DNA_seed | MAPK3 | -0.182584 | STAD | RERF-GC-1B | SCH772984 |
| MVP_top60_DNA_seed | PIK3CA | -0.181396 | STAD | MKN45 | Alpelisib |
| MVP_top60_DNA_seed | MAPK1 | -0.17745 | STAD | TMK-1 | VX-11e |
| MVP_top60_DNA_seed | SRC | -0.177043 | STAD | SNU-1 | Bosutinib |
| MVP_top60_DNA_seed | MAPK1 | -0.175867 | STAD | GCIY | VX-11e |
| MVP_top60_DNA_seed | AKT1 | -0.175351 | STAD | Hs-746T | AZD5363 |
| MVP_top60_DNA_seed | GSK3B | -0.171302 | STAD | HGC-27 | SB216763 |
| MVP_top60_DNA_seed | PIK3CA | -0.170171 | STAD | SNU-1 | Buparlisib |
| MVP_top60_DNA_seed | ERBB2 | -0.169119 | STAD | AGS | Afatinib |
| MVP_top60_DNA_seed | EGFR | -0.169119 | STAD | AGS | Afatinib |
| MVP_top60_DNA_seed | TP53 | -0.164612 | STAD | AGS | MIRA-1 |
| MVP_top60_DNA_seed | MET | -0.164459 | STAD | AGS | Crizotinib |
| MVP_top60_DNA_seed | IGF1R | -0.164212 | STAD | MKN28 | GSK1904529A |
| MVP_top60_DNA_seed | GSK3B | -0.157077 | STAD | RF-48 | SB216763 |
| MVP_top60_DNA_seed | EGFR | -0.154 | STAD | GCIY | Osimertinib |
| MVP_top60_DNA_seed | MAPK1 | -0.148859 | STAD | 23132-87 | Ulixertinib |
| MVP_top60_DNA_seed | MAPK3 | -0.148859 | STAD | 23132-87 | Ulixertinib |
| MVP_top60_DNA_seed | AKT1 | -0.147684 | STAD | SNU-1 | AZD5363 |
| MVP_top60_DNA_seed | ERBB2 | -0.14424 | STAD | 23132-87 | Sapitinib |
| MVP_top60_DNA_seed | ERBB3 | -0.14424 | STAD | 23132-87 | Sapitinib |
| MVP_top60_DNA_seed | EGFR | -0.14424 | STAD | 23132-87 | Sapitinib |
| MVP_top60_DNA_seed | EGFR | -0.143435 | STAD | TGBC11TKB | Osimertinib |
| MVP_top60_DNA_seed | ERBB2 | -0.137624 | STAD | IM-95 | Afatinib |
| MVP_top60_DNA_seed | EGFR | -0.137624 | STAD | IM-95 | Afatinib |
| MVP_top60_DNA_seed | ATM | -0.134883 | STAD | SNU-5 | KU-55933 |
| MVP_top60_DNA_seed | AKT1 | -0.131045 | STAD | SK-GT-2 | GSK2110183B |
| MVP_top60_DNA_seed | AKT1 | -0.131042 | STAD | NUGC-3 | AT13148 |
| MVP_top60_DNA_seed | MAPK1 | -0.1298 | STAD | RERF-GC-1B | Ulixertinib |
| MVP_top60_DNA_seed | MAPK3 | -0.1298 | STAD | RERF-GC-1B | Ulixertinib |
| MVP_top60_DNA_seed | EP300 | -0.129786 | STAD | FU97 | CPI-637 |
| MVP_top60_DNA_seed | AKT1 | -0.124254 | STAD | RERF-GC-1B | Uprosertib |
| MVP_top60_DNA_seed | AKT1 | -0.123406 | STAD | RERF-GC-1B | MK-2206 |
| MVP_top60_DNA_seed | AKT1 | -0.120353 | STAD | MKN45 | AT13148 |
| MVP_top60_DNA_seed | AKT1 | -0.114659 | STAD | GCIY | MK-2206 |
| MVP_top60_DNA_seed | ERBB2 | -0.113922 | STAD | TGBC11TKB | Afatinib |
| MVP_top60_DNA_seed | EGFR | -0.113922 | STAD | TGBC11TKB | Afatinib |
| MVP_top60_DNA_seed | PIK3CA | -0.112712 | STAD | AGS | AZD8186 |
| MVP_top60_DNA_seed | ERBB2 | -0.112354 | STAD | SNU-16 | Sapitinib |
| MVP_top60_DNA_seed | ERBB3 | -0.112354 | STAD | SNU-16 | Sapitinib |
| MVP_top60_DNA_seed | EGFR | -0.112354 | STAD | SNU-16 | Sapitinib |

| | | | | | |
|---|---|---|---|---|---|
| MVP_top60_DNA_seed | EGFR | -0.110913 | STAD | TMK-1 | AZD3759 |
| MVP_top60_DNA_seed | EGFR | -0.110822 | STAD | MKN7 | Gefitinib |
| MVP_top60_DNA_seed | HDAC1 | -0.110771 | STAD | OCUM-1 | Romidepsin |
| MVP_top60_DNA_seed | PIK3CA | -0.110412 | STAD | SK-GT-2 | AZD8186 |
| MVP_top60_DNA_seed | PIK3CA | -0.108094 | STAD | TMK-1 | Buparlisib |
| MVP_top60_DNA_seed | AKT1 | -0.105849 | STAD | FU97 | MK-2206 |
| MVP_top60_DNA_seed | ERBB2 | -0.103653 | STAD | IM-95 | Sapitinib |
| MVP_top60_DNA_seed | ERBB3 | -0.103653 | STAD | IM-95 | Sapitinib |
| MVP_top60_DNA_seed | EGFR | -0.103653 | STAD | IM-95 | Sapitinib |
| MVP_top60_DNA_seed | IGF1R | -0.103205 | STAD | AGS | GSK1904529A |
| MVP_top60_DNA_seed | EGFR | -0.102111 | STAD | IM-95 | Erlotinib |
| MVP_top60_DNA_seed | MAPK1 | -0.100731 | STAD | SK-GT-2 | SCH772984 |
| MVP_top60_DNA_seed | MAPK3 | -0.100731 | STAD | SK-GT-2 | SCH772984 |
| MVP_top60_DNA_seed | ESR1 | -0.100388 | STAD | TGBC11TKB | Tamoxifen |
| MVP_top60_DNA_seed | PIK3CA | -0.100081 | STAD | NCI-N87 | Buparlisib |
| MVP_top60_DNA_seed | ATM | -0.094917 | STAD | MKN28 | KU-55933 |
| MVP_top60_DNA_seed | SRC | -0.092626 | STAD | GCIY | Bosutinib |
| MVP_top60_DNA_seed | IGF1R | -0.092444 | STAD | MKN28 | NVP-ADW742 |
| MVP_top60_DNA_seed | PIK3CA | -0.091567 | STAD | TMK-1 | AZD8186 |
| MVP_top60_DNA_seed | HSP90AA1 | -0.091452 | STAD | SK-GT-2 | Tanespimycin |
| MVP_top60_DNA_seed | AKT1 | -0.086935 | STAD | SNU-5 | Uprosertib |
| MVP_top60_DNA_seed | HSP90AA1 | -0.086544 | STAD | TMK-1 | Elesclomol |
| MVP_top60_DNA_seed | ERBB2 | -0.080522 | STAD | FU97 | Lapatinib |
| MVP_top60_DNA_seed | EGFR | -0.080522 | STAD | FU97 | Lapatinib |
| MVP_top60_DNA_seed | EP300 | -0.078808 | STAD | AGS | SGC-CBP30 |
| MVP_top60_DNA_seed | CREBBP | -0.078808 | STAD | AGS | SGC-CBP30 |
| MVP_top60_DNA_seed | AKT1 | -0.075392 | STAD | SNU-5 | Ipatasertib |
| MVP_top60_DNA_seed | PIK3CA | -0.072562 | STAD | OCUM-1 | Alpelisib |
| MVP_top60_DNA_seed | GSK3B | -0.068422 | STAD | MKN28 | SB216763 |
| MVP_top60_DNA_seed | ATM | -0.068061 | STAD | FU97 | KU-55933 |
| MVP_top60_DNA_seed | MAPK1 | -0.066786 | STAD | SNU-16 | SCH772984 |
| MVP_top60_DNA_seed | MAPK3 | -0.066786 | STAD | SNU-16 | SCH772984 |
| MVP_top60_DNA_seed | IGF1R | -0.066667 | STAD | GCIY | Linsitinib |
| MVP_top60_DNA_seed | ERBB2 | -0.066352 | STAD | MKN7 | Sapitinib |
| MVP_top60_DNA_seed | ERBB3 | -0.066352 | STAD | MKN7 | Sapitinib |
| MVP_top60_DNA_seed | EGFR | -0.066352 | STAD | MKN7 | Sapitinib |
| MVP_top60_DNA_seed | IGF1R | -0.065641 | STAD | TMK-1 | Linsitinib |
| MVP_top60_DNA_seed | TP53 | -0.064001 | STAD | 23132-87 | MIRA-1 |
| MVP_top60_DNA_seed | PIK3CA | -0.061805 | STAD | NUGC-3 | Buparlisib |
| MVP_top60_DNA_seed | ERBB2 | -0.060068 | STAD | SNU-1 | Lapatinib |
| MVP_top60_DNA_seed | EGFR | -0.060068 | STAD | SNU-1 | Lapatinib |

| | | | | | |
|---|---|---|---|---|---|
| MVP_top60_DNA_seed | ERBB2 | -0.059664 | STAD | SNU-1 | Afatinib |
| MVP_top60_DNA_seed | EGFR | -0.059664 | STAD | SNU-1 | Afatinib |
| MVP_top60_DNA_seed | MAPK1 | -0.05529 | STAD | SK-GT-2 | VX-11e |
| MVP_top60_DNA_seed | MAPK1 | -0.053042 | STAD | SNU-1 | Ulixertinib |
| MVP_top60_DNA_seed | MAPK3 | -0.053042 | STAD | SNU-1 | Ulixertinib |
| MVP_top60_DNA_seed | KRAS | -0.051228 | STAD | GCIY | KRAS (G12C) Inhibitor-12 |
| MVP_top60_DNA_seed | MDM2 | -0.039627 | STAD | FU97 | Serdemetan |
| MVP_top60_DNA_seed | HSP90AA1 | -0.037038 | STAD | GCIY | Luminespib |
| MVP_top60_DNA_seed | MDM2 | -0.034759 | STAD | IM-95 | Nutlin-3a (-) |
| MVP_top60_DNA_seed | HDAC1 | -0.034086 | STAD | RERF-GC-1B | Romidepsin |
| MVP_top60_DNA_seed | ERBB2 | -0.033577 | STAD | HGC-27 | Afatinib |
| MVP_top60_DNA_seed | EGFR | -0.033577 | STAD | HGC-27 | Afatinib |
| MVP_top60_DNA_seed | HSP90AA1 | -0.032787 | STAD | MKN7 | Elesclomol |
| MVP_top60_DNA_seed | AKT1 | -0.032066 | STAD | MKN7 | Afuresertib |
| MVP_top60_DNA_seed | MAPK1 | -0.029176 | STAD | NUGC-3 | Ulixertinib |
| MVP_top60_DNA_seed | MAPK3 | -0.029176 | STAD | NUGC-3 | Ulixertinib |
| MVP_top60_DNA_seed | EGFR | -0.028937 | STAD | AGS | AZD3759 |
| MVP_top60_DNA_seed | PIK3CA | -0.023201 | STAD | MKN45 | Buparlisib |
| MVP_top60_DNA_seed | HDAC1 | -0.022067 | STAD | SNU-5 | Romidepsin |
| MVP_top60_DNA_seed | FGFR3 | -0.020568 | STAD | HGC-27 | PD173074 |
| MVP_top60_DNA_seed | ERBB2 | -0.020032 | STAD | AGS | Sapitinib |
| MVP_top60_DNA_seed | ERBB3 | -0.020032 | STAD | AGS | Sapitinib |
| MVP_top60_DNA_seed | EGFR | -0.020032 | STAD | AGS | Sapitinib |
| MVP_top60_DNA_seed | HSP90AA1 | -0.018806 | STAD | 23132-87 | Tanespimycin |
| MVP_top60_DNA_seed | EGFR | -0.017941 | STAD | 23132-87 | Osimertinib |
| MVP_top60_DNA_seed | MET | -0.016684 | STAD | TMK-1 | Crizotinib |
| MVP_top60_DNA_seed | PIK3CA | -0.015433 | STAD | NUGC-3 | Alpelisib |
| MVP_top60_DNA_seed | EGFR | -0.005466 | STAD | RF-48 | AZD3759 |
| MVP_top60_DNA_seed | MAPK1 | 0.001898 | STAD | NUGC-3 | SCH772984 |
| MVP_top60_DNA_seed | MAPK3 | 0.001898 | STAD | NUGC-3 | SCH772984 |
| MVP_top60_DNA_seed | AKT1 | 0.002147 | STAD | SNU-5 | Uprosertib |
| MVP_top60_DNA_seed | ESR1 | 0.002783 | STAD | Hs-746T | GDC0810 |
| MVP_top60_DNA_seed | PIK3CA | 0.005655 | STAD | TGBC11TKB | Buparlisib |
| MVP_top60_DNA_seed | EGFR | 0.00598 | STAD | HGC-27 | Gefitinib |
| MVP_top60_DNA_seed | SRC | 0.007134 | STAD | TMK-1 | Bosutinib |
| MVP_top60_DNA_seed | TP53 | 0.008774 | STAD | IM-95 | MIRA-1 |
| MVP_top60_DNA_seed | AKT1 | 0.012019 | STAD | OCUM-1 | GSK2110183B |
| MVP_top60_DNA_seed | ERBB2 | 0.012472 | STAD | MKN28 | Lapatinib |
| MVP_top60_DNA_seed | EGFR | 0.012472 | STAD | MKN28 | Lapatinib |
| MVP_top60_DNA_seed | HDAC1 | 0.013655 | STAD | Hs-746T | Romidepsin |
| MVP_top60_DNA_seed | AKT1 | 0.013801 | STAD | GCIY | Uprosertib |

| | | | | | |
|---|---|---|---|---|---|
| MVP_top60_DNA_seed | KRAS | 0.015757 | STAD | OCUM-1 | KRAS (G12C) Inhibitor-12 |
| MVP_top60_DNA_seed | SRC | 0.018182 | STAD | NUGC-3 | Bosutinib |
| MVP_top60_DNA_seed | EGFR | 0.019278 | STAD | MKN7 | AZD3759 |
| MVP_top60_DNA_seed | IGF1R | 0.021127 | STAD | RERF-GC-1B | GSK1904529A |
| MVP_top60_DNA_seed | HSP90AA1 | 0.022141 | STAD | MKN28 | Tanespimycin |
| MVP_top60_DNA_seed | MAPK1 | 0.022994 | STAD | 23132-87 | Ulixertinib |
| MVP_top60_DNA_seed | MAPK3 | 0.022994 | STAD | 23132-87 | Ulixertinib |
| MVP_top60_DNA_seed | ESR1 | 0.023539 | STAD | IM-95 | Tamoxifen |
| MVP_top60_DNA_seed | MAPK1 | 0.024982 | STAD | IM-95 | SCH772984 |
| MVP_top60_DNA_seed | MAPK3 | 0.024982 | STAD | IM-95 | SCH772984 |
| MVP_top60_DNA_seed | AKT1 | 0.02593 | STAD | SNU-1 | Uprosertib |
| MVP_top60_DNA_seed | FGFR3 | 0.025966 | STAD | NUGC-3 | PD173074 |
| MVP_top60_DNA_seed | IGF1R | 0.027697 | STAD | AGS | BMS-536924 |
| MVP_top60_DNA_seed | ERBB2 | 0.027895 | STAD | MKN28 | Sapitinib |
| MVP_top60_DNA_seed | ERBB3 | 0.027895 | STAD | MKN28 | Sapitinib |
| MVP_top60_DNA_seed | EGFR | 0.027895 | STAD | MKN28 | Sapitinib |
| MVP_top60_DNA_seed | ESR1 | 0.028829 | STAD | TMK-1 | GDC0810 |
| MVP_top60_DNA_seed | FGFR3 | 0.031612 | STAD | SK-GT-2 | AZD4547 |
| MVP_top60_DNA_seed | HSP90AA1 | 0.033424 | STAD | RF-48 | Tanespimycin |
| MVP_top60_DNA_seed | MAPK1 | 0.033804 | STAD | FU97 | Ulixertinib |
| MVP_top60_DNA_seed | MAPK3 | 0.033804 | STAD | FU97 | Ulixertinib |
| MVP_top60_DNA_seed | PIK3CA | 0.036404 | STAD | MKN7 | AZD8186 |
| MVP_top60_DNA_seed | FGFR3 | 0.038038 | STAD | AGS | AZD4547 |
| MVP_top60_DNA_seed | ERBB2 | 0.03859 | STAD | Hs-746T | Lapatinib |
| MVP_top60_DNA_seed | EGFR | 0.03859 | STAD | Hs-746T | Lapatinib |
| MVP_top60_DNA_seed | EGFR | 0.040425 | STAD | SNU-16 | Erlotinib |
| MVP_top60_DNA_seed | PIK3CA | 0.043728 | STAD | MKN7 | GNE-317 |
| MVP_top60_DNA_seed | ESR1 | 0.044685 | STAD | NUGC-4 | GDC0810 |
| MVP_top60_DNA_seed | IGF1R | 0.04545 | STAD | IM-95 | NVP-ADW742 |
| MVP_top60_DNA_seed | MAPK1 | 0.046051 | STAD | GCIY | Ulixertinib |
| MVP_top60_DNA_seed | MAPK3 | 0.046051 | STAD | GCIY | Ulixertinib |
| MVP_top60_DNA_seed | IGF1R | 0.046795 | STAD | Hs-746T | BMS-754807 |
| MVP_top60_DNA_seed | ERBB2 | 0.046841 | STAD | RERF-GC-1B | Afatinib |
| MVP_top60_DNA_seed | EGFR | 0.046841 | STAD | RERF-GC-1B | Afatinib |
| MVP_top60_DNA_seed | AKT1 | 0.047559 | STAD | Hs-746T | Uprosertib |
| MVP_top60_DNA_seed | EP300 | 0.047649 | STAD | GCIY | SGC-CBP30 |
| MVP_top60_DNA_seed | CREBBP | 0.047649 | STAD | GCIY | SGC-CBP30 |
| MVP_top60_DNA_seed | ATM | 0.0506 | STAD | Hs-746T | KU-55933 |
| MVP_top60_DNA_seed | ERBB2 | 0.050663 | STAD | MKN28 | Afatinib |
| MVP_top60_DNA_seed | EGFR | 0.050663 | STAD | MKN28 | Afatinib |
| MVP_top60_DNA_seed | IGF1R | 0.052342 | STAD | FU97 | BMS-536924 |

| | | | | | |
|---|---|---|---|---|---|
| MVP_top60_DNA_seed | IGF1R | 0.05428 | STAD | Hs-746T | NVP-ADW742 |
| MVP_top60_DNA_seed | AKT1 | 0.057569 | STAD | OCUM-1 | MK-2206 |
| MVP_top60_DNA_seed | EP300 | 0.059794 | STAD | HGC-27 | CPI-637 |
| MVP_top60_DNA_seed | PIK3CA | 0.060791 | STAD | TGBC11TKB | GNE-317 |
| MVP_top60_DNA_seed | AKT1 | 0.061408 | STAD | Hs-746T | Ipatasertib |
| MVP_top60_DNA_seed | PIK3CA | 0.062867 | STAD | GCIY | AZD8186 |
| MVP_top60_DNA_seed | IGF1R | 0.070332 | STAD | SK-GT-2 | BMS-536924 |
| MVP_top60_DNA_seed | PIK3CA | 0.0727 | STAD | RF-48 | AZD8186 |
| MVP_top60_DNA_seed | ERBB2 | 0.073149 | STAD | RERF-GC-1B | Lapatinib |
| MVP_top60_DNA_seed | EGFR | 0.073149 | STAD | RERF-GC-1B | Lapatinib |
| MVP_top60_DNA_seed | MDM2 | 0.074679 | STAD | TGBC11TKB | Serdemetan |
| MVP_top60_DNA_seed | IGF1R | 0.077205 | STAD | 23132-87 | BMS-536924 |
| MVP_top60_DNA_seed | IGF1R | 0.078938 | STAD | GCIY | BMS-754807 |
| MVP_top60_DNA_seed | MDM2 | 0.07984 | STAD | HGC-27 | Nutlin-3a (-) |
| MVP_top60_DNA_seed | AKT1 | 0.089287 | STAD | KATOIII | Uprosertib |
| MVP_top60_DNA_seed | GSK3B | 0.091457 | STAD | IM-95 | SB216763 |
| MVP_top60_DNA_seed | SRC | 0.093718 | STAD | NUGC-4 | Bosutinib |
| MVP_top60_DNA_seed | ESR1 | 0.093965 | STAD | TGBC11TKB | GDC0810 |
| MVP_top60_DNA_seed | FGFR3 | 0.094442 | STAD | TMK-1 | PD173074 |
| MVP_top60_DNA_seed | MAPK1 | 0.097508 | STAD | NUGC-3 | Ulixertinib |
| MVP_top60_DNA_seed | MAPK3 | 0.097508 | STAD | NUGC-3 | Ulixertinib |
| MVP_top60_DNA_seed | IGF1R | 0.099714 | STAD | AGS | Linsitinib |
| MVP_top60_DNA_seed | HDAC1 | 0.107099 | STAD | 23132-87 | Entinostat |
| MVP_top60_DNA_seed | EP300 | 0.107301 | STAD | OCUM-1 | SGC-CBP30 |
| MVP_top60_DNA_seed | CREBBP | 0.107301 | STAD | OCUM-1 | SGC-CBP30 |
| MVP_top60_DNA_seed | EGFR | 0.108621 | STAD | HGC-27 | AZD3759 |
| MVP_top60_DNA_seed | EGFR | 0.109436 | STAD | IM-95 | AZD3759 |
| MVP_top60_DNA_seed | MAPK1 | 0.113167 | STAD | TGBC11TKB | Ulixertinib |
| MVP_top60_DNA_seed | MAPK3 | 0.113167 | STAD | TGBC11TKB | Ulixertinib |
| MVP_top60_DNA_seed | FGFR3 | 0.114731 | STAD | SNU-5 | AZD4547 |
| MVP_top60_DNA_seed | EGFR | 0.116803 | STAD | SNU-5 | Erlotinib |
| MVP_top60_DNA_seed | PIK3CA | 0.116808 | STAD | RF-48 | Alpelisib |
| MVP_top60_DNA_seed | AKT1 | 0.117392 | STAD | RERF-GC-1B | AZD5363 |
| MVP_top60_DNA_seed | HSP90AA1 | 0.118825 | STAD | MKN45 | Luminespib |
| MVP_top60_DNA_seed | MET | 0.123733 | STAD | OCUM-1 | Crizotinib |
| MVP_top60_DNA_seed | AKT1 | 0.126352 | STAD | SNU-1 | MK-2206 |
| MVP_top60_DNA_seed | HDAC1 | 0.130555 | STAD | MKN28 | PCI-34051 |
| MVP_top60_DNA_seed | FGFR3 | 0.13309 | STAD | RF-48 | AZD4547 |
| MVP_top60_DNA_seed | ERBB2 | 0.133165 | STAD | TGBC11TKB | Lapatinib |
| MVP_top60_DNA_seed | EGFR | 0.133165 | STAD | TGBC11TKB | Lapatinib |
| MVP_top60_DNA_seed | AKT1 | 0.136936 | STAD | KATOIII | Uprosertib |

| | | | | | |
|---|---|---|---|---|---|
| MVP_top60_DNA_seed | IGF1R | 0.137153 | STAD | IM-95 | Linsitinib |
| MVP_top60_DNA_seed | AKT1 | 0.1398 | STAD | TGBC11TKB | Uprosertib |
| MVP_top60_DNA_seed | MDM2 | 0.140719 | STAD | GCIY | Serdemetan |
| MVP_top60_DNA_seed | AKT1 | 0.144116 | STAD | RF-48 | Ipatasertib |
| MVP_top60_DNA_seed | HSP90AA1 | 0.145553 | STAD | 23132-87 | Luminespib |
| MVP_top60_DNA_seed | TP53 | 0.145707 | STAD | FU97 | MIRA-1 |
| MVP_top60_DNA_seed | IGF1R | 0.146346 | STAD | OCUM-1 | BMS-754807 |
| MVP_top60_DNA_seed | AKT1 | 0.147023 | STAD | NUGC-3 | Uprosertib |
| MVP_top60_DNA_seed | EP300 | 0.147874 | STAD | SNU-5 | CPI-637 |
| MVP_top60_DNA_seed | AKT1 | 0.147935 | STAD | KATOIII | AZD5363 |
| MVP_top60_DNA_seed | GSK3B | 0.150724 | STAD | SK-GT-2 | SB216763 |
| MVP_top60_DNA_seed | ERBB2 | 0.15512 | STAD | HGC-27 | Sapitinib |
| MVP_top60_DNA_seed | ERBB3 | 0.15512 | STAD | HGC-27 | Sapitinib |
| MVP_top60_DNA_seed | EGFR | 0.15512 | STAD | HGC-27 | Sapitinib |
| MVP_top60_DNA_seed | HDAC1 | 0.162536 | STAD | HGC-27 | PCI-34051 |
| MVP_top60_DNA_seed | TP53 | 0.163433 | STAD | MKN7 | MIRA-1 |
| MVP_top60_DNA_seed | IGF1R | 0.164514 | STAD | NUGC-3 | NVP-ADW742 |
| MVP_top60_DNA_seed | ATM | 0.165988 | STAD | NCI-N87 | KU-55933 |
| MVP_top60_DNA_seed | MAPK1 | 0.166255 | STAD | HGC-27 | Ulixertinib |
| MVP_top60_DNA_seed | MAPK3 | 0.166255 | STAD | HGC-27 | Ulixertinib |
| MVP_top60_DNA_seed | HDAC1 | 0.174872 | STAD | HGC-27 | Romidepsin |
| MVP_top60_DNA_seed | MET | 0.17519 | STAD | NCI-N87 | Savolitinib |
| MVP_top60_DNA_seed | AKT1 | 0.175976 | STAD | RF-48 | Uprosertib |
| MVP_top60_DNA_seed | HSP90AA1 | 0.177348 | STAD | SNU-16 | Luminespib |
| MVP_top60_DNA_seed | IGF1R | 0.178576 | STAD | HGC-27 | Linsitinib |
| MVP_top60_DNA_seed | HSP90AA1 | 0.179068 | STAD | GCIY | Tanespimycin |
| MVP_top60_DNA_seed | MAPK1 | 0.179324 | STAD | RF-48 | SCH772984 |
| MVP_top60_DNA_seed | MAPK3 | 0.179324 | STAD | RF-48 | SCH772984 |
| MVP_top60_DNA_seed | EP300 | 0.182026 | STAD | IM-95 | SGC-CBP30 |
| MVP_top60_DNA_seed | CREBBP | 0.182026 | STAD | IM-95 | SGC-CBP30 |
| MVP_top60_DNA_seed | AKT1 | 0.182445 | STAD | MKN28 | Uprosertib |
| MVP_top60_DNA_seed | EGFR | 0.187375 | STAD | FU97 | Osimertinib |
| MVP_top60_DNA_seed | IGF1R | 0.188258 | STAD | SNU-1 | Linsitinib |
| MVP_top60_DNA_seed | GSK3B | 0.188284 | STAD | SNU-5 | SB216763 |
| MVP_top60_DNA_seed | EP300 | 0.188979 | STAD | TGBC11TKB | CPI-637 |
| MVP_top60_DNA_seed | HDAC1 | 0.189704 | STAD | 23132-87 | PCI-34051 |
| MVP_top60_DNA_seed | MDM2 | 0.193363 | STAD | RERF-GC-1B | Serdemetan |
| MVP_top60_DNA_seed | IGF1R | 0.195355 | STAD | SNU-1 | BMS-536924 |
| MVP_top60_DNA_seed | MET | 0.198369 | STAD | RF-48 | Crizotinib |
| MVP_top60_DNA_seed | EGFR | 0.202177 | STAD | TGBC11TKB | Erlotinib |
| MVP_top60_DNA_seed | EGFR | 0.207298 | STAD | SNU-1 | Gefitinib |

| | | | | | |
|---|---|---|---|---|---|
| MVP_top60_DNA_seed | HSP90AA1 | 0.208561 | STAD | KATOIII | Tanespimycin |
| MVP_top60_DNA_seed | EP300 | 0.209651 | STAD | SNU-1 | CPI-637 |
| MVP_top60_DNA_seed | ESR1 | 0.21113 | STAD | AGS | GDC0810 |
| MVP_top60_DNA_seed | EGFR | 0.21514 | STAD | Hs-746T | Osimertinib |
| MVP_top60_DNA_seed | MDM2 | 0.22093 | STAD | SNU-16 | Nutlin-3a (-) |
| MVP_top60_DNA_seed | MAPK1 | 0.221079 | STAD | NUGC-3 | VX-11e |
| MVP_top60_DNA_seed | MAPK1 | 0.222337 | STAD | NUGC-4 | Ulixertinib |
| MVP_top60_DNA_seed | MAPK3 | 0.222337 | STAD | NUGC-4 | Ulixertinib |
| MVP_top60_DNA_seed | PIK3CA | 0.222733 | STAD | SNU-5 | Alpelisib |
| MVP_top60_DNA_seed | EGFR | 0.225283 | STAD | SNU-5 | AZD3759 |
| MVP_top60_DNA_seed | HDAC1 | 0.22766 | STAD | OCUM-1 | PCI-34051 |
| MVP_top60_DNA_seed | MDM2 | 0.233225 | STAD | SNU-5 | Nutlin-3a (-) |
| MVP_top60_DNA_seed | MDM2 | 0.233994 | STAD | SNU-5 | Serdemetan |
| MVP_top60_DNA_seed | SRC | 0.23633 | STAD | OCUM-1 | Bosutinib |
| MVP_top60_DNA_seed | MDM2 | 0.242162 | STAD | AGS | Serdemetan |
| MVP_top60_DNA_seed | IGF1R | 0.242323 | STAD | 23132-87 | Linsitinib |
| MVP_top60_DNA_seed | AKT1 | 0.242411 | STAD | OCUM-1 | Uprosertib |
| MVP_top60_DNA_seed | ERBB2 | 0.246427 | STAD | MKN7 | Afatinib |
| MVP_top60_DNA_seed | EGFR | 0.246427 | STAD | MKN7 | Afatinib |
| MVP_top60_DNA_seed | PIK3CA | 0.246863 | STAD | RF-48 | GNE-317 |
| MVP_top60_DNA_seed | HSP90AA1 | 0.248015 | STAD | FU97 | Tanespimycin |
| MVP_top60_DNA_seed | ESR1 | 0.248251 | STAD | SNU-1 | GDC0810 |
| MVP_top60_DNA_seed | SRC | 0.250155 | STAD | SK-GT-2 | Bosutinib |
| MVP_top60_DNA_seed | GSK3B | 0.252279 | STAD | MKN45 | SB216763 |
| MVP_top60_DNA_seed | MET | 0.258961 | STAD | 23132-87 | Savolitinib |
| MVP_top60_DNA_seed | HDAC1 | 0.260893 | STAD | NUGC-3 | PCI-34051 |
| MVP_top60_DNA_seed | ERBB2 | 0.261371 | STAD | RF-48 | Afatinib |
| MVP_top60_DNA_seed | EGFR | 0.261371 | STAD | RF-48 | Afatinib |
| MVP_top60_DNA_seed | PIK3CA | 0.261699 | STAD | NUGC-3 | AZD8186 |
| MVP_top60_DNA_seed | FGFR3 | 0.261821 | STAD | MKN28 | PD173074 |
| MVP_top60_DNA_seed | AKT1 | 0.265873 | STAD | NUGC-3 | Afuresertib |
| MVP_top60_DNA_seed | MAPK1 | 0.265918 | STAD | NCI-N87 | VX-11e |
| MVP_top60_DNA_seed | AKT1 | 0.271776 | STAD | MKN28 | AT13148 |
| MVP_top60_DNA_seed | MET | 0.276314 | STAD | MKN28 | Crizotinib |
| MVP_top60_DNA_seed | AKT1 | 0.276999 | STAD | KATOIII | Afuresertib |
| MVP_top60_DNA_seed | EGFR | 0.277814 | STAD | TMK-1 | Gefitinib |
| MVP_top60_DNA_seed | TP53 | 0.278515 | STAD | TMK-1 | MIRA-1 |
| MVP_top60_DNA_seed | HSP90AA1 | 0.287888 | STAD | TMK-1 | Luminespib |
| MVP_top60_DNA_seed | SRC | 0.292812 | STAD | MKN28 | Bosutinib |
| MVP_top60_DNA_seed | ATM | 0.293658 | STAD | NUGC-3 | KU-55933 |
| MVP_top60_DNA_seed | ATM | 0.298397 | STAD | IM-95 | KU-55933 |

| | | | | | |
|---|---|---|---|---|---|
| MVP_top60_DNA_seed | PIK3CA | 0.299203 | STAD | MKN7 | Alpelisib |
| MVP_top60_DNA_seed | HSP90AA1 | 0.299986 | STAD | MKN28 | Luminespib |
| MVP_top60_DNA_seed | MET | 0.301869 | STAD | TGBC11TKB | Crizotinib |
| MVP_top60_DNA_seed | AKT1 | 0.302561 | STAD | TGBC11TKB | Uprosertib |
| MVP_top60_DNA_seed | ERBB2 | 0.305537 | STAD | GCIY | Lapatinib |
| MVP_top60_DNA_seed | EGFR | 0.305537 | STAD | GCIY | Lapatinib |
| MVP_top60_DNA_seed | ESR1 | 0.310481 | STAD | SNU-5 | GDC0810 |
| MVP_top60_DNA_seed | MAPK1 | 0.310979 | STAD | MKN7 | VX-11e |
| MVP_top60_DNA_seed | ERBB2 | 0.312197 | STAD | TMK-1 | Lapatinib |
| MVP_top60_DNA_seed | EGFR | 0.312197 | STAD | TMK-1 | Lapatinib |
| MVP_top60_DNA_seed | IGF1R | 0.316402 | STAD | MKN7 | BMS-754807 |
| MVP_top60_DNA_seed | AKT1 | 0.316851 | STAD | RF-48 | AT13148 |
| MVP_top60_DNA_seed | EP300 | 0.318274 | STAD | RF-48 | CPI-637 |
| MVP_top60_DNA_seed | IGF1R | 0.320863 | STAD | GCIY | GSK1904529A |
| MVP_top60_DNA_seed | HDAC1 | 0.321258 | STAD | SK-GT-2 | PCI-34051 |
| MVP_top60_DNA_seed | AKT1 | 0.324261 | STAD | OCUM-1 | Ipatasertib |
| MVP_top60_DNA_seed | AKT1 | 0.327972 | STAD | NUGC-3 | MK-2206 |
| MVP_top60_DNA_seed | MDM2 | 0.329854 | STAD | RERF-GC-1B | Nutlin-3a (-) |
| MVP_top60_DNA_seed | AKT1 | 0.332001 | STAD | RF-48 | Afuresertib |
| MVP_top60_DNA_seed | HSP90AA1 | 0.340552 | STAD | IM-95 | Luminespib |
| MVP_top60_DNA_seed | MET | 0.343877 | STAD | TMK-1 | Savolitinib |
| MVP_top60_DNA_seed | ESR1 | 0.34605 | STAD | MKN45 | GDC0810 |
| MVP_top60_DNA_seed | TP53 | 0.34861 | STAD | SNU-1 | MIRA-1 |
| MVP_top60_DNA_seed | AKT1 | 0.349796 | STAD | GCIY | Afuresertib |
| MVP_top60_DNA_seed | MAPK1 | 0.358626 | STAD | KATOIII | VX-11e |
| MVP_top60_DNA_seed | AKT1 | 0.359065 | STAD | RF-48 | MK-2206 |
| MVP_top60_DNA_seed | MAPK1 | 0.359989 | STAD | HGC-27 | VX-11e |
| MVP_top60_DNA_seed | ATM | 0.363082 | STAD | SK-GT-2 | KU-55933 |
| MVP_top60_DNA_seed | MAPK1 | 0.368653 | STAD | NCI-N87 | Ulixertinib |
| MVP_top60_DNA_seed | MAPK3 | 0.368653 | STAD | NCI-N87 | Ulixertinib |
| MVP_top60_DNA_seed | AKT1 | 0.368981 | STAD | SNU-1 | Ipatasertib |
| MVP_top60_DNA_seed | PIK3CA | 0.36905 | STAD | MKN28 | Alpelisib |
| MVP_top60_DNA_seed | EGFR | 0.370681 | STAD | MKN28 | Gefitinib |
| MVP_top60_DNA_seed | MAPK1 | 0.37124 | STAD | SNU-1 | SCH772984 |
| MVP_top60_DNA_seed | MAPK3 | 0.37124 | STAD | SNU-1 | SCH772984 |
| MVP_top60_DNA_seed | MAPK1 | 0.374746 | STAD | MKN7 | SCH772984 |
| MVP_top60_DNA_seed | MAPK3 | 0.374746 | STAD | MKN7 | SCH772984 |
| MVP_top60_DNA_seed | AKT1 | 0.375294 | STAD | RF-48 | AZD5363 |
| MVP_top60_DNA_seed | HDAC1 | 0.375703 | STAD | OCUM-1 | Entinostat |
| MVP_top60_DNA_seed | IGF1R | 0.375712 | STAD | SK-GT-2 | GSK1904529A |
| MVP_top60_DNA_seed | FGFR3 | 0.377844 | STAD | Hs-746T | PD173074 |

| | | | | | |
|---|---|---|---|---|---|
| MVP_top60_DNA_seed | IGF1R | 0.378039 | STAD | FU97 | Linsitinib |
| MVP_top60_DNA_seed | MET | 0.381507 | STAD | FU97 | Crizotinib |
| MVP_top60_DNA_seed | TP53 | 0.383155 | STAD | SNU-5 | MIRA-1 |
| MVP_top60_DNA_seed | EGFR | 0.384815 | STAD | SNU-5 | Gefitinib |
| MVP_top60_DNA_seed | IGF1R | 0.387104 | STAD | SNU-5 | Linsitinib |
| MVP_top60_DNA_seed | AKT1 | 0.395153 | STAD | MKN7 | MK-2206 |
| MVP_top60_DNA_seed | ERBB2 | 0.396143 | STAD | MKN45 | Lapatinib |
| MVP_top60_DNA_seed | EGFR | 0.396143 | STAD | MKN45 | Lapatinib |
| MVP_top60_DNA_seed | PIK3CA | 0.396228 | STAD | MKN28 | Buparlisib |
| MVP_top60_DNA_seed | SRC | 0.39658 | STAD | FU97 | Bosutinib |
| MVP_top60_DNA_seed | IGF1R | 0.399759 | STAD | TGBC11TKB | BMS-754807 |
| MVP_top60_DNA_seed | GSK3B | 0.402097 | STAD | TMK-1 | SB216763 |
| MVP_top60_DNA_seed | MAPK1 | 0.402851 | STAD | SK-GT-2 | Ulixertinib |
| MVP_top60_DNA_seed | MAPK3 | 0.402851 | STAD | SK-GT-2 | Ulixertinib |
| MVP_top60_DNA_seed | EGFR | 0.405354 | STAD | RF-48 | Gefitinib |
| MVP_top60_DNA_seed | EP300 | 0.405531 | STAD | Hs-746T | CPI-637 |
| MVP_top60_DNA_seed | TP53 | 0.406069 | STAD | RERF-GC-1B | MIRA-1 |
| MVP_top60_DNA_seed | AKT1 | 0.410688 | STAD | SNU-1 | AT13148 |
| MVP_top60_DNA_seed | MAPK1 | 0.413718 | STAD | SNU-1 | VX-11e |
| MVP_top60_DNA_seed | PIK3CA | 0.415397 | STAD | MKN45 | AZD8186 |
| MVP_top60_DNA_seed | AKT1 | 0.417253 | STAD | RERF-GC-1B | Ipatasertib |
| MVP_top60_DNA_seed | EGFR | 0.417328 | STAD | MKN28 | Osimertinib |
| MVP_top60_DNA_seed | ERBB2 | 0.41743 | STAD | IM-95 | Lapatinib |
| MVP_top60_DNA_seed | EGFR | 0.41743 | STAD | IM-95 | Lapatinib |
| MVP_top60_DNA_seed | MET | 0.424269 | STAD | NUGC-3 | Crizotinib |
| MVP_top60_DNA_seed | MAPK1 | 0.424381 | STAD | FU97 | SCH772984 |
| MVP_top60_DNA_seed | MAPK3 | 0.424381 | STAD | FU97 | SCH772984 |
| MVP_top60_DNA_seed | MAPK1 | 0.425057 | STAD | HGC-27 | SCH772984 |
| MVP_top60_DNA_seed | MAPK3 | 0.425057 | STAD | HGC-27 | SCH772984 |
| MVP_top60_DNA_seed | HDAC1 | 0.429843 | STAD | KATOIII | Entinostat |
| MVP_top60_DNA_seed | TP53 | 0.433549 | STAD | SK-GT-2 | MIRA-1 |
| MVP_top60_DNA_seed | AKT1 | 0.433767 | STAD | MKN7 | AZD5363 |
| MVP_top60_DNA_seed | IGF1R | 0.437526 | STAD | RF-48 | BMS-536924 |
| MVP_top60_DNA_seed | AKT1 | 0.437609 | STAD | KATOIII | Ipatasertib |
| MVP_top60_DNA_seed | FGFR3 | 0.438476 | STAD | FU97 | AZD4547 |
| MVP_top60_DNA_seed | PIK3CA | 0.442209 | STAD | GCIY | Alpelisib |
| MVP_top60_DNA_seed | AKT1 | 0.446665 | STAD | OCUM-1 | Afuresertib |
| MVP_top60_DNA_seed | AKT1 | 0.44744 | STAD | SNU-1 | Afuresertib |
| MVP_top60_DNA_seed | IGF1R | 0.450785 | STAD | RF-48 | BMS-754807 |
| MVP_top60_DNA_seed | AKT1 | 0.454356 | STAD | SNU-5 | MK-2206 |
| MVP_top60_DNA_seed | EP300 | 0.454674 | STAD | SNU-1 | SGC-CBP30 |

| | | | | | |
|---|---|---|---|---|---|
| MVP_top60_DNA_seed | CREBBP | 0.454674 | STAD | SNU-1 | SGC-CBP30 |
| MVP_top60_DNA_seed | KRAS | 0.457661 | STAD | HGC-27 | KRAS (G12C) Inhibitor-12 |
| MVP_top60_DNA_seed | EP300 | 0.461379 | STAD | MKN28 | SGC-CBP30 |
| MVP_top60_DNA_seed | CREBBP | 0.461379 | STAD | MKN28 | SGC-CBP30 |
| MVP_top60_DNA_seed | MAPK1 | 0.468294 | STAD | GCIY | SCH772984 |
| MVP_top60_DNA_seed | MAPK3 | 0.468294 | STAD | GCIY | SCH772984 |
| MVP_top60_DNA_seed | EP300 | 0.471067 | STAD | NUGC-3 | CPI-637 |
| MVP_top60_DNA_seed | AKT1 | 0.471105 | STAD | TMK-1 | MK-2206 |
| MVP_top60_DNA_seed | EGFR | 0.473163 | STAD | MKN7 | Erlotinib |
| MVP_top60_DNA_seed | FGFR3 | 0.474056 | STAD | 23132-87 | PD173074 |
| MVP_top60_DNA_seed | EP300 | 0.476173 | STAD | MKN45 | CPI-637 |
| MVP_top60_DNA_seed | AKT1 | 0.479414 | STAD | SK-GT-2 | Uprosertib |
| MVP_top60_DNA_seed | ERBB2 | 0.479741 | STAD | TGBC11TKB | Sapitinib |
| MVP_top60_DNA_seed | ERBB3 | 0.479741 | STAD | TGBC11TKB | Sapitinib |
| MVP_top60_DNA_seed | EGFR | 0.479741 | STAD | TGBC11TKB | Sapitinib |
| MVP_top60_DNA_seed | ATM | 0.479952 | STAD | 23132-87 | KU-55933 |
| MVP_top60_DNA_seed | FGFR3 | 0.481047 | STAD | IM-95 | PD173074 |
| MVP_top60_DNA_seed | PIK3CA | 0.481446 | STAD | RERF-GC-1B | AZD8186 |
| MVP_top60_DNA_seed | MDM2 | 0.484299 | STAD | NUGC-4 | Nutlin-3a (-) |
| MVP_top60_DNA_seed | EGFR | 0.488501 | STAD | SNU-1 | AZD3759 |
| MVP_top60_DNA_seed | MET | 0.491468 | STAD | NUGC-3 | Savolitinib |
| MVP_top60_DNA_seed | ATM | 0.494666 | STAD | TMK-1 | KU-55933 |
| MVP_top60_DNA_seed | HDAC1 | 0.495248 | STAD | IM-95 | PCI-34051 |
| MVP_top60_DNA_seed | HDAC1 | 0.497644 | STAD | RERF-GC-1B | Entinostat |
| MVP_top60_DNA_seed | GSK3B | 0.499467 | STAD | NUGC-3 | SB216763 |
| MVP_top60_DNA_seed | EGFR | 0.504684 | STAD | 23132-87 | AZD3759 |
| MVP_top60_DNA_seed | EGFR | 0.510394 | STAD | RF-48 | Erlotinib |
| MVP_top60_DNA_seed | ERBB2 | 0.512615 | STAD | KATOIII | Sapitinib |
| MVP_top60_DNA_seed | ERBB3 | 0.512615 | STAD | KATOIII | Sapitinib |
| MVP_top60_DNA_seed | EGFR | 0.512615 | STAD | KATOIII | Sapitinib |
| MVP_top60_DNA_seed | IGF1R | 0.512776 | STAD | NCI-N87 | Linsitinib |
| MVP_top60_DNA_seed | ESR1 | 0.516392 | STAD | MKN28 | GDC0810 |
| MVP_top60_DNA_seed | AKT1 | 0.519589 | STAD | MKN28 | Uprosertib |
| MVP_top60_DNA_seed | FGFR3 | 0.521063 | STAD | OCUM-1 | PD173074 |
| MVP_top60_DNA_seed | IGF1R | 0.529933 | STAD | AGS | BMS-754807 |
| MVP_top60_DNA_seed | AKT1 | 0.52999 | STAD | SNU-1 | GSK2110183B |
| MVP_top60_DNA_seed | MDM2 | 0.53308 | STAD | 23132-87 | Serdemetan |
| MVP_top60_DNA_seed | GSK3B | 0.533731 | STAD | MKN7 | CHIR-99021 |
| MVP_top60_DNA_seed | AKT1 | 0.534085 | STAD | MKN28 | Afuresertib |
| MVP_top60_DNA_seed | AKT1 | 0.535937 | STAD | HSC-39 | Uprosertib |
| MVP_top60_DNA_seed | EP300 | 0.53914 | STAD | SK-GT-2 | SGC-CBP30 |

| | | | | | |
|---|---|---|---|---|---|
| MVP_top60_DNA_seed | CREBBP | 0.53914 | STAD | SK-GT-2 | SGC-CBP30 |
| MVP_top60_DNA_seed | HDAC1 | 0.541107 | STAD | GCIY | Romidepsin |
| MVP_top60_DNA_seed | IGF1R | 0.548114 | STAD | MKN28 | BMS-754807 |
| MVP_top60_DNA_seed | FGFR3 | 0.551204 | STAD | TGBC11TKB | AZD4547 |
| MVP_top60_DNA_seed | MAPK1 | 0.553461 | STAD | IM-95 | VX-11e |
| MVP_top60_DNA_seed | EGFR | 0.553463 | STAD | IM-95 | Gefitinib |
| MVP_top60_DNA_seed | FGFR3 | 0.555995 | STAD | MKN45 | PD173074 |
| MVP_top60_DNA_seed | EGFR | 0.55651 | STAD | RF-48 | Osimertinib |
| MVP_top60_DNA_seed | AKT1 | 0.559525 | STAD | NUGC-3 | Uprosertib |
| MVP_top60_DNA_seed | HDAC1 | 0.564163 | STAD | SNU-1 | Entinostat |
| MVP_top60_DNA_seed | EGFR | 0.571632 | STAD | Hs-746T | Erlotinib |
| MVP_top60_DNA_seed | ESR1 | 0.574937 | STAD | Hs-746T | Tamoxifen |
| MVP_top60_DNA_seed | GSK3B | 0.578226 | STAD | TGBC11TKB | SB216763 |
| MVP_top60_DNA_seed | GSK3B | 0.583817 | STAD | NCI-N87 | SB216763 |
| MVP_top60_DNA_seed | MAPK1 | 0.585739 | STAD | NUGC-4 | Ulixertinib |
| MVP_top60_DNA_seed | MAPK3 | 0.585739 | STAD | NUGC-4 | Ulixertinib |
| MVP_top60_DNA_seed | TP53 | 0.588419 | STAD | TGBC11TKB | MIRA-1 |
| MVP_top60_DNA_seed | IGF1R | 0.589552 | STAD | NUGC-3 | BMS-754807 |
| MVP_top60_DNA_seed | EP300 | 0.590533 | STAD | MKN45 | SGC-CBP30 |
| MVP_top60_DNA_seed | CREBBP | 0.590533 | STAD | MKN45 | SGC-CBP30 |
| MVP_top60_DNA_seed | MDM2 | 0.590792 | STAD | TMK-1 | Nutlin-3a (-) |
| MVP_top60_DNA_seed | EGFR | 0.594909 | STAD | MKN28 | Erlotinib |
| MVP_top60_DNA_seed | EP300 | 0.596401 | STAD | GCIY | CPI-637 |
| MVP_top60_DNA_seed | EGFR | 0.598399 | STAD | Hs-746T | Gefitinib |
| MVP_top60_DNA_seed | PIK3CA | 0.598927 | STAD | SNU-1 | AZD8186 |
| MVP_top60_DNA_seed | IGF1R | 0.598934 | STAD | NUGC-3 | Linsitinib |
| MVP_top60_DNA_seed | EP300 | 0.599162 | STAD | NCI-N87 | SGC-CBP30 |
| MVP_top60_DNA_seed | CREBBP | 0.599162 | STAD | NCI-N87 | SGC-CBP30 |
| MVP_top60_DNA_seed | ERBB2 | 0.601589 | STAD | Hs-746T | Afatinib |
| MVP_top60_DNA_seed | EGFR | 0.601589 | STAD | Hs-746T | Afatinib |
| MVP_top60_DNA_seed | ERBB2 | 0.603182 | STAD | RF-48 | Sapitinib |
| MVP_top60_DNA_seed | ERBB3 | 0.603182 | STAD | RF-48 | Sapitinib |
| MVP_top60_DNA_seed | EGFR | 0.603182 | STAD | RF-48 | Sapitinib |
| MVP_top60_DNA_seed | AKT1 | 0.603229 | STAD | TGBC11TKB | AZD5363 |
| MVP_top60_DNA_seed | EGFR | 0.603392 | STAD | NUGC-4 | Gefitinib |
| MVP_top60_DNA_seed | AKT1 | 0.606153 | STAD | TGBC11TKB | MK-2206 |
| MVP_top60_DNA_seed | HDAC1 | 0.607904 | STAD | HGC-27 | Entinostat |
| MVP_top60_DNA_seed | SRC | 0.612194 | STAD | KATOIII | Bosutinib |
| MVP_top60_DNA_seed | ESR1 | 0.613222 | STAD | SNU-1 | Tamoxifen |
| MVP_top60_DNA_seed | PIK3CA | 0.616749 | STAD | RERF-GC-1B | Buparlisib |
| MVP_top60_DNA_seed | MET | 0.617708 | STAD | KATOIII | Crizotinib |

| | | | | | |
|---|---|---|---|---|---|
| MVP_top60_DNA_seed | AKT1 | 0.618904 | STAD | GCIY | GSK2110183B |
| MVP_top60_DNA_seed | MET | 0.619445 | STAD | SK-GT-2 | Crizotinib |
| MVP_top60_DNA_seed | IGF1R | 0.620162 | STAD | OCUM-1 | NVP-ADW742 |
| MVP_top60_DNA_seed | ESR1 | 0.623097 | STAD | MKN7 | GDC0810 |
| MVP_top60_DNA_seed | AKT1 | 0.626472 | STAD | SNU-5 | Afuresertib |
| MVP_top60_DNA_seed | AKT1 | 0.62767 | STAD | RERF-GC-1B | Afuresertib |
| MVP_top60_DNA_seed | ESR1 | 0.633735 | STAD | MKN45 | Tamoxifen |
| MVP_top60_DNA_seed | IGF1R | 0.635215 | STAD | NCI-N87 | GSK1904529A |
| MVP_top60_DNA_seed | IGF1R | 0.636326 | STAD | MKN7 | BMS-536924 |
| MVP_top60_DNA_seed | MDM2 | 0.642 | STAD | IM-95 | Serdemetan |
| MVP_top60_DNA_seed | MET | 0.642108 | STAD | GCIY | Crizotinib |
| MVP_top60_DNA_seed | AKT1 | 0.647111 | STAD | MKN28 | MK-2206 |
| MVP_top60_DNA_seed | EGFR | 0.648447 | STAD | RERF-GC-1B | Osimertinib |
| MVP_top60_DNA_seed | IGF1R | 0.64852 | STAD | NUGC-3 | GSK1904529A |
| MVP_top60_DNA_seed | AKT1 | 0.651728 | STAD | TGBC11TKB | AT13148 |
| MVP_top60_DNA_seed | EGFR | 0.661869 | STAD | SNU-5 | Osimertinib |
| MVP_top60_DNA_seed | ERBB2 | 0.663193 | STAD | 23132-87 | Lapatinib |
| MVP_top60_DNA_seed | EGFR | 0.663193 | STAD | 23132-87 | Lapatinib |
| MVP_top60_DNA_seed | IGF1R | 0.663443 | STAD | TGBC11TKB | NVP-ADW742 |
| MVP_top60_DNA_seed | ESR1 | 0.663927 | STAD | GCIY | Tamoxifen |
| MVP_top60_DNA_seed | ATM | 0.66448 | STAD | TGBC11TKB | KU-55933 |
| MVP_top60_DNA_seed | HDAC1 | 0.667302 | STAD | SNU-1 | PCI-34051 |
| MVP_top60_DNA_seed | TP53 | 0.67081 | STAD | OCUM-1 | MIRA-1 |
| MVP_top60_DNA_seed | ERBB2 | 0.670885 | STAD | SNU-1 | Sapitinib |
| MVP_top60_DNA_seed | ERBB3 | 0.670885 | STAD | SNU-1 | Sapitinib |
| MVP_top60_DNA_seed | EGFR | 0.670885 | STAD | SNU-1 | Sapitinib |
| MVP_top60_DNA_seed | MAPK1 | 0.672457 | STAD | NUGC-4 | VX-11e |
| MVP_top60_DNA_seed | EP300 | 0.677818 | STAD | KATOIII | SGC-CBP30 |
| MVP_top60_DNA_seed | CREBBP | 0.677818 | STAD | KATOIII | SGC-CBP30 |
| MVP_top60_DNA_seed | SRC | 0.680894 | STAD | 23132-87 | Bosutinib |
| MVP_top60_DNA_seed | KRAS | 0.682097 | STAD | NUGC-4 | (RAS (G12C) Inhibitor-12 |
| MVP_top60_DNA_seed | IGF1R | 0.685379 | STAD | SNU-5 | NVP-ADW742 |
| MVP_top60_DNA_seed | PIK3CA | 0.69062 | STAD | RERF-GC-1B | GNE-317 |
| MVP_top60_DNA_seed | MDM2 | 0.692716 | STAD | SNU-1 | Serdemetan |
| MVP_top60_DNA_seed | MET | 0.699545 | STAD | FU97 | Savolitinib |
| MVP_top60_DNA_seed | FGFR3 | 0.703335 | STAD | RF-48 | PD173074 |
| MVP_top60_DNA_seed | IGF1R | 0.710121 | STAD | TGBC11TKB | GSK1904529A |
| MVP_top60_DNA_seed | AKT1 | 0.71297 | STAD | RERF-GC-1B | AT13148 |
| MVP_top60_DNA_seed | TP53 | 0.719516 | STAD | NUGC-3 | MIRA-1 |
| MVP_top60_DNA_seed | HDAC1 | 0.729204 | STAD | MKN28 | Romidepsin |
| MVP_top60_DNA_seed | PIK3CA | 0.73458 | STAD | TMK-1 | GNE-317 |

| | | | | | |
|---|---|---|---|---|---|
| MVP_top60_DNA_seed | AKT1 | 0.736123 | STAD | FU97 | AT13148 |
| MVP_top60_DNA_seed | HDAC1 | 0.736435 | STAD | NCI-N87 | Entinostat |
| MVP_top60_DNA_seed | HSP90AA1 | 0.743032 | STAD | TGBC11TKB | Luminespib |
| MVP_top60_DNA_seed | IGF1R | 0.743157 | STAD | MKN45 | NVP-ADW742 |
| MVP_top60_DNA_seed | AKT1 | 0.745236 | STAD | MKN28 | AZD5363 |
| MVP_top60_DNA_seed | TP53 | 0.752842 | STAD | NCI-N87 | MIRA-1 |
| MVP_top60_DNA_seed | EP300 | 0.759056 | STAD | NUGC-4 | CPI-637 |
| MVP_top60_DNA_seed | HDAC1 | 0.763636 | STAD | GCIY | PCI-34051 |
| MVP_top60_DNA_seed | MDM2 | 0.76536 | STAD | NCI-N87 | Nutlin-3a (-) |
| MVP_top60_DNA_seed | FGFR3 | 0.770224 | STAD | GCIY | PD173074 |
| MVP_top60_DNA_seed | MAPK1 | 0.770431 | STAD | MKN7 | Ulixertinib |
| MVP_top60_DNA_seed | MAPK3 | 0.770431 | STAD | MKN7 | Ulixertinib |
| MVP_top60_DNA_seed | AKT1 | 0.773986 | STAD | MKN7 | Ipatasertib |
| MVP_top60_DNA_seed | KRAS | 0.778574 | STAD | NUGC-3 | KRAS (G12C) Inhibitor-12 |
| MVP_top60_DNA_seed | EP300 | 0.779691 | STAD | SK-GT-2 | CPI-637 |
| MVP_top60_DNA_seed | IGF1R | 0.785524 | STAD | GCIY | NVP-ADW742 |
| MVP_top60_DNA_seed | HDAC1 | 0.790381 | STAD | MKN45 | Entinostat |
| MVP_top60_DNA_seed | ERBB2 | 0.793279 | STAD | SNU-5 | Afatinib |
| MVP_top60_DNA_seed | EGFR | 0.793279 | STAD | SNU-5 | Afatinib |
| MVP_top60_DNA_seed | PIK3CA | 0.7938 | STAD | SNU-16 | AZD8186 |
| MVP_top60_DNA_seed | HDAC1 | 0.800946 | STAD | Hs-746T | PCI-34051 |
| MVP_top60_DNA_seed | ESR1 | 0.80209 | STAD | NUGC-3 | GDC0810 |
| MVP_top60_DNA_seed | HDAC1 | 0.80635 | STAD | NCI-N87 | PCI-34051 |
| MVP_top60_DNA_seed | AKT1 | 0.806644 | STAD | TGBC11TKB | Ipatasertib |
| MVP_top60_DNA_seed | AR | 0.808352 | STAD | MKN7 | Bicalutamide |
| MVP_top60_DNA_seed | EGFR | 0.81275 | STAD | SNU-1 | Osimertinib |
| MVP_top60_DNA_seed | ERBB2 | 0.817424 | STAD | FU97 | Sapitinib |
| MVP_top60_DNA_seed | ERBB3 | 0.817424 | STAD | FU97 | Sapitinib |
| MVP_top60_DNA_seed | EGFR | 0.817424 | STAD | FU97 | Sapitinib |
| MVP_top60_DNA_seed | IGF1R | 0.817836 | STAD | MKN7 | Linsitinib |
| MVP_top60_DNA_seed | FGFR3 | 0.819547 | STAD | NUGC-3 | AZD4547 |
| MVP_top60_DNA_seed | EP300 | 0.819904 | STAD | IM-95 | CPI-637 |
| MVP_top60_DNA_seed | MDM2 | 0.827733 | STAD | SK-GT-2 | Serdemetan |
| MVP_top60_DNA_seed | IGF1R | 0.829294 | STAD | OCUM-1 | GSK1904529A |
| MVP_top60_DNA_seed | FGFR3 | 0.833458 | STAD | Hs-746T | AZD4547 |
| MVP_top60_DNA_seed | FGFR3 | 0.834795 | STAD | MKN45 | AZD4547 |
| MVP_top60_DNA_seed | AKT1 | 0.836694 | STAD | NUGC-3 | GSK2110183B |
| MVP_top60_DNA_seed | EGFR | 0.837527 | STAD | RERF-GC-1B | AZD3759 |
| MVP_top60_DNA_seed | TP53 | 0.838111 | STAD | MKN28 | MIRA-1 |
| MVP_top60_DNA_seed | IGF1R | 0.849387 | STAD | TGBC11TKB | BMS-536924 |
| MVP_top60_DNA_seed | MDM2 | 0.852634 | STAD | GCIY | Nutlin-3a (-) |

| | | | | | |
|---|---|---|---|---|---|
| MVP_top60_DNA_seed | EGFR | 0.853649 | STAD | AGS | Osimertinib |
| MVP_top60_DNA_seed | PIK3CA | 0.855342 | STAD | SNU-1 | Alpelisib |
| MVP_top60_DNA_seed | PIK3CA | 0.855649 | STAD | NUGC-4 | Alpelisib |
| MVP_top60_DNA_seed | MET | 0.855978 | STAD | GCIY | Savolitinib |
| MVP_top60_DNA_seed | HSP90AA1 | 0.85794 | STAD | TMK-1 | Tanespimycin |
| MVP_top60_DNA_seed | FGFR3 | 0.869434 | STAD | MKN28 | AZD4547 |
| MVP_top60_DNA_seed | MAPK1 | 0.872168 | STAD | FU97 | Ulixertinib |
| MVP_top60_DNA_seed | MAPK3 | 0.872168 | STAD | FU97 | Ulixertinib |
| MVP_top60_DNA_seed | ESR1 | 0.872895 | STAD | NCI-N87 | Tamoxifen |
| MVP_top60_DNA_seed | ERBB2 | 0.876586 | STAD | MKN45 | Afatinib |
| MVP_top60_DNA_seed | EGFR | 0.876586 | STAD | MKN45 | Afatinib |
| MVP_top60_DNA_seed | EP300 | 0.883488 | STAD | KATOIII | CPI-637 |
| MVP_top60_DNA_seed | MAPK1 | 0.883756 | STAD | IM-95 | Ulixertinib |
| MVP_top60_DNA_seed | MAPK3 | 0.883756 | STAD | IM-95 | Ulixertinib |
| MVP_top60_DNA_seed | IGF1R | 0.889232 | STAD | RF-48 | GSK1904529A |
| MVP_top60_DNA_seed | FGFR3 | 0.892925 | STAD | SK-GT-2 | PD173074 |
| MVP_top60_DNA_seed | ATM | 0.89813 | STAD | NUGC-4 | KU-55933 |
| MVP_top60_DNA_seed | EGFR | 0.903246 | STAD | Hs-746T | AZD3759 |
| MVP_top60_DNA_seed | AKT1 | 0.90457 | STAD | TGBC11TKB | Afuresertib |
| MVP_top60_DNA_seed | EP300 | 0.909502 | STAD | 23132-87 | CPI-637 |
| MVP_top60_DNA_seed | KRAS | 0.913606 | STAD | RERF-GC-1B | KRAS (G12C) Inhibitor-12 |
| MVP_top60_DNA_seed | EGFR | 0.916364 | STAD | FU97 | AZD3759 |
| MVP_top60_DNA_seed | IGF1R | 0.91759 | STAD | NCI-N87 | NVP-ADW742 |
| MVP_top60_DNA_seed | PIK3CA | 0.924553 | STAD | MKN28 | AZD8186 |
| MVP_top60_DNA_seed | GSK3B | 0.92934 | STAD | Hs-746T | SB216763 |
| MVP_top60_DNA_seed | FGFR3 | 0.930064 | STAD | SNU-5 | PD173074 |
| MVP_top60_DNA_seed | HDAC1 | 0.932902 | STAD | TMK-1 | PCI-34051 |
| MVP_top60_DNA_seed | FGFR3 | 0.935127 | STAD | OCUM-1 | AZD4547 |
| MVP_top60_DNA_seed | MET | 0.945278 | STAD | NCI-N87 | Crizotinib |
| MVP_top60_DNA_seed | ERBB2 | 0.945989 | STAD | FU97 | Afatinib |
| MVP_top60_DNA_seed | EGFR | 0.945989 | STAD | FU97 | Afatinib |
| MVP_top60_DNA_seed | IGF1R | 0.94638 | STAD | RF-48 | Linsitinib |
| MVP_top60_DNA_seed | ESR1 | 0.958655 | STAD | RF-48 | Tamoxifen |
| MVP_top60_DNA_seed | EGFR | 0.959441 | STAD | OCUM-1 | Gefitinib |
| MVP_top60_DNA_seed | MDM2 | 0.966061 | STAD | SK-GT-2 | Nutlin-3a (-) |
| MVP_top60_DNA_seed | EP300 | 0.967239 | STAD | OCUM-1 | CPI-637 |
| MVP_top60_DNA_seed | HDAC1 | 0.972412 | STAD | SNU-1 | Romidepsin |
| MVP_top60_DNA_seed | HSP90AA1 | 0.973899 | STAD | RERF-GC-1B | Tanespimycin |
| MVP_top60_DNA_seed | PIK3CA | 0.976605 | STAD | OCUM-1 | Buparlisib |
| MVP_top60_DNA_seed | HDAC1 | 0.97777 | STAD | RERF-GC-1B | PCI-34051 |
| MVP_top60_DNA_seed | IGF1R | 0.978083 | STAD | KATOIII | Linsitinib |

| | | | | | |
|---|---|---|---|---|---|
| MVP_top60_DNA_seed | ESR1 | 0.978548 | STAD | MKN7 | Tamoxifen |
| MVP_top60_DNA_seed | FGFR3 | 0.98229 | STAD | TMK-1 | AZD4547 |
| MVP_top60_DNA_seed | MDM2 | 0.986817 | STAD | MKN28 | Nutlin-3a (-) |
| MVP_top60_DNA_seed | AKT1 | 0.98831 | STAD | GCIY | AZD5363 |
| MVP_top60_DNA_seed | MDM2 | 0.996569 | STAD | MKN7 | Serdemetan |
| MVP_top60_DNA_seed | FGFR3 | 0.996848 | STAD | NCI-N87 | AZD4547 |
| MVP_top60_DNA_seed | IGF1R | 1.000706 | STAD | MKN7 | NVP-ADW742 |
| MVP_top60_DNA_seed | EGFR | 1.007633 | STAD | KATOIII | Erlotinib |
| MVP_top60_DNA_seed | MET | 1.010332 | STAD | MKN28 | Savolitinib |
| MVP_top60_DNA_seed | EGFR | 1.033397 | STAD | NCI-N87 | Erlotinib |
| MVP_top60_DNA_seed | TP53 | 1.053476 | STAD | MKN45 | MIRA-1 |
| MVP_top60_DNA_seed | MDM2 | 1.059129 | STAD | MKN28 | Serdemetan |
| MVP_top60_DNA_seed | ERBB2 | 1.059574 | STAD | SNU-5 | Sapitinib |
| MVP_top60_DNA_seed | ERBB3 | 1.059574 | STAD | SNU-5 | Sapitinib |
| MVP_top60_DNA_seed | EGFR | 1.059574 | STAD | SNU-5 | Sapitinib |
| MVP_top60_DNA_seed | AKT1 | 1.060615 | STAD | NUGC-3 | AZD5363 |
| MVP_top60_DNA_seed | EGFR | 1.070197 | STAD | OCUM-1 | AZD3759 |
| MVP_top60_DNA_seed | EP300 | 1.071057 | STAD | RERF-GC-1B | CPI-637 |
| MVP_top60_DNA_seed | EGFR | 1.072267 | STAD | FU97 | Erlotinib |
| MVP_top60_DNA_seed | MDM2 | 1.073453 | STAD | TGBC11TKB | Nutlin-3a (-) |
| MVP_top60_DNA_seed | GSK3B | 1.075772 | STAD | GCIY | SB216763 |
| MVP_top60_DNA_seed | EGFR | 1.079321 | STAD | KATOIII | AZD3759 |
| MVP_top60_DNA_seed | MET | 1.079609 | STAD | 23132-87 | Crizotinib |
| MVP_top60_DNA_seed | PIK3CA | 1.080422 | STAD | OCUM-1 | AZD8186 |
| MVP_top60_DNA_seed | EGFR | 1.080708 | STAD | FU97 | Gefitinib |
| MVP_top60_DNA_seed | ERBB2 | 1.085179 | STAD | SNU-5 | Lapatinib |
| MVP_top60_DNA_seed | EGFR | 1.085179 | STAD | SNU-5 | Lapatinib |
| MVP_top60_DNA_seed | EGFR | 1.087073 | STAD | MKN28 | AZD3759 |
| MVP_top60_DNA_seed | IGF1R | 1.091057 | STAD | MKN7 | GSK1904529A |
| MVP_top60_DNA_seed | ERBB2 | 1.093234 | STAD | OCUM-1 | Sapitinib |
| MVP_top60_DNA_seed | ERBB3 | 1.093234 | STAD | OCUM-1 | Sapitinib |
| MVP_top60_DNA_seed | EGFR | 1.093234 | STAD | OCUM-1 | Sapitinib |
| MVP_top60_DNA_seed | IGF1R | 1.09617 | STAD | SNU-5 | BMS-536924 |
| MVP_top60_DNA_seed | KRAS | 1.096434 | STAD | MKN7 | KRAS (G12C) Inhibitor-12 |
| MVP_top60_DNA_seed | MDM2 | 1.096944 | STAD | Hs-746T | Nutlin-3a (-) |
| MVP_top60_DNA_seed | HDAC1 | 1.109871 | STAD | KATOIII | PCI-34051 |
| MVP_top60_DNA_seed | AKT1 | 1.114398 | STAD | MKN28 | GSK2110183B |
| MVP_top60_DNA_seed | IGF1R | 1.114644 | STAD | MKN45 | GSK1904529A |
| MVP_top60_DNA_seed | AKT1 | 1.143573 | STAD | KATOIII | MK-2206 |
| MVP_top60_DNA_seed | FGFR3 | 1.150755 | STAD | GCIY | AZD4547 |
| MVP_top60_DNA_seed | ESR1 | 1.152807 | STAD | OCUM-1 | Tamoxifen |

| | | | | | |
|---|---|---|---|---|---|
| MVP_top60_DNA_seed | KRAS | 1.159427 | STAD | Hs-746T | KRAS (G12C) Inhibitor-12 |
| MVP_top60_DNA_seed | ESR1 | 1.161804 | STAD | OCUM-1 | GDC0810 |
| MVP_top60_DNA_seed | MDM2 | 1.165518 | STAD | MKN45 | Serdemetan |
| MVP_top60_DNA_seed | SRC | 1.167276 | STAD | MKN7 | Bosutinib |
| MVP_top60_DNA_seed | AKT1 | 1.175589 | STAD | MKN7 | Uprosertib |
| MVP_top60_DNA_seed | AKT1 | 1.177198 | STAD | KATOIII | AT13148 |
| MVP_top60_DNA_seed | IGF1R | 1.185759 | STAD | IM-95 | BMS-536924 |
| MVP_top60_DNA_seed | PIK3CA | 1.187297 | STAD | IM-95 | GNE-317 |
| MVP_top60_DNA_seed | HDAC1 | 1.188913 | STAD | GCIY | Entinostat |
| MVP_top60_DNA_seed | SRC | 1.190962 | STAD | TGBC11TKB | Bosutinib |
| MVP_top60_DNA_seed | AKT1 | 1.20066 | STAD | NUGC-4 | Ipatasertib |
| MVP_top60_DNA_seed | MDM2 | 1.205417 | STAD | KATOIII | Serdemetan |
| MVP_top60_DNA_seed | ESR1 | 1.216674 | STAD | TMK-1 | Tamoxifen |
| MVP_top60_DNA_seed | MDM2 | 1.220894 | STAD | NUGC-3 | Nutlin-3a (-) |
| MVP_top60_DNA_seed | ERBB2 | 1.223624 | STAD | KATOIII | Lapatinib |
| MVP_top60_DNA_seed | EGFR | 1.223624 | STAD | KATOIII | Lapatinib |
| MVP_top60_DNA_seed | PIK3CA | 1.223944 | STAD | MKN7 | Buparlisib |
| MVP_top60_DNA_seed | GSK3B | 1.225564 | STAD | OCUM-1 | SB216763 |
| MVP_top60_DNA_seed | AKT1 | 1.226123 | STAD | MKN7 | AT13148 |
| MVP_top60_DNA_seed | PIK3CA | 1.234249 | STAD | OCUM-1 | GNE-317 |
| MVP_top60_DNA_seed | TP53 | 1.240448 | STAD | NUGC-4 | MIRA-1 |
| MVP_top60_DNA_seed | IGF1R | 1.246206 | STAD | RF-48 | NVP-ADW742 |
| MVP_top60_DNA_seed | EP300 | 1.254091 | STAD | MKN28 | CPI-637 |
| MVP_top60_DNA_seed | ATM | 1.264994 | STAD | OCUM-1 | KU-55933 |
| MVP_top60_DNA_seed | TP53 | 1.27456 | STAD | Hs-746T | MIRA-1 |
| MVP_top60_DNA_seed | EGFR | 1.291515 | STAD | MKN45 | Gefitinib |
| MVP_top60_DNA_seed | MDM2 | 1.294502 | STAD | MKN7 | Nutlin-3a (-) |
| MVP_top60_DNA_seed | HSP90AA1 | 1.299079 | STAD | NUGC-4 | Tanespimycin |
| MVP_top60_DNA_seed | IGF1R | 1.326607 | STAD | SK-GT-2 | Linsitinib |
| MVP_top60_DNA_seed | AKT1 | 1.327374 | STAD | IM-95 | AT13148 |
| MVP_top60_DNA_seed | GSK3B | 1.33164 | STAD | MKN7 | SB216763 |
| MVP_top60_DNA_seed | PIK3CA | 1.33347 | STAD | KATOIII | Alpelisib |
| MVP_top60_DNA_seed | HDAC1 | 1.333627 | STAD | MKN28 | Entinostat |
| MVP_top60_DNA_seed | IGF1R | 1.33705 | STAD | Hs-746T | Linsitinib |
| MVP_top60_DNA_seed | PIK3CA | 1.34511 | STAD | TGBC11TKB | AZD8186 |
| MVP_top60_DNA_seed | SRC | 1.349176 | STAD | MKN45 | Bosutinib |
| MVP_top60_DNA_seed | MDM2 | 1.357746 | STAD | OCUM-1 | Serdemetan |
| MVP_top60_DNA_seed | TP53 | 1.362835 | STAD | KATOIII | MIRA-1 |
| MVP_top60_DNA_seed | MET | 1.363142 | STAD | MKN7 | Crizotinib |
| MVP_top60_DNA_seed | FGFR3 | 1.363982 | STAD | MKN7 | PD173074 |
| MVP_top60_DNA_seed | ATM | 1.369068 | STAD | MKN7 | KU-55933 |

| | | | | | |
|---|---|---|---|---|---|
| MVP_top60_DNA_seed | ATM | 1.379442 | STAD | KATOIII | KU-55933 |
| MVP_top60_DNA_seed | AKT1 | 1.382995 | STAD | GCIY | Ipatasertib |
| MVP_top60_DNA_seed | EGFR | 1.384593 | STAD | MKN45 | AZD3759 |
| MVP_top60_DNA_seed | SRC | 1.419095 | STAD | Hs-746T | Bosutinib |
| MVP_top60_DNA_seed | MET | 1.447934 | STAD | KATOIII | Savolitinib |
| MVP_top60_DNA_seed | KRAS | 1.453123 | STAD | KATOIII | KRAS (G12C) Inhibitor-12 |
| MVP_top60_DNA_seed | PIK3CA | 1.466011 | STAD | Hs-746T | Alpelisib |
| MVP_top60_DNA_seed | AKT1 | 1.470587 | STAD | NUGC-4 | GSK2110183B |
| MVP_top60_DNA_seed | ERBB2 | 1.474767 | STAD | Hs-746T | Sapitinib |
| MVP_top60_DNA_seed | ERBB3 | 1.474767 | STAD | Hs-746T | Sapitinib |
| MVP_top60_DNA_seed | EGFR | 1.474767 | STAD | Hs-746T | Sapitinib |
| MVP_top60_DNA_seed | GSK3B | 1.477037 | STAD | FU97 | SB216763 |
| MVP_top60_DNA_seed | AKT1 | 1.484331 | STAD | NUGC-4 | Uprosertib |
| MVP_top60_DNA_seed | AKT1 | 1.495951 | STAD | Hs-746T | AT13148 |
| MVP_top60_DNA_seed | IGF1R | 1.498209 | STAD | 23132-87 | BMS-754807 |
| MVP_top60_DNA_seed | IGF1R | 1.509833 | STAD | SK-GT-2 | BMS-754807 |
| MVP_top60_DNA_seed | EGFR | 1.531966 | STAD | OCUM-1 | Erlotinib |
| MVP_top60_DNA_seed | HSP90AA1 | 1.542322 | STAD | MKN7 | Luminespib |
| MVP_top60_DNA_seed | MAPK1 | 1.544501 | STAD | RF-48 | VX-11e |
| MVP_top60_DNA_seed | EGFR | 1.561325 | STAD | MKN45 | Erlotinib |
| MVP_top60_DNA_seed | HDAC1 | 1.581781 | STAD | MKN45 | Romidepsin |
| MVP_top60_DNA_seed | EP300 | 1.589912 | STAD | Hs-746T | SGC-CBP30 |
| MVP_top60_DNA_seed | CREBBP | 1.589912 | STAD | Hs-746T | SGC-CBP30 |
| MVP_top60_DNA_seed | MET | 1.613309 | STAD | MKN7 | Savolitinib |
| MVP_top60_DNA_seed | MAPK1 | 1.650232 | STAD | RF-48 | Ulixertinib |
| MVP_top60_DNA_seed | MAPK3 | 1.650232 | STAD | RF-48 | Ulixertinib |
| MVP_top60_DNA_seed | MDM2 | 1.651727 | STAD | KATOIII | Nutlin-3a (-) |
| MVP_top60_DNA_seed | ERBB2 | 1.655498 | STAD | RF-48 | Lapatinib |
| MVP_top60_DNA_seed | EGFR | 1.655498 | STAD | RF-48 | Lapatinib |
| MVP_top60_DNA_seed | AKT1 | 1.659364 | STAD | MKN28 | Ipatasertib |
| MVP_top60_DNA_seed | PIK3CA | 1.678729 | STAD | KATOIII | AZD8186 |
| MVP_top60_DNA_seed | ERBB2 | 1.687067 | STAD | MKN45 | Sapitinib |
| MVP_top60_DNA_seed | ERBB3 | 1.687067 | STAD | MKN45 | Sapitinib |
| MVP_top60_DNA_seed | EGFR | 1.687067 | STAD | MKN45 | Sapitinib |
| MVP_top60_DNA_seed | EGFR | 1.692818 | STAD | KATOIII | Osimertinib |
| MVP_top60_DNA_seed | IGF1R | 1.70497 | STAD | MKN45 | BMS-536924 |
| MVP_top60_DNA_seed | PIK3CA | 1.709535 | STAD | NUGC-4 | AZD8186 |
| MVP_top60_DNA_seed | AKT1 | 1.714454 | STAD | OCUM-1 | AT13148 |
| MVP_top60_DNA_seed | AKT1 | 1.715235 | STAD | NUGC-4 | Uprosertib |
| MVP_top60_DNA_seed | ERBB2 | 1.715738 | STAD | TMK-1 | Afatinib |
| MVP_top60_DNA_seed | EGFR | 1.715738 | STAD | TMK-1 | Afatinib |

| | | | | | |
|---|---|---|---|---|---|
| MVP_top60_DNA_seed | GSK3B | 1.71904 | STAD | KATOIII | SB216763 |
| MVP_top60_DNA_seed | FGFR3 | 1.724345 | STAD | NUGC-4 | AZD4547 |
| MVP_top60_DNA_seed | AKT1 | 1.731918 | STAD | GCIY | AT13148 |
| MVP_top60_DNA_seed | IGF1R | 1.733484 | STAD | Hs-746T | BMS-536924 |
| MVP_top60_DNA_seed | HSP90AA1 | 1.738538 | STAD | KATOIII | Luminespib |
| MVP_top60_DNA_seed | EGFR | 1.74761 | STAD | OCUM-1 | Osimertinib |
| MVP_top60_DNA_seed | ERBB2 | 1.755136 | STAD | OCUM-1 | Afatinib |
| MVP_top60_DNA_seed | EGFR | 1.755136 | STAD | OCUM-1 | Afatinib |
| MVP_top60_DNA_seed | PIK3CA | 1.757653 | STAD | KATOIII | GNE-317 |
| MVP_top60_DNA_seed | EGFR | 1.762082 | STAD | KATOIII | Gefitinib |
| MVP_top60_DNA_seed | HDAC1 | 1.762681 | STAD | NUGC-4 | PCI-34051 |
| MVP_top60_DNA_seed | HSP90AA1 | 1.769519 | STAD | MKN7 | CCT-018159 |
| MVP_top60_DNA_seed | MDM2 | 1.791178 | STAD | NUGC-4 | Serdemetan |
| MVP_top60_DNA_seed | EGFR | 1.841377 | STAD | MKN45 | Osimertinib |
| MVP_top60_DNA_seed | GSK3B | 1.855118 | STAD | NUGC-4 | SB216763 |
| MVP_top60_DNA_seed | IGF1R | 1.880899 | STAD | NUGC-3 | BMS-536924 |
| MVP_top60_DNA_seed | MDM2 | 1.944473 | STAD | Hs-746T | Serdemetan |
| MVP_top60_DNA_seed | HSP90AA1 | 1.952751 | STAD | MKN7 | Tanespimycin |
| MVP_top60_DNA_seed | IGF1R | 1.955467 | STAD | MKN45 | Linsitinib |
| MVP_top60_DNA_seed | PIK3CA | 1.960357 | STAD | NUGC-4 | Buparlisib |
| MVP_top60_DNA_seed | ERBB2 | 1.972143 | STAD | OCUM-1 | Lapatinib |
| MVP_top60_DNA_seed | EGFR | 1.972143 | STAD | OCUM-1 | Lapatinib |
| MVP_top60_DNA_seed | PIK3CA | 1.980186 | STAD | NUGC-4 | GNE-317 |
| MVP_top60_DNA_seed | IGF1R | 1.985715 | STAD | FU97 | BMS-754807 |
| MVP_top60_DNA_seed | IGF1R | 2.046596 | STAD | NUGC-4 | BMS-754807 |
| MVP_top60_DNA_seed | AKT1 | 2.078304 | STAD | NUGC-4 | AT13148 |
| MVP_top60_DNA_seed | AKT1 | 2.082986 | STAD | NUGC-4 | MK-2206 |
| MVP_top60_DNA_seed | ESR1 | 2.096074 | STAD | NUGC-4 | Tamoxifen |
| MVP_top60_DNA_seed | HSP90AA1 | 2.101438 | STAD | NUGC-4 | Luminespib |
| MVP_top60_DNA_seed | IGF1R | 2.113236 | STAD | KATOIII | NVP-ADW742 |
| MVP_top60_DNA_seed | FGFR3 | 2.16447 | STAD | MKN7 | AZD4547 |
| MVP_top60_DNA_seed | IGF1R | 2.229289 | STAD | NUGC-4 | NVP-ADW742 |
| MVP_top60_DNA_seed | FGFR3 | 2.280056 | STAD | NUGC-4 | PD173074 |
| MVP_top60_DNA_seed | PIK3CA | 2.305225 | STAD | KATOIII | Buparlisib |
| MVP_top60_DNA_seed | AKT1 | 2.357672 | STAD | NUGC-4 | Afuresertib |
| MVP_top60_DNA_seed | ERBB2 | 2.393403 | STAD | KATOIII | Afatinib |
| MVP_top60_DNA_seed | EGFR | 2.393403 | STAD | KATOIII | Afatinib |
| MVP_top60_DNA_seed | PIK3CA | 2.561985 | STAD | MKN28 | GNE-317 |
| MVP_top60_DNA_seed | IGF1R | 2.647806 | STAD | NUGC-4 | GSK1904529A |
| MVP_top60_DNA_seed | MET | 2.845476 | STAD | NUGC-4 | Crizotinib |
| MVP_top60_DNA_seed | IGF1R | 2.852806 | STAD | NCI-N87 | BMS-536924 |

| | | | | | |
|---|---|---|---|---|---|
| MVP_top60_DNA_seed | AKT1 | 2.892878 | STAD | NUGC-4 | AZD5363 |
| MVP_top60_DNA_seed | IGF1R | 3.364512 | STAD | NUGC-4 | Linsitinib |
| MVP_top60_DNA_seed | IGF1R | 3.714302 | STAD | NUGC-4 | BMS-536924 |
| MVP_top60_DNA_seed | GSK3B | -4.461638 | THCA | IHH-4 | KIN001-042 |
| MVP_top60_DNA_seed | RET | -3.946255 | THCA | CGTH-W-1 | Motesanib |
| MVP_top60_DNA_seed | BRAF | -3.729294 | THCA | K5 | Dabrafenib |
| MVP_top60_DNA_seed | BRAF | -3.412404 | THCA | BHT-101 | Dabrafenib |
| MVP_top60_DNA_seed | BRAF | -3.190105 | THCA | BHT-101 | PLX-4720 |
| MVP_top60_DNA_seed | BRAF | -3.135102 | THCA | IHH-4 | Dabrafenib |
| MVP_top60_DNA_seed | BRAF | -3.105461 | THCA | HTC-C3 | Dabrafenib |
| MVP_top60_DNA_seed | BRAF | -2.966357 | THCA | CGTH-W-1 | HG6-64-1 |
| MVP_top60_DNA_seed | BRAF | -2.961541 | THCA | K5 | PLX-4720 |
| MVP_top60_DNA_seed | BRAF | -2.905795 | THCA | BHT-101 | PLX-4720 |
| MVP_top60_DNA_seed | BRAF | -2.830382 | THCA | HTC-C3 | PLX-4720 |
| MVP_top60_DNA_seed | BRAF | -2.827309 | THCA | K5 | PLX-4720 |
| MVP_top60_DNA_seed | CHEK2 | -2.727623 | THCA | CGTH-W-1 | AZD7762 |
| MVP_top60_DNA_seed | BRAF | -2.688852 | THCA | HTC-C3 | RAF_9304 |
| MVP_top60_DNA_seed | BRAF | -2.604383 | THCA | BHT-101 | SB590885 |
| MVP_top60_DNA_seed | BRAF | -2.492753 | THCA | HTC-C3 | PLX-4720 |
| MVP_top60_DNA_seed | BRAF | -2.480329 | THCA | B-CPAP | Dabrafenib |
| MVP_top60_DNA_seed | GSK3B | -2.431374 | THCA | B-CPAP | CHIR-99021 |
| MVP_top60_DNA_seed | BRAF | -2.430714 | THCA | K5 | RAF_9304 |
| MVP_top60_DNA_seed | BRAF | -2.371192 | THCA | BHT-101 | RAF_9304 |
| MVP_top60_DNA_seed | GSK3B | -2.310407 | THCA | BHT-101 | CHIR-99021 |
| MVP_top60_DNA_seed | MDM2 | -2.302378 | THCA | K5 | Nutlin-3a (-) |
| MVP_top60_DNA_seed | GSK3B | -2.250906 | THCA | ASH-3 | CHIR-99021 |
| MVP_top60_DNA_seed | GSK3B | -2.250659 | THCA | B-CPAP | CHIR-99021 |
| MVP_top60_DNA_seed | MAPK1 | -2.176373 | THCA | BHT-101 | VX-11e |
| MVP_top60_DNA_seed | BRAF | -2.172551 | THCA | 8305C | Dabrafenib |
| MVP_top60_DNA_seed | HDAC1 | -2.1696 | THCA | CGTH-W-1 | JQ12 |
| MVP_top60_DNA_seed | MDM2 | -2.159461 | THCA | K5 | Serdemetan |
| MVP_top60_DNA_seed | BRAF | -2.151843 | THCA | K5 | AZ628 |
| MVP_top60_DNA_seed | RET | -2.119921 | THCA | HTC-C3 | Motesanib |
| MVP_top60_DNA_seed | GSK3B | -2.047005 | THCA | BHT-101 | SB216763 |
| MVP_top60_DNA_seed | BRAF | -2.037898 | THCA | 8305C | PLX-4720 |
| MVP_top60_DNA_seed | BRAF | -2.027428 | THCA | CGTH-W-1 | AZ628 |
| MVP_top60_DNA_seed | HDAC1 | -1.95383 | THCA | ML-1 | JQ12 |
| MVP_top60_DNA_seed | MDM2 | -1.935515 | THCA | IHH-4 | Nutlin-3a (-) |
| MVP_top60_DNA_seed | BRAF | -1.905405 | THCA | IHH-4 | PLX-4720 |
| MVP_top60_DNA_seed | BRAF | -1.828709 | THCA | TT2609-C02 | HG6-64-1 |
| MVP_top60_DNA_seed | RET | -1.811525 | THCA | TT | Motesanib |

| | | | | | |
|---|---|---|---|---|---|
| MVP_top60_DNA_seed | ATM | -1.797046 | THCA | IHH-4 | KU-60019 |
| MVP_top60_DNA_seed | GSK3B | -1.793115 | THCA | CGTH-W-1 | CHIR-99021 |
| MVP_top60_DNA_seed | BRAF | -1.773559 | THCA | 8305C | PLX-4720 |
| MVP_top60_DNA_seed | PTK2 | -1.758295 | THCA | B-CPAP | PF-562271 |
| MVP_top60_DNA_seed | PTK2 | -1.743982 | THCA | 8305C | PF-562271 |
| MVP_top60_DNA_seed | PTK2 | -1.742409 | THCA | KMH-2 | PF-562271 |
| MVP_top60_DNA_seed | AKT1 | -1.742069 | THCA | BHT-101 | MK-2206 |
| MVP_top60_DNA_seed | AKT2 | -1.742069 | THCA | BHT-101 | MK-2206 |
| MVP_top60_DNA_seed | BRAF | -1.672569 | THCA | IHH-4 | SB590885 |
| MVP_top60_DNA_seed | GSK3B | -1.638434 | THCA | HTC-C3 | KIN001-042 |
| MVP_top60_DNA_seed | HDAC1 | -1.616837 | THCA | TT2609-C02 | JQ12 |
| MVP_top60_DNA_seed | GSK3B | -1.607433 | THCA | CGTH-W-1 | CHIR-99021 |
| MVP_top60_DNA_seed | GSK3B | -1.604268 | THCA | ML-1 | CHIR-99021 |
| MVP_top60_DNA_seed | MDM2 | -1.600604 | THCA | TT | Nutlin-3a (-) |
| MVP_top60_DNA_seed | ATR | -1.591504 | THCA | CGTH-W-1 | QL-VIII-58 |
| MVP_top60_DNA_seed | BRAF | -1.554192 | THCA | B-CPAP | PLX-4720 |
| MVP_top60_DNA_seed | BRAF | -1.542311 | THCA | FTC-133 | PLX-4720 |
| MVP_top60_DNA_seed | GSK3B | -1.535477 | THCA | K5 | CHIR-99021 |
| MVP_top60_DNA_seed | BRAF | -1.51961 | THCA | B-CPAP | PLX-4720 |
| MVP_top60_DNA_seed | ATR | -1.456649 | THCA | CAL-62 | QL-VIII-58 |
| MVP_top60_DNA_seed | PTK2 | -1.455613 | THCA | TT2609-C02 | PF-562271 |
| MVP_top60_DNA_seed | HDAC1 | -1.447506 | THCA | IHH-4 | JQ12 |
| MVP_top60_DNA_seed | ATM | -1.435817 | THCA | IHH-4 | KU-55933 |
| MVP_top60_DNA_seed | BRAF | -1.423171 | THCA | IHH-4 | PLX-4720 |
| MVP_top60_DNA_seed | ATR | -1.418552 | THCA | IHH-4 | AZD6738 |
| MVP_top60_DNA_seed | GSK3B | -1.386729 | THCA | K5 | AZD7969 |
| MVP_top60_DNA_seed | BRAF | -1.362543 | THCA | 8305C | SB590885 |
| MVP_top60_DNA_seed | PTK2 | -1.348531 | THCA | CAL-62 | PF-562271 |
| MVP_top60_DNA_seed | HDAC1 | -1.338739 | THCA | TT2609-C02 | Dacinostat |
| MVP_top60_DNA_seed | AKT1 | -1.335264 | THCA | TT2609-C02 | AKT inhibitor VIII |
| MVP_top60_DNA_seed | AKT2 | -1.335264 | THCA | TT2609-C02 | AKT inhibitor VIII |
| MVP_top60_DNA_seed | AKT3 | -1.335264 | THCA | TT2609-C02 | AKT inhibitor VIII |
| MVP_top60_DNA_seed | GSK3B | -1.329082 | THCA | 8505C | CHIR-99021 |
| MVP_top60_DNA_seed | BRAF | -1.308841 | THCA | IHH-4 | RAF_9304 |
| MVP_top60_DNA_seed | CHEK2 | -1.308715 | THCA | IHH-4 | AZD7762 |
| MVP_top60_DNA_seed | ATM | -1.291053 | THCA | BHT-101 | KU-55933 |
| MVP_top60_DNA_seed | MAPK1 | -1.283128 | THCA | HTC-C3 | VX-11e |
| MVP_top60_DNA_seed | BRAF | -1.282376 | THCA | B-CPAP | HG6-64-1 |
| MVP_top60_DNA_seed | ATR | -1.267009 | THCA | IHH-4 | AZ20 |
| MVP_top60_DNA_seed | ATM | -1.265003 | THCA | K5 | torin2 |
| MVP_top60_DNA_seed | ATR | -1.265003 | THCA | K5 | torin2 |

| | | | | | |
|---|---|---|---|---|---|
| MVP_top60_DNA_seed | AKT1 | -1.262621 | THCA | BHT-101 | AKT inhibitor VIII |
| MVP_top60_DNA_seed | AKT2 | -1.262621 | THCA | BHT-101 | AKT inhibitor VIII |
| MVP_top60_DNA_seed | AKT3 | -1.262621 | THCA | BHT-101 | AKT inhibitor VIII |
| MVP_top60_DNA_seed | ATR | -1.255994 | THCA | CAL-62 | AZD6738 |
| MVP_top60_DNA_seed | GSK3B | -1.239157 | THCA | B-CPAP | AZD7969 |
| MVP_top60_DNA_seed | PTK2 | -1.231651 | THCA | CGTH-W-1 | PF-562271 |
| MVP_top60_DNA_seed | RET | -1.225834 | THCA | TT2609-C02 | Motesanib |
| MVP_top60_DNA_seed | BRAF | -1.224839 | THCA | KMH-2 | SB590885 |
| MVP_top60_DNA_seed | ATM | -1.218818 | THCA | CAL-62 | torin2 |
| MVP_top60_DNA_seed | ATR | -1.218818 | THCA | CAL-62 | torin2 |
| MVP_top60_DNA_seed | MAPK14 | -1.212279 | THCA | IHH-4 | TAK-715 |
| MVP_top60_DNA_seed | ATM | -1.205962 | THCA | 8305C | KU-60019 |
| MVP_top60_DNA_seed | BRAF | -1.201393 | THCA | KMH-2 | Dabrafenib |
| MVP_top60_DNA_seed | GSK3B | -1.192884 | THCA | KMH-2 | SB216763 |
| MVP_top60_DNA_seed | ATM | -1.178386 | THCA | B-CPAP | KU-60019 |
| MVP_top60_DNA_seed | BRAF | -1.173142 | THCA | ML-1 | HG6-64-1 |
| MVP_top60_DNA_seed | BRAF | -1.163381 | THCA | 8305C | HG6-64-1 |
| MVP_top60_DNA_seed | HDAC1 | -1.160062 | THCA | HTC-C3 | JQ12 |
| MVP_top60_DNA_seed | RET | -1.158994 | THCA | CAL-62 | Motesanib |
| MVP_top60_DNA_seed | AKT1 | -1.15598 | THCA | ML-1 | AKT inhibitor VIII |
| MVP_top60_DNA_seed | AKT2 | -1.15598 | THCA | ML-1 | AKT inhibitor VIII |
| MVP_top60_DNA_seed | AKT3 | -1.15598 | THCA | ML-1 | AKT inhibitor VIII |
| MVP_top60_DNA_seed | BRAF | -1.154696 | THCA | ML-1 | PLX-4720 |
| MVP_top60_DNA_seed | HDAC1 | -1.152329 | THCA | CGTH-W-1 | Dacinostat |
| MVP_top60_DNA_seed | AKT1 | -1.135161 | THCA | BHT-101 | BAY AKT1 |
| MVP_top60_DNA_seed | AKT2 | -1.135161 | THCA | BHT-101 | BAY AKT1 |
| MVP_top60_DNA_seed | AKT3 | -1.135161 | THCA | BHT-101 | BAY AKT1 |
| MVP_top60_DNA_seed | GSK3B | -1.131743 | THCA | CAL-62 | AZD7969 |
| MVP_top60_DNA_seed | GSK3B | -1.12833 | THCA | IHH-4 | AZD7969 |
| MVP_top60_DNA_seed | PTK2 | -1.124678 | THCA | IHH-4 | PF-562271 |
| MVP_top60_DNA_seed | PTK2 | -1.110141 | THCA | ASH-3 | PF-562271 |
| MVP_top60_DNA_seed | GSK3B | -1.091751 | THCA | IHH-4 | SB216763 |
| MVP_top60_DNA_seed | ATM | -1.08938 | THCA | 8305C | KU-55933 |
| MVP_top60_DNA_seed | BRAF | -1.088167 | THCA | 8305C | RAF_9304 |
| MVP_top60_DNA_seed | GSK3B | -1.079866 | THCA | 8505C | CHIR-99021 |
| MVP_top60_DNA_seed | AKT1 | -1.066972 | THCA | TT | AKT inhibitor VIII |
| MVP_top60_DNA_seed | AKT2 | -1.066972 | THCA | TT | AKT inhibitor VIII |
| MVP_top60_DNA_seed | AKT3 | -1.066972 | THCA | TT | AKT inhibitor VIII |
| MVP_top60_DNA_seed | GSK3B | -1.054235 | THCA | KMH-2 | CHIR-99021 |
| MVP_top60_DNA_seed | MDM2 | -1.032843 | THCA | IHH-4 | Serdemetan |
| MVP_top60_DNA_seed | BRAF | -1.031488 | THCA | TT2609-C02 | PLX-4720 |

| | | | | | |
|---|---|---|---|---|---|
| MVP_top60_DNA_seed | BRAF | -1.023462 | THCA | IHH-4 | HG6-64-1 |
| MVP_top60_DNA_seed | CHEK2 | -1.013809 | THCA | CAL-62 | AZD7762 |
| MVP_top60_DNA_seed | HDAC1 | -1.009404 | THCA | HTC-C3 | Belinostat |
| MVP_top60_DNA_seed | PTK2 | -1.005164 | THCA | ML-1 | PF-562271 |
| MVP_top60_DNA_seed | GSK3B | -1.001871 | THCA | K5 | CHIR-99021 |
| MVP_top60_DNA_seed | ATR | -0.998455 | THCA | IHH-4 | QL-VIII-58 |
| MVP_top60_DNA_seed | HDAC1 | -0.992644 | THCA | B-CPAP | JQ12 |
| MVP_top60_DNA_seed | PTK2 | -0.983662 | THCA | BHT-101 | PF-562271 |
| MVP_top60_DNA_seed | RET | -0.982642 | THCA | BHT-101 | Motesanib |
| MVP_top60_DNA_seed | ATR | -0.979173 | THCA | RO82-W-1 | QL-VIII-58 |
| MVP_top60_DNA_seed | MAPK1 | -0.950621 | THCA | K5 | FR-180204 |
| MVP_top60_DNA_seed | MAPK3 | -0.950621 | THCA | K5 | FR-180204 |
| MVP_top60_DNA_seed | BRAF | -0.94498 | THCA | CAL-62 | RAF_9304 |
| MVP_top60_DNA_seed | ATM | -0.944882 | THCA | HTC-C3 | CP466722 |
| MVP_top60_DNA_seed | ATM | -0.941654 | THCA | 8505C | KU-60019 |
| MVP_top60_DNA_seed | GSK3B | -0.93908 | THCA | RO82-W-1 | CHIR-99021 |
| MVP_top60_DNA_seed | MAPK1 | -0.933707 | THCA | B-CPAP | VX-11e |
| MVP_top60_DNA_seed | MAPK1 | -0.933083 | THCA | TT2609-C02 | VX-11e |
| MVP_top60_DNA_seed | GSK3B | -0.925468 | THCA | TT2609-C02 | CHIR-99021 |
| MVP_top60_DNA_seed | HDAC1 | -0.92111 | THCA | B-CPAP | CAY10603 |
| MVP_top60_DNA_seed | MAPK1 | -0.915528 | THCA | 8505C | VX-11e |
| MVP_top60_DNA_seed | GSK3B | -0.904701 | THCA | B-CPAP | KIN001-042 |
| MVP_top60_DNA_seed | MAPK1 | -0.899585 | THCA | IHH-4 | VX-11e |
| MVP_top60_DNA_seed | MDM2 | -0.898917 | THCA | FTC-133 | Serdemetan |
| MVP_top60_DNA_seed | ATR | -0.892239 | THCA | HTC-C3 | AZD6738 |
| MVP_top60_DNA_seed | ATM | -0.892202 | THCA | CGTH-W-1 | torin2 |
| MVP_top60_DNA_seed | ATR | -0.892202 | THCA | CGTH-W-1 | torin2 |
| MVP_top60_DNA_seed | HDAC1 | -0.888667 | THCA | 8505C | Dacinostat |
| MVP_top60_DNA_seed | HDAC1 | -0.878096 | THCA | 8505C | JQ12 |
| MVP_top60_DNA_seed | BRAF | -0.870308 | THCA | TT2609-C02 | PLX-4720 |
| MVP_top60_DNA_seed | ATM | -0.868791 | THCA | ML-1 | KU-55933 |
| MVP_top60_DNA_seed | ATR | -0.86763 | THCA | K5 | AZ20 |
| MVP_top60_DNA_seed | HDAC1 | -0.85957 | THCA | BHT-101 | AR-42 |
| MVP_top60_DNA_seed | RET | -0.85325 | THCA | RO82-W-1 | Motesanib |
| MVP_top60_DNA_seed | BRAF | -0.842455 | THCA | HTC-C3 | HG6-64-1 |
| MVP_top60_DNA_seed | ATM | -0.825398 | THCA | KMH-2 | CP466722 |
| MVP_top60_DNA_seed | MDM2 | -0.822725 | THCA | 8505C | Serdemetan |
| MVP_top60_DNA_seed | HDAC1 | -0.817087 | THCA | ML-1 | Dacinostat |
| MVP_top60_DNA_seed | MDM2 | -0.815327 | THCA | BHT-101 | Nutlin-3a (-) |
| MVP_top60_DNA_seed | BRAF | -0.813015 | THCA | HTC-C3 | SB590885 |
| MVP_top60_DNA_seed | GSK3B | -0.803248 | THCA | ASH-3 | CHIR-99021 |

| | | | | | |
|---|---|---|---|---|---|
| MVP_top60_DNA_seed | CHEK2 | -0.802362 | THCA | FTC-133 | AZD7762 |
| MVP_top60_DNA_seed | GSK3B | -0.799404 | THCA | 8305C | CHIR-99021 |
| MVP_top60_DNA_seed | CHEK2 | -0.791571 | THCA | ML-1 | AZD7762 |
| MVP_top60_DNA_seed | RET | -0.791025 | THCA | 8505C | Motesanib |
| MVP_top60_DNA_seed | ATM | -0.773274 | THCA | TT2609-C02 | KU-60019 |
| MVP_top60_DNA_seed | AKT1 | -0.773118 | THCA | FTC-133 | AKT inhibitor VIII |
| MVP_top60_DNA_seed | AKT2 | -0.773118 | THCA | FTC-133 | AKT inhibitor VIII |
| MVP_top60_DNA_seed | AKT3 | -0.773118 | THCA | FTC-133 | AKT inhibitor VIII |
| MVP_top60_DNA_seed | GSK3B | -0.771928 | THCA | RO82-W-1 | AZD7969 |
| MVP_top60_DNA_seed | MDM2 | -0.770772 | THCA | TT2609-C02 | Serdemetan |
| MVP_top60_DNA_seed | HDAC1 | -0.76804 | THCA | B-CPAP | Belinostat |
| MVP_top60_DNA_seed | BRAF | -0.763833 | THCA | B-CPAP | RAF_9304 |
| MVP_top60_DNA_seed | HDAC1 | -0.763123 | THCA | BHT-101 | Belinostat |
| MVP_top60_DNA_seed | BRAF | -0.758091 | THCA | RO82-W-1 | PLX-4720 |
| MVP_top60_DNA_seed | HDAC1 | -0.747378 | THCA | K5 | Entinostat |
| MVP_top60_DNA_seed | ATR | -0.737873 | THCA | ASH-3 | QL-VIII-58 |
| MVP_top60_DNA_seed | HDAC1 | -0.728411 | THCA | KMH-2 | Dacinostat |
| MVP_top60_DNA_seed | BRAF | -0.71913 | THCA | 8505C | Dabrafenib |
| MVP_top60_DNA_seed | GSK3B | -0.718318 | THCA | K5 | SB216763 |
| MVP_top60_DNA_seed | MAPK1 | -0.715936 | THCA | B-CPAP | FR-180204 |
| MVP_top60_DNA_seed | MAPK3 | -0.715936 | THCA | B-CPAP | FR-180204 |
| MVP_top60_DNA_seed | ATM | -0.711223 | THCA | ML-1 | CP466722 |
| MVP_top60_DNA_seed | AKT1 | -0.710688 | THCA | IHH-4 | MK-2206 |
| MVP_top60_DNA_seed | AKT2 | -0.710688 | THCA | IHH-4 | MK-2206 |
| MVP_top60_DNA_seed | HDAC1 | -0.70789 | THCA | 8305C | Dacinostat |
| MVP_top60_DNA_seed | ATM | -0.707837 | THCA | K5 | KU-60019 |
| MVP_top60_DNA_seed | BRAF | -0.706258 | THCA | CAL-62 | Dabrafenib |
| MVP_top60_DNA_seed | HDAC1 | -0.69876 | THCA | HTC-C3 | CAY10603 |
| MVP_top60_DNA_seed | ATR | -0.693007 | THCA | 8505C | QL-VIII-58 |
| MVP_top60_DNA_seed | BRAF | -0.686113 | THCA | TT2609-C02 | SB590885 |
| MVP_top60_DNA_seed | HDAC1 | -0.68595 | THCA | ML-1 | Belinostat |
| MVP_top60_DNA_seed | HDAC1 | -0.683656 | THCA | HTC-C3 | AR-42 |
| MVP_top60_DNA_seed | ATM | -0.674507 | THCA | ASH-3 | KU-60019 |
| MVP_top60_DNA_seed | GSK3B | -0.671053 | THCA | CGTH-W-1 | SB216763 |
| MVP_top60_DNA_seed | HDAC1 | -0.665707 | THCA | BHT-101 | JQ12 |
| MVP_top60_DNA_seed | MAPK1 | -0.650267 | THCA | BHT-101 | FR-180204 |
| MVP_top60_DNA_seed | MAPK3 | -0.650267 | THCA | BHT-101 | FR-180204 |
| MVP_top60_DNA_seed | PTK2 | -0.641648 | THCA | 8505C | PF-562271 |
| MVP_top60_DNA_seed | MAPK14 | -0.638106 | THCA | KMH-2 | TAK-715 |
| MVP_top60_DNA_seed | ATR | -0.636703 | THCA | FTC-133 | AZ20 |
| MVP_top60_DNA_seed | ATM | -0.632848 | THCA | IHH-4 | CP466722 |

| | | | | | |
|---|---|---|---|---|---|
| MVP_top60_DNA_seed | HDAC1 | -0.632105 | THCA | BHT-101 | CAY10603 |
| MVP_top60_DNA_seed | MDM2 | -0.630766 | THCA | ML-1 | Serdemetan |
| MVP_top60_DNA_seed | HDAC1 | -0.613768 | THCA | K5 | JQ12 |
| MVP_top60_DNA_seed | HDAC1 | -0.612377 | THCA | B-CPAP | AR-42 |
| MVP_top60_DNA_seed | BRAF | -0.606979 | THCA | KMH-2 | RAF_9304 |
| MVP_top60_DNA_seed | MAPK1 | -0.594281 | THCA | HTC-C3 | FR-180204 |
| MVP_top60_DNA_seed | MAPK3 | -0.594281 | THCA | HTC-C3 | FR-180204 |
| MVP_top60_DNA_seed | CHEK2 | -0.592674 | THCA | FTC-133 | AZD7762 |
| MVP_top60_DNA_seed | BRAF | -0.5883 | THCA | TT | SB590885 |
| MVP_top60_DNA_seed | PTK2 | -0.587439 | THCA | HTC-C3 | PF-562271 |
| MVP_top60_DNA_seed | HDAC1 | -0.587288 | THCA | ML-1 | CAY10603 |
| MVP_top60_DNA_seed | HDAC1 | -0.585637 | THCA | K5 | Parthenolide |
| MVP_top60_DNA_seed | CHEK2 | -0.575681 | THCA | 8305C | AZD7762 |
| MVP_top60_DNA_seed | GSK3B | -0.564997 | THCA | RO82-W-1 | CHIR-99021 |
| MVP_top60_DNA_seed | AKT1 | -0.56319 | THCA | TT | MK-2206 |
| MVP_top60_DNA_seed | AKT2 | -0.56319 | THCA | TT | MK-2206 |
| MVP_top60_DNA_seed | CHEK2 | -0.555376 | THCA | KMH-2 | AZD7762 |
| MVP_top60_DNA_seed | RET | -0.553464 | THCA | FTC-133 | Motesanib |
| MVP_top60_DNA_seed | HDAC1 | -0.552855 | THCA | TT2609-C02 | Tubastatin A |
| MVP_top60_DNA_seed | HDAC1 | -0.552481 | THCA | B-CPAP | Dacinostat |
| MVP_top60_DNA_seed | RET | -0.550059 | THCA | B-CPAP | Motesanib |
| MVP_top60_DNA_seed | AKT1 | -0.546823 | THCA | ASH-3 | AKT inhibitor VIII |
| MVP_top60_DNA_seed | AKT2 | -0.546823 | THCA | ASH-3 | AKT inhibitor VIII |
| MVP_top60_DNA_seed | AKT3 | -0.546823 | THCA | ASH-3 | AKT inhibitor VIII |
| MVP_top60_DNA_seed | ATM | -0.545485 | THCA | TT2609-C02 | KU-55933 |
| MVP_top60_DNA_seed | CHEK2 | -0.537076 | THCA | KMH-2 | AZD7762 |
| MVP_top60_DNA_seed | GSK3B | -0.535206 | THCA | 8505C | KIN001-042 |
| MVP_top60_DNA_seed | BRAF | -0.53173 | THCA | ASH-3 | RAF_9304 |
| MVP_top60_DNA_seed | CHEK2 | -0.528478 | THCA | IHH-4 | AZD7762 |
| MVP_top60_DNA_seed | GSK3B | -0.525849 | THCA | KMH-2 | CHIR-99021 |
| MVP_top60_DNA_seed | AKT1 | -0.521863 | THCA | CGTH-W-1 | AKT inhibitor VIII |
| MVP_top60_DNA_seed | AKT2 | -0.521863 | THCA | CGTH-W-1 | AKT inhibitor VIII |
| MVP_top60_DNA_seed | AKT3 | -0.521863 | THCA | CGTH-W-1 | AKT inhibitor VIII |
| MVP_top60_DNA_seed | ATM | -0.515829 | THCA | TT | KU-60019 |
| MVP_top60_DNA_seed | CHEK2 | -0.515183 | THCA | 8505C | AZD7762 |
| MVP_top60_DNA_seed | ATM | -0.513165 | THCA | HTC-C3 | KU-60019 |
| MVP_top60_DNA_seed | GSK3B | -0.512483 | THCA | BHT-101 | AZD7969 |
| MVP_top60_DNA_seed | BRAF | -0.489515 | THCA | 8505C | SB590885 |
| MVP_top60_DNA_seed | AKT1 | -0.481397 | THCA | CGTH-W-1 | A-443654 |
| MVP_top60_DNA_seed | AKT2 | -0.481397 | THCA | CGTH-W-1 | A-443654 |
| MVP_top60_DNA_seed | AKT3 | -0.481397 | THCA | CGTH-W-1 | A-443654 |

| | | | | | |
|---|---|---|---|---|---|
| MVP_top60_DNA_seed | ATM | -0.479085 | THCA | BHT-101 | KU-60019 |
| MVP_top60_DNA_seed | HDAC1 | -0.473664 | THCA | TT2609-C02 | Belinostat |
| MVP_top60_DNA_seed | HDAC1 | -0.473279 | THCA | IHH-4 | Belinostat |
| MVP_top60_DNA_seed | GSK3B | -0.467068 | THCA | CAL-62 | CHIR-99021 |
| MVP_top60_DNA_seed | AKT1 | -0.465209 | THCA | IHH-4 | AKT inhibitor VIII |
| MVP_top60_DNA_seed | AKT2 | -0.465209 | THCA | IHH-4 | AKT inhibitor VIII |
| MVP_top60_DNA_seed | AKT3 | -0.465209 | THCA | IHH-4 | AKT inhibitor VIII |
| MVP_top60_DNA_seed | HDAC1 | -0.462103 | THCA | TT2609-C02 | CAY10603 |
| MVP_top60_DNA_seed | ATR | -0.461461 | THCA | FTC-133 | AZD6738 |
| MVP_top60_DNA_seed | MDM2 | -0.460997 | THCA | ML-1 | Nutlin-3a (-) |
| MVP_top60_DNA_seed | MDM2 | -0.460228 | THCA | B-CPAP | Serdemetan |
| MVP_top60_DNA_seed | HDAC1 | -0.458914 | THCA | ML-1 | AR-42 |
| MVP_top60_DNA_seed | AKT1 | -0.454229 | THCA | K5 | BAY AKT1 |
| MVP_top60_DNA_seed | AKT2 | -0.454229 | THCA | K5 | BAY AKT1 |
| MVP_top60_DNA_seed | AKT3 | -0.454229 | THCA | K5 | BAY AKT1 |
| MVP_top60_DNA_seed | MAPK1 | -0.453868 | THCA | CAL-62 | VX-11e |
| MVP_top60_DNA_seed | ATM | -0.452208 | THCA | CGTH-W-1 | KU-55933 |
| MVP_top60_DNA_seed | HDAC1 | -0.443166 | THCA | IHH-4 | AR-42 |
| MVP_top60_DNA_seed | HDAC1 | -0.440462 | THCA | BHT-101 | Tubastatin A |
| MVP_top60_DNA_seed | RET | -0.422726 | THCA | ML-1 | Motesanib |
| MVP_top60_DNA_seed | ATM | -0.421715 | THCA | KMH-2 | KU-60019 |
| MVP_top60_DNA_seed | MAPK1 | -0.414665 | THCA | IHH-4 | FR-180204 |
| MVP_top60_DNA_seed | MAPK3 | -0.414665 | THCA | IHH-4 | FR-180204 |
| MVP_top60_DNA_seed | RET | -0.413693 | THCA | IHH-4 | Motesanib |
| MVP_top60_DNA_seed | ATR | -0.407381 | THCA | CAL-62 | AZ20 |
| MVP_top60_DNA_seed | GSK3B | -0.39953 | THCA | RO82-W-1 | SB216763 |
| MVP_top60_DNA_seed | HDAC1 | -0.398658 | THCA | 8505C | CAY10603 |
| MVP_top60_DNA_seed | HDAC1 | -0.396079 | THCA | TT2609-C02 | AR-42 |
| MVP_top60_DNA_seed | BRAF | -0.392081 | THCA | ASH-3 | Dabrafenib |
| MVP_top60_DNA_seed | ATM | -0.391261 | THCA | ASH-3 | KU-55933 |
| MVP_top60_DNA_seed | MDM2 | -0.386587 | THCA | RO82-W-1 | Nutlin-3a (-) |
| MVP_top60_DNA_seed | ATR | -0.380315 | THCA | B-CPAP | AZ20 |
| MVP_top60_DNA_seed | ATM | -0.374302 | THCA | BHT-101 | CP466722 |
| MVP_top60_DNA_seed | GSK3B | -0.372682 | THCA | IHH-4 | CHIR-99021 |
| MVP_top60_DNA_seed | BRAF | -0.370822 | THCA | K5 | HG6-64-1 |
| MVP_top60_DNA_seed | BRAF | -0.366295 | THCA | CAL-62 | PLX-4720 |
| MVP_top60_DNA_seed | GSK3B | -0.366067 | THCA | B-CPAP | SB216763 |
| MVP_top60_DNA_seed | ATR | -0.352879 | THCA | KMH-2 | AZ20 |
| MVP_top60_DNA_seed | ATR | -0.350383 | THCA | HTC-C3 | AZ20 |
| MVP_top60_DNA_seed | HDAC1 | -0.345252 | THCA | HTC-C3 | Tubastatin A |
| MVP_top60_DNA_seed | HDAC1 | -0.33994 | THCA | KMH-2 | JQ12 |

| | | | | | |
|---|---|---|---|---|---|
| MVP_top60_DNA_seed | HDAC1 | -0.332175 | THCA | CAL-62 | Tubastatin A |
| MVP_top60_DNA_seed | HDAC1 | -0.331273 | THCA | IHH-4 | Dacinostat |
| MVP_top60_DNA_seed | HDAC1 | -0.327175 | THCA | IHH-4 | CAY10603 |
| MVP_top60_DNA_seed | BRAF | -0.324975 | THCA | B-CPAP | SB590885 |
| MVP_top60_DNA_seed | CHEK2 | -0.308969 | THCA | HTC-C3 | AZD7762 |
| MVP_top60_DNA_seed | HDAC1 | -0.303126 | THCA | CAL-62 | Belinostat |
| MVP_top60_DNA_seed | HDAC1 | -0.297678 | THCA | B-CPAP | Tubastatin A |
| MVP_top60_DNA_seed | ATR | -0.297182 | THCA | K5 | AZD6738 |
| MVP_top60_DNA_seed | ATR | -0.292098 | THCA | KMH-2 | AZD6738 |
| MVP_top60_DNA_seed | AKT1 | -0.288646 | THCA | KMH-2 | GSK690693 |
| MVP_top60_DNA_seed | AKT2 | -0.288646 | THCA | KMH-2 | GSK690693 |
| MVP_top60_DNA_seed | AKT3 | -0.288646 | THCA | KMH-2 | GSK690693 |
| MVP_top60_DNA_seed | HDAC1 | -0.287986 | THCA | 8305C | JQ12 |
| MVP_top60_DNA_seed | HDAC1 | -0.281563 | THCA | 8505C | Belinostat |
| MVP_top60_DNA_seed | BRAF | -0.272961 | THCA | 8505C | PLX-4720 |
| MVP_top60_DNA_seed | MDM2 | -0.266613 | THCA | CGTH-W-1 | Serdemetan |
| MVP_top60_DNA_seed | GSK3B | -0.265778 | THCA | HTC-C3 | AZD7969 |
| MVP_top60_DNA_seed | MDM2 | -0.261843 | THCA | TT | Serdemetan |
| MVP_top60_DNA_seed | GSK3B | -0.255147 | THCA | ASH-3 | AZD7969 |
| MVP_top60_DNA_seed | HDAC1 | -0.254665 | THCA | K5 | Dacinostat |
| MVP_top60_DNA_seed | ATR | -0.252201 | THCA | BHT-101 | AZD6738 |
| MVP_top60_DNA_seed | CHEK2 | -0.245498 | THCA | K5 | AZD7762 |
| MVP_top60_DNA_seed | ATR | -0.245353 | THCA | KMH-2 | QL-VIII-58 |
| MVP_top60_DNA_seed | BRAF | -0.242036 | THCA | CAL-62 | HG6-64-1 |
| MVP_top60_DNA_seed | ATM | -0.239267 | THCA | 8505C | KU-55933 |
| MVP_top60_DNA_seed | BRAF | -0.23467 | THCA | 8505C | HG6-64-1 |
| MVP_top60_DNA_seed | ATR | -0.21422 | THCA | B-CPAP | AZD6738 |
| MVP_top60_DNA_seed | ATM | -0.212464 | THCA | 8505C | torin2 |
| MVP_top60_DNA_seed | ATR | -0.212464 | THCA | 8505C | torin2 |
| MVP_top60_DNA_seed | AKT1 | -0.208708 | THCA | B-CPAP | MK-2206 |
| MVP_top60_DNA_seed | AKT2 | -0.208708 | THCA | B-CPAP | MK-2206 |
| MVP_top60_DNA_seed | ATR | -0.202708 | THCA | 8305C | AZD6738 |
| MVP_top60_DNA_seed | ATM | -0.201777 | THCA | ASH-3 | CP466722 |
| MVP_top60_DNA_seed | GSK3B | -0.200796 | THCA | IHH-4 | CHIR-99021 |
| MVP_top60_DNA_seed | MAPK14 | -0.198973 | THCA | B-CPAP | TAK-715 |
| MVP_top60_DNA_seed | HDAC1 | -0.192304 | THCA | BHT-101 | Dacinostat |
| MVP_top60_DNA_seed | BRAF | -0.186451 | THCA | CAL-62 | PLX-4720 |
| MVP_top60_DNA_seed | AKT1 | -0.182832 | THCA | KMH-2 | AKT inhibitor VIII |
| MVP_top60_DNA_seed | AKT2 | -0.182832 | THCA | KMH-2 | AKT inhibitor VIII |
| MVP_top60_DNA_seed | AKT3 | -0.182832 | THCA | KMH-2 | AKT inhibitor VIII |
| MVP_top60_DNA_seed | CHEK2 | -0.174044 | THCA | TT | AZD7762 |

| | | | | | |
|---|---|---|---|---|---|
| MVP_top60_DNA_seed | ATR | -0.173888 | THCA | FTC-133 | QL-VIII-58 |
| MVP_top60_DNA_seed | GSK3B | -0.170817 | THCA | KMH-2 | AZD7969 |
| MVP_top60_DNA_seed | AKT1 | -0.170145 | THCA | IHH-4 | BAY AKT1 |
| MVP_top60_DNA_seed | AKT2 | -0.170145 | THCA | IHH-4 | BAY AKT1 |
| MVP_top60_DNA_seed | AKT3 | -0.170145 | THCA | IHH-4 | BAY AKT1 |
| MVP_top60_DNA_seed | BRAF | -0.164558 | THCA | KMH-2 | PLX-4720 |
| MVP_top60_DNA_seed | BRAF | -0.164224 | THCA | CGTH-W-1 | Dabrafenib |
| MVP_top60_DNA_seed | AKT1 | -0.158883 | THCA | 8505C | MK-2206 |
| MVP_top60_DNA_seed | AKT2 | -0.158883 | THCA | 8505C | MK-2206 |
| MVP_top60_DNA_seed | GSK3B | -0.15719 | THCA | 8505C | AZD7969 |
| MVP_top60_DNA_seed | HDAC1 | -0.156541 | THCA | K5 | AR-42 |
| MVP_top60_DNA_seed | GSK3B | -0.152963 | THCA | TT2609-C02 | CHIR-99021 |
| MVP_top60_DNA_seed | MDM2 | -0.151124 | THCA | TT2609-C02 | Nutlin-3a (-) |
| MVP_top60_DNA_seed | ATR | -0.150256 | THCA | CGTH-W-1 | AZ20 |
| MVP_top60_DNA_seed | ATM | -0.145785 | THCA | FTC-133 | KU-60019 |
| MVP_top60_DNA_seed | CHEK2 | -0.142249 | THCA | ASH-3 | AZD7762 |
| MVP_top60_DNA_seed | MDM2 | -0.141355 | THCA | KMH-2 | Serdemetan |
| MVP_top60_DNA_seed | HDAC1 | -0.133908 | THCA | CAL-62 | Dacinostat |
| MVP_top60_DNA_seed | ATR | -0.131523 | THCA | K5 | QL-VIII-58 |
| MVP_top60_DNA_seed | AKT1 | -0.129599 | THCA | HTC-C3 | AKT inhibitor VIII |
| MVP_top60_DNA_seed | AKT2 | -0.129599 | THCA | HTC-C3 | AKT inhibitor VIII |
| MVP_top60_DNA_seed | AKT3 | -0.129599 | THCA | HTC-C3 | AKT inhibitor VIII |
| MVP_top60_DNA_seed | HDAC1 | -0.11734 | THCA | K5 | CAY10603 |
| MVP_top60_DNA_seed | GSK3B | -0.113085 | THCA | KMH-2 | KIN001-042 |
| MVP_top60_DNA_seed | ATR | -0.107754 | THCA | ML-1 | AZ20 |
| MVP_top60_DNA_seed | HDAC1 | -0.106354 | THCA | CAL-62 | AR-42 |
| MVP_top60_DNA_seed | MAPK14 | -0.099514 | THCA | CAL-62 | TAK-715 |
| MVP_top60_DNA_seed | PTK2 | -0.091913 | THCA | K5 | PF-562271 |
| MVP_top60_DNA_seed | BRAF | -0.085211 | THCA | RO82-W-1 | HG6-64-1 |
| MVP_top60_DNA_seed | ATR | -0.085153 | THCA | 8305C | QL-VIII-58 |
| MVP_top60_DNA_seed | BRAF | -0.081779 | THCA | CGTH-W-1 | PLX-4720 |
| MVP_top60_DNA_seed | HDAC1 | -0.078662 | THCA | 8505C | AR-42 |
| MVP_top60_DNA_seed | GSK3B | -0.073815 | THCA | CAL-62 | CHIR-99021 |
| MVP_top60_DNA_seed | CHEK2 | -0.063791 | THCA | B-CPAP | AZD7762 |
| MVP_top60_DNA_seed | ATM | -0.054763 | THCA | K5 | KU-55933 |
| MVP_top60_DNA_seed | MAPK1 | -0.051215 | THCA | 8305C | FR-180204 |
| MVP_top60_DNA_seed | MAPK3 | -0.051215 | THCA | 8305C | FR-180204 |
| MVP_top60_DNA_seed | AKT1 | -0.042584 | THCA | HTC-C3 | AKT inhibitor VIII |
| MVP_top60_DNA_seed | AKT2 | -0.042584 | THCA | HTC-C3 | AKT inhibitor VIII |
| MVP_top60_DNA_seed | AKT3 | -0.042584 | THCA | HTC-C3 | AKT inhibitor VIII |
| MVP_top60_DNA_seed | BRAF | -0.04043 | THCA | ML-1 | Dabrafenib |

| | | | | | |
|---|---|---|---|---|---|
| MVP_top60_DNA_seed | MAPK1 | -0.036196 | THCA | 8505C | FR-180204 |
| MVP_top60_DNA_seed | MAPK3 | -0.036196 | THCA | 8505C | FR-180204 |
| MVP_top60_DNA_seed | HDAC1 | -0.035633 | THCA | 8505C | Tubastatin A |
| MVP_top60_DNA_seed | ATR | -0.035366 | THCA | BHT-101 | AZ20 |
| MVP_top60_DNA_seed | BRAF | -0.035248 | THCA | KMH-2 | HG6-64-1 |
| MVP_top60_DNA_seed | ATR | -0.030149 | THCA | TT2609-C02 | AZ20 |
| MVP_top60_DNA_seed | GSK3B | -0.0272 | THCA | TT2609-C02 | SB216763 |
| MVP_top60_DNA_seed | AKT1 | -0.025516 | THCA | K5 | A-443654 |
| MVP_top60_DNA_seed | AKT2 | -0.025516 | THCA | K5 | A-443654 |
| MVP_top60_DNA_seed | AKT3 | -0.025516 | THCA | K5 | A-443654 |
| MVP_top60_DNA_seed | AKT1 | -0.024435 | THCA | CGTH-W-1 | MK-2206 |
| MVP_top60_DNA_seed | AKT2 | -0.024435 | THCA | CGTH-W-1 | MK-2206 |
| MVP_top60_DNA_seed | HDAC1 | -0.023376 | THCA | IHH-4 | Tubastatin A |
| MVP_top60_DNA_seed | AKT1 | -0.019583 | THCA | FTC-133 | BAY AKT1 |
| MVP_top60_DNA_seed | AKT2 | -0.019583 | THCA | FTC-133 | BAY AKT1 |
| MVP_top60_DNA_seed | AKT3 | -0.019583 | THCA | FTC-133 | BAY AKT1 |
| MVP_top60_DNA_seed | HDAC1 | -0.016061 | THCA | HTC-C3 | Dacinostat |
| MVP_top60_DNA_seed | CHEK2 | -0.015845 | THCA | K5 | AZD7762 |
| MVP_top60_DNA_seed | CHEK2 | -0.009868 | THCA | CAL-62 | AZD7762 |
| MVP_top60_DNA_seed | HDAC1 | -0.008344 | THCA | CAL-62 | JQ12 |
| MVP_top60_DNA_seed | CHEK2 | -0.005508 | THCA | 8305C | AZD7762 |
| MVP_top60_DNA_seed | HDAC1 | -0.005206 | THCA | RO82-W-1 | CAY10603 |
| MVP_top60_DNA_seed | ATM | -0.001181 | THCA | CAL-62 | KU-60019 |
| MVP_top60_DNA_seed | BRAF | 0.001482 | THCA | 8505C | RAF_9304 |
| MVP_top60_DNA_seed | HDAC1 | 0.003584 | THCA | CGTH-W-1 | Entinostat |
| MVP_top60_DNA_seed | ATM | 0.004693 | THCA | HTC-C3 | KU-55933 |
| MVP_top60_DNA_seed | ATR | 0.007651 | THCA | 8505C | AZ20 |
| MVP_top60_DNA_seed | HDAC1 | 0.009976 | THCA | RO82-W-1 | Belinostat |
| MVP_top60_DNA_seed | AKT1 | 0.012071 | THCA | IHH-4 | AKT inhibitor VIII |
| MVP_top60_DNA_seed | AKT2 | 0.012071 | THCA | IHH-4 | AKT inhibitor VIII |
| MVP_top60_DNA_seed | AKT3 | 0.012071 | THCA | IHH-4 | AKT inhibitor VIII |
| MVP_top60_DNA_seed | HDAC1 | 0.016729 | THCA | TT | JQ12 |
| MVP_top60_DNA_seed | HDAC1 | 0.02288 | THCA | K5 | Belinostat |
| MVP_top60_DNA_seed | ATM | 0.039882 | THCA | K5 | CP466722 |
| MVP_top60_DNA_seed | MDM2 | 0.051386 | THCA | B-CPAP | Nutlin-3a (-) |
| MVP_top60_DNA_seed | MDM2 | 0.065158 | THCA | 8505C | Nutlin-3a (-) |
| MVP_top60_DNA_seed | AKT1 | 0.071723 | THCA | KMH-2 | BAY AKT1 |
| MVP_top60_DNA_seed | AKT2 | 0.071723 | THCA | KMH-2 | BAY AKT1 |
| MVP_top60_DNA_seed | AKT3 | 0.071723 | THCA | KMH-2 | BAY AKT1 |
| MVP_top60_DNA_seed | HDAC1 | 0.073299 | THCA | RO82-W-1 | AR-42 |
| MVP_top60_DNA_seed | AKT1 | 0.074453 | THCA | BHT-101 | AKT inhibitor VIII |

| | | | | | |
|---|---|---|---|---|---|
| MVP_top60_DNA_seed | AKT2 | 0.074453 | THCA | BHT-101 | AKT inhibitor VIII |
| MVP_top60_DNA_seed | AKT3 | 0.074453 | THCA | BHT-101 | AKT inhibitor VIII |
| MVP_top60_DNA_seed | HDAC1 | 0.078315 | THCA | K5 | Tubastatin A |
| MVP_top60_DNA_seed | HDAC1 | 0.093076 | THCA | CGTH-W-1 | AR-42 |
| MVP_top60_DNA_seed | ATM | 0.100739 | THCA | RO82-W-1 | KU-60019 |
| MVP_top60_DNA_seed | ATM | 0.106576 | THCA | CAL-62 | KU-55933 |
| MVP_top60_DNA_seed | RET | 0.108057 | THCA | K5 | Motesanib |
| MVP_top60_DNA_seed | BRAF | 0.111801 | THCA | RO82-W-1 | Dabrafenib |
| MVP_top60_DNA_seed | AKT1 | 0.112808 | THCA | KMH-2 | AKT inhibitor VIII |
| MVP_top60_DNA_seed | AKT2 | 0.112808 | THCA | KMH-2 | AKT inhibitor VIII |
| MVP_top60_DNA_seed | AKT3 | 0.112808 | THCA | KMH-2 | AKT inhibitor VIII |
| MVP_top60_DNA_seed | AKT1 | 0.118602 | THCA | K5 | AKT inhibitor VIII |
| MVP_top60_DNA_seed | AKT2 | 0.118602 | THCA | K5 | AKT inhibitor VIII |
| MVP_top60_DNA_seed | AKT3 | 0.118602 | THCA | K5 | AKT inhibitor VIII |
| MVP_top60_DNA_seed | AKT1 | 0.123679 | THCA | RO82-W-1 | BAY AKT1 |
| MVP_top60_DNA_seed | AKT2 | 0.123679 | THCA | RO82-W-1 | BAY AKT1 |
| MVP_top60_DNA_seed | AKT3 | 0.123679 | THCA | RO82-W-1 | BAY AKT1 |
| MVP_top60_DNA_seed | GSK3B | 0.124593 | THCA | FTC-133 | SB216763 |
| MVP_top60_DNA_seed | AKT1 | 0.128532 | THCA | K5 | GSK690693 |
| MVP_top60_DNA_seed | AKT2 | 0.128532 | THCA | K5 | GSK690693 |
| MVP_top60_DNA_seed | AKT3 | 0.128532 | THCA | K5 | GSK690693 |
| MVP_top60_DNA_seed | BRAF | 0.13551 | THCA | ML-1 | PLX-4720 |
| MVP_top60_DNA_seed | AKT1 | 0.140238 | THCA | TT2609-C02 | MK-2206 |
| MVP_top60_DNA_seed | AKT2 | 0.140238 | THCA | TT2609-C02 | MK-2206 |
| MVP_top60_DNA_seed | CHEK2 | 0.154721 | THCA | HTC-C3 | AZD7762 |
| MVP_top60_DNA_seed | HDAC1 | 0.155716 | THCA | CGTH-W-1 | Belinostat |
| MVP_top60_DNA_seed | BRAF | 0.161866 | THCA | TT | Dabrafenib |
| MVP_top60_DNA_seed | PTK2 | 0.164523 | THCA | RO82-W-1 | PF-562271 |
| MVP_top60_DNA_seed | ATM | 0.164551 | THCA | CGTH-W-1 | KU-60019 |
| MVP_top60_DNA_seed | AKT1 | 0.165025 | THCA | 8305C | MK-2206 |
| MVP_top60_DNA_seed | AKT2 | 0.165025 | THCA | 8305C | MK-2206 |
| MVP_top60_DNA_seed | GSK3B | 0.173615 | THCA | FTC-133 | CHIR-99021 |
| MVP_top60_DNA_seed | AKT1 | 0.180136 | THCA | RO82-W-1 | AKT inhibitor VIII |
| MVP_top60_DNA_seed | AKT2 | 0.180136 | THCA | RO82-W-1 | AKT inhibitor VIII |
| MVP_top60_DNA_seed | AKT3 | 0.180136 | THCA | RO82-W-1 | AKT inhibitor VIII |
| MVP_top60_DNA_seed | MAPK1 | 0.18019 | THCA | TT2609-C02 | FR-180204 |
| MVP_top60_DNA_seed | MAPK3 | 0.18019 | THCA | TT2609-C02 | FR-180204 |
| MVP_top60_DNA_seed | AKT1 | 0.18882 | THCA | 8505C | GSK690693 |
| MVP_top60_DNA_seed | AKT2 | 0.18882 | THCA | 8505C | GSK690693 |
| MVP_top60_DNA_seed | AKT3 | 0.18882 | THCA | 8505C | GSK690693 |
| MVP_top60_DNA_seed | RET | 0.203243 | THCA | KMH-2 | Motesanib |

| | | | | | |
|---|---|---|---|---|---|
| MVP_top60_DNA_seed | AKT1 | 0.215195 | THCA | 8505C | AKT inhibitor VIII |
| MVP_top60_DNA_seed | AKT2 | 0.215195 | THCA | 8505C | AKT inhibitor VIII |
| MVP_top60_DNA_seed | AKT3 | 0.215195 | THCA | 8505C | AKT inhibitor VIII |
| MVP_top60_DNA_seed | CHEK2 | 0.221166 | THCA | TT2609-C02 | AZD7762 |
| MVP_top60_DNA_seed | MAPK1 | 0.222853 | THCA | KMH-2 | FR-180204 |
| MVP_top60_DNA_seed | MAPK3 | 0.222853 | THCA | KMH-2 | FR-180204 |
| MVP_top60_DNA_seed | AKT1 | 0.233858 | THCA | IHH-4 | GSK690693 |
| MVP_top60_DNA_seed | AKT2 | 0.233858 | THCA | IHH-4 | GSK690693 |
| MVP_top60_DNA_seed | AKT3 | 0.233858 | THCA | IHH-4 | GSK690693 |
| MVP_top60_DNA_seed | RET | 0.241375 | THCA | ASH-3 | Motesanib |
| MVP_top60_DNA_seed | BRAF | 0.24194 | THCA | TT2609-C02 | Dabrafenib |
| MVP_top60_DNA_seed | BRAF | 0.241945 | THCA | RO82-W-1 | RAF_9304 |
| MVP_top60_DNA_seed | MAPK1 | 0.249067 | THCA | CAL-62 | FR-180204 |
| MVP_top60_DNA_seed | MAPK3 | 0.249067 | THCA | CAL-62 | FR-180204 |
| MVP_top60_DNA_seed | BRAF | 0.25163 | THCA | 8505C | PLX-4720 |
| MVP_top60_DNA_seed | BRAF | 0.252942 | THCA | TT | RAF_9304 |
| MVP_top60_DNA_seed | HDAC1 | 0.258404 | THCA | KMH-2 | CAY10603 |
| MVP_top60_DNA_seed | MAPK1 | 0.264829 | THCA | CGTH-W-1 | FR-180204 |
| MVP_top60_DNA_seed | MAPK3 | 0.264829 | THCA | CGTH-W-1 | FR-180204 |
| MVP_top60_DNA_seed | GSK3B | 0.265169 | THCA | CAL-62 | KIN001-042 |
| MVP_top60_DNA_seed | HDAC1 | 0.265439 | THCA | CGTH-W-1 | CAY10603 |
| MVP_top60_DNA_seed | ATM | 0.26645 | THCA | ML-1 | KU-60019 |
| MVP_top60_DNA_seed | AKT1 | 0.268555 | THCA | ASH-3 | MK-2206 |
| MVP_top60_DNA_seed | AKT2 | 0.268555 | THCA | ASH-3 | MK-2206 |
| MVP_top60_DNA_seed | HDAC1 | 0.278697 | THCA | FTC-133 | Dacinostat |
| MVP_top60_DNA_seed | MAPK14 | 0.279648 | THCA | BHT-101 | TAK-715 |
| MVP_top60_DNA_seed | MAPK14 | 0.286001 | THCA | 8505C | TAK-715 |
| MVP_top60_DNA_seed | GSK3B | 0.292474 | THCA | BHT-101 | CHIR-99021 |
| MVP_top60_DNA_seed | HDAC1 | 0.29621 | THCA | CAL-62 | CAY10603 |
| MVP_top60_DNA_seed | GSK3B | 0.296564 | THCA | 8305C | AZD7969 |
| MVP_top60_DNA_seed | MAPK1 | 0.298846 | THCA | 8305C | VX-11e |
| MVP_top60_DNA_seed | GSK3B | 0.305956 | THCA | HTC-C3 | CHIR-99021 |
| MVP_top60_DNA_seed | BRAF | 0.307688 | THCA | FTC-133 | SB590885 |
| MVP_top60_DNA_seed | GSK3B | 0.308457 | THCA | BHT-101 | KIN001-042 |
| MVP_top60_DNA_seed | ATM | 0.317324 | THCA | CAL-62 | CP466722 |
| MVP_top60_DNA_seed | AKT1 | 0.318722 | THCA | 8305C | BAY AKT1 |
| MVP_top60_DNA_seed | AKT2 | 0.318722 | THCA | 8305C | BAY AKT1 |
| MVP_top60_DNA_seed | AKT3 | 0.318722 | THCA | 8305C | BAY AKT1 |
| MVP_top60_DNA_seed | CHEK2 | 0.318755 | THCA | B-CPAP | AZD7762 |
| MVP_top60_DNA_seed | GSK3B | 0.32138 | THCA | K5 | KIN001-042 |
| MVP_top60_DNA_seed | BRAF | 0.3238 | THCA | FTC-133 | PLX-4720 |

| | | | | | |
|---|---|---|---|---|---|
| MVP_top60_DNA_seed | MAPK14 | 0.324452 | THCA | HTC-C3 | TAK-715 |
| MVP_top60_DNA_seed | ATM | 0.338034 | THCA | B-CPAP | CP466722 |
| MVP_top60_DNA_seed | ATM | 0.340556 | THCA | CGTH-W-1 | CP466722 |
| MVP_top60_DNA_seed | GSK3B | 0.34145 | THCA | ASH-3 | KIN001-042 |
| MVP_top60_DNA_seed | AKT1 | 0.345478 | THCA | CGTH-W-1 | AKT inhibitor VIII |
| MVP_top60_DNA_seed | AKT2 | 0.345478 | THCA | CGTH-W-1 | AKT inhibitor VIII |
| MVP_top60_DNA_seed | AKT3 | 0.345478 | THCA | CGTH-W-1 | AKT inhibitor VIII |
| MVP_top60_DNA_seed | MDM2 | 0.352277 | THCA | HTC-C3 | Nutlin-3a (-) |
| MVP_top60_DNA_seed | ATM | 0.353416 | THCA | TT | KU-55933 |
| MVP_top60_DNA_seed | MAPK14 | 0.36949 | THCA | ASH-3 | TAK-715 |
| MVP_top60_DNA_seed | ATM | 0.377689 | THCA | 8505C | CP466722 |
| MVP_top60_DNA_seed | BRAF | 0.380782 | THCA | ASH-3 | SB590885 |
| MVP_top60_DNA_seed | ATR | 0.390459 | THCA | ML-1 | AZD6738 |
| MVP_top60_DNA_seed | CHEK2 | 0.391805 | THCA | BHT-101 | AZD7762 |
| MVP_top60_DNA_seed | AKT1 | 0.398747 | THCA | TT | BAY AKT1 |
| MVP_top60_DNA_seed | AKT2 | 0.398747 | THCA | TT | BAY AKT1 |
| MVP_top60_DNA_seed | AKT3 | 0.398747 | THCA | TT | BAY AKT1 |
| MVP_top60_DNA_seed | ATR | 0.402223 | THCA | 8305C | AZ20 |
| MVP_top60_DNA_seed | CHEK2 | 0.413405 | THCA | RO82-W-1 | AZD7762 |
| MVP_top60_DNA_seed | GSK3B | 0.413921 | THCA | CGTH-W-1 | KIN001-042 |
| MVP_top60_DNA_seed | HDAC1 | 0.425525 | THCA | RO82-W-1 | Tubastatin A |
| MVP_top60_DNA_seed | BRAF | 0.427087 | THCA | CGTH-W-1 | PLX-4720 |
| MVP_top60_DNA_seed | HDAC1 | 0.434938 | THCA | ASH-3 | Dacinostat |
| MVP_top60_DNA_seed | CHEK2 | 0.437393 | THCA | 8505C | AZD7762 |
| MVP_top60_DNA_seed | AKT1 | 0.441879 | THCA | B-CPAP | GSK690693 |
| MVP_top60_DNA_seed | AKT2 | 0.441879 | THCA | B-CPAP | GSK690693 |
| MVP_top60_DNA_seed | AKT3 | 0.441879 | THCA | B-CPAP | GSK690693 |
| MVP_top60_DNA_seed | MDM2 | 0.447611 | THCA | CAL-62 | Nutlin-3a (-) |
| MVP_top60_DNA_seed | ATM | 0.448252 | THCA | B-CPAP | KU-55933 |
| MVP_top60_DNA_seed | GSK3B | 0.449242 | THCA | TT | AZD7969 |
| MVP_top60_DNA_seed | ATR | 0.450246 | THCA | ASH-3 | AZ20 |
| MVP_top60_DNA_seed | AKT1 | 0.46 | THCA | FTC-133 | MK-2206 |
| MVP_top60_DNA_seed | AKT2 | 0.46 | THCA | FTC-133 | MK-2206 |
| MVP_top60_DNA_seed | MAPK1 | 0.460452 | THCA | KMH-2 | VX-11e |
| MVP_top60_DNA_seed | AKT1 | 0.464012 | THCA | CAL-62 | AKT inhibitor VIII |
| MVP_top60_DNA_seed | AKT2 | 0.464012 | THCA | CAL-62 | AKT inhibitor VIII |
| MVP_top60_DNA_seed | AKT3 | 0.464012 | THCA | CAL-62 | AKT inhibitor VIII |
| MVP_top60_DNA_seed | MDM2 | 0.466747 | THCA | CGTH-W-1 | Nutlin-3a (-) |
| MVP_top60_DNA_seed | BRAF | 0.47435 | THCA | KMH-2 | PLX-4720 |
| MVP_top60_DNA_seed | BRAF | 0.476929 | THCA | ASH-3 | PLX-4720 |
| MVP_top60_DNA_seed | BRAF | 0.478727 | THCA | FTC-133 | Dabrafenib |

| | | | | | |
|---|---|---|---|---|---|
| MVP_top60_DNA_seed | AKT1 | 0.482172 | THCA | ASH-3 | GSK690693 |
| MVP_top60_DNA_seed | AKT2 | 0.482172 | THCA | ASH-3 | GSK690693 |
| MVP_top60_DNA_seed | AKT3 | 0.482172 | THCA | ASH-3 | GSK690693 |
| MVP_top60_DNA_seed | AKT1 | 0.496458 | THCA | RO82-W-1 | MK-2206 |
| MVP_top60_DNA_seed | AKT2 | 0.496458 | THCA | RO82-W-1 | MK-2206 |
| MVP_top60_DNA_seed | BRAF | 0.503209 | THCA | FTC-133 | RAF_9304 |
| MVP_top60_DNA_seed | MDM2 | 0.50523 | THCA | ASH-3 | Serdemetan |
| MVP_top60_DNA_seed | MDM2 | 0.50823 | THCA | KMH-2 | Nutlin-3a (-) |
| MVP_top60_DNA_seed | MDM2 | 0.511462 | THCA | BHT-101 | Serdemetan |
| MVP_top60_DNA_seed | AKT1 | 0.513617 | THCA | ASH-3 | AKT inhibitor VIII |
| MVP_top60_DNA_seed | AKT2 | 0.513617 | THCA | ASH-3 | AKT inhibitor VIII |
| MVP_top60_DNA_seed | AKT3 | 0.513617 | THCA | ASH-3 | AKT inhibitor VIII |
| MVP_top60_DNA_seed | AKT1 | 0.518521 | THCA | CGTH-W-1 | GSK690693 |
| MVP_top60_DNA_seed | AKT2 | 0.518521 | THCA | CGTH-W-1 | GSK690693 |
| MVP_top60_DNA_seed | AKT3 | 0.518521 | THCA | CGTH-W-1 | GSK690693 |
| MVP_top60_DNA_seed | HDAC1 | 0.530875 | THCA | KMH-2 | Tubastatin A |
| MVP_top60_DNA_seed | ATR | 0.534881 | THCA | CGTH-W-1 | AZD6738 |
| MVP_top60_DNA_seed | AKT1 | 0.537145 | THCA | ASH-3 | BAY AKT1 |
| MVP_top60_DNA_seed | AKT2 | 0.537145 | THCA | ASH-3 | BAY AKT1 |
| MVP_top60_DNA_seed | AKT3 | 0.537145 | THCA | ASH-3 | BAY AKT1 |
| MVP_top60_DNA_seed | MDM2 | 0.540638 | THCA | RO82-W-1 | Serdemetan |
| MVP_top60_DNA_seed | GSK3B | 0.542286 | THCA | HTC-C3 | SB216763 |
| MVP_top60_DNA_seed | BRAF | 0.542429 | THCA | TT2609-C02 | RAF_9304 |
| MVP_top60_DNA_seed | AKT1 | 0.546883 | THCA | HTC-C3 | MK-2206 |
| MVP_top60_DNA_seed | AKT2 | 0.546883 | THCA | HTC-C3 | MK-2206 |
| MVP_top60_DNA_seed | MAPK1 | 0.547965 | THCA | ML-1 | FR-180204 |
| MVP_top60_DNA_seed | MAPK3 | 0.547965 | THCA | ML-1 | FR-180204 |
| MVP_top60_DNA_seed | MDM2 | 0.550663 | THCA | FTC-133 | Nutlin-3a (-) |
| MVP_top60_DNA_seed | MAPK14 | 0.55086 | THCA | TT2609-C02 | TAK-715 |
| MVP_top60_DNA_seed | CHEK2 | 0.551334 | THCA | ASH-3 | AZD7762 |
| MVP_top60_DNA_seed | AKT1 | 0.555869 | THCA | B-CPAP | AKT inhibitor VIII |
| MVP_top60_DNA_seed | AKT2 | 0.555869 | THCA | B-CPAP | AKT inhibitor VIII |
| MVP_top60_DNA_seed | AKT3 | 0.555869 | THCA | B-CPAP | AKT inhibitor VIII |
| MVP_top60_DNA_seed | BRAF | 0.558517 | THCA | ASH-3 | HG6-64-1 |
| MVP_top60_DNA_seed | AKT1 | 0.560545 | THCA | B-CPAP | BAY AKT1 |
| MVP_top60_DNA_seed | AKT2 | 0.560545 | THCA | B-CPAP | BAY AKT1 |
| MVP_top60_DNA_seed | AKT3 | 0.560545 | THCA | B-CPAP | BAY AKT1 |
| MVP_top60_DNA_seed | ATR | 0.573979 | THCA | RO82-W-1 | AZ20 |
| MVP_top60_DNA_seed | HDAC1 | 0.576176 | THCA | FTC-133 | Belinostat |
| MVP_top60_DNA_seed | GSK3B | 0.579491 | THCA | HTC-C3 | CHIR-99021 |
| MVP_top60_DNA_seed | AKT1 | 0.598072 | THCA | TT2609-C02 | AKT inhibitor VIII |

| | | | | | |
|---|---|---|---|---|---|
| MVP_top60_DNA_seed | AKT2 | 0.598072 | THCA | TT2609-C02 | AKT inhibitor VIII |
| MVP_top60_DNA_seed | AKT3 | 0.598072 | THCA | TT2609-C02 | AKT inhibitor VIII |
| MVP_top60_DNA_seed | BRAF | 0.604345 | THCA | BHT-101 | HG6-64-1 |
| MVP_top60_DNA_seed | ATM | 0.604481 | THCA | KMH-2 | KU-55933 |
| MVP_top60_DNA_seed | AKT1 | 0.60813 | THCA | CAL-62 | GSK690693 |
| MVP_top60_DNA_seed | AKT2 | 0.60813 | THCA | CAL-62 | GSK690693 |
| MVP_top60_DNA_seed | AKT3 | 0.60813 | THCA | CAL-62 | GSK690693 |
| MVP_top60_DNA_seed | BRAF | 0.61027 | THCA | CAL-62 | SB590885 |
| MVP_top60_DNA_seed | BRAF | 0.623824 | THCA | RO82-W-1 | PLX-4720 |
| MVP_top60_DNA_seed | AKT1 | 0.630547 | THCA | BHT-101 | GSK690693 |
| MVP_top60_DNA_seed | AKT2 | 0.630547 | THCA | BHT-101 | GSK690693 |
| MVP_top60_DNA_seed | AKT3 | 0.630547 | THCA | BHT-101 | GSK690693 |
| MVP_top60_DNA_seed | AKT1 | 0.632176 | THCA | 8505C | AKT inhibitor VIII |
| MVP_top60_DNA_seed | AKT2 | 0.632176 | THCA | 8505C | AKT inhibitor VIII |
| MVP_top60_DNA_seed | AKT3 | 0.632176 | THCA | 8505C | AKT inhibitor VIII |
| MVP_top60_DNA_seed | MDM2 | 0.641162 | THCA | HTC-C3 | Serdemetan |
| MVP_top60_DNA_seed | AKT1 | 0.642781 | THCA | KMH-2 | MK-2206 |
| MVP_top60_DNA_seed | AKT2 | 0.642781 | THCA | KMH-2 | MK-2206 |
| MVP_top60_DNA_seed | MDM2 | 0.646941 | THCA | 8305C | Serdemetan |
| MVP_top60_DNA_seed | GSK3B | 0.647225 | THCA | ML-1 | KIN001-042 |
| MVP_top60_DNA_seed | AKT1 | 0.647524 | THCA | K5 | AKT inhibitor VIII |
| MVP_top60_DNA_seed | AKT2 | 0.647524 | THCA | K5 | AKT inhibitor VIII |
| MVP_top60_DNA_seed | AKT3 | 0.647524 | THCA | K5 | AKT inhibitor VIII |
| MVP_top60_DNA_seed | BRAF | 0.653543 | THCA | CGTH-W-1 | SB590885 |
| MVP_top60_DNA_seed | GSK3B | 0.657198 | THCA | CAL-62 | SB216763 |
| MVP_top60_DNA_seed | HDAC1 | 0.660448 | THCA | RO82-W-1 | Dacinostat |
| MVP_top60_DNA_seed | CHEK2 | 0.669386 | THCA | RO82-W-1 | AZD7762 |
| MVP_top60_DNA_seed | BRAF | 0.670799 | THCA | TT | HG6-64-1 |
| MVP_top60_DNA_seed | MAPK1 | 0.673209 | THCA | ASH-3 | FR-180204 |
| MVP_top60_DNA_seed | MAPK3 | 0.673209 | THCA | ASH-3 | FR-180204 |
| MVP_top60_DNA_seed | AKT1 | 0.67578 | THCA | TT2609-C02 | GSK690693 |
| MVP_top60_DNA_seed | AKT2 | 0.67578 | THCA | TT2609-C02 | GSK690693 |
| MVP_top60_DNA_seed | AKT3 | 0.67578 | THCA | TT2609-C02 | GSK690693 |
| MVP_top60_DNA_seed | HDAC1 | 0.683032 | THCA | FTC-133 | JQ12 |
| MVP_top60_DNA_seed | ATR | 0.694297 | THCA | 8505C | AZD6738 |
| MVP_top60_DNA_seed | HDAC1 | 0.694364 | THCA | CGTH-W-1 | Tubastatin A |
| MVP_top60_DNA_seed | ATR | 0.700848 | THCA | ASH-3 | AZD6738 |
| MVP_top60_DNA_seed | HDAC1 | 0.707075 | THCA | ASH-3 | JQ12 |
| MVP_top60_DNA_seed | AKT1 | 0.710134 | THCA | FTC-133 | GSK690693 |
| MVP_top60_DNA_seed | AKT2 | 0.710134 | THCA | FTC-133 | GSK690693 |
| MVP_top60_DNA_seed | AKT3 | 0.710134 | THCA | FTC-133 | GSK690693 |

| | | | | | |
|---|---|---|---|---|---|
| MVP_top60_DNA_seed | GSK3B | 0.713104 | THCA | TT2609-C02 | KIN001-042 |
| MVP_top60_DNA_seed | MAPK1 | 0.713488 | THCA | FTC-133 | VX-11e |
| MVP_top60_DNA_seed | HDAC1 | 0.720626 | THCA | ASH-3 | Tubastatin A |
| MVP_top60_DNA_seed | AKT1 | 0.721096 | THCA | HTC-C3 | GSK690693 |
| MVP_top60_DNA_seed | AKT2 | 0.721096 | THCA | HTC-C3 | GSK690693 |
| MVP_top60_DNA_seed | AKT3 | 0.721096 | THCA | HTC-C3 | GSK690693 |
| MVP_top60_DNA_seed | MDM2 | 0.729583 | THCA | CAL-62 | Serdemetan |
| MVP_top60_DNA_seed | HDAC1 | 0.740788 | THCA | FTC-133 | CAY10603 |
| MVP_top60_DNA_seed | MAPK1 | 0.765539 | THCA | K5 | VX-11e |
| MVP_top60_DNA_seed | MAPK14 | 0.766161 | THCA | CGTH-W-1 | TAK-715 |
| MVP_top60_DNA_seed | MAPK1 | 0.775514 | THCA | FTC-133 | FR-180204 |
| MVP_top60_DNA_seed | MAPK3 | 0.775514 | THCA | FTC-133 | FR-180204 |
| MVP_top60_DNA_seed | AKT1 | 0.777805 | THCA | HTC-C3 | BAY AKT1 |
| MVP_top60_DNA_seed | AKT2 | 0.777805 | THCA | HTC-C3 | BAY AKT1 |
| MVP_top60_DNA_seed | AKT3 | 0.777805 | THCA | HTC-C3 | BAY AKT1 |
| MVP_top60_DNA_seed | HDAC1 | 0.778947 | THCA | FTC-133 | AR-42 |
| MVP_top60_DNA_seed | MAPK1 | 0.779093 | THCA | ML-1 | VX-11e |
| MVP_top60_DNA_seed | AKT1 | 0.798693 | THCA | 8305C | AKT inhibitor VIII |
| MVP_top60_DNA_seed | AKT2 | 0.798693 | THCA | 8305C | AKT inhibitor VIII |
| MVP_top60_DNA_seed | AKT3 | 0.798693 | THCA | 8305C | AKT inhibitor VIII |
| MVP_top60_DNA_seed | AKT1 | 0.805927 | THCA | B-CPAP | AKT inhibitor VIII |
| MVP_top60_DNA_seed | AKT2 | 0.805927 | THCA | B-CPAP | AKT inhibitor VIII |
| MVP_top60_DNA_seed | AKT3 | 0.805927 | THCA | B-CPAP | AKT inhibitor VIII |
| MVP_top60_DNA_seed | GSK3B | 0.807173 | THCA | FTC-133 | KIN001-042 |
| MVP_top60_DNA_seed | ATM | 0.815754 | THCA | RO82-W-1 | KU-55933 |
| MVP_top60_DNA_seed | MAPK14 | 0.817884 | THCA | FTC-133 | TAK-715 |
| MVP_top60_DNA_seed | CHEK2 | 0.822093 | THCA | BHT-101 | AZD7762 |
| MVP_top60_DNA_seed | BRAF | 0.831109 | THCA | ASH-3 | PLX-4720 |
| MVP_top60_DNA_seed | RET | 0.846469 | THCA | 8305C | Motesanib |
| MVP_top60_DNA_seed | MDM2 | 0.848292 | THCA | ASH-3 | Nutlin-3a (-) |
| MVP_top60_DNA_seed | MAPK1 | 0.865998 | THCA | CGTH-W-1 | VX-11e |
| MVP_top60_DNA_seed | ATM | 0.876813 | THCA | FTC-133 | KU-55933 |
| MVP_top60_DNA_seed | GSK3B | 0.879238 | THCA | TT2609-C02 | AZD7969 |
| MVP_top60_DNA_seed | HDAC1 | 0.883865 | THCA | FTC-133 | Tubastatin A |
| MVP_top60_DNA_seed | GSK3B | 0.885482 | THCA | 8305C | SB216763 |
| MVP_top60_DNA_seed | MAPK1 | 0.899212 | THCA | ASH-3 | VX-11e |
| MVP_top60_DNA_seed | BRAF | 0.916491 | THCA | TT | PLX-4720 |
| MVP_top60_DNA_seed | HDAC1 | 0.928272 | THCA | CGTH-W-1 | Parthenolide |
| MVP_top60_DNA_seed | GSK3B | 0.934547 | THCA | 8305C | CHIR-99021 |
| MVP_top60_DNA_seed | HDAC1 | 0.942386 | THCA | TT | Dacinostat |
| MVP_top60_DNA_seed | AKT1 | 0.953518 | THCA | CAL-62 | MK-2206 |

| | | | | | |
|---|---|---|---|---|---|
| MVP_top60_DNA_seed | AKT2 | 0.953518 | THCA | CAL-62 | MK-2206 |
| MVP_top60_DNA_seed | AKT1 | 0.984208 | THCA | ML-1 | AKT inhibitor VIII |
| MVP_top60_DNA_seed | AKT2 | 0.984208 | THCA | ML-1 | AKT inhibitor VIII |
| MVP_top60_DNA_seed | AKT3 | 0.984208 | THCA | ML-1 | AKT inhibitor VIII |
| MVP_top60_DNA_seed | ATR | 1.005424 | THCA | RO82-W-1 | AZD6738 |
| MVP_top60_DNA_seed | AKT1 | 1.017811 | THCA | ML-1 | GSK690693 |
| MVP_top60_DNA_seed | AKT2 | 1.017811 | THCA | ML-1 | GSK690693 |
| MVP_top60_DNA_seed | AKT3 | 1.017811 | THCA | ML-1 | GSK690693 |
| MVP_top60_DNA_seed | ATM | 1.023871 | THCA | 8305C | CP466722 |
| MVP_top60_DNA_seed | BRAF | 1.027277 | THCA | RO82-W-1 | SB590885 |
| MVP_top60_DNA_seed | GSK3B | 1.030288 | THCA | ML-1 | SB216763 |
| MVP_top60_DNA_seed | HDAC1 | 1.032466 | THCA | KMH-2 | Belinostat |
| MVP_top60_DNA_seed | HDAC1 | 1.039646 | THCA | ML-1 | Tubastatin A |
| MVP_top60_DNA_seed | CHEK2 | 1.046318 | THCA | TT2609-C02 | AZD7762 |
| MVP_top60_DNA_seed | AKT1 | 1.050863 | THCA | 8305C | GSK690693 |
| MVP_top60_DNA_seed | AKT2 | 1.050863 | THCA | 8305C | GSK690693 |
| MVP_top60_DNA_seed | AKT3 | 1.050863 | THCA | 8305C | GSK690693 |
| MVP_top60_DNA_seed | MAPK14 | 1.05679 | THCA | K5 | TAK-715 |
| MVP_top60_DNA_seed | GSK3B | 1.059125 | THCA | ML-1 | CHIR-99021 |
| MVP_top60_DNA_seed | GSK3B | 1.06425 | THCA | FTC-133 | CHIR-99021 |
| MVP_top60_DNA_seed | AKT1 | 1.085726 | THCA | RO82-W-1 | GSK690693 |
| MVP_top60_DNA_seed | AKT2 | 1.085726 | THCA | RO82-W-1 | GSK690693 |
| MVP_top60_DNA_seed | AKT3 | 1.085726 | THCA | RO82-W-1 | GSK690693 |
| MVP_top60_DNA_seed | AKT1 | 1.106976 | THCA | CAL-62 | AKT inhibitor VIII |
| MVP_top60_DNA_seed | AKT2 | 1.106976 | THCA | CAL-62 | AKT inhibitor VIII |
| MVP_top60_DNA_seed | AKT3 | 1.106976 | THCA | CAL-62 | AKT inhibitor VIII |
| MVP_top60_DNA_seed | ATR | 1.130828 | THCA | BHT-101 | QL-VIII-58 |
| MVP_top60_DNA_seed | MAPK14 | 1.145141 | THCA | TT | TAK-715 |
| MVP_top60_DNA_seed | ATR | 1.146171 | THCA | TT | AZD6738 |
| MVP_top60_DNA_seed | PTK2 | 1.222335 | THCA | FTC-133 | PF-562271 |
| MVP_top60_DNA_seed | GSK3B | 1.240107 | THCA | FTC-133 | AZD7969 |
| MVP_top60_DNA_seed | HDAC1 | 1.262085 | THCA | 8305C | Tubastatin A |
| MVP_top60_DNA_seed | ATR | 1.27277 | THCA | TT2609-C02 | QL-VIII-58 |
| MVP_top60_DNA_seed | AKT1 | 1.295044 | THCA | 8305C | AKT inhibitor VIII |
| MVP_top60_DNA_seed | AKT2 | 1.295044 | THCA | 8305C | AKT inhibitor VIII |
| MVP_top60_DNA_seed | AKT3 | 1.295044 | THCA | 8305C | AKT inhibitor VIII |
| MVP_top60_DNA_seed | MDM2 | 1.296613 | THCA | 8305C | Nutlin-3a (-) |
| MVP_top60_DNA_seed | GSK3B | 1.29885 | THCA | 8305C | KIN001-042 |
| MVP_top60_DNA_seed | BRAF | 1.301277 | THCA | FTC-133 | HG6-64-1 |
| MVP_top60_DNA_seed | MAPK1 | 1.33145 | THCA | RO82-W-1 | VX-11e |
| MVP_top60_DNA_seed | CHEK2 | 1.341407 | THCA | TT | AZD7762 |

| | | | | | |
|---|---|---|---|---|---|
| MVP_top60_DNA_seed | MAPK1 | 1.344383 | THCA | TT | VX-11e |
| MVP_top60_DNA_seed | HDAC1 | 1.344893 | THCA | RO82-W-1 | JQ12 |
| MVP_top60_DNA_seed | BRAF | 1.355515 | THCA | TT | PLX-4720 |
| MVP_top60_DNA_seed | GSK3B | 1.381463 | THCA | RO82-W-1 | KIN001-042 |
| MVP_top60_DNA_seed | AKT1 | 1.413432 | THCA | TT | GSK690693 |
| MVP_top60_DNA_seed | AKT2 | 1.413432 | THCA | TT | GSK690693 |
| MVP_top60_DNA_seed | AKT3 | 1.413432 | THCA | TT | GSK690693 |
| MVP_top60_DNA_seed | MAPK1 | 1.414238 | THCA | RO82-W-1 | FR-180204 |
| MVP_top60_DNA_seed | MAPK3 | 1.414238 | THCA | RO82-W-1 | FR-180204 |
| MVP_top60_DNA_seed | MAPK14 | 1.421002 | THCA | 8305C | TAK-715 |
| MVP_top60_DNA_seed | HDAC1 | 1.509265 | THCA | ASH-3 | CAY10603 |
| MVP_top60_DNA_seed | ATR | 1.538363 | THCA | TT2609-C02 | AZD6738 |
| MVP_top60_DNA_seed | AKT1 | 1.564793 | THCA | TT2609-C02 | BAY AKT1 |
| MVP_top60_DNA_seed | AKT2 | 1.564793 | THCA | TT2609-C02 | BAY AKT1 |
| MVP_top60_DNA_seed | AKT3 | 1.564793 | THCA | TT2609-C02 | BAY AKT1 |
| MVP_top60_DNA_seed | GSK3B | 1.569962 | THCA | TT | CHIR-99021 |
| MVP_top60_DNA_seed | HDAC1 | 1.577116 | THCA | ASH-3 | AR-42 |
| MVP_top60_DNA_seed | AKT1 | 1.58178 | THCA | ML-1 | BAY AKT1 |
| MVP_top60_DNA_seed | AKT2 | 1.58178 | THCA | ML-1 | BAY AKT1 |
| MVP_top60_DNA_seed | AKT3 | 1.58178 | THCA | ML-1 | BAY AKT1 |
| MVP_top60_DNA_seed | MAPK14 | 1.583372 | THCA | ML-1 | TAK-715 |
| MVP_top60_DNA_seed | GSK3B | 1.654439 | THCA | TT | SB216763 |
| MVP_top60_DNA_seed | MAPK14 | 1.68276 | THCA | RO82-W-1 | TAK-715 |
| MVP_top60_DNA_seed | HDAC1 | 1.69053 | THCA | KMH-2 | AR-42 |
| MVP_top60_DNA_seed | HDAC1 | 1.701735 | THCA | TT | Tubastatin A |
| MVP_top60_DNA_seed | GSK3B | 1.715476 | THCA | TT | CHIR-99021 |
| MVP_top60_DNA_seed | AKT1 | 1.752216 | THCA | FTC-133 | AKT inhibitor VIII |
| MVP_top60_DNA_seed | AKT2 | 1.752216 | THCA | FTC-133 | AKT inhibitor VIII |
| MVP_top60_DNA_seed | AKT3 | 1.752216 | THCA | FTC-133 | AKT inhibitor VIII |
| MVP_top60_DNA_seed | ATR | 1.767084 | THCA | TT | AZ20 |
| MVP_top60_DNA_seed | HDAC1 | 1.910143 | THCA | ASH-3 | Belinostat |
| MVP_top60_DNA_seed | AKT1 | 2.049624 | THCA | RO82-W-1 | AKT inhibitor VIII |
| MVP_top60_DNA_seed | AKT2 | 2.049624 | THCA | RO82-W-1 | AKT inhibitor VIII |
| MVP_top60_DNA_seed | AKT3 | 2.049624 | THCA | RO82-W-1 | AKT inhibitor VIII |
| MVP_top60_DNA_seed | ATM | 2.089043 | THCA | TT2609-C02 | CP466722 |
| MVP_top60_DNA_seed | MAPK1 | 2.094196 | THCA | TT | FR-180204 |
| MVP_top60_DNA_seed | MAPK3 | 2.094196 | THCA | TT | FR-180204 |
| MVP_top60_DNA_seed | ATR | 2.108931 | THCA | TT | QL-VIII-58 |
| MVP_top60_DNA_seed | GSK3B | 2.131041 | THCA | TT | KIN001-042 |
| MVP_top60_DNA_seed | PTK2 | 2.198954 | THCA | TT | PF-562271 |
| MVP_top60_DNA_seed | HDAC1 | 2.312474 | THCA | 8305C | AR-42 |

| | | | | | |
|---|---|---|---|---|---|
| MVP_top60_DNA_seed | ATM | 2.326309 | THCA | RO82-W-1 | CP466722 |
| MVP_top60_DNA_seed | AKT1 | 2.457782 | THCA | TT | AKT inhibitor VIII |
| MVP_top60_DNA_seed | AKT2 | 2.457782 | THCA | TT | AKT inhibitor VIII |
| MVP_top60_DNA_seed | AKT3 | 2.457782 | THCA | TT | AKT inhibitor VIII |
| MVP_top60_DNA_seed | ATM | 2.462246 | THCA | FTC-133 | CP466722 |
| MVP_top60_DNA_seed | ATM | 2.506771 | THCA | TT | CP466722 |
| MVP_top60_DNA_seed | HDAC1 | 2.509226 | THCA | TT | AR-42 |
| MVP_top60_DNA_seed | HDAC1 | 2.532809 | THCA | TT | CAY10603 |
| MVP_top60_DNA_seed | HDAC1 | 2.784592 | THCA | TT | Belinostat |
| MVP_top60_DNA_seed | HDAC1 | 2.998205 | THCA | 8305C | CAY10603 |
| MVP_top60_DNA_seed | HDAC1 | 3.113187 | THCA | 8305C | Belinostat |
| MVP_top60_DNA_seed | BRAF | -2.904049 | THCA | K5 | Dabrafenib |
| MVP_top60_DNA_seed | GSK3B | -2.306073 | THCA | B-CPAP | SB216763 |
| MVP_top60_DNA_seed | BRAF | -2.070305 | THCA | BHT-101 | Dabrafenib |
| MVP_top60_DNA_seed | BRAF | -2.008943 | THCA | K5 | PLX-4720 |
| MVP_top60_DNA_seed | MAPK1 | -1.972739 | THCA | KMH-2 | VX-11e |
| MVP_top60_DNA_seed | BRAF | -1.889949 | THCA | HTC-C3 | PLX-4720 |
| MVP_top60_DNA_seed | MAPK1 | -1.76979 | THCA | KMH-2 | SCH772984 |
| MVP_top60_DNA_seed | MAPK3 | -1.76979 | THCA | KMH-2 | SCH772984 |
| MVP_top60_DNA_seed | MAPK1 | -1.757059 | THCA | BHT-101 | VX-11e |
| MVP_top60_DNA_seed | MAPK1 | -1.755432 | THCA | HTC-C3 | Ulixertinib |
| MVP_top60_DNA_seed | MAPK3 | -1.755432 | THCA | HTC-C3 | Ulixertinib |
| MVP_top60_DNA_seed | MAPK1 | -1.730831 | THCA | KMH-2 | Ulixertinib |
| MVP_top60_DNA_seed | MAPK3 | -1.730831 | THCA | KMH-2 | Ulixertinib |
| MVP_top60_DNA_seed | BRAF | -1.699237 | THCA | K5 | SB590885 |
| MVP_top60_DNA_seed | HDAC1 | -1.694317 | THCA | K5 | Romidepsin |
| MVP_top60_DNA_seed | MAPK1 | -1.683262 | THCA | CAL-62 | Ulixertinib |
| MVP_top60_DNA_seed | MAPK3 | -1.683262 | THCA | CAL-62 | Ulixertinib |
| MVP_top60_DNA_seed | MAPK1 | -1.556041 | THCA | BHT-101 | SCH772984 |
| MVP_top60_DNA_seed | MAPK3 | -1.556041 | THCA | BHT-101 | SCH772984 |
| MVP_top60_DNA_seed | BRAF | -1.548644 | THCA | HTC-C3 | Dabrafenib |
| MVP_top60_DNA_seed | MAPK1 | -1.431794 | THCA | IHH-4 | Ulixertinib |
| MVP_top60_DNA_seed | MAPK3 | -1.431794 | THCA | IHH-4 | Ulixertinib |
| MVP_top60_DNA_seed | MAPK1 | -1.365373 | THCA | HTC-C3 | Ulixertinib |
| MVP_top60_DNA_seed | MAPK3 | -1.365373 | THCA | HTC-C3 | Ulixertinib |
| MVP_top60_DNA_seed | BRAF | -1.363286 | THCA | BHT-101 | PLX-4720 |
| MVP_top60_DNA_seed | MAPK1 | -1.343518 | THCA | BHT-101 | Ulixertinib |
| MVP_top60_DNA_seed | MAPK3 | -1.343518 | THCA | BHT-101 | Ulixertinib |
| MVP_top60_DNA_seed | MAPK1 | -1.338722 | THCA | IHH-4 | VX-11e |
| MVP_top60_DNA_seed | BRAF | -1.326271 | THCA | 8305C | Dabrafenib |
| MVP_top60_DNA_seed | MDM2 | -1.311278 | THCA | K5 | Nutlin-3a (-) |

| | | | | | |
|---|---|---|---|---|---|
| MVP_top60_DNA_seed | BRAF | -1.308531 | THCA | IHH-4 | Dabrafenib |
| MVP_top60_DNA_seed | MAPK1 | -1.278527 | THCA | IHH-4 | SCH772984 |
| MVP_top60_DNA_seed | MAPK3 | -1.278527 | THCA | IHH-4 | SCH772984 |
| MVP_top60_DNA_seed | HDAC1 | -1.250755 | THCA | IHH-4 | Romidepsin |
| MVP_top60_DNA_seed | BRAF | -1.24812 | THCA | IHH-4 | PLX-4720 |
| MVP_top60_DNA_seed | AKT1 | -1.24793 | THCA | KMH-2 | AT13148 |
| MVP_top60_DNA_seed | MAPK1 | -1.238702 | THCA | BHT-101 | Ulixertinib |
| MVP_top60_DNA_seed | MAPK3 | -1.238702 | THCA | BHT-101 | Ulixertinib |
| MVP_top60_DNA_seed | BRAF | -1.182661 | THCA | B-CPAP | Dabrafenib |
| MVP_top60_DNA_seed | MDM2 | -1.147367 | THCA | IHH-4 | Nutlin-3a (-) |
| MVP_top60_DNA_seed | RET | -1.146579 | THCA | B-CPAP | Motesanib |
| MVP_top60_DNA_seed | MAPK1 | -1.131306 | THCA | K5 | Ulixertinib |
| MVP_top60_DNA_seed | MAPK3 | -1.131306 | THCA | K5 | Ulixertinib |
| MVP_top60_DNA_seed | MAPK1 | -1.114038 | THCA | HTC-C3 | VX-11e |
| MVP_top60_DNA_seed | AKT1 | -1.105186 | THCA | FTC-133 | AZD5363 |
| MVP_top60_DNA_seed | AKT2 | -1.105186 | THCA | FTC-133 | AZD5363 |
| MVP_top60_DNA_seed | AKT3 | -1.105186 | THCA | FTC-133 | AZD5363 |
| MVP_top60_DNA_seed | CHEK2 | -1.092126 | THCA | CAL-62 | AZD7762 |
| MVP_top60_DNA_seed | GSK3B | -1.089368 | THCA | KMH-2 | SB216763 |
| MVP_top60_DNA_seed | BRAF | -1.081835 | THCA | ML-1 | PLX-4720 |
| MVP_top60_DNA_seed | GSK3B | -1.078602 | THCA | KMH-2 | CHIR-99021 |
| MVP_top60_DNA_seed | HDAC1 | -1.070706 | THCA | KMH-2 | PCI-34051 |
| MVP_top60_DNA_seed | HDAC1 | -1.048756 | THCA | 8505C | PCI-34051 |
| MVP_top60_DNA_seed | USP7 | -1.043553 | THCA | KMH-2 | P22077 |
| MVP_top60_DNA_seed | MAPK1 | -1.038659 | THCA | K5 | SCH772984 |
| MVP_top60_DNA_seed | MAPK3 | -1.038659 | THCA | K5 | SCH772984 |
| MVP_top60_DNA_seed | AKT1 | -1.013336 | THCA | FTC-133 | Afuresertib |
| MVP_top60_DNA_seed | AKT2 | -1.013336 | THCA | FTC-133 | Afuresertib |
| MVP_top60_DNA_seed | AKT3 | -1.013336 | THCA | FTC-133 | Afuresertib |
| MVP_top60_DNA_seed | AKT1 | -1.004826 | THCA | K5 | AT13148 |
| MVP_top60_DNA_seed | ATR | -1.003188 | THCA | CAL-62 | AZD6738 |
| MVP_top60_DNA_seed | USP7 | -0.989122 | THCA | CAL-62 | P22077 |
| MVP_top60_DNA_seed | AKT1 | -0.953548 | THCA | K5 | Afuresertib |
| MVP_top60_DNA_seed | AKT2 | -0.953548 | THCA | K5 | Afuresertib |
| MVP_top60_DNA_seed | AKT3 | -0.953548 | THCA | K5 | Afuresertib |
| MVP_top60_DNA_seed | KRAS | -0.951678 | THCA | IHH-4 | KRAS (G12C) Inhibitor-12 |
| MVP_top60_DNA_seed | KRAS | -0.948296 | THCA | ASH-3 | KRAS (G12C) Inhibitor-12 |
| MVP_top60_DNA_seed | USP7 | -0.893763 | THCA | BHT-101 | P22077 |
| MVP_top60_DNA_seed | GSK3B | -0.862785 | THCA | K5 | SB216763 |
| MVP_top60_DNA_seed | HDAC1 | -0.8621 | THCA | BHT-101 | PCI-34051 |
| MVP_top60_DNA_seed | BRAF | -0.841682 | THCA | KMH-2 | SB590885 |

| | | | | | |
|---|---|---|---|---|---|
| MVP_top60_DNA_seed | ATR | -0.839769 | THCA | IHH-4 | VE821 |
| MVP_top60_DNA_seed | BRAF | -0.811101 | THCA | BHT-101 | SB590885 |
| MVP_top60_DNA_seed | ATR | -0.801901 | THCA | KMH-2 | AZD6738 |
| MVP_top60_DNA_seed | KRAS | -0.791263 | THCA | BHT-101 | KRAS (G12C) Inhibitor-12 |
| MVP_top60_DNA_seed | HDAC1 | -0.787006 | THCA | KMH-2 | Entinostat |
| MVP_top60_DNA_seed | BRAF | -0.779147 | THCA | 8305C | PLX-4720 |
| MVP_top60_DNA_seed | ATR | -0.773189 | THCA | HTC-C3 | AZD6738 |
| MVP_top60_DNA_seed | USP7 | -0.758946 | THCA | HTC-C3 | P22077 |
| MVP_top60_DNA_seed | HDAC1 | -0.744797 | THCA | 8305C | Romidepsin |
| MVP_top60_DNA_seed | BRAF | -0.729044 | THCA | B-CPAP | PLX-4720 |
| MVP_top60_DNA_seed | BRAF | -0.727578 | THCA | KMH-2 | Dabrafenib |
| MVP_top60_DNA_seed | AKT1 | -0.727366 | THCA | K5 | AZD5363 |
| MVP_top60_DNA_seed | AKT2 | -0.727366 | THCA | K5 | AZD5363 |
| MVP_top60_DNA_seed | AKT3 | -0.727366 | THCA | K5 | AZD5363 |
| MVP_top60_DNA_seed | BRAF | -0.724427 | THCA | KMH-2 | PLX-4720 |
| MVP_top60_DNA_seed | MAPK1 | -0.715407 | THCA | IHH-4 | Ulixertinib |
| MVP_top60_DNA_seed | MAPK3 | -0.715407 | THCA | IHH-4 | Ulixertinib |
| MVP_top60_DNA_seed | BRAF | -0.706696 | THCA | 8505C | Dabrafenib |
| MVP_top60_DNA_seed | AKT1 | -0.700062 | THCA | K5 | GSK2110183B |
| MVP_top60_DNA_seed | AKT2 | -0.700062 | THCA | K5 | GSK2110183B |
| MVP_top60_DNA_seed | AKT3 | -0.700062 | THCA | K5 | GSK2110183B |
| MVP_top60_DNA_seed | ATR | -0.698541 | THCA | FTC-133 | AZD6738 |
| MVP_top60_DNA_seed | ATR | -0.687843 | THCA | KMH-2 | VE-822 |
| MVP_top60_DNA_seed | ATM | -0.677234 | THCA | ASH-3 | KU-55933 |
| MVP_top60_DNA_seed | MAPK1 | -0.673281 | THCA | HTC-C3 | SCH772984 |
| MVP_top60_DNA_seed | MAPK3 | -0.673281 | THCA | HTC-C3 | SCH772984 |
| MVP_top60_DNA_seed | MAPK1 | -0.665421 | THCA | K5 | Ulixertinib |
| MVP_top60_DNA_seed | MAPK3 | -0.665421 | THCA | K5 | Ulixertinib |
| MVP_top60_DNA_seed | AKT1 | -0.634041 | THCA | ASH-3 | AT13148 |
| MVP_top60_DNA_seed | AKT1 | -0.626325 | THCA | BHT-101 | AT13148 |
| MVP_top60_DNA_seed | AKT1 | -0.614942 | THCA | FTC-133 | Ipatasertib |
| MVP_top60_DNA_seed | AKT3 | -0.614942 | THCA | FTC-133 | Ipatasertib |
| MVP_top60_DNA_seed | MAPK1 | -0.600139 | THCA | TT | Ulixertinib |
| MVP_top60_DNA_seed | MAPK3 | -0.600139 | THCA | TT | Ulixertinib |
| MVP_top60_DNA_seed | AKT1 | -0.586595 | THCA | K5 | Ipatasertib |
| MVP_top60_DNA_seed | AKT3 | -0.586595 | THCA | K5 | Ipatasertib |
| MVP_top60_DNA_seed | AKT1 | -0.584098 | THCA | K5 | Uprosertib |
| MVP_top60_DNA_seed | AKT2 | -0.584098 | THCA | K5 | Uprosertib |
| MVP_top60_DNA_seed | AKT3 | -0.584098 | THCA | K5 | Uprosertib |
| MVP_top60_DNA_seed | HDAC1 | -0.57334 | THCA | ASH-3 | Romidepsin |
| MVP_top60_DNA_seed | AKT1 | -0.562402 | THCA | IHH-4 | Uprosertib |

| | | | | | |
|---|---|---|---|---|---|
| MVP_top60_DNA_seed | AKT2 | -0.562402 | THCA | IHH-4 | Uprosertib |
| MVP_top60_DNA_seed | AKT3 | -0.562402 | THCA | IHH-4 | Uprosertib |
| MVP_top60_DNA_seed | MAPK1 | -0.549286 | THCA | 8505C | SCH772984 |
| MVP_top60_DNA_seed | MAPK3 | -0.549286 | THCA | 8505C | SCH772984 |
| MVP_top60_DNA_seed | BRAF | -0.507576 | THCA | ML-1 | Dabrafenib |
| MVP_top60_DNA_seed | MDM2 | -0.49863 | THCA | TT | Serdemetan |
| MVP_top60_DNA_seed | MAPK1 | -0.491213 | THCA | TT2609-C02 | SCH772984 |
| MVP_top60_DNA_seed | MAPK3 | -0.491213 | THCA | TT2609-C02 | SCH772984 |
| MVP_top60_DNA_seed | AKT1 | -0.488621 | THCA | ASH-3 | GSK2110183B |
| MVP_top60_DNA_seed | AKT2 | -0.488621 | THCA | ASH-3 | GSK2110183B |
| MVP_top60_DNA_seed | AKT3 | -0.488621 | THCA | ASH-3 | GSK2110183B |
| MVP_top60_DNA_seed | MAPK1 | -0.476449 | THCA | ASH-3 | SCH772984 |
| MVP_top60_DNA_seed | MAPK3 | -0.476449 | THCA | ASH-3 | SCH772984 |
| MVP_top60_DNA_seed | HDAC1 | -0.472437 | THCA | FTC-133 | Romidepsin |
| MVP_top60_DNA_seed | GSK3B | -0.463637 | THCA | BHT-101 | SB216763 |
| MVP_top60_DNA_seed | ATM | -0.461576 | THCA | IHH-4 | KU-55933 |
| MVP_top60_DNA_seed | AKT1 | -0.461444 | THCA | KMH-2 | AZD5363 |
| MVP_top60_DNA_seed | AKT2 | -0.461444 | THCA | KMH-2 | AZD5363 |
| MVP_top60_DNA_seed | AKT3 | -0.461444 | THCA | KMH-2 | AZD5363 |
| MVP_top60_DNA_seed | USP7 | -0.438856 | THCA | FTC-133 | P22077 |
| MVP_top60_DNA_seed | AKT1 | -0.422296 | THCA | FTC-133 | Uprosertib |
| MVP_top60_DNA_seed | AKT2 | -0.422296 | THCA | FTC-133 | Uprosertib |
| MVP_top60_DNA_seed | AKT3 | -0.422296 | THCA | FTC-133 | Uprosertib |
| MVP_top60_DNA_seed | HDAC1 | -0.417698 | THCA | CAL-62 | Romidepsin |
| MVP_top60_DNA_seed | MAPK1 | -0.413029 | THCA | CAL-62 | Ulixertinib |
| MVP_top60_DNA_seed | MAPK3 | -0.413029 | THCA | CAL-62 | Ulixertinib |
| MVP_top60_DNA_seed | AKT1 | -0.408279 | THCA | FTC-133 | GSK2110183B |
| MVP_top60_DNA_seed | AKT2 | -0.408279 | THCA | FTC-133 | GSK2110183B |
| MVP_top60_DNA_seed | AKT3 | -0.408279 | THCA | FTC-133 | GSK2110183B |
| MVP_top60_DNA_seed | CHEK2 | -0.405347 | THCA | FTC-133 | AZD7762 |
| MVP_top60_DNA_seed | ATR | -0.405037 | THCA | BHT-101 | VE-822 |
| MVP_top60_DNA_seed | AKT1 | -0.385085 | THCA | K5 | MK-2206 |
| MVP_top60_DNA_seed | AKT2 | -0.385085 | THCA | K5 | MK-2206 |
| MVP_top60_DNA_seed | AKT1 | -0.384849 | THCA | IHH-4 | Ipatasertib |
| MVP_top60_DNA_seed | AKT3 | -0.384849 | THCA | IHH-4 | Ipatasertib |
| MVP_top60_DNA_seed | BRAF | -0.380654 | THCA | B-CPAP | SB590885 |
| MVP_top60_DNA_seed | BRAF | -0.377407 | THCA | IHH-4 | SB590885 |
| MVP_top60_DNA_seed | AKT1 | -0.372713 | THCA | BHT-101 | MK-2206 |
| MVP_top60_DNA_seed | AKT2 | -0.372713 | THCA | BHT-101 | MK-2206 |
| MVP_top60_DNA_seed | KRAS | -0.36196 | THCA | K5 | KRAS (G12C) Inhibitor-12 |
| MVP_top60_DNA_seed | HDAC1 | -0.358899 | THCA | B-CPAP | Romidepsin |

| | | | | | |
|---|---|---|---|---|---|
| MVP_top60_DNA_seed | ATR | -0.356669 | THCA | K5 | VE821 |
| MVP_top60_DNA_seed | AKT1 | -0.350262 | THCA | B-CPAP | Afuresertib |
| MVP_top60_DNA_seed | AKT2 | -0.350262 | THCA | B-CPAP | Afuresertib |
| MVP_top60_DNA_seed | AKT3 | -0.350262 | THCA | B-CPAP | Afuresertib |
| MVP_top60_DNA_seed | ATR | -0.341487 | THCA | IHH-4 | VE-822 |
| MVP_top60_DNA_seed | MDM2 | -0.328993 | THCA | ML-1 | Serdemetan |
| MVP_top60_DNA_seed | HDAC1 | -0.304496 | THCA | K5 | PCI-34051 |
| MVP_top60_DNA_seed | CHEK2 | -0.303186 | THCA | HTC-C3 | AZD7762 |
| MVP_top60_DNA_seed | MAPK1 | -0.300332 | THCA | ASH-3 | VX-11e |
| MVP_top60_DNA_seed | CHEK2 | -0.296783 | THCA | KMH-2 | AZD7762 |
| MVP_top60_DNA_seed | GSK3B | -0.282 | THCA | IHH-4 | SB216763 |
| MVP_top60_DNA_seed | ATR | -0.270701 | THCA | KMH-2 | VE821 |
| MVP_top60_DNA_seed | AKT1 | -0.265183 | THCA | KMH-2 | MK-2206 |
| MVP_top60_DNA_seed | AKT2 | -0.265183 | THCA | KMH-2 | MK-2206 |
| MVP_top60_DNA_seed | ATR | -0.262717 | THCA | HTC-C3 | VE821 |
| MVP_top60_DNA_seed | AKT1 | -0.259045 | THCA | KMH-2 | Afuresertib |
| MVP_top60_DNA_seed | AKT2 | -0.259045 | THCA | KMH-2 | Afuresertib |
| MVP_top60_DNA_seed | AKT3 | -0.259045 | THCA | KMH-2 | Afuresertib |
| MVP_top60_DNA_seed | AKT1 | -0.254152 | THCA | B-CPAP | AT13148 |
| MVP_top60_DNA_seed | MAPK1 | -0.242485 | THCA | CAL-62 | SCH772984 |
| MVP_top60_DNA_seed | MAPK3 | -0.242485 | THCA | CAL-62 | SCH772984 |
| MVP_top60_DNA_seed | ATR | -0.239914 | THCA | 8305C | VE821 |
| MVP_top60_DNA_seed | AKT1 | -0.238933 | THCA | B-CPAP | Uprosertib |
| MVP_top60_DNA_seed | AKT2 | -0.238933 | THCA | B-CPAP | Uprosertib |
| MVP_top60_DNA_seed | AKT3 | -0.238933 | THCA | B-CPAP | Uprosertib |
| MVP_top60_DNA_seed | BRAF | -0.238721 | THCA | CGTH-W-1 | SB590885 |
| MVP_top60_DNA_seed | HDAC1 | -0.2379 | THCA | FTC-133 | Entinostat |
| MVP_top60_DNA_seed | HDAC1 | -0.232194 | THCA | HTC-C3 | PCI-34051 |
| MVP_top60_DNA_seed | ATR | -0.227652 | THCA | FTC-133 | VE-822 |
| MVP_top60_DNA_seed | CHEK2 | -0.219384 | THCA | FTC-133 | MK-8776 |
| MVP_top60_DNA_seed | HDAC1 | -0.215744 | THCA | IHH-4 | Entinostat |
| MVP_top60_DNA_seed | MAPK1 | -0.203577 | THCA | 8505C | Ulixertinib |
| MVP_top60_DNA_seed | MAPK3 | -0.203577 | THCA | 8505C | Ulixertinib |
| MVP_top60_DNA_seed | MAPK1 | -0.20344 | THCA | ASH-3 | Ulixertinib |
| MVP_top60_DNA_seed | MAPK3 | -0.20344 | THCA | ASH-3 | Ulixertinib |
| MVP_top60_DNA_seed | BRAF | -0.188294 | THCA | CGTH-W-1 | PLX-4720 |
| MVP_top60_DNA_seed | HDAC1 | -0.181671 | THCA | K5 | Entinostat |
| MVP_top60_DNA_seed | BRAF | -0.168751 | THCA | FTC-133 | Dabrafenib |
| MVP_top60_DNA_seed | RET | -0.154526 | THCA | KMH-2 | Motesanib |
| MVP_top60_DNA_seed | MAPK1 | -0.151983 | THCA | FTC-133 | Ulixertinib |
| MVP_top60_DNA_seed | MAPK3 | -0.151983 | THCA | FTC-133 | Ulixertinib |

| | | | | | |
|---|---|---|---|---|---|
| MVP_top60_DNA_seed | MDM2 | -0.150649 | THCA | KMH-2 | Nutlin-3a (-) |
| MVP_top60_DNA_seed | RET | -0.139767 | THCA | IHH-4 | Motesanib |
| MVP_top60_DNA_seed | MAPK1 | -0.132502 | THCA | 8305C | SCH772984 |
| MVP_top60_DNA_seed | MAPK3 | -0.132502 | THCA | 8305C | SCH772984 |
| MVP_top60_DNA_seed | AKT1 | -0.132233 | THCA | CAL-62 | AT13148 |
| MVP_top60_DNA_seed | HDAC1 | -0.130585 | THCA | BHT-101 | Entinostat |
| MVP_top60_DNA_seed | GSK3B | -0.125859 | THCA | TT2609-C02 | SB216763 |
| MVP_top60_DNA_seed | MAPK1 | -0.123184 | THCA | TT2609-C02 | Ulixertinib |
| MVP_top60_DNA_seed | MAPK3 | -0.123184 | THCA | TT2609-C02 | Ulixertinib |
| MVP_top60_DNA_seed | HDAC1 | -0.118586 | THCA | 8505C | Entinostat |
| MVP_top60_DNA_seed | MDM2 | -0.118257 | THCA | K5 | Serdemetan |
| MVP_top60_DNA_seed | ATR | -0.10348 | THCA | 8505C | VE-822 |
| MVP_top60_DNA_seed | CHEK2 | -0.095069 | THCA | IHH-4 | AZD7762 |
| MVP_top60_DNA_seed | MAPK1 | -0.09403 | THCA | ASH-3 | Ulixertinib |
| MVP_top60_DNA_seed | MAPK3 | -0.09403 | THCA | ASH-3 | Ulixertinib |
| MVP_top60_DNA_seed | MAPK1 | -0.089216 | THCA | K5 | VX-11e |
| MVP_top60_DNA_seed | MAPK1 | -0.073967 | THCA | B-CPAP | Ulixertinib |
| MVP_top60_DNA_seed | MAPK3 | -0.073967 | THCA | B-CPAP | Ulixertinib |
| MVP_top60_DNA_seed | HDAC1 | -0.073074 | THCA | HTC-C3 | Entinostat |
| MVP_top60_DNA_seed | AKT1 | -0.070647 | THCA | 8305C | MK-2206 |
| MVP_top60_DNA_seed | AKT2 | -0.070647 | THCA | 8305C | MK-2206 |
| MVP_top60_DNA_seed | MDM2 | -0.065777 | THCA | KMH-2 | Serdemetan |
| MVP_top60_DNA_seed | MAPK1 | -0.065603 | THCA | 8505C | VX-11e |
| MVP_top60_DNA_seed | CHEK2 | -0.062307 | THCA | CGTH-W-1 | MK-8776 |
| MVP_top60_DNA_seed | AKT1 | -0.059484 | THCA | K5 | Uprosertib |
| MVP_top60_DNA_seed | AKT2 | -0.059484 | THCA | K5 | Uprosertib |
| MVP_top60_DNA_seed | AKT3 | -0.059484 | THCA | K5 | Uprosertib |
| MVP_top60_DNA_seed | ATR | -0.056546 | THCA | B-CPAP | AZD6738 |
| MVP_top60_DNA_seed | AKT1 | -0.050801 | THCA | KMH-2 | Ipatasertib |
| MVP_top60_DNA_seed | AKT3 | -0.050801 | THCA | KMH-2 | Ipatasertib |
| MVP_top60_DNA_seed | GSK3B | -0.050169 | THCA | FTC-133 | SB216763 |
| MVP_top60_DNA_seed | MDM2 | -0.045815 | THCA | IHH-4 | Serdemetan |
| MVP_top60_DNA_seed | AKT1 | -0.035112 | THCA | B-CPAP | AZD5363 |
| MVP_top60_DNA_seed | AKT2 | -0.035112 | THCA | B-CPAP | AZD5363 |
| MVP_top60_DNA_seed | AKT3 | -0.035112 | THCA | B-CPAP | AZD5363 |
| MVP_top60_DNA_seed | ATR | -0.033778 | THCA | IHH-4 | AZD6738 |
| MVP_top60_DNA_seed | USP7 | -0.029344 | THCA | K5 | P22077 |
| MVP_top60_DNA_seed | USP7 | -0.022999 | THCA | B-CPAP | P22077 |
| MVP_top60_DNA_seed | MAPK1 | -0.020558 | THCA | B-CPAP | SCH772984 |
| MVP_top60_DNA_seed | MAPK3 | -0.020558 | THCA | B-CPAP | SCH772984 |
| MVP_top60_DNA_seed | USP7 | -0.018396 | THCA | ASH-3 | P22077 |

| | | | | | |
|---|---|---|---|---|---|
| MVP_top60_DNA_seed | AKT1 | -0.012276 | THCA | ASH-3 | Afuresertib |
| MVP_top60_DNA_seed | AKT2 | -0.012276 | THCA | ASH-3 | Afuresertib |
| MVP_top60_DNA_seed | AKT3 | -0.012276 | THCA | ASH-3 | Afuresertib |
| MVP_top60_DNA_seed | GSK3B | -0.005432 | THCA | HTC-C3 | SB216763 |
| MVP_top60_DNA_seed | CHEK2 | -0.003886 | THCA | KMH-2 | MK-8776 |
| MVP_top60_DNA_seed | HDAC1 | -0.000759 | THCA | 8505C | Romidepsin |
| MVP_top60_DNA_seed | AKT1 | 0.006181 | THCA | 8305C | AT13148 |
| MVP_top60_DNA_seed | CHEK2 | 0.012385 | THCA | 8305C | AZD7762 |
| MVP_top60_DNA_seed | AKT1 | 0.022588 | THCA | KMH-2 | Uprosertib |
| MVP_top60_DNA_seed | AKT2 | 0.022588 | THCA | KMH-2 | Uprosertib |
| MVP_top60_DNA_seed | AKT3 | 0.022588 | THCA | KMH-2 | Uprosertib |
| MVP_top60_DNA_seed | CHEK2 | 0.022619 | THCA | CGTH-W-1 | AZD7762 |
| MVP_top60_DNA_seed | CHEK2 | 0.024054 | THCA | HTC-C3 | MK-8776 |
| MVP_top60_DNA_seed | MAPK1 | 0.034688 | THCA | B-CPAP | VX-11e |
| MVP_top60_DNA_seed | CHEK2 | 0.0574 | THCA | IHH-4 | MK-8776 |
| MVP_top60_DNA_seed | MAPK1 | 0.058668 | THCA | B-CPAP | Ulixertinib |
| MVP_top60_DNA_seed | MAPK3 | 0.058668 | THCA | B-CPAP | Ulixertinib |
| MVP_top60_DNA_seed | AKT1 | 0.07147 | THCA | ASH-3 | AZD5363 |
| MVP_top60_DNA_seed | AKT2 | 0.07147 | THCA | ASH-3 | AZD5363 |
| MVP_top60_DNA_seed | AKT3 | 0.07147 | THCA | ASH-3 | AZD5363 |
| MVP_top60_DNA_seed | ATM | 0.082146 | THCA | CGTH-W-1 | KU-55933 |
| MVP_top60_DNA_seed | AKT1 | 0.082997 | THCA | IHH-4 | MK-2206 |
| MVP_top60_DNA_seed | AKT2 | 0.082997 | THCA | IHH-4 | MK-2206 |
| MVP_top60_DNA_seed | AKT1 | 0.088719 | THCA | IHH-4 | Afuresertib |
| MVP_top60_DNA_seed | AKT2 | 0.088719 | THCA | IHH-4 | Afuresertib |
| MVP_top60_DNA_seed | AKT3 | 0.088719 | THCA | IHH-4 | Afuresertib |
| MVP_top60_DNA_seed | HDAC1 | 0.090957 | THCA | ASH-3 | Entinostat |
| MVP_top60_DNA_seed | ATR | 0.093517 | THCA | BHT-101 | AZD6738 |
| MVP_top60_DNA_seed | AKT1 | 0.096746 | THCA | 8305C | Ipatasertib |
| MVP_top60_DNA_seed | AKT3 | 0.096746 | THCA | 8305C | Ipatasertib |
| MVP_top60_DNA_seed | CHEK2 | 0.099675 | THCA | B-CPAP | AZD7762 |
| MVP_top60_DNA_seed | ATM | 0.104186 | THCA | HTC-C3 | KU-55933 |
| MVP_top60_DNA_seed | ATR | 0.105412 | THCA | HTC-C3 | VE-822 |
| MVP_top60_DNA_seed | ATM | 0.11089 | THCA | 8305C | KU-55933 |
| MVP_top60_DNA_seed | AKT1 | 0.116142 | THCA | CAL-62 | Ipatasertib |
| MVP_top60_DNA_seed | AKT3 | 0.116142 | THCA | CAL-62 | Ipatasertib |
| MVP_top60_DNA_seed | RET | 0.121121 | THCA | CAL-62 | Motesanib |
| MVP_top60_DNA_seed | HDAC1 | 0.125716 | THCA | B-CPAP | PCI-34051 |
| MVP_top60_DNA_seed | MAPK1 | 0.134393 | THCA | CAL-62 | VX-11e |
| MVP_top60_DNA_seed | MAPK1 | 0.151567 | THCA | 8505C | Ulixertinib |
| MVP_top60_DNA_seed | MAPK3 | 0.151567 | THCA | 8505C | Ulixertinib |

| | | | | | |
|---|---|---|---|---|---|
| MVP_top60_DNA_seed | GSK3B | 0.157933 | THCA | CGTH-W-1 | SB216763 |
| MVP_top60_DNA_seed | RET | 0.158754 | THCA | ASH-3 | Motesanib |
| MVP_top60_DNA_seed | KRAS | 0.164964 | THCA | KMH-2 | KRAS (G12C) Inhibitor-12 |
| MVP_top60_DNA_seed | ATM | 0.16506 | THCA | KMH-2 | KU-55933 |
| MVP_top60_DNA_seed | KRAS | 0.165603 | THCA | CAL-62 | KRAS (G12C) Inhibitor-12 |
| MVP_top60_DNA_seed | HDAC1 | 0.166609 | THCA | HTC-C3 | Romidepsin |
| MVP_top60_DNA_seed | AKT1 | 0.172248 | THCA | ASH-3 | Ipatasertib |
| MVP_top60_DNA_seed | AKT3 | 0.172248 | THCA | ASH-3 | Ipatasertib |
| MVP_top60_DNA_seed | AKT1 | 0.182526 | THCA | B-CPAP | MK-2206 |
| MVP_top60_DNA_seed | AKT2 | 0.182526 | THCA | B-CPAP | MK-2206 |
| MVP_top60_DNA_seed | AKT1 | 0.193557 | THCA | BHT-101 | Afuresertib |
| MVP_top60_DNA_seed | AKT2 | 0.193557 | THCA | BHT-101 | Afuresertib |
| MVP_top60_DNA_seed | AKT3 | 0.193557 | THCA | BHT-101 | Afuresertib |
| MVP_top60_DNA_seed | HDAC1 | 0.197236 | THCA | 8305C | PCI-34051 |
| MVP_top60_DNA_seed | AKT1 | 0.197588 | THCA | CAL-62 | Afuresertib |
| MVP_top60_DNA_seed | AKT2 | 0.197588 | THCA | CAL-62 | Afuresertib |
| MVP_top60_DNA_seed | AKT3 | 0.197588 | THCA | CAL-62 | Afuresertib |
| MVP_top60_DNA_seed | USP7 | 0.220439 | THCA | TT2609-C02 | P22077 |
| MVP_top60_DNA_seed | ATR | 0.236389 | THCA | K5 | AZD6738 |
| MVP_top60_DNA_seed | CHEK2 | 0.248206 | THCA | ML-1 | MK-8776 |
| MVP_top60_DNA_seed | RET | 0.250691 | THCA | FTC-133 | Motesanib |
| MVP_top60_DNA_seed | ATM | 0.251063 | THCA | K5 | KU-55933 |
| MVP_top60_DNA_seed | RET | 0.25247 | THCA | K5 | Motesanib |
| MVP_top60_DNA_seed | CHEK2 | 0.258476 | THCA | K5 | AZD7762 |
| MVP_top60_DNA_seed | ATM | 0.260663 | THCA | BHT-101 | KU-55933 |
| MVP_top60_DNA_seed | BRAF | 0.268676 | THCA | 8505C | PLX-4720 |
| MVP_top60_DNA_seed | BRAF | 0.269089 | THCA | CAL-62 | SB590885 |
| MVP_top60_DNA_seed | GSK3B | 0.271067 | THCA | CAL-62 | SB216763 |
| MVP_top60_DNA_seed | MDM2 | 0.278969 | THCA | BHT-101 | Serdemetan |
| MVP_top60_DNA_seed | AKT1 | 0.279299 | THCA | 8305C | Afuresertib |
| MVP_top60_DNA_seed | AKT2 | 0.279299 | THCA | 8305C | Afuresertib |
| MVP_top60_DNA_seed | AKT3 | 0.279299 | THCA | 8305C | Afuresertib |
| MVP_top60_DNA_seed | MAPK1 | 0.296411 | THCA | CGTH-W-1 | SCH772984 |
| MVP_top60_DNA_seed | MAPK3 | 0.296411 | THCA | CGTH-W-1 | SCH772984 |
| MVP_top60_DNA_seed | MAPK1 | 0.297869 | THCA | 8305C | VX-11e |
| MVP_top60_DNA_seed | MDM2 | 0.299907 | THCA | FTC-133 | Nutlin-3a (-) |
| MVP_top60_DNA_seed | RET | 0.303606 | THCA | BHT-101 | Motesanib |
| MVP_top60_DNA_seed | AKT1 | 0.309416 | THCA | BHT-101 | Uprosertib |
| MVP_top60_DNA_seed | AKT2 | 0.309416 | THCA | BHT-101 | Uprosertib |
| MVP_top60_DNA_seed | AKT3 | 0.309416 | THCA | BHT-101 | Uprosertib |
| MVP_top60_DNA_seed | ATR | 0.323965 | THCA | K5 | VE-822 |

| | | | | | |
|---|---|---|---|---|---|
| MVP_top60_DNA_seed | AKT1 | 0.346773 | THCA | FTC-133 | MK-2206 |
| MVP_top60_DNA_seed | AKT2 | 0.346773 | THCA | FTC-133 | MK-2206 |
| MVP_top60_DNA_seed | ATR | 0.348762 | THCA | B-CPAP | VE821 |
| MVP_top60_DNA_seed | AKT1 | 0.350825 | THCA | BHT-101 | Ipatasertib |
| MVP_top60_DNA_seed | AKT3 | 0.350825 | THCA | BHT-101 | Ipatasertib |
| MVP_top60_DNA_seed | MAPK1 | 0.357318 | THCA | CGTH-W-1 | VX-11e |
| MVP_top60_DNA_seed | ATR | 0.360264 | THCA | CAL-62 | VE821 |
| MVP_top60_DNA_seed | HDAC1 | 0.367898 | THCA | FTC-133 | PCI-34051 |
| MVP_top60_DNA_seed | RET | 0.368546 | THCA | HTC-C3 | Motesanib |
| MVP_top60_DNA_seed | BRAF | 0.372867 | THCA | FTC-133 | PLX-4720 |
| MVP_top60_DNA_seed | MAPK1 | 0.378 | THCA | RO82-W-1 | Ulixertinib |
| MVP_top60_DNA_seed | MAPK3 | 0.378 | THCA | RO82-W-1 | Ulixertinib |
| MVP_top60_DNA_seed | BRAF | 0.386036 | THCA | ASH-3 | SB590885 |
| MVP_top60_DNA_seed | ATR | 0.386977 | THCA | 8305C | AZD6738 |
| MVP_top60_DNA_seed | CHEK2 | 0.397495 | THCA | TT | MK-8776 |
| MVP_top60_DNA_seed | AKT1 | 0.423141 | THCA | ASH-3 | Uprosertib |
| MVP_top60_DNA_seed | AKT2 | 0.423141 | THCA | ASH-3 | Uprosertib |
| MVP_top60_DNA_seed | AKT3 | 0.423141 | THCA | ASH-3 | Uprosertib |
| MVP_top60_DNA_seed | RET | 0.425459 | THCA | CGTH-W-1 | Motesanib |
| MVP_top60_DNA_seed | CHEK2 | 0.430461 | THCA | CAL-62 | MK-8776 |
| MVP_top60_DNA_seed | KRAS | 0.432497 | THCA | 8505C | KRAS (G12C) Inhibitor-12 |
| MVP_top60_DNA_seed | AKT1 | 0.434285 | THCA | IHH-4 | GSK2110183B |
| MVP_top60_DNA_seed | AKT2 | 0.434285 | THCA | IHH-4 | GSK2110183B |
| MVP_top60_DNA_seed | AKT3 | 0.434285 | THCA | IHH-4 | GSK2110183B |
| MVP_top60_DNA_seed | MDM2 | 0.43624 | THCA | BHT-101 | Nutlin-3a (-) |
| MVP_top60_DNA_seed | HDAC1 | 0.451747 | THCA | BHT-101 | Romidepsin |
| MVP_top60_DNA_seed | CHEK2 | 0.45675 | THCA | 8505C | MK-8776 |
| MVP_top60_DNA_seed | USP7 | 0.459014 | THCA | 8505C | P22077 |
| MVP_top60_DNA_seed | AKT1 | 0.459115 | THCA | BHT-101 | Uprosertib |
| MVP_top60_DNA_seed | AKT2 | 0.459115 | THCA | BHT-101 | Uprosertib |
| MVP_top60_DNA_seed | AKT3 | 0.459115 | THCA | BHT-101 | Uprosertib |
| MVP_top60_DNA_seed | GSK3B | 0.460811 | THCA | ASH-3 | SB216763 |
| MVP_top60_DNA_seed | HDAC1 | 0.467791 | THCA | B-CPAP | Entinostat |
| MVP_top60_DNA_seed | MAPK1 | 0.47052 | THCA | ML-1 | Ulixertinib |
| MVP_top60_DNA_seed | MAPK3 | 0.47052 | THCA | ML-1 | Ulixertinib |
| MVP_top60_DNA_seed | AKT1 | 0.475187 | THCA | BHT-101 | AZD5363 |
| MVP_top60_DNA_seed | AKT2 | 0.475187 | THCA | BHT-101 | AZD5363 |
| MVP_top60_DNA_seed | AKT3 | 0.475187 | THCA | BHT-101 | AZD5363 |
| MVP_top60_DNA_seed | MAPK1 | 0.478319 | THCA | TT2609-C02 | Ulixertinib |
| MVP_top60_DNA_seed | MAPK3 | 0.478319 | THCA | TT2609-C02 | Ulixertinib |
| MVP_top60_DNA_seed | ATM | 0.494355 | THCA | CAL-62 | KU-55933 |

| | | | | | |
|---|---|---|---|---|---|
| MVP_top60_DNA_seed | AKT1 | 0.512286 | THCA | ASH-3 | MK-2206 |
| MVP_top60_DNA_seed | AKT2 | 0.512286 | THCA | ASH-3 | MK-2206 |
| MVP_top60_DNA_seed | AKT1 | 0.517844 | THCA | TT | MK-2206 |
| MVP_top60_DNA_seed | AKT2 | 0.517844 | THCA | TT | MK-2206 |
| MVP_top60_DNA_seed | CHEK2 | 0.519508 | THCA | ML-1 | AZD7762 |
| MVP_top60_DNA_seed | MDM2 | 0.526053 | THCA | FTC-133 | Serdemetan |
| MVP_top60_DNA_seed | BRAF | 0.543479 | THCA | ASH-3 | Dabrafenib |
| MVP_top60_DNA_seed | MAPK1 | 0.543589 | THCA | ML-1 | SCH772984 |
| MVP_top60_DNA_seed | MAPK3 | 0.543589 | THCA | ML-1 | SCH772984 |
| MVP_top60_DNA_seed | MDM2 | 0.547068 | THCA | ASH-3 | Serdemetan |
| MVP_top60_DNA_seed | AKT1 | 0.551077 | THCA | HTC-C3 | Uprosertib |
| MVP_top60_DNA_seed | AKT2 | 0.551077 | THCA | HTC-C3 | Uprosertib |
| MVP_top60_DNA_seed | AKT3 | 0.551077 | THCA | HTC-C3 | Uprosertib |
| MVP_top60_DNA_seed | KRAS | 0.55186 | THCA | HTC-C3 | KRAS (G12C) Inhibitor-12 |
| MVP_top60_DNA_seed | HDAC1 | 0.553659 | THCA | RO82-W-1 | Romidepsin |
| MVP_top60_DNA_seed | CHEK2 | 0.556117 | THCA | ASH-3 | AZD7762 |
| MVP_top60_DNA_seed | ATR | 0.57032 | THCA | ASH-3 | AZD6738 |
| MVP_top60_DNA_seed | HDAC1 | 0.572394 | THCA | ML-1 | PCI-34051 |
| MVP_top60_DNA_seed | BRAF | 0.581894 | THCA | FTC-133 | SB590885 |
| MVP_top60_DNA_seed | AKT1 | 0.587831 | THCA | ML-1 | AT13148 |
| MVP_top60_DNA_seed | BRAF | 0.605339 | THCA | 8305C | SB590885 |
| MVP_top60_DNA_seed | USP7 | 0.605434 | THCA | IHH-4 | P22077 |
| MVP_top60_DNA_seed | AKT1 | 0.605455 | THCA | HTC-C3 | Afuresertib |
| MVP_top60_DNA_seed | AKT2 | 0.605455 | THCA | HTC-C3 | Afuresertib |
| MVP_top60_DNA_seed | AKT3 | 0.605455 | THCA | HTC-C3 | Afuresertib |
| MVP_top60_DNA_seed | MDM2 | 0.606245 | THCA | TT | Nutlin-3a (-) |
| MVP_top60_DNA_seed | AKT1 | 0.611823 | THCA | B-CPAP | Ipatasertib |
| MVP_top60_DNA_seed | AKT3 | 0.611823 | THCA | B-CPAP | Ipatasertib |
| MVP_top60_DNA_seed | BRAF | 0.615266 | THCA | HTC-C3 | SB590885 |
| MVP_top60_DNA_seed | CHEK2 | 0.619692 | THCA | 8305C | MK-8776 |
| MVP_top60_DNA_seed | AKT1 | 0.622402 | THCA | IHH-4 | AT13148 |
| MVP_top60_DNA_seed | MAPK1 | 0.624351 | THCA | CGTH-W-1 | Ulixertinib |
| MVP_top60_DNA_seed | MAPK3 | 0.624351 | THCA | CGTH-W-1 | Ulixertinib |
| MVP_top60_DNA_seed | CHEK2 | 0.626977 | THCA | ASH-3 | MK-8776 |
| MVP_top60_DNA_seed | AKT1 | 0.632152 | THCA | IHH-4 | AZD5363 |
| MVP_top60_DNA_seed | AKT2 | 0.632152 | THCA | IHH-4 | AZD5363 |
| MVP_top60_DNA_seed | AKT3 | 0.632152 | THCA | IHH-4 | AZD5363 |
| MVP_top60_DNA_seed | AKT1 | 0.63469 | THCA | ASH-3 | Uprosertib |
| MVP_top60_DNA_seed | AKT2 | 0.63469 | THCA | ASH-3 | Uprosertib |
| MVP_top60_DNA_seed | AKT3 | 0.63469 | THCA | ASH-3 | Uprosertib |
| MVP_top60_DNA_seed | MAPK1 | 0.636907 | THCA | CGTH-W-1 | Ulixertinib |

| | | | | | |
|---|---|---|---|---|---|
| MVP_top60_DNA_seed | MAPK3 | 0.636907 | THCA | CGTH-W-1 | Ulixertinib |
| MVP_top60_DNA_seed | USP7 | 0.656243 | THCA | CGTH-W-1 | P22077 |
| MVP_top60_DNA_seed | AKT1 | 0.667776 | THCA | 8305C | AZD5363 |
| MVP_top60_DNA_seed | AKT2 | 0.667776 | THCA | 8305C | AZD5363 |
| MVP_top60_DNA_seed | AKT3 | 0.667776 | THCA | 8305C | AZD5363 |
| MVP_top60_DNA_seed | MDM2 | 0.678678 | THCA | CGTH-W-1 | Serdemetan |
| MVP_top60_DNA_seed | GSK3B | 0.680266 | THCA | ML-1 | CHIR-99021 |
| MVP_top60_DNA_seed | HDAC1 | 0.681284 | THCA | IHH-4 | PCI-34051 |
| MVP_top60_DNA_seed | AKT1 | 0.682358 | THCA | IHH-4 | Uprosertib |
| MVP_top60_DNA_seed | AKT2 | 0.682358 | THCA | IHH-4 | Uprosertib |
| MVP_top60_DNA_seed | AKT3 | 0.682358 | THCA | IHH-4 | Uprosertib |
| MVP_top60_DNA_seed | MDM2 | 0.688723 | THCA | B-CPAP | Nutlin-3a (-) |
| MVP_top60_DNA_seed | MDM2 | 0.693233 | THCA | 8505C | Nutlin-3a (-) |
| MVP_top60_DNA_seed | ATM | 0.695164 | THCA | FTC-133 | KU-55933 |
| MVP_top60_DNA_seed | KRAS | 0.703899 | THCA | 8305C | KRAS (G12C) Inhibitor-12 |
| MVP_top60_DNA_seed | MDM2 | 0.705245 | THCA | HTC-C3 | Nutlin-3a (-) |
| MVP_top60_DNA_seed | CHEK2 | 0.711308 | THCA | 8505C | AZD7762 |
| MVP_top60_DNA_seed | ATR | 0.712355 | THCA | B-CPAP | VE-822 |
| MVP_top60_DNA_seed | MAPK1 | 0.712598 | THCA | 8305C | Ulixertinib |
| MVP_top60_DNA_seed | MAPK3 | 0.712598 | THCA | 8305C | Ulixertinib |
| MVP_top60_DNA_seed | AKT1 | 0.71606 | THCA | BHT-101 | GSK2110183B |
| MVP_top60_DNA_seed | AKT2 | 0.71606 | THCA | BHT-101 | GSK2110183B |
| MVP_top60_DNA_seed | AKT3 | 0.71606 | THCA | BHT-101 | GSK2110183B |
| MVP_top60_DNA_seed | MAPK1 | 0.716426 | THCA | TT | Ulixertinib |
| MVP_top60_DNA_seed | MAPK3 | 0.716426 | THCA | TT | Ulixertinib |
| MVP_top60_DNA_seed | ATR | 0.716658 | THCA | ML-1 | AZD6738 |
| MVP_top60_DNA_seed | AKT1 | 0.729228 | THCA | CAL-62 | AZD5363 |
| MVP_top60_DNA_seed | AKT2 | 0.729228 | THCA | CAL-62 | AZD5363 |
| MVP_top60_DNA_seed | AKT3 | 0.729228 | THCA | CAL-62 | AZD5363 |
| MVP_top60_DNA_seed | BRAF | 0.730547 | THCA | CAL-62 | Dabrafenib |
| MVP_top60_DNA_seed | CHEK2 | 0.733719 | THCA | TT2609-C02 | AZD7762 |
| MVP_top60_DNA_seed | RET | 0.739204 | THCA | 8505C | Motesanib |
| MVP_top60_DNA_seed | ATM | 0.756658 | THCA | B-CPAP | KU-55933 |
| MVP_top60_DNA_seed | AKT1 | 0.763665 | THCA | CAL-62 | GSK2110183B |
| MVP_top60_DNA_seed | AKT2 | 0.763665 | THCA | CAL-62 | GSK2110183B |
| MVP_top60_DNA_seed | AKT3 | 0.763665 | THCA | CAL-62 | GSK2110183B |
| MVP_top60_DNA_seed | MDM2 | 0.766016 | THCA | 8305C | Nutlin-3a (-) |
| MVP_top60_DNA_seed | AKT1 | 0.776184 | THCA | HTC-C3 | AZD5363 |
| MVP_top60_DNA_seed | AKT2 | 0.776184 | THCA | HTC-C3 | AZD5363 |
| MVP_top60_DNA_seed | AKT3 | 0.776184 | THCA | HTC-C3 | AZD5363 |
| MVP_top60_DNA_seed | MDM2 | 0.782689 | THCA | CGTH-W-1 | Nutlin-3a (-) |

| | | | | | |
|---|---|---|---|---|---|
| MVP_top60_DNA_seed | MDM2 | 0.78695 | THCA | 8305C | Serdemetan |
| MVP_top60_DNA_seed | MDM2 | 0.793196 | THCA | ASH-3 | Nutlin-3a (-) |
| MVP_top60_DNA_seed | HDAC1 | 0.794174 | THCA | ASH-3 | PCI-34051 |
| MVP_top60_DNA_seed | ATR | 0.804664 | THCA | ML-1 | VE821 |
| MVP_top60_DNA_seed | AKT1 | 0.809369 | THCA | 8505C | Afuresertib |
| MVP_top60_DNA_seed | AKT2 | 0.809369 | THCA | 8505C | Afuresertib |
| MVP_top60_DNA_seed | AKT3 | 0.809369 | THCA | 8505C | Afuresertib |
| MVP_top60_DNA_seed | MDM2 | 0.812724 | THCA | B-CPAP | Serdemetan |
| MVP_top60_DNA_seed | ATR | 0.815436 | THCA | BHT-101 | VE821 |
| MVP_top60_DNA_seed | AKT1 | 0.820935 | THCA | HTC-C3 | MK-2206 |
| MVP_top60_DNA_seed | AKT2 | 0.820935 | THCA | HTC-C3 | MK-2206 |
| MVP_top60_DNA_seed | MDM2 | 0.825721 | THCA | CAL-62 | Nutlin-3a (-) |
| MVP_top60_DNA_seed | AKT1 | 0.833195 | THCA | TT | AZD5363 |
| MVP_top60_DNA_seed | AKT2 | 0.833195 | THCA | TT | AZD5363 |
| MVP_top60_DNA_seed | AKT3 | 0.833195 | THCA | TT | AZD5363 |
| MVP_top60_DNA_seed | BRAF | 0.844063 | THCA | TT2609-C02 | SB590885 |
| MVP_top60_DNA_seed | BRAF | 0.846021 | THCA | CGTH-W-1 | Dabrafenib |
| MVP_top60_DNA_seed | GSK3B | 0.848699 | THCA | 8505C | SB216763 |
| MVP_top60_DNA_seed | CHEK2 | 0.855055 | THCA | BHT-101 | AZD7762 |
| MVP_top60_DNA_seed | MDM2 | 0.860982 | THCA | ML-1 | Nutlin-3a (-) |
| MVP_top60_DNA_seed | KRAS | 0.863473 | THCA | FTC-133 | KRAS (G12C) Inhibitor-12 |
| MVP_top60_DNA_seed | AKT1 | 0.865932 | THCA | HTC-C3 | GSK2110183B |
| MVP_top60_DNA_seed | AKT2 | 0.865932 | THCA | HTC-C3 | GSK2110183B |
| MVP_top60_DNA_seed | AKT3 | 0.865932 | THCA | HTC-C3 | GSK2110183B |
| MVP_top60_DNA_seed | ATR | 0.876637 | THCA | 8505C | AZD6738 |
| MVP_top60_DNA_seed | MAPK1 | 0.877 | THCA | ML-1 | VX-11e |
| MVP_top60_DNA_seed | HDAC1 | 0.88349 | THCA | TT2609-C02 | Entinostat |
| MVP_top60_DNA_seed | MAPK1 | 0.886116 | THCA | FTC-133 | Ulixertinib |
| MVP_top60_DNA_seed | MAPK3 | 0.886116 | THCA | FTC-133 | Ulixertinib |
| MVP_top60_DNA_seed | AKT1 | 0.887738 | THCA | HTC-C3 | Uprosertib |
| MVP_top60_DNA_seed | AKT2 | 0.887738 | THCA | HTC-C3 | Uprosertib |
| MVP_top60_DNA_seed | AKT3 | 0.887738 | THCA | HTC-C3 | Uprosertib |
| MVP_top60_DNA_seed | RET | 0.90392 | THCA | 8305C | Motesanib |
| MVP_top60_DNA_seed | ATR | 0.910131 | THCA | 8305C | VE-822 |
| MVP_top60_DNA_seed | AKT1 | 0.914714 | THCA | TT2609-C02 | AT13148 |
| MVP_top60_DNA_seed | MDM2 | 0.915184 | THCA | HTC-C3 | Serdemetan |
| MVP_top60_DNA_seed | BRAF | 0.919604 | THCA | 8505C | SB590885 |
| MVP_top60_DNA_seed | AKT1 | 0.927061 | THCA | 8505C | Uprosertib |
| MVP_top60_DNA_seed | AKT2 | 0.927061 | THCA | 8505C | Uprosertib |
| MVP_top60_DNA_seed | AKT3 | 0.927061 | THCA | 8505C | Uprosertib |
| MVP_top60_DNA_seed | AKT1 | 0.927763 | THCA | HTC-C3 | AT13148 |

| | | | | | |
|---|---|---|---|---|---|
| MVP_top60_DNA_seed | MAPK1 | 0.931795 | THCA | TT | SCH772984 |
| MVP_top60_DNA_seed | MAPK3 | 0.931795 | THCA | TT | SCH772984 |
| MVP_top60_DNA_seed | ATR | 0.932113 | THCA | TT | VE821 |
| MVP_top60_DNA_seed | BRAF | 0.939749 | THCA | TT2609-C02 | Dabrafenib |
| MVP_top60_DNA_seed | AKT1 | 0.955885 | THCA | CAL-62 | Uprosertib |
| MVP_top60_DNA_seed | AKT2 | 0.955885 | THCA | CAL-62 | Uprosertib |
| MVP_top60_DNA_seed | AKT3 | 0.955885 | THCA | CAL-62 | Uprosertib |
| MVP_top60_DNA_seed | CHEK2 | 0.959975 | THCA | BHT-101 | MK-8776 |
| MVP_top60_DNA_seed | AKT1 | 0.980526 | THCA | CGTH-W-1 | MK-2206 |
| MVP_top60_DNA_seed | AKT2 | 0.980526 | THCA | CGTH-W-1 | MK-2206 |
| MVP_top60_DNA_seed | ATR | 0.98055 | THCA | ASH-3 | VE-822 |
| MVP_top60_DNA_seed | ATR | 0.995768 | THCA | CAL-62 | VE-822 |
| MVP_top60_DNA_seed | GSK3B | 0.998085 | THCA | 8305C | SB216763 |
| MVP_top60_DNA_seed | MAPK1 | 1.002336 | THCA | 8305C | Ulixertinib |
| MVP_top60_DNA_seed | MAPK3 | 1.002336 | THCA | 8305C | Ulixertinib |
| MVP_top60_DNA_seed | HDAC1 | 1.003123 | THCA | CAL-62 | PCI-34051 |
| MVP_top60_DNA_seed | BRAF | 1.008652 | THCA | TT | Dabrafenib |
| MVP_top60_DNA_seed | AKT1 | 1.042761 | THCA | CGTH-W-1 | Uprosertib |
| MVP_top60_DNA_seed | AKT2 | 1.042761 | THCA | CGTH-W-1 | Uprosertib |
| MVP_top60_DNA_seed | AKT3 | 1.042761 | THCA | CGTH-W-1 | Uprosertib |
| MVP_top60_DNA_seed | AKT1 | 1.047035 | THCA | B-CPAP | Uprosertib |
| MVP_top60_DNA_seed | AKT2 | 1.047035 | THCA | B-CPAP | Uprosertib |
| MVP_top60_DNA_seed | AKT3 | 1.047035 | THCA | B-CPAP | Uprosertib |
| MVP_top60_DNA_seed | AKT1 | 1.057772 | THCA | TT | Afuresertib |
| MVP_top60_DNA_seed | AKT2 | 1.057772 | THCA | TT | Afuresertib |
| MVP_top60_DNA_seed | AKT3 | 1.057772 | THCA | TT | Afuresertib |
| MVP_top60_DNA_seed | CHEK2 | 1.062999 | THCA | K5 | MK-8776 |
| MVP_top60_DNA_seed | ATR | 1.066963 | THCA | ASH-3 | VE821 |
| MVP_top60_DNA_seed | BRAF | 1.073604 | THCA | ASH-3 | PLX-4720 |
| MVP_top60_DNA_seed | CHEK2 | 1.080986 | THCA | B-CPAP | MK-8776 |
| MVP_top60_DNA_seed | AKT1 | 1.083041 | THCA | 8505C | MK-2206 |
| MVP_top60_DNA_seed | AKT2 | 1.083041 | THCA | 8505C | MK-2206 |
| MVP_top60_DNA_seed | AKT1 | 1.100716 | THCA | CGTH-W-1 | Afuresertib |
| MVP_top60_DNA_seed | AKT2 | 1.100716 | THCA | CGTH-W-1 | Afuresertib |
| MVP_top60_DNA_seed | AKT3 | 1.100716 | THCA | CGTH-W-1 | Afuresertib |
| MVP_top60_DNA_seed | HDAC1 | 1.103791 | THCA | CAL-62 | Entinostat |
| MVP_top60_DNA_seed | AKT1 | 1.120207 | THCA | 8505C | Ipatasertib |
| MVP_top60_DNA_seed | AKT3 | 1.120207 | THCA | 8505C | Ipatasertib |
| MVP_top60_DNA_seed | CHEK2 | 1.120942 | THCA | TT2609-C02 | MK-8776 |
| MVP_top60_DNA_seed | AKT1 | 1.132721 | THCA | CAL-62 | MK-2206 |
| MVP_top60_DNA_seed | AKT2 | 1.132721 | THCA | CAL-62 | MK-2206 |

| | | | | | |
|---|---|---|---|---|---|
| MVP_top60_DNA_seed | ATR | 1.153743 | THCA | CGTH-W-1 | VE821 |
| MVP_top60_DNA_seed | ATM | 1.184934 | THCA | 8505C | KU-55933 |
| MVP_top60_DNA_seed | AKT1 | 1.186441 | THCA | 8505C | AZD5363 |
| MVP_top60_DNA_seed | AKT2 | 1.186441 | THCA | 8505C | AZD5363 |
| MVP_top60_DNA_seed | AKT3 | 1.186441 | THCA | 8505C | AZD5363 |
| MVP_top60_DNA_seed | RET | 1.194374 | THCA | ML-1 | Motesanib |
| MVP_top60_DNA_seed | AKT1 | 1.201658 | THCA | 8505C | GSK2110183B |
| MVP_top60_DNA_seed | AKT2 | 1.201658 | THCA | 8505C | GSK2110183B |
| MVP_top60_DNA_seed | AKT3 | 1.201658 | THCA | 8505C | GSK2110183B |
| MVP_top60_DNA_seed | AKT1 | 1.211165 | THCA | HTC-C3 | Ipatasertib |
| MVP_top60_DNA_seed | AKT3 | 1.211165 | THCA | HTC-C3 | Ipatasertib |
| MVP_top60_DNA_seed | MDM2 | 1.211927 | THCA | CAL-62 | Serdemetan |
| MVP_top60_DNA_seed | AKT1 | 1.212533 | THCA | B-CPAP | GSK2110183B |
| MVP_top60_DNA_seed | AKT2 | 1.212533 | THCA | B-CPAP | GSK2110183B |
| MVP_top60_DNA_seed | AKT3 | 1.212533 | THCA | B-CPAP | GSK2110183B |
| MVP_top60_DNA_seed | AKT1 | 1.214275 | THCA | TT2609-C02 | Afuresertib |
| MVP_top60_DNA_seed | AKT2 | 1.214275 | THCA | TT2609-C02 | Afuresertib |
| MVP_top60_DNA_seed | AKT3 | 1.214275 | THCA | TT2609-C02 | Afuresertib |
| MVP_top60_DNA_seed | BRAF | 1.232725 | THCA | CAL-62 | PLX-4720 |
| MVP_top60_DNA_seed | KRAS | 1.239146 | THCA | B-CPAP | KRAS (G12C) Inhibitor-12 |
| MVP_top60_DNA_seed | HDAC1 | 1.267906 | THCA | 8305C | Entinostat |
| MVP_top60_DNA_seed | AKT1 | 1.284222 | THCA | CAL-62 | Uprosertib |
| MVP_top60_DNA_seed | AKT2 | 1.284222 | THCA | CAL-62 | Uprosertib |
| MVP_top60_DNA_seed | AKT3 | 1.284222 | THCA | CAL-62 | Uprosertib |
| MVP_top60_DNA_seed | HDAC1 | 1.295316 | THCA | ML-1 | Entinostat |
| MVP_top60_DNA_seed | HDAC1 | 1.295599 | THCA | TT2609-C02 | Romidepsin |
| MVP_top60_DNA_seed | AKT1 | 1.323573 | THCA | CGTH-W-1 | AZD5363 |
| MVP_top60_DNA_seed | AKT2 | 1.323573 | THCA | CGTH-W-1 | AZD5363 |
| MVP_top60_DNA_seed | AKT3 | 1.323573 | THCA | CGTH-W-1 | AZD5363 |
| MVP_top60_DNA_seed | AKT1 | 1.327396 | THCA | 8305C | GSK2110183B |
| MVP_top60_DNA_seed | AKT2 | 1.327396 | THCA | 8305C | GSK2110183B |
| MVP_top60_DNA_seed | AKT3 | 1.327396 | THCA | 8305C | GSK2110183B |
| MVP_top60_DNA_seed | BRAF | 1.327751 | THCA | ML-1 | SB590885 |
| MVP_top60_DNA_seed | AKT1 | 1.342937 | THCA | RO82-W-1 | Uprosertib |
| MVP_top60_DNA_seed | AKT2 | 1.342937 | THCA | RO82-W-1 | Uprosertib |
| MVP_top60_DNA_seed | AKT3 | 1.342937 | THCA | RO82-W-1 | Uprosertib |
| MVP_top60_DNA_seed | GSK3B | 1.382225 | THCA | ML-1 | SB216763 |
| MVP_top60_DNA_seed | USP7 | 1.425846 | THCA | 8305C | P22077 |
| MVP_top60_DNA_seed | AKT1 | 1.444889 | THCA | CGTH-W-1 | Ipatasertib |
| MVP_top60_DNA_seed | AKT3 | 1.444889 | THCA | CGTH-W-1 | Ipatasertib |
| MVP_top60_DNA_seed | ATR | 1.447427 | THCA | ML-1 | VE-822 |

| | | | | | |
|---|---|---|---|---|---|
| MVP_top60_DNA_seed | ATR | 1.463271 | THCA | 8505C | VE821 |
| MVP_top60_DNA_seed | AKT1 | 1.464269 | THCA | RO82-W-1 | AZD5363 |
| MVP_top60_DNA_seed | AKT2 | 1.464269 | THCA | RO82-W-1 | AZD5363 |
| MVP_top60_DNA_seed | AKT3 | 1.464269 | THCA | RO82-W-1 | AZD5363 |
| MVP_top60_DNA_seed | CHEK2 | 1.468932 | THCA | RO82-W-1 | MK-8776 |
| MVP_top60_DNA_seed | HDAC1 | 1.47177 | THCA | TT2609-C02 | PCI-34051 |
| MVP_top60_DNA_seed | CHEK2 | 1.472233 | THCA | TT | AZD7762 |
| MVP_top60_DNA_seed | MDM2 | 1.477213 | THCA | TT2609-C02 | Nutlin-3a (-) |
| MVP_top60_DNA_seed | AKT1 | 1.483532 | THCA | CGTH-W-1 | AT13148 |
| MVP_top60_DNA_seed | BRAF | 1.50131 | THCA | TT | SB590885 |
| MVP_top60_DNA_seed | AKT1 | 1.532642 | THCA | 8305C | Uprosertib |
| MVP_top60_DNA_seed | AKT2 | 1.532642 | THCA | 8305C | Uprosertib |
| MVP_top60_DNA_seed | AKT3 | 1.532642 | THCA | 8305C | Uprosertib |
| MVP_top60_DNA_seed | MDM2 | 1.53329 | THCA | RO82-W-1 | Nutlin-3a (-) |
| MVP_top60_DNA_seed | MAPK1 | 1.53516 | THCA | FTC-133 | SCH772984 |
| MVP_top60_DNA_seed | MAPK3 | 1.53516 | THCA | FTC-133 | SCH772984 |
| MVP_top60_DNA_seed | AKT1 | 1.552267 | THCA | TT | Uprosertib |
| MVP_top60_DNA_seed | AKT2 | 1.552267 | THCA | TT | Uprosertib |
| MVP_top60_DNA_seed | AKT3 | 1.552267 | THCA | TT | Uprosertib |
| MVP_top60_DNA_seed | AKT1 | 1.556382 | THCA | TT | Ipatasertib |
| MVP_top60_DNA_seed | AKT3 | 1.556382 | THCA | TT | Ipatasertib |
| MVP_top60_DNA_seed | ATM | 1.574088 | THCA | ML-1 | KU-55933 |
| MVP_top60_DNA_seed | RET | 1.5804 | THCA | TT2609-C02 | Motesanib |
| MVP_top60_DNA_seed | ATR | 1.592781 | THCA | CGTH-W-1 | AZD6738 |
| MVP_top60_DNA_seed | USP7 | 1.604926 | THCA | ML-1 | P22077 |
| MVP_top60_DNA_seed | AKT1 | 1.605319 | THCA | ML-1 | Ipatasertib |
| MVP_top60_DNA_seed | AKT3 | 1.605319 | THCA | ML-1 | Ipatasertib |
| MVP_top60_DNA_seed | AKT1 | 1.606235 | THCA | TT2609-C02 | GSK2110183B |
| MVP_top60_DNA_seed | AKT2 | 1.606235 | THCA | TT2609-C02 | GSK2110183B |
| MVP_top60_DNA_seed | AKT3 | 1.606235 | THCA | TT2609-C02 | GSK2110183B |
| MVP_top60_DNA_seed | BRAF | 1.632561 | THCA | RO82-W-1 | PLX-4720 |
| MVP_top60_DNA_seed | AKT1 | 1.634191 | THCA | 8505C | Uprosertib |
| MVP_top60_DNA_seed | AKT2 | 1.634191 | THCA | 8505C | Uprosertib |
| MVP_top60_DNA_seed | AKT3 | 1.634191 | THCA | 8505C | Uprosertib |
| MVP_top60_DNA_seed | ATM | 1.660709 | THCA | TT2609-C02 | KU-55933 |
| MVP_top60_DNA_seed | AKT1 | 1.674566 | THCA | ML-1 | AZD5363 |
| MVP_top60_DNA_seed | AKT2 | 1.674566 | THCA | ML-1 | AZD5363 |
| MVP_top60_DNA_seed | AKT3 | 1.674566 | THCA | ML-1 | AZD5363 |
| MVP_top60_DNA_seed | ATR | 1.682319 | THCA | TT2609-C02 | VE-822 |
| MVP_top60_DNA_seed | MDM2 | 1.703892 | THCA | TT2609-C02 | Serdemetan |
| MVP_top60_DNA_seed | AKT1 | 1.713256 | THCA | TT2609-C02 | Ipatasertib |

| | | | | | |
|---|---|---|---|---|---|
| MVP_top60_DNA_seed | AKT3 | 1.713256 | THCA | TT2609-C02 | Ipatasertib |
| MVP_top60_DNA_seed | MDM2 | 1.720719 | THCA | 8505C | Serdemetan |
| MVP_top60_DNA_seed | AKT1 | 1.758311 | THCA | 8505C | AT13148 |
| MVP_top60_DNA_seed | AKT1 | 1.766555 | THCA | 8305C | Uprosertib |
| MVP_top60_DNA_seed | AKT2 | 1.766555 | THCA | 8305C | Uprosertib |
| MVP_top60_DNA_seed | AKT3 | 1.766555 | THCA | 8305C | Uprosertib |
| MVP_top60_DNA_seed | AKT1 | 1.802675 | THCA | TT2609-C02 | Uprosertib |
| MVP_top60_DNA_seed | AKT2 | 1.802675 | THCA | TT2609-C02 | Uprosertib |
| MVP_top60_DNA_seed | AKT3 | 1.802675 | THCA | TT2609-C02 | Uprosertib |
| MVP_top60_DNA_seed | AKT1 | 1.812247 | THCA | ML-1 | MK-2206 |
| MVP_top60_DNA_seed | AKT2 | 1.812247 | THCA | ML-1 | MK-2206 |
| MVP_top60_DNA_seed | RET | 1.819942 | THCA | TT | Motesanib |
| MVP_top60_DNA_seed | HDAC1 | 1.835974 | THCA | RO82-W-1 | PCI-34051 |
| MVP_top60_DNA_seed | ATR | 1.86262 | THCA | TT2609-C02 | AZD6738 |
| MVP_top60_DNA_seed | ATR | 1.864158 | THCA | TT2609-C02 | VE821 |
| MVP_top60_DNA_seed | ATM | 1.880356 | THCA | RO82-W-1 | KU-55933 |
| MVP_top60_DNA_seed | USP7 | 1.888409 | THCA | TT | P22077 |
| MVP_top60_DNA_seed | KRAS | 1.913729 | THCA | ML-1 | KRAS (G12C) Inhibitor-12 |
| MVP_top60_DNA_seed | BRAF | 1.917072 | THCA | TT2609-C02 | PLX-4720 |
| MVP_top60_DNA_seed | AKT1 | 1.933148 | THCA | RO82-W-1 | AT13148 |
| MVP_top60_DNA_seed | AKT1 | 1.960196 | THCA | RO82-W-1 | GSK2110183B |
| MVP_top60_DNA_seed | AKT2 | 1.960196 | THCA | RO82-W-1 | GSK2110183B |
| MVP_top60_DNA_seed | AKT3 | 1.960196 | THCA | RO82-W-1 | GSK2110183B |
| MVP_top60_DNA_seed | AKT1 | 1.96079 | THCA | RO82-W-1 | MK-2206 |
| MVP_top60_DNA_seed | AKT2 | 1.96079 | THCA | RO82-W-1 | MK-2206 |
| MVP_top60_DNA_seed | AKT1 | 1.993555 | THCA | RO82-W-1 | Afuresertib |
| MVP_top60_DNA_seed | AKT2 | 1.993555 | THCA | RO82-W-1 | Afuresertib |
| MVP_top60_DNA_seed | AKT3 | 1.993555 | THCA | RO82-W-1 | Afuresertib |
| MVP_top60_DNA_seed | BRAF | 2.011685 | THCA | TT | PLX-4720 |
| MVP_top60_DNA_seed | AKT1 | 2.025232 | THCA | TT2609-C02 | AZD5363 |
| MVP_top60_DNA_seed | AKT2 | 2.025232 | THCA | TT2609-C02 | AZD5363 |
| MVP_top60_DNA_seed | AKT3 | 2.025232 | THCA | TT2609-C02 | AZD5363 |
| MVP_top60_DNA_seed | BRAF | 2.030313 | THCA | RO82-W-1 | Dabrafenib |
| MVP_top60_DNA_seed | GSK3B | 2.07841 | THCA | TT | SB216763 |
| MVP_top60_DNA_seed | ATM | 2.159676 | THCA | TT | KU-55933 |
| MVP_top60_DNA_seed | KRAS | 2.161661 | THCA | TT2609-C02 | KRAS (G12C) Inhibitor-12 |
| MVP_top60_DNA_seed | MDM2 | 2.187688 | THCA | RO82-W-1 | Serdemetan |
| MVP_top60_DNA_seed | MAPK1 | 2.217249 | THCA | RO82-W-1 | SCH772984 |
| MVP_top60_DNA_seed | MAPK3 | 2.217249 | THCA | RO82-W-1 | SCH772984 |
| MVP_top60_DNA_seed | AKT1 | 2.230667 | THCA | TT2609-C02 | Uprosertib |
| MVP_top60_DNA_seed | AKT2 | 2.230667 | THCA | TT2609-C02 | Uprosertib |

| | | | | | |
|---|---|---|---|---|---|
| MVP_top60_DNA_seed | AKT3 | 2.230667 | THCA | TT2609-C02 | Uprosertib |
| MVP_top60_DNA_seed | MAPK1 | 2.240985 | THCA | TT2609-C02 | VX-11e |
| MVP_top60_DNA_seed | AKT1 | 2.300894 | THCA | ML-1 | Uprosertib |
| MVP_top60_DNA_seed | AKT2 | 2.300894 | THCA | ML-1 | Uprosertib |
| MVP_top60_DNA_seed | AKT3 | 2.300894 | THCA | ML-1 | Uprosertib |
| MVP_top60_DNA_seed | ATR | 2.303272 | THCA | RO82-W-1 | VE-822 |
| MVP_top60_DNA_seed | BRAF | 2.359152 | THCA | RO82-W-1 | SB590885 |
| MVP_top60_DNA_seed | ATR | 2.360714 | THCA | RO82-W-1 | VE821 |
| MVP_top60_DNA_seed | AKT1 | 2.420424 | THCA | TT2609-C02 | MK-2206 |
| MVP_top60_DNA_seed | AKT2 | 2.420424 | THCA | TT2609-C02 | MK-2206 |
| MVP_top60_DNA_seed | AKT1 | 2.431171 | THCA | TT | AT13148 |
| MVP_top60_DNA_seed | AKT1 | 2.510429 | THCA | RO82-W-1 | Ipatasertib |
| MVP_top60_DNA_seed | AKT3 | 2.510429 | THCA | RO82-W-1 | Ipatasertib |
| MVP_top60_DNA_seed | ATR | 2.514321 | THCA | TT | AZD6738 |
| MVP_top60_DNA_seed | MAPK1 | 2.518446 | THCA | TT | VX-11e |
| MVP_top60_DNA_seed | USP7 | 2.522981 | THCA | RO82-W-1 | P22077 |
| MVP_top60_DNA_seed | RET | 2.530202 | THCA | RO82-W-1 | Motesanib |
| MVP_top60_DNA_seed | CHEK2 | 2.539113 | THCA | RO82-W-1 | AZD7762 |
| MVP_top60_DNA_seed | ATR | 2.629672 | THCA | RO82-W-1 | AZD6738 |
| MVP_top60_DNA_seed | MAPK1 | 2.699379 | THCA | RO82-W-1 | VX-11e |
| MVP_top60_DNA_seed | MAPK1 | 2.723591 | THCA | RO82-W-1 | Ulixertinib |
| MVP_top60_DNA_seed | MAPK3 | 2.723591 | THCA | RO82-W-1 | Ulixertinib |
| MVP_top60_DNA_seed | GSK3B | 2.745417 | THCA | RO82-W-1 | SB216763 |
| MVP_top60_DNA_seed | AKT1 | 2.83111 | THCA | RO82-W-1 | Uprosertib |
| MVP_top60_DNA_seed | AKT2 | 2.83111 | THCA | RO82-W-1 | Uprosertib |
| MVP_top60_DNA_seed | AKT3 | 2.83111 | THCA | RO82-W-1 | Uprosertib |
| MVP_top60_DNA_seed | AKT1 | 2.837742 | THCA | ML-1 | Afuresertib |
| MVP_top60_DNA_seed | AKT2 | 2.837742 | THCA | ML-1 | Afuresertib |
| MVP_top60_DNA_seed | AKT3 | 2.837742 | THCA | ML-1 | Afuresertib |
| MVP_top60_DNA_seed | KRAS | 2.906905 | THCA | RO82-W-1 | KRAS (G12C) Inhibitor-12 |
| MVP_top60_DNA_seed | HDAC1 | 3.081271 | THCA | RO82-W-1 | Entinostat |
| MVP_top60_DNA_seed | FGFR2 | -3.839158 | UCEC | MFE-280 | AZD4547 |
| MVP_top60_DNA_seed | FGFR2 | -3.193966 | UCEC | AN3-CA | PD173074 |
| MVP_top60_DNA_seed | FGFR2 | -3.138837 | UCEC | AN3-CA | AZD4547 |
| MVP_top60_DNA_seed | ESR1 | -2.903364 | UCEC | MFE-280 | Tamoxifen |
| MVP_top60_DNA_seed | FGFR2 | -2.831998 | UCEC | AN3-CA | AZD4547 |
| MVP_top60_DNA_seed | JAK1 | -2.681385 | UCEC | AN3-CA | AZD1480 |
| MVP_top60_DNA_seed | JAK1 | -2.574905 | UCEC | MFE-280 | AZD1480 |
| MVP_top60_DNA_seed | AKT1 | -2.540572 | UCEC | COLO-684 | GSK690693 |
| MVP_top60_DNA_seed | PARP1 | -2.445043 | UCEC | HEC-1 | Veliparib |
| MVP_top60_DNA_seed | PARP1 | -2.354548 | UCEC | SNG-M | Veliparib |

| | | | | | |
|---|---|---|---|---|---|
| MVP_top60_DNA_seed | FGFR2 | -2.286659 | UCEC | MFE-296 | AZD4547 |
| MVP_top60_DNA_seed | EGFR | -2.253594 | UCEC | SNG-M | PF-00299804 |
| MVP_top60_DNA_seed | ERBB2 | -2.253594 | UCEC | SNG-M | PF-00299804 |
| MVP_top60_DNA_seed | CDK2 | -2.167338 | UCEC | SNG-M | AZD5438 |
| MVP_top60_DNA_seed | FGFR2 | -2.095604 | UCEC | MFE-280 | FGFR_3831 |
| MVP_top60_DNA_seed | AKT1 | -2.045663 | UCEC | AN3-CA | AKT inhibitor VIII |
| MVP_top60_DNA_seed | AKT1 | -2.045604 | UCEC | MFE-296 | AKT inhibitor VIII |
| MVP_top60_DNA_seed | PARP1 | -2.018407 | UCEC | SNG-M | PARP_9495 |
| MVP_top60_DNA_seed | FGFR2 | -1.92556 | UCEC | MFE-296 | AZD4547 |
| MVP_top60_DNA_seed | FGFR2 | -1.813677 | UCEC | MFE-296 | PD173074 |
| MVP_top60_DNA_seed | EGFR | -1.791614 | UCEC | SNG-M | AZD8931 |
| MVP_top60_DNA_seed | ERBB2 | -1.791614 | UCEC | SNG-M | AZD8931 |
| MVP_top60_DNA_seed | AKT1 | -1.667957 | UCEC | COLO-684 | AKT inhibitor VIII |
| MVP_top60_DNA_seed | MDM2 | -1.665184 | UCEC | SNG-M | Nutlin-3a (-) |
| MVP_top60_DNA_seed | EGFR | -1.6008 | UCEC | SNG-M | CI-1033 |
| MVP_top60_DNA_seed | ERBB2 | -1.6008 | UCEC | SNG-M | CI-1033 |
| MVP_top60_DNA_seed | EGFR | -1.598851 | UCEC | MFE-319 | Cetuximab |
| MVP_top60_DNA_seed | AKT1 | -1.586867 | UCEC | COLO-684 | MK-2206 |
| MVP_top60_DNA_seed | PARP1 | -1.579741 | UCEC | COLO-684 | Olaparib |
| MVP_top60_DNA_seed | EGFR | -1.560385 | UCEC | SNG-M | Pelitinib |
| MVP_top60_DNA_seed | JAK1 | -1.509739 | UCEC | MFE-296 | AZD1480 |
| MVP_top60_DNA_seed | PIK3CA | -1.502812 | UCEC | MFE-280 | PI3Ka_4409 |
| MVP_top60_DNA_seed | PIK3CD | -1.502812 | UCEC | MFE-280 | PI3Ka_4409 |
| MVP_top60_DNA_seed | HSP90AA1 | -1.450464 | UCEC | SNG-M | CCT-018159 |
| MVP_top60_DNA_seed | PARP1 | -1.417198 | UCEC | SNG-M | Talazoparib |
| MVP_top60_DNA_seed | AKT1 | -1.406257 | UCEC | MFE-296 | BAY AKT1 |
| MVP_top60_DNA_seed | PARP1 | -1.405275 | UCEC | HEC-1 | Rucaparib |
| MVP_top60_DNA_seed | AKT1 | -1.390049 | UCEC | SNG-M | GSK690693 |
| MVP_top60_DNA_seed | HSP90AA1 | -1.362448 | UCEC | SNG-M | Luminespib |
| MVP_top60_DNA_seed | PIK3CA | -1.306172 | UCEC | SNG-M | AZD8835 |
| MVP_top60_DNA_seed | PIK3CD | -1.306172 | UCEC | SNG-M | AZD8835 |
| MVP_top60_DNA_seed | EGFR | -1.305625 | UCEC | AN3-CA | CI-1033 |
| MVP_top60_DNA_seed | ERBB2 | -1.305625 | UCEC | AN3-CA | CI-1033 |
| MVP_top60_DNA_seed | ERBB2 | -1.289139 | UCEC | SNG-M | CP724714 |
| MVP_top60_DNA_seed | EGFR | -1.286658 | UCEC | MFE-319 | Afatinib |
| MVP_top60_DNA_seed | ERBB2 | -1.286658 | UCEC | MFE-319 | Afatinib |
| MVP_top60_DNA_seed | HSP90AA1 | -1.280748 | UCEC | AN3-CA | Tanespimycin |
| MVP_top60_DNA_seed | HSP90AA1 | -1.274147 | UCEC | HEC-1 | Tanespimycin |
| MVP_top60_DNA_seed | PARP1 | -1.269042 | UCEC | SNG-M | Olaparib |
| MVP_top60_DNA_seed | AKT1 | -1.264695 | UCEC | MFE-296 | GSK690693 |
| MVP_top60_DNA_seed | AKT1 | -1.262466 | UCEC | AN3-CA | BAY AKT1 |

| | | | | | |
|---|---|---|---|---|---|
| MVP_top60_DNA_seed | AKT1 | -1.253918 | UCEC | COLO-684 | A-443654 |
| MVP_top60_DNA_seed | AR | -1.242198 | UCEC | SNG-M | AZD3514 |
| MVP_top60_DNA_seed | PARP1 | -1.209435 | UCEC | SNG-M | Olaparib |
| MVP_top60_DNA_seed | EGFR | -1.206584 | UCEC | SNG-M | AST-1306 |
| MVP_top60_DNA_seed | PIK3CA | -1.182536 | UCEC | MFE-280 | AZD8835 |
| MVP_top60_DNA_seed | PIK3CD | -1.182536 | UCEC | MFE-280 | AZD8835 |
| MVP_top60_DNA_seed | EGFR | -1.180624 | UCEC | MFE-319 | Gefitinib |
| MVP_top60_DNA_seed | HSP90AA1 | -1.165754 | UCEC | AN3-CA | Luminespib |
| MVP_top60_DNA_seed | AKT1 | -1.155082 | UCEC | MFE-296 | MK-2206 |
| MVP_top60_DNA_seed | SRC | -1.15495 | UCEC | KLE | WH-4-023 |
| MVP_top60_DNA_seed | FGFR2 | -1.154504 | UCEC | MFE-280 | PD173074 |
| MVP_top60_DNA_seed | ATM | -1.138256 | UCEC | MFE-319 | KU-55933 |
| MVP_top60_DNA_seed | AKT1 | -1.136416 | UCEC | MFE-280 | GSK690693 |
| MVP_top60_DNA_seed | PARP1 | -1.134839 | UCEC | MFE-296 | PARP_9482 |
| MVP_top60_DNA_seed | PIK3CD | -1.124603 | UCEC | COLO-684 | Idelalisib |
| MVP_top60_DNA_seed | PARP1 | -1.081834 | UCEC | SNG-M | PARP_9482 |
| MVP_top60_DNA_seed | CDK2 | -1.075432 | UCEC | MFE-296 | AZD5438 |
| MVP_top60_DNA_seed | MDM2 | -1.071433 | UCEC | MFE-296 | Nutlin-3a (-) |
| MVP_top60_DNA_seed | PIK3CA | -1.065015 | UCEC | SNG-M | PI3Ka_4409 |
| MVP_top60_DNA_seed | PIK3CD | -1.065015 | UCEC | SNG-M | PI3Ka_4409 |
| MVP_top60_DNA_seed | EGFR | -1.062848 | UCEC | KLE | AST-1306 |
| MVP_top60_DNA_seed | AR | -1.050465 | UCEC | MFE-319 | AZD3514 |
| MVP_top60_DNA_seed | PARP1 | -1.04306 | UCEC | HEC-1 | Olaparib |
| MVP_top60_DNA_seed | EGFR | -1.038138 | UCEC | MFE-319 | AZD8931 |
| MVP_top60_DNA_seed | ERBB2 | -1.038138 | UCEC | MFE-319 | AZD8931 |
| MVP_top60_DNA_seed | CDK2 | -1.020549 | UCEC | SNG-M | PHA-793887 |
| MVP_top60_DNA_seed | FGFR2 | -1.007139 | UCEC | AN3-CA | FGFR_3831 |
| MVP_top60_DNA_seed | ATM | -0.98435 | UCEC | SNG-M | CP466722 |
| MVP_top60_DNA_seed | PARP1 | -0.983707 | UCEC | AN3-CA | Olaparib |
| MVP_top60_DNA_seed | ATM | -0.967779 | UCEC | MFE-319 | KU-60019 |
| MVP_top60_DNA_seed | PARP1 | -0.965691 | UCEC | MFE-296 | PARP_0108 |
| MVP_top60_DNA_seed | EGFR | -0.965278 | UCEC | MFE-280 | AST-1306 |
| MVP_top60_DNA_seed | HSP90AA1 | -0.942022 | UCEC | HEC-1 | Elesclomol |
| MVP_top60_DNA_seed | ATM | -0.941115 | UCEC | MFE-319 | CP466722 |
| MVP_top60_DNA_seed | AKT1 | -0.938485 | UCEC | MFE-280 | MK-2206 |
| MVP_top60_DNA_seed | SRC | -0.932293 | UCEC | SNG-M | NVP-BHG712 |
| MVP_top60_DNA_seed | EGFR | -0.922846 | UCEC | MFE-280 | CI-1033 |
| MVP_top60_DNA_seed | ERBB2 | -0.922846 | UCEC | MFE-280 | CI-1033 |
| MVP_top60_DNA_seed | ESR1 | -0.92168 | UCEC | AN3-CA | Tamoxifen |
| MVP_top60_DNA_seed | AKT1 | -0.911742 | UCEC | AN3-CA | MK-2206 |
| MVP_top60_DNA_seed | PARP1 | -0.906418 | UCEC | MFE-296 | Talazoparib |

| | | | | | |
|---|---|---|---|---|---|
| MVP_top60_DNA_seed | EGFR | -0.904027 | UCEC | MFE-319 | Afatinib |
| MVP_top60_DNA_seed | ERBB2 | -0.904027 | UCEC | MFE-319 | Afatinib |
| MVP_top60_DNA_seed | PARP1 | -0.89309 | UCEC | COLO-684 | Veliparib |
| MVP_top60_DNA_seed | CDK2 | -0.88684 | UCEC | AN3-CA | AZD5438 |
| MVP_top60_DNA_seed | AKT1 | -0.883088 | UCEC | MFE-319 | AKT inhibitor VIII |
| MVP_top60_DNA_seed | PARP1 | -0.878196 | UCEC | MFE-296 | Olaparib |
| MVP_top60_DNA_seed | PARP1 | -0.877983 | UCEC | MFE-319 | Rucaparib |
| MVP_top60_DNA_seed | HSP90AA1 | -0.867371 | UCEC | COLO-684 | CCT-018159 |
| MVP_top60_DNA_seed | ESR1 | -0.866579 | UCEC | SNG-M | Fulvestrant |
| MVP_top60_DNA_seed | AKT1 | -0.863762 | UCEC | COLO-684 | BAY AKT1 |
| MVP_top60_DNA_seed | PIK3CD | -0.861903 | UCEC | KLE | AZD8186 |
| MVP_top60_DNA_seed | EGFR | -0.856102 | UCEC | HEC-1 | Cetuximab |
| MVP_top60_DNA_seed | PIK3CD | -0.856077 | UCEC | COLO-684 | AZD8186 |
| MVP_top60_DNA_seed | FGFR2 | -0.837374 | UCEC | MFE-296 | FGFR_3831 |
| MVP_top60_DNA_seed | PIK3CD | -0.829826 | UCEC | COLO-684 | IC-87114 |
| MVP_top60_DNA_seed | HSP90AA1 | -0.820435 | UCEC | COLO-684 | Elesclomol |
| MVP_top60_DNA_seed | JAK1 | -0.801292 | UCEC | SNG-M | AZD1480 |
| MVP_top60_DNA_seed | HSP90AA1 | -0.797967 | UCEC | AN3-CA | Elesclomol |
| MVP_top60_DNA_seed | ERBB2 | -0.796951 | UCEC | MFE-280 | CP724714 |
| MVP_top60_DNA_seed | ESR1 | -0.794746 | UCEC | SNG-M | Tamoxifen |
| MVP_top60_DNA_seed | HSP90AA1 | -0.794576 | UCEC | SNG-M | SNX-2112 |
| MVP_top60_DNA_seed | HSP90AA1 | -0.792124 | UCEC | MFE-296 | Tanespimycin |
| MVP_top60_DNA_seed | AR | -0.786412 | UCEC | SNG-M | Bicalutamide |
| MVP_top60_DNA_seed | AKT1 | -0.778096 | UCEC | KLE | AKT inhibitor VIII |
| MVP_top60_DNA_seed | ATM | -0.757481 | UCEC | HEC-1 | KU-60019 |
| MVP_top60_DNA_seed | AKT1 | -0.750188 | UCEC | HEC-1 | AKT inhibitor VIII |
| MVP_top60_DNA_seed | PIK3CD | -0.746986 | UCEC | AN3-CA | AZD8186 |
| MVP_top60_DNA_seed | EGFR | -0.743217 | UCEC | MFE-280 | PF-00299804 |
| MVP_top60_DNA_seed | ERBB2 | -0.743217 | UCEC | MFE-280 | PF-00299804 |
| MVP_top60_DNA_seed | AR | -0.729614 | UCEC | HEC-1 | AZD3514 |
| MVP_top60_DNA_seed | AR | -0.721671 | UCEC | MFE-319 | Bicalutamide |
| MVP_top60_DNA_seed | SRC | -0.697529 | UCEC | SNG-M | Bosutinib |
| MVP_top60_DNA_seed | PIK3CA | -0.695396 | UCEC | AN3-CA | AZD8835 |
| MVP_top60_DNA_seed | PIK3CD | -0.695396 | UCEC | AN3-CA | AZD8835 |
| MVP_top60_DNA_seed | AKT1 | -0.694469 | UCEC | KLE | A-443654 |
| MVP_top60_DNA_seed | AKT1 | -0.686103 | UCEC | MFE-280 | AKT inhibitor VIII |
| MVP_top60_DNA_seed | EP300 | -0.666443 | UCEC | SNG-M | I-CBP112 |
| MVP_top60_DNA_seed | PIK3CD | -0.659503 | UCEC | SNG-M | AZD8186 |
| MVP_top60_DNA_seed | EGFR | -0.650466 | UCEC | KLE | Pelitinib |
| MVP_top60_DNA_seed | HSP90AA1 | -0.6457 | UCEC | COLO-684 | SNX-2112 |
| MVP_top60_DNA_seed | HDAC1 | -0.638229 | UCEC | SNG-M | JQ12 |

| | | | | | |
|---|---|---|---|---|---|
| MVP_top60_DNA_seed | HDAC2 | -0.638229 | UCEC | SNG-M | JQ12 |
| MVP_top60_DNA_seed | EGFR | -0.622425 | UCEC | MFE-280 | CUDC-101 |
| MVP_top60_DNA_seed | ERBB2 | -0.622425 | UCEC | MFE-280 | CUDC-101 |
| MVP_top60_DNA_seed | EGFR | -0.622161 | UCEC | MFE-280 | Pelitinib |
| MVP_top60_DNA_seed | PARP1 | -0.614401 | UCEC | MFE-280 | PARP_9495 |
| MVP_top60_DNA_seed | ATM | -0.608482 | UCEC | COLO-684 | KU-55933 |
| MVP_top60_DNA_seed | HSP90AA1 | -0.598157 | UCEC | HEC-1 | CCT-018159 |
| MVP_top60_DNA_seed | PARP1 | -0.595015 | UCEC | SNG-M | PARP_0108 |
| MVP_top60_DNA_seed | HSP90AA1 | -0.59084 | UCEC | MFE-296 | Luminespib |
| MVP_top60_DNA_seed | ATM | -0.589341 | UCEC | MFE-296 | KU-60019 |
| MVP_top60_DNA_seed | HSP90AA1 | -0.582916 | UCEC | AN3-CA | CCT-018159 |
| MVP_top60_DNA_seed | JAK1 | -0.566585 | UCEC | SNG-M | Ruxolitinib |
| MVP_top60_DNA_seed | ESR1 | -0.559523 | UCEC | MFE-319 | Tamoxifen |
| MVP_top60_DNA_seed | AR | -0.558596 | UCEC | AN3-CA | Bicalutamide |
| MVP_top60_DNA_seed | PIK3CD | -0.555003 | UCEC | MFE-280 | Idelalisib |
| MVP_top60_DNA_seed | JAK1 | -0.550619 | UCEC | SNG-M | JAK1_3715 |
| MVP_top60_DNA_seed | PARP1 | -0.550549 | UCEC | AN3-CA | Olaparib |
| MVP_top60_DNA_seed | PARP1 | -0.546965 | UCEC | MFE-296 | Veliparib |
| MVP_top60_DNA_seed | ATM | -0.540231 | UCEC | AN3-CA | KU-55933 |
| MVP_top60_DNA_seed | HDAC1 | -0.53721 | UCEC | SNG-M | Dacinostat |
| MVP_top60_DNA_seed | ATM | -0.532342 | UCEC | COLO-684 | CP466722 |
| MVP_top60_DNA_seed | AR | -0.529467 | UCEC | HEC-1 | Bicalutamide |
| MVP_top60_DNA_seed | AKT1 | -0.525196 | UCEC | EN | MK-2206 |
| MVP_top60_DNA_seed | HSP90AA1 | -0.523921 | UCEC | EN | Elesclomol |
| MVP_top60_DNA_seed | ESR1 | -0.517665 | UCEC | HEC-1 | Fulvestrant |
| MVP_top60_DNA_seed | PIK3CD | -0.517357 | UCEC | MFE-280 | AZD8186 |
| MVP_top60_DNA_seed | CDK2 | -0.517238 | UCEC | HEC-1 | AZD5438 |
| MVP_top60_DNA_seed | HDAC1 | -0.517153 | UCEC | AN3-CA | Dacinostat |
| MVP_top60_DNA_seed | HSP90AA1 | -0.510537 | UCEC | MFE-319 | Luminespib |
| MVP_top60_DNA_seed | AKT1 | -0.505071 | UCEC | AN3-CA | GSK690693 |
| MVP_top60_DNA_seed | PARP1 | -0.491024 | UCEC | MFE-319 | Veliparib |
| MVP_top60_DNA_seed | AKT1 | -0.483919 | UCEC | MFE-319 | AKT inhibitor VIII |
| MVP_top60_DNA_seed | EGFR | -0.482508 | UCEC | COLO-684 | Gefitinib |
| MVP_top60_DNA_seed | EGFR | -0.473667 | UCEC | SNG-M | Afatinib |
| MVP_top60_DNA_seed | ERBB2 | -0.473667 | UCEC | SNG-M | Afatinib |
| MVP_top60_DNA_seed | AKT1 | -0.469321 | UCEC | EN | GSK690693 |
| MVP_top60_DNA_seed | HSP90AA1 | -0.466099 | UCEC | MFE-319 | Tanespimycin |
| MVP_top60_DNA_seed | SRC | -0.464 | UCEC | KLE | Saracatinib |
| MVP_top60_DNA_seed | ATM | -0.458963 | UCEC | AN3-CA | KU-60019 |
| MVP_top60_DNA_seed | EP300 | -0.453855 | UCEC | KLE | I-CBP112 |
| MVP_top60_DNA_seed | MDM2 | -0.442765 | UCEC | HEC-1 | Nutlin-3a (-) |

| | | | | | |
|---|---|---|---|---|---|
| MVP_top60_DNA_seed | HDAC1 | -0.425759 | UCEC | COLO-684 | CAY10603 |
| MVP_top60_DNA_seed | EGFR | -0.416391 | UCEC | MFE-296 | AZD8931 |
| MVP_top60_DNA_seed | ERBB2 | -0.416391 | UCEC | MFE-296 | AZD8931 |
| MVP_top60_DNA_seed | EGFR | -0.413687 | UCEC | HEC-1 | Afatinib |
| MVP_top60_DNA_seed | ERBB2 | -0.413687 | UCEC | HEC-1 | Afatinib |
| MVP_top60_DNA_seed | CDK2 | -0.407915 | UCEC | COLO-684 | PHA-793887 |
| MVP_top60_DNA_seed | PARP1 | -0.403762 | UCEC | AN3-CA | Veliparib |
| MVP_top60_DNA_seed | AKT1 | -0.400349 | UCEC | KLE | BAY AKT1 |
| MVP_top60_DNA_seed | HSP90AA1 | -0.397375 | UCEC | MFE-296 | Elesclomol |
| MVP_top60_DNA_seed | PIK3CA | -0.392543 | UCEC | AN3-CA | PI3Ka_4409 |
| MVP_top60_DNA_seed | PIK3CD | -0.392543 | UCEC | AN3-CA | PI3Ka_4409 |
| MVP_top60_DNA_seed | AKT1 | -0.381262 | UCEC | SNG-M | AKT inhibitor VIII |
| MVP_top60_DNA_seed | PIK3CD | -0.374953 | UCEC | MFE-280 | IC-87114 |
| MVP_top60_DNA_seed | SRC | -0.374751 | UCEC | MFE-280 | NVP-BHG712 |
| MVP_top60_DNA_seed | AKT1 | -0.366585 | UCEC | AN3-CA | AKT inhibitor VIII |
| MVP_top60_DNA_seed | FGFR2 | -0.355229 | UCEC | SNG-M | FGFR_3831 |
| MVP_top60_DNA_seed | HDAC1 | -0.3448 | UCEC | MFE-280 | CAY10603 |
| MVP_top60_DNA_seed | FGFR2 | -0.342315 | UCEC | COLO-684 | AZD4547 |
| MVP_top60_DNA_seed | HSP90AA1 | -0.333341 | UCEC | MFE-319 | CCT-018159 |
| MVP_top60_DNA_seed | AKT1 | -0.327348 | UCEC | KLE | MK-2206 |
| MVP_top60_DNA_seed | HSP90AA1 | -0.31414 | UCEC | MFE-319 | Elesclomol |
| MVP_top60_DNA_seed | AKT1 | -0.310089 | UCEC | KLE | GSK690693 |
| MVP_top60_DNA_seed | CDK2 | -0.308592 | UCEC | COLO-684 | Seliciclib |
| MVP_top60_DNA_seed | AR | -0.300636 | UCEC | MFE-280 | AZD3514 |
| MVP_top60_DNA_seed | ESR1 | -0.299305 | UCEC | MFE-296 | Tamoxifen |
| MVP_top60_DNA_seed | EGFR | -0.294777 | UCEC | MFE-319 | PF-00299804 |
| MVP_top60_DNA_seed | ERBB2 | -0.294777 | UCEC | MFE-319 | PF-00299804 |
| MVP_top60_DNA_seed | ESR1 | -0.291264 | UCEC | MFE-319 | Fulvestrant |
| MVP_top60_DNA_seed | PARP1 | -0.289161 | UCEC | HEC-1 | Talazoparib |
| MVP_top60_DNA_seed | PARP1 | -0.28234 | UCEC | HEC-1 | PARP_9495 |
| MVP_top60_DNA_seed | HDAC1 | -0.275283 | UCEC | MFE-280 | Belinostat |
| MVP_top60_DNA_seed | EGFR | -0.274487 | UCEC | KLE | Afatinib |
| MVP_top60_DNA_seed | ERBB2 | -0.274487 | UCEC | KLE | Afatinib |
| MVP_top60_DNA_seed | AKT1 | -0.273754 | UCEC | COLO-684 | AKT inhibitor VIII |
| MVP_top60_DNA_seed | EGFR | -0.263096 | UCEC | COLO-684 | CUDC-101 |
| MVP_top60_DNA_seed | ERBB2 | -0.263096 | UCEC | COLO-684 | CUDC-101 |
| MVP_top60_DNA_seed | AKT1 | -0.254252 | UCEC | KLE | AKT inhibitor VIII |
| MVP_top60_DNA_seed | AKT1 | -0.248601 | UCEC | EN | BAY AKT1 |
| MVP_top60_DNA_seed | ATM | -0.244126 | UCEC | HEC-1 | KU-55933 |
| MVP_top60_DNA_seed | HDAC1 | -0.234194 | UCEC | AN3-CA | JQ12 |
| MVP_top60_DNA_seed | HDAC2 | -0.234194 | UCEC | AN3-CA | JQ12 |

| | | | | | |
|---|---|---|---|---|---|
| MVP_top60_DNA_seed | PARP1 | -0.226583 | UCEC | MFE-296 | PARP_9495 |
| MVP_top60_DNA_seed | HDAC1 | -0.226359 | UCEC | HEC-1 | Dacinostat |
| MVP_top60_DNA_seed | MDM2 | -0.22556 | UCEC | MFE-319 | Nutlin-3a (-) |
| MVP_top60_DNA_seed | HDAC1 | -0.221093 | UCEC | COLO-684 | Tubastatin A |
| MVP_top60_DNA_seed | EGFR | -0.213487 | UCEC | KLE | CUDC-101 |
| MVP_top60_DNA_seed | ERBB2 | -0.213487 | UCEC | KLE | CUDC-101 |
| MVP_top60_DNA_seed | HDAC1 | -0.210836 | UCEC | MFE-296 | Dacinostat |
| MVP_top60_DNA_seed | AKT1 | -0.204435 | UCEC | SNG-M | BAY AKT1 |
| MVP_top60_DNA_seed | HDAC1 | -0.195598 | UCEC | EN | Belinostat |
| MVP_top60_DNA_seed | EGFR | -0.190407 | UCEC | KLE | PF-00299804 |
| MVP_top60_DNA_seed | ERBB2 | -0.190407 | UCEC | KLE | PF-00299804 |
| MVP_top60_DNA_seed | HDAC1 | -0.189258 | UCEC | COLO-684 | Dacinostat |
| MVP_top60_DNA_seed | PARP1 | -0.189097 | UCEC | MFE-319 | Olaparib |
| MVP_top60_DNA_seed | ERBB2 | -0.187817 | UCEC | KLE | CP724714 |
| MVP_top60_DNA_seed | PIK3CD | -0.168739 | UCEC | SNG-M | Idelalisib |
| MVP_top60_DNA_seed | PIK3CD | -0.163341 | UCEC | KLE | Idelalisib |
| MVP_top60_DNA_seed | JAK1 | -0.159153 | UCEC | MFE-319 | AZD1480 |
| MVP_top60_DNA_seed | AKT1 | -0.152611 | UCEC | SNG-M | AKT inhibitor VIII |
| MVP_top60_DNA_seed | EGFR | -0.148695 | UCEC | MFE-319 | AST-1306 |
| MVP_top60_DNA_seed | EGFR | -0.13801 | UCEC | SNG-M | Cetuximab |
| MVP_top60_DNA_seed | PIK3CA | -0.137169 | UCEC | COLO-684 | AZD8835 |
| MVP_top60_DNA_seed | PIK3CD | -0.137169 | UCEC | COLO-684 | AZD8835 |
| MVP_top60_DNA_seed | AKT1 | -0.132474 | UCEC | MFE-319 | BAY AKT1 |
| MVP_top60_DNA_seed | AR | -0.128718 | UCEC | MFE-296 | AZD3514 |
| MVP_top60_DNA_seed | CDK2 | -0.128425 | UCEC | MFE-280 | AZD5438 |
| MVP_top60_DNA_seed | HSP90AA1 | -0.125845 | UCEC | MFE-296 | CCT-018159 |
| MVP_top60_DNA_seed | JAK1 | -0.120985 | UCEC | COLO-684 | Ruxolitinib |
| MVP_top60_DNA_seed | EGFR | -0.119893 | UCEC | HEC-1 | AZD8931 |
| MVP_top60_DNA_seed | ERBB2 | -0.119893 | UCEC | HEC-1 | AZD8931 |
| MVP_top60_DNA_seed | HDAC1 | -0.113898 | UCEC | EN | CAY10603 |
| MVP_top60_DNA_seed | EP300 | -0.107449 | UCEC | HEC-1 | I-CBP112 |
| MVP_top60_DNA_seed | MDM2 | -0.103957 | UCEC | HEC-1 | Serdemetan |
| MVP_top60_DNA_seed | JAK1 | -0.103402 | UCEC | HEC-1 | AZD1480 |
| MVP_top60_DNA_seed | HDAC1 | -0.10036 | UCEC | MFE-296 | Tubastatin A |
| MVP_top60_DNA_seed | SRC | -0.099624 | UCEC | COLO-684 | NVP-BHG712 |
| MVP_top60_DNA_seed | ERBB2 | -0.091649 | UCEC | MFE-319 | CP724714 |
| MVP_top60_DNA_seed | PIK3CD | -0.090714 | UCEC | MFE-296 | AZD8186 |
| MVP_top60_DNA_seed | HSP90AA1 | -0.088273 | UCEC | COLO-684 | Luminespib |
| MVP_top60_DNA_seed | HDAC1 | -0.082807 | UCEC | MFE-296 | JQ12 |
| MVP_top60_DNA_seed | HDAC2 | -0.082807 | UCEC | MFE-296 | JQ12 |
| MVP_top60_DNA_seed | MDM2 | -0.079099 | UCEC | SNG-M | Serdemetan |

| | | | | | |
|---|---|---|---|---|---|
| MVP_top60_DNA_seed | SRC | -0.074364 | UCEC | KLE | NVP-BHG712 |
| MVP_top60_DNA_seed | FGFR2 | -0.071005 | UCEC | HEC-1 | AZD4547 |
| MVP_top60_DNA_seed | EGFR | -0.063469 | UCEC | EN | CI-1033 |
| MVP_top60_DNA_seed | ERBB2 | -0.063469 | UCEC | EN | CI-1033 |
| MVP_top60_DNA_seed | FGFR2 | -0.061881 | UCEC | HEC-1 | PD173074 |
| MVP_top60_DNA_seed | ATM | -0.054838 | UCEC | KLE | CP466722 |
| MVP_top60_DNA_seed | HDAC1 | -0.051517 | UCEC | MFE-319 | Dacinostat |
| MVP_top60_DNA_seed | ATM | -0.049526 | UCEC | COLO-684 | KU-60019 |
| MVP_top60_DNA_seed | PARP1 | -0.046973 | UCEC | HEC-1 | Olaparib |
| MVP_top60_DNA_seed | HDAC1 | -0.045742 | UCEC | EN | Tubastatin A |
| MVP_top60_DNA_seed | EGFR | -0.04525 | UCEC | KLE | AZD8931 |
| MVP_top60_DNA_seed | ERBB2 | -0.04525 | UCEC | KLE | AZD8931 |
| MVP_top60_DNA_seed | EGFR | -0.044238 | UCEC | EN | CUDC-101 |
| MVP_top60_DNA_seed | ERBB2 | -0.044238 | UCEC | EN | CUDC-101 |
| MVP_top60_DNA_seed | AKT1 | -0.04317 | UCEC | EN | AKT inhibitor VIII |
| MVP_top60_DNA_seed | JAK1 | -0.029314 | UCEC | MFE-296 | JAK1_3715 |
| MVP_top60_DNA_seed | HDAC1 | -0.026238 | UCEC | HEC-1 | Belinostat |
| MVP_top60_DNA_seed | AKT1 | -0.025501 | UCEC | HEC-1 | MK-2206 |
| MVP_top60_DNA_seed | JAK1 | -0.02547 | UCEC | HEC-1 | JAK1_3715 |
| MVP_top60_DNA_seed | HSP90AA1 | -0.019557 | UCEC | MFE-280 | SNX-2112 |
| MVP_top60_DNA_seed | EP300 | -0.009974 | UCEC | MFE-296 | I-CBP112 |
| MVP_top60_DNA_seed | PIK3CA | -0.004274 | UCEC | KLE | PI3Ka_4409 |
| MVP_top60_DNA_seed | PIK3CD | -0.004274 | UCEC | KLE | PI3Ka_4409 |
| MVP_top60_DNA_seed | PIK3CA | 0.002161 | UCEC | MFE-296 | AZD8835 |
| MVP_top60_DNA_seed | PIK3CD | 0.002161 | UCEC | MFE-296 | AZD8835 |
| MVP_top60_DNA_seed | EGFR | 0.004627 | UCEC | HEC-1 | CI-1033 |
| MVP_top60_DNA_seed | ERBB2 | 0.004627 | UCEC | HEC-1 | CI-1033 |
| MVP_top60_DNA_seed | ESR1 | 0.007057 | UCEC | KLE | Fulvestrant |
| MVP_top60_DNA_seed | PIK3CA | 0.007765 | UCEC | HEC-1 | PI3Ka_4409 |
| MVP_top60_DNA_seed | PIK3CD | 0.007765 | UCEC | HEC-1 | PI3Ka_4409 |
| MVP_top60_DNA_seed | HDAC1 | 0.008111 | UCEC | EN | AR-42 |
| MVP_top60_DNA_seed | EGFR | 0.010594 | UCEC | COLO-684 | CI-1033 |
| MVP_top60_DNA_seed | ERBB2 | 0.010594 | UCEC | COLO-684 | CI-1033 |
| MVP_top60_DNA_seed | EGFR | 0.010679 | UCEC | COLO-684 | Pelitinib |
| MVP_top60_DNA_seed | EGFR | 0.01629 | UCEC | HEC-1 | Gefitinib |
| MVP_top60_DNA_seed | HDAC1 | 0.018214 | UCEC | HEC-1 | AR-42 |
| MVP_top60_DNA_seed | ESR1 | 0.023711 | UCEC | MFE-296 | Fulvestrant |
| MVP_top60_DNA_seed | SRC | 0.026441 | UCEC | EN | NVP-BHG712 |
| MVP_top60_DNA_seed | EGFR | 0.034598 | UCEC | MFE-319 | CI-1033 |
| MVP_top60_DNA_seed | ERBB2 | 0.034598 | UCEC | MFE-319 | CI-1033 |
| MVP_top60_DNA_seed | HDAC1 | 0.035617 | UCEC | MFE-280 | Tubastatin A |

| | | | | | |
|---|---|---|---|---|---|
| MVP_top60_DNA_seed | CDK2 | 0.03767 | UCEC | AN3-CA | PHA-793887 |
| MVP_top60_DNA_seed | ERBB2 | 0.04091 | UCEC | COLO-684 | CP724714 |
| MVP_top60_DNA_seed | HDAC1 | 0.048415 | UCEC | MFE-280 | AR-42 |
| MVP_top60_DNA_seed | EGFR | 0.053618 | UCEC | MFE-296 | AST-1306 |
| MVP_top60_DNA_seed | HDAC1 | 0.058416 | UCEC | MFE-319 | JQ12 |
| MVP_top60_DNA_seed | HDAC2 | 0.058416 | UCEC | MFE-319 | JQ12 |
| MVP_top60_DNA_seed | SRC | 0.0597 | UCEC | HEC-1 | Bosutinib |
| MVP_top60_DNA_seed | EGFR | 0.062381 | UCEC | MFE-296 | CI-1033 |
| MVP_top60_DNA_seed | ERBB2 | 0.062381 | UCEC | MFE-296 | CI-1033 |
| MVP_top60_DNA_seed | JAK1 | 0.063622 | UCEC | MFE-319 | Ruxolitinib |
| MVP_top60_DNA_seed | HDAC1 | 0.06482 | UCEC | KLE | CAY10603 |
| MVP_top60_DNA_seed | HSP90AA1 | 0.070505 | UCEC | COLO-684 | Tanespimycin |
| MVP_top60_DNA_seed | HDAC1 | 0.079084 | UCEC | KLE | Dacinostat |
| MVP_top60_DNA_seed | HSP90AA1 | 0.094957 | UCEC | HEC-1 | Luminespib |
| MVP_top60_DNA_seed | HDAC1 | 0.101031 | UCEC | HEC-1 | JQ12 |
| MVP_top60_DNA_seed | HDAC2 | 0.101031 | UCEC | HEC-1 | JQ12 |
| MVP_top60_DNA_seed | FGFR2 | 0.102188 | UCEC | COLO-684 | PD173074 |
| MVP_top60_DNA_seed | FGFR2 | 0.105396 | UCEC | SNG-M | AZD4547 |
| MVP_top60_DNA_seed | ATM | 0.113108 | UCEC | MFE-296 | KU-55933 |
| MVP_top60_DNA_seed | HDAC1 | 0.126755 | UCEC | KLE | Belinostat |
| MVP_top60_DNA_seed | HDAC1 | 0.128478 | UCEC | HEC-1 | CAY10603 |
| MVP_top60_DNA_seed | SRC | 0.136586 | UCEC | MFE-319 | NVP-BHG712 |
| MVP_top60_DNA_seed | CDK2 | 0.137647 | UCEC | EN | PHA-793887 |
| MVP_top60_DNA_seed | EGFR | 0.149855 | UCEC | HEC-1 | CUDC-101 |
| MVP_top60_DNA_seed | ERBB2 | 0.149855 | UCEC | HEC-1 | CUDC-101 |
| MVP_top60_DNA_seed | HDAC1 | 0.151208 | UCEC | COLO-684 | Belinostat |
| MVP_top60_DNA_seed | PIK3CA | 0.154596 | UCEC | HEC-1 | AZD8835 |
| MVP_top60_DNA_seed | PIK3CD | 0.154596 | UCEC | HEC-1 | AZD8835 |
| MVP_top60_DNA_seed | PIK3CD | 0.167623 | UCEC | MFE-319 | AZD8186 |
| MVP_top60_DNA_seed | PARP1 | 0.171053 | UCEC | COLO-684 | Rucaparib |
| MVP_top60_DNA_seed | PIK3CD | 0.183443 | UCEC | MFE-319 | Idelalisib |
| MVP_top60_DNA_seed | AR | 0.186985 | UCEC | COLO-684 | Bicalutamide |
| MVP_top60_DNA_seed | ATM | 0.192287 | UCEC | EN | CP466722 |
| MVP_top60_DNA_seed | EGFR | 0.207992 | UCEC | MFE-296 | Cetuximab |
| MVP_top60_DNA_seed | EP300 | 0.208765 | UCEC | MFE-319 | I-CBP112 |
| MVP_top60_DNA_seed | EGFR | 0.211331 | UCEC | AN3-CA | AZD8931 |
| MVP_top60_DNA_seed | ERBB2 | 0.211331 | UCEC | AN3-CA | AZD8931 |
| MVP_top60_DNA_seed | PARP1 | 0.213634 | UCEC | MFE-296 | Olaparib |
| MVP_top60_DNA_seed | FGFR2 | 0.220404 | UCEC | EN | FGFR_3831 |
| MVP_top60_DNA_seed | EGFR | 0.221284 | UCEC | AN3-CA | CUDC-101 |
| MVP_top60_DNA_seed | ERBB2 | 0.221284 | UCEC | AN3-CA | CUDC-101 |

| | | | | | |
|---|---|---|---|---|---|
| MVP_top60_DNA_seed | PIK3CD | 0.223966 | UCEC | HEC-1 | IC-87114 |
| MVP_top60_DNA_seed | PARP1 | 0.228563 | UCEC | MFE-319 | Olaparib |
| MVP_top60_DNA_seed | FGFR2 | 0.233763 | UCEC | MFE-319 | PD173074 |
| MVP_top60_DNA_seed | CDK2 | 0.238046 | UCEC | MFE-280 | PHA-793887 |
| MVP_top60_DNA_seed | JAK1 | 0.239651 | UCEC | HEC-1 | Ruxolitinib |
| MVP_top60_DNA_seed | HSP90AA1 | 0.239798 | UCEC | MFE-280 | Tanespimycin |
| MVP_top60_DNA_seed | CDK2 | 0.244684 | UCEC | MFE-296 | PHA-793887 |
| MVP_top60_DNA_seed | FGFR2 | 0.254212 | UCEC | MFE-319 | FGFR_3831 |
| MVP_top60_DNA_seed | EGFR | 0.263839 | UCEC | KLE | CI-1033 |
| MVP_top60_DNA_seed | ERBB2 | 0.263839 | UCEC | KLE | CI-1033 |
| MVP_top60_DNA_seed | AR | 0.274026 | UCEC | MFE-319 | Bicalutamide |
| MVP_top60_DNA_seed | AKT1 | 0.277979 | UCEC | MFE-296 | AKT inhibitor VIII |
| MVP_top60_DNA_seed | ESR1 | 0.282681 | UCEC | HEC-1 | Tamoxifen |
| MVP_top60_DNA_seed | PARP1 | 0.288019 | UCEC | HEC-1 | PARP_0108 |
| MVP_top60_DNA_seed | MDM2 | 0.289948 | UCEC | AN3-CA | Nutlin-3a (-) |
| MVP_top60_DNA_seed | ATM | 0.291618 | UCEC | AN3-CA | CP466722 |
| MVP_top60_DNA_seed | AR | 0.291829 | UCEC | HEC-1 | Bicalutamide |
| MVP_top60_DNA_seed | PARP1 | 0.29662 | UCEC | MFE-280 | PARP_9482 |
| MVP_top60_DNA_seed | ESR1 | 0.299525 | UCEC | MFE-280 | Fulvestrant |
| MVP_top60_DNA_seed | JAK1 | 0.299954 | UCEC | MFE-280 | Ruxolitinib |
| MVP_top60_DNA_seed | FGFR2 | 0.300365 | UCEC | HEC-1 | AZD4547 |
| MVP_top60_DNA_seed | AR | 0.31215 | UCEC | KLE | AZD3514 |
| MVP_top60_DNA_seed | AKT1 | 0.327086 | UCEC | MFE-280 | AKT inhibitor VIII |
| MVP_top60_DNA_seed | AKT1 | 0.327418 | UCEC | MFE-319 | GSK690693 |
| MVP_top60_DNA_seed | ATM | 0.330904 | UCEC | MFE-280 | CP466722 |
| MVP_top60_DNA_seed | PIK3CD | 0.332221 | UCEC | AN3-CA | Idelalisib |
| MVP_top60_DNA_seed | AR | 0.334759 | UCEC | AN3-CA | Bicalutamide |
| MVP_top60_DNA_seed | PIK3CD | 0.335865 | UCEC | MFE-319 | IC-87114 |
| MVP_top60_DNA_seed | ATM | 0.340159 | UCEC | MFE-296 | CP466722 |
| MVP_top60_DNA_seed | AR | 0.342183 | UCEC | KLE | Bicalutamide |
| MVP_top60_DNA_seed | JAK1 | 0.342569 | UCEC | AN3-CA | Ruxolitinib |
| MVP_top60_DNA_seed | FGFR2 | 0.344305 | UCEC | COLO-684 | FGFR_3831 |
| MVP_top60_DNA_seed | PIK3CA | 0.352447 | UCEC | COLO-684 | PI3Ka_4409 |
| MVP_top60_DNA_seed | PIK3CD | 0.352447 | UCEC | COLO-684 | PI3Ka_4409 |
| MVP_top60_DNA_seed | EGFR | 0.35841 | UCEC | AN3-CA | Afatinib |
| MVP_top60_DNA_seed | ERBB2 | 0.35841 | UCEC | AN3-CA | Afatinib |
| MVP_top60_DNA_seed | HDAC1 | 0.361733 | UCEC | KLE | AR-42 |
| MVP_top60_DNA_seed | FGFR2 | 0.369629 | UCEC | MFE-319 | AZD4547 |
| MVP_top60_DNA_seed | EGFR | 0.370158 | UCEC | EN | Cetuximab |
| MVP_top60_DNA_seed | ATM | 0.375809 | UCEC | SNG-M | KU-60019 |
| MVP_top60_DNA_seed | CDK2 | 0.378207 | UCEC | MFE-319 | AZD5438 |

| | | | | | |
|---|---|---|---|---|---|
| MVP_top60_DNA_seed | HDAC1 | 0.382997 | UCEC | AN3-CA | CAY10603 |
| MVP_top60_DNA_seed | PIK3CA | 0.384848 | UCEC | MFE-296 | PI3Ka_4409 |
| MVP_top60_DNA_seed | PIK3CD | 0.384848 | UCEC | MFE-296 | PI3Ka_4409 |
| MVP_top60_DNA_seed | SRC | 0.391633 | UCEC | MFE-319 | Bosutinib |
| MVP_top60_DNA_seed | HDAC1 | 0.395963 | UCEC | AN3-CA | AR-42 |
| MVP_top60_DNA_seed | PARP1 | 0.412603 | UCEC | COLO-684 | Olaparib |
| MVP_top60_DNA_seed | JAK1 | 0.426205 | UCEC | MFE-319 | JAK1_3715 |
| MVP_top60_DNA_seed | MDM2 | 0.426412 | UCEC | EN | Serdemetan |
| MVP_top60_DNA_seed | EGFR | 0.428356 | UCEC | EN | AST-1306 |
| MVP_top60_DNA_seed | HDAC1 | 0.431141 | UCEC | KLE | Tubastatin A |
| MVP_top60_DNA_seed | FGFR2 | 0.441923 | UCEC | SNG-M | PD173074 |
| MVP_top60_DNA_seed | HSP90AA1 | 0.445405 | UCEC | EN | CCT-018159 |
| MVP_top60_DNA_seed | MDM2 | 0.45042 | UCEC | AN3-CA | Serdemetan |
| MVP_top60_DNA_seed | AKT1 | 0.455757 | UCEC | SNG-M | MK-2206 |
| MVP_top60_DNA_seed | HSP90AA1 | 0.457367 | UCEC | SNG-M | Elesclomol |
| MVP_top60_DNA_seed | ATM | 0.458843 | UCEC | SNG-M | KU-55933 |
| MVP_top60_DNA_seed | MDM2 | 0.463917 | UCEC | KLE | Serdemetan |
| MVP_top60_DNA_seed | PARP1 | 0.469077 | UCEC | HEC-1 | PARP_9482 |
| MVP_top60_DNA_seed | EGFR | 0.48369 | UCEC | KLE | Afatinib |
| MVP_top60_DNA_seed | ERBB2 | 0.48369 | UCEC | KLE | Afatinib |
| MVP_top60_DNA_seed | HSP90AA1 | 0.484913 | UCEC | EN | SNX-2112 |
| MVP_top60_DNA_seed | EGFR | 0.486628 | UCEC | AN3-CA | AST-1306 |
| MVP_top60_DNA_seed | ATM | 0.493472 | UCEC | MFE-280 | KU-55933 |
| MVP_top60_DNA_seed | JAK1 | 0.494036 | UCEC | MFE-296 | Ruxolitinib |
| MVP_top60_DNA_seed | PARP1 | 0.49482 | UCEC | EN | PARP_9495 |
| MVP_top60_DNA_seed | PARP1 | 0.503453 | UCEC | MFE-319 | PARP_9482 |
| MVP_top60_DNA_seed | HDAC1 | 0.504916 | UCEC | MFE-280 | Dacinostat |
| MVP_top60_DNA_seed | AR | 0.507807 | UCEC | COLO-684 | AZD3514 |
| MVP_top60_DNA_seed | PIK3CD | 0.508062 | UCEC | AN3-CA | IC-87114 |
| MVP_top60_DNA_seed | SRC | 0.511928 | UCEC | AN3-CA | NVP-BHG712 |
| MVP_top60_DNA_seed | AR | 0.516299 | UCEC | MFE-296 | Bicalutamide |
| MVP_top60_DNA_seed | PARP1 | 0.519955 | UCEC | MFE-319 | PARP_0108 |
| MVP_top60_DNA_seed | JAK1 | 0.52226 | UCEC | MFE-280 | JAK1_3715 |
| MVP_top60_DNA_seed | ESR1 | 0.536283 | UCEC | KLE | Tamoxifen |
| MVP_top60_DNA_seed | HSP90AA1 | 0.537936 | UCEC | KLE | SNX-2112 |
| MVP_top60_DNA_seed | SRC | 0.547493 | UCEC | HEC-1 | NVP-BHG712 |
| MVP_top60_DNA_seed | PARP1 | 0.553506 | UCEC | MFE-319 | Talazoparib |
| MVP_top60_DNA_seed | CDK2 | 0.560813 | UCEC | MFE-319 | PHA-793887 |
| MVP_top60_DNA_seed | PIK3CA | 0.563782 | UCEC | MFE-319 | AZD8835 |
| MVP_top60_DNA_seed | PIK3CD | 0.563782 | UCEC | MFE-319 | AZD8835 |
| MVP_top60_DNA_seed | PARP1 | 0.564015 | UCEC | SNG-M | Rucaparib |

| | | | | | |
|---|---|---|---|---|---|
| MVP_top60_DNA_seed | EP300 | 0.573295 | UCEC | AN3-CA | I-CBP112 |
| MVP_top60_DNA_seed | PARP1 | 0.576186 | UCEC | MFE-319 | PARP_9495 |
| MVP_top60_DNA_seed | EP300 | 0.576443 | UCEC | COLO-684 | I-CBP112 |
| MVP_top60_DNA_seed | HDAC1 | 0.576983 | UCEC | MFE-296 | CAY10603 |
| MVP_top60_DNA_seed | EGFR | 0.582641 | UCEC | AN3-CA | Afatinib |
| MVP_top60_DNA_seed | ERBB2 | 0.582641 | UCEC | AN3-CA | Afatinib |
| MVP_top60_DNA_seed | AR | 0.586072 | UCEC | EN | Bicalutamide |
| MVP_top60_DNA_seed | HDAC1 | 0.587212 | UCEC | KLE | JQ12 |
| MVP_top60_DNA_seed | HDAC2 | 0.587212 | UCEC | KLE | JQ12 |
| MVP_top60_DNA_seed | HSP90AA1 | 0.593566 | UCEC | MFE-280 | Elesclomol |
| MVP_top60_DNA_seed | EGFR | 0.596501 | UCEC | AN3-CA | PF-00299804 |
| MVP_top60_DNA_seed | ERBB2 | 0.596501 | UCEC | AN3-CA | PF-00299804 |
| MVP_top60_DNA_seed | HSP90AA1 | 0.59667 | UCEC | EN | Tanespimycin |
| MVP_top60_DNA_seed | ATM | 0.599189 | UCEC | KLE | KU-60019 |
| MVP_top60_DNA_seed | ERBB2 | 0.6102 | UCEC | HEC-1 | CP724714 |
| MVP_top60_DNA_seed | EGFR | 0.610908 | UCEC | MFE-280 | AZD8931 |
| MVP_top60_DNA_seed | ERBB2 | 0.610908 | UCEC | MFE-280 | AZD8931 |
| MVP_top60_DNA_seed | FGFR2 | 0.612214 | UCEC | EN | AZD4547 |
| MVP_top60_DNA_seed | SRC | 0.616683 | UCEC | MFE-296 | Bosutinib |
| MVP_top60_DNA_seed | HSP90AA1 | 0.624883 | UCEC | MFE-319 | SNX-2112 |
| MVP_top60_DNA_seed | PARP1 | 0.638195 | UCEC | KLE | PARP_9495 |
| MVP_top60_DNA_seed | EGFR | 0.639637 | UCEC | MFE-296 | Afatinib |
| MVP_top60_DNA_seed | ERBB2 | 0.639637 | UCEC | MFE-296 | Afatinib |
| MVP_top60_DNA_seed | FGFR2 | 0.647232 | UCEC | SNG-M | AZD4547 |
| MVP_top60_DNA_seed | JAK1 | 0.650458 | UCEC | EN | AZD1480 |
| MVP_top60_DNA_seed | PARP1 | 0.653298 | UCEC | AN3-CA | Talazoparib |
| MVP_top60_DNA_seed | EGFR | 0.653766 | UCEC | MFE-319 | Pelitinib |
| MVP_top60_DNA_seed | SRC | 0.654226 | UCEC | COLO-684 | Bosutinib |
| MVP_top60_DNA_seed | ERBB2 | 0.654458 | UCEC | MFE-296 | CP724714 |
| MVP_top60_DNA_seed | ERBB2 | 0.655068 | UCEC | AN3-CA | CP724714 |
| MVP_top60_DNA_seed | EGFR | 0.659161 | UCEC | KLE | Gefitinib |
| MVP_top60_DNA_seed | EGFR | 0.663197 | UCEC | AN3-CA | Cetuximab |
| MVP_top60_DNA_seed | PARP1 | 0.665291 | UCEC | AN3-CA | PARP_9495 |
| MVP_top60_DNA_seed | MDM2 | 0.677846 | UCEC | COLO-684 | Nutlin-3a (-) |
| MVP_top60_DNA_seed | EGFR | 0.678647 | UCEC | MFE-296 | Gefitinib |
| MVP_top60_DNA_seed | EGFR | 0.685851 | UCEC | HEC-1 | Afatinib |
| MVP_top60_DNA_seed | ERBB2 | 0.685851 | UCEC | HEC-1 | Afatinib |
| MVP_top60_DNA_seed | HDAC1 | 0.686067 | UCEC | MFE-319 | Tubastatin A |
| MVP_top60_DNA_seed | EGFR | 0.691755 | UCEC | HEC-1 | Pelitinib |
| MVP_top60_DNA_seed | EGFR | 0.699062 | UCEC | COLO-684 | AST-1306 |
| MVP_top60_DNA_seed | HDAC1 | 0.699207 | UCEC | AN3-CA | Belinostat |

| | | | | | |
|---|---|---|---|---|---|
| MVP_top60_DNA_seed | HDAC1 | 0.703336 | UCEC | MFE-296 | Belinostat |
| MVP_top60_DNA_seed | JAK1 | 0.708809 | UCEC | AN3-CA | JAK1_3715 |
| MVP_top60_DNA_seed | PIK3CD | 0.709942 | UCEC | MFE-296 | IC-87114 |
| MVP_top60_DNA_seed | PARP1 | 0.710251 | UCEC | EN | Olaparib |
| MVP_top60_DNA_seed | EGFR | 0.711985 | UCEC | COLO-684 | Lapatinib |
| MVP_top60_DNA_seed | ERBB2 | 0.711985 | UCEC | COLO-684 | Lapatinib |
| MVP_top60_DNA_seed | HSP90AA1 | 0.721388 | UCEC | AN3-CA | SNX-2112 |
| MVP_top60_DNA_seed | HDAC1 | 0.724498 | UCEC | HEC-1 | Tubastatin A |
| MVP_top60_DNA_seed | EGFR | 0.728492 | UCEC | MFE-280 | Gefitinib |
| MVP_top60_DNA_seed | HDAC1 | 0.733862 | UCEC | COLO-684 | AR-42 |
| MVP_top60_DNA_seed | PARP1 | 0.735725 | UCEC | AN3-CA | PARP_0108 |
| MVP_top60_DNA_seed | EGFR | 0.737688 | UCEC | HEC-1 | AST-1306 |
| MVP_top60_DNA_seed | PARP1 | 0.740183 | UCEC | AN3-CA | PARP_9482 |
| MVP_top60_DNA_seed | MDM2 | 0.749433 | UCEC | COLO-684 | Serdemetan |
| MVP_top60_DNA_seed | FGFR2 | 0.75276 | UCEC | COLO-684 | AZD4547 |
| MVP_top60_DNA_seed | ATM | 0.765162 | UCEC | EN | KU-55933 |
| MVP_top60_DNA_seed | CDK2 | 0.769122 | UCEC | KLE | PHA-793887 |
| MVP_top60_DNA_seed | AR | 0.774001 | UCEC | AN3-CA | AZD3514 |
| MVP_top60_DNA_seed | EGFR | 0.77468 | UCEC | SNG-M | Gefitinib |
| MVP_top60_DNA_seed | SRC | 0.776491 | UCEC | COLO-684 | WH-4-023 |
| MVP_top60_DNA_seed | HDAC1 | 0.778957 | UCEC | MFE-280 | JQ12 |
| MVP_top60_DNA_seed | HDAC2 | 0.778957 | UCEC | MFE-280 | JQ12 |
| MVP_top60_DNA_seed | HDAC1 | 0.780934 | UCEC | SNG-M | Tubastatin A |
| MVP_top60_DNA_seed | AKT1 | 0.782152 | UCEC | HEC-1 | GSK690693 |
| MVP_top60_DNA_seed | FGFR2 | 0.786376 | UCEC | MFE-319 | AZD4547 |
| MVP_top60_DNA_seed | PARP1 | 0.787156 | UCEC | MFE-280 | Veliparib |
| MVP_top60_DNA_seed | JAK1 | 0.794613 | UCEC | EN | JAK1_3715 |
| MVP_top60_DNA_seed | AR | 0.804154 | UCEC | MFE-280 | Bicalutamide |
| MVP_top60_DNA_seed | ATM | 0.811403 | UCEC | EN | KU-60019 |
| MVP_top60_DNA_seed | HSP90AA1 | 0.812381 | UCEC | KLE | Luminespib |
| MVP_top60_DNA_seed | EGFR | 0.81677 | UCEC | HEC-1 | PF-00299804 |
| MVP_top60_DNA_seed | ERBB2 | 0.81677 | UCEC | HEC-1 | PF-00299804 |
| MVP_top60_DNA_seed | EGFR | 0.823994 | UCEC | MFE-296 | PF-00299804 |
| MVP_top60_DNA_seed | ERBB2 | 0.823994 | UCEC | MFE-296 | PF-00299804 |
| MVP_top60_DNA_seed | PARP1 | 0.827468 | UCEC | AN3-CA | Rucaparib |
| MVP_top60_DNA_seed | MDM2 | 0.831763 | UCEC | MFE-280 | Nutlin-3a (-) |
| MVP_top60_DNA_seed | PIK3CD | 0.832741 | UCEC | HEC-1 | Idelalisib |
| MVP_top60_DNA_seed | AR | 0.842284 | UCEC | SNG-M | Bicalutamide |
| MVP_top60_DNA_seed | MDM2 | 0.845624 | UCEC | MFE-319 | Serdemetan |
| MVP_top60_DNA_seed | EGFR | 0.847915 | UCEC | EN | PF-00299804 |
| MVP_top60_DNA_seed | ERBB2 | 0.847915 | UCEC | EN | PF-00299804 |

| | | | | | |
|---|---|---|---|---|---|
| MVP_top60_DNA_seed | HDAC1 | 0.851064 | UCEC | COLO-684 | Entinostat |
| MVP_top60_DNA_seed | JAK1 | 0.853787 | UCEC | KLE | Ruxolitinib |
| MVP_top60_DNA_seed | ESR1 | 0.854879 | UCEC | COLO-684 | Tamoxifen |
| MVP_top60_DNA_seed | ERBB2 | 0.876731 | UCEC | EN | CP724714 |
| MVP_top60_DNA_seed | EGFR | 0.884773 | UCEC | KLE | Cetuximab |
| MVP_top60_DNA_seed | JAK1 | 0.884858 | UCEC | COLO-684 | AZD1480 |
| MVP_top60_DNA_seed | EGFR | 0.885137 | UCEC | MFE-280 | Afatinib |
| MVP_top60_DNA_seed | ERBB2 | 0.885137 | UCEC | MFE-280 | Afatinib |
| MVP_top60_DNA_seed | AKT1 | 0.887171 | UCEC | HEC-1 | AKT inhibitor VIII |
| MVP_top60_DNA_seed | HDAC1 | 0.900099 | UCEC | AN3-CA | Tubastatin A |
| MVP_top60_DNA_seed | SRC | 0.90174 | UCEC | AN3-CA | Bosutinib |
| MVP_top60_DNA_seed | FGFR2 | 0.917464 | UCEC | HEC-1 | FGFR_3831 |
| MVP_top60_DNA_seed | AR | 0.917979 | UCEC | MFE-296 | Bicalutamide |
| MVP_top60_DNA_seed | EGFR | 0.919158 | UCEC | COLO-684 | Cetuximab |
| MVP_top60_DNA_seed | PARP1 | 0.924746 | UCEC | MFE-280 | Olaparib |
| MVP_top60_DNA_seed | CDK2 | 0.92558 | UCEC | KLE | AZD5438 |
| MVP_top60_DNA_seed | EGFR | 0.9262 | UCEC | COLO-684 | Erlotinib |
| MVP_top60_DNA_seed | PIK3CD | 0.928035 | UCEC | MFE-296 | Idelalisib |
| MVP_top60_DNA_seed | JAK1 | 0.938225 | UCEC | COLO-684 | JAK1_3715 |
| MVP_top60_DNA_seed | EGFR | 0.960282 | UCEC | COLO-684 | PF-00299804 |
| MVP_top60_DNA_seed | ERBB2 | 0.960282 | UCEC | COLO-684 | PF-00299804 |
| MVP_top60_DNA_seed | PARP1 | 0.964295 | UCEC | KLE | PARP_9482 |
| MVP_top60_DNA_seed | PARP1 | 0.970805 | UCEC | EN | Talazoparib |
| MVP_top60_DNA_seed | ATM | 0.972207 | UCEC | KLE | KU-55933 |
| MVP_top60_DNA_seed | PARP1 | 0.983734 | UCEC | KLE | Olaparib |
| MVP_top60_DNA_seed | EGFR | 0.984534 | UCEC | MFE-280 | Afatinib |
| MVP_top60_DNA_seed | ERBB2 | 0.984534 | UCEC | MFE-280 | Afatinib |
| MVP_top60_DNA_seed | HSP90AA1 | 0.985196 | UCEC | SNG-M | Tanespimycin |
| MVP_top60_DNA_seed | SRC | 0.990642 | UCEC | COLO-684 | Saracatinib |
| MVP_top60_DNA_seed | EGFR | 0.993025 | UCEC | MFE-296 | Afatinib |
| MVP_top60_DNA_seed | ERBB2 | 0.993025 | UCEC | MFE-296 | Afatinib |
| MVP_top60_DNA_seed | AR | 0.99526 | UCEC | COLO-684 | Bicalutamide |
| MVP_top60_DNA_seed | EGFR | 1.003911 | UCEC | COLO-684 | Afatinib |
| MVP_top60_DNA_seed | ERBB2 | 1.003911 | UCEC | COLO-684 | Afatinib |
| MVP_top60_DNA_seed | EGFR | 1.010634 | UCEC | COLO-684 | Afatinib |
| MVP_top60_DNA_seed | ERBB2 | 1.010634 | UCEC | COLO-684 | Afatinib |
| MVP_top60_DNA_seed | FGFR2 | 1.011234 | UCEC | KLE | AZD4547 |
| MVP_top60_DNA_seed | PARP1 | 1.013052 | UCEC | MFE-280 | Olaparib |
| MVP_top60_DNA_seed | PARP1 | 1.019813 | UCEC | MFE-296 | Rucaparib |
| MVP_top60_DNA_seed | HSP90AA1 | 1.020393 | UCEC | MFE-296 | SNX-2112 |
| MVP_top60_DNA_seed | EGFR | 1.02499 | UCEC | AN3-CA | Gefitinib |

| | | | | | |
|---|---|---|---|---|---|
| MVP_top60_DNA_seed | PARP1 | 1.066013 | UCEC | COLO-684 | Talazoparib |
| MVP_top60_DNA_seed | FGFR2 | 1.067797 | UCEC | EN | AZD4547 |
| MVP_top60_DNA_seed | MDM2 | 1.068278 | UCEC | MFE-296 | Serdemetan |
| MVP_top60_DNA_seed | PIK3CD | 1.068292 | UCEC | HEC-1 | AZD8186 |
| MVP_top60_DNA_seed | EP300 | 1.095465 | UCEC | MFE-280 | I-CBP112 |
| MVP_top60_DNA_seed | EGFR | 1.103031 | UCEC | AN3-CA | Pelitinib |
| MVP_top60_DNA_seed | HSP90AA1 | 1.106335 | UCEC | KLE | Elesclomol |
| MVP_top60_DNA_seed | SRC | 1.121127 | UCEC | MFE-296 | NVP-BHG712 |
| MVP_top60_DNA_seed | PARP1 | 1.143687 | UCEC | MFE-280 | Talazoparib |
| MVP_top60_DNA_seed | FGFR2 | 1.159924 | UCEC | EN | PD173074 |
| MVP_top60_DNA_seed | PARP1 | 1.168043 | UCEC | MFE-280 | PARP_0108 |
| MVP_top60_DNA_seed | ESR1 | 1.172355 | UCEC | AN3-CA | Fulvestrant |
| MVP_top60_DNA_seed | HDAC1 | 1.177407 | UCEC | COLO-684 | JQ12 |
| MVP_top60_DNA_seed | HDAC2 | 1.177407 | UCEC | COLO-684 | JQ12 |
| MVP_top60_DNA_seed | PIK3CA | 1.18858 | UCEC | MFE-319 | PI3Ka_4409 |
| MVP_top60_DNA_seed | PIK3CD | 1.18858 | UCEC | MFE-319 | PI3Ka_4409 |
| MVP_top60_DNA_seed | EGFR | 1.188775 | UCEC | EN | Afatinib |
| MVP_top60_DNA_seed | ERBB2 | 1.188775 | UCEC | EN | Afatinib |
| MVP_top60_DNA_seed | PIK3CD | 1.191703 | UCEC | EN | IC-87114 |
| MVP_top60_DNA_seed | HDAC1 | 1.195439 | UCEC | EN | Dacinostat |
| MVP_top60_DNA_seed | PARP1 | 1.213331 | UCEC | COLO-684 | PARP_9495 |
| MVP_top60_DNA_seed | PARP1 | 1.216185 | UCEC | COLO-684 | PARP_0108 |
| MVP_top60_DNA_seed | JAK1 | 1.217774 | UCEC | EN | Ruxolitinib |
| MVP_top60_DNA_seed | EGFR | 1.218919 | UCEC | MFE-296 | Pelitinib |
| MVP_top60_DNA_seed | CDK2 | 1.239535 | UCEC | HEC-1 | PHA-793887 |
| MVP_top60_DNA_seed | PIK3CA | 1.254999 | UCEC | KLE | AZD8835 |
| MVP_top60_DNA_seed | PIK3CD | 1.254999 | UCEC | KLE | AZD8835 |
| MVP_top60_DNA_seed | HDAC1 | 1.269922 | UCEC | MFE-296 | AR-42 |
| MVP_top60_DNA_seed | HSP90AA1 | 1.286752 | UCEC | EN | Luminespib |
| MVP_top60_DNA_seed | PARP1 | 1.299957 | UCEC | COLO-684 | PARP_9482 |
| MVP_top60_DNA_seed | FGFR2 | 1.300353 | UCEC | KLE | FGFR_3831 |
| MVP_top60_DNA_seed | AR | 1.30838 | UCEC | EN | Bicalutamide |
| MVP_top60_DNA_seed | PARP1 | 1.316858 | UCEC | KLE | Rucaparib |
| MVP_top60_DNA_seed | HDAC1 | 1.318465 | UCEC | COLO-684 | Parthenolide |
| MVP_top60_DNA_seed | EGFR | 1.320902 | UCEC | COLO-684 | AZD8931 |
| MVP_top60_DNA_seed | ERBB2 | 1.320902 | UCEC | COLO-684 | AZD8931 |
| MVP_top60_DNA_seed | EGFR | 1.322524 | UCEC | MFE-296 | CUDC-101 |
| MVP_top60_DNA_seed | ERBB2 | 1.322524 | UCEC | MFE-296 | CUDC-101 |
| MVP_top60_DNA_seed | AKT1 | 1.329107 | UCEC | HEC-1 | BAY AKT1 |
| MVP_top60_DNA_seed | PIK3CD | 1.333835 | UCEC | EN | Idelalisib |
| MVP_top60_DNA_seed | HSP90AA1 | 1.344395 | UCEC | MFE-280 | Luminespib |

| | | | | | |
|---|---|---|---|---|---|
| MVP_top60_DNA_seed | ESR1 | 1.360619 | UCEC | EN | Tamoxifen |
| MVP_top60_DNA_seed | PARP1 | 1.375957 | UCEC | MFE-280 | Rucaparib |
| MVP_top60_DNA_seed | HSP90AA1 | 1.386121 | UCEC | KLE | Tanespimycin |
| MVP_top60_DNA_seed | ATM | 1.411894 | UCEC | MFE-280 | KU-60019 |
| MVP_top60_DNA_seed | PARP1 | 1.419227 | UCEC | KLE | Talazoparib |
| MVP_top60_DNA_seed | ATM | 1.419228 | UCEC | HEC-1 | CP466722 |
| MVP_top60_DNA_seed | HDAC1 | 1.427128 | UCEC | MFE-319 | AR-42 |
| MVP_top60_DNA_seed | EGFR | 1.427347 | UCEC | EN | Afatinib |
| MVP_top60_DNA_seed | ERBB2 | 1.427347 | UCEC | EN | Afatinib |
| MVP_top60_DNA_seed | MDM2 | 1.431443 | UCEC | EN | Nutlin-3a (-) |
| MVP_top60_DNA_seed | FGFR2 | 1.445917 | UCEC | MFE-280 | AZD4547 |
| MVP_top60_DNA_seed | EGFR | 1.455239 | UCEC | MFE-319 | CUDC-101 |
| MVP_top60_DNA_seed | ERBB2 | 1.455239 | UCEC | MFE-319 | CUDC-101 |
| MVP_top60_DNA_seed | CDK2 | 1.456808 | UCEC | COLO-684 | AZD5438 |
| MVP_top60_DNA_seed | PARP1 | 1.463433 | UCEC | EN | Olaparib |
| MVP_top60_DNA_seed | PARP1 | 1.467386 | UCEC | EN | Rucaparib |
| MVP_top60_DNA_seed | PIK3CD | 1.47506 | UCEC | KLE | IC-87114 |
| MVP_top60_DNA_seed | EGFR | 1.483304 | UCEC | EN | Pelitinib |
| MVP_top60_DNA_seed | MDM2 | 1.518861 | UCEC | KLE | Nutlin-3a (-) |
| MVP_top60_DNA_seed | SRC | 1.528067 | UCEC | KLE | Bosutinib |
| MVP_top60_DNA_seed | SRC | 1.551759 | UCEC | EN | Bosutinib |
| MVP_top60_DNA_seed | HDAC1 | 1.553633 | UCEC | MFE-319 | CAY10603 |
| MVP_top60_DNA_seed | PARP1 | 1.56297 | UCEC | KLE | PARP_0108 |
| MVP_top60_DNA_seed | PIK3CD | 1.571729 | UCEC | EN | AZD8186 |
| MVP_top60_DNA_seed | HSP90AA1 | 1.606551 | UCEC | HEC-1 | SNX-2112 |
| MVP_top60_DNA_seed | JAK1 | 1.615952 | UCEC | KLE | JAK1_3715 |
| MVP_top60_DNA_seed | FGFR2 | 1.631694 | UCEC | KLE | PD173074 |
| MVP_top60_DNA_seed | PARP1 | 1.65273 | UCEC | KLE | Veliparib |
| MVP_top60_DNA_seed | ESR1 | 1.67379 | UCEC | EN | Fulvestrant |
| MVP_top60_DNA_seed | PARP1 | 1.677433 | UCEC | EN | Veliparib |
| MVP_top60_DNA_seed | HDAC1 | 1.679567 | UCEC | SNG-M | CAY10603 |
| MVP_top60_DNA_seed | ESR1 | 1.726912 | UCEC | COLO-684 | Fulvestrant |
| MVP_top60_DNA_seed | PARP1 | 1.726928 | UCEC | KLE | Olaparib |
| MVP_top60_DNA_seed | PIK3CA | 1.727071 | UCEC | EN | PI3Ka_4409 |
| MVP_top60_DNA_seed | PIK3CD | 1.727071 | UCEC | EN | PI3Ka_4409 |
| MVP_top60_DNA_seed | HDAC1 | 1.744332 | UCEC | MFE-319 | Belinostat |
| MVP_top60_DNA_seed | SRC | 1.746463 | UCEC | MFE-280 | Bosutinib |
| MVP_top60_DNA_seed | HSP90AA1 | 1.808225 | UCEC | KLE | CCT-018159 |
| MVP_top60_DNA_seed | HDAC1 | 1.832348 | UCEC | EN | JQ12 |
| MVP_top60_DNA_seed | HDAC2 | 1.832348 | UCEC | EN | JQ12 |
| MVP_top60_DNA_seed | EGFR | 1.863248 | UCEC | EN | Gefitinib |

| | | | | | |
|---|---|---|---|---|---|
| MVP_top60_DNA_seed | AKT1 | 1.884926 | UCEC | EN | AKT inhibitor VIII |
| MVP_top60_DNA_seed | PARP1 | 1.929522 | UCEC | EN | PARP_9482 |
| MVP_top60_DNA_seed | PARP1 | 2.002474 | UCEC | EN | PARP_0108 |
| MVP_top60_DNA_seed | AR | 2.015584 | UCEC | KLE | Bicalutamide |
| MVP_top60_DNA_seed | FGFR2 | 2.092606 | UCEC | KLE | AZD4547 |
| MVP_top60_DNA_seed | JAK1 | 2.101062 | UCEC | KLE | AZD1480 |
| MVP_top60_DNA_seed | CDK2 | 2.108505 | UCEC | EN | AZD5438 |
| MVP_top60_DNA_seed | HSP90AA1 | 2.13042 | UCEC | MFE-280 | CCT-018159 |
| MVP_top60_DNA_seed | AR | 2.13383 | UCEC | EN | AZD3514 |
| MVP_top60_DNA_seed | MDM2 | 2.430501 | UCEC | MFE-280 | Serdemetan |
| MVP_top60_DNA_seed | EP300 | -1.900587 | UCEC | EN | SGC-CBP30 |
| MVP_top60_DNA_seed | JAK1 | -1.754477 | UCEC | AN3-CA | GSK2276186C |
| MVP_top60_DNA_seed | FGFR2 | -1.681912 | UCEC | AN3-CA | PD173074 |
| MVP_top60_DNA_seed | AKT1 | -1.538701 | UCEC | AN3-CA | GSK2110183B |
| MVP_top60_DNA_seed | AKT1 | -1.461307 | UCEC | COLO-684 | GSK2110183B |
| MVP_top60_DNA_seed | FGFR2 | -1.455099 | UCEC | AN3-CA | AZD4547 |
| MVP_top60_DNA_seed | AKT1 | -1.441089 | UCEC | AN3-CA | Uprosertib |
| MVP_top60_DNA_seed | PIK3CA | -1.433814 | UCEC | MFE-280 | Alpelisib |
| MVP_top60_DNA_seed | JAK1 | -1.401798 | UCEC | AN3-CA | JAK_8517 |
| MVP_top60_DNA_seed | AKT1 | -1.393069 | UCEC | EN | AZD5363 |
| MVP_top60_DNA_seed | AKT1 | -1.363351 | UCEC | AN3-CA | Ipatasertib |
| MVP_top60_DNA_seed | AKT1 | -1.33634 | UCEC | EN | GSK2110183B |
| MVP_top60_DNA_seed | AKT1 | -1.322297 | UCEC | COLO-684 | Uprosertib |
| MVP_top60_DNA_seed | HSP90AA1 | -1.268945 | UCEC | AN3-CA | Tanespimycin |
| MVP_top60_DNA_seed | AKT1 | -1.246177 | UCEC | COLO-684 | Afuresertib |
| MVP_top60_DNA_seed | AKT1 | -1.225323 | UCEC | EN | Afuresertib |
| MVP_top60_DNA_seed | AKT1 | -1.197812 | UCEC | EN | Uprosertib |
| MVP_top60_DNA_seed | AKT1 | -1.173212 | UCEC | AN3-CA | AZD5363 |
| MVP_top60_DNA_seed | CDK2 | -1.137918 | UCEC | SNG-M | AZD5438 |
| MVP_top60_DNA_seed | PIK3CA | -1.129591 | UCEC | AN3-CA | Alpelisib |
| MVP_top60_DNA_seed | PIK3CA | -1.128491 | UCEC | KLE | AZD8186 |
| MVP_top60_DNA_seed | AKT1 | -1.121104 | UCEC | AN3-CA | MK-2206 |
| MVP_top60_DNA_seed | AKT1 | -1.098586 | UCEC | COLO-684 | MK-2206 |
| MVP_top60_DNA_seed | KRAS | -1.093533 | UCEC | SNG-M | KRAS (G12C) Inhibitor-12 |
| MVP_top60_DNA_seed | ATM | -1.082897 | UCEC | MFE-319 | KU-55933 |
| MVP_top60_DNA_seed | AKT1 | -1.07994 | UCEC | MFE-280 | MK-2206 |
| MVP_top60_DNA_seed | AKT1 | -1.069252 | UCEC | AN3-CA | Uprosertib |
| MVP_top60_DNA_seed | PIK3CA | -1.05072 | UCEC | AN3-CA | GNE-317 |
| MVP_top60_DNA_seed | PARP1 | -1.047856 | UCEC | SNG-M | Veliparib |
| MVP_top60_DNA_seed | AKT1 | -1.034058 | UCEC | MFE-280 | Uprosertib |
| MVP_top60_DNA_seed | AKT1 | -1.033309 | UCEC | KLE | MK-2206 |

| | | | | | |
|---|---|---|---|---|---|
| MVP_top60_DNA_seed | AKT1 | -1.029426 | UCEC | EN | Ipatasertib |
| MVP_top60_DNA_seed | PIK3CA | -0.986437 | UCEC | AN3-CA | AZD8186 |
| MVP_top60_DNA_seed | ESR1 | -0.986386 | UCEC | MFE-319 | Tamoxifen |
| MVP_top60_DNA_seed | ATM | -0.980089 | UCEC | AN3-CA | KU-55933 |
| MVP_top60_DNA_seed | ESR1 | -0.97847 | UCEC | COLO-684 | Tamoxifen |
| MVP_top60_DNA_seed | FGFR2 | -0.977111 | UCEC | MFE-280 | AZD4547 |
| MVP_top60_DNA_seed | AKT1 | -0.967928 | UCEC | SNG-M | AT13148 |
| MVP_top60_DNA_seed | JAK1 | -0.957908 | UCEC | COLO-684 | GSK2276186C |
| MVP_top60_DNA_seed | ESR1 | -0.955704 | UCEC | SNG-M | GDC0810 |
| MVP_top60_DNA_seed | AKT1 | -0.948204 | UCEC | COLO-684 | Ipatasertib |
| MVP_top60_DNA_seed | AKT1 | -0.939066 | UCEC | COLO-684 | Uprosertib |
| MVP_top60_DNA_seed | MDM2 | -0.928289 | UCEC | SNG-M | Nutlin-3a (-) |
| MVP_top60_DNA_seed | AKT1 | -0.918815 | UCEC | MFE-296 | Uprosertib |
| MVP_top60_DNA_seed | EP300 | -0.907567 | UCEC | HEC-1 | SGC-CBP30 |
| MVP_top60_DNA_seed | CDK2 | -0.907416 | UCEC | AN3-CA | Dinaciclib |
| MVP_top60_DNA_seed | AKT1 | -0.877355 | UCEC | COLO-684 | AZD5363 |
| MVP_top60_DNA_seed | EP300 | -0.863046 | UCEC | KLE | SGC-CBP30 |
| MVP_top60_DNA_seed | ESR1 | -0.862256 | UCEC | EN | GDC0810 |
| MVP_top60_DNA_seed | CDK2 | -0.859886 | UCEC | MFE-296 | MK-8776 |
| MVP_top60_DNA_seed | EGFR | -0.803748 | UCEC | KLE | Sapitinib |
| MVP_top60_DNA_seed | ERBB2 | -0.803748 | UCEC | KLE | Sapitinib |
| MVP_top60_DNA_seed | PARP1 | -0.802246 | UCEC | SNG-M | Rucaparib |
| MVP_top60_DNA_seed | PIK3CA | -0.799668 | UCEC | AN3-CA | Buparlisib |
| MVP_top60_DNA_seed | PIK3CD | -0.799668 | UCEC | AN3-CA | Buparlisib |
| MVP_top60_DNA_seed | FGFR2 | -0.787283 | UCEC | MFE-296 | AZD4547 |
| MVP_top60_DNA_seed | JAK1 | -0.779399 | UCEC | AN3-CA | Ruxolitinib |
| MVP_top60_DNA_seed | PIK3CA | -0.776155 | UCEC | MFE-319 | GNE-317 |
| MVP_top60_DNA_seed | AKT1 | -0.771595 | UCEC | MFE-280 | Uprosertib |
| MVP_top60_DNA_seed | PARP1 | -0.770478 | UCEC | EN | Veliparib |
| MVP_top60_DNA_seed | EGFR | -0.765043 | UCEC | SNG-M | Sapitinib |
| MVP_top60_DNA_seed | ERBB2 | -0.765043 | UCEC | SNG-M | Sapitinib |
| MVP_top60_DNA_seed | AKT1 | -0.752001 | UCEC | AN3-CA | Afuresertib |
| MVP_top60_DNA_seed | EGFR | -0.740338 | UCEC | MFE-319 | Erlotinib |
| MVP_top60_DNA_seed | AKT1 | -0.693632 | UCEC | MFE-319 | Uprosertib |
| MVP_top60_DNA_seed | PIK3CA | -0.682345 | UCEC | COLO-684 | GNE-317 |
| MVP_top60_DNA_seed | ATM | -0.677619 | UCEC | MFE-280 | KU-55933 |
| MVP_top60_DNA_seed | AKT1 | -0.650013 | UCEC | AN3-CA | AT13148 |
| MVP_top60_DNA_seed | AKT1 | -0.647068 | UCEC | MFE-319 | Ipatasertib |
| MVP_top60_DNA_seed | AKT1 | -0.631172 | UCEC | MFE-319 | AT13148 |
| MVP_top60_DNA_seed | CDK2 | -0.62898 | UCEC | AN3-CA | AZD5438 |
| MVP_top60_DNA_seed | AKT1 | -0.623315 | UCEC | MFE-296 | Afuresertib |

| | | | | | |
|---|---|---|---|---|---|
| MVP_top60_DNA_seed | JAK1 | -0.620585 | UCEC | MFE-280 | JAK_8517 |
| MVP_top60_DNA_seed | CDK2 | -0.618132 | UCEC | SNG-M | Dinaciclib |
| MVP_top60_DNA_seed | EGFR | -0.614033 | UCEC | AN3-CA | Sapitinib |
| MVP_top60_DNA_seed | ERBB2 | -0.614033 | UCEC | AN3-CA | Sapitinib |
| MVP_top60_DNA_seed | AKT1 | -0.609343 | UCEC | MFE-296 | Uprosertib |
| MVP_top60_DNA_seed | TP53 | -0.609038 | UCEC | AN3-CA | MIRA-1 |
| MVP_top60_DNA_seed | EP300 | -0.60117 | UCEC | MFE-296 | SGC-CBP30 |
| MVP_top60_DNA_seed | PARP1 | -0.596758 | UCEC | SNG-M | Niraparib |
| MVP_top60_DNA_seed | AKT1 | -0.595333 | UCEC | EN | Uprosertib |
| MVP_top60_DNA_seed | AKT1 | -0.587729 | UCEC | MFE-280 | GSK2110183B |
| MVP_top60_DNA_seed | ESR1 | -0.580206 | UCEC | MFE-296 | Tamoxifen |
| MVP_top60_DNA_seed | JAK1 | -0.578613 | UCEC | AN3-CA | JAK1_8709 |
| MVP_top60_DNA_seed | HDAC1 | -0.568278 | UCEC | AN3-CA | PCI-34051 |
| MVP_top60_DNA_seed | AKT1 | -0.556221 | UCEC | MFE-280 | Ipatasertib |
| MVP_top60_DNA_seed | PIK3CA | -0.553514 | UCEC | COLO-684 | Alpelisib |
| MVP_top60_DNA_seed | CDK2 | -0.549414 | UCEC | EN | AZD5438 |
| MVP_top60_DNA_seed | AKT1 | -0.526036 | UCEC | EN | MK-2206 |
| MVP_top60_DNA_seed | AKT1 | -0.515101 | UCEC | MFE-319 | Uprosertib |
| MVP_top60_DNA_seed | CDK2 | -0.50745 | UCEC | SNG-M | MK-8776 |
| MVP_top60_DNA_seed | JAK1 | -0.49145 | UCEC | COLO-684 | JAK1_8709 |
| MVP_top60_DNA_seed | HSP90AA1 | -0.485312 | UCEC | MFE-319 | Tanespimycin |
| MVP_top60_DNA_seed | JAK1 | -0.483644 | UCEC | MFE-296 | Ruxolitinib |
| MVP_top60_DNA_seed | ESR1 | -0.474905 | UCEC | MFE-319 | GDC0810 |
| MVP_top60_DNA_seed | HDAC1 | -0.468427 | UCEC | COLO-684 | Romidepsin |
| MVP_top60_DNA_seed | HDAC2 | -0.468427 | UCEC | COLO-684 | Romidepsin |
| MVP_top60_DNA_seed | PARP1 | -0.465023 | UCEC | COLO-684 | Rucaparib |
| MVP_top60_DNA_seed | CDK2 | -0.443393 | UCEC | EN | Dinaciclib |
| MVP_top60_DNA_seed | FGFR2 | -0.441738 | UCEC | MFE-319 | AZD4547 |
| MVP_top60_DNA_seed | EGFR | -0.427167 | UCEC | KLE | Afatinib |
| MVP_top60_DNA_seed | ERBB2 | -0.427167 | UCEC | KLE | Afatinib |
| MVP_top60_DNA_seed | PARP1 | -0.41236 | UCEC | MFE-319 | Rucaparib |
| MVP_top60_DNA_seed | PIK3CA | -0.402583 | UCEC | COLO-684 | AZD8186 |
| MVP_top60_DNA_seed | AKT1 | -0.385786 | UCEC | MFE-296 | Ipatasertib |
| MVP_top60_DNA_seed | HDAC1 | -0.381355 | UCEC | MFE-319 | Entinostat |
| MVP_top60_DNA_seed | AKT1 | -0.375179 | UCEC | MFE-319 | AZD5363 |
| MVP_top60_DNA_seed | KRAS | -0.368617 | UCEC | HEC-1 | KRAS (G12C) Inhibitor-12 |
| MVP_top60_DNA_seed | EP300 | -0.361843 | UCEC | MFE-319 | CPI-637 |
| MVP_top60_DNA_seed | EGFR | -0.344944 | UCEC | MFE-280 | Sapitinib |
| MVP_top60_DNA_seed | ERBB2 | -0.344944 | UCEC | MFE-280 | Sapitinib |
| MVP_top60_DNA_seed | AKT1 | -0.338625 | UCEC | MFE-319 | Afuresertib |
| MVP_top60_DNA_seed | FGFR2 | -0.337154 | UCEC | MFE-280 | PD173074 |

| | | | | | |
|---|---|---|---|---|---|
| MVP_top60_DNA_seed | ESR1 | -0.33711 | UCEC | KLE | GDC0810 |
| MVP_top60_DNA_seed | JAK1 | -0.335557 | UCEC | SNG-M | JAK_8517 |
| MVP_top60_DNA_seed | EP300 | -0.326335 | UCEC | COLO-684 | SGC-CBP30 |
| MVP_top60_DNA_seed | PARP1 | -0.324354 | UCEC | MFE-319 | Olaparib |
| MVP_top60_DNA_seed | PARP1 | -0.32254 | UCEC | AN3-CA | Veliparib |
| MVP_top60_DNA_seed | CDK2 | -0.314826 | UCEC | MFE-319 | MK-8776 |
| MVP_top60_DNA_seed | EP300 | -0.309197 | UCEC | SNG-M | SGC-CBP30 |
| MVP_top60_DNA_seed | HSP90AA1 | -0.30519 | UCEC | AN3-CA | Luminespib |
| MVP_top60_DNA_seed | PIK3CA | -0.300771 | UCEC | KLE | GNE-317 |
| MVP_top60_DNA_seed | PARP1 | -0.290755 | UCEC | COLO-684 | Niraparib |
| MVP_top60_DNA_seed | CDK2 | -0.284026 | UCEC | COLO-684 | AZD5438 |
| MVP_top60_DNA_seed | PIK3CA | -0.277586 | UCEC | EN | Alpelisib |
| MVP_top60_DNA_seed | HDAC1 | -0.272126 | UCEC | MFE-280 | PCI-34051 |
| MVP_top60_DNA_seed | KRAS | -0.264657 | UCEC | EN | KRAS (G12C) Inhibitor-12 |
| MVP_top60_DNA_seed | EGFR | -0.257983 | UCEC | MFE-319 | AZD3759 |
| MVP_top60_DNA_seed | ESR1 | -0.253878 | UCEC | COLO-684 | GDC0810 |
| MVP_top60_DNA_seed | AKT1 | -0.24562 | UCEC | SNG-M | AZD5363 |
| MVP_top60_DNA_seed | AKT1 | -0.239528 | UCEC | MFE-280 | AZD5363 |
| MVP_top60_DNA_seed | HDAC1 | -0.235431 | UCEC | HEC-1 | Romidepsin |
| MVP_top60_DNA_seed | HDAC2 | -0.235431 | UCEC | HEC-1 | Romidepsin |
| MVP_top60_DNA_seed | PIK3CA | -0.22301 | UCEC | MFE-280 | GNE-317 |
| MVP_top60_DNA_seed | PARP1 | -0.221257 | UCEC | EN | Rucaparib |
| MVP_top60_DNA_seed | KRAS | -0.204616 | UCEC | COLO-684 | KRAS (G12C) Inhibitor-12 |
| MVP_top60_DNA_seed | EGFR | -0.201322 | UCEC | KLE | Osimertinib |
| MVP_top60_DNA_seed | HDAC1 | -0.19928 | UCEC | KLE | PCI-34051 |
| MVP_top60_DNA_seed | PARP1 | -0.192794 | UCEC | MFE-296 | Rucaparib |
| MVP_top60_DNA_seed | AKT1 | -0.187275 | UCEC | MFE-280 | Afuresertib |
| MVP_top60_DNA_seed | JAK1 | -0.184399 | UCEC | SNG-M | Ruxolitinib |
| MVP_top60_DNA_seed | HDAC1 | -0.165932 | UCEC | COLO-684 | PCI-34051 |
| MVP_top60_DNA_seed | AKT1 | -0.156282 | UCEC | KLE | Uprosertib |
| MVP_top60_DNA_seed | ATM | -0.154584 | UCEC | MFE-296 | KU-55933 |
| MVP_top60_DNA_seed | EP300 | -0.152548 | UCEC | SNG-M | CPI-637 |
| MVP_top60_DNA_seed | FGFR2 | -0.151758 | UCEC | SNG-M | PD173074 |
| MVP_top60_DNA_seed | MDM2 | -0.141892 | UCEC | AN3-CA | Nutlin-3a (-) |
| MVP_top60_DNA_seed | EGFR | -0.139967 | UCEC | MFE-296 | Erlotinib |
| MVP_top60_DNA_seed | JAK1 | -0.129582 | UCEC | SNG-M | JAK1_8709 |
| MVP_top60_DNA_seed | ATM | -0.127018 | UCEC | SNG-M | KU-55933 |
| MVP_top60_DNA_seed | PIK3CA | -0.117276 | UCEC | MFE-319 | Buparlisib |
| MVP_top60_DNA_seed | PIK3CD | -0.117276 | UCEC | MFE-319 | Buparlisib |
| MVP_top60_DNA_seed | JAK1 | -0.11364 | UCEC | MFE-280 | Ruxolitinib |
| MVP_top60_DNA_seed | CDK2 | -0.109283 | UCEC | MFE-296 | AZD5438 |

| | | | | | |
|---|---|---|---|---|---|
| MVP_top60_DNA_seed | CDK2 | -0.108534 | UCEC | EN | MK-8776 |
| MVP_top60_DNA_seed | JAK1 | -0.095985 | UCEC | MFE-319 | JAK1_8709 |
| MVP_top60_DNA_seed | AKT1 | -0.095928 | UCEC | MFE-296 | AZD5363 |
| MVP_top60_DNA_seed | EGFR | -0.094472 | UCEC | KLE | AZD3759 |
| MVP_top60_DNA_seed | PIK3CA | -0.092167 | UCEC | MFE-280 | AZD8186 |
| MVP_top60_DNA_seed | MDM2 | -0.09068 | UCEC | MFE-296 | Serdemetan |
| MVP_top60_DNA_seed | CDK2 | -0.083845 | UCEC | COLO-684 | MK-8776 |
| MVP_top60_DNA_seed | EGFR | -0.082744 | UCEC | MFE-319 | Gefitinib |
| MVP_top60_DNA_seed | EGFR | -0.082371 | UCEC | AN3-CA | Erlotinib |
| MVP_top60_DNA_seed | MDM2 | -0.068658 | UCEC | MFE-319 | Serdemetan |
| MVP_top60_DNA_seed | EGFR | -0.058552 | UCEC | MFE-319 | Osimertinib |
| MVP_top60_DNA_seed | MDM2 | -0.056338 | UCEC | COLO-684 | Nutlin-3a (-) |
| MVP_top60_DNA_seed | HSP90AA1 | -0.05537 | UCEC | MFE-319 | Luminespib |
| MVP_top60_DNA_seed | PARP1 | -0.053071 | UCEC | MFE-296 | Olaparib |
| MVP_top60_DNA_seed | EP300 | -0.049097 | UCEC | MFE-280 | SGC-CBP30 |
| MVP_top60_DNA_seed | MDM2 | -0.047125 | UCEC | MFE-319 | Nutlin-3a (-) |
| MVP_top60_DNA_seed | ESR1 | -0.046219 | UCEC | MFE-280 | Tamoxifen |
| MVP_top60_DNA_seed | EGFR | -0.044788 | UCEC | SNG-M | AZD3759 |
| MVP_top60_DNA_seed | PIK3CA | -0.035895 | UCEC | COLO-684 | Buparlisib |
| MVP_top60_DNA_seed | PIK3CD | -0.035895 | UCEC | COLO-684 | Buparlisib |
| MVP_top60_DNA_seed | KRAS | -0.031796 | UCEC | AN3-CA | KRAS (G12C) Inhibitor-12 |
| MVP_top60_DNA_seed | EGFR | -0.010186 | UCEC | MFE-280 | Afatinib |
| MVP_top60_DNA_seed | ERBB2 | -0.010186 | UCEC | MFE-280 | Afatinib |
| MVP_top60_DNA_seed | PIK3CA | 0.004545 | UCEC | EN | Buparlisib |
| MVP_top60_DNA_seed | PIK3CD | 0.004545 | UCEC | EN | Buparlisib |
| MVP_top60_DNA_seed | JAK1 | 0.009319 | UCEC | EN | Ruxolitinib |
| MVP_top60_DNA_seed | FGFR2 | 0.011596 | UCEC | MFE-296 | PD173074 |
| MVP_top60_DNA_seed | HSP90AA1 | 0.016781 | UCEC | SNG-M | Luminespib |
| MVP_top60_DNA_seed | AKT1 | 0.028776 | UCEC | SNG-M | Uprosertib |
| MVP_top60_DNA_seed | HSP90AA1 | 0.034726 | UCEC | COLO-684 | Tanespimycin |
| MVP_top60_DNA_seed | ESR1 | 0.048635 | UCEC | SNG-M | Tamoxifen |
| MVP_top60_DNA_seed | EGFR | 0.048945 | UCEC | COLO-684 | Afatinib |
| MVP_top60_DNA_seed | ERBB2 | 0.048945 | UCEC | COLO-684 | Afatinib |
| MVP_top60_DNA_seed | HSP90AA1 | 0.066108 | UCEC | MFE-280 | Tanespimycin |
| MVP_top60_DNA_seed | EGFR | 0.072826 | UCEC | MFE-280 | Lapatinib |
| MVP_top60_DNA_seed | ERBB2 | 0.072826 | UCEC | MFE-280 | Lapatinib |
| MVP_top60_DNA_seed | AKT1 | 0.073645 | UCEC | SNG-M | Ipatasertib |
| MVP_top60_DNA_seed | EGFR | 0.079518 | UCEC | COLO-684 | AZD3759 |
| MVP_top60_DNA_seed | MDM2 | 0.079842 | UCEC | COLO-684 | Serdemetan |
| MVP_top60_DNA_seed | AKT1 | 0.085373 | UCEC | COLO-684 | AT13148 |
| MVP_top60_DNA_seed | PARP1 | 0.096538 | UCEC | AN3-CA | Olaparib |

| | | | | | |
|---|---|---|---|---|---|
| MVP_top60_DNA_seed | EGFR | 0.098884 | UCEC | SNG-M | Afatinib |
| MVP_top60_DNA_seed | ERBB2 | 0.098884 | UCEC | SNG-M | Afatinib |
| MVP_top60_DNA_seed | ESR1 | 0.099888 | UCEC | KLE | Tamoxifen |
| MVP_top60_DNA_seed | EGFR | 0.112023 | UCEC | SNG-M | Osimertinib |
| MVP_top60_DNA_seed | HDAC1 | 0.117694 | UCEC | SNG-M | PCI-34051 |
| MVP_top60_DNA_seed | PIK3CA | 0.121978 | UCEC | KLE | Buparlisib |
| MVP_top60_DNA_seed | PIK3CD | 0.121978 | UCEC | KLE | Buparlisib |
| MVP_top60_DNA_seed | ESR1 | 0.122817 | UCEC | AN3-CA | GDC0810 |
| MVP_top60_DNA_seed | TP53 | 0.125506 | UCEC | SNG-M | MIRA-1 |
| MVP_top60_DNA_seed | CDK2 | 0.12692 | UCEC | MFE-280 | AZD5438 |
| MVP_top60_DNA_seed | MDM2 | 0.127979 | UCEC | MFE-296 | Nutlin-3a (-) |
| MVP_top60_DNA_seed | PARP1 | 0.131189 | UCEC | MFE-280 | Veliparib |
| MVP_top60_DNA_seed | PARP1 | 0.136426 | UCEC | COLO-684 | Veliparib |
| MVP_top60_DNA_seed | PIK3CA | 0.139196 | UCEC | EN | GNE-317 |
| MVP_top60_DNA_seed | AKT1 | 0.143375 | UCEC | EN | AT13148 |
| MVP_top60_DNA_seed | EGFR | 0.147381 | UCEC | SNG-M | Erlotinib |
| MVP_top60_DNA_seed | PARP1 | 0.154685 | UCEC | SNG-M | Olaparib |
| MVP_top60_DNA_seed | TP53 | 0.157663 | UCEC | MFE-296 | MIRA-1 |
| MVP_top60_DNA_seed | PARP1 | 0.160556 | UCEC | MFE-319 | Veliparib |
| MVP_top60_DNA_seed | EGFR | 0.164156 | UCEC | MFE-319 | Lapatinib |
| MVP_top60_DNA_seed | ERBB2 | 0.164156 | UCEC | MFE-319 | Lapatinib |
| MVP_top60_DNA_seed | ESR1 | 0.167485 | UCEC | EN | Tamoxifen |
| MVP_top60_DNA_seed | PARP1 | 0.167759 | UCEC | MFE-280 | Niraparib |
| MVP_top60_DNA_seed | EGFR | 0.168569 | UCEC | KLE | Erlotinib |
| MVP_top60_DNA_seed | TP53 | 0.169217 | UCEC | COLO-684 | MIRA-1 |
| MVP_top60_DNA_seed | EP300 | 0.171773 | UCEC | EN | CPI-637 |
| MVP_top60_DNA_seed | JAK1 | 0.17188 | UCEC | SNG-M | GSK2276186C |
| MVP_top60_DNA_seed | ESR1 | 0.176844 | UCEC | HEC-1 | Tamoxifen |
| MVP_top60_DNA_seed | EGFR | 0.176851 | UCEC | COLO-684 | Lapatinib |
| MVP_top60_DNA_seed | ERBB2 | 0.176851 | UCEC | COLO-684 | Lapatinib |
| MVP_top60_DNA_seed | EGFR | 0.18344 | UCEC | MFE-319 | Sapitinib |
| MVP_top60_DNA_seed | ERBB2 | 0.18344 | UCEC | MFE-319 | Sapitinib |
| MVP_top60_DNA_seed | EP300 | 0.189186 | UCEC | KLE | CPI-637 |
| MVP_top60_DNA_seed | HDAC1 | 0.191042 | UCEC | AN3-CA | Romidepsin |
| MVP_top60_DNA_seed | HDAC2 | 0.191042 | UCEC | AN3-CA | Romidepsin |
| MVP_top60_DNA_seed | EP300 | 0.191117 | UCEC | COLO-684 | CPI-637 |
| MVP_top60_DNA_seed | HDAC1 | 0.194166 | UCEC | MFE-319 | PCI-34051 |
| MVP_top60_DNA_seed | JAK1 | 0.19547 | UCEC | MFE-280 | GSK2276186C |
| MVP_top60_DNA_seed | HDAC1 | 0.198837 | UCEC | SNG-M | Romidepsin |
| MVP_top60_DNA_seed | HDAC2 | 0.198837 | UCEC | SNG-M | Romidepsin |
| MVP_top60_DNA_seed | FGFR2 | 0.204251 | UCEC | KLE | PD173074 |

| | | | | | |
|---|---|---|---|---|---|
| MVP_top60_DNA_seed | EGFR | 0.212163 | UCEC | MFE-296 | AZD3759 |
| MVP_top60_DNA_seed | PIK3CA | 0.230603 | UCEC | SNG-M | Alpelisib |
| MVP_top60_DNA_seed | EP300 | 0.231413 | UCEC | MFE-280 | CPI-637 |
| MVP_top60_DNA_seed | KRAS | 0.232402 | UCEC | MFE-319 | KRAS (G12C) Inhibitor-12 |
| MVP_top60_DNA_seed | PARP1 | 0.236101 | UCEC | MFE-319 | Talazoparib |
| MVP_top60_DNA_seed | JAK1 | 0.239616 | UCEC | COLO-684 | Ruxolitinib |
| MVP_top60_DNA_seed | EP300 | 0.240869 | UCEC | AN3-CA | SGC-CBP30 |
| MVP_top60_DNA_seed | CDK2 | 0.240947 | UCEC | COLO-684 | Dinaciclib |
| MVP_top60_DNA_seed | ESR1 | 0.260571 | UCEC | MFE-280 | GDC0810 |
| MVP_top60_DNA_seed | JAK1 | 0.282599 | UCEC | EN | JAK1_8709 |
| MVP_top60_DNA_seed | PARP1 | 0.285328 | UCEC | SNG-M | Talazoparib |
| MVP_top60_DNA_seed | EGFR | 0.293901 | UCEC | MFE-296 | Osimertinib |
| MVP_top60_DNA_seed | HDAC1 | 0.295173 | UCEC | MFE-296 | PCI-34051 |
| MVP_top60_DNA_seed | EGFR | 0.297354 | UCEC | COLO-684 | Erlotinib |
| MVP_top60_DNA_seed | SRC | 0.300929 | UCEC | SNG-M | Bosutinib |
| MVP_top60_DNA_seed | HDAC1 | 0.304622 | UCEC | MFE-319 | Romidepsin |
| MVP_top60_DNA_seed | HDAC2 | 0.304622 | UCEC | MFE-319 | Romidepsin |
| MVP_top60_DNA_seed | EGFR | 0.311072 | UCEC | HEC-1 | Sapitinib |
| MVP_top60_DNA_seed | ERBB2 | 0.311072 | UCEC | HEC-1 | Sapitinib |
| MVP_top60_DNA_seed | PIK3CA | 0.320685 | UCEC | MFE-319 | AZD8186 |
| MVP_top60_DNA_seed | CDK2 | 0.325627 | UCEC | MFE-319 | Dinaciclib |
| MVP_top60_DNA_seed | AKT1 | 0.336334 | UCEC | SNG-M | GSK2110183B |
| MVP_top60_DNA_seed | CDK2 | 0.337849 | UCEC | AN3-CA | MK-8776 |
| MVP_top60_DNA_seed | HSP90AA1 | 0.34164 | UCEC | COLO-684 | Luminespib |
| MVP_top60_DNA_seed | EGFR | 0.348403 | UCEC | AN3-CA | Gefitinib |
| MVP_top60_DNA_seed | ATM | 0.35817 | UCEC | COLO-684 | KU-55933 |
| MVP_top60_DNA_seed | PARP1 | 0.366463 | UCEC | AN3-CA | Niraparib |
| MVP_top60_DNA_seed | EGFR | 0.371226 | UCEC | AN3-CA | Afatinib |
| MVP_top60_DNA_seed | ERBB2 | 0.371226 | UCEC | AN3-CA | Afatinib |
| MVP_top60_DNA_seed | PIK3CA | 0.376584 | UCEC | SNG-M | GNE-317 |
| MVP_top60_DNA_seed | PARP1 | 0.377331 | UCEC | KLE | Olaparib |
| MVP_top60_DNA_seed | EGFR | 0.380384 | UCEC | KLE | Lapatinib |
| MVP_top60_DNA_seed | ERBB2 | 0.380384 | UCEC | KLE | Lapatinib |
| MVP_top60_DNA_seed | EGFR | 0.384107 | UCEC | SNG-M | Gefitinib |
| MVP_top60_DNA_seed | CDK2 | 0.38513 | UCEC | MFE-280 | Dinaciclib |
| MVP_top60_DNA_seed | ESR1 | 0.389647 | UCEC | HEC-1 | GDC0810 |
| MVP_top60_DNA_seed | AKT1 | 0.393859 | UCEC | MFE-280 | AT13148 |
| MVP_top60_DNA_seed | SRC | 0.394768 | UCEC | COLO-684 | Bosutinib |
| MVP_top60_DNA_seed | EGFR | 0.406535 | UCEC | SNG-M | Lapatinib |
| MVP_top60_DNA_seed | ERBB2 | 0.406535 | UCEC | SNG-M | Lapatinib |
| MVP_top60_DNA_seed | PIK3CA | 0.414572 | UCEC | SNG-M | Buparlisib |

| | | | | | |
|---|---|---|---|---|---|
| MVP_top60_DNA_seed | PIK3CD | 0.414572 | UCEC | SNG-M | Buparlisib |
| MVP_top60_DNA_seed | PARP1 | 0.419411 | UCEC | MFE-296 | Talazoparib |
| MVP_top60_DNA_seed | JAK1 | 0.425695 | UCEC | MFE-319 | Ruxolitinib |
| MVP_top60_DNA_seed | KRAS | 0.426664 | UCEC | MFE-280 | KRAS (G12C) Inhibitor-12 |
| MVP_top60_DNA_seed | ESR1 | 0.42959 | UCEC | MFE-296 | GDC0810 |
| MVP_top60_DNA_seed | EGFR | 0.430831 | UCEC | MFE-319 | Afatinib |
| MVP_top60_DNA_seed | ERBB2 | 0.430831 | UCEC | MFE-319 | Afatinib |
| MVP_top60_DNA_seed | EP300 | 0.432925 | UCEC | MFE-296 | CPI-637 |
| MVP_top60_DNA_seed | EGFR | 0.433326 | UCEC | KLE | Gefitinib |
| MVP_top60_DNA_seed | PARP1 | 0.436597 | UCEC | EN | Olaparib |
| MVP_top60_DNA_seed | AKT1 | 0.437204 | UCEC | MFE-319 | GSK2110183B |
| MVP_top60_DNA_seed | FGFR2 | 0.454732 | UCEC | SNG-M | AZD4547 |
| MVP_top60_DNA_seed | PARP1 | 0.456994 | UCEC | MFE-296 | Niraparib |
| MVP_top60_DNA_seed | PARP1 | 0.469431 | UCEC | MFE-296 | Veliparib |
| MVP_top60_DNA_seed | JAK1 | 0.471991 | UCEC | MFE-296 | GSK2276186C |
| MVP_top60_DNA_seed | EGFR | 0.484221 | UCEC | MFE-296 | Sapitinib |
| MVP_top60_DNA_seed | ERBB2 | 0.484221 | UCEC | MFE-296 | Sapitinib |
| MVP_top60_DNA_seed | PARP1 | 0.485149 | UCEC | AN3-CA | Rucaparib |
| MVP_top60_DNA_seed | PIK3CA | 0.485906 | UCEC | HEC-1 | GNE-317 |
| MVP_top60_DNA_seed | JAK1 | 0.487375 | UCEC | MFE-319 | GSK2276186C |
| MVP_top60_DNA_seed | HSP90AA1 | 0.489232 | UCEC | MFE-296 | Tanespimycin |
| MVP_top60_DNA_seed | HSP90AA1 | 0.493502 | UCEC | MFE-280 | Luminespib |
| MVP_top60_DNA_seed | EGFR | 0.499133 | UCEC | AN3-CA | AZD3759 |
| MVP_top60_DNA_seed | JAK1 | 0.506222 | UCEC | EN | JAK_8517 |
| MVP_top60_DNA_seed | AKT1 | 0.538987 | UCEC | KLE | AZD5363 |
| MVP_top60_DNA_seed | AKT1 | 0.542338 | UCEC | MFE-296 | AT13148 |
| MVP_top60_DNA_seed | PIK3CA | 0.543186 | UCEC | MFE-280 | Buparlisib |
| MVP_top60_DNA_seed | PIK3CD | 0.543186 | UCEC | MFE-280 | Buparlisib |
| MVP_top60_DNA_seed | TP53 | 0.56048 | UCEC | MFE-319 | MIRA-1 |
| MVP_top60_DNA_seed | AKT1 | 0.56375 | UCEC | SNG-M | Afuresertib |
| MVP_top60_DNA_seed | EGFR | 0.564521 | UCEC | MFE-280 | AZD3759 |
| MVP_top60_DNA_seed | JAK1 | 0.578082 | UCEC | MFE-319 | JAK_8517 |
| MVP_top60_DNA_seed | PARP1 | 0.582666 | UCEC | EN | Niraparib |
| MVP_top60_DNA_seed | PARP1 | 0.588089 | UCEC | MFE-280 | Olaparib |
| MVP_top60_DNA_seed | PARP1 | 0.596569 | UCEC | MFE-319 | Niraparib |
| MVP_top60_DNA_seed | HDAC1 | 0.600748 | UCEC | MFE-296 | Romidepsin |
| MVP_top60_DNA_seed | HDAC2 | 0.600748 | UCEC | MFE-296 | Romidepsin |
| MVP_top60_DNA_seed | AKT1 | 0.605984 | UCEC | SNG-M | MK-2206 |
| MVP_top60_DNA_seed | HDAC1 | 0.608743 | UCEC | MFE-280 | Romidepsin |
| MVP_top60_DNA_seed | HDAC2 | 0.608743 | UCEC | MFE-280 | Romidepsin |
| MVP_top60_DNA_seed | FGFR2 | 0.620923 | UCEC | HEC-1 | AZD4547 |

| | | | | | |
|---|---|---|---|---|---|
| MVP_top60_DNA_seed | EGFR | 0.62759 | UCEC | AN3-CA | Lapatinib |
| MVP_top60_DNA_seed | ERBB2 | 0.62759 | UCEC | AN3-CA | Lapatinib |
| MVP_top60_DNA_seed | HDAC1 | 0.633137 | UCEC | HEC-1 | Entinostat |
| MVP_top60_DNA_seed | SRC | 0.650923 | UCEC | AN3-CA | Bosutinib |
| MVP_top60_DNA_seed | SRC | 0.653856 | UCEC | HEC-1 | Bosutinib |
| MVP_top60_DNA_seed | ATM | 0.65422 | UCEC | EN | KU-55933 |
| MVP_top60_DNA_seed | HDAC1 | 0.656557 | UCEC | AN3-CA | Entinostat |
| MVP_top60_DNA_seed | MDM2 | 0.659737 | UCEC | HEC-1 | Nutlin-3a (-) |
| MVP_top60_DNA_seed | EGFR | 0.667135 | UCEC | EN | Afatinib |
| MVP_top60_DNA_seed | ERBB2 | 0.667135 | UCEC | EN | Afatinib |
| MVP_top60_DNA_seed | EGFR | 0.668489 | UCEC | HEC-1 | AZD3759 |
| MVP_top60_DNA_seed | MDM2 | 0.671411 | UCEC | SNG-M | Serdemetan |
| MVP_top60_DNA_seed | EGFR | 0.672078 | UCEC | MFE-296 | Gefitinib |
| MVP_top60_DNA_seed | AKT1 | 0.675819 | UCEC | MFE-296 | GSK2110183B |
| MVP_top60_DNA_seed | AKT1 | 0.692956 | UCEC | HEC-1 | AT13148 |
| MVP_top60_DNA_seed | SRC | 0.694887 | UCEC | MFE-296 | Bosutinib |
| MVP_top60_DNA_seed | EGFR | 0.699991 | UCEC | MFE-280 | Osimertinib |
| MVP_top60_DNA_seed | AKT1 | 0.702212 | UCEC | HEC-1 | AZD5363 |
| MVP_top60_DNA_seed | EP300 | 0.711783 | UCEC | AN3-CA | CPI-637 |
| MVP_top60_DNA_seed | MDM2 | 0.712865 | UCEC | EN | Nutlin-3a (-) |
| MVP_top60_DNA_seed | AKT1 | 0.713039 | UCEC | SNG-M | Uprosertib |
| MVP_top60_DNA_seed | EGFR | 0.716971 | UCEC | AN3-CA | Osimertinib |
| MVP_top60_DNA_seed | EP300 | 0.722915 | UCEC | HEC-1 | CPI-637 |
| MVP_top60_DNA_seed | TP53 | 0.730928 | UCEC | EN | MIRA-1 |
| MVP_top60_DNA_seed | PARP1 | 0.734876 | UCEC | COLO-684 | Talazoparib |
| MVP_top60_DNA_seed | SRC | 0.742165 | UCEC | MFE-319 | Bosutinib |
| MVP_top60_DNA_seed | FGFR2 | 0.750459 | UCEC | COLO-684 | AZD4547 |
| MVP_top60_DNA_seed | PARP1 | 0.754178 | UCEC | HEC-1 | Talazoparib |
| MVP_top60_DNA_seed | CDK2 | 0.756381 | UCEC | MFE-319 | AZD5438 |
| MVP_top60_DNA_seed | PARP1 | 0.764373 | UCEC | MFE-280 | Talazoparib |
| MVP_top60_DNA_seed | HDAC1 | 0.767931 | UCEC | EN | Entinostat |
| MVP_top60_DNA_seed | HDAC1 | 0.790518 | UCEC | HEC-1 | PCI-34051 |
| MVP_top60_DNA_seed | HSP90AA1 | 0.791403 | UCEC | EN | Tanespimycin |
| MVP_top60_DNA_seed | PARP1 | 0.793155 | UCEC | KLE | Talazoparib |
| MVP_top60_DNA_seed | ESR1 | 0.793876 | UCEC | AN3-CA | Tamoxifen |
| MVP_top60_DNA_seed | HSP90AA1 | 0.807676 | UCEC | SNG-M | Tanespimycin |
| MVP_top60_DNA_seed | AKT1 | 0.812305 | UCEC | HEC-1 | GSK2110183B |
| MVP_top60_DNA_seed | CDK2 | 0.814167 | UCEC | HEC-1 | MK-8776 |
| MVP_top60_DNA_seed | PIK3CA | 0.816956 | UCEC | MFE-296 | Buparlisib |
| MVP_top60_DNA_seed | PIK3CD | 0.816956 | UCEC | MFE-296 | Buparlisib |
| MVP_top60_DNA_seed | HDAC1 | 0.817593 | UCEC | EN | PCI-34051 |

| | | | | | |
|---|---|---|---|---|---|
| MVP_top60_DNA_seed | KRAS | 0.824168 | UCEC | KLE | KRAS (G12C) Inhibitor-12 |
| MVP_top60_DNA_seed | HDAC1 | 0.834873 | UCEC | EN | Romidepsin |
| MVP_top60_DNA_seed | HDAC2 | 0.834873 | UCEC | EN | Romidepsin |
| MVP_top60_DNA_seed | JAK1 | 0.841807 | UCEC | MFE-280 | JAK1_8709 |
| MVP_top60_DNA_seed | EP300 | 0.853598 | UCEC | MFE-319 | SGC-CBP30 |
| MVP_top60_DNA_seed | TP53 | 0.860405 | UCEC | MFE-280 | MIRA-1 |
| MVP_top60_DNA_seed | EGFR | 0.863622 | UCEC | EN | Osimertinib |
| MVP_top60_DNA_seed | HDAC1 | 0.864204 | UCEC | MFE-296 | Entinostat |
| MVP_top60_DNA_seed | CDK2 | 0.874853 | UCEC | MFE-280 | MK-8776 |
| MVP_top60_DNA_seed | JAK1 | 0.888952 | UCEC | HEC-1 | GSK2276186C |
| MVP_top60_DNA_seed | ATM | 0.891619 | UCEC | HEC-1 | KU-55933 |
| MVP_top60_DNA_seed | KRAS | 0.892833 | UCEC | MFE-296 | KRAS (G12C) Inhibitor-12 |
| MVP_top60_DNA_seed | JAK1 | 0.896522 | UCEC | EN | GSK2276186C |
| MVP_top60_DNA_seed | EGFR | 0.902998 | UCEC | HEC-1 | Erlotinib |
| MVP_top60_DNA_seed | PIK3CA | 0.916666 | UCEC | HEC-1 | Buparlisib |
| MVP_top60_DNA_seed | PIK3CD | 0.916666 | UCEC | HEC-1 | Buparlisib |
| MVP_top60_DNA_seed | AKT1 | 0.937382 | UCEC | MFE-296 | MK-2206 |
| MVP_top60_DNA_seed | EGFR | 0.938034 | UCEC | EN | AZD3759 |
| MVP_top60_DNA_seed | PIK3CA | 0.94508 | UCEC | EN | AZD8186 |
| MVP_top60_DNA_seed | FGFR2 | 0.946602 | UCEC | EN | AZD4547 |
| MVP_top60_DNA_seed | PIK3CA | 0.953357 | UCEC | MFE-319 | Alpelisib |
| MVP_top60_DNA_seed | HSP90AA1 | 0.954938 | UCEC | MFE-296 | Luminespib |
| MVP_top60_DNA_seed | PARP1 | 0.967666 | UCEC | AN3-CA | Talazoparib |
| MVP_top60_DNA_seed | SRC | 0.974677 | UCEC | KLE | Bosutinib |
| MVP_top60_DNA_seed | AKT1 | 0.976671 | UCEC | KLE | Ipatasertib |
| MVP_top60_DNA_seed | EGFR | 1.004285 | UCEC | EN | Gefitinib |
| MVP_top60_DNA_seed | PARP1 | 1.008732 | UCEC | COLO-684 | Olaparib |
| MVP_top60_DNA_seed | EGFR | 1.017563 | UCEC | EN | Sapitinib |
| MVP_top60_DNA_seed | ERBB2 | 1.017563 | UCEC | EN | Sapitinib |
| MVP_top60_DNA_seed | EGFR | 1.021104 | UCEC | EN | Lapatinib |
| MVP_top60_DNA_seed | ERBB2 | 1.021104 | UCEC | EN | Lapatinib |
| MVP_top60_DNA_seed | PARP1 | 1.021219 | UCEC | HEC-1 | Veliparib |
| MVP_top60_DNA_seed | PIK3CA | 1.02174 | UCEC | HEC-1 | AZD8186 |
| MVP_top60_DNA_seed | AKT1 | 1.043738 | UCEC | HEC-1 | Afuresertib |
| MVP_top60_DNA_seed | PIK3CA | 1.043925 | UCEC | SNG-M | AZD8186 |
| MVP_top60_DNA_seed | EGFR | 1.044043 | UCEC | COLO-684 | Gefitinib |
| MVP_top60_DNA_seed | JAK1 | 1.059244 | UCEC | HEC-1 | Ruxolitinib |
| MVP_top60_DNA_seed | AKT1 | 1.065309 | UCEC | HEC-1 | MK-2206 |
| MVP_top60_DNA_seed | TP53 | 1.069404 | UCEC | HEC-1 | MIRA-1 |
| MVP_top60_DNA_seed | PIK3CA | 1.070358 | UCEC | MFE-296 | Alpelisib |
| MVP_top60_DNA_seed | HDAC1 | 1.08261 | UCEC | MFE-280 | Entinostat |

| | | | | | |
|---|---|---|---|---|---|
| MVP_top60_DNA_seed | MDM2 | 1.084862 | UCEC | KLE | Nutlin-3a (-) |
| MVP_top60_DNA_seed | PIK3CA | 1.09639 | UCEC | MFE-296 | AZD8186 |
| MVP_top60_DNA_seed | JAK1 | 1.109738 | UCEC | MFE-296 | JAK1_8709 |
| MVP_top60_DNA_seed | MDM2 | 1.122706 | UCEC | EN | Serdemetan |
| MVP_top60_DNA_seed | SRC | 1.124856 | UCEC | MFE-280 | Bosutinib |
| MVP_top60_DNA_seed | EGFR | 1.128682 | UCEC | HEC-1 | Osimertinib |
| MVP_top60_DNA_seed | JAK1 | 1.130173 | UCEC | MFE-296 | JAK_8517 |
| MVP_top60_DNA_seed | PARP1 | 1.135546 | UCEC | HEC-1 | Olaparib |
| MVP_top60_DNA_seed | EGFR | 1.136334 | UCEC | HEC-1 | Gefitinib |
| MVP_top60_DNA_seed | MDM2 | 1.180671 | UCEC | MFE-280 | Nutlin-3a (-) |
| MVP_top60_DNA_seed | AKT1 | 1.182879 | UCEC | KLE | Uprosertib |
| MVP_top60_DNA_seed | HDAC1 | 1.183379 | UCEC | SNG-M | Entinostat |
| MVP_top60_DNA_seed | PARP1 | 1.192243 | UCEC | HEC-1 | Niraparib |
| MVP_top60_DNA_seed | PIK3CA | 1.200719 | UCEC | HEC-1 | Alpelisib |
| MVP_top60_DNA_seed | PARP1 | 1.20123 | UCEC | MFE-280 | Rucaparib |
| MVP_top60_DNA_seed | AKT1 | 1.206756 | UCEC | MFE-319 | MK-2206 |
| MVP_top60_DNA_seed | EGFR | 1.228972 | UCEC | EN | Erlotinib |
| MVP_top60_DNA_seed | AKT1 | 1.229678 | UCEC | HEC-1 | Uprosertib |
| MVP_top60_DNA_seed | CDK2 | 1.230365 | UCEC | KLE | Dinaciclib |
| MVP_top60_DNA_seed | FGFR2 | 1.238673 | UCEC | EN | PD173074 |
| MVP_top60_DNA_seed | MDM2 | 1.241378 | UCEC | AN3-CA | Serdemetan |
| MVP_top60_DNA_seed | SRC | 1.241598 | UCEC | EN | Bosutinib |
| MVP_top60_DNA_seed | FGFR2 | 1.252341 | UCEC | KLE | AZD4547 |
| MVP_top60_DNA_seed | AKT1 | 1.264935 | UCEC | HEC-1 | Ipatasertib |
| MVP_top60_DNA_seed | EGFR | 1.265159 | UCEC | HEC-1 | Afatinib |
| MVP_top60_DNA_seed | ERBB2 | 1.265159 | UCEC | HEC-1 | Afatinib |
| MVP_top60_DNA_seed | EGFR | 1.286195 | UCEC | COLO-684 | Sapitinib |
| MVP_top60_DNA_seed | ERBB2 | 1.286195 | UCEC | COLO-684 | Sapitinib |
| MVP_top60_DNA_seed | EGFR | 1.290504 | UCEC | MFE-280 | Gefitinib |
| MVP_top60_DNA_seed | HSP90AA1 | 1.317908 | UCEC | KLE | Luminespib |
| MVP_top60_DNA_seed | CDK2 | 1.317964 | UCEC | KLE | MK-8776 |
| MVP_top60_DNA_seed | PARP1 | 1.352579 | UCEC | KLE | Niraparib |
| MVP_top60_DNA_seed | AKT1 | 1.368983 | UCEC | KLE | Afuresertib |
| MVP_top60_DNA_seed | HSP90AA1 | 1.369409 | UCEC | EN | Luminespib |
| MVP_top60_DNA_seed | EGFR | 1.378763 | UCEC | MFE-296 | Afatinib |
| MVP_top60_DNA_seed | ERBB2 | 1.378763 | UCEC | MFE-296 | Afatinib |
| MVP_top60_DNA_seed | MDM2 | 1.381257 | UCEC | MFE-280 | Serdemetan |
| MVP_top60_DNA_seed | JAK1 | 1.386109 | UCEC | HEC-1 | JAK1_8709 |
| MVP_top60_DNA_seed | HDAC1 | 1.394134 | UCEC | COLO-684 | Entinostat |
| MVP_top60_DNA_seed | HSP90AA1 | 1.40746 | UCEC | KLE | Tanespimycin |
| MVP_top60_DNA_seed | CDK2 | 1.413039 | UCEC | HEC-1 | Dinaciclib |

| | | | | | |
|---|---|---|---|---|---|
| MVP_top60_DNA_seed | PARP1 | 1.414427 | UCEC | EN | Talazoparib |
| MVP_top60_DNA_seed | PARP1 | 1.42695 | UCEC | HEC-1 | Rucaparib |
| MVP_top60_DNA_seed | FGFR2 | 1.456139 | UCEC | MFE-319 | PD173074 |
| MVP_top60_DNA_seed | ATM | 1.460351 | UCEC | KLE | KU-55933 |
| MVP_top60_DNA_seed | AKT1 | 1.480473 | UCEC | KLE | AT13148 |
| MVP_top60_DNA_seed | HDAC1 | 1.500964 | UCEC | KLE | Romidepsin |
| MVP_top60_DNA_seed | HDAC2 | 1.500964 | UCEC | KLE | Romidepsin |
| MVP_top60_DNA_seed | EGFR | 1.524505 | UCEC | MFE-280 | Erlotinib |
| MVP_top60_DNA_seed | CDK2 | 1.552748 | UCEC | MFE-296 | Dinaciclib |
| MVP_top60_DNA_seed | AKT1 | 1.572434 | UCEC | KLE | GSK2110183B |
| MVP_top60_DNA_seed | TP53 | 1.5776 | UCEC | KLE | MIRA-1 |
| MVP_top60_DNA_seed | CDK2 | 1.619899 | UCEC | KLE | AZD5438 |
| MVP_top60_DNA_seed | JAK1 | 1.620162 | UCEC | HEC-1 | JAK_8517 |
| MVP_top60_DNA_seed | EGFR | 1.622469 | UCEC | COLO-684 | Osimertinib |
| MVP_top60_DNA_seed | JAK1 | 1.641106 | UCEC | KLE | Ruxolitinib |
| MVP_top60_DNA_seed | PIK3CA | 1.64178 | UCEC | KLE | Alpelisib |
| MVP_top60_DNA_seed | PIK3CA | 1.651226 | UCEC | MFE-296 | GNE-317 |
| MVP_top60_DNA_seed | JAK1 | 1.666591 | UCEC | KLE | GSK2276186C |
| MVP_top60_DNA_seed | HDAC1 | 1.698881 | UCEC | KLE | Entinostat |
| MVP_top60_DNA_seed | MDM2 | 1.726795 | UCEC | KLE | Serdemetan |
| MVP_top60_DNA_seed | EGFR | 1.72751 | UCEC | MFE-296 | Lapatinib |
| MVP_top60_DNA_seed | ERBB2 | 1.72751 | UCEC | MFE-296 | Lapatinib |
| MVP_top60_DNA_seed | AKT1 | 1.729271 | UCEC | HEC-1 | Uprosertib |
| MVP_top60_DNA_seed | JAK1 | 1.782162 | UCEC | COLO-684 | JAK_8517 |
| MVP_top60_DNA_seed | FGFR2 | 1.811944 | UCEC | COLO-684 | PD173074 |
| MVP_top60_DNA_seed | HSP90AA1 | 1.863317 | UCEC | HEC-1 | Tanespimycin |
| MVP_top60_DNA_seed | FGFR2 | 1.915425 | UCEC | HEC-1 | PD173074 |
| MVP_top60_DNA_seed | MDM2 | 1.915843 | UCEC | HEC-1 | Serdemetan |
| MVP_top60_DNA_seed | EGFR | 1.971861 | UCEC | HEC-1 | Lapatinib |
| MVP_top60_DNA_seed | ERBB2 | 1.971861 | UCEC | HEC-1 | Lapatinib |
| MVP_top60_DNA_seed | HSP90AA1 | 1.980393 | UCEC | HEC-1 | Luminespib |
| MVP_top60_DNA_seed | PARP1 | 2.086899 | UCEC | KLE | Veliparib |
| MVP_top60_DNA_seed | PARP1 | 2.099616 | UCEC | KLE | Rucaparib |
| MVP_top60_DNA_seed | CDK2 | 2.102895 | UCEC | HEC-1 | AZD5438 |
| MVP_top60_DNA_seed | JAK1 | 2.248089 | UCEC | KLE | JAK1_8709 |
| MVP_top60_DNA_seed | JAK1 | 2.426792 | UCEC | KLE | JAK_8517 |
| MVP_top60_DNA_seed | BRAF | -5.08881 | SKCM | G-MEL | SB590885 |
| MVP_top60_DNA_seed | BRAF | -5.015441 | SKCM | IGR-37 | SB590885 |
| MVP_top60_DNA_seed | BRAF | -4.956381 | SKCM | MEL-HO | SB590885 |
| MVP_top60_DNA_seed | BRAF | -4.687931 | SKCM | 451Lu | SB590885 |
| MVP_top60_DNA_seed | BRAF | -4.663005 | SKCM | IGR-37 | PLX-4720 |

| | | | | | |
|---|---|---|---|---|---|
| MVP_top60_DNA_seed | BRAF | -4.608082 | SKCM | C32 | SB590885 |
| MVP_top60_DNA_seed | BRAF | -4.435693 | SKCM | M14 | SB590885 |
| MVP_top60_DNA_seed | BRAF | -4.213756 | SKCM | G-MEL | PLX-4720 |
| MVP_top60_DNA_seed | BRAF | -4.152893 | SKCM | 451Lu | PLX-4720 |
| MVP_top60_DNA_seed | BRAF | -4.105172 | SKCM | C32 | PLX-4720 |
| MVP_top60_DNA_seed | BRAF | -4.095785 | SKCM | MMAc-SF | PLX-4720 |
| MVP_top60_DNA_seed | BRAF | -4.084635 | SKCM | MMAc-SF | Dabrafenib |
| MVP_top60_DNA_seed | BRAF | -4.076641 | SKCM | SH-4 | PLX-4720 |
| MVP_top60_DNA_seed | BRAF | -4.045547 | SKCM | G-MEL | Dabrafenib |
| MVP_top60_DNA_seed | BRAF | -4.036444 | SKCM | IGR-37 | PLX-4720 |
| MVP_top60_DNA_seed | BRAF | -3.995559 | SKCM | COLO-679 | SB590885 |
| MVP_top60_DNA_seed | BRAF | -3.991431 | SKCM | IGR-37 | Dabrafenib |
| MVP_top60_DNA_seed | BRAF | -3.918523 | SKCM | A375 | Dabrafenib |
| MVP_top60_DNA_seed | BRAF | -3.918006 | SKCM | C32 | Dabrafenib |
| MVP_top60_DNA_seed | BRAF | -3.849316 | SKCM | CP50-MEL-B | PLX-4720 |
| MVP_top60_DNA_seed | BRAF | -3.818179 | SKCM | M14 | PLX-4720 |
| MVP_top60_DNA_seed | BRAF | -3.81523 | SKCM | SH-4 | Dabrafenib |
| MVP_top60_DNA_seed | BRAF | -3.800046 | SKCM | G-MEL | PLX-4720 |
| MVP_top60_DNA_seed | BRAF | -3.73954 | SKCM | MEL-HO | Dabrafenib |
| MVP_top60_DNA_seed | BRAF | -3.737306 | SKCM | COLO-679 | PLX-4720 |
| MVP_top60_DNA_seed | BRAF | -3.728287 | SKCM | A375 | PLX-4720 |
| MVP_top60_DNA_seed | BRAF | -3.70433 | SKCM | MEL-HO | PLX-4720 |
| MVP_top60_DNA_seed | BRAF | -3.703903 | SKCM | WM35 | PLX-4720 |
| MVP_top60_DNA_seed | MAP2K1 | -3.701237 | SKCM | IGR-37 | Selumetinib |
| MVP_top60_DNA_seed | BRAF | -3.69593 | SKCM | UACC-62 | PLX-4720 |
| MVP_top60_DNA_seed | BRAF | -3.677782 | SKCM | A375 | SB590885 |
| MVP_top60_DNA_seed | GSK3B | -3.673378 | SKCM | LOXIMVI | CHIR-99021 |
| MVP_top60_DNA_seed | BRAF | -3.661088 | SKCM | A375 | PLX-4720 |
| MVP_top60_DNA_seed | BRAF | -3.642713 | SKCM | MEL-HO | PLX-4720 |
| MVP_top60_DNA_seed | BRAF | -3.633844 | SKCM | M14 | Dabrafenib |
| MVP_top60_DNA_seed | BRAF | -3.630199 | SKCM | HT-144 | SB590885 |
| MVP_top60_DNA_seed | BRAF | -3.621954 | SKCM | CP50-MEL-B | Dabrafenib |
| MVP_top60_DNA_seed | BRAF | -3.59225 | SKCM | WM35 | Dabrafenib |
| MVP_top60_DNA_seed | BRAF | -3.579647 | SKCM | IGR-37 | RAF_9304 |
| MVP_top60_DNA_seed | BRAF | -3.554161 | SKCM | RVH-421 | SB590885 |
| MVP_top60_DNA_seed | BRAF | -3.540122 | SKCM | A101D | PLX-4720 |
| MVP_top60_DNA_seed | BRAF | -3.498877 | SKCM | 451Lu | Dabrafenib |
| MVP_top60_DNA_seed | BRAF | -3.491049 | SKCM | M14 | PLX-4720 |
| MVP_top60_DNA_seed | GSK3B | -3.471882 | SKCM | COLO-829 | CHIR-99021 |
| MVP_top60_DNA_seed | BRAF | -3.453765 | SKCM | UACC-62 | Dabrafenib |
| MVP_top60_DNA_seed | BRAF | -3.406296 | SKCM | SH-4 | PLX-4720 |

| | | | | | |
|---|---|---|---|---|---|
| MVP_top60_DNA_seed | BRAF | -3.399603 | SKCM | UACC-257 | Dabrafenib |
| MVP_top60_DNA_seed | MAP2K1 | -3.377896 | SKCM | M14 | Selumetinib |
| MVP_top60_DNA_seed | BRAF | -3.353063 | SKCM | K2 | PLX-4720 |
| MVP_top60_DNA_seed | BRAF | -3.349799 | SKCM | G-MEL | RAF_9304 |
| MVP_top60_DNA_seed | BRAF | -3.34063 | SKCM | LB2518-MEL | AZ628 |
| MVP_top60_DNA_seed | BRAF | -3.322197 | SKCM | K2 | Dabrafenib |
| MVP_top60_DNA_seed | BRAF | -3.318276 | SKCM | G-361 | Dabrafenib |
| MVP_top60_DNA_seed | BRAF | -3.31556 | SKCM | SK-MEL-28 | Dabrafenib |
| MVP_top60_DNA_seed | BRAF | -3.303363 | SKCM | COLO-829 | SB590885 |
| MVP_top60_DNA_seed | BRAF | -3.299736 | SKCM | COLO-679 | Dabrafenib |
| MVP_top60_DNA_seed | BRAF | -3.295407 | SKCM | C32 | PLX-4720 |
| MVP_top60_DNA_seed | BRAF | -3.28739 | SKCM | SK-MEL-1 | Dabrafenib |
| MVP_top60_DNA_seed | EGFR | -3.276454 | SKCM | CHL-1 | Afatinib |
| MVP_top60_DNA_seed | ERBB2 | -3.276454 | SKCM | CHL-1 | Afatinib |
| MVP_top60_DNA_seed | MAP2K1 | -3.228153 | SKCM | IGR-37 | Refametinib |
| MVP_top60_DNA_seed | JAK3 | -3.223132 | SKCM | CHL-1 | WHI-P97 |
| MVP_top60_DNA_seed | BRAF | -3.203638 | SKCM | LOXIMVI | Dabrafenib |
| MVP_top60_DNA_seed | BRAF | -3.203336 | SKCM | MMAc-SF | PLX-4720 |
| MVP_top60_DNA_seed | MAP2K1 | -3.194339 | SKCM | IGR-37 | Selumetinib |
| MVP_top60_DNA_seed | BRAF | -3.193343 | SKCM | A101D | SB590885 |
| MVP_top60_DNA_seed | BRAF | -3.192999 | SKCM | CP50-MEL-B | PLX-4720 |
| MVP_top60_DNA_seed | BRAF | -3.186265 | SKCM | WM-115 | SB590885 |
| MVP_top60_DNA_seed | BRAF | -3.174985 | SKCM | MZ2-MEL | PLX-4720 |
| MVP_top60_DNA_seed | MAP2K1 | -3.169773 | SKCM | G-MEL | CI-1040 |
| MVP_top60_DNA_seed | BRAF | -3.168905 | SKCM | SH-4 | AZ628 |
| MVP_top60_DNA_seed | BRAF | -3.150297 | SKCM | UACC-257 | PLX-4720 |
| MVP_top60_DNA_seed | BRAF | -3.112875 | SKCM | SK-MEL-28 | PLX-4720 |
| MVP_top60_DNA_seed | EGFR | -3.111399 | SKCM | CHL-1 | Afatinib |
| MVP_top60_DNA_seed | ERBB2 | -3.111399 | SKCM | CHL-1 | Afatinib |
| MVP_top60_DNA_seed | BRAF | -3.095024 | SKCM | SK-MEL-1 | PLX-4720 |
| MVP_top60_DNA_seed | BRAF | -3.083214 | SKCM | A101D | Dabrafenib |
| MVP_top60_DNA_seed | BRAF | -3.079853 | SKCM | HT-144 | AZ628 |
| MVP_top60_DNA_seed | BRAF | -3.067988 | SKCM | SH-4 | SB590885 |
| MVP_top60_DNA_seed | EGFR | -3.062584 | SKCM | WM35 | Afatinib |
| MVP_top60_DNA_seed | ERBB2 | -3.062584 | SKCM | WM35 | Afatinib |
| MVP_top60_DNA_seed | BRAF | -3.059115 | SKCM | WM35 | RAF_9304 |
| MVP_top60_DNA_seed | BRAF | -3.057456 | SKCM | HT-144 | PLX-4720 |
| MVP_top60_DNA_seed | BRAF | -3.057264 | SKCM | RVH-421 | PLX-4720 |
| MVP_top60_DNA_seed | MAP2K1 | -3.051211 | SKCM | G-MEL | Selumetinib |
| MVP_top60_DNA_seed | MAP2K1 | -3.050178 | SKCM | IPC-298 | Selumetinib |
| MVP_top60_DNA_seed | MAP2K1 | -3.048248 | SKCM | IPC-298 | Selumetinib |

| | | | | | |
|---|---|---|---|---|---|
| MVP_top60_DNA_seed | MAP2K1 | -3.047589 | SKCM | IGR-37 | Refametinib |
| MVP_top60_DNA_seed | BRAF | -3.024398 | SKCM | HMV-II | RAF_9304 |
| MVP_top60_DNA_seed | BRAF | -3.018112 | SKCM | WM-115 | PLX-4720 |
| MVP_top60_DNA_seed | PIK3CB | -3.017028 | SKCM | WM793B | AZD6482 |
| MVP_top60_DNA_seed | MAP2K1 | -3.015758 | SKCM | LB2518-MEL | Selumetinib |
| MVP_top60_DNA_seed | BRAF | -3.015636 | SKCM | MEL-HO | RAF_9304 |
| MVP_top60_DNA_seed | EGFR | -3.01504 | SKCM | CHL-1 | PF-00299804 |
| MVP_top60_DNA_seed | ERBB2 | -3.01504 | SKCM | CHL-1 | PF-00299804 |
| MVP_top60_DNA_seed | MAP2K1 | -3.004136 | SKCM | HT-144 | Selumetinib |
| MVP_top60_DNA_seed | BRAF | -2.986544 | SKCM | SK-MEL-24 | Dabrafenib |
| MVP_top60_DNA_seed | MAP2K1 | -2.948451 | SKCM | G-361 | PD0325901 |
| MVP_top60_DNA_seed | MAP2K1 | -2.927875 | SKCM | IGR-37 | PD0325901 |
| MVP_top60_DNA_seed | MAP2K1 | -2.927391 | SKCM | WM35 | Refametinib |
| MVP_top60_DNA_seed | MAP2K1 | -2.924373 | SKCM | CP50-MEL-B | Selumetinib |
| MVP_top60_DNA_seed | MAP2K1 | -2.9208 | SKCM | G-MEL | PD0325901 |
| MVP_top60_DNA_seed | MAP2K1 | -2.91806 | SKCM | MMAc-SF | Refametinib |
| MVP_top60_DNA_seed | BRAF | -2.893544 | SKCM | SK-MEL-28 | PLX-4720 |
| MVP_top60_DNA_seed | BRAF | -2.883344 | SKCM | UACC-257 | SB590885 |
| MVP_top60_DNA_seed | BRAF | -2.869586 | SKCM | MZ7-mel | AZ628 |
| MVP_top60_DNA_seed | BRAF | -2.863258 | SKCM | CP50-MEL-B | SB590885 |
| MVP_top60_DNA_seed | BRAF | -2.857884 | SKCM | HT-144 | Dabrafenib |
| MVP_top60_DNA_seed | MAP2K1 | -2.857076 | SKCM | MEL-HO | Selumetinib |
| MVP_top60_DNA_seed | BRAF | -2.853404 | SKCM | 451Lu | PLX-4720 |
| MVP_top60_DNA_seed | MAP2K1 | -2.8504 | SKCM | MMAc-SF | CI-1040 |
| MVP_top60_DNA_seed | BRAF | -2.849543 | SKCM | SK-MEL-3 | Dabrafenib |
| MVP_top60_DNA_seed | BRAF | -2.834965 | SKCM | COLO-679 | PLX-4720 |
| MVP_top60_DNA_seed | MAP2K1 | -2.8325 | SKCM | IGR-37 | CI-1040 |
| MVP_top60_DNA_seed | BRAF | -2.827968 | SKCM | COLO-783 | Dabrafenib |
| MVP_top60_DNA_seed | BRAF | -2.823639 | SKCM | MZ7-mel | Dabrafenib |
| MVP_top60_DNA_seed | BRAF | -2.822696 | SKCM | COLO-679 | RAF_9304 |
| MVP_top60_DNA_seed | BRAF | -2.813936 | SKCM | COLO-829 | Dabrafenib |
| MVP_top60_DNA_seed | BRAF | -2.716985 | SKCM | A101D | AZ628 |
| MVP_top60_DNA_seed | GSK3B | -2.706632 | SKCM | Hs-940-T | CHIR-99021 |
| MVP_top60_DNA_seed | MAP2K1 | -2.678603 | SKCM | IPC-298 | Refametinib |
| MVP_top60_DNA_seed | MAP2K1 | -2.669169 | SKCM | G-MEL | Refametinib |
| MVP_top60_DNA_seed | MAP2K1 | -2.666692 | SKCM | G-MEL | Refametinib |
| MVP_top60_DNA_seed | MAP2K1 | -2.652613 | SKCM | CP66-MEL | Selumetinib |
| MVP_top60_DNA_seed | MAP2K1 | -2.647937 | SKCM | COLO-679 | Selumetinib |
| MVP_top60_DNA_seed | MAP2K1 | -2.638953 | SKCM | M14 | Refametinib |
| MVP_top60_DNA_seed | MAP2K1 | -2.634295 | SKCM | 451Lu | PD0325901 |
| MVP_top60_DNA_seed | MAP2K1 | -2.625549 | SKCM | M14 | CI-1040 |

| | | | | | |
|---|---|---|---|---|---|
| MVP_top60_DNA_seed | BRAF | -2.621081 | SKCM | COLO-829 | AZ628 |
| MVP_top60_DNA_seed | MAP2K1 | -2.612136 | SKCM | C32 | CI-1040 |
| MVP_top60_DNA_seed | MAP2K1 | -2.599804 | SKCM | MMAc-SF | Selumetinib |
| MVP_top60_DNA_seed | MAP2K1 | -2.596673 | SKCM | IPC-298 | Refametinib |
| MVP_top60_DNA_seed | MAP2K1 | -2.593096 | SKCM | MEL-HO | Selumetinib |
| MVP_top60_DNA_seed | MAP2K1 | -2.592619 | SKCM | MEL-HO | Refametinib |
| MVP_top60_DNA_seed | BRAF | -2.587736 | SKCM | UACC-257 | PLX-4720 |
| MVP_top60_DNA_seed | MAP2K1 | -2.587382 | SKCM | CP50-MEL-B | CI-1040 |
| MVP_top60_DNA_seed | MAP2K1 | -2.571846 | SKCM | K2 | Refametinib |
| MVP_top60_DNA_seed | BRAF | -2.56709 | SKCM | HT-144 | PLX-4720 |
| MVP_top60_DNA_seed | MAP2K1 | -2.56321 | SKCM | SH-4 | Selumetinib |
| MVP_top60_DNA_seed | EGFR | -2.555806 | SKCM | CHL-1 | AZD8931 |
| MVP_top60_DNA_seed | ERBB2 | -2.555806 | SKCM | CHL-1 | AZD8931 |
| MVP_top60_DNA_seed | MAP2K1 | -2.538829 | SKCM | M14 | Selumetinib |
| MVP_top60_DNA_seed | MAPK3 | -2.529404 | SKCM | M14 | FR-180204 |
| MVP_top60_DNA_seed | MAPK1 | -2.529404 | SKCM | M14 | FR-180204 |
| MVP_top60_DNA_seed | BRAF | -2.526483 | SKCM | SK-MEL-1 | SB590885 |
| MVP_top60_DNA_seed | PIK3CB | -2.525415 | SKCM | MEL-HO | AZD8186 |
| MVP_top60_DNA_seed | MAP2K1 | -2.518861 | SKCM | CP50-MEL-B | Refametinib |
| MVP_top60_DNA_seed | BRAF | -2.517707 | SKCM | SH-4 | RAF_9304 |
| MVP_top60_DNA_seed | MAP2K1 | -2.515462 | SKCM | COLO-679 | Refametinib |
| MVP_top60_DNA_seed | MAPK3 | -2.515412 | SKCM | WM35 | FR-180204 |
| MVP_top60_DNA_seed | MAPK1 | -2.515412 | SKCM | WM35 | FR-180204 |
| MVP_top60_DNA_seed | MAP2K1 | -2.513953 | SKCM | K2 | Selumetinib |
| MVP_top60_DNA_seed | PIK3CB | -2.513768 | SKCM | IST-MEL1 | TGX221 |
| MVP_top60_DNA_seed | BRAF | -2.51352 | SKCM | SK-MEL-2 | SB590885 |
| MVP_top60_DNA_seed | MAP2K1 | -2.513502 | SKCM | CP50-MEL-B | Selumetinib |
| MVP_top60_DNA_seed | MAPK1 | -2.512796 | SKCM | MZ7-mel | VX-11e |
| MVP_top60_DNA_seed | MAP2K1 | -2.506865 | SKCM | RVH-421 | Selumetinib |
| MVP_top60_DNA_seed | BRAF | -2.505972 | SKCM | MZ7-mel | RAF_9304 |
| MVP_top60_DNA_seed | MAP2K1 | -2.504149 | SKCM | 451Lu | CI-1040 |
| MVP_top60_DNA_seed | MAP2K1 | -2.503153 | SKCM | MEL-HO | Refametinib |
| MVP_top60_DNA_seed | BRAF | -2.495594 | SKCM | A101D | PLX-4720 |
| MVP_top60_DNA_seed | MAP2K1 | -2.492136 | SKCM | 451Lu | Refametinib |
| MVP_top60_DNA_seed | BRAF | -2.47576 | SKCM | WM278 | PLX-4720 |
| MVP_top60_DNA_seed | MAP2K1 | -2.475082 | SKCM | CP66-MEL | CI-1040 |
| MVP_top60_DNA_seed | BRAF | -2.470813 | SKCM | UACC-257 | AZ628 |
| MVP_top60_DNA_seed | MAP2K1 | -2.467037 | SKCM | M14 | PD0325901 |
| MVP_top60_DNA_seed | MAP2K1 | -2.466572 | SKCM | G-MEL | Trametinib |
| MVP_top60_DNA_seed | MAP2K1 | -2.454051 | SKCM | CP66-MEL | Refametinib |
| MVP_top60_DNA_seed | MAP2K1 | -2.449183 | SKCM | WM-115 | Selumetinib |

| | | | | | |
|---|---|---|---|---|---|
| MVP_top60_DNA_seed | BRAF | -2.437724 | SKCM | SK-MEL-5 | SB590885 |
| MVP_top60_DNA_seed | MAP2K1 | -2.431024 | SKCM | A375 | Selumetinib |
| MVP_top60_DNA_seed | BRAF | -2.43011 | SKCM | C32 | RAF_9304 |
| MVP_top60_DNA_seed | BRAF | -2.428363 | SKCM | M14 | RAF_9304 |
| MVP_top60_DNA_seed | MAP2K1 | -2.425873 | SKCM | M14 | Refametinib |
| MVP_top60_DNA_seed | EGFR | -2.423089 | SKCM | CHL-1 | Gefitinib |
| MVP_top60_DNA_seed | BRAF | -2.415898 | SKCM | LOXIMVI | PLX-4720 |
| MVP_top60_DNA_seed | BRAF | -2.403022 | SKCM | CP50-MEL-B | RAF_9304 |
| MVP_top60_DNA_seed | BRAF | -2.398451 | SKCM | SK-MEL-5 | Dabrafenib |
| MVP_top60_DNA_seed | BRAF | -2.396276 | SKCM | SK-MEL-5 | PLX-4720 |
| MVP_top60_DNA_seed | MAP2K1 | -2.39616 | SKCM | CP66-MEL | PD0325901 |
| MVP_top60_DNA_seed | BRAF | -2.395091 | SKCM | G-361 | PLX-4720 |
| MVP_top60_DNA_seed | ERBB2 | -2.394611 | SKCM | CHL-1 | CP724714 |
| MVP_top60_DNA_seed | MAP2K1 | -2.390166 | SKCM | WM35 | Selumetinib |
| MVP_top60_DNA_seed | BRAF | -2.387781 | SKCM | MMAc-SF | AZ628 |
| MVP_top60_DNA_seed | BRAF | -2.38227 | SKCM | SK-MEL-24 | PLX-4720 |
| MVP_top60_DNA_seed | MAP2K1 | -2.374592 | SKCM | HT-144 | Refametinib |
| MVP_top60_DNA_seed | EGFR | -2.371536 | SKCM | CHL-1 | AST-1306 |
| MVP_top60_DNA_seed | BRAF | -2.369968 | SKCM | MMAc-SF | RAF_9304 |
| MVP_top60_DNA_seed | MAP2K1 | -2.369607 | SKCM | C32 | Refametinib |
| MVP_top60_DNA_seed | PIK3CB | -2.367209 | SKCM | IGR-37 | AZD8186 |
| MVP_top60_DNA_seed | MAPK1 | -2.362173 | SKCM | MEL-HO | VX-11e |
| MVP_top60_DNA_seed | MAP2K1 | -2.358611 | SKCM | A375 | PD0325901 |
| MVP_top60_DNA_seed | MAP2K1 | -2.35771 | SKCM | MMAc-SF | Trametinib |
| MVP_top60_DNA_seed | MAP2K1 | -2.353709 | SKCM | UACC-257 | Selumetinib |
| MVP_top60_DNA_seed | MAP2K1 | -2.350667 | SKCM | IGR-37 | Trametinib |
| MVP_top60_DNA_seed | MAP2K1 | -2.34784 | SKCM | Mewo | Selumetinib |
| MVP_top60_DNA_seed | MAP2K1 | -2.338784 | SKCM | SK-MEL-30 | Selumetinib |
| MVP_top60_DNA_seed | BRAF | -2.338542 | SKCM | MZ7-mel | PLX-4720 |
| MVP_top60_DNA_seed | MAP2K1 | -2.334282 | SKCM | C32 | Trametinib |
| MVP_top60_DNA_seed | MAP2K1 | -2.331378 | SKCM | LB2518-MEL | Selumetinib |
| MVP_top60_DNA_seed | MAP2K1 | -2.322528 | SKCM | COLO-679 | Selumetinib |
| MVP_top60_DNA_seed | EGFR | -2.318534 | SKCM | WM35 | Gefitinib |
| MVP_top60_DNA_seed | EGFR | -2.314327 | SKCM | IGR-1 | PF-00299804 |
| MVP_top60_DNA_seed | ERBB2 | -2.314327 | SKCM | IGR-1 | PF-00299804 |
| MVP_top60_DNA_seed | RAC1 | -2.313797 | SKCM | COLO-792 | EHT-1864 |
| MVP_top60_DNA_seed | MAP2K1 | -2.306366 | SKCM | C32 | Selumetinib |
| MVP_top60_DNA_seed | MAP2K1 | -2.302938 | SKCM | MEL-HO | CI-1040 |
| MVP_top60_DNA_seed | BRAF | -2.300492 | SKCM | A375 | RAF_9304 |
| MVP_top60_DNA_seed | MAPK3 | -2.292976 | SKCM | MZ7-mel | FR-180204 |
| MVP_top60_DNA_seed | MAPK1 | -2.292976 | SKCM | MZ7-mel | FR-180204 |

| | | | | | |
|---|---|---|---|---|---|
| MVP_top60_DNA_seed | MAP2K1 | -2.28899 | SKCM | A375 | Trametinib |
| MVP_top60_DNA_seed | MAPK1 | -2.286097 | SKCM | M14 | VX-11e |
| MVP_top60_DNA_seed | BRAF | -2.282492 | SKCM | RVH-421 | PLX-4720 |
| MVP_top60_DNA_seed | MAP2K1 | -2.281969 | SKCM | C32 | PD0325901 |
| MVP_top60_DNA_seed | MAP2K1 | -2.279729 | SKCM | A375 | CI-1040 |
| MVP_top60_DNA_seed | BRAF | -2.276926 | SKCM | Hs-939-T | PLX-4720 |
| MVP_top60_DNA_seed | MAP2K1 | -2.271759 | SKCM | UACC-257 | Selumetinib |
| MVP_top60_DNA_seed | BRAF | -2.264027 | SKCM | SK-MEL-3 | RAF_9304 |
| MVP_top60_DNA_seed | MAP2K1 | -2.25708 | SKCM | UACC-62 | Selumetinib |
| MVP_top60_DNA_seed | BRAF | -2.248342 | SKCM | LOXIMVI | RAF_9304 |
| MVP_top60_DNA_seed | MAP2K1 | -2.242833 | SKCM | SH-4 | Trametinib |
| MVP_top60_DNA_seed | BRAF | -2.241474 | SKCM | UACC-62 | RAF_9304 |
| MVP_top60_DNA_seed | MAP2K1 | -2.239055 | SKCM | 451Lu | Selumetinib |
| MVP_top60_DNA_seed | BRAF | -2.238509 | SKCM | SK-MEL-5 | PLX-4720 |
| MVP_top60_DNA_seed | MAP2K1 | -2.22715 | SKCM | COLO-679 | CI-1040 |
| MVP_top60_DNA_seed | BRAF | -2.222172 | SKCM | COLO-783 | SB590885 |
| MVP_top60_DNA_seed | BRAF | -2.217851 | SKCM | 451Lu | RAF_9304 |
| MVP_top60_DNA_seed | BRAF | -2.201928 | SKCM | COLO-783 | PLX-4720 |
| MVP_top60_DNA_seed | MAP2K1 | -2.201626 | SKCM | A375 | Refametinib |
| MVP_top60_DNA_seed | MAP2K1 | -2.196316 | SKCM | UACC-62 | Refametinib |
| MVP_top60_DNA_seed | BRAF | -2.195339 | SKCM | COLO-800 | PLX-4720 |
| MVP_top60_DNA_seed | MAP2K1 | -2.186376 | SKCM | CP66-MEL | Refametinib |
| MVP_top60_DNA_seed | MAP2K1 | -2.185201 | SKCM | LOXIMVI | Selumetinib |
| MVP_top60_DNA_seed | BRAF | -2.181887 | SKCM | SK-MEL-3 | PLX-4720 |
| MVP_top60_DNA_seed | MAP2K1 | -2.17642 | SKCM | Hs-939-T | PD0325901 |
| MVP_top60_DNA_seed | BRAF | -2.174501 | SKCM | SK-MEL-3 | SB590885 |
| MVP_top60_DNA_seed | MAP2K1 | -2.173015 | SKCM | WM35 | Selumetinib |
| MVP_top60_DNA_seed | MAP2K1 | -2.165888 | SKCM | CP50-MEL-B | PD0325901 |
| MVP_top60_DNA_seed | MAP2K1 | -2.162171 | SKCM | SK-MEL-28 | Selumetinib |
| MVP_top60_DNA_seed | MAP2K1 | -2.154409 | SKCM | LOXIMVI | CI-1040 |
| MVP_top60_DNA_seed | MAP2K1 | -2.151046 | SKCM | A101D | CI-1040 |
| MVP_top60_DNA_seed | MAP2K1 | -2.148075 | SKCM | CP50-MEL-B | Refametinib |
| MVP_top60_DNA_seed | BRAF | -2.148002 | SKCM | COLO-829 | PLX-4720 |
| MVP_top60_DNA_seed | MAP2K1 | -2.140015 | SKCM | LB2518-MEL | Refametinib |
| MVP_top60_DNA_seed | BRAF | -2.138799 | SKCM | LOXIMVI | SB590885 |
| MVP_top60_DNA_seed | BRAF | -2.13503 | SKCM | K2 | RAF_9304 |
| MVP_top60_DNA_seed | MAP2K1 | -2.132125 | SKCM | WM35 | Trametinib |
| MVP_top60_DNA_seed | BRAF | -2.132021 | SKCM | WM35 | SB590885 |
| MVP_top60_DNA_seed | EGFR | -2.131599 | SKCM | CHL-1 | CI-1033 |
| MVP_top60_DNA_seed | ERBB2 | -2.131599 | SKCM | CHL-1 | CI-1033 |
| MVP_top60_DNA_seed | MAPK1 | -2.130508 | SKCM | A375 | VX-11e |

| | | | | | |
|---|---|---|---|---|---|
| MVP_top60_DNA_seed | BRAF | -2.120201 | SKCM | LOXIMVI | PLX-4720 |
| MVP_top60_DNA_seed | MAP2K1 | -2.119257 | SKCM | M14 | Trametinib |
| MVP_top60_DNA_seed | MAP2K1 | -2.117222 | SKCM | UACC-257 | Refametinib |
| MVP_top60_DNA_seed | MAP2K1 | -2.109091 | SKCM | MEL-HO | Trametinib |
| MVP_top60_DNA_seed | BRAF | -2.107238 | SKCM | RVH-421 | Dabrafenib |
| MVP_top60_DNA_seed | MAP2K1 | -2.104034 | SKCM | HMV-II | Trametinib |
| MVP_top60_DNA_seed | MAP2K1 | -2.089059 | SKCM | IPC-298 | CI-1040 |
| MVP_top60_DNA_seed | MAP2K1 | -2.087151 | SKCM | IPC-298 | Trametinib |
| MVP_top60_DNA_seed | MAP2K1 | -2.079079 | SKCM | SK-MEL-28 | Refametinib |
| MVP_top60_DNA_seed | MAPK1 | -2.076821 | SKCM | SH-4 | VX-11e |
| MVP_top60_DNA_seed | MAPK1 | -2.074456 | SKCM | IGR-37 | VX-11e |
| MVP_top60_DNA_seed | MAP2K1 | -2.07413 | SKCM | MZ7-mel | Refametinib |
| MVP_top60_DNA_seed | EGFR | -2.073096 | SKCM | IGR-1 | AST-1306 |
| MVP_top60_DNA_seed | GSK3B | -2.070672 | SKCM | COLO-783 | CHIR-99021 |
| MVP_top60_DNA_seed | GSK3B | -2.067955 | SKCM | A375 | CHIR-99021 |
| MVP_top60_DNA_seed | MAP2K1 | -2.062088 | SKCM | RVH-421 | Refametinib |
| MVP_top60_DNA_seed | MAP2K1 | -2.061777 | SKCM | SK-MEL-28 | PD0325901 |
| MVP_top60_DNA_seed | MAP2K1 | -2.055525 | SKCM | A375 | Refametinib |
| MVP_top60_DNA_seed | BRAF | -2.053827 | SKCM | WM278 | Dabrafenib |
| MVP_top60_DNA_seed | MAP2K1 | -2.053698 | SKCM | G-361 | Refametinib |
| MVP_top60_DNA_seed | PIK3CB | -2.05145 | SKCM | G-MEL | AZD6482 |
| MVP_top60_DNA_seed | MAP2K1 | -2.049417 | SKCM | K2 | PD0325901 |
| MVP_top60_DNA_seed | MAP2K1 | -2.043975 | SKCM | SK-MEL-28 | CI-1040 |
| MVP_top60_DNA_seed | BRAF | -2.041871 | SKCM | IST-MEL1 | Dabrafenib |
| MVP_top60_DNA_seed | BRAF | -2.036958 | SKCM | G-361 | PLX-4720 |
| MVP_top60_DNA_seed | MAP2K1 | -2.034561 | SKCM | SH-4 | Refametinib |
| MVP_top60_DNA_seed | BRAF | -2.02736 | SKCM | Hs-939-T | PLX-4720 |
| MVP_top60_DNA_seed | MAPK1 | -2.02069 | SKCM | SK-MEL-30 | VX-11e |
| MVP_top60_DNA_seed | GSK3B | -2.02034 | SKCM | WM35 | CHIR-99021 |
| MVP_top60_DNA_seed | MAP2K1 | -2.020034 | SKCM | UACC-257 | Trametinib |
| MVP_top60_DNA_seed | MAPK1 | -2.008555 | SKCM | WM35 | VX-11e |
| MVP_top60_DNA_seed | MAP2K1 | -2.008421 | SKCM | 451Lu | Trametinib |
| MVP_top60_DNA_seed | MAP2K1 | -2.005606 | SKCM | COLO-679 | Trametinib |
| MVP_top60_DNA_seed | MAP2K1 | -2.005588 | SKCM | LB2518-MEL | Trametinib |
| MVP_top60_DNA_seed | BRAF | -2.002465 | SKCM | CP66-MEL | AZ628 |
| MVP_top60_DNA_seed | MAPK1 | -1.994343 | SKCM | HMV-II | VX-11e |
| MVP_top60_DNA_seed | MAP2K1 | -1.990815 | SKCM | LB2518-MEL | Refametinib |
| MVP_top60_DNA_seed | GSK3B | -1.988666 | SKCM | IGR-1 | CHIR-99021 |
| MVP_top60_DNA_seed | MAP2K1 | -1.986639 | SKCM | SK-MEL-30 | Trametinib |
| MVP_top60_DNA_seed | BRAF | -1.983639 | SKCM | HT-144 | RAF_9304 |
| MVP_top60_DNA_seed | MAP2K1 | -1.98157 | SKCM | CP50-MEL-B | Trametinib |

| | | | | | |
|---|---|---|---|---|---|
| MVP_top60_DNA_seed | MAP2K1 | -1.980543 | SKCM | WM-115 | CI-1040 |
| MVP_top60_DNA_seed | MAP2K1 | -1.979524 | SKCM | SK-MEL-2 | Trametinib |
| MVP_top60_DNA_seed | MAPK3 | -1.974832 | SKCM | RVH-421 | FR-180204 |
| MVP_top60_DNA_seed | MAPK1 | -1.974832 | SKCM | RVH-421 | FR-180204 |
| MVP_top60_DNA_seed | GSK3B | -1.974032 | SKCM | RPMI-7951 | CHIR-99021 |
| MVP_top60_DNA_seed | MAP2K1 | -1.973172 | SKCM | C32 | Refametinib |
| MVP_top60_DNA_seed | MAP2K1 | -1.970821 | SKCM | UACC-257 | Refametinib |
| MVP_top60_DNA_seed | BRAF | -1.959714 | SKCM | K2 | SB590885 |
| MVP_top60_DNA_seed | GSK3B | -1.956756 | SKCM | WM278 | CHIR-99021 |
| MVP_top60_DNA_seed | MAP2K1 | -1.953412 | SKCM | SH-4 | CI-1040 |
| MVP_top60_DNA_seed | MAPK3 | -1.953181 | SKCM | CHL-1 | FR-180204 |
| MVP_top60_DNA_seed | MAPK1 | -1.953181 | SKCM | CHL-1 | FR-180204 |
| MVP_top60_DNA_seed | MAP2K1 | -1.946654 | SKCM | SK-MEL-28 | Refametinib |
| MVP_top60_DNA_seed | JAK3 | -1.921974 | SKCM | IST-MEL1 | JAK3_7406 |
| MVP_top60_DNA_seed | MAPK1 | -1.919678 | SKCM | IPC-298 | VX-11e |
| MVP_top60_DNA_seed | MAP2K1 | -1.916767 | SKCM | SK-MEL-2 | Refametinib |
| MVP_top60_DNA_seed | MAP2K1 | -1.913747 | SKCM | SK-MEL-1 | CI-1040 |
| MVP_top60_DNA_seed | PIK3CB | -1.912374 | SKCM | MEL-HO | AZD6482 |
| MVP_top60_DNA_seed | MAP2K1 | -1.898043 | SKCM | MZ7-mel | Selumetinib |
| MVP_top60_DNA_seed | GSK3B | -1.897994 | SKCM | LB2518-MEL | CHIR-99021 |
| MVP_top60_DNA_seed | MAP2K1 | -1.896 | SKCM | UACC-62 | Trametinib |
| MVP_top60_DNA_seed | MAP2K1 | -1.892716 | SKCM | MEL-HO | PD0325901 |
| MVP_top60_DNA_seed | BRAF | -1.891874 | SKCM | Hs-939-T | SB590885 |
| MVP_top60_DNA_seed | MAP2K1 | -1.887865 | SKCM | COLO-679 | Refametinib |
| MVP_top60_DNA_seed | MAP2K1 | -1.869122 | SKCM | SH-4 | PD0325901 |
| MVP_top60_DNA_seed | MAP2K1 | -1.867148 | SKCM | GAK | Trametinib |
| MVP_top60_DNA_seed | BRAF | -1.866642 | SKCM | SK-MEL-24 | RAF_9304 |
| MVP_top60_DNA_seed | MAP2K1 | -1.866616 | SKCM | HT-144 | CI-1040 |
| MVP_top60_DNA_seed | GSK3B | -1.864354 | SKCM | COLO-792 | KIN001-042 |
| MVP_top60_DNA_seed | MAP2K1 | -1.862795 | SKCM | G-361 | Trametinib |
| MVP_top60_DNA_seed | MAP2K1 | -1.861612 | SKCM | LOXIMVI | Selumetinib |
| MVP_top60_DNA_seed | MAP2K1 | -1.861543 | SKCM | 451Lu | Refametinib |
| MVP_top60_DNA_seed | MAP2K1 | -1.858686 | SKCM | HT-144 | PD0325901 |
| MVP_top60_DNA_seed | MAP2K1 | -1.856867 | SKCM | SK-MEL-2 | Selumetinib |
| MVP_top60_DNA_seed | MAP2K1 | -1.856085 | SKCM | SK-MEL-28 | Trametinib |
| MVP_top60_DNA_seed | MAP2K1 | -1.849701 | SKCM | SK-MEL-30 | Refametinib |
| MVP_top60_DNA_seed | MAP2K1 | -1.845179 | SKCM | LOXIMVI | Trametinib |
| MVP_top60_DNA_seed | GSK3B | -1.840426 | SKCM | RPMI-7951 | SB216763 |
| MVP_top60_DNA_seed | MAP2K1 | -1.837892 | SKCM | COLO-679 | PD0325901 |
| MVP_top60_DNA_seed | MAP2K1 | -1.836171 | SKCM | COLO-792 | Selumetinib |
| MVP_top60_DNA_seed | MAP2K1 | -1.83383 | SKCM | SK-MEL-2 | Refametinib |

| | | | | | |
|---|---|---|---|---|---|
| MVP_top60_DNA_seed | PIK3CB | -1.831069 | SKCM | SH-4 | AZD8186 |
| MVP_top60_DNA_seed | MAP2K1 | -1.827852 | SKCM | HMV-II | Selumetinib |
| MVP_top60_DNA_seed | MAP2K1 | -1.819827 | SKCM | IPC-298 | PD0325901 |
| MVP_top60_DNA_seed | MAPK1 | -1.817074 | SKCM | G-MEL | VX-11e |
| MVP_top60_DNA_seed | EGFR | -1.810104 | SKCM | COLO-783 | Cetuximab |
| MVP_top60_DNA_seed | BRAF | -1.803864 | SKCM | SK-MEL-1 | PLX-4720 |
| MVP_top60_DNA_seed | MAP2K1 | -1.7981 | SKCM | K2 | Trametinib |
| MVP_top60_DNA_seed | GSK3B | -1.791539 | SKCM | A375 | SB216763 |
| MVP_top60_DNA_seed | EGFR | -1.784842 | SKCM | HT-144 | AST-1306 |
| MVP_top60_DNA_seed | MAP2K1 | -1.783208 | SKCM | LB2518-MEL | PD0325901 |
| MVP_top60_DNA_seed | GSK3B | -1.781577 | SKCM | LOXIMVI | AZD7969 |
| MVP_top60_DNA_seed | MAP2K1 | -1.781317 | SKCM | CP66-MEL | Trametinib |
| MVP_top60_DNA_seed | GSK3B | -1.781088 | SKCM | A375 | AZD7969 |
| MVP_top60_DNA_seed | JAK3 | -1.778726 | SKCM | COLO-792 | WHI-P97 |
| MVP_top60_DNA_seed | MAP2K1 | -1.777279 | SKCM | SK-MEL-5 | PD0325901 |
| MVP_top60_DNA_seed | MAP2K1 | -1.7771 | SKCM | COLO-792 | Selumetinib |
| MVP_top60_DNA_seed | MAP2K1 | -1.769668 | SKCM | HT-144 | Refametinib |
| MVP_top60_DNA_seed | MAP2K1 | -1.766775 | SKCM | SK-MEL-3 | Selumetinib |
| MVP_top60_DNA_seed | MAPK1 | -1.76119 | SKCM | WM278 | VX-11e |
| MVP_top60_DNA_seed | MAP2K1 | -1.76108 | SKCM | G-361 | Selumetinib |
| MVP_top60_DNA_seed | BRAF | -1.758344 | SKCM | COLO-829 | PLX-4720 |
| MVP_top60_DNA_seed | EGFR | -1.753461 | SKCM | A375 | AST-1306 |
| MVP_top60_DNA_seed | MAP2K1 | -1.748323 | SKCM | LB2518-MEL | CI-1040 |
| MVP_top60_DNA_seed | MAP2K1 | -1.740744 | SKCM | LOXIMVI | Refametinib |
| MVP_top60_DNA_seed | MAP2K1 | -1.73532 | SKCM | HT-144 | Selumetinib |
| MVP_top60_DNA_seed | BRAF | -1.717599 | SKCM | MZ7-mel | SB590885 |
| MVP_top60_DNA_seed | RAC1 | -1.71314 | SKCM | SK-MEL-3 | EHT-1864 |
| MVP_top60_DNA_seed | MAP2K1 | -1.709963 | SKCM | Hs-939-T | CI-1040 |
| MVP_top60_DNA_seed | MAP2K1 | -1.707552 | SKCM | MZ7-mel | Refametinib |
| MVP_top60_DNA_seed | BRAF | -1.70635 | SKCM | IPC-298 | RAF_9304 |
| MVP_top60_DNA_seed | MAP2K1 | -1.705076 | SKCM | MZ7-mel | Trametinib |
| MVP_top60_DNA_seed | BRAF | -1.702459 | SKCM | CHL-1 | Dabrafenib |
| MVP_top60_DNA_seed | MAPK1 | -1.700705 | SKCM | RVH-421 | VX-11e |
| MVP_top60_DNA_seed | MAP2K1 | -1.694938 | SKCM | HT-144 | Trametinib |
| MVP_top60_DNA_seed | MAP2K1 | -1.693962 | SKCM | SH-4 | Refametinib |
| MVP_top60_DNA_seed | PIK3CB | -1.691787 | SKCM | IST-MEL1 | AZD6482 |
| MVP_top60_DNA_seed | BRAF | -1.689875 | SKCM | SK-MEL-3 | PLX-4720 |
| MVP_top60_DNA_seed | GSK3B | -1.678109 | SKCM | G-MEL | AZD7969 |
| MVP_top60_DNA_seed | MAP2K1 | -1.671267 | SKCM | A101D | Refametinib |
| MVP_top60_DNA_seed | EGFR | -1.670705 | SKCM | WM793B | AST-1306 |
| MVP_top60_DNA_seed | JAK1 | -1.668895 | SKCM | IGR-1 | JAK1_3715 |

| | | | | | |
|---|---|---|---|---|---|
| MVP_top60_DNA_seed | MAP2K1 | -1.663295 | SKCM | UACC-257 | PD0325901 |
| MVP_top60_DNA_seed | BRAF | -1.662605 | SKCM | UACC-257 | RAF_9304 |
| MVP_top60_DNA_seed | MAP2K1 | -1.660268 | SKCM | MZ2-MEL | PD0325901 |
| MVP_top60_DNA_seed | EGFR | -1.658819 | SKCM | CHL-1 | Pelitinib |
| MVP_top60_DNA_seed | MAP2K1 | -1.65802 | SKCM | RVH-421 | PD0325901 |
| MVP_top60_DNA_seed | BRAF | -1.656931 | SKCM | WM278 | HG6-64-1 |
| MVP_top60_DNA_seed | GSK3B | -1.653825 | SKCM | A101D | CHIR-99021 |
| MVP_top60_DNA_seed | BRAF | -1.651406 | SKCM | CP66-MEL | PLX-4720 |
| MVP_top60_DNA_seed | RAC1 | -1.650738 | SKCM | UACC-257 | EHT-1864 |
| MVP_top60_DNA_seed | PIK3CB | -1.64861 | SKCM | WM1552C | AZD6482 |
| MVP_top60_DNA_seed | MAP2K1 | -1.645956 | SKCM | K2 | CI-1040 |
| MVP_top60_DNA_seed | EGFR | -1.64431 | SKCM | HT-144 | PF-00299804 |
| MVP_top60_DNA_seed | ERBB2 | -1.64431 | SKCM | HT-144 | PF-00299804 |
| MVP_top60_DNA_seed | MAP2K1 | -1.641504 | SKCM | WM35 | Refametinib |
| MVP_top60_DNA_seed | MAP2K1 | -1.639137 | SKCM | RVH-421 | CI-1040 |
| MVP_top60_DNA_seed | EGFR | -1.638213 | SKCM | A375 | PF-00299804 |
| MVP_top60_DNA_seed | ERBB2 | -1.638213 | SKCM | A375 | PF-00299804 |
| MVP_top60_DNA_seed | GSK3B | -1.6379 | SKCM | WM278 | SB216763 |
| MVP_top60_DNA_seed | MAP2K1 | -1.631961 | SKCM | Hs-939-T | Refametinib |
| MVP_top60_DNA_seed | MAP2K1 | -1.623337 | SKCM | A101D | PD0325901 |
| MVP_top60_DNA_seed | MAP2K1 | -1.612989 | SKCM | SK-MEL-5 | CI-1040 |
| MVP_top60_DNA_seed | MAP2K1 | -1.60837 | SKCM | A101D | Selumetinib |
| MVP_top60_DNA_seed | MAPK1 | -1.603052 | SKCM | LOXIMVI | VX-11e |
| MVP_top60_DNA_seed | BRAF | -1.602716 | SKCM | SK-MEL-30 | RAF_9304 |
| MVP_top60_DNA_seed | BRAF | -1.60183 | SKCM | K2 | PLX-4720 |
| MVP_top60_DNA_seed | EGFR | -1.599725 | SKCM | Hs-940-T | AST-1306 |
| MVP_top60_DNA_seed | BRAF | -1.599522 | SKCM | WM1552C | PLX-4720 |
| MVP_top60_DNA_seed | BRAF | -1.594122 | SKCM | WM278 | RAF_9304 |
| MVP_top60_DNA_seed | JAK2 | -1.591637 | SKCM | SH-4 | AZD1480 |
| MVP_top60_DNA_seed | JAK1 | -1.591637 | SKCM | SH-4 | AZD1480 |
| MVP_top60_DNA_seed | EGFR | -1.587391 | SKCM | SK-MEL-28 | AST-1306 |
| MVP_top60_DNA_seed | PIK3CB | -1.58307 | SKCM | UACC-62 | AZD8186 |
| MVP_top60_DNA_seed | MAP2K1 | -1.580118 | SKCM | MEL-JUSO | PD0325901 |
| MVP_top60_DNA_seed | BRAF | -1.574465 | SKCM | SK-MEL-28 | RAF_9304 |
| MVP_top60_DNA_seed | MAP2K1 | -1.572834 | SKCM | LOXIMVI | Refametinib |
| MVP_top60_DNA_seed | EGFR | -1.570271 | SKCM | WM793B | PF-00299804 |
| MVP_top60_DNA_seed | ERBB2 | -1.570271 | SKCM | WM793B | PF-00299804 |
| MVP_top60_DNA_seed | JAK3 | -1.568135 | SKCM | VMRC-MELG | WHI-P97 |
| MVP_top60_DNA_seed | GSK3B | -1.568001 | SKCM | WM793B | CHIR-99021 |
| MVP_top60_DNA_seed | JAK2 | -1.565359 | SKCM | Hs-940-T | Ruxolitinib |
| MVP_top60_DNA_seed | JAK1 | -1.565359 | SKCM | Hs-940-T | Ruxolitinib |

| | | | | | |
|---|---|---|---|---|---|
| MVP_top60_DNA_seed | MAP2K1 | -1.565193 | SKCM | WM278 | Selumetinib |
| MVP_top60_DNA_seed | BRAF | -1.562431 | SKCM | GAK | RAF_9304 |
| MVP_top60_DNA_seed | MAP2K1 | -1.562055 | SKCM | SK-MEL-24 | Selumetinib |
| MVP_top60_DNA_seed | MAP2K1 | -1.561415 | SKCM | SK-MEL-5 | Refametinib |
| MVP_top60_DNA_seed | BRAF | -1.553998 | SKCM | G-361 | SB590885 |
| MVP_top60_DNA_seed | GSK3B | -1.539519 | SKCM | RPMI-7951 | KIN001-042 |
| MVP_top60_DNA_seed | MAP2K1 | -1.529843 | SKCM | SK-MEL-24 | Trametinib |
| MVP_top60_DNA_seed | MAP2K1 | -1.529393 | SKCM | COLO-792 | Refametinib |
| MVP_top60_DNA_seed | BRAF | -1.526641 | SKCM | MZ2-MEL | PLX-4720 |
| MVP_top60_DNA_seed | MAP2K1 | -1.526606 | SKCM | SH-4 | Selumetinib |
| MVP_top60_DNA_seed | EGFR | -1.523251 | SKCM | WM1552C | AST-1306 |
| MVP_top60_DNA_seed | MAP2K1 | -1.522394 | SKCM | UACC-257 | CI-1040 |
| MVP_top60_DNA_seed | MAPK3 | -1.518145 | SKCM | SH-4 | FR-180204 |
| MVP_top60_DNA_seed | MAPK1 | -1.518145 | SKCM | SH-4 | FR-180204 |
| MVP_top60_DNA_seed | MAP2K1 | -1.518108 | SKCM | WM278 | Refametinib |
| MVP_top60_DNA_seed | MAPK1 | -1.516535 | SKCM | IGR-1 | VX-11e |
| MVP_top60_DNA_seed | MAPK1 | -1.509361 | SKCM | MEL-JUSO | VX-11e |
| MVP_top60_DNA_seed | MAP2K1 | -1.509296 | SKCM | SK-MEL-2 | CI-1040 |
| MVP_top60_DNA_seed | GSK3B | -1.502448 | SKCM | G-361 | CHIR-99021 |
| MVP_top60_DNA_seed | MAP2K1 | -1.489942 | SKCM | G-361 | Refametinib |
| MVP_top60_DNA_seed | BRAF | -1.488305 | SKCM | WM278 | PLX-4720 |
| MVP_top60_DNA_seed | EP300 | -1.485081 | SKCM | UACC-62 | I-CBP112 |
| MVP_top60_DNA_seed | CREBBP | -1.485081 | SKCM | UACC-62 | I-CBP112 |
| MVP_top60_DNA_seed | MAP2K1 | -1.484183 | SKCM | HMV-II | Refametinib |
| MVP_top60_DNA_seed | MAPK1 | -1.482877 | SKCM | MZ2-MEL | VX-11e |
| MVP_top60_DNA_seed | GSK3B | -1.481874 | SKCM | MZ2-MEL | CHIR-99021 |
| MVP_top60_DNA_seed | MAPK1 | -1.481042 | SKCM | 451Lu | VX-11e |
| MVP_top60_DNA_seed | BRAF | -1.478675 | SKCM | COLO-829 | HG6-64-1 |
| MVP_top60_DNA_seed | BRAF | -1.47651 | SKCM | SK-MEL-5 | RAF_9304 |
| MVP_top60_DNA_seed | BRAF | -1.475124 | SKCM | CHL-1 | HG6-64-1 |
| MVP_top60_DNA_seed | MAP2K1 | -1.471093 | SKCM | MZ7-mel | PD0325901 |
| MVP_top60_DNA_seed | MAP2K1 | -1.467845 | SKCM | A101D | Trametinib |
| MVP_top60_DNA_seed | JAK2 | -1.467067 | SKCM | SK-MEL-5 | AZD1480 |
| MVP_top60_DNA_seed | JAK1 | -1.467067 | SKCM | SK-MEL-5 | AZD1480 |
| MVP_top60_DNA_seed | JAK1 | -1.464824 | SKCM | A2058 | JAK1_3715 |
| MVP_top60_DNA_seed | MAPK3 | -1.458562 | SKCM | SK-MEL-24 | FR-180204 |
| MVP_top60_DNA_seed | MAPK1 | -1.458562 | SKCM | SK-MEL-24 | FR-180204 |
| MVP_top60_DNA_seed | BRAF | -1.455948 | SKCM | CHL-1 | PLX-4720 |
| MVP_top60_DNA_seed | MAP2K1 | -1.451773 | SKCM | SK-MEL-2 | PD0325901 |
| MVP_top60_DNA_seed | MAP2K1 | -1.450151 | SKCM | Mewo | PD0325901 |
| MVP_top60_DNA_seed | BRAF | -1.449136 | SKCM | WM793B | SB590885 |

| | | | | | |
|---|---|---|---|---|---|
| MVP_top60_DNA_seed | JAK1 | -1.447103 | SKCM | COLO-800 | JAK1_3715 |
| MVP_top60_DNA_seed | GSK3B | -1.444599 | SKCM | SK-MEL-2 | CHIR-99021 |
| MVP_top60_DNA_seed | BRAF | -1.444524 | SKCM | COLO-783 | PLX-4720 |
| MVP_top60_DNA_seed | GSK3B | -1.434893 | SKCM | VMRC-MELG | CHIR-99021 |
| MVP_top60_DNA_seed | GSK3B | -1.431738 | SKCM | WM-115 | CHIR-99021 |
| MVP_top60_DNA_seed | MAPK3 | -1.430469 | SKCM | MEL-HO | FR-180204 |
| MVP_top60_DNA_seed | MAPK1 | -1.430469 | SKCM | MEL-HO | FR-180204 |
| MVP_top60_DNA_seed | PIK3CB | -1.427322 | SKCM | COLO-829 | TGX221 |
| MVP_top60_DNA_seed | MAPK3 | -1.424598 | SKCM | SK-MEL-3 | FR-180204 |
| MVP_top60_DNA_seed | MAPK1 | -1.424598 | SKCM | SK-MEL-3 | FR-180204 |
| MVP_top60_DNA_seed | MAP2K1 | -1.416787 | SKCM | Hs-939-T | Selumetinib |
| MVP_top60_DNA_seed | MAP2K1 | -1.406705 | SKCM | MZ2-MEL | Refametinib |
| MVP_top60_DNA_seed | EGFR | -1.403681 | SKCM | MMAc-SF | AST-1306 |
| MVP_top60_DNA_seed | BRAF | -1.396823 | SKCM | MZ2-MEL | AZ628 |
| MVP_top60_DNA_seed | MAP2K1 | -1.390146 | SKCM | SK-MEL-5 | Selumetinib |
| MVP_top60_DNA_seed | EGFR | -1.388994 | SKCM | IPC-298 | PF-00299804 |
| MVP_top60_DNA_seed | ERBB2 | -1.388994 | SKCM | IPC-298 | PF-00299804 |
| MVP_top60_DNA_seed | BRAF | -1.385225 | SKCM | IST-MEL1 | PLX-4720 |
| MVP_top60_DNA_seed | GSK3B | -1.378931 | SKCM | MZ2-MEL | KIN001-042 |
| MVP_top60_DNA_seed | MAP2K1 | -1.377907 | SKCM | MEL-JUSO | Selumetinib |
| MVP_top60_DNA_seed | MAPK3 | -1.374109 | SKCM | A375 | FR-180204 |
| MVP_top60_DNA_seed | MAPK1 | -1.374109 | SKCM | A375 | FR-180204 |
| MVP_top60_DNA_seed | MAP2K1 | -1.370362 | SKCM | MEL-JUSO | Trametinib |
| MVP_top60_DNA_seed | GSK3B | -1.36885 | SKCM | WM793B | CHIR-99021 |
| MVP_top60_DNA_seed | EGFR | -1.361279 | SKCM | SK-MEL-28 | PF-00299804 |
| MVP_top60_DNA_seed | ERBB2 | -1.361279 | SKCM | SK-MEL-28 | PF-00299804 |
| MVP_top60_DNA_seed | GSK3B | -1.357193 | SKCM | Hs-940-T | CHIR-99021 |
| MVP_top60_DNA_seed | MAP2K1 | -1.355579 | SKCM | COLO-783 | PD0325901 |
| MVP_top60_DNA_seed | MAP2K1 | -1.353857 | SKCM | LB373-MEL-D | Refametinib |
| MVP_top60_DNA_seed | MAP2K1 | -1.346724 | SKCM | WM-115 | Refametinib |
| MVP_top60_DNA_seed | GSK3B | -1.342862 | SKCM | SK-MEL-2 | CHIR-99021 |
| MVP_top60_DNA_seed | MAP2K1 | -1.342739 | SKCM | COLO-792 | PD0325901 |
| MVP_top60_DNA_seed | MAP2K1 | -1.333917 | SKCM | MZ2-MEL | CI-1040 |
| MVP_top60_DNA_seed | JAK2 | -1.331771 | SKCM | IGR-1 | Ruxolitinib |
| MVP_top60_DNA_seed | JAK1 | -1.331771 | SKCM | IGR-1 | Ruxolitinib |
| MVP_top60_DNA_seed | BRAF | -1.330528 | SKCM | Hs-940-T | PLX-4720 |
| MVP_top60_DNA_seed | BRAF | -1.330121 | SKCM | IST-MEL1 | AZ628 |
| MVP_top60_DNA_seed | GSK3B | -1.328306 | SKCM | A101D | SB216763 |
| MVP_top60_DNA_seed | PIK3CB | -1.328245 | SKCM | UACC-257 | TGX221 |
| MVP_top60_DNA_seed | MAP2K1 | -1.321639 | SKCM | WM35 | CI-1040 |
| MVP_top60_DNA_seed | BRAF | -1.321489 | SKCM | Hs-939-T | RAF_9304 |

| | | | | | |
|---|---|---|---|---|---|
| MVP_top60_DNA_seed | MAPK1 | -1.319316 | SKCM | UACC-257 | VX-11e |
| MVP_top60_DNA_seed | GSK3B | -1.318416 | SKCM | HT-144 | CHIR-99021 |
| MVP_top60_DNA_seed | MAP2K1 | -1.317701 | SKCM | SK-MEL-3 | Refametinib |
| MVP_top60_DNA_seed | BRAF | -1.315079 | SKCM | MZ7-mel | PLX-4720 |
| MVP_top60_DNA_seed | BRAF | -1.31419 | SKCM | SK-MEL-1 | AZ628 |
| MVP_top60_DNA_seed | MAP2K1 | -1.31404 | SKCM | COLO-792 | Refametinib |
| MVP_top60_DNA_seed | MAP2K1 | -1.31079 | SKCM | MEL-JUSO | Refametinib |
| MVP_top60_DNA_seed | EGFR | -1.310732 | SKCM | 451Lu | Cetuximab |
| MVP_top60_DNA_seed | EP300 | -1.308424 | SKCM | COLO-783 | I-CBP112 |
| MVP_top60_DNA_seed | CREBBP | -1.308424 | SKCM | COLO-783 | I-CBP112 |
| MVP_top60_DNA_seed | MAP2K1 | -1.30259 | SKCM | SK-MEL-24 | Refametinib |
| MVP_top60_DNA_seed | EGFR | -1.300864 | SKCM | G-MEL | AST-1306 |
| MVP_top60_DNA_seed | MAP2K1 | -1.299763 | SKCM | GAK | Selumetinib |
| MVP_top60_DNA_seed | EGFR | -1.299268 | SKCM | IGR-37 | AST-1306 |
| MVP_top60_DNA_seed | JAK2 | -1.29807 | SKCM | Hs-940-T | Fedratinib |
| MVP_top60_DNA_seed | MAP2K1 | -1.29773 | SKCM | WM35 | PD0325901 |
| MVP_top60_DNA_seed | RAC1 | -1.297665 | SKCM | COLO-800 | EHT-1864 |
| MVP_top60_DNA_seed | MAP2K1 | -1.293839 | SKCM | LOXIMVI | PD0325901 |
| MVP_top60_DNA_seed | GSK3B | -1.28997 | SKCM | CHL-1 | SB216763 |
| MVP_top60_DNA_seed | GSK3B | -1.289883 | SKCM | M14 | CHIR-99021 |
| MVP_top60_DNA_seed | PIK3CB | -1.283173 | SKCM | GAK | TGX221 |
| MVP_top60_DNA_seed | BRAF | -1.282679 | SKCM | Hs-940-T | PLX-4720 |
| MVP_top60_DNA_seed | MAP2K1 | -1.279876 | SKCM | GAK | Refametinib |
| MVP_top60_DNA_seed | EGFR | -1.276798 | SKCM | SK-MEL-30 | Cetuximab |
| MVP_top60_DNA_seed | BRAF | -1.276418 | SKCM | G-361 | RAF_9304 |
| MVP_top60_DNA_seed | EGFR | -1.276103 | SKCM | WM-115 | Cetuximab |
| MVP_top60_DNA_seed | JAK3 | -1.27429 | SKCM | A375 | JAK3_7406 |
| MVP_top60_DNA_seed | BRAF | -1.272528 | SKCM | HT-144 | HG6-64-1 |
| MVP_top60_DNA_seed | PIK3CB | -1.268474 | SKCM | A101D | AZD6482 |
| MVP_top60_DNA_seed | MAP2K1 | -1.261855 | SKCM | A101D | Refametinib |
| MVP_top60_DNA_seed | MAPK1 | -1.260048 | SKCM | K2 | VX-11e |
| MVP_top60_DNA_seed | MAP2K1 | -1.257974 | SKCM | MZ7-mel | Selumetinib |
| MVP_top60_DNA_seed | MAP2K1 | -1.255989 | SKCM | HMV-II | CI-1040 |
| MVP_top60_DNA_seed | BRAF | -1.255107 | SKCM | WM793B | HG6-64-1 |
| MVP_top60_DNA_seed | BRAF | -1.249927 | SKCM | SK-MEL-2 | RAF_9304 |
| MVP_top60_DNA_seed | MAP2K1 | -1.243072 | SKCM | SK-MEL-1 | Refametinib |
| MVP_top60_DNA_seed | BRAF | -1.239444 | SKCM | IGR-1 | SB590885 |
| MVP_top60_DNA_seed | PIK3CB | -1.23913 | SKCM | G-361 | AZD6482 |
| MVP_top60_DNA_seed | MAP2K1 | -1.237227 | SKCM | COLO-792 | Trametinib |
| MVP_top60_DNA_seed | PIK3CB | -1.237012 | SKCM | CHL-1 | AZD8186 |
| MVP_top60_DNA_seed | MAPK3 | -1.236883 | SKCM | CP50-MEL-B | FR-180204 |

| | | | | | |
|---|---|---|---|---|---|
| MVP_top60_DNA_seed | MAPK1 | -1.236883 | SKCM | CP50-MEL-B | FR-180204 |
| MVP_top60_DNA_seed | EGFR | -1.236066 | SKCM | WM1552C | PF-00299804 |
| MVP_top60_DNA_seed | ERBB2 | -1.236066 | SKCM | WM1552C | PF-00299804 |
| MVP_top60_DNA_seed | PIK3CA | -1.235487 | SKCM | IGR-37 | AZD8835 |
| MVP_top60_DNA_seed | RAC1 | -1.22865 | SKCM | COLO-679 | EHT-1864 |
| MVP_top60_DNA_seed | JAK3 | -1.224283 | SKCM | COLO-800 | JAK3_7406 |
| MVP_top60_DNA_seed | MAP2K1 | -1.220607 | SKCM | LB373-MEL-D | PD0325901 |
| MVP_top60_DNA_seed | MAP2K1 | -1.213926 | SKCM | RVH-421 | Selumetinib |
| MVP_top60_DNA_seed | MAP2K1 | -1.212246 | SKCM | Mewo | Refametinib |
| MVP_top60_DNA_seed | ERBB2 | -1.211877 | SKCM | MEL-HO | CP724714 |
| MVP_top60_DNA_seed | EGFR | -1.209408 | SKCM | IPC-298 | AST-1306 |
| MVP_top60_DNA_seed | JAK1 | -1.204364 | SKCM | RPMI-7951 | JAK1_3715 |
| MVP_top60_DNA_seed | MAP2K1 | -1.204162 | SKCM | SK-MEL-5 | Refametinib |
| MVP_top60_DNA_seed | PIK3CB | -1.198909 | SKCM | IGR-37 | AZD6482 |
| MVP_top60_DNA_seed | BRAF | -1.198875 | SKCM | SK-MEL-24 | PLX-4720 |
| MVP_top60_DNA_seed | MAP2K1 | -1.198422 | SKCM | Mewo | CI-1040 |
| MVP_top60_DNA_seed | MAP2K1 | -1.196346 | SKCM | SK-MEL-1 | Trametinib |
| MVP_top60_DNA_seed | EP300 | -1.194512 | SKCM | 451Lu | I-CBP112 |
| MVP_top60_DNA_seed | CREBBP | -1.194512 | SKCM | 451Lu | I-CBP112 |
| MVP_top60_DNA_seed | EGFR | -1.189494 | SKCM | SK-MEL-1 | Gefitinib |
| MVP_top60_DNA_seed | MAP2K1 | -1.188234 | SKCM | SK-MEL-3 | Trametinib |
| MVP_top60_DNA_seed | BRAF | -1.180196 | SKCM | IPC-298 | PLX-4720 |
| MVP_top60_DNA_seed | EGFR | -1.179985 | SKCM | IGR-1 | CUDC-101 |
| MVP_top60_DNA_seed | ERBB2 | -1.179985 | SKCM | IGR-1 | CUDC-101 |
| MVP_top60_DNA_seed | BRAF | -1.177135 | SKCM | SK-MEL-2 | Dabrafenib |
| MVP_top60_DNA_seed | BRAF | -1.173688 | SKCM | IGR-1 | PLX-4720 |
| MVP_top60_DNA_seed | BRAF | -1.168445 | SKCM | LB373-MEL-D | AZ628 |
| MVP_top60_DNA_seed | BRAF | -1.16803 | SKCM | WM-115 | RAF_9304 |
| MVP_top60_DNA_seed | MAP2K1 | -1.166359 | SKCM | Hs-939-T | Refametinib |
| MVP_top60_DNA_seed | MAP2K1 | -1.160122 | SKCM | WM-115 | PD0325901 |
| MVP_top60_DNA_seed | BRAF | -1.157821 | SKCM | A2058 | PLX-4720 |
| MVP_top60_DNA_seed | MAP2K1 | -1.154576 | SKCM | A101D | Selumetinib |
| MVP_top60_DNA_seed | MAP2K1 | -1.153251 | SKCM | RVH-421 | Refametinib |
| MVP_top60_DNA_seed | MAPK1 | -1.15192 | SKCM | WM-115 | VX-11e |
| MVP_top60_DNA_seed | JAK2 | -1.150077 | SKCM | COLO-800 | AZD1480 |
| MVP_top60_DNA_seed | JAK1 | -1.150077 | SKCM | COLO-800 | AZD1480 |
| MVP_top60_DNA_seed | MAPK1 | -1.146497 | SKCM | SK-MEL-28 | VX-11e |
| MVP_top60_DNA_seed | EP300 | -1.142977 | SKCM | G-MEL | I-CBP112 |
| MVP_top60_DNA_seed | CREBBP | -1.142977 | SKCM | G-MEL | I-CBP112 |
| MVP_top60_DNA_seed | BRAF | -1.142658 | SKCM | RPMI-7951 | PLX-4720 |
| MVP_top60_DNA_seed | MAP2K1 | -1.141438 | SKCM | CHL-1 | Selumetinib |

| | | | | | |
|---|---|---|---|---|---|
| MVP_top60_DNA_seed | EGFR | -1.140637 | SKCM | SK-MEL-2 | CI-1033 |
| MVP_top60_DNA_seed | ERBB2 | -1.140637 | SKCM | SK-MEL-2 | CI-1033 |
| MVP_top60_DNA_seed | BRAF | -1.134287 | SKCM | IST-MEL1 | SB590885 |
| MVP_top60_DNA_seed | JAK3 | -1.132951 | SKCM | G-MEL | JAK3_7406 |
| MVP_top60_DNA_seed | MAP2K1 | -1.132801 | SKCM | GAK | Selumetinib |
| MVP_top60_DNA_seed | MAP2K1 | -1.131501 | SKCM | K2 | Refametinib |
| MVP_top60_DNA_seed | JAK3 | -1.130729 | SKCM | MZ2-MEL | JAK3_7406 |
| MVP_top60_DNA_seed | EGFR | -1.130289 | SKCM | CHL-1 | CUDC-101 |
| MVP_top60_DNA_seed | ERBB2 | -1.130289 | SKCM | CHL-1 | CUDC-101 |
| MVP_top60_DNA_seed | BRAF | -1.129743 | SKCM | WM-115 | PLX-4720 |
| MVP_top60_DNA_seed | MAP2K1 | -1.125812 | SKCM | WM-115 | Trametinib |
| MVP_top60_DNA_seed | MAP2K1 | -1.122865 | SKCM | SK-MEL-3 | Selumetinib |
| MVP_top60_DNA_seed | MAP2K1 | -1.122176 | SKCM | CHL-1 | Refametinib |
| MVP_top60_DNA_seed | BRAF | -1.120565 | SKCM | A101D | HG6-64-1 |
| MVP_top60_DNA_seed | EGFR | -1.120458 | SKCM | MZ7-mel | CUDC-101 |
| MVP_top60_DNA_seed | ERBB2 | -1.120458 | SKCM | MZ7-mel | CUDC-101 |
| MVP_top60_DNA_seed | MAP2K1 | -1.11999 | SKCM | MEL-JUSO | Refametinib |
| MVP_top60_DNA_seed | GSK3B | -1.116764 | SKCM | SK-MEL-2 | AZD7969 |
| MVP_top60_DNA_seed | BRAF | -1.113387 | SKCM | HMV-II | Dabrafenib |
| MVP_top60_DNA_seed | BRAF | -1.113266 | SKCM | SK-MEL-24 | HG6-64-1 |
| MVP_top60_DNA_seed | MAP2K1 | -1.112414 | SKCM | GAK | Refametinib |
| MVP_top60_DNA_seed | MAP2K1 | -1.110318 | SKCM | COLO-783 | Trametinib |
| MVP_top60_DNA_seed | MAPK1 | -1.109459 | SKCM | SK-MEL-24 | VX-11e |
| MVP_top60_DNA_seed | GSK3B | -1.109075 | SKCM | LB373-MEL-D | CHIR-99021 |
| MVP_top60_DNA_seed | EP300 | -1.105961 | SKCM | A101D | I-CBP112 |
| MVP_top60_DNA_seed | CREBBP | -1.105961 | SKCM | A101D | I-CBP112 |
| MVP_top60_DNA_seed | MAPK14 | -1.102839 | SKCM | MEL-HO | TAK-715 |
| MVP_top60_DNA_seed | MAP2K1 | -1.101942 | SKCM | G-361 | CI-1040 |
| MVP_top60_DNA_seed | MAPK14 | -1.101532 | SKCM | A375 | TAK-715 |
| MVP_top60_DNA_seed | EGFR | -1.096994 | SKCM | WM35 | Cetuximab |
| MVP_top60_DNA_seed | EP300 | -1.096713 | SKCM | SK-MEL-5 | I-CBP112 |
| MVP_top60_DNA_seed | CREBBP | -1.096713 | SKCM | SK-MEL-5 | I-CBP112 |
| MVP_top60_DNA_seed | GSK3B | -1.094828 | SKCM | SK-MEL-28 | CHIR-99021 |
| MVP_top60_DNA_seed | JAK2 | -1.093669 | SKCM | MEL-HO | Ruxolitinib |
| MVP_top60_DNA_seed | JAK1 | -1.093669 | SKCM | MEL-HO | Ruxolitinib |
| MVP_top60_DNA_seed | MAPK1 | -1.093413 | SKCM | COLO-679 | VX-11e |
| MVP_top60_DNA_seed | MAP2K1 | -1.087295 | SKCM | SK-MEL-1 | Selumetinib |
| MVP_top60_DNA_seed | MAP2K1 | -1.084948 | SKCM | COLO-829 | CI-1040 |
| MVP_top60_DNA_seed | MAPK14 | -1.083289 | SKCM | SK-MEL-28 | TAK-715 |
| MVP_top60_DNA_seed | BRAF | -1.081893 | SKCM | WM793B | PLX-4720 |
| MVP_top60_DNA_seed | GSK3B | -1.079374 | SKCM | WM-115 | CHIR-99021 |

| | | | | | |
|---|---|---|---|---|---|
| MVP_top60_DNA_seed | MAP2K1 | -1.079286 | SKCM | Mewo | Trametinib |
| MVP_top60_DNA_seed | JAK1 | -1.078596 | SKCM | SK-MEL-28 | JAK1_3715 |
| MVP_top60_DNA_seed | BRAF | -1.076817 | SKCM | MEL-JUSO | RAF_9304 |
| MVP_top60_DNA_seed | MAP2K1 | -1.074468 | SKCM | SK-MEL-3 | PD0325901 |
| MVP_top60_DNA_seed | ERBB2 | -1.070274 | SKCM | G-361 | CP724714 |
| MVP_top60_DNA_seed | BRAF | -1.068949 | SKCM | MEL-HO | HG6-64-1 |
| MVP_top60_DNA_seed | MAPK1 | -1.068862 | SKCM | LB2518-MEL | VX-11e |
| MVP_top60_DNA_seed | MAP2K1 | -1.067865 | SKCM | WM278 | Refametinib |
| MVP_top60_DNA_seed | MAP2K1 | -1.064557 | SKCM | RPMI-7951 | Refametinib |
| MVP_top60_DNA_seed | MAP2K1 | -1.060592 | SKCM | COLO-783 | Selumetinib |
| MVP_top60_DNA_seed | ERBB2 | -1.059511 | SKCM | G-MEL | CP724714 |
| MVP_top60_DNA_seed | BRAF | -1.058739 | SKCM | CP66-MEL | PLX-4720 |
| MVP_top60_DNA_seed | EGFR | -1.05489 | SKCM | K2 | Cetuximab |
| MVP_top60_DNA_seed | GSK3B | -1.054477 | SKCM | SK-MEL-2 | SB216763 |
| MVP_top60_DNA_seed | GSK3B | -1.054359 | SKCM | G-MEL | CHIR-99021 |
| MVP_top60_DNA_seed | MAP2K1 | -1.051019 | SKCM | WM278 | CI-1040 |
| MVP_top60_DNA_seed | MAPK1 | -1.046254 | SKCM | G-361 | VX-11e |
| MVP_top60_DNA_seed | MAP2K1 | -1.044997 | SKCM | COLO-783 | CI-1040 |
| MVP_top60_DNA_seed | MAP2K1 | -1.044662 | SKCM | COLO-829 | Trametinib |
| MVP_top60_DNA_seed | PIK3CB | -1.040105 | SKCM | UACC-62 | AZD6482 |
| MVP_top60_DNA_seed | MAPK3 | -1.038522 | SKCM | VMRC-MELG | FR-180204 |
| MVP_top60_DNA_seed | MAPK1 | -1.038522 | SKCM | VMRC-MELG | FR-180204 |
| MVP_top60_DNA_seed | MAP2K1 | -1.032993 | SKCM | WM-115 | Refametinib |
| MVP_top60_DNA_seed | PIK3CB | -1.031281 | SKCM | Hs-940-T | AZD6482 |
| MVP_top60_DNA_seed | RAC1 | -1.03108 | SKCM | 451Lu | EHT-1864 |
| MVP_top60_DNA_seed | BRAF | -1.030639 | SKCM | 451Lu | HG6-64-1 |
| MVP_top60_DNA_seed | BRAF | -1.030343 | SKCM | IGR-1 | PLX-4720 |
| MVP_top60_DNA_seed | JAK3 | -1.027293 | SKCM | MEL-HO | WHI-P97 |
| MVP_top60_DNA_seed | PIK3CA | -1.021245 | SKCM | WM1552C | PI3Ka_4409 |
| MVP_top60_DNA_seed | JAK3 | -1.019298 | SKCM | CHL-1 | JAK3_7406 |
| MVP_top60_DNA_seed | MAP2K1 | -1.012131 | SKCM | WM1552C | Refametinib |
| MVP_top60_DNA_seed | GSK3B | -1.011961 | SKCM | RPMI-7951 | AZD7969 |
| MVP_top60_DNA_seed | MAP2K1 | -1.009533 | SKCM | SK-MEL-1 | PD0325901 |
| MVP_top60_DNA_seed | BRAF | -1.009434 | SKCM | IST-MEL1 | PLX-4720 |
| MVP_top60_DNA_seed | MAP2K1 | -1.006374 | SKCM | MZ2-MEL | Selumetinib |
| MVP_top60_DNA_seed | EP300 | -1.00634 | SKCM | LB373-MEL-D | I-CBP112 |
| MVP_top60_DNA_seed | CREBBP | -1.00634 | SKCM | LB373-MEL-D | I-CBP112 |
| MVP_top60_DNA_seed | MAP2K1 | -1.005776 | SKCM | RVH-421 | Trametinib |
| MVP_top60_DNA_seed | MAP2K1 | -1.00514 | SKCM | LB373-MEL-D | Selumetinib |
| MVP_top60_DNA_seed | JAK2 | -1.002592 | SKCM | G-MEL | AZD1480 |
| MVP_top60_DNA_seed | JAK1 | -1.002592 | SKCM | G-MEL | AZD1480 |

| | | | | | |
|---|---|---|---|---|---|
| MVP_top60_DNA_seed | BRAF | -0.999764 | SKCM | A2058 | PLX-4720 |
| MVP_top60_DNA_seed | RAC1 | -0.999614 | SKCM | G-361 | EHT-1864 |
| MVP_top60_DNA_seed | MAPK14 | -0.999407 | SKCM | IGR-1 | TAK-715 |
| MVP_top60_DNA_seed | JAK3 | -0.998855 | SKCM | LOXIMVI | JAK3_7406 |
| MVP_top60_DNA_seed | GSK3B | -0.993871 | SKCM | K2 | CHIR-99021 |
| MVP_top60_DNA_seed | JAK3 | -0.993807 | SKCM | MZ2-MEL | WHI-P97 |
| MVP_top60_DNA_seed | BRAF | -0.987726 | SKCM | WM793B | RAF_9304 |
| MVP_top60_DNA_seed | GSK3B | -0.982085 | SKCM | IST-MEL1 | CHIR-99021 |
| MVP_top60_DNA_seed | PIK3CA | -0.97738 | SKCM | MZ2-MEL | AZD8835 |
| MVP_top60_DNA_seed | MAPK1 | -0.974351 | SKCM | CP50-MEL-B | VX-11e |
| MVP_top60_DNA_seed | BRAF | -0.972243 | SKCM | MEL-JUSO | HG6-64-1 |
| MVP_top60_DNA_seed | PIK3CB | -0.969179 | SKCM | COLO-829 | AZD6482 |
| MVP_top60_DNA_seed | PIK3CB | -0.964257 | SKCM | MMAc-SF | TGX221 |
| MVP_top60_DNA_seed | GSK3B | -0.961989 | SKCM | CP50-MEL-B | KIN001-042 |
| MVP_top60_DNA_seed | MAP2K1 | -0.96129 | SKCM | WM-115 | Selumetinib |
| MVP_top60_DNA_seed | EGFR | -0.960091 | SKCM | SK-MEL-30 | AST-1306 |
| MVP_top60_DNA_seed | BRAF | -0.958089 | SKCM | MZ2-MEL | HG6-64-1 |
| MVP_top60_DNA_seed | EP300 | -0.954512 | SKCM | K2 | I-CBP112 |
| MVP_top60_DNA_seed | CREBBP | -0.954512 | SKCM | K2 | I-CBP112 |
| MVP_top60_DNA_seed | GSK3B | -0.950407 | SKCM | WM793B | SB216763 |
| MVP_top60_DNA_seed | GSK3B | -0.949689 | SKCM | COLO-800 | AZD7969 |
| MVP_top60_DNA_seed | MAPK1 | -0.948907 | SKCM | COLO-829 | VX-11e |
| MVP_top60_DNA_seed | MAP2K1 | -0.942242 | SKCM | SK-MEL-1 | Refametinib |
| MVP_top60_DNA_seed | JAK3 | -0.941894 | SKCM | WM793B | JAK3_7406 |
| MVP_top60_DNA_seed | GSK3B | -0.941743 | SKCM | UACC-62 | CHIR-99021 |
| MVP_top60_DNA_seed | MAP2K1 | -0.93234 | SKCM | LB373-MEL-D | Trametinib |
| MVP_top60_DNA_seed | MAP2K1 | -0.929277 | SKCM | COLO-829 | Refametinib |
| MVP_top60_DNA_seed | MAP2K1 | -0.922254 | SKCM | GAK | PD0325901 |
| MVP_top60_DNA_seed | GSK3B | -0.913169 | SKCM | RVH-421 | CHIR-99021 |
| MVP_top60_DNA_seed | BRAF | -0.908717 | SKCM | WM793B | PLX-4720 |
| MVP_top60_DNA_seed | EGFR | -0.908392 | SKCM | SK-MEL-2 | AST-1306 |
| MVP_top60_DNA_seed | MAP2K1 | -0.907106 | SKCM | SK-MEL-1 | Selumetinib |
| MVP_top60_DNA_seed | EGFR | -0.906665 | SKCM | WM278 | PF-00299804 |
| MVP_top60_DNA_seed | ERBB2 | -0.906665 | SKCM | WM278 | PF-00299804 |
| MVP_top60_DNA_seed | BRAF | -0.905572 | SKCM | LB2518-MEL | RAF_9304 |
| MVP_top60_DNA_seed | PIK3CB | -0.902218 | SKCM | MEL-JUSO | AZD8186 |
| MVP_top60_DNA_seed | EGFR | -0.902105 | SKCM | Hs-939-T | AST-1306 |
| MVP_top60_DNA_seed | PIK3CB | -0.898658 | SKCM | WM-115 | AZD8186 |
| MVP_top60_DNA_seed | MAP2K1 | -0.898561 | SKCM | SK-MEL-3 | CI-1040 |
| MVP_top60_DNA_seed | GSK3B | -0.884046 | SKCM | GAK | KIN001-042 |
| MVP_top60_DNA_seed | BRAF | -0.882645 | SKCM | CHL-1 | RAF_9304 |

| | | | | | |
|---|---|---|---|---|---|
| MVP_top60_DNA_seed | PIK3CB | -0.877288 | SKCM | HT-144 | TGX221 |
| MVP_top60_DNA_seed | PIK3CB | -0.877218 | SKCM | CHL-1 | AZD6482 |
| MVP_top60_DNA_seed | MAP2K1 | -0.871811 | SKCM | SK-MEL-2 | Selumetinib |
| MVP_top60_DNA_seed | MAPK1 | -0.869581 | SKCM | SK-MEL-3 | VX-11e |
| MVP_top60_DNA_seed | EGFR | -0.868616 | SKCM | CP50-MEL-B | AST-1306 |
| MVP_top60_DNA_seed | BRAF | -0.86837 | SKCM | COLO-783 | RAF_9304 |
| MVP_top60_DNA_seed | BRAF | -0.866914 | SKCM | IPC-298 | PLX-4720 |
| MVP_top60_DNA_seed | PIK3CA | -0.866719 | SKCM | IGR-37 | PI3Ka_4409 |
| MVP_top60_DNA_seed | BRAF | -0.863686 | SKCM | Mewo | SB590885 |
| MVP_top60_DNA_seed | MAP2K1 | -0.863257 | SKCM | CHL-1 | CI-1040 |
| MVP_top60_DNA_seed | MAP2K1 | -0.857554 | SKCM | WM793B | Trametinib |
| MVP_top60_DNA_seed | EGFR | -0.855871 | SKCM | IGR-1 | CI-1033 |
| MVP_top60_DNA_seed | ERBB2 | -0.855871 | SKCM | IGR-1 | CI-1033 |
| MVP_top60_DNA_seed | EGFR | -0.852579 | SKCM | HT-144 | CI-1033 |
| MVP_top60_DNA_seed | ERBB2 | -0.852579 | SKCM | HT-144 | CI-1033 |
| MVP_top60_DNA_seed | JAK3 | -0.851496 | SKCM | SK-MEL-2 | JAK3_7406 |
| MVP_top60_DNA_seed | BRAF | -0.848532 | SKCM | RVH-421 | RAF_9304 |
| MVP_top60_DNA_seed | EGFR | -0.84652 | SKCM | MEL-HO | CUDC-101 |
| MVP_top60_DNA_seed | ERBB2 | -0.84652 | SKCM | MEL-HO | CUDC-101 |
| MVP_top60_DNA_seed | PIK3CA | -0.84307 | SKCM | SK-MEL-5 | PI3Ka_4409 |
| MVP_top60_DNA_seed | EGFR | -0.838627 | SKCM | SH-4 | PF-00299804 |
| MVP_top60_DNA_seed | ERBB2 | -0.838627 | SKCM | SH-4 | PF-00299804 |
| MVP_top60_DNA_seed | GSK3B | -0.8377 | SKCM | RVH-421 | CHIR-99021 |
| MVP_top60_DNA_seed | MAP2K1 | -0.83733 | SKCM | WM278 | PD0325901 |
| MVP_top60_DNA_seed | EGFR | -0.832601 | SKCM | LOXIMVI | Gefitinib |
| MVP_top60_DNA_seed | EGFR | -0.831839 | SKCM | COLO-800 | CI-1033 |
| MVP_top60_DNA_seed | ERBB2 | -0.831839 | SKCM | COLO-800 | CI-1033 |
| MVP_top60_DNA_seed | MAPK1 | -0.830098 | SKCM | RPMI-7951 | VX-11e |
| MVP_top60_DNA_seed | PIK3CB | -0.82534 | SKCM | A2058 | AZD6482 |
| MVP_top60_DNA_seed | RAC1 | -0.818448 | SKCM | SK-MEL-5 | EHT-1864 |
| MVP_top60_DNA_seed | EGFR | -0.816502 | SKCM | SH-4 | AST-1306 |
| MVP_top60_DNA_seed | BRAF | -0.810236 | SKCM | WM1552C | SB590885 |
| MVP_top60_DNA_seed | MAP2K1 | -0.808551 | SKCM | COLO-792 | CI-1040 |
| MVP_top60_DNA_seed | RAC1 | -0.808184 | SKCM | MEL-JUSO | EHT-1864 |
| MVP_top60_DNA_seed | GSK3B | -0.805912 | SKCM | WM793B | AZD7969 |
| MVP_top60_DNA_seed | JAK3 | -0.805486 | SKCM | RPMI-7951 | JAK3_7406 |
| MVP_top60_DNA_seed | BRAF | -0.804292 | SKCM | G-MEL | HG6-64-1 |
| MVP_top60_DNA_seed | JAK2 | -0.802517 | SKCM | COLO-800 | Ruxolitinib |
| MVP_top60_DNA_seed | JAK1 | -0.802517 | SKCM | COLO-800 | Ruxolitinib |
| MVP_top60_DNA_seed | GSK3B | -0.802042 | SKCM | UACC-257 | SB216763 |
| MVP_top60_DNA_seed | GSK3B | -0.801533 | SKCM | HMV-II | CHIR-99021 |

| | | | | | |
|---|---|---|---|---|---|
| MVP_top60_DNA_seed | BRAF | -0.796694 | SKCM | MZ2-MEL | SB590885 |
| MVP_top60_DNA_seed | MAP2K1 | -0.796059 | SKCM | RPMI-7951 | Selumetinib |
| MVP_top60_DNA_seed | EGFR | -0.794983 | SKCM | Hs-940-T | CI-1033 |
| MVP_top60_DNA_seed | ERBB2 | -0.794983 | SKCM | Hs-940-T | CI-1033 |
| MVP_top60_DNA_seed | GSK3B | -0.792392 | SKCM | CHL-1 | AZD7969 |
| MVP_top60_DNA_seed | EGFR | -0.787508 | SKCM | LB373-MEL-D | Erlotinib |
| MVP_top60_DNA_seed | MAP2K1 | -0.786188 | SKCM | SK-MEL-24 | CI-1040 |
| MVP_top60_DNA_seed | MAPK1 | -0.78475 | SKCM | A101D | VX-11e |
| MVP_top60_DNA_seed | MAP2K1 | -0.781481 | SKCM | WM793B | Selumetinib |
| MVP_top60_DNA_seed | GSK3B | -0.781473 | SKCM | IPC-298 | CHIR-99021 |
| MVP_top60_DNA_seed | BRAF | -0.780405 | SKCM | WM35 | HG6-64-1 |
| MVP_top60_DNA_seed | MAPK3 | -0.778978 | SKCM | IGR-37 | FR-180204 |
| MVP_top60_DNA_seed | MAPK1 | -0.778978 | SKCM | IGR-37 | FR-180204 |
| MVP_top60_DNA_seed | EGFR | -0.777929 | SKCM | IGR-37 | PF-00299804 |
| MVP_top60_DNA_seed | ERBB2 | -0.777929 | SKCM | IGR-37 | PF-00299804 |
| MVP_top60_DNA_seed | BRAF | -0.776445 | SKCM | WM1552C | HG6-64-1 |
| MVP_top60_DNA_seed | BRAF | -0.775535 | SKCM | COLO-679 | HG6-64-1 |
| MVP_top60_DNA_seed | MAPK3 | -0.77544 | SKCM | WM793B | FR-180204 |
| MVP_top60_DNA_seed | MAPK1 | -0.77544 | SKCM | WM793B | FR-180204 |
| MVP_top60_DNA_seed | BRAF | -0.771352 | SKCM | CHL-1 | PLX-4720 |
| MVP_top60_DNA_seed | MAPK1 | -0.771238 | SKCM | GAK | VX-11e |
| MVP_top60_DNA_seed | JAK3 | -0.77088 | SKCM | WM-115 | WHI-P97 |
| MVP_top60_DNA_seed | BRAF | -0.765871 | SKCM | Hs-940-T | HG6-64-1 |
| MVP_top60_DNA_seed | MAP2K1 | -0.762407 | SKCM | CHL-1 | Trametinib |
| MVP_top60_DNA_seed | MAP2K1 | -0.760935 | SKCM | IST-MEL1 | Trametinib |
| MVP_top60_DNA_seed | MAP2K1 | -0.760749 | SKCM | SK-MEL-3 | Refametinib |
| MVP_top60_DNA_seed | PIK3CA | -0.756197 | SKCM | LOXIMVI | AZD8835 |
| MVP_top60_DNA_seed | MAP2K1 | -0.755731 | SKCM | SK-MEL-31 | PD0325901 |
| MVP_top60_DNA_seed | PIK3CA | -0.754921 | SKCM | CHL-1 | PI3Ka_4409 |
| MVP_top60_DNA_seed | JAK3 | -0.754655 | SKCM | IGR-37 | WHI-P97 |
| MVP_top60_DNA_seed | JAK1 | -0.751807 | SKCM | WM1552C | JAK1_3715 |
| MVP_top60_DNA_seed | JAK3 | -0.748661 | SKCM | WM278 | WHI-P97 |
| MVP_top60_DNA_seed | BRAF | -0.748499 | SKCM | WM1552C | Dabrafenib |
| MVP_top60_DNA_seed | MAPK14 | -0.745745 | SKCM | CHL-1 | TAK-715 |
| MVP_top60_DNA_seed | EGFR | -0.74542 | SKCM | HMV-II | AST-1306 |
| MVP_top60_DNA_seed | MAP2K1 | -0.738765 | SKCM | WM1552C | Selumetinib |
| MVP_top60_DNA_seed | EGFR | -0.738126 | SKCM | A375 | CI-1033 |
| MVP_top60_DNA_seed | ERBB2 | -0.738126 | SKCM | A375 | CI-1033 |
| MVP_top60_DNA_seed | BRAF | -0.737577 | SKCM | IGR-1 | HG6-64-1 |
| MVP_top60_DNA_seed | MAP2K1 | -0.73609 | SKCM | COLO-829 | Selumetinib |
| MVP_top60_DNA_seed | JAK2 | -0.733731 | SKCM | MZ2-MEL | Fedratinib |

| | | | | | |
|---|---|---|---|---|---|
| MVP_top60_DNA_seed | MAP2K1 | -0.728614 | SKCM | Hs-939-T | Trametinib |
| MVP_top60_DNA_seed | PIK3CA | -0.725992 | SKCM | MZ2-MEL | PI3Ka_4409 |
| MVP_top60_DNA_seed | MAP2K1 | -0.725903 | SKCM | WM793B | Selumetinib |
| MVP_top60_DNA_seed | RAC1 | -0.722241 | SKCM | G-MEL | EHT-1864 |
| MVP_top60_DNA_seed | GSK3B | -0.715613 | SKCM | IGR-1 | KIN001-042 |
| MVP_top60_DNA_seed | BRAF | -0.708957 | SKCM | A375 | HG6-64-1 |
| MVP_top60_DNA_seed | BRAF | -0.70777 | SKCM | G-361 | HG6-64-1 |
| MVP_top60_DNA_seed | BRAF | -0.702291 | SKCM | SK-MEL-1 | RAF_9304 |
| MVP_top60_DNA_seed | GSK3B | -0.702291 | SKCM | HT-144 | AZD7969 |
| MVP_top60_DNA_seed | GSK3B | -0.70123 | SKCM | IGR-37 | AZD7969 |
| MVP_top60_DNA_seed | ERBB2 | -0.70034 | SKCM | COLO-800 | CP724714 |
| MVP_top60_DNA_seed | JAK2 | -0.697415 | SKCM | COLO-800 | Fedratinib |
| MVP_top60_DNA_seed | MAP2K1 | -0.696318 | SKCM | COLO-829 | PD0325901 |
| MVP_top60_DNA_seed | MAP2K1 | -0.694301 | SKCM | MZ2-MEL | Trametinib |
| MVP_top60_DNA_seed | MAP2K1 | -0.69133 | SKCM | IGR-1 | Selumetinib |
| MVP_top60_DNA_seed | MAPK3 | -0.686521 | SKCM | K2 | FR-180204 |
| MVP_top60_DNA_seed | MAPK1 | -0.686521 | SKCM | K2 | FR-180204 |
| MVP_top60_DNA_seed | MAP2K1 | -0.682964 | SKCM | Mewo | Selumetinib |
| MVP_top60_DNA_seed | MAP2K1 | -0.682786 | SKCM | GAK | CI-1040 |
| MVP_top60_DNA_seed | EGFR | -0.682131 | SKCM | LB2518-MEL | Erlotinib |
| MVP_top60_DNA_seed | PIK3CB | -0.681486 | SKCM | MZ2-MEL | TGX221 |
| MVP_top60_DNA_seed | GSK3B | -0.677446 | SKCM | IGR-1 | AZD7969 |
| MVP_top60_DNA_seed | MAPK14 | -0.676829 | SKCM | G-361 | TAK-715 |
| MVP_top60_DNA_seed | EGFR | -0.676541 | SKCM | WM-115 | CI-1033 |
| MVP_top60_DNA_seed | ERBB2 | -0.676541 | SKCM | WM-115 | CI-1033 |
| MVP_top60_DNA_seed | EGFR | -0.674901 | SKCM | HT-144 | Gefitinib |
| MVP_top60_DNA_seed | GSK3B | -0.672593 | SKCM | WM278 | AZD7969 |
| MVP_top60_DNA_seed | BRAF | -0.671997 | SKCM | IGR-1 | RAF_9304 |
| MVP_top60_DNA_seed | EGFR | -0.67064 | SKCM | CP66-MEL | Erlotinib |
| MVP_top60_DNA_seed | JAK1 | -0.669322 | SKCM | A375 | JAK1_3715 |
| MVP_top60_DNA_seed | PIK3CB | -0.666834 | SKCM | SK-MEL-30 | AZD6482 |
| MVP_top60_DNA_seed | EP300 | -0.666374 | SKCM | CHL-1 | I-CBP112 |
| MVP_top60_DNA_seed | CREBBP | -0.666374 | SKCM | CHL-1 | I-CBP112 |
| MVP_top60_DNA_seed | PIK3CB | -0.660735 | SKCM | MZ7-mel | TGX221 |
| MVP_top60_DNA_seed | PIK3CA | -0.659856 | SKCM | CP50-MEL-B | AZD8835 |
| MVP_top60_DNA_seed | GSK3B | -0.658038 | SKCM | Hs-939-T | CHIR-99021 |
| MVP_top60_DNA_seed | EGFR | -0.657636 | SKCM | VMRC-MELG | CI-1033 |
| MVP_top60_DNA_seed | ERBB2 | -0.657636 | SKCM | VMRC-MELG | CI-1033 |
| MVP_top60_DNA_seed | MAP2K1 | -0.655199 | SKCM | COLO-783 | Refametinib |
| MVP_top60_DNA_seed | GSK3B | -0.654866 | SKCM | LOXIMVI | CHIR-99021 |
| MVP_top60_DNA_seed | PIK3CA | -0.652599 | SKCM | WM35 | PI3Ka_4409 |

| | | | | | |
|---|---|---|---|---|---|
| MVP_top60_DNA_seed | EGFR | -0.651564 | SKCM | WM35 | CUDC-101 |
| MVP_top60_DNA_seed | ERBB2 | -0.651564 | SKCM | WM35 | CUDC-101 |
| MVP_top60_DNA_seed | EGFR | -0.650264 | SKCM | COLO-800 | AZD8931 |
| MVP_top60_DNA_seed | ERBB2 | -0.650264 | SKCM | COLO-800 | AZD8931 |
| MVP_top60_DNA_seed | GSK3B | -0.650253 | SKCM | WM35 | SB216763 |
| MVP_top60_DNA_seed | PIK3CB | -0.648506 | SKCM | RPMI-7951 | AZD8186 |
| MVP_top60_DNA_seed | GSK3B | -0.64783 | SKCM | SK-MEL-1 | SB216763 |
| MVP_top60_DNA_seed | MAP2K1 | -0.6468 | SKCM | WM1552C | Refametinib |
| MVP_top60_DNA_seed | MAP2K1 | -0.646312 | SKCM | LB373-MEL-D | Refametinib |
| MVP_top60_DNA_seed | PIK3CB | -0.645899 | SKCM | SK-MEL-24 | AZD8186 |
| MVP_top60_DNA_seed | PIK3CA | -0.643715 | SKCM | A375 | AZD8835 |
| MVP_top60_DNA_seed | EGFR | -0.639842 | SKCM | CP50-MEL-B | CI-1033 |
| MVP_top60_DNA_seed | ERBB2 | -0.639842 | SKCM | CP50-MEL-B | CI-1033 |
| MVP_top60_DNA_seed | MAP2K1 | -0.633481 | SKCM | CHL-1 | PD0325901 |
| MVP_top60_DNA_seed | PIK3CB | -0.631622 | SKCM | Hs-939-T | AZD8186 |
| MVP_top60_DNA_seed | JAK3 | -0.630929 | SKCM | IPC-298 | JAK3_7406 |
| MVP_top60_DNA_seed | RAC1 | -0.630922 | SKCM | IGR-1 | EHT-1864 |
| MVP_top60_DNA_seed | JAK3 | -0.629105 | SKCM | WM-115 | JAK3_7406 |
| MVP_top60_DNA_seed | PIK3CB | -0.625255 | SKCM | MMAc-SF | AZD8186 |
| MVP_top60_DNA_seed | PIK3CA | -0.624959 | SKCM | WM278 | AZD8835 |
| MVP_top60_DNA_seed | JAK2 | -0.623615 | SKCM | G-MEL | Ruxolitinib |
| MVP_top60_DNA_seed | JAK1 | -0.623615 | SKCM | G-MEL | Ruxolitinib |
| MVP_top60_DNA_seed | MAP2K1 | -0.617233 | SKCM | MZ2-MEL | Refametinib |
| MVP_top60_DNA_seed | JAK2 | -0.617084 | SKCM | CHL-1 | AZD1480 |
| MVP_top60_DNA_seed | JAK1 | -0.617084 | SKCM | CHL-1 | AZD1480 |
| MVP_top60_DNA_seed | BRAF | -0.616135 | SKCM | SH-4 | HG6-64-1 |
| MVP_top60_DNA_seed | EP300 | -0.615514 | SKCM | A375 | I-CBP112 |
| MVP_top60_DNA_seed | CREBBP | -0.615514 | SKCM | A375 | I-CBP112 |
| MVP_top60_DNA_seed | PIK3CB | -0.612334 | SKCM | SK-MEL-5 | AZD8186 |
| MVP_top60_DNA_seed | MAP2K1 | -0.611945 | SKCM | IST-MEL1 | Refametinib |
| MVP_top60_DNA_seed | EGFR | -0.611113 | SKCM | WM1552C | Pelitinib |
| MVP_top60_DNA_seed | ERBB2 | -0.607891 | SKCM | RVH-421 | CP724714 |
| MVP_top60_DNA_seed | BRAF | -0.602072 | SKCM | CP66-MEL | RAF_9304 |
| MVP_top60_DNA_seed | EGFR | -0.601274 | SKCM | IGR-37 | CUDC-101 |
| MVP_top60_DNA_seed | ERBB2 | -0.601274 | SKCM | IGR-37 | CUDC-101 |
| MVP_top60_DNA_seed | RAC1 | -0.59942 | SKCM | CHL-1 | EHT-1864 |
| MVP_top60_DNA_seed | JAK2 | -0.598138 | SKCM | G-MEL | Fedratinib |
| MVP_top60_DNA_seed | GSK3B | -0.590376 | SKCM | UACC-62 | SB216763 |
| MVP_top60_DNA_seed | BRAF | -0.58618 | SKCM | IST-MEL1 | RAF_9304 |
| MVP_top60_DNA_seed | MAPK3 | -0.584876 | SKCM | SK-MEL-28 | FR-180204 |
| MVP_top60_DNA_seed | MAPK1 | -0.584876 | SKCM | SK-MEL-28 | FR-180204 |

| | | | | | |
|---|---|---|---|---|---|
| MVP_top60_DNA_seed | PIK3CB | -0.583447 | SKCM | MEL-JUSO | AZD6482 |
| MVP_top60_DNA_seed | EGFR | -0.578407 | SKCM | SK-MEL-30 | Gefitinib |
| MVP_top60_DNA_seed | GSK3B | -0.577617 | SKCM | HT-144 | SB216763 |
| MVP_top60_DNA_seed | ERBB2 | -0.577286 | SKCM | COLO-792 | CP724714 |
| MVP_top60_DNA_seed | GSK3B | -0.57551 | SKCM | IST-MEL1 | AZD7969 |
| MVP_top60_DNA_seed | BRAF | -0.570504 | SKCM | COLO-829 | RAF_9304 |
| MVP_top60_DNA_seed | JAK2 | -0.570438 | SKCM | WM35 | Fedratinib |
| MVP_top60_DNA_seed | GSK3B | -0.568087 | SKCM | A101D | AZD7969 |
| MVP_top60_DNA_seed | EGFR | -0.567456 | SKCM | LB2518-MEL | Cetuximab |
| MVP_top60_DNA_seed | JAK2 | -0.565948 | SKCM | WM1552C | Fedratinib |
| MVP_top60_DNA_seed | MAP2K1 | -0.564998 | SKCM | IST-MEL1 | Refametinib |
| MVP_top60_DNA_seed | EGFR | -0.56468 | SKCM | WM793B | CI-1033 |
| MVP_top60_DNA_seed | ERBB2 | -0.56468 | SKCM | WM793B | CI-1033 |
| MVP_top60_DNA_seed | EGFR | -0.564462 | SKCM | WM1552C | AZD8931 |
| MVP_top60_DNA_seed | ERBB2 | -0.564462 | SKCM | WM1552C | AZD8931 |
| MVP_top60_DNA_seed | EGFR | -0.564152 | SKCM | HMV-II | PF-00299804 |
| MVP_top60_DNA_seed | ERBB2 | -0.564152 | SKCM | HMV-II | PF-00299804 |
| MVP_top60_DNA_seed | JAK2 | -0.564098 | SKCM | IGR-37 | Ruxolitinib |
| MVP_top60_DNA_seed | JAK1 | -0.564098 | SKCM | IGR-37 | Ruxolitinib |
| MVP_top60_DNA_seed | JAK1 | -0.558541 | SKCM | SK-MEL-5 | JAK1_3715 |
| MVP_top60_DNA_seed | EP300 | -0.558152 | SKCM | IPC-298 | I-CBP112 |
| MVP_top60_DNA_seed | CREBBP | -0.558152 | SKCM | IPC-298 | I-CBP112 |
| MVP_top60_DNA_seed | PIK3CA | -0.557003 | SKCM | A101D | AZD8835 |
| MVP_top60_DNA_seed | EGFR | -0.549936 | SKCM | Hs-940-T | PF-00299804 |
| MVP_top60_DNA_seed | ERBB2 | -0.549936 | SKCM | Hs-940-T | PF-00299804 |
| MVP_top60_DNA_seed | GSK3B | -0.549176 | SKCM | IPC-298 | AZD7969 |
| MVP_top60_DNA_seed | MAP2K1 | -0.548201 | SKCM | IST-MEL1 | Selumetinib |
| MVP_top60_DNA_seed | MAP2K1 | -0.547888 | SKCM | Mewo | Refametinib |
| MVP_top60_DNA_seed | EGFR | -0.545229 | SKCM | MEL-JUSO | AST-1306 |
| MVP_top60_DNA_seed | ERBB2 | -0.544097 | SKCM | RPMI-7951 | CP724714 |
| MVP_top60_DNA_seed | EGFR | -0.544053 | SKCM | MEL-JUSO | CUDC-101 |
| MVP_top60_DNA_seed | ERBB2 | -0.544053 | SKCM | MEL-JUSO | CUDC-101 |
| MVP_top60_DNA_seed | GSK3B | -0.542302 | SKCM | WM1552C | AZD7969 |
| MVP_top60_DNA_seed | PIK3CB | -0.541072 | SKCM | GAK | AZD6482 |
| MVP_top60_DNA_seed | GSK3B | -0.539395 | SKCM | M14 | AZD7969 |
| MVP_top60_DNA_seed | BRAF | -0.535898 | SKCM | GAK | PLX-4720 |
| MVP_top60_DNA_seed | BRAF | -0.533949 | SKCM | UACC-62 | HG6-64-1 |
| MVP_top60_DNA_seed | EGFR | -0.532414 | SKCM | MZ2-MEL | CI-1033 |
| MVP_top60_DNA_seed | ERBB2 | -0.532414 | SKCM | MZ2-MEL | CI-1033 |
| MVP_top60_DNA_seed | PIK3CB | -0.530839 | SKCM | RVH-421 | AZD8186 |
| MVP_top60_DNA_seed | GSK3B | -0.530378 | SKCM | RVH-421 | AZD7969 |

| | | | | | |
|---|---|---|---|---|---|
| MVP_top60_DNA_seed | EGFR | -0.530363 | SKCM | MEL-HO | CI-1033 |
| MVP_top60_DNA_seed | ERBB2 | -0.530363 | SKCM | MEL-HO | CI-1033 |
| MVP_top60_DNA_seed | BRAF | -0.529819 | SKCM | A2058 | HG6-64-1 |
| MVP_top60_DNA_seed | JAK2 | -0.528847 | SKCM | WM793B | Ruxolitinib |
| MVP_top60_DNA_seed | JAK1 | -0.528847 | SKCM | WM793B | Ruxolitinib |
| MVP_top60_DNA_seed | EGFR | -0.527596 | SKCM | 451Lu | AST-1306 |
| MVP_top60_DNA_seed | JAK1 | -0.526451 | SKCM | WM278 | JAK1_3715 |
| MVP_top60_DNA_seed | EGFR | -0.525105 | SKCM | IST-MEL1 | Lapatinib |
| MVP_top60_DNA_seed | ERBB2 | -0.525105 | SKCM | IST-MEL1 | Lapatinib |
| MVP_top60_DNA_seed | PIK3CB | -0.520935 | SKCM | VMRC-MELG | AZD6482 |
| MVP_top60_DNA_seed | GSK3B | -0.518214 | SKCM | M14 | CHIR-99021 |
| MVP_top60_DNA_seed | MAP2K1 | -0.513358 | SKCM | COLO-783 | Refametinib |
| MVP_top60_DNA_seed | EGFR | -0.513153 | SKCM | MEL-JUSO | CI-1033 |
| MVP_top60_DNA_seed | ERBB2 | -0.513153 | SKCM | MEL-JUSO | CI-1033 |
| MVP_top60_DNA_seed | BRAF | -0.510743 | SKCM | LOXIMVI | HG6-64-1 |
| MVP_top60_DNA_seed | GSK3B | -0.507326 | SKCM | A375 | KIN001-042 |
| MVP_top60_DNA_seed | GSK3B | -0.507006 | SKCM | IST-MEL1 | SB216763 |
| MVP_top60_DNA_seed | PIK3CA | -0.506539 | SKCM | CP66-MEL | PI3Ka_4409 |
| MVP_top60_DNA_seed | GSK3B | -0.50292 | SKCM | SK-MEL-31 | CHIR-99021 |
| MVP_top60_DNA_seed | EGFR | -0.50192 | SKCM | RPMI-7951 | CI-1033 |
| MVP_top60_DNA_seed | ERBB2 | -0.50192 | SKCM | RPMI-7951 | CI-1033 |
| MVP_top60_DNA_seed | RAC1 | -0.501876 | SKCM | MMAc-SF | EHT-1864 |
| MVP_top60_DNA_seed | JAK3 | -0.50023 | SKCM | SK-MEL-3 | WHI-P97 |
| MVP_top60_DNA_seed | MAP2K1 | -0.499054 | SKCM | A2058 | CI-1040 |
| MVP_top60_DNA_seed | GSK3B | -0.497615 | SKCM | SK-MEL-24 | CHIR-99021 |
| MVP_top60_DNA_seed | JAK3 | -0.497518 | SKCM | HT-144 | JAK3_7406 |
| MVP_top60_DNA_seed | BRAF | -0.497117 | SKCM | WM1552C | PLX-4720 |
| MVP_top60_DNA_seed | JAK3 | -0.494452 | SKCM | SK-MEL-24 | WHI-P97 |
| MVP_top60_DNA_seed | MAP2K1 | -0.494067 | SKCM | WM793B | Refametinib |
| MVP_top60_DNA_seed | EGFR | -0.490487 | SKCM | WM-115 | AZD8931 |
| MVP_top60_DNA_seed | ERBB2 | -0.490487 | SKCM | WM-115 | AZD8931 |
| MVP_top60_DNA_seed | EGFR | -0.487032 | SKCM | SK-MEL-28 | Gefitinib |
| MVP_top60_DNA_seed | MAP2K1 | -0.486681 | SKCM | MEL-JUSO | CI-1040 |
| MVP_top60_DNA_seed | EGFR | -0.482712 | SKCM | COLO-792 | PF-00299804 |
| MVP_top60_DNA_seed | ERBB2 | -0.482712 | SKCM | COLO-792 | PF-00299804 |
| MVP_top60_DNA_seed | EGFR | -0.480452 | SKCM | M14 | AZD8931 |
| MVP_top60_DNA_seed | ERBB2 | -0.480452 | SKCM | M14 | AZD8931 |
| MVP_top60_DNA_seed | EP300 | -0.480391 | SKCM | G-361 | I-CBP112 |
| MVP_top60_DNA_seed | CREBBP | -0.480391 | SKCM | G-361 | I-CBP112 |
| MVP_top60_DNA_seed | PIK3CB | -0.480336 | SKCM | Hs-939-T | AZD6482 |
| MVP_top60_DNA_seed | MAP2K1 | -0.479487 | SKCM | RPMI-7951 | CI-1040 |

| | | | | | |
|---|---|---|---|---|---|
| MVP_top60_DNA_seed | PIK3CA | -0.474166 | SKCM | MEL-JUSO | PI3Ka_4409 |
| MVP_top60_DNA_seed | JAK3 | -0.473522 | SKCM | M14 | WHI-P97 |
| MVP_top60_DNA_seed | BRAF | -0.472887 | SKCM | RPMI-7951 | RAF_9304 |
| MVP_top60_DNA_seed | GSK3B | -0.4727 | SKCM | G-MEL | SB216763 |
| MVP_top60_DNA_seed | EGFR | -0.471621 | SKCM | WM1552C | CI-1033 |
| MVP_top60_DNA_seed | ERBB2 | -0.471621 | SKCM | WM1552C | CI-1033 |
| MVP_top60_DNA_seed | JAK2 | -0.470848 | SKCM | WM793B | Fedratinib |
| MVP_top60_DNA_seed | BRAF | -0.465994 | SKCM | COLO-800 | SB590885 |
| MVP_top60_DNA_seed | EGFR | -0.465357 | SKCM | MMAc-SF | CI-1033 |
| MVP_top60_DNA_seed | ERBB2 | -0.465357 | SKCM | MMAc-SF | CI-1033 |
| MVP_top60_DNA_seed | PIK3CB | -0.463855 | SKCM | RVH-421 | AZD6482 |
| MVP_top60_DNA_seed | EGFR | -0.46317 | SKCM | LOXIMVI | CI-1033 |
| MVP_top60_DNA_seed | ERBB2 | -0.46317 | SKCM | LOXIMVI | CI-1033 |
| MVP_top60_DNA_seed | GSK3B | -0.463015 | SKCM | WM-115 | SB216763 |
| MVP_top60_DNA_seed | MAPK14 | -0.460674 | SKCM | COLO-800 | TAK-715 |
| MVP_top60_DNA_seed | MAP2K1 | -0.460138 | SKCM | WM793B | CI-1040 |
| MVP_top60_DNA_seed | EP300 | -0.4575 | SKCM | COLO-800 | I-CBP112 |
| MVP_top60_DNA_seed | CREBBP | -0.4575 | SKCM | COLO-800 | I-CBP112 |
| MVP_top60_DNA_seed | GSK3B | -0.454862 | SKCM | COLO-829 | CHIR-99021 |
| MVP_top60_DNA_seed | JAK2 | -0.454 | SKCM | COLO-783 | Ruxolitinib |
| MVP_top60_DNA_seed | JAK1 | -0.454 | SKCM | COLO-783 | Ruxolitinib |
| MVP_top60_DNA_seed | MAPK1 | -0.453836 | SKCM | C32 | VX-11e |
| MVP_top60_DNA_seed | MAPK14 | -0.452975 | SKCM | SK-MEL-30 | TAK-715 |
| MVP_top60_DNA_seed | BRAF | -0.451403 | SKCM | LB2518-MEL | PLX-4720 |
| MVP_top60_DNA_seed | ERBB2 | -0.449899 | SKCM | UACC-62 | CP724714 |
| MVP_top60_DNA_seed | JAK2 | -0.448156 | SKCM | CHL-1 | Fedratinib |
| MVP_top60_DNA_seed | MAP2K1 | -0.445853 | SKCM | CHL-1 | Selumetinib |
| MVP_top60_DNA_seed | BRAF | -0.442684 | SKCM | IPC-298 | SB590885 |
| MVP_top60_DNA_seed | EGFR | -0.442627 | SKCM | LB373-MEL-D | CUDC-101 |
| MVP_top60_DNA_seed | ERBB2 | -0.442627 | SKCM | LB373-MEL-D | CUDC-101 |
| MVP_top60_DNA_seed | EGFR | -0.441464 | SKCM | RVH-421 | CI-1033 |
| MVP_top60_DNA_seed | ERBB2 | -0.441464 | SKCM | RVH-421 | CI-1033 |
| MVP_top60_DNA_seed | EGFR | -0.441121 | SKCM | COLO-792 | CI-1033 |
| MVP_top60_DNA_seed | ERBB2 | -0.441121 | SKCM | COLO-792 | CI-1033 |
| MVP_top60_DNA_seed | RAC1 | -0.440633 | SKCM | IPC-298 | EHT-1864 |
| MVP_top60_DNA_seed | EGFR | -0.440367 | SKCM | G-MEL | PF-00299804 |
| MVP_top60_DNA_seed | ERBB2 | -0.440367 | SKCM | G-MEL | PF-00299804 |
| MVP_top60_DNA_seed | GSK3B | -0.43923 | SKCM | UACC-62 | KIN001-042 |
| MVP_top60_DNA_seed | PIK3CA | -0.438869 | SKCM | CP66-MEL | AZD8835 |
| MVP_top60_DNA_seed | PIK3CB | -0.438789 | SKCM | HT-144 | AZD6482 |
| MVP_top60_DNA_seed | PIK3CA | -0.430033 | SKCM | COLO-800 | AZD8835 |

| | | | | | |
|---|---|---|---|---|---|
| MVP_top60_DNA_seed | ERBB2 | -0.430015 | SKCM | HMV-II | CP724714 |
| MVP_top60_DNA_seed | EGFR | -0.429583 | SKCM | RPMI-7951 | CUDC-101 |
| MVP_top60_DNA_seed | ERBB2 | -0.429583 | SKCM | RPMI-7951 | CUDC-101 |
| MVP_top60_DNA_seed | JAK1 | -0.429369 | SKCM | CP50-MEL-B | JAK1_3715 |
| MVP_top60_DNA_seed | GSK3B | -0.426548 | SKCM | MZ2-MEL | CHIR-99021 |
| MVP_top60_DNA_seed | EGFR | -0.425797 | SKCM | IPC-298 | CUDC-101 |
| MVP_top60_DNA_seed | ERBB2 | -0.425797 | SKCM | IPC-298 | CUDC-101 |
| MVP_top60_DNA_seed | JAK2 | -0.425156 | SKCM | RPMI-7951 | Ruxolitinib |
| MVP_top60_DNA_seed | JAK1 | -0.425156 | SKCM | RPMI-7951 | Ruxolitinib |
| MVP_top60_DNA_seed | PIK3CA | -0.423997 | SKCM | MMAc-SF | AZD8835 |
| MVP_top60_DNA_seed | BRAF | -0.422768 | SKCM | GAK | Dabrafenib |
| MVP_top60_DNA_seed | EGFR | -0.415218 | SKCM | G-MEL | CUDC-101 |
| MVP_top60_DNA_seed | ERBB2 | -0.415218 | SKCM | G-MEL | CUDC-101 |
| MVP_top60_DNA_seed | GSK3B | -0.414158 | SKCM | WM35 | CHIR-99021 |
| MVP_top60_DNA_seed | MAPK14 | -0.410912 | SKCM | A2058 | TAK-715 |
| MVP_top60_DNA_seed | EGFR | -0.409922 | SKCM | UACC-257 | CUDC-101 |
| MVP_top60_DNA_seed | ERBB2 | -0.409922 | SKCM | UACC-257 | CUDC-101 |
| MVP_top60_DNA_seed | MAP2K1 | -0.407203 | SKCM | CHL-1 | Refametinib |
| MVP_top60_DNA_seed | MAP2K1 | -0.40705 | SKCM | MZ2-MEL | Selumetinib |
| MVP_top60_DNA_seed | PIK3CA | -0.405591 | SKCM | LOXIMVI | PI3Ka_4409 |
| MVP_top60_DNA_seed | EGFR | -0.404402 | SKCM | G-MEL | Pelitinib |
| MVP_top60_DNA_seed | PIK3CB | -0.401356 | SKCM | RPMI-7951 | AZD6482 |
| MVP_top60_DNA_seed | MAPK14 | -0.397734 | SKCM | G-MEL | TAK-715 |
| MVP_top60_DNA_seed | MAP2K1 | -0.394374 | SKCM | A2058 | Refametinib |
| MVP_top60_DNA_seed | BRAF | -0.392957 | SKCM | MMAc-SF | HG6-64-1 |
| MVP_top60_DNA_seed | EGFR | -0.392421 | SKCM | A2058 | Gefitinib |
| MVP_top60_DNA_seed | EGFR | -0.391622 | SKCM | RPMI-7951 | Pelitinib |
| MVP_top60_DNA_seed | PIK3CB | -0.390679 | SKCM | SK-MEL-28 | AZD8186 |
| MVP_top60_DNA_seed | PIK3CB | -0.389284 | SKCM | COLO-800 | AZD6482 |
| MVP_top60_DNA_seed | JAK3 | -0.38814 | SKCM | IGR-37 | JAK3_7406 |
| MVP_top60_DNA_seed | EGFR | -0.386596 | SKCM | CP50-MEL-B | AZD8931 |
| MVP_top60_DNA_seed | ERBB2 | -0.386596 | SKCM | CP50-MEL-B | AZD8931 |
| MVP_top60_DNA_seed | MAPK3 | -0.386377 | SKCM | WM278 | FR-180204 |
| MVP_top60_DNA_seed | MAPK1 | -0.386377 | SKCM | WM278 | FR-180204 |
| MVP_top60_DNA_seed | MAP2K1 | -0.383425 | SKCM | WM278 | Trametinib |
| MVP_top60_DNA_seed | JAK3 | -0.383094 | SKCM | SH-4 | WHI-P97 |
| MVP_top60_DNA_seed | PIK3CB | -0.381252 | SKCM | MMAc-SF | AZD6482 |
| MVP_top60_DNA_seed | PIK3CA | -0.381221 | SKCM | A375 | PI3Ka_4409 |
| MVP_top60_DNA_seed | BRAF | -0.380382 | SKCM | WM1552C | RAF_9304 |
| MVP_top60_DNA_seed | EGFR | -0.379561 | SKCM | SK-MEL-30 | PF-00299804 |
| MVP_top60_DNA_seed | ERBB2 | -0.379561 | SKCM | SK-MEL-30 | PF-00299804 |

| | | | | | |
|---|---|---|---|---|---|
| MVP_top60_DNA_seed | EGFR | -0.379043 | SKCM | WM35 | CI-1033 |
| MVP_top60_DNA_seed | ERBB2 | -0.379043 | SKCM | WM35 | CI-1033 |
| MVP_top60_DNA_seed | JAK2 | -0.376916 | SKCM | A375 | AZD1480 |
| MVP_top60_DNA_seed | JAK1 | -0.376916 | SKCM | A375 | AZD1480 |
| MVP_top60_DNA_seed | GSK3B | -0.376807 | SKCM | LB2518-MEL | KIN001-042 |
| MVP_top60_DNA_seed | MAP2K1 | -0.37593 | SKCM | WM1552C | Selumetinib |
| MVP_top60_DNA_seed | GSK3B | -0.3742 | SKCM | MZ7-mel | CHIR-99021 |
| MVP_top60_DNA_seed | PIK3CB | -0.37287 | SKCM | COLO-800 | TGX221 |
| MVP_top60_DNA_seed | EGFR | -0.372566 | SKCM | G-361 | CI-1033 |
| MVP_top60_DNA_seed | ERBB2 | -0.372566 | SKCM | G-361 | CI-1033 |
| MVP_top60_DNA_seed | EGFR | -0.370918 | SKCM | SK-MEL-30 | CUDC-101 |
| MVP_top60_DNA_seed | ERBB2 | -0.370918 | SKCM | SK-MEL-30 | CUDC-101 |
| MVP_top60_DNA_seed | EGFR | -0.370362 | SKCM | COLO-800 | Afatinib |
| MVP_top60_DNA_seed | ERBB2 | -0.370362 | SKCM | COLO-800 | Afatinib |
| MVP_top60_DNA_seed | RAC1 | -0.370289 | SKCM | HT-144 | EHT-1864 |
| MVP_top60_DNA_seed | EGFR | -0.367712 | SKCM | COLO-792 | AZD8931 |
| MVP_top60_DNA_seed | ERBB2 | -0.367712 | SKCM | COLO-792 | AZD8931 |
| MVP_top60_DNA_seed | BRAF | -0.364594 | SKCM | CHL-1 | SB590885 |
| MVP_top60_DNA_seed | EGFR | -0.363577 | SKCM | IPC-298 | CI-1033 |
| MVP_top60_DNA_seed | ERBB2 | -0.363577 | SKCM | IPC-298 | CI-1033 |
| MVP_top60_DNA_seed | GSK3B | -0.360956 | SKCM | HT-144 | KIN001-042 |
| MVP_top60_DNA_seed | EGFR | -0.360864 | SKCM | G-361 | CUDC-101 |
| MVP_top60_DNA_seed | ERBB2 | -0.360864 | SKCM | G-361 | CUDC-101 |
| MVP_top60_DNA_seed | GSK3B | -0.360542 | SKCM | G-MEL | KIN001-042 |
| MVP_top60_DNA_seed | BRAF | -0.358696 | SKCM | SK-MEL-2 | AZ628 |
| MVP_top60_DNA_seed | GSK3B | -0.355236 | SKCM | LB373-MEL-D | CHIR-99021 |
| MVP_top60_DNA_seed | RAC1 | -0.355009 | SKCM | MZ7-mel | EHT-1864 |
| MVP_top60_DNA_seed | BRAF | -0.353964 | SKCM | RPMI-7951 | PLX-4720 |
| MVP_top60_DNA_seed | GSK3B | -0.352691 | SKCM | HMV-II | CHIR-99021 |
| MVP_top60_DNA_seed | PIK3CB | -0.352103 | SKCM | WM35 | AZD6482 |
| MVP_top60_DNA_seed | ERBB2 | -0.351846 | SKCM | SK-MEL-30 | CP724714 |
| MVP_top60_DNA_seed | EGFR | -0.349126 | SKCM | SK-MEL-5 | AST-1306 |
| MVP_top60_DNA_seed | EP300 | -0.348877 | SKCM | COLO-792 | I-CBP112 |
| MVP_top60_DNA_seed | CREBBP | -0.348877 | SKCM | COLO-792 | I-CBP112 |
| MVP_top60_DNA_seed | EGFR | -0.347237 | SKCM | 451Lu | CI-1033 |
| MVP_top60_DNA_seed | ERBB2 | -0.347237 | SKCM | 451Lu | CI-1033 |
| MVP_top60_DNA_seed | EGFR | -0.346576 | SKCM | IST-MEL1 | Pelitinib |
| MVP_top60_DNA_seed | MAP2K1 | -0.343696 | SKCM | WM793B | Refametinib |
| MVP_top60_DNA_seed | PIK3CB | -0.339081 | SKCM | COLO-679 | AZD6482 |
| MVP_top60_DNA_seed | MAP2K1 | -0.339081 | SKCM | RPMI-7951 | PD0325901 |
| MVP_top60_DNA_seed | JAK3 | -0.33865 | SKCM | WM278 | JAK3_7406 |

| | | | | | |
|---|---|---|---|---|---|
| MVP_top60_DNA_seed | JAK2 | -0.338024 | SKCM | RPMI-7951 | AZD1480 |
| MVP_top60_DNA_seed | JAK1 | -0.338024 | SKCM | RPMI-7951 | AZD1480 |
| MVP_top60_DNA_seed | MAPK1 | -0.33737 | SKCM | COLO-792 | VX-11e |
| MVP_top60_DNA_seed | MAPK3 | -0.336262 | SKCM | IGR-1 | FR-180204 |
| MVP_top60_DNA_seed | MAPK1 | -0.336262 | SKCM | IGR-1 | FR-180204 |
| MVP_top60_DNA_seed | MAPK1 | -0.336191 | SKCM | MMAc-SF | VX-11e |
| MVP_top60_DNA_seed | MAP2K1 | -0.336097 | SKCM | IGR-1 | PD0325901 |
| MVP_top60_DNA_seed | EGFR | -0.33441 | SKCM | SK-MEL-5 | CI-1033 |
| MVP_top60_DNA_seed | ERBB2 | -0.33441 | SKCM | SK-MEL-5 | CI-1033 |
| MVP_top60_DNA_seed | PIK3CB | -0.326295 | SKCM | SK-MEL-3 | AZD6482 |
| MVP_top60_DNA_seed | RAC1 | -0.326137 | SKCM | WM35 | EHT-1864 |
| MVP_top60_DNA_seed | MAPK1 | -0.325887 | SKCM | LB373-MEL-D | VX-11e |
| MVP_top60_DNA_seed | JAK2 | -0.324456 | SKCM | G-361 | Ruxolitinib |
| MVP_top60_DNA_seed | JAK1 | -0.324456 | SKCM | G-361 | Ruxolitinib |
| MVP_top60_DNA_seed | BRAF | -0.323334 | SKCM | SK-MEL-2 | PLX-4720 |
| MVP_top60_DNA_seed | JAK2 | -0.323243 | SKCM | A375 | Ruxolitinib |
| MVP_top60_DNA_seed | JAK1 | -0.323243 | SKCM | A375 | Ruxolitinib |
| MVP_top60_DNA_seed | MAPK14 | -0.323233 | SKCM | WM1552C | TAK-715 |
| MVP_top60_DNA_seed | GSK3B | -0.323086 | SKCM | CP50-MEL-B | SB216763 |
| MVP_top60_DNA_seed | EGFR | -0.321244 | SKCM | MZ2-MEL | Lapatinib |
| MVP_top60_DNA_seed | ERBB2 | -0.321244 | SKCM | MZ2-MEL | Lapatinib |
| MVP_top60_DNA_seed | EP300 | -0.32106 | SKCM | SK-MEL-31 | I-CBP112 |
| MVP_top60_DNA_seed | CREBBP | -0.32106 | SKCM | SK-MEL-31 | I-CBP112 |
| MVP_top60_DNA_seed | ERBB2 | -0.320426 | SKCM | SK-MEL-5 | CP724714 |
| MVP_top60_DNA_seed | GSK3B | -0.320271 | SKCM | CP50-MEL-B | AZD7969 |
| MVP_top60_DNA_seed | EGFR | -0.320226 | SKCM | MZ7-mel | Afatinib |
| MVP_top60_DNA_seed | ERBB2 | -0.320226 | SKCM | MZ7-mel | Afatinib |
| MVP_top60_DNA_seed | MAP2K1 | -0.320145 | SKCM | WM1552C | PD0325901 |
| MVP_top60_DNA_seed | PIK3CB | -0.319984 | SKCM | WM278 | AZD8186 |
| MVP_top60_DNA_seed | JAK2 | -0.319652 | SKCM | A101D | Ruxolitinib |
| MVP_top60_DNA_seed | JAK1 | -0.319652 | SKCM | A101D | Ruxolitinib |
| MVP_top60_DNA_seed | MAPK14 | -0.317123 | SKCM | WM793B | TAK-715 |
| MVP_top60_DNA_seed | EGFR | -0.317034 | SKCM | IGR-1 | AZD8931 |
| MVP_top60_DNA_seed | ERBB2 | -0.317034 | SKCM | IGR-1 | AZD8931 |
| MVP_top60_DNA_seed | EGFR | -0.316332 | SKCM | M14 | Cetuximab |
| MVP_top60_DNA_seed | MAP2K1 | -0.316288 | SKCM | A2058 | PD0325901 |
| MVP_top60_DNA_seed | MAPK14 | -0.314378 | SKCM | MEL-JUSO | TAK-715 |
| MVP_top60_DNA_seed | GSK3B | -0.313618 | SKCM | G-361 | AZD7969 |
| MVP_top60_DNA_seed | JAK3 | -0.313523 | SKCM | HMV-II | JAK3_7406 |
| MVP_top60_DNA_seed | EGFR | -0.311848 | SKCM | Hs-939-T | PF-00299804 |
| MVP_top60_DNA_seed | ERBB2 | -0.311848 | SKCM | Hs-939-T | PF-00299804 |

| | | | | | |
|---|---|---|---|---|---|
| MVP_top60_DNA_seed | EGFR | -0.309552 | SKCM | G-361 | Pelitinib |
| MVP_top60_DNA_seed | GSK3B | -0.309295 | SKCM | GAK | CHIR-99021 |
| MVP_top60_DNA_seed | EP300 | -0.308918 | SKCM | MEL-HO | I-CBP112 |
| MVP_top60_DNA_seed | CREBBP | -0.308918 | SKCM | MEL-HO | I-CBP112 |
| MVP_top60_DNA_seed | EGFR | -0.307665 | SKCM | SK-MEL-2 | AZD8931 |
| MVP_top60_DNA_seed | ERBB2 | -0.307665 | SKCM | SK-MEL-2 | AZD8931 |
| MVP_top60_DNA_seed | JAK3 | -0.305524 | SKCM | MEL-HO | JAK3_7406 |
| MVP_top60_DNA_seed | RAC1 | -0.3024 | SKCM | A2058 | EHT-1864 |
| MVP_top60_DNA_seed | RAC1 | -0.29905 | SKCM | CP50-MEL-B | EHT-1864 |
| MVP_top60_DNA_seed | JAK1 | -0.298521 | SKCM | G-MEL | JAK1_3715 |
| MVP_top60_DNA_seed | PIK3CA | -0.297439 | SKCM | SK-MEL-28 | AZD8835 |
| MVP_top60_DNA_seed | EGFR | -0.296121 | SKCM | COLO-792 | Pelitinib |
| MVP_top60_DNA_seed | EGFR | -0.295701 | SKCM | SK-MEL-30 | CI-1033 |
| MVP_top60_DNA_seed | ERBB2 | -0.295701 | SKCM | SK-MEL-30 | CI-1033 |
| MVP_top60_DNA_seed | EGFR | -0.29538 | SKCM | RVH-421 | AZD8931 |
| MVP_top60_DNA_seed | ERBB2 | -0.29538 | SKCM | RVH-421 | AZD8931 |
| MVP_top60_DNA_seed | BRAF | -0.294952 | SKCM | IST-MEL1 | HG6-64-1 |
| MVP_top60_DNA_seed | EGFR | -0.294249 | SKCM | CP50-MEL-B | CUDC-101 |
| MVP_top60_DNA_seed | ERBB2 | -0.294249 | SKCM | CP50-MEL-B | CUDC-101 |
| MVP_top60_DNA_seed | GSK3B | -0.294019 | SKCM | UACC-62 | AZD7969 |
| MVP_top60_DNA_seed | EGFR | -0.293632 | SKCM | WM793B | Pelitinib |
| MVP_top60_DNA_seed | MAPK14 | -0.292121 | SKCM | LB2518-MEL | TAK-715 |
| MVP_top60_DNA_seed | GSK3B | -0.291181 | SKCM | Hs-940-T | AZD7969 |
| MVP_top60_DNA_seed | GSK3B | -0.289291 | SKCM | A2058 | CHIR-99021 |
| MVP_top60_DNA_seed | JAK3 | -0.288069 | SKCM | CP50-MEL-B | WHI-P97 |
| MVP_top60_DNA_seed | EGFR | -0.286717 | SKCM | COLO-792 | AST-1306 |
| MVP_top60_DNA_seed | EGFR | -0.28634 | SKCM | COLO-800 | AST-1306 |
| MVP_top60_DNA_seed | PIK3CB | -0.281185 | SKCM | WM-115 | AZD6482 |
| MVP_top60_DNA_seed | MAPK14 | -0.280167 | SKCM | IGR-37 | TAK-715 |
| MVP_top60_DNA_seed | MAP2K1 | -0.275669 | SKCM | IGR-1 | Refametinib |
| MVP_top60_DNA_seed | EGFR | -0.274439 | SKCM | WM793B | AZD8931 |
| MVP_top60_DNA_seed | ERBB2 | -0.274439 | SKCM | WM793B | AZD8931 |
| MVP_top60_DNA_seed | BRAF | -0.270534 | SKCM | Mewo | PLX-4720 |
| MVP_top60_DNA_seed | PIK3CA | -0.269608 | SKCM | MMAc-SF | PI3Ka_4409 |
| MVP_top60_DNA_seed | BRAF | -0.268966 | SKCM | LOXIMVI | AZ628 |
| MVP_top60_DNA_seed | JAK3 | -0.268612 | SKCM | SK-MEL-28 | WHI-P97 |
| MVP_top60_DNA_seed | MAP2K1 | -0.268427 | SKCM | SK-MEL-30 | CI-1040 |
| MVP_top60_DNA_seed | EGFR | -0.267293 | SKCM | WM278 | CI-1033 |
| MVP_top60_DNA_seed | ERBB2 | -0.267293 | SKCM | WM278 | CI-1033 |
| MVP_top60_DNA_seed | PIK3CB | -0.266975 | SKCM | M14 | AZD8186 |
| MVP_top60_DNA_seed | BRAF | -0.265707 | SKCM | IGR-37 | HG6-64-1 |

| | | | | | |
|---|---|---|---|---|---|
| MVP_top60_DNA_seed | GSK3B | -0.261297 | SKCM | COLO-792 | AZD7969 |
| MVP_top60_DNA_seed | MAPK1 | -0.256067 | SKCM | SK-MEL-5 | VX-11e |
| MVP_top60_DNA_seed | BRAF | -0.252395 | SKCM | SK-MEL-31 | PLX-4720 |
| MVP_top60_DNA_seed | JAK2 | -0.252248 | SKCM | MZ7-mel | Ruxolitinib |
| MVP_top60_DNA_seed | JAK1 | -0.252248 | SKCM | MZ7-mel | Ruxolitinib |
| MVP_top60_DNA_seed | JAK2 | -0.25137 | SKCM | COLO-792 | Fedratinib |
| MVP_top60_DNA_seed | JAK1 | -0.246353 | SKCM | CHL-1 | JAK1_3715 |
| MVP_top60_DNA_seed | EGFR | -0.245525 | SKCM | A375 | Cetuximab |
| MVP_top60_DNA_seed | BRAF | -0.238589 | SKCM | MZ7-mel | HG6-64-1 |
| MVP_top60_DNA_seed | GSK3B | -0.23742 | SKCM | VMRC-MELG | CHIR-99021 |
| MVP_top60_DNA_seed | JAK2 | -0.236823 | SKCM | RPMI-7951 | Fedratinib |
| MVP_top60_DNA_seed | MAPK14 | -0.235986 | SKCM | MZ7-mel | TAK-715 |
| MVP_top60_DNA_seed | MAPK3 | -0.235795 | SKCM | Hs-939-T | FR-180204 |
| MVP_top60_DNA_seed | MAPK1 | -0.235795 | SKCM | Hs-939-T | FR-180204 |
| MVP_top60_DNA_seed | PIK3CB | -0.235691 | SKCM | MZ2-MEL | AZD8186 |
| MVP_top60_DNA_seed | RAC1 | -0.233321 | SKCM | SK-MEL-24 | EHT-1864 |
| MVP_top60_DNA_seed | GSK3B | -0.23156 | SKCM | A101D | CHIR-99021 |
| MVP_top60_DNA_seed | MAPK1 | -0.231054 | SKCM | SK-MEL-1 | VX-11e |
| MVP_top60_DNA_seed | EGFR | -0.229651 | SKCM | WM278 | AZD8931 |
| MVP_top60_DNA_seed | ERBB2 | -0.229651 | SKCM | WM278 | AZD8931 |
| MVP_top60_DNA_seed | JAK3 | -0.227461 | SKCM | 451Lu | JAK3_7406 |
| MVP_top60_DNA_seed | EGFR | -0.225172 | SKCM | UACC-62 | AST-1306 |
| MVP_top60_DNA_seed | EGFR | -0.222205 | SKCM | HT-144 | AZD8931 |
| MVP_top60_DNA_seed | ERBB2 | -0.222205 | SKCM | HT-144 | AZD8931 |
| MVP_top60_DNA_seed | PIK3CB | -0.219821 | SKCM | G-MEL | AZD8186 |
| MVP_top60_DNA_seed | JAK3 | -0.219721 | SKCM | COLO-792 | JAK3_7406 |
| MVP_top60_DNA_seed | JAK3 | -0.219126 | SKCM | SK-MEL-5 | WHI-P97 |
| MVP_top60_DNA_seed | MAP2K1 | -0.217637 | SKCM | CP66-MEL | Selumetinib |
| MVP_top60_DNA_seed | EGFR | -0.217168 | SKCM | SH-4 | Afatinib |
| MVP_top60_DNA_seed | ERBB2 | -0.217168 | SKCM | SH-4 | Afatinib |
| MVP_top60_DNA_seed | JAK3 | -0.216046 | SKCM | Mewo | JAK3_7406 |
| MVP_top60_DNA_seed | MAP2K1 | -0.215944 | SKCM | WM793B | PD0325901 |
| MVP_top60_DNA_seed | BRAF | -0.215587 | SKCM | A2058 | SB590885 |
| MVP_top60_DNA_seed | JAK3 | -0.214583 | SKCM | 451Lu | WHI-P97 |
| MVP_top60_DNA_seed | GSK3B | -0.21426 | SKCM | COLO-679 | KIN001-042 |
| MVP_top60_DNA_seed | GSK3B | -0.213921 | SKCM | Hs-939-T | CHIR-99021 |
| MVP_top60_DNA_seed | EGFR | -0.213332 | SKCM | G-MEL | AZD8931 |
| MVP_top60_DNA_seed | ERBB2 | -0.213332 | SKCM | G-MEL | AZD8931 |
| MVP_top60_DNA_seed | PIK3CA | -0.211812 | SKCM | WM35 | AZD8835 |
| MVP_top60_DNA_seed | JAK3 | -0.210689 | SKCM | WM35 | JAK3_7406 |
| MVP_top60_DNA_seed | GSK3B | -0.208767 | SKCM | CP50-MEL-B | CHIR-99021 |

| | | | | | |
|---|---|---|---|---|---|
| MVP_top60_DNA_seed | EGFR | -0.207708 | SKCM | IGR-37 | AZD8931 |
| MVP_top60_DNA_seed | ERBB2 | -0.207708 | SKCM | IGR-37 | AZD8931 |
| MVP_top60_DNA_seed | JAK1 | -0.206573 | SKCM | M14 | JAK1_3715 |
| MVP_top60_DNA_seed | PIK3CB | -0.20524 | SKCM | LB2518-MEL | TGX221 |
| MVP_top60_DNA_seed | EGFR | -0.204859 | SKCM | COLO-783 | CI-1033 |
| MVP_top60_DNA_seed | ERBB2 | -0.204859 | SKCM | COLO-783 | CI-1033 |
| MVP_top60_DNA_seed | JAK2 | -0.202334 | SKCM | SK-MEL-3 | Fedratinib |
| MVP_top60_DNA_seed | GSK3B | -0.201612 | SKCM | MEL-JUSO | KIN001-042 |
| MVP_top60_DNA_seed | GSK3B | -0.201284 | SKCM | 451Lu | KIN001-042 |
| MVP_top60_DNA_seed | PIK3CA | -0.200966 | SKCM | CHL-1 | AZD8835 |
| MVP_top60_DNA_seed | EGFR | -0.200707 | SKCM | SK-MEL-28 | CI-1033 |
| MVP_top60_DNA_seed | ERBB2 | -0.200707 | SKCM | SK-MEL-28 | CI-1033 |
| MVP_top60_DNA_seed | GSK3B | -0.198815 | SKCM | MEL-JUSO | CHIR-99021 |
| MVP_top60_DNA_seed | EGFR | -0.197291 | SKCM | IST-MEL1 | CI-1033 |
| MVP_top60_DNA_seed | ERBB2 | -0.197291 | SKCM | IST-MEL1 | CI-1033 |
| MVP_top60_DNA_seed | GSK3B | -0.193688 | SKCM | WM35 | AZD7969 |
| MVP_top60_DNA_seed | PIK3CB | -0.192519 | SKCM | MZ2-MEL | AZD6482 |
| MVP_top60_DNA_seed | GSK3B | -0.190764 | SKCM | COLO-792 | CHIR-99021 |
| MVP_top60_DNA_seed | MAPK1 | -0.190243 | SKCM | WM793B | VX-11e |
| MVP_top60_DNA_seed | JAK3 | -0.18995 | SKCM | UACC-62 | JAK3_7406 |
| MVP_top60_DNA_seed | ERBB2 | -0.188969 | SKCM | WM-115 | CP724714 |
| MVP_top60_DNA_seed | MAP2K1 | -0.188685 | SKCM | IST-MEL1 | Selumetinib |
| MVP_top60_DNA_seed | GSK3B | -0.188522 | SKCM | C32 | SB216763 |
| MVP_top60_DNA_seed | EGFR | -0.187997 | SKCM | A375 | AZD8931 |
| MVP_top60_DNA_seed | ERBB2 | -0.187997 | SKCM | A375 | AZD8931 |
| MVP_top60_DNA_seed | EGFR | -0.18671 | SKCM | MZ2-MEL | Pelitinib |
| MVP_top60_DNA_seed | MAPK14 | -0.185915 | SKCM | WM35 | TAK-715 |
| MVP_top60_DNA_seed | EGFR | -0.185774 | SKCM | A375 | CUDC-101 |
| MVP_top60_DNA_seed | ERBB2 | -0.185774 | SKCM | A375 | CUDC-101 |
| MVP_top60_DNA_seed | ERBB2 | -0.183884 | SKCM | MZ2-MEL | CP724714 |
| MVP_top60_DNA_seed | JAK3 | -0.182646 | SKCM | G-361 | WHI-P97 |
| MVP_top60_DNA_seed | JAK3 | -0.182139 | SKCM | Hs-940-T | JAK3_7406 |
| MVP_top60_DNA_seed | PIK3CA | -0.180798 | SKCM | CP50-MEL-B | PI3Ka_4409 |
| MVP_top60_DNA_seed | MAPK1 | -0.18069 | SKCM | CHL-1 | VX-11e |
| MVP_top60_DNA_seed | MAPK14 | -0.178057 | SKCM | SK-MEL-2 | TAK-715 |
| MVP_top60_DNA_seed | ERBB2 | -0.176427 | SKCM | IST-MEL1 | CP724714 |
| MVP_top60_DNA_seed | MAP2K1 | -0.176097 | SKCM | LB373-MEL-D | CI-1040 |
| MVP_top60_DNA_seed | EGFR | -0.174362 | SKCM | IST-MEL1 | CUDC-101 |
| MVP_top60_DNA_seed | ERBB2 | -0.174362 | SKCM | IST-MEL1 | CUDC-101 |
| MVP_top60_DNA_seed | JAK3 | -0.173527 | SKCM | G-MEL | WHI-P97 |
| MVP_top60_DNA_seed | EGFR | -0.173036 | SKCM | A101D | Erlotinib |

| | | | | | |
|---|---|---|---|---|---|
| MVP_top60_DNA_seed | EGFR | -0.173005 | SKCM | MEL-JUSO | Pelitinib |
| MVP_top60_DNA_seed | EGFR | -0.172073 | SKCM | IPC-298 | Pelitinib |
| MVP_top60_DNA_seed | ERBB2 | -0.171244 | SKCM | LB2518-MEL | CP724714 |
| MVP_top60_DNA_seed | EP300 | -0.171229 | SKCM | MZ2-MEL | I-CBP112 |
| MVP_top60_DNA_seed | CREBBP | -0.171229 | SKCM | MZ2-MEL | I-CBP112 |
| MVP_top60_DNA_seed | EGFR | -0.170547 | SKCM | LOXIMVI | CUDC-101 |
| MVP_top60_DNA_seed | ERBB2 | -0.170547 | SKCM | LOXIMVI | CUDC-101 |
| MVP_top60_DNA_seed | PIK3CB | -0.167329 | SKCM | A101D | AZD8186 |
| MVP_top60_DNA_seed | JAK2 | -0.164319 | SKCM | MEL-JUSO | Fedratinib |
| MVP_top60_DNA_seed | RAC1 | -0.163316 | SKCM | IGR-37 | EHT-1864 |
| MVP_top60_DNA_seed | PIK3CB | -0.161957 | SKCM | LOXIMVI | AZD8186 |
| MVP_top60_DNA_seed | JAK2 | -0.156514 | SKCM | IGR-37 | AZD1480 |
| MVP_top60_DNA_seed | JAK1 | -0.156514 | SKCM | IGR-37 | AZD1480 |
| MVP_top60_DNA_seed | BRAF | -0.15607 | SKCM | SK-MEL-30 | PLX-4720 |
| MVP_top60_DNA_seed | BRAF | -0.153906 | SKCM | Mewo | RAF_9304 |
| MVP_top60_DNA_seed | EGFR | -0.152462 | SKCM | A101D | Gefitinib |
| MVP_top60_DNA_seed | EGFR | -0.151869 | SKCM | WM35 | AZD8931 |
| MVP_top60_DNA_seed | ERBB2 | -0.151869 | SKCM | WM35 | AZD8931 |
| MVP_top60_DNA_seed | MAP2K1 | -0.148224 | SKCM | WM1552C | Trametinib |
| MVP_top60_DNA_seed | EGFR | -0.147358 | SKCM | IGR-37 | CI-1033 |
| MVP_top60_DNA_seed | ERBB2 | -0.147358 | SKCM | IGR-37 | CI-1033 |
| MVP_top60_DNA_seed | EGFR | -0.146577 | SKCM | SK-MEL-3 | CUDC-101 |
| MVP_top60_DNA_seed | ERBB2 | -0.146577 | SKCM | SK-MEL-3 | CUDC-101 |
| MVP_top60_DNA_seed | EGFR | -0.145121 | SKCM | WM793B | Afatinib |
| MVP_top60_DNA_seed | ERBB2 | -0.145121 | SKCM | WM793B | Afatinib |
| MVP_top60_DNA_seed | BRAF | -0.144491 | SKCM | MEL-JUSO | Dabrafenib |
| MVP_top60_DNA_seed | EGFR | -0.140632 | SKCM | CP66-MEL | AZD8931 |
| MVP_top60_DNA_seed | ERBB2 | -0.140632 | SKCM | CP66-MEL | AZD8931 |
| MVP_top60_DNA_seed | GSK3B | -0.138707 | SKCM | C32 | CHIR-99021 |
| MVP_top60_DNA_seed | PIK3CA | -0.136964 | SKCM | VMRC-MELG | AZD8835 |
| MVP_top60_DNA_seed | EP300 | -0.136069 | SKCM | MMAc-SF | I-CBP112 |
| MVP_top60_DNA_seed | CREBBP | -0.136069 | SKCM | MMAc-SF | I-CBP112 |
| MVP_top60_DNA_seed | BRAF | -0.136066 | SKCM | Mewo | Dabrafenib |
| MVP_top60_DNA_seed | PIK3CA | -0.13518 | SKCM | COLO-792 | PI3Ka_4409 |
| MVP_top60_DNA_seed | PIK3CB | -0.135005 | SKCM | CP50-MEL-B | AZD8186 |
| MVP_top60_DNA_seed | MAPK14 | -0.134218 | SKCM | RVH-421 | TAK-715 |
| MVP_top60_DNA_seed | PIK3CB | -0.13419 | SKCM | UACC-257 | AZD8186 |
| MVP_top60_DNA_seed | MAPK1 | -0.130489 | SKCM | COLO-783 | VX-11e |
| MVP_top60_DNA_seed | MAP2K1 | -0.128272 | SKCM | G-MEL | Selumetinib |
| MVP_top60_DNA_seed | EGFR | -0.126179 | SKCM | SK-MEL-5 | PF-00299804 |
| MVP_top60_DNA_seed | ERBB2 | -0.126179 | SKCM | SK-MEL-5 | PF-00299804 |

| | | | | | |
|---|---|---|---|---|---|
| MVP_top60_DNA_seed | EGFR | -0.125841 | SKCM | Mewo | Afatinib |
| MVP_top60_DNA_seed | ERBB2 | -0.125841 | SKCM | Mewo | Afatinib |
| MVP_top60_DNA_seed | BRAF | -0.125291 | SKCM | SK-MEL-30 | HG6-64-1 |
| MVP_top60_DNA_seed | EGFR | -0.122956 | SKCM | A375 | Gefitinib |
| MVP_top60_DNA_seed | EGFR | -0.122886 | SKCM | MMAc-SF | Cetuximab |
| MVP_top60_DNA_seed | GSK3B | -0.122804 | SKCM | WM1552C | KIN001-042 |
| MVP_top60_DNA_seed | JAK3 | -0.122038 | SKCM | IGR-1 | JAK3_7406 |
| MVP_top60_DNA_seed | BRAF | -0.119272 | SKCM | SK-MEL-3 | HG6-64-1 |
| MVP_top60_DNA_seed | GSK3B | -0.118239 | SKCM | MZ7-mel | AZD7969 |
| MVP_top60_DNA_seed | PIK3CA | -0.115885 | SKCM | RPMI-7951 | AZD8835 |
| MVP_top60_DNA_seed | BRAF | -0.114366 | SKCM | K2 | HG6-64-1 |
| MVP_top60_DNA_seed | EGFR | -0.111436 | SKCM | A101D | CI-1033 |
| MVP_top60_DNA_seed | ERBB2 | -0.111436 | SKCM | A101D | CI-1033 |
| MVP_top60_DNA_seed | EGFR | -0.110702 | SKCM | SK-MEL-24 | CUDC-101 |
| MVP_top60_DNA_seed | ERBB2 | -0.110702 | SKCM | SK-MEL-24 | CUDC-101 |
| MVP_top60_DNA_seed | MAPK14 | -0.103464 | SKCM | RPMI-7951 | TAK-715 |
| MVP_top60_DNA_seed | JAK3 | -0.101634 | SKCM | LB373-MEL-D | WHI-P97 |
| MVP_top60_DNA_seed | EGFR | -0.10139 | SKCM | K2 | AZD8931 |
| MVP_top60_DNA_seed | ERBB2 | -0.10139 | SKCM | K2 | AZD8931 |
| MVP_top60_DNA_seed | BRAF | -0.101109 | SKCM | RVH-421 | HG6-64-1 |
| MVP_top60_DNA_seed | JAK1 | -0.101012 | SKCM | SK-MEL-30 | JAK1_3715 |
| MVP_top60_DNA_seed | BRAF | -0.09878 | SKCM | COLO-800 | AZ628 |
| MVP_top60_DNA_seed | GSK3B | -0.094549 | SKCM | MMAc-SF | AZD7969 |
| MVP_top60_DNA_seed | GSK3B | -0.094499 | SKCM | RVH-421 | SB216763 |
| MVP_top60_DNA_seed | BRAF | -0.090442 | SKCM | SK-MEL-2 | HG6-64-1 |
| MVP_top60_DNA_seed | JAK2 | -0.086567 | SKCM | SK-MEL-30 | Ruxolitinib |
| MVP_top60_DNA_seed | JAK1 | -0.086567 | SKCM | SK-MEL-30 | Ruxolitinib |
| MVP_top60_DNA_seed | MAP2K1 | -0.086068 | SKCM | SK-MEL-24 | Refametinib |
| MVP_top60_DNA_seed | EGFR | -0.085566 | SKCM | WM278 | Cetuximab |
| MVP_top60_DNA_seed | GSK3B | -0.085408 | SKCM | G-361 | CHIR-99021 |
| MVP_top60_DNA_seed | JAK2 | -0.084006 | SKCM | SK-MEL-24 | AZD1480 |
| MVP_top60_DNA_seed | JAK1 | -0.084006 | SKCM | SK-MEL-24 | AZD1480 |
| MVP_top60_DNA_seed | GSK3B | -0.083296 | SKCM | LB2518-MEL | CHIR-99021 |
| MVP_top60_DNA_seed | ERBB2 | -0.082256 | SKCM | CP50-MEL-B | CP724714 |
| MVP_top60_DNA_seed | PIK3CA | -0.081863 | SKCM | SH-4 | PI3Ka_4409 |
| MVP_top60_DNA_seed | GSK3B | -0.079565 | SKCM | MEL-HO | AZD7969 |
| MVP_top60_DNA_seed | PIK3CB | -0.077212 | SKCM | LB373-MEL-D | TGX221 |
| MVP_top60_DNA_seed | GSK3B | -0.076943 | SKCM | HT-144 | CHIR-99021 |
| MVP_top60_DNA_seed | EGFR | -0.076764 | SKCM | COLO-800 | Afatinib |
| MVP_top60_DNA_seed | ERBB2 | -0.076764 | SKCM | COLO-800 | Afatinib |
| MVP_top60_DNA_seed | JAK3 | -0.075694 | SKCM | WM1552C | JAK3_7406 |

| | | | | | |
|---|---|---|---|---|---|
| MVP_top60_DNA_seed | EGFR | -0.068891 | SKCM | Hs-940-T | Pelitinib |
| MVP_top60_DNA_seed | GSK3B | -0.066472 | SKCM | K2 | AZD7969 |
| MVP_top60_DNA_seed | EGFR | -0.066242 | SKCM | WM793B | Gefitinib |
| MVP_top60_DNA_seed | PIK3CB | -0.062717 | SKCM | K2 | AZD6482 |
| MVP_top60_DNA_seed | GSK3B | -0.061903 | SKCM | COLO-800 | SB216763 |
| MVP_top60_DNA_seed | JAK2 | -0.0617 | SKCM | LOXIMVI | Fedratinib |
| MVP_top60_DNA_seed | EGFR | -0.059757 | SKCM | LB373-MEL-D | CI-1033 |
| MVP_top60_DNA_seed | ERBB2 | -0.059757 | SKCM | LB373-MEL-D | CI-1033 |
| MVP_top60_DNA_seed | GSK3B | -0.057959 | SKCM | IST-MEL1 | CHIR-99021 |
| MVP_top60_DNA_seed | EGFR | -0.057603 | SKCM | WM-115 | Pelitinib |
| MVP_top60_DNA_seed | MAPK3 | -0.054392 | SKCM | MZ2-MEL | FR-180204 |
| MVP_top60_DNA_seed | MAPK1 | -0.054392 | SKCM | MZ2-MEL | FR-180204 |
| MVP_top60_DNA_seed | EGFR | -0.052738 | SKCM | UACC-62 | Cetuximab |
| MVP_top60_DNA_seed | EGFR | -0.049093 | SKCM | WM1552C | Cetuximab |
| MVP_top60_DNA_seed | EP300 | -0.047877 | SKCM | SK-MEL-30 | I-CBP112 |
| MVP_top60_DNA_seed | CREBBP | -0.047877 | SKCM | SK-MEL-30 | I-CBP112 |
| MVP_top60_DNA_seed | JAK2 | -0.04735 | SKCM | WM35 | AZD1480 |
| MVP_top60_DNA_seed | JAK1 | -0.04735 | SKCM | WM35 | AZD1480 |
| MVP_top60_DNA_seed | EGFR | -0.047302 | SKCM | COLO-792 | Gefitinib |
| MVP_top60_DNA_seed | JAK1 | -0.047105 | SKCM | SK-MEL-2 | JAK1_3715 |
| MVP_top60_DNA_seed | MAPK3 | -0.045436 | SKCM | G-361 | FR-180204 |
| MVP_top60_DNA_seed | MAPK1 | -0.045436 | SKCM | G-361 | FR-180204 |
| MVP_top60_DNA_seed | GSK3B | -0.043819 | SKCM | COLO-792 | CHIR-99021 |
| MVP_top60_DNA_seed | JAK2 | -0.0411 | SKCM | A2058 | AZD1480 |
| MVP_top60_DNA_seed | JAK1 | -0.0411 | SKCM | A2058 | AZD1480 |
| MVP_top60_DNA_seed | EGFR | -0.040736 | SKCM | RPMI-7951 | AZD8931 |
| MVP_top60_DNA_seed | ERBB2 | -0.040736 | SKCM | RPMI-7951 | AZD8931 |
| MVP_top60_DNA_seed | EGFR | -0.037757 | SKCM | RPMI-7951 | Afatinib |
| MVP_top60_DNA_seed | ERBB2 | -0.037757 | SKCM | RPMI-7951 | Afatinib |
| MVP_top60_DNA_seed | EGFR | -0.034548 | SKCM | COLO-679 | CI-1033 |
| MVP_top60_DNA_seed | ERBB2 | -0.034548 | SKCM | COLO-679 | CI-1033 |
| MVP_top60_DNA_seed | ERBB2 | -0.034134 | SKCM | WM35 | CP724714 |
| MVP_top60_DNA_seed | EP300 | -0.030394 | SKCM | IGR-37 | I-CBP112 |
| MVP_top60_DNA_seed | CREBBP | -0.030394 | SKCM | IGR-37 | I-CBP112 |
| MVP_top60_DNA_seed | JAK2 | -0.028996 | SKCM | LB2518-MEL | Ruxolitinib |
| MVP_top60_DNA_seed | JAK1 | -0.028996 | SKCM | LB2518-MEL | Ruxolitinib |
| MVP_top60_DNA_seed | MAPK3 | -0.0289 | SKCM | MEL-JUSO | FR-180204 |
| MVP_top60_DNA_seed | MAPK1 | -0.0289 | SKCM | MEL-JUSO | FR-180204 |
| MVP_top60_DNA_seed | JAK3 | -0.028536 | SKCM | MMAc-SF | JAK3_7406 |
| MVP_top60_DNA_seed | PIK3CA | -0.028411 | SKCM | SK-MEL-2 | AZD8835 |
| MVP_top60_DNA_seed | GSK3B | -0.028297 | SKCM | HMV-II | AZD7969 |

| | | | | | |
|---|---|---|---|---|---|
| MVP_top60_DNA_seed | JAK2 | -0.028134 | SKCM | CP50-MEL-B | AZD1480 |
| MVP_top60_DNA_seed | JAK1 | -0.028134 | SKCM | CP50-MEL-B | AZD1480 |
| MVP_top60_DNA_seed | PIK3CB | -0.027069 | SKCM | HT-144 | AZD8186 |
| MVP_top60_DNA_seed | EGFR | -0.023404 | SKCM | UACC-62 | Afatinib |
| MVP_top60_DNA_seed | ERBB2 | -0.023404 | SKCM | UACC-62 | Afatinib |
| MVP_top60_DNA_seed | MAPK3 | -0.023127 | SKCM | RPMI-7951 | FR-180204 |
| MVP_top60_DNA_seed | MAPK1 | -0.023127 | SKCM | RPMI-7951 | FR-180204 |
| MVP_top60_DNA_seed | JAK2 | -0.023095 | SKCM | SK-MEL-30 | AZD1480 |
| MVP_top60_DNA_seed | JAK1 | -0.023095 | SKCM | SK-MEL-30 | AZD1480 |
| MVP_top60_DNA_seed | MAPK1 | -0.021313 | SKCM | Hs-939-T | VX-11e |
| MVP_top60_DNA_seed | PIK3CA | -0.021001 | SKCM | Hs-939-T | AZD8835 |
| MVP_top60_DNA_seed | EGFR | -0.019612 | SKCM | SK-MEL-3 | Cetuximab |
| MVP_top60_DNA_seed | JAK2 | -0.018733 | SKCM | MEL-JUSO | AZD1480 |
| MVP_top60_DNA_seed | JAK1 | -0.018733 | SKCM | MEL-JUSO | AZD1480 |
| MVP_top60_DNA_seed | BRAF | -0.017785 | SKCM | IPC-298 | HG6-64-1 |
| MVP_top60_DNA_seed | BRAF | -0.01767 | SKCM | RPMI-7951 | HG6-64-1 |
| MVP_top60_DNA_seed | BRAF | -0.015164 | SKCM | SK-MEL-30 | Dabrafenib |
| MVP_top60_DNA_seed | PIK3CB | -0.014213 | SKCM | SK-MEL-28 | AZD6482 |
| MVP_top60_DNA_seed | EGFR | -0.013156 | SKCM | SK-MEL-24 | Afatinib |
| MVP_top60_DNA_seed | ERBB2 | -0.013156 | SKCM | SK-MEL-24 | Afatinib |
| MVP_top60_DNA_seed | EGFR | -0.012512 | SKCM | LOXIMVI | AZD8931 |
| MVP_top60_DNA_seed | ERBB2 | -0.012512 | SKCM | LOXIMVI | AZD8931 |
| MVP_top60_DNA_seed | JAK2 | -0.008939 | SKCM | WM-115 | Fedratinib |
| MVP_top60_DNA_seed | EGFR | -0.006357 | SKCM | MMAc-SF | Gefitinib |
| MVP_top60_DNA_seed | BRAF | -0.00588 | SKCM | SK-MEL-31 | PLX-4720 |
| MVP_top60_DNA_seed | MAPK3 | -0.001024 | SKCM | COLO-792 | FR-180204 |
| MVP_top60_DNA_seed | MAPK1 | -0.001024 | SKCM | COLO-792 | FR-180204 |
| MVP_top60_DNA_seed | PIK3CA | 0.000655 | SKCM | COLO-800 | PI3Ka_4409 |
| MVP_top60_DNA_seed | MAPK1 | 0.003478 | SKCM | CP66-MEL | VX-11e |
| MVP_top60_DNA_seed | GSK3B | 0.003975 | SKCM | MZ7-mel | SB216763 |
| MVP_top60_DNA_seed | EGFR | 0.004444 | SKCM | UACC-62 | CI-1033 |
| MVP_top60_DNA_seed | ERBB2 | 0.004444 | SKCM | UACC-62 | CI-1033 |
| MVP_top60_DNA_seed | GSK3B | 0.005792 | SKCM | Hs-939-T | KIN001-042 |
| MVP_top60_DNA_seed | GSK3B | 0.006256 | SKCM | IPC-298 | CHIR-99021 |
| MVP_top60_DNA_seed | EGFR | 0.009579 | SKCM | SK-MEL-28 | Cetuximab |
| MVP_top60_DNA_seed | JAK2 | 0.01141 | SKCM | LOXIMVI | AZD1480 |
| MVP_top60_DNA_seed | JAK1 | 0.01141 | SKCM | LOXIMVI | AZD1480 |
| MVP_top60_DNA_seed | BRAF | 0.014112 | SKCM | MEL-JUSO | PLX-4720 |
| MVP_top60_DNA_seed | PIK3CA | 0.015623 | SKCM | UACC-62 | PI3Ka_4409 |
| MVP_top60_DNA_seed | EGFR | 0.016412 | SKCM | HMV-II | CUDC-101 |
| MVP_top60_DNA_seed | ERBB2 | 0.016412 | SKCM | HMV-II | CUDC-101 |

| | | | | | |
|---|---|---|---|---|---|
| MVP_top60_DNA_seed | BRAF | 0.018719 | SKCM | Hs-939-T | HG6-64-1 |
| MVP_top60_DNA_seed | EGFR | 0.019999 | SKCM | COLO-829 | CUDC-101 |
| MVP_top60_DNA_seed | ERBB2 | 0.019999 | SKCM | COLO-829 | CUDC-101 |
| MVP_top60_DNA_seed | EGFR | 0.022951 | SKCM | UACC-62 | AZD8931 |
| MVP_top60_DNA_seed | ERBB2 | 0.022951 | SKCM | UACC-62 | AZD8931 |
| MVP_top60_DNA_seed | EGFR | 0.02437 | SKCM | WM1552C | CUDC-101 |
| MVP_top60_DNA_seed | ERBB2 | 0.02437 | SKCM | WM1552C | CUDC-101 |
| MVP_top60_DNA_seed | JAK1 | 0.028197 | SKCM | K2 | JAK1_3715 |
| MVP_top60_DNA_seed | MAPK14 | 0.030713 | SKCM | HMV-II | TAK-715 |
| MVP_top60_DNA_seed | PIK3CA | 0.032344 | SKCM | G-MEL | AZD8835 |
| MVP_top60_DNA_seed | EGFR | 0.032773 | SKCM | MZ2-MEL | CUDC-101 |
| MVP_top60_DNA_seed | ERBB2 | 0.032773 | SKCM | MZ2-MEL | CUDC-101 |
| MVP_top60_DNA_seed | EP300 | 0.03314 | SKCM | WM278 | I-CBP112 |
| MVP_top60_DNA_seed | CREBBP | 0.03314 | SKCM | WM278 | I-CBP112 |
| MVP_top60_DNA_seed | MAPK14 | 0.034032 | SKCM | IPC-298 | TAK-715 |
| MVP_top60_DNA_seed | EGFR | 0.034665 | SKCM | MZ2-MEL | Afatinib |
| MVP_top60_DNA_seed | ERBB2 | 0.034665 | SKCM | MZ2-MEL | Afatinib |
| MVP_top60_DNA_seed | GSK3B | 0.035894 | SKCM | SK-MEL-30 | AZD7969 |
| MVP_top60_DNA_seed | EGFR | 0.036762 | SKCM | MEL-HO | Cetuximab |
| MVP_top60_DNA_seed | JAK2 | 0.037214 | SKCM | WM1552C | AZD1480 |
| MVP_top60_DNA_seed | JAK1 | 0.037214 | SKCM | WM1552C | AZD1480 |
| MVP_top60_DNA_seed | ERBB2 | 0.037849 | SKCM | MEL-JUSO | CP724714 |
| MVP_top60_DNA_seed | EGFR | 0.040065 | SKCM | VMRC-MELG | AST-1306 |
| MVP_top60_DNA_seed | EGFR | 0.04018 | SKCM | SK-MEL-2 | Cetuximab |
| MVP_top60_DNA_seed | EGFR | 0.041102 | SKCM | WM793B | Cetuximab |
| MVP_top60_DNA_seed | GSK3B | 0.042082 | SKCM | K2 | CHIR-99021 |
| MVP_top60_DNA_seed | EGFR | 0.04248 | SKCM | IST-MEL1 | AST-1306 |
| MVP_top60_DNA_seed | EGFR | 0.042729 | SKCM | VMRC-MELG | CUDC-101 |
| MVP_top60_DNA_seed | ERBB2 | 0.042729 | SKCM | VMRC-MELG | CUDC-101 |
| MVP_top60_DNA_seed | JAK1 | 0.04285 | SKCM | COLO-792 | JAK1_3715 |
| MVP_top60_DNA_seed | PIK3CA | 0.04308 | SKCM | UACC-62 | AZD8835 |
| MVP_top60_DNA_seed | JAK2 | 0.047793 | SKCM | IPC-298 | Ruxolitinib |
| MVP_top60_DNA_seed | JAK1 | 0.047793 | SKCM | IPC-298 | Ruxolitinib |
| MVP_top60_DNA_seed | EGFR | 0.048002 | SKCM | WM-115 | CUDC-101 |
| MVP_top60_DNA_seed | ERBB2 | 0.048002 | SKCM | WM-115 | CUDC-101 |
| MVP_top60_DNA_seed | GSK3B | 0.053296 | SKCM | IGR-1 | SB216763 |
| MVP_top60_DNA_seed | JAK2 | 0.053741 | SKCM | CP50-MEL-B | Ruxolitinib |
| MVP_top60_DNA_seed | JAK1 | 0.053741 | SKCM | CP50-MEL-B | Ruxolitinib |
| MVP_top60_DNA_seed | JAK2 | 0.054798 | SKCM | SK-MEL-2 | Fedratinib |
| MVP_top60_DNA_seed | PIK3CA | 0.055744 | SKCM | MEL-JUSO | AZD8835 |
| MVP_top60_DNA_seed | EGFR | 0.057033 | SKCM | A2058 | CI-1033 |

| | | | | | |
|---|---|---|---|---|---|
| MVP_top60_DNA_seed | ERBB2 | 0.057033 | SKCM | A2058 | CI-1033 |
| MVP_top60_DNA_seed | GSK3B | 0.05738 | SKCM | COLO-783 | KIN001-042 |
| MVP_top60_DNA_seed | EGFR | 0.058295 | SKCM | G-MEL | CI-1033 |
| MVP_top60_DNA_seed | ERBB2 | 0.058295 | SKCM | G-MEL | CI-1033 |
| MVP_top60_DNA_seed | MAP2K1 | 0.059009 | SKCM | Hs-940-T | PD0325901 |
| MVP_top60_DNA_seed | PIK3CB | 0.062967 | SKCM | VMRC-MELG | AZD8186 |
| MVP_top60_DNA_seed | EGFR | 0.063432 | SKCM | COLO-679 | CUDC-101 |
| MVP_top60_DNA_seed | ERBB2 | 0.063432 | SKCM | COLO-679 | CUDC-101 |
| MVP_top60_DNA_seed | EGFR | 0.065531 | SKCM | Mewo | Gefitinib |
| MVP_top60_DNA_seed | JAK2 | 0.066049 | SKCM | GAK | Ruxolitinib |
| MVP_top60_DNA_seed | JAK1 | 0.066049 | SKCM | GAK | Ruxolitinib |
| MVP_top60_DNA_seed | GSK3B | 0.067833 | SKCM | CHL-1 | CHIR-99021 |
| MVP_top60_DNA_seed | BRAF | 0.068592 | SKCM | GAK | PLX-4720 |
| MVP_top60_DNA_seed | PIK3CA | 0.072663 | SKCM | G-MEL | PI3Ka_4409 |
| MVP_top60_DNA_seed | EGFR | 0.074263 | SKCM | CP50-MEL-B | Afatinib |
| MVP_top60_DNA_seed | ERBB2 | 0.074263 | SKCM | CP50-MEL-B | Afatinib |
| MVP_top60_DNA_seed | JAK3 | 0.080821 | SKCM | A375 | WHI-P97 |
| MVP_top60_DNA_seed | PIK3CA | 0.081584 | SKCM | WM278 | PI3Ka_4409 |
| MVP_top60_DNA_seed | MAPK3 | 0.082672 | SKCM | 451Lu | FR-180204 |
| MVP_top60_DNA_seed | MAPK1 | 0.082672 | SKCM | 451Lu | FR-180204 |
| MVP_top60_DNA_seed | GSK3B | 0.084015 | SKCM | WM-115 | AZD7969 |
| MVP_top60_DNA_seed | EGFR | 0.085975 | SKCM | SK-MEL-24 | AST-1306 |
| MVP_top60_DNA_seed | EGFR | 0.086201 | SKCM | 451Lu | CUDC-101 |
| MVP_top60_DNA_seed | ERBB2 | 0.086201 | SKCM | 451Lu | CUDC-101 |
| MVP_top60_DNA_seed | JAK2 | 0.08679 | SKCM | WM1552C | Ruxolitinib |
| MVP_top60_DNA_seed | JAK1 | 0.08679 | SKCM | WM1552C | Ruxolitinib |
| MVP_top60_DNA_seed | PIK3CA | 0.087879 | SKCM | SK-MEL-5 | AZD8835 |
| MVP_top60_DNA_seed | JAK2 | 0.088163 | SKCM | CHL-1 | Ruxolitinib |
| MVP_top60_DNA_seed | JAK1 | 0.088163 | SKCM | CHL-1 | Ruxolitinib |
| MVP_top60_DNA_seed | EGFR | 0.091479 | SKCM | RVH-421 | Afatinib |
| MVP_top60_DNA_seed | ERBB2 | 0.091479 | SKCM | RVH-421 | Afatinib |
| MVP_top60_DNA_seed | EGFR | 0.094495 | SKCM | SK-MEL-28 | Afatinib |
| MVP_top60_DNA_seed | ERBB2 | 0.094495 | SKCM | SK-MEL-28 | Afatinib |
| MVP_top60_DNA_seed | GSK3B | 0.097448 | SKCM | GAK | SB216763 |
| MVP_top60_DNA_seed | EGFR | 0.098939 | SKCM | RVH-421 | AST-1306 |
| MVP_top60_DNA_seed | EGFR | 0.099646 | SKCM | HT-144 | Erlotinib |
| MVP_top60_DNA_seed | MAP2K1 | 0.103799 | SKCM | IST-MEL1 | PD0325901 |
| MVP_top60_DNA_seed | EGFR | 0.106169 | SKCM | SK-MEL-1 | CI-1033 |
| MVP_top60_DNA_seed | ERBB2 | 0.106169 | SKCM | SK-MEL-1 | CI-1033 |
| MVP_top60_DNA_seed | EGFR | 0.107474 | SKCM | IGR-1 | Pelitinib |
| MVP_top60_DNA_seed | JAK3 | 0.109569 | SKCM | K2 | JAK3_7406 |

| | | | | | |
|---|---|---|---|---|---|
| MVP_top60_DNA_seed | GSK3B | 0.110323 | SKCM | WM278 | CHIR-99021 |
| MVP_top60_DNA_seed | EGFR | 0.110446 | SKCM | SK-MEL-24 | CI-1033 |
| MVP_top60_DNA_seed | ERBB2 | 0.110446 | SKCM | SK-MEL-24 | CI-1033 |
| MVP_top60_DNA_seed | PIK3CA | 0.112372 | SKCM | SK-MEL-28 | PI3Ka_4409 |
| MVP_top60_DNA_seed | EGFR | 0.1137 | SKCM | COLO-679 | Pelitinib |
| MVP_top60_DNA_seed | EGFR | 0.113957 | SKCM | MZ7-mel | CI-1033 |
| MVP_top60_DNA_seed | ERBB2 | 0.113957 | SKCM | MZ7-mel | CI-1033 |
| MVP_top60_DNA_seed | EP300 | 0.118328 | SKCM | GAK | I-CBP112 |
| MVP_top60_DNA_seed | CREBBP | 0.118328 | SKCM | GAK | I-CBP112 |
| MVP_top60_DNA_seed | BRAF | 0.121078 | SKCM | COLO-800 | RAF_9304 |
| MVP_top60_DNA_seed | EGFR | 0.121956 | SKCM | MMAc-SF | Erlotinib |
| MVP_top60_DNA_seed | MAP2K1 | 0.124152 | SKCM | SK-MEL-24 | PD0325901 |
| MVP_top60_DNA_seed | EGFR | 0.124332 | SKCM | IST-MEL1 | AZD8931 |
| MVP_top60_DNA_seed | ERBB2 | 0.124332 | SKCM | IST-MEL1 | AZD8931 |
| MVP_top60_DNA_seed | EGFR | 0.124348 | SKCM | Hs-939-T | CI-1033 |
| MVP_top60_DNA_seed | ERBB2 | 0.124348 | SKCM | Hs-939-T | CI-1033 |
| MVP_top60_DNA_seed | EGFR | 0.12482 | SKCM | A101D | Lapatinib |
| MVP_top60_DNA_seed | ERBB2 | 0.12482 | SKCM | A101D | Lapatinib |
| MVP_top60_DNA_seed | EGFR | 0.126005 | SKCM | SK-MEL-3 | Pelitinib |
| MVP_top60_DNA_seed | PIK3CB | 0.126396 | SKCM | 451Lu | AZD6482 |
| MVP_top60_DNA_seed | JAK3 | 0.127225 | SKCM | A101D | JAK3_7406 |
| MVP_top60_DNA_seed | PIK3CB | 0.129303 | SKCM | COLO-679 | AZD8186 |
| MVP_top60_DNA_seed | PIK3CB | 0.131985 | SKCM | G-361 | AZD8186 |
| MVP_top60_DNA_seed | GSK3B | 0.132032 | SKCM | CP66-MEL | CHIR-99021 |
| MVP_top60_DNA_seed | JAK2 | 0.136503 | SKCM | Hs-939-T | Ruxolitinib |
| MVP_top60_DNA_seed | JAK1 | 0.136503 | SKCM | Hs-939-T | Ruxolitinib |
| MVP_top60_DNA_seed | EGFR | 0.137262 | SKCM | LB2518-MEL | CI-1033 |
| MVP_top60_DNA_seed | ERBB2 | 0.137262 | SKCM | LB2518-MEL | CI-1033 |
| MVP_top60_DNA_seed | MAPK14 | 0.138088 | SKCM | CP66-MEL | TAK-715 |
| MVP_top60_DNA_seed | EGFR | 0.138142 | SKCM | A101D | CUDC-101 |
| MVP_top60_DNA_seed | ERBB2 | 0.138142 | SKCM | A101D | CUDC-101 |
| MVP_top60_DNA_seed | GSK3B | 0.142809 | SKCM | WM278 | KIN001-042 |
| MVP_top60_DNA_seed | EGFR | 0.143011 | SKCM | Mewo | CUDC-101 |
| MVP_top60_DNA_seed | ERBB2 | 0.143011 | SKCM | Mewo | CUDC-101 |
| MVP_top60_DNA_seed | JAK2 | 0.145026 | SKCM | SK-MEL-28 | Ruxolitinib |
| MVP_top60_DNA_seed | JAK1 | 0.145026 | SKCM | SK-MEL-28 | Ruxolitinib |
| MVP_top60_DNA_seed | PIK3CA | 0.145198 | SKCM | IPC-298 | AZD8835 |
| MVP_top60_DNA_seed | JAK3 | 0.145254 | SKCM | CP66-MEL | WHI-P97 |
| MVP_top60_DNA_seed | EP300 | 0.147125 | SKCM | HT-144 | I-CBP112 |
| MVP_top60_DNA_seed | CREBBP | 0.147125 | SKCM | HT-144 | I-CBP112 |
| MVP_top60_DNA_seed | PIK3CA | 0.147579 | SKCM | COLO-783 | AZD8835 |

| | | | | | |
|---|---|---|---|---|---|
| MVP_top60_DNA_seed | EGFR | 0.148049 | SKCM | Hs-939-T | AZD8931 |
| MVP_top60_DNA_seed | ERBB2 | 0.148049 | SKCM | Hs-939-T | AZD8931 |
| MVP_top60_DNA_seed | PIK3CA | 0.150437 | SKCM | SK-MEL-30 | AZD8835 |
| MVP_top60_DNA_seed | GSK3B | 0.150837 | SKCM | SK-MEL-24 | AZD7969 |
| MVP_top60_DNA_seed | JAK1 | 0.152169 | SKCM | COLO-829 | JAK1_3715 |
| MVP_top60_DNA_seed | EGFR | 0.153418 | SKCM | Hs-940-T | CUDC-101 |
| MVP_top60_DNA_seed | ERBB2 | 0.153418 | SKCM | Hs-940-T | CUDC-101 |
| MVP_top60_DNA_seed | ERBB2 | 0.154712 | SKCM | A2058 | CP724714 |
| MVP_top60_DNA_seed | JAK2 | 0.157888 | SKCM | IPC-298 | Fedratinib |
| MVP_top60_DNA_seed | BRAF | 0.159752 | SKCM | RPMI-7951 | SB590885 |
| MVP_top60_DNA_seed | EGFR | 0.16271 | SKCM | IGR-37 | Cetuximab |
| MVP_top60_DNA_seed | EGFR | 0.162901 | SKCM | MZ7-mel | Cetuximab |
| MVP_top60_DNA_seed | JAK2 | 0.162969 | SKCM | LOXIMVI | Ruxolitinib |
| MVP_top60_DNA_seed | JAK1 | 0.162969 | SKCM | LOXIMVI | Ruxolitinib |
| MVP_top60_DNA_seed | EGFR | 0.163576 | SKCM | IGR-1 | Gefitinib |
| MVP_top60_DNA_seed | EGFR | 0.165312 | SKCM | 451Lu | Afatinib |
| MVP_top60_DNA_seed | ERBB2 | 0.165312 | SKCM | 451Lu | Afatinib |
| MVP_top60_DNA_seed | MAPK3 | 0.166359 | SKCM | SK-MEL-30 | FR-180204 |
| MVP_top60_DNA_seed | MAPK1 | 0.166359 | SKCM | SK-MEL-30 | FR-180204 |
| MVP_top60_DNA_seed | EGFR | 0.166618 | SKCM | HMV-II | Pelitinib |
| MVP_top60_DNA_seed | GSK3B | 0.167651 | SKCM | WM-115 | KIN001-042 |
| MVP_top60_DNA_seed | GSK3B | 0.169129 | SKCM | CP66-MEL | AZD7969 |
| MVP_top60_DNA_seed | EGFR | 0.170393 | SKCM | G-MEL | Gefitinib |
| MVP_top60_DNA_seed | EGFR | 0.171889 | SKCM | GAK | AZD8931 |
| MVP_top60_DNA_seed | ERBB2 | 0.171889 | SKCM | GAK | AZD8931 |
| MVP_top60_DNA_seed | JAK1 | 0.172456 | SKCM | GAK | JAK1_3715 |
| MVP_top60_DNA_seed | EGFR | 0.173249 | SKCM | MZ2-MEL | AZD8931 |
| MVP_top60_DNA_seed | ERBB2 | 0.173249 | SKCM | MZ2-MEL | AZD8931 |
| MVP_top60_DNA_seed | JAK3 | 0.17726 | SKCM | MMAc-SF | WHI-P97 |
| MVP_top60_DNA_seed | PIK3CA | 0.178081 | SKCM | COLO-792 | AZD8835 |
| MVP_top60_DNA_seed | GSK3B | 0.178664 | SKCM | 451Lu | CHIR-99021 |
| MVP_top60_DNA_seed | PIK3CB | 0.179471 | SKCM | UACC-257 | AZD6482 |
| MVP_top60_DNA_seed | BRAF | 0.180461 | SKCM | HMV-II | PLX-4720 |
| MVP_top60_DNA_seed | BRAF | 0.18138 | SKCM | COLO-792 | RAF_9304 |
| MVP_top60_DNA_seed | EGFR | 0.181672 | SKCM | SK-MEL-5 | Afatinib |
| MVP_top60_DNA_seed | ERBB2 | 0.181672 | SKCM | SK-MEL-5 | Afatinib |
| MVP_top60_DNA_seed | GSK3B | 0.184618 | SKCM | SK-MEL-28 | CHIR-99021 |
| MVP_top60_DNA_seed | PIK3CB | 0.185201 | SKCM | COLO-800 | AZD8186 |
| MVP_top60_DNA_seed | EGFR | 0.185864 | SKCM | SK-MEL-24 | AZD8931 |
| MVP_top60_DNA_seed | ERBB2 | 0.185864 | SKCM | SK-MEL-24 | AZD8931 |
| MVP_top60_DNA_seed | MAP2K1 | 0.186669 | SKCM | IST-MEL1 | CI-1040 |

| | | | | | |
|---|---|---|---|---|---|
| MVP_top60_DNA_seed | JAK3 | 0.189839 | SKCM | GAK | JAK3_7406 |
| MVP_top60_DNA_seed | BRAF | 0.190607 | SKCM | VMRC-MELG | Dabrafenib |
| MVP_top60_DNA_seed | ERBB2 | 0.190635 | SKCM | GAK | CP724714 |
| MVP_top60_DNA_seed | MAPK3 | 0.191635 | SKCM | A101D | FR-180204 |
| MVP_top60_DNA_seed | MAPK1 | 0.191635 | SKCM | A101D | FR-180204 |
| MVP_top60_DNA_seed | GSK3B | 0.192808 | SKCM | SK-MEL-5 | AZD7969 |
| MVP_top60_DNA_seed | ERBB2 | 0.193116 | SKCM | IGR-1 | CP724714 |
| MVP_top60_DNA_seed | MAP2K1 | 0.193465 | SKCM | SK-MEL-31 | Refametinib |
| MVP_top60_DNA_seed | MAPK3 | 0.194506 | SKCM | IPC-298 | FR-180204 |
| MVP_top60_DNA_seed | MAPK1 | 0.194506 | SKCM | IPC-298 | FR-180204 |
| MVP_top60_DNA_seed | MAPK3 | 0.196218 | SKCM | MMAc-SF | FR-180204 |
| MVP_top60_DNA_seed | MAPK1 | 0.196218 | SKCM | MMAc-SF | FR-180204 |
| MVP_top60_DNA_seed | MAPK3 | 0.196356 | SKCM | SK-MEL-5 | FR-180204 |
| MVP_top60_DNA_seed | MAPK1 | 0.196356 | SKCM | SK-MEL-5 | FR-180204 |
| MVP_top60_DNA_seed | GSK3B | 0.196604 | SKCM | LOXIMVI | SB216763 |
| MVP_top60_DNA_seed | MAPK14 | 0.197589 | SKCM | MZ2-MEL | TAK-715 |
| MVP_top60_DNA_seed | GSK3B | 0.199348 | SKCM | SK-MEL-28 | KIN001-042 |
| MVP_top60_DNA_seed | EGFR | 0.200532 | SKCM | LB2518-MEL | CUDC-101 |
| MVP_top60_DNA_seed | ERBB2 | 0.200532 | SKCM | LB2518-MEL | CUDC-101 |
| MVP_top60_DNA_seed | GSK3B | 0.202093 | SKCM | IGR-37 | KIN001-042 |
| MVP_top60_DNA_seed | PIK3CB | 0.203194 | SKCM | HMV-II | AZD6482 |
| MVP_top60_DNA_seed | EGFR | 0.203439 | SKCM | M14 | CUDC-101 |
| MVP_top60_DNA_seed | ERBB2 | 0.203439 | SKCM | M14 | CUDC-101 |
| MVP_top60_DNA_seed | ERBB2 | 0.206964 | SKCM | WM793B | CP724714 |
| MVP_top60_DNA_seed | MAPK14 | 0.207087 | SKCM | GAK | TAK-715 |
| MVP_top60_DNA_seed | RAC1 | 0.20856 | SKCM | A375 | EHT-1864 |
| MVP_top60_DNA_seed | EGFR | 0.208626 | SKCM | SK-MEL-28 | CUDC-101 |
| MVP_top60_DNA_seed | ERBB2 | 0.208626 | SKCM | SK-MEL-28 | CUDC-101 |
| MVP_top60_DNA_seed | ERBB2 | 0.209909 | SKCM | A375 | CP724714 |
| MVP_top60_DNA_seed | PIK3CB | 0.210994 | SKCM | LB373-MEL-D | AZD8186 |
| MVP_top60_DNA_seed | EGFR | 0.211728 | SKCM | WM35 | AST-1306 |
| MVP_top60_DNA_seed | EGFR | 0.21218 | SKCM | RVH-421 | Pelitinib |
| MVP_top60_DNA_seed | EP300 | 0.212457 | SKCM | WM35 | I-CBP112 |
| MVP_top60_DNA_seed | CREBBP | 0.212457 | SKCM | WM35 | I-CBP112 |
| MVP_top60_DNA_seed | EGFR | 0.213213 | SKCM | G-361 | AZD8931 |
| MVP_top60_DNA_seed | ERBB2 | 0.213213 | SKCM | G-361 | AZD8931 |
| MVP_top60_DNA_seed | GSK3B | 0.213846 | SKCM | IGR-37 | CHIR-99021 |
| MVP_top60_DNA_seed | JAK2 | 0.215227 | SKCM | UACC-62 | Ruxolitinib |
| MVP_top60_DNA_seed | JAK1 | 0.215227 | SKCM | UACC-62 | Ruxolitinib |
| MVP_top60_DNA_seed | GSK3B | 0.215233 | SKCM | LOXIMVI | KIN001-042 |
| MVP_top60_DNA_seed | JAK3 | 0.215762 | SKCM | RVH-421 | JAK3_7406 |

| | | | | | |
|---|---|---|---|---|---|
| MVP_top60_DNA_seed | PIK3CB | 0.21615 | SKCM | Mewo | AZD8186 |
| MVP_top60_DNA_seed | JAK3 | 0.219626 | SKCM | WM793B | WHI-P97 |
| MVP_top60_DNA_seed | JAK3 | 0.221459 | SKCM | MEL-JUSO | JAK3_7406 |
| MVP_top60_DNA_seed | EGFR | 0.227019 | SKCM | SK-MEL-30 | AZD8931 |
| MVP_top60_DNA_seed | ERBB2 | 0.227019 | SKCM | SK-MEL-30 | AZD8931 |
| MVP_top60_DNA_seed | EGFR | 0.228299 | SKCM | COLO-800 | PF-00299804 |
| MVP_top60_DNA_seed | ERBB2 | 0.228299 | SKCM | COLO-800 | PF-00299804 |
| MVP_top60_DNA_seed | BRAF | 0.230979 | SKCM | UACC-62 | SB590885 |
| MVP_top60_DNA_seed | JAK2 | 0.231049 | SKCM | LB373-MEL-D | Ruxolitinib |
| MVP_top60_DNA_seed | JAK1 | 0.231049 | SKCM | LB373-MEL-D | Ruxolitinib |
| MVP_top60_DNA_seed | BRAF | 0.234865 | SKCM | CP66-MEL | Dabrafenib |
| MVP_top60_DNA_seed | MAPK3 | 0.235171 | SKCM | LOXIMVI | FR-180204 |
| MVP_top60_DNA_seed | MAPK1 | 0.235171 | SKCM | LOXIMVI | FR-180204 |
| MVP_top60_DNA_seed | EGFR | 0.235244 | SKCM | C32 | AZD8931 |
| MVP_top60_DNA_seed | ERBB2 | 0.235244 | SKCM | C32 | AZD8931 |
| MVP_top60_DNA_seed | EGFR | 0.237912 | SKCM | COLO-679 | Afatinib |
| MVP_top60_DNA_seed | ERBB2 | 0.237912 | SKCM | COLO-679 | Afatinib |
| MVP_top60_DNA_seed | MAPK1 | 0.238125 | SKCM | HT-144 | VX-11e |
| MVP_top60_DNA_seed | BRAF | 0.239094 | SKCM | MEL-JUSO | PLX-4720 |
| MVP_top60_DNA_seed | ERBB2 | 0.239462 | SKCM | IPC-298 | CP724714 |
| MVP_top60_DNA_seed | EGFR | 0.239757 | SKCM | M14 | CI-1033 |
| MVP_top60_DNA_seed | ERBB2 | 0.239757 | SKCM | M14 | CI-1033 |
| MVP_top60_DNA_seed | EGFR | 0.239953 | SKCM | COLO-783 | CUDC-101 |
| MVP_top60_DNA_seed | ERBB2 | 0.239953 | SKCM | COLO-783 | CUDC-101 |
| MVP_top60_DNA_seed | JAK2 | 0.240869 | SKCM | IGR-1 | AZD1480 |
| MVP_top60_DNA_seed | JAK1 | 0.240869 | SKCM | IGR-1 | AZD1480 |
| MVP_top60_DNA_seed | BRAF | 0.245239 | SKCM | COLO-800 | Dabrafenib |
| MVP_top60_DNA_seed | PIK3CA | 0.247149 | SKCM | K2 | AZD8835 |
| MVP_top60_DNA_seed | EGFR | 0.247297 | SKCM | SK-MEL-24 | PF-00299804 |
| MVP_top60_DNA_seed | ERBB2 | 0.247297 | SKCM | SK-MEL-24 | PF-00299804 |
| MVP_top60_DNA_seed | EP300 | 0.251101 | SKCM | Mewo | I-CBP112 |
| MVP_top60_DNA_seed | CREBBP | 0.251101 | SKCM | Mewo | I-CBP112 |
| MVP_top60_DNA_seed | PIK3CA | 0.251621 | SKCM | COLO-783 | PI3Ka_4409 |
| MVP_top60_DNA_seed | PIK3CB | 0.252797 | SKCM | MZ7-mel | AZD6482 |
| MVP_top60_DNA_seed | PIK3CB | 0.253068 | SKCM | GAK | AZD8186 |
| MVP_top60_DNA_seed | JAK3 | 0.254328 | SKCM | K2 | WHI-P97 |
| MVP_top60_DNA_seed | BRAF | 0.255124 | SKCM | M14 | HG6-64-1 |
| MVP_top60_DNA_seed | PIK3CB | 0.256453 | SKCM | SH-4 | AZD6482 |
| MVP_top60_DNA_seed | GSK3B | 0.259105 | SKCM | CHL-1 | KIN001-042 |
| MVP_top60_DNA_seed | PIK3CB | 0.261421 | SKCM | LOXIMVI | AZD6482 |
| MVP_top60_DNA_seed | RAC1 | 0.261736 | SKCM | COLO-783 | EHT-1864 |

| | | | | | |
|---|---|---|---|---|---|
| MVP_top60_DNA_seed | PIK3CA | 0.262914 | SKCM | GAK | AZD8835 |
| MVP_top60_DNA_seed | MAP2K1 | 0.263919 | SKCM | WM1552C | CI-1040 |
| MVP_top60_DNA_seed | EGFR | 0.264668 | SKCM | UACC-62 | PF-00299804 |
| MVP_top60_DNA_seed | ERBB2 | 0.264668 | SKCM | UACC-62 | PF-00299804 |
| MVP_top60_DNA_seed | EGFR | 0.264782 | SKCM | G-MEL | Afatinib |
| MVP_top60_DNA_seed | ERBB2 | 0.264782 | SKCM | G-MEL | Afatinib |
| MVP_top60_DNA_seed | EGFR | 0.266299 | SKCM | UACC-257 | Afatinib |
| MVP_top60_DNA_seed | ERBB2 | 0.266299 | SKCM | UACC-257 | Afatinib |
| MVP_top60_DNA_seed | EGFR | 0.266752 | SKCM | WM793B | CUDC-101 |
| MVP_top60_DNA_seed | ERBB2 | 0.266752 | SKCM | WM793B | CUDC-101 |
| MVP_top60_DNA_seed | ERBB2 | 0.267128 | SKCM | MZ7-mel | CP724714 |
| MVP_top60_DNA_seed | JAK3 | 0.268889 | SKCM | MZ7-mel | WHI-P97 |
| MVP_top60_DNA_seed | JAK1 | 0.26913 | SKCM | LOXIMVI | JAK1_3715 |
| MVP_top60_DNA_seed | GSK3B | 0.269995 | SKCM | WM1552C | SB216763 |
| MVP_top60_DNA_seed | JAK1 | 0.270253 | SKCM | IPC-298 | JAK1_3715 |
| MVP_top60_DNA_seed | BRAF | 0.274802 | SKCM | LB2518-MEL | HG6-64-1 |
| MVP_top60_DNA_seed | BRAF | 0.275418 | SKCM | UACC-257 | HG6-64-1 |
| MVP_top60_DNA_seed | EGFR | 0.275562 | SKCM | A375 | Afatinib |
| MVP_top60_DNA_seed | ERBB2 | 0.275562 | SKCM | A375 | Afatinib |
| MVP_top60_DNA_seed | JAK1 | 0.276476 | SKCM | IGR-37 | JAK1_3715 |
| MVP_top60_DNA_seed | GSK3B | 0.280973 | SKCM | SK-MEL-2 | KIN001-042 |
| MVP_top60_DNA_seed | EGFR | 0.284083 | SKCM | MMAc-SF | Lapatinib |
| MVP_top60_DNA_seed | ERBB2 | 0.284083 | SKCM | MMAc-SF | Lapatinib |
| MVP_top60_DNA_seed | EGFR | 0.284996 | SKCM | MEL-JUSO | AZD8931 |
| MVP_top60_DNA_seed | ERBB2 | 0.284996 | SKCM | MEL-JUSO | AZD8931 |
| MVP_top60_DNA_seed | EGFR | 0.285941 | SKCM | CP66-MEL | AST-1306 |
| MVP_top60_DNA_seed | MAPK14 | 0.286178 | SKCM | MMAc-SF | TAK-715 |
| MVP_top60_DNA_seed | PIK3CB | 0.287747 | SKCM | IPC-298 | AZD6482 |
| MVP_top60_DNA_seed | JAK2 | 0.289102 | SKCM | WM35 | Ruxolitinib |
| MVP_top60_DNA_seed | JAK1 | 0.289102 | SKCM | WM35 | Ruxolitinib |
| MVP_top60_DNA_seed | PIK3CB | 0.290468 | SKCM | A375 | AZD6482 |
| MVP_top60_DNA_seed | GSK3B | 0.291853 | SKCM | G-MEL | CHIR-99021 |
| MVP_top60_DNA_seed | EGFR | 0.292097 | SKCM | A101D | Cetuximab |
| MVP_top60_DNA_seed | PIK3CB | 0.295922 | SKCM | A375 | AZD8186 |
| MVP_top60_DNA_seed | JAK3 | 0.297088 | SKCM | COLO-679 | WHI-P97 |
| MVP_top60_DNA_seed | EGFR | 0.298381 | SKCM | MMAc-SF | CUDC-101 |
| MVP_top60_DNA_seed | ERBB2 | 0.298381 | SKCM | MMAc-SF | CUDC-101 |
| MVP_top60_DNA_seed | MAPK3 | 0.298886 | SKCM | WM-115 | FR-180204 |
| MVP_top60_DNA_seed | MAPK1 | 0.298886 | SKCM | WM-115 | FR-180204 |
| MVP_top60_DNA_seed | GSK3B | 0.300106 | SKCM | IPC-298 | KIN001-042 |
| MVP_top60_DNA_seed | GSK3B | 0.300958 | SKCM | 451Lu | CHIR-99021 |

| | | | | | |
|---|---|---|---|---|---|
| MVP_top60_DNA_seed | EGFR | 0.301374 | SKCM | COLO-829 | AZD8931 |
| MVP_top60_DNA_seed | ERBB2 | 0.301374 | SKCM | COLO-829 | AZD8931 |
| MVP_top60_DNA_seed | EGFR | 0.30306 | SKCM | SK-MEL-30 | Pelitinib |
| MVP_top60_DNA_seed | EGFR | 0.308371 | SKCM | GAK | CUDC-101 |
| MVP_top60_DNA_seed | ERBB2 | 0.308371 | SKCM | GAK | CUDC-101 |
| MVP_top60_DNA_seed | EGFR | 0.309951 | SKCM | COLO-800 | Pelitinib |
| MVP_top60_DNA_seed | JAK3 | 0.31063 | SKCM | IPC-298 | WHI-P97 |
| MVP_top60_DNA_seed | GSK3B | 0.311609 | SKCM | MEL-HO | KIN001-042 |
| MVP_top60_DNA_seed | EGFR | 0.31194 | SKCM | WM35 | Afatinib |
| MVP_top60_DNA_seed | ERBB2 | 0.31194 | SKCM | WM35 | Afatinib |
| MVP_top60_DNA_seed | MAP2K1 | 0.313057 | SKCM | UACC-62 | CI-1040 |
| MVP_top60_DNA_seed | EGFR | 0.313785 | SKCM | SK-MEL-24 | Afatinib |
| MVP_top60_DNA_seed | ERBB2 | 0.313785 | SKCM | SK-MEL-24 | Afatinib |
| MVP_top60_DNA_seed | EGFR | 0.31489 | SKCM | UACC-62 | CUDC-101 |
| MVP_top60_DNA_seed | ERBB2 | 0.31489 | SKCM | UACC-62 | CUDC-101 |
| MVP_top60_DNA_seed | EGFR | 0.315836 | SKCM | MEL-HO | Gefitinib |
| MVP_top60_DNA_seed | GSK3B | 0.316066 | SKCM | SK-MEL-5 | CHIR-99021 |
| MVP_top60_DNA_seed | MAPK3 | 0.31678 | SKCM | WM1552C | FR-180204 |
| MVP_top60_DNA_seed | MAPK1 | 0.31678 | SKCM | WM1552C | FR-180204 |
| MVP_top60_DNA_seed | MAPK14 | 0.317484 | SKCM | HT-144 | TAK-715 |
| MVP_top60_DNA_seed | MAPK14 | 0.317509 | SKCM | SK-MEL-24 | TAK-715 |
| MVP_top60_DNA_seed | MAPK14 | 0.319729 | SKCM | LOXIMVI | TAK-715 |
| MVP_top60_DNA_seed | RAC1 | 0.320038 | SKCM | SK-MEL-30 | EHT-1864 |
| MVP_top60_DNA_seed | EGFR | 0.32188 | SKCM | COLO-800 | Cetuximab |
| MVP_top60_DNA_seed | EGFR | 0.322135 | SKCM | UACC-62 | Afatinib |
| MVP_top60_DNA_seed | ERBB2 | 0.322135 | SKCM | UACC-62 | Afatinib |
| MVP_top60_DNA_seed | EGFR | 0.322736 | SKCM | CHL-1 | Cetuximab |
| MVP_top60_DNA_seed | EGFR | 0.324552 | SKCM | G-MEL | Cetuximab |
| MVP_top60_DNA_seed | EGFR | 0.325625 | SKCM | MMAc-SF | AZD8931 |
| MVP_top60_DNA_seed | ERBB2 | 0.325625 | SKCM | MMAc-SF | AZD8931 |
| MVP_top60_DNA_seed | JAK2 | 0.32789 | SKCM | SK-MEL-24 | Ruxolitinib |
| MVP_top60_DNA_seed | JAK1 | 0.32789 | SKCM | SK-MEL-24 | Ruxolitinib |
| MVP_top60_DNA_seed | BRAF | 0.330204 | SKCM | LB373-MEL-D | RAF_9304 |
| MVP_top60_DNA_seed | GSK3B | 0.330303 | SKCM | COLO-800 | KIN001-042 |
| MVP_top60_DNA_seed | EP300 | 0.330755 | SKCM | SK-MEL-28 | I-CBP112 |
| MVP_top60_DNA_seed | CREBBP | 0.330755 | SKCM | SK-MEL-28 | I-CBP112 |
| MVP_top60_DNA_seed | EGFR | 0.331135 | SKCM | COLO-829 | CI-1033 |
| MVP_top60_DNA_seed | ERBB2 | 0.331135 | SKCM | COLO-829 | CI-1033 |
| MVP_top60_DNA_seed | EGFR | 0.332906 | SKCM | M14 | AST-1306 |
| MVP_top60_DNA_seed | JAK2 | 0.332941 | SKCM | 451Lu | Ruxolitinib |
| MVP_top60_DNA_seed | JAK1 | 0.332941 | SKCM | 451Lu | Ruxolitinib |

| | | | | | |
|---|---|---|---|---|---|
| MVP_top60_DNA_seed | EGFR | 0.333509 | SKCM | RVH-421 | PF-00299804 |
| MVP_top60_DNA_seed | ERBB2 | 0.333509 | SKCM | RVH-421 | PF-00299804 |
| MVP_top60_DNA_seed | MAP2K1 | 0.334916 | SKCM | A375 | Selumetinib |
| MVP_top60_DNA_seed | EGFR | 0.335071 | SKCM | LB373-MEL-D | Cetuximab |
| MVP_top60_DNA_seed | RAC1 | 0.337059 | SKCM | COLO-829 | EHT-1864 |
| MVP_top60_DNA_seed | BRAF | 0.338068 | SKCM | GAK | HG6-64-1 |
| MVP_top60_DNA_seed | MAPK3 | 0.338205 | SKCM | UACC-257 | FR-180204 |
| MVP_top60_DNA_seed | MAPK1 | 0.338205 | SKCM | UACC-257 | FR-180204 |
| MVP_top60_DNA_seed | EGFR | 0.34158 | SKCM | A375 | Afatinib |
| MVP_top60_DNA_seed | ERBB2 | 0.34158 | SKCM | A375 | Afatinib |
| MVP_top60_DNA_seed | JAK1 | 0.341929 | SKCM | G-361 | JAK1_3715 |
| MVP_top60_DNA_seed | GSK3B | 0.343936 | SKCM | GAK | CHIR-99021 |
| MVP_top60_DNA_seed | EGFR | 0.344439 | SKCM | SK-MEL-5 | Cetuximab |
| MVP_top60_DNA_seed | PIK3CA | 0.34451 | SKCM | VMRC-MELG | PI3Ka_4409 |
| MVP_top60_DNA_seed | JAK3 | 0.346155 | SKCM | IGR-1 | WHI-P97 |
| MVP_top60_DNA_seed | PIK3CA | 0.346387 | SKCM | Hs-939-T | PI3Ka_4409 |
| MVP_top60_DNA_seed | JAK2 | 0.350074 | SKCM | SK-MEL-2 | AZD1480 |
| MVP_top60_DNA_seed | JAK1 | 0.350074 | SKCM | SK-MEL-2 | AZD1480 |
| MVP_top60_DNA_seed | GSK3B | 0.35045 | SKCM | MEL-JUSO | CHIR-99021 |
| MVP_top60_DNA_seed | ERBB2 | 0.350671 | SKCM | UACC-257 | CP724714 |
| MVP_top60_DNA_seed | MAPK3 | 0.3516 | SKCM | COLO-800 | FR-180204 |
| MVP_top60_DNA_seed | MAPK1 | 0.3516 | SKCM | COLO-800 | FR-180204 |
| MVP_top60_DNA_seed | EGFR | 0.3526 | SKCM | MZ7-mel | Pelitinib |
| MVP_top60_DNA_seed | BRAF | 0.35273 | SKCM | VMRC-MELG | PLX-4720 |
| MVP_top60_DNA_seed | JAK2 | 0.352746 | SKCM | SK-MEL-2 | Ruxolitinib |
| MVP_top60_DNA_seed | JAK1 | 0.352746 | SKCM | SK-MEL-2 | Ruxolitinib |
| MVP_top60_DNA_seed | MAPK3 | 0.355914 | SKCM | LB373-MEL-D | FR-180204 |
| MVP_top60_DNA_seed | MAPK1 | 0.355914 | SKCM | LB373-MEL-D | FR-180204 |
| MVP_top60_DNA_seed | EGFR | 0.358205 | SKCM | LB373-MEL-D | Lapatinib |
| MVP_top60_DNA_seed | ERBB2 | 0.358205 | SKCM | LB373-MEL-D | Lapatinib |
| MVP_top60_DNA_seed | BRAF | 0.360966 | SKCM | GAK | SB590885 |
| MVP_top60_DNA_seed | JAK2 | 0.361728 | SKCM | COLO-679 | AZD1480 |
| MVP_top60_DNA_seed | JAK1 | 0.361728 | SKCM | COLO-679 | AZD1480 |
| MVP_top60_DNA_seed | GSK3B | 0.361882 | SKCM | G-361 | KIN001-042 |
| MVP_top60_DNA_seed | GSK3B | 0.364915 | SKCM | MZ2-MEL | AZD7969 |
| MVP_top60_DNA_seed | PIK3CA | 0.365091 | SKCM | COLO-679 | AZD8835 |
| MVP_top60_DNA_seed | JAK2 | 0.370717 | SKCM | A2058 | Ruxolitinib |
| MVP_top60_DNA_seed | JAK1 | 0.370717 | SKCM | A2058 | Ruxolitinib |
| MVP_top60_DNA_seed | GSK3B | 0.372402 | SKCM | LB2518-MEL | AZD7969 |
| MVP_top60_DNA_seed | GSK3B | 0.372437 | SKCM | 451Lu | AZD7969 |
| MVP_top60_DNA_seed | EGFR | 0.372709 | SKCM | IST-MEL1 | Cetuximab |

| | | | | | |
|---|---|---|---|---|---|
| MVP_top60_DNA_seed | JAK1 | 0.373226 | SKCM | Hs-939-T | JAK1_3715 |
| MVP_top60_DNA_seed | GSK3B | 0.373247 | SKCM | WM1552C | CHIR-99021 |
| MVP_top60_DNA_seed | EGFR | 0.373976 | SKCM | A2058 | Pelitinib |
| MVP_top60_DNA_seed | EGFR | 0.374181 | SKCM | IPC-298 | Afatinib |
| MVP_top60_DNA_seed | ERBB2 | 0.374181 | SKCM | IPC-298 | Afatinib |
| MVP_top60_DNA_seed | JAK2 | 0.375624 | SKCM | MEL-JUSO | Ruxolitinib |
| MVP_top60_DNA_seed | JAK1 | 0.375624 | SKCM | MEL-JUSO | Ruxolitinib |
| MVP_top60_DNA_seed | EGFR | 0.376165 | SKCM | WM35 | Pelitinib |
| MVP_top60_DNA_seed | GSK3B | 0.376573 | SKCM | LB2518-MEL | SB216763 |
| MVP_top60_DNA_seed | EGFR | 0.37769 | SKCM | UACC-257 | CI-1033 |
| MVP_top60_DNA_seed | ERBB2 | 0.37769 | SKCM | UACC-257 | CI-1033 |
| MVP_top60_DNA_seed | JAK2 | 0.38187 | SKCM | SK-MEL-5 | Fedratinib |
| MVP_top60_DNA_seed | BRAF | 0.383017 | SKCM | CP66-MEL | SB590885 |
| MVP_top60_DNA_seed | GSK3B | 0.383951 | SKCM | SK-MEL-3 | AZD7969 |
| MVP_top60_DNA_seed | PIK3CA | 0.38519 | SKCM | SK-MEL-24 | PI3Ka_4409 |
| MVP_top60_DNA_seed | EGFR | 0.386726 | SKCM | WM-115 | Gefitinib |
| MVP_top60_DNA_seed | EP300 | 0.387078 | SKCM | SK-MEL-3 | I-CBP112 |
| MVP_top60_DNA_seed | CREBBP | 0.387078 | SKCM | SK-MEL-3 | I-CBP112 |
| MVP_top60_DNA_seed | EGFR | 0.388118 | SKCM | K2 | CUDC-101 |
| MVP_top60_DNA_seed | ERBB2 | 0.388118 | SKCM | K2 | CUDC-101 |
| MVP_top60_DNA_seed | EP300 | 0.389509 | SKCM | MEL-JUSO | I-CBP112 |
| MVP_top60_DNA_seed | CREBBP | 0.389509 | SKCM | MEL-JUSO | I-CBP112 |
| MVP_top60_DNA_seed | EGFR | 0.390158 | SKCM | IGR-1 | Afatinib |
| MVP_top60_DNA_seed | ERBB2 | 0.390158 | SKCM | IGR-1 | Afatinib |
| MVP_top60_DNA_seed | ERBB2 | 0.390216 | SKCM | Mewo | CP724714 |
| MVP_top60_DNA_seed | MAPK1 | 0.391463 | SKCM | COLO-800 | VX-11e |
| MVP_top60_DNA_seed | JAK3 | 0.392113 | SKCM | Hs-939-T | JAK3_7406 |
| MVP_top60_DNA_seed | EGFR | 0.39245 | SKCM | SK-MEL-1 | CUDC-101 |
| MVP_top60_DNA_seed | ERBB2 | 0.39245 | SKCM | SK-MEL-1 | CUDC-101 |
| MVP_top60_DNA_seed | GSK3B | 0.393067 | SKCM | A2058 | KIN001-042 |
| MVP_top60_DNA_seed | JAK1 | 0.393124 | SKCM | MEL-HO | JAK1_3715 |
| MVP_top60_DNA_seed | GSK3B | 0.395163 | SKCM | CP50-MEL-B | CHIR-99021 |
| MVP_top60_DNA_seed | MAPK1 | 0.395779 | SKCM | UACC-62 | VX-11e |
| MVP_top60_DNA_seed | EGFR | 0.39634 | SKCM | COLO-829 | Afatinib |
| MVP_top60_DNA_seed | ERBB2 | 0.39634 | SKCM | COLO-829 | Afatinib |
| MVP_top60_DNA_seed | EGFR | 0.397707 | SKCM | IST-MEL1 | Afatinib |
| MVP_top60_DNA_seed | ERBB2 | 0.397707 | SKCM | IST-MEL1 | Afatinib |
| MVP_top60_DNA_seed | EGFR | 0.398834 | SKCM | MZ2-MEL | Erlotinib |
| MVP_top60_DNA_seed | EGFR | 0.399002 | SKCM | G-361 | Cetuximab |
| MVP_top60_DNA_seed | EGFR | 0.399239 | SKCM | IST-MEL1 | Erlotinib |
| MVP_top60_DNA_seed | JAK2 | 0.399592 | SKCM | UACC-62 | AZD1480 |

| | | | | | |
|---|---|---|---|---|---|
| MVP_top60_DNA_seed | JAK1 | 0.399592 | SKCM | UACC-62 | AZD1480 |
| MVP_top60_DNA_seed | PIK3CB | 0.400103 | SKCM | LOXIMVI | TGX221 |
| MVP_top60_DNA_seed | MAP2K1 | 0.400818 | SKCM | VMRC-MELG | Selumetinib |
| MVP_top60_DNA_seed | EGFR | 0.401702 | SKCM | COLO-829 | Erlotinib |
| MVP_top60_DNA_seed | JAK2 | 0.401942 | SKCM | VMRC-MELG | Ruxolitinib |
| MVP_top60_DNA_seed | JAK1 | 0.401942 | SKCM | VMRC-MELG | Ruxolitinib |
| MVP_top60_DNA_seed | ERBB2 | 0.402774 | SKCM | IGR-37 | CP724714 |
| MVP_top60_DNA_seed | MAP2K1 | 0.402807 | SKCM | IGR-1 | CI-1040 |
| MVP_top60_DNA_seed | JAK2 | 0.406964 | SKCM | WM278 | Ruxolitinib |
| MVP_top60_DNA_seed | JAK1 | 0.406964 | SKCM | WM278 | Ruxolitinib |
| MVP_top60_DNA_seed | PIK3CB | 0.406997 | SKCM | WM793B | AZD8186 |
| MVP_top60_DNA_seed | EGFR | 0.408484 | SKCM | Mewo | Afatinib |
| MVP_top60_DNA_seed | ERBB2 | 0.408484 | SKCM | Mewo | Afatinib |
| MVP_top60_DNA_seed | EP300 | 0.408749 | SKCM | WM793B | I-CBP112 |
| MVP_top60_DNA_seed | CREBBP | 0.408749 | SKCM | WM793B | I-CBP112 |
| MVP_top60_DNA_seed | ERBB2 | 0.410049 | SKCM | COLO-679 | CP724714 |
| MVP_top60_DNA_seed | JAK1 | 0.412961 | SKCM | WM35 | JAK1_3715 |
| MVP_top60_DNA_seed | JAK3 | 0.413134 | SKCM | COLO-800 | WHI-P97 |
| MVP_top60_DNA_seed | PIK3CB | 0.413152 | SKCM | COLO-783 | AZD8186 |
| MVP_top60_DNA_seed | JAK3 | 0.4168 | SKCM | WM35 | WHI-P97 |
| MVP_top60_DNA_seed | MAPK3 | 0.418198 | SKCM | CP66-MEL | FR-180204 |
| MVP_top60_DNA_seed | MAPK1 | 0.418198 | SKCM | CP66-MEL | FR-180204 |
| MVP_top60_DNA_seed | MAP2K1 | 0.420119 | SKCM | SK-MEL-5 | Selumetinib |
| MVP_top60_DNA_seed | PIK3CA | 0.421412 | SKCM | RPMI-7951 | PI3Ka_4409 |
| MVP_top60_DNA_seed | EGFR | 0.424515 | SKCM | WM-115 | AST-1306 |
| MVP_top60_DNA_seed | GSK3B | 0.424729 | SKCM | MMAc-SF | CHIR-99021 |
| MVP_top60_DNA_seed | GSK3B | 0.427201 | SKCM | HMV-II | SB216763 |
| MVP_top60_DNA_seed | MAP2K1 | 0.429075 | SKCM | UACC-62 | PD0325901 |
| MVP_top60_DNA_seed | EP300 | 0.429507 | SKCM | MZ7-mel | I-CBP112 |
| MVP_top60_DNA_seed | CREBBP | 0.429507 | SKCM | MZ7-mel | I-CBP112 |
| MVP_top60_DNA_seed | ERBB2 | 0.431181 | SKCM | LB373-MEL-D | CP724714 |
| MVP_top60_DNA_seed | GSK3B | 0.43161 | SKCM | CP66-MEL | CHIR-99021 |
| MVP_top60_DNA_seed | EGFR | 0.431681 | SKCM | COLO-800 | Gefitinib |
| MVP_top60_DNA_seed | MAPK3 | 0.431685 | SKCM | HMV-II | FR-180204 |
| MVP_top60_DNA_seed | MAPK1 | 0.431685 | SKCM | HMV-II | FR-180204 |
| MVP_top60_DNA_seed | PIK3CA | 0.431706 | SKCM | M14 | AZD8835 |
| MVP_top60_DNA_seed | EGFR | 0.431817 | SKCM | WM-115 | Afatinib |
| MVP_top60_DNA_seed | ERBB2 | 0.431817 | SKCM | WM-115 | Afatinib |
| MVP_top60_DNA_seed | PIK3CA | 0.433131 | SKCM | A101D | PI3Ka_4409 |
| MVP_top60_DNA_seed | JAK3 | 0.433289 | SKCM | C32 | JAK3_7406 |
| MVP_top60_DNA_seed | JAK2 | 0.433802 | SKCM | SK-MEL-3 | Ruxolitinib |

| | | | | | |
|---|---|---|---|---|---|
| MVP_top60_DNA_seed | JAK1 | 0.433802 | SKCM | SK-MEL-3 | Ruxolitinib |
| MVP_top60_DNA_seed | EGFR | 0.434658 | SKCM | Hs-939-T | Cetuximab |
| MVP_top60_DNA_seed | EGFR | 0.435358 | SKCM | LB2518-MEL | AZD8931 |
| MVP_top60_DNA_seed | ERBB2 | 0.435358 | SKCM | LB2518-MEL | AZD8931 |
| MVP_top60_DNA_seed | MAPK3 | 0.435451 | SKCM | UACC-62 | FR-180204 |
| MVP_top60_DNA_seed | MAPK1 | 0.435451 | SKCM | UACC-62 | FR-180204 |
| MVP_top60_DNA_seed | EGFR | 0.437109 | SKCM | SK-MEL-28 | Afatinib |
| MVP_top60_DNA_seed | ERBB2 | 0.437109 | SKCM | SK-MEL-28 | Afatinib |
| MVP_top60_DNA_seed | PIK3CB | 0.438878 | SKCM | WM35 | AZD8186 |
| MVP_top60_DNA_seed | JAK3 | 0.43916 | SKCM | Hs-939-T | WHI-P97 |
| MVP_top60_DNA_seed | EGFR | 0.440139 | SKCM | CP50-MEL-B | Afatinib |
| MVP_top60_DNA_seed | ERBB2 | 0.440139 | SKCM | CP50-MEL-B | Afatinib |
| MVP_top60_DNA_seed | EGFR | 0.440518 | SKCM | IPC-298 | Cetuximab |
| MVP_top60_DNA_seed | BRAF | 0.442011 | SKCM | LB2518-MEL | Dabrafenib |
| MVP_top60_DNA_seed | MAPK14 | 0.443417 | SKCM | SK-MEL-5 | TAK-715 |
| MVP_top60_DNA_seed | ERBB2 | 0.443803 | SKCM | 451Lu | CP724714 |
| MVP_top60_DNA_seed | EGFR | 0.444075 | SKCM | COLO-783 | Afatinib |
| MVP_top60_DNA_seed | ERBB2 | 0.444075 | SKCM | COLO-783 | Afatinib |
| MVP_top60_DNA_seed | GSK3B | 0.44547 | SKCM | A2058 | SB216763 |
| MVP_top60_DNA_seed | BRAF | 0.446103 | SKCM | MZ2-MEL | RAF_9304 |
| MVP_top60_DNA_seed | EGFR | 0.446428 | SKCM | SK-MEL-2 | CUDC-101 |
| MVP_top60_DNA_seed | ERBB2 | 0.446428 | SKCM | SK-MEL-2 | CUDC-101 |
| MVP_top60_DNA_seed | EGFR | 0.446479 | SKCM | SK-MEL-24 | Gefitinib |
| MVP_top60_DNA_seed | EGFR | 0.447658 | SKCM | SH-4 | Pelitinib |
| MVP_top60_DNA_seed | EGFR | 0.449563 | SKCM | COLO-679 | Afatinib |
| MVP_top60_DNA_seed | ERBB2 | 0.449563 | SKCM | COLO-679 | Afatinib |
| MVP_top60_DNA_seed | JAK2 | 0.450871 | SKCM | SH-4 | Ruxolitinib |
| MVP_top60_DNA_seed | JAK1 | 0.450871 | SKCM | SH-4 | Ruxolitinib |
| MVP_top60_DNA_seed | JAK2 | 0.452047 | SKCM | IST-MEL1 | Fedratinib |
| MVP_top60_DNA_seed | GSK3B | 0.453885 | SKCM | Mewo | AZD7969 |
| MVP_top60_DNA_seed | JAK2 | 0.454631 | SKCM | M14 | Ruxolitinib |
| MVP_top60_DNA_seed | JAK1 | 0.454631 | SKCM | M14 | Ruxolitinib |
| MVP_top60_DNA_seed | EGFR | 0.455025 | SKCM | UACC-62 | Gefitinib |
| MVP_top60_DNA_seed | EGFR | 0.456115 | SKCM | WM793B | Afatinib |
| MVP_top60_DNA_seed | ERBB2 | 0.456115 | SKCM | WM793B | Afatinib |
| MVP_top60_DNA_seed | EGFR | 0.45673 | SKCM | LB373-MEL-D | AZD8931 |
| MVP_top60_DNA_seed | ERBB2 | 0.45673 | SKCM | LB373-MEL-D | AZD8931 |
| MVP_top60_DNA_seed | EGFR | 0.457838 | SKCM | WM1552C | Afatinib |
| MVP_top60_DNA_seed | ERBB2 | 0.457838 | SKCM | WM1552C | Afatinib |
| MVP_top60_DNA_seed | EGFR | 0.45797 | SKCM | GAK | Afatinib |
| MVP_top60_DNA_seed | ERBB2 | 0.45797 | SKCM | GAK | Afatinib |

| | | | | | |
|---|---|---|---|---|---|
| MVP_top60_DNA_seed | EGFR | 0.459173 | SKCM | GAK | Afatinib |
| MVP_top60_DNA_seed | ERBB2 | 0.459173 | SKCM | GAK | Afatinib |
| MVP_top60_DNA_seed | JAK3 | 0.460874 | SKCM | SK-MEL-5 | JAK3_7406 |
| MVP_top60_DNA_seed | EGFR | 0.461306 | SKCM | WM278 | CUDC-101 |
| MVP_top60_DNA_seed | ERBB2 | 0.461306 | SKCM | WM278 | CUDC-101 |
| MVP_top60_DNA_seed | JAK1 | 0.461983 | SKCM | MMAc-SF | JAK1_3715 |
| MVP_top60_DNA_seed | GSK3B | 0.46219 | SKCM | MEL-JUSO | AZD7969 |
| MVP_top60_DNA_seed | JAK2 | 0.463748 | SKCM | SK-MEL-30 | Fedratinib |
| MVP_top60_DNA_seed | EGFR | 0.465082 | SKCM | MEL-JUSO | Cetuximab |
| MVP_top60_DNA_seed | PIK3CA | 0.46513 | SKCM | HT-144 | PI3Ka_4409 |
| MVP_top60_DNA_seed | PIK3CB | 0.465448 | SKCM | IGR-1 | AZD6482 |
| MVP_top60_DNA_seed | ERBB2 | 0.468154 | SKCM | LOXIMVI | CP724714 |
| MVP_top60_DNA_seed | GSK3B | 0.468274 | SKCM | Hs-939-T | AZD7969 |
| MVP_top60_DNA_seed | PIK3CA | 0.46856 | SKCM | HT-144 | AZD8835 |
| MVP_top60_DNA_seed | GSK3B | 0.468683 | SKCM | MEL-HO | SB216763 |
| MVP_top60_DNA_seed | EGFR | 0.469675 | SKCM | COLO-800 | Lapatinib |
| MVP_top60_DNA_seed | ERBB2 | 0.469675 | SKCM | COLO-800 | Lapatinib |
| MVP_top60_DNA_seed | BRAF | 0.470943 | SKCM | SK-MEL-30 | PLX-4720 |
| MVP_top60_DNA_seed | PIK3CB | 0.472322 | SKCM | SH-4 | TGX221 |
| MVP_top60_DNA_seed | EGFR | 0.472352 | SKCM | LOXIMVI | Afatinib |
| MVP_top60_DNA_seed | ERBB2 | 0.472352 | SKCM | LOXIMVI | Afatinib |
| MVP_top60_DNA_seed | JAK1 | 0.473687 | SKCM | HT-144 | JAK1_3715 |
| MVP_top60_DNA_seed | GSK3B | 0.473835 | SKCM | COLO-829 | SB216763 |
| MVP_top60_DNA_seed | MAPK14 | 0.474203 | SKCM | WM278 | TAK-715 |
| MVP_top60_DNA_seed | ERBB2 | 0.477233 | SKCM | CP66-MEL | CP724714 |
| MVP_top60_DNA_seed | EGFR | 0.479116 | SKCM | IST-MEL1 | PF-00299804 |
| MVP_top60_DNA_seed | ERBB2 | 0.479116 | SKCM | IST-MEL1 | PF-00299804 |
| MVP_top60_DNA_seed | JAK2 | 0.479298 | SKCM | HT-144 | AZD1480 |
| MVP_top60_DNA_seed | JAK1 | 0.479298 | SKCM | HT-144 | AZD1480 |
| MVP_top60_DNA_seed | MAPK14 | 0.480199 | SKCM | COLO-783 | TAK-715 |
| MVP_top60_DNA_seed | PIK3CA | 0.480823 | SKCM | Hs-940-T | PI3Ka_4409 |
| MVP_top60_DNA_seed | PIK3CA | 0.481285 | SKCM | WM-115 | AZD8835 |
| MVP_top60_DNA_seed | EGFR | 0.481705 | SKCM | A2058 | Afatinib |
| MVP_top60_DNA_seed | ERBB2 | 0.481705 | SKCM | A2058 | Afatinib |
| MVP_top60_DNA_seed | EGFR | 0.483348 | SKCM | Mewo | Cetuximab |
| MVP_top60_DNA_seed | JAK3 | 0.48347 | SKCM | M14 | JAK3_7406 |
| MVP_top60_DNA_seed | BRAF | 0.485086 | SKCM | UACC-62 | PLX-4720 |
| MVP_top60_DNA_seed | GSK3B | 0.485184 | SKCM | WM35 | KIN001-042 |
| MVP_top60_DNA_seed | JAK2 | 0.485243 | SKCM | A2058 | Fedratinib |
| MVP_top60_DNA_seed | BRAF | 0.486652 | SKCM | SK-MEL-31 | RAF_9304 |
| MVP_top60_DNA_seed | PIK3CA | 0.487149 | SKCM | IST-MEL1 | PI3Ka_4409 |

| | | | | | |
|---|---|---|---|---|---|
| MVP_top60_DNA_seed | JAK3 | 0.487686 | SKCM | UACC-62 | WHI-P97 |
| MVP_top60_DNA_seed | MAPK14 | 0.488149 | SKCM | A101D | TAK-715 |
| MVP_top60_DNA_seed | PIK3CB | 0.488687 | SKCM | IST-MEL1 | AZD8186 |
| MVP_top60_DNA_seed | PIK3CB | 0.489583 | SKCM | COLO-792 | AZD6482 |
| MVP_top60_DNA_seed | EGFR | 0.4898 | SKCM | MZ7-mel | Lapatinib |
| MVP_top60_DNA_seed | ERBB2 | 0.4898 | SKCM | MZ7-mel | Lapatinib |
| MVP_top60_DNA_seed | JAK3 | 0.490351 | SKCM | COLO-783 | WHI-P97 |
| MVP_top60_DNA_seed | EGFR | 0.49151 | SKCM | GAK | CI-1033 |
| MVP_top60_DNA_seed | ERBB2 | 0.49151 | SKCM | GAK | CI-1033 |
| MVP_top60_DNA_seed | JAK2 | 0.491686 | SKCM | IST-MEL1 | AZD1480 |
| MVP_top60_DNA_seed | JAK1 | 0.491686 | SKCM | IST-MEL1 | AZD1480 |
| MVP_top60_DNA_seed | EGFR | 0.492689 | SKCM | CP50-MEL-B | Cetuximab |
| MVP_top60_DNA_seed | EGFR | 0.494801 | SKCM | COLO-800 | Erlotinib |
| MVP_top60_DNA_seed | EGFR | 0.495033 | SKCM | LOXIMVI | Afatinib |
| MVP_top60_DNA_seed | ERBB2 | 0.495033 | SKCM | LOXIMVI | Afatinib |
| MVP_top60_DNA_seed | EGFR | 0.495981 | SKCM | 451Lu | PF-00299804 |
| MVP_top60_DNA_seed | ERBB2 | 0.495981 | SKCM | 451Lu | PF-00299804 |
| MVP_top60_DNA_seed | EGFR | 0.501834 | SKCM | SK-MEL-2 | PF-00299804 |
| MVP_top60_DNA_seed | ERBB2 | 0.501834 | SKCM | SK-MEL-2 | PF-00299804 |
| MVP_top60_DNA_seed | GSK3B | 0.504961 | SKCM | SK-MEL-30 | KIN001-042 |
| MVP_top60_DNA_seed | BRAF | 0.506954 | SKCM | SK-MEL-31 | Dabrafenib |
| MVP_top60_DNA_seed | EGFR | 0.507061 | SKCM | A2058 | PF-00299804 |
| MVP_top60_DNA_seed | ERBB2 | 0.507061 | SKCM | A2058 | PF-00299804 |
| MVP_top60_DNA_seed | RAC1 | 0.507125 | SKCM | RPMI-7951 | EHT-1864 |
| MVP_top60_DNA_seed | JAK1 | 0.510789 | SKCM | UACC-62 | JAK1_3715 |
| MVP_top60_DNA_seed | GSK3B | 0.511381 | SKCM | COLO-829 | AZD7969 |
| MVP_top60_DNA_seed | EGFR | 0.51165 | SKCM | GAK | Cetuximab |
| MVP_top60_DNA_seed | EGFR | 0.512215 | SKCM | Hs-939-T | CUDC-101 |
| MVP_top60_DNA_seed | ERBB2 | 0.512215 | SKCM | Hs-939-T | CUDC-101 |
| MVP_top60_DNA_seed | EGFR | 0.514103 | SKCM | SK-MEL-3 | Afatinib |
| MVP_top60_DNA_seed | ERBB2 | 0.514103 | SKCM | SK-MEL-3 | Afatinib |
| MVP_top60_DNA_seed | JAK2 | 0.51656 | SKCM | COLO-792 | Ruxolitinib |
| MVP_top60_DNA_seed | JAK1 | 0.51656 | SKCM | COLO-792 | Ruxolitinib |
| MVP_top60_DNA_seed | JAK2 | 0.521375 | SKCM | M14 | AZD1480 |
| MVP_top60_DNA_seed | JAK1 | 0.521375 | SKCM | M14 | AZD1480 |
| MVP_top60_DNA_seed | GSK3B | 0.522814 | SKCM | VMRC-MELG | KIN001-042 |
| MVP_top60_DNA_seed | EGFR | 0.523699 | SKCM | HMV-II | Gefitinib |
| MVP_top60_DNA_seed | JAK2 | 0.523797 | SKCM | COLO-679 | Fedratinib |
| MVP_top60_DNA_seed | JAK3 | 0.524044 | SKCM | A101D | WHI-P97 |
| MVP_top60_DNA_seed | ERBB2 | 0.525428 | SKCM | Hs-940-T | CP724714 |
| MVP_top60_DNA_seed | PIK3CA | 0.52545 | SKCM | WM1552C | AZD8835 |

| | | | | | |
|---|---|---|---|---|---|
| MVP_top60_DNA_seed | EGFR | 0.525647 | SKCM | SK-MEL-3 | CI-1033 |
| MVP_top60_DNA_seed | ERBB2 | 0.525647 | SKCM | SK-MEL-3 | CI-1033 |
| MVP_top60_DNA_seed | EGFR | 0.527763 | SKCM | MEL-HO | Afatinib |
| MVP_top60_DNA_seed | ERBB2 | 0.527763 | SKCM | MEL-HO | Afatinib |
| MVP_top60_DNA_seed | EGFR | 0.534032 | SKCM | HT-144 | Cetuximab |
| MVP_top60_DNA_seed | EGFR | 0.534068 | SKCM | COLO-783 | AZD8931 |
| MVP_top60_DNA_seed | ERBB2 | 0.534068 | SKCM | COLO-783 | AZD8931 |
| MVP_top60_DNA_seed | MAPK3 | 0.535143 | SKCM | LB2518-MEL | FR-180204 |
| MVP_top60_DNA_seed | MAPK1 | 0.535143 | SKCM | LB2518-MEL | FR-180204 |
| MVP_top60_DNA_seed | EP300 | 0.53581 | SKCM | LB2518-MEL | I-CBP112 |
| MVP_top60_DNA_seed | CREBBP | 0.53581 | SKCM | LB2518-MEL | I-CBP112 |
| MVP_top60_DNA_seed | GSK3B | 0.535902 | SKCM | Hs-940-T | KIN001-042 |
| MVP_top60_DNA_seed | PIK3CB | 0.538693 | SKCM | SK-MEL-3 | AZD8186 |
| MVP_top60_DNA_seed | GSK3B | 0.541863 | SKCM | VMRC-MELG | SB216763 |
| MVP_top60_DNA_seed | MAPK14 | 0.542667 | SKCM | K2 | TAK-715 |
| MVP_top60_DNA_seed | PIK3CB | 0.543492 | SKCM | C32 | AZD8186 |
| MVP_top60_DNA_seed | EGFR | 0.543783 | SKCM | RPMI-7951 | Gefitinib |
| MVP_top60_DNA_seed | EGFR | 0.543802 | SKCM | GAK | AST-1306 |
| MVP_top60_DNA_seed | GSK3B | 0.54384 | SKCM | SK-MEL-28 | SB216763 |
| MVP_top60_DNA_seed | RAC1 | 0.54423 | SKCM | Hs-940-T | EHT-1864 |
| MVP_top60_DNA_seed | MAPK3 | 0.546475 | SKCM | HT-144 | FR-180204 |
| MVP_top60_DNA_seed | MAPK1 | 0.546475 | SKCM | HT-144 | FR-180204 |
| MVP_top60_DNA_seed | MAPK3 | 0.546888 | SKCM | G-MEL | FR-180204 |
| MVP_top60_DNA_seed | MAPK1 | 0.546888 | SKCM | G-MEL | FR-180204 |
| MVP_top60_DNA_seed | EGFR | 0.547837 | SKCM | UACC-257 | Erlotinib |
| MVP_top60_DNA_seed | GSK3B | 0.548427 | SKCM | Mewo | CHIR-99021 |
| MVP_top60_DNA_seed | PIK3CB | 0.548615 | SKCM | COLO-792 | AZD8186 |
| MVP_top60_DNA_seed | MAP2K1 | 0.551901 | SKCM | Hs-940-T | Refametinib |
| MVP_top60_DNA_seed | JAK2 | 0.552114 | SKCM | UACC-257 | Fedratinib |
| MVP_top60_DNA_seed | MAPK14 | 0.552582 | SKCM | CP50-MEL-B | TAK-715 |
| MVP_top60_DNA_seed | PIK3CA | 0.55454 | SKCM | IST-MEL1 | AZD8835 |
| MVP_top60_DNA_seed | MAP2K1 | 0.55615 | SKCM | Hs-940-T | Trametinib |
| MVP_top60_DNA_seed | PIK3CB | 0.557446 | SKCM | CP66-MEL | TGX221 |
| MVP_top60_DNA_seed | JAK1 | 0.557981 | SKCM | VMRC-MELG | JAK1_3715 |
| MVP_top60_DNA_seed | MAPK3 | 0.558855 | SKCM | GAK | FR-180204 |
| MVP_top60_DNA_seed | MAPK1 | 0.558855 | SKCM | GAK | FR-180204 |
| MVP_top60_DNA_seed | GSK3B | 0.559362 | SKCM | IPC-298 | SB216763 |
| MVP_top60_DNA_seed | PIK3CB | 0.55941 | SKCM | MZ7-mel | AZD8186 |
| MVP_top60_DNA_seed | EGFR | 0.562658 | SKCM | SK-MEL-2 | Lapatinib |
| MVP_top60_DNA_seed | ERBB2 | 0.562658 | SKCM | SK-MEL-2 | Lapatinib |
| MVP_top60_DNA_seed | JAK1 | 0.564582 | SKCM | MEL-JUSO | JAK1_3715 |

| | | | | | |
|---|---|---|---|---|---|
| MVP_top60_DNA_seed | BRAF | 0.565339 | SKCM | MEL-JUSO | SB590885 |
| MVP_top60_DNA_seed | EGFR | 0.566298 | SKCM | SK-MEL-5 | Pelitinib |
| MVP_top60_DNA_seed | ERBB2 | 0.566444 | SKCM | SK-MEL-2 | CP724714 |
| MVP_top60_DNA_seed | EGFR | 0.567874 | SKCM | SK-MEL-3 | Afatinib |
| MVP_top60_DNA_seed | ERBB2 | 0.567874 | SKCM | SK-MEL-3 | Afatinib |
| MVP_top60_DNA_seed | MAPK14 | 0.568379 | SKCM | WM-115 | TAK-715 |
| MVP_top60_DNA_seed | GSK3B | 0.569193 | SKCM | Mewo | CHIR-99021 |
| MVP_top60_DNA_seed | EGFR | 0.569963 | SKCM | MEL-JUSO | Afatinib |
| MVP_top60_DNA_seed | ERBB2 | 0.569963 | SKCM | MEL-JUSO | Afatinib |
| MVP_top60_DNA_seed | EP300 | 0.570962 | SKCM | WM1552C | I-CBP112 |
| MVP_top60_DNA_seed | CREBBP | 0.570962 | SKCM | WM1552C | I-CBP112 |
| MVP_top60_DNA_seed | EGFR | 0.570975 | SKCM | WM1552C | Afatinib |
| MVP_top60_DNA_seed | ERBB2 | 0.570975 | SKCM | WM1552C | Afatinib |
| MVP_top60_DNA_seed | PIK3CB | 0.571824 | SKCM | IPC-298 | AZD8186 |
| MVP_top60_DNA_seed | PIK3CA | 0.572253 | SKCM | LB373-MEL-D | PI3Ka_4409 |
| MVP_top60_DNA_seed | RAC1 | 0.573872 | SKCM | UACC-62 | EHT-1864 |
| MVP_top60_DNA_seed | EGFR | 0.574059 | SKCM | MEL-HO | AZD8931 |
| MVP_top60_DNA_seed | ERBB2 | 0.574059 | SKCM | MEL-HO | AZD8931 |
| MVP_top60_DNA_seed | JAK2 | 0.574652 | SKCM | COLO-679 | Ruxolitinib |
| MVP_top60_DNA_seed | JAK1 | 0.574652 | SKCM | COLO-679 | Ruxolitinib |
| MVP_top60_DNA_seed | EGFR | 0.57484 | SKCM | SH-4 | Cetuximab |
| MVP_top60_DNA_seed | EP300 | 0.577581 | SKCM | COLO-679 | I-CBP112 |
| MVP_top60_DNA_seed | CREBBP | 0.577581 | SKCM | COLO-679 | I-CBP112 |
| MVP_top60_DNA_seed | JAK2 | 0.579281 | SKCM | UACC-257 | AZD1480 |
| MVP_top60_DNA_seed | JAK1 | 0.579281 | SKCM | UACC-257 | AZD1480 |
| MVP_top60_DNA_seed | PIK3CA | 0.581204 | SKCM | SK-MEL-2 | PI3Ka_4409 |
| MVP_top60_DNA_seed | EGFR | 0.582695 | SKCM | CP66-MEL | CI-1033 |
| MVP_top60_DNA_seed | ERBB2 | 0.582695 | SKCM | CP66-MEL | CI-1033 |
| MVP_top60_DNA_seed | GSK3B | 0.58384 | SKCM | COLO-679 | CHIR-99021 |
| MVP_top60_DNA_seed | EGFR | 0.584232 | SKCM | CP66-MEL | Afatinib |
| MVP_top60_DNA_seed | ERBB2 | 0.584232 | SKCM | CP66-MEL | Afatinib |
| MVP_top60_DNA_seed | MAP2K1 | 0.584286 | SKCM | COLO-800 | PD0325901 |
| MVP_top60_DNA_seed | MAPK14 | 0.585292 | SKCM | IST-MEL1 | TAK-715 |
| MVP_top60_DNA_seed | PIK3CA | 0.586003 | SKCM | G-361 | AZD8835 |
| MVP_top60_DNA_seed | EGFR | 0.586823 | SKCM | SK-MEL-30 | Afatinib |
| MVP_top60_DNA_seed | ERBB2 | 0.586823 | SKCM | SK-MEL-30 | Afatinib |
| MVP_top60_DNA_seed | JAK1 | 0.586826 | SKCM | WM793B | JAK1_3715 |
| MVP_top60_DNA_seed | EP300 | 0.587365 | SKCM | RVH-421 | I-CBP112 |
| MVP_top60_DNA_seed | CREBBP | 0.587365 | SKCM | RVH-421 | I-CBP112 |
| MVP_top60_DNA_seed | JAK1 | 0.589131 | SKCM | LB2518-MEL | JAK1_3715 |
| MVP_top60_DNA_seed | MAP2K1 | 0.590632 | SKCM | Hs-939-T | Selumetinib |

| | | | | | |
|---|---|---|---|---|---|
| MVP_top60_DNA_seed | ERBB2 | 0.593058 | SKCM | COLO-783 | CP724714 |
| MVP_top60_DNA_seed | RAC1 | 0.593284 | SKCM | Mewo | EHT-1864 |
| MVP_top60_DNA_seed | EGFR | 0.594846 | SKCM | CP50-MEL-B | Gefitinib |
| MVP_top60_DNA_seed | JAK3 | 0.596317 | SKCM | LB2518-MEL | WHI-P97 |
| MVP_top60_DNA_seed | GSK3B | 0.597031 | SKCM | COLO-679 | AZD7969 |
| MVP_top60_DNA_seed | PIK3CA | 0.597341 | SKCM | LB373-MEL-D | AZD8835 |
| MVP_top60_DNA_seed | RAC1 | 0.598339 | SKCM | GAK | EHT-1864 |
| MVP_top60_DNA_seed | GSK3B | 0.598977 | SKCM | C32 | CHIR-99021 |
| MVP_top60_DNA_seed | PIK3CB | 0.598997 | SKCM | LB2518-MEL | AZD6482 |
| MVP_top60_DNA_seed | ERBB2 | 0.601289 | SKCM | HT-144 | CP724714 |
| MVP_top60_DNA_seed | PIK3CB | 0.601895 | SKCM | SK-MEL-31 | AZD8186 |
| MVP_top60_DNA_seed | GSK3B | 0.601896 | SKCM | G-361 | SB216763 |
| MVP_top60_DNA_seed | JAK2 | 0.603083 | SKCM | UACC-257 | Ruxolitinib |
| MVP_top60_DNA_seed | JAK1 | 0.603083 | SKCM | UACC-257 | Ruxolitinib |
| MVP_top60_DNA_seed | EGFR | 0.603662 | SKCM | SH-4 | CI-1033 |
| MVP_top60_DNA_seed | ERBB2 | 0.603662 | SKCM | SH-4 | CI-1033 |
| MVP_top60_DNA_seed | EGFR | 0.60462 | SKCM | MZ2-MEL | Cetuximab |
| MVP_top60_DNA_seed | JAK3 | 0.60831 | SKCM | GAK | WHI-P97 |
| MVP_top60_DNA_seed | EGFR | 0.609679 | SKCM | SK-MEL-5 | AZD8931 |
| MVP_top60_DNA_seed | ERBB2 | 0.609679 | SKCM | SK-MEL-5 | AZD8931 |
| MVP_top60_DNA_seed | EGFR | 0.612472 | SKCM | HT-144 | Afatinib |
| MVP_top60_DNA_seed | ERBB2 | 0.612472 | SKCM | HT-144 | Afatinib |
| MVP_top60_DNA_seed | GSK3B | 0.613301 | SKCM | COLO-783 | CHIR-99021 |
| MVP_top60_DNA_seed | MAP2K1 | 0.614561 | SKCM | UACC-62 | Selumetinib |
| MVP_top60_DNA_seed | MAPK1 | 0.615385 | SKCM | A2058 | VX-11e |
| MVP_top60_DNA_seed | EGFR | 0.616905 | SKCM | G-361 | Afatinib |
| MVP_top60_DNA_seed | ERBB2 | 0.616905 | SKCM | G-361 | Afatinib |
| MVP_top60_DNA_seed | MAPK14 | 0.61736 | SKCM | LB373-MEL-D | TAK-715 |
| MVP_top60_DNA_seed | GSK3B | 0.617896 | SKCM | MMAc-SF | CHIR-99021 |
| MVP_top60_DNA_seed | EGFR | 0.61831 | SKCM | MEL-JUSO | PF-00299804 |
| MVP_top60_DNA_seed | ERBB2 | 0.61831 | SKCM | MEL-JUSO | PF-00299804 |
| MVP_top60_DNA_seed | JAK1 | 0.618461 | SKCM | 451Lu | JAK1_3715 |
| MVP_top60_DNA_seed | ERBB2 | 0.618487 | SKCM | SK-MEL-28 | CP724714 |
| MVP_top60_DNA_seed | BRAF | 0.618568 | SKCM | WM35 | PLX-4720 |
| MVP_top60_DNA_seed | JAK3 | 0.621584 | SKCM | LOXIMVI | WHI-P97 |
| MVP_top60_DNA_seed | EGFR | 0.623351 | SKCM | LB2518-MEL | Gefitinib |
| MVP_top60_DNA_seed | PIK3CA | 0.623542 | SKCM | COLO-829 | PI3Ka_4409 |
| MVP_top60_DNA_seed | EGFR | 0.624935 | SKCM | 451Lu | AZD8931 |
| MVP_top60_DNA_seed | ERBB2 | 0.624935 | SKCM | 451Lu | AZD8931 |
| MVP_top60_DNA_seed | GSK3B | 0.625297 | SKCM | MMAc-SF | KIN001-042 |
| MVP_top60_DNA_seed | ERBB2 | 0.626943 | SKCM | VMRC-MELG | CP724714 |

| | | | | | |
|---|---|---|---|---|---|
| MVP_top60_DNA_seed | BRAF | 0.629503 | SKCM | LB2518-MEL | PLX-4720 |
| MVP_top60_DNA_seed | JAK2 | 0.630545 | SKCM | HMV-II | Fedratinib |
| MVP_top60_DNA_seed | ERBB2 | 0.630667 | SKCM | SK-MEL-3 | CP724714 |
| MVP_top60_DNA_seed | EGFR | 0.631147 | SKCM | IGR-37 | Gefitinib |
| MVP_top60_DNA_seed | EGFR | 0.631194 | SKCM | A2058 | CUDC-101 |
| MVP_top60_DNA_seed | ERBB2 | 0.631194 | SKCM | A2058 | CUDC-101 |
| MVP_top60_DNA_seed | EGFR | 0.63595 | SKCM | VMRC-MELG | PF-00299804 |
| MVP_top60_DNA_seed | ERBB2 | 0.63595 | SKCM | VMRC-MELG | PF-00299804 |
| MVP_top60_DNA_seed | EGFR | 0.637122 | SKCM | IGR-1 | Afatinib |
| MVP_top60_DNA_seed | ERBB2 | 0.637122 | SKCM | IGR-1 | Afatinib |
| MVP_top60_DNA_seed | EGFR | 0.638238 | SKCM | GAK | Gefitinib |
| MVP_top60_DNA_seed | BRAF | 0.638491 | SKCM | LB2518-MEL | SB590885 |
| MVP_top60_DNA_seed | ERBB2 | 0.638722 | SKCM | WM1552C | CP724714 |
| MVP_top60_DNA_seed | EGFR | 0.640062 | SKCM | MMAc-SF | Afatinib |
| MVP_top60_DNA_seed | ERBB2 | 0.640062 | SKCM | MMAc-SF | Afatinib |
| MVP_top60_DNA_seed | MAP2K1 | 0.64058 | SKCM | SK-MEL-31 | Trametinib |
| MVP_top60_DNA_seed | JAK3 | 0.641213 | SKCM | UACC-257 | JAK3_7406 |
| MVP_top60_DNA_seed | EGFR | 0.641938 | SKCM | WM278 | Afatinib |
| MVP_top60_DNA_seed | ERBB2 | 0.641938 | SKCM | WM278 | Afatinib |
| MVP_top60_DNA_seed | EGFR | 0.643052 | SKCM | RPMI-7951 | Afatinib |
| MVP_top60_DNA_seed | ERBB2 | 0.643052 | SKCM | RPMI-7951 | Afatinib |
| MVP_top60_DNA_seed | EGFR | 0.643892 | SKCM | COLO-783 | Gefitinib |
| MVP_top60_DNA_seed | PIK3CA | 0.64409 | SKCM | MEL-HO | AZD8835 |
| MVP_top60_DNA_seed | EGFR | 0.645149 | SKCM | Hs-940-T | AZD8931 |
| MVP_top60_DNA_seed | ERBB2 | 0.645149 | SKCM | Hs-940-T | AZD8931 |
| MVP_top60_DNA_seed | EGFR | 0.645295 | SKCM | MMAc-SF | Afatinib |
| MVP_top60_DNA_seed | ERBB2 | 0.645295 | SKCM | MMAc-SF | Afatinib |
| MVP_top60_DNA_seed | RAC1 | 0.645978 | SKCM | MZ2-MEL | EHT-1864 |
| MVP_top60_DNA_seed | EGFR | 0.651447 | SKCM | MZ7-mel | AZD8931 |
| MVP_top60_DNA_seed | ERBB2 | 0.651447 | SKCM | MZ7-mel | AZD8931 |
| MVP_top60_DNA_seed | MAPK14 | 0.652022 | SKCM | SK-MEL-3 | TAK-715 |
| MVP_top60_DNA_seed | EGFR | 0.654172 | SKCM | MEL-HO | PF-00299804 |
| MVP_top60_DNA_seed | ERBB2 | 0.654172 | SKCM | MEL-HO | PF-00299804 |
| MVP_top60_DNA_seed | BRAF | 0.654869 | SKCM | SK-MEL-2 | PLX-4720 |
| MVP_top60_DNA_seed | GSK3B | 0.655006 | SKCM | MEL-JUSO | SB216763 |
| MVP_top60_DNA_seed | JAK2 | 0.656624 | SKCM | GAK | AZD1480 |
| MVP_top60_DNA_seed | JAK1 | 0.656624 | SKCM | GAK | AZD1480 |
| MVP_top60_DNA_seed | PIK3CA | 0.659296 | SKCM | HMV-II | AZD8835 |
| MVP_top60_DNA_seed | GSK3B | 0.659708 | SKCM | LB373-MEL-D | AZD7969 |
| MVP_top60_DNA_seed | JAK2 | 0.659795 | SKCM | G-361 | AZD1480 |
| MVP_top60_DNA_seed | JAK1 | 0.659795 | SKCM | G-361 | AZD1480 |

| | | | | | |
|---|---|---|---|---|---|
| MVP_top60_DNA_seed | JAK2 | 0.66088 | SKCM | WM-115 | AZD1480 |
| MVP_top60_DNA_seed | JAK1 | 0.66088 | SKCM | WM-115 | AZD1480 |
| MVP_top60_DNA_seed | EGFR | 0.66162 | SKCM | Mewo | AZD8931 |
| MVP_top60_DNA_seed | ERBB2 | 0.66162 | SKCM | Mewo | AZD8931 |
| MVP_top60_DNA_seed | EGFR | 0.667105 | SKCM | MZ2-MEL | AST-1306 |
| MVP_top60_DNA_seed | GSK3B | 0.668496 | SKCM | SK-MEL-30 | CHIR-99021 |
| MVP_top60_DNA_seed | GSK3B | 0.66907 | SKCM | WM793B | KIN001-042 |
| MVP_top60_DNA_seed | JAK1 | 0.671466 | SKCM | MZ7-mel | JAK1_3715 |
| MVP_top60_DNA_seed | JAK3 | 0.673295 | SKCM | MEL-JUSO | WHI-P97 |
| MVP_top60_DNA_seed | JAK2 | 0.676421 | SKCM | CP66-MEL | Ruxolitinib |
| MVP_top60_DNA_seed | JAK1 | 0.676421 | SKCM | CP66-MEL | Ruxolitinib |
| MVP_top60_DNA_seed | EGFR | 0.678114 | SKCM | Mewo | CI-1033 |
| MVP_top60_DNA_seed | ERBB2 | 0.678114 | SKCM | Mewo | CI-1033 |
| MVP_top60_DNA_seed | JAK3 | 0.67963 | SKCM | SK-MEL-28 | JAK3_7406 |
| MVP_top60_DNA_seed | RAC1 | 0.680653 | SKCM | VMRC-MELG | EHT-1864 |
| MVP_top60_DNA_seed | EGFR | 0.682074 | SKCM | CP66-MEL | CUDC-101 |
| MVP_top60_DNA_seed | ERBB2 | 0.682074 | SKCM | CP66-MEL | CUDC-101 |
| MVP_top60_DNA_seed | EP300 | 0.682265 | SKCM | IST-MEL1 | I-CBP112 |
| MVP_top60_DNA_seed | CREBBP | 0.682265 | SKCM | IST-MEL1 | I-CBP112 |
| MVP_top60_DNA_seed | MAP2K1 | 0.682674 | SKCM | COLO-800 | Selumetinib |
| MVP_top60_DNA_seed | EGFR | 0.683053 | SKCM | UACC-257 | Pelitinib |
| MVP_top60_DNA_seed | JAK1 | 0.683984 | SKCM | MZ2-MEL | JAK1_3715 |
| MVP_top60_DNA_seed | EGFR | 0.687609 | SKCM | LOXIMVI | Lapatinib |
| MVP_top60_DNA_seed | ERBB2 | 0.687609 | SKCM | LOXIMVI | Lapatinib |
| MVP_top60_DNA_seed | EGFR | 0.689386 | SKCM | IGR-37 | Afatinib |
| MVP_top60_DNA_seed | ERBB2 | 0.689386 | SKCM | IGR-37 | Afatinib |
| MVP_top60_DNA_seed | EGFR | 0.689842 | SKCM | MEL-HO | Afatinib |
| MVP_top60_DNA_seed | ERBB2 | 0.689842 | SKCM | MEL-HO | Afatinib |
| MVP_top60_DNA_seed | GSK3B | 0.690723 | SKCM | MEL-HO | CHIR-99021 |
| MVP_top60_DNA_seed | EGFR | 0.691074 | SKCM | MMAc-SF | Pelitinib |
| MVP_top60_DNA_seed | JAK2 | 0.691463 | SKCM | MZ2-MEL | AZD1480 |
| MVP_top60_DNA_seed | JAK1 | 0.691463 | SKCM | MZ2-MEL | AZD1480 |
| MVP_top60_DNA_seed | MAP2K1 | 0.693983 | SKCM | COLO-800 | Refametinib |
| MVP_top60_DNA_seed | EGFR | 0.694764 | SKCM | HMV-II | Cetuximab |
| MVP_top60_DNA_seed | EGFR | 0.695905 | SKCM | MEL-HO | AST-1306 |
| MVP_top60_DNA_seed | JAK1 | 0.698535 | SKCM | COLO-783 | JAK1_3715 |
| MVP_top60_DNA_seed | EGFR | 0.699276 | SKCM | LOXIMVI | Erlotinib |
| MVP_top60_DNA_seed | ERBB2 | 0.702671 | SKCM | SH-4 | CP724714 |
| MVP_top60_DNA_seed | PIK3CA | 0.704266 | SKCM | WM-115 | PI3Ka_4409 |
| MVP_top60_DNA_seed | BRAF | 0.705163 | SKCM | COLO-792 | Dabrafenib |
| MVP_top60_DNA_seed | JAK3 | 0.705186 | SKCM | RVH-421 | WHI-P97 |

| | | | | | |
|---|---|---|---|---|---|
| MVP_top60_DNA_seed | EP300 | 0.705227 | SKCM | CP66-MEL | I-CBP112 |
| MVP_top60_DNA_seed | CREBBP | 0.705227 | SKCM | CP66-MEL | I-CBP112 |
| MVP_top60_DNA_seed | PIK3CB | 0.705402 | SKCM | CP66-MEL | AZD8186 |
| MVP_top60_DNA_seed | JAK3 | 0.70557 | SKCM | COLO-829 | JAK3_7406 |
| MVP_top60_DNA_seed | JAK1 | 0.706061 | SKCM | WM-115 | JAK1_3715 |
| MVP_top60_DNA_seed | MAP2K1 | 0.706158 | SKCM | COLO-800 | Selumetinib |
| MVP_top60_DNA_seed | MAPK3 | 0.706193 | SKCM | A2058 | FR-180204 |
| MVP_top60_DNA_seed | MAPK1 | 0.706193 | SKCM | A2058 | FR-180204 |
| MVP_top60_DNA_seed | EGFR | 0.706641 | SKCM | IGR-37 | Afatinib |
| MVP_top60_DNA_seed | ERBB2 | 0.706641 | SKCM | IGR-37 | Afatinib |
| MVP_top60_DNA_seed | MAP2K1 | 0.707052 | SKCM | SK-MEL-31 | Selumetinib |
| MVP_top60_DNA_seed | PIK3CA | 0.708756 | SKCM | SK-MEL-30 | PI3Ka_4409 |
| MVP_top60_DNA_seed | EGFR | 0.709784 | SKCM | LB2518-MEL | Afatinib |
| MVP_top60_DNA_seed | ERBB2 | 0.709784 | SKCM | LB2518-MEL | Afatinib |
| MVP_top60_DNA_seed | RAC1 | 0.709871 | SKCM | HMV-II | EHT-1864 |
| MVP_top60_DNA_seed | GSK3B | 0.709999 | SKCM | CHL-1 | CHIR-99021 |
| MVP_top60_DNA_seed | JAK2 | 0.711795 | SKCM | Mewo | AZD1480 |
| MVP_top60_DNA_seed | JAK1 | 0.711795 | SKCM | Mewo | AZD1480 |
| MVP_top60_DNA_seed | EP300 | 0.715596 | SKCM | LOXIMVI | I-CBP112 |
| MVP_top60_DNA_seed | CREBBP | 0.715596 | SKCM | LOXIMVI | I-CBP112 |
| MVP_top60_DNA_seed | MAPK14 | 0.716419 | SKCM | COLO-679 | TAK-715 |
| MVP_top60_DNA_seed | PIK3CB | 0.724816 | SKCM | WM1552C | AZD8186 |
| MVP_top60_DNA_seed | MAPK14 | 0.726607 | SKCM | SH-4 | TAK-715 |
| MVP_top60_DNA_seed | EGFR | 0.727819 | SKCM | SK-MEL-31 | Afatinib |
| MVP_top60_DNA_seed | ERBB2 | 0.727819 | SKCM | SK-MEL-31 | Afatinib |
| MVP_top60_DNA_seed | MAP2K1 | 0.728592 | SKCM | COLO-800 | Refametinib |
| MVP_top60_DNA_seed | GSK3B | 0.728681 | SKCM | A101D | KIN001-042 |
| MVP_top60_DNA_seed | EGFR | 0.731418 | SKCM | SH-4 | Gefitinib |
| MVP_top60_DNA_seed | EGFR | 0.732965 | SKCM | A2058 | Afatinib |
| MVP_top60_DNA_seed | ERBB2 | 0.732965 | SKCM | A2058 | Afatinib |
| MVP_top60_DNA_seed | MAP2K1 | 0.736299 | SKCM | LB373-MEL-D | Selumetinib |
| MVP_top60_DNA_seed | GSK3B | 0.739065 | SKCM | SK-MEL-28 | AZD7969 |
| MVP_top60_DNA_seed | EGFR | 0.74223 | SKCM | G-361 | Afatinib |
| MVP_top60_DNA_seed | ERBB2 | 0.74223 | SKCM | G-361 | Afatinib |
| MVP_top60_DNA_seed | PIK3CB | 0.745734 | SKCM | CP66-MEL | AZD6482 |
| MVP_top60_DNA_seed | GSK3B | 0.746546 | SKCM | MZ2-MEL | SB216763 |
| MVP_top60_DNA_seed | EGFR | 0.746867 | SKCM | MEL-JUSO | Afatinib |
| MVP_top60_DNA_seed | ERBB2 | 0.746867 | SKCM | MEL-JUSO | Afatinib |
| MVP_top60_DNA_seed | GSK3B | 0.748508 | SKCM | SK-MEL-30 | SB216763 |
| MVP_top60_DNA_seed | PIK3CB | 0.749765 | SKCM | COLO-829 | AZD8186 |
| MVP_top60_DNA_seed | GSK3B | 0.750299 | SKCM | UACC-257 | CHIR-99021 |

| | | | | | |
|---|---|---|---|---|---|
| MVP_top60_DNA_seed | EGFR | 0.75036 | SKCM | SK-MEL-28 | AZD8931 |
| MVP_top60_DNA_seed | ERBB2 | 0.75036 | SKCM | SK-MEL-28 | AZD8931 |
| MVP_top60_DNA_seed | EGFR | 0.751616 | SKCM | LB2518-MEL | Lapatinib |
| MVP_top60_DNA_seed | ERBB2 | 0.751616 | SKCM | LB2518-MEL | Lapatinib |
| MVP_top60_DNA_seed | BRAF | 0.756074 | SKCM | COLO-800 | PLX-4720 |
| MVP_top60_DNA_seed | EGFR | 0.758069 | SKCM | LB373-MEL-D | Pelitinib |
| MVP_top60_DNA_seed | EGFR | 0.759235 | SKCM | A101D | Afatinib |
| MVP_top60_DNA_seed | ERBB2 | 0.759235 | SKCM | A101D | Afatinib |
| MVP_top60_DNA_seed | JAK3 | 0.762546 | SKCM | RPMI-7951 | WHI-P97 |
| MVP_top60_DNA_seed | EGFR | 0.763765 | SKCM | GAK | PF-00299804 |
| MVP_top60_DNA_seed | ERBB2 | 0.763765 | SKCM | GAK | PF-00299804 |
| MVP_top60_DNA_seed | PIK3CA | 0.764658 | SKCM | GAK | PI3Ka_4409 |
| MVP_top60_DNA_seed | PIK3CA | 0.764852 | SKCM | RVH-421 | AZD8835 |
| MVP_top60_DNA_seed | ERBB2 | 0.768325 | SKCM | WM278 | CP724714 |
| MVP_top60_DNA_seed | EGFR | 0.768637 | SKCM | C32 | CI-1033 |
| MVP_top60_DNA_seed | ERBB2 | 0.768637 | SKCM | C32 | CI-1033 |
| MVP_top60_DNA_seed | GSK3B | 0.770462 | SKCM | SK-MEL-1 | KIN001-042 |
| MVP_top60_DNA_seed | EGFR | 0.770837 | SKCM | COLO-783 | AST-1306 |
| MVP_top60_DNA_seed | BRAF | 0.771435 | SKCM | VMRC-MELG | HG6-64-1 |
| MVP_top60_DNA_seed | GSK3B | 0.772486 | SKCM | RVH-421 | KIN001-042 |
| MVP_top60_DNA_seed | JAK3 | 0.772891 | SKCM | HT-144 | WHI-P97 |
| MVP_top60_DNA_seed | PIK3CA | 0.776731 | SKCM | MEL-HO | PI3Ka_4409 |
| MVP_top60_DNA_seed | EGFR | 0.777348 | SKCM | HT-144 | Afatinib |
| MVP_top60_DNA_seed | ERBB2 | 0.777348 | SKCM | HT-144 | Afatinib |
| MVP_top60_DNA_seed | EGFR | 0.778714 | SKCM | SK-MEL-5 | Afatinib |
| MVP_top60_DNA_seed | ERBB2 | 0.778714 | SKCM | SK-MEL-5 | Afatinib |
| MVP_top60_DNA_seed | JAK1 | 0.779461 | SKCM | IST-MEL1 | JAK1_3715 |
| MVP_top60_DNA_seed | EGFR | 0.780946 | SKCM | MZ7-mel | Gefitinib |
| MVP_top60_DNA_seed | BRAF | 0.782026 | SKCM | HMV-II | PLX-4720 |
| MVP_top60_DNA_seed | BRAF | 0.782456 | SKCM | LB373-MEL-D | SB590885 |
| MVP_top60_DNA_seed | JAK2 | 0.783927 | SKCM | COLO-783 | Fedratinib |
| MVP_top60_DNA_seed | EGFR | 0.785492 | SKCM | SK-MEL-2 | Erlotinib |
| MVP_top60_DNA_seed | PIK3CA | 0.785887 | SKCM | LB2518-MEL | PI3Ka_4409 |
| MVP_top60_DNA_seed | MAPK3 | 0.786765 | SKCM | COLO-829 | FR-180204 |
| MVP_top60_DNA_seed | MAPK1 | 0.786765 | SKCM | COLO-829 | FR-180204 |
| MVP_top60_DNA_seed | JAK2 | 0.788722 | SKCM | SH-4 | Fedratinib |
| MVP_top60_DNA_seed | EGFR | 0.78986 | SKCM | M14 | Afatinib |
| MVP_top60_DNA_seed | ERBB2 | 0.78986 | SKCM | M14 | Afatinib |
| MVP_top60_DNA_seed | GSK3B | 0.790506 | SKCM | COLO-800 | CHIR-99021 |
| MVP_top60_DNA_seed | EGFR | 0.790999 | SKCM | G-MEL | Afatinib |
| MVP_top60_DNA_seed | ERBB2 | 0.790999 | SKCM | G-MEL | Afatinib |

| | | | | | |
|---|---|---|---|---|---|
| MVP_top60_DNA_seed | EGFR | 0.792309 | SKCM | C32 | Cetuximab |
| MVP_top60_DNA_seed | EGFR | 0.793004 | SKCM | UACC-257 | Cetuximab |
| MVP_top60_DNA_seed | EGFR | 0.793766 | SKCM | LOXIMVI | Pelitinib |
| MVP_top60_DNA_seed | EP300 | 0.794663 | SKCM | Hs-939-T | I-CBP112 |
| MVP_top60_DNA_seed | CREBBP | 0.794663 | SKCM | Hs-939-T | I-CBP112 |
| MVP_top60_DNA_seed | GSK3B | 0.795522 | SKCM | COLO-783 | AZD7969 |
| MVP_top60_DNA_seed | EGFR | 0.796695 | SKCM | MZ2-MEL | Afatinib |
| MVP_top60_DNA_seed | ERBB2 | 0.796695 | SKCM | MZ2-MEL | Afatinib |
| MVP_top60_DNA_seed | GSK3B | 0.798402 | SKCM | COLO-679 | CHIR-99021 |
| MVP_top60_DNA_seed | EP300 | 0.798959 | SKCM | UACC-257 | I-CBP112 |
| MVP_top60_DNA_seed | CREBBP | 0.798959 | SKCM | UACC-257 | I-CBP112 |
| MVP_top60_DNA_seed | JAK2 | 0.800155 | SKCM | A101D | AZD1480 |
| MVP_top60_DNA_seed | JAK1 | 0.800155 | SKCM | A101D | AZD1480 |
| MVP_top60_DNA_seed | MAPK1 | 0.802306 | SKCM | IST-MEL1 | VX-11e |
| MVP_top60_DNA_seed | EGFR | 0.803362 | SKCM | HT-144 | Pelitinib |
| MVP_top60_DNA_seed | MAPK14 | 0.809264 | SKCM | 451Lu | TAK-715 |
| MVP_top60_DNA_seed | JAK1 | 0.811352 | SKCM | SK-MEL-3 | JAK1_3715 |
| MVP_top60_DNA_seed | EGFR | 0.812237 | SKCM | IPC-298 | Afatinib |
| MVP_top60_DNA_seed | ERBB2 | 0.812237 | SKCM | IPC-298 | Afatinib |
| MVP_top60_DNA_seed | EGFR | 0.813178 | SKCM | CP50-MEL-B | PF-00299804 |
| MVP_top60_DNA_seed | ERBB2 | 0.813178 | SKCM | CP50-MEL-B | PF-00299804 |
| MVP_top60_DNA_seed | EGFR | 0.815161 | SKCM | 451Lu | Pelitinib |
| MVP_top60_DNA_seed | BRAF | 0.815956 | SKCM | Hs-940-T | SB590885 |
| MVP_top60_DNA_seed | BRAF | 0.816054 | SKCM | WM-115 | HG6-64-1 |
| MVP_top60_DNA_seed | JAK3 | 0.818348 | SKCM | SK-MEL-24 | JAK3_7406 |
| MVP_top60_DNA_seed | EGFR | 0.818939 | SKCM | C32 | Gefitinib |
| MVP_top60_DNA_seed | JAK1 | 0.820835 | SKCM | UACC-257 | JAK1_3715 |
| MVP_top60_DNA_seed | RAC1 | 0.821186 | SKCM | WM1552C | EHT-1864 |
| MVP_top60_DNA_seed | PIK3CB | 0.82157 | SKCM | M14 | AZD6482 |
| MVP_top60_DNA_seed | MAP2K1 | 0.821712 | SKCM | SK-MEL-31 | CI-1040 |
| MVP_top60_DNA_seed | EGFR | 0.822053 | SKCM | LOXIMVI | Cetuximab |
| MVP_top60_DNA_seed | MAPK14 | 0.823169 | SKCM | UACC-257 | TAK-715 |
| MVP_top60_DNA_seed | EGFR | 0.824454 | SKCM | COLO-783 | PF-00299804 |
| MVP_top60_DNA_seed | ERBB2 | 0.824454 | SKCM | COLO-783 | PF-00299804 |
| MVP_top60_DNA_seed | EGFR | 0.825889 | SKCM | IPC-298 | Gefitinib |
| MVP_top60_DNA_seed | JAK2 | 0.826696 | SKCM | VMRC-MELG | Fedratinib |
| MVP_top60_DNA_seed | JAK1 | 0.828731 | SKCM | COLO-679 | JAK1_3715 |
| MVP_top60_DNA_seed | PIK3CA | 0.829178 | SKCM | UACC-257 | PI3Ka_4409 |
| MVP_top60_DNA_seed | PIK3CB | 0.830953 | SKCM | K2 | AZD8186 |
| MVP_top60_DNA_seed | PIK3CA | 0.833258 | SKCM | Hs-940-T | AZD8835 |
| MVP_top60_DNA_seed | JAK2 | 0.83563 | SKCM | CP66-MEL | AZD1480 |

| | | | | | |
|---|---|---|---|---|---|
| MVP_top60_DNA_seed | JAK1 | 0.83563 | SKCM | CP66-MEL | AZD1480 |
| MVP_top60_DNA_seed | GSK3B | 0.841182 | SKCM | SK-MEL-3 | CHIR-99021 |
| MVP_top60_DNA_seed | MAP2K1 | 0.842718 | SKCM | VMRC-MELG | Trametinib |
| MVP_top60_DNA_seed | EGFR | 0.845172 | SKCM | COLO-783 | Afatinib |
| MVP_top60_DNA_seed | ERBB2 | 0.845172 | SKCM | COLO-783 | Afatinib |
| MVP_top60_DNA_seed | GSK3B | 0.845273 | SKCM | MMAc-SF | SB216763 |
| MVP_top60_DNA_seed | MAPK14 | 0.846411 | SKCM | M14 | TAK-715 |
| MVP_top60_DNA_seed | GSK3B | 0.846742 | SKCM | COLO-829 | KIN001-042 |
| MVP_top60_DNA_seed | MAPK14 | 0.847603 | SKCM | COLO-792 | TAK-715 |
| MVP_top60_DNA_seed | EGFR | 0.848005 | SKCM | SH-4 | Afatinib |
| MVP_top60_DNA_seed | ERBB2 | 0.848005 | SKCM | SH-4 | Afatinib |
| MVP_top60_DNA_seed | EGFR | 0.848194 | SKCM | IST-MEL1 | Afatinib |
| MVP_top60_DNA_seed | ERBB2 | 0.848194 | SKCM | IST-MEL1 | Afatinib |
| MVP_top60_DNA_seed | EGFR | 0.848638 | SKCM | UACC-62 | Pelitinib |
| MVP_top60_DNA_seed | JAK3 | 0.849888 | SKCM | SK-MEL-2 | WHI-P97 |
| MVP_top60_DNA_seed | JAK3 | 0.851734 | SKCM | SK-MEL-30 | WHI-P97 |
| MVP_top60_DNA_seed | MAPK3 | 0.857905 | SKCM | SK-MEL-2 | FR-180204 |
| MVP_top60_DNA_seed | MAPK1 | 0.857905 | SKCM | SK-MEL-2 | FR-180204 |
| MVP_top60_DNA_seed | MAPK3 | 0.859335 | SKCM | COLO-679 | FR-180204 |
| MVP_top60_DNA_seed | MAPK1 | 0.859335 | SKCM | COLO-679 | FR-180204 |
| MVP_top60_DNA_seed | BRAF | 0.860431 | SKCM | COLO-792 | SB590885 |
| MVP_top60_DNA_seed | EGFR | 0.861429 | SKCM | COLO-679 | Cetuximab |
| MVP_top60_DNA_seed | EGFR | 0.865228 | SKCM | C32 | Afatinib |
| MVP_top60_DNA_seed | ERBB2 | 0.865228 | SKCM | C32 | Afatinib |
| MVP_top60_DNA_seed | GSK3B | 0.86529 | SKCM | SK-MEL-3 | SB216763 |
| MVP_top60_DNA_seed | JAK2 | 0.866256 | SKCM | MMAc-SF | Ruxolitinib |
| MVP_top60_DNA_seed | JAK1 | 0.866256 | SKCM | MMAc-SF | Ruxolitinib |
| MVP_top60_DNA_seed | JAK2 | 0.867259 | SKCM | MMAc-SF | Fedratinib |
| MVP_top60_DNA_seed | EGFR | 0.867546 | SKCM | 451Lu | Gefitinib |
| MVP_top60_DNA_seed | GSK3B | 0.868466 | SKCM | COLO-679 | SB216763 |
| MVP_top60_DNA_seed | ERBB2 | 0.87099 | SKCM | MMAc-SF | CP724714 |
| MVP_top60_DNA_seed | GSK3B | 0.87142 | SKCM | IGR-37 | SB216763 |
| MVP_top60_DNA_seed | EGFR | 0.872116 | SKCM | G-361 | Gefitinib |
| MVP_top60_DNA_seed | JAK2 | 0.875845 | SKCM | HMV-II | Ruxolitinib |
| MVP_top60_DNA_seed | JAK1 | 0.875845 | SKCM | HMV-II | Ruxolitinib |
| MVP_top60_DNA_seed | JAK2 | 0.875945 | SKCM | IPC-298 | AZD1480 |
| MVP_top60_DNA_seed | JAK1 | 0.875945 | SKCM | IPC-298 | AZD1480 |
| MVP_top60_DNA_seed | EGFR | 0.876982 | SKCM | UACC-257 | Lapatinib |
| MVP_top60_DNA_seed | ERBB2 | 0.876982 | SKCM | UACC-257 | Lapatinib |
| MVP_top60_DNA_seed | EGFR | 0.877694 | SKCM | CP50-MEL-B | Pelitinib |
| MVP_top60_DNA_seed | GSK3B | 0.878338 | SKCM | SH-4 | SB216763 |

| | | | | | |
|---|---|---|---|---|---|
| MVP_top60_DNA_seed | JAK3 | 0.878943 | SKCM | COLO-679 | JAK3_7406 |
| MVP_top60_DNA_seed | GSK3B | 0.880047 | SKCM | GAK | AZD7969 |
| MVP_top60_DNA_seed | MAP2K1 | 0.880147 | SKCM | G-361 | Selumetinib |
| MVP_top60_DNA_seed | EGFR | 0.881447 | SKCM | WM35 | PF-00299804 |
| MVP_top60_DNA_seed | ERBB2 | 0.881447 | SKCM | WM35 | PF-00299804 |
| MVP_top60_DNA_seed | PIK3CB | 0.883237 | SKCM | Mewo | AZD6482 |
| MVP_top60_DNA_seed | JAK2 | 0.884354 | SKCM | MZ2-MEL | Ruxolitinib |
| MVP_top60_DNA_seed | JAK1 | 0.884354 | SKCM | MZ2-MEL | Ruxolitinib |
| MVP_top60_DNA_seed | JAK2 | 0.884759 | SKCM | VMRC-MELG | AZD1480 |
| MVP_top60_DNA_seed | JAK1 | 0.884759 | SKCM | VMRC-MELG | AZD1480 |
| MVP_top60_DNA_seed | PIK3CB | 0.88546 | SKCM | Hs-940-T | AZD8186 |
| MVP_top60_DNA_seed | EGFR | 0.886029 | SKCM | MZ7-mel | Erlotinib |
| MVP_top60_DNA_seed | JAK1 | 0.887048 | SKCM | CP66-MEL | JAK1_3715 |
| MVP_top60_DNA_seed | EGFR | 0.887535 | SKCM | CP66-MEL | Pelitinib |
| MVP_top60_DNA_seed | EGFR | 0.890048 | SKCM | UACC-257 | Afatinib |
| MVP_top60_DNA_seed | ERBB2 | 0.890048 | SKCM | UACC-257 | Afatinib |
| MVP_top60_DNA_seed | JAK3 | 0.892044 | SKCM | IST-MEL1 | WHI-P97 |
| MVP_top60_DNA_seed | EGFR | 0.893321 | SKCM | HMV-II | Afatinib |
| MVP_top60_DNA_seed | ERBB2 | 0.893321 | SKCM | HMV-II | Afatinib |
| MVP_top60_DNA_seed | EGFR | 0.893451 | SKCM | SK-MEL-2 | Afatinib |
| MVP_top60_DNA_seed | ERBB2 | 0.893451 | SKCM | SK-MEL-2 | Afatinib |
| MVP_top60_DNA_seed | JAK2 | 0.893925 | SKCM | LB373-MEL-D | AZD1480 |
| MVP_top60_DNA_seed | JAK1 | 0.893925 | SKCM | LB373-MEL-D | AZD1480 |
| MVP_top60_DNA_seed | JAK1 | 0.896392 | SKCM | A101D | JAK1_3715 |
| MVP_top60_DNA_seed | JAK3 | 0.902247 | SKCM | A2058 | WHI-P97 |
| MVP_top60_DNA_seed | EGFR | 0.90743 | SKCM | VMRC-MELG | AZD8931 |
| MVP_top60_DNA_seed | ERBB2 | 0.90743 | SKCM | VMRC-MELG | AZD8931 |
| MVP_top60_DNA_seed | EGFR | 0.910416 | SKCM | WM278 | Gefitinib |
| MVP_top60_DNA_seed | MAP2K1 | 0.912968 | SKCM | COLO-800 | Trametinib |
| MVP_top60_DNA_seed | EGFR | 0.914664 | SKCM | COLO-792 | Afatinib |
| MVP_top60_DNA_seed | ERBB2 | 0.914664 | SKCM | COLO-792 | Afatinib |
| MVP_top60_DNA_seed | GSK3B | 0.914781 | SKCM | SK-MEL-3 | KIN001-042 |
| MVP_top60_DNA_seed | EGFR | 0.918529 | SKCM | Hs-940-T | Cetuximab |
| MVP_top60_DNA_seed | EGFR | 0.919349 | SKCM | MZ7-mel | Afatinib |
| MVP_top60_DNA_seed | ERBB2 | 0.919349 | SKCM | MZ7-mel | Afatinib |
| MVP_top60_DNA_seed | EP300 | 0.919546 | SKCM | M14 | I-CBP112 |
| MVP_top60_DNA_seed | CREBBP | 0.919546 | SKCM | M14 | I-CBP112 |
| MVP_top60_DNA_seed | EGFR | 0.920251 | SKCM | MEL-JUSO | Gefitinib |
| MVP_top60_DNA_seed | EGFR | 0.922633 | SKCM | SK-MEL-30 | Afatinib |
| MVP_top60_DNA_seed | ERBB2 | 0.922633 | SKCM | SK-MEL-30 | Afatinib |
| MVP_top60_DNA_seed | EGFR | 0.922897 | SKCM | COLO-679 | AST-1306 |

| | | | | | |
|---|---|---|---|---|---|
| MVP_top60_DNA_seed | EGFR | 0.922962 | SKCM | LB373-MEL-D | Afatinib |
| MVP_top60_DNA_seed | ERBB2 | 0.922962 | SKCM | LB373-MEL-D | Afatinib |
| MVP_top60_DNA_seed | EGFR | 0.923362 | SKCM | CP66-MEL | Lapatinib |
| MVP_top60_DNA_seed | ERBB2 | 0.923362 | SKCM | CP66-MEL | Lapatinib |
| MVP_top60_DNA_seed | EGFR | 0.923486 | SKCM | LB2518-MEL | Afatinib |
| MVP_top60_DNA_seed | ERBB2 | 0.923486 | SKCM | LB2518-MEL | Afatinib |
| MVP_top60_DNA_seed | EGFR | 0.923565 | SKCM | SK-MEL-1 | Afatinib |
| MVP_top60_DNA_seed | ERBB2 | 0.923565 | SKCM | SK-MEL-1 | Afatinib |
| MVP_top60_DNA_seed | RAC1 | 0.925655 | SKCM | LOXIMVI | EHT-1864 |
| MVP_top60_DNA_seed | EGFR | 0.929069 | SKCM | COLO-792 | Cetuximab |
| MVP_top60_DNA_seed | JAK2 | 0.931665 | SKCM | HMV-II | AZD1480 |
| MVP_top60_DNA_seed | JAK1 | 0.931665 | SKCM | HMV-II | AZD1480 |
| MVP_top60_DNA_seed | MAP2K1 | 0.932151 | SKCM | COLO-800 | CI-1040 |
| MVP_top60_DNA_seed | EGFR | 0.934027 | SKCM | MMAc-SF | PF-00299804 |
| MVP_top60_DNA_seed | ERBB2 | 0.934027 | SKCM | MMAc-SF | PF-00299804 |
| MVP_top60_DNA_seed | EGFR | 0.934783 | SKCM | CP66-MEL | Cetuximab |
| MVP_top60_DNA_seed | JAK2 | 0.936043 | SKCM | IST-MEL1 | Ruxolitinib |
| MVP_top60_DNA_seed | JAK1 | 0.936043 | SKCM | IST-MEL1 | Ruxolitinib |
| MVP_top60_DNA_seed | EGFR | 0.940286 | SKCM | VMRC-MELG | Afatinib |
| MVP_top60_DNA_seed | ERBB2 | 0.940286 | SKCM | VMRC-MELG | Afatinib |
| MVP_top60_DNA_seed | EGFR | 0.942069 | SKCM | Hs-939-T | Afatinib |
| MVP_top60_DNA_seed | ERBB2 | 0.942069 | SKCM | Hs-939-T | Afatinib |
| MVP_top60_DNA_seed | EGFR | 0.942841 | SKCM | CP66-MEL | Afatinib |
| MVP_top60_DNA_seed | ERBB2 | 0.942841 | SKCM | CP66-MEL | Afatinib |
| MVP_top60_DNA_seed | EGFR | 0.943501 | SKCM | CP66-MEL | PF-00299804 |
| MVP_top60_DNA_seed | ERBB2 | 0.943501 | SKCM | CP66-MEL | PF-00299804 |
| MVP_top60_DNA_seed | BRAF | 0.943639 | SKCM | COLO-792 | HG6-64-1 |
| MVP_top60_DNA_seed | PIK3CA | 0.944455 | SKCM | IPC-298 | PI3Ka_4409 |
| MVP_top60_DNA_seed | JAK3 | 0.944763 | SKCM | UACC-257 | WHI-P97 |
| MVP_top60_DNA_seed | EGFR | 0.945617 | SKCM | COLO-829 | PF-00299804 |
| MVP_top60_DNA_seed | ERBB2 | 0.945617 | SKCM | COLO-829 | PF-00299804 |
| MVP_top60_DNA_seed | EGFR | 0.946064 | SKCM | A375 | Pelitinib |
| MVP_top60_DNA_seed | PIK3CA | 0.947339 | SKCM | COLO-829 | AZD8835 |
| MVP_top60_DNA_seed | GSK3B | 0.948368 | SKCM | M14 | SB216763 |
| MVP_top60_DNA_seed | PIK3CB | 0.948559 | SKCM | SK-MEL-5 | AZD6482 |
| MVP_top60_DNA_seed | MAP2K1 | 0.94932 | SKCM | 451Lu | Selumetinib |
| MVP_top60_DNA_seed | JAK3 | 0.950635 | SKCM | G-361 | JAK3_7406 |
| MVP_top60_DNA_seed | JAK2 | 0.952955 | SKCM | Hs-940-T | AZD1480 |
| MVP_top60_DNA_seed | JAK1 | 0.952955 | SKCM | Hs-940-T | AZD1480 |
| MVP_top60_DNA_seed | PIK3CA | 0.957619 | SKCM | RVH-421 | PI3Ka_4409 |
| MVP_top60_DNA_seed | EGFR | 0.961034 | SKCM | SH-4 | Lapatinib |

| | | | | | |
|---|---|---|---|---|---|
| MVP_top60_DNA_seed | ERBB2 | 0.961034 | SKCM | SH-4 | Lapatinib |
| MVP_top60_DNA_seed | EGFR | 0.961426 | SKCM | VMRC-MELG | Afatinib |
| MVP_top60_DNA_seed | ERBB2 | 0.961426 | SKCM | VMRC-MELG | Afatinib |
| MVP_top60_DNA_seed | JAK2 | 0.962168 | SKCM | LB2518-MEL | AZD1480 |
| MVP_top60_DNA_seed | JAK1 | 0.962168 | SKCM | LB2518-MEL | AZD1480 |
| MVP_top60_DNA_seed | GSK3B | 0.963819 | SKCM | 451Lu | SB216763 |
| MVP_top60_DNA_seed | GSK3B | 0.964782 | SKCM | CP66-MEL | KIN001-042 |
| MVP_top60_DNA_seed | GSK3B | 0.965947 | SKCM | SK-MEL-5 | KIN001-042 |
| MVP_top60_DNA_seed | GSK3B | 0.966071 | SKCM | SK-MEL-30 | CHIR-99021 |
| MVP_top60_DNA_seed | JAK2 | 0.967635 | SKCM | WM278 | AZD1480 |
| MVP_top60_DNA_seed | JAK1 | 0.967635 | SKCM | WM278 | AZD1480 |
| MVP_top60_DNA_seed | EGFR | 0.969129 | SKCM | UACC-257 | AZD8931 |
| MVP_top60_DNA_seed | ERBB2 | 0.969129 | SKCM | UACC-257 | AZD8931 |
| MVP_top60_DNA_seed | EGFR | 0.971626 | SKCM | WM278 | Afatinib |
| MVP_top60_DNA_seed | ERBB2 | 0.971626 | SKCM | WM278 | Afatinib |
| MVP_top60_DNA_seed | RAC1 | 0.974053 | SKCM | WM278 | EHT-1864 |
| MVP_top60_DNA_seed | EGFR | 0.974766 | SKCM | SK-MEL-3 | AST-1306 |
| MVP_top60_DNA_seed | PIK3CB | 0.975312 | SKCM | IGR-1 | AZD8186 |
| MVP_top60_DNA_seed | EGFR | 0.976929 | SKCM | LB373-MEL-D | PF-00299804 |
| MVP_top60_DNA_seed | ERBB2 | 0.976929 | SKCM | LB373-MEL-D | PF-00299804 |
| MVP_top60_DNA_seed | EGFR | 0.978261 | SKCM | Hs-939-T | Afatinib |
| MVP_top60_DNA_seed | ERBB2 | 0.978261 | SKCM | Hs-939-T | Afatinib |
| MVP_top60_DNA_seed | RAC1 | 0.980407 | SKCM | RVH-421 | EHT-1864 |
| MVP_top60_DNA_seed | PIK3CA | 0.98103 | SKCM | UACC-257 | AZD8835 |
| MVP_top60_DNA_seed | GSK3B | 0.981428 | SKCM | IST-MEL1 | KIN001-042 |
| MVP_top60_DNA_seed | MAP2K1 | 0.982265 | SKCM | Hs-940-T | Refametinib |
| MVP_top60_DNA_seed | GSK3B | 0.982428 | SKCM | IGR-37 | CHIR-99021 |
| MVP_top60_DNA_seed | EGFR | 0.982618 | SKCM | A101D | AST-1306 |
| MVP_top60_DNA_seed | MAPK14 | 0.983255 | SKCM | SK-MEL-1 | TAK-715 |
| MVP_top60_DNA_seed | ERBB2 | 0.984208 | SKCM | M14 | CP724714 |
| MVP_top60_DNA_seed | EP300 | 0.9855 | SKCM | Hs-940-T | I-CBP112 |
| MVP_top60_DNA_seed | CREBBP | 0.9855 | SKCM | Hs-940-T | I-CBP112 |
| MVP_top60_DNA_seed | PIK3CA | 0.986184 | SKCM | WM793B | PI3Ka_4409 |
| MVP_top60_DNA_seed | PIK3CA | 0.9869 | SKCM | G-361 | PI3Ka_4409 |
| MVP_top60_DNA_seed | EGFR | 0.992908 | SKCM | M14 | Gefitinib |
| MVP_top60_DNA_seed | JAK2 | 0.995517 | SKCM | HT-144 | Ruxolitinib |
| MVP_top60_DNA_seed | JAK1 | 0.995517 | SKCM | HT-144 | Ruxolitinib |
| MVP_top60_DNA_seed | GSK3B | 0.996608 | SKCM | COLO-792 | SB216763 |
| MVP_top60_DNA_seed | BRAF | 0.998149 | SKCM | LB373-MEL-D | PLX-4720 |
| MVP_top60_DNA_seed | GSK3B | 0.998218 | SKCM | C32 | AZD7969 |
| MVP_top60_DNA_seed | EGFR | 0.998237 | SKCM | M14 | PF-00299804 |

| | | | | | |
|---|---|---|---|---|---|
| MVP_top60_DNA_seed | ERBB2 | 0.998237 | SKCM | M14 | PF-00299804 |
| MVP_top60_DNA_seed | JAK2 | 0.999348 | SKCM | COLO-783 | AZD1480 |
| MVP_top60_DNA_seed | JAK1 | 0.999348 | SKCM | COLO-783 | AZD1480 |
| MVP_top60_DNA_seed | JAK2 | 0.999507 | SKCM | MEL-HO | AZD1480 |
| MVP_top60_DNA_seed | JAK1 | 0.999507 | SKCM | MEL-HO | AZD1480 |
| MVP_top60_DNA_seed | EP300 | 0.999597 | SKCM | SK-MEL-24 | I-CBP112 |
| MVP_top60_DNA_seed | CREBBP | 0.999597 | SKCM | SK-MEL-24 | I-CBP112 |
| MVP_top60_DNA_seed | MAPK14 | 1.002789 | SKCM | UACC-62 | TAK-715 |
| MVP_top60_DNA_seed | JAK3 | 1.004954 | SKCM | WM1552C | WHI-P97 |
| MVP_top60_DNA_seed | JAK2 | 1.005373 | SKCM | COLO-829 | Fedratinib |
| MVP_top60_DNA_seed | PIK3CA | 1.007116 | SKCM | LB2518-MEL | AZD8835 |
| MVP_top60_DNA_seed | MAP2K1 | 1.007286 | SKCM | SK-MEL-31 | Refametinib |
| MVP_top60_DNA_seed | EP300 | 1.009256 | SKCM | WM-115 | I-CBP112 |
| MVP_top60_DNA_seed | CREBBP | 1.009256 | SKCM | WM-115 | I-CBP112 |
| MVP_top60_DNA_seed | EGFR | 1.012803 | SKCM | MEL-HO | Pelitinib |
| MVP_top60_DNA_seed | EGFR | 1.013235 | SKCM | COLO-679 | AZD8931 |
| MVP_top60_DNA_seed | ERBB2 | 1.013235 | SKCM | COLO-679 | AZD8931 |
| MVP_top60_DNA_seed | EGFR | 1.013365 | SKCM | COLO-829 | Afatinib |
| MVP_top60_DNA_seed | ERBB2 | 1.013365 | SKCM | COLO-829 | Afatinib |
| MVP_top60_DNA_seed | EGFR | 1.015465 | SKCM | SK-MEL-24 | Cetuximab |
| MVP_top60_DNA_seed | GSK3B | 1.016058 | SKCM | Mewo | SB216763 |
| MVP_top60_DNA_seed | EGFR | 1.016586 | SKCM | SK-MEL-3 | PF-00299804 |
| MVP_top60_DNA_seed | ERBB2 | 1.016586 | SKCM | SK-MEL-3 | PF-00299804 |
| MVP_top60_DNA_seed | PIK3CB | 1.017804 | SKCM | SK-MEL-2 | TGX221 |
| MVP_top60_DNA_seed | PIK3CB | 1.019176 | SKCM | HMV-II | AZD8186 |
| MVP_top60_DNA_seed | RAC1 | 1.021037 | SKCM | SH-4 | EHT-1864 |
| MVP_top60_DNA_seed | EGFR | 1.021926 | SKCM | Hs-940-T | Afatinib |
| MVP_top60_DNA_seed | ERBB2 | 1.021926 | SKCM | Hs-940-T | Afatinib |
| MVP_top60_DNA_seed | ERBB2 | 1.024466 | SKCM | K2 | CP724714 |
| MVP_top60_DNA_seed | JAK2 | 1.025889 | SKCM | Hs-939-T | AZD1480 |
| MVP_top60_DNA_seed | JAK1 | 1.025889 | SKCM | Hs-939-T | AZD1480 |
| MVP_top60_DNA_seed | PIK3CA | 1.026061 | SKCM | COLO-679 | PI3Ka_4409 |
| MVP_top60_DNA_seed | MAP2K1 | 1.026864 | SKCM | MEL-JUSO | Selumetinib |
| MVP_top60_DNA_seed | MAPK1 | 1.027535 | SKCM | VMRC-MELG | VX-11e |
| MVP_top60_DNA_seed | GSK3B | 1.027538 | SKCM | MZ7-mel | CHIR-99021 |
| MVP_top60_DNA_seed | EGFR | 1.032616 | SKCM | SK-MEL-3 | Gefitinib |
| MVP_top60_DNA_seed | GSK3B | 1.034535 | SKCM | UACC-257 | CHIR-99021 |
| MVP_top60_DNA_seed | PIK3CB | 1.036028 | SKCM | SK-MEL-2 | AZD8186 |
| MVP_top60_DNA_seed | EGFR | 1.036927 | SKCM | Mewo | Pelitinib |
| MVP_top60_DNA_seed | JAK1 | 1.037635 | SKCM | SK-MEL-24 | JAK1_3715 |
| MVP_top60_DNA_seed | GSK3B | 1.038074 | SKCM | M14 | KIN001-042 |

| | | | | | |
|---|---|---|---|---|---|
| MVP_top60_DNA_seed | EGFR | 1.040129 | SKCM | SK-MEL-2 | Afatinib |
| MVP_top60_DNA_seed | ERBB2 | 1.040129 | SKCM | SK-MEL-2 | Afatinib |
| MVP_top60_DNA_seed | EGFR | 1.040253 | SKCM | COLO-679 | PF-00299804 |
| MVP_top60_DNA_seed | ERBB2 | 1.040253 | SKCM | COLO-679 | PF-00299804 |
| MVP_top60_DNA_seed | MAPK14 | 1.040332 | SKCM | VMRC-MELG | TAK-715 |
| MVP_top60_DNA_seed | BRAF | 1.043476 | SKCM | COLO-792 | PLX-4720 |
| MVP_top60_DNA_seed | EGFR | 1.044719 | SKCM | WM1552C | Gefitinib |
| MVP_top60_DNA_seed | EGFR | 1.045475 | SKCM | RVH-421 | Afatinib |
| MVP_top60_DNA_seed | ERBB2 | 1.045475 | SKCM | RVH-421 | Afatinib |
| MVP_top60_DNA_seed | EGFR | 1.046442 | SKCM | MZ2-MEL | Gefitinib |
| MVP_top60_DNA_seed | EGFR | 1.046447 | SKCM | RVH-421 | CUDC-101 |
| MVP_top60_DNA_seed | ERBB2 | 1.046447 | SKCM | RVH-421 | CUDC-101 |
| MVP_top60_DNA_seed | ERBB2 | 1.046743 | SKCM | Hs-939-T | CP724714 |
| MVP_top60_DNA_seed | MAPK3 | 1.047308 | SKCM | IST-MEL1 | FR-180204 |
| MVP_top60_DNA_seed | MAPK1 | 1.047308 | SKCM | IST-MEL1 | FR-180204 |
| MVP_top60_DNA_seed | JAK3 | 1.047356 | SKCM | COLO-829 | WHI-P97 |
| MVP_top60_DNA_seed | GSK3B | 1.047706 | SKCM | UACC-257 | KIN001-042 |
| MVP_top60_DNA_seed | EGFR | 1.048105 | SKCM | SK-MEL-2 | Gefitinib |
| MVP_top60_DNA_seed | PIK3CA | 1.050329 | SKCM | C32 | AZD8835 |
| MVP_top60_DNA_seed | GSK3B | 1.051851 | SKCM | SK-MEL-1 | CHIR-99021 |
| MVP_top60_DNA_seed | EGFR | 1.054901 | SKCM | MZ2-MEL | PF-00299804 |
| MVP_top60_DNA_seed | ERBB2 | 1.054901 | SKCM | MZ2-MEL | PF-00299804 |
| MVP_top60_DNA_seed | PIK3CA | 1.055867 | SKCM | Mewo | AZD8835 |
| MVP_top60_DNA_seed | GSK3B | 1.057367 | SKCM | WM1552C | CHIR-99021 |
| MVP_top60_DNA_seed | EGFR | 1.061062 | SKCM | SK-MEL-1 | AST-1306 |
| MVP_top60_DNA_seed | MAPK3 | 1.061089 | SKCM | Hs-940-T | FR-180204 |
| MVP_top60_DNA_seed | MAPK1 | 1.061089 | SKCM | Hs-940-T | FR-180204 |
| MVP_top60_DNA_seed | EGFR | 1.062187 | SKCM | UACC-257 | PF-00299804 |
| MVP_top60_DNA_seed | ERBB2 | 1.062187 | SKCM | UACC-257 | PF-00299804 |
| MVP_top60_DNA_seed | EGFR | 1.062942 | SKCM | SK-MEL-1 | Afatinib |
| MVP_top60_DNA_seed | ERBB2 | 1.062942 | SKCM | SK-MEL-1 | Afatinib |
| MVP_top60_DNA_seed | JAK3 | 1.063065 | SKCM | SK-MEL-1 | WHI-P97 |
| MVP_top60_DNA_seed | RAC1 | 1.063418 | SKCM | IST-MEL1 | EHT-1864 |
| MVP_top60_DNA_seed | MAPK1 | 1.063438 | SKCM | SK-MEL-2 | VX-11e |
| MVP_top60_DNA_seed | MAPK1 | 1.06497 | SKCM | WM1552C | VX-11e |
| MVP_top60_DNA_seed | EGFR | 1.066917 | SKCM | SK-MEL-2 | Pelitinib |
| MVP_top60_DNA_seed | GSK3B | 1.067096 | SKCM | SH-4 | AZD7969 |
| MVP_top60_DNA_seed | BRAF | 1.068896 | SKCM | LB373-MEL-D | Dabrafenib |
| MVP_top60_DNA_seed | EGFR | 1.06901 | SKCM | M14 | Afatinib |
| MVP_top60_DNA_seed | ERBB2 | 1.06901 | SKCM | M14 | Afatinib |
| MVP_top60_DNA_seed | EGFR | 1.069397 | SKCM | IGR-37 | Pelitinib |

| | | | | | |
|---|---|---|---|---|---|
| MVP_top60_DNA_seed | EGFR | 1.070372 | SKCM | COLO-792 | Afatinib |
| MVP_top60_DNA_seed | ERBB2 | 1.070372 | SKCM | COLO-792 | Afatinib |
| MVP_top60_DNA_seed | JAK1 | 1.070663 | SKCM | Hs-940-T | JAK1_3715 |
| MVP_top60_DNA_seed | PIK3CA | 1.072282 | SKCM | IGR-1 | AZD8835 |
| MVP_top60_DNA_seed | EGFR | 1.076596 | SKCM | SK-MEL-5 | CUDC-101 |
| MVP_top60_DNA_seed | ERBB2 | 1.076596 | SKCM | SK-MEL-5 | CUDC-101 |
| MVP_top60_DNA_seed | EGFR | 1.078157 | SKCM | Hs-940-T | Afatinib |
| MVP_top60_DNA_seed | ERBB2 | 1.078157 | SKCM | Hs-940-T | Afatinib |
| MVP_top60_DNA_seed | EGFR | 1.081332 | SKCM | Mewo | PF-00299804 |
| MVP_top60_DNA_seed | ERBB2 | 1.081332 | SKCM | Mewo | PF-00299804 |
| MVP_top60_DNA_seed | RAC1 | 1.085875 | SKCM | LB373-MEL-D | EHT-1864 |
| MVP_top60_DNA_seed | EGFR | 1.086101 | SKCM | COLO-829 | AST-1306 |
| MVP_top60_DNA_seed | JAK3 | 1.086539 | SKCM | SH-4 | JAK3_7406 |
| MVP_top60_DNA_seed | RAC1 | 1.086619 | SKCM | SK-MEL-31 | EHT-1864 |
| MVP_top60_DNA_seed | EGFR | 1.08827 | SKCM | A101D | Afatinib |
| MVP_top60_DNA_seed | ERBB2 | 1.08827 | SKCM | A101D | Afatinib |
| MVP_top60_DNA_seed | BRAF | 1.089648 | SKCM | Mewo | PLX-4720 |
| MVP_top60_DNA_seed | EGFR | 1.093038 | SKCM | 451Lu | Afatinib |
| MVP_top60_DNA_seed | ERBB2 | 1.093038 | SKCM | 451Lu | Afatinib |
| MVP_top60_DNA_seed | PIK3CA | 1.093163 | SKCM | K2 | PI3Ka_4409 |
| MVP_top60_DNA_seed | GSK3B | 1.093202 | SKCM | SK-MEL-24 | CHIR-99021 |
| MVP_top60_DNA_seed | EGFR | 1.094504 | SKCM | WM-115 | Afatinib |
| MVP_top60_DNA_seed | ERBB2 | 1.094504 | SKCM | WM-115 | Afatinib |
| MVP_top60_DNA_seed | MAP2K1 | 1.095962 | SKCM | COLO-783 | Selumetinib |
| MVP_top60_DNA_seed | RAC1 | 1.097133 | SKCM | Hs-939-T | EHT-1864 |
| MVP_top60_DNA_seed | EGFR | 1.101788 | SKCM | COLO-679 | Gefitinib |
| MVP_top60_DNA_seed | EGFR | 1.10512 | SKCM | WM-115 | PF-00299804 |
| MVP_top60_DNA_seed | ERBB2 | 1.10512 | SKCM | WM-115 | PF-00299804 |
| MVP_top60_DNA_seed | JAK2 | 1.106618 | SKCM | WM793B | AZD1480 |
| MVP_top60_DNA_seed | JAK1 | 1.106618 | SKCM | WM793B | AZD1480 |
| MVP_top60_DNA_seed | JAK3 | 1.106955 | SKCM | CP50-MEL-B | JAK3_7406 |
| MVP_top60_DNA_seed | PIK3CA | 1.107456 | SKCM | SK-MEL-31 | PI3Ka_4409 |
| MVP_top60_DNA_seed | JAK1 | 1.109337 | SKCM | LB373-MEL-D | JAK1_3715 |
| MVP_top60_DNA_seed | GSK3B | 1.112397 | SKCM | SK-MEL-3 | CHIR-99021 |
| MVP_top60_DNA_seed | GSK3B | 1.115281 | SKCM | SK-MEL-1 | CHIR-99021 |
| MVP_top60_DNA_seed | EGFR | 1.118624 | SKCM | VMRC-MELG | Gefitinib |
| MVP_top60_DNA_seed | EGFR | 1.120784 | SKCM | COLO-829 | Pelitinib |
| MVP_top60_DNA_seed | JAK2 | 1.121636 | SKCM | RVH-421 | AZD1480 |
| MVP_top60_DNA_seed | JAK1 | 1.121636 | SKCM | RVH-421 | AZD1480 |
| MVP_top60_DNA_seed | RAC1 | 1.122353 | SKCM | WM793B | EHT-1864 |
| MVP_top60_DNA_seed | BRAF | 1.124 | SKCM | LB373-MEL-D | PLX-4720 |

| | | | | | |
|---|---|---|---|---|---|
| MVP_top60_DNA_seed | GSK3B | 1.124228 | SKCM | Mewo | KIN001-042 |
| MVP_top60_DNA_seed | ERBB2 | 1.125232 | SKCM | COLO-829 | CP724714 |
| MVP_top60_DNA_seed | EGFR | 1.132474 | SKCM | SK-MEL-28 | Pelitinib |
| MVP_top60_DNA_seed | EGFR | 1.13379 | SKCM | SK-MEL-1 | Erlotinib |
| MVP_top60_DNA_seed | JAK2 | 1.134434 | SKCM | COLO-829 | Ruxolitinib |
| MVP_top60_DNA_seed | JAK1 | 1.134434 | SKCM | COLO-829 | Ruxolitinib |
| MVP_top60_DNA_seed | RAC1 | 1.138323 | SKCM | SK-MEL-1 | EHT-1864 |
| MVP_top60_DNA_seed | EGFR | 1.138329 | SKCM | VMRC-MELG | Pelitinib |
| MVP_top60_DNA_seed | EGFR | 1.142121 | SKCM | SH-4 | CUDC-101 |
| MVP_top60_DNA_seed | ERBB2 | 1.142121 | SKCM | SH-4 | CUDC-101 |
| MVP_top60_DNA_seed | PIK3CA | 1.142659 | SKCM | SK-MEL-3 | AZD8835 |
| MVP_top60_DNA_seed | RAC1 | 1.146071 | SKCM | MEL-HO | EHT-1864 |
| MVP_top60_DNA_seed | EGFR | 1.146903 | SKCM | SH-4 | Erlotinib |
| MVP_top60_DNA_seed | BRAF | 1.147562 | SKCM | SK-MEL-28 | HG6-64-1 |
| MVP_top60_DNA_seed | EGFR | 1.150279 | SKCM | RVH-421 | Gefitinib |
| MVP_top60_DNA_seed | PIK3CB | 1.150866 | SKCM | SK-MEL-2 | AZD6482 |
| MVP_top60_DNA_seed | EGFR | 1.152199 | SKCM | LB373-MEL-D | Afatinib |
| MVP_top60_DNA_seed | ERBB2 | 1.152199 | SKCM | LB373-MEL-D | Afatinib |
| MVP_top60_DNA_seed | EGFR | 1.153038 | SKCM | UACC-257 | AST-1306 |
| MVP_top60_DNA_seed | EGFR | 1.153375 | SKCM | Mewo | AST-1306 |
| MVP_top60_DNA_seed | EGFR | 1.15892 | SKCM | GAK | Pelitinib |
| MVP_top60_DNA_seed | GSK3B | 1.159498 | SKCM | SK-MEL-24 | SB216763 |
| MVP_top60_DNA_seed | MAPK14 | 1.160767 | SKCM | Hs-939-T | TAK-715 |
| MVP_top60_DNA_seed | JAK3 | 1.162134 | SKCM | CP66-MEL | JAK3_7406 |
| MVP_top60_DNA_seed | JAK2 | 1.163446 | SKCM | 451Lu | Fedratinib |
| MVP_top60_DNA_seed | EGFR | 1.166361 | SKCM | A101D | PF-00299804 |
| MVP_top60_DNA_seed | ERBB2 | 1.166361 | SKCM | A101D | PF-00299804 |
| MVP_top60_DNA_seed | EGFR | 1.167407 | SKCM | LB373-MEL-D | Gefitinib |
| MVP_top60_DNA_seed | JAK2 | 1.168945 | SKCM | Mewo | Ruxolitinib |
| MVP_top60_DNA_seed | JAK1 | 1.168945 | SKCM | Mewo | Ruxolitinib |
| MVP_top60_DNA_seed | EGFR | 1.170785 | SKCM | SK-MEL-31 | Afatinib |
| MVP_top60_DNA_seed | ERBB2 | 1.170785 | SKCM | SK-MEL-31 | Afatinib |
| MVP_top60_DNA_seed | PIK3CB | 1.178165 | SKCM | SK-MEL-1 | TGX221 |
| MVP_top60_DNA_seed | BRAF | 1.178902 | SKCM | CP66-MEL | HG6-64-1 |
| MVP_top60_DNA_seed | JAK1 | 1.179134 | SKCM | Mewo | JAK1_3715 |
| MVP_top60_DNA_seed | EGFR | 1.180522 | SKCM | RVH-421 | Cetuximab |
| MVP_top60_DNA_seed | JAK2 | 1.181388 | SKCM | COLO-792 | AZD1480 |
| MVP_top60_DNA_seed | JAK1 | 1.181388 | SKCM | COLO-792 | AZD1480 |
| MVP_top60_DNA_seed | RAC1 | 1.185137 | SKCM | SK-MEL-28 | EHT-1864 |
| MVP_top60_DNA_seed | MAP2K1 | 1.187273 | SKCM | VMRC-MELG | CI-1040 |
| MVP_top60_DNA_seed | ERBB2 | 1.18889 | SKCM | SK-MEL-24 | CP724714 |

| | | | | | |
|---|---|---|---|---|---|
| MVP_top60_DNA_seed | JAK2 | 1.19011 | SKCM | C32 | AZD1480 |
| MVP_top60_DNA_seed | JAK1 | 1.19011 | SKCM | C32 | AZD1480 |
| MVP_top60_DNA_seed | EP300 | 1.190512 | SKCM | SK-MEL-1 | I-CBP112 |
| MVP_top60_DNA_seed | CREBBP | 1.190512 | SKCM | SK-MEL-1 | I-CBP112 |
| MVP_top60_DNA_seed | EGFR | 1.194281 | SKCM | CP66-MEL | Gefitinib |
| MVP_top60_DNA_seed | EGFR | 1.195582 | SKCM | IST-MEL1 | Gefitinib |
| MVP_top60_DNA_seed | EGFR | 1.197052 | SKCM | HMV-II | CI-1033 |
| MVP_top60_DNA_seed | ERBB2 | 1.197052 | SKCM | HMV-II | CI-1033 |
| MVP_top60_DNA_seed | ERBB2 | 1.200658 | SKCM | A101D | CP724714 |
| MVP_top60_DNA_seed | GSK3B | 1.204424 | SKCM | SK-MEL-31 | SB216763 |
| MVP_top60_DNA_seed | EGFR | 1.204607 | SKCM | COLO-829 | Lapatinib |
| MVP_top60_DNA_seed | ERBB2 | 1.204607 | SKCM | COLO-829 | Lapatinib |
| MVP_top60_DNA_seed | EGFR | 1.206715 | SKCM | UACC-257 | Gefitinib |
| MVP_top60_DNA_seed | MAP2K1 | 1.211663 | SKCM | K2 | Selumetinib |
| MVP_top60_DNA_seed | BRAF | 1.212192 | SKCM | SK-MEL-31 | SB590885 |
| MVP_top60_DNA_seed | PIK3CB | 1.213254 | SKCM | COLO-783 | AZD6482 |
| MVP_top60_DNA_seed | JAK1 | 1.214509 | SKCM | RVH-421 | JAK1_3715 |
| MVP_top60_DNA_seed | JAK1 | 1.221813 | SKCM | SH-4 | JAK1_3715 |
| MVP_top60_DNA_seed | PIK3CB | 1.22336 | SKCM | CP50-MEL-B | AZD6482 |
| MVP_top60_DNA_seed | EGFR | 1.225501 | SKCM | SK-MEL-1 | PF-00299804 |
| MVP_top60_DNA_seed | ERBB2 | 1.225501 | SKCM | SK-MEL-1 | PF-00299804 |
| MVP_top60_DNA_seed | JAK2 | 1.22636 | SKCM | SK-MEL-5 | Ruxolitinib |
| MVP_top60_DNA_seed | JAK1 | 1.22636 | SKCM | SK-MEL-5 | Ruxolitinib |
| MVP_top60_DNA_seed | GSK3B | 1.234077 | SKCM | SK-MEL-5 | CHIR-99021 |
| MVP_top60_DNA_seed | JAK3 | 1.234559 | SKCM | SK-MEL-30 | JAK3_7406 |
| MVP_top60_DNA_seed | EGFR | 1.234995 | SKCM | HT-144 | Lapatinib |
| MVP_top60_DNA_seed | ERBB2 | 1.234995 | SKCM | HT-144 | Lapatinib |
| MVP_top60_DNA_seed | EP300 | 1.237815 | SKCM | SH-4 | I-CBP112 |
| MVP_top60_DNA_seed | CREBBP | 1.237815 | SKCM | SH-4 | I-CBP112 |
| MVP_top60_DNA_seed | GSK3B | 1.25337 | SKCM | SK-MEL-24 | KIN001-042 |
| MVP_top60_DNA_seed | BRAF | 1.265911 | SKCM | COLO-800 | HG6-64-1 |
| MVP_top60_DNA_seed | GSK3B | 1.269864 | SKCM | MZ7-mel | KIN001-042 |
| MVP_top60_DNA_seed | GSK3B | 1.273936 | SKCM | HMV-II | KIN001-042 |
| MVP_top60_DNA_seed | EGFR | 1.275745 | SKCM | SK-MEL-5 | Gefitinib |
| MVP_top60_DNA_seed | JAK3 | 1.276418 | SKCM | HMV-II | WHI-P97 |
| MVP_top60_DNA_seed | JAK3 | 1.277838 | SKCM | COLO-783 | JAK3_7406 |
| MVP_top60_DNA_seed | EGFR | 1.27967 | SKCM | K2 | Afatinib |
| MVP_top60_DNA_seed | ERBB2 | 1.27967 | SKCM | K2 | Afatinib |
| MVP_top60_DNA_seed | EGFR | 1.279848 | SKCM | K2 | Pelitinib |
| MVP_top60_DNA_seed | JAK2 | 1.281345 | SKCM | SK-MEL-28 | AZD1480 |
| MVP_top60_DNA_seed | JAK1 | 1.281345 | SKCM | SK-MEL-28 | AZD1480 |

| | | | | | |
|---|---|---|---|---|---|
| MVP_top60_DNA_seed | PIK3CA | 1.284251 | SKCM | SH-4 | AZD8835 |
| MVP_top60_DNA_seed | MAPK3 | 1.287575 | SKCM | COLO-783 | FR-180204 |
| MVP_top60_DNA_seed | MAPK1 | 1.287575 | SKCM | COLO-783 | FR-180204 |
| MVP_top60_DNA_seed | PIK3CA | 1.28839 | SKCM | HMV-II | PI3Ka_4409 |
| MVP_top60_DNA_seed | EGFR | 1.290145 | SKCM | K2 | Afatinib |
| MVP_top60_DNA_seed | ERBB2 | 1.290145 | SKCM | K2 | Afatinib |
| MVP_top60_DNA_seed | EGFR | 1.294183 | SKCM | WM278 | Pelitinib |
| MVP_top60_DNA_seed | MAP2K1 | 1.295891 | SKCM | Hs-940-T | Selumetinib |
| MVP_top60_DNA_seed | EGFR | 1.311322 | SKCM | SK-MEL-1 | Lapatinib |
| MVP_top60_DNA_seed | ERBB2 | 1.311322 | SKCM | SK-MEL-1 | Lapatinib |
| MVP_top60_DNA_seed | BRAF | 1.312416 | SKCM | CP50-MEL-B | HG6-64-1 |
| MVP_top60_DNA_seed | EGFR | 1.313934 | SKCM | LB2518-MEL | Pelitinib |
| MVP_top60_DNA_seed | EGFR | 1.318079 | SKCM | COLO-829 | Gefitinib |
| MVP_top60_DNA_seed | EGFR | 1.325689 | SKCM | SK-MEL-31 | Cetuximab |
| MVP_top60_DNA_seed | RAC1 | 1.326086 | SKCM | M14 | EHT-1864 |
| MVP_top60_DNA_seed | JAK2 | 1.330389 | SKCM | 451Lu | AZD1480 |
| MVP_top60_DNA_seed | JAK1 | 1.330389 | SKCM | 451Lu | AZD1480 |
| MVP_top60_DNA_seed | JAK2 | 1.333807 | SKCM | IGR-1 | Fedratinib |
| MVP_top60_DNA_seed | MAPK1 | 1.334394 | SKCM | Hs-940-T | VX-11e |
| MVP_top60_DNA_seed | JAK3 | 1.335269 | SKCM | MZ7-mel | JAK3_7406 |
| MVP_top60_DNA_seed | EGFR | 1.335774 | SKCM | Hs-940-T | Gefitinib |
| MVP_top60_DNA_seed | JAK3 | 1.341284 | SKCM | LB2518-MEL | JAK3_7406 |
| MVP_top60_DNA_seed | MAPK3 | 1.345607 | SKCM | Mewo | FR-180204 |
| MVP_top60_DNA_seed | MAPK1 | 1.345607 | SKCM | Mewo | FR-180204 |
| MVP_top60_DNA_seed | JAK2 | 1.346871 | SKCM | SK-MEL-28 | Fedratinib |
| MVP_top60_DNA_seed | JAK2 | 1.354362 | SKCM | WM-115 | Ruxolitinib |
| MVP_top60_DNA_seed | JAK1 | 1.354362 | SKCM | WM-115 | Ruxolitinib |
| MVP_top60_DNA_seed | MAP2K1 | 1.357069 | SKCM | Hs-940-T | Selumetinib |
| MVP_top60_DNA_seed | JAK3 | 1.365613 | SKCM | VMRC-MELG | JAK3_7406 |
| MVP_top60_DNA_seed | JAK2 | 1.365697 | SKCM | K2 | AZD1480 |
| MVP_top60_DNA_seed | JAK1 | 1.365697 | SKCM | K2 | AZD1480 |
| MVP_top60_DNA_seed | PIK3CB | 1.366846 | SKCM | 451Lu | AZD8186 |
| MVP_top60_DNA_seed | JAK2 | 1.367418 | SKCM | MMAc-SF | AZD1480 |
| MVP_top60_DNA_seed | JAK1 | 1.367418 | SKCM | MMAc-SF | AZD1480 |
| MVP_top60_DNA_seed | JAK3 | 1.367967 | SKCM | Hs-940-T | WHI-P97 |
| MVP_top60_DNA_seed | EGFR | 1.370375 | SKCM | COLO-783 | Pelitinib |
| MVP_top60_DNA_seed | JAK2 | 1.371258 | SKCM | RVH-421 | Ruxolitinib |
| MVP_top60_DNA_seed | JAK1 | 1.371258 | SKCM | RVH-421 | Ruxolitinib |
| MVP_top60_DNA_seed | JAK2 | 1.371339 | SKCM | A375 | Fedratinib |
| MVP_top60_DNA_seed | JAK3 | 1.371968 | SKCM | SK-MEL-3 | JAK3_7406 |
| MVP_top60_DNA_seed | PIK3CA | 1.3723 | SKCM | MZ7-mel | AZD8835 |

| | | | | | |
|---|---|---|---|---|---|
| MVP_top60_DNA_seed | JAK2 | 1.375359 | SKCM | COLO-829 | AZD1480 |
| MVP_top60_DNA_seed | JAK1 | 1.375359 | SKCM | COLO-829 | AZD1480 |
| MVP_top60_DNA_seed | BRAF | 1.37581 | SKCM | COLO-792 | PLX-4720 |
| MVP_top60_DNA_seed | MAP2K1 | 1.380776 | SKCM | UACC-62 | Refametinib |
| MVP_top60_DNA_seed | PIK3CB | 1.398336 | SKCM | SK-MEL-30 | AZD8186 |
| MVP_top60_DNA_seed | MAPK1 | 1.407199 | SKCM | Mewo | VX-11e |
| MVP_top60_DNA_seed | BRAF | 1.413755 | SKCM | COLO-783 | HG6-64-1 |
| MVP_top60_DNA_seed | PIK3CA | 1.415069 | SKCM | Mewo | PI3Ka_4409 |
| MVP_top60_DNA_seed | ERBB2 | 1.419058 | SKCM | SK-MEL-1 | CP724714 |
| MVP_top60_DNA_seed | JAK1 | 1.427257 | SKCM | SK-MEL-31 | JAK1_3715 |
| MVP_top60_DNA_seed | EGFR | 1.427887 | SKCM | SH-4 | AZD8931 |
| MVP_top60_DNA_seed | ERBB2 | 1.427887 | SKCM | SH-4 | AZD8931 |
| MVP_top60_DNA_seed | EGFR | 1.431424 | SKCM | M14 | Pelitinib |
| MVP_top60_DNA_seed | GSK3B | 1.436343 | SKCM | SH-4 | CHIR-99021 |
| MVP_top60_DNA_seed | RAC1 | 1.436512 | SKCM | A101D | EHT-1864 |
| MVP_top60_DNA_seed | JAK2 | 1.438633 | SKCM | GAK | Fedratinib |
| MVP_top60_DNA_seed | RAC1 | 1.441465 | SKCM | LB2518-MEL | EHT-1864 |
| MVP_top60_DNA_seed | PIK3CA | 1.44298 | SKCM | C32 | PI3Ka_4409 |
| MVP_top60_DNA_seed | MAPK3 | 1.449872 | SKCM | SK-MEL-1 | FR-180204 |
| MVP_top60_DNA_seed | MAPK1 | 1.449872 | SKCM | SK-MEL-1 | FR-180204 |
| MVP_top60_DNA_seed | MAPK14 | 1.453469 | SKCM | C32 | TAK-715 |
| MVP_top60_DNA_seed | JAK3 | 1.469016 | SKCM | Mewo | WHI-P97 |
| MVP_top60_DNA_seed | EGFR | 1.472916 | SKCM | Hs-939-T | Pelitinib |
| MVP_top60_DNA_seed | PIK3CA | 1.474416 | SKCM | 451Lu | AZD8835 |
| MVP_top60_DNA_seed | JAK2 | 1.477958 | SKCM | K2 | Ruxolitinib |
| MVP_top60_DNA_seed | JAK1 | 1.477958 | SKCM | K2 | Ruxolitinib |
| MVP_top60_DNA_seed | JAK2 | 1.478885 | SKCM | SK-MEL-31 | AZD1480 |
| MVP_top60_DNA_seed | JAK1 | 1.478885 | SKCM | SK-MEL-31 | AZD1480 |
| MVP_top60_DNA_seed | GSK3B | 1.47968 | SKCM | SK-MEL-1 | AZD7969 |
| MVP_top60_DNA_seed | PIK3CB | 1.481121 | SKCM | SK-MEL-24 | AZD6482 |
| MVP_top60_DNA_seed | GSK3B | 1.482048 | SKCM | SH-4 | CHIR-99021 |
| MVP_top60_DNA_seed | RAC1 | 1.483558 | SKCM | SK-MEL-2 | EHT-1864 |
| MVP_top60_DNA_seed | EGFR | 1.483567 | SKCM | C32 | Afatinib |
| MVP_top60_DNA_seed | ERBB2 | 1.483567 | SKCM | C32 | Afatinib |
| MVP_top60_DNA_seed | JAK1 | 1.48462 | SKCM | HMV-II | JAK1_3715 |
| MVP_top60_DNA_seed | JAK2 | 1.488585 | SKCM | IGR-37 | Fedratinib |
| MVP_top60_DNA_seed | EGFR | 1.489599 | SKCM | A101D | Pelitinib |
| MVP_top60_DNA_seed | JAK2 | 1.491411 | SKCM | MZ7-mel | Fedratinib |
| MVP_top60_DNA_seed | EGFR | 1.491517 | SKCM | SK-MEL-31 | Gefitinib |
| MVP_top60_DNA_seed | EGFR | 1.499703 | SKCM | HMV-II | Afatinib |
| MVP_top60_DNA_seed | ERBB2 | 1.499703 | SKCM | HMV-II | Afatinib |

| | | | | | |
|---|---|---|---|---|---|
| MVP_top60_DNA_seed | EGFR | 1.517543 | SKCM | SK-MEL-1 | Cetuximab |
| MVP_top60_DNA_seed | RAC1 | 1.525037 | SKCM | WM-115 | EHT-1864 |
| MVP_top60_DNA_seed | JAK2 | 1.525742 | SKCM | MEL-HO | Fedratinib |
| MVP_top60_DNA_seed | BRAF | 1.525955 | SKCM | SK-MEL-1 | HG6-64-1 |
| MVP_top60_DNA_seed | JAK2 | 1.526855 | SKCM | CP50-MEL-B | Fedratinib |
| MVP_top60_DNA_seed | JAK2 | 1.52697 | SKCM | G-361 | Fedratinib |
| MVP_top60_DNA_seed | PIK3CA | 1.530646 | SKCM | SK-MEL-24 | AZD8835 |
| MVP_top60_DNA_seed | EGFR | 1.535219 | SKCM | SK-MEL-24 | Pelitinib |
| MVP_top60_DNA_seed | PIK3CA | 1.536904 | SKCM | MZ7-mel | PI3Ka_4409 |
| MVP_top60_DNA_seed | BRAF | 1.537083 | SKCM | SK-MEL-5 | HG6-64-1 |
| MVP_top60_DNA_seed | JAK1 | 1.537257 | SKCM | SK-MEL-1 | JAK1_3715 |
| MVP_top60_DNA_seed | PIK3CA | 1.542196 | SKCM | IGR-1 | PI3Ka_4409 |
| MVP_top60_DNA_seed | EGFR | 1.545233 | SKCM | Hs-939-T | Gefitinib |
| MVP_top60_DNA_seed | MAP2K1 | 1.554564 | SKCM | SK-MEL-31 | Selumetinib |
| MVP_top60_DNA_seed | EGFR | 1.559655 | SKCM | COLO-800 | CUDC-101 |
| MVP_top60_DNA_seed | ERBB2 | 1.559655 | SKCM | COLO-800 | CUDC-101 |
| MVP_top60_DNA_seed | JAK2 | 1.564051 | SKCM | UACC-62 | Fedratinib |
| MVP_top60_DNA_seed | GSK3B | 1.568573 | SKCM | SH-4 | KIN001-042 |
| MVP_top60_DNA_seed | EP300 | 1.577734 | SKCM | HMV-II | I-CBP112 |
| MVP_top60_DNA_seed | CREBBP | 1.577734 | SKCM | HMV-II | I-CBP112 |
| MVP_top60_DNA_seed | GSK3B | 1.589223 | SKCM | K2 | SB216763 |
| MVP_top60_DNA_seed | GSK3B | 1.59054 | SKCM | UACC-257 | AZD7969 |
| MVP_top60_DNA_seed | JAK2 | 1.60257 | SKCM | SK-MEL-1 | Ruxolitinib |
| MVP_top60_DNA_seed | JAK1 | 1.60257 | SKCM | SK-MEL-1 | Ruxolitinib |
| MVP_top60_DNA_seed | JAK2 | 1.607064 | SKCM | Mewo | Fedratinib |
| MVP_top60_DNA_seed | EGFR | 1.609626 | SKCM | SK-MEL-1 | Pelitinib |
| MVP_top60_DNA_seed | EP300 | 1.612256 | SKCM | C32 | I-CBP112 |
| MVP_top60_DNA_seed | CREBBP | 1.612256 | SKCM | C32 | I-CBP112 |
| MVP_top60_DNA_seed | MAPK14 | 1.650103 | SKCM | Hs-940-T | TAK-715 |
| MVP_top60_DNA_seed | EGFR | 1.650493 | SKCM | K2 | Gefitinib |
| MVP_top60_DNA_seed | RAC1 | 1.659207 | SKCM | C32 | EHT-1864 |
| MVP_top60_DNA_seed | JAK3 | 1.671423 | SKCM | SK-MEL-1 | JAK3_7406 |
| MVP_top60_DNA_seed | JAK2 | 1.687395 | SKCM | MZ7-mel | AZD1480 |
| MVP_top60_DNA_seed | JAK1 | 1.687395 | SKCM | MZ7-mel | AZD1480 |
| MVP_top60_DNA_seed | JAK2 | 1.689867 | SKCM | C32 | Ruxolitinib |
| MVP_top60_DNA_seed | JAK1 | 1.689867 | SKCM | C32 | Ruxolitinib |
| MVP_top60_DNA_seed | JAK2 | 1.69737 | SKCM | SK-MEL-3 | AZD1480 |
| MVP_top60_DNA_seed | JAK1 | 1.69737 | SKCM | SK-MEL-3 | AZD1480 |
| MVP_top60_DNA_seed | PIK3CA | 1.698605 | SKCM | SK-MEL-1 | PI3Ka_4409 |
| MVP_top60_DNA_seed | RAC1 | 1.717789 | SKCM | K2 | EHT-1864 |
| MVP_top60_DNA_seed | JAK2 | 1.730113 | SKCM | LB2518-MEL | Fedratinib |

| | | | | | |
|---|---|---|---|---|---|
| MVP_top60_DNA_seed | EGFR | 1.730119 | SKCM | C32 | CUDC-101 |
| MVP_top60_DNA_seed | ERBB2 | 1.730119 | SKCM | C32 | CUDC-101 |
| MVP_top60_DNA_seed | GSK3B | 1.732361 | SKCM | SK-MEL-31 | AZD7969 |
| MVP_top60_DNA_seed | JAK3 | 1.735496 | SKCM | LB373-MEL-D | JAK3_7406 |
| MVP_top60_DNA_seed | MAP2K1 | 1.738289 | SKCM | MZ7-mel | CI-1040 |
| MVP_top60_DNA_seed | PIK3CA | 1.74875 | SKCM | WM793B | AZD8835 |
| MVP_top60_DNA_seed | JAK3 | 1.756478 | SKCM | SK-MEL-31 | JAK3_7406 |
| MVP_top60_DNA_seed | JAK2 | 1.766061 | SKCM | CP66-MEL | Fedratinib |
| MVP_top60_DNA_seed | JAK2 | 1.769527 | SKCM | RVH-421 | Fedratinib |
| MVP_top60_DNA_seed | EGFR | 1.780036 | SKCM | SK-MEL-3 | AZD8931 |
| MVP_top60_DNA_seed | ERBB2 | 1.780036 | SKCM | SK-MEL-3 | AZD8931 |
| MVP_top60_DNA_seed | JAK2 | 1.82084 | SKCM | WM278 | Fedratinib |
| MVP_top60_DNA_seed | PIK3CA | 1.822467 | SKCM | M14 | PI3Ka_4409 |
| MVP_top60_DNA_seed | MAPK14 | 1.832376 | SKCM | Mewo | TAK-715 |
| MVP_top60_DNA_seed | JAK2 | 1.833976 | SKCM | HT-144 | Fedratinib |
| MVP_top60_DNA_seed | JAK2 | 1.853529 | SKCM | M14 | Fedratinib |
| MVP_top60_DNA_seed | BRAF | 1.855514 | SKCM | Mewo | HG6-64-1 |
| MVP_top60_DNA_seed | PIK3CA | 1.864362 | SKCM | SK-MEL-31 | AZD8835 |
| MVP_top60_DNA_seed | PIK3CB | 1.87633 | SKCM | SK-MEL-1 | AZD8186 |
| MVP_top60_DNA_seed | JAK2 | 1.884593 | SKCM | Hs-939-T | Fedratinib |
| MVP_top60_DNA_seed | GSK3B | 1.885757 | SKCM | VMRC-MELG | AZD7969 |
| MVP_top60_DNA_seed | EGFR | 1.887109 | SKCM | C32 | Pelitinib |
| MVP_top60_DNA_seed | RAC1 | 1.89529 | SKCM | CP66-MEL | EHT-1864 |
| MVP_top60_DNA_seed | JAK2 | 1.900155 | SKCM | K2 | Fedratinib |
| MVP_top60_DNA_seed | PIK3CB | 1.902261 | SKCM | SK-MEL-1 | AZD6482 |
| MVP_top60_DNA_seed | MAPK14 | 1.904226 | SKCM | COLO-829 | TAK-715 |
| MVP_top60_DNA_seed | EGFR | 1.928823 | SKCM | IPC-298 | AZD8931 |
| MVP_top60_DNA_seed | ERBB2 | 1.928823 | SKCM | IPC-298 | AZD8931 |
| MVP_top60_DNA_seed | PIK3CA | 1.946575 | SKCM | 451Lu | PI3Ka_4409 |
| MVP_top60_DNA_seed | JAK2 | 1.950246 | SKCM | LB373-MEL-D | Fedratinib |
| MVP_top60_DNA_seed | JAK2 | 1.964268 | SKCM | C32 | Fedratinib |
| MVP_top60_DNA_seed | JAK2 | 1.971767 | SKCM | SK-MEL-24 | Fedratinib |
| MVP_top60_DNA_seed | JAK2 | 1.979197 | SKCM | A101D | Fedratinib |
| MVP_top60_DNA_seed | ERBB2 | 1.982941 | SKCM | C32 | CP724714 |
| MVP_top60_DNA_seed | EP300 | 1.986073 | SKCM | SK-MEL-2 | I-CBP112 |
| MVP_top60_DNA_seed | CREBBP | 1.986073 | SKCM | SK-MEL-2 | I-CBP112 |
| MVP_top60_DNA_seed | EGFR | 2.025321 | SKCM | C32 | PF-00299804 |
| MVP_top60_DNA_seed | ERBB2 | 2.025321 | SKCM | C32 | PF-00299804 |
| MVP_top60_DNA_seed | PIK3CB | 2.029319 | SKCM | C32 | AZD6482 |
| MVP_top60_DNA_seed | PIK3CA | 2.039954 | SKCM | SK-MEL-1 | AZD8835 |
| MVP_top60_DNA_seed | BRAF | 2.054559 | SKCM | C32 | HG6-64-1 |

| | | | | | |
|---|---|---|---|---|---|
| MVP_top60_DNA_seed | BRAF | 2.055568 | SKCM | VMRC-MELG | RAF_9304 |
| MVP_top60_DNA_seed | JAK2 | 2.085036 | SKCM | SK-MEL-1 | Fedratinib |
| MVP_top60_DNA_seed | BRAF | 2.099185 | SKCM | Hs-940-T | RAF_9304 |
| MVP_top60_DNA_seed | JAK3 | 2.137641 | SKCM | C32 | WHI-P97 |
| MVP_top60_DNA_seed | EGFR | 2.179433 | SKCM | SK-MEL-31 | AZD8931 |
| MVP_top60_DNA_seed | ERBB2 | 2.179433 | SKCM | SK-MEL-31 | AZD8931 |
| MVP_top60_DNA_seed | GSK3B | 2.216102 | SKCM | C32 | KIN001-042 |
| MVP_top60_DNA_seed | MAPK3 | 2.277363 | SKCM | C32 | FR-180204 |
| MVP_top60_DNA_seed | MAPK1 | 2.277363 | SKCM | C32 | FR-180204 |
| MVP_top60_DNA_seed | EGFR | 2.298868 | SKCM | SK-MEL-1 | AZD8931 |
| MVP_top60_DNA_seed | ERBB2 | 2.298868 | SKCM | SK-MEL-1 | AZD8931 |
| MVP_top60_DNA_seed | JAK2 | 2.332877 | SKCM | SK-MEL-1 | AZD1480 |
| MVP_top60_DNA_seed | JAK1 | 2.332877 | SKCM | SK-MEL-1 | AZD1480 |
| MVP_top60_DNA_seed | EGFR | 2.551463 | SKCM | HMV-II | AZD8931 |
| MVP_top60_DNA_seed | ERBB2 | 2.551463 | SKCM | HMV-II | AZD8931 |
| MVP_top60_DNA_seed | EGFR | 2.640632 | SKCM | HT-144 | CUDC-101 |
| MVP_top60_DNA_seed | ERBB2 | 2.640632 | SKCM | HT-144 | CUDC-101 |
| MVP_top60_DNA_seed | JAK1 | 3.031742 | SKCM | C32 | JAK1_3715 |
| MVP_top60_DNA_seed | BRAF | -4.749342 | SKCM | G-MEL | Dabrafenib |
| MVP_top60_DNA_seed | BRAF | -4.690012 | SKCM | SK-MEL-28 | Dabrafenib |
| MVP_top60_DNA_seed | BRAF | -4.659915 | SKCM | CP50-MEL-B | Dabrafenib |
| MVP_top60_DNA_seed | BRAF | -4.254281 | SKCM | SK-MEL-28 | PLX-4720 |
| MVP_top60_DNA_seed | BRAF | -4.215455 | SKCM | M14 | Dabrafenib |
| MVP_top60_DNA_seed | BRAF | -4.160715 | SKCM | A375 | Dabrafenib |
| MVP_top60_DNA_seed | BRAF | -4.107077 | SKCM | C32 | Dabrafenib |
| MVP_top60_DNA_seed | BRAF | -4.090654 | SKCM | CP50-MEL-B | PLX-4720 |
| MVP_top60_DNA_seed | BRAF | -4.02731 | SKCM | IGR-37 | Dabrafenib |
| MVP_top60_DNA_seed | BRAF | -3.992912 | SKCM | G-MEL | SB590885 |
| MVP_top60_DNA_seed | BRAF | -3.991288 | SKCM | SH-4 | Dabrafenib |
| MVP_top60_DNA_seed | BRAF | -3.955507 | SKCM | UACC-62 | Dabrafenib |
| MVP_top60_DNA_seed | BRAF | -3.95452 | SKCM | HT-144 | Dabrafenib |
| MVP_top60_DNA_seed | BRAF | -3.747167 | SKCM | UACC-62 | SB590885 |
| MVP_top60_DNA_seed | BRAF | -3.732422 | SKCM | G-MEL | PLX-4720 |
| MVP_top60_DNA_seed | BRAF | -3.731334 | SKCM | WM35 | Dabrafenib |
| MVP_top60_DNA_seed | BRAF | -3.708066 | SKCM | MEL-HO | Dabrafenib |
| MVP_top60_DNA_seed | BRAF | -3.698163 | SKCM | K2 | Dabrafenib |
| MVP_top60_DNA_seed | BRAF | -3.6342 | SKCM | Hs-939-T | PLX-4720 |
| MVP_top60_DNA_seed | MAP2K1 | -3.552189 | SKCM | CP50-MEL-B | Selumetinib |
| MVP_top60_DNA_seed | BRAF | -3.492056 | SKCM | CP50-MEL-B | SB590885 |
| MVP_top60_DNA_seed | BRAF | -3.489487 | SKCM | Hs-939-T | Dabrafenib |
| MVP_top60_DNA_seed | BRAF | -3.482924 | SKCM | COLO-679 | Dabrafenib |

| | | | | | |
|---|---|---|---|---|---|
| MVP_top60_DNA_seed | MAP2K1 | -3.482251 | SKCM | IPC-298 | Selumetinib |
| MVP_top60_DNA_seed | BRAF | -3.47547 | SKCM | COLO-800 | Dabrafenib |
| MVP_top60_DNA_seed | MAP2K1 | -3.43862 | SKCM | G-MEL | Selumetinib |
| MVP_top60_DNA_seed | BRAF | -3.422786 | SKCM | IGR-37 | SB590885 |
| MVP_top60_DNA_seed | BRAF | -3.375367 | SKCM | MEL-HO | SB590885 |
| MVP_top60_DNA_seed | BRAF | -3.350935 | SKCM | 451Lu | Dabrafenib |
| MVP_top60_DNA_seed | BRAF | -3.282869 | SKCM | COLO-679 | SB590885 |
| MVP_top60_DNA_seed | BRAF | -3.279193 | SKCM | MZ7-mel | Dabrafenib |
| MVP_top60_DNA_seed | BRAF | -3.205137 | SKCM | WM278 | SB590885 |
| MVP_top60_DNA_seed | BRAF | -3.189075 | SKCM | UACC-62 | PLX-4720 |
| MVP_top60_DNA_seed | MAP2K1 | -3.155879 | SKCM | G-MEL | PD0325901 |
| MVP_top60_DNA_seed | BRAF | -3.145685 | SKCM | MEL-HO | PLX-4720 |
| MVP_top60_DNA_seed | BRAF | -3.139668 | SKCM | 451Lu | PLX-4720 |
| MVP_top60_DNA_seed | MAP2K1 | -3.110759 | SKCM | G-MEL | Trametinib |
| MVP_top60_DNA_seed | MAP2K1 | -3.097058 | SKCM | G-MEL | Selumetinib |
| MVP_top60_DNA_seed | BRAF | -3.092362 | SKCM | M14 | PLX-4720 |
| MVP_top60_DNA_seed | MAP2K1 | -3.088878 | SKCM | IGR-37 | Selumetinib |
| MVP_top60_DNA_seed | BRAF | -3.076376 | SKCM | IGR-37 | PLX-4720 |
| MVP_top60_DNA_seed | MAP2K1 | -2.993739 | SKCM | CP50-MEL-B | PD0325901 |
| MVP_top60_DNA_seed | BRAF | -2.982128 | SKCM | COLO-800 | PLX-4720 |
| MVP_top60_DNA_seed | MAP2K1 | -2.944054 | SKCM | K2 | Selumetinib |
| MVP_top60_DNA_seed | MAP2K1 | -2.93872 | SKCM | M14 | Trametinib |
| MVP_top60_DNA_seed | BRAF | -2.934823 | SKCM | A375 | PLX-4720 |
| MVP_top60_DNA_seed | MAP2K1 | -2.922682 | SKCM | CP50-MEL-B | Refametinib |
| MVP_top60_DNA_seed | MAP2K1 | -2.916285 | SKCM | CP66-MEL | Selumetinib |
| MVP_top60_DNA_seed | BRAF | -2.899576 | SKCM | K2 | PLX-4720 |
| MVP_top60_DNA_seed | MAP2K1 | -2.85862 | SKCM | CP50-MEL-B | Trametinib |
| MVP_top60_DNA_seed | BRAF | -2.855013 | SKCM | A101D | Dabrafenib |
| MVP_top60_DNA_seed | MAP2K1 | -2.841852 | SKCM | M14 | Selumetinib |
| MVP_top60_DNA_seed | MAPK3 | -2.835554 | SKCM | SK-MEL-28 | Ulixertinib |
| MVP_top60_DNA_seed | MAPK1 | -2.835554 | SKCM | SK-MEL-28 | Ulixertinib |
| MVP_top60_DNA_seed | BRAF | -2.832276 | SKCM | C32 | PLX-4720 |
| MVP_top60_DNA_seed | MAP2K1 | -2.831211 | SKCM | MZ7-mel | Selumetinib |
| MVP_top60_DNA_seed | MAP2K1 | -2.822679 | SKCM | IPC-298 | PD0325901 |
| MVP_top60_DNA_seed | BRAF | -2.812637 | SKCM | HT-144 | PLX-4720 |
| MVP_top60_DNA_seed | BRAF | -2.78915 | SKCM | WM35 | SB590885 |
| MVP_top60_DNA_seed | BRAF | -2.780108 | SKCM | UACC-257 | SB590885 |
| MVP_top60_DNA_seed | MAPK3 | -2.773647 | SKCM | G-MEL | Ulixertinib |
| MVP_top60_DNA_seed | MAPK1 | -2.773647 | SKCM | G-MEL | Ulixertinib |
| MVP_top60_DNA_seed | MAP2K1 | -2.772513 | SKCM | SK-MEL-28 | Selumetinib |
| MVP_top60_DNA_seed | MAP2K1 | -2.753236 | SKCM | SH-4 | Trametinib |

| | | | | | |
|---|---|---|---|---|---|
| MVP_top60_DNA_seed | MAP2K1 | -2.751778 | SKCM | UACC-62 | Selumetinib |
| MVP_top60_DNA_seed | MAP2K1 | -2.748112 | SKCM | UACC-62 | PD0325901 |
| MVP_top60_DNA_seed | BRAF | -2.742669 | SKCM | COLO-679 | PLX-4720 |
| MVP_top60_DNA_seed | MAP2K1 | -2.732855 | SKCM | M14 | PD0325901 |
| MVP_top60_DNA_seed | BRAF | -2.722951 | SKCM | A375 | SB590885 |
| MVP_top60_DNA_seed | MAP2K1 | -2.722853 | SKCM | G-MEL | Refametinib |
| MVP_top60_DNA_seed | MAP2K1 | -2.722648 | SKCM | 451Lu | Selumetinib |
| MVP_top60_DNA_seed | MAP2K1 | -2.712123 | SKCM | IGR-37 | PD0325901 |
| MVP_top60_DNA_seed | BRAF | -2.709563 | SKCM | SH-4 | PLX-4720 |
| MVP_top60_DNA_seed | BRAF | -2.686826 | SKCM | SK-MEL-1 | Dabrafenib |
| MVP_top60_DNA_seed | BRAF | -2.682732 | SKCM | K2 | SB590885 |
| MVP_top60_DNA_seed | MAP2K1 | -2.672794 | SKCM | COLO-679 | Trametinib |
| MVP_top60_DNA_seed | MAPK3 | -2.667315 | SKCM | CP50-MEL-B | SCH772984 |
| MVP_top60_DNA_seed | MAPK1 | -2.667315 | SKCM | CP50-MEL-B | SCH772984 |
| MVP_top60_DNA_seed | MAP2K1 | -2.657585 | SKCM | MEL-HO | Selumetinib |
| MVP_top60_DNA_seed | MAP2K1 | -2.655351 | SKCM | IPC-298 | Refametinib |
| MVP_top60_DNA_seed | MAP2K1 | -2.650814 | SKCM | WM35 | Trametinib |
| MVP_top60_DNA_seed | MAP2K1 | -2.647445 | SKCM | UACC-62 | Refametinib |
| MVP_top60_DNA_seed | BRAF | -2.646627 | SKCM | G-361 | Dabrafenib |
| MVP_top60_DNA_seed | MAP2K1 | -2.644394 | SKCM | SK-MEL-28 | PD0325901 |
| MVP_top60_DNA_seed | MAPK3 | -2.63665 | SKCM | MZ7-mel | SCH772984 |
| MVP_top60_DNA_seed | MAPK1 | -2.63665 | SKCM | MZ7-mel | SCH772984 |
| MVP_top60_DNA_seed | MAP2K1 | -2.634892 | SKCM | SH-4 | PD0325901 |
| MVP_top60_DNA_seed | MAP2K1 | -2.632691 | SKCM | UACC-62 | Trametinib |
| MVP_top60_DNA_seed | MAP2K1 | -2.630649 | SKCM | C32 | PD0325901 |
| MVP_top60_DNA_seed | BRAF | -2.627807 | SKCM | 451Lu | SB590885 |
| MVP_top60_DNA_seed | MAP2K1 | -2.624875 | SKCM | LB2518-MEL | Selumetinib |
| MVP_top60_DNA_seed | BRAF | -2.620304 | SKCM | HT-144 | SB590885 |
| MVP_top60_DNA_seed | MAP2K1 | -2.605855 | SKCM | SH-4 | Refametinib |
| MVP_top60_DNA_seed | BRAF | -2.605377 | SKCM | MMAc-SF | PLX-4720 |
| MVP_top60_DNA_seed | MAP2K1 | -2.599717 | SKCM | IPC-298 | Trametinib |
| MVP_top60_DNA_seed | MAPK3 | -2.59758 | SKCM | CP50-MEL-B | Ulixertinib |
| MVP_top60_DNA_seed | MAPK1 | -2.59758 | SKCM | CP50-MEL-B | Ulixertinib |
| MVP_top60_DNA_seed | MAP2K1 | -2.597146 | SKCM | LB2518-MEL | PD0325901 |
| MVP_top60_DNA_seed | MAP2K1 | -2.594749 | SKCM | IGR-37 | Trametinib |
| MVP_top60_DNA_seed | MAP2K1 | -2.594502 | SKCM | IGR-37 | Refametinib |
| MVP_top60_DNA_seed | MAP2K1 | -2.589407 | SKCM | K2 | PD0325901 |
| MVP_top60_DNA_seed | BRAF | -2.57747 | SKCM | COLO-783 | SB590885 |
| MVP_top60_DNA_seed | MAP2K1 | -2.571909 | SKCM | COLO-679 | Selumetinib |
| MVP_top60_DNA_seed | BRAF | -2.567505 | SKCM | COLO-783 | PLX-4720 |
| MVP_top60_DNA_seed | MAP2K1 | -2.550407 | SKCM | SH-4 | Selumetinib |

| | | | | | |
|---|---|---|---|---|---|
| MVP_top60_DNA_seed | BRAF | -2.548212 | SKCM | SK-MEL-1 | SB590885 |
| MVP_top60_DNA_seed | BRAF | -2.545772 | SKCM | A101D | SB590885 |
| MVP_top60_DNA_seed | BRAF | -2.543506 | SKCM | Hs-939-T | SB590885 |
| MVP_top60_DNA_seed | MAP2K1 | -2.530352 | SKCM | Hs-939-T | Selumetinib |
| MVP_top60_DNA_seed | MAP2K1 | -2.51119 | SKCM | C32 | Selumetinib |
| MVP_top60_DNA_seed | MAP2K1 | -2.505585 | SKCM | K2 | Trametinib |
| MVP_top60_DNA_seed | MAPK3 | -2.50447 | SKCM | M14 | SCH772984 |
| MVP_top60_DNA_seed | MAPK1 | -2.50447 | SKCM | M14 | SCH772984 |
| MVP_top60_DNA_seed | MAP2K1 | -2.494199 | SKCM | COLO-783 | Selumetinib |
| MVP_top60_DNA_seed | MAP2K1 | -2.473138 | SKCM | SH-4 | Selumetinib |
| MVP_top60_DNA_seed | MAP2K1 | -2.451865 | SKCM | COLO-679 | Refametinib |
| MVP_top60_DNA_seed | MAP2K1 | -2.444091 | SKCM | C32 | Selumetinib |
| MVP_top60_DNA_seed | MAP2K1 | -2.436441 | SKCM | C32 | Trametinib |
| MVP_top60_DNA_seed | MAP2K1 | -2.433307 | SKCM | COLO-679 | Selumetinib |
| MVP_top60_DNA_seed | MAP2K1 | -2.431327 | SKCM | CP66-MEL | Selumetinib |
| MVP_top60_DNA_seed | MAP2K1 | -2.426451 | SKCM | CP66-MEL | PD0325901 |
| MVP_top60_DNA_seed | MAP2K1 | -2.411795 | SKCM | COLO-679 | PD0325901 |
| MVP_top60_DNA_seed | BRAF | -2.399939 | SKCM | COLO-829 | PLX-4720 |
| MVP_top60_DNA_seed | MAPK1 | -2.377244 | SKCM | SH-4 | VX-11e |
| MVP_top60_DNA_seed | MAPK3 | -2.376194 | SKCM | WM35 | SCH772984 |
| MVP_top60_DNA_seed | MAPK1 | -2.376194 | SKCM | WM35 | SCH772984 |
| MVP_top60_DNA_seed | MAPK3 | -2.372189 | SKCM | HT-144 | SCH772984 |
| MVP_top60_DNA_seed | MAPK1 | -2.372189 | SKCM | HT-144 | SCH772984 |
| MVP_top60_DNA_seed | MAP2K1 | -2.366541 | SKCM | A375 | PD0325901 |
| MVP_top60_DNA_seed | BRAF | -2.355924 | SKCM | SK-MEL-28 | SB590885 |
| MVP_top60_DNA_seed | MAP2K1 | -2.35392 | SKCM | WM35 | Selumetinib |
| MVP_top60_DNA_seed | MAP2K1 | -2.347397 | SKCM | MEL-HO | PD0325901 |
| MVP_top60_DNA_seed | MAP2K1 | -2.347301 | SKCM | A375 | Selumetinib |
| MVP_top60_DNA_seed | MAP2K1 | -2.344159 | SKCM | IGR-37 | Selumetinib |
| MVP_top60_DNA_seed | MAP2K1 | -2.34291 | SKCM | SK-MEL-30 | Selumetinib |
| MVP_top60_DNA_seed | MAP2K1 | -2.335528 | SKCM | HT-144 | Trametinib |
| MVP_top60_DNA_seed | MAPK3 | -2.333343 | SKCM | G-MEL | SCH772984 |
| MVP_top60_DNA_seed | MAPK1 | -2.333343 | SKCM | G-MEL | SCH772984 |
| MVP_top60_DNA_seed | MAP2K1 | -2.328341 | SKCM | LB2518-MEL | Refametinib |
| MVP_top60_DNA_seed | MAP2K1 | -2.326105 | SKCM | SK-MEL-1 | PD0325901 |
| MVP_top60_DNA_seed | MAP2K1 | -2.325255 | SKCM | SK-MEL-28 | Trametinib |
| MVP_top60_DNA_seed | MAPK3 | -2.322594 | SKCM | SH-4 | SCH772984 |
| MVP_top60_DNA_seed | MAPK1 | -2.322594 | SKCM | SH-4 | SCH772984 |
| MVP_top60_DNA_seed | MAPK3 | -2.319386 | SKCM | M14 | Ulixertinib |
| MVP_top60_DNA_seed | MAPK1 | -2.319386 | SKCM | M14 | Ulixertinib |
| MVP_top60_DNA_seed | BRAF | -2.315428 | SKCM | UACC-257 | Dabrafenib |

| | | | | | |
|---|---|---|---|---|---|
| MVP_top60_DNA_seed | MAP2K1 | -2.312432 | SKCM | MEL-HO | Trametinib |
| MVP_top60_DNA_seed | BRAF | -2.298401 | SKCM | COLO-800 | SB590885 |
| MVP_top60_DNA_seed | MAPK3 | -2.280169 | SKCM | COLO-679 | SCH772984 |
| MVP_top60_DNA_seed | MAPK1 | -2.280169 | SKCM | COLO-679 | SCH772984 |
| MVP_top60_DNA_seed | MAPK3 | -2.274379 | SKCM | K2 | SCH772984 |
| MVP_top60_DNA_seed | MAPK1 | -2.274379 | SKCM | K2 | SCH772984 |
| MVP_top60_DNA_seed | BRAF | -2.273836 | SKCM | WM793B | SB590885 |
| MVP_top60_DNA_seed | RAC1 | -2.273696 | SKCM | A2058 | EHT-1864 |
| MVP_top60_DNA_seed | MAP2K1 | -2.270637 | SKCM | COLO-800 | PD0325901 |
| MVP_top60_DNA_seed | MAPK3 | -2.267008 | SKCM | SH-4 | Ulixertinib |
| MVP_top60_DNA_seed | MAPK1 | -2.267008 | SKCM | SH-4 | Ulixertinib |
| MVP_top60_DNA_seed | MAP2K1 | -2.262342 | SKCM | UACC-62 | Selumetinib |
| MVP_top60_DNA_seed | MAP2K1 | -2.256774 | SKCM | MZ7-mel | Refametinib |
| MVP_top60_DNA_seed | MAP2K1 | -2.253501 | SKCM | A375 | Trametinib |
| MVP_top60_DNA_seed | BRAF | -2.248185 | SKCM | MZ7-mel | PLX-4720 |
| MVP_top60_DNA_seed | BRAF | -2.24625 | SKCM | SH-4 | SB590885 |
| MVP_top60_DNA_seed | MAP2K1 | -2.24311 | SKCM | COLO-800 | Trametinib |
| MVP_top60_DNA_seed | MAP2K1 | -2.241345 | SKCM | HMV-II | Selumetinib |
| MVP_top60_DNA_seed | MAPK3 | -2.239943 | SKCM | SK-MEL-28 | SCH772984 |
| MVP_top60_DNA_seed | MAPK1 | -2.239943 | SKCM | SK-MEL-28 | SCH772984 |
| MVP_top60_DNA_seed | MAP2K1 | -2.236189 | SKCM | C32 | Refametinib |
| MVP_top60_DNA_seed | MAPK3 | -2.235109 | SKCM | MEL-HO | SCH772984 |
| MVP_top60_DNA_seed | MAPK1 | -2.235109 | SKCM | MEL-HO | SCH772984 |
| MVP_top60_DNA_seed | MAPK3 | -2.232363 | SKCM | COLO-783 | SCH772984 |
| MVP_top60_DNA_seed | MAPK1 | -2.232363 | SKCM | COLO-783 | SCH772984 |
| MVP_top60_DNA_seed | MAP2K1 | -2.230612 | SKCM | WM278 | Selumetinib |
| MVP_top60_DNA_seed | MAP2K1 | -2.229385 | SKCM | MZ7-mel | PD0325901 |
| MVP_top60_DNA_seed | MAP2K1 | -2.220539 | SKCM | MEL-HO | Refametinib |
| MVP_top60_DNA_seed | MAPK3 | -2.213335 | SKCM | UACC-62 | Ulixertinib |
| MVP_top60_DNA_seed | MAPK1 | -2.213335 | SKCM | UACC-62 | Ulixertinib |
| MVP_top60_DNA_seed | MAP2K1 | -2.211931 | SKCM | HMV-II | Trametinib |
| MVP_top60_DNA_seed | MAPK3 | -2.21103 | SKCM | IPC-298 | SCH772984 |
| MVP_top60_DNA_seed | MAPK1 | -2.21103 | SKCM | IPC-298 | SCH772984 |
| MVP_top60_DNA_seed | MAP2K1 | -2.189513 | SKCM | 451Lu | PD0325901 |
| MVP_top60_DNA_seed | BRAF | -2.184988 | SKCM | WM278 | PLX-4720 |
| MVP_top60_DNA_seed | MAP2K1 | -2.178323 | SKCM | HT-144 | Selumetinib |
| MVP_top60_DNA_seed | MAP2K1 | -2.174601 | SKCM | MZ7-mel | Trametinib |
| MVP_top60_DNA_seed | BRAF | -2.166504 | SKCM | C32 | SB590885 |
| MVP_top60_DNA_seed | MAP2K1 | -2.163624 | SKCM | WM35 | PD0325901 |
| MVP_top60_DNA_seed | MAP2K1 | -2.152862 | SKCM | UACC-257 | Selumetinib |
| MVP_top60_DNA_seed | MAPK3 | -2.152619 | SKCM | IPC-298 | Ulixertinib |

| | | | | | |
|---|---|---|---|---|---|
| MVP_top60_DNA_seed | MAPK1 | -2.152619 | SKCM | IPC-298 | Ulixertinib |
| MVP_top60_DNA_seed | MAP2K1 | -2.147712 | SKCM | MZ2-MEL | Selumetinib |
| MVP_top60_DNA_seed | MAP2K1 | -2.147026 | SKCM | WM35 | Refametinib |
| MVP_top60_DNA_seed | MAPK3 | -2.137663 | SKCM | COLO-800 | SCH772984 |
| MVP_top60_DNA_seed | MAPK1 | -2.137663 | SKCM | COLO-800 | SCH772984 |
| MVP_top60_DNA_seed | MAPK3 | -2.135038 | SKCM | UACC-62 | SCH772984 |
| MVP_top60_DNA_seed | MAPK1 | -2.135038 | SKCM | UACC-62 | SCH772984 |
| MVP_top60_DNA_seed | MAP2K1 | -2.128967 | SKCM | COLO-800 | Selumetinib |
| MVP_top60_DNA_seed | MAP2K1 | -2.11581 | SKCM | K2 | Refametinib |
| MVP_top60_DNA_seed | GSK3B | -2.113951 | SKCM | SK-MEL-2 | SB216763 |
| MVP_top60_DNA_seed | MAPK1 | -2.110834 | SKCM | CP50-MEL-B | VX-11e |
| MVP_top60_DNA_seed | MAP2K1 | -2.107213 | SKCM | A375 | Refametinib |
| MVP_top60_DNA_seed | MAP2K1 | -2.097859 | SKCM | HT-144 | PD0325901 |
| MVP_top60_DNA_seed | BRAF | -2.097784 | SKCM | SK-MEL-24 | Dabrafenib |
| MVP_top60_DNA_seed | MAPK1 | -2.09221 | SKCM | HT-144 | VX-11e |
| MVP_top60_DNA_seed | MAPK3 | -2.091357 | SKCM | SK-MEL-30 | SCH772984 |
| MVP_top60_DNA_seed | MAPK1 | -2.091357 | SKCM | SK-MEL-30 | SCH772984 |
| MVP_top60_DNA_seed | MAPK1 | -2.091052 | SKCM | LOXIMVI | VX-11e |
| MVP_top60_DNA_seed | MAP2K1 | -2.086895 | SKCM | A375 | Selumetinib |
| MVP_top60_DNA_seed | MAPK3 | -2.080631 | SKCM | C32 | SCH772984 |
| MVP_top60_DNA_seed | MAPK1 | -2.080631 | SKCM | C32 | SCH772984 |
| MVP_top60_DNA_seed | BRAF | -2.079743 | SKCM | G-361 | PLX-4720 |
| MVP_top60_DNA_seed | MAP2K1 | -2.053889 | SKCM | Hs-939-T | PD0325901 |
| MVP_top60_DNA_seed | MAPK1 | -2.051063 | SKCM | G-MEL | VX-11e |
| MVP_top60_DNA_seed | BRAF | -2.048532 | SKCM | G-361 | SB590885 |
| MVP_top60_DNA_seed | MAPK3 | -2.033086 | SKCM | MEL-HO | Ulixertinib |
| MVP_top60_DNA_seed | MAPK1 | -2.033086 | SKCM | MEL-HO | Ulixertinib |
| MVP_top60_DNA_seed | MAP2K1 | -2.016321 | SKCM | SK-MEL-1 | Trametinib |
| MVP_top60_DNA_seed | GSK3B | -2.011794 | SKCM | COLO-829 | SB216763 |
| MVP_top60_DNA_seed | MAP2K1 | -1.995084 | SKCM | 451Lu | Refametinib |
| MVP_top60_DNA_seed | MAPK3 | -1.985991 | SKCM | SK-MEL-30 | Ulixertinib |
| MVP_top60_DNA_seed | MAPK1 | -1.985991 | SKCM | SK-MEL-30 | Ulixertinib |
| MVP_top60_DNA_seed | MAP2K1 | -1.984689 | SKCM | SK-MEL-1 | Selumetinib |
| MVP_top60_DNA_seed | BRAF | -1.979924 | SKCM | SK-MEL-24 | SB590885 |
| MVP_top60_DNA_seed | MAP2K1 | -1.976511 | SKCM | HT-144 | Refametinib |
| MVP_top60_DNA_seed | MAP2K1 | -1.970819 | SKCM | 451Lu | Trametinib |
| MVP_top60_DNA_seed | MAP2K1 | -1.962531 | SKCM | HT-144 | Selumetinib |
| MVP_top60_DNA_seed | MAPK3 | -1.951521 | SKCM | K2 | Ulixertinib |
| MVP_top60_DNA_seed | MAPK1 | -1.951521 | SKCM | K2 | Ulixertinib |
| MVP_top60_DNA_seed | MAPK1 | -1.950415 | SKCM | COLO-800 | VX-11e |
| MVP_top60_DNA_seed | MAPK3 | -1.943518 | SKCM | COLO-783 | Ulixertinib |

| | | | | | |
|---|---|---|---|---|---|
| MVP_top60_DNA_seed | MAPK1 | -1.943518 | SKCM | COLO-783 | Ulixertinib |
| MVP_top60_DNA_seed | BRAF | -1.937656 | SKCM | A101D | PLX-4720 |
| MVP_top60_DNA_seed | MAP2K1 | -1.929417 | SKCM | UACC-257 | Refametinib |
| MVP_top60_DNA_seed | MAPK3 | -1.922601 | SKCM | HMV-II | SCH772984 |
| MVP_top60_DNA_seed | MAPK1 | -1.922601 | SKCM | HMV-II | SCH772984 |
| MVP_top60_DNA_seed | MAP2K1 | -1.910581 | SKCM | UACC-257 | PD0325901 |
| MVP_top60_DNA_seed | MAP2K1 | -1.908154 | SKCM | MMAc-SF | PD0325901 |
| MVP_top60_DNA_seed | MAP2K1 | -1.908098 | SKCM | SK-MEL-30 | Trametinib |
| MVP_top60_DNA_seed | BRAF | -1.905691 | SKCM | SK-MEL-3 | Dabrafenib |
| MVP_top60_DNA_seed | BRAF | -1.905041 | SKCM | Hs-940-T | Dabrafenib |
| MVP_top60_DNA_seed | MAPK1 | -1.902392 | SKCM | M14 | VX-11e |
| MVP_top60_DNA_seed | MAP2K1 | -1.890006 | SKCM | COLO-800 | Refametinib |
| MVP_top60_DNA_seed | MAP2K1 | -1.880715 | SKCM | Hs-939-T | Trametinib |
| MVP_top60_DNA_seed | MAPK3 | -1.880169 | SKCM | 451Lu | SCH772984 |
| MVP_top60_DNA_seed | MAPK1 | -1.880169 | SKCM | 451Lu | SCH772984 |
| MVP_top60_DNA_seed | MAP2K1 | -1.877505 | SKCM | G-361 | Selumetinib |
| MVP_top60_DNA_seed | MAPK3 | -1.867353 | SKCM | COLO-679 | Ulixertinib |
| MVP_top60_DNA_seed | MAPK1 | -1.867353 | SKCM | COLO-679 | Ulixertinib |
| MVP_top60_DNA_seed | MAPK3 | -1.86715 | SKCM | LOXIMVI | SCH772984 |
| MVP_top60_DNA_seed | MAPK1 | -1.86715 | SKCM | LOXIMVI | SCH772984 |
| MVP_top60_DNA_seed | MAP2K1 | -1.862247 | SKCM | SK-MEL-30 | Refametinib |
| MVP_top60_DNA_seed | MAPK3 | -1.853978 | SKCM | A375 | SCH772984 |
| MVP_top60_DNA_seed | MAPK1 | -1.853978 | SKCM | A375 | SCH772984 |
| MVP_top60_DNA_seed | MAP2K1 | -1.84896 | SKCM | HMV-II | PD0325901 |
| MVP_top60_DNA_seed | MAPK3 | -1.840384 | SKCM | M14 | Ulixertinib |
| MVP_top60_DNA_seed | MAPK1 | -1.840384 | SKCM | M14 | Ulixertinib |
| MVP_top60_DNA_seed | MAPK1 | -1.835075 | SKCM | SK-MEL-28 | VX-11e |
| MVP_top60_DNA_seed | MAP2K1 | -1.820607 | SKCM | CP66-MEL | Trametinib |
| MVP_top60_DNA_seed | BRAF | -1.817332 | SKCM | MMAc-SF | Dabrafenib |
| MVP_top60_DNA_seed | EGFR | -1.815562 | SKCM | CHL-1 | Sapitinib |
| MVP_top60_DNA_seed | ERBB2 | -1.815562 | SKCM | CHL-1 | Sapitinib |
| MVP_top60_DNA_seed | MAP2K1 | -1.814267 | SKCM | COLO-783 | PD0325901 |
| MVP_top60_DNA_seed | MAP2K1 | -1.810216 | SKCM | UACC-257 | Selumetinib |
| MVP_top60_DNA_seed | MAP2K1 | -1.802946 | SKCM | G-361 | PD0325901 |
| MVP_top60_DNA_seed | GSK3B | -1.786498 | SKCM | COLO-792 | SB216763 |
| MVP_top60_DNA_seed | MAPK3 | -1.777688 | SKCM | IGR-37 | SCH772984 |
| MVP_top60_DNA_seed | MAPK1 | -1.777688 | SKCM | IGR-37 | SCH772984 |
| MVP_top60_DNA_seed | MAP2K1 | -1.777163 | SKCM | HMV-II | Selumetinib |
| MVP_top60_DNA_seed | MAPK3 | -1.770214 | SKCM | MZ7-mel | Ulixertinib |
| MVP_top60_DNA_seed | MAPK1 | -1.770214 | SKCM | MZ7-mel | Ulixertinib |
| MVP_top60_DNA_seed | TP53 | -1.767456 | SKCM | RPMI-7951 | MIRA-1 |

| | | | | | |
|---|---|---|---|---|---|
| MVP_top60_DNA_seed | MAP2K1 | -1.765577 | SKCM | G-361 | Trametinib |
| MVP_top60_DNA_seed | MAPK1 | -1.754816 | SKCM | HMV-II | VX-11e |
| MVP_top60_DNA_seed | BRAF | -1.753284 | SKCM | SK-MEL-24 | PLX-4720 |
| MVP_top60_DNA_seed | MAP2K1 | -1.74861 | SKCM | CP66-MEL | Refametinib |
| MVP_top60_DNA_seed | BRAF | -1.747297 | SKCM | WM278 | Dabrafenib |
| MVP_top60_DNA_seed | MAP2K1 | -1.746935 | SKCM | COLO-783 | Refametinib |
| MVP_top60_DNA_seed | MAP2K1 | -1.745859 | SKCM | LB2518-MEL | Trametinib |
| MVP_top60_DNA_seed | MAP2K1 | -1.743801 | SKCM | SK-MEL-28 | Refametinib |
| MVP_top60_DNA_seed | MAPK3 | -1.740016 | SKCM | HMV-II | Ulixertinib |
| MVP_top60_DNA_seed | MAPK1 | -1.740016 | SKCM | HMV-II | Ulixertinib |
| MVP_top60_DNA_seed | MAPK3 | -1.737904 | SKCM | IGR-37 | Ulixertinib |
| MVP_top60_DNA_seed | MAPK1 | -1.737904 | SKCM | IGR-37 | Ulixertinib |
| MVP_top60_DNA_seed | MAP2K1 | -1.732732 | SKCM | MZ2-MEL | Refametinib |
| MVP_top60_DNA_seed | MAPK3 | -1.730609 | SKCM | LB2518-MEL | SCH772984 |
| MVP_top60_DNA_seed | MAPK1 | -1.730609 | SKCM | LB2518-MEL | SCH772984 |
| MVP_top60_DNA_seed | MAP2K1 | -1.727084 | SKCM | SK-MEL-30 | PD0325901 |
| MVP_top60_DNA_seed | MAP2K1 | -1.724833 | SKCM | MMAc-SF | Selumetinib |
| MVP_top60_DNA_seed | MAPK3 | -1.724164 | SKCM | WM35 | Ulixertinib |
| MVP_top60_DNA_seed | MAPK1 | -1.724164 | SKCM | WM35 | Ulixertinib |
| MVP_top60_DNA_seed | MAPK3 | -1.720185 | SKCM | A375 | Ulixertinib |
| MVP_top60_DNA_seed | MAPK1 | -1.720185 | SKCM | A375 | Ulixertinib |
| MVP_top60_DNA_seed | MAPK3 | -1.710426 | SKCM | HT-144 | Ulixertinib |
| MVP_top60_DNA_seed | MAPK1 | -1.710426 | SKCM | HT-144 | Ulixertinib |
| MVP_top60_DNA_seed | MAP2K1 | -1.709681 | SKCM | Hs-939-T | Refametinib |
| MVP_top60_DNA_seed | MAP2K1 | -1.708774 | SKCM | MMAc-SF | Trametinib |
| MVP_top60_DNA_seed | MAP2K1 | -1.696473 | SKCM | SK-MEL-2 | Trametinib |
| MVP_top60_DNA_seed | EP300 | -1.695592 | SKCM | RPMI-7951 | SGC-CBP30 |
| MVP_top60_DNA_seed | CREBBP | -1.695592 | SKCM | RPMI-7951 | SGC-CBP30 |
| MVP_top60_DNA_seed | MAPK3 | -1.685959 | SKCM | COLO-800 | Ulixertinib |
| MVP_top60_DNA_seed | MAPK1 | -1.685959 | SKCM | COLO-800 | Ulixertinib |
| MVP_top60_DNA_seed | BRAF | -1.685814 | SKCM | WM-115 | Dabrafenib |
| MVP_top60_DNA_seed | MAPK1 | -1.684838 | SKCM | A375 | VX-11e |
| MVP_top60_DNA_seed | MAP2K1 | -1.684224 | SKCM | GAK | PD0325901 |
| MVP_top60_DNA_seed | MAP2K1 | -1.68328 | SKCM | WM278 | PD0325901 |
| MVP_top60_DNA_seed | MAPK1 | -1.671378 | SKCM | IPC-298 | VX-11e |
| MVP_top60_DNA_seed | EGFR | -1.662963 | SKCM | CHL-1 | Osimertinib |
| MVP_top60_DNA_seed | MAP2K1 | -1.657146 | SKCM | GAK | Trametinib |
| MVP_top60_DNA_seed | MAPK1 | -1.655541 | SKCM | IGR-37 | VX-11e |
| MVP_top60_DNA_seed | BRAF | -1.654505 | SKCM | UACC-257 | PLX-4720 |
| MVP_top60_DNA_seed | MAPK3 | -1.652784 | SKCM | Hs-939-T | SCH772984 |
| MVP_top60_DNA_seed | MAPK1 | -1.652784 | SKCM | Hs-939-T | SCH772984 |

| | | | | | |
|---|---|---|---|---|---|
| MVP_top60_DNA_seed | BRAF | -1.632477 | SKCM | WM-115 | PLX-4720 |
| MVP_top60_DNA_seed | PIK3CB | -1.632018 | SKCM | VMRC-MELG | AZD6482 |
| MVP_top60_DNA_seed | MAPK1 | -1.631803 | SKCM | COLO-679 | VX-11e |
| MVP_top60_DNA_seed | BRAF | -1.627317 | SKCM | SK-MEL-30 | SB590885 |
| MVP_top60_DNA_seed | MAP2K1 | -1.624005 | SKCM | HMV-II | Refametinib |
| MVP_top60_DNA_seed | MAP2K1 | -1.622216 | SKCM | WM278 | Refametinib |
| MVP_top60_DNA_seed | MAP2K1 | -1.620029 | SKCM | G-361 | Selumetinib |
| MVP_top60_DNA_seed | MAP2K1 | -1.617176 | SKCM | MZ2-MEL | Trametinib |
| MVP_top60_DNA_seed | MAP2K1 | -1.616435 | SKCM | SK-MEL-1 | Refametinib |
| MVP_top60_DNA_seed | MAPK3 | -1.616261 | SKCM | HMV-II | Ulixertinib |
| MVP_top60_DNA_seed | MAPK1 | -1.616261 | SKCM | HMV-II | Ulixertinib |
| MVP_top60_DNA_seed | MAP2K1 | -1.611897 | SKCM | COLO-783 | Trametinib |
| MVP_top60_DNA_seed | MAP2K1 | -1.608514 | SKCM | UACC-257 | Trametinib |
| MVP_top60_DNA_seed | EGFR | -1.602795 | SKCM | VMRC-MELG | AZD3759 |
| MVP_top60_DNA_seed | MAP2K1 | -1.596118 | SKCM | MZ2-MEL | PD0325901 |
| MVP_top60_DNA_seed | MAPK3 | -1.589228 | SKCM | G-361 | SCH772984 |
| MVP_top60_DNA_seed | MAPK1 | -1.589228 | SKCM | G-361 | SCH772984 |
| MVP_top60_DNA_seed | MAP2K1 | -1.586711 | SKCM | GAK | Selumetinib |
| MVP_top60_DNA_seed | EGFR | -1.584782 | SKCM | MMAc-SF | AZD3759 |
| MVP_top60_DNA_seed | BRAF | -1.576498 | SKCM | MZ7-mel | SB590885 |
| MVP_top60_DNA_seed | BRAF | -1.575211 | SKCM | LOXIMVI | Dabrafenib |
| MVP_top60_DNA_seed | MAP2K1 | -1.568594 | SKCM | COLO-792 | Trametinib |
| MVP_top60_DNA_seed | MAPK1 | -1.561034 | SKCM | MEL-HO | VX-11e |
| MVP_top60_DNA_seed | MAP2K1 | -1.557991 | SKCM | COLO-829 | Selumetinib |
| MVP_top60_DNA_seed | MAP2K1 | -1.553764 | SKCM | LOXIMVI | Refametinib |
| MVP_top60_DNA_seed | EGFR | -1.548683 | SKCM | CHL-1 | Afatinib |
| MVP_top60_DNA_seed | ERBB2 | -1.548683 | SKCM | CHL-1 | Afatinib |
| MVP_top60_DNA_seed | MAP2K1 | -1.542425 | SKCM | A101D | Selumetinib |
| MVP_top60_DNA_seed | MAPK3 | -1.542131 | SKCM | HT-144 | Ulixertinib |
| MVP_top60_DNA_seed | MAPK1 | -1.542131 | SKCM | HT-144 | Ulixertinib |
| MVP_top60_DNA_seed | MAP2K1 | -1.540587 | SKCM | COLO-829 | PD0325901 |
| MVP_top60_DNA_seed | MAPK3 | -1.534728 | SKCM | MZ2-MEL | SCH772984 |
| MVP_top60_DNA_seed | MAPK1 | -1.534728 | SKCM | MZ2-MEL | SCH772984 |
| MVP_top60_DNA_seed | MAP2K1 | -1.532257 | SKCM | WM278 | Selumetinib |
| MVP_top60_DNA_seed | MAPK3 | -1.522021 | SKCM | UACC-257 | SCH772984 |
| MVP_top60_DNA_seed | MAPK1 | -1.522021 | SKCM | UACC-257 | SCH772984 |
| MVP_top60_DNA_seed | MAP2K1 | -1.521655 | SKCM | LOXIMVI | Selumetinib |
| MVP_top60_DNA_seed | MAP2K1 | -1.507703 | SKCM | WM-115 | Selumetinib |
| MVP_top60_DNA_seed | MAPK3 | -1.507638 | SKCM | UACC-257 | Ulixertinib |
| MVP_top60_DNA_seed | MAPK1 | -1.507638 | SKCM | UACC-257 | Ulixertinib |
| MVP_top60_DNA_seed | BRAF | -1.503855 | SKCM | SK-MEL-3 | PLX-4720 |

| | | | | | |
|---|---|---|---|---|---|
| MVP_top60_DNA_seed | MAP2K1 | -1.502616 | SKCM | GAK | Refametinib |
| MVP_top60_DNA_seed | BRAF | -1.488247 | SKCM | LOXIMVI | SB590885 |
| MVP_top60_DNA_seed | MAPK1 | -1.485598 | SKCM | Hs-939-T | VX-11e |
| MVP_top60_DNA_seed | GSK3B | -1.483469 | SKCM | RPMI-7951 | SB216763 |
| MVP_top60_DNA_seed | PIK3CA | -1.467881 | SKCM | VMRC-MELG | AZD8186 |
| MVP_top60_DNA_seed | PIK3CB | -1.467881 | SKCM | VMRC-MELG | AZD8186 |
| MVP_top60_DNA_seed | MAP2K1 | -1.465839 | SKCM | GAK | Selumetinib |
| MVP_top60_DNA_seed | MAPK3 | -1.461793 | SKCM | Hs-939-T | Ulixertinib |
| MVP_top60_DNA_seed | MAPK1 | -1.461793 | SKCM | Hs-939-T | Ulixertinib |
| MVP_top60_DNA_seed | MAP2K1 | -1.456416 | SKCM | G-361 | Refametinib |
| MVP_top60_DNA_seed | BRAF | -1.450751 | SKCM | WM-115 | SB590885 |
| MVP_top60_DNA_seed | MAP2K1 | -1.440086 | SKCM | LB373-MEL-D | Selumetinib |
| MVP_top60_DNA_seed | MAP2K1 | -1.438143 | SKCM | LOXIMVI | Trametinib |
| MVP_top60_DNA_seed | MAP2K1 | -1.436527 | SKCM | A101D | PD0325901 |
| MVP_top60_DNA_seed | RAC1 | -1.429039 | SKCM | CHL-1 | EHT-1864 |
| MVP_top60_DNA_seed | PIK3CA | -1.424087 | SKCM | VMRC-MELG | Alpelisib |
| MVP_top60_DNA_seed | MAP2K1 | -1.421525 | SKCM | COLO-829 | Trametinib |
| MVP_top60_DNA_seed | MAPK1 | -1.418107 | SKCM | UACC-62 | VX-11e |
| MVP_top60_DNA_seed | GSK3B | -1.406919 | SKCM | Hs-940-T | CHIR-99021 |
| MVP_top60_DNA_seed | PIK3CA | -1.395357 | SKCM | RVH-421 | AZD8186 |
| MVP_top60_DNA_seed | PIK3CB | -1.395357 | SKCM | RVH-421 | AZD8186 |
| MVP_top60_DNA_seed | MAPK1 | -1.39383 | SKCM | SK-MEL-30 | VX-11e |
| MVP_top60_DNA_seed | MAPK3 | -1.391638 | SKCM | C32 | Ulixertinib |
| MVP_top60_DNA_seed | MAPK1 | -1.391638 | SKCM | C32 | Ulixertinib |
| MVP_top60_DNA_seed | BRAF | -1.386463 | SKCM | WM793B | PLX-4720 |
| MVP_top60_DNA_seed | MAPK3 | -1.385684 | SKCM | COLO-792 | Ulixertinib |
| MVP_top60_DNA_seed | MAPK1 | -1.385684 | SKCM | COLO-792 | Ulixertinib |
| MVP_top60_DNA_seed | MAPK3 | -1.380537 | SKCM | CP66-MEL | Ulixertinib |
| MVP_top60_DNA_seed | MAPK1 | -1.380537 | SKCM | CP66-MEL | Ulixertinib |
| MVP_top60_DNA_seed | MAPK3 | -1.380269 | SKCM | UACC-62 | Ulixertinib |
| MVP_top60_DNA_seed | MAPK1 | -1.380269 | SKCM | UACC-62 | Ulixertinib |
| MVP_top60_DNA_seed | BRAF | -1.375744 | SKCM | SK-MEL-5 | SB590885 |
| MVP_top60_DNA_seed | BRAF | -1.374494 | SKCM | WM35 | PLX-4720 |
| MVP_top60_DNA_seed | MAPK3 | -1.370251 | SKCM | LOXIMVI | Ulixertinib |
| MVP_top60_DNA_seed | MAPK1 | -1.370251 | SKCM | LOXIMVI | Ulixertinib |
| MVP_top60_DNA_seed | MAPK1 | -1.368883 | SKCM | MZ7-mel | VX-11e |
| MVP_top60_DNA_seed | MAPK1 | -1.364872 | SKCM | GAK | VX-11e |
| MVP_top60_DNA_seed | MAP2K1 | -1.358394 | SKCM | LOXIMVI | PD0325901 |
| MVP_top60_DNA_seed | MAPK3 | -1.352662 | SKCM | COLO-792 | SCH772984 |
| MVP_top60_DNA_seed | MAPK1 | -1.352662 | SKCM | COLO-792 | SCH772984 |
| MVP_top60_DNA_seed | MAPK3 | -1.342497 | SKCM | CP66-MEL | SCH772984 |

| | | | | | |
|---|---|---|---|---|---|
| MVP_top60_DNA_seed | MAPK1 | -1.342497 | SKCM | CP66-MEL | SCH772984 |
| MVP_top60_DNA_seed | MAPK3 | -1.331774 | SKCM | COLO-679 | Ulixertinib |
| MVP_top60_DNA_seed | MAPK1 | -1.331774 | SKCM | COLO-679 | Ulixertinib |
| MVP_top60_DNA_seed | TP53 | -1.331377 | SKCM | A101D | MIRA-1 |
| MVP_top60_DNA_seed | MAPK1 | -1.329965 | SKCM | WM35 | VX-11e |
| MVP_top60_DNA_seed | MAP2K1 | -1.319481 | SKCM | SK-MEL-5 | Selumetinib |
| MVP_top60_DNA_seed | PIK3CA | -1.318423 | SKCM | WM-115 | AZD8186 |
| MVP_top60_DNA_seed | PIK3CB | -1.318423 | SKCM | WM-115 | AZD8186 |
| MVP_top60_DNA_seed | MAPK3 | -1.3128 | SKCM | WM35 | Ulixertinib |
| MVP_top60_DNA_seed | MAPK1 | -1.3128 | SKCM | WM35 | Ulixertinib |
| MVP_top60_DNA_seed | MAP2K1 | -1.309084 | SKCM | SK-MEL-2 | PD0325901 |
| MVP_top60_DNA_seed | JAK1 | -1.306934 | SKCM | LOXIMVI | JAK1_8709 |
| MVP_top60_DNA_seed | MAPK3 | -1.303818 | SKCM | UACC-257 | Ulixertinib |
| MVP_top60_DNA_seed | MAPK1 | -1.303818 | SKCM | UACC-257 | Ulixertinib |
| MVP_top60_DNA_seed | MAP2K1 | -1.302205 | SKCM | LOXIMVI | Selumetinib |
| MVP_top60_DNA_seed | MAPK3 | -1.301603 | SKCM | COLO-829 | SCH772984 |
| MVP_top60_DNA_seed | MAPK1 | -1.301603 | SKCM | COLO-829 | SCH772984 |
| MVP_top60_DNA_seed | MAP2K1 | -1.286703 | SKCM | A101D | Refametinib |
| MVP_top60_DNA_seed | GSK3B | -1.285156 | SKCM | IGR-1 | SB216763 |
| MVP_top60_DNA_seed | BRAF | -1.284334 | SKCM | Hs-940-T | PLX-4720 |
| MVP_top60_DNA_seed | EP300 | -1.284136 | SKCM | CHL-1 | CPI-637 |
| MVP_top60_DNA_seed | PIK3CA | -1.281923 | SKCM | SK-MEL-3 | Alpelisib |
| MVP_top60_DNA_seed | MAP2K1 | -1.279752 | SKCM | A101D | Selumetinib |
| MVP_top60_DNA_seed | BRAF | -1.274919 | SKCM | IPC-298 | PLX-4720 |
| MVP_top60_DNA_seed | BRAF | -1.272359 | SKCM | LOXIMVI | PLX-4720 |
| MVP_top60_DNA_seed | MAPK3 | -1.271775 | SKCM | A375 | Ulixertinib |
| MVP_top60_DNA_seed | MAPK1 | -1.271775 | SKCM | A375 | Ulixertinib |
| MVP_top60_DNA_seed | EP300 | -1.266887 | SKCM | A2058 | SGC-CBP30 |
| MVP_top60_DNA_seed | CREBBP | -1.266887 | SKCM | A2058 | SGC-CBP30 |
| MVP_top60_DNA_seed | TP53 | -1.265636 | SKCM | Hs-940-T | MIRA-1 |
| MVP_top60_DNA_seed | BRAF | -1.259437 | SKCM | SK-MEL-5 | Dabrafenib |
| MVP_top60_DNA_seed | MAP2K1 | -1.255731 | SKCM | SK-MEL-5 | Selumetinib |
| MVP_top60_DNA_seed | MAPK3 | -1.253686 | SKCM | LB2518-MEL | Ulixertinib |
| MVP_top60_DNA_seed | MAPK1 | -1.253686 | SKCM | LB2518-MEL | Ulixertinib |
| MVP_top60_DNA_seed | PIK3CA | -1.252788 | SKCM | UACC-62 | AZD8186 |
| MVP_top60_DNA_seed | PIK3CB | -1.252788 | SKCM | UACC-62 | AZD8186 |
| MVP_top60_DNA_seed | BRAF | -1.251223 | SKCM | SK-MEL-3 | SB590885 |
| MVP_top60_DNA_seed | MAPK3 | -1.248906 | SKCM | WM278 | SCH772984 |
| MVP_top60_DNA_seed | MAPK1 | -1.248906 | SKCM | WM278 | SCH772984 |
| MVP_top60_DNA_seed | BRAF | -1.245366 | SKCM | COLO-829 | SB590885 |
| MVP_top60_DNA_seed | MAPK1 | -1.243315 | SKCM | 451Lu | VX-11e |

| | | | | | |
|---|---|---|---|---|---|
| MVP_top60_DNA_seed | MAPK3 | -1.239768 | SKCM | MZ2-MEL | Ulixertinib |
| MVP_top60_DNA_seed | MAPK1 | -1.239768 | SKCM | MZ2-MEL | Ulixertinib |
| MVP_top60_DNA_seed | MAPK1 | -1.238462 | SKCM | LB2518-MEL | VX-11e |
| MVP_top60_DNA_seed | MAPK3 | -1.236355 | SKCM | SK-MEL-1 | Ulixertinib |
| MVP_top60_DNA_seed | MAPK1 | -1.236355 | SKCM | SK-MEL-1 | Ulixertinib |
| MVP_top60_DNA_seed | MAP2K1 | -1.223028 | SKCM | WM278 | Trametinib |
| MVP_top60_DNA_seed | MAPK3 | -1.221961 | SKCM | CHL-1 | Ulixertinib |
| MVP_top60_DNA_seed | MAPK1 | -1.221961 | SKCM | CHL-1 | Ulixertinib |
| MVP_top60_DNA_seed | BRAF | -1.218829 | SKCM | IGR-1 | Dabrafenib |
| MVP_top60_DNA_seed | JAK1 | -1.20923 | SKCM | GAK | JAK1_8709 |
| MVP_top60_DNA_seed | MAP2K1 | -1.19757 | SKCM | SK-MEL-2 | Selumetinib |
| MVP_top60_DNA_seed | BRAF | -1.194543 | SKCM | MMAc-SF | SB590885 |
| MVP_top60_DNA_seed | RAC1 | -1.186301 | SKCM | GAK | EHT-1864 |
| MVP_top60_DNA_seed | MAPK1 | -1.185257 | SKCM | K2 | VX-11e |
| MVP_top60_DNA_seed | MAPK3 | -1.17053 | SKCM | SH-4 | Ulixertinib |
| MVP_top60_DNA_seed | MAPK1 | -1.17053 | SKCM | SH-4 | Ulixertinib |
| MVP_top60_DNA_seed | GSK3B | -1.17005 | SKCM | A375 | CHIR-99021 |
| MVP_top60_DNA_seed | MAPK1 | -1.166888 | SKCM | A101D | VX-11e |
| MVP_top60_DNA_seed | MAP2K1 | -1.165649 | SKCM | A101D | Trametinib |
| MVP_top60_DNA_seed | MAPK1 | -1.157729 | SKCM | COLO-783 | VX-11e |
| MVP_top60_DNA_seed | MAP2K1 | -1.147955 | SKCM | SK-MEL-5 | PD0325901 |
| MVP_top60_DNA_seed | MAPK3 | -1.145851 | SKCM | 451Lu | Ulixertinib |
| MVP_top60_DNA_seed | MAPK1 | -1.145851 | SKCM | 451Lu | Ulixertinib |
| MVP_top60_DNA_seed | MAP2K1 | -1.141238 | SKCM | SK-MEL-24 | Selumetinib |
| MVP_top60_DNA_seed | BRAF | -1.140318 | SKCM | SK-MEL-1 | PLX-4720 |
| MVP_top60_DNA_seed | BRAF | -1.131181 | SKCM | IPC-298 | SB590885 |
| MVP_top60_DNA_seed | MAPK3 | -1.127055 | SKCM | WM278 | Ulixertinib |
| MVP_top60_DNA_seed | MAPK1 | -1.127055 | SKCM | WM278 | Ulixertinib |
| MVP_top60_DNA_seed | MAPK1 | -1.126336 | SKCM | G-361 | VX-11e |
| MVP_top60_DNA_seed | MAPK1 | -1.122269 | SKCM | SK-MEL-2 | VX-11e |
| MVP_top60_DNA_seed | MAP2K1 | -1.117758 | SKCM | SK-MEL-24 | PD0325901 |
| MVP_top60_DNA_seed | MAPK3 | -1.108123 | SKCM | GAK | SCH772984 |
| MVP_top60_DNA_seed | MAPK1 | -1.108123 | SKCM | GAK | SCH772984 |
| MVP_top60_DNA_seed | MAP2K1 | -1.107378 | SKCM | COLO-792 | Selumetinib |
| MVP_top60_DNA_seed | MAP2K1 | -1.093217 | SKCM | SK-MEL-2 | Refametinib |
| MVP_top60_DNA_seed | BRAF | -1.07447 | SKCM | IPC-298 | Dabrafenib |
| MVP_top60_DNA_seed | BRAF | -1.072549 | SKCM | SK-MEL-5 | PLX-4720 |
| MVP_top60_DNA_seed | MAPK3 | -1.067409 | SKCM | SK-MEL-2 | SCH772984 |
| MVP_top60_DNA_seed | MAPK1 | -1.067409 | SKCM | SK-MEL-2 | SCH772984 |
| MVP_top60_DNA_seed | RAC1 | -1.063166 | SKCM | G-361 | EHT-1864 |
| MVP_top60_DNA_seed | MAP2K1 | -1.060798 | SKCM | SK-MEL-24 | Selumetinib |

| | | | | | |
|---|---|---|---|---|---|
| MVP_top60_DNA_seed | JAK1 | -1.057658 | SKCM | G-MEL | JAK1_8709 |
| MVP_top60_DNA_seed | MAP2K1 | -1.056982 | SKCM | LB373-MEL-D | Refametinib |
| MVP_top60_DNA_seed | MAPK1 | -1.054945 | SKCM | MZ2-MEL | VX-11e |
| MVP_top60_DNA_seed | PIK3CA | -1.054124 | SKCM | LOXIMVI | GNE-317 |
| MVP_top60_DNA_seed | EGFR | -1.05132 | SKCM | SK-MEL-31 | AZD3759 |
| MVP_top60_DNA_seed | EGFR | -1.040913 | SKCM | MZ7-mel | Lapatinib |
| MVP_top60_DNA_seed | ERBB2 | -1.040913 | SKCM | MZ7-mel | Lapatinib |
| MVP_top60_DNA_seed | GSK3B | -1.04075 | SKCM | SK-MEL-31 | CHIR-99021 |
| MVP_top60_DNA_seed | PIK3CA | -1.040469 | SKCM | MEL-HO | AZD8186 |
| MVP_top60_DNA_seed | PIK3CB | -1.040469 | SKCM | MEL-HO | AZD8186 |
| MVP_top60_DNA_seed | JAK2 | -1.040296 | SKCM | RPMI-7951 | GSK2276186C |
| MVP_top60_DNA_seed | JAK1 | -1.040296 | SKCM | RPMI-7951 | GSK2276186C |
| MVP_top60_DNA_seed | JAK3 | -1.040296 | SKCM | RPMI-7951 | GSK2276186C |
| MVP_top60_DNA_seed | MAPK3 | -1.04005 | SKCM | LOXIMVI | Ulixertinib |
| MVP_top60_DNA_seed | MAPK1 | -1.04005 | SKCM | LOXIMVI | Ulixertinib |
| MVP_top60_DNA_seed | PIK3CA | -1.034067 | SKCM | MZ2-MEL | GNE-317 |
| MVP_top60_DNA_seed | MAP2K1 | -1.033895 | SKCM | COLO-829 | Selumetinib |
| MVP_top60_DNA_seed | MAPK3 | -1.032181 | SKCM | IGR-37 | Ulixertinib |
| MVP_top60_DNA_seed | MAPK1 | -1.032181 | SKCM | IGR-37 | Ulixertinib |
| MVP_top60_DNA_seed | BRAF | -1.03042 | SKCM | IST-MEL1 | PLX-4720 |
| MVP_top60_DNA_seed | EGFR | -1.024676 | SKCM | G-MEL | Lapatinib |
| MVP_top60_DNA_seed | ERBB2 | -1.024676 | SKCM | G-MEL | Lapatinib |
| MVP_top60_DNA_seed | MAP2K1 | -1.021741 | SKCM | COLO-792 | PD0325901 |
| MVP_top60_DNA_seed | MAPK3 | -1.021393 | SKCM | A101D | Ulixertinib |
| MVP_top60_DNA_seed | MAPK1 | -1.021393 | SKCM | A101D | Ulixertinib |
| MVP_top60_DNA_seed | MAP2K1 | -1.018123 | SKCM | COLO-829 | Refametinib |
| MVP_top60_DNA_seed | JAK2 | -1.016901 | SKCM | LOXIMVI | AZ960 |
| MVP_top60_DNA_seed | JAK3 | -1.016901 | SKCM | LOXIMVI | AZ960 |
| MVP_top60_DNA_seed | MAP2K1 | -1.01671 | SKCM | SK-MEL-2 | Selumetinib |
| MVP_top60_DNA_seed | TP53 | -1.016494 | SKCM | LB2518-MEL | MIRA-1 |
| MVP_top60_DNA_seed | JAK2 | -1.013193 | SKCM | IGR-37 | GSK2276186C |
| MVP_top60_DNA_seed | JAK1 | -1.013193 | SKCM | IGR-37 | GSK2276186C |
| MVP_top60_DNA_seed | JAK3 | -1.013193 | SKCM | IGR-37 | GSK2276186C |
| MVP_top60_DNA_seed | KRAS | -1.008418 | SKCM | A375 | KRAS (G12C) Inhibitor-12 |
| MVP_top60_DNA_seed | TP53 | -0.997342 | SKCM | COLO-783 | MIRA-1 |
| MVP_top60_DNA_seed | JAK1 | -0.996746 | SKCM | HMV-II | JAK1_8709 |
| MVP_top60_DNA_seed | MAPK1 | -0.995952 | SKCM | C32 | VX-11e |
| MVP_top60_DNA_seed | MAPK3 | -0.982674 | SKCM | G-361 | Ulixertinib |
| MVP_top60_DNA_seed | MAPK1 | -0.982674 | SKCM | G-361 | Ulixertinib |
| MVP_top60_DNA_seed | TP53 | -0.97468 | SKCM | SK-MEL-2 | MIRA-1 |
| MVP_top60_DNA_seed | MAPK3 | -0.970724 | SKCM | SK-MEL-1 | SCH772984 |

| | | | | | |
|---|---|---|---|---|---|
| MVP_top60_DNA_seed | MAPK1 | -0.970724 | SKCM | SK-MEL-1 | SCH772984 |
| MVP_top60_DNA_seed | BRAF | -0.968986 | SKCM | RVH-421 | SB590885 |
| MVP_top60_DNA_seed | TP53 | -0.967579 | SKCM | WM793B | MIRA-1 |
| MVP_top60_DNA_seed | KRAS | -0.965095 | SKCM | MZ7-mel | KRAS (G12C) Inhibitor-12 |
| MVP_top60_DNA_seed | MAP2K1 | -0.963515 | SKCM | WM-115 | Refametinib |
| MVP_top60_DNA_seed | PIK3CA | -0.960085 | SKCM | RPMI-7951 | AZD8186 |
| MVP_top60_DNA_seed | PIK3CB | -0.960085 | SKCM | RPMI-7951 | AZD8186 |
| MVP_top60_DNA_seed | PIK3CA | -0.955818 | SKCM | G-MEL | GNE-317 |
| MVP_top60_DNA_seed | MAPK3 | -0.955664 | SKCM | A101D | SCH772984 |
| MVP_top60_DNA_seed | MAPK1 | -0.955664 | SKCM | A101D | SCH772984 |
| MVP_top60_DNA_seed | MAP2K1 | -0.951242 | SKCM | LB373-MEL-D | PD0325901 |
| MVP_top60_DNA_seed | JAK2 | -0.94921 | SKCM | Hs-940-T | AZ960 |
| MVP_top60_DNA_seed | JAK3 | -0.94921 | SKCM | Hs-940-T | AZ960 |
| MVP_top60_DNA_seed | GSK3B | -0.945432 | SKCM | COLO-800 | CHIR-99021 |
| MVP_top60_DNA_seed | BRAF | -0.945144 | SKCM | RVH-421 | PLX-4720 |
| MVP_top60_DNA_seed | PIK3CA | -0.942673 | SKCM | COLO-783 | Alpelisib |
| MVP_top60_DNA_seed | PIK3CA | -0.941343 | SKCM | Hs-939-T | AZD8186 |
| MVP_top60_DNA_seed | PIK3CB | -0.941343 | SKCM | Hs-939-T | AZD8186 |
| MVP_top60_DNA_seed | JAK2 | -0.938618 | SKCM | WM1552C | AZ960 |
| MVP_top60_DNA_seed | JAK3 | -0.938618 | SKCM | WM1552C | AZ960 |
| MVP_top60_DNA_seed | BRAF | -0.935767 | SKCM | WM1552C | SB590885 |
| MVP_top60_DNA_seed | BRAF | -0.933673 | SKCM | IGR-1 | PLX-4720 |
| MVP_top60_DNA_seed | BRAF | -0.929291 | SKCM | MZ2-MEL | PLX-4720 |
| MVP_top60_DNA_seed | MAPK3 | -0.926908 | SKCM | WM793B | Ulixertinib |
| MVP_top60_DNA_seed | MAPK1 | -0.926908 | SKCM | WM793B | Ulixertinib |
| MVP_top60_DNA_seed | MAP2K1 | -0.908236 | SKCM | MMAc-SF | Refametinib |
| MVP_top60_DNA_seed | KRAS | -0.905436 | SKCM | LOXIMVI | KRAS (G12C) Inhibitor-12 |
| MVP_top60_DNA_seed | MAP2K1 | -0.905264 | SKCM | WM793B | PD0325901 |
| MVP_top60_DNA_seed | MAP2K1 | -0.902761 | SKCM | SK-MEL-5 | Trametinib |
| MVP_top60_DNA_seed | JAK2 | -0.89771 | SKCM | G-361 | Ruxolitinib |
| MVP_top60_DNA_seed | JAK1 | -0.89771 | SKCM | G-361 | Ruxolitinib |
| MVP_top60_DNA_seed | JAK2 | -0.897429 | SKCM | G-MEL | GSK2276186C |
| MVP_top60_DNA_seed | JAK1 | -0.897429 | SKCM | G-MEL | GSK2276186C |
| MVP_top60_DNA_seed | JAK3 | -0.897429 | SKCM | G-MEL | GSK2276186C |
| MVP_top60_DNA_seed | MAP2K1 | -0.895494 | SKCM | SK-MEL-24 | Refametinib |
| MVP_top60_DNA_seed | MAP2K1 | -0.893167 | SKCM | SK-MEL-24 | Trametinib |
| MVP_top60_DNA_seed | PIK3CA | -0.888834 | SKCM | A101D | Buparlisib |
| MVP_top60_DNA_seed | PIK3CB | -0.888834 | SKCM | A101D | Buparlisib |
| MVP_top60_DNA_seed | MAP2K1 | -0.886267 | SKCM | SK-MEL-3 | Selumetinib |
| MVP_top60_DNA_seed | MAP2K1 | -0.884787 | SKCM | RVH-421 | Trametinib |
| MVP_top60_DNA_seed | PIK3CA | -0.880495 | SKCM | MZ2-MEL | AZD8186 |

| | | | | | |
|---|---|---|---|---|---|
| MVP_top60_DNA_seed | PIK3CB | -0.880495 | SKCM | MZ2-MEL | AZD8186 |
| MVP_top60_DNA_seed | EGFR | -0.876347 | SKCM | G-MEL | AZD3759 |
| MVP_top60_DNA_seed | MAPK3 | -0.87575 | SKCM | GAK | Ulixertinib |
| MVP_top60_DNA_seed | MAPK1 | -0.87575 | SKCM | GAK | Ulixertinib |
| MVP_top60_DNA_seed | PIK3CA | -0.873981 | SKCM | A101D | AZD8186 |
| MVP_top60_DNA_seed | PIK3CB | -0.873981 | SKCM | A101D | AZD8186 |
| MVP_top60_DNA_seed | MAP2K1 | -0.869167 | SKCM | WM-115 | Trametinib |
| MVP_top60_DNA_seed | EP300 | -0.86724 | SKCM | LOXIMVI | CPI-637 |
| MVP_top60_DNA_seed | PIK3CA | -0.860194 | SKCM | COLO-792 | AZD8186 |
| MVP_top60_DNA_seed | PIK3CB | -0.860194 | SKCM | COLO-792 | AZD8186 |
| MVP_top60_DNA_seed | EGFR | -0.856941 | SKCM | COLO-783 | Sapitinib |
| MVP_top60_DNA_seed | ERBB2 | -0.856941 | SKCM | COLO-783 | Sapitinib |
| MVP_top60_DNA_seed | PIK3CB | -0.854753 | SKCM | COLO-800 | AZD6482 |
| MVP_top60_DNA_seed | EGFR | -0.853889 | SKCM | VMRC-MELG | Gefitinib |
| MVP_top60_DNA_seed | MAPK3 | -0.853039 | SKCM | SK-MEL-5 | Ulixertinib |
| MVP_top60_DNA_seed | MAPK1 | -0.853039 | SKCM | SK-MEL-5 | Ulixertinib |
| MVP_top60_DNA_seed | BRAF | -0.852645 | SKCM | HMV-II | Dabrafenib |
| MVP_top60_DNA_seed | RAC1 | -0.85098 | SKCM | COLO-792 | EHT-1864 |
| MVP_top60_DNA_seed | MAPK3 | -0.847357 | SKCM | G-361 | Ulixertinib |
| MVP_top60_DNA_seed | MAPK1 | -0.847357 | SKCM | G-361 | Ulixertinib |
| MVP_top60_DNA_seed | MAPK3 | -0.839088 | SKCM | SK-MEL-1 | Ulixertinib |
| MVP_top60_DNA_seed | MAPK1 | -0.839088 | SKCM | SK-MEL-1 | Ulixertinib |
| MVP_top60_DNA_seed | EGFR | -0.832043 | SKCM | CHL-1 | Lapatinib |
| MVP_top60_DNA_seed | ERBB2 | -0.832043 | SKCM | CHL-1 | Lapatinib |
| MVP_top60_DNA_seed | MAP2K1 | -0.827992 | SKCM | WM793B | Selumetinib |
| MVP_top60_DNA_seed | EGFR | -0.826339 | SKCM | M14 | Erlotinib |
| MVP_top60_DNA_seed | EGFR | -0.825865 | SKCM | HT-144 | Lapatinib |
| MVP_top60_DNA_seed | ERBB2 | -0.825865 | SKCM | HT-144 | Lapatinib |
| MVP_top60_DNA_seed | BRAF | -0.823021 | SKCM | WM1552C | PLX-4720 |
| MVP_top60_DNA_seed | PIK3CA | -0.81823 | SKCM | COLO-792 | GNE-317 |
| MVP_top60_DNA_seed | MAP2K1 | -0.813982 | SKCM | COLO-792 | Refametinib |
| MVP_top60_DNA_seed | EGFR | -0.811936 | SKCM | IGR-1 | Lapatinib |
| MVP_top60_DNA_seed | ERBB2 | -0.811936 | SKCM | IGR-1 | Lapatinib |
| MVP_top60_DNA_seed | MAP2K1 | -0.809314 | SKCM | WM793B | Selumetinib |
| MVP_top60_DNA_seed | KRAS | -0.796789 | SKCM | SK-MEL-3 | KRAS (G12C) Inhibitor-12 |
| MVP_top60_DNA_seed | RAC1 | -0.791529 | SKCM | SK-MEL-5 | EHT-1864 |
| MVP_top60_DNA_seed | RAC1 | -0.7898 | SKCM | LB373-MEL-D | EHT-1864 |
| MVP_top60_DNA_seed | MAPK3 | -0.788466 | SKCM | RVH-421 | Ulixertinib |
| MVP_top60_DNA_seed | MAPK1 | -0.788466 | SKCM | RVH-421 | Ulixertinib |
| MVP_top60_DNA_seed | EGFR | -0.78481 | SKCM | SK-MEL-28 | Erlotinib |
| MVP_top60_DNA_seed | MAPK1 | -0.784418 | SKCM | COLO-792 | VX-11e |

| | | | | | |
|---|---|---|---|---|---|
| MVP_top60_DNA_seed | GSK3B | -0.779651 | SKCM | A101D | SB216763 |
| MVP_top60_DNA_seed | BRAF | -0.776845 | SKCM | SK-MEL-31 | PLX-4720 |
| MVP_top60_DNA_seed | KRAS | -0.775947 | SKCM | G-MEL | KRAS (G12C) Inhibitor-12 |
| MVP_top60_DNA_seed | EGFR | -0.775846 | SKCM | A2058 | Lapatinib |
| MVP_top60_DNA_seed | ERBB2 | -0.775846 | SKCM | A2058 | Lapatinib |
| MVP_top60_DNA_seed | MAP2K1 | -0.774937 | SKCM | WM-115 | PD0325901 |
| MVP_top60_DNA_seed | EP300 | -0.774582 | SKCM | M14 | SGC-CBP30 |
| MVP_top60_DNA_seed | CREBBP | -0.774582 | SKCM | M14 | SGC-CBP30 |
| MVP_top60_DNA_seed | KRAS | -0.773535 | SKCM | IGR-37 | KRAS (G12C) Inhibitor-12 |
| MVP_top60_DNA_seed | EGFR | -0.772249 | SKCM | MEL-HO | Lapatinib |
| MVP_top60_DNA_seed | ERBB2 | -0.772249 | SKCM | MEL-HO | Lapatinib |
| MVP_top60_DNA_seed | PIK3CA | -0.769652 | SKCM | SK-MEL-3 | GNE-317 |
| MVP_top60_DNA_seed | BRAF | -0.76818 | SKCM | RPMI-7951 | PLX-4720 |
| MVP_top60_DNA_seed | MAP2K1 | -0.766914 | SKCM | WM793B | Refametinib |
| MVP_top60_DNA_seed | BRAF | -0.762441 | SKCM | A2058 | SB590885 |
| MVP_top60_DNA_seed | JAK1 | -0.761378 | SKCM | G-361 | JAK1_8709 |
| MVP_top60_DNA_seed | EGFR | -0.757467 | SKCM | G-361 | Lapatinib |
| MVP_top60_DNA_seed | ERBB2 | -0.757467 | SKCM | G-361 | Lapatinib |
| MVP_top60_DNA_seed | JAK2 | -0.755143 | SKCM | VMRC-MELG | AZ960 |
| MVP_top60_DNA_seed | JAK3 | -0.755143 | SKCM | VMRC-MELG | AZ960 |
| MVP_top60_DNA_seed | MAPK3 | -0.750108 | SKCM | MMAc-SF | SCH772984 |
| MVP_top60_DNA_seed | MAPK1 | -0.750108 | SKCM | MMAc-SF | SCH772984 |
| MVP_top60_DNA_seed | KRAS | -0.747761 | SKCM | 451Lu | KRAS (G12C) Inhibitor-12 |
| MVP_top60_DNA_seed | PIK3CA | -0.745578 | SKCM | A2058 | GNE-317 |
| MVP_top60_DNA_seed | PIK3CA | -0.743391 | SKCM | RPMI-7951 | GNE-317 |
| MVP_top60_DNA_seed | RAC1 | -0.740063 | SKCM | VMRC-MELG | EHT-1864 |
| MVP_top60_DNA_seed | MAPK3 | -0.73676 | SKCM | GAK | Ulixertinib |
| MVP_top60_DNA_seed | MAPK1 | -0.73676 | SKCM | GAK | Ulixertinib |
| MVP_top60_DNA_seed | EP300 | -0.736198 | SKCM | GAK | SGC-CBP30 |
| MVP_top60_DNA_seed | CREBBP | -0.736198 | SKCM | GAK | SGC-CBP30 |
| MVP_top60_DNA_seed | MAP2K1 | -0.735611 | SKCM | WM1552C | Refametinib |
| MVP_top60_DNA_seed | PIK3CB | -0.728569 | SKCM | SK-MEL-3 | AZD6482 |
| MVP_top60_DNA_seed | MAP2K1 | -0.727231 | SKCM | LB373-MEL-D | Trametinib |
| MVP_top60_DNA_seed | MAP2K1 | -0.726046 | SKCM | A2058 | Selumetinib |
| MVP_top60_DNA_seed | MAPK1 | -0.724548 | SKCM | CP66-MEL | VX-11e |
| MVP_top60_DNA_seed | MAP2K1 | -0.722341 | SKCM | A2058 | Trametinib |
| MVP_top60_DNA_seed | PIK3CA | -0.720784 | SKCM | Hs-939-T | GNE-317 |
| MVP_top60_DNA_seed | MAPK3 | -0.71937 | SKCM | WM278 | Ulixertinib |
| MVP_top60_DNA_seed | MAPK1 | -0.71937 | SKCM | WM278 | Ulixertinib |
| MVP_top60_DNA_seed | EGFR | -0.717264 | SKCM | COLO-800 | AZD3759 |
| MVP_top60_DNA_seed | EGFR | -0.713509 | SKCM | RVH-421 | AZD3759 |

| | | | | | |
|---|---|---|---|---|---|
| MVP_top60_DNA_seed | JAK2 | -0.708722 | SKCM | WM1552C | JAK_8517 |
| MVP_top60_DNA_seed | JAK1 | -0.708722 | SKCM | WM1552C | JAK_8517 |
| MVP_top60_DNA_seed | MAP2K1 | -0.707181 | SKCM | WM1552C | PD0325901 |
| MVP_top60_DNA_seed | JAK1 | -0.705485 | SKCM | SH-4 | JAK1_8709 |
| MVP_top60_DNA_seed | MAPK3 | -0.704927 | SKCM | SK-MEL-2 | Ulixertinib |
| MVP_top60_DNA_seed | MAPK1 | -0.704927 | SKCM | SK-MEL-2 | Ulixertinib |
| MVP_top60_DNA_seed | PIK3CA | -0.700045 | SKCM | SK-MEL-31 | AZD8186 |
| MVP_top60_DNA_seed | PIK3CB | -0.700045 | SKCM | SK-MEL-31 | AZD8186 |
| MVP_top60_DNA_seed | PIK3CA | -0.699129 | SKCM | IGR-1 | Buparlisib |
| MVP_top60_DNA_seed | PIK3CB | -0.699129 | SKCM | IGR-1 | Buparlisib |
| MVP_top60_DNA_seed | BRAF | -0.697813 | SKCM | HMV-II | PLX-4720 |
| MVP_top60_DNA_seed | PIK3CA | -0.697586 | SKCM | A2058 | Buparlisib |
| MVP_top60_DNA_seed | PIK3CB | -0.697586 | SKCM | A2058 | Buparlisib |
| MVP_top60_DNA_seed | MAPK1 | -0.696316 | SKCM | UACC-257 | VX-11e |
| MVP_top60_DNA_seed | PIK3CB | -0.695016 | SKCM | CP50-MEL-B | AZD6482 |
| MVP_top60_DNA_seed | JAK2 | -0.694875 | SKCM | COLO-792 | GSK2276186C |
| MVP_top60_DNA_seed | JAK1 | -0.694875 | SKCM | COLO-792 | GSK2276186C |
| MVP_top60_DNA_seed | JAK3 | -0.694875 | SKCM | COLO-792 | GSK2276186C |
| MVP_top60_DNA_seed | EGFR | -0.687625 | SKCM | CHL-1 | Gefitinib |
| MVP_top60_DNA_seed | MAPK3 | -0.687222 | SKCM | LB373-MEL-D | SCH772984 |
| MVP_top60_DNA_seed | MAPK1 | -0.687222 | SKCM | LB373-MEL-D | SCH772984 |
| MVP_top60_DNA_seed | MAPK3 | -0.686141 | SKCM | A2058 | Ulixertinib |
| MVP_top60_DNA_seed | MAPK1 | -0.686141 | SKCM | A2058 | Ulixertinib |
| MVP_top60_DNA_seed | MAPK3 | -0.682248 | SKCM | SK-MEL-3 | SCH772984 |
| MVP_top60_DNA_seed | MAPK1 | -0.682248 | SKCM | SK-MEL-3 | SCH772984 |
| MVP_top60_DNA_seed | PIK3CA | -0.680681 | SKCM | RPMI-7951 | Buparlisib |
| MVP_top60_DNA_seed | PIK3CB | -0.680681 | SKCM | RPMI-7951 | Buparlisib |
| MVP_top60_DNA_seed | PIK3CA | -0.680434 | SKCM | CP50-MEL-B | AZD8186 |
| MVP_top60_DNA_seed | PIK3CB | -0.680434 | SKCM | CP50-MEL-B | AZD8186 |
| MVP_top60_DNA_seed | MAP2K1 | -0.677515 | SKCM | RPMI-7951 | Refametinib |
| MVP_top60_DNA_seed | JAK2 | -0.675472 | SKCM | LOXIMVI | JAK_8517 |
| MVP_top60_DNA_seed | JAK1 | -0.675472 | SKCM | LOXIMVI | JAK_8517 |
| MVP_top60_DNA_seed | PIK3CA | -0.674592 | SKCM | COLO-792 | Alpelisib |
| MVP_top60_DNA_seed | MAP2K1 | -0.671116 | SKCM | COLO-792 | Selumetinib |
| MVP_top60_DNA_seed | EGFR | -0.669948 | SKCM | WM793B | Erlotinib |
| MVP_top60_DNA_seed | MAP2K1 | -0.664635 | SKCM | MEL-JUSO | Trametinib |
| MVP_top60_DNA_seed | PIK3CA | -0.662107 | SKCM | M14 | GNE-317 |
| MVP_top60_DNA_seed | EP300 | -0.661976 | SKCM | A375 | SGC-CBP30 |
| MVP_top60_DNA_seed | CREBBP | -0.661976 | SKCM | A375 | SGC-CBP30 |
| MVP_top60_DNA_seed | MAPK3 | -0.658801 | SKCM | SK-MEL-2 | Ulixertinib |
| MVP_top60_DNA_seed | MAPK1 | -0.658801 | SKCM | SK-MEL-2 | Ulixertinib |

| | | | | | |
|---|---|---|---|---|---|
| MVP_top60_DNA_seed | MAPK3 | -0.657367 | SKCM | A2058 | SCH772984 |
| MVP_top60_DNA_seed | MAPK1 | -0.657367 | SKCM | A2058 | SCH772984 |
| MVP_top60_DNA_seed | MAP2K1 | -0.656007 | SKCM | WM793B | Trametinib |
| MVP_top60_DNA_seed | BRAF | -0.653236 | SKCM | VMRC-MELG | PLX-4720 |
| MVP_top60_DNA_seed | MAP2K1 | -0.652109 | SKCM | MEL-JUSO | PD0325901 |
| MVP_top60_DNA_seed | MAPK1 | -0.651711 | SKCM | SK-MEL-1 | VX-11e |
| MVP_top60_DNA_seed | PIK3CB | -0.651334 | SKCM | Hs-939-T | AZD6482 |
| MVP_top60_DNA_seed | MAPK3 | -0.648679 | SKCM | C32 | Ulixertinib |
| MVP_top60_DNA_seed | MAPK1 | -0.648679 | SKCM | C32 | Ulixertinib |
| MVP_top60_DNA_seed | EGFR | -0.641067 | SKCM | MZ2-MEL | Lapatinib |
| MVP_top60_DNA_seed | ERBB2 | -0.641067 | SKCM | MZ2-MEL | Lapatinib |
| MVP_top60_DNA_seed | EGFR | -0.639191 | SKCM | LB2518-MEL | Gefitinib |
| MVP_top60_DNA_seed | RAC1 | -0.634977 | SKCM | A375 | EHT-1864 |
| MVP_top60_DNA_seed | PIK3CA | -0.633191 | SKCM | IST-MEL1 | GNE-317 |
| MVP_top60_DNA_seed | EGFR | -0.632257 | SKCM | CP50-MEL-B | AZD3759 |
| MVP_top60_DNA_seed | EGFR | -0.628944 | SKCM | A101D | Lapatinib |
| MVP_top60_DNA_seed | ERBB2 | -0.628944 | SKCM | A101D | Lapatinib |
| MVP_top60_DNA_seed | BRAF | -0.628599 | SKCM | WM1552C | Dabrafenib |
| MVP_top60_DNA_seed | EGFR | -0.627314 | SKCM | G-MEL | Gefitinib |
| MVP_top60_DNA_seed | MAP2K1 | -0.622689 | SKCM | WM-115 | Selumetinib |
| MVP_top60_DNA_seed | BRAF | -0.622453 | SKCM | IGR-1 | SB590885 |
| MVP_top60_DNA_seed | MAPK1 | -0.622204 | SKCM | RVH-421 | VX-11e |
| MVP_top60_DNA_seed | PIK3CA | -0.621649 | SKCM | G-MEL | Buparlisib |
| MVP_top60_DNA_seed | PIK3CB | -0.621649 | SKCM | G-MEL | Buparlisib |
| MVP_top60_DNA_seed | EGFR | -0.619783 | SKCM | A101D | Erlotinib |
| MVP_top60_DNA_seed | PIK3CA | -0.61919 | SKCM | WM793B | GNE-317 |
| MVP_top60_DNA_seed | JAK2 | -0.617061 | SKCM | HT-144 | GSK2276186C |
| MVP_top60_DNA_seed | JAK1 | -0.617061 | SKCM | HT-144 | GSK2276186C |
| MVP_top60_DNA_seed | JAK3 | -0.617061 | SKCM | HT-144 | GSK2276186C |
| MVP_top60_DNA_seed | MAP2K1 | -0.616397 | SKCM | A2058 | PD0325901 |
| MVP_top60_DNA_seed | MAPK3 | -0.61165 | SKCM | SK-MEL-5 | SCH772984 |
| MVP_top60_DNA_seed | MAPK1 | -0.61165 | SKCM | SK-MEL-5 | SCH772984 |
| MVP_top60_DNA_seed | KRAS | -0.611209 | SKCM | RPMI-7951 | KRAS (G12C) Inhibitor-12 |
| MVP_top60_DNA_seed | BRAF | -0.607267 | SKCM | RVH-421 | Dabrafenib |
| MVP_top60_DNA_seed | PIK3CA | -0.606888 | SKCM | LOXIMVI | Buparlisib |
| MVP_top60_DNA_seed | PIK3CB | -0.606888 | SKCM | LOXIMVI | Buparlisib |
| MVP_top60_DNA_seed | MAPK3 | -0.605249 | SKCM | IGR-1 | Ulixertinib |
| MVP_top60_DNA_seed | MAPK1 | -0.605249 | SKCM | IGR-1 | Ulixertinib |
| MVP_top60_DNA_seed | MAPK1 | -0.596518 | SKCM | IST-MEL1 | VX-11e |
| MVP_top60_DNA_seed | PIK3CA | -0.58903 | SKCM | COLO-792 | Buparlisib |
| MVP_top60_DNA_seed | PIK3CB | -0.58903 | SKCM | COLO-792 | Buparlisib |

| | | | | | |
|---|---|---|---|---|---|
| MVP_top60_DNA_seed | EGFR | -0.587305 | SKCM | CHL-1 | Erlotinib |
| MVP_top60_DNA_seed | MAP2K1 | -0.586865 | SKCM | IST-MEL1 | Refametinib |
| MVP_top60_DNA_seed | MAPK3 | -0.583555 | SKCM | MEL-JUSO | SCH772984 |
| MVP_top60_DNA_seed | MAPK1 | -0.583555 | SKCM | MEL-JUSO | SCH772984 |
| MVP_top60_DNA_seed | TP53 | -0.579629 | SKCM | VMRC-MELG | MIRA-1 |
| MVP_top60_DNA_seed | KRAS | -0.577844 | SKCM | A101D | KRAS (G12C) Inhibitor-12 |
| MVP_top60_DNA_seed | BRAF | -0.574058 | SKCM | A2058 | PLX-4720 |
| MVP_top60_DNA_seed | PIK3CB | -0.572845 | SKCM | RVH-421 | AZD6482 |
| MVP_top60_DNA_seed | MAP2K1 | -0.572735 | SKCM | MEL-JUSO | Selumetinib |
| MVP_top60_DNA_seed | PIK3CA | -0.566861 | SKCM | VMRC-MELG | GNE-317 |
| MVP_top60_DNA_seed | EP300 | -0.566823 | SKCM | RPMI-7951 | CPI-637 |
| MVP_top60_DNA_seed | PIK3CB | -0.566223 | SKCM | MZ2-MEL | AZD6482 |
| MVP_top60_DNA_seed | MAP2K1 | -0.565017 | SKCM | SK-MEL-5 | Refametinib |
| MVP_top60_DNA_seed | TP53 | -0.56118 | SKCM | UACC-62 | MIRA-1 |
| MVP_top60_DNA_seed | PIK3CA | -0.560586 | SKCM | WM1552C | GNE-317 |
| MVP_top60_DNA_seed | BRAF | -0.559475 | SKCM | CP66-MEL | Dabrafenib |
| MVP_top60_DNA_seed | PIK3CB | -0.556943 | SKCM | MEL-HO | AZD6482 |
| MVP_top60_DNA_seed | PIK3CA | -0.553493 | SKCM | IGR-1 | GNE-317 |
| MVP_top60_DNA_seed | PIK3CA | -0.552016 | SKCM | LOXIMVI | Alpelisib |
| MVP_top60_DNA_seed | MAP2K1 | -0.549103 | SKCM | Mewo | PD0325901 |
| MVP_top60_DNA_seed | EGFR | -0.546952 | SKCM | RPMI-7951 | Lapatinib |
| MVP_top60_DNA_seed | ERBB2 | -0.546952 | SKCM | RPMI-7951 | Lapatinib |
| MVP_top60_DNA_seed | MAP2K1 | -0.540763 | SKCM | RVH-421 | Selumetinib |
| MVP_top60_DNA_seed | PIK3CA | -0.53816 | SKCM | WM35 | Buparlisib |
| MVP_top60_DNA_seed | PIK3CB | -0.53816 | SKCM | WM35 | Buparlisib |
| MVP_top60_DNA_seed | EGFR | -0.537724 | SKCM | A2058 | Erlotinib |
| MVP_top60_DNA_seed | PIK3CA | -0.536385 | SKCM | A101D | GNE-317 |
| MVP_top60_DNA_seed | PIK3CA | -0.52192 | SKCM | SK-MEL-28 | AZD8186 |
| MVP_top60_DNA_seed | PIK3CB | -0.52192 | SKCM | SK-MEL-28 | AZD8186 |
| MVP_top60_DNA_seed | MAPK1 | -0.520634 | SKCM | WM278 | VX-11e |
| MVP_top60_DNA_seed | EP300 | -0.518872 | SKCM | HMV-II | CPI-637 |
| MVP_top60_DNA_seed | PIK3CA | -0.517301 | SKCM | IST-MEL1 | AZD8186 |
| MVP_top60_DNA_seed | PIK3CB | -0.517301 | SKCM | IST-MEL1 | AZD8186 |
| MVP_top60_DNA_seed | EGFR | -0.512094 | SKCM | MZ7-mel | Erlotinib |
| MVP_top60_DNA_seed | MAP2K1 | -0.50802 | SKCM | RVH-421 | PD0325901 |
| MVP_top60_DNA_seed | PIK3CA | -0.505146 | SKCM | K2 | GNE-317 |
| MVP_top60_DNA_seed | PIK3CA | -0.505137 | SKCM | MZ2-MEL | Alpelisib |
| MVP_top60_DNA_seed | MAP2K1 | -0.500171 | SKCM | IGR-1 | PD0325901 |
| MVP_top60_DNA_seed | MAP2K1 | -0.499261 | SKCM | Mewo | Trametinib |
| MVP_top60_DNA_seed | EGFR | -0.49865 | SKCM | SK-MEL-24 | Lapatinib |
| MVP_top60_DNA_seed | ERBB2 | -0.49865 | SKCM | SK-MEL-24 | Lapatinib |

| | | | | | |
|---|---|---|---|---|---|
| MVP_top60_DNA_seed | MAP2K1 | -0.488036 | SKCM | MEL-JUSO | Refametinib |
| MVP_top60_DNA_seed | BRAF | -0.481898 | SKCM | WM793B | Dabrafenib |
| MVP_top60_DNA_seed | TP53 | -0.477698 | SKCM | LOXIMVI | MIRA-1 |
| MVP_top60_DNA_seed | MAPK3 | -0.475015 | SKCM | WM-115 | SCH772984 |
| MVP_top60_DNA_seed | MAPK1 | -0.475015 | SKCM | WM-115 | SCH772984 |
| MVP_top60_DNA_seed | MAPK1 | -0.473626 | SKCM | WM-115 | VX-11e |
| MVP_top60_DNA_seed | KRAS | -0.467291 | SKCM | MZ2-MEL | KRAS (G12C) Inhibitor-12 |
| MVP_top60_DNA_seed | EGFR | -0.465269 | SKCM | SK-MEL-30 | Lapatinib |
| MVP_top60_DNA_seed | ERBB2 | -0.465269 | SKCM | SK-MEL-30 | Lapatinib |
| MVP_top60_DNA_seed | EGFR | -0.459988 | SKCM | IPC-298 | Lapatinib |
| MVP_top60_DNA_seed | ERBB2 | -0.459988 | SKCM | IPC-298 | Lapatinib |
| MVP_top60_DNA_seed | BRAF | -0.459969 | SKCM | MZ2-MEL | Dabrafenib |
| MVP_top60_DNA_seed | EP300 | -0.459491 | SKCM | LOXIMVI | SGC-CBP30 |
| MVP_top60_DNA_seed | CREBBP | -0.459491 | SKCM | LOXIMVI | SGC-CBP30 |
| MVP_top60_DNA_seed | JAK2 | -0.45853 | SKCM | RPMI-7951 | AZ960 |
| MVP_top60_DNA_seed | JAK3 | -0.45853 | SKCM | RPMI-7951 | AZ960 |
| MVP_top60_DNA_seed | GSK3B | -0.454342 | SKCM | VMRC-MELG | CHIR-99021 |
| MVP_top60_DNA_seed | PIK3CA | -0.453205 | SKCM | CP50-MEL-B | GNE-317 |
| MVP_top60_DNA_seed | EGFR | -0.452742 | SKCM | 451Lu | Gefitinib |
| MVP_top60_DNA_seed | MAP2K1 | -0.450742 | SKCM | IST-MEL1 | Trametinib |
| MVP_top60_DNA_seed | EGFR | -0.450467 | SKCM | IPC-298 | Gefitinib |
| MVP_top60_DNA_seed | EGFR | -0.445828 | SKCM | SK-MEL-31 | Lapatinib |
| MVP_top60_DNA_seed | ERBB2 | -0.445828 | SKCM | SK-MEL-31 | Lapatinib |
| MVP_top60_DNA_seed | MAPK3 | -0.445413 | SKCM | A101D | Ulixertinib |
| MVP_top60_DNA_seed | MAPK1 | -0.445413 | SKCM | A101D | Ulixertinib |
| MVP_top60_DNA_seed | GSK3B | -0.445243 | SKCM | G-MEL | SB216763 |
| MVP_top60_DNA_seed | EP300 | -0.443542 | SKCM | SK-MEL-2 | SGC-CBP30 |
| MVP_top60_DNA_seed | CREBBP | -0.443542 | SKCM | SK-MEL-2 | SGC-CBP30 |
| MVP_top60_DNA_seed | EGFR | -0.441195 | SKCM | MZ2-MEL | Osimertinib |
| MVP_top60_DNA_seed | EGFR | -0.440978 | SKCM | LOXIMVI | Lapatinib |
| MVP_top60_DNA_seed | ERBB2 | -0.440978 | SKCM | LOXIMVI | Lapatinib |
| MVP_top60_DNA_seed | EGFR | -0.435271 | SKCM | LB373-MEL-D | AZD3759 |
| MVP_top60_DNA_seed | PIK3CA | -0.434494 | SKCM | Hs-939-T | Alpelisib |
| MVP_top60_DNA_seed | MAPK3 | -0.434298 | SKCM | WM793B | SCH772984 |
| MVP_top60_DNA_seed | MAPK1 | -0.434298 | SKCM | WM793B | SCH772984 |
| MVP_top60_DNA_seed | PIK3CA | -0.429787 | SKCM | M14 | Buparlisib |
| MVP_top60_DNA_seed | PIK3CB | -0.429787 | SKCM | M14 | Buparlisib |
| MVP_top60_DNA_seed | EP300 | -0.429412 | SKCM | WM1552C | CPI-637 |
| MVP_top60_DNA_seed | KRAS | -0.428581 | SKCM | WM35 | KRAS (G12C) Inhibitor-12 |
| MVP_top60_DNA_seed | EGFR | -0.428439 | SKCM | MEL-HO | Osimertinib |
| MVP_top60_DNA_seed | EGFR | -0.428003 | SKCM | Hs-939-T | AZD3759 |

| | | | | | |
|---|---|---|---|---|---|
| MVP_top60_DNA_seed | GSK3B | -0.427819 | SKCM | SK-MEL-24 | SB216763 |
| MVP_top60_DNA_seed | EGFR | -0.423604 | SKCM | IGR-37 | Gefitinib |
| MVP_top60_DNA_seed | MAPK3 | -0.422719 | SKCM | COLO-829 | Ulixertinib |
| MVP_top60_DNA_seed | MAPK1 | -0.422719 | SKCM | COLO-829 | Ulixertinib |
| MVP_top60_DNA_seed | PIK3CA | -0.421331 | SKCM | UACC-62 | GNE-317 |
| MVP_top60_DNA_seed | MAP2K1 | -0.419675 | SKCM | RPMI-7951 | PD0325901 |
| MVP_top60_DNA_seed | EGFR | -0.418733 | SKCM | HMV-II | Lapatinib |
| MVP_top60_DNA_seed | ERBB2 | -0.418733 | SKCM | HMV-II | Lapatinib |
| MVP_top60_DNA_seed | MAP2K1 | -0.418607 | SKCM | IGR-1 | Selumetinib |
| MVP_top60_DNA_seed | KRAS | -0.418495 | SKCM | GAK | KRAS (G12C) Inhibitor-12 |
| MVP_top60_DNA_seed | MAPK3 | -0.418319 | SKCM | A2058 | Ulixertinib |
| MVP_top60_DNA_seed | MAPK1 | -0.418319 | SKCM | A2058 | Ulixertinib |
| MVP_top60_DNA_seed | EGFR | -0.413794 | SKCM | MZ2-MEL | Gefitinib |
| MVP_top60_DNA_seed | TP53 | -0.413487 | SKCM | COLO-792 | MIRA-1 |
| MVP_top60_DNA_seed | BRAF | -0.412815 | SKCM | SK-MEL-31 | Dabrafenib |
| MVP_top60_DNA_seed | MAPK3 | -0.409308 | SKCM | WM1552C | SCH772984 |
| MVP_top60_DNA_seed | MAPK1 | -0.409308 | SKCM | WM1552C | SCH772984 |
| MVP_top60_DNA_seed | MAPK3 | -0.406204 | SKCM | COLO-829 | Ulixertinib |
| MVP_top60_DNA_seed | MAPK1 | -0.406204 | SKCM | COLO-829 | Ulixertinib |
| MVP_top60_DNA_seed | MAPK3 | -0.402047 | SKCM | IST-MEL1 | SCH772984 |
| MVP_top60_DNA_seed | MAPK1 | -0.402047 | SKCM | IST-MEL1 | SCH772984 |
| MVP_top60_DNA_seed | KRAS | -0.401995 | SKCM | Hs-939-T | KRAS (G12C) Inhibitor-12 |
| MVP_top60_DNA_seed | JAK2 | -0.400978 | SKCM | RPMI-7951 | Ruxolitinib |
| MVP_top60_DNA_seed | JAK1 | -0.400978 | SKCM | RPMI-7951 | Ruxolitinib |
| MVP_top60_DNA_seed | GSK3B | -0.400684 | SKCM | HT-144 | SB216763 |
| MVP_top60_DNA_seed | MAP2K1 | -0.399974 | SKCM | IST-MEL1 | PD0325901 |
| MVP_top60_DNA_seed | PIK3CB | -0.396645 | SKCM | SK-MEL-28 | AZD6482 |
| MVP_top60_DNA_seed | MAP2K1 | -0.396032 | SKCM | IGR-1 | Selumetinib |
| MVP_top60_DNA_seed | JAK1 | -0.395275 | SKCM | A2058 | JAK1_8709 |
| MVP_top60_DNA_seed | TP53 | -0.393712 | SKCM | G-MEL | MIRA-1 |
| MVP_top60_DNA_seed | BRAF | -0.386309 | SKCM | IST-MEL1 | Dabrafenib |
| MVP_top60_DNA_seed | MAPK3 | -0.383415 | SKCM | COLO-792 | Ulixertinib |
| MVP_top60_DNA_seed | MAPK1 | -0.383415 | SKCM | COLO-792 | Ulixertinib |
| MVP_top60_DNA_seed | BRAF | -0.382748 | SKCM | IST-MEL1 | SB590885 |
| MVP_top60_DNA_seed | MAPK1 | -0.38252 | SKCM | WM1552C | VX-11e |
| MVP_top60_DNA_seed | MAP2K1 | -0.379252 | SKCM | WM1552C | Selumetinib |
| MVP_top60_DNA_seed | MAP2K1 | -0.378472 | SKCM | RPMI-7951 | Trametinib |
| MVP_top60_DNA_seed | RAC1 | -0.376188 | SKCM | RVH-421 | EHT-1864 |
| MVP_top60_DNA_seed | KRAS | -0.37097 | SKCM | K2 | KRAS (G12C) Inhibitor-12 |
| MVP_top60_DNA_seed | JAK1 | -0.368955 | SKCM | A375 | JAK1_8709 |
| MVP_top60_DNA_seed | JAK2 | -0.36829 | SKCM | RPMI-7951 | JAK_8517 |

| | | | | | |
|---|---|---|---|---|---|
| MVP_top60_DNA_seed | JAK1 | -0.36829 | SKCM | RPMI-7951 | JAK_8517 |
| MVP_top60_DNA_seed | EP300 | -0.363712 | SKCM | WM35 | CPI-637 |
| MVP_top60_DNA_seed | EP300 | -0.362274 | SKCM | A375 | CPI-637 |
| MVP_top60_DNA_seed | EP300 | -0.36223 | SKCM | SK-MEL-2 | CPI-637 |
| MVP_top60_DNA_seed | RAC1 | -0.360817 | SKCM | MZ2-MEL | EHT-1864 |
| MVP_top60_DNA_seed | JAK2 | -0.360009 | SKCM | MZ2-MEL | AZ960 |
| MVP_top60_DNA_seed | JAK3 | -0.360009 | SKCM | MZ2-MEL | AZ960 |
| MVP_top60_DNA_seed | PIK3CA | -0.35848 | SKCM | CHL-1 | AZD8186 |
| MVP_top60_DNA_seed | PIK3CB | -0.35848 | SKCM | CHL-1 | AZD8186 |
| MVP_top60_DNA_seed | MAPK1 | -0.353345 | SKCM | RPMI-7951 | VX-11e |
| MVP_top60_DNA_seed | PIK3CA | -0.351758 | SKCM | G-361 | Buparlisib |
| MVP_top60_DNA_seed | PIK3CB | -0.351758 | SKCM | G-361 | Buparlisib |
| MVP_top60_DNA_seed | EGFR | -0.350458 | SKCM | MEL-HO | AZD3759 |
| MVP_top60_DNA_seed | EP300 | -0.350454 | SKCM | COLO-792 | CPI-637 |
| MVP_top60_DNA_seed | MAP2K1 | -0.348584 | SKCM | SK-MEL-3 | Refametinib |
| MVP_top60_DNA_seed | EGFR | -0.341096 | SKCM | LB2518-MEL | AZD3759 |
| MVP_top60_DNA_seed | MAP2K1 | -0.338905 | SKCM | WM1552C | Trametinib |
| MVP_top60_DNA_seed | MAPK3 | -0.338675 | SKCM | MMAc-SF | Ulixertinib |
| MVP_top60_DNA_seed | MAPK1 | -0.338675 | SKCM | MMAc-SF | Ulixertinib |
| MVP_top60_DNA_seed | EP300 | -0.336471 | SKCM | M14 | CPI-637 |
| MVP_top60_DNA_seed | MAPK3 | -0.335965 | SKCM | Mewo | SCH772984 |
| MVP_top60_DNA_seed | MAPK1 | -0.335965 | SKCM | Mewo | SCH772984 |
| MVP_top60_DNA_seed | MAPK3 | -0.33526 | SKCM | SK-MEL-31 | SCH772984 |
| MVP_top60_DNA_seed | MAPK1 | -0.33526 | SKCM | SK-MEL-31 | SCH772984 |
| MVP_top60_DNA_seed | EP300 | -0.331273 | SKCM | SH-4 | CPI-637 |
| MVP_top60_DNA_seed | MAPK1 | -0.330688 | SKCM | MMAc-SF | VX-11e |
| MVP_top60_DNA_seed | MAPK3 | -0.32933 | SKCM | RPMI-7951 | Ulixertinib |
| MVP_top60_DNA_seed | MAPK1 | -0.32933 | SKCM | RPMI-7951 | Ulixertinib |
| MVP_top60_DNA_seed | PIK3CA | -0.328448 | SKCM | MEL-HO | GNE-317 |
| MVP_top60_DNA_seed | MAPK1 | -0.324355 | SKCM | SK-MEL-24 | VX-11e |
| MVP_top60_DNA_seed | GSK3B | -0.323983 | SKCM | WM278 | SB216763 |
| MVP_top60_DNA_seed | MAP2K1 | -0.322888 | SKCM | WM1552C | Selumetinib |
| MVP_top60_DNA_seed | MAPK3 | -0.320114 | SKCM | SK-MEL-24 | SCH772984 |
| MVP_top60_DNA_seed | MAPK1 | -0.320114 | SKCM | SK-MEL-24 | SCH772984 |
| MVP_top60_DNA_seed | PIK3CA | -0.31446 | SKCM | WM793B | Buparlisib |
| MVP_top60_DNA_seed | PIK3CB | -0.31446 | SKCM | WM793B | Buparlisib |
| MVP_top60_DNA_seed | JAK1 | -0.312221 | SKCM | IGR-1 | JAK1_8709 |
| MVP_top60_DNA_seed | EGFR | -0.31107 | SKCM | SK-MEL-3 | Osimertinib |
| MVP_top60_DNA_seed | GSK3B | -0.310129 | SKCM | GAK | SB216763 |
| MVP_top60_DNA_seed | EGFR | -0.30995 | SKCM | MMAc-SF | Sapitinib |
| MVP_top60_DNA_seed | ERBB2 | -0.30995 | SKCM | MMAc-SF | Sapitinib |

| | | | | | |
|---|---|---|---|---|---|
| MVP_top60_DNA_seed | MAPK3 | -0.304207 | SKCM | LB373-MEL-D | Ulixertinib |
| MVP_top60_DNA_seed | MAPK1 | -0.304207 | SKCM | LB373-MEL-D | Ulixertinib |
| MVP_top60_DNA_seed | EGFR | -0.299195 | SKCM | G-MEL | Erlotinib |
| MVP_top60_DNA_seed | PIK3CA | -0.298096 | SKCM | CP50-MEL-B | Alpelisib |
| MVP_top60_DNA_seed | MAPK3 | -0.295464 | SKCM | WM1552C | Ulixertinib |
| MVP_top60_DNA_seed | MAPK1 | -0.295464 | SKCM | WM1552C | Ulixertinib |
| MVP_top60_DNA_seed | EGFR | -0.295048 | SKCM | MZ2-MEL | Erlotinib |
| MVP_top60_DNA_seed | JAK1 | -0.294437 | SKCM | IGR-37 | JAK1_8709 |
| MVP_top60_DNA_seed | EGFR | -0.292146 | SKCM | HT-144 | Erlotinib |
| MVP_top60_DNA_seed | MAP2K1 | -0.287518 | SKCM | A2058 | Selumetinib |
| MVP_top60_DNA_seed | PIK3CA | -0.287231 | SKCM | IPC-298 | GNE-317 |
| MVP_top60_DNA_seed | PIK3CA | -0.28483 | SKCM | G-361 | AZD8186 |
| MVP_top60_DNA_seed | PIK3CB | -0.28483 | SKCM | G-361 | AZD8186 |
| MVP_top60_DNA_seed | PIK3CA | -0.284829 | SKCM | MZ7-mel | GNE-317 |
| MVP_top60_DNA_seed | JAK2 | -0.284451 | SKCM | A375 | AZ960 |
| MVP_top60_DNA_seed | JAK3 | -0.284451 | SKCM | A375 | AZ960 |
| MVP_top60_DNA_seed | EGFR | -0.28326 | SKCM | RVH-421 | Osimertinib |
| MVP_top60_DNA_seed | EGFR | -0.283083 | SKCM | IPC-298 | Erlotinib |
| MVP_top60_DNA_seed | EGFR | -0.282564 | SKCM | COLO-800 | Lapatinib |
| MVP_top60_DNA_seed | ERBB2 | -0.282564 | SKCM | COLO-800 | Lapatinib |
| MVP_top60_DNA_seed | RAC1 | -0.282021 | SKCM | RPMI-7951 | EHT-1864 |
| MVP_top60_DNA_seed | PIK3CA | -0.281224 | SKCM | Hs-940-T | GNE-317 |
| MVP_top60_DNA_seed | JAK2 | -0.2808 | SKCM | SK-MEL-3 | AZ960 |
| MVP_top60_DNA_seed | JAK3 | -0.2808 | SKCM | SK-MEL-3 | AZ960 |
| MVP_top60_DNA_seed | PIK3CA | -0.279915 | SKCM | COLO-783 | AZD8186 |
| MVP_top60_DNA_seed | PIK3CB | -0.279915 | SKCM | COLO-783 | AZD8186 |
| MVP_top60_DNA_seed | TP53 | -0.276219 | SKCM | MZ7-mel | MIRA-1 |
| MVP_top60_DNA_seed | KRAS | -0.275201 | SKCM | HT-144 | KRAS (G12C) Inhibitor-12 |
| MVP_top60_DNA_seed | GSK3B | -0.273287 | SKCM | IPC-298 | CHIR-99021 |
| MVP_top60_DNA_seed | EGFR | -0.271108 | SKCM | M14 | Lapatinib |
| MVP_top60_DNA_seed | ERBB2 | -0.271108 | SKCM | M14 | Lapatinib |
| MVP_top60_DNA_seed | EGFR | -0.268619 | SKCM | G-361 | Erlotinib |
| MVP_top60_DNA_seed | PIK3CA | -0.268411 | SKCM | IPC-298 | Alpelisib |
| MVP_top60_DNA_seed | MAP2K1 | -0.268235 | SKCM | SK-MEL-31 | Trametinib |
| MVP_top60_DNA_seed | EGFR | -0.265331 | SKCM | K2 | AZD3759 |
| MVP_top60_DNA_seed | JAK2 | -0.263861 | SKCM | IGR-1 | GSK2276186C |
| MVP_top60_DNA_seed | JAK1 | -0.263861 | SKCM | IGR-1 | GSK2276186C |
| MVP_top60_DNA_seed | JAK3 | -0.263861 | SKCM | IGR-1 | GSK2276186C |
| MVP_top60_DNA_seed | MAP2K1 | -0.263046 | SKCM | SK-MEL-3 | Trametinib |
| MVP_top60_DNA_seed | EGFR | -0.257829 | SKCM | K2 | Gefitinib |
| MVP_top60_DNA_seed | PIK3CA | -0.255322 | SKCM | A375 | GNE-317 |

| | | | | | |
|---|---|---|---|---|---|
| MVP_top60_DNA_seed | JAK2 | -0.249924 | SKCM | A375 | JAK_8517 |
| MVP_top60_DNA_seed | JAK1 | -0.249924 | SKCM | A375 | JAK_8517 |
| MVP_top60_DNA_seed | EP300 | -0.24831 | SKCM | G-361 | SGC-CBP30 |
| MVP_top60_DNA_seed | CREBBP | -0.24831 | SKCM | G-361 | SGC-CBP30 |
| MVP_top60_DNA_seed | EGFR | -0.248287 | SKCM | A2058 | AZD3759 |
| MVP_top60_DNA_seed | MAPK3 | -0.247218 | SKCM | RPMI-7951 | Ulixertinib |
| MVP_top60_DNA_seed | MAPK1 | -0.247218 | SKCM | RPMI-7951 | Ulixertinib |
| MVP_top60_DNA_seed | PIK3CA | -0.246085 | SKCM | A375 | Buparlisib |
| MVP_top60_DNA_seed | PIK3CB | -0.246085 | SKCM | A375 | Buparlisib |
| MVP_top60_DNA_seed | KRAS | -0.245119 | SKCM | CP50-MEL-B | KRAS (G12C) Inhibitor-12 |
| MVP_top60_DNA_seed | EGFR | -0.243053 | SKCM | LOXIMVI | Afatinib |
| MVP_top60_DNA_seed | ERBB2 | -0.243053 | SKCM | LOXIMVI | Afatinib |
| MVP_top60_DNA_seed | JAK2 | -0.239072 | SKCM | G-MEL | Ruxolitinib |
| MVP_top60_DNA_seed | JAK1 | -0.239072 | SKCM | G-MEL | Ruxolitinib |
| MVP_top60_DNA_seed | PIK3CA | -0.23327 | SKCM | LOXIMVI | AZD8186 |
| MVP_top60_DNA_seed | PIK3CB | -0.23327 | SKCM | LOXIMVI | AZD8186 |
| MVP_top60_DNA_seed | PIK3CA | -0.228887 | SKCM | HT-144 | Buparlisib |
| MVP_top60_DNA_seed | PIK3CB | -0.228887 | SKCM | HT-144 | Buparlisib |
| MVP_top60_DNA_seed | GSK3B | -0.228441 | SKCM | WM793B | SB216763 |
| MVP_top60_DNA_seed | PIK3CA | -0.22819 | SKCM | A2058 | AZD8186 |
| MVP_top60_DNA_seed | PIK3CB | -0.22819 | SKCM | A2058 | AZD8186 |
| MVP_top60_DNA_seed | KRAS | -0.221303 | SKCM | LB2518-MEL | KRAS (G12C) Inhibitor-12 |
| MVP_top60_DNA_seed | MAPK1 | -0.217783 | SKCM | LB373-MEL-D | VX-11e |
| MVP_top60_DNA_seed | PIK3CB | -0.216908 | SKCM | IPC-298 | AZD6482 |
| MVP_top60_DNA_seed | GSK3B | -0.21539 | SKCM | A375 | SB216763 |
| MVP_top60_DNA_seed | PIK3CA | -0.213641 | SKCM | SK-MEL-31 | Alpelisib |
| MVP_top60_DNA_seed | JAK1 | -0.212821 | SKCM | CP66-MEL | JAK1_8709 |
| MVP_top60_DNA_seed | PIK3CA | -0.211404 | SKCM | SH-4 | GNE-317 |
| MVP_top60_DNA_seed | MAPK1 | -0.209195 | SKCM | SK-MEL-5 | VX-11e |
| MVP_top60_DNA_seed | GSK3B | -0.209064 | SKCM | Hs-939-T | CHIR-99021 |
| MVP_top60_DNA_seed | JAK1 | -0.203869 | SKCM | A101D | JAK1_8709 |
| MVP_top60_DNA_seed | MAPK3 | -0.203175 | SKCM | IST-MEL1 | Ulixertinib |
| MVP_top60_DNA_seed | MAPK1 | -0.203175 | SKCM | IST-MEL1 | Ulixertinib |
| MVP_top60_DNA_seed | EP300 | -0.202401 | SKCM | G-MEL | SGC-CBP30 |
| MVP_top60_DNA_seed | CREBBP | -0.202401 | SKCM | G-MEL | SGC-CBP30 |
| MVP_top60_DNA_seed | MAPK1 | -0.199164 | SKCM | SK-MEL-3 | VX-11e |
| MVP_top60_DNA_seed | EP300 | -0.197599 | SKCM | IGR-1 | CPI-637 |
| MVP_top60_DNA_seed | JAK2 | -0.195617 | SKCM | WM35 | GSK2276186C |
| MVP_top60_DNA_seed | JAK1 | -0.195617 | SKCM | WM35 | GSK2276186C |
| MVP_top60_DNA_seed | JAK3 | -0.195617 | SKCM | WM35 | GSK2276186C |
| MVP_top60_DNA_seed | EGFR | -0.195533 | SKCM | COLO-783 | Lapatinib |

| | | | | | |
|---|---|---|---|---|---|
| MVP_top60_DNA_seed | ERBB2 | -0.195533 | SKCM | COLO-783 | Lapatinib |
| MVP_top60_DNA_seed | EP300 | -0.195031 | SKCM | CHL-1 | SGC-CBP30 |
| MVP_top60_DNA_seed | CREBBP | -0.195031 | SKCM | CHL-1 | SGC-CBP30 |
| MVP_top60_DNA_seed | JAK2 | -0.19302 | SKCM | CHL-1 | JAK_8517 |
| MVP_top60_DNA_seed | JAK1 | -0.19302 | SKCM | CHL-1 | JAK_8517 |
| MVP_top60_DNA_seed | PIK3CA | -0.192537 | SKCM | MZ7-mel | Buparlisib |
| MVP_top60_DNA_seed | PIK3CB | -0.192537 | SKCM | MZ7-mel | Buparlisib |
| MVP_top60_DNA_seed | JAK2 | -0.189358 | SKCM | SK-MEL-31 | AZ960 |
| MVP_top60_DNA_seed | JAK3 | -0.189358 | SKCM | SK-MEL-31 | AZ960 |
| MVP_top60_DNA_seed | PIK3CA | -0.189018 | SKCM | SK-MEL-3 | AZD8186 |
| MVP_top60_DNA_seed | PIK3CB | -0.189018 | SKCM | SK-MEL-3 | AZD8186 |
| MVP_top60_DNA_seed | MAPK3 | -0.18678 | SKCM | IGR-1 | SCH772984 |
| MVP_top60_DNA_seed | MAPK1 | -0.18678 | SKCM | IGR-1 | SCH772984 |
| MVP_top60_DNA_seed | MAP2K1 | -0.185374 | SKCM | RPMI-7951 | Selumetinib |
| MVP_top60_DNA_seed | MAP2K1 | -0.184495 | SKCM | IST-MEL1 | Selumetinib |
| MVP_top60_DNA_seed | MAPK3 | -0.183633 | SKCM | RVH-421 | SCH772984 |
| MVP_top60_DNA_seed | MAPK1 | -0.183633 | SKCM | RVH-421 | SCH772984 |
| MVP_top60_DNA_seed | TP53 | -0.183487 | SKCM | CP50-MEL-B | MIRA-1 |
| MVP_top60_DNA_seed | KRAS | -0.182803 | SKCM | CHL-1 | KRAS (G12C) Inhibitor-12 |
| MVP_top60_DNA_seed | BRAF | -0.179315 | SKCM | RPMI-7951 | SB590885 |
| MVP_top60_DNA_seed | RAC1 | -0.178073 | SKCM | G-MEL | EHT-1864 |
| MVP_top60_DNA_seed | PIK3CA | -0.174179 | SKCM | SK-MEL-28 | Alpelisib |
| MVP_top60_DNA_seed | EGFR | -0.171398 | SKCM | COLO-792 | Afatinib |
| MVP_top60_DNA_seed | ERBB2 | -0.171398 | SKCM | COLO-792 | Afatinib |
| MVP_top60_DNA_seed | EGFR | -0.171083 | SKCM | CHL-1 | AZD3759 |
| MVP_top60_DNA_seed | TP53 | -0.16971 | SKCM | GAK | MIRA-1 |
| MVP_top60_DNA_seed | MAP2K1 | -0.169666 | SKCM | IGR-1 | Trametinib |
| MVP_top60_DNA_seed | PIK3CA | -0.168681 | SKCM | CP50-MEL-B | Buparlisib |
| MVP_top60_DNA_seed | PIK3CB | -0.168681 | SKCM | CP50-MEL-B | Buparlisib |
| MVP_top60_DNA_seed | RAC1 | -0.16659 | SKCM | 451Lu | EHT-1864 |
| MVP_top60_DNA_seed | BRAF | -0.166111 | SKCM | CP66-MEL | PLX-4720 |
| MVP_top60_DNA_seed | BRAF | -0.164214 | SKCM | SK-MEL-30 | PLX-4720 |
| MVP_top60_DNA_seed | PIK3CA | -0.161486 | SKCM | IGR-37 | Buparlisib |
| MVP_top60_DNA_seed | PIK3CB | -0.161486 | SKCM | IGR-37 | Buparlisib |
| MVP_top60_DNA_seed | PIK3CA | -0.160859 | SKCM | MZ2-MEL | Buparlisib |
| MVP_top60_DNA_seed | PIK3CB | -0.160859 | SKCM | MZ2-MEL | Buparlisib |
| MVP_top60_DNA_seed | EGFR | -0.158544 | SKCM | SK-MEL-2 | Osimertinib |
| MVP_top60_DNA_seed | MAPK3 | -0.156118 | SKCM | RPMI-7951 | SCH772984 |
| MVP_top60_DNA_seed | MAPK1 | -0.156118 | SKCM | RPMI-7951 | SCH772984 |
| MVP_top60_DNA_seed | JAK1 | -0.154623 | SKCM | WM793B | JAK1_8709 |
| MVP_top60_DNA_seed | EGFR | -0.153244 | SKCM | RPMI-7951 | AZD3759 |

| | | | | | |
|---|---|---|---|---|---|
| MVP_top60_DNA_seed | RAC1 | -0.152638 | SKCM | SK-MEL-1 | EHT-1864 |
| MVP_top60_DNA_seed | EGFR | -0.150442 | SKCM | WM793B | Lapatinib |
| MVP_top60_DNA_seed | ERBB2 | -0.150442 | SKCM | WM793B | Lapatinib |
| MVP_top60_DNA_seed | PIK3CB | -0.148627 | SKCM | LB373-MEL-D | AZD6482 |
| MVP_top60_DNA_seed | EGFR | -0.147476 | SKCM | Hs-939-T | Gefitinib |
| MVP_top60_DNA_seed | JAK2 | -0.146899 | SKCM | A2058 | AZ960 |
| MVP_top60_DNA_seed | JAK3 | -0.146899 | SKCM | A2058 | AZ960 |
| MVP_top60_DNA_seed | GSK3B | -0.144339 | SKCM | COLO-783 | SB216763 |
| MVP_top60_DNA_seed | MAP2K1 | -0.144089 | SKCM | SK-MEL-3 | PD0325901 |
| MVP_top60_DNA_seed | EGFR | -0.142593 | SKCM | MZ7-mel | AZD3759 |
| MVP_top60_DNA_seed | MAP2K1 | -0.141688 | SKCM | Mewo | Refametinib |
| MVP_top60_DNA_seed | RAC1 | -0.140162 | SKCM | LOXIMVI | EHT-1864 |
| MVP_top60_DNA_seed | EGFR | -0.134208 | SKCM | SK-MEL-3 | AZD3759 |
| MVP_top60_DNA_seed | PIK3CA | -0.133698 | SKCM | WM35 | GNE-317 |
| MVP_top60_DNA_seed | GSK3B | -0.13348 | SKCM | A2058 | SB216763 |
| MVP_top60_DNA_seed | EGFR | -0.132535 | SKCM | G-MEL | Afatinib |
| MVP_top60_DNA_seed | ERBB2 | -0.132535 | SKCM | G-MEL | Afatinib |
| MVP_top60_DNA_seed | EP300 | -0.131705 | SKCM | A2058 | CPI-637 |
| MVP_top60_DNA_seed | MAP2K1 | -0.129817 | SKCM | MEL-JUSO | Selumetinib |
| MVP_top60_DNA_seed | TP53 | -0.127838 | SKCM | HT-144 | MIRA-1 |
| MVP_top60_DNA_seed | EP300 | -0.126332 | SKCM | Mewo | CPI-637 |
| MVP_top60_DNA_seed | JAK2 | -0.122481 | SKCM | MZ7-mel | Ruxolitinib |
| MVP_top60_DNA_seed | JAK1 | -0.122481 | SKCM | MZ7-mel | Ruxolitinib |
| MVP_top60_DNA_seed | PIK3CA | -0.117395 | SKCM | Hs-940-T | AZD8186 |
| MVP_top60_DNA_seed | PIK3CB | -0.117395 | SKCM | Hs-940-T | AZD8186 |
| MVP_top60_DNA_seed | PIK3CB | -0.115671 | SKCM | COLO-783 | AZD6482 |
| MVP_top60_DNA_seed | JAK2 | -0.114814 | SKCM | SK-MEL-28 | AZ960 |
| MVP_top60_DNA_seed | JAK3 | -0.114814 | SKCM | SK-MEL-28 | AZ960 |
| MVP_top60_DNA_seed | EGFR | -0.110379 | SKCM | K2 | Sapitinib |
| MVP_top60_DNA_seed | ERBB2 | -0.110379 | SKCM | K2 | Sapitinib |
| MVP_top60_DNA_seed | PIK3CA | -0.108529 | SKCM | COLO-800 | AZD8186 |
| MVP_top60_DNA_seed | PIK3CB | -0.108529 | SKCM | COLO-800 | AZD8186 |
| MVP_top60_DNA_seed | TP53 | -0.106484 | SKCM | SK-MEL-1 | MIRA-1 |
| MVP_top60_DNA_seed | JAK1 | -0.103914 | SKCM | COLO-792 | JAK1_8709 |
| MVP_top60_DNA_seed | JAK2 | -0.103862 | SKCM | GAK | Ruxolitinib |
| MVP_top60_DNA_seed | JAK1 | -0.103862 | SKCM | GAK | Ruxolitinib |
| MVP_top60_DNA_seed | JAK1 | -0.103284 | SKCM | WM1552C | JAK1_8709 |
| MVP_top60_DNA_seed | EGFR | -0.099835 | SKCM | UACC-62 | AZD3759 |
| MVP_top60_DNA_seed | GSK3B | -0.094627 | SKCM | SK-MEL-3 | CHIR-99021 |
| MVP_top60_DNA_seed | TP53 | -0.094193 | SKCM | SK-MEL-3 | MIRA-1 |
| MVP_top60_DNA_seed | JAK2 | -0.091901 | SKCM | CP50-MEL-B | AZ960 |

| | | | | | |
|---|---|---|---|---|---|
| MVP_top60_DNA_seed | JAK3 | -0.091901 | SKCM | CP50-MEL-B | AZ960 |
| MVP_top60_DNA_seed | JAK2 | -0.091732 | SKCM | IGR-37 | Ruxolitinib |
| MVP_top60_DNA_seed | JAK1 | -0.091732 | SKCM | IGR-37 | Ruxolitinib |
| MVP_top60_DNA_seed | EP300 | -0.09123 | SKCM | IGR-1 | SGC-CBP30 |
| MVP_top60_DNA_seed | CREBBP | -0.09123 | SKCM | IGR-1 | SGC-CBP30 |
| MVP_top60_DNA_seed | JAK2 | -0.090218 | SKCM | SK-MEL-3 | Ruxolitinib |
| MVP_top60_DNA_seed | JAK1 | -0.090218 | SKCM | SK-MEL-3 | Ruxolitinib |
| MVP_top60_DNA_seed | EGFR | -0.090215 | SKCM | HT-144 | Afatinib |
| MVP_top60_DNA_seed | ERBB2 | -0.090215 | SKCM | HT-144 | Afatinib |
| MVP_top60_DNA_seed | EGFR | -0.083537 | SKCM | WM35 | Erlotinib |
| MVP_top60_DNA_seed | PIK3CA | -0.08291 | SKCM | CHL-1 | Buparlisib |
| MVP_top60_DNA_seed | PIK3CB | -0.08291 | SKCM | CHL-1 | Buparlisib |
| MVP_top60_DNA_seed | EGFR | -0.082897 | SKCM | COLO-783 | Afatinib |
| MVP_top60_DNA_seed | ERBB2 | -0.082897 | SKCM | COLO-783 | Afatinib |
| MVP_top60_DNA_seed | EP300 | -0.082687 | SKCM | HT-144 | CPI-637 |
| MVP_top60_DNA_seed | PIK3CA | -0.081702 | SKCM | WM1552C | Alpelisib |
| MVP_top60_DNA_seed | PIK3CA | -0.079356 | SKCM | IGR-37 | AZD8186 |
| MVP_top60_DNA_seed | PIK3CB | -0.079356 | SKCM | IGR-37 | AZD8186 |
| MVP_top60_DNA_seed | JAK1 | -0.077965 | SKCM | M14 | JAK1_8709 |
| MVP_top60_DNA_seed | EGFR | -0.077558 | SKCM | IGR-1 | Erlotinib |
| MVP_top60_DNA_seed | EGFR | -0.077388 | SKCM | WM793B | Gefitinib |
| MVP_top60_DNA_seed | RAC1 | -0.077293 | SKCM | COLO-783 | EHT-1864 |
| MVP_top60_DNA_seed | PIK3CA | -0.074229 | SKCM | COLO-829 | AZD8186 |
| MVP_top60_DNA_seed | PIK3CB | -0.074229 | SKCM | COLO-829 | AZD8186 |
| MVP_top60_DNA_seed | EGFR | -0.071801 | SKCM | SK-MEL-1 | Lapatinib |
| MVP_top60_DNA_seed | ERBB2 | -0.071801 | SKCM | SK-MEL-1 | Lapatinib |
| MVP_top60_DNA_seed | PIK3CA | -0.071301 | SKCM | RPMI-7951 | Alpelisib |
| MVP_top60_DNA_seed | JAK2 | -0.067333 | SKCM | IPC-298 | AZ960 |
| MVP_top60_DNA_seed | JAK3 | -0.067333 | SKCM | IPC-298 | AZ960 |
| MVP_top60_DNA_seed | PIK3CA | -0.066392 | SKCM | MMAc-SF | AZD8186 |
| MVP_top60_DNA_seed | PIK3CB | -0.066392 | SKCM | MMAc-SF | AZD8186 |
| MVP_top60_DNA_seed | PIK3CA | -0.065515 | SKCM | Mewo | AZD8186 |
| MVP_top60_DNA_seed | PIK3CB | -0.065515 | SKCM | Mewo | AZD8186 |
| MVP_top60_DNA_seed | EGFR | -0.062139 | SKCM | MEL-HO | Sapitinib |
| MVP_top60_DNA_seed | ERBB2 | -0.062139 | SKCM | MEL-HO | Sapitinib |
| MVP_top60_DNA_seed | EGFR | -0.060891 | SKCM | WM35 | Gefitinib |
| MVP_top60_DNA_seed | TP53 | -0.059965 | SKCM | A2058 | MIRA-1 |
| MVP_top60_DNA_seed | EGFR | -0.05584 | SKCM | HT-144 | Gefitinib |
| MVP_top60_DNA_seed | EGFR | -0.055629 | SKCM | WM793B | Afatinib |
| MVP_top60_DNA_seed | ERBB2 | -0.055629 | SKCM | WM793B | Afatinib |
| MVP_top60_DNA_seed | EGFR | -0.054488 | SKCM | SK-MEL-2 | Afatinib |

| | | | | | |
|---|---|---|---|---|---|
| MVP_top60_DNA_seed | ERBB2 | -0.054488 | SKCM | SK-MEL-2 | Afatinib |
| MVP_top60_DNA_seed | EGFR | -0.052825 | SKCM | A2058 | Osimertinib |
| MVP_top60_DNA_seed | EGFR | -0.05033 | SKCM | K2 | Erlotinib |
| MVP_top60_DNA_seed | EGFR | -0.049933 | SKCM | Hs-940-T | Sapitinib |
| MVP_top60_DNA_seed | ERBB2 | -0.049933 | SKCM | Hs-940-T | Sapitinib |
| MVP_top60_DNA_seed | EGFR | -0.047674 | SKCM | MZ2-MEL | Afatinib |
| MVP_top60_DNA_seed | ERBB2 | -0.047674 | SKCM | MZ2-MEL | Afatinib |
| MVP_top60_DNA_seed | KRAS | -0.046265 | SKCM | A2058 | KRAS (G12C) Inhibitor-12 |
| MVP_top60_DNA_seed | EGFR | -0.044376 | SKCM | Hs-939-T | Osimertinib |
| MVP_top60_DNA_seed | BRAF | -0.043138 | SKCM | GAK | Dabrafenib |
| MVP_top60_DNA_seed | PIK3CB | -0.041491 | SKCM | G-MEL | AZD6482 |
| MVP_top60_DNA_seed | EP300 | -0.0411 | SKCM | A101D | SGC-CBP30 |
| MVP_top60_DNA_seed | CREBBP | -0.0411 | SKCM | A101D | SGC-CBP30 |
| MVP_top60_DNA_seed | KRAS | -0.037128 | SKCM | WM793B | KRAS (G12C) Inhibitor-12 |
| MVP_top60_DNA_seed | MAPK1 | -0.031787 | SKCM | WM793B | VX-11e |
| MVP_top60_DNA_seed | JAK2 | -0.029534 | SKCM | K2 | Ruxolitinib |
| MVP_top60_DNA_seed | JAK1 | -0.029534 | SKCM | K2 | Ruxolitinib |
| MVP_top60_DNA_seed | TP53 | -0.028765 | SKCM | LB373-MEL-D | MIRA-1 |
| MVP_top60_DNA_seed | EGFR | -0.0275 | SKCM | IPC-298 | Afatinib |
| MVP_top60_DNA_seed | ERBB2 | -0.0275 | SKCM | IPC-298 | Afatinib |
| MVP_top60_DNA_seed | PIK3CA | -0.025635 | SKCM | HMV-II | Alpelisib |
| MVP_top60_DNA_seed | MAPK3 | -0.020683 | SKCM | MEL-JUSO | Ulixertinib |
| MVP_top60_DNA_seed | MAPK1 | -0.020683 | SKCM | MEL-JUSO | Ulixertinib |
| MVP_top60_DNA_seed | PIK3CA | -0.019752 | SKCM | M14 | AZD8186 |
| MVP_top60_DNA_seed | PIK3CB | -0.019752 | SKCM | M14 | AZD8186 |
| MVP_top60_DNA_seed | EGFR | -0.019005 | SKCM | A101D | Afatinib |
| MVP_top60_DNA_seed | ERBB2 | -0.019005 | SKCM | A101D | Afatinib |
| MVP_top60_DNA_seed | JAK2 | -0.018538 | SKCM | A101D | AZ960 |
| MVP_top60_DNA_seed | JAK3 | -0.018538 | SKCM | A101D | AZ960 |
| MVP_top60_DNA_seed | EP300 | -0.01711 | SKCM | GAK | CPI-637 |
| MVP_top60_DNA_seed | BRAF | -0.013826 | SKCM | GAK | SB590885 |
| MVP_top60_DNA_seed | JAK2 | -0.011375 | SKCM | IGR-1 | Ruxolitinib |
| MVP_top60_DNA_seed | JAK1 | -0.011375 | SKCM | IGR-1 | Ruxolitinib |
| MVP_top60_DNA_seed | JAK2 | -0.008755 | SKCM | Hs-939-T | AZ960 |
| MVP_top60_DNA_seed | JAK3 | -0.008755 | SKCM | Hs-939-T | AZ960 |
| MVP_top60_DNA_seed | GSK3B | -0.007065 | SKCM | LB2518-MEL | CHIR-99021 |
| MVP_top60_DNA_seed | TP53 | -0.006167 | SKCM | A375 | MIRA-1 |
| MVP_top60_DNA_seed | EGFR | -0.003563 | SKCM | MMAc-SF | Gefitinib |
| MVP_top60_DNA_seed | MAPK1 | -0.001335 | SKCM | A2058 | VX-11e |
| MVP_top60_DNA_seed | PIK3CA | -0.000659 | SKCM | CP66-MEL | Alpelisib |
| MVP_top60_DNA_seed | EP300 | 0.001357 | SKCM | G-361 | CPI-637 |

| | | | | | |
|---|---|---|---|---|---|
| MVP_top60_DNA_seed | JAK2 | 0.00361 | SKCM | G-MEL | JAK_8517 |
| MVP_top60_DNA_seed | JAK1 | 0.00361 | SKCM | G-MEL | JAK_8517 |
| MVP_top60_DNA_seed | EGFR | 0.003688 | SKCM | SK-MEL-30 | Afatinib |
| MVP_top60_DNA_seed | ERBB2 | 0.003688 | SKCM | SK-MEL-30 | Afatinib |
| MVP_top60_DNA_seed | EGFR | 0.005899 | SKCM | COLO-783 | AZD3759 |
| MVP_top60_DNA_seed | RAC1 | 0.007366 | SKCM | SK-MEL-3 | EHT-1864 |
| MVP_top60_DNA_seed | PIK3CB | 0.010484 | SKCM | SK-MEL-5 | AZD6482 |
| MVP_top60_DNA_seed | EGFR | 0.012753 | SKCM | 451Lu | Sapitinib |
| MVP_top60_DNA_seed | ERBB2 | 0.012753 | SKCM | 451Lu | Sapitinib |
| MVP_top60_DNA_seed | MAP2K1 | 0.016805 | SKCM | SK-MEL-31 | PD0325901 |
| MVP_top60_DNA_seed | EP300 | 0.017343 | SKCM | A101D | CPI-637 |
| MVP_top60_DNA_seed | EGFR | 0.021691 | SKCM | MZ7-mel | Sapitinib |
| MVP_top60_DNA_seed | ERBB2 | 0.021691 | SKCM | MZ7-mel | Sapitinib |
| MVP_top60_DNA_seed | JAK1 | 0.025245 | SKCM | UACC-62 | JAK1_8709 |
| MVP_top60_DNA_seed | JAK2 | 0.027389 | SKCM | HT-144 | AZ960 |
| MVP_top60_DNA_seed | JAK3 | 0.027389 | SKCM | HT-144 | AZ960 |
| MVP_top60_DNA_seed | BRAF | 0.029126 | SKCM | MEL-JUSO | Dabrafenib |
| MVP_top60_DNA_seed | JAK2 | 0.031986 | SKCM | HMV-II | GSK2276186C |
| MVP_top60_DNA_seed | JAK1 | 0.031986 | SKCM | HMV-II | GSK2276186C |
| MVP_top60_DNA_seed | JAK3 | 0.031986 | SKCM | HMV-II | GSK2276186C |
| MVP_top60_DNA_seed | MAPK3 | 0.034761 | SKCM | WM-115 | Ulixertinib |
| MVP_top60_DNA_seed | MAPK1 | 0.034761 | SKCM | WM-115 | Ulixertinib |
| MVP_top60_DNA_seed | PIK3CA | 0.035518 | SKCM | WM1552C | Buparlisib |
| MVP_top60_DNA_seed | PIK3CB | 0.035518 | SKCM | WM1552C | Buparlisib |
| MVP_top60_DNA_seed | EGFR | 0.036171 | SKCM | RPMI-7951 | Erlotinib |
| MVP_top60_DNA_seed | PIK3CA | 0.036636 | SKCM | IPC-298 | AZD8186 |
| MVP_top60_DNA_seed | PIK3CB | 0.036636 | SKCM | IPC-298 | AZD8186 |
| MVP_top60_DNA_seed | KRAS | 0.040256 | SKCM | SK-MEL-30 | KRAS (G12C) Inhibitor-12 |
| MVP_top60_DNA_seed | TP53 | 0.041386 | SKCM | IPC-298 | MIRA-1 |
| MVP_top60_DNA_seed | GSK3B | 0.043218 | SKCM | IPC-298 | SB216763 |
| MVP_top60_DNA_seed | BRAF | 0.044274 | SKCM | A2058 | Dabrafenib |
| MVP_top60_DNA_seed | PIK3CA | 0.045792 | SKCM | SK-MEL-28 | Buparlisib |
| MVP_top60_DNA_seed | PIK3CB | 0.045792 | SKCM | SK-MEL-28 | Buparlisib |
| MVP_top60_DNA_seed | GSK3B | 0.046015 | SKCM | WM35 | SB216763 |
| MVP_top60_DNA_seed | GSK3B | 0.048129 | SKCM | LOXIMVI | SB216763 |
| MVP_top60_DNA_seed | GSK3B | 0.048143 | SKCM | MZ2-MEL | CHIR-99021 |
| MVP_top60_DNA_seed | EGFR | 0.0488 | SKCM | CP66-MEL | AZD3759 |
| MVP_top60_DNA_seed | PIK3CA | 0.055844 | SKCM | SK-MEL-5 | GNE-317 |
| MVP_top60_DNA_seed | EP300 | 0.058441 | SKCM | WM35 | SGC-CBP30 |
| MVP_top60_DNA_seed | CREBBP | 0.058441 | SKCM | WM35 | SGC-CBP30 |
| MVP_top60_DNA_seed | EGFR | 0.059311 | SKCM | WM1552C | Afatinib |

| | | | | | |
|---|---|---|---|---|---|
| MVP_top60_DNA_seed | ERBB2 | 0.059311 | SKCM | WM1552C | Afatinib |
| MVP_top60_DNA_seed | PIK3CA | 0.062335 | SKCM | COLO-800 | Alpelisib |
| MVP_top60_DNA_seed | PIK3CA | 0.064294 | SKCM | Mewo | GNE-317 |
| MVP_top60_DNA_seed | EGFR | 0.066855 | SKCM | Hs-940-T | Osimertinib |
| MVP_top60_DNA_seed | MAPK3 | 0.069574 | SKCM | Mewo | Ulixertinib |
| MVP_top60_DNA_seed | MAPK1 | 0.069574 | SKCM | Mewo | Ulixertinib |
| MVP_top60_DNA_seed | EGFR | 0.071364 | SKCM | SK-MEL-30 | Gefitinib |
| MVP_top60_DNA_seed | EGFR | 0.071874 | SKCM | G-361 | Gefitinib |
| MVP_top60_DNA_seed | PIK3CA | 0.073796 | SKCM | GAK | GNE-317 |
| MVP_top60_DNA_seed | JAK2 | 0.073894 | SKCM | COLO-792 | Ruxolitinib |
| MVP_top60_DNA_seed | JAK1 | 0.073894 | SKCM | COLO-792 | Ruxolitinib |
| MVP_top60_DNA_seed | PIK3CA | 0.074412 | SKCM | UACC-62 | Buparlisib |
| MVP_top60_DNA_seed | PIK3CB | 0.074412 | SKCM | UACC-62 | Buparlisib |
| MVP_top60_DNA_seed | MAP2K1 | 0.074685 | SKCM | RVH-421 | Refametinib |
| MVP_top60_DNA_seed | MAPK1 | 0.075238 | SKCM | Hs-940-T | VX-11e |
| MVP_top60_DNA_seed | GSK3B | 0.078978 | SKCM | G-MEL | CHIR-99021 |
| MVP_top60_DNA_seed | PIK3CA | 0.080747 | SKCM | SK-MEL-30 | Alpelisib |
| MVP_top60_DNA_seed | TP53 | 0.080939 | SKCM | K2 | MIRA-1 |
| MVP_top60_DNA_seed | MAPK3 | 0.081584 | SKCM | SK-MEL-3 | Ulixertinib |
| MVP_top60_DNA_seed | MAPK1 | 0.081584 | SKCM | SK-MEL-3 | Ulixertinib |
| MVP_top60_DNA_seed | TP53 | 0.084145 | SKCM | G-361 | MIRA-1 |
| MVP_top60_DNA_seed | JAK2 | 0.091093 | SKCM | M14 | GSK2276186C |
| MVP_top60_DNA_seed | JAK1 | 0.091093 | SKCM | M14 | GSK2276186C |
| MVP_top60_DNA_seed | JAK3 | 0.091093 | SKCM | M14 | GSK2276186C |
| MVP_top60_DNA_seed | EGFR | 0.095754 | SKCM | COLO-679 | AZD3759 |
| MVP_top60_DNA_seed | PIK3CA | 0.096106 | SKCM | G-MEL | Alpelisib |
| MVP_top60_DNA_seed | JAK2 | 0.096298 | SKCM | CHL-1 | GSK2276186C |
| MVP_top60_DNA_seed | JAK1 | 0.096298 | SKCM | CHL-1 | GSK2276186C |
| MVP_top60_DNA_seed | JAK3 | 0.096298 | SKCM | CHL-1 | GSK2276186C |
| MVP_top60_DNA_seed | MAPK3 | 0.096877 | SKCM | WM1552C | Ulixertinib |
| MVP_top60_DNA_seed | MAPK1 | 0.096877 | SKCM | WM1552C | Ulixertinib |
| MVP_top60_DNA_seed | JAK2 | 0.098041 | SKCM | LOXIMVI | Ruxolitinib |
| MVP_top60_DNA_seed | JAK1 | 0.098041 | SKCM | LOXIMVI | Ruxolitinib |
| MVP_top60_DNA_seed | EGFR | 0.098488 | SKCM | MEL-HO | Gefitinib |
| MVP_top60_DNA_seed | EGFR | 0.101064 | SKCM | LB373-MEL-D | Osimertinib |
| MVP_top60_DNA_seed | PIK3CA | 0.102219 | SKCM | WM-115 | GNE-317 |
| MVP_top60_DNA_seed | MAP2K1 | 0.106321 | SKCM | Hs-940-T | Refametinib |
| MVP_top60_DNA_seed | EGFR | 0.10794 | SKCM | UACC-257 | Erlotinib |
| MVP_top60_DNA_seed | MAP2K1 | 0.111556 | SKCM | SK-MEL-31 | Selumetinib |
| MVP_top60_DNA_seed | RAC1 | 0.11165 | SKCM | SH-4 | EHT-1864 |
| MVP_top60_DNA_seed | EGFR | 0.113388 | SKCM | RPMI-7951 | Afatinib |

| | | | | | |
|---|---|---|---|---|---|
| MVP_top60_DNA_seed | ERBB2 | 0.113388 | SKCM | RPMI-7951 | Afatinib |
| MVP_top60_DNA_seed | EGFR | 0.115678 | SKCM | SK-MEL-5 | AZD3759 |
| MVP_top60_DNA_seed | EGFR | 0.115799 | SKCM | A375 | Afatinib |
| MVP_top60_DNA_seed | ERBB2 | 0.115799 | SKCM | A375 | Afatinib |
| MVP_top60_DNA_seed | JAK2 | 0.116162 | SKCM | LB373-MEL-D | Ruxolitinib |
| MVP_top60_DNA_seed | JAK1 | 0.116162 | SKCM | LB373-MEL-D | Ruxolitinib |
| MVP_top60_DNA_seed | EGFR | 0.119524 | SKCM | M14 | Afatinib |
| MVP_top60_DNA_seed | ERBB2 | 0.119524 | SKCM | M14 | Afatinib |
| MVP_top60_DNA_seed | MAP2K1 | 0.121085 | SKCM | Mewo | Selumetinib |
| MVP_top60_DNA_seed | MAPK3 | 0.121438 | SKCM | MEL-JUSO | Ulixertinib |
| MVP_top60_DNA_seed | MAPK1 | 0.121438 | SKCM | MEL-JUSO | Ulixertinib |
| MVP_top60_DNA_seed | MAPK1 | 0.121472 | SKCM | IGR-1 | VX-11e |
| MVP_top60_DNA_seed | GSK3B | 0.121998 | SKCM | Hs-940-T | SB216763 |
| MVP_top60_DNA_seed | GSK3B | 0.126261 | SKCM | CP50-MEL-B | SB216763 |
| MVP_top60_DNA_seed | EGFR | 0.12921 | SKCM | CP50-MEL-B | Lapatinib |
| MVP_top60_DNA_seed | ERBB2 | 0.12921 | SKCM | CP50-MEL-B | Lapatinib |
| MVP_top60_DNA_seed | EGFR | 0.136586 | SKCM | SK-MEL-2 | Gefitinib |
| MVP_top60_DNA_seed | KRAS | 0.145376 | SKCM | M14 | KRAS (G12C) Inhibitor-12 |
| MVP_top60_DNA_seed | JAK2 | 0.14657 | SKCM | MEL-JUSO | Ruxolitinib |
| MVP_top60_DNA_seed | JAK1 | 0.14657 | SKCM | MEL-JUSO | Ruxolitinib |
| MVP_top60_DNA_seed | EGFR | 0.14659 | SKCM | RVH-421 | Gefitinib |
| MVP_top60_DNA_seed | EGFR | 0.146851 | SKCM | COLO-800 | Erlotinib |
| MVP_top60_DNA_seed | MAPK1 | 0.152108 | SKCM | COLO-829 | VX-11e |
| MVP_top60_DNA_seed | EGFR | 0.152939 | SKCM | HMV-II | Afatinib |
| MVP_top60_DNA_seed | ERBB2 | 0.152939 | SKCM | HMV-II | Afatinib |
| MVP_top60_DNA_seed | BRAF | 0.157413 | SKCM | HMV-II | SB590885 |
| MVP_top60_DNA_seed | PIK3CA | 0.157819 | SKCM | SK-MEL-5 | Buparlisib |
| MVP_top60_DNA_seed | PIK3CB | 0.157819 | SKCM | SK-MEL-5 | Buparlisib |
| MVP_top60_DNA_seed | PIK3CA | 0.159039 | SKCM | MEL-HO | Alpelisib |
| MVP_top60_DNA_seed | JAK2 | 0.160386 | SKCM | MZ7-mel | AZ960 |
| MVP_top60_DNA_seed | JAK3 | 0.160386 | SKCM | MZ7-mel | AZ960 |
| MVP_top60_DNA_seed | EGFR | 0.166333 | SKCM | SK-MEL-3 | Erlotinib |
| MVP_top60_DNA_seed | JAK2 | 0.174019 | SKCM | G-MEL | AZ960 |
| MVP_top60_DNA_seed | JAK3 | 0.174019 | SKCM | G-MEL | AZ960 |
| MVP_top60_DNA_seed | TP53 | 0.175197 | SKCM | CHL-1 | MIRA-1 |
| MVP_top60_DNA_seed | BRAF | 0.175232 | SKCM | RPMI-7951 | Dabrafenib |
| MVP_top60_DNA_seed | PIK3CA | 0.175685 | SKCM | WM35 | AZD8186 |
| MVP_top60_DNA_seed | PIK3CB | 0.175685 | SKCM | WM35 | AZD8186 |
| MVP_top60_DNA_seed | EP300 | 0.175691 | SKCM | WM278 | SGC-CBP30 |
| MVP_top60_DNA_seed | CREBBP | 0.175691 | SKCM | WM278 | SGC-CBP30 |
| MVP_top60_DNA_seed | EGFR | 0.175844 | SKCM | COLO-783 | Erlotinib |

| | | | | | |
|---|---|---|---|---|---|
| MVP_top60_DNA_seed | TP53 | 0.176308 | SKCM | SK-MEL-5 | MIRA-1 |
| MVP_top60_DNA_seed | JAK2 | 0.177627 | SKCM | SK-MEL-2 | Ruxolitinib |
| MVP_top60_DNA_seed | JAK1 | 0.177627 | SKCM | SK-MEL-2 | Ruxolitinib |
| MVP_top60_DNA_seed | RAC1 | 0.181429 | SKCM | MEL-HO | EHT-1864 |
| MVP_top60_DNA_seed | EGFR | 0.182272 | SKCM | GAK | Lapatinib |
| MVP_top60_DNA_seed | ERBB2 | 0.182272 | SKCM | GAK | Lapatinib |
| MVP_top60_DNA_seed | PIK3CB | 0.182945 | SKCM | MMAc-SF | AZD6482 |
| MVP_top60_DNA_seed | PIK3CA | 0.183003 | SKCM | LB2518-MEL | Alpelisib |
| MVP_top60_DNA_seed | PIK3CA | 0.183227 | SKCM | 451Lu | GNE-317 |
| MVP_top60_DNA_seed | PIK3CA | 0.185117 | SKCM | WM-115 | Alpelisib |
| MVP_top60_DNA_seed | EGFR | 0.191118 | SKCM | MZ2-MEL | AZD3759 |
| MVP_top60_DNA_seed | EGFR | 0.191811 | SKCM | GAK | Afatinib |
| MVP_top60_DNA_seed | ERBB2 | 0.191811 | SKCM | GAK | Afatinib |
| MVP_top60_DNA_seed | KRAS | 0.194455 | SKCM | MMAc-SF | KRAS (G12C) Inhibitor-12 |
| MVP_top60_DNA_seed | EGFR | 0.195187 | SKCM | COLO-800 | Sapitinib |
| MVP_top60_DNA_seed | ERBB2 | 0.195187 | SKCM | COLO-800 | Sapitinib |
| MVP_top60_DNA_seed | BRAF | 0.198774 | SKCM | SK-MEL-30 | Dabrafenib |
| MVP_top60_DNA_seed | GSK3B | 0.20011 | SKCM | Mewo | SB216763 |
| MVP_top60_DNA_seed | EGFR | 0.200129 | SKCM | CP50-MEL-B | Osimertinib |
| MVP_top60_DNA_seed | TP53 | 0.200981 | SKCM | IGR-37 | MIRA-1 |
| MVP_top60_DNA_seed | MAPK3 | 0.201706 | SKCM | CHL-1 | Ulixertinib |
| MVP_top60_DNA_seed | MAPK1 | 0.201706 | SKCM | CHL-1 | Ulixertinib |
| MVP_top60_DNA_seed | EGFR | 0.202885 | SKCM | COLO-800 | Gefitinib |
| MVP_top60_DNA_seed | EGFR | 0.203125 | SKCM | CP50-MEL-B | Sapitinib |
| MVP_top60_DNA_seed | ERBB2 | 0.203125 | SKCM | CP50-MEL-B | Sapitinib |
| MVP_top60_DNA_seed | TP53 | 0.206601 | SKCM | MZ2-MEL | MIRA-1 |
| MVP_top60_DNA_seed | EGFR | 0.20733 | SKCM | IPC-298 | Sapitinib |
| MVP_top60_DNA_seed | ERBB2 | 0.20733 | SKCM | IPC-298 | Sapitinib |
| MVP_top60_DNA_seed | RAC1 | 0.207672 | SKCM | CP50-MEL-B | EHT-1864 |
| MVP_top60_DNA_seed | RAC1 | 0.209983 | SKCM | LB2518-MEL | EHT-1864 |
| MVP_top60_DNA_seed | BRAF | 0.210061 | SKCM | CP66-MEL | SB590885 |
| MVP_top60_DNA_seed | JAK2 | 0.210157 | SKCM | WM-115 | JAK_8517 |
| MVP_top60_DNA_seed | JAK1 | 0.210157 | SKCM | WM-115 | JAK_8517 |
| MVP_top60_DNA_seed | EGFR | 0.210765 | SKCM | IGR-37 | Erlotinib |
| MVP_top60_DNA_seed | MAP2K1 | 0.213134 | SKCM | RPMI-7951 | Selumetinib |
| MVP_top60_DNA_seed | JAK2 | 0.214618 | SKCM | WM793B | AZ960 |
| MVP_top60_DNA_seed | JAK3 | 0.214618 | SKCM | WM793B | AZ960 |
| MVP_top60_DNA_seed | TP53 | 0.215063 | SKCM | CP66-MEL | MIRA-1 |
| MVP_top60_DNA_seed | TP53 | 0.217295 | SKCM | WM-115 | MIRA-1 |
| MVP_top60_DNA_seed | EP300 | 0.218545 | SKCM | WM793B | CPI-637 |
| MVP_top60_DNA_seed | EGFR | 0.221161 | SKCM | IGR-1 | Osimertinib |

| | | | | | |
|---|---|---|---|---|---|
| MVP_top60_DNA_seed | GSK3B | 0.222235 | SKCM | IGR-37 | SB216763 |
| MVP_top60_DNA_seed | RAC1 | 0.223018 | SKCM | HMV-II | EHT-1864 |
| MVP_top60_DNA_seed | TP53 | 0.223794 | SKCM | MMAc-SF | MIRA-1 |
| MVP_top60_DNA_seed | JAK2 | 0.229838 | SKCM | C32 | AZ960 |
| MVP_top60_DNA_seed | JAK3 | 0.229838 | SKCM | C32 | AZ960 |
| MVP_top60_DNA_seed | EP300 | 0.230762 | SKCM | MEL-JUSO | CPI-637 |
| MVP_top60_DNA_seed | JAK2 | 0.231771 | SKCM | HMV-II | JAK_8517 |
| MVP_top60_DNA_seed | JAK1 | 0.231771 | SKCM | HMV-II | JAK_8517 |
| MVP_top60_DNA_seed | PIK3CA | 0.232165 | SKCM | K2 | Buparlisib |
| MVP_top60_DNA_seed | PIK3CB | 0.232165 | SKCM | K2 | Buparlisib |
| MVP_top60_DNA_seed | JAK2 | 0.232279 | SKCM | A2058 | JAK_8517 |
| MVP_top60_DNA_seed | JAK1 | 0.232279 | SKCM | A2058 | JAK_8517 |
| MVP_top60_DNA_seed | PIK3CA | 0.239679 | SKCM | Mewo | Alpelisib |
| MVP_top60_DNA_seed | EGFR | 0.241952 | SKCM | SK-MEL-2 | Sapitinib |
| MVP_top60_DNA_seed | ERBB2 | 0.241952 | SKCM | SK-MEL-2 | Sapitinib |
| MVP_top60_DNA_seed | BRAF | 0.242275 | SKCM | CHL-1 | PLX-4720 |
| MVP_top60_DNA_seed | PIK3CA | 0.24436 | SKCM | G-361 | GNE-317 |
| MVP_top60_DNA_seed | KRAS | 0.244946 | SKCM | CP66-MEL | KRAS (G12C) Inhibitor-12 |
| MVP_top60_DNA_seed | MAP2K1 | 0.245532 | SKCM | IGR-1 | Refametinib |
| MVP_top60_DNA_seed | JAK2 | 0.245643 | SKCM | WM1552C | GSK2276186C |
| MVP_top60_DNA_seed | JAK1 | 0.245643 | SKCM | WM1552C | GSK2276186C |
| MVP_top60_DNA_seed | JAK3 | 0.245643 | SKCM | WM1552C | GSK2276186C |
| MVP_top60_DNA_seed | EGFR | 0.249155 | SKCM | A375 | Lapatinib |
| MVP_top60_DNA_seed | ERBB2 | 0.249155 | SKCM | A375 | Lapatinib |
| MVP_top60_DNA_seed | MAP2K1 | 0.250616 | SKCM | A2058 | Refametinib |
| MVP_top60_DNA_seed | EGFR | 0.250646 | SKCM | GAK | Erlotinib |
| MVP_top60_DNA_seed | JAK2 | 0.257195 | SKCM | Hs-940-T | Ruxolitinib |
| MVP_top60_DNA_seed | JAK1 | 0.257195 | SKCM | Hs-940-T | Ruxolitinib |
| MVP_top60_DNA_seed | EGFR | 0.257767 | SKCM | COLO-792 | Sapitinib |
| MVP_top60_DNA_seed | ERBB2 | 0.257767 | SKCM | COLO-792 | Sapitinib |
| MVP_top60_DNA_seed | GSK3B | 0.261822 | SKCM | UACC-257 | SB216763 |
| MVP_top60_DNA_seed | EGFR | 0.264484 | SKCM | IST-MEL1 | AZD3759 |
| MVP_top60_DNA_seed | JAK2 | 0.264563 | SKCM | M14 | JAK_8517 |
| MVP_top60_DNA_seed | JAK1 | 0.264563 | SKCM | M14 | JAK_8517 |
| MVP_top60_DNA_seed | PIK3CA | 0.265176 | SKCM | K2 | AZD8186 |
| MVP_top60_DNA_seed | PIK3CB | 0.265176 | SKCM | K2 | AZD8186 |
| MVP_top60_DNA_seed | EGFR | 0.267128 | SKCM | SK-MEL-28 | AZD3759 |
| MVP_top60_DNA_seed | MAPK1 | 0.268702 | SKCM | Mewo | VX-11e |
| MVP_top60_DNA_seed | JAK2 | 0.26913 | SKCM | A2058 | Ruxolitinib |
| MVP_top60_DNA_seed | JAK1 | 0.26913 | SKCM | A2058 | Ruxolitinib |
| MVP_top60_DNA_seed | JAK2 | 0.2694 | SKCM | A2058 | GSK2276186C |

| | | | | | |
|---|---|---|---|---|---|
| MVP_top60_DNA_seed | JAK1 | 0.2694 | SKCM | A2058 | GSK2276186C |
| MVP_top60_DNA_seed | JAK3 | 0.2694 | SKCM | A2058 | GSK2276186C |
| MVP_top60_DNA_seed | BRAF | 0.27152 | SKCM | MZ2-MEL | SB590885 |
| MVP_top60_DNA_seed | EGFR | 0.271639 | SKCM | SK-MEL-2 | AZD3759 |
| MVP_top60_DNA_seed | EGFR | 0.272235 | SKCM | Hs-940-T | AZD3759 |
| MVP_top60_DNA_seed | EP300 | 0.275427 | SKCM | COLO-792 | SGC-CBP30 |
| MVP_top60_DNA_seed | CREBBP | 0.275427 | SKCM | COLO-792 | SGC-CBP30 |
| MVP_top60_DNA_seed | JAK1 | 0.27738 | SKCM | COLO-679 | JAK1_8709 |
| MVP_top60_DNA_seed | MAPK3 | 0.278168 | SKCM | IST-MEL1 | Ulixertinib |
| MVP_top60_DNA_seed | MAPK1 | 0.278168 | SKCM | IST-MEL1 | Ulixertinib |
| MVP_top60_DNA_seed | TP53 | 0.279229 | SKCM | MEL-JUSO | MIRA-1 |
| MVP_top60_DNA_seed | EGFR | 0.280209 | SKCM | RPMI-7951 | Sapitinib |
| MVP_top60_DNA_seed | ERBB2 | 0.280209 | SKCM | RPMI-7951 | Sapitinib |
| MVP_top60_DNA_seed | EGFR | 0.281626 | SKCM | MZ7-mel | Osimertinib |
| MVP_top60_DNA_seed | EP300 | 0.282277 | SKCM | HT-144 | SGC-CBP30 |
| MVP_top60_DNA_seed | CREBBP | 0.282277 | SKCM | HT-144 | SGC-CBP30 |
| MVP_top60_DNA_seed | PIK3CA | 0.282331 | SKCM | WM793B | AZD8186 |
| MVP_top60_DNA_seed | PIK3CB | 0.282331 | SKCM | WM793B | AZD8186 |
| MVP_top60_DNA_seed | TP53 | 0.284184 | SKCM | MEL-HO | MIRA-1 |
| MVP_top60_DNA_seed | MAP2K1 | 0.285974 | SKCM | SK-MEL-31 | Refametinib |
| MVP_top60_DNA_seed | EGFR | 0.288361 | SKCM | G-361 | Afatinib |
| MVP_top60_DNA_seed | ERBB2 | 0.288361 | SKCM | G-361 | Afatinib |
| MVP_top60_DNA_seed | EGFR | 0.290214 | SKCM | M14 | AZD3759 |
| MVP_top60_DNA_seed | RAC1 | 0.290532 | SKCM | WM35 | EHT-1864 |
| MVP_top60_DNA_seed | GSK3B | 0.29401 | SKCM | MZ2-MEL | SB216763 |
| MVP_top60_DNA_seed | PIK3CA | 0.29742 | SKCM | K2 | Alpelisib |
| MVP_top60_DNA_seed | MAPK3 | 0.297665 | SKCM | SK-MEL-24 | Ulixertinib |
| MVP_top60_DNA_seed | MAPK1 | 0.297665 | SKCM | SK-MEL-24 | Ulixertinib |
| MVP_top60_DNA_seed | EGFR | 0.300122 | SKCM | COLO-783 | Osimertinib |
| MVP_top60_DNA_seed | EP300 | 0.301714 | SKCM | UACC-62 | SGC-CBP30 |
| MVP_top60_DNA_seed | CREBBP | 0.301714 | SKCM | UACC-62 | SGC-CBP30 |
| MVP_top60_DNA_seed | MAP2K1 | 0.303538 | SKCM | Hs-940-T | Trametinib |
| MVP_top60_DNA_seed | GSK3B | 0.305863 | SKCM | MZ7-mel | SB216763 |
| MVP_top60_DNA_seed | JAK2 | 0.306608 | SKCM | SK-MEL-5 | Ruxolitinib |
| MVP_top60_DNA_seed | JAK1 | 0.306608 | SKCM | SK-MEL-5 | Ruxolitinib |
| MVP_top60_DNA_seed | EGFR | 0.310323 | SKCM | A2058 | Gefitinib |
| MVP_top60_DNA_seed | PIK3CA | 0.312216 | SKCM | SK-MEL-5 | AZD8186 |
| MVP_top60_DNA_seed | PIK3CB | 0.312216 | SKCM | SK-MEL-5 | AZD8186 |
| MVP_top60_DNA_seed | MAPK3 | 0.316161 | SKCM | Hs-940-T | SCH772984 |
| MVP_top60_DNA_seed | MAPK1 | 0.316161 | SKCM | Hs-940-T | SCH772984 |
| MVP_top60_DNA_seed | PIK3CA | 0.316613 | SKCM | SK-MEL-31 | GNE-317 |

| | | | | | |
|---|---|---|---|---|---|
| MVP_top60_DNA_seed | PIK3CA | 0.320473 | SKCM | SK-MEL-31 | Buparlisib |
| MVP_top60_DNA_seed | PIK3CB | 0.320473 | SKCM | SK-MEL-31 | Buparlisib |
| MVP_top60_DNA_seed | TP53 | 0.321996 | SKCM | COLO-800 | MIRA-1 |
| MVP_top60_DNA_seed | RAC1 | 0.323875 | SKCM | IGR-1 | EHT-1864 |
| MVP_top60_DNA_seed | JAK2 | 0.326891 | SKCM | GAK | JAK_8517 |
| MVP_top60_DNA_seed | JAK1 | 0.326891 | SKCM | GAK | JAK_8517 |
| MVP_top60_DNA_seed | MAPK3 | 0.328077 | SKCM | SK-MEL-31 | Ulixertinib |
| MVP_top60_DNA_seed | MAPK1 | 0.328077 | SKCM | SK-MEL-31 | Ulixertinib |
| MVP_top60_DNA_seed | PIK3CA | 0.328341 | SKCM | GAK | Buparlisib |
| MVP_top60_DNA_seed | PIK3CB | 0.328341 | SKCM | GAK | Buparlisib |
| MVP_top60_DNA_seed | MAPK3 | 0.328501 | SKCM | WM-115 | Ulixertinib |
| MVP_top60_DNA_seed | MAPK1 | 0.328501 | SKCM | WM-115 | Ulixertinib |
| MVP_top60_DNA_seed | EGFR | 0.333267 | SKCM | MZ7-mel | Afatinib |
| MVP_top60_DNA_seed | ERBB2 | 0.333267 | SKCM | MZ7-mel | Afatinib |
| MVP_top60_DNA_seed | EGFR | 0.334253 | SKCM | IGR-1 | Afatinib |
| MVP_top60_DNA_seed | ERBB2 | 0.334253 | SKCM | IGR-1 | Afatinib |
| MVP_top60_DNA_seed | PIK3CA | 0.336373 | SKCM | WM35 | Alpelisib |
| MVP_top60_DNA_seed | JAK2 | 0.34386 | SKCM | GAK | GSK2276186C |
| MVP_top60_DNA_seed | JAK1 | 0.34386 | SKCM | GAK | GSK2276186C |
| MVP_top60_DNA_seed | JAK3 | 0.34386 | SKCM | GAK | GSK2276186C |
| MVP_top60_DNA_seed | GSK3B | 0.346183 | SKCM | MEL-JUSO | SB216763 |
| MVP_top60_DNA_seed | PIK3CA | 0.346439 | SKCM | HMV-II | Buparlisib |
| MVP_top60_DNA_seed | PIK3CB | 0.346439 | SKCM | HMV-II | Buparlisib |
| MVP_top60_DNA_seed | JAK2 | 0.34654 | SKCM | SK-MEL-24 | JAK_8517 |
| MVP_top60_DNA_seed | JAK1 | 0.34654 | SKCM | SK-MEL-24 | JAK_8517 |
| MVP_top60_DNA_seed | EGFR | 0.349734 | SKCM | SK-MEL-3 | Lapatinib |
| MVP_top60_DNA_seed | ERBB2 | 0.349734 | SKCM | SK-MEL-3 | Lapatinib |
| MVP_top60_DNA_seed | TP53 | 0.350432 | SKCM | M14 | MIRA-1 |
| MVP_top60_DNA_seed | EGFR | 0.351546 | SKCM | SK-MEL-5 | Gefitinib |
| MVP_top60_DNA_seed | PIK3CA | 0.353332 | SKCM | COLO-783 | Buparlisib |
| MVP_top60_DNA_seed | PIK3CB | 0.353332 | SKCM | COLO-783 | Buparlisib |
| MVP_top60_DNA_seed | JAK2 | 0.358305 | SKCM | CHL-1 | AZ960 |
| MVP_top60_DNA_seed | JAK3 | 0.358305 | SKCM | CHL-1 | AZ960 |
| MVP_top60_DNA_seed | JAK2 | 0.359308 | SKCM | WM-115 | Ruxolitinib |
| MVP_top60_DNA_seed | JAK1 | 0.359308 | SKCM | WM-115 | Ruxolitinib |
| MVP_top60_DNA_seed | EGFR | 0.35933 | SKCM | VMRC-MELG | Lapatinib |
| MVP_top60_DNA_seed | ERBB2 | 0.35933 | SKCM | VMRC-MELG | Lapatinib |
| MVP_top60_DNA_seed | EGFR | 0.35963 | SKCM | MZ2-MEL | Sapitinib |
| MVP_top60_DNA_seed | ERBB2 | 0.35963 | SKCM | MZ2-MEL | Sapitinib |
| MVP_top60_DNA_seed | JAK2 | 0.363853 | SKCM | SH-4 | JAK_8517 |
| MVP_top60_DNA_seed | JAK1 | 0.363853 | SKCM | SH-4 | JAK_8517 |

| | | | | | |
|---|---|---|---|---|---|
| MVP_top60_DNA_seed | MAP2K1 | 0.36456 | SKCM | CHL-1 | Refametinib |
| MVP_top60_DNA_seed | EP300 | 0.364944 | SKCM | SK-MEL-24 | CPI-637 |
| MVP_top60_DNA_seed | EGFR | 0.366821 | SKCM | CP50-MEL-B | Gefitinib |
| MVP_top60_DNA_seed | EGFR | 0.367166 | SKCM | VMRC-MELG | Osimertinib |
| MVP_top60_DNA_seed | JAK2 | 0.370307 | SKCM | MEL-HO | Ruxolitinib |
| MVP_top60_DNA_seed | JAK1 | 0.370307 | SKCM | MEL-HO | Ruxolitinib |
| MVP_top60_DNA_seed | EGFR | 0.371425 | SKCM | G-MEL | Sapitinib |
| MVP_top60_DNA_seed | ERBB2 | 0.371425 | SKCM | G-MEL | Sapitinib |
| MVP_top60_DNA_seed | EGFR | 0.372438 | SKCM | SK-MEL-31 | Gefitinib |
| MVP_top60_DNA_seed | KRAS | 0.372782 | SKCM | COLO-792 | [illegible]RAS (G12C) Inhibitor-12 |
| MVP_top60_DNA_seed | EGFR | 0.372829 | SKCM | 451Lu | AZD3759 |
| MVP_top60_DNA_seed | EGFR | 0.373082 | SKCM | WM1552C | Osimertinib |
| MVP_top60_DNA_seed | JAK2 | 0.373591 | SKCM | M14 | Ruxolitinib |
| MVP_top60_DNA_seed | JAK1 | 0.373591 | SKCM | M14 | Ruxolitinib |
| MVP_top60_DNA_seed | JAK2 | 0.375111 | SKCM | COLO-783 | AZ960 |
| MVP_top60_DNA_seed | JAK3 | 0.375111 | SKCM | COLO-783 | AZ960 |
| MVP_top60_DNA_seed | PIK3CA | 0.378165 | SKCM | WM1552C | AZD8186 |
| MVP_top60_DNA_seed | PIK3CB | 0.378165 | SKCM | WM1552C | AZD8186 |
| MVP_top60_DNA_seed | GSK3B | 0.379017 | SKCM | LB373-MEL-D | CHIR-99021 |
| MVP_top60_DNA_seed | JAK2 | 0.381347 | SKCM | RVH-421 | Ruxolitinib |
| MVP_top60_DNA_seed | JAK1 | 0.381347 | SKCM | RVH-421 | Ruxolitinib |
| MVP_top60_DNA_seed | GSK3B | 0.382529 | SKCM | COLO-800 | SB216763 |
| MVP_top60_DNA_seed | RAC1 | 0.383732 | SKCM | IGR-37 | EHT-1864 |
| MVP_top60_DNA_seed | EGFR | 0.386483 | SKCM | RPMI-7951 | Gefitinib |
| MVP_top60_DNA_seed | EGFR | 0.387325 | SKCM | HMV-II | Osimertinib |
| MVP_top60_DNA_seed | EP300 | 0.388207 | SKCM | SK-MEL-1 | CPI-637 |
| MVP_top60_DNA_seed | GSK3B | 0.388618 | SKCM | MEL-HO | SB216763 |
| MVP_top60_DNA_seed | EGFR | 0.391283 | SKCM | UACC-62 | Afatinib |
| MVP_top60_DNA_seed | ERBB2 | 0.391283 | SKCM | UACC-62 | Afatinib |
| MVP_top60_DNA_seed | TP53 | 0.392118 | SKCM | SH-4 | MIRA-1 |
| MVP_top60_DNA_seed | PIK3CA | 0.394201 | SKCM | MZ7-mel | AZD8186 |
| MVP_top60_DNA_seed | PIK3CB | 0.394201 | SKCM | MZ7-mel | AZD8186 |
| MVP_top60_DNA_seed | EGFR | 0.396428 | SKCM | A2058 | Sapitinib |
| MVP_top60_DNA_seed | ERBB2 | 0.396428 | SKCM | A2058 | Sapitinib |
| MVP_top60_DNA_seed | EGFR | 0.397606 | SKCM | Hs-940-T | Erlotinib |
| MVP_top60_DNA_seed | MAPK1 | 0.399792 | SKCM | MEL-JUSO | VX-11e |
| MVP_top60_DNA_seed | BRAF | 0.399824 | SKCM | LB373-MEL-D | PLX-4720 |
| MVP_top60_DNA_seed | GSK3B | 0.400502 | SKCM | UACC-62 | SB216763 |
| MVP_top60_DNA_seed | EP300 | 0.400701 | SKCM | COLO-679 | CPI-637 |
| MVP_top60_DNA_seed | EGFR | 0.402448 | SKCM | HT-144 | Sapitinib |
| MVP_top60_DNA_seed | ERBB2 | 0.402448 | SKCM | HT-144 | Sapitinib |

| | | | | | |
|---|---|---|---|---|---|
| MVP_top60_DNA_seed | KRAS | 0.402783 | SKCM | HMV-II | KRAS (G12C) Inhibitor-12 |
| MVP_top60_DNA_seed | EGFR | 0.403779 | SKCM | M14 | Gefitinib |
| MVP_top60_DNA_seed | EGFR | 0.404043 | SKCM | G-MEL | Osimertinib |
| MVP_top60_DNA_seed | JAK2 | 0.406572 | SKCM | CP66-MEL | Ruxolitinib |
| MVP_top60_DNA_seed | JAK1 | 0.406572 | SKCM | CP66-MEL | Ruxolitinib |
| MVP_top60_DNA_seed | EGFR | 0.407007 | SKCM | A375 | AZD3759 |
| MVP_top60_DNA_seed | EP300 | 0.408314 | SKCM | IGR-37 | SGC-CBP30 |
| MVP_top60_DNA_seed | CREBBP | 0.408314 | SKCM | IGR-37 | SGC-CBP30 |
| MVP_top60_DNA_seed | EGFR | 0.408686 | SKCM | IST-MEL1 | Sapitinib |
| MVP_top60_DNA_seed | ERBB2 | 0.408686 | SKCM | IST-MEL1 | Sapitinib |
| MVP_top60_DNA_seed | PIK3CA | 0.409335 | SKCM | MMAc-SF | Alpelisib |
| MVP_top60_DNA_seed | GSK3B | 0.409533 | SKCM | G-361 | SB216763 |
| MVP_top60_DNA_seed | KRAS | 0.409653 | SKCM | COLO-800 | KRAS (G12C) Inhibitor-12 |
| MVP_top60_DNA_seed | PIK3CA | 0.410784 | SKCM | LB373-MEL-D | Alpelisib |
| MVP_top60_DNA_seed | EGFR | 0.411518 | SKCM | UACC-62 | Erlotinib |
| MVP_top60_DNA_seed | MAP2K1 | 0.412265 | SKCM | CHL-1 | Trametinib |
| MVP_top60_DNA_seed | EP300 | 0.412924 | SKCM | WM1552C | SGC-CBP30 |
| MVP_top60_DNA_seed | CREBBP | 0.412924 | SKCM | WM1552C | SGC-CBP30 |
| MVP_top60_DNA_seed | EGFR | 0.416432 | SKCM | WM35 | Afatinib |
| MVP_top60_DNA_seed | ERBB2 | 0.416432 | SKCM | WM35 | Afatinib |
| MVP_top60_DNA_seed | JAK2 | 0.417163 | SKCM | IGR-37 | JAK_8517 |
| MVP_top60_DNA_seed | JAK1 | 0.417163 | SKCM | IGR-37 | JAK_8517 |
| MVP_top60_DNA_seed | KRAS | 0.418826 | SKCM | C32 | KRAS (G12C) Inhibitor-12 |
| MVP_top60_DNA_seed | PIK3CA | 0.421542 | SKCM | SK-MEL-5 | Alpelisib |
| MVP_top60_DNA_seed | PIK3CA | 0.421563 | SKCM | VMRC-MELG | Buparlisib |
| MVP_top60_DNA_seed | PIK3CB | 0.421563 | SKCM | VMRC-MELG | Buparlisib |
| MVP_top60_DNA_seed | EGFR | 0.42221 | SKCM | SK-MEL-30 | Erlotinib |
| MVP_top60_DNA_seed | TP53 | 0.423116 | SKCM | IGR-1 | MIRA-1 |
| MVP_top60_DNA_seed | BRAF | 0.424186 | SKCM | CHL-1 | Dabrafenib |
| MVP_top60_DNA_seed | KRAS | 0.424505 | SKCM | COLO-679 | KRAS (G12C) Inhibitor-12 |
| MVP_top60_DNA_seed | EGFR | 0.425883 | SKCM | 451Lu | Lapatinib |
| MVP_top60_DNA_seed | ERBB2 | 0.425883 | SKCM | 451Lu | Lapatinib |
| MVP_top60_DNA_seed | EGFR | 0.427537 | SKCM | A375 | Erlotinib |
| MVP_top60_DNA_seed | JAK2 | 0.427747 | SKCM | SK-MEL-2 | GSK2276186C |
| MVP_top60_DNA_seed | JAK1 | 0.427747 | SKCM | SK-MEL-2 | GSK2276186C |
| MVP_top60_DNA_seed | JAK3 | 0.427747 | SKCM | SK-MEL-2 | GSK2276186C |
| MVP_top60_DNA_seed | EGFR | 0.430707 | SKCM | M14 | Osimertinib |
| MVP_top60_DNA_seed | EGFR | 0.432331 | SKCM | A375 | Osimertinib |
| MVP_top60_DNA_seed | JAK2 | 0.432457 | SKCM | LOXIMVI | GSK2276186C |
| MVP_top60_DNA_seed | JAK1 | 0.432457 | SKCM | LOXIMVI | GSK2276186C |
| MVP_top60_DNA_seed | JAK3 | 0.432457 | SKCM | LOXIMVI | GSK2276186C |

| | | | | | |
|---|---|---|---|---|---|
| MVP_top60_DNA_seed | EGFR | 0.43669 | SKCM | IGR-1 | AZD3759 |
| MVP_top60_DNA_seed | GSK3B | 0.436915 | SKCM | LB373-MEL-D | SB216763 |
| MVP_top60_DNA_seed | GSK3B | 0.43789 | SKCM | COLO-783 | CHIR-99021 |
| MVP_top60_DNA_seed | EGFR | 0.43819 | SKCM | SK-MEL-24 | Afatinib |
| MVP_top60_DNA_seed | ERBB2 | 0.43819 | SKCM | SK-MEL-24 | Afatinib |
| MVP_top60_DNA_seed | BRAF | 0.438214 | SKCM | COLO-792 | SB590885 |
| MVP_top60_DNA_seed | BRAF | 0.439929 | SKCM | VMRC-MELG | Dabrafenib |
| MVP_top60_DNA_seed | PIK3CA | 0.441297 | SKCM | SK-MEL-3 | Buparlisib |
| MVP_top60_DNA_seed | PIK3CB | 0.441297 | SKCM | SK-MEL-3 | Buparlisib |
| MVP_top60_DNA_seed | EGFR | 0.442109 | SKCM | A101D | AZD3759 |
| MVP_top60_DNA_seed | BRAF | 0.44273 | SKCM | LB373-MEL-D | SB590885 |
| MVP_top60_DNA_seed | BRAF | 0.444993 | SKCM | CHL-1 | SB590885 |
| MVP_top60_DNA_seed | EGFR | 0.451935 | SKCM | IGR-37 | AZD3759 |
| MVP_top60_DNA_seed | JAK2 | 0.45516 | SKCM | CP66-MEL | JAK_8517 |
| MVP_top60_DNA_seed | JAK1 | 0.45516 | SKCM | CP66-MEL | JAK_8517 |
| MVP_top60_DNA_seed | JAK2 | 0.455984 | SKCM | A101D | Ruxolitinib |
| MVP_top60_DNA_seed | JAK1 | 0.455984 | SKCM | A101D | Ruxolitinib |
| MVP_top60_DNA_seed | GSK3B | 0.456509 | SKCM | SK-MEL-3 | SB216763 |
| MVP_top60_DNA_seed | EGFR | 0.459402 | SKCM | IPC-298 | AZD3759 |
| MVP_top60_DNA_seed | JAK2 | 0.46187 | SKCM | HT-144 | Ruxolitinib |
| MVP_top60_DNA_seed | JAK1 | 0.46187 | SKCM | HT-144 | Ruxolitinib |
| MVP_top60_DNA_seed | PIK3CA | 0.462099 | SKCM | SK-MEL-28 | GNE-317 |
| MVP_top60_DNA_seed | JAK2 | 0.464327 | SKCM | CP50-MEL-B | Ruxolitinib |
| MVP_top60_DNA_seed | JAK1 | 0.464327 | SKCM | CP50-MEL-B | Ruxolitinib |
| MVP_top60_DNA_seed | EGFR | 0.464474 | SKCM | MMAc-SF | Osimertinib |
| MVP_top60_DNA_seed | PIK3CA | 0.465968 | SKCM | SK-MEL-2 | Buparlisib |
| MVP_top60_DNA_seed | PIK3CB | 0.465968 | SKCM | SK-MEL-2 | Buparlisib |
| MVP_top60_DNA_seed | EGFR | 0.466799 | SKCM | A2058 | Afatinib |
| MVP_top60_DNA_seed | ERBB2 | 0.466799 | SKCM | A2058 | Afatinib |
| MVP_top60_DNA_seed | TP53 | 0.467687 | SKCM | 451Lu | MIRA-1 |
| MVP_top60_DNA_seed | EP300 | 0.467908 | SKCM | SK-MEL-24 | SGC-CBP30 |
| MVP_top60_DNA_seed | CREBBP | 0.467908 | SKCM | SK-MEL-24 | SGC-CBP30 |
| MVP_top60_DNA_seed | EGFR | 0.468246 | SKCM | 451Lu | Afatinib |
| MVP_top60_DNA_seed | ERBB2 | 0.468246 | SKCM | 451Lu | Afatinib |
| MVP_top60_DNA_seed | MAP2K1 | 0.474571 | SKCM | CHL-1 | PD0325901 |
| MVP_top60_DNA_seed | EP300 | 0.474735 | SKCM | COLO-679 | SGC-CBP30 |
| MVP_top60_DNA_seed | CREBBP | 0.474735 | SKCM | COLO-679 | SGC-CBP30 |
| MVP_top60_DNA_seed | TP53 | 0.474866 | SKCM | SK-MEL-24 | MIRA-1 |
| MVP_top60_DNA_seed | MAPK3 | 0.478294 | SKCM | SK-MEL-24 | Ulixertinib |
| MVP_top60_DNA_seed | MAPK1 | 0.478294 | SKCM | SK-MEL-24 | Ulixertinib |
| MVP_top60_DNA_seed | GSK3B | 0.479737 | SKCM | CP50-MEL-B | CHIR-99021 |

| | | | | | |
|---|---|---|---|---|---|
| MVP_top60_DNA_seed | EGFR | 0.481266 | SKCM | IGR-1 | Gefitinib |
| MVP_top60_DNA_seed | JAK1 | 0.481821 | SKCM | CHL-1 | JAK1_8709 |
| MVP_top60_DNA_seed | RAC1 | 0.482432 | SKCM | Hs-939-T | EHT-1864 |
| MVP_top60_DNA_seed | BRAF | 0.482456 | SKCM | LB2518-MEL | Dabrafenib |
| MVP_top60_DNA_seed | EGFR | 0.485239 | SKCM | MZ7-mel | Gefitinib |
| MVP_top60_DNA_seed | EGFR | 0.48606 | SKCM | Hs-940-T | Afatinib |
| MVP_top60_DNA_seed | ERBB2 | 0.48606 | SKCM | Hs-940-T | Afatinib |
| MVP_top60_DNA_seed | RAC1 | 0.488109 | SKCM | MEL-JUSO | EHT-1864 |
| MVP_top60_DNA_seed | EGFR | 0.488422 | SKCM | SK-MEL-3 | Gefitinib |
| MVP_top60_DNA_seed | TP53 | 0.489754 | SKCM | WM1552C | MIRA-1 |
| MVP_top60_DNA_seed | PIK3CA | 0.489892 | SKCM | A375 | AZD8186 |
| MVP_top60_DNA_seed | PIK3CB | 0.489892 | SKCM | A375 | AZD8186 |
| MVP_top60_DNA_seed | GSK3B | 0.498177 | SKCM | CHL-1 | SB216763 |
| MVP_top60_DNA_seed | PIK3CA | 0.498458 | SKCM | IGR-1 | AZD8186 |
| MVP_top60_DNA_seed | PIK3CB | 0.498458 | SKCM | IGR-1 | AZD8186 |
| MVP_top60_DNA_seed | KRAS | 0.499277 | SKCM | WM1552C | KRAS (G12C) Inhibitor-12 |
| MVP_top60_DNA_seed | EGFR | 0.50202 | SKCM | G-361 | AZD3759 |
| MVP_top60_DNA_seed | GSK3B | 0.503989 | SKCM | SK-MEL-30 | SB216763 |
| MVP_top60_DNA_seed | KRAS | 0.507161 | SKCM | UACC-257 | KRAS (G12C) Inhibitor-12 |
| MVP_top60_DNA_seed | GSK3B | 0.507924 | SKCM | 451Lu | SB216763 |
| MVP_top60_DNA_seed | EGFR | 0.507952 | SKCM | RPMI-7951 | Osimertinib |
| MVP_top60_DNA_seed | PIK3CA | 0.509077 | SKCM | COLO-800 | GNE-317 |
| MVP_top60_DNA_seed | JAK2 | 0.509772 | SKCM | HMV-II | AZ960 |
| MVP_top60_DNA_seed | JAK3 | 0.509772 | SKCM | HMV-II | AZ960 |
| MVP_top60_DNA_seed | PIK3CA | 0.511301 | SKCM | WM-115 | Buparlisib |
| MVP_top60_DNA_seed | PIK3CB | 0.511301 | SKCM | WM-115 | Buparlisib |
| MVP_top60_DNA_seed | EGFR | 0.512705 | SKCM | SK-MEL-31 | Osimertinib |
| MVP_top60_DNA_seed | PIK3CA | 0.514761 | SKCM | MMAc-SF | GNE-317 |
| MVP_top60_DNA_seed | TP53 | 0.5162 | SKCM | RVH-421 | MIRA-1 |
| MVP_top60_DNA_seed | EGFR | 0.516755 | SKCM | G-361 | Osimertinib |
| MVP_top60_DNA_seed | JAK2 | 0.516939 | SKCM | G-361 | AZ960 |
| MVP_top60_DNA_seed | JAK3 | 0.516939 | SKCM | G-361 | AZ960 |
| MVP_top60_DNA_seed | EGFR | 0.517757 | SKCM | WM278 | Erlotinib |
| MVP_top60_DNA_seed | JAK2 | 0.518407 | SKCM | COLO-792 | AZ960 |
| MVP_top60_DNA_seed | JAK3 | 0.518407 | SKCM | COLO-792 | AZ960 |
| MVP_top60_DNA_seed | EGFR | 0.518583 | SKCM | IST-MEL1 | Osimertinib |
| MVP_top60_DNA_seed | EP300 | 0.520379 | SKCM | C32 | CPI-637 |
| MVP_top60_DNA_seed | PIK3CA | 0.520866 | SKCM | LB2518-MEL | Buparlisib |
| MVP_top60_DNA_seed | PIK3CB | 0.520866 | SKCM | LB2518-MEL | Buparlisib |
| MVP_top60_DNA_seed | EGFR | 0.521671 | SKCM | LOXIMVI | Sapitinib |
| MVP_top60_DNA_seed | ERBB2 | 0.521671 | SKCM | LOXIMVI | Sapitinib |

| | | | | | |
|---|---|---|---|---|---|
| MVP_top60_DNA_seed | EGFR | 0.523511 | SKCM | SH-4 | Gefitinib |
| MVP_top60_DNA_seed | TP53 | 0.524747 | SKCM | SK-MEL-30 | MIRA-1 |
| MVP_top60_DNA_seed | EGFR | 0.525339 | SKCM | SK-MEL-30 | Osimertinib |
| MVP_top60_DNA_seed | PIK3CA | 0.5262 | SKCM | HT-144 | GNE-317 |
| MVP_top60_DNA_seed | EGFR | 0.527778 | SKCM | COLO-792 | Erlotinib |
| MVP_top60_DNA_seed | EGFR | 0.529028 | SKCM | A101D | Gefitinib |
| MVP_top60_DNA_seed | JAK2 | 0.529891 | SKCM | MEL-JUSO | GSK2276186C |
| MVP_top60_DNA_seed | JAK1 | 0.529891 | SKCM | MEL-JUSO | GSK2276186C |
| MVP_top60_DNA_seed | JAK3 | 0.529891 | SKCM | MEL-JUSO | GSK2276186C |
| MVP_top60_DNA_seed | EGFR | 0.531213 | SKCM | VMRC-MELG | Erlotinib |
| MVP_top60_DNA_seed | EGFR | 0.536154 | SKCM | LOXIMVI | Gefitinib |
| MVP_top60_DNA_seed | KRAS | 0.536214 | SKCM | UACC-62 | KRAS (G12C) Inhibitor-12 |
| MVP_top60_DNA_seed | EGFR | 0.538793 | SKCM | COLO-679 | Gefitinib |
| MVP_top60_DNA_seed | TP53 | 0.538996 | SKCM | IST-MEL1 | MIRA-1 |
| MVP_top60_DNA_seed | EGFR | 0.539527 | SKCM | GAK | Gefitinib |
| MVP_top60_DNA_seed | PIK3CB | 0.539832 | SKCM | MZ7-mel | AZD6482 |
| MVP_top60_DNA_seed | TP53 | 0.540632 | SKCM | SK-MEL-28 | MIRA-1 |
| MVP_top60_DNA_seed | PIK3CA | 0.541673 | SKCM | A375 | Alpelisib |
| MVP_top60_DNA_seed | KRAS | 0.542594 | SKCM | IGR-1 | KRAS (G12C) Inhibitor-12 |
| MVP_top60_DNA_seed | EGFR | 0.543049 | SKCM | WM1552C | Lapatinib |
| MVP_top60_DNA_seed | ERBB2 | 0.543049 | SKCM | WM1552C | Lapatinib |
| MVP_top60_DNA_seed | TP53 | 0.54756 | SKCM | WM35 | MIRA-1 |
| MVP_top60_DNA_seed | PIK3CA | 0.548437 | SKCM | IST-MEL1 | Buparlisib |
| MVP_top60_DNA_seed | PIK3CB | 0.548437 | SKCM | IST-MEL1 | Buparlisib |
| MVP_top60_DNA_seed | PIK3CA | 0.549245 | SKCM | MEL-HO | Buparlisib |
| MVP_top60_DNA_seed | PIK3CB | 0.549245 | SKCM | MEL-HO | Buparlisib |
| MVP_top60_DNA_seed | EGFR | 0.549555 | SKCM | SK-MEL-28 | Gefitinib |
| MVP_top60_DNA_seed | PIK3CA | 0.551292 | SKCM | A101D | Alpelisib |
| MVP_top60_DNA_seed | JAK2 | 0.551417 | SKCM | G-361 | GSK2276186C |
| MVP_top60_DNA_seed | JAK1 | 0.551417 | SKCM | G-361 | GSK2276186C |
| MVP_top60_DNA_seed | JAK3 | 0.551417 | SKCM | G-361 | GSK2276186C |
| MVP_top60_DNA_seed | GSK3B | 0.552143 | SKCM | VMRC-MELG | SB216763 |
| MVP_top60_DNA_seed | EGFR | 0.553275 | SKCM | MEL-JUSO | Lapatinib |
| MVP_top60_DNA_seed | ERBB2 | 0.553275 | SKCM | MEL-JUSO | Lapatinib |
| MVP_top60_DNA_seed | KRAS | 0.553725 | SKCM | SH-4 | KRAS (G12C) Inhibitor-12 |
| MVP_top60_DNA_seed | EGFR | 0.553763 | SKCM | IST-MEL1 | Afatinib |
| MVP_top60_DNA_seed | ERBB2 | 0.553763 | SKCM | IST-MEL1 | Afatinib |
| MVP_top60_DNA_seed | JAK2 | 0.557694 | SKCM | C32 | JAK_8517 |
| MVP_top60_DNA_seed | JAK1 | 0.557694 | SKCM | C32 | JAK_8517 |
| MVP_top60_DNA_seed | BRAF | 0.559152 | SKCM | MEL-JUSO | SB590885 |
| MVP_top60_DNA_seed | BRAF | 0.560395 | SKCM | SK-MEL-31 | SB590885 |

| | | | | | |
|---|---|---|---|---|---|
| MVP_top60_DNA_seed | PIK3CB | 0.562014 | SKCM | SK-MEL-31 | AZD6482 |
| MVP_top60_DNA_seed | JAK2 | 0.562235 | SKCM | COLO-783 | Ruxolitinib |
| MVP_top60_DNA_seed | JAK1 | 0.562235 | SKCM | COLO-783 | Ruxolitinib |
| MVP_top60_DNA_seed | JAK2 | 0.563558 | SKCM | SK-MEL-5 | AZ960 |
| MVP_top60_DNA_seed | JAK3 | 0.563558 | SKCM | SK-MEL-5 | AZ960 |
| MVP_top60_DNA_seed | TP53 | 0.564183 | SKCM | Hs-939-T | MIRA-1 |
| MVP_top60_DNA_seed | EGFR | 0.565187 | SKCM | LOXIMVI | Erlotinib |
| MVP_top60_DNA_seed | GSK3B | 0.565746 | SKCM | SH-4 | SB216763 |
| MVP_top60_DNA_seed | JAK2 | 0.567543 | SKCM | HMV-II | Ruxolitinib |
| MVP_top60_DNA_seed | JAK1 | 0.567543 | SKCM | HMV-II | Ruxolitinib |
| MVP_top60_DNA_seed | JAK2 | 0.568033 | SKCM | GAK | AZ960 |
| MVP_top60_DNA_seed | JAK3 | 0.568033 | SKCM | GAK | AZ960 |
| MVP_top60_DNA_seed | GSK3B | 0.568881 | SKCM | Hs-939-T | SB216763 |
| MVP_top60_DNA_seed | JAK2 | 0.569101 | SKCM | HT-144 | JAK_8517 |
| MVP_top60_DNA_seed | JAK1 | 0.569101 | SKCM | HT-144 | JAK_8517 |
| MVP_top60_DNA_seed | PIK3CA | 0.56948 | SKCM | CHL-1 | GNE-317 |
| MVP_top60_DNA_seed | PIK3CA | 0.570138 | SKCM | CP66-MEL | AZD8186 |
| MVP_top60_DNA_seed | PIK3CB | 0.570138 | SKCM | CP66-MEL | AZD8186 |
| MVP_top60_DNA_seed | MAPK3 | 0.570653 | SKCM | IGR-1 | Ulixertinib |
| MVP_top60_DNA_seed | MAPK1 | 0.570653 | SKCM | IGR-1 | Ulixertinib |
| MVP_top60_DNA_seed | JAK2 | 0.571364 | SKCM | SH-4 | GSK2276186C |
| MVP_top60_DNA_seed | JAK1 | 0.571364 | SKCM | SH-4 | GSK2276186C |
| MVP_top60_DNA_seed | JAK3 | 0.571364 | SKCM | SH-4 | GSK2276186C |
| MVP_top60_DNA_seed | PIK3CA | 0.572854 | SKCM | COLO-829 | GNE-317 |
| MVP_top60_DNA_seed | EGFR | 0.57286 | SKCM | LB2518-MEL | Sapitinib |
| MVP_top60_DNA_seed | ERBB2 | 0.57286 | SKCM | LB2518-MEL | Sapitinib |
| MVP_top60_DNA_seed | EGFR | 0.573034 | SKCM | Hs-939-T | Sapitinib |
| MVP_top60_DNA_seed | ERBB2 | 0.573034 | SKCM | Hs-939-T | Sapitinib |
| MVP_top60_DNA_seed | GSK3B | 0.57439 | SKCM | WM1552C | SB216763 |
| MVP_top60_DNA_seed | GSK3B | 0.574964 | SKCM | MZ7-mel | CHIR-99021 |
| MVP_top60_DNA_seed | BRAF | 0.575202 | SKCM | LB2518-MEL | PLX-4720 |
| MVP_top60_DNA_seed | EGFR | 0.577258 | SKCM | IGR-1 | Sapitinib |
| MVP_top60_DNA_seed | ERBB2 | 0.577258 | SKCM | IGR-1 | Sapitinib |
| MVP_top60_DNA_seed | PIK3CA | 0.578106 | SKCM | UACC-62 | Alpelisib |
| MVP_top60_DNA_seed | PIK3CA | 0.578894 | SKCM | COLO-783 | GNE-317 |
| MVP_top60_DNA_seed | EGFR | 0.580685 | SKCM | UACC-62 | Osimertinib |
| MVP_top60_DNA_seed | EGFR | 0.581337 | SKCM | HMV-II | Gefitinib |
| MVP_top60_DNA_seed | EGFR | 0.584795 | SKCM | WM1552C | AZD3759 |
| MVP_top60_DNA_seed | EGFR | 0.58527 | SKCM | COLO-792 | Gefitinib |
| MVP_top60_DNA_seed | PIK3CA | 0.588974 | SKCM | HT-144 | AZD8186 |
| MVP_top60_DNA_seed | PIK3CB | 0.588974 | SKCM | HT-144 | AZD8186 |

| | | | | | |
|---|---|---|---|---|---|
| MVP_top60_DNA_seed | EGFR | 0.589614 | SKCM | LB373-MEL-D | Sapitinib |
| MVP_top60_DNA_seed | ERBB2 | 0.589614 | SKCM | LB373-MEL-D | Sapitinib |
| MVP_top60_DNA_seed | EGFR | 0.593757 | SKCM | IGR-37 | Afatinib |
| MVP_top60_DNA_seed | ERBB2 | 0.593757 | SKCM | IGR-37 | Afatinib |
| MVP_top60_DNA_seed | JAK2 | 0.594036 | SKCM | MZ2-MEL | Ruxolitinib |
| MVP_top60_DNA_seed | JAK1 | 0.594036 | SKCM | MZ2-MEL | Ruxolitinib |
| MVP_top60_DNA_seed | PIK3CA | 0.59447 | SKCM | C32 | AZD8186 |
| MVP_top60_DNA_seed | PIK3CB | 0.59447 | SKCM | C32 | AZD8186 |
| MVP_top60_DNA_seed | EGFR | 0.595228 | SKCM | C32 | Osimertinib |
| MVP_top60_DNA_seed | MAP2K1 | 0.595664 | SKCM | VMRC-MELG | Trametinib |
| MVP_top60_DNA_seed | PIK3CB | 0.598265 | SKCM | CP66-MEL | AZD6482 |
| MVP_top60_DNA_seed | GSK3B | 0.603489 | SKCM | RVH-421 | SB216763 |
| MVP_top60_DNA_seed | JAK2 | 0.605874 | SKCM | WM793B | GSK2276186C |
| MVP_top60_DNA_seed | JAK1 | 0.605874 | SKCM | WM793B | GSK2276186C |
| MVP_top60_DNA_seed | JAK3 | 0.605874 | SKCM | WM793B | GSK2276186C |
| MVP_top60_DNA_seed | EP300 | 0.606174 | SKCM | C32 | SGC-CBP30 |
| MVP_top60_DNA_seed | CREBBP | 0.606174 | SKCM | C32 | SGC-CBP30 |
| MVP_top60_DNA_seed | JAK2 | 0.607446 | SKCM | IPC-298 | Ruxolitinib |
| MVP_top60_DNA_seed | JAK1 | 0.607446 | SKCM | IPC-298 | Ruxolitinib |
| MVP_top60_DNA_seed | RAC1 | 0.610505 | SKCM | SK-MEL-24 | EHT-1864 |
| MVP_top60_DNA_seed | KRAS | 0.610514 | SKCM | LB373-MEL-D | KRAS (G12C) Inhibitor-12 |
| MVP_top60_DNA_seed | EGFR | 0.611829 | SKCM | GAK | AZD3759 |
| MVP_top60_DNA_seed | JAK2 | 0.613504 | SKCM | WM1552C | Ruxolitinib |
| MVP_top60_DNA_seed | JAK1 | 0.613504 | SKCM | WM1552C | Ruxolitinib |
| MVP_top60_DNA_seed | JAK2 | 0.613546 | SKCM | WM793B | Ruxolitinib |
| MVP_top60_DNA_seed | JAK1 | 0.613546 | SKCM | WM793B | Ruxolitinib |
| MVP_top60_DNA_seed | EGFR | 0.615822 | SKCM | Mewo | Afatinib |
| MVP_top60_DNA_seed | ERBB2 | 0.615822 | SKCM | Mewo | Afatinib |
| MVP_top60_DNA_seed | JAK2 | 0.618755 | SKCM | COLO-800 | AZ960 |
| MVP_top60_DNA_seed | JAK3 | 0.618755 | SKCM | COLO-800 | AZ960 |
| MVP_top60_DNA_seed | PIK3CA | 0.619007 | SKCM | COLO-829 | Buparlisib |
| MVP_top60_DNA_seed | PIK3CB | 0.619007 | SKCM | COLO-829 | Buparlisib |
| MVP_top60_DNA_seed | RAC1 | 0.619903 | SKCM | C32 | EHT-1864 |
| MVP_top60_DNA_seed | EGFR | 0.620045 | SKCM | HMV-II | Erlotinib |
| MVP_top60_DNA_seed | EGFR | 0.620279 | SKCM | UACC-62 | Lapatinib |
| MVP_top60_DNA_seed | ERBB2 | 0.620279 | SKCM | UACC-62 | Lapatinib |
| MVP_top60_DNA_seed | PIK3CB | 0.62532 | SKCM | Hs-940-T | AZD6482 |
| MVP_top60_DNA_seed | EGFR | 0.625928 | SKCM | LOXIMVI | AZD3759 |
| MVP_top60_DNA_seed | JAK1 | 0.627207 | SKCM | RPMI-7951 | JAK1_8709 |
| MVP_top60_DNA_seed | GSK3B | 0.62951 | SKCM | SK-MEL-31 | SB216763 |
| MVP_top60_DNA_seed | PIK3CA | 0.633578 | SKCM | WM278 | AZD8186 |

| | | | | | |
|---|---|---|---|---|---|
| MVP_top60_DNA_seed | PIK3CB | 0.633578 | SKCM | WM278 | AZD8186 |
| MVP_top60_DNA_seed | EGFR | 0.637513 | SKCM | LB373-MEL-D | Gefitinib |
| MVP_top60_DNA_seed | EGFR | 0.64397 | SKCM | MEL-JUSO | Osimertinib |
| MVP_top60_DNA_seed | GSK3B | 0.644912 | SKCM | MEL-HO | CHIR-99021 |
| MVP_top60_DNA_seed | JAK2 | 0.645442 | SKCM | SK-MEL-28 | Ruxolitinib |
| MVP_top60_DNA_seed | JAK1 | 0.645442 | SKCM | SK-MEL-28 | Ruxolitinib |
| MVP_top60_DNA_seed | EGFR | 0.645789 | SKCM | K2 | Afatinib |
| MVP_top60_DNA_seed | ERBB2 | 0.645789 | SKCM | K2 | Afatinib |
| MVP_top60_DNA_seed | GSK3B | 0.645962 | SKCM | K2 | SB216763 |
| MVP_top60_DNA_seed | JAK2 | 0.649376 | SKCM | UACC-62 | GSK2276186C |
| MVP_top60_DNA_seed | JAK1 | 0.649376 | SKCM | UACC-62 | GSK2276186C |
| MVP_top60_DNA_seed | JAK3 | 0.649376 | SKCM | UACC-62 | GSK2276186C |
| MVP_top60_DNA_seed | EGFR | 0.651209 | SKCM | WM793B | Osimertinib |
| MVP_top60_DNA_seed | PIK3CA | 0.653437 | SKCM | Hs-940-T | Alpelisib |
| MVP_top60_DNA_seed | JAK1 | 0.654334 | SKCM | HT-144 | JAK1_8709 |
| MVP_top60_DNA_seed | EGFR | 0.654901 | SKCM | C32 | AZD3759 |
| MVP_top60_DNA_seed | EGFR | 0.655421 | SKCM | Hs-940-T | Lapatinib |
| MVP_top60_DNA_seed | ERBB2 | 0.655421 | SKCM | Hs-940-T | Lapatinib |
| MVP_top60_DNA_seed | BRAF | 0.655727 | SKCM | SK-MEL-2 | Dabrafenib |
| MVP_top60_DNA_seed | PIK3CA | 0.656623 | SKCM | MEL-JUSO | Buparlisib |
| MVP_top60_DNA_seed | PIK3CB | 0.656623 | SKCM | MEL-JUSO | Buparlisib |
| MVP_top60_DNA_seed | KRAS | 0.656681 | SKCM | WM-115 | KRAS (G12C) Inhibitor-12 |
| MVP_top60_DNA_seed | GSK3B | 0.657264 | SKCM | WM-115 | SB216763 |
| MVP_top60_DNA_seed | PIK3CB | 0.660065 | SKCM | A375 | AZD6482 |
| MVP_top60_DNA_seed | PIK3CA | 0.660509 | SKCM | Mewo | Buparlisib |
| MVP_top60_DNA_seed | PIK3CB | 0.660509 | SKCM | Mewo | Buparlisib |
| MVP_top60_DNA_seed | EGFR | 0.661518 | SKCM | SH-4 | Afatinib |
| MVP_top60_DNA_seed | ERBB2 | 0.661518 | SKCM | SH-4 | Afatinib |
| MVP_top60_DNA_seed | RAC1 | 0.662249 | SKCM | Hs-940-T | EHT-1864 |
| MVP_top60_DNA_seed | JAK1 | 0.662934 | SKCM | MEL-JUSO | JAK1_8709 |
| MVP_top60_DNA_seed | EGFR | 0.666682 | SKCM | WM793B | Sapitinib |
| MVP_top60_DNA_seed | ERBB2 | 0.666682 | SKCM | WM793B | Sapitinib |
| MVP_top60_DNA_seed | EGFR | 0.66869 | SKCM | SK-MEL-3 | Sapitinib |
| MVP_top60_DNA_seed | ERBB2 | 0.66869 | SKCM | SK-MEL-3 | Sapitinib |
| MVP_top60_DNA_seed | JAK2 | 0.669572 | SKCM | COLO-829 | JAK_8517 |
| MVP_top60_DNA_seed | JAK1 | 0.669572 | SKCM | COLO-829 | JAK_8517 |
| MVP_top60_DNA_seed | RAC1 | 0.675353 | SKCM | A101D | EHT-1864 |
| MVP_top60_DNA_seed | PIK3CA | 0.67602 | SKCM | LB2518-MEL | AZD8186 |
| MVP_top60_DNA_seed | PIK3CB | 0.67602 | SKCM | LB2518-MEL | AZD8186 |
| MVP_top60_DNA_seed | PIK3CA | 0.676899 | SKCM | HT-144 | Alpelisib |
| MVP_top60_DNA_seed | JAK2 | 0.677824 | SKCM | CHL-1 | Ruxolitinib |

| | | | | | |
|---|---|---|---|---|---|
| MVP_top60_DNA_seed | JAK1 | 0.677824 | SKCM | CHL-1 | Ruxolitinib |
| MVP_top60_DNA_seed | EGFR | 0.67819 | SKCM | C32 | Gefitinib |
| MVP_top60_DNA_seed | EGFR | 0.682922 | SKCM | SK-MEL-5 | Osimertinib |
| MVP_top60_DNA_seed | EGFR | 0.682936 | SKCM | COLO-792 | AZD3759 |
| MVP_top60_DNA_seed | EP300 | 0.684497 | SKCM | WM793B | SGC-CBP30 |
| MVP_top60_DNA_seed | CREBBP | 0.684497 | SKCM | WM793B | SGC-CBP30 |
| MVP_top60_DNA_seed | EGFR | 0.685112 | SKCM | WM1552C | Sapitinib |
| MVP_top60_DNA_seed | ERBB2 | 0.685112 | SKCM | WM1552C | Sapitinib |
| MVP_top60_DNA_seed | EP300 | 0.686904 | SKCM | IGR-37 | CPI-637 |
| MVP_top60_DNA_seed | GSK3B | 0.687446 | SKCM | SK-MEL-28 | SB216763 |
| MVP_top60_DNA_seed | JAK1 | 0.696613 | SKCM | SK-MEL-24 | JAK1_8709 |
| MVP_top60_DNA_seed | EGFR | 0.697721 | SKCM | MMAc-SF | Afatinib |
| MVP_top60_DNA_seed | ERBB2 | 0.697721 | SKCM | MMAc-SF | Afatinib |
| MVP_top60_DNA_seed | JAK2 | 0.697956 | SKCM | WM-115 | GSK2276186C |
| MVP_top60_DNA_seed | JAK1 | 0.697956 | SKCM | WM-115 | GSK2276186C |
| MVP_top60_DNA_seed | JAK3 | 0.697956 | SKCM | WM-115 | GSK2276186C |
| MVP_top60_DNA_seed | PIK3CA | 0.700021 | SKCM | CP66-MEL | GNE-317 |
| MVP_top60_DNA_seed | EGFR | 0.700241 | SKCM | IPC-298 | Osimertinib |
| MVP_top60_DNA_seed | EGFR | 0.700608 | SKCM | WM35 | AZD3759 |
| MVP_top60_DNA_seed | TP53 | 0.709167 | SKCM | SK-MEL-31 | MIRA-1 |
| MVP_top60_DNA_seed | PIK3CA | 0.710813 | SKCM | IPC-298 | Buparlisib |
| MVP_top60_DNA_seed | PIK3CB | 0.710813 | SKCM | IPC-298 | Buparlisib |
| MVP_top60_DNA_seed | EGFR | 0.712463 | SKCM | SK-MEL-30 | AZD3759 |
| MVP_top60_DNA_seed | TP53 | 0.712729 | SKCM | COLO-679 | MIRA-1 |
| MVP_top60_DNA_seed | EGFR | 0.714786 | SKCM | Mewo | Osimertinib |
| MVP_top60_DNA_seed | EGFR | 0.715953 | SKCM | SK-MEL-28 | Osimertinib |
| MVP_top60_DNA_seed | JAK2 | 0.719888 | SKCM | COLO-800 | Ruxolitinib |
| MVP_top60_DNA_seed | JAK1 | 0.719888 | SKCM | COLO-800 | Ruxolitinib |
| MVP_top60_DNA_seed | BRAF | 0.722336 | SKCM | Hs-940-T | SB590885 |
| MVP_top60_DNA_seed | BRAF | 0.723359 | SKCM | LB373-MEL-D | Dabrafenib |
| MVP_top60_DNA_seed | EGFR | 0.726966 | SKCM | IST-MEL1 | Lapatinib |
| MVP_top60_DNA_seed | ERBB2 | 0.726966 | SKCM | IST-MEL1 | Lapatinib |
| MVP_top60_DNA_seed | EGFR | 0.72829 | SKCM | GAK | Sapitinib |
| MVP_top60_DNA_seed | ERBB2 | 0.72829 | SKCM | GAK | Sapitinib |
| MVP_top60_DNA_seed | EGFR | 0.729397 | SKCM | CP50-MEL-B | Erlotinib |
| MVP_top60_DNA_seed | EGFR | 0.732229 | SKCM | SK-MEL-31 | Sapitinib |
| MVP_top60_DNA_seed | ERBB2 | 0.732229 | SKCM | SK-MEL-31 | Sapitinib |
| MVP_top60_DNA_seed | PIK3CB | 0.733432 | SKCM | K2 | AZD6482 |
| MVP_top60_DNA_seed | EGFR | 0.736973 | SKCM | UACC-62 | Gefitinib |
| MVP_top60_DNA_seed | EGFR | 0.737205 | SKCM | COLO-829 | Erlotinib |
| MVP_top60_DNA_seed | BRAF | 0.739716 | SKCM | COLO-792 | Dabrafenib |

| | | | | | |
|---|---|---|---|---|---|
| MVP_top60_DNA_seed | EGFR | 0.740084 | SKCM | COLO-783 | Gefitinib |
| MVP_top60_DNA_seed | PIK3CB | 0.746147 | SKCM | LB2518-MEL | AZD6482 |
| MVP_top60_DNA_seed | JAK2 | 0.748003 | SKCM | SK-MEL-30 | Ruxolitinib |
| MVP_top60_DNA_seed | JAK1 | 0.748003 | SKCM | SK-MEL-30 | Ruxolitinib |
| MVP_top60_DNA_seed | EGFR | 0.750619 | SKCM | WM35 | Lapatinib |
| MVP_top60_DNA_seed | ERBB2 | 0.750619 | SKCM | WM35 | Lapatinib |
| MVP_top60_DNA_seed | PIK3CA | 0.750704 | SKCM | RVH-421 | GNE-317 |
| MVP_top60_DNA_seed | EGFR | 0.754043 | SKCM | RVH-421 | Erlotinib |
| MVP_top60_DNA_seed | GSK3B | 0.755558 | SKCM | LB2518-MEL | SB216763 |
| MVP_top60_DNA_seed | EGFR | 0.757018 | SKCM | SK-MEL-28 | Sapitinib |
| MVP_top60_DNA_seed | ERBB2 | 0.757018 | SKCM | SK-MEL-28 | Sapitinib |
| MVP_top60_DNA_seed | PIK3CA | 0.75716 | SKCM | MZ7-mel | Alpelisib |
| MVP_top60_DNA_seed | EGFR | 0.757715 | SKCM | COLO-679 | Afatinib |
| MVP_top60_DNA_seed | ERBB2 | 0.757715 | SKCM | COLO-679 | Afatinib |
| MVP_top60_DNA_seed | EGFR | 0.75801 | SKCM | SK-MEL-2 | Erlotinib |
| MVP_top60_DNA_seed | BRAF | 0.761686 | SKCM | COLO-792 | PLX-4720 |
| MVP_top60_DNA_seed | BRAF | 0.763247 | SKCM | LB2518-MEL | SB590885 |
| MVP_top60_DNA_seed | EGFR | 0.764134 | SKCM | WM278 | Sapitinib |
| MVP_top60_DNA_seed | ERBB2 | 0.764134 | SKCM | WM278 | Sapitinib |
| MVP_top60_DNA_seed | EGFR | 0.765275 | SKCM | LB2518-MEL | Afatinib |
| MVP_top60_DNA_seed | ERBB2 | 0.765275 | SKCM | LB2518-MEL | Afatinib |
| MVP_top60_DNA_seed | EGFR | 0.765961 | SKCM | A375 | Sapitinib |
| MVP_top60_DNA_seed | ERBB2 | 0.765961 | SKCM | A375 | Sapitinib |
| MVP_top60_DNA_seed | EGFR | 0.767246 | SKCM | Hs-939-T | Lapatinib |
| MVP_top60_DNA_seed | ERBB2 | 0.767246 | SKCM | Hs-939-T | Lapatinib |
| MVP_top60_DNA_seed | EGFR | 0.767687 | SKCM | Hs-939-T | Afatinib |
| MVP_top60_DNA_seed | ERBB2 | 0.767687 | SKCM | Hs-939-T | Afatinib |
| MVP_top60_DNA_seed | PIK3CA | 0.770767 | SKCM | G-MEL | AZD8186 |
| MVP_top60_DNA_seed | PIK3CB | 0.770767 | SKCM | G-MEL | AZD8186 |
| MVP_top60_DNA_seed | RAC1 | 0.771116 | SKCM | COLO-800 | EHT-1864 |
| MVP_top60_DNA_seed | EGFR | 0.772437 | SKCM | WM35 | Osimertinib |
| MVP_top60_DNA_seed | RAC1 | 0.776763 | SKCM | K2 | EHT-1864 |
| MVP_top60_DNA_seed | GSK3B | 0.780825 | SKCM | HMV-II | SB216763 |
| MVP_top60_DNA_seed | EGFR | 0.782648 | SKCM | Mewo | Sapitinib |
| MVP_top60_DNA_seed | ERBB2 | 0.782648 | SKCM | Mewo | Sapitinib |
| MVP_top60_DNA_seed | JAK2 | 0.783651 | SKCM | 451Lu | Ruxolitinib |
| MVP_top60_DNA_seed | JAK1 | 0.783651 | SKCM | 451Lu | Ruxolitinib |
| MVP_top60_DNA_seed | EGFR | 0.784935 | SKCM | SK-MEL-30 | Sapitinib |
| MVP_top60_DNA_seed | ERBB2 | 0.784935 | SKCM | SK-MEL-30 | Sapitinib |
| MVP_top60_DNA_seed | PIK3CA | 0.786839 | SKCM | COLO-679 | AZD8186 |
| MVP_top60_DNA_seed | PIK3CB | 0.786839 | SKCM | COLO-679 | AZD8186 |

| | | | | | |
|---|---|---|---|---|---|
| MVP_top60_DNA_seed | EP300 | 0.791185 | SKCM | WM-115 | CPI-637 |
| MVP_top60_DNA_seed | BRAF | 0.795289 | SKCM | Mewo | SB590885 |
| MVP_top60_DNA_seed | EGFR | 0.798261 | SKCM | MEL-HO | Erlotinib |
| MVP_top60_DNA_seed | EGFR | 0.800242 | SKCM | COLO-792 | Osimertinib |
| MVP_top60_DNA_seed | PIK3CA | 0.804235 | SKCM | SK-MEL-24 | AZD8186 |
| MVP_top60_DNA_seed | PIK3CB | 0.804235 | SKCM | SK-MEL-24 | AZD8186 |
| MVP_top60_DNA_seed | MAP2K1 | 0.804564 | SKCM | VMRC-MELG | Refametinib |
| MVP_top60_DNA_seed | GSK3B | 0.807504 | SKCM | MMAc-SF | SB216763 |
| MVP_top60_DNA_seed | RAC1 | 0.810253 | SKCM | IPC-298 | EHT-1864 |
| MVP_top60_DNA_seed | EGFR | 0.81364 | SKCM | HMV-II | Sapitinib |
| MVP_top60_DNA_seed | ERBB2 | 0.81364 | SKCM | HMV-II | Sapitinib |
| MVP_top60_DNA_seed | EGFR | 0.817569 | SKCM | MEL-JUSO | Gefitinib |
| MVP_top60_DNA_seed | RAC1 | 0.818102 | SKCM | UACC-62 | EHT-1864 |
| MVP_top60_DNA_seed | JAK1 | 0.818932 | SKCM | UACC-257 | JAK1_8709 |
| MVP_top60_DNA_seed | PIK3CA | 0.819066 | SKCM | C32 | Buparlisib |
| MVP_top60_DNA_seed | PIK3CB | 0.819066 | SKCM | C32 | Buparlisib |
| MVP_top60_DNA_seed | RAC1 | 0.820885 | SKCM | WM278 | EHT-1864 |
| MVP_top60_DNA_seed | EGFR | 0.821258 | SKCM | SK-MEL-24 | Gefitinib |
| MVP_top60_DNA_seed | JAK2 | 0.82342 | SKCM | SK-MEL-30 | AZ960 |
| MVP_top60_DNA_seed | JAK3 | 0.82342 | SKCM | SK-MEL-30 | AZ960 |
| MVP_top60_DNA_seed | EGFR | 0.823614 | SKCM | SH-4 | Lapatinib |
| MVP_top60_DNA_seed | ERBB2 | 0.823614 | SKCM | SH-4 | Lapatinib |
| MVP_top60_DNA_seed | TP53 | 0.825158 | SKCM | Mewo | MIRA-1 |
| MVP_top60_DNA_seed | PIK3CA | 0.825477 | SKCM | 451Lu | Buparlisib |
| MVP_top60_DNA_seed | PIK3CB | 0.825477 | SKCM | 451Lu | Buparlisib |
| MVP_top60_DNA_seed | JAK2 | 0.827487 | SKCM | IGR-1 | AZ960 |
| MVP_top60_DNA_seed | JAK3 | 0.827487 | SKCM | IGR-1 | AZ960 |
| MVP_top60_DNA_seed | JAK2 | 0.828434 | SKCM | COLO-829 | AZ960 |
| MVP_top60_DNA_seed | JAK3 | 0.828434 | SKCM | COLO-829 | AZ960 |
| MVP_top60_DNA_seed | EGFR | 0.83096 | SKCM | 451Lu | Erlotinib |
| MVP_top60_DNA_seed | JAK2 | 0.834962 | SKCM | UACC-62 | Ruxolitinib |
| MVP_top60_DNA_seed | JAK1 | 0.834962 | SKCM | UACC-62 | Ruxolitinib |
| MVP_top60_DNA_seed | MAP2K1 | 0.835508 | SKCM | Hs-940-T | Selumetinib |
| MVP_top60_DNA_seed | GSK3B | 0.838607 | SKCM | RVH-421 | CHIR-99021 |
| MVP_top60_DNA_seed | MAPK3 | 0.838767 | SKCM | WM793B | Ulixertinib |
| MVP_top60_DNA_seed | MAPK1 | 0.838767 | SKCM | WM793B | Ulixertinib |
| MVP_top60_DNA_seed | PIK3CA | 0.839797 | SKCM | MEL-JUSO | GNE-317 |
| MVP_top60_DNA_seed | PIK3CA | 0.842326 | SKCM | Hs-939-T | Buparlisib |
| MVP_top60_DNA_seed | PIK3CB | 0.842326 | SKCM | Hs-939-T | Buparlisib |
| MVP_top60_DNA_seed | KRAS | 0.842525 | SKCM | COLO-783 | KRAS (G12C) Inhibitor-12 |
| MVP_top60_DNA_seed | EGFR | 0.845577 | SKCM | HT-144 | AZD3759 |

| | | | | | |
|---|---|---|---|---|---|
| MVP_top60_DNA_seed | EGFR | 0.847965 | SKCM | GAK | Osimertinib |
| MVP_top60_DNA_seed | JAK2 | 0.849224 | SKCM | SK-MEL-24 | AZ960 |
| MVP_top60_DNA_seed | JAK3 | 0.849224 | SKCM | SK-MEL-24 | AZ960 |
| MVP_top60_DNA_seed | EP300 | 0.850135 | SKCM | UACC-257 | SGC-CBP30 |
| MVP_top60_DNA_seed | CREBBP | 0.850135 | SKCM | UACC-257 | SGC-CBP30 |
| MVP_top60_DNA_seed | GSK3B | 0.858204 | SKCM | SK-MEL-28 | CHIR-99021 |
| MVP_top60_DNA_seed | PIK3CA | 0.860411 | SKCM | UACC-257 | Buparlisib |
| MVP_top60_DNA_seed | PIK3CB | 0.860411 | SKCM | UACC-257 | Buparlisib |
| MVP_top60_DNA_seed | JAK2 | 0.862125 | SKCM | SK-MEL-5 | GSK2276186C |
| MVP_top60_DNA_seed | JAK1 | 0.862125 | SKCM | SK-MEL-5 | GSK2276186C |
| MVP_top60_DNA_seed | JAK3 | 0.862125 | SKCM | SK-MEL-5 | GSK2276186C |
| MVP_top60_DNA_seed | EGFR | 0.862604 | SKCM | WM793B | AZD3759 |
| MVP_top60_DNA_seed | EGFR | 0.864198 | SKCM | WM-115 | Afatinib |
| MVP_top60_DNA_seed | ERBB2 | 0.864198 | SKCM | WM-115 | Afatinib |
| MVP_top60_DNA_seed | JAK2 | 0.866354 | SKCM | SH-4 | Ruxolitinib |
| MVP_top60_DNA_seed | JAK1 | 0.866354 | SKCM | SH-4 | Ruxolitinib |
| MVP_top60_DNA_seed | KRAS | 0.868749 | SKCM | WM278 | (RAS (G12C) Inhibitor-12 |
| MVP_top60_DNA_seed | BRAF | 0.871041 | SKCM | GAK | PLX-4720 |
| MVP_top60_DNA_seed | RAC1 | 0.872865 | SKCM | UACC-257 | EHT-1864 |
| MVP_top60_DNA_seed | EGFR | 0.874341 | SKCM | UACC-257 | Afatinib |
| MVP_top60_DNA_seed | ERBB2 | 0.874341 | SKCM | UACC-257 | Afatinib |
| MVP_top60_DNA_seed | EGFR | 0.875152 | SKCM | MEL-JUSO | Afatinib |
| MVP_top60_DNA_seed | ERBB2 | 0.875152 | SKCM | MEL-JUSO | Afatinib |
| MVP_top60_DNA_seed | GSK3B | 0.882116 | SKCM | SK-MEL-1 | SB216763 |
| MVP_top60_DNA_seed | KRAS | 0.883548 | SKCM | MEL-HO | (RAS (G12C) Inhibitor-12 |
| MVP_top60_DNA_seed | EGFR | 0.884458 | SKCM | COLO-800 | Osimertinib |
| MVP_top60_DNA_seed | JAK2 | 0.8851 | SKCM | WM793B | JAK_8517 |
| MVP_top60_DNA_seed | JAK1 | 0.8851 | SKCM | WM793B | JAK_8517 |
| MVP_top60_DNA_seed | EGFR | 0.886769 | SKCM | SK-MEL-1 | Afatinib |
| MVP_top60_DNA_seed | ERBB2 | 0.886769 | SKCM | SK-MEL-1 | Afatinib |
| MVP_top60_DNA_seed | EGFR | 0.886845 | SKCM | IGR-37 | Osimertinib |
| MVP_top60_DNA_seed | JAK2 | 0.887147 | SKCM | SK-MEL-5 | JAK_8517 |
| MVP_top60_DNA_seed | JAK1 | 0.887147 | SKCM | SK-MEL-5 | JAK_8517 |
| MVP_top60_DNA_seed | PIK3CA | 0.888832 | SKCM | COLO-800 | Buparlisib |
| MVP_top60_DNA_seed | PIK3CB | 0.888832 | SKCM | COLO-800 | Buparlisib |
| MVP_top60_DNA_seed | JAK2 | 0.889279 | SKCM | A101D | GSK2276186C |
| MVP_top60_DNA_seed | JAK1 | 0.889279 | SKCM | A101D | GSK2276186C |
| MVP_top60_DNA_seed | JAK3 | 0.889279 | SKCM | A101D | GSK2276186C |
| MVP_top60_DNA_seed | PIK3CA | 0.891461 | SKCM | HMV-II | GNE-317 |
| MVP_top60_DNA_seed | KRAS | 0.893332 | SKCM | RVH-421 | (RAS (G12C) Inhibitor-12 |
| MVP_top60_DNA_seed | JAK2 | 0.893859 | SKCM | Hs-939-T | Ruxolitinib |

| | | | | | |
|---|---|---|---|---|---|
| MVP_top60_DNA_seed | JAK1 | 0.893859 | SKCM | Hs-939-T | Ruxolitinib |
| MVP_top60_DNA_seed | PIK3CA | 0.894951 | SKCM | WM793B | Alpelisib |
| MVP_top60_DNA_seed | PIK3CA | 0.895952 | SKCM | LB373-MEL-D | AZD8186 |
| MVP_top60_DNA_seed | PIK3CB | 0.895952 | SKCM | LB373-MEL-D | AZD8186 |
| MVP_top60_DNA_seed | BRAF | 0.896578 | SKCM | VMRC-MELG | SB590885 |
| MVP_top60_DNA_seed | EGFR | 0.900733 | SKCM | VMRC-MELG | Sapitinib |
| MVP_top60_DNA_seed | ERBB2 | 0.900733 | SKCM | VMRC-MELG | Sapitinib |
| MVP_top60_DNA_seed | PIK3CA | 0.901935 | SKCM | SH-4 | Buparlisib |
| MVP_top60_DNA_seed | PIK3CB | 0.901935 | SKCM | SH-4 | Buparlisib |
| MVP_top60_DNA_seed | PIK3CB | 0.904049 | SKCM | SK-MEL-30 | AZD6482 |
| MVP_top60_DNA_seed | GSK3B | 0.906579 | SKCM | COLO-679 | SB216763 |
| MVP_top60_DNA_seed | EGFR | 0.910035 | SKCM | LB373-MEL-D | Lapatinib |
| MVP_top60_DNA_seed | ERBB2 | 0.910035 | SKCM | LB373-MEL-D | Lapatinib |
| MVP_top60_DNA_seed | JAK2 | 0.912121 | SKCM | WM278 | Ruxolitinib |
| MVP_top60_DNA_seed | JAK1 | 0.912121 | SKCM | WM278 | Ruxolitinib |
| MVP_top60_DNA_seed | GSK3B | 0.913865 | SKCM | IST-MEL1 | SB216763 |
| MVP_top60_DNA_seed | MAPK3 | 0.917389 | SKCM | Mewo | Ulixertinib |
| MVP_top60_DNA_seed | MAPK1 | 0.917389 | SKCM | Mewo | Ulixertinib |
| MVP_top60_DNA_seed | EGFR | 0.918875 | SKCM | UACC-257 | Gefitinib |
| MVP_top60_DNA_seed | MAP2K1 | 0.919046 | SKCM | CHL-1 | Selumetinib |
| MVP_top60_DNA_seed | PIK3CA | 0.920461 | SKCM | IST-MEL1 | Alpelisib |
| MVP_top60_DNA_seed | EP300 | 0.922848 | SKCM | UACC-257 | CPI-637 |
| MVP_top60_DNA_seed | JAK1 | 0.923864 | SKCM | COLO-829 | JAK1_8709 |
| MVP_top60_DNA_seed | PIK3CA | 0.925931 | SKCM | 451Lu | Alpelisib |
| MVP_top60_DNA_seed | PIK3CA | 0.925994 | SKCM | C32 | GNE-317 |
| MVP_top60_DNA_seed | EGFR | 0.926423 | SKCM | WM278 | Afatinib |
| MVP_top60_DNA_seed | ERBB2 | 0.926423 | SKCM | WM278 | Afatinib |
| MVP_top60_DNA_seed | EGFR | 0.927565 | SKCM | LOXIMVI | Osimertinib |
| MVP_top60_DNA_seed | MAP2K1 | 0.928572 | SKCM | CHL-1 | Selumetinib |
| MVP_top60_DNA_seed | EGFR | 0.929968 | SKCM | RVH-421 | Sapitinib |
| MVP_top60_DNA_seed | ERBB2 | 0.929968 | SKCM | RVH-421 | Sapitinib |
| MVP_top60_DNA_seed | GSK3B | 0.931034 | SKCM | K2 | CHIR-99021 |
| MVP_top60_DNA_seed | EGFR | 0.933272 | SKCM | CP50-MEL-B | Afatinib |
| MVP_top60_DNA_seed | ERBB2 | 0.933272 | SKCM | CP50-MEL-B | Afatinib |
| MVP_top60_DNA_seed | EGFR | 0.937696 | SKCM | A375 | Gefitinib |
| MVP_top60_DNA_seed | EP300 | 0.941294 | SKCM | SK-MEL-5 | SGC-CBP30 |
| MVP_top60_DNA_seed | CREBBP | 0.941294 | SKCM | SK-MEL-5 | SGC-CBP30 |
| MVP_top60_DNA_seed | PIK3CA | 0.946555 | SKCM | WM278 | GNE-317 |
| MVP_top60_DNA_seed | JAK2 | 0.948523 | SKCM | COLO-679 | Ruxolitinib |
| MVP_top60_DNA_seed | JAK1 | 0.948523 | SKCM | COLO-679 | Ruxolitinib |
| MVP_top60_DNA_seed | EGFR | 0.949198 | SKCM | WM1552C | Gefitinib |

| | | | | | |
|---|---|---|---|---|---|
| MVP_top60_DNA_seed | GSK3B | 0.949821 | SKCM | MMAc-SF | CHIR-99021 |
| MVP_top60_DNA_seed | EP300 | 0.95196 | SKCM | UACC-62 | CPI-637 |
| MVP_top60_DNA_seed | EGFR | 0.953239 | SKCM | WM1552C | Erlotinib |
| MVP_top60_DNA_seed | KRAS | 0.954843 | SKCM | IPC-298 | KRAS (G12C) Inhibitor-12 |
| MVP_top60_DNA_seed | EGFR | 0.954851 | SKCM | LB2518-MEL | Erlotinib |
| MVP_top60_DNA_seed | MAPK1 | 0.959157 | SKCM | VMRC-MELG | VX-11e |
| MVP_top60_DNA_seed | TP53 | 0.961851 | SKCM | WM278 | MIRA-1 |
| MVP_top60_DNA_seed | PIK3CA | 0.962901 | SKCM | WM278 | Buparlisib |
| MVP_top60_DNA_seed | PIK3CB | 0.962901 | SKCM | WM278 | Buparlisib |
| MVP_top60_DNA_seed | BRAF | 0.963493 | SKCM | Mewo | Dabrafenib |
| MVP_top60_DNA_seed | JAK2 | 0.963759 | SKCM | COLO-679 | GSK2276186C |
| MVP_top60_DNA_seed | JAK1 | 0.963759 | SKCM | COLO-679 | GSK2276186C |
| MVP_top60_DNA_seed | JAK3 | 0.963759 | SKCM | COLO-679 | GSK2276186C |
| MVP_top60_DNA_seed | GSK3B | 0.964521 | SKCM | 451Lu | CHIR-99021 |
| MVP_top60_DNA_seed | RAC1 | 0.966156 | SKCM | HT-144 | EHT-1864 |
| MVP_top60_DNA_seed | TP53 | 0.966158 | SKCM | HMV-II | MIRA-1 |
| MVP_top60_DNA_seed | JAK2 | 0.968569 | SKCM | M14 | AZ960 |
| MVP_top60_DNA_seed | JAK3 | 0.968569 | SKCM | M14 | AZ960 |
| MVP_top60_DNA_seed | EGFR | 0.969151 | SKCM | RVH-421 | Lapatinib |
| MVP_top60_DNA_seed | ERBB2 | 0.969151 | SKCM | RVH-421 | Lapatinib |
| MVP_top60_DNA_seed | EGFR | 0.971481 | SKCM | WM278 | Lapatinib |
| MVP_top60_DNA_seed | ERBB2 | 0.971481 | SKCM | WM278 | Lapatinib |
| MVP_top60_DNA_seed | EP300 | 0.972035 | SKCM | MEL-JUSO | SGC-CBP30 |
| MVP_top60_DNA_seed | CREBBP | 0.972035 | SKCM | MEL-JUSO | SGC-CBP30 |
| MVP_top60_DNA_seed | JAK2 | 0.972086 | SKCM | WM278 | GSK2276186C |
| MVP_top60_DNA_seed | JAK1 | 0.972086 | SKCM | WM278 | GSK2276186C |
| MVP_top60_DNA_seed | JAK3 | 0.972086 | SKCM | WM278 | GSK2276186C |
| MVP_top60_DNA_seed | EGFR | 0.972547 | SKCM | SK-MEL-24 | Sapitinib |
| MVP_top60_DNA_seed | ERBB2 | 0.972547 | SKCM | SK-MEL-24 | Sapitinib |
| MVP_top60_DNA_seed | RAC1 | 0.972712 | SKCM | CP66-MEL | EHT-1864 |
| MVP_top60_DNA_seed | EGFR | 0.973787 | SKCM | LB2518-MEL | Osimertinib |
| MVP_top60_DNA_seed | EGFR | 0.974526 | SKCM | Mewo | Lapatinib |
| MVP_top60_DNA_seed | ERBB2 | 0.974526 | SKCM | Mewo | Lapatinib |
| MVP_top60_DNA_seed | EGFR | 0.980719 | SKCM | SK-MEL-2 | Lapatinib |
| MVP_top60_DNA_seed | ERBB2 | 0.980719 | SKCM | SK-MEL-2 | Lapatinib |
| MVP_top60_DNA_seed | GSK3B | 0.981154 | SKCM | CP66-MEL | SB216763 |
| MVP_top60_DNA_seed | PIK3CA | 0.986257 | SKCM | SK-MEL-24 | Buparlisib |
| MVP_top60_DNA_seed | PIK3CB | 0.986257 | SKCM | SK-MEL-24 | Buparlisib |
| MVP_top60_DNA_seed | EGFR | 0.990778 | SKCM | COLO-829 | Osimertinib |
| MVP_top60_DNA_seed | PIK3CB | 0.99256 | SKCM | C32 | AZD6482 |
| MVP_top60_DNA_seed | PIK3CA | 0.996864 | SKCM | IGR-37 | Alpelisib |

| | | | | | |
|---|---|---|---|---|---|
| MVP_top60_DNA_seed | PIK3CA | 1.004055 | SKCM | A2058 | Alpelisib |
| MVP_top60_DNA_seed | JAK2 | 1.004215 | SKCM | IGR-37 | AZ960 |
| MVP_top60_DNA_seed | JAK3 | 1.004215 | SKCM | IGR-37 | AZ960 |
| MVP_top60_DNA_seed | EGFR | 1.005806 | SKCM | WM35 | Sapitinib |
| MVP_top60_DNA_seed | ERBB2 | 1.005806 | SKCM | WM35 | Sapitinib |
| MVP_top60_DNA_seed | GSK3B | 1.006581 | SKCM | SK-MEL-30 | CHIR-99021 |
| MVP_top60_DNA_seed | TP53 | 1.007744 | SKCM | UACC-257 | MIRA-1 |
| MVP_top60_DNA_seed | RAC1 | 1.009965 | SKCM | WM793B | EHT-1864 |
| MVP_top60_DNA_seed | JAK2 | 1.010674 | SKCM | WM-115 | AZ960 |
| MVP_top60_DNA_seed | JAK3 | 1.010674 | SKCM | WM-115 | AZ960 |
| MVP_top60_DNA_seed | EP300 | 1.011005 | SKCM | WM-115 | SGC-CBP30 |
| MVP_top60_DNA_seed | CREBBP | 1.011005 | SKCM | WM-115 | SGC-CBP30 |
| MVP_top60_DNA_seed | EGFR | 1.013053 | SKCM | MEL-JUSO | Sapitinib |
| MVP_top60_DNA_seed | ERBB2 | 1.013053 | SKCM | MEL-JUSO | Sapitinib |
| MVP_top60_DNA_seed | EGFR | 1.014451 | SKCM | Hs-940-T | Gefitinib |
| MVP_top60_DNA_seed | EGFR | 1.017328 | SKCM | IST-MEL1 | Erlotinib |
| MVP_top60_DNA_seed | EGFR | 1.018487 | SKCM | A101D | Sapitinib |
| MVP_top60_DNA_seed | ERBB2 | 1.018487 | SKCM | A101D | Sapitinib |
| MVP_top60_DNA_seed | EGFR | 1.021341 | SKCM | SH-4 | AZD3759 |
| MVP_top60_DNA_seed | PIK3CA | 1.021909 | SKCM | SK-MEL-30 | AZD8186 |
| MVP_top60_DNA_seed | PIK3CB | 1.021909 | SKCM | SK-MEL-30 | AZD8186 |
| MVP_top60_DNA_seed | EGFR | 1.027463 | SKCM | SK-MEL-5 | Lapatinib |
| MVP_top60_DNA_seed | ERBB2 | 1.027463 | SKCM | SK-MEL-5 | Lapatinib |
| MVP_top60_DNA_seed | JAK2 | 1.027981 | SKCM | MMAc-SF | Ruxolitinib |
| MVP_top60_DNA_seed | JAK1 | 1.027981 | SKCM | MMAc-SF | Ruxolitinib |
| MVP_top60_DNA_seed | EGFR | 1.035697 | SKCM | COLO-800 | Afatinib |
| MVP_top60_DNA_seed | ERBB2 | 1.035697 | SKCM | COLO-800 | Afatinib |
| MVP_top60_DNA_seed | EGFR | 1.036356 | SKCM | WM278 | Gefitinib |
| MVP_top60_DNA_seed | KRAS | 1.036729 | SKCM | Hs-940-T | (RAS (G12C) Inhibitor-12 |
| MVP_top60_DNA_seed | PIK3CA | 1.038299 | SKCM | COLO-679 | Alpelisib |
| MVP_top60_DNA_seed | EGFR | 1.038611 | SKCM | SH-4 | Erlotinib |
| MVP_top60_DNA_seed | JAK1 | 1.039807 | SKCM | SK-MEL-2 | JAK1_8709 |
| MVP_top60_DNA_seed | JAK2 | 1.039967 | SKCM | A375 | Ruxolitinib |
| MVP_top60_DNA_seed | JAK1 | 1.039967 | SKCM | A375 | Ruxolitinib |
| MVP_top60_DNA_seed | JAK2 | 1.047851 | SKCM | VMRC-MELG | Ruxolitinib |
| MVP_top60_DNA_seed | JAK1 | 1.047851 | SKCM | VMRC-MELG | Ruxolitinib |
| MVP_top60_DNA_seed | JAK2 | 1.050188 | SKCM | A101D | JAK_8517 |
| MVP_top60_DNA_seed | JAK1 | 1.050188 | SKCM | A101D | JAK_8517 |
| MVP_top60_DNA_seed | EGFR | 1.051417 | SKCM | SK-MEL-24 | Osimertinib |
| MVP_top60_DNA_seed | BRAF | 1.051503 | SKCM | MEL-JUSO | PLX-4720 |
| MVP_top60_DNA_seed | JAK1 | 1.054588 | SKCM | SK-MEL-5 | JAK1_8709 |

| | | | | | |
|---|---|---|---|---|---|
| MVP_top60_DNA_seed | EP300 | 1.056834 | SKCM | SH-4 | SGC-CBP30 |
| MVP_top60_DNA_seed | CREBBP | 1.056834 | SKCM | SH-4 | SGC-CBP30 |
| MVP_top60_DNA_seed | PIK3CA | 1.056957 | SKCM | SK-MEL-30 | GNE-317 |
| MVP_top60_DNA_seed | BRAF | 1.057173 | SKCM | SK-MEL-2 | PLX-4720 |
| MVP_top60_DNA_seed | MAPK3 | 1.059564 | SKCM | Hs-940-T | Ulixertinib |
| MVP_top60_DNA_seed | MAPK1 | 1.059564 | SKCM | Hs-940-T | Ulixertinib |
| MVP_top60_DNA_seed | EGFR | 1.062173 | SKCM | IGR-37 | Lapatinib |
| MVP_top60_DNA_seed | ERBB2 | 1.062173 | SKCM | IGR-37 | Lapatinib |
| MVP_top60_DNA_seed | JAK2 | 1.062431 | SKCM | LB2518-MEL | Ruxolitinib |
| MVP_top60_DNA_seed | JAK1 | 1.062431 | SKCM | LB2518-MEL | Ruxolitinib |
| MVP_top60_DNA_seed | KRAS | 1.063899 | SKCM | SK-MEL-2 | KRAS (G12C) Inhibitor-12 |
| MVP_top60_DNA_seed | JAK2 | 1.065807 | SKCM | SK-MEL-24 | Ruxolitinib |
| MVP_top60_DNA_seed | JAK1 | 1.065807 | SKCM | SK-MEL-24 | Ruxolitinib |
| MVP_top60_DNA_seed | JAK2 | 1.073601 | SKCM | SK-MEL-24 | GSK2276186C |
| MVP_top60_DNA_seed | JAK1 | 1.073601 | SKCM | SK-MEL-24 | GSK2276186C |
| MVP_top60_DNA_seed | JAK3 | 1.073601 | SKCM | SK-MEL-24 | GSK2276186C |
| MVP_top60_DNA_seed | RAC1 | 1.077505 | SKCM | SK-MEL-31 | EHT-1864 |
| MVP_top60_DNA_seed | EGFR | 1.080004 | SKCM | SK-MEL-31 | Erlotinib |
| MVP_top60_DNA_seed | EGFR | 1.083379 | SKCM | CP66-MEL | Sapitinib |
| MVP_top60_DNA_seed | ERBB2 | 1.083379 | SKCM | CP66-MEL | Sapitinib |
| MVP_top60_DNA_seed | EGFR | 1.084471 | SKCM | VMRC-MELG | Afatinib |
| MVP_top60_DNA_seed | ERBB2 | 1.084471 | SKCM | VMRC-MELG | Afatinib |
| MVP_top60_DNA_seed | EP300 | 1.084473 | SKCM | WM278 | CPI-637 |
| MVP_top60_DNA_seed | EGFR | 1.08512 | SKCM | COLO-792 | Lapatinib |
| MVP_top60_DNA_seed | ERBB2 | 1.08512 | SKCM | COLO-792 | Lapatinib |
| MVP_top60_DNA_seed | KRAS | 1.085705 | SKCM | G-361 | KRAS (G12C) Inhibitor-12 |
| MVP_top60_DNA_seed | EGFR | 1.087395 | SKCM | COLO-679 | Erlotinib |
| MVP_top60_DNA_seed | EGFR | 1.087606 | SKCM | SK-MEL-24 | AZD3759 |
| MVP_top60_DNA_seed | EP300 | 1.090602 | SKCM | HMV-II | SGC-CBP30 |
| MVP_top60_DNA_seed | CREBBP | 1.090602 | SKCM | HMV-II | SGC-CBP30 |
| MVP_top60_DNA_seed | EGFR | 1.090893 | SKCM | M14 | Sapitinib |
| MVP_top60_DNA_seed | ERBB2 | 1.090893 | SKCM | M14 | Sapitinib |
| MVP_top60_DNA_seed | JAK2 | 1.091822 | SKCM | CP66-MEL | GSK2276186C |
| MVP_top60_DNA_seed | JAK1 | 1.091822 | SKCM | CP66-MEL | GSK2276186C |
| MVP_top60_DNA_seed | JAK3 | 1.091822 | SKCM | CP66-MEL | GSK2276186C |
| MVP_top60_DNA_seed | EGFR | 1.098437 | SKCM | MEL-HO | Afatinib |
| MVP_top60_DNA_seed | ERBB2 | 1.098437 | SKCM | MEL-HO | Afatinib |
| MVP_top60_DNA_seed | EGFR | 1.106463 | SKCM | HMV-II | AZD3759 |
| MVP_top60_DNA_seed | TP53 | 1.109707 | SKCM | C32 | MIRA-1 |
| MVP_top60_DNA_seed | EGFR | 1.111222 | SKCM | CP66-MEL | Osimertinib |
| MVP_top60_DNA_seed | EGFR | 1.112053 | SKCM | IGR-37 | Sapitinib |

| | | | | | |
|---|---|---|---|---|---|
| MVP_top60_DNA_seed | ERBB2 | 1.112053 | SKCM | IGR-37 | Sapitinib |
| MVP_top60_DNA_seed | PIK3CA | 1.112307 | SKCM | SK-MEL-2 | AZD8186 |
| MVP_top60_DNA_seed | PIK3CB | 1.112307 | SKCM | SK-MEL-2 | AZD8186 |
| MVP_top60_DNA_seed | MAP2K1 | 1.114929 | SKCM | Hs-940-T | PD0325901 |
| MVP_top60_DNA_seed | EGFR | 1.11494 | SKCM | HT-144 | Osimertinib |
| MVP_top60_DNA_seed | EGFR | 1.116299 | SKCM | WM-115 | Osimertinib |
| MVP_top60_DNA_seed | EGFR | 1.119081 | SKCM | G-361 | Sapitinib |
| MVP_top60_DNA_seed | ERBB2 | 1.119081 | SKCM | G-361 | Sapitinib |
| MVP_top60_DNA_seed | BRAF | 1.130583 | SKCM | Mewo | PLX-4720 |
| MVP_top60_DNA_seed | EGFR | 1.130687 | SKCM | UACC-62 | Sapitinib |
| MVP_top60_DNA_seed | ERBB2 | 1.130687 | SKCM | UACC-62 | Sapitinib |
| MVP_top60_DNA_seed | EGFR | 1.133111 | SKCM | SK-MEL-5 | Sapitinib |
| MVP_top60_DNA_seed | ERBB2 | 1.133111 | SKCM | SK-MEL-5 | Sapitinib |
| MVP_top60_DNA_seed | JAK2 | 1.134791 | SKCM | UACC-257 | Ruxolitinib |
| MVP_top60_DNA_seed | JAK1 | 1.134791 | SKCM | UACC-257 | Ruxolitinib |
| MVP_top60_DNA_seed | JAK1 | 1.136733 | SKCM | C32 | JAK1_8709 |
| MVP_top60_DNA_seed | GSK3B | 1.140743 | SKCM | SK-MEL-5 | SB216763 |
| MVP_top60_DNA_seed | EGFR | 1.147106 | SKCM | IST-MEL1 | Gefitinib |
| MVP_top60_DNA_seed | EGFR | 1.147337 | SKCM | LB2518-MEL | Lapatinib |
| MVP_top60_DNA_seed | ERBB2 | 1.147337 | SKCM | LB2518-MEL | Lapatinib |
| MVP_top60_DNA_seed | EGFR | 1.156175 | SKCM | COLO-829 | Sapitinib |
| MVP_top60_DNA_seed | ERBB2 | 1.156175 | SKCM | COLO-829 | Sapitinib |
| MVP_top60_DNA_seed | KRAS | 1.15728 | SKCM | MEL-JUSO | KRAS (G12C) Inhibitor-12 |
| MVP_top60_DNA_seed | EP300 | 1.163479 | SKCM | COLO-829 | CPI-637 |
| MVP_top60_DNA_seed | EGFR | 1.164823 | SKCM | MEL-JUSO | AZD3759 |
| MVP_top60_DNA_seed | RAC1 | 1.168165 | SKCM | MMAc-SF | EHT-1864 |
| MVP_top60_DNA_seed | EGFR | 1.169139 | SKCM | K2 | Lapatinib |
| MVP_top60_DNA_seed | ERBB2 | 1.169139 | SKCM | K2 | Lapatinib |
| MVP_top60_DNA_seed | GSK3B | 1.172897 | SKCM | CP66-MEL | CHIR-99021 |
| MVP_top60_DNA_seed | EGFR | 1.174203 | SKCM | SK-MEL-24 | Erlotinib |
| MVP_top60_DNA_seed | EGFR | 1.180923 | SKCM | WM-115 | Gefitinib |
| MVP_top60_DNA_seed | PIK3CA | 1.182168 | SKCM | G-361 | Alpelisib |
| MVP_top60_DNA_seed | GSK3B | 1.182195 | SKCM | SK-MEL-5 | CHIR-99021 |
| MVP_top60_DNA_seed | EGFR | 1.187206 | SKCM | UACC-257 | Sapitinib |
| MVP_top60_DNA_seed | ERBB2 | 1.187206 | SKCM | UACC-257 | Sapitinib |
| MVP_top60_DNA_seed | PIK3CA | 1.188893 | SKCM | SK-MEL-24 | Alpelisib |
| MVP_top60_DNA_seed | JAK2 | 1.190031 | SKCM | UACC-257 | GSK2276186C |
| MVP_top60_DNA_seed | JAK1 | 1.190031 | SKCM | UACC-257 | GSK2276186C |
| MVP_top60_DNA_seed | JAK3 | 1.190031 | SKCM | UACC-257 | GSK2276186C |
| MVP_top60_DNA_seed | PIK3CA | 1.196751 | SKCM | LB2518-MEL | GNE-317 |
| MVP_top60_DNA_seed | JAK2 | 1.204733 | SKCM | MEL-HO | AZ960 |

| | | | | | |
|---|---|---|---|---|---|
| MVP_top60_DNA_seed | JAK3 | 1.204733 | SKCM | MEL-HO | AZ960 |
| MVP_top60_DNA_seed | PIK3CA | 1.205041 | SKCM | Hs-940-T | Buparlisib |
| MVP_top60_DNA_seed | PIK3CB | 1.205041 | SKCM | Hs-940-T | Buparlisib |
| MVP_top60_DNA_seed | EGFR | 1.206287 | SKCM | SK-MEL-3 | Afatinib |
| MVP_top60_DNA_seed | ERBB2 | 1.206287 | SKCM | SK-MEL-3 | Afatinib |
| MVP_top60_DNA_seed | JAK2 | 1.209597 | SKCM | SK-MEL-31 | Ruxolitinib |
| MVP_top60_DNA_seed | JAK1 | 1.209597 | SKCM | SK-MEL-31 | Ruxolitinib |
| MVP_top60_DNA_seed | JAK2 | 1.217689 | SKCM | LB373-MEL-D | AZ960 |
| MVP_top60_DNA_seed | JAK3 | 1.217689 | SKCM | LB373-MEL-D | AZ960 |
| MVP_top60_DNA_seed | EGFR | 1.219044 | SKCM | C32 | Sapitinib |
| MVP_top60_DNA_seed | ERBB2 | 1.219044 | SKCM | C32 | Sapitinib |
| MVP_top60_DNA_seed | EGFR | 1.227077 | SKCM | SH-4 | Sapitinib |
| MVP_top60_DNA_seed | ERBB2 | 1.227077 | SKCM | SH-4 | Sapitinib |
| MVP_top60_DNA_seed | EGFR | 1.228315 | SKCM | 451Lu | Osimertinib |
| MVP_top60_DNA_seed | TP53 | 1.231431 | SKCM | COLO-829 | MIRA-1 |
| MVP_top60_DNA_seed | EGFR | 1.233595 | SKCM | SK-MEL-1 | Gefitinib |
| MVP_top60_DNA_seed | KRAS | 1.238492 | SKCM | SK-MEL-28 | KRAS (G12C) Inhibitor-12 |
| MVP_top60_DNA_seed | PIK3CA | 1.241056 | SKCM | SK-MEL-2 | Alpelisib |
| MVP_top60_DNA_seed | EGFR | 1.243248 | SKCM | CP66-MEL | Gefitinib |
| MVP_top60_DNA_seed | EP300 | 1.244164 | SKCM | IST-MEL1 | CPI-637 |
| MVP_top60_DNA_seed | EGFR | 1.250757 | SKCM | WM-115 | AZD3759 |
| MVP_top60_DNA_seed | EGFR | 1.254805 | SKCM | COLO-679 | Osimertinib |
| MVP_top60_DNA_seed | JAK2 | 1.257225 | SKCM | COLO-829 | Ruxolitinib |
| MVP_top60_DNA_seed | JAK1 | 1.257225 | SKCM | COLO-829 | Ruxolitinib |
| MVP_top60_DNA_seed | PIK3CA | 1.259859 | SKCM | GAK | AZD8186 |
| MVP_top60_DNA_seed | PIK3CB | 1.259859 | SKCM | GAK | AZD8186 |
| MVP_top60_DNA_seed | EGFR | 1.260759 | SKCM | MEL-JUSO | Erlotinib |
| MVP_top60_DNA_seed | PIK3CA | 1.262605 | SKCM | GAK | Alpelisib |
| MVP_top60_DNA_seed | PIK3CA | 1.26523 | SKCM | C32 | Alpelisib |
| MVP_top60_DNA_seed | EGFR | 1.265311 | SKCM | WM278 | Osimertinib |
| MVP_top60_DNA_seed | PIK3CA | 1.270803 | SKCM | UACC-257 | GNE-317 |
| MVP_top60_DNA_seed | PIK3CA | 1.282817 | SKCM | SK-MEL-30 | Buparlisib |
| MVP_top60_DNA_seed | PIK3CB | 1.282817 | SKCM | SK-MEL-30 | Buparlisib |
| MVP_top60_DNA_seed | PIK3CA | 1.284462 | SKCM | IGR-1 | Alpelisib |
| MVP_top60_DNA_seed | GSK3B | 1.285965 | SKCM | C32 | SB216763 |
| MVP_top60_DNA_seed | PIK3CA | 1.292984 | SKCM | UACC-257 | AZD8186 |
| MVP_top60_DNA_seed | PIK3CB | 1.292984 | SKCM | UACC-257 | AZD8186 |
| MVP_top60_DNA_seed | PIK3CA | 1.297477 | SKCM | WM278 | Alpelisib |
| MVP_top60_DNA_seed | PIK3CA | 1.301907 | SKCM | CP66-MEL | Buparlisib |
| MVP_top60_DNA_seed | PIK3CB | 1.301907 | SKCM | CP66-MEL | Buparlisib |
| MVP_top60_DNA_seed | PIK3CB | 1.313708 | SKCM | 451Lu | AZD6482 |

| | | | | | |
|---|---|---|---|---|---|
| MVP_top60_DNA_seed | JAK2 | 1.315728 | SKCM | C32 | Ruxolitinib |
| MVP_top60_DNA_seed | JAK1 | 1.315728 | SKCM | C32 | Ruxolitinib |
| MVP_top60_DNA_seed | PIK3CA | 1.317568 | SKCM | MEL-JUSO | Alpelisib |
| MVP_top60_DNA_seed | EGFR | 1.320099 | SKCM | A101D | Osimertinib |
| MVP_top60_DNA_seed | PIK3CA | 1.327032 | SKCM | CHL-1 | Alpelisib |
| MVP_top60_DNA_seed | EGFR | 1.329269 | SKCM | COLO-829 | Gefitinib |
| MVP_top60_DNA_seed | MAPK3 | 1.335314 | SKCM | CHL-1 | SCH772984 |
| MVP_top60_DNA_seed | MAPK1 | 1.335314 | SKCM | CHL-1 | SCH772984 |
| MVP_top60_DNA_seed | EGFR | 1.338378 | SKCM | COLO-829 | AZD3759 |
| MVP_top60_DNA_seed | PIK3CA | 1.359436 | SKCM | SH-4 | Alpelisib |
| MVP_top60_DNA_seed | EGFR | 1.360292 | SKCM | C32 | Afatinib |
| MVP_top60_DNA_seed | ERBB2 | 1.360292 | SKCM | C32 | Afatinib |
| MVP_top60_DNA_seed | EGFR | 1.371081 | SKCM | COLO-679 | Sapitinib |
| MVP_top60_DNA_seed | ERBB2 | 1.371081 | SKCM | COLO-679 | Sapitinib |
| MVP_top60_DNA_seed | EGFR | 1.374559 | SKCM | SK-MEL-28 | Lapatinib |
| MVP_top60_DNA_seed | ERBB2 | 1.374559 | SKCM | SK-MEL-28 | Lapatinib |
| MVP_top60_DNA_seed | EGFR | 1.374726 | SKCM | SK-MEL-5 | Afatinib |
| MVP_top60_DNA_seed | ERBB2 | 1.374726 | SKCM | SK-MEL-5 | Afatinib |
| MVP_top60_DNA_seed | PIK3CA | 1.376799 | SKCM | COLO-679 | Buparlisib |
| MVP_top60_DNA_seed | PIK3CB | 1.376799 | SKCM | COLO-679 | Buparlisib |
| MVP_top60_DNA_seed | PIK3CA | 1.388759 | SKCM | SK-MEL-2 | GNE-317 |
| MVP_top60_DNA_seed | EGFR | 1.39581 | SKCM | C32 | Erlotinib |
| MVP_top60_DNA_seed | PIK3CA | 1.396392 | SKCM | SK-MEL-1 | Alpelisib |
| MVP_top60_DNA_seed | JAK2 | 1.413361 | SKCM | UACC-62 | JAK_8517 |
| MVP_top60_DNA_seed | JAK1 | 1.413361 | SKCM | UACC-62 | JAK_8517 |
| MVP_top60_DNA_seed | EGFR | 1.419716 | SKCM | MMAc-SF | Lapatinib |
| MVP_top60_DNA_seed | ERBB2 | 1.419716 | SKCM | MMAc-SF | Lapatinib |
| MVP_top60_DNA_seed | EGFR | 1.420693 | SKCM | UACC-257 | Osimertinib |
| MVP_top60_DNA_seed | EGFR | 1.420713 | SKCM | C32 | Lapatinib |
| MVP_top60_DNA_seed | ERBB2 | 1.420713 | SKCM | C32 | Lapatinib |
| MVP_top60_DNA_seed | PIK3CA | 1.423375 | SKCM | RVH-421 | Alpelisib |
| MVP_top60_DNA_seed | EGFR | 1.428762 | SKCM | CP66-MEL | Erlotinib |
| MVP_top60_DNA_seed | PIK3CA | 1.429829 | SKCM | M14 | Alpelisib |
| MVP_top60_DNA_seed | JAK2 | 1.430394 | SKCM | SK-MEL-2 | JAK_8517 |
| MVP_top60_DNA_seed | JAK1 | 1.430394 | SKCM | SK-MEL-2 | JAK_8517 |
| MVP_top60_DNA_seed | PIK3CA | 1.434345 | SKCM | HMV-II | AZD8186 |
| MVP_top60_DNA_seed | PIK3CB | 1.434345 | SKCM | HMV-II | AZD8186 |
| MVP_top60_DNA_seed | JAK2 | 1.438792 | SKCM | A375 | GSK2276186C |
| MVP_top60_DNA_seed | JAK1 | 1.438792 | SKCM | A375 | GSK2276186C |
| MVP_top60_DNA_seed | JAK3 | 1.438792 | SKCM | A375 | GSK2276186C |
| MVP_top60_DNA_seed | EGFR | 1.439869 | SKCM | SK-MEL-1 | Erlotinib |

| | | | | | |
|---|---|---|---|---|---|
| MVP_top60_DNA_seed | JAK2 | 1.445097 | SKCM | IGR-1 | JAK_8517 |
| MVP_top60_DNA_seed | JAK1 | 1.445097 | SKCM | IGR-1 | JAK_8517 |
| MVP_top60_DNA_seed | EGFR | 1.466311 | SKCM | WM-115 | Sapitinib |
| MVP_top60_DNA_seed | ERBB2 | 1.466311 | SKCM | WM-115 | Sapitinib |
| MVP_top60_DNA_seed | EGFR | 1.475105 | SKCM | Mewo | Gefitinib |
| MVP_top60_DNA_seed | EGFR | 1.477272 | SKCM | Hs-939-T | Erlotinib |
| MVP_top60_DNA_seed | EGFR | 1.479434 | SKCM | LB373-MEL-D | Erlotinib |
| MVP_top60_DNA_seed | EGFR | 1.480658 | SKCM | Mewo | Erlotinib |
| MVP_top60_DNA_seed | JAK2 | 1.489382 | SKCM | COLO-679 | JAK_8517 |
| MVP_top60_DNA_seed | JAK1 | 1.489382 | SKCM | COLO-679 | JAK_8517 |
| MVP_top60_DNA_seed | PIK3CA | 1.490068 | SKCM | 451Lu | AZD8186 |
| MVP_top60_DNA_seed | PIK3CB | 1.490068 | SKCM | 451Lu | AZD8186 |
| MVP_top60_DNA_seed | JAK2 | 1.490198 | SKCM | SH-4 | AZ960 |
| MVP_top60_DNA_seed | JAK3 | 1.490198 | SKCM | SH-4 | AZ960 |
| MVP_top60_DNA_seed | EGFR | 1.501959 | SKCM | SK-MEL-5 | Erlotinib |
| MVP_top60_DNA_seed | JAK2 | 1.506296 | SKCM | COLO-792 | JAK_8517 |
| MVP_top60_DNA_seed | JAK1 | 1.506296 | SKCM | COLO-792 | JAK_8517 |
| MVP_top60_DNA_seed | JAK2 | 1.506951 | SKCM | MEL-JUSO | JAK_8517 |
| MVP_top60_DNA_seed | JAK1 | 1.506951 | SKCM | MEL-JUSO | JAK_8517 |
| MVP_top60_DNA_seed | JAK2 | 1.51467 | SKCM | MEL-JUSO | AZ960 |
| MVP_top60_DNA_seed | JAK3 | 1.51467 | SKCM | MEL-JUSO | AZ960 |
| MVP_top60_DNA_seed | EGFR | 1.516604 | SKCM | WM-115 | Lapatinib |
| MVP_top60_DNA_seed | ERBB2 | 1.516604 | SKCM | WM-115 | Lapatinib |
| MVP_top60_DNA_seed | JAK1 | 1.517785 | SKCM | WM278 | JAK1_8709 |
| MVP_top60_DNA_seed | JAK2 | 1.518779 | SKCM | 451Lu | AZ960 |
| MVP_top60_DNA_seed | JAK3 | 1.518779 | SKCM | 451Lu | AZ960 |
| MVP_top60_DNA_seed | PIK3CA | 1.51955 | SKCM | MMAc-SF | Buparlisib |
| MVP_top60_DNA_seed | PIK3CB | 1.51955 | SKCM | MMAc-SF | Buparlisib |
| MVP_top60_DNA_seed | EGFR | 1.521958 | SKCM | WM278 | AZD3759 |
| MVP_top60_DNA_seed | RAC1 | 1.522295 | SKCM | Mewo | EHT-1864 |
| MVP_top60_DNA_seed | KRAS | 1.524405 | SKCM | SK-MEL-31 | (RAS (G12C) Inhibitor-12 |
| MVP_top60_DNA_seed | MAPK1 | 1.525721 | SKCM | CHL-1 | VX-11e |
| MVP_top60_DNA_seed | JAK2 | 1.529703 | SKCM | G-361 | JAK_8517 |
| MVP_top60_DNA_seed | JAK1 | 1.529703 | SKCM | G-361 | JAK_8517 |
| MVP_top60_DNA_seed | RAC1 | 1.532025 | SKCM | IST-MEL1 | EHT-1864 |
| MVP_top60_DNA_seed | EP300 | 1.54458 | SKCM | CP66-MEL | SGC-CBP30 |
| MVP_top60_DNA_seed | CREBBP | 1.54458 | SKCM | CP66-MEL | SGC-CBP30 |
| MVP_top60_DNA_seed | RAC1 | 1.557631 | SKCM | COLO-679 | EHT-1864 |
| MVP_top60_DNA_seed | EGFR | 1.563734 | SKCM | WM-115 | Erlotinib |
| MVP_top60_DNA_seed | KRAS | 1.564083 | SKCM | SK-MEL-24 | (RAS (G12C) Inhibitor-12 |
| MVP_top60_DNA_seed | KRAS | 1.564907 | SKCM | SK-MEL-5 | (RAS (G12C) Inhibitor-12 |

| | | | | | |
|---|---|---|---|---|---|
| MVP_top60_DNA_seed | RAC1 | 1.566175 | SKCM | SK-MEL-30 | EHT-1864 |
| MVP_top60_DNA_seed | EGFR | 1.574995 | SKCM | K2 | Osimertinib |
| MVP_top60_DNA_seed | EGFR | 1.592959 | SKCM | Mewo | AZD3759 |
| MVP_top60_DNA_seed | EGFR | 1.602122 | SKCM | SK-MEL-1 | Sapitinib |
| MVP_top60_DNA_seed | ERBB2 | 1.602122 | SKCM | SK-MEL-1 | Sapitinib |
| MVP_top60_DNA_seed | PIK3CA | 1.602884 | SKCM | RVH-421 | Buparlisib |
| MVP_top60_DNA_seed | PIK3CB | 1.602884 | SKCM | RVH-421 | Buparlisib |
| MVP_top60_DNA_seed | JAK2 | 1.603299 | SKCM | WM278 | AZ960 |
| MVP_top60_DNA_seed | JAK3 | 1.603299 | SKCM | WM278 | AZ960 |
| MVP_top60_DNA_seed | EGFR | 1.606884 | SKCM | SH-4 | Osimertinib |
| MVP_top60_DNA_seed | EGFR | 1.619955 | SKCM | UACC-257 | AZD3759 |
| MVP_top60_DNA_seed | JAK2 | 1.620572 | SKCM | WM278 | JAK_8517 |
| MVP_top60_DNA_seed | JAK1 | 1.620572 | SKCM | WM278 | JAK_8517 |
| MVP_top60_DNA_seed | KRAS | 1.62419 | SKCM | VMRC-MELG | KRAS (G12C) Inhibitor-12 |
| MVP_top60_DNA_seed | JAK2 | 1.647043 | SKCM | UACC-62 | AZ960 |
| MVP_top60_DNA_seed | JAK3 | 1.647043 | SKCM | UACC-62 | AZ960 |
| MVP_top60_DNA_seed | EGFR | 1.649471 | SKCM | LB373-MEL-D | Afatinib |
| MVP_top60_DNA_seed | ERBB2 | 1.649471 | SKCM | LB373-MEL-D | Afatinib |
| MVP_top60_DNA_seed | MAP2K1 | 1.666415 | SKCM | VMRC-MELG | Selumetinib |
| MVP_top60_DNA_seed | RAC1 | 1.674037 | SKCM | MZ7-mel | EHT-1864 |
| MVP_top60_DNA_seed | EGFR | 1.687124 | SKCM | SK-MEL-1 | AZD3759 |
| MVP_top60_DNA_seed | JAK2 | 1.706009 | SKCM | C32 | GSK2276186C |
| MVP_top60_DNA_seed | JAK1 | 1.706009 | SKCM | C32 | GSK2276186C |
| MVP_top60_DNA_seed | JAK3 | 1.706009 | SKCM | C32 | GSK2276186C |
| MVP_top60_DNA_seed | MAPK3 | 1.706016 | SKCM | VMRC-MELG | SCH772984 |
| MVP_top60_DNA_seed | MAPK1 | 1.706016 | SKCM | VMRC-MELG | SCH772984 |
| MVP_top60_DNA_seed | JAK2 | 1.716178 | SKCM | COLO-679 | AZ960 |
| MVP_top60_DNA_seed | JAK3 | 1.716178 | SKCM | COLO-679 | AZ960 |
| MVP_top60_DNA_seed | PIK3CA | 1.724842 | SKCM | UACC-257 | Alpelisib |
| MVP_top60_DNA_seed | PIK3CA | 1.732286 | SKCM | MEL-JUSO | AZD8186 |
| MVP_top60_DNA_seed | PIK3CB | 1.732286 | SKCM | MEL-JUSO | AZD8186 |
| MVP_top60_DNA_seed | RAC1 | 1.737224 | SKCM | COLO-829 | EHT-1864 |
| MVP_top60_DNA_seed | EGFR | 1.753932 | SKCM | SK-MEL-1 | Osimertinib |
| MVP_top60_DNA_seed | JAK1 | 1.780004 | SKCM | WM-115 | JAK1_8709 |
| MVP_top60_DNA_seed | GSK3B | 1.781897 | SKCM | C32 | CHIR-99021 |
| MVP_top60_DNA_seed | PIK3CA | 1.78337 | SKCM | SH-4 | AZD8186 |
| MVP_top60_DNA_seed | PIK3CB | 1.78337 | SKCM | SH-4 | AZD8186 |
| MVP_top60_DNA_seed | JAK2 | 1.786633 | SKCM | LB2518-MEL | AZ960 |
| MVP_top60_DNA_seed | JAK3 | 1.786633 | SKCM | LB2518-MEL | AZ960 |
| MVP_top60_DNA_seed | EGFR | 1.788596 | SKCM | UACC-257 | Lapatinib |
| MVP_top60_DNA_seed | ERBB2 | 1.788596 | SKCM | UACC-257 | Lapatinib |

| | | | | | |
|---|---|---|---|---|---|
| MVP_top60_DNA_seed | PIK3CA | 1.794828 | SKCM | SK-MEL-1 | GNE-317 |
| MVP_top60_DNA_seed | PIK3CA | 1.804611 | SKCM | LB373-MEL-D | GNE-317 |
| MVP_top60_DNA_seed | JAK2 | 1.809204 | SKCM | UACC-257 | JAK_8517 |
| MVP_top60_DNA_seed | JAK1 | 1.809204 | SKCM | UACC-257 | JAK_8517 |
| MVP_top60_DNA_seed | PIK3CA | 1.821961 | SKCM | COLO-679 | GNE-317 |
| MVP_top60_DNA_seed | JAK2 | 1.823875 | SKCM | COLO-829 | GSK2276186C |
| MVP_top60_DNA_seed | JAK1 | 1.823875 | SKCM | COLO-829 | GSK2276186C |
| MVP_top60_DNA_seed | JAK3 | 1.823875 | SKCM | COLO-829 | GSK2276186C |
| MVP_top60_DNA_seed | EGFR | 1.846431 | SKCM | CP66-MEL | Afatinib |
| MVP_top60_DNA_seed | ERBB2 | 1.846431 | SKCM | CP66-MEL | Afatinib |
| MVP_top60_DNA_seed | EGFR | 1.881867 | SKCM | MMAc-SF | Erlotinib |
| MVP_top60_DNA_seed | EGFR | 1.925021 | SKCM | CP66-MEL | Lapatinib |
| MVP_top60_DNA_seed | ERBB2 | 1.925021 | SKCM | CP66-MEL | Lapatinib |
| MVP_top60_DNA_seed | JAK2 | 1.933417 | SKCM | K2 | AZ960 |
| MVP_top60_DNA_seed | JAK3 | 1.933417 | SKCM | K2 | AZ960 |
| MVP_top60_DNA_seed | MAP2K1 | 1.947151 | SKCM | VMRC-MELG | PD0325901 |
| MVP_top60_DNA_seed | EGFR | 1.951589 | SKCM | COLO-829 | Afatinib |
| MVP_top60_DNA_seed | ERBB2 | 1.951589 | SKCM | COLO-829 | Afatinib |
| MVP_top60_DNA_seed | PIK3CA | 1.951942 | SKCM | SK-MEL-1 | Buparlisib |
| MVP_top60_DNA_seed | PIK3CB | 1.951942 | SKCM | SK-MEL-1 | Buparlisib |
| MVP_top60_DNA_seed | RAC1 | 1.972615 | SKCM | SK-MEL-28 | EHT-1864 |
| MVP_top60_DNA_seed | EGFR | 1.974493 | SKCM | SK-MEL-31 | Afatinib |
| MVP_top60_DNA_seed | ERBB2 | 1.974493 | SKCM | SK-MEL-31 | Afatinib |
| MVP_top60_DNA_seed | RAC1 | 2.034286 | SKCM | WM1552C | EHT-1864 |
| MVP_top60_DNA_seed | RAC1 | 2.066773 | SKCM | WM-115 | EHT-1864 |
| MVP_top60_DNA_seed | PIK3CA | 2.074385 | SKCM | COLO-829 | Alpelisib |
| MVP_top60_DNA_seed | JAK2 | 2.074618 | SKCM | SK-MEL-2 | AZ960 |
| MVP_top60_DNA_seed | JAK3 | 2.074618 | SKCM | SK-MEL-2 | AZ960 |
| MVP_top60_DNA_seed | JAK2 | 2.078961 | SKCM | MMAc-SF | AZ960 |
| MVP_top60_DNA_seed | JAK3 | 2.078961 | SKCM | MMAc-SF | AZ960 |
| MVP_top60_DNA_seed | MAPK1 | 2.10026 | SKCM | SK-MEL-31 | VX-11e |
| MVP_top60_DNA_seed | EGFR | 2.102406 | SKCM | RVH-421 | Afatinib |
| MVP_top60_DNA_seed | ERBB2 | 2.102406 | SKCM | RVH-421 | Afatinib |
| MVP_top60_DNA_seed | JAK2 | 2.138103 | SKCM | RVH-421 | AZ960 |
| MVP_top60_DNA_seed | JAK3 | 2.138103 | SKCM | RVH-421 | AZ960 |
| MVP_top60_DNA_seed | PIK3CA | 2.175623 | SKCM | SK-MEL-1 | AZD8186 |
| MVP_top60_DNA_seed | PIK3CB | 2.175623 | SKCM | SK-MEL-1 | AZD8186 |
| MVP_top60_DNA_seed | KRAS | 2.191249 | SKCM | COLO-829 | KRAS (G12C) Inhibitor-12 |
| MVP_top60_DNA_seed | EP300 | 2.229299 | SKCM | COLO-829 | SGC-CBP30 |
| MVP_top60_DNA_seed | CREBBP | 2.229299 | SKCM | COLO-829 | SGC-CBP30 |
| MVP_top60_DNA_seed | EGFR | 2.29687 | SKCM | COLO-679 | Lapatinib |

| | | | | | |
|---|---|---|---|---|---|
| MVP_top60_DNA_seed | ERBB2 | 2.29687 | SKCM | COLO-679 | Lapatinib |
| MVP_top60_DNA_seed | EGFR | 2.321365 | SKCM | COLO-829 | Lapatinib |
| MVP_top60_DNA_seed | ERBB2 | 2.321365 | SKCM | COLO-829 | Lapatinib |
| MVP_top60_DNA_seed | PIK3CA | 2.327058 | SKCM | LB373-MEL-D | Buparlisib |
| MVP_top60_DNA_seed | PIK3CB | 2.327058 | SKCM | LB373-MEL-D | Buparlisib |
| MVP_top60_DNA_seed | MAPK3 | 2.333303 | SKCM | VMRC-MELG | Ulixertinib |
| MVP_top60_DNA_seed | MAPK1 | 2.333303 | SKCM | VMRC-MELG | Ulixertinib |
| MVP_top60_DNA_seed | JAK2 | 2.347762 | SKCM | CP66-MEL | AZ960 |
| MVP_top60_DNA_seed | JAK3 | 2.347762 | SKCM | CP66-MEL | AZ960 |
| MVP_top60_DNA_seed | JAK2 | 2.369911 | SKCM | UACC-257 | AZ960 |
| MVP_top60_DNA_seed | JAK3 | 2.369911 | SKCM | UACC-257 | AZ960 |
| MVP_top60_DNA_seed | EGFR | 2.437064 | SKCM | SK-MEL-28 | Afatinib |
| MVP_top60_DNA_seed | ERBB2 | 2.437064 | SKCM | SK-MEL-28 | Afatinib |
| MVP_top60_DNA_seed | PIK3CA | 2.893731 | SKCM | SK-MEL-24 | GNE-317 |
| top60_altered_DNA_seed | EP300 | -2.606148 | BLCA | BFTC-905 | I-CBP112 |
| top60_altered_DNA_seed | PIK3CA | -2.363975 | BLCA | CAL-29 | AZD8835 |
| top60_altered_DNA_seed | PIK3CA | -2.256468 | BLCA | CAL-29 | PI3Ka_4409 |
| top60_altered_DNA_seed | PIK3CA | -1.581022 | BLCA | BFTC-905 | AZD8835 |
| top60_altered_DNA_seed | PIK3CA | -1.503629 | BLCA | BFTC-905 | PI3Ka_4409 |
| top60_altered_DNA_seed | EP300 | -1.304192 | BLCA | SW780 | I-CBP112 |
| top60_altered_DNA_seed | PIK3CA | -1.067843 | BLCA | HT-1197 | AZD8835 |
| top60_altered_DNA_seed | PIK3CA | -0.997563 | BLCA | RT-112 | PI3Ka_4409 |
| top60_altered_DNA_seed | EP300 | -0.631944 | BLCA | LB831-BLC | I-CBP112 |
| top60_altered_DNA_seed | PIK3CA | -0.625709 | BLCA | RT-112 | AZD8835 |
| top60_altered_DNA_seed | PIK3CA | -0.57933 | BLCA | TCCSUP | AZD8835 |
| top60_altered_DNA_seed | PIK3CA | -0.521185 | BLCA | SW1710 | AZD8835 |
| top60_altered_DNA_seed | PIK3CA | -0.475212 | BLCA | 5637 | PI3Ka_4409 |
| top60_altered_DNA_seed | PIK3CA | -0.367211 | BLCA | 639-V | AZD8835 |
| top60_altered_DNA_seed | PIK3CA | -0.302125 | BLCA | SW780 | PI3Ka_4409 |
| top60_altered_DNA_seed | PIK3CA | -0.295805 | BLCA | LB831-BLC | AZD8835 |
| top60_altered_DNA_seed | PIK3CA | -0.227711 | BLCA | HT-1197 | PI3Ka_4409 |
| top60_altered_DNA_seed | PIK3CA | -0.226616 | BLCA | 647-V | PI3Ka_4409 |
| top60_altered_DNA_seed | EP300 | -0.156056 | BLCA | J82 | I-CBP112 |
| top60_altered_DNA_seed | EP300 | -0.150236 | BLCA | 639-V | I-CBP112 |
| top60_altered_DNA_seed | EP300 | -0.115471 | BLCA | 5637 | I-CBP112 |
| top60_altered_DNA_seed | EP300 | -0.056293 | BLCA | UM-UC-3 | I-CBP112 |
| top60_altered_DNA_seed | EP300 | -0.031533 | BLCA | SW1710 | I-CBP112 |
| top60_altered_DNA_seed | PIK3CA | 0.049539 | BLCA | SW1710 | PI3Ka_4409 |
| top60_altered_DNA_seed | PIK3CA | 0.050942 | BLCA | J82 | PI3Ka_4409 |
| top60_altered_DNA_seed | EP300 | 0.10791 | BLCA | HT-1197 | I-CBP112 |
| top60_altered_DNA_seed | PIK3CA | 0.120483 | BLCA | DSH1 | AZD8835 |

| | | | | | |
|---|---|---|---|---|---|
| top60_altered_DNA_seed | PIK3CA | 0.123523 | BLCA | 5637 | AZD8835 |
| top60_altered_DNA_seed | EP300 | 0.212048 | BLCA | DSH1 | I-CBP112 |
| top60_altered_DNA_seed | PIK3CA | 0.212412 | BLCA | DSH1 | PI3Ka_4409 |
| top60_altered_DNA_seed | PIK3CA | 0.217414 | BLCA | T24 | AZD8835 |
| top60_altered_DNA_seed | PIK3CA | 0.230402 | BLCA | 639-V | PI3Ka_4409 |
| top60_altered_DNA_seed | PIK3CA | 0.306539 | BLCA | UM-UC-3 | AZD8835 |
| top60_altered_DNA_seed | EP300 | 0.351883 | BLCA | CAL-29 | I-CBP112 |
| top60_altered_DNA_seed | PIK3CA | 0.378449 | BLCA | 647-V | AZD8835 |
| top60_altered_DNA_seed | PIK3CA | 0.379359 | BLCA | LB831-BLC | PI3Ka_4409 |
| top60_altered_DNA_seed | EP300 | 0.410601 | BLCA | T24 | I-CBP112 |
| top60_altered_DNA_seed | EP300 | 0.458008 | BLCA | 647-V | I-CBP112 |
| top60_altered_DNA_seed | EP300 | 0.480541 | BLCA | RT-112 | I-CBP112 |
| top60_altered_DNA_seed | PIK3CA | 0.485958 | BLCA | J82 | AZD8835 |
| top60_altered_DNA_seed | EP300 | 0.653108 | BLCA | RT4 | I-CBP112 |
| top60_altered_DNA_seed | PIK3CA | 0.670531 | BLCA | HT-1376 | AZD8835 |
| top60_altered_DNA_seed | PIK3CA | 0.758114 | BLCA | KU-19-19 | AZD8835 |
| top60_altered_DNA_seed | PIK3CA | 0.811732 | BLCA | UM-UC-3 | PI3Ka_4409 |
| top60_altered_DNA_seed | EP300 | 0.818554 | BLCA | TCCSUP | I-CBP112 |
| top60_altered_DNA_seed | PIK3CA | 0.831832 | BLCA | HT-1376 | PI3Ka_4409 |
| top60_altered_DNA_seed | PIK3CA | 0.840047 | BLCA | RT4 | AZD8835 |
| top60_altered_DNA_seed | PIK3CA | 0.936587 | BLCA | SW780 | AZD8835 |
| top60_altered_DNA_seed | PIK3CA | 0.983085 | BLCA | T24 | PI3Ka_4409 |
| top60_altered_DNA_seed | PIK3CA | 1.110391 | BLCA | KU-19-19 | PI3Ka_4409 |
| top60_altered_DNA_seed | EP300 | 1.592007 | BLCA | HT-1376 | I-CBP112 |
| top60_altered_DNA_seed | PIK3CA | 1.846686 | BLCA | RT4 | PI3Ka_4409 |
| top60_altered_DNA_seed | PIK3CA | -1.984079 | BLCA | CAL-29 | Alpelisib |
| top60_altered_DNA_seed | TP53 | -1.730724 | BLCA | BFTC-905 | MIRA-1 |
| top60_altered_DNA_seed | EP300 | -1.208234 | BLCA | BFTC-905 | SGC-CBP30 |
| top60_altered_DNA_seed | PIK3CA | -1.147597 | BLCA | CAL-29 | GNE-317 |
| top60_altered_DNA_seed | PIK3CA | -1.094133 | BLCA | CAL-29 | Buparlisib |
| top60_altered_DNA_seed | EP300 | -1.03655 | BLCA | BFTC-905 | CPI-637 |
| top60_altered_DNA_seed | EP300 | -0.963364 | BLCA | RT4 | SGC-CBP30 |
| top60_altered_DNA_seed | PIK3CA | -0.884619 | BLCA | CAL-29 | AZD8186 |
| top60_altered_DNA_seed | EP300 | -0.879263 | BLCA | SW1710 | SGC-CBP30 |
| top60_altered_DNA_seed | PIK3CA | -0.837054 | BLCA | BFTC-905 | Buparlisib |
| top60_altered_DNA_seed | PIK3CA | -0.799839 | BLCA | BFTC-905 | Alpelisib |
| top60_altered_DNA_seed | PIK3CA | -0.777686 | BLCA | 5637 | Buparlisib |
| top60_altered_DNA_seed | PIK3CA | -0.69325 | BLCA | HT-1197 | Buparlisib |
| top60_altered_DNA_seed | PIK3CA | -0.670616 | BLCA | RT-112 | Alpelisib |
| top60_altered_DNA_seed | EP300 | -0.567635 | BLCA | KU-19-19 | SGC-CBP30 |
| top60_altered_DNA_seed | PIK3CA | -0.562267 | BLCA | DSH1 | GNE-317 |

| | | | | | |
|---|---|---|---|---|---|
| top60_altered_DNA_seed | PIK3CA | -0.533608 | BLCA | HT-1197 | GNE-317 |
| top60_altered_DNA_seed | PIK3CA | -0.481732 | BLCA | KU-19-19 | Buparlisib |
| top60_altered_DNA_seed | PIK3CA | -0.476776 | BLCA | KU-19-19 | GNE-317 |
| top60_altered_DNA_seed | EP300 | -0.459899 | BLCA | DSH1 | SGC-CBP30 |
| top60_altered_DNA_seed | EP300 | -0.45414 | BLCA | SW780 | SGC-CBP30 |
| top60_altered_DNA_seed | PIK3CA | -0.335329 | BLCA | 647-V | AZD8186 |
| top60_altered_DNA_seed | EP300 | -0.327979 | BLCA | KU-19-19 | CPI-637 |
| top60_altered_DNA_seed | PIK3CA | -0.311178 | BLCA | 639-V | AZD8186 |
| top60_altered_DNA_seed | PIK3CA | -0.297558 | BLCA | 639-V | Buparlisib |
| top60_altered_DNA_seed | PIK3CA | -0.28342 | BLCA | RT-112 | GNE-317 |
| top60_altered_DNA_seed | EP300 | -0.281952 | BLCA | 639-V | CPI-637 |
| top60_altered_DNA_seed | EP300 | -0.273802 | BLCA | J82 | SGC-CBP30 |
| top60_altered_DNA_seed | PIK3CA | -0.273018 | BLCA | BFTC-905 | GNE-317 |
| top60_altered_DNA_seed | PIK3CA | -0.270785 | BLCA | TCCSUP | GNE-317 |
| top60_altered_DNA_seed | EP300 | -0.250773 | BLCA | TCCSUP | SGC-CBP30 |
| top60_altered_DNA_seed | PIK3CA | -0.22432 | BLCA | UM-UC-3 | GNE-317 |
| top60_altered_DNA_seed | TP53 | -0.214167 | BLCA | 639-V | MIRA-1 |
| top60_altered_DNA_seed | PIK3CA | -0.202548 | BLCA | HT-1197 | AZD8186 |
| top60_altered_DNA_seed | EP300 | -0.180177 | BLCA | DSH1 | CPI-637 |
| top60_altered_DNA_seed | PIK3CA | -0.167433 | BLCA | RT-112 | Buparlisib |
| top60_altered_DNA_seed | PIK3CA | -0.141735 | BLCA | UM-UC-3 | AZD8186 |
| top60_altered_DNA_seed | TP53 | -0.130501 | BLCA | RT-112 | MIRA-1 |
| top60_altered_DNA_seed | PIK3CA | -0.11969 | BLCA | 647-V | GNE-317 |
| top60_altered_DNA_seed | PIK3CA | -0.110066 | BLCA | TCCSUP | Buparlisib |
| top60_altered_DNA_seed | PIK3CA | -0.101739 | BLCA | BFTC-905 | AZD8186 |
| top60_altered_DNA_seed | PIK3CA | -0.087925 | BLCA | LB831-BLC | Alpelisib |
| top60_altered_DNA_seed | PIK3CA | -0.070269 | BLCA | DSH1 | Buparlisib |
| top60_altered_DNA_seed | PIK3CA | -0.013456 | BLCA | 647-V | Buparlisib |
| top60_altered_DNA_seed | PIK3CA | -0.010391 | BLCA | 5637 | GNE-317 |
| top60_altered_DNA_seed | PIK3CA | -0.005178 | BLCA | HT-1197 | Alpelisib |
| top60_altered_DNA_seed | EP300 | 0.003175 | BLCA | 639-V | SGC-CBP30 |
| top60_altered_DNA_seed | EP300 | 0.007679 | BLCA | RT4 | CPI-637 |
| top60_altered_DNA_seed | PIK3CA | 0.010687 | BLCA | 5637 | Alpelisib |
| top60_altered_DNA_seed | PIK3CA | 0.019742 | BLCA | UM-UC-3 | Buparlisib |
| top60_altered_DNA_seed | TP53 | 0.039981 | BLCA | KU-19-19 | MIRA-1 |
| top60_altered_DNA_seed | PIK3CA | 0.070222 | BLCA | DSH1 | Alpelisib |
| top60_altered_DNA_seed | PIK3CA | 0.077521 | BLCA | 639-V | GNE-317 |
| top60_altered_DNA_seed | TP53 | 0.082077 | BLCA | CAL-29 | MIRA-1 |
| top60_altered_DNA_seed | PIK3CA | 0.095867 | BLCA | SW1710 | Buparlisib |
| top60_altered_DNA_seed | EP300 | 0.126142 | BLCA | 5637 | SGC-CBP30 |
| top60_altered_DNA_seed | EP300 | 0.129522 | BLCA | SW1710 | CPI-637 |

| | | | | | |
|---|---|---|---|---|---|
| top60_altered_DNA_seed | EP300 | 0.159683 | BLCA | CAL-29 | SGC-CBP30 |
| top60_altered_DNA_seed | EP300 | 0.162684 | BLCA | HT-1376 | CPI-637 |
| top60_altered_DNA_seed | EP300 | 0.163524 | BLCA | 647-V | CPI-637 |
| top60_altered_DNA_seed | PIK3CA | 0.16847 | BLCA | SW780 | GNE-317 |
| top60_altered_DNA_seed | TP53 | 0.170108 | BLCA | UM-UC-3 | MIRA-1 |
| top60_altered_DNA_seed | TP53 | 0.209842 | BLCA | DSH1 | MIRA-1 |
| top60_altered_DNA_seed | EP300 | 0.219325 | BLCA | LB831-BLC | CPI-637 |
| top60_altered_DNA_seed | PIK3CA | 0.220832 | BLCA | SW1710 | GNE-317 |
| top60_altered_DNA_seed | PIK3CA | 0.255993 | BLCA | LB831-BLC | GNE-317 |
| top60_altered_DNA_seed | PIK3CA | 0.336452 | BLCA | LB831-BLC | AZD8186 |
| top60_altered_DNA_seed | EP300 | 0.343881 | BLCA | SW780 | CPI-637 |
| top60_altered_DNA_seed | PIK3CA | 0.349232 | BLCA | SW1710 | AZD8186 |
| top60_altered_DNA_seed | PIK3CA | 0.380056 | BLCA | DSH1 | AZD8186 |
| top60_altered_DNA_seed | TP53 | 0.382708 | BLCA | 647-V | MIRA-1 |
| top60_altered_DNA_seed | PIK3CA | 0.404888 | BLCA | 5637 | AZD8186 |
| top60_altered_DNA_seed | TP53 | 0.417231 | BLCA | SW1710 | MIRA-1 |
| top60_altered_DNA_seed | PIK3CA | 0.435335 | BLCA | HT-1376 | AZD8186 |
| top60_altered_DNA_seed | PIK3CA | 0.44544 | BLCA | SW780 | Buparlisib |
| top60_altered_DNA_seed | EP300 | 0.457723 | BLCA | CAL-29 | CPI-637 |
| top60_altered_DNA_seed | EP300 | 0.474652 | BLCA | 5637 | CPI-637 |
| top60_altered_DNA_seed | TP53 | 0.480846 | BLCA | T24 | MIRA-1 |
| top60_altered_DNA_seed | PIK3CA | 0.491344 | BLCA | RT4 | Buparlisib |
| top60_altered_DNA_seed | PIK3CA | 0.505059 | BLCA | TCCSUP | Alpelisib |
| top60_altered_DNA_seed | TP53 | 0.520183 | BLCA | SW780 | MIRA-1 |
| top60_altered_DNA_seed | TP53 | 0.520968 | BLCA | RT4 | MIRA-1 |
| top60_altered_DNA_seed | EP300 | 0.543802 | BLCA | RT-112 | SGC-CBP30 |
| top60_altered_DNA_seed | PIK3CA | 0.557464 | BLCA | RT-112 | AZD8186 |
| top60_altered_DNA_seed | EP300 | 0.572369 | BLCA | J82 | CPI-637 |
| top60_altered_DNA_seed | PIK3CA | 0.599658 | BLCA | 639-V | Alpelisib |
| top60_altered_DNA_seed | PIK3CA | 0.61902 | BLCA | LB831-BLC | Buparlisib |
| top60_altered_DNA_seed | TP53 | 0.670544 | BLCA | 5637 | MIRA-1 |
| top60_altered_DNA_seed | PIK3CA | 0.672557 | BLCA | TCCSUP | AZD8186 |
| top60_altered_DNA_seed | TP53 | 0.706531 | BLCA | HT-1197 | MIRA-1 |
| top60_altered_DNA_seed | TP53 | 0.720144 | BLCA | TCCSUP | MIRA-1 |
| top60_altered_DNA_seed | TP53 | 0.736355 | BLCA | LB831-BLC | MIRA-1 |
| top60_altered_DNA_seed | EP300 | 0.736911 | BLCA | 647-V | SGC-CBP30 |
| top60_altered_DNA_seed | PIK3CA | 0.770447 | BLCA | RT4 | Alpelisib |
| top60_altered_DNA_seed | EP300 | 0.772096 | BLCA | LB831-BLC | SGC-CBP30 |
| top60_altered_DNA_seed | TP53 | 0.778608 | BLCA | J82 | MIRA-1 |
| top60_altered_DNA_seed | PIK3CA | 0.807117 | BLCA | T24 | AZD8186 |
| top60_altered_DNA_seed | PIK3CA | 0.864754 | BLCA | KU-19-19 | AZD8186 |

| | | | | | |
|---|---|---|---|---|---|
| top60_altered_DNA_seed | EP300 | 0.883701 | BLCA | RT-112 | CPI-637 |
| top60_altered_DNA_seed | PIK3CA | 0.933356 | BLCA | T24 | GNE-317 |
| top60_altered_DNA_seed | EP300 | 0.95824 | BLCA | TCCSUP | CPI-637 |
| top60_altered_DNA_seed | TP53 | 0.988377 | BLCA | HT-1376 | MIRA-1 |
| top60_altered_DNA_seed | EP300 | 1.009242 | BLCA | UM-UC-3 | CPI-637 |
| top60_altered_DNA_seed | PIK3CA | 1.024332 | BLCA | J82 | AZD8186 |
| top60_altered_DNA_seed | PIK3CA | 1.077254 | BLCA | KU-19-19 | Alpelisib |
| top60_altered_DNA_seed | PIK3CA | 1.103866 | BLCA | T24 | Alpelisib |
| top60_altered_DNA_seed | PIK3CA | 1.110764 | BLCA | RT4 | GNE-317 |
| top60_altered_DNA_seed | PIK3CA | 1.13296 | BLCA | SW1710 | Alpelisib |
| top60_altered_DNA_seed | PIK3CA | 1.261823 | BLCA | 647-V | Alpelisib |
| top60_altered_DNA_seed | PIK3CA | 1.29491 | BLCA | UM-UC-3 | Alpelisib |
| top60_altered_DNA_seed | PIK3CA | 1.309147 | BLCA | J82 | Alpelisib |
| top60_altered_DNA_seed | PIK3CA | 1.405059 | BLCA | SW780 | Alpelisib |
| top60_altered_DNA_seed | EP300 | 1.51147 | BLCA | HT-1197 | SGC-CBP30 |
| top60_altered_DNA_seed | PIK3CA | 1.680588 | BLCA | T24 | Buparlisib |
| top60_altered_DNA_seed | PIK3CA | 1.706332 | BLCA | RT4 | AZD8186 |
| top60_altered_DNA_seed | PIK3CA | 1.715415 | BLCA | HT-1376 | GNE-317 |
| top60_altered_DNA_seed | PIK3CA | 1.716928 | BLCA | SW780 | AZD8186 |
| top60_altered_DNA_seed | PIK3CA | 1.756588 | BLCA | HT-1376 | Alpelisib |
| top60_altered_DNA_seed | PIK3CA | 1.951983 | BLCA | J82 | Buparlisib |
| top60_altered_DNA_seed | PIK3CA | 2.716316 | BLCA | HT-1376 | Buparlisib |
| top60_altered_DNA_seed | PIK3CA | 2.971117 | BLCA | J82 | GNE-317 |
| top60_altered_DNA_seed | PIK3CA | -2.548715 | BRCA | OCUB-M | PI3Ka_4409 |
| top60_altered_DNA_seed | PIK3CA | -2.357662 | BRCA | MDA-MB-453 | PI3Ka_4409 |
| top60_altered_DNA_seed | PIK3CA | -2.077815 | BRCA | MDA-MB-453 | AZD8835 |
| top60_altered_DNA_seed | PIK3CA | -2.033158 | BRCA | HCC202 | PI3Ka_4409 |
| top60_altered_DNA_seed | PIK3CA | -2.01571 | BRCA | MDA-MB-361 | PI3Ka_4409 |
| top60_altered_DNA_seed | PIK3CA | -1.993881 | BRCA | MDA-MB-361 | AZD8835 |
| top60_altered_DNA_seed | PIK3CA | -1.923421 | BRCA | HCC1954 | PI3Ka_4409 |
| top60_altered_DNA_seed | PIK3CA | -1.888956 | BRCA | MDA-MB-330 | AZD8835 |
| top60_altered_DNA_seed | PIK3CA | -1.819452 | BRCA | OCUB-M | AZD8835 |
| top60_altered_DNA_seed | PIK3CA | -1.72137 | BRCA | HCC1599 | PI3Ka_4409 |
| top60_altered_DNA_seed | PIK3CA | -1.437506 | BRCA | JIMT-1 | AZD8835 |
| top60_altered_DNA_seed | PIK3CA | -1.311004 | BRCA | EFM-19 | PI3Ka_4409 |
| top60_altered_DNA_seed | PIK3CA | -1.288787 | BRCA | EFM-192A | PI3Ka_4409 |
| top60_altered_DNA_seed | PIK3CA | -1.199455 | BRCA | BT-20 | AZD8835 |
| top60_altered_DNA_seed | PIK3CA | -1.194497 | BRCA | EFM-19 | AZD8835 |
| top60_altered_DNA_seed | PIK3CA | -1.156943 | BRCA | HCC1599 | AZD8835 |
| top60_altered_DNA_seed | PIK3CA | -1.1376 | BRCA | UACC-893 | PI3Ka_4409 |
| top60_altered_DNA_seed | PIK3CA | -1.130069 | BRCA | T47D | PI3Ka_4409 |

| | | | | | |
|---|---|---|---|---|---|
| top60_altered_DNA_seed | PIK3CA | -1.12156 | BRCA | MCF7 | AZD8835 |
| top60_altered_DNA_seed | PIK3CA | -1.072273 | BRCA | JIMT-1 | PI3Ka_4409 |
| top60_altered_DNA_seed | PIK3CA | -0.962684 | BRCA | EVSA-T | PI3Ka_4409 |
| top60_altered_DNA_seed | PIK3CA | -0.937617 | BRCA | HCC1954 | AZD8835 |
| top60_altered_DNA_seed | PIK3CA | -0.895769 | BRCA | BT-20 | PI3Ka_4409 |
| top60_altered_DNA_seed | PIK3CA | -0.8937 | BRCA | BT-474 | PI3Ka_4409 |
| top60_altered_DNA_seed | PIK3CA | -0.852086 | BRCA | UACC-893 | AZD8835 |
| top60_altered_DNA_seed | PIK3CA | -0.832895 | BRCA | MFM-223 | AZD8835 |
| top60_altered_DNA_seed | PIK3CA | -0.787269 | BRCA | MCF7 | PI3Ka_4409 |
| top60_altered_DNA_seed | PIK3CA | -0.782566 | BRCA | EFM-192A | AZD8835 |
| top60_altered_DNA_seed | PIK3CA | -0.778535 | BRCA | BT-474 | AZD8835 |
| top60_altered_DNA_seed | PIK3CA | -0.767928 | BRCA | HCC202 | AZD8835 |
| top60_altered_DNA_seed | PIK3CA | -0.574712 | BRCA | HCC1187 | AZD8835 |
| top60_altered_DNA_seed | PIK3CA | -0.573288 | BRCA | T47D | AZD8835 |
| top60_altered_DNA_seed | PIK3CA | -0.568687 | BRCA | MFM-223 | PI3Ka_4409 |
| top60_altered_DNA_seed | PIK3CA | -0.565849 | BRCA | HCC1187 | PI3Ka_4409 |
| top60_altered_DNA_seed | PIK3CA | -0.453369 | BRCA | MDA-MB-330 | PI3Ka_4409 |
| top60_altered_DNA_seed | PIK3CA | -0.43744 | BRCA | EVSA-T | AZD8835 |
| top60_altered_DNA_seed | PIK3CA | -0.297693 | BRCA | MDA-MB-175-VII | AZD8835 |
| top60_altered_DNA_seed | PIK3CA | -0.24184 | BRCA | AU565 | AZD8835 |
| top60_altered_DNA_seed | PIK3CA | -0.234688 | BRCA | MDA-MB-175-VII | PI3Ka_4409 |
| top60_altered_DNA_seed | PIK3CA | -0.185494 | BRCA | HCC1569 | PI3Ka_4409 |
| top60_altered_DNA_seed | PIK3CA | -0.182922 | BRCA | CAL-51 | PI3Ka_4409 |
| top60_altered_DNA_seed | PIK3CA | -0.171098 | BRCA | CAL-120 | AZD8835 |
| top60_altered_DNA_seed | PIK3CA | -0.14942 | BRCA | CAL-85-1 | PI3Ka_4409 |
| top60_altered_DNA_seed | PIK3CA | -0.149209 | BRCA | AU565 | PI3Ka_4409 |
| top60_altered_DNA_seed | PIK3CA | -0.111097 | BRCA | HCC2218 | PI3Ka_4409 |
| top60_altered_DNA_seed | PIK3CA | -0.057415 | BRCA | HCC70 | PI3Ka_4409 |
| top60_altered_DNA_seed | PIK3CA | -0.021242 | BRCA | HCC1143 | AZD8835 |
| top60_altered_DNA_seed | PIK3CA | 0.017213 | BRCA | HCC2157 | AZD8835 |
| top60_altered_DNA_seed | PIK3CA | 0.04396 | BRCA | HCC1143 | PI3Ka_4409 |
| top60_altered_DNA_seed | PIK3CA | 0.062903 | BRCA | MDA-MB-415 | PI3Ka_4409 |
| top60_altered_DNA_seed | PIK3CA | 0.070522 | BRCA | HCC2157 | PI3Ka_4409 |
| top60_altered_DNA_seed | PIK3CA | 0.077335 | BRCA | COLO-824 | AZD8835 |
| top60_altered_DNA_seed | PIK3CA | 0.089588 | BRCA | BT-549 | PI3Ka_4409 |
| top60_altered_DNA_seed | PIK3CA | 0.096349 | BRCA | HCC1569 | AZD8835 |
| top60_altered_DNA_seed | PIK3CA | 0.140748 | BRCA | COLO-824 | PI3Ka_4409 |
| top60_altered_DNA_seed | PIK3CA | 0.195146 | BRCA | CAL-120 | PI3Ka_4409 |
| top60_altered_DNA_seed | PIK3CA | 0.20947 | BRCA | DU-4475 | AZD8835 |
| top60_altered_DNA_seed | PIK3CA | 0.263913 | BRCA | Hs-578-T | PI3Ka_4409 |
| top60_altered_DNA_seed | PIK3CA | 0.273674 | BRCA | MDA-MB-231 | AZD8835 |

| | | | | | |
|---|---|---|---|---|---|
| top60_altered_DNA_seed | PIK3CA | 0.357184 | BRCA | BT-483 | PI3Ka_4409 |
| top60_altered_DNA_seed | PIK3CA | 0.494257 | BRCA | HDQ-P1 | PI3Ka_4409 |
| top60_altered_DNA_seed | PIK3CA | 0.513118 | BRCA | HCC2218 | AZD8835 |
| top60_altered_DNA_seed | PIK3CA | 0.51515 | BRCA | DU-4475 | PI3Ka_4409 |
| top60_altered_DNA_seed | PIK3CA | 0.551516 | BRCA | HCC1419 | AZD8835 |
| top60_altered_DNA_seed | PIK3CA | 0.586171 | BRCA | CAL-85-1 | AZD8835 |
| top60_altered_DNA_seed | PIK3CA | 0.605227 | BRCA | HDQ-P1 | AZD8835 |
| top60_altered_DNA_seed | PIK3CA | 0.650719 | BRCA | HCC70 | AZD8835 |
| top60_altered_DNA_seed | PIK3CA | 0.674489 | BRCA | BT-549 | AZD8835 |
| top60_altered_DNA_seed | PIK3CA | 0.676875 | BRCA | HCC1419 | PI3Ka_4409 |
| top60_altered_DNA_seed | PIK3CA | 0.754134 | BRCA | MDA-MB-468 | PI3Ka_4409 |
| top60_altered_DNA_seed | PIK3CA | 0.936188 | BRCA | MDA-MB-415 | AZD8835 |
| top60_altered_DNA_seed | PIK3CA | 0.972751 | BRCA | Hs-578-T | AZD8835 |
| top60_altered_DNA_seed | PIK3CA | 0.987493 | BRCA | HCC1395 | AZD8835 |
| top60_altered_DNA_seed | PIK3CA | 0.997241 | BRCA | ZR-75-30 | AZD8835 |
| top60_altered_DNA_seed | PIK3CA | 1.012949 | BRCA | HCC38 | PI3Ka_4409 |
| top60_altered_DNA_seed | PIK3CA | 1.082358 | BRCA | HCC38 | AZD8835 |
| top60_altered_DNA_seed | PIK3CA | 1.15106 | BRCA | BT-483 | AZD8835 |
| top60_altered_DNA_seed | PIK3CA | 1.15663 | BRCA | MDA-MB-468 | AZD8835 |
| top60_altered_DNA_seed | PIK3CA | 1.175306 | BRCA | CAL-51 | AZD8835 |
| top60_altered_DNA_seed | PIK3CA | 1.217205 | BRCA | HCC1937 | PI3Ka_4409 |
| top60_altered_DNA_seed | PIK3CA | 1.260121 | BRCA | MDA-MB-231 | PI3Ka_4409 |
| top60_altered_DNA_seed | PIK3CA | 1.317085 | BRCA | MDA-MB-157 | AZD8835 |
| top60_altered_DNA_seed | PIK3CA | 1.320549 | BRCA | HCC1428 | AZD8835 |
| top60_altered_DNA_seed | PIK3CA | 1.344327 | BRCA | MDA-MB-436 | AZD8835 |
| top60_altered_DNA_seed | PIK3CA | 1.386546 | BRCA | HCC1428 | PI3Ka_4409 |
| top60_altered_DNA_seed | PIK3CA | 1.450804 | BRCA | ZR-75-30 | PI3Ka_4409 |
| top60_altered_DNA_seed | PIK3CA | 1.479792 | BRCA | HCC1937 | AZD8835 |
| top60_altered_DNA_seed | PIK3CA | 1.482499 | BRCA | MDA-MB-157 | PI3Ka_4409 |
| top60_altered_DNA_seed | PIK3CA | 1.682452 | BRCA | HCC1500 | AZD8835 |
| top60_altered_DNA_seed | PIK3CA | 1.886279 | BRCA | MRK-nu-1 | PI3Ka_4409 |
| top60_altered_DNA_seed | PIK3CA | 1.903586 | BRCA | MRK-nu-1 | AZD8835 |
| top60_altered_DNA_seed | PIK3CA | 1.962095 | BRCA | HCC1500 | PI3Ka_4409 |
| top60_altered_DNA_seed | PIK3CA | 2.047064 | BRCA | HCC1395 | PI3Ka_4409 |
| top60_altered_DNA_seed | PIK3CA | 2.180415 | BRCA | CAMA-1 | AZD8835 |
| top60_altered_DNA_seed | PIK3CA | 2.290532 | BRCA | CAMA-1 | PI3Ka_4409 |
| top60_altered_DNA_seed | PIK3CA | 2.332789 | BRCA | MDA-MB-436 | PI3Ka_4409 |
| top60_altered_DNA_seed | PIK3CA | -4.087508 | BRCA | YMB-1-E | AZD8186 |
| top60_altered_DNA_seed | PIK3CA | -3.846124 | BRCA | EVSA-T | AZD8186 |
| top60_altered_DNA_seed | PIK3CA | -3.378294 | BRCA | MDA-MB-415 | AZD8186 |
| top60_altered_DNA_seed | PIK3CA | -2.478395 | BRCA | MDA-MB-361 | Alpelisib |

| | | | | | |
|---|---|---|---|---|---|
| top60_altered_DNA_seed | PIK3CA | -2.467028 | BRCA | MDA-MB-330 | Alpelisib |
| top60_altered_DNA_seed | PIK3CA | -2.349952 | BRCA | OCUB-M | Alpelisib |
| top60_altered_DNA_seed | TP53 | -2.331311 | BRCA | HCC1599 | MIRA-1 |
| top60_altered_DNA_seed | PIK3CA | -2.200339 | BRCA | UACC-893 | Alpelisib |
| top60_altered_DNA_seed | PIK3CA | -2.198675 | BRCA | MDA-MB-453 | Alpelisib |
| top60_altered_DNA_seed | PIK3CA | -2.109527 | BRCA | HCC70 | AZD8186 |
| top60_altered_DNA_seed | TP53 | -2.017852 | BRCA | MRK-nu-1 | MIRA-1 |
| top60_altered_DNA_seed | PIK3CA | -1.979001 | BRCA | HCC2157 | AZD8186 |
| top60_altered_DNA_seed | PIK3CA | -1.88702 | BRCA | MDA-MB-330 | AZD8186 |
| top60_altered_DNA_seed | PIK3CA | -1.882454 | BRCA | CAMA-1 | AZD8186 |
| top60_altered_DNA_seed | PIK3CA | -1.729419 | BRCA | CAL-148 | Alpelisib |
| top60_altered_DNA_seed | PIK3CA | -1.675579 | BRCA | BT-20 | Alpelisib |
| top60_altered_DNA_seed | PIK3CA | -1.673389 | BRCA | OCUB-M | GNE-317 |
| top60_altered_DNA_seed | PIK3CA | -1.607008 | BRCA | EFM-19 | Alpelisib |
| top60_altered_DNA_seed | PIK3CA | -1.544994 | BRCA | MRK-nu-1 | GNE-317 |
| top60_altered_DNA_seed | PIK3CA | -1.452375 | BRCA | MDA-MB-415 | GNE-317 |
| top60_altered_DNA_seed | PIK3CA | -1.424528 | BRCA | HCC2157 | GNE-317 |
| top60_altered_DNA_seed | PIK3CA | -1.401092 | BRCA | HCC1954 | Alpelisib |
| top60_altered_DNA_seed | PIK3CA | -1.397942 | BRCA | MDA-MB-175-VII | GNE-317 |
| top60_altered_DNA_seed | PIK3CA | -1.37047 | BRCA | MDA-MB-361 | GNE-317 |
| top60_altered_DNA_seed | PIK3CA | -1.319474 | BRCA | EVSA-T | GNE-317 |
| top60_altered_DNA_seed | PIK3CA | -1.305722 | BRCA | T47D | Alpelisib |
| top60_altered_DNA_seed | PIK3CA | -1.30157 | BRCA | HDQ-P1 | GNE-317 |
| top60_altered_DNA_seed | PIK3CA | -1.298818 | BRCA | HCC2157 | Buparlisib |
| top60_altered_DNA_seed | PIK3CA | -1.282233 | BRCA | MDA-MB-330 | GNE-317 |
| top60_altered_DNA_seed | PIK3CA | -1.243567 | BRCA | HDQ-P1 | AZD8186 |
| top60_altered_DNA_seed | PIK3CA | -1.240029 | BRCA | EFM-19 | GNE-317 |
| top60_altered_DNA_seed | PIK3CA | -1.236731 | BRCA | CAL-51 | GNE-317 |
| top60_altered_DNA_seed | PIK3CA | -1.223717 | BRCA | EVSA-T | Buparlisib |
| top60_altered_DNA_seed | PIK3CA | -1.217177 | BRCA | BT-474 | Alpelisib |
| top60_altered_DNA_seed | PIK3CA | -1.207284 | BRCA | UACC-893 | GNE-317 |
| top60_altered_DNA_seed | TP53 | -1.185478 | BRCA | HCC2157 | MIRA-1 |
| top60_altered_DNA_seed | TP53 | -1.155661 | BRCA | HCC1419 | MIRA-1 |
| top60_altered_DNA_seed | PIK3CA | -1.128218 | BRCA | MDA-MB-175-VII | Alpelisib |
| top60_altered_DNA_seed | PIK3CA | -1.098943 | BRCA | CAL-148 | GNE-317 |
| top60_altered_DNA_seed | PIK3CA | -1.078618 | BRCA | HCC1395 | AZD8186 |
| top60_altered_DNA_seed | PIK3CA | -1.076969 | BRCA | CAMA-1 | GNE-317 |
| top60_altered_DNA_seed | PIK3CA | -1.074375 | BRCA | T47D | GNE-317 |
| top60_altered_DNA_seed | PIK3CA | -1.073624 | BRCA | HCC1599 | Alpelisib |
| top60_altered_DNA_seed | PIK3CA | -1.026325 | BRCA | CAL-51 | Buparlisib |
| top60_altered_DNA_seed | TP53 | -1.02495 | BRCA | OCUB-M | MIRA-1 |

| | | | | | |
|---|---|---|---|---|---|
| top60_altered_DNA_seed | PIK3CA | -1.022631 | BRCA | EFM-19 | AZD8186 |
| top60_altered_DNA_seed | PIK3CA | -1.019348 | BRCA | MDA-MB-361 | Buparlisib |
| top60_altered_DNA_seed | PIK3CA | -1.018997 | BRCA | MDA-MB-175-VII | Buparlisib |
| top60_altered_DNA_seed | PIK3CA | -1.016139 | BRCA | HCC1569 | Alpelisib |
| top60_altered_DNA_seed | PIK3CA | -0.932406 | BRCA | HDQ-P1 | Buparlisib |
| top60_altered_DNA_seed | PIK3CA | -0.931641 | BRCA | AU565 | Alpelisib |
| top60_altered_DNA_seed | PIK3CA | -0.927964 | BRCA | T47D | AZD8186 |
| top60_altered_DNA_seed | TP53 | -0.920382 | BRCA | MDA-MB-415 | MIRA-1 |
| top60_altered_DNA_seed | PIK3CA | -0.917627 | BRCA | CAL-120 | GNE-317 |
| top60_altered_DNA_seed | PIK3CA | -0.892798 | BRCA | T47D | Buparlisib |
| top60_altered_DNA_seed | PIK3CA | -0.882716 | BRCA | AU565 | AZD8186 |
| top60_altered_DNA_seed | PIK3CA | -0.880401 | BRCA | Hs-578-T | AZD8186 |
| top60_altered_DNA_seed | PIK3CA | -0.853556 | BRCA | MCF7 | Alpelisib |
| top60_altered_DNA_seed | PIK3CA | -0.850261 | BRCA | EFM-192A | Alpelisib |
| top60_altered_DNA_seed | PIK3CA | -0.840747 | BRCA | MDA-MB-361 | AZD8186 |
| top60_altered_DNA_seed | PIK3CA | -0.819784 | BRCA | COLO-824 | AZD8186 |
| top60_altered_DNA_seed | PIK3CA | -0.809161 | BRCA | MRK-nu-1 | Buparlisib |
| top60_altered_DNA_seed | PIK3CA | -0.772239 | BRCA | OCUB-M | Buparlisib |
| top60_altered_DNA_seed | PIK3CA | -0.745503 | BRCA | HCC1143 | AZD8186 |
| top60_altered_DNA_seed | PIK3CA | -0.743736 | BRCA | Hs-578-T | Buparlisib |
| top60_altered_DNA_seed | PIK3CA | -0.741553 | BRCA | MDA-MB-468 | AZD8186 |
| top60_altered_DNA_seed | PIK3CA | -0.73116 | BRCA | JIMT-1 | GNE-317 |
| top60_altered_DNA_seed | PIK3CA | -0.730409 | BRCA | Hs-578-T | Alpelisib |
| top60_altered_DNA_seed | PIK3CA | -0.704781 | BRCA | CAL-148 | AZD8186 |
| top60_altered_DNA_seed | PIK3CA | -0.702855 | BRCA | HCC1395 | Buparlisib |
| top60_altered_DNA_seed | PIK3CA | -0.691065 | BRCA | EFM-192A | GNE-317 |
| top60_altered_DNA_seed | PIK3CA | -0.67704 | BRCA | UACC-812 | AZD8186 |
| top60_altered_DNA_seed | PIK3CA | -0.660445 | BRCA | EFM-192A | Buparlisib |
| top60_altered_DNA_seed | PIK3CA | -0.653864 | BRCA | AU565 | GNE-317 |
| top60_altered_DNA_seed | PIK3CA | -0.652526 | BRCA | MDA-MB-330 | Buparlisib |
| top60_altered_DNA_seed | PIK3CA | -0.652473 | BRCA | UACC-812 | Alpelisib |
| top60_altered_DNA_seed | TP53 | -0.647937 | BRCA | EVSA-T | MIRA-1 |
| top60_altered_DNA_seed | PIK3CA | -0.607739 | BRCA | Hs-578-T | GNE-317 |
| top60_altered_DNA_seed | PIK3CA | -0.601721 | BRCA | COLO-824 | Alpelisib |
| top60_altered_DNA_seed | PIK3CA | -0.596498 | BRCA | DU-4475 | Buparlisib |
| top60_altered_DNA_seed | PIK3CA | -0.595752 | BRCA | HCC1143 | Buparlisib |
| top60_altered_DNA_seed | PIK3CA | -0.592111 | BRCA | MDA-MB-175-VII | AZD8186 |
| top60_altered_DNA_seed | PIK3CA | -0.57039 | BRCA | HCC1569 | AZD8186 |
| top60_altered_DNA_seed | PIK3CA | -0.568248 | BRCA | CAL-148 | Buparlisib |
| top60_altered_DNA_seed | TP53 | -0.527752 | BRCA | HCC70 | MIRA-1 |
| top60_altered_DNA_seed | PIK3CA | -0.515105 | BRCA | AU565 | Buparlisib |

| | | | | | |
|---|---|---|---|---|---|
| top60_altered_DNA_seed | PIK3CA | -0.514386 | BRCA | HCC1143 | GNE-317 |
| top60_altered_DNA_seed | PIK3CA | -0.503639 | BRCA | HCC70 | GNE-317 |
| top60_altered_DNA_seed | TP53 | -0.499832 | BRCA | CAL-148 | MIRA-1 |
| top60_altered_DNA_seed | PIK3CA | -0.486361 | BRCA | HCC1954 | Buparlisib |
| top60_altered_DNA_seed | PIK3CA | -0.464579 | BRCA | MDA-MB-231 | GNE-317 |
| top60_altered_DNA_seed | TP53 | -0.454454 | BRCA | DU-4475 | MIRA-1 |
| top60_altered_DNA_seed | PIK3CA | -0.441574 | BRCA | COLO-824 | Buparlisib |
| top60_altered_DNA_seed | PIK3CA | -0.441497 | BRCA | HCC1143 | Alpelisib |
| top60_altered_DNA_seed | PIK3CA | -0.438056 | BRCA | HCC1954 | GNE-317 |
| top60_altered_DNA_seed | PIK3CA | -0.43752 | BRCA | MCF7 | Buparlisib |
| top60_altered_DNA_seed | PIK3CA | -0.436057 | BRCA | HDQ-P1 | Alpelisib |
| top60_altered_DNA_seed | PIK3CA | -0.413514 | BRCA | EFM-192A | AZD8186 |
| top60_altered_DNA_seed | PIK3CA | -0.410153 | BRCA | YMB-1-E | GNE-317 |
| top60_altered_DNA_seed | PIK3CA | -0.402635 | BRCA | EFM-19 | Buparlisib |
| top60_altered_DNA_seed | TP53 | -0.401514 | BRCA | AU565 | MIRA-1 |
| top60_altered_DNA_seed | PIK3CA | -0.372092 | BRCA | ZR-75-30 | Alpelisib |
| top60_altered_DNA_seed | PIK3CA | -0.37003 | BRCA | HCC1599 | Buparlisib |
| top60_altered_DNA_seed | PIK3CA | -0.332338 | BRCA | HCC70 | Buparlisib |
| top60_altered_DNA_seed | PIK3CA | -0.32986 | BRCA | MFM-223 | Alpelisib |
| top60_altered_DNA_seed | PIK3CA | -0.328263 | BRCA | CAL-85-1 | GNE-317 |
| top60_altered_DNA_seed | PIK3CA | -0.321277 | BRCA | MDA-MB-453 | Buparlisib |
| top60_altered_DNA_seed | PIK3CA | -0.320262 | BRCA | BT-20 | Buparlisib |
| top60_altered_DNA_seed | PIK3CA | -0.312548 | BRCA | BT-20 | GNE-317 |
| top60_altered_DNA_seed | PIK3CA | -0.261162 | BRCA | JIMT-1 | Alpelisib |
| top60_altered_DNA_seed | PIK3CA | -0.238073 | BRCA | CAL-85-1 | Alpelisib |
| top60_altered_DNA_seed | PIK3CA | -0.212837 | BRCA | HCC1937 | AZD8186 |
| top60_altered_DNA_seed | PIK3CA | -0.208771 | BRCA | MFM-223 | GNE-317 |
| top60_altered_DNA_seed | PIK3CA | -0.208619 | BRCA | HCC1569 | GNE-317 |
| top60_altered_DNA_seed | TP53 | -0.205086 | BRCA | HCC1187 | MIRA-1 |
| top60_altered_DNA_seed | TP53 | -0.201608 | BRCA | HCC1500 | MIRA-1 |
| top60_altered_DNA_seed | PIK3CA | -0.175846 | BRCA | UACC-812 | GNE-317 |
| top60_altered_DNA_seed | PIK3CA | -0.175299 | BRCA | UACC-893 | AZD8186 |
| top60_altered_DNA_seed | PIK3CA | -0.168908 | BRCA | MDA-MB-453 | AZD8186 |
| top60_altered_DNA_seed | TP53 | -0.167296 | BRCA | BT-20 | MIRA-1 |
| top60_altered_DNA_seed | TP53 | -0.164861 | BRCA | MDA-MB-453 | MIRA-1 |
| top60_altered_DNA_seed | PIK3CA | -0.147878 | BRCA | MDA-MB-468 | GNE-317 |
| top60_altered_DNA_seed | PIK3CA | -0.142764 | BRCA | CAL-51 | Alpelisib |
| top60_altered_DNA_seed | PIK3CA | -0.132617 | BRCA | JIMT-1 | Buparlisib |
| top60_altered_DNA_seed | PIK3CA | -0.13028 | BRCA | YMB-1-E | Buparlisib |
| top60_altered_DNA_seed | TP53 | -0.128637 | BRCA | HCC2218 | MIRA-1 |
| top60_altered_DNA_seed | PIK3CA | -0.122867 | BRCA | HCC1187 | AZD8186 |

| | | | | | |
|---|---|---|---|---|---|
| top60_altered_DNA_seed | PIK3CA | -0.105804 | BRCA | MDA-MB-453 | GNE-317 |
| top60_altered_DNA_seed | PIK3CA | -0.096342 | BRCA | HCC2218 | Alpelisib |
| top60_altered_DNA_seed | PIK3CA | -0.09236 | BRCA | HCC70 | Alpelisib |
| top60_altered_DNA_seed | PIK3CA | -0.086838 | BRCA | DU-4475 | GNE-317 |
| top60_altered_DNA_seed | PIK3CA | -0.077314 | BRCA | HCC2157 | Alpelisib |
| top60_altered_DNA_seed | PIK3CA | -0.064745 | BRCA | MDA-MB-415 | Buparlisib |
| top60_altered_DNA_seed | PIK3CA | -0.014331 | BRCA | BT-549 | GNE-317 |
| top60_altered_DNA_seed | PIK3CA | -0.006998 | BRCA | MCF7 | AZD8186 |
| top60_altered_DNA_seed | PIK3CA | -0.003606 | BRCA | COLO-824 | GNE-317 |
| top60_altered_DNA_seed | PIK3CA | -0.003436 | BRCA | EVSA-T | Alpelisib |
| top60_altered_DNA_seed | TP53 | 0.010507 | BRCA | MDA-MB-330 | MIRA-1 |
| top60_altered_DNA_seed | PIK3CA | 0.031282 | BRCA | HCC1954 | AZD8186 |
| top60_altered_DNA_seed | PIK3CA | 0.044886 | BRCA | BT-549 | AZD8186 |
| top60_altered_DNA_seed | PIK3CA | 0.053041 | BRCA | CAMA-1 | Buparlisib |
| top60_altered_DNA_seed | PIK3CA | 0.05419 | BRCA | MDA-MB-468 | Buparlisib |
| top60_altered_DNA_seed | TP53 | 0.067588 | BRCA | CAL-85-1 | MIRA-1 |
| top60_altered_DNA_seed | PIK3CA | 0.072001 | BRCA | HCC1395 | Alpelisib |
| top60_altered_DNA_seed | TP53 | 0.079871 | BRCA | CAMA-1 | MIRA-1 |
| top60_altered_DNA_seed | TP53 | 0.09638 | BRCA | HDQ-P1 | MIRA-1 |
| top60_altered_DNA_seed | PIK3CA | 0.103107 | BRCA | UACC-893 | Buparlisib |
| top60_altered_DNA_seed | TP53 | 0.112399 | BRCA | HCC1569 | MIRA-1 |
| top60_altered_DNA_seed | TP53 | 0.124896 | BRCA | Hs-578-T | MIRA-1 |
| top60_altered_DNA_seed | TP53 | 0.129468 | BRCA | JIMT-1 | MIRA-1 |
| top60_altered_DNA_seed | TP53 | 0.15232 | BRCA | MDA-MB-231 | MIRA-1 |
| top60_altered_DNA_seed | PIK3CA | 0.163942 | BRCA | CAL-51 | AZD8186 |
| top60_altered_DNA_seed | PIK3CA | 0.19012 | BRCA | MDA-MB-436 | AZD8186 |
| top60_altered_DNA_seed | PIK3CA | 0.213926 | BRCA | MDA-MB-157 | Buparlisib |
| top60_altered_DNA_seed | TP53 | 0.221206 | BRCA | UACC-812 | MIRA-1 |
| top60_altered_DNA_seed | PIK3CA | 0.227552 | BRCA | CAL-85-1 | Buparlisib |
| top60_altered_DNA_seed | TP53 | 0.238802 | BRCA | HCC1954 | MIRA-1 |
| top60_altered_DNA_seed | TP53 | 0.246462 | BRCA | MDA-MB-361 | MIRA-1 |
| top60_altered_DNA_seed | PIK3CA | 0.268441 | BRCA | YMB-1-E | Alpelisib |
| top60_altered_DNA_seed | PIK3CA | 0.273017 | BRCA | BT-483 | Alpelisib |
| top60_altered_DNA_seed | PIK3CA | 0.274275 | BRCA | CAL-120 | Buparlisib |
| top60_altered_DNA_seed | PIK3CA | 0.280119 | BRCA | DU-4475 | Alpelisib |
| top60_altered_DNA_seed | PIK3CA | 0.286683 | BRCA | BT-474 | AZD8186 |
| top60_altered_DNA_seed | TP53 | 0.294139 | BRCA | MDA-MB-157 | MIRA-1 |
| top60_altered_DNA_seed | TP53 | 0.311105 | BRCA | UACC-893 | MIRA-1 |
| top60_altered_DNA_seed | PIK3CA | 0.315274 | BRCA | DU-4475 | AZD8186 |
| top60_altered_DNA_seed | PIK3CA | 0.319258 | BRCA | HCC1395 | GNE-317 |
| top60_altered_DNA_seed | PIK3CA | 0.342549 | BRCA | HCC1599 | GNE-317 |

| | | | | | |
|---|---|---|---|---|---|
| top60_altered_DNA_seed | PIK3CA | 0.349377 | BRCA | HCC38 | GNE-317 |
| top60_altered_DNA_seed | PIK3CA | 0.358545 | BRCA | HCC2218 | AZD8186 |
| top60_altered_DNA_seed | PIK3CA | 0.373971 | BRCA | BT-20 | AZD8186 |
| top60_altered_DNA_seed | TP53 | 0.389817 | BRCA | CAL-51 | MIRA-1 |
| top60_altered_DNA_seed | TP53 | 0.393963 | BRCA | YMB-1-E | MIRA-1 |
| top60_altered_DNA_seed | PIK3CA | 0.396797 | BRCA | OCUB-M | AZD8186 |
| top60_altered_DNA_seed | PIK3CA | 0.405435 | BRCA | UACC-812 | Buparlisib |
| top60_altered_DNA_seed | PIK3CA | 0.422393 | BRCA | HCC1419 | AZD8186 |
| top60_altered_DNA_seed | TP53 | 0.453655 | BRCA | MDA-MB-468 | MIRA-1 |
| top60_altered_DNA_seed | PIK3CA | 0.459333 | BRCA | BT-549 | Buparlisib |
| top60_altered_DNA_seed | PIK3CA | 0.4689 | BRCA | MDA-MB-415 | Alpelisib |
| top60_altered_DNA_seed | PIK3CA | 0.487822 | BRCA | CAL-85-1 | AZD8186 |
| top60_altered_DNA_seed | TP53 | 0.499196 | BRCA | T47D | MIRA-1 |
| top60_altered_DNA_seed | PIK3CA | 0.501662 | BRCA | HCC1937 | Alpelisib |
| top60_altered_DNA_seed | PIK3CA | 0.522996 | BRCA | MDA-MB-157 | GNE-317 |
| top60_altered_DNA_seed | PIK3CA | 0.530461 | BRCA | HCC38 | Alpelisib |
| top60_altered_DNA_seed | PIK3CA | 0.533957 | BRCA | HCC1419 | Alpelisib |
| top60_altered_DNA_seed | PIK3CA | 0.535915 | BRCA | HCC38 | AZD8186 |
| top60_altered_DNA_seed | PIK3CA | 0.540628 | BRCA | JIMT-1 | AZD8186 |
| top60_altered_DNA_seed | PIK3CA | 0.541288 | BRCA | BT-549 | Alpelisib |
| top60_altered_DNA_seed | TP53 | 0.557608 | BRCA | ZR-75-30 | MIRA-1 |
| top60_altered_DNA_seed | PIK3CA | 0.561574 | BRCA | MFM-223 | Buparlisib |
| top60_altered_DNA_seed | PIK3CA | 0.56285 | BRCA | MDA-MB-231 | Buparlisib |
| top60_altered_DNA_seed | TP53 | 0.597125 | BRCA | COLO-824 | MIRA-1 |
| top60_altered_DNA_seed | PIK3CA | 0.60026 | BRCA | ZR-75-30 | AZD8186 |
| top60_altered_DNA_seed | PIK3CA | 0.600468 | BRCA | MCF7 | GNE-317 |
| top60_altered_DNA_seed | PIK3CA | 0.604254 | BRCA | MDA-MB-157 | Alpelisib |
| top60_altered_DNA_seed | PIK3CA | 0.611107 | BRCA | CAMA-1 | Alpelisib |
| top60_altered_DNA_seed | TP53 | 0.612158 | BRCA | EFM-19 | MIRA-1 |
| top60_altered_DNA_seed | PIK3CA | 0.618825 | BRCA | HCC1187 | Buparlisib |
| top60_altered_DNA_seed | TP53 | 0.671396 | BRCA | EFM-192A | MIRA-1 |
| top60_altered_DNA_seed | PIK3CA | 0.678447 | BRCA | MDA-MB-231 | Alpelisib |
| top60_altered_DNA_seed | PIK3CA | 0.685429 | BRCA | MRK-nu-1 | Alpelisib |
| top60_altered_DNA_seed | PIK3CA | 0.706951 | BRCA | MDA-MB-468 | Alpelisib |
| top60_altered_DNA_seed | TP53 | 0.756287 | BRCA | HCC38 | MIRA-1 |
| top60_altered_DNA_seed | TP53 | 0.759119 | BRCA | HCC1395 | MIRA-1 |
| top60_altered_DNA_seed | PIK3CA | 0.776656 | BRCA | HCC1937 | GNE-317 |
| top60_altered_DNA_seed | PIK3CA | 0.776945 | BRCA | HCC1187 | Alpelisib |
| top60_altered_DNA_seed | PIK3CA | 0.857775 | BRCA | HCC2218 | Buparlisib |
| top60_altered_DNA_seed | TP53 | 0.860765 | BRCA | MCF7 | MIRA-1 |
| top60_altered_DNA_seed | PIK3CA | 0.8721 | BRCA | HCC1599 | AZD8186 |

| | | | | | |
|---|---|---|---|---|---|
| top60_altered_DNA_seed | PIK3CA | 0.901775 | BRCA | BT-474 | GNE-317 |
| top60_altered_DNA_seed | PIK3CA | 0.911682 | BRCA | MRK-nu-1 | AZD8186 |
| top60_altered_DNA_seed | PIK3CA | 0.919742 | BRCA | HCC1569 | Buparlisib |
| top60_altered_DNA_seed | PIK3CA | 0.9207 | BRCA | MFM-223 | AZD8186 |
| top60_altered_DNA_seed | PIK3CA | 0.951276 | BRCA | MDA-MB-436 | Alpelisib |
| top60_altered_DNA_seed | TP53 | 1.036629 | BRCA | HCC1937 | MIRA-1 |
| top60_altered_DNA_seed | PIK3CA | 1.040492 | BRCA | HCC38 | Buparlisib |
| top60_altered_DNA_seed | PIK3CA | 1.045803 | BRCA | MDA-MB-231 | AZD8186 |
| top60_altered_DNA_seed | PIK3CA | 1.071977 | BRCA | MDA-MB-436 | GNE-317 |
| top60_altered_DNA_seed | PIK3CA | 1.098639 | BRCA | HCC1419 | Buparlisib |
| top60_altered_DNA_seed | TP53 | 1.099032 | BRCA | BT-549 | MIRA-1 |
| top60_altered_DNA_seed | PIK3CA | 1.099167 | BRCA | BT-474 | Buparlisib |
| top60_altered_DNA_seed | PIK3CA | 1.100603 | BRCA | ZR-75-30 | GNE-317 |
| top60_altered_DNA_seed | PIK3CA | 1.140993 | BRCA | HCC1937 | Buparlisib |
| top60_altered_DNA_seed | TP53 | 1.142254 | BRCA | HCC1143 | MIRA-1 |
| top60_altered_DNA_seed | PIK3CA | 1.149817 | BRCA | HCC1500 | Alpelisib |
| top60_altered_DNA_seed | PIK3CA | 1.15996 | BRCA | MDA-MB-157 | AZD8186 |
| top60_altered_DNA_seed | PIK3CA | 1.160204 | BRCA | MDA-MB-436 | Buparlisib |
| top60_altered_DNA_seed | TP53 | 1.167902 | BRCA | MFM-223 | MIRA-1 |
| top60_altered_DNA_seed | TP53 | 1.204967 | BRCA | MDA-MB-436 | MIRA-1 |
| top60_altered_DNA_seed | TP53 | 1.22708 | BRCA | CAL-120 | MIRA-1 |
| top60_altered_DNA_seed | PIK3CA | 1.246289 | BRCA | BT-483 | GNE-317 |
| top60_altered_DNA_seed | PIK3CA | 1.275595 | BRCA | CAL-120 | AZD8186 |
| top60_altered_DNA_seed | TP53 | 1.29189 | BRCA | BT-474 | MIRA-1 |
| top60_altered_DNA_seed | TP53 | 1.444911 | BRCA | HCC1428 | MIRA-1 |
| top60_altered_DNA_seed | PIK3CA | 1.481992 | BRCA | HCC1500 | Buparlisib |
| top60_altered_DNA_seed | PIK3CA | 1.553813 | BRCA | HCC1187 | GNE-317 |
| top60_altered_DNA_seed | PIK3CA | 1.582609 | BRCA | HCC1419 | GNE-317 |
| top60_altered_DNA_seed | PIK3CA | 1.601177 | BRCA | HCC1500 | AZD8186 |
| top60_altered_DNA_seed | PIK3CA | 1.70168 | BRCA | ZR-75-30 | Buparlisib |
| top60_altered_DNA_seed | TP53 | 1.829054 | BRCA | MDA-MB-175-VII | MIRA-1 |
| top60_altered_DNA_seed | TP53 | 1.908261 | BRCA | BT-483 | MIRA-1 |
| top60_altered_DNA_seed | PIK3CA | 1.92054 | BRCA | HCC1500 | GNE-317 |
| top60_altered_DNA_seed | PIK3CA | 2.052789 | BRCA | CAL-120 | Alpelisib |
| top60_altered_DNA_seed | PIK3CA | 2.131534 | BRCA | BT-483 | Buparlisib |
| top60_altered_DNA_seed | PIK3CA | 2.322717 | BRCA | HCC1428 | GNE-317 |
| top60_altered_DNA_seed | PIK3CA | 2.57169 | BRCA | HCC1428 | Buparlisib |
| top60_altered_DNA_seed | PIK3CA | 2.676332 | BRCA | HCC1428 | AZD8186 |
| top60_altered_DNA_seed | PIK3CA | 2.896574 | BRCA | HCC1428 | Alpelisib |
| top60_altered_DNA_seed | PIK3CA | 2.896804 | BRCA | BT-483 | AZD8186 |
| top60_altered_DNA_seed | PIK3CA | -2.885395 | COADREAD | SNU-407 | AZD8835 |

| | | | | | |
|---|---|---|---|---|---|
| top60_altered_DNA_seed | ATM | -2.26219 | COADREAD | LoVo | KU-55933 |
| top60_altered_DNA_seed | PIK3CA | -1.570969 | COADREAD | CaR-1 | PI3Ka_4409 |
| top60_altered_DNA_seed | ATM | -1.540048 | COADREAD | LS-411N | KU-60019 |
| top60_altered_DNA_seed | PIK3CA | -1.527487 | COADREAD | T84 | AZD8835 |
| top60_altered_DNA_seed | PIK3CA | -1.479836 | COADREAD | SW48 | AZD8835 |
| top60_altered_DNA_seed | ATM | -1.289227 | COADREAD | HCT-116 | KU-60019 |
| top60_altered_DNA_seed | ATM | -1.191718 | COADREAD | MDST8 | torin2 |
| top60_altered_DNA_seed | PIK3CA | -1.178649 | COADREAD | SK-CO-1 | AZD8835 |
| top60_altered_DNA_seed | ATM | -1.134419 | COADREAD | NCI-H716 | KU-60019 |
| top60_altered_DNA_seed | ATM | -1.124209 | COADREAD | SNU-175 | CP466722 |
| top60_altered_DNA_seed | PIK3CA | -1.086516 | COADREAD | GP5d | AZD8835 |
| top60_altered_DNA_seed | PIK3CA | -1.084276 | COADREAD | DiFi | PI3Ka_4409 |
| top60_altered_DNA_seed | PIK3CA | -1.081583 | COADREAD | SNU-407 | PI3Ka_4409 |
| top60_altered_DNA_seed | PIK3CA | -1.060381 | COADREAD | NCI-H747 | AZD8835 |
| top60_altered_DNA_seed | ATM | -1.042063 | COADREAD | NCI-H716 | KU-55933 |
| top60_altered_DNA_seed | PIK3CA | -1.013241 | COADREAD | HCT-116 | PI3Ka_4409 |
| top60_altered_DNA_seed | ATM | -1.012839 | COADREAD | SNU-C5 | KU-55933 |
| top60_altered_DNA_seed | PIK3CA | -0.984215 | COADREAD | GP5d | PI3Ka_4409 |
| top60_altered_DNA_seed | ATM | -0.98135 | COADREAD | KM12 | KU-60019 |
| top60_altered_DNA_seed | PIK3CA | -0.928663 | COADREAD | HCT-116 | AZD8835 |
| top60_altered_DNA_seed | ATM | -0.890736 | COADREAD | LoVo | KU-60019 |
| top60_altered_DNA_seed | PIK3CA | -0.869761 | COADREAD | RKO | AZD8835 |
| top60_altered_DNA_seed | PIK3CA | -0.844898 | COADREAD | SNU-C1 | AZD8835 |
| top60_altered_DNA_seed | PIK3CA | -0.843751 | COADREAD | SW48 | PI3Ka_4409 |
| top60_altered_DNA_seed | ATM | -0.825948 | COADREAD | CL-11 | KU-60019 |
| top60_altered_DNA_seed | ATM | -0.790348 | COADREAD | HCT-116 | torin2 |
| top60_altered_DNA_seed | ATM | -0.790196 | COADREAD | RKO | KU-60019 |
| top60_altered_DNA_seed | ATM | -0.776531 | COADREAD | SNU-81 | CP466722 |
| top60_altered_DNA_seed | PIK3CA | -0.72489 | COADREAD | COLO-320-HSR | AZD8835 |
| top60_altered_DNA_seed | PIK3CA | -0.715688 | COADREAD | RKO | PI3Ka_4409 |
| top60_altered_DNA_seed | ATM | -0.709043 | COADREAD | HCT-116 | CP466722 |
| top60_altered_DNA_seed | ATM | -0.698908 | COADREAD | LS-180 | CP466722 |
| top60_altered_DNA_seed | PIK3CA | -0.694177 | COADREAD | COLO-205 | PI3Ka_4409 |
| top60_altered_DNA_seed | ATM | -0.673183 | COADREAD | SNU-81 | KU-55933 |
| top60_altered_DNA_seed | ATM | -0.635834 | COADREAD | CW-2 | KU-60019 |
| top60_altered_DNA_seed | PIK3CA | -0.627517 | COADREAD | DiFi | AZD8835 |
| top60_altered_DNA_seed | PIK3CA | -0.624684 | COADREAD | SNU-C5 | AZD8835 |
| top60_altered_DNA_seed | ATM | -0.61809 | COADREAD | NCI-H747 | CP466722 |
| top60_altered_DNA_seed | PIK3CA | -0.61323 | COADREAD | SK-CO-1 | PI3Ka_4409 |
| top60_altered_DNA_seed | ATM | -0.587908 | COADREAD | CCK-81 | KU-60019 |
| top60_altered_DNA_seed | PIK3CA | -0.586716 | COADREAD | SNU-C5 | PI3Ka_4409 |

| | | | | | |
|---|---|---|---|---|---|
| top60_altered_DNA_seed | ATM | -0.584066 | COADREAD | COLO-320-HSR | KU-60019 |
| top60_altered_DNA_seed | PIK3CA | -0.572921 | COADREAD | SNU-C1 | PI3Ka_4409 |
| top60_altered_DNA_seed | PIK3CA | -0.542571 | COADREAD | LoVo | AZD8835 |
| top60_altered_DNA_seed | PIK3CA | -0.539297 | COADREAD | NCI-H747 | PI3Ka_4409 |
| top60_altered_DNA_seed | PIK3CA | -0.534642 | COADREAD | HT-29 | AZD8835 |
| top60_altered_DNA_seed | PIK3CA | -0.521684 | COADREAD | NCI-H716 | PI3Ka_4409 |
| top60_altered_DNA_seed | ATM | -0.519922 | COADREAD | CL-11 | KU-55933 |
| top60_altered_DNA_seed | ATM | -0.518202 | COADREAD | SK-CO-1 | KU-55933 |
| top60_altered_DNA_seed | ATM | -0.511225 | COADREAD | T84 | CP466722 |
| top60_altered_DNA_seed | ATM | -0.510789 | COADREAD | SNU-1040 | CP466722 |
| top60_altered_DNA_seed | PIK3CA | -0.474138 | COADREAD | SW948 | PI3Ka_4409 |
| top60_altered_DNA_seed | ATM | -0.463427 | COADREAD | COLO-320-HSR | CP466722 |
| top60_altered_DNA_seed | ATM | -0.461765 | COADREAD | CCK-81 | CP466722 |
| top60_altered_DNA_seed | PIK3CA | -0.433547 | COADREAD | NCI-H508 | PI3Ka_4409 |
| top60_altered_DNA_seed | ATM | -0.432508 | COADREAD | HCT-15 | CP466722 |
| top60_altered_DNA_seed | ATM | -0.432033 | COADREAD | SNU-81 | KU-60019 |
| top60_altered_DNA_seed | ATM | -0.414445 | COADREAD | KM12 | CP466722 |
| top60_altered_DNA_seed | ATM | -0.38166 | COADREAD | COLO-205 | CP466722 |
| top60_altered_DNA_seed | PIK3CA | -0.36886 | COADREAD | LS-513 | PI3Ka_4409 |
| top60_altered_DNA_seed | ATM | -0.360689 | COADREAD | HCT-15 | KU-60019 |
| top60_altered_DNA_seed | ATM | -0.353132 | COADREAD | RKO | CP466722 |
| top60_altered_DNA_seed | ATM | -0.353111 | COADREAD | SW620 | CP466722 |
| top60_altered_DNA_seed | PIK3CA | -0.347499 | COADREAD | COLO-320-HSR | PI3Ka_4409 |
| top60_altered_DNA_seed | PIK3CA | -0.344309 | COADREAD | NCI-H716 | AZD8835 |
| top60_altered_DNA_seed | PIK3CA | -0.336371 | COADREAD | SW1463 | PI3Ka_4409 |
| top60_altered_DNA_seed | PIK3CA | -0.319794 | COADREAD | COLO-205 | AZD8835 |
| top60_altered_DNA_seed | PIK3CA | -0.318476 | COADREAD | MDST8 | AZD8835 |
| top60_altered_DNA_seed | ATM | -0.283284 | COADREAD | SW48 | KU-55933 |
| top60_altered_DNA_seed | ATM | -0.277802 | COADREAD | MDST8 | KU-60019 |
| top60_altered_DNA_seed | ATM | -0.273401 | COADREAD | SK-CO-1 | KU-60019 |
| top60_altered_DNA_seed | PIK3CA | -0.2684 | COADREAD | CCK-81 | AZD8835 |
| top60_altered_DNA_seed | ATM | -0.243581 | COADREAD | LS-411N | CP466722 |
| top60_altered_DNA_seed | PIK3CA | -0.191277 | COADREAD | LS-180 | AZD8835 |
| top60_altered_DNA_seed | PIK3CA | -0.182657 | COADREAD | LoVo | PI3Ka_4409 |
| top60_altered_DNA_seed | ATM | -0.182176 | COADREAD | SW837 | CP466722 |
| top60_altered_DNA_seed | ATM | -0.173003 | COADREAD | LoVo | torin2 |
| top60_altered_DNA_seed | PIK3CA | -0.168674 | COADREAD | HCT-15 | AZD8835 |
| top60_altered_DNA_seed | ATM | -0.151994 | COADREAD | HCT-15 | KU-55933 |
| top60_altered_DNA_seed | PIK3CA | -0.130352 | COADREAD | T84 | PI3Ka_4409 |
| top60_altered_DNA_seed | PIK3CA | -0.118668 | COADREAD | NCI-H508 | AZD8835 |
| top60_altered_DNA_seed | PIK3CA | -0.113655 | COADREAD | HT-115 | PI3Ka_4409 |

| | | | | | |
|---|---|---|---|---|---|
| top60_altered_DNA_seed | PIK3CA | -0.099566 | COADREAD | LS-513 | AZD8835 |
| top60_altered_DNA_seed | PIK3CA | -0.092356 | COADREAD | HT-115 | AZD8835 |
| top60_altered_DNA_seed | PIK3CA | -0.086547 | COADREAD | LS-180 | PI3Ka_4409 |
| top60_altered_DNA_seed | PIK3CA | -0.077205 | COADREAD | CCK-81 | PI3Ka_4409 |
| top60_altered_DNA_seed | ATM | -0.067091 | COADREAD | SNU-407 | CP466722 |
| top60_altered_DNA_seed | ATM | -0.057984 | COADREAD | SW48 | torin2 |
| top60_altered_DNA_seed | ATM | -0.037897 | COADREAD | LS-180 | KU-55933 |
| top60_altered_DNA_seed | ATM | -0.035626 | COADREAD | GP5d | CP466722 |
| top60_altered_DNA_seed | ATM | -0.012139 | COADREAD | RKO | KU-55933 |
| top60_altered_DNA_seed | PIK3CA | 0.022367 | COADREAD | SNU-81 | AZD8835 |
| top60_altered_DNA_seed | PIK3CA | 0.028046 | COADREAD | CL-11 | PI3Ka_4409 |
| top60_altered_DNA_seed | PIK3CA | 0.037754 | COADREAD | LS-123 | PI3Ka_4409 |
| top60_altered_DNA_seed | PIK3CA | 0.038105 | COADREAD | SW837 | PI3Ka_4409 |
| top60_altered_DNA_seed | PIK3CA | 0.039911 | COADREAD | SW948 | AZD8835 |
| top60_altered_DNA_seed | PIK3CA | 0.057106 | COADREAD | HT-29 | PI3Ka_4409 |
| top60_altered_DNA_seed | ATM | 0.062083 | COADREAD | SW620 | KU-60019 |
| top60_altered_DNA_seed | PIK3CA | 0.064424 | COADREAD | LS-1034 | PI3Ka_4409 |
| top60_altered_DNA_seed | PIK3CA | 0.078169 | COADREAD | SNU-1040 | PI3Ka_4409 |
| top60_altered_DNA_seed | ATM | 0.079666 | COADREAD | HT-29 | CP466722 |
| top60_altered_DNA_seed | ATM | 0.083235 | COADREAD | C2BBe1 | KU-60019 |
| top60_altered_DNA_seed | ATM | 0.090307 | COADREAD | CL-11 | torin2 |
| top60_altered_DNA_seed | ATM | 0.114673 | COADREAD | GP5d | torin2 |
| top60_altered_DNA_seed | ATM | 0.125947 | COADREAD | HT-115 | torin2 |
| top60_altered_DNA_seed | PIK3CA | 0.148652 | COADREAD | LS-1034 | AZD8835 |
| top60_altered_DNA_seed | PIK3CA | 0.151439 | COADREAD | HT55 | PI3Ka_4409 |
| top60_altered_DNA_seed | PIK3CA | 0.165612 | COADREAD | CL-11 | AZD8835 |
| top60_altered_DNA_seed | ATM | 0.166745 | COADREAD | HCC2998 | CP466722 |
| top60_altered_DNA_seed | ATM | 0.191918 | COADREAD | HCT-116 | KU-55933 |
| top60_altered_DNA_seed | PIK3CA | 0.200779 | COADREAD | LS-123 | AZD8835 |
| top60_altered_DNA_seed | ATM | 0.220984 | COADREAD | CaR-1 | KU-55933 |
| top60_altered_DNA_seed | ATM | 0.232157 | COADREAD | LS-123 | CP466722 |
| top60_altered_DNA_seed | PIK3CA | 0.265577 | COADREAD | HT55 | AZD8835 |
| top60_altered_DNA_seed | ATM | 0.267102 | COADREAD | COLO-320-HSR | KU-55933 |
| top60_altered_DNA_seed | PIK3CA | 0.271829 | COADREAD | SW1463 | AZD8835 |
| top60_altered_DNA_seed | ATM | 0.279929 | COADREAD | SNU-61 | CP466722 |
| top60_altered_DNA_seed | PIK3CA | 0.286264 | COADREAD | SNU-C2B | PI3Ka_4409 |
| top60_altered_DNA_seed | ATM | 0.287877 | COADREAD | GP5d | KU-60019 |
| top60_altered_DNA_seed | ATM | 0.289545 | COADREAD | NCI-H508 | KU-55933 |
| top60_altered_DNA_seed | ATM | 0.299476 | COADREAD | LS-1034 | KU-60019 |
| top60_altered_DNA_seed | PIK3CA | 0.315191 | COADREAD | HCC2998 | PI3Ka_4409 |
| top60_altered_DNA_seed | PIK3CA | 0.320958 | COADREAD | HCT-15 | PI3Ka_4409 |

| | | | | | |
|---|---|---|---|---|---|
| top60_altered_DNA_seed | PIK3CA | 0.331059 | COADREAD | SW1417 | PI3Ka_4409 |
| top60_altered_DNA_seed | ATM | 0.340794 | COADREAD | SW48 | CP466722 |
| top60_altered_DNA_seed | ATM | 0.345977 | COADREAD | NCI-H747 | KU-60019 |
| top60_altered_DNA_seed | ATM | 0.356093 | COADREAD | HT-115 | KU-60019 |
| top60_altered_DNA_seed | ATM | 0.364793 | COADREAD | NCI-H716 | CP466722 |
| top60_altered_DNA_seed | ATM | 0.366626 | COADREAD | LS-513 | CP466722 |
| top60_altered_DNA_seed | ATM | 0.374839 | COADREAD | LS-1034 | CP466722 |
| top60_altered_DNA_seed | ATM | 0.38181 | COADREAD | C2BBe1 | torin2 |
| top60_altered_DNA_seed | ATM | 0.382052 | COADREAD | CaR-1 | torin2 |
| top60_altered_DNA_seed | PIK3CA | 0.388994 | COADREAD | SNU-81 | PI3Ka_4409 |
| top60_altered_DNA_seed | ATM | 0.395235 | COADREAD | HCT-15 | torin2 |
| top60_altered_DNA_seed | ATM | 0.397829 | COADREAD | HT-115 | KU-55933 |
| top60_altered_DNA_seed | ATM | 0.403881 | COADREAD | C2BBe1 | KU-55933 |
| top60_altered_DNA_seed | ATM | 0.404426 | COADREAD | MDST8 | KU-55933 |
| top60_altered_DNA_seed | PIK3CA | 0.416232 | COADREAD | MDST8 | PI3Ka_4409 |
| top60_altered_DNA_seed | ATM | 0.450843 | COADREAD | SW620 | torin2 |
| top60_altered_DNA_seed | ATM | 0.459969 | COADREAD | SW837 | KU-60019 |
| top60_altered_DNA_seed | PIK3CA | 0.465554 | COADREAD | SW837 | AZD8835 |
| top60_altered_DNA_seed | ATM | 0.532861 | COADREAD | COLO-205 | KU-55933 |
| top60_altered_DNA_seed | ATM | 0.540154 | COADREAD | CaR-1 | CP466722 |
| top60_altered_DNA_seed | ATM | 0.556719 | COADREAD | SW48 | KU-60019 |
| top60_altered_DNA_seed | ATM | 0.564396 | COADREAD | SNU-C2B | torin2 |
| top60_altered_DNA_seed | PIK3CA | 0.59458 | COADREAD | SNU-C2B | AZD8835 |
| top60_altered_DNA_seed | ATM | 0.652341 | COADREAD | SNU-C2B | KU-60019 |
| top60_altered_DNA_seed | ATM | 0.654055 | COADREAD | SNU-407 | KU-55933 |
| top60_altered_DNA_seed | ATM | 0.677636 | COADREAD | SW1463 | KU-55933 |
| top60_altered_DNA_seed | PIK3CA | 0.686876 | COADREAD | HCC2998 | AZD8835 |
| top60_altered_DNA_seed | ATM | 0.705629 | COADREAD | SW1417 | CP466722 |
| top60_altered_DNA_seed | ATM | 0.713882 | COADREAD | CaR-1 | KU-60019 |
| top60_altered_DNA_seed | PIK3CA | 0.737672 | COADREAD | CW-2 | AZD8835 |
| top60_altered_DNA_seed | ATM | 0.742282 | COADREAD | HCC2998 | KU-60019 |
| top60_altered_DNA_seed | PIK3CA | 0.746074 | COADREAD | RCM-1 | AZD8835 |
| top60_altered_DNA_seed | ATM | 0.772567 | COADREAD | GP5d | KU-55933 |
| top60_altered_DNA_seed | ATM | 0.776763 | COADREAD | HCC2998 | KU-55933 |
| top60_altered_DNA_seed | ATM | 0.782786 | COADREAD | LoVo | CP466722 |
| top60_altered_DNA_seed | ATM | 0.796034 | COADREAD | SNU-C2B | CP466722 |
| top60_altered_DNA_seed | ATM | 0.806017 | COADREAD | SW1417 | KU-60019 |
| top60_altered_DNA_seed | ATM | 0.810344 | COADREAD | LS-123 | KU-60019 |
| top60_altered_DNA_seed | ATM | 0.811286 | COADREAD | LS-411N | KU-55933 |
| top60_altered_DNA_seed | ATM | 0.840684 | COADREAD | SNU-C5 | KU-60019 |
| top60_altered_DNA_seed | ATM | 0.84323 | COADREAD | CCK-81 | KU-55933 |

| | | | | | |
|---|---|---|---|---|---|
| top60_altered_DNA_seed | ATM | 0.853584 | COADREAD | C2BBe1 | CP466722 |
| top60_altered_DNA_seed | ATM | 0.871659 | COADREAD | LS-1034 | KU-55933 |
| top60_altered_DNA_seed | PIK3CA | 0.898392 | COADREAD | SW620 | AZD8835 |
| top60_altered_DNA_seed | ATM | 0.90616 | COADREAD | KM12 | KU-55933 |
| top60_altered_DNA_seed | PIK3CA | 0.918071 | COADREAD | SW1116 | PI3Ka_4409 |
| top60_altered_DNA_seed | ATM | 0.975841 | COADREAD | DiFi | KU-55933 |
| top60_altered_DNA_seed | ATM | 0.979241 | COADREAD | SNU-C1 | KU-55933 |
| top60_altered_DNA_seed | ATM | 1.00267 | COADREAD | HT-115 | CP466722 |
| top60_altered_DNA_seed | PIK3CA | 1.012971 | COADREAD | COLO-678 | AZD8835 |
| top60_altered_DNA_seed | ATM | 1.01836 | COADREAD | DiFi | KU-60019 |
| top60_altered_DNA_seed | PIK3CA | 1.026844 | COADREAD | CaR-1 | AZD8835 |
| top60_altered_DNA_seed | ATM | 1.112672 | COADREAD | RCM-1 | torin2 |
| top60_altered_DNA_seed | ATM | 1.149447 | COADREAD | HT-29 | KU-55933 |
| top60_altered_DNA_seed | ATM | 1.170588 | COADREAD | SW1417 | KU-55933 |
| top60_altered_DNA_seed | ATM | 1.192943 | COADREAD | COLO-678 | KU-60019 |
| top60_altered_DNA_seed | ATM | 1.199291 | COADREAD | CL-11 | CP466722 |
| top60_altered_DNA_seed | PIK3CA | 1.211845 | COADREAD | C2BBe1 | AZD8835 |
| top60_altered_DNA_seed | ATM | 1.224194 | COADREAD | SNU-C5 | CP466722 |
| top60_altered_DNA_seed | PIK3CA | 1.228973 | COADREAD | SW1417 | AZD8835 |
| top60_altered_DNA_seed | ATM | 1.232944 | COADREAD | SNU-407 | KU-60019 |
| top60_altered_DNA_seed | ATM | 1.24555 | COADREAD | T84 | torin2 |
| top60_altered_DNA_seed | ATM | 1.280746 | COADREAD | SW948 | CP466722 |
| top60_altered_DNA_seed | ATM | 1.283206 | COADREAD | T84 | KU-55933 |
| top60_altered_DNA_seed | PIK3CA | 1.290498 | COADREAD | RCM-1 | PI3Ka_4409 |
| top60_altered_DNA_seed | ATM | 1.300873 | COADREAD | COLO-205 | KU-60019 |
| top60_altered_DNA_seed | ATM | 1.330659 | COADREAD | RCM-1 | CP466722 |
| top60_altered_DNA_seed | PIK3CA | 1.336968 | COADREAD | LS-411N | AZD8835 |
| top60_altered_DNA_seed | ATM | 1.362202 | COADREAD | SNU-C1 | KU-60019 |
| top60_altered_DNA_seed | PIK3CA | 1.377467 | COADREAD | CW-2 | PI3Ka_4409 |
| top60_altered_DNA_seed | ATM | 1.385027 | COADREAD | NCI-H747 | KU-55933 |
| top60_altered_DNA_seed | ATM | 1.389542 | COADREAD | SW1116 | CP466722 |
| top60_altered_DNA_seed | ATM | 1.398281 | COADREAD | CW-2 | torin2 |
| top60_altered_DNA_seed | ATM | 1.398285 | COADREAD | SNU-C1 | CP466722 |
| top60_altered_DNA_seed | ATM | 1.416189 | COADREAD | CW-2 | KU-55933 |
| top60_altered_DNA_seed | ATM | 1.417218 | COADREAD | HT55 | KU-55933 |
| top60_altered_DNA_seed | ATM | 1.45201 | COADREAD | SW620 | KU-55933 |
| top60_altered_DNA_seed | ATM | 1.501923 | COADREAD | HT55 | KU-60019 |
| top60_altered_DNA_seed | ATM | 1.50684 | COADREAD | RCM-1 | KU-60019 |
| top60_altered_DNA_seed | PIK3CA | 1.564331 | COADREAD | SNU-61 | AZD8835 |
| top60_altered_DNA_seed | PIK3CA | 1.574111 | COADREAD | SW1116 | AZD8835 |
| top60_altered_DNA_seed | ATM | 1.598014 | COADREAD | RCM-1 | KU-55933 |

| | | | | | |
|---|---|---|---|---|---|
| top60_altered_DNA_seed | PIK3CA | 1.61439 | COADREAD | COLO-678 | PI3Ka_4409 |
| top60_altered_DNA_seed | ATM | 1.635518 | COADREAD | CW-2 | CP466722 |
| top60_altered_DNA_seed | PIK3CA | 1.673282 | COADREAD | SW620 | PI3Ka_4409 |
| top60_altered_DNA_seed | ATM | 1.693951 | COADREAD | SW1463 | CP466722 |
| top60_altered_DNA_seed | ATM | 1.695339 | COADREAD | MDST8 | CP466722 |
| top60_altered_DNA_seed | ATM | 1.724064 | COADREAD | SNU-C2B | KU-55933 |
| top60_altered_DNA_seed | PIK3CA | 1.773409 | COADREAD | LS-411N | PI3Ka_4409 |
| top60_altered_DNA_seed | ATM | 1.782405 | COADREAD | LS-513 | KU-55933 |
| top60_altered_DNA_seed | ATM | 1.796726 | COADREAD | COLO-678 | KU-55933 |
| top60_altered_DNA_seed | PIK3CA | 1.799843 | COADREAD | SNU-61 | PI3Ka_4409 |
| top60_altered_DNA_seed | PIK3CA | 1.826066 | COADREAD | KM12 | AZD8835 |
| top60_altered_DNA_seed | PIK3CA | 1.869052 | COADREAD | C2BBe1 | PI3Ka_4409 |
| top60_altered_DNA_seed | PIK3CA | 1.928324 | COADREAD | SNU-1040 | AZD8835 |
| top60_altered_DNA_seed | ATM | 1.9308 | COADREAD | SW837 | KU-55933 |
| top60_altered_DNA_seed | ATM | 2.012473 | COADREAD | COLO-678 | CP466722 |
| top60_altered_DNA_seed | ATM | 2.01646 | COADREAD | T84 | KU-60019 |
| top60_altered_DNA_seed | PIK3CA | 2.09473 | COADREAD | KM12 | PI3Ka_4409 |
| top60_altered_DNA_seed | ATM | 2.112754 | COADREAD | LS-123 | KU-55933 |
| top60_altered_DNA_seed | ATM | 2.116676 | COADREAD | HT55 | torin2 |
| top60_altered_DNA_seed | ATM | 2.148201 | COADREAD | HT-29 | KU-60019 |
| top60_altered_DNA_seed | ATM | 2.230647 | COADREAD | SW948 | KU-55933 |
| top60_altered_DNA_seed | ATM | 2.230884 | COADREAD | LS-513 | KU-60019 |
| top60_altered_DNA_seed | ATM | 2.296439 | COADREAD | KM12 | torin2 |
| top60_altered_DNA_seed | ATM | 2.299113 | COADREAD | SW948 | KU-60019 |
| top60_altered_DNA_seed | ATM | 2.384769 | COADREAD | LS-180 | KU-60019 |
| top60_altered_DNA_seed | ATM | 2.446635 | COADREAD | SNU-61 | KU-55933 |
| top60_altered_DNA_seed | ATM | 2.534752 | COADREAD | LS-180 | torin2 |
| top60_altered_DNA_seed | ATM | 2.624244 | COADREAD | HT-29 | torin2 |
| top60_altered_DNA_seed | ATM | 2.62843 | COADREAD | HT55 | CP466722 |
| top60_altered_DNA_seed | ATM | 2.713114 | COADREAD | SNU-61 | KU-60019 |
| top60_altered_DNA_seed | ATM | 2.716752 | COADREAD | SW1116 | KU-60019 |
| top60_altered_DNA_seed | ATM | 2.752819 | COADREAD | SW1116 | KU-55933 |
| top60_altered_DNA_seed | ATM | 3.090585 | COADREAD | SW1463 | KU-60019 |
| top60_altered_DNA_seed | ATM | 3.099592 | COADREAD | SW948 | torin2 |
| top60_altered_DNA_seed | ATM | 3.150686 | COADREAD | NCI-H508 | KU-60019 |
| top60_altered_DNA_seed | PIK3CA | -2.306434 | COADREAD | C2BBe1 | AZD8186 |
| top60_altered_DNA_seed | PIK3CA | -2.162493 | COADREAD | NCI-H508 | Alpelisib |
| top60_altered_DNA_seed | PIK3CA | -2.120441 | COADREAD | LoVo | AZD8186 |
| top60_altered_DNA_seed | ATM | -1.525765 | COADREAD | SK-CO-1 | KU-55933 |
| top60_altered_DNA_seed | PIK3CA | -1.501377 | COADREAD | GP5d | Alpelisib |
| top60_altered_DNA_seed | ATM | -1.481905 | COADREAD | RKO | KU-55933 |

| | | | | | |
|---|---|---|---|---|---|
| top60_altered_DNA_seed | KRAS | -1.426956 | COADREAD | KM12 | KRAS (G12C) Inhibitor-12 |
| top60_altered_DNA_seed | ATM | -1.418079 | COADREAD | CCK-81 | KU-55933 |
| top60_altered_DNA_seed | ATM | -1.3741 | COADREAD | LoVo | KU-55933 |
| top60_altered_DNA_seed | ATM | -1.312324 | COADREAD | T84 | KU-55933 |
| top60_altered_DNA_seed | KRAS | -1.301712 | COADREAD | HCT-15 | KRAS (G12C) Inhibitor-12 |
| top60_altered_DNA_seed | PIK3CA | -1.201292 | COADREAD | SW48 | AZD8186 |
| top60_altered_DNA_seed | PIK3CA | -1.197354 | COADREAD | HCT-15 | GNE-317 |
| top60_altered_DNA_seed | PIK3CA | -1.158287 | COADREAD | NCI-H508 | AZD8186 |
| top60_altered_DNA_seed | PIK3CA | -1.125492 | COADREAD | SNU-81 | AZD8186 |
| top60_altered_DNA_seed | PIK3CA | -1.117837 | COADREAD | SW48 | GNE-317 |
| top60_altered_DNA_seed | PIK3CA | -1.104252 | COADREAD | C2BBe1 | GNE-317 |
| top60_altered_DNA_seed | TP53 | -1.101466 | COADREAD | COLO-320-HSR | MIRA-1 |
| top60_altered_DNA_seed | PIK3CA | -1.052704 | COADREAD | SW48 | Alpelisib |
| top60_altered_DNA_seed | PIK3CA | -1.051702 | COADREAD | COLO-320-HSR | Buparlisib |
| top60_altered_DNA_seed | PIK3CA | -1.051678 | COADREAD | T84 | Alpelisib |
| top60_altered_DNA_seed | PIK3CA | -1.026602 | COADREAD | HCT-15 | Alpelisib |
| top60_altered_DNA_seed | ATM | -1.019802 | COADREAD | HCC2998 | KU-55933 |
| top60_altered_DNA_seed | PIK3CA | -1.008135 | COADREAD | DiFi | Alpelisib |
| top60_altered_DNA_seed | PIK3CA | -1.004211 | COADREAD | SK-CO-1 | AZD8186 |
| top60_altered_DNA_seed | PIK3CA | -0.993072 | COADREAD | MDST8 | GNE-317 |
| top60_altered_DNA_seed | PIK3CA | -0.977944 | COADREAD | SK-CO-1 | Buparlisib |
| top60_altered_DNA_seed | PIK3CA | -0.958464 | COADREAD | NCI-H716 | AZD8186 |
| top60_altered_DNA_seed | KRAS | -0.948158 | COADREAD | HT55 | KRAS (G12C) Inhibitor-12 |
| top60_altered_DNA_seed | PIK3CA | -0.935631 | COADREAD | NCI-H747 | Alpelisib |
| top60_altered_DNA_seed | PIK3CA | -0.924875 | COADREAD | CaR-1 | GNE-317 |
| top60_altered_DNA_seed | PIK3CA | -0.911292 | COADREAD | COLO-205 | GNE-317 |
| top60_altered_DNA_seed | TP53 | -0.910323 | COADREAD | COLO-205 | MIRA-1 |
| top60_altered_DNA_seed | PIK3CA | -0.897857 | COADREAD | HT55 | Alpelisib |
| top60_altered_DNA_seed | PIK3CA | -0.889066 | COADREAD | CCK-81 | Alpelisib |
| top60_altered_DNA_seed | PIK3CA | -0.83054 | COADREAD | HT-29 | Alpelisib |
| top60_altered_DNA_seed | PIK3CA | -0.827186 | COADREAD | HCT-15 | Buparlisib |
| top60_altered_DNA_seed | PIK3CA | -0.826032 | COADREAD | SK-CO-1 | GNE-317 |
| top60_altered_DNA_seed | PIK3CA | -0.80982 | COADREAD | GP5d | Buparlisib |
| top60_altered_DNA_seed | PIK3CA | -0.808737 | COADREAD | COLO-320-HSR | Alpelisib |
| top60_altered_DNA_seed | PIK3CA | -0.807439 | COADREAD | MDST8 | Buparlisib |
| top60_altered_DNA_seed | PIK3CA | -0.797717 | COADREAD | NCI-H747 | Buparlisib |
| top60_altered_DNA_seed | PIK3CA | -0.795901 | COADREAD | NCI-H747 | GNE-317 |
| top60_altered_DNA_seed | PIK3CA | -0.783071 | COADREAD | SW948 | Alpelisib |
| top60_altered_DNA_seed | PIK3CA | -0.772557 | COADREAD | LoVo | Buparlisib |
| top60_altered_DNA_seed | PIK3CA | -0.765258 | COADREAD | COLO-205 | Buparlisib |
| top60_altered_DNA_seed | PIK3CA | -0.758564 | COADREAD | SNU-175 | Buparlisib |

| | | | | | |
|---|---|---|---|---|---|
| top60_altered_DNA_seed | PIK3CA | -0.755599 | COADREAD | SW48 | Buparlisib |
| top60_altered_DNA_seed | PIK3CA | -0.753669 | COADREAD | LoVo | GNE-317 |
| top60_altered_DNA_seed | TP53 | -0.749332 | COADREAD | LS-513 | MIRA-1 |
| top60_altered_DNA_seed | PIK3CA | -0.735045 | COADREAD | SW948 | GNE-317 |
| top60_altered_DNA_seed | PIK3CA | -0.732431 | COADREAD | C2BBe1 | Buparlisib |
| top60_altered_DNA_seed | PIK3CA | -0.728024 | COADREAD | KM12 | Buparlisib |
| top60_altered_DNA_seed | PIK3CA | -0.698622 | COADREAD | COLO-320-HSR | GNE-317 |
| top60_altered_DNA_seed | PIK3CA | -0.691121 | COADREAD | DiFi | GNE-317 |
| top60_altered_DNA_seed | ATM | -0.679532 | COADREAD | SNU-C5 | KU-55933 |
| top60_altered_DNA_seed | PIK3CA | -0.661258 | COADREAD | DiFi | Buparlisib |
| top60_altered_DNA_seed | ATM | -0.660589 | COADREAD | MDST8 | KU-55933 |
| top60_altered_DNA_seed | KRAS | -0.652878 | COADREAD | HCC2998 | KRAS (G12C) Inhibitor-12 |
| top60_altered_DNA_seed | ATM | -0.620347 | COADREAD | SW837 | KU-55933 |
| top60_altered_DNA_seed | PIK3CA | -0.60258 | COADREAD | SK-CO-1 | Alpelisib |
| top60_altered_DNA_seed | PIK3CA | -0.602487 | COADREAD | SNU-C5 | Buparlisib |
| top60_altered_DNA_seed | PIK3CA | -0.594705 | COADREAD | HCT-116 | Buparlisib |
| top60_altered_DNA_seed | PIK3CA | -0.551992 | COADREAD | COLO-320-HSR | AZD8186 |
| top60_altered_DNA_seed | PIK3CA | -0.527602 | COADREAD | HT-115 | Buparlisib |
| top60_altered_DNA_seed | PIK3CA | -0.508767 | COADREAD | HCT-116 | Alpelisib |
| top60_altered_DNA_seed | TP53 | -0.502326 | COADREAD | GP5d | MIRA-1 |
| top60_altered_DNA_seed | PIK3CA | -0.484905 | COADREAD | CCK-81 | Buparlisib |
| top60_altered_DNA_seed | PIK3CA | -0.48204 | COADREAD | CaR-1 | AZD8186 |
| top60_altered_DNA_seed | PIK3CA | -0.478742 | COADREAD | NCI-H747 | AZD8186 |
| top60_altered_DNA_seed | KRAS | -0.460015 | COADREAD | GP5d | KRAS (G12C) Inhibitor-12 |
| top60_altered_DNA_seed | KRAS | -0.452548 | COADREAD | LoVo | KRAS (G12C) Inhibitor-12 |
| top60_altered_DNA_seed | ATM | -0.441003 | COADREAD | CaR-1 | KU-55933 |
| top60_altered_DNA_seed | KRAS | -0.43604 | COADREAD | SW620 | KRAS (G12C) Inhibitor-12 |
| top60_altered_DNA_seed | PIK3CA | -0.432524 | COADREAD | SNU-407 | Buparlisib |
| top60_altered_DNA_seed | TP53 | -0.424729 | COADREAD | DiFi | MIRA-1 |
| top60_altered_DNA_seed | PIK3CA | -0.42315 | COADREAD | RKO | Alpelisib |
| top60_altered_DNA_seed | PIK3CA | -0.408004 | COADREAD | HCT-116 | GNE-317 |
| top60_altered_DNA_seed | PIK3CA | -0.39723 | COADREAD | SNU-407 | Alpelisib |
| top60_altered_DNA_seed | ATM | -0.391766 | COADREAD | C2BBe1 | KU-55933 |
| top60_altered_DNA_seed | PIK3CA | -0.389519 | COADREAD | LS-180 | GNE-317 |
| top60_altered_DNA_seed | PIK3CA | -0.380122 | COADREAD | COLO-205 | AZD8186 |
| top60_altered_DNA_seed | KRAS | -0.379246 | COADREAD | CW-2 | KRAS (G12C) Inhibitor-12 |
| top60_altered_DNA_seed | ATM | -0.378328 | COADREAD | SNU-C1 | KU-55933 |
| top60_altered_DNA_seed | TP53 | -0.365414 | COADREAD | CW-2 | MIRA-1 |
| top60_altered_DNA_seed | TP53 | -0.364305 | COADREAD | HT55 | MIRA-1 |
| top60_altered_DNA_seed | TP53 | -0.337806 | COADREAD | CCK-81 | MIRA-1 |
| top60_altered_DNA_seed | PIK3CA | -0.337026 | COADREAD | COLO-205 | Alpelisib |

| | | | | | |
|---|---|---|---|---|---|
| top60_altered_DNA_seed | KRAS | -0.334025 | COADREAD | RKO | KRAS (G12C) Inhibitor-12 |
| top60_altered_DNA_seed | PIK3CA | -0.321428 | COADREAD | CaR-1 | Alpelisib |
| top60_altered_DNA_seed | KRAS | -0.297829 | COADREAD | SW48 | KRAS (G12C) Inhibitor-12 |
| top60_altered_DNA_seed | PIK3CA | -0.290618 | COADREAD | LS-411N | GNE-317 |
| top60_altered_DNA_seed | ATM | -0.278066 | COADREAD | CW-2 | KU-55933 |
| top60_altered_DNA_seed | KRAS | -0.259001 | COADREAD | NCI-H716 | KRAS (G12C) Inhibitor-12 |
| top60_altered_DNA_seed | KRAS | -0.249931 | COADREAD | SNU-C5 | KRAS (G12C) Inhibitor-12 |
| top60_altered_DNA_seed | PIK3CA | -0.24948 | COADREAD | KM12 | GNE-317 |
| top60_altered_DNA_seed | PIK3CA | -0.233439 | COADREAD | CaR-1 | Buparlisib |
| top60_altered_DNA_seed | PIK3CA | -0.227941 | COADREAD | LS-123 | GNE-317 |
| top60_altered_DNA_seed | KRAS | -0.211738 | COADREAD | LS-513 | KRAS (G12C) Inhibitor-12 |
| top60_altered_DNA_seed | PIK3CA | -0.194916 | COADREAD | HT-29 | GNE-317 |
| top60_altered_DNA_seed | PIK3CA | -0.192095 | COADREAD | LS-513 | Alpelisib |
| top60_altered_DNA_seed | TP53 | -0.191951 | COADREAD | MDST8 | MIRA-1 |
| top60_altered_DNA_seed | PIK3CA | -0.190287 | COADREAD | HCC2998 | Buparlisib |
| top60_altered_DNA_seed | ATM | -0.186846 | COADREAD | NCI-H716 | KU-55933 |
| top60_altered_DNA_seed | ATM | -0.184286 | COADREAD | HT55 | KU-55933 |
| top60_altered_DNA_seed | PIK3CA | -0.172189 | COADREAD | LS-180 | Alpelisib |
| top60_altered_DNA_seed | PIK3CA | -0.170209 | COADREAD | SNU-407 | AZD8186 |
| top60_altered_DNA_seed | PIK3CA | -0.164333 | COADREAD | HT-29 | AZD8186 |
| top60_altered_DNA_seed | PIK3CA | -0.158809 | COADREAD | HT-115 | Alpelisib |
| top60_altered_DNA_seed | PIK3CA | -0.156407 | COADREAD | MDST8 | AZD8186 |
| top60_altered_DNA_seed | PIK3CA | -0.155478 | COADREAD | SW948 | Buparlisib |
| top60_altered_DNA_seed | PIK3CA | -0.13814 | COADREAD | CW-2 | Alpelisib |
| top60_altered_DNA_seed | PIK3CA | -0.136585 | COADREAD | LS-123 | AZD8186 |
| top60_altered_DNA_seed | ATM | -0.125136 | COADREAD | KM12 | KU-55933 |
| top60_altered_DNA_seed | PIK3CA | -0.115279 | COADREAD | SNU-C5 | Alpelisib |
| top60_altered_DNA_seed | PIK3CA | -0.115211 | COADREAD | LoVo | Alpelisib |
| top60_altered_DNA_seed | PIK3CA | -0.09687 | COADREAD | HT55 | AZD8186 |
| top60_altered_DNA_seed | PIK3CA | -0.085083 | COADREAD | T84 | AZD8186 |
| top60_altered_DNA_seed | PIK3CA | -0.080345 | COADREAD | HCC2998 | GNE-317 |
| top60_altered_DNA_seed | PIK3CA | -0.072293 | COADREAD | RCM-1 | Alpelisib |
| top60_altered_DNA_seed | PIK3CA | -0.072201 | COADREAD | GP5d | GNE-317 |
| top60_altered_DNA_seed | KRAS | -0.072151 | COADREAD | NCI-H747 | KRAS (G12C) Inhibitor-12 |
| top60_altered_DNA_seed | KRAS | -0.069158 | COADREAD | HT-29 | KRAS (G12C) Inhibitor-12 |
| top60_altered_DNA_seed | PIK3CA | -0.066991 | COADREAD | CW-2 | Buparlisib |
| top60_altered_DNA_seed | PIK3CA | -0.060572 | COADREAD | SW1463 | GNE-317 |
| top60_altered_DNA_seed | ATM | -0.059558 | COADREAD | HT-115 | KU-55933 |
| top60_altered_DNA_seed | PIK3CA | -0.058033 | COADREAD | HCC2998 | AZD8186 |
| top60_altered_DNA_seed | KRAS | -0.055151 | COADREAD | HCT-116 | KRAS (G12C) Inhibitor-12 |
| top60_altered_DNA_seed | TP53 | -0.052856 | COADREAD | HCT-116 | MIRA-1 |

| | | | | | |
|---|---|---|---|---|---|
| top60_altered_DNA_seed | ATM | -0.0517 | COADREAD | GP5d | KU-55933 |
| top60_altered_DNA_seed | KRAS | -0.034769 | COADREAD | SNU-1040 | KRAS (G12C) Inhibitor-12 |
| top60_altered_DNA_seed | PIK3CA | -0.027099 | COADREAD | RKO | Buparlisib |
| top60_altered_DNA_seed | PIK3CA | -0.026078 | COADREAD | SNU-C5 | AZD8186 |
| top60_altered_DNA_seed | PIK3CA | -0.019827 | COADREAD | NCI-H508 | Buparlisib |
| top60_altered_DNA_seed | PIK3CA | -0.000728 | COADREAD | SNU-407 | GNE-317 |
| top60_altered_DNA_seed | PIK3CA | 0.002855 | COADREAD | SNU-175 | AZD8186 |
| top60_altered_DNA_seed | PIK3CA | 0.006593 | COADREAD | SNU-175 | GNE-317 |
| top60_altered_DNA_seed | PIK3CA | 0.0087 | COADREAD | HT-29 | Buparlisib |
| top60_altered_DNA_seed | PIK3CA | 0.017624 | COADREAD | CCK-81 | AZD8186 |
| top60_altered_DNA_seed | ATM | 0.018433 | COADREAD | COLO-320-HSR | KU-55933 |
| top60_altered_DNA_seed | TP53 | 0.031989 | COADREAD | HCC2998 | MIRA-1 |
| top60_altered_DNA_seed | KRAS | 0.053306 | COADREAD | LS-411N | KRAS (G12C) Inhibitor-12 |
| top60_altered_DNA_seed | PIK3CA | 0.06485 | COADREAD | DiFi | AZD8186 |
| top60_altered_DNA_seed | TP53 | 0.065601 | COADREAD | SK-CO-1 | MIRA-1 |
| top60_altered_DNA_seed | PIK3CA | 0.067512 | COADREAD | SNU-175 | Alpelisib |
| top60_altered_DNA_seed | PIK3CA | 0.085727 | COADREAD | GP5d | AZD8186 |
| top60_altered_DNA_seed | PIK3CA | 0.087524 | COADREAD | SW837 | GNE-317 |
| top60_altered_DNA_seed | TP53 | 0.089289 | COADREAD | HT-115 | MIRA-1 |
| top60_altered_DNA_seed | KRAS | 0.104894 | COADREAD | SNU-175 | KRAS (G12C) Inhibitor-12 |
| top60_altered_DNA_seed | TP53 | 0.111455 | COADREAD | SNU-175 | MIRA-1 |
| top60_altered_DNA_seed | PIK3CA | 0.117066 | COADREAD | HCC2998 | Alpelisib |
| top60_altered_DNA_seed | PIK3CA | 0.140505 | COADREAD | T84 | Buparlisib |
| top60_altered_DNA_seed | PIK3CA | 0.152698 | COADREAD | HT55 | Buparlisib |
| top60_altered_DNA_seed | PIK3CA | 0.152922 | COADREAD | SNU-C5 | GNE-317 |
| top60_altered_DNA_seed | PIK3CA | 0.153117 | COADREAD | NCI-H508 | GNE-317 |
| top60_altered_DNA_seed | KRAS | 0.172869 | COADREAD | SNU-407 | KRAS (G12C) Inhibitor-12 |
| top60_altered_DNA_seed | PIK3CA | 0.178998 | COADREAD | LS-180 | AZD8186 |
| top60_altered_DNA_seed | ATM | 0.195901 | COADREAD | LS-1034 | KU-55933 |
| top60_altered_DNA_seed | TP53 | 0.205136 | COADREAD | T84 | MIRA-1 |
| top60_altered_DNA_seed | KRAS | 0.207918 | COADREAD | SK-CO-1 | KRAS (G12C) Inhibitor-12 |
| top60_altered_DNA_seed | PIK3CA | 0.220829 | COADREAD | SNU-C1 | GNE-317 |
| top60_altered_DNA_seed | PIK3CA | 0.22992 | COADREAD | HCT-15 | AZD8186 |
| top60_altered_DNA_seed | ATM | 0.233929 | COADREAD | SW1463 | KU-55933 |
| top60_altered_DNA_seed | PIK3CA | 0.24029 | COADREAD | SW620 | GNE-317 |
| top60_altered_DNA_seed | TP53 | 0.240471 | COADREAD | LS-180 | MIRA-1 |
| top60_altered_DNA_seed | TP53 | 0.240883 | COADREAD | SW48 | MIRA-1 |
| top60_altered_DNA_seed | PIK3CA | 0.246012 | COADREAD | HT55 | GNE-317 |
| top60_altered_DNA_seed | ATM | 0.257987 | COADREAD | HCT-116 | KU-55933 |
| top60_altered_DNA_seed | TP53 | 0.268349 | COADREAD | SNU-81 | MIRA-1 |
| top60_altered_DNA_seed | TP53 | 0.268778 | COADREAD | RKO | MIRA-1 |

| | | | | | |
|---|---|---|---|---|---|
| top60_altered_DNA_seed | ATM | 0.269069 | COADREAD | SNU-81 | KU-55933 |
| top60_altered_DNA_seed | TP53 | 0.280323 | COADREAD | HCT-15 | MIRA-1 |
| top60_altered_DNA_seed | PIK3CA | 0.310936 | COADREAD | T84 | GNE-317 |
| top60_altered_DNA_seed | ATM | 0.339776 | COADREAD | SNU-175 | KU-55933 |
| top60_altered_DNA_seed | PIK3CA | 0.348843 | COADREAD | SNU-81 | GNE-317 |
| top60_altered_DNA_seed | PIK3CA | 0.364046 | COADREAD | LS-123 | Buparlisib |
| top60_altered_DNA_seed | ATM | 0.380863 | COADREAD | CL-11 | KU-55933 |
| top60_altered_DNA_seed | PIK3CA | 0.381011 | COADREAD | SW620 | Buparlisib |
| top60_altered_DNA_seed | TP53 | 0.382935 | COADREAD | CaR-1 | MIRA-1 |
| top60_altered_DNA_seed | PIK3CA | 0.389427 | COADREAD | NCI-H716 | Buparlisib |
| top60_altered_DNA_seed | KRAS | 0.394184 | COADREAD | DiFi | KRAS (G12C) Inhibitor-12 |
| top60_altered_DNA_seed | TP53 | 0.396854 | COADREAD | NCI-H508 | MIRA-1 |
| top60_altered_DNA_seed | PIK3CA | 0.409103 | COADREAD | SW837 | AZD8186 |
| top60_altered_DNA_seed | PIK3CA | 0.411115 | COADREAD | HT-115 | GNE-317 |
| top60_altered_DNA_seed | PIK3CA | 0.414408 | COADREAD | CCK-81 | GNE-317 |
| top60_altered_DNA_seed | PIK3CA | 0.425256 | COADREAD | SW1463 | AZD8186 |
| top60_altered_DNA_seed | PIK3CA | 0.426361 | COADREAD | HCT-116 | AZD8186 |
| top60_altered_DNA_seed | ATM | 0.433564 | COADREAD | HCT-15 | KU-55933 |
| top60_altered_DNA_seed | TP53 | 0.436215 | COADREAD | KM12 | MIRA-1 |
| top60_altered_DNA_seed | PIK3CA | 0.447664 | COADREAD | LS-180 | Buparlisib |
| top60_altered_DNA_seed | PIK3CA | 0.45689 | COADREAD | LS-411N | Buparlisib |
| top60_altered_DNA_seed | PIK3CA | 0.460474 | COADREAD | SNU-C1 | Alpelisib |
| top60_altered_DNA_seed | TP53 | 0.484948 | COADREAD | LS-411N | MIRA-1 |
| top60_altered_DNA_seed | PIK3CA | 0.504079 | COADREAD | SNU-81 | Buparlisib |
| top60_altered_DNA_seed | TP53 | 0.519311 | COADREAD | LoVo | MIRA-1 |
| top60_altered_DNA_seed | PIK3CA | 0.522049 | COADREAD | C2BBe1 | Alpelisib |
| top60_altered_DNA_seed | ATM | 0.522708 | COADREAD | DiFi | KU-55933 |
| top60_altered_DNA_seed | PIK3CA | 0.523605 | COADREAD | SW1417 | AZD8186 |
| top60_altered_DNA_seed | PIK3CA | 0.523866 | COADREAD | SNU-C2B | GNE-317 |
| top60_altered_DNA_seed | PIK3CA | 0.546419 | COADREAD | SW1116 | Alpelisib |
| top60_altered_DNA_seed | KRAS | 0.556824 | COADREAD | CaR-1 | KRAS (G12C) Inhibitor-12 |
| top60_altered_DNA_seed | PIK3CA | 0.57669 | COADREAD | LS-123 | Alpelisib |
| top60_altered_DNA_seed | ATM | 0.59262 | COADREAD | NCI-H747 | KU-55933 |
| top60_altered_DNA_seed | ATM | 0.596974 | COADREAD | SNU-407 | KU-55933 |
| top60_altered_DNA_seed | PIK3CA | 0.603429 | COADREAD | NCI-H716 | Alpelisib |
| top60_altered_DNA_seed | TP53 | 0.611766 | COADREAD | LS-1034 | MIRA-1 |
| top60_altered_DNA_seed | ATM | 0.629547 | COADREAD | NCI-H508 | KU-55933 |
| top60_altered_DNA_seed | PIK3CA | 0.645643 | COADREAD | CL-11 | Alpelisib |
| top60_altered_DNA_seed | TP53 | 0.646022 | COADREAD | SW1463 | MIRA-1 |
| top60_altered_DNA_seed | ATM | 0.651683 | COADREAD | LS-513 | KU-55933 |
| top60_altered_DNA_seed | PIK3CA | 0.651935 | COADREAD | HT-115 | AZD8186 |

| | | | | | |
|---|---|---|---|---|---|
| top60_altered_DNA_seed | ATM | 0.668588 | COADREAD | COLO-205 | KU-55933 |
| top60_altered_DNA_seed | KRAS | 0.679288 | COADREAD | SW948 | KRAS (G12C) Inhibitor-12 |
| top60_altered_DNA_seed | PIK3CA | 0.683489 | COADREAD | CL-11 | AZD8186 |
| top60_altered_DNA_seed | KRAS | 0.687276 | COADREAD | HT-115 | KRAS (G12C) Inhibitor-12 |
| top60_altered_DNA_seed | ATM | 0.694096 | COADREAD | SW948 | KU-55933 |
| top60_altered_DNA_seed | ATM | 0.704125 | COADREAD | HT-29 | KU-55933 |
| top60_altered_DNA_seed | PIK3CA | 0.716953 | COADREAD | RCM-1 | AZD8186 |
| top60_altered_DNA_seed | PIK3CA | 0.732518 | COADREAD | CL-11 | Buparlisib |
| top60_altered_DNA_seed | ATM | 0.740156 | COADREAD | SW48 | KU-55933 |
| top60_altered_DNA_seed | PIK3CA | 0.743821 | COADREAD | RKO | AZD8186 |
| top60_altered_DNA_seed | TP53 | 0.752339 | COADREAD | CL-11 | MIRA-1 |
| top60_altered_DNA_seed | PIK3CA | 0.769725 | COADREAD | SW620 | Alpelisib |
| top60_altered_DNA_seed | PIK3CA | 0.794784 | COADREAD | SNU-C2B | Buparlisib |
| top60_altered_DNA_seed | ATM | 0.804114 | COADREAD | RCM-1 | KU-55933 |
| top60_altered_DNA_seed | ATM | 0.808598 | COADREAD | SW1116 | KU-55933 |
| top60_altered_DNA_seed | PIK3CA | 0.811062 | COADREAD | SW1417 | GNE-317 |
| top60_altered_DNA_seed | PIK3CA | 0.843262 | COADREAD | SNU-61 | Alpelisib |
| top60_altered_DNA_seed | PIK3CA | 0.846979 | COADREAD | SNU-C1 | Buparlisib |
| top60_altered_DNA_seed | PIK3CA | 0.849273 | COADREAD | SW837 | Buparlisib |
| top60_altered_DNA_seed | KRAS | 0.850839 | COADREAD | COLO-320-HSR | KRAS (G12C) Inhibitor-12 |
| top60_altered_DNA_seed | PIK3CA | 0.87246 | COADREAD | SW948 | AZD8186 |
| top60_altered_DNA_seed | TP53 | 0.872614 | COADREAD | HT-29 | MIRA-1 |
| top60_altered_DNA_seed | PIK3CA | 0.87469 | COADREAD | SW1463 | Alpelisib |
| top60_altered_DNA_seed | TP53 | 0.877092 | COADREAD | SW948 | MIRA-1 |
| top60_altered_DNA_seed | TP53 | 0.892991 | COADREAD | SNU-407 | MIRA-1 |
| top60_altered_DNA_seed | PIK3CA | 0.893689 | COADREAD | LS-1034 | Alpelisib |
| top60_altered_DNA_seed | TP53 | 0.920579 | COADREAD | SNU-61 | MIRA-1 |
| top60_altered_DNA_seed | ATM | 0.966502 | COADREAD | SNU-61 | KU-55933 |
| top60_altered_DNA_seed | PIK3CA | 0.97764 | COADREAD | SW1417 | Alpelisib |
| top60_altered_DNA_seed | PIK3CA | 0.98394 | COADREAD | SNU-61 | Buparlisib |
| top60_altered_DNA_seed | KRAS | 0.989445 | COADREAD | CL-11 | KRAS (G12C) Inhibitor-12 |
| top60_altered_DNA_seed | ATM | 1.018111 | COADREAD | LS-411N | KU-55933 |
| top60_altered_DNA_seed | ATM | 1.028655 | COADREAD | LS-180 | KU-55933 |
| top60_altered_DNA_seed | PIK3CA | 1.029105 | COADREAD | MDST8 | Alpelisib |
| top60_altered_DNA_seed | TP53 | 1.043725 | COADREAD | SNU-C5 | MIRA-1 |
| top60_altered_DNA_seed | TP53 | 1.048788 | COADREAD | SNU-C1 | MIRA-1 |
| top60_altered_DNA_seed | KRAS | 1.051334 | COADREAD | CCK-81 | KRAS (G12C) Inhibitor-12 |
| top60_altered_DNA_seed | KRAS | 1.053421 | COADREAD | MDST8 | KRAS (G12C) Inhibitor-12 |
| top60_altered_DNA_seed | PIK3CA | 1.054854 | COADREAD | SNU-C2B | Alpelisib |
| top60_altered_DNA_seed | PIK3CA | 1.068569 | COADREAD | SNU-C2B | AZD8186 |
| top60_altered_DNA_seed | TP53 | 1.072246 | COADREAD | SW837 | MIRA-1 |

| | | | | | |
|---|---|---|---|---|---|
| top60_altered_DNA_seed | TP53 | 1.083472 | COADREAD | NCI-H716 | MIRA-1 |
| top60_altered_DNA_seed | TP53 | 1.084915 | COADREAD | SW620 | MIRA-1 |
| top60_altered_DNA_seed | KRAS | 1.118692 | COADREAD | SW837 | KRAS (G12C) Inhibitor-12 |
| top60_altered_DNA_seed | KRAS | 1.119802 | COADREAD | LS-1034 | KRAS (G12C) Inhibitor-12 |
| top60_altered_DNA_seed | KRAS | 1.127681 | COADREAD | SNU-61 | KRAS (G12C) Inhibitor-12 |
| top60_altered_DNA_seed | KRAS | 1.130554 | COADREAD | NCI-H508 | KRAS (G12C) Inhibitor-12 |
| top60_altered_DNA_seed | PIK3CA | 1.142895 | COADREAD | LS-1034 | GNE-317 |
| top60_altered_DNA_seed | PIK3CA | 1.160921 | COADREAD | SNU-1040 | Alpelisib |
| top60_altered_DNA_seed | KRAS | 1.198722 | COADREAD | C2BBe1 | KRAS (G12C) Inhibitor-12 |
| top60_altered_DNA_seed | KRAS | 1.199411 | COADREAD | COLO-205 | KRAS (G12C) Inhibitor-12 |
| top60_altered_DNA_seed | KRAS | 1.200548 | COADREAD | T84 | KRAS (G12C) Inhibitor-12 |
| top60_altered_DNA_seed | PIK3CA | 1.217937 | COADREAD | NCI-H716 | GNE-317 |
| top60_altered_DNA_seed | KRAS | 1.230805 | COADREAD | SNU-81 | KRAS (G12C) Inhibitor-12 |
| top60_altered_DNA_seed | TP53 | 1.236705 | COADREAD | LS-123 | MIRA-1 |
| top60_altered_DNA_seed | KRAS | 1.240603 | COADREAD | SNU-C1 | KRAS (G12C) Inhibitor-12 |
| top60_altered_DNA_seed | TP53 | 1.244492 | COADREAD | SW1116 | MIRA-1 |
| top60_altered_DNA_seed | PIK3CA | 1.245301 | COADREAD | CW-2 | AZD8186 |
| top60_altered_DNA_seed | PIK3CA | 1.253307 | COADREAD | RCM-1 | GNE-317 |
| top60_altered_DNA_seed | PIK3CA | 1.257325 | COADREAD | SNU-61 | AZD8186 |
| top60_altered_DNA_seed | PIK3CA | 1.270247 | COADREAD | SW1463 | Buparlisib |
| top60_altered_DNA_seed | KRAS | 1.278234 | COADREAD | RCM-1 | KRAS (G12C) Inhibitor-12 |
| top60_altered_DNA_seed | KRAS | 1.302596 | COADREAD | LS-180 | KRAS (G12C) Inhibitor-12 |
| top60_altered_DNA_seed | ATM | 1.357096 | COADREAD | SW1417 | KU-55933 |
| top60_altered_DNA_seed | PIK3CA | 1.368682 | COADREAD | CL-11 | GNE-317 |
| top60_altered_DNA_seed | TP53 | 1.369924 | COADREAD | RCM-1 | MIRA-1 |
| top60_altered_DNA_seed | PIK3CA | 1.369951 | COADREAD | LS-411N | Alpelisib |
| top60_altered_DNA_seed | PIK3CA | 1.385943 | COADREAD | SW620 | AZD8186 |
| top60_altered_DNA_seed | PIK3CA | 1.393056 | COADREAD | SW1417 | Buparlisib |
| top60_altered_DNA_seed | PIK3CA | 1.412744 | COADREAD | SNU-81 | Alpelisib |
| top60_altered_DNA_seed | KRAS | 1.442541 | COADREAD | SW1463 | KRAS (G12C) Inhibitor-12 |
| top60_altered_DNA_seed | KRAS | 1.460753 | COADREAD | SNU-C2B | KRAS (G12C) Inhibitor-12 |
| top60_altered_DNA_seed | PIK3CA | 1.474853 | COADREAD | SW837 | Alpelisib |
| top60_altered_DNA_seed | PIK3CA | 1.489188 | COADREAD | KM12 | Alpelisib |
| top60_altered_DNA_seed | PIK3CA | 1.493804 | COADREAD | RKO | GNE-317 |
| top60_altered_DNA_seed | PIK3CA | 1.499855 | COADREAD | LS-1034 | Buparlisib |
| top60_altered_DNA_seed | PIK3CA | 1.548009 | COADREAD | LS-411N | AZD8186 |
| top60_altered_DNA_seed | PIK3CA | 1.603978 | COADREAD | SNU-C1 | AZD8186 |
| top60_altered_DNA_seed | KRAS | 1.604202 | COADREAD | SW1116 | KRAS (G12C) Inhibitor-12 |
| top60_altered_DNA_seed | ATM | 1.637887 | COADREAD | LS-123 | KU-55933 |
| top60_altered_DNA_seed | PIK3CA | 1.652514 | COADREAD | LS-1034 | AZD8186 |
| top60_altered_DNA_seed | TP53 | 1.663562 | COADREAD | C2BBe1 | MIRA-1 |

| | | | | | |
|---|---|---|---|---|---|
| top60_altered_DNA_seed | PIK3CA | 1.670726 | COADREAD | KM12 | AZD8186 |
| top60_altered_DNA_seed | TP53 | 1.727207 | COADREAD | NCI-H747 | MIRA-1 |
| top60_altered_DNA_seed | ATM | 1.762167 | COADREAD | SNU-C2B | KU-55933 |
| top60_altered_DNA_seed | PIK3CA | 1.771889 | COADREAD | SW1116 | Buparlisib |
| top60_altered_DNA_seed | PIK3CA | 1.863039 | COADREAD | CW-2 | GNE-317 |
| top60_altered_DNA_seed | PIK3CA | 1.876554 | COADREAD | SW1116 | GNE-317 |
| top60_altered_DNA_seed | PIK3CA | 1.907945 | COADREAD | COLO-678 | AZD8186 |
| top60_altered_DNA_seed | PIK3CA | 1.98122 | COADREAD | SW1116 | AZD8186 |
| top60_altered_DNA_seed | KRAS | 2.035954 | COADREAD | LS-123 | KRAS (G12C) Inhibitor-12 |
| top60_altered_DNA_seed | TP53 | 2.051736 | COADREAD | SNU-C2B | MIRA-1 |
| top60_altered_DNA_seed | PIK3CA | 2.296675 | COADREAD | RCM-1 | Buparlisib |
| top60_altered_DNA_seed | ATM | 2.380035 | COADREAD | SW620 | KU-55933 |
| top60_altered_DNA_seed | KRAS | 2.416192 | COADREAD | SW1417 | KRAS (G12C) Inhibitor-12 |
| top60_altered_DNA_seed | TP53 | 2.502555 | COADREAD | SW1417 | MIRA-1 |
| top60_altered_DNA_seed | PIK3CA | 2.581429 | COADREAD | COLO-678 | Alpelisib |
| top60_altered_DNA_seed | KRAS | 2.680078 | COADREAD | COLO-678 | KRAS (G12C) Inhibitor-12 |
| top60_altered_DNA_seed | PIK3CA | 2.855048 | COADREAD | SNU-61 | GNE-317 |
| top60_altered_DNA_seed | ATM | 2.874474 | COADREAD | COLO-678 | KU-55933 |
| top60_altered_DNA_seed | TP53 | 3.209116 | COADREAD | COLO-678 | MIRA-1 |
| top60_altered_DNA_seed | PIK3CA | 3.578788 | COADREAD | COLO-678 | Buparlisib |
| top60_altered_DNA_seed | PIK3CA | 3.798474 | COADREAD | COLO-678 | GNE-317 |
| top60_altered_DNA_seed | PDGFRA | -3.050319 | GBM | D-247MG | Amuvatinib |
| top60_altered_DNA_seed | KIT | -3.050319 | GBM | D-247MG | Amuvatinib |
| top60_altered_DNA_seed | PDGFRA | -2.850486 | GBM | D-247MG | Masitinib |
| top60_altered_DNA_seed | KIT | -2.850486 | GBM | D-247MG | Masitinib |
| top60_altered_DNA_seed | KIT | -2.660137 | GBM | 42-MG-BA | Pazopanib |
| top60_altered_DNA_seed | MDM4 | -2.500866 | GBM | SF268 | NSC-207895 |
| top60_altered_DNA_seed | PDGFRA | -2.358508 | GBM | KS-1 | Amuvatinib |
| top60_altered_DNA_seed | KIT | -2.358508 | GBM | KS-1 | Amuvatinib |
| top60_altered_DNA_seed | KIT | -2.272548 | GBM | YH-13 | Motesanib |
| top60_altered_DNA_seed | KIT | -2.085568 | GBM | DK-MG | OSI-930 |
| top60_altered_DNA_seed | KIT | -1.79081 | GBM | 42-MG-BA | Motesanib |
| top60_altered_DNA_seed | KIT | -1.727975 | GBM | SW1088 | Motesanib |
| top60_altered_DNA_seed | PDGFRA | -1.687812 | GBM | AM-38 | Amuvatinib |
| top60_altered_DNA_seed | KIT | -1.687812 | GBM | AM-38 | Amuvatinib |
| top60_altered_DNA_seed | KIT | -1.65303 | GBM | SNB75 | Pazopanib |
| top60_altered_DNA_seed | CDK4 | -1.652208 | GBM | D-263MG | Palbociclib |
| top60_altered_DNA_seed | KIT | -1.571127 | GBM | SF126 | Pazopanib |
| top60_altered_DNA_seed | KIT | -1.559146 | GBM | D-247MG | OSI-930 |
| top60_altered_DNA_seed | EGFR | -1.546145 | GBM | KS-1 | Erlotinib |
| top60_altered_DNA_seed | KIT | -1.524173 | GBM | M059J | Pazopanib |

| | | | | | |
|---|---|---|---|---|---|
| top60_altered_DNA_seed | KIT | -1.471504 | GBM | D-263MG | Motesanib |
| top60_altered_DNA_seed | MDM4 | -1.454502 | GBM | D-423MG | NSC-207895 |
| top60_altered_DNA_seed | CDK4 | -1.453012 | GBM | U251 | Palbociclib |
| top60_altered_DNA_seed | KIT | -1.412645 | GBM | YH-13 | Pazopanib |
| top60_altered_DNA_seed | KIT | -1.383928 | GBM | A172 | Pazopanib |
| top60_altered_DNA_seed | KIT | -1.356714 | GBM | YH-13 | Axitinib |
| top60_altered_DNA_seed | KIT | -1.349951 | GBM | D-566MG | Axitinib |
| top60_altered_DNA_seed | KIT | -1.348018 | GBM | LN-18 | Motesanib |
| top60_altered_DNA_seed | KIT | -1.240521 | GBM | D-392MG | Pazopanib |
| top60_altered_DNA_seed | KIT | -1.216698 | GBM | DK-MG | Pazopanib |
| top60_altered_DNA_seed | EGFR | -1.189085 | GBM | U251 | CUDC-101 |
| top60_altered_DNA_seed | KIT | -1.170538 | GBM | 8-MG-BA | Sorafenib |
| top60_altered_DNA_seed | EGFR | -1.16436 | GBM | DK-MG | Pelitinib |
| top60_altered_DNA_seed | KIT | -1.161978 | GBM | GB-1 | Pazopanib |
| top60_altered_DNA_seed | KIT | -1.154051 | GBM | AM-38 | Motesanib |
| top60_altered_DNA_seed | CDK4 | -1.153023 | GBM | YH-13 | Palbociclib |
| top60_altered_DNA_seed | CDK4 | -1.122478 | GBM | KS-1 | CGP-082996 |
| top60_altered_DNA_seed | PDGFRA | -1.114523 | GBM | DK-MG | Masitinib |
| top60_altered_DNA_seed | KIT | -1.114523 | GBM | DK-MG | Masitinib |
| top60_altered_DNA_seed | CDK4 | -1.097164 | GBM | GB-1 | CGP-082996 |
| top60_altered_DNA_seed | MDM4 | -1.0886 | GBM | GB-1 | NSC-207895 |
| top60_altered_DNA_seed | KIT | -1.085279 | GBM | 8-MG-BA | OSI-930 |
| top60_altered_DNA_seed | KIT | -1.051823 | GBM | D-392MG | Sorafenib |
| top60_altered_DNA_seed | KIT | -1.044783 | GBM | CAS-1 | Pazopanib |
| top60_altered_DNA_seed | KIT | -1.044413 | GBM | A172 | Axitinib |
| top60_altered_DNA_seed | MDM4 | -1.03399 | GBM | D-566MG | NSC-207895 |
| top60_altered_DNA_seed | EGFR | -1.028082 | GBM | Becker | Erlotinib |
| top60_altered_DNA_seed | CDK4 | -1.011535 | GBM | SF268 | Palbociclib |
| top60_altered_DNA_seed | KIT | -1.008269 | GBM | D-247MG | Sorafenib |
| top60_altered_DNA_seed | KIT | -1.000371 | GBM | U251 | OSI-930 |
| top60_altered_DNA_seed | KIT | -0.979923 | GBM | DK-MG | Axitinib |
| top60_altered_DNA_seed | MDM4 | -0.95628 | GBM | SK-MG-1 | NSC-207895 |
| top60_altered_DNA_seed | EGFR | -0.954878 | GBM | U-87-MG | Afatinib |
| top60_altered_DNA_seed | KIT | -0.949293 | GBM | SF126 | Axitinib |
| top60_altered_DNA_seed | KIT | -0.948321 | GBM | D-392MG | Axitinib |
| top60_altered_DNA_seed | PDGFRA | -0.941443 | GBM | D-423MG | Amuvatinib |
| top60_altered_DNA_seed | KIT | -0.941443 | GBM | D-423MG | Amuvatinib |
| top60_altered_DNA_seed | EGFR | -0.941235 | GBM | KS-1 | Lapatinib |
| top60_altered_DNA_seed | CDK4 | -0.93056 | GBM | YKG-1 | Palbociclib |
| top60_altered_DNA_seed | EGFR | -0.924194 | GBM | SF268 | CUDC-101 |
| top60_altered_DNA_seed | EGFR | -0.919244 | GBM | A172 | Cetuximab |

| | | | | | |
|---|---|---|---|---|---|
| top60_altered_DNA_seed | KIT | -0.914208 | GBM | D-566MG | Pazopanib |
| top60_altered_DNA_seed | CDK4 | -0.9049 | GBM | Becker | Palbociclib |
| top60_altered_DNA_seed | KIT | -0.904738 | GBM | T98G | Pazopanib |
| top60_altered_DNA_seed | KIT | -0.900213 | GBM | SW1088 | Pazopanib |
| top60_altered_DNA_seed | CDK4 | -0.881495 | GBM | T98G | Palbociclib |
| top60_altered_DNA_seed | CDK4 | -0.874117 | GBM | SW1088 | Palbociclib |
| top60_altered_DNA_seed | EGFR | -0.859529 | GBM | GB-1 | CUDC-101 |
| top60_altered_DNA_seed | KIT | -0.843251 | GBM | DK-MG | Motesanib |
| top60_altered_DNA_seed | PDGFRA | -0.835034 | GBM | U251 | Amuvatinib |
| top60_altered_DNA_seed | KIT | -0.835034 | GBM | U251 | Amuvatinib |
| top60_altered_DNA_seed | KIT | -0.810695 | GBM | KS-1 | Pazopanib |
| top60_altered_DNA_seed | KIT | -0.802047 | GBM | SF126 | Motesanib |
| top60_altered_DNA_seed | KIT | -0.779205 | GBM | LN-405 | Pazopanib |
| top60_altered_DNA_seed | KIT | -0.777983 | GBM | SF126 | Sorafenib |
| top60_altered_DNA_seed | CDK4 | -0.764434 | GBM | SF295 | Palbociclib |
| top60_altered_DNA_seed | EGFR | -0.747323 | GBM | Becker | AZD8931 |
| top60_altered_DNA_seed | CDK4 | -0.730447 | GBM | D-423MG | Palbociclib |
| top60_altered_DNA_seed | EGFR | -0.730374 | GBM | D-542MG | Cetuximab |
| top60_altered_DNA_seed | EGFR | -0.724267 | GBM | KS-1 | CI-1033 |
| top60_altered_DNA_seed | CDK4 | -0.720499 | GBM | D-247MG | Palbociclib |
| top60_altered_DNA_seed | PDGFRA | -0.712378 | GBM | U251 | Masitinib |
| top60_altered_DNA_seed | KIT | -0.712378 | GBM | U251 | Masitinib |
| top60_altered_DNA_seed | CDK4 | -0.709831 | GBM | DK-MG | Palbociclib |
| top60_altered_DNA_seed | EGFR | -0.704283 | GBM | YH-13 | AZD8931 |
| top60_altered_DNA_seed | CDK4 | -0.680791 | GBM | SNB75 | CGP-082996 |
| top60_altered_DNA_seed | EGFR | -0.671917 | GBM | U-87-MG | Cetuximab |
| top60_altered_DNA_seed | KIT | -0.671466 | GBM | U-118-MG | Pazopanib |
| top60_altered_DNA_seed | EGFR | -0.652341 | GBM | SW1088 | Gefitinib |
| top60_altered_DNA_seed | PDGFRA | -0.646932 | GBM | SK-MG-1 | Masitinib |
| top60_altered_DNA_seed | KIT | -0.646932 | GBM | SK-MG-1 | Masitinib |
| top60_altered_DNA_seed | EGFR | -0.645346 | GBM | 8-MG-BA | Pelitinib |
| top60_altered_DNA_seed | KIT | -0.637122 | GBM | SF268 | Pazopanib |
| top60_altered_DNA_seed | KIT | -0.633538 | GBM | 8-MG-BA | Axitinib |
| top60_altered_DNA_seed | EGFR | -0.630226 | GBM | D-566MG | CUDC-101 |
| top60_altered_DNA_seed | PDGFRA | -0.624878 | GBM | 8-MG-BA | Masitinib |
| top60_altered_DNA_seed | KIT | -0.624878 | GBM | 8-MG-BA | Masitinib |
| top60_altered_DNA_seed | KIT | -0.622387 | GBM | SF295 | Axitinib |
| top60_altered_DNA_seed | EGFR | -0.622044 | GBM | YKG-1 | Afatinib |
| top60_altered_DNA_seed | EGFR | -0.621896 | GBM | D-423MG | CUDC-101 |
| top60_altered_DNA_seed | CDK4 | -0.589324 | GBM | GB-1 | Palbociclib |
| top60_altered_DNA_seed | MDM4 | -0.583662 | GBM | AM-38 | NSC-207895 |

| | | | | | |
|---|---|---|---|---|---|
| top60_altered_DNA_seed | EGFR | -0.55904 | GBM | SF295 | Afatinib |
| top60_altered_DNA_seed | EGFR | -0.556108 | GBM | KS-1 | Pelitinib |
| top60_altered_DNA_seed | PDGFRA | -0.548783 | GBM | D-566MG | Masitinib |
| top60_altered_DNA_seed | KIT | -0.548783 | GBM | D-566MG | Masitinib |
| top60_altered_DNA_seed | EGFR | -0.539459 | GBM | D-542MG | Erlotinib |
| top60_altered_DNA_seed | KIT | -0.537768 | GBM | LN-405 | Motesanib |
| top60_altered_DNA_seed | EGFR | -0.53299 | GBM | CAS-1 | Pelitinib |
| top60_altered_DNA_seed | KIT | -0.529625 | GBM | D-247MG | Motesanib |
| top60_altered_DNA_seed | EGFR | -0.528168 | GBM | YH-13 | AST-1306 |
| top60_altered_DNA_seed | CDK4 | -0.522497 | GBM | KS-1 | Palbociclib |
| top60_altered_DNA_seed | EGFR | -0.521341 | GBM | D-392MG | Lapatinib |
| top60_altered_DNA_seed | KIT | -0.518883 | GBM | LNZTA3WT4 | Axitinib |
| top60_altered_DNA_seed | PDGFRA | -0.516043 | GBM | YKG-1 | Masitinib |
| top60_altered_DNA_seed | KIT | -0.516043 | GBM | YKG-1 | Masitinib |
| top60_altered_DNA_seed | EGFR | -0.501998 | GBM | A172 | Gefitinib |
| top60_altered_DNA_seed | EGFR | -0.498351 | GBM | GB-1 | Cetuximab |
| top60_altered_DNA_seed | CDK4 | -0.488739 | GBM | SK-MG-1 | Palbociclib |
| top60_altered_DNA_seed | KIT | -0.476714 | GBM | D-542MG | Motesanib |
| top60_altered_DNA_seed | KIT | -0.467782 | GBM | D-423MG | Axitinib |
| top60_altered_DNA_seed | KIT | -0.464692 | GBM | U251 | Motesanib |
| top60_altered_DNA_seed | EGFR | -0.464147 | GBM | DK-MG | CI-1033 |
| top60_altered_DNA_seed | EGFR | -0.463366 | GBM | SF268 | Cetuximab |
| top60_altered_DNA_seed | PDGFRA | -0.456348 | GBM | A172 | Masitinib |
| top60_altered_DNA_seed | KIT | -0.456348 | GBM | A172 | Masitinib |
| top60_altered_DNA_seed | KIT | -0.450407 | GBM | SF268 | Axitinib |
| top60_altered_DNA_seed | KIT | -0.433233 | GBM | T98G | Axitinib |
| top60_altered_DNA_seed | EGFR | -0.42457 | GBM | Becker | Lapatinib |
| top60_altered_DNA_seed | EGFR | -0.42128 | GBM | D-247MG | Lapatinib |
| top60_altered_DNA_seed | KIT | -0.414221 | GBM | D-542MG | Sorafenib |
| top60_altered_DNA_seed | KIT | -0.409304 | GBM | D-392MG | Motesanib |
| top60_altered_DNA_seed | KIT | -0.408316 | GBM | 42-MG-BA | Axitinib |
| top60_altered_DNA_seed | EGFR | -0.40314 | GBM | YH-13 | Afatinib |
| top60_altered_DNA_seed | KIT | -0.402132 | GBM | SK-MG-1 | Motesanib |
| top60_altered_DNA_seed | EGFR | -0.398414 | GBM | SF126 | Cetuximab |
| top60_altered_DNA_seed | EGFR | -0.397437 | GBM | D-247MG | CUDC-101 |
| top60_altered_DNA_seed | CDK4 | -0.396022 | GBM | LN-18 | Palbociclib |
| top60_altered_DNA_seed | KIT | -0.393166 | GBM | D-263MG | Axitinib |
| top60_altered_DNA_seed | EGFR | -0.392801 | GBM | U251 | AZD8931 |
| top60_altered_DNA_seed | CDK4 | -0.384323 | GBM | 8-MG-BA | CGP-082996 |
| top60_altered_DNA_seed | EGFR | -0.383351 | GBM | SF295 | Gefitinib |
| top60_altered_DNA_seed | EGFR | -0.378426 | GBM | KS-1 | CUDC-101 |

| | | | | | |
|---|---|---|---|---|---|
| top60_altered_DNA_seed | KIT | -0.377473 | GBM | SF295 | Motesanib |
| top60_altered_DNA_seed | KIT | -0.374604 | GBM | D-542MG | OSI-930 |
| top60_altered_DNA_seed | EGFR | -0.373229 | GBM | D-566MG | Gefitinib |
| top60_altered_DNA_seed | KIT | -0.371311 | GBM | YKG-1 | OSI-930 |
| top60_altered_DNA_seed | EGFR | -0.366672 | GBM | SK-MG-1 | AZD8931 |
| top60_altered_DNA_seed | EGFR | -0.36401 | GBM | T98G | Cetuximab |
| top60_altered_DNA_seed | KIT | -0.359892 | GBM | Becker | Pazopanib |
| top60_altered_DNA_seed | EGFR | -0.357658 | GBM | SF268 | AZD8931 |
| top60_altered_DNA_seed | KIT | -0.356805 | GBM | D-423MG | Pazopanib |
| top60_altered_DNA_seed | MDM4 | -0.352334 | GBM | YKG-1 | NSC-207895 |
| top60_altered_DNA_seed | KIT | -0.347847 | GBM | T98G | Motesanib |
| top60_altered_DNA_seed | PDGFRA | -0.345152 | GBM | SF268 | Masitinib |
| top60_altered_DNA_seed | KIT | -0.345152 | GBM | SF268 | Masitinib |
| top60_altered_DNA_seed | EGFR | -0.341445 | GBM | D-542MG | AZD8931 |
| top60_altered_DNA_seed | KIT | -0.339833 | GBM | SW1088 | Axitinib |
| top60_altered_DNA_seed | EGFR | -0.335944 | GBM | D-566MG | Afatinib |
| top60_altered_DNA_seed | KIT | -0.333278 | GBM | A172 | OSI-930 |
| top60_altered_DNA_seed | EGFR | -0.331832 | GBM | AM-38 | Pelitinib |
| top60_altered_DNA_seed | PDGFRA | -0.317828 | GBM | D-423MG | Masitinib |
| top60_altered_DNA_seed | KIT | -0.317828 | GBM | D-423MG | Masitinib |
| top60_altered_DNA_seed | PDGFRA | -0.311239 | GBM | AM-38 | Masitinib |
| top60_altered_DNA_seed | KIT | -0.311239 | GBM | AM-38 | Masitinib |
| top60_altered_DNA_seed | EGFR | -0.309137 | GBM | 8-MG-BA | Gefitinib |
| top60_altered_DNA_seed | CDK4 | -0.302098 | GBM | D-566MG | Palbociclib |
| top60_altered_DNA_seed | PDGFRA | -0.299402 | GBM | YH-13 | Masitinib |
| top60_altered_DNA_seed | KIT | -0.299402 | GBM | YH-13 | Masitinib |
| top60_altered_DNA_seed | EGFR | -0.297176 | GBM | D-392MG | Erlotinib |
| top60_altered_DNA_seed | PDGFRA | -0.296604 | GBM | SF295 | Masitinib |
| top60_altered_DNA_seed | KIT | -0.296604 | GBM | SF295 | Masitinib |
| top60_altered_DNA_seed | PDGFRA | -0.294946 | GBM | LNZTA3WT4 | Amuvatinib |
| top60_altered_DNA_seed | KIT | -0.294946 | GBM | LNZTA3WT4 | Amuvatinib |
| top60_altered_DNA_seed | EGFR | -0.290201 | GBM | SF268 | Lapatinib |
| top60_altered_DNA_seed | EGFR | -0.283792 | GBM | U-87-MG | Afatinib |
| top60_altered_DNA_seed | CDK4 | -0.283675 | GBM | SF268 | CGP-082996 |
| top60_altered_DNA_seed | EGFR | -0.277359 | GBM | D-247MG | Erlotinib |
| top60_altered_DNA_seed | EGFR | -0.27486 | GBM | KS-1 | AZD8931 |
| top60_altered_DNA_seed | PDGFRA | -0.27437 | GBM | SF126 | Masitinib |
| top60_altered_DNA_seed | KIT | -0.27437 | GBM | SF126 | Masitinib |
| top60_altered_DNA_seed | EGFR | -0.270597 | GBM | KS-1 | PF-00299804 |
| top60_altered_DNA_seed | EGFR | -0.269198 | GBM | SF126 | AZD8931 |
| top60_altered_DNA_seed | EGFR | -0.259102 | GBM | D-542MG | Lapatinib |

| | | | | | |
|---|---|---|---|---|---|
| top60_altered_DNA_seed | EGFR | -0.252138 | GBM | SNB75 | AZD8931 |
| top60_altered_DNA_seed | PDGFRA | -0.250136 | GBM | GB-1 | Amuvatinib |
| top60_altered_DNA_seed | KIT | -0.250136 | GBM | GB-1 | Amuvatinib |
| top60_altered_DNA_seed | EGFR | -0.249357 | GBM | 8-MG-BA | CUDC-101 |
| top60_altered_DNA_seed | EGFR | -0.233569 | GBM | DBTRG-05MG | Pelitinib |
| top60_altered_DNA_seed | KIT | -0.229462 | GBM | SF268 | Motesanib |
| top60_altered_DNA_seed | KIT | -0.226997 | GBM | 8-MG-BA | Pazopanib |
| top60_altered_DNA_seed | PDGFRA | -0.226488 | GBM | KS-1 | Masitinib |
| top60_altered_DNA_seed | KIT | -0.226488 | GBM | KS-1 | Masitinib |
| top60_altered_DNA_seed | CDK4 | -0.222332 | GBM | U-118-MG | Palbociclib |
| top60_altered_DNA_seed | EGFR | -0.218535 | GBM | D-423MG | Pelitinib |
| top60_altered_DNA_seed | KIT | -0.218461 | GBM | Becker | Axitinib |
| top60_altered_DNA_seed | EGFR | -0.217538 | GBM | SK-MG-1 | Cetuximab |
| top60_altered_DNA_seed | KIT | -0.217188 | GBM | D-392MG | OSI-930 |
| top60_altered_DNA_seed | KIT | -0.216375 | GBM | GB-1 | Sorafenib |
| top60_altered_DNA_seed | EGFR | -0.208843 | GBM | LN-229 | Afatinib |
| top60_altered_DNA_seed | EGFR | -0.207617 | GBM | LN-405 | Gefitinib |
| top60_altered_DNA_seed | KIT | -0.198885 | GBM | LNZTA3WT4 | OSI-930 |
| top60_altered_DNA_seed | EGFR | -0.192046 | GBM | GB-1 | AST-1306 |
| top60_altered_DNA_seed | EGFR | -0.192031 | GBM | U-87-MG | AZD8931 |
| top60_altered_DNA_seed | EGFR | -0.190368 | GBM | KS-1 | Afatinib |
| top60_altered_DNA_seed | EGFR | -0.186879 | GBM | YKG-1 | CI-1033 |
| top60_altered_DNA_seed | EGFR | -0.180533 | GBM | KS-1 | Afatinib |
| top60_altered_DNA_seed | EGFR | -0.179381 | GBM | DBTRG-05MG | Gefitinib |
| top60_altered_DNA_seed | KIT | -0.173465 | GBM | LN-18 | Axitinib |
| top60_altered_DNA_seed | EGFR | -0.171645 | GBM | U251 | Pelitinib |
| top60_altered_DNA_seed | KIT | -0.169749 | GBM | U251 | Pazopanib |
| top60_altered_DNA_seed | MDM4 | -0.163512 | GBM | DK-MG | NSC-207895 |
| top60_altered_DNA_seed | EGFR | -0.158119 | GBM | D-542MG | CUDC-101 |
| top60_altered_DNA_seed | EGFR | -0.157381 | GBM | 8-MG-BA | Cetuximab |
| top60_altered_DNA_seed | KIT | -0.152528 | GBM | LNZTA3WT4 | Pazopanib |
| top60_altered_DNA_seed | KIT | -0.151015 | GBM | SK-MG-1 | Axitinib |
| top60_altered_DNA_seed | EGFR | -0.146351 | GBM | A172 | CUDC-101 |
| top60_altered_DNA_seed | EGFR | -0.143505 | GBM | GB-1 | CI-1033 |
| top60_altered_DNA_seed | EGFR | -0.142824 | GBM | YKG-1 | Cetuximab |
| top60_altered_DNA_seed | EGFR | -0.13723 | GBM | DK-MG | PF-00299804 |
| top60_altered_DNA_seed | CDK4 | -0.130802 | GBM | AM-38 | Palbociclib |
| top60_altered_DNA_seed | PDGFRA | -0.12509 | GBM | U-87-MG | Masitinib |
| top60_altered_DNA_seed | KIT | -0.12509 | GBM | U-87-MG | Masitinib |
| top60_altered_DNA_seed | EGFR | -0.119082 | GBM | T98G | AZD8931 |
| top60_altered_DNA_seed | KIT | -0.118261 | GBM | LN-18 | Pazopanib |

| | | | | | |
|---|---|---|---|---|---|
| top60_altered_DNA_seed | EGFR | -0.116551 | GBM | SF126 | CI-1033 |
| top60_altered_DNA_seed | PDGFRA | -0.110574 | GBM | CAS-1 | Amuvatinib |
| top60_altered_DNA_seed | KIT | -0.110574 | GBM | CAS-1 | Amuvatinib |
| top60_altered_DNA_seed | EGFR | -0.108835 | GBM | M059J | AZD8931 |
| top60_altered_DNA_seed | PDGFRA | -0.107572 | GBM | SF268 | Amuvatinib |
| top60_altered_DNA_seed | KIT | -0.107572 | GBM | SF268 | Amuvatinib |
| top60_altered_DNA_seed | EGFR | -0.095753 | GBM | CAS-1 | AST-1306 |
| top60_altered_DNA_seed | EGFR | -0.085668 | GBM | AM-38 | Gefitinib |
| top60_altered_DNA_seed | KIT | -0.079933 | GBM | D-566MG | OSI-930 |
| top60_altered_DNA_seed | EGFR | -0.078794 | GBM | M059J | CUDC-101 |
| top60_altered_DNA_seed | CDK4 | -0.07251 | GBM | D-392MG | Palbociclib |
| top60_altered_DNA_seed | EGFR | -0.057445 | GBM | AM-38 | AST-1306 |
| top60_altered_DNA_seed | KIT | -0.052292 | GBM | Becker | Sorafenib |
| top60_altered_DNA_seed | EGFR | -0.046977 | GBM | AM-38 | CUDC-101 |
| top60_altered_DNA_seed | EGFR | -0.038738 | GBM | D-423MG | Cetuximab |
| top60_altered_DNA_seed | KIT | -0.035361 | GBM | LN-18 | OSI-930 |
| top60_altered_DNA_seed | EGFR | -0.033848 | GBM | YH-13 | CI-1033 |
| top60_altered_DNA_seed | EGFR | -0.029853 | GBM | 42-MG-BA | CUDC-101 |
| top60_altered_DNA_seed | EGFR | -0.024533 | GBM | KS-1 | AST-1306 |
| top60_altered_DNA_seed | KIT | -0.021645 | GBM | SF295 | OSI-930 |
| top60_altered_DNA_seed | EGFR | -0.021607 | GBM | 8-MG-BA | Erlotinib |
| top60_altered_DNA_seed | KIT | -0.016623 | GBM | U-118-MG | OSI-930 |
| top60_altered_DNA_seed | EGFR | -0.013575 | GBM | SF126 | AST-1306 |
| top60_altered_DNA_seed | EGFR | -0.011909 | GBM | YKG-1 | PF-00299804 |
| top60_altered_DNA_seed | EGFR | -0.010827 | GBM | LN-229 | AST-1306 |
| top60_altered_DNA_seed | PDGFRA | -0.009646 | GBM | LN-18 | Masitinib |
| top60_altered_DNA_seed | KIT | -0.009646 | GBM | LN-18 | Masitinib |
| top60_altered_DNA_seed | CDK4 | -0.000924 | GBM | LNZTA3WT4 | Palbociclib |
| top60_altered_DNA_seed | PDGFRA | -0.000135 | GBM | 8-MG-BA | Amuvatinib |
| top60_altered_DNA_seed | KIT | -0.000135 | GBM | 8-MG-BA | Amuvatinib |
| top60_altered_DNA_seed | PDGFRA | -0.000123 | GBM | D-542MG | Amuvatinib |
| top60_altered_DNA_seed | KIT | -0.000123 | GBM | D-542MG | Amuvatinib |
| top60_altered_DNA_seed | KIT | 0.006691 | GBM | CAS-1 | Axitinib |
| top60_altered_DNA_seed | EGFR | 0.006771 | GBM | SNB75 | Lapatinib |
| top60_altered_DNA_seed | EGFR | 0.007005 | GBM | YKG-1 | Pelitinib |
| top60_altered_DNA_seed | PDGFRA | 0.009774 | GBM | LN-405 | Masitinib |
| top60_altered_DNA_seed | KIT | 0.009774 | GBM | LN-405 | Masitinib |
| top60_altered_DNA_seed | EGFR | 0.012495 | GBM | U251 | Cetuximab |
| top60_altered_DNA_seed | EGFR | 0.01419 | GBM | U251 | PF-00299804 |
| top60_altered_DNA_seed | EGFR | 0.01544 | GBM | 8-MG-BA | CI-1033 |
| top60_altered_DNA_seed | EGFR | 0.020324 | GBM | D-423MG | Afatinib |

| | | | | | |
|---|---|---|---|---|---|
| top60_altered_DNA_seed | KIT | 0.022968 | GBM | Becker | OSI-930 |
| top60_altered_DNA_seed | EGFR | 0.029027 | GBM | 8-MG-BA | Lapatinib |
| top60_altered_DNA_seed | EGFR | 0.032605 | GBM | LN-18 | AST-1306 |
| top60_altered_DNA_seed | EGFR | 0.037946 | GBM | LNZTA3WT4 | AST-1306 |
| top60_altered_DNA_seed | KIT | 0.039707 | GBM | A172 | Motesanib |
| top60_altered_DNA_seed | EGFR | 0.04348 | GBM | D-423MG | PF-00299804 |
| top60_altered_DNA_seed | EGFR | 0.044373 | GBM | U251 | AST-1306 |
| top60_altered_DNA_seed | EGFR | 0.044568 | GBM | D-263MG | AST-1306 |
| top60_altered_DNA_seed | EGFR | 0.045027 | GBM | LN-229 | CI-1033 |
| top60_altered_DNA_seed | PDGFRA | 0.047682 | GBM | A172 | Amuvatinib |
| top60_altered_DNA_seed | KIT | 0.047682 | GBM | A172 | Amuvatinib |
| top60_altered_DNA_seed | EGFR | 0.04783 | GBM | SK-MG-1 | CUDC-101 |
| top60_altered_DNA_seed | EGFR | 0.053342 | GBM | LN-18 | Cetuximab |
| top60_altered_DNA_seed | EGFR | 0.055296 | GBM | LN-18 | AZD8931 |
| top60_altered_DNA_seed | KIT | 0.058221 | GBM | D-566MG | Motesanib |
| top60_altered_DNA_seed | KIT | 0.059381 | GBM | LNZTA3WT4 | Motesanib |
| top60_altered_DNA_seed | EGFR | 0.066144 | GBM | SF295 | AZD8931 |
| top60_altered_DNA_seed | KIT | 0.066404 | GBM | LN-405 | OSI-930 |
| top60_altered_DNA_seed | EGFR | 0.067477 | GBM | LN-18 | PF-00299804 |
| top60_altered_DNA_seed | KIT | 0.072607 | GBM | CAS-1 | OSI-930 |
| top60_altered_DNA_seed | EGFR | 0.078088 | GBM | 8-MG-BA | Afatinib |
| top60_altered_DNA_seed | KIT | 0.083062 | GBM | T98G | OSI-930 |
| top60_altered_DNA_seed | EGFR | 0.083535 | GBM | U251 | CI-1033 |
| top60_altered_DNA_seed | PDGFRA | 0.085676 | GBM | YKG-1 | Amuvatinib |
| top60_altered_DNA_seed | KIT | 0.085676 | GBM | YKG-1 | Amuvatinib |
| top60_altered_DNA_seed | EGFR | 0.087284 | GBM | 8-MG-BA | AST-1306 |
| top60_altered_DNA_seed | KIT | 0.09315 | GBM | 8-MG-BA | Imatinib |
| top60_altered_DNA_seed | CDK4 | 0.09599 | GBM | DBTRG-05MG | Palbociclib |
| top60_altered_DNA_seed | EGFR | 0.100039 | GBM | SK-MG-1 | PF-00299804 |
| top60_altered_DNA_seed | EGFR | 0.100493 | GBM | D-566MG | CI-1033 |
| top60_altered_DNA_seed | EGFR | 0.101348 | GBM | YKG-1 | AZD8931 |
| top60_altered_DNA_seed | EGFR | 0.101937 | GBM | LN-229 | AZD8931 |
| top60_altered_DNA_seed | KIT | 0.1034 | GBM | SK-MG-1 | Pazopanib |
| top60_altered_DNA_seed | KIT | 0.103442 | GBM | U251 | Axitinib |
| top60_altered_DNA_seed | KIT | 0.1047 | GBM | YKG-1 | Axitinib |
| top60_altered_DNA_seed | EGFR | 0.106911 | GBM | D-542MG | AST-1306 |
| top60_altered_DNA_seed | EGFR | 0.109626 | GBM | LN-229 | PF-00299804 |
| top60_altered_DNA_seed | KIT | 0.115877 | GBM | KS-1 | Sorafenib |
| top60_altered_DNA_seed | KIT | 0.117192 | GBM | SW1088 | OSI-930 |
| top60_altered_DNA_seed | EGFR | 0.120889 | GBM | AM-38 | CI-1033 |
| top60_altered_DNA_seed | EGFR | 0.122608 | GBM | 42-MG-BA | CI-1033 |

| | | | | | |
|---|---|---|---|---|---|
| top60_altered_DNA_seed | EGFR | 0.122964 | GBM | YH-13 | Pelitinib |
| top60_altered_DNA_seed | EGFR | 0.126075 | GBM | D-247MG | AZD8931 |
| top60_altered_DNA_seed | EGFR | 0.129469 | GBM | LN-229 | Afatinib |
| top60_altered_DNA_seed | PDGFRA | 0.132873 | GBM | Becker | Amuvatinib |
| top60_altered_DNA_seed | KIT | 0.132873 | GBM | Becker | Amuvatinib |
| top60_altered_DNA_seed | EGFR | 0.133898 | GBM | 8-MG-BA | PF-00299804 |
| top60_altered_DNA_seed | EGFR | 0.134567 | GBM | D-423MG | AZD8931 |
| top60_altered_DNA_seed | KIT | 0.138209 | GBM | 8-MG-BA | Motesanib |
| top60_altered_DNA_seed | KIT | 0.139832 | GBM | SF126 | Imatinib |
| top60_altered_DNA_seed | EGFR | 0.140854 | GBM | D-566MG | Pelitinib |
| top60_altered_DNA_seed | KIT | 0.141263 | GBM | D-423MG | OSI-930 |
| top60_altered_DNA_seed | EGFR | 0.156294 | GBM | D-566MG | AST-1306 |
| top60_altered_DNA_seed | EGFR | 0.160378 | GBM | AM-38 | PF-00299804 |
| top60_altered_DNA_seed | EGFR | 0.160936 | GBM | Becker | Pelitinib |
| top60_altered_DNA_seed | EGFR | 0.165266 | GBM | U-87-MG | AST-1306 |
| top60_altered_DNA_seed | EGFR | 0.167431 | GBM | SNB75 | AST-1306 |
| top60_altered_DNA_seed | EGFR | 0.167731 | GBM | DBTRG-05MG | CUDC-101 |
| top60_altered_DNA_seed | EGFR | 0.171606 | GBM | D-392MG | Cetuximab |
| top60_altered_DNA_seed | PDGFRA | 0.176476 | GBM | DBTRG-05MG | Masitinib |
| top60_altered_DNA_seed | KIT | 0.176476 | GBM | DBTRG-05MG | Masitinib |
| top60_altered_DNA_seed | EGFR | 0.176952 | GBM | SF126 | Erlotinib |
| top60_altered_DNA_seed | EGFR | 0.185456 | GBM | A172 | AZD8931 |
| top60_altered_DNA_seed | EGFR | 0.188649 | GBM | SF295 | CUDC-101 |
| top60_altered_DNA_seed | EGFR | 0.193134 | GBM | YH-13 | CUDC-101 |
| top60_altered_DNA_seed | KIT | 0.19325 | GBM | Becker | Imatinib |
| top60_altered_DNA_seed | KIT | 0.201687 | GBM | D-263MG | Sorafenib |
| top60_altered_DNA_seed | EGFR | 0.203313 | GBM | LN-229 | CUDC-101 |
| top60_altered_DNA_seed | KIT | 0.203491 | GBM | U-87-MG | Imatinib |
| top60_altered_DNA_seed | EGFR | 0.207045 | GBM | GB-1 | Lapatinib |
| top60_altered_DNA_seed | KIT | 0.213831 | GBM | DBTRG-05MG | Axitinib |
| top60_altered_DNA_seed | EGFR | 0.216187 | GBM | SK-MG-1 | AST-1306 |
| top60_altered_DNA_seed | EGFR | 0.22013 | GBM | YKG-1 | AST-1306 |
| top60_altered_DNA_seed | EGFR | 0.220175 | GBM | LN-229 | Gefitinib |
| top60_altered_DNA_seed | PDGFRA | 0.225253 | GBM | M059J | Masitinib |
| top60_altered_DNA_seed | KIT | 0.225253 | GBM | M059J | Masitinib |
| top60_altered_DNA_seed | KIT | 0.229827 | GBM | D-247MG | Imatinib |
| top60_altered_DNA_seed | EGFR | 0.231418 | GBM | GB-1 | Afatinib |
| top60_altered_DNA_seed | EGFR | 0.239334 | GBM | D-263MG | Erlotinib |
| top60_altered_DNA_seed | KIT | 0.240315 | GBM | DBTRG-05MG | Pazopanib |
| top60_altered_DNA_seed | EGFR | 0.241185 | GBM | YH-13 | Gefitinib |
| top60_altered_DNA_seed | MDM4 | 0.241462 | GBM | KS-1 | NSC-207895 |

| | | | | | |
|---|---|---|---|---|---|
| top60_altered_DNA_seed | KIT | 0.243139 | GBM | D-542MG | Axitinib |
| top60_altered_DNA_seed | KIT | 0.243608 | GBM | 42-MG-BA | OSI-930 |
| top60_altered_DNA_seed | KIT | 0.245897 | GBM | U-87-MG | Motesanib |
| top60_altered_DNA_seed | EGFR | 0.248324 | GBM | LN-18 | CI-1033 |
| top60_altered_DNA_seed | CDK4 | 0.248666 | GBM | SF126 | CGP-082996 |
| top60_altered_DNA_seed | EGFR | 0.255388 | GBM | Becker | Cetuximab |
| top60_altered_DNA_seed | EGFR | 0.25594 | GBM | SF295 | Cetuximab |
| top60_altered_DNA_seed | EGFR | 0.261073 | GBM | KS-1 | Gefitinib |
| top60_altered_DNA_seed | CDK4 | 0.267169 | GBM | D-247MG | CGP-082996 |
| top60_altered_DNA_seed | EGFR | 0.269698 | GBM | LN-18 | Afatinib |
| top60_altered_DNA_seed | EGFR | 0.274048 | GBM | SF126 | CUDC-101 |
| top60_altered_DNA_seed | EGFR | 0.276723 | GBM | DBTRG-05MG | AZD8931 |
| top60_altered_DNA_seed | EGFR | 0.276843 | GBM | SW1088 | Afatinib |
| top60_altered_DNA_seed | EGFR | 0.279396 | GBM | 8-MG-BA | AZD8931 |
| top60_altered_DNA_seed | EGFR | 0.282952 | GBM | CAS-1 | PF-00299804 |
| top60_altered_DNA_seed | KIT | 0.285659 | GBM | KS-1 | Axitinib |
| top60_altered_DNA_seed | EGFR | 0.286063 | GBM | SNB75 | Afatinib |
| top60_altered_DNA_seed | EGFR | 0.287584 | GBM | D-263MG | AZD8931 |
| top60_altered_DNA_seed | EGFR | 0.290069 | GBM | U251 | Gefitinib |
| top60_altered_DNA_seed | EGFR | 0.292605 | GBM | D-247MG | AST-1306 |
| top60_altered_DNA_seed | EGFR | 0.294226 | GBM | GB-1 | Pelitinib |
| top60_altered_DNA_seed | KIT | 0.300276 | GBM | AM-38 | Imatinib |
| top60_altered_DNA_seed | EGFR | 0.3009 | GBM | Becker | Afatinib |
| top60_altered_DNA_seed | EGFR | 0.306633 | GBM | LN-18 | Gefitinib |
| top60_altered_DNA_seed | EGFR | 0.309742 | GBM | D-392MG | CUDC-101 |
| top60_altered_DNA_seed | EGFR | 0.310051 | GBM | LN-18 | Afatinib |
| top60_altered_DNA_seed | EGFR | 0.311172 | GBM | M059J | CI-1033 |
| top60_altered_DNA_seed | KIT | 0.313197 | GBM | D-247MG | Axitinib |
| top60_altered_DNA_seed | EGFR | 0.313652 | GBM | CAS-1 | CI-1033 |
| top60_altered_DNA_seed | EGFR | 0.315478 | GBM | KS-1 | Cetuximab |
| top60_altered_DNA_seed | EGFR | 0.317128 | GBM | GB-1 | PF-00299804 |
| top60_altered_DNA_seed | MDM4 | 0.32474 | GBM | T98G | NSC-207895 |
| top60_altered_DNA_seed | EGFR | 0.324796 | GBM | U251 | Afatinib |
| top60_altered_DNA_seed | EGFR | 0.32485 | GBM | D-566MG | Afatinib |
| top60_altered_DNA_seed | EGFR | 0.325862 | GBM | D-263MG | Lapatinib |
| top60_altered_DNA_seed | EGFR | 0.328818 | GBM | D-423MG | CI-1033 |
| top60_altered_DNA_seed | PDGFRA | 0.331779 | GBM | T98G | Amuvatinib |
| top60_altered_DNA_seed | KIT | 0.331779 | GBM | T98G | Amuvatinib |
| top60_altered_DNA_seed | MDM4 | 0.334359 | GBM | SF295 | NSC-207895 |
| top60_altered_DNA_seed | KIT | 0.336955 | GBM | SF268 | Sorafenib |
| top60_altered_DNA_seed | EGFR | 0.34101 | GBM | DK-MG | AST-1306 |

| | | | | | |
|---|---|---|---|---|---|
| top60_altered_DNA_seed | KIT | 0.348106 | GBM | SNB75 | Motesanib |
| top60_altered_DNA_seed | EGFR | 0.349775 | GBM | T98G | Afatinib |
| top60_altered_DNA_seed | PDGFRA | 0.353126 | GBM | D-392MG | Masitinib |
| top60_altered_DNA_seed | KIT | 0.353126 | GBM | D-392MG | Masitinib |
| top60_altered_DNA_seed | KIT | 0.358007 | GBM | U-118-MG | Axitinib |
| top60_altered_DNA_seed | EGFR | 0.359435 | GBM | T98G | Pelitinib |
| top60_altered_DNA_seed | PDGFRA | 0.361447 | GBM | T98G | Masitinib |
| top60_altered_DNA_seed | KIT | 0.361447 | GBM | T98G | Masitinib |
| top60_altered_DNA_seed | EGFR | 0.361453 | GBM | D-392MG | AST-1306 |
| top60_altered_DNA_seed | KIT | 0.362003 | GBM | D-263MG | OSI-930 |
| top60_altered_DNA_seed | EGFR | 0.362003 | GBM | U251 | Afatinib |
| top60_altered_DNA_seed | KIT | 0.362144 | GBM | DBTRG-05MG | Motesanib |
| top60_altered_DNA_seed | KIT | 0.363008 | GBM | KS-1 | Motesanib |
| top60_altered_DNA_seed | EGFR | 0.365374 | GBM | M059J | Cetuximab |
| top60_altered_DNA_seed | EGFR | 0.36579 | GBM | SF126 | PF-00299804 |
| top60_altered_DNA_seed | EGFR | 0.368807 | GBM | D-566MG | AZD8931 |
| top60_altered_DNA_seed | EGFR | 0.374295 | GBM | YH-13 | PF-00299804 |
| top60_altered_DNA_seed | KIT | 0.375078 | GBM | U-118-MG | Motesanib |
| top60_altered_DNA_seed | EGFR | 0.376762 | GBM | U-118-MG | CUDC-101 |
| top60_altered_DNA_seed | EGFR | 0.376843 | GBM | M059J | Gefitinib |
| top60_altered_DNA_seed | EGFR | 0.379354 | GBM | 42-MG-BA | Cetuximab |
| top60_altered_DNA_seed | EGFR | 0.39038 | GBM | A172 | Pelitinib |
| top60_altered_DNA_seed | EGFR | 0.390957 | GBM | D-392MG | Gefitinib |
| top60_altered_DNA_seed | KIT | 0.39282 | GBM | M059J | Axitinib |
| top60_altered_DNA_seed | CDK4 | 0.398742 | GBM | U-87-MG | Palbociclib |
| top60_altered_DNA_seed | EGFR | 0.3991 | GBM | D-392MG | AZD8931 |
| top60_altered_DNA_seed | EGFR | 0.399554 | GBM | M059J | AST-1306 |
| top60_altered_DNA_seed | EGFR | 0.403308 | GBM | SF268 | Pelitinib |
| top60_altered_DNA_seed | PDGFRA | 0.408351 | GBM | D-392MG | Amuvatinib |
| top60_altered_DNA_seed | KIT | 0.408351 | GBM | D-392MG | Amuvatinib |
| top60_altered_DNA_seed | EGFR | 0.41024 | GBM | 8-MG-BA | Afatinib |
| top60_altered_DNA_seed | MDM4 | 0.421423 | GBM | U251 | NSC-207895 |
| top60_altered_DNA_seed | KIT | 0.422007 | GBM | LN-229 | Pazopanib |
| top60_altered_DNA_seed | EGFR | 0.427033 | GBM | SK-MG-1 | Afatinib |
| top60_altered_DNA_seed | EGFR | 0.433777 | GBM | SK-MG-1 | Afatinib |
| top60_altered_DNA_seed | EGFR | 0.436719 | GBM | SF268 | Afatinib |
| top60_altered_DNA_seed | PDGFRA | 0.440862 | GBM | SW1088 | Masitinib |
| top60_altered_DNA_seed | KIT | 0.440862 | GBM | SW1088 | Masitinib |
| top60_altered_DNA_seed | EGFR | 0.45681 | GBM | LNZTA3WT4 | CI-1033 |
| top60_altered_DNA_seed | KIT | 0.458569 | GBM | M059J | Motesanib |
| top60_altered_DNA_seed | EGFR | 0.462306 | GBM | D-423MG | AST-1306 |

| | | | | | |
|---|---|---|---|---|---|
| top60_altered_DNA_seed | KIT | 0.462812 | GBM | CAS-1 | Imatinib |
| top60_altered_DNA_seed | PDGFRA | 0.466775 | GBM | SF295 | Amuvatinib |
| top60_altered_DNA_seed | KIT | 0.466775 | GBM | SF295 | Amuvatinib |
| top60_altered_DNA_seed | EGFR | 0.468353 | GBM | DK-MG | CUDC-101 |
| top60_altered_DNA_seed | EGFR | 0.46971 | GBM | DK-MG | Cetuximab |
| top60_altered_DNA_seed | EGFR | 0.474493 | GBM | 42-MG-BA | PF-00299804 |
| top60_altered_DNA_seed | EGFR | 0.476711 | GBM | SNB75 | CUDC-101 |
| top60_altered_DNA_seed | EGFR | 0.480983 | GBM | LN-229 | Cetuximab |
| top60_altered_DNA_seed | EGFR | 0.481954 | GBM | LN-405 | AST-1306 |
| top60_altered_DNA_seed | EGFR | 0.485131 | GBM | SW1088 | AST-1306 |
| top60_altered_DNA_seed | MDM4 | 0.489949 | GBM | 8-MG-BA | NSC-207895 |
| top60_altered_DNA_seed | KIT | 0.497011 | GBM | DBTRG-05MG | OSI-930 |
| top60_altered_DNA_seed | EGFR | 0.498474 | GBM | D-542MG | PF-00299804 |
| top60_altered_DNA_seed | KIT | 0.499232 | GBM | U-87-MG | Sorafenib |
| top60_altered_DNA_seed | EGFR | 0.500836 | GBM | LN-229 | Pelitinib |
| top60_altered_DNA_seed | EGFR | 0.51882 | GBM | U-118-MG | Gefitinib |
| top60_altered_DNA_seed | KIT | 0.520885 | GBM | D-263MG | Pazopanib |
| top60_altered_DNA_seed | EGFR | 0.525061 | GBM | YKG-1 | Gefitinib |
| top60_altered_DNA_seed | PDGFRA | 0.525591 | GBM | D-566MG | Amuvatinib |
| top60_altered_DNA_seed | KIT | 0.525591 | GBM | D-566MG | Amuvatinib |
| top60_altered_DNA_seed | EGFR | 0.528014 | GBM | D-263MG | Afatinib |
| top60_altered_DNA_seed | EGFR | 0.530853 | GBM | D-423MG | Gefitinib |
| top60_altered_DNA_seed | KIT | 0.534042 | GBM | SF268 | Imatinib |
| top60_altered_DNA_seed | EGFR | 0.534881 | GBM | SF295 | Afatinib |
| top60_altered_DNA_seed | EGFR | 0.535548 | GBM | A172 | Afatinib |
| top60_altered_DNA_seed | KIT | 0.53667 | GBM | LN-229 | Motesanib |
| top60_altered_DNA_seed | CDK4 | 0.537366 | GBM | SF126 | Palbociclib |
| top60_altered_DNA_seed | EGFR | 0.545301 | GBM | LN-18 | CUDC-101 |
| top60_altered_DNA_seed | PDGFRA | 0.545993 | GBM | LN-18 | Amuvatinib |
| top60_altered_DNA_seed | KIT | 0.545993 | GBM | LN-18 | Amuvatinib |
| top60_altered_DNA_seed | EGFR | 0.546202 | GBM | Becker | Afatinib |
| top60_altered_DNA_seed | EGFR | 0.548579 | GBM | SW1088 | AZD8931 |
| top60_altered_DNA_seed | PDGFRA | 0.551922 | GBM | CAS-1 | Masitinib |
| top60_altered_DNA_seed | KIT | 0.551922 | GBM | CAS-1 | Masitinib |
| top60_altered_DNA_seed | PDGFRA | 0.556644 | GBM | U-118-MG | Masitinib |
| top60_altered_DNA_seed | KIT | 0.556644 | GBM | U-118-MG | Masitinib |
| top60_altered_DNA_seed | EGFR | 0.557431 | GBM | SF126 | Pelitinib |
| top60_altered_DNA_seed | KIT | 0.55828 | GBM | GB-1 | Axitinib |
| top60_altered_DNA_seed | EGFR | 0.562046 | GBM | DBTRG-05MG | AST-1306 |
| top60_altered_DNA_seed | PDGFRA | 0.562627 | GBM | YH-13 | Amuvatinib |
| top60_altered_DNA_seed | KIT | 0.562627 | GBM | YH-13 | Amuvatinib |

| | | | | | |
|---|---|---|---|---|---|
| top60_altered_DNA_seed | EGFR | 0.57615 | GBM | D-263MG | CI-1033 |
| top60_altered_DNA_seed | EGFR | 0.578175 | GBM | 42-MG-BA | Afatinib |
| top60_altered_DNA_seed | EGFR | 0.584384 | GBM | D-566MG | PF-00299804 |
| top60_altered_DNA_seed | EGFR | 0.588596 | GBM | T98G | Afatinib |
| top60_altered_DNA_seed | EGFR | 0.59153 | GBM | SK-MG-1 | CI-1033 |
| top60_altered_DNA_seed | EGFR | 0.592515 | GBM | D-263MG | PF-00299804 |
| top60_altered_DNA_seed | EGFR | 0.593466 | GBM | GB-1 | Afatinib |
| top60_altered_DNA_seed | EGFR | 0.596205 | GBM | LN-405 | Cetuximab |
| top60_altered_DNA_seed | KIT | 0.59876 | GBM | AM-38 | OSI-930 |
| top60_altered_DNA_seed | EGFR | 0.599575 | GBM | D-263MG | Cetuximab |
| top60_altered_DNA_seed | EGFR | 0.603259 | GBM | Becker | AST-1306 |
| top60_altered_DNA_seed | PDGFRA | 0.604991 | GBM | 42-MG-BA | Masitinib |
| top60_altered_DNA_seed | KIT | 0.604991 | GBM | 42-MG-BA | Masitinib |
| top60_altered_DNA_seed | PDGFRA | 0.608412 | GBM | U-118-MG | Amuvatinib |
| top60_altered_DNA_seed | KIT | 0.608412 | GBM | U-118-MG | Amuvatinib |
| top60_altered_DNA_seed | EGFR | 0.611763 | GBM | DK-MG | Afatinib |
| top60_altered_DNA_seed | EGFR | 0.613775 | GBM | T98G | Gefitinib |
| top60_altered_DNA_seed | PDGFRA | 0.619007 | GBM | SF126 | Amuvatinib |
| top60_altered_DNA_seed | KIT | 0.619007 | GBM | SF126 | Amuvatinib |
| top60_altered_DNA_seed | EGFR | 0.620391 | GBM | 42-MG-BA | AST-1306 |
| top60_altered_DNA_seed | EGFR | 0.62599 | GBM | SF268 | PF-00299804 |
| top60_altered_DNA_seed | KIT | 0.626929 | GBM | GB-1 | Imatinib |
| top60_altered_DNA_seed | EGFR | 0.627451 | GBM | A172 | Afatinib |
| top60_altered_DNA_seed | KIT | 0.633945 | GBM | SNB75 | Axitinib |
| top60_altered_DNA_seed | EGFR | 0.636266 | GBM | LNZTA3WT4 | Cetuximab |
| top60_altered_DNA_seed | EGFR | 0.639196 | GBM | D-392MG | PF-00299804 |
| top60_altered_DNA_seed | EGFR | 0.643621 | GBM | Becker | CI-1033 |
| top60_altered_DNA_seed | EGFR | 0.645694 | GBM | CAS-1 | CUDC-101 |
| top60_altered_DNA_seed | EGFR | 0.648066 | GBM | U-118-MG | PF-00299804 |
| top60_altered_DNA_seed | EGFR | 0.648765 | GBM | D-263MG | Gefitinib |
| top60_altered_DNA_seed | EGFR | 0.651567 | GBM | D-247MG | Cetuximab |
| top60_altered_DNA_seed | EGFR | 0.654301 | GBM | SF295 | PF-00299804 |
| top60_altered_DNA_seed | MDM4 | 0.658592 | GBM | LN-18 | NSC-207895 |
| top60_altered_DNA_seed | EGFR | 0.660387 | GBM | U-87-MG | PF-00299804 |
| top60_altered_DNA_seed | KIT | 0.660892 | GBM | SK-MG-1 | OSI-930 |
| top60_altered_DNA_seed | EGFR | 0.661416 | GBM | 42-MG-BA | Gefitinib |
| top60_altered_DNA_seed | EGFR | 0.666765 | GBM | CAS-1 | Lapatinib |
| top60_altered_DNA_seed | MDM4 | 0.674033 | GBM | D-247MG | NSC-207895 |
| top60_altered_DNA_seed | EGFR | 0.674141 | GBM | A172 | AST-1306 |
| top60_altered_DNA_seed | EGFR | 0.675234 | GBM | M059J | PF-00299804 |
| top60_altered_DNA_seed | EGFR | 0.677006 | GBM | D-263MG | Afatinib |

| | | | | | |
|---|---|---|---|---|---|
| top60_altered_DNA_seed | EGFR | 0.6825 | GBM | D-392MG | Afatinib |
| top60_altered_DNA_seed | EGFR | 0.686034 | GBM | U-87-MG | CUDC-101 |
| top60_altered_DNA_seed | EGFR | 0.689103 | GBM | M059J | Afatinib |
| top60_altered_DNA_seed | EGFR | 0.695419 | GBM | U-118-MG | AZD8931 |
| top60_altered_DNA_seed | EGFR | 0.695518 | GBM | Becker | Gefitinib |
| top60_altered_DNA_seed | EGFR | 0.698169 | GBM | 42-MG-BA | Afatinib |
| top60_altered_DNA_seed | EGFR | 0.698915 | GBM | 42-MG-BA | AZD8931 |
| top60_altered_DNA_seed | EGFR | 0.699406 | GBM | YKG-1 | Afatinib |
| top60_altered_DNA_seed | EGFR | 0.699493 | GBM | CAS-1 | Cetuximab |
| top60_altered_DNA_seed | EGFR | 0.699883 | GBM | SK-MG-1 | Pelitinib |
| top60_altered_DNA_seed | PDGFRA | 0.701246 | GBM | GB-1 | Masitinib |
| top60_altered_DNA_seed | KIT | 0.701246 | GBM | GB-1 | Masitinib |
| top60_altered_DNA_seed | PDGFRA | 0.706838 | GBM | D-263MG | Amuvatinib |
| top60_altered_DNA_seed | KIT | 0.706838 | GBM | D-263MG | Amuvatinib |
| top60_altered_DNA_seed | EGFR | 0.708314 | GBM | T98G | AST-1306 |
| top60_altered_DNA_seed | EGFR | 0.709652 | GBM | SW1088 | Pelitinib |
| top60_altered_DNA_seed | EGFR | 0.715007 | GBM | U-87-MG | CI-1033 |
| top60_altered_DNA_seed | KIT | 0.715535 | GBM | YKG-1 | Motesanib |
| top60_altered_DNA_seed | KIT | 0.715967 | GBM | SF126 | OSI-930 |
| top60_altered_DNA_seed | KIT | 0.716577 | GBM | GB-1 | OSI-930 |
| top60_altered_DNA_seed | EGFR | 0.716797 | GBM | YH-13 | Afatinib |
| top60_altered_DNA_seed | EGFR | 0.730349 | GBM | 42-MG-BA | Pelitinib |
| top60_altered_DNA_seed | EGFR | 0.732461 | GBM | SF126 | Gefitinib |
| top60_altered_DNA_seed | EGFR | 0.741936 | GBM | D-392MG | CI-1033 |
| top60_altered_DNA_seed | PDGFRA | 0.742057 | GBM | DBTRG-05MG | Amuvatinib |
| top60_altered_DNA_seed | KIT | 0.742057 | GBM | DBTRG-05MG | Amuvatinib |
| top60_altered_DNA_seed | EGFR | 0.748635 | GBM | AM-38 | Lapatinib |
| top60_altered_DNA_seed | MDM4 | 0.749432 | GBM | LNZTA3WT4 | NSC-207895 |
| top60_altered_DNA_seed | EGFR | 0.753517 | GBM | SF126 | Afatinib |
| top60_altered_DNA_seed | EGFR | 0.755547 | GBM | U-87-MG | Lapatinib |
| top60_altered_DNA_seed | KIT | 0.766075 | GBM | Becker | Motesanib |
| top60_altered_DNA_seed | EGFR | 0.774173 | GBM | SF295 | Pelitinib |
| top60_altered_DNA_seed | EGFR | 0.775129 | GBM | AM-38 | Afatinib |
| top60_altered_DNA_seed | EGFR | 0.778509 | GBM | D-392MG | Pelitinib |
| top60_altered_DNA_seed | EGFR | 0.780786 | GBM | AM-38 | Erlotinib |
| top60_altered_DNA_seed | EGFR | 0.782544 | GBM | LN-405 | CUDC-101 |
| top60_altered_DNA_seed | EGFR | 0.782733 | GBM | SNB75 | Cetuximab |
| top60_altered_DNA_seed | EGFR | 0.785877 | GBM | SF126 | Lapatinib |
| top60_altered_DNA_seed | EGFR | 0.786171 | GBM | LN-405 | AZD8931 |
| top60_altered_DNA_seed | KIT | 0.786733 | GBM | D-263MG | Imatinib |
| top60_altered_DNA_seed | MDM4 | 0.79203 | GBM | Becker | NSC-207895 |

| | | | | | |
|---|---|---|---|---|---|
| top60_altered_DNA_seed | EGFR | 0.792512 | GBM | LN-405 | Afatinib |
| top60_altered_DNA_seed | KIT | 0.795167 | GBM | YH-13 | OSI-930 |
| top60_altered_DNA_seed | KIT | 0.795643 | GBM | LN-229 | OSI-930 |
| top60_altered_DNA_seed | EGFR | 0.798119 | GBM | SF126 | Afatinib |
| top60_altered_DNA_seed | EGFR | 0.799122 | GBM | U-87-MG | Gefitinib |
| top60_altered_DNA_seed | EGFR | 0.803099 | GBM | GB-1 | AZD8931 |
| top60_altered_DNA_seed | EGFR | 0.81115 | GBM | SF295 | AST-1306 |
| top60_altered_DNA_seed | EGFR | 0.813821 | GBM | LNZTA3WT4 | PF-00299804 |
| top60_altered_DNA_seed | KIT | 0.813889 | GBM | D-423MG | Motesanib |
| top60_altered_DNA_seed | KIT | 0.820614 | GBM | AM-38 | Axitinib |
| top60_altered_DNA_seed | MDM4 | 0.821518 | GBM | A172 | NSC-207895 |
| top60_altered_DNA_seed | EGFR | 0.824329 | GBM | SF268 | Erlotinib |
| top60_altered_DNA_seed | KIT | 0.829619 | GBM | M059J | OSI-930 |
| top60_altered_DNA_seed | EGFR | 0.848985 | GBM | T98G | PF-00299804 |
| top60_altered_DNA_seed | EGFR | 0.85078 | GBM | D-423MG | Afatinib |
| top60_altered_DNA_seed | EGFR | 0.852691 | GBM | LNZTA3WT4 | Afatinib |
| top60_altered_DNA_seed | EGFR | 0.854566 | GBM | GB-1 | Gefitinib |
| top60_altered_DNA_seed | KIT | 0.856831 | GBM | KS-1 | Imatinib |
| top60_altered_DNA_seed | PDGFRA | 0.861017 | GBM | SNB75 | Masitinib |
| top60_altered_DNA_seed | KIT | 0.861017 | GBM | SNB75 | Masitinib |
| top60_altered_DNA_seed | MDM4 | 0.866497 | GBM | M059J | NSC-207895 |
| top60_altered_DNA_seed | EGFR | 0.868661 | GBM | M059J | Afatinib |
| top60_altered_DNA_seed | PDGFRA | 0.869844 | GBM | LNZTA3WT4 | Masitinib |
| top60_altered_DNA_seed | KIT | 0.869844 | GBM | LNZTA3WT4 | Masitinib |
| top60_altered_DNA_seed | EGFR | 0.871061 | GBM | SK-MG-1 | Gefitinib |
| top60_altered_DNA_seed | EGFR | 0.875322 | GBM | LNZTA3WT4 | Afatinib |
| top60_altered_DNA_seed | KIT | 0.886004 | GBM | AM-38 | Sorafenib |
| top60_altered_DNA_seed | EGFR | 0.893863 | GBM | SF268 | AST-1306 |
| top60_altered_DNA_seed | PDGFRA | 0.896973 | GBM | 42-MG-BA | Amuvatinib |
| top60_altered_DNA_seed | KIT | 0.896973 | GBM | 42-MG-BA | Amuvatinib |
| top60_altered_DNA_seed | EGFR | 0.905008 | GBM | CAS-1 | AZD8931 |
| top60_altered_DNA_seed | EGFR | 0.905612 | GBM | LNZTA3WT4 | Pelitinib |
| top60_altered_DNA_seed | KIT | 0.907527 | GBM | GB-1 | Motesanib |
| top60_altered_DNA_seed | EGFR | 0.907562 | GBM | U-118-MG | AST-1306 |
| top60_altered_DNA_seed | PDGFRA | 0.907839 | GBM | D-542MG | Masitinib |
| top60_altered_DNA_seed | KIT | 0.907839 | GBM | D-542MG | Masitinib |
| top60_altered_DNA_seed | EGFR | 0.911172 | GBM | Becker | CUDC-101 |
| top60_altered_DNA_seed | EGFR | 0.918206 | GBM | LN-405 | PF-00299804 |
| top60_altered_DNA_seed | EGFR | 0.922185 | GBM | A172 | PF-00299804 |
| top60_altered_DNA_seed | EGFR | 0.922255 | GBM | SNB75 | PF-00299804 |
| top60_altered_DNA_seed | EGFR | 0.928215 | GBM | SW1088 | PF-00299804 |

| | | | | | |
|---|---|---|---|---|---|
| top60_altered_DNA_seed | EGFR | 0.930239 | GBM | SF268 | Afatinib |
| top60_altered_DNA_seed | EGFR | 0.930559 | GBM | DBTRG-05MG | Cetuximab |
| top60_altered_DNA_seed | EGFR | 0.934676 | GBM | D-247MG | Gefitinib |
| top60_altered_DNA_seed | KIT | 0.94659 | GBM | SF268 | OSI-930 |
| top60_altered_DNA_seed | EGFR | 0.948982 | GBM | DBTRG-05MG | PF-00299804 |
| top60_altered_DNA_seed | EGFR | 0.950111 | GBM | LN-18 | Pelitinib |
| top60_altered_DNA_seed | EGFR | 0.950739 | GBM | D-263MG | CUDC-101 |
| top60_altered_DNA_seed | EGFR | 0.963154 | GBM | D-247MG | PF-00299804 |
| top60_altered_DNA_seed | EGFR | 0.966872 | GBM | LN-405 | Afatinib |
| top60_altered_DNA_seed | EGFR | 0.973124 | GBM | GB-1 | Erlotinib |
| top60_altered_DNA_seed | CDK4 | 0.97739 | GBM | 42-MG-BA | Palbociclib |
| top60_altered_DNA_seed | PDGFRA | 0.9916 | GBM | M059J | Amuvatinib |
| top60_altered_DNA_seed | KIT | 0.9916 | GBM | M059J | Amuvatinib |
| top60_altered_DNA_seed | EGFR | 0.997767 | GBM | U-118-MG | Cetuximab |
| top60_altered_DNA_seed | KIT | 1.001311 | GBM | SF295 | Pazopanib |
| top60_altered_DNA_seed | MDM4 | 1.002031 | GBM | SF126 | NSC-207895 |
| top60_altered_DNA_seed | EGFR | 1.007573 | GBM | DK-MG | Gefitinib |
| top60_altered_DNA_seed | EGFR | 1.013694 | GBM | U-118-MG | Afatinib |
| top60_altered_DNA_seed | KIT | 1.017944 | GBM | SNB75 | Imatinib |
| top60_altered_DNA_seed | MDM4 | 1.01938 | GBM | LN-229 | NSC-207895 |
| top60_altered_DNA_seed | EGFR | 1.023486 | GBM | DBTRG-05MG | Afatinib |
| top60_altered_DNA_seed | EGFR | 1.028719 | GBM | D-392MG | Afatinib |
| top60_altered_DNA_seed | EGFR | 1.030523 | GBM | SW1088 | Afatinib |
| top60_altered_DNA_seed | EGFR | 1.034087 | GBM | Becker | PF-00299804 |
| top60_altered_DNA_seed | KIT | 1.040731 | GBM | KS-1 | OSI-930 |
| top60_altered_DNA_seed | EGFR | 1.041416 | GBM | DBTRG-05MG | Afatinib |
| top60_altered_DNA_seed | EGFR | 1.041725 | GBM | SF268 | Gefitinib |
| top60_altered_DNA_seed | EGFR | 1.047505 | GBM | CAS-1 | Erlotinib |
| top60_altered_DNA_seed | MDM4 | 1.049135 | GBM | YH-13 | NSC-207895 |
| top60_altered_DNA_seed | CDK4 | 1.050188 | GBM | 8-MG-BA | Palbociclib |
| top60_altered_DNA_seed | EGFR | 1.051861 | GBM | D-542MG | Afatinib |
| top60_altered_DNA_seed | PDGFRA | 1.054111 | GBM | LN-405 | Amuvatinib |
| top60_altered_DNA_seed | KIT | 1.054111 | GBM | LN-405 | Amuvatinib |
| top60_altered_DNA_seed | EGFR | 1.060722 | GBM | DK-MG | Afatinib |
| top60_altered_DNA_seed | MDM4 | 1.06462 | GBM | LN-405 | NSC-207895 |
| top60_altered_DNA_seed | EGFR | 1.06613 | GBM | D-542MG | Afatinib |
| top60_altered_DNA_seed | EGFR | 1.074244 | GBM | AM-38 | Cetuximab |
| top60_altered_DNA_seed | CDK4 | 1.080758 | GBM | CAS-1 | Palbociclib |
| top60_altered_DNA_seed | EGFR | 1.084594 | GBM | CAS-1 | Afatinib |
| top60_altered_DNA_seed | EGFR | 1.088397 | GBM | SNB75 | Pelitinib |
| top60_altered_DNA_seed | EGFR | 1.091108 | GBM | M059J | Pelitinib |

| | | | | | |
|---|---|---|---|---|---|
| top60_altered_DNA_seed | EGFR | 1.092916 | GBM | T98G | CUDC-101 |
| top60_altered_DNA_seed | EGFR | 1.093055 | GBM | SW1088 | Cetuximab |
| top60_altered_DNA_seed | EGFR | 1.093282 | GBM | D-542MG | CI-1033 |
| top60_altered_DNA_seed | MDM4 | 1.111887 | GBM | 42-MG-BA | NSC-207895 |
| top60_altered_DNA_seed | EGFR | 1.114354 | GBM | SNB75 | CI-1033 |
| top60_altered_DNA_seed | EGFR | 1.11661 | GBM | LNZTA3WT4 | Gefitinib |
| top60_altered_DNA_seed | EGFR | 1.123878 | GBM | AM-38 | Afatinib |
| top60_altered_DNA_seed | CDK4 | 1.124655 | GBM | U-87-MG | CGP-082996 |
| top60_altered_DNA_seed | EGFR | 1.128897 | GBM | D-247MG | Afatinib |
| top60_altered_DNA_seed | KIT | 1.129922 | GBM | AM-38 | Pazopanib |
| top60_altered_DNA_seed | KIT | 1.13374 | GBM | U-87-MG | Axitinib |
| top60_altered_DNA_seed | EGFR | 1.164147 | GBM | LN-405 | Pelitinib |
| top60_altered_DNA_seed | EGFR | 1.169363 | GBM | D-247MG | Afatinib |
| top60_altered_DNA_seed | KIT | 1.183016 | GBM | LN-229 | Axitinib |
| top60_altered_DNA_seed | EGFR | 1.183565 | GBM | D-542MG | Gefitinib |
| top60_altered_DNA_seed | KIT | 1.192739 | GBM | U-87-MG | Pazopanib |
| top60_altered_DNA_seed | EGFR | 1.217418 | GBM | U-87-MG | Erlotinib |
| top60_altered_DNA_seed | CDK4 | 1.232426 | GBM | D-263MG | CGP-082996 |
| top60_altered_DNA_seed | EGFR | 1.234474 | GBM | CAS-1 | Afatinib |
| top60_altered_DNA_seed | EGFR | 1.248653 | GBM | SNB75 | Afatinib |
| top60_altered_DNA_seed | CDK4 | 1.249184 | GBM | AM-38 | CGP-082996 |
| top60_altered_DNA_seed | EGFR | 1.261068 | GBM | U-118-MG | Afatinib |
| top60_altered_DNA_seed | EGFR | 1.280577 | GBM | CAS-1 | Gefitinib |
| top60_altered_DNA_seed | PDGFRA | 1.280743 | GBM | LN-229 | Amuvatinib |
| top60_altered_DNA_seed | KIT | 1.280743 | GBM | LN-229 | Amuvatinib |
| top60_altered_DNA_seed | MDM4 | 1.283486 | GBM | D-263MG | NSC-207895 |
| top60_altered_DNA_seed | KIT | 1.286309 | GBM | CAS-1 | Motesanib |
| top60_altered_DNA_seed | KIT | 1.294529 | GBM | U-87-MG | OSI-930 |
| top60_altered_DNA_seed | MDM4 | 1.315538 | GBM | DBTRG-05MG | NSC-207895 |
| top60_altered_DNA_seed | PDGFRA | 1.317396 | GBM | Becker | Masitinib |
| top60_altered_DNA_seed | KIT | 1.317396 | GBM | Becker | Masitinib |
| top60_altered_DNA_seed | EGFR | 1.318872 | GBM | DK-MG | AZD8931 |
| top60_altered_DNA_seed | EGFR | 1.321979 | GBM | SNB75 | Erlotinib |
| top60_altered_DNA_seed | PDGFRA | 1.330843 | GBM | SW1088 | Amuvatinib |
| top60_altered_DNA_seed | KIT | 1.330843 | GBM | SW1088 | Amuvatinib |
| top60_altered_DNA_seed | PDGFRA | 1.340726 | GBM | SNB75 | Amuvatinib |
| top60_altered_DNA_seed | KIT | 1.340726 | GBM | SNB75 | Amuvatinib |
| top60_altered_DNA_seed | PDGFRA | 1.356297 | GBM | DK-MG | Amuvatinib |
| top60_altered_DNA_seed | KIT | 1.356297 | GBM | DK-MG | Amuvatinib |
| top60_altered_DNA_seed | CDK4 | 1.357365 | GBM | LN-405 | Palbociclib |
| top60_altered_DNA_seed | EGFR | 1.364496 | GBM | U-118-MG | Pelitinib |

| | | | | | |
|---|---|---|---|---|---|
| top60_altered_DNA_seed | MDM4 | 1.370385 | GBM | D-542MG | NSC-207895 |
| top60_altered_DNA_seed | MDM4 | 1.378786 | GBM | SW1088 | NSC-207895 |
| top60_altered_DNA_seed | CDK4 | 1.396558 | GBM | Becker | CGP-082996 |
| top60_altered_DNA_seed | EGFR | 1.39991 | GBM | U-87-MG | Pelitinib |
| top60_altered_DNA_seed | EGFR | 1.433478 | GBM | SF268 | CI-1033 |
| top60_altered_DNA_seed | EGFR | 1.439725 | GBM | D-263MG | Pelitinib |
| top60_altered_DNA_seed | KIT | 1.475745 | GBM | SNB75 | OSI-930 |
| top60_altered_DNA_seed | MDM4 | 1.477133 | GBM | U-87-MG | NSC-207895 |
| top60_altered_DNA_seed | PDGFRA | 1.501964 | GBM | U-87-MG | Amuvatinib |
| top60_altered_DNA_seed | KIT | 1.501964 | GBM | U-87-MG | Amuvatinib |
| top60_altered_DNA_seed | MDM4 | 1.509498 | GBM | U-118-MG | NSC-207895 |
| top60_altered_DNA_seed | EGFR | 1.516762 | GBM | D-247MG | Pelitinib |
| top60_altered_DNA_seed | EGFR | 1.520523 | GBM | SNB75 | Gefitinib |
| top60_altered_DNA_seed | PDGFRA | 1.533048 | GBM | LN-229 | Masitinib |
| top60_altered_DNA_seed | KIT | 1.533048 | GBM | LN-229 | Masitinib |
| top60_altered_DNA_seed | MDM4 | 1.533469 | GBM | CAS-1 | NSC-207895 |
| top60_altered_DNA_seed | EGFR | 1.541826 | GBM | SW1088 | CI-1033 |
| top60_altered_DNA_seed | EGFR | 1.544437 | GBM | U-118-MG | CI-1033 |
| top60_altered_DNA_seed | EGFR | 1.553352 | GBM | D-542MG | Pelitinib |
| top60_altered_DNA_seed | CDK4 | 1.554434 | GBM | M059J | Palbociclib |
| top60_altered_DNA_seed | EGFR | 1.571145 | GBM | D-247MG | CI-1033 |
| top60_altered_DNA_seed | KIT | 1.599698 | GBM | LN-405 | Axitinib |
| top60_altered_DNA_seed | MDM4 | 1.62402 | GBM | SNB75 | NSC-207895 |
| top60_altered_DNA_seed | KIT | 1.670904 | GBM | CAS-1 | Sorafenib |
| top60_altered_DNA_seed | EGFR | 1.672108 | GBM | AM-38 | AZD8931 |
| top60_altered_DNA_seed | CDK4 | 1.678389 | GBM | D-542MG | Palbociclib |
| top60_altered_DNA_seed | CDK4 | 1.67935 | GBM | CAS-1 | CGP-082996 |
| top60_altered_DNA_seed | MDM4 | 1.716764 | GBM | D-392MG | NSC-207895 |
| top60_altered_DNA_seed | PDGFRA | 1.747289 | GBM | D-263MG | Masitinib |
| top60_altered_DNA_seed | KIT | 1.747289 | GBM | D-263MG | Masitinib |
| top60_altered_DNA_seed | EGFR | 1.79721 | GBM | SF295 | CI-1033 |
| top60_altered_DNA_seed | EGFR | 1.824225 | GBM | A172 | CI-1033 |
| top60_altered_DNA_seed | EGFR | 1.873966 | GBM | T98G | CI-1033 |
| top60_altered_DNA_seed | EGFR | 1.886643 | GBM | LNZTA3WT4 | AZD8931 |
| top60_altered_DNA_seed | EGFR | 1.894947 | GBM | DBTRG-05MG | CI-1033 |
| top60_altered_DNA_seed | EGFR | 1.929988 | GBM | LN-405 | CI-1033 |
| top60_altered_DNA_seed | EGFR | 1.939787 | GBM | SW1088 | CUDC-101 |
| top60_altered_DNA_seed | CDK4 | 1.965836 | GBM | SNB75 | Palbociclib |
| top60_altered_DNA_seed | EGFR | 2.206699 | GBM | LNZTA3WT4 | CUDC-101 |
| top60_altered_DNA_seed | KIT | 2.210286 | GBM | SNB75 | Sorafenib |
| top60_altered_DNA_seed | KIT | -2.444714 | GBM | 42-MG-BA | Motesanib |

| | | | | | |
|---|---|---|---|---|---|
| top60_altered_DNA_seed | KIT | -1.42003 | GBM | D-245MG | Axitinib |
| top60_altered_DNA_seed | CDK4 | -1.409729 | GBM | D-245MG | Ribociclib |
| top60_altered_DNA_seed | KIT | -1.403938 | GBM | DK-MG | Motesanib |
| top60_altered_DNA_seed | KIT | -1.177564 | GBM | LN-405 | Motesanib |
| top60_altered_DNA_seed | KIT | -1.097832 | GBM | D-423MG | Sorafenib |
| top60_altered_DNA_seed | TP53 | -1.090076 | GBM | LNZTA3WT4 | MIRA-1 |
| top60_altered_DNA_seed | EGFR | -1.068129 | GBM | D-423MG | AZD3759 |
| top60_altered_DNA_seed | CDK4 | -1.023605 | GBM | YH-13 | Ribociclib |
| top60_altered_DNA_seed | CDK4 | -1.003435 | GBM | A172 | Ribociclib |
| top60_altered_DNA_seed | TP53 | -0.962093 | GBM | D-566MG | MIRA-1 |
| top60_altered_DNA_seed | EGFR | -0.844443 | GBM | LN-229 | Erlotinib |
| top60_altered_DNA_seed | KIT | -0.806644 | GBM | 42-MG-BA | Axitinib |
| top60_altered_DNA_seed | EGFR | -0.783918 | GBM | 8-MG-BA | Lapatinib |
| top60_altered_DNA_seed | KIT | -0.768842 | GBM | M059J | Axitinib |
| top60_altered_DNA_seed | KIT | -0.756471 | GBM | A172 | Axitinib |
| top60_altered_DNA_seed | EGFR | -0.74154 | GBM | D-566MG | Lapatinib |
| top60_altered_DNA_seed | KIT | -0.71489 | GBM | SW1088 | Axitinib |
| top60_altered_DNA_seed | KIT | -0.702457 | GBM | D-566MG | Sorafenib |
| top60_altered_DNA_seed | EGFR | -0.69609 | GBM | LN-18 | Lapatinib |
| top60_altered_DNA_seed | EGFR | -0.690795 | GBM | SF126 | Osimertinib |
| top60_altered_DNA_seed | KIT | -0.678608 | GBM | DK-MG | Axitinib |
| top60_altered_DNA_seed | CDK4 | -0.676501 | GBM | LNZTA3WT4 | Palbociclib |
| top60_altered_DNA_seed | KIT | -0.674652 | GBM | 8-MG-BA | Sorafenib |
| top60_altered_DNA_seed | CDK4 | -0.663907 | GBM | DK-MG | Ribociclib |
| top60_altered_DNA_seed | KIT | -0.660411 | GBM | LN-18 | Sorafenib |
| top60_altered_DNA_seed | EGFR | -0.641273 | GBM | D-423MG | Lapatinib |
| top60_altered_DNA_seed | KIT | -0.634731 | GBM | D-392MG | Axitinib |
| top60_altered_DNA_seed | KIT | -0.615117 | GBM | SF126 | Axitinib |
| top60_altered_DNA_seed | KIT | -0.600814 | GBM | 42-MG-BA | Sorafenib |
| top60_altered_DNA_seed | KIT | -0.588784 | GBM | D-263MG | Axitinib |
| top60_altered_DNA_seed | KIT | -0.579761 | GBM | SF295 | Sorafenib |
| top60_altered_DNA_seed | TP53 | -0.55726 | GBM | SF295 | MIRA-1 |
| top60_altered_DNA_seed | KIT | -0.545149 | GBM | D-423MG | Axitinib |
| top60_altered_DNA_seed | EGFR | -0.514164 | GBM | SF126 | Lapatinib |
| top60_altered_DNA_seed | CDK4 | -0.506462 | GBM | D-263MG | Ribociclib |
| top60_altered_DNA_seed | CDK4 | -0.498988 | GBM | A172 | Palbociclib |
| top60_altered_DNA_seed | TP53 | -0.487817 | GBM | GB-1 | MIRA-1 |
| top60_altered_DNA_seed | KIT | -0.485328 | GBM | CAS-1 | Axitinib |
| top60_altered_DNA_seed | EGFR | -0.478769 | GBM | SF295 | AZD3759 |
| top60_altered_DNA_seed | EGFR | -0.470561 | GBM | LN-18 | Erlotinib |
| top60_altered_DNA_seed | KIT | -0.456512 | GBM | LNZTA3WT4 | Axitinib |

| | | | | | |
|---|---|---|---|---|---|
| top60_altered_DNA_seed | EGFR | -0.439711 | GBM | 8-MG-BA | Gefitinib |
| top60_altered_DNA_seed | KIT | -0.422334 | GBM | AM-38 | Sorafenib |
| top60_altered_DNA_seed | EGFR | -0.409178 | GBM | YKG-1 | Sapitinib |
| top60_altered_DNA_seed | TP53 | -0.396213 | GBM | D-423MG | MIRA-1 |
| top60_altered_DNA_seed | EGFR | -0.393865 | GBM | GB-1 | Lapatinib |
| top60_altered_DNA_seed | KIT | -0.385714 | GBM | D-566MG | Motesanib |
| top60_altered_DNA_seed | EGFR | -0.377618 | GBM | YKG-1 | Lapatinib |
| top60_altered_DNA_seed | KIT | -0.37585 | GBM | A172 | Sorafenib |
| top60_altered_DNA_seed | CDK4 | -0.373544 | GBM | LNZTA3WT4 | Ribociclib |
| top60_altered_DNA_seed | CDK4 | -0.369037 | GBM | Becker | Palbociclib |
| top60_altered_DNA_seed | EGFR | -0.358775 | GBM | SNB75 | AZD3759 |
| top60_altered_DNA_seed | EGFR | -0.356905 | GBM | LN-229 | Sapitinib |
| top60_altered_DNA_seed | CDK4 | -0.355037 | GBM | YH-13 | Palbociclib |
| top60_altered_DNA_seed | EGFR | -0.339388 | GBM | GB-1 | Erlotinib |
| top60_altered_DNA_seed | KIT | -0.334249 | GBM | SF126 | Sorafenib |
| top60_altered_DNA_seed | EGFR | -0.334036 | GBM | YKG-1 | AZD3759 |
| top60_altered_DNA_seed | KIT | -0.333345 | GBM | D-247MG | Motesanib |
| top60_altered_DNA_seed | TP53 | -0.319202 | GBM | KS-1 | MIRA-1 |
| top60_altered_DNA_seed | EGFR | -0.317899 | GBM | D-423MG | Osimertinib |
| top60_altered_DNA_seed | TP53 | -0.309237 | GBM | LN-18 | MIRA-1 |
| top60_altered_DNA_seed | TP53 | -0.290196 | GBM | D-247MG | MIRA-1 |
| top60_altered_DNA_seed | KIT | -0.281704 | GBM | D-247MG | Axitinib |
| top60_altered_DNA_seed | CDK4 | -0.271687 | GBM | 42-MG-BA | Ribociclib |
| top60_altered_DNA_seed | EGFR | -0.26647 | GBM | DK-MG | Osimertinib |
| top60_altered_DNA_seed | CDK4 | -0.262916 | GBM | D-263MG | Palbociclib |
| top60_altered_DNA_seed | KIT | -0.259552 | GBM | 8-MG-BA | Motesanib |
| top60_altered_DNA_seed | EGFR | -0.258488 | GBM | SF126 | Gefitinib |
| top60_altered_DNA_seed | EGFR | -0.252165 | GBM | U-87-MG | Erlotinib |
| top60_altered_DNA_seed | TP53 | -0.246912 | GBM | A172 | MIRA-1 |
| top60_altered_DNA_seed | TP53 | -0.241901 | GBM | YH-13 | MIRA-1 |
| top60_altered_DNA_seed | EGFR | -0.235615 | GBM | LN-229 | Osimertinib |
| top60_altered_DNA_seed | CDK4 | -0.231495 | GBM | D-566MG | Palbociclib |
| top60_altered_DNA_seed | EGFR | -0.23003 | GBM | AM-38 | Erlotinib |
| top60_altered_DNA_seed | EGFR | -0.225836 | GBM | 42-MG-BA | Lapatinib |
| top60_altered_DNA_seed | KIT | -0.212426 | GBM | YKG-1 | Sorafenib |
| top60_altered_DNA_seed | CDK4 | -0.208943 | GBM | SF295 | Palbociclib |
| top60_altered_DNA_seed | KIT | -0.208197 | GBM | D-245MG | Motesanib |
| top60_altered_DNA_seed | TP53 | -0.206846 | GBM | 8-MG-BA | MIRA-1 |
| top60_altered_DNA_seed | EGFR | -0.19335 | GBM | D-245MG | Sapitinib |
| top60_altered_DNA_seed | CDK4 | -0.181636 | GBM | YKG-1 | Palbociclib |
| top60_altered_DNA_seed | EGFR | -0.17646 | GBM | SW1088 | Sapitinib |

| | | | | | |
|---|---|---|---|---|---|
| top60_altered_DNA_seed | TP53 | -0.170488 | GBM | 42-MG-BA | MIRA-1 |
| top60_altered_DNA_seed | KIT | -0.17028 | GBM | D-566MG | Axitinib |
| top60_altered_DNA_seed | TP53 | -0.162285 | GBM | D-263MG | MIRA-1 |
| top60_altered_DNA_seed | KIT | -0.149323 | GBM | D-423MG | Motesanib |
| top60_altered_DNA_seed | EGFR | -0.138111 | GBM | 42-MG-BA | Osimertinib |
| top60_altered_DNA_seed | CDK4 | -0.137955 | GBM | AM-38 | Palbociclib |
| top60_altered_DNA_seed | KIT | -0.128427 | GBM | SF295 | Axitinib |
| top60_altered_DNA_seed | EGFR | -0.12271 | GBM | 8-MG-BA | Erlotinib |
| top60_altered_DNA_seed | CDK4 | -0.12219 | GBM | D-566MG | Ribociclib |
| top60_altered_DNA_seed | EGFR | -0.116719 | GBM | A172 | Afatinib |
| top60_altered_DNA_seed | CDK4 | -0.102853 | GBM | D-247MG | Palbociclib |
| top60_altered_DNA_seed | KIT | -0.09975 | GBM | SF126 | Motesanib |
| top60_altered_DNA_seed | EGFR | -0.0997 | GBM | CAS-1 | Lapatinib |
| top60_altered_DNA_seed | CDK4 | -0.086184 | GBM | KS-1 | Ribociclib |
| top60_altered_DNA_seed | EGFR | -0.082638 | GBM | SNB75 | Osimertinib |
| top60_altered_DNA_seed | KIT | -0.076167 | GBM | D-542MG | Sorafenib |
| top60_altered_DNA_seed | EGFR | -0.06101 | GBM | LN-229 | Gefitinib |
| top60_altered_DNA_seed | EGFR | -0.059066 | GBM | 8-MG-BA | Sapitinib |
| top60_altered_DNA_seed | EGFR | -0.057422 | GBM | A172 | Gefitinib |
| top60_altered_DNA_seed | EGFR | -0.055395 | GBM | D-542MG | Lapatinib |
| top60_altered_DNA_seed | CDK4 | -0.053038 | GBM | U251 | Palbociclib |
| top60_altered_DNA_seed | EGFR | -0.052 | GBM | 42-MG-BA | Afatinib |
| top60_altered_DNA_seed | EGFR | -0.050785 | GBM | A172 | Osimertinib |
| top60_altered_DNA_seed | EGFR | -0.042264 | GBM | D-566MG | Erlotinib |
| top60_altered_DNA_seed | TP53 | -0.03173 | GBM | M059J | MIRA-1 |
| top60_altered_DNA_seed | EGFR | -0.0309 | GBM | SF295 | Osimertinib |
| top60_altered_DNA_seed | CDK4 | -0.030002 | GBM | U-87-MG | Palbociclib |
| top60_altered_DNA_seed | CDK4 | -0.021426 | GBM | 8-MG-BA | Palbociclib |
| top60_altered_DNA_seed | EGFR | -0.015156 | GBM | U-87-MG | AZD3759 |
| top60_altered_DNA_seed | EGFR | -0.014364 | GBM | YH-13 | Erlotinib |
| top60_altered_DNA_seed | CDK4 | -0.012447 | GBM | 8-MG-BA | Ribociclib |
| top60_altered_DNA_seed | EGFR | -0.011375 | GBM | YKG-1 | Osimertinib |
| top60_altered_DNA_seed | EGFR | -0.007482 | GBM | YH-13 | Osimertinib |
| top60_altered_DNA_seed | EGFR | -0.005381 | GBM | SF126 | Sapitinib |
| top60_altered_DNA_seed | KIT | -0.001062 | GBM | U-118-MG | Sorafenib |
| top60_altered_DNA_seed | EGFR | 0.002614 | GBM | M059J | Erlotinib |
| top60_altered_DNA_seed | EGFR | 0.004394 | GBM | U251 | Erlotinib |
| top60_altered_DNA_seed | TP53 | 0.007679 | GBM | T98G | MIRA-1 |
| top60_altered_DNA_seed | EGFR | 0.008988 | GBM | KS-1 | Sapitinib |
| top60_altered_DNA_seed | EGFR | 0.010215 | GBM | SF268 | Erlotinib |
| top60_altered_DNA_seed | KIT | 0.010475 | GBM | YH-13 | Motesanib |

| | | | | | |
|---|---|---|---|---|---|
| top60_altered_DNA_seed | EGFR | 0.027993 | GBM | A172 | Lapatinib |
| top60_altered_DNA_seed | EGFR | 0.029904 | GBM | YKG-1 | Gefitinib |
| top60_altered_DNA_seed | EGFR | 0.030789 | GBM | 8-MG-BA | Afatinib |
| top60_altered_DNA_seed | CDK4 | 0.031313 | GBM | SW1088 | Palbociclib |
| top60_altered_DNA_seed | EGFR | 0.035055 | GBM | LN-18 | Sapitinib |
| top60_altered_DNA_seed | EGFR | 0.036472 | GBM | DK-MG | Erlotinib |
| top60_altered_DNA_seed | KIT | 0.060232 | GBM | U251 | Sorafenib |
| top60_altered_DNA_seed | KIT | 0.060283 | GBM | LN-18 | Axitinib |
| top60_altered_DNA_seed | TP53 | 0.063766 | GBM | SNB75 | MIRA-1 |
| top60_altered_DNA_seed | EGFR | 0.065361 | GBM | D-566MG | Gefitinib |
| top60_altered_DNA_seed | EGFR | 0.067748 | GBM | LNZTA3WT4 | Afatinib |
| top60_altered_DNA_seed | CDK4 | 0.076872 | GBM | D-423MG | Palbociclib |
| top60_altered_DNA_seed | CDK4 | 0.080189 | GBM | D-245MG | Palbociclib |
| top60_altered_DNA_seed | TP53 | 0.090294 | GBM | SF126 | MIRA-1 |
| top60_altered_DNA_seed | EGFR | 0.091768 | GBM | U-118-MG | AZD3759 |
| top60_altered_DNA_seed | EGFR | 0.100732 | GBM | SF295 | Lapatinib |
| top60_altered_DNA_seed | EGFR | 0.110759 | GBM | U-118-MG | Osimertinib |
| top60_altered_DNA_seed | EGFR | 0.114846 | GBM | SK-MG-1 | Afatinib |
| top60_altered_DNA_seed | EGFR | 0.115143 | GBM | DK-MG | Sapitinib |
| top60_altered_DNA_seed | EGFR | 0.118834 | GBM | D-542MG | Osimertinib |
| top60_altered_DNA_seed | KIT | 0.122009 | GBM | LN-18 | Motesanib |
| top60_altered_DNA_seed | KIT | 0.122163 | GBM | GB-1 | Sorafenib |
| top60_altered_DNA_seed | EGFR | 0.124166 | GBM | SF126 | AZD3759 |
| top60_altered_DNA_seed | EGFR | 0.12491 | GBM | AM-38 | AZD3759 |
| top60_altered_DNA_seed | CDK4 | 0.128484 | GBM | SF268 | Ribociclib |
| top60_altered_DNA_seed | KIT | 0.129903 | GBM | LNZTA3WT4 | Motesanib |
| top60_altered_DNA_seed | EGFR | 0.132447 | GBM | D-566MG | AZD3759 |
| top60_altered_DNA_seed | EGFR | 0.134693 | GBM | LNZTA3WT4 | Lapatinib |
| top60_altered_DNA_seed | EGFR | 0.135413 | GBM | T98G | Gefitinib |
| top60_altered_DNA_seed | KIT | 0.139652 | GBM | T98G | Motesanib |
| top60_altered_DNA_seed | KIT | 0.140043 | GBM | D-245MG | Sorafenib |
| top60_altered_DNA_seed | KIT | 0.146053 | GBM | A172 | Motesanib |
| top60_altered_DNA_seed | EGFR | 0.148283 | GBM | DK-MG | Afatinib |
| top60_altered_DNA_seed | EGFR | 0.149432 | GBM | D-263MG | AZD3759 |
| top60_altered_DNA_seed | CDK4 | 0.152862 | GBM | SF126 | Ribociclib |
| top60_altered_DNA_seed | CDK4 | 0.169151 | GBM | D-423MG | Ribociclib |
| top60_altered_DNA_seed | CDK4 | 0.169831 | GBM | LN-18 | Ribociclib |
| top60_altered_DNA_seed | EGFR | 0.17414 | GBM | U-118-MG | Erlotinib |
| top60_altered_DNA_seed | EGFR | 0.175123 | GBM | D-423MG | Gefitinib |
| top60_altered_DNA_seed | EGFR | 0.177191 | GBM | SF268 | Afatinib |
| top60_altered_DNA_seed | EGFR | 0.180797 | GBM | A172 | AZD3759 |

| | | | | | |
|---|---|---|---|---|---|
| top60_altered_DNA_seed | EGFR | 0.181391 | GBM | A172 | Erlotinib |
| top60_altered_DNA_seed | EGFR | 0.183624 | GBM | D-245MG | Afatinib |
| top60_altered_DNA_seed | CDK4 | 0.18412 | GBM | DK-MG | Palbociclib |
| top60_altered_DNA_seed | KIT | 0.188521 | GBM | AM-38 | Axitinib |
| top60_altered_DNA_seed | EGFR | 0.192616 | GBM | D-245MG | Osimertinib |
| top60_altered_DNA_seed | EGFR | 0.192717 | GBM | SF268 | Gefitinib |
| top60_altered_DNA_seed | CDK4 | 0.195795 | GBM | U-118-MG | Palbociclib |
| top60_altered_DNA_seed | EGFR | 0.197386 | GBM | D-247MG | Osimertinib |
| top60_altered_DNA_seed | KIT | 0.197784 | GBM | AM-38 | Motesanib |
| top60_altered_DNA_seed | EGFR | 0.200155 | GBM | AM-38 | Lapatinib |
| top60_altered_DNA_seed | EGFR | 0.209204 | GBM | U251 | AZD3759 |
| top60_altered_DNA_seed | EGFR | 0.20971 | GBM | YH-13 | Afatinib |
| top60_altered_DNA_seed | KIT | 0.211268 | GBM | T98G | Axitinib |
| top60_altered_DNA_seed | KIT | 0.21285 | GBM | YH-13 | Axitinib |
| top60_altered_DNA_seed | EGFR | 0.225471 | GBM | M059J | Afatinib |
| top60_altered_DNA_seed | EGFR | 0.232195 | GBM | M059J | Lapatinib |
| top60_altered_DNA_seed | EGFR | 0.233071 | GBM | DK-MG | AZD3759 |
| top60_altered_DNA_seed | KIT | 0.233351 | GBM | SF268 | Axitinib |
| top60_altered_DNA_seed | KIT | 0.235526 | GBM | SK-MG-1 | Axitinib |
| top60_altered_DNA_seed | EGFR | 0.237068 | GBM | U-87-MG | Gefitinib |
| top60_altered_DNA_seed | EGFR | 0.241009 | GBM | U-87-MG | Sapitinib |
| top60_altered_DNA_seed | KIT | 0.243335 | GBM | SF268 | Sorafenib |
| top60_altered_DNA_seed | KIT | 0.248351 | GBM | U251 | Axitinib |
| top60_altered_DNA_seed | EGFR | 0.255549 | GBM | LN-229 | AZD3759 |
| top60_altered_DNA_seed | CDK4 | 0.261538 | GBM | SF295 | Ribociclib |
| top60_altered_DNA_seed | KIT | 0.262318 | GBM | CAS-1 | Motesanib |
| top60_altered_DNA_seed | TP53 | 0.264647 | GBM | YKG-1 | MIRA-1 |
| top60_altered_DNA_seed | KIT | 0.264698 | GBM | Becker | Axitinib |
| top60_altered_DNA_seed | EGFR | 0.269102 | GBM | 42-MG-BA | Sapitinib |
| top60_altered_DNA_seed | EGFR | 0.27457 | GBM | SW1088 | Erlotinib |
| top60_altered_DNA_seed | EGFR | 0.275207 | GBM | D-542MG | Gefitinib |
| top60_altered_DNA_seed | EGFR | 0.280485 | GBM | Becker | AZD3759 |
| top60_altered_DNA_seed | CDK4 | 0.290798 | GBM | U-87-MG | Ribociclib |
| top60_altered_DNA_seed | EGFR | 0.293897 | GBM | SW1088 | Osimertinib |
| top60_altered_DNA_seed | EGFR | 0.294884 | GBM | T98G | Lapatinib |
| top60_altered_DNA_seed | EGFR | 0.296246 | GBM | CAS-1 | Sapitinib |
| top60_altered_DNA_seed | KIT | 0.296414 | GBM | KS-1 | Axitinib |
| top60_altered_DNA_seed | TP53 | 0.299196 | GBM | AM-38 | MIRA-1 |
| top60_altered_DNA_seed | CDK4 | 0.305782 | GBM | CAS-1 | Palbociclib |
| top60_altered_DNA_seed | KIT | 0.310924 | GBM | YKG-1 | Motesanib |
| top60_altered_DNA_seed | TP53 | 0.3112 | GBM | D-542MG | MIRA-1 |

| | | | | | |
|---|---|---|---|---|---|
| top60_altered_DNA_seed | TP53 | 0.313675 | GBM | Becker | MIRA-1 |
| top60_altered_DNA_seed | KIT | 0.321616 | GBM | SF295 | Motesanib |
| top60_altered_DNA_seed | KIT | 0.326282 | GBM | DK-MG | Sorafenib |
| top60_altered_DNA_seed | EGFR | 0.327969 | GBM | D-423MG | Erlotinib |
| top60_altered_DNA_seed | EGFR | 0.334174 | GBM | 8-MG-BA | AZD3759 |
| top60_altered_DNA_seed | EGFR | 0.334651 | GBM | D-423MG | Afatinib |
| top60_altered_DNA_seed | EGFR | 0.337738 | GBM | D-566MG | Osimertinib |
| top60_altered_DNA_seed | EGFR | 0.364198 | GBM | SF268 | Sapitinib |
| top60_altered_DNA_seed | EGFR | 0.364904 | GBM | D-247MG | Sapitinib |
| top60_altered_DNA_seed | EGFR | 0.366709 | GBM | AM-38 | Gefitinib |
| top60_altered_DNA_seed | CDK4 | 0.375127 | GBM | D-392MG | Ribociclib |
| top60_altered_DNA_seed | KIT | 0.376224 | GBM | T98G | Sorafenib |
| top60_altered_DNA_seed | EGFR | 0.37903 | GBM | SF295 | Gefitinib |
| top60_altered_DNA_seed | EGFR | 0.379162 | GBM | YKG-1 | Erlotinib |
| top60_altered_DNA_seed | EGFR | 0.382277 | GBM | D-247MG | Lapatinib |
| top60_altered_DNA_seed | EGFR | 0.384935 | GBM | 42-MG-BA | AZD3759 |
| top60_altered_DNA_seed | CDK4 | 0.385871 | GBM | SF126 | Palbociclib |
| top60_altered_DNA_seed | EGFR | 0.3876 | GBM | KS-1 | Afatinib |
| top60_altered_DNA_seed | EGFR | 0.393013 | GBM | YH-13 | Sapitinib |
| top60_altered_DNA_seed | EGFR | 0.393487 | GBM | SK-MG-1 | Osimertinib |
| top60_altered_DNA_seed | CDK4 | 0.394151 | GBM | D-542MG | Ribociclib |
| top60_altered_DNA_seed | KIT | 0.395221 | GBM | YKG-1 | Axitinib |
| top60_altered_DNA_seed | KIT | 0.396427 | GBM | U251 | Motesanib |
| top60_altered_DNA_seed | EGFR | 0.400711 | GBM | D-245MG | Gefitinib |
| top60_altered_DNA_seed | EGFR | 0.402606 | GBM | SK-MG-1 | Erlotinib |
| top60_altered_DNA_seed | KIT | 0.406433 | GBM | D-263MG | Motesanib |
| top60_altered_DNA_seed | CDK4 | 0.410973 | GBM | Becker | Ribociclib |
| top60_altered_DNA_seed | EGFR | 0.412974 | GBM | D-566MG | Sapitinib |
| top60_altered_DNA_seed | EGFR | 0.417341 | GBM | U-118-MG | Gefitinib |
| top60_altered_DNA_seed | EGFR | 0.431461 | GBM | LNZTA3WT4 | AZD3759 |
| top60_altered_DNA_seed | KIT | 0.433319 | GBM | SF268 | Motesanib |
| top60_altered_DNA_seed | KIT | 0.442759 | GBM | SK-MG-1 | Motesanib |
| top60_altered_DNA_seed | EGFR | 0.447714 | GBM | D-423MG | Sapitinib |
| top60_altered_DNA_seed | KIT | 0.452088 | GBM | Becker | Motesanib |
| top60_altered_DNA_seed | EGFR | 0.455249 | GBM | M059J | AZD3759 |
| top60_altered_DNA_seed | TP53 | 0.47018 | GBM | D-245MG | MIRA-1 |
| top60_altered_DNA_seed | KIT | 0.47121 | GBM | D-263MG | Sorafenib |
| top60_altered_DNA_seed | CDK4 | 0.472011 | GBM | D-392MG | Palbociclib |
| top60_altered_DNA_seed | EGFR | 0.475655 | GBM | A172 | Sapitinib |
| top60_altered_DNA_seed | CDK4 | 0.480546 | GBM | SK-MG-1 | Palbociclib |
| top60_altered_DNA_seed | EGFR | 0.4854 | GBM | KS-1 | Erlotinib |

| | | | | | |
|---|---|---|---|---|---|
| top60_altered_DNA_seed | TP53 | 0.49485 | GBM | U251 | MIRA-1 |
| top60_altered_DNA_seed | EGFR | 0.503098 | GBM | D-263MG | Gefitinib |
| top60_altered_DNA_seed | CDK4 | 0.505729 | GBM | KS-1 | Palbociclib |
| top60_altered_DNA_seed | EGFR | 0.506116 | GBM | LN-229 | Afatinib |
| top60_altered_DNA_seed | EGFR | 0.509287 | GBM | U-87-MG | Osimertinib |
| top60_altered_DNA_seed | EGFR | 0.510367 | GBM | SNB75 | Gefitinib |
| top60_altered_DNA_seed | EGFR | 0.514688 | GBM | SW1088 | Afatinib |
| top60_altered_DNA_seed | CDK4 | 0.515567 | GBM | T98G | Palbociclib |
| top60_altered_DNA_seed | KIT | 0.518554 | GBM | KS-1 | Motesanib |
| top60_altered_DNA_seed | EGFR | 0.520009 | GBM | LN-405 | Gefitinib |
| top60_altered_DNA_seed | EGFR | 0.526224 | GBM | D-542MG | Afatinib |
| top60_altered_DNA_seed | CDK4 | 0.527481 | GBM | AM-38 | Ribociclib |
| top60_altered_DNA_seed | EGFR | 0.530148 | GBM | D-542MG | Sapitinib |
| top60_altered_DNA_seed | KIT | 0.530999 | GBM | SNB75 | Sorafenib |
| top60_altered_DNA_seed | CDK4 | 0.533868 | GBM | LN-18 | Palbociclib |
| top60_altered_DNA_seed | EGFR | 0.537031 | GBM | 42-MG-BA | Erlotinib |
| top60_altered_DNA_seed | CDK4 | 0.537833 | GBM | D-247MG | Ribociclib |
| top60_altered_DNA_seed | EGFR | 0.546951 | GBM | DK-MG | Lapatinib |
| top60_altered_DNA_seed | EGFR | 0.547239 | GBM | SF268 | Lapatinib |
| top60_altered_DNA_seed | CDK4 | 0.557772 | GBM | SK-MG-1 | Ribociclib |
| top60_altered_DNA_seed | KIT | 0.562065 | GBM | M059J | Motesanib |
| top60_altered_DNA_seed | EGFR | 0.56236 | GBM | 8-MG-BA | Osimertinib |
| top60_altered_DNA_seed | EGFR | 0.575883 | GBM | D-542MG | AZD3759 |
| top60_altered_DNA_seed | EGFR | 0.579175 | GBM | YH-13 | Lapatinib |
| top60_altered_DNA_seed | EGFR | 0.587528 | GBM | 42-MG-BA | Gefitinib |
| top60_altered_DNA_seed | EGFR | 0.590257 | GBM | M059J | Sapitinib |
| top60_altered_DNA_seed | EGFR | 0.595524 | GBM | SK-MG-1 | AZD3759 |
| top60_altered_DNA_seed | EGFR | 0.595763 | GBM | D-542MG | Erlotinib |
| top60_altered_DNA_seed | EGFR | 0.595809 | GBM | D-245MG | Lapatinib |
| top60_altered_DNA_seed | KIT | 0.600619 | GBM | U-87-MG | Sorafenib |
| top60_altered_DNA_seed | CDK4 | 0.605292 | GBM | GB-1 | Palbociclib |
| top60_altered_DNA_seed | KIT | 0.611181 | GBM | SNB75 | Motesanib |
| top60_altered_DNA_seed | CDK4 | 0.61874 | GBM | YKG-1 | Ribociclib |
| top60_altered_DNA_seed | EGFR | 0.621027 | GBM | LN-18 | Gefitinib |
| top60_altered_DNA_seed | KIT | 0.621581 | GBM | 8-MG-BA | Axitinib |
| top60_altered_DNA_seed | KIT | 0.62534 | GBM | Becker | Sorafenib |
| top60_altered_DNA_seed | EGFR | 0.627379 | GBM | Becker | Osimertinib |
| top60_altered_DNA_seed | EGFR | 0.628974 | GBM | GB-1 | Sapitinib |
| top60_altered_DNA_seed | EGFR | 0.629459 | GBM | Becker | Afatinib |
| top60_altered_DNA_seed | EGFR | 0.635474 | GBM | D-247MG | Afatinib |
| top60_altered_DNA_seed | EGFR | 0.639898 | GBM | T98G | AZD3759 |

| | | | | | |
|---|---|---|---|---|---|
| top60_altered_DNA_seed | KIT | 0.643224 | GBM | U-87-MG | Axitinib |
| top60_altered_DNA_seed | KIT | 0.643827 | GBM | YH-13 | Sorafenib |
| top60_altered_DNA_seed | TP53 | 0.645376 | GBM | LN-229 | MIRA-1 |
| top60_altered_DNA_seed | KIT | 0.647223 | GBM | D-542MG | Motesanib |
| top60_altered_DNA_seed | TP53 | 0.648846 | GBM | U-118-MG | MIRA-1 |
| top60_altered_DNA_seed | EGFR | 0.662326 | GBM | YH-13 | AZD3759 |
| top60_altered_DNA_seed | CDK4 | 0.663048 | GBM | SW1088 | Ribociclib |
| top60_altered_DNA_seed | EGFR | 0.666117 | GBM | DBTRG-05MG | Erlotinib |
| top60_altered_DNA_seed | EGFR | 0.667379 | GBM | AM-38 | Osimertinib |
| top60_altered_DNA_seed | TP53 | 0.673706 | GBM | D-392MG | MIRA-1 |
| top60_altered_DNA_seed | EGFR | 0.686248 | GBM | AM-38 | Afatinib |
| top60_altered_DNA_seed | EGFR | 0.688348 | GBM | CAS-1 | Osimertinib |
| top60_altered_DNA_seed | CDK4 | 0.691742 | GBM | M059J | Ribociclib |
| top60_altered_DNA_seed | EGFR | 0.702379 | GBM | D-392MG | Osimertinib |
| top60_altered_DNA_seed | CDK4 | 0.703697 | GBM | U251 | Ribociclib |
| top60_altered_DNA_seed | CDK4 | 0.704074 | GBM | SNB75 | Ribociclib |
| top60_altered_DNA_seed | EGFR | 0.706663 | GBM | M059J | Osimertinib |
| top60_altered_DNA_seed | TP53 | 0.714854 | GBM | U-87-MG | MIRA-1 |
| top60_altered_DNA_seed | KIT | 0.716923 | GBM | D-392MG | Motesanib |
| top60_altered_DNA_seed | EGFR | 0.71768 | GBM | T98G | Afatinib |
| top60_altered_DNA_seed | EGFR | 0.722859 | GBM | SW1088 | Lapatinib |
| top60_altered_DNA_seed | EGFR | 0.7261 | GBM | Becker | Gefitinib |
| top60_altered_DNA_seed | KIT | 0.726755 | GBM | LN-405 | Axitinib |
| top60_altered_DNA_seed | EGFR | 0.738527 | GBM | LN-229 | Lapatinib |
| top60_altered_DNA_seed | CDK4 | 0.744679 | GBM | 42-MG-BA | Palbociclib |
| top60_altered_DNA_seed | EGFR | 0.746715 | GBM | CAS-1 | Gefitinib |
| top60_altered_DNA_seed | EGFR | 0.758851 | GBM | SK-MG-1 | Lapatinib |
| top60_altered_DNA_seed | KIT | 0.759812 | GBM | LN-229 | Sorafenib |
| top60_altered_DNA_seed | EGFR | 0.760432 | GBM | D-392MG | AZD3759 |
| top60_altered_DNA_seed | EGFR | 0.780566 | GBM | D-392MG | Afatinib |
| top60_altered_DNA_seed | EGFR | 0.787238 | GBM | SW1088 | AZD3759 |
| top60_altered_DNA_seed | CDK4 | 0.792326 | GBM | SNB75 | Palbociclib |
| top60_altered_DNA_seed | EGFR | 0.803738 | GBM | CAS-1 | Afatinib |
| top60_altered_DNA_seed | KIT | 0.807835 | GBM | M059J | Sorafenib |
| top60_altered_DNA_seed | EGFR | 0.809603 | GBM | LN-405 | Erlotinib |
| top60_altered_DNA_seed | EGFR | 0.810487 | GBM | U251 | Gefitinib |
| top60_altered_DNA_seed | EGFR | 0.811865 | GBM | SNB75 | Sapitinib |
| top60_altered_DNA_seed | EGFR | 0.820737 | GBM | D-245MG | AZD3759 |
| top60_altered_DNA_seed | EGFR | 0.822151 | GBM | D-247MG | Gefitinib |
| top60_altered_DNA_seed | EGFR | 0.823413 | GBM | KS-1 | Osimertinib |
| top60_altered_DNA_seed | KIT | 0.825736 | GBM | SW1088 | Motesanib |

| | | | | | |
|---|---|---|---|---|---|
| top60_altered_DNA_seed | CDK4 | 0.825781 | GBM | T98G | Ribociclib |
| top60_altered_DNA_seed | KIT | 0.835293 | GBM | U-118-MG | Axitinib |
| top60_altered_DNA_seed | EGFR | 0.835318 | GBM | YH-13 | Gefitinib |
| top60_altered_DNA_seed | EGFR | 0.849635 | GBM | T98G | Sapitinib |
| top60_altered_DNA_seed | TP53 | 0.851192 | GBM | SK-MG-1 | MIRA-1 |
| top60_altered_DNA_seed | EGFR | 0.860964 | GBM | SF295 | Sapitinib |
| top60_altered_DNA_seed | EGFR | 0.871233 | GBM | LN-405 | AZD3759 |
| top60_altered_DNA_seed | TP53 | 0.871374 | GBM | SF268 | MIRA-1 |
| top60_altered_DNA_seed | EGFR | 0.87195 | GBM | D-263MG | Erlotinib |
| top60_altered_DNA_seed | KIT | 0.880972 | GBM | SK-MG-1 | Sorafenib |
| top60_altered_DNA_seed | EGFR | 0.884814 | GBM | D-392MG | Gefitinib |
| top60_altered_DNA_seed | EGFR | 0.884914 | GBM | KS-1 | Gefitinib |
| top60_altered_DNA_seed | KIT | 0.886867 | GBM | GB-1 | Motesanib |
| top60_altered_DNA_seed | EGFR | 0.888304 | GBM | SK-MG-1 | Sapitinib |
| top60_altered_DNA_seed | KIT | 0.894385 | GBM | LN-229 | Motesanib |
| top60_altered_DNA_seed | EGFR | 0.895519 | GBM | SF126 | Erlotinib |
| top60_altered_DNA_seed | CDK4 | 0.899147 | GBM | SF268 | Palbociclib |
| top60_altered_DNA_seed | EGFR | 0.908304 | GBM | SF268 | AZD3759 |
| top60_altered_DNA_seed | EGFR | 0.910745 | GBM | LNZTA3WT4 | Sapitinib |
| top60_altered_DNA_seed | KIT | 0.914464 | GBM | SW1088 | Sorafenib |
| top60_altered_DNA_seed | CDK4 | 0.917941 | GBM | LN-229 | Ribociclib |
| top60_altered_DNA_seed | EGFR | 0.931245 | GBM | U251 | Osimertinib |
| top60_altered_DNA_seed | EGFR | 0.931524 | GBM | SF126 | Afatinib |
| top60_altered_DNA_seed | EGFR | 0.932418 | GBM | CAS-1 | Erlotinib |
| top60_altered_DNA_seed | EGFR | 0.938563 | GBM | T98G | Erlotinib |
| top60_altered_DNA_seed | EGFR | 0.941274 | GBM | D-566MG | Afatinib |
| top60_altered_DNA_seed | KIT | 0.942809 | GBM | U-118-MG | Motesanib |
| top60_altered_DNA_seed | EGFR | 0.945968 | GBM | D-247MG | Erlotinib |
| top60_altered_DNA_seed | EGFR | 0.949568 | GBM | U-118-MG | Afatinib |
| top60_altered_DNA_seed | EGFR | 0.955973 | GBM | T98G | Osimertinib |
| top60_altered_DNA_seed | EGFR | 0.956074 | GBM | D-263MG | Sapitinib |
| top60_altered_DNA_seed | KIT | 0.958155 | GBM | D-542MG | Axitinib |
| top60_altered_DNA_seed | KIT | 0.958709 | GBM | LN-229 | Axitinib |
| top60_altered_DNA_seed | KIT | 0.959321 | GBM | D-247MG | Sorafenib |
| top60_altered_DNA_seed | EGFR | 0.965185 | GBM | SNB75 | Lapatinib |
| top60_altered_DNA_seed | EGFR | 0.972104 | GBM | LN-18 | AZD3759 |
| top60_altered_DNA_seed | EGFR | 0.973538 | GBM | SF295 | Afatinib |
| top60_altered_DNA_seed | TP53 | 0.975512 | GBM | DK-MG | MIRA-1 |
| top60_altered_DNA_seed | EGFR | 0.979684 | GBM | SK-MG-1 | Gefitinib |
| top60_altered_DNA_seed | EGFR | 0.980727 | GBM | LN-405 | Sapitinib |
| top60_altered_DNA_seed | EGFR | 0.986548 | GBM | D-263MG | Osimertinib |

| | | | | | |
|---|---|---|---|---|---|
| top60_altered_DNA_seed | EGFR | 0.9873 | GBM | D-263MG | Afatinib |
| top60_altered_DNA_seed | EGFR | 0.991396 | GBM | M059J | Gefitinib |
| top60_altered_DNA_seed | EGFR | 1.000318 | GBM | YKG-1 | Afatinib |
| top60_altered_DNA_seed | CDK4 | 1.00091 | GBM | LN-405 | Ribociclib |
| top60_altered_DNA_seed | EGFR | 1.023988 | GBM | Becker | Sapitinib |
| top60_altered_DNA_seed | EGFR | 1.027722 | GBM | U251 | Afatinib |
| top60_altered_DNA_seed | EGFR | 1.039273 | GBM | LN-18 | Afatinib |
| top60_altered_DNA_seed | EGFR | 1.039408 | GBM | KS-1 | AZD3759 |
| top60_altered_DNA_seed | EGFR | 1.04507 | GBM | LN-18 | Osimertinib |
| top60_altered_DNA_seed | EGFR | 1.049049 | GBM | D-247MG | AZD3759 |
| top60_altered_DNA_seed | KIT | 1.05376 | GBM | GB-1 | Axitinib |
| top60_altered_DNA_seed | EGFR | 1.056207 | GBM | SW1088 | Gefitinib |
| top60_altered_DNA_seed | EGFR | 1.063211 | GBM | GB-1 | Gefitinib |
| top60_altered_DNA_seed | KIT | 1.065315 | GBM | LN-405 | Sorafenib |
| top60_altered_DNA_seed | KIT | 1.080951 | GBM | SNB75 | Axitinib |
| top60_altered_DNA_seed | EGFR | 1.086889 | GBM | GB-1 | Afatinib |
| top60_altered_DNA_seed | EGFR | 1.088469 | GBM | Becker | Erlotinib |
| top60_altered_DNA_seed | KIT | 1.095995 | GBM | CAS-1 | Sorafenib |
| top60_altered_DNA_seed | TP53 | 1.104613 | GBM | LN-405 | MIRA-1 |
| top60_altered_DNA_seed | EGFR | 1.128677 | GBM | AM-38 | Sapitinib |
| top60_altered_DNA_seed | EGFR | 1.130392 | GBM | D-392MG | Erlotinib |
| top60_altered_DNA_seed | EGFR | 1.14068 | GBM | D-392MG | Sapitinib |
| top60_altered_DNA_seed | CDK4 | 1.157249 | GBM | U-118-MG | Ribociclib |
| top60_altered_DNA_seed | CDK4 | 1.178543 | GBM | M059J | Palbociclib |
| top60_altered_DNA_seed | CDK4 | 1.180443 | GBM | CAS-1 | Ribociclib |
| top60_altered_DNA_seed | EGFR | 1.180972 | GBM | U251 | Lapatinib |
| top60_altered_DNA_seed | TP53 | 1.185029 | GBM | SW1088 | MIRA-1 |
| top60_altered_DNA_seed | EGFR | 1.186611 | GBM | SF268 | Osimertinib |
| top60_altered_DNA_seed | EGFR | 1.187698 | GBM | DBTRG-05MG | Osimertinib |
| top60_altered_DNA_seed | EGFR | 1.201006 | GBM | LN-405 | Osimertinib |
| top60_altered_DNA_seed | EGFR | 1.222906 | GBM | U-118-MG | Lapatinib |
| top60_altered_DNA_seed | EGFR | 1.234905 | GBM | DK-MG | Gefitinib |
| top60_altered_DNA_seed | EGFR | 1.247878 | GBM | CAS-1 | AZD3759 |
| top60_altered_DNA_seed | TP53 | 1.249454 | GBM | CAS-1 | MIRA-1 |
| top60_altered_DNA_seed | EGFR | 1.260517 | GBM | SNB75 | Erlotinib |
| top60_altered_DNA_seed | CDK4 | 1.263292 | GBM | LN-405 | Palbociclib |
| top60_altered_DNA_seed | EGFR | 1.283699 | GBM | D-263MG | Lapatinib |
| top60_altered_DNA_seed | EGFR | 1.288037 | GBM | D-245MG | Erlotinib |
| top60_altered_DNA_seed | CDK4 | 1.299051 | GBM | D-542MG | Palbociclib |
| top60_altered_DNA_seed | CDK4 | 1.331047 | GBM | GB-1 | Ribociclib |
| top60_altered_DNA_seed | EGFR | 1.375176 | GBM | SNB75 | Afatinib |

| | | | | | |
|---|---|---|---|---|---|
| top60_altered_DNA_seed | EGFR | 1.378183 | GBM | DBTRG-05MG | AZD3759 |
| top60_altered_DNA_seed | EGFR | 1.39833 | GBM | GB-1 | AZD3759 |
| top60_altered_DNA_seed | EGFR | 1.401945 | GBM | U251 | Sapitinib |
| top60_altered_DNA_seed | EGFR | 1.425539 | GBM | U-87-MG | Afatinib |
| top60_altered_DNA_seed | EGFR | 1.430242 | GBM | U-87-MG | Lapatinib |
| top60_altered_DNA_seed | EGFR | 1.446664 | GBM | DBTRG-05MG | Afatinib |
| top60_altered_DNA_seed | EGFR | 1.4674 | GBM | Becker | Lapatinib |
| top60_altered_DNA_seed | CDK4 | 1.489772 | GBM | LN-229 | Palbociclib |
| top60_altered_DNA_seed | EGFR | 1.52125 | GBM | LN-405 | Afatinib |
| top60_altered_DNA_seed | EGFR | 1.562033 | GBM | KS-1 | Lapatinib |
| top60_altered_DNA_seed | CDK4 | 1.691322 | GBM | DBTRG-05MG | Ribociclib |
| top60_altered_DNA_seed | EGFR | 1.716475 | GBM | SF295 | Erlotinib |
| top60_altered_DNA_seed | EGFR | 1.747985 | GBM | LN-405 | Lapatinib |
| top60_altered_DNA_seed | EGFR | 1.750427 | GBM | U-118-MG | Sapitinib |
| top60_altered_DNA_seed | KIT | 1.751138 | GBM | DBTRG-05MG | Sorafenib |
| top60_altered_DNA_seed | EGFR | 1.767129 | GBM | DBTRG-05MG | Sapitinib |
| top60_altered_DNA_seed | EGFR | 1.805704 | GBM | GB-1 | Osimertinib |
| top60_altered_DNA_seed | KIT | 1.822663 | GBM | KS-1 | Sorafenib |
| top60_altered_DNA_seed | KIT | 1.831669 | GBM | DBTRG-05MG | Motesanib |
| top60_altered_DNA_seed | CDK4 | 2.065252 | GBM | DBTRG-05MG | Palbociclib |
| top60_altered_DNA_seed | EGFR | 2.079235 | GBM | D-392MG | Lapatinib |
| top60_altered_DNA_seed | EGFR | 2.094157 | GBM | DBTRG-05MG | Gefitinib |
| top60_altered_DNA_seed | TP53 | 2.125063 | GBM | DBTRG-05MG | MIRA-1 |
| top60_altered_DNA_seed | KIT | 2.201168 | GBM | D-392MG | Sorafenib |
| top60_altered_DNA_seed | KIT | 2.251505 | GBM | DBTRG-05MG | Axitinib |
| top60_altered_DNA_seed | EGFR | 2.576791 | GBM | DBTRG-05MG | Lapatinib |
| top60_altered_DNA_seed | PIK3CA | -2.592722 | HNSC | CAL-33 | AZD8835 |
| top60_altered_DNA_seed | PIK3CA | -2.355176 | HNSC | JHU-029 | PI3Ka_4409 |
| top60_altered_DNA_seed | PIK3CA | -1.907966 | HNSC | BB49-HNC | AZD8835 |
| top60_altered_DNA_seed | PIK3CA | -1.848721 | HNSC | JHU-029 | AZD8835 |
| top60_altered_DNA_seed | PIK3CA | -1.837295 | HNSC | HSC-2 | PI3Ka_4409 |
| top60_altered_DNA_seed | PIK3CA | -1.796513 | HNSC | CAL-33 | PI3Ka_4409 |
| top60_altered_DNA_seed | PIK3CA | -1.785323 | HNSC | DOK | AZD8835 |
| top60_altered_DNA_seed | PIK3CA | -1.782516 | HNSC | HSC-3 | PI3Ka_4409 |
| top60_altered_DNA_seed | PIK3CA | -1.678015 | HNSC | SCC-9 | PI3Ka_4409 |
| top60_altered_DNA_seed | PIK3CA | -1.492015 | HNSC | DOK | PI3Ka_4409 |
| top60_altered_DNA_seed | PIK3CA | -1.487289 | HNSC | BICR10 | AZD8835 |
| top60_altered_DNA_seed | PIK3CA | -1.463342 | HNSC | BICR22 | PI3Ka_4409 |
| top60_altered_DNA_seed | PIK3CA | -1.423077 | HNSC | HSC-4 | PI3Ka_4409 |
| top60_altered_DNA_seed | PIK3CA | -1.387447 | HNSC | BB49-HNC | PI3Ka_4409 |
| top60_altered_DNA_seed | PIK3CA | -1.367361 | HNSC | SCC-25 | PI3Ka_4409 |

| | | | | | |
|---|---|---|---|---|---|
| top60_altered_DNA_seed | PIK3CA | -1.298999 | HNSC | HSC-2 | AZD8835 |
| top60_altered_DNA_seed | PIK3CA | -1.199689 | HNSC | SAT | AZD8835 |
| top60_altered_DNA_seed | PIK3CA | -1.172892 | HNSC | SCC-9 | AZD8835 |
| top60_altered_DNA_seed | PIK3CA | -1.164822 | HNSC | CAL-27 | PI3Ka_4409 |
| top60_altered_DNA_seed | PIK3CA | -1.158928 | HNSC | CAL-27 | AZD8835 |
| top60_altered_DNA_seed | PIK3CA | -1.155628 | HNSC | PE-CA-PJ15 | PI3Ka_4409 |
| top60_altered_DNA_seed | PIK3CA | -1.127244 | HNSC | HSC-4 | AZD8835 |
| top60_altered_DNA_seed | PIK3CA | -1.008638 | HNSC | BB30-HNC | PI3Ka_4409 |
| top60_altered_DNA_seed | PIK3CA | -1.001532 | HNSC | BICR10 | PI3Ka_4409 |
| top60_altered_DNA_seed | PIK3CA | -1.000452 | HNSC | JHU-011 | PI3Ka_4409 |
| top60_altered_DNA_seed | PIK3CA | -0.994369 | HNSC | SAT | PI3Ka_4409 |
| top60_altered_DNA_seed | PIK3CA | -0.973117 | HNSC | SCC-15 | PI3Ka_4409 |
| top60_altered_DNA_seed | PIK3CA | -0.955939 | HNSC | Detroit562 | PI3Ka_4409 |
| top60_altered_DNA_seed | PIK3CA | -0.916753 | HNSC | JHU-022 | PI3Ka_4409 |
| top60_altered_DNA_seed | PIK3CA | -0.908083 | HNSC | SCC-15 | AZD8835 |
| top60_altered_DNA_seed | PIK3CA | -0.897997 | HNSC | PE-CA-PJ15 | AZD8835 |
| top60_altered_DNA_seed | PIK3CA | -0.889663 | HNSC | KOSC-2 | PI3Ka_4409 |
| top60_altered_DNA_seed | PIK3CA | -0.881566 | HNSC | BHY | PI3Ka_4409 |
| top60_altered_DNA_seed | PIK3CA | -0.863537 | HNSC | BICR78 | AZD8835 |
| top60_altered_DNA_seed | PIK3CA | -0.830883 | HNSC | BB30-HNC | AZD8835 |
| top60_altered_DNA_seed | PIK3CA | -0.815559 | HNSC | HO-1-N-1 | PI3Ka_4409 |
| top60_altered_DNA_seed | PIK3CA | -0.815453 | HNSC | Detroit562 | AZD8835 |
| top60_altered_DNA_seed | PIK3CA | -0.80159 | HNSC | OSC-20 | PI3Ka_4409 |
| top60_altered_DNA_seed | PIK3CA | -0.712964 | HNSC | LB771-HNC | AZD8835 |
| top60_altered_DNA_seed | PIK3CA | -0.676735 | HNSC | SCC-25 | AZD8835 |
| top60_altered_DNA_seed | PIK3CA | -0.673846 | HNSC | KON | PI3Ka_4409 |
| top60_altered_DNA_seed | PIK3CA | -0.65167 | HNSC | HSC-3 | AZD8835 |
| top60_altered_DNA_seed | PIK3CA | -0.612883 | HNSC | Ca9-22 | AZD8835 |
| top60_altered_DNA_seed | PIK3CA | -0.604195 | HNSC | HO-1-N-1 | AZD8835 |
| top60_altered_DNA_seed | PIK3CA | -0.602938 | HNSC | SKN-3 | PI3Ka_4409 |
| top60_altered_DNA_seed | PIK3CA | -0.54341 | HNSC | KOSC-2 | AZD8835 |
| top60_altered_DNA_seed | PIK3CA | -0.516948 | HNSC | OSC-19 | PI3Ka_4409 |
| top60_altered_DNA_seed | PIK3CA | -0.515743 | HNSC | BICR22 | AZD8835 |
| top60_altered_DNA_seed | PIK3CA | -0.480643 | HNSC | OSC-20 | AZD8835 |
| top60_altered_DNA_seed | PIK3CA | -0.444205 | HNSC | Ca9-22 | PI3Ka_4409 |
| top60_altered_DNA_seed | PIK3CA | -0.44101 | HNSC | KON | AZD8835 |
| top60_altered_DNA_seed | PIK3CA | -0.346995 | HNSC | LB771-HNC | PI3Ka_4409 |
| top60_altered_DNA_seed | PIK3CA | -0.339834 | HNSC | JHU-011 | AZD8835 |
| top60_altered_DNA_seed | PIK3CA | -0.31855 | HNSC | BICR78 | PI3Ka_4409 |
| top60_altered_DNA_seed | PIK3CA | -0.308458 | HNSC | OSC-19 | AZD8835 |
| top60_altered_DNA_seed | PIK3CA | -0.308287 | HNSC | PCI-4B | AZD8835 |

| | | | | | |
|---|---|---|---|---|---|
| top60_altered_DNA_seed | PIK3CA | -0.286122 | HNSC | BHY | AZD8835 |
| top60_altered_DNA_seed | PIK3CA | -0.269128 | HNSC | HO-1-u-1 | AZD8835 |
| top60_altered_DNA_seed | PIK3CA | -0.237089 | HNSC | FADU | PI3Ka_4409 |
| top60_altered_DNA_seed | PIK3CA | -0.197253 | HNSC | PCI-4B | PI3Ka_4409 |
| top60_altered_DNA_seed | PIK3CA | -0.188754 | HNSC | PCI-15A | PI3Ka_4409 |
| top60_altered_DNA_seed | PIK3CA | -0.18721 | HNSC | PCI-6A | AZD8835 |
| top60_altered_DNA_seed | PIK3CA | -0.179975 | HNSC | PCI-15A | AZD8835 |
| top60_altered_DNA_seed | PIK3CA | -0.168298 | HNSC | PCI-38 | PI3Ka_4409 |
| top60_altered_DNA_seed | PIK3CA | -0.126852 | HNSC | PCI-6A | PI3Ka_4409 |
| top60_altered_DNA_seed | PIK3CA | -0.094636 | HNSC | SKN-3 | AZD8835 |
| top60_altered_DNA_seed | PIK3CA | -0.073796 | HNSC | PCI-38 | AZD8835 |
| top60_altered_DNA_seed | PIK3CA | -0.063626 | HNSC | SCC-4 | PI3Ka_4409 |
| top60_altered_DNA_seed | PIK3CA | -0.030052 | HNSC | HO-1-u-1 | PI3Ka_4409 |
| top60_altered_DNA_seed | PIK3CA | -0.005308 | HNSC | FADU | AZD8835 |
| top60_altered_DNA_seed | PIK3CA | 0.127621 | HNSC | SAS | AZD8835 |
| top60_altered_DNA_seed | PIK3CA | 0.236136 | HNSC | SCC-4 | AZD8835 |
| top60_altered_DNA_seed | PIK3CA | 0.333339 | HNSC | SCC90 | AZD8835 |
| top60_altered_DNA_seed | PIK3CA | 0.58502 | HNSC | PCI-30 | AZD8835 |
| top60_altered_DNA_seed | PIK3CA | 0.854607 | HNSC | PCI-30 | PI3Ka_4409 |
| top60_altered_DNA_seed | PIK3CA | 1.534807 | HNSC | SAS | PI3Ka_4409 |
| top60_altered_DNA_seed | PIK3CA | 1.761379 | HNSC | SCC90 | PI3Ka_4409 |
| top60_altered_DNA_seed | PIK3CA | -2.249485 | HNSC | DOK | Alpelisib |
| top60_altered_DNA_seed | PIK3CA | -2.022601 | HNSC | BB49-HNC | AZD8186 |
| top60_altered_DNA_seed | PIK3CA | -2.021446 | HNSC | BB49-HNC | Alpelisib |
| top60_altered_DNA_seed | PIK3CA | -1.98395 | HNSC | BB49-HNC | Buparlisib |
| top60_altered_DNA_seed | PIK3CA | -1.76856 | HNSC | HSC-2 | Alpelisib |
| top60_altered_DNA_seed | PIK3CA | -1.763759 | HNSC | JHU-011 | Alpelisib |
| top60_altered_DNA_seed | PIK3CA | -1.722505 | HNSC | CAL-33 | Alpelisib |
| top60_altered_DNA_seed | TP53 | -1.685645 | HNSC | HO-1-N-1 | MIRA-1 |
| top60_altered_DNA_seed | PIK3CA | -1.575222 | HNSC | BB49-HNC | GNE-317 |
| top60_altered_DNA_seed | PIK3CA | -1.51221 | HNSC | HO-1-u-1 | GNE-317 |
| top60_altered_DNA_seed | PIK3CA | -1.367034 | HNSC | BB30-HNC | Alpelisib |
| top60_altered_DNA_seed | PIK3CA | -1.341142 | HNSC | DOK | AZD8186 |
| top60_altered_DNA_seed | PIK3CA | -1.314742 | HNSC | HO-1-u-1 | Alpelisib |
| top60_altered_DNA_seed | PIK3CA | -1.29089 | HNSC | DOK | Buparlisib |
| top60_altered_DNA_seed | PIK3CA | -1.206075 | HNSC | PCI-15A | Alpelisib |
| top60_altered_DNA_seed | PIK3CA | -1.194457 | HNSC | JHU-029 | Alpelisib |
| top60_altered_DNA_seed | PIK3CA | -1.186851 | HNSC | BICR10 | Buparlisib |
| top60_altered_DNA_seed | PIK3CA | -1.147633 | HNSC | BICR22 | Alpelisib |
| top60_altered_DNA_seed | PIK3CA | -1.144714 | HNSC | PCI-15A | GNE-317 |
| top60_altered_DNA_seed | PIK3CA | -1.115337 | HNSC | Detroit562 | GNE-317 |

| | | | | | |
|---|---|---|---|---|---|
| top60_altered_DNA_seed | PIK3CA | -1.10622 | HNSC | SAS | GNE-317 |
| top60_altered_DNA_seed | PIK3CA | -1.044558 | HNSC | HO-1-N-1 | Buparlisib |
| top60_altered_DNA_seed | PIK3CA | -0.983473 | HNSC | HO-1-u-1 | Buparlisib |
| top60_altered_DNA_seed | PIK3CA | -0.980902 | HNSC | HSC-3 | GNE-317 |
| top60_altered_DNA_seed | PIK3CA | -0.973561 | HNSC | KON | GNE-317 |
| top60_altered_DNA_seed | PIK3CA | -0.970934 | HNSC | JHU-011 | GNE-317 |
| top60_altered_DNA_seed | PIK3CA | -0.934206 | HNSC | CAL-27 | Buparlisib |
| top60_altered_DNA_seed | PIK3CA | -0.934102 | HNSC | KON | Alpelisib |
| top60_altered_DNA_seed | PIK3CA | -0.911513 | HNSC | BICR10 | GNE-317 |
| top60_altered_DNA_seed | PIK3CA | -0.902106 | HNSC | KON | AZD8186 |
| top60_altered_DNA_seed | PIK3CA | -0.884112 | HNSC | Detroit562 | Alpelisib |
| top60_altered_DNA_seed | PIK3CA | -0.866948 | HNSC | PCI-6A | Alpelisib |
| top60_altered_DNA_seed | PIK3CA | -0.807013 | HNSC | SAT | Alpelisib |
| top60_altered_DNA_seed | TP53 | -0.78828 | HNSC | OSC-19 | MIRA-1 |
| top60_altered_DNA_seed | TP53 | -0.78149 | HNSC | PCI-15A | MIRA-1 |
| top60_altered_DNA_seed | PIK3CA | -0.780878 | HNSC | Detroit562 | Buparlisib |
| top60_altered_DNA_seed | PIK3CA | -0.760683 | HNSC | JHU-029 | AZD8186 |
| top60_altered_DNA_seed | PIK3CA | -0.755008 | HNSC | JHU-011 | Buparlisib |
| top60_altered_DNA_seed | PIK3CA | -0.753698 | HNSC | HO-1-N-1 | Alpelisib |
| top60_altered_DNA_seed | PIK3CA | -0.713635 | HNSC | BB30-HNC | AZD8186 |
| top60_altered_DNA_seed | PIK3CA | -0.693506 | HNSC | SCC-9 | Alpelisib |
| top60_altered_DNA_seed | PIK3CA | -0.669504 | HNSC | BICR10 | Alpelisib |
| top60_altered_DNA_seed | PIK3CA | -0.66092 | HNSC | HO-1-N-1 | AZD8186 |
| top60_altered_DNA_seed | PIK3CA | -0.633116 | HNSC | HSC-2 | Buparlisib |
| top60_altered_DNA_seed | PIK3CA | -0.632378 | HNSC | HSC-3 | Alpelisib |
| top60_altered_DNA_seed | PIK3CA | -0.629658 | HNSC | Detroit562 | AZD8186 |
| top60_altered_DNA_seed | PIK3CA | -0.609705 | HNSC | SAS | Alpelisib |
| top60_altered_DNA_seed | PIK3CA | -0.596823 | HNSC | BB30-HNC | Buparlisib |
| top60_altered_DNA_seed | PIK3CA | -0.58603 | HNSC | SAS | AZD8186 |
| top60_altered_DNA_seed | PIK3CA | -0.577704 | HNSC | HSC-3 | Buparlisib |
| top60_altered_DNA_seed | TP53 | -0.558305 | HNSC | BHY | MIRA-1 |
| top60_altered_DNA_seed | PIK3CA | -0.549105 | HNSC | BICR22 | Buparlisib |
| top60_altered_DNA_seed | PIK3CA | -0.536458 | HNSC | Ca9-22 | Alpelisib |
| top60_altered_DNA_seed | PIK3CA | -0.508338 | HNSC | BB30-HNC | GNE-317 |
| top60_altered_DNA_seed | PIK3CA | -0.485251 | HNSC | CAL-27 | AZD8186 |
| top60_altered_DNA_seed | TP53 | -0.473581 | HNSC | BB30-HNC | MIRA-1 |
| top60_altered_DNA_seed | PIK3CA | -0.465323 | HNSC | JHU-022 | GNE-317 |
| top60_altered_DNA_seed | PIK3CA | -0.46297 | HNSC | SCC-4 | Alpelisib |
| top60_altered_DNA_seed | PIK3CA | -0.437463 | HNSC | OSC-19 | GNE-317 |
| top60_altered_DNA_seed | PIK3CA | -0.432018 | HNSC | OSC-19 | AZD8186 |
| top60_altered_DNA_seed | PIK3CA | -0.41911 | HNSC | HO-1-N-1 | GNE-317 |

| | | | | | |
|---|---|---|---|---|---|
| top60_altered_DNA_seed | PIK3CA | -0.415452 | HNSC | KOSC-2 | Alpelisib |
| top60_altered_DNA_seed | PIK3CA | -0.409117 | HNSC | PCI-4B | Alpelisib |
| top60_altered_DNA_seed | TP53 | -0.395709 | HNSC | BICR31 | MIRA-1 |
| top60_altered_DNA_seed | PIK3CA | -0.391237 | HNSC | LB771-HNC | Alpelisib |
| top60_altered_DNA_seed | PIK3CA | -0.388996 | HNSC | JHU-029 | Buparlisib |
| top60_altered_DNA_seed | PIK3CA | -0.385124 | HNSC | BICR22 | GNE-317 |
| top60_altered_DNA_seed | PIK3CA | -0.383097 | HNSC | BICR31 | Alpelisib |
| top60_altered_DNA_seed | TP53 | -0.372726 | HNSC | JHU-011 | MIRA-1 |
| top60_altered_DNA_seed | PIK3CA | -0.352846 | HNSC | PCI-15A | Buparlisib |
| top60_altered_DNA_seed | PIK3CA | -0.352723 | HNSC | HSC-2 | AZD8186 |
| top60_altered_DNA_seed | PIK3CA | -0.345583 | HNSC | SCC-25 | Alpelisib |
| top60_altered_DNA_seed | PIK3CA | -0.337761 | HNSC | JHU-022 | Alpelisib |
| top60_altered_DNA_seed | PIK3CA | -0.32346 | HNSC | JHU-011 | AZD8186 |
| top60_altered_DNA_seed | PIK3CA | -0.310817 | HNSC | BICR10 | AZD8186 |
| top60_altered_DNA_seed | PIK3CA | -0.297602 | HNSC | SCC-15 | AZD8186 |
| top60_altered_DNA_seed | TP53 | -0.291126 | HNSC | HO-1-u-1 | MIRA-1 |
| top60_altered_DNA_seed | PIK3CA | -0.290049 | HNSC | PCI-15A | AZD8186 |
| top60_altered_DNA_seed | PIK3CA | -0.280666 | HNSC | KOSC-2 | Buparlisib |
| top60_altered_DNA_seed | PIK3CA | -0.270935 | HNSC | Ca9-22 | Buparlisib |
| top60_altered_DNA_seed | PIK3CA | -0.26389 | HNSC | CAL-33 | GNE-317 |
| top60_altered_DNA_seed | PIK3CA | -0.26291 | HNSC | OSC-19 | Alpelisib |
| top60_altered_DNA_seed | PIK3CA | -0.260552 | HNSC | SCC-15 | Buparlisib |
| top60_altered_DNA_seed | PIK3CA | -0.251431 | HNSC | KON | Buparlisib |
| top60_altered_DNA_seed | PIK3CA | -0.245717 | HNSC | PE-CA-PJ15 | Alpelisib |
| top60_altered_DNA_seed | PIK3CA | -0.24413 | HNSC | CAL-33 | AZD8186 |
| top60_altered_DNA_seed | PIK3CA | -0.234926 | HNSC | HSC-4 | AZD8186 |
| top60_altered_DNA_seed | PIK3CA | -0.226396 | HNSC | CAL-33 | Buparlisib |
| top60_altered_DNA_seed | TP53 | -0.225344 | HNSC | HSC-2 | MIRA-1 |
| top60_altered_DNA_seed | PIK3CA | -0.223246 | HNSC | SAS | Buparlisib |
| top60_altered_DNA_seed | TP53 | -0.219795 | HNSC | PCI-4B | MIRA-1 |
| top60_altered_DNA_seed | PIK3CA | -0.188249 | HNSC | FADU | Alpelisib |
| top60_altered_DNA_seed | TP53 | -0.184433 | HNSC | SCC-25 | MIRA-1 |
| top60_altered_DNA_seed | PIK3CA | -0.16585 | HNSC | SCC-9 | AZD8186 |
| top60_altered_DNA_seed | TP53 | -0.158759 | HNSC | DOK | MIRA-1 |
| top60_altered_DNA_seed | PIK3CA | -0.155686 | HNSC | JHU-022 | AZD8186 |
| top60_altered_DNA_seed | TP53 | -0.128942 | HNSC | KON | MIRA-1 |
| top60_altered_DNA_seed | PIK3CA | -0.113985 | HNSC | SAT | GNE-317 |
| top60_altered_DNA_seed | TP53 | -0.09071 | HNSC | JHU-029 | MIRA-1 |
| top60_altered_DNA_seed | TP53 | -0.081288 | HNSC | SAS | MIRA-1 |
| top60_altered_DNA_seed | TP53 | -0.077083 | HNSC | PCI-6A | MIRA-1 |
| top60_altered_DNA_seed | PIK3CA | -0.070107 | HNSC | PCI-30 | GNE-317 |

| | | | | | |
|---|---|---|---|---|---|
| top60_altered_DNA_seed | PIK3CA | -0.048748 | HNSC | CAL-27 | Alpelisib |
| top60_altered_DNA_seed | TP53 | -0.048623 | HNSC | FADU | MIRA-1 |
| top60_altered_DNA_seed | TP53 | -0.0398 | HNSC | KOSC-2 | MIRA-1 |
| top60_altered_DNA_seed | PIK3CA | -0.032409 | HNSC | SCC-15 | GNE-317 |
| top60_altered_DNA_seed | PIK3CA | -0.028222 | HNSC | OSC-19 | Buparlisib |
| top60_altered_DNA_seed | PIK3CA | -0.021695 | HNSC | FADU | Buparlisib |
| top60_altered_DNA_seed | PIK3CA | -0.011501 | HNSC | PE-CA-PJ15 | AZD8186 |
| top60_altered_DNA_seed | PIK3CA | 0.00364 | HNSC | HO-1-u-1 | AZD8186 |
| top60_altered_DNA_seed | PIK3CA | 0.017505 | HNSC | BHY | AZD8186 |
| top60_altered_DNA_seed | PIK3CA | 0.019557 | HNSC | LB771-HNC | Buparlisib |
| top60_altered_DNA_seed | TP53 | 0.023865 | HNSC | BICR22 | MIRA-1 |
| top60_altered_DNA_seed | TP53 | 0.031459 | HNSC | CAL-33 | MIRA-1 |
| top60_altered_DNA_seed | PIK3CA | 0.035033 | HNSC | PE-CA-PJ15 | GNE-317 |
| top60_altered_DNA_seed | TP53 | 0.04735 | HNSC | PE-CA-PJ15 | MIRA-1 |
| top60_altered_DNA_seed | TP53 | 0.050948 | HNSC | HSC-3 | MIRA-1 |
| top60_altered_DNA_seed | PIK3CA | 0.067968 | HNSC | SAT | AZD8186 |
| top60_altered_DNA_seed | PIK3CA | 0.08018 | HNSC | JHU-029 | GNE-317 |
| top60_altered_DNA_seed | PIK3CA | 0.107142 | HNSC | PCI-6A | Buparlisib |
| top60_altered_DNA_seed | PIK3CA | 0.189775 | HNSC | SCC-15 | Alpelisib |
| top60_altered_DNA_seed | PIK3CA | 0.205448 | HNSC | BICR31 | AZD8186 |
| top60_altered_DNA_seed | TP53 | 0.225644 | HNSC | Ca9-22 | MIRA-1 |
| top60_altered_DNA_seed | PIK3CA | 0.246362 | HNSC | JHU-022 | Buparlisib |
| top60_altered_DNA_seed | PIK3CA | 0.266267 | HNSC | HSC-3 | AZD8186 |
| top60_altered_DNA_seed | PIK3CA | 0.27247 | HNSC | BHY | Alpelisib |
| top60_altered_DNA_seed | PIK3CA | 0.27338 | HNSC | PCI-30 | AZD8186 |
| top60_altered_DNA_seed | PIK3CA | 0.274282 | HNSC | PCI-6A | AZD8186 |
| top60_altered_DNA_seed | PIK3CA | 0.275438 | HNSC | CAL-27 | GNE-317 |
| top60_altered_DNA_seed | PIK3CA | 0.279707 | HNSC | FADU | GNE-317 |
| top60_altered_DNA_seed | TP53 | 0.283242 | HNSC | Detroit562 | MIRA-1 |
| top60_altered_DNA_seed | TP53 | 0.317842 | HNSC | CAL-27 | MIRA-1 |
| top60_altered_DNA_seed | PIK3CA | 0.338833 | HNSC | PCI-6A | GNE-317 |
| top60_altered_DNA_seed | PIK3CA | 0.35721 | HNSC | LB771-HNC | GNE-317 |
| top60_altered_DNA_seed | PIK3CA | 0.365457 | HNSC | Ca9-22 | GNE-317 |
| top60_altered_DNA_seed | TP53 | 0.371511 | HNSC | PCI-30 | MIRA-1 |
| top60_altered_DNA_seed | PIK3CA | 0.388751 | HNSC | PCI-4B | Buparlisib |
| top60_altered_DNA_seed | PIK3CA | 0.389747 | HNSC | PCI-30 | Alpelisib |
| top60_altered_DNA_seed | PIK3CA | 0.39866 | HNSC | BICR78 | Alpelisib |
| top60_altered_DNA_seed | PIK3CA | 0.402136 | HNSC | BICR22 | AZD8186 |
| top60_altered_DNA_seed | TP53 | 0.411201 | HNSC | SCC-9 | MIRA-1 |
| top60_altered_DNA_seed | PIK3CA | 0.412275 | HNSC | BICR31 | Buparlisib |
| top60_altered_DNA_seed | PIK3CA | 0.438892 | HNSC | KOSC-2 | GNE-317 |

| | | | | | |
|---|---|---|---|---|---|
| top60_altered_DNA_seed | PIK3CA | 0.461071 | HNSC | SCC-4 | Buparlisib |
| top60_altered_DNA_seed | TP53 | 0.463248 | HNSC | OSC-20 | MIRA-1 |
| top60_altered_DNA_seed | TP53 | 0.483098 | HNSC | SCC-15 | MIRA-1 |
| top60_altered_DNA_seed | PIK3CA | 0.483358 | HNSC | OSC-20 | Buparlisib |
| top60_altered_DNA_seed | TP53 | 0.51814 | HNSC | BB49-HNC | MIRA-1 |
| top60_altered_DNA_seed | PIK3CA | 0.531023 | HNSC | Ca9-22 | AZD8186 |
| top60_altered_DNA_seed | PIK3CA | 0.555687 | HNSC | HSC-4 | GNE-317 |
| top60_altered_DNA_seed | PIK3CA | 0.569282 | HNSC | SCC-4 | AZD8186 |
| top60_altered_DNA_seed | PIK3CA | 0.587562 | HNSC | FADU | AZD8186 |
| top60_altered_DNA_seed | TP53 | 0.619898 | HNSC | HSC-4 | MIRA-1 |
| top60_altered_DNA_seed | PIK3CA | 0.634428 | HNSC | SCC-9 | Buparlisib |
| top60_altered_DNA_seed | PIK3CA | 0.655852 | HNSC | SCC-4 | GNE-317 |
| top60_altered_DNA_seed | PIK3CA | 0.675844 | HNSC | KOSC-2 | AZD8186 |
| top60_altered_DNA_seed | PIK3CA | 0.685895 | HNSC | PE-CA-PJ15 | Buparlisib |
| top60_altered_DNA_seed | TP53 | 0.692683 | HNSC | BICR10 | MIRA-1 |
| top60_altered_DNA_seed | TP53 | 0.696963 | HNSC | BICR78 | MIRA-1 |
| top60_altered_DNA_seed | PIK3CA | 0.713261 | HNSC | OSC-20 | AZD8186 |
| top60_altered_DNA_seed | TP53 | 0.737087 | HNSC | SKN-3 | MIRA-1 |
| top60_altered_DNA_seed | PIK3CA | 0.76117 | HNSC | HSC-4 | Buparlisib |
| top60_altered_DNA_seed | PIK3CA | 0.770498 | HNSC | BHY | GNE-317 |
| top60_altered_DNA_seed | PIK3CA | 0.77162 | HNSC | OSC-20 | Alpelisib |
| top60_altered_DNA_seed | TP53 | 0.79274 | HNSC | SCC-4 | MIRA-1 |
| top60_altered_DNA_seed | PIK3CA | 0.833152 | HNSC | SKN-3 | AZD8186 |
| top60_altered_DNA_seed | PIK3CA | 0.85678 | HNSC | PCI-4B | GNE-317 |
| top60_altered_DNA_seed | PIK3CA | 0.898326 | HNSC | PCI-30 | Buparlisib |
| top60_altered_DNA_seed | PIK3CA | 0.917451 | HNSC | BICR31 | GNE-317 |
| top60_altered_DNA_seed | PIK3CA | 0.968149 | HNSC | SKN-3 | GNE-317 |
| top60_altered_DNA_seed | PIK3CA | 0.980543 | HNSC | LB771-HNC | AZD8186 |
| top60_altered_DNA_seed | PIK3CA | 0.993353 | HNSC | PCI-4B | AZD8186 |
| top60_altered_DNA_seed | TP53 | 1.029272 | HNSC | SAT | MIRA-1 |
| top60_altered_DNA_seed | PIK3CA | 1.047838 | HNSC | BICR78 | Buparlisib |
| top60_altered_DNA_seed | PIK3CA | 1.048236 | HNSC | SKN-3 | Buparlisib |
| top60_altered_DNA_seed | PIK3CA | 1.05214 | HNSC | SCC90 | AZD8186 |
| top60_altered_DNA_seed | PIK3CA | 1.080515 | HNSC | BICR78 | AZD8186 |
| top60_altered_DNA_seed | PIK3CA | 1.084815 | HNSC | SAT | Buparlisib |
| top60_altered_DNA_seed | PIK3CA | 1.086612 | HNSC | OSC-20 | GNE-317 |
| top60_altered_DNA_seed | TP53 | 1.097963 | HNSC | SCC90 | MIRA-1 |
| top60_altered_DNA_seed | PIK3CA | 1.103864 | HNSC | SCC-9 | GNE-317 |
| top60_altered_DNA_seed | PIK3CA | 1.113835 | HNSC | HSC-4 | Alpelisib |
| top60_altered_DNA_seed | TP53 | 1.159736 | HNSC | JHU-022 | MIRA-1 |
| top60_altered_DNA_seed | TP53 | 1.159898 | HNSC | LB771-HNC | MIRA-1 |

| | | | | | |
|---|---|---|---|---|---|
| top60_altered_DNA_seed | PIK3CA | 1.171475 | HNSC | SKN-3 | Alpelisib |
| top60_altered_DNA_seed | PIK3CA | 1.205565 | HNSC | BICR78 | GNE-317 |
| top60_altered_DNA_seed | PIK3CA | 1.278254 | HNSC | SCC90 | Alpelisib |
| top60_altered_DNA_seed | PIK3CA | 1.349365 | HNSC | SCC-25 | AZD8186 |
| top60_altered_DNA_seed | PIK3CA | 1.68388 | HNSC | BHY | Buparlisib |
| top60_altered_DNA_seed | PIK3CA | 1.696895 | HNSC | SCC90 | GNE-317 |
| top60_altered_DNA_seed | PIK3CA | 1.801314 | HNSC | SCC90 | Buparlisib |
| top60_altered_DNA_seed | PIK3CA | 1.832461 | HNSC | SCC-25 | GNE-317 |
| top60_altered_DNA_seed | PIK3CA | 3.472935 | HNSC | SCC-25 | Buparlisib |
| top60_altered_DNA_seed | FGFR4 | -2.4591 | KIRC | U031 | FGFR_0939 |
| top60_altered_DNA_seed | FGFR4 | -1.657996 | KIRC | RCC-JW | FGFR_0939 |
| top60_altered_DNA_seed | FGFR4 | -1.281831 | KIRC | HA7-RCC | FGFR_0939 |
| top60_altered_DNA_seed | FGFR4 | -0.9546 | KIRC | LB1047-RCC | FGFR_0939 |
| top60_altered_DNA_seed | FGFR4 | -0.919515 | KIRC | RCC-AB | FGFR_0939 |
| top60_altered_DNA_seed | FGFR4 | -0.892812 | KIRC | 786-0 | FGFR_0939 |
| top60_altered_DNA_seed | FGFR4 | -0.875187 | KIRC | A704 | FGFR_0939 |
| top60_altered_DNA_seed | FGFR4 | -0.775996 | KIRC | NCC010 | FGFR_3831 |
| top60_altered_DNA_seed | FGFR4 | -0.749345 | KIRC | BFTC-909 | FGFR_0939 |
| top60_altered_DNA_seed | FGFR4 | -0.73585 | KIRC | RCC10RGB | FGFR_0939 |
| top60_altered_DNA_seed | FGFR4 | -0.713748 | KIRC | A704 | FGFR_3831 |
| top60_altered_DNA_seed | FGFR4 | -0.655322 | KIRC | RCC-FG2 | FGFR_0939 |
| top60_altered_DNA_seed | FGFR4 | -0.65324 | KIRC | RXF393 | FGFR_3831 |
| top60_altered_DNA_seed | FGFR4 | -0.652829 | KIRC | LB996-RCC | FGFR_0939 |
| top60_altered_DNA_seed | FGFR4 | -0.609385 | KIRC | RCC-JF | FGFR_0939 |
| top60_altered_DNA_seed | FGFR4 | -0.494367 | KIRC | 769-P | FGFR_0939 |
| top60_altered_DNA_seed | FGFR4 | -0.486649 | KIRC | NCC021 | FGFR_0939 |
| top60_altered_DNA_seed | FGFR4 | -0.461804 | KIRC | SN12C | FGFR_0939 |
| top60_altered_DNA_seed | FGFR4 | -0.453204 | KIRC | LB2241-RCC | FGFR_3831 |
| top60_altered_DNA_seed | FGFR4 | -0.405782 | KIRC | BFTC-909 | FGFR_3831 |
| top60_altered_DNA_seed | FGFR4 | -0.374645 | KIRC | HA7-RCC | FGFR_3831 |
| top60_altered_DNA_seed | FGFR4 | -0.3568 | KIRC | LB1047-RCC | FGFR_3831 |
| top60_altered_DNA_seed | FGFR4 | -0.302501 | KIRC | NCC010 | FGFR_0939 |
| top60_altered_DNA_seed | FGFR4 | -0.296539 | KIRC | LB2241-RCC | FGFR_0939 |
| top60_altered_DNA_seed | FGFR4 | -0.262488 | KIRC | 769-P | FGFR_3831 |
| top60_altered_DNA_seed | FGFR4 | -0.261901 | KIRC | KMRC-20 | FGFR_0939 |
| top60_altered_DNA_seed | FGFR4 | -0.258943 | KIRC | RCC-JW | FGFR_3831 |
| top60_altered_DNA_seed | FGFR4 | -0.248438 | KIRC | VMRC-RCZ | FGFR_0939 |
| top60_altered_DNA_seed | FGFR4 | -0.241968 | KIRC | RCC-MF | FGFR_0939 |
| top60_altered_DNA_seed | FGFR4 | -0.219742 | KIRC | RXF393 | FGFR_0939 |
| top60_altered_DNA_seed | FGFR4 | -0.199676 | KIRC | SW156 | FGFR_0939 |
| top60_altered_DNA_seed | FGFR4 | -0.116518 | KIRC | A498 | FGFR_3831 |

| | | | | | |
|---|---|---|---|---|---|
| top60_altered_DNA_seed | FGFR4 | -0.112227 | KIRC | 786-0 | FGFR_3831 |
| top60_altered_DNA_seed | FGFR4 | -0.109773 | KIRC | BB65-RCC | FGFR_0939 |
| top60_altered_DNA_seed | FGFR4 | -0.080865 | KIRC | SN12C | FGFR_3831 |
| top60_altered_DNA_seed | FGFR4 | -0.052158 | KIRC | KMRC-1 | FGFR_0939 |
| top60_altered_DNA_seed | FGFR4 | -0.044434 | KIRC | LB996-RCC | FGFR_3831 |
| top60_altered_DNA_seed | FGFR4 | 0.022847 | KIRC | RCC-AB | FGFR_3831 |
| top60_altered_DNA_seed | FGFR4 | 0.035524 | KIRC | CAKI-1 | FGFR_0939 |
| top60_altered_DNA_seed | FGFR4 | 0.06514 | KIRC | ACHN | FGFR_3831 |
| top60_altered_DNA_seed | FGFR4 | 0.088982 | KIRC | RCC-ER | FGFR_0939 |
| top60_altered_DNA_seed | FGFR4 | 0.152314 | KIRC | RCC-JF | FGFR_3831 |
| top60_altered_DNA_seed | FGFR4 | 0.175746 | KIRC | NCC021 | FGFR_3831 |
| top60_altered_DNA_seed | FGFR4 | 0.242805 | KIRC | CAKI-1 | FGFR_3831 |
| top60_altered_DNA_seed | FGFR4 | 0.253419 | KIRC | TK10 | FGFR_0939 |
| top60_altered_DNA_seed | FGFR4 | 0.284177 | KIRC | KMRC-20 | FGFR_3831 |
| top60_altered_DNA_seed | FGFR4 | 0.386611 | KIRC | KMRC-1 | FGFR_3831 |
| top60_altered_DNA_seed | FGFR4 | 0.407777 | KIRC | CAL-54 | FGFR_3831 |
| top60_altered_DNA_seed | FGFR4 | 0.476178 | KIRC | BB65-RCC | FGFR_3831 |
| top60_altered_DNA_seed | FGFR4 | 0.522441 | KIRC | A498 | FGFR_0939 |
| top60_altered_DNA_seed | FGFR4 | 0.558079 | KIRC | U031 | FGFR_3831 |
| top60_altered_DNA_seed | FGFR4 | 0.621739 | KIRC | VMRC-RCZ | FGFR_3831 |
| top60_altered_DNA_seed | FGFR4 | 0.668249 | KIRC | OS-RC-2 | FGFR_0939 |
| top60_altered_DNA_seed | FGFR4 | 0.736576 | KIRC | ACHN | FGFR_0939 |
| top60_altered_DNA_seed | FGFR4 | 0.824612 | KIRC | TK10 | FGFR_3831 |
| top60_altered_DNA_seed | FGFR4 | 0.932604 | KIRC | CAL-54 | FGFR_0939 |
| top60_altered_DNA_seed | FGFR4 | 1.151913 | KIRC | SW156 | FGFR_3831 |
| top60_altered_DNA_seed | FGFR4 | 1.15443 | KIRC | OS-RC-2 | FGFR_3831 |
| top60_altered_DNA_seed | FGFR4 | 1.190802 | KIRC | RCC-FG2 | FGFR_3831 |
| top60_altered_DNA_seed | FGFR4 | 1.559144 | KIRC | RCC-MF | FGFR_3831 |
| top60_altered_DNA_seed | FGFR4 | 1.7371 | KIRC | RCC-ER | FGFR_3831 |
| top60_altered_DNA_seed | FGFR4 | 2.045145 | KIRC | RCC10RGB | FGFR_3831 |
| top60_altered_DNA_seed | HSF1 | -2.260876 | LGG | H4 | CCT245232 |
| top60_altered_DNA_seed | EGFR | -1.781339 | LGG | D-502MG | Lapatinib |
| top60_altered_DNA_seed | EGFR | -1.665939 | LGG | D-502MG | Erlotinib |
| top60_altered_DNA_seed | HSF1 | -1.643141 | LGG | MOG-G-UVW | CCT245232 |
| top60_altered_DNA_seed | HSF1 | -1.451036 | LGG | MOG-G-CCM | CCT245467 |
| top60_altered_DNA_seed | EGFR | -1.349885 | LGG | NMC-G1 | Erlotinib |
| top60_altered_DNA_seed | EGFR | -1.348828 | LGG | MOG-G-CCM | Cetuximab |
| top60_altered_DNA_seed | EGFR | -1.161462 | LGG | GI-1 | Lapatinib |
| top60_altered_DNA_seed | EGFR | -1.130372 | LGG | NMC-G1 | Lapatinib |
| top60_altered_DNA_seed | HSF1 | -0.792226 | LGG | GAMG | CCT245232 |
| top60_altered_DNA_seed | EGFR | -0.738873 | LGG | D-336MG | Lapatinib |

| | | | | | |
|---|---|---|---|---|---|
| top60_altered_DNA_seed | EGFR | -0.710063 | LGG | Hs-683 | Cetuximab |
| top60_altered_DNA_seed | PIK3CA | -0.698952 | LGG | H4 | AZD8835 |
| top60_altered_DNA_seed | PIK3CA | -0.68251 | LGG | GAMG | AZD8835 |
| top60_altered_DNA_seed | EGFR | -0.674168 | LGG | GAMG | Gefitinib |
| top60_altered_DNA_seed | EGFR | -0.636895 | LGG | GI-1 | CUDC-101 |
| top60_altered_DNA_seed | EGFR | -0.631103 | LGG | GAMG | Afatinib |
| top60_altered_DNA_seed | EGFR | -0.629785 | LGG | SF539 | Lapatinib |
| top60_altered_DNA_seed | EGFR | -0.627747 | LGG | NMC-G1 | Cetuximab |
| top60_altered_DNA_seed | EGFR | -0.597681 | LGG | no-11 | Erlotinib |
| top60_altered_DNA_seed | HSF1 | -0.594091 | LGG | Hs-683 | CCT245232 |
| top60_altered_DNA_seed | HSF1 | -0.58396 | LGG | SF539 | CCT245232 |
| top60_altered_DNA_seed | HSF1 | -0.567149 | LGG | GI-1 | CCT245232 |
| top60_altered_DNA_seed | EGFR | -0.566105 | LGG | GAMG | CUDC-101 |
| top60_altered_DNA_seed | EGFR | -0.541582 | LGG | KALS-1 | AZD8931 |
| top60_altered_DNA_seed | EGFR | -0.538609 | LGG | GI-1 | Cetuximab |
| top60_altered_DNA_seed | PIK3CA | -0.534211 | LGG | GI-1 | PI3Ka_4409 |
| top60_altered_DNA_seed | EGFR | -0.496939 | LGG | D-502MG | Afatinib |
| top60_altered_DNA_seed | EGFR | -0.494186 | LGG | MOG-G-CCM | CUDC-101 |
| top60_altered_DNA_seed | EGFR | -0.47902 | LGG | MOG-G-CCM | Pelitinib |
| top60_altered_DNA_seed | EGFR | -0.47516 | LGG | KINGS-1 | CUDC-101 |
| top60_altered_DNA_seed | EGFR | -0.468754 | LGG | KNS-42 | Lapatinib |
| top60_altered_DNA_seed | HSF1 | -0.467839 | LGG | KALS-1 | CCT245467 |
| top60_altered_DNA_seed | EGFR | -0.460047 | LGG | MOG-G-UVW | Afatinib |
| top60_altered_DNA_seed | EGFR | -0.456558 | LGG | MOG-G-UVW | CUDC-101 |
| top60_altered_DNA_seed | HSF1 | -0.431512 | LGG | D-336MG | CCT245232 |
| top60_altered_DNA_seed | EGFR | -0.424855 | LGG | D-502MG | Pelitinib |
| top60_altered_DNA_seed | EGFR | -0.416291 | LGG | H4 | AZD8931 |
| top60_altered_DNA_seed | HSF1 | -0.414379 | LGG | NMC-G1 | CCT245232 |
| top60_altered_DNA_seed | PIK3CA | -0.411424 | LGG | D-336MG | PI3Ka_4409 |
| top60_altered_DNA_seed | PIK3CA | -0.367911 | LGG | D-336MG | AZD8835 |
| top60_altered_DNA_seed | PIK3CA | -0.365229 | LGG | GI-1 | AZD8835 |
| top60_altered_DNA_seed | EGFR | -0.350702 | LGG | SF539 | AZD8931 |
| top60_altered_DNA_seed | HSF1 | -0.340787 | LGG | Hs-683 | CCT245467 |
| top60_altered_DNA_seed | EGFR | -0.340037 | LGG | D-336MG | CI-1033 |
| top60_altered_DNA_seed | HSF1 | -0.33906 | LGG | SW1783 | CCT245467 |
| top60_altered_DNA_seed | EGFR | -0.335629 | LGG | D-336MG | AST-1306 |
| top60_altered_DNA_seed | EGFR | -0.327108 | LGG | NMC-G1 | Pelitinib |
| top60_altered_DNA_seed | EGFR | -0.323543 | LGG | D-336MG | PF-00299804 |
| top60_altered_DNA_seed | EGFR | -0.317082 | LGG | MOG-G-UVW | Cetuximab |
| top60_altered_DNA_seed | EGFR | -0.306079 | LGG | SW1783 | AST-1306 |
| top60_altered_DNA_seed | EGFR | -0.289355 | LGG | MOG-G-CCM | CI-1033 |

| | | | | | |
|---|---|---|---|---|---|
| top60_altered_DNA_seed | EGFR | -0.278218 | LGG | KALS-1 | Erlotinib |
| top60_altered_DNA_seed | EGFR | -0.258231 | LGG | D-502MG | Cetuximab |
| top60_altered_DNA_seed | EGFR | -0.256302 | LGG | KNS-42 | Erlotinib |
| top60_altered_DNA_seed | EGFR | -0.254777 | LGG | SF539 | CI-1033 |
| top60_altered_DNA_seed | PIK3CA | -0.239207 | LGG | GAMG | PI3Ka_4409 |
| top60_altered_DNA_seed | EGFR | -0.229225 | LGG | GI-1 | AST-1306 |
| top60_altered_DNA_seed | EGFR | -0.226739 | LGG | SF539 | Afatinib |
| top60_altered_DNA_seed | EGFR | -0.220696 | LGG | SF539 | Pelitinib |
| top60_altered_DNA_seed | EGFR | -0.210579 | LGG | GAMG | Pelitinib |
| top60_altered_DNA_seed | EGFR | -0.208433 | LGG | MOG-G-CCM | AST-1306 |
| top60_altered_DNA_seed | EGFR | -0.20048 | LGG | D-502MG | CUDC-101 |
| top60_altered_DNA_seed | PIK3CA | -0.196107 | LGG | H4 | PI3Ka_4409 |
| top60_altered_DNA_seed | EGFR | -0.192387 | LGG | GAMG | AST-1306 |
| top60_altered_DNA_seed | EGFR | -0.180613 | LGG | MOG-G-UVW | AZD8931 |
| top60_altered_DNA_seed | EGFR | -0.170201 | LGG | GI-1 | AZD8931 |
| top60_altered_DNA_seed | EGFR | -0.1693 | LGG | D-336MG | Pelitinib |
| top60_altered_DNA_seed | EGFR | -0.163842 | LGG | H4 | Pelitinib |
| top60_altered_DNA_seed | EGFR | -0.116983 | LGG | KNS-81-FD | Lapatinib |
| top60_altered_DNA_seed | HSF1 | -0.110421 | LGG | KALS-1 | CCT245232 |
| top60_altered_DNA_seed | HSF1 | -0.106538 | LGG | SF539 | CCT245467 |
| top60_altered_DNA_seed | EGFR | -0.102049 | LGG | MOG-G-CCM | PF-00299804 |
| top60_altered_DNA_seed | PIK3CA | -0.101891 | LGG | MOG-G-UVW | AZD8835 |
| top60_altered_DNA_seed | EGFR | -0.099033 | LGG | no-11 | Pelitinib |
| top60_altered_DNA_seed | EGFR | -0.096522 | LGG | GAMG | Cetuximab |
| top60_altered_DNA_seed | EGFR | -0.093728 | LGG | GI-1 | CI-1033 |
| top60_altered_DNA_seed | PIK3CA | -0.088347 | LGG | NMC-G1 | AZD8835 |
| top60_altered_DNA_seed | EGFR | -0.086896 | LGG | no-11 | Afatinib |
| top60_altered_DNA_seed | EGFR | -0.081845 | LGG | SW1783 | CI-1033 |
| top60_altered_DNA_seed | EGFR | -0.062129 | LGG | GAMG | PF-00299804 |
| top60_altered_DNA_seed | EGFR | -0.05156 | LGG | GAMG | CI-1033 |
| top60_altered_DNA_seed | EGFR | -0.044553 | LGG | GI-1 | PF-00299804 |
| top60_altered_DNA_seed | EGFR | -0.043042 | LGG | SF539 | Erlotinib |
| top60_altered_DNA_seed | EGFR | -0.038619 | LGG | H4 | Afatinib |
| top60_altered_DNA_seed | EGFR | -0.033075 | LGG | NMC-G1 | PF-00299804 |
| top60_altered_DNA_seed | EGFR | -0.016892 | LGG | KNS-81-FD | CUDC-101 |
| top60_altered_DNA_seed | HSF1 | -0.013066 | LGG | no-10 | CCT245232 |
| top60_altered_DNA_seed | EGFR | -0.012796 | LGG | no-10 | Lapatinib |
| top60_altered_DNA_seed | PIK3CA | -0.001245 | LGG | SW1783 | PI3Ka_4409 |
| top60_altered_DNA_seed | EGFR | 0.001096 | LGG | D-336MG | AZD8931 |
| top60_altered_DNA_seed | HSF1 | 0.001849 | LGG | KINGS-1 | CCT245232 |
| top60_altered_DNA_seed | EGFR | 0.011479 | LGG | NMC-G1 | AZD8931 |

| | | | | | |
|---|---|---|---|---|---|
| top60_altered_DNA_seed | EGFR | 0.016618 | LGG | SW1783 | PF-00299804 |
| top60_altered_DNA_seed | HSF1 | 0.016783 | LGG | D-502MG | CCT245232 |
| top60_altered_DNA_seed | EGFR | 0.017897 | LGG | GAMG | AZD8931 |
| top60_altered_DNA_seed | EGFR | 0.026012 | LGG | SF539 | PF-00299804 |
| top60_altered_DNA_seed | EGFR | 0.028518 | LGG | SF539 | Afatinib |
| top60_altered_DNA_seed | EGFR | 0.037276 | LGG | KNS-42 | AZD8931 |
| top60_altered_DNA_seed | EGFR | 0.039318 | LGG | KALS-1 | CUDC-101 |
| top60_altered_DNA_seed | EGFR | 0.039762 | LGG | no-10 | AZD8931 |
| top60_altered_DNA_seed | EGFR | 0.044501 | LGG | SF539 | AST-1306 |
| top60_altered_DNA_seed | EGFR | 0.045337 | LGG | KINGS-1 | Afatinib |
| top60_altered_DNA_seed | HSF1 | 0.051369 | LGG | GI-1 | CCT245467 |
| top60_altered_DNA_seed | EGFR | 0.054718 | LGG | MOG-G-CCM | AZD8931 |
| top60_altered_DNA_seed | EGFR | 0.056061 | LGG | Hs-683 | AZD8931 |
| top60_altered_DNA_seed | PIK3CA | 0.060692 | LGG | SF539 | AZD8835 |
| top60_altered_DNA_seed | EGFR | 0.061681 | LGG | D-336MG | Afatinib |
| top60_altered_DNA_seed | PIK3CA | 0.064094 | LGG | SF539 | PI3Ka_4409 |
| top60_altered_DNA_seed | EGFR | 0.070173 | LGG | GAMG | Afatinib |
| top60_altered_DNA_seed | EGFR | 0.083334 | LGG | D-502MG | CI-1033 |
| top60_altered_DNA_seed | EGFR | 0.103801 | LGG | no-10 | CUDC-101 |
| top60_altered_DNA_seed | EGFR | 0.109636 | LGG | SW1783 | Afatinib |
| top60_altered_DNA_seed | EGFR | 0.111063 | LGG | H4 | PF-00299804 |
| top60_altered_DNA_seed | EGFR | 0.113206 | LGG | SW1783 | Gefitinib |
| top60_altered_DNA_seed | EGFR | 0.115724 | LGG | SW1783 | AZD8931 |
| top60_altered_DNA_seed | PIK3CA | 0.124961 | LGG | MOG-G-CCM | PI3Ka_4409 |
| top60_altered_DNA_seed | EGFR | 0.130562 | LGG | GI-1 | Pelitinib |
| top60_altered_DNA_seed | EGFR | 0.130695 | LGG | H4 | Cetuximab |
| top60_altered_DNA_seed | EGFR | 0.144527 | LGG | KNS-42 | CUDC-101 |
| top60_altered_DNA_seed | EGFR | 0.155229 | LGG | SF539 | Cetuximab |
| top60_altered_DNA_seed | PIK3CA | 0.157451 | LGG | KINGS-1 | PI3Ka_4409 |
| top60_altered_DNA_seed | EGFR | 0.171176 | LGG | no-10 | Cetuximab |
| top60_altered_DNA_seed | EGFR | 0.172969 | LGG | GI-1 | Afatinib |
| top60_altered_DNA_seed | EGFR | 0.183783 | LGG | GI-1 | Gefitinib |
| top60_altered_DNA_seed | EGFR | 0.213495 | LGG | NMC-G1 | CUDC-101 |
| top60_altered_DNA_seed | EGFR | 0.222707 | LGG | no-10 | Afatinib |
| top60_altered_DNA_seed | EGFR | 0.222872 | LGG | MOG-G-UVW | Gefitinib |
| top60_altered_DNA_seed | EGFR | 0.230755 | LGG | no-11 | Cetuximab |
| top60_altered_DNA_seed | PIK3CA | 0.234733 | LGG | MOG-G-UVW | PI3Ka_4409 |
| top60_altered_DNA_seed | EGFR | 0.240156 | LGG | Hs-683 | AST-1306 |
| top60_altered_DNA_seed | EGFR | 0.246354 | LGG | H4 | CI-1033 |
| top60_altered_DNA_seed | EGFR | 0.258762 | LGG | KINGS-1 | PF-00299804 |
| top60_altered_DNA_seed | EGFR | 0.266573 | LGG | GI-1 | Erlotinib |

| | | | | | |
|---|---|---|---|---|---|
| top60_altered_DNA_seed | HSF1 | 0.268307 | LGG | GAMG | CCT245467 |
| top60_altered_DNA_seed | EGFR | 0.278956 | LGG | NMC-G1 | Afatinib |
| top60_altered_DNA_seed | EGFR | 0.28175 | LGG | H4 | AST-1306 |
| top60_altered_DNA_seed | PIK3CA | 0.282967 | LGG | D-502MG | PI3Ka_4409 |
| top60_altered_DNA_seed | EGFR | 0.284077 | LGG | NMC-G1 | CI-1033 |
| top60_altered_DNA_seed | EGFR | 0.289358 | LGG | KINGS-1 | AZD8931 |
| top60_altered_DNA_seed | HSF1 | 0.306013 | LGG | KNS-81-FD | CCT245232 |
| top60_altered_DNA_seed | PIK3CA | 0.332913 | LGG | KALS-1 | PI3Ka_4409 |
| top60_altered_DNA_seed | EGFR | 0.349894 | LGG | SW1783 | Pelitinib |
| top60_altered_DNA_seed | EGFR | 0.356587 | LGG | MOG-G-CCM | Gefitinib |
| top60_altered_DNA_seed | EGFR | 0.393673 | LGG | no-10 | Afatinib |
| top60_altered_DNA_seed | EGFR | 0.395668 | LGG | D-502MG | AST-1306 |
| top60_altered_DNA_seed | EGFR | 0.409953 | LGG | KALS-1 | Lapatinib |
| top60_altered_DNA_seed | HSF1 | 0.422368 | LGG | NMC-G1 | CCT245467 |
| top60_altered_DNA_seed | EGFR | 0.424023 | LGG | Hs-683 | Afatinib |
| top60_altered_DNA_seed | EGFR | 0.424677 | LGG | SF539 | Gefitinib |
| top60_altered_DNA_seed | EGFR | 0.434767 | LGG | GI-1 | Afatinib |
| top60_altered_DNA_seed | EGFR | 0.443061 | LGG | D-336MG | Gefitinib |
| top60_altered_DNA_seed | EGFR | 0.456989 | LGG | D-502MG | Afatinib |
| top60_altered_DNA_seed | HSF1 | 0.471667 | LGG | SW1783 | CCT245232 |
| top60_altered_DNA_seed | EGFR | 0.478771 | LGG | KNS-42 | Afatinib |
| top60_altered_DNA_seed | EGFR | 0.486965 | LGG | D-502MG | PF-00299804 |
| top60_altered_DNA_seed | EGFR | 0.490698 | LGG | D-336MG | Erlotinib |
| top60_altered_DNA_seed | HSF1 | 0.505731 | LGG | KINGS-1 | CCT245467 |
| top60_altered_DNA_seed | EGFR | 0.507585 | LGG | MOG-G-UVW | Afatinib |
| top60_altered_DNA_seed | EGFR | 0.508347 | LGG | KNS-81-FD | Erlotinib |
| top60_altered_DNA_seed | PIK3CA | 0.515561 | LGG | D-502MG | AZD8835 |
| top60_altered_DNA_seed | EGFR | 0.516424 | LGG | H4 | Afatinib |
| top60_altered_DNA_seed | EGFR | 0.519947 | LGG | no-10 | Pelitinib |
| top60_altered_DNA_seed | EGFR | 0.525044 | LGG | H4 | Gefitinib |
| top60_altered_DNA_seed | EGFR | 0.526874 | LGG | no-10 | AST-1306 |
| top60_altered_DNA_seed | EGFR | 0.532922 | LGG | MOG-G-UVW | Pelitinib |
| top60_altered_DNA_seed | EGFR | 0.533552 | LGG | SW1783 | Afatinib |
| top60_altered_DNA_seed | EGFR | 0.539288 | LGG | Hs-683 | Pelitinib |
| top60_altered_DNA_seed | EGFR | 0.540151 | LGG | D-336MG | Afatinib |
| top60_altered_DNA_seed | EGFR | 0.564708 | LGG | D-336MG | Cetuximab |
| top60_altered_DNA_seed | EGFR | 0.589554 | LGG | no-11 | CUDC-101 |
| top60_altered_DNA_seed | EGFR | 0.596731 | LGG | no-10 | CI-1033 |
| top60_altered_DNA_seed | EGFR | 0.597685 | LGG | D-502MG | AZD8931 |
| top60_altered_DNA_seed | EGFR | 0.624476 | LGG | SW1783 | CUDC-101 |
| top60_altered_DNA_seed | EGFR | 0.628077 | LGG | MOG-G-UVW | AST-1306 |

| | | | | | |
|---|---|---|---|---|---|
| top60_altered_DNA_seed | HSF1 | 0.632688 | LGG | no-11 | CCT245232 |
| top60_altered_DNA_seed | PIK3CA | 0.63559 | LGG | MOG-G-CCM | AZD8835 |
| top60_altered_DNA_seed | EGFR | 0.646058 | LGG | MOG-G-UVW | CI-1033 |
| top60_altered_DNA_seed | EGFR | 0.679651 | LGG | no-11 | CI-1033 |
| top60_altered_DNA_seed | EGFR | 0.688353 | LGG | MOG-G-UVW | PF-00299804 |
| top60_altered_DNA_seed | PIK3CA | 0.703455 | LGG | KINGS-1 | AZD8835 |
| top60_altered_DNA_seed | EGFR | 0.71433 | LGG | no-10 | Erlotinib |
| top60_altered_DNA_seed | EGFR | 0.716901 | LGG | KALS-1 | Pelitinib |
| top60_altered_DNA_seed | EGFR | 0.741187 | LGG | Hs-683 | Afatinib |
| top60_altered_DNA_seed | EGFR | 0.767565 | LGG | NMC-G1 | AST-1306 |
| top60_altered_DNA_seed | PIK3CA | 0.768108 | LGG | SW1783 | AZD8835 |
| top60_altered_DNA_seed | HSF1 | 0.769501 | LGG | KNS-42 | CCT245232 |
| top60_altered_DNA_seed | EGFR | 0.774391 | LGG | KNS-42 | Cetuximab |
| top60_altered_DNA_seed | EGFR | 0.782337 | LGG | no-11 | Gefitinib |
| top60_altered_DNA_seed | PIK3CA | 0.783126 | LGG | no-10 | PI3Ka_4409 |
| top60_altered_DNA_seed | EGFR | 0.789883 | LGG | KNS-42 | Gefitinib |
| top60_altered_DNA_seed | EGFR | 0.790636 | LGG | no-11 | PF-00299804 |
| top60_altered_DNA_seed | EGFR | 0.796652 | LGG | MOG-G-CCM | Afatinib |
| top60_altered_DNA_seed | EGFR | 0.818872 | LGG | KNS-81-FD | Afatinib |
| top60_altered_DNA_seed | EGFR | 0.821511 | LGG | NMC-G1 | Afatinib |
| top60_altered_DNA_seed | EGFR | 0.832433 | LGG | KINGS-1 | Erlotinib |
| top60_altered_DNA_seed | EGFR | 0.847427 | LGG | KALS-1 | Afatinib |
| top60_altered_DNA_seed | EGFR | 0.851345 | LGG | Hs-683 | Gefitinib |
| top60_altered_DNA_seed | PIK3CA | 0.851969 | LGG | NMC-G1 | PI3Ka_4409 |
| top60_altered_DNA_seed | EGFR | 0.853546 | LGG | Hs-683 | CI-1033 |
| top60_altered_DNA_seed | EGFR | 0.86105 | LGG | KALS-1 | PF-00299804 |
| top60_altered_DNA_seed | EGFR | 0.868104 | LGG | KALS-1 | Gefitinib |
| top60_altered_DNA_seed | EGFR | 0.870005 | LGG | KINGS-1 | Cetuximab |
| top60_altered_DNA_seed | EGFR | 0.874775 | LGG | no-10 | PF-00299804 |
| top60_altered_DNA_seed | EGFR | 0.881212 | LGG | Hs-683 | PF-00299804 |
| top60_altered_DNA_seed | PIK3CA | 0.897285 | LGG | Hs-683 | AZD8835 |
| top60_altered_DNA_seed | PIK3CA | 0.908864 | LGG | KNS-42 | PI3Ka_4409 |
| top60_altered_DNA_seed | EGFR | 0.916778 | LGG | MOG-G-CCM | Afatinib |
| top60_altered_DNA_seed | HSF1 | 0.923046 | LGG | MOG-G-CCM | CCT245232 |
| top60_altered_DNA_seed | HSF1 | 0.930243 | LGG | D-502MG | CCT245467 |
| top60_altered_DNA_seed | EGFR | 0.936563 | LGG | KNS-81-FD | AST-1306 |
| top60_altered_DNA_seed | EGFR | 0.949388 | LGG | KNS-81-FD | Gefitinib |
| top60_altered_DNA_seed | PIK3CA | 0.952488 | LGG | KALS-1 | AZD8835 |
| top60_altered_DNA_seed | EGFR | 0.966507 | LGG | NMC-G1 | Gefitinib |
| top60_altered_DNA_seed | PIK3CA | 0.990872 | LGG | KNS-42 | AZD8835 |
| top60_altered_DNA_seed | EGFR | 0.998428 | LGG | no-11 | Afatinib |

| | | | | | |
|---|---|---|---|---|---|
| top60_altered_DNA_seed | EGFR | 1.003117 | LGG | KNS-42 | Afatinib |
| top60_altered_DNA_seed | HSF1 | 1.016141 | LGG | KNS-42 | CCT245467 |
| top60_altered_DNA_seed | HSF1 | 1.016795 | LGG | no-10 | CCT245467 |
| top60_altered_DNA_seed | EGFR | 1.020504 | LGG | KINGS-1 | Afatinib |
| top60_altered_DNA_seed | EGFR | 1.026118 | LGG | KINGS-1 | CI-1033 |
| top60_altered_DNA_seed | PIK3CA | 1.0522 | LGG | no-10 | AZD8835 |
| top60_altered_DNA_seed | EGFR | 1.061562 | LGG | KALS-1 | Afatinib |
| top60_altered_DNA_seed | EGFR | 1.061812 | LGG | Hs-683 | CUDC-101 |
| top60_altered_DNA_seed | PIK3CA | 1.106367 | LGG | no-11 | AZD8835 |
| top60_altered_DNA_seed | EGFR | 1.117272 | LGG | H4 | CUDC-101 |
| top60_altered_DNA_seed | EGFR | 1.126234 | LGG | no-10 | Gefitinib |
| top60_altered_DNA_seed | EGFR | 1.128556 | LGG | KINGS-1 | Pelitinib |
| top60_altered_DNA_seed | HSF1 | 1.143047 | LGG | D-336MG | CCT245467 |
| top60_altered_DNA_seed | EGFR | 1.157726 | LGG | KNS-42 | PF-00299804 |
| top60_altered_DNA_seed | HSF1 | 1.181149 | LGG | MOG-G-UVW | CCT245467 |
| top60_altered_DNA_seed | EGFR | 1.190454 | LGG | KINGS-1 | AST-1306 |
| top60_altered_DNA_seed | EGFR | 1.208594 | LGG | KALS-1 | Cetuximab |
| top60_altered_DNA_seed | EGFR | 1.214041 | LGG | no-11 | AZD8931 |
| top60_altered_DNA_seed | EGFR | 1.230171 | LGG | KNS-81-FD | Pelitinib |
| top60_altered_DNA_seed | PIK3CA | 1.259339 | LGG | KNS-81-FD | AZD8835 |
| top60_altered_DNA_seed | EGFR | 1.262328 | LGG | KALS-1 | AST-1306 |
| top60_altered_DNA_seed | EGFR | 1.282193 | LGG | KNS-81-FD | PF-00299804 |
| top60_altered_DNA_seed | HSF1 | 1.287093 | LGG | H4 | CCT245467 |
| top60_altered_DNA_seed | EGFR | 1.289915 | LGG | no-11 | Lapatinib |
| top60_altered_DNA_seed | EGFR | 1.291035 | LGG | KNS-81-FD | Afatinib |
| top60_altered_DNA_seed | EGFR | 1.295543 | LGG | KINGS-1 | Gefitinib |
| top60_altered_DNA_seed | EGFR | 1.336917 | LGG | D-502MG | Gefitinib |
| top60_altered_DNA_seed | HSF1 | 1.353296 | LGG | KNS-81-FD | CCT245467 |
| top60_altered_DNA_seed | PIK3CA | 1.372272 | LGG | Hs-683 | PI3Ka_4409 |
| top60_altered_DNA_seed | EGFR | 1.379924 | LGG | KINGS-1 | Lapatinib |
| top60_altered_DNA_seed | EGFR | 1.431837 | LGG | KNS-42 | Pelitinib |
| top60_altered_DNA_seed | EGFR | 1.52666 | LGG | no-11 | AST-1306 |
| top60_altered_DNA_seed | EGFR | 1.686139 | LGG | KALS-1 | CI-1033 |
| top60_altered_DNA_seed | PIK3CA | 1.693859 | LGG | KNS-81-FD | PI3Ka_4409 |
| top60_altered_DNA_seed | EGFR | 1.742203 | LGG | KNS-81-FD | Cetuximab |
| top60_altered_DNA_seed | HSF1 | 1.999872 | LGG | no-11 | CCT245467 |
| top60_altered_DNA_seed | EGFR | 2.352454 | LGG | KNS-42 | AST-1306 |
| top60_altered_DNA_seed | EGFR | 2.464475 | LGG | KNS-81-FD | CI-1033 |
| top60_altered_DNA_seed | EGFR | 2.524426 | LGG | KNS-81-FD | AZD8931 |
| top60_altered_DNA_seed | EGFR | 2.592692 | LGG | KNS-42 | CI-1033 |
| top60_altered_DNA_seed | PIK3CA | -1.687761 | LGG | D-336MG | AZD8186 |

| | | | | | |
|---|---|---|---|---|---|
| top60_altered_DNA_seed | PIK3CA | -1.512682 | LGG | MOG-G-CCM | AZD8186 |
| top60_altered_DNA_seed | PIK3CA | -1.232974 | LGG | SW1783 | GNE-317 |
| top60_altered_DNA_seed | EGFR | -1.141818 | LGG | Hs-683 | Gefitinib |
| top60_altered_DNA_seed | PIK3CA | -1.121296 | LGG | SW1783 | AZD8186 |
| top60_altered_DNA_seed | EGFR | -1.089063 | LGG | GI-1 | Gefitinib |
| top60_altered_DNA_seed | PIK3CA | -1.060958 | LGG | MOG-G-UVW | GNE-317 |
| top60_altered_DNA_seed | PIK3CA | -1.020332 | LGG | H4 | AZD8186 |
| top60_altered_DNA_seed | PIK3CA | -0.981862 | LGG | GAMG | AZD8186 |
| top60_altered_DNA_seed | EGFR | -0.979745 | LGG | Hs-683 | AZD3759 |
| top60_altered_DNA_seed | PIK3CA | -0.967746 | LGG | D-336MG | Alpelisib |
| top60_altered_DNA_seed | TP53 | -0.959389 | LGG | MOG-G-UVW | MIRA-1 |
| top60_altered_DNA_seed | PIK3CA | -0.940227 | LGG | Hs-683 | AZD8186 |
| top60_altered_DNA_seed | TP53 | -0.810133 | LGG | SW1783 | MIRA-1 |
| top60_altered_DNA_seed | PIK3CA | -0.773063 | LGG | D-336MG | Buparlisib |
| top60_altered_DNA_seed | EGFR | -0.768215 | LGG | D-336MG | AZD3759 |
| top60_altered_DNA_seed | PIK3CA | -0.688327 | LGG | MOG-G-UVW | Buparlisib |
| top60_altered_DNA_seed | PIK3CA | -0.676352 | LGG | GI-1 | Buparlisib |
| top60_altered_DNA_seed | PIK3CA | -0.662721 | LGG | MOG-G-UVW | AZD8186 |
| top60_altered_DNA_seed | PIK3CA | -0.648271 | LGG | H4 | Buparlisib |
| top60_altered_DNA_seed | PIK3CA | -0.646293 | LGG | MOG-G-CCM | GNE-317 |
| top60_altered_DNA_seed | EGFR | -0.613204 | LGG | Hs-683 | Lapatinib |
| top60_altered_DNA_seed | PIK3CA | -0.594503 | LGG | MOG-G-CCM | Buparlisib |
| top60_altered_DNA_seed | PIK3CA | -0.577715 | LGG | SF539 | Buparlisib |
| top60_altered_DNA_seed | EGFR | -0.56726 | LGG | SF539 | Erlotinib |
| top60_altered_DNA_seed | EGFR | -0.56455 | LGG | D-502MG | AZD3759 |
| top60_altered_DNA_seed | PIK3CA | -0.563741 | LGG | Hs-683 | Buparlisib |
| top60_altered_DNA_seed | EGFR | -0.556105 | LGG | D-336MG | Lapatinib |
| top60_altered_DNA_seed | EGFR | -0.540131 | LGG | D-336MG | Osimertinib |
| top60_altered_DNA_seed | EGFR | -0.517011 | LGG | H4 | Erlotinib |
| top60_altered_DNA_seed | EGFR | -0.479491 | LGG | SF539 | Lapatinib |
| top60_altered_DNA_seed | PIK3CA | -0.477739 | LGG | Hs-683 | Alpelisib |
| top60_altered_DNA_seed | TP53 | -0.460786 | LGG | no-11 | MIRA-1 |
| top60_altered_DNA_seed | TP53 | -0.442532 | LGG | D-502MG | MIRA-1 |
| top60_altered_DNA_seed | PIK3CA | -0.417037 | LGG | SW1783 | Buparlisib |
| top60_altered_DNA_seed | EGFR | -0.410901 | LGG | MOG-G-UVW | AZD3759 |
| top60_altered_DNA_seed | IDH1 | -0.356654 | LGG | Hs-683 | AGI-5198 |
| top60_altered_DNA_seed | PIK3CA | -0.352235 | LGG | D-502MG | Alpelisib |
| top60_altered_DNA_seed | PIK3CA | -0.333807 | LGG | KALS-1 | AZD8186 |
| top60_altered_DNA_seed | EGFR | -0.311257 | LGG | D-502MG | Lapatinib |
| top60_altered_DNA_seed | EGFR | -0.307399 | LGG | no-10 | AZD3759 |
| top60_altered_DNA_seed | PIK3CA | -0.285899 | LGG | SF539 | AZD8186 |

| | | | | | |
|---|---|---|---|---|---|
| top60_altered_DNA_seed | PIK3CA | -0.209957 | LGG | H4 | Alpelisib |
| top60_altered_DNA_seed | PIK3CA | -0.196285 | LGG | GI-1 | AZD8186 |
| top60_altered_DNA_seed | PIK3CA | -0.186453 | LGG | GAMG | GNE-317 |
| top60_altered_DNA_seed | EGFR | -0.179159 | LGG | H4 | Afatinib |
| top60_altered_DNA_seed | TP53 | -0.172212 | LGG | GI-1 | MIRA-1 |
| top60_altered_DNA_seed | IDH1 | -0.15915 | LGG | GI-1 | AGI-5198 |
| top60_altered_DNA_seed | PIK3CA | -0.155749 | LGG | KINGS-1 | AZD8186 |
| top60_altered_DNA_seed | IDH1 | -0.155203 | LGG | SF539 | AGI-5198 |
| top60_altered_DNA_seed | EGFR | -0.149404 | LGG | no-10 | Lapatinib |
| top60_altered_DNA_seed | EGFR | -0.1312 | LGG | D-502MG | Sapitinib |
| top60_altered_DNA_seed | PIK3CA | -0.130225 | LGG | KINGS-1 | Alpelisib |
| top60_altered_DNA_seed | EGFR | -0.121743 | LGG | H4 | Sapitinib |
| top60_altered_DNA_seed | PIK3CA | -0.112874 | LGG | KNS-81-FD | Alpelisib |
| top60_altered_DNA_seed | PIK3CA | -0.112242 | LGG | GAMG | Buparlisib |
| top60_altered_DNA_seed | IDH1 | -0.104133 | LGG | KNS-81-FD | AGI-5198 |
| top60_altered_DNA_seed | EGFR | -0.093755 | LGG | SF539 | Osimertinib |
| top60_altered_DNA_seed | EGFR | -0.086449 | LGG | GAMG | Afatinib |
| top60_altered_DNA_seed | PIK3CA | -0.066356 | LGG | D-502MG | AZD8186 |
| top60_altered_DNA_seed | EGFR | -0.048049 | LGG | no-10 | Sapitinib |
| top60_altered_DNA_seed | EGFR | -0.045606 | LGG | GI-1 | Erlotinib |
| top60_altered_DNA_seed | EGFR | -0.03322 | LGG | SF539 | Afatinib |
| top60_altered_DNA_seed | TP53 | -0.030854 | LGG | D-336MG | MIRA-1 |
| top60_altered_DNA_seed | PIK3CA | -0.017429 | LGG | SF539 | GNE-317 |
| top60_altered_DNA_seed | EGFR | -0.006992 | LGG | KNS-81-FD | Gefitinib |
| top60_altered_DNA_seed | EGFR | 0.008738 | LGG | GI-1 | Afatinib |
| top60_altered_DNA_seed | EGFR | 0.019251 | LGG | SF539 | Gefitinib |
| top60_altered_DNA_seed | PIK3CA | 0.031366 | LGG | GI-1 | Alpelisib |
| top60_altered_DNA_seed | TP53 | 0.040713 | LGG | KNS-42 | MIRA-1 |
| top60_altered_DNA_seed | PIK3CA | 0.042034 | LGG | KNS-81-FD | GNE-317 |
| top60_altered_DNA_seed | PIK3CA | 0.043355 | LGG | SF539 | Alpelisib |
| top60_altered_DNA_seed | TP53 | 0.070502 | LGG | SF539 | MIRA-1 |
| top60_altered_DNA_seed | IDH1 | 0.078202 | LGG | H4 | AGI-5198 |
| top60_altered_DNA_seed | EGFR | 0.080258 | LGG | SW1783 | Osimertinib |
| top60_altered_DNA_seed | TP53 | 0.100083 | LGG | Hs-683 | MIRA-1 |
| top60_altered_DNA_seed | TP53 | 0.117569 | LGG | no-10 | MIRA-1 |
| top60_altered_DNA_seed | IDH1 | 0.139609 | LGG | MOG-G-UVW | AGI-5198 |
| top60_altered_DNA_seed | EGFR | 0.140876 | LGG | SW1783 | Afatinib |
| top60_altered_DNA_seed | PIK3CA | 0.142299 | LGG | MOG-G-CCM | Alpelisib |
| top60_altered_DNA_seed | TP53 | 0.147656 | LGG | H4 | MIRA-1 |
| top60_altered_DNA_seed | EGFR | 0.151751 | LGG | MOG-G-UVW | Gefitinib |
| top60_altered_DNA_seed | EGFR | 0.153475 | LGG | MOG-G-UVW | Lapatinib |

| | | | | | |
|---|---|---|---|---|---|
| top60_altered_DNA_seed | EGFR | 0.160787 | LGG | GAMG | AZD3759 |
| top60_altered_DNA_seed | EGFR | 0.161313 | LGG | MOG-G-UVW | Afatinib |
| top60_altered_DNA_seed | EGFR | 0.164179 | LGG | H4 | Gefitinib |
| top60_altered_DNA_seed | EGFR | 0.165006 | LGG | SF539 | Sapitinib |
| top60_altered_DNA_seed | EGFR | 0.175121 | LGG | MOG-G-CCM | Lapatinib |
| top60_altered_DNA_seed | TP53 | 0.179887 | LGG | GAMG | MIRA-1 |
| top60_altered_DNA_seed | PIK3CA | 0.183884 | LGG | KINGS-1 | GNE-317 |
| top60_altered_DNA_seed | EGFR | 0.186305 | LGG | D-336MG | Sapitinib |
| top60_altered_DNA_seed | PIK3CA | 0.191285 | LGG | no-10 | AZD8186 |
| top60_altered_DNA_seed | PIK3CA | 0.192235 | LGG | Hs-683 | GNE-317 |
| top60_altered_DNA_seed | EGFR | 0.216492 | LGG | SW1783 | Gefitinib |
| top60_altered_DNA_seed | EGFR | 0.246484 | LGG | H4 | AZD3759 |
| top60_altered_DNA_seed | PIK3CA | 0.267498 | LGG | no-11 | AZD8186 |
| top60_altered_DNA_seed | EGFR | 0.270554 | LGG | SF539 | AZD3759 |
| top60_altered_DNA_seed | PIK3CA | 0.274647 | LGG | GAMG | Alpelisib |
| top60_altered_DNA_seed | EGFR | 0.279487 | LGG | H4 | Lapatinib |
| top60_altered_DNA_seed | PIK3CA | 0.301169 | LGG | KALS-1 | GNE-317 |
| top60_altered_DNA_seed | EGFR | 0.309872 | LGG | KNS-42 | AZD3759 |
| top60_altered_DNA_seed | EGFR | 0.334534 | LGG | D-336MG | Gefitinib |
| top60_altered_DNA_seed | EGFR | 0.345412 | LGG | no-10 | Afatinib |
| top60_altered_DNA_seed | EGFR | 0.34689 | LGG | GI-1 | Sapitinib |
| top60_altered_DNA_seed | EGFR | 0.354282 | LGG | D-502MG | Erlotinib |
| top60_altered_DNA_seed | EGFR | 0.360726 | LGG | GAMG | Gefitinib |
| top60_altered_DNA_seed | EGFR | 0.372986 | LGG | KNS-81-FD | AZD3759 |
| top60_altered_DNA_seed | EGFR | 0.386058 | LGG | Hs-683 | Sapitinib |
| top60_altered_DNA_seed | EGFR | 0.397765 | LGG | D-502MG | Osimertinib |
| top60_altered_DNA_seed | EGFR | 0.40411 | LGG | SW1783 | Sapitinib |
| top60_altered_DNA_seed | EGFR | 0.411407 | LGG | GAMG | Lapatinib |
| top60_altered_DNA_seed | EGFR | 0.419301 | LGG | SW1783 | AZD3759 |
| top60_altered_DNA_seed | IDH1 | 0.419509 | LGG | no-10 | AGI-5198 |
| top60_altered_DNA_seed | PIK3CA | 0.432726 | LGG | MOG-G-UVW | Alpelisib |
| top60_altered_DNA_seed | EGFR | 0.440839 | LGG | H4 | Osimertinib |
| top60_altered_DNA_seed | EGFR | 0.44375 | LGG | GI-1 | Osimertinib |
| top60_altered_DNA_seed | EGFR | 0.465577 | LGG | KALS-1 | Afatinib |
| top60_altered_DNA_seed | PIK3CA | 0.477954 | LGG | SW1783 | Alpelisib |
| top60_altered_DNA_seed | EGFR | 0.480667 | LGG | Hs-683 | Erlotinib |
| top60_altered_DNA_seed | EGFR | 0.490183 | LGG | D-502MG | Afatinib |
| top60_altered_DNA_seed | PIK3CA | 0.490219 | LGG | KINGS-1 | Buparlisib |
| top60_altered_DNA_seed | EGFR | 0.498904 | LGG | Hs-683 | Osimertinib |
| top60_altered_DNA_seed | EGFR | 0.505316 | LGG | no-11 | AZD3759 |
| top60_altered_DNA_seed | IDH1 | 0.520709 | LGG | MOG-G-CCM | AGI-5198 |

| | | | | | |
|---|---|---|---|---|---|
| top60_altered_DNA_seed | EGFR | 0.531287 | LGG | MOG-G-CCM | Afatinib |
| top60_altered_DNA_seed | PIK3CA | 0.534214 | LGG | KNS-81-FD | AZD8186 |
| top60_altered_DNA_seed | PIK3CA | 0.535369 | LGG | D-336MG | GNE-317 |
| top60_altered_DNA_seed | EGFR | 0.53921 | LGG | KALS-1 | Lapatinib |
| top60_altered_DNA_seed | EGFR | 0.539901 | LGG | D-336MG | Afatinib |
| top60_altered_DNA_seed | EGFR | 0.561184 | LGG | GAMG | Sapitinib |
| top60_altered_DNA_seed | EGFR | 0.571943 | LGG | KNS-42 | Gefitinib |
| top60_altered_DNA_seed | EGFR | 0.576865 | LGG | GI-1 | AZD3759 |
| top60_altered_DNA_seed | PIK3CA | 0.586555 | LGG | KALS-1 | Buparlisib |
| top60_altered_DNA_seed | EGFR | 0.586737 | LGG | D-336MG | Erlotinib |
| top60_altered_DNA_seed | PIK3CA | 0.590988 | LGG | no-11 | Alpelisib |
| top60_altered_DNA_seed | PIK3CA | 0.611713 | LGG | no-10 | Alpelisib |
| top60_altered_DNA_seed | EGFR | 0.614726 | LGG | SW1783 | Lapatinib |
| top60_altered_DNA_seed | IDH1 | 0.617818 | LGG | KNS-42 | AGI-5198 |
| top60_altered_DNA_seed | EGFR | 0.643642 | LGG | GI-1 | Lapatinib |
| top60_altered_DNA_seed | EGFR | 0.680258 | LGG | KNS-81-FD | Lapatinib |
| top60_altered_DNA_seed | EGFR | 0.692155 | LGG | no-11 | Erlotinib |
| top60_altered_DNA_seed | EGFR | 0.722897 | LGG | MOG-G-UVW | Erlotinib |
| top60_altered_DNA_seed | TP53 | 0.723695 | LGG | KALS-1 | MIRA-1 |
| top60_altered_DNA_seed | PIK3CA | 0.727532 | LGG | KALS-1 | Alpelisib |
| top60_altered_DNA_seed | EGFR | 0.742559 | LGG | MOG-G-UVW | Osimertinib |
| top60_altered_DNA_seed | EGFR | 0.771658 | LGG | D-502MG | Gefitinib |
| top60_altered_DNA_seed | TP53 | 0.786286 | LGG | KNS-81-FD | MIRA-1 |
| top60_altered_DNA_seed | EGFR | 0.797162 | LGG | no-10 | Gefitinib |
| top60_altered_DNA_seed | PIK3CA | 0.802433 | LGG | D-502MG | GNE-317 |
| top60_altered_DNA_seed | EGFR | 0.803018 | LGG | no-11 | Osimertinib |
| top60_altered_DNA_seed | IDH1 | 0.807931 | LGG | D-502MG | AGI-5198 |
| top60_altered_DNA_seed | IDH1 | 0.814015 | LGG | D-336MG | AGI-5198 |
| top60_altered_DNA_seed | EGFR | 0.814218 | LGG | KALS-1 | Sapitinib |
| top60_altered_DNA_seed | EGFR | 0.875275 | LGG | MOG-G-UVW | Sapitinib |
| top60_altered_DNA_seed | EGFR | 0.883305 | LGG | SW1783 | Erlotinib |
| top60_altered_DNA_seed | PIK3CA | 0.898276 | LGG | D-502MG | Buparlisib |
| top60_altered_DNA_seed | EGFR | 0.901063 | LGG | KNS-81-FD | Afatinib |
| top60_altered_DNA_seed | EGFR | 0.908559 | LGG | KNS-81-FD | Sapitinib |
| top60_altered_DNA_seed | EGFR | 0.912961 | LGG | KNS-42 | Erlotinib |
| top60_altered_DNA_seed | EGFR | 0.925221 | LGG | MOG-G-CCM | Sapitinib |
| top60_altered_DNA_seed | PIK3CA | 0.93336 | LGG | KNS-42 | Alpelisib |
| top60_altered_DNA_seed | EGFR | 0.939677 | LGG | KINGS-1 | Afatinib |
| top60_altered_DNA_seed | EGFR | 0.951583 | LGG | KNS-81-FD | Osimertinib |
| top60_altered_DNA_seed | IDH1 | 0.955434 | LGG | SW1783 | AGI-5198 |
| top60_altered_DNA_seed | EGFR | 0.959616 | LGG | KALS-1 | Gefitinib |

| | | | | | |
|---|---|---|---|---|---|
| top60_altered_DNA_seed | EGFR | 0.980044 | LGG | KNS-42 | Lapatinib |
| top60_altered_DNA_seed | EGFR | 0.999901 | LGG | KALS-1 | Osimertinib |
| top60_altered_DNA_seed | EGFR | 1.019898 | LGG | no-10 | Erlotinib |
| top60_altered_DNA_seed | EGFR | 1.02098 | LGG | KALS-1 | AZD3759 |
| top60_altered_DNA_seed | PIK3CA | 1.072966 | LGG | NMC-G1 | AZD8186 |
| top60_altered_DNA_seed | EGFR | 1.076377 | LGG | MOG-G-CCM | Osimertinib |
| top60_altered_DNA_seed | EGFR | 1.115263 | LGG | KALS-1 | Erlotinib |
| top60_altered_DNA_seed | EGFR | 1.121947 | LGG | no-10 | Osimertinib |
| top60_altered_DNA_seed | IDH1 | 1.137279 | LGG | GAMG | AGI-5198 |
| top60_altered_DNA_seed | EGFR | 1.148711 | LGG | KNS-42 | Osimertinib |
| top60_altered_DNA_seed | EGFR | 1.151089 | LGG | KNS-42 | Afatinib |
| top60_altered_DNA_seed | EGFR | 1.154416 | LGG | Hs-683 | Afatinib |
| top60_altered_DNA_seed | EGFR | 1.163253 | LGG | MOG-G-CCM | Erlotinib |
| top60_altered_DNA_seed | PIK3CA | 1.184239 | LGG | no-11 | GNE-317 |
| top60_altered_DNA_seed | EGFR | 1.209865 | LGG | KNS-42 | Sapitinib |
| top60_altered_DNA_seed | IDH1 | 1.226974 | LGG | KINGS-1 | AGI-5198 |
| top60_altered_DNA_seed | EGFR | 1.232984 | LGG | GAMG | Erlotinib |
| top60_altered_DNA_seed | PIK3CA | 1.28262 | LGG | KNS-42 | AZD8186 |
| top60_altered_DNA_seed | EGFR | 1.300919 | LGG | KINGS-1 | Sapitinib |
| top60_altered_DNA_seed | EGFR | 1.307215 | LGG | no-11 | Gefitinib |
| top60_altered_DNA_seed | TP53 | 1.333023 | LGG | MOG-G-CCM | MIRA-1 |
| top60_altered_DNA_seed | TP53 | 1.334479 | LGG | KINGS-1 | MIRA-1 |
| top60_altered_DNA_seed | EGFR | 1.335166 | LGG | KINGS-1 | AZD3759 |
| top60_altered_DNA_seed | PIK3CA | 1.365099 | LGG | no-10 | Buparlisib |
| top60_altered_DNA_seed | EGFR | 1.386507 | LGG | GAMG | Osimertinib |
| top60_altered_DNA_seed | EGFR | 1.414827 | LGG | MOG-G-CCM | Gefitinib |
| top60_altered_DNA_seed | EGFR | 1.443705 | LGG | MOG-G-CCM | AZD3759 |
| top60_altered_DNA_seed | PIK3CA | 1.445666 | LGG | no-10 | GNE-317 |
| top60_altered_DNA_seed | EGFR | 1.51225 | LGG | no-11 | Sapitinib |
| top60_altered_DNA_seed | EGFR | 1.539234 | LGG | NMC-G1 | Erlotinib |
| top60_altered_DNA_seed | PIK3CA | 1.565584 | LGG | NMC-G1 | GNE-317 |
| top60_altered_DNA_seed | EGFR | 1.566406 | LGG | KINGS-1 | Osimertinib |
| top60_altered_DNA_seed | IDH1 | 1.678283 | LGG | no-11 | AGI-5198 |
| top60_altered_DNA_seed | EGFR | 1.69048 | LGG | no-11 | Lapatinib |
| top60_altered_DNA_seed | IDH1 | 1.695498 | LGG | KALS-1 | AGI-5198 |
| top60_altered_DNA_seed | PIK3CA | 1.705865 | LGG | KNS-42 | GNE-317 |
| top60_altered_DNA_seed | PIK3CA | 1.7939 | LGG | KNS-81-FD | Buparlisib |
| top60_altered_DNA_seed | EGFR | 1.79458 | LGG | KINGS-1 | Lapatinib |
| top60_altered_DNA_seed | EGFR | 1.857789 | LGG | KINGS-1 | Gefitinib |
| top60_altered_DNA_seed | PIK3CA | 1.861603 | LGG | KNS-42 | Buparlisib |
| top60_altered_DNA_seed | EGFR | 1.87069 | LGG | NMC-G1 | Osimertinib |

| | | | | | |
|---|---|---|---|---|---|
| top60_altered_DNA_seed | EGFR | 1.941697 | LGG | KINGS-1 | Erlotinib |
| top60_altered_DNA_seed | IDH1 | 2.050594 | LGG | NMC-G1 | AGI-5198 |
| top60_altered_DNA_seed | EGFR | 2.072975 | LGG | NMC-G1 | Sapitinib |
| top60_altered_DNA_seed | EGFR | 2.10689 | LGG | KNS-81-FD | Erlotinib |
| top60_altered_DNA_seed | PIK3CA | 2.257283 | LGG | no-11 | Buparlisib |
| top60_altered_DNA_seed | EGFR | 2.263084 | LGG | NMC-G1 | Gefitinib |
| top60_altered_DNA_seed | TP53 | 2.315779 | LGG | NMC-G1 | MIRA-1 |
| top60_altered_DNA_seed | EGFR | 2.325145 | LGG | NMC-G1 | AZD3759 |
| top60_altered_DNA_seed | PIK3CA | 2.538988 | LGG | NMC-G1 | Alpelisib |
| top60_altered_DNA_seed | EGFR | 2.718584 | LGG | NMC-G1 | Lapatinib |
| top60_altered_DNA_seed | EGFR | 2.856642 | LGG | no-11 | Afatinib |
| top60_altered_DNA_seed | EGFR | 2.954802 | LGG | NMC-G1 | Afatinib |
| top60_altered_DNA_seed | PIK3CA | 5.209719 | LGG | NMC-G1 | Buparlisib |
| top60_altered_DNA_seed | KRAS | -1.609231 | LUAD | EMC-BAC-1 | KRAS (G12C) Inhibitor-12 |
| top60_altered_DNA_seed | KRAS | -1.223262 | LUAD | NCI-H2122 | KRAS (G12C) Inhibitor-12 |
| top60_altered_DNA_seed | KRAS | -1.128535 | LUAD | NCI-H1975 | KRAS (G12C) Inhibitor-12 |
| top60_altered_DNA_seed | TP53 | -1.049009 | LUAD | NCI-H1734 | MIRA-1 |
| top60_altered_DNA_seed | KRAS | -0.907714 | LUAD | HCC-78 | KRAS (G12C) Inhibitor-12 |
| top60_altered_DNA_seed | KRAS | -0.868207 | LUAD | LC-2-ad | KRAS (G12C) Inhibitor-12 |
| top60_altered_DNA_seed | KRAS | -0.854987 | LUAD | NCI-H1792 | KRAS (G12C) Inhibitor-12 |
| top60_altered_DNA_seed | KRAS | -0.825506 | LUAD | NCI-H1437 | KRAS (G12C) Inhibitor-12 |
| top60_altered_DNA_seed | KRAS | -0.781584 | LUAD | NCI-H1993 | KRAS (G12C) Inhibitor-12 |
| top60_altered_DNA_seed | KRAS | -0.75525 | LUAD | NCI-H23 | KRAS (G12C) Inhibitor-12 |
| top60_altered_DNA_seed | TP53 | -0.730298 | LUAD | NCI-H1435 | MIRA-1 |
| top60_altered_DNA_seed | TP53 | -0.664239 | LUAD | NCI-H1792 | MIRA-1 |
| top60_altered_DNA_seed | KRAS | -0.608473 | LUAD | PC-14 | KRAS (G12C) Inhibitor-12 |
| top60_altered_DNA_seed | KRAS | -0.576165 | LUAD | A549 | KRAS (G12C) Inhibitor-12 |
| top60_altered_DNA_seed | KRAS | -0.523116 | LUAD | NCI-H650 | KRAS (G12C) Inhibitor-12 |
| top60_altered_DNA_seed | KRAS | -0.455088 | LUAD | EMC-BAC-2 | KRAS (G12C) Inhibitor-12 |
| top60_altered_DNA_seed | KRAS | -0.414707 | LUAD | NCI-H1734 | KRAS (G12C) Inhibitor-12 |
| top60_altered_DNA_seed | KRAS | -0.360798 | LUAD | NCI-H1781 | KRAS (G12C) Inhibitor-12 |
| top60_altered_DNA_seed | KRAS | -0.347633 | LUAD | RERF-LC-MS | KRAS (G12C) Inhibitor-12 |
| top60_altered_DNA_seed | KRAS | -0.29333 | LUAD | HCC-44 | KRAS (G12C) Inhibitor-12 |
| top60_altered_DNA_seed | TP53 | -0.266147 | LUAD | NCI-H2030 | MIRA-1 |
| top60_altered_DNA_seed | KRAS | -0.263595 | LUAD | LXF-289 | KRAS (G12C) Inhibitor-12 |
| top60_altered_DNA_seed | TP53 | -0.25003 | LUAD | NCI-H2122 | MIRA-1 |
| top60_altered_DNA_seed | KRAS | -0.239604 | LUAD | NCI-H322M | KRAS (G12C) Inhibitor-12 |
| top60_altered_DNA_seed | TP53 | -0.228242 | LUAD | LC-2-ad | MIRA-1 |
| top60_altered_DNA_seed | TP53 | -0.13766 | LUAD | RERF-LC-MS | MIRA-1 |
| top60_altered_DNA_seed | KRAS | -0.117287 | LUAD | NCI-H838 | KRAS (G12C) Inhibitor-12 |
| top60_altered_DNA_seed | TP53 | -0.070238 | LUAD | LXF-289 | MIRA-1 |

| | | | | | |
|---|---|---|---|---|---|
| top60_altered_DNA_seed | TP53 | -0.063293 | LUAD | NCI-H1437 | MIRA-1 |
| top60_altered_DNA_seed | KRAS | -0.036458 | LUAD | NCI-H2023 | KRAS (G12C) Inhibitor-12 |
| top60_altered_DNA_seed | TP53 | -0.021429 | LUAD | NCI-H522 | MIRA-1 |
| top60_altered_DNA_seed | TP53 | -0.021345 | LUAD | RERF-LC-KJ | MIRA-1 |
| top60_altered_DNA_seed | TP53 | -0.020545 | LUAD | NCI-H1651 | MIRA-1 |
| top60_altered_DNA_seed | KRAS | 0.013546 | LUAD | NCI-H3122 | KRAS (G12C) Inhibitor-12 |
| top60_altered_DNA_seed | KRAS | 0.100027 | LUAD | NCI-H1651 | KRAS (G12C) Inhibitor-12 |
| top60_altered_DNA_seed | KRAS | 0.105467 | LUAD | H3255 | KRAS (G12C) Inhibitor-12 |
| top60_altered_DNA_seed | TP53 | 0.109622 | LUAD | NCI-H2009 | MIRA-1 |
| top60_altered_DNA_seed | TP53 | 0.122093 | LUAD | VMRC-LCD | MIRA-1 |
| top60_altered_DNA_seed | KRAS | 0.179106 | LUAD | HOP-62 | KRAS (G12C) Inhibitor-12 |
| top60_altered_DNA_seed | TP53 | 0.184186 | LUAD | NCI-H838 | MIRA-1 |
| top60_altered_DNA_seed | TP53 | 0.194821 | LUAD | NCI-H1648 | MIRA-1 |
| top60_altered_DNA_seed | KRAS | 0.20407 | LUAD | ABC-1 | KRAS (G12C) Inhibitor-12 |
| top60_altered_DNA_seed | KRAS | 0.257163 | LUAD | RERF-LC-KJ | KRAS (G12C) Inhibitor-12 |
| top60_altered_DNA_seed | KRAS | 0.265622 | LUAD | NCI-H1666 | KRAS (G12C) Inhibitor-12 |
| top60_altered_DNA_seed | TP53 | 0.271202 | LUAD | NCI-H2085 | MIRA-1 |
| top60_altered_DNA_seed | TP53 | 0.273078 | LUAD | NCI-H23 | MIRA-1 |
| top60_altered_DNA_seed | TP53 | 0.366569 | LUAD | EMC-BAC-1 | MIRA-1 |
| top60_altered_DNA_seed | TP53 | 0.439079 | LUAD | NCI-H650 | MIRA-1 |
| top60_altered_DNA_seed | TP53 | 0.444214 | LUAD | NCI-H3122 | MIRA-1 |
| top60_altered_DNA_seed | KRAS | 0.480611 | LUAD | VMRC-LCD | KRAS (G12C) Inhibitor-12 |
| top60_altered_DNA_seed | KRAS | 0.497351 | LUAD | Calu-6 | KRAS (G12C) Inhibitor-12 |
| top60_altered_DNA_seed | TP53 | 0.50233 | LUAD | HCC-44 | MIRA-1 |
| top60_altered_DNA_seed | TP53 | 0.510946 | LUAD | NCI-H1993 | MIRA-1 |
| top60_altered_DNA_seed | TP53 | 0.529901 | LUAD | NCI-H1573 | MIRA-1 |
| top60_altered_DNA_seed | TP53 | 0.546422 | LUAD | NCI-H1781 | MIRA-1 |
| top60_altered_DNA_seed | TP53 | 0.556919 | LUAD | ABC-1 | MIRA-1 |
| top60_altered_DNA_seed | KRAS | 0.557187 | LUAD | NCI-H1563 | KRAS (G12C) Inhibitor-12 |
| top60_altered_DNA_seed | TP53 | 0.60034 | LUAD | NCI-H1755 | MIRA-1 |
| top60_altered_DNA_seed | KRAS | 0.638042 | LUAD | NCI-H522 | KRAS (G12C) Inhibitor-12 |
| top60_altered_DNA_seed | KRAS | 0.63857 | LUAD | NCI-H2087 | KRAS (G12C) Inhibitor-12 |
| top60_altered_DNA_seed | KRAS | 0.664545 | LUAD | NCI-H1944 | KRAS (G12C) Inhibitor-12 |
| top60_altered_DNA_seed | TP53 | 0.690013 | LUAD | NCI-H1944 | MIRA-1 |
| top60_altered_DNA_seed | TP53 | 0.718681 | LUAD | EMC-BAC-2 | MIRA-1 |
| top60_altered_DNA_seed | TP53 | 0.734693 | LUAD | NCI-H1693 | MIRA-1 |
| top60_altered_DNA_seed | TP53 | 0.738409 | LUAD | NCI-H2023 | MIRA-1 |
| top60_altered_DNA_seed | TP53 | 0.758629 | LUAD | NCI-H1563 | MIRA-1 |
| top60_altered_DNA_seed | TP53 | 0.774438 | LUAD | HCC-78 | MIRA-1 |
| top60_altered_DNA_seed | TP53 | 0.813338 | LUAD | PC-3_[JPC-3] | MIRA-1 |
| top60_altered_DNA_seed | TP53 | 0.848045 | LUAD | NCI-H1623 | MIRA-1 |

| | | | | | |
|---|---|---|---|---|---|
| top60_altered_DNA_seed | KRAS | 0.849262 | LUAD | NCI-H1435 | KRAS (G12C) Inhibitor-12 |
| top60_altered_DNA_seed | KRAS | 0.86381 | LUAD | NCI-H358 | KRAS (G12C) Inhibitor-12 |
| top60_altered_DNA_seed | TP53 | 0.877456 | LUAD | NCI-H2342 | MIRA-1 |
| top60_altered_DNA_seed | KRAS | 0.886349 | LUAD | NCI-H1623 | KRAS (G12C) Inhibitor-12 |
| top60_altered_DNA_seed | TP53 | 0.914092 | LUAD | Calu-6 | MIRA-1 |
| top60_altered_DNA_seed | TP53 | 0.92166 | LUAD | NCI-H1975 | MIRA-1 |
| top60_altered_DNA_seed | KRAS | 0.936083 | LUAD | NCI-H2030 | KRAS (G12C) Inhibitor-12 |
| top60_altered_DNA_seed | KRAS | 0.946933 | LUAD | NCI-H2085 | KRAS (G12C) Inhibitor-12 |
| top60_altered_DNA_seed | KRAS | 0.969335 | LUAD | HCC-827 | KRAS (G12C) Inhibitor-12 |
| top60_altered_DNA_seed | TP53 | 0.977542 | LUAD | NCI-H322M | MIRA-1 |
| top60_altered_DNA_seed | TP53 | 0.986453 | LUAD | PC-14 | MIRA-1 |
| top60_altered_DNA_seed | KRAS | 0.98968 | LUAD | Calu-3 | KRAS (G12C) Inhibitor-12 |
| top60_altered_DNA_seed | TP53 | 1.010753 | LUAD | HOP-62 | MIRA-1 |
| top60_altered_DNA_seed | TP53 | 1.015556 | LUAD | NCI-H358 | MIRA-1 |
| top60_altered_DNA_seed | KRAS | 1.033303 | LUAD | EKVX | KRAS (G12C) Inhibitor-12 |
| top60_altered_DNA_seed | TP53 | 1.043648 | LUAD | SK-LU-1 | MIRA-1 |
| top60_altered_DNA_seed | TP53 | 1.057105 | LUAD | A549 | MIRA-1 |
| top60_altered_DNA_seed | KRAS | 1.058704 | LUAD | NCI-H1355 | KRAS (G12C) Inhibitor-12 |
| top60_altered_DNA_seed | TP53 | 1.060177 | LUAD | NCI-H2087 | MIRA-1 |
| top60_altered_DNA_seed | TP53 | 1.071087 | LUAD | NCI-H2291 | MIRA-1 |
| top60_altered_DNA_seed | TP53 | 1.075039 | LUAD | Calu-3 | MIRA-1 |
| top60_altered_DNA_seed | TP53 | 1.144447 | LUAD | NCI-H1838 | MIRA-1 |
| top60_altered_DNA_seed | TP53 | 1.162837 | LUAD | H3255 | MIRA-1 |
| top60_altered_DNA_seed | TP53 | 1.172228 | LUAD | NCI-H1355 | MIRA-1 |
| top60_altered_DNA_seed | KRAS | 1.209619 | LUAD | NCI-H2291 | KRAS (G12C) Inhibitor-12 |
| top60_altered_DNA_seed | TP53 | 1.221488 | LUAD | COR-L105 | MIRA-1 |
| top60_altered_DNA_seed | KRAS | 1.250427 | LUAD | NCI-H1793 | KRAS (G12C) Inhibitor-12 |
| top60_altered_DNA_seed | KRAS | 1.30271 | LUAD | NCI-H1755 | KRAS (G12C) Inhibitor-12 |
| top60_altered_DNA_seed | TP53 | 1.320283 | LUAD | EKVX | MIRA-1 |
| top60_altered_DNA_seed | TP53 | 1.342672 | LUAD | SW1573 | MIRA-1 |
| top60_altered_DNA_seed | KRAS | 1.453596 | LUAD | NCI-H1693 | KRAS (G12C) Inhibitor-12 |
| top60_altered_DNA_seed | TP53 | 1.479916 | LUAD | NCI-H2228 | MIRA-1 |
| top60_altered_DNA_seed | TP53 | 1.497721 | LUAD | NCI-H1666 | MIRA-1 |
| top60_altered_DNA_seed | TP53 | 1.500118 | LUAD | NCI-H1650 | MIRA-1 |
| top60_altered_DNA_seed | KRAS | 1.630854 | LUAD | COR-L105 | KRAS (G12C) Inhibitor-12 |
| top60_altered_DNA_seed | TP53 | 1.691954 | LUAD | NCI-H1568 | MIRA-1 |
| top60_altered_DNA_seed | KRAS | 1.703993 | LUAD | NCI-H1650 | KRAS (G12C) Inhibitor-12 |
| top60_altered_DNA_seed | TP53 | 1.763346 | LUAD | NCI-H1793 | MIRA-1 |
| top60_altered_DNA_seed | TP53 | 1.824734 | LUAD | HCC-827 | MIRA-1 |
| top60_altered_DNA_seed | KRAS | 1.826049 | LUAD | NCI-H2405 | KRAS (G12C) Inhibitor-12 |
| top60_altered_DNA_seed | TP53 | 1.91054 | LUAD | NCI-H2347 | MIRA-1 |

| | | | | | |
|---|---|---|---|---|---|
| top60_altered_DNA_seed | KRAS | 1.93085 | LUAD | NCI-H1568 | KRAS (G12C) Inhibitor-12 |
| top60_altered_DNA_seed | TP53 | 1.932986 | LUAD | NCI-H441 | MIRA-1 |
| top60_altered_DNA_seed | KRAS | 2.042551 | LUAD | NCI-H2228 | KRAS (G12C) Inhibitor-12 |
| top60_altered_DNA_seed | KRAS | 2.098972 | LUAD | SW1573 | KRAS (G12C) Inhibitor-12 |
| top60_altered_DNA_seed | KRAS | 2.199916 | LUAD | SK-LU-1 | KRAS (G12C) Inhibitor-12 |
| top60_altered_DNA_seed | TP53 | 2.464529 | LUAD | NCI-H2405 | MIRA-1 |
| top60_altered_DNA_seed | KRAS | 2.603151 | LUAD | NCI-H441 | KRAS (G12C) Inhibitor-12 |
| top60_altered_DNA_seed | PIK3CA | -1.665707 | LUSC | SK-MES-1 | PI3Ka_4409 |
| top60_altered_DNA_seed | PIK3CA | -1.469562 | LUSC | RERF-LC-Sq1 | PI3Ka_4409 |
| top60_altered_DNA_seed | PIK3CA | -1.37393 | LUSC | SK-MES-1 | AZD8835 |
| top60_altered_DNA_seed | PIK3CA | -1.09166 | LUSC | RERF-LC-Sq1 | AZD8835 |
| top60_altered_DNA_seed | PIK3CA | -1.001354 | LUSC | EPLC-272H | PI3Ka_4409 |
| top60_altered_DNA_seed | PIK3CA | -0.834848 | LUSC | EPLC-272H | AZD8835 |
| top60_altered_DNA_seed | PIK3CA | -0.809777 | LUSC | LOU-NH91 | AZD8835 |
| top60_altered_DNA_seed | PIK3CA | -0.633699 | LUSC | EBC-1 | PI3Ka_4409 |
| top60_altered_DNA_seed | PIK3CA | -0.455436 | LUSC | KNS-62 | PI3Ka_4409 |
| top60_altered_DNA_seed | PIK3CA | -0.250907 | LUSC | LK-2 | AZD8835 |
| top60_altered_DNA_seed | PIK3CA | -0.247775 | LUSC | HCC-15 | PI3Ka_4409 |
| top60_altered_DNA_seed | PIK3CA | -0.233333 | LUSC | NCI-H226 | AZD8835 |
| top60_altered_DNA_seed | PIK3CA | -0.221388 | LUSC | NCI-H1869 | PI3Ka_4409 |
| top60_altered_DNA_seed | PIK3CA | -0.202814 | LUSC | HARA | AZD8835 |
| top60_altered_DNA_seed | PIK3CA | -0.155866 | LUSC | HARA | PI3Ka_4409 |
| top60_altered_DNA_seed | PIK3CA | -0.032854 | LUSC | NCI-H2170 | PI3Ka_4409 |
| top60_altered_DNA_seed | PIK3CA | 0.041565 | LUSC | LOU-NH91 | PI3Ka_4409 |
| top60_altered_DNA_seed | PIK3CA | 0.050047 | LUSC | HCC-15 | AZD8835 |
| top60_altered_DNA_seed | PIK3CA | 0.109597 | LUSC | LC-1-sq | AZD8835 |
| top60_altered_DNA_seed | PIK3CA | 0.293316 | LUSC | NCI-H2170 | AZD8835 |
| top60_altered_DNA_seed | PIK3CA | 0.321961 | LUSC | LC-1-sq | PI3Ka_4409 |
| top60_altered_DNA_seed | PIK3CA | 0.33785 | LUSC | LK-2 | PI3Ka_4409 |
| top60_altered_DNA_seed | PIK3CA | 0.846248 | LUSC | EBC-1 | AZD8835 |
| top60_altered_DNA_seed | PIK3CA | 1.193603 | LUSC | NCI-H226 | PI3Ka_4409 |
| top60_altered_DNA_seed | PIK3CA | 1.349994 | LUSC | NCI-H520 | AZD8835 |
| top60_altered_DNA_seed | PIK3CA | 1.576912 | LUSC | NCI-H1869 | AZD8835 |
| top60_altered_DNA_seed | PIK3CA | 1.590637 | LUSC | NCI-H520 | PI3Ka_4409 |
| top60_altered_DNA_seed | PIK3CA | 2.274945 | LUSC | KNS-62 | AZD8835 |
| top60_altered_DNA_seed | PIK3CA | -1.338512 | LUSC | RERF-LC-Sq1 | Alpelisib |
| top60_altered_DNA_seed | PIK3CA | -1.220521 | LUSC | EPLC-272H | Alpelisib |
| top60_altered_DNA_seed | PIK3CA | -1.145638 | LUSC | EPLC-272H | Buparlisib |
| top60_altered_DNA_seed | PIK3CA | -1.121422 | LUSC | NCI-H2170 | Buparlisib |
| top60_altered_DNA_seed | PIK3CA | -1.08911 | LUSC | EPLC-272H | AZD8186 |
| top60_altered_DNA_seed | PIK3CA | -1.036936 | LUSC | EPLC-272H | GNE-317 |

| | | | | | |
|---|---|---|---|---|---|
| top60_altered_DNA_seed | PIK3CA | -0.955533 | LUSC | HARA | Buparlisib |
| top60_altered_DNA_seed | PIK3CA | -0.913577 | LUSC | HARA | Alpelisib |
| top60_altered_DNA_seed | PIK3CA | -0.830417 | LUSC | HARA | GNE-317 |
| top60_altered_DNA_seed | PIK3CA | -0.822014 | LUSC | EBC-1 | AZD8186 |
| top60_altered_DNA_seed | PIK3CA | -0.786418 | LUSC | RERF-LC-Sq1 | GNE-317 |
| top60_altered_DNA_seed | PIK3CA | -0.781629 | LUSC | NCI-H2170 | Alpelisib |
| top60_altered_DNA_seed | PIK3CA | -0.729955 | LUSC | EBC-1 | GNE-317 |
| top60_altered_DNA_seed | PIK3CA | -0.713796 | LUSC | NCI-H520 | AZD8186 |
| top60_altered_DNA_seed | PIK3CA | -0.531937 | LUSC | HCC-15 | Buparlisib |
| top60_altered_DNA_seed | PIK3CA | -0.461466 | LUSC | NCI-H2170 | AZD8186 |
| top60_altered_DNA_seed | PIK3CA | -0.33911 | LUSC | NCI-H520 | GNE-317 |
| top60_altered_DNA_seed | PIK3CA | -0.313509 | LUSC | LOU-NH91 | Alpelisib |
| top60_altered_DNA_seed | PIK3CA | -0.290278 | LUSC | RERF-LC-Sq1 | Buparlisib |
| top60_altered_DNA_seed | PIK3CA | -0.287494 | LUSC | LOU-NH91 | GNE-317 |
| top60_altered_DNA_seed | PIK3CA | -0.283547 | LUSC | LK-2 | GNE-317 |
| top60_altered_DNA_seed | PIK3CA | -0.280465 | LUSC | LOU-NH91 | AZD8186 |
| top60_altered_DNA_seed | PIK3CA | -0.27608 | LUSC | NCI-H1869 | AZD8186 |
| top60_altered_DNA_seed | PIK3CA | -0.270405 | LUSC | LK-2 | Buparlisib |
| top60_altered_DNA_seed | PIK3CA | -0.210267 | LUSC | EBC-1 | Buparlisib |
| top60_altered_DNA_seed | PIK3CA | -0.086508 | LUSC | LC-1-sq | Alpelisib |
| top60_altered_DNA_seed | PIK3CA | -0.039436 | LUSC | LOU-NH91 | Buparlisib |
| top60_altered_DNA_seed | TP53 | -0.002669 | LUSC | NCI-H1869 | MIRA-1 |
| top60_altered_DNA_seed | TP53 | 0.006589 | LUSC | NCI-H2170 | MIRA-1 |
| top60_altered_DNA_seed | PIK3CA | 0.035824 | LUSC | SK-MES-1 | AZD8186 |
| top60_altered_DNA_seed | TP53 | 0.085711 | LUSC | HCC-15 | MIRA-1 |
| top60_altered_DNA_seed | TP53 | 0.117814 | LUSC | NCI-H520 | MIRA-1 |
| top60_altered_DNA_seed | TP53 | 0.127705 | LUSC | HARA | MIRA-1 |
| top60_altered_DNA_seed | TP53 | 0.175425 | LUSC | LC-1-sq | MIRA-1 |
| top60_altered_DNA_seed | PIK3CA | 0.179874 | LUSC | NCI-H520 | Buparlisib |
| top60_altered_DNA_seed | TP53 | 0.215474 | LUSC | LK-2 | MIRA-1 |
| top60_altered_DNA_seed | PIK3CA | 0.328737 | LUSC | SK-MES-1 | GNE-317 |
| top60_altered_DNA_seed | PIK3CA | 0.332925 | LUSC | HARA | AZD8186 |
| top60_altered_DNA_seed | PIK3CA | 0.357542 | LUSC | NCI-H1869 | GNE-317 |
| top60_altered_DNA_seed | TP53 | 0.371749 | LUSC | EPLC-272H | MIRA-1 |
| top60_altered_DNA_seed | PIK3CA | 0.384334 | LUSC | RERF-LC-Sq1 | AZD8186 |
| top60_altered_DNA_seed | PIK3CA | 0.493276 | LUSC | LC-1-sq | Buparlisib |
| top60_altered_DNA_seed | PIK3CA | 0.528927 | LUSC | HCC-15 | GNE-317 |
| top60_altered_DNA_seed | PIK3CA | 0.568534 | LUSC | KNS-62 | Alpelisib |
| top60_altered_DNA_seed | TP53 | 0.593917 | LUSC | RERF-LC-Sq1 | MIRA-1 |
| top60_altered_DNA_seed | PIK3CA | 0.621815 | LUSC | KNS-62 | GNE-317 |
| top60_altered_DNA_seed | PIK3CA | 0.637805 | LUSC | NCI-H520 | Alpelisib |

| | | | | | |
|---|---|---|---|---|---|
| top60_altered_DNA_seed | PIK3CA | 0.638856 | LUSC | SK-MES-1 | Buparlisib |
| top60_altered_DNA_seed | PIK3CA | 0.654019 | LUSC | KNS-62 | Buparlisib |
| top60_altered_DNA_seed | TP53 | 0.691211 | LUSC | KNS-62 | MIRA-1 |
| top60_altered_DNA_seed | TP53 | 0.717299 | LUSC | LOU-NH91 | MIRA-1 |
| top60_altered_DNA_seed | PIK3CA | 0.724967 | LUSC | LC-1-sq | AZD8186 |
| top60_altered_DNA_seed | PIK3CA | 0.786867 | LUSC | LC-1-sq | GNE-317 |
| top60_altered_DNA_seed | PIK3CA | 0.810293 | LUSC | NCI-H1869 | Alpelisib |
| top60_altered_DNA_seed | PIK3CA | 0.833017 | LUSC | HCC-15 | AZD8186 |
| top60_altered_DNA_seed | TP53 | 0.841218 | LUSC | EBC-1 | MIRA-1 |
| top60_altered_DNA_seed | TP53 | 0.849479 | LUSC | SK-MES-1 | MIRA-1 |
| top60_altered_DNA_seed | PIK3CA | 0.906604 | LUSC | LK-2 | AZD8186 |
| top60_altered_DNA_seed | PIK3CA | 0.914289 | LUSC | EBC-1 | Alpelisib |
| top60_altered_DNA_seed | PIK3CA | 1.049427 | LUSC | NCI-H226 | Alpelisib |
| top60_altered_DNA_seed | PIK3CA | 1.066896 | LUSC | KNS-62 | AZD8186 |
| top60_altered_DNA_seed | PIK3CA | 1.11521 | LUSC | HCC-15 | Alpelisib |
| top60_altered_DNA_seed | PIK3CA | 1.146915 | LUSC | SK-MES-1 | Alpelisib |
| top60_altered_DNA_seed | PIK3CA | 1.175026 | LUSC | LK-2 | Alpelisib |
| top60_altered_DNA_seed | PIK3CA | 1.288456 | LUSC | NCI-H1869 | Buparlisib |
| top60_altered_DNA_seed | TP53 | 1.773584 | LUSC | NCI-H226 | MIRA-1 |
| top60_altered_DNA_seed | PIK3CA | 2.103956 | LUSC | NCI-H226 | AZD8186 |
| top60_altered_DNA_seed | PIK3CA | 2.336999 | LUSC | NCI-H226 | GNE-317 |
| top60_altered_DNA_seed | PIK3CA | 2.454696 | LUSC | NCI-H226 | Buparlisib |
| top60_altered_DNA_seed | PTK2 | -1.808301 | OV | IGROV-1 | PF-562271 |
| top60_altered_DNA_seed | PTK2 | -1.766052 | OV | OVTOKO | PF-562271 |
| top60_altered_DNA_seed | PTK2 | -1.466293 | OV | OVCAR-5 | PF-562271 |
| top60_altered_DNA_seed | PTK2 | -1.010748 | OV | TOV-112D | PF-562271 |
| top60_altered_DNA_seed | PTK2 | -0.473914 | OV | OVMIU | PF-562271 |
| top60_altered_DNA_seed | PTK2 | -0.409326 | OV | OC-314 | PF-562271 |
| top60_altered_DNA_seed | PTK2 | -0.369307 | OV | OVCAR-3 | PF-562271 |
| top60_altered_DNA_seed | PTK2 | -0.353918 | OV | SK-OV-3 | PF-562271 |
| top60_altered_DNA_seed | PTK2 | 0.042812 | OV | TOV-21G | PF-562271 |
| top60_altered_DNA_seed | PTK2 | 0.071001 | OV | OVCAR-4 | PF-562271 |
| top60_altered_DNA_seed | PTK2 | 0.1039 | OV | OVISE | PF-562271 |
| top60_altered_DNA_seed | PTK2 | 0.115421 | OV | OVCAR-8 | PF-562271 |
| top60_altered_DNA_seed | PTK2 | 0.130294 | OV | SW626 | PF-562271 |
| top60_altered_DNA_seed | PTK2 | 0.151912 | OV | EFO-21 | PF-562271 |
| top60_altered_DNA_seed | PTK2 | 0.403877 | OV | FU-OV-1 | PF-562271 |
| top60_altered_DNA_seed | PTK2 | 0.711747 | OV | Caov-3 | PF-562271 |
| top60_altered_DNA_seed | PTK2 | 1.025922 | OV | OAW-42 | PF-562271 |
| top60_altered_DNA_seed | PTK2 | 1.132375 | OV | OV-90 | PF-562271 |
| top60_altered_DNA_seed | PTK2 | 1.234536 | OV | OAW-28 | PF-562271 |

| | | | | | |
|---|---|---|---|---|---|
| top60_altered_DNA_seed | PTK2 | 2.324968 | OV | Caov-4 | PF-562271 |
| top60_altered_DNA_seed | PTK2 | 2.985457 | OV | OVKATE | PF-562271 |
| top60_altered_DNA_seed | TP53 | -0.014816 | PRAD | PWR-1E | MIRA-1 |
| top60_altered_DNA_seed | TP53 | 0.243205 | PRAD | VCaP | MIRA-1 |
| top60_altered_DNA_seed | TP53 | 0.687703 | PRAD | LNCaP-Clone-FGC | MIRA-1 |
| top60_altered_DNA_seed | TP53 | 0.706283 | PRAD | 22RV1 | MIRA-1 |
| top60_altered_DNA_seed | TP53 | 0.955249 | PRAD | DU-145 | MIRA-1 |
| top60_altered_DNA_seed | TP53 | 1.463099 | PRAD | PC-3 | MIRA-1 |
| top60_altered_DNA_seed | ERBB2 | -5.506139 | STAD | NCI-N87 | CP724714 |
| top60_altered_DNA_seed | ERBB2 | -4.174126 | STAD | Hs-746T | CP724714 |
| top60_altered_DNA_seed | ERBB2 | -3.997646 | STAD | SK-GT-2 | Afatinib |
| top60_altered_DNA_seed | ERBB2 | -3.708978 | STAD | SK-GT-2 | Afatinib |
| top60_altered_DNA_seed | ERBB2 | -3.579769 | STAD | NUGC-4 | Afatinib |
| top60_altered_DNA_seed | ERBB2 | -3.19964 | STAD | NCI-N87 | Afatinib |
| top60_altered_DNA_seed | ERBB2 | -3.012876 | STAD | NUGC-4 | AZD8931 |
| top60_altered_DNA_seed | ERBB2 | -2.840983 | STAD | NCI-N87 | Afatinib |
| top60_altered_DNA_seed | PIK3CA | -2.507956 | STAD | IM-95 | AZD8835 |
| top60_altered_DNA_seed | ERBB2 | -2.458722 | STAD | NUGC-4 | PF-00299804 |
| top60_altered_DNA_seed | ERBB2 | -2.202897 | STAD | SK-GT-2 | AZD8931 |
| top60_altered_DNA_seed | ERBB2 | -2.197683 | STAD | TGBC11TKB | CP724714 |
| top60_altered_DNA_seed | ERBB2 | -2.084813 | STAD | AGS | PF-00299804 |
| top60_altered_DNA_seed | PIK3CA | -2.011361 | STAD | AGS | AZD8835 |
| top60_altered_DNA_seed | ERBB2 | -1.999928 | STAD | NUGC-3 | Afatinib |
| top60_altered_DNA_seed | ERBB2 | -1.907664 | STAD | GCIY | CI-1033 |
| top60_altered_DNA_seed | ERBB2 | -1.847543 | STAD | NCI-N87 | AZD8931 |
| top60_altered_DNA_seed | ERBB2 | -1.746646 | STAD | NUGC-3 | Afatinib |
| top60_altered_DNA_seed | ERBB2 | -1.727366 | STAD | GCIY | PF-00299804 |
| top60_altered_DNA_seed | ERBB2 | -1.633241 | STAD | IM-95 | AZD8931 |
| top60_altered_DNA_seed | PIK3CA | -1.601479 | STAD | AGS | PI3Ka_4409 |
| top60_altered_DNA_seed | ERBB2 | -1.527312 | STAD | NUGC-4 | CI-1033 |
| top60_altered_DNA_seed | PIK3CA | -1.49039 | STAD | HSC-39 | PI3Ka_4409 |
| top60_altered_DNA_seed | ERBB2 | -1.456809 | STAD | HSC-39 | CI-1033 |
| top60_altered_DNA_seed | LRRK2 | -1.352028 | STAD | MKN28 | XMD11-85h |
| top60_altered_DNA_seed | ERBB2 | -1.309811 | STAD | RF-48 | CUDC-101 |
| top60_altered_DNA_seed | ERBB2 | -1.29334 | STAD | AGS | CI-1033 |
| top60_altered_DNA_seed | LRRK2 | -1.276783 | STAD | MKN28 | XMD11-50 |
| top60_altered_DNA_seed | ERBB2 | -1.248274 | STAD | GCIY | Afatinib |
| top60_altered_DNA_seed | LRRK2 | -1.240374 | STAD | AGS | XMD11-85h |
| top60_altered_DNA_seed | ERBB2 | -1.180036 | STAD | NUGC-4 | CUDC-101 |
| top60_altered_DNA_seed | ERBB2 | -1.09391 | STAD | NCI-N87 | CUDC-101 |
| top60_altered_DNA_seed | ERBB2 | -1.08654 | STAD | SNU-1 | PF-00299804 |

| | | | | | |
|---|---|---|---|---|---|
| top60_altered_DNA_seed | LRRK2 | -1.058802 | STAD | AGS | XMD11-50 |
| top60_altered_DNA_seed | LRRK2 | -1.057558 | STAD | SNU-16 | XMD11-85h |
| top60_altered_DNA_seed | ERBB2 | -1.040137 | STAD | NUGC-4 | Afatinib |
| top60_altered_DNA_seed | LRRK2 | -1.022759 | STAD | NUGC-4 | XMD11-85h |
| top60_altered_DNA_seed | LRRK2 | -1.019998 | STAD | SNU-16 | XMD11-50 |
| top60_altered_DNA_seed | ERBB2 | -1.019476 | STAD | NUGC-3 | AZD8931 |
| top60_altered_DNA_seed | ERBB2 | -0.995298 | STAD | AGS | AZD8931 |
| top60_altered_DNA_seed | ERBB2 | -0.968535 | STAD | HSC-39 | PF-00299804 |
| top60_altered_DNA_seed | PIK3CA | -0.955002 | STAD | HSC-39 | AZD8835 |
| top60_altered_DNA_seed | LRRK2 | -0.944716 | STAD | HGC-27 | XMD11-85h |
| top60_altered_DNA_seed | ERBB2 | -0.940647 | STAD | SNU-1 | CI-1033 |
| top60_altered_DNA_seed | ERBB2 | -0.930647 | STAD | RERF-GC-1B | CP724714 |
| top60_altered_DNA_seed | ERBB2 | -0.910739 | STAD | GCIY | Lapatinib |
| top60_altered_DNA_seed | ERBB2 | -0.876224 | STAD | SNU-1 | CP724714 |
| top60_altered_DNA_seed | ERBB2 | -0.845644 | STAD | HGC-27 | CP724714 |
| top60_altered_DNA_seed | ERBB2 | -0.839761 | STAD | RERF-GC-1B | Afatinib |
| top60_altered_DNA_seed | PIK3CA | -0.771988 | STAD | NCI-N87 | PI3Ka_4409 |
| top60_altered_DNA_seed | ERBB2 | -0.75817 | STAD | MKN7 | Afatinib |
| top60_altered_DNA_seed | ERBB2 | -0.747106 | STAD | RF-48 | Afatinib |
| top60_altered_DNA_seed | PIK3CA | -0.739382 | STAD | SNU-5 | PI3Ka_4409 |
| top60_altered_DNA_seed | ERBB2 | -0.687259 | STAD | TMK-1 | Afatinib |
| top60_altered_DNA_seed | ERBB2 | -0.66765 | STAD | 23132-87 | CUDC-101 |
| top60_altered_DNA_seed | ERBB2 | -0.650746 | STAD | IM-95 | CP724714 |
| top60_altered_DNA_seed | ERBB2 | -0.648542 | STAD | GCIY | CP724714 |
| top60_altered_DNA_seed | ERBB2 | -0.605117 | STAD | IM-95 | CI-1033 |
| top60_altered_DNA_seed | ERBB2 | -0.602136 | STAD | MKN28 | AZD8931 |
| top60_altered_DNA_seed | PIK3CA | -0.58183 | STAD | 23132-87 | AZD8835 |
| top60_altered_DNA_seed | ERBB2 | -0.576329 | STAD | Hs-746T | CUDC-101 |
| top60_altered_DNA_seed | ERBB2 | -0.576186 | STAD | KATOIII | PF-00299804 |
| top60_altered_DNA_seed | PIK3CA | -0.544564 | STAD | NCI-N87 | AZD8835 |
| top60_altered_DNA_seed | ERBB2 | -0.53759 | STAD | RERF-GC-1B | CI-1033 |
| top60_altered_DNA_seed | ERBB2 | -0.507793 | STAD | NUGC-4 | CP724714 |
| top60_altered_DNA_seed | ERBB2 | -0.490537 | STAD | SNU-1 | CUDC-101 |
| top60_altered_DNA_seed | ERBB2 | -0.458125 | STAD | AGS | CUDC-101 |
| top60_altered_DNA_seed | ERBB2 | -0.457756 | STAD | MKN28 | CI-1033 |
| top60_altered_DNA_seed | ERBB2 | -0.428087 | STAD | KATOIII | CI-1033 |
| top60_altered_DNA_seed | ERBB2 | -0.423921 | STAD | MKN7 | Afatinib |
| top60_altered_DNA_seed | PIK3CA | -0.406898 | STAD | TMK-1 | PI3Ka_4409 |
| top60_altered_DNA_seed | ERBB2 | -0.406648 | STAD | FU97 | CI-1033 |
| top60_altered_DNA_seed | PIK3CA | -0.366374 | STAD | FU97 | PI3Ka_4409 |
| top60_altered_DNA_seed | ERBB2 | -0.314995 | STAD | MKN7 | PF-00299804 |

| | | | | | |
|---|---|---|---|---|---|
| top60_altered_DNA_seed | ERBB2 | -0.314747 | STAD | IM-95 | CUDC-101 |
| top60_altered_DNA_seed | ERBB2 | -0.284447 | STAD | 23132-87 | AZD8931 |
| top60_altered_DNA_seed | LRRK2 | -0.258105 | STAD | TMK-1 | XMD11-50 |
| top60_altered_DNA_seed | ERBB2 | -0.24229 | STAD | RERF-GC-1B | AZD8931 |
| top60_altered_DNA_seed | LRRK2 | -0.241996 | STAD | TMK-1 | XMD11-85h |
| top60_altered_DNA_seed | ERBB2 | -0.23349 | STAD | SNU-16 | AZD8931 |
| top60_altered_DNA_seed | ERBB2 | -0.21364 | STAD | TGBC11TKB | CI-1033 |
| top60_altered_DNA_seed | ERBB2 | -0.200245 | STAD | NUGC-3 | CUDC-101 |
| top60_altered_DNA_seed | ERBB2 | -0.193304 | STAD | RERF-GC-1B | PF-00299804 |
| top60_altered_DNA_seed | PIK3CA | -0.182956 | STAD | RERF-GC-1B | PI3Ka_4409 |
| top60_altered_DNA_seed | ERBB2 | -0.175941 | STAD | MKN7 | CI-1033 |
| top60_altered_DNA_seed | PIK3CA | -0.166059 | STAD | FU97 | AZD8835 |
| top60_altered_DNA_seed | ERBB2 | -0.160746 | STAD | HGC-27 | CUDC-101 |
| top60_altered_DNA_seed | ERBB2 | -0.144342 | STAD | IM-95 | Afatinib |
| top60_altered_DNA_seed | ERBB2 | -0.138867 | STAD | AGS | CP724714 |
| top60_altered_DNA_seed | ERBB2 | -0.134016 | STAD | FU97 | CP724714 |
| top60_altered_DNA_seed | PIK3CA | -0.130699 | STAD | 23132-87 | PI3Ka_4409 |
| top60_altered_DNA_seed | PIK3CA | -0.128391 | STAD | RERF-GC-1B | AZD8835 |
| top60_altered_DNA_seed | ERBB2 | -0.128348 | STAD | TGBC11TKB | PF-00299804 |
| top60_altered_DNA_seed | ERBB2 | -0.125099 | STAD | HGC-27 | CI-1033 |
| top60_altered_DNA_seed | ERBB2 | -0.120265 | STAD | GCIY | CUDC-101 |
| top60_altered_DNA_seed | ERBB2 | -0.120054 | STAD | HSC-39 | Lapatinib |
| top60_altered_DNA_seed | ERBB2 | -0.093925 | STAD | NUGC-3 | CI-1033 |
| top60_altered_DNA_seed | ERBB2 | -0.065151 | STAD | AGS | Afatinib |
| top60_altered_DNA_seed | ERBB2 | -0.05119 | STAD | 23132-87 | CI-1033 |
| top60_altered_DNA_seed | PIK3CA | -0.050562 | STAD | Hs-746T | PI3Ka_4409 |
| top60_altered_DNA_seed | ERBB2 | -0.047554 | STAD | HSC-39 | AZD8931 |
| top60_altered_DNA_seed | ERBB2 | -0.032571 | STAD | HSC-39 | CP724714 |
| top60_altered_DNA_seed | ERBB2 | -0.020592 | STAD | NUGC-3 | CP724714 |
| top60_altered_DNA_seed | ERBB2 | -0.000572 | STAD | TMK-1 | Afatinib |
| top60_altered_DNA_seed | ERBB2 | 0.004358 | STAD | Hs-746T | CI-1033 |
| top60_altered_DNA_seed | ERBB2 | 0.007664 | STAD | HSC-39 | Afatinib |
| top60_altered_DNA_seed | PIK3CA | 0.020711 | STAD | IM-95 | PI3Ka_4409 |
| top60_altered_DNA_seed | ERBB2 | 0.037018 | STAD | HSC-39 | Afatinib |
| top60_altered_DNA_seed | LRRK2 | 0.05224 | STAD | KATOIII | XMD11-85h |
| top60_altered_DNA_seed | LRRK2 | 0.068953 | STAD | NCI-N87 | XMD11-50 |
| top60_altered_DNA_seed | PIK3CA | 0.073779 | STAD | SNU-16 | AZD8835 |
| top60_altered_DNA_seed | ERBB2 | 0.094896 | STAD | FU97 | CUDC-101 |
| top60_altered_DNA_seed | PIK3CA | 0.121784 | STAD | TGBC11TKB | AZD8835 |
| top60_altered_DNA_seed | PIK3CA | 0.128448 | STAD | NUGC-4 | AZD8835 |
| top60_altered_DNA_seed | PIK3CA | 0.137472 | STAD | MKN28 | AZD8835 |

| | | | | | |
|---|---|---|---|---|---|
| top60_altered_DNA_seed | ERBB2 | 0.138652 | STAD | SNU-16 | CP724714 |
| top60_altered_DNA_seed | ERBB2 | 0.146955 | STAD | MKN7 | CP724714 |
| top60_altered_DNA_seed | ERBB2 | 0.160058 | STAD | MKN7 | AZD8931 |
| top60_altered_DNA_seed | ERBB2 | 0.167947 | STAD | HGC-27 | Afatinib |
| top60_altered_DNA_seed | PIK3CA | 0.178054 | STAD | SK-GT-2 | PI3Ka_4409 |
| top60_altered_DNA_seed | ERBB2 | 0.191637 | STAD | IM-95 | PF-00299804 |
| top60_altered_DNA_seed | PIK3CA | 0.203951 | STAD | SNU-5 | AZD8835 |
| top60_altered_DNA_seed | LRRK2 | 0.219075 | STAD | SK-GT-2 | XMD11-85h |
| top60_altered_DNA_seed | PIK3CA | 0.249282 | STAD | KATOIII | AZD8835 |
| top60_altered_DNA_seed | ERBB2 | 0.256146 | STAD | SNU-5 | Afatinib |
| top60_altered_DNA_seed | ERBB2 | 0.268978 | STAD | SNU-1 | Afatinib |
| top60_altered_DNA_seed | ERBB2 | 0.279114 | STAD | TGBC11TKB | Afatinib |
| top60_altered_DNA_seed | ERBB2 | 0.28185 | STAD | SNU-16 | Lapatinib |
| top60_altered_DNA_seed | ERBB2 | 0.305071 | STAD | SNU-5 | AZD8931 |
| top60_altered_DNA_seed | ERBB2 | 0.314203 | STAD | MKN28 | CP724714 |
| top60_altered_DNA_seed | ERBB2 | 0.340594 | STAD | HGC-27 | Afatinib |
| top60_altered_DNA_seed | ERBB2 | 0.347888 | STAD | NUGC-3 | PF-00299804 |
| top60_altered_DNA_seed | LRRK2 | 0.349183 | STAD | HSC-39 | XMD11-85h |
| top60_altered_DNA_seed | ERBB2 | 0.381472 | STAD | FU97 | AZD8931 |
| top60_altered_DNA_seed | ERBB2 | 0.394715 | STAD | Hs-746T | PF-00299804 |
| top60_altered_DNA_seed | ERBB2 | 0.424051 | STAD | SNU-5 | Afatinib |
| top60_altered_DNA_seed | ERBB2 | 0.42897 | STAD | MKN45 | CI-1033 |
| top60_altered_DNA_seed | ERBB2 | 0.45463 | STAD | RERF-GC-1B | Afatinib |
| top60_altered_DNA_seed | LRRK2 | 0.455366 | STAD | NCI-N87 | XMD11-85h |
| top60_altered_DNA_seed | ERBB2 | 0.462993 | STAD | Hs-746T | AZD8931 |
| top60_altered_DNA_seed | LRRK2 | 0.463826 | STAD | KATOIII | XMD11-50 |
| top60_altered_DNA_seed | ERBB2 | 0.48105 | STAD | MKN28 | CUDC-101 |
| top60_altered_DNA_seed | ERBB2 | 0.489956 | STAD | 23132-87 | Afatinib |
| top60_altered_DNA_seed | ERBB2 | 0.496494 | STAD | KATOIII | AZD8931 |
| top60_altered_DNA_seed | PIK3CA | 0.500928 | STAD | MKN45 | PI3Ka_4409 |
| top60_altered_DNA_seed | ERBB2 | 0.507041 | STAD | 23132-87 | Afatinib |
| top60_altered_DNA_seed | PIK3CA | 0.511737 | STAD | MKN28 | PI3Ka_4409 |
| top60_altered_DNA_seed | ERBB2 | 0.514996 | STAD | RF-48 | CI-1033 |
| top60_altered_DNA_seed | PIK3CA | 0.519231 | STAD | NUGC-4 | PI3Ka_4409 |
| top60_altered_DNA_seed | ERBB2 | 0.522888 | STAD | MKN7 | CUDC-101 |
| top60_altered_DNA_seed | PIK3CA | 0.523535 | STAD | SNU-16 | PI3Ka_4409 |
| top60_altered_DNA_seed | ERBB2 | 0.536215 | STAD | 23132-87 | CP724714 |
| top60_altered_DNA_seed | PIK3CA | 0.554986 | STAD | KATOIII | PI3Ka_4409 |
| top60_altered_DNA_seed | ERBB2 | 0.563994 | STAD | OCUM-1 | AZD8931 |
| top60_altered_DNA_seed | ERBB2 | 0.564898 | STAD | 23132-87 | PF-00299804 |
| top60_altered_DNA_seed | ERBB2 | 0.569323 | STAD | FU97 | Afatinib |

| | | | | | |
|---|---|---|---|---|---|
| top60_altered_DNA_seed | PIK3CA | 0.572245 | STAD | OCUM-1 | PI3Ka_4409 |
| top60_altered_DNA_seed | ERBB2 | 0.577607 | STAD | KATOIII | Afatinib |
| top60_altered_DNA_seed | PIK3CA | 0.589523 | STAD | MKN7 | AZD8835 |
| top60_altered_DNA_seed | ERBB2 | 0.624443 | STAD | MKN45 | Afatinib |
| top60_altered_DNA_seed | ERBB2 | 0.628557 | STAD | TGBC11TKB | CUDC-101 |
| top60_altered_DNA_seed | PIK3CA | 0.634969 | STAD | NUGC-3 | AZD8835 |
| top60_altered_DNA_seed | ERBB2 | 0.65275 | STAD | TMK-1 | CUDC-101 |
| top60_altered_DNA_seed | ERBB2 | 0.673728 | STAD | SNU-16 | Afatinib |
| top60_altered_DNA_seed | ERBB2 | 0.674507 | STAD | OCUM-1 | CP724714 |
| top60_altered_DNA_seed | ERBB2 | 0.68628 | STAD | MKN28 | Afatinib |
| top60_altered_DNA_seed | PIK3CA | 0.712438 | STAD | NUGC-3 | PI3Ka_4409 |
| top60_altered_DNA_seed | ERBB2 | 0.713247 | STAD | RF-48 | AZD8931 |
| top60_altered_DNA_seed | ERBB2 | 0.725846 | STAD | SNU-5 | Lapatinib |
| top60_altered_DNA_seed | PIK3CA | 0.732466 | STAD | TGBC11TKB | PI3Ka_4409 |
| top60_altered_DNA_seed | ERBB2 | 0.741339 | STAD | KATOIII | Afatinib |
| top60_altered_DNA_seed | ERBB2 | 0.753648 | STAD | FU97 | Afatinib |
| top60_altered_DNA_seed | ERBB2 | 0.754714 | STAD | TGBC11TKB | Afatinib |
| top60_altered_DNA_seed | PIK3CA | 0.758768 | STAD | SK-GT-2 | AZD8835 |
| top60_altered_DNA_seed | ERBB2 | 0.760857 | STAD | HGC-27 | PF-00299804 |
| top60_altered_DNA_seed | PIK3CA | 0.800803 | STAD | MKN7 | PI3Ka_4409 |
| top60_altered_DNA_seed | ERBB2 | 0.81431 | STAD | MKN28 | PF-00299804 |
| top60_altered_DNA_seed | ERBB2 | 0.846034 | STAD | FU97 | PF-00299804 |
| top60_altered_DNA_seed | ERBB2 | 0.911389 | STAD | TMK-1 | CP724714 |
| top60_altered_DNA_seed | ERBB2 | 0.918981 | STAD | RF-48 | Lapatinib |
| top60_altered_DNA_seed | PIK3CA | 0.952346 | STAD | RF-48 | AZD8835 |
| top60_altered_DNA_seed | ERBB2 | 0.952632 | STAD | TGBC11TKB | AZD8931 |
| top60_altered_DNA_seed | ERBB2 | 0.953741 | STAD | Hs-746T | Afatinib |
| top60_altered_DNA_seed | ERBB2 | 0.95716 | STAD | MKN45 | PF-00299804 |
| top60_altered_DNA_seed | ERBB2 | 0.957312 | STAD | MKN45 | AZD8931 |
| top60_altered_DNA_seed | ERBB2 | 0.961597 | STAD | Hs-746T | Afatinib |
| top60_altered_DNA_seed | ERBB2 | 0.963799 | STAD | OCUM-1 | Afatinib |
| top60_altered_DNA_seed | ERBB2 | 0.9741 | STAD | SNU-16 | Afatinib |
| top60_altered_DNA_seed | ERBB2 | 0.981788 | STAD | RF-48 | CP724714 |
| top60_altered_DNA_seed | PIK3CA | 1.028821 | STAD | OCUM-1 | AZD8835 |
| top60_altered_DNA_seed | ERBB2 | 1.030123 | STAD | MKN45 | Afatinib |
| top60_altered_DNA_seed | ERBB2 | 1.039935 | STAD | SNU-1 | Lapatinib |
| top60_altered_DNA_seed | ERBB2 | 1.077495 | STAD | OCUM-1 | Afatinib |
| top60_altered_DNA_seed | ERBB2 | 1.095849 | STAD | RF-48 | PF-00299804 |
| top60_altered_DNA_seed | ERBB2 | 1.106125 | STAD | RF-48 | Afatinib |
| top60_altered_DNA_seed | ERBB2 | 1.116749 | STAD | SNU-5 | CUDC-101 |
| top60_altered_DNA_seed | LRRK2 | 1.123016 | STAD | IM-95 | XMD11-85h |

| | | | | | |
|---|---|---|---|---|---|
| top60_altered_DNA_seed | PIK3CA | 1.134076 | STAD | MKN45 | AZD8835 |
| top60_altered_DNA_seed | PIK3CA | 1.154174 | STAD | RF-48 | PI3Ka_4409 |
| top60_altered_DNA_seed | ERBB2 | 1.157487 | STAD | SNU-1 | Afatinib |
| top60_altered_DNA_seed | PIK3CA | 1.214622 | STAD | Hs-746T | AZD8835 |
| top60_altered_DNA_seed | ERBB2 | 1.235015 | STAD | SNU-16 | CUDC-101 |
| top60_altered_DNA_seed | ERBB2 | 1.254838 | STAD | SNU-5 | CP724714 |
| top60_altered_DNA_seed | ERBB2 | 1.306524 | STAD | MKN45 | CP724714 |
| top60_altered_DNA_seed | LRRK2 | 1.32848 | STAD | MKN7 | XMD11-85h |
| top60_altered_DNA_seed | ERBB2 | 1.395564 | STAD | MKN45 | CUDC-101 |
| top60_altered_DNA_seed | ERBB2 | 1.538234 | STAD | SNU-16 | CI-1033 |
| top60_altered_DNA_seed | LRRK2 | 1.543215 | STAD | SNU-5 | XMD11-85h |
| top60_altered_DNA_seed | ERBB2 | 1.742186 | STAD | RERF-GC-1B | CUDC-101 |
| top60_altered_DNA_seed | ERBB2 | 1.870352 | STAD | SNU-16 | PF-00299804 |
| top60_altered_DNA_seed | PIK3CA | 1.925783 | STAD | SNU-1 | PI3Ka_4409 |
| top60_altered_DNA_seed | PIK3CA | 2.177344 | STAD | SNU-1 | AZD8835 |
| top60_altered_DNA_seed | ERBB2 | 2.219959 | STAD | OCUM-1 | CUDC-101 |
| top60_altered_DNA_seed | ERBB2 | 2.327314 | STAD | SNU-1 | AZD8931 |
| top60_altered_DNA_seed | ERBB2 | 2.343604 | STAD | OCUM-1 | CI-1033 |
| top60_altered_DNA_seed | ERBB2 | 2.62529 | STAD | SNU-5 | CI-1033 |
| top60_altered_DNA_seed | ERBB2 | 3.33718 | STAD | SNU-5 | PF-00299804 |
| top60_altered_DNA_seed | ERBB2 | 3.35161 | STAD | OCUM-1 | PF-00299804 |
| top60_altered_DNA_seed | ERBB2 | -4.964046 | STAD | NCI-N87 | Afatinib |
| top60_altered_DNA_seed | ERBB2 | -4.104673 | STAD | SK-GT-2 | Afatinib |
| top60_altered_DNA_seed | ERBB2 | -3.70213 | STAD | NCI-N87 | Lapatinib |
| top60_altered_DNA_seed | ERBB2 | -3.096553 | STAD | NCI-N87 | Sapitinib |
| top60_altered_DNA_seed | ERBB2 | -2.940765 | STAD | SK-GT-2 | Sapitinib |
| top60_altered_DNA_seed | ERBB2 | -2.431962 | STAD | SK-GT-2 | Lapatinib |
| top60_altered_DNA_seed | PIK3CA | -2.195662 | STAD | NCI-N87 | Alpelisib |
| top60_altered_DNA_seed | PIK3CA | -2.178419 | STAD | HGC-27 | Alpelisib |
| top60_altered_DNA_seed | PIK3CA | -2.002491 | STAD | HSC-39 | Alpelisib |
| top60_altered_DNA_seed | PIK3CA | -1.889177 | STAD | HGC-27 | Buparlisib |
| top60_altered_DNA_seed | PIK3CA | -1.879573 | STAD | 23132-87 | AZD8186 |
| top60_altered_DNA_seed | PIK3CA | -1.873533 | STAD | HGC-27 | GNE-317 |
| top60_altered_DNA_seed | LRRK2 | -1.849874 | STAD | HSC-39 | GSK2578215A |
| top60_altered_DNA_seed | PIK3CA | -1.806595 | STAD | AGS | Alpelisib |
| top60_altered_DNA_seed | ERBB2 | -1.699082 | STAD | NUGC-3 | Sapitinib |
| top60_altered_DNA_seed | PIK3CA | -1.690563 | STAD | HGC-27 | AZD8186 |
| top60_altered_DNA_seed | PIK3CA | -1.661051 | STAD | FU97 | Alpelisib |
| top60_altered_DNA_seed | PIK3CA | -1.513459 | STAD | HSC-39 | GNE-317 |
| top60_altered_DNA_seed | ERBB2 | -1.340055 | STAD | NUGC-3 | Afatinib |
| top60_altered_DNA_seed | PIK3CA | -1.255553 | STAD | 23132-87 | Buparlisib |

| | | | | | |
|---|---|---|---|---|---|
| top60_altered_DNA_seed | ERBB2 | -1.18579 | STAD | NUGC-3 | Lapatinib |
| top60_altered_DNA_seed | PIK3CA | -1.166136 | STAD | 23132-87 | GNE-317 |
| top60_altered_DNA_seed | ERBB2 | -1.149545 | STAD | GCIY | Afatinib |
| top60_altered_DNA_seed | LRRK2 | -1.144119 | STAD | SNU-16 | GSK2578215A |
| top60_altered_DNA_seed | ERBB2 | -1.120754 | STAD | GCIY | Sapitinib |
| top60_altered_DNA_seed | PIK3CA | -1.107574 | STAD | HSC-39 | Buparlisib |
| top60_altered_DNA_seed | PIK3CA | -1.096937 | STAD | IM-95 | Alpelisib |
| top60_altered_DNA_seed | PIK3CA | -1.074511 | STAD | 23132-87 | Alpelisib |
| top60_altered_DNA_seed | ERBB2 | -1.0707 | STAD | NUGC-4 | Sapitinib |
| top60_altered_DNA_seed | PIK3CA | -1.064823 | STAD | FU97 | GNE-317 |
| top60_altered_DNA_seed | ERBB2 | -1.064414 | STAD | HGC-27 | Lapatinib |
| top60_altered_DNA_seed | PIK3CA | -1.052359 | STAD | SNU-16 | Buparlisib |
| top60_altered_DNA_seed | ERBB2 | -0.992582 | STAD | HSC-39 | Lapatinib |
| top60_altered_DNA_seed | TP53 | -0.964362 | STAD | SNU-16 | MIRA-1 |
| top60_altered_DNA_seed | ERBB2 | -0.961978 | STAD | RERF-GC-1B | Sapitinib |
| top60_altered_DNA_seed | PIK3CA | -0.94861 | STAD | HSC-39 | AZD8186 |
| top60_altered_DNA_seed | LRRK2 | -0.914916 | STAD | MKN7 | GSK2578215A |
| top60_altered_DNA_seed | PIK3CA | -0.895344 | STAD | FU97 | Buparlisib |
| top60_altered_DNA_seed | PIK3CA | -0.860398 | STAD | AGS | GNE-317 |
| top60_altered_DNA_seed | PIK3CA | -0.842493 | STAD | NCI-N87 | AZD8186 |
| top60_altered_DNA_seed | PIK3CA | -0.832559 | STAD | GCIY | GNE-317 |
| top60_altered_DNA_seed | PIK3CA | -0.829642 | STAD | SNU-5 | GNE-317 |
| top60_altered_DNA_seed | ERBB2 | -0.814856 | STAD | SNU-16 | Lapatinib |
| top60_altered_DNA_seed | PIK3CA | -0.814266 | STAD | SNU-5 | AZD8186 |
| top60_altered_DNA_seed | PIK3CA | -0.791431 | STAD | AGS | Buparlisib |
| top60_altered_DNA_seed | TP53 | -0.79102 | STAD | HSC-39 | MIRA-1 |
| top60_altered_DNA_seed | PIK3CA | -0.790039 | STAD | NUGC-3 | GNE-317 |
| top60_altered_DNA_seed | PIK3CA | -0.770271 | STAD | FU97 | AZD8186 |
| top60_altered_DNA_seed | PIK3CA | -0.768064 | STAD | Hs-746T | GNE-317 |
| top60_altered_DNA_seed | ERBB2 | -0.729675 | STAD | NUGC-4 | Afatinib |
| top60_altered_DNA_seed | ERBB2 | -0.724525 | STAD | MKN7 | Lapatinib |
| top60_altered_DNA_seed | TP53 | -0.720545 | STAD | RF-48 | MIRA-1 |
| top60_altered_DNA_seed | PIK3CA | -0.640799 | STAD | MKN45 | GNE-317 |
| top60_altered_DNA_seed | PIK3CA | -0.636908 | STAD | SNU-5 | Buparlisib |
| top60_altered_DNA_seed | PIK3CA | -0.633687 | STAD | SK-GT-2 | Buparlisib |
| top60_altered_DNA_seed | ERBB2 | -0.628949 | STAD | HSC-39 | Sapitinib |
| top60_altered_DNA_seed | PIK3CA | -0.603623 | STAD | SK-GT-2 | Alpelisib |
| top60_altered_DNA_seed | ERBB2 | -0.596463 | STAD | TMK-1 | Sapitinib |
| top60_altered_DNA_seed | LRRK2 | -0.587554 | STAD | SNU-1 | GSK2578215A |
| top60_altered_DNA_seed | TP53 | -0.578312 | STAD | GCIY | MIRA-1 |
| top60_altered_DNA_seed | PIK3CA | -0.553995 | STAD | IM-95 | Buparlisib |

| | | | | | |
|---|---|---|---|---|---|
| top60_altered_DNA_seed | ERBB2 | -0.544321 | STAD | SNU-16 | Afatinib |
| top60_altered_DNA_seed | PIK3CA | -0.536074 | STAD | SK-GT-2 | GNE-317 |
| top60_altered_DNA_seed | PIK3CA | -0.502607 | STAD | RERF-GC-1B | Alpelisib |
| top60_altered_DNA_seed | PIK3CA | -0.494304 | STAD | TMK-1 | Alpelisib |
| top60_altered_DNA_seed | PIK3CA | -0.492705 | STAD | RF-48 | Buparlisib |
| top60_altered_DNA_seed | LRRK2 | -0.461563 | STAD | RERF-GC-1B | GSK2578215A |
| top60_altered_DNA_seed | PIK3CA | -0.458624 | STAD | NCI-N87 | GNE-317 |
| top60_altered_DNA_seed | LRRK2 | -0.452884 | STAD | HGC-27 | GSK2578215A |
| top60_altered_DNA_seed | LRRK2 | -0.435433 | STAD | NUGC-4 | GSK2578215A |
| top60_altered_DNA_seed | PIK3CA | -0.432008 | STAD | IM-95 | AZD8186 |
| top60_altered_DNA_seed | LRRK2 | -0.395166 | STAD | IM-95 | GSK2578215A |
| top60_altered_DNA_seed | PIK3CA | -0.392938 | STAD | Hs-746T | Buparlisib |
| top60_altered_DNA_seed | PIK3CA | -0.359993 | STAD | Hs-746T | AZD8186 |
| top60_altered_DNA_seed | ERBB2 | -0.336204 | STAD | HSC-39 | Afatinib |
| top60_altered_DNA_seed | PIK3CA | -0.33418 | STAD | SNU-16 | Alpelisib |
| top60_altered_DNA_seed | LRRK2 | -0.315705 | STAD | NUGC-3 | GSK2578215A |
| top60_altered_DNA_seed | ERBB2 | -0.293342 | STAD | 23132-87 | Afatinib |
| top60_altered_DNA_seed | ERBB2 | -0.289654 | STAD | NUGC-4 | Lapatinib |
| top60_altered_DNA_seed | TP53 | -0.272887 | STAD | HGC-27 | MIRA-1 |
| top60_altered_DNA_seed | PIK3CA | -0.263263 | STAD | GCIY | Buparlisib |
| top60_altered_DNA_seed | LRRK2 | -0.2261 | STAD | FU97 | GSK2578215A |
| top60_altered_DNA_seed | PIK3CA | -0.191993 | STAD | TGBC11TKB | Alpelisib |
| top60_altered_DNA_seed | ERBB2 | -0.189838 | STAD | AGS | Lapatinib |
| top60_altered_DNA_seed | PIK3CA | -0.189207 | STAD | SNU-16 | GNE-317 |
| top60_altered_DNA_seed | PIK3CA | -0.181396 | STAD | MKN45 | Alpelisib |
| top60_altered_DNA_seed | PIK3CA | -0.170171 | STAD | SNU-1 | Buparlisib |
| top60_altered_DNA_seed | ERBB2 | -0.169119 | STAD | AGS | Afatinib |
| top60_altered_DNA_seed | TP53 | -0.164612 | STAD | AGS | MIRA-1 |
| top60_altered_DNA_seed | ERBB2 | -0.14424 | STAD | 23132-87 | Sapitinib |
| top60_altered_DNA_seed | ERBB2 | -0.137624 | STAD | IM-95 | Afatinib |
| top60_altered_DNA_seed | ERBB2 | -0.113922 | STAD | TGBC11TKB | Afatinib |
| top60_altered_DNA_seed | PIK3CA | -0.112712 | STAD | AGS | AZD8186 |
| top60_altered_DNA_seed | ERBB2 | -0.112354 | STAD | SNU-16 | Sapitinib |
| top60_altered_DNA_seed | PIK3CA | -0.110412 | STAD | SK-GT-2 | AZD8186 |
| top60_altered_DNA_seed | PIK3CA | -0.108094 | STAD | TMK-1 | Buparlisib |
| top60_altered_DNA_seed | ERBB2 | -0.103653 | STAD | IM-95 | Sapitinib |
| top60_altered_DNA_seed | PIK3CA | -0.100081 | STAD | NCI-N87 | Buparlisib |
| top60_altered_DNA_seed | PIK3CA | -0.091567 | STAD | TMK-1 | AZD8186 |
| top60_altered_DNA_seed | ERBB2 | -0.080522 | STAD | FU97 | Lapatinib |
| top60_altered_DNA_seed | PIK3CA | -0.072562 | STAD | OCUM-1 | Alpelisib |
| top60_altered_DNA_seed | ERBB2 | -0.066352 | STAD | MKN7 | Sapitinib |

| | | | | | |
|---|---|---|---|---|---|
| top60_altered_DNA_seed | TP53 | -0.064001 | STAD | 23132-87 | MIRA-1 |
| top60_altered_DNA_seed | PIK3CA | -0.061805 | STAD | NUGC-3 | Buparlisib |
| top60_altered_DNA_seed | ERBB2 | -0.060068 | STAD | SNU-1 | Lapatinib |
| top60_altered_DNA_seed | ERBB2 | -0.059664 | STAD | SNU-1 | Afatinib |
| top60_altered_DNA_seed | ERBB2 | -0.033577 | STAD | HGC-27 | Afatinib |
| top60_altered_DNA_seed | PIK3CA | -0.023201 | STAD | MKN45 | Buparlisib |
| top60_altered_DNA_seed | ERBB2 | -0.020032 | STAD | AGS | Sapitinib |
| top60_altered_DNA_seed | PIK3CA | -0.015433 | STAD | NUGC-3 | Alpelisib |
| top60_altered_DNA_seed | PIK3CA | 0.005655 | STAD | TGBC11TKB | Buparlisib |
| top60_altered_DNA_seed | TP53 | 0.008774 | STAD | IM-95 | MIRA-1 |
| top60_altered_DNA_seed | ERBB2 | 0.012472 | STAD | MKN28 | Lapatinib |
| top60_altered_DNA_seed | ERBB2 | 0.027895 | STAD | MKN28 | Sapitinib |
| top60_altered_DNA_seed | PIK3CA | 0.036404 | STAD | MKN7 | AZD8186 |
| top60_altered_DNA_seed | ERBB2 | 0.03859 | STAD | Hs-746T | Lapatinib |
| top60_altered_DNA_seed | PIK3CA | 0.043728 | STAD | MKN7 | GNE-317 |
| top60_altered_DNA_seed | ERBB2 | 0.046841 | STAD | RERF-GC-1B | Afatinib |
| top60_altered_DNA_seed | ERBB2 | 0.050663 | STAD | MKN28 | Afatinib |
| top60_altered_DNA_seed | PIK3CA | 0.060791 | STAD | TGBC11TKB | GNE-317 |
| top60_altered_DNA_seed | PIK3CA | 0.062867 | STAD | GCIY | AZD8186 |
| top60_altered_DNA_seed | PIK3CA | 0.0727 | STAD | RF-48 | AZD8186 |
| top60_altered_DNA_seed | ERBB2 | 0.073149 | STAD | RERF-GC-1B | Lapatinib |
| top60_altered_DNA_seed | LRRK2 | 0.108515 | STAD | Hs-746T | GSK2578215A |
| top60_altered_DNA_seed | PIK3CA | 0.116808 | STAD | RF-48 | Alpelisib |
| top60_altered_DNA_seed | LRRK2 | 0.131607 | STAD | TGBC11TKB | GSK2578215A |
| top60_altered_DNA_seed | ERBB2 | 0.133165 | STAD | TGBC11TKB | Lapatinib |
| top60_altered_DNA_seed | TP53 | 0.145707 | STAD | FU97 | MIRA-1 |
| top60_altered_DNA_seed | LRRK2 | 0.148303 | STAD | 23132-87 | GSK2578215A |
| top60_altered_DNA_seed | ERBB2 | 0.15512 | STAD | HGC-27 | Sapitinib |
| top60_altered_DNA_seed | TP53 | 0.163433 | STAD | MKN7 | MIRA-1 |
| top60_altered_DNA_seed | LRRK2 | 0.18397 | STAD | AGS | GSK2578215A |
| top60_altered_DNA_seed | PIK3CA | 0.222733 | STAD | SNU-5 | Alpelisib |
| top60_altered_DNA_seed | ERBB2 | 0.246427 | STAD | MKN7 | Afatinib |
| top60_altered_DNA_seed | PIK3CA | 0.246863 | STAD | RF-48 | GNE-317 |
| top60_altered_DNA_seed | ERBB2 | 0.261371 | STAD | RF-48 | Afatinib |
| top60_altered_DNA_seed | PIK3CA | 0.261699 | STAD | NUGC-3 | AZD8186 |
| top60_altered_DNA_seed | LRRK2 | 0.264798 | STAD | MKN45 | GSK2578215A |
| top60_altered_DNA_seed | TP53 | 0.278515 | STAD | TMK-1 | MIRA-1 |
| top60_altered_DNA_seed | PIK3CA | 0.299203 | STAD | MKN7 | Alpelisib |
| top60_altered_DNA_seed | LRRK2 | 0.300579 | STAD | TMK-1 | GSK2578215A |
| top60_altered_DNA_seed | ERBB2 | 0.305537 | STAD | GCIY | Lapatinib |
| top60_altered_DNA_seed | ERBB2 | 0.312197 | STAD | TMK-1 | Lapatinib |

| | | | | | |
|---|---|---|---|---|---|
| top60_altered_DNA_seed | LRRK2 | 0.326649 | STAD | RF-48 | GSK2578215A |
| top60_altered_DNA_seed | LRRK2 | 0.342437 | STAD | SNU-5 | GSK2578215A |
| top60_altered_DNA_seed | TP53 | 0.34861 | STAD | SNU-1 | MIRA-1 |
| top60_altered_DNA_seed | PIK3CA | 0.36905 | STAD | MKN28 | Alpelisib |
| top60_altered_DNA_seed | TP53 | 0.383155 | STAD | SNU-5 | MIRA-1 |
| top60_altered_DNA_seed | ERBB2 | 0.396143 | STAD | MKN45 | Lapatinib |
| top60_altered_DNA_seed | PIK3CA | 0.396228 | STAD | MKN28 | Buparlisib |
| top60_altered_DNA_seed | LRRK2 | 0.398734 | STAD | SK-GT-2 | GSK2578215A |
| top60_altered_DNA_seed | TP53 | 0.406069 | STAD | RERF-GC-1B | MIRA-1 |
| top60_altered_DNA_seed | PIK3CA | 0.415397 | STAD | MKN45 | AZD8186 |
| top60_altered_DNA_seed | ERBB2 | 0.41743 | STAD | IM-95 | Lapatinib |
| top60_altered_DNA_seed | TP53 | 0.433549 | STAD | SK-GT-2 | MIRA-1 |
| top60_altered_DNA_seed | PIK3CA | 0.442209 | STAD | GCIY | Alpelisib |
| top60_altered_DNA_seed | ERBB2 | 0.479741 | STAD | TGBC11TKB | Sapitinib |
| top60_altered_DNA_seed | PIK3CA | 0.481446 | STAD | RERF-GC-1B | AZD8186 |
| top60_altered_DNA_seed | ERBB2 | 0.512615 | STAD | KATOIII | Sapitinib |
| top60_altered_DNA_seed | LRRK2 | 0.517656 | STAD | KATOIII | GSK2578215A |
| top60_altered_DNA_seed | TP53 | 0.588419 | STAD | TGBC11TKB | MIRA-1 |
| top60_altered_DNA_seed | PIK3CA | 0.598927 | STAD | SNU-1 | AZD8186 |
| top60_altered_DNA_seed | ERBB2 | 0.601589 | STAD | Hs-746T | Afatinib |
| top60_altered_DNA_seed | ERBB2 | 0.603182 | STAD | RF-48 | Sapitinib |
| top60_altered_DNA_seed | PIK3CA | 0.616749 | STAD | RERF-GC-1B | Buparlisib |
| top60_altered_DNA_seed | ERBB2 | 0.663193 | STAD | 23132-87 | Lapatinib |
| top60_altered_DNA_seed | TP53 | 0.67081 | STAD | OCUM-1 | MIRA-1 |
| top60_altered_DNA_seed | ERBB2 | 0.670885 | STAD | SNU-1 | Sapitinib |
| top60_altered_DNA_seed | PIK3CA | 0.69062 | STAD | RERF-GC-1B | GNE-317 |
| top60_altered_DNA_seed | TP53 | 0.719516 | STAD | NUGC-3 | MIRA-1 |
| top60_altered_DNA_seed | PIK3CA | 0.73458 | STAD | TMK-1 | GNE-317 |
| top60_altered_DNA_seed | TP53 | 0.752842 | STAD | NCI-N87 | MIRA-1 |
| top60_altered_DNA_seed | ERBB2 | 0.793279 | STAD | SNU-5 | Afatinib |
| top60_altered_DNA_seed | PIK3CA | 0.7938 | STAD | SNU-16 | AZD8186 |
| top60_altered_DNA_seed | ERBB2 | 0.817424 | STAD | FU97 | Sapitinib |
| top60_altered_DNA_seed | TP53 | 0.838111 | STAD | MKN28 | MIRA-1 |
| top60_altered_DNA_seed | LRRK2 | 0.84813 | STAD | GCIY | GSK2578215A |
| top60_altered_DNA_seed | PIK3CA | 0.855342 | STAD | SNU-1 | Alpelisib |
| top60_altered_DNA_seed | PIK3CA | 0.855649 | STAD | NUGC-4 | Alpelisib |
| top60_altered_DNA_seed | ERBB2 | 0.876586 | STAD | MKN45 | Afatinib |
| top60_altered_DNA_seed | LRRK2 | 0.876706 | STAD | MKN28 | GSK2578215A |
| top60_altered_DNA_seed | PIK3CA | 0.924553 | STAD | MKN28 | AZD8186 |
| top60_altered_DNA_seed | ERBB2 | 0.945989 | STAD | FU97 | Afatinib |
| top60_altered_DNA_seed | PIK3CA | 0.976605 | STAD | OCUM-1 | Buparlisib |

| | | | | | |
|---|---|---|---|---|---|
| top60_altered_DNA_seed | LRRK2 | 0.994491 | STAD | NCI-N87 | GSK2578215A |
| top60_altered_DNA_seed | TP53 | 1.053476 | STAD | MKN45 | MIRA-1 |
| top60_altered_DNA_seed | ERBB2 | 1.059574 | STAD | SNU-5 | Sapitinib |
| top60_altered_DNA_seed | PIK3CA | 1.080422 | STAD | OCUM-1 | AZD8186 |
| top60_altered_DNA_seed | ERBB2 | 1.085179 | STAD | SNU-5 | Lapatinib |
| top60_altered_DNA_seed | ERBB2 | 1.093234 | STAD | OCUM-1 | Sapitinib |
| top60_altered_DNA_seed | LRRK2 | 1.185292 | STAD | OCUM-1 | GSK2578215A |
| top60_altered_DNA_seed | PIK3CA | 1.187297 | STAD | IM-95 | GNE-317 |
| top60_altered_DNA_seed | ERBB2 | 1.223624 | STAD | KATOIII | Lapatinib |
| top60_altered_DNA_seed | PIK3CA | 1.223944 | STAD | MKN7 | Buparlisib |
| top60_altered_DNA_seed | PIK3CA | 1.234249 | STAD | OCUM-1 | GNE-317 |
| top60_altered_DNA_seed | TP53 | 1.240448 | STAD | NUGC-4 | MIRA-1 |
| top60_altered_DNA_seed | TP53 | 1.27456 | STAD | Hs-746T | MIRA-1 |
| top60_altered_DNA_seed | PIK3CA | 1.33347 | STAD | KATOIII | Alpelisib |
| top60_altered_DNA_seed | PIK3CA | 1.34511 | STAD | TGBC11TKB | AZD8186 |
| top60_altered_DNA_seed | TP53 | 1.362835 | STAD | KATOIII | MIRA-1 |
| top60_altered_DNA_seed | PIK3CA | 1.466011 | STAD | Hs-746T | Alpelisib |
| top60_altered_DNA_seed | ERBB2 | 1.474767 | STAD | Hs-746T | Sapitinib |
| top60_altered_DNA_seed | ERBB2 | 1.655498 | STAD | RF-48 | Lapatinib |
| top60_altered_DNA_seed | PIK3CA | 1.678729 | STAD | KATOIII | AZD8186 |
| top60_altered_DNA_seed | ERBB2 | 1.687067 | STAD | MKN45 | Sapitinib |
| top60_altered_DNA_seed | PIK3CA | 1.709535 | STAD | NUGC-4 | AZD8186 |
| top60_altered_DNA_seed | ERBB2 | 1.715738 | STAD | TMK-1 | Afatinib |
| top60_altered_DNA_seed | ERBB2 | 1.755136 | STAD | OCUM-1 | Afatinib |
| top60_altered_DNA_seed | PIK3CA | 1.757653 | STAD | KATOIII | GNE-317 |
| top60_altered_DNA_seed | PIK3CA | 1.960357 | STAD | NUGC-4 | Buparlisib |
| top60_altered_DNA_seed | ERBB2 | 1.972143 | STAD | OCUM-1 | Lapatinib |
| top60_altered_DNA_seed | PIK3CA | 1.980186 | STAD | NUGC-4 | GNE-317 |
| top60_altered_DNA_seed | PIK3CA | 2.305225 | STAD | KATOIII | Buparlisib |
| top60_altered_DNA_seed | ERBB2 | 2.393403 | STAD | KATOIII | Afatinib |
| top60_altered_DNA_seed | PIK3CA | 2.561985 | STAD | MKN28 | GNE-317 |
| top60_altered_DNA_seed | RET | -3.946255 | THCA | CGTH-W-1 | Motesanib |
| top60_altered_DNA_seed | BRAF | -3.729294 | THCA | K5 | Dabrafenib |
| top60_altered_DNA_seed | BRAF | -3.412404 | THCA | BHT-101 | Dabrafenib |
| top60_altered_DNA_seed | BRAF | -3.190105 | THCA | BHT-101 | PLX-4720 |
| top60_altered_DNA_seed | BRAF | -3.135102 | THCA | IHH-4 | Dabrafenib |
| top60_altered_DNA_seed | BRAF | -3.105461 | THCA | HTC-C3 | Dabrafenib |
| top60_altered_DNA_seed | BRAF | -2.966357 | THCA | CGTH-W-1 | HG6-64-1 |
| top60_altered_DNA_seed | BRAF | -2.961541 | THCA | K5 | PLX-4720 |
| top60_altered_DNA_seed | BRAF | -2.905795 | THCA | BHT-101 | PLX-4720 |
| top60_altered_DNA_seed | BRAF | -2.830382 | THCA | HTC-C3 | PLX-4720 |

| | | | | | |
|---|---|---|---|---|---|
| top60_altered_DNA_seed | BRAF | -2.827309 | THCA | K5 | PLX-4720 |
| top60_altered_DNA_seed | BRAF | -2.688852 | THCA | HTC-C3 | RAF_9304 |
| top60_altered_DNA_seed | BRAF | -2.604383 | THCA | BHT-101 | SB590885 |
| top60_altered_DNA_seed | BRAF | -2.492753 | THCA | HTC-C3 | PLX-4720 |
| top60_altered_DNA_seed | BRAF | -2.480329 | THCA | B-CPAP | Dabrafenib |
| top60_altered_DNA_seed | BRAF | -2.430714 | THCA | K5 | RAF_9304 |
| top60_altered_DNA_seed | NTRK1 | -2.403245 | THCA | BHT-101 | GW441756 |
| top60_altered_DNA_seed | BRAF | -2.371192 | THCA | BHT-101 | RAF_9304 |
| top60_altered_DNA_seed | BRAF | -2.172551 | THCA | 8305C | Dabrafenib |
| top60_altered_DNA_seed | BRAF | -2.151843 | THCA | K5 | AZ628 |
| top60_altered_DNA_seed | RET | -2.119921 | THCA | HTC-C3 | Motesanib |
| top60_altered_DNA_seed | BRAF | -2.037898 | THCA | 8305C | PLX-4720 |
| top60_altered_DNA_seed | BRAF | -2.027428 | THCA | CGTH-W-1 | AZ628 |
| top60_altered_DNA_seed | BRAF | -1.905405 | THCA | IHH-4 | PLX-4720 |
| top60_altered_DNA_seed | BRAF | -1.828709 | THCA | TT2609-C02 | HG6-64-1 |
| top60_altered_DNA_seed | RET | -1.811525 | THCA | TT | Motesanib |
| top60_altered_DNA_seed | ATM | -1.797046 | THCA | IHH-4 | KU-60019 |
| top60_altered_DNA_seed | BRAF | -1.773559 | THCA | 8305C | PLX-4720 |
| top60_altered_DNA_seed | BRAF | -1.672569 | THCA | IHH-4 | SB590885 |
| top60_altered_DNA_seed | BRAF | -1.554192 | THCA | B-CPAP | PLX-4720 |
| top60_altered_DNA_seed | BRAF | -1.542311 | THCA | FTC-133 | PLX-4720 |
| top60_altered_DNA_seed | BRAF | -1.51961 | THCA | B-CPAP | PLX-4720 |
| top60_altered_DNA_seed | ATM | -1.435817 | THCA | IHH-4 | KU-55933 |
| top60_altered_DNA_seed | BRAF | -1.423171 | THCA | IHH-4 | PLX-4720 |
| top60_altered_DNA_seed | NTRK1 | -1.374276 | THCA | KMH-2 | AZD1332 |
| top60_altered_DNA_seed | NTRK3 | -1.374276 | THCA | KMH-2 | AZD1332 |
| top60_altered_DNA_seed | BRAF | -1.362543 | THCA | 8305C | SB590885 |
| top60_altered_DNA_seed | BRAF | -1.308841 | THCA | IHH-4 | RAF_9304 |
| top60_altered_DNA_seed | ATM | -1.291053 | THCA | BHT-101 | KU-55933 |
| top60_altered_DNA_seed | BRAF | -1.282376 | THCA | B-CPAP | HG6-64-1 |
| top60_altered_DNA_seed | ATM | -1.265003 | THCA | K5 | torin2 |
| top60_altered_DNA_seed | NTRK1 | -1.240635 | THCA | CGTH-W-1 | GW441756 |
| top60_altered_DNA_seed | RET | -1.225834 | THCA | TT2609-C02 | Motesanib |
| top60_altered_DNA_seed | BRAF | -1.224839 | THCA | KMH-2 | SB590885 |
| top60_altered_DNA_seed | ATM | -1.218818 | THCA | CAL-62 | torin2 |
| top60_altered_DNA_seed | ATM | -1.205962 | THCA | 8305C | KU-60019 |
| top60_altered_DNA_seed | BRAF | -1.201393 | THCA | KMH-2 | Dabrafenib |
| top60_altered_DNA_seed | ATM | -1.178386 | THCA | B-CPAP | KU-60019 |
| top60_altered_DNA_seed | BRAF | -1.173142 | THCA | ML-1 | HG6-64-1 |
| top60_altered_DNA_seed | BRAF | -1.163381 | THCA | 8305C | HG6-64-1 |
| top60_altered_DNA_seed | RET | -1.158994 | THCA | CAL-62 | Motesanib |

| | | | | | |
|---|---|---|---|---|---|
| top60_altered_DNA_seed | BRAF | -1.154696 | THCA | ML-1 | PLX-4720 |
| top60_altered_DNA_seed | NTRK1 | -1.147626 | THCA | RO82-W-1 | AZD1332 |
| top60_altered_DNA_seed | NTRK3 | -1.147626 | THCA | RO82-W-1 | AZD1332 |
| top60_altered_DNA_seed | ATM | -1.08938 | THCA | 8305C | KU-55933 |
| top60_altered_DNA_seed | BRAF | -1.088167 | THCA | 8305C | RAF_9304 |
| top60_altered_DNA_seed | BRAF | -1.031488 | THCA | TT2609-C02 | PLX-4720 |
| top60_altered_DNA_seed | BRAF | -1.023462 | THCA | IHH-4 | HG6-64-1 |
| top60_altered_DNA_seed | NTRK1 | -1.022147 | THCA | CAL-62 | AZD1332 |
| top60_altered_DNA_seed | NTRK3 | -1.022147 | THCA | CAL-62 | AZD1332 |
| top60_altered_DNA_seed | RET | -0.982642 | THCA | BHT-101 | Motesanib |
| top60_altered_DNA_seed | NTRK1 | -0.977355 | THCA | IHH-4 | AZD1332 |
| top60_altered_DNA_seed | NTRK3 | -0.977355 | THCA | IHH-4 | AZD1332 |
| top60_altered_DNA_seed | BRAF | -0.94498 | THCA | CAL-62 | RAF_9304 |
| top60_altered_DNA_seed | ATM | -0.944882 | THCA | HTC-C3 | CP466722 |
| top60_altered_DNA_seed | ATM | -0.941654 | THCA | 8505C | KU-60019 |
| top60_altered_DNA_seed | ATM | -0.892202 | THCA | CGTH-W-1 | torin2 |
| top60_altered_DNA_seed | NTRK1 | -0.891329 | THCA | ML-1 | GW441756 |
| top60_altered_DNA_seed | NTRK1 | -0.874411 | THCA | IHH-4 | GW441756 |
| top60_altered_DNA_seed | BRAF | -0.870308 | THCA | TT2609-C02 | PLX-4720 |
| top60_altered_DNA_seed | ATM | -0.868791 | THCA | ML-1 | KU-55933 |
| top60_altered_DNA_seed | RET | -0.85325 | THCA | RO82-W-1 | Motesanib |
| top60_altered_DNA_seed | BRAF | -0.842455 | THCA | HTC-C3 | HG6-64-1 |
| top60_altered_DNA_seed | ATM | -0.825398 | THCA | KMH-2 | CP466722 |
| top60_altered_DNA_seed | BRAF | -0.813015 | THCA | HTC-C3 | SB590885 |
| top60_altered_DNA_seed | RET | -0.791025 | THCA | 8505C | Motesanib |
| top60_altered_DNA_seed | ATM | -0.773274 | THCA | TT2609-C02 | KU-60019 |
| top60_altered_DNA_seed | BRAF | -0.763833 | THCA | B-CPAP | RAF_9304 |
| top60_altered_DNA_seed | BRAF | -0.758091 | THCA | RO82-W-1 | PLX-4720 |
| top60_altered_DNA_seed | BRAF | -0.71913 | THCA | 8505C | Dabrafenib |
| top60_altered_DNA_seed | ATM | -0.711223 | THCA | ML-1 | CP466722 |
| top60_altered_DNA_seed | ATM | -0.707837 | THCA | K5 | KU-60019 |
| top60_altered_DNA_seed | BRAF | -0.706258 | THCA | CAL-62 | Dabrafenib |
| top60_altered_DNA_seed | BRAF | -0.686113 | THCA | TT2609-C02 | SB590885 |
| top60_altered_DNA_seed | ATM | -0.674507 | THCA | ASH-3 | KU-60019 |
| top60_altered_DNA_seed | ATM | -0.632848 | THCA | IHH-4 | CP466722 |
| top60_altered_DNA_seed | BRAF | -0.606979 | THCA | KMH-2 | RAF_9304 |
| top60_altered_DNA_seed | NTRK1 | -0.59068 | THCA | K5 | AZD1332 |
| top60_altered_DNA_seed | NTRK3 | -0.59068 | THCA | K5 | AZD1332 |
| top60_altered_DNA_seed | BRAF | -0.5883 | THCA | TT | SB590885 |
| top60_altered_DNA_seed | RET | -0.553464 | THCA | FTC-133 | Motesanib |
| top60_altered_DNA_seed | RET | -0.550059 | THCA | B-CPAP | Motesanib |

| | | | | | |
|---|---|---|---|---|---|
| top60_altered_DNA_seed | ATM | -0.545485 | THCA | TT2609-C02 | KU-55933 |
| top60_altered_DNA_seed | BRAF | -0.53173 | THCA | ASH-3 | RAF_9304 |
| top60_altered_DNA_seed | NTRK1 | -0.531148 | THCA | BHT-101 | AZD1332 |
| top60_altered_DNA_seed | NTRK3 | -0.531148 | THCA | BHT-101 | AZD1332 |
| top60_altered_DNA_seed | NTRK1 | -0.518844 | THCA | B-CPAP | AZD1332 |
| top60_altered_DNA_seed | NTRK3 | -0.518844 | THCA | B-CPAP | AZD1332 |
| top60_altered_DNA_seed | ATM | -0.515829 | THCA | TT | KU-60019 |
| top60_altered_DNA_seed | ATM | -0.513165 | THCA | HTC-C3 | KU-60019 |
| top60_altered_DNA_seed | NTRK1 | -0.508802 | THCA | TT | AZD1332 |
| top60_altered_DNA_seed | NTRK3 | -0.508802 | THCA | TT | AZD1332 |
| top60_altered_DNA_seed | BRAF | -0.489515 | THCA | 8505C | SB590885 |
| top60_altered_DNA_seed | ATM | -0.479085 | THCA | BHT-101 | KU-60019 |
| top60_altered_DNA_seed | ATM | -0.452208 | THCA | CGTH-W-1 | KU-55933 |
| top60_altered_DNA_seed | NTRK1 | -0.450033 | THCA | KMH-2 | GW441756 |
| top60_altered_DNA_seed | RET | -0.422726 | THCA | ML-1 | Motesanib |
| top60_altered_DNA_seed | ATM | -0.421715 | THCA | KMH-2 | KU-60019 |
| top60_altered_DNA_seed | NTRK1 | -0.418248 | THCA | HTC-C3 | AZD1332 |
| top60_altered_DNA_seed | NTRK3 | -0.418248 | THCA | HTC-C3 | AZD1332 |
| top60_altered_DNA_seed | RET | -0.413693 | THCA | IHH-4 | Motesanib |
| top60_altered_DNA_seed | BRAF | -0.392081 | THCA | ASH-3 | Dabrafenib |
| top60_altered_DNA_seed | ATM | -0.391261 | THCA | ASH-3 | KU-55933 |
| top60_altered_DNA_seed | NTRK1 | -0.378228 | THCA | TT | GW441756 |
| top60_altered_DNA_seed | ATM | -0.374302 | THCA | BHT-101 | CP466722 |
| top60_altered_DNA_seed | BRAF | -0.370822 | THCA | K5 | HG6-64-1 |
| top60_altered_DNA_seed | BRAF | -0.366295 | THCA | CAL-62 | PLX-4720 |
| top60_altered_DNA_seed | BRAF | -0.324975 | THCA | B-CPAP | SB590885 |
| top60_altered_DNA_seed | BRAF | -0.272961 | THCA | 8505C | PLX-4720 |
| top60_altered_DNA_seed | BRAF | -0.242036 | THCA | CAL-62 | HG6-64-1 |
| top60_altered_DNA_seed | ATM | -0.239267 | THCA | 8505C | KU-55933 |
| top60_altered_DNA_seed | BRAF | -0.23467 | THCA | 8505C | HG6-64-1 |
| top60_altered_DNA_seed | ATM | -0.212464 | THCA | 8505C | torin2 |
| top60_altered_DNA_seed | ATM | -0.201777 | THCA | ASH-3 | CP466722 |
| top60_altered_DNA_seed | BRAF | -0.186451 | THCA | CAL-62 | PLX-4720 |
| top60_altered_DNA_seed | BRAF | -0.164558 | THCA | KMH-2 | PLX-4720 |
| top60_altered_DNA_seed | BRAF | -0.164224 | THCA | CGTH-W-1 | Dabrafenib |
| top60_altered_DNA_seed | ATM | -0.145785 | THCA | FTC-133 | KU-60019 |
| top60_altered_DNA_seed | BRAF | -0.085211 | THCA | RO82-W-1 | HG6-64-1 |
| top60_altered_DNA_seed | BRAF | -0.081779 | THCA | CGTH-W-1 | PLX-4720 |
| top60_altered_DNA_seed | NTRK1 | -0.079601 | THCA | ASH-3 | AZD1332 |
| top60_altered_DNA_seed | NTRK3 | -0.079601 | THCA | ASH-3 | AZD1332 |
| top60_altered_DNA_seed | ATM | -0.054763 | THCA | K5 | KU-55933 |

| | | | | | |
|---|---|---|---|---|---|
| top60_altered_DNA_seed | BRAF | -0.04043 | THCA | ML-1 | Dabrafenib |
| top60_altered_DNA_seed | BRAF | -0.035248 | THCA | KMH-2 | HG6-64-1 |
| top60_altered_DNA_seed | NTRK1 | -0.001777 | THCA | B-CPAP | GW441756 |
| top60_altered_DNA_seed | ATM | -0.001181 | THCA | CAL-62 | KU-60019 |
| top60_altered_DNA_seed | BRAF | 0.001482 | THCA | 8505C | RAF_9304 |
| top60_altered_DNA_seed | ATM | 0.004693 | THCA | HTC-C3 | KU-55933 |
| top60_altered_DNA_seed | NTRK1 | 0.018944 | THCA | 8505C | GW441756 |
| top60_altered_DNA_seed | NTRK1 | 0.034081 | THCA | 8505C | AZD1332 |
| top60_altered_DNA_seed | NTRK3 | 0.034081 | THCA | 8505C | AZD1332 |
| top60_altered_DNA_seed | ATM | 0.039882 | THCA | K5 | CP466722 |
| top60_altered_DNA_seed | ATM | 0.100739 | THCA | RO82-W-1 | KU-60019 |
| top60_altered_DNA_seed | ATM | 0.106576 | THCA | CAL-62 | KU-55933 |
| top60_altered_DNA_seed | RET | 0.108057 | THCA | K5 | Motesanib |
| top60_altered_DNA_seed | BRAF | 0.111801 | THCA | RO82-W-1 | Dabrafenib |
| top60_altered_DNA_seed | BRAF | 0.13551 | THCA | ML-1 | PLX-4720 |
| top60_altered_DNA_seed | NTRK1 | 0.159006 | THCA | CAL-62 | GW441756 |
| top60_altered_DNA_seed | BRAF | 0.161866 | THCA | TT | Dabrafenib |
| top60_altered_DNA_seed | ATM | 0.164551 | THCA | CGTH-W-1 | KU-60019 |
| top60_altered_DNA_seed | RET | 0.203243 | THCA | KMH-2 | Motesanib |
| top60_altered_DNA_seed | RET | 0.241375 | THCA | ASH-3 | Motesanib |
| top60_altered_DNA_seed | BRAF | 0.24194 | THCA | TT2609-C02 | Dabrafenib |
| top60_altered_DNA_seed | BRAF | 0.241945 | THCA | RO82-W-1 | RAF_9304 |
| top60_altered_DNA_seed | BRAF | 0.25163 | THCA | 8505C | PLX-4720 |
| top60_altered_DNA_seed | BRAF | 0.252942 | THCA | TT | RAF_9304 |
| top60_altered_DNA_seed | ATM | 0.26645 | THCA | ML-1 | KU-60019 |
| top60_altered_DNA_seed | NTRK1 | 0.286499 | THCA | 8305C | AZD1332 |
| top60_altered_DNA_seed | NTRK3 | 0.286499 | THCA | 8305C | AZD1332 |
| top60_altered_DNA_seed | BRAF | 0.307688 | THCA | FTC-133 | SB590885 |
| top60_altered_DNA_seed | ATM | 0.317324 | THCA | CAL-62 | CP466722 |
| top60_altered_DNA_seed | BRAF | 0.3238 | THCA | FTC-133 | PLX-4720 |
| top60_altered_DNA_seed | NTRK1 | 0.333757 | THCA | K5 | GW441756 |
| top60_altered_DNA_seed | ATM | 0.338034 | THCA | B-CPAP | CP466722 |
| top60_altered_DNA_seed | ATM | 0.340556 | THCA | CGTH-W-1 | CP466722 |
| top60_altered_DNA_seed | ATM | 0.353416 | THCA | TT | KU-55933 |
| top60_altered_DNA_seed | ATM | 0.377689 | THCA | 8505C | CP466722 |
| top60_altered_DNA_seed | BRAF | 0.380782 | THCA | ASH-3 | SB590885 |
| top60_altered_DNA_seed | BRAF | 0.427087 | THCA | CGTH-W-1 | PLX-4720 |
| top60_altered_DNA_seed | NTRK1 | 0.43456 | THCA | FTC-133 | AZD1332 |
| top60_altered_DNA_seed | NTRK3 | 0.43456 | THCA | FTC-133 | AZD1332 |
| top60_altered_DNA_seed | ATM | 0.448252 | THCA | B-CPAP | KU-55933 |
| top60_altered_DNA_seed | BRAF | 0.47435 | THCA | KMH-2 | PLX-4720 |

| | | | | | |
|---|---|---|---|---|---|
| top60_altered_DNA_seed | BRAF | 0.476929 | THCA | ASH-3 | PLX-4720 |
| top60_altered_DNA_seed | BRAF | 0.478727 | THCA | FTC-133 | Dabrafenib |
| top60_altered_DNA_seed | BRAF | 0.503209 | THCA | FTC-133 | RAF_9304 |
| top60_altered_DNA_seed | BRAF | 0.542429 | THCA | TT2609-C02 | RAF_9304 |
| top60_altered_DNA_seed | BRAF | 0.558517 | THCA | ASH-3 | HG6-64-1 |
| top60_altered_DNA_seed | NTRK1 | 0.560261 | THCA | TT2609-C02 | GW441756 |
| top60_altered_DNA_seed | BRAF | 0.604345 | THCA | BHT-101 | HG6-64-1 |
| top60_altered_DNA_seed | ATM | 0.604481 | THCA | KMH-2 | KU-55933 |
| top60_altered_DNA_seed | BRAF | 0.61027 | THCA | CAL-62 | SB590885 |
| top60_altered_DNA_seed | BRAF | 0.623824 | THCA | RO82-W-1 | PLX-4720 |
| top60_altered_DNA_seed | BRAF | 0.653543 | THCA | CGTH-W-1 | SB590885 |
| top60_altered_DNA_seed | BRAF | 0.670799 | THCA | TT | HG6-64-1 |
| top60_altered_DNA_seed | NTRK1 | 0.810941 | THCA | TT2609-C02 | AZD1332 |
| top60_altered_DNA_seed | NTRK3 | 0.810941 | THCA | TT2609-C02 | AZD1332 |
| top60_altered_DNA_seed | ATM | 0.815754 | THCA | RO82-W-1 | KU-55933 |
| top60_altered_DNA_seed | BRAF | 0.831109 | THCA | ASH-3 | PLX-4720 |
| top60_altered_DNA_seed | RET | 0.846469 | THCA | 8305C | Motesanib |
| top60_altered_DNA_seed | NTRK1 | 0.851058 | THCA | FTC-133 | GW441756 |
| top60_altered_DNA_seed | NTRK1 | 0.87567 | THCA | HTC-C3 | GW441756 |
| top60_altered_DNA_seed | ATM | 0.876813 | THCA | FTC-133 | KU-55933 |
| top60_altered_DNA_seed | BRAF | 0.916491 | THCA | TT | PLX-4720 |
| top60_altered_DNA_seed | ATM | 1.023871 | THCA | 8305C | CP466722 |
| top60_altered_DNA_seed | BRAF | 1.027277 | THCA | RO82-W-1 | SB590885 |
| top60_altered_DNA_seed | NTRK1 | 1.139921 | THCA | 8305C | GW441756 |
| top60_altered_DNA_seed | NTRK1 | 1.29246 | THCA | RO82-W-1 | GW441756 |
| top60_altered_DNA_seed | BRAF | 1.301277 | THCA | FTC-133 | HG6-64-1 |
| top60_altered_DNA_seed | BRAF | 1.355515 | THCA | TT | PLX-4720 |
| top60_altered_DNA_seed | ATM | 2.089043 | THCA | TT2609-C02 | CP466722 |
| top60_altered_DNA_seed | ATM | 2.326309 | THCA | RO82-W-1 | CP466722 |
| top60_altered_DNA_seed | ATM | 2.462246 | THCA | FTC-133 | CP466722 |
| top60_altered_DNA_seed | ATM | 2.506771 | THCA | TT | CP466722 |
| top60_altered_DNA_seed | BRAF | -2.904049 | THCA | K5 | Dabrafenib |
| top60_altered_DNA_seed | BRAF | -2.070305 | THCA | BHT-101 | Dabrafenib |
| top60_altered_DNA_seed | BRAF | -2.008943 | THCA | K5 | PLX-4720 |
| top60_altered_DNA_seed | BRAF | -1.889949 | THCA | HTC-C3 | PLX-4720 |
| top60_altered_DNA_seed | BRAF | -1.699237 | THCA | K5 | SB590885 |
| top60_altered_DNA_seed | BRAF | -1.548644 | THCA | HTC-C3 | Dabrafenib |
| top60_altered_DNA_seed | BRAF | -1.363286 | THCA | BHT-101 | PLX-4720 |
| top60_altered_DNA_seed | BRAF | -1.326271 | THCA | 8305C | Dabrafenib |
| top60_altered_DNA_seed | BRAF | -1.308531 | THCA | IHH-4 | Dabrafenib |
| top60_altered_DNA_seed | BRAF | -1.24812 | THCA | IHH-4 | PLX-4720 |

| | | | | | |
|---|---|---|---|---|---|
| top60_altered_DNA_seed | BRAF | -1.182661 | THCA | B-CPAP | Dabrafenib |
| top60_altered_DNA_seed | RET | -1.146579 | THCA | B-CPAP | Motesanib |
| top60_altered_DNA_seed | BRAF | -1.081835 | THCA | ML-1 | PLX-4720 |
| top60_altered_DNA_seed | BRAF | -0.841682 | THCA | KMH-2 | SB590885 |
| top60_altered_DNA_seed | BRAF | -0.811101 | THCA | BHT-101 | SB590885 |
| top60_altered_DNA_seed | BRAF | -0.779147 | THCA | 8305C | PLX-4720 |
| top60_altered_DNA_seed | EHMT1 | -0.759624 | THCA | K5 | A-366 |
| top60_altered_DNA_seed | EHMT1 | -0.743298 | THCA | CAL-62 | A-366 |
| top60_altered_DNA_seed | BRAF | -0.729044 | THCA | B-CPAP | PLX-4720 |
| top60_altered_DNA_seed | BRAF | -0.727578 | THCA | KMH-2 | Dabrafenib |
| top60_altered_DNA_seed | BRAF | -0.724427 | THCA | KMH-2 | PLX-4720 |
| top60_altered_DNA_seed | BRAF | -0.706696 | THCA | 8505C | Dabrafenib |
| top60_altered_DNA_seed | ATM | -0.677234 | THCA | ASH-3 | KU-55933 |
| top60_altered_DNA_seed | EHMT1 | -0.653728 | THCA | CGTH-W-1 | A-366 |
| top60_altered_DNA_seed | NTRK1 | -0.540702 | THCA | FTC-133 | AZD1332 |
| top60_altered_DNA_seed | NTRK3 | -0.540702 | THCA | FTC-133 | AZD1332 |
| top60_altered_DNA_seed | BRAF | -0.507576 | THCA | ML-1 | Dabrafenib |
| top60_altered_DNA_seed | NTRK1 | -0.501919 | THCA | K5 | AZD1332 |
| top60_altered_DNA_seed | NTRK3 | -0.501919 | THCA | K5 | AZD1332 |
| top60_altered_DNA_seed | NTRK1 | -0.48447 | THCA | B-CPAP | GW441756 |
| top60_altered_DNA_seed | NTRK1 | -0.468359 | THCA | B-CPAP | AZD1332 |
| top60_altered_DNA_seed | NTRK3 | -0.468359 | THCA | B-CPAP | AZD1332 |
| top60_altered_DNA_seed | ATM | -0.461576 | THCA | IHH-4 | KU-55933 |
| top60_altered_DNA_seed | NTRK1 | -0.443973 | THCA | KMH-2 | GW441756 |
| top60_altered_DNA_seed | EHMT1 | -0.400559 | THCA | IHH-4 | A-366 |
| top60_altered_DNA_seed | NTRK1 | -0.388949 | THCA | IHH-4 | GW441756 |
| top60_altered_DNA_seed | NTRK1 | -0.387193 | THCA | CAL-62 | GW441756 |
| top60_altered_DNA_seed | BRAF | -0.380654 | THCA | B-CPAP | SB590885 |
| top60_altered_DNA_seed | BRAF | -0.377407 | THCA | IHH-4 | SB590885 |
| top60_altered_DNA_seed | NTRK1 | -0.362272 | THCA | ASH-3 | AZD1332 |
| top60_altered_DNA_seed | NTRK3 | -0.362272 | THCA | ASH-3 | AZD1332 |
| top60_altered_DNA_seed | NTRK1 | -0.35082 | THCA | FTC-133 | GW441756 |
| top60_altered_DNA_seed | NTRK1 | -0.316476 | THCA | HTC-C3 | GW441756 |
| top60_altered_DNA_seed | BRAF | -0.238721 | THCA | CGTH-W-1 | SB590885 |
| top60_altered_DNA_seed | NTRK1 | -0.237606 | THCA | ASH-3 | GW441756 |
| top60_altered_DNA_seed | BRAF | -0.188294 | THCA | CGTH-W-1 | PLX-4720 |
| top60_altered_DNA_seed | BRAF | -0.168751 | THCA | FTC-133 | Dabrafenib |
| top60_altered_DNA_seed | RET | -0.154526 | THCA | KMH-2 | Motesanib |
| top60_altered_DNA_seed | RET | -0.139767 | THCA | IHH-4 | Motesanib |
| top60_altered_DNA_seed | EHMT1 | -0.107764 | THCA | ASH-3 | A-366 |
| top60_altered_DNA_seed | NTRK1 | -0.056998 | THCA | BHT-101 | AZD1332 |

| | | | | | |
|---|---|---|---|---|---|
| top60_altered_DNA_seed | NTRK3 | -0.056998 | THCA | BHT-101 | AZD1332 |
| top60_altered_DNA_seed | NTRK1 | -0.043358 | THCA | BHT-101 | GW441756 |
| top60_altered_DNA_seed | EHMT1 | -0.026972 | THCA | B-CPAP | A-366 |
| top60_altered_DNA_seed | ATM | 0.082146 | THCA | CGTH-W-1 | KU-55933 |
| top60_altered_DNA_seed | ATM | 0.104186 | THCA | HTC-C3 | KU-55933 |
| top60_altered_DNA_seed | ATM | 0.11089 | THCA | 8305C | KU-55933 |
| top60_altered_DNA_seed | RET | 0.121121 | THCA | CAL-62 | Motesanib |
| top60_altered_DNA_seed | RET | 0.158754 | THCA | ASH-3 | Motesanib |
| top60_altered_DNA_seed | ATM | 0.16506 | THCA | KMH-2 | KU-55933 |
| top60_altered_DNA_seed | EHMT1 | 0.230574 | THCA | 8505C | A-366 |
| top60_altered_DNA_seed | RET | 0.250691 | THCA | FTC-133 | Motesanib |
| top60_altered_DNA_seed | ATM | 0.251063 | THCA | K5 | KU-55933 |
| top60_altered_DNA_seed | RET | 0.25247 | THCA | K5 | Motesanib |
| top60_altered_DNA_seed | ATM | 0.260663 | THCA | BHT-101 | KU-55933 |
| top60_altered_DNA_seed | BRAF | 0.268676 | THCA | 8505C | PLX-4720 |
| top60_altered_DNA_seed | BRAF | 0.269089 | THCA | CAL-62 | SB590885 |
| top60_altered_DNA_seed | NTRK1 | 0.29298 | THCA | CAL-62 | AZD1332 |
| top60_altered_DNA_seed | NTRK3 | 0.29298 | THCA | CAL-62 | AZD1332 |
| top60_altered_DNA_seed | RET | 0.303606 | THCA | BHT-101 | Motesanib |
| top60_altered_DNA_seed | NTRK1 | 0.330976 | THCA | 8505C | GW441756 |
| top60_altered_DNA_seed | NTRK1 | 0.345098 | THCA | HTC-C3 | AZD1332 |
| top60_altered_DNA_seed | NTRK3 | 0.345098 | THCA | HTC-C3 | AZD1332 |
| top60_altered_DNA_seed | EHMT1 | 0.351451 | THCA | BHT-101 | A-366 |
| top60_altered_DNA_seed | RET | 0.368546 | THCA | HTC-C3 | Motesanib |
| top60_altered_DNA_seed | BRAF | 0.372867 | THCA | FTC-133 | PLX-4720 |
| top60_altered_DNA_seed | BRAF | 0.386036 | THCA | ASH-3 | SB590885 |
| top60_altered_DNA_seed | NTRK1 | 0.418265 | THCA | CGTH-W-1 | GW441756 |
| top60_altered_DNA_seed | RET | 0.425459 | THCA | CGTH-W-1 | Motesanib |
| top60_altered_DNA_seed | ATM | 0.494355 | THCA | CAL-62 | KU-55933 |
| top60_altered_DNA_seed | BRAF | 0.543479 | THCA | ASH-3 | Dabrafenib |
| top60_altered_DNA_seed | BRAF | 0.581894 | THCA | FTC-133 | SB590885 |
| top60_altered_DNA_seed | EHMT1 | 0.601777 | THCA | 8305C | A-366 |
| top60_altered_DNA_seed | BRAF | 0.605339 | THCA | 8305C | SB590885 |
| top60_altered_DNA_seed | BRAF | 0.615266 | THCA | HTC-C3 | SB590885 |
| top60_altered_DNA_seed | NTRK1 | 0.693841 | THCA | K5 | GW441756 |
| top60_altered_DNA_seed | ATM | 0.695164 | THCA | FTC-133 | KU-55933 |
| top60_altered_DNA_seed | BRAF | 0.730547 | THCA | CAL-62 | Dabrafenib |
| top60_altered_DNA_seed | RET | 0.739204 | THCA | 8505C | Motesanib |
| top60_altered_DNA_seed | ATM | 0.756658 | THCA | B-CPAP | KU-55933 |
| top60_altered_DNA_seed | EHMT1 | 0.801091 | THCA | HTC-C3 | A-366 |
| top60_altered_DNA_seed | BRAF | 0.844063 | THCA | TT2609-C02 | SB590885 |

| | | | | | |
|---|---|---|---|---|---|
| top60_altered_DNA_seed | BRAF | 0.846021 | THCA | CGTH-W-1 | Dabrafenib |
| top60_altered_DNA_seed | NTRK1 | 0.86336 | THCA | TT2609-C02 | GW441756 |
| top60_altered_DNA_seed | EHMT1 | 0.898935 | THCA | FTC-133 | A-366 |
| top60_altered_DNA_seed | RET | 0.90392 | THCA | 8305C | Motesanib |
| top60_altered_DNA_seed | BRAF | 0.919604 | THCA | 8505C | SB590885 |
| top60_altered_DNA_seed | NTRK1 | 0.920197 | THCA | IHH-4 | AZD1332 |
| top60_altered_DNA_seed | NTRK3 | 0.920197 | THCA | IHH-4 | AZD1332 |
| top60_altered_DNA_seed | BRAF | 0.939749 | THCA | TT2609-C02 | Dabrafenib |
| top60_altered_DNA_seed | BRAF | 1.008652 | THCA | TT | Dabrafenib |
| top60_altered_DNA_seed | BRAF | 1.073604 | THCA | ASH-3 | PLX-4720 |
| top60_altered_DNA_seed | NTRK1 | 1.090459 | THCA | TT | GW441756 |
| top60_altered_DNA_seed | ATM | 1.184934 | THCA | 8505C | KU-55933 |
| top60_altered_DNA_seed | RET | 1.194374 | THCA | ML-1 | Motesanib |
| top60_altered_DNA_seed | BRAF | 1.232725 | THCA | CAL-62 | PLX-4720 |
| top60_altered_DNA_seed | BRAF | 1.327751 | THCA | ML-1 | SB590885 |
| top60_altered_DNA_seed | NTRK1 | 1.359431 | THCA | 8305C | GW441756 |
| top60_altered_DNA_seed | BRAF | 1.50131 | THCA | TT | SB590885 |
| top60_altered_DNA_seed | EHMT1 | 1.504114 | THCA | TT2609-C02 | A-366 |
| top60_altered_DNA_seed | NTRK1 | 1.529692 | THCA | TT2609-C02 | AZD1332 |
| top60_altered_DNA_seed | NTRK3 | 1.529692 | THCA | TT2609-C02 | AZD1332 |
| top60_altered_DNA_seed | ATM | 1.574088 | THCA | ML-1 | KU-55933 |
| top60_altered_DNA_seed | RET | 1.5804 | THCA | TT2609-C02 | Motesanib |
| top60_altered_DNA_seed | NTRK1 | 1.59525 | THCA | 8305C | AZD1332 |
| top60_altered_DNA_seed | NTRK3 | 1.59525 | THCA | 8305C | AZD1332 |
| top60_altered_DNA_seed | BRAF | 1.632561 | THCA | RO82-W-1 | PLX-4720 |
| top60_altered_DNA_seed | NTRK1 | 1.654662 | THCA | 8505C | AZD1332 |
| top60_altered_DNA_seed | NTRK3 | 1.654662 | THCA | 8505C | AZD1332 |
| top60_altered_DNA_seed | ATM | 1.660709 | THCA | TT2609-C02 | KU-55933 |
| top60_altered_DNA_seed | RET | 1.819942 | THCA | TT | Motesanib |
| top60_altered_DNA_seed | ATM | 1.880356 | THCA | RO82-W-1 | KU-55933 |
| top60_altered_DNA_seed | BRAF | 1.917072 | THCA | TT2609-C02 | PLX-4720 |
| top60_altered_DNA_seed | NTRK1 | 1.918867 | THCA | ML-1 | GW441756 |
| top60_altered_DNA_seed | BRAF | 2.011685 | THCA | TT | PLX-4720 |
| top60_altered_DNA_seed | BRAF | 2.030313 | THCA | RO82-W-1 | Dabrafenib |
| top60_altered_DNA_seed | EHMT1 | 2.122321 | THCA | TT | A-366 |
| top60_altered_DNA_seed | ATM | 2.159676 | THCA | TT | KU-55933 |
| top60_altered_DNA_seed | BRAF | 2.359152 | THCA | RO82-W-1 | SB590885 |
| top60_altered_DNA_seed | NTRK1 | 2.516265 | THCA | RO82-W-1 | AZD1332 |
| top60_altered_DNA_seed | NTRK3 | 2.516265 | THCA | RO82-W-1 | AZD1332 |
| top60_altered_DNA_seed | RET | 2.530202 | THCA | RO82-W-1 | Motesanib |
| top60_altered_DNA_seed | NTRK1 | 2.881846 | THCA | RO82-W-1 | GW441756 |

| | | | | | |
|---|---|---|---|---|---|
| top60_altered_DNA_seed | EHMT1 | 3.201287 | THCA | RO82-W-1 | A-366 |
| top60_altered_DNA_seed | PIK3CA | -1.502812 | UCEC | MFE-280 | PI3Ka_4409 |
| top60_altered_DNA_seed | PIK3CA | -1.306172 | UCEC | SNG-M | AZD8835 |
| top60_altered_DNA_seed | PIK3CA | -1.182536 | UCEC | MFE-280 | AZD8835 |
| top60_altered_DNA_seed | ATM | -1.138256 | UCEC | MFE-319 | KU-55933 |
| top60_altered_DNA_seed | PIK3CA | -1.065015 | UCEC | SNG-M | PI3Ka_4409 |
| top60_altered_DNA_seed | ATM | -0.98435 | UCEC | SNG-M | CP466722 |
| top60_altered_DNA_seed | ATM | -0.967779 | UCEC | MFE-319 | KU-60019 |
| top60_altered_DNA_seed | ATM | -0.941115 | UCEC | MFE-319 | CP466722 |
| top60_altered_DNA_seed | ATM | -0.757481 | UCEC | HEC-1 | KU-60019 |
| top60_altered_DNA_seed | PIK3CA | -0.695396 | UCEC | AN3-CA | AZD8835 |
| top60_altered_DNA_seed | ATM | -0.608482 | UCEC | COLO-684 | KU-55933 |
| top60_altered_DNA_seed | ATM | -0.589341 | UCEC | MFE-296 | KU-60019 |
| top60_altered_DNA_seed | ATM | -0.540231 | UCEC | AN3-CA | KU-55933 |
| top60_altered_DNA_seed | ATM | -0.532342 | UCEC | COLO-684 | CP466722 |
| top60_altered_DNA_seed | ATM | -0.458963 | UCEC | AN3-CA | KU-60019 |
| top60_altered_DNA_seed | PIK3CA | -0.392543 | UCEC | AN3-CA | PI3Ka_4409 |
| top60_altered_DNA_seed | ATM | -0.244126 | UCEC | HEC-1 | KU-55933 |
| top60_altered_DNA_seed | PIK3CA | -0.137169 | UCEC | COLO-684 | AZD8835 |
| top60_altered_DNA_seed | ATM | -0.054838 | UCEC | KLE | CP466722 |
| top60_altered_DNA_seed | ATM | -0.049526 | UCEC | COLO-684 | KU-60019 |
| top60_altered_DNA_seed | PIK3CA | -0.004274 | UCEC | KLE | PI3Ka_4409 |
| top60_altered_DNA_seed | PIK3CA | 0.002161 | UCEC | MFE-296 | AZD8835 |
| top60_altered_DNA_seed | PIK3CA | 0.007765 | UCEC | HEC-1 | PI3Ka_4409 |
| top60_altered_DNA_seed | ATM | 0.113108 | UCEC | MFE-296 | KU-55933 |
| top60_altered_DNA_seed | PIK3CA | 0.154596 | UCEC | HEC-1 | AZD8835 |
| top60_altered_DNA_seed | ATM | 0.192287 | UCEC | EN | CP466722 |
| top60_altered_DNA_seed | ATM | 0.291618 | UCEC | AN3-CA | CP466722 |
| top60_altered_DNA_seed | ATM | 0.330904 | UCEC | MFE-280 | CP466722 |
| top60_altered_DNA_seed | ATM | 0.340159 | UCEC | MFE-296 | CP466722 |
| top60_altered_DNA_seed | PIK3CA | 0.352447 | UCEC | COLO-684 | PI3Ka_4409 |
| top60_altered_DNA_seed | ATM | 0.375809 | UCEC | SNG-M | KU-60019 |
| top60_altered_DNA_seed | PIK3CA | 0.384848 | UCEC | MFE-296 | PI3Ka_4409 |
| top60_altered_DNA_seed | ATM | 0.458843 | UCEC | SNG-M | KU-55933 |
| top60_altered_DNA_seed | ATM | 0.493472 | UCEC | MFE-280 | KU-55933 |
| top60_altered_DNA_seed | PIK3CA | 0.563782 | UCEC | MFE-319 | AZD8835 |
| top60_altered_DNA_seed | ATM | 0.599189 | UCEC | KLE | KU-60019 |
| top60_altered_DNA_seed | ATM | 0.765162 | UCEC | EN | KU-55933 |
| top60_altered_DNA_seed | ATM | 0.811403 | UCEC | EN | KU-60019 |
| top60_altered_DNA_seed | ATM | 0.972207 | UCEC | KLE | KU-55933 |
| top60_altered_DNA_seed | PIK3CA | 1.18858 | UCEC | MFE-319 | PI3Ka_4409 |

| | | | | | |
|---|---|---|---|---|---|
| top60_altered_DNA_seed | PIK3CA | 1.254999 | UCEC | KLE | AZD8835 |
| top60_altered_DNA_seed | ATM | 1.411894 | UCEC | MFE-280 | KU-60019 |
| top60_altered_DNA_seed | ATM | 1.419228 | UCEC | HEC-1 | CP466722 |
| top60_altered_DNA_seed | PIK3CA | 1.727071 | UCEC | EN | PI3Ka_4409 |
| top60_altered_DNA_seed | PIK3CA | -1.433814 | UCEC | MFE-280 | Alpelisib |
| top60_altered_DNA_seed | TAF1 | -1.274367 | UCEC | MFE-319 | TAF1_5496 |
| top60_altered_DNA_seed | PIK3CA | -1.129591 | UCEC | AN3-CA | Alpelisib |
| top60_altered_DNA_seed | PIK3CA | -1.128491 | UCEC | KLE | AZD8186 |
| top60_altered_DNA_seed | KRAS | -1.093533 | UCEC | SNG-M | KRAS (G12C) Inhibitor-12 |
| top60_altered_DNA_seed | TAF1 | -1.08519 | UCEC | HEC-1 | TAF1_5496 |
| top60_altered_DNA_seed | ATM | -1.082897 | UCEC | MFE-319 | KU-55933 |
| top60_altered_DNA_seed | PIK3CA | -1.05072 | UCEC | AN3-CA | GNE-317 |
| top60_altered_DNA_seed | PIK3CA | -0.986437 | UCEC | AN3-CA | AZD8186 |
| top60_altered_DNA_seed | ATM | -0.980089 | UCEC | AN3-CA | KU-55933 |
| top60_altered_DNA_seed | PIK3CA | -0.799668 | UCEC | AN3-CA | Buparlisib |
| top60_altered_DNA_seed | PIK3CA | -0.776155 | UCEC | MFE-319 | GNE-317 |
| top60_altered_DNA_seed | TAF1 | -0.711839 | UCEC | MFE-296 | TAF1_5496 |
| top60_altered_DNA_seed | PIK3CA | -0.682345 | UCEC | COLO-684 | GNE-317 |
| top60_altered_DNA_seed | ATM | -0.677619 | UCEC | MFE-280 | KU-55933 |
| top60_altered_DNA_seed | TP53 | -0.609038 | UCEC | AN3-CA | MIRA-1 |
| top60_altered_DNA_seed | PIK3CA | -0.553514 | UCEC | COLO-684 | Alpelisib |
| top60_altered_DNA_seed | PIK3CA | -0.402583 | UCEC | COLO-684 | AZD8186 |
| top60_altered_DNA_seed | KRAS | -0.368617 | UCEC | HEC-1 | KRAS (G12C) Inhibitor-12 |
| top60_altered_DNA_seed | TAF1 | -0.337792 | UCEC | MFE-280 | TAF1_5496 |
| top60_altered_DNA_seed | PIK3CA | -0.300771 | UCEC | KLE | GNE-317 |
| top60_altered_DNA_seed | PIK3CA | -0.277586 | UCEC | EN | Alpelisib |
| top60_altered_DNA_seed | KRAS | -0.264657 | UCEC | EN | KRAS (G12C) Inhibitor-12 |
| top60_altered_DNA_seed | PIK3CA | -0.22301 | UCEC | MFE-280 | GNE-317 |
| top60_altered_DNA_seed | KRAS | -0.204616 | UCEC | COLO-684 | KRAS (G12C) Inhibitor-12 |
| top60_altered_DNA_seed | ATM | -0.154584 | UCEC | MFE-296 | KU-55933 |
| top60_altered_DNA_seed | ATM | -0.127018 | UCEC | SNG-M | KU-55933 |
| top60_altered_DNA_seed | PIK3CA | -0.117276 | UCEC | MFE-319 | Buparlisib |
| top60_altered_DNA_seed | PIK3CA | -0.092167 | UCEC | MFE-280 | AZD8186 |
| top60_altered_DNA_seed | PIK3CA | -0.035895 | UCEC | COLO-684 | Buparlisib |
| top60_altered_DNA_seed | KRAS | -0.031796 | UCEC | AN3-CA | KRAS (G12C) Inhibitor-12 |
| top60_altered_DNA_seed | PIK3CA | 0.004545 | UCEC | EN | Buparlisib |
| top60_altered_DNA_seed | PIK3CA | 0.121978 | UCEC | KLE | Buparlisib |
| top60_altered_DNA_seed | TP53 | 0.125506 | UCEC | SNG-M | MIRA-1 |
| top60_altered_DNA_seed | PIK3CA | 0.139196 | UCEC | EN | GNE-317 |
| top60_altered_DNA_seed | TP53 | 0.157663 | UCEC | MFE-296 | MIRA-1 |
| top60_altered_DNA_seed | TAF1 | 0.166665 | UCEC | COLO-684 | TAF1_5496 |

| | | | | | |
|---|---|---|---|---|---|
| top60_altered_DNA_seed | TP53 | 0.169217 | UCEC | COLO-684 | MIRA-1 |
| top60_altered_DNA_seed | TAF1 | 0.193506 | UCEC | AN3-CA | TAF1_5496 |
| top60_altered_DNA_seed | PIK3CA | 0.230603 | UCEC | SNG-M | Alpelisib |
| top60_altered_DNA_seed | KRAS | 0.232402 | UCEC | MFE-319 | KRAS (G12C) Inhibitor-12 |
| top60_altered_DNA_seed | TAF1 | 0.258555 | UCEC | SNG-M | TAF1_5496 |
| top60_altered_DNA_seed | PIK3CA | 0.320685 | UCEC | MFE-319 | AZD8186 |
| top60_altered_DNA_seed | ATM | 0.35817 | UCEC | COLO-684 | KU-55933 |
| top60_altered_DNA_seed | PIK3CA | 0.376584 | UCEC | SNG-M | GNE-317 |
| top60_altered_DNA_seed | PIK3CA | 0.414572 | UCEC | SNG-M | Buparlisib |
| top60_altered_DNA_seed | KRAS | 0.426664 | UCEC | MFE-280 | KRAS (G12C) Inhibitor-12 |
| top60_altered_DNA_seed | TAF1 | 0.442137 | UCEC | EN | TAF1_5496 |
| top60_altered_DNA_seed | PIK3CA | 0.485906 | UCEC | HEC-1 | GNE-317 |
| top60_altered_DNA_seed | PIK3CA | 0.543186 | UCEC | MFE-280 | Buparlisib |
| top60_altered_DNA_seed | TP53 | 0.56048 | UCEC | MFE-319 | MIRA-1 |
| top60_altered_DNA_seed | ATM | 0.65422 | UCEC | EN | KU-55933 |
| top60_altered_DNA_seed | TP53 | 0.730928 | UCEC | EN | MIRA-1 |
| top60_altered_DNA_seed | PIK3CA | 0.816956 | UCEC | MFE-296 | Buparlisib |
| top60_altered_DNA_seed | KRAS | 0.824168 | UCEC | KLE | KRAS (G12C) Inhibitor-12 |
| top60_altered_DNA_seed | TP53 | 0.860405 | UCEC | MFE-280 | MIRA-1 |
| top60_altered_DNA_seed | ATM | 0.891619 | UCEC | HEC-1 | KU-55933 |
| top60_altered_DNA_seed | KRAS | 0.892833 | UCEC | MFE-296 | KRAS (G12C) Inhibitor-12 |
| top60_altered_DNA_seed | PIK3CA | 0.916666 | UCEC | HEC-1 | Buparlisib |
| top60_altered_DNA_seed | PIK3CA | 0.94508 | UCEC | EN | AZD8186 |
| top60_altered_DNA_seed | PIK3CA | 0.953357 | UCEC | MFE-319 | Alpelisib |
| top60_altered_DNA_seed | PIK3CA | 1.02174 | UCEC | HEC-1 | AZD8186 |
| top60_altered_DNA_seed | PIK3CA | 1.043925 | UCEC | SNG-M | AZD8186 |
| top60_altered_DNA_seed | TP53 | 1.069404 | UCEC | HEC-1 | MIRA-1 |
| top60_altered_DNA_seed | PIK3CA | 1.070358 | UCEC | MFE-296 | Alpelisib |
| top60_altered_DNA_seed | PIK3CA | 1.09639 | UCEC | MFE-296 | AZD8186 |
| top60_altered_DNA_seed | TAF1 | 1.123393 | UCEC | KLE | TAF1_5496 |
| top60_altered_DNA_seed | PIK3CA | 1.200719 | UCEC | HEC-1 | Alpelisib |
| top60_altered_DNA_seed | ATM | 1.460351 | UCEC | KLE | KU-55933 |
| top60_altered_DNA_seed | TP53 | 1.5776 | UCEC | KLE | MIRA-1 |
| top60_altered_DNA_seed | PIK3CA | 1.64178 | UCEC | KLE | Alpelisib |
| top60_altered_DNA_seed | PIK3CA | 1.651226 | UCEC | MFE-296 | GNE-317 |
| top60_altered_DNA_seed | BRAF | -5.08881 | SKCM | G-MEL | SB590885 |
| top60_altered_DNA_seed | BRAF | -5.015441 | SKCM | IGR-37 | SB590885 |
| top60_altered_DNA_seed | BRAF | -4.956381 | SKCM | MEL-HO | SB590885 |
| top60_altered_DNA_seed | BRAF | -4.687931 | SKCM | 451Lu | SB590885 |
| top60_altered_DNA_seed | BRAF | -4.663005 | SKCM | IGR-37 | PLX-4720 |
| top60_altered_DNA_seed | BRAF | -4.608082 | SKCM | C32 | SB590885 |

| | | | | | |
|---|---|---|---|---|---|
| top60_altered_DNA_seed | BRAF | -4.435693 | SKCM | M14 | SB590885 |
| top60_altered_DNA_seed | BRAF | -4.213756 | SKCM | G-MEL | PLX-4720 |
| top60_altered_DNA_seed | BRAF | -4.152893 | SKCM | 451Lu | PLX-4720 |
| top60_altered_DNA_seed | BRAF | -4.105172 | SKCM | C32 | PLX-4720 |
| top60_altered_DNA_seed | BRAF | -4.095785 | SKCM | MMAc-SF | PLX-4720 |
| top60_altered_DNA_seed | BRAF | -4.084635 | SKCM | MMAc-SF | Dabrafenib |
| top60_altered_DNA_seed | BRAF | -4.076641 | SKCM | SH-4 | PLX-4720 |
| top60_altered_DNA_seed | BRAF | -4.045547 | SKCM | G-MEL | Dabrafenib |
| top60_altered_DNA_seed | BRAF | -4.036444 | SKCM | IGR-37 | PLX-4720 |
| top60_altered_DNA_seed | BRAF | -3.995559 | SKCM | COLO-679 | SB590885 |
| top60_altered_DNA_seed | BRAF | -3.991431 | SKCM | IGR-37 | Dabrafenib |
| top60_altered_DNA_seed | BRAF | -3.918523 | SKCM | A375 | Dabrafenib |
| top60_altered_DNA_seed | BRAF | -3.918006 | SKCM | C32 | Dabrafenib |
| top60_altered_DNA_seed | BRAF | -3.849316 | SKCM | CP50-MEL-B | PLX-4720 |
| top60_altered_DNA_seed | BRAF | -3.818179 | SKCM | M14 | PLX-4720 |
| top60_altered_DNA_seed | BRAF | -3.81523 | SKCM | SH-4 | Dabrafenib |
| top60_altered_DNA_seed | BRAF | -3.800046 | SKCM | G-MEL | PLX-4720 |
| top60_altered_DNA_seed | BRAF | -3.73954 | SKCM | MEL-HO | Dabrafenib |
| top60_altered_DNA_seed | BRAF | -3.737306 | SKCM | COLO-679 | PLX-4720 |
| top60_altered_DNA_seed | BRAF | -3.728287 | SKCM | A375 | PLX-4720 |
| top60_altered_DNA_seed | BRAF | -3.70433 | SKCM | MEL-HO | PLX-4720 |
| top60_altered_DNA_seed | BRAF | -3.703903 | SKCM | WM35 | PLX-4720 |
| top60_altered_DNA_seed | BRAF | -3.69593 | SKCM | UACC-62 | PLX-4720 |
| top60_altered_DNA_seed | BRAF | -3.677782 | SKCM | A375 | SB590885 |
| top60_altered_DNA_seed | BRAF | -3.661088 | SKCM | A375 | PLX-4720 |
| top60_altered_DNA_seed | BRAF | -3.642713 | SKCM | MEL-HO | PLX-4720 |
| top60_altered_DNA_seed | BRAF | -3.633844 | SKCM | M14 | Dabrafenib |
| top60_altered_DNA_seed | BRAF | -3.630199 | SKCM | HT-144 | SB590885 |
| top60_altered_DNA_seed | BRAF | -3.621954 | SKCM | CP50-MEL-B | Dabrafenib |
| top60_altered_DNA_seed | BRAF | -3.59225 | SKCM | WM35 | Dabrafenib |
| top60_altered_DNA_seed | BRAF | -3.579647 | SKCM | IGR-37 | RAF_9304 |
| top60_altered_DNA_seed | BRAF | -3.554161 | SKCM | RVH-421 | SB590885 |
| top60_altered_DNA_seed | BRAF | -3.540122 | SKCM | A101D | PLX-4720 |
| top60_altered_DNA_seed | BRAF | -3.498877 | SKCM | 451Lu | Dabrafenib |
| top60_altered_DNA_seed | BRAF | -3.491049 | SKCM | M14 | PLX-4720 |
| top60_altered_DNA_seed | BRAF | -3.453765 | SKCM | UACC-62 | Dabrafenib |
| top60_altered_DNA_seed | BRAF | -3.406296 | SKCM | SH-4 | PLX-4720 |
| top60_altered_DNA_seed | BRAF | -3.399603 | SKCM | UACC-257 | Dabrafenib |
| top60_altered_DNA_seed | BRAF | -3.353063 | SKCM | K2 | PLX-4720 |
| top60_altered_DNA_seed | BRAF | -3.349799 | SKCM | G-MEL | RAF_9304 |
| top60_altered_DNA_seed | BRAF | -3.34063 | SKCM | LB2518-MEL | AZ628 |

| | | | | | |
|---|---|---|---|---|---|
| top60_altered_DNA_seed | BRAF | -3.322197 | SKCM | K2 | Dabrafenib |
| top60_altered_DNA_seed | BRAF | -3.318276 | SKCM | G-361 | Dabrafenib |
| top60_altered_DNA_seed | BRAF | -3.31556 | SKCM | SK-MEL-28 | Dabrafenib |
| top60_altered_DNA_seed | BRAF | -3.303363 | SKCM | COLO-829 | SB590885 |
| top60_altered_DNA_seed | BRAF | -3.299736 | SKCM | COLO-679 | Dabrafenib |
| top60_altered_DNA_seed | BRAF | -3.295407 | SKCM | C32 | PLX-4720 |
| top60_altered_DNA_seed | BRAF | -3.28739 | SKCM | SK-MEL-1 | Dabrafenib |
| top60_altered_DNA_seed | BRAF | -3.203638 | SKCM | LOXIMVI | Dabrafenib |
| top60_altered_DNA_seed | BRAF | -3.203336 | SKCM | MMAc-SF | PLX-4720 |
| top60_altered_DNA_seed | BRAF | -3.193343 | SKCM | A101D | SB590885 |
| top60_altered_DNA_seed | BRAF | -3.192999 | SKCM | CP50-MEL-B | PLX-4720 |
| top60_altered_DNA_seed | BRAF | -3.186265 | SKCM | WM-115 | SB590885 |
| top60_altered_DNA_seed | BRAF | -3.174985 | SKCM | MZ2-MEL | PLX-4720 |
| top60_altered_DNA_seed | BRAF | -3.168905 | SKCM | SH-4 | AZ628 |
| top60_altered_DNA_seed | BRAF | -3.150297 | SKCM | UACC-257 | PLX-4720 |
| top60_altered_DNA_seed | BRAF | -3.112875 | SKCM | SK-MEL-28 | PLX-4720 |
| top60_altered_DNA_seed | BRAF | -3.095024 | SKCM | SK-MEL-1 | PLX-4720 |
| top60_altered_DNA_seed | BRAF | -3.083214 | SKCM | A101D | Dabrafenib |
| top60_altered_DNA_seed | BRAF | -3.079853 | SKCM | HT-144 | AZ628 |
| top60_altered_DNA_seed | BRAF | -3.067988 | SKCM | SH-4 | SB590885 |
| top60_altered_DNA_seed | BRAF | -3.059115 | SKCM | WM35 | RAF_9304 |
| top60_altered_DNA_seed | BRAF | -3.057456 | SKCM | HT-144 | PLX-4720 |
| top60_altered_DNA_seed | BRAF | -3.057264 | SKCM | RVH-421 | PLX-4720 |
| top60_altered_DNA_seed | BRAF | -3.024398 | SKCM | HMV-II | RAF_9304 |
| top60_altered_DNA_seed | BRAF | -3.018112 | SKCM | WM-115 | PLX-4720 |
| top60_altered_DNA_seed | BRAF | -3.015636 | SKCM | MEL-HO | RAF_9304 |
| top60_altered_DNA_seed | BRAF | -2.986544 | SKCM | SK-MEL-24 | Dabrafenib |
| top60_altered_DNA_seed | BRAF | -2.893544 | SKCM | SK-MEL-28 | PLX-4720 |
| top60_altered_DNA_seed | BRAF | -2.883344 | SKCM | UACC-257 | SB590885 |
| top60_altered_DNA_seed | BRAF | -2.869586 | SKCM | MZ7-mel | AZ628 |
| top60_altered_DNA_seed | BRAF | -2.863258 | SKCM | CP50-MEL-B | SB590885 |
| top60_altered_DNA_seed | BRAF | -2.857884 | SKCM | HT-144 | Dabrafenib |
| top60_altered_DNA_seed | BRAF | -2.853404 | SKCM | 451Lu | PLX-4720 |
| top60_altered_DNA_seed | BRAF | -2.849543 | SKCM | SK-MEL-3 | Dabrafenib |
| top60_altered_DNA_seed | BRAF | -2.834965 | SKCM | COLO-679 | PLX-4720 |
| top60_altered_DNA_seed | BRAF | -2.827968 | SKCM | COLO-783 | Dabrafenib |
| top60_altered_DNA_seed | BRAF | -2.823639 | SKCM | MZ7-mel | Dabrafenib |
| top60_altered_DNA_seed | BRAF | -2.822696 | SKCM | COLO-679 | RAF_9304 |
| top60_altered_DNA_seed | BRAF | -2.813936 | SKCM | COLO-829 | Dabrafenib |
| top60_altered_DNA_seed | BRAF | -2.716985 | SKCM | A101D | AZ628 |
| top60_altered_DNA_seed | BRAF | -2.621081 | SKCM | COLO-829 | AZ628 |

| | | | | | |
|---|---|---|---|---|---|
| top60_altered_DNA_seed | BRAF | -2.587736 | SKCM | UACC-257 | PLX-4720 |
| top60_altered_DNA_seed | BRAF | -2.56709 | SKCM | HT-144 | PLX-4720 |
| top60_altered_DNA_seed | BRAF | -2.526483 | SKCM | SK-MEL-1 | SB590885 |
| top60_altered_DNA_seed | BRAF | -2.517707 | SKCM | SH-4 | RAF_9304 |
| top60_altered_DNA_seed | BRAF | -2.51352 | SKCM | SK-MEL-2 | SB590885 |
| top60_altered_DNA_seed | BRAF | -2.505972 | SKCM | MZ7-mel | RAF_9304 |
| top60_altered_DNA_seed | BRAF | -2.495594 | SKCM | A101D | PLX-4720 |
| top60_altered_DNA_seed | BRAF | -2.47576 | SKCM | WM278 | PLX-4720 |
| top60_altered_DNA_seed | BRAF | -2.470813 | SKCM | UACC-257 | AZ628 |
| top60_altered_DNA_seed | BRAF | -2.437724 | SKCM | SK-MEL-5 | SB590885 |
| top60_altered_DNA_seed | BRAF | -2.43011 | SKCM | C32 | RAF_9304 |
| top60_altered_DNA_seed | BRAF | -2.428363 | SKCM | M14 | RAF_9304 |
| top60_altered_DNA_seed | BRAF | -2.415898 | SKCM | LOXIMVI | PLX-4720 |
| top60_altered_DNA_seed | BRAF | -2.403022 | SKCM | CP50-MEL-B | RAF_9304 |
| top60_altered_DNA_seed | BRAF | -2.398451 | SKCM | SK-MEL-5 | Dabrafenib |
| top60_altered_DNA_seed | BRAF | -2.396276 | SKCM | SK-MEL-5 | PLX-4720 |
| top60_altered_DNA_seed | BRAF | -2.395091 | SKCM | G-361 | PLX-4720 |
| top60_altered_DNA_seed | BRAF | -2.387781 | SKCM | MMAc-SF | AZ628 |
| top60_altered_DNA_seed | BRAF | -2.38227 | SKCM | SK-MEL-24 | PLX-4720 |
| top60_altered_DNA_seed | BRAF | -2.369968 | SKCM | MMAc-SF | RAF_9304 |
| top60_altered_DNA_seed | BRAF | -2.338542 | SKCM | MZ7-mel | PLX-4720 |
| top60_altered_DNA_seed | BRAF | -2.300492 | SKCM | A375 | RAF_9304 |
| top60_altered_DNA_seed | BRAF | -2.282492 | SKCM | RVH-421 | PLX-4720 |
| top60_altered_DNA_seed | BRAF | -2.276926 | SKCM | Hs-939-T | PLX-4720 |
| top60_altered_DNA_seed | BRAF | -2.264027 | SKCM | SK-MEL-3 | RAF_9304 |
| top60_altered_DNA_seed | BRAF | -2.248342 | SKCM | LOXIMVI | RAF_9304 |
| top60_altered_DNA_seed | BRAF | -2.241474 | SKCM | UACC-62 | RAF_9304 |
| top60_altered_DNA_seed | BRAF | -2.238509 | SKCM | SK-MEL-5 | PLX-4720 |
| top60_altered_DNA_seed | BRAF | -2.222172 | SKCM | COLO-783 | SB590885 |
| top60_altered_DNA_seed | BRAF | -2.217851 | SKCM | 451Lu | RAF_9304 |
| top60_altered_DNA_seed | BRAF | -2.201928 | SKCM | COLO-783 | PLX-4720 |
| top60_altered_DNA_seed | BRAF | -2.195339 | SKCM | COLO-800 | PLX-4720 |
| top60_altered_DNA_seed | BRAF | -2.181887 | SKCM | SK-MEL-3 | PLX-4720 |
| top60_altered_DNA_seed | BRAF | -2.174501 | SKCM | SK-MEL-3 | SB590885 |
| top60_altered_DNA_seed | BRAF | -2.148002 | SKCM | COLO-829 | PLX-4720 |
| top60_altered_DNA_seed | BRAF | -2.138799 | SKCM | LOXIMVI | SB590885 |
| top60_altered_DNA_seed | BRAF | -2.13503 | SKCM | K2 | RAF_9304 |
| top60_altered_DNA_seed | BRAF | -2.132021 | SKCM | WM35 | SB590885 |
| top60_altered_DNA_seed | BRAF | -2.120201 | SKCM | LOXIMVI | PLX-4720 |
| top60_altered_DNA_seed | BRAF | -2.107238 | SKCM | RVH-421 | Dabrafenib |
| top60_altered_DNA_seed | BRAF | -2.053827 | SKCM | WM278 | Dabrafenib |

| | | | | | |
|---|---|---|---|---|---|
| top60_altered_DNA_seed | BRAF | -2.041871 | SKCM | IST-MEL1 | Dabrafenib |
| top60_altered_DNA_seed | BRAF | -2.036958 | SKCM | G-361 | PLX-4720 |
| top60_altered_DNA_seed | BRAF | -2.02736 | SKCM | Hs-939-T | PLX-4720 |
| top60_altered_DNA_seed | BRAF | -2.002465 | SKCM | CP66-MEL | AZ628 |
| top60_altered_DNA_seed | BRAF | -1.983639 | SKCM | HT-144 | RAF_9304 |
| top60_altered_DNA_seed | BRAF | -1.959714 | SKCM | K2 | SB590885 |
| top60_altered_DNA_seed | BRAF | -1.891874 | SKCM | Hs-939-T | SB590885 |
| top60_altered_DNA_seed | BRAF | -1.866642 | SKCM | SK-MEL-24 | RAF_9304 |
| top60_altered_DNA_seed | BRAF | -1.803864 | SKCM | SK-MEL-1 | PLX-4720 |
| top60_altered_DNA_seed | BRAF | -1.758344 | SKCM | COLO-829 | PLX-4720 |
| top60_altered_DNA_seed | BRAF | -1.717599 | SKCM | MZ7-mel | SB590885 |
| top60_altered_DNA_seed | BRAF | -1.70635 | SKCM | IPC-298 | RAF_9304 |
| top60_altered_DNA_seed | BRAF | -1.702459 | SKCM | CHL-1 | Dabrafenib |
| top60_altered_DNA_seed | BRAF | -1.689875 | SKCM | SK-MEL-3 | PLX-4720 |
| top60_altered_DNA_seed | BRAF | -1.662605 | SKCM | UACC-257 | RAF_9304 |
| top60_altered_DNA_seed | BRAF | -1.656931 | SKCM | WM278 | HG6-64-1 |
| top60_altered_DNA_seed | BRAF | -1.651406 | SKCM | CP66-MEL | PLX-4720 |
| top60_altered_DNA_seed | BRAF | -1.602716 | SKCM | SK-MEL-30 | RAF_9304 |
| top60_altered_DNA_seed | BRAF | -1.60183 | SKCM | K2 | PLX-4720 |
| top60_altered_DNA_seed | BRAF | -1.599522 | SKCM | WM1552C | PLX-4720 |
| top60_altered_DNA_seed | BRAF | -1.594122 | SKCM | WM278 | RAF_9304 |
| top60_altered_DNA_seed | BRAF | -1.574465 | SKCM | SK-MEL-28 | RAF_9304 |
| top60_altered_DNA_seed | BRAF | -1.562431 | SKCM | GAK | RAF_9304 |
| top60_altered_DNA_seed | BRAF | -1.553998 | SKCM | G-361 | SB590885 |
| top60_altered_DNA_seed | BRAF | -1.526641 | SKCM | MZ2-MEL | PLX-4720 |
| top60_altered_DNA_seed | BRAF | -1.488305 | SKCM | WM278 | PLX-4720 |
| top60_altered_DNA_seed | BRAF | -1.478675 | SKCM | COLO-829 | HG6-64-1 |
| top60_altered_DNA_seed | BRAF | -1.47651 | SKCM | SK-MEL-5 | RAF_9304 |
| top60_altered_DNA_seed | BRAF | -1.475124 | SKCM | CHL-1 | HG6-64-1 |
| top60_altered_DNA_seed | BRAF | -1.455948 | SKCM | CHL-1 | PLX-4720 |
| top60_altered_DNA_seed | BRAF | -1.449136 | SKCM | WM793B | SB590885 |
| top60_altered_DNA_seed | BRAF | -1.444524 | SKCM | COLO-783 | PLX-4720 |
| top60_altered_DNA_seed | BRAF | -1.396823 | SKCM | MZ2-MEL | AZ628 |
| top60_altered_DNA_seed | BRAF | -1.385225 | SKCM | IST-MEL1 | PLX-4720 |
| top60_altered_DNA_seed | BRAF | -1.330528 | SKCM | Hs-940-T | PLX-4720 |
| top60_altered_DNA_seed | BRAF | -1.330121 | SKCM | IST-MEL1 | AZ628 |
| top60_altered_DNA_seed | BRAF | -1.321489 | SKCM | Hs-939-T | RAF_9304 |
| top60_altered_DNA_seed | BRAF | -1.315079 | SKCM | MZ7-mel | PLX-4720 |
| top60_altered_DNA_seed | BRAF | -1.31419 | SKCM | SK-MEL-1 | AZ628 |
| top60_altered_DNA_seed | BRAF | -1.282679 | SKCM | Hs-940-T | PLX-4720 |
| top60_altered_DNA_seed | BRAF | -1.276418 | SKCM | G-361 | RAF_9304 |

| | | | | | |
|---|---|---|---|---|---|
| top60_altered_DNA_seed | BRAF | -1.272528 | SKCM | HT-144 | HG6-64-1 |
| top60_altered_DNA_seed | BRAF | -1.255107 | SKCM | WM793B | HG6-64-1 |
| top60_altered_DNA_seed | BRAF | -1.249927 | SKCM | SK-MEL-2 | RAF_9304 |
| top60_altered_DNA_seed | BRAF | -1.239444 | SKCM | IGR-1 | SB590885 |
| top60_altered_DNA_seed | BRAF | -1.198875 | SKCM | SK-MEL-24 | PLX-4720 |
| top60_altered_DNA_seed | BRAF | -1.180196 | SKCM | IPC-298 | PLX-4720 |
| top60_altered_DNA_seed | BRAF | -1.177135 | SKCM | SK-MEL-2 | Dabrafenib |
| top60_altered_DNA_seed | BRAF | -1.173688 | SKCM | IGR-1 | PLX-4720 |
| top60_altered_DNA_seed | BRAF | -1.168445 | SKCM | LB373-MEL-D | AZ628 |
| top60_altered_DNA_seed | BRAF | -1.16803 | SKCM | WM-115 | RAF_9304 |
| top60_altered_DNA_seed | BRAF | -1.157821 | SKCM | A2058 | PLX-4720 |
| top60_altered_DNA_seed | BRAF | -1.142658 | SKCM | RPMI-7951 | PLX-4720 |
| top60_altered_DNA_seed | BRAF | -1.134287 | SKCM | IST-MEL1 | SB590885 |
| top60_altered_DNA_seed | BRAF | -1.129743 | SKCM | WM-115 | PLX-4720 |
| top60_altered_DNA_seed | BRAF | -1.120565 | SKCM | A101D | HG6-64-1 |
| top60_altered_DNA_seed | BRAF | -1.113387 | SKCM | HMV-II | Dabrafenib |
| top60_altered_DNA_seed | BRAF | -1.113266 | SKCM | SK-MEL-24 | HG6-64-1 |
| top60_altered_DNA_seed | BRAF | -1.081893 | SKCM | WM793B | PLX-4720 |
| top60_altered_DNA_seed | BRAF | -1.076817 | SKCM | MEL-JUSO | RAF_9304 |
| top60_altered_DNA_seed | BRAF | -1.068949 | SKCM | MEL-HO | HG6-64-1 |
| top60_altered_DNA_seed | BRAF | -1.058739 | SKCM | CP66-MEL | PLX-4720 |
| top60_altered_DNA_seed | BRAF | -1.030639 | SKCM | 451Lu | HG6-64-1 |
| top60_altered_DNA_seed | BRAF | -1.030343 | SKCM | IGR-1 | PLX-4720 |
| top60_altered_DNA_seed | BRAF | -1.009434 | SKCM | IST-MEL1 | PLX-4720 |
| top60_altered_DNA_seed | BRAF | -0.999764 | SKCM | A2058 | PLX-4720 |
| top60_altered_DNA_seed | BRAF | -0.987726 | SKCM | WM793B | RAF_9304 |
| top60_altered_DNA_seed | BRAF | -0.972243 | SKCM | MEL-JUSO | HG6-64-1 |
| top60_altered_DNA_seed | BRAF | -0.958089 | SKCM | MZ2-MEL | HG6-64-1 |
| top60_altered_DNA_seed | BRAF | -0.908717 | SKCM | WM793B | PLX-4720 |
| top60_altered_DNA_seed | BRAF | -0.905572 | SKCM | LB2518-MEL | RAF_9304 |
| top60_altered_DNA_seed | BRAF | -0.882645 | SKCM | CHL-1 | RAF_9304 |
| top60_altered_DNA_seed | BRAF | -0.86837 | SKCM | COLO-783 | RAF_9304 |
| top60_altered_DNA_seed | BRAF | -0.866914 | SKCM | IPC-298 | PLX-4720 |
| top60_altered_DNA_seed | BRAF | -0.863686 | SKCM | Mewo | SB590885 |
| top60_altered_DNA_seed | BRAF | -0.848532 | SKCM | RVH-421 | RAF_9304 |
| top60_altered_DNA_seed | BRAF | -0.810236 | SKCM | WM1552C | SB590885 |
| top60_altered_DNA_seed | BRAF | -0.804292 | SKCM | G-MEL | HG6-64-1 |
| top60_altered_DNA_seed | BRAF | -0.796694 | SKCM | MZ2-MEL | SB590885 |
| top60_altered_DNA_seed | BRAF | -0.780405 | SKCM | WM35 | HG6-64-1 |
| top60_altered_DNA_seed | BRAF | -0.776445 | SKCM | WM1552C | HG6-64-1 |
| top60_altered_DNA_seed | BRAF | -0.775535 | SKCM | COLO-679 | HG6-64-1 |

| | | | | | |
|---|---|---|---|---|---|
| top60_altered_DNA_seed | BRAF | -0.771352 | SKCM | CHL-1 | PLX-4720 |
| top60_altered_DNA_seed | BRAF | -0.765871 | SKCM | Hs-940-T | HG6-64-1 |
| top60_altered_DNA_seed | BRAF | -0.748499 | SKCM | WM1552C | Dabrafenib |
| top60_altered_DNA_seed | BRAF | -0.737577 | SKCM | IGR-1 | HG6-64-1 |
| top60_altered_DNA_seed | BRAF | -0.708957 | SKCM | A375 | HG6-64-1 |
| top60_altered_DNA_seed | BRAF | -0.70777 | SKCM | G-361 | HG6-64-1 |
| top60_altered_DNA_seed | BRAF | -0.702291 | SKCM | SK-MEL-1 | RAF_9304 |
| top60_altered_DNA_seed | BRAF | -0.671997 | SKCM | IGR-1 | RAF_9304 |
| top60_altered_DNA_seed | BRAF | -0.616135 | SKCM | SH-4 | HG6-64-1 |
| top60_altered_DNA_seed | BRAF | -0.602072 | SKCM | CP66-MEL | RAF_9304 |
| top60_altered_DNA_seed | BRAF | -0.58618 | SKCM | IST-MEL1 | RAF_9304 |
| top60_altered_DNA_seed | BRAF | -0.570504 | SKCM | COLO-829 | RAF_9304 |
| top60_altered_DNA_seed | BRAF | -0.535898 | SKCM | GAK | PLX-4720 |
| top60_altered_DNA_seed | BRAF | -0.533949 | SKCM | UACC-62 | HG6-64-1 |
| top60_altered_DNA_seed | BRAF | -0.529819 | SKCM | A2058 | HG6-64-1 |
| top60_altered_DNA_seed | BRAF | -0.510743 | SKCM | LOXIMVI | HG6-64-1 |
| top60_altered_DNA_seed | BRAF | -0.497117 | SKCM | WM1552C | PLX-4720 |
| top60_altered_DNA_seed | BRAF | -0.472887 | SKCM | RPMI-7951 | RAF_9304 |
| top60_altered_DNA_seed | BRAF | -0.465994 | SKCM | COLO-800 | SB590885 |
| top60_altered_DNA_seed | BRAF | -0.451403 | SKCM | LB2518-MEL | PLX-4720 |
| top60_altered_DNA_seed | BRAF | -0.442684 | SKCM | IPC-298 | SB590885 |
| top60_altered_DNA_seed | BRAF | -0.422768 | SKCM | GAK | Dabrafenib |
| top60_altered_DNA_seed | BRAF | -0.392957 | SKCM | MMAc-SF | HG6-64-1 |
| top60_altered_DNA_seed | BRAF | -0.380382 | SKCM | WM1552C | RAF_9304 |
| top60_altered_DNA_seed | BRAF | -0.364594 | SKCM | CHL-1 | SB590885 |
| top60_altered_DNA_seed | BRAF | -0.358696 | SKCM | SK-MEL-2 | AZ628 |
| top60_altered_DNA_seed | BRAF | -0.353964 | SKCM | RPMI-7951 | PLX-4720 |
| top60_altered_DNA_seed | BRAF | -0.323334 | SKCM | SK-MEL-2 | PLX-4720 |
| top60_altered_DNA_seed | BRAF | -0.294952 | SKCM | IST-MEL1 | HG6-64-1 |
| top60_altered_DNA_seed | BRAF | -0.270534 | SKCM | Mewo | PLX-4720 |
| top60_altered_DNA_seed | BRAF | -0.268966 | SKCM | LOXIMVI | AZ628 |
| top60_altered_DNA_seed | BRAF | -0.265707 | SKCM | IGR-37 | HG6-64-1 |
| top60_altered_DNA_seed | BRAF | -0.252395 | SKCM | SK-MEL-31 | PLX-4720 |
| top60_altered_DNA_seed | BRAF | -0.238589 | SKCM | MZ7-mel | HG6-64-1 |
| top60_altered_DNA_seed | BRAF | -0.215587 | SKCM | A2058 | SB590885 |
| top60_altered_DNA_seed | BRAF | -0.15607 | SKCM | SK-MEL-30 | PLX-4720 |
| top60_altered_DNA_seed | BRAF | -0.153906 | SKCM | Mewo | RAF_9304 |
| top60_altered_DNA_seed | BRAF | -0.144491 | SKCM | MEL-JUSO | Dabrafenib |
| top60_altered_DNA_seed | BRAF | -0.136066 | SKCM | Mewo | Dabrafenib |
| top60_altered_DNA_seed | BRAF | -0.125291 | SKCM | SK-MEL-30 | HG6-64-1 |
| top60_altered_DNA_seed | BRAF | -0.119272 | SKCM | SK-MEL-3 | HG6-64-1 |

| | | | | | |
|---|---|---|---|---|---|
| top60_altered_DNA_seed | BRAF | -0.114366 | SKCM | K2 | HG6-64-1 |
| top60_altered_DNA_seed | BRAF | -0.101109 | SKCM | RVH-421 | HG6-64-1 |
| top60_altered_DNA_seed | BRAF | -0.09878 | SKCM | COLO-800 | AZ628 |
| top60_altered_DNA_seed | BRAF | -0.090442 | SKCM | SK-MEL-2 | HG6-64-1 |
| top60_altered_DNA_seed | BRAF | -0.017785 | SKCM | IPC-298 | HG6-64-1 |
| top60_altered_DNA_seed | BRAF | -0.01767 | SKCM | RPMI-7951 | HG6-64-1 |
| top60_altered_DNA_seed | BRAF | -0.015164 | SKCM | SK-MEL-30 | Dabrafenib |
| top60_altered_DNA_seed | BRAF | -0.00588 | SKCM | SK-MEL-31 | PLX-4720 |
| top60_altered_DNA_seed | BRAF | 0.014112 | SKCM | MEL-JUSO | PLX-4720 |
| top60_altered_DNA_seed | BRAF | 0.018719 | SKCM | Hs-939-T | HG6-64-1 |
| top60_altered_DNA_seed | BRAF | 0.068592 | SKCM | GAK | PLX-4720 |
| top60_altered_DNA_seed | BRAF | 0.121078 | SKCM | COLO-800 | RAF_9304 |
| top60_altered_DNA_seed | BRAF | 0.159752 | SKCM | RPMI-7951 | SB590885 |
| top60_altered_DNA_seed | BRAF | 0.180461 | SKCM | HMV-II | PLX-4720 |
| top60_altered_DNA_seed | BRAF | 0.18138 | SKCM | COLO-792 | RAF_9304 |
| top60_altered_DNA_seed | BRAF | 0.190607 | SKCM | VMRC-MELG | Dabrafenib |
| top60_altered_DNA_seed | BRAF | 0.230979 | SKCM | UACC-62 | SB590885 |
| top60_altered_DNA_seed | BRAF | 0.234865 | SKCM | CP66-MEL | Dabrafenib |
| top60_altered_DNA_seed | BRAF | 0.239094 | SKCM | MEL-JUSO | PLX-4720 |
| top60_altered_DNA_seed | BRAF | 0.245239 | SKCM | COLO-800 | Dabrafenib |
| top60_altered_DNA_seed | BRAF | 0.255124 | SKCM | M14 | HG6-64-1 |
| top60_altered_DNA_seed | BRAF | 0.274802 | SKCM | LB2518-MEL | HG6-64-1 |
| top60_altered_DNA_seed | BRAF | 0.275418 | SKCM | UACC-257 | HG6-64-1 |
| top60_altered_DNA_seed | BRAF | 0.330204 | SKCM | LB373-MEL-D | RAF_9304 |
| top60_altered_DNA_seed | BRAF | 0.338068 | SKCM | GAK | HG6-64-1 |
| top60_altered_DNA_seed | BRAF | 0.35273 | SKCM | VMRC-MELG | PLX-4720 |
| top60_altered_DNA_seed | BRAF | 0.360966 | SKCM | GAK | SB590885 |
| top60_altered_DNA_seed | BRAF | 0.383017 | SKCM | CP66-MEL | SB590885 |
| top60_altered_DNA_seed | BRAF | 0.442011 | SKCM | LB2518-MEL | Dabrafenib |
| top60_altered_DNA_seed | BRAF | 0.446103 | SKCM | MZ2-MEL | RAF_9304 |
| top60_altered_DNA_seed | BRAF | 0.470943 | SKCM | SK-MEL-30 | PLX-4720 |
| top60_altered_DNA_seed | BRAF | 0.485086 | SKCM | UACC-62 | PLX-4720 |
| top60_altered_DNA_seed | BRAF | 0.486652 | SKCM | SK-MEL-31 | RAF_9304 |
| top60_altered_DNA_seed | BRAF | 0.506954 | SKCM | SK-MEL-31 | Dabrafenib |
| top60_altered_DNA_seed | BRAF | 0.565339 | SKCM | MEL-JUSO | SB590885 |
| top60_altered_DNA_seed | BRAF | 0.618568 | SKCM | WM35 | PLX-4720 |
| top60_altered_DNA_seed | BRAF | 0.629503 | SKCM | LB2518-MEL | PLX-4720 |
| top60_altered_DNA_seed | BRAF | 0.638491 | SKCM | LB2518-MEL | SB590885 |
| top60_altered_DNA_seed | BRAF | 0.654869 | SKCM | SK-MEL-2 | PLX-4720 |
| top60_altered_DNA_seed | BRAF | 0.705163 | SKCM | COLO-792 | Dabrafenib |
| top60_altered_DNA_seed | BRAF | 0.756074 | SKCM | COLO-800 | PLX-4720 |

| | | | | | |
|---|---|---|---|---|---|
| top60_altered_DNA_seed | BRAF | 0.771435 | SKCM | VMRC-MELG | HG6-64-1 |
| top60_altered_DNA_seed | BRAF | 0.782026 | SKCM | HMV-II | PLX-4720 |
| top60_altered_DNA_seed | BRAF | 0.782456 | SKCM | LB373-MEL-D | SB590885 |
| top60_altered_DNA_seed | BRAF | 0.815956 | SKCM | Hs-940-T | SB590885 |
| top60_altered_DNA_seed | BRAF | 0.816054 | SKCM | WM-115 | HG6-64-1 |
| top60_altered_DNA_seed | BRAF | 0.860431 | SKCM | COLO-792 | SB590885 |
| top60_altered_DNA_seed | BRAF | 0.943639 | SKCM | COLO-792 | HG6-64-1 |
| top60_altered_DNA_seed | BRAF | 0.998149 | SKCM | LB373-MEL-D | PLX-4720 |
| top60_altered_DNA_seed | BRAF | 1.043476 | SKCM | COLO-792 | PLX-4720 |
| top60_altered_DNA_seed | BRAF | 1.068896 | SKCM | LB373-MEL-D | Dabrafenib |
| top60_altered_DNA_seed | BRAF | 1.089648 | SKCM | Mewo | PLX-4720 |
| top60_altered_DNA_seed | BRAF | 1.124 | SKCM | LB373-MEL-D | PLX-4720 |
| top60_altered_DNA_seed | BRAF | 1.147562 | SKCM | SK-MEL-28 | HG6-64-1 |
| top60_altered_DNA_seed | BRAF | 1.178902 | SKCM | CP66-MEL | HG6-64-1 |
| top60_altered_DNA_seed | BRAF | 1.212192 | SKCM | SK-MEL-31 | SB590885 |
| top60_altered_DNA_seed | BRAF | 1.265911 | SKCM | COLO-800 | HG6-64-1 |
| top60_altered_DNA_seed | BRAF | 1.312416 | SKCM | CP50-MEL-B | HG6-64-1 |
| top60_altered_DNA_seed | BRAF | 1.37581 | SKCM | COLO-792 | PLX-4720 |
| top60_altered_DNA_seed | BRAF | 1.413755 | SKCM | COLO-783 | HG6-64-1 |
| top60_altered_DNA_seed | BRAF | 1.525955 | SKCM | SK-MEL-1 | HG6-64-1 |
| top60_altered_DNA_seed | BRAF | 1.537083 | SKCM | SK-MEL-5 | HG6-64-1 |
| top60_altered_DNA_seed | BRAF | 1.855514 | SKCM | Mewo | HG6-64-1 |
| top60_altered_DNA_seed | BRAF | 2.054559 | SKCM | C32 | HG6-64-1 |
| top60_altered_DNA_seed | BRAF | 2.055568 | SKCM | VMRC-MELG | RAF_9304 |
| top60_altered_DNA_seed | BRAF | 2.099185 | SKCM | Hs-940-T | RAF_9304 |
| top60_altered_DNA_seed | BRAF | -4.749342 | SKCM | G-MEL | Dabrafenib |
| top60_altered_DNA_seed | BRAF | -4.690012 | SKCM | SK-MEL-28 | Dabrafenib |
| top60_altered_DNA_seed | BRAF | -4.659915 | SKCM | CP50-MEL-B | Dabrafenib |
| top60_altered_DNA_seed | BRAF | -4.254281 | SKCM | SK-MEL-28 | PLX-4720 |
| top60_altered_DNA_seed | BRAF | -4.215455 | SKCM | M14 | Dabrafenib |
| top60_altered_DNA_seed | BRAF | -4.160715 | SKCM | A375 | Dabrafenib |
| top60_altered_DNA_seed | BRAF | -4.107077 | SKCM | C32 | Dabrafenib |
| top60_altered_DNA_seed | BRAF | -4.090654 | SKCM | CP50-MEL-B | PLX-4720 |
| top60_altered_DNA_seed | BRAF | -4.02731 | SKCM | IGR-37 | Dabrafenib |
| top60_altered_DNA_seed | BRAF | -3.992912 | SKCM | G-MEL | SB590885 |
| top60_altered_DNA_seed | BRAF | -3.991288 | SKCM | SH-4 | Dabrafenib |
| top60_altered_DNA_seed | BRAF | -3.955507 | SKCM | UACC-62 | Dabrafenib |
| top60_altered_DNA_seed | BRAF | -3.95452 | SKCM | HT-144 | Dabrafenib |
| top60_altered_DNA_seed | BRAF | -3.747167 | SKCM | UACC-62 | SB590885 |
| top60_altered_DNA_seed | BRAF | -3.732422 | SKCM | G-MEL | PLX-4720 |
| top60_altered_DNA_seed | BRAF | -3.731334 | SKCM | WM35 | Dabrafenib |

| | | | | | |
|---|---|---|---|---|---|
| top60_altered_DNA_seed | BRAF | -3.708066 | SKCM | MEL-HO | Dabrafenib |
| top60_altered_DNA_seed | BRAF | -3.698163 | SKCM | K2 | Dabrafenib |
| top60_altered_DNA_seed | BRAF | -3.6342 | SKCM | Hs-939-T | PLX-4720 |
| top60_altered_DNA_seed | BRAF | -3.492056 | SKCM | CP50-MEL-B | SB590885 |
| top60_altered_DNA_seed | BRAF | -3.489487 | SKCM | Hs-939-T | Dabrafenib |
| top60_altered_DNA_seed | BRAF | -3.482924 | SKCM | COLO-679 | Dabrafenib |
| top60_altered_DNA_seed | BRAF | -3.47547 | SKCM | COLO-800 | Dabrafenib |
| top60_altered_DNA_seed | BRAF | -3.422786 | SKCM | IGR-37 | SB590885 |
| top60_altered_DNA_seed | BRAF | -3.375367 | SKCM | MEL-HO | SB590885 |
| top60_altered_DNA_seed | BRAF | -3.350935 | SKCM | 451Lu | Dabrafenib |
| top60_altered_DNA_seed | BRAF | -3.282869 | SKCM | COLO-679 | SB590885 |
| top60_altered_DNA_seed | BRAF | -3.279193 | SKCM | MZ7-mel | Dabrafenib |
| top60_altered_DNA_seed | BRAF | -3.205137 | SKCM | WM278 | SB590885 |
| top60_altered_DNA_seed | BRAF | -3.189075 | SKCM | UACC-62 | PLX-4720 |
| top60_altered_DNA_seed | BRAF | -3.145685 | SKCM | MEL-HO | PLX-4720 |
| top60_altered_DNA_seed | BRAF | -3.139668 | SKCM | 451Lu | PLX-4720 |
| top60_altered_DNA_seed | BRAF | -3.092362 | SKCM | M14 | PLX-4720 |
| top60_altered_DNA_seed | BRAF | -3.076376 | SKCM | IGR-37 | PLX-4720 |
| top60_altered_DNA_seed | BRAF | -2.982128 | SKCM | COLO-800 | PLX-4720 |
| top60_altered_DNA_seed | BRAF | -2.934823 | SKCM | A375 | PLX-4720 |
| top60_altered_DNA_seed | BRAF | -2.899576 | SKCM | K2 | PLX-4720 |
| top60_altered_DNA_seed | BRAF | -2.855013 | SKCM | A101D | Dabrafenib |
| top60_altered_DNA_seed | BRAF | -2.832276 | SKCM | C32 | PLX-4720 |
| top60_altered_DNA_seed | BRAF | -2.812637 | SKCM | HT-144 | PLX-4720 |
| top60_altered_DNA_seed | BRAF | -2.78915 | SKCM | WM35 | SB590885 |
| top60_altered_DNA_seed | BRAF | -2.780108 | SKCM | UACC-257 | SB590885 |
| top60_altered_DNA_seed | BRAF | -2.742669 | SKCM | COLO-679 | PLX-4720 |
| top60_altered_DNA_seed | BRAF | -2.722951 | SKCM | A375 | SB590885 |
| top60_altered_DNA_seed | BRAF | -2.709563 | SKCM | SH-4 | PLX-4720 |
| top60_altered_DNA_seed | BRAF | -2.686826 | SKCM | SK-MEL-1 | Dabrafenib |
| top60_altered_DNA_seed | BRAF | -2.682732 | SKCM | K2 | SB590885 |
| top60_altered_DNA_seed | BRAF | -2.646627 | SKCM | G-361 | Dabrafenib |
| top60_altered_DNA_seed | BRAF | -2.627807 | SKCM | 451Lu | SB590885 |
| top60_altered_DNA_seed | BRAF | -2.620304 | SKCM | HT-144 | SB590885 |
| top60_altered_DNA_seed | BRAF | -2.605377 | SKCM | MMAc-SF | PLX-4720 |
| top60_altered_DNA_seed | BRAF | -2.57747 | SKCM | COLO-783 | SB590885 |
| top60_altered_DNA_seed | BRAF | -2.567505 | SKCM | COLO-783 | PLX-4720 |
| top60_altered_DNA_seed | BRAF | -2.548212 | SKCM | SK-MEL-1 | SB590885 |
| top60_altered_DNA_seed | BRAF | -2.545772 | SKCM | A101D | SB590885 |
| top60_altered_DNA_seed | BRAF | -2.543506 | SKCM | Hs-939-T | SB590885 |
| top60_altered_DNA_seed | BRAF | -2.399939 | SKCM | COLO-829 | PLX-4720 |

| | | | | | |
|---|---|---|---|---|---|
| top60_altered_DNA_seed | BRAF | -2.355924 | SKCM | SK-MEL-28 | SB590885 |
| top60_altered_DNA_seed | BRAF | -2.315428 | SKCM | UACC-257 | Dabrafenib |
| top60_altered_DNA_seed | BRAF | -2.298401 | SKCM | COLO-800 | SB590885 |
| top60_altered_DNA_seed | BRAF | -2.273836 | SKCM | WM793B | SB590885 |
| top60_altered_DNA_seed | BRAF | -2.248185 | SKCM | MZ7-mel | PLX-4720 |
| top60_altered_DNA_seed | BRAF | -2.24625 | SKCM | SH-4 | SB590885 |
| top60_altered_DNA_seed | BRAF | -2.184988 | SKCM | WM278 | PLX-4720 |
| top60_altered_DNA_seed | BRAF | -2.166504 | SKCM | C32 | SB590885 |
| top60_altered_DNA_seed | BRAF | -2.097784 | SKCM | SK-MEL-24 | Dabrafenib |
| top60_altered_DNA_seed | BRAF | -2.079743 | SKCM | G-361 | PLX-4720 |
| top60_altered_DNA_seed | BRAF | -2.048532 | SKCM | G-361 | SB590885 |
| top60_altered_DNA_seed | BRAF | -1.979924 | SKCM | SK-MEL-24 | SB590885 |
| top60_altered_DNA_seed | BRAF | -1.937656 | SKCM | A101D | PLX-4720 |
| top60_altered_DNA_seed | BRAF | -1.905691 | SKCM | SK-MEL-3 | Dabrafenib |
| top60_altered_DNA_seed | BRAF | -1.905041 | SKCM | Hs-940-T | Dabrafenib |
| top60_altered_DNA_seed | BRAF | -1.817332 | SKCM | MMAc-SF | Dabrafenib |
| top60_altered_DNA_seed | BRAF | -1.753284 | SKCM | SK-MEL-24 | PLX-4720 |
| top60_altered_DNA_seed | BRAF | -1.747297 | SKCM | WM278 | Dabrafenib |
| top60_altered_DNA_seed | BRAF | -1.685814 | SKCM | WM-115 | Dabrafenib |
| top60_altered_DNA_seed | BRAF | -1.654505 | SKCM | UACC-257 | PLX-4720 |
| top60_altered_DNA_seed | BRAF | -1.632477 | SKCM | WM-115 | PLX-4720 |
| top60_altered_DNA_seed | BRAF | -1.627317 | SKCM | SK-MEL-30 | SB590885 |
| top60_altered_DNA_seed | BRAF | -1.576498 | SKCM | MZ7-mel | SB590885 |
| top60_altered_DNA_seed | BRAF | -1.575211 | SKCM | LOXIMVI | Dabrafenib |
| top60_altered_DNA_seed | BRAF | -1.503855 | SKCM | SK-MEL-3 | PLX-4720 |
| top60_altered_DNA_seed | BRAF | -1.488247 | SKCM | LOXIMVI | SB590885 |
| top60_altered_DNA_seed | BRAF | -1.450751 | SKCM | WM-115 | SB590885 |
| top60_altered_DNA_seed | BRAF | -1.386463 | SKCM | WM793B | PLX-4720 |
| top60_altered_DNA_seed | BRAF | -1.375744 | SKCM | SK-MEL-5 | SB590885 |
| top60_altered_DNA_seed | BRAF | -1.374494 | SKCM | WM35 | PLX-4720 |
| top60_altered_DNA_seed | BRAF | -1.284334 | SKCM | Hs-940-T | PLX-4720 |
| top60_altered_DNA_seed | BRAF | -1.274919 | SKCM | IPC-298 | PLX-4720 |
| top60_altered_DNA_seed | BRAF | -1.272359 | SKCM | LOXIMVI | PLX-4720 |
| top60_altered_DNA_seed | BRAF | -1.259437 | SKCM | SK-MEL-5 | Dabrafenib |
| top60_altered_DNA_seed | BRAF | -1.251223 | SKCM | SK-MEL-3 | SB590885 |
| top60_altered_DNA_seed | BRAF | -1.245366 | SKCM | COLO-829 | SB590885 |
| top60_altered_DNA_seed | BRAF | -1.218829 | SKCM | IGR-1 | Dabrafenib |
| top60_altered_DNA_seed | BRAF | -1.194543 | SKCM | MMAc-SF | SB590885 |
| top60_altered_DNA_seed | BRAF | -1.140318 | SKCM | SK-MEL-1 | PLX-4720 |
| top60_altered_DNA_seed | BRAF | -1.131181 | SKCM | IPC-298 | SB590885 |
| top60_altered_DNA_seed | BRAF | -1.07447 | SKCM | IPC-298 | Dabrafenib |

| | | | | | |
|---|---|---|---|---|---|
| top60_altered_DNA_seed | BRAF | -1.072549 | SKCM | SK-MEL-5 | PLX-4720 |
| top60_altered_DNA_seed | BRAF | -1.03042 | SKCM | IST-MEL1 | PLX-4720 |
| top60_altered_DNA_seed | BRAF | -0.968986 | SKCM | RVH-421 | SB590885 |
| top60_altered_DNA_seed | BRAF | -0.945144 | SKCM | RVH-421 | PLX-4720 |
| top60_altered_DNA_seed | BRAF | -0.935767 | SKCM | WM1552C | SB590885 |
| top60_altered_DNA_seed | BRAF | -0.933673 | SKCM | IGR-1 | PLX-4720 |
| top60_altered_DNA_seed | BRAF | -0.929291 | SKCM | MZ2-MEL | PLX-4720 |
| top60_altered_DNA_seed | BRAF | -0.852645 | SKCM | HMV-II | Dabrafenib |
| top60_altered_DNA_seed | BRAF | -0.823021 | SKCM | WM1552C | PLX-4720 |
| top60_altered_DNA_seed | BRAF | -0.776845 | SKCM | SK-MEL-31 | PLX-4720 |
| top60_altered_DNA_seed | BRAF | -0.76818 | SKCM | RPMI-7951 | PLX-4720 |
| top60_altered_DNA_seed | BRAF | -0.762441 | SKCM | A2058 | SB590885 |
| top60_altered_DNA_seed | BRAF | -0.697813 | SKCM | HMV-II | PLX-4720 |
| top60_altered_DNA_seed | BRAF | -0.653236 | SKCM | VMRC-MELG | PLX-4720 |
| top60_altered_DNA_seed | BRAF | -0.628599 | SKCM | WM1552C | Dabrafenib |
| top60_altered_DNA_seed | BRAF | -0.622453 | SKCM | IGR-1 | SB590885 |
| top60_altered_DNA_seed | BRAF | -0.607267 | SKCM | RVH-421 | Dabrafenib |
| top60_altered_DNA_seed | BRAF | -0.574058 | SKCM | A2058 | PLX-4720 |
| top60_altered_DNA_seed | BRAF | -0.559475 | SKCM | CP66-MEL | Dabrafenib |
| top60_altered_DNA_seed | BRAF | -0.481898 | SKCM | WM793B | Dabrafenib |
| top60_altered_DNA_seed | BRAF | -0.459969 | SKCM | MZ2-MEL | Dabrafenib |
| top60_altered_DNA_seed | BRAF | -0.412815 | SKCM | SK-MEL-31 | Dabrafenib |
| top60_altered_DNA_seed | BRAF | -0.386309 | SKCM | IST-MEL1 | Dabrafenib |
| top60_altered_DNA_seed | BRAF | -0.382748 | SKCM | IST-MEL1 | SB590885 |
| top60_altered_DNA_seed | BRAF | -0.179315 | SKCM | RPMI-7951 | SB590885 |
| top60_altered_DNA_seed | BRAF | -0.166111 | SKCM | CP66-MEL | PLX-4720 |
| top60_altered_DNA_seed | BRAF | -0.164214 | SKCM | SK-MEL-30 | PLX-4720 |
| top60_altered_DNA_seed | BRAF | -0.043138 | SKCM | GAK | Dabrafenib |
| top60_altered_DNA_seed | BRAF | -0.013826 | SKCM | GAK | SB590885 |
| top60_altered_DNA_seed | BRAF | 0.029126 | SKCM | MEL-JUSO | Dabrafenib |
| top60_altered_DNA_seed | BRAF | 0.044274 | SKCM | A2058 | Dabrafenib |
| top60_altered_DNA_seed | BRAF | 0.157413 | SKCM | HMV-II | SB590885 |
| top60_altered_DNA_seed | BRAF | 0.175232 | SKCM | RPMI-7951 | Dabrafenib |
| top60_altered_DNA_seed | BRAF | 0.198774 | SKCM | SK-MEL-30 | Dabrafenib |
| top60_altered_DNA_seed | BRAF | 0.210061 | SKCM | CP66-MEL | SB590885 |
| top60_altered_DNA_seed | BRAF | 0.242275 | SKCM | CHL-1 | PLX-4720 |
| top60_altered_DNA_seed | BRAF | 0.27152 | SKCM | MZ2-MEL | SB590885 |
| top60_altered_DNA_seed | BRAF | 0.399824 | SKCM | LB373-MEL-D | PLX-4720 |
| top60_altered_DNA_seed | BRAF | 0.424186 | SKCM | CHL-1 | Dabrafenib |
| top60_altered_DNA_seed | BRAF | 0.438214 | SKCM | COLO-792 | SB590885 |
| top60_altered_DNA_seed | BRAF | 0.439929 | SKCM | VMRC-MELG | Dabrafenib |

| | | | | | |
|---|---|---|---|---|---|
| top60_altered_DNA_seed | BRAF | 0.44273 | SKCM | LB373-MEL-D | SB590885 |
| top60_altered_DNA_seed | BRAF | 0.444993 | SKCM | CHL-1 | SB590885 |
| top60_altered_DNA_seed | BRAF | 0.482456 | SKCM | LB2518-MEL | Dabrafenib |
| top60_altered_DNA_seed | BRAF | 0.559152 | SKCM | MEL-JUSO | SB590885 |
| top60_altered_DNA_seed | BRAF | 0.560395 | SKCM | SK-MEL-31 | SB590885 |
| top60_altered_DNA_seed | BRAF | 0.575202 | SKCM | LB2518-MEL | PLX-4720 |
| top60_altered_DNA_seed | BRAF | 0.655727 | SKCM | SK-MEL-2 | Dabrafenib |
| top60_altered_DNA_seed | BRAF | 0.722336 | SKCM | Hs-940-T | SB590885 |
| top60_altered_DNA_seed | BRAF | 0.723359 | SKCM | LB373-MEL-D | Dabrafenib |
| top60_altered_DNA_seed | BRAF | 0.739716 | SKCM | COLO-792 | Dabrafenib |
| top60_altered_DNA_seed | BRAF | 0.761686 | SKCM | COLO-792 | PLX-4720 |
| top60_altered_DNA_seed | BRAF | 0.763247 | SKCM | LB2518-MEL | SB590885 |
| top60_altered_DNA_seed | BRAF | 0.795289 | SKCM | Mewo | SB590885 |
| top60_altered_DNA_seed | BRAF | 0.871041 | SKCM | GAK | PLX-4720 |
| top60_altered_DNA_seed | BRAF | 0.896578 | SKCM | VMRC-MELG | SB590885 |
| top60_altered_DNA_seed | BRAF | 0.963493 | SKCM | Mewo | Dabrafenib |
| top60_altered_DNA_seed | BRAF | 1.051503 | SKCM | MEL-JUSO | PLX-4720 |
| top60_altered_DNA_seed | BRAF | 1.057173 | SKCM | SK-MEL-2 | PLX-4720 |
| top60_altered_DNA_seed | BRAF | 1.130583 | SKCM | Mewo | PLX-4720 |
