## Supplementary material for "Probabilistic graph-based model uncovers previously unseen druggable vulnerabilities in major solid cancers": supp table 4

| Tumor type | Method | GDSC target count | GDSC target count with low z-scored LN_IC50 | Percent of top 20 genes with low z-scored LN_IC50 (%) | GDSC targets among top 20 with low z-scored LN_IC50 |
|---|---|---|---|---|---|
| **BLCA** | **$A_3D_3$a's MVP** | **9** | **8** | **40** | **CREBBP,EP300,ERBB2,ERBB3,ESR1,FGFR3,PIK3CA,TP53** |
| BLCA | Top altered in seed | 0 | 0 | 0 | |
| **BRCA** | **$A_3D_3$a's MVP** | **10** | **10** | **50** | **AKT1,CDK2,EGFR,EP300,ERBB2,ESR1,PIK3CA,PLK1,SRC,TP53** |
| BRCA | Top altered in seed | 1 | 1 | 5 | PLK1 |
| **COADREAD** | **$A_3D_3$a's MVP** | **8** | **6** | **30** | **AKT1,ATM,HSP90AA1,PIK3CA,PIK3CB,SRC** |
| COADREAD | Top altered in seed | 0 | 0 | 0 | |
| **GBM** | **$A_3D_3$a's MVP** | **6** | **5** | **25** | **EGFR,PDGFRA,PIK3CA,PIK3CB,PIK3CD** |
| GBM | Top altered in seed | 1 | 1 | 5 | EGFR |
| **KIRC** | **$A_3D_3$a's MVP** | **11** | **10** | **50** | **AR,ATM,EP300,ESR1,FGFR4,HDAC1,HSP90AA1,IGF1R,PIK3CA, SMARCA4** |
| KIRC | Top altered in seed | 0 | 0 | 0 | |
| **LUAD** | **$A_3D_3$a's MVP** | **10** | **9** | **45** | **AKT1,ATM,EGFR,HDAC1,HSP90AA1,KRAS,MET,PIK3CA,SRC** |
| LUAD | Top altered in seed | 1 | 0 | 0 | |
| **LUSC** | **$A_3D_3$a's MVP** | **5** | **4** | **20** | **AR,CREBBP,HDAC1,PIK3CA** |
| LUSC | Top altered in seed | 1 | 0 | 0 | |
| **PRAD** | **$A_3D_3$a's MVP** | **8** | **6** | **30** | **AR,ATM,EP300,ESR1,GSK3B,PIK3CA** |
| PRAD | Top altered in seed | 1 | 0 | 0 | |
| **STAD** | **$A_3D_3$a's MVP** | **11** | **9** | **45** | **AR,ATM,EP300,ERBB2,ERBB3,ERBB4,ESR1,PIK3CA,SRC** |
| STAD | Top altered in seed | 1 | 0 | 0 | |
| **THCA** | **$A_3D_3$a's MVP** | **9** | **7** | **35** | **AKT1,AKT2,ATM,BRAF,GSK3B,MAPK1,MAPK3** |
| THCA | Top altered in seed | 3 | 2 | 10 | BRAF,RET |
| **UCEC** | **$A_3D_3$a's MVP** | **8** | **5** | **25** | **AKT3,EGFR,EP300,ESR1,PIK3CA** |
| UCEC | Top altered in seed | 1 | 1 | 5 | AKT3 |
