## Supplementary material for "Probabilistic graph-based model uncovers previously unseen druggable vulnerabilities in major solid cancers": supp table 5

| Target | Disease | oncoKB | oncoKB_level | canSAR | canSAR_type | MVP_top60 | Top60_altered |
|---|---|---|---|---|---|---|---|
| CD274 | BLCA | No | | Yes | targetted | No | No |
| EGFR | BLCA | No | | Yes | targetted | No | No |
| ERBB2 | BLCA | No | | Yes | targetted | Yes | No |
| ERCC2 | BLCA | Yes | [3] | No | | No | No |
| ESR1 | BLCA | No | | Yes | targetted | Yes | No |
| ESR2 | BLCA | No | | Yes | targetted | No | No |
| FGFR1 | BLCA | No | | Yes | targetted | No | No |
| FGFR2 | BLCA | Yes | [1] | Yes | targetted | No | No |
| FGFR3 | BLCA | Yes | [1, 3] | Yes | targetted | Yes | No |
| FGFR4 | BLCA | No | | Yes | targetted | No | No |
| HDAC1 | BLCA | No | | Yes | targetted | Yes | No |
| HDAC2 | BLCA | No | | Yes | targetted | No | No |
| HDAC3 | BLCA | No | | Yes | targetted | No | No |
| HDAC6 | BLCA | No | | Yes | targetted | No | No |
| KDR | BLCA | No | | Yes | targetted | No | No |
| MTOR | BLCA | Yes | [3] | No | | No | No |
| PDCD1 | BLCA | No | | Yes | targetted | No | No |
| TACSTD2 | BLCA | No | | Yes | targetted | No | No |
| TOP1 | BLCA | No | | Yes | targetted | No | No |
| AKT1 | BRCA | Yes | [3] | No | | Yes | No |
| BRCA1 | BRCA | Yes | [3] | No | | Yes | No |
| BRCA2 | BRCA | Yes | [3] | No | | No | No |
| CCND1 | BRCA | No | | Yes | targetted | Yes | Yes |
| CDK4 | BRCA | No | | Yes | targetted | No | No |
| CDK6 | BRCA | No | | Yes | targetted | No | No |
| CYP19A1 | BRCA | No | | Yes | targetted | No | No |
| EGFR | BRCA | No | | Yes | targetted | Yes | No |
| ERBB2 | BRCA | Yes | [1, 3] | Yes | targetted | Yes | No |
| ESR1 | BRCA | Yes | [1, 3] | Yes | targetted | Yes | No |
| ESR2 | BRCA | No | | Yes | targetted | No | No |
| FKBP1A | BRCA | No | | Yes | targetted | No | No |
| GNRHR | BRCA | No | | Yes | targetted | No | No |
| MTOR | BRCA | No | | Yes | targetted | No | No |
| PARP1 | BRCA | No | | Yes | targetted | No | No |
| PARP2 | BRCA | No | | Yes | targetted | No | No |
| PARP3 | BRCA | No | | Yes | targetted | No | No |
| PDCD1 | BRCA | No | | Yes | targetted | No | No |
| PIK3CA | BRCA | Yes | [1, 2] | Yes | targetted | Yes | Yes |
| TACSTD2 | BRCA | No | | Yes | targetted | No | No |
| TNFSF11 | BRCA | No | | Yes | targetted | No | No |
| TOP1 | BRCA | No | | Yes | targetted | No | No |
| ABL1 | COADREAD | No | | Yes | targetted | Yes | No |
| BCR | COADREAD | No | | Yes | targetted | No | No |
| BRAF | COADREAD | Yes | [1, 2] | Yes | targetted | Yes | No |
| CTLA4 | COADREAD | No | | Yes | targetted | No | No |
| EGFR | COADREAD | No | | Yes | targetted | No | No |
| ERBB2 | COADREAD | Yes | [1, 2] | Yes | targetted | No | No |
| FKBP1A | COADREAD | No | | Yes | targetted | No | No |
| KDR | COADREAD | No | | Yes | targetted | No | No |
| KIT | COADREAD | No | | Yes | targetted | No | No |
| KRAS | COADREAD | Yes | [1, 3] | No | | Yes | Yes |
| MTOR | COADREAD | No | | Yes | targetted | No | No |

| | | | | | | | |
|---|---|---|---|---|---|---|---|
| NRAS | COADREAD | Yes | [1] | No | | Yes | No |
| PDCD1 | COADREAD | No | | Yes | targetted | No | No |
| PDGFRA | COADREAD | No | | Yes | targetted | No | No |
| PDGFRB | COADREAD | No | | Yes | targetted | No | No |
| RAF1 | COADREAD | No | | Yes | targetted | No | No |
| TLR7 | COADREAD | No | | Yes | targetted | No | No |
| VEGFA | COADREAD | No | | Yes | targetted | No | No |
| VEGFB | COADREAD | No | | Yes | targetted | No | No |
| EGFR | HNSC | No | | Yes | targetted | Yes | No |
| HRAS | HNSC | Yes | [3] | No | | Yes | No |
| PDCD1 | HNSC | No | | Yes | targetted | No | No |
| CD274 | KIRC | No | | Yes | targetted | No | No |
| CTLA4 | KIRC | No | | Yes | targetted | No | No |
| EPAS1 | KIRC | No | | Yes | targetted | No | No |
| FKBP1A | KIRC | No | | Yes | targetted | No | No |
| FLT1 | KIRC | No | | Yes | targetted | Yes | No |
| FLT4 | KIRC | No | | Yes | targetted | Yes | No |
| IL2RA | KIRC | No | | Yes | targetted | No | No |
| KDR | KIRC | No | | Yes | targetted | No | No |
| MTOR | KIRC | Yes | [3] | Yes | targetted | Yes | No |
| PDCD1 | KIRC | No | | Yes | targetted | No | No |
| VEGFA | KIRC | No | | Yes | targetted | No | No |
| VEGFB | KIRC | No | | Yes | targetted | No | No |
| FKBP1A | LGG | No | | Yes | targetted | No | No |
| IDH1 | LGG | Yes | [3] | No | | Yes | Yes |
| MTOR | LGG | No | | Yes | targetted | No | No |
| VEGFA | LGG | No | | Yes | targetted | No | No |
| VEGFB | LGG | No | | Yes | targetted | No | No |
| ALK | LUAD | Yes | [1] | Yes | targetted | No | No |
| ARAF | LUAD | Yes | [3] | No | | No | No |
| BRAF | LUAD | Yes | [1] | No | | Yes | No |
| CCDC6 | LUAD | No | | Yes | targetted | No | No |
| CD274 | LUAD | No | | Yes | targetted | No | No |
| CTLA4 | LUAD | No | | Yes | targetted | No | No |
| DHFR | LUAD | No | | Yes | targetted | No | No |
| EGFR | LUAD | Yes | [1, 2, 3] | Yes | targetted | Yes | No |
| EML4 | LUAD | No | | Yes | targetted | No | No |
| ERBB2 | LUAD | Yes | [1, 2, 3] | Yes | targetted | Yes | No |
| ERBB4 | LUAD | No | | Yes | targetted | No | No |
| FKBP1A | LUAD | No | | Yes | targetted | No | No |
| GART | LUAD | No | | Yes | targetted | No | No |
| KDR | LUAD | No | | Yes | targetted | No | No |
| KIF5B | LUAD | No | | Yes | targetted | No | No |
| KRAS | LUAD | Yes | [1] | Yes | targetted | Yes | Yes |
| MAP2K1 | LUAD | Yes | [3] | Yes | targetted | No | No |
| MAP2K2 | LUAD | No | | Yes | targetted | No | No |
| MET | LUAD | Yes | [1, 2, 3] | Yes | targetted | Yes | No |
| MTOR | LUAD | No | | Yes | targetted | No | No |
| NRG1 | LUAD | Yes | [3] | No | | No | No |
| PDCD1 | LUAD | No | | Yes | targetted | No | No |
| RET | LUAD | Yes | [1, 2, 3] | Yes | targetted | No | No |
| ROS1 | LUAD | Yes | [1, 2, 3] | Yes | targetted | No | No |
| THYA | LUAD | No | | Yes | targetted | No | No |

| | | | | | | | |
|---|---|---|---|---|---|---|---|
| TYMS | LUAD | No | | Yes | targetted | No | No |
| VEGFA | LUAD | No | | Yes | targetted | No | No |
| VEGFB | LUAD | No | | Yes | targetted | No | No |
| AKT1 | OV | Yes | [3] | No | | No | No |
| BRAF | OV | Yes | [3] | No | | No | No |
| BRCA1 | OV | Yes | [1] | No | | Yes | No |
| BRCA2 | OV | Yes | [1] | No | | No | No |
| PARP1 | OV | No | | Yes | targetted | No | No |
| PARP2 | OV | No | | Yes | targetted | No | No |
| PARP3 | OV | No | | Yes | targetted | No | No |
| VEGFA | OV | No | | Yes | targetted | No | No |
| VEGFB | OV | No | | Yes | targetted | No | No |
| AR | PRAD | No | | Yes | targetted | Yes | No |
| ATM | PRAD | Yes | [1] | No | | Yes | No |
| BARD1 | PRAD | Yes | [1] | No | | No | No |
| BRCA1 | PRAD | Yes | [1] | No | | No | No |
| BRCA2 | PRAD | Yes | [1] | No | | Yes | No |
| BRIP1 | PRAD | Yes | [1] | No | | No | No |
| CDK12 | PRAD | Yes | [1] | No | | No | No |
| CHEK1 | PRAD | Yes | [1] | No | | No | No |
| CHEK2 | PRAD | Yes | [1] | No | | No | No |
| FANCL | PRAD | Yes | [1] | No | | No | No |
| GNRHR | PRAD | No | | Yes | targetted | No | No |
| PALB2 | PRAD | Yes | [1] | No | | No | No |
| PARP1 | PRAD | No | | Yes | targetted | Yes | No |
| PARP2 | PRAD | No | | Yes | targetted | No | No |
| PARP3 | PRAD | No | | Yes | targetted | No | No |
| RAD51B | PRAD | Yes | [1] | No | | No | No |
| RAD51C | PRAD | Yes | [1] | No | | No | No |
| RAD51D | PRAD | Yes | [1] | No | | No | No |
| RAD54L | PRAD | Yes | [1] | No | | No | No |
| TNFSF11 | PRAD | No | | Yes | targetted | No | No |
| BRAF | SKCM | Yes | [1, 2, 3] | Yes | targetted | Yes | Yes |
| CD274 | SKCM | No | | Yes | targetted | No | No |
| CTLA4 | SKCM | No | | Yes | targetted | No | No |
| IL2RA | SKCM | No | | Yes | targetted | No | No |
| KIT | SKCM | Yes | [2] | No | | No | No |
| MAP2K1 | SKCM | Yes | [3] | Yes | targetted | Yes | No |
| MAP2K2 | SKCM | No | | Yes | targetted | No | No |
| MAP3K20 | SKCM | No | | Yes | targetted | No | No |
| NRAS | SKCM | Yes | [3] | No | | Yes | Yes |
| PDCD1 | SKCM | No | | Yes | targetted | No | No |
| RAF1 | SKCM | No | | Yes | targetted | No | No |
| TLR7 | SKCM | No | | Yes | targetted | No | No |
| CCDC6 | THCA | No | | Yes | targetted | Yes | Yes |
| CRBN | THCA | No | | Yes | targetted | No | No |
| FLT1 | THCA | No | | Yes | targetted | No | No |
| FLT4 | THCA | No | | Yes | targetted | No | No |
| KDR | THCA | No | | Yes | targetted | No | No |
| KIF5B | THCA | No | | Yes | targetted | No | No |
| MAP2K1 | THCA | No | | Yes | targetted | No | No |
| MAP2K2 | THCA | No | | Yes | targetted | No | No |
| MS4A1 | THCA | No | | Yes | targetted | No | No |

| | | | | | | | |
|---|---|---|---|---|---|---|---|
| NRAS | THCA | Yes | [3] | No | | Yes | Yes |
| PIK3CA | THCA | No | | Yes | targetted | No | No |
| PIK3CD | THCA | No | | Yes | targetted | No | No |
| RET | THCA | Yes | [1] | Yes | targetted | Yes | Yes |
| AKT1 | UCEC | Yes | [3] | No | | Yes | No |
| ERBB2 | UCEC | Yes | [2] | No | | Yes | No |
| FLT1 | UCEC | No | | Yes | targetted | No | No |
| FLT4 | UCEC | No | | Yes | targetted | No | No |
| KDR | UCEC | No | | Yes | targetted | No | No |
| PDCD1 | UCEC | No | | Yes | targetted | No | No |
