## Supplementary material for "Probabilistic graph-based model uncovers previously unseen druggable vulnerabilities in major solid cancers": supp table 6

| Tumor type | Clinically relevant genes | Initial score | MVP pre-rank | MVP post-rank | Network size | Normalized degree | MVP rank change |
|---|---|---|---|---|---|---|---|
| COADREAD | ABL1 | 0 | 42 | 42 | 632 | 0.105 | 0 |
| UCEC | AKT1 | 0 | 34 | 34 | 644 | 0.154 | 0 |
| PRAD | AR | 0 | 15 | 15 | 638 | 0.127 | 0 |
| BRCA | EGFR | 0 | 16 | 16 | 638 | 0.294 | 0 |
| HNSC | EGFR | 0 | 14 | 14 | 636 | 0.268 | 0 |
| LUAD | ERBB2 | 0 | 13 | 13 | 630 | 0.192 | 0 |
| UCEC | ERBB2 | 0 | 45 | 45 | 644 | 0.134 | 0 |
| BLCA | ESR1 | 0 | 18 | 18 | 643 | 0.316 | 0 |
| BRCA | ESR1 | 0 | 17 | 17 | 638 | 0.281 | 0 |
| BLCA | HDAC1 | 0 | 56 | 56 | 643 | 0.195 | 0 |
| THCA | NRAS | 0.075 | 3 | 3 | 633 | 0.252 | 0 |
| PRAD | PARP1 | 0 | 35 | 35 | 638 | 0.072 | 0 |
| OV | BRCA1 | 0.009 | 5 | 6 | 633 | 0.383 | 1 |
| LUAD | EGFR | 0.123 | 2 | 3 | 630 | 0.388 | 1 |
| KIRC | FLT1 | 0.013 | 17 | 18 | 637 | 0.131 | 1 |
| PRAD | ATM | 0.041 | 5 | 7 | 638 | 0.192 | 2 |
| COADREAD | NRAS | 0.036 | 7 | 9 | 632 | 0.228 | 2 |
| BRCA | BRCA1 | 0.02 | 6 | 9 | 638 | 0.364 | 3 |
| COADREAD | KRAS | 0.277 | 2 | 9 | 632 | 0.233 | 7 |
| LUAD | MET | 0.039 | 15 | 23 | 630 | 0.146 | 8 |
| BRCA | ERBB2 | 0.022 | 14 | 24 | 638 | 0.223 | 10 |
| BRCA | AKT1 | 0.023 | 11 | 22 | 638 | 0.254 | 11 |
| PRAD | BRCA2 | 0.016 | 52 | 66 | 638 | 0.058 | 14 |
| KIRC | FLT4 | 0.079 | 9 | 23 | 637 | 0.118 | 14 |
| LUAD | KRAS | 0.295 | 3 | 19 | 630 | 0.148 | 16 |
| BLCA | ERBB2 | 0.118 | 7 | 27 | 643 | 0.268 | 20 |
| SKCM | MAP2K1 | 0.055 | 45 | 65 | 626 | 0.094 | 20 |
| KIRC | MTOR | 0.066 | 15 | 35 | 637 | 0.097 | 20 |
| THCA | CCDC6 | 0.043 | 58 | 79 | 633 | 0.041 | 21 |
| THCA | RET | 0.067 | 20 | 43 | 633 | 0.066 | 23 |
| SKCM | NRAS | 0.306 | 2 | 26 | 626 | 0.166 | 24 |
| HNSC | HRAS | 0.058 | 19 | 51 | 636 | 0.167 | 32 |
| LUAD | BRAF | 0.08 | 45 | 81 | 630 | 0.064 | 36 |
| BRCA | PIK3CA | 0.258 | 2 | 42 | 638 | 0.179 | 40 |
| COADREAD | BRAF | 0.09 | 41 | 86 | 632 | 0.067 | 45 |
| BLCA | FGFR3 | 0.145 | 10 | 60 | 643 | 0.195 | 50 |
| BRCA | CCND1 | 0.156 | 4 | 56 | 638 | 0.16 | 52 |
| SKCM | BRAF | 0.51 | 4 | 95 | 626 | 0.074 | 91 |
| LGG | IDH1 | 0.773 | 34 | 341 | 640 | 0.02 | 307 |
| OV | TOP1MT | 0.277 | 10 | 581 | 633 | 0.005 | 571 |
