## Supplementary material for "Probabilistic graph-based model uncovers previously unseen druggable vulnerabilities in major solid cancers": supp table 7

| Type | cancer_type | top_index | top_genes | depmap_essential | common_essential |
|---|---|---|---|---|---|
| DNA as seed | BLCA | 1 | TP53 | 0 | 0 |
| DNA as seed | BLCA | 2 | EP300 | 1 | 0 |
| DNA as seed | BLCA | 3 | PIK3CA | 1 | 0 |
| DNA as seed | BLCA | 4 | CDKN2A | 0 | 0 |
| DNA as seed | BLCA | 5 | CREBBP | 1 | 0 |
| DNA as seed | BLCA | 6 | RB1 | 0 | 0 |
| DNA as seed | BLCA | 7 | ERBB2 | 0 | 0 |
| DNA as seed | BLCA | 8 | YWHAZ | 0 | 0 |
| DNA as seed | BLCA | 9 | CDKN1A | 0 | 0 |
| DNA as seed | BLCA | 10 | FGFR3 | 0 | 0 |
| DNA as seed | BLCA | 11 | ERBB3 | 0 | 0 |
| DNA as seed | BLCA | 12 | ATM | 0 | 0 |
| DNA as seed | BLCA | 13 | ARID1A | 0 | 0 |
| DNA as seed | BLCA | 14 | FCGR2A | 0 | 0 |
| DNA as seed | BLCA | 15 | USF1 | 0 | 0 |
| DNA as seed | BLCA | 16 | E2F3 | 1 | 0 |
| DNA as seed | BLCA | 17 | FCER1G | 0 | 0 |
| DNA as seed | BLCA | 18 | ESR1 | 0 | 0 |
| DNA as seed | BLCA | 19 | FCGR3A | 0 | 0 |
| DNA as seed | BLCA | 20 | NCOR1 | 0 | 0 |
| DNA as seed | BLCA | 21 | MYC | 1 | 0 |
| DNA as seed | BLCA | 22 | HRAS | 1 | 0 |
| DNA as seed | BLCA | 23 | GRB2 | 1 | 1 |
| DNA as seed | BLCA | 24 | CDKN2B | 0 | 0 |
| DNA as seed | BLCA | 25 | PIK3R1 | 0 | 0 |
| DNA as seed | BLCA | 26 | SP1 | 0 | 0 |
| DNA as seed | BLCA | 27 | SHC1 | 0 | 0 |
| DNA as seed | BLCA | 28 | FCGR2B | 0 | 0 |
| DNA as seed | BLCA | 29 | RELA | 0 | 0 |
| DNA as seed | BLCA | 30 | CUL1 | 1 | 1 |
| DNA as seed | BLCA | 31 | PIK3R2 | 0 | 0 |
| DNA as seed | BLCA | 32 | FBXW7 | 0 | 0 |
| DNA as seed | BLCA | 33 | UBC | 1 | 0 |
| DNA as seed | BLCA | 34 | AR | 0 | 0 |
| DNA as seed | BLCA | 35 | CDK2 | 1 | 0 |
| DNA as seed | BLCA | 36 | RXRA | 0 | 0 |
| DNA as seed | BLCA | 37 | CTNNB1 | 1 | 0 |
| DNA as seed | BLCA | 38 | KRAS | 1 | 0 |
| DNA as seed | BLCA | 39 | BRCA1 | 0 | 1 |
| DNA as seed | BLCA | 40 | SRC | 0 | 0 |
| DNA as seed | BLCA | 41 | MYH9 | 1 | 0 |
| DNA as seed | BLCA | 42 | ELAVL1 | 0 | 0 |
| DNA as seed | BLCA | 43 | MAPK1 | 0 | 0 |
| DNA as seed | BLCA | 44 | CBL | 0 | 0 |
| DNA as seed | BLCA | 45 | HSP90AA1 | 0 | 0 |
| DNA as seed | BLCA | 46 | MAPK3 | 0 | 0 |
| DNA as seed | BLCA | 47 | RAC1 | 1 | 1 |
| DNA as seed | BLCA | 48 | PLCG1 | 0 | 0 |
| DNA as seed | BLCA | 49 | PIK3CB | 0 | 0 |
| DNA as seed | BLCA | 50 | PTEN | 0 | 0 |
| DNA as seed | BLCA | 51 | JUN | 0 | 0 |
| DNA as seed | BLCA | 52 | PABPC1 | 1 | 0 |
| DNA as seed | BLCA | 53 | HSPA6 | 0 | 0 |
| DNA as seed | BLCA | 54 | SOS1 | 0 | 0 |
| DNA as seed | BLCA | 55 | NFKB1 | 0 | 0 |

| | | | | | |
|---|---|---|---|---|---|
| DNA as seed | BLCA | 56 | HDAC1 | 0 | 0 |
| DNA as seed | BLCA | 57 | LYN | 0 | 0 |
| DNA as seed | BLCA | 58 | KDM6A | 0 | 0 |
| DNA as seed | BLCA | 59 | EPHA2 | 0 | 0 |
| DNA as seed | BLCA | 60 | SMAD3 | 0 | 0 |
| DNA as seed | BLCA | 61 | HIF1A | 0 | 0 |
| DNA as seed | BLCA | 62 | STAT3 | 0 | 0 |
| DNA as seed | BLCA | 63 | CRK | 0 | 0 |
| DNA as seed | BLCA | 64 | AKT1 | 0 | 0 |
| DNA as seed | BLCA | 65 | E2F1 | 0 | 0 |
| DNA as seed | BLCA | 66 | RHOA | 1 | 1 |
| DNA as seed | BLCA | 67 | SYK | 0 | 0 |
| DNA as seed | BLCA | 68 | USP21 | 0 | 0 |
| DNA as seed | BLCA | 69 | MET | 0 | 0 |
| DNA as seed | BLCA | 70 | UBB | 0 | 0 |
| DNA as seed | BLCA | 71 | PPARG | 1 | 0 |
| DNA as seed | BLCA | 72 | PRKCA | 0 | 0 |
| DNA as seed | BLCA | 73 | SOX4 | 0 | 0 |
| DNA as seed | BLCA | 74 | FOS | 0 | 0 |
| DNA as seed | BLCA | 75 | MDM2 | 1 | 0 |
| DNA as seed | BLCA | 76 | SPTAN1 | 0 | 0 |
| DNA as seed | BLCA | 77 | PLCG2 | 0 | 0 |
| DNA as seed | BLCA | 78 | GSK3B | 0 | 0 |
| DNA as seed | BLCA | 79 | SMAD2 | 0 | 0 |
| DNA as seed | BLCA | 80 | MAPK14 | 0 | 0 |
| DNA as seed | BLCA | 81 | VAV1 | 0 | 0 |
| DNA as seed | BLCA | 82 | CDH1 | 0 | 0 |
| DNA as seed | BLCA | 83 | HSPA8 | 1 | 1 |
| DNA as seed | BLCA | 84 | HSP90AB1 | 0 | 0 |
| DNA as seed | BLCA | 85 | CCND1 | 1 | 0 |
| DNA as seed | BLCA | 86 | MAPK8 | 0 | 0 |
| DNA as seed | BLCA | 87 | HDAC3 | 1 | 0 |
| DNA as seed | BLCA | 88 | TRRAP | 1 | 1 |
| DNA as seed | BLCA | 89 | RPS27A | 1 | 0 |
| DNA as seed | BLCA | 90 | GAB2 | 0 | 0 |
| DNA as seed | BLCA | 91 | FYN | 0 | 0 |
| DNA as seed | BLCA | 92 | SMARCA4 | 0 | 0 |
| DNA as seed | BLCA | 93 | ACTB | 1 | 0 |
| DNA as seed | BLCA | 94 | UBE2I | 1 | 1 |
| DNA as seed | BLCA | 95 | VAV2 | 0 | 0 |
| DNA as seed | BLCA | 96 | VAV3 | 0 | 0 |
| DNA as seed | BLCA | 97 | PCNA | 1 | 1 |
| DNA as seed | BLCA | 98 | KLF5 | 1 | 0 |
| DNA as seed | BLCA | 99 | NCOA3 | 0 | 0 |
| DNA as seed | BLCA | 100 | NFE2L2 | 0 | 0 |
| DNA as seed | BRCA | 1 | TP53 | 0 | 0 |
| DNA as seed | BRCA | 2 | PIK3CA | 1 | 0 |
| DNA as seed | BRCA | 3 | MYC | 1 | 0 |
| DNA as seed | BRCA | 4 | CCND1 | 1 | 0 |
| DNA as seed | BRCA | 5 | CDH1 | 0 | 0 |
| DNA as seed | BRCA | 6 | BRCA1 | 1 | 1 |
| DNA as seed | BRCA | 7 | FGF3 | 0 | 0 |
| DNA as seed | BRCA | 8 | FGF4 | 0 | 0 |
| DNA as seed | BRCA | 9 | FGF19 | 0 | 0 |
| DNA as seed | BRCA | 10 | CTTN | 0 | 0 |
| DNA as seed | BRCA | 11 | AKT1 | 1 | 0 |

| | | | | | |
|---|---|---|---|---|---|
| DNA as seed | BRCA | 12 | RAD21 | 1 | 0 |
| DNA as seed | BRCA | 13 | PIK3R1 | 0 | 0 |
| DNA as seed | BRCA | 14 | ERBB2 | 1 | 0 |
| DNA as seed | BRCA | 15 | PTEN | 0 | 0 |
| DNA as seed | BRCA | 16 | EGFR | 1 | 0 |
| DNA as seed | BRCA | 17 | ESR1 | 1 | 0 |
| DNA as seed | BRCA | 18 | RB1 | 0 | 0 |
| DNA as seed | BRCA | 19 | UBC | 1 | 0 |
| DNA as seed | BRCA | 20 | EP300 | 1 | 0 |
| DNA as seed | BRCA | 21 | MAP3K1 | 0 | 0 |
| DNA as seed | BRCA | 22 | GRB2 | 1 | 1 |
| DNA as seed | BRCA | 23 | SP1 | 0 | 0 |
| DNA as seed | BRCA | 24 | SRC | 0 | 0 |
| DNA as seed | BRCA | 25 | NCOR1 | 0 | 0 |
| DNA as seed | BRCA | 26 | ATM | 0 | 0 |
| DNA as seed | BRCA | 27 | RUNX1 | 1 | 0 |
| DNA as seed | BRCA | 28 | HDAC1 | 0 | 0 |
| DNA as seed | BRCA | 29 | RELA | 1 | 0 |
| DNA as seed | BRCA | 30 | CDK2 | 1 | 0 |
| DNA as seed | BRCA | 31 | HSP90AA1 | 0 | 0 |
| DNA as seed | BRCA | 32 | AR | 0 | 0 |
| DNA as seed | BRCA | 33 | CTNNB1 | 1 | 0 |
| DNA as seed | BRCA | 34 | MAPK3 | 0 | 0 |
| DNA as seed | BRCA | 35 | ERBB3 | 1 | 0 |
| DNA as seed | BRCA | 36 | MAPK1 | 0 | 0 |
| DNA as seed | BRCA | 37 | CREBBP | 0 | 0 |
| DNA as seed | BRCA | 38 | GATA3 | 1 | 0 |
| DNA as seed | BRCA | 39 | STAT3 | 0 | 0 |
| DNA as seed | BRCA | 40 | CDKN1A | 0 | 0 |
| DNA as seed | BRCA | 41 | MDM2 | 1 | 0 |
| DNA as seed | BRCA | 42 | JUN | 0 | 0 |
| DNA as seed | BRCA | 43 | NFKB1 | 0 | 0 |
| DNA as seed | BRCA | 44 | CASP8 | 0 | 0 |
| DNA as seed | BRCA | 45 | MYH9 | 1 | 0 |
| DNA as seed | BRCA | 46 | HSP90AB1 | 0 | 0 |
| DNA as seed | BRCA | 47 | FBXW7 | 1 | 0 |
| DNA as seed | BRCA | 48 | SMAD3 | 0 | 0 |
| DNA as seed | BRCA | 49 | PIK3R2 | 0 | 0 |
| DNA as seed | BRCA | 50 | CDK1 | 1 | 1 |
| DNA as seed | BRCA | 51 | HDAC2 | 1 | 0 |
| DNA as seed | BRCA | 52 | NPM1 | 0 | 0 |
| DNA as seed | BRCA | 53 | HIF1A | 0 | 0 |
| DNA as seed | BRCA | 54 | SHC1 | 0 | 0 |
| DNA as seed | BRCA | 55 | FADD | 1 | 0 |
| DNA as seed | BRCA | 56 | FOXA1 | 1 | 0 |
| DNA as seed | BRCA | 57 | FOS | 0 | 0 |
| DNA as seed | BRCA | 58 | ATAD2 | 1 | 0 |
| DNA as seed | BRCA | 59 | UBE2I | 1 | 1 |
| DNA as seed | BRCA | 60 | CBL | 0 | 0 |
| DNA as seed | BRCA | 61 | RPS27A | 1 | 0 |
| DNA as seed | BRCA | 62 | HSPA8 | 1 | 1 |
| DNA as seed | BRCA | 63 | SIRT7 | 0 | 0 |
| DNA as seed | BRCA | 64 | UBB | 0 | 0 |
| DNA as seed | BRCA | 65 | E2F1 | 1 | 0 |
| DNA as seed | BRCA | 66 | PLCG1 | 0 | 0 |
| DNA as seed | BRCA | 67 | HDAC3 | 1 | 0 |

| | | | | | |
|---|---|---|---|---|---|
| DNA as seed | BRCA | 68 | ABL1 | 0 | 0 |
| DNA as seed | BRCA | 69 | l2\|H3C2\|H3C3 | 0 | 0 |
| DNA as seed | BRCA | 70 | CRK | 0 | 0 |
| DNA as seed | BRCA | 71 | SMAD2 | 0 | 0 |
| DNA as seed | BRCA | 72 | ASAP1 | 0 | 0 |
| DNA as seed | BRCA | 73 | H3-4 | 0 | 0 |
| DNA as seed | BRCA | 74 | HRAS | 0 | 0 |
| DNA as seed | BRCA | 75 | CHD4 | 1 | 0 |
| DNA as seed | BRCA | 76 | PARP1 | 0 | 0 |
| DNA as seed | BRCA | 77 | SHANK2 | 0 | 0 |
| DNA as seed | BRCA | 78 | CREB1 | 0 | 0 |
| DNA as seed | BRCA | 79 | PPARG | 0 | 0 |
| DNA as seed | BRCA | 80 | EGR1 | 0 | 0 |
| DNA as seed | BRCA | 81 | FYN | 0 | 0 |
| DNA as seed | BRCA | 82 | SNW1 | 1 | 1 |
| DNA as seed | BRCA | 83 | PML | 0 | 0 |
| DNA as seed | BRCA | 84 | MET | 0 | 0 |
| DNA as seed | BRCA | 85 | SIRT1 | 0 | 0 |
| DNA as seed | BRCA | 86 | BRCA2 | 1 | 1 |
| DNA as seed | BRCA | 87 | 14\|H4C15\|H4 | 0 | 0 |
| DNA as seed | BRCA | 88 | ERBB4 | 0 | 0 |
| DNA as seed | BRCA | 89 | LMNA | 0 | 0 |
| DNA as seed | BRCA | 90 | UBA52 | 1 | 0 |
| DNA as seed | BRCA | 91 | NRAS | 0 | 0 |
| DNA as seed | BRCA | 92 | KAT2B | 0 | 0 |
| DNA as seed | BRCA | 93 | KRAS | 1 | 0 |
| DNA as seed | BRCA | 94 | EZH2 | 0 | 0 |
| DNA as seed | BRCA | 95 | FLNA | 0 | 0 |
| DNA as seed | BRCA | 96 | NOTCH1 | 0 | 0 |
| DNA as seed | BRCA | 97 | SMARCA4 | 1 | 0 |
| DNA as seed | BRCA | 98 | IGF1R | 0 | 0 |
| DNA as seed | BRCA | 99 | SMAD4 | 0 | 0 |
| DNA as seed | BRCA | 100 | H2AX | 1 | 0 |
| DNA as seed | COADREAD | 1 | TP53 | 0 | 0 |
| DNA as seed | COADREAD | 2 | KRAS | 1 | 0 |
| DNA as seed | COADREAD | 3 | APC | 1 | 0 |
| DNA as seed | COADREAD | 4 | PIK3CA | 1 | 0 |
| DNA as seed | COADREAD | 5 | CTNNB1 | 1 | 0 |
| DNA as seed | COADREAD | 6 | PIK3R1 | 0 | 0 |
| DNA as seed | COADREAD | 7 | NRAS | 0 | 0 |
| DNA as seed | COADREAD | 8 | ATM | 0 | 0 |
| DNA as seed | COADREAD | 9 | SMAD4 | 1 | 0 |
| DNA as seed | COADREAD | 10 | SRC | 0 | 0 |
| DNA as seed | COADREAD | 11 | PTEN | 0 | 0 |
| DNA as seed | COADREAD | 12 | ESR1 | 0 | 0 |
| DNA as seed | COADREAD | 13 | FBXW7 | 0 | 0 |
| DNA as seed | COADREAD | 14 | WWOX | 0 | 0 |
| DNA as seed | COADREAD | 15 | PIK3CB | 0 | 0 |
| DNA as seed | COADREAD | 16 | ELAVL1 | 1 | 0 |
| DNA as seed | COADREAD | 17 | MAPK1 | 1 | 0 |
| DNA as seed | COADREAD | 18 | PRKDC | 0 | 0 |
| DNA as seed | COADREAD | 19 | TCF7L2 | 1 | 0 |
| DNA as seed | COADREAD | 20 | SMAD3 | 0 | 0 |
| DNA as seed | COADREAD | 21 | HSP90AA1 | 0 | 0 |
| DNA as seed | COADREAD | 22 | AKT1 | 0 | 0 |
| DNA as seed | COADREAD | 23 | CDH1 | 1 | 0 |

| | | | | | |
|---|---|---|---|---|---|
| DNA as seed | COADREAD | 24 | ITCH | 0 | 0 |
| DNA as seed | COADREAD | 25 | PIK3R3 | 0 | 0 |
| DNA as seed | COADREAD | 26 | SHC1 | 0 | 0 |
| DNA as seed | COADREAD | 27 | FYN | 0 | 0 |
| DNA as seed | COADREAD | 28 | CDK2 | 1 | 0 |
| DNA as seed | COADREAD | 29 | PIK3CD | 0 | 0 |
| DNA as seed | COADREAD | 30 | SMAD2 | 0 | 0 |
| DNA as seed | COADREAD | 31 | AR | 0 | 0 |
| DNA as seed | COADREAD | 32 | CSNK2A1 | 0 | 0 |
| DNA as seed | COADREAD | 33 | GSK3B | 0 | 0 |
| DNA as seed | COADREAD | 34 | CREBBP | 1 | 0 |
| DNA as seed | COADREAD | 35 | JUN | 1 | 0 |
| DNA as seed | COADREAD | 36 | YWHAZ | 1 | 0 |
| DNA as seed | COADREAD | 37 | HSP90AB1 | 0 | 0 |
| DNA as seed | COADREAD | 38 | RPS27A | 1 | 0 |
| DNA as seed | COADREAD | 39 | HCK | 0 | 0 |
| DNA as seed | COADREAD | 40 | HLA-B | 0 | 0 |
| DNA as seed | COADREAD | 41 | BRAF | 1 | 0 |
| DNA as seed | COADREAD | 42 | ABL1 | 0 | 0 |
| DNA as seed | COADREAD | 43 | MAPK8 | 0 | 0 |
| DNA as seed | COADREAD | 44 | BCL2L1 | 1 | 1 |
| DNA as seed | COADREAD | 45 | ERBB3 | 0 | 0 |
| DNA as seed | COADREAD | 46 | UBE2I | 1 | 1 |
| DNA as seed | COADREAD | 47 | B2M | 0 | 0 |
| DNA as seed | COADREAD | 48 | MAPK14 | 0 | 0 |
| DNA as seed | COADREAD | 49 | CDC37 | 1 | 1 |
| DNA as seed | COADREAD | 50 | IGF1R | 1 | 0 |
| DNA as seed | COADREAD | 51 | CRK | 0 | 0 |
| DNA as seed | COADREAD | 52 | MET | 0 | 0 |
| DNA as seed | COADREAD | 53 | ERBB4 | 0 | 0 |
| DNA as seed | COADREAD | 54 | CREB1 | 0 | 0 |
| DNA as seed | COADREAD | 55 | HIF1A | 0 | 0 |
| DNA as seed | COADREAD | 56 | PTK2 | 1 | 0 |
| DNA as seed | COADREAD | 57 | IKBKB | 0 | 0 |
| DNA as seed | COADREAD | 58 | NPM1 | 0 | 0 |
| DNA as seed | COADREAD | 59 | EWSR1 | 1 | 1 |
| DNA as seed | COADREAD | 60 | FOS | 0 | 0 |
| DNA as seed | COADREAD | 61 | IRS1 | 0 | 0 |
| DNA as seed | COADREAD | 62 | PARP1 | 0 | 0 |
| DNA as seed | COADREAD | 63 | BCL2 | 0 | 0 |
| DNA as seed | COADREAD | 64 | SMAD1 | 0 | 0 |
| DNA as seed | COADREAD | 65 | FGFR2 | 0 | 0 |
| DNA as seed | COADREAD | 66 | RBM39 | 1 | 1 |
| DNA as seed | COADREAD | 67 | CDC42 | 1 | 0 |
| DNA as seed | COADREAD | 68 | BMPR2 | 1 | 0 |
| DNA as seed | COADREAD | 69 | NOTCH1 | 0 | 0 |
| DNA as seed | COADREAD | 70 | NCK1 | 0 | 0 |
| DNA as seed | COADREAD | 71 | BTRC | 0 | 0 |
| DNA as seed | COADREAD | 72 | PCBP1 | 1 | 0 |
| DNA as seed | COADREAD | 73 | CDKN2A | 0 | 0 |
| DNA as seed | COADREAD | 74 | MAPK9 | 0 | 0 |
| DNA as seed | COADREAD | 75 | YWHAQ | 0 | 0 |
| DNA as seed | COADREAD | 76 | RRAS2 | 0 | 0 |
| DNA as seed | COADREAD | 77 | FOXO3 | 0 | 0 |
| DNA as seed | COADREAD | 78 | CHUK | 0 | 0 |
| DNA as seed | COADREAD | 79 | CASP3 | 0 | 0 |

| | | | | | |
|---|---|---|---|---|---|
| DNA as seed | COADREAD | 80 | XRCC6 | 1 | 1 |
| DNA as seed | COADREAD | 81 | HSPA1B | 0 | 0 |
| DNA as seed | COADREAD | 82 | FUS | 0 | 0 |
| DNA as seed | COADREAD | 83 | KIT | 0 | 0 |
| DNA as seed | COADREAD | 84 | EZH2 | 0 | 0 |
| DNA as seed | COADREAD | 85 | CSNK2B | 1 | 1 |
| DNA as seed | COADREAD | 86 | FGFR4 | 0 | 0 |
| DNA as seed | COADREAD | 87 | PPP2CB | 0 | 0 |
| DNA as seed | COADREAD | 88 | CASP8 | 0 | 0 |
| DNA as seed | COADREAD | 89 | NCOA6 | 1 | 0 |
| DNA as seed | COADREAD | 90 | TRIM25 | 0 | 0 |
| DNA as seed | COADREAD | 91 | CAMK2A | 0 | 0 |
| DNA as seed | COADREAD | 92 | ID1 | 0 | 0 |
| DNA as seed | COADREAD | 93 | FLT1 | 0 | 0 |
| DNA as seed | COADREAD | 94 | TGFBR1 | 0 | 0 |
| DNA as seed | COADREAD | 95 | HSPA1A | 0 | 0 |
| DNA as seed | COADREAD | 96 | MAPK10 | 0 | 0 |
| DNA as seed | COADREAD | 97 | MAP1LC3A | 0 | 0 |
| DNA as seed | COADREAD | 98 | FLT3 | 0 | 0 |
| DNA as seed | COADREAD | 99 | LEF1 | 0 | 0 |
| DNA as seed | COADREAD | 100 | CHEK1 | 1 | 1 |
| DNA as seed | GBM | 1 | EGFR | 0 | 0 |
| DNA as seed | GBM | 2 | TP53 | 0 | 0 |
| DNA as seed | GBM | 3 | IFNA5 | 0 | 0 |
| DNA as seed | GBM | 4 | PTEN | 0 | 0 |
| DNA as seed | GBM | 5 | CDKN2A | 0 | 0 |
| DNA as seed | GBM | 6 | IFNA2 | 0 | 0 |
| DNA as seed | GBM | 7 | PIK3R1 | 0 | 0 |
| DNA as seed | GBM | 8 | IFNA8 | 0 | 0 |
| DNA as seed | GBM | 9 | IFNA6 | 0 | 0 |
| DNA as seed | GBM | 10 | IFNA4 | 0 | 0 |
| DNA as seed | GBM | 11 | IFNA14 | 0 | 0 |
| DNA as seed | GBM | 12 | PDGFRA | 1 | 0 |
| DNA as seed | GBM | 13 | IFNA21 | 0 | 0 |
| DNA as seed | GBM | 14 | PIK3CA | 1 | 0 |
| DNA as seed | GBM | 15 | IFNA1 | 0 | 0 |
| DNA as seed | GBM | 16 | IFNA17 | 0 | 0 |
| DNA as seed | GBM | 17 | IFNE | 0 | 0 |
| DNA as seed | GBM | 18 | IFNA7 | 0 | 0 |
| DNA as seed | GBM | 19 | IFNA13 | 0 | 0 |
| DNA as seed | GBM | 20 | IFNA10 | 0 | 0 |
| DNA as seed | GBM | 21 | IFNA16 | 0 | 0 |
| DNA as seed | GBM | 22 | IFNB1 | 0 | 0 |
| DNA as seed | GBM | 23 | IFNW1 | 0 | 0 |
| DNA as seed | GBM | 24 | PIK3CB | 0 | 0 |
| DNA as seed | GBM | 25 | PTPN11 | 1 | 0 |
| DNA as seed | GBM | 26 | PIK3R2 | 0 | 0 |
| DNA as seed | GBM | 27 | JAK2 | 0 | 0 |
| DNA as seed | GBM | 28 | RELA | 1 | 0 |
| DNA as seed | GBM | 29 | JAK1 | 0 | 0 |
| DNA as seed | GBM | 30 | PIK3CD | 0 | 0 |
| DNA as seed | GBM | 31 | PIK3R3 | 0 | 0 |
| DNA as seed | GBM | 32 | CDKN2B | 0 | 0 |
| DNA as seed | GBM | 33 | JAK3 | 0 | 0 |
| DNA as seed | GBM | 34 | PDGFRB | 0 | 0 |
| DNA as seed | GBM | 35 | NFKB1 | 0 | 0 |

| | | | | | |
|---|---|---|---|---|---|
| DNA as seed | GBM | 36 | PLCG1 | 0 | 0 |
| DNA as seed | GBM | 37 | TYK2 | 0 | 0 |
| DNA as seed | GBM | 38 | STAT1 | 0 | 0 |
| DNA as seed | GBM | 39 | SHC1 | 0 | 0 |
| DNA as seed | GBM | 40 | CDK4 | 1 | 0 |
| DNA as seed | GBM | 41 | PTPN6 | 0 | 0 |
| DNA as seed | GBM | 42 | STAT3 | 0 | 0 |
| DNA as seed | GBM | 43 | SP1 | 0 | 0 |
| DNA as seed | GBM | 44 | RB1 | 0 | 0 |
| DNA as seed | GBM | 45 | CSF2RB | 0 | 0 |
| DNA as seed | GBM | 46 | IL2RG | 0 | 0 |
| DNA as seed | GBM | 47 | IFNGR1 | 0 | 0 |
| DNA as seed | GBM | 48 | IL6ST | 0 | 0 |
| DNA as seed | GBM | 49 | HRAS | 0 | 0 |
| DNA as seed | GBM | 50 | IL2RB | 0 | 0 |
| DNA as seed | GBM | 51 | PDGFB | 0 | 0 |
| DNA as seed | GBM | 52 | CTNNB1 | 0 | 0 |
| DNA as seed | GBM | 53 | EP300 | 0 | 0 |
| DNA as seed | GBM | 54 | IL7R | 0 | 0 |
| DNA as seed | GBM | 55 | SOCS1 | 0 | 0 |
| DNA as seed | GBM | 56 | EPOR | 0 | 0 |
| DNA as seed | GBM | 57 | IL3RA | 0 | 0 |
| DNA as seed | GBM | 58 | IL2RA | 0 | 0 |
| DNA as seed | GBM | 59 | PDGFA | 0 | 0 |
| DNA as seed | GBM | 60 | IFNGR2 | 0 | 0 |
| DNA as seed | GBM | 61 | EGF | 0 | 0 |
| DNA as seed | GBM | 62 | CBL | 0 | 0 |
| DNA as seed | GBM | 63 | GHR | 0 | 0 |
| DNA as seed | GBM | 64 | CSF2RA | 0 | 0 |
| DNA as seed | GBM | 65 | IFNAR1 | 0 | 0 |
| DNA as seed | GBM | 66 | IRF3 | 0 | 0 |
| DNA as seed | GBM | 67 | IFNAR2 | 0 | 0 |
| DNA as seed | GBM | 68 | IL5RA | 0 | 0 |
| DNA as seed | GBM | 69 | IL4R | 0 | 0 |
| DNA as seed | GBM | 70 | SOCS3 | 1 | 0 |
| DNA as seed | GBM | 71 | PTPN1 | 0 | 0 |
| DNA as seed | GBM | 72 | LANCL2 | 0 | 0 |
| DNA as seed | GBM | 73 | LIFR | 0 | 0 |
| DNA as seed | GBM | 74 | IL12RB1 | 0 | 0 |
| DNA as seed | GBM | 75 | IL6R | 0 | 0 |
| DNA as seed | GBM | 76 | NRAS | 0 | 0 |
| DNA as seed | GBM | 77 | LEPR | 0 | 0 |
| DNA as seed | GBM | 78 | PLCG2 | 0 | 0 |
| DNA as seed | GBM | 79 | PRLR | 0 | 0 |
| DNA as seed | GBM | 80 | IL13RA2 | 0 | 0 |
| DNA as seed | GBM | 81 | IL20RB | 0 | 0 |
| DNA as seed | GBM | 82 | IL12RB2 | 0 | 0 |
| DNA as seed | GBM | 83 | STAT2 | 0 | 0 |
| DNA as seed | GBM | 84 | IL20RA | 0 | 0 |
| DNA as seed | GBM | 85 | IRF7 | 0 | 0 |
| DNA as seed | GBM | 86 | MET | 0 | 0 |
| DNA as seed | GBM | 87 | CSF3R | 0 | 0 |
| DNA as seed | GBM | 88 | IL15RA | 0 | 0 |
| DNA as seed | GBM | 89 | IL22RA1 | 0 | 0 |
| DNA as seed | GBM | 90 | IL10RB | 0 | 0 |
| DNA as seed | GBM | 91 | AKT1 | 0 | 0 |

| | | | | | |
|---|---|---|---|---|---|
| DNA as seed | GBM | 92 | IL2 | 0 | 0 |
| DNA as seed | GBM | 93 | MPL | 0 | 0 |
| DNA as seed | GBM | 94 | LYN | 0 | 0 |
| DNA as seed | GBM | 95 | IL23R | 0 | 0 |
| DNA as seed | GBM | 96 | IL21R | 0 | 0 |
| DNA as seed | GBM | 97 | IL10RA | 0 | 0 |
| DNA as seed | GBM | 98 | CREBBP | 0 | 0 |
| DNA as seed | GBM | 99 | KRAS | 1 | 0 |
| DNA as seed | GBM | 100 | IL3 | 0 | 0 |
| DNA as seed | HNSC | 1 | TP53 | 0 | 0 |
| DNA as seed | HNSC | 2 | EP300 | 0 | 0 |
| DNA as seed | HNSC | 3 | CCND1 | 1 | 0 |
| DNA as seed | HNSC | 4 | CDKN2A | 0 | 0 |
| DNA as seed | HNSC | 5 | CREBBP | 0 | 0 |
| DNA as seed | HNSC | 6 | PIK3CA | 0 | 0 |
| DNA as seed | HNSC | 7 | NOTCH1 | 0 | 0 |
| DNA as seed | HNSC | 8 | MYC | 1 | 0 |
| DNA as seed | HNSC | 9 | NTRK1 | 0 | 0 |
| DNA as seed | HNSC | 10 | ESR1 | 0 | 0 |
| DNA as seed | HNSC | 11 | TP63 | 1 | 0 |
| DNA as seed | HNSC | 12 | CUL3 | 0 | 0 |
| DNA as seed | HNSC | 13 | UBC | 0 | 0 |
| DNA as seed | HNSC | 14 | EGFR | 1 | 0 |
| DNA as seed | HNSC | 15 | CTNNB1 | 1 | 0 |
| DNA as seed | HNSC | 16 | HUWE1 | 1 | 0 |
| DNA as seed | HNSC | 17 | BRCA1 | 0 | 1 |
| DNA as seed | HNSC | 18 | RB1 | 0 | 0 |
| DNA as seed | HNSC | 19 | HRAS | 0 | 0 |
| DNA as seed | HNSC | 20 | RELA | 0 | 0 |
| DNA as seed | HNSC | 21 | SOX2 | 1 | 0 |
| DNA as seed | HNSC | 22 | HDAC1 | 0 | 0 |
| DNA as seed | HNSC | 23 | MAPK1 | 0 | 0 |
| DNA as seed | HNSC | 24 | SMAD3 | 0 | 0 |
| DNA as seed | HNSC | 25 | GRB2 | 1 | 1 |
| DNA as seed | HNSC | 26 | SRC | 0 | 0 |
| DNA as seed | HNSC | 27 | SMARCA4 | 0 | 0 |
| DNA as seed | HNSC | 28 | SP1 | 0 | 0 |
| DNA as seed | HNSC | 29 | CASP8 | 0 | 0 |
| DNA as seed | HNSC | 30 | CDK2 | 0 | 0 |
| DNA as seed | HNSC | 31 | FBXW7 | 0 | 0 |
| DNA as seed | HNSC | 32 | SMAD4 | 0 | 0 |
| DNA as seed | HNSC | 33 | CDKN1A | 0 | 0 |
| DNA as seed | HNSC | 34 | NCOR1 | 0 | 0 |
| DNA as seed | HNSC | 35 | AR | 0 | 0 |
| DNA as seed | HNSC | 36 | HSP90AA1 | 0 | 0 |
| DNA as seed | HNSC | 37 | MAPK3 | 0 | 0 |
| DNA as seed | HNSC | 38 | FGF4 | 0 | 0 |
| DNA as seed | HNSC | 39 | MDM2 | 1 | 0 |
| DNA as seed | HNSC | 40 | FGF3 | 0 | 0 |
| DNA as seed | HNSC | 41 | STAT3 | 0 | 0 |
| DNA as seed | HNSC | 42 | SMAD2 | 0 | 0 |
| DNA as seed | HNSC | 43 | PTEN | 0 | 0 |
| DNA as seed | HNSC | 44 | CDH1 | 0 | 0 |
| DNA as seed | HNSC | 45 | AKT1 | 0 | 0 |
| DNA as seed | HNSC | 46 | ACTL6A | 1 | 1 |
| DNA as seed | HNSC | 47 | FXR1 | 0 | 0 |

| | | | | | |
|---|---|---|---|---|---|
| DNA as seed | HNSC | 48 | RAC1 | 0 | 1 |
| DNA as seed | HNSC | 49 | PIK3R1 | 0 | 0 |
| DNA as seed | HNSC | 50 | CTTN | 0 | 0 |
| DNA as seed | HNSC | 51 | AP2M1 | 1 | 0 |
| DNA as seed | HNSC | 52 | FGF19 | 0 | 0 |
| DNA as seed | HNSC | 53 | NFKB1 | 0 | 0 |
| DNA as seed | HNSC | 54 | HDAC3 | 1 | 0 |
| DNA as seed | HNSC | 55 | PPARG | 0 | 0 |
| DNA as seed | HNSC | 56 | SNW1 | 1 | 1 |
| DNA as seed | HNSC | 57 | HIF1A | 0 | 0 |
| DNA as seed | HNSC | 58 | UBB | 0 | 0 |
| DNA as seed | HNSC | 59 | PIK3R2 | 0 | 0 |
| DNA as seed | HNSC | 60 | CDKN2B | 0 | 0 |
| DNA as seed | HNSC | 61 | ACTB | 1 | 0 |
| DNA as seed | HNSC | 62 | HSP90AB1 | 0 | 0 |
| DNA as seed | HNSC | 63 | NPM1 | 0 | 0 |
| DNA as seed | HNSC | 64 | PRKCA | 0 | 0 |
| DNA as seed | HNSC | 65 | GSK3B | 0 | 0 |
| DNA as seed | HNSC | 66 | MYH9 | 1 | 0 |
| DNA as seed | HNSC | 67 | RPS27A | 1 | 0 |
| DNA as seed | HNSC | 68 | SHC1 | 0 | 0 |
| DNA as seed | HNSC | 69 | UBE2I | 1 | 1 |
| DNA as seed | HNSC | 70 | ERBB2 | 0 | 0 |
| DNA as seed | HNSC | 71 | HDAC2 | 0 | 0 |
| DNA as seed | HNSC | 72 | MAPK8 | 0 | 0 |
| DNA as seed | HNSC | 73 | TBL1XR1 | 1 | 0 |
| DNA as seed | HNSC | 74 | NCOR2 | 0 | 0 |
| DNA as seed | HNSC | 75 | TGFBR2 | 0 | 0 |
| DNA as seed | HNSC | 76 | FOS | 0 | 0 |
| DNA as seed | HNSC | 77 | RHOA | 0 | 1 |
| DNA as seed | HNSC | 78 | KAT2B | 0 | 0 |
| DNA as seed | HNSC | 79 | RXRA | 0 | 0 |
| DNA as seed | HNSC | 80 | STAT1 | 0 | 0 |
| DNA as seed | HNSC | 81 | EWSR1 | 1 | 1 |
| DNA as seed | HNSC | 82 | PML | 0 | 0 |
| DNA as seed | HNSC | 83 | L2\|H3C2\|H3C[illegible] | 0 | 0 |
| DNA as seed | HNSC | 84 | NFE2L2 | 1 | 0 |
| DNA as seed | HNSC | 85 | PIK3CB | 0 | 0 |
| DNA as seed | HNSC | 86 | ATM | 0 | 0 |
| DNA as seed | HNSC | 87 | SIRT1 | 0 | 0 |
| DNA as seed | HNSC | 88 | EZH2 | 0 | 0 |
| DNA as seed | HNSC | 89 | DCUN1D1 | 0 | 0 |
| DNA as seed | HNSC | 90 | UBA52 | 1 | 0 |
| DNA as seed | HNSC | 91 | CBL | 0 | 0 |
| DNA as seed | HNSC | 92 | NCOA3 | 0 | 0 |
| DNA as seed | HNSC | 93 | PARP1 | 0 | 0 |
| DNA as seed | HNSC | 94 | PLCG1 | 0 | 0 |
| DNA as seed | HNSC | 95 | PIK3R3 | 0 | 0 |
| DNA as seed | HNSC | 96 | HLA-B | 0 | 0 |
| DNA as seed | HNSC | 97 | FADD | 0 | 0 |
| DNA as seed | HNSC | 98 | ABL1 | 0 | 0 |
| DNA as seed | HNSC | 99 | CASP3 | 0 | 0 |
| DNA as seed | HNSC | 100 | SIN3A | 1 | 0 |
| DNA as seed | KIRC | 1 | VHL | 1 | 0 |
| DNA as seed | KIRC | 2 | TP53 | 0 | 0 |
| DNA as seed | KIRC | 3 | PTEN | 0 | 0 |

| | | | | | |
|---|---|---|---|---|---|
| DNA as seed | KIRC | 4 | ATM | 0 | 0 |
| DNA as seed | KIRC | 5 | PBRM1 | 0 | 0 |
| DNA as seed | KIRC | 6 | ESR1 | 0 | 0 |
| DNA as seed | KIRC | 7 | PIK3CA | 0 | 0 |
| DNA as seed | KIRC | 8 | FGFR4 | 0 | 0 |
| DNA as seed | KIRC | 9 | FLT4 | 0 | 0 |
| DNA as seed | KIRC | 10 | SMARCA4 | 0 | 0 |
| DNA as seed | KIRC | 11 | ACTB | 1 | 0 |
| DNA as seed | KIRC | 12 | DBN1 | 0 | 0 |
| DNA as seed | KIRC | 13 | CTNNB1 | 0 | 0 |
| DNA as seed | KIRC | 14 | IGF1R | 0 | 0 |
| DNA as seed | KIRC | 15 | MTOR | 1 | 1 |
| DNA as seed | KIRC | 16 | EP300 | 0 | 0 |
| DNA as seed | KIRC | 17 | FLT1 | 0 | 0 |
| DNA as seed | KIRC | 18 | HSP90AA1 | 0 | 0 |
| DNA as seed | KIRC | 19 | HDAC1 | 0 | 0 |
| DNA as seed | KIRC | 20 | PDLIM7 | 0 | 0 |
| DNA as seed | KIRC | 21 | SIRT7 | 0 | 0 |
| DNA as seed | KIRC | 22 | HIF1A | 0 | 0 |
| DNA as seed | KIRC | 23 | AR | 0 | 0 |
| DNA as seed | KIRC | 24 | CSNK2A1 | 0 | 0 |
| DNA as seed | KIRC | 25 | SHC1 | 0 | 0 |
| DNA as seed | KIRC | 26 | NXF1 | 1 | 0 |
| DNA as seed | KIRC | 27 | SPTAN1 | 0 | 0 |
| DNA as seed | KIRC | 28 | HSP90AB1 | 0 | 0 |
| DNA as seed | KIRC | 29 | ACTG1 | 1 | 0 |
| DNA as seed | KIRC | 30 | FLNA | 0 | 0 |
| DNA as seed | KIRC | 31 | NF2 | 0 | 0 |
| DNA as seed | KIRC | 32 | l2\|H3C2\|H3C3 | 0 | 0 |
| DNA as seed | KIRC | 33 | RNF2 | 0 | 0 |
| DNA as seed | KIRC | 34 | OBSL1 | 0 | 0 |
| DNA as seed | KIRC | 35 | SNW1 | 1 | 1 |
| DNA as seed | KIRC | 36 | MOV10 | 0 | 0 |
| DNA as seed | KIRC | 37 | MYH9 | 1 | 0 |
| DNA as seed | KIRC | 38 | PIK3R3 | 0 | 0 |
| DNA as seed | KIRC | 39 | SMAD3 | 0 | 0 |
| DNA as seed | KIRC | 40 | CBL | 0 | 0 |
| DNA as seed | KIRC | 41 | E2F1 | 0 | 0 |
| DNA as seed | KIRC | 42 | PARP1 | 0 | 0 |
| DNA as seed | KIRC | 43 | LIMA1 | 0 | 0 |
| DNA as seed | KIRC | 44 | PIK3CD | 0 | 0 |
| DNA as seed | KIRC | 45 | STAT1 | 0 | 0 |
| DNA as seed | KIRC | 46 | PPARG | 0 | 0 |
| DNA as seed | KIRC | 47 | MAX | 1 | 1 |
| DNA as seed | KIRC | 48 | 14\|H4C15\|H4 | 0 | 0 |
| DNA as seed | KIRC | 49 | RPA1 | 1 | 1 |
| DNA as seed | KIRC | 50 | HUWE1 | 1 | 0 |
| DNA as seed | KIRC | 51 | RPA2 | 1 | 1 |
| DNA as seed | KIRC | 52 | ABL1 | 0 | 0 |
| DNA as seed | KIRC | 53 | H3-4 | 0 | 0 |
| DNA as seed | KIRC | 54 | PTK2 | 1 | 0 |
| DNA as seed | KIRC | 55 | HSPA8 | 1 | 1 |
| DNA as seed | KIRC | 56 | SMARCC1 | 0 | 0 |
| DNA as seed | KIRC | 57 | CRK | 1 | 0 |
| DNA as seed | KIRC | 58 | UIMC1 | 0 | 0 |
| DNA as seed | KIRC | 59 | FAF2 | 1 | 0 |

| | | | | | |
|---|---|---|---|---|---|
| DNA as seed | KIRC | 60 | SMARCC2 | 0 | 0 |
| DNA as seed | KIRC | 61 | GSK3B | 0 | 0 |
| DNA as seed | KIRC | 62 | EED | 0 | 0 |
| DNA as seed | KIRC | 63 | KAT2B | 0 | 0 |
| DNA as seed | KIRC | 64 | HNRNPAB | 0 | 0 |
| DNA as seed | KIRC | 65 | SMARCA2 | 0 | 0 |
| DNA as seed | KIRC | 66 | DDB1 | 1 | 1 |
| DNA as seed | KIRC | 67 | CDC42 | 1 | 0 |
| DNA as seed | KIRC | 68 | SPTBN1 | 0 | 0 |
| DNA as seed | KIRC | 69 | SMURF1 | 0 | 0 |
| DNA as seed | KIRC | 70 | HSPA4 | 0 | 0 |
| DNA as seed | KIRC | 71 | ARID1A | 0 | 0 |
| DNA as seed | KIRC | 72 | POLR2A | 0 | 0 |
| DNA as seed | KIRC | 73 | NOTCH1 | 0 | 0 |
| DNA as seed | KIRC | 74 | PRKCZ | 0 | 0 |
| DNA as seed | KIRC | 75 | ATR | 1 | 1 |
| DNA as seed | KIRC | 76 | SIN3A | 1 | 0 |
| DNA as seed | KIRC | 77 | SMARCB1 | 1 | 1 |
| DNA as seed | KIRC | 78 | CAPZA2 | 0 | 0 |
| DNA as seed | KIRC | 79 | TERF1 | 1 | 0 |
| DNA as seed | KIRC | 80 | ROCK1 | 0 | 0 |
| DNA as seed | KIRC | 81 | BAP1 | 1 | 0 |
| DNA as seed | KIRC | 82 | IKBKB | 0 | 0 |
| DNA as seed | KIRC | 83 | MYO1C | 0 | 0 |
| DNA as seed | KIRC | 84 | EPAS1 | 0 | 0 |
| DNA as seed | KIRC | 85 | PRPF8 | 1 | 1 |
| DNA as seed | KIRC | 86 | PKM | 1 | 0 |
| DNA as seed | KIRC | 87 | SHMT2 | 0 | 0 |
| DNA as seed | KIRC | 88 | NR3C1 | 0 | 0 |
| DNA as seed | KIRC | 89 | SQSTM1 | 0 | 0 |
| DNA as seed | KIRC | 90 | UBE2D1 | 0 | 0 |
| DNA as seed | KIRC | 91 | SKP2 | 1 | 0 |
| DNA as seed | KIRC | 92 | WDR5 | 1 | 1 |
| DNA as seed | KIRC | 93 | FBXW7 | 0 | 0 |
| DNA as seed | KIRC | 94 | UBE2D3 | 1 | 0 |
| DNA as seed | KIRC | 95 | FOXO3 | 0 | 0 |
| DNA as seed | KIRC | 96 | IRS1 | 0 | 0 |
| DNA as seed | KIRC | 97 | VEGFA | 0 | 0 |
| DNA as seed | KIRC | 98 | AKT2 | 0 | 0 |
| DNA as seed | KIRC | 99 | FOXO1 | 0 | 0 |
| DNA as seed | KIRC | 100 | NSD1 | 0 | 0 |
| DNA as seed | LGG | 1 | TP53 | 0 | 0 |
| DNA as seed | LGG | 2 | EGFR | 0 | 0 |
| DNA as seed | LGG | 3 | PIK3R1 | 0 | 0 |
| DNA as seed | LGG | 4 | PIK3CA | 0 | 0 |
| DNA as seed | LGG | 5 | NTRK1 | 0 | 0 |
| DNA as seed | LGG | 6 | RELA | 1 | 0 |
| DNA as seed | LGG | 7 | MYC | 1 | 0 |
| DNA as seed | LGG | 8 | PTPN11 | 1 | 0 |
| DNA as seed | LGG | 9 | MAPK1 | 0 | 0 |
| DNA as seed | LGG | 10 | CDKN2A | 0 | 0 |
| DNA as seed | LGG | 11 | SHC1 | 0 | 0 |
| DNA as seed | LGG | 12 | PTEN | 0 | 0 |
| DNA as seed | LGG | 13 | NFKB1 | 0 | 0 |
| DNA as seed | LGG | 14 | HSP90AA1 | 0 | 0 |
| DNA as seed | LGG | 15 | NRAS | 0 | 0 |

| | | | | | |
|---|---|---|---|---|---|
| DNA as seed | LGG | 16 | MAPK3 | 0 | 0 |
| DNA as seed | LGG | 17 | SP1 | 0 | 0 |
| DNA as seed | LGG | 18 | HRAS | 0 | 0 |
| DNA as seed | LGG | 19 | HSF1 | 0 | 0 |
| DNA as seed | LGG | 20 | JAK2 | 0 | 0 |
| DNA as seed | LGG | 21 | SMARCA4 | 0 | 0 |
| DNA as seed | LGG | 22 | AR | 0 | 0 |
| DNA as seed | LGG | 23 | PIK3R3 | 0 | 0 |
| DNA as seed | LGG | 24 | RB1 | 0 | 0 |
| DNA as seed | LGG | 25 | PIK3CD | 0 | 0 |
| DNA as seed | LGG | 26 | STAT1 | 0 | 0 |
| DNA as seed | LGG | 27 | CDK2 | 1 | 0 |
| DNA as seed | LGG | 28 | HDAC1 | 0 | 0 |
| DNA as seed | LGG | 29 | JAK1 | 0 | 0 |
| DNA as seed | LGG | 30 | JAK3 | 0 | 0 |
| DNA as seed | LGG | 31 | IFNA5 | 0 | 0 |
| DNA as seed | LGG | 32 | CREBBP | 0 | 0 |
| DNA as seed | LGG | 33 | IFNA8 | 0 | 0 |
| DNA as seed | LGG | 34 | IDH1 | 0 | 0 |
| DNA as seed | LGG | 35 | IFNA2 | 0 | 0 |
| DNA as seed | LGG | 36 | MDM2 | 1 | 0 |
| DNA as seed | LGG | 37 | CDKN1A | 0 | 0 |
| DNA as seed | LGG | 38 | PTPN6 | 0 | 0 |
| DNA as seed | LGG | 39 | IFNA6 | 0 | 0 |
| DNA as seed | LGG | 40 | PDGFRB | 0 | 0 |
| DNA as seed | LGG | 41 | TYK2 | 0 | 0 |
| DNA as seed | LGG | 42 | HIF1A | 0 | 0 |
| DNA as seed | LGG | 43 | PDGFB | 0 | 0 |
| DNA as seed | LGG | 44 | HSPA8 | 1 | 1 |
| DNA as seed | LGG | 45 | FOS | 0 | 0 |
| DNA as seed | LGG | 46 | EGF | 0 | 0 |
| DNA as seed | LGG | 47 | CDH1 | 0 | 0 |
| DNA as seed | LGG | 48 | NOTCH1 | 0 | 0 |
| DNA as seed | LGG | 49 | SMAD3 | 0 | 0 |
| DNA as seed | LGG | 50 | CSF2 | 0 | 0 |
| DNA as seed | LGG | 51 | IGF1R | 0 | 0 |
| DNA as seed | LGG | 52 | IFNA14 | 0 | 0 |
| DNA as seed | LGG | 53 | PDGFA | 0 | 0 |
| DNA as seed | LGG | 54 | PDGFRA | 0 | 0 |
| DNA as seed | LGG | 55 | IFNA4 | 0 | 0 |
| DNA as seed | LGG | 56 | IL3 | 0 | 0 |
| DNA as seed | LGG | 57 | MET | 0 | 0 |
| DNA as seed | LGG | 58 | IFNA1 | 0 | 0 |
| DNA as seed | LGG | 59 | CUL4B | 0 | 0 |
| DNA as seed | LGG | 60 | IFNA13 | 0 | 0 |
| DNA as seed | LGG | 61 | NR3C1 | 0 | 0 |
| DNA as seed | LGG | 62 | MAPK8 | 0 | 0 |
| DNA as seed | LGG | 63 | PPARG | 0 | 0 |
| DNA as seed | LGG | 64 | IL5 | 0 | 0 |
| DNA as seed | LGG | 65 | ATRX | 0 | 1 |
| DNA as seed | LGG | 66 | SIRT1 | 0 | 0 |
| DNA as seed | LGG | 67 | HCFC1 | 1 | 1 |
| DNA as seed | LGG | 68 | EZH2 | 0 | 0 |
| DNA as seed | LGG | 69 | PML | 0 | 0 |
| DNA as seed | LGG | 70 | IFNE | 0 | 0 |
| DNA as seed | LGG | 71 | SMAD2 | 0 | 0 |

| | | | | | |
|---|---|---|---|---|---|
| DNA as seed | LGG | 72 | MAPK14 | 0 | 0 |
| DNA as seed | LGG | 73 | CCND1 | 1 | 0 |
| DNA as seed | LGG | 74 | ERBB4 | 0 | 0 |
| DNA as seed | LGG | 75 | IRS1 | 0 | 0 |
| DNA as seed | LGG | 76 | HDAC3 | 1 | 0 |
| DNA as seed | LGG | 77 | EED | 0 | 0 |
| DNA as seed | LGG | 78 | FOXO3 | 0 | 0 |
| DNA as seed | LGG | 79 | PTK2 | 0 | 0 |
| DNA as seed | LGG | 80 | FOXO1 | 0 | 0 |
| DNA as seed | LGG | 81 | SIRT7 | 0 | 0 |
| DNA as seed | LGG | 82 | CDK4 | 1 | 0 |
| DNA as seed | LGG | 83 | SOCS1 | 0 | 0 |
| DNA as seed | LGG | 84 | USP7 | 1 | 0 |
| DNA as seed | LGG | 85 | STAT2 | 0 | 0 |
| DNA as seed | LGG | 86 | SNW1 | 1 | 1 |
| DNA as seed | LGG | 87 | MAPK9 | 0 | 0 |
| DNA as seed | LGG | 88 | CRKL | 1 | 0 |
| DNA as seed | LGG | 89 | SOCS3 | 1 | 0 |
| DNA as seed | LGG | 90 | HSPA4 | 0 | 0 |
| DNA as seed | LGG | 91 | NCL | 1 | 1 |
| DNA as seed | LGG | 92 | STUB1 | 0 | 0 |
| DNA as seed | LGG | 93 | SMARCB1 | 1 | 1 |
| DNA as seed | LGG | 94 | CDC5L | 1 | 1 |
| DNA as seed | LGG | 95 | PTPN1 | 0 | 0 |
| DNA as seed | LGG | 96 | IL7R | 0 | 0 |
| DNA as seed | LGG | 97 | SMARCC1 | 0 | 0 |
| DNA as seed | LGG | 98 | EPHA2 | 0 | 0 |
| DNA as seed | LGG | 99 | RUNX1 | 0 | 0 |
| DNA as seed | LGG | 100 | ARID1A | 0 | 0 |
| DNA as seed | LUAD | 1 | TP53 | 0 | 0 |
| DNA as seed | LUAD | 2 | EGFR | 1 | 0 |
| DNA as seed | LUAD | 3 | KRAS | 1 | 0 |
| DNA as seed | LUAD | 4 | CTNNB1 | 1 | 0 |
| DNA as seed | LUAD | 5 | CDKN2A | 0 | 0 |
| DNA as seed | LUAD | 6 | ATM | 0 | 0 |
| DNA as seed | LUAD | 7 | MYC | 1 | 0 |
| DNA as seed | LUAD | 8 | PIK3CA | 1 | 0 |
| DNA as seed | LUAD | 9 | SRC | 0 | 0 |
| DNA as seed | LUAD | 10 | RB1 | 0 | 0 |
| DNA as seed | LUAD | 11 | SMARCA4 | 0 | 0 |
| DNA as seed | LUAD | 12 | STK11 | 0 | 0 |
| DNA as seed | LUAD | 13 | ERBB2 | 0 | 0 |
| DNA as seed | LUAD | 14 | SP1 | 0 | 0 |
| DNA as seed | LUAD | 15 | MET | 0 | 0 |
| DNA as seed | LUAD | 16 | TERT | 0 | 0 |
| DNA as seed | LUAD | 17 | HSP90AA1 | 0 | 0 |
| DNA as seed | LUAD | 18 | MAPK1 | 0 | 0 |
| DNA as seed | LUAD | 19 | HRAS | 0 | 0 |
| DNA as seed | LUAD | 20 | SPTA1 | 0 | 0 |
| DNA as seed | LUAD | 21 | APC | 0 | 0 |
| DNA as seed | LUAD | 22 | NFKBIA | 0 | 0 |
| DNA as seed | LUAD | 23 | CDKN1A | 0 | 0 |
| DNA as seed | LUAD | 24 | CREBBP | 0 | 0 |
| DNA as seed | LUAD | 25 | NRAS | 1 | 0 |
| DNA as seed | LUAD | 26 | HDAC1 | 0 | 0 |
| DNA as seed | LUAD | 27 | AKT1 | 0 | 0 |

| | | | | | |
|---|---|---|---|---|---|
| DNA as seed | LUAD | 28 | SMAD3 | 0 | 0 |
| DNA as seed | LUAD | 29 | STAT3 | 1 | 0 |
| DNA as seed | LUAD | 30 | MDM2 | 1 | 0 |
| DNA as seed | LUAD | 31 | PRKCA | 0 | 0 |
| DNA as seed | LUAD | 32 | NOTCH1 | 0 | 0 |
| DNA as seed | LUAD | 33 | ERBB3 | 0 | 0 |
| DNA as seed | LUAD | 34 | SMAD2 | 0 | 0 |
| DNA as seed | LUAD | 35 | NF1 | 0 | 0 |
| DNA as seed | LUAD | 36 | GSK3B | 0 | 0 |
| DNA as seed | LUAD | 37 | NOTCH4 | 0 | 0 |
| DNA as seed | LUAD | 38 | HIF1A | 0 | 0 |
| DNA as seed | LUAD | 39 | NOTCH2 | 0 | 0 |
| DNA as seed | LUAD | 40 | YWHAZ | 1 | 0 |
| DNA as seed | LUAD | 41 | AURKA | 1 | 0 |
| DNA as seed | LUAD | 42 | CCND1 | 1 | 0 |
| DNA as seed | LUAD | 43 | PTK2 | 1 | 0 |
| DNA as seed | LUAD | 44 | IGF1R | 1 | 0 |
| DNA as seed | LUAD | 45 | BRAF | 0 | 0 |
| DNA as seed | LUAD | 46 | FGFR1 | 0 | 0 |
| DNA as seed | LUAD | 47 | A2M | 0 | 0 |
| DNA as seed | LUAD | 48 | CDC37 | 1 | 1 |
| DNA as seed | LUAD | 49 | KEAP1 | 1 | 0 |
| DNA as seed | LUAD | 50 | EZH2 | 0 | 0 |
| DNA as seed | LUAD | 51 | PML | 0 | 0 |
| DNA as seed | LUAD | 52 | H3-4 | 0 | 0 |
| DNA as seed | LUAD | 53 | PPARG | 0 | 0 |
| DNA as seed | LUAD | 54 | L1CAM | 0 | 0 |
| DNA as seed | LUAD | 55 | ABL1 | 0 | 0 |
| DNA as seed | LUAD | 56 | PTEN | 0 | 0 |
| DNA as seed | LUAD | 57 | PDCD6 | 0 | 0 |
| DNA as seed | LUAD | 58 | CDKN2B | 0 | 0 |
| DNA as seed | LUAD | 59 | KAT2B | 0 | 0 |
| DNA as seed | LUAD | 60 | FBXO6 | 0 | 0 |
| DNA as seed | LUAD | 61 | PIK3R3 | 0 | 0 |
| DNA as seed | LUAD | 62 | HSPA1A | 0 | 0 |
| DNA as seed | LUAD | 63 | YWHAQ | 0 | 0 |
| DNA as seed | LUAD | 64 | ARID1A | 0 | 0 |
| DNA as seed | LUAD | 65 | ITGB1 | 1 | 0 |
| DNA as seed | LUAD | 66 | YAP1 | 1 | 0 |
| DNA as seed | LUAD | 67 | FOXO3 | 0 | 0 |
| DNA as seed | LUAD | 68 | HSPA4 | 0 | 0 |
| DNA as seed | LUAD | 69 | COL3A1 | 0 | 0 |
| DNA as seed | LUAD | 70 | FOXO1 | 0 | 0 |
| DNA as seed | LUAD | 71 | HSPA1B | 0 | 0 |
| DNA as seed | LUAD | 72 | VCAN | 0 | 0 |
| DNA as seed | LUAD | 73 | BCL2 | 0 | 0 |
| DNA as seed | LUAD | 74 | CDK4 | 1 | 0 |
| DNA as seed | LUAD | 75 | ATR | 1 | 1 |
| DNA as seed | LUAD | 76 | VEGFA | 0 | 0 |
| DNA as seed | LUAD | 77 | HSPA5 | 1 | 1 |
| DNA as seed | LUAD | 78 | SQSTM1 | 0 | 0 |
| DNA as seed | LUAD | 79 | CASP3 | 0 | 0 |
| DNA as seed | LUAD | 80 | H2AX | 1 | 0 |
| DNA as seed | LUAD | 81 | VDR | 0 | 0 |
| DNA as seed | LUAD | 82 | MIPOL1 | 0 | 0 |
| DNA as seed | LUAD | 83 | IRS1 | 0 | 0 |

| | | | | | |
|---|---|---|---|---|---|
| DNA as seed | LUAD | 84 | BTRC | 0 | 0 |
| DNA as seed | LUAD | 85 | CHUK | 0 | 0 |
| DNA as seed | LUAD | 86 | SP3 | 0 | 0 |
| DNA as seed | LUAD | 87 | SMARCB1 | 1 | 1 |
| DNA as seed | LUAD | 88 | PRKCZ | 0 | 0 |
| DNA as seed | LUAD | 89 | SKP1 | 1 | 0 |
| DNA as seed | LUAD | 90 | CTBP1 | 0 | 0 |
| DNA as seed | LUAD | 91 | WT1 | 0 | 0 |
| DNA as seed | LUAD | 92 | MBIP | 0 | 0 |
| DNA as seed | LUAD | 93 | YY1 | 1 | 1 |
| DNA as seed | LUAD | 94 | CDK5 | 0 | 0 |
| DNA as seed | LUAD | 95 | SDHA | 0 | 1 |
| DNA as seed | LUAD | 96 | NFATC2 | 0 | 0 |
| DNA as seed | LUAD | 97 | TFAP2A | 0 | 0 |
| DNA as seed | LUAD | 98 | CD44 | 0 | 0 |
| DNA as seed | LUAD | 99 | TCF7L2 | 0 | 0 |
| DNA as seed | LUAD | 100 | DDB1 | 1 | 1 |
| DNA as seed | LUSC | 1 | TP53 | 0 | 0 |
| DNA as seed | LUSC | 2 | CUL3 | 1 | 0 |
| DNA as seed | LUSC | 3 | SOX2 | 1 | 0 |
| DNA as seed | LUSC | 4 | PIK3CA | 1 | 0 |
| DNA as seed | LUSC | 5 | FXR1 | 0 | 0 |
| DNA as seed | LUSC | 6 | AP2M1 | 1 | 0 |
| DNA as seed | LUSC | 7 | CREBBP | 1 | 0 |
| DNA as seed | LUSC | 8 | DCUN1D1 | 0 | 0 |
| DNA as seed | LUSC | 9 | FN1 | 0 | 0 |
| DNA as seed | LUSC | 10 | ACTL6A | 1 | 1 |
| DNA as seed | LUSC | 11 | PTEN | 0 | 0 |
| DNA as seed | LUSC | 12 | UBC | 1 | 0 |
| DNA as seed | LUSC | 13 | ESR1 | 0 | 0 |
| DNA as seed | LUSC | 14 | EGFR | 1 | 0 |
| DNA as seed | LUSC | 15 | CDKN2A | 0 | 0 |
| DNA as seed | LUSC | 16 | RELA | 0 | 0 |
| DNA as seed | LUSC | 17 | RB1 | 0 | 0 |
| DNA as seed | LUSC | 18 | CTNNB1 | 0 | 0 |
| DNA as seed | LUSC | 19 | NOTCH1 | 0 | 0 |
| DNA as seed | LUSC | 20 | SP1 | 0 | 0 |
| DNA as seed | LUSC | 21 | HDAC1 | 0 | 0 |
| DNA as seed | LUSC | 22 | GRB2 | 1 | 1 |
| DNA as seed | LUSC | 23 | DVL3 | 0 | 0 |
| DNA as seed | LUSC | 24 | AR | 0 | 0 |
| DNA as seed | LUSC | 25 | TBL1XR1 | 0 | 0 |
| DNA as seed | LUSC | 26 | NFE2L2 | 1 | 0 |
| DNA as seed | LUSC | 27 | CSNK2A1 | 0 | 0 |
| DNA as seed | LUSC | 28 | SNW1 | 1 | 1 |
| DNA as seed | LUSC | 29 | PRKCA | 0 | 0 |
| DNA as seed | LUSC | 30 | HIF1A | 0 | 0 |
| DNA as seed | LUSC | 31 | PLD1 | 0 | 0 |
| DNA as seed | LUSC | 32 | FBXW7 | 0 | 0 |
| DNA as seed | LUSC | 33 | GSK3B | 0 | 0 |
| DNA as seed | LUSC | 34 | HDAC3 | 1 | 0 |
| DNA as seed | LUSC | 35 | PPARG | 0 | 0 |
| DNA as seed | LUSC | 36 | HDAC2 | 0 | 0 |
| DNA as seed | LUSC | 37 | SMARCA4 | 0 | 0 |
| DNA as seed | LUSC | 38 | GNB4 | 0 | 0 |
| DNA as seed | LUSC | 39 | ZMAT3 | 0 | 0 |

| | | | | | |
|---|---|---|---|---|---|
| DNA as seed | LUSC | 40 | HSPA8 | 1 | 1 |
| DNA as seed | LUSC | 41 | KAT2B | 0 | 0 |
| DNA as seed | LUSC | 42 | CCND1 | 1 | 0 |
| DNA as seed | LUSC | 43 | RXRA | 0 | 0 |
| DNA as seed | LUSC | 44 | PML | 0 | 0 |
| DNA as seed | LUSC | 45 | CBL | 0 | 0 |
| DNA as seed | LUSC | 46 | BMI1 | 0 | 0 |
| DNA as seed | LUSC | 47 | RBX1 | 1 | 1 |
| DNA as seed | LUSC | 48 | PIK3R3 | 0 | 0 |
| DNA as seed | LUSC | 49 | IKBKB | 0 | 0 |
| DNA as seed | LUSC | 50 | NF1 | 0 | 0 |
| DNA as seed | LUSC | 51 | RAC1 | 1 | 1 |
| DNA as seed | LUSC | 52 | NCOR1 | 0 | 0 |
| DNA as seed | LUSC | 53 | SMARCA2 | 0 | 0 |
| DNA as seed | LUSC | 54 | EZH2 | 0 | 0 |
| DNA as seed | LUSC | 55 | INSR | 0 | 0 |
| DNA as seed | LUSC | 56 | TNFSF10 | 0 | 0 |
| DNA as seed | LUSC | 57 | KEAP1 | 1 | 0 |
| DNA as seed | LUSC | 58 | HRAS | 0 | 0 |
| DNA as seed | LUSC | 59 | PRKCD | 0 | 0 |
| DNA as seed | LUSC | 60 | ILK | 1 | 0 |
| DNA as seed | LUSC | 61 | NFATC1 | 0 | 0 |
| DNA as seed | LUSC | 62 | ARID1A | 0 | 0 |
| DNA as seed | LUSC | 63 | NCOR2 | 0 | 0 |
| DNA as seed | LUSC | 64 | SMARCD1 | 0 | 0 |
| DNA as seed | LUSC | 65 | SMARCB1 | 1 | 1 |
| DNA as seed | LUSC | 66 | DVL2 | 0 | 0 |
| DNA as seed | LUSC | 67 | RASA1 | 0 | 0 |
| DNA as seed | LUSC | 68 | PRKCB | 0 | 0 |
| DNA as seed | LUSC | 69 | RARA | 0 | 0 |
| DNA as seed | LUSC | 70 | SQSTM1 | 0 | 0 |
| DNA as seed | LUSC | 71 | YY1 | 1 | 1 |
| DNA as seed | LUSC | 72 | HSPA1B | 0 | 0 |
| DNA as seed | LUSC | 73 | SMARCC1 | 0 | 0 |
| DNA as seed | LUSC | 74 | TYK2 | 0 | 0 |
| DNA as seed | LUSC | 75 | PRKCZ | 0 | 0 |
| DNA as seed | LUSC | 76 | MET | 0 | 0 |
| DNA as seed | LUSC | 77 | KLF5 | 1 | 0 |
| DNA as seed | LUSC | 78 | TBP | 1 | 0 |
| DNA as seed | LUSC | 79 | WDR5 | 1 | 1 |
| DNA as seed | LUSC | 80 | NCL | 0 | 1 |
| DNA as seed | LUSC | 81 | IRS1 | 0 | 0 |
| DNA as seed | LUSC | 82 | RUNX1 | 0 | 0 |
| DNA as seed | LUSC | 83 | TUBB | 1 | 1 |
| DNA as seed | LUSC | 84 | SKP1 | 1 | 0 |
| DNA as seed | LUSC | 85 | CCNB1 | 1 | 0 |
| DNA as seed | LUSC | 86 | SMARCC2 | 0 | 0 |
| DNA as seed | LUSC | 87 | IRF1 | 0 | 0 |
| DNA as seed | LUSC | 88 | RBPJ | 0 | 0 |
| DNA as seed | LUSC | 89 | CAV1 | 0 | 0 |
| DNA as seed | LUSC | 90 | SUMO1 | 0 | 0 |
| DNA as seed | LUSC | 91 | RUVBL1 | 1 | 1 |
| DNA as seed | LUSC | 92 | RUVBL2 | 1 | 1 |
| DNA as seed | LUSC | 93 | TUBA1C | 0 | 0 |
| DNA as seed | LUSC | 94 | CASP3 | 0 | 0 |
| DNA as seed | LUSC | 95 | SREBF1 | 1 | 0 |

| | | | | | |
|---|---|---|---|---|---|
| DNA as seed | LUSC | 96 | CASP8 | 0 | 0 |
| DNA as seed | LUSC | 97 | FOXO3 | 0 | 0 |
| DNA as seed | LUSC | 98 | MED1 | 1 | 1 |
| DNA as seed | LUSC | 99 | FLNA | 0 | 0 |
| DNA as seed | LUSC | 100 | TRIM28 | 0 | 0 |
| DNA as seed | OV | 1 | MYC | 1 | 0 |
| DNA as seed | OV | 2 | TP53 | 0 | 0 |
| DNA as seed | OV | 3 | PTK2 | 1 | 0 |
| DNA as seed | OV | 4 | PLEC | 0 | 0 |
| DNA as seed | OV | 5 | BRCA1 | 0 | 1 |
| DNA as seed | OV | 6 | NDRG1 | 0 | 0 |
| DNA as seed | OV | 7 | EPPK1 | 0 | 0 |
| DNA as seed | OV | 8 | EEF1D | 0 | 0 |
| DNA as seed | OV | 9 | CDK2 | 1 | 0 |
| DNA as seed | OV | 10 | TOP1MT | 0 | 0 |
| DNA as seed | OV | 11 | ST3GAL1 | 0 | 0 |
| DNA as seed | OV | 12 | MECOM | 0 | 0 |
| DNA as seed | OV | 13 | EGFR | 1 | 0 |
| DNA as seed | OV | 14 | SCRIB | 0 | 0 |
| DNA as seed | OV | 15 | RB1 | 0 | 0 |
| DNA as seed | OV | 16 | ASAP1 | 0 | 0 |
| DNA as seed | OV | 17 | PUF60 | 1 | 0 |
| DNA as seed | OV | 18 | SP1 | 0 | 0 |
| DNA as seed | OV | 19 | CDK1 | 1 | 1 |
| DNA as seed | OV | 20 | PCNA | 1 | 1 |
| DNA as seed | OV | 21 | GRB2 | 1 | 1 |
| DNA as seed | OV | 22 | MCM2 | 1 | 1 |
| DNA as seed | OV | 23 | CREBBP | 0 | 0 |
| DNA as seed | OV | 24 | SNW1 | 1 | 1 |
| DNA as seed | OV | 25 | CDH1 | 0 | 0 |
| DNA as seed | OV | 26 | CUL7 | 0 | 0 |
| DNA as seed | OV | 27 | OBSL1 | 0 | 0 |
| DNA as seed | OV | 28 | MAPK1 | 0 | 0 |
| DNA as seed | OV | 29 | CDKN1B | 0 | 0 |
| DNA as seed | OV | 30 | CCDC8 | 0 | 0 |
| DNA as seed | OV | 31 | MAPK3 | 0 | 0 |
| DNA as seed | OV | 32 | CDC5L | 1 | 1 |
| DNA as seed | OV | 33 | E2F1 | 0 | 0 |
| DNA as seed | OV | 34 | CSNK2A1 | 0 | 0 |
| DNA as seed | OV | 35 | FOS | 0 | 0 |
| DNA as seed | OV | 36 | UBE2I | 1 | 1 |
| DNA as seed | OV | 37 | ERBB2 | 1 | 0 |
| DNA as seed | OV | 38 | PRKCA | 0 | 0 |
| DNA as seed | OV | 39 | SMC3 | 1 | 1 |
| DNA as seed | OV | 40 | HDAC2 | 0 | 0 |
| DNA as seed | OV | 41 | ABL1 | 0 | 0 |
| DNA as seed | OV | 42 | PIK3R1 | 0 | 0 |
| DNA as seed | OV | 43 | VCP | 1 | 1 |
| DNA as seed | OV | 44 | PLK1 | 1 | 1 |
| DNA as seed | OV | 45 | YWHAG | 0 | 0 |
| DNA as seed | OV | 46 | SMAD2 | 0 | 0 |
| DNA as seed | OV | 47 | H3-4 | 0 | 0 |
| DNA as seed | OV | 48 | CCNB1 | 1 | 0 |
| DNA as seed | OV | 49 | PPP1CA | 1 | 0 |
| DNA as seed | OV | 50 | SIRT1 | 0 | 0 |
| DNA as seed | OV | 51 | PIK3CA | 1 | 0 |

| | | | | | |
|---|---|---|---|---|---|
| DNA as seed | OV | 52 | ATR | 1 | 1 |
| DNA as seed | OV | 53 | KHDRBS3 | 0 | 0 |
| DNA as seed | OV | 54 | EXOSC4 | 1 | 0 |
| DNA as seed | OV | 55 | STAT5A | 0 | 0 |
| DNA as seed | OV | 56 | PPP1CC | 0 | 0 |
| DNA as seed | OV | 57 | NCL | 1 | 1 |
| DNA as seed | OV | 58 | ACTG1 | 1 | 0 |
| DNA as seed | OV | 59 | SHC1 | 0 | 0 |
| DNA as seed | OV | 60 | DYNC1H1 | 1 | 1 |
| DNA as seed | OV | 61 | PRKDC | 0 | 0 |
| DNA as seed | OV | 62 | IQGAP1 | 0 | 0 |
| DNA as seed | OV | 63 | POLR2A | 0 | 0 |
| DNA as seed | OV | 64 | CDK4 | 1 | 0 |
| DNA as seed | OV | 65 | XRCC5 | 1 | 0 |
| DNA as seed | OV | 66 | KAT2B | 0 | 0 |
| DNA as seed | OV | 67 | HSPA4 | 0 | 0 |
| DNA as seed | OV | 68 | HIF1A | 0 | 0 |
| DNA as seed | OV | 69 | SQSTM1 | 0 | 0 |
| DNA as seed | OV | 70 | CBL | 0 | 0 |
| DNA as seed | OV | 71 | PPP1CB | 1 | 0 |
| DNA as seed | OV | 72 | RBL1 | 0 | 0 |
| DNA as seed | OV | 73 | CRK | 1 | 0 |
| DNA as seed | OV | 74 | ZC3H3 | 1 | 0 |
| DNA as seed | OV | 75 | PPARG | 0 | 0 |
| DNA as seed | OV | 76 | AHNAK | 0 | 0 |
| DNA as seed | OV | 77 | TP73 | 0 | 0 |
| DNA as seed | OV | 78 | FOXO3 | 0 | 0 |
| DNA as seed | OV | 79 | YY1 | 1 | 1 |
| DNA as seed | OV | 80 | PRKCD | 0 | 0 |
| DNA as seed | OV | 81 | ETS1 | 0 | 0 |
| DNA as seed | OV | 82 | PKM | 1 | 0 |
| DNA as seed | OV | 83 | TOP2A | 1 | 1 |
| DNA as seed | OV | 84 | IFI16 | 0 | 0 |
| DNA as seed | OV | 85 | DDB1 | 1 | 1 |
| DNA as seed | OV | 86 | PXN | 0 | 0 |
| DNA as seed | OV | 87 | SMARCA4 | 1 | 0 |
| DNA as seed | OV | 88 | PIK3R3 | 0 | 0 |
| DNA as seed | OV | 89 | KCNQ3 | 0 | 0 |
| DNA as seed | OV | 90 | DDX5 | 1 | 0 |
| DNA as seed | OV | 91 | CCNE1 | 1 | 0 |
| DNA as seed | OV | 92 | APC | 0 | 0 |
| DNA as seed | OV | 93 | E2F4 | 0 | 0 |
| DNA as seed | OV | 94 | PIN1 | 0 | 0 |
| DNA as seed | OV | 95 | EEF1G | 1 | 0 |
| DNA as seed | OV | 96 | BCR | 0 | 0 |
| DNA as seed | OV | 97 | GAPDH | 1 | 1 |
| DNA as seed | OV | 98 | TP53BP1 | 0 | 0 |
| DNA as seed | OV | 99 | RAD51 | 1 | 1 |
| DNA as seed | OV | 100 | E2F3 | 1 | 0 |
| DNA as seed | PRAD | 1 | TP53 | 0 | 0 |
| DNA as seed | PRAD | 2 | ERG | 1 | 0 |
| DNA as seed | PRAD | 3 | PTEN | 0 | 0 |
| DNA as seed | PRAD | 4 | CTNNB1 | 0 | 0 |
| DNA as seed | PRAD | 5 | ATM | 0 | 0 |
| DNA as seed | PRAD | 6 | MYC | 1 | 0 |
| DNA as seed | PRAD | 7 | HSPA8 | 0 | 1 |

| | | | | | |
|---|---|---|---|---|---|
| DNA as seed | PRAD | 8 | ESR1 | 0 | 0 |
| DNA as seed | PRAD | 9 | EP300 | 1 | 0 |
| DNA as seed | PRAD | 10 | SNW1 | 1 | 1 |
| DNA as seed | PRAD | 11 | PIK3CA | 0 | 0 |
| DNA as seed | PRAD | 12 | SP1 | 0 | 0 |
| DNA as seed | PRAD | 13 | HRAS | 0 | 0 |
| DNA as seed | PRAD | 14 | HDAC1 | 0 | 0 |
| DNA as seed | PRAD | 15 | AR | 0 | 0 |
| DNA as seed | PRAD | 16 | FOXA1 | 1 | 0 |
| DNA as seed | PRAD | 17 | MAPK3 | 0 | 0 |
| DNA as seed | PRAD | 18 | GSK3B | 0 | 0 |
| DNA as seed | PRAD | 19 | SMAD3 | 0 | 0 |
| DNA as seed | PRAD | 20 | CDKN1B | 0 | 0 |
| DNA as seed | PRAD | 21 | APC | 1 | 0 |
| DNA as seed | PRAD | 22 | UBB | 0 | 0 |
| DNA as seed | PRAD | 23 | FOS | 0 | 0 |
| DNA as seed | PRAD | 24 | YWHAZ | 1 | 0 |
| DNA as seed | PRAD | 25 | ETS2 | 0 | 0 |
| DNA as seed | PRAD | 26 | RB1 | 0 | 0 |
| DNA as seed | PRAD | 27 | ABL1 | 0 | 0 |
| DNA as seed | PRAD | 28 | SPOP | 0 | 0 |
| DNA as seed | PRAD | 29 | TMPRSS2 | 0 | 0 |
| DNA as seed | PRAD | 30 | TP53BP1 | 0 | 0 |
| DNA as seed | PRAD | 31 | HIF1A | 0 | 0 |
| DNA as seed | PRAD | 32 | ETS1 | 0 | 0 |
| DNA as seed | PRAD | 33 | KPNA3 | 0 | 0 |
| DNA as seed | PRAD | 34 | CDK4 | 1 | 0 |
| DNA as seed | PRAD | 35 | PARP1 | 0 | 0 |
| DNA as seed | PRAD | 36 | CBL | 0 | 0 |
| DNA as seed | PRAD | 37 | RPA2 | 1 | 1 |
| DNA as seed | PRAD | 38 | KAT2B | 0 | 0 |
| DNA as seed | PRAD | 39 | ATR | 1 | 1 |
| DNA as seed | PRAD | 40 | MAPK8 | 0 | 0 |
| DNA as seed | PRAD | 41 | ESR2 | 0 | 0 |
| DNA as seed | PRAD | 42 | NCOA3 | 0 | 0 |
| DNA as seed | PRAD | 43 | USP7 | 1 | 0 |
| DNA as seed | PRAD | 44 | SKP1 | 1 | 0 |
| DNA as seed | PRAD | 45 | NOTCH1 | 0 | 0 |
| DNA as seed | PRAD | 46 | SUMO1 | 0 | 0 |
| DNA as seed | PRAD | 47 | FLNA | 0 | 0 |
| DNA as seed | PRAD | 48 | BMI1 | 1 | 0 |
| DNA as seed | PRAD | 49 | BTRC | 0 | 0 |
| DNA as seed | PRAD | 50 | DDB1 | 1 | 1 |
| DNA as seed | PRAD | 51 | SUMO2 | 1 | 0 |
| DNA as seed | PRAD | 52 | BRCA2 | 0 | 1 |
| DNA as seed | PRAD | 53 | PRKDC | 0 | 0 |
| DNA as seed | PRAD | 54 | BRAF | 0 | 0 |
| DNA as seed | PRAD | 55 | FBXW7 | 0 | 0 |
| DNA as seed | PRAD | 56 | PRKCZ | 0 | 0 |
| DNA as seed | PRAD | 57 | WDR5 | 1 | 1 |
| DNA as seed | PRAD | 58 | CASP8 | 0 | 0 |
| DNA as seed | PRAD | 59 | PTK2 | 0 | 0 |
| DNA as seed | PRAD | 60 | EZH2 | 0 | 0 |
| DNA as seed | PRAD | 61 | PGR | 0 | 0 |
| DNA as seed | PRAD | 62 | JAK2 | 0 | 0 |
| DNA as seed | PRAD | 63 | FOXO3 | 0 | 0 |

| | | | | | |
|---|---|---|---|---|---|
| DNA as seed | PRAD | 64 | SMARCA4 | 0 | 0 |
| DNA as seed | PRAD | 65 | PIAS1 | 1 | 0 |
| DNA as seed | PRAD | 66 | PRKCD | 0 | 0 |
| DNA as seed | PRAD | 67 | TP73 | 0 | 0 |
| DNA as seed | PRAD | 68 | RAD51 | 1 | 1 |
| DNA as seed | PRAD | 69 | PRKAA1 | 0 | 0 |
| DNA as seed | PRAD | 70 | UBR5 | 0 | 0 |
| DNA as seed | PRAD | 71 | COPS6 | 1 | 1 |
| DNA as seed | PRAD | 72 | NCOA1 | 0 | 0 |
| DNA as seed | PRAD | 73 | LYN | 0 | 0 |
| DNA as seed | PRAD | 74 | DAXX | 0 | 0 |
| DNA as seed | PRAD | 75 | IGF1R | 1 | 0 |
| DNA as seed | PRAD | 76 | NANOG | 0 | 0 |
| DNA as seed | PRAD | 77 | TERT | 0 | 0 |
| DNA as seed | PRAD | 78 | MYB | 0 | 0 |
| DNA as seed | PRAD | 79 | NCOA2 | 0 | 0 |
| DNA as seed | PRAD | 80 | TCF7L2 | 0 | 0 |
| DNA as seed | PRAD | 81 | IRS1 | 0 | 0 |
| DNA as seed | PRAD | 82 | USF1 | 0 | 0 |
| DNA as seed | PRAD | 83 | RBX1 | 1 | 1 |
| DNA as seed | PRAD | 84 | DDX5 | 1 | 0 |
| DNA as seed | PRAD | 85 | TGFBR2 | 0 | 0 |
| DNA as seed | PRAD | 86 | CAMK2A | 0 | 0 |
| DNA as seed | PRAD | 87 | PDGFRB | 0 | 0 |
| DNA as seed | PRAD | 88 | UBE3A | 0 | 0 |
| DNA as seed | PRAD | 89 | MAP2K1 | 0 | 0 |
| DNA as seed | PRAD | 90 | LAMA3 | 0 | 0 |
| DNA as seed | PRAD | 91 | BLM | 0 | 1 |
| DNA as seed | PRAD | 92 | STK11 | 0 | 0 |
| DNA as seed | PRAD | 93 | MMP2 | 0 | 0 |
| DNA as seed | PRAD | 94 | FGFR2 | 0 | 0 |
| DNA as seed | PRAD | 95 | CSNK1A1 | 1 | 1 |
| DNA as seed | PRAD | 96 | CCNA2 | 1 | 1 |
| DNA as seed | PRAD | 97 | WT1 | 0 | 0 |
| DNA as seed | PRAD | 98 | DNMT1 | 0 | 1 |
| DNA as seed | PRAD | 99 | UBE2D2 | 0 | 0 |
| DNA as seed | PRAD | 100 | CD44 | 0 | 0 |
| DNA as seed | STAD | 1 | TP53 | 0 | 0 |
| DNA as seed | STAD | 2 | MYC | 1 | 0 |
| DNA as seed | STAD | 3 | ERBB2 | 0 | 0 |
| DNA as seed | STAD | 4 | WWOX | 0 | 0 |
| DNA as seed | STAD | 5 | CTNNB1 | 1 | 0 |
| DNA as seed | STAD | 6 | CDH1 | 0 | 0 |
| DNA as seed | STAD | 7 | ERBB4 | 0 | 0 |
| DNA as seed | STAD | 8 | PIK3CA | 1 | 0 |
| DNA as seed | STAD | 9 | ERBB3 | 0 | 0 |
| DNA as seed | STAD | 10 | EGFR | 1 | 0 |
| DNA as seed | STAD | 11 | ATM | 0 | 0 |
| DNA as seed | STAD | 12 | PTEN | 0 | 0 |
| DNA as seed | STAD | 13 | CDKN2A | 0 | 0 |
| DNA as seed | STAD | 14 | SMAD4 | 0 | 0 |
| DNA as seed | STAD | 15 | KRAS | 1 | 0 |
| DNA as seed | STAD | 16 | ESR1 | 0 | 0 |
| DNA as seed | STAD | 17 | ARID1A | 0 | 0 |
| DNA as seed | STAD | 18 | FBXW7 | 0 | 0 |
| DNA as seed | STAD | 19 | SRC | 0 | 0 |

| | | | | | |
|---|---|---|---|---|---|
| DNA as seed | STAD | 20 | SP1 | 0 | 0 |
| DNA as seed | STAD | 21 | APC | 0 | 0 |
| DNA as seed | STAD | 22 | EP300 | 0 | 0 |
| DNA as seed | STAD | 23 | UBC | 1 | 0 |
| DNA as seed | STAD | 24 | AR | 0 | 0 |
| DNA as seed | STAD | 25 | CDKN1A | 0 | 0 |
| DNA as seed | STAD | 26 | SMAD3 | 0 | 0 |
| DNA as seed | STAD | 27 | RARA | 0 | 0 |
| DNA as seed | STAD | 28 | MAPK1 | 1 | 0 |
| DNA as seed | STAD | 29 | MET | 1 | 0 |
| DNA as seed | STAD | 30 | MAPK3 | 0 | 0 |
| DNA as seed | STAD | 31 | MDM2 | 1 | 0 |
| DNA as seed | STAD | 32 | PIK3R1 | 0 | 0 |
| DNA as seed | STAD | 33 | FGFR3 | 0 | 0 |
| DNA as seed | STAD | 34 | CREBBP | 1 | 0 |
| DNA as seed | STAD | 35 | IGF1R | 1 | 0 |
| DNA as seed | STAD | 36 | RHOA | 0 | 1 |
| DNA as seed | STAD | 37 | SMAD2 | 0 | 0 |
| DNA as seed | STAD | 38 | HSP90AA1 | 1 | 0 |
| DNA as seed | STAD | 39 | HDAC1 | 1 | 0 |
| DNA as seed | STAD | 40 | GSK3B | 0 | 0 |
| DNA as seed | STAD | 41 | HRAS | 0 | 0 |
| DNA as seed | STAD | 42 | AKT1 | 1 | 0 |
| DNA as seed | STAD | 43 | UBB | 0 | 0 |
| DNA as seed | STAD | 44 | HIF1A | 0 | 0 |
| DNA as seed | STAD | 45 | NRAS | 0 | 0 |
| DNA as seed | STAD | 46 | CCNE1 | 1 | 0 |
| DNA as seed | STAD | 47 | SHC1 | 0 | 0 |
| DNA as seed | STAD | 48 | RPS27A | 1 | 0 |
| DNA as seed | STAD | 49 | THRA | 0 | 0 |
| DNA as seed | STAD | 50 | CBL | 0 | 0 |
| DNA as seed | STAD | 51 | E2F1 | 0 | 0 |
| DNA as seed | STAD | 52 | ABL1 | 0 | 0 |
| DNA as seed | STAD | 53 | PDGFRA | 1 | 0 |
| DNA as seed | STAD | 54 | UBR5 | 0 | 0 |
| DNA as seed | STAD | 55 | PPARG | 0 | 0 |
| DNA as seed | STAD | 56 | CCND1 | 1 | 0 |
| DNA as seed | STAD | 57 | SNW1 | 1 | 1 |
| DNA as seed | STAD | 58 | PCNA | 1 | 1 |
| DNA as seed | STAD | 59 | PIK3R2 | 0 | 0 |
| DNA as seed | STAD | 60 | ACTB | 1 | 0 |
| DNA as seed | STAD | 61 | FYN | 0 | 0 |
| DNA as seed | STAD | 62 | HLA-B | 0 | 0 |
| DNA as seed | STAD | 63 | HSPA8 | 1 | 1 |
| DNA as seed | STAD | 64 | MAPK8 | 0 | 0 |
| DNA as seed | STAD | 65 | CTNND1 | 0 | 0 |
| DNA as seed | STAD | 66 | EWSR1 | 1 | 1 |
| DNA as seed | STAD | 67 | HSP90AB1 | 0 | 0 |
| DNA as seed | STAD | 68 | SPTA1 | 0 | 0 |
| DNA as seed | STAD | 69 | UBA52 | 1 | 0 |
| DNA as seed | STAD | 70 | HDAC3 | 1 | 0 |
| DNA as seed | STAD | 71 | PARP1 | 0 | 0 |
| DNA as seed | STAD | 72 | GRB7 | 0 | 0 |
| DNA as seed | STAD | 73 | PTK2 | 1 | 0 |
| DNA as seed | STAD | 74 | EZH2 | 0 | 0 |
| DNA as seed | STAD | 75 | HUWE1 | 1 | 0 |

| | | | | | |
|---|---|---|---|---|---|
| DNA as seed | STAD | 76 | CDC42 | 1 | 0 |
| DNA as seed | STAD | 77 | SMAD1 | 0 | 0 |
| DNA as seed | STAD | 78 | NOTCH1 | 0 | 0 |
| DNA as seed | STAD | 79 | CDC6 | 1 | 1 |
| DNA as seed | STAD | 80 | CRK | 0 | 0 |
| DNA as seed | STAD | 81 | PLK1 | 1 | 1 |
| DNA as seed | STAD | 82 | BTRC | 0 | 0 |
| DNA as seed | STAD | 83 | SMARCA4 | 1 | 0 |
| DNA as seed | STAD | 84 | FOXO3 | 0 | 0 |
| DNA as seed | STAD | 85 | NCOR1 | 0 | 0 |
| DNA as seed | STAD | 86 | NCOA3 | 0 | 0 |
| DNA as seed | STAD | 87 | CDKN1B | 0 | 0 |
| DNA as seed | STAD | 88 | CDK4 | 1 | 0 |
| DNA as seed | STAD | 89 | PML | 0 | 0 |
| DNA as seed | STAD | 90 | BMI1 | 0 | 0 |
| DNA as seed | STAD | 91 | SQSTM1 | 0 | 0 |
| DNA as seed | STAD | 92 | KAT2B | 0 | 0 |
| DNA as seed | STAD | 93 | SKP1 | 1 | 0 |
| DNA as seed | STAD | 94 | FLNA | 0 | 0 |
| DNA as seed | STAD | 95 | NCOA1 | 0 | 0 |
| DNA as seed | STAD | 96 | YY1 | 1 | 1 |
| DNA as seed | STAD | 97 | CDKN2B | 0 | 0 |
| DNA as seed | STAD | 98 | VEGFA | 0 | 0 |
| DNA as seed | STAD | 99 | CALM3 | 0 | 0 |
| DNA as seed | STAD | 100 | IQGAP1 | 0 | 0 |
| DNA as seed | THCA | 1 | BRAF | 0 | 0 |
| DNA as seed | THCA | 2 | HRAS | 0 | 0 |
| DNA as seed | THCA | 3 | NRAS | 0 | 0 |
| DNA as seed | THCA | 4 | AKT1 | 0 | 0 |
| DNA as seed | THCA | 5 | KRAS | 0 | 0 |
| DNA as seed | THCA | 6 | MAPK1 | 0 | 0 |
| DNA as seed | THCA | 7 | TRAF2 | 0 | 0 |
| DNA as seed | THCA | 8 | MAPK3 | 0 | 0 |
| DNA as seed | THCA | 9 | ATM | 0 | 0 |
| DNA as seed | THCA | 10 | PIK3R1 | 0 | 0 |
| DNA as seed | THCA | 11 | MAPK14 | 0 | 0 |
| DNA as seed | THCA | 12 | TSC1 | 0 | 0 |
| DNA as seed | THCA | 13 | MAP3K3 | 0 | 0 |
| DNA as seed | THCA | 14 | YWHAZ | 0 | 0 |
| DNA as seed | THCA | 15 | SHC1 | 0 | 0 |
| DNA as seed | THCA | 16 | IKBKG | 0 | 0 |
| DNA as seed | THCA | 17 | NOTCH1 | 0 | 0 |
| DNA as seed | THCA | 18 | GSK3B | 0 | 0 |
| DNA as seed | THCA | 19 | AKT2 | 0 | 0 |
| DNA as seed | THCA | 20 | RET | 0 | 0 |
| DNA as seed | THCA | 21 | AKT3 | 0 | 0 |
| DNA as seed | THCA | 22 | RAF1 | 0 | 0 |
| DNA as seed | THCA | 23 | MDM2 | 0 | 0 |
| DNA as seed | THCA | 24 | YWHAG | 0 | 0 |
| DNA as seed | THCA | 25 | CDC37 | 0 | 1 |
| DNA as seed | THCA | 26 | [illegible]2\|H3C2\|H3C[illegible] | 0 | 0 |
| DNA as seed | THCA | 27 | HDAC1 | 0 | 0 |
| DNA as seed | THCA | 28 | PPP2CB | 0 | 0 |
| DNA as seed | THCA | 29 | CREB1 | 0 | 0 |
| DNA as seed | THCA | 30 | CBL | 0 | 0 |
| DNA as seed | THCA | 31 | YWHAE | 0 | 0 |

| | | | | | |
|---|---|---|---|---|---|
| DNA as seed | THCA | 32 | MAPK9 | 0 | 0 |
| DNA as seed | THCA | 33 | WDR5 | 1 | 1 |
| DNA as seed | THCA | 34 | IKBKB | 0 | 0 |
| DNA as seed | THCA | 35 | PRKAA1 | 0 | 0 |
| DNA as seed | THCA | 36 | YWHAB | 0 | 0 |
| DNA as seed | THCA | 37 | RXRA | 0 | 0 |
| DNA as seed | THCA | 38 | CALM3 | 0 | 0 |
| DNA as seed | THCA | 39 | CHEK2 | 0 | 0 |
| DNA as seed | THCA | 40 | RB1 | 0 | 0 |
| DNA as seed | THCA | 41 | MAP3K1 | 0 | 0 |
| DNA as seed | THCA | 42 | CALM1 | 0 | 0 |
| DNA as seed | THCA | 43 | CALM2 | 0 | 0 |
| DNA as seed | THCA | 44 | PTK2 | 0 | 0 |
| DNA as seed | THCA | 45 | CHUK | 0 | 0 |
| DNA as seed | THCA | 46 | VHL | 1 | 0 |
| DNA as seed | THCA | 47 | PRKDC | 0 | 0 |
| DNA as seed | THCA | 48 | GNAI2 | 0 | 0 |
| DNA as seed | THCA | 49 | GNB2 | 0 | 0 |
| DNA as seed | THCA | 50 | PKM | 0 | 0 |
| DNA as seed | THCA | 51 | FOXO3 | 0 | 0 |
| DNA as seed | THCA | 52 | IL7R | 0 | 0 |
| DNA as seed | THCA | 53 | USP7 | 0 | 0 |
| DNA as seed | THCA | 54 | KAT2B | 0 | 0 |
| DNA as seed | THCA | 55 | RPS6KA3 | 0 | 0 |
| DNA as seed | THCA | 56 | ATR | 0 | 1 |
| DNA as seed | THCA | 57 | POU5F1 | 0 | 0 |
| DNA as seed | THCA | 58 | CCDC6 | 0 | 0 |
| DNA as seed | THCA | 59 | FBXW7 | 0 | 0 |
| DNA as seed | THCA | 60 | RPS6KA1 | 0 | 0 |
| DNA as seed | THCA | 61 | CDKN2A | 0 | 0 |
| DNA as seed | THCA | 62 | IRS1 | 0 | 0 |
| DNA as seed | THCA | 63 | TBC1D7 | 0 | 0 |
| DNA as seed | THCA | 64 | GNAI3 | 0 | 0 |
| DNA as seed | THCA | 65 | PDPK1 | 1 | 1 |
| DNA as seed | THCA | 66 | RPTOR | 0 | 1 |
| DNA as seed | THCA | 67 | PIK3R5 | 0 | 0 |
| DNA as seed | THCA | 68 | PIK3CG | 0 | 0 |
| DNA as seed | THCA | 69 | GNB4 | 0 | 0 |
| DNA as seed | THCA | 70 | MAP3K5 | 0 | 0 |
| DNA as seed | THCA | 71 | CDKN1B | 0 | 0 |
| DNA as seed | THCA | 72 | CAMK4 | 0 | 0 |
| DNA as seed | THCA | 73 | TRAF3 | 0 | 0 |
| DNA as seed | THCA | 74 | GNB3 | 0 | 0 |
| DNA as seed | THCA | 75 | DLC1 | 0 | 0 |
| DNA as seed | THCA | 76 | CACNA1B | 0 | 0 |
| DNA as seed | THCA | 77 | BIRC3 | 0 | 0 |
| DNA as seed | THCA | 78 | BCL10 | 0 | 0 |
| DNA as seed | THCA | 79 | SEC16A | 0 | 1 |
| DNA as seed | THCA | 80 | ODF2 | 0 | 0 |
| DNA as seed | THCA | 81 | FRS2 | 0 | 0 |
| DNA as seed | THCA | 82 | RASGRF1 | 0 | 0 |
| DNA as seed | THCA | 83 | NANOG | 0 | 0 |
| DNA as seed | THCA | 84 | PIK3R6 | 0 | 0 |
| DNA as seed | THCA | 85 | SPTA1 | 0 | 0 |
| DNA as seed | THCA | 86 | SMAD1 | 0 | 0 |
| DNA as seed | THCA | 87 | TGFB1 | 0 | 0 |

| | | | | | |
|---|---|---|---|---|---|
| DNA as seed | THCA | 88 | BIRC2 | 0 | 0 |
| DNA as seed | THCA | 89 | NEDD4L | 0 | 0 |
| DNA as seed | THCA | 90 | NR4A1 | 0 | 0 |
| DNA as seed | THCA | 91 | GNG5 | 0 | 0 |
| DNA as seed | THCA | 92 | GNG4 | 0 | 0 |
| DNA as seed | THCA | 93 | TERF1 | 0 | 0 |
| DNA as seed | THCA | 94 | BCL6 | 0 | 0 |
| DNA as seed | THCA | 95 | RPS6KA6 | 0 | 0 |
| DNA as seed | THCA | 96 | RPS6KA2 | 0 | 0 |
| DNA as seed | THCA | 97 | USP9X | 0 | 0 |
| DNA as seed | THCA | 98 | VBP1 | 0 | 0 |
| DNA as seed | THCA | 99 | USP20 | 0 | 0 |
| DNA as seed | THCA | 100 | FOXO4 | 0 | 0 |
| DNA as seed | UCEC | 1 | PTEN | 0 | 0 |
| DNA as seed | UCEC | 2 | TP53 | 0 | 0 |
| DNA as seed | UCEC | 3 | PIK3CA | 1 | 0 |
| DNA as seed | UCEC | 4 | PIK3R1 | 0 | 0 |
| DNA as seed | UCEC | 5 | CTNNB1 | 0 | 0 |
| DNA as seed | UCEC | 6 | MYC | 1 | 0 |
| DNA as seed | UCEC | 7 | EP300 | 0 | 0 |
| DNA as seed | UCEC | 8 | ATM | 0 | 0 |
| DNA as seed | UCEC | 9 | KRAS | 1 | 0 |
| DNA as seed | UCEC | 10 | FBXW7 | 0 | 0 |
| DNA as seed | UCEC | 11 | PRKDC | 0 | 0 |
| DNA as seed | UCEC | 12 | EGFR | 0 | 0 |
| DNA as seed | UCEC | 13 | ARID1A | 0 | 0 |
| DNA as seed | UCEC | 14 | LMNA | 0 | 0 |
| DNA as seed | UCEC | 15 | UBC | 1 | 0 |
| DNA as seed | UCEC | 16 | PPP2R1A | 1 | 1 |
| DNA as seed | UCEC | 17 | ESR1 | 0 | 0 |
| DNA as seed | UCEC | 18 | FGFR2 | 1 | 0 |
| DNA as seed | UCEC | 19 | CHD4 | 1 | 0 |
| DNA as seed | UCEC | 20 | PIK3R2 | 0 | 0 |
| DNA as seed | UCEC | 21 | BRCA1 | 0 | 1 |
| DNA as seed | UCEC | 22 | ELAVL1 | 0 | 0 |
| DNA as seed | UCEC | 23 | GRB2 | 1 | 1 |
| DNA as seed | UCEC | 24 | SRC | 0 | 0 |
| DNA as seed | UCEC | 25 | JAK1 | 0 | 0 |
| DNA as seed | UCEC | 26 | PIK3R3 | 0 | 0 |
| DNA as seed | UCEC | 27 | HDAC1 | 1 | 0 |
| DNA as seed | UCEC | 28 | CDK2 | 1 | 0 |
| DNA as seed | UCEC | 29 | HSP90AA1 | 0 | 0 |
| DNA as seed | UCEC | 30 | PLCG1 | 0 | 0 |
| DNA as seed | UCEC | 31 | AR | 0 | 0 |
| DNA as seed | UCEC | 32 | MDM2 | 1 | 0 |
| DNA as seed | UCEC | 33 | PIK3CD | 0 | 0 |
| DNA as seed | UCEC | 34 | AKT1 | 0 | 0 |
| DNA as seed | UCEC | 35 | INPPL1 | 0 | 0 |
| DNA as seed | UCEC | 36 | CHD3 | 0 | 0 |
| DNA as seed | UCEC | 37 | STAT3 | 0 | 0 |
| DNA as seed | UCEC | 38 | SMAD3 | 0 | 0 |
| DNA as seed | UCEC | 39 | MAP3K1 | 0 | 0 |
| DNA as seed | UCEC | 40 | PRKCA | 0 | 0 |
| DNA as seed | UCEC | 41 | HDAC2 | 0 | 0 |
| DNA as seed | UCEC | 42 | SHC1 | 0 | 0 |
| DNA as seed | UCEC | 43 | NPM1 | 0 | 0 |

| | | | | | |
|---|---|---|---|---|---|
| DNA as seed | UCEC | 44 | PIP5K1A | 1 | 0 |
| DNA as seed | UCEC | 45 | ERBB2 | 0 | 0 |
| DNA as seed | UCEC | 46 | YWHAZ | 0 | 0 |
| DNA as seed | UCEC | 47 | PARP1 | 0 | 0 |
| DNA as seed | UCEC | 48 | CBL | 0 | 0 |
| DNA as seed | UCEC | 49 | ABL1 | 0 | 0 |
| DNA as seed | UCEC | 50 | UBE2I | 1 | 1 |
| DNA as seed | UCEC | 51 | RNF2 | 0 | 0 |
| DNA as seed | UCEC | 52 | CTCF | 1 | 1 |
| DNA as seed | UCEC | 53 | EED | 0 | 0 |
| DNA as seed | UCEC | 54 | PLCG2 | 0 | 0 |
| DNA as seed | UCEC | 55 | HIF1A | 0 | 0 |
| DNA as seed | UCEC | 56 | PCNA | 1 | 1 |
| DNA as seed | UCEC | 57 | SMAD2 | 0 | 0 |
| DNA as seed | UCEC | 58 | CCNE1 | 1 | 0 |
| DNA as seed | UCEC | 59 | ECT2 | 1 | 1 |
| DNA as seed | UCEC | 60 | CREB1 | 0 | 0 |
| DNA as seed | UCEC | 61 | E2F1 | 0 | 0 |
| DNA as seed | UCEC | 62 | FOS | 0 | 0 |
| DNA as seed | UCEC | 63 | SNW1 | 1 | 1 |
| DNA as seed | UCEC | 64 | NF1 | 0 | 0 |
| DNA as seed | UCEC | 65 | MAPK8 | 0 | 0 |
| DNA as seed | UCEC | 66 | RPA1 | 1 | 1 |
| DNA as seed | UCEC | 67 | MET | 0 | 0 |
| DNA as seed | UCEC | 68 | XRCC6 | 1 | 1 |
| DNA as seed | UCEC | 69 | SIRT7 | 0 | 0 |
| DNA as seed | UCEC | 70 | PPARG | 0 | 0 |
| DNA as seed | UCEC | 71 | SIRT1 | 0 | 0 |
| DNA as seed | UCEC | 72 | ZMYM2 | 0 | 0 |
| DNA as seed | UCEC | 73 | XRCC5 | 1 | 0 |
| DNA as seed | UCEC | 74 | ERBB3 | 0 | 0 |
| DNA as seed | UCEC | 75 | RPA2 | 1 | 1 |
| DNA as seed | UCEC | 76 | PRKN | 0 | 0 |
| DNA as seed | UCEC | 77 | H3-4 | 0 | 0 |
| DNA as seed | UCEC | 78 | RAC1 | 0 | 1 |
| DNA as seed | UCEC | 79 | MECOM | 0 | 0 |
| DNA as seed | UCEC | 80 | SMURF1 | 0 | 0 |
| DNA as seed | UCEC | 81 | PML | 0 | 0 |
| DNA as seed | UCEC | 82 | PLK1 | 1 | 1 |
| DNA as seed | UCEC | 83 | PRKCD | 0 | 0 |
| DNA as seed | UCEC | 84 | VCAM1 | 0 | 0 |
| DNA as seed | UCEC | 85 | PI4KB | 0 | 0 |
| DNA as seed | UCEC | 86 | YAP1 | 0 | 0 |
| DNA as seed | UCEC | 87 | PDGFRB | 0 | 0 |
| DNA as seed | UCEC | 88 | NR3C1 | 0 | 0 |
| DNA as seed | UCEC | 89 | PDGFRA | 0 | 0 |
| DNA as seed | UCEC | 90 | FOXO3 | 0 | 0 |
| DNA as seed | UCEC | 91 | USP7 | 1 | 0 |
| DNA as seed | UCEC | 92 | SMARCA4 | 1 | 0 |
| DNA as seed | UCEC | 93 | FGFR1 | 0 | 0 |
| DNA as seed | UCEC | 94 | CDC42 | 1 | 0 |
| DNA as seed | UCEC | 95 | BMI1 | 0 | 0 |
| DNA as seed | UCEC | 96 | PRKCZ | 0 | 0 |
| DNA as seed | UCEC | 97 | RPA3 | 1 | 0 |
| DNA as seed | UCEC | 98 | YY1 | 1 | 1 |
| DNA as seed | UCEC | 99 | TFRC | 1 | 0 |

| | | | | | |
|---|---|---|---|---|---|
| DNA as seed | UCEC | 100 | CCDC8 | 0 | 0 |
| DNA as seed | SKCM | 1 | TP53 | 0 | 0 |
| DNA as seed | SKCM | 2 | NRAS | 1 | 0 |
| DNA as seed | SKCM | 3 | CDKN2A | 0 | 0 |
| DNA as seed | SKCM | 4 | BRAF | 1 | 0 |
| DNA as seed | SKCM | 5 | EGFR | 0 | 0 |
| DNA as seed | SKCM | 6 | PTEN | 0 | 0 |
| DNA as seed | SKCM | 7 | RAC1 | 1 | 1 |
| DNA as seed | SKCM | 8 | PLCB4 | 0 | 0 |
| DNA as seed | SKCM | 9 | TP63 | 0 | 0 |
| DNA as seed | SKCM | 10 | CCND1 | 1 | 0 |
| DNA as seed | SKCM | 11 | GRB2 | 0 | 1 |
| DNA as seed | SKCM | 12 | RELA | 0 | 0 |
| DNA as seed | SKCM | 13 | MAPK3 | 0 | 0 |
| DNA as seed | SKCM | 14 | MAPK1 | 1 | 0 |
| DNA as seed | SKCM | 15 | EP300 | 0 | 0 |
| DNA as seed | SKCM | 16 | IFNA2 | 0 | 0 |
| DNA as seed | SKCM | 17 | IFNA5 | 0 | 0 |
| DNA as seed | SKCM | 18 | IFNA14 | 0 | 0 |
| DNA as seed | SKCM | 19 | CTNNB1 | 0 | 0 |
| DNA as seed | SKCM | 20 | NFKB1 | 0 | 0 |
| DNA as seed | SKCM | 21 | IFNA21 | 0 | 0 |
| DNA as seed | SKCM | 22 | IFNB1 | 0 | 0 |
| DNA as seed | SKCM | 23 | IFNA8 | 0 | 0 |
| DNA as seed | SKCM | 24 | IFNA4 | 0 | 0 |
| DNA as seed | SKCM | 25 | HRAS | 1 | 0 |
| DNA as seed | SKCM | 26 | IFNA6 | 0 | 0 |
| DNA as seed | SKCM | 27 | CDKN1A | 0 | 0 |
| DNA as seed | SKCM | 28 | JAK2 | 0 | 0 |
| DNA as seed | SKCM | 29 | JAK1 | 0 | 0 |
| DNA as seed | SKCM | 30 | PIK3R2 | 0 | 0 |
| DNA as seed | SKCM | 31 | DCC | 0 | 0 |
| DNA as seed | SKCM | 32 | STAT1 | 0 | 0 |
| DNA as seed | SKCM | 33 | PIK3CA | 0 | 0 |
| DNA as seed | SKCM | 34 | IFNA7 | 0 | 0 |
| DNA as seed | SKCM | 35 | IFNA17 | 0 | 0 |
| DNA as seed | SKCM | 36 | PLCG1 | 0 | 0 |
| DNA as seed | SKCM | 37 | IFNA10 | 0 | 0 |
| DNA as seed | SKCM | 38 | KRAS | 0 | 0 |
| DNA as seed | SKCM | 39 | PTPN11 | 0 | 0 |
| DNA as seed | SKCM | 40 | CREBBP | 0 | 0 |
| DNA as seed | SKCM | 41 | IFNA16 | 0 | 0 |
| DNA as seed | SKCM | 42 | JAK3 | 0 | 0 |
| DNA as seed | SKCM | 43 | PIK3CB | 0 | 0 |
| DNA as seed | SKCM | 44 | MAPK8 | 0 | 0 |
| DNA as seed | SKCM | 45 | MAP2K1 | 0 | 0 |
| DNA as seed | SKCM | 46 | PIK3R3 | 0 | 0 |
| DNA as seed | SKCM | 47 | IFNA1 | 0 | 0 |
| DNA as seed | SKCM | 48 | MAPK14 | 0 | 0 |
| DNA as seed | SKCM | 49 | ERBB2 | 0 | 0 |
| DNA as seed | SKCM | 50 | IFNA13 | 0 | 0 |
| DNA as seed | SKCM | 51 | NF1 | 0 | 0 |
| DNA as seed | SKCM | 52 | GSK3B | 0 | 0 |
| DNA as seed | SKCM | 53 | IFNE | 0 | 0 |
| DNA as seed | SKCM | 54 | SMAD3 | 0 | 0 |
| DNA as seed | SKCM | 55 | TYK2 | 0 | 0 |

| | | | | | |
|---|---|---|---|---|---|
| DNA as seed | SKCM | 56 | YWHAZ | 0 | 0 |
| DNA as seed | SKCM | 57 | PTPN6 | 0 | 0 |
| DNA as seed | SKCM | 58 | IFNW1 | 0 | 0 |
| DNA as seed | SKCM | 59 | FGF3 | 0 | 0 |
| DNA as seed | SKCM | 60 | FOS | 0 | 0 |
| DNA as seed | SKCM | 61 | HIF1A | 0 | 0 |
| DNA as seed | SKCM | 62 | TERT | 0 | 0 |
| DNA as seed | SKCM | 63 | CDKN2B | 0 | 0 |
| DNA as seed | SKCM | 64 | TRRAP | 1 | 1 |
| DNA as seed | SKCM | 65 | PDGFRB | 0 | 0 |
| DNA as seed | SKCM | 66 | FGF19 | 0 | 0 |
| DNA as seed | SKCM | 67 | MAPK9 | 0 | 0 |
| DNA as seed | SKCM | 68 | FGF4 | 0 | 0 |
| DNA as seed | SKCM | 69 | CREB1 | 0 | 0 |
| DNA as seed | SKCM | 70 | DSP | 0 | 0 |
| DNA as seed | SKCM | 71 | PTK2 | 1 | 0 |
| DNA as seed | SKCM | 72 | STAT2 | 0 | 0 |
| DNA as seed | SKCM | 73 | PDGFRA | 0 | 0 |
| DNA as seed | SKCM | 74 | YWHAB | 0 | 0 |
| DNA as seed | SKCM | 75 | PRKCD | 0 | 0 |
| DNA as seed | SKCM | 76 | RPS27A | 1 | 0 |
| DNA as seed | SKCM | 77 | MAPK13 | 0 | 0 |
| DNA as seed | SKCM | 78 | SOCS3 | 1 | 0 |
| DNA as seed | SKCM | 79 | SOCS1 | 0 | 0 |
| DNA as seed | SKCM | 80 | SOS1 | 0 | 0 |
| DNA as seed | SKCM | 81 | MAPK10 | 0 | 0 |
| DNA as seed | SKCM | 82 | APOB | 0 | 0 |
| DNA as seed | SKCM | 83 | TP73 | 0 | 0 |
| DNA as seed | SKCM | 84 | ACTB | 0 | 0 |
| DNA as seed | SKCM | 85 | CALM3 | 0 | 0 |
| DNA as seed | SKCM | 86 | KAT2B | 0 | 0 |
| DNA as seed | SKCM | 87 | MET | 0 | 0 |
| DNA as seed | SKCM | 88 | MAPK11 | 0 | 0 |
| DNA as seed | SKCM | 89 | PTPN1 | 0 | 0 |
| DNA as seed | SKCM | 90 | MITF | 0 | 0 |
| DNA as seed | SKCM | 91 | IFNGR1 | 0 | 0 |
| DNA as seed | SKCM | 92 | PRKCE | 0 | 0 |
| DNA as seed | SKCM | 93 | PRKCZ | 0 | 0 |
| DNA as seed | SKCM | 94 | IGF1R | 1 | 0 |
| DNA as seed | SKCM | 95 | CSF2RB | 0 | 0 |
| DNA as seed | SKCM | 96 | PML | 0 | 0 |
| DNA as seed | SKCM | 97 | NFATC1 | 0 | 0 |
| DNA as seed | SKCM | 98 | IRF3 | 0 | 0 |
| DNA as seed | SKCM | 99 | PTPRB | 0 | 0 |
| DNA as seed | SKCM | 100 | FOXO3 | 0 | 0 |
| DNA+RNA as seed | BLCA | 1 | TP53 | 0 | 0 |
| DNA+RNA as seed | BLCA | 2 | EP300 | 1 | 0 |
| DNA+RNA as seed | BLCA | 3 | NR4A1 | 0 | 0 |
| DNA+RNA as seed | BLCA | 4 | PIK3CA | 1 | 0 |
| DNA+RNA as seed | BLCA | 5 | CREBBP | 1 | 0 |
| DNA+RNA as seed | BLCA | 6 | ERBB2 | 0 | 0 |
| DNA+RNA as seed | BLCA | 7 | CDKN1A | 0 | 0 |
| DNA+RNA as seed | BLCA | 8 | ESR1 | 0 | 0 |
| DNA+RNA as seed | BLCA | 9 | RB1 | 0 | 0 |
| DNA+RNA as seed | BLCA | 10 | TERT | 0 | 0 |
| DNA+RNA as seed | BLCA | 11 | MYC | 1 | 0 |

| | | | | | |
|---|---|---|---|---|---|
| DNA+RNA as seed | BLCA | 12 | CDKN2A | 0 | 0 |
| DNA+RNA as seed | BLCA | 13 | YWHAZ | 0 | 0 |
| DNA+RNA as seed | BLCA | 14 | SP1 | 0 | 0 |
| DNA+RNA as seed | BLCA | 15 | PIK3R1 | 0 | 0 |
| DNA+RNA as seed | BLCA | 16 | FGFR3 | 0 | 0 |
| DNA+RNA as seed | BLCA | 17 | PIK3R2 | 0 | 0 |
| DNA+RNA as seed | BLCA | 18 | SHC1 | 0 | 0 |
| DNA+RNA as seed | BLCA | 19 | RELA | 0 | 0 |
| DNA+RNA as seed | BLCA | 20 | ERBB3 | 0 | 0 |
| DNA+RNA as seed | BLCA | 21 | MAPK1 | 0 | 0 |
| DNA+RNA as seed | BLCA | 22 | JUN | 0 | 0 |
| DNA+RNA as seed | BLCA | 23 | UBC | 1 | 0 |
| DNA+RNA as seed | BLCA | 24 | SRC | 0 | 0 |
| DNA+RNA as seed | BLCA | 25 | HRAS | 1 | 0 |
| DNA+RNA as seed | BLCA | 26 | CTNNB1 | 1 | 0 |
| DNA+RNA as seed | BLCA | 27 | AR | 0 | 0 |
| DNA+RNA as seed | BLCA | 28 | CBL | 0 | 0 |
| DNA+RNA as seed | BLCA | 29 | MAPK3 | 0 | 0 |
| DNA+RNA as seed | BLCA | 30 | RXRA | 0 | 0 |
| DNA+RNA as seed | BLCA | 31 | PLCG1 | 0 | 0 |
| DNA+RNA as seed | BLCA | 32 | RAC1 | 1 | 1 |
| DNA+RNA as seed | BLCA | 33 | CDK2 | 1 | 0 |
| DNA+RNA as seed | BLCA | 34 | NCOR1 | 0 | 0 |
| DNA+RNA as seed | BLCA | 35 | SOS1 | 0 | 0 |
| DNA+RNA as seed | BLCA | 36 | SMAD3 | 0 | 0 |
| DNA+RNA as seed | BLCA | 37 | FCGR2A | 0 | 0 |
| DNA+RNA as seed | BLCA | 38 | HDAC1 | 0 | 0 |
| DNA+RNA as seed | BLCA | 39 | NFKB1 | 0 | 0 |
| DNA+RNA as seed | BLCA | 40 | ATM | 0 | 0 |
| DNA+RNA as seed | BLCA | 41 | STAT3 | 0 | 0 |
| DNA+RNA as seed | BLCA | 42 | HSP90AA1 | 0 | 0 |
| DNA+RNA as seed | BLCA | 43 | FHL1 | 0 | 0 |
| DNA+RNA as seed | BLCA | 44 | CUL1 | 1 | 1 |
| DNA+RNA as seed | BLCA | 45 | FCGR3A | 0 | 0 |
| DNA+RNA as seed | BLCA | 46 | KRAS | 1 | 0 |
| DNA+RNA as seed | BLCA | 47 | PPARG | 1 | 0 |
| DNA+RNA as seed | BLCA | 48 | GSK3B | 0 | 0 |
| DNA+RNA as seed | BLCA | 49 | AKT1 | 0 | 0 |
| DNA+RNA as seed | BLCA | 50 | FCER1G | 0 | 0 |
| DNA+RNA as seed | BLCA | 51 | LYN | 0 | 0 |
| DNA+RNA as seed | BLCA | 52 | CRK | 0 | 0 |
| DNA+RNA as seed | BLCA | 53 | HIF1A | 0 | 0 |
| DNA+RNA as seed | BLCA | 54 | FBXW7 | 0 | 0 |
| DNA+RNA as seed | BLCA | 55 | SYK | 0 | 0 |
| DNA+RNA as seed | BLCA | 56 | E2F1 | 0 | 0 |
| DNA+RNA as seed | BLCA | 57 | RHOA | 1 | 1 |
| DNA+RNA as seed | BLCA | 58 | SMAD2 | 0 | 0 |
| DNA+RNA as seed | BLCA | 59 | H2BC9 | 1 | 0 |
| DNA+RNA as seed | BLCA | 60 | PRKCA | 0 | 0 |
| DNA+RNA as seed | BLCA | 61 | FCGR2B | 0 | 0 |
| DNA+RNA as seed | BLCA | 62 | MET | 0 | 0 |
| DNA+RNA as seed | BLCA | 63 | PLCG2 | 0 | 0 |
| DNA+RNA as seed | BLCA | 64 | HDAC3 | 1 | 0 |
| DNA+RNA as seed | BLCA | 65 | PTEN | 0 | 0 |
| DNA+RNA as seed | BLCA | 66 | MYH9 | 1 | 0 |
| DNA+RNA as seed | BLCA | 67 | VAV1 | 0 | 0 |

| | | | | | |
|---|---|---|---|---|---|
| DNA+RNA as seed | BLCA | 68 | UBB | 0 | 0 |
| DNA+RNA as seed | BLCA | 69 | USF1 | 0 | 0 |
| DNA+RNA as seed | BLCA | 70 | MAPK14 | 0 | 0 |
| DNA+RNA as seed | BLCA | 71 | CDH1 | 0 | 0 |
| DNA+RNA as seed | BLCA | 72 | NCOR2 | 0 | 0 |
| DNA+RNA as seed | BLCA | 73 | MDM2 | 1 | 0 |
| DNA+RNA as seed | BLCA | 74 | HSP90AB1 | 0 | 0 |
| DNA+RNA as seed | BLCA | 75 | CCND1 | 1 | 0 |
| DNA+RNA as seed | BLCA | 76 | NCOA3 | 0 | 0 |
| DNA+RNA as seed | BLCA | 77 | MAPK8 | 0 | 0 |
| DNA+RNA as seed | BLCA | 78 | RPS27A | 1 | 0 |
| DNA+RNA as seed | BLCA | 79 | VAV2 | 0 | 0 |
| DNA+RNA as seed | BLCA | 80 | PCNA | 1 | 1 |
| DNA+RNA as seed | BLCA | 81 | MCM2 | 1 | 1 |
| DNA+RNA as seed | BLCA | 82 | EPHA2 | 0 | 0 |
| DNA+RNA as seed | BLCA | 83 | HSPA8 | 1 | 1 |
| DNA+RNA as seed | BLCA | 84 | NR3C1 | 0 | 0 |
| DNA+RNA as seed | BLCA | 85 | VAV3 | 0 | 0 |
| DNA+RNA as seed | BLCA | 86 | ABL1 | 0 | 0 |
| DNA+RNA as seed | BLCA | 87 | NCOA1 | 0 | 0 |
| DNA+RNA as seed | BLCA | 88 | ACTB | 1 | 0 |
| DNA+RNA as seed | BLCA | 89 | FYN | 0 | 0 |
| DNA+RNA as seed | BLCA | 90 | KAT2B | 0 | 0 |
| DNA+RNA as seed | BLCA | 91 | PARP1 | 0 | 0 |
| DNA+RNA as seed | BLCA | 92 | MED1 | 1 | 1 |
| DNA+RNA as seed | BLCA | 93 | CDK4 | 1 | 0 |
| DNA+RNA as seed | BLCA | 94 | CDKN1B | 0 | 0 |
| DNA+RNA as seed | BLCA | 95 | IGHG1 | 0 | 0 |
| DNA+RNA as seed | BLCA | 96 | E2F3 | 1 | 0 |
| DNA+RNA as seed | BLCA | 97 | PTPN11 | 1 | 0 |
| DNA+RNA as seed | BLCA | 98 | UBA52 | 1 | 0 |
| DNA+RNA as seed | BLCA | 99 | EZH2 | 0 | 0 |
| DNA+RNA as seed | BLCA | 100 | PPARGC1A | 0 | 0 |
| DNA+RNA as seed | BRCA | 1 | TP53 | 0 | 0 |
| DNA+RNA as seed | BRCA | 2 | PLK1 | 1 | 1 |
| DNA+RNA as seed | BRCA | 3 | MYC | 1 | 0 |
| DNA+RNA as seed | BRCA | 4 | BRCA1 | 1 | 1 |
| DNA+RNA as seed | BRCA | 5 | CDC25C | 0 | 0 |
| DNA+RNA as seed | BRCA | 6 | PIK3CA | 1 | 0 |
| DNA+RNA as seed | BRCA | 7 | EGFR | 1 | 0 |
| DNA+RNA as seed | BRCA | 8 | MMP1 | 0 | 0 |
| DNA+RNA as seed | BRCA | 9 | CDH1 | 0 | 0 |
| DNA+RNA as seed | BRCA | 10 | SP1 | 0 | 0 |
| DNA+RNA as seed | BRCA | 11 | ESR1 | 1 | 0 |
| DNA+RNA as seed | BRCA | 12 | PIK3R1 | 0 | 0 |
| DNA+RNA as seed | BRCA | 13 | EP300 | 1 | 0 |
| DNA+RNA as seed | BRCA | 14 | CCND1 | 1 | 0 |
| DNA+RNA as seed | BRCA | 15 | AKT1 | 1 | 0 |
| DNA+RNA as seed | BRCA | 16 | SPRY2 | 0 | 0 |
| DNA+RNA as seed | BRCA | 17 | RELA | 1 | 0 |
| DNA+RNA as seed | BRCA | 18 | CDK2 | 1 | 0 |
| DNA+RNA as seed | BRCA | 19 | ERBB2 | 1 | 0 |
| DNA+RNA as seed | BRCA | 20 | SRC | 0 | 0 |
| DNA+RNA as seed | BRCA | 21 | CTNNB1 | 1 | 0 |
| DNA+RNA as seed | BRCA | 22 | UBC | 1 | 0 |
| DNA+RNA as seed | BRCA | 23 | HSP90AA1 | 0 | 0 |

| | | | | | |
|---|---|---|---|---|---|
| DNA+RNA as seed | BRCA | 24 | JUN | 0 | 0 |
| DNA+RNA as seed | BRCA | 25 | GRB2 | 1 | 1 |
| DNA+RNA as seed | BRCA | 26 | CDK1 | 1 | 1 |
| DNA+RNA as seed | BRCA | 27 | MAPK3 | 0 | 0 |
| DNA+RNA as seed | BRCA | 28 | STAT3 | 0 | 0 |
| DNA+RNA as seed | BRCA | 29 | MAPK1 | 0 | 0 |
| DNA+RNA as seed | BRCA | 30 | TPX2 | 1 | 1 |
| DNA+RNA as seed | BRCA | 31 | PTEN | 0 | 0 |
| DNA+RNA as seed | BRCA | 32 | RB1 | 0 | 0 |
| DNA+RNA as seed | BRCA | 33 | NFKB1 | 0 | 0 |
| DNA+RNA as seed | BRCA | 34 | AR | 0 | 0 |
| DNA+RNA as seed | BRCA | 35 | ATM | 0 | 0 |
| DNA+RNA as seed | BRCA | 36 | HDAC1 | 0 | 0 |
| DNA+RNA as seed | BRCA | 37 | CDKN1A | 0 | 0 |
| DNA+RNA as seed | BRCA | 38 | SORBS1 | 0 | 0 |
| DNA+RNA as seed | BRCA | 39 | VEGFD | 0 | 0 |
| DNA+RNA as seed | BRCA | 40 | MDM2 | 1 | 0 |
| DNA+RNA as seed | BRCA | 41 | SMAD3 | 0 | 0 |
| DNA+RNA as seed | BRCA | 42 | MMP13 | 0 | 0 |
| DNA+RNA as seed | BRCA | 43 | HIF1A | 0 | 0 |
| DNA+RNA as seed | BRCA | 44 | FOS | 0 | 0 |
| DNA+RNA as seed | BRCA | 45 | ERBB3 | 1 | 0 |
| DNA+RNA as seed | BRCA | 46 | RAD21 | 1 | 0 |
| DNA+RNA as seed | BRCA | 47 | SHC1 | 0 | 0 |
| DNA+RNA as seed | BRCA | 48 | HSP90AB1 | 0 | 0 |
| DNA+RNA as seed | BRCA | 49 | CSNK2A1 | 0 | 0 |
| DNA+RNA as seed | BRCA | 50 | UBB | 0 | 0 |
| DNA+RNA as seed | BRCA | 51 | HDAC2 | 1 | 0 |
| DNA+RNA as seed | BRCA | 52 | HRAS | 0 | 0 |
| DNA+RNA as seed | BRCA | 53 | CBL | 0 | 0 |
| DNA+RNA as seed | BRCA | 54 | MAPK8 | 0 | 0 |
| DNA+RNA as seed | BRCA | 55 | RPS27A | 1 | 0 |
| DNA+RNA as seed | BRCA | 56 | SPC24 | 1 | 1 |
| DNA+RNA as seed | BRCA | 57 | NPM1 | 0 | 0 |
| DNA+RNA as seed | BRCA | 58 | MAPK14 | 0 | 0 |
| DNA+RNA as seed | BRCA | 59 | PPARG | 0 | 0 |
| DNA+RNA as seed | BRCA | 60 | SMAD2 | 0 | 0 |
| DNA+RNA as seed | BRCA | 61 | CREB1 | 0 | 0 |
| DNA+RNA as seed | BRCA | 62 | PARP1 | 0 | 0 |
| DNA+RNA as seed | BRCA | 63 | ABL1 | 0 | 0 |
| DNA+RNA as seed | BRCA | 64 | HSPA8 | 1 | 1 |
| DNA+RNA as seed | BRCA | 65 | CRK | 0 | 0 |
| DNA+RNA as seed | BRCA | 66 | FGF3 | 0 | 0 |
| DNA+RNA as seed | BRCA | 67 | NCOR1 | 0 | 0 |
| DNA+RNA as seed | BRCA | 68 | NR3C1 | 0 | 0 |
| DNA+RNA as seed | BRCA | 69 | CASP8 | 0 | 0 |
| DNA+RNA as seed | BRCA | 70 | SNW1 | 1 | 1 |
| DNA+RNA as seed | BRCA | 71 | CCNB1 | 1 | 0 |
| DNA+RNA as seed | BRCA | 72 | RUNX1 | 1 | 0 |
| DNA+RNA as seed | BRCA | 73 | MYH9 | 1 | 0 |
| DNA+RNA as seed | BRCA | 74 | IKBKG | 0 | 0 |
| DNA+RNA as seed | BRCA | 75 | PML | 0 | 0 |
| DNA+RNA as seed | BRCA | 76 | PKMYT1 | 1 | 0 |
| DNA+RNA as seed | BRCA | 77 | HDAC3 | 1 | 0 |
| DNA+RNA as seed | BRCA | 78 | ERBB4 | 0 | 0 |
| DNA+RNA as seed | BRCA | 79 | PPP1CC | 0 | 0 |

| | | | | | |
|---|---|---|---|---|---|
| DNA+RNA as seed | BRCA | 80 | 14\|H4C15\|H4 | 0 | 0 |
| DNA+RNA as seed | BRCA | 81 | H3-4 | 0 | 0 |
| DNA+RNA as seed | BRCA | 82 | RPA2 | 1 | 1 |
| DNA+RNA as seed | BRCA | 83 | RPA1 | 1 | 1 |
| DNA+RNA as seed | BRCA | 84 | LMNA | 0 | 0 |
| DNA+RNA as seed | BRCA | 85 | PPP2CA | 1 | 1 |
| DNA+RNA as seed | BRCA | 86 | SIRT1 | 0 | 0 |
| DNA+RNA as seed | BRCA | 87 | FBXW7 | 1 | 0 |
| DNA+RNA as seed | BRCA | 88 | ETS1 | 0 | 0 |
| DNA+RNA as seed | BRCA | 89 | CDK4 | 1 | 0 |
| DNA+RNA as seed | BRCA | 90 | UBA52 | 1 | 0 |
| DNA+RNA as seed | BRCA | 91 | PLCG1 | 0 | 0 |
| DNA+RNA as seed | BRCA | 92 | VEGFA | 0 | 0 |
| DNA+RNA as seed | BRCA | 93 | KRAS | 1 | 0 |
| DNA+RNA as seed | BRCA | 94 | FGFR3 | 0 | 0 |
| DNA+RNA as seed | BRCA | 95 | FGF4 | 0 | 0 |
| DNA+RNA as seed | BRCA | 96 | NOTCH1 | 0 | 0 |
| DNA+RNA as seed | BRCA | 97 | H2AX | 1 | 0 |
| DNA+RNA as seed | BRCA | 98 | NRAS | 0 | 0 |
| DNA+RNA as seed | BRCA | 99 | MAP3K1 | 0 | 0 |
| DNA+RNA as seed | BRCA | 100 | PPP1CA | 1 | 0 |
| DNA+RNA as seed | COADREAD | 1 | TP53 | 0 | 0 |
| DNA+RNA as seed | COADREAD | 2 | CTNNB1 | 1 | 0 |
| DNA+RNA as seed | COADREAD | 3 | KRAS | 1 | 0 |
| DNA+RNA as seed | COADREAD | 4 | PIK3CA | 1 | 0 |
| DNA+RNA as seed | COADREAD | 5 | PIK3R1 | 0 | 0 |
| DNA+RNA as seed | COADREAD | 6 | NRAS | 0 | 0 |
| DNA+RNA as seed | COADREAD | 7 | SRC | 0 | 0 |
| DNA+RNA as seed | COADREAD | 8 | APC | 1 | 0 |
| DNA+RNA as seed | COADREAD | 9 | ATM | 0 | 0 |
| DNA+RNA as seed | COADREAD | 10 | PIK3CB | 0 | 0 |
| DNA+RNA as seed | COADREAD | 11 | SMAD4 | 1 | 0 |
| DNA+RNA as seed | COADREAD | 12 | MMP7 | 0 | 0 |
| DNA+RNA as seed | COADREAD | 13 | CDH1 | 1 | 0 |
| DNA+RNA as seed | COADREAD | 14 | SMAD3 | 0 | 0 |
| DNA+RNA as seed | COADREAD | 15 | PIK3R3 | 0 | 0 |
| DNA+RNA as seed | COADREAD | 16 | HSP90AA1 | 0 | 0 |
| DNA+RNA as seed | COADREAD | 17 | PTEN | 0 | 0 |
| DNA+RNA as seed | COADREAD | 18 | AKT1 | 0 | 0 |
| DNA+RNA as seed | COADREAD | 19 | SMAD2 | 0 | 0 |
| DNA+RNA as seed | COADREAD | 20 | STAT3 | 0 | 0 |
| DNA+RNA as seed | COADREAD | 21 | PIK3CD | 0 | 0 |
| DNA+RNA as seed | COADREAD | 22 | FYN | 0 | 0 |
| DNA+RNA as seed | COADREAD | 23 | SHC1 | 0 | 0 |
| DNA+RNA as seed | COADREAD | 24 | JUN | 1 | 0 |
| DNA+RNA as seed | COADREAD | 25 | FBXW7 | 0 | 0 |
| DNA+RNA as seed | COADREAD | 26 | WWOX | 0 | 0 |
| DNA+RNA as seed | COADREAD | 27 | AR | 0 | 0 |
| DNA+RNA as seed | COADREAD | 28 | HSP90AB1 | 0 | 0 |
| DNA+RNA as seed | COADREAD | 29 | GSK3B | 0 | 0 |
| DNA+RNA as seed | COADREAD | 30 | TCF7L2 | 1 | 0 |
| DNA+RNA as seed | COADREAD | 31 | MMP1 | 0 | 0 |
| DNA+RNA as seed | COADREAD | 32 | CSNK2A1 | 0 | 0 |
| DNA+RNA as seed | COADREAD | 33 | PRKDC | 0 | 0 |
| DNA+RNA as seed | COADREAD | 34 | CDK2 | 1 | 0 |
| DNA+RNA as seed | COADREAD | 35 | YWHAZ | 1 | 0 |

| | | | | | |
|---|---|---|---|---|---|
| DNA+RNA as seed | COADREAD | 36 | ERBB3 | 0 | 0 |
| DNA+RNA as seed | COADREAD | 37 | ITCH | 0 | 0 |
| DNA+RNA as seed | COADREAD | 38 | HLA-B | 0 | 0 |
| DNA+RNA as seed | COADREAD | 39 | MAPK8 | 0 | 0 |
| DNA+RNA as seed | COADREAD | 40 | IGF1R | 1 | 0 |
| DNA+RNA as seed | COADREAD | 41 | MET | 0 | 0 |
| DNA+RNA as seed | COADREAD | 42 | CRK | 0 | 0 |
| DNA+RNA as seed | COADREAD | 43 | CDC37 | 1 | 1 |
| DNA+RNA as seed | COADREAD | 44 | KLK6 | 0 | 0 |
| DNA+RNA as seed | COADREAD | 45 | CREB1 | 0 | 0 |
| DNA+RNA as seed | COADREAD | 46 | FOS | 0 | 0 |
| DNA+RNA as seed | COADREAD | 47 | ABL1 | 0 | 0 |
| DNA+RNA as seed | COADREAD | 48 | ERBB4 | 0 | 0 |
| DNA+RNA as seed | COADREAD | 49 | B2M | 0 | 0 |
| DNA+RNA as seed | COADREAD | 50 | HCK | 0 | 0 |
| DNA+RNA as seed | COADREAD | 51 | HIF1A | 0 | 0 |
| DNA+RNA as seed | COADREAD | 52 | PTK2 | 1 | 0 |
| DNA+RNA as seed | COADREAD | 53 | BCL2 | 0 | 0 |
| DNA+RNA as seed | COADREAD | 54 | SIM2 | 0 | 0 |
| DNA+RNA as seed | COADREAD | 55 | BCL2L1 | 1 | 1 |
| DNA+RNA as seed | COADREAD | 56 | IRS1 | 0 | 0 |
| DNA+RNA as seed | COADREAD | 57 | IKBKB | 0 | 0 |
| DNA+RNA as seed | COADREAD | 58 | FOXO3 | 0 | 0 |
| DNA+RNA as seed | COADREAD | 59 | BTRC | 0 | 0 |
| DNA+RNA as seed | COADREAD | 60 | SMAD1 | 0 | 0 |
| DNA+RNA as seed | COADREAD | 61 | PPARG | 0 | 0 |
| DNA+RNA as seed | COADREAD | 62 | PDGFRA | 0 | 0 |
| DNA+RNA as seed | COADREAD | 63 | FGFR2 | 0 | 0 |
| DNA+RNA as seed | COADREAD | 64 | STK11 | 0 | 0 |
| DNA+RNA as seed | COADREAD | 65 | NOTCH1 | 0 | 0 |
| DNA+RNA as seed | COADREAD | 66 | NCK1 | 0 | 0 |
| DNA+RNA as seed | COADREAD | 67 | CASP3 | 0 | 0 |
| DNA+RNA as seed | COADREAD | 68 | CDKN2A | 0 | 0 |
| DNA+RNA as seed | COADREAD | 69 | XRCC6 | 1 | 1 |
| DNA+RNA as seed | COADREAD | 70 | EWSR1 | 1 | 1 |
| DNA+RNA as seed | COADREAD | 71 | YWHAQ | 0 | 0 |
| DNA+RNA as seed | COADREAD | 72 | PPP2CB | 0 | 0 |
| DNA+RNA as seed | COADREAD | 73 | PARP1 | 0 | 0 |
| DNA+RNA as seed | COADREAD | 74 | BRAF | 1 | 0 |
| DNA+RNA as seed | COADREAD | 75 | HSPA1A | 0 | 0 |
| DNA+RNA as seed | COADREAD | 76 | RRAS2 | 0 | 0 |
| DNA+RNA as seed | COADREAD | 77 | FLT3 | 0 | 0 |
| DNA+RNA as seed | COADREAD | 78 | MAPK9 | 0 | 0 |
| DNA+RNA as seed | COADREAD | 79 | BMPR2 | 1 | 0 |
| DNA+RNA as seed | COADREAD | 80 | FGFR4 | 0 | 0 |
| DNA+RNA as seed | COADREAD | 81 | LEF1 | 0 | 0 |
| DNA+RNA as seed | COADREAD | 82 | FLT1 | 0 | 0 |
| DNA+RNA as seed | COADREAD | 83 | FUS | 0 | 0 |
| DNA+RNA as seed | COADREAD | 84 | CHUK | 0 | 0 |
| DNA+RNA as seed | COADREAD | 85 | EZH2 | 0 | 0 |
| DNA+RNA as seed | COADREAD | 86 | CASP8 | 0 | 0 |
| DNA+RNA as seed | COADREAD | 87 | CDH3 | 0 | 0 |
| DNA+RNA as seed | COADREAD | 88 | PDPK1 | 1 | 1 |
| DNA+RNA as seed | COADREAD | 89 | TP63 | 0 | 0 |
| DNA+RNA as seed | COADREAD | 90 | WNT2 | 0 | 0 |
| DNA+RNA as seed | COADREAD | 91 | BTK | 0 | 0 |

| | | | | | |
|---|---|---|---|---|---|
| DNA+RNA as seed | COADREAD | 92 | MMP2 | 0 | 0 |
| DNA+RNA as seed | COADREAD | 93 | CSNK2B | 1 | 1 |
| DNA+RNA as seed | COADREAD | 94 | NGF | 0 | 0 |
| DNA+RNA as seed | COADREAD | 95 | SMURF2 | 0 | 0 |
| DNA+RNA as seed | COADREAD | 96 | ITGB1 | 1 | 0 |
| DNA+RNA as seed | COADREAD | 97 | NR4A1 | 0 | 0 |
| DNA+RNA as seed | COADREAD | 98 | TP73 | 0 | 0 |
| DNA+RNA as seed | COADREAD | 99 | FRS3 | 0 | 0 |
| DNA+RNA as seed | COADREAD | 100 | DVL2 | 0 | 0 |
| DNA+RNA as seed | GBM | 1 | EGFR | 0 | 0 |
| DNA+RNA as seed | GBM | 2 | PIK3R1 | 0 | 0 |
| DNA+RNA as seed | GBM | 3 | PIK3CA | 1 | 0 |
| DNA+RNA as seed | GBM | 4 | TP53 | 0 | 0 |
| DNA+RNA as seed | GBM | 5 | PIK3CB | 0 | 0 |
| DNA+RNA as seed | GBM | 6 | PTEN | 0 | 0 |
| DNA+RNA as seed | GBM | 7 | PDGFRA | 1 | 0 |
| DNA+RNA as seed | GBM | 8 | CDKN2A | 0 | 0 |
| DNA+RNA as seed | GBM | 9 | PIK3R3 | 0 | 0 |
| DNA+RNA as seed | GBM | 10 | PIK3CD | 0 | 0 |
| DNA+RNA as seed | GBM | 11 | PTPN11 | 1 | 0 |
| DNA+RNA as seed | GBM | 12 | IFNA5 | 0 | 0 |
| DNA+RNA as seed | GBM | 13 | IFNA2 | 0 | 0 |
| DNA+RNA as seed | GBM | 14 | IFNA8 | 0 | 0 |
| DNA+RNA as seed | GBM | 15 | IFNA14 | 0 | 0 |
| DNA+RNA as seed | GBM | 16 | IFNA21 | 0 | 0 |
| DNA+RNA as seed | GBM | 17 | IFNA4 | 0 | 0 |
| DNA+RNA as seed | GBM | 18 | IFNA6 | 0 | 0 |
| DNA+RNA as seed | GBM | 19 | PLCG1 | 0 | 0 |
| DNA+RNA as seed | GBM | 20 | SHC1 | 0 | 0 |
| DNA+RNA as seed | GBM | 21 | RELA | 1 | 0 |
| DNA+RNA as seed | GBM | 22 | JAK2 | 0 | 0 |
| DNA+RNA as seed | GBM | 23 | IFNA7 | 0 | 0 |
| DNA+RNA as seed | GBM | 24 | IFNA17 | 0 | 0 |
| DNA+RNA as seed | GBM | 25 | JAK1 | 0 | 0 |
| DNA+RNA as seed | GBM | 26 | IFNA10 | 0 | 0 |
| DNA+RNA as seed | GBM | 27 | IFNB1 | 0 | 0 |
| DNA+RNA as seed | GBM | 28 | PDGFRB | 0 | 0 |
| DNA+RNA as seed | GBM | 29 | IFNA16 | 0 | 0 |
| DNA+RNA as seed | GBM | 30 | IFNA1 | 0 | 0 |
| DNA+RNA as seed | GBM | 31 | JAK3 | 0 | 0 |
| DNA+RNA as seed | GBM | 32 | IFNA13 | 0 | 0 |
| DNA+RNA as seed | GBM | 33 | NFKB1 | 0 | 0 |
| DNA+RNA as seed | GBM | 34 | STAT3 | 0 | 0 |
| DNA+RNA as seed | GBM | 35 | IFNE | 0 | 0 |
| DNA+RNA as seed | GBM | 36 | STAT1 | 0 | 0 |
| DNA+RNA as seed | GBM | 37 | HRAS | 0 | 0 |
| DNA+RNA as seed | GBM | 38 | TYK2 | 0 | 0 |
| DNA+RNA as seed | GBM | 39 | SP1 | 0 | 0 |
| DNA+RNA as seed | GBM | 40 | PTPN6 | 0 | 0 |
| DNA+RNA as seed | GBM | 41 | IFNW1 | 0 | 0 |
| DNA+RNA as seed | GBM | 42 | CTNNB1 | 0 | 0 |
| DNA+RNA as seed | GBM | 43 | MET | 0 | 0 |
| DNA+RNA as seed | GBM | 44 | EP300 | 0 | 0 |
| DNA+RNA as seed | GBM | 45 | PDGFB | 0 | 0 |
| DNA+RNA as seed | GBM | 46 | NRAS | 0 | 0 |
| DNA+RNA as seed | GBM | 47 | ERBB3 | 0 | 0 |

| | | | | | |
|---|---|---|---|---|---|
| DNA+RNA as seed | GBM | 48 | IGF1R | 0 | 0 |
| DNA+RNA as seed | GBM | 49 | PDGFA | 0 | 0 |
| DNA+RNA as seed | GBM | 50 | EGF | 0 | 0 |
| DNA+RNA as seed | GBM | 51 | RB1 | 0 | 0 |
| DNA+RNA as seed | GBM | 52 | INSR | 0 | 0 |
| DNA+RNA as seed | GBM | 53 | KRAS | 1 | 0 |
| DNA+RNA as seed | GBM | 54 | KIT | 0 | 0 |
| DNA+RNA as seed | GBM | 55 | PLCG2 | 0 | 0 |
| DNA+RNA as seed | GBM | 56 | CDK4 | 1 | 0 |
| DNA+RNA as seed | GBM | 57 | CREBBP | 0 | 0 |
| DNA+RNA as seed | GBM | 58 | AKT1 | 0 | 0 |
| DNA+RNA as seed | GBM | 59 | CRK | 0 | 0 |
| DNA+RNA as seed | GBM | 60 | LYN | 0 | 0 |
| DNA+RNA as seed | GBM | 61 | IL2 | 0 | 0 |
| DNA+RNA as seed | GBM | 62 | SOCS1 | 0 | 0 |
| DNA+RNA as seed | GBM | 63 | STAT5A | 0 | 0 |
| DNA+RNA as seed | GBM | 64 | IL3 | 0 | 0 |
| DNA+RNA as seed | GBM | 65 | CSF2RB | 0 | 0 |
| DNA+RNA as seed | GBM | 66 | PTPN1 | 0 | 0 |
| DNA+RNA as seed | GBM | 67 | IL5 | 0 | 0 |
| DNA+RNA as seed | GBM | 68 | CSF2 | 0 | 0 |
| DNA+RNA as seed | GBM | 69 | IL2RG | 0 | 0 |
| DNA+RNA as seed | GBM | 70 | SOCS3 | 1 | 0 |
| DNA+RNA as seed | GBM | 71 | IL2RB | 0 | 0 |
| DNA+RNA as seed | GBM | 72 | IFNGR1 | 0 | 0 |
| DNA+RNA as seed | GBM | 73 | IL7R | 0 | 0 |
| DNA+RNA as seed | GBM | 74 | IL6ST | 0 | 0 |
| DNA+RNA as seed | GBM | 75 | CRKL | 1 | 0 |
| DNA+RNA as seed | GBM | 76 | EPOR | 0 | 0 |
| DNA+RNA as seed | GBM | 77 | ERBB4 | 0 | 0 |
| DNA+RNA as seed | GBM | 78 | EPO | 0 | 0 |
| DNA+RNA as seed | GBM | 79 | IL2RA | 0 | 0 |
| DNA+RNA as seed | GBM | 80 | IL3RA | 0 | 0 |
| DNA+RNA as seed | GBM | 81 | IFNGR2 | 0 | 0 |
| DNA+RNA as seed | GBM | 82 | CDKN2B | 0 | 0 |
| DNA+RNA as seed | GBM | 83 | IRS1 | 0 | 0 |
| DNA+RNA as seed | GBM | 84 | PLCB2 | 0 | 0 |
| DNA+RNA as seed | GBM | 85 | GHR | 0 | 0 |
| DNA+RNA as seed | GBM | 86 | IL5RA | 0 | 0 |
| DNA+RNA as seed | GBM | 87 | MAPK8 | 0 | 0 |
| DNA+RNA as seed | GBM | 88 | FLT1 | 0 | 0 |
| DNA+RNA as seed | GBM | 89 | CSF2RA | 0 | 0 |
| DNA+RNA as seed | GBM | 90 | SMAD3 | 0 | 0 |
| DNA+RNA as seed | GBM | 91 | IL24 | 0 | 0 |
| DNA+RNA as seed | GBM | 92 | IFNAR1 | 0 | 0 |
| DNA+RNA as seed | GBM | 93 | PRKCD | 0 | 0 |
| DNA+RNA as seed | GBM | 94 | PLCB3 | 0 | 0 |
| DNA+RNA as seed | GBM | 95 | PLCB1 | 0 | 0 |
| DNA+RNA as seed | GBM | 96 | STAT2 | 0 | 0 |
| DNA+RNA as seed | GBM | 97 | GAB1 | 0 | 0 |
| DNA+RNA as seed | GBM | 98 | IFNAR2 | 0 | 0 |
| DNA+RNA as seed | GBM | 99 | IRF3 | 0 | 0 |
| DNA+RNA as seed | GBM | 100 | PLCB4 | 0 | 0 |
| DNA+RNA as seed | KIRC | 1 | TP53 | 0 | 0 |
| DNA+RNA as seed | KIRC | 2 | VHL | 1 | 0 |
| DNA+RNA as seed | KIRC | 3 | PTEN | 0 | 0 |

| | | | | | |
|---|---|---|---|---|---|
| DNA+RNA as seed | KIRC | 4 | PIK3CA | 0 | 0 |
| DNA+RNA as seed | KIRC | 5 | ATM | 0 | 0 |
| DNA+RNA as seed | KIRC | 6 | ESR1 | 0 | 0 |
| DNA+RNA as seed | KIRC | 7 | SPTBN2 | 0 | 0 |
| DNA+RNA as seed | KIRC | 8 | SP1 | 0 | 0 |
| DNA+RNA as seed | KIRC | 9 | IGF1R | 0 | 0 |
| DNA+RNA as seed | KIRC | 10 | ACTB | 1 | 0 |
| DNA+RNA as seed | KIRC | 11 | FGFR4 | 0 | 0 |
| DNA+RNA as seed | KIRC | 12 | FLT4 | 0 | 0 |
| DNA+RNA as seed | KIRC | 13 | FLT1 | 0 | 0 |
| DNA+RNA as seed | KIRC | 14 | EP300 | 0 | 0 |
| DNA+RNA as seed | KIRC | 15 | HSP90AA1 | 0 | 0 |
| DNA+RNA as seed | KIRC | 16 | SMARCA4 | 0 | 0 |
| DNA+RNA as seed | KIRC | 17 | HIF1A | 0 | 0 |
| DNA+RNA as seed | KIRC | 18 | AR | 0 | 0 |
| DNA+RNA as seed | KIRC | 19 | DBN1 | 0 | 0 |
| DNA+RNA as seed | KIRC | 20 | HDAC1 | 0 | 0 |
| DNA+RNA as seed | KIRC | 21 | SIRT7 | 0 | 0 |
| DNA+RNA as seed | KIRC | 22 | CSNK2A1 | 0 | 0 |
| DNA+RNA as seed | KIRC | 23 | ACTG1 | 1 | 0 |
| DNA+RNA as seed | KIRC | 24 | SPTAN1 | 0 | 0 |
| DNA+RNA as seed | KIRC | 25 | FLNA | 0 | 0 |
| DNA+RNA as seed | KIRC | 26 | MTOR | 1 | 1 |
| DNA+RNA as seed | KIRC | 27 | HSP90AB1 | 0 | 0 |
| DNA+RNA as seed | KIRC | 28 | MYH9 | 1 | 0 |
| DNA+RNA as seed | KIRC | 29 | PIK3R3 | 0 | 0 |
| DNA+RNA as seed | KIRC | 30 | PDLIM7 | 0 | 0 |
| DNA+RNA as seed | KIRC | 31 | IQGAP1 | 0 | 0 |
| DNA+RNA as seed | KIRC | 32 | SNW1 | 1 | 1 |
| DNA+RNA as seed | KIRC | 33 | CA9 | 0 | 0 |
| DNA+RNA as seed | KIRC | 34 | PIK3CD | 0 | 0 |
| DNA+RNA as seed | KIRC | 35 | PPP1CB | 1 | 0 |
| DNA+RNA as seed | KIRC | 36 | OBSL1 | 0 | 0 |
| DNA+RNA as seed | KIRC | 37 | LIMA1 | 0 | 0 |
| DNA+RNA as seed | KIRC | 38 | NF2 | 0 | 0 |
| DNA+RNA as seed | KIRC | 39 | PPARG | 0 | 0 |
| DNA+RNA as seed | KIRC | 40 | MAX | 1 | 1 |
| DNA+RNA as seed | KIRC | 41 | PTK2 | 1 | 0 |
| DNA+RNA as seed | KIRC | 42 | HSPA8 | 1 | 1 |
| DNA+RNA as seed | KIRC | 43 | RPA2 | 1 | 1 |
| DNA+RNA as seed | KIRC | 44 | CRK | 1 | 0 |
| DNA+RNA as seed | KIRC | 45 | E2F1 | 0 | 0 |
| DNA+RNA as seed | KIRC | 46 | SPTBN1 | 0 | 0 |
| DNA+RNA as seed | KIRC | 47 | SMARCC2 | 0 | 0 |
| DNA+RNA as seed | KIRC | 48 | SMARCC1 | 0 | 0 |
| DNA+RNA as seed | KIRC | 49 | SMARCA2 | 0 | 0 |
| DNA+RNA as seed | KIRC | 50 | CAPZA2 | 0 | 0 |
| DNA+RNA as seed | KIRC | 51 | KISS1R | 0 | 0 |
| DNA+RNA as seed | KIRC | 52 | EED | 0 | 0 |
| DNA+RNA as seed | KIRC | 53 | FAF2 | 1 | 0 |
| DNA+RNA as seed | KIRC | 54 | MYO1C | 0 | 0 |
| DNA+RNA as seed | KIRC | 55 | EPAS1 | 0 | 0 |
| DNA+RNA as seed | KIRC | 56 | SMARCB1 | 1 | 1 |
| DNA+RNA as seed | KIRC | 57 | DDB1 | 1 | 1 |
| DNA+RNA as seed | KIRC | 58 | POLR2A | 0 | 0 |
| DNA+RNA as seed | KIRC | 59 | PBRM1 | 0 | 0 |

| | | | | | |
|---|---|---|---|---|---|
| DNA+RNA as seed | KIRC | 60 | PRKCZ | 0 | 0 |
| DNA+RNA as seed | KIRC | 61 | ROCK1 | 0 | 0 |
| DNA+RNA as seed | KIRC | 62 | H3-4 | 0 | 0 |
| DNA+RNA as seed | KIRC | 63 | KAT2B | 0 | 0 |
| DNA+RNA as seed | KIRC | 64 | SMURF1 | 0 | 0 |
| DNA+RNA as seed | KIRC | 65 | ATR | 1 | 1 |
| DNA+RNA as seed | KIRC | 66 | TERF1 | 1 | 0 |
| DNA+RNA as seed | KIRC | 67 | FBXW7 | 0 | 0 |
| DNA+RNA as seed | KIRC | 68 | ARID1A | 0 | 0 |
| DNA+RNA as seed | KIRC | 69 | MUC15 | 0 | 0 |
| DNA+RNA as seed | KIRC | 70 | NOTCH1 | 0 | 0 |
| DNA+RNA as seed | KIRC | 71 | SQSTM1 | 0 | 0 |
| DNA+RNA as seed | KIRC | 72 | NR3C1 | 0 | 0 |
| DNA+RNA as seed | KIRC | 73 | IKBKB | 0 | 0 |
| DNA+RNA as seed | KIRC | 74 | CD44 | 0 | 0 |
| DNA+RNA as seed | KIRC | 75 | TFAP2A | 0 | 0 |
| DNA+RNA as seed | KIRC | 76 | SKP2 | 1 | 0 |
| DNA+RNA as seed | KIRC | 77 | SHMT2 | 0 | 0 |
| DNA+RNA as seed | KIRC | 78 | PKM | 1 | 0 |
| DNA+RNA as seed | KIRC | 79 | PIP4K2A | 0 | 0 |
| DNA+RNA as seed | KIRC | 80 | IRS1 | 0 | 0 |
| DNA+RNA as seed | KIRC | 81 | NEDD4 | 0 | 0 |
| DNA+RNA as seed | KIRC | 82 | UBE2D3 | 1 | 0 |
| DNA+RNA as seed | KIRC | 83 | ITGB1 | 0 | 0 |
| DNA+RNA as seed | KIRC | 84 | ANXA2 | 0 | 0 |
| DNA+RNA as seed | KIRC | 85 | PLCB1 | 0 | 0 |
| DNA+RNA as seed | KIRC | 86 | VEGFA | 0 | 0 |
| DNA+RNA as seed | KIRC | 87 | WDR5 | 1 | 1 |
| DNA+RNA as seed | KIRC | 88 | SREBF1 | 0 | 0 |
| DNA+RNA as seed | KIRC | 89 | PCK1 | 0 | 0 |
| DNA+RNA as seed | KIRC | 90 | BAP1 | 1 | 0 |
| DNA+RNA as seed | KIRC | 91 | PRPF8 | 1 | 1 |
| DNA+RNA as seed | KIRC | 92 | UIMC1 | 0 | 0 |
| DNA+RNA as seed | KIRC | 93 | UBA1 | 1 | 1 |
| DNA+RNA as seed | KIRC | 94 | SVIL | 0 | 0 |
| DNA+RNA as seed | KIRC | 95 | PIP5K1A | 1 | 0 |
| DNA+RNA as seed | KIRC | 96 | INPP5E | 0 | 0 |
| DNA+RNA as seed | KIRC | 97 | MUC17 | 0 | 0 |
| DNA+RNA as seed | KIRC | 98 | BCL7C | 0 | 0 |
| DNA+RNA as seed | KIRC | 99 | PRKCI | 0 | 0 |
| DNA+RNA as seed | KIRC | 100 | PLCB3 | 0 | 0 |
| DNA+RNA as seed | LUAD | 1 | TP53 | 0 | 0 |
| DNA+RNA as seed | LUAD | 2 | EGFR | 1 | 0 |
| DNA+RNA as seed | LUAD | 3 | CTNNB1 | 1 | 0 |
| DNA+RNA as seed | LUAD | 4 | KRAS | 1 | 0 |
| DNA+RNA as seed | LUAD | 5 | CDKN2A | 0 | 0 |
| DNA+RNA as seed | LUAD | 6 | RELA | 1 | 0 |
| DNA+RNA as seed | LUAD | 7 | ATM | 0 | 0 |
| DNA+RNA as seed | LUAD | 8 | SRC | 0 | 0 |
| DNA+RNA as seed | LUAD | 9 | SP1 | 0 | 0 |
| DNA+RNA as seed | LUAD | 10 | PIK3CA | 1 | 0 |
| DNA+RNA as seed | LUAD | 11 | HSP90AA1 | 0 | 0 |
| DNA+RNA as seed | LUAD | 12 | RB1 | 0 | 0 |
| DNA+RNA as seed | LUAD | 13 | JUN | 0 | 0 |
| DNA+RNA as seed | LUAD | 14 | NFKB1 | 0 | 0 |
| DNA+RNA as seed | LUAD | 15 | MET | 0 | 0 |

| | | | | | |
|---|---|---|---|---|---|
| DNA+RNA as seed | LUAD | 16 | AKT1 | 0 | 0 |
| DNA+RNA as seed | LUAD | 17 | HDAC1 | 0 | 0 |
| DNA+RNA as seed | LUAD | 18 | HRAS | 0 | 0 |
| DNA+RNA as seed | LUAD | 19 | CDKN1A | 0 | 0 |
| DNA+RNA as seed | LUAD | 20 | NRAS | 1 | 0 |
| DNA+RNA as seed | LUAD | 21 | MDM2 | 1 | 0 |
| DNA+RNA as seed | LUAD | 22 | PITX2 | 0 | 0 |
| DNA+RNA as seed | LUAD | 23 | SMAD3 | 0 | 0 |
| DNA+RNA as seed | LUAD | 24 | EEF1A2 | 0 | 0 |
| DNA+RNA as seed | LUAD | 25 | APC | 0 | 0 |
| DNA+RNA as seed | LUAD | 26 | PRKCA | 0 | 0 |
| DNA+RNA as seed | LUAD | 27 | SMARCA4 | 0 | 0 |
| DNA+RNA as seed | LUAD | 28 | STK11 | 0 | 0 |
| DNA+RNA as seed | LUAD | 29 | NFKBIA | 0 | 0 |
| DNA+RNA as seed | LUAD | 30 | NOTCH1 | 0 | 0 |
| DNA+RNA as seed | LUAD | 31 | TERT | 0 | 0 |
| DNA+RNA as seed | LUAD | 32 | SMAD2 | 0 | 0 |
| DNA+RNA as seed | LUAD | 33 | HIF1A | 0 | 0 |
| DNA+RNA as seed | LUAD | 34 | TFF1 | 0 | 0 |
| DNA+RNA as seed | LUAD | 35 | CCND1 | 1 | 0 |
| DNA+RNA as seed | LUAD | 36 | AURKA | 1 | 0 |
| DNA+RNA as seed | LUAD | 37 | IGF1R | 1 | 0 |
| DNA+RNA as seed | LUAD | 38 | YWHAZ | 1 | 0 |
| DNA+RNA as seed | LUAD | 39 | FGFR1 | 0 | 0 |
| DNA+RNA as seed | LUAD | 40 | PTK2 | 1 | 0 |
| DNA+RNA as seed | LUAD | 41 | MAPK14 | 0 | 0 |
| DNA+RNA as seed | LUAD | 42 | PPARG | 0 | 0 |
| DNA+RNA as seed | LUAD | 43 | CDC37 | 1 | 1 |
| DNA+RNA as seed | LUAD | 44 | PML | 0 | 0 |
| DNA+RNA as seed | LUAD | 45 | A2M | 0 | 0 |
| DNA+RNA as seed | LUAD | 46 | NOTCH2 | 0 | 0 |
| DNA+RNA as seed | LUAD | 47 | MAPK8 | 0 | 0 |
| DNA+RNA as seed | LUAD | 48 | ABL1 | 0 | 0 |
| DNA+RNA as seed | LUAD | 49 | EZH2 | 0 | 0 |
| DNA+RNA as seed | LUAD | 50 | SPTA1 | 0 | 0 |
| DNA+RNA as seed | LUAD | 51 | NOTCH4 | 0 | 0 |
| DNA+RNA as seed | LUAD | 52 | PIK3R3 | 0 | 0 |
| DNA+RNA as seed | LUAD | 53 | ETS1 | 0 | 0 |
| DNA+RNA as seed | LUAD | 54 | PTEN | 0 | 0 |
| DNA+RNA as seed | LUAD | 55 | KAT2B | 0 | 0 |
| DNA+RNA as seed | LUAD | 56 | FLT4 | 0 | 0 |
| DNA+RNA as seed | LUAD | 57 | H3-4 | 0 | 0 |
| DNA+RNA as seed | LUAD | 58 | SP3 | 0 | 0 |
| DNA+RNA as seed | LUAD | 59 | SQSTM1 | 0 | 0 |
| DNA+RNA as seed | LUAD | 60 | BRAF | 0 | 0 |
| DNA+RNA as seed | LUAD | 61 | FOXO1 | 0 | 0 |
| DNA+RNA as seed | LUAD | 62 | HSPA4 | 0 | 0 |
| DNA+RNA as seed | LUAD | 63 | BTRC | 0 | 0 |
| DNA+RNA as seed | LUAD | 64 | FOXO3 | 0 | 0 |
| DNA+RNA as seed | LUAD | 65 | COL3A1 | 0 | 0 |
| DNA+RNA as seed | LUAD | 66 | HSPA5 | 1 | 1 |
| DNA+RNA as seed | LUAD | 67 | CASP3 | 0 | 0 |
| DNA+RNA as seed | LUAD | 68 | YWHAQ | 0 | 0 |
| DNA+RNA as seed | LUAD | 69 | VDR | 0 | 0 |
| DNA+RNA as seed | LUAD | 70 | TFAP2A | 0 | 0 |
| DNA+RNA as seed | LUAD | 71 | HSPA1A | 0 | 0 |

| | | | | | |
|---|---|---|---|---|---|
| DNA+RNA as seed | LUAD | 72 | CHUK | 0 | 0 |
| DNA+RNA as seed | LUAD | 73 | ATR | 1 | 1 |
| DNA+RNA as seed | LUAD | 74 | VEGFA | 0 | 0 |
| DNA+RNA as seed | LUAD | 75 | MMP13 | 0 | 0 |
| DNA+RNA as seed | LUAD | 76 | WT1 | 0 | 0 |
| DNA+RNA as seed | LUAD | 77 | IRS1 | 0 | 0 |
| DNA+RNA as seed | LUAD | 78 | CTNNA1 | 0 | 0 |
| DNA+RNA as seed | LUAD | 79 | ITGB1 | 1 | 0 |
| DNA+RNA as seed | LUAD | 80 | AGER | 0 | 0 |
| DNA+RNA as seed | LUAD | 81 | MMP2 | 0 | 0 |
| DNA+RNA as seed | LUAD | 82 | COL11A1 | 0 | 0 |
| DNA+RNA as seed | LUAD | 83 | PRKCZ | 0 | 0 |
| DNA+RNA as seed | LUAD | 84 | NFKB2 | 0 | 0 |
| DNA+RNA as seed | LUAD | 85 | SMARCB1 | 1 | 1 |
| DNA+RNA as seed | LUAD | 86 | TP63 | 0 | 0 |
| DNA+RNA as seed | LUAD | 87 | CA9 | 0 | 0 |
| DNA+RNA as seed | LUAD | 88 | SKP1 | 1 | 0 |
| DNA+RNA as seed | LUAD | 89 | MAPK11 | 0 | 0 |
| DNA+RNA as seed | LUAD | 90 | TCF7L2 | 0 | 0 |
| DNA+RNA as seed | LUAD | 91 | KEAP1 | 1 | 0 |
| DNA+RNA as seed | LUAD | 92 | WWOX | 0 | 0 |
| DNA+RNA as seed | LUAD | 93 | CD44 | 0 | 0 |
| DNA+RNA as seed | LUAD | 94 | FAS | 0 | 0 |
| DNA+RNA as seed | LUAD | 95 | CTBP1 | 0 | 0 |
| DNA+RNA as seed | LUAD | 96 | HTR3A | 0 | 0 |
| DNA+RNA as seed | LUAD | 97 | CDK5 | 0 | 0 |
| DNA+RNA as seed | LUAD | 98 | TGFB1 | 0 | 0 |
| DNA+RNA as seed | LUAD | 99 | LEF1 | 0 | 0 |
| DNA+RNA as seed | LUAD | 100 | ARID1A | 0 | 0 |
| DNA+RNA as seed | LUSC | 1 | TP53 | 0 | 0 |
| DNA+RNA as seed | LUSC | 2 | CUL3 | 1 | 0 |
| DNA+RNA as seed | LUSC | 3 | CREBBP | 1 | 0 |
| DNA+RNA as seed | LUSC | 4 | SOX2 | 1 | 0 |
| DNA+RNA as seed | LUSC | 5 | SERPINB5 | 0 | 0 |
| DNA+RNA as seed | LUSC | 6 | FN1 | 0 | 0 |
| DNA+RNA as seed | LUSC | 7 | PIK3CA | 1 | 0 |
| DNA+RNA as seed | LUSC | 8 | DCUN1D1 | 0 | 0 |
| DNA+RNA as seed | LUSC | 9 | FXR1 | 0 | 0 |
| DNA+RNA as seed | LUSC | 10 | AP2M1 | 1 | 0 |
| DNA+RNA as seed | LUSC | 11 | HDAC1 | 0 | 0 |
| DNA+RNA as seed | LUSC | 12 | PTEN | 0 | 0 |
| DNA+RNA as seed | LUSC | 13 | RB1 | 0 | 0 |
| DNA+RNA as seed | LUSC | 14 | CTNNB1 | 0 | 0 |
| DNA+RNA as seed | LUSC | 15 | NOTCH1 | 0 | 0 |
| DNA+RNA as seed | LUSC | 16 | GRB2 | 1 | 1 |
| DNA+RNA as seed | LUSC | 17 | CDKN2A | 0 | 0 |
| DNA+RNA as seed | LUSC | 18 | AR | 0 | 0 |
| DNA+RNA as seed | LUSC | 19 | CSNK2A1 | 0 | 0 |
| DNA+RNA as seed | LUSC | 20 | CALML3 | 0 | 0 |
| DNA+RNA as seed | LUSC | 21 | FOXE1 | 0 | 0 |
| DNA+RNA as seed | LUSC | 22 | SNW1 | 1 | 1 |
| DNA+RNA as seed | LUSC | 23 | PRKCA | 0 | 0 |
| DNA+RNA as seed | LUSC | 24 | NFE2L2 | 1 | 0 |
| DNA+RNA as seed | LUSC | 25 | ACTL6A | 1 | 1 |
| DNA+RNA as seed | LUSC | 26 | DVL3 | 0 | 0 |
| DNA+RNA as seed | LUSC | 27 | FBXW7 | 0 | 0 |

| | | | | | |
|---|---|---|---|---|---|
| DNA+RNA as seed | LUSC | 28 | HSP90AB1 | 1 | 0 |
| DNA+RNA as seed | LUSC | 29 | HOXD13 | 0 | 0 |
| DNA+RNA as seed | LUSC | 30 | SMARCA4 | 0 | 0 |
| DNA+RNA as seed | LUSC | 31 | HDAC3 | 1 | 0 |
| DNA+RNA as seed | LUSC | 32 | SMAD2 | 0 | 0 |
| DNA+RNA as seed | LUSC | 33 | CREB1 | 0 | 0 |
| DNA+RNA as seed | LUSC | 34 | HIF1A | 0 | 0 |
| DNA+RNA as seed | LUSC | 35 | PPARG | 0 | 0 |
| DNA+RNA as seed | LUSC | 36 | TBL1XR1 | 0 | 0 |
| DNA+RNA as seed | LUSC | 37 | KAT2B | 0 | 0 |
| DNA+RNA as seed | LUSC | 38 | HSPA8 | 1 | 1 |
| DNA+RNA as seed | LUSC | 39 | CCND1 | 1 | 0 |
| DNA+RNA as seed | LUSC | 40 | RBX1 | 1 | 1 |
| DNA+RNA as seed | LUSC | 41 | PML | 0 | 0 |
| DNA+RNA as seed | LUSC | 42 | PIK3R3 | 0 | 0 |
| DNA+RNA as seed | LUSC | 43 | RBPJ | 0 | 0 |
| DNA+RNA as seed | LUSC | 44 | NCOR1 | 0 | 0 |
| DNA+RNA as seed | LUSC | 45 | EZH2 | 0 | 0 |
| DNA+RNA as seed | LUSC | 46 | HRAS | 0 | 0 |
| DNA+RNA as seed | LUSC | 47 | POU6F2 | 0 | 0 |
| DNA+RNA as seed | LUSC | 48 | PLD1 | 0 | 0 |
| DNA+RNA as seed | LUSC | 49 | GNB4 | 0 | 0 |
| DNA+RNA as seed | LUSC | 50 | PRKCB | 0 | 0 |
| DNA+RNA as seed | LUSC | 51 | NFATC1 | 0 | 0 |
| DNA+RNA as seed | LUSC | 52 | IKBKB | 0 | 0 |
| DNA+RNA as seed | LUSC | 53 | SMARCB1 | 1 | 1 |
| DNA+RNA as seed | LUSC | 54 | KRAS | 1 | 0 |
| DNA+RNA as seed | LUSC | 55 | PRAME | 0 | 0 |
| DNA+RNA as seed | LUSC | 56 | TP73 | 0 | 0 |
| DNA+RNA as seed | LUSC | 57 | PRKCD | 0 | 0 |
| DNA+RNA as seed | LUSC | 58 | RASA1 | 0 | 0 |
| DNA+RNA as seed | LUSC | 59 | RARA | 0 | 0 |
| DNA+RNA as seed | LUSC | 60 | SMARCA2 | 0 | 0 |
| DNA+RNA as seed | LUSC | 61 | TBP | 1 | 0 |
| DNA+RNA as seed | LUSC | 62 | SMARCC1 | 0 | 0 |
| DNA+RNA as seed | LUSC | 63 | YY1 | 1 | 1 |
| DNA+RNA as seed | LUSC | 64 | RUNX1 | 0 | 0 |
| DNA+RNA as seed | LUSC | 65 | ARID1A | 0 | 0 |
| DNA+RNA as seed | LUSC | 66 | SMARCD1 | 0 | 0 |
| DNA+RNA as seed | LUSC | 67 | DVL2 | 0 | 0 |
| DNA+RNA as seed | LUSC | 68 | NCOR2 | 0 | 0 |
| DNA+RNA as seed | LUSC | 69 | SUMO1 | 0 | 0 |
| DNA+RNA as seed | LUSC | 70 | NF1 | 0 | 0 |
| DNA+RNA as seed | LUSC | 71 | HOXC13 | 0 | 0 |
| DNA+RNA as seed | LUSC | 72 | FOXO1 | 0 | 0 |
| DNA+RNA as seed | LUSC | 73 | SMARCC2 | 0 | 0 |
| DNA+RNA as seed | LUSC | 74 | CASP8 | 0 | 0 |
| DNA+RNA as seed | LUSC | 75 | XRCC6 | 1 | 1 |
| DNA+RNA as seed | LUSC | 76 | INSR | 0 | 0 |
| DNA+RNA as seed | LUSC | 77 | KEAP1 | 1 | 0 |
| DNA+RNA as seed | LUSC | 78 | ETS1 | 0 | 0 |
| DNA+RNA as seed | LUSC | 79 | AURKB | 1 | 0 |
| DNA+RNA as seed | LUSC | 80 | SKP1 | 1 | 0 |
| DNA+RNA as seed | LUSC | 81 | PRKAA2 | 0 | 0 |
| DNA+RNA as seed | LUSC | 82 | HTR2C | 0 | 0 |
| DNA+RNA as seed | LUSC | 83 | KLF5 | 1 | 0 |

| | | | | | |
|---|---|---|---|---|---|
| DNA+RNA as seed | LUSC | 84 | H2BC21 | 0 | 0 |
| DNA+RNA as seed | LUSC | 85 | SPI1 | 0 | 0 |
| DNA+RNA as seed | LUSC | 86 | CEBPA | 0 | 0 |
| DNA+RNA as seed | LUSC | 87 | MET | 0 | 0 |
| DNA+RNA as seed | LUSC | 88 | MYOD1 | 0 | 0 |
| DNA+RNA as seed | LUSC | 89 | IRS1 | 0 | 0 |
| DNA+RNA as seed | LUSC | 90 | WDR5 | 1 | 1 |
| DNA+RNA as seed | LUSC | 91 | CAV1 | 0 | 0 |
| DNA+RNA as seed | LUSC | 92 | HMGA1 | 1 | 0 |
| DNA+RNA as seed | LUSC | 93 | FOXK2 | 0 | 0 |
| DNA+RNA as seed | LUSC | 94 | RHOA | 1 | 1 |
| DNA+RNA as seed | LUSC | 95 | RACK1 | 1 | 0 |
| DNA+RNA as seed | LUSC | 96 | CAMK2G | 0 | 0 |
| DNA+RNA as seed | LUSC | 97 | COPS3 | 1 | 1 |
| DNA+RNA as seed | LUSC | 98 | VEGFA | 0 | 0 |
| DNA+RNA as seed | LUSC | 99 | SP3 | 0 | 0 |
| DNA+RNA as seed | LUSC | 100 | COPS2 | 1 | 1 |
| DNA+RNA as seed | PRAD | 1 | TP53 | 0 | 0 |
| DNA+RNA as seed | PRAD | 2 | CTNNB1 | 0 | 0 |
| DNA+RNA as seed | PRAD | 3 | PTEN | 0 | 0 |
| DNA+RNA as seed | PRAD | 4 | ATM | 0 | 0 |
| DNA+RNA as seed | PRAD | 5 | MYC | 1 | 0 |
| DNA+RNA as seed | PRAD | 6 | HSPA8 | 0 | 1 |
| DNA+RNA as seed | PRAD | 7 | ERG | 1 | 0 |
| DNA+RNA as seed | PRAD | 8 | ESR1 | 0 | 0 |
| DNA+RNA as seed | PRAD | 9 | EP300 | 1 | 0 |
| DNA+RNA as seed | PRAD | 10 | PIK3CA | 0 | 0 |
| DNA+RNA as seed | PRAD | 11 | SNW1 | 1 | 1 |
| DNA+RNA as seed | PRAD | 12 | HDAC1 | 0 | 0 |
| DNA+RNA as seed | PRAD | 13 | SP1 | 0 | 0 |
| DNA+RNA as seed | PRAD | 14 | HRAS | 0 | 0 |
| DNA+RNA as seed | PRAD | 15 | AR | 0 | 0 |
| DNA+RNA as seed | PRAD | 16 | FOXA1 | 1 | 0 |
| DNA+RNA as seed | PRAD | 17 | SMAD3 | 0 | 0 |
| DNA+RNA as seed | PRAD | 18 | GSK3B | 0 | 0 |
| DNA+RNA as seed | PRAD | 19 | CDKN1B | 0 | 0 |
| DNA+RNA as seed | PRAD | 20 | APC | 1 | 0 |
| DNA+RNA as seed | PRAD | 21 | UBB | 0 | 0 |
| DNA+RNA as seed | PRAD | 22 | YWHAZ | 1 | 0 |
| DNA+RNA as seed | PRAD | 23 | CREB1 | 0 | 0 |
| DNA+RNA as seed | PRAD | 24 | RB1 | 0 | 0 |
| DNA+RNA as seed | PRAD | 25 | ABL1 | 0 | 0 |
| DNA+RNA as seed | PRAD | 26 | TP53BP1 | 0 | 0 |
| DNA+RNA as seed | PRAD | 27 | ETS1 | 0 | 0 |
| DNA+RNA as seed | PRAD | 28 | ETS2 | 0 | 0 |
| DNA+RNA as seed | PRAD | 29 | SPOP | 0 | 0 |
| DNA+RNA as seed | PRAD | 30 | XRCC6 | 1 | 1 |
| DNA+RNA as seed | PRAD | 31 | HIF1A | 0 | 0 |
| DNA+RNA as seed | PRAD | 32 | ATR | 1 | 1 |
| DNA+RNA as seed | PRAD | 33 | RPA2 | 1 | 1 |
| DNA+RNA as seed | PRAD | 34 | PARP1 | 0 | 0 |
| DNA+RNA as seed | PRAD | 35 | CDK4 | 1 | 0 |
| DNA+RNA as seed | PRAD | 36 | KAT2B | 0 | 0 |
| DNA+RNA as seed | PRAD | 37 | PRKDC | 0 | 0 |
| DNA+RNA as seed | PRAD | 38 | NOTCH1 | 0 | 0 |
| DNA+RNA as seed | PRAD | 39 | ESR2 | 0 | 0 |

| | | | | | |
|---|---|---|---|---|---|
| DNA+RNA as seed | PRAD | 40 | CBL | 0 | 0 |
| DNA+RNA as seed | PRAD | 41 | NCOA3 | 0 | 0 |
| DNA+RNA as seed | PRAD | 42 | MAPK8 | 0 | 0 |
| DNA+RNA as seed | PRAD | 43 | SKP1 | 1 | 0 |
| DNA+RNA as seed | PRAD | 44 | EZH2 | 0 | 0 |
| DNA+RNA as seed | PRAD | 45 | CASP8 | 0 | 0 |
| DNA+RNA as seed | PRAD | 46 | FLNA | 0 | 0 |
| DNA+RNA as seed | PRAD | 47 | KPNA3 | 0 | 0 |
| DNA+RNA as seed | PRAD | 48 | FBXW7 | 0 | 0 |
| DNA+RNA as seed | PRAD | 49 | BRCA2 | 0 | 1 |
| DNA+RNA as seed | PRAD | 50 | SMARCA4 | 0 | 0 |
| DNA+RNA as seed | PRAD | 51 | POU5F1 | 0 | 0 |
| DNA+RNA as seed | PRAD | 52 | SUMO1 | 0 | 0 |
| DNA+RNA as seed | PRAD | 53 | DDB1 | 1 | 1 |
| DNA+RNA as seed | PRAD | 54 | WDR5 | 1 | 1 |
| DNA+RNA as seed | PRAD | 55 | SUMO2 | 1 | 0 |
| DNA+RNA as seed | PRAD | 56 | PRKCZ | 0 | 0 |
| DNA+RNA as seed | PRAD | 57 | PRKCD | 0 | 0 |
| DNA+RNA as seed | PRAD | 58 | TCF7L2 | 0 | 0 |
| DNA+RNA as seed | PRAD | 59 | BMI1 | 1 | 0 |
| DNA+RNA as seed | PRAD | 60 | PGR | 0 | 0 |
| DNA+RNA as seed | PRAD | 61 | PIAS1 | 1 | 0 |
| DNA+RNA as seed | PRAD | 62 | RAD51 | 1 | 1 |
| DNA+RNA as seed | PRAD | 63 | PTK2 | 0 | 0 |
| DNA+RNA as seed | PRAD | 64 | BRAF | 0 | 0 |
| DNA+RNA as seed | PRAD | 65 | JAK2 | 0 | 0 |
| DNA+RNA as seed | PRAD | 66 | FOXO3 | 0 | 0 |
| DNA+RNA as seed | PRAD | 67 | NANOG | 0 | 0 |
| DNA+RNA as seed | PRAD | 68 | TERT | 0 | 0 |
| DNA+RNA as seed | PRAD | 69 | UBR5 | 0 | 0 |
| DNA+RNA as seed | PRAD | 70 | USF1 | 0 | 0 |
| DNA+RNA as seed | PRAD | 71 | PRKAA1 | 0 | 0 |
| DNA+RNA as seed | PRAD | 72 | IGF1R | 1 | 0 |
| DNA+RNA as seed | PRAD | 73 | RBX1 | 1 | 1 |
| DNA+RNA as seed | PRAD | 74 | NCOA1 | 0 | 0 |
| DNA+RNA as seed | PRAD | 75 | DAXX | 0 | 0 |
| DNA+RNA as seed | PRAD | 76 | NCOA2 | 0 | 0 |
| DNA+RNA as seed | PRAD | 77 | PDGFRB | 0 | 0 |
| DNA+RNA as seed | PRAD | 78 | LYN | 0 | 0 |
| DNA+RNA as seed | PRAD | 79 | GLI2 | 0 | 0 |
| DNA+RNA as seed | PRAD | 80 | MYB | 0 | 0 |
| DNA+RNA as seed | PRAD | 81 | TGFBR2 | 0 | 0 |
| DNA+RNA as seed | PRAD | 82 | FGFR2 | 0 | 0 |
| DNA+RNA as seed | PRAD | 83 | COPS6 | 1 | 1 |
| DNA+RNA as seed | PRAD | 84 | UBE3A | 0 | 0 |
| DNA+RNA as seed | PRAD | 85 | IRS1 | 0 | 0 |
| DNA+RNA as seed | PRAD | 86 | CCNA2 | 1 | 1 |
| DNA+RNA as seed | PRAD | 87 | MMP2 | 0 | 0 |
| DNA+RNA as seed | PRAD | 88 | RBPJ | 0 | 0 |
| DNA+RNA as seed | PRAD | 89 | CD44 | 0 | 0 |
| DNA+RNA as seed | PRAD | 90 | BLM | 0 | 1 |
| DNA+RNA as seed | PRAD | 91 | WT1 | 0 | 0 |
| DNA+RNA as seed | PRAD | 92 | DDX5 | 1 | 0 |
| DNA+RNA as seed | PRAD | 93 | STK11 | 0 | 0 |
| DNA+RNA as seed | PRAD | 94 | ITGB3 | 0 | 0 |
| DNA+RNA as seed | PRAD | 95 | LAMA3 | 0 | 0 |

| | | | | | |
|---|---|---|---|---|---|
| DNA+RNA as seed | PRAD | 96 | BRD4 | 1 | 1 |
| DNA+RNA as seed | PRAD | 97 | CSNK1A1 | 1 | 1 |
| DNA+RNA as seed | PRAD | 98 | UBE2D2 | 0 | 0 |
| DNA+RNA as seed | PRAD | 99 | E2F3 | 1 | 0 |
| DNA+RNA as seed | PRAD | 100 | FOXM1 | 0 | 0 |
| DNA+RNA as seed | STAD | 1 | TP53 | 0 | 0 |
| DNA+RNA as seed | STAD | 2 | MYC | 1 | 0 |
| DNA+RNA as seed | STAD | 3 | ERBB2 | 0 | 0 |
| DNA+RNA as seed | STAD | 4 | CTNNB1 | 1 | 0 |
| DNA+RNA as seed | STAD | 5 | CDH1 | 0 | 0 |
| DNA+RNA as seed | STAD | 6 | ERBB3 | 0 | 0 |
| DNA+RNA as seed | STAD | 7 | ERBB4 | 0 | 0 |
| DNA+RNA as seed | STAD | 8 | ESR1 | 0 | 0 |
| DNA+RNA as seed | STAD | 9 | PIK3CA | 1 | 0 |
| DNA+RNA as seed | STAD | 10 | SMAD4 | 0 | 0 |
| DNA+RNA as seed | STAD | 11 | SP1 | 0 | 0 |
| DNA+RNA as seed | STAD | 12 | SRC | 0 | 0 |
| DNA+RNA as seed | STAD | 13 | EP300 | 0 | 0 |
| DNA+RNA as seed | STAD | 14 | ATM | 0 | 0 |
| DNA+RNA as seed | STAD | 15 | UBC | 1 | 0 |
| DNA+RNA as seed | STAD | 16 | AR | 0 | 0 |
| DNA+RNA as seed | STAD | 17 | WWOX | 0 | 0 |
| DNA+RNA as seed | STAD | 18 | SMAD3 | 0 | 0 |
| DNA+RNA as seed | STAD | 19 | PTEN | 0 | 0 |
| DNA+RNA as seed | STAD | 20 | KRAS | 1 | 0 |
| DNA+RNA as seed | STAD | 21 | CDKN2A | 0 | 0 |
| DNA+RNA as seed | STAD | 22 | STAT3 | 0 | 0 |
| DNA+RNA as seed | STAD | 23 | MAPK1 | 1 | 0 |
| DNA+RNA as seed | STAD | 24 | CDKN1A | 0 | 0 |
| DNA+RNA as seed | STAD | 25 | MAPK3 | 0 | 0 |
| DNA+RNA as seed | STAD | 26 | PIK3R1 | 0 | 0 |
| DNA+RNA as seed | STAD | 27 | CREBBP | 1 | 0 |
| DNA+RNA as seed | STAD | 28 | MET | 1 | 0 |
| DNA+RNA as seed | STAD | 29 | FGFR3 | 0 | 0 |
| DNA+RNA as seed | STAD | 30 | SMAD2 | 0 | 0 |
| DNA+RNA as seed | STAD | 31 | HDAC1 | 1 | 0 |
| DNA+RNA as seed | STAD | 32 | HSP90AA1 | 1 | 0 |
| DNA+RNA as seed | STAD | 33 | FBXW7 | 0 | 0 |
| DNA+RNA as seed | STAD | 34 | HRAS | 0 | 0 |
| DNA+RNA as seed | STAD | 35 | GSK3B | 0 | 0 |
| DNA+RNA as seed | STAD | 36 | MDM2 | 1 | 0 |
| DNA+RNA as seed | STAD | 37 | HIF1A | 0 | 0 |
| DNA+RNA as seed | STAD | 38 | APC | 0 | 0 |
| DNA+RNA as seed | STAD | 39 | NRAS | 0 | 0 |
| DNA+RNA as seed | STAD | 40 | RARA | 0 | 0 |
| DNA+RNA as seed | STAD | 41 | E2F1 | 0 | 0 |
| DNA+RNA as seed | STAD | 42 | RHOA | 0 | 1 |
| DNA+RNA as seed | STAD | 43 | SHC1 | 0 | 0 |
| DNA+RNA as seed | STAD | 44 | CCND1 | 1 | 0 |
| DNA+RNA as seed | STAD | 45 | PPARG | 0 | 0 |
| DNA+RNA as seed | STAD | 46 | ABL1 | 0 | 0 |
| DNA+RNA as seed | STAD | 47 | RPS27A | 1 | 0 |
| DNA+RNA as seed | STAD | 48 | CBL | 0 | 0 |
| DNA+RNA as seed | STAD | 49 | HSP90AB1 | 0 | 0 |
| DNA+RNA as seed | STAD | 50 | MAPK8 | 0 | 0 |
| DNA+RNA as seed | STAD | 51 | PIK3R2 | 0 | 0 |

| | | | | | |
|---|---|---|---|---|---|
| DNA+RNA as seed | STAD | 52 | HDAC3 | 1 | 0 |
| DNA+RNA as seed | STAD | 53 | UBA52 | 1 | 0 |
| DNA+RNA as seed | STAD | 54 | PARP1 | 0 | 0 |
| DNA+RNA as seed | STAD | 55 | FYN | 0 | 0 |
| DNA+RNA as seed | STAD | 56 | SMAD1 | 0 | 0 |
| DNA+RNA as seed | STAD | 57 | THRA | 0 | 0 |
| DNA+RNA as seed | STAD | 58 | PTK2 | 1 | 0 |
| DNA+RNA as seed | STAD | 59 | EZH2 | 0 | 0 |
| DNA+RNA as seed | STAD | 60 | CTNND1 | 0 | 0 |
| DNA+RNA as seed | STAD | 61 | CDKN1B | 0 | 0 |
| DNA+RNA as seed | STAD | 62 | NOTCH1 | 0 | 0 |
| DNA+RNA as seed | STAD | 63 | PML | 0 | 0 |
| DNA+RNA as seed | STAD | 64 | NCOA3 | 0 | 0 |
| DNA+RNA as seed | STAD | 65 | CCNE1 | 1 | 0 |
| DNA+RNA as seed | STAD | 66 | NCOR1 | 0 | 0 |
| DNA+RNA as seed | STAD | 67 | HLA-B | 0 | 0 |
| DNA+RNA as seed | STAD | 68 | CDC42 | 1 | 0 |
| DNA+RNA as seed | STAD | 69 | YY1 | 1 | 1 |
| DNA+RNA as seed | STAD | 70 | MAPK9 | 0 | 0 |
| DNA+RNA as seed | STAD | 71 | BTRC | 0 | 0 |
| DNA+RNA as seed | STAD | 72 | FOXO1 | 0 | 0 |
| DNA+RNA as seed | STAD | 73 | NCOA1 | 0 | 0 |
| DNA+RNA as seed | STAD | 74 | FOXO3 | 0 | 0 |
| DNA+RNA as seed | STAD | 75 | CDK4 | 1 | 0 |
| DNA+RNA as seed | STAD | 76 | HUWE1 | 1 | 0 |
| DNA+RNA as seed | STAD | 77 | KAT2B | 0 | 0 |
| DNA+RNA as seed | STAD | 78 | PIN1 | 0 | 0 |
| DNA+RNA as seed | STAD | 79 | SKP1 | 1 | 0 |
| DNA+RNA as seed | STAD | 80 | SMARCA4 | 1 | 0 |
| DNA+RNA as seed | STAD | 81 | NCOR2 | 0 | 0 |
| DNA+RNA as seed | STAD | 82 | JUP | 0 | 0 |
| DNA+RNA as seed | STAD | 83 | MED1 | 1 | 1 |
| DNA+RNA as seed | STAD | 84 | CDK8 | 0 | 0 |
| DNA+RNA as seed | STAD | 85 | ACTG1 | 1 | 0 |
| DNA+RNA as seed | STAD | 86 | FLNA | 0 | 0 |
| DNA+RNA as seed | STAD | 87 | CTNNA1 | 0 | 0 |
| DNA+RNA as seed | STAD | 88 | VEGFA | 0 | 0 |
| DNA+RNA as seed | STAD | 89 | ARID1A | 0 | 0 |
| DNA+RNA as seed | STAD | 90 | TGFBR2 | 0 | 0 |
| DNA+RNA as seed | STAD | 91 | MAPK10 | 0 | 0 |
| DNA+RNA as seed | STAD | 92 | PKM | 1 | 0 |
| DNA+RNA as seed | STAD | 93 | THRB | 0 | 0 |
| DNA+RNA as seed | STAD | 94 | IQGAP1 | 0 | 0 |
| DNA+RNA as seed | STAD | 95 | TP63 | 0 | 0 |
| DNA+RNA as seed | STAD | 96 | TCF7L2 | 1 | 0 |
| DNA+RNA as seed | STAD | 97 | BMI1 | 0 | 0 |
| DNA+RNA as seed | STAD | 98 | HOXC11 | 0 | 0 |
| DNA+RNA as seed | STAD | 99 | PIWIL1 | 0 | 0 |
| DNA+RNA as seed | STAD | 100 | SREBF1 | 0 | 0 |
| DNA+RNA as seed | THCA | 1 | HRAS | 0 | 0 |
| DNA+RNA as seed | THCA | 2 | NRAS | 0 | 0 |
| DNA+RNA as seed | THCA | 3 | KRAS | 0 | 0 |
| DNA+RNA as seed | THCA | 4 | AKT1 | 0 | 0 |
| DNA+RNA as seed | THCA | 5 | BRAF | 0 | 0 |
| DNA+RNA as seed | THCA | 6 | TRAF2 | 0 | 0 |
| DNA+RNA as seed | THCA | 7 | MAPK1 | 0 | 0 |

| | | | | | |
|---|---|---|---|---|---|
| DNA+RNA as seed | THCA | 8 | MAPK3 | 0 | 0 |
| DNA+RNA as seed | THCA | 9 | ATM | 0 | 0 |
| DNA+RNA as seed | THCA | 10 | PIK3R1 | 0 | 0 |
| DNA+RNA as seed | THCA | 11 | PRKCB | 0 | 0 |
| DNA+RNA as seed | THCA | 12 | TSC1 | 0 | 0 |
| DNA+RNA as seed | THCA | 13 | SHC1 | 0 | 0 |
| DNA+RNA as seed | THCA | 14 | GSK3B | 0 | 0 |
| DNA+RNA as seed | THCA | 15 | AKT2 | 0 | 0 |
| DNA+RNA as seed | THCA | 16 | AKT3 | 0 | 0 |
| DNA+RNA as seed | THCA | 17 | NOTCH1 | 0 | 0 |
| DNA+RNA as seed | THCA | 18 | MAP3K3 | 0 | 0 |
| DNA+RNA as seed | THCA | 19 | IKBKG | 0 | 0 |
| DNA+RNA as seed | THCA | 20 | RAF1 | 0 | 0 |
| DNA+RNA as seed | THCA | 21 | RET | 0 | 0 |
| DNA+RNA as seed | THCA | 22 | PPP2CB | 0 | 0 |
| DNA+RNA as seed | THCA | 23 | CREB1 | 0 | 0 |
| DNA+RNA as seed | THCA | 24 | MAPK9 | 0 | 0 |
| DNA+RNA as seed | THCA | 25 | WDR5 | 1 | 1 |
| DNA+RNA as seed | THCA | 26 | PRKAA1 | 0 | 0 |
| DNA+RNA as seed | THCA | 27 | CBL | 0 | 0 |
| DNA+RNA as seed | THCA | 28 | HDAC1 | 0 | 0 |
| DNA+RNA as seed | THCA | 29 | IKBKB | 0 | 0 |
| DNA+RNA as seed | THCA | 30 | RB1 | 0 | 0 |
| DNA+RNA as seed | THCA | 31 | CALM3 | 0 | 0 |
| DNA+RNA as seed | THCA | 32 | CHEK2 | 0 | 0 |
| DNA+RNA as seed | THCA | 33 | RXRA | 0 | 0 |
| DNA+RNA as seed | THCA | 34 | PTK2 | 0 | 0 |
| DNA+RNA as seed | THCA | 35 | CALM2 | 0 | 0 |
| DNA+RNA as seed | THCA | 36 | YWHAB | 0 | 0 |
| DNA+RNA as seed | THCA | 37 | VHL | 1 | 0 |
| DNA+RNA as seed | THCA | 38 | YWHAE | 0 | 0 |
| DNA+RNA as seed | THCA | 39 | CHUK | 0 | 0 |
| DNA+RNA as seed | THCA | 40 | NBPF19 | 0 | 0 |
| DNA+RNA as seed | THCA | 41 | MAP3K1 | 0 | 0 |
| DNA+RNA as seed | THCA | 42 | GNB2 | 0 | 0 |
| DNA+RNA as seed | THCA | 43 | GNAI2 | 0 | 0 |
| DNA+RNA as seed | THCA | 44 | RPS6KA1 | 0 | 0 |
| DNA+RNA as seed | THCA | 45 | RPS6KA3 | 0 | 0 |
| DNA+RNA as seed | THCA | 46 | PRKDC | 0 | 0 |
| DNA+RNA as seed | THCA | 47 | IRS1 | 0 | 0 |
| DNA+RNA as seed | THCA | 48 | IL7R | 0 | 0 |
| DNA+RNA as seed | THCA | 49 | PDPK1 | 1 | 1 |
| DNA+RNA as seed | THCA | 50 | FOXO3 | 0 | 0 |
| DNA+RNA as seed | THCA | 51 | FRS3 | 0 | 0 |
| DNA+RNA as seed | THCA | 52 | CCDC6 | 0 | 0 |
| DNA+RNA as seed | THCA | 53 | PKM | 0 | 0 |
| DNA+RNA as seed | THCA | 54 | FBXW7 | 0 | 0 |
| DNA+RNA as seed | THCA | 55 | DLC1 | 0 | 0 |
| DNA+RNA as seed | THCA | 56 | CDKN1B | 0 | 0 |
| DNA+RNA as seed | THCA | 57 | FRS2 | 0 | 0 |
| DNA+RNA as seed | THCA | 58 | PIK3CG | 0 | 0 |
| DNA+RNA as seed | THCA | 59 | CDKN2A | 0 | 0 |
| DNA+RNA as seed | THCA | 60 | TBC1D7 | 0 | 0 |
| DNA+RNA as seed | THCA | 61 | USP7 | 0 | 0 |
| DNA+RNA as seed | THCA | 62 | CAMK4 | 0 | 0 |
| DNA+RNA as seed | THCA | 63 | SMAD1 | 0 | 0 |

| | | | | | |
|---|---|---|---|---|---|
| DNA+RNA as seed | THCA | 64 | GNB4 | 0 | 0 |
| DNA+RNA as seed | THCA | 65 | GNB3 | 0 | 0 |
| DNA+RNA as seed | THCA | 66 | ODF2 | 0 | 0 |
| DNA+RNA as seed | THCA | 67 | ATR | 0 | 1 |
| DNA+RNA as seed | THCA | 68 | CACNA1B | 0 | 0 |
| DNA+RNA as seed | THCA | 69 | KAT2B | 0 | 0 |
| DNA+RNA as seed | THCA | 70 | PIK3R5 | 0 | 0 |
| DNA+RNA as seed | THCA | 71 | RPTOR | 0 | 1 |
| DNA+RNA as seed | THCA | 72 | TGFB1 | 0 | 0 |
| DNA+RNA as seed | THCA | 73 | BCL10 | 0 | 0 |
| DNA+RNA as seed | THCA | 74 | BIRC3 | 0 | 0 |
| DNA+RNA as seed | THCA | 75 | SPTA1 | 0 | 0 |
| DNA+RNA as seed | THCA | 76 | RASGRF1 | 0 | 0 |
| DNA+RNA as seed | THCA | 77 | PIK3R6 | 0 | 0 |
| DNA+RNA as seed | THCA | 78 | TTF1 | 0 | 1 |
| DNA+RNA as seed | THCA | 79 | BIRC2 | 0 | 0 |
| DNA+RNA as seed | THCA | 80 | RPS6KA6 | 0 | 0 |
| DNA+RNA as seed | THCA | 81 | RASGRF2 | 0 | 0 |
| DNA+RNA as seed | THCA | 82 | GNG5 | 0 | 0 |
| DNA+RNA as seed | THCA | 83 | GNG4 | 0 | 0 |
| DNA+RNA as seed | THCA | 84 | VBP1 | 0 | 0 |
| DNA+RNA as seed | THCA | 85 | NEDD4L | 0 | 0 |
| DNA+RNA as seed | THCA | 86 | MAP3K5 | 0 | 0 |
| DNA+RNA as seed | THCA | 87 | RPS6KA2 | 0 | 0 |
| DNA+RNA as seed | THCA | 88 | GNB5 | 0 | 0 |
| DNA+RNA as seed | THCA | 89 | GNG12 | 0 | 0 |
| DNA+RNA as seed | THCA | 90 | GNG3 | 0 | 0 |
| DNA+RNA as seed | THCA | 91 | LZTS2 | 0 | 0 |
| DNA+RNA as seed | THCA | 92 | BCL6 | 0 | 0 |
| DNA+RNA as seed | THCA | 93 | CREB5 | 0 | 0 |
| DNA+RNA as seed | THCA | 94 | FCN1 | 0 | 0 |
| DNA+RNA as seed | THCA | 95 | GNG8 | 0 | 0 |
| DNA+RNA as seed | THCA | 96 | GNGT2 | 0 | 0 |
| DNA+RNA as seed | THCA | 97 | GNG11 | 0 | 0 |
| DNA+RNA as seed | THCA | 98 | GNG10 | 0 | 0 |
| DNA+RNA as seed | THCA | 99 | SMARCA4 | 0 | 0 |
| DNA+RNA as seed | THCA | 100 | PPP1R3A | 0 | 0 |
| DNA+RNA as seed | UCEC | 1 | TP53 | 0 | 0 |
| DNA+RNA as seed | UCEC | 2 | PTEN | 0 | 0 |
| DNA+RNA as seed | UCEC | 3 | AKT3 | 0 | 0 |
| DNA+RNA as seed | UCEC | 4 | TFAP2A | 0 | 0 |
| DNA+RNA as seed | UCEC | 5 | PIK3CA | 1 | 0 |
| DNA+RNA as seed | UCEC | 6 | CAV1 | 0 | 0 |
| DNA+RNA as seed | UCEC | 7 | PIK3R1 | 0 | 0 |
| DNA+RNA as seed | UCEC | 8 | CTNNB1 | 0 | 0 |
| DNA+RNA as seed | UCEC | 9 | MYC | 1 | 0 |
| DNA+RNA as seed | UCEC | 10 | CAMK2A | 0 | 0 |
| DNA+RNA as seed | UCEC | 11 | EP300 | 0 | 0 |
| DNA+RNA as seed | UCEC | 12 | EGFR | 0 | 0 |
| DNA+RNA as seed | UCEC | 13 | GNGT1 | 0 | 0 |
| DNA+RNA as seed | UCEC | 14 | ATM | 0 | 0 |
| DNA+RNA as seed | UCEC | 15 | FHL1 | 0 | 0 |
| DNA+RNA as seed | UCEC | 16 | ESR1 | 0 | 0 |
| DNA+RNA as seed | UCEC | 17 | PIK3R2 | 0 | 0 |
| DNA+RNA as seed | UCEC | 18 | KRAS | 1 | 0 |
| DNA+RNA as seed | UCEC | 19 | UBC | 1 | 0 |

| | | | | | |
|---|---|---|---|---|---|
| DNA+RNA as seed | UCEC | 20 | SP1 | 0 | 0 |
| DNA+RNA as seed | UCEC | 21 | MAPK1 | 0 | 0 |
| DNA+RNA as seed | UCEC | 22 | SRC | 0 | 0 |
| DNA+RNA as seed | UCEC | 23 | BRCA1 | 0 | 1 |
| DNA+RNA as seed | UCEC | 24 | MAPK3 | 0 | 0 |
| DNA+RNA as seed | UCEC | 25 | PRKCA | 0 | 0 |
| DNA+RNA as seed | UCEC | 26 | AKT1 | 0 | 0 |
| DNA+RNA as seed | UCEC | 27 | PPP2R1A | 1 | 1 |
| DNA+RNA as seed | UCEC | 28 | PIK3R3 | 0 | 0 |
| DNA+RNA as seed | UCEC | 29 | PIK3CB | 0 | 0 |
| DNA+RNA as seed | UCEC | 30 | HSP90AA1 | 0 | 0 |
| DNA+RNA as seed | UCEC | 31 | LMNA | 0 | 0 |
| DNA+RNA as seed | UCEC | 32 | PLCG1 | 0 | 0 |
| DNA+RNA as seed | UCEC | 33 | PRKDC | 0 | 0 |
| DNA+RNA as seed | UCEC | 34 | STAT3 | 0 | 0 |
| DNA+RNA as seed | UCEC | 35 | AR | 0 | 0 |
| DNA+RNA as seed | UCEC | 36 | PIK3CD | 0 | 0 |
| DNA+RNA as seed | UCEC | 37 | MDM2 | 1 | 0 |
| DNA+RNA as seed | UCEC | 38 | HDAC1 | 1 | 0 |
| DNA+RNA as seed | UCEC | 39 | SMAD3 | 0 | 0 |
| DNA+RNA as seed | UCEC | 40 | MYH11 | 0 | 0 |
| DNA+RNA as seed | UCEC | 41 | JUN | 0 | 0 |
| DNA+RNA as seed | UCEC | 42 | NFKB1 | 0 | 0 |
| DNA+RNA as seed | UCEC | 43 | ERBB2 | 0 | 0 |
| DNA+RNA as seed | UCEC | 44 | HRAS | 0 | 0 |
| DNA+RNA as seed | UCEC | 45 | FBXW7 | 0 | 0 |
| DNA+RNA as seed | UCEC | 46 | NRAS | 0 | 0 |
| DNA+RNA as seed | UCEC | 47 | CSNK2A1 | 0 | 0 |
| DNA+RNA as seed | UCEC | 48 | PPP2CA | 1 | 1 |
| DNA+RNA as seed | UCEC | 49 | FGFR2 | 1 | 0 |
| DNA+RNA as seed | UCEC | 50 | SHC1 | 0 | 0 |
| DNA+RNA as seed | UCEC | 51 | JAK1 | 0 | 0 |
| DNA+RNA as seed | UCEC | 52 | HSP90AB1 | 0 | 0 |
| DNA+RNA as seed | UCEC | 53 | NPM1 | 0 | 0 |
| DNA+RNA as seed | UCEC | 54 | CREB1 | 0 | 0 |
| DNA+RNA as seed | UCEC | 55 | HDAC2 | 0 | 0 |
| DNA+RNA as seed | UCEC | 56 | MAP3K1 | 0 | 0 |
| DNA+RNA as seed | UCEC | 57 | HIF1A | 0 | 0 |
| DNA+RNA as seed | UCEC | 58 | SMAD2 | 0 | 0 |
| DNA+RNA as seed | UCEC | 59 | ABL1 | 0 | 0 |
| DNA+RNA as seed | UCEC | 60 | PLCG2 | 0 | 0 |
| DNA+RNA as seed | UCEC | 61 | CBL | 0 | 0 |
| DNA+RNA as seed | UCEC | 62 | PARP1 | 0 | 0 |
| DNA+RNA as seed | UCEC | 63 | MAPK14 | 0 | 0 |
| DNA+RNA as seed | UCEC | 64 | MAPK8 | 0 | 0 |
| DNA+RNA as seed | UCEC | 65 | EED | 0 | 0 |
| DNA+RNA as seed | UCEC | 66 | FGFR3 | 0 | 0 |
| DNA+RNA as seed | UCEC | 67 | PRKCD | 0 | 0 |
| DNA+RNA as seed | UCEC | 68 | RAC1 | 0 | 1 |
| DNA+RNA as seed | UCEC | 69 | RB1 | 0 | 0 |
| DNA+RNA as seed | UCEC | 70 | CHD4 | 1 | 0 |
| DNA+RNA as seed | UCEC | 71 | SMAD4 | 0 | 0 |
| DNA+RNA as seed | UCEC | 72 | PCNA | 1 | 1 |
| DNA+RNA as seed | UCEC | 73 | RHOA | 1 | 1 |
| DNA+RNA as seed | UCEC | 74 | HSPA8 | 1 | 1 |
| DNA+RNA as seed | UCEC | 75 | PPARG | 0 | 0 |

| | | | | | |
|---|---|---|---|---|---|
| DNA+RNA as seed | UCEC | 76 | PRKN | 0 | 0 |
| DNA+RNA as seed | UCEC | 77 | MET | 0 | 0 |
| DNA+RNA as seed | UCEC | 78 | BCL2 | 0 | 0 |
| DNA+RNA as seed | UCEC | 79 | SMURF1 | 0 | 0 |
| DNA+RNA as seed | UCEC | 80 | E2F1 | 0 | 0 |
| DNA+RNA as seed | UCEC | 81 | IGF1R | 0 | 0 |
| DNA+RNA as seed | UCEC | 82 | PRKCZ | 0 | 0 |
| DNA+RNA as seed | UCEC | 83 | PIP5K1A | 1 | 0 |
| DNA+RNA as seed | UCEC | 84 | CALM3 | 0 | 0 |
| DNA+RNA as seed | UCEC | 85 | FOXO3 | 0 | 0 |
| DNA+RNA as seed | UCEC | 86 | RPA1 | 1 | 1 |
| DNA+RNA as seed | UCEC | 87 | YY1 | 1 | 1 |
| DNA+RNA as seed | UCEC | 88 | NR3C1 | 0 | 0 |
| DNA+RNA as seed | UCEC | 89 | PPP1R12B | 0 | 0 |
| DNA+RNA as seed | UCEC | 90 | PLK1 | 1 | 1 |
| DNA+RNA as seed | UCEC | 91 | PPP2CB | 0 | 0 |
| DNA+RNA as seed | UCEC | 92 | NOTCH1 | 0 | 0 |
| DNA+RNA as seed | UCEC | 93 | RPA2 | 1 | 1 |
| DNA+RNA as seed | UCEC | 94 | PLCB1 | 0 | 0 |
| DNA+RNA as seed | UCEC | 95 | PML | 0 | 0 |
| DNA+RNA as seed | UCEC | 96 | SIRT1 | 0 | 0 |
| DNA+RNA as seed | UCEC | 97 | RAF1 | 0 | 0 |
| DNA+RNA as seed | UCEC | 98 | XRCC6 | 1 | 1 |
| DNA+RNA as seed | UCEC | 99 | PLCB2 | 0 | 0 |
| DNA+RNA as seed | UCEC | 100 | TIMP3 | 0 | 0 |
