## Supplementary material for "Probabilistic graph-based model uncovers previously unseen druggable vulnerabilities in major solid cancers": supp table 8

| cancer_type | top_index | top_genes | depmap_essential | common_essential | GDSC_target |
|---|---|---|---|---|---|
| CESC | 1 | PIK3CA | 1 | 0 | 1 |
| CESC | 2 | EP300 | 0 | 0 | 0 |
| CESC | 3 | TP53 | 0 | 0 | 0 |
| CESC | 4 | CREBBP | 1 | 0 | 0 |
| CESC | 5 | TP63 | 1 | 0 | 0 |
| CESC | 6 | MAPK1 | 0 | 0 | 1 |
| CESC | 7 | ERBB2 | 0 | 0 | 1 |
| CESC | 8 | FBXW7 | 0 | 0 | 0 |
| CESC | 9 | RB1 | 0 | 0 | 0 |
| CESC | 10 | PTEN | 0 | 0 | 0 |
| CESC | 11 | PRKCI | 0 | 0 | 0 |
| CESC | 12 | SMAD4 | 0 | 0 | 0 |
| CESC | 13 | PLD1 | 0 | 0 | 0 |
| CESC | 14 | MYC | 1 | 0 | 0 |
| CESC | 15 | ESR1 | 0 | 0 | 1 |
| CESC | 16 | KRAS | 1 | 0 | 0 |
| CESC | 17 | ECT2 | 1 | 1 | 0 |
| CESC | 18 | ERBB3 | 0 | 0 | 1 |
| CESC | 19 | HUWE1 | 1 | 0 | 0 |
| CESC | 20 | GNB4 | 0 | 0 | 0 |
| CESC | 21 | SP1 | 0 | 0 | 0 |
| CESC | 22 | ACTB | 1 | 0 | 0 |
| CESC | 23 | UBC | 1 | 0 | 0 |
| CESC | 24 | RELA | 0 | 0 | 0 |
| CESC | 25 | CTNNB1 | 0 | 0 | 0 |
| CESC | 26 | SRC | 0 | 0 | 1 |
| CESC | 27 | HES1 | 0 | 0 | 0 |
| CESC | 28 | MECOM | 0 | 0 | 0 |
| CESC | 29 | SMAD3 | 0 | 0 | 0 |
| CESC | 30 | HDAC1 | 0 | 0 | 0 |
| CESC | 31 | CASP8 | 0 | 0 | 0 |
| CESC | 32 | PIK3R1 | 0 | 0 | 0 |
| CESC | 33 | STAT3 | 0 | 0 | 0 |
| CESC | 34 | ACTL6A | 1 | 1 | 0 |
| CESC | 35 | SKIL | 0 | 0 | 0 |
| CESC | 36 | DLG1 | 0 | 0 | 0 |
| CESC | 37 | NFKB1 | 0 | 0 | 0 |
| CESC | 38 | TBL1XR1 | 1 | 0 | 0 |
| CESC | 39 | SMAD2 | 0 | 0 | 0 |
| CESC | 40 | CDK2 | 1 | 0 | 1 |
| CESC | 41 | PRKCA | 0 | 0 | 0 |
| CESC | 42 | CDKN1A | 0 | 0 | 0 |
| CESC | 43 | AR | 0 | 0 | 1 |
| CESC | 44 | CDH1 | 0 | 0 | 0 |
| CESC | 45 | HIF1A | 0 | 0 | 0 |
| CESC | 46 | HRAS | 0 | 0 | 0 |
| CESC | 47 | TNIK | 0 | 0 | 0 |
| CESC | 48 | HLA-B | 0 | 0 | 0 |
| CESC | 49 | PIK3R2 | 0 | 0 | 0 |
| CESC | 50 | NFE2L2 | 0 | 0 | 0 |
| CESC | 51 | PPARG | 0 | 0 | 0 |
| CESC | 52 | STAT1 | 0 | 0 | 0 |

| | | | | | |
|---|---|---|---|---|---|
| CESC | 53 | AKT1 | 0 | 0 | 1 |
| CESC | 54 | NRAS | 0 | 0 | 0 |
| CESC | 55 | MAPK14 | 0 | 0 | 0 |
| CESC | 56 | YWHAZ | 0 | 0 | 0 |
| CESC | 57 | UBB | 0 | 0 | 0 |
| CESC | 58 | FOS | 0 | 0 | 0 |
| CESC | 59 | MAPK8 | 0 | 0 | 0 |
| CESC | 60 | NCOR1 | 0 | 0 | 0 |
| ESCA | 1 | TP53 | 0 | 0 | 0 |
| ESCA | 2 | MYC | 1 | 0 | 0 |
| ESCA | 3 | CCND1 | 1 | 0 | 0 |
| ESCA | 4 | CDKN2A | 0 | 0 | 0 |
| ESCA | 5 | WWOX | 0 | 0 | 0 |
| ESCA | 6 | HTT | 0 | 0 | 0 |
| ESCA | 7 | TP63 | 0 | 0 | 0 |
| ESCA | 8 | RB1 | 0 | 0 | 0 |
| ESCA | 9 | EP300 | 0 | 0 | 0 |
| ESCA | 10 | SMAD4 | 0 | 0 | 0 |
| ESCA | 11 | EGFR | 1 | 0 | 1 |
| ESCA | 12 | ESR1 | 0 | 0 | 1 |
| ESCA | 13 | ERBB2 | 0 | 0 | 1 |
| ESCA | 14 | CTNNB1 | 0 | 0 | 0 |
| ESCA | 15 | HDAC1 | 0 | 0 | 1 |
| ESCA | 16 | UBC | 1 | 0 | 0 |
| ESCA | 17 | SP1 | 0 | 0 | 0 |
| ESCA | 18 | SMAD3 | 0 | 0 | 0 |
| ESCA | 19 | CREBBP | 0 | 0 | 0 |
| ESCA | 20 | SMARCA4 | 0 | 0 | 1 |
| ESCA | 21 | NOTCH1 | 0 | 0 | 0 |
| ESCA | 22 | CDKN1A | 0 | 0 | 0 |
| ESCA | 23 | BRCA1 | 0 | 1 | 0 |
| ESCA | 24 | JUN | 0 | 0 | 0 |
| ESCA | 25 | AR | 0 | 0 | 1 |
| ESCA | 26 | FGF3 | 0 | 0 | 0 |
| ESCA | 27 | SMAD2 | 0 | 0 | 0 |
| ESCA | 28 | PIK3CA | 1 | 0 | 1 |
| ESCA | 29 | STAT3 | 0 | 0 | 0 |
| ESCA | 30 | MAPK1 | 0 | 0 | 1 |
| ESCA | 31 | MAPK3 | 0 | 0 | 1 |
| ESCA | 32 | FBXW7 | 0 | 0 | 0 |
| ESCA | 33 | PRKDC | 0 | 0 | 0 |
| ESCA | 34 | MDM2 | 0 | 0 | 0 |
| ESCA | 35 | GRB2 | 1 | 1 | 0 |
| ESCA | 36 | CUL3 | 1 | 0 | 0 |
| ESCA | 37 | UBE2I | 1 | 1 | 0 |
| ESCA | 38 | FGF4 | 0 | 0 | 0 |
| ESCA | 39 | HIF1A | 0 | 0 | 0 |
| ESCA | 40 | CDKN2B | 0 | 0 | 0 |
| ESCA | 41 | CSNK2A1 | 0 | 0 | 0 |
| ESCA | 42 | NFKB1 | 0 | 0 | 0 |
| ESCA | 43 | FGF19 | 0 | 0 | 0 |
| ESCA | 44 | CDH1 | 0 | 0 | 0 |
| ESCA | 45 | HDAC2 | 0 | 0 | 0 |

| | | | | | |
|---|---|---|---|---|---|
| ESCA | 46 | NFE2L2 | 0 | 0 | 0 |
| ESCA | 47 | PML | 0 | 0 | 0 |
| ESCA | 48 | TP73 | 0 | 0 | 0 |
| ESCA | 49 | PPARG | 0 | 0 | 0 |
| ESCA | 50 | PIK3R1 | 0 | 0 | 0 |
| ESCA | 51 | MAPK8 | 0 | 0 | 0 |
| ESCA | 52 | TGFBR2 | 0 | 0 | 0 |
| ESCA | 53 | PARP1 | 0 | 0 | 1 |
| ESCA | 54 | ATM | 0 | 0 | 1 |
| ESCA | 55 | SIRT1 | 0 | 0 | 0 |
| ESCA | 56 | HDAC3 | 1 | 0 | 0 |
| ESCA | 57 | CTTN | 0 | 0 | 0 |
| ESCA | 58 | RPS27A | 1 | 0 | 0 |
| ESCA | 59 | UBB | 0 | 0 | 0 |
| ESCA | 60 | HDAC4 | 0 | 0 | 0 |
| LIHC | 1 | TP53 | 0 | 0 | 0 |
| LIHC | 2 | CTNNB1 | 1 | 0 | 0 |
| LIHC | 3 | MYC | 1 | 0 | 0 |
| LIHC | 4 | RB1 | 0 | 0 | 0 |
| LIHC | 5 | MUC1 | 0 | 0 | 0 |
| LIHC | 6 | EP300 | 1 | 0 | 1 |
| LIHC | 7 | NTRK1 | 0 | 0 | 1 |
| LIHC | 8 | SMARCA4 | 0 | 0 | 1 |
| LIHC | 9 | MCL1 | 1 | 0 | 1 |
| LIHC | 10 | ESR1 | 0 | 0 | 1 |
| LIHC | 11 | PRKDC | 0 | 0 | 0 |
| LIHC | 12 | AXIN1 | 0 | 0 | 0 |
| LIHC | 13 | BRCA1 | 0 | 1 | 0 |
| LIHC | 14 | CDKN1A | 0 | 0 | 0 |
| LIHC | 15 | STAT3 | 0 | 0 | 0 |
| LIHC | 16 | CDKN2A | 0 | 0 | 0 |
| LIHC | 17 | PIK3CA | 0 | 0 | 1 |
| LIHC | 18 | SMAD3 | 0 | 0 | 0 |
| LIHC | 19 | HSP90AA1 | 0 | 0 | 0 |
| LIHC | 20 | GRB2 | 1 | 1 | 0 |
| LIHC | 21 | AR | 0 | 0 | 1 |
| LIHC | 22 | EEF1A1 | 0 | 0 | 0 |
| LIHC | 23 | GSK3B | 0 | 0 | 1 |
| LIHC | 24 | ZHX1 | 0 | 0 | 0 |
| LIHC | 25 | ALB | 0 | 0 | 0 |
| LIHC | 26 | ARID1A | 0 | 0 | 0 |
| LIHC | 27 | ACTB | 1 | 0 | 0 |
| LIHC | 28 | HSPA8 | 1 | 1 | 0 |
| LIHC | 29 | PCNA | 1 | 1 | 0 |
| LIHC | 30 | SMAD2 | 0 | 0 | 0 |
| LIHC | 31 | HIF1A | 0 | 0 | 0 |
| LIHC | 32 | CUL1 | 1 | 1 | 0 |
| LIHC | 33 | CDH1 | 0 | 0 | 0 |
| LIHC | 34 | CCND1 | 1 | 0 | 0 |
| LIHC | 35 | CREB1 | 0 | 0 | 0 |
| LIHC | 36 | NFE2L2 | 1 | 0 | 0 |
| LIHC | 37 | ABL1 | 0 | 0 | 0 |
| LIHC | 38 | ATM | 0 | 0 | 1 |

| | | | | | |
|---|---|---|---|---|---|
| LIHC | 39 | FOS | 0 | 0 | 0 |
| LIHC | 40 | [illegible]2\|H3C2\|H3C[illegible] | 0 | 0 | 0 |
| LIHC | 41 | TSC2 | 0 | 0 | 0 |
| LIHC | 42 | ERBB2 | 0 | 0 | 1 |
| LIHC | 43 | PPARG | 0 | 0 | 0 |
| LIHC | 44 | NPM1 | 0 | 0 | 0 |
| LIHC | 45 | CDKN1B | 0 | 0 | 0 |
| LIHC | 46 | EFNA1 | 0 | 0 | 0 |
| LIHC | 47 | NR3C1 | 0 | 0 | 0 |
| LIHC | 48 | EFNA4 | 0 | 0 | 0 |
| LIHC | 49 | FYN | 0 | 0 | 0 |
| LIHC | 50 | SIRT1 | 0 | 0 | 1 |
| LIHC | 51 | EZH2 | 0 | 0 | 0 |
| LIHC | 52 | HSPA1B | 0 | 0 | 0 |
| LIHC | 53 | PML | 0 | 0 | 0 |
| LIHC | 54 | EFNA3 | 0 | 0 | 0 |
| LIHC | 55 | PPP1CA | 1 | 0 | 0 |
| LIHC | 56 | BCL2 | 0 | 0 | 1 |
| LIHC | 57 | RPS6KA3 | 0 | 0 | 0 |
| LIHC | 58 | PPP1CC | 0 | 0 | 0 |
| LIHC | 59 | CBL | 0 | 0 | 0 |
| LIHC | 60 | SMARCB1 | 1 | 1 | 0 |
| PAAD | 1 | TP53 | 0 | 0 | 0 |
| PAAD | 2 | KRAS | 1 | 0 | 0 |
| PAAD | 3 | MYC | 1 | 0 | 0 |
| PAAD | 4 | SMAD4 | 0 | 0 | 0 |
| PAAD | 5 | CDKN2A | 0 | 0 | 0 |
| PAAD | 6 | ATM | 0 | 0 | 1 |
| PAAD | 7 | EGFR | 1 | 0 | 1 |
| PAAD | 8 | IFNA5 | 0 | 0 | 0 |
| PAAD | 9 | IFNA2 | 0 | 0 | 0 |
| PAAD | 10 | RELA | 1 | 0 | 0 |
| PAAD | 11 | IFNA8 | 0 | 0 | 0 |
| PAAD | 12 | IFNA6 | 0 | 0 | 0 |
| PAAD | 13 | IFNA14 | 0 | 0 | 0 |
| PAAD | 14 | SMAD3 | 0 | 0 | 0 |
| PAAD | 15 | IFNA4 | 0 | 0 | 0 |
| PAAD | 16 | EP300 | 0 | 0 | 0 |
| PAAD | 17 | SMAD2 | 0 | 0 | 0 |
| PAAD | 18 | IFNA7 | 0 | 0 | 0 |
| PAAD | 19 | IFNA17 | 0 | 0 | 0 |
| PAAD | 20 | IFNA10 | 0 | 0 | 0 |
| PAAD | 21 | TGFBR1 | 0 | 0 | 1 |
| PAAD | 22 | IFNA16 | 0 | 0 | 0 |
| PAAD | 23 | NFKB1 | 0 | 0 | 0 |
| PAAD | 24 | MAPK1 | 1 | 0 | 1 |
| PAAD | 25 | HSP90AA1 | 0 | 0 | 1 |
| PAAD | 26 | IFNA1 | 0 | 0 | 0 |
| PAAD | 27 | IFNA13 | 0 | 0 | 0 |
| PAAD | 28 | JAK2 | 0 | 0 | 1 |
| PAAD | 29 | TGFBR2 | 0 | 0 | 0 |
| PAAD | 30 | MAPK3 | 0 | 0 | 1 |
| PAAD | 31 | CDKN1A | 0 | 0 | 0 |

| | | | | | |
|---|---|---|---|---|---|
| PAAD | 32 | JAK1 | 0 | 0 | 1 |
| PAAD | 33 | STAT1 | 0 | 0 | 0 |
| PAAD | 34 | IFNE | 0 | 0 | 0 |
| PAAD | 35 | AKT1 | 0 | 0 | 1 |
| PAAD | 36 | JAK3 | 0 | 0 | 1 |
| PAAD | 37 | TYK2 | 0 | 0 | 0 |
| PAAD | 38 | FBXW7 | 0 | 0 | 0 |
| PAAD | 39 | CDKN2B | 0 | 0 | 0 |
| PAAD | 40 | AR | 0 | 0 | 0 |
| PAAD | 41 | GSK3B | 0 | 0 | 1 |
| PAAD | 42 | STK11 | 0 | 0 | 0 |
| PAAD | 43 | SQSTM1 | 0 | 0 | 0 |
| PAAD | 44 | MAPK14 | 0 | 0 | 0 |
| PAAD | 45 | IFNA21 | 0 | 0 | 0 |
| PAAD | 46 | HECW2 | 0 | 0 | 0 |
| PAAD | 47 | UBB | 0 | 0 | 0 |
| PAAD | 48 | MAPK8 | 0 | 0 | 0 |
| PAAD | 49 | PTPN11 | 1 | 0 | 0 |
| PAAD | 50 | SMAD1 | 0 | 0 | 0 |
| PAAD | 51 | IL24 | 0 | 0 | 0 |
| PAAD | 52 | HIF1A | 0 | 0 | 0 |
| PAAD | 53 | RPS27A | 1 | 0 | 0 |
| PAAD | 54 | PPP2CA | 1 | 1 | 0 |
| PAAD | 55 | SMURF1 | 0 | 0 | 0 |
| PAAD | 56 | STAT2 | 0 | 0 | 0 |
| PAAD | 57 | PTPN6 | 0 | 0 | 0 |
| PAAD | 58 | SOCS3 | 0 | 0 | 0 |
| PAAD | 59 | IKBKB | 0 | 0 | 0 |
| PAAD | 60 | PTPN1 | 0 | 0 | 0 |
