## Supplementary material for "Probabilistic graph-based model uncovers previously unseen druggable vulnerabilities in major solid cancers": supp table 10

| gene | main_category | sub_category | accessible_primary | accessible_secondary | indications_with_nonsense_mutation (>2% patients) | indications_with_copynumber deletion (>2% patients) | indications_with_low_RNA_expression | indications_with_high_RNA_expression | among_MVP_top20 | Depmap_dependency | GDSC_target | overlap_indication_with_Depmap | overlap_indication_with_GDSC | Depmap_hit_as_least_one_indication | GDSC_hit_as_least_one_indication |
|---|---|---|---|---|---|---|---|---|---|---|---|---|---|---|---|
| ACTB | D | 1 | Yes | Yes | | | | GBM | KIRC | BLCA,BRCA,COADREAD,GBM,HNSC,KIRC,LGG,LUAD,LUSC,OV,STAD,UCEC | | KIRC | | 1 | 0 |
| ACTL6A | D | 1 | Yes | No | | | | BLCA,GBM,LUSC | LUSC | BLCA,BRCA,COADREAD,GBM,HNSC,KIRC,LGG,LUAD,LUSC,OV,PRAD,SKCM,STAD,THCA,UCEC | | LUSC | | 1 | 0 |
| AKT1 | B | 1 | Yes | No | | | | | BRCA,COADREAD,LUAD,THCA | BRCA,STAD | BLCA,BRCA,COADREAD,GBM,HNSC,KIRC,LGG,LUAD,LUSC,OV,PRAD,SKCM,STAD,THCA,UCEC | BRCA | BRCA,COADREAD,LUAD,THCA | 1 | 1 |
| AKT2 | B | 1 | Yes | No | | | | | THCA | | BLCA,BRCA,COADREAD,GBM,HNSC,KIRC,LGG,LUAD,LUSC,OV,PRAD,SKCM,STAD,THCA,UCEC | | THCA | 0 | 1 |
| AKT3 | C | 1 | Yes | No | | | BLCA,COADREAD,GBM,UCEC | | THCA,UCEC | | BLCA,BRCA,COADREAD,GBM,HNSC,KIRC,LGG,LUAD,LUSC,OV,PRAD,SKCM,STAD,UCEC | | UCEC | 0 | 1 |
| AP2M1 | D | 1 | Yes | No | | | | LUSC | LUSC | BLCA,BRCA,COADREAD,GBM,HNSC,KIRC,LUAD,LUSC,OV,STAD | | LUSC | | 1 | 0 |
| APC | C | 1 | Yes | No | COADREAD,STAD,UCEC | COADREAD,OV,PRAD,STAD | GBM | | COADREAD,PRAD | COADREAD,PRAD | | COADREAD,PRAD | | 1 | 0 |
| AR | A | 1 | Yes | Yes | | LUSC | BLCA,COADREAD,LUSC,STAD,UCEC | GBM | KIRC,LUSC,PRAD,STAD | | BLCA,BRCA,COADREAD,GBM,HNSC,KIRC,LUAD,LUSC,OV,PRAD,SKCM,STAD,THCA | | KIRC,LUSC,PRAD,STAD | 0 | 1 |
| ARID1A | | | No | No | BLCA,LUAD,STAD,UCEC | STAD | | | BLCA,STAD,UCEC | | | | | 0 | 0 |
| ASAP1 | | | No | No | | | | KIRC | OV | | | | | 0 | 0 |
| ATM | B | 1 | Yes | Yes | BLCA,COADREAD,UCEC | SKCM | | | BLCA,COADREAD,KIRC,LUAD,PRAD,STAD,THCA,UCEC | | BLCA,BRCA,COADREAD,GBM,HNSC,KIRC,LGG,LUAD,LUSC,OV,PRAD,SKCM,STAD,THCA | | BLCA,COADREAD,KIRC,LUAD,PRAD,STAD,THCA | 0 | 1 |
| BRAF | A | 1 | Yes | Yes | | | GBM | | SKCM,THCA | BRCA,COADREAD,GBM,OV,SKCM | BLCA,BRCA,COADREAD,GBM,HNSC,KIRC,LGG,LUAD,LUSC,OV,SKCM,STAD,THCA,UCEC | SKCM | SKCM,THCA | 1 | 1 |
| BRCA1 | | | Yes | No | | | | BLCA,BRCA,COADREAD,GBM,LUAD,LUSC,STAD,UCEC | BRCA,HNSC,OV | BRCA,GBM | | BRCA | | 1 | 0 |
| CALML3 | | | No | No | | | KIRC,PRAD | LUAD,LUSC | LUSC | | | | | 0 | 0 |
| CAMK2A | C | 2 | Yes | No | | | BLCA,BRCA,COADREAD,GBM,KIRC,LUAD,LUSC,PRAD,STAD,UCEC | | UCEC | | | | | 0 | 0 |
| CAV1 | | | No | No | | | BLCA,BRCA,COADREAD,LUAD,LUSC,PRAD,STAD,UCEC | GBM,KIRC | UCEC | | | | | 0 | 0 |
| CCND1 | A | 1 | Yes | No | | | | BRCA,COADREAD,GBM,KIRC,THCA | BRCA,HNSC,SKCM | BLCA,BRCA,COADREAD,GBM,HNSC,KIRC,LGG,LUAD,LUSC,OV,PRAD,SKCM,STAD,UCEC | | BRCA,HNSC,SKCM | | 1 | 0 |
| CDC25C | D | 2 | Yes | No | | | | BLCA,BRCA,COADREAD,GBM,KIRC,LUAD,LUSC,PRAD,STAD,UCEC | BRCA | | | | | 0 | 0 |
| CDH1 | | | Yes | No | BRCA | OV,PRAD | KIRC | LUAD,LUSC,UCEC | BRCA,COADREAD,STAD | COADREAD | | COADREAD | | 1 | 0 |
| CDK1 | B | 1 | Yes | No | | | | BLCA,BRCA,COADREAD,GBM,LUAD,LUSC,PRAD,STAD,UCEC | OV | BLCA,BRCA,COADREAD,GBM,HNSC,KIRC,LGG,LUAD,LUSC,OV,PRAD,SKCM,STAD,THCA,UCEC | BLCA,BRCA,COADREAD,GBM,HNSC,KIRC,LGG,LUAD,LUSC,OV,SKCM,STAD,THCA,UCEC | OV | OV | 1 | 1 |
| CDK2 | B | 1 | Yes | Yes | | | | BLCA,COADREAD,GBM | BRCA,OV | BLCA,BRCA,COADREAD,GBM,KIRC,LGG,LUAD,LUSC,OV,PRAD,STAD,UCEC | BLCA,BRCA,COADREAD,GBM,HNSC,KIRC,LGG,LUAD,LUSC,OV,STAD,THCA,UCEC | BRCA,OV | BRCA,OV | 1 | 1 |
| CDKN1A | | | No | No | | | BLCA,STAD | GBM,THCA | BLCA,LUAD | | | | | 0 | 0 |
| CDKN1B | | | No | No | | PRAD | UCEC | | PRAD | | | | | 0 | 0 |
| CDKN2A | | | No | No | HNSC,LUSC,SKCM | BLCA,BRCA,GBM,HNSC,KIRC,LGG,LUAD,LUSC,OV,STAD,SKCM | | BLCA,BRCA,COADREAD,GBM,KIRC,LUAD,LUSC,PRAD,STAD,THCA,UCEC | BLCA,GBM,HNSC,LGG,LUAD,LUSC,SKCM,STAD | | | | | 0 | 0 |
| CHD4 | | | Yes | No | | | | BRCA,THCA | UCEC | BLCA,BRCA,COADREAD,GBM,HNSC,KIRC,LGG,LUAD,LUSC,OV,PRAD,SKCM,STAD,THCA,UCEC | | UCEC | | 1 | 0 |
| CREBBP | B | 1 | Yes | Yes | BLCA | BLCA,OV | | | BLCA,HNSC,LUSC | BLCA,COADREAD,LUSC,STAD,UCEC | BLCA,BRCA,COADREAD,HNSC,KIRC,LGG,LUAD,LUSC,OV,SKCM,STAD,THCA,UCEC | BLCA,LUSC | BLCA,HNSC,LUSC | 1 | 1 |
| CSNK2A1 | C | 2 | Yes | Yes | | | | | LUSC | | | | | 0 | 0 |
| CTNNB1 | C | 1 | Yes | No | | KIRC | | | COADREAD,HNSC,KIRC,LUAD,LUSC,PRAD,SKCM,STAD,UCEC | BLCA,BRCA,COADREAD,HNSC,LUAD,STAD | | COADREAD,HNSC,LUAD,STAD | | 1 | 0 |
| CTTN | | | No | No | | | | | BRCA | | | | | 0 | 0 |
| CUL3 | | | No | No | | BLCA,HNSC | | | HNSC,LUSC | BLCA,BRCA,COADREAD,GBM,KIRC,LGG,LUAD,LUSC,OV,PRAD,STAD,UCEC | | LUSC | | 1 | 0 |
| DBN1 | | | No | No | | | | LUAD,LUSC | KIRC | | | | | 0 | 0 |
| DCUN1D1 | C | 2 | Yes | Yes | | | | LUSC | LUSC | | | | | 0 | 0 |
| E2F3 | | | No | No | | | | BLCA,BRCA,LUAD,LUSC,STAD,UCEC | BLCA | BLCA,GBM,LGG,LUAD,OV,PRAD | | BLCA | | 1 | 0 |
| EEF1D | | | No | No | | | | | OV | | | | | 0 | 0 |
| EGFR | A | 1 | Yes | Yes | | | | GBM,KIRC,LUSC | BRCA,GBM,HNSC,LGG,LUAD,LUSC,OV,SKCM,STAD,UCEC | BLCA,BRCA,COADREAD,HNSC,LUAD,LUSC,OV,PRAD,STAD | BLCA,BRCA,COADREAD,GBM,HNSC,KIRC,LGG,LUAD,LUSC,OV,PRAD,SKCM,STAD,THCA,UCEC | BRCA,HNSC,LUAD,LUSC,OV,STAD | BRCA,GBM,HNSC,LGG,LUAD,LUSC,OV,SKCM,STAD,UCEC | 1 | 1 |
| ELAVL1 | C | 1 | Yes | No | | | | | COADREAD | COADREAD | | COADREAD | | 1 | 0 |
| EP300 | B | 1 | Yes | Yes | BLCA,UCEC | | | | BLCA,BRCA,HNSC,KIRC,PRAD,SKCM,STAD,UCEC | BLCA,BRCA,PRAD | BLCA,BRCA,COADREAD,HNSC,KIRC,LGG,LUAD,LUSC,OV,PRAD,SKCM,STAD,THCA,UCEC | BLCA,BRCA,PRAD | BLCA,BRCA,HNSC,KIRC,PRAD,SKCM,STAD,UCEC | 1 | 1 |
| EPPK1 | | | No | No | | | | BRCA,LUSC,THCA,UCEC | OV | | | | | 0 | 0 |
| ERBB2 | A | 1 | Yes | Yes | | | KIRC | BLCA,BRCA,GBM,LUAD,STAD,UCEC | BLCA,BRCA,LUAD,STAD | BRCA,COADREAD,GBM,LUSC,OV | BLCA,BRCA,COADREAD,GBM,HNSC,KIRC,LGG,LUAD,LUSC,OV,PRAD,SKCM,STAD,UCEC | BRCA | BLCA,BRCA,LUAD,STAD | 1 | 1 |
| ERBB3 | A | 1 | Yes | Yes | | | GBM | BLCA,BRCA,THCA,UCEC | BLCA,STAD | BRCA | BLCA,BRCA,COADREAD,HNSC,KIRC,LUAD,LUSC,OV,SKCM,STAD,UCEC | | BLCA,STAD | 0 | 1 |
| ERBB4 | C | 1 | Yes | No | | BLCA,LUSC | COADREAD,GBM,KIRC,LUAD,LUSC,PRAD,THCA | | STAD | | BLCA,BRCA,COADREAD,HNSC,KIRC,LUAD,OV,SKCM,STAD,THCA,UCEC | | STAD | 0 | 1 |
| ERG | | | No | No | | PRAD | BRCA,LUAD,LUSC,UCEC | PRAD | PRAD | PRAD | | PRAD | | 1 | 0 |
| ESR1 | A | 1 | Yes | Yes | | | BLCA | BRCA,THCA | BLCA,BRCA,COADREAD,HNSC,KIRC,LUSC,PRAD,STAD,UCEC | BRCA | BLCA,BRCA,COADREAD,GBM,HNSC,KIRC,LGG,LUAD,LUSC,OV,PRAD,SKCM,STAD,THCA,UCEC | BRCA | BLCA,BRCA,COADREAD,HNSC,KIRC,LUSC,PRAD,STAD,UCEC | 1 | 1 |
| FBXW7 | D | 2 | Yes | No | BLCA,COADREAD,UCEC | | GBM | | COADREAD,STAD,UCEC | BRCA | | | | 0 | 0 |
| FCER1G | | | No | No | | | LUAD,LUSC | GBM,KIRC,STAD,THCA,UCEC | BLCA | | | | | 0 | 0 |
| FCGR2A | | | Yes | No | | | LUSC | GBM,KIRC,STAD,THCA | BLCA | | | | | 0 | 0 |
| FCGR3A | | | Yes | No | | | LUAD,LUSC | BLCA,BRCA,COADREAD,GBM,KIRC,STAD,THCA,UCEC | BLCA | | | | | 0 | 0 |
| FGF19 | | | No | No | | | | COADREAD,GBM,LUAD,LUSC,STAD,THCA,UCEC | BRCA | | | | | 0 | 0 |
| FGF3 | | | No | No | | | | COADREAD,GBM,STAD | BRCA | | | | | 0 | 0 |
| FGF4 | | | No | No | | | | BRCA,COADREAD,STAD,THCA,UCEC | BRCA | | | | | 0 | 0 |
| FGFR2 | A | 1 | Yes | No | | | COADREAD,GBM,KIRC,LUAD,PRAD,THCA | | UCEC | UCEC | BLCA,BRCA,COADREAD,GBM,HNSC,KIRC,LGG,LUSC,OV,PRAD,SKCM,STAD,THCA,UCEC | UCEC | UCEC | 1 | 1 |
| FGFR3 | A | 1 | Yes | No | | | | BRCA,LUSC,THCA,UCEC | BLCA | | BLCA,BRCA,COADREAD,GBM,HNSC,KIRC,LGG,LUSC,OV,PRAD,SKCM,STAD,THCA,UCEC | | BLCA | 0 | 1 |
| FGFR4 | A | 1 | Yes | No | | | LUAD,LUSC | BRCA,COADREAD,GBM,STAD,UCEC | KIRC | BRCA | BLCA,BRCA,COADREAD,GBM,HNSC,KIRC,LGG,LUAD,LUSC,OV,PRAD,SKCM,STAD,UCEC | | KIRC | 0 | 1 |
| FHL1 | | | No | No | | | BLCA,BRCA,COADREAD,LUAD,LUSC,PRAD,STAD,THCA,UCEC | GBM,KIRC | UCEC | | | | | 0 | 0 |
| FLT1 | A | 2 | Yes | No | | | LUSC,UCEC | KIRC | KIRC | | | | | 0 | 0 |
| FLT4 | A | 2 | Yes | No | | | BRCA,LUAD,LUSC,UCEC | GBM,KIRC | KIRC | | | | | 0 | 0 |
| FN1 | | | Yes | Yes | | | PRAD | BRCA,GBM,KIRC,THCA | LUSC | | | | | 0 | 0 |
| FOXA1 | | | No | No | | | | BRCA,GBM,LUAD,THCA,UCEC | PRAD | BRCA,COADREAD,LUAD,PRAD,STAD | | PRAD | | 1 | 0 |
| FXR1 | | | No | No | | | | LUSC | LUSC | | | | | 0 | 0 |
| GNGT1 | | | No | No | | | | BLCA,BRCA,COADREAD,GBM,KIRC,LUAD,LUSC,STAD,UCEC | UCEC | | | | | 0 | 0 |

| | | | | | | | | | | | | | | | |
|---|---|---|---|---|---|---|---|---|---|---|---|---|---|---|---|
| GRB2 | C | 1 | Yes | Yes | | | | | LUSC,SKCM | BLCA,BRCA,COADREAD,GBM,HNSC,KIRC,LGG,LUAD,LUSC,OV,PRAD,STAD,UCEC | | LUSC | | 1 | 0 |
| GSK3B | B | 1 | Yes | No | | | | | PRAD,THCA | | BLCA,BRCA,COADREAD,GBM,HNSC,KIRC,LGG,LUAD,LUSC,OV,PRAD,SKCM,STAD,THCA,UCEC | | PRAD,THCA | 0 | 1 |
| HDAC1 | A | 1 | Yes | No | | | | GBM,LUSC | KIRC,LUAD,LUSC,PRAD | COADREAD,STAD,UCEC | BLCA,BRCA,COADREAD,GBM,HNSC,KIRC,LGG,LUAD,LUSC,OV,SKCM,STAD,THCA | | KIRC,LUAD,LUSC | 0 | 1 |
| HIF1A | C | 2 | No | No | | | KIRC | GBM | KIRC | | | | | 0 | 0 |
| HRAS | C | 2 | Yes | Yes | | LGG | | LUSC,UCEC | HNSC,LGG,LUAD,PRAD,THCA | BLCA,SKCM | | | | 0 | 0 |
| HSF1 | B | 1 | No | No | | | | | LGG | BLCA,BRCA,COADREAD,HNSC,KIRC,LUAD,OV,STAD | BRCA,COADREAD,GBM,HNSC,KIRC,LGG,LUAD,LUSC,OV,SKCM,STAD,THCA,UCEC | | LGG | 0 | 1 |
| HSP90AA1 | B | 1 | Yes | No | | | | STAD | COADREAD,KIRC,LGG,LUAD | STAD | BLCA,BRCA,COADREAD,GBM,HNSC,KIRC,LGG,LUAD,LUSC,OV,SKCM,STAD,THCA | | COADREAD,KIRC,LGG,LUAD | 0 | 1 |
| HSPA8 | D | 2 | Yes | Yes | | SKCM | | | PRAD | BLCA,BRCA,COADREAD,GBM,HNSC,KIRC,LGG,LUAD,LUSC,OV,SKCM,STAD,UCEC | | | | 0 | 0 |
| HUWE1 | D | 1 | Yes | No | UCEC | | | | HNSC | BLCA,BRCA,COADREAD,GBM,HNSC,KIRC,LGG,LUAD,LUSC,OV,PRAD,SKCM,STAD,UCEC | | HNSC | | 1 | 0 |
| IFNA1 | | | No | No | | BLCA,GBM,HNSC,LGG,LUAD,LUSC,OV,STAD,SKCM | | GBM,THCA | GBM | | | | | 0 | 0 |
| IFNA10 | | | No | No | | BLCA,GBM,HNSC,LGG,LUAD,LUSC,STAD,SKCM | | | GBM | | | | | 0 | 0 |
| IFNA13 | | | No | No | | BLCA,GBM,HNSC,LGG,LUAD,LUSC,OV,STAD,SKCM | GBM | | GBM | | | | | 0 | 0 |
| IFNA14 | | | No | No | | BLCA,GBM,HNSC,LGG,LUAD,LUSC,STAD,SKCM | KIRC | | GBM,SKCM | | | | | 0 | 0 |
| IFNA17 | | | No | No | | BLCA,GBM,HNSC,LGG,LUAD,LUSC,STAD,SKCM | | | GBM | | | | | 0 | 0 |
| IFNA2 | D | 2 | Yes | No | | BLCA,GBM,HNSC,LGG,LUAD,LUSC,OV,STAD,SKCM | | | GBM,SKCM | | | | | 0 | 0 |
| IFNA21 | | | No | No | | BLCA,GBM,HNSC,LGG,LUAD,LUSC,STAD,SKCM | GBM | | GBM | | | | | 0 | 0 |
| IFNA4 | | | No | No | | BLCA,GBM,HNSC,LGG,LUAD,LUSC,STAD,SKCM | | | GBM | | | | | 0 | 0 |
| IFNA5 | | | No | No | | BLCA,GBM,HNSC,LGG,LUAD,LUSC,STAD,SKCM | LUSC | | GBM,SKCM | | | | | 0 | 0 |
| IFNA6 | | | No | No | | BLCA,GBM,HNSC,LGG,LUAD,LUSC,STAD,SKCM | GBM | | GBM | | | | | 0 | 0 |
| IFNA7 | | | No | No | | BLCA,GBM,HNSC,LGG,LUAD,LUSC,STAD,SKCM | | | GBM | | | | | 0 | 0 |
| IFNA8 | | | Yes | No | | BLCA,GBM,HNSC,LGG,LUAD,LUSC,OV,STAD,SKCM | | | GBM | | | | | 0 | 0 |
| IFNE | | | No | No | | BLCA,GBM,HNSC,LGG,LUAD,LUSC,OV,STAD,SKCM | PRAD | COADREAD,KIRC,LUAD,LUSC,STAD,THCA | GBM | | | | | 0 | 0 |
| IGF1R | A | 1 | Yes | Yes | | | | BRCA,LUSC | KIRC | BLCA,COADREAD,HNSC,LUAD,OV,PRAD,SKCM,STAD | BLCA,BRCA,COADREAD,GBM,HNSC,KIRC,LUAD,OV,PRAD,SKCM,STAD,THCA | | KIRC | 0 | 1 |
| IKBKG | C | 2 | No | No | | | | | THCA | | | | | 0 | 0 |
| JAK2 | A | 1 | Yes | Yes | UCEC | LUSC,STAD | UCEC | | LGG | | BLCA,BRCA,COADREAD,GBM,HNSC,KIRC,LGG,LUAD,LUSC,PRAD,SKCM,STAD,UCEC | | LGG | 0 | 1 |
| JUN | C | 2 | No | No | | | BLCA,BRCA,LUAD,LUSC,STAD,THCA,UCEC | | LUAD | COADREAD,GBM,HNSC,LGG | | | | 0 | 0 |
| KRAS | A | 1 | Yes | Yes | | | | STAD | COADREAD,LUAD,STAD,THCA,UCEC | BLCA,BRCA,COADREAD,GBM,HNSC,LUAD,LUSC,OV,STAD,UCEC | GBM,LUAD,PRAD | COADREAD,LUAD,STAD,UCEC | LUAD | 1 | 1 |
| LMNA | | | No | Yes | | | | | UCEC | GBM,HNSC | | | | 0 | 0 |
| MAP3K3 | C | 2 | No | No | | | LUAD,LUSC,UCEC | | THCA | | | | | 0 | 0 |
| MAPK1 | B | 1 | Yes | Yes | | | GBM | | COADREAD,LGG,LUAD,SKCM,THCA | COADREAD,GBM,SKCM,STAD | BLCA,BRCA,COADREAD,GBM,HNSC,KIRC,LUAD,LUSC,OV,PRAD,SKCM,STAD,THCA,UCEC | COADREAD,SKCM | COADREAD,LUAD,SKCM,THCA | 1 | 1 |
| MAPK14 | B | 2 | Yes | Yes | | | | | THCA | | COADREAD,LUAD,OV,STAD | | | 0 | 0 |
| MAPK3 | B | 1 | Yes | No | | | GBM | | LGG,PRAD,SKCM,THCA | | BLCA,BRCA,COADREAD,GBM,HNSC,KIRC,LUAD,LUSC,OV,SKCM,STAD,THCA,UCEC | | SKCM,THCA | 0 | 1 |
| MECOM | | | No | No | | | BRCA,KIRC,PRAD | GBM,THCA,UCEC | OV | | | | | 0 | 0 |
| MET | A | 1 | Yes | No | | | BRCA | COADREAD,KIRC,LUAD,STAD,THCA,UCEC | LUAD | STAD | BRCA,COADREAD,GBM,HNSC,KIRC,LGG,LUAD,LUSC,OV,SKCM,STAD | | LUAD | 0 | 1 |
| MMP1 | A | 2 | Yes | No | | | | BLCA,BRCA,COADREAD,GBM,KIRC,LUAD,LUSC,STAD,THCA,UCEC | BRCA | | | | | 0 | 0 |
| MMP7 | C | 2 | Yes | No | | | | BLCA,COADREAD,GBM,LUAD,LUSC,STAD,THCA,UCEC | COADREAD | | | | | 0 | 0 |
| MTOR | B | 1 | Yes | No | | | KIRC | | KIRC | BLCA,BRCA,COADREAD,GBM,HNSC,KIRC,LGG,LUAD,LUSC,OV,PRAD,SKCM,STAD,THCA,UCEC | BLCA,BRCA,COADREAD,GBM,HNSC,KIRC,LGG,LUAD,LUSC,OV,PRAD,SKCM,STAD,THCA,UCEC | KIRC | KIRC | 1 | 1 |
| MYC | C | 1 | Yes | No | | | BLCA | COADREAD,GBM,KIRC | BLCA,BRCA,HNSC,LGG,LUAD,OV,PRAD,STAD,UCEC | BLCA,BRCA,COADREAD,GBM,HNSC,KIRC,LGG,LUAD,LUSC,OV,PRAD,SKCM,STAD,THCA,UCEC | | BLCA,BRCA,HNSC,LGG,LUAD,OV,PRAD,STAD,UCEC | | 1 | 0 |
| NCOR1 | C | 2 | No | No | | | | | BLCA | | | | | 0 | 0 |
| NDRG1 | | | No | No | | | COADREAD | KIRC,LUSC | OV | | | | | 0 | 0 |
| NFKB1 | B | 2 | No | No | | | | | LGG,LUAD,SKCM | | | | | 0 | 0 |
| NOTCH1 | B | 2 | Yes | No | HNSC,LUSC | | | | HNSC,LUSC,THCA | | | | | 0 | 0 |
| NR4A1 | C | 2 | Yes | No | | | BLCA,BRCA,LUAD,LUSC,STAD,THCA,UCEC | | BLCA | | | | | 0 | 0 |
| NRAS | C | 1 | Yes | Yes | | HNSC | | GBM | COADREAD,LGG,LUAD,SKCM,THCA | BLCA,LUAD,OV,SKCM | | LUAD,SKCM | | 1 | 0 |
| NTRK1 | A | 1 | Yes | Yes | | | KIRC,PRAD | GBM,THCA | HNSC,LGG | | BLCA,BRCA,COADREAD,GBM,HNSC,KIRC,LGG,LUAD,LUSC,OV,PRAD,SKCM,STAD,THCA,UCEC | | HNSC,LGG | 0 | 1 |
| PBRM1 | C | 2 | Yes | Yes | KIRC,UCEC | KIRC | | | KIRC | | | | | 0 | 0 |
| PCNA | B | 1 | Yes | Yes | | | | BLCA,BRCA,COADREAD,GBM,LUAD,LUSC,STAD,UCEC | OV | BLCA,BRCA,COADREAD,GBM,HNSC,KIRC,LGG,LUAD,LUSC,OV,PRAD,SKCM,STAD,THCA,UCEC | | OV | | 1 | 0 |
| PDGFRA | A | 1 | Yes | No | | | BLCA,BRCA,COADREAD,KIRC,THCA,UCEC | GBM | GBM | GBM,HNSC,STAD | BLCA,BRCA,COADREAD,GBM,HNSC,KIRC,LUAD,OV,SKCM,STAD,THCA | GBM | GBM | 1 | 1 |
| PDLIM7 | | | No | No | | | BLCA,STAD,UCEC | GBM,KIRC,THCA | KIRC | | | | | 0 | 0 |
| PIK3CA | A | 1 | Yes | Yes | | | | | BLCA,BRCA,COADREAD,GBM,HNSC,KIRC,LGG,LUAD,LUSC,PRAD,STAD,UCEC | BLCA,BRCA,COADREAD,GBM,LUAD,LUSC,OV,STAD,UCEC | BLCA,BRCA,COADREAD,GBM,HNSC,KIRC,LGG,LUAD,LUSC,OV,PRAD,STAD,THCA,UCEC | BLCA,BRCA,COADREAD,GBM,LUAD,LUSC,STAD,UCEC | BLCA,BRCA,COADREAD,GBM,HNSC,KIRC,LGG,LUAD,LUSC,PRAD,STAD,UCEC | 1 | 1 |
| PIK3CB | B | 1 | No | No | | | GBM | | COADREAD,GBM | BRCA,KIRC | BLCA,BRCA,COADREAD,GBM,HNSC,KIRC,LGG,LUAD,LUSC,OV,PRAD,SKCM,STAD | | COADREAD,GBM | 0 | 1 |
| PIK3CD | A | 1 | Yes | Yes | | | | | GBM | | BLCA,BRCA,COADREAD,GBM,HNSC,KIRC,LGG,LUAD,LUSC,OV,PRAD,SKCM,STAD,THCA,UCEC | | GBM | 0 | 1 |
| PIK3R1 | D | 2 | Yes | No | UCEC | OV,PRAD | BLCA,BRCA,LUAD | | BLCA,BRCA,COADREAD,GBM,LGG,THCA,UCEC | | | | | 0 | 0 |
| PIK3R2 | C | 2 | No | No | | | | COADREAD,KIRC,LUSC | BLCA,UCEC | | | | | 0 | 0 |
| PIK3R3 | | | No | No | | | | UCEC | COADREAD,GBM | | | | | 0 | 0 |
| PLCB4 | | | No | No | SKCM | | BLCA | COADREAD,KIRC | SKCM | | | | | 0 | 0 |
| PLCG1 | | | No | No | | | | COADREAD | GBM | | | | | 0 | 0 |
| PLEC | | | Yes | No | | | | | OV | | | | | 0 | 0 |
| PLK1 | B | 1 | Yes | No | | | | BLCA,BRCA,COADREAD,GBM,KIRC,LUAD,LUSC,PRAD,STAD,UCEC | BRCA | BLCA,BRCA,COADREAD,GBM,HNSC,KIRC,LGG,LUAD,LUSC,OV,PRAD,SKCM,STAD,THCA,UCEC | BLCA,BRCA,COADREAD,HNSC,KIRC,LGG,LUAD,LUSC,STAD,THCA | BRCA | BRCA | 1 | 1 |
| PPP2R1A | D | 1 | Yes | Yes | | LGG | | | UCEC | BLCA,BRCA,COADREAD,GBM,HNSC,KIRC,LGG,LUAD,LUSC,OV,PRAD,SKCM,STAD,UCEC | | UCEC | | 1 | 0 |
| PRKCB | B | 2 | Yes | No | | | BLCA,COADREAD,GBM,LUSC,PRAD,STAD,UCEC | | THCA | | | | | 0 | 0 |
| PRKDC | B | 2 | Yes | No | UCEC | | | COADREAD,LUAD,LUSC,STAD | COADREAD,UCEC | | | | | 0 | 0 |
| PTEN | D | 2 | Yes | No | GBM,LUSC,UCEC | BLCA,BRCA,COADREAD,GBM,HNSC,LUSC,OV,PRAD,STAD,UCEC,SKCM | | | BRCA,COADREAD,GBM,KIRC,LGG,LUSC,PRAD,SKCM,STAD,UCEC | | | | | 0 | 0 |
| PTK2 | B | 1 | Yes | No | | | GBM | | OV | BRCA,COADREAD,GBM,KIRC,LUAD,LUSC,OV,SKCM,STAD | BLCA,BRCA,COADREAD,GBM,HNSC,OV,SKCM,STAD,THCA | OV | OV | 1 | 1 |
| PTPN11 | C | 1 | Yes | Yes | | | | | GBM,LGG | BLCA,BRCA,COADREAD,GBM,HNSC,KIRC,LGG,LUAD,LUSC,OV,PRAD,STAD,UCEC | | GBM,LGG | | 1 | 0 |

| | | | | | | | | | | | | | | | |
|---|---|---|---|---|---|---|---|---|---|---|---|---|---|---|---|
| PUF60 | D | 1 | Yes | No | | | | | OV | BLCA,BRCA,COADREAD,GBM,HNSC,KIRC,LGG,LUAD,LUSC,OV,PRAD,SKCM,STAD,THCA,UCEC | | OV | | 1 | 0 |
| RAC1 | D | 1 | Yes | Yes | | | | | SKCM | BLCA,BRCA,COADREAD,GBM,KIRC,LGG,LUAD,LUSC,OV,SKCM,STAD | BLCA,BRCA,COADREAD,GBM,HNSC,KIRC,LUAD,OV,SKCM,STAD,UCEC | SKCM | SKCM | 1 | 1 |
| RAD21 | | | Yes | No | | | | BRCA | BRCA | BLCA,BRCA,COADREAD,GBM,HNSC,KIRC,LGG,LUAD,LUSC,OV,PRAD,SKCM,STAD,THCA,UCEC | | BRCA | | 1 | 0 |
| RAF1 | A | 2 | Yes | Yes | | KIRC | | | THCA | COADREAD,LUSC,SKCM | | | | 0 | 0 |
| RB1 | B | 2 | Yes | No | BLCA,GBM,LUSC,UCEC,SKCM | BLCA,BRCA,GBM,LUSC,OV,PRAD | | GBM | BLCA,BRCA,HNSC,LUAD,LUSC,OV | | | | | 0 | 0 |
| RELA | | | Yes | No | | | | GBM | BLCA,BRCA,HNSC,LGG,LUAD,LUSC,SKCM | BRCA,GBM,LGG,LUAD,OV | | BRCA,LGG,LUAD | | 1 | 0 |
| RET | A | 1 | Yes | No | | | | BRCA,LUAD,LUSC,PRAD,THCA,UCEC | THCA | | BLCA,BRCA,COADREAD,GBM,HNSC,KIRC,LUAD,LUSC,OV,PRAD,SKCM,STAD,THCA,UCEC | | THCA | 0 | 1 |
| SCRIB | | | Yes | No | | | | BRCA,LUAD,LUSC,UCEC | OV | | | | | 0 | 0 |
| SERPINB5 | | | No | No | | HNSC,STAD | GBM,PRAD | COADREAD,LUAD,LUSC,STAD,THCA,UCEC | LUSC | | | | | 0 | 0 |
| SHC1 | D | 2 | Yes | No | | | | GBM,KIRC | BLCA,GBM,LGG,THCA | | | | | 0 | 0 |
| SMAD2 | | | Yes | No | | | | | COADREAD | | | | | 0 | 0 |
| SMAD3 | | | Yes | No | | COADREAD | | | COADREAD,PRAD,STAD | | | | | 0 | 0 |
| SMAD4 | D | 1 | Yes | No | | COADREAD,HNSC,STAD | | | COADREAD,STAD | COADREAD | | COADREAD | | 1 | 0 |
| SMARCA4 | B | 1 | Yes | Yes | LUAD | | | BRCA,UCEC | KIRC,LUAD | BRCA,COADREAD,OV,STAD,UCEC | BRCA,COADREAD,GBM,HNSC,KIRC,LUAD,LUSC,OV,SKCM,UCEC | | KIRC,LUAD | 0 | 1 |
| SNW1 | | | No | No | | | | | PRAD | BLCA,BRCA,COADREAD,GBM,HNSC,KIRC,LGG,LUAD,LUSC,OV,PRAD,SKCM,STAD,THCA,UCEC | | PRAD | | 1 | 0 |
| SOX2 | | | No | No | | | | BLCA,BRCA,GBM,LUAD,LUSC,UCEC | LUSC | COADREAD,GBM,HNSC,LGG,LUAD,LUSC | | LUSC | | 1 | 0 |
| SP1 | | | No | No | | | | | BLCA,BRCA,KIRC,LGG,LUAD,LUSC,OV,PRAD,STAD,UCEC | | | | | 0 | 0 |
| SPRY2 | | | No | No | | PRAD | BLCA,BRCA | GBM | BRCA | | | | | 0 | 0 |
| SPTA1 | | | No | No | LUAD,UCEC | | BRCA,UCEC | GBM,KIRC,STAD | LUAD | | | | | 0 | 0 |
| SPTBN2 | | | No | No | | | GBM,KIRC | BRCA,COADREAD,LUAD,LUSC,PRAD,THCA,UCEC | KIRC | | | | | 0 | 0 |
| SRC | A | 1 | Yes | Yes | | | | | BRCA,COADREAD,LUAD,STAD | | BLCA,BRCA,COADREAD,GBM,HNSC,KIRC,LGG,LUAD,LUSC,OV,SKCM,STAD,THCA | | BRCA,COADREAD,LUAD,STAD | 0 | 1 |
| ST3GAL1 | | | No | No | | | | | OV | | | | | 0 | 0 |
| STAT3 | C | 2 | Yes | No | | | | GBM | COADREAD | LUAD | | | | 0 | 0 |
| STK11 | D | 2 | Yes | No | LUAD | OV | | | LUAD | BLCA,KIRC | | | | 0 | 0 |
| TCF7L2 | | | No | No | | | | | COADREAD | COADREAD,STAD | | COADREAD | | 1 | 0 |
| TERT | B | 2 | Yes | No | | | | BLCA,BRCA,COADREAD,GBM,KIRC,LUAD,LUSC,PRAD,STAD,UCEC | BLCA,LUAD | | BRCA,COADREAD,HNSC,LGG,PRAD,STAD | | | 0 | 0 |
| TFAP2A | | | No | No | | | KIRC | BLCA,BRCA,COADREAD,GBM,LUAD,LUSC,STAD,UCEC | UCEC | SKCM | | | | 0 | 0 |
| TOP1MT | | | No | No | | | | COADREAD,LUSC,STAD | OV | | | | | 0 | 0 |
| TP53 | B | 1 | Yes | No | BLCA,BRCA,COADREAD,GBM,HNSC,LGG,LUAD,LUSC,STAD,UCEC,SKCM | PRAD | | GBM | BLCA,BRCA,COADREAD,GBM,HNSC,KIRC,LGG,LUAD,LUSC,OV,PRAD,SKCM,STAD,UCEC | | BLCA,BRCA,HNSC,SKCM | | BLCA,BRCA,HNSC,SKCM | 0 | 1 |
| TP63 | D | 1 | Yes | No | | | BRCA,KIRC,PRAD | GBM,LUAD,LUSC,THCA,UCEC | HNSC,SKCM | BLCA,HNSC,LUSC | | HNSC | | 1 | 0 |
| TRAF2 | | | No | No | | | | BLCA,BRCA,KIRC,LUAD,UCEC | THCA | BLCA,BRCA,LGG,LUSC,OV,STAD | | | | 0 | 0 |
| TSC1 | | | No | No | BLCA | | GBM | | THCA | COADREAD | | | | 0 | 0 |
| UBC | D | 1 | Yes | Yes | | | | | BRCA,HNSC,LUSC,STAD,UCEC | BLCA,BRCA,COADREAD,GBM,KIRC,LGG,LUAD,LUSC,OV,PRAD,SKCM,STAD,UCEC | | BRCA,LUSC,STAD,UCEC | | 1 | 0 |
| USF1 | | | No | No | | | | | BLCA | | | | | 0 | 0 |
| VHL | C | 1 | Yes | No | KIRC | KIRC | | BLCA | KIRC | BLCA,BRCA,COADREAD,GBM,HNSC,KIRC,LGG,LUAD,LUSC,OV,PRAD,SKCM,STAD,THCA,UCEC | | KIRC | | 1 | 0 |
| WWOX | | | No | No | | BLCA,BRCA,COADREAD,HNSC,LUAD,LUSC,OV,PRAD,STAD,UCEC | | | COADREAD,STAD | | | | | 0 | 0 |
| YWHAZ | D | 2 | Yes | No | | | | LUSC | BLCA,THCA | BRCA,COADREAD,GBM,HNSC,KIRC,LGG,LUAD,LUSC,PRAD | | | | 0 | 0 |
