## Supplementary material for "Probabilistic graph-based model uncovers previously unseen druggable vulnerabilities in major solid cancers": supp table 11

| Cancer | Gene | Source | CNV_Amp | CNV_Del | final_rank | Sample_size |
|---|---|---|---|---|---|---|
| LGG | TP53 | Mutation | | | 1 | 510 |
| LGG | EGFR | ['CNV', 'Mutation'] | 39 | 0 | 2 | 510 |
| LGG | PIK3R1 | Mutation | | | 3 | 510 |
| LGG | PIK3CA | Mutation | | | 4 | 510 |
| LGG | NTRK1 | first neighbor | | | 5 | 510 |
| LGG | RELA | first neighbor | | | 6 | 510 |
| LGG | MYC | first neighbor | | | 7 | 510 |
| LGG | PTPN11 | Mutation | | | 8 | 510 |
| LGG | MAPK1 | first neighbor | | | 9 | 510 |
| LGG | CDKN2A | CNV | 0 | 55 | 10 | 510 |
| LGG | SHC1 | first neighbor | | | 11 | 510 |
| LGG | PTEN | Mutation | | | 12 | 510 |
| LGG | NFKB1 | first neighbor | | | 13 | 510 |
| LGG | HSP90AA1 | first neighbor | | | 14 | 510 |
| LGG | NRAS | Mutation | | | 15 | 510 |
| LGG | MAPK3 | first neighbor | | | 16 | 510 |
| LGG | SP1 | first neighbor | | | 17 | 510 |
| LGG | HRAS | first neighbor | | | 18 | 510 |
| LGG | HSF1 | CNV | 18 | 8 | 19 | 510 |
| LGG | JAK2 | first neighbor | | | 20 | 510 |
| LGG | SMARCA4 | Mutation | | | 21 | 510 |
| LGG | AR | first neighbor | | | 22 | 510 |
| LGG | PIK3R3 | first neighbor | | | 23 | 510 |
| LGG | RB1 | Mutation | | | 24 | 510 |
| LGG | PIK3CD | first neighbor | | | 25 | 510 |
| LGG | STAT1 | first neighbor | | | 26 | 510 |
| LGG | CDK2 | first neighbor | | | 27 | 510 |
| LGG | HDAC1 | first neighbor | | | 28 | 510 |
| LGG | JAK1 | first neighbor | | | 29 | 510 |
| LGG | JAK3 | first neighbor | | | 30 | 510 |
| LGG | IFNA5 | CNV | 0 | 26 | 31 | 510 |
| LGG | CREBBP | first neighbor | | | 32 | 510 |
| LGG | IFNA8 | CNV | 0 | 27 | 33 | 510 |
| LGG | IDH1 | Mutation | | | 34 | 510 |
| LGG | IFNA2 | CNV | 0 | 26 | 35 | 510 |
| LGG | MDM2 | first neighbor | | | 36 | 510 |
| LGG | CDKN1A | first neighbor | | | 37 | 510 |
| LGG | PTPN6 | first neighbor | | | 38 | 510 |
| LGG | IFNA6 | CNV | 0 | 26 | 39 | 510 |
| LGG | PDGFRB | first neighbor | | | 40 | 510 |
| LGG | TYK2 | first neighbor | | | 41 | 510 |
| LGG | HIF1A | first neighbor | | | 42 | 510 |
| LGG | PDGFB | first neighbor | | | 43 | 510 |
| LGG | HSPA8 | first neighbor | | | 44 | 510 |
| LGG | FOS | first neighbor | | | 45 | 510 |
| LGG | EGF | first neighbor | | | 46 | 510 |
| LGG | CDH1 | first neighbor | | | 47 | 510 |
| LGG | NOTCH1 | Mutation | | | 48 | 510 |
| LGG | SMAD3 | first neighbor | | | 49 | 510 |
| LGG | CSF2 | first neighbor | | | 50 | 510 |
| LGG | IGF1R | first neighbor | | | 51 | 510 |
| LGG | IFNA14 | first neighbor | | | 52 | 510 |
| LGG | PDGFA | first neighbor | | | 53 | 510 |

| | | | | | | |
|---|---|---|---|---|---|---|
| LGG | PDGFRA | first neighbor | | | 54 | 510 |
| LGG | IFNA4 | first neighbor | | | 55 | 510 |
| LGG | IL3 | first neighbor | | | 56 | 510 |
| LGG | MET | first neighbor | | | 57 | 510 |
| LGG | IFNA1 | CNV | 0 | 28 | 58 | 510 |
| LGG | CUL4B | Mutation | | | 59 | 510 |
| LGG | IFNA13 | CNV | 0 | 26 | 60 | 510 |
| LGG | NR3C1 | first neighbor | | | 61 | 510 |
| LGG | MAPK8 | first neighbor | | | 62 | 510 |
| LGG | PPARG | first neighbor | | | 63 | 510 |
| LGG | IL5 | first neighbor | | | 64 | 510 |
| LGG | ATRX | Mutation | | | 65 | 510 |
| LGG | SIRT1 | first neighbor | | | 66 | 510 |
| LGG | HCFC1 | Mutation | | | 67 | 510 |
| LGG | EZH2 | first neighbor | | | 68 | 510 |
| LGG | PML | first neighbor | | | 69 | 510 |
| LGG | IFNE | CNV | 0 | 28 | 70 | 510 |
| LGG | SMAD2 | first neighbor | | | 71 | 510 |
| LGG | MAPK14 | first neighbor | | | 72 | 510 |
| LGG | CCND1 | first neighbor | | | 73 | 510 |
| LGG | ERBB4 | first neighbor | | | 74 | 510 |
| LGG | IRS1 | first neighbor | | | 75 | 510 |
| LGG | HDAC3 | first neighbor | | | 76 | 510 |
| LGG | EED | first neighbor | | | 77 | 510 |
| LGG | FOXO3 | first neighbor | | | 78 | 510 |
| LGG | PTK2 | first neighbor | | | 79 | 510 |
| LGG | FOXO1 | first neighbor | | | 80 | 510 |
| LGG | SIRT7 | first neighbor | | | 81 | 510 |
| LGG | CDK4 | first neighbor | | | 82 | 510 |
| LGG | SOCS1 | first neighbor | | | 83 | 510 |
| LGG | USP7 | first neighbor | | | 84 | 510 |
| LGG | STAT2 | first neighbor | | | 85 | 510 |
| LGG | SNW1 | first neighbor | | | 86 | 510 |
| LGG | MAPK9 | first neighbor | | | 87 | 510 |
| LGG | CRKL | first neighbor | | | 88 | 510 |
| LGG | SOCS3 | first neighbor | | | 89 | 510 |
| LGG | HSPA4 | first neighbor | | | 90 | 510 |
| LGG | NCL | first neighbor | | | 91 | 510 |
| LGG | STUB1 | first neighbor | | | 92 | 510 |
| LGG | SMARCB1 | first neighbor | | | 93 | 510 |
| LGG | CDC5L | first neighbor | | | 94 | 510 |
| LGG | PTPN1 | first neighbor | | | 95 | 510 |
| LGG | IL7R | first neighbor | | | 96 | 510 |
| LGG | SMARCC1 | first neighbor | | | 97 | 510 |
| LGG | EPHA2 | first neighbor | | | 98 | 510 |
| LGG | RUNX1 | first neighbor | | | 99 | 510 |
| LGG | ARID1A | Mutation | | | 100 | 510 |
| HNSC | TP53 | Mutation | | | 1 | 517 |
| HNSC | EP300 | Mutation | | | 2 | 517 |
| HNSC | CCND1 | CNV | 120 | 0 | 3 | 517 |
| HNSC | CDKN2A | ['CNV', 'Mutation'] | 0 | 159 | 4 | 517 |
| HNSC | CREBBP | Mutation | | | 5 | 517 |
| HNSC | PIK3CA | ['CNV', 'Mutation'] | 81 | 0 | 6 | 517 |
| HNSC | NOTCH1 | Mutation | | | 7 | 517 |

| | | | | | | |
|---|---|---|---|---|---|---|
| HNSC | MYC | first neighbor | | | 8 | 517 |
| HNSC | NTRK1 | first neighbor | | | 9 | 517 |
| HNSC | ESR1 | first neighbor | | | 10 | 517 |
| HNSC | TP63 | CNV | 83 | 0 | 11 | 517 |
| HNSC | CUL3 | Mutation | | | 12 | 517 |
| HNSC | UBC | first neighbor | | | 13 | 517 |
| HNSC | EGFR | first neighbor | | | 14 | 517 |
| HNSC | CTNNB1 | first neighbor | | | 15 | 517 |
| HNSC | HUWE1 | Mutation | | | 16 | 517 |
| HNSC | BRCA1 | first neighbor | | | 17 | 517 |
| HNSC | RB1 | Mutation | | | 18 | 517 |
| HNSC | HRAS | Mutation | | | 19 | 517 |
| HNSC | RELA | first neighbor | | | 20 | 517 |
| HNSC | SOX2 | CNV | 81 | 0 | 21 | 517 |
| HNSC | HDAC1 | first neighbor | | | 22 | 517 |
| HNSC | MAPK1 | first neighbor | | | 23 | 517 |
| HNSC | SMAD3 | first neighbor | | | 24 | 517 |
| HNSC | GRB2 | first neighbor | | | 25 | 517 |
| HNSC | SRC | first neighbor | | | 26 | 517 |
| HNSC | SMARCA4 | Mutation | | | 27 | 517 |
| HNSC | SP1 | first neighbor | | | 28 | 517 |
| HNSC | CASP8 | Mutation | | | 29 | 517 |
| HNSC | CDK2 | first neighbor | | | 30 | 517 |
| HNSC | FBXW7 | Mutation | | | 31 | 517 |
| HNSC | SMAD4 | Mutation | | | 32 | 517 |
| HNSC | CDKN1A | first neighbor | | | 33 | 517 |
| HNSC | NCOR1 | Mutation | | | 34 | 517 |
| HNSC | AR | first neighbor | | | 35 | 517 |
| HNSC | HSP90AA1 | first neighbor | | | 36 | 517 |
| HNSC | MAPK3 | first neighbor | | | 37 | 517 |
| HNSC | FGF4 | CNV | 123 | 0 | 38 | 517 |
| HNSC | MDM2 | first neighbor | | | 39 | 517 |
| HNSC | FGF3 | CNV | 124 | 0 | 40 | 517 |
| HNSC | STAT3 | first neighbor | | | 41 | 517 |
| HNSC | SMAD2 | first neighbor | | | 42 | 517 |
| HNSC | PTEN | Mutation | | | 43 | 517 |
| HNSC | CDH1 | first neighbor | | | 44 | 517 |
| HNSC | AKT1 | first neighbor | | | 45 | 517 |
| HNSC | ACTL6A | CNV | 79 | 0 | 46 | 517 |
| HNSC | FXR1 | CNV | 79 | 0 | 47 | 517 |
| HNSC | RAC1 | Mutation | | | 48 | 517 |
| HNSC | PIK3R1 | first neighbor | | | 49 | 517 |
| HNSC | CTTN | CNV | 126 | 0 | 50 | 517 |
| HNSC | AP2M1 | CNV | 78 | 0 | 51 | 517 |
| HNSC | FGF19 | CNV | 121 | 0 | 52 | 517 |
| HNSC | NFKB1 | first neighbor | | | 53 | 517 |
| HNSC | HDAC3 | first neighbor | | | 54 | 517 |
| HNSC | PPARG | first neighbor | | | 55 | 517 |
| HNSC | SNW1 | first neighbor | | | 56 | 517 |
| HNSC | HIF1A | first neighbor | | | 57 | 517 |
| HNSC | UBB | first neighbor | | | 58 | 517 |
| HNSC | PIK3R2 | first neighbor | | | 59 | 517 |
| HNSC | CDKN2B | CNV | 0 | 144 | 60 | 517 |
| HNSC | ACTB | first neighbor | | | 61 | 517 |

| | | | | | |
|---|---|---|---|---|---|
| HNSC | HSP90AB1 | first neighbor | | | 62 | 517 |
| HNSC | NPM1 | first neighbor | | | 63 | 517 |
| HNSC | PRKCA | first neighbor | | | 64 | 517 |
| HNSC | GSK3B | first neighbor | | | 65 | 517 |
| HNSC | MYH9 | Mutation | | | 66 | 517 |
| HNSC | RPS27A | first neighbor | | | 67 | 517 |
| HNSC | SHC1 | first neighbor | | | 68 | 517 |
| HNSC | UBE2I | first neighbor | | | 69 | 517 |
| HNSC | ERBB2 | first neighbor | | | 70 | 517 |
| HNSC | HDAC2 | first neighbor | | | 71 | 517 |
| HNSC | MAPK8 | first neighbor | | | 72 | 517 |
| HNSC | TBL1XR1 | CNV | 78 | 0 | 73 | 517 |
| HNSC | NCOR2 | first neighbor | | | 74 | 517 |
| HNSC | TGFBR2 | Mutation | | | 75 | 517 |
| HNSC | FOS | first neighbor | | | 76 | 517 |
| HNSC | RHOA | Mutation | | | 77 | 517 |
| HNSC | KAT2B | first neighbor | | | 78 | 517 |
| HNSC | RXRA | first neighbor | | | 79 | 517 |
| HNSC | STAT1 | first neighbor | | | 80 | 517 |
| HNSC | EWSR1 | first neighbor | | | 81 | 517 |
| HNSC | PML | first neighbor | | | 82 | 517 |
| HNSC | H3C1\|H3C | first neighbor | | | 83 | 517 |
| HNSC | NFE2L2 | Mutation | | | 84 | 517 |
| HNSC | PIK3CB | first neighbor | | | 85 | 517 |
| HNSC | ATM | first neighbor | | | 86 | 517 |
| HNSC | SIRT1 | first neighbor | | | 87 | 517 |
| HNSC | EZH2 | first neighbor | | | 88 | 517 |
| HNSC | DCUN1D1 | CNV | 80 | 0 | 89 | 517 |
| HNSC | UBA52 | first neighbor | | | 90 | 517 |
| HNSC | CBL | first neighbor | | | 91 | 517 |
| HNSC | NCOA3 | first neighbor | | | 92 | 517 |
| HNSC | PARP1 | first neighbor | | | 93 | 517 |
| HNSC | PLCG1 | first neighbor | | | 94 | 517 |
| HNSC | PIK3R3 | first neighbor | | | 95 | 517 |
| HNSC | HLA-B | Mutation | | | 96 | 517 |
| HNSC | FADD | CNV | 127 | 0 | 97 | 517 |
| HNSC | ABL1 | first neighbor | | | 98 | 517 |
| HNSC | CASP3 | first neighbor | | | 99 | 517 |
| HNSC | SIN3A | first neighbor | | | 100 | 517 |
| THCA | BRAF | ['Mutation', 'SV'] | | | 1 | 494 |
| THCA | HRAS | Mutation | | | 2 | 494 |
| THCA | NRAS | Mutation | | | 3 | 494 |
| THCA | AKT1 | Mutation | | | 4 | 494 |
| THCA | KRAS | first neighbor | | | 5 | 494 |
| THCA | MAPK1 | first neighbor | | | 6 | 494 |
| THCA | TRAF2 | CNV | 1 | 4 | 7 | 494 |
| THCA | MAPK3 | first neighbor | | | 8 | 494 |
| THCA | ATM | Mutation | | | 9 | 494 |
| THCA | PIK3R1 | first neighbor | | | 10 | 494 |
| THCA | MAPK14 | first neighbor | | | 11 | 494 |
| THCA | TSC1 | CNV | 1 | 4 | 12 | 494 |
| THCA | MAP3K3 | Mutation | | | 13 | 494 |
| THCA | YWHAZ | first neighbor | | | 14 | 494 |
| THCA | SHC1 | first neighbor | | | 15 | 494 |

| | | | | | | |
|---|---|---|---|---|---|---|
| THCA | IKBKG | first neighbor | | | 16 | 494 |
| THCA | NOTCH1 | CNV | 2 | 4 | 17 | 494 |
| THCA | GSK3B | first neighbor | | | 18 | 494 |
| THCA | AKT2 | first neighbor | | | 19 | 494 |
| THCA | RET | SV | | | 20 | 494 |
| THCA | AKT3 | first neighbor | | | 21 | 494 |
| THCA | RAF1 | first neighbor | | | 22 | 494 |
| THCA | MDM2 | first neighbor | | | 23 | 494 |
| THCA | YWHAG | first neighbor | | | 24 | 494 |
| THCA | CDC37 | first neighbor | | | 25 | 494 |
| THCA | H3C1\|H3C | first neighbor | | | 26 | 494 |
| THCA | HDAC1 | first neighbor | | | 27 | 494 |
| THCA | PPP2CB | first neighbor | | | 28 | 494 |
| THCA | CREB1 | first neighbor | | | 29 | 494 |
| THCA | CBL | first neighbor | | | 30 | 494 |
| THCA | YWHAE | first neighbor | | | 31 | 494 |
| THCA | MAPK9 | first neighbor | | | 32 | 494 |
| THCA | WDR5 | CNV | 1 | 4 | 33 | 494 |
| THCA | IKBKB | first neighbor | | | 34 | 494 |
| THCA | PRKAA1 | first neighbor | | | 35 | 494 |
| THCA | YWHAB | first neighbor | | | 36 | 494 |
| THCA | RXRA | CNV | 1 | 4 | 37 | 494 |
| THCA | CALM3 | first neighbor | | | 38 | 494 |
| THCA | CHEK2 | Mutation | | | 39 | 494 |
| THCA | RB1 | first neighbor | | | 40 | 494 |
| THCA | MAP3K1 | first neighbor | | | 41 | 494 |
| THCA | CALM1 | first neighbor | | | 42 | 494 |
| THCA | CALM2 | first neighbor | | | 43 | 494 |
| THCA | PTK2 | first neighbor | | | 44 | 494 |
| THCA | CHUK | first neighbor | | | 45 | 494 |
| THCA | VHL | first neighbor | | | 46 | 494 |
| THCA | PRKDC | first neighbor | | | 47 | 494 |
| THCA | GNAI2 | first neighbor | | | 48 | 494 |
| THCA | GNB2 | first neighbor | | | 49 | 494 |
| THCA | PKM | first neighbor | | | 50 | 494 |
| THCA | FOXO3 | first neighbor | | | 51 | 494 |
| THCA | IL7R | Mutation | | | 52 | 494 |
| THCA | USP7 | first neighbor | | | 53 | 494 |
| THCA | KAT2B | first neighbor | | | 54 | 494 |
| THCA | RPS6KA3 | first neighbor | | | 55 | 494 |
| THCA | ATR | first neighbor | | | 56 | 494 |
| THCA | POU5F1 | first neighbor | | | 57 | 494 |
| THCA | CCDC6 | ['CNV', 'SV'] | 0 | 6 | 58 | 494 |
| THCA | FBXW7 | first neighbor | | | 59 | 494 |
| THCA | RPS6KA1 | first neighbor | | | 60 | 494 |
| THCA | CDKN2A | first neighbor | | | 61 | 494 |
| THCA | IRS1 | first neighbor | | | 62 | 494 |
| THCA | TBC1D7 | Mutation | | | 63 | 494 |
| THCA | GNAI3 | first neighbor | | | 64 | 494 |
| THCA | PDPK1 | first neighbor | | | 65 | 494 |
| THCA | RPTOR | first neighbor | | | 66 | 494 |
| THCA | PIK3R5 | Mutation | | | 67 | 494 |
| THCA | PIK3CG | first neighbor | | | 68 | 494 |
| THCA | GNB4 | first neighbor | | | 69 | 494 |

| | | | | | | |
|---|---|---|---|---|---|---|
| THCA | MAP3K5 | first neighbor | | | 70 | 494 |
| THCA | CDKN1B | first neighbor | | | 71 | 494 |
| THCA | CAMK4 | first neighbor | | | 72 | 494 |
| THCA | TRAF3 | first neighbor | | | 73 | 494 |
| THCA | GNB3 | first neighbor | | | 74 | 494 |
| THCA | DLC1 | Mutation | | | 75 | 494 |
| THCA | CACNA1B | CNV | 3 | 5 | 76 | 494 |
| THCA | BIRC3 | first neighbor | | | 77 | 494 |
| THCA | BCL10 | first neighbor | | | 78 | 494 |
| THCA | SEC16A | CNV | 2 | 4 | 79 | 494 |
| THCA | ODF2 | CNV | 1 | 4 | 80 | 494 |
| THCA | FRS2 | first neighbor | | | 81 | 494 |
| THCA | RASGRF1 | first neighbor | | | 82 | 494 |
| THCA | NANOG | first neighbor | | | 83 | 494 |
| THCA | PIK3R6 | first neighbor | | | 84 | 494 |
| THCA | SPTA1 | Mutation | | | 85 | 494 |
| THCA | SMAD1 | first neighbor | | | 86 | 494 |
| THCA | TGFB1 | first neighbor | | | 87 | 494 |
| THCA | BIRC2 | first neighbor | | | 88 | 494 |
| THCA | NEDD4L | first neighbor | | | 89 | 494 |
| THCA | NR4A1 | first neighbor | | | 90 | 494 |
| THCA | GNG5 | first neighbor | | | 91 | 494 |
| THCA | GNG4 | first neighbor | | | 92 | 494 |
| THCA | TERF1 | first neighbor | | | 93 | 494 |
| THCA | BCL6 | first neighbor | | | 94 | 494 |
| THCA | RPS6KA6 | first neighbor | | | 95 | 494 |
| THCA | RPS6KA2 | first neighbor | | | 96 | 494 |
| THCA | USP9X | Mutation | | | 97 | 494 |
| THCA | VBP1 | CNV | 2 | 3 | 98 | 494 |
| THCA | USP20 | CNV | 1 | 4 | 99 | 494 |
| THCA | FOXO4 | first neighbor | | | 100 | 494 |
| PRAD | TP53 | Mutation | | | 1 | 489 |
| PRAD | ERG | ['CNV', 'SV'] | 1 | 49 | 2 | 489 |
| PRAD | PTEN | ['CNV', 'Mutation'] | 1 | 85 | 3 | 489 |
| PRAD | CTNNB1 | Mutation | | | 4 | 489 |
| PRAD | ATM | Mutation | | | 5 | 489 |
| PRAD | MYC | first neighbor | | | 6 | 489 |
| PRAD | HSPA8 | Mutation | | | 7 | 489 |
| PRAD | ESR1 | first neighbor | | | 8 | 489 |
| PRAD | EP300 | first neighbor | | | 9 | 489 |
| PRAD | SNW1 | Mutation | | | 10 | 489 |
| PRAD | PIK3CA | Mutation | | | 11 | 489 |
| PRAD | SP1 | first neighbor | | | 12 | 489 |
| PRAD | HRAS | Mutation | | | 13 | 489 |
| PRAD | HDAC1 | first neighbor | | | 14 | 489 |
| PRAD | AR | first neighbor | | | 15 | 489 |
| PRAD | FOXA1 | Mutation | | | 16 | 489 |
| PRAD | MAPK3 | first neighbor | | | 17 | 489 |
| PRAD | GSK3B | first neighbor | | | 18 | 489 |
| PRAD | SMAD3 | first neighbor | | | 19 | 489 |
| PRAD | CDKN1B | Mutation | | | 20 | 489 |
| PRAD | APC | Mutation | | | 21 | 489 |
| PRAD | UBB | first neighbor | | | 22 | 489 |
| PRAD | FOS | first neighbor | | | 23 | 489 |

| | | | | | | |
|---|---|---|---|---|---|---|
| PRAD | YWHAZ | first neighbor | | | 24 | 489 |
| PRAD | ETS2 | CNV | 1 | 46 | 25 | 489 |
| PRAD | RB1 | first neighbor | | | 26 | 489 |
| PRAD | ABL1 | first neighbor | | | 27 | 489 |
| PRAD | SPOP | Mutation | | | 28 | 489 |
| PRAD | TMPRSS2 | ['CNV', 'SV'] | 1 | 58 | 29 | 489 |
| PRAD | TP53BP1 | Mutation | | | 30 | 489 |
| PRAD | HIF1A | first neighbor | | | 31 | 489 |
| PRAD | ETS1 | first neighbor | | | 32 | 489 |
| PRAD | KPNA3 | CNV | 1 | 48 | 33 | 489 |
| PRAD | CDK4 | first neighbor | | | 34 | 489 |
| PRAD | PARP1 | first neighbor | | | 35 | 489 |
| PRAD | CBL | first neighbor | | | 36 | 489 |
| PRAD | RPA2 | first neighbor | | | 37 | 489 |
| PRAD | KAT2B | first neighbor | | | 38 | 489 |
| PRAD | ATR | first neighbor | | | 39 | 489 |
| PRAD | MAPK8 | first neighbor | | | 40 | 489 |
| PRAD | ESR2 | first neighbor | | | 41 | 489 |
| PRAD | NCOA3 | first neighbor | | | 42 | 489 |
| PRAD | USP7 | first neighbor | | | 43 | 489 |
| PRAD | SKP1 | first neighbor | | | 44 | 489 |
| PRAD | NOTCH1 | first neighbor | | | 45 | 489 |
| PRAD | SUMO1 | first neighbor | | | 46 | 489 |
| PRAD | FLNA | first neighbor | | | 47 | 489 |
| PRAD | BMI1 | first neighbor | | | 48 | 489 |
| PRAD | BTRC | first neighbor | | | 49 | 489 |
| PRAD | DDB1 | first neighbor | | | 50 | 489 |
| PRAD | SUMO2 | first neighbor | | | 51 | 489 |
| PRAD | BRCA2 | Mutation | | | 52 | 489 |
| PRAD | PRKDC | first neighbor | | | 53 | 489 |
| PRAD | BRAF | Mutation | | | 54 | 489 |
| PRAD | FBXW7 | first neighbor | | | 55 | 489 |
| PRAD | PRKCZ | first neighbor | | | 56 | 489 |
| PRAD | WDR5 | first neighbor | | | 57 | 489 |
| PRAD | CASP8 | first neighbor | | | 58 | 489 |
| PRAD | PTK2 | first neighbor | | | 59 | 489 |
| PRAD | EZH2 | first neighbor | | | 60 | 489 |
| PRAD | PGR | first neighbor | | | 61 | 489 |
| PRAD | JAK2 | first neighbor | | | 62 | 489 |
| PRAD | FOXO3 | first neighbor | | | 63 | 489 |
| PRAD | SMARCA4 | first neighbor | | | 64 | 489 |
| PRAD | PIAS1 | first neighbor | | | 65 | 489 |
| PRAD | PRKCD | first neighbor | | | 66 | 489 |
| PRAD | TP73 | first neighbor | | | 67 | 489 |
| PRAD | RAD51 | first neighbor | | | 68 | 489 |
| PRAD | PRKAA1 | first neighbor | | | 69 | 489 |
| PRAD | UBR5 | first neighbor | | | 70 | 489 |
| PRAD | COPS6 | first neighbor | | | 71 | 489 |
| PRAD | NCOA1 | first neighbor | | | 72 | 489 |
| PRAD | LYN | first neighbor | | | 73 | 489 |
| PRAD | DAXX | first neighbor | | | 74 | 489 |
| PRAD | IGF1R | first neighbor | | | 75 | 489 |
| PRAD | NANOG | first neighbor | | | 76 | 489 |
| PRAD | TERT | first neighbor | | | 77 | 489 |

| | | | | | | |
|---|---|---|---|---|---|---|
| PRAD | MYB | first neighbor | | | 78 | 489 |
| PRAD | NCOA2 | first neighbor | | | 79 | 489 |
| PRAD | TCF7L2 | first neighbor | | | 80 | 489 |
| PRAD | IRS1 | first neighbor | | | 81 | 489 |
| PRAD | USF1 | first neighbor | | | 82 | 489 |
| PRAD | RBX1 | first neighbor | | | 83 | 489 |
| PRAD | DDX5 | first neighbor | | | 84 | 489 |
| PRAD | TGFBR2 | first neighbor | | | 85 | 489 |
| PRAD | CAMK2A | first neighbor | | | 86 | 489 |
| PRAD | PDGFRB | first neighbor | | | 87 | 489 |
| PRAD | UBE3A | first neighbor | | | 88 | 489 |
| PRAD | MAP2K1 | first neighbor | | | 89 | 489 |
| PRAD | LAMA3 | Mutation | | | 90 | 489 |
| PRAD | BLM | first neighbor | | | 91 | 489 |
| PRAD | STK11 | first neighbor | | | 92 | 489 |
| PRAD | MMP2 | first neighbor | | | 93 | 489 |
| PRAD | FGFR2 | first neighbor | | | 94 | 489 |
| PRAD | CSNK1A1 | first neighbor | | | 95 | 489 |
| PRAD | CCNA2 | first neighbor | | | 96 | 489 |
| PRAD | WT1 | first neighbor | | | 97 | 489 |
| PRAD | DNMT1 | first neighbor | | | 98 | 489 |
| PRAD | UBE2D2 | first neighbor | | | 99 | 489 |
| PRAD | CD44 | first neighbor | | | 100 | 489 |
| SKCM | TP53 | Mutation | | | 1 | 363 |
| SKCM | NRAS | Mutation | | | 2 | 363 |
| SKCM | CDKN2A | ['CNV', 'Mutation'] | 1 | 112 | 3 | 363 |
| SKCM | BRAF | Mutation | | | 4 | 363 |
| SKCM | EGFR | first neighbor | | | 5 | 363 |
| SKCM | PTEN | ['CNV', 'Mutation'] | 0 | 28 | 6 | 363 |
| SKCM | RAC1 | Mutation | | | 7 | 363 |
| SKCM | PLCB4 | Mutation | | | 8 | 363 |
| SKCM | TP63 | Mutation | | | 9 | 363 |
| SKCM | CCND1 | CNV | 23 | 1 | 10 | 363 |
| SKCM | GRB2 | first neighbor | | | 11 | 363 |
| SKCM | RELA | first neighbor | | | 12 | 363 |
| SKCM | MAPK3 | first neighbor | | | 13 | 363 |
| SKCM | MAPK1 | first neighbor | | | 14 | 363 |
| SKCM | EP300 | first neighbor | | | 15 | 363 |
| SKCM | IFNA2 | CNV | 1 | 34 | 16 | 363 |
| SKCM | IFNA5 | CNV | 1 | 29 | 17 | 363 |
| SKCM | IFNA14 | CNV | 1 | 30 | 18 | 363 |
| SKCM | CTNNB1 | first neighbor | | | 19 | 363 |
| SKCM | NFKB1 | first neighbor | | | 20 | 363 |
| SKCM | IFNA21 | CNV | 1 | 28 | 21 | 363 |
| SKCM | IFNB1 | CNV | 1 | 27 | 22 | 363 |
| SKCM | IFNA8 | CNV | 1 | 34 | 23 | 363 |
| SKCM | IFNA4 | CNV | 1 | 29 | 24 | 363 |
| SKCM | HRAS | first neighbor | | | 25 | 363 |
| SKCM | IFNA6 | CNV | 1 | 32 | 26 | 363 |
| SKCM | CDKN1A | first neighbor | | | 27 | 363 |
| SKCM | JAK2 | first neighbor | | | 28 | 363 |
| SKCM | JAK1 | first neighbor | | | 29 | 363 |
| SKCM | PIK3R2 | first neighbor | | | 30 | 363 |
| SKCM | DCC | Mutation | | | 31 | 363 |

| | | | | | | |
|---|---|---|---|---|---|---|
| SKCM | STAT1 | first neighbor | | | 32 | 363 |
| SKCM | PIK3CA | first neighbor | | | 33 | 363 |
| SKCM | IFNA7 | CNV | 1 | 29 | 34 | 363 |
| SKCM | IFNA17 | CNV | 1 | 30 | 35 | 363 |
| SKCM | PLCG1 | first neighbor | | | 36 | 363 |
| SKCM | IFNA10 | CNV | 1 | 29 | 37 | 363 |
| SKCM | KRAS | first neighbor | | | 38 | 363 |
| SKCM | PTPN11 | first neighbor | | | 39 | 363 |
| SKCM | CREBBP | first neighbor | | | 40 | 363 |
| SKCM | IFNA16 | CNV | 1 | 29 | 41 | 363 |
| SKCM | JAK3 | first neighbor | | | 42 | 363 |
| SKCM | PIK3CB | first neighbor | | | 43 | 363 |
| SKCM | MAPK8 | first neighbor | | | 44 | 363 |
| SKCM | MAP2K1 | Mutation | | | 45 | 363 |
| SKCM | PIK3R3 | first neighbor | | | 46 | 363 |
| SKCM | IFNA1 | CNV | 1 | 34 | 47 | 363 |
| SKCM | MAPK14 | first neighbor | | | 48 | 363 |
| SKCM | ERBB2 | first neighbor | | | 49 | 363 |
| SKCM | IFNA13 | CNV | 1 | 32 | 50 | 363 |
| SKCM | NF1 | Mutation | | | 51 | 363 |
| SKCM | GSK3B | first neighbor | | | 52 | 363 |
| SKCM | IFNE | CNV | 1 | 36 | 53 | 363 |
| SKCM | SMAD3 | first neighbor | | | 54 | 363 |
| SKCM | TYK2 | first neighbor | | | 55 | 363 |
| SKCM | YWHAZ | first neighbor | | | 56 | 363 |
| SKCM | PTPN6 | first neighbor | | | 57 | 363 |
| SKCM | IFNW1 | CNV | 1 | 27 | 58 | 363 |
| SKCM | FGF3 | CNV | 21 | 1 | 59 | 363 |
| SKCM | FOS | first neighbor | | | 60 | 363 |
| SKCM | HIF1A | first neighbor | | | 61 | 363 |
| SKCM | TERT | CNV | 19 | 3 | 62 | 363 |
| SKCM | CDKN2B | CNV | 2 | 108 | 63 | 363 |
| SKCM | TRRAP | Mutation | | | 64 | 363 |
| SKCM | PDGFRB | first neighbor | | | 65 | 363 |
| SKCM | FGF19 | CNV | 20 | 1 | 66 | 363 |
| SKCM | MAPK9 | first neighbor | | | 67 | 363 |
| SKCM | FGF4 | CNV | 20 | 1 | 68 | 363 |
| SKCM | CREB1 | first neighbor | | | 69 | 363 |
| SKCM | DSP | Mutation | | | 70 | 363 |
| SKCM | PTK2 | first neighbor | | | 71 | 363 |
| SKCM | STAT2 | first neighbor | | | 72 | 363 |
| SKCM | PDGFRA | first neighbor | | | 73 | 363 |
| SKCM | YWHAB | first neighbor | | | 74 | 363 |
| SKCM | PRKCD | first neighbor | | | 75 | 363 |
| SKCM | RPS27A | first neighbor | | | 76 | 363 |
| SKCM | MAPK13 | first neighbor | | | 77 | 363 |
| SKCM | SOCS3 | first neighbor | | | 78 | 363 |
| SKCM | SOCS1 | first neighbor | | | 79 | 363 |
| SKCM | SOS1 | first neighbor | | | 80 | 363 |
| SKCM | MAPK10 | first neighbor | | | 81 | 363 |
| SKCM | APOB | Mutation | | | 82 | 363 |
| SKCM | TP73 | first neighbor | | | 83 | 363 |
| SKCM | ACTB | first neighbor | | | 84 | 363 |
| SKCM | CALM3 | first neighbor | | | 85 | 363 |

| | | | | | | |
|---|---|---|---|---|---|---|
| SKCM | KAT2B | first neighbor | | | 86 | 363 |
| SKCM | MET | first neighbor | | | 87 | 363 |
| SKCM | MAPK11 | first neighbor | | | 88 | 363 |
| SKCM | PTPN1 | first neighbor | | | 89 | 363 |
| SKCM | MITF | CNV | 24 | 0 | 90 | 363 |
| SKCM | IFNGR1 | first neighbor | | | 91 | 363 |
| SKCM | PRKCE | first neighbor | | | 92 | 363 |
| SKCM | PRKCZ | first neighbor | | | 93 | 363 |
| SKCM | IGF1R | first neighbor | | | 94 | 363 |
| SKCM | CSF2RB | first neighbor | | | 95 | 363 |
| SKCM | PML | first neighbor | | | 96 | 363 |
| SKCM | NFATC1 | first neighbor | | | 97 | 363 |
| SKCM | IRF3 | first neighbor | | | 98 | 363 |
| SKCM | PTPRB | Mutation | | | 99 | 363 |
| SKCM | FOXO3 | first neighbor | | | 100 | 363 |
| KIRC | VHL | Mutation | | | 1 | 468 |
| KIRC | TP53 | Mutation | | | 2 | 468 |
| KIRC | PTEN | Mutation | | | 3 | 468 |
| KIRC | ATM | Mutation | | | 4 | 468 |
| KIRC | PBRM1 | Mutation | | | 5 | 468 |
| KIRC | ESR1 | first neighbor | | | 6 | 468 |
| KIRC | PIK3CA | Mutation | | | 7 | 468 |
| KIRC | FGFR4 | CNV | 38 | 0 | 8 | 468 |
| KIRC | FLT4 | CNV | 37 | 0 | 9 | 468 |
| KIRC | SMARCA4 | Mutation | | | 10 | 468 |
| KIRC | ACTB | first neighbor | | | 11 | 468 |
| KIRC | DBN1 | CNV | 38 | 0 | 12 | 468 |
| KIRC | CTNNB1 | first neighbor | | | 13 | 468 |
| KIRC | IGF1R | first neighbor | | | 14 | 468 |
| KIRC | MTOR | Mutation | | | 15 | 468 |
| KIRC | EP300 | first neighbor | | | 16 | 468 |
| KIRC | FLT1 | Mutation | | | 17 | 468 |
| KIRC | HSP90AA1 | first neighbor | | | 18 | 468 |
| KIRC | HDAC1 | first neighbor | | | 19 | 468 |
| KIRC | PDLIM7 | CNV | 38 | 0 | 20 | 468 |
| KIRC | SIRT7 | first neighbor | | | 21 | 468 |
| KIRC | HIF1A | first neighbor | | | 22 | 468 |
| KIRC | AR | first neighbor | | | 23 | 468 |
| KIRC | CSNK2A1 | first neighbor | | | 24 | 468 |
| KIRC | SHC1 | first neighbor | | | 25 | 468 |
| KIRC | NXF1 | first neighbor | | | 26 | 468 |
| KIRC | SPTAN1 | Mutation | | | 27 | 468 |
| KIRC | HSP90AB1 | first neighbor | | | 28 | 468 |
| KIRC | ACTG1 | first neighbor | | | 29 | 468 |
| KIRC | FLNA | first neighbor | | | 30 | 468 |
| KIRC | NF2 | Mutation | | | 31 | 468 |
| KIRC | H3C1\|H3C | first neighbor | | | 32 | 468 |
| KIRC | RNF2 | first neighbor | | | 33 | 468 |
| KIRC | OBSL1 | first neighbor | | | 34 | 468 |
| KIRC | SNW1 | first neighbor | | | 35 | 468 |
| KIRC | MOV10 | first neighbor | | | 36 | 468 |
| KIRC | MYH9 | first neighbor | | | 37 | 468 |
| KIRC | PIK3R3 | first neighbor | | | 38 | 468 |
| KIRC | SMAD3 | first neighbor | | | 39 | 468 |

| | | | | | | |
|---|---|---|---|---|---|---|
| KIRC | CBL | first neighbor | | | 40 | 468 |
| KIRC | E2F1 | first neighbor | | | 41 | 468 |
| KIRC | PARP1 | first neighbor | | | 42 | 468 |
| KIRC | LIMA1 | first neighbor | | | 43 | 468 |
| KIRC | PIK3CD | first neighbor | | | 44 | 468 |
| KIRC | STAT1 | first neighbor | | | 45 | 468 |
| KIRC | PPARG | first neighbor | | | 46 | 468 |
| KIRC | MAX | Mutation | | | 47 | 468 |
| KIRC | H4-16\|H4C | first neighbor | | | 48 | 468 |
| KIRC | RPA1 | first neighbor | | | 49 | 468 |
| KIRC | HUWE1 | first neighbor | | | 50 | 468 |
| KIRC | RPA2 | first neighbor | | | 51 | 468 |
| KIRC | ABL1 | first neighbor | | | 52 | 468 |
| KIRC | H3-4 | first neighbor | | | 53 | 468 |
| KIRC | PTK2 | first neighbor | | | 54 | 468 |
| KIRC | HSPA8 | first neighbor | | | 55 | 468 |
| KIRC | SMARCC1 | first neighbor | | | 56 | 468 |
| KIRC | CRK | first neighbor | | | 57 | 468 |
| KIRC | UIMC1 | CNV | 38 | 0 | 58 | 468 |
| KIRC | FAF2 | CNV | 37 | 0 | 59 | 468 |
| KIRC | SMARCC2 | first neighbor | | | 60 | 468 |
| KIRC | GSK3B | first neighbor | | | 61 | 468 |
| KIRC | EED | first neighbor | | | 62 | 468 |
| KIRC | KAT2B | first neighbor | | | 63 | 468 |
| KIRC | HNRNPAB | CNV | 37 | 0 | 64 | 468 |
| KIRC | SMARCA2 | first neighbor | | | 65 | 468 |
| KIRC | DDB1 | first neighbor | | | 66 | 468 |
| KIRC | CDC42 | first neighbor | | | 67 | 468 |
| KIRC | SPTBN1 | first neighbor | | | 68 | 468 |
| KIRC | SMURF1 | first neighbor | | | 69 | 468 |
| KIRC | HSPA4 | first neighbor | | | 70 | 468 |
| KIRC | ARID1A | Mutation | | | 71 | 468 |
| KIRC | POLR2A | first neighbor | | | 72 | 468 |
| KIRC | NOTCH1 | first neighbor | | | 73 | 468 |
| KIRC | PRKCZ | first neighbor | | | 74 | 468 |
| KIRC | ATR | first neighbor | | | 75 | 468 |
| KIRC | SIN3A | first neighbor | | | 76 | 468 |
| KIRC | SMARCB1 | first neighbor | | | 77 | 468 |
| KIRC | CAPZA2 | first neighbor | | | 78 | 468 |
| KIRC | TERF1 | first neighbor | | | 79 | 468 |
| KIRC | ROCK1 | Mutation | | | 80 | 468 |
| KIRC | BAP1 | Mutation | | | 81 | 468 |
| KIRC | IKBKB | first neighbor | | | 82 | 468 |
| KIRC | MYO1C | first neighbor | | | 83 | 468 |
| KIRC | EPAS1 | first neighbor | | | 84 | 468 |
| KIRC | PRPF8 | Mutation | | | 85 | 468 |
| KIRC | PKM | first neighbor | | | 86 | 468 |
| KIRC | SHMT2 | first neighbor | | | 87 | 468 |
| KIRC | NR3C1 | first neighbor | | | 88 | 468 |
| KIRC | SQSTM1 | first neighbor | | | 89 | 468 |
| KIRC | UBE2D1 | first neighbor | | | 90 | 468 |
| KIRC | SKP2 | first neighbor | | | 91 | 468 |
| KIRC | WDR5 | first neighbor | | | 92 | 468 |
| KIRC | FBXW7 | first neighbor | | | 93 | 468 |

| | | | | | | |
|---|---|---|---|---|---|---|
| KIRC | UBE2D3 | first neighbor | | | 94 | 468 |
| KIRC | FOXO3 | first neighbor | | | 95 | 468 |
| KIRC | IRS1 | first neighbor | | | 96 | 468 |
| KIRC | VEGFA | first neighbor | | | 97 | 468 |
| KIRC | AKT2 | first neighbor | | | 98 | 468 |
| KIRC | FOXO1 | first neighbor | | | 99 | 468 |
| KIRC | NSD1 | CNV | 38 | 0 | 100 | 468 |
| COADREAD | TP53 | Mutation | | | 1 | 581 |
| COADREAD | KRAS | Mutation | | | 2 | 581 |
| COADREAD | APC | Mutation | | | 3 | 581 |
| COADREAD | PIK3CA | Mutation | | | 4 | 581 |
| COADREAD | CTNNB1 | Mutation | | | 5 | 581 |
| COADREAD | PIK3R1 | Mutation | | | 6 | 581 |
| COADREAD | NRAS | Mutation | | | 7 | 581 |
| COADREAD | ATM | Mutation | | | 8 | 581 |
| COADREAD | SMAD4 | Mutation | | | 9 | 581 |
| COADREAD | SRC | first neighbor | | | 10 | 581 |
| COADREAD | PTEN | Mutation | | | 11 | 581 |
| COADREAD | ESR1 | first neighbor | | | 12 | 581 |
| COADREAD | FBXW7 | Mutation | | | 13 | 581 |
| COADREAD | WWOX | CNV | 3 | 65 | 14 | 581 |
| COADREAD | PIK3CB | first neighbor | | | 15 | 581 |
| COADREAD | ELAVL1 | first neighbor | | | 16 | 581 |
| COADREAD | MAPK1 | first neighbor | | | 17 | 581 |
| COADREAD | PRKDC | Mutation | | | 18 | 581 |
| COADREAD | TCF7L2 | Mutation | | | 19 | 581 |
| COADREAD | SMAD3 | first neighbor | | | 20 | 581 |
| COADREAD | HSP90AA1 | first neighbor | | | 21 | 581 |
| COADREAD | AKT1 | first neighbor | | | 22 | 581 |
| COADREAD | CDH1 | first neighbor | | | 23 | 581 |
| COADREAD | ITCH | CNV | 51 | 0 | 24 | 581 |
| COADREAD | PIK3R3 | first neighbor | | | 25 | 581 |
| COADREAD | SHC1 | first neighbor | | | 26 | 581 |
| COADREAD | FYN | first neighbor | | | 27 | 581 |
| COADREAD | CDK2 | first neighbor | | | 28 | 581 |
| COADREAD | PIK3CD | first neighbor | | | 29 | 581 |
| COADREAD | SMAD2 | first neighbor | | | 30 | 581 |
| COADREAD | AR | first neighbor | | | 31 | 581 |
| COADREAD | CSNK2A1 | first neighbor | | | 32 | 581 |
| COADREAD | GSK3B | first neighbor | | | 33 | 581 |
| COADREAD | CREBBP | first neighbor | | | 34 | 581 |
| COADREAD | JUN | first neighbor | | | 35 | 581 |
| COADREAD | YWHAZ | first neighbor | | | 36 | 581 |
| COADREAD | HSP90AB1 | first neighbor | | | 37 | 581 |
| COADREAD | RPS27A | first neighbor | | | 38 | 581 |
| COADREAD | HCK | CNV | 54 | 0 | 39 | 581 |
| COADREAD | HLA-B | Mutation | | | 40 | 581 |
| COADREAD | BRAF | Mutation | | | 41 | 581 |
| COADREAD | ABL1 | first neighbor | | | 42 | 581 |
| COADREAD | MAPK8 | first neighbor | | | 43 | 581 |
| COADREAD | BCL2L1 | CNV | 50 | 0 | 44 | 581 |
| COADREAD | ERBB3 | first neighbor | | | 45 | 581 |
| COADREAD | UBE2I | first neighbor | | | 46 | 581 |
| COADREAD | B2M | Mutation | | | 47 | 581 |

| | | | | | | |
|---|---|---|---|---|---|---|
| COADREAD | MAPK14 | first neighbor | | | 48 | 581 |
| COADREAD | CDC37 | first neighbor | | | 49 | 581 |
| COADREAD | IGF1R | first neighbor | | | 50 | 581 |
| COADREAD | CRK | first neighbor | | | 51 | 581 |
| COADREAD | MET | first neighbor | | | 52 | 581 |
| COADREAD | ERBB4 | first neighbor | | | 53 | 581 |
| COADREAD | CREB1 | first neighbor | | | 54 | 581 |
| COADREAD | HIF1A | first neighbor | | | 55 | 581 |
| COADREAD | PTK2 | first neighbor | | | 56 | 581 |
| COADREAD | IKBKB | first neighbor | | | 57 | 581 |
| COADREAD | NPM1 | first neighbor | | | 58 | 581 |
| COADREAD | EWSR1 | first neighbor | | | 59 | 581 |
| COADREAD | FOS | first neighbor | | | 60 | 581 |
| COADREAD | IRS1 | first neighbor | | | 61 | 581 |
| COADREAD | PARP1 | first neighbor | | | 62 | 581 |
| COADREAD | BCL2 | first neighbor | | | 63 | 581 |
| COADREAD | SMAD1 | first neighbor | | | 64 | 581 |
| COADREAD | FGFR2 | first neighbor | | | 65 | 581 |
| COADREAD | RBM39 | CNV | 49 | 1 | 66 | 581 |
| COADREAD | CDC42 | first neighbor | | | 67 | 581 |
| COADREAD | BMPR2 | Mutation | | | 68 | 581 |
| COADREAD | NOTCH1 | first neighbor | | | 69 | 581 |
| COADREAD | NCK1 | first neighbor | | | 70 | 581 |
| COADREAD | BTRC | first neighbor | | | 71 | 581 |
| COADREAD | PCBP1 | Mutation | | | 72 | 581 |
| COADREAD | CDKN2A | first neighbor | | | 73 | 581 |
| COADREAD | MAPK9 | first neighbor | | | 74 | 581 |
| COADREAD | YWHAQ | first neighbor | | | 75 | 581 |
| COADREAD | RRAS2 | first neighbor | | | 76 | 581 |
| COADREAD | FOXO3 | first neighbor | | | 77 | 581 |
| COADREAD | CHUK | first neighbor | | | 78 | 581 |
| COADREAD | CASP3 | first neighbor | | | 79 | 581 |
| COADREAD | XRCC6 | first neighbor | | | 80 | 581 |
| COADREAD | HSPA1B | first neighbor | | | 81 | 581 |
| COADREAD | FUS | first neighbor | | | 82 | 581 |
| COADREAD | KIT | first neighbor | | | 83 | 581 |
| COADREAD | EZH2 | first neighbor | | | 84 | 581 |
| COADREAD | CSNK2B | first neighbor | | | 85 | 581 |
| COADREAD | FGFR4 | first neighbor | | | 86 | 581 |
| COADREAD | PPP2CB | first neighbor | | | 87 | 581 |
| COADREAD | CASP8 | first neighbor | | | 88 | 581 |
| COADREAD | NCOA6 | CNV | 49 | 0 | 89 | 581 |
| COADREAD | TRIM25 | first neighbor | | | 90 | 581 |
| COADREAD | CAMK2A | first neighbor | | | 91 | 581 |
| COADREAD | ID1 | CNV | 51 | 0 | 92 | 581 |
| COADREAD | FLT1 | first neighbor | | | 93 | 581 |
| COADREAD | TGFBR1 | first neighbor | | | 94 | 581 |
| COADREAD | HSPA1A | first neighbor | | | 95 | 581 |
| COADREAD | MAPK10 | first neighbor | | | 96 | 581 |
| COADREAD | MAP1LC3A | CNV | 50 | 0 | 97 | 581 |
| COADREAD | FLT3 | first neighbor | | | 98 | 581 |
| COADREAD | LEF1 | first neighbor | | | 99 | 581 |
| COADREAD | CHEK1 | first neighbor | | | 100 | 581 |
| LUSC | TP53 | Mutation | | | 1 | 469 |

| | | | | | | |
|---|---|---|---|---|---|---|
| LUSC | CUL3 | Mutation | | | 2 | 469 |
| LUSC | SOX2 | CNV | 194 | 0 | 3 | 469 |
| LUSC | PIK3CA | ['CNV', 'Mutation'] | 184 | 0 | 4 | 469 |
| LUSC | FXR1 | CNV | 188 | 0 | 5 | 469 |
| LUSC | AP2M1 | CNV | 190 | 0 | 6 | 469 |
| LUSC | CREBBP | Mutation | | | 7 | 469 |
| LUSC | DCUN1D1 | CNV | 195 | 0 | 8 | 469 |
| LUSC | FN1 | Mutation | | | 9 | 469 |
| LUSC | ACTL6A | CNV | 182 | 0 | 10 | 469 |
| LUSC | PTEN | Mutation | | | 11 | 469 |
| LUSC | UBC | first neighbor | | | 12 | 469 |
| LUSC | ESR1 | first neighbor | | | 13 | 469 |
| LUSC | EGFR | first neighbor | | | 14 | 469 |
| LUSC | CDKN2A | Mutation | | | 15 | 469 |
| LUSC | RELA | first neighbor | | | 16 | 469 |
| LUSC | RB1 | Mutation | | | 17 | 469 |
| LUSC | CTNNB1 | first neighbor | | | 18 | 469 |
| LUSC | NOTCH1 | Mutation | | | 19 | 469 |
| LUSC | SP1 | first neighbor | | | 20 | 469 |
| LUSC | HDAC1 | first neighbor | | | 21 | 469 |
| LUSC | GRB2 | first neighbor | | | 22 | 469 |
| LUSC | DVL3 | CNV | 190 | 0 | 23 | 469 |
| LUSC | AR | first neighbor | | | 24 | 469 |
| LUSC | TBL1XR1 | CNV | 179 | 0 | 25 | 469 |
| LUSC | NFE2L2 | Mutation | | | 26 | 469 |
| LUSC | CSNK2A1 | first neighbor | | | 27 | 469 |
| LUSC | SNW1 | first neighbor | | | 28 | 469 |
| LUSC | PRKCA | first neighbor | | | 29 | 469 |
| LUSC | HIF1A | first neighbor | | | 30 | 469 |
| LUSC | PLD1 | CNV | 179 | 0 | 31 | 469 |
| LUSC | FBXW7 | Mutation | | | 32 | 469 |
| LUSC | GSK3B | first neighbor | | | 33 | 469 |
| LUSC | HDAC3 | first neighbor | | | 34 | 469 |
| LUSC | PPARG | first neighbor | | | 35 | 469 |
| LUSC | HDAC2 | first neighbor | | | 36 | 469 |
| LUSC | SMARCA4 | first neighbor | | | 37 | 469 |
| LUSC | GNB4 | CNV | 183 | 0 | 38 | 469 |
| LUSC | ZMAT3 | CNV | 183 | 0 | 39 | 469 |
| LUSC | HSPA8 | first neighbor | | | 40 | 469 |
| LUSC | KAT2B | first neighbor | | | 41 | 469 |
| LUSC | CCND1 | first neighbor | | | 42 | 469 |
| LUSC | RXRA | first neighbor | | | 43 | 469 |
| LUSC | PML | first neighbor | | | 44 | 469 |
| LUSC | CBL | first neighbor | | | 45 | 469 |
| LUSC | BMI1 | first neighbor | | | 46 | 469 |
| LUSC | RBX1 | first neighbor | | | 47 | 469 |
| LUSC | PIK3R3 | first neighbor | | | 48 | 469 |
| LUSC | IKBKB | first neighbor | | | 49 | 469 |
| LUSC | NF1 | Mutation | | | 50 | 469 |
| LUSC | RAC1 | first neighbor | | | 51 | 469 |
| LUSC | NCOR1 | first neighbor | | | 52 | 469 |
| LUSC | SMARCA2 | first neighbor | | | 53 | 469 |
| LUSC | EZH2 | first neighbor | | | 54 | 469 |
| LUSC | INSR | Mutation | | | 55 | 469 |

| | | | | | | |
|---|---|---|---|---|---|---|
| LUSC | TNFSF10 | CNV | 179 | 0 | 56 | 469 |
| LUSC | KEAP1 | Mutation | | | 57 | 469 |
| LUSC | HRAS | first neighbor | | | 58 | 469 |
| LUSC | PRKCD | first neighbor | | | 59 | 469 |
| LUSC | ILK | first neighbor | | | 60 | 469 |
| LUSC | NFATC1 | first neighbor | | | 61 | 469 |
| LUSC | ARID1A | Mutation | | | 62 | 469 |
| LUSC | NCOR2 | first neighbor | | | 63 | 469 |
| LUSC | SMARCD1 | first neighbor | | | 64 | 469 |
| LUSC | SMARCB1 | first neighbor | | | 65 | 469 |
| LUSC | DVL2 | first neighbor | | | 66 | 469 |
| LUSC | RASA1 | Mutation | | | 67 | 469 |
| LUSC | PRKCB | first neighbor | | | 68 | 469 |
| LUSC | RARA | first neighbor | | | 69 | 469 |
| LUSC | SQSTM1 | first neighbor | | | 70 | 469 |
| LUSC | YY1 | first neighbor | | | 71 | 469 |
| LUSC | HSPA1B | first neighbor | | | 72 | 469 |
| LUSC | SMARCC1 | first neighbor | | | 73 | 469 |
| LUSC | TYK2 | Mutation | | | 74 | 469 |
| LUSC | PRKCZ | first neighbor | | | 75 | 469 |
| LUSC | MET | first neighbor | | | 76 | 469 |
| LUSC | KLF5 | Mutation | | | 77 | 469 |
| LUSC | TBP | first neighbor | | | 78 | 469 |
| LUSC | WDR5 | first neighbor | | | 79 | 469 |
| LUSC | NCL | first neighbor | | | 80 | 469 |
| LUSC | IRS1 | first neighbor | | | 81 | 469 |
| LUSC | RUNX1 | first neighbor | | | 82 | 469 |
| LUSC | TUBB | first neighbor | | | 83 | 469 |
| LUSC | SKP1 | first neighbor | | | 84 | 469 |
| LUSC | CCNB1 | first neighbor | | | 85 | 469 |
| LUSC | SMARCC2 | first neighbor | | | 86 | 469 |
| LUSC | IRF1 | first neighbor | | | 87 | 469 |
| LUSC | RBPJ | first neighbor | | | 88 | 469 |
| LUSC | CAV1 | first neighbor | | | 89 | 469 |
| LUSC | SUMO1 | first neighbor | | | 90 | 469 |
| LUSC | RUVBL1 | first neighbor | | | 91 | 469 |
| LUSC | RUVBL2 | first neighbor | | | 92 | 469 |
| LUSC | TUBA1C | first neighbor | | | 93 | 469 |
| LUSC | CASP3 | first neighbor | | | 94 | 469 |
| LUSC | SREBF1 | first neighbor | | | 95 | 469 |
| LUSC | CASP8 | first neighbor | | | 96 | 469 |
| LUSC | FOXO3 | first neighbor | | | 97 | 469 |
| LUSC | MED1 | first neighbor | | | 98 | 469 |
| LUSC | FLNA | first neighbor | | | 99 | 469 |
| LUSC | TRIM28 | first neighbor | | | 100 | 469 |
| LUAD | TP53 | Mutation | | | 1 | 502 |
| LUAD | EGFR | Mutation | | | 2 | 502 |
| LUAD | KRAS | Mutation | | | 3 | 502 |
| LUAD | CTNNB1 | Mutation | | | 4 | 502 |
| LUAD | CDKN2A | ['CNV', 'Mutation'] | 0 | 86 | 5 | 502 |
| LUAD | ATM | Mutation | | | 6 | 502 |
| LUAD | MYC | first neighbor | | | 7 | 502 |
| LUAD | PIK3CA | Mutation | | | 8 | 502 |
| LUAD | SRC | first neighbor | | | 9 | 502 |

| | | | | | | |
|---|---|---|---|---|---|---|
| LUAD | RB1 | Mutation | | | 10 | 502 |
| LUAD | SMARCA4 | Mutation | | | 11 | 502 |
| LUAD | STK11 | Mutation | | | 12 | 502 |
| LUAD | ERBB2 | first neighbor | | | 13 | 502 |
| LUAD | SP1 | first neighbor | | | 14 | 502 |
| LUAD | MET | Mutation | | | 15 | 502 |
| LUAD | TERT | CNV | 66 | 0 | 16 | 502 |
| LUAD | HSP90AA1 | first neighbor | | | 17 | 502 |
| LUAD | MAPK1 | first neighbor | | | 18 | 502 |
| LUAD | HRAS | first neighbor | | | 19 | 502 |
| LUAD | SPTA1 | Mutation | | | 20 | 502 |
| LUAD | APC | Mutation | | | 21 | 502 |
| LUAD | NFKBIA | CNV | 57 | 1 | 22 | 502 |
| LUAD | CDKN1A | first neighbor | | | 23 | 502 |
| LUAD | CREBBP | first neighbor | | | 24 | 502 |
| LUAD | NRAS | first neighbor | | | 25 | 502 |
| LUAD | HDAC1 | first neighbor | | | 26 | 502 |
| LUAD | AKT1 | first neighbor | | | 27 | 502 |
| LUAD | SMAD3 | first neighbor | | | 28 | 502 |
| LUAD | STAT3 | first neighbor | | | 29 | 502 |
| LUAD | MDM2 | first neighbor | | | 30 | 502 |
| LUAD | PRKCA | first neighbor | | | 31 | 502 |
| LUAD | NOTCH1 | first neighbor | | | 32 | 502 |
| LUAD | ERBB3 | first neighbor | | | 33 | 502 |
| LUAD | SMAD2 | first neighbor | | | 34 | 502 |
| LUAD | NF1 | Mutation | | | 35 | 502 |
| LUAD | GSK3B | first neighbor | | | 36 | 502 |
| LUAD | NOTCH4 | Mutation | | | 37 | 502 |
| LUAD | HIF1A | first neighbor | | | 38 | 502 |
| LUAD | NOTCH2 | CNV | 50 | 7 | 39 | 502 |
| LUAD | YWHAZ | first neighbor | | | 40 | 502 |
| LUAD | AURKA | first neighbor | | | 41 | 502 |
| LUAD | CCND1 | first neighbor | | | 42 | 502 |
| LUAD | PTK2 | first neighbor | | | 43 | 502 |
| LUAD | IGF1R | first neighbor | | | 44 | 502 |
| LUAD | BRAF | Mutation | | | 45 | 502 |
| LUAD | FGFR1 | first neighbor | | | 46 | 502 |
| LUAD | A2M | Mutation | | | 47 | 502 |
| LUAD | CDC37 | first neighbor | | | 48 | 502 |
| LUAD | KEAP1 | Mutation | | | 49 | 502 |
| LUAD | EZH2 | first neighbor | | | 50 | 502 |
| LUAD | PML | first neighbor | | | 51 | 502 |
| LUAD | H3-4 | first neighbor | | | 52 | 502 |
| LUAD | PPARG | first neighbor | | | 53 | 502 |
| LUAD | L1CAM | Mutation | | | 54 | 502 |
| LUAD | ABL1 | first neighbor | | | 55 | 502 |
| LUAD | PTEN | first neighbor | | | 56 | 502 |
| LUAD | PDCD6 | CNV | 66 | 0 | 57 | 502 |
| LUAD | CDKN2B | CNV | 0 | 84 | 58 | 502 |
| LUAD | KAT2B | first neighbor | | | 59 | 502 |
| LUAD | FBXO6 | first neighbor | | | 60 | 502 |
| LUAD | PIK3R3 | first neighbor | | | 61 | 502 |
| LUAD | HSPA1A | first neighbor | | | 62 | 502 |
| LUAD | YWHAQ | first neighbor | | | 63 | 502 |

| | | | | | | |
|---|---|---|---|---|---|---|
| LUAD | ARID1A | Mutation | | | 64 | 502 |
| LUAD | ITGB1 | first neighbor | | | 65 | 502 |
| LUAD | YAP1 | first neighbor | | | 66 | 502 |
| LUAD | FOXO3 | first neighbor | | | 67 | 502 |
| LUAD | HSPA4 | first neighbor | | | 68 | 502 |
| LUAD | COL3A1 | Mutation | | | 69 | 502 |
| LUAD | FOXO1 | first neighbor | | | 70 | 502 |
| LUAD | HSPA1B | first neighbor | | | 71 | 502 |
| LUAD | VCAN | Mutation | | | 72 | 502 |
| LUAD | BCL2 | first neighbor | | | 73 | 502 |
| LUAD | CDK4 | first neighbor | | | 74 | 502 |
| LUAD | ATR | first neighbor | | | 75 | 502 |
| LUAD | VEGFA | first neighbor | | | 76 | 502 |
| LUAD | HSPA5 | first neighbor | | | 77 | 502 |
| LUAD | SQSTM1 | first neighbor | | | 78 | 502 |
| LUAD | CASP3 | first neighbor | | | 79 | 502 |
| LUAD | H2AX | first neighbor | | | 80 | 502 |
| LUAD | VDR | first neighbor | | | 81 | 502 |
| LUAD | MIPOL1 | CNV | 57 | 0 | 82 | 502 |
| LUAD | IRS1 | first neighbor | | | 83 | 502 |
| LUAD | BTRC | first neighbor | | | 84 | 502 |
| LUAD | CHUK | first neighbor | | | 85 | 502 |
| LUAD | SP3 | first neighbor | | | 86 | 502 |
| LUAD | SMARCB1 | first neighbor | | | 87 | 502 |
| LUAD | PRKCZ | first neighbor | | | 88 | 502 |
| LUAD | SKP1 | first neighbor | | | 89 | 502 |
| LUAD | CTBP1 | first neighbor | | | 90 | 502 |
| LUAD | WT1 | first neighbor | | | 91 | 502 |
| LUAD | MBIP | CNV | 67 | 0 | 92 | 502 |
| LUAD | YY1 | first neighbor | | | 93 | 502 |
| LUAD | CDK5 | first neighbor | | | 94 | 502 |
| LUAD | SDHA | CNV | 66 | 0 | 95 | 502 |
| LUAD | NFATC2 | first neighbor | | | 96 | 502 |
| LUAD | TFAP2A | first neighbor | | | 97 | 502 |
| LUAD | CD44 | first neighbor | | | 98 | 502 |
| LUAD | TCF7L2 | first neighbor | | | 99 | 502 |
| LUAD | DDB1 | first neighbor | | | 100 | 502 |
| OV | MYC | CNV | 190 | 0 | 1 | 564 |
| OV | TP53 | Mutation | | | 2 | 564 |
| OV | PTK2 | CNV | 159 | 2 | 3 | 564 |
| OV | PLEC | CNV | 155 | 1 | 4 | 564 |
| OV | BRCA1 | Mutation | | | 5 | 564 |
| OV | NDRG1 | CNV | 165 | 1 | 6 | 564 |
| OV | EPPK1 | CNV | 155 | 1 | 7 | 564 |
| OV | EEF1D | CNV | 156 | 1 | 8 | 564 |
| OV | CDK2 | first neighbor | | | 9 | 564 |
| OV | TOP1MT | CNV | 155 | 1 | 10 | 564 |
| OV | ST3GAL1 | CNV | 165 | 1 | 11 | 564 |
| OV | MECOM | CNV | 159 | 0 | 12 | 564 |
| OV | EGFR | first neighbor | | | 13 | 564 |
| OV | SCRIB | CNV | 155 | 1 | 14 | 564 |
| OV | RB1 | Mutation | | | 15 | 564 |
| OV | ASAP1 | CNV | 182 | 1 | 16 | 564 |
| OV | PUF60 | CNV | 155 | 1 | 17 | 564 |

| | | | | | | |
|---|---|---|---|---|---|---|
| OV | SP1 | first neighbor | | | 18 | 564 |
| OV | CDK1 | first neighbor | | | 19 | 564 |
| OV | PCNA | first neighbor | | | 20 | 564 |
| OV | GRB2 | first neighbor | | | 21 | 564 |
| OV | MCM2 | first neighbor | | | 22 | 564 |
| OV | CREBBP | first neighbor | | | 23 | 564 |
| OV | SNW1 | first neighbor | | | 24 | 564 |
| OV | CDH1 | first neighbor | | | 25 | 564 |
| OV | CUL7 | first neighbor | | | 26 | 564 |
| OV | OBSL1 | first neighbor | | | 27 | 564 |
| OV | MAPK1 | first neighbor | | | 28 | 564 |
| OV | CDKN1B | Mutation | | | 29 | 564 |
| OV | CCDC8 | first neighbor | | | 30 | 564 |
| OV | MAPK3 | first neighbor | | | 31 | 564 |
| OV | CDC5L | first neighbor | | | 32 | 564 |
| OV | E2F1 | first neighbor | | | 33 | 564 |
| OV | CSNK2A1 | first neighbor | | | 34 | 564 |
| OV | FOS | first neighbor | | | 35 | 564 |
| OV | UBE2I | first neighbor | | | 36 | 564 |
| OV | ERBB2 | first neighbor | | | 37 | 564 |
| OV | PRKCA | first neighbor | | | 38 | 564 |
| OV | SMC3 | Mutation | | | 39 | 564 |
| OV | HDAC2 | first neighbor | | | 40 | 564 |
| OV | ABL1 | first neighbor | | | 41 | 564 |
| OV | PIK3R1 | first neighbor | | | 42 | 564 |
| OV | VCP | first neighbor | | | 43 | 564 |
| OV | PLK1 | first neighbor | | | 44 | 564 |
| OV | YWHAG | first neighbor | | | 45 | 564 |
| OV | SMAD2 | first neighbor | | | 46 | 564 |
| OV | H3-4 | first neighbor | | | 47 | 564 |
| OV | CCNB1 | first neighbor | | | 48 | 564 |
| OV | PPP1CA | first neighbor | | | 49 | 564 |
| OV | SIRT1 | first neighbor | | | 50 | 564 |
| OV | PIK3CA | Mutation | | | 51 | 564 |
| OV | ATR | first neighbor | | | 52 | 564 |
| OV | KHDRBS3 | CNV | 162 | 1 | 53 | 564 |
| OV | EXOSC4 | CNV | 155 | 1 | 54 | 564 |
| OV | STAT5A | first neighbor | | | 55 | 564 |
| OV | PPP1CC | first neighbor | | | 56 | 564 |
| OV | NCL | first neighbor | | | 57 | 564 |
| OV | ACTG1 | first neighbor | | | 58 | 564 |
| OV | SHC1 | first neighbor | | | 59 | 564 |
| OV | DYNC1H1 | Mutation | | | 60 | 564 |
| OV | PRKDC | first neighbor | | | 61 | 564 |
| OV | IQGAP1 | first neighbor | | | 62 | 564 |
| OV | POLR2A | first neighbor | | | 63 | 564 |
| OV | CDK4 | first neighbor | | | 64 | 564 |
| OV | XRCC5 | first neighbor | | | 65 | 564 |
| OV | KAT2B | first neighbor | | | 66 | 564 |
| OV | HSPA4 | first neighbor | | | 67 | 564 |
| OV | HIF1A | first neighbor | | | 68 | 564 |
| OV | SQSTM1 | first neighbor | | | 69 | 564 |
| OV | CBL | first neighbor | | | 70 | 564 |
| OV | PPP1CB | first neighbor | | | 71 | 564 |

| | | | | | | |
|---|---|---|---|---|---|---|
| OV | RBL1 | first neighbor | | | 72 | 564 |
| OV | CRK | first neighbor | | | 73 | 564 |
| OV | ZC3H3 | CNV | 157 | 1 | 74 | 564 |
| OV | PPARG | first neighbor | | | 75 | 564 |
| OV | AHNAK | Mutation | | | 76 | 564 |
| OV | TP73 | first neighbor | | | 77 | 564 |
| OV | FOXO3 | first neighbor | | | 78 | 564 |
| OV | YY1 | first neighbor | | | 79 | 564 |
| OV | PRKCD | first neighbor | | | 80 | 564 |
| OV | ETS1 | first neighbor | | | 81 | 564 |
| OV | PKM | first neighbor | | | 82 | 564 |
| OV | TOP2A | Mutation | | | 83 | 564 |
| OV | IFI16 | first neighbor | | | 84 | 564 |
| OV | DDB1 | first neighbor | | | 85 | 564 |
| OV | PXN | first neighbor | | | 86 | 564 |
| OV | SMARCA4 | first neighbor | | | 87 | 564 |
| OV | PIK3R3 | first neighbor | | | 88 | 564 |
| OV | KCNQ3 | CNV | 168 | 1 | 89 | 564 |
| OV | DDX5 | first neighbor | | | 90 | 564 |
| OV | CCNE1 | first neighbor | | | 91 | 564 |
| OV | APC | first neighbor | | | 92 | 564 |
| OV | E2F4 | first neighbor | | | 93 | 564 |
| OV | PIN1 | first neighbor | | | 94 | 564 |
| OV | EEF1G | first neighbor | | | 95 | 564 |
| OV | BCR | Mutation | | | 96 | 564 |
| OV | GAPDH | first neighbor | | | 97 | 564 |
| OV | TP53BP1 | first neighbor | | | 98 | 564 |
| OV | RAD51 | first neighbor | | | 99 | 564 |
| OV | E2F3 | first neighbor | | | 100 | 564 |
| GBM | EGFR | ['CNV', 'Mutation', 'SV'] | 179 | 0 | 1 | 378 |
| GBM | TP53 | Mutation | | | 2 | 378 |
| GBM | IFNA5 | CNV | 0 | 94 | 3 | 378 |
| GBM | PTEN | Mutation | | | 4 | 378 |
| GBM | CDKN2A | CNV | 0 | 213 | 5 | 378 |
| GBM | IFNA2 | CNV | 0 | 102 | 6 | 378 |
| GBM | PIK3R1 | Mutation | | | 7 | 378 |
| GBM | IFNA8 | CNV | 0 | 104 | 8 | 378 |
| GBM | IFNA6 | CNV | 0 | 98 | 9 | 378 |
| GBM | IFNA4 | CNV | 0 | 87 | 10 | 378 |
| GBM | IFNA14 | CNV | 0 | 90 | 11 | 378 |
| GBM | PDGFRA | ['CNV', 'Mutation'] | 55 | 0 | 12 | 378 |
| GBM | IFNA21 | CNV | 0 | 85 | 13 | 378 |
| GBM | PIK3CA | Mutation | | | 14 | 378 |
| GBM | IFNA1 | CNV | 0 | 108 | 15 | 378 |
| GBM | IFNA17 | CNV | 0 | 88 | 16 | 378 |
| GBM | IFNE | CNV | 0 | 111 | 17 | 378 |
| GBM | IFNA7 | CNV | 0 | 86 | 18 | 378 |
| GBM | IFNA13 | CNV | 0 | 102 | 19 | 378 |
| GBM | IFNA10 | CNV | 0 | 86 | 20 | 378 |
| GBM | IFNA16 | CNV | 0 | 86 | 21 | 378 |
| GBM | IFNB1 | CNV | 0 | 70 | 22 | 378 |
| GBM | IFNW1 | CNV | 0 | 84 | 23 | 378 |
| GBM | PIK3CB | Mutation | | | 24 | 378 |
| GBM | PTPN11 | Mutation | | | 25 | 378 |

| | | | | | | |
|---|---|---|---|---|---|---|
| GBM | PIK3R2 | first neighbor | | | 26 | 378 |
| GBM | JAK2 | first neighbor | | | 27 | 378 |
| GBM | RELA | first neighbor | | | 28 | 378 |
| GBM | JAK1 | first neighbor | | | 29 | 378 |
| GBM | PIK3CD | first neighbor | | | 30 | 378 |
| GBM | PIK3R3 | first neighbor | | | 31 | 378 |
| GBM | CDKN2B | CNV | 0 | 208 | 32 | 378 |
| GBM | JAK3 | first neighbor | | | 33 | 378 |
| GBM | PDGFRB | first neighbor | | | 34 | 378 |
| GBM | NFKB1 | first neighbor | | | 35 | 378 |
| GBM | PLCG1 | first neighbor | | | 36 | 378 |
| GBM | TYK2 | first neighbor | | | 37 | 378 |
| GBM | STAT1 | first neighbor | | | 38 | 378 |
| GBM | SHC1 | first neighbor | | | 39 | 378 |
| GBM | CDK4 | CNV | 60 | 0 | 40 | 378 |
| GBM | PTPN6 | first neighbor | | | 41 | 378 |
| GBM | STAT3 | first neighbor | | | 42 | 378 |
| GBM | SP1 | first neighbor | | | 43 | 378 |
| GBM | RB1 | Mutation | | | 44 | 378 |
| GBM | CSF2RB | first neighbor | | | 45 | 378 |
| GBM | IL2RG | first neighbor | | | 46 | 378 |
| GBM | IFNGR1 | first neighbor | | | 47 | 378 |
| GBM | IL6ST | first neighbor | | | 48 | 378 |
| GBM | HRAS | first neighbor | | | 49 | 378 |
| GBM | IL2RB | first neighbor | | | 50 | 378 |
| GBM | PDGFB | first neighbor | | | 51 | 378 |
| GBM | CTNNB1 | first neighbor | | | 52 | 378 |
| GBM | EP300 | first neighbor | | | 53 | 378 |
| GBM | IL7R | first neighbor | | | 54 | 378 |
| GBM | SOCS1 | first neighbor | | | 55 | 378 |
| GBM | EPOR | first neighbor | | | 56 | 378 |
| GBM | IL3RA | first neighbor | | | 57 | 378 |
| GBM | IL2RA | first neighbor | | | 58 | 378 |
| GBM | PDGFA | first neighbor | | | 59 | 378 |
| GBM | IFNGR2 | first neighbor | | | 60 | 378 |
| GBM | EGF | first neighbor | | | 61 | 378 |
| GBM | CBL | first neighbor | | | 62 | 378 |
| GBM | GHR | first neighbor | | | 63 | 378 |
| GBM | CSF2RA | first neighbor | | | 64 | 378 |
| GBM | IFNAR1 | first neighbor | | | 65 | 378 |
| GBM | IRF3 | first neighbor | | | 66 | 378 |
| GBM | IFNAR2 | first neighbor | | | 67 | 378 |
| GBM | IL5RA | first neighbor | | | 68 | 378 |
| GBM | IL4R | first neighbor | | | 69 | 378 |
| GBM | SOCS3 | first neighbor | | | 70 | 378 |
| GBM | PTPN1 | first neighbor | | | 71 | 378 |
| GBM | LANCL2 | CNV | 115 | 0 | 72 | 378 |
| GBM | LIFR | first neighbor | | | 73 | 378 |
| GBM | IL12RB1 | first neighbor | | | 74 | 378 |
| GBM | IL6R | first neighbor | | | 75 | 378 |
| GBM | NRAS | first neighbor | | | 76 | 378 |
| GBM | LEPR | first neighbor | | | 77 | 378 |
| GBM | PLCG2 | first neighbor | | | 78 | 378 |
| GBM | PRLR | first neighbor | | | 79 | 378 |

| | | | | | | |
|---|---|---|---|---|---|---|
| GBM | IL13RA2 | first neighbor | | | 80 | 378 |
| GBM | IL20RB | first neighbor | | | 81 | 378 |
| GBM | IL12RB2 | first neighbor | | | 82 | 378 |
| GBM | STAT2 | first neighbor | | | 83 | 378 |
| GBM | IL20RA | first neighbor | | | 84 | 378 |
| GBM | IRF7 | first neighbor | | | 85 | 378 |
| GBM | MET | first neighbor | | | 86 | 378 |
| GBM | CSF3R | first neighbor | | | 87 | 378 |
| GBM | IL15RA | first neighbor | | | 88 | 378 |
| GBM | IL22RA1 | first neighbor | | | 89 | 378 |
| GBM | IL10RB | first neighbor | | | 90 | 378 |
| GBM | AKT1 | first neighbor | | | 91 | 378 |
| GBM | IL2 | first neighbor | | | 92 | 378 |
| GBM | MPL | first neighbor | | | 93 | 378 |
| GBM | LYN | first neighbor | | | 94 | 378 |
| GBM | IL23R | first neighbor | | | 95 | 378 |
| GBM | IL21R | first neighbor | | | 96 | 378 |
| GBM | IL10RA | first neighbor | | | 97 | 378 |
| GBM | CREBBP | first neighbor | | | 98 | 378 |
| GBM | KRAS | first neighbor | | | 99 | 378 |
| GBM | IL3 | first neighbor | | | 100 | 378 |
| STAD | TP53 | Mutation | | | 1 | 436 |
| STAD | MYC | CNV | 53 | 0 | 2 | 436 |
| STAD | ERBB2 | CNV | 58 | 0 | 3 | 436 |
| STAD | WWOX | CNV | 7 | 139 | 4 | 436 |
| STAD | CTNNB1 | Mutation | | | 5 | 436 |
| STAD | CDH1 | Mutation | | | 6 | 436 |
| STAD | ERBB4 | Mutation | | | 7 | 436 |
| STAD | PIK3CA | Mutation | | | 8 | 436 |
| STAD | ERBB3 | Mutation | | | 9 | 436 |
| STAD | EGFR | first neighbor | | | 10 | 436 |
| STAD | ATM | Mutation | | | 11 | 436 |
| STAD | PTEN | Mutation | | | 12 | 436 |
| STAD | CDKN2A | CNV | 1 | 50 | 13 | 436 |
| STAD | SMAD4 | Mutation | | | 14 | 436 |
| STAD | KRAS | Mutation | | | 15 | 436 |
| STAD | ESR1 | first neighbor | | | 16 | 436 |
| STAD | ARID1A | Mutation | | | 17 | 436 |
| STAD | FBXW7 | Mutation | | | 18 | 436 |
| STAD | SRC | first neighbor | | | 19 | 436 |
| STAD | SP1 | first neighbor | | | 20 | 436 |
| STAD | APC | Mutation | | | 21 | 436 |
| STAD | EP300 | first neighbor | | | 22 | 436 |
| STAD | UBC | first neighbor | | | 23 | 436 |
| STAD | AR | first neighbor | | | 24 | 436 |
| STAD | CDKN1A | first neighbor | | | 25 | 436 |
| STAD | SMAD3 | first neighbor | | | 26 | 436 |
| STAD | RARA | CNV | 37 | 1 | 27 | 436 |
| STAD | MAPK1 | first neighbor | | | 28 | 436 |
| STAD | MET | first neighbor | | | 29 | 436 |
| STAD | MAPK3 | first neighbor | | | 30 | 436 |
| STAD | MDM2 | first neighbor | | | 31 | 436 |
| STAD | PIK3R1 | first neighbor | | | 32 | 436 |
| STAD | FGFR3 | first neighbor | | | 33 | 436 |

| | | | | | |
|---|---|---|---|---|---|
| STAD | CREBBP | first neighbor | | | 34 | 436 |
| STAD | IGF1R | first neighbor | | | 35 | 436 |
| STAD | RHOA | Mutation | | | 36 | 436 |
| STAD | SMAD2 | first neighbor | | | 37 | 436 |
| STAD | HSP90AA1 | first neighbor | | | 38 | 436 |
| STAD | HDAC1 | first neighbor | | | 39 | 436 |
| STAD | GSK3B | first neighbor | | | 40 | 436 |
| STAD | HRAS | first neighbor | | | 41 | 436 |
| STAD | AKT1 | first neighbor | | | 42 | 436 |
| STAD | UBB | first neighbor | | | 43 | 436 |
| STAD | HIF1A | first neighbor | | | 44 | 436 |
| STAD | NRAS | first neighbor | | | 45 | 436 |
| STAD | CCNE1 | CNV | 49 | 1 | 46 | 436 |
| STAD | SHC1 | first neighbor | | | 47 | 436 |
| STAD | RPS27A | first neighbor | | | 48 | 436 |
| STAD | THRA | CNV | 38 | 1 | 49 | 436 |
| STAD | CBL | first neighbor | | | 50 | 436 |
| STAD | E2F1 | first neighbor | | | 51 | 436 |
| STAD | ABL1 | first neighbor | | | 52 | 436 |
| STAD | PDGFRA | first neighbor | | | 53 | 436 |
| STAD | UBR5 | Mutation | | | 54 | 436 |
| STAD | PPARG | first neighbor | | | 55 | 436 |
| STAD | CCND1 | first neighbor | | | 56 | 436 |
| STAD | SNW1 | first neighbor | | | 57 | 436 |
| STAD | PCNA | first neighbor | | | 58 | 436 |
| STAD | PIK3R2 | first neighbor | | | 59 | 436 |
| STAD | ACTB | first neighbor | | | 60 | 436 |
| STAD | FYN | first neighbor | | | 61 | 436 |
| STAD | HLA-B | Mutation | | | 62 | 436 |
| STAD | HSPA8 | first neighbor | | | 63 | 436 |
| STAD | MAPK8 | first neighbor | | | 64 | 436 |
| STAD | CTNND1 | Mutation | | | 65 | 436 |
| STAD | EWSR1 | first neighbor | | | 66 | 436 |
| STAD | HSP90AB1 | first neighbor | | | 67 | 436 |
| STAD | SPTA1 | Mutation | | | 68 | 436 |
| STAD | UBA52 | first neighbor | | | 69 | 436 |
| STAD | HDAC3 | first neighbor | | | 70 | 436 |
| STAD | PARP1 | first neighbor | | | 71 | 436 |
| STAD | GRB7 | CNV | 58 | 1 | 72 | 436 |
| STAD | PTK2 | first neighbor | | | 73 | 436 |
| STAD | EZH2 | first neighbor | | | 74 | 436 |
| STAD | HUWE1 | first neighbor | | | 75 | 436 |
| STAD | CDC42 | first neighbor | | | 76 | 436 |
| STAD | SMAD1 | first neighbor | | | 77 | 436 |
| STAD | NOTCH1 | first neighbor | | | 78 | 436 |
| STAD | CDC6 | CNV | 37 | 1 | 79 | 436 |
| STAD | CRK | first neighbor | | | 80 | 436 |
| STAD | PLK1 | first neighbor | | | 81 | 436 |
| STAD | BTRC | first neighbor | | | 82 | 436 |
| STAD | SMARCA4 | first neighbor | | | 83 | 436 |
| STAD | FOXO3 | first neighbor | | | 84 | 436 |
| STAD | NCOR1 | first neighbor | | | 85 | 436 |
| STAD | NCOA3 | first neighbor | | | 86 | 436 |
| STAD | CDKN1B | first neighbor | | | 87 | 436 |

| | | | | | | |
|---|---|---|---|---|---|---|
| STAD | CDK4 | first neighbor | | | 88 | 436 |
| STAD | PML | first neighbor | | | 89 | 436 |
| STAD | BMI1 | first neighbor | | | 90 | 436 |
| STAD | SQSTM1 | first neighbor | | | 91 | 436 |
| STAD | KAT2B | first neighbor | | | 92 | 436 |
| STAD | SKP1 | first neighbor | | | 93 | 436 |
| STAD | FLNA | first neighbor | | | 94 | 436 |
| STAD | NCOA1 | first neighbor | | | 95 | 436 |
| STAD | YY1 | first neighbor | | | 96 | 436 |
| STAD | CDKN2B | CNV | 1 | 48 | 97 | 436 |
| STAD | VEGFA | first neighbor | | | 98 | 436 |
| STAD | CALM3 | first neighbor | | | 99 | 436 |
| STAD | IQGAP1 | first neighbor | | | 100 | 436 |
| BLCA | TP53 | Mutation | | | 1 | 408 |
| BLCA | EP300 | Mutation | | | 2 | 408 |
| BLCA | PIK3CA | Mutation | | | 3 | 408 |
| BLCA | CDKN2A | ['CNV', 'Mutation'] | 1 | 130 | 4 | 408 |
| BLCA | CREBBP | Mutation | | | 5 | 408 |
| BLCA | RB1 | Mutation | | | 6 | 408 |
| BLCA | ERBB2 | Mutation | | | 7 | 408 |
| BLCA | YWHAZ | CNV | 65 | 0 | 8 | 408 |
| BLCA | CDKN1A | Mutation | | | 9 | 408 |
| BLCA | FGFR3 | Mutation | | | 10 | 408 |
| BLCA | ERBB3 | Mutation | | | 11 | 408 |
| BLCA | ATM | Mutation | | | 12 | 408 |
| BLCA | ARID1A | Mutation | | | 13 | 408 |
| BLCA | FCGR2A | CNV | 60 | 0 | 14 | 408 |
| BLCA | USF1 | CNV | 69 | 0 | 15 | 408 |
| BLCA | E2F3 | CNV | 64 | 1 | 16 | 408 |
| BLCA | FCER1G | CNV | 66 | 0 | 17 | 408 |
| BLCA | ESR1 | first neighbor | | | 18 | 408 |
| BLCA | FCGR3A | CNV | 60 | 0 | 19 | 408 |
| BLCA | NCOR1 | Mutation | | | 20 | 408 |
| BLCA | MYC | first neighbor | | | 21 | 408 |
| BLCA | HRAS | Mutation | | | 22 | 408 |
| BLCA | GRB2 | first neighbor | | | 23 | 408 |
| BLCA | CDKN2B | CNV | 1 | 128 | 24 | 408 |
| BLCA | PIK3R1 | first neighbor | | | 25 | 408 |
| BLCA | SP1 | first neighbor | | | 26 | 408 |
| BLCA | SHC1 | first neighbor | | | 27 | 408 |
| BLCA | FCGR2B | CNV | 60 | 0 | 28 | 408 |
| BLCA | RELA | first neighbor | | | 29 | 408 |
| BLCA | CUL1 | Mutation | | | 30 | 408 |
| BLCA | PIK3R2 | first neighbor | | | 31 | 408 |
| BLCA | FBXW7 | Mutation | | | 32 | 408 |
| BLCA | UBC | first neighbor | | | 33 | 408 |
| BLCA | AR | first neighbor | | | 34 | 408 |
| BLCA | CDK2 | first neighbor | | | 35 | 408 |
| BLCA | RXRA | Mutation | | | 36 | 408 |
| BLCA | CTNNB1 | first neighbor | | | 37 | 408 |
| BLCA | KRAS | Mutation | | | 38 | 408 |
| BLCA | BRCA1 | first neighbor | | | 39 | 408 |
| BLCA | SRC | first neighbor | | | 40 | 408 |
| BLCA | MYH9 | Mutation | | | 41 | 408 |

| | | | | | | |
|---|---|---|---|---|---|---|
| BLCA | ELAVL1 | first neighbor | | | 42 | 408 |
| BLCA | MAPK1 | first neighbor | | | 43 | 408 |
| BLCA | CBL | first neighbor | | | 44 | 408 |
| BLCA | HSP90AA1 | first neighbor | | | 45 | 408 |
| BLCA | MAPK3 | first neighbor | | | 46 | 408 |
| BLCA | RAC1 | first neighbor | | | 47 | 408 |
| BLCA | PLCG1 | first neighbor | | | 48 | 408 |
| BLCA | PIK3CB | first neighbor | | | 49 | 408 |
| BLCA | PTEN | Mutation | | | 50 | 408 |
| BLCA | JUN | first neighbor | | | 51 | 408 |
| BLCA | PABPC1 | CNV | 63 | 0 | 52 | 408 |
| BLCA | HSPA6 | CNV | 60 | 0 | 53 | 408 |
| BLCA | SOS1 | first neighbor | | | 54 | 408 |
| BLCA | NFKB1 | first neighbor | | | 55 | 408 |
| BLCA | HDAC1 | first neighbor | | | 56 | 408 |
| BLCA | LYN | first neighbor | | | 57 | 408 |
| BLCA | KDM6A | Mutation | | | 58 | 408 |
| BLCA | EPHA2 | Mutation | | | 59 | 408 |
| BLCA | SMAD3 | first neighbor | | | 60 | 408 |
| BLCA | HIF1A | first neighbor | | | 61 | 408 |
| BLCA | STAT3 | first neighbor | | | 62 | 408 |
| BLCA | CRK | first neighbor | | | 63 | 408 |
| BLCA | AKT1 | first neighbor | | | 64 | 408 |
| BLCA | E2F1 | first neighbor | | | 65 | 408 |
| BLCA | RHOA | Mutation | | | 66 | 408 |
| BLCA | SYK | first neighbor | | | 67 | 408 |
| BLCA | USP21 | CNV | 66 | 0 | 68 | 408 |
| BLCA | MET | first neighbor | | | 69 | 408 |
| BLCA | UBB | first neighbor | | | 70 | 408 |
| BLCA | PPARG | first neighbor | | | 71 | 408 |
| BLCA | PRKCA | first neighbor | | | 72 | 408 |
| BLCA | SOX4 | CNV | 64 | 1 | 73 | 408 |
| BLCA | FOS | first neighbor | | | 74 | 408 |
| BLCA | MDM2 | first neighbor | | | 75 | 408 |
| BLCA | SPTAN1 | Mutation | | | 76 | 408 |
| BLCA | PLCG2 | first neighbor | | | 77 | 408 |
| BLCA | GSK3B | first neighbor | | | 78 | 408 |
| BLCA | SMAD2 | first neighbor | | | 79 | 408 |
| BLCA | MAPK14 | first neighbor | | | 80 | 408 |
| BLCA | VAV1 | first neighbor | | | 81 | 408 |
| BLCA | CDH1 | first neighbor | | | 82 | 408 |
| BLCA | HSPA8 | first neighbor | | | 83 | 408 |
| BLCA | HSP90AB1 | first neighbor | | | 84 | 408 |
| BLCA | CCND1 | first neighbor | | | 85 | 408 |
| BLCA | MAPK8 | first neighbor | | | 86 | 408 |
| BLCA | HDAC3 | first neighbor | | | 87 | 408 |
| BLCA | TRRAP | Mutation | | | 88 | 408 |
| BLCA | RPS27A | first neighbor | | | 89 | 408 |
| BLCA | GAB2 | first neighbor | | | 90 | 408 |
| BLCA | FYN | first neighbor | | | 91 | 408 |
| BLCA | SMARCA4 | first neighbor | | | 92 | 408 |
| BLCA | ACTB | first neighbor | | | 93 | 408 |
| BLCA | UBE2I | first neighbor | | | 94 | 408 |
| BLCA | VAV2 | first neighbor | | | 95 | 408 |

| | | | | | | |
|---|---|---|---|---|---|---|
| BLCA | VAV3 | first neighbor | | | 96 | 408 |
| BLCA | PCNA | first neighbor | | | 97 | 408 |
| BLCA | KLF5 | Mutation | | | 98 | 408 |
| BLCA | NCOA3 | first neighbor | | | 99 | 408 |
| BLCA | NFE2L2 | Mutation | | | 100 | 408 |
| UCEC | PTEN | Mutation | | | 1 | 408 |
| UCEC | TP53 | Mutation | | | 2 | 408 |
| UCEC | PIK3CA | ['CNV', 'Mutation'] | 35 | 1 | 3 | 408 |
| UCEC | PIK3R1 | Mutation | | | 4 | 408 |
| UCEC | CTNNB1 | Mutation | | | 5 | 408 |
| UCEC | MYC | CNV | 45 | 0 | 6 | 408 |
| UCEC | EP300 | Mutation | | | 7 | 408 |
| UCEC | ATM | Mutation | | | 8 | 408 |
| UCEC | KRAS | Mutation | | | 9 | 408 |
| UCEC | FBXW7 | Mutation | | | 10 | 408 |
| UCEC | PRKDC | Mutation | | | 11 | 408 |
| UCEC | EGFR | first neighbor | | | 12 | 408 |
| UCEC | ARID1A | Mutation | | | 13 | 408 |
| UCEC | LMNA | CNV | 39 | 0 | 14 | 408 |
| UCEC | UBC | first neighbor | | | 15 | 408 |
| UCEC | PPP2R1A | Mutation | | | 16 | 408 |
| UCEC | ESR1 | first neighbor | | | 17 | 408 |
| UCEC | FGFR2 | Mutation | | | 18 | 408 |
| UCEC | CHD4 | Mutation | | | 19 | 408 |
| UCEC | PIK3R2 | first neighbor | | | 20 | 408 |
| UCEC | BRCA1 | first neighbor | | | 21 | 408 |
| UCEC | ELAVL1 | first neighbor | | | 22 | 408 |
| UCEC | GRB2 | first neighbor | | | 23 | 408 |
| UCEC | SRC | first neighbor | | | 24 | 408 |
| UCEC | JAK1 | Mutation | | | 25 | 408 |
| UCEC | PIK3R3 | first neighbor | | | 26 | 408 |
| UCEC | HDAC1 | first neighbor | | | 27 | 408 |
| UCEC | CDK2 | first neighbor | | | 28 | 408 |
| UCEC | HSP90AA1 | first neighbor | | | 29 | 408 |
| UCEC | PLCG1 | first neighbor | | | 30 | 408 |
| UCEC | AR | first neighbor | | | 31 | 408 |
| UCEC | MDM2 | first neighbor | | | 32 | 408 |
| UCEC | PIK3CD | first neighbor | | | 33 | 408 |
| UCEC | AKT1 | first neighbor | | | 34 | 408 |
| UCEC | INPPL1 | Mutation | | | 35 | 408 |
| UCEC | CHD3 | Mutation | | | 36 | 408 |
| UCEC | STAT3 | first neighbor | | | 37 | 408 |
| UCEC | SMAD3 | first neighbor | | | 38 | 408 |
| UCEC | MAP3K1 | Mutation | | | 39 | 408 |
| UCEC | PRKCA | first neighbor | | | 40 | 408 |
| UCEC | HDAC2 | first neighbor | | | 41 | 408 |
| UCEC | SHC1 | first neighbor | | | 42 | 408 |
| UCEC | NPM1 | first neighbor | | | 43 | 408 |
| UCEC | PIP5K1A | CNV | 36 | 0 | 44 | 408 |
| UCEC | ERBB2 | first neighbor | | | 45 | 408 |
| UCEC | YWHAZ | first neighbor | | | 46 | 408 |
| UCEC | PARP1 | first neighbor | | | 47 | 408 |
| UCEC | CBL | first neighbor | | | 48 | 408 |
| UCEC | ABL1 | first neighbor | | | 49 | 408 |

| | | | | | | |
|---|---|---|---|---|---|---|
| UCEC | UBE2I | first neighbor | | | 50 | 408 |
| UCEC | RNF2 | first neighbor | | | 51 | 408 |
| UCEC | CTCF | Mutation | | | 52 | 408 |
| UCEC | EED | first neighbor | | | 53 | 408 |
| UCEC | PLCG2 | first neighbor | | | 54 | 408 |
| UCEC | HIF1A | first neighbor | | | 55 | 408 |
| UCEC | PCNA | first neighbor | | | 56 | 408 |
| UCEC | SMAD2 | first neighbor | | | 57 | 408 |
| UCEC | CCNE1 | CNV | 40 | 0 | 58 | 408 |
| UCEC | ECT2 | CNV | 37 | 1 | 59 | 408 |
| UCEC | CREB1 | first neighbor | | | 60 | 408 |
| UCEC | E2F1 | first neighbor | | | 61 | 408 |
| UCEC | FOS | first neighbor | | | 62 | 408 |
| UCEC | SNW1 | first neighbor | | | 63 | 408 |
| UCEC | NF1 | Mutation | | | 64 | 408 |
| UCEC | MAPK8 | first neighbor | | | 65 | 408 |
| UCEC | RPA1 | first neighbor | | | 66 | 408 |
| UCEC | MET | first neighbor | | | 67 | 408 |
| UCEC | XRCC6 | first neighbor | | | 68 | 408 |
| UCEC | SIRT7 | first neighbor | | | 69 | 408 |
| UCEC | PPARG | first neighbor | | | 70 | 408 |
| UCEC | SIRT1 | first neighbor | | | 71 | 408 |
| UCEC | ZMYM2 | Mutation | | | 72 | 408 |
| UCEC | XRCC5 | first neighbor | | | 73 | 408 |
| UCEC | ERBB3 | first neighbor | | | 74 | 408 |
| UCEC | RPA2 | first neighbor | | | 75 | 408 |
| UCEC | PRKN | first neighbor | | | 76 | 408 |
| UCEC | H3-4 | first neighbor | | | 77 | 408 |
| UCEC | RAC1 | first neighbor | | | 78 | 408 |
| UCEC | MECOM | CNV | 62 | 0 | 79 | 408 |
| UCEC | SMURF1 | first neighbor | | | 80 | 408 |
| UCEC | PML | first neighbor | | | 81 | 408 |
| UCEC | PLK1 | first neighbor | | | 82 | 408 |
| UCEC | PRKCD | first neighbor | | | 83 | 408 |
| UCEC | VCAM1 | first neighbor | | | 84 | 408 |
| UCEC | PI4KB | CNV | 38 | 0 | 85 | 408 |
| UCEC | YAP1 | first neighbor | | | 86 | 408 |
| UCEC | PDGFRB | first neighbor | | | 87 | 408 |
| UCEC | NR3C1 | first neighbor | | | 88 | 408 |
| UCEC | PDGFRA | first neighbor | | | 89 | 408 |
| UCEC | FOXO3 | first neighbor | | | 90 | 408 |
| UCEC | USP7 | first neighbor | | | 91 | 408 |
| UCEC | SMARCA4 | first neighbor | | | 92 | 408 |
| UCEC | FGFR1 | first neighbor | | | 93 | 408 |
| UCEC | CDC42 | first neighbor | | | 94 | 408 |
| UCEC | BMI1 | first neighbor | | | 95 | 408 |
| UCEC | PRKCZ | first neighbor | | | 96 | 408 |
| UCEC | RPA3 | first neighbor | | | 97 | 408 |
| UCEC | YY1 | first neighbor | | | 98 | 408 |
| UCEC | TFRC | CNV | 36 | 0 | 99 | 408 |
| UCEC | CCDC8 | first neighbor | | | 100 | 408 |
| BRCA | TP53 | Mutation | | | 1 | 1034 |
| BRCA | PIK3CA | Mutation | | | 2 | 1034 |
| BRCA | MYC | CNV | 161 | 0 | 3 | 1034 |

| | | | | | | |
|---|---|---|---|---|---|---|
| BRCA | CCND1 | CNV | 161 | 0 | 4 | 1034 |
| BRCA | CDH1 | Mutation | | | 5 | 1034 |
| BRCA | BRCA1 | Mutation | | | 6 | 1034 |
| BRCA | FGF3 | CNV | 154 | 0 | 7 | 1034 |
| BRCA | FGF4 | CNV | 155 | 0 | 8 | 1034 |
| BRCA | FGF19 | CNV | 156 | 0 | 9 | 1034 |
| BRCA | CTTN | CNV | 145 | 0 | 10 | 1034 |
| BRCA | AKT1 | Mutation | | | 11 | 1034 |
| BRCA | RAD21 | CNV | 141 | 1 | 12 | 1034 |
| BRCA | PIK3R1 | Mutation | | | 13 | 1034 |
| BRCA | ERBB2 | Mutation | | | 14 | 1034 |
| BRCA | PTEN | Mutation | | | 15 | 1034 |
| BRCA | EGFR | first neighbor | | | 16 | 1034 |
| BRCA | ESR1 | first neighbor | | | 17 | 1034 |
| BRCA | RB1 | Mutation | | | 18 | 1034 |
| BRCA | UBC | first neighbor | | | 19 | 1034 |
| BRCA | EP300 | first neighbor | | | 20 | 1034 |
| BRCA | MAP3K1 | Mutation | | | 21 | 1034 |
| BRCA | GRB2 | first neighbor | | | 22 | 1034 |
| BRCA | SP1 | first neighbor | | | 23 | 1034 |
| BRCA | SRC | first neighbor | | | 24 | 1034 |
| BRCA | NCOR1 | Mutation | | | 25 | 1034 |
| BRCA | ATM | Mutation | | | 26 | 1034 |
| BRCA | RUNX1 | Mutation | | | 27 | 1034 |
| BRCA | HDAC1 | first neighbor | | | 28 | 1034 |
| BRCA | RELA | first neighbor | | | 29 | 1034 |
| BRCA | CDK2 | first neighbor | | | 30 | 1034 |
| BRCA | HSP90AA1 | first neighbor | | | 31 | 1034 |
| BRCA | AR | first neighbor | | | 32 | 1034 |
| BRCA | CTNNB1 | first neighbor | | | 33 | 1034 |
| BRCA | MAPK3 | first neighbor | | | 34 | 1034 |
| BRCA | ERBB3 | Mutation | | | 35 | 1034 |
| BRCA | MAPK1 | first neighbor | | | 36 | 1034 |
| BRCA | CREBBP | first neighbor | | | 37 | 1034 |
| BRCA | GATA3 | Mutation | | | 38 | 1034 |
| BRCA | STAT3 | first neighbor | | | 39 | 1034 |
| BRCA | CDKN1A | first neighbor | | | 40 | 1034 |
| BRCA | MDM2 | first neighbor | | | 41 | 1034 |
| BRCA | JUN | first neighbor | | | 42 | 1034 |
| BRCA | NFKB1 | first neighbor | | | 43 | 1034 |
| BRCA | CASP8 | Mutation | | | 44 | 1034 |
| BRCA | MYH9 | Mutation | | | 45 | 1034 |
| BRCA | HSP90AB1 | first neighbor | | | 46 | 1034 |
| BRCA | FBXW7 | Mutation | | | 47 | 1034 |
| BRCA | SMAD3 | first neighbor | | | 48 | 1034 |
| BRCA | PIK3R2 | first neighbor | | | 49 | 1034 |
| BRCA | CDK1 | first neighbor | | | 50 | 1034 |
| BRCA | HDAC2 | first neighbor | | | 51 | 1034 |
| BRCA | NPM1 | first neighbor | | | 52 | 1034 |
| BRCA | HIF1A | first neighbor | | | 53 | 1034 |
| BRCA | SHC1 | first neighbor | | | 54 | 1034 |
| BRCA | FADD | CNV | 149 | 0 | 55 | 1034 |
| BRCA | FOXA1 | Mutation | | | 56 | 1034 |
| BRCA | FOS | first neighbor | | | 57 | 1034 |

| | | | | | | |
|---|---|---|---|---|---|---|
| BRCA | ATAD2 | CNV | 134 | 1 | 58 | 1034 |
| BRCA | UBE2I | first neighbor | | | 59 | 1034 |
| BRCA | CBL | first neighbor | | | 60 | 1034 |
| BRCA | RPS27A | first neighbor | | | 61 | 1034 |
| BRCA | HSPA8 | first neighbor | | | 62 | 1034 |
| BRCA | SIRT7 | first neighbor | | | 63 | 1034 |
| BRCA | UBB | first neighbor | | | 64 | 1034 |
| BRCA | E2F1 | first neighbor | | | 65 | 1034 |
| BRCA | PLCG1 | first neighbor | | | 66 | 1034 |
| BRCA | HDAC3 | first neighbor | | | 67 | 1034 |
| BRCA | ABL1 | first neighbor | | | 68 | 1034 |
| BRCA | H3C1\|H3C | first neighbor | | | 69 | 1034 |
| BRCA | CRK | first neighbor | | | 70 | 1034 |
| BRCA | SMAD2 | first neighbor | | | 71 | 1034 |
| BRCA | ASAP1 | CNV | 133 | 2 | 72 | 1034 |
| BRCA | H3-4 | first neighbor | | | 73 | 1034 |
| BRCA | HRAS | first neighbor | | | 74 | 1034 |
| BRCA | CHD4 | Mutation | | | 75 | 1034 |
| BRCA | PARP1 | first neighbor | | | 76 | 1034 |
| BRCA | SHANK2 | ['CNV', 'SV'] | 144 | 0 | 77 | 1034 |
| BRCA | CREB1 | first neighbor | | | 78 | 1034 |
| BRCA | PPARG | first neighbor | | | 79 | 1034 |
| BRCA | EGR1 | first neighbor | | | 80 | 1034 |
| BRCA | FYN | first neighbor | | | 81 | 1034 |
| BRCA | SNW1 | first neighbor | | | 82 | 1034 |
| BRCA | PML | first neighbor | | | 83 | 1034 |
| BRCA | MET | first neighbor | | | 84 | 1034 |
| BRCA | SIRT1 | first neighbor | | | 85 | 1034 |
| BRCA | BRCA2 | Mutation | | | 86 | 1034 |
| BRCA | H4-16\|H4C | first neighbor | | | 87 | 1034 |
| BRCA | ERBB4 | first neighbor | | | 88 | 1034 |
| BRCA | LMNA | first neighbor | | | 89 | 1034 |
| BRCA | UBA52 | first neighbor | | | 90 | 1034 |
| BRCA | NRAS | first neighbor | | | 91 | 1034 |
| BRCA | KAT2B | first neighbor | | | 92 | 1034 |
| BRCA | KRAS | first neighbor | | | 93 | 1034 |
| BRCA | EZH2 | first neighbor | | | 94 | 1034 |
| BRCA | FLNA | first neighbor | | | 95 | 1034 |
| BRCA | NOTCH1 | first neighbor | | | 96 | 1034 |
| BRCA | SMARCA4 | first neighbor | | | 97 | 1034 |
| BRCA | IGF1R | first neighbor | | | 98 | 1034 |
| BRCA | SMAD4 | first neighbor | | | 99 | 1034 |
| BRCA | H2AX | first neighbor | | | 100 | 1034 |
