## supplementary information for "Probabilistic graph-based model uncovers previously unseen druggable vulnerabilities in major solid cancers"

### Handling study bias in model development and optimization

To characterize study bias, we calculated the Spearman correlation between the ranking of a given gene returned by each model and the binary logarithm of the number of publications for that gene. For the Markov chain model (MCm), we also compared how the values of α, the cooperativity factor (Fig. S3A), and $W_m$, the weight parameter of the self-loop (Fig. S3B), related to study bias.

Across indications, we found that the absolute correlation between the rank of the top 20 genes and the number of publications did not show clear differences across models, but this absolute correlation is significantly reduced for the Markov chain model compared to Page Rank or diffusion models when the top 100 genes are considered for networks derived from DNA data only (Fig. S3C) and reduced from DNA and RNA-seq data (Fig. S2E).

Compared to other models, the MCm is less sensitive to study bias, since the Spearman correlation of the MCm is -0.33 ± -0.15, while that of the Page Rank model is -0.46 ± 0.09, Personalized Page Rank model is -0.39 ± 0.17, and Raw Diffusion model is -0.47 ± 0.14. This is well exemplified by the correlations shown in Fig. S3D for the application of the models to the bladder cancer (BLCA) DNA data; the ranking returned by the MCm implementation of $A_3D_3a$'s MVP displays the weakest correlation, r = -0.23, to number of publications of all models considered.

### Commentary on cell lines as model systems

The high-throughput genetic perturbation data made available through DepMap[1] comprise an invaluable resource for understanding genetic dependencies by highlighting genes required for cell growth. Given DepMap's rigorous and standardized assay conditions, dependencies reported can be reliably regarded as true.

However, we expect that additional dependencies exist beyond those identified in DepMap. As an illustration, AR, the key driver gene in prostate cancer, has an average Chronos score of -0.16 and thus, is not reported as a dependency in that disease by DepMap. This can be partly attributed to the challenge of growing some cancer cell lines, like those from prostate cancers. Since DepMap's standardized assay time (~72 hours) was optimized for "average" cell lines, dependencies in slowly growing cell lines are more difficult to capture within the timeframe of the assay.

Finally, cancer cell lines, like any model system, are not necessarily representative of the disease itself. As an example, we reviewed the prostate cancer cell lines included in DepMap and found that their origins[2] ranged from a serially propagated xenograft to a benign prostatic hyperplasia. Despite the described limitations, we rely upon public databases like DepMap as they are invaluable, large-scale experimental datasets for contextualization of our results.

Interestingly, although we explicitly developed $A_3D_3a$'s MVP *only* using patient data and without use of cell line data, it is able to crystallize key vulnerability information beyond what is obvious. As a test, we compared it a recent analysis that included cell line data to identify cell line genetic dependencies[3] (Fig. S16). $A_3D_3a$'s MVP, despite not including any cell line data as input, performed equally at identifying genetic dependencies for the same cell lines. Moreover, it was significantly better at identifying pharmacological dependencies! This further highlights the value of cooperativity to uncover true biological signal.

### List of Supplementary Tables provided as files

Results come from models generated with DNA data only unless otherwise noted for each file.

1. Cohort sizes across indications
2. Alteration type for DNA seed genes
3. Z-scored ln(IC50) values for top 60 genes across indications and models
4. GDSC validation of top 20 ranked genes from model using DNA and RNA data
5. Clinically relevant genes from oncoKB and canSAR
6. Setting initial scores of clinically relevant genes to zero
7. Top 100 ranked genes with DepMap information
8. DepMap and GDSC information for top 60 genes across indications in test set
9. Druggability, DepMap, and GDSC information for top 20 genes across indications
10. Top 20 genes across indications annotated with accessibility of cysteine, lysine, and tyrosine residues; loss information; and relevant indications
11. Copy number alterations in top 100 genes by indication
